# Genome-wide identification of microsatellite markers and their application in genetic studies of wolfberry (*Lycium barbarum*)

**DOI:** 10.1101/2022.05.25.493367

**Authors:** Yue Yin, Xiaoya Qin, Jianhua Zhao, Wei An, Yanlong Li, Yunfang Fan, Yajun Wang, Youlong Cao

**Author notes:** **Corresponding author:** National Wolfberry Engineering Research Center, Ningxia Academy of Agriculture and Forestry Sciences, Yinchuan 750021, Ningxia, China.

## Abstract

The wolfberry (*Lycium barbarum*) has been an important traditional Chinese food and medicine for hundreds of years. However, few microsatellite markers have been developed for assessing the genetic diversity and genetic linkage maps among *Lycium* species in genetic research. In the present study, a total of 397,558 microsatellite loci were identified from whole wolfberry genome sequences with an overall density of 212.1 Mb/SSRs. Dinucleotide microsatellites were the most abundant type, representing 58.26% of the total microsatellite loci, followed by tri-, penta-, hexa-, and tetranucleotide. The AT motifs were the most abundant of all nucleotide repeat motifs, accounting for 44.52%. A set of 600 primer pairs were synthesized and preliminarily screened among four wolfberry accessions. Of these, 277 primer pairs were selected as polymorphic markers. In total, 22 highly polymorphic SSR markers were selected for the characterization of genetic diversity among 37 wolfberry accessions. A total of 323 alleles were detected with an average of 14.7 alleles per marker. The polymorphic information content (PIC) values ranged from 0.612 to 0.911. The dendrogram revealed that the phylogenetic relationships among these wolfberry accessions were consistent with their genetic background. Thus, the large number of genome-wide microsatellite markers developed from the wolfberry genome provides a valuable resource for genetic diversity, genetic map construction, quantitative trait locus (QTL) mapping, and marker-assisted selection breeding in *Lycium* species.

## Introduction

The wolfberry (*Lycium barbarum*), also called the goji berry, is a deciduous shrub plant in the Solanaceae family. The adult tree is 1.5-2 m tall and its fruits are 1.5-1.9 cm long, bright orange-red ellipsoid berries (Figure 1). The fruits, leaves, and roots are rich in polysaccharides, carotenoids, flavonoids, and vitamins, which are responsible for anti-aging, immuno-enhancement, and radio-resistance effects (Hempel et al. 2017; Skenderidis et al. 2019; Gao et al. 2019). Wolfberries have been extensively cultivated in China and are consumed globally (Wetters et al. 2018). Therefore, there is an urgent demand for breeding new wolfberry cultivars with high levels of secondary metabolites. However, the traditional breeding method based only on general appearance and agronomic performance is time consuming and financially intensive. With the development of molecular breeding technology, marker-assisted selection (MAS) is an effective tool for precision plant breeding in crops (Ma et al. 2020). DNA-based markers have enormous potential for improving the efficiency and precision of conventional plant breeding via MAS breeding.

**Figure 1.**
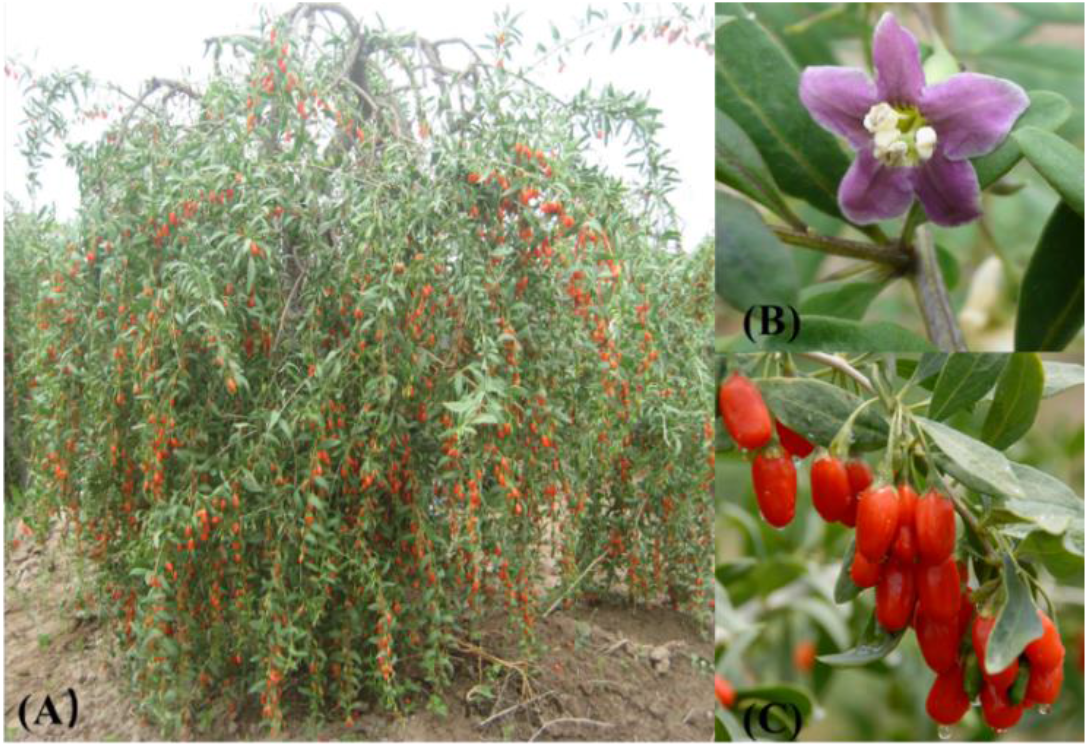
Adult tree (A), flowers (B), and fruits (C) of *Lycium* barbarum ‘Ningqi 1’

Microsatellites, or simple sequence repeats (SSRs), are motifs comprising one to six nucleotides that are found in both the coding and non-coding regions of all prokaryotic and eukaryotic genomes (Zane et al. 2002). Due to their co-dominant inheritance, high reproducibility, and abundance in the genome (Kalia et al. 2011), SSRs serve as useful and powerful markers for genetic studies, including studies of genetic diversity and population structure as well as in genetic map construction, comparative genomics, and molecular marker-assistant breeding for various plants species (Hipparagi et al. 2017; Ouni et al. 2020; GU et al. 2018; Han et al. 2019; Tan et al. 2018).

With the rapid development of high-throughput sequencing technology, an increasing number of plant genomes have been sequenced and assembled using PacBio, Hi-C, and Bio Nano Technology (Kersey 2019; Chen et al. 2019). These whole-genome DNA sequences are valuable resources for SSR development. Genome-wide identification of microsatellite markers have been investigated in many plant species, such as the Chinese jujube, watermelon, tea, eggplant, tobacco, pear, and peanut (Xiao et al. 2015; Zhu et al. 2016; Liu et al. 2018; Portis et al. 2018; Wang et al. 2018; Xue et al. 2018; Lu et al. 2019). In the recent past, there have been only a few reports for the wolfberry plant detailing the development of microsatellite markers using motif-enriched libraries (Kwon et al. 2009; Chen and Zhong 2014) and transcriptome sequences (Chen et al. 2017a; Chen et al. 2017b). Thus, the number of publicly available SSRs for wolfberry is still insufficient for genetic studies, such as high-density linkage map construction, quantitative trait locus mapping analyses, and evolutionary studies.

In the present study, we obtained whole genome sequences from the National Wolfberry Engineering Research Center using Illumina and PacBio sequencing platforms (manuscript under preparation). We performed genome-wide detection of microsatellite sequences and analyzed the frequency distribution of microsatellite motifs in the genome. The SSR markers developed from *Lycium* barbarum were used for genetic studies. This is the first report of large-scale generation of genomic SSR markers in the wolfberry plant. The newly developed SSR markers in L. barbarum provide a valuable dataset for further studies in QTL mapping, genetic resource conservation, and evolutional analysis.

## Materials and Methods

### Plant materials and DNA extraction

A total of 37 wolfberry accessions, planted in the Wolfberry Germplasm Resources Garden in Yinchuan (38°38’49”N, 106°9’10”E), National Wolfberry Engineering Research Center, Ningxia Academy of Agriculture and Forestry Sciences, were used in this study (Table 1).

**Table 1.** The 37 wolfberry accessions used in this study.

| Code | Name | Species | Type | Fruit color |
| --- | --- | --- | --- | --- |
| L1 | Damaye | *L. barbarum* | Landrace | Red |
| L2 | Xiaomaye | *L. barbarum* | Landrace | Red |
| C3 | Ningqi 2 | *L. barbarum* | Cultivar | Red |
| C4 | Ningqi 3 | *L. barbarum* | Cultivar | Red |
| C5 | Ningqi 4 | *L. barbarum* | Cultivar | Red |
| C6 | Ningqi 5 | *L. barbarum* | Cultivar | Red |
| C7 | Ningqi 6 | *L. barbarum* | Cultivar | Red |
| C8 | Ningqi 7 | *L. barbarum* | Cultivar | Red |
| C9 | Ningnongqi 1 | *L. barbarum* | Cultivar | Red |
| C10 | Ningnongqi 2 | *L. barbarum* | Cultivar | Red |
| C11 | Ningnongqi 3 | *L. barbarum* | Cultivar | Red |
| C12 | Ningnongqi 9 | *L. barbarum* | Cultivar | Red |
| C13 | Mengqi 1 | *L. barbarum* | Cultivar | Red |
| C14 | Tianjing 3 | Unknown | Cultivar | Red |
| C15 | Tianjing 5 | Unknown | Cultivar | Red |
| H16 | 9001 | Hybrid | New line | Red |
| W17 | BG-01 | *L. ruthenicum* | Wild | White |
| W18 | HGB | Unknown | Wild | Yellow |
| W19 | W-12-30 | Unknown | Wild | Yellow |
| W20 | W-12-27 | Unknown | Wild | Yellow |
| W21 | W-11-15 | Unknown | Wild | Yellow |
| W22 | W-13-30 | Unknown | Wild | Yellow |
| W23 | W-13-26 | Unknown | Wild | Yellow |
| W24 | W-12-26 | Unknown | Wild | Yellow |
| W25 | NXHG | *L. barbarum* var. *auranticarpum* | Wild | Yellow |
| W26 | DHG | *L. barbarum* | Wild | Yellow |
| W27 | BF-01 | *L. chinense* var. *potaninii* | Wild | Red |
| W28 | YN-01 | *L. yunnanense* | Wild | Red |
| W29 | LM-01 | *L. ruthenicum* | Wild | Black |
| W30 | YN-14 | Unknown | Wild | Red |
| W31 | YN | Unknown | Wild | Red |
| W32 | QH-13-02 | Unknown | Wild | Black |
| W33 | Mansheng | Unknown | Wild | Red |
| W34 | QH-13-05 | Unknown | Wild | Purple |
| W35 | HB-09 | Unknown | Wild | Red |
| W36 | SC-09 | Unknown | Wild | Red |
| W37 | Changji | Unknown | Wild | Purple |

Total genomic DNA was extracted using a Plant Genomic DNA Kit (Tiangen, Beijing, China). DNA concentrations and quality were detected using a Nanodrop spectrophotometer (Nanodrop products, Wilmington, DE, USA) and 1% agarose gel electrophoresis. The final concentrations were adjusted by 10 ng/ μL and stored at ⌝-20 °C until use.

### SSR identification and primer design

Wolfberry genomic sequencing was performed by our lab (unpublished data). The MISA (MIcroSAtellite identification) (http://pgrc.ipk-gatersleben.de/misa/) tool was used to identify microsatellite loci in the sequence database. Sequences were considered as potential SSRs only if their repeat unit was based on 2–6 bp. The minimum SSR length criteria were defined as six, five, four, three, and three reiterations for di-, tri-, tetra-, penta-, and hexanucleotide, respectively. Subsequently, the SSR primers were designed using BatchPrimer3 interface modules (You et al. 2008). Primers were designed under the following parameters: product size from 100 to 500 bp, primer size from 18 to 23 bp with an optional size of 21 bp, annealing temperatures from 50 to 60 °C (optimally 55 °C), GC content from 40% to 60% (optimally 50%). All primers were synthesized by Shanghai Generay Biotech Co., Ltd (Shanghai, China).

### Functional annotation of genome-wide SSR markers

Go annotation was performed by searching the flanking region against the GenBank database using Blast2Go software (Conesa and Gotz, 2008). The GO categorization results were expressed as three independent hierarchies for biological process, cellular component and molecular function.

### SSR markers polymorphism

A total of 600 SSR markers were designed and synthesized, and four wolfberry accessions were selected for polymorphism screening. Polymerase chain reaction (PCR) reactions were conducted in 15 μL volumes containing 5 ng/μL DNA, 1× PCR mix, 0.25 μM M13 primer, 0.06 μM forward primer, 0.25 μM reverse primer, and 2.6 μL ddH_2_O. The M13 universal primer was labeled with 6-FAM (blue) and HEX (Green) fluorescent tags (Generay, Shanghai, China). PCR amplifications were performed using a T100 Thermal Cycler (Bio-Rad, USA) under the following conditions: denaturation at 95 °C for 5 min; followed by 25 cycles of 95 °C for 30 s, 55 °C for 30 s, and 72 °C for 30 s; 15 cycles of 95 °C for 30 s, 53 °C for 30 s, 72 °C for 30 s, and a final step at 72 °C for 5 min. PCR products were separated by capillary electrophoresis with an ABI37370 DNA analyzer (Applied Biosystems).

### Data analysis

DNA fragments were obtained and analyzed by Genemapper^®^ Software v4.0 (Applied Biosystems, Foster City, CA, USA). The number of alleles (Na), expected heterozygosity (He), observed heterozygosity (Ho), and polymorphic information content (PIC) were calculated via the Power Marker v3.25 (Liu and Muse 2005). The polygenetic tree was constructed based on Nei’s coefficient distances using the unweighted pair group method with arithmetic (UPGMA) method and viewed using MEGA 6.06 (Tamura et al. 2013). The confidence level of the branch support was evaluated by a bootstrap analysis with 1000 replicates. Population structure was analyzed using the mixed model within STRUCTURE 2.3.4 (Pritchard et al. 2000). The K value, which indicates population number in the admixture model, ranged from 1 to 10, with 10 iterations and the burn-in MCMC set at 10,000 and 100,000, respectively. The STRUCTURE HARVESTER software tool was used to determine the optimum number of the K (Earl 2012).

## Results

### Frequency, distribution, and characterization of microsatellite in the wolfberry genome

In the present study, 1,873,983,257 bp available wolfberry genome sequences were searched for SSR loci. A total of 397,558 SSRs comprising di-to hexanucleotides were identified with an average frequency of ~212.1 SSRs/Mb (Table S1). Dinucleotide was the most abundant type, with a proportion of 58.26%, followed by tri- (16.48%), penta- (15.07%), and hexanucleotide (6.77%). Tetranucleotide was the lowest repeat type, accounting for 3.42% (Figure 2A). Based on the length of the repeat motif, a total of 132,943 (33.44%) microsatellites were grouped into the long and hyper variable class I (⩾20 bp) type, while the remaining 264,615 (66.56%) microsatellites were grouped into variable class II (12-19 bp) type (Figure 2B).

**Figure 2.**
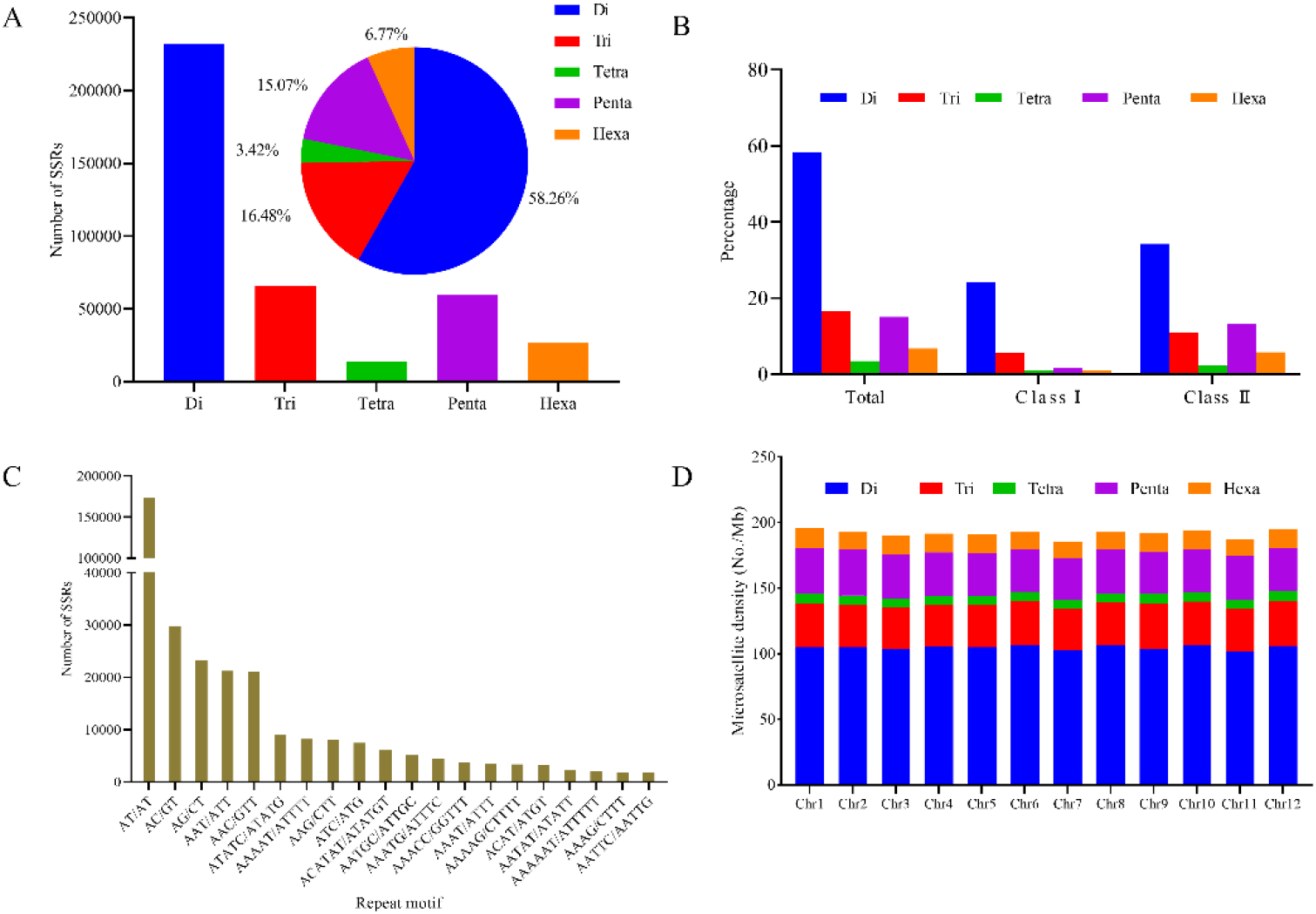
Frequency and distribution of microsatellites identified in the wolfberry genome. (A) Numbers of microsatellites with different motif types. (B) Percentage of long and hypervariable class I (≥20bp) and variable class II (12-19 bp) microsatellites in the genome. (C) The top 20 most frequently occurring SSR motifs in the wolfberry genome. (D) The distribution of SSR repeats on different chromosomes.

A total of 463 types of SSR motifs were identified in the wolfberry genome. The repeat number of all these motifs varied from 2 to 1675. The majority of these motifs belonged within the top 20 most frequently occurring motifs. The percentages of penta- and hexanucleotides with three repeats were much higher than other types. For all five SSR motif types, the occurrence decreased with an increase in motif length (Figure S1). Among the SSR motif types, the AT/AT motif of dinucleotide was the most common motif in the entire wolfberry genome, accounting for 44.52%, followed by AC/GT and AG/CT, which accounted for 7.63% and 5.98%, respectively (Figure 2C). For each type of SSR motif, from di-to hexanucleotide, the richest motifs were AT (76.41%), ATT (33.28%), ATTT (29.22%), ATTTT (14.68%) and GTTTTT (4.78%), respectively.

Furthermore, we analyzed SSR distribution on each chromosome, which showed that the largest number of SSRs were present on chromosome 4 (28,360) and chromosome 2 (28,122) (Table S2), while chromosome 12 (20,315) contained the lowest number of SSRs. A positive correlation was observed between chromosome length and the number of SSRs for each chromosome (R2 = 0.9785, p < 0.01) (Figure S2). In addition, the average SSR density was 191.7 SSRs/Mb, ranging from 185.7 SSRs/Mb on chromosome 7 to 195.4 SSRs/Mb on chromosome 1 (Figure 2D, Table S2).

### Functional annotation

We analyzed the microsatellite distribution in both coding and non-coding regions of the genome. Microsatellites were mainly located in non-coding regions (313,038, 99.28%), whereas approximately 0.72% (2266) of the microsatellites were located in coding regions (Figure 3A). For the microsatellites in coding regions, the functional annotation of SSR loci was performed via Blast2Go analysis. A total of 401 genes were assigned to a molecular functional category (Figure 3B). Binding (54.1%) was the most dominant group followed by catalytic activity (31.7%) and transcription regulator activity (8.0%). Cellular process (23.3%) was the most enriched group based on the annotations of biological process category (Figure 3C). With regard to the cellular component, 23.9% of the sequences were assigned to the cell, followed by organelle (19.1%), membrane (8.9%), and organelle (8.2%) (Figure 3D).

**Figure 3.**
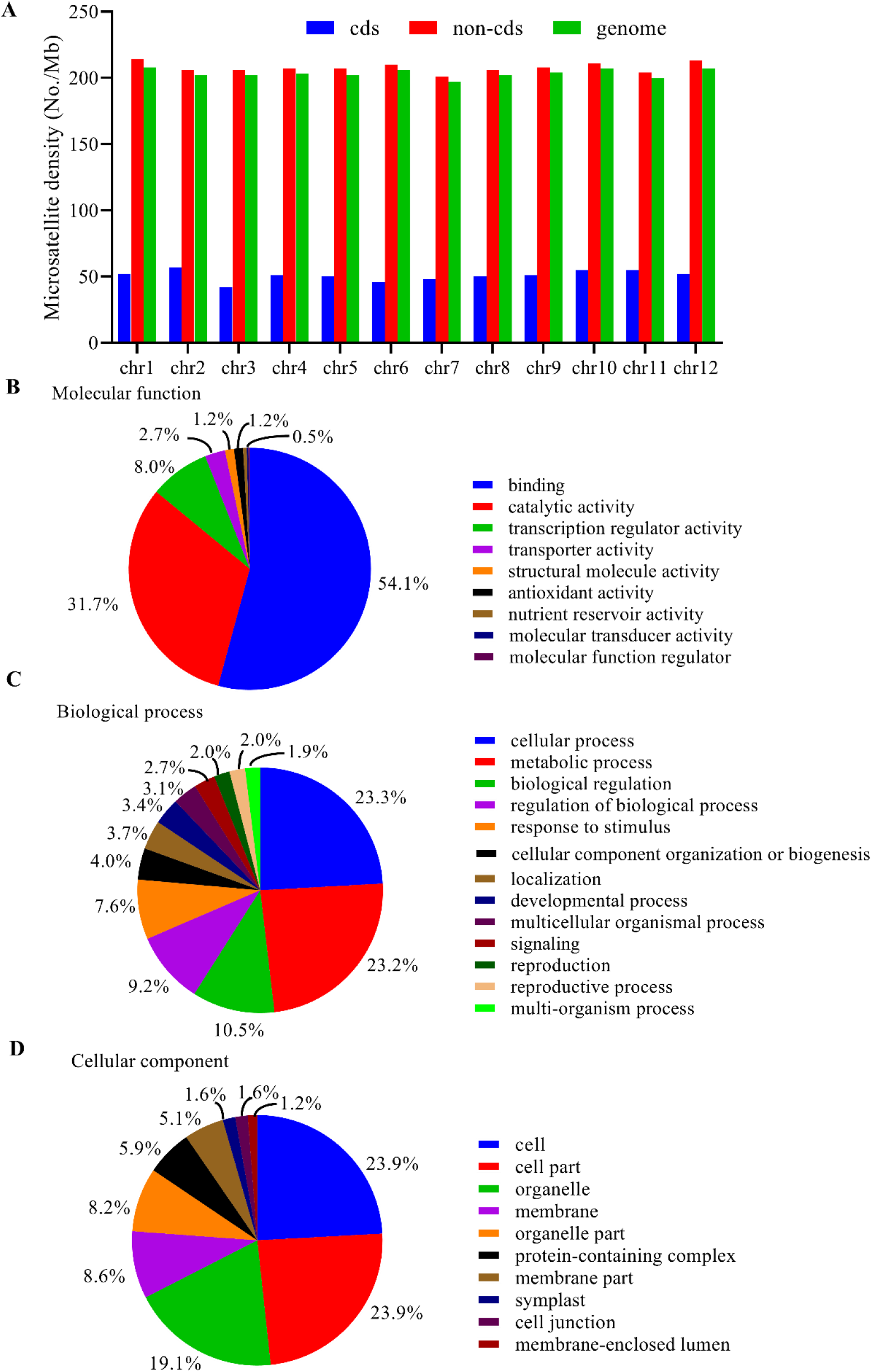
Distribution of microsatellites in coding regions (cds) in the wolfberry genome. (A) Microsatellite density in cds and non-cds on each chromosome. Breakdown of categories (B) molecular function, (C) biological process, and (D) cellular component.

### SSR marker development and polymorphism

The flanking sequences of 397,558 SSRs were used to design suitable forward and reverse primer pairs. A total of 42,141 SSR markers were successfully developed on 12 chromosomes, which accounted for 10.6% of all identified SSRs. Among these markers, 600 SSR markers (Table S3) were randomly selected to screen polymorphism on four genotypes with significant phenotypic variation. Of them, 277 (46.17%) were polymorphic among four accessions, while most of them (61.01%) were trinucleotide, followed by di- (34.66%) and penta- (3.61%). To identify and validate an appropriate set of SSR markers for characterizing L. barbarum germplasm collections, 22 highly informative SSR markers were selected for polymorphism analysis. The 22 selected SSR markers could be detected in 323 alleles, representing an average of 14.7 alleles per marker. The observed heterozygosity ranged from 0.162 to 1.000 and expected heterozygosity from 0.638 to 0.917. The PIC value for each locus varied from 0.612 to 0.911 with a mean value of 0.832 (Table 2).

**Table 2.**
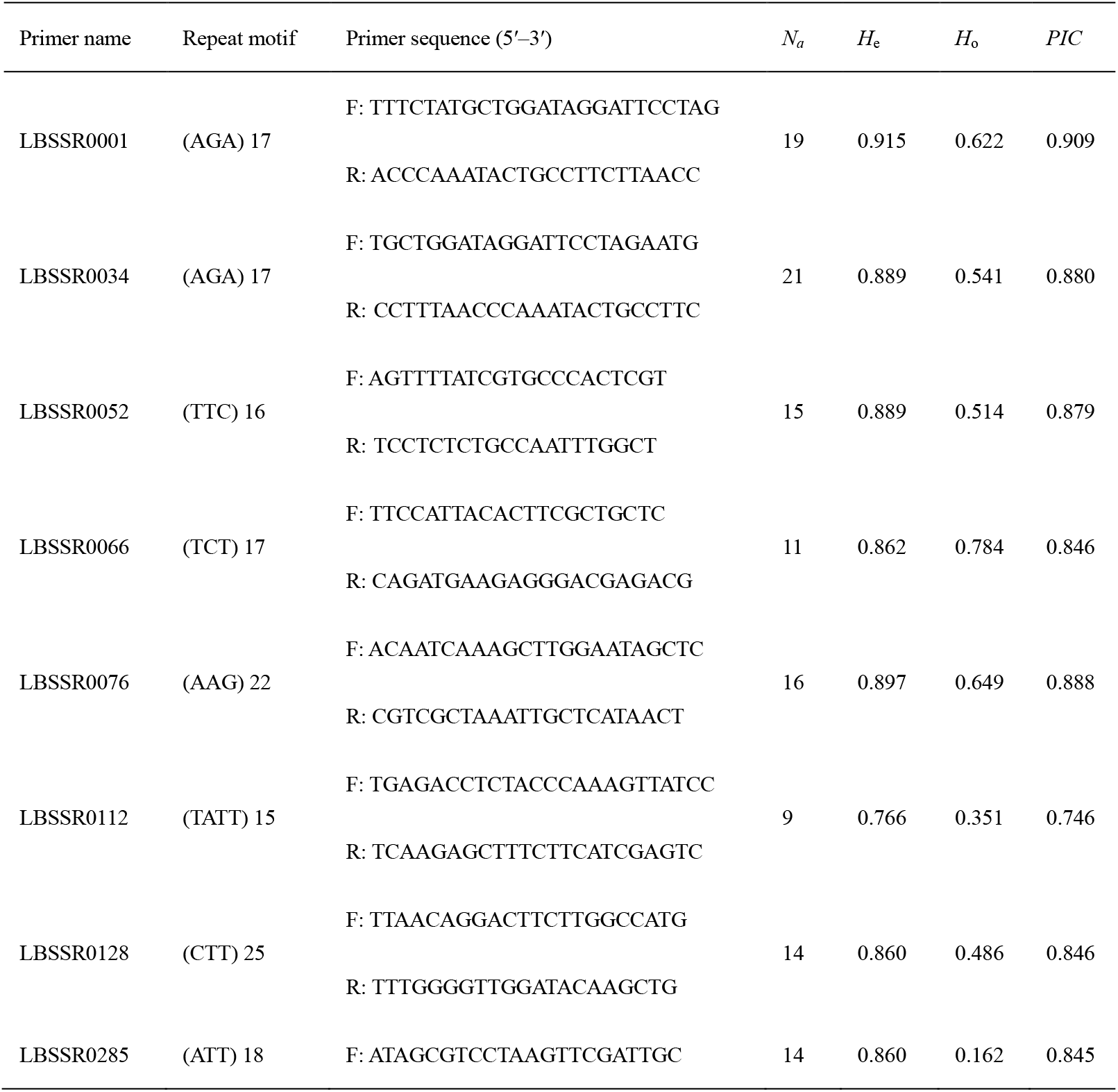
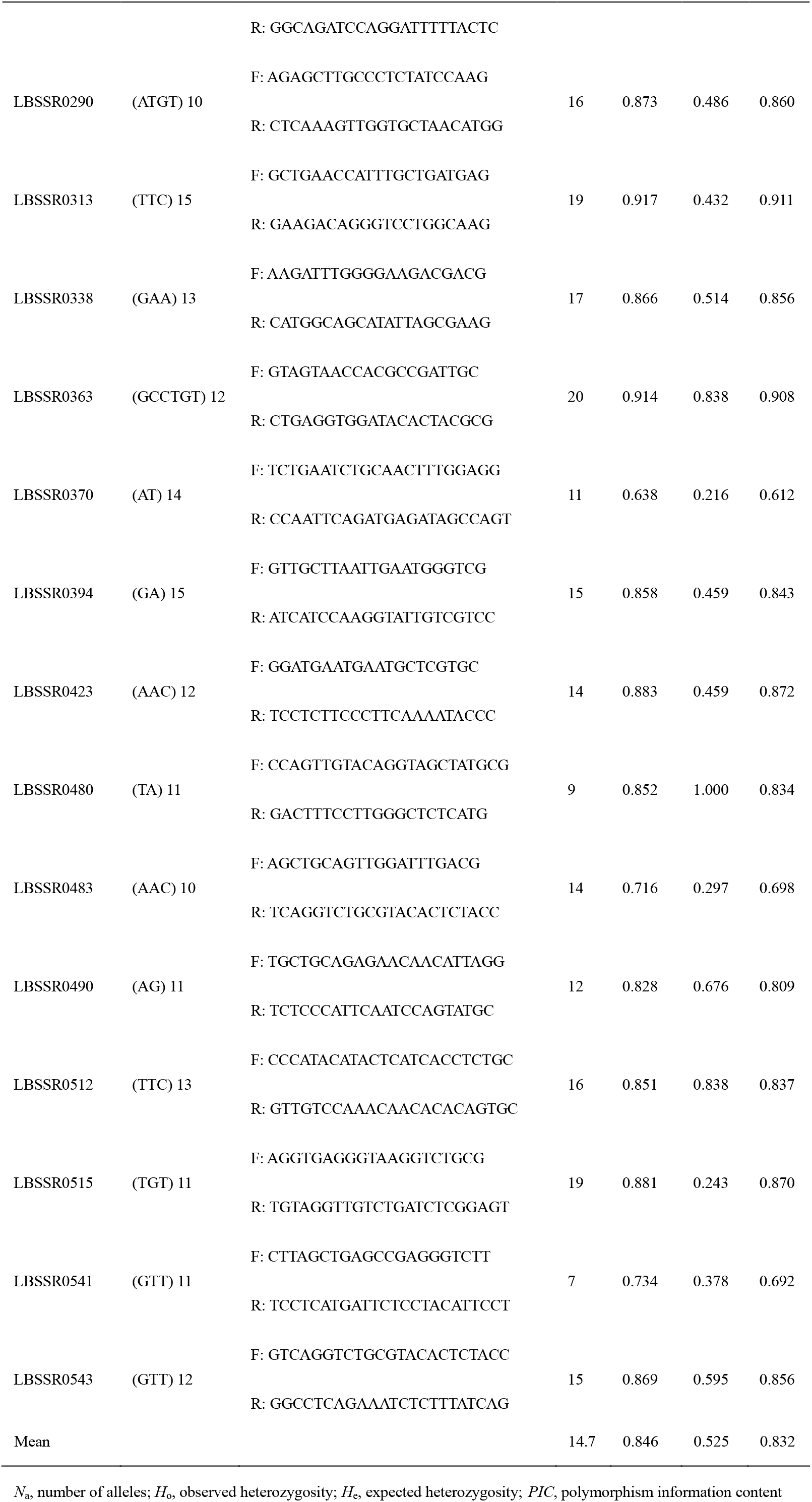
The 22 polymorphic SSR markers selected for validation among 37 wolfberry accessions.

### Genetic diversity and population structure analysis

We analyzed the genetic structure of 37 individuals based on the 22 highly informative SSR markers using STRUCTURE software. According to Evanno (Evanno et al., 2005), the maximum K represents the optional number of clusters and K = 2 was the largest value (Figure 4). A total of 37 wolfberry accessions clustered into two groups (Figure 5). The first group contained 22 wolfberry accessions (red block), 20 of which belonged to the L. barbarum species (cultivar type). The other two accessions were wild type. The second group included 15 accessions, mainly from wild type accessions except for two cultivars: C14 (Tianjing 3) and C15 (Tianjing 5). Phylogenetic trees were constructed for the 37 wolfberry accessions using the UPGMA method (Figure 6). These wolfberry samples also clustered into two groups. One corresponded to the cultivar type and the other to wild type accessions. The results correspond with the conclusions of STRUCTURE analysis.

**Figure 4.**
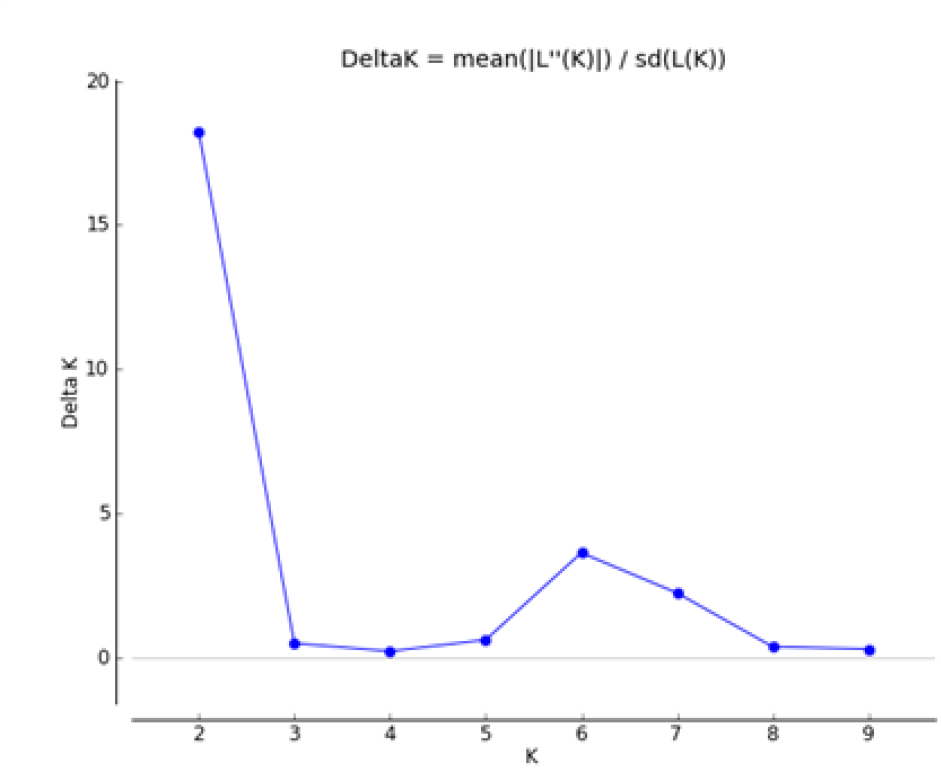
Estimated ΔK values for a given K in the structure analysis.

**Figure 5.**
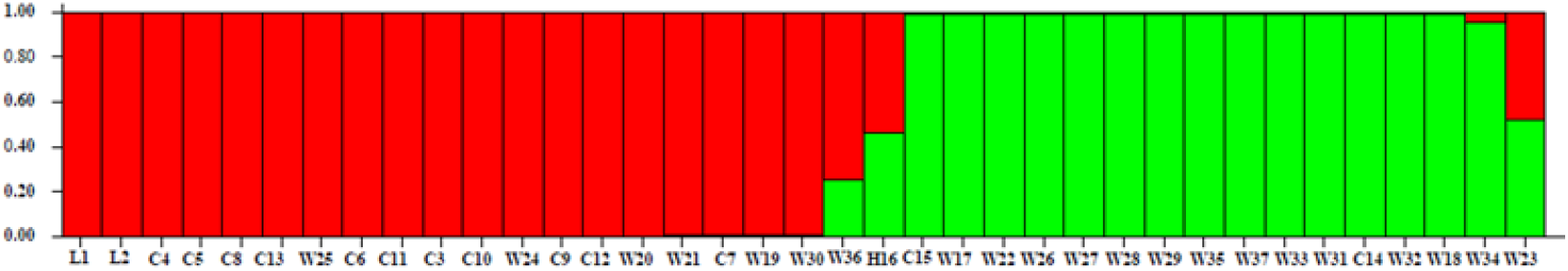
Population structure of 37 wolfberry accessions. Clusters are indicated by different colors. Samples code are the same as given in Table1.

**Figure 6.**
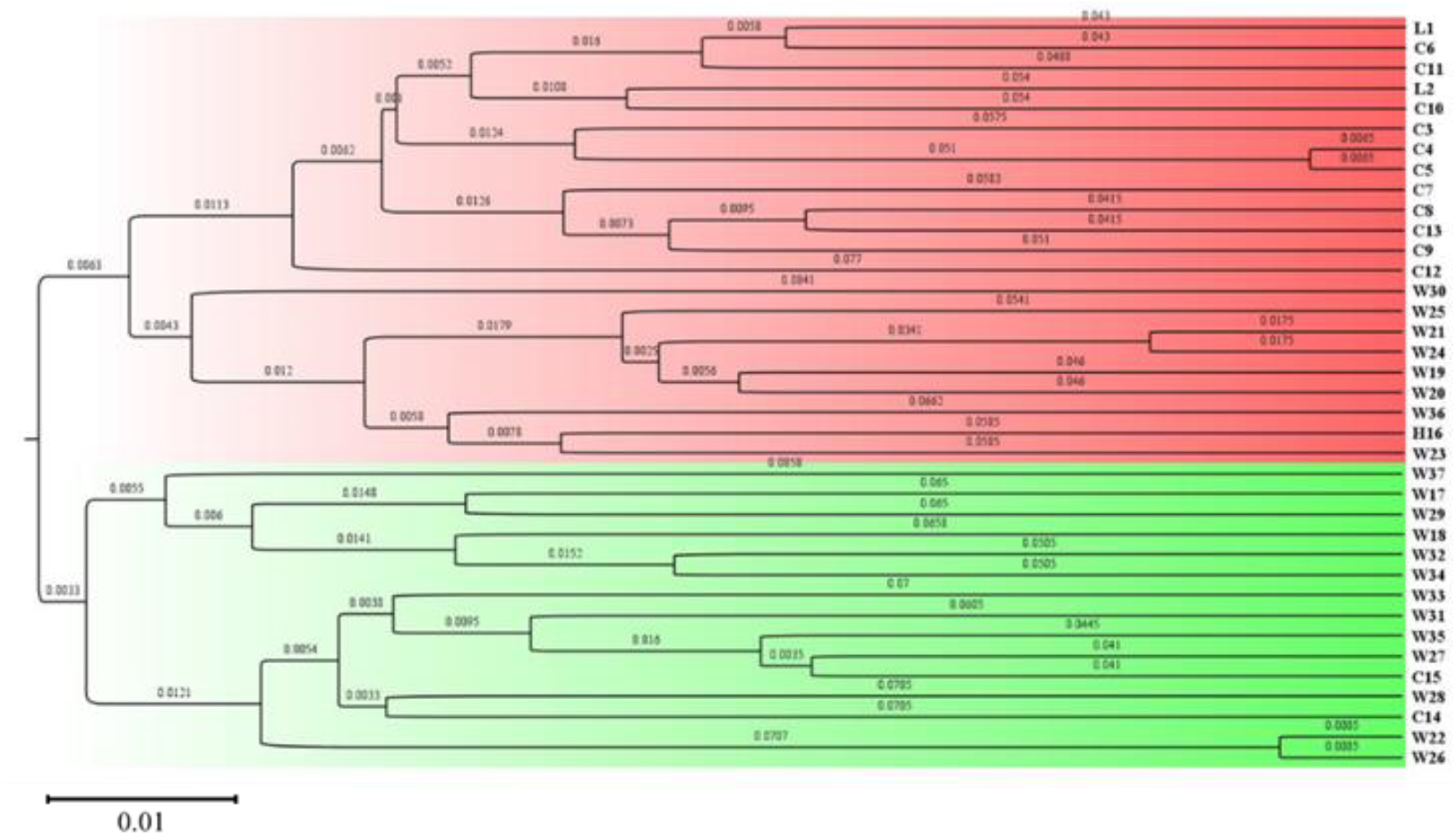
The UPGMA phylogenetic trees of 37 accessions based on 22 SSR markers. Sample codes are the same as given in Table 1.

### Suitability of SSR Markers for Genetic Mapping

SSR markers are effective tools for genetic map construction. To determine the suitability of the newly developed SSR markers, an F1 population derived from W25 (male) × W27 (female) was constructed and used for validation. As a cross population (CP), four different alleles may be segregated, and five segregation types codes were detected. A total of 236 SSR markers were used to performed segregation types. Of these, 106 SSR markers were suitable for genetic map construction of this population, which consisted of 59, 19, 15, 9, and 4 SSR markers belonging to nn × np, lm × ll, ab × cd, ef × eg, and hk × hk segregation types, respectively (Figure 7).

**Figure 7.**
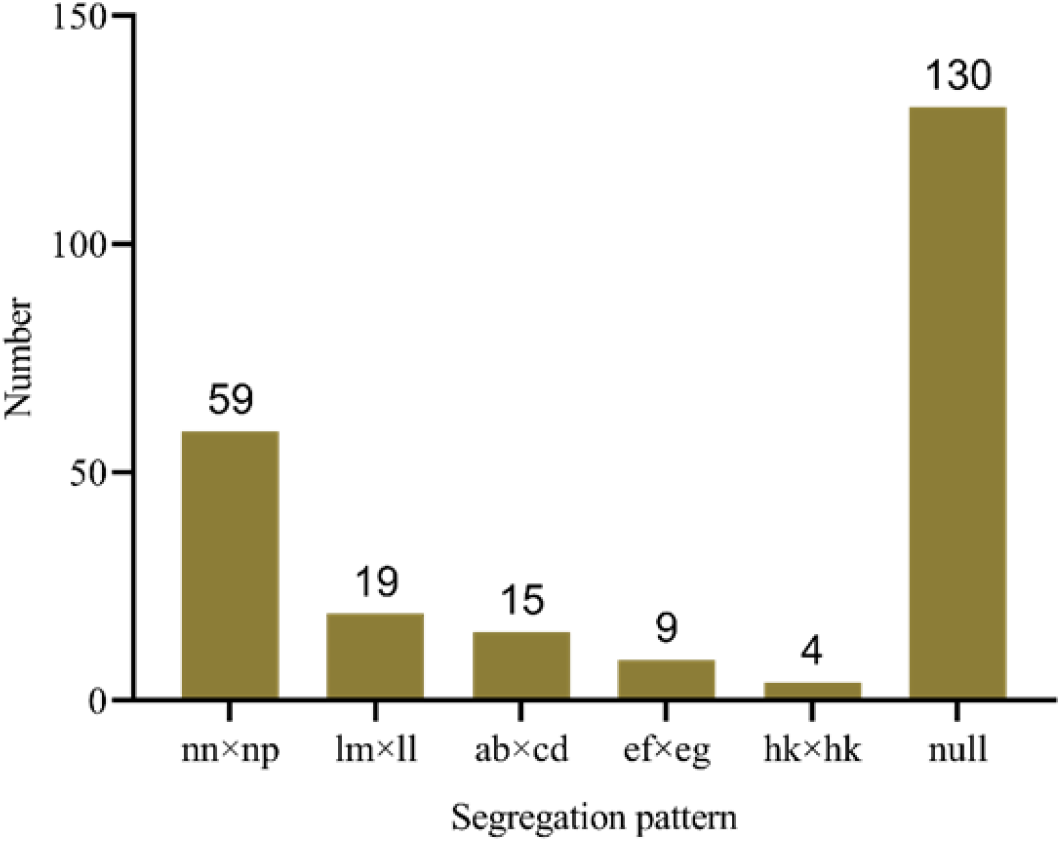
Number of SSR markers in each of the five segregation patterns.

## Discussion

SSR molecular markers are highly polymorphic, co-dominant, reproducible, and easy to use. They are widely used for genetic diversity of wolfberry germplasm resources (Kwon et al., 2009; Zhao et al., 2010). However, the wolfberry genome has not yet been fully deciphered, and the development of SSR markers is only at the level of ESTs. To the best of our knowledge, this is the first study to describe use of the genomic data of cultivated wolfberry species for the analysis of relevant SSR information such as their genome distribution frequency, motif types, and sequence characteristics, providing important data support for further wolfberry genome genetic research and molecular marker development.

### Frequency, distribution, and characterization of microsatellite in the wolfberry genome

Whole-genome sequences have provided useful resources for microsatellite identification and large-scale development of microsatellite markers in various kinds of plant species, such as jujube, watermelon, tea, eggplant, tobacco, pear, and peanut (Liu et al., 2018; Lu et al., 2019; Portis et al., 2018; Wang et al., 2018; Xiao et al., 2015; Xue et al., 2018; Zhu et al., 2016). In the current study, we obtained the high-quality assembly whole genome sequences for *Lycium* barbarum ‘Ningqi 1’ based on high-throughput sequencing data. The genome size of wolfberry was estimated to be 1.8 Gb. As a result, a total of 397,558 microsatellite loci were identified from 783,061 sequences with a density of 212.1 SSRs/Mb. The number of microsatellites and their density as identified in our study was higher than for watermelon (111 SSRs/Mb) and eggplant (125.5 SSRs/Mb), while lower than jujube (387 SSRs/Mb) and tea (217 SSRs/Mb). Previous studies have reported a positive correlation between chromosome length and SSR density in plants (Lu et al., 2019; Portis et al., 2018). Our findings were in accordance with this general trend (Figure S2). However, some studies have found no correlation between chromosome size and SSR density (Biswas et al., 2014; Zhu et al., 2016). These differences could be due to the variation in search criteria, genome size, and bioinformatics software tools used for microsatellite identification (Biswas et al., 2014; Wang et al., 2018; Xue et al., 2018; Zhu et al., 2016).

We found that dinucleotides and trinucleotides were the most abundant type of SSR repeat motif, which together accounted for 74.74% of all SSRs. Pentanucleotides, tetranucleotides, and hexanucleotides were less abundant, together accounting for 25.26% (Figure 2A) in the wolfberry genome. Similar results have been reported for several other plant species (Cheng et al., 2016; Kumari et al., 2019; Liu et al., 2018; Pandey et al., 2013). Previous studies have reported that tetranucleotide, pentanucleotide, and hexanucleotide repeats are the most common repeat in the cucumber (Cavagnaro et al., 2010), peanut (Lu et al., 2019), and Gossypium (Wang et al., 2015) genomes, respectively. Toth et al. (Tóth et al., 2000) reported that having a majority of repeat motifs that are short indicates a relatively high level of evolution. Harr et al. (Harr and Schlötterer, 2000) reported that the majority of long repeat motifs have a relatively low mutation frequency or relatively short evolution time. Our results therefore further indicate that wolfberry is at a relatively high level in terms of evolution and has a similar nucleotide motif composition as most dicot plant species.

In a given SSR type, there may be significant differences in the frequency of specific repeat bases. In the wolfberry genome, the base composition of SSR motifs was strongly biased toward A and T. For example, AT/AT, AC/GT, and AG/CT motifs were the most abundant, while CG/GC motifs were very rare in dinucleotide repeats. Similarly, AAT/ATT and ATATC/ATATG motifs were the most abundant in trinucleotide and tetranucleotide repeats, respectively, but the percentages of motifs with lower content of GC repeats were extremely low. Similar results have been reported in a number of plant species (Biswas et al., 2014; Cavagnaro et al., 2010; Han et al., 2015; Xiao et al., 2015). It is noteworthy that the nature of the most abundant dinucleotide repeat sequences of differs among plant species. For example, AG/CT is most abundant in tea (Liu et al., 2018), foxtail millet (Pandey et al., 2013), and Chinese spring wheat (Han et al., 2015), whereas AT/TA is most abundant in watermelon (Zhu et al., 2016), black pepper (Kumari et al., 2019), and cucumber (Cavagnaro et al., 2010).

### SSR marker development and polymorphism analysis

With the rapid development of next-generation whole-genome sequencing technologies, it is becoming increasingly the case for more species that large numbers of SSR markers are being characterized that cover their entire genome. For *Lycium* species, only a few genomic SSR markers have been developed and are based on BAC libraries (Chen and Zhong, 2014; Kwon et al., 2009). In the present study, we first developed SSR markers using the whole genome sequence for wolfberry. Only di- to hexanucleotides repeats were only considered for SSR marker development. In this size range, 397,558 SSRs were identified in the wolfberry genome, but only 42,141 (10.6%) loci were suitable for SSR marker development. The efficiency of SSR marker suitable for development in wolfberry was higher than for peanut (11.69%) (Lu et al., 2019) but lower than sweet orange (62.76%) (Biswas et al., 2014).

Among these new development markers, 600 SSR markers were randomly selected to screen for polymorphism using four accessions with significant phenotypic differences. Through marker screening, the 277 SSR markers indicated high polymorphism. The ratio of polymorphism was 46.17%. The PCR product size was as expected. In addition, polymorphic information content (PIC) was considered an important index for measuring polymorphism (Hipparagi et al., 2017; Luo et al., 2018). According to Botstein (Botstein et al., 1980), PIC values greater than 0.5 are indicative of highly polymorphic loci, while a PIC value between 0.25 and 0.5 indicates moderate polymorphism, and PIC values lower than 0.25 indicate low polymorphism. In the present study, PIC values ranged from 0.612 to 0.911 with an average of 0.832, meaning that the loci display high polymorphism. Remarkably, the PIC value for most SSR markers were greater than those reported in previous studies for both EST-SSR markers (Chen et al., 2017b) and genomic SSR markers (Kwon et al., 2009). This is probably because the genome sequence contains introns and exons, while ESTs only correspond to exons. Another alternative explanation is that different species and different numbers of plant materials were used in previous studies. Overall, the PIC values calculated for all SSR markers were greater than 0.5, indicating that these newly developed SSR markers will be very useful for programming in the genetic diversity analysis and genetic linkage construction of the *Lycium* species.

### Application in genetic diversity and genetic map construction

Microsatellites are an ideal marker for genetic diversity analysis, mapping construction, and other genetic research because of their abundance, reproducibility, and high polymorphism. As far as we are aware, there are few studies that have reported diversity analysis using SSR markers. Wei-Guo Zhao et al. (Zhao et al., 2010) reported the development of SSR markers for the study of genetic variation in L. chinense accessions. In the present study, 22 highly polymorphic SSR markers were used for genetic diversity and population structure of 37 wolfberry accessions. The value for genetic diversity (heterozygosity) of wolfberry was determined as 0.846 in the present study. The clustering results revealed that 37 wolfberry accessions fell into two groups that were consistent with the genetic background and source of the material. Group I was represented by the cultivar type (*L. barbarum*) while the other group was mainly wild type. Similar results have been observed in previous studies (Chen et al., 2017a; Zhao et al., 2010). The genetic diversity results further illustrate that test materials had a close genetic relationship because the existing cultivars of *Lycium* barbarum were mainly obtained by group selection.

We previously obtained an F1 population derived from interspecific hybridization of two specific cultivars of two *Lycium* species, namely *L. barbarum* var. auranticarpum (W25) and L. chinense var. potaninii (W27). We selected four F1 hybrids together with two parents to initially assess the applicability of the 236 SSR markers for linkage map construction. The results showed that a total of 166 SSR markers corresponding to well-amplified fragments that reflect the polymorphism in length and heterozygosity between the two parents. Moreover, segregated genotypes were observed among the different F1 individuals. According to the results from 236 SSR markers, the percentage of polymorphism between male and female parents among the SSR loci was 70.3%. Finally, genotype data for 106 SSR markers confirmed cross pollinators (CP) population and were applicable to genetic map construction for this F1 population. Thus, having a greater number of SSRs from the whole-genome sequence of wolfberry will be useful in advancing wolfberry plant genetic breeding programs.

## Acknowledgments

This study was supported by the Natural Science Foundation of Ningxia (2020AAC03284), the Whole Industry Chain Innovation Demonstration project of Ningxia Academy of Agriculture and Forestry Sciences (YES-16-0405), the Third Batch of Ningxia Youth Talents Supporting Program (TJGC2018022), the Special Foundation for Agricultural Breeding of the Ningxia Hui Autonomous Region (2013NYYZ0101), and the All-China Federation of Trade Unions Staff Innovation Grant Program (2018300002). The wolfberry genome sequence was provided by National Wolfberry Engineering Research Center, Ningxia Academy of Agriculture and Forestry Sciences.

