## Supplementary material for "Genome-wide identification of microsatellite markers and their application in genetic studies of wolfberry (*Lycium barbarum*)": Supplementary Figure S1.docx

Percentage of total SSRs

| Repeat number | Hexa | Penta | Tetra | Tri | Di | Total |
| --- | --- | --- | --- | --- | --- | --- |
| 3 | 5.78 | 13.38 | 0 | 0 | 0 | 19.16 |
| 4 | 0.75 | 1.27 | 2.37 | 0 | 0 | 4.39 |
| 5 | 0.15 | 0.26 | 0.6 | 7.5 | 0 | 8.51 |
| 6 | 0.05 | 0.08 | 0.22 | 3.28 | 14.96 | 18.59 |
| 7 | 0.02 | 0.03 | 0.11 | 1.43 | 8.9 | 10.49 |
| 8 | 0.01 | 0.01 | 0.05 | 0.82 | 6.04 | 6.93 |
| 9 | 0.01 | 0.01 | 0.03 | 0.55 | 4.36 | 4.96 |
| 10 | 0 | 0.01 | 0.02 | 0.42 | 3.38 | 3.83 |
| 11 | 0 | 0 | 0.01 | 0.35 | 2.72 | 3.08 |
| 12 | 0 | 0 | 0.01 | 0.28 | 2.12 | 2.41 |
| 13 | 0 | 0 | 0 | 0.24 | 1.84 | 2.08 |
| 14 | 0 | 0 | 0 | 0.24 | 1.59 | 1.83 |
| 15 | 0 | 0 | 0 | 0.2 | 1.45 | 1.65 |
| 16 | 0 | 0 | 0 | 0.16 | 1.25 | 1.41 |
| 17 | 0 | 0 | 0 | 0.15 | 1.05 | 1.2 |
| 18 | 0 | 0 | 0 | 0.14 | 0.87 | 1.01 |
| 19 | 0 | 0 | 0 | 0.12 | 0.75 | 0.87 |
| 20 | 0 | 0 | 0 | 0.11 | 0.65 | 0.76 |
| 21 | 0 | 0 | 0 | 0.09 | 0.57 | 0.66 |
| 22 | 0 | 0 | 0 | 0.08 | 0.51 | 0.59 |
| 23 | 0 | 0 | 0 | 0.07 | 0.49 | 0.56 |
| 24 | 0 | 0 | 0 | 0.04 | 0.43 | 0.47 |
| 25 | 0 | 0 | 0 | 0.03 | 0.39 | 0.42 |
| 26 | 0 | 0 | 0 | 0.03 | 0.38 | 0.41 |
| 27 | 0 | 0 | 0 | 0.02 | 0.36 | 0.38 |
| 28 | 0 | 0 | 0 | 0.02 | 0.34 | 0.36 |
| 29 | 0 | 0 | 0 | 0.01 | 0.33 | 0.34 |
| 30 | 0 | 0 | 0 | 0.01 | 0.31 | 0.32 |
| ＞30 | 0 | 0 | 0 | 0.07 | 2.24 | 2.31 |

**Figure S1** Percentage distribution of SSRs with different motif types and repeat numbers. The vertical axis showed the abundance of SSRs with different repeat numbers. Motif types were indicated in different colors.
