## Supplementary material for "Genome-wide identification of microsatellite markers and their application in genetic studies of wolfberry (*Lycium barbarum*)": Supplementary Table S4.pdf

Table S4 Details of 42141 microsatellites markers mapped on 12 chromosommes of goji berry

| Marker ID | SSR type | SSR motif | FORWARD PRIMER (5'-3') | REVERSE PRIMER (5'-3') | size (bp) |
|---|---|---|---|---|---|
| chr1:175896-177911 | p3 | (ATA)5 | CAAAACCTGTTCTTGGGGAA | CCGCTCCCTTTCTCTTTCTT | 285 |
| chr1:235572-237596 | p2 | (TC)6 | GGGCCAACAAACAAAAAGAA | CCGAATGATTTTGCCTTCAT | 174 |
| chr1:235614-237653 | p2 | (TC)6 | GGGCCAACAAACAAAAAGAA | CCGAATGATTTTGCCTTCAT | 132 |
| chr1:275654-277669 | p5 | (TTGGT)3 | TCATTCTTTTTGGGACGGAC | TCAACCCTGGAGGTAAGGTG | 204 |
| chr1:309499-311511 | p2 | (AT)6 | GCAGGGAAGGTAACCAATCA | AGAGAATTGCGTGGTAACGG | 108 |
| chr1:339475-341490 | p5 | (AGTCA)3 | CCAGTTGCCTTGCTTCTTTC | GCATCCACAACTTCTTGGGT | 186 |
| chr1:339475-341490 | p5 | (TGAAC)3 | CCAGTTGCCTTGCTTCTTTC | GCATCCACAACTTCTTGGGT | 186 |
| chr1:409464-411492 | p2 | (AT)14 | TATCCAGTGGCAAGGTGTCA | ATGTAGCAGCCTTGTTGGCT | 123 |
| chr1:409464-411492 | p3 | (CAA)5 | TATCCAGTGGCAAGGTGTCA | ATGTAGCAGCCTTGTTGGCT | 123 |
| chr1:412238-414262 | p6 | (CCCCAA)4 | CATGTTTTTCTGCATGCCAC | TAGATTTCACCGGCAACTCC | 298 |
| chr1:437493-439508 | p3 | (GAA)5 | GCTTTCGATGGGATAGTGGA | GCTTGGGGGTTGTTGTTAGA | 239 |
| chr1:443153-445167 | p2 | (AT)7 | AGCTCAAACCCTGCCTTGTA | GTTTGGGCTGTAGTGCGTTT | 248 |
| chr1:443153-445167 | p5 | (GAAAA)3 | AGCTCAAACCCTGCCTTGTA | GTTTGGGCTGTAGTGCGTTT | 248 |
| chr1:454929-456944 | p5 | (TGATA)3 | ATAGGCGAAAAGAGCAACGA | ATTTTCAGATTGGCGTTTGC | 213 |
| chr1:889591-891603 | p2 | (CT)6 | CCATTGACACGTTATTTGCG | AGGGTGTGCAAATGGCTAAG | 202 |
| chr1:932299-934320 | p3 | (GAA)7 | CAAGGCCGACACAGAACATA | GAATTGCTCAAGGGTGCATT | 126 |
| chr1:939888-941900 | p2 | (TA)6 | ACAATTTTTGCGCAGGTTTC | TCGACCTTGAGAATCCAACC | 211 |
| chr1:939888-941900 | p2 | (AT)9 | ACAATTTTTGCGCAGGTTTC | TCGACCTTGAGAATCCAACC | 211 |
| chr1:978698-980713 | p5 | (TTGGT)3 | ACCATAGCCATGAAAGCCTG | ACCGACCAAGAGAGCTTCAA | 253 |
| chr1:978698-980713 | p2 | (TA)10 | ACCATAGCCATGAAAGCCTG | ACCGACCAAGAGAGCTTCAA | 253 |
| chr1:1005351-1007366 | p5 | (CTTTT)3 | GCTCGGACACATGCAAGTAA | TCCACAACAACCTTTCATGG | 227 |
| chr1:1005351-1007366 | p2 | (AT)6 | GCTCGGACACATGCAAGTAA | TCCACAACAACCTTTCATGG | 227 |
| chr1:1006347-1008359 | p5 | (CTTTT)3 | ACATTCAGTGTATGCACGCC | AATAAGTGTGGCGTTAGCGG | 230 |
| chr1:1006347-1008359 | p5 | (AAACC)3 | ACATTCAGTGTATGCACGCC | AATAAGTGTGGCGTTAGCGG | 230 |
| chr1:1006347-1008359 | p2 | (TA)18 | ACATTCAGTGTATGCACGCC | AATAAGTGTGGCGTTAGCGG | 230 |
| chr1:1013103-1015115 | p5 | (CTTTT)3 | GGCGGAACTAGAGGACGTAA | TAAAAATGTGGCCCGTTAGC | 262 |
| chr1:1013103-1015115 | p2 | (AT)6 | GGCGGAACTAGAGGACGTAA | TAAAAATGTGGCCCGTTAGC | 262 |
| chr1:1022832-1024854 | p2 | (AT)11 | AATCCTCCTAAATGGGTGGG | ATTTTGACGATCTCGTTGGC | 150 |
| chr1:1022832-1024854 | p2 | (AT)6 | AATCCTCCTAAATGGGTGGG | ATTTTGACGATCTCGTTGGC | 150 |
| chr1:1092223-1094259 | p6 | (ACTTGG)6 | CTTCCAGGCACTTCTTCTCG | TTTTGGAGCACTTTTGGACC | 147 |
| chr1:1108928-1110943 | p3 | (TCT)7 | CAATTTGCCTCCAACATCCT | AGCCACTGCAGAAAAGGAAA | 184 |
| chr1:1108928-1110943 | p3 | (TCT)5 | CAATTTGCCTCCAACATCCT | AGCCACTGCAGAAAAGGAAA | 184 |
| chr1:1248837-1250855 | p6 | (ATCTAT)3 | TCAGCGTCAAAGCTCTTCAA | ACACCAGTATTCCCCTTCCC | 265 |
| chr1:1248837-1250855 | p3 | (AAT)6 | TCAGCGTCAAAGCTCTTCAA | ACACCAGTATTCCCCTTCCC | 265 |
| chr1:1337415-1339431 | p2 | (AT)8 | GCTGAGCGAAAAGAATGGAC | ACAAATATCGGACCGGAACA | 180 |

| Marker ID | SSR type | SSR motif | FORWARD PRIMER (5'-3') | REVERSE PRIMER (5'-3') | size (bp) |
|---|---|---|---|---|---|
| chr1:1337415-1339431 | p2 | (AT)8 | GCTGAGCGAAAAGAATGGAC | ACAAATATCGGACCGGAACA | 180 |
| chr1:1337415-1339431 | p2 | (AT)8 | GCTGAGCGAAAAGAATGGAC | ACAAATATCGGACCGGAACA | 180 |
| chr1:1337415-1339431 | p2 | (AT)8 | GCTGAGCGAAAAGAATGGAC | ACAAATATCGGACCGGAACA | 180 |
| chr1:1337415-1339431 | p2 | (AT)8 | GCTGAGCGAAAAGAATGGAC | ACAAATATCGGACCGGAACA | 180 |
| chr1:1337415-1339431 | p3 | (AGC)6 | GCTGAGCGAAAAGAATGGAC | ACAAATATCGGACCGGAACA | 180 |
| chr1:1398015-1400040 | p5 | (AAACA)5 | GGGACCAAAAACCTCGAAAT | GTAGGTGACGGTCGGAATGT | 172 |
| chr1:1414660-1416684 | p3 | (AAT)8 | CAGTTACCCGGAAAACGAGA | AATCCAATAAGCTTCCCGCT | 296 |
| chr1:1437460-1439475 | p6 | (AAAAAG)4 | TCGCTAAGCTTGTCCGAAAT | CTCCAAAGCAAACCCAACTC | 300 |
| chr1:1515262-1517274 | p2 | (AT)6 | TACCAGCCCATTTGAAGGAC | AGTGCTTTGACCAAATTGCC | 275 |
| chr1:1650362-1652377 | p5 | (GTTTG)3 | ACGCCAAGTCTAGTCGAGGA | CCATGGCATTTGAACACTTG | 293 |
| chr1:1684498-1686519 | p3 | (GTA)7 | TGTATTCAGCCCAAAGGGAC | GAAATGGGCAGAATTTTCCA | 158 |
| chr1:1729926-1731944 | p3 | (GCT)6 | CGGGTTGAGCTAAATCCAAA | AAGCTAGCCCCTTCAAAAGC | 246 |
| chr1:1786029-1788044 | p5 | (AAAAT)3 | CAAGCAGAGCAACTCTTCCC | GTTGTTGGCAGTGCAGTTTG | 118 |
| chr1:1786029-1788044 | p5 | (AAACC)3 | CAAGCAGAGCAACTCTTCCC | GTTGTTGGCAGTGCAGTTTG | 118 |
| chr1:1786029-1788044 | p5 | (AAAAT)3 | CAAGCAGAGCAACTCTTCCC | GTTGTTGGCAGTGCAGTTTG | 118 |
| chr1:1786564-1788579 | p5 | (AAAAT)3 | TGATATTTGTTGTGCCTCGG | TCCGAAATTGAAAAACCCAA | 172 |
| chr1:1786564-1788579 | p5 | (AAACC)3 | TGATATTTGTTGTGCCTCGG | TCCGAAATTGAAAAACCCAA | 172 |
| chr1:1786564-1788579 | p5 | (AAAAT)3 | TGATATTTGTTGTGCCTCGG | TCCGAAATTGAAAAACCCAA | 172 |
| chr1:1817483-1819509 | p2 | (TA)7 | CAGACGACATCTTCTTGCCA | CGGAAGAGGTTCGGATGATA | 284 |
| chr1:1841735-1843750 | p5 | (CCTCT)3 | CAAAGGTCCCCAGATTGAAA | GTGGGACAAAATTGGGTGTC | 116 |
| chr1:1878239-1880257 | p6 | (TATTTT)3 | TTGAAGAAGACCGGGTTGAC | ATTTAGGGGAAAATGTCGGG | 138 |
| chr1:1904777-1906795 | p2 | (AT)9 | TCTGCTCGAATGATGACTGC | GGGAGCTGTCACATCACCTT | 119 |
| chr1:1922064-1924096 | p2 | (AT)16 | AGCATTTCGTGGAACAATCC | CTAATTTCGAGTCGGAGCCA | 186 |
| chr1:1938282-1940315 | p3 | (ATG)11 | TGCATTGAAACAAAATCGGA | AAGGCCAAGTTCACCACAAC | 191 |
| chr1:1954124-1956139 | p5 | (ACCAA)3 | CGAACGAAGTGGGAGGTTTA | CGAAGCAATTGGTTTGGTTT | 106 |
| chr1:1954124-1956139 | p5 | (AAACC)3 | CGAACGAAGTGGGAGGTTTA | CGAAGCAATTGGTTTGGTTT | 106 |
| chr1:2001649-2003685 | p3 | (TTA)12 | CCAGTCGTCATGCAAAGATG | TTGTTTCTCGCATGTTCTCG | 264 |
| chr1:2022637-2024652 | p5 | (GTTCG)3 | TATACGCCTTGCAGCCTCTT | GACCAAATCATCCCGTTGAC | 127 |
| chr1:2022637-2024652 | p5 | (AAAAG)3 | TATACGCCTTGCAGCCTCTT | GACCAAATCATCCCGTTGAC | 127 |
| chr1:2052385-2054401 | p4 | (TTCC)4 | TGGTGGTGAATGGTGATTGT | TATGACAGCAATGCCCAAAA | 235 |
| chr1:2056364-2058400 | p3 | (GAA)6 | TGCAAGAAAGCAAGCAAGAA | TGTTGGATTGAGAGCTGCTG | 230 |
| chr1:2061757-2063777 | p2 | (TA)8 | CCCATTAGTGGGAGCAAATG | TGTCAATGAATCCATCAGGC | 149 |
| chr1:2209095-2211122 | p3 | (ACA)9 | AACCTCCAGCTTAGCGTTCA | CAGCTCTTGGCCCATTTTTA | 137 |
| chr1:2241078-2243099 | p5 | (TTGTC)3 | CCACCGAAACTTAAAGGCAA | CGCATCTTGTCTTCCACTGA | 196 |
| chr1:2241078-2243099 | p3 | (TTG)7 | CCACCGAAACTTAAAGGCAA | CGCATCTTGTCTTCCACTGA | 196 |
| chr1:2245791-2247806 | p5 | (AAATG)3 | ATCCCCACATACCTTTGACG | GCGTTGTTGTGAATGTTTGG | 253 |

| Marker ID | SSR type | SSR motif | FORWARD PRIMER (5'-3') | REVERSE PRIMER (5'-3') | size (bp) |
|---|---|---|---|---|---|
| chr1:2276371-2278385 | p2 | (TA)7 | AATGAAATCGTGGGTTCTCG | ATCCGACCATCTAGGCACAG | 214 |
| chr1:2284229-2286243 | p6 | (TATATG)3 | GGGCTTCCCAGATTGGTATT | CTTTAGCGCTTATTCGCCAC | 226 |
| chr1:2284229-2286243 | p2 | (TA)7 | GGGCTTCCCAGATTGGTATT | CTTTAGCGCTTATTCGCCAC | 226 |
| chr1:2312857-2314872 | p5 | (AAAAT)3 | GTCGCCAAAGGCTCATCTAC | GGGTAAGCTGTTTCCCATCA | 227 |
| chr1:2332942-2334962 | p5 | (AAATA)4 | GGATGGAATCTCCAGAACGA | TTTGTGGCTGGAATTTGACA | 146 |
| chr1:2372730-2374748 | p6 | (TAAAAA)3 | AGCCTACCAGCATTTGCAGT | AATACCCCACTTCCAGGACC | 148 |
| chr1:2402299-2404339 | p5 | (GAAAA)3 | TTTGCCCCTACTTCCATGAG | TACGGCTATGTCCAGCCTTT | 284 |
| chr1:2443267-2445301 | p2 | (AT)8 | CGTCGGTATGGGAGAGGTAA | ATGTTGGCTTGCATGGATTT | 222 |
| chr1:2443267-2445301 | p5 | (TATTT)3 | CGTCGGTATGGGAGAGGTAA | ATGTTGGCTTGCATGGATTT | 222 |
| chr1:2443310-2445326 | p2 | (AT)8 | AAAAATCCATGCAAGCCAAC | GCCAAAGGCAAAGGGATATT | 275 |
| chr1:2443310-2445326 | p5 | (TATTT)3 | AAAAATCCATGCAAGCCAAC | GCCAAAGGCAAAGGGATATT | 275 |
| chr1:2500010-2502024 | p2 | (TC)7 | CCTATGCCCATTTGCTGATT | ATTTCGTGGTCAAATGAGGC | 276 |
| chr1:2542820-2544836 | p3 | (GTT)8 | CACGTTGTGGGATTTCACTG | TCCAATATGGCCAGTCCTTC | 278 |
| chr1:2542838-2544850 | p3 | (GTT)8 | CACGTTGTGGGATTTCACTG | TCCAATATGGCCAGTCCTTC | 260 |
| chr1:2798858-2800870 | p2 | (GA)6 | CTGTGAGTGTGCGAACTGCT | GCCACCTCATCACCTCCTAA | 125 |
| chr1:2798858-2800870 | p2 | (AC)6 | CTGTGAGTGTGCGAACTGCT | GCCACCTCATCACCTCCTAA | 125 |
| chr1:2896503-2898515 | p2 | (TA)6 | CCCCCTACAAACAACACCAT | GTGTATTGTCGTATGCCCCC | 193 |
| chr1:2896503-2898515 | p2 | (TA)8 | CCCCCTACAAACAACACCAT | GTGTATTGTCGTATGCCCCC | 193 |
| chr1:2916789-2918804 | p6 | (CCACCC)3 | TTTGGATGGTGGTTTACCGT | GGTTAGTGGTAGGGTGGGGT | 219 |
| chr1:2916789-2918804 | p5 | (TACAA)3 | TTTGGATGGTGGTTTACCGT | GGTTAGTGGTAGGGTGGGGT | 219 |
| chr1:2916789-2918804 | p2 | (TA)16 | TTTGGATGGTGGTTTACCGT | GGTTAGTGGTAGGGTGGGGT | 219 |
| chr1:2959242-2961257 | p6 | (CCACCC)3 | TTTGGATGGTGGTTTACCGT | GGTTAGTGGTAGGGTGGGGT | 219 |
| chr1:2959242-2961257 | p5 | (TACAA)3 | TTTGGATGGTGGTTTACCGT | GGTTAGTGGTAGGGTGGGGT | 219 |
| chr1:2959242-2961257 | p2 | (TA)16 | TTTGGATGGTGGTTTACCGT | GGTTAGTGGTAGGGTGGGGT | 219 |
| chr1:3077647-3079665 | p3 | (TTG)6 | GGGGTATTTTGGCCACTTTT | AACCCAGCTCATGAGGTCAC | 177 |
| chr1:3078955-3080970 | p5 | (TTCGG)3 | CTTGGGCCTAAGGATGTGAA | TTAAATAACCGACCGAACCG | 124 |
| chr1:3078955-3080970 | p2 | (AT)12 | CTTGGGCCTAAGGATGTGAA | TTAAATAACCGACCGAACCG | 124 |
| chr1:3079559-3081625 | p5 | (TTCGG)3 | CCAATCAACTAAAGGCCCAA | CCGAACCTAACCATACCGAA | 263 |
| chr1:3079559-3081625 | p2 | (AT)12 | CCAATCAACTAAAGGCCCAA | CCGAACCTAACCATACCGAA | 263 |
| chr1:3182099-3184177 | p2 | (TA)11 | CTTAAAAGTCCCTGCGCTTG | GGTATTGGAGAAGGGCAACA | 184 |
| chr1:3274503-3276515 | p2 | (TA)6 | CGTCAAAAGATGGGAGAGGA | CACGTCGGCCCTGTACTTAT | 202 |
| chr1:3325357-3327369 | p2 | (TA)6 | TAGCAAAAAGGGTTTGTCGG | ACCATGCTAAGACGTCCACC | 288 |
| chr1:3325357-3327369 | p2 | (AT)8 | TAGCAAAAAGGGTTTGTCGG | ACCATGCTAAGACGTCCACC | 288 |
| chr1:3325877-3327893 | p2 | (TA)6 | AACCAAAACTCAGAAGCGGA | TTTCGTTAATTTCCCCCACA | 111 |
| chr1:3325877-3327893 | p2 | (AT)8 | AACCAAAACTCAGAAGCGGA | TTTCGTTAATTTCCCCCACA | 111 |
| chr1:3348761-3350797 | p3 | (ATT)12 | TTGCCACACCAATTTGACTC | TATTTTAGCCACAATGGCCC | 118 |

| Marker ID | SSR type | SSR motif | FORWARD PRIMER (5'-3') | REVERSE PRIMER (5'-3') | size (bp) |
|---|---|---|---|---|---|
| chr1:3406061-3408079 | p2 | (AT)9 | CGAGAAGATGGTGCACAAGA | TGATACGCATTTTGGCTGTC | 253 |
| chr1:3464503-3466518 | p5 | (AACCG)3 | TTCATCTGCTTCATTGACCG | CAACGTAGATGGGCCAGATT | 293 |
| chr1:3464531-3466546 | p5 | (AACCG)3 | TTCATCTGCTTCATTGACCG | CAACGTAGATGGGCCAGATT | 265 |
| chr1:3484626-3486680 | p3 | (ATT)18 | TTCAACTTTCGAAACAGGGG | AGTTCGCTGCCGAAGAAATA | 255 |
| chr1:3484626-3486680 | p2 | (CT)6 | TTCAACTTTCGAAACAGGGG | AGTTCGCTGCCGAAGAAATA | 255 |
| chr1:3485815-3487843 | p2 | (CT)6 | TTTCGGACCAATCCACATTT | CGAGTGGGGAACAGAGAGAG | 121 |
| chr1:3485815-3487843 | p2 | (AT)14 | TTTCGGACCAATCCACATTT | CGAGTGGGGAACAGAGAGAG | 121 |
| chr1:3623394-3625420 | p2 | (AT)13 | TGCAACAGCATGTCTTCACA | AAACGGACACACTTTCCCTG | 249 |
| chr1:3726190-3728214 | p3 | (AAT)8 | GGCTGACAGCTCCAGACTTC | TCGTTCATGATTTCATCCCA | 249 |
| chr1:3751462-3753477 | p5 | (AAAAT)3 | CGAGGAAGCTGATTGGAAAG | ATGCTGAAGGATGACCCAAC | 129 |
| chr1:3789075-3791090 | p3 | (AGA)5 | AGAAGGCATGGACAATGGAG | CCATGCAGCCCAAGATTATT | 291 |
| chr1:3809488-3811503 | p3 | (ACC)5 | CTTCTGCGCATTTTTCATCA | TCGAAACACAAAGATCGACG | 114 |
| chr1:3845308-3847323 | p5 | (TGCTT)3 | GCATGAAGGAGTGGTTGGAT | CAATAGCCATCGCCCATACT | 115 |
| chr1:3880880-3882896 | p2 | (AT)8 | ATGCTGCCATGCTCTAGTGA | GATAACGGACGACGAAGCAT | 263 |
| chr1:3880880-3882896 | p2 | (TA)8 | ATGCTGCCATGCTCTAGTGA | GATAACGGACGACGAAGCAT | 263 |
| chr1:3880880-3882896 | p5 | (TTTGT)3 | ATGCTGCCATGCTCTAGTGA | GATAACGGACGACGAAGCAT | 263 |
| chr1:4079952-4081968 | p4 | (AATA)4 | AGTCTAGCTGCAAAGCCTGC | GACAGCTAACGGCTTTGGTC | 109 |
| chr1:4144487-4146502 | p3 | (GTT)5 | CCAAGTGTAAGCCCTCCAAA | CGGCCTATGGTTGGAAAGTA | 299 |
| chr1:4146508-4148532 | p2 | (AT)12 | ATTTTTACCGAGGGGTACGG | TGTGAGACAAGAGGTGCTCG | 172 |
| chr1:4146508-4148532 | p2 | (AT)10 | ATTTTTACCGAGGGGTACGG | TGTGAGACAAGAGGTGCTCG | 172 |
| chr1:4146686-4148706 | p2 | (AT)12 | GGACGATACCGTCATTGCTT | TTGAGAGCGCTCCAGTTCTT | 298 |
| chr1:4146686-4148706 | p2 | (AT)10 | GGACGATACCGTCATTGCTT | TTGAGAGCGCTCCAGTTCTT | 298 |
| chr1:4160507-4162519 | p2 | (AT)6 | TGGTATAGTTCAGGGACGCC | CCGATAGAGCCTGTGGACAT | 136 |
| chr1:4226428-4228458 | p2 | (AT)15 | TTCAATTTCCTTTTGCCTGC | GTGAAAATCCGTTCCCAAGA | 291 |
| chr1:4230599-4232615 | p4 | (CTAT)4 | TGAGGTTGTATTGGTGGGGT | AACGTGGTTGTCAAAGCTCC | 161 |
| chr1:4235678-4237690 | p2 | (TG)6 | TAAAGCGTGGAATGTTCGTG | TCGCATGTTATTCCCAGTCA | 133 |
| chr1:4241641-4243656 | p3 | (ATA)8 | TGTCACGGGATTCAAAATGA | TCCGAGGCTACTTTTGTTGG | 253 |
| chr1:4241641-4243656 | p5 | (TATAT)3 | TGTCACGGGATTCAAAATGA | TCCGAGGCTACTTTTGTTGG | 253 |
| chr1:4252276-4254300 | p2 | (AT)12 | GGGAATAAACACAGGCCCTT | CAGCACTCATCACGTTGGTT | 263 |
| chr1:4291342-4293357 | p3 | (TTG)5 | GCTACTGCAACGACACCGTA | TTGGTGGATCGAAGTGACAA | 119 |
| chr1:4376403-4378421 | p6 | (AAATCG)3 | GAATAACGGCGACCCAAGTA | TGAGGCCCAAATCGACTATC | 201 |
| chr1:4387027-4389051 | p2 | (TG)7 | CATCACTTCGCTGGGATTTT | TGGTCGGCTTTCTTCAACTT | 299 |
| chr1:4387027-4389051 | p2 | (AT)6 | CATCACTTCGCTGGGATTTT | TGGTCGGCTTTCTTCAACTT | 299 |
| chr1:4387027-4389051 | p2 | (TA)12 | CATCACTTCGCTGGGATTTT | TGGTCGGCTTTCTTCAACTT | 299 |
| chr1:4412429-4414447 | p2 | (AT)9 | TGGTGGGTTTAGAGGAGTGG | AACAACTTGCCTCCGATCAC | 244 |
| chr1:4442274-4444289 | p5 | (TTTTA)3 | ACGCCGTCACATTTGTTTCT | AGCTCCAGCTATGCGTAGGA | 179 |

| Marker ID | SSR type | SSR motif | FORWARD PRIMER (5'-3') | REVERSE PRIMER (5'-3') | size (bp) |
|---|---|---|---|---|---|
| chr1:4507260-4509272 | p2 | (GA)6 | ATGACATGGTTCTGCCTTCC | GATCACACACCATCACAGCC | 189 |
| chr1:4514162-4516182 | p5 | (TTACT)4 | AAAGGAAGCGGAGTGAGTGA | TGCTCTTTGATTGCTTGTGC | 144 |
| chr1:4514162-4516182 | p2 | (TA)6 | AAAGGAAGCGGAGTGAGTGA | TGCTCTTTGATTGCTTGTGC | 144 |
| chr1:4514162-4516182 | p3 | (ACA)6 | AAAGGAAGCGGAGTGAGTGA | TGCTCTTTGATTGCTTGTGC | 144 |
| chr1:4585617-4587647 | p6 | (CACCTC)5 | GAACCTTTGTACCTTGCCCA | CACAACCGGGAAACACTCTT | 191 |
| chr1:4585617-4587647 | p2 | (AG)6 | GAACCTTTGTACCTTGCCCA | CACAACCGGGAAACACTCTT | 191 |
| chr1:4586346-4588358 | p6 | (CACCTC)5 | CACCTCCAGTCCACTCACCT | ACTGGTGGTAGTGGTGAGGC | 109 |
| chr1:4586346-4588358 | p2 | (AG)6 | CACCTCCAGTCCACTCACCT | ACTGGTGGTAGTGGTGAGGC | 109 |
| chr1:4597221-4599242 | p3 | (TGT)7 | TTTTGCAGGGATATCGGAAG | ATTCTTCAAGCAGCAGGCAT | 120 |
| chr1:4692214-4694232 | p6 | (TTTTTG)3 | GGTTGCTTCTGCATGGAAGT | CTTCCTCTTCTACCCCCACC | 230 |
| chr1:4781770-4783806 | p3 | (ACA)5 | AATGCTCCATTCTTTGCACC | TACAAAGCGGCCATTAGGTC | 299 |
| chr1:4781804-4783822 | p3 | (ACA)5 | AATGCTCCATTCTTTGCACC | TACAAAGCGGCCATTAGGTC | 265 |
| chr1:4785742-4787758 | p2 | (TA)8 | AATGAACCATAACCGCTTCC | CATGCAATGTCACTTCACCC | 148 |
| chr1:4838353-4840367 | p2 | (AT)7 | CAGCACGAATGGTTCAGCTA | TAAGGGGTGCTCGGTACAAG | 208 |
| chr1:4931160-4933172 | p5 | (TTGTC)3 | TCAGTGGAAGACAAGATGCG | GGTCATTTCCTCCGTAAGCA | 266 |
| chr1:4931160-4933172 | p2 | (AT)6 | TCAGTGGAAGACAAGATGCG | GGTCATTTCCTCCGTAAGCA | 266 |
| chr1:4974796-4976810 | p2 | (AT)7 | TGGGGACCCTGTAAAAATGA | GCCAAAGATAGAGCAAACCG | 263 |
| chr1:4974796-4976810 | p2 | (TA)10 | TGGGGACCCTGTAAAAATGA | GCCAAAGATAGAGCAAACCG | 263 |
| chr1:4974861-4976885 | p2 | (AT)7 | TGGGGACCCTGTAAAAATGA | GCCAAAGATAGAGCAAACCG | 198 |
| chr1:4974861-4976885 | p2 | (TA)10 | TGGGGACCCTGTAAAAATGA | GCCAAAGATAGAGCAAACCG | 198 |
| chr1:4995642-4997657 | p5 | (AACCG)3 | AAATGCGCAGGGAGAGTAGA | CGGTTCGGTTTGTTGAAGTT | 253 |
| chr1:4995642-4997657 | p5 | (GAACC)3 | AAATGCGCAGGGAGAGTAGA | CGGTTCGGTTTGTTGAAGTT | 253 |
| chr1:5105774-5107790 | p2 | (TA)21 | CCCATTAGTGGGAGCAAATG | AAATGACATGGCCGAATAGC | 195 |
| chr1:5105774-5107790 | p2 | (TA)8 | CCCATTAGTGGGAGCAAATG | AAATGACATGGCCGAATAGC | 195 |
| chr1:5133245-5135260 | p5 | (TTTAT)3 | ACATTTCCAAGCCAATTCCA | TTCCATTGCAACAAGGATCA | 183 |
| chr1:5184771-5186811 | p2 | (TA)20 | GCAACAACAAACGACCAAAA | CAAGGGGAATAAATGAGCCA | 274 |
| chr1:5244717-5246757 | p2 | (TA)20 | GCAACAACAAACGACCAAAA | CAAGGGGAATAAATGAGCCA | 274 |
| chr1:5313818-5315830 | p2 | (TA)6 | ATCAGGTCCAGAGGTCATGC | ATGACGAAATTGCCCTTCTG | 229 |
| chr1:5313818-5315830 | p6 | (TGTTAT)3 | ATCAGGTCCAGAGGTCATGC | ATGACGAAATTGCCCTTCTG | 229 |
| chr1:5389572-5391593 | p3 | (ACA)7 | AAACTTTAGGCGAAGGCGAT | ATGTGATGACACGAGCCAAA | 198 |
| chr1:5398257-5400285 | p2 | (TA)14 | AGGCTACATCCAACATTGCC | TAACATCCTACGCCAAACCC | 201 |
| chr1:5455436-5457458 | p2 | (AG)11 | TAAGGAAAGGCGTTGGTCAC | TCTCAATTGGAGCCAGCTTT | 135 |
| chr1:5465108-5467126 | p3 | (GTT)6 | ATCCTCTCCGATAGCCCACT | ATCCTCTCCGATAGCCCACT | 287 |
| chr1:5505434-5507448 | p2 | (AC)7 | ATGTGCAACTTGGTCATGGA | AGAACTCGTGCCAAGCAGAT | 190 |
| chr1:5558455-5560470 | p5 | (GGGTG)3 | GTTGGAAAAGCGACAACCAT | GGACATTTCGGCATTCACTT | 147 |
| chr1:5558455-5560470 | p6 | (AAACCG)3 | GTTGGAAAAGCGACAACCAT | GGACATTTCGGCATTCACTT | 147 |

| Marker ID | SSR type | SSR motif | FORWARD PRIMER (5'-3') | REVERSE PRIMER (5'-3') | size (bp) |
|---|---|---|---|---|---|
| chr1:5578091-5580129 | p4 | (TTAT)4 | TTGCCGTAAACATGGAGACA | CTGTCCAGGATGCACTGAGA | 182 |
| chr1:5578091-5580129 | p2 | (AG)19 | TTGCCGTAAACATGGAGACA | CTGTCCAGGATGCACTGAGA | 182 |
| chr1:5578091-5580129 | p2 | (AT)7 | TTGCCGTAAACATGGAGACA | CTGTCCAGGATGCACTGAGA | 182 |
| chr1:5581604-5583642 | p4 | (TTAT)4 | TTGCCGTAAACATGGAGACA | CTGTCCAGGATGCACTGAGA | 182 |
| chr1:5581604-5583642 | p2 | (AG)19 | TTGCCGTAAACATGGAGACA | CTGTCCAGGATGCACTGAGA | 182 |
| chr1:5581604-5583642 | p2 | (AT)7 | TTGCCGTAAACATGGAGACA | CTGTCCAGGATGCACTGAGA | 182 |
| chr1:5596670-5598688 | p3 | (CAA)6 | TCCTTTCCCCATCTTCCTCT | TTCAAGGTGAGACACGATGC | 186 |
| chr1:5663942-5665956 | p2 | (TA)7 | GTTTTCCAGGTTTGGTGGTG | CTCGCCAAACGCCTACTAAG | 126 |
| chr1:5746822-5748842 | p2 | (AT)10 | CCGAGACTCTGTGGGGATTA | GACCGTCCCAGAAAAACAAA | 137 |
| chr1:5834618-5836633 | p3 | (TAA)5 | CCAAGTCATCAACTGCGCTA | CGTTCACGACCACACTTGTC | 269 |
| chr1:5857700-5859721 | p3 | (TGT)7 | GGCTTTAAAAGGGGACAAGC | GTAAACCCCGCTCCTGTGTA | 189 |
| chr1:5871647-5873659 | p2 | (CA)6 | CGTACAGTTGGGTCTCGGTT | AAGGCAGCCATCACCAATAG | 174 |
| chr1:5908214-5910226 | p2 | (AC)6 | TGTTCACGTCATTGGGCTAA | CTGTCCGCAGCACAAATAGA | 157 |
| chr1:5983591-5985617 | p2 | (AT)10 | ATCCATCGATCACCAAGGAG | AGAGGTGCTGTGGAGATGCT | 148 |
| chr1:5983591-5985617 | p2 | (TA)13 | ATCCATCGATCACCAAGGAG | AGAGGTGCTGTGGAGATGCT | 148 |
| chr1:6026057-6028073 | p6 | (TTTCGG)3 | CAAAAGCAAAAGAGGGCTTG | ACCGAAACCGAACTTTGATG | 109 |
| chr1:6026057-6028073 | p4 | (GAAA)4 | CAAAAGCAAAAGAGGGCTTG | ACCGAAACCGAACTTTGATG | 109 |
| chr1:6086568-6088583 | p3 | (AAG)5 | GGGCAGGTGGGTTAATTTTT | GCGTAGACAGCCCTTTCATC | 211 |
| chr1:6086568-6088583 | p3 | (GAT)5 | GGGCAGGTGGGTTAATTTTT | GCGTAGACAGCCCTTTCATC | 211 |
| chr1:6086568-6088583 | p3 | (ATA)5 | GGGCAGGTGGGTTAATTTTT | GCGTAGACAGCCCTTTCATC | 211 |
| chr1:6145972-6147984 | p2 | (TA)6 | AAGAGAAGATTGACCGCATCA | TCGGAAAAGGCGTACCTAGA | 236 |
| chr1:6145972-6147984 | p2 | (TA)6 | AAGAGAAGATTGACCGCATCA | TCGGAAAAGGCGTACCTAGA | 236 |
| chr1:6145972-6147984 | p6 | (ATCTAT)3 | AAGAGAAGATTGACCGCATCA | TCGGAAAAGGCGTACCTAGA | 236 |
| chr1:6523515-6525530 | p5 | (AAAAT)3 | GATTGCATGGCTTCCTGTTT | ATTAATCGACATGCACGCAA | 204 |
| chr1:6546450-6548468 | p2 | (AT)7 | ATACATTTTGAAGCGGCGAC | GTGTCATTTGAACCCCTCGT | 117 |
| chr1:6546450-6548468 | p2 | (AT)9 | ATACATTTTGAAGCGGCGAC | GTGTCATTTGAACCCCTCGT | 117 |
| chr1:6546450-6548468 | p2 | (AT)6 | ATACATTTTGAAGCGGCGAC | GTGTCATTTGAACCCCTCGT | 117 |
| chr1:6579569-6581584 | p5 | (TTTTC)3 | GCGACACATGAGCAGGTAGA | GTAGCATCGTGGGCTTTGTT | 190 |
| chr1:6670664-6672679 | p5 | (CTCTT)3 | TTACGTTTGCAGACAGGCAC | CAAGCACCTCGTAGGCTCTC | 145 |
| chr1:6674066-6676084 | p6 | (ATTTTT)3 | GCCCCCTTTAAGCAAAGAAG | CTATTTTGCGCCACCAAAGT | 127 |
| chr1:6789151-6791169 | p3 | (AAT)6 | TTCGAGATTAACGGCTGCTT | CCCAGGTCCTTGATTTTGAA | 253 |
| chr1:6882428-6884464 | p2 | (AG)18 | CTAACTCCGGTGAAGCGGTA | TGGCTGAATGGCCTTATTTC | 134 |
| chr1:6896564-6898598 | p2 | (TA)6 | GGGTTTAGACCCATGCTTCA | GATGTGGCGAGTTCCACTTT | 275 |
| chr1:6897432-6899458 | p2 | (TA)6 | GGGCCCCATACTAAATGACA | AGGGACAACGGACAATGAAG | 210 |
| chr1:6897432-6899458 | p5 | (TTTAT)4 | GGGCCCCATACTAAATGACA | AGGGACAACGGACAATGAAG | 210 |
| chr1:6897456-6899477 | p2 | (TA)6 | AGACCCATGCTTCATTGTCC | TTTGGATGTGGTGAGTTCCA | 258 |

| Marker ID | SSR type | SSR motif | FORWARD PRIMER (5'-3') | REVERSE PRIMER (5'-3') | size (bp) |
|---|---|---|---|---|---|
| chr1:6897456-6899477 | p5 | (TTTAT)4 | AGACCCATGCTTCATTGTCC | TTTGGATGTGGTGAGTTCCA | 258 |
| chr1:7005462-7007484 | p2 | (AT)11 | CTTCCTGCCAAGTTGACACA | CCAATGCAGGAATTGGAGAT | 209 |
| chr1:7005462-7007484 | p6 | (AAAGAT)3 | CTTCCTGCCAAGTTGACACA | CCAATGCAGGAATTGGAGAT | 209 |
| chr1:7011252-7013267 | p5 | (ACTCC)3 | AATGATGAAAGCAAGCCGAT | ATCATTCATCCCGACCTGAG | 149 |
| chr1:7024227-7026241 | p2 | (TA)7 | ATCGATGGCCATTTTGAGTC | GCCTTTGGTAAAAATGGGGT | 120 |
| chr1:7024227-7026241 | p2 | (AT)9 | ATCGATGGCCATTTTGAGTC | GCCTTTGGTAAAAATGGGGT | 120 |
| chr1:7041147-7043165 | p6 | (TTCCTT)3 | AGATTTCGCAGTCGCTCAAT | TTCACGAGATCAAAAGCACG | 168 |
| chr1:7065368-7067438 | p2 | (TA)35 | CCAAAAGGGTGGACCTCATA | TTTTCCCCATGAGTTTGGAG | 256 |
| chr1:7065368-7067438 | p2 | (AT)7 | CCAAAAGGGTGGACCTCATA | TTTTCCCCATGAGTTTGGAG | 256 |
| chr1:7116881-7118893 | p2 | (TA)6 | AACGGTCTCCCTCACACAAC | CGGATTGTCCCAAAAGACAT | 176 |
| chr1:7186863-7188881 | p2 | (GA)9 | TTCAAGGTGGACCCTTCAAC | AAGCTTTGGTTCTCAGGCAA | 208 |
| chr1:7186863-7188881 | p2 | (TC)6 | TTCAAGGTGGACCCTTCAAC | AAGCTTTGGTTCTCAGGCAA | 208 |
| chr1:7204872-7206886 | p6 | (ATTTTT)3 | AGCAATGAAACCATCCCTTG | GATGATCGGAAAAGCCACAT | 291 |
| chr1:7290496-7292516 | p2 | (AT)10 | TGATTCCGCGTCTGTGATTA | GACGTGGAATGCTTGAGACA | 167 |
| chr1:7330385-7332407 | p2 | (TA)10 | TTGTTCCACGAAATGCAAAA | TGTTTCCACGAAATGCAAAA | 151 |
| chr1:7370016-7372040 | p2 | (TA)12 | TTGTGACGCGAGAGCATAAC | AACAATGATGAGGCTTCGCT | 210 |
| chr1:7415425-7417437 | p2 | (CT)6 | CAAAGTTAGGCCTCAGGCAG | TCCTTCAAAGGCACTGGTCT | 160 |
| chr1:7495467-7497482 | p3 | (TTA)5 | TCAGGATAGTTTGCTTGGGG | AGGGAGGAAGAGAATCGAGG | 228 |
| chr1:7499194-7501208 | p2 | (AT)7 | GGAAAATGAACCCTCGGACT | GCGTCCTATTGGGAACAAAA | 157 |
| chr1:7529663-7531678 | p5 | (AAAAT)3 | TGAGGCTTCAAATACCCCTG | GGTGGCCATAGAAAGTCCAA | 278 |
| chr1:7534002-7536020 | p6 | (TTTTAT)3 | TGCACACTCTGTGGTGTTGA | GCCCACGAACACTAGCATTT | 280 |
| chr1:7555805-7557823 | p2 | (TA)9 | GGTTGTGAATTTGGACCCAC | CCAAATTCTTGAACTCCCGA | 254 |
| chr1:7555805-7557823 | p2 | (AT)8 | GGTTGTGAATTTGGACCCAC | CCAAATTCTTGAACTCCCGA | 254 |
| chr1:7566946-7568960 | p2 | (TA)7 | CCACCCACCTCCAAACATAC | GACCCATAGCTAGCTCGACG | 265 |
| chr1:7615290-7617306 | p2 | (AT)8 | TTTTGTTATTCCTTTCGCCG | CATGTAAGTCATGCATCGGG | 278 |
| chr1:7790528-7792552 | p3 | (AAG)8 | TGCAACGGGAAAGATAGGAC | TGTGTGTTTCCTGGGATGAA | 285 |
| chr1:7790528-7792552 | p5 | (AAACT)3 | TGCAACGGGAAAGATAGGAC | TGTGTGTTTCCTGGGATGAA | 285 |
| chr1:7793650-7795665 | p3 | (CAA)5 | GCTCCAAAATCAGGGTTTCA | GGCACTTGGGGAAGATGATA | 279 |
| chr1:7816224-7818238 | p2 | (TA)7 | CGAGAGGCCTGTGTAAAAGC | CACGAAAGAAATGTGGCCTT | 204 |
| chr1:7865423-7867437 | p2 | (GA)7 | ATTAATAGTGGTGCTGCGGG | TTCCGCTATTGGAAGATTCG | 107 |
| chr1:7887298-7889316 | p2 | (TA)6 | TAATGTTCAAAAGGGTGGGC | TTTAAGGGACAACGGACGAC | 272 |
| chr1:7887298-7889316 | p6 | (ATAAAA)3 | TAATGTTCAAAAGGGTGGGC | TTTAAGGGACAACGGACGAC | 272 |
| chr1:7887298-7889316 | p2 | (TA)7 | TAATGTTCAAAAGGGTGGGC | TTTAAGGGACAACGGACGAC | 272 |
| chr1:7895862-7897898 | p3 | (TCT)12 | CCAGGAGCAGAAGAGGTGTC | TCCTGCTTGAGGGTTCAGTT | 297 |
| chr1:7905746-7907766 | p2 | (TA)10 | GCCTAAGGTGTTGCACCAAT | TATCTCCAGCCACAACATGC | 233 |
| chr1:7938237-7940249 | p2 | (TA)6 | CATCCCAGCGAATGGTAATC | ACTCCCCCTTGTTGAGTCCT | 155 |

| Marker ID | SSR type | SSR motif | FORWARD PRIMER (5'-3') | REVERSE PRIMER (5'-3') | size (bp) |
|---|---|---|---|---|---|
| chr1:7956552-7958612 | p3 | (TAT)20 | TTCCTTTTGGTTTTGATGGG | ATGTCTCGGTCTCGCAACTT | 241 |
| chr1:8023900-8025915 | p4 | (ATAA)4 | GATAGCATACCCCTGTGCGT | TAGGGTTTGATCACCGGAAG | 236 |
| chr1:8023900-8025915 | p3 | (AGT)5 | GATAGCATACCCCTGTGCGT | TAGGGTTTGATCACCGGAAG | 236 |
| chr1:8023900-8025915 | p5 | (TTTTA)4 | GATAGCATACCCCTGTGCGT | TAGGGTTTGATCACCGGAAG | 236 |
| chr1:8029683-8031697 | p2 | (AT)7 | CCCATTAGTGGGAGCAAATG | ATAAATGGCATGGCGGAATA | 140 |
| chr1:8083270-8085288 | p3 | (GTT)5 | GGAAGTTACTCCGGTGTCCA | CCAGCAAGGGGCTACTACTG | 130 |
| chr1:8083270-8085288 | p3 | (TTA)6 | GGAAGTTACTCCGGTGTCCA | CCAGCAAGGGGCTACTACTG | 130 |
| chr1:8098100-8100118 | p3 | (TAT)6 | TTGCATAAACATCGACGCAT | GGAAGCCGAGTGTCGAGTAG | 131 |
| chr1:8098100-8100118 | p3 | (TTA)6 | TTGCATAAACATCGACGCAT | GGAAGCCGAGTGTCGAGTAG | 131 |
| chr1:8186733-8188747 | p2 | (TG)7 | AGAAATCGGCATTGTTGCTC | TGACCAGTGGCAGAGTCAAG | 141 |
| chr1:8212581-8214593 | p2 | (TC)6 | TGAACTTGTTGGATTGCAGG | CCAATTGCTCATTGCTGCTA | 255 |
| chr1:8231867-8233879 | p2 | (TC)6 | TGAACTTGTTGGATTGCAGG | CCAATTGCTCATTGCTGCTA | 255 |
| chr1:8274583-8276607 | p3 | (TAA)8 | AATGGAAATAGGTTGGCACG | AATATTCTTTGGCGACGGTG | 159 |
| chr1:8350283-8352299 | p3 | (AGA)6 | GCAATCCCCACTCTTTTTGA | GTTCCACTTTCTGTTGCCGT | 223 |
| chr1:8350283-8352299 | p2 | (AT)8 | GCAATCCCCACTCTTTTTGA | GTTCCACTTTCTGTTGCCGT | 223 |
| chr1:8380695-8382716 | p3 | (AAT)7 | ATTCAGACCAACACCCGAAG | TCGCTCAAGCCATTCTTTTT | 164 |
| chr1:8384417-8386438 | p3 | (AAT)7 | ATTCAGACCAACACCCGAAG | TCGCTCAAGCCATTCTTTTT | 164 |
| chr1:8537662-8539706 | p2 | (TA)14 | CAACATTGCCATCAACAAGG | GCTTCCACTTTTGCCAACAT | 132 |
| chr1:8537662-8539706 | p4 | (AGAA)4 | CAACATTGCCATCAACAAGG | GCTTCCACTTTTGCCAACAT | 132 |
| chr1:8607880-8609898 | p3 | (TGT)6 | GTTGTACGGGGCAGAGTGTT | ATCTGTGCCTCTCCTCCTCA | 289 |
| chr1:8619542-8621560 | p3 | (AAT)6 | TGTTCCAGGGGTACCAGAAG | CACGAGATCTTCATTTGCGA | 165 |
| chr1:8637109-8639130 | p3 | (TAA)10 | AAAGGAAGAAGAAGAGGCGG | ATTTTGACTTACCGGACCCC | 154 |
| chr1:8637109-8639130 | p3 | (ATA)7 | AAAGGAAGAAGAAGAGGCGG | ATTTTGACTTACCGGACCCC | 154 |
| chr1:8637109-8639130 | p3 | (TTG)5 | AAAGGAAGAAGAAGAGGCGG | ATTTTGACTTACCGGACCCC | 154 |
| chr1:8637109-8639130 | p2 | (AT)13 | AAAGGAAGAAGAAGAGGCGG | ATTTTGACTTACCGGACCCC | 154 |
| chr1:8641683-8643698 | p5 | (ATGGT)3 | TCCGTCGGTTTTATCCTGAG | TCCCTTAAGTTCGCCAAATG | 251 |
| chr1:8749320-8751334 | p2 | (AT)7 | CAGCAATTCGTAGGGGAGAA | AGGTTGGGTCGTGACAAAAG | 153 |
| chr1:8749320-8751334 | p2 | (AT)7 | CAGCAATTCGTAGGGGAGAA | AGGTTGGGTCGTGACAAAAG | 153 |
| chr1:8920339-8922357 | p6 | (TAAAAA)3 | TGGTAAATGATTGATGGGGG | TGGTCGTGGTTTGAAATCTG | 186 |
| chr1:8920339-8922357 | p2 | (AT)9 | TGGTAAATGATTGATGGGGG | TGGTCGTGGTTTGAAATCTG | 186 |
| chr1:8975260-8977282 | p2 | (AT)11 | CTACTCGCAAAAGAGGGTCG | CATGTAAGTCATGCATCGGG | 299 |
| chr1:8975260-8977282 | p3 | (GAT)6 | CTACTCGCAAAAGAGGGTCG | CATGTAAGTCATGCATCGGG | 299 |
| chr1:9042408-9044426 | p3 | (TGT)6 | GCTTTTGGAGTGCTGTGTGA | ACCCCGGATATCTTCATTCC | 214 |
| chr1:9074842-9076857 | p5 | (TTTAT)3 | CTCTCCCCATTTTTCCTTCC | GGGGTTTATGCCTCCAAAAT | 103 |
| chr1:9087854-9089866 | p2 | (TA)6 | CCGGGTTAGTTCGGATTTTT | CAAGAAAACCCAAGGTCGAA | 264 |
| chr1:9201183-9203209 | p2 | (AT)13 | CTCCGTTTCCCTTTTTCCTC | TTTTCAGGTTCATCCTTCGG | 219 |

| Marker ID | SSR type | SSR motif | FORWARD PRIMER (5'-3') | REVERSE PRIMER (5'-3') | size (bp) |
|---|---|---|---|---|---|
| chr1:9225917-9227933 | p4 | (TATT)4 | TCGCGAAAGTTCTGCCTTAT | ACTGTCCTTCTTTCGGGGTT | 184 |
| chr1:9264256-9266268 | p2 | (TC)6 | AAATGGTGGTACCCTCCTCC | ACTAGCTATGTTGGCACCCG | 216 |
| chr1:9489302-9491317 | p3 | (TTA)5 | AGGCGGTAATGGTGGTAATG | TATACCCCCGTGTTATCGGA | 214 |
| chr1:9489302-9491317 | p4 | (GAAA)6 | AGGCGGTAATGGTGGTAATG | TATACCCCCGTGTTATCGGA | 214 |
| chr1:9578885-9580900 | p5 | (TATTT)3 | GAAGTTGGGCTTTTGGATGA | TTGTCTGTTGCAAAAGGTCG | 275 |
| chr1:9578885-9580900 | p5 | (TAAAA)3 | GAAGTTGGGCTTTTGGATGA | TTGTCTGTTGCAAAAGGTCG | 275 |
| chr1:9673011-9675023 | p6 | (AAACGG)3 | CTCCGATGCTAAACCTCCAA | CAGTTCCGGTTCCGATTTTA | 171 |
| chr1:9673011-9675023 | p2 | (AT)6 | CTCCGATGCTAAACCTCCAA | CAGTTCCGGTTCCGATTTTA | 171 |
| chr1:9695976-9697992 | p2 | (TA)8 | GCTGCGGAATCAAGATTTTC | GGATCCACACTCGTCGAAAG | 193 |
| chr1:9853275-9855287 | p2 | (TA)6 | CTACTGCAGTGCCGATGTGT | ATGTCACCTTGGCTGGTCTC | 157 |
| chr1:10053775-10055789 | p2 | (TA)7 | TAGTTCTGTGAAGGCACCCC | AATGGAGCAATGCAACATCA | 252 |
| chr1:10067707-10069727 | p2 | (CT)10 | TTGCCAACAGTCAGATCGAG | GCAAAGAATGGCCAACAAAT | 289 |
| chr1:10131468-10133496 | p2 | (TA)14 | GGGCCAAAGATGTGGAGATA | AGGCTTGAAAGGGATTTGGT | 200 |
| chr1:10264146-10266170 | p6 | (CCGGTT)4 | CATTCTGCAATTCCCCATTC | GAGGCAACAATCCAACATCC | 288 |
| chr1:10264146-10266170 | p6 | (CCGGTT)4 | CATTCTGCAATTCCCCATTC | GAGGCAACAATCCAACATCC | 288 |
| chr1:10264146-10266170 | p6 | (CCGGTT)4 | CATTCTGCAATTCCCCATTC | GAGGCAACAATCCAACATCC | 288 |
| chr1:10264146-10266170 | p6 | (CCGGTT)4 | CATTCTGCAATTCCCCATTC | GAGGCAACAATCCAACATCC | 288 |
| chr1:10264146-10266170 | p6 | (CCGGTT)4 | CATTCTGCAATTCCCCATTC | GAGGCAACAATCCAACATCC | 288 |
| chr1:10264378-10266402 | p6 | (CCGGTT)4 | ATTGGGCAACGTGCTACTTC | TGCTGCAGACATGTGAGGTT | 200 |
| chr1:10264378-10266402 | p6 | (CCGGTT)4 | ATTGGGCAACGTGCTACTTC | TGCTGCAGACATGTGAGGTT | 200 |
| chr1:10264378-10266402 | p6 | (CCGGTT)4 | ATTGGGCAACGTGCTACTTC | TGCTGCAGACATGTGAGGTT | 200 |
| chr1:10264378-10266402 | p6 | (CCGGTT)4 | ATTGGGCAACGTGCTACTTC | TGCTGCAGACATGTGAGGTT | 200 |
| chr1:10264378-10266402 | p6 | (CCGGTT)4 | ATTGGGCAACGTGCTACTTC | TGCTGCAGACATGTGAGGTT | 200 |
| chr1:10264378-10266402 | p6 | (CCGGTT)4 | ATTGGGCAACGTGCTACTTC | TGCTGCAGACATGTGAGGTT | 200 |
| chr1:10320641-10322657 | p2 | (AT)8 | GAGTTTGGATGTGGTGGGTT | CCATGCGTATTCGTAGCAAA | 119 |
| chr1:10320641-10322657 | p2 | (TA)9 | GAGTTTGGATGTGGTGGGTT | CCATGCGTATTCGTAGCAAA | 119 |
| chr1:10396211-10398226 | p5 | (TTGGT)3 | AGTGGTTTGGTTTGACTCGG | GTAATGTCGGGTTGGTTTGG | 277 |
| chr1:10396251-10398266 | p5 | (TTGGT)3 | AGTGGTTTGGTTTGACTCGG | GTAATGTCGGGTTGGTTTGG | 237 |
| chr1:10417507-10419535 | p4 | (AAAG)7 | GAACCCGGCCCTCTAGTAAG | GGTTCTGTTATTGTGCGCCT | 178 |
| chr1:10431748-10433760 | p2 | (CT)6 | TGTTGGTTCTTGCCACTCTG | CAAATCTTGCTTCCAAAGCC | 194 |
| chr1:10449478-10451490 | p2 | (GA)6 | CGATCAATCCAGCTTTCGAT | TTGGTTGCAACACTCGGATA | 108 |
| chr1:10488574-10490612 | p2 | (CT)19 | GCGATTCCCATGTTTCAGAT | AAAACATGGCATGACGACAA | 276 |
| chr1:10622443-10624458 | p5 | (TTATT)3 | AACCCCCTGCTCAGAAAAAT | CGCTTAATAAGGTGGCCAAA | 292 |
| chr1:10622443-10624458 | p2 | (TA)9 | AACCCCCTGCTCAGAAAAAT | CGCTTAATAAGGTGGCCAAA | 292 |
| chr1:10623156-10625174 | p5 | (TTATT)3 | TGGTAGAGCCCTGTAATGCC | TTTCGAAACCGACCTATTGC | 226 |
| chr1:10623156-10625174 | p2 | (TA)9 | TGGTAGAGCCCTGTAATGCC | TTTCGAAACCGACCTATTGC | 226 |

| Marker ID | SSR type | SSR motif | FORWARD PRIMER (5'-3') | REVERSE PRIMER (5'-3') | size (bp) |
|---|---|---|---|---|---|
| chr1:10656049-10658088 | p2 | (GA)6 | GATTTGGACCTACAAGCGGA | GCGTGGTCGATTTAAGGAAA | 296 |
| chr1:10656049-10658088 | p3 | (ATA)13 | GATTTGGACCTACAAGCGGA | GCGTGGTCGATTTAAGGAAA | 296 |
| chr1:10656049-10658088 | p5 | (TTATT)3 | GATTTGGACCTACAAGCGGA | GCGTGGTCGATTTAAGGAAA | 296 |
| chr1:10786154-10788184 | p4 | (CAAT)4 | GGGGTCTGGAGAGGGTAGAG | CCCAAATGGAAGAGGATTCA | 125 |
| chr1:10786154-10788184 | p6 | (TCTCTT)5 | GGGGTCTGGAGAGGGTAGAG | CCCAAATGGAAGAGGATTCA | 125 |
| chr1:10855929-10857941 | p2 | (AG)6 | TCACAATCATATCGGCTGGA | CCATTGCCCTTCAATGAGAT | 117 |
| chr1:10969848-10971860 | p2 | (AT)9 | CAAAAAGGCAAGGGCAATAA | GATCACGAACTGCCCATTTT | 180 |
| chr1:11016893-11018911 | p6 | (GCAGTA)3 | TCTCCGCTGCCTTGTTATCT | TCAAGTTGCGGCAGTGATAG | 281 |
| chr1:11136696-11138726 | p5 | (AAAAT)3 | CGCGTAGACCTCAAATCCAT | CGTCTCCTTGTGCTCCTTTT | 144 |
| chr1:11136844-11138856 | p5 | (AAAAT)3 | AAAGGAGCACAAGGAGACGA | TTTTGATCCAAACCCAAACA | 196 |
| chr1:11136872-11138887 | p5 | (AAAAT)3 | AAAGGAGCACAAGGAGACGA | TTTTGATCCAAACCCAAACA | 168 |
| chr1:11136882-11138896 | p5 | (AAAAT)3 | AAAGGAGCACAAGGAGACGA | TTTTGATCCAAACCCAAACA | 158 |
| chr1:11200595-11202613 | p3 | (TGT)6 | GGAGGATCACGATCCATTTG | TCTTGTCCTCCACAGAGGCT | 217 |
| chr1:11271235-11273249 | p2 | (GA)7 | AATAATTTCGCAAACGCCAG | ATTGAATCCCCTGCCTTTCT | 134 |
| chr1:11271235-11273249 | p2 | (AC)7 | AATAATTTCGCAAACGCCAG | ATTGAATCCCCTGCCTTTCT | 134 |
| chr1:11272123-11274137 | p2 | (GA)7 | ATGAGCTTGTTTGGATTGGC | AGCCTGTTTGGATTGGCTTA | 248 |
| chr1:11272123-11274137 | p2 | (AC)7 | ATGAGCTTGTTTGGATTGGC | AGCCTGTTTGGATTGGCTTA | 248 |
| chr1:11392613-11394625 | p2 | (AT)6 | TGTGGCATGCCTGAATTTTA | GCATCGGGCTAGTCCATTTA | 177 |
| chr1:11393412-11395424 | p2 | (AT)6 | TCCACAATGACTTGGCTTGA | TTTGGGGGAAAATATTGCTG | 256 |
| chr1:11393430-11395474 | p2 | (AT)6 | TCCACAATGACTTGGCTTGA | TTTGGGGGAAAATATTGCTG | 238 |
| chr1:11393476-11395488 | p2 | (AT)6 | TCCACAATGACTTGGCTTGA | TTTGGGGGAAAATATTGCTG | 192 |
| chr1:11425325-11427351 | p2 | (AT)13 | CCTGGGGATCCTGAAAAGTT | GCAAATGATGCTTCCAGGTT | 176 |
| chr1:11425325-11427351 | p6 | (GATGAG)3 | CCTGGGGATCCTGAAAAGTT | GCAAATGATGCTTCCAGGTT | 176 |
| chr1:11518296-11520314 | p6 | (TGAAGG)3 | ACAGATTGGAGGGAGTGGTG | CATTCATGCTTGCTTCTCCA | 282 |
| chr1:11518296-11520314 | p3 | (TAA)5 | ACAGATTGGAGGGAGTGGTG | CATTCATGCTTGCTTCTCCA | 282 |
| chr1:11573397-11575411 | p2 | (AC)7 | TAGAGAGCCAAGAAAGGCCA | TCGAGAAGAAACTCGCTGGT | 291 |
| chr1:11619787-11621799 | p2 | (TA)6 | TATTTTTGGTTCCGACTCCG | CTGGGGAAACTGAAAAGCAG | 254 |
| chr1:11906081-11908105 | p3 | (AAT)8 | CCGAACTCGAAAAGAGTTGC | ACATGTGCATCCAATGTGCT | 267 |
| chr1:11942055-11944070 | p5 | (AAGAA)3 | CCATGTCTTCAGCACTCCAA | CCTTCCCCAACAGACCAGTA | 175 |
| chr1:11998916-12000998 | p2 | (TA)41 | TTGCTGTCCCAACATTTTCA | CACAGGTCCATTTCAAGCCT | 276 |
| chr1:12015055-12017067 | p2 | (TG)8 | CGTGGTCAACGAGGGATACT | ATTCGGTCGCTATCATTTGC | 243 |
| chr1:12015055-12017067 | p2 | (AT)6 | CGTGGTCAACGAGGGATACT | ATTCGGTCGCTATCATTTGC | 243 |
| chr1:12015055-12017067 | p3 | (GTT)8 | CGTGGTCAACGAGGGATACT | ATTCGGTCGCTATCATTTGC | 243 |
| chr1:12017270-12019290 | p5 | (GGGCC)3 | TCCCTGCTCTTCTTCTTCCA | CTTGTGGCCCACGTTAAAAT | 212 |
| chr1:12017270-12019290 | p2 | (TA)10 | TCCCTGCTCTTCTTCTTCCA | CTTGTGGCCCACGTTAAAAT | 212 |
| chr1:12119455-12121470 | p3 | (AAT)5 | GAGGAAGCATCACTTTTGGC | CAGACTGAATCTGTTGGCGA | 295 |

| Marker ID | SSR type | SSR motif | FORWARD PRIMER (5'-3') | REVERSE PRIMER (5'-3') | size (bp) |
|---|---|---|---|---|---|
| chr1:12155140-12157158 | p2 | (AG)9 | CCAGACTCCACTTATGCGGT | ATTTGCGAAAAGCCGATATG | 273 |
| chr1:12155140-12157158 | p3 | (ATC)5 | CCAGACTCCACTTATGCGGT | ATTTGCGAAAAGCCGATATG | 273 |
| chr1:12168942-12170958 | p3 | (ACA)5 | TGTGGAGGCAGAATTTGACA | AATCCAGATAATTCGCACCG | 284 |
| chr1:12168942-12170958 | p4 | (AAAT)4 | TGTGGAGGCAGAATTTGACA | AATCCAGATAATTCGCACCG | 284 |
| chr1:12229331-12231355 | p3 | (GTT)8 | GACAAAAAGGGAACGACCAA | ACGTTAACACAGGCTCCAGG | 192 |
| chr1:12291792-12293831 | p3 | (TAT)13 | GCAACAGCACAAGCAAAGAA | CGATCTCCACTCCGTCATTT | 253 |
| chr1:12305428-12307440 | p2 | (AG)6 | TTTTCCAGGCACCCAGATAG | GAAGGAAACTGCTGACCTGC | 266 |
| chr1:12313191-12315211 | p2 | (AC)10 | AAAAGCCACAGCCTTTAGCA | GCACTGAAGAATCACCCGAT | 184 |
| chr1:12364912-12366926 | p2 | (AT)7 | CAAGGGAGTACAAGGGTGGA | AGATTGCTGGTGTGCAAGAA | 174 |
| chr1:12379497-12381509 | p2 | (TA)8 | TCATAACTTTGCCCTCGGAC | ATGCCAATTTGGGGAAAAAT | 268 |
| chr1:12379497-12381509 | p2 | (TA)7 | TCATAACTTTGCCCTCGGAC | ATGCCAATTTGGGGAAAAAT | 268 |
| chr1:12379523-12381547 | p2 | (TA)7 | TCATAACTTTGCCCTCGGAC | ATGCCAATTTGGGGAAAAAT | 242 |
| chr1:12379643-12381657 | p2 | (TA)7 | TCATAACTTTGCCCTCGGAC | ATGCCAATTTGGGGAAAAAT | 122 |
| chr1:12385605-12387635 | p2 | (TA)14 | GAAGAACGTGCTACACCCGT | TCCAGTTCCTTGATATCCGC | 140 |
| chr1:12385893-12387921 | p2 | (TA)14 | AAGTATCAACCGTGGCTTCG | CCCTCCTCCGCTTCTAAGTT | 110 |
| chr1:12407595-12409611 | p2 | (TA)8 | GCATTTAGCCGAGGAAGACA | TCTATGTTTTGGTTGCGTCG | 150 |
| chr1:12448764-12450779 | p5 | (AATAA)3 | GTTTGGAAAGCCACCTGGTA | ACACTTGTGCAACCAAACGA | 195 |
| chr1:12463267-12465279 | p2 | (AT)8 | GGAGGTGAGCCACCATTAGA | TTGCCCATAAACCCCATAAA | 298 |
| chr1:12463267-12465279 | p2 | (TA)8 | GGAGGTGAGCCACCATTAGA | TTGCCCATAAACCCCATAAA | 298 |
| chr1:12511756-12513774 | p6 | (ATTTTT)3 | CAGGCTGTCCACATTTAGCA | TCTGAATGGATCAAGAGGGG | 225 |
| chr1:12545439-12547455 | p4 | (TAAA)4 | CAAGCTGGGGCTATGAAAAA | GCAAAAGCTTCAAGAATGGC | 176 |
| chr1:12670982-12673000 | p3 | (TTG)6 | ATGCGGAAGAATATCGTTGG | TGGGGTTATGGGTATGGCTA | 239 |
| chr1:12670982-12673000 | p6 | (CCATGA)3 | ATGCGGAAGAATATCGTTGG | TGGGGTTATGGGTATGGCTA | 239 |
| chr1:12689001-12691027 | p2 | (AT)7 | GTCCGTTTTACGCGTTTTGT | CCCCGAGATTTTTACGTTGA | 278 |
| chr1:12689001-12691027 | p2 | (TA)13 | GTCCGTTTTACGCGTTTTGT | CCCCGAGATTTTTACGTTGA | 278 |
| chr1:12689152-12691174 | p2 | (TA)13 | GTCCGTTTTACGCGTTTTGT | CCCCGAGATTTTTACGTTGA | 127 |
| chr1:12701090-12703105 | p3 | (TGT)5 | CAAAGCAAGAACCAACAGCA | GGTTGTCTTCATTGGCGTCT | 133 |
| chr1:12740186-12742212 | p2 | (AT)7 | GTCCGTTTTACGCGTTTTGT | CCCCGAGATTTTTACGTTGA | 278 |
| chr1:12740186-12742212 | p2 | (TA)13 | GTCCGTTTTACGCGTTTTGT | CCCCGAGATTTTTACGTTGA | 278 |
| chr1:12740337-12742359 | p2 | (TA)13 | GTCCGTTTTACGCGTTTTGT | CCCCGAGATTTTTACGTTGA | 127 |
| chr1:12752275-12754290 | p3 | (TGT)5 | CAAAGCAAGAACCAACAGCA | GGTTGTCTTCATTGGCGTCT | 133 |
| chr1:12781263-12783283 | p2 | (TA)10 | AGGCATGAGTCCATCCAAAC | TGGGCTGTTTAATATTGGGC | 170 |
| chr1:12815561-12817576 | p3 | (ACA)5 | ACCATCAGGGATTGTTTCCA | CAAAAGGGGTGGTTCTTCAA | 120 |
| chr1:12866976-12868992 | p2 | (TG)8 | TAGAAGATGACCAATCCCGC | CATCACTACGGTCGGGCTAT | 176 |
| chr1:12897132-12899164 | p2 | (AT)16 | AGTCGTATGGGACACAAGCC | TGTACGGAGCACTCCACTTG | 236 |
| chr1:12953608-12955624 | p2 | (AT)8 | CATCCTAGGAAGGGTGGGTT | TAGGAGCAGCAAGGGAGTGT | 112 |

| Marker ID | SSR type | SSR motif | FORWARD PRIMER (5'-3') | REVERSE PRIMER (5'-3') | size (bp) |
|---|---|---|---|---|---|
| chr1:12953608-12955624 | p2 | (AT)7 | CATCCTAGGAAGGGTGGGTT | TAGGAGCAGCAAGGGAGTGT | 112 |
| chr1:12953608-12955624 | p5 | (TTTAT)4 | CATCCTAGGAAGGGTGGGTT | TAGGAGCAGCAAGGGAGTGT | 112 |
| chr1:13010474-13012492 | p2 | (AC)9 | CAAAAACGCACAGCAGAAAA | CACCGTCGAATGAAACACAG | 237 |
| chr1:13012764-13014779 | p5 | (TTGGT)3 | AGTAAACCGACAAACCGCAC | CAAACCAATTATGGTCGGGT | 191 |
| chr1:13012804-13014819 | p5 | (TTGGT)3 | AGTAAACCGACAAACCGCAC | CAAACCAATTATGGTCGGGT | 151 |
| chr1:13042896-13044911 | p5 | (CCCCA)3 | TCTCGTGTGCAAACCAACTT | ATGCTAACTTGCCAAGAGGG | 209 |
| chr1:13148439-13150451 | p2 | (TA)6 | TGAATACACCGCATTTCTCG | CAACTCCACCAATCATGCAC | 240 |
| chr1:13148439-13150451 | p2 | (TA)11 | TGAATACACCGCATTTCTCG | CAACTCCACCAATCATGCAC | 240 |
| chr1:13154659-13156671 | p2 | (GC)6 | TTGGCCTACCCCTACCTCTT | AAGACAAGATGCGGGAAATG | 229 |
| chr1:13338807-13340819 | p2 | (AT)6 | TTTGGTGTGCGACAATTCAT | TTTTTCGTGGGCATTAAAGG | 241 |
| chr1:13343004-13345022 | p6 | (TGGAGT)3 | CACACCCTTTGCCTCTCTTC | AGTCAATGTCCGTGTGGTGA | 284 |
| chr1:13343004-13345022 | p2 | (TA)9 | CACACCCTTTGCCTCTCTTC | AGTCAATGTCCGTGTGGTGA | 284 |
| chr1:13343004-13345022 | p2 | (TA)21 | CACACCCTTTGCCTCTCTTC | AGTCAATGTCCGTGTGGTGA | 284 |
| chr1:13359313-13361373 | p2 | (AT)30 | CGCCTTCCTAATATCCAGCA | AATCAGCAAATGACAAGGGG | 209 |
| chr1:13464092-13466124 | p2 | (TA)16 | TATGGCATCTTTGGGAAAGG | AACGTTGCCTCCGAACATAC | 245 |
| chr1:13477944-13479959 | p3 | (ATA)5 | TTCAGAGGATTTTTGGTCCG | CTTTCCCTCCTTCTTCTCCG | 117 |
| chr1:13578772-13580786 | p2 | (TA)7 | TCGAATAGGGCATCTTGCTT | CTCGTTTGTCTTCCGTGTGA | 231 |
| chr1:13836352-13838372 | p5 | (TTGGT)4 | CCGATAACGGTGCAAAAAGT | CCGATAACGGTGCAAAAAGT | 231 |
| chr1:13836352-13838372 | p5 | (TTGGT)3 | CCGATAACGGTGCAAAAAGT | CCGATAACGGTGCAAAAAGT | 231 |
| chr1:13852670-13854716 | p2 | (TG)7 | AAACACAGCCGACAATGACA | CCTCATTTCCCTTCCCCTTA | 232 |
| chr1:13852670-13854716 | p2 | (AT)23 | AAACACAGCCGACAATGACA | CCTCATTTCCCTTCCCCTTA | 232 |
| chr1:13852670-13854716 | p2 | (TG)7 | AAACACAGCCGACAATGACA | CCTCATTTCCCTTCCCCTTA | 232 |
| chr1:13852670-13854716 | p2 | (AT)24 | AAACACAGCCGACAATGACA | CCTCATTTCCCTTCCCCTTA | 232 |
| chr1:13859992-13862007 | p5 | (CATAG)3 | TGAGAGAGGAGTGGGATTGG | AAAACGACAGACAAGGGGTG | 168 |
| chr1:13890674-13892686 | p2 | (AT)7 | GTTTCGGAGAGCGCATTTAG | CTCTTCCTCTTCGCCTTCCT | 259 |
| chr1:13890674-13892686 | p5 | (TCCGG)3 | GTTTCGGAGAGCGCATTTAG | CTCTTCCTCTTCGCCTTCCT | 259 |
| chr1:13917264-13919279 | p5 | (TTGGT)3 | GGTACCCCTAAAAGTCCCCA | TTGGCCGAAGGATAGCTAAA | 240 |
| chr1:13917264-13919279 | p5 | (TTGGT)3 | GGTACCCCTAAAAGTCCCCA | TTGGCCGAAGGATAGCTAAA | 240 |
| chr1:14020669-14022725 | p2 | (AT)28 | CCATCTAGCTGAGTTTGGGC | CATAAACAGCAGGCTCACCA | 265 |
| chr1:14020669-14022725 | p2 | (AT)7 | CCATCTAGCTGAGTTTGGGC | CATAAACAGCAGGCTCACCA | 265 |
| chr1:14064174-14066198 | p2 | (AT)12 | ACCAGGGAGTTTGCAATCAG | GTTTTGCTGGGATCGTCAAT | 143 |
| chr1:14082871-14084889 | p6 | (GTGGGC)3 | CAGTCCGTCGATTTTTGCTT | TTTCTCTGCACAGTGTTGGC | 108 |
| chr1:14197577-14199589 | p2 | (AT)6 | GCCAATGATTTTCCTCCAGA | CAAGTCTTTGGGGTGATCGT | 175 |
| chr1:14306912-14308947 | p2 | (TG)8 | AAATAAAGGGGTGAGTGGGG | TTTTGCTACAGTGACACGGC | 210 |
| chr1:14445146-14447161 | p5 | (TTGTT)3 | AAAAAGGGCAATTGATGCAG | CCCAATTCCAATGTAGGGTG | 184 |
| chr1:14543346-14545361 | p3 | (GTT)5 | CTTCTTAAACCTACCCGCCC | GGTATGCCATTTCCCATGTC | 174 |

| Marker ID | SSR type | SSR motif | FORWARD PRIMER (5'-3') | REVERSE PRIMER (5'-3') | size (bp) |
|---|---|---|---|---|---|
| chr1:14543346-14545361 | p3 | (ACA)7 | CTTCTTAAACCTACCCGCCC | GGTATGCCATTTCCCATGTC | 174 |
| chr1:14592533-14594561 | p2 | (TA)6 | TCAAGTCCAAAAGCGTGATG | TCGTTAGCGTGAAGGGGATA | 298 |
| chr1:14592533-14594561 | p2 | (AT)14 | TCAAGTCCAAAAGCGTGATG | TCGTTAGCGTGAAGGGGATA | 298 |
| chr1:14628732-14630748 | p4 | (AATA)4 | GGTGGAGGGGACAGAGTCTT | TATGGCCAAACACAACTCCA | 227 |
| chr1:14633414-14635468 | p3 | (TAT)5 | TGGTTAGCCAATCCCTTGAG | TGCGTGCAATTACACCTGTT | 118 |
| chr1:14655773-14657787 | p2 | (AT)7 | ACCAGCTACCCCTATCCTGC | CAGGACGAGAATTTTGGGAA | 113 |
| chr1:14684427-14686442 | p5 | (AAGAA)3 | ATGGTGGCAAGAATGAGACC | CCCACCATGTAAGTGGAACC | 180 |
| chr1:14694892-14696910 | p6 | (TTTTAT)3 | AATTTTTCGTGGCAAAAAGC | CACTGCGGCAAATAATCCTT | 114 |
| chr1:14713850-14715865 | p5 | (AAAAT)3 | AATTTTCCATCCCAAGTCCC | GTTGACATGCTTCTCGGGTT | 180 |
| chr1:14713850-14715865 | p3 | (CTT)17 | AATTTTCCATCCCAAGTCCC | GTTGACATGCTTCTCGGGTT | 180 |
| chr1:14720320-14722335 | p5 | (CAATA)3 | CGATGTCCGAAGATCCAGTT | GGAAGGGTTCTCCTACCGAG | 164 |
| chr1:14720320-14722335 | p2 | (AT)14 | CGATGTCCGAAGATCCAGTT | GGAAGGGTTCTCCTACCGAG | 164 |
| chr1:14826382-14828402 | p4 | (TTTC)4 | AACCACCCGAGAACTTGTTG | TGCAGTGTTCGGCTTGTTAG | 208 |
| chr1:14826382-14828402 | p2 | (CT)6 | AACCACCCGAGAACTTGTTG | TGCAGTGTTCGGCTTGTTAG | 208 |
| chr1:14826382-14828402 | p2 | (TC)10 | AACCACCCGAGAACTTGTTG | TGCAGTGTTCGGCTTGTTAG | 208 |
| chr1:14899804-14901819 | p2 | (AC)7 | TAATCATTGTCGGCCTGTGA | AGATGTCACCCTCGCGTATC | 152 |
| chr1:14899804-14901819 | p3 | (TGC)5 | TAATCATTGTCGGCCTGTGA | AGATGTCACCCTCGCGTATC | 152 |
| chr1:14921667-14923700 | p5 | (ACCCC)3 | GCCCAAATATACCCCTCGTT | TGAATTGGGCTCCACCTTAG | 192 |
| chr1:15171099-15173114 | p3 | (AGT)5 | GGTTTCCCTTCGTTATGCAA | ACTGAAACTCCGGCAACATC | 276 |
| chr1:15175216-15177231 | p5 | (ATTTT)3 | CCCTAATCAAAGGGAACCGT | TCCGAATAATCCGGTAGCAC | 101 |
| chr1:15238308-15240320 | p2 | (AG)6 | ACCGGGCACTCAAGAATATG | GCTACCAACCGCGGTATAAG | 210 |
| chr1:15319840-15321866 | p3 | (TTA)9 | TGGATGAAACCCATGGAAGT | GGCCGGATCAGATATTGAGA | 195 |
| chr1:15319840-15321866 | p2 | (TA)13 | TGGATGAAACCCATGGAAGT | GGCCGGATCAGATATTGAGA | 195 |
| chr1:15328517-15330533 | p4 | (TATT)4 | TCTTTTGCTATTCCCCTCCA | AAGTTGGGACGTTCCTTGAA | 287 |
| chr1:15364807-15366822 | p3 | (TTG)5 | TGCAGGATGGATGAAATGAA | ATTCGCATTTCCACGACTTC | 139 |
| chr1:15372177-15374199 | p6 | (ATTTTT)3 | TAAATGGGTACCGTTCCGAG | TTGCTTATGCTTTTCCCCAG | 121 |
| chr1:15372177-15374199 | p2 | (AT)11 | TAAATGGGTACCGTTCCGAG | TTGCTTATGCTTTTCCCCAG | 121 |
| chr1:15427382-15429396 | p2 | (AT)7 | TCCAGCACGCAATAGAACAC | GCGCACTCTGAAATAGGAGC | 287 |
| chr1:15472838-15474856 | p3 | (TAT)5 | ATTTTGTTTCCCTTTTGGGG | ATCGAATGAAACGACACCGT | 194 |
| chr1:15472838-15474856 | p3 | (TCT)5 | ATTTTGTTTCCCTTTTGGGG | ATCGAATGAAACGACACCGT | 194 |
| chr1:15472838-15474856 | p6 | (CGGGTT)3 | ATTTTGTTTCCCTTTTGGGG | ATCGAATGAAACGACACCGT | 194 |
| chr1:15483439-15485467 | p2 | (AC)7 | ATGGTGGAAGAAATGCAAGG | GTACTGGCAAGCTAAAGCCG | 181 |
| chr1:15483439-15485467 | p3 | (ACA)5 | ATGGTGGAAGAAATGCAAGG | GTACTGGCAAGCTAAAGCCG | 181 |
| chr1:15483439-15485467 | p2 | (AT)18 | ATGGTGGAAGAAATGCAAGG | GTACTGGCAAGCTAAAGCCG | 181 |
| chr1:15484374-15486410 | p3 | (ACA)5 | GTGAGGCTCGTGGTAAGCTC | CTGGTGGGACTACAATGGGT | 257 |
| chr1:15484498-15486522 | p3 | (ACA)5 | GTGAGGCTCGTGGTAAGCTC | CTGGTGGGACTACAATGGGT | 133 |

| Marker ID | SSR type | SSR motif | FORWARD PRIMER (5'-3') | REVERSE PRIMER (5'-3') | size (bp) |
|---|---|---|---|---|---|
| chr1:15526135-15528157 | p2 | (AC)11 | ATGGAAACCTGTTCCACGAG | AGCCTATTCGCCTTTTGTGA | 243 |
| chr1:15533923-15535945 | p2 | (CT)11 | TAGATCGCACAAAGCACGTC | AAGAATTGGGACATTGTGGC | 151 |
| chr1:15594001-15596029 | p5 | (ATAAA)3 | GTGTAGCCACGACGACAATG | ATTTTATGAGGCCCACAACG | 110 |
| chr1:15594001-15596029 | p2 | (AT)14 | GTGTAGCCACGACGACAATG | ATTTTATGAGGCCCACAACG | 110 |
| chr1:15629951-15632011 | p3 | (AAT)20 | CTCGAGTTTCCAAGGTGAGC | AACCCTCATCCCAACAATGA | 159 |
| chr1:15666474-15668489 | p3 | (TGT)5 | GCTCTTGGGGAGTAGCACTG | CCAGAGGCATTGACAAAGGT | 141 |
| chr1:15703020-15705045 | p3 | (TTA)5 | TGGTCATCGGAGGTGTCATA | GATCGCCACATTAATGGCTT | 272 |
| chr1:15703020-15705045 | p2 | (AT)6 | TGGTCATCGGAGGTGTCATA | GATCGCCACATTAATGGCTT | 272 |
| chr1:15746532-15748548 | p2 | (TA)8 | AATTTTTCCCCTTCGGCTAA | TATGTCAAGTGGGGGAGGAG | 232 |
| chr1:15746532-15748548 | p2 | (TA)17 | AATTTTTCCCCTTCGGCTAA | TATGTCAAGTGGGGGAGGAG | 232 |
| chr1:15750769-15752791 | p2 | (AT)11 | CCTGTCCATGAAAAATGCCT | GGCAATAAATTCTGGGACGA | 145 |
| chr1:15775150-15777184 | p2 | (AC)6 | TTCTTTTTGGGCAAAGATGC | CAGTTTGTAACGGCAAGGGT | 184 |
| chr1:15875025-15877037 | p2 | (AT)6 | ATTTGGGGGAAAAAGACCAC | CAGGCAAAGTGTCCAAGTCA | 172 |
| chr1:16015177-16017189 | p2 | (AT)6 | GGAGGAGCATACTTTGCCTG | CAGAGGCCCATGGAATCTAA | 273 |
| chr1:16111086-16113098 | p2 | (TA)6 | GATGTTTCGGCCTCACTGTT | CAAACCTTTGGAAAGCAAGC | 103 |
| chr1:16218264-16220296 | p5 | (AAACC)3 | CGGATCTGGTGTTCACCTTT | AGGGCCAAATATACCCTTCG | 193 |
| chr1:16218264-16220296 | p4 | (AAAT)8 | CGGATCTGGTGTTCACCTTT | AGGGCCAAATATACCCTTCG | 193 |
| chr1:16218264-16220296 | p2 | (TA)32 | CGGATCTGGTGTTCACCTTT | AGGGCCAAATATACCCTTCG | 193 |
| chr1:16244129-16246141 | p2 | (TA)6 | ACAGCTGGGTGTTTTGCTTT | TTTGCTGACTTGCTGTGTCC | 121 |
| chr1:16360988-16363002 | p2 | (TA)7 | GGATGGAGGACCCTATCCAT | AGCAAAACCTTTCCCTTGCT | 193 |
| chr1:16365939-16367951 | p2 | (TA)6 | AAGTGGAACCCACCACATCT | ATATGGGGCCCACAATTTTT | 259 |
| chr1:16370990-16373002 | p5 | (AAGAG)3 | GAGAATCGGGGAATGTTTGA | GTTTCTACCCAGCCCTTTCC | 248 |
| chr1:16370990-16373002 | p2 | (AT)6 | GAGAATCGGGGAATGTTTGA | GTTTCTACCCAGCCCTTTCC | 248 |
| chr1:16370990-16373002 | p2 | (AT)11 | GAGAATCGGGGAATGTTTGA | GTTTCTACCCAGCCCTTTCC | 248 |
| chr1:16372886-16374920 | p2 | (AT)11 | CCCAGTGGAAGCAAATGAAA | ATGACATGGGGGAATAGCAA | 224 |
| chr1:16372966-16374980 | p2 | (TA)10 | CCCAGTGGAAGCAAATGAAA | ATGACATGGGGGAATAGCAA | 144 |
| chr1:16392134-16394152 | p3 | (TCA)5 | TTTCTTCATTAAGCGCACCC | GGAAATTAATCCGGCCATTT | 273 |
| chr1:16392134-16394152 | p6 | (CCGGTT)3 | TTTCTTCATTAAGCGCACCC | GGAAATTAATCCGGCCATTT | 273 |
| chr1:16431294-16433316 | p2 | (AT)11 | GGGCCAATTATCCCATTTTT | ATGATATGCCCTCAGATCGC | 161 |
| chr1:16458457-16460473 | p4 | (AAAG)4 | GACGGGACACAAACGAAAAT | AAGGCATGTGGAGGAATGAG | 179 |
| chr1:16458457-16460473 | p5 | (AAGGC)3 | GACGGGACACAAACGAAAAT | AAGGCATGTGGAGGAATGAG | 179 |
| chr1:16462013-16464028 | p5 | (GTTTT)3 | TTGGGACTTGGCTACCAAAC | TCCAACTTGCCTGTTAAGGG | 242 |
| chr1:16495531-16497545 | p2 | (CT)7 | CACCACAAGATTGGCTTCCT | AATTATTTTGGACCGGAGGG | 170 |
| chr1:16529702-16531717 | p3 | (TGT)5 | TGGTCACATGGCAACAACTT | TTGCCCAAAATGGAGAAGTC | 146 |
| chr1:16539070-16541086 | p4 | (TTTA)4 | GGGGCAAAGGAAACTTAACC | GGTGTTTCTGAAAAGGGCAA | 189 |
| chr1:16659325-16661337 | p2 | (TG)6 | CTCCGGACTTCCCATAACAA | TTTGACAGAAGCAAAAGCCA | 271 |

| Marker ID | SSR type | SSR motif | FORWARD PRIMER (5'-3') | REVERSE PRIMER (5'-3') | size (bp) |
|---|---|---|---|---|---|
| chr1:16659325-16661337 | p2 | (AT)9 | CTCCGGACTTCCCATAACAA | TTTGACAGAAGCAAAAGCCA | 271 |
| chr1:16757370-16759385 | p5 | (AAATA)3 | TTGCTGCATTTCAAACTTGG | TATTGAACCCCATCCCGATA | 208 |
| chr1:16757370-16759385 | p5 | (ATAAA)3 | TTGCTGCATTTCAAACTTGG | TATTGAACCCCATCCCGATA | 208 |
| chr1:16949096-16951114 | p3 | (TGT)6 | CCACCAAAACTTAAGGGCAA | CTAATCCTGTCACGCCTGGT | 231 |
| chr1:17027803-17029823 | p5 | (TTCTC)4 | TGAAACAAATCTTGCCACCA | AACCGTCCTTCGATTCTCCT | 242 |
| chr1:17043746-17045770 | p3 | (GAT)8 | GCGCTTTATTTGGCACTTTC | ATTTGTCCAGGGGTTAAGGG | 140 |
| chr1:17047221-17049263 | p2 | (TA)21 | TTGCCAAGGTTCATTTCTCC | CTTTGCTTTTTGCTTTTGGG | 196 |
| chr1:17080294-17082308 | p2 | (TA)7 | AAACCCCAAATCCAAGGAAC | CCGAAAACCCAAAGCTCATA | 163 |
| chr1:17086092-17088107 | p5 | (TTTCC)3 | ATATTTGGAGGTCAAGGGGG | TCAATAGCACGCAACTCTGC | 271 |
| chr1:17111143-17113158 | p2 | (TA)7 | CACCCTCTGCCAAGGATAAC | AGTTATGCAGGGTCGTTTGG | 229 |
| chr1:17111143-17113158 | p3 | (AAT)5 | CACCCTCTGCCAAGGATAAC | AGTTATGCAGGGTCGTTTGG | 229 |
| chr1:17111143-17113158 | p3 | (TAA)5 | CACCCTCTGCCAAGGATAAC | AGTTATGCAGGGTCGTTTGG | 229 |
| chr1:17111143-17113158 | p3 | (TGA)6 | CACCCTCTGCCAAGGATAAC | AGTTATGCAGGGTCGTTTGG | 229 |
| chr1:17111269-17113284 | p2 | (TA)7 | CACCCTCTGCCAAGGATAAC | AGTTATGCAGGGTCGTTTGG | 103 |
| chr1:17111269-17113284 | p3 | (AAT)5 | CACCCTCTGCCAAGGATAAC | AGTTATGCAGGGTCGTTTGG | 103 |
| chr1:17111269-17113284 | p3 | (TAA)5 | CACCCTCTGCCAAGGATAAC | AGTTATGCAGGGTCGTTTGG | 103 |
| chr1:17111269-17113284 | p3 | (TGA)6 | CACCCTCTGCCAAGGATAAC | AGTTATGCAGGGTCGTTTGG | 103 |
| chr1:17298298-17300312 | p2 | (GA)7 | ACTCAGGGAGGAGCAACTGA | GAGCAGGATCCCATTGCTTA | 100 |
| chr1:17323847-17325865 | p2 | (AT)9 | ACAAAAGCCTTGCCTTGAGA | CGGCCATACAGAGCATTTTT | 176 |
| chr1:17345307-17347322 | p5 | (AAAAT)3 | CTCTTGAGGGGAGTATTGCG | TTGCCATTGCCAAGTTTGTA | 277 |
| chr1:17345307-17347322 | p5 | (ACCGA)3 | CTCTTGAGGGGAGTATTGCG | TTGCCATTGCCAAGTTTGTA | 277 |
| chr1:17530307-17532323 | p2 | (TA)8 | CATGTTGGGAGAGAAACGGT | ACCTTGGTGGTCCACTTCTG | 182 |
| chr1:17530307-17532323 | p2 | (AT)9 | CATGTTGGGAGAGAAACGGT | ACCTTGGTGGTCCACTTCTG | 182 |
| chr1:17563195-17565215 | p2 | (AC)10 | TATCAGGCCTTAGCCCGTAA | CAAGTCAGAAGGGGGAACAA | 213 |
| chr1:17604156-17606168 | p2 | (GA)6 | GAGTACACTGCTGGGGCAAT | CTGTTGGGATCCTCCTGTGT | 127 |
| chr1:17604156-17606168 | p5 | (CAAAA)3 | GAGTACACTGCTGGGGCAAT | CTGTTGGGATCCTCCTGTGT | 127 |
| chr1:17807566-17809581 | p5 | (TTTTC)3 | GAGAAATGGCGGTGAGAGAG | GTCTCAATGAAGGCAACCGT | 259 |
| chr1:17908402-17910422 | p2 | (AT)6 | TCCCACCACTTCATCCAAAT | GCTAAGGATGTGGACGGGTA | 140 |
| chr1:17908562-17910574 | p2 | (AT)6 | TACCCGTCCACATCCTTAGC | TAGAGGGGGTAAGTGGGGTC | 212 |
| chr1:17940445-17942460 | p5 | (CAAAC)3 | AGTTGCAGCAAATGCTGATG | GTTTTAGGACGGAGGGGGTA | 270 |
| chr1:17989469-17991484 | p3 | (TTA)5 | CCCTTCTCAATGATGGGCTA | TGCTGCTGTTATGGGATTTG | 236 |
| chr1:18008962-18010977 | p5 | (AAGGA)3 | CGCTATAACGGTGGACAGGT | CCCTAGATGCTTGTTGGGAA | 297 |
| chr1:18052181-18054211 | p2 | (GA)15 | AAAGGAAAATCAACGGACCC | CGGTGTTGTCCACATTCAAC | 160 |
| chr1:18052181-18054211 | p4 | (TAGT)4 | AAAGGAAAATCAACGGACCC | CGGTGTTGTCCACATTCAAC | 160 |
| chr1:18054155-18056170 | p3 | (TTA)6 | TCAGATGTTCAGGCTGATGC | CATGCATTCCTCATTTGCTG | 268 |
| chr1:18054328-18056346 | p3 | (TTA)6 | GCAAGAGCTGAGATTCCAGG | CATGCATTCCTCATTTGCTG | 113 |

| Marker ID | SSR type | SSR motif | FORWARD PRIMER (5'-3') | REVERSE PRIMER (5'-3') | size (bp) |
|---|---|---|---|---|---|
| chr1:18115027-18117051 | p2 | (AT)12 | ACCAGACTCCACGTACCCAC | CGTACCTGCATCGTGAAATG | 276 |
| chr1:18180821-18182836 | p3 | (ACA)5 | AAAGCAAAGAGGGAGAAGCC | GCTCTTACACGTGGCAAACA | 200 |
| chr1:18256747-18258771 | p3 | (TTG)8 | CAATCGTGGTCTCGGAAAGT | AATTAGTCCGCCTCCTTGCT | 296 |
| chr1:18256747-18258771 | p2 | (TA)8 | CAATCGTGGTCTCGGAAAGT | AATTAGTCCGCCTCCTTGCT | 296 |
| chr1:18256886-18258902 | p3 | (TTG)8 | CAATCGTGGTCTCGGAAAGT | AATTAGTCCGCCTCCTTGCT | 157 |
| chr1:18256886-18258902 | p2 | (TA)8 | CAATCGTGGTCTCGGAAAGT | AATTAGTCCGCCTCCTTGCT | 157 |
| chr1:18256886-18258902 | p5 | (AAAAT)4 | CAATCGTGGTCTCGGAAAGT | AATTAGTCCGCCTCCTTGCT | 157 |
| chr1:18278152-18280170 | p2 | (CA)7 | CCCAGAATTTAACTCGGCAA | TCGAATGGCTTAAAATTCCG | 174 |
| chr1:18353300-18355312 | p2 | (AT)6 | CGTGCACTTGCCAAAGTTTA | CCCAGTGTCATTCTTGCTGA | 260 |
| chr1:18415021-18417033 | p2 | (AT)6 | CGTGCACTTGCCAAAGTTTA | CCCAGTGTCATTCTTGCTGA | 260 |
| chr1:18611244-18613276 | p2 | (AT)16 | GCGGAACACGAATCAAAGAT | TGGGGATTTATGGAAGGTGA | 112 |
| chr1:18658474-18660486 | p2 | (TA)6 | ACGCAACCATCACAAACGTA | GTACTCGCTTAATCCGAGCG | 248 |
| chr1:18804655-18806682 | p2 | (TA)7 | CCGTAGGATGAGAATCGCTC | CAGTCTTTGCCAGGGAACAT | 298 |
| chr1:18895315-18897331 | p2 | (TA)8 | AGATCTATGCGGACAAACCG | CCGCAGAAGTGAAATTGACA | 229 |
| chr1:18895315-18897331 | p2 | (TA)10 | AGATCTATGCGGACAAACCG | CCGCAGAAGTGAAATTGACA | 229 |
| chr1:18895590-18897602 | p2 | (TA)8 | TGTCAATTTCACTTCTGCGG | CGTGTGTCAAGACTCCGAAA | 129 |
| chr1:18895590-18897602 | p2 | (TA)10 | TGTCAATTTCACTTCTGCGG | CGTGTGTCAAGACTCCGAAA | 129 |
| chr1:18898121-18900159 | p2 | (TA)11 | ACAAACTTACTTCGGGCACC | CGTGTGTCAAGACTCCGAAA | 226 |
| chr1:18898121-18900159 | p2 | (AT)6 | ACAAACTTACTTCGGGCACC | CGTGTGTCAAGACTCCGAAA | 226 |
| chr1:18898121-18900159 | p2 | (AT)19 | ACAAACTTACTTCGGGCACC | CGTGTGTCAAGACTCCGAAA | 226 |
| chr1:18898121-18900159 | p2 | (TA)6 | ACAAACTTACTTCGGGCACC | CGTGTGTCAAGACTCCGAAA | 226 |
| chr1:18900367-18902381 | p2 | (TA)9 | TGTCAATTTCACTTCTGCGG | CGTGTGTCAAGACTCCGAAA | 300 |
| chr1:18900367-18902381 | p2 | (AT)7 | TGTCAATTTCACTTCTGCGG | CGTGTGTCAAGACTCCGAAA | 300 |
| chr1:18900367-18902381 | p2 | (TA)8 | TGTCAATTTCACTTCTGCGG | CGTGTGTCAAGACTCCGAAA | 300 |
| chr1:18900382-18902418 | p2 | (TA)9 | TGTCAATTTCACTTCTGCGG | CGTGTGTCAAGACTCCGAAA | 285 |
| chr1:18900382-18902418 | p2 | (AT)7 | TGTCAATTTCACTTCTGCGG | CGTGTGTCAAGACTCCGAAA | 285 |
| chr1:18900382-18902418 | p2 | (TA)8 | TGTCAATTTCACTTCTGCGG | CGTGTGTCAAGACTCCGAAA | 285 |
| chr1:18962993-18965019 | p2 | (AT)13 | ACCCACCATCTCCAAACTCA | TTGGAGTCGACCCTTTATGG | 145 |
| chr1:19124934-19126949 | p3 | (GAA)5 | GCTAGTTCCTCCAAGGGGAC | GCCATACCATTTTTCGCACT | 105 |
| chr1:19169670-19171684 | p2 | (AT)7 | TTTTGGTTGCAACTTGGTCA | AGAAGTCTCTCCCATTCCCC | 159 |
| chr1:19185923-19187937 | p2 | (AT)7 | TTTTGGTTGCAACTTGGTCA | AGAAGTCTCTCCCATTCCCC | 159 |
| chr1:19250748-19252780 | p2 | (TA)16 | TTTGAGCTGTTGCTTCGTTG | AAACATAACAGCCCAGTGGC | 268 |
| chr1:19287925-19289940 | p5 | (TTGGT)3 | ACTCCCTGATTTTCGGTGTG | CTTCTTGGTTGCCTCAGCTC | 163 |
| chr1:19320084-19322108 | p2 | (AT)6 | TTTTTCCCCATTGAGTTTGG | GTCGTCCGTTGTCCCTTAAA | 300 |
| chr1:19320121-19322141 | p2 | (AT)6 | TTTTTCCCCATTGAGTTTGG | GTCGTCCGTTGTCCCTTAAA | 263 |
| chr1:19375079-19377097 | p2 | (AT)8 | CAAGGGGTGTCAATCGAATC | GGGATTGGATGAAAAAGGGT | 243 |

| Marker ID | SSR type | SSR motif | FORWARD PRIMER (5'-3') | REVERSE PRIMER (5'-3') | size (bp) |
|---|---|---|---|---|---|
| chr1:19375141-19377161 | p2 | (AT)8 | CAAGGGGTGTCAATCGAATC | GGGATTGGATGAAAAAGGGT | 181 |
| chr1:19375201-19377219 | p2 | (AT)8 | CAAGGGGTGTCAATCGAATC | GGGATTGGATGAAAAAGGGT | 121 |
| chr1:19387480-19389506 | p2 | (TA)13 | TTAGAATTCATCGGTTCGGC | TCTCACTTGGGTTTTGGTCC | 190 |
| chr1:19390222-19392238 | p2 | (TA)8 | CTTTTGTGGCTGGAGTGGTT | TCGCGTGTATTTGACGTTGT | 219 |
| chr1:19401112-19403130 | p6 | (AAGAAA)3 | TAATGCACAAGTGGAACCCA | AAACAAACGGCCCCTAAACT | 287 |
| chr1:19408256-19410274 | p6 | (ATGATC)3 | GCCAATGGAGGTTTTGAAGA | CTTCCCCTTGAGATGGTTGA | 122 |
| chr1:19451912-19453926 | p2 | (TA)7 | TACTTGGTTTTGTCTCCGCC | CAAACCAACGACCCCTTAGA | 280 |
| chr1:19538678-19540692 | p2 | (AG)6 | TGTCGGCAACAACATCAAAT | CACTTTGGGGTTTCTGCACT | 261 |
| chr1:19538678-19540692 | p2 | (TA)7 | TGTCGGCAACAACATCAAAT | CACTTTGGGGTTTCTGCACT | 261 |
| chr1:19614491-19616503 | p2 | (AT)6 | GGGAAAATGTGGGGTTCTTT | CATAGCCCACATTCATGCAC | 242 |
| chr1:19682932-19684990 | p2 | (AT)29 | AAATATCAACCGTGGCTTCG | CTCCGATCTAATCCAATCGC | 193 |
| chr1:19712798-19714830 | p2 | (TA)16 | ATTGTTGCCAGATTTCGACC | TTTTTCCAGATTTTCGACCG | 131 |
| chr1:19801871-19803886 | p5 | (TTTTC)3 | CCTGTGATCATGGCTCAGAA | GTGTTCGCCTTATGCGGTAT | 121 |
| chr1:19801871-19803886 | p3 | (AGA)5 | CCTGTGATCATGGCTCAGAA | GTGTTCGCCTTATGCGGTAT | 121 |
| chr1:19848636-19850651 | p3 | (TGC)5 | CAGGGAAAATTTGAAGGCAA | AAACCGTACCCAACCATTGA | 281 |
| chr1:19848636-19850651 | p6 | (GTGTTG)3 | CAGGGAAAATTTGAAGGCAA | AAACCGTACCCAACCATTGA | 281 |
| chr1:19848636-19850651 | p3 | (TTG)7 | CAGGGAAAATTTGAAGGCAA | AAACCGTACCCAACCATTGA | 281 |
| chr1:19849087-19851105 | p3 | (TGC)5 | TACAAAAGCTGACGTGTGGG | TGGAAGAAGTGCGAGGAAGT | 227 |
| chr1:19849087-19851105 | p6 | (GTGTTG)3 | TACAAAAGCTGACGTGTGGG | TGGAAGAAGTGCGAGGAAGT | 227 |
| chr1:19849087-19851105 | p3 | (TTG)7 | TACAAAAGCTGACGTGTGGG | TGGAAGAAGTGCGAGGAAGT | 227 |
| chr1:19849335-19851356 | p3 | (TGC)5 | CCCTCGTCTATAGGTGCTGC | TTTACAATCGCAAAAACCCC | 287 |
| chr1:19849335-19851356 | p6 | (GTGTTG)3 | CCCTCGTCTATAGGTGCTGC | TTTACAATCGCAAAAACCCC | 287 |
| chr1:19849335-19851356 | p3 | (TTG)7 | CCCTCGTCTATAGGTGCTGC | TTTACAATCGCAAAAACCCC | 287 |
| chr1:19863500-19865515 | p3 | (ACA)5 | GACGTCTTGAGGAGGCAAAG | GTGCTATGCTCTCCTGGCTC | 159 |
| chr1:19869111-19871126 | p5 | (TTTTG)3 | GACGACCTTTGGTTCTTCCA | GGAACTCAAAATCGTCGGAA | 126 |
| chr1:20050430-20052484 | p2 | (AT)27 | TACCCGGTCTTTTGTTTTCG | AGCATCTGTTGTCACACCCA | 189 |
| chr1:20073436-20075450 | p2 | (TG)7 | ACCCCAAATTGGAAAACTCC | TGGCCGTCTAATGAGGAATC | 242 |
| chr1:20247622-20249637 | p3 | (TTG)5 | TCTTTCCTCTCTTCACCCCA | TCAGTACGATCAGCCACAGG | 178 |
| chr1:20274136-20276151 | p3 | (TCA)5 | AGCACCATCAACCTCACTCC | TTTTATTGGCCCATTTTGGA | 150 |
| chr1:20274923-20276941 | p3 | (TCA)5 | GTAGATGCACACATTGCGCT | CCGACCCGATTTGTACCTAA | 256 |
| chr1:20274933-20276948 | p3 | (TCA)5 | GTAGATGCACACATTGCGCT | CCGACCCGATTTGTACCTAA | 246 |
| chr1:20418898-20420916 | p2 | (AT)12 | GACAACGGACGACGAAGAAT | AGGTGGGCCCCATACTAAAC | 210 |
| chr1:20418898-20420916 | p2 | (AT)11 | GACAACGGACGACGAAGAAT | AGGTGGGCCCCATACTAAAC | 210 |
| chr1:20418898-20420916 | p2 | (TA)9 | GACAACGGACGACGAAGAAT | AGGTGGGCCCCATACTAAAC | 210 |
| chr1:20435993-20438011 | p6 | (AGCACC)3 | AGAATTCTCCGGGACCAAGT | CGATCAACCCTAGCTCTTCG | 121 |
| chr1:20508579-20510594 | p5 | (CTTGT)3 | CCTCCTGAAACGCTGAACTC | CTCTCTTGGGACTCCACCTG | 102 |

| Marker ID | SSR type | SSR motif | FORWARD PRIMER (5'-3') | REVERSE PRIMER (5'-3') | size (bp) |
|---|---|---|---|---|---|
| chr1:20577604-20579616 | p2 | (AT)6 | AGATTTTTGCGGGGATTTCT | CAGTGAGGTGCAAGCTTCTG | 104 |
| chr1:20713588-20715603 | p5 | (AACCC)3 | ATTTTTGGGCAATTAGCACG | CTTTTCCGGCACAGTTTTGT | 126 |
| chr1:20713588-20715603 | p5 | (TTGTT)3 | ATTTTTGGGCAATTAGCACG | CTTTTCCGGCACAGTTTTGT | 126 |
| chr1:20754324-20756342 | p2 | (TA)9 | CGGTTGGGATGACTTCACTT | GGCGGATATAGATTTTGCCA | 188 |
| chr1:20796831-20798843 | p2 | (CT)6 | CCAAGGGGTCAATGGATATG | TTTGGTGGATGCGAATATCA | 230 |
| chr1:20837606-20839621 | p5 | (CCACC)3 | CTTACCCCATGTTTGGATGG | TGGTGGAAGGTGGGGTATAA | 186 |
| chr1:20837627-20839642 | p5 | (CCACC)3 | CTTACCCCATGTTTGGATGG | TGGTGGAAGGTGGGGTATAA | 165 |
| chr1:20957782-20959797 | p5 | (TTTGG)3 | TCAAATTTGGAAAAGTGCCC | ATTGGTTTTTCACCAAACCG | 271 |
| chr1:21032735-21034751 | p2 | (TA)8 | CACCTGATTCTGGGTGGAAC | AAGGGAACACCTTGGGAGAT | 241 |
| chr1:21041378-21043393 | p5 | (TTGAT)3 | TACGTGATGAAAGCGCAAAG | TCCTCCGCTTAAAATGATGG | 271 |
| chr1:21056221-21058235 | p2 | (TC)7 | TTTGATTCCGGACCCAATTA | CAGGAGCAAAAGTGCAATGA | 299 |
| chr1:21130271-21132285 | p2 | (AT)14 | ATTAAAATTGGGGGTAGCGG | GTTCATTTGGAGGAGTCGGA | 246 |
| chr1:21130271-21132285 | p2 | (TA)7 | ATTAAAATTGGGGGTAGCGG | GTTCATTTGGAGGAGTCGGA | 246 |
| chr1:21149578-21151594 | p2 | (AT)8 | GAATTTGTTCCGCATGGTTT | AACGGCCTTCTTATCCGTTT | 220 |
| chr1:21230147-21232173 | p2 | (TA)13 | TCCAACAGCAACCAAACAAC | TTTGAGCTTACTGATGGGGG | 143 |
| chr1:21352945-21354989 | p4 | (TTTA)11 | AACGTCGGACATCCTTTCAC | TGCCAACCGTCTCTTTTCTT | 115 |
| chr1:21352945-21354989 | p2 | (TA)21 | AACGTCGGACATCCTTTCAC | TGCCAACCGTCTCTTTTCTT | 115 |
| chr1:21373254-21375269 | p5 | (TAAAA)3 | GTAGCACGAGGCCATTGTTT | CATGCAGCCTCAATTCTCAA | 193 |
| chr1:21377962-21379976 | p2 | (TA)7 | ACCTTTACGGCCGACCTATT | TTGTAGCAGTGACGTAGCGG | 206 |
| chr1:21382537-21384552 | p3 | (AAT)5 | GAGGGATAACAAACGAGGCA | TCCCTGCACACAATATCCAA | 297 |
| chr1:21492494-21494510 | p3 | (ATT)5 | CATGACCTTGGCACAGATTG | TTCGAGAAAAATGGATTCCG | 221 |
| chr1:21492494-21494510 | p2 | (TC)8 | CATGACCTTGGCACAGATTG | TTCGAGAAAAATGGATTCCG | 221 |
| chr1:21661769-21663795 | p2 | (TA)8 | TCCACCAACTACTGCCATCA | CCACTGGTGTTTGTCAATGC | 234 |
| chr1:21679828-21681850 | p5 | (ATTTT)3 | CAACGGCCACTTAAGGTGTT | TTTGATTTAGCTGGGGTTGG | 223 |
| chr1:21687134-21689149 | p5 | (TTTTA)3 | GTGAACAAAAGCCAACAGCA | ATCGAAGCTTCCACCAAAGA | 258 |
| chr1:21724769-21726793 | p4 | (TTTA)6 | TCGTGAGCCGTTATGAAAGA | ACAAGAAACGCAAATCACCC | 273 |
| chr1:21742845-21744929 | p2 | (TA)9 | ACATGCAGAGAGGTATGGGC | TTGCCTCGAAAAGGAGCTAA | 151 |
| chr1:21761836-21763851 | p5 | (AAAAG)3 | TCATACACACGCTTTTGGGA | GCAGATGCAGATTTTGCTGA | 120 |
| chr1:21768908-21770926 | p6 | (AAAATT)3 | CGATGGATAAAATCGAGGGA | ACAAAACCGACGGACGTTAG | 269 |
| chr1:21975814-21977832 | p2 | (TA)6 | TGGCCCCATAGCACTATTTT | ATACGCACGTGACAAATGGA | 142 |
| chr1:21975814-21977832 | p2 | (AT)6 | TGGCCCCATAGCACTATTTT | ATACGCACGTGACAAATGGA | 142 |
| chr1:22077535-22079550 | p5 | (AAAAT)3 | ATTTGGCACGTTAGACTGGC | TAAACAGGATATCGCGAGGC | 241 |
| chr1:22086991-22089003 | p2 | (AT)6 | GGTTGCCTAGAACCACGGTA | GTCTTTTTCTCTTGGCGGTG | 288 |
| chr1:22086991-22089003 | p3 | (TTG)5 | GGTTGCCTAGAACCACGGTA | GTCTTTTTCTCTTGGCGGTG | 288 |
| chr1:22127768-22129786 | p6 | (AAAAGG)3 | GTATGTTGCATCTGCATGGC | GCTTCCTTTTCCTGCCTTCT | 146 |
| chr1:22201025-22203041 | p2 | (AT)8 | ACCCCACTTTGCACTTCAAC | CAATCGCTGCTATCACTGGA | 216 |

| Marker ID | SSR type | SSR motif | FORWARD PRIMER (5'-3') | REVERSE PRIMER (5'-3') | size (bp) |
|---|---|---|---|---|---|
| chr1:22257272-22259290 | p2 | (TA)9 | GATCAGGTGCAATCAACGAA | TAATCACTCCAAACCGGACC | 224 |
| chr1:22257272-22259290 | p2 | (TA)7 | GATCAGGTGCAATCAACGAA | TAATCACTCCAAACCGGACC | 224 |
| chr1:22257272-22259290 | p6 | (CCGGTT)3 | GATCAGGTGCAATCAACGAA | TAATCACTCCAAACCGGACC | 224 |
| chr1:22485972-22487987 | p5 | (TTGGT)3 | GGCTTGACTTGGTTTGGTGT | GTAATGTCGGGTTGGTTTGG | 203 |
| chr1:22485972-22487987 | p5 | (TTGGT)3 | GGCTTGACTTGGTTTGGTGT | GTAATGTCGGGTTGGTTTGG | 203 |
| chr1:22492561-22494575 | p2 | (AT)7 | AAAGTCGGTCTCACCACCAC | TTGGTGCAGGATGAATGAAA | 199 |
| chr1:22536814-22538832 | p6 | (TCTTTC)3 | CGAGAAGATGGCAACAGTGA | TTGAAAAGGGTAATGCCTGC | 206 |
| chr1:22595170-22597188 | p6 | (ATAATG)3 | GTAGTTGGAGGCTTGCCTTG | TTGGCCAATTTCAATCAACA | 135 |
| chr1:22611200-22613266 | p2 | (AT)33 | AGCGCCTTTGTTTCTTCAAA | TGATGGAGAAGATGCAGACG | 131 |
| chr1:22679850-22681870 | p5 | (TGGTT)4 | TATTTGCTTGCCTCCACCTT | TAGCACGTCCAAAACATCCA | 286 |
| chr1:22705233-22707289 | p2 | (AT)28 | GCATGACTTTTTGTTGGCCT | CACCGTAAAAGCAGGGAAAA | 297 |
| chr1:22887695-22889713 | p6 | (AAGAAA)3 | CTAAGCCTCAGCAACCAACC | ATTCGTCCCATCAAATGAGC | 293 |
| chr1:22898027-22900042 | p5 | (TTTGG)3 | ACTGGCTCATCATTTCCGAC | ATAAGGGGGCTGAGTCCATC | 132 |
| chr1:23029337-23031358 | p3 | (CAA)7 | CCCCTTCCCCATAAAATCAT | CTTCTTAAGTCCCGAAGGGG | 266 |
| chr1:23123252-23125276 | p3 | (ATA)8 | CTGGTTTTGCAAGGCAATTT | ACACCCACTCCTCTGACACC | 136 |
| chr1:23131717-23133755 | p3 | (ATT)17 | GCCACCCACAAAGACTGATT | GCCCTCTCATTCCCTTTCTC | 197 |
| chr1:23131717-23133755 | p2 | (AT)19 | GCCACCCACAAAGACTGATT | GCCCTCTCATTCCCTTTCTC | 197 |
| chr1:23307017-23309069 | p3 | (TCT)10 | CGCGTGTAACATATTGTGGG | CGTGCAAGACGAGATAGCAA | 223 |
| chr1:23307017-23309069 | p5 | (ATACT)3 | CGCGTGTAACATATTGTGGG | CGTGCAAGACGAGATAGCAA | 223 |
| chr1:23446541-23448553 | p2 | (TA)6 | ACACCCAGGACCACGTCTAC | TAGGTGGTTGAAAAGGTGCC | 114 |
| chr1:23446541-23448553 | p2 | (TA)6 | ACACCCAGGACCACGTCTAC | TAGGTGGTTGAAAAGGTGCC | 114 |
| chr1:23446712-23448724 | p2 | (TA)6 | GGCACCTTTTCAACCACCTA | TGTGTCAAGACTCCGAAACG | 140 |
| chr1:23446712-23448724 | p2 | (TA)6 | GGCACCTTTTCAACCACCTA | TGTGTCAAGACTCCGAAACG | 140 |
| chr1:23460292-23462304 | p2 | (TA)6 | TTGCTCCACGCCCTACTACT | ACCATGCTAAGACGTCCACC | 123 |
| chr1:23460292-23462304 | p2 | (AT)7 | TTGCTCCACGCCCTACTACT | ACCATGCTAAGACGTCCACC | 123 |
| chr1:23460441-23462455 | p2 | (TA)6 | AAGTATCAACCGTGGCTTCG | TCCGCTTCTGAGTTTTGGTT | 211 |
| chr1:23460441-23462455 | p2 | (AT)7 | AAGTATCAACCGTGGCTTCG | TCCGCTTCTGAGTTTTGGTT | 211 |
| chr1:23479483-23481495 | p2 | (TA)6 | TTGCTCCACGCCCTACTACT | ACCATGCTAAGACGTCCACC | 123 |
| chr1:23479483-23481495 | p2 | (AT)7 | TTGCTCCACGCCCTACTACT | ACCATGCTAAGACGTCCACC | 123 |
| chr1:23479632-23481646 | p2 | (TA)6 | AAGTATCAACCGTGGCTTCG | TCCGCTTCTGAGTTTTGGTT | 211 |
| chr1:23479632-23481646 | p2 | (AT)7 | AAGTATCAACCGTGGCTTCG | TCCGCTTCTGAGTTTTGGTT | 211 |
| chr1:23542292-23544310 | p2 | (CA)9 | ATGTCATGTGGCAGCCATTA | ATGCCAAAAATTCGCAAGAG | 103 |
| chr1:23567184-23569212 | p2 | (AT)14 | CAAAACCACGGGTGTCTTCT | TGGCATTTCTTTTAGGGTCG | 207 |
| chr1:23591409-23593423 | p2 | (AC)7 | GGTAAATCACCAAGAGCCCA | CTTGGAACGTCGGTGAAGAT | 100 |
| chr1:23660986-23663020 | p2 | (AT)17 | ATCCCATAGCCTAGCCGAAT | AGCCCACCATTCCTTCTCTT | 162 |
| chr1:23683452-23685467 | p5 | (TAAAA)3 | AGCCTCGCCCCATATTAGAT | TTATGGCGACCAACCAAAAT | 135 |

| Marker ID | SSR type | SSR motif | FORWARD PRIMER (5'-3') | REVERSE PRIMER (5'-3') | size (bp) |
|---|---|---|---|---|---|
| chr1:23700490-23702506 | p2 | (AT)8 | AAGGGGGTCTGATTTCCAAC | CTATACTCACACCCGGCGAT | 102 |
| chr1:23916781-23918796 | p5 | (AAAAG)3 | CCGGAACAGGAAAAACTTCA | ATTCCGGCATAATTTCGTGA | 279 |
| chr1:23961994-23964006 | p2 | (TA)6 | ATAGGCATAGGCCAAAAGGG | TGCCTTAAAACGCGAAAACT | 282 |
| chr1:24109810-24111825 | p3 | (TGT)5 | GGCCAGAACTTGATTCTCCA | CTCACCCCCTACTTCAACCA | 136 |
| chr1:24193822-24195846 | p2 | (AT)12 | AGAGTAGTTGGCCATGCGTT | ACACGCCTAGGGAAAAGGAT | 150 |
| chr1:24338051-24340071 | p2 | (TA)10 | AATCCGCCCCATACAACATA | CCCAGGTCATCCAGAAGAAA | 157 |
| chr1:24353737-24355759 | p2 | (TA)6 | CCAAAGTTCCCGTAAAGCAA | GTGCCCAACACATTCATCAC | 232 |
| chr1:24353737-24355759 | p2 | (AT)11 | CCAAAGTTCCCGTAAAGCAA | GTGCCCAACACATTCATCAC | 232 |
| chr1:24353737-24355759 | p2 | (TA)8 | CCAAAGTTCCCGTAAAGCAA | GTGCCCAACACATTCATCAC | 232 |
| chr1:24368838-24370856 | p6 | (AAAACC)3 | CTTGCTCGTACGTCGTCAAA | GGGGGTGATGTAACCCTTTC | 231 |
| chr1:24378321-24380336 | p5 | (AATGA)3 | GCTTCTCGTCATCGGAGAAC | GAAAGGCTAACCCTTCGCTT | 110 |
| chr1:24452753-24454771 | p2 | (AT)9 | GGCCACTGGTTTATTTCCTG | GCCTTTTGAATCACGGGTAA | 242 |
| chr1:24490133-24492151 | p2 | (AT)9 | GGCCACTGGTTTATTTCCTG | GCCTTTTGAATCACGGGTAA | 242 |
| chr1:24553830-24555850 | p5 | (AAACC)3 | AGTCGAAAACGGAGCAAGAA | CGACCATGAACACCCCTAAT | 268 |
| chr1:24562405-24564431 | p2 | (TA)6 | CTTCGCGCGGTATGAATAAT | ATTAAGCGAGACCGGAGGAT | 118 |
| chr1:24671681-24673717 | p2 | (TA)18 | GTTGAAGACTGCGTGACAGC | TGGCCCAAACTTTTAGATGG | 287 |
| chr1:24679188-24681200 | p2 | (TA)6 | GAAAGCCGAATGCTCTTCAC | AAACCAAAGAAAACCCGTCC | 175 |
| chr1:24679188-24681200 | p2 | (TC)9 | GAAAGCCGAATGCTCTTCAC | AAACCAAAGAAAACCCGTCC | 175 |
| chr1:24711208-24713220 | p2 | (CT)6 | TCGGGGAAATTGCTAAACAG | GAAATCTTCTGGAACGCTGC | 130 |
| chr1:24711208-24713220 | p2 | (TA)6 | TCGGGGAAATTGCTAAACAG | GAAATCTTCTGGAACGCTGC | 130 |
| chr1:24779273-24781291 | p6 | (ATTTTT)3 | GTCCAAACCATTGATACCCG | TGGTTGGGATTTTTCGTTGT | 199 |
| chr1:24784929-24786944 | p3 | (CTC)5 | GGTGGTACGACACTCGGACT | AAACAGATTGATCGTGGGCT | 126 |
| chr1:24784929-24786944 | p3 | (AAG)9 | GGTGGTACGACACTCGGACT | AAACAGATTGATCGTGGGCT | 126 |
| chr1:24894728-24896748 | p2 | (GA)10 | AATTTGCCCCTACCGTTACC | ACCTCATCCAAATGTCAGGC | 222 |
| chr1:24919877-24921895 | p6 | (AATGTA)3 | ATGGGTGTCAGAAAAGTCGC | GTCGTTTGGAGGGATTTTGA | 128 |
| chr1:24951473-24953491 | p6 | (TTTTTA)3 | GATGGAGCTTGCTCAACACA | TGATGGCATTACCCAGTGAA | 104 |
| chr1:25002189-25004204 | p3 | (GTT)5 | CCGTATTTGGACAACCCAAC | AACGAGCATCGGGATCTATG | 204 |
| chr1:25002189-25004204 | p3 | (GTT)5 | CCGTATTTGGACAACCCAAC | AACGAGCATCGGGATCTATG | 204 |
| chr1:25002189-25004204 | p3 | (GTT)6 | CCGTATTTGGACAACCCAAC | AACGAGCATCGGGATCTATG | 204 |
| chr1:25007411-25009429 | p3 | (TGT)6 | CGCCAAAACTTAAAGGCAAG | TTCCGCGACTTTCATCTTCT | 213 |
| chr1:25073474-25075509 | p2 | (AT)6 | GCCGAAGAATAAGGGAAAGG | CGGAGACCAACAAGAACGAT | 222 |
| chr1:25154019-25156051 | p2 | (AT)16 | CCGAAGGATGAACCTGAAAA | ATCACACCCATCTCAGGAGG | 204 |
| chr1:25161599-25163613 | p2 | (TA)7 | TTTTCAAGAAAACCCATCGC | AACATGGCGTCAGGAAGAAC | 221 |
| chr1:25164861-25166876 | p3 | (GAA)5 | TGACGCACTTAATCTTTGCG | GGTGGTGGGTGAGAGGATAA | 123 |
| chr1:25164861-25166876 | p2 | (AT)17 | TGACGCACTTAATCTTTGCG | GGTGGTGGGTGAGAGGATAA | 123 |
| chr1:25272180-25274242 | p2 | (TA)31 | GAGAATGTGCGTGGTTGAGA | TGAACGGATTTGTTGCACAT | 201 |

| Marker ID | SSR type | SSR motif | FORWARD PRIMER (5'-3') | REVERSE PRIMER (5'-3') | size (bp) |
|---|---|---|---|---|---|
| chr1:25272180-25274242 | p6 | (GAAAAA)3 | GAGAATGTGCGTGGTTGAGA | TGAACGGATTTGTTGCACAT | 201 |
| chr1:25272180-25274242 | p2 | (TA)7 | GAGAATGTGCGTGGTTGAGA | TGAACGGATTTGTTGCACAT | 201 |
| chr1:25398387-25400402 | p5 | (TAAAA)3 | CAAGTGCTCCTCCTTTTTCG | ATTGGGTCTATTTCGAGGGG | 270 |
| chr1:25410810-25412825 | p4 | (AATT)4 | CCACTGTTTTTGCTCGTTCA | GCGCAATAGGACCAAACTGT | 233 |
| chr1:25410810-25412825 | p5 | (ATTTT)3 | CCACTGTTTTTGCTCGTTCA | GCGCAATAGGACCAAACTGT | 233 |
| chr1:25410810-25412825 | p5 | (AAAAT)3 | CCACTGTTTTTGCTCGTTCA | GCGCAATAGGACCAAACTGT | 233 |
| chr1:25422489-25424507 | p6 | (CTGCTA)3 | GGGCCATGATCAGAAGATGT | GGCCAACGCGAGATAAAATA | 225 |
| chr1:25425620-25427636 | p2 | (GA)8 | AGCCTCCAACTTACCCACCT | ATTGTGGTTTTTGGGAGTGG | 115 |
| chr1:25436535-25438550 | p5 | (TTTTC)3 | TTCCTTTGTTTTGACGGAGG | TGCATGAAACAAGGGACAAA | 292 |
| chr1:25465453-25467465 | p2 | (AT)6 | TTGCCCACTTCTGTCCTCTT | TGATCTTTTCACGTGCTTCG | 262 |
| chr1:25500439-25502454 | p3 | (TCA)5 | ATATTACCGGGCAGAAGCCT | ATATCTCGGGCGTAGGTGTG | 162 |
| chr1:25515827-25517860 | p3 | (CTT)11 | TCCTCTTCCTCCTCCTCCTC | TGCATGAAAATCGGGTTGTA | 287 |
| chr1:25516001-25518022 | p3 | (CTT)11 | TCCTCTTCCTCCTCCTCCTC | TGCATGAAAATCGGGTTGTA | 113 |
| chr1:25529861-25531879 | p2 | (TA)9 | TAAGAACGGAGTCAGCAGCA | TGCAGTAAGACACCGAGTGC | 289 |
| chr1:25593507-25595527 | p4 | (ATAG)5 | TGTTCACCAAATCTTTGCCA | TCACTCCCCCTGATGAAAAC | 284 |
| chr1:25594534-25596552 | p6 | (ATTCAG)3 | ATGAATGTGCCAAGTCAGCA | CTTTGTGGGTAGGGAAAACG | 142 |
| chr1:25635420-25637432 | p2 | (TA)6 | GGCACAAAAGGAGAGAACCA | TGCAAGGCTTCCAAGAAAGT | 183 |
| chr1:25754691-25756709 | p3 | (ATC)6 | TTTTTCCAGATGAACTCGGG | AGGGAGTTCGATGTTGTTGG | 165 |
| chr1:25754691-25756709 | p2 | (TC)6 | TTTTTCCAGATGAACTCGGG | AGGGAGTTCGATGTTGTTGG | 165 |
| chr1:25769348-25771363 | p3 | (TAG)5 | ATTTTCACCCCCACTTTTCC | TCAAGCGAATCTTGTGGCTA | 149 |
| chr1:25955325-25957337 | p2 | (AT)6 | ATTTGTGGGCTGTTTTGAGG | AACGGCTTACTTTCAAGCCA | 285 |
| chr1:26004171-26006186 | p2 | (AG)7 | GGGAATTTGAAGCAACTGGA | CCGGTCTTGAATCGGTTAGA | 217 |
| chr1:26004171-26006186 | p3 | (TAA)6 | GGGAATTTGAAGCAACTGGA | CCGGTCTTGAATCGGTTAGA | 217 |
| chr1:26004171-26006186 | p5 | (TTTTA)3 | GGGAATTTGAAGCAACTGGA | CCGGTCTTGAATCGGTTAGA | 217 |
| chr1:26017381-26019396 | p2 | (AT)18 | AAGGGCATTTGCATAAATCG | CCCAGAACCTCAGGTATCCA | 205 |
| chr1:26017381-26019396 | p5 | (TTTTA)3 | AAGGGCATTTGCATAAATCG | CCCAGAACCTCAGGTATCCA | 205 |
| chr1:26017381-26019396 | p5 | (AATTT)3 | AAGGGCATTTGCATAAATCG | CCCAGAACCTCAGGTATCCA | 205 |
| chr1:26039164-26041190 | p2 | (AT)13 | TGATCACTCCGTTTAGTGCG | TTGATGGTGTTTTGGGTCAG | 266 |
| chr1:26054003-26056023 | p5 | (TTATT)4 | AGTAGGCGTTTGGATATGCG | GTTGGTAAATGGTTGGTGGG | 197 |
| chr1:26055009-26057027 | p5 | (TATTT)3 | CAGCCCTTCTAGCTGACCAC | ATCTTGGTGCACTGGTGTGA | 248 |
| chr1:26055009-26057027 | p6 | (ATATTA)3 | CAGCCCTTCTAGCTGACCAC | ATCTTGGTGCACTGGTGTGA | 248 |
| chr1:26072237-26074255 | p3 | (TAT)6 | TGTGTTTGCCTTGCTATTCG | CACGGTGGCTACTCTTCCTC | 151 |
| chr1:26076801-26078825 | p2 | (CT)12 | AACCCCTTGGCTATACCTGG | GTGGGGAAGGGGTGTTAAGT | 268 |
| chr1:26124791-26126806 | p5 | (ATATA)3 | GATGCGCACTATGCTAACCA | GTTTGGCTTGTCGGCTGTAT | 113 |
| chr1:26175570-26177598 | p2 | (AT)6 | CCTGATTTCGGAGAGTGCAT | ACTTCCTCTTCGCCTTCCTC | 278 |
| chr1:26175570-26177598 | p2 | (GA)6 | CCTGATTTCGGAGAGTGCAT | ACTTCCTCTTCGCCTTCCTC | 278 |

| Marker ID | SSR type | SSR motif | FORWARD PRIMER (5'-3') | REVERSE PRIMER (5'-3') | size (bp) |
|---|---|---|---|---|---|
| chr1:26223018-26225033 | p5 | (TTAGA)3 | CTGCGTAAAAATCGAACCGT | GCGAAATGCCAATTTAAGGA | 276 |
| chr1:26238833-26240853 | p5 | (GAGAG)4 | TGCTGTAAGTTGCTGCTGCT | TCCCTCAAAATACAGCCACC | 139 |
| chr1:26295445-26297463 | p6 | (GGGGGT)3 | CTGTCCAAATCCTCCCAAAA | TTCTCAGGTAGGTCGAGGGA | 151 |
| chr1:26307120-26309138 | p6 | (GACAGA)3 | TCACCGACGGTAAAAGAACC | ACTTCACGCGCTTAGGTGTT | 220 |
| chr1:26318844-26320860 | p2 | (AT)6 | CAACCGTGGCTTCGTAGAGT | TCCGACCTAATCCAATCTCG | 237 |
| chr1:26318844-26320860 | p5 | (AATAA)3 | CAACCGTGGCTTCGTAGAGT | TCCGACCTAATCCAATCTCG | 237 |
| chr1:26321523-26323538 | p2 | (TA)6 | TACTCGGGCAAGGTATGAGG | GTGGGTACGTGGAGTCTGGT | 151 |
| chr1:26321523-26323538 | p3 | (TTG)5 | TACTCGGGCAAGGTATGAGG | GTGGGTACGTGGAGTCTGGT | 151 |
| chr1:26386680-26388704 | p2 | (AT)6 | AAAGGTGAACCACCCTCTCC | TATTATCGGGTGGCACGTTT | 101 |
| chr1:26420185-26422205 | p2 | (AT)10 | CATGTTGCTTTTGCATGGTT | GCTATTCCGCCATGTCATTT | 127 |
| chr1:26499645-26501660 | p5 | (AAACA)3 | AGGTTACCATGTCACCTCGG | ACGGACATTTAAGCATTCGG | 231 |
| chr1:26499645-26501660 | p3 | (TTG)5 | AGGTTACCATGTCACCTCGG | ACGGACATTTAAGCATTCGG | 231 |
| chr1:26522234-26524250 | p4 | (AAAT)4 | GCCTCTTTCCTAGTGGTCCC | TTCACGTGAGTTTGGACTGC | 152 |
| chr1:26577630-26579642 | p2 | (TG)6 | CATCACCTCGAACACCAATG | GTTGGTGCTCCTTTAGCTGC | 117 |
| chr1:26629109-26631127 | p6 | (TAATTT)3 | CAAGTCTGCCCATAAGGCAT | CTGAGTTTGAACTCCCGCTC | 101 |
| chr1:26630890-26632918 | p2 | (AT)14 | CGCCATTGAAGAGTGCAGTA | TAGCGAGCGAGGACCTTAAC | 294 |
| chr1:26630890-26632918 | p2 | (TA)13 | CGCCATTGAAGAGTGCAGTA | TAGCGAGCGAGGACCTTAAC | 294 |
| chr1:26664963-26666978 | p3 | (CAA)5 | CCGTCACACATCACAACACA | CTATGATATCGTTCGCCGGT | 115 |
| chr1:26808772-26810832 | p3 | (TTA)20 | TTCGGAGCACTTTACAACCC | CAAACTAGGCTTCACGCACA | 209 |
| chr1:26847975-26849993 | p2 | (AT)9 | CTGGGCATTCTTGATCTGGT | TTTTCAGCGTGTGTATTGCC | 160 |
| chr1:26926264-26928279 | p5 | (GTTTG)3 | ATGCAGCAATGACACCAAAA | TGAACCAAAACACCCTCTCC | 163 |
| chr1:26940010-26942024 | p2 | (TC)7 | CAAGTGGCATTTCAAGCAAA | GTGGGAAAGGATTGACGAGA | 226 |
| chr1:26940010-26942024 | p2 | (TA)8 | CAAGTGGCATTTCAAGCAAA | GTGGGAAAGGATTGACGAGA | 226 |
| chr1:26978321-26980341 | p2 | (AT)6 | ATTAGGTCATCTGCATGGGG | CTCGCCTATCTTTAAGGGGG | 249 |
| chr1:26978321-26980341 | p2 | (TA)10 | ATTAGGTCATCTGCATGGGG | CTCGCCTATCTTTAAGGGGG | 249 |
| chr1:26978321-26980341 | p2 | (TC)7 | ATTAGGTCATCTGCATGGGG | CTCGCCTATCTTTAAGGGGG | 249 |
| chr1:26985462-26987474 | p2 | (TA)6 | GCTCCCATGTAGGGCAATAA | GCACAGAACTGGGGACTGTT | 109 |
| chr1:26995995-26998013 | p6 | (AAAGCA)3 | CGGAGATGAAAGCAGGCTAC | GAAGGAGTCAAAGGTGCCAG | 169 |
| chr1:26995995-26998013 | p2 | (AT)20 | CGGAGATGAAAGCAGGCTAC | GAAGGAGTCAAAGGTGCCAG | 169 |
| chr1:27065418-27067438 | p3 | (CTA)5 | TCCCTCTCCCATTTCTCCTT | GCCGGTGCAAGATATTGTTT | 170 |
| chr1:27065418-27067438 | p2 | (TA)10 | TCCCTCTCCCATTTCTCCTT | GCCGGTGCAAGATATTGTTT | 170 |
| chr1:27073425-27075440 | p5 | (ATATT)3 | TGGAACAGCCTCACCACATA | GCACCACCACCTTCGTTACT | 170 |
| chr1:27170790-27172805 | p3 | (CTT)5 | AAGACCACCCCAAAAGAAGG | CAGTGGGAGACTGGGAGTGT | 102 |
| chr1:27187838-27189856 | p3 | (TTG)6 | GCCCACACGTCAAGATTTTT | GAGTTTGCCCAAATTCCAAA | 211 |
| chr1:27334007-27336022 | p5 | (TAAAA)3 | CATGAGCATTTCACACCAGG | CCGTACATGTCAGAACGGTG | 121 |
| chr1:27372587-27374617 | p2 | (AT)15 | TCGCCTTGAGGTTCAAAGTT | GACAGAAGCTGCGGGTAAAC | 128 |

| Marker ID | SSR type | SSR motif | FORWARD PRIMER (5'-3') | REVERSE PRIMER (5'-3') | size (bp) |
|---|---|---|---|---|---|
| chr1:27372587-27374617 | p2 | (AT)8 | TCGCCTTGAGGTTCAAAGTT | GACAGAAGCTGCGGGTAAAC | 128 |
| chr1:27393875-27395890 | p5 | (ACCCG)3 | GTGACTCATCCACTTCCCGT | AGTTTTGCCTTATGGGCCTT | 117 |
| chr1:27493939-27495951 | p2 | (AT)6 | CAACTCAACCCAGCCCTAAA | AACTTAAAGGGCTGACCCGT | 190 |
| chr1:27529136-27531152 | p2 | (TA)8 | AAATGCTACACCCGTTGCTC | CTAAATGCACTTTCCGAGGC | 233 |
| chr1:27530152-27532173 | p3 | (ATT)7 | TTTAGTCGTCCTCACAGGGG | TGGCTATTGAGACCCACTGC | 281 |
| chr1:27572330-27574348 | p3 | (AAT)6 | TTGTCCATGTTTTACCCCGT | TATCGTGTCCCGTGAGCATA | 108 |
| chr1:27595860-27597875 | p5 | (AATAA)3 | TTGCGCCTCTTTTCATCTTT | AATTGCGGCTAAGTGTGCTT | 118 |
| chr1:27705661-27707701 | p2 | (AT)7 | TGGTATTCAGAGGCCAATCC | GCTTGGCGCAGGTAAATTAG | 150 |
| chr1:27705661-27707701 | p2 | (AT)20 | TGGTATTCAGAGGCCAATCC | GCTTGGCGCAGGTAAATTAG | 150 |
| chr1:27705661-27707701 | p3 | (TTG)8 | TGGTATTCAGAGGCCAATCC | GCTTGGCGCAGGTAAATTAG | 150 |
| chr1:27717906-27719921 | p3 | (GTT)5 | GGTGTCGTTCTCCAGTTGGT | AGCCGGTCTTGGATTACCTT | 111 |
| chr1:27727510-27729524 | p2 | (TA)7 | GGCCATCTTTATGGTTGTGC | AGTATGGGGCTCACCCTTTT | 126 |
| chr1:27727510-27729524 | p2 | (TA)29 | GGCCATCTTTATGGTTGTGC | AGTATGGGGCTCACCCTTTT | 126 |
| chr1:27729783-27731801 | p2 | (TA)9 | TTCCCAAAGTATCCCCATCA | AAATGACATGGCGGAATAGC | 161 |
| chr1:27738014-27740032 | p2 | (TA)9 | TTCCCAAAGTATCCCCATCA | AAATGACATGGCGGAATAGC | 161 |
| chr1:27782617-27784629 | p2 | (TG)6 | TGATGTGTTACGGTTTCCGA | ACGAACACACCGTCAACGTA | 264 |
| chr1:27870893-27872907 | p2 | (AT)7 | ATTTGCCTTGTGCGGATTAC | CATATTGCGTGGCTTCACAC | 300 |
| chr1:27967689-27969704 | p5 | (TAAAT)3 | CCAAGGTTTGAAAAGACCCA | GATGCACGAGCACTATGAGC | 237 |
| chr1:28020402-28022414 | p2 | (TA)6 | GGGGAAATGATTGCTTCAAA | AATACGGGTCGCAAACGTAG | 281 |
| chr1:28049270-28051292 | p2 | (AT)11 | CAACGGAGGAAGGCATAAAA | TGCCCTCAACATTTGCATTA | 164 |
| chr1:28064446-28066514 | p2 | (AT)34 | CTCCGCCATCTTAATTTTGG | AAGTTTACGCGGTGGACATC | 121 |
| chr1:28104652-28106674 | p2 | (AT)11 | CAACGGAGGAAGGCATAAAA | TGCCCTCAACATTTGCATTA | 164 |
| chr1:28119828-28121896 | p2 | (AT)34 | CTCCGCCATCTTAATTTTGG | AAGTTTACGCGGTGGACATC | 121 |
| chr1:28137383-28139398 | p3 | (TTG)5 | AAAAGGAAAGTCGCGGAAAT | CTTTGCCTCTCTTGAAACCG | 300 |
| chr1:28137383-28139398 | p2 | (TG)11 | AAAAGGAAAGTCGCGGAAAT | CTTTGCCTCTCTTGAAACCG | 300 |
| chr1:28146974-28148994 | p4 | (TCCT)5 | AGAGGCTGTCTGGTTCTCCA | AAATCCTCTGGGGCAATTCT | 165 |
| chr1:28162639-28164663 | p3 | (AGG)8 | CCTAGTCGGGTCGTGACACT | TAAGATTTGAACGTTCCGGG | 114 |
| chr1:28162639-28164663 | p3 | (TCT)10 | CCTAGTCGGGTCGTGACACT | TAAGATTTGAACGTTCCGGG | 114 |
| chr1:28183497-28185513 | p4 | (CATT)4 | ATAGTCTCGCGAAGACGGAA | GGGGTGATCTATCCACCAGA | 181 |
| chr1:28231639-28233657 | p6 | (TATATT)3 | CTTACGCGCTGAAGGAGAGT | GGTATCACAGTGTCCCAGGC | 220 |
| chr1:28231639-28233657 | p2 | (TA)9 | CTTACGCGCTGAAGGAGAGT | GGTATCACAGTGTCCCAGGC | 220 |
| chr1:28367725-28369740 | p5 | (AAAAG)3 | ACCAGTGATGCACGATTTGA | TTAACCCCGAAATTGGGAA | 187 |
| chr1:28489219-28491273 | p3 | (TAA)18 | ACCCCTCAACACTGAGATGG | CCTTCCTCCATCAGGATTCA | 195 |
| chr1:28494617-28496661 | p2 | (TA)22 | TGTTCGTGTTTCAGCGTCTC | CCGAGAAATTCAGGCACAAT | 181 |
| chr1:28667754-28669784 | p5 | (TACAA)3 | CCTGTTTGGATGGTGGTTTC | TCACCAAATCCGTGGTTACA | 230 |
| chr1:28667754-28669784 | p6 | (AAAAAT)5 | CCTGTTTGGATGGTGGTTTC | TCACCAAATCCGTGGTTACA | 230 |

| Marker ID | SSR type | SSR motif | FORWARD PRIMER (5'-3') | REVERSE PRIMER (5'-3') | size (bp) |
|---|---|---|---|---|---|
| chr1:28821981-28824021 | p2 | (AG)9 | CGAACCGTCCAATGAAGAAT | AAAGACTGCACCACTTTGGG | 289 |
| chr1:29019754-29021768 | p2 | (TG)7 | AAAGGAAAGCAACTGCGAGA | TTTGTTTGCGTGTATTGGGA | 244 |
| chr1:29096536-29098551 | p5 | (AATAA)3 | AAGAAGTCCCTGCTGCTGAA | CGACACAACTTTGTCAGCGT | 187 |
| chr1:29096536-29098551 | p5 | (AATTA)4 | AAGAAGTCCCTGCTGCTGAA | CGACACAACTTTGTCAGCGT | 187 |
| chr1:29096536-29098551 | p2 | (AT)6 | AAGAAGTCCCTGCTGCTGAA | CGACACAACTTTGTCAGCGT | 187 |
| chr1:29205001-29207016 | p6 | (CTGAAA)3 | GAATTAGCCTTCAATGGGCA | TAGAAAAGGGACCAAATGCG | 182 |
| chr1:29205001-29207016 | p5 | (ATAAT)3 | GAATTAGCCTTCAATGGGCA | TAGAAAAGGGACCAAATGCG | 182 |
| chr1:29239817-29241880 | p3 | (TAA)17 | CGAGTTGAAAATCATGCCCT | AACAACAATAGTCGACCCCG | 201 |
| chr1:29246479-29248503 | p2 | (TA)12 | CCCAAAAAATTCCAACACACC | AATTTTGGAGCCAATCCCTT | 154 |
| chr1:29411798-29413814 | p2 | (AT)8 | TTGATGCTAACACCAGCAGC | AGCAATGCAGAAAGCCAAAT | 266 |
| chr1:29453206-29455226 | p2 | (AT)10 | CATGCTCATGCTCCTGTCAT | CAGAACCGGTTAAATCGGAA | 242 |
| chr1:29453206-29455226 | p2 | (AT)6 | CATGCTCATGCTCCTGTCAT | CAGAACCGGTTAAATCGGAA | 242 |
| chr1:29453460-29455472 | p2 | (AT)10 | TAGCATTAAGCCCGTCAACC | GGTCTCAAACCGGTAAACCA | 151 |
| chr1:29453460-29455472 | p2 | (AT)6 | TAGCATTAAGCCCGTCAACC | GGTCTCAAACCGGTAAACCA | 151 |
| chr1:29458475-29460493 | p3 | (TCA)5 | CCTCATGCCCTAAAAGTGGA | GGTATTGTTAGGGGTGGGCT | 150 |
| chr1:29458475-29460493 | p6 | (TTTCGA)3 | CCTCATGCCCTAAAAGTGGA | GGTATTGTTAGGGGTGGGCT | 150 |
| chr1:29517771-29519819 | p3 | (AAT)16 | TTTAGGTTTTTCGCGGTTTG | CAAACCCACTCATCCCTGTT | 228 |
| chr1:29532479-29534543 | p2 | (TA)6 | GCCATCATTATTTAGCCGGA | GCATGCACGAAGTCTCAAAA | 269 |
| chr1:29532479-29534543 | p2 | (TA)32 | GCCATCATTATTTAGCCGGA | GCATGCACGAAGTCTCAAAA | 269 |
| chr1:29621479-29623493 | p2 | (TA)7 | ATGTGCGCAATAGGGGTTAC | CCCTAAAACGACGATCCAAA | 177 |
| chr1:29634094-29636108 | p2 | (AC)7 | GTGAGGCTCAGCAAAGGAAG | GTCCTGCTGGTTTCCCATAA | 283 |
| chr1:29654111-29656125 | p2 | (AC)7 | GTGAGGCTCAGCAAAGGAAG | GTCCTGCTGGTTTCCCATAA | 283 |
| chr1:29673880-29675894 | p2 | (AT)7 | GTTGCTCCACGCCCTACTAA | CGAAGCCACGGTTGATACTT | 190 |
| chr1:29674286-29676298 | p2 | (AT)7 | TTGGATTAGATCGGAGCGAC | TCATATTCGGCCATGGAGAT | 133 |
| chr1:29677850-29679868 | p6 | (CCCCCA)3 | CAGCCATAGTCGCATAGCAA | AAGGGCGTGAGGAAAAGATT | 294 |
| chr1:29677850-29679868 | p3 | (ATT)6 | CAGCCATAGTCGCATAGCAA | AAGGGCGTGAGGAAAAGATT | 294 |
| chr1:29677850-29679868 | p2 | (TG)6 | CAGCCATAGTCGCATAGCAA | AAGGGCGTGAGGAAAAGATT | 294 |
| chr1:29766680-29768695 | p5 | (AAATA)3 | TCCGAGCCTATTTTGCAGTT | CACAAGACCTAGCAGGAGCC | 186 |
| chr1:29766680-29768695 | p5 | (AAATA)3 | TCCGAGCCTATTTTGCAGTT | CACAAGACCTAGCAGGAGCC | 186 |
| chr1:29766680-29768695 | p5 | (TTTAT)3 | TCCGAGCCTATTTTGCAGTT | CACAAGACCTAGCAGGAGCC | 186 |
| chr1:29766680-29768695 | p5 | (TTTAT)4 | TCCGAGCCTATTTTGCAGTT | CACAAGACCTAGCAGGAGCC | 186 |
| chr1:29910165-29912185 | p2 | (TA)16 | GGATGAAGAAATTGGCTGGA | ATTCCAGCTACAGCACCACC | 234 |
| chr1:29910165-29912185 | p2 | (TA)10 | GGATGAAGAAATTGGCTGGA | ATTCCAGCTACAGCACCACC | 234 |
| chr1:30011462-30013483 | p2 | (AG)10 | GGGGATCTCTCGGCTAAAAC | ACTTCCCTCTGTTCCCCACT | 237 |
| chr1:30011486-30013531 | p2 | (AG)10 | GGGGATCTCTCGGCTAAAAC | ACTTCCCTCTGTTCCCCACT | 213 |
| chr1:30185160-30187178 | p6 | (TGTGTA)3 | CACATGTTCCCTCCACACAG | GAAATGTGAGGTGGGGTTTG | 207 |

| Marker ID | SSR type | SSR motif | FORWARD PRIMER (5'-3') | REVERSE PRIMER (5'-3') | size (bp) |
|---|---|---|---|---|---|
| chr1:30218902-30220956 | p3 | (ATA)18 | GCCCAGGTGGATTGTCTTTA | TTTTTGATTTTCCGGTTGGT | 211 |
| chr1:30218902-30220956 | p3 | (GAA)5 | GCCCAGGTGGATTGTCTTTA | TTTTTGATTTTCCGGTTGGT | 211 |
| chr1:30326570-30328585 | p3 | (TAT)5 | ATTGTGGAGCCCAAACAGAG | TCAACTTTTGACCCGAAACC | 106 |
| chr1:30404158-30406173 | p5 | (ACAAC)3 | TGCTGACGTGTACGAGGATT | TTCTGGTCACCACCGTGTAA | 192 |
| chr1:30463334-30465346 | p2 | (TA)7 | GTTGCTCCACGCCCTACTAC | CGAAGCCACGGTTGATACTT | 205 |
| chr1:30463361-30465373 | p2 | (TA)7 | GTTGCTCCACGCCCTACTAC | CGAAGCCACGGTTGATACTT | 178 |
| chr1:30463402-30465414 | p2 | (TA)7 | GTTGCTCCACGCCCTACTAC | CGAAGCCACGGTTGATACTT | 137 |
| chr1:30465631-30467645 | p6 | (CCGGTT)6 | CACGACACTCGATATGGTGC | TTTAAAACCGGTAAACGGGA | 291 |
| chr1:30550916-30552930 | p2 | (AT)7 | GGACATGCCAAAAAGAAGGA | TCGGGATGGCCTAATTACAG | 218 |
| chr1:30550916-30552930 | p2 | (TC)7 | GGACATGCCAAAAAGAAGGA | TCGGGATGGCCTAATTACAG | 218 |
| chr1:30626626-30628649 | p2 | (TG)6 | GTGGGGGCCATTATAACCTT | AATAGTTTGGCGTGGTGGTC | 230 |
| chr1:30643485-30645500 | p5 | (TGGCA)3 | TTCACATTTCCTTCCCTTGG | TGTATCGTGTCCTGATCCCA | 108 |
| chr1:30717362-30719386 | p3 | (TAA)8 | ATGCGCCACAATTACATCAA | TTGAATGAAACTCCCTTCGG | 110 |
| chr1:30776207-30778227 | p2 | (AT)7 | GAAATGCTGCAACCGATTTT | TCACGTTGATTACGATGGGA | 108 |
| chr1:30776207-30778227 | p5 | (TATAT)3 | GAAATGCTGCAACCGATTTT | TCACGTTGATTACGATGGGA | 108 |
| chr1:30795238-30797256 | p6 | (AATTTT)3 | TAATGTCGGGTTGGTTTGGT | CCCGACTACTGGTTTGGTGT | 227 |
| chr1:30795238-30797256 | p3 | (ACA)5 | TAATGTCGGGTTGGTTTGGT | CCCGACTACTGGTTTGGTGT | 227 |
| chr1:30815451-30817467 | p2 | (TA)8 | TCCTAAAGCGGATAGGAGCA | CTGAACAATATGGCTCGGGT | 112 |
| chr1:30815451-30817467 | p2 | (AT)13 | TCCTAAAGCGGATAGGAGCA | CTGAACAATATGGCTCGGGT | 112 |
| chr1:30844060-30846076 | p4 | (AAAT)4 | CCAAGAGTGTGCTTGCATGT | ACGCGTACCCAGAGACTAGC | 131 |
| chr1:30853832-30855847 | p5 | (TAAAA)3 | TCCCTAGCGCGAGACTCTTA | GAGATTCCTCGCACGCTTAC | 137 |
| chr1:30853832-30855847 | p2 | (AT)7 | TCCCTAGCGCGAGACTCTTA | GAGATTCCTCGCACGCTTAC | 137 |
| chr1:30854833-30856865 | p5 | (AAAAT)3 | CTCCTCATCTTCTGCGGTTC | TTCAGCAAGCCATGTGATTC | 296 |
| chr1:30854833-30856865 | p2 | (AT)16 | CTCCTCATCTTCTGCGGTTC | TTCAGCAAGCCATGTGATTC | 296 |
| chr1:30907413-30909428 | p5 | (GACGA)3 | CTTAGCTTTGGTGGGCAGAG | AAGAAGCCTAAGGGGGTTCA | 270 |
| chr1:30916299-30918320 | p3 | (GTT)7 | GTGATGGGTGTTCTAGCCGT | ATCAGTCCGCAACGCTTACT | 166 |
| chr1:30937227-30939305 | p3 | (AGA)26 | GATTCAAACCCAGAACCTCG | CAAGTCAACCGCTACCCATT | 187 |
| chr1:31049722-31051738 | p2 | (TA)11 | TCATTAAGTCAATCGCGTGC | TGACATGGCGGAATAACAAA | 227 |
| chr1:31049722-31051738 | p2 | (AT)8 | TCATTAAGTCAATCGCGTGC | TGACATGGCGGAATAACAAA | 227 |
| chr1:31078323-31080341 | p6 | (AAATGG)3 | GGAGGAATGTGTCCCAAGAA | GAAATGGTGGCGAGATCTGT | 265 |
| chr1:31107264-31109279 | p5 | (CCCAA)3 | TTCATGACTTGAGGGGGTTC | ATACCGGGGGTATTTTAGCG | 263 |
| chr1:31146128-31148146 | p6 | (AAATGG)3 | GGAGGAATGTGTCCCAAGAA | GAAATGGTGGCGAGATCTGT | 265 |
| chr1:31192225-31194237 | p2 | (TG)6 | ATCCGCTCCATTTAAACACG | AAGCGATCAACTTGTTTCCG | 250 |
| chr1:31195305-31197357 | p2 | (AG)26 | CTACCCCTTAAATTGCCGGT | AAGGCCTAACATTTGCACGA | 249 |
| chr1:31357422-31359438 | p2 | (AT)15 | CATACAGAAGCCGCTAAGGC | CTAAGCGGTCCCATTTGCTA | 197 |
| chr1:31394205-31396244 | p3 | (TCT)13 | AGTCGGCCAAGAATATGACG | AACAGATGTTTCCGCAGCTT | 168 |

| Marker ID | SSR type | SSR motif | FORWARD PRIMER (5'-3') | REVERSE PRIMER (5'-3') | size (bp) |
|---|---|---|---|---|---|
| chr1:31408493-31410515 | p2 | (TA)7 | AGTGCTAAACGACCAAACGG | AACCCCAAGTGCCTTTTCAT | 237 |
| chr1:31408493-31410515 | p2 | (TA)11 | AGTGCTAAACGACCAAACGG | AACCCCAAGTGCCTTTTCAT | 237 |
| chr1:31408493-31410515 | p3 | (CTA)5 | AGTGCTAAACGACCAAACGG | AACCCCAAGTGCCTTTTCAT | 237 |
| chr1:31408493-31410515 | p2 | (AC)8 | AGTGCTAAACGACCAAACGG | AACCCCAAGTGCCTTTTCAT | 237 |
| chr1:31408493-31410515 | p3 | (CTA)6 | AGTGCTAAACGACCAAACGG | AACCCCAAGTGCCTTTTCAT | 237 |
| chr1:31409439-31411457 | p2 | (TA)11 | GCCACTTGGCATCAAATTCT | CATTATTCACGCGCCACTT | 126 |
| chr1:31409439-31411457 | p3 | (CTA)5 | GCCACTTGGCATCAAATTCT | CATTATTCACGCGCCACTT | 126 |
| chr1:31409439-31411457 | p2 | (AC)8 | GCCACTTGGCATCAAATTCT | CATTATTCACGCGCCACTT | 126 |
| chr1:31409439-31411457 | p3 | (CTA)6 | GCCACTTGGCATCAAATTCT | CATTATTCACGCGCCACTT | 126 |
| chr1:31409439-31411457 | p2 | (TA)9 | GCCACTTGGCATCAAATTCT | CATTATTCACGCGCCACTT | 126 |
| chr1:31493538-31495556 | p3 | (TAA)6 | TTTCAAAACGAGACGAGCAA | CGTTCTTGGCTTTCCCACTA | 151 |
| chr1:31493538-31495556 | p3 | (ATA)6 | TTTCAAAACGAGACGAGCAA | CGTTCTTGGCTTTCCCACTA | 151 |
| chr1:31600295-31602349 | p3 | (AAT)18 | AAACCAAGCGACTGCATTTT | TGGGGTGAAGGTTTCTTTTG | 153 |
| chr1:31698091-31700103 | p2 | (AT)6 | TGCTAGCTGAACGATCCCTT | CAGAAGGGCAAAGCTACGAC | 215 |
| chr1:31719508-31721523 | p5 | (AAATT)3 | TTAAACGAAAGCCGCCTAGA | ACCAGAAGCCGTCTAACCAA | 134 |
| chr1:31719508-31721523 | p3 | (TTA)5 | TTAAACGAAAGCCGCCTAGA | ACCAGAAGCCGTCTAACCAA | 134 |
| chr1:31829437-31831455 | p6 | (TTTTTA)3 | GTTCTGAGTCGTTGGGTCGT | CCAACAATGAATGTGTGGGA | 122 |
| chr1:31829437-31831455 | p2 | (AT)14 | GTTCTGAGTCGTTGGGTCGT | CCAACAATGAATGTGTGGGA | 122 |
| chr1:31830293-31832321 | p6 | (TTTTTA)3 | TGCTTTTAAAGTCCCGGAAA | AGTATGGGACGCATCGAATC | 192 |
| chr1:31830293-31832321 | p2 | (AT)14 | TGCTTTTAAAGTCCCGGAAA | AGTATGGGACGCATCGAATC | 192 |
| chr1:31830293-31832321 | p2 | (TA)15 | TGCTTTTAAAGTCCCGGAAA | AGTATGGGACGCATCGAATC | 192 |
| chr1:31830955-31832985 | p2 | (AT)14 | GGTGGCGACTCCTTAAAACA | GGGGTCCATTTTGTTTTTCC | 254 |
| chr1:31830955-31832985 | p2 | (TA)15 | GGTGGCGACTCCTTAAAACA | GGGGTCCATTTTGTTTTTCC | 254 |
| chr1:31830955-31832985 | p2 | (TA)29 | GGTGGCGACTCCTTAAAACA | GGGGTCCATTTTGTTTTTCC | 254 |
| chr1:31862663-31864681 | p6 | (GTCCAT)3 | AAAACAAGCCCGACAACAAC | CTCAAGGACTCCACCCATGT | 239 |
| chr1:31892219-31894233 | p2 | (AT)7 | ATTCTGCTTGAAGGGCAAAA | CCAAATAAATTGCGGGCTAA | 281 |
| chr1:31945146-31947164 | p2 | (TA)9 | CCCCACTCAACTTCTTCCAA | TCATGATGATTCAGCAAGCC | 169 |
| chr1:31945146-31947164 | p5 | (ATAGA)3 | CCCCACTCAACTTCTTCCAA | TCATGATGATTCAGCAAGCC | 169 |
| chr1:31953503-31955518 | p3 | (ATA)5 | GGTTTCCCTCATTTTGCTGA | ACCATTTCTTGCACTCGGTC | 217 |
| chr1:31953503-31955518 | p3 | (TAA)7 | GGTTTCCCTCATTTTGCTGA | ACCATTTCTTGCACTCGGTC | 217 |
| chr1:31976311-31978341 | p2 | (AT)15 | AATGCAGTCAGCAATTTCCC | CAGATCAAACCCCTGACGAT | 165 |
| chr1:32022630-32024650 | p4 | (TTTA)5 | CATGGACGATGGAATTGTGA | CGGATGGTAGAGCGAAGAAC | 172 |
| chr1:32082888-32084903 | p3 | (AAG)5 | AGTTTGAATCTCGGAAGCGA | TGTTGCAAAAGCTCGTCATC | 166 |
| chr1:32173233-32175248 | p3 | (ATA)5 | AGTGGTGTAGTGGAGGGTGC | CGACCCGTCTTTATCCAAAA | 200 |
| chr1:32173233-32175248 | p3 | (ATA)13 | AGTGGTGTAGTGGAGGGTGC | CGACCCGTCTTTATCCAAAA | 200 |
| chr1:32256207-32258231 | p6 | (CCGAAG)4 | AACATTCTCTGCTGCCATCC | AACTCCCTCTGCAACTGGAA | 167 |

| Marker ID | SSR type | SSR motif | FORWARD PRIMER (5'-3') | REVERSE PRIMER (5'-3') | size (bp) |
|---|---|---|---|---|---|
| chr1:32304400-32306416 | p5 | (ACACA)3 | TCACACAACCACATTTGCCT | TTATTCCCCGGAACCTCTCT | 259 |
| chr1:32304426-32306444 | p5 | (ACACA)3 | TCACACAACCACATTTGCCT | TTATTCCCCGGAACCTCTCT | 233 |
| chr1:32363404-32365419 | p6 | (AAGATC)3 | TTGGGAAAATTAAGCATCCG | CCGAATTAGTAAGCTCCCCC | 139 |
| chr1:32363404-32365419 | p5 | (AGAAA)3 | TTGGGAAAATTAAGCATCCG | CCGAATTAGTAAGCTCCCCC | 139 |
| chr1:32365086-32367098 | p2 | (TA)6 | GCGTGTCAAAACCAAACCTT | TGGGTTGCAAGGAGGTTATC | 260 |
| chr1:32382245-32384269 | p2 | (CA)6 | TCTCTTTGCCACCCTTCTGT | CTTGAAATCGTTGTTCCGGT | 297 |
| chr1:32382245-32384269 | p6 | (AAGCCA)4 | TCTCTTTGCCACCCTTCTGT | CTTGAAATCGTTGTTCCGGT | 297 |
| chr1:32382245-32384269 | p3 | (TGT)7 | TCTCTTTGCCACCCTTCTGT | CTTGAAATCGTTGTTCCGGT | 297 |
| chr1:32383114-32385135 | p6 | (AAGCCA)4 | AATAAGCCAAAGCCAAAGCC | CGTGCTAGGGCATCTGGTAT | 128 |
| chr1:32383114-32385135 | p3 | (TGT)7 | AATAAGCCAAAGCCAAAGCC | CGTGCTAGGGCATCTGGTAT | 128 |
| chr1:32406634-32408667 | p3 | (CTT)11 | AAGGCAGAATTTGCATGACC | CCGTTTTCTCAGCGTGTGTA | 115 |
| chr1:32411176-32413188 | p2 | (TA)8 | AAAACAGTGACGGCAAGGTC | TGGGCGGTGTAGCTTTAATC | 300 |
| chr1:32411176-32413188 | p2 | (TA)6 | AAAACAGTGACGGCAAGGTC | TGGGCGGTGTAGCTTTAATC | 300 |
| chr1:32423727-32425743 | p6 | (CATGAG)3 | TACTGAGCAGGTGCGTTGTC | GAGATCAGACTCCACGCTCC | 241 |
| chr1:32423727-32425743 | p4 | (TACA)4 | TACTGAGCAGGTGCGTTGTC | GAGATCAGACTCCACGCTCC | 241 |
| chr1:32477005-32479023 | p3 | (ATG)6 | CTGGTAAACCAGAACCGGAA | AGAGGGGGAGGAAAATGAAA | 252 |
| chr1:32483845-32485860 | p5 | (AAAAT)3 | TGGTCAAACAAACAGCCAAA | GCAAAACTCTCCCCATTTCA | 111 |
| chr1:32624864-32626879 | p3 | (TTA)5 | GAGAACCCTTGAGTCGTGGA | CCATATTTTGCTGGGCAGTT | 177 |
| chr1:32624880-32626898 | p3 | (TTA)5 | GAGAACCCTTGAGTCGTGGA | CCATATTTTGCTGGGCAGTT | 161 |
| chr1:32726968-32729000 | p2 | (TA)16 | CACCCAACCCCGTATTTATG | AGCTATCACGGGATCACCAG | 203 |
| chr1:32833982-32835998 | p4 | (TACA)4 | GTGGTATCCGAGGGTGTCAT | GAAGAGCCTCACCAATGAGC | 152 |
| chr1:32854681-32856699 | p6 | (TTTTAT)3 | TCCTACCCATCCCTAAACCC | TCCCAGTGTGTCTCACCAAA | 227 |
| chr1:32911157-32913169 | p2 | (AT)6 | AATCACGGGCTCATACGTTC | GGACCTCCTCCTTGTTGGAT | 218 |
| chr1:32911157-32913169 | p2 | (TC)6 | AATCACGGGCTCATACGTTC | GGACCTCCTCCTTGTTGGAT | 218 |
| chr1:32987463-32989478 | p5 | (CATTG)3 | CATTGGTTTGATCGTTCGTG | ACCATGGTGACAAGCTCACA | 236 |
| chr1:33047302-33049344 | p2 | (AT)14 | CCCAGGGAAAGGTAAGGTGT | ACCACGCCCTTTTGTCTATG | 268 |
| chr1:33047302-33049344 | p2 | (AT)21 | CCCAGGGAAAGGTAAGGTGT | ACCACGCCCTTTTGTCTATG | 268 |
| chr1:33087934-33089946 | p2 | (TA)6 | GGGCCGTAATAGGTCCGTAG | CAGCCTATACACACGCATCG | 114 |
| chr1:33105501-33107527 | p2 | (AT)13 | CCTTGGACAACCTAGGCTCA | TCAAGCAGGATTGAGGCTTT | 226 |
| chr1:33130856-33132871 | p5 | (ATTTT)3 | CCTGCATGTCTGTGTTCACC | TTTTTCTCGGTTTGATTCGG | 202 |
| chr1:33130856-33132871 | p5 | (TTGGT)3 | CCTGCATGTCTGTGTTCACC | TTTTTCTCGGTTTGATTCGG | 202 |
| chr1:33142424-33144439 | p5 | (TTGGT)3 | CCTGCATGTCTGTGTTCACC | TTTTTCTCGGTTTGATTCGG | 210 |
| chr1:33142424-33144439 | p5 | (ATTTT)3 | CCTGCATGTCTGTGTTCACC | TTTTTCTCGGTTTGATTCGG | 210 |
| chr1:33246381-33248399 | p2 | (TA)9 | CCGGAAAGCAGAGAGTTGAC | TTTCGAGTCGTGGGACTCTT | 183 |
| chr1:33258021-33260037 | p4 | (ATTT)4 | GCAAACTTCCAGCCATGATT | CACAAACAAAATTGCCCAGA | 106 |
| chr1:33703335-33705350 | p5 | (ATAAA)3 | TCAAATGCAAACCAGACCAA | CCGTGACAGCGTATGTTTGT | 238 |

| Marker ID | SSR type | SSR motif | FORWARD PRIMER (5'-3') | REVERSE PRIMER (5'-3') | size (bp) |
|---|---|---|---|---|---|
| chr1:33717704-33719722 | p2 | (TC)9 | ATGTTTTCTGTACGGGCGAC | TAATGTTGTCACGACCCAGC | 174 |
| chr1:33759893-33761965 | p6 | (CATCAC)3 | TAATTCCACAAGCCCACCTC | CCACATGCAGTCGTTAATGG | 130 |
| chr1:33762431-33765040 | p6 | (TAAAAA)3 | CTTTACATCCCTTGCCCAAA | GCAAAGCCAAGGTTGATGAT | 255 |
| chr1:33913671-33915683 | p2 | (TA)6 | TTGCTCCACGCCCTACTACT | ACCATGCTAAGACGTCCACC | 187 |
| chr1:33913671-33915683 | p2 | (TA)12 | TTGCTCCACGCCCTACTACT | ACCATGCTAAGACGTCCACC | 187 |
| chr1:33913905-33915923 | p2 | (TA)6 | AAGTATCAACCGTGGCTTCG | TTTCGTTAATTTCCCCCACA | 282 |
| chr1:33913905-33915923 | p2 | (TA)12 | AAGTATCAACCGTGGCTTCG | TTTCGTTAATTTCCCCCACA | 282 |
| chr1:33995485-33997503 | p6 | (ATGTTC)3 | TTTTCACATTCTCCAAGCCC | CCACACGTGGATAGAAACCC | 236 |
| chr1:33995485-33997503 | p2 | (TA)60 | TTTTCACATTCTCCAAGCCC | CCACACGTGGATAGAAACCC | 236 |
| chr1:34012930-34014945 | p5 | (TGAAC)3 | GAGCCTCCTTGGGAATTAGG | TCAATTGATCCCTCTCTGCC | 106 |
| chr1:34037563-34039579 | p2 | (TG)8 | GCTAGTGGCCAAACCAAAAG | GTGTCGGGATCTGGTCCTTA | 150 |
| chr1:34047857-34049872 | p3 | (TTG)5 | AGAAGATGAAAGTCGCGGAA | ATTTCTTCGGCCTCCCTTTA | 300 |
| chr1:34080957-34082975 | p2 | (TA)14 | GGACAATGTCTAAACCCCGA | ATGTCCCCTTTTTATTGCCC | 300 |
| chr1:34080957-34082975 | p2 | (AT)9 | GGACAATGTCTAAACCCCGA | ATGTCCCCTTTTTATTGCCC | 300 |
| chr1:34095960-34097975 | p3 | (TTA)18 | CAGGTTCATTTGCGTGCTTA | CTGGGATATCCACATCCACC | 208 |
| chr1:34095960-34097975 | p5 | (ACCCG)3 | CAGGTTCATTTGCGTGCTTA | CTGGGATATCCACATCCACC | 208 |
| chr1:34176316-34178336 | p2 | (GA)10 | AAACCGGTGGTCATTGTTGT | GGGCAGATGGCTGAATTAGA | 177 |
| chr1:34178086-34180106 | p2 | (GA)10 | CAAAAATGGCAGAATGAGGG | TGCAGCAGCATTTTAACCAG | 181 |
| chr1:34263489-34265504 | p5 | (AAGCC)3 | AGAAAATTGGGCAAACAACG | TAGCCTACGGGCAAATCAAC | 248 |
| chr1:34350585-34352597 | p2 | (AT)6 | TTGGAACCACCACAAGTTCA | ATCTTCAACGACACTTGCCC | 229 |
| chr1:34350585-34352597 | p2 | (AT)6 | TTGGAACCACCACAAGTTCA | ATCTTCAACGACACTTGCCC | 229 |
| chr1:34404115-34406130 | p5 | (TTTCT)3 | TGTGCCTACATATGGGGGAT | ATTCTGCCACATCATCACGA | 284 |
| chr1:34430864-34432876 | p2 | (TA)7 | CCACCGAACCGACTTTTAGA | CAACGCCCAAGGTAGAAAAA | 211 |
| chr1:34430864-34432876 | p2 | (TA)6 | CCACCGAACCGACTTTTAGA | CAACGCCCAAGGTAGAAAAA | 211 |
| chr1:34433328-34435346 | p3 | (TTG)6 | CCAAGATTCGAACCTGCACT | TCAGAATTTTAAGACGGCGG | 168 |
| chr1:34545315-34547341 | p2 | (AT)13 | GCTTGGCTATCGAAGTGGAG | CTCGATCTCCTTGTCTTCCG | 124 |
| chr1:34548223-34550247 | p2 | (CT)12 | ACACCATACACACGGCTCAA | ACCCACTATTTGGAGGTCCC | 216 |
| chr1:34612806-34614821 | p3 | (TTC)5 | CAGTGCGTTTGAAGGGAAAT | TGCACAATTCAGGAAATGGA | 233 |
| chr1:34665432-34667446 | p2 | (AT)7 | GCGGATATCGAACCAGAAAA | AGCTATCGAACCTGGGGACT | 113 |
| chr1:34675304-34677322 | p2 | (TA)9 | CGTGCCACGTATCAAATGAC | TTTTGGCTTTGTGAAGACCC | 159 |
| chr1:34675304-34677322 | p2 | (AT)6 | CGTGCCACGTATCAAATGAC | TTTTGGCTTTGTGAAGACCC | 159 |
| chr1:34726104-34728122 | p6 | (AACCGG)3 | ATCGCCGATTACTACCGTTG | TGAGGAATGCTAAATGCACG | 182 |
| chr1:34726104-34728122 | p2 | (AT)7 | ATCGCCGATTACTACCGTTG | TGAGGAATGCTAAATGCACG | 182 |
| chr1:34726104-34728122 | p2 | (AT)7 | ATCGCCGATTACTACCGTTG | TGAGGAATGCTAAATGCACG | 182 |
| chr1:34729892-34731904 | p2 | (TG)6 | TAAGCCATAAGGACGGCAAC | AATTGCAACGTTCAACCTCC | 240 |
| chr1:34759925-34761937 | p2 | (AG)6 | TTGACGGCTCTCCCAGTAGT | GGGCCGTACAACATCTGACT | 111 |

| Marker ID | SSR type | SSR motif | FORWARD PRIMER (5'-3') | REVERSE PRIMER (5'-3') | size (bp) |
|---|---|---|---|---|---|
| chr1:34787413-34789428 | p5 | (TTGGT)3 | GTGGTTTGATTTGCCTTGGT | GTAATGTCGGGTTGGTTTGG | 188 |
| chr1:34787413-34789428 | p2 | (TA)8 | GTGGTTTGATTTGCCTTGGT | GTAATGTCGGGTTGGTTTGG | 188 |
| chr1:34833933-34835987 | p2 | (AT)27 | GCATCTTGGTCATTTCGGAC | TGCTGAGATAAGCAAGGGGT | 295 |
| chr1:35000808-35002828 | p6 | (TTTCTT)3 | TCCGTTGATTCACGCAATAA | TGTGAAATTGCACGGTTCAT | 282 |
| chr1:35000808-35002828 | p2 | (TA)8 | TCCGTTGATTCACGCAATAA | TGTGAAATTGCACGGTTCAT | 282 |
| chr1:35000916-35002932 | p6 | (TTTCTT)3 | TCCGTTGATTCACGCAATAA | TGTGAAATTGCACGGTTCAT | 174 |
| chr1:35000916-35002932 | p2 | (TA)8 | TCCGTTGATTCACGCAATAA | TGTGAAATTGCACGGTTCAT | 174 |
| chr1:35102170-35104185 | p3 | (AAC)5 | TGCTTTAACCGTCACGAATG | CATCTAGCCGTTTCCAGAGC | 250 |
| chr1:35133207-35135225 | p2 | (AT)28 | ATTTCGCTGGGATTGTAACG | AGCTCATCATGCACTTCACG | 286 |
| chr1:35133207-35135225 | p2 | (AT)7 | ATTTCGCTGGGATTGTAACG | AGCTCATCATGCACTTCACG | 286 |
| chr1:35133231-35135247 | p2 | (AT)28 | ATTTCGCTGGGATTGTAACG | AGCTCATCATGCACTTCACG | 262 |
| chr1:35133231-35135247 | p2 | (AT)7 | ATTTCGCTGGGATTGTAACG | AGCTCATCATGCACTTCACG | 262 |
| chr1:35133246-35135264 | p2 | (AT)28 | ATTTCGCTGGGATTGTAACG | AGCTCATCATGCACTTCACG | 247 |
| chr1:35133246-35135264 | p2 | (AT)7 | ATTTCGCTGGGATTGTAACG | AGCTCATCATGCACTTCACG | 247 |
| chr1:35152704-35154716 | p2 | (AT)6 | GGGATGGCTGAAGTGTTGTT | AAAGCAAAGTCGTCGAAGGA | 262 |
| chr1:35203082-35205100 | p6 | (ACTGAA)3 | TTCTCTCGCAGGCATTTTCT | CCAATGGGTCGCTCTTATGT | 222 |
| chr1:35310730-35312748 | p2 | (AT)9 | TTTAAATGTAGCCGTTGGGC | GGGGGATGGGTTAAAGTGTT | 297 |
| chr1:35317161-35319176 | p5 | (AAAAT)3 | TCCTTTTTGGTCGTCATTCC | TTTGGATGTGGTGAGTTCCA | 270 |
| chr1:35351662-35353676 | p2 | (GT)7 | CCCGTATGTCTCTGCTAGGC | ATTATGAATTCGCCACTCGG | 239 |
| chr1:35358394-35360409 | p5 | (TTTTA)3 | CATCTTTTGCCGAAGCTACC | TAGCTGTGCATTTGACCCAC | 245 |
| chr1:35367227-35369249 | p2 | (AT)11 | TCTCAGAAGCCCTGACCAGT | CGGGCTAGGAGGTGATGATA | 121 |
| chr1:35367227-35369249 | p2 | (AT)29 | TCTCAGAAGCCCTGACCAGT | CGGGCTAGGAGGTGATGATA | 121 |
| chr1:35369346-35371384 | p2 | (AT)19 | CTCGATCACTTGCCAAACAA | TGAATTTACAGGGGAGGTGC | 231 |
| chr1:35410022-35412036 | p2 | (CT)7 | TCTGTTCATGGATGAGGCAA | TCGGTGAGCAATGGTGATAA | 223 |
| chr1:35476320-35478335 | p5 | (GATAT)3 | GCAAGCTTCACTCAACACCA | GCCGAATTTGGCCACTATAA | 247 |
| chr1:35540333-35542363 | p2 | (TA)18 | AAGTACCCCGGATTTTGGAC | CTAGGGAAATGGCTGCATTG | 113 |
| chr1:35557602-35559620 | p6 | (TTTTTC)3 | CGATCGGTTCTTTCCACATT | TCCGATGATGACGACGTAAA | 218 |
| chr1:35625903-35627941 | p2 | (TC)19 | TAGGCGTCGTGGTTTATTCC | GATGAAAGAATTTTCGGGCA | 116 |
| chr1:35628774-35630786 | p2 | (TA)6 | TTTCGGTTTGTTTGAGGGTC | ATGCCCAAAACCACACTAGG | 205 |
| chr1:35636495-35638510 | p5 | (TAGTT)3 | CCTGCTCCTCTCTTCCCTTT | GAATTCAACTCGCGACCATT | 208 |
| chr1:35639429-35641477 | p2 | (TG)10 | TTACCACGTGCGTGTCAATC | GACATGGCGGAATAGCAAAT | 245 |
| chr1:35639429-35641477 | p2 | (TA)10 | TTACCACGTGCGTGTCAATC | GACATGGCGGAATAGCAAAT | 245 |
| chr1:35642208-35644280 | p2 | (AT)16 | AAATCCTTTTTGGTCCCACC | TTGGCGTTTTTAGTATGGGG | 202 |
| chr1:35642290-35644368 | p2 | (AT)16 | AAATCCTTTTTGGTCCCACC | TTGGCGTTTTTAGTATGGGG | 120 |
| chr1:35659264-35661286 | p2 | (AT)11 | GGACTACCCGGAGCATGTTA | ACACCATGAACCACCAACCT | 238 |
| chr1:35699290-35701302 | p5 | (CACCC)5 | ATTCCACGCTGTTTGTTTCC | AGCGAAAATGCTTTCTTGGA | 111 |

| Marker ID | SSR type | SSR motif | FORWARD PRIMER (5'-3') | REVERSE PRIMER (5'-3') | size (bp) |
|---|---|---|---|---|---|
| chr1:35699290-35701302 | p2 | (AT)6 | ATTCCACGCTGTTTGTTTCC | AGCGAAAATGCTTTCTTGGA | 111 |
| chr1:35699290-35701302 | p2 | (GA)8 | ATTCCACGCTGTTTGTTTCC | AGCGAAAATGCTTTCTTGGA | 111 |
| chr1:35699290-35701302 | p2 | (GA)8 | ATTCCACGCTGTTTGTTTCC | AGCGAAAATGCTTTCTTGGA | 111 |
| chr1:35777871-35779883 | p2 | (TA)6 | AAAAAGGGGAACCCAACATC | ACGACGAAGCATGGACATTT | 290 |
| chr1:35777871-35779883 | p2 | (TA)17 | AAAAAGGGGAACCCAACATC | ACGACGAAGCATGGACATTT | 290 |
| chr1:35778646-35780680 | p2 | (TA)6 | GGGCCCCATACTAAACAACA | TTTTTAGGGACAACGGACGA | 276 |
| chr1:35778646-35780680 | p2 | (TA)17 | GGGCCCCATACTAAACAACA | TTTTTAGGGACAACGGACGA | 276 |
| chr1:35852992-35855006 | p2 | (AT)7 | CGATGTGTGACATCCTCGTC | GGTCCCTTTATGTTACGCGA | 111 |
| chr1:35886975-35888989 | p2 | (AT)7 | CGATGTGTGACATCCTCGTC | GGTCCCTTTATGTTACGCGA | 111 |
| chr1:36051286-36053301 | p5 | (AAATG)3 | TTGAGGGGATTTGCTATTGG | TCCTTCAGCGCGTAAGAGAT | 230 |
| chr1:36066795-36068813 | p6 | (AGAAAA)3 | GCTGCAAGACAAATCGAGGT | CCCAATATTCACCGGCTCTA | 108 |
| chr1:36068659-36070677 | p6 | (TGTTGC)3 | AGGAGGTGTCCCTTGTGATG | GCGGGCTCTAAAATCAACAG | 241 |
| chr1:36115167-36117191 | p4 | (AAAG)6 | TTTTGCCCTTTTCCCTTTTT | TACTGGAAAGGGTGTCCCAG | 230 |
| chr1:36115167-36117191 | p6 | (GGGGGT)4 | TTTTGCCCTTTTCCCTTTTT | TACTGGAAAGGGTGTCCCAG | 230 |
| chr1:36125414-36127426 | p2 | (CT)6 | CCGACGCCTTCATTACATTT | CGGACTGCTAGTGCTAAGGC | 169 |
| chr1:36128172-36130186 | p2 | (CT)7 | CCGACGCCTTCATTACATTT | AGGCTGGACGACAGGATATG | 169 |
| chr1:36132261-36134276 | p3 | (CAA)5 | CAACCTGGGAGTAACCTGGA | ATCCTAGCTAAGCCGGAAGC | 135 |
| chr1:36132261-36134276 | p2 | (CT)6 | CAACCTGGGAGTAACCTGGA | ATCCTAGCTAAGCCGGAAGC | 135 |
| chr1:36135656-36137668 | p2 | (CT)6 | CCGACGCCTTCATTACATTT | CGGACTGCTAGTGCTAAGGC | 169 |
| chr1:36138414-36140428 | p2 | (CT)7 | CCGACGCCTTCATTACATTT | AGGCTGGACGACAGGATATG | 169 |
| chr1:36142503-36144518 | p3 | (CAA)5 | CAACCTGGGAGTAACCTGGA | ATCCTAGCTAAGCCGGAAGC | 135 |
| chr1:36142503-36144518 | p2 | (CT)6 | CAACCTGGGAGTAACCTGGA | ATCCTAGCTAAGCCGGAAGC | 135 |
| chr1:36185695-36187710 | p5 | (ATATA)3 | CAATTCTGTCAGCTTTGCCA | CTTTCCAGGGATTGTGGTGT | 159 |
| chr1:36226007-36228041 | p2 | (TA)11 | TAGGCATTTCCGACCAACTC | TGGTTGCTGCTTCTTTTGTG | 113 |
| chr1:36239317-36241329 | p2 | (AT)6 | TGAGCCGGTAAATGCTATCC | GGAGGCTTGGTACATCCTCA | 115 |
| chr1:36239317-36241329 | p4 | (CATA)4 | TGAGCCGGTAAATGCTATCC | GGAGGCTTGGTACATCCTCA | 115 |
| chr1:36315047-36317067 | p5 | (TGCAT)3 | GGGTCCTCTGCAGGTTATGA | TTCAACAACCCCACCAAAAT | 221 |
| chr1:36315047-36317067 | p4 | (TATT)5 | GGGTCCTCTGCAGGTTATGA | TTCAACAACCCCACCAAAAT | 221 |
| chr1:36315047-36317067 | p4 | (ATAG)5 | GGGTCCTCTGCAGGTTATGA | TTCAACAACCCCACCAAAAT | 221 |
| chr1:36342083-36344097 | p2 | (TA)8 | CGGGGTCTTGTCTTGGTAAA | ATACCCACCCCCTTGTTTGT | 295 |
| chr1:36342099-36344113 | p2 | (TA)8 | CGGGGTCTTGTCTTGGTAAA | ATACCCACCCCCTTGTTTGT | 279 |
| chr1:36342115-36344129 | p2 | (TA)8 | CGGGGTCTTGTCTTGGTAAA | ATACCCACCCCCTTGTTTGT | 263 |
| chr1:36342131-36344145 | p2 | (TA)8 | CGGGGTCTTGTCTTGGTAAA | ATACCCACCCCCTTGTTTGT | 247 |
| chr1:36342147-36344161 | p2 | (TA)8 | CGGGGTCTTGTCTTGGTAAA | ATACCCACCCCCTTGTTTGT | 231 |
| chr1:36342163-36344177 | p2 | (TA)8 | CGGGGTCTTGTCTTGGTAAA | ATACCCACCCCCTTGTTTGT | 215 |
| chr1:36342179-36344193 | p2 | (TA)8 | CGGGGTCTTGTCTTGGTAAA | ATACCCACCCCCTTGTTTGT | 199 |

| Marker ID | SSR type | SSR motif | FORWARD PRIMER (5'-3') | REVERSE PRIMER (5'-3') | size (bp) |
|---|---|---|---|---|---|
| chr1:36342195-36344209 | p2 | (TA)8 | CGGGGTCTTGTCTTGGTAAA | ATACCCACCCCCTTGTTTGT | 183 |
| chr1:36342211-36344225 | p2 | (TA)8 | CGGGGTCTTGTCTTGGTAAA | ATACCCACCCCCTTGTTTGT | 167 |
| chr1:36342227-36344241 | p2 | (TA)8 | CGGGGTCTTGTCTTGGTAAA | ATACCCACCCCCTTGTTTGT | 151 |
| chr1:36368462-36370476 | p2 | (TA)8 | CGGGGTCTTGTCTTGGTAAA | ATACCCACCCCCTTGTTTGT | 295 |
| chr1:36368478-36370492 | p2 | (TA)8 | CGGGGTCTTGTCTTGGTAAA | ATACCCACCCCCTTGTTTGT | 279 |
| chr1:36368494-36370508 | p2 | (TA)8 | CGGGGTCTTGTCTTGGTAAA | ATACCCACCCCCTTGTTTGT | 263 |
| chr1:36368510-36370524 | p2 | (TA)8 | CGGGGTCTTGTCTTGGTAAA | ATACCCACCCCCTTGTTTGT | 247 |
| chr1:36368526-36370540 | p2 | (TA)8 | CGGGGTCTTGTCTTGGTAAA | ATACCCACCCCCTTGTTTGT | 231 |
| chr1:36368542-36370556 | p2 | (TA)8 | CGGGGTCTTGTCTTGGTAAA | ATACCCACCCCCTTGTTTGT | 215 |
| chr1:36368558-36370572 | p2 | (TA)8 | CGGGGTCTTGTCTTGGTAAA | ATACCCACCCCCTTGTTTGT | 199 |
| chr1:36368574-36370588 | p2 | (TA)8 | CGGGGTCTTGTCTTGGTAAA | ATACCCACCCCCTTGTTTGT | 183 |
| chr1:36368590-36370604 | p2 | (TA)8 | CGGGGTCTTGTCTTGGTAAA | ATACCCACCCCCTTGTTTGT | 167 |
| chr1:36368606-36370620 | p2 | (TA)8 | CGGGGTCTTGTCTTGGTAAA | ATACCCACCCCCTTGTTTGT | 151 |
| chr1:36404441-36406463 | p2 | (AT)11 | ATGATTCATGCATCCGGTTT | AAGTGGGAAGCCCTCAAGTT | 296 |
| chr1:36404441-36406463 | p2 | (AT)38 | ATGATTCATGCATCCGGTTT | AAGTGGGAAGCCCTCAAGTT | 296 |
| chr1:36490357-36492381 | p6 | (GGATTT)4 | CAATGTTCGGGAACAAGGAT | ACCCTTAACCGTTCTTGGCT | 290 |
| chr1:36493644-36495658 | p6 | (AAAAAT)3 | AGCTCGATGCACCGTCTTAT | AAAGCAGATCGAGGAGGGAT | 194 |
| chr1:36493644-36495658 | p2 | (TA)7 | AGCTCGATGCACCGTCTTAT | AAAGCAGATCGAGGAGGGAT | 194 |
| chr1:36493644-36495658 | p6 | (GAAGGT)3 | AGCTCGATGCACCGTCTTAT | AAAGCAGATCGAGGAGGGAT | 194 |
| chr1:36563367-36565383 | p4 | (AAAG)4 | AATGTGTTGTTTTCGGGAGC | GCCACAAATTGGATATTCGG | 143 |
| chr1:36622737-36624775 | p2 | (AT)19 | GACTGCAACGTCCCAAAAAT | AGCAAGTGAATACGATCCGC | 119 |
| chr1:36622737-36624775 | p2 | (TA)7 | GACTGCAACGTCCCAAAAAT | AGCAAGTGAATACGATCCGC | 119 |
| chr1:36622737-36624775 | p2 | (TA)6 | GACTGCAACGTCCCAAAAAT | AGCAAGTGAATACGATCCGC | 119 |
| chr1:36871064-36873082 | p6 | (CCGGTT)3 | CATTAAGCCCACCCCTAACA | TCCGGCAATTTGGTACTTTC | 207 |
| chr1:36881827-36883845 | p2 | (AT)9 | CGTTTCGGAGTCTTGACACA | AGCGCAGTAAAATAGCGAGC | 271 |
| chr1:36881827-36883845 | p5 | (CATAT)3 | CGTTTCGGAGTCTTGACACA | AGCGCAGTAAAATAGCGAGC | 271 |
| chr1:36890028-36892052 | p2 | (GA)12 | TCAGTGAGGCGCCTTTTAGT | GCCCAACTACTCTGGGATCA | 101 |
| chr1:36901153-36903167 | p2 | (TA)7 | AGGTGGGCCCCATACTAAAC | TTTTTCCCCATGAGTTTGGA | 275 |
| chr1:36918317-36920357 | p2 | (TA)20 | CCCCAATCGGTTGATTACAC | CTTCGTAGCTTGAGGCTTGC | 249 |
| chr1:36984458-36986470 | p2 | (AT)9 | TTTTGTGGATGTTCCAACGA | TTCCACGAAATGCAAATCAA | 227 |
| chr1:36984473-36986519 | p2 | (AT)9 | TTTTGTGGATGTTCCAACGA | TTCCACGAAATGCAAATCAA | 212 |
| chr1:36984784-36986798 | p2 | (AT)9 | TTCAAAAATTGAAACCGCAA | ACACAAACTACCGCAAGGCT | 213 |
| chr1:36984811-36986833 | p2 | (AT)9 | TTCAAAAATTGAAACCGCAA | ACACAAACTACCGCAAGGCT | 186 |
| chr1:37164928-37166943 | p5 | (ATTTC)3 | AAGCATGCGATCCAGATACC | ATAACAATGCCTCACAGGGC | 183 |
| chr1:37188972-37190984 | p2 | (TA)7 | TTACAGACGTCCCGGATAGG | TTTCTTTTTCATGGTTCGGG | 104 |
| chr1:37188972-37190984 | p5 | (ATATA)3 | TTACAGACGTCCCGGATAGG | TTTCTTTTTCATGGTTCGGG | 104 |

| Marker ID | SSR type | SSR motif | FORWARD PRIMER (5'-3') | REVERSE PRIMER (5'-3') | size (bp) |
|---|---|---|---|---|---|
| chr1:37188972-37190984 | p2 | (AT)6 | TTACAGACGTCCCGGATAGG | TTTCTTTTTCATGGTTCGGG | 104 |
| chr1:37248944-37250958 | p2 | (TA)7 | CACCCCAATACCTTCTTCCA | AAGCGCTCTTTGGAATTTGA | 155 |
| chr1:37259294-37261324 | p2 | (TA)15 | GATACGGAGCACAAACAGCA | AAAAATGGAAACGGACCAGA | 228 |
| chr1:37259417-37261429 | p2 | (TA)15 | GATACGGAGCACAAACAGCA | AAAAATGGAAACGGACCAGA | 105 |
| chr1:37315184-37317199 | p5 | (TAAAA)3 | GTCAAATGTTCTTGCCCGAT | GCAAGTTATCTCCGCTCCAC | 187 |
| chr1:37381391-37383409 | p2 | (AT)9 | GGGCTATCCGGATTGGTATT | AAAGACGCCAACAACTACGG | 239 |
| chr1:37413866-37415882 | p2 | (TA)8 | CAGGTGGGTTCACTTGGAGT | CATATTCCCTGTGGGATTCG | 236 |
| chr1:37488981-37490996 | p3 | (TCA)5 | TGGTGCAAAGATGGAATGAA | ATGTGGATCCAGAACTTGCC | 107 |
| chr1:37511073-37513085 | p2 | (TA)6 | TTGATTCCGATTCGGTCTTC | CCATATTCCTGTGGGATTCG | 249 |
| chr1:37555912-37557927 | p5 | (ATTTA)3 | CGACTTCGCTTCGTATCACA | CAAAATCGCGCCATAGAAAT | 159 |
| chr1:37690945-37692987 | p2 | (AT)8 | CATGAGGCATAACATGAGCG | TCGTATAGCCATTTCGGCTC | 297 |
| chr1:37690945-37692987 | p2 | (TA)21 | CATGAGGCATAACATGAGCG | TCGTATAGCCATTTCGGCTC | 297 |
| chr1:37772408-37774428 | p2 | (GA)10 | AAGATGGTATGCCACTTGCC | CGATGCCTTGTCCTGTCTTT | 225 |
| chr1:37849268-37851282 | p2 | (AT)7 | AAGACTTGCCTTGCCACAAC | TGCCAGGACATGTGAACAAT | 115 |
| chr1:37849268-37851282 | p2 | (TA)7 | AAGACTTGCCTTGCCACAAC | TGCCAGGACATGTGAACAAT | 115 |
| chr1:38017865-38019880 | p3 | (CAA)5 | GCCTTATTGGCATCAAAGGA | TTGAATGCTTGGGAAGAAGG | 126 |
| chr1:38065455-38067470 | p5 | (TCGGT)3 | ATGCCCTTCTCCCAAAAAGT | ATTCCTTCGCCATTCCTTCT | 106 |
| chr1:38075322-38077334 | p2 | (TA)6 | CGTCAGTTGCCCTTCTTGAT | AAAACCCAGCAAAAGGAAGA | 143 |
| chr1:38086168-38088182 | p2 | (AT)7 | AGCTCCGAAGGCAGTCTGTA | ATGTTTGCCTTTGGGCTATG | 228 |
| chr1:38154983-38157004 | p3 | (ATA)7 | ATAACGTGGTAGTGGAGCGG | CCGCTCTTCCCTCTCTTCTT | 225 |
| chr1:38186542-38188564 | p2 | (AT)11 | TGCATGCCCCTTAAGAAATC | TTCCCATCTTATCCTCTCCGT | 210 |
| chr1:38210546-38212564 | p6 | (CCGGTT)3 | GTGATTGTGCAGGGCCTTAT | GTGGTGCCGGTAGTAGTGGT | 139 |
| chr1:38231129-38233153 | p3 | (GGA)5 | GGTGGTTCGCTCTGTCAAAT | TGCAGCTTGCAACTGAGTCT | 132 |
| chr1:38231129-38233153 | p2 | (AT)12 | GGTGGTTCGCTCTGTCAAAT | TGCAGCTTGCAACTGAGTCT | 132 |
| chr1:38271049-38273117 | p2 | (TA)34 | ACCTTGCCATTGACACTTCC | AATCCTGTTTGCGGTTGAAG | 143 |
| chr1:38271049-38273117 | p2 | (TA)6 | ACCTTGCCATTGACACTTCC | AATCCTGTTTGCGGTTGAAG | 143 |
| chr1:38271049-38273117 | p2 | (AT)6 | ACCTTGCCATTGACACTTCC | AATCCTGTTTGCGGTTGAAG | 143 |
| chr1:38459011-38461056 | p3 | (AAG)6 | AAACTTCCCAAACCGTTCCT | TAGGCGAGGGTGAAGAAGAA | 164 |
| chr1:38459011-38461056 | p3 | (TTG)15 | AAACTTCCCAAACCGTTCCT | TAGGCGAGGGTGAAGAAGAA | 164 |
| chr1:38522832-38524852 | p5 | (AAATA)4 | TTTGGTGTTTCCATGCTTGA | CCACCGTTACAGTAGCGACA | 297 |
| chr1:38551593-38553608 | p3 | (GAA)5 | CGGAGCGGGAATATGATAAA | GGAGAAAGCGACTTGACTGC | 272 |
| chr1:38551593-38553608 | p5 | (AATAA)3 | CGGAGCGGGAATATGATAAA | GGAGAAAGCGACTTGACTGC | 272 |
| chr1:38588355-38590369 | p2 | (TA)6 | GGGCATACGCCGTCTAGTAA | TGAGATGGTGGGTTTCCACT | 217 |
| chr1:38588355-38590369 | p2 | (TA)7 | GGGCATACGCCGTCTAGTAA | TGAGATGGTGGGTTTCCACT | 217 |
| chr1:38588355-38590369 | p6 | (TTTTTA)3 | GGGCATACGCCGTCTAGTAA | TGAGATGGTGGGTTTCCACT | 217 |
| chr1:38588355-38590369 | p2 | (TA)7 | GGGCATACGCCGTCTAGTAA | TGAGATGGTGGGTTTCCACT | 217 |

| Marker ID | SSR type | SSR motif | FORWARD PRIMER (5'-3') | REVERSE PRIMER (5'-3') | size (bp) |
|---|---|---|---|---|---|
| chr1:38599031-38601095 | p2 | (TA)32 | GGTCCTGTCCCGCTTATGTA | TCCGTCATGCTAATCCCATT | 289 |
| chr1:38604283-38606307 | p3 | (TAT)8 | AAAAACCCTAGCCTCCAAGC | TTGCAAAAAGGGGACAGAAC | 254 |
| chr1:38634492-38636520 | p2 | (GT)6 | AAGGCACAACCTCGAAGAGA | TTTAAGGGTCTGGGGGATTC | 131 |
| chr1:38634492-38636520 | p5 | (AAAAT)3 | AAGGCACAACCTCGAAGAGA | TTTAAGGGTCTGGGGGATTC | 131 |
| chr1:38741937-38743969 | p2 | (GA)16 | CTCGCGGGATGCTATTATGT | CGCCGCTCGTTCTATTACTC | 145 |
| chr1:38741937-38743969 | p2 | (TG)7 | CTCGCGGGATGCTATTATGT | CGCCGCTCGTTCTATTACTC | 145 |
| chr1:38752377-38754391 | p2 | (TC)7 | AAGCTCGGGAGGATAGGTGT | TGGAGCATGAATCAAATGGA | 246 |
| chr1:38778392-38780462 | p2 | (AT)6 | TGTCAAGCGACACTCTTTCG | TCCATTTTGACAGTGGCTGA | 115 |
| chr1:38783957-38785969 | p2 | (CA)6 | CTAAAGGCAATTTGAACGGC | GTGAAAATGGTTGGGGTTTG | 256 |
| chr1:38790955-38792977 | p2 | (AT)11 | TTGATGAGCTAGGCTCGGTT | GGTAGCACAACCCAAGGAGA | 172 |
| chr1:38790955-38792977 | p4 | (TTAA)5 | TTGATGAGCTAGGCTCGGTT | GGTAGCACAACCCAAGGAGA | 172 |
| chr1:38791716-38793736 | p2 | (AT)11 | ACCCTGGGGGTGTCATTATT | ACGACCCATTTTGGCATAAG | 203 |
| chr1:38791716-38793736 | p4 | (TTAA)5 | ACCCTGGGGGTGTCATTATT | ACGACCCATTTTGGCATAAG | 203 |
| chr1:38915584-38917602 | p5 | (GGGTT)3 | ATGGATGAACCATTTCCCAA | TGGTTAAGTAGGCGTTTGGC | 198 |
| chr1:38915584-38917602 | p3 | (AAG)5 | ATGGATGAACCATTTCCCAA | TGGTTAAGTAGGCGTTTGGC | 198 |
| chr1:38915584-38917602 | p2 | (AT)9 | ATGGATGAACCATTTCCCAA | TGGTTAAGTAGGCGTTTGGC | 198 |
| chr1:38915584-38917602 | p3 | (AAT)7 | ATGGATGAACCATTTCCCAA | TGGTTAAGTAGGCGTTTGGC | 198 |
| chr1:38963165-38965203 | p2 | (TA)19 | TATTCCTCTCTTTGGGCGTG | CACCACCCCATTATCCTCAC | 249 |
| chr1:38969403-38971418 | p3 | (ACA)5 | GCGTGTACGCATTCTTTTGA | TTCTCGCTTCATTGACCCTT | 122 |
| chr1:39065852-39067867 | p5 | (AAAGG)3 | TATTTTTGGGCGAATTCTGG | TAACCAGGCCTGATACCGTC | 162 |
| chr1:39068982-39070994 | p2 | (AT)6 | CCGAATTACTGAACCCGAAA | GGGCCAAAATTAGGTGTCAA | 158 |
| chr1:39144627-39146661 | p2 | (TA)17 | TTCCCAAAGTATCCCCATCA | GACATGGCGGAATAGCAAAT | 268 |
| chr1:39145795-39147837 | p2 | (TA)21 | ATCCAAGTTCCTTTCCCCAG | GATGGGCTTATGGCTTTTGA | 156 |
| chr1:39201351-39203437 | p2 | (TA)6 | CCCTAGGGAAGGTTTTACGC | CTCGAGAGAAATCCCGAGTG | 157 |
| chr1:39201351-39203437 | p6 | (TCCTTC)3 | CCCTAGGGAAGGTTTTACGC | CTCGAGAGAAATCCCGAGTG | 157 |
| chr1:39245913-39247928 | p3 | (TAA)5 | CCATTCGCAGCCTTGATATT | GCGAAGTTTGTGGAAAGAGC | 136 |
| chr1:39270947-39272959 | p5 | (ATTTT)3 | TGTTGATTTCCCTCCCATGT | ACGTGCCAGTTCTTCTTGCT | 146 |
| chr1:39270947-39272959 | p2 | (AC)6 | TGTTGATTTCCCTCCCATGT | ACGTGCCAGTTCTTCTTGCT | 146 |
| chr1:39380650-39382668 | p3 | (TTG)6 | ATCCTGCATTTTCGAACGTC | ATTGTCCTTTTGGAGGGCTT | 166 |
| chr1:39415892-39417914 | p2 | (TA)8 | AGAACTGTTAGGCGAGCCCT | TTGTTCGCAGTTTGCTATCG | 128 |
| chr1:39415892-39417914 | p2 | (AT)11 | AGAACTGTTAGGCGAGCCCT | TTGTTCGCAGTTTGCTATCG | 128 |
| chr1:39416053-39418113 | p2 | (TA)8 | CTACGGACGAACTTTCTGGC | AAGCAATGGTTCACATGCAA | 216 |
| chr1:39416053-39418113 | p2 | (AT)11 | CTACGGACGAACTTTCTGGC | AAGCAATGGTTCACATGCAA | 216 |
| chr1:39416163-39418175 | p2 | (TA)8 | CTACGGACGAACTTTCTGGC | AAGCAATGGTTCACATGCAA | 106 |
| chr1:39416163-39418175 | p2 | (AT)11 | CTACGGACGAACTTTCTGGC | AAGCAATGGTTCACATGCAA | 106 |
| chr1:39476384-39478441 | p3 | (GTA)5 | TTTTACGGTGGGATTTGCTC | TTACGGTCCCATTTTACGGA | 290 |

| Marker ID | SSR type | SSR motif | FORWARD PRIMER (5'-3') | REVERSE PRIMER (5'-3') | size (bp) |
|---|---|---|---|---|---|
| chr1:39506657-39508677 | p4 | (AAAT)5 | AGCAGGTTCCCAGGAAAGAT | CAACGTGGACGTAGCTCTCA | 271 |
| chr1:39509983-39511998 | p5 | (TTTAT)3 | AACGAATGATAGGCAGGTGG | AAACATCATAGACTGGCGGC | 201 |
| chr1:39538350-39540365 | p3 | (TAC)5 | TGCGTTAATGAAAAATGGCA | TCGGAAAAGGCTCAGGTATG | 215 |
| chr1:39628601-39630619 | p3 | (GAG)6 | AGGGTTATTAAGGGGAGGGG | GAGCGCACTTTCTGAACACA | 269 |
| chr1:39661699-39663711 | p2 | (AC)6 | GCTTTCGTGCATCCAATTTT | GCTTGTTTGTTGGTTGGGTT | 176 |
| chr1:39661699-39663711 | p2 | (AT)6 | GCTTTCGTGCATCCAATTTT | GCTTGTTTGTTGGTTGGGTT | 176 |
| chr1:39662220-39664232 | p2 | (AC)6 | AAACAGCCCGCAACAATTAC | TGGACTGTTTTGCTTGTCCA | 226 |
| chr1:39662220-39664232 | p2 | (AT)6 | AAACAGCCCGCAACAATTAC | TGGACTGTTTTGCTTGTCCA | 226 |
| chr1:39679846-39681866 | p5 | (TACAA)4 | ATGGTGGTTTCTCGTGGTTC | GAGTGGTAGAGTGGGGGTGA | 221 |
| chr1:39679846-39681866 | p2 | (AT)8 | ATGGTGGTTTCTCGTGGTTC | GAGTGGTAGAGTGGGGGTGA | 221 |
| chr1:39848468-39850483 | p5 | (AAAAC)3 | GGATCCAACGTATGCTCCAT | CTTTGTCCAGTCAAACCCGT | 127 |
| chr1:39907433-39909447 | p2 | (AT)7 | AACACATCCAATGCCTCACA | TTGTCCCTCCGAAAGGTATG | 151 |
| chr1:39912542-39914569 | p3 | (TTA)9 | AGTCCAAGTCAAATCCACCG | GGCTAACAGAGCAGAAACGC | 262 |
| chr1:39974084-39976100 | p2 | (AT)8 | TCGGCAATATTTTAGGGTCG | CTGGGCACTATGGTCGAAAT | 208 |
| chr1:39989640-39991652 | p2 | (AG)6 | TTGAGTCTACCCGTTTTGCC | CGGAAAAGATGAAGCAGAGC | 186 |
| chr1:40062827-40064841 | p2 | (AT)7 | AACAAATCTCCGCTGGACAC | GATAGGGGAGTGGGGAATGT | 112 |
| chr1:40062827-40064841 | p2 | (AT)12 | AACAAATCTCCGCTGGACAC | GATAGGGGAGTGGGGAATGT | 112 |
| chr1:40074584-40076599 | p3 | (AAG)11 | CATGTCGATATGCCAAATGC | TCCTTCTCTCCTTCAGCAGC | 182 |
| chr1:40074584-40076599 | p5 | (TTAAA)3 | CATGTCGATATGCCAAATGC | TCCTTCTCTCCTTCAGCAGC | 182 |
| chr1:40074584-40076599 | p5 | (AATAA)3 | CATGTCGATATGCCAAATGC | TCCTTCTCTCCTTCAGCAGC | 182 |
| chr1:40078571-40080583 | p2 | (AT)6 | CCGTTGTTACTCCATTGGCT | TCGAGACTCGGTGTGTAACG | 106 |
| chr1:40087398-40089413 | p5 | (TGAAT)3 | TCCAAGTTAAGGTGGGCTTG | TTGTTCCAACAACCACTCCA | 256 |
| chr1:40087398-40089413 | p6 | (TATATG)3 | TCCAAGTTAAGGTGGGCTTG | TTGTTCCAACAACCACTCCA | 256 |
| chr1:40253403-40255476 | p2 | (AT)8 | TTGCTGCAGATCAGGATGAC | TCCCACTTGTCTTCAAACCC | 148 |
| chr1:40253403-40255476 | p2 | (AT)8 | TTGCTGCAGATCAGGATGAC | TCCCACTTGTCTTCAAACCC | 148 |
| chr1:40253403-40255476 | p2 | (AT)31 | TTGCTGCAGATCAGGATGAC | TCCCACTTGTCTTCAAACCC | 148 |
| chr1:40253831-40255857 | p2 | (AT)8 | ATATGCGTTTTTCCCATTGC | ATTGGATCCGGGGTTAAGAC | 247 |
| chr1:40253831-40255857 | p2 | (AT)8 | ATATGCGTTTTTCCCATTGC | ATTGGATCCGGGGTTAAGAC | 247 |
| chr1:40253831-40255857 | p2 | (AT)31 | ATATGCGTTTTTCCCATTGC | ATTGGATCCGGGGTTAAGAC | 247 |
| chr1:40253906-40255968 | p2 | (AT)8 | ATATGCGTTTTTCCCATTGC | ATTGGATCCGGGGTTAAGAC | 172 |
| chr1:40253906-40255968 | p2 | (AT)8 | ATATGCGTTTTTCCCATTGC | ATTGGATCCGGGGTTAAGAC | 172 |
| chr1:40253906-40255968 | p2 | (AT)31 | ATATGCGTTTTTCCCATTGC | ATTGGATCCGGGGTTAAGAC | 172 |
| chr1:40323074-40325090 | p2 | (AT)8 | GCTTTGGGGGTAATTCCATT | AGAAAGCTCGAGAGCAGTGG | 262 |
| chr1:40428730-40430796 | p3 | (ATA)22 | ATCAATTTCCAGGGGGTTTC | TCGGTTGATTCGTAAGAGGG | 203 |
| chr1:40428730-40430796 | p2 | (TA)21 | ATCAATTTCCAGGGGGTTTC | TCGGTTGATTCGTAAGAGGG | 203 |
| chr1:40455210-40457248 | p2 | (TA)19 | TGGCATGGAAGAGTCCTACC | TGTTCAACAAGCCAACAAGG | 162 |

| Marker ID | SSR type | SSR motif | FORWARD PRIMER (5'-3') | REVERSE PRIMER (5'-3') | size (bp) |
|---|---|---|---|---|---|
| chr1:40489579-40491594 | p2 | (AT)6 | TTTCAAACCACCCAACCTTC | ATGGTTGTGGCCGAATAGAG | 141 |
| chr1:40489579-40491594 | p3 | (ACA)5 | TTTCAAACCACCCAACCTTC | ATGGTTGTGGCCGAATAGAG | 141 |
| chr1:40489893-40491911 | p2 | (AT)6 | ATGCCTTCAATTTTTGGTGC | AGTTCTTGAAGGGCCATGTG | 183 |
| chr1:40489893-40491911 | p3 | (ACA)5 | ATGCCTTCAATTTTTGGTGC | AGTTCTTGAAGGGCCATGTG | 183 |
| chr1:40519236-40521251 | p3 | (AGA)11 | AGTCCCCAATGATGAAGCTG | TGGAGCCATTAATTTCCTCG | 271 |
| chr1:40519236-40521251 | p5 | (AATTA)3 | AGTCCCCAATGATGAAGCTG | TGGAGCCATTAATTTCCTCG | 271 |
| chr1:40519236-40521251 | p5 | (TTTAT)3 | AGTCCCCAATGATGAAGCTG | TGGAGCCATTAATTTCCTCG | 271 |
| chr1:40519236-40521251 | p5 | (TTTAT)4 | AGTCCCCAATGATGAAGCTG | TGGAGCCATTAATTTCCTCG | 271 |
| chr1:40582448-40584463 | p5 | (AAAAC)3 | GCTTTGAACGGGAAACAAAA | AAATGGTTCACAGGCACCTC | 113 |
| chr1:40582448-40584463 | p6 | (CAGCAA)3 | GCTTTGAACGGGAAACAAAA | AAATGGTTCACAGGCACCTC | 113 |
| chr1:40597047-40599113 | p2 | (AT)33 | CCCAACGCCTCATTCTACAT | CAAAGTGATCACCATGAGCG | 132 |
| chr1:40885307-40887327 | p2 | (TC)10 | TTCCCTTCAAATTGGTGGTC | TTTGCATAAGGCCCAAATTC | 250 |
| chr1:40898764-40900782 | p6 | (ATTTTT)3 | AGTGCTTTGACCAAATTGCC | TTAAACACCAGCGCATTTGA | 185 |
| chr1:40960293-40962311 | p2 | (AT)9 | TATTTGGCTCACGCAGATGA | ATCATTTGGCTACACAGCCC | 223 |
| chr1:40970324-40972342 | p6 | (ATTTTT)3 | AGTGCTTTGACCAAATTGCC | TTAAACACCAGCGCATTTGA | 185 |
| chr1:41036358-41038373 | p3 | (ATA)5 | AACCCCCAAACATCAAACAA | GGCTTCGACATCCAGTTCAT | 247 |
| chr1:41036358-41038373 | p2 | (AT)8 | AACCCCCAAACATCAAACAA | GGCTTCGACATCCAGTTCAT | 247 |
| chr1:41036488-41038512 | p3 | (ATA)5 | AACCCCCAAACATCAAACAA | GGCTTCGACATCCAGTTCAT | 117 |
| chr1:41036488-41038512 | p2 | (AT)21 | AACCCCCAAACATCAAACAA | GGCTTCGACATCCAGTTCAT | 117 |
| chr1:41174289-41176304 | p5 | (TTTTG)3 | AGCTTGGTGCACGAGAGAAT | GATCCTCATTGTTTGGGTGG | 101 |
| chr1:41481257-41483273 | p4 | (ATTT)4 | ATGTGCAAACCTTGGAGCTT | GTGACCCATTTAGCCCCTTT | 160 |
| chr1:41535243-41537259 | p4 | (ATTT)4 | ATGTGCAAACCTTGGAGCTT | GTGACCCATTTAGCCCCTTT | 160 |
| chr1:41580829-41582841 | p2 | (TA)6 | CGACGGCATTTGATTTAGGT | GTATGTTGTTGTTGTGGCCG | 252 |
| chr1:41609595-41611607 | p2 | (TA)6 | CGACGGCATTTGATTTAGGT | GTATGTTGTTGTTGTGGCCG | 252 |
| chr1:41672337-41674369 | p2 | (TA)16 | TCATGGAATCGAGGGTTCTC | ATGTTTGGCCATCTAGGCAC | 243 |
| chr1:41706919-41708937 | p6 | (CATATA)3 | GGGAGGGTAAAGTGTACGCA | CGCATGTGGTAGGAGTCAGA | 116 |
| chr1:41766320-41768338 | p6 | (TATATG)3 | CGGATTGATCCGAAACTTGT | TGAGGATTGCACGAACAGAG | 118 |
| chr1:41768462-41770480 | p6 | (TATATG)3 | CGGATTGATCCGAAACTTGT | TGAGGATTGCACGAACAGAG | 118 |
| chr1:41801040-41803058 | p5 | (TGAAT)3 | TTAGATTCTCGGGGATGGTG | GCTTAATTGTCCAAGCTCGC | 276 |
| chr1:41801040-41803058 | p6 | (TATATG)3 | TTAGATTCTCGGGGATGGTG | GCTTAATTGTCCAAGCTCGC | 276 |
| chr1:41842842-41844856 | p3 | (TTG)6 | CTCCCCCAAAAATGCTTGTA | AACCCAATTTCCAAAATCCC | 276 |
| chr1:41885439-41887453 | p2 | (TA)7 | ATCGTCCTTCTGTACCGTCG | TGACAATTTCCCTTCCTTCG | 210 |
| chr1:41887491-41889503 | p2 | (AT)6 | CTTCACCCAAAACCTTCCAA | TGGAAGTGAGAAATGCGTTG | 177 |
| chr1:41893070-41895082 | p2 | (AG)6 | GGGGGCTTCACCTTAAAATC | GCCAAGATATGAGGGACCAA | 233 |
| chr1:41921227-41923242 | p3 | (TTG)5 | TCGTTGCTGTAAGCCAAGTG | TTTTGACCCCTCGAAGTCAG | 214 |
| chr1:41934069-41936096 | p3 | (AAT)9 | CAAACACTCCGCTAGTGCAA | TCTGTAGACAGCTGGCATGG | 180 |

| Marker ID | SSR type | SSR motif | FORWARD PRIMER (5'-3') | REVERSE PRIMER (5'-3') | size (bp) |
|---|---|---|---|---|---|
| chr1:42183871-42185883 | p2 | (TA)6 | CGTTCGCAAGGTAGGTAAGG | ACCCGGATATCCTCTATGGG | 198 |
| chr1:42264089-42266101 | p2 | (AT)6 | ATGATGGAATGGAGGGTTCA | TATCGTATAGCTGCCCGCTT | 184 |
| chr1:42264144-42266159 | p2 | (AT)6 | ATGATGGAATGGAGGGTTCA | TATCGTATAGCTGCCCGCTT | 129 |
| chr1:42266371-42268437 | p2 | (AT)8 | GGCAGCATTACACCGTACCT | AGGAGGGTCAGTACGAGCAA | 215 |
| chr1:42266371-42268437 | p2 | (AT)7 | GGCAGCATTACACCGTACCT | AGGAGGGTCAGTACGAGCAA | 215 |
| chr1:42266439-42268451 | p2 | (AT)8 | GGCAGCATTACACCGTACCT | AGGAGGGTCAGTACGAGCAA | 147 |
| chr1:42266439-42268451 | p2 | (AT)7 | GGCAGCATTACACCGTACCT | AGGAGGGTCAGTACGAGCAA | 147 |
| chr1:42324338-42326350 | p2 | (TA)6 | GGAGTCTCGGAATCTGCAAC | GATGAGAGCCAAGTGGAAGC | 272 |
| chr1:42345912-42347932 | p2 | (AT)10 | CTTTCAGGGTGCTGGATCAT | GGCCACCAAGAATGAAAAGA | 111 |
| chr1:42441506-42443542 | p5 | (TTTTG)3 | TTTCCCCCAAATTACCCTTC | CTTCCAAGATTACCCCCGAT | 206 |
| chr1:42441506-42443542 | p3 | (TTA)12 | TTTCCCCCAAATTACCCTTC | CTTCCAAGATTACCCCCGAT | 206 |
| chr1:42471969-42473983 | p3 | (CAA)5 | ATCAAGTCTTGGGCTCTCCA | TGTCGAAGGTTGGAGGTTTC | 195 |
| chr1:42471969-42473983 | p2 | (AC)6 | ATCAAGTCTTGGGCTCTCCA | TGTCGAAGGTTGGAGGTTTC | 195 |
| chr1:42471969-42473983 | p2 | (TC)7 | ATCAAGTCTTGGGCTCTCCA | TGTCGAAGGTTGGAGGTTTC | 195 |
| chr1:42482391-42484409 | p2 | (GA)9 | CGGTGTTTTCAATTCAGGCT | CTGCTGAGCTGCTGAAACTG | 129 |
| chr1:42591684-42593698 | p2 | (CA)7 | TTATGCTTGATGGGGTAGGG | TTCCCCTCAAATTGTTGGTC | 199 |
| chr1:42591684-42593698 | p2 | (AT)18 | TTATGCTTGATGGGGTAGGG | TTCCCCTCAAATTGTTGGTC | 199 |
| chr1:42695456-42697486 | p3 | (TAT)10 | TCATGAATCATCGGCCTACA | AGTTCGCTGCCGAAGAAATA | 239 |
| chr1:42702925-42704937 | p2 | (AT)6 | TCACTCTGAATCGCATCCTG | TTAGGCGTTCGCTTTCTGAT | 157 |
| chr1:42702925-42704937 | p2 | (AT)13 | TCACTCTGAATCGCATCCTG | TTAGGCGTTCGCTTTCTGAT | 157 |
| chr1:42714664-42716679 | p3 | (AAT)5 | TTGCTTCAGCAGGTACTCCA | CCAGATCGCTTCAACTGTGA | 185 |
| chr1:42714664-42716679 | p3 | (TAT)5 | TTGCTTCAGCAGGTACTCCA | CCAGATCGCTTCAACTGTGA | 185 |
| chr1:42729269-42731283 | p2 | (AT)7 | GACTCGGCATTCTTTTCGAG | ATCAACGGACAAACACCCAT | 297 |
| chr1:42729363-42731379 | p2 | (AT)7 | GACTCGGCATTCTTTTCGAG | ATCAACGGACAAACACCCAT | 203 |
| chr1:42729363-42731379 | p2 | (AT)10 | GACTCGGCATTCTTTTCGAG | ATCAACGGACAAACACCCAT | 203 |
| chr1:42730322-42732342 | p2 | (AT)8 | TGTTTCCACGAAATGCAAAA | TTGTTCCACGAAAATGCAAA | 137 |
| chr1:42735655-42737675 | p2 | (TA)10 | CTCGCCTAATCCAAAATTCG | AAATAAAGTGCAGCGGATGG | 145 |
| chr1:42865264-42867279 | p3 | (TTG)5 | CCACCGAAACTTAAAGGCAA | CGCATCTTGTCTTCCACTGA | 202 |
| chr1:42865264-42867279 | p3 | (TGT)6 | CCACCGAAACTTAAAGGCAA | CGCATCTTGTCTTCCACTGA | 202 |
| chr1:42876122-42878154 | p3 | (TAG)6 | TCACCTTTTCTCCGAAATGC | CAATCCTTCCTCCTCCACAA | 285 |
| chr1:42876179-42878241 | p3 | (TAG)6 | TCACCTTTTCTCCGAAATGC | CAATCCTTCCTCCTCCACAA | 228 |
| chr1:42888708-42890722 | p2 | (TA)7 | CACAGGCATTGAGCAACACT | GCTGATGGTGATGCAGAGAA | 116 |
| chr1:42935607-42937643 | p2 | (AT)18 | TGCGTAACTGACAATGCCTC | ATGCCATAGAGGCGTACAGG | 267 |
| chr1:43032843-43034855 | p2 | (AT)6 | CAAAGTGGAGAAATCGGGAG | AAAAATACATTTTGGCCCCC | 144 |
| chr1:43079937-43081952 | p5 | (GCATT)3 | TCCAATCCTTCCGACACTTC | ACGTTCGCCAAGCCTAAGTA | 158 |
| chr1:43108922-43110937 | p3 | (TTG)5 | TGAAAAATGTTGATGCGGAA | ATGGCCTTCTTTTGTTGTGC | 233 |

| Marker ID | SSR type | SSR motif | FORWARD PRIMER (5'-3') | REVERSE PRIMER (5'-3') | size (bp) |
|---|---|---|---|---|---|
| chr1:43201638-43203656 | p2 | (TC)9 | AATCGAACGAACAACCGAAC | GTGGTTGAGCCCCTATTTCA | 265 |
| chr1:43239333-43241361 | p4 | (TTTA)7 | AACCCTGCTTAGGCGAAAAT | ATACCCCGAGGTTCTGCTTT | 167 |
| chr1:43245539-43247555 | p2 | (TA)8 | ATATGTGTGCGGGGCTTTAG | AGTTTCCAACGCAACAAACC | 296 |
| chr1:43258988-43261006 | p6 | (CCCCCT)3 | CCAGATATCTTGGGCGTTGT | TGGTACAACTGTGCGCTAGG | 215 |
| chr1:43263075-43265093 | p6 | (GGGGAA)3 | TCCAAGTTCAAATTCCCAGG | TCCTTGCTGCACAATGTCTC | 101 |
| chr1:43290197-43292212 | p5 | (CCAAT)3 | CAATGATCTTGCCGATGATG | TCAGTGACTGGGGGTAGGTC | 241 |
| chr1:43360258-43362273 | p2 | (AT)13 | AAGTTTCGACTAAGGGGCGT | AAGTACATCACCTCTCGCGG | 281 |
| chr1:43360258-43362273 | p5 | (AAAAT)3 | AAGTTTCGACTAAGGGGCGT | AAGTACATCACCTCTCGCGG | 281 |
| chr1:43360258-43362273 | p5 | (AAAAT)3 | AAGTTTCGACTAAGGGGCGT | AAGTACATCACCTCTCGCGG | 281 |
| chr1:43368963-43370978 | p2 | (AT)7 | AAAGACGGTAGAAAGCGCAA | GTAGGCCCAATGCTCATGTT | 275 |
| chr1:43368963-43370978 | p5 | (CATAT)3 | AAAGACGGTAGAAAGCGCAA | GTAGGCCCAATGCTCATGTT | 275 |
| chr1:43414168-43416183 | p5 | (TATAA)3 | ACCACCATCTGAGCGAGACT | TCTGCCTCTCAATGTTGTCG | 286 |
| chr1:43428658-43430673 | p5 | (CCCCA)3 | GTTTGACTCGGCACGAAGTT | TTTTCCAATTCCAAGGCAAG | 259 |
| chr1:43592507-43594527 | p2 | (TA)10 | TGGCAGAGGGACATGACATA | TACCGGAGTAACGGGAAGTG | 168 |
| chr1:43693972-43695988 | p4 | (TTGA)4 | GCTAATCGTCGTCATCAGCA | ATCCCCAGGGCCTAGTCTAA | 110 |
| chr1:43730260-43732278 | p6 | (CAAAAA)3 | CTGATTTCTACTGAGCCCGC | CATGCTCAACAATGGACTGG | 114 |
| chr1:43760802-43762820 | p6 | (CAAAAA)3 | CTGATTTCTACTGAGCCCGC | CATGCTCAACAATGGACTGG | 114 |
| chr1:43936281-43938299 | p6 | (TCAGTT)3 | TCGTGCCCACTAAGTGTTTG | CAGCCTTTGAGCGTGATACA | 283 |
| chr1:44247100-44249115 | p5 | (CCCCA)3 | GGCCTGTAGGATAAAAGCCC | TCCAATTCCCCATTTTTCAA | 241 |
| chr1:44259784-44261812 | p2 | (TA)14 | TTTTTAAACCCGCTCCATCA | TACTCCCAGTTGGGGTTTTG | 254 |
| chr1:44295484-44297510 | p2 | (AT)13 | CAGTGAACAACAACATCCCG | CGGGTACTGGCTATGGTGAT | 144 |
| chr1:44295484-44297510 | p2 | (TA)7 | CAGTGAACAACAACATCCCG | CGGGTACTGGCTATGGTGAT | 144 |
| chr1:44326446-44328462 | p2 | (TA)8 | GAGGATTCTGGAATTTGGCA | TTTCGGCCCAGTTAAATACG | 149 |
| chr1:44340688-44342706 | p3 | (CAA)6 | ACCGGCGAACGATATCATAG | GAAAAGCTCGGATTGCTTTG | 259 |
| chr1:44436594-44438606 | p2 | (AG)6 | CTCGTGGTCTGAGGTTCGTT | CGCAAACAAGAAATGGAACA | 102 |
| chr1:44474346-44476362 | p2 | (AT)6 | GCTCCAGTGGCAATCCTAAC | TCATTTGCAAAATGTCTCGG | 290 |
| chr1:44480048-44482112 | p3 | (TTG)5 | CCCCTTCTTCTTCAAGTCCC | AGGCCAAAGCTTCCATTTCT | 145 |
| chr1:44480048-44482112 | p2 | (AT)32 | CCCCTTCTTCTTCAAGTCCC | AGGCCAAAGCTTCCATTTCT | 145 |
| chr1:44512785-44514801 | p2 | (AT)8 | TGCGTGGACCATGAAACTAA | CTAAATGGCTGGGCAGGATA | 143 |
| chr1:44512785-44514801 | p2 | (AT)17 | TGCGTGGACCATGAAACTAA | CTAAATGGCTGGGCAGGATA | 143 |
| chr1:44562222-44564242 | p2 | (AT)10 | AGCTGTCCATTGCAGTGTTG | AGCTTTCACTTGGGCTCTCA | 230 |
| chr1:44562222-44564242 | p2 | (AT)15 | AGCTGTCCATTGCAGTGTTG | AGCTTTCACTTGGGCTCTCA | 230 |
| chr1:44570464-44572478 | p5 | (ATATC)3 | TTATCCGACATGCCTCTTCC | CCCAGCCATAGGAAATGAAA | 272 |
| chr1:44570464-44572478 | p2 | (TA)7 | TTATCCGACATGCCTCTTCC | CCCAGCCATAGGAAATGAAA | 272 |
| chr1:44570464-44572478 | p2 | (AT)7 | TTATCCGACATGCCTCTTCC | CCCAGCCATAGGAAATGAAA | 272 |
| chr1:44608732-44610746 | p2 | (AT)7 | GAACACGGCTCGTAGACACA | TCGGCATGTTTGTTGATGTT | 117 |

| Marker ID | SSR type | SSR motif | FORWARD PRIMER (5'-3') | REVERSE PRIMER (5'-3') | size (bp) |
|---|---|---|---|---|---|
| chr1:44636167-44638207 | p2 | (AT)20 | CTCAGACGAGAAAGGGATGG | AAAAGCTTTGGAGCCAGACA | 299 |
| chr1:44707613-44709677 | p2 | (AT)8 | ACCGATCCATGGTCTAGTGC | CCATGTCCTGGACGATCTCT | 265 |
| chr1:44707613-44709677 | p2 | (AT)32 | ACCGATCCATGGTCTAGTGC | CCATGTCCTGGACGATCTCT | 265 |
| chr1:44884636-44886648 | p2 | (AT)6 | TTGAGGGATCTTCAATTGGG | TCAATGGCAACAAGAATGGA | 163 |
| chr1:44939777-44941793 | p4 | (TTCT)4 | TGTTTGTCCATTTGTTGGGA | AAGGGCAAACATTGCGTATC | 246 |
| chr1:45017088-45019142 | p2 | (AT)27 | GCTATAGCGACGTAGCGACC | CCCTTGAGAGGCTTTTGTTG | 240 |
| chr1:45058927-45060943 | p2 | (AC)8 | GAAAAGGGTGGAAACGATGA | CCATATCCAAACAGCCCATC | 205 |
| chr1:45068216-45070230 | p5 | (AATGC)3 | CGAGATGTGCTACCTGCGTA | ACTTCATCTTGATGCCCAGC | 173 |
| chr1:45112310-45114352 | p2 | (TA)11 | TATCATTCGAACCCCCAGAC | ATTTGCCGATTTTGTGTTCA | 188 |
| chr1:45112310-45114352 | p2 | (AT)21 | TATCATTCGAACCCCCAGAC | ATTTGCCGATTTTGTGTTCA | 188 |
| chr1:45208859-45210875 | p2 | (CT)8 | CTTATGGGTGTTCCATTCGG | GGCCAGTGTTTGGAACCTTA | 137 |
| chr1:45251431-45253469 | p2 | (AT)6 | TAGGATTGGCAAGTTCCACC | CGTACCCTCCTGCTCGTAAG | 249 |
| chr1:45251471-45253529 | p2 | (AT)6 | TAGGATTGGCAAGTTCCACC | CGTACCCTCCTGCTCGTAAG | 209 |
| chr1:45254017-45256032 | p5 | (AAATG)3 | TAGCGTTCGCTTGTTCTCCT | AGCGTTGGGAGTTGATATGG | 299 |
| chr1:45291576-45293591 | p3 | (ATA)5 | ACCTGGAGCACGAACTCTGT | CCACTCACGGCTATTTGGAT | 175 |
| chr1:45291576-45293591 | p5 | (ATGCA)3 | ACCTGGAGCACGAACTCTGT | CCACTCACGGCTATTTGGAT | 175 |
| chr1:45300199-45302217 | p5 | (AAAAG)3 | GCTAGTAGTATGCCCGCCTG | TGCTCGTCGTTTTTGTCTTG | 134 |
| chr1:45300199-45302217 | p3 | (AAG)11 | GCTAGTAGTATGCCCGCCTG | TGCTCGTCGTTTTTGTCTTG | 134 |
| chr1:45300785-45302818 | p3 | (AAG)11 | CACCATGCCAGAAGGTTTTT | CCGTTGACGAGTGTTTTCCT | 163 |
| chr1:45318273-45320285 | p2 | (AT)6 | TACTGGCCGTATTTCGGTTC | TTGGGAAATCTTCCTTGCTG | 100 |
| chr1:45438813-45440831 | p3 | (TTG)6 | TCAAAAGCAACATCGCACTC | GGCCTGGAGGTAGCTAAACC | 274 |
| chr1:45440035-45442050 | p5 | (TTTGT)3 | TTCTCCAGTCCGTGAAATCC | TAGGGGTCTTGCTATGGTGC | 170 |
| chr1:45495244-45497259 | p5 | (AAAGG)3 | GGATTTCGGATGGAGATTGA | ACTGTCTCTTAGACCGCCGA | 188 |
| chr1:45585382-45587394 | p6 | (GGAAGA)3 | TTGGACGTCTTAGCATGGTG | CCAAAGCCACGGTTGATACT | 165 |
| chr1:45585382-45587394 | p2 | (AT)6 | TTGGACGTCTTAGCATGGTG | CCAAAGCCACGGTTGATACT | 165 |
| chr1:45695234-45697249 | p3 | (TCC)5 | TTTTCCCGGCTTAAATCCTT | CCTCTGTTTGGTACCCTGGA | 240 |
| chr1:45695234-45697249 | p3 | (CTC)5 | TTTTCCCGGCTTAAATCCTT | CCTCTGTTTGGTACCCTGGA | 240 |
| chr1:45712366-45714382 | p2 | (AT)8 | TACGGATCCAGGCAATAACC | TCATCCTTCCGACTACCCTG | 260 |
| chr1:45712458-45714470 | p2 | (AT)8 | TACGGATCCAGGCAATAACC | TCATCCTTCCGACTACCCTG | 168 |
| chr1:45757138-45759158 | p2 | (GT)10 | TTTGAATTGGGGTGGAGTTC | GGGTAGGGAAGGAGAAATGG | 239 |
| chr1:45760400-45762416 | p2 | (AT)8 | AACCAACGAAAAACCCACAC | CAACACGAATTGCACCAAAG | 174 |
| chr1:45794125-45796140 | p5 | (TTTCC)3 | TCTAATTTTGCCCTTGACGG | ACTTTTCTCAATGCCTCCGA | 163 |
| chr1:45794125-45796140 | p3 | (GAA)5 | TCTAATTTTGCCCTTGACGG | ACTTTTCTCAATGCCTCCGA | 163 |
| chr1:45823431-45825446 | p5 | (ATGAT)3 | GATGCTGCCCAAATTTCTGT | GACAATTTCAAACGCGGATT | 114 |
| chr1:45839460-45841484 | p3 | (TTG)8 | TTTTGGACTGATGAATGGCA | TTCCGCGACTTTCATCTTCT | 120 |
| chr1:45919646-45921660 | p2 | (TC)7 | CTGCTTTCGTCAGCCTATCC | CGAAGCTAACCCAGAAGTCG | 282 |

| Marker ID | SSR type | SSR motif | FORWARD PRIMER (5'-3') | REVERSE PRIMER (5'-3') | size (bp) |
|---|---|---|---|---|---|
| chr1:46040789-46042815 | p2 | (AT)13 | AGCCCCACTTTATTCGAGGT | ATTTCACCGAACACTACGGG | 224 |
| chr1:46157787-46159807 | p5 | (GCATT)4 | CGTAGGGGACAGTGGTGAGT | ATTTCAGACCAGCCCAACAC | 226 |
| chr1:46175925-46177945 | p5 | (GCATT)4 | CGTAGGGGACAGTGGTGAGT | ATTTCAGACCAGCCCAACAC | 226 |
| chr1:46264746-46266784 | p2 | (AT)9 | GTACGAAAAGAGCCGAACCA | GGCTTGGACTCACACCGTAT | 230 |
| chr1:46300450-46302465 | p5 | (CCCAA)3 | GCGCCAAATTGTTTACCCTA | CGGTGCCCTCCATAATTAGA | 295 |
| chr1:46637604-46639648 | p2 | (AT)22 | CAGCAGAATCTCCAGAAGGC | TTTGTTGCAGCAAGGAAGTG | 193 |
| chr1:46656276-46658294 | p6 | (CATATA)3 | ATCATAGCAAATGGAACCGC | CACGTGAGTGACCCAACAAC | 198 |
| chr1:46742929-46744944 | p5 | (CATAG)3 | ACCGAGCAGCCCTTACCTAT | TGTGTATGTATCCTCGGGCA | 260 |
| chr1:46792319-46794331 | p2 | (AC)6 | TGATTCAGCTTTTAACGGGG | CTGGAGGTCTGTGATGGGAT | 192 |
| chr1:46810720-46812738 | p6 | (ATTTTT)3 | CTAACCTCTCACACCTCCGC | ACTTTGTTGTACGGGGCAAG | 171 |
| chr1:46853766-46855781 | p3 | (GTT)5 | GTACCGCTGCTGCTTCTTCT | GCAAAAACCCCAAAAAGTCA | 285 |
| chr1:46923264-46925279 | p3 | (TGC)5 | TCGGGAGGGCTACTTGTATG | TTCGAGAAAAATGGATTCCG | 236 |
| chr1:46931815-46933897 | p2 | (TA)8 | AGCTCATGCCTTTCCTTCAA | GGTATCTCCATTCAGGCCAA | 183 |
| chr1:46931815-46933897 | p5 | (ATTTT)3 | AGCTCATGCCTTTCCTTCAA | GGTATCTCCATTCAGGCCAA | 183 |
| chr1:46933393-46935421 | p2 | (TA)8 | CACGGTGTTTAATATGGGGC | AAAGTGGAACCCACCACATC | 245 |
| chr1:46933393-46935421 | p5 | (ATTTT)3 | CACGGTGTTTAATATGGGGC | AAAGTGGAACCCACCACATC | 245 |
| chr1:46933393-46935421 | p5 | (ATTTT)3 | CACGGTGTTTAATATGGGGC | AAAGTGGAACCCACCACATC | 245 |
| chr1:46933393-46935421 | p2 | (AT)14 | CACGGTGTTTAATATGGGGC | AAAGTGGAACCCACCACATC | 245 |
| chr1:46962403-46964431 | p2 | (TA)14 | GTCTCCAAATGGCCGATAGA | AAGGATGACAAAATCTGCCG | 290 |
| chr1:46978595-46980631 | p2 | (TA)18 | GGTAGCGATTAGCAAGCACC | TCTGGCCAATTTTTCCTTTG | 113 |
| chr1:47008751-47010763 | p2 | (AT)6 | AAGTATCAACCGTGGCTTCG | TTGCTCCGCTTCTGAGTTTT | 105 |
| chr1:47008751-47010763 | p2 | (TC)6 | AAGTATCAACCGTGGCTTCG | TTGCTCCGCTTCTGAGTTTT | 105 |
| chr1:47043951-47045966 | p5 | (ATTTT)3 | AAACGTTGGAACTTGATCGG | AAAGATTCACCCTCCCCTGT | 200 |
| chr1:47045340-47047510 | p2 | (AT)85 | TTCAGCGAATGGTTTGATTG | TGAAGCTGAAAAACCCGACT | 273 |
| chr1:47045340-47047510 | p2 | (AT)10 | TTCAGCGAATGGTTTGATTG | TGAAGCTGAAAAACCCGACT | 273 |
| chr1:47174872-47176887 | p2 | (AT)12 | GACGGAAAGCAGAATTCGAG | TTAAAGACCCATGGGGATGA | 295 |
| chr1:47174872-47176887 | p5 | (TACAA)3 | GACGGAAAGCAGAATTCGAG | TTAAAGACCCATGGGGATGA | 295 |
| chr1:47231633-47233699 | p2 | (TA)33 | CAGAAGGGATATTGCCTCCA | CCCCAAGACAACCTACTCCA | 294 |
| chr1:47244204-47246231 | p5 | (ATTTT)3 | GCCATTGAAAGACAATCCGT | GTAGGAAGAGGAGAACGGGG | 242 |
| chr1:47244204-47246231 | p5 | (TTCTC)5 | GCCATTGAAAGACAATCCGT | GTAGGAAGAGGAGAACGGGG | 242 |
| chr1:47244252-47246279 | p5 | (ATTTT)3 | GCCATTGAAAGACAATCCGT | GGTAGGAAGAGGAGAACGGG | 194 |
| chr1:47244252-47246279 | p5 | (TTCTC)5 | GCCATTGAAAGACAATCCGT | GGTAGGAAGAGGAGAACGGG | 194 |
| chr1:47244312-47246339 | p5 | (ATTTT)3 | GCCATTGAAAGACAATCCGT | GGTAGGAAGAGGAGAACGGG | 134 |
| chr1:47244312-47246339 | p5 | (TTCTC)5 | GCCATTGAAAGACAATCCGT | GGTAGGAAGAGGAGAACGGG | 134 |
| chr1:47256893-47258908 | p2 | (AT)8 | TACGCCTCGCCCAATACTTA | TTGCCATGTCGATCCTAACA | 134 |
| chr1:47256893-47258908 | p5 | (AAAAT)3 | TACGCCTCGCCCAATACTTA | TTGCCATGTCGATCCTAACA | 134 |

| Marker ID | SSR type | SSR motif | FORWARD PRIMER (5'-3') | REVERSE PRIMER (5'-3') | size (bp) |
|---|---|---|---|---|---|
| chr1:47291297-47293312 | p2 | (AT)8 | TACGCCTCGCCCAATACTTA | TTGCCATGTCGATCCTAACA | 134 |
| chr1:47291297-47293312 | p5 | (AAAAT)3 | TACGCCTCGCCCAATACTTA | TTGCCATGTCGATCCTAACA | 134 |
| chr1:47350938-47352953 | p5 | (AAAAT)3 | GTTGTGTGGGTTGTTAGGGG | CTTTGCCACCCACTACCACT | 159 |
| chr1:47541477-47543537 | p3 | (ATA)20 | CTTACCACGGAGCATCCATT | ATATTCGGCGAAGTTTGTGG | 125 |
| chr1:47546505-47548523 | p3 | (CTT)6 | TTCGCTTTGTTGGGTTTCTC | TTGAAGAGAGAGATGGCGGT | 181 |
| chr1:47614289-47616304 | p5 | (CATTG)3 | TCGCTTGTTGTCCTGATTTG | CACAAGTACCTCAACGGGGT | 266 |
| chr1:47625978-47627994 | p2 | (TA)12 | ATGTTAGCTCCGCAGTTGCT | CAAGGTGCACCAGCTGATTA | 137 |
| chr1:47625978-47627994 | p2 | (AT)8 | ATGTTAGCTCCGCAGTTGCT | CAAGGTGCACCAGCTGATTA | 137 |
| chr1:47753341-47755356 | p5 | (AAAAT)3 | CTGCTTATTCCCTTCTCCCC | ATCATCTACGCCATTCTCCG | 157 |
| chr1:47858526-47860552 | p2 | (TA)13 | GAAGGTTTCCTTCTCCCACC | TGATGTTCCCACACGTTGAT | 109 |
| chr1:47860799-47862831 | p2 | (TC)16 | TTACGCTTAACGGTTTGGCT | TTTTACGGAGTGCAGAGGCT | 177 |
| chr1:47999787-48001802 | p3 | (GCA)5 | CGGTGAAATTACGCAAGGAT | CTCTTGTCTCGGACTCGGAC | 160 |
| chr1:48012532-48014608 | p6 | (AGTGAT)3 | TCAGGCAGCGATAACTCCTT | TCCTTGCTCCGATCTCTGAT | 208 |
| chr1:48132229-48134259 | p2 | (CT)6 | CAATGATTTCCTCGGCACTT | ACACTCTCCGTCGTTCCATC | 186 |
| chr1:48144545-48146560 | p5 | (TTAAT)3 | GAACAAGTCGGGAATTGGAA | TCCAGGACCATCAAAACTCC | 118 |
| chr1:48186831-48188865 | p2 | (AT)8 | ATTGCAAATGGCCAAAGAAG | AGCACCAACCAAAGATCCAC | 230 |
| chr1:48186831-48188865 | p2 | (TA)17 | ATTGCAAATGGCCAAAGAAG | AGCACCAACCAAAGATCCAC | 230 |
| chr1:48219008-48221020 | p2 | (TG)6 | TGGCCACTGACTTAAGGGAC | TTCGTCTCGAGCTTTTCCAT | 104 |
| chr1:48219008-48221020 | p3 | (TGT)5 | TGGCCACTGACTTAAGGGAC | TTCGTCTCGAGCTTTTCCAT | 104 |
| chr1:48342879-48344894 | p5 | (CCAAA)3 | CCAAGTTGGGTTGACTCGAT | CCATCCAAGCCTGAACACTT | 192 |
| chr1:48385582-48387596 | p2 | (TG)7 | GACCTTGTTTCTATCCGCCA | ACAGGAGAGCTAACCTGCCA | 131 |
| chr1:48385582-48387596 | p2 | (AC)6 | GACCTTGTTTCTATCCGCCA | ACAGGAGAGCTAACCTGCCA | 131 |
| chr1:48390785-48392845 | p2 | (AT)30 | TTGCTCCACGCCCTACTACT | CGGATGACTCGCCTTGTATT | 106 |
| chr1:48462080-48464100 | p2 | (TG)8 | CGCGTTCACTGTTAGAACCA | CATCTTCTTACTCCGCTGGC | 112 |
| chr1:48462871-48464887 | p2 | (TG)8 | GACACCTTGCACGCAGAATA | GTCTGGCCTTGTTGGTCATT | 192 |
| chr1:48531522-48533537 | p5 | (AACAT)3 | TAAACCATGGCCTCGAAAAG | TCAATCTCTTCCTCTTCGCC | 193 |
| chr1:48623233-48625249 | p5 | (TTTGT)3 | CAAGCGCATGTAGACCAAAA | GATTGTTTCACGAAATGCGA | 249 |
| chr1:48623233-48625249 | p2 | (TA)6 | CAAGCGCATGTAGACCAAAA | GATTGTTTCACGAAATGCGA | 249 |
| chr1:48629237-48631253 | p2 | (AT)8 | TCCTCAACCAATCCTTCACC | CAAAAATGCAGCCACAGAGA | 143 |
| chr1:48681786-48683801 | p3 | (TGT)5 | GGGCCTACATTGTGAATGCT | CAAAATTTAAAGGCCACCCA | 292 |
| chr1:48688473-48690488 | p5 | (TTTTA)3 | TGCCCTTTACTTTTCCGTTG | GCCAGCAATATTGGCATCTT | 290 |
| chr1:48696580-48698600 | p4 | (TCTT)4 | CACACCACAACATTCCCAAG | ATGAAATCATGCCCCAAAAA | 123 |
| chr1:48696580-48698600 | p5 | (TTTAG)4 | CACACCACAACATTCCCAAG | ATGAAATCATGCCCCAAAAA | 123 |
| chr1:48718506-48720522 | p2 | (AT)8 | TGCACAAGTGATAAGGTCCG | TGTTGTGACGAGGCAAATGT | 284 |
| chr1:48872626-48874642 | p2 | (TA)8 | GAACCTCTCTCGCCACTTTG | GGAGAACCAGTTCGGATTCA | 275 |
| chr1:48885303-48887315 | p2 | (AT)6 | GAAGGAAATCATGCCTTGGA | AAGGCTCGTCCCTTTCTTTC | 140 |

| Marker ID | SSR type | SSR motif | FORWARD PRIMER (5'-3') | REVERSE PRIMER (5'-3') | size (bp) |
|---|---|---|---|---|---|
| chr1:48898323-48900338 | p3 | (TCA)5 | CGGTAGCTCGATAGCACTCC | CAGGTTCACAGACAGCCAGA | 155 |
| chr1:48898323-48900338 | p2 | (AT)7 | CGGTAGCTCGATAGCACTCC | CAGGTTCACAGACAGCCAGA | 155 |
| chr1:48952469-48954491 | p2 | (AT)11 | CATGACAAGGGCACACAAAC | CTCAGTGGTTAGCCCCGATA | 193 |
| chr1:49043189-49045204 | p2 | (AG)6 | GTCGATGGGAGATCGGAGTA | CCGTCACTTGAAAATTGCCT | 175 |
| chr1:49043189-49045204 | p5 | (AAATA)3 | GTCGATGGGAGATCGGAGTA | CCGTCACTTGAAAATTGCCT | 175 |
| chr1:49045563-49047578 | p5 | (ACAAA)3 | ACCATATGCAGCAAAGGAGG | CATTTTAGCCTTGCCTGCTC | 135 |
| chr1:49059872-49061894 | p2 | (AT)11 | TGTTCTTTTGTCACGACCCA | TGTAGTGGCAGCCTTGTCAG | 166 |
| chr1:49059872-49061894 | p2 | (AT)9 | TGTTCTTTTGTCACGACCCA | TGTAGTGGCAGCCTTGTCAG | 166 |
| chr1:49105671-49107697 | p2 | (TA)7 | AGGACGAATGTTCAGCGAGT | CATCATATACCGTACCCGGC | 137 |
| chr1:49136119-49138137 | p6 | (CCAATG)3 | ACAATTGGCCGTCTACCAAG | CGGACATGCCGAAATAAACT | 136 |
| chr1:49154644-49156659 | p5 | (AAAAT)3 | GTTTGATTGGACGCGAAGTT | TGTTATGTTGCCAGGGGAAT | 228 |
| chr1:49209633-49211645 | p2 | (TA)6 | TGAACTTGGCATGGCATAAA | TCCTGAATCCCTCCACAGAC | 280 |
| chr1:49310783-49312798 | p5 | (TTCTT)3 | CGATTTTGAAGGGCGTTTTA | AAGTGCTGATCCCAATGACC | 154 |
| chr1:49495891-49497903 | p2 | (TA)6 | CAGACAACCCCTAGGACGAA | GCAAGCCGACGTACCATTAT | 189 |
| chr1:49552584-49554680 | p2 | (AT)48 | TTGTCTACCTGTGGGCATGA | TGATTATCACACCGATCCGA | 124 |
| chr1:49576607-49578623 | p4 | (AAAT)4 | TGGTTACTGGCTCGTGTTTG | CTAAGCAGCAACCCTCGAAC | 186 |
| chr1:49606879-49608894 | p2 | (TA)16 | GGTTGGATATTCCCTTGGGT | AGCACCAACAATGGGTAAGC | 210 |
| chr1:49609739-49611765 | p2 | (AT)13 | GAGAGGTTGGGCATGTTGTT | GCATCTGAGACCTCTGCCTC | 181 |
| chr1:49628940-49630955 | p3 | (TTA)5 | AAGATGGTGATGATGGAGGC | CTCTCGACAGTCATGACCCA | 220 |
| chr1:49633664-49635676 | p2 | (AT)6 | AGGTGCCACATTATCTTGCC | CCCTCAAGTTGGACCAAAAA | 271 |
| chr1:49753798-49755818 | p2 | (TA)10 | TACCTTCACACCATGCCTGA | CGGTGTAATTGGTGCAAGTG | 284 |
| chr1:49842368-49844382 | p2 | (TG)7 | TCACAAATATAGGCCGAGGG | ATCATAGGGTTGCGGATGAG | 235 |
| chr1:49894416-49896428 | p2 | (AT)6 | AGTTCTAGCCACTCCCCGAT | ACCATACCGAAGGGAAAAGG | 199 |
| chr1:49931970-49933982 | p2 | (TA)6 | GGGGTGTAAGCCCAGAAAAT | TTCGAGGCACACCACATAAA | 279 |
| chr1:49958508-49960532 | p2 | (GA)6 | CGACGATGAGAGAAAGAGGG | TTACCGAACCCCAAATCAAA | 297 |
| chr1:49983850-49985864 | p2 | (TA)7 | CGGTCCGTCATTTCAATTTT | AGGTATGGCTCCAGCAAATG | 258 |
| chr1:49983850-49985864 | p3 | (CAT)6 | CGGTCCGTCATTTCAATTTT | AGGTATGGCTCCAGCAAATG | 258 |
| chr1:49984047-49986065 | p2 | (TA)7 | CATTTGCTGGAGCCATACCT | CCCGTAGAAAACCGGTTACA | 155 |
| chr1:49984047-49986065 | p3 | (CAT)6 | CATTTGCTGGAGCCATACCT | CCCGTAGAAAACCGGTTACA | 155 |
| chr1:50121446-50123458 | p2 | (AC)6 | TATCAAACGGGACTTCGGAC | GGCTTGCCCTTTCTTTCTTT | 292 |
| chr1:50123501-50125521 | p5 | (AAATA)4 | GGAGCATGAGGAACTGAAGC | CATCAACAACAACCTCACCG | 168 |
| chr1:50123501-50125521 | p5 | (AATAA)3 | GGAGCATGAGGAACTGAAGC | CATCAACAACAACCTCACCG | 168 |
| chr1:50123501-50125521 | p5 | (TTTAT)3 | GGAGCATGAGGAACTGAAGC | CATCAACAACAACCTCACCG | 168 |
| chr1:50215575-50217593 | p6 | (CATATA)3 | AAATGTCCGCAGCTTCTTGT | GCAGTTCAGTCCTTGGGTGT | 129 |
| chr1:50215575-50217593 | p3 | (TCA)5 | AAATGTCCGCAGCTTCTTGT | GCAGTTCAGTCCTTGGGTGT | 129 |
| chr1:50217679-50219694 | p5 | (GAAAT)3 | ATACGGCCAGTATGTTTGGC | TTGCCAAGAGAAGGCAAAGT | 282 |

| Marker ID | SSR type | SSR motif | FORWARD PRIMER (5'-3') | REVERSE PRIMER (5'-3') | size (bp) |
|---|---|---|---|---|---|
| chr1:50217704-50219719 | p5 | (GAAAT)3 | ATACGGCCAGTATGTTTGGC | TTGCCAAGAGAAGGCAAAGT | 257 |
| chr1:50409396-50411410 | p2 | (AT)7 | TAGCAAGTCAAGGCTGAGCA | CATATCGTTATGCGGTGTCG | 276 |
| chr1:50463943-50465959 | p2 | (AT)6 | CGTGGGCGCATATTTTTACT | GGAACTCCGGTACATAGCGA | 180 |
| chr1:50463943-50465959 | p2 | (TA)8 | CGTGGGCGCATATTTTTACT | GGAACTCCGGTACATAGCGA | 180 |
| chr1:50482491-50484503 | p2 | (TA)6 | CCCTCTTCGCGAAACAGTAG | AAGGACGAGGGGCTATCCTA | 241 |
| chr1:50541437-50543455 | p2 | (TA)9 | TGAGTCCATTTCAACTTGCG | GGATGCCTTACCTCGTGTGT | 223 |
| chr1:50694826-50696844 | p6 | (ATATGT)3 | ATGTTGGATATGAGCCTCGG | GTCGGGTCCTATTTATGCGA | 163 |
| chr1:50694826-50696844 | p2 | (AT)6 | ATGTTGGATATGAGCCTCGG | GTCGGGTCCTATTTATGCGA | 163 |
| chr1:50716198-50718216 | p2 | (TA)8 | TTGCCTCATAAGGCGGATAC | TAAATGGATAGGGCCATTCG | 221 |
| chr1:50716198-50718216 | p2 | (TG)23 | TTGCCTCATAAGGCGGATAC | TAAATGGATAGGGCCATTCG | 221 |
| chr1:50716198-50718216 | p3 | (ATC)6 | TTGCCTCATAAGGCGGATAC | TAAATGGATAGGGCCATTCG | 221 |
| chr1:50735596-50737608 | p2 | (CA)6 | GCTTTTCAATTCGTCTTCGC | TTCTCCCTCTCCACATGACC | 222 |
| chr1:50800532-50802547 | p3 | (ATT)5 | AAAATGCCTAAAATTGCCCC | AGGAAGGAAGGGCTGGTAAA | 193 |
| chr1:50895687-50897705 | p3 | (TTG)6 | GCCCACACGTCAAGATTTTT | TTTATTTCTGGGCGTTGTCC | 243 |
| chr1:50940974-50942992 | p3 | (TTG)6 | AGTGTCGTGAAGGTTGGAGG | CACCGTGTACTGTTCATCCG | 112 |
| chr1:50956576-50958640 | p5 | (TATGA)3 | TGTTAGTCTCCCCTCACGCT | CGGAGTGACATAAGGTCGGT | 287 |
| chr1:50956576-50958640 | p2 | (AT)32 | TGTTAGTCTCCCCTCACGCT | CGGAGTGACATAAGGTCGGT | 287 |
| chr1:51197153-51199168 | p5 | (TCATA)3 | TTGCAGACAACCCCTAGGAC | TGTCACGCGTTCCATATGAT | 293 |
| chr1:51204235-51206261 | p2 | (TA)9 | GTTTGGGATATTGTGTGGGC | AGGCGACGTTTGCTCTTTTA | 211 |
| chr1:51204235-51206261 | p2 | (AT)8 | GTTTGGGATATTGTGTGGGC | AGGCGACGTTTGCTCTTTTA | 211 |
| chr1:51272271-51274287 | p4 | (TGTA)4 | TTTGATTCACCGAGTCCCTC | ACCGTACCCAGCCCTCTAGT | 252 |
| chr1:51370296-51372320 | p3 | (TTG)8 | CGTATTGGAGCAGGGTGAAT | CATCTCATTTCCGGCACTTT | 181 |
| chr1:51386344-51388371 | p3 | (TCT)9 | CAAGTGGTCGCTCCTCTTTC | TGGATGGTTCATCCGGTTAT | 284 |
| chr1:51431364-51433378 | p2 | (TA)7 | AGTAAGCGAGCGAGTTCCAA | TCATCATCTATAAGCCGCCC | 174 |
| chr1:51431364-51433378 | p2 | (TA)7 | AGTAAGCGAGCGAGTTCCAA | TCATCATCTATAAGCCGCCC | 174 |
| chr1:51518404-51520418 | p2 | (GT)7 | TATTGGAAGCACGAACCCTC | ATATATGTTCAGCGAGGGCG | 207 |
| chr1:51871634-51873649 | p5 | (ACACA)3 | TACATACCTTTGCCCGCTTC | GAATTCTCGGGGTTGTTGAA | 299 |
| chr1:51886529-51888545 | p4 | (GAAA)4 | GTCACGCGACAAGCTTAACA | CAAGGGCAACCAAGCATAAT | 165 |
| chr1:51901928-51903943 | p2 | (TA)6 | GAGGATGACGTTTGCTGCTA | GAATTGTTGCCAAGTGGCTT | 236 |
| chr1:51901928-51903943 | p2 | (TA)7 | GAGGATGACGTTTGCTGCTA | GAATTGTTGCCAAGTGGCTT | 236 |
| chr1:51901928-51903943 | p3 | (TAA)5 | GAGGATGACGTTTGCTGCTA | GAATTGTTGCCAAGTGGCTT | 236 |
| chr1:51901928-51903943 | p3 | (CTA)8 | GAGGATGACGTTTGCTGCTA | GAATTGTTGCCAAGTGGCTT | 236 |
| chr1:51901928-51903943 | p2 | (TA)6 | GAGGATGACGTTTGCTGCTA | GAATTGTTGCCAAGTGGCTT | 236 |
| chr1:51901928-51903943 | p3 | (CTA)6 | GAGGATGACGTTTGCTGCTA | GAATTGTTGCCAAGTGGCTT | 236 |
| chr1:51901928-51903943 | p2 | (AT)10 | GAGGATGACGTTTGCTGCTA | GAATTGTTGCCAAGTGGCTT | 236 |
| chr1:51901974-51904000 | p2 | (TA)6 | GAGGATGACGTTTGCTGCTA | GAATTGTTGCCAAGTGGCTT | 190 |

| Marker ID | SSR type | SSR motif | FORWARD PRIMER (5'-3') | REVERSE PRIMER (5'-3') | size (bp) |
|---|---|---|---|---|---|
| chr1:51901974-51904000 | p2 | (TA)7 | GAGGATGACGTTTGCTGCTA | GAATTGTTGCCAAGTGGCTT | 190 |
| chr1:51901974-51904000 | p3 | (TAA)5 | GAGGATGACGTTTGCTGCTA | GAATTGTTGCCAAGTGGCTT | 190 |
| chr1:51901974-51904000 | p3 | (CTA)8 | GAGGATGACGTTTGCTGCTA | GAATTGTTGCCAAGTGGCTT | 190 |
| chr1:51901974-51904000 | p2 | (TA)6 | GAGGATGACGTTTGCTGCTA | GAATTGTTGCCAAGTGGCTT | 190 |
| chr1:51901974-51904000 | p3 | (CTA)6 | GAGGATGACGTTTGCTGCTA | GAATTGTTGCCAAGTGGCTT | 190 |
| chr1:51901974-51904000 | p2 | (AT)10 | GAGGATGACGTTTGCTGCTA | GAATTGTTGCCAAGTGGCTT | 190 |
| chr1:51902017-51904039 | p2 | (TA)6 | GAGGATGACGTTTGCTGCTA | GAATTGTTGCCAAGTGGCTT | 147 |
| chr1:51902017-51904039 | p2 | (TA)7 | GAGGATGACGTTTGCTGCTA | GAATTGTTGCCAAGTGGCTT | 147 |
| chr1:51902017-51904039 | p3 | (TAA)5 | GAGGATGACGTTTGCTGCTA | GAATTGTTGCCAAGTGGCTT | 147 |
| chr1:51902017-51904039 | p3 | (CTA)8 | GAGGATGACGTTTGCTGCTA | GAATTGTTGCCAAGTGGCTT | 147 |
| chr1:51902017-51904039 | p2 | (TA)6 | GAGGATGACGTTTGCTGCTA | GAATTGTTGCCAAGTGGCTT | 147 |
| chr1:51902017-51904039 | p3 | (CTA)6 | GAGGATGACGTTTGCTGCTA | GAATTGTTGCCAAGTGGCTT | 147 |
| chr1:51902017-51904039 | p2 | (AT)10 | GAGGATGACGTTTGCTGCTA | GAATTGTTGCCAAGTGGCTT | 147 |
| chr1:51902017-51904039 | p6 | (ATCTAT)3 | GAGGATGACGTTTGCTGCTA | GAATTGTTGCCAAGTGGCTT | 147 |
| chr1:51902169-51904193 | p2 | (TA)6 | AGCCAAGTGGCGTAATATGAA | TGTTGCAAAGTGGCTTGTTC | 282 |
| chr1:51902169-51904193 | p2 | (TA)7 | AGCCAAGTGGCGTAATATGAA | TGTTGCAAAGTGGCTTGTTC | 282 |
| chr1:51902169-51904193 | p3 | (TAA)5 | AGCCAAGTGGCGTAATATGAA | TGTTGCAAAGTGGCTTGTTC | 282 |
| chr1:51902169-51904193 | p3 | (CTA)8 | AGCCAAGTGGCGTAATATGAA | TGTTGCAAAGTGGCTTGTTC | 282 |
| chr1:51902169-51904193 | p2 | (TA)6 | AGCCAAGTGGCGTAATATGAA | TGTTGCAAAGTGGCTTGTTC | 282 |
| chr1:51902169-51904193 | p3 | (CTA)6 | AGCCAAGTGGCGTAATATGAA | TGTTGCAAAGTGGCTTGTTC | 282 |
| chr1:51902169-51904193 | p2 | (AT)10 | AGCCAAGTGGCGTAATATGAA | TGTTGCAAAGTGGCTTGTTC | 282 |
| chr1:51902169-51904193 | p6 | (ATCTAT)3 | AGCCAAGTGGCGTAATATGAA | TGTTGCAAAGTGGCTTGTTC | 282 |
| chr1:51903185-51905200 | p2 | (TA)6 | GAACAAGCAACTTGGCAACA | TTGTTCATTTACACCACTTGGC | 244 |
| chr1:51903185-51905200 | p3 | (CTA)6 | GAACAAGCAACTTGGCAACA | TTGTTCATTTACACCACTTGGC | 244 |
| chr1:51903185-51905200 | p2 | (AT)10 | GAACAAGCAACTTGGCAACA | TTGTTCATTTACACCACTTGGC | 244 |
| chr1:51903185-51905200 | p6 | (ATCTAT)3 | GAACAAGCAACTTGGCAACA | TTGTTCATTTACACCACTTGGC | 244 |
| chr1:51903185-51905200 | p3 | (TAA)5 | GAACAAGCAACTTGGCAACA | TTGTTCATTTACACCACTTGGC | 244 |
| chr1:51903185-51905200 | p3 | (ATA)5 | GAACAAGCAACTTGGCAACA | TTGTTCATTTACACCACTTGGC | 244 |
| chr1:51903185-51905200 | p3 | (CTA)6 | GAACAAGCAACTTGGCAACA | TTGTTCATTTACACCACTTGGC | 244 |
| chr1:51903610-51905625 | p2 | (AT)10 | TGAACAAGCCACTTAGCAACA | GGCTGTGTTCATATTACGCC | 281 |
| chr1:51903610-51905625 | p6 | (ATCTAT)3 | TGAACAAGCCACTTAGCAACA | GGCTGTGTTCATATTACGCC | 281 |
| chr1:51903610-51905625 | p3 | (TAA)5 | TGAACAAGCCACTTAGCAACA | GGCTGTGTTCATATTACGCC | 281 |
| chr1:51903610-51905625 | p3 | (ATA)5 | TGAACAAGCCACTTAGCAACA | GGCTGTGTTCATATTACGCC | 281 |
| chr1:51903610-51905625 | p3 | (CTA)6 | TGAACAAGCCACTTAGCAACA | GGCTGTGTTCATATTACGCC | 281 |
| chr1:51903610-51905625 | p3 | (TAA)6 | TGAACAAGCCACTTAGCAACA | GGCTGTGTTCATATTACGCC | 281 |
| chr1:51903610-51905625 | p3 | (TAA)10 | TGAACAAGCCACTTAGCAACA | GGCTGTGTTCATATTACGCC | 281 |

| Marker ID | SSR type | SSR motif | FORWARD PRIMER (5'-3') | REVERSE PRIMER (5'-3') | size (bp) |
|---|---|---|---|---|---|
| chr1:51959200-51961212 | p2 | (AT)6 | TCTTCAACTCCCTGACGCTT | AACGTGGCCATCAAGAAGAC | 126 |
| chr1:51970451-51972463 | p2 | (TA)6 | ACGTGTGCGAACATCATTGT | GGATTACGGACGGATTCTCA | 216 |
| chr1:51975773-51977788 | p3 | (ACA)5 | ACAAACAAAAACGTGCACCA | CGCAAACCTTGTCCAAAAAT | 136 |
| chr1:51978493-51980508 | p5 | (ATGCA)3 | TCCATAGGCTCGGTTGATTC | TAGTTTCGTTGGGTTGAGGG | 186 |
| chr1:51994735-51996756 | p3 | (TAT)7 | GTGGATGTCCCAATGTTTCC | AGGATCAGAAAAGGGGCTGT | 237 |
| chr1:52012785-52014805 | p4 | (TGTA)5 | GTTCAGATCCCTCTGGGTGA | TTTAGCATTACCCGTGGAGG | 256 |
| chr1:52139062-52141077 | p5 | (TAAAC)3 | CCACTACAGCGGTCCATTTT | GGTGAGTCCGTAGCGTGTTT | 274 |
| chr1:52148854-52150898 | p4 | (GAAA)11 | GTCAACAGAGGAGACCCAGC | CCAGTTACAGGGCTGGTGAT | 275 |
| chr1:52214528-52216546 | p6 | (TCAGTT)3 | TGCGTGATGGAGAATGATGT | AATTCCAGCCTTTGAGCGTA | 171 |
| chr1:52231839-52233854 | p3 | (TCT)5 | ACATCCTTCACGTCCAGACC | CTCTCTCCGGTATCATCCCA | 233 |
| chr1:52249910-52251925 | p5 | (TCATA)3 | TCATGTCACGACCCGTCTAA | GTGGCGTAGATGCCTATGGT | 266 |
| chr1:52388796-52390812 | p2 | (GA)8 | GGGGCAAATTTGATCCTTTT | AGTTGTTGCTGGCTGTCCTT | 100 |
| chr1:52397725-52399743 | p3 | (AAC)6 | ACGAGCGGAGTAGTGAGGAA | TATGTTTTTCATAGGCGGGC | 298 |
| chr1:52411739-52413753 | p2 | (TG)7 | AAGATTTTGAGCGAGGGGAT | GACTAGCTCTGGGTCCACCA | 134 |
| chr1:52422465-52424479 | p2 | (TG)7 | AAGATTTTGAGCGAGGGGAT | GACTAGCTCTGGGTCCACCA | 134 |
| chr1:52447981-52449996 | p5 | (CAGAA)3 | TCAATATCTTTCAGCGCACG | TTCCTCAGGGTTTTGGTCAG | 285 |
| chr1:52447981-52449996 | p3 | (TGT)5 | TCAATATCTTTCAGCGCACG | TTCCTCAGGGTTTTGGTCAG | 285 |
| chr1:52472747-52474759 | p2 | (AT)6 | CTGGCAGGACTGGTAAAAGC | AGCACATGGTTAAGGCCAAC | 174 |
| chr1:52631729-52633763 | p2 | (AT)17 | TGCAACCTTGTCGAGAATTG | GATTGTGAGGAGGAAACCCA | 222 |
| chr1:52642256-52644274 | p6 | (CTGAAA)3 | TTGGACCTTGAATTTCCTGC | GAAACCGTGTCCAGTGAGGT | 155 |
| chr1:52650382-52652414 | p2 | (AT)16 | ATGGAAATCAAGCAAGGCAC | CTATGAAACCCTAGCACCGC | 247 |
| chr1:52717405-52719420 | p5 | (AAATA)3 | ACGAGCATGCCTGGAATATC | GTATGCGTGGACCAGAACCT | 169 |
| chr1:52723006-52725038 | p2 | (AT)16 | GAGGAATTCAAGTGGGTGGA | CTTCTTTGGCGAGTTCAAGC | 262 |
| chr1:52807416-52809428 | p2 | (AT)6 | TTTGAGCGATTGGTTTGTTG | GAGCCCCACCAACAATTTTA | 122 |
| chr1:52833275-52835290 | p5 | (TGGTT)3 | ACAAAGCAAACGGACAAACC | ACCAAACCAAACCAGACCAA | 150 |
| chr1:52931863-52933941 | p3 | (TAT)26 | ACAGCCACAACCTCAAAACC | TGAAGAGGTGGGAACTGGTC | 148 |
| chr1:52931863-52933941 | p5 | (TTCTC)3 | ACAGCCACAACCTCAAAACC | TGAAGAGGTGGGAACTGGTC | 148 |
| chr1:52945636-52947652 | p5 | (TTCTC)3 | TTGCGGACAAATTCACAAAA | GATGGGCTGGTCTGAAATGT | 154 |
| chr1:52945636-52947652 | p2 | (AG)8 | TTGCGGACAAATTCACAAAA | GATGGGCTGGTCTGAAATGT | 154 |
| chr1:52947843-52949858 | p5 | (TTGGT)3 | CAGGCGAAACTCGTTAGAGG | CTCCACCCAATTTTAAGCCA | 121 |
| chr1:53023616-53025630 | p2 | (AG)7 | CCTACATACCTCGGCGAAAA | CATTTGGTTTAACGGCGACT | 101 |
| chr1:53053107-53055121 | p2 | (TA)7 | AAAATCACGCCACATCAACA | GGGGTGGTCAGTTGAAAGAA | 270 |
| chr1:53054216-53056234 | p6 | (TGGTGC)3 | AAAAAGGCTCAAAAGGGGAA | ATGCCGATGTGTGTGTGAGT | 104 |
| chr1:53116380-53118398 | p2 | (TA)9 | CGAGCTTGGAAGGATAAACG | CACCATCATCATGCTCAACC | 195 |
| chr1:53204344-53206356 | p2 | (TC)6 | TGCTATCCCCGTCACTTTCT | AAGGTTGTTTGTGGGTTTCG | 142 |
| chr1:53279551-53281575 | p2 | (AT)12 | GCCATATCCTTAGTCCGCAA | TTCGGTATTTCACGTGTCCA | 210 |

| Marker ID | SSR type | SSR motif | FORWARD PRIMER (5'-3') | REVERSE PRIMER (5'-3') | size (bp) |
|---|---|---|---|---|---|
| chr1:53291240-53293256 | p2 | (AT)8 | TCGACAACAATTCAACGCAT | TTTGGAAGAGGGGAAATGTG | 212 |
| chr1:53291240-53293256 | p2 | (TA)7 | TCGACAACAATTCAACGCAT | TTTGGAAGAGGGGAAATGTG | 212 |
| chr1:53381033-53383047 | p3 | (CAA)5 | GTGAGCTAAGCGGGTGAAAG | CTCGAAAATATCCCCGTGAA | 127 |
| chr1:53381033-53383047 | p2 | (CT)7 | GTGAGCTAAGCGGGTGAAAG | CTCGAAAATATCCCCGTGAA | 127 |
| chr1:53551583-53553598 | p5 | (CTTTG)3 | AATCCCCAGCAAAGAAAGGT | CCAATCCAAGGGTTGTTGAT | 236 |
| chr1:53607359-53609375 | p3 | (TGT)5 | GCCATGATCAACTCGGCTAT | GAAATTCACGGTGGAGTCGT | 226 |
| chr1:53607359-53609375 | p4 | (AAAC)4 | GCCATGATCAACTCGGCTAT | GAAATTCACGGTGGAGTCGT | 226 |
| chr1:53656825-53658917 | p2 | (AT)6 | TTGTTACTTGGACAAGGCCC | TTTCCAGAGCATCCCTTGTT | 292 |
| chr1:53656919-53658941 | p2 | (AT)6 | TTGTTACTTGGACAAGGCCC | TTTCCAGAGCATCCCTTGTT | 198 |
| chr1:53656943-53658965 | p2 | (AT)6 | TTGTTACTTGGACAAGGCCC | TTTCCAGAGCATCCCTTGTT | 174 |
| chr1:53656967-53658989 | p2 | (AT)6 | TTGTTACTTGGACAAGGCCC | TTTCCAGAGCATCCCTTGTT | 150 |
| chr1:53656991-53659013 | p2 | (AT)6 | TTGTTACTTGGACAAGGCCC | TTTCCAGAGCATCCCTTGTT | 126 |
| chr1:53657015-53659037 | p2 | (AT)6 | TTGTTACTTGGACAAGGCCC | TTTCCAGAGCATCCCTTGTT | 102 |
| chr1:53657183-53659205 | p2 | (AT)6 | AAACAAGGGATGCTCTGGAA | CTCTCCCCACATGCAAGAAT | 170 |
| chr1:53657207-53659229 | p2 | (AT)6 | AAACAAGGGATGCTCTGGAA | CTCTCCCCACATGCAAGAAT | 146 |
| chr1:53657231-53659253 | p2 | (AT)6 | AAACAAGGGATGCTCTGGAA | CTCTCCCCACATGCAAGAAT | 122 |
| chr1:53657255-53659277 | p2 | (AT)6 | ATTCTTGCATGTGGGGAGAG | TACCCCGGCAAAGTATGAGA | 242 |
| chr1:53657279-53659301 | p2 | (AT)6 | ATTCTTGCATGTGGGGAGAG | TACCCCGGCAAAGTATGAGA | 218 |
| chr1:53657303-53659325 | p2 | (AT)6 | ATTCTTGCATGTGGGGAGAG | TACCCCGGCAAAGTATGAGA | 194 |
| chr1:53657327-53659349 | p2 | (AT)6 | ATTCTTGCATGTGGGGAGAG | TACCCCGGCAAAGTATGAGA | 170 |
| chr1:53657351-53659373 | p2 | (AT)6 | ATTCTTGCATGTGGGGAGAG | TACCCCGGCAAAGTATGAGA | 146 |
| chr1:53717334-53719350 | p4 | (TAAT)4 | TACCCCTGCCGTTAACAAAG | ATGAAATCAGGGCTGAGTCG | 290 |
| chr1:53958392-53960407 | p3 | (TAT)5 | TTGAGTGACGTGAGAGGTCG | TACTCCGGACGGAAATATCG | 232 |
| chr1:53990107-53992143 | p2 | (TA)18 | TATGAATGCGATTTCGCTGA | AAAGGAAATTGGTTTTGGGG | 156 |
| chr1:54149144-54151162 | p6 | (GAAGTT)3 | TTATCCCATGCGTACGTTGA | ACAGCTTGGCCTTGTCTTGT | 192 |
| chr1:54191287-54193325 | p5 | (AAATG)3 | ATGTCATGCAGCTTCCTGTG | CACCTAGTGACCCTCGTGGT | 244 |
| chr1:54191379-54193394 | p5 | (AAATG)3 | ATGTCATGCAGCTTCCTGTG | CACCTAGTGACCCTCGTGGT | 152 |
| chr1:54238019-54240034 | p5 | (AATAC)3 | TGGACACAGCTCAAGAATGG | CCGTGGTTACAAAAATTGGG | 267 |
| chr1:54238019-54240034 | p2 | (TA)16 | TGGACACAGCTCAAGAATGG | CCGTGGTTACAAAAATTGGG | 267 |
| chr1:54505984-54508018 | p6 | (ATACAT)3 | TGAACAGCCTCGATCACTTG | CCGCCCAATTTAGAAGAACA | 286 |
| chr1:54546871-54548885 | p2 | (AT)9 | TCTGGTTGTCTTCCCTGTCC | CACTGATGTGGGCCTATCCT | 202 |
| chr1:54546871-54548885 | p2 | (TA)7 | TCTGGTTGTCTTCCCTGTCC | CACTGATGTGGGCCTATCCT | 202 |
| chr1:54553784-54555796 | p2 | (AT)6 | CCTCTAGCATGCTCCACCTC | TGATCGATCCCACAGTTTGA | 131 |
| chr1:54553784-54555796 | p2 | (TA)6 | CCTCTAGCATGCTCCACCTC | TGATCGATCCCACAGTTTGA | 131 |
| chr1:54571978-54573993 | p5 | (GCTTA)3 | CTATTGCCCCAAAATCCCTT | TGGTTCTACCTTGTTTGGGC | 293 |
| chr1:54588453-54590475 | p2 | (AT)11 | TATTTGGTTCGCGTAGGCTC | GAGTGCACGCATTTCACACT | 263 |

| Marker ID | SSR type | SSR motif | FORWARD PRIMER (5'-3') | REVERSE PRIMER (5'-3') | size (bp) |
|---|---|---|---|---|---|
| chr1:54588453-54590475 | p2 | (TA)9 | TATTTGGTTCGCGTAGGCTC | GAGTGCACGCATTTCACACT | 263 |
| chr1:54809551-54811563 | p4 | (TATG)5 | TGTTAAAAGGTGCCTACGGG | CTCATACCTGGCCCTACCAA | 214 |
| chr1:54809551-54811563 | p2 | (TG)6 | TGTTAAAAGGTGCCTACGGG | CTCATACCTGGCCCTACCAA | 214 |
| chr1:54883224-54885252 | p4 | (GAAA)7 | GAGGCAAATTTGTTGCCCTA | CCTAGGCTAGTGGGAGGACC | 227 |
| chr1:55080333-55082378 | p3 | (CTT)15 | CTCCCAAATCCGAAGATTCA | CACCAAAAACACCCAAAACC | 209 |
| chr1:55086788-55088833 | p3 | (CTT)15 | CTCCCAAATCCGAAGATTCA | CACCAAAAACACCCAAAACC | 209 |
| chr1:55088893-55090933 | p2 | (TA)20 | TATGGGTCTTCAATGGCCTC | GGTGAAGCTCGTCCTCAGAC | 288 |
| chr1:55088893-55090933 | p2 | (TA)8 | TATGGGTCTTCAATGGCCTC | GGTGAAGCTCGTCCTCAGAC | 288 |
| chr1:55104624-55106642 | p6 | (CATATA)3 | AATGTACAAGGCCGACAAGG | GAGCTCCATCTCATTCGGAG | 280 |
| chr1:55190044-55192112 | p2 | (TA)7 | TGGTTGTTGGTGGTGAGTGT | GTGTGCTGGCAACCTAGTGA | 108 |
| chr1:55190118-55192140 | p2 | (TA)7 | CTCACTAGGTTGCCAGCACA | CAAGGAAGGTCAAAAGGCAG | 127 |
| chr1:55227588-55229600 | p2 | (TA)6 | TACTAGGCAATTTGGGCCAC | ATGGATGGGCCTTCATCATA | 229 |
| chr1:55243390-55245411 | p3 | (TCA)7 | CCTTCCGAGTGGTCATTTGT | AGCCCACATTGAGGTAGTGG | 169 |
| chr1:55331445-55333465 | p3 | (TGT)6 | GAACACTGGAATATCGGGGA | CACTTCCAAAACAGGCCAAT | 163 |
| chr1:55331445-55333465 | p2 | (GT)10 | GAACACTGGAATATCGGGGA | CACTTCCAAAACAGGCCAAT | 163 |
| chr1:55331445-55333465 | p2 | (AT)6 | GAACACTGGAATATCGGGGA | CACTTCCAAAACAGGCCAAT | 163 |
| chr1:55331445-55333465 | p2 | (AT)6 | GAACACTGGAATATCGGGGA | CACTTCCAAAACAGGCCAAT | 163 |
| chr1:55337508-55339520 | p2 | (GA)6 | TAGGTGGAAGAGCTCGCAAT | AACATTCCTTCCCCCTTTTG | 110 |
| chr1:55337508-55339520 | p3 | (GTT)6 | TAGGTGGAAGAGCTCGCAAT | AACATTCCTTCCCCCTTTTG | 110 |
| chr1:55366928-55368943 | p3 | (CAC)5 | TAAACGTGAAAAATGGGGGA | ATCCCCATTCAGAAGTGCAG | 258 |
| chr1:55377010-55379025 | p3 | (TTA)5 | TACAGTTCAGAGATCCGGGG | TTCATGGTCTTGGGAAGGAG | 188 |
| chr1:55502594-55504606 | p2 | (TA)6 | CGGGGAGTCAGTACGGATAA | ATGATGGGATGCCATAGAGG | 105 |
| chr1:55619611-55621626 | p5 | (ATCAT)3 | GTAGGGGTAAACCGAGCACA | CTGTCCCGCCATATGATTCT | 176 |
| chr1:55626590-55628605 | p5 | (ATCAT)3 | GTAGGGGTAAACCGAGCACA | CGGAGCCCATACTTTTGGTA | 178 |
| chr1:55678241-55680253 | p2 | (TC)6 | GCAAAGGCCCTATCAAATCA | GGGTGTTACATCATCCCCAC | 194 |
| chr1:55759855-55761867 | p2 | (AT)6 | AATTGGGCCCTTTTTCATTC | ACAATAAGGAAAATCCGGGG | 131 |
| chr1:55769979-55771991 | p2 | (CA)6 | ACATGCCATCCTAGTGGCTC | CGAGGGATCGAGATTTGGTA | 157 |
| chr1:55818431-55820467 | p2 | (AT)18 | CCCTTACGTGATCAATGCCT | AAGGTACATCACAGCCCTGG | 215 |
| chr1:55869904-55871916 | p2 | (CA)6 | CCATACCCTGAACTGTGCCT | AGGAAAGTGAGTCCCAGGGT | 251 |
| chr1:55877859-55879877 | p6 | (TGTAAG)3 | TAACCCTGACGCCTTGGTAG | ACTCGACCATACAGGATCGG | 159 |
| chr1:55954695-55956710 | p5 | (TTCAT)3 | ACGTACACGAATCACGCAAG | ACCCGGAGTATCATGACCAA | 250 |
| chr1:56005084-56007096 | p2 | (GT)6 | GGCCTGTTGGTACCTGAAAA | CCTCGTTTTGTAATTCCCGA | 282 |
| chr1:56064322-56066337 | p3 | (CAA)5 | ATGTACGCGTATACCTGCCC | AATGTTTCCCACACTGAGCC | 134 |
| chr1:56064322-56066337 | p2 | (AC)6 | ATGTACGCGTATACCTGCCC | AATGTTTCCCACACTGAGCC | 134 |
| chr1:56204839-56206857 | p2 | (TA)9 | GGCTCTCCCTTGTGTTTGAG | CAGTCTCTCTACCCCGCAAG | 199 |
| chr1:56204839-56206857 | p2 | (AG)8 | GGCTCTCCCTTGTGTTTGAG | CAGTCTCTCTACCCCGCAAG | 199 |

| Marker ID | SSR type | SSR motif | FORWARD PRIMER (5'-3') | REVERSE PRIMER (5'-3') | size (bp) |
|---|---|---|---|---|---|
| chr1:56204839-56206857 | p2 | (AT)16 | GGCTCTCCCTTGTGTTTGAG | CAGTCTCTCTACCCCGCAAG | 199 |
| chr1:56319868-56321883 | p6 | (ATACAT)3 | CGCTTAGACGCAAATCACAA | CCTTGGAGTGCTCCATTGTT | 214 |
| chr1:56319868-56321883 | p3 | (CAT)5 | CGCTTAGACGCAAATCACAA | CCTTGGAGTGCTCCATTGTT | 214 |
| chr1:56361592-56363607 | p5 | (ATATT)3 | TTAGCCAGGGTCTCGTCTGT | ATGTACCCGATCCAATCCAA | 266 |
| chr1:56375675-56377695 | p4 | (AGAA)5 | CCGTCTTGTGTGCCTCTGTA | ACCAACAGGTTCAGGACTGG | 117 |
| chr1:56395729-56397744 | p5 | (TTTCA)3 | TGATAAGGGGTGCTCGGTAG | CCTCTTGGCTTTTGATGCTC | 196 |
| chr1:56507145-56509163 | p6 | (AGGTAG)3 | TTATTCATGCCTCCGAGACC | TTCAAGAACCTGTCCGCTCT | 153 |
| chr1:56583275-56585287 | p2 | (AT)6 | GATCTGTGTATGGGGATGGC | ATTAATACAGATCCCGGCCC | 114 |
| chr1:56583275-56585287 | p3 | (TAA)5 | GATCTGTGTATGGGGATGGC | ATTAATACAGATCCCGGCCC | 114 |
| chr1:56643081-56645093 | p2 | (AT)6 | GGAAGATTGCGGATTGTGTT | TTAACGTCCCCGAGAATCAC | 142 |
| chr1:56658378-56660390 | p2 | (GT)6 | GGATGTCATGCACACACACA | TTTCCAATTGTGCGAAAACA | 156 |
| chr1:56665948-56667960 | p2 | (TG)6 | GGACTTCACGGTGTTCCTGT | ATCCCAGCTCCCTCTGAAAT | 116 |
| chr1:56790264-56792279 | p3 | (CAT)5 | AGTGTCACGACCCGATTAGG | TGAGTGGTGCTTGGTGGTTA | 270 |
| chr1:56790264-56792279 | p2 | (AT)6 | AGTGTCACGACCCGATTAGG | TGAGTGGTGCTTGGTGGTTA | 270 |
| chr1:56909260-56911276 | p2 | (AT)8 | AAGGCAAGTGACGATATGGC | TGCTTCCTTCTGTTAGGCGT | 210 |
| chr1:56953507-56955535 | p2 | (AT)14 | TTTTAAGCGAGCTTTTCGGA | ATTTGGAAAAACGTTGGCAG | 230 |
| chr1:56968866-56970881 | p5 | (ATTTC)3 | AAGAAATGACATGGGAAGCG | ATAACAATGCCTCACAGGGC | 187 |
| chr1:57283449-57285464 | p3 | (TCA)5 | TGTCGCATTGACAAATGGTT | CTTGGAACCAAAGGAAACGA | 107 |
| chr1:57287194-57289206 | p2 | (AT)6 | CTCCCGGGTGAACAGAATAA | TGAGATTTCTGGAGATGGGG | 243 |
| chr1:57287194-57289206 | p3 | (TTG)5 | CTCCCGGGTGAACAGAATAA | TGAGATTTCTGGAGATGGGG | 243 |
| chr1:57287640-57289655 | p2 | (AT)6 | GGATGCTAAGCGAATCGAAG | TGCACTGCCTCAACTTCAAC | 248 |
| chr1:57287640-57289655 | p3 | (TTG)5 | GGATGCTAAGCGAATCGAAG | TGCACTGCCTCAACTTCAAC | 248 |
| chr1:57366133-57368145 | p2 | (CA)6 | CGAATCGGGCAAAAAGATTA | TTCTATCGCTCTCGCAGGTT | 193 |
| chr1:57422394-57424409 | p5 | (AAAGG)3 | GGAGACCATAAAAGGCCACA | ACTCGGGCTTCGATATCCTT | 183 |
| chr1:57435716-57437734 | p3 | (TTG)6 | TGCCACCAAAACTTAAAGGC | GCTTTCCGCATCTTGTCTTC | 188 |
| chr1:57512027-57514041 | p3 | (CAT)5 | TCCCCAACTATCCCCTCTTT | CACGAGCCGAACCCTATAAA | 171 |
| chr1:57574453-57576469 | p2 | (TA)8 | CGTGCTAGCTGAAAGATCCC | AGAAGGGCAAAGCTACGACA | 214 |
| chr1:57603995-57606011 | p2 | (AT)6 | AACGGATACGCACCAAAAAG | TTATGCGATAACACGGGACA | 113 |
| chr1:57603995-57606011 | p2 | (GT)8 | AACGGATACGCACCAAAAAG | TTATGCGATAACACGGGACA | 113 |
| chr1:57699398-57701418 | p3 | (TCA)5 | TATTGCTCGCTCGTTCTCCT | CAAAGGAAATGCTGCCAAAT | 298 |
| chr1:57699398-57701418 | p5 | (AAATG)4 | TATTGCTCGCTCGTTCTCCT | CAAAGGAAATGCTGCCAAAT | 298 |
| chr1:57767101-57769119 | p6 | (ACCAGC)3 | ACCGACTATATGAGGTGGCG | GGAGGCAGAAAATGCTCAAG | 204 |
| chr1:57879938-57881952 | p2 | (CA)7 | ACAGAATATTCCCCGAGGCT | GGAGGAAACGGGAGTTAAGC | 210 |
| chr1:57939735-57941747 | p3 | (AAC)5 | GGAGTGGGTCGCCTTACATA | TGTGATGATGTTGGGCTGTT | 259 |
| chr1:57939735-57941747 | p2 | (AC)6 | GGAGTGGGTCGCCTTACATA | TGTGATGATGTTGGGCTGTT | 259 |
| chr1:58032769-58034801 | p2 | (AT)15 | GGCGATGCTCGATCATATTT | ATAATCCACTGCGCCAACTC | 238 |

| Marker ID | SSR type | SSR motif | FORWARD PRIMER (5'-3') | REVERSE PRIMER (5'-3') | size (bp) |
|---|---|---|---|---|---|
| chr1:58032769-58034801 | p2 | (TA)16 | GGCGATGCTCGATCATATTT | ATAATCCACTGCGCCAACTC | 238 |
| chr1:58058240-58060284 | p2 | (AT)8 | CGGCTTGGTCTGTTTAGAGG | ATATAAAAACGCACGCCGAC | 174 |
| chr1:58166505-58168517 | p2 | (AT)6 | CTTGCTCTCACACAGTCCCA | GCTCTCGTTATCGTCCCTTG | 272 |
| chr1:58166505-58168517 | p2 | (TA)6 | CTTGCTCTCACACAGTCCCA | GCTCTCGTTATCGTCCCTTG | 272 |
| chr1:58213137-58215152 | p5 | (ACAAT)3 | CTCCTATTCCTGCGAACGAG | TATTCATGCGGTCCACAAAA | 179 |
| chr1:58244904-58246936 | p5 | (ATAAA)3 | CTGTCATCGAGGAACCCCTA | CAGGCTTGCAAACGATGTAA | 222 |
| chr1:58244904-58246936 | p2 | (AT)16 | CTGTCATCGAGGAACCCCTA | CAGGCTTGCAAACGATGTAA | 222 |
| chr1:58256844-58258859 | p3 | (CTG)5 | ACAGGAAGCTAGCCAGTCCA | CACCCGAACCTATAGAGCCA | 248 |
| chr1:58290290-58292305 | p5 | (GAAAT)3 | ACGTATAGGTGCCCAGTTCG | CATTTGAGGAGAACGTGCAA | 218 |
| chr1:58315409-58317424 | p3 | (TCA)5 | ATAAGTCAATGGCACCTCCG | CCGTTTTCTTTGGATGCTGT | 178 |
| chr1:58317942-58319958 | p2 | (TA)8 | GGACCACGAGGGTCATTAGA | CCACGATGGCCTTGTAGACT | 215 |
| chr1:58566574-58568598 | p2 | (AT)12 | TCATGGCTCTATCCTCCACC | AGTTACTAGGGCCCATGCCT | 256 |
| chr1:58601356-58603374 | p6 | (CATATC)3 | CACATCTAGGGGAGCCACAT | GTCCTGTCCCGCCTATATGA | 284 |
| chr1:58617498-58619513 | p5 | (TGCAT)3 | TAAGGTGCGGATTCATGTCA | GTTCCGCATGTGTGTTTGTC | 298 |
| chr1:58617498-58619513 | p4 | (AACA)4 | TAAGGTGCGGATTCATGTCA | GTTCCGCATGTGTGTTTGTC | 298 |
| chr1:58618180-58620196 | p5 | (TGCAT)3 | TAAAAACCCCACATTGGCAT | ACTGGCCAACTGTCCCATAC | 121 |
| chr1:58618180-58620196 | p4 | (AACA)4 | TAAAAACCCCACATTGGCAT | ACTGGCCAACTGTCCCATAC | 121 |
| chr1:58668530-58670548 | p6 | (TTTTCA)3 | CATCTGCAGAAAGGCATTCA | CGCTTGGAACATTTCATTCA | 162 |
| chr1:58813544-58815556 | p2 | (AT)6 | CATTAGCAGCGTCACCGTAA | GACAGGTGGACGTGATCCTT | 178 |
| chr1:58835365-58837380 | p2 | (AT)6 | AAGGATGCTCTCCAACATGG | ATTCGGCCAACACCATACAT | 259 |
| chr1:58835365-58837380 | p5 | (TGAAT)3 | AAGGATGCTCTCCAACATGG | ATTCGGCCAACACCATACAT | 259 |
| chr1:58961218-58963234 | p2 | (AT)8 | AGAGCTCGAGACCTGAGTCG | GGTCATGCCAAAAAGGAAGA | 114 |
| chr1:58972047-58974059 | p2 | (CA)6 | GGACATGCCAAAAGAAGGAA | GCATTGTTGATGTTGTTGGG | 256 |
| chr1:59016099-59018141 | p3 | (TAA)14 | CCTCTGAAGTCCAGCACACA | GTTTGGACTGCATCACATGG | 152 |
| chr1:59157837-59159851 | p3 | (TTG)7 | TTATGAAGGGGGTGGATTGA | ATAACCTTCGGAAAATCGGG | 233 |
| chr1:59157837-59159851 | p2 | (TA)7 | TTATGAAGGGGGTGGATTGA | ATAACCTTCGGAAAATCGGG | 233 |
| chr1:59195854-59197872 | p3 | (GTT)6 | CATCGTATTGGTGCAGGATG | CATTTCGGCCACTTTCATCT | 151 |
| chr1:59221786-59223802 | p4 | (GTAA)4 | CGTGGAAGTTGAACCGAAAT | GTAGCAAGAACCATGCCGTT | 281 |
| chr1:59252757-59254772 | p5 | (CCACA)3 | GTCAAACGGAGCAATAGGGA | AGCCTCGGGGACAACTCTAT | 139 |
| chr1:59267442-59269460 | p6 | (TATATG)3 | GATTCTCGGGGATGGGTATT | AGGGCGAGCCTTACATACCT | 246 |
| chr1:59412389-59414427 | p2 | (TA)19 | TTCATCCCAGATTGGTAGCC | AATCTCACAATCTGTCCCGC | 234 |
| chr1:59412389-59414427 | p6 | (ATGAAT)3 | TTCATCCCAGATTGGTAGCC | AATCTCACAATCTGTCCCGC | 234 |
| chr1:59421329-59423343 | p2 | (TA)7 | ATAGATTGGGACGGAGGAGG | ATTGGCCAAACATCACATCA | 204 |
| chr1:59461370-59463394 | p2 | (TA)12 | AAGCATGATTCCGGTCTGTC | TAAATCCATCTCGAATCGCC | 156 |
| chr1:59489966-59491986 | p2 | (AT)10 | GACAATTTCAAACGCGGATT | GCCCAAATTTCTGTAGGCAA | 183 |
| chr1:59536732-59538750 | p3 | (AAC)6 | ATGCAATATGGAGGCCAAAG | GACATATTGCCTGGCCACTT | 227 |

| Marker ID | SSR type | SSR motif | FORWARD PRIMER (5'-3') | REVERSE PRIMER (5'-3') | size (bp) |
|---|---|---|---|---|---|
| chr1:59592286-59594298 | p2 | (AT)6 | CCGACGAGGCTATCAAAAAG | TTGTGGAAGCTGCGTCATAC | 107 |
| chr1:59752666-59754686 | p3 | (GTT)9 | GGAAAACGTCGTGAGGGATA | TCGGGTAGCATCTCTCCTGT | 299 |
| chr1:59757827-59759848 | p3 | (GTT)7 | AGGAGGACCCTTAAGCCAAA | ATGAGGGTCCACGACCTATG | 238 |
| chr1:59757827-59759848 | p2 | (TA)6 | AGGAGGACCCTTAAGCCAAA | ATGAGGGTCCACGACCTATG | 238 |
| chr1:59861548-59863568 | p2 | (TA)10 | ATGGCAATTCCTTCAAGGTG | GCCCCATAGGCTCAGTGTAA | 249 |
| chr1:59864091-59866113 | p2 | (TA)11 | ATGGCAATTCCTTCAAGGTG | GCCCCATAGGCTCAGTGTAA | 249 |
| chr1:59905704-59907722 | p3 | (ATA)6 | CGAGCCCAATCATACCAACT | GCTTAGGTCACCGACACGAT | 140 |
| chr1:59943253-59945271 | p3 | (ATA)6 | CGAGCCCAATCATACCAACT | GCTTAGGTCACCGACACGAT | 140 |
| chr1:59985746-59987762 | p2 | (AT)8 | TCACACAATTCATCGGCATT | TTGGGAGGAAAGGAAATGTG | 183 |
| chr1:60030258-60032273 | p5 | (CTTTT)3 | ATCGGATACTCGTCTGCTGG | GATGGGTTCGTAGCAGGTGT | 213 |
| chr1:60107298-60109310 | p2 | (AT)6 | TATGCAAGCTGACAAGGCTG | CCTGTCCCGCCATATGATAC | 237 |
| chr1:60126350-60128365 | p3 | (ACA)5 | AAGCTACCCCGAAATTCGAT | TCAGTGGATATAGCCCTCGG | 252 |
| chr1:60187044-60189056 | p2 | (AT)6 | CGGAAGTTGTCAAGAAAGGC | CCCCAATCCTTATGGAACCT | 290 |
| chr1:60217547-60219561 | p2 | (AC)6 | AAAACATCACCATTCTCGGC | AGTTGTGGCCGTGTGTTGTA | 225 |
| chr1:60217547-60219561 | p2 | (TA)34 | AAAACATCACCATTCTCGGC | AGTTGTGGCCGTGTGTTGTA | 225 |
| chr1:60217547-60219561 | p2 | (TA)7 | AAAACATCACCATTCTCGGC | AGTTGTGGCCGTGTGTTGTA | 225 |
| chr1:60313350-60315364 | p3 | (AAC)5 | AACTTGTCGAACCACCCATC | GATGTAGTGTTGTGGCCGTG | 188 |
| chr1:60313350-60315364 | p2 | (TA)7 | AACTTGTCGAACCACCCATC | GATGTAGTGTTGTGGCCGTG | 188 |
| chr1:60332649-60334685 | p2 | (TA)18 | CCTTCATGGTGAGGCAGAAT | GTATACGCTGCCCGCTTTAG | 254 |
| chr1:60372458-60374474 | p4 | (TATG)4 | ATTCTCTCTTTGTGGGTGCG | AACATGTTCCAGGGCAAGTC | 172 |
| chr1:60412079-60414125 | p2 | (TA)14 | CAGATTGTGCAGAGTTCGGA | CAAGGGCATACCCGACTAGA | 110 |
| chr1:60412079-60414125 | p2 | (TA)11 | CAGATTGTGCAGAGTTCGGA | CAAGGGCATACCCGACTAGA | 110 |
| chr1:60422268-60424289 | p2 | (AG)6 | CCAAGTGTAAGCCCTCCAAA | CATTCACATAACAGGCCACG | 130 |
| chr1:60422268-60424289 | p3 | (TTG)7 | CCAAGTGTAAGCCCTCCAAA | CATTCACATAACAGGCCACG | 130 |
| chr1:60422268-60424289 | p3 | (TTG)7 | CCAAGTGTAAGCCCTCCAAA | CATTCACATAACAGGCCACG | 130 |
| chr1:60422268-60424289 | p5 | (TATAT)3 | CCAAGTGTAAGCCCTCCAAA | CATTCACATAACAGGCCACG | 130 |
| chr1:60442531-60444547 | p4 | (ACAT)4 | CGGCAGCATTTCCTCTGTAT | GGATGCTTCCGACGTATGTT | 231 |
| chr1:60511298-60513360 | p2 | (TA)31 | CCAGGTGGTTGAAGGAAGAA | ATTCAATCCCCCAAAACTCC | 292 |
| chr1:60609275-60611290 | p5 | (TATAT)3 | TGGATTGGGTAAAATGGGAA | CGAACAATCCAACAACCCTT | 242 |
| chr1:60652722-60654742 | p2 | (AT)10 | GGACGCTCGAAAAATATGGA | CGCCAACAGTCTTTTTGTGA | 121 |
| chr1:60652722-60654742 | p5 | (TGAAT)3 | GGACGCTCGAAAAATATGGA | CGCCAACAGTCTTTTTGTGA | 121 |
| chr1:60796930-60798948 | p2 | (TA)9 | TGGCAATGTCAGCTTGTAGC | CTTGCTGCCCAAAAACACTT | 285 |
| chr1:60896898-60898913 | p5 | (TTATG)3 | TAGATTCTGGGTGAAACGGG | TACACCAAGGACATGCCAAA | 169 |
| chr1:60963283-60965298 | p5 | (AATAT)3 | CTCTAGTTTGCGAACCTCCG | GCCCGAGTATCAACCGAGTA | 298 |
| chr1:61041778-61043792 | p2 | (AT)7 | CGACAAGGCTATCACACGAA | CCCTTTTTGCTCAGAGTTGC | 251 |
| chr1:61080902-61082920 | p6 | (TATTGT)3 | TTTGCATGTGGGATTCGTTA | TTTCCTTGATTCACTTGCCC | 147 |

| Marker ID | SSR type | SSR motif | FORWARD PRIMER (5'-3') | REVERSE PRIMER (5'-3') | size (bp) |
|---|---|---|---|---|---|
| chr1:61136066-61138080 | p2 | (TA)7 | GGGGATGAAATGTGGAAATG | ACGACCCGTAAAATGTGTCC | 122 |
| chr1:61197992-61200006 | p2 | (AT)7 | CGGTCTTCGTTAGGACTTGC | CTGGCAAAGCTAAAAGTGGC | 275 |
| chr1:61247066-61249081 | p5 | (TGAAT)3 | AGCCTAGGGCCATGAAGAAT | ACCACACCACCTCTATTCGG | 293 |
| chr1:61254781-61256799 | p3 | (CAA)6 | GGACTCGGCGAACGATATAA | TCAGATTGACGCGAAACTTG | 231 |
| chr1:61262567-61264597 | p5 | (ATACA)6 | CCTCGCAACATCAAGAATCA | TTAGCTCCGAGTGCCTTCAT | 254 |
| chr1:61262567-61264597 | p3 | (CAT)5 | CCTCGCAACATCAAGAATCA | TTAGCTCCGAGTGCCTTCAT | 254 |
| chr1:61265251-61267269 | p2 | (AG)9 | TGATGAATGGTTTGTTGCGT | TTTTGGGTCAACTTTGGAGG | 134 |
| chr1:61298471-61300485 | p2 | (TA)7 | ACAGAAATTTGGGCAGCATC | ATTGGATTGGAAGTGCGAAG | 279 |
| chr1:61389164-61391182 | p2 | (AT)6 | GCCGTGACTAGTGTCCGAAT | TTTGGAGTTGCGTGTAGTGC | 189 |
| chr1:61389164-61391182 | p6 | (CATAAA)3 | GCCGTGACTAGTGTCCGAAT | TTTGGAGTTGCGTGTAGTGC | 189 |
| chr1:61450238-61452256 | p2 | (AT)7 | TGAGAACATACTCCGATGCG | CATACTGACGTCCCTTTGCC | 197 |
| chr1:61450238-61452256 | p6 | (TTATCA)3 | TGAGAACATACTCCGATGCG | CATACTGACGTCCCTTTGCC | 197 |
| chr1:61472153-61474169 | p2 | (TA)8 | TAACCTTGAGCCTTGGTTGG | GCACTTACTTGCCGTGCATA | 194 |
| chr1:61476942-61478954 | p2 | (TG)6 | AAAGGTGCCTACGGGCTATT | TCATACCTGGCCCTACCAAG | 222 |
| chr1:61509543-61511555 | p2 | (CA)6 | CATGCCAAGAGAAGAGAGGG | GCATTGTTGGGTTGTTGTTG | 254 |
| chr1:61531741-61533756 | p5 | (ACACA)3 | ACACGGCCAACTTAGACCAC | TCATGGAGTGGGTGTGTTGT | 199 |
| chr1:61546508-61548548 | p2 | (AT)20 | CAGAAGCGCAAAACACAAAA | GAAGGCAATGGCATGAATTT | 178 |
| chr1:61555930-61557942 | p5 | (AAGAA)3 | CAACCGAAAACCCAAAAGAA | TTTTTCAAGCCCAAACATCC | 295 |
| chr1:61555930-61557942 | p2 | (GT)6 | CAACCGAAAACCCAAAAGAA | TTTTTCAAGCCCAAACATCC | 295 |
| chr1:61593506-61595518 | p2 | (GT)6 | CCGACGTCCACAGGTAATCT | CTTCGTCCCCGAATGTTAAA | 193 |
| chr1:61610156-61612170 | p2 | (AT)7 | ATTTATGCAAGCGACAAGGG | TCCGCCAGCTTAGAACACTT | 294 |
| chr1:61610156-61612170 | p2 | (AT)7 | ATTTATGCAAGCGACAAGGG | TCCGCCAGCTTAGAACACTT | 294 |
| chr1:61623336-61625350 | p2 | (AT)7 | GTCAAGTTTTCCCGGATTGA | GAACGCGAGATAAGAGGTGC | 228 |
| chr1:61623336-61625350 | p6 | (CATATA)3 | GTCAAGTTTTCCCGGATTGA | GAACGCGAGATAAGAGGTGC | 228 |
| chr1:61630487-61632505 | p6 | (CATATA)3 | GCACCTCTTATCTCGCGTTC | ATCTTGTTCATCCCTGTCGC | 199 |
| chr1:61630487-61632505 | p3 | (CAA)5 | GCACCTCTTATCTCGCGTTC | ATCTTGTTCATCCCTGTCGC | 199 |
| chr1:61695652-61697670 | p3 | (TAA)6 | GTTGTTCATGACGTCGGATG | GTCCTGGGAGTTTTTGGTCA | 143 |
| chr1:61716043-61718061 | p6 | (CATATA)3 | ACATCCGTTTTCTTTGGACG | CATAAGCACCTCCGGACAAT | 117 |
| chr1:61839893-61841911 | p6 | (TATATG)3 | GTTGATGATTTGGCCGAGTT | TTAAGTCTCGGCTTTCCCAA | 156 |
| chr1:61839893-61841911 | p6 | (TATATG)3 | GTTGATGATTTGGCCGAGTT | TTAAGTCTCGGCTTTCCCAA | 156 |
| chr1:61848551-61850566 | p5 | (AAAAT)3 | ACACTTCCTGCCATACAGGG | GCTAGAAGATATGCTGCGGG | 154 |
| chr1:61888155-61890167 | p2 | (AT)6 | TTGAATCTGCGACAGAGGTG | ATTGTAATCGGCTCCACCAG | 283 |
| chr1:61892662-61894674 | p2 | (AT)6 | CCTTAGACCGACCTGCTCTG | GGCTCCACCAGTCGATATGT | 271 |
| chr1:61959955-61961967 | p2 | (GT)6 | TGTTGATGCGGAAATATGGA | GCCATGGTTGTTGTTCTCCT | 298 |
| chr1:62157981-62159996 | p6 | (ATATGT)3 | TATATGGGGAAAGGGGAAGG | TCACATTTACAGACGCCGAG | 292 |
| chr1:62157981-62159996 | p5 | (GAAAA)3 | TATATGGGGAAAGGGGAAGG | TCACATTTACAGACGCCGAG | 292 |

| Marker ID | SSR type | SSR motif | FORWARD PRIMER (5'-3') | REVERSE PRIMER (5'-3') | size (bp) |
|---|---|---|---|---|---|
| chr1:62175513-62177541 | p5 | (CAATG)3 | TCTACACCCCGAAAAGGATG | CGACATGGGAATGGTAAAGG | 116 |
| chr1:62175513-62177541 | p2 | (AT)14 | TCTACACCCCGAAAAGGATG | CGACATGGGAATGGTAAAGG | 116 |
| chr1:62234976-62236994 | p2 | (AT)9 | TCCCGAAGTAGTATGACGCC | GCCCAATTGCTGGCTAAATA | 261 |
| chr1:62368790-62370804 | p2 | (TG)7 | GAAATTTGGGGGCAAAAAGT | CACTCTTGGCCGTTTTTCTC | 114 |
| chr1:62405977-62408009 | p2 | (TA)10 | GCCATCTTTCTTTCGGACAA | TGTGTTGCAAGGCACTAAGC | 143 |
| chr1:62405977-62408009 | p3 | (ATA)7 | GCCATCTTTCTTTCGGACAA | TGTGTTGCAAGGCACTAAGC | 143 |
| chr1:62424388-62426408 | p5 | (TCATA)4 | TTTCAGATCGTGGTGCTCAG | ACACCTTCGCCCATACAGAG | 123 |
| chr1:62443594-62445610 | p2 | (AT)14 | CGAATGCGAAAATGATGATG | ACATAAGGTCGAGAACCCCC | 131 |
| chr1:62443594-62445610 | p2 | (AT)8 | CGAATGCGAAAATGATGATG | ACATAAGGTCGAGAACCCCC | 131 |
| chr1:62460012-62462058 | p2 | (AT)23 | CGAATGCGAAAATGATGATG | TGACAGAAGGTCGAGAACCC | 184 |
| chr1:62486949-62488961 | p2 | (TG)6 | TGATGTGTTACGGTTTCCGA | ACAAACACACCGTTATCGCA | 257 |
| chr1:62527458-62529504 | p2 | (TA)13 | GCCTCTGTACTACGGGTGGA | GTGCCTCATCAGGGCTAAAA | 256 |
| chr1:62527458-62529504 | p2 | (TA)23 | GCCTCTGTACTACGGGTGGA | GTGCCTCATCAGGGCTAAAA | 256 |
| chr1:62556121-62558139 | p2 | (AT)9 | TGTGAAATCGTGGGTTGAAA | GCCTAACCCGAATTTCCAAT | 180 |
| chr1:62562709-62564721 | p2 | (AT)6 | ACGAGTTAACCCTACCGGCT | ATCTGCCGATCAAGGTATGG | 205 |
| chr1:62564558-62566573 | p5 | (ATATA)3 | TGTGAAAATCGTGGGTTGAA | CCTCCTCCTGTCACACGAAT | 191 |
| chr1:62581618-62583632 | p2 | (AT)7 | TGTACCAGCATCATAGGCCA | TGGGTCGTGACACATTCACT | 198 |
| chr1:62581618-62583632 | p2 | (CA)7 | TGTACCAGCATCATAGGCCA | TGGGTCGTGACACATTCACT | 198 |
| chr1:62581618-62583632 | p3 | (ATC)5 | TGTACCAGCATCATAGGCCA | TGGGTCGTGACACATTCACT | 198 |
| chr1:62645650-62647690 | p2 | (TA)20 | GAGCCCCTAGGATGGCTAAC | ACATATCAAGGGCTCGATGG | 196 |
| chr1:62645650-62647690 | p2 | (TA)16 | GAGCCCCTAGGATGGCTAAC | ACATATCAAGGGCTCGATGG | 196 |
| chr1:62679128-62681248 | p2 | (TC)7 | TACATACCTTGCCCGCTTTC | CTCGGGGATGGTAGCATTTA | 251 |
| chr1:62884191-62886206 | p5 | (TGAAT)3 | TCGTTCGCTTGTTCTCCTTT | TGTGAATATTTTTGCGAGCG | 297 |
| chr1:62905882-62907897 | p5 | (TGAAT)3 | TCGTTCGCTTGTTCTCCTTT | TGTGAATATTTTTGCGAGCG | 297 |
| chr1:62922321-62924337 | p4 | (TGAT)4 | TAGACAGGGAGGTGGTCAGG | TGAATGGAAGCTGCACAGAC | 165 |
| chr1:62933772-62935787 | p5 | (GAAAT)3 | GGTGGCGGAGGTGTTATAGA | CCCTATTTCGATTCTTGCCA | 146 |
| chr1:62947127-62949141 | p2 | (TA)7 | TTGACCGTGAGTATGGGACA | AACATTTAGACGGCGTTTCG | 187 |
| chr1:63006633-63008648 | p5 | (ATGCA)3 | ACCTTAGTTGGCGAACCCTT | GACCAGATGCCTTGGGTAGA | 129 |
| chr1:63021205-63023219 | p2 | (AC)7 | GCGCGAGGAGTCATAATAGC | TGCTGGTTTAAGGGGAGTTG | 216 |
| chr1:63035252-63037267 | p5 | (TGAAT)3 | TCGTTCGCTTGTTCTCCTTT | GTGCTGCCCAATTTCCTCTA | 297 |
| chr1:63247829-63249841 | p2 | (AC)6 | ATTATCCCCGAGGCTCAAGT | GATGATTCTTGGGGGTGCTA | 110 |
| chr1:63441876-63443896 | p2 | (AT)14 | ACACAACATGCGGAAGAACA | GCAGTGGTCAGTATGCTCCA | 106 |
| chr1:63450000-63452016 | p2 | (AT)8 | GTCGTAGATCTTGCGAAGCC | TCACAAGGATCATGCCAAAA | 110 |
| chr1:63461451-63463471 | p5 | (CATAT)4 | ACATACCTTTCGGATGTCGC | GACATACGTGTGGGTTGTGC | 260 |
| chr1:63618394-63620412 | p5 | (TTAAA)3 | CCGATTGTCACATTTTCCCT | GGATGTGTTTTCTGGGCTGT | 200 |
| chr1:63618394-63620412 | p6 | (ACCAAT)3 | CCGATTGTCACATTTTCCCT | GGATGTGTTTTCTGGGCTGT | 200 |

| Marker ID | SSR type | SSR motif | FORWARD PRIMER (5'-3') | REVERSE PRIMER (5'-3') | size (bp) |
|---|---|---|---|---|---|
| chr1:63618394-63620412 | p2 | (TA)20 | CCGATTGTCACATTTTCCCT | GGATGTGTTTTCTGGGCTGT | 200 |
| chr1:63678159-63680171 | p2 | (AT)6 | AAAAATTGTGGCAACTTCGC | ACTGCACAGCCTCCTCCTTA | 118 |
| chr1:63678159-63680171 | p5 | (ATCAG)5 | AAAAATTGTGGCAACTTCGC | ACTGCACAGCCTCCTCCTTA | 118 |
| chr1:63706167-63708217 | p5 | (TGTTT)3 | ACATGGATGATAGGCCCAAA | GAACGTGCAAAATACGGGAT | 133 |
| chr1:63719064-63721079 | p3 | (GAT)5 | AAACACGAATGGGACGAAAG | TTCCCGTTTCCAAATGCTAC | 166 |
| chr1:63747982-63750008 | p2 | (TG)6 | CCGCTGTATATTGGGCCTTA | CCAAAATTGCTATGGGCTGT | 271 |
| chr1:63804635-63806647 | p2 | (AG)6 | TCCGTCTGACTCTGAACACG | GATTCGTAACTGGGATCGGA | 174 |
| chr1:63852665-63854680 | p2 | (TG)6 | AAGGGTGTGATGAGATTCGG | TCCAGGTCGGATGGTAAGTC | 189 |
| chr1:63852665-63854680 | p3 | (GTT)5 | AAGGGTGTGATGAGATTCGG | TCCAGGTCGGATGGTAAGTC | 189 |
| chr1:63900277-63902289 | p2 | (AT)6 | TCCCCTAGAGTCACGGCTAA | GGCCCTGTCCCTTCATATTT | 254 |
| chr1:63932482-63934502 | p5 | (AAAAT)4 | ACGTATCGGGAATCATGAGC | CATTGAAGGAGAACGTGCAA | 171 |
| chr1:64027659-64029675 | p2 | (AT)8 | AAGAGGGTGCTTCATTGGTG | AGTGTGTCGGGCAACTTAGG | 173 |
| chr1:64088738-64090754 | p2 | (TC)8 | TGGTCTCAAAAATCTTGGGC | TGTTGGTTAGAATTTCGGGC | 258 |
| chr1:64184921-64186939 | p3 | (CAA)6 | CGTACCTGGCCATATCGACT | GTTGGGTACGGATGTTACCG | 146 |
| chr1:64308138-64310156 | p6 | (TGTAAG)3 | TTGGTAGGGCCAGGTAGATG | AGCATACCCGACCATACAGG | 161 |
| chr1:64308138-64310156 | p2 | (TA)6 | TTGGTAGGGCCAGGTAGATG | AGCATACCCGACCATACAGG | 161 |
| chr1:64394940-64396960 | p2 | (TA)19 | AGGGTAATTGGAAATTCGGG | TTAGCGACTATTCGCAACCC | 161 |
| chr1:64455952-64457968 | p4 | (TTCT)4 | TTTGGAAGTGTTCCGACTCC | AGGCGGTATCGAGGTATGTG | 130 |
| chr1:64519739-64521777 | p2 | (AT)20 | CCGAACTACAGCTACCCTGC | ATGTGCGAGCATGAAGTCAG | 130 |
| chr1:64533295-64535315 | p4 | (AGAA)5 | CCGTCTTGTGTGCCTCTGTA | ACCAACAGGTTCAGGACTGG | 117 |
| chr1:64549594-64551610 | p4 | (AATT)4 | TATATGCGCCTAACCCGAAC | TCGATTTGGATTGGAAAAGG | 160 |
| chr1:64591192-64593210 | p6 | (TATATG)3 | TGGCCAAGTTAAATTCTCGG | TACCTTGCCCGTTTTCTACG | 126 |
| chr1:64703266-64705281 | p5 | (CATAT)3 | GCAAGGAATCATAGTCGGGA | TAAGGGGTGCTCGGTACAAG | 136 |
| chr1:64732081-64734093 | p2 | (AT)8 | GGTTGACATTCCGAGCTCAT | AATAGCCCGTAGGCACCTTT | 102 |
| chr1:64732081-64734093 | p3 | (ATG)5 | GGTTGACATTCCGAGCTCAT | AATAGCCCGTAGGCACCTTT | 102 |
| chr1:64732081-64734093 | p2 | (TG)6 | GGTTGACATTCCGAGCTCAT | AATAGCCCGTAGGCACCTTT | 102 |
| chr1:64741235-64743249 | p2 | (AT)7 | TCGAGGTCTTACGGATTTCG | CGAGAATGGTGAGTGGGAAT | 255 |
| chr1:64783304-64785316 | p2 | (GA)6 | AGGGAGGCTACAAAACAGCA | GTGTCACGCTCAGGCTATGA | 238 |
| chr1:64786269-64788281 | p2 | (AT)6 | TCCTTTAACGATCCGTCCTG | GATCGTCCAATAGAGCCCAA | 251 |
| chr1:64786269-64788281 | p2 | (AT)7 | TCCTTTAACGATCCGTCCTG | GATCGTCCAATAGAGCCCAA | 251 |
| chr1:64831559-64833575 | p2 | (TA)8 | TTTCGATTCCAATTTCGTCC | AGCTAAGTCCGGGGATGTTT | 186 |
| chr1:64878003-64880030 | p3 | (ATC)9 | GCCAGTGTAGACCACCGATT | GTGTACTCTGCCTGGCCATT | 105 |
| chr1:64918893-64920908 | p3 | (GTA)5 | CGACTGGACTAGTTGGAGCC | TGGTTTCAATTGGGGGAATA | 121 |
| chr1:64921021-64923036 | p5 | (ATGCA)3 | ATCCTTCTGCATTGGACACC | TATTGTTCCCCATTTCCCAA | 215 |
| chr1:65018397-65020412 | p5 | (ATCAC)3 | AGCCAAAAGCTCACCTTGAA | TGTGGGTGGAAATCCTTGAT | 264 |
| chr1:65072635-65074657 | p2 | (AT)11 | TGCCTTGTAGCCTCCTGTTT | CATCACGACTCATCGGATTG | 180 |

| Marker ID | SSR type | SSR motif | FORWARD PRIMER (5'-3') | REVERSE PRIMER (5'-3') | size (bp) |
|---|---|---|---|---|---|
| chr1:65079929-65081941 | p5 | (TGAAT)4 | TGGTAATTGTCACGACCCAA | GGCCTTGTCGGCCTATATTT | 268 |
| chr1:65079947-65081969 | p5 | (TGAAT)4 | TGGTAATTGTCACGACCCAA | GGCCTTGTCGGCCTATATTT | 250 |
| chr1:65085209-65087224 | p2 | (AT)7 | TTCTTGAATTCTTGTGGGGG | TTGGTCCCTAGAACTGGTCG | 210 |
| chr1:65085209-65087224 | p2 | (AT)8 | TTCTTGAATTCTTGTGGGGG | TTGGTCCCTAGAACTGGTCG | 210 |
| chr1:65085209-65087224 | p5 | (TGTGT)3 | TTCTTGAATTCTTGTGGGGG | TTGGTCCCTAGAACTGGTCG | 210 |
| chr1:65119971-65121991 | p5 | (TGAAT)4 | CAAAACAAAAACATGTGGCG | CATCTTTGATCCTCCGCAGT | 163 |
| chr1:65119971-65121991 | p6 | (TATATG)3 | CAAAACAAAAACATGTGGCG | CATCTTTGATCCTCCGCAGT | 163 |
| chr1:65184043-65186058 | p2 | (AT)27 | TTTGGTGGGATTGCTAGGAG | GGCCCATGTGTCCATATTTC | 120 |
| chr1:65184043-65186058 | p3 | (TCA)5 | TTTGGTGGGATTGCTAGGAG | GGCCCATGTGTCCATATTTC | 120 |
| chr1:65193337-65195361 | p2 | (TA)12 | TGAGGGAAAGAGATCCATGC | GTTGGCCTGATTTGGAAGAA | 274 |
| chr1:65285011-65287029 | p2 | (AT)9 | CGTCTCGGCCAAGTCTTTAC | TCTTCAGGCTCGGACGTTAT | 151 |
| chr1:65285011-65287029 | p2 | (AT)8 | CGTCTCGGCCAAGTCTTTAC | TCTTCAGGCTCGGACGTTAT | 151 |
| chr1:65285011-65287029 | p2 | (AT)9 | CGTCTCGGCCAAGTCTTTAC | TCTTCAGGCTCGGACGTTAT | 151 |
| chr1:65407391-65409423 | p2 | (TA)8 | ATCCATGCAAGGAGGAACAG | TAATTTTGGGGGTTGTGCAT | 156 |
| chr1:65416141-65418155 | p2 | (CA)7 | AGAGTGTGCGGTGGAGACTT | ATGCCATGCTATGCAAATGA | 146 |
| chr1:65423369-65425384 | p5 | (CATAT)3 | ACTTGGTTTCATAGGCGACG | TGAATACGAGCACGAAGTGG | 266 |
| chr1:65432555-65434579 | p2 | (AT)6 | AAATGCCATGTAGCCTCCTG | CCCTAGTCGGGTCGTTACAA | 294 |
| chr1:65432583-65434601 | p2 | (AT)6 | AAATGCCATGTAGCCTCCTG | CCCTAGTCGGGTCGTTACAA | 266 |
| chr1:65503439-65505513 | p2 | (TA)7 | ACGATGGAGCTAGTCAACGG | ACAGGCTCGGTTCTTGCTAA | 105 |
| chr1:65519728-65521748 | p2 | (TA)10 | GGCAAGCTGAACGTACCATT | ACGGTGTAATCATAAGCCGC | 163 |
| chr1:65524924-65526960 | p2 | (AT)18 | TTTAGCCCATCTTGACCCAG | AATTTCCGATCAAGGACACG | 120 |
| chr1:65530743-65532758 | p5 | (AAGCG)3 | TTGGGGGTTTTAATCATGGA | AAACCAAGCTCCTTCGTCAA | 129 |
| chr1:65622629-65624641 | p2 | (CT)6 | CCCGATTTTTCCCATCCTAT | GGATCAAGCCTGTCTCCTTG | 123 |
| chr1:65703715-65705737 | p2 | (TA)11 | TTTCGCAAATAATTCTCGGG | ACGACGATGTTGGGAAAGAG | 179 |
| chr1:65703715-65705737 | p2 | (TA)9 | TTTCGCAAATAATTCTCGGG | ACGACGATGTTGGGAAAGAG | 179 |
| chr1:65703896-65705914 | p2 | (TA)11 | CTCTTTCCCAACATCGTCGT | CGAACCCTGATAGCCATTGT | 116 |
| chr1:65703896-65705914 | p2 | (TA)9 | CTCTTTCCCAACATCGTCGT | CGAACCCTGATAGCCATTGT | 116 |
| chr1:65755941-65757956 | p5 | (GTCCT)3 | GTCCCTTCTTTTCCAAAGGC | AGGCGAGGGTATCAACAATG | 149 |
| chr1:65895811-65897823 | p2 | (AT)6 | TTCCATGAGCCATTTTGTGA | TTCGTACGTTCGGAAAATCC | 287 |
| chr1:65895861-65897873 | p2 | (AT)6 | TTCCATGAGCCATTTTGTGA | TTCGTACGTTCGGAAAATCC | 237 |
| chr1:65954465-65956479 | p2 | (TA)7 | GGAGGTAGAGTCGCTGATGC | GGACGAATGGAAGGGAATTT | 116 |
| chr1:65961906-65963918 | p2 | (AT)6 | CCACCTGAGCACCCTCTTAG | GTTGTCCAATTGGGACTCGT | 151 |
| chr1:65968845-65970857 | p2 | (TG)6 | GTGAGGCGTGGTTGGTATTT | TCCAAATAGTCGGCGAAATC | 258 |
| chr1:65975317-65977341 | p2 | (AT)12 | GTTGACTTCCCTTTTTGGCA | GAGCATGGCAAGAACTGTGA | 152 |
| chr1:66067385-66069397 | p2 | (AT)6 | GGACGCGAACAATTCAAAAT | CTAACAGCCACACTGACGGA | 203 |
| chr1:66088769-66090825 | p2 | (AT)28 | CCTTCTGTCACCCCGATAGA | CTGGGGTACCATCTCATGCT | 289 |

| Marker ID | SSR type | SSR motif | FORWARD PRIMER (5'-3') | REVERSE PRIMER (5'-3') | size (bp) |
|---|---|---|---|---|---|
| chr1:66100185-66102200 | p5 | (ATTTC)3 | CCGACTATGGTGGCTGAGAT | GGGAAATATCCATCCCGTCT | 265 |
| chr1:66102954-66104969 | p5 | (AAATG)3 | CCTCACATACCTTGCCGTTT | TGCCCAAATTTTCCAAGAAC | 292 |
| chr1:66189194-66191209 | p3 | (ACA)5 | CTCACAGCCAAACAGCACAT | GGGGCTGTTGTAAGAATGGA | 281 |
| chr1:66276641-66278665 | p4 | (TTAT)6 | CTCATTGTGTTCGCAAAGGA | GTCAAAACCAAGTCTGGGGA | 161 |
| chr1:66406225-66408237 | p2 | (AT)6 | GGTTTCGGAGGCATGAATAA | TAACTCCAATCTCCCGGTTG | 217 |
| chr1:66580171-66582189 | p2 | (AT)9 | GCGTGCAACACACCATAAAC | ATTATGCAAATGCCAGAGGG | 163 |
| chr1:66609256-66611268 | p2 | (TA)6 | GTCGTTCCGGTAGTGCGTAT | GGGAACTTTTGATCCCGATT | 102 |
| chr1:66609256-66611268 | p2 | (AG)6 | GTCGTTCCGGTAGTGCGTAT | GGGAACTTTTGATCCCGATT | 102 |
| chr1:66705423-66707438 | p3 | (ATC)5 | AGCCCACAAGGCTACAAATG | GGCCTGTTCCATCCTTATGA | 105 |
| chr1:66734416-66736428 | p2 | (AT)6 | GTTGCGTTGCTCTTTCTTCC | GACCCCCTGGGAAATGTACT | 102 |
| chr1:66826629-66828641 | p2 | (CA)6 | GGTCATGCCAAAAAGGAAGA | TGAATTTACAGGGGAGGTGC | 229 |
| chr1:66826629-66828641 | p2 | (AT)8 | GGTCATGCCAAAAAGGAAGA | TGAATTTACAGGGGAGGTGC | 229 |
| chr1:66937184-66939206 | p2 | (AT)34 | TCACATTTACGGTTGCAGGA | TTGTGTGTAGGGTTCGTGCT | 113 |
| chr1:66937184-66939206 | p2 | (TA)11 | TCACATTTACGGTTGCAGGA | TTGTGTGTAGGGTTCGTGCT | 113 |
| chr1:67002739-67004751 | p2 | (AC)6 | TTGGAGTGCTCCCTTGTTCT | GGCCGTCATAGCAAATAGGA | 281 |
| chr1:67044826-67046841 | p2 | (AT)18 | ACCGTACCTGGCCCTCTAGT | TCGTGCATGCCTCAATACAT | 222 |
| chr1:67044826-67046841 | p2 | (AT)7 | ACCGTACCTGGCCCTCTAGT | TCGTGCATGCCTCAATACAT | 222 |
| chr1:67044826-67046841 | p3 | (ACA)5 | ACCGTACCTGGCCCTCTAGT | TCGTGCATGCCTCAATACAT | 222 |
| chr1:67044826-67046841 | p2 | (CT)7 | ACCGTACCTGGCCCTCTAGT | TCGTGCATGCCTCAATACAT | 222 |
| chr1:67059766-67061820 | p5 | (TATCA)3 | GGCGCCCAATTACAAGTAGA | CCCAGCCATAGGAAATGAAA | 101 |
| chr1:67059766-67061820 | p2 | (AT)7 | GGCGCCCAATTACAAGTAGA | CCCAGCCATAGGAAATGAAA | 101 |
| chr1:67059766-67061820 | p3 | (ACA)5 | GGCGCCCAATTACAAGTAGA | CCCAGCCATAGGAAATGAAA | 101 |
| chr1:67086977-67088992 | p3 | (TAT)5 | ATACTCCGACGCGGTATCAC | TTTGCAGGAATGTTGTCCAG | 277 |
| chr1:67094953-67097007 | p5 | (TATCA)3 | GGCGCCCAATTACAAGTAGA | CCCAGCCATAGGAAATGAAA | 101 |
| chr1:67094953-67097007 | p2 | (AT)7 | GGCGCCCAATTACAAGTAGA | CCCAGCCATAGGAAATGAAA | 101 |
| chr1:67094953-67097007 | p3 | (ACA)5 | GGCGCCCAATTACAAGTAGA | CCCAGCCATAGGAAATGAAA | 101 |
| chr1:67122164-67124179 | p3 | (TAT)5 | ATACTCCGACGCGGTATCAC | TTTGCAGGAATGTTGTCCAG | 277 |
| chr1:67445681-67447701 | p2 | (AT)10 | AGGGGGTCAGTACGAAAGGT | GTATGCTGTGTTGTGGCACC | 251 |
| chr1:67488382-67490406 | p6 | (TATATG)4 | TTGATGTTGTTGGCACGATT | ATTTAGGCAGCATCTCCCCT | 114 |
| chr1:67498212-67500226 | p2 | (AT)7 | ACTGTGCATGCCTGTCTACG | ATCGAGAACGCTCCAGTTGT | 248 |
| chr1:67530580-67532658 | p2 | (AT)39 | ATCTCGAAACAGCCTGCACT | ATGGGTTGTTTGGAGCATGT | 158 |
| chr1:67530580-67532658 | p2 | (AT)6 | ATCTCGAAACAGCCTGCACT | ATGGGTTGTTTGGAGCATGT | 158 |
| chr1:67530850-67532862 | p2 | (AT)39 | TCCAATTCATGTCCCAACAA | AGAATTGGATGCAAGGAACG | 231 |
| chr1:67530850-67532862 | p2 | (AT)6 | TCCAATTCATGTCCCAACAA | AGAATTGGATGCAAGGAACG | 231 |
| chr1:67572210-67574242 | p2 | (TA)21 | CCGGGTAAGGTAAGCGTGTA | CGTAACCGGCCACTCATTAT | 174 |
| chr1:67572210-67574242 | p2 | (AT)16 | CCGGGTAAGGTAAGCGTGTA | CGTAACCGGCCACTCATTAT | 174 |

| Marker ID | SSR type | SSR motif | FORWARD PRIMER (5'-3') | REVERSE PRIMER (5'-3') | size (bp) |
|---|---|---|---|---|---|
| chr1:67582637-67584649 | p2 | (AT)13 | ATACGAGTGTCCAGTTCGGG | GGAGGCATCGCTGTAAACAT | 150 |
| chr1:67582637-67584649 | p2 | (CA)6 | ATACGAGTGTCCAGTTCGGG | GGAGGCATCGCTGTAAACAT | 150 |
| chr1:67692537-67694559 | p2 | (AT)11 | AGTGGCGGTGTAGCAGAACT | CTATGCCTCCAAAGAAAGCG | 300 |
| chr1:67692537-67694559 | p2 | (AT)11 | AGTGGCGGTGTAGCAGAACT | CTATGCCTCCAAAGAAAGCG | 300 |
| chr1:67773795-67775807 | p2 | (AT)6 | ATGTTGGATATGAGCCTCGG | GTCGGGTCCTATTTATGCGA | 148 |
| chr1:67773795-67775807 | p6 | (TATATG)4 | ATGTTGGATATGAGCCTCGG | GTCGGGTCCTATTTATGCGA | 148 |
| chr1:67813886-67815910 | p2 | (AT)12 | TAATCCCTGCATTACCCGAA | CCTAGCTGGGTCGTGACAAT | 140 |
| chr1:67924064-67926078 | p2 | (GT)7 | ACCACACCAACCATTACCGT | CAACCTTGACCTTCGCTCAT | 242 |
| chr1:67937906-67939950 | p2 | (AT)22 | CGTAGAGTCTGCCAAGGGTC | GCGAGCTCTTCTGCTGAAAT | 162 |
| chr1:68005184-68007226 | p3 | (AAT)14 | AACCCTTTCAACGAACATCG | TAGGGTTCCCTTCCCATTTC | 297 |
| chr1:68005184-68007226 | p2 | (GC)6 | AACCCTTTCAACGAACATCG | TAGGGTTCCCTTCCCATTTC | 297 |
| chr1:68020254-68022275 | p3 | (AAT)17 | GCTGATGAAGGAAATGGGAA | AATTCAACCAGCCCAACATC | 124 |
| chr1:68020254-68022275 | p3 | (CAT)7 | GCTGATGAAGGAAATGGGAA | AATTCAACCAGCCCAACATC | 124 |
| chr1:68160400-68162418 | p6 | (CATATA)3 | TATGCCCGCATTTACCTCTC | CGTTGGCGTGTATTTTGTTG | 267 |
| chr1:68191874-68193902 | p2 | (TA)14 | CCTGCCTCAGTCCAGAAGAG | TGTCCCAAACAATGGGATTT | 141 |
| chr1:68191874-68193902 | p5 | (TCATA)3 | CCTGCCTCAGTCCAGAAGAG | TGTCCCAAACAATGGGATTT | 141 |
| chr1:68215832-68217844 | p2 | (AT)6 | CGAGCCAGATATGCACAGAA | CATTTCCCCGAGCTTGTTTA | 129 |
| chr1:68286706-68288726 | p2 | (AT)10 | CTAGCTGCCCTATCACTCGG | CGAATGTAGAAGAGGGCGAG | 216 |
| chr1:68317498-68319518 | p2 | (TA)6 | GGAACCCTAGACTCCCCAAG | CTCCGATTCATCCCTCAAAA | 118 |
| chr1:68317498-68319518 | p2 | (AC)10 | GGAACCCTAGACTCCCCAAG | CTCCGATTCATCCCTCAAAA | 118 |
| chr1:68345419-68347455 | p5 | (GAAAT)3 | AAAAAGGGGTTGAAAGGGAA | GAACAAGTCGAGGCGAGAAC | 265 |
| chr1:68345559-68347577 | p5 | (GAAAT)3 | AAAAAGGGGTTGAAAGGGAA | GAACAAGTCGAGGCGAGAAC | 125 |
| chr1:68345559-68347577 | p3 | (TAA)6 | AAAAAGGGGTTGAAAGGGAA | GAACAAGTCGAGGCGAGAAC | 125 |
| chr1:68345686-68347701 | p5 | (GAAAT)3 | GTTCTCGCCTCGACTTGTTC | AGGAGGCAAGGCATACGATA | 256 |
| chr1:68345686-68347701 | p3 | (TAA)6 | GTTCTCGCCTCGACTTGTTC | AGGAGGCAAGGCATACGATA | 256 |
| chr1:68417692-68419707 | p5 | (ACATA)3 | CTAGCCAAGGAATTGAAGCG | GCTCCGGTTGATACATGCTT | 199 |
| chr1:68456315-68458330 | p5 | (ATCAT)3 | TTTCGGTGTCCCTTTACTCG | GCTCGGGAATTAGGATAGGC | 143 |
| chr1:68474428-68476442 | p2 | (AC)7 | CTGTTCTGTGGATCGGGTTT | GCCGAAATTTTTGTAGGCAA | 167 |
| chr1:68492012-68494042 | p2 | (TA)15 | GATTAGGTCTTCGGGCATGA | TAACAGGGCAACAAACCACA | 288 |
| chr1:68530295-68532310 | p3 | (CAA)5 | TGGGAACTTCTTCCCCTTTT | ACGAACCCAGCAAATAAACG | 117 |
| chr1:68555061-68557076 | p3 | (CAG)5 | GATAGGACCCTGAAGGAGGC | AGTCTCTGAAGCCCTGACCA | 254 |
| chr1:68571797-68573809 | p2 | (AG)6 | AGCGATAATTGGAGGGGTCT | TCTCACATCCGATTCCCTTC | 111 |
| chr1:68594127-68596139 | p2 | (AG)6 | TGCAAAAGCAGAAGAAGGGT | ATAAGCATCGGGTCAAGACG | 141 |
| chr1:68676775-68678787 | p2 | (AC)6 | TTTCGAGGCTCAAATCCAAT | TAGATTTTGGACGAAACGGG | 229 |
| chr1:68682952-68684968 | p2 | (AC)8 | CGGATTTCGAGAAGCAAAAC | CTGAGCGAAACGGGACTTAG | 197 |
| chr1:68762138-68764150 | p2 | (TA)6 | CTCCCTCACTTTCCCAAACA | GAGGTGCTATTCGTGGTGGT | 100 |

| Marker ID | SSR type | SSR motif | FORWARD PRIMER (5'-3') | REVERSE PRIMER (5'-3') | size (bp) |
|---|---|---|---|---|---|
| chr1:68775731-68777747 | p2 | (TA)8 | GATGGGCTACAGGGCATTTA | TTTTCCCTTGTACGGCATTC | 300 |
| chr1:68778712-68780724 | p2 | (AT)6 | CAAGAGCATCCGAGACAACA | TGTTGAAACACCGACCTTCA | 286 |
| chr1:68872104-68874174 | p2 | (AT)35 | GCCTAACAGAAGGAAGCGTG | CCTCTGGGCTGCTAATTGTC | 201 |
| chr1:68872104-68874174 | p2 | (AT)18 | GCCTAACAGAAGGAAGCGTG | CCTCTGGGCTGCTAATTGTC | 201 |
| chr1:68896960-68898978 | p3 | (CAA)6 | GGCCGTAGCACAGTGGTAAT | GGTCTACGACCAGTCGGGTA | 183 |
| chr1:68917546-68919564 | p2 | (AT)9 | CGCAGCATCTGTCGAATAAA | GGCTCGTAGACGCAGGTTAG | 104 |
| chr1:68996908-68998932 | p6 | (CAGCCA)4 | GAGCAACCCTGTCCACTGAT | TGCGTCATTAGTTGAGGCTG | 194 |
| chr1:69018481-69020496 | p5 | (TTTCA)3 | GCCTCGGCAATATGAGATGT | CCACAAGAAGGGTCGTCAGT | 108 |
| chr1:69045131-69047143 | p6 | (CATATA)3 | GTTGATGATTTGGCCGAGTT | TCACAAGGATCATGCCAAAA | 255 |
| chr1:69045229-69047247 | p6 | (CATATA)3 | GTTGATGATTTGGCCGAGTT | TTCACAAGGATCATGCCAAA | 157 |
| chr1:69060137-69062149 | p2 | (AT)6 | ATGCAACAAGGCAATGACAA | TTGTAGTGTGGTGGCCGATA | 141 |
| chr1:69065281-69067293 | p2 | (AT)6 | CACCTTACCCGAAATCTCCA | TGAAATCGTTGTGGTGTCGT | 109 |
| chr1:69155179-69157194 | p5 | (TCGGT)3 | TACACTTGGCCTTATTCCGC | CCATGGGTTCCGTGATATTC | 260 |
| chr1:69182745-69184757 | p2 | (CA)6 | CGCTCTCGCCAAAACTAAAC | CAATCAGTCTGTCGACGCAT | 154 |
| chr1:69194014-69196026 | p2 | (GT)6 | AGAAATGACATGGGGAGCAG | ATGGTGGGGAGTCTCTCTT | 108 |
| chr1:69203574-69205592 | p6 | (TATATG)3 | TTTACAAGGGAAATGCTGCC | GCTACGCTTATTCGTCCGAG | 184 |
| chr1:69229691-69231706 | p5 | (AAACC)3 | GACTACCATTTTGAGGGGCA | TTGGTTTGGTTTTGGTGGTT | 122 |
| chr1:69283947-69285962 | p3 | (TGT)5 | GAAGGTGCCACCGAAACTTA | AGGCTACCCCAACCTTGTCT | 276 |
| chr1:69319540-69321558 | p6 | (CCTATT)3 | AAACAAGGGCATCAACTTGG | TTGTGGTTGATGAAAAGGCA | 202 |
| chr1:69372494-69374512 | p3 | (GTT)6 | ACCCTACTACAACATGGCCG | TGGCTGTGTGAAATTCTTGC | 231 |
| chr1:69373985-69376019 | p2 | (TA)17 | ATGATTATGGCACCGAGTCC | TACCTGCGGGTATGAAAAGC | 167 |
| chr1:69373985-69376019 | p4 | (AAAG)6 | ATGATTATGGCACCGAGTCC | TACCTGCGGGTATGAAAAGC | 167 |
| chr1:69373985-69376019 | p3 | (CAA)6 | ATGATTATGGCACCGAGTCC | TACCTGCGGGTATGAAAAGC | 167 |
| chr1:69393459-69395479 | p2 | (AT)10 | ACCAAATCCTCATTCGTTCG | GCCGCCTAAAGATCTTCCTT | 276 |
| chr1:69445092-69447116 | p2 | (AG)12 | GTCTTGCTCTTGACCTTCCG | GCCTTTGGAATCACCTCAAA | 256 |
| chr1:69558412-69560430 | p5 | (TTTTG)3 | CATAAAACTCTGGCGGCTTC | GCAGGAAAGTGTTCCAGCTC | 263 |
| chr1:69558412-69560430 | p6 | (TTAACA)3 | CATAAAACTCTGGCGGCTTC | GCAGGAAAGTGTTCCAGCTC | 263 |
| chr1:69643228-69645246 | p3 | (ATC)6 | TTGACCAAGCACCCCTATTC | TAGCCTTGTCGGCTCTGATT | 182 |
| chr1:69700634-69702652 | p4 | (CATA)4 | TTACCCCTAGCGGATCAGTG | TCGTTCCTTCGAGGTGAGAT | 291 |
| chr1:69700634-69702652 | p3 | (CAA)6 | TTACCCCTAGCGGATCAGTG | TCGTTCCTTCGAGGTGAGAT | 291 |
| chr1:69838267-69840282 | p5 | (ATATG)3 | AATAGGGGTGAATTGGGAGC | TCAAGTCCCAAACGTCAACA | 130 |
| chr1:69838267-69840282 | p2 | (CT)7 | AATAGGGGTGAATTGGGAGC | TCAAGTCCCAAACGTCAACA | 130 |
| chr1:69857988-69860003 | p3 | (GAT)5 | TTCCTCCAAAGTACGGGATG | GAGGCCCAAGTTATGGGAAT | 130 |
| chr1:69874847-69876862 | p3 | (TGA)5 | TTTTTGCCTCCAAAGTACGG | ATCGGTGTTAAGGGGAGCTT | 117 |
| chr1:69887849-69889861 | p2 | (AT)6 | TCTGGGATCGTTGTTGCATA | ACATCTCGGTAGCACAACCC | 259 |
| chr1:69989804-69991824 | p5 | (TCATT)4 | GAAACAACAACCGGGAAGAA | ATTTTGCCAGAGTTTCCCCT | 173 |

| Marker ID | SSR type | SSR motif | FORWARD PRIMER (5'-3') | REVERSE PRIMER (5'-3') | size (bp) |
|---|---|---|---|---|---|
| chr1:69989804-69991824 | p3 | (ATG)5 | GAAACAACAACCGGGAAGAA | ATTTTGCCAGAGTTTCCCCT | 173 |
| chr1:70304234-70306252 | p2 | (AT)9 | GGATGGCAATGTTATACCCG | TGTGATTGATTGAGGACCCA | 270 |
| chr1:70326145-70328175 | p2 | (AT)6 | ACTTTTGTCATGACCCAGCC | TGTTCTGTCGTGGTAGCAGC | 295 |
| chr1:70326145-70328175 | p2 | (AT)15 | ACTTTTGTCATGACCCAGCC | TGTTCTGTCGTGGTAGCAGC | 295 |
| chr1:70394726-70396738 | p2 | (AT)6 | TGGTATGGCTGAAAGAGCCT | TGTTTGCTGTGCATCTAGCC | 173 |
| chr1:70433469-70435505 | p2 | (AT)18 | AGCGTCCACAAACAAAAAGG | TCATGGCTGGGTATGTGAGA | 300 |
| chr1:70433469-70435505 | p2 | (AT)7 | AGCGTCCACAAACAAAAAGG | TCATGGCTGGGTATGTGAGA | 300 |
| chr1:70608517-70610533 | p5 | (TATGA)3 | AAATGTGCTGCCCAATTTTC | TTCTTTCCGAGCTCACGATT | 286 |
| chr1:70644111-70646125 | p2 | (TA)7 | AATACCGCCACATCTTGGAG | GCCCGTGATTTCATGAGTTT | 140 |
| chr1:70644111-70646125 | p3 | (ATC)5 | AATACCGCCACATCTTGGAG | GCCCGTGATTTCATGAGTTT | 140 |
| chr1:70788636-70790648 | p2 | (AG)6 | TTCCTTAATCCCGCATTTTG | CCATTAGCCAAGCACCATTT | 235 |
| chr1:70788650-70790665 | p2 | (AG)6 | TTCCTTAATCCCGCATTTTG | CCATTAGCCAAGCACCATTT | 221 |
| chr1:70870219-70872239 | p2 | (AT)7 | TGGACCGGTAGGGTTATGAA | CTCCACTTTATCGAGCTGCC | 231 |
| chr1:70870219-70872239 | p2 | (AT)6 | TGGACCGGTAGGGTTATGAA | CTCCACTTTATCGAGCTGCC | 231 |
| chr1:70870219-70872239 | p2 | (TA)10 | TGGACCGGTAGGGTTATGAA | CTCCACTTTATCGAGCTGCC | 231 |
| chr1:70870219-70872239 | p3 | (TCA)5 | TGGACCGGTAGGGTTATGAA | CTCCACTTTATCGAGCTGCC | 231 |
| chr1:70906036-70908048 | p2 | (TG)6 | TCTTATGAAGAGCAGCCCGT | TCGAAATTACCTGAGGGCAC | 131 |
| chr1:70906036-70908048 | p3 | (TTG)5 | TCTTATGAAGAGCAGCCCGT | TCGAAATTACCTGAGGGCAC | 131 |
| chr1:70906643-70908658 | p2 | (TG)6 | AGTCCCTTTGCAACTTGGTG | CCCACTTTGCAAACTTCCAT | 177 |
| chr1:70906643-70908658 | p3 | (TTG)5 | AGTCCCTTTGCAACTTGGTG | CCCACTTTGCAAACTTCCAT | 177 |
| chr1:70966582-70968614 | p2 | (AT)16 | CTAGCTCCACGGACTCGTTC | GGGCTGAGAGGTGATGATGT | 154 |
| chr1:70973795-70975810 | p5 | (GATAT)3 | GTGGATGTCCCAATGTTTCC | GAGGAATCAGAAAAAGGGGC | 225 |
| chr1:70973795-70975810 | p3 | (TAT)5 | GTGGATGTCCCAATGTTTCC | GAGGAATCAGAAAAAGGGGC | 225 |
| chr1:70977923-70979993 | p2 | (AT)35 | TAAACAGGGGAGATGTTGCC | TTGAAAGAAATGGACGAGCC | 138 |
| chr1:70996828-70998840 | p2 | (TA)6 | TGTAAGTGGGGTATGGGGAA | TACTGAAAATGCCTTTGGGG | 234 |
| chr1:71032160-71034184 | p5 | (TGAAT)3 | CCAAAATTTCGGCAGATCAT | AACATACCTGCTCCACGTCC | 237 |
| chr1:71032186-71034204 | p5 | (TGAAT)3 | CCAAAATTTCGGCAGATCAT | AACATACCTGCTCCACGTCC | 211 |
| chr1:71058293-71060315 | p2 | (GT)11 | ACTTGGGGGCTTTTTATGCT | TTTGAAGGAACATTCTGGGG | 278 |
| chr1:71147240-71149258 | p3 | (ACA)6 | CATGGTTCAAAATGCTGTGC | GTATCTGTACCTGCGGGCAT | 196 |
| chr1:71222021-71224035 | p2 | (CT)7 | GTAACTCGATCGTGGGGAGA | CAACCGCAAAACTCAAACAA | 225 |
| chr1:71222021-71224035 | p2 | (GT)6 | GTAACTCGATCGTGGGGAGA | CAACCGCAAAACTCAAACAA | 225 |
| chr1:71222021-71224035 | p2 | (GT)7 | GTAACTCGATCGTGGGGAGA | CAACCGCAAAACTCAAACAA | 225 |
| chr1:71254371-71256397 | p2 | (TA)13 | AATGAAATCGAGGGATCACG | ATTTGGCCATGGTGTATCGT | 246 |
| chr1:71303857-71305869 | p2 | (TA)6 | TGTGGGCTGTTTTGAAATGA | GAACTCAACAATCCCCGAGA | 134 |
| chr1:71363590-71365602 | p2 | (TA)6 | GTCCAATTTCCCACAACACC | GTGCCCGATTGCTTGTAAAT | 105 |
| chr1:71426682-71428694 | p2 | (TA)6 | TCATCGCATCTCCTCATACG | GTGCAAAAGGAAGGATTCCA | 216 |

| Marker ID | SSR type | SSR motif | FORWARD PRIMER (5'-3') | REVERSE PRIMER (5'-3') | size (bp) |
|---|---|---|---|---|---|
| chr1:71540218-71542234 | p4 | (TTTC)4 | GCTCTACCATCGGGCAATAA | CTGCAGGGAAGAAGACAAGC | 145 |
| chr1:71602568-71604580 | p2 | (TA)12 | CTCACGACCAACACCAACAC | ATAAGAGCATCGTCATCGGG | 197 |
| chr1:71602568-71604580 | p2 | (AT)6 | CTCACGACCAACACCAACAC | ATAAGAGCATCGTCATCGGG | 197 |
| chr1:71784478-71786492 | p2 | (TG)7 | AAAGGTGCCTACGGGCTATT | CTCATACCTGGCCCTACCAA | 173 |
| chr1:71870179-71872203 | p2 | (TA)6 | ATGTATCGGGCTTCACGTTC | TGTCTACCTGCTGACTTGCG | 250 |
| chr1:71870179-71872203 | p2 | (TA)12 | ATGTATCGGGCTTCACGTTC | TGTCTACCTGCTGACTTGCG | 250 |
| chr1:71957180-71959194 | p2 | (AT)7 | TTGAGAAATGAGTCCCGACC | TGGTCCAAGAATAGAACCCG | 285 |
| chr1:72001661-72003713 | p2 | (AT)26 | AATGTACATAAGCCGACGGG | CAGGGACTACATAGCGGAGC | 258 |
| chr1:72030843-72032857 | p2 | (AT)7 | CCAACTCCACGGATTCAGTT | CTCCAACTCCAATTCCTCCA | 172 |
| chr1:72030843-72032857 | p2 | (AT)7 | CCAACTCCACGGATTCAGTT | CTCCAACTCCAATTCCTCCA | 172 |
| chr1:72036740-72038764 | p2 | (TA)12 | GGTCCATCGCCAAGACTTAC | GACTGAGCAAGGGATTGGAG | 117 |
| chr1:72088818-72090856 | p2 | (TA)13 | TAAATTTGGGCAGCATCTCC | GGGGTTGGTAGCTCGTGTAG | 111 |
| chr1:72088818-72090856 | p2 | (AT)13 | TAAATTTGGGCAGCATCTCC | GGGGTTGGTAGCTCGTGTAG | 111 |
| chr1:72306858-72308873 | p5 | (TGATA)3 | CGTGCATGCCTCAATACATC | AACTCGGTGAATATGGTCGC | 293 |
| chr1:72317789-72319803 | p2 | (TA)7 | CATTTTTCCTAGATGCCCCA | GGGTTCGTGACAAAAGTGGT | 102 |
| chr1:72434653-72436668 | p5 | (GTTTA)3 | TAAAAATGTGGGCTTGGACC | TGCTAAGACATTGGGGAAGG | 188 |
| chr1:72434653-72436668 | p2 | (AT)15 | TAAAAATGTGGGCTTGGACC | TGCTAAGACATTGGGGAAGG | 188 |
| chr1:72526589-72528604 | p3 | (ACA)5 | TGAGCAAGCAAATGCAAATC | TTCCGATCTTTGGAACATCC | 255 |
| chr1:72614415-72616430 | p5 | (TTCAT)3 | ATTTCAGCCGATGGTATTCG | TGTCCATTGCAATTTGGCTA | 179 |
| chr1:72647125-72649140 | p5 | (GTGTT)3 | GGAAAGTTAAGGGCGAGACC | CAACCCACTGCATCATAACG | 134 |
| chr1:72648565-72650583 | p6 | (ATATGA)3 | TTGCAACTTGAATGAGTCGC | TCGGGTCCTATTTATGCGAG | 155 |
| chr1:72754379-72756391 | p2 | (AT)6 | CTCAGTGACCTTTTTGGGGA | ACACTACGAACCTCCGTTGG | 185 |
| chr1:72906713-72908729 | p2 | (TG)8 | CACCTACTCCGAGCCTTCTG | CATAATGAATGGCACGAACG | 127 |
| chr1:73057411-73059429 | p2 | (AT)6 | AGCCCTTCGGCTATCAAAAT | ATGCCCACAGGTAGACAAGG | 209 |
| chr1:73057411-73059429 | p2 | (TA)9 | AGCCCTTCGGCTATCAAAAT | ATGCCCACAGGTAGACAAGG | 209 |
| chr1:73057411-73059429 | p2 | (TA)23 | AGCCCTTCGGCTATCAAAAT | ATGCCCACAGGTAGACAAGG | 209 |
| chr1:73057736-73059782 | p2 | (TA)9 | TGCGTAATTTCTAGGGGTGG | ATGATCACAACTAGCGTGCG | 261 |
| chr1:73057736-73059782 | p2 | (TA)23 | TGCGTAATTTCTAGGGGTGG | ATGATCACAACTAGCGTGCG | 261 |
| chr1:73062493-73064537 | p2 | (TA)22 | GTTGTTTTCCAAAATGCCGT | TTTTGCATCATTTGCGTCAT | 123 |
| chr1:73098237-73100253 | p2 | (TG)6 | AAGGCAGGTAAGCGCTATGA | ATATGGCTCCGAACGAAATG | 151 |
| chr1:73098237-73100253 | p2 | (AT)8 | AAGGCAGGTAAGCGCTATGA | ATATGGCTCCGAACGAAATG | 151 |
| chr1:73143182-73145206 | p2 | (AT)12 | GCGAACCCCATTAGTTCTCA | AAGCCGACGTACCATTATGC | 234 |
| chr1:73283064-73285076 | p2 | (AC)6 | ATAGGTTTTTGCAGTTGCGG | CTCTTGGGCTGTCTCCTGTC | 158 |
| chr1:73393171-73395189 | p6 | (TCCACA)3 | TCCACATCCACAAAAATCCA | AATTTTGGGGCATAGACGTG | 298 |
| chr1:73396405-73398417 | p2 | (AT)6 | TGTCAGGTGTTCATCCGGTA | CTTGCGGGTCGTATACTGGT | 193 |
| chr1:73396405-73398417 | p3 | (CAA)6 | TGTCAGGTGTTCATCCGGTA | CTTGCGGGTCGTATACTGGT | 193 |

| Marker ID | SSR type | SSR motif | FORWARD PRIMER (5'-3') | REVERSE PRIMER (5'-3') | size (bp) |
|---|---|---|---|---|---|
| chr1:73403597-73405641 | p2 | (AT)22 | CTCGAGCAAGGTTCCAACTC | CTATTTTGGGTGGTCGTCGT | 185 |
| chr1:73463365-73465380 | p3 | (CAT)5 | GACCCTTACTTAGGGCCGAG | CATCAGCCGATGTGTCATTC | 139 |
| chr1:73602250-73604265 | p3 | (GCT)5 | GCCTTGGAGCTCTGATTGAC | AGTGGCTCTTCAGCCAGGTA | 100 |
| chr1:73647805-73649820 | p2 | (TG)7 | GAATGTTCTTGAAGGGCCAA | TCTCTTGCAAGCTTTGTCCA | 149 |
| chr1:73647805-73649820 | p3 | (GTT)5 | GAATGTTCTTGAAGGGCCAA | TCTCTTGCAAGCTTTGTCCA | 149 |
| chr1:73722163-73724183 | p2 | (AT)10 | ATTACCGTGATCTCAACCGC | TGAGTCCGAAAGGCATTACC | 168 |
| chr1:73765315-73767375 | p2 | (TA)30 | TCGTTTGGATTTTGAGGGAC | CACCCCGGAAGTCAAGTAAA | 214 |
| chr1:73765315-73767375 | p2 | (AT)28 | TCGTTTGGATTTTGAGGGAC | CACCCCGGAAGTCAAGTAAA | 214 |
| chr1:73773046-73775088 | p3 | (GAA)14 | CGGGGCCTCTTTGTATTTTT | AAGATGACGGGTGAAGATCG | 247 |
| chr1:73773046-73775088 | p3 | (GAA)14 | CGGGGCCTCTTTGTATTTTT | AAGATGACGGGTGAAGATCG | 247 |
| chr1:73956438-73958456 | p6 | (TATATG)3 | TTTTGGCATGATCCTTGTGA | CGAACGTGCTAGCATTTGAA | 200 |
| chr1:73970119-73972131 | p2 | (AT)6 | CTTTGGCAAGCACAGATTCA | GCTAATCGGTTTGGGTCGTA | 252 |
| chr1:73970459-73972525 | p2 | (AT)6 | GAATGCCATTATACACGGGG | GTACCGCCAGGTTTCACTGT | 221 |
| chr1:73970560-73972602 | p2 | (AT)6 | GAATGCCATTATACACGGGG | GTACCGCCAGGTTTCACTGT | 120 |
| chr1:74079349-74081365 | p2 | (AT)8 | TGCTCGGCTTATATGCCTTT | ATGTATAGGACGTGCTGCCC | 266 |
| chr1:74094580-74096595 | p2 | (TA)19 | TGCAAACCGAGCTTACCTCT | ATGAATCGGACAACACCTCC | 190 |
| chr1:74094580-74096595 | p2 | (TA)16 | TGCAAACCGAGCTTACCTCT | ATGAATCGGACAACACCTCC | 190 |
| chr1:74094607-74096631 | p2 | (TA)19 | TGCAAACCGAGCTTACCTCT | ATGAATCGGACAACACCTCC | 163 |
| chr1:74094607-74096631 | p2 | (TA)16 | TGCAAACCGAGCTTACCTCT | ATGAATCGGACAACACCTCC | 163 |
| chr1:74207754-74209770 | p2 | (TA)8 | CGTGCGATAGAAGCAAGTCA | AGCCTGGTTCCCCTTGTACT | 125 |
| chr1:74212554-74214572 | p2 | (TA)9 | GGAATGCGAACCATCAGAAT | AGGTATACGGACCGGGTTTC | 264 |
| chr1:74212554-74214572 | p2 | (AT)9 | GGAATGCGAACCATCAGAAT | AGGTATACGGACCGGGTTTC | 264 |
| chr1:74215732-74217748 | p2 | (TA)8 | CCGGATGGCTAGAATGTGAT | TTTGAATCCTCCAACAAGCC | 263 |
| chr1:74224800-74226814 | p5 | (TGATC)3 | CAGGGCCAGGATCTGTATGT | ACCCATACGGACGTACCAAA | 225 |
| chr1:74224800-74226814 | p2 | (TA)7 | CAGGGCCAGGATCTGTATGT | ACCCATACGGACGTACCAAA | 225 |
| chr1:74233485-74235515 | p2 | (AT)15 | CCTCACAAATTTGGGCCTTA | TGACTTCAAGGGGCAAAAAC | 144 |
| chr1:74276078-74278096 | p6 | (ATATGT)3 | ATGTTGGATATGAGCCTCGG | TCGGGTCCTATTTATGCGAG | 236 |
| chr1:74424903-74426919 | p4 | (TCTG)4 | AAACCGCTCCAATTCCTTCT | AGGACCAACTCGAGGCTACA | 134 |
| chr1:74439431-74441455 | p6 | (TATATG)4 | TATGGCCGAGTTGGATTCTC | GACATCGGAATCATGCCTTT | 258 |
| chr1:74459614-74461638 | p6 | (TATATG)4 | TATGGCCGAGTTGGATTCTC | GACATCGGAATCATGCCTTT | 258 |
| chr1:74492867-74494882 | p3 | (ATG)5 | TCCATGCTAAACCCAAGAGG | TCTTCCTTCAACCAAATGGC | 146 |
| chr1:74612918-74614934 | p2 | (AC)8 | GGAAAACTTTAACGTGCGGA | TTGGAATAACAAAAAGGGCG | 175 |
| chr1:74617271-74619283 | p2 | (AT)6 | CTGTCAAGGTGGGAAATGGT | ACCCCTCAATGAGTTTGCAC | 295 |
| chr1:74623079-74625103 | p6 | (TATTCA)4 | TATGTCACGACCCAACTGGA | GATAGCCTCGTCGGCTTATG | 299 |
| chr1:74652634-74654649 | p3 | (AAC)5 | ACATATCCAAGGCTCGATGG | AGAGCCGCTATGATGGCTAA | 223 |
| chr1:74729280-74731298 | p2 | (AT)6 | CACAGCATAACACCGGAATG | TTTTCATAGGCGGCTCAGTT | 215 |

| Marker ID | SSR type | SSR motif | FORWARD PRIMER (5'-3') | REVERSE PRIMER (5'-3') | size (bp) |
|---|---|---|---|---|---|
| chr1:74729280-74731298 | p3 | (CAA)6 | CACAGCATAACACCGGAATG | TTTTCATAGGCGGCTCAGTT | 215 |
| chr1:74754142-74756157 | p3 | (AAC)5 | CACGGTGTGGTCACATTAGC | ATTGGAGCTTACCGGACCTT | 115 |
| chr1:74754142-74756157 | p2 | (CT)10 | CACGGTGTGGTCACATTAGC | ATTGGAGCTTACCGGACCTT | 115 |
| chr1:74870749-74872767 | p2 | (AT)9 | AATTTCCCTTTTTGCCCCTA | TGTATACACCTGCGTGGCAT | 117 |
| chr1:74870749-74872767 | p2 | (TA)7 | AATTTCCCTTTTTGCCCCTA | TGTATACACCTGCGTGGCAT | 117 |
| chr1:74938478-74940502 | p6 | (TATATG)4 | GCGAGCTTGGACAATTAAGC | GAATGCCCTTCGGAAACATA | 300 |
| chr1:74938478-74940502 | p4 | (TTAC)4 | GCGAGCTTGGACAATTAAGC | GAATGCCCTTCGGAAACATA | 300 |
| chr1:74985194-74987208 | p2 | (GT)7 | TTTTGTTAGCCTTGTCGGCT | TTATGCGAAAACATGGGACA | 114 |
| chr1:74993276-74995294 | p2 | (GT)9 | TAATTTAGCATTCGGGGACG | GCTCGTTTTGTAATTCCCGA | 293 |
| chr1:75038723-75040737 | p2 | (TA)7 | CACTGTAGCTAACCCGCCTC | CTACACCAGCCAATGGGAGT | 167 |
| chr1:75041958-75043973 | p2 | (GT)6 | CGTCTCTTTTGCTAGGGCAC | TATAATCCGGCACGGCTATC | 105 |
| chr1:75041958-75043973 | p5 | (ATTTC)3 | CGTCTCTTTTGCTAGGGCAC | TATAATCCGGCACGGCTATC | 105 |
| chr1:75047308-75049320 | p2 | (CA)6 | AACGACCATTTTGAGTTCCG | CACAAATGATCCGCATGTTC | 163 |
| chr1:75077768-75079782 | p2 | (AT)7 | AAAAGCCTTAGGGGTCCTCA | ACAAGTGACCGAGGCCATAC | 154 |
| chr1:75121502-75123516 | p2 | (CT)7 | TCCAATCAACCCACATGCTA | GGGCGTGTCGTTATTCTTGT | 168 |
| chr1:75188638-75190650 | p2 | (AT)6 | TGATATGTTTCCGAATGGCA | TTCGGGTCAATCCGACTTAG | 231 |
| chr1:75191884-75193900 | p2 | (AT)8 | ACTTTTGTCACAGCCCAACC | TTTATCGGGTAGCCTTGTCG | 296 |
| chr1:75191884-75193900 | p2 | (AT)8 | ACTTTTGTCACAGCCCAACC | TTTATCGGGTAGCCTTGTCG | 296 |
| chr1:75191884-75193900 | p3 | (CAA)5 | ACTTTTGTCACAGCCCAACC | TTTATCGGGTAGCCTTGTCG | 296 |
| chr1:75264029-75266041 | p2 | (TG)6 | TCGATGCTGGTTCTGCTATG | TCTCCGTACAGTCCCCAAAC | 186 |
| chr1:75264029-75266041 | p2 | (TG)6 | TCGATGCTGGTTCTGCTATG | TCTCCGTACAGTCCCCAAAC | 186 |
| chr1:75276831-75278845 | p2 | (AT)7 | AGTTTACGGACCGTCCCTCT | TTGCAACTGCCTAAGTGTCG | 155 |
| chr1:75332750-75334766 | p2 | (TA)8 | GTAGACTTGCGGGCTCAGAC | CGGATGGGAATCATACCTTG | 194 |
| chr1:75332750-75334766 | p2 | (AT)8 | GTAGACTTGCGGGCTCAGAC | CGGATGGGAATCATACCTTG | 194 |
| chr1:75443020-75445032 | p2 | (TA)6 | CAAATTCGGACTCATTCGGT | TTTGGGCCTCAACTTTGTTC | 127 |
| chr1:75443020-75445032 | p3 | (GTT)5 | CAAATTCGGACTCATTCGGT | TTTGGGCCTCAACTTTGTTC | 127 |
| chr1:75463986-75466000 | p2 | (AT)7 | ACTGTTACGACCCGAACTGG | TGTGATGTAGCCTTGTCGGA | 161 |
| chr1:75572811-75574823 | p2 | (AT)6 | GCGCATTGCTGACACTTAAA | GGGGATGCTAACAGCAAAAA | 168 |
| chr1:75580396-75582411 | p3 | (AGA)5 | AATTCCCCAACCTTCCAAAC | ATTAAACCATAACGCGCTGC | 157 |
| chr1:75700561-75702576 | p3 | (ATG)5 | AAAAGGAGTTTTGGGGCATT | GCAGCATTTCCCCTATTCAA | 295 |
| chr1:75728055-75730067 | p2 | (AT)6 | GCGCATTTTGTTTCCTTGAT | GACCCGCAAGGTTTTATTGA | 103 |
| chr1:75869860-75871872 | p6 | (CATATA)3 | AAAGTGACCCCACTGGACAC | AAGTTCCATGTTACGCCCAG | 280 |
| chr1:75931534-75933549 | p5 | (CATAT)3 | TGGCCCTAGCACACTTAACC | AGGCTGTTAAGACCCCGATT | 103 |
| chr1:76004336-76006348 | p2 | (AT)6 | GGCCCGTAAACTGAGTCGTA | TGAACAACAGTCGGGGTGTA | 141 |
| chr1:76065855-76067867 | p2 | (AG)6 | AGGCCATTTAGGGATTTGCT | TGCCTCACAGAGCATACGAC | 149 |
| chr1:76115236-76117248 | p2 | (TA)6 | GAAGGTCCATCGGAAAGACA | CCCTTTCCCGTCCTTTCTAC | 236 |

| Marker ID | SSR type | SSR motif | FORWARD PRIMER (5'-3') | REVERSE PRIMER (5'-3') | size (bp) |
|---|---|---|---|---|---|
| chr1:76133114-76135129 | p5 | (TCAAT)3 | TTTGTGCATCCTTGTGTCGT | AGCCATACAACGAAATTGCC | 222 |
| chr1:76151098-76153110 | p2 | (TG)6 | CGAGCTTATAATGCCGGGTA | CGACCTCATAGGATCGGTGT | 281 |
| chr1:76180901-76182916 | p5 | (TGTCC)3 | TGCACTGTCAAATCTCCTGC | CCGCTATTTGATTGATGGCT | 213 |
| chr1:76225799-76227819 | p3 | (ATG)5 | CCGATTTTCATTGGCTCATT | AGCTTTTTGGGACCAAGGAT | 180 |
| chr1:76225799-76227819 | p2 | (AT)8 | CCGATTTTCATTGGCTCATT | AGCTTTTTGGGACCAAGGAT | 180 |
| chr1:76225814-76227832 | p3 | (ATG)5 | CCGATTTTCATTGGCTCATT | AGCTTTTTGGGACCAAGGAT | 165 |
| chr1:76225814-76227832 | p2 | (AT)8 | CCGATTTTCATTGGCTCATT | AGCTTTTTGGGACCAAGGAT | 165 |
| chr1:76252917-76254932 | p5 | (ACATT)3 | TGGCTTAGCCGTAGAGTCGT | TATACGCTACCTCATCCGGC | 120 |
| chr1:76342235-76344247 | p2 | (TA)6 | CGATATTCGCTGCGTACAGA | AACGATGTTTCCTTTCCACG | 290 |
| chr1:76345841-76347859 | p5 | (TGAAT)3 | TTACAGGGGAGATGCTACCG | CCAGCTCGCTATCGCTAATC | 195 |
| chr1:76345841-76347859 | p6 | (TATATG)3 | TTACAGGGGAGATGCTACCG | CCAGCTCGCTATCGCTAATC | 195 |
| chr1:76478730-76480745 | p5 | (ATTTC)3 | CGTGGCCAAATTTTTATGCT | ATTTCGCCAGAGTTTTCCCT | 295 |
| chr1:76503957-76505975 | p2 | (GT)9 | TGTGCCTTTTGGTCATGAAA | CCGTTTTCTCCATGCTCATT | 284 |
| chr1:76503957-76505975 | p5 | (GTGTT)3 | TGTGCCTTTTGGTCATGAAA | CCGTTTTCTCCATGCTCATT | 284 |
| chr1:76526829-76528844 | p5 | (AAAGA)3 | GGCAAGTTGAGTTTTTGGGA | TCAATGGCCACTCACTCAAG | 110 |
| chr1:76569321-76571333 | p2 | (AG)6 | TGGGTGTTTTGAAGGTGTCA | CTCTTCCGTCACCTGGACAT | 121 |
| chr1:76789668-76791683 | p3 | (TTG)5 | AACAAGTGTGACGACGGTGA | CCACGTGGATGTGTGTCAAT | 208 |
| chr1:76789668-76791683 | p2 | (TA)6 | AACAAGTGTGACGACGGTGA | CCACGTGGATGTGTGTCAAT | 208 |
| chr1:76804559-76806571 | p2 | (AT)7 | TGCAAAAACTCCAAAGGGTC | AGCCTTGTCAGCTTGCGTAT | 221 |
| chr1:76804559-76806571 | p5 | (AAAAG)3 | TGCAAAAACTCCAAAGGGTC | AGCCTTGTCAGCTTGCGTAT | 221 |
| chr1:76804572-76806594 | p2 | (AT)7 | TGCAAAAACTCCAAAGGGTC | AGCCTTGTCAGCTTGCGTAT | 208 |
| chr1:76804572-76806594 | p5 | (AAAAG)3 | TGCAAAAACTCCAAAGGGTC | AGCCTTGTCAGCTTGCGTAT | 208 |
| chr1:77101504-77103519 | p3 | (CAA)5 | CGATGAGGCAATCAAAGACA | TCACGTCATACTTTCGAGCG | 256 |
| chr1:77143101-77145116 | p3 | (TGT)5 | TGACCCTTCCCAAGTGTTTC | ACCAGAACCACATCTTTGGC | 140 |
| chr1:77154479-77156494 | p3 | (CAA)5 | ATCTGGATGCCAAAGAATGC | GCCACTTGGTCAGTGAGACA | 185 |
| chr1:77169978-77171993 | p2 | (AT)10 | CGAGGCCATAATCTGGACAT | GTGTGTGTGTGTAGGACGGG | 160 |
| chr1:77169978-77171993 | p5 | (AAAAT)3 | CGAGGCCATAATCTGGACAT | GTGTGTGTGTGTAGGACGGG | 160 |
| chr1:77192901-77194929 | p2 | (TA)14 | GTGTGGGCTGTTTGGAATCT | TGCTGCCCAAAATATGTCAG | 300 |
| chr1:77196036-77198054 | p6 | (ATGTAT)3 | CTCCGAGACTCTTTCGGTTG | AAAACGGGGTTCATTTCACA | 173 |
| chr1:77198074-77200086 | p2 | (TA)14 | TTAGCACAGCCCATGTGAAG | CCCAGTTCGAGGAAAATTGA | 178 |
| chr1:77198074-77200086 | p2 | (AT)6 | TTAGCACAGCCCATGTGAAG | CCCAGTTCGAGGAAAATTGA | 178 |
| chr1:77265402-77267420 | p2 | (TA)14 | TATCCAAGGCAAGAAGGTGG | AGTCGAGGTCAGGCACTCAT | 119 |
| chr1:77275854-77277866 | p2 | (AG)6 | GGAAGGAATCATGCTTTGGA | TGCCGCCCAATTTACTTTAG | 188 |
| chr1:77351776-77353790 | p2 | (AT)7 | ACAACCATCCGGGAATGTAA | CGTATGCGGTTGATTGAATG | 107 |
| chr1:77383314-77385329 | p3 | (AGC)5 | CCGCCACATGTAGAGGAGAT | ACTCACGGGCTCTCGAACTA | 165 |
| chr1:77407333-77409345 | p2 | (AT)10 | AGTGCACCACACACCGATAA | GGGGTGCTCGGTAGTCAATA | 129 |

| Marker ID | SSR type | SSR motif | FORWARD PRIMER (5'-3') | REVERSE PRIMER (5'-3') | size (bp) |
|---|---|---|---|---|---|
| chr1:77407333-77409345 | p2 | (AT)6 | AGTGCACCACACACCGATAA | GGGGTGCTCGGTAGTCAATA | 129 |
| chr1:77416704-77418716 | p2 | (AG)6 | GGGAAGTCTCATCGACTCCA | TTGAGTGACTGTGACGAGGC | 234 |
| chr1:77511947-77513968 | p2 | (AT)6 | TGACCGTGTGACATGTCCTT | GGTCACATGCCCCTTTATGT | 247 |
| chr1:77511947-77513968 | p3 | (GTT)7 | TGACCGTGTGACATGTCCTT | GGTCACATGCCCCTTTATGT | 247 |
| chr1:77511947-77513968 | p5 | (TGAAT)4 | TGACCGTGTGACATGTCCTT | GGTCACATGCCCCTTTATGT | 247 |
| chr1:77512496-77514516 | p3 | (GTT)7 | CATGAGTTATGTGCGGCTTG | TGTCATACCAAATCAGCCCA | 178 |
| chr1:77512496-77514516 | p5 | (TGAAT)4 | CATGAGTTATGTGCGGCTTG | TGTCATACCAAATCAGCCCA | 178 |
| chr1:77517487-77519507 | p3 | (GTT)6 | GATGTTATGTGTGTTGGCCG | TGTCATACCAAATCAGCCCA | 262 |
| chr1:77517487-77519507 | p5 | (TGAAT)4 | GATGTTATGTGTGTTGGCCG | TGTCATACCAAATCAGCCCA | 262 |
| chr1:77537272-77539284 | p2 | (AG)6 | GAAACGAAAAGGGTTCACGA | AAGGGGAAGGCTCAATCACT | 247 |
| chr1:77783888-77785900 | p2 | (AT)6 | AGGTTTTGTCCCGGAGAACT | TCTGTTACACTCCCCTTGCC | 193 |
| chr1:77857344-77859362 | p3 | (TTG)6 | TAAAATGGACAGCCAGAGGG | ATGGGGATGAGAGAATGGTG | 247 |
| chr1:77973018-77975030 | p2 | (AT)6 | GATGAGCCGTGACGAGTACA | GTAGCGGCCTAATCAGCTTG | 294 |
| chr1:77979913-77981945 | p2 | (AT)16 | TATGGCTGACTTCACACCCA | CCAGGTGACAGCATAGCTCA | 277 |
| chr1:77979913-77981945 | p2 | (AT)19 | TATGGCTGACTTCACACCCA | CCAGGTGACAGCATAGCTCA | 277 |
| chr1:77979913-77981945 | p3 | (ATC)7 | TATGGCTGACTTCACACCCA | CCAGGTGACAGCATAGCTCA | 277 |
| chr1:77980081-77982119 | p2 | (AT)16 | TATGGCTGACTTCACACCCA | CCAGGTGACAGCATAGCTCA | 109 |
| chr1:77980081-77982119 | p2 | (AT)19 | TATGGCTGACTTCACACCCA | CCAGGTGACAGCATAGCTCA | 109 |
| chr1:77980081-77982119 | p3 | (ATC)7 | TATGGCTGACTTCACACCCA | CCAGGTGACAGCATAGCTCA | 109 |
| chr1:77980354-77982375 | p2 | (AT)16 | AAAACAAGTGCCAACAAGGC | CTCCCCTCAGTCGCAGTTAG | 171 |
| chr1:77980354-77982375 | p2 | (AT)19 | AAAACAAGTGCCAACAAGGC | CTCCCCTCAGTCGCAGTTAG | 171 |
| chr1:77980354-77982375 | p3 | (ATC)7 | AAAACAAGTGCCAACAAGGC | CTCCCCTCAGTCGCAGTTAG | 171 |
| chr1:78074446-78076462 | p2 | (AT)8 | ACAAAATTCGACGGCATTTC | TGTTGGTTGGGTTGTTTTGA | 134 |
| chr1:78083430-78085456 | p2 | (AT)13 | CTCGAACTACCAGCTCCCTG | AGGTTCAGAGTGAGAGCCCA | 195 |
| chr1:78177642-78179666 | p6 | (CATATA)4 | GATGGAGCCGCTTACAAGAC | TGGCAACCTACATTTCCTCC | 189 |
| chr1:78337609-78339625 | p4 | (TGTA)4 | GCTCCAATTCCTTCAGCAAG | ACCAACTCGAGGCGACATAC | 289 |
| chr1:78381952-78383967 | p3 | (GTT)5 | ACGTTTTCGGGTGATCTGAC | AATGTGGAGGAGAAGGCTGA | 173 |
| chr1:78381952-78383967 | p2 | (TA)7 | ACGTTTTCGGGTGATCTGAC | AATGTGGAGGAGAAGGCTGA | 173 |
| chr1:78406848-78408864 | p2 | (GA)6 | CGGATTAGTCGTAAGCGGAG | CTTTGATTTGGCTCATGGGT | 241 |
| chr1:78406848-78408864 | p4 | (TATT)4 | CGGATTAGTCGTAAGCGGAG | CTTTGATTTGGCTCATGGGT | 241 |
| chr1:78413107-78415122 | p3 | (GTT)5 | ACGTTTTCGGGTGATCTGAC | AATGTGGAGGAGAAGGCTGA | 173 |
| chr1:78413107-78415122 | p2 | (TA)7 | ACGTTTTCGGGTGATCTGAC | AATGTGGAGGAGAAGGCTGA | 173 |
| chr1:78435047-78437065 | p6 | (TATATG)3 | TTAGATTCTCGGGGATGGTG | CTTGGCCTTCCCAACATCTA | 191 |
| chr1:78500686-78502701 | p5 | (TGATA)3 | GACCGTTTGTAGGTTTCGGA | TGGGACAAGGGCAAGTTTAG | 182 |
| chr1:78539585-78541799 | p2 | (AT)107 | TGGTGATTGGGTGTTCTTGA | GCTCATACATTTGCGAAGCA | 184 |
| chr1:78541790-78543802 | p2 | (AT)6 | CGGAAGTGGCATATGTGTTG | ATCAATACCTGCAGCCTTCG | 248 |

| Marker ID | SSR type | SSR motif | FORWARD PRIMER (5'-3') | REVERSE PRIMER (5'-3') | size (bp) |
|---|---|---|---|---|---|
| chr1:78547600-78549615 | p3 | (GTT)5 | ACGTTTTCGGGTGATCTGAC | CAGGTCCACGACCTATGGTT | 226 |
| chr1:78642925-78644970 | p2 | (AT)26 | AAGTCCGCAAACACAAAACC | CCATGGTGTCCATGTAGCAG | 272 |
| chr1:78642925-78644970 | p3 | (TTA)15 | AAGTCCGCAAACACAAAACC | CCATGGTGTCCATGTAGCAG | 272 |
| chr1:78775489-78777501 | p2 | (TA)6 | GCTAGCATTGGAGAGGAACG | TCGTTGCGTATCAAGAATCG | 260 |
| chr1:78775489-78777501 | p2 | (AT)6 | GCTAGCATTGGAGAGGAACG | TCGTTGCGTATCAAGAATCG | 260 |
| chr1:78856224-78858242 | p3 | (TTG)6 | AAGCCCTCCAAAAGGATGAT | TTTGGCAGCTTGGAGTCTTT | 184 |
| chr1:78856224-78858242 | p3 | (ATG)5 | AAGCCCTCCAAAAGGATGAT | TTTGGCAGCTTGGAGTCTTT | 184 |
| chr1:78899509-78901521 | p2 | (AT)6 | GAATAGAGTTGTCGCCGAGG | AGCGTCTCCAGACGTGTTCT | 136 |
| chr1:78956381-78958399 | p4 | (CATA)5 | GTCCATCATATCCGTACCCG | TAACACCAAGTCCCGAAAGG | 198 |
| chr1:78956381-78958399 | p3 | (CAA)6 | GTCCATCATATCCGTACCCG | TAACACCAAGTCCCGAAAGG | 198 |
| chr1:78956381-78958399 | p2 | (AT)7 | GTCCATCATATCCGTACCCG | TAACACCAAGTCCCGAAAGG | 198 |
| chr1:78988001-78990013 | p2 | (TA)6 | GACAGTGTCTCGAACTGGCA | CCGTCCTTTGCCTTATGAAA | 100 |
| chr1:78990624-78992640 | p2 | (TA)8 | ACGTTTGATGACCTTAGGCG | GCAAAAAGTTGAAGCCTTGC | 259 |
| chr1:78990624-78992640 | p5 | (TGAAT)5 | ACGTTTGATGACCTTAGGCG | GCAAAAAGTTGAAGCCTTGC | 259 |
| chr1:78991358-78993383 | p2 | (TA)8 | GGAGCAAACCAAGTGGTCAT | TTTGGGACAGGATTTCAAGG | 150 |
| chr1:78991358-78993383 | p5 | (TGAAT)5 | GGAGCAAACCAAGTGGTCAT | TTTGGGACAGGATTTCAAGG | 150 |
| chr1:79139358-79141390 | p2 | (AT)16 | GGTGCGTAATCGGGTTTCTA | AGGAGAAAAGTGTGCGAGGA | 149 |
| chr1:79139358-79141390 | p2 | (TG)7 | GGTGCGTAATCGGGTTTCTA | AGGAGAAAAGTGTGCGAGGA | 149 |
| chr1:79205807-79207845 | p2 | (AT)19 | TGCCTGTGTCTGAGACCTTG | CCTTTGGTCAAGGGAATGAA | 130 |
| chr1:79205807-79207845 | p3 | (ACA)5 | TGCCTGTGTCTGAGACCTTG | CCTTTGGTCAAGGGAATGAA | 130 |
| chr1:79217126-79219138 | p2 | (TA)6 | TCTGTAGGGTTGGCAAGGTC | ACCTGACGATCCAGTATGGC | 253 |
| chr1:79233721-79235733 | p2 | (AT)6 | CAGTCTCAGAAGCCCTGACC | GGTGATGATGTGGACAGACG | 163 |
| chr1:79233721-79235733 | p4 | (TAAA)4 | CAGTCTCAGAAGCCCTGACC | GGTGATGATGTGGACAGACG | 163 |
| chr1:79310341-79312355 | p2 | (AT)7 | TCCCGATAACCTGTCTCCTG | TGTTCGTCTGTAGTGGCAGC | 298 |
| chr1:79431877-79433945 | p2 | (AT)30 | TGCATTATGTGGATAGCCGA | ACTGTCACGACCCGACTAGG | 160 |
| chr1:79431877-79433945 | p2 | (TA)34 | TGCATTATGTGGATAGCCGA | ACTGTCACGACCCGACTAGG | 160 |
| chr1:79440885-79442903 | p2 | (TA)9 | CCAATGAGCCAAGGATCTGT | AAAACAACCTACGCCACCAC | 282 |
| chr1:79488330-79490352 | p2 | (AC)11 | AAGAGGGAGAACTTCCCCAA | ATATGCTCGCAAACCGTCTC | 113 |
| chr1:79511140-79513155 | p3 | (AGT)5 | GTGCTCTCTGGGCTATGTCC | TAATCTCGTCCCGCCATTAC | 168 |
| chr1:79551173-79553191 | p2 | (AT)9 | CCACGTCCCGGAGTCTATAA | CCATTGACACAGTTGGTTGC | 151 |
| chr1:79649181-79651201 | p5 | (CATTG)3 | GGTTTGGCTATTTGGGGTTT | TTCCTCTTTGGAGAGCCTCA | 172 |
| chr1:79697695-79699707 | p2 | (TA)6 | GATTCATGCTCCGAGGGTAA | AGATCGATTGTTGACAGCCC | 152 |
| chr1:79697695-79699707 | p5 | (TGAAT)3 | GATTCATGCTCCGAGGGTAA | AGATCGATTGTTGACAGCCC | 152 |
| chr1:79698576-79700591 | p2 | (TA)6 | CGAACCTAGGACATGGAGGA | TTATGGATTTGTGCCAACGA | 232 |
| chr1:79698576-79700591 | p5 | (TGAAT)3 | CGAACCTAGGACATGGAGGA | TTATGGATTTGTGCCAACGA | 232 |
| chr1:79759239-79761253 | p2 | (TA)7 | ATATGAAAGCCGGAAAGCCT | TCCCGTCATATTTCACCGTT | 143 |

| Marker ID | SSR type | SSR motif | FORWARD PRIMER (5'-3') | REVERSE PRIMER (5'-3') | size (bp) |
|---|---|---|---|---|---|
| chr1:79759239-79761253 | p2 | (TA)7 | ATATGAAAGCCGGAAAGCCT | TCCCGTCATATTTCACCGTT | 143 |
| chr1:79823426-79825448 | p2 | (TA)11 | CTCGCAAAAGAACTCCAAGG | GTTCGTCTGTAGTGGCAGCA | 197 |
| chr1:79823426-79825448 | p3 | (ACA)7 | CTCGCAAAAGAACTCCAAGG | GTTCGTCTGTAGTGGCAGCA | 197 |
| chr1:79842968-79844980 | p2 | (AT)7 | GCCTAAGATGGACACCCGTA | ATGTGCGTCTGTAGTGGCAG | 178 |
| chr1:79842968-79844980 | p2 | (AT)6 | GCCTAAGATGGACACCCGTA | ATGTGCGTCTGTAGTGGCAG | 178 |
| chr1:79954554-79956566 | p2 | (TA)6 | TGGTTTCACGATTTCCCATT | TGCGCTCATGCATTAATCTC | 100 |
| chr1:79984001-79986013 | p2 | (CA)6 | TTCGGACGCTTACTTGCTTT | AACGAAACGGGATTTGAGTG | 133 |
| chr1:80036502-80038538 | p2 | (TA)18 | GTAATCGGTTTCTACCGCCA | CTAATGGTTGGTTCGGAGGA | 159 |
| chr1:80166807-80168822 | p2 | (CA)8 | CCAACATTTCCACCAAAACC | GGCTGATTTGGAAGCCAATA | 254 |
| chr1:80166807-80168822 | p3 | (TCT)5 | CCAACATTTCCACCAAAACC | GGCTGATTTGGAAGCCAATA | 254 |
| chr1:80227517-80229541 | p2 | (TA)12 | GGTGGCGGCTTATAGATGAA | TGGTGTTGCCCAGCTATGTA | 124 |
| chr1:80258941-80260955 | p3 | (AAC)5 | GGCCGAGCCTAATACATTCA | GTTGTTGTTTGGAGCCGAGT | 109 |
| chr1:80258941-80260955 | p2 | (AC)7 | GGCCGAGCCTAATACATTCA | GTTGTTGTTTGGAGCCGAGT | 109 |
| chr1:80330715-80332743 | p2 | (AT)14 | GATGCGTAATCGGGTTCCTA | GGAGGAAGAAACCCAAGGAG | 161 |
| chr1:80330715-80332743 | p2 | (TA)47 | GATGCGTAATCGGGTTCCTA | GGAGGAAGAAACCCAAGGAG | 161 |
| chr1:80333151-80335169 | p6 | (CATATA)3 | AGCGAACCCCTTTAGCTCTC | ACAGAGTCATGGGTAAGCGG | 289 |
| chr1:80344072-80346090 | p3 | (GTT)6 | CGTTTTCGGGTGATCTGAGT | CATGAAGGCCCACGAACTAT | 218 |
| chr1:80355238-80357256 | p3 | (GTT)6 | ACGTTTTCGGGTGATCTGAG | AAGCCCACGAACTATGATGG | 231 |
| chr1:80381786-80383798 | p2 | (TG)6 | TGTGACGGAACAATAACCGA | ACGGGGTATAACATCCTCCC | 124 |
| chr1:80381786-80383798 | p2 | (TA)6 | TGTGACGGAACAATAACCGA | ACGGGGTATAACATCCTCCC | 124 |
| chr1:80442702-80444717 | p3 | (ACA)5 | ACGAACAGTTTGTTGCGTTG | GACCTTGGAACCAAAGGACA | 190 |
| chr1:80454067-80456082 | p5 | (TGATA)3 | CCGATTCTTCTTTGGCATGT | GGGGGCATTCCATTAATCTT | 200 |
| chr1:80510322-80512337 | p5 | (TTTCA)3 | GGAACTCGTATGGAAGCCAA | CGCACCCTTCGTCATTCTAT | 293 |
| chr1:80559788-80561800 | p2 | (CT)6 | TTCCTAAGTGTTCCATGCCC | GAGCAGGTGTTACGTCCCAT | 209 |
| chr1:80661857-80663872 | p5 | (ATTTC)3 | GTGCCTTGGGAACCAAACTA | CGAGGTGATTGGGTGAGATT | 173 |
| chr1:80711656-80713680 | p2 | (TA)12 | ATATGGACGGGATAGGACCC | TGTTCATGCCCACAGGTAGA | 252 |
| chr1:80718981-80720995 | p2 | (AG)7 | ATTGAAAATTTGGCCGAGTG | ACATACCTGCTCTGCGTCCT | 128 |
| chr1:80768916-80770928 | p5 | (ATTCA)5 | GGATCATGCCTTTGGAAGAA | TTCCCGGATTGTTGTTGATT | 190 |
| chr1:80768916-80770928 | p2 | (CA)6 | GGATCATGCCTTTGGAAGAA | TTCCCGGATTGTTGTTGATT | 190 |
| chr1:80768916-80770928 | p2 | (AT)10 | GGATCATGCCTTTGGAAGAA | TTCCCGGATTGTTGTTGATT | 190 |
| chr1:80842448-80844466 | p3 | (CAA)6 | ACGAGCGGAGTAGTGAGGAA | CCTCAGAAAAGCTCGGATTG | 289 |
| chr1:80855565-80857589 | p6 | (TATATG)4 | TCACCTCCCTGTCTACCTGG | TTTCCAAGCTCGTTATCGCT | 211 |
| chr1:80893619-80895633 | p2 | (AT)7 | TAGCTCCTTCCACAACCCAC | CAACATCGAATCCCAGTCCT | 211 |
| chr1:80971247-80973262 | p2 | (AG)7 | CCATAGGTCTTGGGCCTGTA | TGAGGAGGAGAAGGCTGAAA | 266 |
| chr1:80971247-80973262 | p3 | (TGT)5 | CCATAGGTCTTGGGCCTGTA | TGAGGAGGAGAAGGCTGAAA | 266 |
| chr1:80985764-80987779 | p5 | (TTTTG)3 | GACGGCGTTTCGTTTATGTT | CCGTTTGAAATGTTGTGCAG | 129 |

| Marker ID | SSR type | SSR motif | FORWARD PRIMER (5'-3') | REVERSE PRIMER (5'-3') | size (bp) |
|---|---|---|---|---|---|
| chr1:81000599-81002614 | p5 | (GAAAA)3 | TACGCGTCCGTTCCTAATTC | CCCCACATAATCATCGAACC | 224 |
| chr1:81004425-81006440 | p2 | (AG)6 | CTGACTTTGGGAGGCCATAA | TGAGGAGGAGAAGGCTGAAA | 186 |
| chr1:81004425-81006440 | p3 | (GTT)5 | CTGACTTTGGGAGGCCATAA | TGAGGAGGAGAAGGCTGAAA | 186 |
| chr1:81136796-81138824 | p2 | (AT)6 | ACTCAGGTGTTGGAACCTCG | AGTTCTGCTACACCGCCACT | 167 |
| chr1:81136796-81138824 | p2 | (AT)14 | ACTCAGGTGTTGGAACCTCG | AGTTCTGCTACACCGCCACT | 167 |
| chr1:81166143-81168159 | p4 | (ATAA)4 | GCTTAAGAAACCGATCGCAG | GATCCGGAGCCCATATTTTT | 242 |
| chr1:81167318-81169330 | p2 | (AC)6 | TACCTCGATCGCTCGCTAAT | TTGGCCGAGTTAAATTCTCG | 277 |
| chr1:81175384-81177396 | p2 | (AC)6 | CAAAGACATTTGGCCATCCT | TTACAGGGGAGATGCTACCG | 163 |
| chr1:81389656-81391718 | p3 | (TTG)5 | ATTCGGAAGATAGGCGAGGT | CCCAGCTCATTTCCTCTCTG | 289 |
| chr1:81546450-81548462 | p2 | (AC)6 | ACTAGTCATGGCTCATCGGG | ATCACAGCTTTGCTTTGCCT | 229 |
| chr1:81689374-81691392 | p6 | (CATATA)3 | TACTGACCCCGGTTCTTTTG | GCCGCTTACAGACTGACACA | 224 |
| chr1:81709494-81711506 | p2 | (TA)6 | GGGTAGTGCCATGCTCTCTC | GCCAAACAATGAGCCAAGAT | 147 |
| chr1:81771130-81773210 | p2 | (TA)10 | TTCACGACCTTCTTTCACCC | AACGGTGGAATCGTATCGTC | 177 |
| chr1:81771130-81773210 | p2 | (TA)40 | TTCACGACCTTCTTTCACCC | AACGGTGGAATCGTATCGTC | 177 |
| chr1:81915776-81917788 | p2 | (AT)6 | GGTGGGGTCCACACATAGAG | CACCCGGAAACGTTTTCTTA | 180 |
| chr1:81953837-81955852 | p2 | (AT)8 | TTTATGGGGACACGCATTTT | AATAAGTTTTGTGGCCGTCG | 235 |
| chr1:81953837-81955852 | p5 | (AAATA)3 | TTTATGGGGACACGCATTTT | AATAAGTTTTGTGGCCGTCG | 235 |
| chr1:81980574-81982589 | p5 | (AAGAG)3 | CATTATGAGGCGAGGTTGGT | ATAGACCGTCCGAACACAGG | 206 |
| chr1:81981634-81983658 | p3 | (TTA)8 | GAGCCGGCAGAACAAGATAG | ATTCGGGAAACATCCAATGA | 249 |
| chr1:82002559-82004591 | p2 | (TA)7 | TCACTACTCTGCTCGTGCGT | GGCCGCCATAACATATCATC | 276 |
| chr1:82002559-82004591 | p2 | (TA)6 | TCACTACTCTGCTCGTGCGT | GGCCGCCATAACATATCATC | 276 |
| chr1:82002593-82004625 | p2 | (TA)7 | TCACTACTCTGCTCGTGCGT | GGCCGCCATAACATATCATC | 242 |
| chr1:82002593-82004625 | p2 | (TA)6 | TCACTACTCTGCTCGTGCGT | GGCCGCCATAACATATCATC | 242 |
| chr1:82002627-82004659 | p2 | (TA)7 | TCACTACTCTGCTCGTGCGT | GGCCGCCATAACATATCATC | 208 |
| chr1:82002627-82004659 | p2 | (TA)6 | TCACTACTCTGCTCGTGCGT | GGCCGCCATAACATATCATC | 208 |
| chr1:82002661-82004693 | p2 | (TA)7 | TCACTACTCTGCTCGTGCGT | GGCCGCCATAACATATCATC | 174 |
| chr1:82002661-82004693 | p2 | (TA)6 | TCACTACTCTGCTCGTGCGT | GGCCGCCATAACATATCATC | 174 |
| chr1:82002695-82004727 | p2 | (TA)7 | TCACTACTCTGCTCGTGCGT | GGCCGCCATAACATATCATC | 140 |
| chr1:82002695-82004727 | p2 | (TA)6 | TCACTACTCTGCTCGTGCGT | GGCCGCCATAACATATCATC | 140 |
| chr1:82002729-82004761 | p2 | (TA)7 | TCACTACTCTGCTCGTGCGT | GGCCGCCATAACATATCATC | 106 |
| chr1:82002729-82004761 | p2 | (TA)6 | TCACTACTCTGCTCGTGCGT | GGCCGCCATAACATATCATC | 106 |
| chr1:82145722-82147756 | p2 | (AT)17 | CCAAAAGAGCCCATTTTTCA | CTCAACTCCCTCCAAAACCA | 121 |
| chr1:82243192-82245216 | p2 | (TA)7 | CACCTACGTGGCATCAAATG | GCATGTATGGTATCCGCCTT | 181 |
| chr1:82243192-82245216 | p4 | (CATA)6 | CACCTACGTGGCATCAAATG | GCATGTATGGTATCCGCCTT | 181 |
| chr1:82243192-82245216 | p3 | (CAA)6 | CACCTACGTGGCATCAAATG | GCATGTATGGTATCCGCCTT | 181 |
| chr1:82285277-82287292 | p3 | (ACA)5 | TAAAACAGGCCCTTGTCCAC | GAAATGGTTTGTGGGCTGTT | 276 |

| Marker ID | SSR type | SSR motif | FORWARD PRIMER (5'-3') | REVERSE PRIMER (5'-3') | size (bp) |
|---|---|---|---|---|---|
| chr1:82285277-82287292 | p3 | (ACA)5 | TAAAACAGGCCCTTGTCCAC | GAAATGGTTTGTGGGCTGTT | 276 |
| chr1:82292660-82294674 | p2 | (AT)7 | ACCGCTTTCTCTTGGAACCT | ATCTGGCCTAGACCTTGGGT | 128 |
| chr1:82292660-82294674 | p2 | (TA)7 | ACCGCTTTCTCTTGGAACCT | ATCTGGCCTAGACCTTGGGT | 128 |
| chr1:82406913-82408931 | p6 | (CATATA)3 | AAGGCTGCCACTATGAAACG | GTTCCGGAGCCTAGCTCTTT | 281 |
| chr1:82447876-82449890 | p2 | (GT)7 | ATCGGGATGAGATGACTTGG | AAGTCTTATGATCGTGGCGG | 102 |
| chr1:82538535-82540555 | p5 | (TTCAT)4 | GCTGCCTGAGAACAATCCAT | GGCCAATTGCATTTTGATCT | 285 |
| chr1:82576492-82578508 | p4 | (CAGA)4 | ACTTCTGTCACGACCCAACC | GTACGCGTGCCGTATCTGTA | 221 |
| chr1:82581243-82583277 | p2 | (TA)6 | CCTGGACCTGTCAATCCCTA | TGAGACACCTGTCCAAGCAG | 292 |
| chr1:82581243-82583277 | p2 | (AT)17 | CCTGGACCTGTCAATCCCTA | TGAGACACCTGTCCAAGCAG | 292 |
| chr1:82581243-82583277 | p2 | (AT)11 | CCTGGACCTGTCAATCCCTA | TGAGACACCTGTCCAAGCAG | 292 |
| chr1:82581243-82583277 | p2 | (TA)7 | CCTGGACCTGTCAATCCCTA | TGAGACACCTGTCCAAGCAG | 292 |
| chr1:82582057-82584071 | p2 | (AT)17 | CGAGCTGACTAGGCCGTAAC | CGAAGGACCTCTTCAGTTGG | 241 |
| chr1:82582057-82584071 | p2 | (AT)11 | CGAGCTGACTAGGCCGTAAC | CGAAGGACCTCTTCAGTTGG | 241 |
| chr1:82582057-82584071 | p2 | (TA)7 | CGAGCTGACTAGGCCGTAAC | CGAAGGACCTCTTCAGTTGG | 241 |
| chr1:82763484-82765500 | p5 | (GCATT)3 | TGTTACATGGCCGATTCTGA | ACATCTCGGTAGCACAACCC | 215 |
| chr1:82763484-82765500 | p2 | (TA)8 | TGTTACATGGCCGATTCTGA | ACATCTCGGTAGCACAACCC | 215 |
| chr1:82847185-82849201 | p5 | (GCATT)3 | TGTTACATGGCCGATTCTGA | ACATCTCGGTAGCACAACCC | 215 |
| chr1:82847185-82849201 | p2 | (TA)8 | TGTTACATGGCCGATTCTGA | ACATCTCGGTAGCACAACCC | 215 |
| chr1:82933086-82935101 | p5 | (CCACA)3 | GTCAAACGGAGCAATAGGGA | CGATGACCCTAAACCCTCAA | 136 |
| chr1:82991326-82993348 | p2 | (AT)11 | AATTCACACGGCTACAAGGG | GAAGTCGGGACTTTTGACCA | 147 |
| chr1:82991569-82993581 | p2 | (AT)11 | GGGACGTAACATTCACCACC | GGTCCAACATCAACCCAAAC | 127 |
| chr1:82991583-82993619 | p2 | (AT)11 | GGGACGTAACATTCACCACC | GGTCCAACATCAACCCAAAC | 113 |
| chr1:83110674-83112688 | p2 | (TA)7 | AAACAATCGTGCCAACAACA | ATTTGGAAGGTGTTGGGTGA | 156 |
| chr1:83110674-83112688 | p2 | (TA)6 | AAACAATCGTGCCAACAACA | ATTTGGAAGGTGTTGGGTGA | 156 |
| chr1:83126292-83128322 | p2 | (TA)15 | GCAACTCCAACCATTTTCGT | GGCTGTGGTTTTTGCATTTT | 162 |
| chr1:83134907-83136965 | p2 | (AT)29 | CATAATCGAGGCGGAAACAT | AGGAACCCAATGGATTGACA | 147 |
| chr1:83153784-83155800 | p4 | (AGAA)4 | GGAGCAATAAGGTTTTGGCA | GTTTCCATAGGTGGGGTCCT | 184 |
| chr1:83153784-83155800 | p5 | (TTGAA)4 | GGAGCAATAAGGTTTTGGCA | GTTTCCATAGGTGGGGTCCT | 184 |
| chr1:83173739-83175751 | p2 | (GT)6 | TCTTGAATTGGGTCTCGTCC | TCCCATTTTGTAATTCCCGA | 254 |
| chr1:83173739-83175751 | p5 | (ATTCA)3 | TCTTGAATTGGGTCTCGTCC | TCCCATTTTGTAATTCCCGA | 254 |
| chr1:83213462-83215476 | p2 | (AG)7 | ATCGTCAAAGAAAACGGACG | CAGTTACAGATTGCTCCGCA | 188 |
| chr1:83220147-83222161 | p2 | (AT)7 | ACAAATTCCTGCCAAACAGG | CCGTGGAGTTGGCATCTTAT | 128 |
| chr1:83220147-83222161 | p2 | (AT)6 | ACAAATTCCTGCCAAACAGG | CCGTGGAGTTGGCATCTTAT | 128 |
| chr1:83228383-83230398 | p3 | (ACA)5 | AACTGACCGTAGAACGGTGG | AGGGGATATTTGGATCAGGC | 188 |
| chr1:83234865-83236903 | p3 | (ATC)5 | CCATTTCTAAGTGGGCCAAA | ATATACTTGGGCATGGCGAG | 255 |
| chr1:83234865-83236903 | p3 | (AAC)6 | CCATTTCTAAGTGGGCCAAA | ATATACTTGGGCATGGCGAG | 255 |

| Marker ID | SSR type | SSR motif | FORWARD PRIMER (5'-3') | REVERSE PRIMER (5'-3') | size (bp) |
|---|---|---|---|---|---|
| chr1:83234941-83236981 | p3 | (ATC)5 | CCATTTCTAAGTGGGCCAAA | ATATACTTGGGCATGGCGAG | 179 |
| chr1:83234941-83236981 | p3 | (AAC)6 | CCATTTCTAAGTGGGCCAAA | ATATACTTGGGCATGGCGAG | 179 |
| chr1:83272583-83274598 | p5 | (TCATA)3 | TATCCTCGTTCATCGAACCC | ATGGAATGAGAATTCAGGCG | 103 |
| chr1:83403563-83405578 | p5 | (ATGTC)3 | CCATTGTCGGAGGTGCTTAT | GCTCAGTTAGGGCCATCAAA | 147 |
| chr1:83420258-83422273 | p5 | (CAATT)3 | AAGGTCGGTAACCTCCGATT | CCATAACGAATATGTCGGGG | 208 |
| chr1:83498306-83500321 | p5 | (TCATT)3 | AAACAAAGCGTAGCTTCGGA | ATCATGAACCCCTTCTGTCG | 195 |
| chr1:83517625-83519637 | p2 | (TA)6 | AGGGAAGCATACACAATGCC | GTTTTGTCTGCCTGGTCCAT | 289 |
| chr1:83538120-83540135 | p3 | (ATC)5 | ATTATCCCCGAATCCTTTGC | CGCTCTATTTGTTTCGGAGC | 140 |
| chr1:83545957-83547973 | p2 | (AT)8 | ATATGACCCGACAAAGCTGC | ACATATGTGTATGGCGGGGT | 124 |
| chr1:83573919-83575933 | p2 | (AG)7 | GCCTCACATACCTTCGTCGT | AAGGAAGTCCTGCCCAATTT | 295 |
| chr1:83578800-83580814 | p2 | (AT)7 | CTTCATCCGGTTTAGACCCA | CAACAAGTACCGAGGCCATT | 168 |
| chr1:83612581-83614633 | p2 | (AT)26 | CCTAGCATGTCCCCTTTTCA | TATATGTTGGCTTGTGGGCA | 118 |
| chr1:83612581-83614633 | p2 | (TA)6 | CCTAGCATGTCCCCTTTTCA | TATATGTTGGCTTGTGGGCA | 118 |
| chr1:83625616-83627634 | p6 | (ATCCGT)3 | ACTCCACCGATTCAGTGTCC | TAGGGTGTGTCTACGAGCCC | 184 |
| chr1:83680721-83682739 | p6 | (CATATA)3 | CCGTTTGGCGAAGTAAATGT | TGGGACAATCATTGGGAACT | 130 |
| chr1:83747754-83749772 | p3 | (GAT)6 | CCGATCATCAAAGGGGTAAA | AAACGGCAGAAAGCGTTAGA | 297 |
| chr1:83839280-83841292 | p2 | (AT)6 | GACCTCACGTGAACCCTCAT | GAGGAGGAAGAAACCCAAGG | 114 |
| chr1:83898815-83900847 | p2 | (AT)30 | AAAATGATACGCAAGCCGAC | ACTGTCCCGTCATATTTCGC | 251 |
| chr1:83898815-83900847 | p2 | (TA)16 | AAAATGATACGCAAGCCGAC | ACTGTCCCGTCATATTTCGC | 251 |
| chr1:84014611-84016623 | p2 | (TA)6 | GCGACCACGATTTCCATAAT | TTTCCATCTTTCCCGTATGC | 210 |
| chr1:84020908-84022938 | p2 | (TA)15 | GCACCAGCAAATGAAGTCAA | CATGGATCTTCTTCCCGTGT | 257 |
| chr1:84146921-84148941 | p4 | (ATGA)5 | CTACCCCTTCAGTGTTCGGA | CACTGGTGATCTTGGGACCT | 288 |
| chr1:84198130-84200192 | p2 | (AT)31 | TTTGGAAAGGCTTACCGTTG | TTTCGCCTATCGGATGAAAC | 109 |
| chr1:84212338-84214356 | p6 | (TATATG)3 | GCCGCAATTTCGGTAGAATA | GCCACGCCTTATTAGCTACG | 103 |
| chr1:84229725-84231739 | p2 | (AT)7 | TCAAGAAATCACGGTCCCTC | GCTATGCGAGACGCAATACA | 169 |
| chr1:84241296-84243314 | p6 | (GTGTAT)3 | TTGAGTTGGACACGTGTGGT | TCACAGGGCCAACACATTTA | 136 |
| chr1:84313134-84315148 | p2 | (AT)7 | GCAAGTGCAAGTGAACGGTA | ATCCCCCGATCTTTTTATGG | 153 |
| chr1:84332360-84334374 | p2 | (AT)6 | AACATCGTGTGGGGAAATGT | CAGCCATCTTCCCCTGTTTA | 159 |
| chr1:84332360-84334374 | p2 | (TA)6 | AACATCGTGTGGGGAAATGT | CAGCCATCTTCCCCTGTTTA | 159 |
| chr1:84332400-84334414 | p2 | (AT)6 | AACATCGTGTGGGGAAATGT | CAGCCATCTTCCCCTGTTTA | 119 |
| chr1:84332400-84334414 | p2 | (TA)6 | AACATCGTGTGGGGAAATGT | CAGCCATCTTCCCCTGTTTA | 119 |
| chr1:84429620-84431635 | p5 | (AAAAC)3 | AAATCCTCACCTCACATGGC | TGGGCCTGAGCTTGAGTAGT | 295 |
| chr1:84442183-84444198 | p3 | (GAA)5 | CAAGGTGATGTACTGGGCCT | TGTCCCGGTGGTTAATGAAT | 166 |
| chr1:84530979-84532997 | p6 | (GTGGAC)3 | AAGGGCACATAATCGTGGAG | CCTAATTCACGTCCCGTTGT | 274 |
| chr1:84530979-84532997 | p6 | (GTGGAC)3 | AAGGGCACATAATCGTGGAG | CCTAATTCACGTCCCGTTGT | 274 |
| chr1:84530979-84532997 | p6 | (GTGGAT)3 | AAGGGCACATAATCGTGGAG | CCTAATTCACGTCCCGTTGT | 274 |

| Marker ID | SSR type | SSR motif | FORWARD PRIMER (5'-3') | REVERSE PRIMER (5'-3') | size (bp) |
|---|---|---|---|---|---|
| chr1:84531204-84533222 | p6 | (GTGGAC)3 | TCACAACGGGACGTGAATTA | CCACAATATCCACCATGCAG | 268 |
| chr1:84531204-84533222 | p6 | (GTGGAC)3 | TCACAACGGGACGTGAATTA | CCACAATATCCACCATGCAG | 268 |
| chr1:84531204-84533222 | p6 | (GTGGAT)3 | TCACAACGGGACGTGAATTA | CCACAATATCCACCATGCAG | 268 |
| chr1:84648337-84650351 | p2 | (AT)7 | GGGCCAGGTAAGGTATGTGA | TCCAGAGTAACGCTGCAATG | 267 |
| chr1:84906990-84909026 | p3 | (GTT)5 | TGGATAAACATCGTGTGGGA | TTTTACCAAAATTCGGGCAG | 178 |
| chr1:84906990-84909026 | p3 | (ATG)5 | TGGATAAACATCGTGTGGGA | TTTTACCAAAATTCGGGCAG | 178 |
| chr1:84966590-84968605 | p5 | (GCATT)3 | AAAGGCAAGGCAAAGAGTGA | TAAGGCAATATCCAACCCGA | 104 |
| chr1:84966590-84968605 | p2 | (AT)44 | AAAGGCAAGGCAAAGAGTGA | TAAGGCAATATCCAACCCGA | 104 |
| chr1:85057103-85059117 | p2 | (AT)7 | ATGTGTTACAGATGCCGCAA | TTTCTTCATTCCGTTCCACC | 158 |
| chr1:85057103-85059117 | p2 | (AT)6 | ATGTGTTACAGATGCCGCAA | TTTCTTCATTCCGTTCCACC | 158 |
| chr1:85130368-85132398 | p5 | (CATAT)6 | GGATGATCCGCTGGAGTTTA | TCTTTCGTGCGCTACATCTG | 233 |
| chr1:85244954-85246969 | p5 | (CCGAA)3 | TCTCCTCCCATCTTTCCCTT | ACCTGTGTCCATTTGGGGTA | 283 |
| chr1:85244954-85246969 | p5 | (AACCG)3 | TCTCCTCCCATCTTTCCCTT | ACCTGTGTCCATTTGGGGTA | 283 |
| chr1:85261322-85263337 | p5 | (TTTTG)3 | GTCGTACCCTGTGAAGGCAT | TTTAGGGAACGAGAACCCCT | 190 |
| chr1:85265249-85267291 | p3 | (TTA)14 | GGCGTTCCATGTCACCTTAT | TGTCTCATACCTTCCCCTCG | 267 |
| chr1:85290571-85292589 | p3 | (CAA)6 | ACGAGCGGAGTAGTGAGGAA | CCTCAGAAAAGCTCGGATTG | 282 |
| chr1:85316943-85318961 | p6 | (GACGAA)3 | GTCGAAGCTCCAACGAGAAC | TGTGTGTGCTTGTGTCCAGA | 157 |
| chr1:85316966-85318978 | p6 | (GACGAA)3 | GTCGAAGCTCCAACGAGAAC | TGTGTGTGCTTGTGTCCAGA | 134 |
| chr1:85343401-85345489 | p5 | (CAATG)3 | TCTTCTCCTCTGCTTCCCAA | ATTCCATGTTTCGATGCTCC | 222 |
| chr1:85364537-85366553 | p2 | (AT)8 | TTTTGGTCGAGCATCAGTTG | TTTCATTTTTGTGGGGGAAA | 238 |
| chr1:85461471-85463501 | p2 | (TA)13 | AAAGGTGCCTTTGATGATGG | CAAACGCCACCAATCTTTTT | 183 |
| chr1:85461471-85463501 | p2 | (TA)15 | AAAGGTGCCTTTGATGATGG | CAAACGCCACCAATCTTTTT | 183 |
| chr1:85469758-85471774 | p4 | (TTTC)4 | ACGGTTACTTTGGCGATGAC | TGCACTCAAAAGAATGCAGC | 232 |
| chr1:85488730-85490742 | p2 | (AT)13 | CTGGAGCCAAATCAAACGAT | ACTTTGGCCGAATTTCCTTT | 230 |
| chr1:85488730-85490742 | p2 | (AT)6 | CTGGAGCCAAATCAAACGAT | ACTTTGGCCGAATTTCCTTT | 230 |
| chr1:85512433-85514448 | p5 | (AGAAA)3 | GCCATTGTCAGGCTTGTTTT | CAGAGGGGTGTGCTCAATTT | 182 |
| chr1:85530867-85532882 | p2 | (TA)6 | ATATTGGGCCAAAGGCTTCT | CCACATTTGGACATTGCTTG | 111 |
| chr1:85530867-85532882 | p5 | (GAAAT)3 | ATATTGGGCCAAAGGCTTCT | CCACATTTGGACATTGCTTG | 111 |
| chr1:85607298-85609314 | p4 | (ACAT)4 | ATTTTGTGAGAGATTGCCCG | CGCCGTATCTGTATTGGCTT | 141 |
| chr1:85667730-85669745 | p5 | (AAGTT)3 | GGTACATTGCCCGCTACAGT | GTTTTTGAGGCCTTGGATGA | 140 |
| chr1:85741193-85743205 | p2 | (TA)6 | ACACTTGCGATAAGAACGGG | AGATCGCTGAGTTTGTGGCT | 188 |
| chr1:85771588-85773602 | p2 | (TA)7 | TTGTGCAAAGGGTTGGTGTA | AAGTTGGGTAGCATTTCCCC | 245 |
| chr1:85815672-85817704 | p2 | (TA)16 | GAAAGCCTTAGCGACACCTG | GTGTTGGTGTTGGTCGTGAG | 155 |
| chr1:85815672-85817704 | p3 | (AAC)5 | GAAAGCCTTAGCGACACCTG | GTGTTGGTGTTGGTCGTGAG | 155 |
| chr1:86072328-86074343 | p3 | (CAA)5 | ATGGGACTCGGTGAATGAAG | ACTAGAGGGCCAGGTACGGT | 109 |
| chr1:86233218-86235250 | p2 | (AT)16 | GACAGGCACTCGGAGCTAAC | GCGTCCCTTCTGTTTGTGAT | 209 |

| Marker ID | SSR type | SSR motif | FORWARD PRIMER (5'-3') | REVERSE PRIMER (5'-3') | size (bp) |
|---|---|---|---|---|---|
| chr1:86275712-86277732 | p2 | (TA)10 | CTAGAGTGGCATGACCAGCA | CCCAATTCAGACGAGATGGT | 296 |
| chr1:86333270-86335282 | p2 | (CA)6 | CTGAACAAACTCGCACGAAA | ATCATGACCGTGATTTGGGT | 156 |
| chr1:86357429-86359471 | p2 | (TA)10 | AGGTGCGCTTTCATGAAGTT | TCTCGACAAGCTGAGGGAGT | 233 |
| chr1:86357429-86359471 | p2 | (TA)14 | AGGTGCGCTTTCATGAAGTT | TCTCGACAAGCTGAGGGAGT | 233 |
| chr1:86422068-86424083 | p5 | (TCACA)3 | GCGCACAACACACCATAAAC | ACGAGCGGTGCTCAGTAGTT | 276 |
| chr1:86431037-86433055 | p2 | (AT)9 | ACCCAAGTGAAGGGGAAACT | TTCCCACATTCTTGACCCAT | 139 |
| chr1:86459883-86461895 | p2 | (AT)6 | GATGTCAAGAACTCGGGGAA | TTTGGGCCTCAACTTTGTTC | 245 |
| chr1:86749897-86751909 | p2 | (TA)6 | TTGGAGAGCTCCATTGTTCC | TACGCAAGATGACAAGGCTG | 168 |
| chr1:86769003-86771021 | p6 | (CATATA)3 | GACCCTTACTTAGGGCCGAG | CAGTCAGCCGATGTGTCATT | 117 |
| chr1:86780719-86782731 | p2 | (AT)6 | GCCTGTTATGCCCAAACTGT | CAACCTACCTGGATAGCCGA | 143 |
| chr1:86838238-86840253 | p5 | (TAATA)3 | TCTTAAACTCCCACGGCATC | GACACGACCCGAAGACATTT | 129 |
| chr1:86908273-86910313 | p2 | (TA)26 | TTTGACCCTTCCCCTAGCTT | GATCGTGACATTTCCGACCT | 109 |
| chr1:86908273-86910313 | p2 | (AT)20 | TTTGACCCTTCCCCTAGCTT | GATCGTGACATTTCCGACCT | 109 |
| chr1:86925810-86927825 | p2 | (AT)7 | TTGTGGCCACTTAGCTTTCC | AATACACAGGAGGTGCAGGG | 100 |
| chr1:86926170-86928184 | p2 | (AT)7 | AGCCGACAAGGCAACTAAAA | TCTTTGGATGCTGCGTTATG | 174 |
| chr1:86986933-86988955 | p2 | (AT)11 | CAAGCCGAGTAAACAGAGCC | TTTCGGAGCCTCACTTCTGT | 149 |
| chr1:87097267-87099289 | p2 | (AT)11 | GTTGCAAAATCTTCCCCAAA | TTAAGGTGAACATGAGGCCC | 143 |
| chr1:87184893-87186945 | p6 | (TATTCG)3 | ATAAGCCGACGAGGCTAACA | GTCCTGTCCCGCCTATATGA | 202 |
| chr1:87199266-87201294 | p2 | (AT)14 | TGGAGAGTCCTCGGAAGCTA | TGATTTTGTGGATTGACCGA | 166 |
| chr1:87202184-87204199 | p5 | (AAAGG)3 | TATAGCGATCACCCTACCGC | GATTGTCCCCCATGAATCAC | 300 |
| chr1:87210530-87212542 | p2 | (TG)6 | TGTAGAGGTGGGTCCCAGAG | TTCCAAATTTGCCCTTTGTC | 114 |
| chr1:87449843-87451857 | p5 | (TATTT)3 | AGAGGGATCCTCCCGAAGTA | TGCTCTTAACTTCCCGCAAT | 201 |
| chr1:87449843-87451857 | p2 | (AG)7 | AGAGGGATCCTCCCGAAGTA | TGCTCTTAACTTCCCGCAAT | 201 |
| chr1:87494187-87496217 | p2 | (TA)15 | TTTGGGGGATTACAGGTCAA | TCAGAGGAGCCTAGAGCAGC | 209 |
| chr1:87528229-87530254 | p3 | (TTC)5 | AACAATGCAGAGGAAAACGC | TGTGCATATGGTTTCGCACT | 237 |
| chr1:87584446-87586482 | p2 | (TA)18 | CCGTGGTACAGGTGGTCTCT | CCAGTTAGTAGCCGCTCAGG | 161 |
| chr1:87584446-87586482 | p2 | (TA)7 | CCGTGGTACAGGTGGTCTCT | CCAGTTAGTAGCCGCTCAGG | 161 |
| chr1:87596943-87598958 | p5 | (AAAAT)3 | CGGAAACTTGATGCAATCCT | GGGTTCGTCCACCCTAAAAT | 144 |
| chr1:87633520-87635532 | p2 | (TA)6 | ACAAAATATGCAAGCCGGAC | TTGTTCCGGAGCTTGAGTTT | 242 |
| chr1:87633520-87635532 | p3 | (CAA)5 | ACAAAATATGCAAGCCGGAC | TTGTTCCGGAGCTTGAGTTT | 242 |
| chr1:87764779-87766809 | p2 | (AT)15 | ACCACCTTCCATCTTGCATC | AGCAATGAGTTTCCCTGGTG | 260 |
| chr1:87765350-87767432 | p2 | (AT)15 | GGCACGGCTCATCATACTTT | GAGGCTCGTAGACAGTTGGG | 300 |
| chr1:87828084-87830096 | p2 | (TC)6 | CCGGTGTTTACTCCCTCTCA | TTGGAGAATCGAAGGCAGTT | 266 |
| chr1:87984359-87986381 | p2 | (TA)11 | TTAGCCCGAAATCACAATCC | TTTGAGCATGTTGTGGTCGT | 104 |
| chr1:88013275-88015290 | p5 | (TATTT)3 | TGACCAAAATTGAGGTCGAAA | ATTGCCACGATTGTCATCAA | 163 |
| chr1:88013275-88015290 | p2 | (AT)12 | TGACCAAAATTGAGGTCGAAA | ATTGCCACGATTGTCATCAA | 163 |

| Marker ID | SSR type | SSR motif | FORWARD PRIMER (5'-3') | REVERSE PRIMER (5'-3') | size (bp) |
|---|---|---|---|---|---|
| chr1:88016488-88018503 | p5 | (AACCG)3 | TTGATGAACAAAAGGGGGAG | TTTCGGTATTTATGCCCAGC | 194 |
| chr1:88016488-88018503 | p5 | (AACCG)3 | TTGATGAACAAAAGGGGGAG | TTTCGGTATTTATGCCCAGC | 194 |
| chr1:88272400-88274420 | p2 | (AT)14 | GCCACCGATTTCGATTTTTA | GAAAACCCCTTCTTTTTCCG | 240 |
| chr1:88272400-88274420 | p6 | (AACCGG)3 | GCCACCGATTTCGATTTTTA | GAAAACCCCTTCTTTTTCCG | 240 |
| chr1:88272400-88274420 | p2 | (TA)10 | GCCACCGATTTCGATTTTTA | GAAAACCCCTTCTTTTTCCG | 240 |
| chr1:88272400-88274420 | p2 | (TA)6 | GCCACCGATTTCGATTTTTA | GAAAACCCCTTCTTTTTCCG | 240 |
| chr1:88272400-88274420 | p2 | (AT)10 | GCCACCGATTTCGATTTTTA | GAAAACCCCTTCTTTTTCCG | 240 |
| chr1:88396925-88398941 | p2 | (AT)11 | TCCAGATAACCTGTCCCCTG | TGTGCAATGGATTTGTTCGT | 288 |
| chr1:88396925-88398941 | p2 | (AT)8 | TCCAGATAACCTGTCCCCTG | TGTGCAATGGATTTGTTCGT | 288 |
| chr1:88396925-88398941 | p3 | (ACA)5 | TCCAGATAACCTGTCCCCTG | TGTGCAATGGATTTGTTCGT | 288 |
| chr1:88440638-88442656 | p3 | (TTG)6 | GACCCTTGAGCCAAATCAAA | CTTATCCCGACGTCACCTGT | 182 |
| chr1:88474217-88476235 | p3 | (CAA)6 | ACATCACAACAAAGGCCACA | GCACAGTTGACAGGGAGTGA | 178 |
| chr1:88479139-88481154 | p5 | (ATGAT)3 | GATCGGGACTTGAGTTTGGA | GGTCGTAATTGTGCTCGGAT | 174 |
| chr1:88519781-88521796 | p5 | (TTTTA)3 | AAACCGTTCGTCGTACCTTG | GTAATCCACCCCCATTTCCT | 195 |
| chr1:88522997-88525012 | p5 | (TTGCA)3 | AGAAGGGAAGGAGTTTGGGA | ACCCACTTGATGATCCTTGC | 144 |
| chr1:88620311-88622323 | p2 | (AT)6 | AAGAGACAACCTCGCCTCAA | GTATGGTTAGCTTGGCGCAT | 258 |
| chr1:88656728-88658743 | p5 | (ATTGC)3 | TGGTGTATCCGCTGTTGGTA | ATGAGAGAATGCGTGCAATG | 288 |
| chr1:88656728-88658743 | p5 | (TATAT)3 | TGGTGTATCCGCTGTTGGTA | ATGAGAGAATGCGTGCAATG | 288 |
| chr1:88657285-88659300 | p5 | (ATTGC)3 | TAGTCAGGGTCCCGTCTGTC | GCAATGCAATGCAATAATGG | 281 |
| chr1:88657285-88659300 | p5 | (TATAT)3 | TAGTCAGGGTCCCGTCTGTC | GCAATGCAATGCAATAATGG | 281 |
| chr1:88676018-88678038 | p2 | (TA)10 | TCCGGTAGGATTTGACTTGG | ACGGAGTTGGTATGCGGTAG | 128 |
| chr1:88858975-88860993 | p2 | (AT)6 | ATATATACCGTACCCGGCCC | GTGAGCGAGCGATGAACATA | 198 |
| chr1:88858975-88860993 | p3 | (CAA)6 | ATATATACCGTACCCGGCCC | GTGAGCGAGCGATGAACATA | 198 |
| chr1:88909382-88911559 | p3 | (ATG)6 | TCGAAGCTGTGAGTGTTTGG | AACAACCCGTTTGGTCGATA | 262 |
| chr1:89050049-89052061 | p2 | (GT)6 | AATCCGTAGCTTTCTGGGGT | CTTTGGCCCAATTCTTGAAA | 287 |
| chr1:89050049-89052061 | p2 | (GT)6 | AATCCGTAGCTTTCTGGGGT | CTTTGGCCCAATTCTTGAAA | 287 |
| chr1:89050049-89052061 | p3 | (TGT)5 | AATCCGTAGCTTTCTGGGGT | CTTTGGCCCAATTCTTGAAA | 287 |
| chr1:89050190-89052202 | p2 | (GT)6 | AATCCGTAGCTTTCTGGGGT | CTTTGGCCCAATTCTTGAAA | 146 |
| chr1:89050190-89052202 | p2 | (GT)6 | AATCCGTAGCTTTCTGGGGT | CTTTGGCCCAATTCTTGAAA | 146 |
| chr1:89050190-89052202 | p3 | (TGT)5 | AATCCGTAGCTTTCTGGGGT | CTTTGGCCCAATTCTTGAAA | 146 |
| chr1:89138030-89140048 | p6 | (ACACTC)3 | AACTCGATGGACCGTCTTTG | CAAAAAGACGGGAAGGTTGA | 205 |
| chr1:89155847-89157877 | p2 | (TA)15 | TGGATGGCGAGAATGTGATA | ATCTCCAGCAAAACCGATTG | 263 |
| chr1:89168228-89170246 | p6 | (ATATGT)3 | ATGTTGGATATGAGCCTCGG | GTCGGGTCCTATTTATGCGA | 226 |
| chr1:89206550-89208564 | p2 | (AT)7 | TGTCCGAGGCATGTATCAAA | TCATTACCTTGAACCCTCCG | 280 |
| chr1:89275076-89277094 | p6 | (TTTTAT)3 | TTACTCCTAGCCCTCCCACC | TGCACTGTGGCAAATAATCC | 115 |
| chr1:89446820-89448834 | p2 | (TA)7 | ACGACCTGGTTTACGGTCTG | TCCAGTGGTCGTATATGGCA | 295 |

| Marker ID | SSR type | SSR motif | FORWARD PRIMER (5'-3') | REVERSE PRIMER (5'-3') | size (bp) |
|---|---|---|---|---|---|
| chr1:89452125-89454143 | p6 | (ATACAT)3 | CACGGTTGGGTTTAGATGCT | TCGTGTTATTTGGTGCATGG | 235 |
| chr1:89475819-89477831 | p2 | (AT)6 | CGCAGGCCCCATATAAGTAA | CATGCAGAAGATTGAAGCGA | 122 |
| chr1:89475819-89477831 | p2 | (TA)6 | CGCAGGCCCCATATAAGTAA | CATGCAGAAGATTGAAGCGA | 122 |
| chr1:89490198-89492210 | p2 | (AT)6 | TATCAAATCCGGTAGGCTGG | TCCTTTGTGAGGGAGGTTTG | 131 |
| chr1:89662060-89664075 | p5 | (GGTGC)3 | GAGGCGAGTGTTCGAGGTAG | GGGCAAGAACCTCATTTTCA | 219 |
| chr1:89662060-89664075 | p5 | (GGTGC)3 | GAGGCGAGTGTTCGAGGTAG | GGGCAAGAACCTCATTTTCA | 219 |
| chr1:89843126-89845144 | p3 | (ATT)6 | TCCTAGGCCCCTCAAGGTAT | CCCTTCAACTTCCCATTCAA | 185 |
| chr1:89909808-89911820 | p2 | (TA)9 | GTCCAGACCAGATTTTGGGA | TATCCTCTTGTGGGCAAACC | 140 |
| chr1:89909808-89911820 | p2 | (AT)6 | GTCCAGACCAGATTTTGGGA | TATCCTCTTGTGGGCAAACC | 140 |
| chr1:89966550-89968584 | p2 | (TG)7 | CGCATTGTGTTTTTCATTGG | ACATAAGGTCGAGAACCCCC | 104 |
| chr1:90047442-90049454 | p2 | (AC)6 | CAGCTTGTAGACTCCCTGCC | TTTGGAAAAATCCACGCTTC | 184 |
| chr1:90063704-90065722 | p2 | (AT)9 | GCACATATGAGGCCCCTAAA | GCAACTTTGGAGCCAATTTC | 148 |
| chr1:90071649-90073669 | p2 | (TA)10 | CCCAACTAATTTGGCATTGG | TTGCCACCAACAGGTCAGTA | 287 |
| chr1:90071649-90073669 | p2 | (AT)8 | CCCAACTAATTTGGCATTGG | TTGCCACCAACAGGTCAGTA | 287 |
| chr1:90081362-90083428 | p2 | (TA)33 | GAACCCACGACCTATGGATG | TTCGGGTAACCTGACTTTGG | 192 |
| chr1:90081362-90083428 | p5 | (TTTCA)3 | GAACCCACGACCTATGGATG | TTCGGGTAACCTGACTTTGG | 192 |
| chr1:90094775-90096789 | p2 | (AC)7 | GCCCTTCACGAATTGTGTTT | GCCACTAGAATGGCTAACGC | 134 |
| chr1:90220329-90222341 | p2 | (TA)6 | CGCACTCGGAGCTAACCTAC | GGTACCTTGTCGGCTTGTGT | 220 |
| chr1:90234192-90236204 | p2 | (CA)6 | CCCTCACCCTTTTCAAATCA | GCTTGGTTGTTGTTTGGGTT | 296 |
| chr1:90234192-90236204 | p2 | (CA)6 | CCCTCACCCTTTTCAAATCA | GCTTGGTTGTTGTTTGGGTT | 296 |
| chr1:90268717-90270729 | p2 | (AT)6 | GGACGCTTTTCCTTAGGGTC | CCCACAAAGGGTAATCGAGA | 129 |
| chr1:90268717-90270729 | p2 | (AT)6 | GGACGCTTTTCCTTAGGGTC | CCCACAAAGGGTAATCGAGA | 129 |
| chr1:90281533-90283551 | p6 | (TATATG)3 | TTGGCCGAGTTAAATTCTCG | GGGATCATGCCTTGAAAGAA | 208 |
| chr1:90334134-90336146 | p2 | (TA)6 | GATTTGGGGAATGACATTGG | AGGAACTAGCGGGGGTAAAA | 144 |
| chr1:90346147-90348162 | p3 | (ATG)5 | CGCAACAAACCTCTCATCAA | GACCTAGAATTTTCGCGCAC | 109 |
| chr1:90420063-90422099 | p3 | (TGA)12 | GCTGGCCAGACCTCTACAAC | ACGTAGGGTTTTGCATGGAG | 179 |
| chr1:90465284-90467298 | p2 | (TA)7 | CCAACTAGTGGGGCTGTGAT | TAGGCTCAACATGCTTTCCC | 210 |
| chr1:90465284-90467298 | p2 | (TG)8 | CCAACTAGTGGGGCTGTGAT | TAGGCTCAACATGCTTTCCC | 210 |
| chr1:90495574-90497589 | p3 | (AGA)5 | CTGGTGCAGACTTTGCGTTA | TCGCTCCTAAATGGATCACC | 159 |
| chr1:90495574-90497589 | p2 | (GT)12 | CTGGTGCAGACTTTGCGTTA | TCGCTCCTAAATGGATCACC | 159 |
| chr1:90729364-90731379 | p5 | (ATCAT)3 | ACAACCATGCAAAAACACCA | CAGCAGACAGGTGGAGTTGA | 174 |
| chr1:90832460-90834520 | p2 | (AT)23 | AAACCACACTTTACGGGTCG | CCGTATACTGCCATCGTCCT | 122 |
| chr1:90832460-90834520 | p2 | (AC)7 | AAACCACACTTTACGGGTCG | CCGTATACTGCCATCGTCCT | 122 |
| chr1:90832460-90834520 | p2 | (AT)30 | AAACCACACTTTACGGGTCG | CCGTATACTGCCATCGTCCT | 122 |
| chr1:90846471-90848485 | p2 | (AT)7 | GTTGTCCAATTGGGACTCGT | CGGCCATATAAGCTTGGTGT | 273 |
| chr1:90846471-90848485 | p2 | (GT)7 | GTTGTCCAATTGGGACTCGT | CGGCCATATAAGCTTGGTGT | 273 |

| Marker ID | SSR type | SSR motif | FORWARD PRIMER (5'-3') | REVERSE PRIMER (5'-3') | size (bp) |
|---|---|---|---|---|---|
| chr1:90846471-90848485 | p5 | (AATTG)5 | GTTGTCCAATTGGGACTCGT | CGGCCATATAAGCTTGGTGT | 273 |
| chr1:90848152-90850194 | p5 | (AATTG)5 | ATTGGAAGGTGTTGGATGGA | GAGAATTCAAATCGGCCAAA | 126 |
| chr1:90848152-90850194 | p6 | (TATATG)7 | ATTGGAAGGTGTTGGATGGA | GAGAATTCAAATCGGCCAAA | 126 |
| chr1:90848152-90850194 | p6 | (TATATG)3 | ATTGGAAGGTGTTGGATGGA | GAGAATTCAAATCGGCCAAA | 126 |
| chr1:90848152-90850194 | p2 | (TA)12 | ATTGGAAGGTGTTGGATGGA | GAGAATTCAAATCGGCCAAA | 126 |
| chr1:90868490-90870502 | p2 | (AC)6 | CCAAACTCACTTGCCGTTTT | AGAGGCGAGTTGAATCCTCA | 197 |
| chr1:90891559-90893574 | p5 | (TCTTT)3 | GGTGGTCTTGTGGCCTTAAA | CCATCCCAAAACTTCCTTCA | 143 |
| chr1:90891559-90893574 | p3 | (TGA)5 | GGTGGTCTTGTGGCCTTAAA | CCATCCCAAAACTTCCTTCA | 143 |
| chr1:90891559-90893574 | p2 | (TA)20 | GGTGGTCTTGTGGCCTTAAA | CCATCCCAAAACTTCCTTCA | 143 |
| chr1:90901823-90903837 | p2 | (AT)7 | GCATTTTCAGAGCATGAGCA | CTTCCACTTCGATTTGGCTC | 143 |
| chr1:90932424-90934464 | p2 | (AT)20 | TGTTACACCCCTCGTTTTCC | CGATGATGATTGGTGTCTGC | 226 |
| chr1:90943132-90945148 | p2 | (TA)8 | GTTTTCCGCTTTTTGGTTCA | AGTCGGGTCCCATTCATACA | 167 |
| chr1:91078577-91080589 | p4 | (TATG)5 | GATAACACCGAGTCCCGAAA | AATGTTATCTTACCCGGCCC | 214 |
| chr1:91078577-91080589 | p2 | (AT)6 | GATAACACCGAGTCCCGAAA | AATGTTATCTTACCCGGCCC | 214 |
| chr1:91172597-91174612 | p5 | (CATAA)3 | CCGTCTCCCATAATCTCGAA | GAAACAGCGACATCATCCCT | 191 |
| chr1:91246443-91248458 | p5 | (AATTG)3 | ACGAGAAATTCGCAAAATGG | CATGGCTCATGAAAGTGTGG | 253 |
| chr1:91246443-91248458 | p6 | (TATATG)3 | ACGAGAAATTCGCAAAATGG | CATGGCTCATGAAAGTGTGG | 253 |
| chr1:91257663-91259678 | p5 | (TATGA)3 | TTAGAATCTCGGGGATGGTG | CCGCGCCTTACTAGCTACAC | 191 |
| chr1:91265127-91267145 | p6 | (TATATG)3 | CAACAATCCACCCAGCTTTT | GTTCTCCGCTGTCGAGGTAG | 230 |
| chr1:91265127-91267145 | p5 | (TATGA)3 | CAACAATCCACCCAGCTTTT | GTTCTCCGCTGTCGAGGTAG | 230 |
| chr1:91309914-91311932 | p3 | (ACA)6 | CTTTCCATCATTTGCCCACT | TGCGACGTGAATTTGGAATA | 225 |
| chr1:91366085-91368097 | p2 | (AG)6 | TGGCCAAGGAGTTAAAATGG | GCTCCGGTTGATACATGCTT | 137 |
| chr1:91399795-91401815 | p5 | (ATGAT)4 | AGATGTATGGCCTCGGTCAC | GATTTAAGAAATGCCGCCAA | 106 |
| chr1:91410437-91412455 | p2 | (AT)12 | TGTACCGAGCACCCCTTATC | TGGTTGTCCCTGTCGATGTA | 146 |
| chr1:91410437-91412455 | p6 | (CATATA)3 | TGTACCGAGCACCCCTTATC | TGGTTGTCCCTGTCGATGTA | 146 |
| chr1:91427403-91429431 | p2 | (TG)6 | CCGTACTGACACCCGTTTCT | GTACCAAAGTAGGCACCGGA | 109 |
| chr1:91427403-91429431 | p2 | (TG)14 | CCGTACTGACACCCGTTTCT | GTACCAAAGTAGGCACCGGA | 109 |
| chr1:91452668-91454680 | p2 | (AG)6 | AGAGGAATGTGCGGTATTCG | CTAGCCACGCCATGGTTACT | 189 |
| chr1:91493394-91495412 | p6 | (CATATA)3 | CCCGACCCTTACTTAGAGCC | TTATAACCGGCTCCATCAGC | 109 |
| chr1:91543083-91545097 | p2 | (AT)7 | TGTGCAAGCTGTCCAAAGTC | GCCCAAAAACACTTATCCCA | 230 |
| chr1:91554657-91556671 | p2 | (TA)7 | TGTGCAAGCTGACCAAAGTC | TACTTGCATGCCCAAAAACA | 198 |
| chr1:91581229-91583273 | p2 | (AT)22 | CCCGAGATGGTGAAGAAGAA | TCAAATCCATCCCCAATCAT | 281 |
| chr1:91595358-91597372 | p2 | (AT)7 | AAAGGAACCTCTGCCCAAAT | CTTCACATGGGCTGTGCTAA | 127 |
| chr1:91600346-91602361 | p5 | (CATAT)3 | TCATCTTTGTTTTCCAGGGC | GTGTGATTTGGAATGCATCG | 179 |
| chr1:91662882-91664900 | p3 | (AAC)6 | ACGTGGTGCTTTTCCATAGG | TTGGCGAGAGATGAAAAGGT | 128 |
| chr1:91662882-91664900 | p2 | (CT)6 | ACGTGGTGCTTTTCCATAGG | TTGGCGAGAGATGAAAAGGT | 128 |

| Marker ID | SSR type | SSR motif | FORWARD PRIMER (5'-3') | REVERSE PRIMER (5'-3') | size (bp) |
|---|---|---|---|---|---|
| chr1:91741583-91743595 | p2 | (CA)6 | ATGCCGAAGGAAAGAAAGGT | GGTGCGCATTAATGTTGTTG | 233 |
| chr1:91790532-91792544 | p2 | (AC)6 | AATTCAAATTCCCGGTTTCC | TTGGCCGAGTTAAATTCTCG | 197 |
| chr1:91811404-91813428 | p6 | (CATATA)4 | TCACTTGGACTCGCATTCTG | CCATCCCGCCTAATGTAGAA | 143 |
| chr1:91844208-91846228 | p2 | (AT)10 | CTATCTACACCTGCGGGCAT | ATATGAGTGTCCAGCTCGGG | 101 |
| chr1:91844208-91846228 | p3 | (ACA)5 | CTATCTACACCTGCGGGCAT | ATATGAGTGTCCAGCTCGGG | 101 |
| chr1:92174418-92176482 | p2 | (TA)32 | GAGCCTCAGGGACAACTCTG | CGGTTCCCATTTATGCAAGT | 233 |
| chr1:92204381-92206399 | p2 | (TA)9 | TTCAGCTCGGCCAAATTATC | AATGCCGTTGAATTGTGTGA | 100 |
| chr1:92320040-92322054 | p2 | (TG)7 | TAGCCAGTACCAGGGGTTTG | GGGGTAGGAGAATGCCAAAT | 158 |
| chr1:92374807-92376819 | p2 | (AT)6 | CCTTGTTGTTGGGCTTGAGT | ATTAGGGGAGCAAGAAGGGA | 217 |
| chr1:92378856-92380871 | p5 | (AATGC)3 | TGCAAATTACTGGCTAGGGG | TGCACACAATAGGGATTCCA | 164 |
| chr1:92414522-92416540 | p6 | (TCCATA)3 | ACGAAATTTCCCATCGTGAG | GCTTGTGGCCTGTAGACTCC | 190 |
| chr1:92431883-92433901 | p3 | (TGT)6 | AGTGAAGGGATCGACCAATG | GTCCTTGCCTCATATCCCAA | 109 |
| chr1:92431883-92433901 | p2 | (GT)6 | AGTGAAGGGATCGACCAATG | GTCCTTGCCTCATATCCCAA | 109 |
| chr1:92442188-92444216 | p2 | (AG)14 | ACGTCTACGCGGAAGAGGTA | CTCTCCCTTTCATTTCGCAG | 208 |
| chr1:92442188-92444216 | p3 | (TGT)5 | ACGTCTACGCGGAAGAGGTA | CTCTCCCTTTCATTTCGCAG | 208 |
| chr1:92448675-92450687 | p2 | (AT)6 | CATATCGGTGGAGGAGCATT | TCTTTGGGTCAAAATCGGAG | 114 |
| chr1:92448675-92450687 | p4 | (CATA)6 | CATATCGGTGGAGGAGCATT | TCTTTGGGTCAAAATCGGAG | 114 |
| chr1:92487774-92489789 | p2 | (TA)7 | GTGTACATACATGGGCACGG | AGCCGACAAGGCTAAAGACA | 150 |
| chr1:92487774-92489789 | p5 | (ATCAT)3 | GTGTACATACATGGGCACGG | AGCCGACAAGGCTAAAGACA | 150 |
| chr1:92550372-92552390 | p6 | (CATATA)3 | ATTTCGCTAGGGTGTGATGG | TCAGTTCGGTAAATGGCTCC | 110 |
| chr1:92687060-92689075 | p5 | (TGATA)3 | CTTTTGATTCGGAGCCCATA | AACAGACGGACCGCATTAAG | 107 |
| chr1:92687060-92689075 | p2 | (AT)6 | CTTTTGATTCGGAGCCCATA | AACAGACGGACCGCATTAAG | 107 |
| chr1:92860298-92862328 | p2 | (TA)13 | TCCATAATACAATCGGGGGA | GGTGTCTACCCATGCCATTC | 267 |
| chr1:92860298-92862328 | p2 | (TA)15 | TCCATAATACAATCGGGGGA | GGTGTCTACCCATGCCATTC | 267 |
| chr1:92927911-92929941 | p2 | (TA)13 | TCCATAATACAATCGGGGGA | GGTGTCTACCCATGCCATTC | 267 |
| chr1:92927911-92929941 | p2 | (TA)15 | TCCATAATACAATCGGGGGA | GGTGTCTACCCATGCCATTC | 267 |
| chr1:92969294-92971309 | p3 | (ATC)5 | TCGTATGAGCGGGATAGGAC | TCTTGTTTGTGGACGCTGAG | 259 |
| chr1:92979922-92981934 | p2 | (AT)6 | CACCAAGTTGCAAAGGGACT | AGGGCCCCATTAATTTTGAC | 207 |
| chr1:93150433-93152451 | p6 | (CATATA)3 | ACTCACGGCCAACACATACA | TCAGCAGCAGTCAGTACGCT | 233 |
| chr1:93157488-93159506 | p6 | (CATATA)3 | ACTCACGGCCAACACATACA | TCAGCAGCAGTCAGTACGCT | 233 |
| chr1:93212312-93214330 | p3 | (TTG)6 | TTTGGAAGCCACATGGTGTA | TTCTTGAATCGTCCCTTTGG | 101 |
| chr1:93287936-93289948 | p2 | (AT)6 | ATCTTCCGTACTGACCCCCT | CCCAAAATATGGCTCCTGAA | 256 |
| chr1:93312013-93314031 | p3 | (TTG)6 | CTTGCACCAAAACCCAAACT | CTCCGCACATCAAGAAACCT | 173 |
| chr1:93457161-93459183 | p2 | (TA)11 | CTACCCGCCGTTTTATCGTA | CATGAGGCTACAGGGCATTT | 107 |
| chr1:93465414-93467432 | p6 | (TATGAA)3 | ATGAATCACCTGGTTGGAGC | ATGGCTGGCGTACAACCTAC | 142 |
| chr1:93560616-93562631 | p5 | (TGATA)3 | GAAACGGGAATTGAGTTGGA | GGGCAATCCCATATCCTCTT | 196 |

| Marker ID | SSR type | SSR motif | FORWARD PRIMER (5'-3') | REVERSE PRIMER (5'-3') | size (bp) |
|---|---|---|---|---|---|
| chr1:93604948-93606963 | p3 | (TTA)6 | CGGGTCCTGTTTATGCAAGT | AAACCCCATTCTCCGAAATC | 187 |
| chr1:93648183-93650211 | p2 | (AT)14 | TGAACCTTGGGTGGAAACTC | CCCTTTTCACTCCCCTTCTC | 217 |
| chr1:93681673-93683688 | p5 | (TATCG)3 | ATGCTTTGAAGATATGGCGG | TTGGATTTTCCCGGATGATA | 239 |
| chr1:93681673-93683688 | p2 | (TA)8 | ATGCTTTGAAGATATGGCGG | TTGGATTTTCCCGGATGATA | 239 |
| chr1:93897702-93899720 | p2 | (AT)9 | AAACTCATGAAATCACGGGC | CACCGCCCAATTTAGAAGAA | 281 |
| chr1:93978379-93980394 | p5 | (AAATG)3 | TTGGTCATGTGCTTGGTCAT | TAATGCTCCCCACTCCAATC | 137 |
| chr1:94047012-94049026 | p2 | (AT)7 | GGGCATCTAGGGATTGTTGA | GCCCTTGTGACCTTTCCATA | 278 |
| chr1:94047012-94049026 | p2 | (TA)31 | GGGCATCTAGGGATTGTTGA | GCCCTTGTGACCTTTCCATA | 278 |
| chr1:94047187-94049249 | p2 | (AT)7 | GGGCATCTAGGGATTGTTGA | GCCCTTGTGACCTTTCCATA | 103 |
| chr1:94047187-94049249 | p2 | (TA)31 | GGGCATCTAGGGATTGTTGA | GCCCTTGTGACCTTTCCATA | 103 |
| chr1:94063106-94065120 | p2 | (TA)7 | ACTGAAGAGCGAGGCACAAT | GCAAAAGGGATTCACGGATA | 103 |
| chr1:94364606-94366621 | p2 | (AT)6 | GTGCTCCTCTTATTCCGCAG | ATCAGAAGCCGACAAGGCTA | 100 |
| chr1:94364606-94366621 | p3 | (AAC)5 | GTGCTCCTCTTATTCCGCAG | ATCAGAAGCCGACAAGGCTA | 100 |
| chr1:94364606-94366621 | p2 | (TC)8 | GTGCTCCTCTTATTCCGCAG | ATCAGAAGCCGACAAGGCTA | 100 |
| chr1:94393388-94395406 | p2 | (AT)6 | CCGGAGCCTACATTTTGGTA | ATCAGAAGCCGACAAGGCTA | 115 |
| chr1:94393388-94395406 | p3 | (AAC)6 | CCGGAGCCTACATTTTGGTA | ATCAGAAGCCGACAAGGCTA | 115 |
| chr1:94393388-94395406 | p2 | (TC)6 | CCGGAGCCTACATTTTGGTA | ATCAGAAGCCGACAAGGCTA | 115 |
| chr1:94412735-94414750 | p2 | (AT)6 | CCGGAGCCTACATTTTGGTA | ATCAGAAGCCGACAAGGCTA | 116 |
| chr1:94412735-94414750 | p3 | (AAC)5 | CCGGAGCCTACATTTTGGTA | ATCAGAAGCCGACAAGGCTA | 116 |
| chr1:94412735-94414750 | p2 | (TC)10 | CCGGAGCCTACATTTTGGTA | ATCAGAAGCCGACAAGGCTA | 116 |
| chr1:94454068-94456086 | p6 | (AGTGCT)3 | AGAATCAAGAAGCCGAACGA | AACTCAGGGGCACGTTTATG | 159 |
| chr1:94558517-94560551 | p2 | (AT)17 | CAACAACACATGCCACAACA | GACGTTCGGAGGACAATCAT | 122 |
| chr1:94563777-94565792 | p2 | (GA)6 | AATCAACCAAGGAGGAGGGT | TCACAATACCAACGGACGAA | 280 |
| chr1:94563777-94565792 | p3 | (TTG)5 | AATCAACCAAGGAGGAGGGT | TCACAATACCAACGGACGAA | 280 |
| chr1:94563777-94565792 | p5 | (TGATA)5 | AATCAACCAAGGAGGAGGGT | TCACAATACCAACGGACGAA | 280 |
| chr1:94567333-94569345 | p2 | (GA)6 | ACAAGTGGTTCCCAGCAGTC | TCTCTCCATTCCCGAACATC | 212 |
| chr1:94644996-94647010 | p2 | (TA)7 | GCTTATGCCGTCCATGCTAT | CCACTGATGGGCTTTTTCAT | 106 |
| chr1:94710253-94712268 | p5 | (ATAAA)3 | CATATTGCGTTTGTCGTTGG | ATAAAATCGAGGGGGAAGGA | 143 |
| chr1:94710253-94712268 | p2 | (AT)9 | CATATTGCGTTTGTCGTTGG | ATAAAATCGAGGGGGAAGGA | 143 |
| chr1:94725775-94727791 | p2 | (AT)14 | CTATCACTACCCGCAGGCAT | TTGGCCACTTTGTCAGTGAG | 154 |
| chr1:94725775-94727791 | p2 | (AT)8 | CTATCACTACCCGCAGGCAT | TTGGCCACTTTGTCAGTGAG | 154 |
| chr1:94777843-94779855 | p2 | (AT)6 | TCCGGCCACTGTAACTTTTC | GTTCATCCGAGGGTGTGTCT | 120 |
| chr1:94788591-94790603 | p2 | (GT)6 | CCAGTCTCCAGGACCGAATA | ACATAGCCGGTCTAACGTGG | 183 |
| chr1:95083288-95085314 | p2 | (TA)13 | AAACTTTCCAACACAACCCG | GTGGGATGATTTGGAGGCTA | 253 |
| chr1:95083288-95085314 | p5 | (TTTTC)3 | AAACTTTCCAACACAACCCG | GTGGGATGATTTGGAGGCTA | 253 |
| chr1:95083440-95085455 | p2 | (TA)13 | AAACTTTCCAACACAACCCG | GTGGGATGATTTGGAGGCTA | 101 |

| Marker ID | SSR type | SSR motif | FORWARD PRIMER (5'-3') | REVERSE PRIMER (5'-3') | size (bp) |
|---|---|---|---|---|---|
| chr1:95083440-95085455 | p5 | (TTTTC)3 | AAACTTTCCAACACAACCCG | GTGGGATGATTTGGAGGCTA | 101 |
| chr1:95175087-95177101 | p2 | (CT)7 | ATAGGGCACAATTTGCCAAC | TCCGACGTATGTTGGAATGA | 153 |
| chr1:95211872-95213900 | p2 | (AT)8 | AGATCAAGCATCAAATCCGC | TTCGAATTCCATCCATAGCC | 239 |
| chr1:95214364-95216384 | p2 | (TA)10 | TTTGGCCGAGTTTGATTCTC | TTACATACCTTGCCCGCTTC | 285 |
| chr1:95316565-95318583 | p2 | (AT)9 | GGACACGACTTTGCAGACAA | TTGGCTCTGAGTATGGGGTC | 183 |
| chr1:95384824-95386839 | p3 | (AAC)5 | AACTCGGCCAAATCATCAAC | GAAATGTCGAGCGGATTGTT | 178 |
| chr1:95384824-95386839 | p5 | (TTCTT)3 | AACTCGGCCAAATCATCAAC | GAAATGTCGAGCGGATTGTT | 178 |
| chr1:95392335-95394383 | p2 | (AT)8 | AGCGGAGATGGCTTGAGTTA | GGACGAACAAAAATCCCAGA | 268 |
| chr1:95392394-95394408 | p2 | (AT)8 | AGCGGAGATGGCTTGAGTTA | GGACGAACAAAAATCCCAGA | 209 |
| chr1:95392995-95395011 | p2 | (AT)8 | TGACAGGCTGACGAAATCAG | ATGACTGATGTTTTTGGGGC | 290 |
| chr1:95417925-95419940 | p3 | (AAC)5 | AACTCGGCCAAATCATCAAC | GAAATGTCGAGCGGATTGTT | 178 |
| chr1:95417925-95419940 | p5 | (TTCTT)3 | AACTCGGCCAAATCATCAAC | GAAATGTCGAGCGGATTGTT | 178 |
| chr1:95448069-95450085 | p2 | (AT)8 | GATTGAAGCTGGACAACGGT | TCACATGCCCAACACAAACT | 245 |
| chr1:95465636-95467650 | p2 | (TA)6 | CGCGGCATTTCTAAAATCAT | CCCTATTACGGCCTGACAAA | 282 |
| chr1:95465636-95467650 | p2 | (AT)7 | CGCGGCATTTCTAAAATCAT | CCCTATTACGGCCTGACAAA | 282 |
| chr1:95508600-95510636 | p2 | (AT)18 | CAATTCCCGTTTCCTTCAAA | GTTGATTGTTTTGGCCGAGT | 185 |
| chr1:95534429-95536441 | p2 | (AC)6 | CGGAGTCGTACTCAAGGCTC | ATCTCGGGGAGGCTTATTGT | 209 |
| chr1:95561636-95563652 | p2 | (AT)12 | CGCGAGAACAGAAACATCAA | GCTATTCGATCGGACCATGT | 138 |
| chr1:95597914-95599938 | p2 | (AT)12 | ATTTCACTTCCAGTGTCGGC | TGGGATGGACGGAGTAATTC | 123 |
| chr1:95608826-95610844 | p2 | (AC)6 | GAGAATTCAACTCGGCCAAA | TTTGAATGTTGTGCGGGTTA | 185 |
| chr1:95608826-95610844 | p6 | (TCTTTT)3 | GAGAATTCAACTCGGCCAAA | TTTGAATGTTGTGCGGGTTA | 185 |
| chr1:95706725-95708745 | p2 | (AT)10 | ATTCATTCGGCTTTGACCTG | CATCCAGTTCCTCCACTGGT | 237 |
| chr1:95767104-95769128 | p3 | (GAT)8 | AATTTTAACCGGAACCGGAC | GGGGGCCAAAAATTAAGAAA | 213 |
| chr1:95776938-95778950 | p2 | (GT)6 | CACCCAATTTCCTAGCGTGT | CAGAGGTTACATCCCTCCCA | 255 |
| chr1:95794057-95796077 | p2 | (TA)6 | TGGGCCATTAAGGTAGGGTT | ATAGGGATAGGTGCGTGTCG | 225 |
| chr1:95798822-95800837 | p5 | (AATAC)3 | GTGGGGAGTCTTGTCTGGAA | TGTTGTGAATGTGGCCTGTT | 229 |
| chr1:95841027-95843045 | p2 | (GT)9 | ATGAAAAGGACAAGATGCGG | TTGTACGAAAACACGGGACA | 241 |
| chr1:95841027-95843045 | p3 | (GTT)5 | ATGAAAAGGACAAGATGCGG | TTGTACGAAAACACGGGACA | 241 |
| chr1:95855908-95857920 | p2 | (TA)6 | CAGTCCTAGGAGGGACCACA | TTTGGTGAACCAGGTGAACA | 126 |
| chr1:96013853-96015873 | p2 | (AT)10 | AAGCAATGAGGTAGCCTGGA | CGCCCTTCACATATTCAGGT | 212 |
| chr1:96131772-96133784 | p2 | (TA)6 | GATTGGGGACCCAGGTAAGT | AGTTTTGCCAACATCGAACC | 175 |
| chr1:96159725-96161737 | p2 | (CT)6 | CTCAAAGCTTCCTTTCACCG | ACTTGCCTTCAAAATGCCAC | 286 |
| chr1:96172348-96174360 | p2 | (AG)6 | CCGTTTGGAATAAAGTGCGT | TCCGACATGGCTTCTTTACC | 261 |
| chr1:96487871-96489886 | p5 | (TTCAT)3 | TTAGCGATCTGCCCTGCTAT | TGAACGAAAAATTTCCCGAG | 220 |
| chr1:96515189-96517204 | p5 | (TTACT)3 | GTCCATCAGGCTTCTGGTGT | TGGTAGGGAGACGACCTTTG | 270 |
| chr1:96519899-96521913 | p2 | (AT)7 | GCACTTTACGCCTTTCCAAG | GTTGATGTTTTGGCCGAGTT | 135 |

| Marker ID | SSR type | SSR motif | FORWARD PRIMER (5'-3') | REVERSE PRIMER (5'-3') | size (bp) |
|---|---|---|---|---|---|
| chr1:96544374-96546394 | p5 | (CAAAA)4 | TTTGCGAATTTCCCTTTTTG | GTCGTGGCAAAGTTAAGGGA | 204 |
| chr1:96586322-96588342 | p5 | (CAAAA)4 | TTTGCGAATTTCCCTTTTTG | GTCGTGGCAAAGTTAAGGGA | 204 |
| chr1:96656555-96658569 | p2 | (AT)7 | ACCCGAATTCTTCCAAATCC | GCCGTGTGGTATGTATGCTG | 233 |
| chr1:96756187-96758205 | p2 | (AT)13 | AGGGACTTGGTGAACAATGC | GTGAGCGAACGACGAACATA | 226 |
| chr1:96756187-96758205 | p3 | (CAA)6 | AGGGACTTGGTGAACAATGC | GTGAGCGAACGACGAACATA | 226 |
| chr1:96766296-96768311 | p5 | (ATTTC)3 | AAGAAATGACATGGGAAGCG | TTAAGTCGTCGGGGATTCTG | 144 |
| chr1:96772232-96774247 | p5 | (AATAA)3 | ACGCCATAGACCAACCAAAG | TACCCCTTTGAAATTGAGCG | 239 |
| chr1:96863106-96865121 | p3 | (TGT)5 | ACGTTTTCGGGTGATCTGAC | TAACCCACTGACCTTCCTGC | 218 |
| chr1:96886021-96888039 | p6 | (ATACAT)3 | TAAGTACAAGGCCGACGAGG | GCAGAGCTCCATTTCTTTCG | 211 |
| chr1:97014908-97016920 | p2 | (AT)6 | CCGACAAGGCTACACCATCT | TCCGTACTAACTCCCCGTTG | 285 |
| chr1:97019314-97021342 | p2 | (AT)14 | AACGCATCCCTTACCCTTTT | CCCCCTCAGGAAAAGAACTC | 191 |
| chr1:97041016-97043030 | p2 | (TA)7 | GCCCCTAGAAAAATCCCAAG | CCCAGAGTTCTACGGGTCAA | 183 |
| chr1:97159546-97161562 | p2 | (AT)8 | CGACCACCCTTCAAGACAAT | TGTCGAATAAGGCCAAGGAC | 101 |
| chr1:97196283-97198295 | p2 | (TA)6 | CCAACTCCACGGATTCAGTT | CGGGTCAACAAATACCGAGT | 137 |
| chr1:97209662-97211677 | p5 | (ATATG)3 | CCACGAAAACAACATACCCC | GAGAGGGAGGAGAGCCCTTA | 183 |
| chr1:97215852-97217872 | p4 | (AGAA)5 | ATCTTTTGGCAATCCACTCG | GAACATGTGGATCATTTGCG | 100 |
| chr1:97255624-97257656 | p2 | (TA)16 | CTTCACCTCGCGTGTACAGA | TTTCTGCCAGTTCCCTGAGT | 191 |
| chr1:97326191-97328251 | p2 | (AT)30 | CCAACTCCACGGATTCAGTT | CAAGTACCAAAGCCCGTGAT | 164 |
| chr1:97326191-97328251 | p2 | (TA)7 | CCAACTCCACGGATTCAGTT | CAAGTACCAAAGCCCGTGAT | 164 |
| chr1:97326369-97328383 | p2 | (AT)30 | ATCACGGGCTTTGGTACTTG | CTCCAGTTCCAATTCCTCCA | 113 |
| chr1:97326369-97328383 | p2 | (TA)7 | ATCACGGGCTTTGGTACTTG | CTCCAGTTCCAATTCCTCCA | 113 |
| chr1:97537934-97539972 | p2 | (TA)19 | CCCAATGCTGATGAGTTGTG | TATTGGTACGACGGGGTGTT | 124 |
| chr1:97593344-97595356 | p2 | (AG)6 | GACAGATGAGGCCACCCTTA | CTCCACTCCTCCCCTACCTC | 130 |
| chr1:97683354-97685369 | p3 | (ATG)5 | GAATCACTGCCCAACATCCT | TTTCGCATATTTTGGAACCC | 108 |
| chr1:97713256-97715268 | p2 | (TA)6 | ACGCACAACGCTTTTCTCTT | TTTGAGGGCTGGTAATTTGC | 241 |
| chr1:97758516-97760530 | p2 | (AT)7 | ATTGCGTACCAAAGACAGGG | TTCCGTGTCATGGGATTTCT | 296 |
| chr1:97810388-97812428 | p2 | (AT)20 | TGGGAACACAAGTTGGATGA | TTAGGAATGGGCTTGGTGTC | 243 |
| chr1:97810388-97812428 | p2 | (AT)19 | TGGGAACACAAGTTGGATGA | TTAGGAATGGGCTTGGTGTC | 243 |
| chr1:97861887-97863919 | p2 | (TA)16 | ATCACGACCTTACGTCACCC | TGTGTATATCTGCCCGACCA | 289 |
| chr1:97895681-97897696 | p5 | (TGTGT)3 | TGTGGCTGCAAACCTAGATG | TTGTAATTGTCACGACCCGA | 175 |
| chr1:97908676-97910691 | p5 | (ATATC)3 | TGCTCCGATCTCTGTCACAC | GTGTTATATGTCCGTCCCCG | 204 |
| chr1:97978967-97981055 | p2 | (TA)31 | TTGGGCTCATTTTGAAGGTC | AAATGGATCCCCTCCTCATT | 283 |
| chr1:98049627-98051642 | p5 | (TGTTG)3 | AATTGCGGGAAGACAACAAG | AGCCCACAAAGAAGTGGATG | 290 |
| chr1:98106926-98108944 | p6 | (CCCCAC)3 | CGGAGTTAACGCTAAGCTGG | ATCATCGTCAATCTCCCTGC | 145 |
| chr1:98106926-98108944 | p3 | (GTT)5 | CGGAGTTAACGCTAAGCTGG | ATCATCGTCAATCTCCCTGC | 145 |
| chr1:98148919-98150934 | p2 | (AT)6 | GGGCCATAGTAGCTCGAGTG | TGGGTTGTGACAAGTTGGAA | 100 |

| Marker ID | SSR type | SSR motif | FORWARD PRIMER (5'-3') | REVERSE PRIMER (5'-3') | size (bp) |
|---|---|---|---|---|---|
| chr1:98148919-98150934 | p2 | (AT)11 | GGGCCATAGTAGCTCGAGTG | TGGGTTGTGACAAGTTGGAA | 100 |
| chr1:98158737-98160757 | p5 | (AATGC)3 | ACAGACAGAAGAGACCCGGA | AGTCCCGAAATCGTGAATTG | 130 |
| chr1:98158737-98160757 | p2 | (AG)10 | ACAGACAGAAGAGACCCGGA | AGTCCCGAAATCGTGAATTG | 130 |
| chr1:98234194-98236206 | p2 | (CT)6 | TGCTTCAAATGCTCCCTTCT | CTTGCGTACCAACGGAAGAT | 162 |
| chr1:98294918-98296936 | p5 | (CCCAC)3 | TTTGATGCACCGCATGTTAT | CCTAAGGTCGGAACCATTCA | 104 |
| chr1:98294918-98296936 | p3 | (ACC)6 | TTTGATGCACCGCATGTTAT | CCTAAGGTCGGAACCATTCA | 104 |
| chr1:98294918-98296936 | p3 | (CAC)9 | TTTGATGCACCGCATGTTAT | CCTAAGGTCGGAACCATTCA | 104 |
| chr1:98348375-98350399 | p2 | (AC)12 | GGTCATGCCAAAAAGGAAGA | GTTGATGATTTTGGCCGAGT | 248 |
| chr1:98348375-98350399 | p2 | (CT)7 | GGTCATGCCAAAAAGGAAGA | GTTGATGATTTTGGCCGAGT | 248 |
| chr1:98366973-98369012 | p2 | (AT)47 | TTCTACTGCCCATCCGTACC | CCTTGTCCAAAATCCCCTTT | 228 |
| chr1:98366973-98369012 | p2 | (AT)13 | TTCTACTGCCCATCCGTACC | CCTTGTCCAAAATCCCCTTT | 228 |
| chr1:98367013-98369064 | p2 | (AT)47 | TTCTACTGCCCATCCGTACC | CCTTGTCCAAAATCCCCTTT | 188 |
| chr1:98367013-98369064 | p2 | (AT)13 | TTCTACTGCCCATCCGTACC | CCTTGTCCAAAATCCCCTTT | 188 |
| chr1:98404858-98406909 | p3 | (ATT)17 | CCAGCAGCACCACTAGATGA | GGGGTACCTAAGGAGCTTGG | 187 |
| chr1:98429636-98431660 | p2 | (CT)6 | GGACTCGGCGAACGATATAA | ATACTTGTCCGTGGTCTCCG | 251 |
| chr1:98448412-98450430 | p6 | (TCATCG)3 | GGCAAACAAAAGGAAATCCA | AGAGCTCCACTTCATTCGGA | 112 |
| chr1:98448412-98450430 | p5 | (GAAAT)3 | GGCAAACAAAAGGAAATCCA | AGAGCTCCACTTCATTCGGA | 112 |
| chr1:98475710-98477725 | p3 | (TCA)5 | TTTACCACCTTCACCCTTGC | CGGAGGTGCCATTGATTTAT | 100 |
| chr1:98599698-98601710 | p2 | (AT)6 | AAATTTCATCCCCCAAGACC | GGTCAGGCCCATTTCCTTAT | 205 |
| chr1:98637209-98639224 | p5 | (TTTTA)3 | ATTATGAAGTGGAGCAGCGG | GGTCGGACCATCTCAACACT | 209 |
| chr1:98744692-98746712 | p5 | (AAATA)4 | CCAAAATGCCCAGAAGACTC | TTATAAGGGCATCTGGGTCG | 128 |
| chr1:98744692-98746712 | p5 | (TTTAT)4 | CCAAAATGCCCAGAAGACTC | TTATAAGGGCATCTGGGTCG | 128 |
| chr1:98748592-98750606 | p2 | (TA)7 | CTTGTACCGAGCACCCCTTA | TCCGGAGCCTTATTACATCG | 164 |
| chr1:98768459-98770477 | p6 | (CATATC)3 | ACTTTTGTCACGACCCAACC | AGGTATTGTTGCCCAATGGA | 198 |
| chr1:98904777-98906797 | p2 | (AT)10 | ATCCCGTTTCCTCCAAAATC | GTTGACATTTTGGCCGAGTT | 186 |
| chr1:98945314-98947326 | p2 | (TA)6 | TCATGGCTCCGAAAATAAGG | CAGAATTCGATCCCCTTCAA | 290 |
| chr1:99029487-99031502 | p3 | (AAT)5 | TGTTGTTCGGCTGTGTTAGC | CATGACATAACACGCAAGCC | 222 |
| chr1:99068075-99070093 | p6 | (TTTTTC)3 | ATGTCGTACCCTGTGAAGGC | TTCCCTGGTTGCATTTTAGG | 166 |
| chr1:99070106-99072121 | p5 | (TCGGT)3 | ATGGTCGTTAGGAGTGTCCG | AGCGACGAGAAATTTGGATG | 285 |
| chr1:99118406-99120422 | p2 | (TA)8 | GAGAGGTGCCACATCACCTT | AACAGGTGCAACCGAAAAAG | 181 |
| chr1:99140191-99142207 | p2 | (AT)8 | GGATAATTTGCGACCGTGAT | AACGGCAGAGAAAACGTGTC | 173 |
| chr1:99140191-99142207 | p2 | (AT)6 | GGATAATTTGCGACCGTGAT | AACGGCAGAGAAAACGTGTC | 173 |
| chr1:99140191-99142207 | p2 | (AT)14 | GGATAATTTGCGACCGTGAT | AACGGCAGAGAAAACGTGTC | 173 |
| chr1:99152033-99154051 | p6 | (CAAAAA)3 | CCATTTTTGCGACCTTTGTT | ACCAAGGCTCCTTCCAAGTT | 196 |
| chr1:99234877-99236889 | p2 | (AT)6 | GACAATTTCAAACGCGGATT | GATGCTGCCCAAATTTCTGT | 181 |
| chr1:99243655-99245670 | p5 | (AATTC)3 | AACAAGTTTCGGTCAATCCG | CGATGATGCCAAAGATGAGA | 151 |

| Marker ID | SSR type | SSR motif | FORWARD PRIMER (5'-3') | REVERSE PRIMER (5'-3') | size (bp) |
|---|---|---|---|---|---|
| chr1:99243655-99245670 | p5 | (TCCAT)3 | AACAAGTTTCGGTCAATCCG | CGATGATGCCAAAGATGAGA | 151 |
| chr1:99286537-99288564 | p3 | (ATA)9 | CTTCTGTCACACCCCAACCT | GTTGTTGGGTTCTTCTGCGT | 274 |
| chr1:99310805-99312819 | p2 | (TA)6 | AAACCCGAAGGAAAGGAAAA | ACGATAGACCCACGAACTGG | 266 |
| chr1:99310805-99312819 | p2 | (AT)7 | AAACCCGAAGGAAAGGAAAA | ACGATAGACCCACGAACTGG | 266 |
| chr1:99380949-99382963 | p2 | (TA)7 | GATTTGTGGCACATTCAACG | CTTTGCTTTTTGCTTTTGGG | 235 |
| chr1:99380949-99382963 | p2 | (TA)12 | GATTTGTGGCACATTCAACG | CTTTGCTTTTTGCTTTTGGG | 235 |
| chr1:99381938-99383976 | p2 | (TA)7 | TATAACAAGGACTTGGGGCG | CTAATGGGTCGCGGTTATGT | 146 |
| chr1:99381938-99383976 | p2 | (TA)19 | TATAACAAGGACTTGGGGCG | CTAATGGGTCGCGGTTATGT | 146 |
| chr1:99381938-99383976 | p2 | (TA)13 | TATAACAAGGACTTGGGGCG | CTAATGGGTCGCGGTTATGT | 146 |
| chr1:99402658-99404672 | p2 | (GC)7 | GAATCCAGGAAACGTCAGGA | CTTGATCTTTGGGCGTTAGG | 149 |
| chr1:99406292-99408310 | p2 | (TA)9 | AACAGACAAGGGAAAAGGCA | TTTCCAAGGCGTTTCTGTCT | 112 |
| chr1:99560134-99562152 | p6 | (GAATAT)3 | ATGGGTGTGAGGATGTTGGT | GCATTTCCCCTATTTCGTCA | 255 |
| chr1:99647986-99650001 | p2 | (AT)8 | CCCCTATATACGAGCCGACA | TTCATGCCCACAGGTAAACA | 268 |
| chr1:99647986-99650001 | p5 | (AATGC)3 | CCCCTATATACGAGCCGACA | TTCATGCCCACAGGTAAACA | 268 |
| chr1:99699121-99701136 | p3 | (ATC)5 | ACAGGAATCCATGAAGACGG | TCATGCCTGCAGGTAGACAG | 248 |
| chr1:99714241-99716255 | p2 | (TA)7 | TTGGGGTGAATTTTTGGTGT | ATGCAGCAGAAACGCAGTAA | 141 |
| chr1:99742268-99744280 | p2 | (TA)6 | AAAAGACGCCTATCGGAGGT | CTTCCGGTCAAGGGTCAATA | 144 |
| chr1:99847539-99849591 | p2 | (AT)26 | GTGTAGCCACTCCCCAATGT | CCAAAGGGAAAAGGTTGTCA | 262 |
| chr1:99874698-99876734 | p2 | (AT)7 | TAGCCGACAAGGCCAAATAC | GGAGCTCAATTTCATTCGGA | 289 |
| chr1:99874698-99876734 | p2 | (AT)18 | TAGCCGACAAGGCCAAATAC | GGAGCTCAATTTCATTCGGA | 289 |
| chr1:99942544-99944562 | p6 | (TATTTT)3 | CAACTTCGGCTGGATTGTTT | ATGAATTTTGGGGGTGATGA | 127 |
| chr1:99985404-99987422 | p6 | (TTTTTA)3 | AATTTGGGTGAACAGCAAGG | TGACAATCCATCCAAAAGCA | 284 |
| chr1:100046384-100048399 | p2 | (TG)7 | CCTCCCACCAGCAATAGGTA | AAGTGGGCAAGAACCATGAG | 264 |
| chr1:100046384-100048399 | p3 | (TCT)5 | CCTCCCACCAGCAATAGGTA | AAGTGGGCAAGAACCATGAG | 264 |
| chr1:100182983-100185013 | p2 | (TA)15 | GGACCGTCATTTAGCTCCAA | TAAGTTGGGGGTGGTGATGT | 179 |
| chr1:100225772-100227786 | p2 | (GT)7 | TGATAGGCTTGAAAATGGGG | GCCATTTCACCTCAGTGTCC | 223 |
| chr1:100242428-100244446 | p3 | (ACA)6 | GGGATGCTGTTCCTCCTGTA | GAACAGAGAGCCGTTTTTGC | 197 |
| chr1:100431163-100433175 | p2 | (AC)6 | ATAGATCCGCATCAAAACCG | TTAGCGTAGAAAGCGGGAAA | 129 |
| chr1:100489195-100491209 | p2 | (AG)7 | ATCTTCGTGTTTCCAGGGTG | GGAAAATTCAACGGGTCAGA | 215 |
| chr1:100522684-100524699 | p3 | (ACA)5 | GGAACTAGGTCGATGGGGAT | ACACAAACGCCATTCAACAA | 238 |
| chr1:100566575-100568593 | p6 | (TCCACA)3 | TTGCTGCGAGCTTAACTCAA | GGCCTAAACTTCCTCAACCC | 233 |
| chr1:100676128-100678140 | p2 | (AT)6 | ACCAAATTTCGATCACTCGC | ATGTTGGGTTGGTTGTTGGT | 235 |
| chr1:100676128-100678140 | p2 | (TC)6 | ACCAAATTTCGATCACTCGC | ATGTTGGGTTGGTTGTTGGT | 235 |
| chr1:100678077-100680115 | p2 | (TA)19 | CTTTCACCCTTGGGAAACAA | GGCCAACATTTGTAGTGGCT | 182 |
| chr1:100809382-100811406 | p2 | (AT)12 | TGTGTCAAGACTCCGAAACG | CGGACACCTTTTCAACACCT | 180 |
| chr1:100809382-100811406 | p2 | (TA)6 | TGTGTCAAGACTCCGAAACG | CGGACACCTTTTCAACACCT | 180 |

| Marker ID | SSR type | SSR motif | FORWARD PRIMER (5'-3') | REVERSE PRIMER (5'-3') | size (bp) |
|---|---|---|---|---|---|
| chr1:100809665-100811711 | p2 | (AT)12 | CCCCGAGATTTTTACGTTGA | GTTTGGCTGAGCCGATTTTA | 167 |
| chr1:100809665-100811711 | p2 | (TA)6 | CCCCGAGATTTTTACGTTGA | GTTTGGCTGAGCCGATTTTA | 167 |
| chr1:100857489-100859555 | p2 | (AT)33 | AGAACGATGGCTCTTGGAGA | TGGATCCACCCAGTTCATCT | 129 |
| chr1:100896531-100898546 | p5 | (TCATT)3 | CAAGAAGCCAACCAATCCAT | GTCGATTTAGAGTCGAGCCG | 269 |
| chr1:100898108-100900120 | p2 | (TA)6 | CGACCAAGGTATGTGAGGCT | TCTATCGGGGTGACGTAAGG | 158 |
| chr1:100931941-100933956 | p5 | (ATATA)3 | CAGACAACCCCTAGGACGAA | AAGGGTTTGCTCAGCTTGAA | 116 |
| chr1:100931941-100933956 | p3 | (CAA)6 | CAGACAACCCCTAGGACGAA | AAGGGTTTGCTCAGCTTGAA | 116 |
| chr1:100943821-100945841 | p5 | (ATTGC)3 | GTGATTCTGATGAGGCCGAT | AATTGCTCACATGGGGAGTC | 134 |
| chr1:101044223-101046238 | p5 | (TCATT)3 | CGCCCTTGCAAGAGTAAAAC | CATGGAATTGGGGTTTGAAG | 238 |
| chr1:101130294-101132310 | p4 | (CAGA)4 | ATAGGCTGCAACAATCCACC | GTATTTTCTGTGCGCAGGGT | 257 |
| chr1:101166013-101168028 | p5 | (CTAGT)3 | CCGAATAGCCATCAAGGAAA | AAGAAAGGAGGGATGACGGT | 238 |
| chr1:101261816-101263831 | p5 | (TCATT)3 | GGGGTGTGTTTTAATGGTGG | AAAACATGCCAAATAAGCGG | 288 |
| chr1:101303151-101305166 | p5 | (TTTAA)3 | TTCTTCGCCATTCCATTGTT | CCCACAGTTTTCCTCCGTTA | 199 |
| chr1:101350003-101352018 | p3 | (ATC)5 | GCCCACGAGGCTACAAAATA | CCTGTCCCGTCGTTATGATT | 102 |
| chr1:101497483-101499495 | p2 | (AG)6 | ACGATGCAGACCACATACCA | CTTTAGGTTTCAACGCTGGC | 147 |
| chr1:101512118-101514140 | p2 | (AC)6 | CGGCCAAACACACTACACAC | ATTAGTTTGGCACGAGGACG | 179 |
| chr1:101512118-101514140 | p2 | (AG)11 | CGGCCAAACACACTACACAC | ATTAGTTTGGCACGAGGACG | 179 |
| chr1:101668311-101670331 | p2 | (AT)10 | AGTACGCTCGGCTTGTCTGT | CACGGTATTCCTTCTGCCAT | 101 |
| chr1:101730929-101732949 | p2 | (TA)10 | TCCCGATAGATGATGTGCAA | GTTTGCGATTCGTCTTGGAT | 162 |
| chr1:101759644-101761672 | p2 | (GT)6 | ACCATCCTATGATGCTTGCC | GCTTACAAGGCATGATGCAA | 250 |
| chr1:101785175-101787187 | p2 | (TA)6 | AGGAACGACAGCAGAAGCAT | TCAAGGAAACCAAGGTGGAC | 157 |
| chr1:101940168-101942180 | p2 | (AT)7 | GTGCAATAGTGCAGGCAAAA | ATCAAGTTCCACCCCATCAA | 106 |
| chr1:101940168-101942180 | p2 | (CA)6 | GTGCAATAGTGCAGGCAAAA | ATCAAGTTCCACCCCATCAA | 106 |
| chr1:101973172-101975190 | p2 | (TA)9 | TAAACAGGGAAAATGCTGCC | TCCCGTCTTGCAACTCTCTT | 173 |
| chr1:102016170-102018188 | p6 | (ATCCAC)3 | GGGTGCCTGAACTGGAAATA | GTGGAAGTGGACGTGGATCT | 191 |
| chr1:102123387-102125402 | p5 | (TTATT)3 | CAGTCTTGGGCAATCCTCAT | GATGTGGGACTCAACACCCT | 226 |
| chr1:102147642-102149657 | p3 | (AAT)5 | TCAGGTTCCAGAGCTCGTTT | CTTTGTCACCCGCTCTTCTC | 153 |
| chr1:102147642-102149657 | p5 | (AAAAG)3 | TCAGGTTCCAGAGCTCGTTT | CTTTGTCACCCGCTCTTCTC | 153 |
| chr1:102202062-102204077 | p2 | (GA)6 | CTCAGGCTGATAACCTTCGG | GATGTTATTCCCGAGGACGA | 160 |
| chr1:102202062-102204077 | p3 | (ACA)5 | CTCAGGCTGATAACCTTCGG | GATGTTATTCCCGAGGACGA | 160 |
| chr1:102285752-102287764 | p5 | (TTCTT)3 | AGCGTATATTTGGGCGACAG | ATCGTGCAAAACACGTTTGA | 175 |
| chr1:102285752-102287764 | p2 | (TA)6 | AGCGTATATTTGGGCGACAG | ATCGTGCAAAACACGTTTGA | 175 |
| chr1:102285752-102287764 | p2 | (AT)6 | AGCGTATATTTGGGCGACAG | ATCGTGCAAAACACGTTTGA | 175 |
| chr1:102456183-102458198 | p5 | (ATTGC)3 | TAGGGCATTGACTCCGTAGG | GCAATGCAATGCAATAATGG | 250 |
| chr1:102456183-102458198 | p5 | (TTATA)3 | TAGGGCATTGACTCCGTAGG | GCAATGCAATGCAATAATGG | 250 |
| chr1:102554573-102556591 | p3 | (TTG)6 | ATATCGTATTGCGGCGAATC | TCCATAAACATCCCACACGA | 265 |

| Marker ID | SSR type | SSR motif | FORWARD PRIMER (5'-3') | REVERSE PRIMER (5'-3') | size (bp) |
|---|---|---|---|---|---|
| chr1:102554573-102556591 | p3 | (ATG)6 | ATATCGTATTGCGGCGAATC | TCCATAAACATCCCACACGA | 265 |
| chr1:102571232-102573304 | p2 | (TA)11 | TTCCCGCTTTGAACTACAGG | CCTTAGCGCTCTTTCCCTTT | 230 |
| chr1:102610222-102612284 | p2 | (AT)31 | AAACTCATGAAATCACGGGC | CCGCCCAATTTAGAAGAACA | 272 |
| chr1:102610222-102612284 | p2 | (AT)6 | AAACTCATGAAATCACGGGC | CCGCCCAATTTAGAAGAACA | 272 |
| chr1:102610508-102612520 | p2 | (AT)31 | TTTCCACCCGTCTTTTCATC | GTTTATGGTGTGTTGCGTGC | 251 |
| chr1:102610508-102612520 | p2 | (AT)6 | TTTCCACCCGTCTTTTCATC | GTTTATGGTGTGTTGCGTGC | 251 |
| chr1:102653119-102655135 | p2 | (TA)8 | GCGGCTTATGATTACACCGT | TCTCCCTGTCTACCTGTGGG | 234 |
| chr1:102696865-102698880 | p5 | (AAACT)3 | TTGTTGTCCCCAGTTTCTCC | AAATGGTCACGACCCAACTC | 101 |
| chr1:102696865-102698880 | p4 | (AATA)4 | TTGTTGTCCCCAGTTTCTCC | AAATGGTCACGACCCAACTC | 101 |
| chr1:102720156-102722168 | p2 | (AT)6 | GCACTAGCACATGAAGCCAA | ACATGGATCTTCTTCCCGTG | 152 |
| chr1:102722129-102724147 | p6 | (ATATGT)3 | GCCTCGGGAAAAACTCTACC | ACAATAAGGAAAATCCGGGG | 170 |
| chr1:102924591-102926603 | p2 | (TG)6 | GGGCTATAATGATACCGCGA | AGTGTGCCCGACCATTTAAG | 283 |
| chr1:102994908-102996922 | p2 | (AT)30 | GTCGGCCCCATACTAAACAA | CCCATGAGTTTGGAGATGGT | 272 |
| chr1:102994908-102996922 | p2 | (TA)7 | GTCGGCCCCATACTAAACAA | CCCATGAGTTTGGAGATGGT | 272 |
| chr1:102994908-102996922 | p2 | (TA)10 | GTCGGCCCCATACTAAACAA | CCCATGAGTTTGGAGATGGT | 272 |
| chr1:103232483-103234498 | p5 | (TTGGT)3 | CGATTTTGTTATGGTCCGCT | GTAATGTCGGGTTGGTTTGG | 127 |
| chr1:103309500-103311530 | p2 | (AG)7 | TTGGGGAGGTCTCTACCCTT | GGTGGAAGGTGTTGAAGGAA | 181 |
| chr1:103309500-103311530 | p3 | (TTC)10 | TTGGGGAGGTCTCTACCCTT | GGTGGAAGGTGTTGAAGGAA | 181 |
| chr1:103309500-103311530 | p3 | (TTC)6 | TTGGGGAGGTCTCTACCCTT | GGTGGAAGGTGTTGAAGGAA | 181 |
| chr1:103391354-103393370 | p2 | (AT)8 | AAGGGATTCGATCCCATCTT | GATTTCTTGGTGCTCCGGTA | 229 |
| chr1:103396955-103398971 | p2 | (GA)8 | GAGCTTCAGCAGCCTCAAGT | CCAGATTTGGCAAGAAAGGA | 120 |
| chr1:103423678-103425690 | p2 | (AT)6 | CCGATAGATTGGCGGAAATA | TCCGGAGCTCATATTTTTGG | 115 |
| chr1:103468943-103470958 | p3 | (TAT)5 | GGTAACTAAAGGGGGTCCCA | CTAATGCCCTGAGCAACACA | 210 |
| chr1:103502620-103504650 | p2 | (AG)15 | TCTTTTCGCTTCCTTTTGGA | TATGCGCAAAAGCCCTAGAT | 108 |
| chr1:103534343-103536367 | p2 | (TG)12 | TGGGTGATTTGAAATGGGTT | CCAATCAGTGAATGTCGTGG | 208 |
| chr1:103534343-103536367 | p2 | (TA)32 | TGGGTGATTTGAAATGGGTT | CCAATCAGTGAATGTCGTGG | 208 |
| chr1:103545361-103547376 | p5 | (GTATT)3 | GGCCTGTAAAAGCCCCTAAC | TTTTGGCAATTTCTGAAGGG | 260 |
| chr1:103596441-103598461 | p5 | (TTTGT)4 | GGTGGATGCATCTTCTCCAT | TGAGGAAGGGAAGCAGTTGT | 125 |
| chr1:103629313-103631328 | p2 | (GA)8 | TCTCTTTTGCAAACCTGCCT | GAAAACCAACATTTCCCCCT | 113 |
| chr1:103723922-103725980 | p2 | (AT)29 | TGTCACCCATCCCCATAAAT | GTACCGACCCATTCAGTCGT | 160 |
| chr1:103750073-103752093 | p2 | (TA)7 | GCCAAGCAGATTCTACGGTT | TAAACAACACGTGGTCCGAA | 149 |
| chr1:103750073-103752093 | p2 | (TA)10 | GCCAAGCAGATTCTACGGTT | TAAACAACACGTGGTCCGAA | 149 |
| chr1:103759902-103761920 | p3 | (TTC)6 | TCAATTCACGCACCACAGAT | TTTGTAAGCCACCGTTCTCC | 136 |
| chr1:103778620-103780635 | p6 | (TGTGTT)3 | GTTTGTGTTTGGTGGGCTTT | CCCCTCATCCAGTGTCTGAT | 289 |
| chr1:103778620-103780635 | p5 | (TTTCT)3 | GTTTGTGTTTGGTGGGCTTT | CCCCTCATCCAGTGTCTGAT | 289 |
| chr1:103966356-103968390 | p2 | (AT)17 | GCAAAAACATGTTGACGGTG | ATTCCTAGTCGGGTACGCCT | 111 |

| Marker ID | SSR type | SSR motif | FORWARD PRIMER (5'-3') | REVERSE PRIMER (5'-3') | size (bp) |
|---|---|---|---|---|---|
| chr1:103992643-103994659 | p2 | (CA)8 | CATGCTGAAATGCTTCCTCA | CAACAATTCCTTCACGGGAG | 256 |
| chr1:104079934-104081948 | p2 | (AG)7 | ACAGCTCCCAAACACCAAAC | TAAATCCGAGACCTCGATGC | 275 |
| chr1:104091997-104094009 | p2 | (AT)53 | GAATATGGCGACTTAGGGCA | CGACCTAGGTGTTGTGAGCA | 178 |
| chr1:104092010-104094074 | p2 | (AT)53 | GAATATGGCGACTTAGGGCA | CGACCTAGGTGTTGTGAGCA | 165 |
| chr1:104111255-104113267 | p2 | (AT)6 | AAAGCTGGTGGCAAAGCTAA | ATTTGGGAGAAAGGGCAAGT | 187 |
| chr1:104111255-104113267 | p2 | (TA)59 | AAAGCTGGTGGCAAAGCTAA | ATTTGGGAGAAAGGGCAAGT | 187 |
| chr1:104144446-104146467 | p3 | (TTG)7 | AGTTTCCAACGCAACCAAAC | AACTACGTCCGCAGATCACC | 158 |
| chr1:104147989-104150003 | p2 | (AC)6 | TTCTGGTCCGCTGTAGCTTT | TTCGTAGCGATAGAGCCCAT | 172 |
| chr1:104147989-104150003 | p2 | (TA)7 | TTCTGGTCCGCTGTAGCTTT | TTCGTAGCGATAGAGCCCAT | 172 |
| chr1:104147989-104150003 | p3 | (CAT)10 | TTCTGGTCCGCTGTAGCTTT | TTCGTAGCGATAGAGCCCAT | 172 |
| chr1:104208379-104210394 | p5 | (ATAGT)3 | TGGGAGTGAGTTTAGGGGTG | GTCAAGGGGTCGGTACTTCA | 159 |
| chr1:104263450-104265465 | p5 | (TGGCT)3 | GAACCAGGGACGGAGTACAA | ATGGGGATTTGAAAGGAACC | 253 |
| chr1:104297815-104299827 | p2 | (AT)6 | GTGTATGGCCTCGGTCACTT | CAGAAGGGCAAAGCTACGAC | 298 |
| chr1:104486050-104488065 | p5 | (TTTGA)3 | TACTGTGACGGTTCGCTTTG | AAACCCTTCGAAACCGACTT | 162 |
| chr1:104488704-104490718 | p2 | (AC)7 | CACAGTCATGGACAAGTGGG | CAAATGTCTGCCCAGTCTGA | 243 |
| chr1:104498253-104500268 | p5 | (AAAGA)3 | AAGCCCATCCTACCTGACCT | AGAGCAGTGTGGATCGTGTG | 189 |
| chr1:104531158-104533173 | p3 | (TTA)5 | AATTCCTCCCAAAGGCTGTT | CCACTTTATCGCCCCAGTTA | 271 |
| chr1:104621742-104623754 | p2 | (AG)6 | TGGGAGAACCAACAACAACA | TGAGGAAACCCCACATCTTC | 236 |
| chr1:104627144-104629170 | p2 | (TA)8 | TTGGACGTCTTAGCATGGTG | GCCTTCCTCCGCTTCTAAGT | 104 |
| chr1:104725605-104727619 | p2 | (AT)7 | GGCAAATAATGGACCGAAGA | GTGGGAAGTGAGGGCAATTA | 296 |
| chr1:104725605-104727619 | p4 | (TTTG)4 | GGCAAATAATGGACCGAAGA | GTGGGAAGTGAGGGCAATTA | 296 |
| chr1:104855815-104857830 | p5 | (AAAAT)3 | GTGCAGCATGTTTGATTTGG | GAAGAGTGGTTTGCCAGAGC | 178 |
| chr1:104907378-104909390 | p2 | (TA)6 | ACATTGTGTTTTGTGCTCCG | GTGGAACTCAGACACTGCCA | 150 |
| chr1:104907378-104909390 | p2 | (TA)7 | ACATTGTGTTTTGTGCTCCG | GTGGAACTCAGACACTGCCA | 150 |
| chr1:104907378-104909390 | p2 | (TA)6 | ACATTGTGTTTTGTGCTCCG | GTGGAACTCAGACACTGCCA | 150 |
| chr1:104907378-104909390 | p2 | (GT)8 | ACATTGTGTTTTGTGCTCCG | GTGGAACTCAGACACTGCCA | 150 |
| chr1:104907378-104909390 | p2 | (AT)7 | ACATTGTGTTTTGTGCTCCG | GTGGAACTCAGACACTGCCA | 150 |
| chr1:104914803-104916815 | p2 | (AT)6 | TTTGGAAAATGCAAGAACCC | GGACTAAGGTTGTCGTTCCG | 127 |
| chr1:104914803-104916815 | p4 | (ATAC)4 | TTTGGAAAATGCAAGAACCC | GGACTAAGGTTGTCGTTCCG | 127 |
| chr1:104915692-104917706 | p2 | (AT)6 | GGTATGTCGCTCCGTTGATT | TCAAAAACAAAGTTCCCCCA | 289 |
| chr1:104915692-104917706 | p4 | (ATAC)4 | GGTATGTCGCTCCGTTGATT | TCAAAAACAAAGTTCCCCCA | 289 |
| chr1:104915692-104917706 | p2 | (AT)6 | GGTATGTCGCTCCGTTGATT | TCAAAAACAAAGTTCCCCCA | 289 |
| chr1:104915708-104917722 | p2 | (AT)6 | GGTATGTCGCTCCGTTGATT | TCAAAAACAAAGTTCCCCCA | 273 |
| chr1:104915708-104917722 | p4 | (ATAC)4 | GGTATGTCGCTCCGTTGATT | TCAAAAACAAAGTTCCCCCA | 273 |
| chr1:104915708-104917722 | p2 | (AT)6 | GGTATGTCGCTCCGTTGATT | TCAAAAACAAAGTTCCCCCA | 273 |
| chr1:104915755-104917769 | p2 | (AT)6 | GGTATGTCGCTCCGTTGATT | TCAAAAACAAAGTTCCCCCA | 226 |

| Marker ID | SSR type | SSR motif | FORWARD PRIMER (5'-3') | REVERSE PRIMER (5'-3') | size (bp) |
|---|---|---|---|---|---|
| chr1:104915755-104917769 | p4 | (ATAC)4 | GGTATGTCGCTCCGTTGATT | TCAAAAACAAAGTTCCCCCA | 226 |
| chr1:104915755-104917769 | p2 | (AT)6 | GGTATGTCGCTCCGTTGATT | TCAAAAACAAAGTTCCCCCA | 226 |
| chr1:105121607-105123622 | p6 | (ATCTAT)3 | ATGCAATGGGTTCACACAGA | AAAGTGCTGCCACTGAAGGT | 248 |
| chr1:105121607-105123622 | p5 | (AAATA)3 | ATGCAATGGGTTCACACAGA | AAAGTGCTGCCACTGAAGGT | 248 |
| chr1:105252093-105254108 | p5 | (AAAGG)3 | GAGAAATCTGGCTCTGGTGC | TGAAAACCCTTCCAAAATCG | 242 |
| chr1:105252093-105254108 | p5 | (ATTGC)3 | GAGAAATCTGGCTCTGGTGC | TGAAAACCCTTCCAAAATCG | 242 |
| chr1:105283217-105285253 | p2 | (AT)18 | TTCAATTATGGCCCCGTTTA | CAATTTCGCAAAATCAGCCT | 242 |
| chr1:105283217-105285253 | p2 | (AT)6 | TTCAATTATGGCCCCGTTTA | CAATTTCGCAAAATCAGCCT | 242 |
| chr1:105295896-105297912 | p5 | (AACTA)3 | GATTAGGAAGCCTTTGTGCG | AAAGCCATTCTGGGGTTTCT | 192 |
| chr1:105295896-105297912 | p4 | (TAAA)4 | GATTAGGAAGCCTTTGTGCG | AAAGCCATTCTGGGGTTTCT | 192 |
| chr1:105410692-105412704 | p2 | (GT)6 | CTCGAGGCAGACAACATGAA | ATTGGTTGGGTGGTTTTGAA | 287 |
| chr1:105410692-105412704 | p2 | (TA)8 | CTCGAGGCAGACAACATGAA | ATTGGTTGGGTGGTTTTGAA | 287 |
| chr1:105411482-105413498 | p2 | (GT)6 | AGGCACGGAACACGAACTAT | GTTGTTTAGTATGGGGCCGA | 153 |
| chr1:105411482-105413498 | p2 | (TA)8 | AGGCACGGAACACGAACTAT | GTTGTTTAGTATGGGGCCGA | 153 |
| chr1:105413649-105415667 | p3 | (ATA)16 | GTTGTCGGAACCTCCAAAGA | TATTTGGTGCGATGTGCTTC | 113 |
| chr1:105463689-105465729 | p2 | (AC)9 | CCCATCTCATTCGAGGTGTT | GCCAACGAGGCTAACTCAAC | 291 |
| chr1:105463739-105465759 | p2 | (AT)8 | CCCATCTCATTCGAGGTGTT | GCCAACGAGGCTAACTCAAC | 241 |
| chr1:105463739-105465759 | p2 | (AC)9 | CCCATCTCATTCGAGGTGTT | GCCAACGAGGCTAACTCAAC | 241 |
| chr1:105463763-105465779 | p2 | (AC)9 | CCCATCTCATTCGAGGTGTT | GCCAACGAGGCTAACTCAAC | 217 |
| chr1:105503869-105505884 | p2 | (AT)6 | CGTCCGCTAACCTTGAAGAC | TGAAATAGCCAAACCGAACC | 183 |
| chr1:105503869-105505884 | p5 | (GTTCG)3 | CGTCCGCTAACCTTGAAGAC | TGAAATAGCCAAACCGAACC | 183 |
| chr1:105539385-105541401 | p4 | (TTGT)4 | TGTTCACGGCTATTTCCACA | AGGTTCCATTGAAACGTTGG | 218 |
| chr1:105612552-105614570 | p2 | (AT)9 | TGTTTTCCTGCACAGATCCA | GGCACCATCCCATATCAATC | 262 |
| chr1:105628892-105630926 | p5 | (TTAAT)3 | CAAGGATGCTTCAATCCACA | CGTGGCTCTTGTTTCATTCA | 109 |
| chr1:105664819-105666833 | p2 | (TA)12 | TACCTTCGGGAGATGTACGG | TTCTATGGCCAAACGCCTAC | 168 |
| chr1:105664819-105666833 | p2 | (TG)7 | TACCTTCGGGAGATGTACGG | TTCTATGGCCAAACGCCTAC | 168 |
| chr1:105710154-105712169 | p2 | (GT)6 | ACATTTCAATTGGGTCGGAG | TCTCTTCTTCCTCGCCTTCA | 104 |
| chr1:105710154-105712169 | p3 | (ACA)5 | ACATTTCAATTGGGTCGGAG | TCTCTTCTTCCTCGCCTTCA | 104 |
| chr1:105724416-105726438 | p2 | (CA)6 | TTGGATCAAAGATGGAAGGC | ATTTAAAAATGGCCCAAGCC | 297 |
| chr1:105724416-105726438 | p4 | (TTTC)4 | TTGGATCAAAGATGGAAGGC | ATTTAAAAATGGCCCAAGCC | 297 |
| chr1:105724416-105726438 | p2 | (CT)11 | TTGGATCAAAGATGGAAGGC | ATTTAAAAATGGCCCAAGCC | 297 |
| chr1:105831942-105833972 | p2 | (AG)15 | ATTCGGAAGATAGGCGAGGT | CCTTACCTGACGGTCCAAAA | 168 |
| chr1:105831942-105833972 | p2 | (TG)8 | ATTCGGAAGATAGGCGAGGT | CCTTACCTGACGGTCCAAAA | 168 |
| chr1:105831942-105833972 | p3 | (TTG)5 | ATTCGGAAGATAGGCGAGGT | CCTTACCTGACGGTCCAAAA | 168 |
| chr1:105869744-105871758 | p2 | (GT)7 | AGCGGAGAACGAGATGAAGA | TTACAAGCATTTTGGGGGAG | 115 |
| chr1:105869744-105871758 | p3 | (TTG)5 | AGCGGAGAACGAGATGAAGA | TTACAAGCATTTTGGGGGAG | 115 |

| Marker ID | SSR type | SSR motif | FORWARD PRIMER (5'-3') | REVERSE PRIMER (5'-3') | size (bp) |
|---|---|---|---|---|---|
| chr1:105909333-105911351 | p2 | (TG)6 | GGCCTTTTATGCAAATGGAA | TCCGAAATCTCCAACCTTTG | 280 |
| chr1:105909333-105911351 | p2 | (AT)9 | GGCCTTTTATGCAAATGGAA | TCCGAAATCTCCAACCTTTG | 280 |
| chr1:106006057-106008069 | p2 | (TG)6 | TTTGAAACAGTGGGGAGACC | CCGGAACACACTTTCCAGAT | 281 |
| chr1:106071385-106073400 | p5 | (ACCGA)4 | AACACCACGTACCTTCAGGC | CATTTGAAAACGGGCTCTGT | 208 |
| chr1:106071385-106073400 | p5 | (ACCGA)3 | AACACCACGTACCTTCAGGC | CATTTGAAAACGGGCTCTGT | 208 |
| chr1:106071385-106073400 | p6 | (TACCGA)3 | AACACCACGTACCTTCAGGC | CATTTGAAAACGGGCTCTGT | 208 |
| chr1:106084139-106086155 | p2 | (AC)8 | CCATTGTGCAGTCAATTTGG | CTCTTCTTCGGTCAGCAAGG | 165 |
| chr1:106084139-106086155 | p2 | (AT)10 | CCATTGTGCAGTCAATTTGG | CTCTTCTTCGGTCAGCAAGG | 165 |
| chr1:106109992-106112010 | p3 | (TGT)6 | CGTCATGGCCCTATCAGTTT | AACACTCTGCCCCGTACAAC | 176 |
| chr1:106228350-106230370 | p4 | (TTAT)5 | ATTCAGATGGGATGCCAAAG | GATTTTGTAAGGCTGGGCAA | 189 |
| chr1:106566504-106568518 | p2 | (TA)7 | TGAACCCCACAAGGAATAGC | GGCCGGTATCCTAAGTCTCC | 286 |
| chr1:106647195-106649210 | p5 | (AATGC)3 | GCCCCAATGGTAGAGTGAAA | GACTTAAGGGAAGGGTTCGC | 192 |
| chr1:106663188-106665202 | p2 | (AT)7 | ATGCTTGGGCTTGTTTTGTC | AATCAAGTAAAGGGTGGGGG | 242 |
| chr1:106668052-106670070 | p6 | (CCATCT)3 | GGCAGGGTTTTCAAAATTCA | AACCCCCAGAAGTCTGCTTT | 205 |
| chr1:106674894-106676934 | p2 | (AT)6 | CAATCGAAACTCCAACAGCA | GCATTTGCCGGCTATTATGT | 158 |
| chr1:106690103-106692167 | p2 | (AT)8 | GCTTTATAAGTGCCCCTCCC | GTGTCCTCCCATAGCCAAAA | 209 |
| chr1:106716059-106718077 | p2 | (TC)9 | TTGAATGCCCAACAGAGACA | CGGGGACCATCGAGTTACTA | 126 |
| chr1:106759412-106761430 | p6 | (CAACTC)3 | CCCTAACGGACGAGTGATGT | TCTTGGTAAGCACCCTCCAC | 227 |
| chr1:106790899-106792914 | p5 | (CATAG)3 | TGTACCGAGCACCCCTTATC | ACAGTACTCGGCCCCTTTCT | 249 |
| chr1:106804883-106806898 | p5 | (AAAAG)3 | AGGCCTCACAAACACAAACC | CTTCCGGTCGTTATGGAAAA | 172 |
| chr1:106823554-106825586 | p2 | (AT)16 | TGTCCTGCAGCCTCCTATCT | TCATATATGGGCCACCCTGT | 178 |
| chr1:106843190-106845224 | p2 | (AT)17 | GGAAAGATTGGAGGGTCACA | CAACGTGGACGTAGCTCTCA | 289 |
| chr1:106843190-106845224 | p2 | (TA)6 | GGAAAGATTGGAGGGTCACA | CAACGTGGACGTAGCTCTCA | 289 |
| chr1:106936383-106938398 | p5 | (TCTTT)3 | GGGAAGCATACACAATGCCT | ATGTTTTGTCTGCCTGGTCC | 265 |
| chr1:106936383-106938398 | p6 | (TTTTCC)3 | GGGAAGCATACACAATGCCT | ATGTTTTGTCTGCCTGGTCC | 265 |
| chr1:106963136-106965151 | p3 | (TGT)5 | TCATTTTGACCCGATGTTGA | CCCAAGGTGGAGAGTTGGTA | 274 |
| chr1:107057034-107059058 | p2 | (TA)12 | CGTATTCGGTAGAACGGGAA | GTGTCCAGTGGGGTCACTTT | 165 |
| chr1:107059367-107061408 | p2 | (AT)10 | TGTTACAATTCGCAAGGCAA | GGGCAAGTCCAAAACTCCTT | 182 |
| chr1:107072071-107074083 | p2 | (AT)6 | TAATTGACCTCCCCCAACTG | ATGACACGGAACCAGGACTC | 187 |
| chr1:107085150-107087162 | p2 | (AT)6 | TAATTGACCTCCCCCAACTG | ATGACACGGAACCAGGACTC | 187 |
| chr1:107262282-107264297 | p3 | (ATG)5 | CCTCCAAAGTTGACCCAAAA | ATCGGAGATGTCCCATGAAG | 215 |
| chr1:107337038-107339056 | p3 | (CAA)6 | ATCTTCCGTACTGACCCCCT | GCTCCTGAACAAAGGAGCAC | 153 |
| chr1:107358979-107360994 | p5 | (ATTAT)3 | CGTAGATTTGGGGCGTAAAA | TTCTTGATGATGGGAGAGGG | 289 |
| chr1:107372431-107374443 | p2 | (AT)6 | AAGATAGGCTGTCAGGGCAA | TGGATGAGGAAATGAGGAGG | 184 |
| chr1:107398383-107400398 | p3 | (TAA)5 | ATTCCTGCAAGCATTGATCC | AAAAATGAGGTCATCGGTGC | 156 |
| chr1:107413562-107415607 | p3 | (TGT)15 | TATGGAGATCCGTACCCGAG | TCGACAGCGTAAAACGACAG | 288 |

| Marker ID | SSR type | SSR motif | FORWARD PRIMER (5'-3') | REVERSE PRIMER (5'-3') | size (bp) |
|---|---|---|---|---|---|
| chr1:107423757-107425779 | p2 | (AT)11 | TCTTCGGACCTCATGGAATC | ACTTGTGGGTATGGGTGGAA | 161 |
| chr1:107620403-107622417 | p2 | (CA)7 | TGAGGCATGTATCAATCGGA | AGGCACAGTTCAGGGTATGG | 107 |
| chr1:107750627-107752653 | p2 | (AT)13 | GCCACACTTTTACCCTTCCA | AACCATAGCGTGCCTCAAAC | 221 |
| chr1:107913757-107915773 | p2 | (TA)8 | TACGGACCGTCCTTCATAGC | GCTTAAGTTGGGGGTGTTGA | 108 |
| chr1:107955049-107957118 | p5 | (TAATT)3 | GACAAAGCCGTCGGTATGAT | TAGACCCGCAATTGATTCGT | 132 |
| chr1:107955201-107957237 | p5 | (TAATT)3 | ACGAATCAATTGCGGGTCTA | ATTCACCTGCAGGGTTTTCA | 227 |
| chr1:107955252-107957282 | p5 | (TAATT)3 | ACGAATCAATTGCGGGTCTA | ATTCACCTGCAGGGTTTTCA | 176 |
| chr1:107995561-107997579 | p6 | (GTGGGT)3 | TCGAGCCAGACTCGTAGGAT | GCTGTAGAAAGGACGGGACA | 241 |
| chr1:108035717-108037735 | p3 | (TTG)6 | GAAGGTACAAGCCATGGGAA | ATGGAATCTTCCCCAACTCC | 135 |
| chr1:108035717-108037735 | p2 | (TA)14 | GAAGGTACAAGCCATGGGAA | ATGGAATCTTCCCCAACTCC | 135 |
| chr1:108053762-108055778 | p5 | (CCCAC)3 | ACCACTAGCTTCACCCCTCC | GTGTTGGTAGTGAGTGGGGG | 130 |
| chr1:108053762-108055778 | p2 | (TC)8 | ACCACTAGCTTCACCCCTCC | GTGTTGGTAGTGAGTGGGGG | 130 |
| chr1:108077909-108080007 | p2 | (TA)6 | ACATGAGGACGGGACAAGAC | TGTTCATGCCCACAGGTAGA | 273 |
| chr1:108077909-108080007 | p2 | (AT)45 | ACATGAGGACGGGACAAGAC | TGTTCATGCCCACAGGTAGA | 273 |
| chr1:108077909-108080007 | p2 | (AT)49 | ACATGAGGACGGGACAAGAC | TGTTCATGCCCACAGGTAGA | 273 |
| chr1:108090629-108092727 | p2 | (AT)7 | ACCAGCCGACAAGGCTACTA | GGAGCACTCCACTTTCTTCG | 252 |
| chr1:108227571-108229591 | p2 | (AT)10 | CTTCCTGAATCTCCCAACCA | CACATCCCGTTGCAGTTATG | 198 |
| chr1:108227571-108229591 | p5 | (AATGC)3 | CTTCCTGAATCTCCCAACCA | CACATCCCGTTGCAGTTATG | 198 |
| chr1:108310754-108312790 | p2 | (AT)18 | ACTCGCAACGGTACTTTCGT | TGCGGTTTCTGACTTGTTTG | 278 |
| chr1:108433827-108435842 | p5 | (TTAAA)3 | GCGTTAGAGGCAGAATTTCG | CCCTATTTCGGGGTGATCTT | 145 |
| chr1:108436702-108438730 | p2 | (TA)8 | ACCTCCCCGAAGAGTTGATT | TAACACGTGTCAAATGGGGA | 189 |
| chr1:108467945-108469960 | p3 | (GAT)5 | CAAAATAACCCCCACGTTTG | GCTCTTGTCCACATCAGCAA | 185 |
| chr1:108508151-108510173 | p5 | (TCTTT)3 | GGATCGGAGAAAGAAAAGGG | TAATGAGACACCGGGGAGAG | 251 |
| chr1:108508210-108510228 | p5 | (TCTTT)3 | GGATCGGAGAAAGAAAAGGG | TAATGAGACACCGGGGAGAG | 192 |
| chr1:108523200-108525215 | p3 | (TAT)10 | TTCGGGTCCAAACTTCTGTC | CAAACATTCCCGATTCTCGT | 275 |
| chr1:108523200-108525215 | p5 | (AAGAG)3 | TTCGGGTCCAAACTTCTGTC | CAAACATTCCCGATTCTCGT | 275 |
| chr1:108523226-108525241 | p3 | (TAT)10 | TTCGGGTCCAAACTTCTGTC | CAAACATTCCCGATTCTCGT | 249 |
| chr1:108523226-108525241 | p5 | (AAGAG)3 | TTCGGGTCCAAACTTCTGTC | CAAACATTCCCGATTCTCGT | 249 |
| chr1:108523298-108525331 | p3 | (TAT)10 | TTCGGGTCCAAACTTCTGTC | CAAACATTCCCGATTCTCGT | 177 |
| chr1:108523298-108525331 | p5 | (AAGAG)3 | TTCGGGTCCAAACTTCTGTC | CAAACATTCCCGATTCTCGT | 177 |
| chr1:108523346-108525403 | p3 | (TAT)10 | TTCGGGTCCAAACTTCTGTC | CAAACATTCCCGATTCTCGT | 129 |
| chr1:108523346-108525403 | p5 | (AAGAG)3 | TTCGGGTCCAAACTTCTGTC | CAAACATTCCCGATTCTCGT | 129 |
| chr1:108536161-108538176 | p5 | (ATAAA)3 | GTTGGGTTGGAGTTGGTGAG | GGATGTGTTCGGTATGGAGG | 157 |
| chr1:108639959-108641975 | p2 | (GT)8 | TGGATTGGTTGGCTTCTTTC | CCGTCTTTTGTAATTCCCGA | 264 |
| chr1:108688130-108690160 | p3 | (ATA)10 | GGAGAAAGAGGAACGGGAAC | TACCCCTCTGACACCTCCAC | 258 |
| chr1:108763055-108765069 | p2 | (AT)7 | ACCTCTCGTTGGCACACTCT | GCGGCTCGGTAATGAAAATA | 210 |

| Marker ID | SSR type | SSR motif | FORWARD PRIMER (5'-3') | REVERSE PRIMER (5'-3') | size (bp) |
|---|---|---|---|---|---|
| chr1:108781452-108783476 | p3 | (CTT)8 | ACTTACCATCAGTGCCCAGG | AAACTTGGTTGATCGTTGCC | 160 |
| chr1:108781452-108783476 | p3 | (CTT)6 | ACTTACCATCAGTGCCCAGG | AAACTTGGTTGATCGTTGCC | 160 |
| chr1:108802499-108804511 | p2 | (TA)12 | ACACCTTTCACTGAGTCCCG | GATGTATAACGTGTCCCCCG | 208 |
| chr1:108802499-108804511 | p2 | (AT)6 | ACACCTTTCACTGAGTCCCG | GATGTATAACGTGTCCCCCG | 208 |
| chr1:108807414-108809429 | p3 | (ATT)5 | AACGGCAAAATTTGAAAACG | TGGAATTGAGACGGAAGGAG | 162 |
| chr1:108916162-108918178 | p4 | (ATAA)4 | GCATCATTTCCATTTCGTCC | CTCGCCGATCCCAATTACTA | 160 |
| chr1:108916162-108918178 | p2 | (AT)10 | GCATCATTTCCATTTCGTCC | CTCGCCGATCCCAATTACTA | 160 |
| chr1:108917961-108920001 | p2 | (TA)20 | CCAACCCCCTCCATTTTTAT | ATTAATGCACACAGCAGCCA | 256 |
| chr1:108957469-108959484 | p5 | (AAAAC)3 | CCTTACCTTTCGCCCTTTCT | ATTTCAACCATTGGAGACGC | 113 |
| chr1:109087361-109089376 | p5 | (ATGCA)3 | CATGCTTGGACAACTGGATG | TGAATTGAGGGTTCGCCTAC | 196 |
| chr1:109175644-109177658 | p2 | (AT)7 | ACCCAAGTAGCCATGAGTGG | TGGGATCTTTGGGGATGTAA | 194 |
| chr1:109182125-109184153 | p2 | (AT)14 | ACAGGTACGCGAGCTCCTAA | GTTTCGTCGCGTGGATCTAT | 297 |
| chr1:109302900-109304930 | p2 | (AT)15 | AATTCTGGAACCGTTGTTGC | ATGAATGCAATGCTATGCCA | 195 |
| chr1:109400323-109402347 | p5 | (CATTG)3 | TTGGTCCCGAAATCATTGTT | GACTCGCGAATCCTAAGTGC | 234 |
| chr1:109400323-109402347 | p4 | (ATAC)6 | TTGGTCCCGAAATCATTGTT | GACTCGCGAATCCTAAGTGC | 234 |
| chr1:109520741-109522753 | p2 | (GT)6 | AATCACGTCACCCACACAGA | AACTTAGAAGGATGGGCCGT | 218 |
| chr1:109574637-109576653 | p4 | (TTTG)4 | GCCGAATTTCATGGAGAAAA | GCAAAAGAAGGAAAGGGGAC | 137 |
| chr1:109773099-109775114 | p2 | (TA)6 | TATCATTGGCCACCTCAACA | CGATTTCGCTTCAAGTCACA | 217 |
| chr1:109773099-109775114 | p3 | (TTC)5 | TATCATTGGCCACCTCAACA | CGATTTCGCTTCAAGTCACA | 217 |
| chr1:109788040-109790052 | p2 | (AT)6 | CATAAAGGGTACGCCCTGAA | CTCGGCATATGGACGTTTTT | 258 |
| chr1:109788040-109790052 | p2 | (GA)6 | CATAAAGGGTACGCCCTGAA | CTCGGCATATGGACGTTTTT | 258 |
| chr1:109788040-109790052 | p2 | (GA)6 | CATAAAGGGTACGCCCTGAA | CTCGGCATATGGACGTTTTT | 258 |
| chr1:109788174-109790186 | p2 | (AT)6 | CATAAAGGGTACGCCCTGAA | CTCGGCATATGGACGTTTTT | 124 |
| chr1:109788174-109790186 | p2 | (GA)6 | CATAAAGGGTACGCCCTGAA | CTCGGCATATGGACGTTTTT | 124 |
| chr1:109788174-109790186 | p2 | (GA)6 | CATAAAGGGTACGCCCTGAA | CTCGGCATATGGACGTTTTT | 124 |
| chr1:109851912-109853928 | p2 | (TC)6 | CCGCATATTCCGATCCATAC | TTCTTCTTTGGACCACCACC | 117 |
| chr1:109851912-109853928 | p3 | (AAT)7 | CCGCATATTCCGATCCATAC | TTCTTCTTTGGACCACCACC | 117 |
| chr1:109851912-109853928 | p2 | (TA)8 | CCGCATATTCCGATCCATAC | TTCTTCTTTGGACCACCACC | 117 |
| chr1:109894891-109896906 | p5 | (TGACT)3 | GGCTTGTTACACCTCGGAAA | CCTTGCCTTCTTCAACGAAC | 127 |
| chr1:109922553-109924568 | p5 | (AAAAT)3 | GGGTTTGTGGAAATTTGTGG | GCACCTTTTACCCTCCATCA | 240 |
| chr1:110044915-110046930 | p3 | (ACA)5 | TTGATGATCATGGTGTGGCT | TAGCAAAAGTAGGCGAGGGA | 219 |
| chr1:110060927-110062942 | p5 | (AAATA)3 | ACTGCAGCCCAGACAGAGAT | AAGGTCTGATGAAACACGGC | 151 |
| chr1:110141607-110143622 | p3 | (ACA)5 | TTGATGATCATGGTGTGGCT | TAGCAAAAGTAGGCGAGGGA | 219 |
| chr1:110157619-110159634 | p5 | (AAATA)3 | ACTGCAGCCCAGACAGAGAT | AAGGTCTGATGAAACACGGC | 151 |
| chr1:110221018-110223048 | p2 | (TA)15 | TTTCGGCGATCCAACTTATC | TTGCTCGGAGATTATACGGG | 179 |
| chr1:110287884-110289896 | p2 | (AT)6 | AACAACGCATGCCATATTCA | GATACCCATTTCGGCGACTA | 192 |

| Marker ID | SSR type | SSR motif | FORWARD PRIMER (5'-3') | REVERSE PRIMER (5'-3') | size (bp) |
|---|---|---|---|---|---|
| chr1:110287884-110289896 | p2 | (AT)7 | AACAACGCATGCCATATTCA | GATACCCATTTCGGCGACTA | 192 |
| chr1:110375510-110377538 | p2 | (TA)14 | AACAAAATCCACCCCTTCCT | AGGACGCTCCAACTCTAGCA | 280 |
| chr1:110414235-110416251 | p2 | (AT)8 | TACTCATGACTCCGCTCGTG | TCATAACACAGCATAGCGCC | 214 |
| chr1:110463949-110465969 | p2 | (TA)10 | CCCACTGACATCCATATCCC | GTTTTTGGGGTGGCTTTCTT | 249 |
| chr1:110478691-110480729 | p2 | (AT)19 | TAGTGGCATGGTGCATTTGT | TTCCAATCAAAACATGGCAA | 199 |
| chr1:110523128-110525144 | p2 | (TC)8 | TCAGTCCGCAATATTCACCA | GGGGTGTGATTGGGTTAATG | 284 |
| chr1:110526598-110528610 | p2 | (AT)6 | AACGTAAAGCGGCAGAAAAA | GTGCGTTGGTTAAAAAGGGA | 115 |
| chr1:110526598-110528610 | p5 | (GTGGG)3 | AACGTAAAGCGGCAGAAAAA | GTGCGTTGGTTAAAAAGGGA | 115 |
| chr1:110589500-110591516 | p4 | (TTTA)4 | ATCTATGCGGACAAACCGTC | CCCCAGGATTTTTACGTTGA | 267 |
| chr1:110622884-110624914 | p2 | (AT)15 | GTGATGACCAAGCACCATTG | TCTGGCCAATTTTTCCTTTG | 121 |
| chr1:110714833-110716848 | p5 | (CTTTT)3 | GCACTCAATCTTTTGCCCAT | TGGTATGGGCGTCTTGATTT | 134 |
| chr1:110717310-110719338 | p4 | (AAAT)7 | TCGGTTGGACAATGATCAGA | CATTCCAAGGCCAACTTGAT | 257 |
| chr1:110909239-110911264 | p5 | (ATCAT)5 | AGGCACGATGAAAATTCGAC | ACTGCCATCCAACCAGTCTC | 156 |
| chr1:110909239-110911264 | p6 | (CATATA)3 | AGGCACGATGAAAATTCGAC | ACTGCCATCCAACCAGTCTC | 156 |
| chr1:111040887-111042934 | p2 | (GT)7 | ATGGATTTTGAATTGGCAGC | GATCCGCACTCTCCACTAGC | 110 |
| chr1:111045074-111047096 | p2 | (TA)11 | AAGATTGGGATATGGGAGGG | ACGAAAATCAGGCAGCACTT | 281 |
| chr1:111045074-111047096 | p2 | (AT)14 | AAGATTGGGATATGGGAGGG | ACGAAAATCAGGCAGCACTT | 281 |
| chr1:111068302-111070316 | p2 | (TA)6 | AAAAATTGAAACCGCAGACG | CGCCGATTAAAACCCATTTA | 252 |
| chr1:111068302-111070316 | p2 | (AT)6 | AAAAATTGAAACCGCAGACG | CGCCGATTAAAACCCATTTA | 252 |
| chr1:111068302-111070316 | p2 | (TA)10 | AAAAATTGAAACCGCAGACG | CGCCGATTAAAACCCATTTA | 252 |
| chr1:111068302-111070316 | p2 | (AT)7 | AAAAATTGAAACCGCAGACG | CGCCGATTAAAACCCATTTA | 252 |
| chr1:111089711-111091743 | p2 | (TA)16 | GAGCAGCAATAGCACCCTTC | ACCATTGAACCACCACCATT | 218 |
| chr1:111089711-111091743 | p2 | (CT)7 | GAGCAGCAATAGCACCCTTC | ACCATTGAACCACCACCATT | 218 |
| chr1:111101907-111103922 | p3 | (ATG)5 | GTTTGGTGTTGGAGTTGGCT | TTTCCGTAAAGAAGCCGAAA | 105 |
| chr1:111134303-111136323 | p2 | (AT)10 | CCCCCTTGATAACCTCCAGT | GACGCTAGTACAAGAGCCCG | 115 |
| chr1:111154181-111156197 | p2 | (AT)22 | GTCCTGTCCCGTCCTTATGA | GGTACCCGGAGCTACCCTAC | 201 |
| chr1:111154198-111156213 | p2 | (AT)22 | GTCCTGTCCCGTCCTTATGA | GGTACCCGGAGCTACCCTAC | 184 |
| chr1:111166311-111168326 | p5 | (TTATT)3 | ACCACAAATATGAGGGGCAA | AGTAGTCGTTTGGTACGCGG | 169 |
| chr1:111339622-111341634 | p2 | (AT)6 | AACTCCATGACCCTGCAAAG | TTGGCCAATTAAAAGGTTGG | 190 |
| chr1:111375488-111377506 | p6 | (CCGGTT)3 | TCATTAAGCGCACCCCTAAC | TCTAGCCGCCTTCACTTTGT | 255 |
| chr1:111447868-111449882 | p2 | (TA)7 | CACAAATGTGGGTCACTTGC | AATTGCACGGTTTGTCCTTC | 269 |
| chr1:111487062-111489088 | p2 | (AT)13 | CAAAACACCGAAACCGAAAT | TTGTTTCCGTTAGACCCTCG | 134 |
| chr1:111513318-111515358 | p5 | (TAAAA)3 | GGATGGCGTTGTAAATGACC | TTGACACCCTCAATTCCACA | 128 |
| chr1:111513318-111515358 | p2 | (AT)20 | GGATGGCGTTGTAAATGACC | TTGACACCCTCAATTCCACA | 128 |
| chr1:111582613-111584631 | p3 | (AAT)6 | CAAGGTTTTAGGCGTTTTCC | GTGCCTGTTTGGTCATGAGA | 211 |
| chr1:111597312-111599328 | p2 | (AG)8 | GCCTATTTCAACCGGCAGTA | CCCCCTACCTCGCTATTGTT | 216 |

| Marker ID | SSR type | SSR motif | FORWARD PRIMER (5'-3') | REVERSE PRIMER (5'-3') | size (bp) |
|---|---|---|---|---|---|
| chr1:111630710-111632722 | p2 | (TA)6 | ATTTCCCATCATGCGTTTTC | CCCTTTTTCATCCTGTCCAA | 177 |
| chr1:111723976-111725994 | p2 | (AT)9 | TATTTGGCTCACGCAGATGA | ATCATTTGGCTACACAGCCC | 223 |
| chr1:111806976-111808994 | p6 | (GTGGAC)4 | GGTGGACGTTGTTCCACTCT | TTGATCCCTCGAGAAGTGCT | 206 |
| chr1:111806976-111808994 | p6 | (GTGGAC)3 | GGTGGACGTTGTTCCACTCT | TTGATCCCTCGAGAAGTGCT | 206 |
| chr1:111806976-111808994 | p6 | (GTGGAC)5 | GGTGGACGTTGTTCCACTCT | TTGATCCCTCGAGAAGTGCT | 206 |
| chr1:111807204-111809234 | p6 | (GTGGAC)4 | CAGGATCGAAACTTGCATCA | TTCGACGTCCACAATATCCA | 233 |
| chr1:111807204-111809234 | p6 | (GTGGAC)3 | CAGGATCGAAACTTGCATCA | TTCGACGTCCACAATATCCA | 233 |
| chr1:111807204-111809234 | p6 | (GTGGAC)5 | CAGGATCGAAACTTGCATCA | TTCGACGTCCACAATATCCA | 233 |
| chr1:111947717-111949732 | p5 | (ATCAT)3 | ACCTCGAAGAACTCCAAGCA | ACATTAGTCACGGCCTACGG | 109 |
| chr1:111962662-111964710 | p3 | (ACA)16 | GCACGGGTCTTTGACTTGAT | TTGCGGCTGAAAATCTCTTT | 208 |
| chr1:111985105-111987120 | p5 | (ATTTC)3 | CGCATTTAATTCAGCCCCTA | ATAACAATGCCTCACAGGGC | 206 |
| chr1:111992850-111994865 | p5 | (ATTTC)3 | GACATCCAAGTACCCGCATT | ATAACAATGCCTCACAGGGC | 192 |
| chr1:112000197-112002212 | p5 | (ATTTC)3 | GACATCCAAGTACCCGCATT | ATAACAATGCCTCACAGGGC | 192 |
| chr1:112072340-112074356 | p2 | (TA)7 | GTGGAATTCACCCTCTCCAA | CGAATACAAAGCCAGGCACT | 277 |
| chr1:112072340-112074356 | p2 | (CT)8 | GTGGAATTCACCCTCTCCAA | CGAATACAAAGCCAGGCACT | 277 |
| chr1:112133866-112135878 | p3 | (AAT)5 | GCCAAAGTTTTCCTGCACTC | TTTAGGGGCCACAGTAAGGA | 168 |
| chr1:112133866-112135878 | p2 | (AT)6 | GCCAAAGTTTTCCTGCACTC | TTTAGGGGCCACAGTAAGGA | 168 |
| chr1:112322905-112324961 | p2 | (TA)8 | TCCATTAAAGGACTGACCCG | CCGATGACGAGGAGAGAGAG | 192 |
| chr1:112322905-112324961 | p2 | (AT)8 | TCCATTAAAGGACTGACCCG | CCGATGACGAGGAGAGAGAG | 192 |
| chr1:112322905-112324961 | p2 | (AT)28 | TCCATTAAAGGACTGACCCG | CCGATGACGAGGAGAGAGAG | 192 |
| chr1:112322905-112324961 | p2 | (TA)7 | TCCATTAAAGGACTGACCCG | CCGATGACGAGGAGAGAGAG | 192 |
| chr1:112326108-112328160 | p2 | (AT)26 | ATAACGCCATCAAGCCTCTG | CGACCTTACGTCACTCCGAT | 275 |
| chr1:112355186-112357198 | p5 | (ATAGC)3 | AAGTCGACTGGGGGAGAAGT | GACGCGATATGCTACGACAA | 248 |
| chr1:112355197-112357239 | p5 | (ATAGC)3 | AAGTCGACTGGGGGAGAAGT | GACGCGATATGCTACGACAA | 237 |
| chr1:112355252-112357268 | p5 | (ATAGC)3 | AAGTCGACTGGGGGAGAAGT | GACGCGATATGCTACGACAA | 182 |
| chr1:112355270-112357282 | p5 | (ATAGC)3 | AAGTCGACTGGGGGAGAAGT | GACGCGATATGCTACGACAA | 164 |
| chr1:112370324-112372344 | p4 | (TTAA)5 | TAGGACAAGTCATCATGGCG | TCATCACGAGAATGGGTTGA | 107 |
| chr1:112388546-112390574 | p2 | (GA)6 | ACCGTAAATGTGGTGGAAGC | TCCGAATCGAAGCTCAAAAT | 164 |
| chr1:112388546-112390574 | p5 | (TATAA)3 | ACCGTAAATGTGGTGGAAGC | TCCGAATCGAAGCTCAAAAT | 164 |
| chr1:112412883-112414898 | p3 | (TTC)5 | AGTGGTAGGGTTTTGAGGGG | TAGGGTGGAATCAAAGGTGC | 129 |
| chr1:112477789-112479803 | p2 | (AC)7 | CAATCTAGACGGCATGAGCA | CTCGGAGATACTGAGTCCGC | 184 |
| chr1:112483694-112485709 | p5 | (AATGC)3 | TGACACCTGCAGAATTGAGG | TCTTTCCGCTTGCTTGATTT | 205 |
| chr1:112598334-112600349 | p2 | (AT)9 | CGGTAGGTCTGTCCGTCATT | CCGACAACTCCTCACAGACA | 117 |
| chr1:112605390-112607408 | p2 | (AT)7 | AGGCTGCGTCTAAACGATGT | TGATACTGCAAGCACAAGGC | 262 |
| chr1:112605390-112607408 | p2 | (AT)6 | AGGCTGCGTCTAAACGATGT | TGATACTGCAAGCACAAGGC | 262 |
| chr1:112605390-112607408 | p2 | (TA)9 | AGGCTGCGTCTAAACGATGT | TGATACTGCAAGCACAAGGC | 262 |

| Marker ID | SSR type | SSR motif | FORWARD PRIMER (5'-3') | REVERSE PRIMER (5'-3') | size (bp) |
|---|---|---|---|---|---|
| chr1:112826092-112828104 | p2 | (TA)6 | TACGGACCATCCTTTTGCTC | TGTTGATGTGCCGTGAATTT | 174 |
| chr1:112908077-112910137 | p6 | (ATTGGG)3 | CCTCTCCCAAACCATTCAAA | TTTTCGCCCATCTTATCTCG | 169 |
| chr1:112977516-112979552 | p3 | (ATT)12 | CTGCCTCAACAAGATGTCCA | TTTTGAAGCTTGTGCCAGTG | 299 |
| chr1:112977516-112979552 | p5 | (ATAAA)3 | CTGCCTCAACAAGATGTCCA | TTTTGAAGCTTGTGCCAGTG | 299 |
| chr1:113038408-113040420 | p2 | (CT)6 | TTTGGTTGCTTCTACCCAGG | TCAATCAGGCAACAAATGGA | 287 |
| chr1:113093702-113095718 | p4 | (CTAT)4 | ATTGTCTCTTCCCCCACCTC | GTTTGGCACGTGTCTCTTCA | 243 |
| chr1:113118090-113120108 | p3 | (AAG)6 | AAGCCCCCTAAGGTCGATAA | TACATGTCATCGGCGAGTGT | 104 |
| chr1:113224749-113226770 | p3 | (AAT)7 | CTCCACTTCATGCACCAAAA | TGGTTGTTTCCTTGTCACCA | 183 |
| chr1:113271082-113273100 | p3 | (TAA)6 | TTTTATGGGGTTTATGGGCA | GGCACCTGACGAGCTTCTAC | 155 |
| chr1:113271082-113273100 | p5 | (AAGAG)3 | TTTTATGGGGTTTATGGGCA | GGCACCTGACGAGCTTCTAC | 155 |
| chr1:113271905-113273944 | p3 | (TAA)6 | CGGGTCCAAACTTCTGTCAT | CAAACATTCCCCATTCTCGT | 227 |
| chr1:113271905-113273944 | p5 | (AAGAG)3 | CGGGTCCAAACTTCTGTCAT | CAAACATTCCCCATTCTCGT | 227 |
| chr1:113300777-113302789 | p2 | (AT)21 | GTAGATGAAGCCGAACTCGC | CGACAAGCCCTTTTTGCTAC | 118 |
| chr1:113301058-113303100 | p2 | (AT)21 | TGTGCATTCACGATTGGATT | ACCATTCGTTAATTTCCCCC | 252 |
| chr1:113301058-113303100 | p2 | (AT)22 | TGTGCATTCACGATTGGATT | ACCATTCGTTAATTTCCCCC | 252 |
| chr1:113305270-113307282 | p3 | (CAA)5 | TTCCCGTTTCGTCAAAAATC | GTTGTTGGTGTTGTTGCTGG | 153 |
| chr1:113305270-113307282 | p2 | (AC)6 | TTCCCGTTTCGTCAAAAATC | GTTGTTGGTGTTGTTGCTGG | 153 |
| chr1:113307386-113309406 | p5 | (AAATC)4 | TTTTATGGGGTTTATGGGCA | TTGTCACGACCCAAATCTGA | 281 |
| chr1:113325551-113327563 | p3 | (CAA)5 | TTCCCGTTTCGTCAAAAATC | GTTGTTGGTGTTGTTGCTGG | 153 |
| chr1:113325551-113327563 | p2 | (AC)6 | TTCCCGTTTCGTCAAAAATC | GTTGTTGGTGTTGTTGCTGG | 153 |
| chr1:113327667-113329687 | p5 | (AAATC)4 | TTTTATGGGGTTTATGGGCA | TTGTCACGACCCAAATCTGA | 281 |
| chr1:113357819-113359835 | p2 | (TA)6 | ATATCACATGCAAGCCGACA | TTTATCCAGGGGTACGACGA | 289 |
| chr1:113357819-113359835 | p2 | (AT)8 | ATATCACATGCAAGCCGACA | TTTATCCAGGGGTACGACGA | 289 |
| chr1:113366144-113368176 | p2 | (TC)16 | CCCGCCAAAATGAGAAGATA | CAAAAGGCAGATCAATCGGT | 285 |
| chr1:113366144-113368176 | p2 | (TC)16 | CCCGCCAAAATGAGAAGATA | CAAAAGGCAGATCAATCGGT | 285 |
| chr1:113386475-113388490 | p3 | (ACA)5 | CTTGCCATGGGATGAATCTT | TTGTTTCTCCCCCTCCTTTT | 226 |
| chr1:113469134-113471152 | p3 | (TTG)6 | AGACCCTCCAAAAGGACGAT | GGTCATAACGTTTGATCCCG | 249 |
| chr1:113599621-113601639 | p5 | (GTTCG)5 | CACACACAGTCATGGGGAAG | AAGGCTTCAGGTAGGGGAGA | 269 |
| chr1:113599621-113601639 | p6 | (AAAATC)3 | CACACACAGTCATGGGGAAG | AAGGCTTCAGGTAGGGGAGA | 269 |
| chr1:113689357-113691372 | p3 | (ACA)5 | CCTTTGTTTGGATCGGTTTC | TGATTTCACTGGAGCGTGTT | 130 |
| chr1:113743700-113745712 | p2 | (TC)6 | CTTGTGGTGACCGTTCAATG | AGCCCCACAGTCCTTTTCTT | 156 |
| chr1:113785095-113787125 | p2 | (TA)7 | AGGAAAGGTACGGAGAGGGA | TTGCCTCCAATCAACAATGA | 252 |
| chr1:113837030-113839044 | p2 | (TA)7 | CAATGACCATGCCTGAAGTG | GGCTAAATGCACTTTCCGAG | 124 |
| chr1:113857387-113859405 | p3 | (ACA)6 | TCAGATTGCTTGTGAGGTGC | GCGTGGATTCTTCATTCTCC | 259 |
| chr1:113894131-113896146 | p3 | (AAG)5 | ATCTACGGTCATGTCCTCGG | TGAGGATATTCGGGACAAGG | 154 |
| chr1:113899489-113901507 | p6 | (AAAAAT)3 | TCTATAAGCAGCTCGGCCAT | CAGACGGTGGTTACAACGTG | 234 |

| Marker ID | SSR type | SSR motif | FORWARD PRIMER (5'-3') | REVERSE PRIMER (5'-3') | size (bp) |
|---|---|---|---|---|---|
| chr1:113899489-113901507 | p2 | (AT)33 | TCTATAAGCAGCTCGGCCAT | CAGACGGTGGTTACAACGTG | 234 |
| chr1:114037979-114040001 | p2 | (AT)11 | CTTTGGAACTTGGGTTGGAA | CGGAGCCTGTCATACTCCTC | 130 |
| chr1:114038387-114040405 | p2 | (AT)11 | CCCACAATGTCCATAAGCCT | GAGTTATCGAGGAGCAACCG | 190 |
| chr1:114038400-114040462 | p2 | (AT)11 | CCCACAATGTCCATAAGCCT | GAGTTATCGAGGAGCAACCG | 177 |
| chr1:114046588-114048603 | p5 | (AAATA)3 | GCACAGGTTCCTTGCTTGAT | AGTTTCAGCATGACGCACAC | 237 |
| chr1:114053160-114055175 | p5 | (TTTTA)3 | TATGCTGGGGCCTGACTAAT | GGAAAATAATGGGGCTTGTTC | 129 |
| chr1:114053160-114055175 | p5 | (TTTTA)3 | TATGCTGGGGCCTGACTAAT | GGAAAATAATGGGGCTTGTTC | 129 |
| chr1:114053160-114055175 | p5 | (TTTTA)3 | TATGCTGGGGCCTGACTAAT | GGAAAATAATGGGGCTTGTTC | 129 |
| chr1:114066913-114068927 | p2 | (AT)6 | ACTTAGAAGCGGAGGAAGGC | GGTTAATTTTCCGCCATTCA | 270 |
| chr1:114066931-114068953 | p2 | (AT)6 | ACTTAGAAGCGGAGGAAGGC | GGTTAATTTTCCGCCATTCA | 252 |
| chr1:114066960-114068976 | p2 | (AT)6 | ACTTAGAAGCGGAGGAAGGC | GGTTAATTTTCCGCCATTCA | 223 |
| chr1:114169046-114171064 | p6 | (TCAGTT)3 | ACCAGTAATGGAGATTGCGG | AATTCCAGCCTTTGAGCGTA | 257 |
| chr1:114173082-114175098 | p2 | (AT)8 | TGGAGTATGAGGAACGAGGC | TATGCCCCTCCAAAGTTGAC | 196 |
| chr1:114433382-114435410 | p2 | (AT)14 | ACAAACCAAATGATGTGCCA | TAACACCCTCGTTGAAAGGG | 245 |
| chr1:114450637-114452653 | p6 | (CATGAG)3 | GCAACCGGAAGAAACAACAT | TTAGCTGATCGGGCAGAGAT | 259 |
| chr1:114450637-114452653 | p4 | (ATAC)4 | GCAACCGGAAGAAACAACAT | TTAGCTGATCGGGCAGAGAT | 259 |
| chr1:114509213-114511249 | p2 | (AG)7 | GCGGAAAAAGAAAAGGGAAG | TCCTAAACGAATACAGGCGG | 256 |
| chr1:114509258-114511276 | p2 | (AG)7 | GCGGAAAAAGAAAAGGGAAG | TCCTAAACGAATACAGGCGG | 211 |
| chr1:114509303-114511321 | p2 | (AG)7 | GCGGAAAAAGAAAAGGGAAG | TCCTAAACGAATACAGGCGG | 166 |
| chr1:114509318-114511366 | p2 | (AG)7 | GCGGAAAAAGAAAAGGGAAG | TCCTAAACGAATACAGGCGG | 151 |
| chr1:114661753-114663771 | p6 | (AAAATA)3 | GCTGGCTATTGTTCGGAGAC | ACTGGCACTTGTGGGAAAAC | 163 |
| chr1:114661753-114663771 | p5 | (CAAAC)3 | GCTGGCTATTGTTCGGAGAC | ACTGGCACTTGTGGGAAAAC | 163 |
| chr1:114699963-114701981 | p6 | (TCTTTT)3 | CCATTCTCAATGGTTCGTCA | TCAGCTAGCGAATCAAGGGT | 230 |
| chr1:114720039-114722057 | p6 | (TATTTT)3 | TCAAAACGTGCTTGTGGAAG | ACCTTAATGCGATGGGTCTG | 155 |
| chr1:114720039-114722057 | p6 | (TTTAAA)3 | TCAAAACGTGCTTGTGGAAG | ACCTTAATGCGATGGGTCTG | 155 |
| chr1:114720039-114722057 | p2 | (TA)11 | TCAAAACGTGCTTGTGGAAG | ACCTTAATGCGATGGGTCTG | 155 |
| chr1:114755079-114757103 | p3 | (GTT)8 | GTGATGGGTGTTCTAGCCGT | ATCAGTCCGCAACGCTTACT | 277 |
| chr1:114772434-114774488 | p2 | (AT)27 | GAGGAGAGGCACAGATGCTC | TCTACCCTAATCCTCGCCCT | 163 |
| chr1:114774777-114776792 | p4 | (TTAT)4 | TTGCAGTCACGAGGACCATA | TAGTGCCCTTCATTCAGGCT | 107 |
| chr1:114774777-114776792 | p3 | (AGT)5 | TTGCAGTCACGAGGACCATA | TAGTGCCCTTCATTCAGGCT | 107 |
| chr1:114793916-114795931 | p3 | (ATT)5 | AGCACCGAATCAATTTGTCC | AACAACATGCACGAAACGAA | 288 |
| chr1:114793916-114795931 | p2 | (TA)7 | AGCACCGAATCAATTTGTCC | AACAACATGCACGAAACGAA | 288 |
| chr1:114818805-114820844 | p3 | (ACA)13 | CCCAAACCATTCATCAATCC | TATTGGCCCCAAAGTCTCAG | 214 |
| chr1:114872950-114875014 | p2 | (AT)32 | TTCATCAACTCAGTCACGCC | TTATTTTAGACCGCGGCATT | 231 |
| chr1:115031510-115033530 | p4 | (AAGA)5 | CGCACCTCGACTAATTCCAT | GATTGCGAACCATGAGGTTT | 172 |
| chr1:115078380-115080392 | p2 | (AC)6 | AAGCTTGGCAAAACAGGCTA | CAGAGAAGCCGAAAAGCAAC | 288 |

| Marker ID | SSR type | SSR motif | FORWARD PRIMER (5'-3') | REVERSE PRIMER (5'-3') | size (bp) |
|---|---|---|---|---|---|
| chr1:115093607-115095625 | p2 | (TA)9 | ATGGCGTTATGTTGGGATGT | CATGTAAGTCATGCATCGGG | 284 |
| chr1:115093607-115095625 | p6 | (TATATG)3 | ATGGCGTTATGTTGGGATGT | CATGTAAGTCATGCATCGGG | 284 |
| chr1:115310294-115312339 | p3 | (TAA)15 | TTTCAGTCGCCCTCTCACTT | CGAGTTTGCCTCCTCTCATC | 179 |
| chr1:115310294-115312339 | p2 | (AT)6 | TTTCAGTCGCCCTCTCACTT | CGAGTTTGCCTCCTCTCATC | 179 |
| chr1:115335843-115337909 | p4 | (AATA)4 | ATAACAAGGCCAAAAATGCG | GAAGTCGCCGGAAAATAACA | 135 |
| chr1:115335843-115337909 | p5 | (CGGGT)3 | ATAACAAGGCCAAAAATGCG | GAAGTCGCCGGAAAATAACA | 135 |
| chr1:115372151-115374167 | p4 | (AAAT)4 | GCTCCATTCAAAGCACAACA | AATGCAATACAAGCCGAACC | 286 |
| chr1:115403321-115405339 | p2 | (AT)9 | TTACATGTTGGGGTTTTCCC | TTGCGGCAAATGACATAGTT | 113 |
| chr1:115414042-115416060 | p2 | (TA)9 | CCCCTCCAACTCATGAAAAA | TTGATGGGGATACTTTGGGA | 178 |
| chr1:115464528-115466554 | p2 | (AT)13 | GGCTAATGCTACCACCTCCA | TTGCAATTGCCACAATCAAT | 296 |
| chr1:115566762-115568784 | p2 | (CT)11 | TTTACGAGGATGGGGACTTG | ACTGGACATCTTGACCTGGC | 159 |
| chr1:115600527-115602551 | p5 | (ATTGC)4 | GCACGCACTCTCATTGTCAT | TTCCGGAACTAACATCGGAC | 111 |
| chr1:115742735-115744750 | p3 | (ACA)5 | TCCGTTAGAAAGAAGCGGAA | CGGGGAATATGTGTAATGGC | 166 |
| chr1:115742735-115744750 | p3 | (AAC)5 | TCCGTTAGAAAGAAGCGGAA | CGGGGAATATGTGTAATGGC | 166 |
| chr1:115777936-115779956 | p2 | (TA)10 | TTGGAGTCCACCTTGGTTTC | GCATATCAAATTTTGGCGCT | 225 |
| chr1:115941989-115944005 | p2 | (AG)8 | TCTGGGTCCTCAGCTTCAGT | TTTTTACGAAAATCGCCCAG | 151 |
| chr1:115941989-115944005 | p3 | (GTT)5 | TCTGGGTCCTCAGCTTCAGT | TTTTTACGAAAATCGCCCAG | 151 |
| chr1:115951874-115953886 | p3 | (TAT)5 | CGACCCTATGTCACCTCGAT | TGCCATCTGGCTATGGTGTA | 288 |
| chr1:115951874-115953886 | p2 | (TA)6 | CGACCCTATGTCACCTCGAT | TGCCATCTGGCTATGGTGTA | 288 |
| chr1:115957867-115959881 | p2 | (AT)7 | CCCAAAACAAAACCCACTTC | ACAAGTTTAAAACGGGGCTG | 114 |
| chr1:115978690-115980706 | p2 | (TA)8 | GGCCCACAAGTCGAGATTTA | TTTTGCCCCTAGATCACACC | 204 |
| chr1:115996326-115998344 | p6 | (TAAAAA)3 | GGATTGGGTTCAAGAGTGGA | CCCATGTTCCAAAAAGGTTG | 266 |
| chr1:116079302-116081352 | p5 | (ATAAA)3 | CGATGAAAACGGTGTGTTTG | TTGTTGTGTCAAATGGGGAA | 234 |
| chr1:116079302-116081352 | p2 | (AT)25 | CGATGAAAACGGTGTGTTTG | TTGTTGTGTCAAATGGGGAA | 234 |
| chr1:116100387-116102417 | p2 | (TA)15 | CAGAAAGGGCCCAACATAAA | ATGGGTATGACGAAGCCTTG | 300 |
| chr1:116100387-116102417 | p3 | (CAA)5 | CAGAAAGGGCCCAACATAAA | ATGGGTATGACGAAGCCTTG | 300 |
| chr1:116244183-116246207 | p3 | (ATT)8 | ATTCATGCCCGCAGATAGAC | GCCATGCCCCATATATGAAC | 131 |
| chr1:116338817-116340839 | p2 | (AC)11 | CTCTGGTGAACTCGGACCAT | TTCTTGATCTGGATCCCTCG | 103 |
| chr1:116338817-116340839 | p2 | (AT)13 | CTCTGGTGAACTCGGACCAT | TTCTTGATCTGGATCCCTCG | 103 |
| chr1:116347743-116349758 | p3 | (AAG)5 | CCAGCTTTGGGGATAAAACA | GGAACAACGGATAGTTGGGA | 284 |
| chr1:116395691-116397709 | p3 | (CAA)6 | ATTGTCCGGAGGTGCTTATG | TTGTGAGACCATCTGACCCA | 183 |
| chr1:116412650-116414662 | p2 | (AC)6 | AAAGGAATTGGAGCGGTTTT | ATACAGAGGAAATGCTGCCG | 210 |
| chr1:116471650-116473665 | p5 | (AATGC)3 | ATTAGGGATGGGAAGGGTTG | CGGCATAACACAGGAGAGGT | 165 |
| chr1:116503015-116505041 | p2 | (AT)13 | TGACATCCTGCTCAACAAGC | GCAGGTCAGAGGGTAATGGA | 148 |
| chr1:116503015-116505041 | p2 | (AT)8 | TGACATCCTGCTCAACAAGC | GCAGGTCAGAGGGTAATGGA | 148 |
| chr1:116512942-116514962 | p5 | (CATAT)4 | GCTATAGTCGGTCGGGCATA | TTTCGGGACTCGGTGTTATC | 206 |

| Marker ID | SSR type | SSR motif | FORWARD PRIMER (5'-3') | REVERSE PRIMER (5'-3') | size (bp) |
|---|---|---|---|---|---|
| chr1:116542895-116544910 | p3 | (ACA)6 | GGGTTCGAACATGGATATGG | CCGCAATTAAGCCCAAATAA | 293 |
| chr1:116542895-116544910 | p3 | (TTA)5 | GGGTTCGAACATGGATATGG | CCGCAATTAAGCCCAAATAA | 293 |
| chr1:116753578-116755594 | p2 | (AT)6 | AGCATAAATGTCACACGCCA | AAGTGGCTCTGCAAGACGAT | 192 |
| chr1:116753578-116755594 | p2 | (AT)6 | AGCATAAATGTCACACGCCA | AAGTGGCTCTGCAAGACGAT | 192 |
| chr1:116753595-116755611 | p2 | (AT)6 | AGCATAAATGTCACACGCCA | AAGTGGCTCTGCAAGACGAT | 175 |
| chr1:116753595-116755611 | p2 | (AT)6 | AGCATAAATGTCACACGCCA | AAGTGGCTCTGCAAGACGAT | 175 |
| chr1:116753615-116755629 | p2 | (AT)6 | AGCATAAATGTCACACGCCA | AAGTGGCTCTGCAAGACGAT | 155 |
| chr1:116753615-116755629 | p2 | (AT)6 | AGCATAAATGTCACACGCCA | AAGTGGCTCTGCAAGACGAT | 155 |
| chr1:116753630-116755660 | p2 | (AT)6 | AGCATAAATGTCACACGCCA | AAGTGGCTCTGCAAGACGAT | 140 |
| chr1:116753630-116755660 | p2 | (AT)6 | AGCATAAATGTCACACGCCA | AAGTGGCTCTGCAAGACGAT | 140 |
| chr1:116814636-116816648 | p2 | (TA)6 | GAAGCGCATTTTCAAACCAT | TCAACGTACACACGAACGGT | 110 |
| chr1:117076115-117078130 | p5 | (AATAA)3 | GCACATGTTCGTCATTTTCG | CTGCGTCTGTGCATTCAACT | 208 |
| chr1:117274146-117276381 | p3 | (TTG)6 | TCTAGTTGGTCGTCGCACAG | TCAAGGGGTGAGCAAAAATC | 249 |
| chr1:117274146-117276381 | p5 | (GATAT)3 | TCTAGTTGGTCGTCGCACAG | TCAAGGGGTGAGCAAAAATC | 249 |
| chr1:117274146-117276381 | p6 | (TATATG)3 | TCTAGTTGGTCGTCGCACAG | TCAAGGGGTGAGCAAAAATC | 249 |
| chr1:117274146-117276381 | p5 | (ATGAT)47 | TCTAGTTGGTCGTCGCACAG | TCAAGGGGTGAGCAAAAATC | 249 |
| chr1:117400419-117402431 | p3 | (GTT)6 | CTCTACCCTCCTCAGACCCC | TCGCACATCATGGGTAGTGT | 278 |
| chr1:117400419-117402431 | p2 | (AG)6 | CTCTACCCTCCTCAGACCCC | TCGCACATCATGGGTAGTGT | 278 |
| chr1:117532732-117534748 | p4 | (AAAT)4 | TGCGTTATACCCTTCAACCC | ATCAAATGGATATCCGAGCG | 249 |
| chr1:117539422-117541434 | p2 | (TA)6 | CTCCAGTTCAGAGGGTGAGC | CTGCCAAAGAAAAGCAGAGG | 119 |
| chr1:117622346-117624374 | p2 | (TA)17 | TTCCTTCTGTTCATGGGTCC | TCCCTATCCCATATAGGCCC | 144 |
| chr1:117622346-117624374 | p2 | (TG)6 | TTCCTTCTGTTCATGGGTCC | TCCCTATCCCATATAGGCCC | 144 |
| chr1:117622346-117624374 | p2 | (TA)14 | TTCCTTCTGTTCATGGGTCC | TCCCTATCCCATATAGGCCC | 144 |
| chr1:117622346-117624374 | p3 | (GTT)5 | TTCCTTCTGTTCATGGGTCC | TCCCTATCCCATATAGGCCC | 144 |
| chr1:117845772-117847784 | p2 | (AG)6 | CACCGGATTACCAAACAAGG | AATCGTGTTTCTGTTTCCCG | 123 |
| chr1:117866272-117868287 | p5 | (TATCA)3 | TCCAACACAACAACCAAGGA | AGCCAGTGAGTGGAGCAAAT | 208 |
| chr1:117896969-117898987 | p6 | (TTTATT)3 | GCTCGAATTTCGGCTCAGTA | GCCAAAAATACCCCCAGATT | 123 |
| chr1:117936056-117938074 | p6 | (TTGGAG)3 | ATCGGTGACAGACGAAAACC | CGGTTAATTAGTTTGGGGCA | 192 |
| chr1:117936056-117938074 | p6 | (AGTTGG)3 | ATCGGTGACAGACGAAAACC | CGGTTAATTAGTTTGGGGCA | 192 |
| chr1:117956935-117958953 | p2 | (TA)7 | ATTTTAGGATTTGACGGGGC | ATATGGTGGACCAAAACCCA | 128 |
| chr1:117964519-117966534 | p3 | (CAA)5 | TTGCAAAATCACCAACTCCA | CTTGTTCCTTCAAGGTGGGA | 160 |
| chr1:118009292-118011355 | p2 | (AG)11 | TAGGCGTGGGATTTCAAAAG | ACGGACAGAGGTTGTTCACC | 224 |
| chr1:118009292-118011355 | p3 | (ATA)21 | TAGGCGTGGGATTTCAAAAG | ACGGACAGAGGTTGTTCACC | 224 |
| chr1:118028948-118030966 | p6 | (TTCATC)3 | ACGTAGGGATGCATGAGGAG | AAGAGTTTGTTTTGGCGTGG | 196 |
| chr1:118155146-118157164 | p2 | (AT)9 | AGTTGAAGCAGCCACCTCTC | CGCTAGCGAATAATGAAGGC | 249 |
| chr1:118175356-118177384 | p2 | (AT)7 | GGATTCCTTGGACTGCAGAA | ATTCGAATCAACCCAGTCCA | 237 |

| Marker ID | SSR type | SSR motif | FORWARD PRIMER (5'-3') | REVERSE PRIMER (5'-3') | size (bp) |
|---|---|---|---|---|---|
| chr1:118191554-118193568 | p2 | (AC)7 | CAATGGGAAATGGGAACTTG | ACCAAAAAGAACCAGCCTCA | 138 |
| chr1:118258254-118260269 | p5 | (GATAT)3 | TGATGATTCAGTGCGAAAGC | TTTGTCACTGCAGCCTGAAC | 166 |
| chr1:118431531-118433546 | p5 | (AAAAT)3 | TAAGAGGCAGGTGGTCATCC | TGGGAGAGCCAGTCTTCAGT | 258 |
| chr1:118458083-118460101 | p2 | (AT)11 | GCCAAGTAATATGGGTCGGA | AAGAGGGACTAACGCTGCAA | 265 |
| chr1:118458083-118460101 | p2 | (TA)9 | GCCAAGTAATATGGGTCGGA | AAGAGGGACTAACGCTGCAA | 265 |
| chr1:118458083-118460101 | p3 | (AAC)5 | GCCAAGTAATATGGGTCGGA | AAGAGGGACTAACGCTGCAA | 265 |
| chr1:118471369-118473383 | p2 | (TC)7 | AAGCAGGTCGTGGAATCAAG | AGCAGCCTCAAAACCAAAAA | 121 |
| chr1:118514732-118516746 | p3 | (AAG)5 | GGTCTTGGCTAATTCTCCCC | GCATTGGAATTTCAGGCATT | 213 |
| chr1:118514732-118516746 | p2 | (TA)7 | GGTCTTGGCTAATTCTCCCC | GCATTGGAATTTCAGGCATT | 213 |
| chr1:118514732-118516746 | p2 | (TA)10 | GGTCTTGGCTAATTCTCCCC | GCATTGGAATTTCAGGCATT | 213 |
| chr1:118619417-118621432 | p3 | (ACA)5 | GCCCGAGACTCAGTGAAAAG | ACGTTCCTCGCTATGGCTTA | 207 |
| chr1:118629552-118631570 | p6 | (CGGCAT)3 | TCGCAGAGTTGACACTGGAC | CCCAAAACAGGCAGCATAAT | 157 |
| chr1:118845630-118847648 | p6 | (ATATAG)3 | TTTGAATTGCACATGCTTTACC | TCGACAAGTTTTATGGCTCTGA | 180 |
| chr1:118845630-118847648 | p3 | (TAT)6 | TTTGAATTGCACATGCTTTACC | TCGACAAGTTTTATGGCTCTGA | 180 |
| chr1:118845630-118847648 | p3 | (AGT)6 | TTTGAATTGCACATGCTTTACC | TCGACAAGTTTTATGGCTCTGA | 180 |
| chr1:118845649-118847670 | p6 | (ATATAG)3 | TTTGAATTGCACATGCTTTACC | TCGACAAGTTTTATGGCTCTGA | 161 |
| chr1:118845649-118847670 | p3 | (TAT)6 | TTTGAATTGCACATGCTTTACC | TCGACAAGTTTTATGGCTCTGA | 161 |
| chr1:118845649-118847670 | p3 | (AGT)6 | TTTGAATTGCACATGCTTTACC | TCGACAAGTTTTATGGCTCTGA | 161 |
| chr1:118845672-118847687 | p6 | (ATATAG)3 | TTTGAATTGCACATGCTTTACC | TCGACAAGTTTTATGGCTCTGA | 138 |
| chr1:118845672-118847687 | p3 | (TAT)6 | TTTGAATTGCACATGCTTTACC | TCGACAAGTTTTATGGCTCTGA | 138 |
| chr1:118845672-118847687 | p3 | (AGT)6 | TTTGAATTGCACATGCTTTACC | TCGACAAGTTTTATGGCTCTGA | 138 |
| chr1:118855608-118857620 | p2 | (AT)6 | TGGGTTGGGTGATAAATGGT | CTGACGATGATTCAGGCAGA | 178 |
| chr1:118990972-118992990 | p2 | (AT)9 | AGCTTTGGCATGCTTGAAGT | GGAGCGTGACACTTCTGTGA | 163 |
| chr1:118992973-118994991 | p6 | (TTTTTA)4 | AGCACAGAGCTCCGTTTGAT | ACAGTTTTCGGGTACTTGCG | 166 |
| chr1:118992973-118994991 | p6 | (TTTTTA)3 | AGCACAGAGCTCCGTTTGAT | ACAGTTTTCGGGTACTTGCG | 166 |
| chr1:118992973-118994991 | p3 | (AAG)15 | AGCACAGAGCTCCGTTTGAT | ACAGTTTTCGGGTACTTGCG | 166 |
| chr1:119057152-119059172 | p2 | (TA)31 | ATGACTCGTTGCATTTGTCG | CTTTTCAAAATTCTGCGCCT | 118 |
| chr1:119057152-119059172 | p2 | (TA)10 | ATGACTCGTTGCATTTGTCG | CTTTTCAAAATTCTGCGCCT | 118 |
| chr1:119057308-119059326 | p2 | (TA)31 | GGCGCAGAATTTTGAAAAGA | TACAAATTCAAAGGCCCCC | 176 |
| chr1:119057308-119059326 | p2 | (TA)10 | GGCGCAGAATTTTGAAAAGA | TACAAATTCAAAGGCCCCC | 176 |
| chr1:119057341-119059399 | p2 | (TA)31 | GGCGCAGAATTTTGAAAAGA | TACAAATTCAAAGGCCCCC | 143 |
| chr1:119057341-119059399 | p2 | (TA)10 | GGCGCAGAATTTTGAAAAGA | TACAAATTCAAAGGCCCCC | 143 |
| chr1:119061066-119063080 | p2 | (AT)6 | TACCTGAGGTTCTGGGTTCG | TGCCCCATAAGGCATAGTTC | 163 |
| chr1:119061066-119063080 | p2 | (TA)7 | TACCTGAGGTTCTGGGTTCG | TGCCCCATAAGGCATAGTTC | 163 |
| chr1:119142980-119144995 | p5 | (TCATA)3 | GTGAGCAAGTTTGGCTCCTC | GACACTTGATGGAAGACGCA | 131 |
| chr1:119264331-119266353 | p2 | (TA)10 | AAGATTGGAGCGACACAACC | TTTCGTTAATTTCCCCCACA | 152 |

| Marker ID | SSR type | SSR motif | FORWARD PRIMER (5'-3') | REVERSE PRIMER (5'-3') | size (bp) |
|---|---|---|---|---|---|
| chr1:119264331-119266353 | p3 | (TGT)5 | AAGATTGGAGCGACACAACC | TTTCGTTAATTTCCCCCACA | 152 |
| chr1:119264331-119266353 | p2 | (AT)11 | AAGATTGGAGCGACACAACC | TTTCGTTAATTTCCCCCACA | 152 |
| chr1:119303485-119305500 | p5 | (AAAAT)3 | ACTGACTGCTGTGGCATCTG | GCTGTGAACCTCTGGGTAGC | 194 |
| chr1:119387635-119389650 | p5 | (AAAAT)3 | ATTTTAGGGTGGACGAACCC | TTGTCCGCCGTTAATAGGAC | 254 |
| chr1:119432256-119434304 | p3 | (ATT)16 | GTGGAAGCTTGTCTGGAAGC | TAATGCAGCATCACCGAGAG | 291 |
| chr1:119434735-119436783 | p3 | (ATT)16 | GTGGAAGCTTGTCTGGAAGC | TAATGCAGCATCACCGAGAG | 291 |
| chr1:119451202-119453244 | p2 | (AC)11 | CCTCGACATCAGCTTCAACA | ATTGGTCTCCTCACACGGAC | 129 |
| chr1:119451202-119453244 | p2 | (AC)7 | CCTCGACATCAGCTTCAACA | ATTGGTCTCCTCACACGGAC | 129 |
| chr1:119451202-119453244 | p6 | (TAGTGT)3 | CCTCGACATCAGCTTCAACA | ATTGGTCTCCTCACACGGAC | 129 |
| chr1:119452115-119454133 | p2 | (AC)7 | CACAGAACAAATCGGCAATG | CAGTTCTGTCTTTCTGCCCC | 229 |
| chr1:119452115-119454133 | p6 | (TAGTGT)3 | CACAGAACAAATCGGCAATG | CAGTTCTGTCTTTCTGCCCC | 229 |
| chr1:119466823-119468835 | p2 | (TA)6 | CATATTATGGGGTTGTGGGG | TTTTTCCTTTCCCCAGTCCT | 126 |
| chr1:119466823-119468835 | p5 | (AAAAT)4 | CATATTATGGGGTTGTGGGG | TTTTTCCTTTCCCCAGTCCT | 126 |
| chr1:119530399-119532423 | p4 | (TTCT)6 | TTGGTCATGATAAGCGCAAG | ATACGCAAAAGTTCGGATGG | 203 |
| chr1:119532360-119534376 | p4 | (GAAA)4 | GCTTTACCACCATGCCAAGT | TCAATTTCAATTTTCCCCCA | 154 |
| chr1:119534953-119536968 | p2 | (AG)8 | ACCATAGGTTGTGGGACTGC | TTCTGGGTTTGTTGGTGTGA | 116 |
| chr1:119534953-119536968 | p3 | (TTG)5 | ACCATAGGTTGTGGGACTGC | TTCTGGGTTTGTTGGTGTGA | 116 |
| chr1:119537846-119539864 | p3 | (GTT)6 | TACCCATGCCATTTAAGGGA | CTCTTCATGAGCTTCCCGAG | 152 |
| chr1:119544791-119546812 | p3 | (ACA)7 | ATGCCAAAAGAAGGGAAGGT | CCACCAATTGACAGACATGC | 177 |
| chr1:119544791-119546812 | p2 | (TC)7 | ATGCCAAAAGAAGGGAAGGT | CCACCAATTGACAGACATGC | 177 |
| chr1:119585794-119587809 | p2 | (CT)7 | ACTCCGTGGATGAACTGCTC | CCTTGTCAGCTTGCGTATCA | 142 |
| chr1:119585815-119587831 | p2 | (CT)7 | ACTCCGTGGATGAACTGCTC | CCTTGTCAGCTTGCGTATCA | 121 |
| chr1:119685254-119687275 | p5 | (AAATT)3 | GTGGGCAAATGTGTGAACAG | GGGGTTGTTTTGTTTTGCAG | 178 |
| chr1:119685254-119687275 | p3 | (GAA)7 | GTGGGCAAATGTGTGAACAG | GGGGTTGTTTTGTTTTGCAG | 178 |
| chr1:119712053-119714067 | p2 | (AT)7 | CGAAAACGTTAAGGCTACGG | TTTCGTTCGCATTCAAATCA | 101 |
| chr1:119714705-119716717 | p2 | (TA)6 | CAGGTAAGCAGGGGTACGAA | ATCATAGCCACACAGGGCTC | 202 |
| chr1:119714705-119716717 | p2 | (TA)6 | CAGGTAAGCAGGGGTACGAA | ATCATAGCCACACAGGGCTC | 202 |
| chr1:119714705-119716717 | p2 | (TA)6 | CAGGTAAGCAGGGGTACGAA | ATCATAGCCACACAGGGCTC | 202 |
| chr1:119714705-119716717 | p2 | (TA)6 | CAGGTAAGCAGGGGTACGAA | ATCATAGCCACACAGGGCTC | 202 |
| chr1:119714705-119716717 | p2 | (TA)6 | CAGGTAAGCAGGGGTACGAA | ATCATAGCCACACAGGGCTC | 202 |
| chr1:119714705-119716717 | p2 | (TA)6 | CAGGTAAGCAGGGGTACGAA | ATCATAGCCACACAGGGCTC | 202 |
| chr1:119714705-119716717 | p2 | (TA)6 | CAGGTAAGCAGGGGTACGAA | ATCATAGCCACACAGGGCTC | 202 |
| chr1:119714705-119716717 | p2 | (TA)6 | CAGGTAAGCAGGGGTACGAA | ATCATAGCCACACAGGGCTC | 202 |
| chr1:119714705-119716717 | p2 | (TA)6 | CAGGTAAGCAGGGGTACGAA | ATCATAGCCACACAGGGCTC | 202 |
| chr1:119714705-119716717 | p2 | (TA)6 | CAGGTAAGCAGGGGTACGAA | ATCATAGCCACACAGGGCTC | 202 |
| chr1:119714705-119716717 | p2 | (TA)6 | CAGGTAAGCAGGGGTACGAA | ATCATAGCCACACAGGGCTC | 202 |

| Marker ID | SSR type | SSR motif | FORWARD PRIMER (5'-3') | REVERSE PRIMER (5'-3') | size (bp) |
|---|---|---|---|---|---|
| chr1:119714705-119716717 | p2 | (TA)6 | CAGGTAAGCAGGGGTACGAA | ATCATAGCCACACAGGGCTC | 202 |
| chr1:119714705-119716717 | p2 | (TA)6 | CAGGTAAGCAGGGGTACGAA | ATCATAGCCACACAGGGCTC | 202 |
| chr1:119714705-119716717 | p2 | (TA)6 | CAGGTAAGCAGGGGTACGAA | ATCATAGCCACACAGGGCTC | 202 |
| chr1:119714945-119716957 | p2 | (TA)6 | AATATTGGGAGCCCTGTGTG | GGCCCTTTAATGAGGGACTC | 121 |
| chr1:119714945-119716957 | p2 | (TA)6 | AATATTGGGAGCCCTGTGTG | GGCCCTTTAATGAGGGACTC | 121 |
| chr1:119714945-119716957 | p2 | (TA)6 | AATATTGGGAGCCCTGTGTG | GGCCCTTTAATGAGGGACTC | 121 |
| chr1:119714945-119716957 | p2 | (TA)6 | AATATTGGGAGCCCTGTGTG | GGCCCTTTAATGAGGGACTC | 121 |
| chr1:119714945-119716957 | p2 | (TA)6 | AATATTGGGAGCCCTGTGTG | GGCCCTTTAATGAGGGACTC | 121 |
| chr1:119714945-119716957 | p2 | (TA)6 | AATATTGGGAGCCCTGTGTG | GGCCCTTTAATGAGGGACTC | 121 |
| chr1:119714945-119716957 | p2 | (TA)6 | AATATTGGGAGCCCTGTGTG | GGCCCTTTAATGAGGGACTC | 121 |
| chr1:119714945-119716957 | p2 | (TA)6 | AATATTGGGAGCCCTGTGTG | GGCCCTTTAATGAGGGACTC | 121 |
| chr1:119714945-119716957 | p2 | (TA)6 | AATATTGGGAGCCCTGTGTG | GGCCCTTTAATGAGGGACTC | 121 |
| chr1:119714945-119716957 | p2 | (TA)6 | AATATTGGGAGCCCTGTGTG | GGCCCTTTAATGAGGGACTC | 121 |
| chr1:119714945-119716957 | p2 | (TA)6 | AATATTGGGAGCCCTGTGTG | GGCCCTTTAATGAGGGACTC | 121 |
| chr1:119714945-119716957 | p2 | (TA)6 | AATATTGGGAGCCCTGTGTG | GGCCCTTTAATGAGGGACTC | 121 |
| chr1:119714945-119716957 | p2 | (TA)6 | AATATTGGGAGCCCTGTGTG | GGCCCTTTAATGAGGGACTC | 121 |
| chr1:119714945-119716957 | p2 | (TA)6 | AATATTGGGAGCCCTGTGTG | GGCCCTTTAATGAGGGACTC | 121 |
| chr1:119714945-119716957 | p2 | (TA)6 | AATATTGGGAGCCCTGTGTG | GGCCCTTTAATGAGGGACTC | 121 |
| chr1:119714945-119716957 | p2 | (TA)6 | AATATTGGGAGCCCTGTGTG | GGCCCTTTAATGAGGGACTC | 121 |
| chr1:119801760-119803775 | p5 | (TTTGA)3 | CTGCATTTTTGTTGGGGTTT | CCGAGTTTTCTCAACAAGGC | 254 |
| chr1:119820250-119822274 | p3 | (TTA)8 | TTGTGTATCGGGCATGGATA | GTTTCTGTCGACCGACCAAT | 160 |
| chr1:119835681-119837701 | p2 | (TA)10 | CAATCTCACTCTCGCTTCCC | AGATTGATGAGGATGTCGGG | 294 |
| chr1:119891595-119893618 | p2 | (TA)6 | TCCTTTGTTCCGTGAAAACC | GTATGTAGGCGAAGGGCAAA | 297 |
| chr1:119891974-119893986 | p2 | (TA)6 | TAAGTTTTGCCCTTCGCCTA | TGGCCCTCAAAATGGTAGTC | 144 |
| chr1:119904141-119906156 | p2 | (TA)6 | ACATAGTCATCGGGAGGGTG | CGCTGCATTTATCCCTCTTC | 246 |
| chr1:119904141-119906156 | p3 | (TAC)5 | ACATAGTCATCGGGAGGGTG | CGCTGCATTTATCCCTCTTC | 246 |
| chr1:119932631-119934646 | p3 | (TAA)5 | CATGTATCTGGAGGGGCACT | GACGAACACCCAACCAAAGT | 140 |
| chr1:119943965-119945977 | p2 | (TA)6 | CATCGCCTGAAAATTCACCT | GTTGGCGTAGGTGGTCTGTT | 203 |
| chr1:120018925-120020940 | p3 | (TTC)5 | TGGGTTAAATAATGGAGCGG | CGGCCATTAAAGTTTTCCAA | 212 |
| chr1:120018925-120020940 | p3 | (AAC)5 | TGGGTTAAATAATGGAGCGG | CGGCCATTAAAGTTTTCCAA | 212 |
| chr1:120018925-120020940 | p3 | (GAA)7 | TGGGTTAAATAATGGAGCGG | CGGCCATTAAAGTTTTCCAA | 212 |
| chr1:120018925-120020940 | p3 | (CAC)5 | TGGGTTAAATAATGGAGCGG | CGGCCATTAAAGTTTTCCAA | 212 |
| chr1:120022359-120024383 | p6 | (GTGTTG)4 | CAGGCCCAAAGAATTAGCAG | AATTGGGCTTTGATGTGAGG | 115 |
| chr1:120022359-120024383 | p3 | (TTG)7 | CAGGCCCAAAGAATTAGCAG | AATTGGGCTTTGATGTGAGG | 115 |
| chr1:120022359-120024383 | p3 | (TTA)10 | CAGGCCCAAAGAATTAGCAG | AATTGGGCTTTGATGTGAGG | 115 |
| chr1:120100233-120102248 | p5 | (ACCGG)3 | GGACTTGGAATGAGCCAAAA | CTGCATCATCATCCAACAGG | 184 |

| Marker ID | SSR type | SSR motif | FORWARD PRIMER (5'-3') | REVERSE PRIMER (5'-3') | size (bp) |
|---|---|---|---|---|---|
| chr1:120100233-120102248 | p2 | (TA)21 | GGACTTGGAATGAGCCAAAA | CTGCATCATCATCCAACAGG | 184 |
| chr1:120195882-120197897 | p2 | (AC)8 | AGTAAACCGACAAACCGCAC | GCTAAAACCAAACCAACCCA | 168 |
| chr1:120195882-120197897 | p5 | (TTGGT)3 | AGTAAACCGACAAACCGCAC | GCTAAAACCAAACCAACCCA | 168 |
| chr1:120341802-120343820 | p3 | (TTA)11 | TTAAAAGAGCTTGCGCCATT | GCCAACATCCGGAACTCTAA | 106 |
| chr1:120341802-120343820 | p6 | (ATTCTA)3 | TTAAAAGAGCTTGCGCCATT | GCCAACATCCGGAACTCTAA | 106 |
| chr1:120341802-120343820 | p2 | (TA)7 | TTAAAAGAGCTTGCGCCATT | GCCAACATCCGGAACTCTAA | 106 |
| chr1:120692435-120694451 | p4 | (AAAT)4 | CACCCACCTACCTTACCCCT | TGTTCGCTAGTGATGGTGGA | 210 |
| chr1:120731397-120733412 | p5 | (ATTTT)3 | TTAAATGTCGTGCCCAGTCA | CGTGCCAATTCAACAACAAC | 270 |
| chr1:120837225-120839246 | p3 | (AAT)7 | GTGGGTCCACCTCCTACTCA | ACATGTGGATCCCCACCTTA | 113 |
| chr1:120878096-120880120 | p2 | (AT)12 | AAAGTGGACCACGCAGAAGT | GCTTAGCACGGGCCTTTAAT | 195 |
| chr1:120938562-120940574 | p2 | (AT)6 | AAAATGTTTCACGCCACGAT | TATTACATGCGGGTGTCGTC | 247 |
| chr1:120938562-120940574 | p5 | (AAAAT)3 | AAAATGTTTCACGCCACGAT | TATTACATGCGGGTGTCGTC | 247 |
| chr1:120938599-120940623 | p2 | (AT)6 | AAAATGTTTCACGCCACGAT | TATTACATGCGGGTGTCGTC | 210 |
| chr1:120938599-120940623 | p5 | (AAAAT)3 | AAAATGTTTCACGCCACGAT | TATTACATGCGGGTGTCGTC | 210 |
| chr1:121014432-121016466 | p2 | (AT)17 | AGCCAACGAGGCTAACAAAA | CAGGGACTACATAGCGGAGC | 286 |
| chr1:121098677-121100702 | p5 | (TACAA)5 | CCGAGATTCAAACTGCATGA | CACCGCTGCTGCTGTATCTA | 209 |
| chr1:121102532-121104556 | p2 | (TA)7 | CTTGACGCCAAAAGTGGTTT | CGAACTATTGACGAGTGGCA | 239 |
| chr1:121102552-121104567 | p2 | (TA)7 | CTTGACGCCAAAAGTGGTTT | CGAACTATTGACGAGTGGCA | 219 |
| chr1:121144415-121146439 | p3 | (AAT)8 | AGCAATGGGTCTTATGCCAG | GCCTGACCGTACATCCAAAG | 219 |
| chr1:121268846-121270861 | p3 | (TCT)5 | CCTGACTGCGACATCACTTG | GGTAGCAGAAAATGGGACGA | 298 |
| chr1:121323905-121325926 | p3 | (TAA)7 | GGGAGAAGATGAAGACGCAG | ATTTTGACTTACCGGACCCC | 290 |
| chr1:121324034-121326094 | p3 | (TAA)7 | GGGAGAAGATGAAGACGCAG | ATTTTGACTTACCGGACCCC | 161 |
| chr1:121347706-121349738 | p6 | (TTGGAG)3 | ATTGGAACGGAGCAACTGAC | ACGCCGAGCTCATACTCACT | 256 |
| chr1:121347789-121349815 | p6 | (TTGGAG)3 | ATTGGAACGGAGCAACTGAC | ACGCCGAGCTCATACTCACT | 173 |
| chr1:121348713-121350731 | p6 | (TTGGAG)3 | ATCCTCAGCCCCACCTAAAT | AAGCTTGGTCAAATCGGCTA | 236 |
| chr1:121398955-121400970 | p5 | (AATAA)3 | TGGCCTACGTATCCTCATCC | CGCACCTGATTCTCATCAAA | 186 |
| chr1:121412091-121414107 | p4 | (AAGA)4 | CAAGTCTGCCCATAAGGCAT | TTCACGCAGAAATCATGAGG | 122 |
| chr1:121414580-121416624 | p2 | (TA)22 | TACACCTACCCGCTTCATCC | TATCTTTAGGGGCTCGGCTT | 188 |
| chr1:121447223-121449238 | p3 | (AAC)5 | AAATGCGCAAATGAAGTTCC | TGTGAGCTGTTTCTATGCCG | 189 |
| chr1:121513953-121515995 | p5 | (AAAAG)3 | CGCGATCACATGAAAAGAGA | CATCTTCATTTGAGCTGCCA | 150 |
| chr1:121513953-121515995 | p2 | (AT)21 | CGCGATCACATGAAAAGAGA | CATCTTCATTTGAGCTGCCA | 150 |
| chr1:121513953-121515995 | p2 | (AT)6 | CGCGATCACATGAAAAGAGA | CATCTTCATTTGAGCTGCCA | 150 |
| chr1:121545601-121547613 | p2 | (AT)6 | GACATGGCGGAATAGCAAAT | TGTCACGTGCGTATCAATCC | 162 |
| chr1:121545601-121547613 | p2 | (TA)6 | GACATGGCGGAATAGCAAAT | TGTCACGTGCGTATCAATCC | 162 |
| chr1:121568417-121570432 | p5 | (AATTA)3 | TGGACCTTATGCGTTGTGAA | AATGGGAAAGGTTAAACGGG | 160 |
| chr1:121704915-121706930 | p2 | (AT)19 | CTAAATATACCCCCGCCGTT | GTATATGGAGGGCATGGTGG | 233 |

| Marker ID | SSR type | SSR motif | FORWARD PRIMER (5'-3') | REVERSE PRIMER (5'-3') | size (bp) |
|---|---|---|---|---|---|
| chr1:121704915-121706930 | p5 | (AAAAT)3 | CTAAATATACCCCCGCCGTT | GTATATGGAGGGCATGGTGG | 233 |
| chr1:121713620-121715674 | p3 | (CTT)18 | CACATGGCATTTTCTTGTGC | TAAGCGAATTTCTCTGCCCA | 109 |
| chr1:121790587-121792602 | p5 | (ATTGC)3 | TATCGATATTGAGGCCCAGC | TCCGAACTTCATACCCGAAG | 221 |
| chr1:121811415-121813427 | p2 | (TA)6 | ATTGACCAAAACGGTTCCAA | GTGGGGGAGGGTTAAAGTGT | 243 |
| chr1:121862350-121864362 | p2 | (AT)6 | TAACACTTAACCCCTCCCCC | TTGCCGGACTAATTTCCAAC | 154 |
| chr1:121862350-121864362 | p2 | (TA)6 | TAACACTTAACCCCTCCCCC | TTGCCGGACTAATTTCCAAC | 154 |
| chr1:121935182-121937198 | p4 | (ACAT)4 | GGCCACGACTGGTATCCTAA | TTAGCCTCATCGGCTTGTCT | 187 |
| chr1:121971887-121973905 | p6 | (TATTTT)3 | GGTTAGACCCTGGACCCCTA | TGTAGGCCTAAATCCCACCA | 235 |
| chr1:122001411-122003429 | p3 | (AAT)6 | TGCGAGAGGCGTTAAAAACT | GCGGGATAATGGAGGATTTT | 188 |
| chr1:122001411-122003429 | p5 | (AAAGG)5 | TGCGAGAGGCGTTAAAAACT | GCGGGATAATGGAGGATTTT | 188 |
| chr1:122069398-122071413 | p3 | (CGG)5 | ATGACCGGATTGGTTATGGA | ATCCGATTTCGAAGGGATCT | 196 |
| chr1:122156695-122158721 | p2 | (CA)13 | TCCTTTGGAGATCTTGTGCC | CCAGGTAAGCCCTGATGGTA | 209 |
| chr1:122199276-122201300 | p6 | (CATATT)4 | GTTTGCTCACCAATCCGAGT | GGGTCGTGACAATATCGCTT | 124 |
| chr1:122266737-122268752 | p3 | (TAA)5 | CTATGCTCGAACACGAACGA | CCAAGGATCTGGAACCAAAA | 243 |
| chr1:122513969-122515987 | p6 | (AACCCT)3 | ATTCCTGGTTCCGACTCCTT | CTCGAAGGAGAATAAGGGGG | 262 |
| chr1:122513969-122515987 | p2 | (AT)8 | ATTCCTGGTTCCGACTCCTT | CTCGAAGGAGAATAAGGGGG | 262 |
| chr1:122514112-122516128 | p6 | (AACCCT)3 | ATTCCTGGTTCCGACTCCTT | CTCGAAGGAGAATAAGGGGG | 119 |
| chr1:122514112-122516128 | p2 | (AT)8 | ATTCCTGGTTCCGACTCCTT | CTCGAAGGAGAATAAGGGGG | 119 |
| chr1:122544345-122546363 | p3 | (GAA)6 | CCACTGACATTGGTTGTTCG | GATTTGCTTTCGGGTGAGAA | 246 |
| chr1:122641918-122643933 | p3 | (TGT)5 | TGCCACCAAAACTTAAAGGC | TGCCACTCCCACCTTATCTC | 239 |
| chr1:122659098-122661113 | p5 | (TAAAA)3 | ACCGAATGGGAACATCAATC | GAAAAGGAATGAGGGCAACA | 287 |
| chr1:122697025-122699047 | p2 | (AT)8 | TATGTATGTGTGTGCGCGTG | TCTTCTTTAACCACACCCCG | 183 |
| chr1:122697025-122699047 | p2 | (AT)11 | TATGTATGTGTGTGCGCGTG | TCTTCTTTAACCACACCCCG | 183 |
| chr1:122728257-122730272 | p5 | (ACCAA)3 | GGACCGGGTTAGTTTGGATT | AAAACCCGAGAAAACCGAAT | 208 |
| chr1:122728257-122730272 | p5 | (AAACC)3 | GGACCGGGTTAGTTTGGATT | AAAACCCGAGAAAACCGAAT | 208 |
| chr1:122728257-122730272 | p2 | (TA)7 | GGACCGGGTTAGTTTGGATT | AAAACCCGAGAAAACCGAAT | 208 |
| chr1:122742891-122744903 | p2 | (AT)6 | CGTGACCCACACTTCAACAC | CTTGCTGACCGCGATATTTT | 192 |
| chr1:122742891-122744903 | p2 | (TA)8 | CGTGACCCACACTTCAACAC | CTTGCTGACCGCGATATTTT | 192 |
| chr1:122742944-122744962 | p2 | (AT)6 | CGTGACCCACACTTCAACAC | ACTTGCTGACCGCGATATTT | 139 |
| chr1:122742944-122744962 | p2 | (TA)8 | CGTGACCCACACTTCAACAC | ACTTGCTGACCGCGATATTT | 139 |
| chr1:122822459-122824480 | p3 | (AGA)7 | CCTCCCACTAGTTCGTCCAA | TGGGGTGGGCATTTTAGTAG | 181 |
| chr1:122854882-122856898 | p4 | (TTAA)4 | CCTCCGAAGTCCATGAGGTA | TGCTGGTCGAAGTTCATTTG | 216 |
| chr1:122874577-122876589 | p2 | (GA)6 | CACAAACTGTTGCCTTCCCT | GATTCCAGCAGATAACCGGA | 150 |
| chr1:122921336-122923351 | p5 | (CAAAT)3 | ACGTCAAAAGCTAACGGTGC | TGAATCACCTCCCCAACTTC | 233 |
| chr1:122937785-122939809 | p3 | (ATT)8 | AAACAGAGGGAGTGCATTGG | GCAACTAGCGGAGGCTAAAA | 232 |
| chr1:122937785-122939809 | p2 | (AT)15 | AAACAGAGGGAGTGCATTGG | GCAACTAGCGGAGGCTAAAA | 232 |

| Marker ID | SSR type | SSR motif | FORWARD PRIMER (5'-3') | REVERSE PRIMER (5'-3') | size (bp) |
|---|---|---|---|---|---|
| chr1:122956937-122958949 | p2 | (TA)6 | ACTGAATGGGGATTGAGCAC | CGCTTGAAGGGTTCTGGTAA | 275 |
| chr1:123051300-123053354 | p2 | (AT)27 | AAGGGCATAAGCCTCACAGA | GTTCACCGCGTAAAGGTGAT | 291 |
| chr1:123080643-123082661 | p2 | (TA)9 | AGGACATCAGGGACAAGGTG | TAAAACCGTCCAAAGCCAAC | 285 |
| chr1:123095806-123097832 | p3 | (AAC)7 | CTTGGGTGCTTATCTCTGGC | TTCCATTGCCTCGACCTTAC | 263 |
| chr1:123095806-123097832 | p2 | (AG)13 | CTTGGGTGCTTATCTCTGGC | TTCCATTGCCTCGACCTTAC | 263 |
| chr1:123121760-123123778 | p6 | (ATCACC)3 | GGAGGTGATTAGGCAGGACA | ATGAAAAGCGCCCAATTATG | 157 |
| chr1:123129506-123131532 | p4 | (ACAT)4 | GAGCCATTTCCGATAGTTCG | TTTGCTTTGGACCCACTTTC | 101 |
| chr1:123141547-123143562 | p6 | (CCCCCA)3 | TTTCACGACTCATCACCCAA | CGCATGACGCCCTATTATTT | 277 |
| chr1:123141547-123143562 | p5 | (TTTTA)3 | TTTCACGACTCATCACCCAA | CGCATGACGCCCTATTATTT | 277 |
| chr1:123141547-123143562 | p2 | (AT)8 | TTTCACGACTCATCACCCAA | CGCATGACGCCCTATTATTT | 277 |
| chr1:123147958-123149976 | p3 | (TAA)6 | AGGGCGAATTAGGGTTTGTT | ACTTCACTTTGTTCCCCACG | 106 |
| chr1:123192974-123194989 | p5 | (TTTAA)3 | GGTAGGCGAGACTGAGATGG | CGACTTCCACACCTTCCAAT | 104 |
| chr1:123216916-123218930 | p2 | (TA)7 | CTGGATCCGGCATACACTTT | TGCCTTATGGGAAGACTTGG | 229 |
| chr1:123216916-123218930 | p2 | (TA)9 | CTGGATCCGGCATACACTTT | TGCCTTATGGGAAGACTTGG | 229 |
| chr1:123216916-123218930 | p3 | (ACA)5 | CTGGATCCGGCATACACTTT | TGCCTTATGGGAAGACTTGG | 229 |
| chr1:123317095-123319109 | p2 | (AT)7 | TTCTAATACATCGGTGGGGC | CCCTCGTGTTTAGTTGCCAT | 189 |
| chr1:123317095-123319109 | p3 | (ACA)6 | TTCTAATACATCGGTGGGGC | CCCTCGTGTTTAGTTGCCAT | 189 |
| chr1:123389550-123391570 | p2 | (TA)10 | CAACGTGTGGACTGATTTGG | TATTTCTTGGAAATTGGGCG | 253 |
| chr1:123405651-123407669 | p2 | (AT)9 | TAAAGGTTGTGGGGGATGAG | TCGACATCTCATCGACTTGC | 216 |
| chr1:123417444-123419459 | p4 | (AAAT)6 | GGATTCACCCATCACTGCTT | TGAAATCGAACCTGAAACCC | 194 |
| chr1:123417444-123419459 | p5 | (AACCG)3 | GGATTCACCCATCACTGCTT | TGAAATCGAACCTGAAACCC | 194 |
| chr1:123433726-123435801 | p3 | (ATA)25 | TAAAGGAGCCAGACCCGATA | TGGATGCCGACACTAATTCA | 219 |
| chr1:123433726-123435801 | p5 | (GGATT)3 | TAAAGGAGCCAGACCCGATA | TGGATGCCGACACTAATTCA | 219 |
| chr1:123449971-123451991 | p2 | (TA)10 | GTCGTTGAGGTTCCGTTTGT | GGAACGCGGACATAGATGAT | 245 |
| chr1:123464790-123466808 | p6 | (AAAAAG)3 | GGAGCATCAGAGAAACGAGG | CCCAAACCCAAATTAACCCT | 192 |
| chr1:123469136-123471154 | p6 | (TCCAAT)3 | GGGGGCATAAACCAAAAACT | AGCACTCAGGATGGCACTTT | 276 |
| chr1:123546567-123548579 | p2 | (TA)6 | TGCTCCACGCCCTACTACTT | CGAAGCCACGGTTGATACTT | 192 |
| chr1:123546567-123548579 | p2 | (AT)6 | TGCTCCACGCCCTACTACTT | CGAAGCCACGGTTGATACTT | 192 |
| chr1:123547673-123549691 | p2 | (AT)6 | CGGTTGACAGGCCTAGTTGT | TATCAAGGGACCGGTGAAAC | 216 |
| chr1:123547673-123549691 | p6 | (CCCCCA)3 | CGGTTGACAGGCCTAGTTGT | TATCAAGGGACCGGTGAAAC | 216 |
| chr1:123592631-123594685 | p3 | (TTA)18 | GAGCCAAAATACAAGGGCAG | CAAAAACAACGGCATTTCCT | 219 |
| chr1:123611010-123613030 | p5 | (TGGGG)4 | ATAAAGCCCCAAAACCTGCT | AGATCTCTCCGTCTGCTCCA | 116 |
| chr1:123774667-123776700 | p3 | (AAT)11 | GCCAACGGACAAAGTTCATT | GCATGGCTTATTTGGGAAAA | 270 |
| chr1:123810538-123812553 | p5 | (TTTGA)3 | GGACACAATTCATGCACCAG | ACGGGATGATGGGAGTATGA | 213 |
| chr1:123928757-123930769 | p2 | (TA)6 | TGGCCCCTCAGTTTATTTGT | TTGAGAAGTTTAATGGCCCG | 104 |
| chr1:123931945-123933960 | p5 | (AAAAT)3 | CAACTTTCGCCAGTTTTGGT | TTTTTGAGCAGCTTTGGTCC | 126 |

| Marker ID | SSR type | SSR motif | FORWARD PRIMER (5'-3') | REVERSE PRIMER (5'-3') | size (bp) |
|---|---|---|---|---|---|
| chr1:123983517-123985532 | p5 | (CAAAC)3 | TAGTGGCTGCACCATTTTTG | AATTGAGGGCAAAATGCAAG | 117 |
| chr1:124064701-124066716 | p3 | (GTT)5 | TTCCCTTTGACTGCATAGCC | CGGAAAGGTTTCTTTCCTCC | 296 |
| chr1:124079787-124081805 | p3 | (AAG)6 | CTATTTCAGCCGGAGATGGA | GACCCCAGCCAACAAAAATA | 105 |
| chr1:124109519-124111531 | p2 | (AT)6 | AGGAACTGGTCCCTTGTGTG | AATTCATCTGTGGCACCTCC | 294 |
| chr1:124121611-124123629 | p2 | (TA)11 | GGTGGGCCCCATACTAAACT | GTTCGGAGATGGTGAGTTCC | 138 |
| chr1:124121611-124123629 | p2 | (TA)9 | GGTGGGCCCCATACTAAACT | GTTCGGAGATGGTGAGTTCC | 138 |
| chr1:124209088-124211124 | p2 | (TA)18 | TGTGAAGAGGAGGTGTGTCG | ATCACCTCCCCGAGTTCTTT | 213 |
| chr1:124222624-124224660 | p5 | (CCGGT)3 | AACACTTTAACCCATCCCCC | ACAAAGTGAAGGCGGCTAGA | 221 |
| chr1:124229211-124231249 | p2 | (TA)19 | TTTTAAACGAACCAGACCCG | CCCCCATTACAAAAGTGGTG | 251 |
| chr1:124229211-124231249 | p3 | (TAC)6 | TTTTAAACGAACCAGACCCG | CCCCCATTACAAAAGTGGTG | 251 |
| chr1:124309138-124311156 | p2 | (AT)9 | ATCGGATTTGTAGATTGCGG | CCCCCTGCTAACTTTTGACA | 167 |
| chr1:124316995-124319011 | p5 | (TATAT)3 | TGTCAGATAGGGTGGGTTCA | GATAACGGACGACGAAGCAT | 287 |
| chr1:124316995-124319011 | p3 | (TAT)5 | TGTCAGATAGGGTGGGTTCA | GATAACGGACGACGAAGCAT | 287 |
| chr1:124316995-124319011 | p2 | (TA)8 | TGTCAGATAGGGTGGGTTCA | GATAACGGACGACGAAGCAT | 287 |
| chr1:124316995-124319011 | p2 | (AT)19 | TGTCAGATAGGGTGGGTTCA | GATAACGGACGACGAAGCAT | 287 |
| chr1:124328457-124330472 | p5 | (AGAGA)3 | ATGAAATGTGCTTTCTGGGC | TTGCAACAATCCTGCCTACA | 223 |
| chr1:124348356-124350407 | p2 | (TA)12 | TGATACGCTCATTACGTGTGG | TGGCAGTCAGAGTGACCAAG | 105 |
| chr1:124348356-124350407 | p3 | (AAT)6 | TGATACGCTCATTACGTGTGG | TGGCAGTCAGAGTGACCAAG | 105 |
| chr1:124419628-124421640 | p2 | (TA)6 | TCCGGCTTTAGTTTGCAAGT | CAATTTCTCTCTCGGCCTTG | 278 |
| chr1:124419628-124421640 | p2 | (TA)7 | TCCGGCTTTAGTTTGCAAGT | CAATTTCTCTCTCGGCCTTG | 278 |
| chr1:124423571-124425583 | p2 | (AT)6 | ATGTGACGTACCATGACCGA | TTCATCGAGCCACAAAACAA | 242 |
| chr1:124535752-124537764 | p2 | (TA)7 | GGGGAAAGGAGAAGAATTGG | TTGCACCTGCACCTTCTACA | 299 |
| chr1:124535752-124537764 | p2 | (TG)6 | GGGGAAAGGAGAAGAATTGG | TTGCACCTGCACCTTCTACA | 299 |
| chr1:124598803-124600828 | p5 | (CCGAA)3 | GTTTCCACCCCTGAGACGTA | GATTTTCAGCTTTGGCTTCG | 225 |
| chr1:124600293-124602308 | p5 | (CCGAA)3 | TATGAAAAACCGACCGAACC | AAAAAGACCGCCCTAATCGT | 223 |
| chr1:124600293-124602308 | p5 | (AACCG)3 | TATGAAAAACCGACCGAACC | AAAAAGACCGCCCTAATCGT | 223 |
| chr1:124708849-124710869 | p2 | (TA)10 | GTCAAGAGGTGCAAAAAGCC | TTGTGGTGTGTAGCGGACAT | 126 |
| chr1:124708849-124710869 | p2 | (TA)9 | GTCAAGAGGTGCAAAAAGCC | TTGTGGTGTGTAGCGGACAT | 126 |
| chr1:124743717-124745732 | p3 | (ACA)5 | TAGCCTTGAAAAGGGCAAGA | CAATTGTAGGGAAAACGGGA | 120 |
| chr1:124743717-124745732 | p3 | (CAA)5 | TAGCCTTGAAAAGGGCAAGA | CAATTGTAGGGAAAACGGGA | 120 |
| chr1:124765441-124767465 | p6 | (GATTGT)4 | GTCCAACCAAACTGCCTTGT | GAGCCCCATTTCAAGAATGA | 299 |
| chr1:124945290-124947310 | p5 | (TTTTA)4 | ATTCGTCCTCATCGATCCAC | CATCACTACACACGCTCGCT | 267 |
| chr1:124945290-124947310 | p2 | (AT)11 | ATTCGTCCTCATCGATCCAC | CATCACTACACACGCTCGCT | 267 |
| chr1:124945604-124947626 | p5 | (TTTTA)4 | GCGAGCGTGTGTAGTGATGT | TTCATTTGAACTCCCTTCGC | 107 |
| chr1:124945604-124947626 | p2 | (AT)11 | GCGAGCGTGTGTAGTGATGT | TTCATTTGAACTCCCTTCGC | 107 |
| chr1:124977377-124979393 | p2 | (TA)8 | AAATGACAATCAAGCGGACC | ATTTTGGTCCTTTTCTCGCA | 132 |

| Marker ID | SSR type | SSR motif | FORWARD PRIMER (5'-3') | REVERSE PRIMER (5'-3') | size (bp) |
|---|---|---|---|---|---|
| chr1:125153050-125155064 | p2 | (AT)7 | TTTGGAGGCCTGAAAGTTTG | GCAGGCATGTGTATGTGTCC | 205 |
| chr1:125169120-125171138 | p6 | (AAAAGA)3 | TTGTTCAGCAGCATGAGGAG | ATTGGGAAAAATGTCCACCA | 194 |
| chr1:125304603-125306627 | p2 | (TA)12 | CTGGGTGAATTGGACGTCTT | TCCGCTTCTGAGTTTTGGTT | 247 |
| chr1:125304603-125306627 | p2 | (TA)13 | CTGGGTGAATTGGACGTCTT | TCCGCTTCTGAGTTTTGGTT | 247 |
| chr1:125304603-125306627 | p2 | (AT)6 | CTGGGTGAATTGGACGTCTT | TCCGCTTCTGAGTTTTGGTT | 247 |
| chr1:125304849-125306885 | p2 | (TA)12 | CGCATTTAGCCAAGGAAGAC | TCCGCTTCTGAGTTTTGGTT | 117 |
| chr1:125304849-125306885 | p2 | (TA)13 | CGCATTTAGCCAAGGAAGAC | TCCGCTTCTGAGTTTTGGTT | 117 |
| chr1:125304849-125306885 | p2 | (AT)6 | CGCATTTAGCCAAGGAAGAC | TCCGCTTCTGAGTTTTGGTT | 117 |
| chr1:125330256-125332276 | p5 | (CTTTT)4 | GTCAAGCAAAACATGAGGCA | ATTCGACCACTTTTTGGCAC | 146 |
| chr1:125448618-125450636 | p3 | (ACA)6 | CCACCGCCTTTATCACAACT | TGCATCATGTCCACCCTAAA | 155 |
| chr1:125510719-125512731 | p2 | (AC)8 | TGAACCTTCTGCCTCATTCC | ATATGCCCTGTGGATTCAGC | 120 |
| chr1:125510719-125512731 | p2 | (AC)9 | TGAACCTTCTGCCTCATTCC | ATATGCCCTGTGGATTCAGC | 120 |
| chr1:125510719-125512731 | p2 | (AC)6 | TGAACCTTCTGCCTCATTCC | ATATGCCCTGTGGATTCAGC | 120 |
| chr1:125510719-125512731 | p2 | (AC)6 | TGAACCTTCTGCCTCATTCC | ATATGCCCTGTGGATTCAGC | 120 |
| chr1:125510719-125512731 | p2 | (AT)9 | TGAACCTTCTGCCTCATTCC | ATATGCCCTGTGGATTCAGC | 120 |
| chr1:125510719-125512731 | p2 | (CA)6 | TGAACCTTCTGCCTCATTCC | ATATGCCCTGTGGATTCAGC | 120 |
| chr1:125569884-125571928 | p2 | (AC)6 | AAGCAACAACTAGCAGCCGT | ACCTGGAAAAATCCCCATTC | 105 |
| chr1:125591465-125593485 | p2 | (AT)10 | ATCTGTTTGGAGAACGTGCC | ATTCACATTTGGCTGCATCA | 219 |
| chr1:125591465-125593485 | p2 | (AT)8 | ATCTGTTTGGAGAACGTGCC | ATTCACATTTGGCTGCATCA | 219 |
| chr1:125591791-125593807 | p2 | (AT)10 | GCAACCAAAACTTAGAGGCG | TTTCGTTAATTTCCCCCACA | 288 |
| chr1:125591791-125593807 | p2 | (AT)8 | GCAACCAAAACTTAGAGGCG | TTTCGTTAATTTCCCCCACA | 288 |
| chr1:125595749-125597779 | p2 | (AT)15 | AGGACTGTCCCCAAAGGACT | AGAAATTCGAATGAACCCCC | 111 |
| chr1:125628657-125630683 | p2 | (TA)13 | TCCGGATGTTACCATGTTCC | TCCATCAACCCAAATCCACT | 183 |
| chr1:125689413-125691425 | p2 | (TA)6 | TAGTTCCCGAACTGGACACC | TGGCTTGCATATGACACGAT | 201 |
| chr1:125738098-125740116 | p2 | (AG)8 | TTGGAATGAATGGAAGGCAT | AATAGGGGATTTTGGGTTCG | 295 |
| chr1:125738098-125740116 | p2 | (AT)9 | TTGGAATGAATGGAAGGCAT | AATAGGGGATTTTGGGTTCG | 295 |
| chr1:125756155-125758241 | p2 | (AC)9 | GCGGAAGTGAAGGTGATGAT | ACAACATTTTCTGCGGGAAC | 228 |
| chr1:125781107-125783122 | p2 | (AT)7 | CACCCCTACCTGGCAATATG | GTGTGTTGCTGTTACGTGGG | 244 |
| chr1:125781107-125783122 | p2 | (AT)7 | CACCCCTACCTGGCAATATG | GTGTGTTGCTGTTACGTGGG | 244 |
| chr1:125781107-125783122 | p5 | (ACCGG)3 | CACCCCTACCTGGCAATATG | GTGTGTTGCTGTTACGTGGG | 244 |
| chr1:125807530-125809548 | p3 | (ATT)6 | GGCAGGATCACAAGGGTCTA | GAGCCCTCAAATTGAAGCAG | 176 |
| chr1:125890852-125892878 | p2 | (AC)7 | ACCAAAGGGAAAGTGTGTCG | TTCGATCCACTGTTTCTCCC | 166 |
| chr1:125890852-125892878 | p2 | (AT)9 | ACCAAAGGGAAAGTGTGTCG | TTCGATCCACTGTTTCTCCC | 166 |
| chr1:125914427-125916449 | p5 | (TGTTT)3 | TGAGTTCTGGTGTCTTGGGA | ATCAAGTCGATGTTGACCCC | 267 |
| chr1:125927261-125929276 | p3 | (CAA)5 | CCATTCGGTGTCTTCCAGTT | ACATCACTCCTTCGGGTTTG | 168 |
| chr1:125960351-125962363 | p2 | (GT)6 | GACGATTTTGCTGAGAAGCC | ACCCTGCGCATTGTCTTAAC | 138 |

| Marker ID | SSR type | SSR motif | FORWARD PRIMER (5'-3') | REVERSE PRIMER (5'-3') | size (bp) |
|---|---|---|---|---|---|
| chr1:125960351-125962363 | p5 | (ATAAA)3 | GACGATTTTGCTGAGAAGCC | ACCCTGCGCATTGTCTTAAC | 138 |
| chr1:125971126-125973146 | p2 | (TC)10 | TTGTCGGGCCTTATTTTGAG | AGGAGTATGACGGGCTCTGA | 204 |
| chr1:125978409-125980424 | p3 | (CAA)5 | CCACTCCATTGTTCCGTTCT | AACAGCTTCACCCCTTTGTG | 145 |
| chr1:126006394-126008412 | p2 | (TC)9 | CAATTCAACCATGTGGGGTT | AATTAGGTTGGGCAATTCCA | 203 |
| chr1:126013711-126015723 | p2 | (AT)6 | CTGGAACACATCCAGCAATG | GTGATCTGCAGGTGACATGG | 181 |
| chr1:126013711-126015723 | p2 | (TA)9 | CTGGAACACATCCAGCAATG | GTGATCTGCAGGTGACATGG | 181 |
| chr1:126025094-126027112 | p3 | (AAT)6 | TTTTCCTTTCGTCGGTGTTT | AAAGGAAAGAATCGCGACAA | 271 |
| chr1:126025094-126027112 | p2 | (TA)44 | TTTTCCTTTCGTCGGTGTTT | AAAGGAAAGAATCGCGACAA | 271 |
| chr1:126030868-126032884 | p2 | (CA)8 | CGGGAATGGCTACAGAGGTA | TAACGGACCATAAAGACGCC | 232 |
| chr1:126051590-126053608 | p6 | (GTTGTG)3 | GCAATATGCTTGTTGGAGCA | CACAAATTCCAATTTTGCCC | 126 |
| chr1:126146414-126148432 | p6 | (CGTCCC)3 | GCCCCCATATCCTTCAATTT | CATTCCGAGGGACAACCTTA | 128 |
| chr1:126148256-126150283 | p3 | (CAA)7 | GCACTCCAAGCTAGCCAATC | AGTTAAATTCTCGGGGTGGC | 202 |
| chr1:126215819-126217869 | p2 | (TA)25 | TCGGGGTAAACAAGATGGAG | CCGGTTTTGTCGTGAAAGAT | 193 |
| chr1:126226250-126228268 | p6 | (CATGAG)3 | AACCGGCCCTTAGCAGTTAT | TGTGGTTCACTCGGTCACAT | 270 |
| chr1:126242594-126244609 | p5 | (TTTTA)3 | AAAATCACGCCACATCAACA | AAAAGAAATTGGGGTGGTCC | 267 |
| chr1:126255164-126257178 | p2 | (AT)7 | AGCCGATAGGTGATGACGAC | TTCTAAATGCAACCGAACCC | 156 |
| chr1:126289749-126291765 | p2 | (TA)8 | ACTCGGCTCCAAACATCAAC | CCGAATGGAATTGGAGTGAT | 101 |
| chr1:126292402-126294458 | p2 | (TA)28 | TTAATATCCCGTGGGAACCA | GGCATTGGGAGCTCATTAGA | 170 |
| chr1:126331978-126333990 | p2 | (AT)6 | ACCAGCTTCAAAACCACCAG | ATTCAAGTATGCCTGGGTGC | 153 |
| chr1:126382158-126384173 | p5 | (AATAA)3 | ACGCCTACTTAGGACCCGTT | GAACCGTTTGGCCATAAGAA | 152 |
| chr1:126476059-126478100 | p2 | (GT)7 | TCAATCACACCGAGGAATCA | TTCCTGAGGGTGCTACTTGC | 151 |
| chr1:126489078-126491126 | p2 | (TA)10 | TGACCCAACCCATTTTTAGC | GAGAGAGGTGTCAAATGGGC | 242 |
| chr1:126489127-126491175 | p2 | (TA)10 | TGACCCAACCCATTTTTAGC | GAGAGAGGTGTCAAATGGGC | 193 |
| chr1:126620634-126622649 | p3 | (TTA)5 | TGAGAATAGCCAACCCCAAC | ATCACCCCAAAAAGGGAATC | 163 |
| chr1:126627309-126629324 | p5 | (AAAAT)3 | GGTTTTACATCCCCATGCAC | ATGCATCCACAAAATGCTGA | 102 |
| chr1:126635182-126637194 | p2 | (TG)6 | GTGCAGATTCAAGGTCCCAT | ACCATCAGATCAGCGAAAGG | 265 |
| chr1:126658727-126660743 | p2 | (CG)8 | CTAACGTGCGGGCTTAGTTC | TCACCCTCATGACGTTTTCA | 278 |
| chr1:126658727-126660743 | p3 | (AAT)11 | CTAACGTGCGGGCTTAGTTC | TCACCCTCATGACGTTTTCA | 278 |
| chr1:126739078-126741096 | p6 | (TATTTT)3 | GCCCGAAGATCAATTTCAAG | CCGGTTTGGACTTATGGAGA | 172 |
| chr1:126739078-126741096 | p2 | (TA)6 | GCCCGAAGATCAATTTCAAG | CCGGTTTGGACTTATGGAGA | 172 |
| chr1:126739265-126741277 | p6 | (TATTTT)3 | CGGTCCGTTATGTAGGCACT | GAATGAAGACTCCCAATGGC | 107 |
| chr1:126739265-126741277 | p2 | (TA)6 | CGGTCCGTTATGTAGGCACT | GAATGAAGACTCCCAATGGC | 107 |
| chr11:100767-102782 | p5 | (TTTCT)3 | GTGGTGCCAGGAACTGATTT | AAGAATGGGTCAGCCAAATG | 248 |
| chr11:134245-136260 | p3 | (CAA)5 | GGCTCAGTGTGGGAAACATT | CCCCATTGGATTTGTTTGTC | 209 |
| chr11:137561-139576 | p3 | (ATT)5 | ACAATTTGGTTTGAGAGGCG | CACGCAAAGCTTGACACCTA | 128 |
| chr11:169174-171186 | p5 | (TTTTG)3 | CCTGTTTGCTCACTAACGCA | AGAACTGTTAGGCGAGCCCT | 245 |

| Marker ID | SSR type | SSR motif | FORWARD PRIMER (5'-3') | REVERSE PRIMER (5'-3') | size (bp) |
|---|---|---|---|---|---|
| chr11:169187-171229 | p5 | (TTTTG)3 | CCTGTTTGCTCACTAACGCA | AGAACTGTTAGGCGAGCCCT | 232 |
| chr11:248917-250935 | p2 | (AT)9 | AAGCTGGCAAGAGGTTTCAA | GAAGCCTGAAGGATGGAACA | 144 |
| chr11:273653-275665 | p2 | (TC)6 | TTTTTAATCCCAACGCGAAG | TTGCAAGTCCTACATGCCAG | 124 |
| chr11:390571-392591 | p5 | (AAGAA)4 | AAGAAGAGTGGCAGAGCGAG | GGCCTATTGGTTGGTCTTCA | 232 |
| chr11:403821-405839 | p6 | (AAAATC)3 | CTCCGTCCAAGTTCATGGTT | TTGAGTATGGCCAAGGAAGG | 144 |
| chr11:438946-440970 | p3 | (TTA)8 | CCGAACACCCTCGATAAAAA | TAAGAATCGAACCTGGGCAC | 286 |
| chr11:501891-503906 | p3 | (GCT)7 | AGCAGAGGCGTAGACAGCTT | AAACGAAACGAAGCAAAACG | 119 |
| chr11:501891-503906 | p5 | (ATTTT)3 | AGCAGAGGCGTAGACAGCTT | AAACGAAACGAAGCAAAACG | 119 |
| chr11:501891-503906 | p2 | (AT)6 | AGCAGAGGCGTAGACAGCTT | AAACGAAACGAAGCAAAACG | 119 |
| chr11:501891-503906 | p2 | (AT)9 | AGCAGAGGCGTAGACAGCTT | AAACGAAACGAAGCAAAACG | 119 |
| chr11:501891-503906 | p2 | (AT)8 | AGCAGAGGCGTAGACAGCTT | AAACGAAACGAAGCAAAACG | 119 |
| chr11:537407-539422 | p5 | (CGCAC)3 | ACTGTTTTCAAATGCCCTGC | TTCGCTGCCGAAGAAATAGT | 206 |
| chr11:537424-539463 | p5 | (CGCAC)3 | ACTGTTTTCAAATGCCCTGC | TTCGCTGCCGAAGAAATAGT | 189 |
| chr11:614419-616434 | p5 | (TATTT)3 | TTTGTTCTTTTGTTGCACGC | ACAATTGGCTTACGTGTCCC | 175 |
| chr11:631111-633127 | p2 | (AT)8 | TAAGAGTCCGGAAGCCAAAA | ATTCCGGCTATGAACAGTGC | 282 |
| chr11:631111-633127 | p2 | (CA)7 | TAAGAGTCCGGAAGCCAAAA | ATTCCGGCTATGAACAGTGC | 282 |
| chr11:655683-657698 | p5 | (AATTA)3 | CAAATTCTGCAAAGGGCAAT | TCAAACATGCAAGTTGCTCC | 180 |
| chr11:695273-697288 | p3 | (ACA)5 | AGCGAACTGCAGCCATAACT | GAATTTGGACTGGCAAGGAA | 154 |
| chr11:771973-774003 | p2 | (AT)15 | TTCCCCAAGTATCTTTCCCC | GTGCATCACGGTGAGATTTG | 190 |
| chr11:774867-776891 | p4 | (TTGT)6 | AACGACCCCCTAACCAATTC | TGCCAAAAGATGCAGATGAC | 127 |
| chr11:774867-776891 | p2 | (AT)8 | AACGACCCCCTAACCAATTC | TGCCAAAAGATGCAGATGAC | 127 |
| chr11:778738-780786 | p2 | (AT)11 | TGGCAGAAGAGGAGGAGAAA | TAGGCCGGTGATTAGACTGG | 136 |
| chr11:807100-809115 | p2 | (AT)14 | AGTTGCACCGAAGGATATGG | TACATGGCCTCGAAGGAAAC | 140 |
| chr11:807100-809115 | p5 | (AAAAG)3 | AGTTGCACCGAAGGATATGG | TACATGGCCTCGAAGGAAAC | 140 |
| chr11:925034-927056 | p2 | (TA)16 | TTCCCAAATTATCCCCATCA | AAATGACATGGCGGAATAGC | 159 |
| chr11:1009548-1011566 | p6 | (TATATG)3 | GTCTAGCTTGGAGATTGCGG | ATGCTCACACGTTCAAGTGC | 145 |
| chr11:1009548-1011566 | p2 | (AT)9 | GTCTAGCTTGGAGATTGCGG | ATGCTCACACGTTCAAGTGC | 145 |
| chr11:1015830-1017842 | p2 | (TA)6 | ATTCGTGAAAAATTCGACGC | CTTATCGACGGTCCAAAGGA | 232 |
| chr11:1189801-1191825 | p2 | (CT)12 | GTGTCGGTGAGGACAAGGAT | TCACCTCACACCCAATTCAA | 211 |
| chr11:1189801-1191825 | p2 | (AT)38 | GTGTCGGTGAGGACAAGGAT | TCACCTCACACCCAATTCAA | 211 |
| chr11:1216483-1218495 | p2 | (AT)6 | ATTAAGAGGCGCGACAAAAA | AAACGCATTTGGGTTACCAG | 292 |
| chr11:1216483-1218495 | p2 | (TA)6 | ATTAAGAGGCGCGACAAAAA | AAACGCATTTGGGTTACCAG | 292 |
| chr11:1216649-1218661 | p2 | (AT)6 | AATTAAGAGGCGCGACAAAA | AAACGCATTTGGGTTACCAG | 125 |
| chr11:1216649-1218661 | p2 | (TA)6 | AATTAAGAGGCGCGACAAAA | AAACGCATTTGGGTTACCAG | 125 |
| chr11:1241172-1243194 | p2 | (AT)11 | GGGACCCCATGTAACACATC | GAAAGAGACAGCAAGCCAGG | 238 |
| chr11:1251354-1253390 | p3 | (ATT)12 | AGGCTTTTGTTTTGCTGGAA | TGTAGGAAGGCTCACGAGGT | 292 |

| Marker ID | SSR type | SSR motif | FORWARD PRIMER (5'-3') | REVERSE PRIMER (5'-3') | size (bp) |
|---|---|---|---|---|---|
| chr11:1328177-1330193 | p2 | (TA)8 | CACTTGACCCCGTATTTTGG | TTTTTGTCCTTCAAAACGGC | 274 |
| chr11:1353357-1355369 | p2 | (AT)11 | GTGTACATGAGGCCGTTGTG | GTCAACTTGTCGGCTTCCAT | 141 |
| chr11:1353357-1355369 | p2 | (TG)6 | GTGTACATGAGGCCGTTGTG | GTCAACTTGTCGGCTTCCAT | 141 |
| chr11:1375970-1377986 | p2 | (GT)8 | TTGGTGCCCTCAACACAATA | GTTGCTGTCATGTGACCTGG | 227 |
| chr11:1561302-1563326 | p3 | (TAT)8 | CCAAGGGGGAGTGTTATGAA | GTTTGCCCCTCAACCTACAA | 148 |
| chr11:1618471-1620483 | p2 | (TA)6 | AATCCTTGCGGTGAAAAAGA | TTGCATTGTGATATTCCCGA | 247 |
| chr11:1618471-1620483 | p2 | (AT)6 | AATCCTTGCGGTGAAAAAGA | TTGCATTGTGATATTCCCGA | 247 |
| chr11:1634348-1636372 | p2 | (TA)12 | TCTAGCCGCCTTCACTTTGT | CTACTACCGGCACCACGTCT | 222 |
| chr11:1634348-1636372 | p2 | (AT)6 | TCTAGCCGCCTTCACTTTGT | CTACTACCGGCACCACGTCT | 222 |
| chr11:1715720-1717738 | p2 | (TA)7 | GTATGTCAGGGTGCCCAAGT | TGATCAATGAAACGGGTTGA | 165 |
| chr11:1715720-1717738 | p6 | (TACATA)3 | GTATGTCAGGGTGCCCAAGT | TGATCAATGAAACGGGTTGA | 165 |
| chr11:1745149-1747171 | p2 | (AT)11 | TTGAATCCTCCAAAACTGCC | TAATGGGCCTACAAAACCCA | 256 |
| chr11:1755523-1757541 | p6 | (AAAAAG)3 | ACTGCAAGTGATCCTTCTCCA | AATACGCCGTGAAGACGAAG | 192 |
| chr11:1795092-1797106 | p2 | (TA)7 | CGATGGATAAAGCAAAGGGA | GAAGGGATGAGTCAAGCGAC | 295 |
| chr11:1972745-1974763 | p6 | (CTTTTT)3 | CACTTCCCGTTACTCCGGTA | AAACCGGACTGGCTATGATG | 214 |
| chr11:2009104-2011138 | p5 | (TCTGA)3 | GATTGCTCCTGCATTGTTGA | ATATTCCTGTTTCGCCATCG | 261 |
| chr11:2009148-2011182 | p5 | (TCTGA)3 | GATTGCTCCTGCATTGTTGA | ATATTCCTGTTTCGCCATCG | 217 |
| chr11:2009192-2011226 | p5 | (TCTGA)3 | GATTGCTCCTGCATTGTTGA | ATATTCCTGTTTCGCCATCG | 173 |
| chr11:2009236-2011270 | p5 | (TCTGA)3 | GATTGCTCCTGCATTGTTGA | ATATTCCTGTTTCGCCATCG | 129 |
| chr11:2009500-2011534 | p5 | (TCTGA)3 | AAAACAGAGGCCTGAACGTG | CTTCGAACTCCTGCTTCGTC | 290 |
| chr11:2009544-2011578 | p5 | (TCTGA)3 | AAAACAGAGGCCTGAACGTG | CTTCGAACTCCTGCTTCGTC | 246 |
| chr11:2009588-2011622 | p5 | (TCTGA)3 | AAAACAGAGGCCTGAACGTG | CTTCGAACTCCTGCTTCGTC | 202 |
| chr11:2009632-2011666 | p5 | (TCTGA)3 | AAAACAGAGGCCTGAACGTG | CTTCGAACTCCTGCTTCGTC | 158 |
| chr11:2009676-2011710 | p5 | (TCTGA)3 | AAAACAGAGGCCTGAACGTG | CTTCGAACTCCTGCTTCGTC | 114 |
| chr11:2016546-2018586 | p2 | (AT)20 | CCACCTCATTCGAAGGGTTA | CCTTCCTAGTCCCCCAGAAC | 248 |
| chr11:2037762-2039780 | p3 | (CTT)6 | TCCTCGCTGATATTTGTCCC | TTTTTGGATTGATGATGGCA | 287 |
| chr11:2228721-2230739 | p6 | (CAATTT)3 | TGGACCTCGCCACTTATTTT | TTCGTTATTTGTTTGCCGGT | 186 |
| chr11:2237330-2239384 | p2 | (TA)6 | CACGAAAATCATGGGCTTCT | ACGATCCTTGTCCTACGGTG | 262 |
| chr11:2237330-2239384 | p3 | (ATA)18 | CACGAAAATCATGGGCTTCT | ACGATCCTTGTCCTACGGTG | 262 |
| chr11:2257864-2259882 | p2 | (TA)9 | AGTATTGGTCTGGTCAGGCG | GCGAATTGCCCTACCAGTTA | 178 |
| chr11:2263332-2265350 | p6 | (ATAAAA)3 | GGGATCCAATAAAGCAGCAG | AATGTAGCAGGAAGCATGGG | 145 |
| chr11:2274260-2276276 | p2 | (TA)7 | CACAAATAAATTGGGGTGGC | CGTGCAATAGACCCTTGTGA | 262 |
| chr11:2274260-2276276 | p4 | (ATTT)4 | CACAAATAAATTGGGGTGGC | CGTGCAATAGACCCTTGTGA | 262 |
| chr11:2274260-2276276 | p4 | (ATTT)4 | CACAAATAAATTGGGGTGGC | CGTGCAATAGACCCTTGTGA | 262 |
| chr11:2274260-2276276 | p4 | (ATTT)4 | CACAAATAAATTGGGGTGGC | CGTGCAATAGACCCTTGTGA | 262 |
| chr11:2274260-2276276 | p4 | (ATTT)4 | CACAAATAAATTGGGGTGGC | CGTGCAATAGACCCTTGTGA | 262 |

| Marker ID | SSR type | SSR motif | FORWARD PRIMER (5'-3') | REVERSE PRIMER (5'-3') | size (bp) |
|---|---|---|---|---|---|
| chr11:2274260-2276276 | p4 | (ATTT)4 | CACAAATAAATTGGGGTGGC | CGTGCAATAGACCCTTGTGA | 262 |
| chr11:2325092-2327107 | p5 | (TAAAA)3 | ATCTACGATCAATCGCCCAC | AAAGCCTATTTTTCCCCCAA | 245 |
| chr11:2361141-2363153 | p2 | (TA)6 | GGTCACGTGTTCTTGAGGGT | TTGGCTAAACGGATTGAACC | 209 |
| chr11:2361141-2363153 | p2 | (AT)8 | GGTCACGTGTTCTTGAGGGT | TTGGCTAAACGGATTGAACC | 209 |
| chr11:2487361-2489407 | p2 | (TA)8 | CCCACCATCTCCAAACTCAC | GGTTGGGTTGTGCTCCTAAA | 296 |
| chr11:2487361-2489407 | p2 | (TA)23 | CCCACCATCTCCAAACTCAC | GGTTGGGTTGTGCTCCTAAA | 296 |
| chr11:2487361-2489407 | p2 | (AT)10 | CCCACCATCTCCAAACTCAC | GGTTGGGTTGTGCTCCTAAA | 296 |
| chr11:2488290-2490310 | p2 | (TA)23 | TTACCACGTGCGTGTCAATC | AAATGACATGGCGGAATAGC | 232 |
| chr11:2488290-2490310 | p2 | (AT)10 | TTACCACGTGCGTGTCAATC | AAATGACATGGCGGAATAGC | 232 |
| chr11:2732051-2734117 | p3 | (AAT)22 | TCGATTTACCCTCGAACACC | GCTCCCTCCACTCACTTCTG | 178 |
| chr11:2854385-2856405 | p2 | (AG)10 | TTCCACATGAACTCAGCCAA | CGCGTCCTTATTCTCAGCTC | 260 |
| chr11:2863772-2865799 | p2 | (AG)9 | CCACGGTTTGGACTGTTTTT | GTTTGGAAATGAAGCCGTGT | 285 |
| chr11:2977244-2979266 | p2 | (TA)11 | TTTCCCATGGAAGCCATTAG | CCTTTGAGCCACCGTCTCTA | 108 |
| chr11:2977244-2979266 | p4 | (ATTT)4 | TTTCCCATGGAAGCCATTAG | CCTTTGAGCCACCGTCTCTA | 108 |
| chr11:2998343-3000370 | p5 | (ATTAA)3 | TGTGCAGTTTCTTCGTTTGG | AAGGACATTCCAGCGAATTG | 131 |
| chr11:3033560-3035576 | p5 | (ACCGG)3 | CACCCGTTAACAAAAATGGC | TGGGTTAATGGTTCGGGTTA | 156 |
| chr11:3033560-3035576 | p2 | (TA)8 | CACCCGTTAACAAAAATGGC | TGGGTTAATGGTTCGGGTTA | 156 |
| chr11:3058532-3060544 | p2 | (AT)6 | CTATTTCCATCCGCTAGCCA | GCCTGATTTCTGTAGCAGCC | 259 |
| chr11:3124257-3126279 | p2 | (TA)9 | AAACAAAAAGGGAGTGGCCT | TGACAAGTGTTGGCGTCATT | 143 |
| chr11:3124257-3126279 | p2 | (AT)11 | AAACAAAAAGGGAGTGGCCT | TGACAAGTGTTGGCGTCATT | 143 |
| chr11:3168110-3170125 | p5 | (AATTA)3 | GGCCGCGTTACTTAAAATGA | ACGGTCACGCTTCAAGAGAT | 247 |
| chr11:3192581-3194599 | p6 | (TAAAAA)3 | CCCAACTTGTCGGTTTCCTA | AGCCAAGAAAGAGATGCCAA | 206 |
| chr11:3328542-3330556 | p2 | (AG)7 | TCGCTCCATCAACCTCTTCT | GCCTAGTTGTGCTCGTGTGA | 169 |
| chr11:3328542-3330556 | p3 | (TTG)5 | TCGCTCCATCAACCTCTTCT | GCCTAGTTGTGCTCGTGTGA | 169 |
| chr11:3413742-3415757 | p3 | (AGA)5 | ATCTCTCACACCTCCGCACT | TACTTTTGAAAATTGGGCGG | 231 |
| chr11:3463618-3465632 | p2 | (AG)7 | TCGCTCCATCAACCTCTTCT | GCCTAGTTGTGCTCGTGTGA | 169 |
| chr11:3463618-3465632 | p3 | (TTG)5 | TCGCTCCATCAACCTCTTCT | GCCTAGTTGTGCTCGTGTGA | 169 |
| chr11:3548818-3550833 | p3 | (AGA)5 | ATCTCTCACACCTCCGCACT | TACTTTTGAAAATTGGGCGG | 231 |
| chr11:3630408-3632422 | p2 | (TA)13 | AAGTATCAACCGTGGCTTCG | CCCCCACATCATATTCGTTC | 247 |
| chr11:3630429-3632443 | p2 | (TA)13 | AAGTATCAACCGTGGCTTCG | CCCCCACATCATATTCGTTC | 226 |
| chr11:3630467-3632530 | p2 | (TA)13 | AAGTATCAACCGTGGCTTCG | CCCCCACATCATATTCGTTC | 188 |
| chr11:3630602-3632632 | p2 | (TA)13 | AAAACTCAGAAGCGGAGCAA | TTCCCCCACATCATATTCGT | 191 |
| chr11:3630643-3632657 | p2 | (TA)13 | AAAACTCAGAAGCGGAGCAA | TTCCCCCACATCATATTCGT | 150 |
| chr11:3646064-3648082 | p6 | (TTGCTT)3 | GGATGAAATGGAGGCTCGTA | TCCCTCATCTTGTCCTCCAC | 191 |
| chr11:3646064-3648082 | p3 | (TGT)6 | GGATGAAATGGAGGCTCGTA | TCCCTCATCTTGTCCTCCAC | 191 |
| chr11:3661962-3663980 | p2 | (TA)9 | TCTCAATGCCCGTTAACTCC | TCAGGGCTAAGAAGAGCGAA | 199 |

| Marker ID | SSR type | SSR motif | FORWARD PRIMER (5'-3') | REVERSE PRIMER (5'-3') | size (bp) |
|---|---|---|---|---|---|
| chr11:3669112-3671127 | p5 | (TTTTA)4 | ACCCCCTTCTTCCTATCCCT | TGAAATTGATCACGTGGGTG | 252 |
| chr11:3705644-3707656 | p2 | (AT)6 | CCAATTTACGACCTGCCAGT | ACGTTTGATGACCTTAGGCG | 131 |
| chr11:3829938-3831953 | p5 | (AACCG)4 | TTGACTCGCCTCTTTTGGAT | CGGTTTGTTGAAGTTTCGGT | 268 |
| chr11:3829938-3831953 | p5 | (AACCG)3 | TTGACTCGCCTCTTTTGGAT | CGGTTTGTTGAAGTTTCGGT | 268 |
| chr11:3874843-3876864 | p3 | (CAT)7 | TTGGCAAGCATACAATCTGG | GCTTGTCGTCAAGTAAGCCC | 284 |
| chr11:3924415-3926430 | p5 | (TTGTA)3 | TGTGGGAGAATCAAGAACCC | CCCACGAGATTGTTGAGGTT | 239 |
| chr11:3941912-3943952 | p2 | (AT)20 | TCCCAAAGTATCCCCATCAA | AAATGACATGGCGGAATAGC | 235 |
| chr11:4000896-4002936 | p2 | (AT)20 | TCCCAAAGTATCCCCATCAA | AAATGACATGGCGGAATAGC | 235 |
| chr11:4046799-4048815 | p4 | (TATT)4 | TTTGGGCCATAATCCAACTC | AAGTCACCACCAAGATTCGG | 165 |
| chr11:4077375-4079390 | p5 | (AAATC)3 | GCTTCTTTGACACTTCCCCA | TTTAACCCAAAACCGATCCA | 173 |
| chr11:4093989-4096004 | p5 | (AAATC)3 | GCTTCTTTGACACTTCCCCA | TTTAACCCAAAACCGATCCA | 173 |
| chr11:4105231-4107251 | p2 | (TA)10 | CCAGCTAGGACAGGCTTCAG | TTGTCTTCTGGGCGATTTTC | 102 |
| chr11:4165994-4168009 | p5 | (AACCC)3 | TTTGGCCCATGTTTGAATTT | TTGAAGCTTTTCTGTCGCCT | 204 |
| chr11:4261512-4263590 | p2 | (AT)39 | GGACTTGGGAGCTGTTTTGA | TATGGGGTACATGTGCAACG | 168 |
| chr11:4264164-4266198 | p2 | (TA)17 | ATTGGACAAATTTGCAAGGG | TTGAACCGCGTATGGATGTA | 175 |
| chr11:4264164-4266198 | p4 | (TTTC)4 | ATTGGACAAATTTGCAAGGG | TTGAACCGCGTATGGATGTA | 175 |
| chr11:4378734-4380754 | p5 | (TTGTT)4 | GACCAAACCCAAAACGAGAA | TGCATTGACGAAGATTGGAG | 189 |
| chr11:4588366-4590386 | p2 | (AT)6 | GCAAGAAAAGGAAATGCTGC | GTCCTTGGTCCCTGGTTTTT | 280 |
| chr11:4588366-4590386 | p2 | (AT)10 | GCAAGAAAAGGAAATGCTGC | GTCCTTGGTCCCTGGTTTTT | 280 |
| chr11:4722940-4724962 | p2 | (TA)11 | GTTGATCCAGTTGGTGCCTT | GCTGCTCGTCATCTCTGTCA | 101 |
| chr11:4867031-4869047 | p6 | (GGCATG)3 | ACCGCCCATACCCTAAATTC | ACCTCACAAACGTTGCCTTC | 217 |
| chr11:4867031-4869047 | p4 | (ATAA)4 | ACCGCCCATACCCTAAATTC | ACCTCACAAACGTTGCCTTC | 217 |
| chr11:4894714-4896760 | p2 | (CG)7 | ATCTTGGTAAAAGCGGCAGA | GCGCGTGTGTATTTGACAGT | 272 |
| chr11:4942227-4944241 | p2 | (AT)7 | GTCGGGAACCACCTGACTTA | TTTTGTAGCTCCACCGGTTC | 207 |
| chr11:4956345-4958384 | p3 | (AAT)13 | AACAACCCCCACCTTCTTTT | ATATGGCAACCCCTGTTTCA | 152 |
| chr11:5009067-5011079 | p2 | (AT)6 | AAGTATCAACCGTGGCTTCG | TCCGCTTCTGAGTTTTGGTT | 127 |
| chr11:5009154-5011166 | p2 | (AT)6 | AAAACTCAGAAGCGGAGCAA | TTTCGTTAATTTCCCCCACA | 178 |
| chr11:5044175-5046235 | p2 | (AT)30 | AAATCCGTATCCGGGCTAGT | CCCTAACGGTCGACTGATGT | 218 |
| chr11:5059085-5061100 | p5 | (ATTTT)3 | CAAGCATTTTTGGGCTTCAT | CCCTTTTGGTACTTTTGGCA | 214 |
| chr11:5132604-5134622 | p2 | (AG)9 | TAATAACCGGGTGCTCCAAG | TGCCCATTGACACACAGATT | 235 |
| chr11:5157702-5159720 | p2 | (TA)9 | CCTCGCTGCTTAATGGACTC | CCCAGAACCTCAGGTATCCA | 256 |
| chr11:5189763-5191779 | p2 | (AT)8 | TTGTGCGCGCTTTGATATAC | TTCAATTCTCTGAATCCCCG | 240 |
| chr11:5242888-5244903 | p5 | (TTTCT)3 | TCACACTAGCAAAGGTTGCG | ATGACTTCGTTTTCACGGCT | 260 |
| chr11:5255681-5257712 | p2 | (AG)8 | CCAGTCGAAACAGAGCAACA | TTGTTTTCCGGAGGTTTCTG | 111 |
| chr11:5533106-5535131 | p5 | (ATAAA)5 | CAAAACGTTACAGGCGGATT | TGGGATACAAATCAGTGGCA | 128 |
| chr11:5562756-5564771 | p5 | (TTGTA)3 | ATTTTGCGGAGCTCTGTGAT | AAGCGTGGAGTGGTGGATAC | 191 |

| Marker ID | SSR type | SSR motif | FORWARD PRIMER (5'-3') | REVERSE PRIMER (5'-3') | size (bp) |
|---|---|---|---|---|---|
| chr11:5562756-5564771 | p5 | (GTGGG)3 | ATTTTGCGGAGCTCTGTGAT | AAGCGTGGAGTGGTGGATAC | 191 |
| chr11:5742683-5744697 | p2 | (TA)7 | CCTGTTTTTAGTGGCGGAAA | CAGGGAGACATACGCTGACA | 217 |
| chr11:5928590-5930605 | p5 | (AAAAT)3 | GGTATAATCGCGTTTGGGAG | GTGAGATATTGCCGGGAGAA | 252 |
| chr11:5936298-5938344 | p2 | (AT)23 | AAAGTGGGCAGAGATGATGG | CTTTACGGGGCAGTGACATT | 274 |
| chr11:5943622-5945636 | p2 | (AT)7 | GTGAGTTGCTGAACCATCCA | GATGACTGGAGGATTTCCGA | 113 |
| chr11:5943622-5945636 | p2 | (TA)7 | GTGAGTTGCTGAACCATCCA | GATGACTGGAGGATTTCCGA | 113 |
| chr11:5943839-5945853 | p2 | (AT)7 | TTCATCAAACTCATGGCTGC | TAATGGTTGGAAGGTACCCG | 218 |
| chr11:5943839-5945853 | p2 | (TA)7 | TTCATCAAACTCATGGCTGC | TAATGGTTGGAAGGTACCCG | 218 |
| chr11:5958553-5960568 | p2 | (AT)9 | CTTTGTGGGATCTGGACCAT | GTGGCAACAAAAGGACGAAT | 277 |
| chr11:5958553-5960568 | p5 | (AACTC)3 | CTTTGTGGGATCTGGACCAT | GTGGCAACAAAAGGACGAAT | 277 |
| chr11:6074237-6076249 | p5 | (TAAAT)3 | TGAATAGATCCGCCTTCTGG | TGGCAATGTCCCTCAAAAAT | 260 |
| chr11:6074250-6076266 | p5 | (TAAAT)3 | TGAATAGATCCGCCTTCTGG | TGGCAATGTCCCTCAAAAAT | 247 |
| chr11:6097675-6099689 | p5 | (AATGC)3 | GCAGTGCCTCAAAGACATGA | TTTTGCCGGAGATTGAGAGT | 215 |
| chr11:6097675-6099689 | p2 | (AT)7 | GCAGTGCCTCAAAGACATGA | TTTTGCCGGAGATTGAGAGT | 215 |
| chr11:6107632-6109650 | p3 | (TGT)6 | GACCGGTGCAAATAGAGGAA | TCACTGCTTCTTCACCAACG | 220 |
| chr11:6146807-6148822 | p2 | (TA)9 | AAAAGGCGATTGTTGACACC | GAACCATACCGTACCATGCC | 147 |
| chr11:6148408-6150422 | p2 | (TA)23 | AATTCGGGCAACATCTGAAC | CGGGCATAAGCCGTCTAGTA | 141 |
| chr11:6148408-6150422 | p2 | (AT)7 | AATTCGGGCAACATCTGAAC | CGGGCATAAGCCGTCTAGTA | 141 |
| chr11:6150281-6152301 | p5 | (TTTAT)4 | ATGTGGCAATGAATCCATCA | CAAAAAGTGGAACCCACCAC | 274 |
| chr11:6210102-6212117 | p3 | (CAT)5 | ACAAGCCAACGAGGCTAAAA | ACATATGGGCATGGTGGAGT | 282 |
| chr11:6229766-6231784 | p3 | (GAT)6 | TGGGGGAAAATTGAAATTGA | GGGCCATTTTGCTACTGAAA | 168 |
| chr11:6246416-6248428 | p2 | (AC)6 | CTTTGGTGTGCTGGCATAGA | CGGTTTGTCGGTTTCCTTTA | 208 |
| chr11:6248787-6250879 | p2 | (TA)19 | CAAGGCAGCCCAAATACAAT | GAACAGGGACATCGGAAAAA | 176 |
| chr11:6248787-6250879 | p2 | (AT)20 | CAAGGCAGCCCAAATACAAT | GAACAGGGACATCGGAAAAA | 176 |
| chr11:6273117-6275135 | p3 | (TCA)6 | GGGGGTTGTGTTTCATCATC | GTTAGGGGTGCGCTTAATGA | 131 |
| chr11:6273117-6275135 | p6 | (CCGGTT)3 | GGGGGTTGTGTTTCATCATC | GTTAGGGGTGCGCTTAATGA | 131 |
| chr11:6518554-6520580 | p2 | (TA)7 | TCACGCTGTGGCTTATTCAG | TCCAGCCTATCTTGTGCTCC | 120 |
| chr11:6518554-6520580 | p2 | (TA)7 | TCACGCTGTGGCTTATTCAG | TCCAGCCTATCTTGTGCTCC | 120 |
| chr11:6520468-6522480 | p2 | (TA)7 | ATCTAAAGTGTTGGCCCGTG | GCACACCAGTGAAAAAGCCT | 102 |
| chr11:6520468-6522480 | p2 | (TG)6 | ATCTAAAGTGTTGGCCCGTG | GCACACCAGTGAAAAAGCCT | 102 |
| chr11:6680754-6682768 | p2 | (TA)9 | TGCTACACCTGTTGCTCCAC | TCCACATTTGGCTGCATAAA | 176 |
| chr11:6680769-6682781 | p2 | (TA)9 | TGCTACACCTGTTGCTCCAC | TCCACATTTGGCTGCATAAA | 161 |
| chr11:6680783-6682803 | p2 | (TA)9 | TGCTACACCTGTTGCTCCAC | TCCACATTTGGCTGCATAAA | 147 |
| chr11:6680801-6682813 | p2 | (TA)9 | TGCTACACCTGTTGCTCCAC | TCCACATTTGGCTGCATAAA | 129 |
| chr11:6693612-6695628 | p4 | (TTGT)4 | AGTGGTGGTGGTGGTATGGT | GTCAAAAATGGCACGTGATG | 104 |
| chr11:6693612-6695628 | p5 | (ACCAA)3 | AGTGGTGGTGGTGGTATGGT | GTCAAAAATGGCACGTGATG | 104 |

| Marker ID | SSR type | SSR motif | FORWARD PRIMER (5'-3') | REVERSE PRIMER (5'-3') | size (bp) |
|---|---|---|---|---|---|
| chr11:6803018-6805033 | p5 | (TATTT)3 | CAAAGGGAAAGGGAGGAAAC | GGGCGGGGGAGTATTATCTA | 198 |
| chr11:6803018-6805033 | p5 | (ACCAA)3 | CAAAGGGAAAGGGAGGAAAC | GGGCGGGGGAGTATTATCTA | 198 |
| chr11:6833278-6835292 | p2 | (CT)7 | CCGGGGTCAATTGAGAGTAA | AGGGACAAAATTGGGTGTCA | 286 |
| chr11:6903141-6905159 | p6 | (CAACTC)3 | TTAGGGGTAATGAGGTGGCA | GCCAAACACCCACGAAGTAT | 115 |
| chr11:6903141-6905159 | p2 | (AT)9 | TTAGGGGTAATGAGGTGGCA | GCCAAACACCCACGAAGTAT | 115 |
| chr11:6903141-6905159 | p2 | (TC)7 | TTAGGGGTAATGAGGTGGCA | GCCAAACACCCACGAAGTAT | 115 |
| chr11:6994967-6996981 | p2 | (TA)7 | TTACCCCGTACCTCTGTTGG | TTTCTGTGCATGGTGGGTTA | 139 |
| chr11:7100510-7102528 | p3 | (TTC)6 | GTTTATCGATTTGGTGCGGT | TTCACTCAACTGGTGCCATC | 257 |
| chr11:7126864-7128884 | p5 | (AAGAG)4 | TCACCCAAATCCTCCAGAAG | GCTAAGGATGTGGACGGGTA | 172 |
| chr11:7135932-7137947 | p5 | (AAATA)3 | TTTCAGATTCCGTTTCCGTC | GGCCGTACAGTCTTCGTCAT | 289 |
| chr11:7161854-7163872 | p3 | (TGA)6 | CAGAACCGGTAAAACCGAAA | GCCATTTTTGCAAATGGACT | 224 |
| chr11:7176377-7178395 | p3 | (TGA)6 | CAGAACCGGTAAAACCGAAA | GCCATTTTTGCAAATGGACT | 224 |
| chr11:7288220-7290250 | p2 | (TA)15 | TTTTGGGCGGTTTTGTTAAG | CACGTGAAAAGTGCCACATC | 245 |
| chr11:7288220-7290250 | p2 | (AT)7 | TTTTGGGCGGTTTTGTTAAG | CACGTGAAAAGTGCCACATC | 245 |
| chr11:7288220-7290250 | p2 | (AT)12 | TTTTGGGCGGTTTTGTTAAG | CACGTGAAAAGTGCCACATC | 245 |
| chr11:7288220-7290250 | p4 | (TTTA)4 | TTTTGGGCGGTTTTGTTAAG | CACGTGAAAAGTGCCACATC | 245 |
| chr11:7348593-7350623 | p2 | (TA)15 | GTTTTTGGGCGGTTTTGTTA | GTACCCAGCGACGGAGATTA | 243 |
| chr11:7348593-7350623 | p2 | (AT)7 | GTTTTTGGGCGGTTTTGTTA | GTACCCAGCGACGGAGATTA | 243 |
| chr11:7348593-7350623 | p2 | (AT)12 | GTTTTTGGGCGGTTTTGTTA | GTACCCAGCGACGGAGATTA | 243 |
| chr11:7348593-7350623 | p2 | (AT)33 | GTTTTTGGGCGGTTTTGTTA | GTACCCAGCGACGGAGATTA | 243 |
| chr11:7352265-7354283 | p3 | (TCA)6 | GGGGGTTGTGTTTCATCATC | AAACGCGGTGTCGGTAGTAG | 174 |
| chr11:7352265-7354283 | p6 | (CCGGTT)3 | GGGGGTTGTGTTTCATCATC | AAACGCGGTGTCGGTAGTAG | 174 |
| chr11:7352265-7354283 | p5 | (ATTGT)3 | GGGGGTTGTGTTTCATCATC | AAACGCGGTGTCGGTAGTAG | 174 |
| chr11:7390305-7392331 | p5 | (ACCGG)3 | GCTTTTGCGCCTTTGAGTAG | TTTCTTGGACTAGCAGCCGT | 232 |
| chr11:7390305-7392331 | p2 | (TA)6 | GCTTTTGCGCCTTTGAGTAG | TTTCTTGGACTAGCAGCCGT | 232 |
| chr11:7390305-7392331 | p2 | (TA)13 | GCTTTTGCGCCTTTGAGTAG | TTTCTTGGACTAGCAGCCGT | 232 |
| chr11:7390305-7392331 | p2 | (TA)10 | GCTTTTGCGCCTTTGAGTAG | TTTCTTGGACTAGCAGCCGT | 232 |
| chr11:7390305-7392331 | p2 | (AC)22 | GCTTTTGCGCCTTTGAGTAG | TTTCTTGGACTAGCAGCCGT | 232 |
| chr11:7459660-7461684 | p4 | (AAAC)6 | AATTAGACCAGCGCAGGAAA | ACGTGAGAAAGCCACGAAGT | 115 |
| chr11:7493619-7495631 | p2 | (TA)6 | GCCACGTTTCAGTAGTGGGT | GTATGGAATGGAGAAGGCGA | 164 |
| chr11:7509103-7511118 | p5 | (ACAAA)3 | TCAGTTCATCCCCGTTTTTC | ATTTCCGCTCTCTCCACTCA | 167 |
| chr11:7516315-7518327 | p2 | (TA)7 | TTGGGTTTGTTTGAAGAGCA | CCGCATTTTACTTAGCGCAT | 116 |
| chr11:7516315-7518327 | p2 | (TA)6 | TTGGGTTTGTTTGAAGAGCA | CCGCATTTTACTTAGCGCAT | 116 |
| chr11:7516315-7518327 | p2 | (TA)6 | TTGGGTTTGTTTGAAGAGCA | CCGCATTTTACTTAGCGCAT | 116 |
| chr11:7567987-7570071 | p2 | (AG)6 | AGCCTGTTCTTTTTGGGGAT | TTCGTTTCCTCCCTTTTCCT | 229 |
| chr11:7567987-7570071 | p3 | (AAT)28 | AGCCTGTTCTTTTTGGGGAT | TTCGTTTCCTCCCTTTTCCT | 229 |

| Marker ID | SSR type | SSR motif | FORWARD PRIMER (5'-3') | REVERSE PRIMER (5'-3') | size (bp) |
|---|---|---|---|---|---|
| chr11:7602340-7604355 | p5 | (TCTTC)3 | AAATTGGGAGTGTCGTCAGG | CTTTATTTCCATCTCCGCCA | 290 |
| chr11:7602340-7604355 | p6 | (AGGCCT)3 | AAATTGGGAGTGTCGTCAGG | CTTTATTTCCATCTCCGCCA | 290 |
| chr11:7654441-7656457 | p4 | (AGTG)4 | CAATGAAGGATGGCGAATTT | AACTTCCTTCCCCTGCATCT | 221 |
| chr11:7662508-7664526 | p6 | (TAGAAA)3 | GTTTAATATGGAGCCCGCAA | CCGAAATTACCCTTGAACGA | 195 |
| chr11:7700924-7702966 | p2 | (TA)21 | GGCCAAAACTCCAACTTCAA | TGCAGAACGACAATTTGGAA | 209 |
| chr11:7700924-7702966 | p2 | (AT)13 | GGCCAAAACTCCAACTTCAA | TGCAGAACGACAATTTGGAA | 209 |
| chr11:7802255-7804270 | p3 | (TAA)5 | TATTGCGTCCTTCCTTCACC | TTGCATGTTCCTCAGTGACC | 165 |
| chr11:7802255-7804270 | p5 | (TATAA)3 | TATTGCGTCCTTCCTTCACC | TTGCATGTTCCTCAGTGACC | 165 |
| chr11:7802287-7804299 | p5 | (TATAA)3 | TATTGCGTCCTTCCTTCACC | TTGCATGTTCCTCAGTGACC | 133 |
| chr11:7884383-7886401 | p6 | (TTTTTA)3 | TGCCCAAAATTTTCCTTCTG | TAAAGTTGGGGACGGTCAAA | 263 |
| chr11:7889359-7891374 | p3 | (TGG)5 | TCCTGCACCAATACGTGTGT | CCCAGGTCATCCAAAAGAAA | 118 |
| chr11:7889359-7891374 | p3 | (TGG)5 | TCCTGCACCAATACGTGTGT | CCCAGGTCATCCAAAAGAAA | 118 |
| chr11:7980289-7982304 | p5 | (TTTTA)3 | TCAAGTGGGTTGTAGGGACC | CAAGCGTGCGGTCTTTTACT | 166 |
| chr11:8002620-8004642 | p2 | (AT)11 | AAAGAAAGTGGAACCCACCA | TTTTTCCCGTGATTTTGGAG | 286 |
| chr11:8018680-8020692 | p2 | (TC)6 | GTGGGTGTGGTATGAGTCCC | CACCCTCCACCCCTAATTTT | 135 |
| chr11:8039762-8041776 | p5 | (ATATC)4 | GTTATGCACGCGGATAAGGT | GCTGGGGCATACTAGCCATA | 126 |
| chr11:8156377-8158398 | p3 | (ATT)7 | TCGATGATTCCTATCGAGCA | TTGAGCGAGCTCCCACTATT | 125 |
| chr11:8190388-8192402 | p3 | (CAA)5 | AAAATACGATTGGGCTCACG | TTTGACCCGGCAATAGAATC | 288 |
| chr11:8190388-8192402 | p2 | (AT)7 | AAAATACGATTGGGCTCACG | TTTGACCCGGCAATAGAATC | 288 |
| chr11:8195062-8197076 | p2 | (AT)7 | CGCCAATCCTGAATGTTTTT | AGCCATCTCAAGTTTGGTGC | 224 |
| chr11:8197948-8199968 | p2 | (AT)10 | CTGATGAACGCTGCATTTGT | GCAACAGGACACAATGGTTG | 227 |
| chr11:8221489-8223537 | p3 | (AAT)16 | TTTTCGTGTGGAATCGTCAA | GCGAAGTTTGTGGAAAGAGC | 193 |
| chr11:8302677-8304691 | p2 | (AT)7 | AGTGCAGGCTTTGTATTCGG | TGGTGGGATCCATTTTTGTT | 106 |
| chr11:8302677-8304691 | p2 | (TA)16 | AGTGCAGGCTTTGTATTCGG | TGGTGGGATCCATTTTTGTT | 106 |
| chr11:8401323-8403345 | p2 | (AT)14 | TGATTTTCACCCCCTTTTTG | GAGATTGCCGTCCTTCCATA | 193 |
| chr11:8401356-8403370 | p2 | (AT)14 | TGATTTTCACCCCCTTTTTG | GAGATTGCCGTCCTTCCATA | 160 |
| chr11:8495936-8497951 | p3 | (TAA)5 | TCAAGTTGCGGCAGTGATAG | TCTCCGCTGCCTTGTTATCT | 220 |
| chr11:8521159-8523175 | p4 | (AAAC)4 | GCACGGGAGGAACTAATGAA | AACATGGCAAACCAAGGAAG | 171 |
| chr11:8654061-8656076 | p3 | (AAT)5 | GAAGAAGGAGGGAAAGGGTG | CATTTTCCTCCATACCCACG | 229 |
| chr11:8654061-8656076 | p3 | (AAT)13 | GAAGAAGGAGGGAAAGGGTG | CATTTTCCTCCATACCCACG | 229 |
| chr11:8654061-8656076 | p5 | (AAAAC)3 | GAAGAAGGAGGGAAAGGGTG | CATTTTCCTCCATACCCACG | 229 |
| chr11:8700404-8702419 | p3 | (ACA)5 | TCTGTTGAGTCCGAAAAGGG | GTTGCCTAAGTGTTCTCGGC | 159 |
| chr11:8902102-8904117 | p3 | (TGT)5 | TCACCTACCCCTAGGCCTTT | TCAGGGAGGGAAACATCAAG | 196 |
| chr11:8909016-8911032 | p4 | (ATAA)4 | GTCATAACCTGCATGGGACC | ATCCTCGTCCGAGGTTCTTT | 277 |
| chr11:8929709-8931739 | p2 | (AT)15 | GCGCCCAAAATAACGTAAAA | CAAAACAATTGGCTGTGTGG | 116 |
| chr11:9027143-9029191 | p2 | (AT)24 | CAAGAGGACACAGTACGCCA | CATGTGTGGGAACACGAAAG | 108 |

| Marker ID | SSR type | SSR motif | FORWARD PRIMER (5'-3') | REVERSE PRIMER (5'-3') | size (bp) |
|---|---|---|---|---|---|
| chr11:9027143-9029191 | p5 | (GGTTT)3 | CAAGAGGACACAGTACGCCA | CATGTGTGGGAACACGAAAG | 108 |
| chr11:9027143-9029191 | p2 | (TA)6 | CAAGAGGACACAGTACGCCA | CATGTGTGGGAACACGAAAG | 108 |
| chr11:9190521-9192536 | p3 | (ATT)5 | AACGACTTCGTTTGGACACC | TCTCGGACTAAAACGCCACT | 117 |
| chr11:9210990-9213005 | p3 | (CAA)5 | GCTTTGTTCATTGCATGTGG | AGCAGGGCATCTCCTCACTA | 184 |
| chr11:9257366-9259426 | p2 | (AT)30 | TTTAAGCCCATCCAAACAGG | AGAGCACTCAAAGTCCGAGC | 185 |
| chr11:9278816-9280830 | p2 | (TA)7 | AAAAACAAAAGCAAAGCCGA | AAAAGGGAAAGCTAGCAGCC | 122 |
| chr11:9345570-9347620 | p2 | (TA)9 | TGGGCGATTCTTTGATAACC | CGGTTTTGGTTTTGTAACCG | 266 |
| chr11:9345570-9347620 | p2 | (AT)10 | TGGGCGATTCTTTGATAACC | CGGTTTTGGTTTTGTAACCG | 266 |
| chr11:9345658-9347670 | p2 | (TA)9 | TGGGCGATTCTTTGATAACC | CGGTTTTGGTTTTGTAACCG | 178 |
| chr11:9345658-9347670 | p2 | (AT)10 | TGGGCGATTCTTTGATAACC | CGGTTTTGGTTTTGTAACCG | 178 |
| chr11:9345679-9347729 | p2 | (TA)9 | GGGCGATTCTTTGATAACCA | CGGTTTTGGTTTTGTAACCG | 158 |
| chr11:9345679-9347729 | p2 | (AT)10 | GGGCGATTCTTTGATAACCA | CGGTTTTGGTTTTGTAACCG | 158 |
| chr11:9385545-9387563 | p6 | (GTAAAA)3 | GATGCTTGAGGCTGAACACA | CGTCCTTTCTTGGGAATGAA | 237 |
| chr11:9463267-9465285 | p5 | (ATTTT)3 | ACCCTTCCCATTTAACCCAC | GGTGGTGGTTTTGCTCTGTT | 229 |
| chr11:9463267-9465285 | p6 | (GGAGAT)3 | ACCCTTCCCATTTAACCCAC | GGTGGTGGTTTTGCTCTGTT | 229 |
| chr11:9586881-9588907 | p2 | (TA)7 | AAATGACATGGCGGAATAGC | ATGATGGATTCATTGCCACA | 269 |
| chr11:9681151-9683167 | p2 | (AT)8 | AAGTATCAACCGTGGCTTCG | TTGCTCCGCTTCTGAGTTTT | 105 |
| chr11:9721635-9723659 | p4 | (TCTT)6 | CTCTTTCTGCCACTATGCCC | ACTCGAGCAGAAGCCTTCAC | 179 |
| chr11:9747060-9749078 | p6 | (AAAATA)3 | ACAAAGGTAGACATTGCCCG | ACCTCACTCCTCAGCCTTCA | 287 |
| chr11:9821923-9823941 | p2 | (AT)9 | AAAGCCAGAGCCCAACTACA | CCTCTTTCATGTTTGCGGTT | 226 |
| chr11:9828492-9830510 | p6 | (CTATAT)3 | AGGATGCGACGTCCTTTCTA | GAAATACGTAGGCCCCCAAT | 174 |
| chr11:9888658-9890672 | p3 | (GAA)5 | GCAAACCCCTCTGTCACATT | GCCCAAATGTCTGCAATTTT | 226 |
| chr11:9888761-9890776 | p3 | (GAA)5 | GCAAACCCCTCTGTCACATT | GCCCAAATGTCTGCAATTTT | 123 |
| chr11:9926986-9929001 | p3 | (TAA)5 | CCCTCCCTTGTTGTTGTTGT | TCCGCTGTTCAGTCCTTTCT | 213 |
| chr11:9926986-9929001 | p4 | (ATTT)4 | CCCTCCCTTGTTGTTGTTGT | TCCGCTGTTCAGTCCTTTCT | 213 |
| chr11:9940913-9942931 | p6 | (TAGTAA)3 | TGCTTGCTGTCATCCTGTTC | CAATTACATGCCCAGGGAGT | 166 |
| chr11:10027992-10030004 | p2 | (TA)8 | GCCCCCATTAGTCAACAAGA | TGTGTTAATTTGGGGTCCGT | 244 |
| chr11:10027992-10030004 | p2 | (TA)9 | GCCCCCATTAGTCAACAAGA | TGTGTTAATTTGGGGTCCGT | 244 |
| chr11:10028012-10030028 | p2 | (TA)8 | TGGGCCCCAATATTAAACAG | CGTGAGTTTTGGATGTGGTG | 111 |
| chr11:10028012-10030028 | p2 | (TA)9 | TGGGCCCCAATATTAAACAG | CGTGAGTTTTGGATGTGGTG | 111 |
| chr11:10092443-10094459 | p4 | (TAAA)4 | TTTTGGCGCTACACACAAAG | TCGAAGAAAAGGGGGAAGTT | 100 |
| chr11:10125059-10127075 | p4 | (AATG)4 | CCATTTGGCTCATTTATGCC | CACCTTACCCACCCACAATC | 180 |
| chr11:10139585-10141600 | p3 | (ACA)5 | TCTGTTGAGTCCGAAAAGGG | TATCCAACCAGCGCAATGTA | 184 |
| chr11:10139585-10141600 | p3 | (ATT)5 | TCTGTTGAGTCCGAAAAGGG | TATCCAACCAGCGCAATGTA | 184 |
| chr11:10254696-10256710 | p2 | (TA)8 | GGTGGTGGCAGAACTAGGAA | TCACCTTGCCAAGATTAGGG | 163 |
| chr11:10254696-10256710 | p2 | (TC)7 | GGTGGTGGCAGAACTAGGAA | TCACCTTGCCAAGATTAGGG | 163 |

| Marker ID | SSR type | SSR motif | FORWARD PRIMER (5'-3') | REVERSE PRIMER (5'-3') | size (bp) |
|---|---|---|---|---|---|
| chr11:10309161-10311176 | p5 | (CTTTT)3 | AATTCCCTTGCTGTTATGCG | GTGTCGGTTAACACCCCAAC | 134 |
| chr11:10355156-10357172 | p4 | (AATA)4 | ACCAATTCCAGCAGCCATAC | AGCTTCTTGTCCAGCGTGTT | 271 |
| chr11:10424537-10426549 | p2 | (AT)6 | GCACTCTTGCTTATAGCGGC | TGAATGGAAACACGATCCAA | 167 |
| chr11:10424537-10426549 | p2 | (AT)8 | GCACTCTTGCTTATAGCGGC | TGAATGGAAACACGATCCAA | 167 |
| chr11:10424537-10426549 | p2 | (AT)7 | GCACTCTTGCTTATAGCGGC | TGAATGGAAACACGATCCAA | 167 |
| chr11:10425404-10427418 | p2 | (AT)6 | GCTTCGAATACAAAATCGGC | GCTATTCCGCCATGTCATTT | 217 |
| chr11:10425404-10427418 | p2 | (AT)8 | GCTTCGAATACAAAATCGGC | GCTATTCCGCCATGTCATTT | 217 |
| chr11:10425404-10427418 | p2 | (AT)7 | GCTTCGAATACAAAATCGGC | GCTATTCCGCCATGTCATTT | 217 |
| chr11:10479410-10481436 | p2 | (AT)13 | GTCCACGATGAGGAGGAAAA | ACATGAGGATGAGAGTGGGG | 107 |
| chr11:10495288-10497312 | p3 | (TTA)8 | CTACATTTAGGCCGAGCGAG | AAGTTTTCGCACGGATTGTC | 190 |
| chr11:10621077-10623092 | p5 | (TTTCT)3 | TTAACTTGTGCGCTGCATTC | CAAAAGTCGTGGCTGCAATA | 155 |
| chr11:10665066-10667082 | p2 | (AT)8 | TGGTTACCGCAAGTTCCTTC | GGTCTAACTTGGGATTGGCA | 181 |
| chr11:10717260-10719276 | p4 | (ATTT)4 | ACATGAACAAGCACGCTGAG | GTGATCCTCGAGTTTCTCGG | 299 |
| chr11:10722258-10724273 | p5 | (TTTAA)3 | TTGAACTGGGGGTTCTCAAC | GGGACATCCGACCTACAAAA | 123 |
| chr11:10743091-10745105 | p2 | (AG)7 | GTTCAACTTGCACCAAGCAA | GTAAGCGTGAGAGCCATTCC | 158 |
| chr11:10775134-10777149 | p5 | (GTTCG)3 | GCAAAACTGGAAGAATGGGA | GTCCCCAGCAAACCTTTGTA | 268 |
| chr11:10836154-10838170 | p5 | (TCGGT)3 | AATTTAGGGGCTGCAAAGGT | CCGAACCGAAATTTCAAAAA | 258 |
| chr11:10836171-10838189 | p5 | (TCGGT)3 | AATTTAGGGGCTGCAAAGGT | CCGAACCGAAATTTCAAAAA | 241 |
| chr11:10836190-10838206 | p5 | (TCGGT)3 | AATTTAGGGGCTGCAAAGGT | CCGAACCGAAATTTCAAAAA | 222 |
| chr11:10836207-10838225 | p5 | (TCGGT)3 | AATTTAGGGGCTGCAAAGGT | CCGAACCGAAATTTCAAAAA | 205 |
| chr11:10836226-10838242 | p5 | (TCGGT)3 | AATTTAGGGGCTGCAAAGGT | CCGAACCGAAATTTCAAAAA | 186 |
| chr11:10836243-10838261 | p5 | (TCGGT)3 | AATTTAGGGGCTGCAAAGGT | CCGAACCGAAATTTCAAAAA | 169 |
| chr11:10836262-10838278 | p5 | (TCGGT)3 | AATTTAGGGGCTGCAAAGGT | CCGAACCGAAATTTCAAAAA | 150 |
| chr11:10836279-10838297 | p5 | (TCGGT)3 | AATTTAGGGGCTGCAAAGGT | CCGAACCGAAATTTCAAAAA | 133 |
| chr11:10836298-10838314 | p5 | (TCGGT)3 | AATTTAGGGGCTGCAAAGGT | CCGAACCGAAATTTCAAAAA | 114 |
| chr11:10875699-10877715 | p5 | (TCGGT)3 | AATTTAGGGGCTGCAAAGGT | CCGAACCGAAATTTCAAAAA | 258 |
| chr11:10875716-10877734 | p5 | (TCGGT)3 | AATTTAGGGGCTGCAAAGGT | CCGAACCGAAATTTCAAAAA | 241 |
| chr11:10875735-10877751 | p5 | (TCGGT)3 | AATTTAGGGGCTGCAAAGGT | CCGAACCGAAATTTCAAAAA | 222 |
| chr11:10875752-10877770 | p5 | (TCGGT)3 | AATTTAGGGGCTGCAAAGGT | CCGAACCGAAATTTCAAAAA | 205 |
| chr11:10875771-10877787 | p5 | (TCGGT)3 | AATTTAGGGGCTGCAAAGGT | CCGAACCGAAATTTCAAAAA | 186 |
| chr11:10875788-10877806 | p5 | (TCGGT)3 | AATTTAGGGGCTGCAAAGGT | CCGAACCGAAATTTCAAAAA | 169 |
| chr11:10875807-10877823 | p5 | (TCGGT)3 | AATTTAGGGGCTGCAAAGGT | CCGAACCGAAATTTCAAAAA | 150 |
| chr11:10875824-10877842 | p5 | (TCGGT)3 | AATTTAGGGGCTGCAAAGGT | CCGAACCGAAATTTCAAAAA | 133 |
| chr11:10875843-10877859 | p5 | (TCGGT)3 | AATTTAGGGGCTGCAAAGGT | CCGAACCGAAATTTCAAAAA | 114 |
| chr11:10946844-10948859 | p3 | (TAT)5 | GTGAGGATGACGAGCCTAGC | TCATAGCGCATCGTCTGAAC | 111 |
| chr11:10949130-10951178 | p2 | (TA)12 | CAGCCTCCCTTTGTTTACCA | CGGTTGCCAGGTTTAGTTGT | 246 |

| Marker ID | SSR type | SSR motif | FORWARD PRIMER (5'-3') | REVERSE PRIMER (5'-3') | size (bp) |
|---|---|---|---|---|---|
| chr11:10949130-10951178 | p2 | (TA)24 | CAGCCTCCCTTTGTTTACCA | CGGTTGCCAGGTTTAGTTGT | 246 |
| chr11:11032844-11034859 | p2 | (TA)19 | TGACGATGATGTTCTCCGAA | CCCAAAAGTCGTGGCTCTAA | 300 |
| chr11:11032844-11034859 | p5 | (TATTT)3 | TGACGATGATGTTCTCCGAA | CCCAAAAGTCGTGGCTCTAA | 300 |
| chr11:11038119-11040133 | p2 | (TA)7 | CCCCAATTAATCCAGCTTCA | GATGGGTTGCGAACTCACTT | 186 |
| chr11:11180440-11182467 | p2 | (TA)8 | AAATACAGAAGCCCCAGGGT | TTGCAAAAGTGACTTGCCTG | 244 |
| chr11:11209983-11212005 | p3 | (GTT)6 | ATGGATGGTTTCAGACGAGG | CGCAAGCGAAACAGATAACA | 138 |
| chr11:11209983-11212005 | p2 | (AT)11 | ATGGATGGTTTCAGACGAGG | CGCAAGCGAAACAGATAACA | 138 |
| chr11:11228758-11230776 | p6 | (TTTTTG)3 | CAAAACCAGCTCCATCCACT | CAACCCAGATCGCCTCTTAG | 244 |
| chr11:11228758-11230776 | p2 | (TA)13 | CAAAACCAGCTCCATCCACT | CAACCCAGATCGCCTCTTAG | 244 |
| chr11:11231489-11233507 | p6 | (TTTTTG)3 | TACCAAAACCAGCTCCATCC | CAACCCAGATCGCCTCTTAG | 241 |
| chr11:11231489-11233507 | p2 | (TA)14 | TACCAAAACCAGCTCCATCC | CAACCCAGATCGCCTCTTAG | 241 |
| chr11:11258519-11260531 | p2 | (AT)8 | GGCGTATTCCGTTGATCATT | TTGTCGACGAAAGTGCAAAA | 113 |
| chr11:11258519-11260531 | p2 | (TA)9 | GGCGTATTCCGTTGATCATT | TTGTCGACGAAAGTGCAAAA | 113 |
| chr11:11258519-11260531 | p2 | (TA)11 | GGCGTATTCCGTTGATCATT | TTGTCGACGAAAGTGCAAAA | 113 |
| chr11:11258532-11260548 | p2 | (AT)8 | GGCGTATTCCGTTGATCATT | TTGTCGACGAAAGTGCAAAA | 100 |
| chr11:11258532-11260548 | p2 | (TA)9 | GGCGTATTCCGTTGATCATT | TTGTCGACGAAAGTGCAAAA | 100 |
| chr11:11258532-11260548 | p2 | (TA)11 | GGCGTATTCCGTTGATCATT | TTGTCGACGAAAGTGCAAAA | 100 |
| chr11:11259501-11261523 | p2 | (TA)11 | GAAACCGCAGACCCAGTAAA | CCGAGATCTATGCACCATCA | 237 |
| chr11:11263804-11265820 | p2 | (TG)8 | GACGGATTCTGAAAATGGGA | GAGAGGGCTTGGTCACAGAG | 141 |
| chr11:11278748-11280763 | p2 | (TA)8 | CCGGCTATGAAACGAAAGAA | ACTTCCCTCTGTTCCCCACT | 245 |
| chr11:11278850-11280892 | p2 | (TA)8 | CCGGCTATGAAACGAAAGAA | ACTTCCCTCTGTTCCCCACT | 143 |
| chr11:11587235-11589253 | p3 | (GAT)8 | CCTGGCCAACTATTCAGCAT | ACTTTGCCCAACGCATTTAC | 268 |
| chr11:11587235-11589253 | p3 | (AAG)6 | CCTGGCCAACTATTCAGCAT | ACTTTGCCCAACGCATTTAC | 268 |
| chr11:11747249-11749261 | p3 | (CAA)5 | CCCCTTGCCTTAGCTCTCTT | TGCAGCTCTGATACCCAATG | 265 |
| chr11:11747249-11749261 | p2 | (TG)6 | CCCCTTGCCTTAGCTCTCTT | TGCAGCTCTGATACCCAATG | 265 |
| chr11:11761344-11763370 | p2 | (AT)13 | GAAGATTACTGCTGGCTCGG | AATCGTCCTTGTATGCCGTC | 198 |
| chr11:11781212-11783232 | p5 | (GGGGT)4 | CCCATCAGATTCTCCGACAT | TATCGATTTTTGGCACGTCA | 185 |
| chr11:11781212-11783232 | p5 | (GGGGT)4 | CCCATCAGATTCTCCGACAT | TATCGATTTTTGGCACGTCA | 185 |
| chr11:11797832-11799858 | p2 | (TA)13 | ATATGTGCATGCCATTTCCA | GCTTGGTTGAACTACGCCAT | 100 |
| chr11:11853328-11855343 | p5 | (AAAAG)3 | GGAGCAATGCCATTACCTGT | TGATCAGATTTTCACGCTGC | 281 |
| chr11:11915854-11917869 | p2 | (AT)8 | TTTCGTGACACAAATCCGAA | CGTGAAACATGCAACGAAAT | 278 |
| chr11:11916006-11918022 | p2 | (AT)8 | TTCGTGACACAAATCCGAAA | CGTGAAACATGCAACGAAAT | 127 |
| chr11:12088038-12090056 | p6 | (GTGGGG)3 | CACCCTCGTACTTGCCGTAT | AATTGGGTTAAGGGGACACC | 229 |
| chr11:12093294-12095324 | p2 | (TA)15 | GAAATCTGCCCACGTTGACT | ATTTGGACATCTGCAAAGGC | 152 |
| chr11:12119373-12121393 | p2 | (AT)10 | GTAGCTGCATCAAACGACGA | TCAAAATGGAAGTGATGCCA | 279 |
| chr11:12136521-12138573 | p2 | (AT)26 | TTGAGTGGACCATGCCAATA | ATGGGAGAAAGTCGAGCAGA | 189 |

| Marker ID | SSR type | SSR motif | FORWARD PRIMER (5'-3') | REVERSE PRIMER (5'-3') | size (bp) |
|---|---|---|---|---|---|
| chr11:12229636-12231650 | p2 | (AT)9 | ATTGGACCAAGAGCACGAGT | TTGCCATCAACGTGGTTTTA | 297 |
| chr11:12229691-12231727 | p2 | (AT)9 | ATTGGACCAAGAGCACGAGT | TTGCCATCAACGTGGTTTTA | 242 |
| chr11:12265947-12267965 | p6 | (TTTTTG)3 | GCCAGGGCATTTGAAAATAA | ATTTGGCCCTGTGTATGCTC | 156 |
| chr11:12313561-12315576 | p3 | (AGA)5 | ATGTGCAAACCTTGGAGCTT | TTTGCAATAATGGTGACCCA | 293 |
| chr11:12313561-12315576 | p2 | (TA)27 | ATGTGCAAACCTTGGAGCTT | TTTGCAATAATGGTGACCCA | 293 |
| chr11:12317846-12319861 | p6 | (CCCCAA)3 | CGAGGACAAAGGGTGTGTTT | TGGGAGTAGGGTGTGGGTAG | 188 |
| chr11:12317846-12319861 | p5 | (GGGGT)3 | CGAGGACAAAGGGTGTGTTT | TGGGAGTAGGGTGTGGGTAG | 188 |
| chr11:12368893-12370919 | p2 | (TA)13 | CTTGAGCAACTCGGTGATGA | GGAATTCCGGCGAAATAAAT | 185 |
| chr11:12560531-12562549 | p3 | (AAG)6 | AGCATCCAATTTCTGTTCCG | TCAGGCCCTTGAAATATTGG | 147 |
| chr11:12560531-12562549 | p5 | (ATTTT)3 | AGCATCCAATTTCTGTTCCG | TCAGGCCCTTGAAATATTGG | 147 |
| chr11:12560531-12562549 | p5 | (TAAAG)3 | AGCATCCAATTTCTGTTCCG | TCAGGCCCTTGAAATATTGG | 147 |
| chr11:12560531-12562549 | p2 | (TA)7 | AGCATCCAATTTCTGTTCCG | TCAGGCCCTTGAAATATTGG | 147 |
| chr11:12621186-12623204 | p6 | (ACTATC)3 | AAAAATTATGCAGCCGTTGG | GGCACCCCCACTACAAAGTA | 169 |
| chr11:12640877-12642892 | p5 | (TTTTA)3 | TGAGCAGTTCACAGAAGGGA | TTTCATGTCTTATTGCCCCC | 220 |
| chr11:12642861-12644885 | p6 | (GGGGTG)4 | GGGTGTGTTTGGTATGGAGG | TGGTGTGGTATGAGGTGGTG | 108 |
| chr11:12697959-12699971 | p3 | (GTT)7 | CCAAAAGCAGCCAAAAAGAG | TCTTTGGCCTACCCCTACCT | 234 |
| chr11:12697959-12699971 | p2 | (TA)6 | CCAAAAGCAGCCAAAAAGAG | TCTTTGGCCTACCCCTACCT | 234 |
| chr11:12723717-12725729 | p2 | (AG)6 | CGTCATTAGCATCAATTGCG | CACCCGATGCGACTAAGTTT | 213 |
| chr11:12723717-12725729 | p5 | (TTTCT)3 | CGTCATTAGCATCAATTGCG | CACCCGATGCGACTAAGTTT | 213 |
| chr11:12790094-12792109 | p3 | (TTA)5 | GTTTTGTTACACCCCGCACT | TTCCCATGGCTTGTACCTTC | 210 |
| chr11:12790094-12792109 | p3 | (TTG)5 | GTTTTGTTACACCCCGCACT | TTCCCATGGCTTGTACCTTC | 210 |
| chr11:12824562-12826576 | p2 | (AC)7 | CTTTCGAGGAGCAGAAATGC | TCATAGGGGAAAGTGCTGCT | 185 |
| chr11:12951770-12953784 | p3 | (GAT)5 | GAGGGAGCGTTTTCCTCTTT | GGCCATCTTGAAATGCTGTT | 130 |
| chr11:12951770-12953784 | p2 | (AT)7 | GAGGGAGCGTTTTCCTCTTT | GGCCATCTTGAAATGCTGTT | 130 |
| chr11:13130423-13132438 | p5 | (CATAT)3 | TACATGCACTCTGACTCGGC | CGGGAAATAGCGCTTTGATA | 259 |
| chr11:13243454-13245476 | p2 | (CT)11 | TAATGATGTTGCTGCTTGCC | TCTGCAGGGAAGGTTATGCT | 261 |
| chr11:13479101-13481121 | p2 | (AT)10 | AAGTATCAACCGTGGCTTCG | TTGCTCCGCTTCTGAGTTTT | 175 |
| chr11:13557742-13559758 | p2 | (TC)8 | CAAGGTTGAAGCTCTTTCCG | TTAACCGGTGAGCATTGACA | 275 |
| chr11:13680277-13682292 | p5 | (AAAAC)3 | TTATTTGCATGCCCAACTCA | TTATTAGTGGCGGGTCCAAC | 229 |
| chr11:13796534-13798600 | p3 | (ATA)22 | TGGATTAGGGTTTTCGTTCG | TCTCTCCTCCATCTCCCTCA | 299 |
| chr11:13809307-13811322 | p5 | (TCTTG)3 | CCTTGAGAGGCTCTGTGTCC | TCATGAACAGCCAACGGTAA | 283 |
| chr11:13809307-13811322 | p3 | (ACA)5 | CCTTGAGAGGCTCTGTGTCC | TCATGAACAGCCAACGGTAA | 283 |
| chr11:13818545-13820560 | p5 | (AAACC)3 | GGTCGTCAATCGTGCTACCT | CCTACCTAGTGCCACCCAAA | 120 |
| chr11:13845703-13847718 | p5 | (CGGGT)3 | ATTTGGATTCACACCGCATT | GGAACGTTTATGGTGGGATG | 146 |
| chr11:13870677-13872692 | p3 | (TGT)5 | GTTGTACGGGATGGAGTGCT | ATATGCCCGAACCATCTCAG | 258 |
| chr11:13896336-13898357 | p3 | (TGT)7 | GCTTCCTATCACGCAAGGAG | GGGACTGGGGGTATTGAGAT | 151 |

| Marker ID | SSR type | SSR motif | FORWARD PRIMER (5'-3') | REVERSE PRIMER (5'-3') | size (bp) |
|---|---|---|---|---|---|
| chr11:13914794-13916808 | p2 | (TA)7 | CTTTTCCCCCTCCTCTTTTG | CCATGGATGGGTCTTCATTC | 140 |
| chr11:13975063-13977075 | p2 | (TA)6 | TGCCAGCTACCTCAACACAC | GTTTCTTTTCTGGCAGGTCG | 240 |
| chr11:13975063-13977075 | p2 | (AT)6 | TGCCAGCTACCTCAACACAC | GTTTCTTTTCTGGCAGGTCG | 240 |
| chr11:14019498-14021516 | p6 | (CATGAG)3 | GCGTATTCGGTAGAATGGGA | CGCGCCACACATATAAACAG | 183 |
| chr11:14032412-14034424 | p3 | (TTA)6 | AGGCTAAAATTGCCAGAGCA | GAATTGACTCCCGAAAAGCA | 106 |
| chr11:14032412-14034424 | p2 | (AT)9 | AGGCTAAAATTGCCAGAGCA | GAATTGACTCCCGAAAAGCA | 106 |
| chr11:14032412-14034424 | p2 | (TA)6 | AGGCTAAAATTGCCAGAGCA | GAATTGACTCCCGAAAAGCA | 106 |
| chr11:14088066-14090081 | p5 | (ATTTG)3 | ACGGGTTTGGCATAACTTTG | GAACCCCCACTCAGTCAAAA | 198 |
| chr11:14100524-14102539 | p2 | (TA)6 | CCATAATCATTGTGTCCCCC | AATTGCAGGCCACGTTTAAG | 218 |
| chr11:14100524-14102539 | p3 | (GTA)5 | CCATAATCATTGTGTCCCCC | AATTGCAGGCCACGTTTAAG | 218 |
| chr11:14100524-14102539 | p3 | (TTA)5 | CCATAATCATTGTGTCCCCC | AATTGCAGGCCACGTTTAAG | 218 |
| chr11:14100524-14102539 | p2 | (AT)33 | CCATAATCATTGTGTCCCCC | AATTGCAGGCCACGTTTAAG | 218 |
| chr11:14183271-14185283 | p2 | (AT)6 | TCCGCAATTACACCATGCTA | CACAACTGGTACACCGCATC | 215 |
| chr11:14183271-14185283 | p2 | (AT)8 | TCCGCAATTACACCATGCTA | CACAACTGGTACACCGCATC | 215 |
| chr11:14252882-14254896 | p2 | (TG)7 | CATCAAGAAAAAGCCCCAAA | TTTCACCAAGGGGCATCTAC | 272 |
| chr11:14254895-14256913 | p2 | (AC)7 | CTGGCTGATTGAGACATGGA | GTCATTGGTGTCCTGACGTG | 117 |
| chr11:14254895-14256913 | p6 | (TTTAAT)3 | CTGGCTGATTGAGACATGGA | GTCATTGGTGTCCTGACGTG | 117 |
| chr11:14419107-14421129 | p2 | (TA)11 | TTCCTTCGGGATGATCACAT | TCAAAAGGGACGAGGTGAAG | 210 |
| chr11:14471631-14473665 | p2 | (TA)17 | GGCTGCTTTCAGAAAACCAA | CATCTGGCAGAGTTGACGAA | 182 |
| chr11:14479222-14481237 | p2 | (AC)6 | GAACGGCGAAATATTCCAAA | CACTTGTGAAAAGGCACGAA | 232 |
| chr11:14479222-14481237 | p5 | (CGGCG)3 | GAACGGCGAAATATTCCAAA | CACTTGTGAAAAGGCACGAA | 232 |
| chr11:14479222-14481237 | p4 | (AAAT)4 | GAACGGCGAAATATTCCAAA | CACTTGTGAAAAGGCACGAA | 232 |
| chr11:14606296-14608326 | p3 | (TGT)10 | CTGCAGCAGAGATGAAGCAC | AGCTTAATTCAAGGAGCGCA | 187 |
| chr11:14680892-14682904 | p2 | (GT)6 | GGCTACCTCACGGGTTGATA | TTTTCCTCTCCCCCTCCTAA | 285 |
| chr11:14695135-14697161 | p2 | (AT)13 | TTGCTTGTTTTTCTCCCACC | AAAATTGAACTGCTGCCACC | 278 |
| chr11:14695135-14697161 | p2 | (TA)9 | TTGCTTGTTTTTCTCCCACC | AAAATTGAACTGCTGCCACC | 278 |
| chr11:14957099-14959119 | p2 | (CT)10 | TATGCCCCAAGGACACTTTC | AAAACCCCATGGTTCTCCTC | 277 |
| chr11:15031122-15033140 | p2 | (AT)9 | AAATCCTCCCCAAACCATTC | TTTCTCTCCCTTCCCCTCTC | 102 |
| chr11:15179176-15181190 | p2 | (GA)7 | GGGGCTGTACTATTTTGCCA | TGCTGATTGTTTTGTTGGGA | 188 |
| chr11:15267108-15269130 | p2 | (TA)11 | TAATGCCAGTGGGTCTCCAT | GGGCACTAAAAGGGTGATGA | 165 |
| chr11:15267556-15269576 | p2 | (TA)11 | GGCTCTATTTTCTGGCGAAC | TTGTGATCCAAACCCAAACA | 173 |
| chr11:15294144-15296160 | p2 | (AT)8 | TGGAAGTGGTCTTGGAAAGG | CCTTACATCCTTGTCGCCAT | 273 |
| chr11:15294144-15296160 | p3 | (TTG)5 | TGGAAGTGGTCTTGGAAAGG | CCTTACATCCTTGTCGCCAT | 273 |
| chr11:15356011-15358027 | p4 | (TTAA)4 | CGGATTCTAGGGTTTCCACA | CACGAAAACTCACCCCAAAT | 194 |
| chr11:15356011-15358027 | p4 | (TTAA)4 | CGGATTCTAGGGTTTCCACA | CACGAAAACTCACCCCAAAT | 194 |
| chr11:15378279-15380291 | p2 | (TG)6 | GTTGCATAGGGGTCCTTCAA | GAAGAAAATATGGGCTGCCA | 246 |

| Marker ID | SSR type | SSR motif | FORWARD PRIMER (5'-3') | REVERSE PRIMER (5'-3') | size (bp) |
|---|---|---|---|---|---|
| chr11:15378279-15380291 | p5 | (TTCAT)3 | GTTGCATAGGGGTCCTTCAA | GAAGAAAATATGGGCTGCCA | 246 |
| chr11:15387333-15389375 | p3 | (ATA)14 | ACACTGTGTTTCGCTGCTTG | TTGGGAGGAGAATGTGAAGG | 146 |
| chr11:15387333-15389375 | p2 | (TA)10 | ACACTGTGTTTCGCTGCTTG | TTGGGAGGAGAATGTGAAGG | 146 |
| chr11:15388615-15390653 | p2 | (TA)10 | CGGTTTATGGCACTGTTTCA | CTTTCTCACATCCCAGCCAT | 166 |
| chr11:15388615-15390653 | p2 | (TA)19 | CGGTTTATGGCACTGTTTCA | CTTTCTCACATCCCAGCCAT | 166 |
| chr11:15388615-15390653 | p2 | (AT)6 | CGGTTTATGGCACTGTTTCA | CTTTCTCACATCCCAGCCAT | 166 |
| chr11:15421435-15423453 | p6 | (TAGATC)3 | GGCACAGAAAAGAAGCCTTG | ATGGCCTAGCTGTTCTTGGA | 142 |
| chr11:15437666-15439712 | p2 | (TA)23 | TTGGCCAGAAAGTTATCGCT | GACCAGTTCCCACCTCTTCA | 172 |
| chr11:15517266-15519281 | p5 | (TTAAA)3 | TGAACAAATCAAGCTTTGCG | CACGGAACTGTTGAGCAAAT | 273 |
| chr11:15517266-15519281 | p2 | (AT)6 | TGAACAAATCAAGCTTTGCG | CACGGAACTGTTGAGCAAAT | 273 |
| chr11:15532861-15534881 | p2 | (CT)10 | TGGTGGCCTGGACTTCTTAC | CCCCTTCCCCCTATGAATAA | 177 |
| chr11:15532861-15534881 | p2 | (AT)16 | TGGTGGCCTGGACTTCTTAC | CCCCTTCCCCCTATGAATAA | 177 |
| chr11:15544708-15546724 | p2 | (AG)8 | TTTCTGGCACATGGAATGAA | TGGCCATCTTCTTAAAACGG | 274 |
| chr11:15588530-15590546 | p2 | (TA)8 | TTTAAGTTTTGGCATTCGCC | ATTTCCCTTCGTTAGGGGTG | 171 |
| chr11:15672603-15674619 | p2 | (GA)8 | TAGAGTCGTCTCATGGCGTG | AGAGGAAACGAAGGGGAGAA | 292 |
| chr11:15741266-15743300 | p2 | (TA)17 | CAGCAGTTGTTCGTCCAAAA | CAGCAGTTGTTCGTCCAAAA | 178 |
| chr11:15748378-15750398 | p5 | (TGGCT)4 | AAGTTTTGGGCACGATTCAC | TGAAAGAAACCATCATGCCA | 196 |
| chr11:15796897-15798912 | p5 | (AAACC)3 | GGTTTGAACCCAGAACCTCA | ATTCCAACTCGGGCCTAACT | 184 |
| chr11:15796937-15798952 | p5 | (AAACC)3 | GGTTTGAACCCAGAACCTCA | ATTCCAACTCGGGCCTAACT | 144 |
| chr11:15819669-15821684 | p5 | (CTTTG)3 | CTTGAAGGTGCCCACAATCT | ACCGATGTAACGGAGTTTGC | 160 |
| chr11:15819669-15821684 | p5 | (CTTTG)3 | CTTGAAGGTGCCCACAATCT | ACCGATGTAACGGAGTTTGC | 160 |
| chr11:15965188-15967206 | p2 | (TA)9 | GAACCAAAGAGCCGTGAAAA | TTTTGGACATAAAAACCGCC | 288 |
| chr11:15967882-15969894 | p2 | (AT)6 | GACAAAGGCGGAATAGCAAA | TTGCCACATGCGTTATCAAT | 270 |
| chr11:16002013-16004025 | p2 | (TA)6 | TATTGCCTACACCCGAGTCC | TTGCAAGGAAAAATCAAGGG | 196 |
| chr11:16002013-16004025 | p2 | (CT)6 | TATTGCCTACACCCGAGTCC | TTGCAAGGAAAAATCAAGGG | 196 |
| chr11:16057855-16059871 | p4 | (TATT)4 | TCAACCCAAATCACCCATTT | AAATACACACCGGAGCTTGG | 258 |
| chr11:16057855-16059871 | p4 | (TATT)4 | TCAACCCAAATCACCCATTT | AAATACACACCGGAGCTTGG | 258 |
| chr11:16265419-16267434 | p4 | (CCAC)4 | TTGATTCCCACTTTCCTTGC | AAAGGCTGACGTGGAAGAGA | 159 |
| chr11:16265419-16267434 | p5 | (TTTAA)3 | TTGATTCCCACTTTCCTTGC | AAAGGCTGACGTGGAAGAGA | 159 |
| chr11:16289785-16291811 | p2 | (TC)7 | GTTAGCGGTCTAGGGGGTTC | CACCACCAAGGCAGATTACC | 183 |
| chr11:16289785-16291811 | p2 | (TA)13 | GTTAGCGGTCTAGGGGGTTC | CACCACCAAGGCAGATTACC | 183 |
| chr11:16410872-16412887 | p5 | (GGTTT)3 | ACGAGTCTGTATCCGGGTTG | GATGAGCAAATCCAGCACAA | 208 |
| chr11:16410872-16412887 | p5 | (TTGGT)3 | ACGAGTCTGTATCCGGGTTG | GATGAGCAAATCCAGCACAA | 208 |
| chr11:16448856-16450871 | p3 | (ATA)5 | TCCCGCCTTATGATTCTGTC | CTGGACACCCATACGTACCC | 210 |
| chr11:16473382-16475397 | p3 | (TAA)5 | CGTTGATTCGGGTTTCTTGT | TGTCCCCCACTTGTCTCTCT | 182 |
| chr11:16473382-16475397 | p3 | (AAT)5 | CGTTGATTCGGGTTTCTTGT | TGTCCCCCACTTGTCTCTCT | 182 |

| Marker ID | SSR type | SSR motif | FORWARD PRIMER (5'-3') | REVERSE PRIMER (5'-3') | size (bp) |
|---|---|---|---|---|---|
| chr11:16473382-16475397 | p5 | (GGGTC)4 | CGTTGATTCGGGTTTCTTGT | TGTCCCCCACTTGTCTCTCT | 182 |
| chr11:16490037-16492059 | p2 | (CT)11 | CTCTCATTGCAAACAAGCCA | CTTTGATTCCGAGCCTTCTG | 264 |
| chr11:16521237-16523255 | p6 | (TTTTCT)3 | TGCATAAAATTGGGGTCGAT | CTCTTGCATGCCAACTTGAA | 165 |
| chr11:16538552-16540568 | p2 | (AT)8 | AAGGAAAGCCTCATGCTTCA | CTCTGATCACCGCAATAGCA | 300 |
| chr11:16565539-16567554 | p5 | (TTAGT)3 | CAAGGATTCGGTTTTGAGGA | CGAACAAACACTTCTTGCGA | 217 |
| chr11:16565539-16567554 | p2 | (CT)6 | CAAGGATTCGGTTTTGAGGA | CGAACAAACACTTCTTGCGA | 217 |
| chr11:16585527-16587545 | p6 | (AAAATT)3 | CCAAACAATGTTGGCAAATG | GTTAGCCAGGATGCGCTTAG | 278 |
| chr11:16642962-16644977 | p5 | (TAAAC)3 | TGCCAAAAAGGAATTTGGAC | TTTTGCAATATCCGTGCGTA | 167 |
| chr11:16642962-16644977 | p5 | (GAGAG)5 | TGCCAAAAAGGAATTTGGAC | TTTTGCAATATCCGTGCGTA | 167 |
| chr11:16643887-16645912 | p5 | (TAAAC)3 | CAGAACAGAGCTGAACGCAG | TCCGATATACACACCGGGTT | 113 |
| chr11:16643887-16645912 | p5 | (GAGAG)5 | CAGAACAGAGCTGAACGCAG | TCCGATATACACACCGGGTT | 113 |
| chr11:16702159-16704191 | p2 | (AT)16 | TCATTGACCACTAGGCCACA | TGCCACCCATGAAATCTACA | 120 |
| chr11:16710311-16712350 | p3 | (TTG)13 | TTCCCTCCCTACAACTCCCT | TTGAAGAAGAAGCAGCAGCA | 135 |
| chr11:16728789-16730804 | p5 | (TTTAG)3 | CCATAACCCGGTCAGACCTA | AGACAAGATTAAGCGGGGGT | 193 |
| chr11:16757490-16759504 | p2 | (TA)7 | TCACCTGGTTCACCAAAACA | CATATTCCCTGTGGGATTCG | 275 |
| chr11:16757490-16759504 | p2 | (TC)6 | TCACCTGGTTCACCAAAACA | CATATTCCCTGTGGGATTCG | 275 |
| chr11:16798228-16800242 | p2 | (TA)7 | ATGCTGCAATTCAGCTGCTA | AGGAAAAACTCATGATGGCG | 289 |
| chr11:16798326-16800361 | p2 | (TA)7 | ATGCTGCAATTCAGCTGCTA | AGGAAAAACTCATGATGGCG | 191 |
| chr11:16873933-16875948 | p5 | (AAACC)3 | ACCGACCAAGAGAGCTTCAA | GATTGGCTTGCCATTCTGTT | 180 |
| chr11:16907063-16909085 | p2 | (AT)11 | CAACCGTTTGGACAACAATG | TAGCCGTCCACTTTGGGTAG | 106 |
| chr11:16916058-16918073 | p5 | (AAAAT)3 | GGCAATGTCGGAAACAGTCT | ATGAAATTTGAAGCGGGTTG | 217 |
| chr11:17101777-17103789 | p2 | (TA)6 | AAGAGGGACGGAAGAAGGAG | TTCCCAACCCAAATTCTCTG | 171 |
| chr11:17101777-17103789 | p2 | (AT)9 | AAGAGGGACGGAAGAAGGAG | TTCCCAACCCAAATTCTCTG | 171 |
| chr11:17101777-17103789 | p2 | (TA)8 | AAGAGGGACGGAAGAAGGAG | TTCCCAACCCAAATTCTCTG | 171 |
| chr11:17101966-17103978 | p2 | (TA)6 | GTTTAAACAAAGGGGTGGGG | GGACTGCTTGCTTAACCGAC | 164 |
| chr11:17101966-17103978 | p2 | (AT)9 | GTTTAAACAAAGGGGTGGGG | GGACTGCTTGCTTAACCGAC | 164 |
| chr11:17101966-17103978 | p2 | (TA)8 | GTTTAAACAAAGGGGTGGGG | GGACTGCTTGCTTAACCGAC | 164 |
| chr11:17101966-17103978 | p2 | (TA)8 | GTTTAAACAAAGGGGTGGGG | GGACTGCTTGCTTAACCGAC | 164 |
| chr11:17102009-17104027 | p2 | (TA)6 | GTTTAAACAAAGGGGTGGGG | GGACTGCTTGCTTAACCGAC | 121 |
| chr11:17102009-17104027 | p2 | (AT)9 | GTTTAAACAAAGGGGTGGGG | GGACTGCTTGCTTAACCGAC | 121 |
| chr11:17102009-17104027 | p2 | (TA)8 | GTTTAAACAAAGGGGTGGGG | GGACTGCTTGCTTAACCGAC | 121 |
| chr11:17102009-17104027 | p2 | (TA)8 | GTTTAAACAAAGGGGTGGGG | GGACTGCTTGCTTAACCGAC | 121 |
| chr11:17131286-17133301 | p3 | (AAG)5 | CCTCTCCGCCCCATATTACT | CTTGCCACTTTCCTTTTTGC | 198 |
| chr11:17141851-17143867 | p6 | (ATTTTT)3 | TGAAATAGGACTCTGCCGCT | AGCAATCGCCTATTACCACG | 131 |
| chr11:17141851-17143867 | p4 | (TTTA)4 | TGAAATAGGACTCTGCCGCT | AGCAATCGCCTATTACCACG | 131 |
| chr11:17170253-17172268 | p5 | (TTTAT)3 | GATCAGGCCGTACACCACTT | TTGTGGTGGAGAAAAGAGGG | 220 |

| Marker ID | SSR type | SSR motif | FORWARD PRIMER (5'-3') | REVERSE PRIMER (5'-3') | size (bp) |
|---|---|---|---|---|---|
| chr11:17245030-17247045 | p5 | (GTTCG)3 | CATGTGATCAGGAGGTCACG | TTGCATCCAATTTACCCCAT | 216 |
| chr11:17245030-17247045 | p5 | (GTTCG)3 | CATGTGATCAGGAGGTCACG | TTGCATCCAATTTACCCCAT | 216 |
| chr11:17247068-17249088 | p2 | (AT)10 | TCAATTTCTGGTCAATGGCA | AATGTGTTTTGCAAACGTGG | 295 |
| chr11:17272831-17274846 | p5 | (CAAAC)3 | CGGTTTGGTTGTAGTCGGTT | TTTTGTGTTCGGCAATGAAA | 264 |
| chr11:17272831-17274846 | p5 | (AAACC)3 | CGGTTTGGTTGTAGTCGGTT | TTTTGTGTTCGGCAATGAAA | 264 |
| chr11:17291093-17293107 | p2 | (CT)7 | CGTTGGTTTTAGGGAGGTCA | TCTCATGCGAACACCAAAAG | 258 |
| chr11:17348478-17350505 | p3 | (TGT)9 | CACCCACATCAATGCAAGAG | ATCATTGAACCACCCCAAAA | 261 |
| chr11:17362537-17364552 | p5 | (TGATT)3 | CAGAGTTGCCTCTAGGACGG | CGGTACCTGTGCTGGTAGGT | 267 |
| chr11:17364328-17366343 | p5 | (CTTTT)3 | AGATTTTGTTGAATTGCCGC | CTGCTGATTATGGGCCATCT | 291 |
| chr11:17364328-17366343 | p5 | (TGATT)3 | AGATTTTGTTGAATTGCCGC | CTGCTGATTATGGGCCATCT | 291 |
| chr11:17375535-17377547 | p2 | (CT)6 | CTATCGAAATGTCGTCGGGT | CCGTAGGAGGATGTTCCAAA | 182 |
| chr11:17397277-17399292 | p5 | (AATAA)3 | ACCCTTATCATGCGGAACTG | ACCATGACCCAATGGCTAAG | 259 |
| chr11:17397277-17399292 | p5 | (AATAA)3 | ACCCTTATCATGCGGAACTG | ACCATGACCCAATGGCTAAG | 259 |
| chr11:17409564-17411580 | p4 | (TTAG)4 | GAATTCAAATCCGCGACCTA | CACCAGAACACCAAAGCTGA | 179 |
| chr11:17425447-17427465 | p6 | (AATTCA)3 | TATGGAGCCTACAGTTGCGA | AGGGAAGGACCTAATGGTGG | 229 |
| chr11:17452550-17454562 | p6 | (CCCACC)3 | ACGCAAATGCATGAGGTGTA | TGTGCTTTAAGAGTCCCGCT | 168 |
| chr11:17524406-17526422 | p2 | (TC)8 | CTGGGGGATTTCAGTTGCTA | TCGTTCCTCTCTGCTGGAAT | 259 |
| chr11:17569177-17571216 | p2 | (TG)6 | ATTTAGCGGACCCAGGATTT | TGTGCCCTCTCTATTGGCTT | 159 |
| chr11:17569177-17571216 | p3 | (AAT)13 | ATTTAGCGGACCCAGGATTT | TGTGCCCTCTCTATTGGCTT | 159 |
| chr11:17579473-17581489 | p3 | (GCG)6 | AAATTCATCAACCCCACCAA | TGCTCTTCAACACGCTATGG | 297 |
| chr11:17579509-17581537 | p3 | (GCG)6 | AAATTCATCAACCCCACCAA | TGCTCTTCAACACGCTATGG | 261 |
| chr11:17586250-17588268 | p2 | (AT)6 | AAATTGTGGCCCCTCTTCTT | ACGAACGTCAGGCAATGAAT | 297 |
| chr11:17586250-17588268 | p2 | (TA)6 | AAATTGTGGCCCCTCTTCTT | ACGAACGTCAGGCAATGAAT | 297 |
| chr11:17586273-17588293 | p5 | (ATTAT)3 | AAATTGTGGCCCCTCTTCTT | ACGAACGTCAGGCAATGAAT | 274 |
| chr11:17586273-17588293 | p2 | (AT)6 | AAATTGTGGCCCCTCTTCTT | ACGAACGTCAGGCAATGAAT | 274 |
| chr11:17586273-17588293 | p2 | (TA)8 | AAATTGTGGCCCCTCTTCTT | ACGAACGTCAGGCAATGAAT | 274 |
| chr11:17613277-17615289 | p2 | (CT)6 | TAGCGCATTTCATCATGAGC | GTTTGGACGGTAAATGTGGG | 232 |
| chr11:17667764-17669784 | p5 | (GGGCC)4 | GTATTGGAGGGTGGGAGGAT | GGCCTATGGGTGTGTTTGAT | 106 |
| chr11:17696572-17698587 | p5 | (CAAAA)3 | AGGATGCATGAACAACCACA | GGATAGTAGGGCAGCAACCA | 275 |
| chr11:18078600-18080615 | p5 | (CCATT)3 | TAAGTTAGTTCGCGCCGTTT | TGCAGATCAGCAAATGCTTC | 286 |
| chr11:18135819-18137840 | p3 | (ACA)7 | TTGTAAGGGCTGCCTTCATC | GGACGCCAGTGGTACCTTTA | 207 |
| chr11:18222845-18224860 | p5 | (TGCTT)3 | TGCGAGTTTCCGCTAGACTT | ACACATTTGGCACAATCGAA | 124 |
| chr11:18259963-18261978 | p5 | (TGCTT)3 | TGCGAGTTTCCGCTAGACTT | ACACATTTGGCACAATCGAA | 124 |
| chr11:18410325-18412343 | p6 | (CCAGCT)3 | CTTCGATGGTCTCGATGGAT | TGCCCTTTCTTCTGTGCTTT | 102 |
| chr11:18413633-18415657 | p2 | (TA)12 | TCGCCACCTAGCATTTAAGG | GCACATCGCCTCGTATAGGT | 246 |
| chr11:18506061-18508079 | p6 | (AAAATA)3 | ATATATGGGTCCAAACGCCA | ATGCATATGTGGGCCCTTTA | 251 |

| Marker ID | SSR type | SSR motif | FORWARD PRIMER (5'-3') | REVERSE PRIMER (5'-3') | size (bp) |
|---|---|---|---|---|---|
| chr11:18585006-18587072 | p2 | (AT)33 | GATGCAATTACAAGCAGCGA | ATTCATCTGGTCATGCCGTT | 275 |
| chr11:18616999-18619013 | p2 | (GA)7 | TTGTTTGTGGACACTAGCGG | GCCAAACATCCGCCTATAAA | 260 |
| chr11:18616999-18619013 | p5 | (ACCAA)3 | TTGTTTGTGGACACTAGCGG | GCCAAACATCCGCCTATAAA | 260 |
| chr11:18671355-18673370 | p5 | (TTAGG)3 | TCCACATAACCAGGTGCAAA | GGTAGCTCTTAACACGCCCA | 283 |
| chr11:18686585-18688603 | p6 | (ACCGAA)3 | GGTTTGGATTTTGGTGTTGG | CCTTTCCCAATTTACCCGAT | 283 |
| chr11:18756824-18758862 | p2 | (AT)19 | TTCCCCACTTTCGATTTTTG | TACCCTTTAGGCTCCGGACT | 174 |
| chr11:18756824-18758862 | p2 | (AT)7 | TTCCCCACTTTCGATTTTTG | TACCCTTTAGGCTCCGGACT | 174 |
| chr11:18765413-18767425 | p2 | (AT)6 | GTATGGCCTCGGTCACTTGT | TGCGATAGAGCCGTGTGTAG | 300 |
| chr11:18986754-18988774 | p5 | (GGTTT)4 | ACGGATATGCTCCTTGTTCG | TATGTGGCGGGGGAATATAA | 188 |
| chr11:19034103-19036117 | p2 | (CT)7 | GTGTTCAAAGGTAGGGCAGG | GCCAAAAGAGTTGATGGGAA | 142 |
| chr11:19066636-19068648 | p2 | (GA)6 | AAAAGCGACAGGTGCATTCT | TCTTCCTTTTTGGTTCCCCT | 257 |
| chr11:19122533-19124547 | p2 | (TC)7 | TGTGGCACATTAGATTGGGA | AAGGAACAAAGATGGGGCTT | 172 |
| chr11:19212201-19214221 | p4 | (TATT)5 | AGGGCCTACATTTCACGATG | CGGCACACCTCACTGACTAA | 131 |
| chr11:19217442-19219460 | p6 | (TTCAGT)3 | TTTTCCGATCTCCCTTTCCT | ACTTCCACACGTTGTCACCA | 116 |
| chr11:19286574-19288592 | p2 | (TA)9 | CGTGGTCAACTGGGGATACT | CTGAACAACCAGCCACTGAA | 237 |
| chr11:19361689-19363711 | p2 | (AT)11 | TAGATGCTCCTGGCCAGACT | AATCGACGATGCATTTCACA | 297 |
| chr11:19376846-19378861 | p5 | (AAAAT)3 | TTTGTTGTCAGCATACCCCA | GCATATTGAACCCCAGCACT | 159 |
| chr11:19381868-19383892 | p2 | (AT)12 | AGAAACAGGTTCATGGTCGG | TCCTTTCCCCAACTCCTCTT | 243 |
| chr11:19395304-19397328 | p2 | (TA)17 | GGGACAAATCCAACAAATGG | TGAAGGCTCACGAGTAGCAA | 138 |
| chr11:19395304-19397328 | p4 | (TTAA)6 | GGGACAAATCCAACAAATGG | TGAAGGCTCACGAGTAGCAA | 138 |
| chr11:19441578-19443634 | p2 | (TA)28 | ATCATCACCTCCACCACCAT | CCATTTGTTGGATTTGTCCC | 166 |
| chr11:19451724-19453742 | p2 | (TA)9 | AACAATGTCTCTTCGACGGC | CACTGTTGCACACCGAAATC | 146 |
| chr11:19477563-19479581 | p2 | (AT)10 | AAGTATCAACCGTGGCTTCG | CGCTCCGATCTAATCCAAAC | 281 |
| chr11:19477604-19479616 | p2 | (AT)10 | AAGTATCAACCGTGGCTTCG | CGCTCCGATCTAATCCAAAC | 240 |
| chr11:19477699-19479711 | p2 | (AT)10 | AAGTATCAACCGTGGCTTCG | CGCTCCGATCTAATCCAAAC | 145 |
| chr11:19523039-19525055 | p4 | (TTAA)4 | GCTCTGCCAGGCATGTATTT | ATTAATCCCTCACGATCCCC | 157 |
| chr11:19532283-19534301 | p6 | (ATATGA)3 | ACACAGTCGAAGTGGGGAAC | CCCTCTTCGTCTCCACCATA | 114 |
| chr11:19713508-19715526 | p6 | (AACCGG)3 | GATGCAAGAGGCCCAAATTA | TTCCATGCTGAAGGAAAAGC | 278 |
| chr11:19713508-19715526 | p2 | (TA)9 | GATGCAAGAGGCCCAAATTA | TTCCATGCTGAAGGAAAAGC | 278 |
| chr11:19713508-19715526 | p5 | (ATAGT)5 | GATGCAAGAGGCCCAAATTA | TTCCATGCTGAAGGAAAAGC | 278 |
| chr11:19713508-19715526 | p2 | (TA)7 | GATGCAAGAGGCCCAAATTA | TTCCATGCTGAAGGAAAAGC | 278 |
| chr11:19713508-19715526 | p2 | (AT)8 | GATGCAAGAGGCCCAAATTA | TTCCATGCTGAAGGAAAAGC | 278 |
| chr11:19758551-19760569 | p3 | (ATT)6 | TAGTTGGGGGCTGTTTTGAC | GACTTGAAGAAAATCGGGCA | 221 |
| chr11:19781317-19783351 | p4 | (AATA)4 | GACCACAAAGTACCTGGCGT | GCGTGATCTCTTGGGTTCAT | 246 |
| chr11:19781317-19783351 | p6 | (TTACTA)3 | GACCACAAAGTACCTGGCGT | GCGTGATCTCTTGGGTTCAT | 246 |
| chr11:19883755-19885835 | p2 | (TA)40 | ACTTGTTTGGCCAATTCCAC | TGACTAGTGGTGCTTCCCCT | 184 |

| Marker ID | SSR type | SSR motif | FORWARD PRIMER (5'-3') | REVERSE PRIMER (5'-3') | size (bp) |
|---|---|---|---|---|---|
| chr11:19939134-19941149 | p3 | (TGT)5 | CCTATTCCAAATTCTGGGCA | CAAATGCTGAATCCTCCGTT | 237 |
| chr11:19946112-19948127 | p3 | (GTT)6 | CACGATGTGGGATTTCACTG | CCAAGGGGCTTCAACATCTA | 230 |
| chr11:19946112-19948127 | p3 | (AGA)5 | CACGATGTGGGATTTCACTG | CCAAGGGGCTTCAACATCTA | 230 |
| chr11:19986606-19988621 | p3 | (AAC)5 | CCTCGGGGAAGTTACTGTCA | ATCCACCACCAGAGAACAGG | 175 |
| chr11:20002260-20004275 | p2 | (AT)6 | GAAAAGCATGAATTGGGGAA | GGCTGCTTCAATGGTGATTT | 124 |
| chr11:20014931-20016946 | p5 | (GTTGA)3 | GCGAGTGTTTGATTGCTTCA | GCAGCCTCATCACCCTGTAT | 193 |
| chr11:20258941-20260963 | p2 | (AT)11 | TGTTGTCGTGCACACTTTGA | CCGTATCCCCGTCACATATC | 282 |
| chr11:20258941-20260963 | p5 | (GGGGT)3 | TGTTGTCGTGCACACTTTGA | CCGTATCCCCGTCACATATC | 282 |
| chr11:20347913-20349928 | p3 | (AAT)12 | GGCTGATGAGAATGGAAGGA | TTCATCCTGGCACAATCAAA | 176 |
| chr11:20347913-20349928 | p3 | (TTC)5 | GGCTGATGAGAATGGAAGGA | TTCATCCTGGCACAATCAAA | 176 |
| chr11:20503550-20505584 | p2 | (AT)17 | TTCACTCCCCATTCGTTCTC | CGGCCGGATCATTTCTAATA | 169 |
| chr11:20564773-20566791 | p2 | (AC)9 | ATTCCGGGGAATAGGTTGTC | TTGGAATAACAAAAAGGGCG | 187 |
| chr11:20566268-20568286 | p6 | (CCCCAC)3 | TCCATAATTCCACGGGTCAT | CGCATTTTTGTACAATCGGA | 192 |
| chr11:20570386-20572402 | p4 | (AACA)4 | CATGAGCTATTGCTGGGGTT | GGTAACATGCACGCAATGAC | 191 |
| chr11:20570386-20572402 | p5 | (ATAAC)3 | CATGAGCTATTGCTGGGGTT | GGTAACATGCACGCAATGAC | 191 |
| chr11:20663490-20665505 | p5 | (AAAAG)3 | CACCACTCCACATACCACCA | ACCAGTCTAGGGGCTGGTTT | 157 |
| chr11:20663490-20665505 | p3 | (AAG)6 | CACCACTCCACATACCACCA | ACCAGTCTAGGGGCTGGTTT | 157 |
| chr11:20677251-20679272 | p3 | (TTG)7 | TTTTTATTCGGCAAAAACGG | AAATGTGCTTCATTCCACCC | 253 |
| chr11:20724338-20726358 | p2 | (TA)10 | TTCCATTGCTATGTGCTTCG | CAAAGTCTAGGCCCCCTACC | 283 |
| chr11:20780897-20782943 | p2 | (TA)23 | TTTTATTGAGCTGGGGTTCG | ACCCTTCTTTTGGGTGGTCT | 249 |
| chr11:20841124-20843142 | p2 | (AT)9 | TATCACAAGGGGACCTCGAC | CAACCGCCCTAGTTTGACAT | 189 |
| chr11:20841124-20843142 | p6 | (TTATTT)3 | TATCACAAGGGGACCTCGAC | CAACCGCCCTAGTTTGACAT | 189 |
| chr11:20860972-20863005 | p3 | (CAA)11 | ATGTGGGCTGTTCGGTAGAC | AATGCGCCGTTTTAATTCTG | 117 |
| chr11:20867643-20869659 | p2 | (CA)8 | TAGTTGGAAATGCAGAGGGG | GAGTTCGAATCCCCACATTG | 177 |
| chr11:20883602-20885616 | p4 | (AAAT)4 | CGGAAACCAAATTGCGTACT | TTACACTCCCCGACCTCAAG | 129 |
| chr11:20883602-20885616 | p2 | (AT)7 | CGGAAACCAAATTGCGTACT | TTACACTCCCCGACCTCAAG | 129 |
| chr11:20883602-20885616 | p3 | (CTT)6 | CGGAAACCAAATTGCGTACT | TTACACTCCCCGACCTCAAG | 129 |
| chr11:20894001-20896019 | p3 | (TGT)6 | GGTCGGTTACCACCGACTTA | TGAAGCGCTTTGAAGGAAAT | 151 |
| chr11:20912727-20914742 | p3 | (AAT)5 | CCTAGGTGGCTCTAATCCCC | GAGTGCCGTTTCGAACTCTC | 260 |
| chr11:20918084-20920102 | p6 | (AAAAAC)3 | GTTGCTCCTGATTGGTCCAT | CGTCACCTGGTGTCCCTACT | 244 |
| chr11:20989480-20991498 | p2 | (AT)9 | TCCCTCATCTTGTCCTCCAC | GTGGTTGTGAGACCGACCTT | 292 |
| chr11:21023207-21025222 | p3 | (TTG)5 | ACTACGACCCCGTATGCTTG | ATGCCCAAACCATCTCAGTC | 215 |
| chr11:21031527-21033554 | p3 | (TGT)9 | GCGAAATCTGCAAGAGGAAC | CTGGGGAAGAGCATAAGCAA | 193 |
| chr11:21031527-21033554 | p3 | (TGT)6 | GCGAAATCTGCAAGAGGAAC | CTGGGGAAGAGCATAAGCAA | 193 |
| chr11:21058803-21060819 | p4 | (AAAT)4 | AGAGGGCTAAGGGTGTTCGT | ATCCAATGACGGATTCAGGA | 161 |
| chr11:21238001-21240022 | p2 | (TC)6 | TCTCGCGTCTCTCTCTCTCC | ACCGCAATCATAGGTTCCAC | 206 |

| Marker ID | SSR type | SSR motif | FORWARD PRIMER (5'-3') | REVERSE PRIMER (5'-3') | size (bp) |
|---|---|---|---|---|---|
| chr11:21238001-21240022 | p3 | (GTT)7 | TCTCGCGTCTCTCTCTCTCC | ACCGCAATCATAGGTTCCAC | 206 |
| chr11:21253047-21255068 | p3 | (TGT)7 | GATGTCACGCATCGTATTGG | ATCCTATCTCGCCTGGTGTG | 132 |
| chr11:21390936-21392951 | p4 | (ATAA)4 | AATTTGGGCTCATTTGATGG | TGACCCGTCCAATTTTTCAT | 104 |
| chr11:21390936-21392951 | p3 | (TTC)5 | AATTTGGGCTCATTTGATGG | TGACCCGTCCAATTTTTCAT | 104 |
| chr11:21503920-21505936 | p4 | (CATA)4 | TGAAATTTTTGGAAGGTCGG | GGTTTGCACTTTGACCCAGT | 103 |
| chr11:21503920-21505936 | p2 | (AT)97 | TGAAATTTTTGGAAGGTCGG | GGTTTGCACTTTGACCCAGT | 103 |
| chr11:21532425-21534437 | p3 | (TGT)6 | CCACGAGGTCAAGGATGTTT | TCGTTGCCTGTCTCTTTCCT | 202 |
| chr11:21532425-21534437 | p2 | (TA)6 | CCACGAGGTCAAGGATGTTT | TCGTTGCCTGTCTCTTTCCT | 202 |
| chr11:21644229-21646244 | p5 | (TATTA)3 | CCGTTGCATGTAAAGCAAAA | CAGAAGGTAGAGCCATTGCC | 280 |
| chr11:21687936-21689954 | p6 | (CTCTAC)3 | CTGCGTCCTGATGAGTCAAA | ACGTCATGAACAACATCGGA | 215 |
| chr11:21790235-21792250 | p2 | (TA)7 | GAATCCAGATCTGACCGGAA | GCTTTTGTCGTCCTTTGCAT | 211 |
| chr11:21790235-21792250 | p5 | (AAACC)3 | GAATCCAGATCTGACCGGAA | GCTTTTGTCGTCCTTTGCAT | 211 |
| chr11:21790273-21792288 | p2 | (TA)7 | GAATCCAGATCTGACCGGAA | GCTTTTGTCGTCCTTTGCAT | 173 |
| chr11:21790273-21792288 | p5 | (AAACC)3 | GAATCCAGATCTGACCGGAA | GCTTTTGTCGTCCTTTGCAT | 173 |
| chr11:21791044-21793059 | p5 | (AAACC)3 | CTAGGGGTGTACATGGACCG | CCGACTTTTGTTGGTTTGGT | 149 |
| chr11:21868201-21870213 | p2 | (TA)6 | TATCATTTGCCATCTTCCCC | TAAAAACGGTGCTGGGTAGC | 104 |
| chr11:21868730-21870742 | p2 | (TA)6 | CGTAAGCGTGGTCGATAGGT | GCACAATCCCAACCAATTCT | 100 |
| chr11:21879569-21881581 | p2 | (AT)6 | TCCCGTCTTTCATGTTCCTC | ATGCCAAAAATTTGCTCTGG | 136 |
| chr11:21976857-21978875 | p6 | (AAGAAA)3 | ACGAGATTTGGATGACTGGG | GGTCAAATCGCTTCCAAAAA | 143 |
| chr11:21991082-21993097 | p3 | (AAC)5 | CAGCCCTTCCTCCTCTCTCT | GAATAGTGTCGCCCATTCGT | 199 |
| chr11:22043768-22045783 | p6 | (AACCGG)3 | ATCCGTAAAGGTGTCAACCG | ATTGGGGGCCAAAAATTAAG | 129 |
| chr11:22043768-22045783 | p3 | (GAT)5 | ATCCGTAAAGGTGTCAACCG | ATTGGGGGCCAAAAATTAAG | 129 |
| chr11:22048527-22050539 | p2 | (TA)6 | AAGTATCAACCGTGGCTTCG | GTCGCTCCGATCTAATCCAA | 286 |
| chr11:22048527-22050539 | p2 | (AT)8 | AAGTATCAACCGTGGCTTCG | GTCGCTCCGATCTAATCCAA | 286 |
| chr11:22068340-22070355 | p6 | (GGGGTG)3 | GGAGCACTCCACTTTCTTCG | AGCCGGCAAAGCTAACAGTA | 213 |
| chr11:22068340-22070355 | p5 | (GGGGT)3 | GGAGCACTCCACTTTCTTCG | AGCCGGCAAAGCTAACAGTA | 213 |
| chr11:22091627-22093642 | p6 | (AACCGG)3 | ATCCGTAAAGGTGTCAACCG | ATTGGGGGCCAAAAATTAAG | 129 |
| chr11:22091627-22093642 | p3 | (GAT)5 | ATCCGTAAAGGTGTCAACCG | ATTGGGGGCCAAAAATTAAG | 129 |
| chr11:22096386-22098398 | p2 | (TA)6 | AAGTATCAACCGTGGCTTCG | GTCGCTCCGATCTAATCCAA | 286 |
| chr11:22096386-22098398 | p2 | (AT)8 | AAGTATCAACCGTGGCTTCG | GTCGCTCCGATCTAATCCAA | 286 |
| chr11:22116199-22118214 | p6 | (GGGGTG)3 | GGAGCACTCCACTTTCTTCG | AGCCGGCAAAGCTAACAGTA | 213 |
| chr11:22116199-22118214 | p5 | (GGGGT)3 | GGAGCACTCCACTTTCTTCG | AGCCGGCAAAGCTAACAGTA | 213 |
| chr11:22129282-22131300 | p3 | (CTT)6 | CTCCACCTTTTTGGGGGTAT | GATTGGCATTGGGAAGAAAA | 152 |
| chr11:22152286-22154302 | p4 | (ACAA)5 | CGACGAGCAGGGTAAACTGT | CAATTGCGCTGATGTAATGG | 296 |
| chr11:22152286-22154302 | p2 | (GT)6 | CGACGAGCAGGGTAAACTGT | CAATTGCGCTGATGTAATGG | 296 |
| chr11:22152300-22154315 | p4 | (ACAA)5 | CGACGAGCAGGGTAAACTGT | CAATTGCGCTGATGTAATGG | 282 |

| Marker ID | SSR type | SSR motif | FORWARD PRIMER (5'-3') | REVERSE PRIMER (5'-3') | size (bp) |
|---|---|---|---|---|---|
| chr11:22152300-22154315 | p2 | (GT)6 | CGACGAGCAGGGTAAACTGT | CAATTGCGCTGATGTAATGG | 282 |
| chr11:22349364-22351382 | p3 | (GTG)6 | TGGAAAAGCGTGTCAGTCAG | GACACTGCTTGGTCGGATTT | 138 |
| chr11:22349364-22351382 | p5 | (TTTTA)3 | TGGAAAAGCGTGTCAGTCAG | GACACTGCTTGGTCGGATTT | 138 |
| chr11:22447411-22449426 | p5 | (TAAAA)3 | TTTCTTTCGGGTGCTGCTAT | CCTTGAAGGAAAGAAATTTGGA | 150 |
| chr11:22828722-22830758 | p2 | (AT)18 | CGCCTCCTCAAGAAACAAAG | CTTATGAGAGCCGTTTTGGC | 160 |
| chr11:22837293-22839309 | p4 | (ATAA)4 | CCTGCCATCAAGTTCCAAAT | AAATGCGAAAACATTCCAGC | 122 |
| chr11:22849487-22851555 | p2 | (TA)34 | GTGGTACATTCGACGCCTTT | GGATGGTCCGTCAATCAATC | 162 |
| chr11:22863735-22865786 | p3 | (CAA)6 | TCCAATCTGATTCGCAAGAA | ATCAATTCCTCCGTGACGAT | 281 |
| chr11:22863799-22865832 | p3 | (CAA)6 | TCCAATCTGATTCGCAAGAA | ATCAATTCCTCCGTGACGAT | 217 |
| chr11:22863836-22865852 | p3 | (CAA)6 | TCCAATCTGATTCGCAAGAA | ATCAATTCCTCCGTGACGAT | 180 |
| chr11:22863850-22865919 | p3 | (CAA)6 | TCCAATCTGATTCGCAAGAA | ATCAATTCCTCCGTGACGAT | 166 |
| chr11:22881111-22883127 | p4 | (TCTT)4 | TCCTACCCATCCCTAAACCC | CCAAATGAAGAGAAATGCCC | 221 |
| chr11:22904107-22906170 | p3 | (TAT)21 | CGGCTTAGAAAGCCTCATTG | AGTTCGCTGCCGAAGAAATA | 109 |
| chr11:22904107-22906170 | p5 | (TTCTC)3 | CGGCTTAGAAAGCCTCATTG | AGTTCGCTGCCGAAGAAATA | 109 |
| chr11:22904107-22906170 | p6 | (CTTTTT)3 | CGGCTTAGAAAGCCTCATTG | AGTTCGCTGCCGAAGAAATA | 109 |
| chr11:22942960-22944992 | p2 | (TA)10 | CCAAAGCATTACGGAAGGAA | TAAAGAAATGGCCACAAGCC | 107 |
| chr11:22942960-22944992 | p2 | (AT)16 | CCAAAGCATTACGGAAGGAA | TAAAGAAATGGCCACAAGCC | 107 |
| chr11:22942960-22944992 | p2 | (AC)7 | CCAAAGCATTACGGAAGGAA | TAAAGAAATGGCCACAAGCC | 107 |
| chr11:22996072-22998086 | p2 | (AT)7 | TGGGCCCCATACTAAACAAC | TTTAAGGGACAACGGACGAC | 153 |
| chr11:22996072-22998086 | p2 | (AT)31 | TGGGCCCCATACTAAACAAC | TTTAAGGGACAACGGACGAC | 153 |
| chr11:22996072-22998086 | p2 | (AT)16 | TGGGCCCCATACTAAACAAC | TTTAAGGGACAACGGACGAC | 153 |
| chr11:23002521-23004537 | p4 | (ATGT)4 | AGGTCATCGGACGAGTTTTG | CAACGCAAGTCGTGTTGACT | 180 |
| chr11:23005022-23007034 | p2 | (TA)6 | GTACTAAGAGGGCCGGGTTC | AATCGGAACCCATACAGACG | 186 |
| chr11:23086077-23088093 | p2 | (AT)12 | CCGAATATCCAATTTGTGGC | GGACTGACCTTCCCTCCCTA | 262 |
| chr11:23086077-23088093 | p4 | (AAGA)4 | CCGAATATCCAATTTGTGGC | GGACTGACCTTCCCTCCCTA | 262 |
| chr11:23108912-23110926 | p2 | (GT)7 | GGCTTGTGGTGGAAGTTGTT | TCGCAAATAACATTGCCAAA | 115 |
| chr11:23222228-23224249 | p3 | (GTT)7 | GACCAGCCATTCTGGACAGT | GCAACAACGTAACCCGCTAT | 216 |
| chr11:23230815-23232831 | p4 | (AAGA)4 | ACATCCATCGTGAAAAAGCC | AGCCAACTGATAATGCAGGG | 245 |
| chr11:23236894-23238912 | p6 | (TTTTTC)3 | TTGCAGGGTTTTTCAAGGAG | ATCACTGGCCGAAGCTAGAA | 264 |
| chr11:23281561-23283575 | p2 | (AT)7 | ACCTCACCTTATGCCCACAC | TCTTTTACAGGTGGTTGCCC | 163 |
| chr11:23286693-23288705 | p2 | (TA)6 | AATGAATCAGAAGGCAACGC | GATTCTCGCCATGCGTTATT | 128 |
| chr11:23405900-23407914 | p2 | (AT)7 | TTTGATGTGCCATTGACGTT | CCACTCCGCGTAATCAAACT | 146 |
| chr11:23425987-23428005 | p6 | (CTTTTT)3 | CGAAAATTAACCGAACCGAA | GCTGCTTTTTCTTCCGAATG | 245 |
| chr11:23446813-23448825 | p2 | (AT)6 | AGGACTTTTTGGTGTGACGC | TTTTCCTTATCCGGTGATCG | 151 |
| chr11:23499405-23501423 | p3 | (AAT)6 | AGCCGATAAAGCTGCAAAGA | TTGGTGTTTGGACTTGTGGA | 273 |
| chr11:23524732-23526762 | p4 | (TCTT)4 | GTATTCTCTTTCCCCCAGGC | TCTTGTTCCCTTCACTTGGG | 280 |

| Marker ID | SSR type | SSR motif | FORWARD PRIMER (5'-3') | REVERSE PRIMER (5'-3') | size (bp) |
|---|---|---|---|---|---|
| chr11:23525712-23527728 | p4 | (TCTT)4 | AACAAGTTGGATTAGGGCGA | CCAAGGGTGTGCAGGTTACT | 200 |
| chr11:23555750-23557765 | p5 | (TTGGT)3 | GATTGCCCTTATTTTGGGGT | ACAAGAGTATGCTGGGTCCG | 164 |
| chr11:23569971-23571986 | p5 | (TGGGT)3 | TAATGCATATGCGGCACCTA | TATATGAGCGAGAGGCTGGG | 123 |
| chr11:23569971-23571986 | p2 | (AT)7 | TAATGCATATGCGGCACCTA | TATATGAGCGAGAGGCTGGG | 123 |
| chr11:23572766-23574784 | p5 | (AAAAT)4 | GCACTCTTTTTGGGACAAGC | CAACGTACCGAACTCGAACC | 247 |
| chr11:23572766-23574784 | p6 | (TTTTAT)3 | GCACTCTTTTTGGGACAAGC | CAACGTACCGAACTCGAACC | 247 |
| chr11:23652132-23654144 | p2 | (GT)6 | TGCAGAGTTCGGTGATTACG | TACTCCGGGAGGGTTCTCTT | 135 |
| chr11:23678682-23680700 | p2 | (AT)38 | TGGTTGGGCCTTTATCACAT | ACACGGACGACGAAAGTAGC | 213 |
| chr11:23678682-23680700 | p6 | (ATAAAA)3 | TGGTTGGGCCTTTATCACAT | ACACGGACGACGAAAGTAGC | 213 |
| chr11:23678682-23680700 | p2 | (TA)10 | TGGTTGGGCCTTTATCACAT | ACACGGACGACGAAAGTAGC | 213 |
| chr11:23678682-23680700 | p6 | (ATAAAA)3 | TGGTTGGGCCTTTATCACAT | ACACGGACGACGAAAGTAGC | 213 |
| chr11:23688193-23690208 | p5 | (TTGGT)3 | AGTAAACCGACAAACGGCAC | ACCAAACCAATTATGGTCGG | 125 |
| chr11:23688193-23690208 | p5 | (TTGGT)3 | AGTAAACCGACAAACGGCAC | ACCAAACCAATTATGGTCGG | 125 |
| chr11:23691500-23693516 | p4 | (TAAT)4 | TCAACCATGCCAGGTCTACA | ACATCGGGCCTAGACCTTTT | 111 |
| chr11:23724921-23726933 | p2 | (AT)6 | GCGAATCTCTAATCCCACCA | TAGGAACTAAGTCCCGCACG | 169 |
| chr11:23724935-23726965 | p2 | (AT)6 | GCGAATCTCTAATCCCACCA | TAGGAACTAAGTCCCGCACG | 155 |
| chr11:23742718-23744736 | p3 | (TAA)6 | TTTTTGAGAGGGGTGGTTGA | CCCTGTTTTGCACTGGTCTT | 299 |
| chr11:23772586-23775084 | p2 | (TA)249 | TGTCTCCGCATCTCATCTTG | AAGAGATCGTGGGCTAGCAA | 114 |
| chr11:23816044-23818059 | p5 | (CAATT)3 | AGTGTTTGGTTTGGGGTGAA | TGGAGGGAAAAGATTTGGTG | 192 |
| chr11:23816044-23818059 | p3 | (TCT)11 | AGTGTTTGGTTTGGGGTGAA | TGGAGGGAAAAGATTTGGTG | 192 |
| chr11:23822541-23824562 | p2 | (TA)18 | CGCAACTGCTAAGATGCAAA | GGTGTAAAATGCCTACCCCA | 276 |
| chr11:23822541-23824562 | p3 | (ATT)7 | CGCAACTGCTAAGATGCAAA | GGTGTAAAATGCCTACCCCA | 276 |
| chr11:23935531-23937558 | p3 | (AAC)9 | CACTCGAAATCTCACCAGCA | TTTTTACAGGGGCATGAAGG | 287 |
| chr11:23955000-23957012 | p2 | (TA)25 | ATTTGGTAACGGTTGGTGGA | GAATCAGAATCCCCGTTGAA | 263 |
| chr11:23955000-23957012 | p2 | (AC)6 | ATTTGGTAACGGTTGGTGGA | GAATCAGAATCCCCGTTGAA | 263 |
| chr11:23955000-23957012 | p2 | (AC)6 | ATTTGGTAACGGTTGGTGGA | GAATCAGAATCCCCGTTGAA | 263 |
| chr11:24009263-24011281 | p3 | (CAA)6 | GGACCTTGGGCTACTCTTCC | TTAGCTACTCGGGGCTTCAA | 199 |
| chr11:24009263-24011281 | p2 | (CA)6 | GGACCTTGGGCTACTCTTCC | TTAGCTACTCGGGGCTTCAA | 199 |
| chr11:24076889-24078903 | p2 | (TC)7 | TCCTCGTTTAATGGGCCTAA | TACGTGAACTCGTTGCTTGC | 248 |
| chr11:24112169-24114184 | p5 | (TATTT)3 | ACTCATTTCCTTCGGTTCCC | GGACACGCACAAAGAACTCA | 143 |
| chr11:24143581-24145596 | p5 | (TTACT)3 | TTCAGTTTTTGGGTTTTGGC | TTAATCGCCACCCAAAGAAG | 162 |
| chr11:24144700-24146721 | p3 | (TTG)7 | GCTGCTGCTGCTTCTTCATT | AATTGCAAAAACCCCAACAA | 272 |
| chr11:24171927-24173942 | p5 | (CAAAC)3 | CTTTGGGGCAATACTTGCAT | GTAGTCGAGACCCTAGCCCC | 280 |
| chr11:24264462-24266476 | p2 | (AT)7 | GCATTCTGGAATTTGGCAAT | TCCGTTTGACCCCCATAATA | 101 |
| chr11:24408862-24410877 | p3 | (ATG)5 | GCACCGCTGGAACTTTTTAG | TCCAACTCCAACTTCCCAAG | 189 |
| chr11:24515977-24517989 | p2 | (TA)6 | TGGATGATTTGTGGGATGTG | CAAAATCATGTCATCCGCTG | 217 |

| Marker ID | SSR type | SSR motif | FORWARD PRIMER (5'-3') | REVERSE PRIMER (5'-3') | size (bp) |
|---|---|---|---|---|---|
| chr11:24515977-24517989 | p2 | (AT)7 | TGGATGATTTGTGGGATGTG | CAAAATCATGTCATCCGCTG | 217 |
| chr11:24572505-24574520 | p3 | (ACA)5 | AGATGATGGCTTCCGAATTG | TGTGCAAATGGCTAAGTGGA | 203 |
| chr11:24577524-24579539 | p5 | (AACAA)3 | CTATCATTTCCGACCCTCCA | TCCCATTTAGCACCAAGTCC | 167 |
| chr11:24577524-24579539 | p5 | (CCCAA)4 | CTATCATTTCCGACCCTCCA | TCCCATTTAGCACCAAGTCC | 167 |
| chr11:24607597-24609617 | p2 | (TA)10 | CAATTGATCTGAGCCGTTGA | TCAGGGGCATTTTTGTTCTT | 195 |
| chr11:24621744-24623768 | p5 | (AAAAT)3 | CCCCATATTAAACACCGTGC | TGTCAATGTGGGATACCGTG | 230 |
| chr11:24621771-24623793 | p5 | (AAAAT)3 | CCCCATATTAAACACCGTGC | TGTCAATGTGGGATACCGTG | 203 |
| chr11:24621801-24623813 | p5 | (AAAAT)3 | CCCCATATTAAACACCGTGC | TGTCAATGTGGGATACCGTG | 173 |
| chr11:24639088-24641104 | p4 | (TTAA)4 | TTGGGGCATTAAAACCACTC | TGAGTCAATGAGCAACCGTC | 110 |
| chr11:24642492-24644506 | p2 | (AC)7 | CAGAATATTGAAGGGGGCAA | TTCATGTCACCCAGTTTCCA | 192 |
| chr11:24777054-24779072 | p2 | (AT)11 | AGGCTTATTCGGGCAAAAGT | TGCCGGTCTTTAATTTCTGG | 137 |
| chr11:24777054-24779072 | p6 | (ATCTAT)3 | AGGCTTATTCGGGCAAAAGT | TGCCGGTCTTTAATTTCTGG | 137 |
| chr11:24807563-24809609 | p2 | (TA)23 | AACAACTGACGGAACCAACC | TTTTGTTAGCCTTGTCGGCT | 213 |
| chr11:24822046-24824058 | p2 | (AT)6 | GCCGATCGAACTGCTTATTC | TTTACCGGTACAGAGTGGGC | 259 |
| chr11:24822196-24824212 | p2 | (AT)6 | GCCGATCGAACTGCTTATTC | TTTACCGGTACAGAGTGGGC | 109 |
| chr11:24822297-24824309 | p2 | (AT)6 | ACCATTGCCCACTCTGTACC | CGTCCACTTGGGGAAAACTA | 236 |
| chr11:24955124-24957136 | p2 | (TC)6 | CCGAAGACGGTAATTCTCCA | GGTGTCAACACCGCATACAC | 128 |
| chr11:24981876-24983894 | p2 | (TA)9 | CAGGATTTGACACGTGTTGG | GGCCATAAAATGGTGGACAG | 151 |
| chr11:25039610-25041630 | p2 | (TA)10 | AAGTGTGCCGAAAAATGTCC | GCCCGCATAAGGCATAACTA | 197 |
| chr11:25054260-25056272 | p2 | (TA)6 | AGGCCTGCTTTAACGAAACA | AATCTGAAGTGTTGGGCCTG | 196 |
| chr11:25070478-25072494 | p6 | (TTTTTG)3 | CGTTACGGGTCAAGCGTTAT | GCTAGCCGTAAATTCCCCTC | 148 |
| chr11:25070478-25072494 | p4 | (CCTG)4 | CGTTACGGGTCAAGCGTTAT | GCTAGCCGTAAATTCCCCTC | 148 |
| chr11:25099124-25101138 | p2 | (AT)7 | TACGGGCACTCGTCATCATA | TTTTGCTAGCCTTGTTGGCT | 217 |
| chr11:25151992-25154013 | p2 | (AG)13 | CGATCAAAACCAAGGGAGAA | TGAAAACAGCAAAATCGCAG | 137 |
| chr11:25152016-25154043 | p2 | (AG)13 | CGATCAAAACCAAGGGAGAA | TGAAAACAGCAAAATCGCAG | 113 |
| chr11:25312307-25314322 | p5 | (ACCAA)3 | TCGAAATGGAGAGTTTGGGT | TGGCATTCCAACAACCACTA | 198 |
| chr11:25403126-25405141 | p5 | (ATATA)3 | ATGAAATTTCTCATGCGGGT | CGGCTGGTATCAAATGAAGC | 122 |
| chr11:25412165-25414183 | p3 | (TAT)6 | ACCCGCTACGTACCAAATGA | CACCTTCTTCATTGCCCATT | 211 |
| chr11:25412165-25414183 | p5 | (ATTTT)3 | ACCCGCTACGTACCAAATGA | CACCTTCTTCATTGCCCATT | 211 |
| chr11:25441414-25443432 | p2 | (AT)9 | TTAAAATATGGTGCCCGTCC | CGTCCGTTGTCCCTTAAAAA | 277 |
| chr11:25441414-25443432 | p2 | (AT)9 | TTAAAATATGGTGCCCGTCC | CGTCCGTTGTCCCTTAAAAA | 277 |
| chr11:25441414-25443432 | p2 | (AT)6 | TTAAAATATGGTGCCCGTCC | CGTCCGTTGTCCCTTAAAAA | 277 |
| chr11:25676166-25678181 | p5 | (ATATG)3 | AACCATTGTAGGTGACCCCA | TTTCTTTAGGACCGCTCGAA | 224 |
| chr11:25676166-25678181 | p2 | (TA)8 | AACCATTGTAGGTGACCCCA | TTTCTTTAGGACCGCTCGAA | 224 |
| chr11:25944861-25946879 | p6 | (TTCAGT)3 | CTGGGAATTGGGTGAAGAAA | CATGGCGACACATTCTGAAC | 268 |
| chr11:26188935-26190953 | p6 | (GCAGTA)3 | AATTGTGGCAGTTTCCCTTG | TCAAGTTGCGGCAGTGATAG | 142 |

| Marker ID | SSR type | SSR motif | FORWARD PRIMER (5'-3') | REVERSE PRIMER (5'-3') | size (bp) |
|---|---|---|---|---|---|
| chr11:26200587-26202611 | p6 | (GGGGAC)4 | CCGTTCATATGTGCCTGTTG | ATAGCATTGTTTGGCCTTGG | 232 |
| chr11:26200587-26202611 | p3 | (TTC)12 | CCGTTCATATGTGCCTGTTG | ATAGCATTGTTTGGCCTTGG | 232 |
| chr11:26241726-26243806 | p5 | (AAACC)3 | ATCACTAGGGGTGCACTTGG | ACCCGACTTTTGTTGGTTTG | 212 |
| chr11:26241804-26243828 | p5 | (AAACC)3 | ATCACTAGGGGTGCACTTGG | ACCCGACTTTTGTTGGTTTG | 134 |
| chr11:26253418-26255430 | p2 | (TG)6 | AAAGGTGCCTACGGGCTATT | ACTTAGCCAACCAAGGCTCA | 255 |
| chr11:26261043-26263055 | p2 | (TG)6 | ATGATGCCAAAGGTGCCTAC | CGACCATAAGGTTCGGTGTT | 247 |
| chr11:26387590-26389608 | p5 | (TTATT)3 | GCAATGCTTCAGTCTTGCAG | ACTTACTGCAATGCTCATCGG | 117 |
| chr11:26415649-26417667 | p3 | (ATT)6 | CTTGGCACGCAACTCACTTA | GCATTGATGTGGCAATAACG | 287 |
| chr11:26415649-26417667 | p3 | (GAA)5 | CTTGGCACGCAACTCACTTA | GCATTGATGTGGCAATAACG | 287 |
| chr11:26415649-26417667 | p3 | (GAA)5 | CTTGGCACGCAACTCACTTA | GCATTGATGTGGCAATAACG | 287 |
| chr11:26425513-26427525 | p2 | (AT)11 | AGCAGCGTTTTCGTCTAGGA | CCCAAAAACTTCACCTCCAA | 168 |
| chr11:26425513-26427525 | p2 | (AT)6 | AGCAGCGTTTTCGTCTAGGA | CCCAAAAACTTCACCTCCAA | 168 |
| chr11:26425513-26427525 | p2 | (AT)8 | AGCAGCGTTTTCGTCTAGGA | CCCAAAAACTTCACCTCCAA | 168 |
| chr11:26425513-26427525 | p2 | (AT)10 | AGCAGCGTTTTCGTCTAGGA | CCCAAAAACTTCACCTCCAA | 168 |
| chr11:26425513-26427525 | p2 | (AT)8 | AGCAGCGTTTTCGTCTAGGA | CCCAAAAACTTCACCTCCAA | 168 |
| chr11:26462664-26464682 | p3 | (ATT)6 | CTTGGCACGCAACTCACTTA | GCATTGATGTGGCAATAACG | 287 |
| chr11:26462664-26464682 | p3 | (GAA)5 | CTTGGCACGCAACTCACTTA | GCATTGATGTGGCAATAACG | 287 |
| chr11:26462664-26464682 | p3 | (GAA)5 | CTTGGCACGCAACTCACTTA | GCATTGATGTGGCAATAACG | 287 |
| chr11:26472528-26474540 | p2 | (AT)11 | AGCAGCGTTTTCGTCTAGGA | CCCAAAAACTTCACCTCCAA | 168 |
| chr11:26472528-26474540 | p2 | (AT)6 | AGCAGCGTTTTCGTCTAGGA | CCCAAAAACTTCACCTCCAA | 168 |
| chr11:26472528-26474540 | p2 | (AT)8 | AGCAGCGTTTTCGTCTAGGA | CCCAAAAACTTCACCTCCAA | 168 |
| chr11:26472528-26474540 | p2 | (AT)10 | AGCAGCGTTTTCGTCTAGGA | CCCAAAAACTTCACCTCCAA | 168 |
| chr11:26472528-26474540 | p2 | (AT)8 | AGCAGCGTTTTCGTCTAGGA | CCCAAAAACTTCACCTCCAA | 168 |
| chr11:26521172-26523200 | p2 | (AT)14 | ATGCGATTATGACACGTCCA | TCGGTTAGGGACTGTTTTGG | 280 |
| chr11:26643230-26645242 | p2 | (AG)6 | GCAGTCCAAAGCCAACATTT | TGAACAGCCCTATCCCTACG | 204 |
| chr11:26654596-26656611 | p2 | (AT)13 | TGCTGAACTGCAAGTTGTCC | ACGTCTTCCGTTGTATTGGG | 197 |
| chr11:26654596-26656611 | p3 | (ATC)5 | TGCTGAACTGCAAGTTGTCC | ACGTCTTCCGTTGTATTGGG | 197 |
| chr11:26654596-26656611 | p2 | (TA)6 | TGCTGAACTGCAAGTTGTCC | ACGTCTTCCGTTGTATTGGG | 197 |
| chr11:26707381-26709395 | p5 | (TGGGG)4 | GCGCCCTGATTAATGACCTA | ATGGCTTATGCTTTGGCAAC | 135 |
| chr11:26707381-26709395 | p2 | (TA)7 | GCGCCCTGATTAATGACCTA | ATGGCTTATGCTTTGGCAAC | 135 |
| chr11:26707381-26709395 | p2 | (TA)8 | GCGCCCTGATTAATGACCTA | ATGGCTTATGCTTTGGCAAC | 135 |
| chr11:26708022-26710038 | p2 | (TA)7 | GCCGTAAATTAAAGGTGGCA | TTTAACCACTCCAAACCGGA | 188 |
| chr11:26708022-26710038 | p2 | (TA)8 | GCCGTAAATTAAAGGTGGCA | TTTAACCACTCCAAACCGGA | 188 |
| chr11:26733156-26735171 | p5 | (TTCGG)3 | TCCTCTCAAATCAGCAACCC | CGAACTTGCAAAAACCGAAT | 101 |
| chr11:26733156-26735171 | p5 | (TTGGT)3 | TCCTCTCAAATCAGCAACCC | CGAACTTGCAAAAACCGAAT | 101 |
| chr11:26858080-26860095 | p5 | (TTTTG)3 | CCATTTGTATAACGGCGAGG | AACTAGGGAAAAGGGGCAAA | 177 |

| Marker ID | SSR type | SSR motif | FORWARD PRIMER (5'-3') | REVERSE PRIMER (5'-3') | size (bp) |
|---|---|---|---|---|---|
| chr11:27027820-27029835 | p5 | (GGTGG)3 | TGAACCCAAGACTTTCCACC | CTGAATTGCGCTTTGATTGA | 239 |
| chr11:27140121-27142139 | p6 | (TTGAAT)3 | AGCCAAGTGGCAGGCTAATA | CAATTTGCAGGAAAAAGGGA | 118 |
| chr11:27150597-27152612 | p5 | (AGGAA)3 | CAAAGAGCAAAGGCTGATCC | AACGACCCTGCTCTGTAGGA | 222 |
| chr11:27295292-27297310 | p6 | (GTGGAT)3 | TTTAGCTTGCGCTAAACGGT | TGATCAATCGAGAAATGCCA | 149 |
| chr11:27335087-27337099 | p2 | (AT)6 | ATGGCCTAACGGGTCCTAAG | CAGCGATTTCCAAATGATGA | 142 |
| chr11:27463411-27465429 | p6 | (ATCTTC)3 | GCTTCTGCATCAGTGGACAA | CAGCTTGTTGCTCACGACAT | 269 |
| chr11:27466879-27468915 | p2 | (AT)18 | TGAAAGTTGTCGGTCCATGA | AAGTGTGGACGGGTGGAATA | 278 |
| chr11:27479719-27481734 | p5 | (ACACA)3 | TGTTTGGCCATGAAAATCAA | TAAGTGAGGTGCGTTTTCCC | 300 |
| chr11:27479719-27481734 | p2 | (TC)6 | TGTTTGGCCATGAAAATCAA | TAAGTGAGGTGCGTTTTCCC | 300 |
| chr11:27482582-27484598 | p4 | (AAGA)4 | AACCACTGCATGTGTTCCAA | TGAGGCACAGATCATTTCCA | 106 |
| chr11:27519178-27521194 | p4 | (AGTG)4 | TGTCATGGCCAGTTGGAATA | CGCTCTCCTATAAAGCTGCG | 186 |
| chr11:27524073-27526087 | p2 | (TA)7 | CCGGTCAATCTTGACGTCTT | AAGTTTGTCCGCCGTTACAC | 114 |
| chr11:27550889-27552905 | p2 | (AG)8 | TCGCATAAGGTGTCCAATGA | ATTTTGAGAGCGGATGGTTG | 167 |
| chr11:27550889-27552905 | p2 | (AG)9 | TCGCATAAGGTGTCCAATGA | ATTTTGAGAGCGGATGGTTG | 167 |
| chr11:27564188-27566203 | p5 | (ATGAA)3 | TAGCCAATGAGGTGCTGACA | CCTGAACACGTGGAACCTTT | 206 |
| chr11:27734644-27736656 | p2 | (AT)6 | TTTTCAACGAACACGCACTC | TTGTATCCAAACAGGCACCA | 212 |
| chr11:27809229-27811247 | p2 | (GA)9 | ATGCCCGAAAAACCTCTTCT | GGGGGAATTGAATCCTGTTT | 220 |
| chr11:27968489-27970505 | p2 | (AT)8 | ATTTTTGGCTTGGCAAAGTG | CTTTAAAGCTTTGATGCCGC | 219 |
| chr11:27971460-27973476 | p6 | (TCCAAA)3 | AGACGCCAAACAAACCAATC | GGTGTACAGTACGCACCCCT | 123 |
| chr11:27971460-27973476 | p4 | (TTTG)4 | AGACGCCAAACAAACCAATC | GGTGTACAGTACGCACCCCT | 123 |
| chr11:28002087-28004099 | p2 | (AT)6 | TCCCCGCCCTTTTATTCTAT | TTGTTGCAGCAAACCTGAAG | 195 |
| chr11:28090752-28092782 | p2 | (CA)7 | AATGCACCGTACAATCCACA | AAGCAGAGGAAGGCATACGA | 116 |
| chr11:28180016-28182036 | p5 | (TTCTT)4 | CGGGTTGCCTCAATAACTGT | CGCAGATGAAACCAACAGAA | 150 |
| chr11:28199190-28201205 | p5 | (AAACC)3 | GAGTGTACATGGACCGGGTT | ACTCGAGCAAACCGAAAGAA | 139 |
| chr11:28199190-28201205 | p5 | (AAACC)3 | GAGTGTACATGGACCGGGTT | ACTCGAGCAAACCGAAAGAA | 139 |
| chr11:28327306-28329322 | p2 | (TC)7 | ACGGCAAGAATATCATCGGT | AGCCAAAAATGAAAATTGCG | 138 |
| chr11:28327306-28329322 | p4 | (TTTG)4 | ACGGCAAGAATATCATCGGT | AGCCAAAAATGAAAATTGCG | 138 |
| chr11:28429062-28431116 | p2 | (AT)27 | ATCACTGCTGCTTGTCATGG | ACGCTTGGGACAATTTCAAC | 212 |
| chr11:28454354-28456366 | p2 | (GT)6 | GGCGCAAAATCTGTTTTCAT | CGAAGCCTGATTTCGTTCTC | 100 |
| chr11:28475688-28477704 | p4 | (TTTC)4 | GATTCCGAGAATTGCCTTGA | CCTACTTCCTGCCCACACAT | 183 |
| chr11:28571820-28573832 | p2 | (AT)6 | CCTCCTTGGGTTTCTTCCTC | AAATTAACACGGTTGCCTCG | 298 |
| chr11:28596964-28598976 | p2 | (AT)6 | CCTCCTTGGGTTTCTTCCTC | AAATTAACACGGTTGCCTCG | 298 |
| chr11:28887808-28889820 | p2 | (TA)6 | TTTAGCTTTGGACACAGGGG | CTCCTTTTCTCCCACCTTCC | 180 |
| chr11:29187512-29189532 | p5 | (CCCCA)4 | TGATGACCCTCTTCCACTCC | GTCAAGGTGGCTATGGGAAA | 218 |
| chr11:29290182-29292203 | p3 | (GTT)7 | ACAAGCCCCCATTTTCTTCT | CCTTGTCCCGAATATCCTCA | 237 |
| chr11:29303474-29305495 | p5 | (ACCCG)3 | TGCATTAGCATCAGCAAAGG | TAGTTCGCTGCCGAAGAAAT | 229 |

| Marker ID | SSR type | SSR motif | FORWARD PRIMER (5'-3') | REVERSE PRIMER (5'-3') | size (bp) |
|---|---|---|---|---|---|
| chr11:29303497-29305530 | p5 | (ACCCG)3 | TGCATTAGCATCAGCAAAGG | TAGTTCGCTGCCGAAGAAAT | 206 |
| chr11:29389098-29391120 | p2 | (AT)11 | TTGTCTTGGAGTTGCTGTCG | ATGAGCCCAAAGGACATACG | 126 |
| chr11:29411045-29413060 | p3 | (TCT)5 | CAACCCATTTTCTTCCTCCA | CCGCTCCATCTTTTCCATAA | 271 |
| chr11:29411045-29413060 | p3 | (GAA)5 | CAACCCATTTTCTTCCTCCA | CCGCTCCATCTTTTCCATAA | 271 |
| chr11:29456921-29458936 | p3 | (GAA)5 | TCATTTCGCCACCTATTTCC | TTTGATTGTTGCCACGTGAT | 206 |
| chr11:29467550-29469564 | p2 | (AT)7 | GGCAATTCCAGGACAACCTA | AGCGGGAACTAACGAAGACA | 279 |
| chr11:29500414-29502429 | p3 | (ACA)5 | TCCCTTCAGGTGTGTTCCTC | TTTTGGTTACTCTGGCACCC | 257 |
| chr11:29532410-29534425 | p3 | (ACA)5 | GGCCAAGCAAGAGAAGTGAC | TAGTCCATGACCCGTTGACC | 240 |
| chr11:29538293-29540308 | p5 | (CACCC)3 | TACCCGTGGTTCATTCCATT | GGGTGAGTGGTAAGGTGTGG | 268 |
| chr11:29538293-29540308 | p2 | (AT)8 | TACCCGTGGTTCATTCCATT | GGGTGAGTGGTAAGGTGTGG | 268 |
| chr11:29538293-29540308 | p5 | (TACAA)3 | TACCCGTGGTTCATTCCATT | GGGTGAGTGGTAAGGTGTGG | 268 |
| chr11:29551774-29553786 | p2 | (AT)6 | CAGCAATTCCCAGTGGTCTT | TCAATTGGAGATTGTGGCAA | 204 |
| chr11:29606041-29608055 | p2 | (AT)7 | CCAGTGGTGAAGGGTCAAGT | TTGTGTGTTTGAGTCTGGGC | 229 |
| chr11:29627740-29629754 | p5 | (CAAAC)4 | TTCGGTTTGAATCGGTTTTT | TGGCAAGTTTTGTGGCAATA | 255 |
| chr11:29627740-29629754 | p2 | (AT)7 | TTCGGTTTGAATCGGTTTTT | TGGCAAGTTTTGTGGCAATA | 255 |
| chr11:29645599-29647641 | p2 | (TA)21 | GGAAGGGTTAGTCCGGAGAC | TAGGCGACGTCAGTTGAGTG | 115 |
| chr11:29780339-29782354 | p3 | (CAG)5 | CACCCTATGTTGCTCGGATT | GGGGCATCTCAAAATGAAAA | 131 |
| chr11:29780339-29782354 | p3 | (CAG)5 | CACCCTATGTTGCTCGGATT | GGGGCATCTCAAAATGAAAA | 131 |
| chr11:29780960-29782975 | p3 | (CAG)5 | AATTGCAGCATGAAAAGGCT | ATCACCCATGGAAGAGCAAG | 257 |
| chr11:29780960-29782975 | p3 | (CAG)5 | AATTGCAGCATGAAAAGGCT | ATCACCCATGGAAGAGCAAG | 257 |
| chr11:29843189-29845209 | p4 | (ACCC)5 | TCAACAACTTCCACATTGGC | TTGGGGCAAGTCAAAACTTC | 194 |
| chr11:29843189-29845209 | p2 | (AG)6 | TCAACAACTTCCACATTGGC | TTGGGGCAAGTCAAAACTTC | 194 |
| chr11:29859629-29861644 | p3 | (GTT)5 | GAAATGGAGGCTTGCTTCTG | CACAGATTCCACTCCCACCT | 219 |
| chr11:29958045-29960060 | p3 | (TTG)5 | AAACAAGGTGCGACCAAAAC | AACGTTTTCATCTCGGCAAC | 105 |
| chr11:29981019-29983034 | p5 | (ATTAA)3 | GACGGCTCGTTCAGATAAGC | CATGGATGGACACGACTGAG | 279 |
| chr11:30001987-30004002 | p3 | (TAT)5 | GCTCTCTTTCCCTTTTGGCT | TCTCCAGGACTGCTTCGATT | 206 |
| chr11:30036848-30038864 | p2 | (TA)6 | TGGGCCCCATACTAAACAAC | CCCATGAGTTTGGAGATGGT | 260 |
| chr11:30036848-30038864 | p2 | (TA)8 | TGGGCCCCATACTAAACAAC | CCCATGAGTTTGGAGATGGT | 260 |
| chr11:30130087-30132102 | p5 | (TTTTA)3 | CCAATCTTGAATGTGGCCTT | CCTTCCGCTCTGTAACCAAA | 104 |
| chr11:30172041-30174075 | p2 | (TC)17 | AATTGCACGGATTGTCCTTC | TTTCCCCTCTGCTAGGGTTT | 283 |
| chr11:30307303-30309318 | p2 | (AT)9 | ATTAATTGGGCTCAAATGCG | CGAACCTGGGGACTTTTGTA | 201 |
| chr11:30307303-30309318 | p5 | (AAAAT)3 | ATTAATTGGGCTCAAATGCG | CGAACCTGGGGACTTTTGTA | 201 |
| chr11:30345879-30347894 | p3 | (TGA)5 | TAAAATCGGAACTGGAACCG | GGGTGGGGATGGTTTAAAGT | 133 |
| chr11:30361341-30363356 | p3 | (AAT)5 | TGGGTCGTGACAAGTTGGTA | GTTCGCGCAAGAATTTAAGG | 184 |
| chr11:30376037-30378079 | p2 | (AT)21 | CCGAAGAAGGTAATTTCCCC | AAGCAGAGCCCAACTCGATA | 146 |
| chr11:30403722-30405742 | p2 | (GA)10 | CACCGTCGATGTTTGTTTTG | TTTTCCCGCAAAGTGTTTTC | 281 |

| Marker ID | SSR type | SSR motif | FORWARD PRIMER (5'-3') | REVERSE PRIMER (5'-3') | size (bp) |
|---|---|---|---|---|---|
| chr11:30410499-30412513 | p2 | (TA)7 | TTGGAGTCCACCTTGGTTTC | GTATCAAATTTTTGCGCCGT | 223 |
| chr11:30512961-30514975 | p2 | (AT)7 | TGCCAAAAGAAAAAGGGTTG | TTGGGATGATTGTTGGGATT | 253 |
| chr11:30514905-30516917 | p2 | (AT)40 | TCGGAGGAATGAAATTGTCC | AAACGGGCAACATTTCACAT | 259 |
| chr11:30514905-30516917 | p2 | (AT)6 | TCGGAGGAATGAAATTGTCC | AAACGGGCAACATTTCACAT | 259 |
| chr11:30671172-30673270 | p2 | (AT)49 | GCTGCCCGGTGATAAATAAA | GCTTAGTCCAGCCAGAGTGG | 189 |
| chr11:30806197-30808221 | p3 | (TCT)8 | TCTTTTGTTTGTGGACGCTG | CGGGACAGGACTTCTCCATA | 191 |
| chr11:30806197-30808221 | p3 | (GAA)5 | TCTTTTGTTTGTGGACGCTG | CGGGACAGGACTTCTCCATA | 191 |
| chr11:30853527-30855543 | p2 | (AT)25 | TGGCGTGCTTCATAAAAGTG | AGCTTTGTCTTCGGTTTCCA | 139 |
| chr11:30853527-30855543 | p2 | (TA)8 | TGGCGTGCTTCATAAAAGTG | AGCTTTGTCTTCGGTTTCCA | 139 |
| chr11:30994248-30996266 | p3 | (TTG)6 | GAGGGGCAATTTTAGCACAA | ATGAGCAGTTTGTTGCGTTG | 291 |
| chr11:30994248-30996266 | p3 | (ATG)5 | GAGGGGCAATTTTAGCACAA | ATGAGCAGTTTGTTGCGTTG | 291 |
| chr11:31046527-31048539 | p2 | (GT)6 | CAAAACTCCCCCTTTCCTTC | TTTCAGTCTGCGTTACGGTG | 297 |
| chr11:31083263-31085278 | p5 | (TAAAA)3 | TTTCCGGGTATGGGTATGAA | TTCGGTTCAAATATCCCTGC | 296 |
| chr11:31106473-31108488 | p3 | (ACA)5 | TCTATAATATCGCTCGCGCC | CATATCCATGTTCGAACCCC | 219 |
| chr11:31106473-31108488 | p3 | (ACA)5 | TCTATAATATCGCTCGCGCC | CATATCCATGTTCGAACCCC | 219 |
| chr11:31364478-31366520 | p3 | (TAT)14 | GCCAAAGGCATCATTCTCAT | GTTCGCTGCCGAAGAAATAG | 171 |
| chr11:31364478-31366520 | p2 | (TC)6 | GCCAAAGGCATCATTCTCAT | GTTCGCTGCCGAAGAAATAG | 171 |
| chr11:31472877-31474891 | p2 | (TA)7 | GGCGGCTAAAAGTCTCATTG | CAAGTCTAAAAGCCTTGCCG | 206 |
| chr11:31518595-31520607 | p3 | (GAT)5 | ATCCATTTCATTCCCAACCA | TTCATGCATCCTCATCCTCA | 218 |
| chr11:31518595-31520607 | p2 | (TC)6 | ATCCATTTCATTCCCAACCA | TTCATGCATCCTCATCCTCA | 218 |
| chr11:31518595-31520607 | p5 | (TCGGT)3 | ATCCATTTCATTCCCAACCA | TTCATGCATCCTCATCCTCA | 218 |
| chr11:31568101-31570116 | p5 | (TATTA)3 | CCATCACATCCTAGCGGAAT | TCCTAGATGCTCGTAGCCGT | 196 |
| chr11:31576171-31578191 | p5 | (TCTTT)4 | TCGAGTTTGAGTTCGTCGTG | AGGAAACGACCCTGTGATTG | 272 |
| chr11:31621176-31623191 | p3 | (AAT)5 | ACGGACGTGTAAGTTGCCTT | GAAGCTGTCCCCTAAGCCTC | 204 |
| chr11:31641685-31643721 | p3 | (TTC)12 | ATTGTCACGTAGGCACCACA | TTGCCCTCGGTTAGTCAAAG | 161 |
| chr11:31641685-31643721 | p3 | (TTC)6 | ATTGTCACGTAGGCACCACA | TTGCCCTCGGTTAGTCAAAG | 161 |
| chr11:31651396-31653468 | p3 | (TTC)24 | TTTACCATGAAAGCCCAAGG | AAAGCACAAGGAGGCACTGT | 120 |
| chr11:31656591-31658613 | p2 | (TA)11 | AATTTGGATCCTGCAAGGTG | AACCTTCAACCTGGGCTTTT | 293 |
| chr11:31721911-31723925 | p2 | (TC)7 | TGCTCCATCCAATAACCCTC | AAATGGAGAATTTTGTGCCG | 205 |
| chr11:31775343-31777359 | p2 | (TA)8 | TTGGCTCTTGCTTGTTGATG | TCTGGCCAATTTTTCCTTTG | 212 |
| chr11:31793051-31795079 | p2 | (TA)14 | TGCTTGAGGACAAGCAAAGA | GCTTATCGACGCTTCAAAGG | 188 |
| chr11:31806193-31808208 | p5 | (TACAA)3 | TGAAACCGTGTTTGTTTCCA | ACCACGGAAAAATCACCAAA | 247 |
| chr11:31850544-31852568 | p2 | (TA)12 | CTCTGCAAGCCGATGTATCA | AACCCGCTGGTTCTCCTTAT | 132 |
| chr11:31933372-31935387 | p2 | (AG)7 | GATTTGCTGCCATGAAACCT | TCATCCGACCAAAGGCTAAC | 206 |
| chr11:31933449-31935464 | p2 | (AG)7 | GATTTGCTGCCATGAAACCT | TCATCCGACCAAAGGCTAAC | 129 |
| chr11:31933777-31935791 | p2 | (AG)7 | GTCGTCGTTGTCGTCGTCTA | CCTCTGGGGTTTCTGTTTCA | 240 |

| Marker ID | SSR type | SSR motif | FORWARD PRIMER (5'-3') | REVERSE PRIMER (5'-3') | size (bp) |
|---|---|---|---|---|---|
| chr11:31975679-31977703 | p6 | (AAAAGA)4 | AACAAAGCACCACAAAAGGC | GGCGACGATCGACTTGTATT | 135 |
| chr11:32082396-32084410 | p2 | (AT)7 | CATGATCTCAGGGAGGGAAA | CCCTCACGTTTTGCTTCATT | 168 |
| chr11:32082396-32084410 | p6 | (GAGCGT)3 | CATGATCTCAGGGAGGGAAA | CCCTCACGTTTTGCTTCATT | 168 |
| chr11:32136978-32139002 | p3 | (ACA)8 | TTTAGCAACAGCAGGACACG | TGGCCATGTATGGGCTTAAT | 285 |
| chr11:32146644-32148664 | p2 | (AT)10 | TTTCTTTGCCATTGTCGTGA | GGAAATGCCAAAAGGAAAAA | 273 |
| chr11:32181770-32183782 | p2 | (AT)6 | TTCGGATCCAAACTGAGAGC | TGCAAAGACGAGGGAGATTT | 101 |
| chr11:32204464-32206480 | p2 | (AG)8 | TCTTTTCGCTTCCTTTTGGA | CTCACTCAAACCCCGGTAAA | 113 |
| chr11:32253435-32255447 | p2 | (CA)6 | AAATGCACGACATGGTTCAA | TGTGTAAGGAGCCACCCTTC | 300 |
| chr11:32278922-32280943 | p3 | (TTA)7 | CAAGTCTGCCCATAAGGCAT | AAATTGCAAGGCAAGGTTTG | 120 |
| chr11:32278922-32280943 | p2 | (AT)12 | CAAGTCTGCCCATAAGGCAT | AAATTGCAAGGCAAGGTTTG | 120 |
| chr11:32352672-32354687 | p3 | (TTC)5 | GTGTTCGCCGCTTTATGAGT | CGTACGATCGGAACCAAAAT | 145 |
| chr11:32358855-32360870 | p5 | (TTGGT)3 | CGAGAAAATCCGAGGTTGAA | GGCCAGGAAGGTTTGATTTT | 111 |
| chr11:32358855-32360870 | p5 | (TTTGG)3 | CGAGAAAATCCGAGGTTGAA | GGCCAGGAAGGTTTGATTTT | 111 |
| chr11:32358855-32360870 | p6 | (ATTTTA)3 | CGAGAAAATCCGAGGTTGAA | GGCCAGGAAGGTTTGATTTT | 111 |
| chr11:32384198-32386213 | p5 | (GCATT)3 | GGCAAGGCAAATGATTGAGT | AGCATACAACAAGGCCCAAC | 136 |
| chr11:32436372-32438394 | p2 | (TA)11 | GGAACCCAAATCATGTGGTC | TCTCTCCATTTTCCCTGTGG | 274 |
| chr11:32459646-32461710 | p2 | (CA)6 | AAAAATGGCGAGATGACGAG | GTGTGTGGAAATTGTGCTGC | 240 |
| chr11:32459646-32461710 | p2 | (AT)32 | AAAAATGGCGAGATGACGAG | GTGTGTGGAAATTGTGCTGC | 240 |
| chr11:32540442-32542457 | p5 | (TTAGT)3 | CGGCCCACTAAAGCATTAAA | AAATTGGAAATGAGCGTTGG | 282 |
| chr11:32574373-32576389 | p2 | (TA)8 | GGGACCAAGCAAATCAAAGA | GATGAGCGGTTTATTGCGTT | 121 |
| chr11:32574373-32576389 | p2 | (TA)17 | GGGACCAAGCAAATCAAAGA | GATGAGCGGTTTATTGCGTT | 121 |
| chr11:32722768-32724784 | p2 | (AC)8 | GAGTTGCTTCAAGAGCTGGG | CAAGCTTCTGGCACAAAACA | 144 |
| chr11:32727287-32729302 | p5 | (ACCCC)3 | CCTCCATACCGAACACATCC | GATTCGGACATGAGGCAAAT | 219 |
| chr11:32727287-32729302 | p3 | (CAC)5 | CCTCCATACCGAACACATCC | GATTCGGACATGAGGCAAAT | 219 |
| chr11:32733260-32735275 | p5 | (CTTTT)3 | CGAGGAATATCACGAACCGT | GACATGCCCATTGTGAAGTG | 142 |
| chr11:32733260-32735275 | p2 | (CA)6 | CGAGGAATATCACGAACCGT | GACATGCCCATTGTGAAGTG | 142 |
| chr11:32750149-32752161 | p6 | (AATAAA)3 | TATCTCACCAAGCACACCCA | CATCGGGATCTGGAGAAAAA | 129 |
| chr11:32750149-32752161 | p2 | (CT)6 | TATCTCACCAAGCACACCCA | CATCGGGATCTGGAGAAAAA | 129 |
| chr11:32832957-32834972 | p5 | (AACCC)3 | TCACGTGGGGGATTGTTAGT | TTGGGCAACTTCAAAACTCC | 199 |
| chr11:32884867-32886901 | p2 | (CT)10 | GCCATGGCTGTAGCTCTCTC | ATTGGATGGACCCCACTACA | 168 |
| chr11:32884867-32886901 | p2 | (AT)17 | GCCATGGCTGTAGCTCTCTC | ATTGGATGGACCCCACTACA | 168 |
| chr11:32886934-32888949 | p3 | (AAC)5 | TGTTCAGCAACTGCCTTCTG | ACTGACAATCAATCCCCCAA | 223 |
| chr11:32995416-32997431 | p5 | (AAAAT)3 | CCATGGTACCCCGTCAGTAT | TCCAATCCTCTATGATGCCC | 124 |
| chr11:33015645-33017705 | p3 | (TAT)20 | TCTTGAGGGAATGGAAATGC | AAGGCACGAAGTGGCTCTAA | 139 |
| chr11:33027218-33029294 | p2 | (AT)38 | GGTTAATGTTGCGATGGCTT | TTGCCTATCTACCATGCCGT | 278 |
| chr11:33027218-33029294 | p2 | (AT)7 | GGTTAATGTTGCGATGGCTT | TTGCCTATCTACCATGCCGT | 278 |

| Marker ID | SSR type | SSR motif | FORWARD PRIMER (5'-3') | REVERSE PRIMER (5'-3') | size (bp) |
|---|---|---|---|---|---|
| chr11:33135775-33137791 | p2 | (TA)12 | GTCGTCCGTTGTCCCTTAAA | CCATGTGTTTGGAGATGGTG | 244 |
| chr11:33135775-33137791 | p3 | (ATA)24 | GTCGTCCGTTGTCCCTTAAA | CCATGTGTTTGGAGATGGTG | 244 |
| chr11:33135775-33137791 | p2 | (TA)8 | GTCGTCCGTTGTCCCTTAAA | CCATGTGTTTGGAGATGGTG | 244 |
| chr11:33135775-33137791 | p2 | (AT)31 | GTCGTCCGTTGTCCCTTAAA | CCATGTGTTTGGAGATGGTG | 244 |
| chr11:33143058-33145070 | p2 | (AT)6 | GAAGGGGCCATCCTTATTGT | TATTTGAACCGTGCCGTGTA | 235 |
| chr11:33187084-33189100 | p4 | (TTCT)4 | TTCAAGAAAGCTGTTGCCCT | CATCCCACTTCTGTCCTGGT | 262 |
| chr11:33313153-33315169 | p4 | (TTTC)4 | GCGCTTTGAAACCTTAATCG | CCCATCCAAACAGGCTCTTA | 279 |
| chr11:33407889-33409905 | p2 | (CA)8 | TGGGAGGAATTATGAGTCGG | AGGGCTAGAGGAAAAGGCAG | 188 |
| chr11:33476690-33478730 | p2 | (AT)20 | TTAGGCCTCCTTTGGGTTTT | TACCTGAGGAGCCCATTTTG | 109 |
| chr11:33507872-33509890 | p3 | (GAT)6 | CACCTAACCTTTGCGTTGGT | TGTGGCATCTATGTGGCAGT | 166 |
| chr11:33513712-33515727 | p3 | (GAT)5 | ACACCTAACCTTTGCGTTGG | TGTGGCATCTATGTGGCAGT | 165 |
| chr11:33589427-33591475 | p2 | (AT)6 | TCCATTTATCCAGGAGCGAC | CTTCGTGAGTCGGTCAACAA | 140 |
| chr11:33589427-33591475 | p2 | (AT)24 | TCCATTTATCCAGGAGCGAC | CTTCGTGAGTCGGTCAACAA | 140 |
| chr11:33595430-33597478 | p2 | (AT)6 | AGCCCATTACTCCCGCTATT | TCCATGTTGATCACTTCCCA | 121 |
| chr11:33595430-33597478 | p2 | (AT)24 | AGCCCATTACTCCCGCTATT | TCCATGTTGATCACTTCCCA | 121 |
| chr11:33595430-33597478 | p2 | (AT)8 | AGCCCATTACTCCCGCTATT | TCCATGTTGATCACTTCCCA | 121 |
| chr11:33670496-33672510 | p2 | (AT)7 | ATCGTTTTCCACTGGTTTCG | TGAAGCTCATAAACAGGGGG | 195 |
| chr11:33670496-33672510 | p6 | (CATATA)3 | ATCGTTTTCCACTGGTTTCG | TGAAGCTCATAAACAGGGGG | 195 |
| chr11:33683045-33685059 | p2 | (TA)8 | TTTGATAGCCGAAGGGCTTA | TCCGAGGACTGAACTGCTCT | 168 |
| chr11:33683045-33685059 | p2 | (TA)7 | TTTGATAGCCGAAGGGCTTA | TCCGAGGACTGAACTGCTCT | 168 |
| chr11:33689270-33691346 | p5 | (ACATA)3 | GTAGACAGGGAGGTGACCCA | AGACATATGGACGGGACAGG | 246 |
| chr11:33689347-33691395 | p5 | (ACATA)3 | GTAGACAGGGAGGTGACCCA | AGACATATGGACGGGACAGG | 169 |
| chr11:33850034-33852100 | p2 | (TA)33 | ACCCGCTGTTGGTATTTCTG | CATCGGTCATCGAACTCTCA | 271 |
| chr11:33861039-33863054 | p5 | (TTCGG)3 | AGACCAATACGGTGCGAATC | GAACTAATTCGGTGCGGTGT | 207 |
| chr11:33918198-33920218 | p5 | (TATTT)4 | TCGAGGGCAATAGCTCAACT | GCTAAAAGAGTGACGAGGCG | 300 |
| chr11:33976003-33978027 | p5 | (TTCTT)5 | TACAGCACGCTTGTTTTTGC | CCTCGGCTTGACACATTTTT | 278 |
| chr11:34043624-34045642 | p3 | (TGT)6 | TGTCTCATCTCCTCCCATCC | TGGTCCAAGTTGGGATCATT | 194 |
| chr11:34133033-34135051 | p6 | (TTTTTA)3 | CGCGTAGAGTAAAATTCGCC | AAAGTTGGTATCCGTCGTGC | 183 |
| chr11:34427326-34429344 | p3 | (CAA)6 | TACTGCCGAGAAAGGGATGT | TTCCTTCGTCAAATTCAGGC | 178 |
| chr11:34427326-34429344 | p5 | (TTTAT)3 | TACTGCCGAGAAAGGGATGT | TTCCTTCGTCAAATTCAGGC | 178 |
| chr11:34428059-34430074 | p3 | (CAA)6 | CAACAACCCAGTGAAATCCC | AATGACTCCCGGAAACAGC | 279 |
| chr11:34428059-34430074 | p5 | (TTTAT)3 | CAACAACCCAGTGAAATCCC | AATGACTCCCGGAAACAGC | 279 |
| chr11:34428224-34430257 | p3 | (CAA)6 | CAACAACCCAGTGAAATCCC | AATGACTCCCGGAAACAGC | 114 |
| chr11:34428224-34430257 | p5 | (TTTAT)3 | CAACAACCCAGTGAAATCCC | AATGACTCCCGGAAACAGC | 114 |
| chr11:34511973-34514007 | p5 | (TTGGT)3 | TCCATGTAAACCCCTACCCA | GCAACAAATACACGCGAAAA | 101 |
| chr11:34531674-34533695 | p3 | (AAT)7 | GATCCGAAACAGAACCCAGA | TTTTAGCCGAGAAATCCCCT | 193 |

| Marker ID | SSR type | SSR motif | FORWARD PRIMER (5'-3') | REVERSE PRIMER (5'-3') | size (bp) |
|---|---|---|---|---|---|
| chr11:34609997-34612012 | p5 | (CATTT)3 | AAAGAAGTCGCTTCGGTCAA | GCCACACACGACTAAAAGCA | 172 |
| chr11:34673926-34675938 | p2 | (TG)6 | CGGATGAGAAGGTGTTGGAT | TCCTCCCAATAGTACTGCCG | 292 |
| chr11:34772746-34774762 | p2 | (AT)8 | ACATGCGCATCAATCCATTA | AAATGACATGGCGGAATAGC | 116 |
| chr11:34772746-34774762 | p2 | (AT)6 | ACATGCGCATCAATCCATTA | AAATGACATGGCGGAATAGC | 116 |
| chr11:34805420-34807435 | p3 | (TGT)5 | AAAGGGTTATGACACGCCAC | TTCTTCACCGACCTTGCTCT | 169 |
| chr11:34910815-34912833 | p2 | (AT)9 | GGGCACCCTCTTAGCTTACC | TGTTGGGCCCATATTTTTGT | 258 |
| chr11:34910815-34912833 | p3 | (TAT)22 | GGGCACCCTCTTAGCTTACC | TGTTGGGCCCATATTTTTGT | 258 |
| chr11:34940014-34942032 | p6 | (AAAAAT)3 | TGCCCAAAATCATCCTCATT | GGCCATTGTCCACTGAGTTT | 204 |
| chr11:34955249-34957267 | p2 | (AT)9 | GCTTATGCATATGGGGCAGT | TCAAGTCCCCGAAGACAATC | 145 |
| chr11:34981137-34983157 | p2 | (AT)10 | AAATGTCCGCAGCTTCTTGT | GGCCGTGACAGAAGTGGTAT | 137 |
| chr11:35003964-35005976 | p2 | (CG)6 | ATAACCCCCACGCATTAACA | ACTCTTCCATTTGGCAGTCG | 193 |
| chr11:35041153-35043167 | p2 | (TC)7 | CACCTGGAGCTTTGTCTTCC | CACTCACGGATGCAGAAGAA | 106 |
| chr11:35041153-35043167 | p5 | (TTTGT)3 | CACCTGGAGCTTTGTCTTCC | CACTCACGGATGCAGAAGAA | 106 |
| chr11:35041153-35043167 | p4 | (TTTG)4 | CACCTGGAGCTTTGTCTTCC | CACTCACGGATGCAGAAGAA | 106 |
| chr11:35041920-35043936 | p2 | (TC)7 | CGTTCCTCTCTCTCCATTGC | AACTCACAAACCCCCACAAG | 296 |
| chr11:35041920-35043936 | p5 | (TTTGT)3 | CGTTCCTCTCTCTCCATTGC | AACTCACAAACCCCCACAAG | 296 |
| chr11:35041920-35043936 | p4 | (TTTG)4 | CGTTCCTCTCTCTCCATTGC | AACTCACAAACCCCCACAAG | 296 |
| chr11:35041920-35043936 | p3 | (GAA)5 | CGTTCCTCTCTCTCCATTGC | AACTCACAAACCCCCACAAG | 296 |
| chr11:35081532-35083558 | p5 | (ACCGG)3 | TGGTGACGGTTTGTCTTTCA | CTGCAATCATCATGGAGGTG | 150 |
| chr11:35091669-35093684 | p3 | (AAC)5 | TTAGGCAATCCAATGAAGCC | TTGGCAAATGAACAGCTCAC | 267 |
| chr11:35097099-35099114 | p5 | (ATTTT)3 | GCCTGGGAGTCCAACAATTA | AGGTGAGTGGGTTAGCCCTT | 124 |
| chr11:35215257-35217275 | p6 | (ACACCC)3 | GCAACTCTACCCACTTCCCA | AACTGGATTGGACATGAGCC | 118 |
| chr11:35368866-35370928 | p2 | (TA)31 | GGGTTCATAACCCCAATGTG | CAACAAGCCAGTCATAGGCA | 221 |
| chr11:35368866-35370928 | p2 | (TA)6 | GGGTTCATAACCCCAATGTG | CAACAAGCCAGTCATAGGCA | 221 |
| chr11:35368866-35370928 | p2 | (TA)7 | GGGTTCATAACCCCAATGTG | CAACAAGCCAGTCATAGGCA | 221 |
| chr11:35369995-35372007 | p2 | (TA)6 | TACCCAACTTAAACTGCCCG | TCAAACCGGTAAATCGGAAC | 125 |
| chr11:35369995-35372007 | p2 | (TA)7 | TACCCAACTTAAACTGCCCG | TCAAACCGGTAAATCGGAAC | 125 |
| chr11:35369995-35372007 | p2 | (TA)6 | TACCCAACTTAAACTGCCCG | TCAAACCGGTAAATCGGAAC | 125 |
| chr11:35383185-35385221 | p2 | (AT)13 | TTCTCCTACCAACCACCGAC | CACTTATAAGCAGGAGGCCG | 156 |
| chr11:35383185-35385221 | p2 | (TA)18 | TTCTCCTACCAACCACCGAC | CACTTATAAGCAGGAGGCCG | 156 |
| chr11:35419756-35421774 | p6 | (AGAGGA)3 | TTTGCTTCAGAGGGGAACAC | TGAGTCCTGGGCTAAGTGCT | 293 |
| chr11:35442270-35444294 | p3 | (TCA)5 | CGAGTTCTAGGCGGTTTGTC | GGGGTGTGCTTAATGAGGAA | 124 |
| chr11:35442270-35444294 | p6 | (CCGGTT)4 | CGAGTTCTAGGCGGTTTGTC | GGGGTGTGCTTAATGAGGAA | 124 |
| chr11:35517671-35519689 | p3 | (AAT)6 | CCAAATCACCGCCCTATCTA | GCAAGAAGCAAAAGTCGTCC | 146 |
| chr11:35552117-35554132 | p3 | (ACA)5 | CTTGCAAAATGGCCAAAGAT | TTTGCAAAACTGACCAGTGC | 288 |
| chr11:35775010-35777028 | p3 | (CAA)6 | CACAGCATAACACCGGAATG | TTCGGGTCTATACTCGTCCG | 270 |

| Marker ID | SSR type | SSR motif | FORWARD PRIMER (5'-3') | REVERSE PRIMER (5'-3') | size (bp) |
|---|---|---|---|---|---|
| chr11:35822321-35824337 | p2 | (AT)8 | TAGACACGTGGCCGATGTTA | AGCCATATCTGGTGCAATCC | 240 |
| chr11:35889222-35891240 | p2 | (CA)9 | GCCCATTCTGTGGTTTCACT | AATGTTATCCTGCGCCAAAG | 100 |
| chr11:35940907-35942929 | p2 | (AT)11 | TGCATTCCTACCTCAAGGCT | TCCGAGAACTTTTTGCTTCG | 176 |
| chr11:35970030-35972042 | p2 | (TA)6 | TTGACGACGACATTGATGGT | TTAACCAGTGGGTCGGAAAG | 137 |
| chr11:35986807-35988822 | p5 | (TATAA)3 | GACAGACGGCCCACATTAGT | AGAGAGAAGGGAGAGTCGGC | 195 |
| chr11:36193118-36195133 | p3 | (CAA)5 | GGACCAAAACTAGACGCCAA | GATACTTGATGAAGCCGGGA | 275 |
| chr11:36196015-36198033 | p3 | (TAA)10 | TAGGTTACCGAATTGACGCC | TTAGGTGACTGAACCGCCTT | 123 |
| chr11:36212519-36214534 | p5 | (TATTT)3 | GCGTACTTCCGCCAAATAAA | AATGTATCCGTGTGCACCAA | 172 |
| chr11:36212519-36214534 | p6 | (ATTTCT)3 | GCGTACTTCCGCCAAATAAA | AATGTATCCGTGTGCACCAA | 172 |
| chr11:36353275-36355289 | p2 | (TA)7 | ATGGCTTATGGTGAGTTGGC | AGGCCAATAGCGGAAAGAAT | 296 |
| chr11:36470783-36472798 | p5 | (AAATC)3 | CTTTGGTTTCCTGTTGGCAT | GACGAAAGAAGGCAAGCAAG | 279 |
| chr11:36513317-36515347 | p5 | (AATTC)6 | ATCGGAGTGACGTAAGGTCG | AGCCAGGCATGAGAGAGTGT | 212 |
| chr11:36513317-36515347 | p5 | (ACACA)3 | ATCGGAGTGACGTAAGGTCG | AGCCAGGCATGAGAGAGTGT | 212 |
| chr11:36513317-36515347 | p2 | (TA)8 | ATCGGAGTGACGTAAGGTCG | AGCCAGGCATGAGAGAGTGT | 212 |
| chr11:36575993-36578009 | p2 | (AT)8 | TGGGATGGAAAGCAAACTTC | CCCTTCGGCAGAAAATTACA | 152 |
| chr11:36747290-36749328 | p2 | (TA)19 | CAACTGGCTGAACGACTGAA | CTCTGAGTGCGTCCACTTGA | 183 |
| chr11:36781700-36783712 | p2 | (TA)6 | CTTGGGTTGGGAAGTTGAAA | GGCTCATCAGTTTGGGTCAT | 100 |
| chr11:36825816-36827831 | p5 | (AACCG)3 | GGTGTGGAGGTCGAGGATTA | CCCTGGCTCAAGTAAAGCAG | 201 |
| chr11:36825816-36827831 | p5 | (AACCG)3 | GGTGTGGAGGTCGAGGATTA | CCCTGGCTCAAGTAAAGCAG | 201 |
| chr11:36863286-36865306 | p4 | (GGTG)5 | CAAATCTTGAATTGTGGCCC | CAGGGTGTTTGATGATGTGC | 106 |
| chr11:37033453-37035469 | p2 | (AT)8 | GTGAAGTGTGCTCCTCGTCA | CTATCGAGACTGACCCCTCG | 156 |
| chr11:37054104-37056124 | p2 | (TC)10 | AAAATCGCTATTTGCCCAAC | AATTCAGTGTGCCGGGTAAC | 127 |
| chr11:37079914-37081929 | p5 | (TTATT)3 | GCAGGTTCACTTTGGGATGT | GCTTCCCACCTACCACTTCA | 146 |
| chr11:37200058-37202070 | p2 | (TA)6 | CCTATTGGTGCAACCAACCT | GCAGAAGAGACAACCCTTGC | 145 |
| chr11:37208327-37210349 | p2 | (TA)11 | AGAGTGCCACTGTGCATACG | ATCCAACCCCGAGAACTCTT | 143 |
| chr11:37234650-37236668 | p6 | (ACATAT)3 | TCGGGTGTTCTTAATGCTCC | TCAATGGCAACAAGAATGGA | 154 |
| chr11:37234650-37236668 | p3 | (CAA)6 | TCGGGTGTTCTTAATGCTCC | TCAATGGCAACAAGAATGGA | 154 |
| chr11:37321981-37323993 | p2 | (TA)7 | GAGAGCGCCATTTTAGCAAG | TGTAAGTTTTGGTTGCGTCG | 204 |
| chr11:37321981-37323993 | p2 | (AT)6 | GAGAGCGCCATTTTAGCAAG | TGTAAGTTTTGGTTGCGTCG | 204 |
| chr11:37352942-37354960 | p2 | (AG)6 | TCTGAAGGAGAAAGGACCGA | TCCTAAACCATTCGACCCAG | 284 |
| chr11:37352962-37355010 | p2 | (AG)6 | TCTGAAGGAGAAAGGACCGA | TCCTAAACCATTCGACCCAG | 264 |
| chr11:37361023-37363038 | p5 | (ATGCA)3 | ATTGTTGACCGGATGGTGAT | CTTTTGAGTTGAGGGTTCGC | 247 |
| chr11:37396437-37398452 | p3 | (ATA)5 | TTCAGGAACCTCCCAACAAG | TATAGCCCCGCATTTAGGTG | 139 |
| chr11:37396437-37398452 | p2 | (AT)22 | TTCAGGAACCTCCCAACAAG | TATAGCCCCGCATTTAGGTG | 139 |
| chr11:37520900-37522912 | p5 | (ATTTC)3 | CAAGAACGTGCTCAACCAAA | TGATGTGGAAGTGATCGGAA | 177 |
| chr11:37520900-37522912 | p2 | (CT)6 | CAAGAACGTGCTCAACCAAA | TGATGTGGAAGTGATCGGAA | 177 |

| Marker ID | SSR type | SSR motif | FORWARD PRIMER (5'-3') | REVERSE PRIMER (5'-3') | size (bp) |
|---|---|---|---|---|---|
| chr11:37561479-37563503 | p2 | (TA)12 | CAGGAAAGGCCCAACATAGA | TTCTGATAGCCTTGTCGGCT | 193 |
| chr11:37659297-37661333 | p2 | (TA)12 | CTTCGCGCGGTATGAATAAT | ACAACATGGTATCGGCAACA | 100 |
| chr11:37672918-37674933 | p5 | (GTGCG)3 | GCCGTGTGTAGTTCTGGGAT | CTATTTCTTCGGCAGCGAAC | 277 |
| chr11:37694219-37696243 | p2 | (AT)12 | GTTTGGAAAGCCACCTGGTA | TGAGATGACATGCCAAAGGA | 239 |
| chr11:37713714-37715746 | p2 | (AT)8 | GAATCTCTTTGTGGCGCTTC | TTATTTTCCATGAAAGCCCG | 271 |
| chr11:37713714-37715746 | p2 | (AT)16 | GAATCTCTTTGTGGCGCTTC | TTATTTTCCATGAAAGCCCG | 271 |
| chr11:37713714-37715746 | p2 | (AT)6 | GAATCTCTTTGTGGCGCTTC | TTATTTTCCATGAAAGCCCG | 271 |
| chr11:37872390-37874411 | p3 | (AAC)7 | CGACCCGTTGACAGGTTTAT | TCTGGACCTACTCGAGGCAT | 187 |
| chr11:38202323-38204335 | p2 | (AG)6 | ACCAGGTGGTGCTGAAACTC | TTAGGATCTGAATCCCGTGG | 152 |
| chr11:38227094-38229112 | p3 | (AAG)6 | AGCACTAGGATTGCGGAAGA | GTGCTAAGCCTTAATTGGCG | 272 |
| chr11:38408632-38410650 | p2 | (TA)9 | GGGTAGGATCAGAGAAGGGG | TTCTACACCGACAACCACCA | 102 |
| chr11:38451038-38453056 | p2 | (TA)9 | GGGTAGGATCAGAGAAGGGG | TTCTACACCGACAACCACCA | 102 |
| chr11:38464628-38466640 | p2 | (AC)6 | AAACTGGATAGGGTGGGGTC | TCAACTGTGGGAGCTTTGTG | 237 |
| chr11:38472958-38474974 | p4 | (CATA)4 | TTGTAGGAACCGAACGAACC | GAGGCTTTGCAGACAGTTCC | 121 |
| chr11:38494538-38496554 | p2 | (AT)8 | CCAAGCCTAAAGTCGACGAG | TTGTGGGGTTGGGTGTTTAT | 174 |
| chr11:38494538-38496554 | p2 | (TA)6 | CCAAGCCTAAAGTCGACGAG | TTGTGGGGTTGGGTGTTTAT | 174 |
| chr11:38501031-38503069 | p2 | (AT)15 | AATACGCGTTGGGACGTAAC | TTAATGAAATGGGACCAGGC | 100 |
| chr11:38501031-38503069 | p2 | (CT)19 | AATACGCGTTGGGACGTAAC | TTAATGAAATGGGACCAGGC | 100 |
| chr11:38557554-38559566 | p2 | (TA)6 | TTTTGAACCTCGGAGCATCT | TTTGGAGGGTTTCCACTTTG | 151 |
| chr11:38560410-38562424 | p2 | (AT)7 | GGCCTATGTGTGCCAAGTCT | TACGTACCCGATACCCCAAA | 177 |
| chr11:38565690-38567708 | p6 | (TTCCTA)3 | CGCGACGGTATATTCGGTAT | GATCACGGAGGTTTGCAGTT | 286 |
| chr11:38597454-38599466 | p2 | (AG)6 | CAAAGAGGCCGTGATTGATT | TACATCTTAGTCGCCCCTGG | 179 |
| chr11:38597454-38599466 | p5 | (TTCTT)3 | CAAAGAGGCCGTGATTGATT | TACATCTTAGTCGCCCCTGG | 179 |
| chr11:38659573-38661617 | p2 | (TA)22 | GCGGATCGTATTCACTTGCT | GTTCAACAACCACCTTCGCT | 210 |
| chr11:38659573-38661617 | p2 | (TA)6 | GCGGATCGTATTCACTTGCT | GTTCAACAACCACCTTCGCT | 210 |
| chr11:38659573-38661617 | p2 | (TA)7 | GCGGATCGTATTCACTTGCT | GTTCAACAACCACCTTCGCT | 210 |
| chr11:38659763-38661783 | p2 | (TA)22 | CTTAGTGGCGAGGATGCAGT | GTTCAACAACCACCTTCGCT | 102 |
| chr11:38659763-38661783 | p2 | (TA)6 | CTTAGTGGCGAGGATGCAGT | GTTCAACAACCACCTTCGCT | 102 |
| chr11:38659763-38661783 | p2 | (TA)7 | CTTAGTGGCGAGGATGCAGT | GTTCAACAACCACCTTCGCT | 102 |
| chr11:38681164-38683179 | p3 | (TCA)5 | CATCGTGACGACCTTTTCCT | TCTTACCGCTGATTTGGGTC | 242 |
| chr11:38681164-38683179 | p5 | (TGAAA)3 | CATCGTGACGACCTTTTCCT | TCTTACCGCTGATTTGGGTC | 242 |
| chr11:38723201-38725233 | p4 | (ATTT)8 | ATGTTAGTCCCGGAGTCGTG | CCCAAGGTGACTCACGAAGT | 183 |
| chr11:38805187-38807209 | p2 | (AT)11 | TGGATAATTGTCACGACCCA | TGATGTTGCCTTGTCAGCTC | 216 |
| chr11:38830768-38832790 | p2 | (AT)11 | TGGATAATTGTCACGACCCA | TGATGTTGCCTTGTCAGCTC | 216 |
| chr11:38881422-38883448 | p2 | (TC)13 | AGAACTACACGCGGCTTGAT | GATGAAAGAATTTTCGGGCA | 114 |
| chr11:38932478-38934498 | p4 | (TATT)5 | GAGAGAGGTTTCCCTTTCGG | AATTCTCTCGCACCAACCAC | 151 |

| Marker ID | SSR type | SSR motif | FORWARD PRIMER (5'-3') | REVERSE PRIMER (5'-3') | size (bp) |
|---|---|---|---|---|---|
| chr11:38977537-38979552 | p2 | (AT)7 | CTGCCCGGAACACTGTAAAT | AACCAGCGTCGACTTATTGG | 229 |
| chr11:38977554-38979590 | p2 | (AT)7 | CTGCCCGGAACACTGTAAAT | AACCAGCGTCGACTTATTGG | 212 |
| chr11:38982188-38984203 | p5 | (TTTAT)3 | CGTGTGCTTCTTGAGTTGGA | TCGCTGAAAGCAAATTGTTG | 130 |
| chr11:39031647-39033662 | p3 | (ACA)5 | TTGTAACCAGAGGCGAATCC | TAAACCACCGAGCCAACTTC | 100 |
| chr11:39102211-39104244 | p3 | (GAA)5 | TGCTGATGTGGATAAGAGCG | ATGTGGCATCTATGTGGCAG | 249 |
| chr11:39130470-39132485 | p5 | (GGGTC)3 | TTTTGATATTGCCGTCGTGA | TGGGGATTTCCATCCACTTA | 299 |
| chr11:39155027-39157039 | p3 | (TAT)5 | AACCTCTTCCTCCGTCCATT | CACCTTTACCTTCCAACCCA | 295 |
| chr11:39155027-39157039 | p2 | (AG)22 | AACCTCTTCCTCCGTCCATT | CACCTTTACCTTCCAACCCA | 295 |
| chr11:39155069-39157089 | p3 | (TAT)5 | AACCTCTTCCTCCGTCCATT | CACCTTTACCTTCCAACCCA | 253 |
| chr11:39155069-39157089 | p2 | (AG)22 | AACCTCTTCCTCCGTCCATT | CACCTTTACCTTCCAACCCA | 253 |
| chr11:39157856-39159874 | p6 | (ATATGT)3 | TTCGGATGTATCTCCGGGTA | ACTTTTGGCCGATGAAGAAA | 243 |
| chr11:39374173-39376185 | p2 | (CT)6 | TCTTCTTTTCCCCGGATTCT | GAAACCAGTTGGACATGGCT | 226 |
| chr11:39434221-39436260 | p3 | (GAA)13 | TAACCAGTTGCACAATGGGA | TGGCAATGGTAGTGGTGAAA | 242 |
| chr11:39511415-39513429 | p2 | (AT)7 | AAGTATCAACCGTGGCTTCG | TTGCTCCGCTTCTGAGTTTT | 126 |
| chr11:39525219-39527237 | p5 | (TCATG)3 | ATCGGAGTGACGTAAGGTCG | CGATGCCTTATGACCTGGTT | 261 |
| chr11:39525219-39527237 | p3 | (CAA)6 | ATCGGAGTGACGTAAGGTCG | CGATGCCTTATGACCTGGTT | 261 |
| chr11:39528494-39530512 | p6 | (TGAAGA)3 | TGACTTGGTTTGGTGTTGGA | GACTGCCACATCAGCAAAAA | 120 |
| chr11:39528494-39530512 | p5 | (TTGGT)3 | TGACTTGGTTTGGTGTTGGA | GACTGCCACATCAGCAAAAA | 120 |
| chr11:39552770-39554788 | p3 | (TAA)6 | AAGGTTGGCCATAAACACCA | GCCACAATTGGAGAACAGGT | 254 |
| chr11:39748392-39750407 | p3 | (TTG)5 | AAGCCCTCCAAAAGGATGAT | TTTGGCAGCTTGGAGTCTTT | 189 |
| chr11:39748392-39750407 | p4 | (TATG)4 | AAGCCCTCCAAAAGGATGAT | TTTGGCAGCTTGGAGTCTTT | 189 |
| chr11:39808861-39810875 | p2 | (AT)7 | GCACAGTCGGACCCTATCTC | GGCAAGAGGGTGTCCAATAA | 271 |
| chr11:39823083-39825105 | p5 | (TTATA)3 | GACAAAGCTCATTTGAGGGG | TGGCCATTGAAAACAATTCA | 254 |
| chr11:39850423-39852438 | p5 | (TTTTG)3 | CAGAGCAATCACCATAGGCA | ACGGCGTGGGTACATATCAT | 123 |
| chr11:39960550-39962571 | p3 | (AAT)7 | CTCTGTTCCGATTTCCCTCA | AATGAAGTTGTAAACGGCGG | 236 |
| chr11:40070142-40072166 | p2 | (TA)12 | TATGCGAATGGACGAAATCA | TCAACTCCCAAACGTCAACA | 221 |
| chr11:40110892-40112910 | p2 | (TA)9 | ACGGGGACCGTTATATTTCC | TAAGGCCACCAGTGCTTTCT | 174 |
| chr11:40110892-40112910 | p2 | (AT)8 | ACGGGGACCGTTATATTTCC | TAAGGCCACCAGTGCTTTCT | 174 |
| chr11:40110892-40112910 | p2 | (AT)6 | ACGGGGACCGTTATATTTCC | TAAGGCCACCAGTGCTTTCT | 174 |
| chr11:40124624-40126638 | p2 | (AC)7 | TTTCATTATCCATTCCCCCA | TCACCCCGTACTACCGTTTC | 242 |
| chr11:40124624-40126638 | p3 | (ACA)5 | TTTCATTATCCATTCCCCCA | TCACCCCGTACTACCGTTTC | 242 |
| chr11:40174554-40176574 | p5 | (TTCAT)4 | TGTGGAGAAACAACAACCGA | ATTTCGCCAGAGTTTTCCCT | 141 |
| chr11:40253731-40255752 | p3 | (AAT)7 | ATCCCGAATTAGGGTTTTCG | TCCCTCGTCTCTCTTCTCCA | 280 |
| chr11:40290193-40292213 | p5 | (TGATG)4 | ACCACCAGTCATGTTGCTCA | TCATGGGATCACCTGTGCTA | 158 |
| chr11:40358298-40360313 | p5 | (ATTGC)3 | CCCTCTCCGAGCATTGATTA | TTGACGAATTCCCGAAAAAG | 125 |
| chr11:40381012-40383048 | p3 | (ATT)12 | ATATTTCAGTGCAGGCACCC | ACACTTGCAACCCCTGTTTC | 189 |

| Marker ID | SSR type | SSR motif | FORWARD PRIMER (5'-3') | REVERSE PRIMER (5'-3') | size (bp) |
|---|---|---|---|---|---|
| chr11:40432786-40434804 | p6 | (TATATG)3 | TAAACAGGGAAAATGCTGCC | GCTACGCTTATTCGTCCGAG | 208 |
| chr11:40477732-40479759 | p3 | (ATT)9 | TTTGAGGTCGAAACTCCAGG | CCAGCCTCATACGGAAACAT | 140 |
| chr11:40574117-40576132 | p3 | (TTA)5 | ATTTTTGCATACTCACCCGC | TTGCCAAATATGGCGAAGAT | 139 |
| chr11:40683638-40685658 | p2 | (TA)10 | ACGAGCTTCAACAGCGAAAT | AGCCATCAAGGCTACGAAAA | 121 |
| chr11:40768362-40770398 | p2 | (TA)21 | ATCTTTGGCACGAATCTTGG | AAATGTCGTAACCTCGGACG | 267 |
| chr11:40768362-40770398 | p2 | (AC)6 | ATCTTTGGCACGAATCTTGG | AAATGTCGTAACCTCGGACG | 267 |
| chr11:40768414-40770442 | p2 | (TA)21 | ATCTTTGGCACGAATCTTGG | AAATGTCGTAACCTCGGACG | 215 |
| chr11:40768414-40770442 | p2 | (AC)6 | ATCTTTGGCACGAATCTTGG | AAATGTCGTAACCTCGGACG | 215 |
| chr11:40831612-40833630 | p2 | (TA)31 | GCCATTAAGGCTCGGTGTTA | GGGCGTGACAATTGACTTCT | 290 |
| chr11:40831612-40833630 | p6 | (AACTGA)3 | GCCATTAAGGCTCGGTGTTA | GGGCGTGACAATTGACTTCT | 290 |
| chr11:40869400-40871418 | p3 | (TTG)6 | CGGAGGGTCTGACACAGATT | CAATAGCCCAAGGTGGAAAA | 241 |
| chr11:40869400-40871418 | p6 | (ATTTTT)3 | CGGAGGGTCTGACACAGATT | CAATAGCCCAAGGTGGAAAA | 241 |
| chr11:40949951-40951966 | p5 | (AATTA)3 | TAGTCGAAGTGCACAGTGGC | AACTGCCAGGTCGTATTCCA | 294 |
| chr11:40956506-40958520 | p2 | (AT)7 | ATCCAGCGTTTGATAATGGG | CACCACTTCCGCTCTTCTTC | 200 |
| chr11:41027791-41029830 | p3 | (TGC)6 | TGAAGCTTGGGTATTTTCGG | CAATCAAACGGCAAGTAGCA | 127 |
| chr11:41091770-41093833 | p3 | (AAT)21 | AATCCAAACAGATGCCAAGG | CGCTCCCCTCTCTCTTTCTT | 199 |
| chr11:41142386-41144404 | p2 | (AT)9 | CTGGAGGTTGCTTTCAGTCC | CTGGTGGGCCTATCCAGTTA | 243 |
| chr11:41203648-41205704 | p2 | (AT)28 | AAATGAACCCCAACTGCTTC | CATGCAATGTCACTTCACCC | 297 |
| chr11:41203648-41205704 | p2 | (AT)10 | AAATGAACCCCAACTGCTTC | CATGCAATGTCACTTCACCC | 297 |
| chr11:41258415-41260430 | p5 | (TAAAC)3 | TTGGCAACCAACTCAAAACA | GTTTTTGACCATGCCGTCTT | 211 |
| chr11:41277398-41279418 | p2 | (TA)10 | AGTTGGTGCAACATTTTCCC | TTCACACAGGCTGGAATGAG | 103 |
| chr11:41314332-41316347 | p2 | (TA)13 | GAAATTGCTGATGCGTGAAA | CATTATCGGCGGAGACATTT | 292 |
| chr11:41314332-41316347 | p5 | (CGGCC)3 | GAAATTGCTGATGCGTGAAA | CATTATCGGCGGAGACATTT | 292 |
| chr11:41339031-41341046 | p5 | (ACAAC)3 | TGGAATATAGCCTTGTCGGC | TGAACCCTCCCCAGTGTTAG | 165 |
| chr11:41385689-41387709 | p2 | (AT)10 | TTCCTCCAAAGTGACGAACC | TATCATGGAAAGAGTCGGGG | 128 |
| chr11:41438167-41440191 | p2 | (TA)12 | GAACAACGGAGCTCTCCAAG | CGTACTGACCCCTTTCTTCG | 145 |
| chr11:41481214-41483228 | p2 | (TA)7 | TTTGGTGCCTATTTCAAGCC | GGGGTGTGACACATATTCCC | 116 |
| chr11:41481214-41483228 | p3 | (ACA)5 | TTTGGTGCCTATTTCAAGCC | GGGGTGTGACACATATTCCC | 116 |
| chr11:41519839-41521854 | p5 | (ACCGT)3 | ATGGCGGCAGAGACTGATAG | CCACTCCTCCATCATTACCG | 186 |
| chr11:41667854-41669905 | p3 | (CTT)17 | GGCGAACTCAATACCAGAGC | TGTCAACAACTGCTCCTTCG | 281 |
| chr11:41675780-41677795 | p5 | (AAAGT)3 | CCGAACGTGTTGCTCAGTTA | GAATCAGCTTTCGACCTTCG | 199 |
| chr11:41701828-41703854 | p2 | (TA)8 | GGTTCAAAAATCAACCGTGG | GTCGCTCCGATCTAATCCAA | 141 |
| chr11:41701828-41703854 | p2 | (AT)13 | GGTTCAAAAATCAACCGTGG | GTCGCTCCGATCTAATCCAA | 141 |
| chr11:41825547-41827561 | p2 | (TA)7 | ATCCACCCGAGAGTTGTACG | GGCACGTGTTATTGAGAGCA | 126 |
| chr11:41826589-41828607 | p2 | (TG)9 | TACAACCTCGAATAACCGGC | GGAGACCGAAGAAGAAAGGG | 236 |
| chr11:41891395-41893410 | p5 | (ATTTT)3 | AGGATGTTGCACCTGTGTGA | AATATCCGCTTACCCAACCC | 228 |

| Marker ID | SSR type | SSR motif | FORWARD PRIMER (5'-3') | REVERSE PRIMER (5'-3') | size (bp) |
|---|---|---|---|---|---|
| chr11:41891395-41893410 | p5 | (TTATT)3 | AGGATGTTGCACCTGTGTGA | AATATCCGCTTACCCAACCC | 228 |
| chr11:41916539-41918557 | p6 | (ACGTGG)3 | GTTCCCCGATATGCACAAGT | TGCATGAAGAGTGGAACAGC | 214 |
| chr11:41932109-41934169 | p2 | (AT)7 | CAGTTGAACCCCAACCTTGT | AATATTGGGGGACTTTTGCC | 180 |
| chr11:41932178-41934190 | p2 | (AT)7 | CAGTTGAACCCCAACCTTGT | AATATTGGGGGACTTTTGCC | 111 |
| chr11:42076306-42078318 | p2 | (AT)6 | CGCATGCTCATGCTTATGTT | TGATGTTTCTGTATGGGCGA | 210 |
| chr11:42099261-42101309 | p2 | (TA)24 | TGGATAGGATGCAATTTCCC | TCCTCACGGGACATCAAAAT | 176 |
| chr11:42141798-42143816 | p2 | (TA)9 | TGGCAAGGGTTTTAATGAGG | AGCATTGTTATGCTTTCCCG | 150 |
| chr11:42175809-42177824 | p3 | (AAT)5 | GATCGCCATATAGCCACGTT | AACTGGGGCTCACCACATAG | 167 |
| chr11:42206031-42208075 | p2 | (AT)22 | AGTTGACTTTGCAGGGATGG | GATGTTGAAACGTGTGACGG | 255 |
| chr11:42338446-42340462 | p2 | (AT)8 | AAGTATCAACCGTGGCTTCG | TCCGCTTCTGAGTTTTGGTT | 105 |
| chr11:42407519-42409539 | p5 | (GTTCG)4 | CGTGTATTTGGTCCGTTTCC | CCAGGTACCAGAATCCCTCA | 180 |
| chr11:42410451-42412507 | p5 | (GCATT)3 | CGTGAGATGGACGATGTTTG | ACATCTCGGTAGCACAACCC | 217 |
| chr11:42410451-42412507 | p2 | (TA)28 | CGTGAGATGGACGATGTTTG | ACATCTCGGTAGCACAACCC | 217 |
| chr11:42483895-42485915 | p5 | (TTGCA)4 | TGCTTGAAAGGGGTTGAATC | TCCTTTTCAAATGGCGAAAC | 145 |
| chr11:42489886-42491924 | p2 | (AT)19 | TCCATAATGATATCGGGGGA | TGCCAAAAGGTCATGGTGTA | 183 |
| chr11:42702567-42704581 | p2 | (AT)7 | GCTTCCCTTGGATTGATTGA | GGTCGTGACAACATCCCTTT | 118 |
| chr11:42865684-42867702 | p2 | (AT)9 | TTAGATCAAGTGGGGCTTGG | CGGCAAGCATATTCAGGATT | 251 |
| chr11:42887153-42889193 | p2 | (AT)6 | GCACAAAACTCCGTGCCTAT | CTAATCATCATCCCGCGTCT | 272 |
| chr11:42887153-42889193 | p2 | (AT)10 | GCACAAAACTCCGTGCCTAT | CTAATCATCATCCCGCGTCT | 272 |
| chr11:42887202-42889228 | p2 | (AT)6 | GCACAAAACTCCGTGCCTAT | CTAATCATCATCCCGCGTCT | 223 |
| chr11:42887202-42889228 | p2 | (AT)10 | GCACAAAACTCCGTGCCTAT | CTAATCATCATCCCGCGTCT | 223 |
| chr11:42968807-42970823 | p4 | (ATAA)4 | CGAAAAACACTTTCGCCAAT | GATGGCCAAACGCCTACTAA | 266 |
| chr11:43039843-43041858 | p3 | (ATC)5 | AGGTTCGGATCCTGGAAGTT | AATACTCCCTCCGCCTCAAT | 176 |
| chr11:43092357-43094384 | p5 | (ATGCA)4 | TCTGGAATCACTCCTCGGAC | GAGCTCGTGGTCTGAGGTTC | 228 |
| chr11:43092357-43094384 | p3 | (CAA)9 | TCTGGAATCACTCCTCGGAC | GAGCTCGTGGTCTGAGGTTC | 228 |
| chr11:43152253-43154268 | p5 | (AATAA)3 | ATGGCAGAGGTGGTTAGTGG | ACTTTTTATCACCGGCTCCA | 104 |
| chr11:43155801-43157816 | p5 | (AATAA)3 | ATGGCAGAGGTGGTTAGTGG | ACTTTTTATCACCGGCTCCA | 104 |
| chr11:43178674-43180694 | p2 | (AT)7 | GATGAATTCGGCGGTAGAAA | TTAGCCTCGTCGGCTTATGT | 111 |
| chr11:43255981-43258037 | p2 | (TA)28 | TTGTGGTTGACAGGTGGAAA | ATTATGCTTGAGGGCAATGG | 128 |
| chr11:43255981-43258037 | p4 | (ATTT)4 | TTGTGGTTGACAGGTGGAAA | ATTATGCTTGAGGGCAATGG | 128 |
| chr11:43345022-43347037 | p5 | (TTCTC)3 | CCCCAATAGATCCTCTGCAA | AGCGAAAGGACATTGACGAC | 203 |
| chr11:43384025-43386040 | p3 | (CAT)5 | ACTGAATAAAAGGGTGGCCC | GCCGGCTTACATTGTTGTTT | 283 |
| chr11:43414198-43416216 | p6 | (TAACCA)3 | CGGGGTCACTGTAGCATTTT | ACTCGTCGATAGGACATCGG | 153 |
| chr11:43415832-43417847 | p5 | (AATGC)3 | TCCGTGACCTCTTCGAGTTT | GGTGGAAAAGGTAAGTGCCA | 279 |
| chr11:43419809-43421824 | p3 | (CAA)5 | AACATTAGGGCATCCACGAG | CCGATTTCATTGCTTTGGTT | 177 |
| chr11:43427683-43429698 | p3 | (CAA)5 | GGCTCAGTGTGGGAAACATT | GCCATTCGATATGCCTCATT | 271 |

| Marker ID | SSR type | SSR motif | FORWARD PRIMER (5'-3') | REVERSE PRIMER (5'-3') | size (bp) |
|---|---|---|---|---|---|
| chr11:43509521-43511537 | p2 | (CA)8 | GCCTGACCCGACTTTGAATA | CGTTTTCGGGTGATCTGACT | 286 |
| chr11:43514872-43516887 | p5 | (AATGC)3 | TGGGCACCTACCTTCTCAAC | GGTCATGGGGAATTGTATCG | 146 |
| chr11:43816710-43818725 | p5 | (TTAAG)3 | TAGTACACCGAGTCCAGCCC | GGGTTAGCAAAGGGAAAAGG | 297 |
| chr11:43825021-43827036 | p3 | (AAG)5 | TCCCGAGTGTGTCCTAGTCC | CCCCAAGGAAGGTTTTAAGC | 202 |
| chr11:43849703-43851727 | p3 | (GTT)8 | TGGACTTCCTCTTCCTCCCT | CGGCACTGATAAGGACCAAT | 144 |
| chr11:43849703-43851727 | p5 | (TAAAA)3 | TGGACTTCCTCTTCCTCCCT | CGGCACTGATAAGGACCAAT | 144 |
| chr11:43927771-43929789 | p6 | (CCCCCA)3 | CGAATTGGTTAAGGCTTCCA | CTCATTCGACCCGGTTTAGA | 255 |
| chr11:44102786-44104801 | p5 | (CAAAC)3 | ACGCCCAACCCTCATACATA | AAGTAGCGTGGAAGCATCGT | 189 |
| chr11:44117418-44119439 | p3 | (GTT)7 | TATTGGGGTAGGGTGGATGA | CTATCGCTCCTAGTGTGCCC | 145 |
| chr11:44266934-44268946 | p2 | (AT)6 | CCTCCGAATCGTGATAGGAA | ATTCGACAAAGGCAACAAGG | 125 |
| chr11:44393423-44395435 | p2 | (TA)6 | ACTATCGCAGTCTCGCAGGT | TATGAACCTGTCGCGGTGTA | 143 |
| chr11:44435259-44437273 | p2 | (AG)7 | CCAAGGGAATCAAGAACCAA | TTTACCCAGTATGGCGAAGC | 195 |
| chr11:44506103-44508118 | p3 | (AAT)5 | TGCCTGCCATCTTACAAACA | TCCCCAGAAGAGAAGCAAGA | 269 |
| chr11:44507276-44509296 | p2 | (AT)10 | TTGACATCGACTGGCTTTTG | TCAAGGACCGGTTGATGAAT | 273 |
| chr11:44514477-44516492 | p3 | (ATG)5 | TTAAATCGCCTCCAAGATGC | CGGGTGTCGATTTCAAGTTT | 183 |
| chr11:44514477-44516492 | p5 | (GATAC)3 | TTAAATCGCCTCCAAGATGC | CGGGTGTCGATTTCAAGTTT | 183 |
| chr11:44621545-44623560 | p3 | (TAA)5 | AAATGGTTCGACCACTCTGG | AAGCCGTGAACAGCTTGTCT | 268 |
| chr11:44660210-44662238 | p2 | (AT)14 | CGTTTTCTTTTTCTGGTCGC | AGTGAGGCGAAAGTCTCGAA | 129 |
| chr11:44710696-44712708 | p2 | (TC)6 | ACTAGAACTTGCCCGGTGTG | TGGAAGATTTTTCCGAGGTG | 278 |
| chr11:44723613-44725645 | p2 | (TA)16 | TGTCATTTCTTTCATGGCGA | AGCAGGCTTGCAAATGAGAT | 167 |
| chr11:44723613-44725645 | p3 | (AAT)5 | TGTCATTTCTTTCATGGCGA | AGCAGGCTTGCAAATGAGAT | 167 |
| chr11:44732797-44734835 | p2 | (AT)19 | AAACAAGAGCACCAACCACC | GGAGAGAGGTTTCCCTTTCG | 276 |
| chr11:44761606-44763634 | p2 | (AT)14 | CGTTTTCTTTTTCTGGTCGC | AGTGAGGCGAAAGTCTCGAA | 129 |
| chr11:45021063-45023078 | p5 | (TACAA)3 | TGACTATTCAGGGGTACGGC | TTTCATTTGCGTTCCCTCTT | 215 |
| chr11:45112790-45114814 | p4 | (AATA)6 | TTTATGGCCCTCTTCATTGG | AACCGTGACACCGGAGATAG | 204 |
| chr11:45357381-45359393 | p2 | (TG)6 | CACGACGTTCCAAATGTACG | CATTCCAGTCAACCCGAACT | 237 |
| chr11:45372347-45374371 | p2 | (AG)12 | GCCAGGTTGCCTACGAATTA | CTCACCTGACGATCCGAAAT | 191 |
| chr11:45513371-45515441 | p6 | (ATCATT)3 | AACGTCCACAAGAAAGGACG | GGTATCCGCCTTATGAAGCA | 285 |
| chr11:45513371-45515441 | p6 | (ATCATT)3 | AACGTCCACAAGAAAGGACG | GGTATCCGCCTTATGAAGCA | 285 |
| chr11:45513675-45515693 | p6 | (ATCATT)3 | TAATAGTGCTGCGGAACGTG | TCTCGCGTCACTTCTCTCAA | 189 |
| chr11:45513675-45515693 | p6 | (ATCATT)3 | TAATAGTGCTGCGGAACGTG | TCTCGCGTCACTTCTCTCAA | 189 |
| chr11:45514102-45516118 | p2 | (TA)16 | CTCACCTTTCCCCATACCCT | CGACCTTACGTCACTCCGAT | 133 |
| chr11:45514102-45516118 | p6 | (ATCATT)3 | CTCACCTTTCCCCATACCCT | CGACCTTACGTCACTCCGAT | 133 |
| chr11:45514102-45516118 | p6 | (ATCATT)3 | CTCACCTTTCCCCATACCCT | CGACCTTACGTCACTCCGAT | 133 |
| chr11:45514120-45516134 | p2 | (TA)7 | CTCACCTTTCCCCATACCCT | CGACCTTACGTCACTCCGAT | 115 |
| chr11:45514120-45516134 | p6 | (ATCATT)3 | CTCACCTTTCCCCATACCCT | CGACCTTACGTCACTCCGAT | 115 |

| Marker ID | SSR type | SSR motif | FORWARD PRIMER (5'-3') | REVERSE PRIMER (5'-3') | size (bp) |
|---|---|---|---|---|---|
| chr11:45514120-45516134 | p6 | (ATCATT)3 | CTCACCTTTCCCCATACCCT | CGACCTTACGTCACTCCGAT | 115 |
| chr11:45520182-45522220 | p2 | (AT)19 | TTTGGATGTGTTCTTTGGCA | GATAGCATGCGAGGATGGTT | 215 |
| chr11:45576133-45578149 | p6 | (TATTCA)3 | GTAGCTCCGGATGAAATGGA | CGTCCTTGTGTTTGTGGATG | 280 |
| chr11:45576133-45578149 | p2 | (AT)8 | GTAGCTCCGGATGAAATGGA | CGTCCTTGTGTTTGTGGATG | 280 |
| chr11:45578880-45580894 | p2 | (TA)7 | AGTTGTCGGATCACCAAAGC | TTTCACGCTTTACATGCTCG | 232 |
| chr11:45635731-45637794 | p3 | (GAA)21 | TCCTGTCGTGCTCAGCTTTA | ATTCACCGTCGAAGGAACTG | 188 |
| chr11:45644226-45646238 | p6 | (TTTTCT)3 | CCCCGCCTATTTTCTTTTTC | CATGCATCAAGGGGAAGTTT | 196 |
| chr11:45644226-45646238 | p2 | (GA)6 | CCCCGCCTATTTTCTTTTTC | CATGCATCAAGGGGAAGTTT | 196 |
| chr11:45644226-45646238 | p2 | (AT)11 | CCCCGCCTATTTTCTTTTTC | CATGCATCAAGGGGAAGTTT | 196 |
| chr11:45719339-45721355 | p2 | (AT)8 | AGTTGGCAGCTCCGTTTCTA | TCTCCTCCCTAATGGTGTGG | 180 |
| chr11:45744288-45746303 | p5 | (TATCG)3 | GTAAAATTTGAGGCGGTGGA | TTCAATTCCAAGGTTACGCC | 189 |
| chr11:45744288-45746303 | p2 | (AT)6 | GTAAAATTTGAGGCGGTGGA | TTCAATTCCAAGGTTACGCC | 189 |
| chr11:45775565-45777583 | p6 | (ATTATG)3 | TGGTAAAGCCTTGAGGGAGA | CGGTCCGTATAAGTGTCCGT | 211 |
| chr11:46007773-46009785 | p2 | (TA)6 | TTCTCTAGCTCCCGGTTCAA | CGCTGGTTTCTGTTCCGTAT | 127 |
| chr11:46040757-46042769 | p3 | (CAA)5 | AGAACACTTTCCGAGGCTCA | TTGGCCGAGTTAAATTCTCG | 104 |
| chr11:46040757-46042769 | p2 | (AC)6 | AGAACACTTTCCGAGGCTCA | TTGGCCGAGTTAAATTCTCG | 104 |
| chr11:46063120-46065135 | p2 | (AT)7 | TGTAGTCTTGCCCTTTTGGG | TTTCAGAGCAGTTCCGTCCT | 103 |
| chr11:46063120-46065135 | p3 | (CAT)5 | TGTAGTCTTGCCCTTTTGGG | TTTCAGAGCAGTTCCGTCCT | 103 |
| chr11:46376408-46378420 | p2 | (AT)6 | GTGTAGTGGGCGGCTATGAT | GGGCACTAGTCATGGCCTAA | 204 |
| chr11:46385013-46387025 | p2 | (AT)6 | ATGGGCCATGACTAGTGACC | CGTCTTAGATGTCCCCGTGT | 268 |
| chr11:46385013-46387025 | p3 | (CAA)5 | ATGGGCCATGACTAGTGACC | CGTCTTAGATGTCCCCGTGT | 268 |
| chr11:46390268-46392340 | p2 | (CT)8 | CCCTCACCAAAGCCAGTAAA | AATGCTACAGTGACCCGACC | 222 |
| chr11:46495748-46497763 | p5 | (GATAT)3 | TTTTGCAACCCACAATACGA | TTGAAAGTGGTGATGGGTCA | 182 |
| chr11:46514440-46516460 | p5 | (ATATG)3 | CTTCAAGGTGGGGCAGAATA | AGGCGCGGTGTAATATCATC | 296 |
| chr11:46514440-46516460 | p2 | (TA)10 | CTTCAAGGTGGGGCAGAATA | AGGCGCGGTGTAATATCATC | 296 |
| chr11:46608505-46610517 | p2 | (AT)6 | CGGAAAAACCTCCTAGACCC | TGAGGTCGGGATCAAAAGTC | 130 |
| chr11:46657753-46659771 | p2 | (AT)9 | GTCCCTTTCTTCAAATGGCA | TTGTGCTCTACCGGGGTAAC | 156 |
| chr11:46663518-46665532 | p2 | (CT)7 | GAATTGAAATTGCCTTCGGA | CTCGATTATTGTTCCGCCAT | 158 |
| chr11:46665046-46667070 | p2 | (AT)12 | ACGACCAAGGTATGTGAGGC | ATCATTTGTGCTCTACCGGG | 176 |
| chr11:46754297-46756321 | p2 | (AT)12 | GTTGGGGCTGAGTGACCTAC | AACACCCTGGACCTTGACTG | 136 |
| chr11:46756467-46758482 | p5 | (AACCG)3 | TGTCTTGTGTTTGCTAGCGG | ATCTTGGCGGTCAATTCAAG | 271 |
| chr11:46926038-46928134 | p2 | (TA)48 | GCACCGGATTTCATCAGAGT | TTCACCGGTCTAGTCTGCCT | 177 |
| chr11:47006708-47008723 | p5 | (AGTAT)3 | TTAATGCATTTTCCCGAAGG | TTTCGGGCATGTTTAGGTGT | 182 |
| chr11:47006708-47008723 | p3 | (TTA)10 | TTAATGCATTTTCCCGAAGG | TTTCGGGCATGTTTAGGTGT | 182 |
| chr11:47006708-47008723 | p2 | (GA)6 | TTAATGCATTTTCCCGAAGG | TTTCGGGCATGTTTAGGTGT | 182 |
| chr11:47096656-47098676 | p2 | (TA)10 | ACGTGACTGAGCGTGCTAGA | ACCTGAGGCCACATATCCTG | 217 |

| Marker ID | SSR type | SSR motif | FORWARD PRIMER (5'-3') | REVERSE PRIMER (5'-3') | size (bp) |
|---|---|---|---|---|---|
| chr11:47195053-47197071 | p3 | (CAA)6 | ACTCCCTTCGGAGCAATTTT | ACGAACCCAGCAAATAAACG | 198 |
| chr11:47206225-47208240 | p3 | (TTG)5 | TGGAGACCGTATGATGCAAA | GGCCCAAGGGTCTTTCTTAG | 169 |
| chr11:47420201-47422217 | p2 | (AC)8 | TCCCAATTTCGTCCAGAATC | GAGTCGAGCTATGTCTCGGG | 197 |
| chr11:47455161-47457173 | p2 | (TA)6 | GTATGGCCTCGGTCACTTGT | GCAAAGCTACGACGAAAAGG | 284 |
| chr11:47545496-47547508 | p2 | (AT)6 | GAATGCGGACTGTTTGGAAT | GCGCTTATTCGTCACACAAA | 127 |
| chr11:47545496-47547508 | p2 | (AT)7 | GAATGCGGACTGTTTGGAAT | GCGCTTATTCGTCACACAAA | 127 |
| chr11:47621015-47623033 | p2 | (AT)9 | GCAGGTTGAGGTAGAGGCAG | GGATCACCAACCGGAGTAGA | 217 |
| chr11:47647235-47649250 | p2 | (TA)14 | AAGCGACGGTACGAAAGATG | AACACCAACAATGCCTCACA | 211 |
| chr11:47647235-47649250 | p5 | (TTTAT)3 | AAGCGACGGTACGAAAGATG | AACACCAACAATGCCTCACA | 211 |
| chr11:47653959-47655971 | p2 | (AT)6 | AACTTGGGCGGCTTAATCTT | TGTAGGGGGTGCCACTACTC | 122 |
| chr11:47653959-47655971 | p5 | (AACAT)3 | AACTTGGGCGGCTTAATCTT | TGTAGGGGGTGCCACTACTC | 122 |
| chr11:47654503-47656518 | p2 | (AT)6 | AGCTCGTCATACCGATCCAC | TGGGAAAATTGTAAGTGCCC | 227 |
| chr11:47654503-47656518 | p5 | (AACAT)3 | AGCTCGTCATACCGATCCAC | TGGGAAAATTGTAAGTGCCC | 227 |
| chr11:47713042-47715084 | p2 | (AT)21 | TGGCCTCCTTCTTTCTCTGA | AGGGATGTCTATGAGCCGTG | 157 |
| chr11:47750664-47752680 | p2 | (TA)8 | CCCACTAGCGAGGAAACAAG | CTGCGGGTGTAACATCCTTT | 152 |
| chr11:47809751-47811763 | p2 | (GT)6 | ATGAGTTCTCAGGCCTCCCT | GGGCTACGAATTATGCCAGA | 177 |
| chr11:47967939-47969971 | p2 | (AT)16 | ACATCCGTTTTCTTTGGACG | TAATCAAGCACCTCCGGAAC | 254 |
| chr11:47983620-47985632 | p2 | (TC)6 | CTGCACGACCAGCAATCTTA | ACCCGTATAGAACGCATTGG | 272 |
| chr11:47983620-47985632 | p3 | (TTG)9 | CTGCACGACCAGCAATCTTA | ACCCGTATAGAACGCATTGG | 272 |
| chr11:47983620-47985632 | p3 | (TGT)5 | CTGCACGACCAGCAATCTTA | ACCCGTATAGAACGCATTGG | 272 |
| chr11:48068163-48070183 | p2 | (AT)10 | GGGTCGTGACAGGAGGATTA | GCATATCGAGCCCCATCTAA | 165 |
| chr11:48140796-48142811 | p3 | (ATC)5 | ATGGCCACAGGAAAAACAAG | AGGCAGATATCCTCCTCCGT | 133 |
| chr11:48190549-48192561 | p2 | (TC)6 | TCGGCCAAAATACAACAACA | GTGCGGATTATTTGAAGCGT | 168 |
| chr11:48228508-48230529 | p3 | (CTA)7 | TCTTGGGCAATATGAAAGGC | CTCAGCTCAGGATTCCCAAC | 180 |
| chr11:48228508-48230529 | p2 | (AT)8 | TCTTGGGCAATATGAAAGGC | CTCAGCTCAGGATTCCCAAC | 180 |
| chr11:48228508-48230529 | p3 | (ATA)6 | TCTTGGGCAATATGAAAGGC | CTCAGCTCAGGATTCCCAAC | 180 |
| chr11:48228707-48230722 | p3 | (CTA)7 | ATGTTGGGAATCCTGAGCTG | TGTTGTCCAAGTGGCTTGTT | 174 |
| chr11:48228707-48230722 | p2 | (AT)8 | ATGTTGGGAATCCTGAGCTG | TGTTGTCCAAGTGGCTTGTT | 174 |
| chr11:48228707-48230722 | p3 | (ATA)6 | ATGTTGGGAATCCTGAGCTG | TGTTGTCCAAGTGGCTTGTT | 174 |
| chr11:48228707-48230722 | p3 | (TAA)5 | ATGTTGGGAATCCTGAGCTG | TGTTGTCCAAGTGGCTTGTT | 174 |
| chr11:48228707-48230722 | p3 | (ATA)5 | ATGTTGGGAATCCTGAGCTG | TGTTGTCCAAGTGGCTTGTT | 174 |
| chr11:48228749-48230789 | p3 | (CTA)7 | ATGTTGGGAATCCTGAGCTG | TGTTGTCCAAGTGGCTTGTT | 132 |
| chr11:48228749-48230789 | p2 | (AT)8 | ATGTTGGGAATCCTGAGCTG | TGTTGTCCAAGTGGCTTGTT | 132 |
| chr11:48228749-48230789 | p3 | (ATA)6 | ATGTTGGGAATCCTGAGCTG | TGTTGTCCAAGTGGCTTGTT | 132 |
| chr11:48228749-48230789 | p3 | (TAA)5 | ATGTTGGGAATCCTGAGCTG | TGTTGTCCAAGTGGCTTGTT | 132 |
| chr11:48228749-48230789 | p3 | (ATA)5 | ATGTTGGGAATCCTGAGCTG | TGTTGTCCAAGTGGCTTGTT | 132 |

| Marker ID | SSR type | SSR motif | FORWARD PRIMER (5'-3') | REVERSE PRIMER (5'-3') | size (bp) |
|---|---|---|---|---|---|
| chr11:48393883-48395899 | p3 | (AAC)7 | ACCAGGAGAATGTCACGACC | TGTTTCCAGTTTTCGGTTCC | 245 |
| chr11:48393911-48395927 | p3 | (AAC)7 | ACCAGGAGAATGTCACGACC | TGTTTCCAGTTTTCGGTTCC | 217 |
| chr11:48393934-48395946 | p3 | (AAC)7 | ACCAGGAGAATGTCACGACC | TGTTTCCAGTTTTCGGTTCC | 194 |
| chr11:48395344-48397358 | p3 | (AAC)7 | CATACCTTTCGGTCGCCTTA | TGAGCGTATGACATTGAGCC | 198 |
| chr11:48395344-48397358 | p2 | (AG)7 | CATACCTTTCGGTCGCCTTA | TGAGCGTATGACATTGAGCC | 198 |
| chr11:48398882-48400918 | p2 | (AT)8 | AACAGGAAGAAGCGTGGAAA | TGTCGACTTTCCCTGAATCC | 172 |
| chr11:48398882-48400918 | p2 | (TA)10 | AACAGGAAGAAGCGTGGAAA | TGTCGACTTTCCCTGAATCC | 172 |
| chr11:48417533-48419545 | p5 | (ATATC)3 | CCGTACTCGCAACTCAGTCA | ATGATATTGGCCCTGTGAGG | 166 |
| chr11:48417533-48419545 | p2 | (AT)6 | CCGTACTCGCAACTCAGTCA | ATGATATTGGCCCTGTGAGG | 166 |
| chr11:48542270-48544303 | p3 | (AAC)11 | ACATCACAACAAAGGCCACA | AGGCATAGTTGACAGGGGTG | 119 |
| chr11:48623543-48625567 | p6 | (CTTACA)4 | TGTCGTTCCCGTTCTAGTCC | CTGGCTCGGCTGAGATAAAG | 100 |
| chr11:48639501-48641527 | p2 | (GA)6 | TGGGTTCTCATTAGGGTTGC | TGTTCCCAACAACCAAACAA | 294 |
| chr11:48766729-48768749 | p2 | (TA)10 | ACCAAAAATCCCAACACCAA | TTTGACCATTTTTCCCTTGC | 118 |
| chr11:48782560-48784572 | p2 | (AT)6 | CAGCATACGGCCGAATTTAT | GTCGGCAAAACACTTTTGGT | 252 |
| chr11:48782560-48784572 | p2 | (CA)7 | CAGCATACGGCCGAATTTAT | GTCGGCAAAACACTTTTGGT | 252 |
| chr11:48788925-48790937 | p2 | (AT)6 | CAGCATACGGCCGAATTTAT | GTCGGCAAAACACTTTTGGT | 252 |
| chr11:48788925-48790937 | p2 | (CA)7 | CAGCATACGGCCGAATTTAT | GTCGGCAAAACACTTTTGGT | 252 |
| chr11:48799747-48801759 | p2 | (CT)6 | TGGGTTGCTGTGACAGGTTA | GCCACACACCCATTTTATCC | 278 |
| chr11:48801276-48803310 | p2 | (AT)17 | AAGGCTTGTCCCTTTCCTTC | CCCTTTAAACCCGGAATCAT | 179 |
| chr11:48801276-48803310 | p2 | (TA)6 | AAGGCTTGTCCCTTTCCTTC | CCCTTTAAACCCGGAATCAT | 179 |
| chr11:48944452-48946470 | p5 | (ATTCA)4 | AAAGGGACGAGCCCTAACAT | GAAATTTCGGGCAGGTCATA | 181 |
| chr11:48944452-48946470 | p2 | (CA)9 | AAAGGGACGAGCCCTAACAT | GAAATTTCGGGCAGGTCATA | 181 |
| chr11:48944452-48946470 | p2 | (AT)9 | AAAGGGACGAGCCCTAACAT | GAAATTTCGGGCAGGTCATA | 181 |
| chr11:48995083-48997098 | p3 | (TTG)5 | ACTGCCGAATCGTTAACCTG | CGTATTTTCAGCGATGGGTT | 284 |
| chr11:48995083-48997098 | p3 | (ATG)5 | ACTGCCGAATCGTTAACCTG | CGTATTTTCAGCGATGGGTT | 284 |
| chr11:49131775-49133787 | p2 | (TG)6 | TACTCTGCTGCTGAGCCTGA | CCCGATGTCTTAGCACCAAT | 216 |
| chr11:49163540-49165568 | p4 | (TAAT)7 | GCAACGCTCCAGGGATATTA | TGCATTTCGAGTTGTTCTGC | 284 |
| chr11:49191249-49193270 | p3 | (AAC)7 | GGCGACATATAGTGCTCGGT | TTATTGCCTGGCCTATGGTC | 234 |
| chr11:49254198-49256210 | p2 | (TA)6 | ACCATGCTAGATCGGCAAGT | AATGGCGTCTTTGCTTGAGT | 177 |
| chr11:49264714-49266729 | p5 | (ATATA)3 | CCTCCACCGGTATAGCAAGA | ACGTGGTCTCCCATATCTGC | 116 |
| chr11:49281075-49283090 | p2 | (TG)6 | GTGTGTGTGAAAATTTGGCG | GTTATGTCACGACCCAACCC | 205 |
| chr11:49281075-49283090 | p5 | (TTTTG)3 | GTGTGTGTGAAAATTTGGCG | GTTATGTCACGACCCAACCC | 205 |
| chr11:49402824-49404842 | p3 | (AAC)6 | CGACGAACGAAGTCATAGCA | AGCAATGACCTATGAACGGG | 156 |
| chr11:49481957-49483971 | p2 | (CA)7 | TACATACCTTGCCCGCTTTC | TTGGCGAGTTAAATTCTCGG | 279 |
| chr11:49548206-49550221 | p3 | (TTG)5 | ACAACTCAAGCCAATTTCGG | ACCAAGTGCTCCTTCTTCCA | 216 |
| chr11:49559298-49561313 | p3 | (TTG)5 | ACAACTCAAGCCAATTTCGG | ACCAAGTGCTCCTTCTTCCA | 216 |

| Marker ID | SSR type | SSR motif | FORWARD PRIMER (5'-3') | REVERSE PRIMER (5'-3') | size (bp) |
|---|---|---|---|---|---|
| chr11:49584586-49586601 | p3 | (TTG)5 | TGAAAAATGTTGATGCGGAA | AATTTAGCACATGGGCTTGG | 235 |
| chr11:49649577-49651589 | p2 | (TA)6 | GGCGGGCAGCTTATACATTA | TCCTCCCTGTCTACCTGTGG | 177 |
| chr11:49665835-49667853 | p3 | (ATA)6 | CCAAGTGTAAGCCCTCCAAA | ATAGCAAGGGAAATGGGCTT | 226 |
| chr11:49690640-49692655 | p3 | (ACA)5 | ATGGCACAGCATACTCCACA | AATGTTTCCCACACTGAGCC | 109 |
| chr11:49719858-49721880 | p2 | (AT)11 | CCGGAGTCTATAGGATGCCA | GCTCAAGCACAACGACAAAA | 146 |
| chr11:49800289-49802345 | p2 | (AT)28 | CTTACCTGCATGTGGTGGTG | ATCACGACTCCTTGGTTTGG | 229 |
| chr11:49800289-49802345 | p2 | (TA)6 | CTTACCTGCATGTGGTGGTG | ATCACGACTCCTTGGTTTGG | 229 |
| chr11:49800289-49802345 | p2 | (AT)7 | CTTACCTGCATGTGGTGGTG | ATCACGACTCCTTGGTTTGG | 229 |
| chr11:49800671-49802685 | p2 | (AT)28 | AAGATAACGGTGCAACCCAG | GCAGGTTCAGGTAGTCAGGC | 260 |
| chr11:49800671-49802685 | p2 | (TA)6 | AAGATAACGGTGCAACCCAG | GCAGGTTCAGGTAGTCAGGC | 260 |
| chr11:49800671-49802685 | p2 | (AT)7 | AAGATAACGGTGCAACCCAG | GCAGGTTCAGGTAGTCAGGC | 260 |
| chr11:49800671-49802685 | p3 | (CAA)5 | AAGATAACGGTGCAACCCAG | GCAGGTTCAGGTAGTCAGGC | 260 |
| chr11:49834891-49836903 | p2 | (AT)6 | ATCGGGGTACTTTGAAGCCT | CACATGCCTATCACACCAGG | 131 |
| chr11:49851502-49853516 | p2 | (AT)7 | AGAATCGGCATACAAATGGC | GACTTGCGTCTAAGCGGTTC | 297 |
| chr11:49869601-49871616 | p5 | (AGCAA)3 | GCAAAACAGACAGAATCGGG | CAAAATCTGTTTGCCCTGCT | 287 |
| chr11:49927554-49929566 | p2 | (GT)6 | TTCAGTTCTGTTGGGGGAAC | TCCTCAATATTTCCGCATCC | 114 |
| chr11:49963622-49965637 | p3 | (GAA)5 | CAAGGTGATGTACTGGGCCT | TGTCCCGGTGGTTAATGAAT | 154 |
| chr11:49975002-49977017 | p3 | (GAA)5 | CAAGGTGATGTACTGGGCCT | TGTCCCGGTGGTTAATGAAT | 154 |
| chr11:50033919-50035931 | p2 | (AT)6 | AGGGGGTGCGAGATTTTACT | AATAATAATGGCCCCTTCGG | 284 |
| chr11:50067116-50069128 | p2 | (AT)6 | AGGGGGTGCGAGATTTTACT | AATAATAATGGCCCCTTCGG | 284 |
| chr11:50130383-50132401 | p2 | (TA)9 | GACGTTCGAAAATGCAGGAT | CCCTCTCAACTCGGTGACAT | 272 |
| chr11:50166906-50168934 | p2 | (TA)14 | TTTAGGCTCGTGCTCAAGGT | GGGCCACCATAATGTTCAGT | 181 |
| chr11:50166906-50168934 | p3 | (CAA)6 | TTTAGGCTCGTGCTCAAGGT | GGGCCACCATAATGTTCAGT | 181 |
| chr11:50168420-50170448 | p3 | (CAA)6 | CAAGTTTCCCCACTAACCGA | CTTCTGGAATCCTCCACCAA | 256 |
| chr11:50168420-50170448 | p2 | (TA)14 | CAAGTTTCCCCACTAACCGA | CTTCTGGAATCCTCCACCAA | 256 |
| chr11:50168420-50170448 | p2 | (AT)23 | CAAGTTTCCCCACTAACCGA | CTTCTGGAATCCTCCACCAA | 256 |
| chr11:50174016-50176031 | p5 | (CCCCA)3 | CATCCATCGGAAAGGGTAGA | ACAGGAGCCATGAACCACTC | 131 |
| chr11:50174016-50176031 | p3 | (CTT)13 | CATCCATCGGAAAGGGTAGA | ACAGGAGCCATGAACCACTC | 131 |
| chr11:50388176-50390194 | p6 | (GAGTTG)3 | GGTCGTGTGAGATCATGGTG | AGCCAAAGACTTTCTTGCGA | 155 |
| chr11:50433053-50435071 | p6 | (TTAGTG)3 | TTTCCTTGATTCACTTGCCC | TGGGTCGTGACAGATTTGAA | 210 |
| chr11:50433053-50435071 | p3 | (ATC)5 | TTTCCTTGATTCACTTGCCC | TGGGTCGTGACAGATTTGAA | 210 |
| chr11:50433808-50435823 | p6 | (TTAGTG)3 | GAAGGGGACGTCAGTACGAA | GTTGCACGTTCTGCAACACT | 106 |
| chr11:50433808-50435823 | p3 | (ATC)5 | GAAGGGGACGTCAGTACGAA | GTTGCACGTTCTGCAACACT | 106 |
| chr11:50698816-50700831 | p5 | (TGCAA)3 | AAATGGAGCTGATCGTACCG | GGTAGGTGTCCGCATGACTT | 122 |
| chr11:50807339-50809353 | p2 | (AT)7 | GCCGACTAGGCCGATACATA | TCCACTTGATCCGGAGTTTC | 289 |
| chr11:50819491-50821503 | p2 | (AT)6 | CCCGAGGAAAGTCTTAACCC | AAGCAGGATATGTGGGATGC | 259 |

| Marker ID | SSR type | SSR motif | FORWARD PRIMER (5'-3') | REVERSE PRIMER (5'-3') | size (bp) |
|---|---|---|---|---|---|
| chr11:50831835-50833850 | p3 | (AAC)5 | AATGGAGCTCTCTCCAAGCA | ATTTTCGCGTACTGACCCAC | 147 |
| chr11:50835083-50837098 | p5 | (AGAAA)3 | CCAACCTTAACGATCCTCCA | AAAAAGGCAGCGAGAAATGA | 221 |
| chr11:50838164-50840180 | p2 | (AC)8 | TTCCTGCTTTCCTCCTCAAA | ATACTGACACGGTTGCCCTC | 264 |
| chr11:50867149-50869164 | p5 | (GCATT)3 | TGACTATTTGACCTTCGGGG | CATTCCATTTTCGGAGTGCT | 214 |
| chr11:50945524-50947536 | p2 | (AG)6 | GCACTTGGTCACCCATATCC | GCGAAAATTTCTGGGGTGTA | 233 |
| chr11:50945524-50947536 | p5 | (TATAT)4 | GCACTTGGTCACCCATATCC | GCGAAAATTTCTGGGGTGTA | 233 |
| chr11:51001489-51003507 | p6 | (TATATG)3 | TGTTGTTGTTGTGGTTGGCT | AATCACCACATCGGGATCAT | 189 |
| chr11:51006582-51008632 | p2 | (TA)25 | TTCTCCAACCCCTGTTATCG | CATTCCTCCATCAGATGCCT | 201 |
| chr11:51336993-51339011 | p5 | (AAGAG)3 | CAGTTCACGCCTGTTCTTCA | AAACCCTCGAATTCGTACCC | 270 |
| chr11:51336993-51339011 | p3 | (AAT)19 | CAGTTCACGCCTGTTCTTCA | AAACCCTCGAATTCGTACCC | 270 |
| chr11:51337015-51339117 | p5 | (AAGAG)3 | CAGTTCACGCCTGTTCTTCA | AAACCCTCGAATTCGTACCC | 248 |
| chr11:51337015-51339117 | p3 | (AAT)19 | CAGTTCACGCCTGTTCTTCA | AAACCCTCGAATTCGTACCC | 248 |
| chr11:51443104-51445138 | p2 | (TA)17 | CCCCATAGTGTCCTGCACTT | CCAGGGTTGCAAGTGCTAAT | 293 |
| chr11:51485656-51487677 | p3 | (TGA)7 | AAATGGTGTCCAGTTTTCGC | TTAACGTTCACCTCCCAAGC | 217 |
| chr11:51485656-51487677 | p2 | (TA)16 | AAATGGTGTCCAGTTTTCGC | TTAACGTTCACCTCCCAAGC | 217 |
| chr11:51485700-51487722 | p3 | (TGA)7 | AAATGGTGTCCAGTTTTCGC | TTAACGTTCACCTCCCAAGC | 173 |
| chr11:51485700-51487722 | p2 | (TA)16 | AAATGGTGTCCAGTTTTCGC | TTAACGTTCACCTCCCAAGC | 173 |
| chr11:51524440-51526455 | p5 | (ATTTC)3 | CGTAAATCACCATGTTCCCC | GAGAAGGGGTGAAGTGTGGA | 127 |
| chr11:51582056-51584078 | p3 | (TCA)5 | AGGGATGAGCCCAAGAAGAT | GGCCCTGTCTGACCTTGTTA | 283 |
| chr11:51582056-51584078 | p2 | (TA)11 | AGGGATGAGCCCAAGAAGAT | GGCCCTGTCTGACCTTGTTA | 283 |
| chr11:51793012-51795026 | p2 | (TA)7 | TCCCCCATGCACAATAGAAT | GCAGCAGGAGTTAGTGGTCC | 166 |
| chr11:51843168-51845186 | p2 | (TA)9 | GACCAAGTCTGGCCCTATCA | ACGGGCATGTATGATGGATT | 250 |
| chr11:51910847-51912862 | p3 | (CAT)5 | TATCATTAGGCAACGCCACA | GGGACTGCATAGAGGAGCTG | 248 |
| chr11:51955050-51957082 | p2 | (AT)16 | TTGCCAATTCATTCAGGTCA | AGGCCATTTGCTGACAGACT | 214 |
| chr11:52019067-52021087 | p4 | (GAAA)5 | CCCGGTAGAACTGAATCCAA | AATGGGGATAGTGGAGACCC | 243 |
| chr11:52041597-52043627 | p2 | (TA)15 | GCCCAACGGGATTTTAAGTT | ATCTCCCTTGGATTTTGGCT | 146 |
| chr11:52041597-52043627 | p2 | (AT)12 | GCCCAACGGGATTTTAAGTT | ATCTCCCTTGGATTTTGGCT | 146 |
| chr11:52119027-52121042 | p3 | (TGG)5 | AAGCCCTCCAAAAGGATGAT | ACCCCTCATAGGTTGGTTCC | 190 |
| chr11:52123794-52125809 | p3 | (ATC)5 | TCTGAGCAACTGCACGTAGG | ACGTCCTTCTTGTGGTACGG | 189 |
| chr11:52289278-52291296 | p2 | (TA)15 | ACTGCTTAGCTCGGGTGAAA | GGTGAGGTCTGAGAACCGAG | 163 |
| chr11:52338316-52340337 | p3 | (GTT)7 | CGAAGAACTGTTCGGAGGAG | TGCTGCCTTGCACAATAGAC | 298 |
| chr11:52338316-52340337 | p2 | (CT)6 | CGAAGAACTGTTCGGAGGAG | TGCTGCCTTGCACAATAGAC | 298 |
| chr11:52350339-52352363 | p2 | (TA)14 | GGTGGACTCAATCCCTCCTT | ATCCATCTCAACTGGCCAAC | 262 |
| chr11:52350339-52352363 | p3 | (GTT)8 | GGTGGACTCAATCCCTCCTT | ATCCATCTCAACTGGCCAAC | 262 |
| chr11:52453046-52455062 | p4 | (CATA)4 | TCAATTGCTCCTAAATGCCC | GGTTGGGTCGTGACAAAAGT | 236 |
| chr11:52453046-52455062 | p3 | (CAT)5 | TCAATTGCTCCTAAATGCCC | GGTTGGGTCGTGACAAAAGT | 236 |

| Marker ID | SSR type | SSR motif | FORWARD PRIMER (5'-3') | REVERSE PRIMER (5'-3') | size (bp) |
|---|---|---|---|---|---|
| chr11:52517986-52520004 | p6 | (TAAAAA)3 | TGCAATACATAAGGCCACCA | ACGCACCATCCATATTGTGA | 105 |
| chr11:52550072-52552087 | p5 | (GAAAA)3 | AACCTGGTGGATGAAACAGC | AGGTCGGTTGACACAAAACC | 241 |
| chr11:52615486-52617500 | p2 | (TA)7 | GCAGAATACTTTTGCGAGGC | TGGAAAGGCGTAAAGAGCAT | 222 |
| chr11:52621879-52623895 | p2 | (TC)8 | CACACAACCACCATTGAAGC | TGAACTAACCCCTCAAACGG | 149 |
| chr11:52623780-52625798 | p6 | (AAAAAG)3 | TGGTGGATGTATCTCGCTTG | ATTTCCGAGTTGATTGGGTG | 204 |
| chr11:52623780-52625798 | p6 | (AAAAAG)3 | TGGTGGATGTATCTCGCTTG | ATTTCCGAGTTGATTGGGTG | 204 |
| chr11:52668696-52670717 | p3 | (TTG)7 | CCACCGAAACTTAAAGGCAA | CATTTCCGCCACTTTCATCT | 202 |
| chr11:52668696-52670717 | p2 | (TA)6 | CCACCGAAACTTAAAGGCAA | CATTTCCGCCACTTTCATCT | 202 |
| chr11:52720815-52722845 | p2 | (AT)10 | GGCATCGCTGAAAACGTAAT | TACAAATAGGAGGTTGCGGG | 131 |
| chr11:52720815-52722845 | p2 | (TA)15 | GGCATCGCTGAAAACGTAAT | TACAAATAGGAGGTTGCGGG | 131 |
| chr11:52850834-52852854 | p5 | (AAAAT)4 | TGGCCTTTAAGTGAGGGATG | CGGTGTCATGGAAAATGTTG | 159 |
| chr11:52860684-52862699 | p5 | (ATTGC)3 | ATAGCATGTGAACCTTCGGG | TACCGAATCACAAGCACCAA | 174 |
| chr11:52879179-52881218 | p3 | (TAT)13 | TACTGACCTGCACTTGCTGC | ATGCTCACCCTGGAAACTTG | 176 |
| chr11:52879179-52881218 | p3 | (TAT)5 | TACTGACCTGCACTTGCTGC | ATGCTCACCCTGGAAACTTG | 176 |
| chr11:52879179-52881218 | p4 | (AATA)5 | TACTGACCTGCACTTGCTGC | ATGCTCACCCTGGAAACTTG | 176 |
| chr11:52906434-52908449 | p5 | (AAACC)3 | CTTTGGGGCAGTACTTGCAT | GACCCTAGCCTTGGGGATAA | 278 |
| chr11:52918778-52920802 | p6 | (ATGTAT)4 | ACCTTGAAATTCGCACCAAC | AGTGAGGGTTGATCACTGGG | 141 |
| chr11:52946586-52948660 | p2 | (AT)13 | TGAGGGAAAAGCAATCGAAC | GTGATACCCCGATCTCATGG | 235 |
| chr11:52946685-52948769 | p2 | (AT)13 | TGAGGGAAAAGCAATCGAAC | GTGATACCCCGATCTCATGG | 136 |
| chr11:52985421-52987433 | p2 | (TA)10 | CCCGAGTTGGACACTCGTAT | CCTTGTCGGCTTGCATATTT | 265 |
| chr11:52985421-52987433 | p2 | (AT)6 | CCCGAGTTGGACACTCGTAT | CCTTGTCGGCTTGCATATTT | 265 |
| chr11:53090864-53092876 | p2 | (AT)14 | TTTTTGAAAATGGTCTCGCC | CGTAGGCCAGGCAAAATAAA | 112 |
| chr11:53133211-53135223 | p2 | (AT)6 | AATGAAATCGAGGGTTCACG | CGTCCGGAGTACCACTTCAT | 177 |
| chr11:53242773-53244788 | p5 | (TGATA)3 | GGTCTTGGGGAAAACTCCAT | ATTAGGGGCCTTTTGGTGTC | 264 |
| chr11:53254841-53256883 | p2 | (AT)21 | GTGGCTTGTTGCCCTGTTAT | CGTCATTCTTCCTTTCTCCG | 248 |
| chr11:53274732-53276747 | p2 | (AT)13 | ACACATCAACCGTCCATTCA | CGTAGATACGCATGTGTGGG | 216 |
| chr11:53274732-53276747 | p2 | (AT)6 | ACACATCAACCGTCCATTCA | CGTAGATACGCATGTGTGGG | 216 |
| chr11:53274732-53276747 | p3 | (TTA)5 | ACACATCAACCGTCCATTCA | CGTAGATACGCATGTGTGGG | 216 |
| chr11:53274732-53276747 | p3 | (TCA)5 | ACACATCAACCGTCCATTCA | CGTAGATACGCATGTGTGGG | 216 |
| chr11:53275030-53277045 | p2 | (AT)13 | AATGAGTCAGTGGCACCTCC | CCATTTTCTTTGGACGCTGT | 180 |
| chr11:53275030-53277045 | p2 | (AT)6 | AATGAGTCAGTGGCACCTCC | CCATTTTCTTTGGACGCTGT | 180 |
| chr11:53275030-53277045 | p3 | (TTA)5 | AATGAGTCAGTGGCACCTCC | CCATTTTCTTTGGACGCTGT | 180 |
| chr11:53275030-53277045 | p3 | (TCA)5 | AATGAGTCAGTGGCACCTCC | CCATTTTCTTTGGACGCTGT | 180 |
| chr11:53277771-53279786 | p3 | (ATC)5 | TCATGCCCCAAATATCCCTA | TCTAATCCTTAGGAGCGCCA | 139 |
| chr11:53296018-53298033 | p3 | (TGT)5 | TTGGGTCAACTTTAGAGGCG | GGTTTGGTGCGTTGAAAACT | 233 |
| chr11:53332894-53334929 | p5 | (ATTCA)7 | TTTCCCACCTGTGAAGGAAC | TGACAACGGTGACGTCAAAT | 180 |

| Marker ID | SSR type | SSR motif | FORWARD PRIMER (5'-3') | REVERSE PRIMER (5'-3') | size (bp) |
|---|---|---|---|---|---|
| chr11:53332894-53334929 | p2 | (TA)8 | TTTCCCACCTGTGAAGGAAC | TGACAACGGTGACGTCAAAT | 180 |
| chr11:53349616-53351628 | p2 | (AG)6 | CTCCTCTTCGGGGTCTTCTT | GCTCGAAAGAAACTGCAACC | 178 |
| chr11:53357346-53359362 | p4 | (ATTT)4 | TCGGTTTCTCGAGGTAATGG | TTTGGAGGGTGGTTGAAAAG | 288 |
| chr11:53408565-53410580 | p5 | (ATTGC)3 | ACCGTGGATTGAGTGGTAGC | ATCTACTATCACCACGCGGC | 216 |
| chr11:53424563-53426578 | p3 | (TGT)5 | ACGTTTTCGGATGACCTGAC | GGTTTCACCGACTTCCTCAA | 244 |
| chr11:53435389-53437409 | p5 | (TGATA)4 | TCAGAGGCATTTTGGAAAGG | TCCACACCTGAAGGCCTAAC | 183 |
| chr11:53440350-53442365 | p5 | (TTTCA)3 | CCTTTCACGTCAATGGGACT | TTATCCTGACAAACACGGCA | 204 |
| chr11:53481437-53483452 | p5 | (AGGAC)3 | TGGGACATCAACTACCCACA | CACTATTCCGGAAGTGCCAT | 179 |
| chr11:53481437-53483452 | p3 | (ATC)5 | TGGGACATCAACTACCCACA | CACTATTCCGGAAGTGCCAT | 179 |
| chr11:53484362-53486377 | p5 | (AGGAC)3 | TGGGACATCAACTACCCACA | CACTATTCCGGAAGTGCCAT | 179 |
| chr11:53484362-53486377 | p3 | (ATC)5 | TGGGACATCAACTACCCACA | CACTATTCCGGAAGTGCCAT | 179 |
| chr11:53517539-53519557 | p3 | (GAA)6 | GGTACCTTGTCAATGTGGGG | TTGGTCTGCCTTTAACACCC | 192 |
| chr11:53525984-53528002 | p2 | (CA)9 | TCGAAACCATTCGATGTGAA | GATCATTGGGACCAGCACTT | 233 |
| chr11:53648508-53650523 | p5 | (CGGGT)3 | TTTCATCTTCTGAACGTGCG | ACACATGAGACTGGCATGGA | 172 |
| chr11:53746362-53748382 | p6 | (TTCAGT)3 | ATAGCCCAGTCTCGGTCTCA | GTGCACAAAAACTTTGGCCT | 230 |
| chr11:53746362-53748382 | p5 | (GAGGA)4 | ATAGCCCAGTCTCGGTCTCA | GTGCACAAAAACTTTGGCCT | 230 |
| chr11:53751618-53753660 | p2 | (AT)21 | GATCGAGTTCTTATGCCCCA | CGTGCGATAGAGCTGAGTTG | 279 |
| chr11:53751618-53753660 | p2 | (AT)13 | GATCGAGTTCTTATGCCCCA | CGTGCGATAGAGCTGAGTTG | 279 |
| chr11:53768953-53770999 | p2 | (AG)8 | TGAATTGCTGCTGTACTCGG | TTAAAAGGAGGTCGGGTGTG | 285 |
| chr11:53769001-53771013 | p2 | (AG)8 | TGAATTGCTGCTGTACTCGG | TTAAAAGGAGGTCGGGTGTG | 237 |
| chr11:53769134-53771150 | p2 | (AG)8 | TGAATTGCTGCTGTACTCGG | TTAAAAGGAGGTCGGGTGTG | 104 |
| chr11:53839967-53842001 | p2 | (TA)17 | GACGTAGGGCCTGTCACAAT | ACCAGTTTGAGAACGGATCG | 266 |
| chr11:53902015-53904031 | p2 | (AT)19 | TAGTGAGACCATGCAACCCA | TGTTTTCTTTGGACGCTGTG | 213 |
| chr11:53902033-53904053 | p2 | (AT)19 | TAGTGAGACCATGCAACCCA | TGTTTTCTTTGGACGCTGTG | 195 |
| chr11:53902065-53904085 | p2 | (AT)19 | TAGTGAGACCATGCAACCCA | TGTTTTCTTTGGACGCTGTG | 163 |
| chr11:53902097-53904117 | p2 | (AT)19 | TAGTGAGACCATGCAACCCA | TGTTTTCTTTGGACGCTGTG | 131 |
| chr11:53902257-53904277 | p2 | (AT)19 | ATAAATTTGGATCGCCTCCC | CCTCCCTTAAACGTTACGCA | 142 |
| chr11:53902289-53904309 | p2 | (AT)19 | CACAGCGTCCAAAGAAAACA | CCTCCCTTAAACGTTACGCA | 150 |
| chr11:53902321-53904387 | p2 | (AT)19 | CACAGCGTCCAAAGAAAACA | CCTCCCTTAAACGTTACGCA | 118 |
| chr11:53929360-53931388 | p2 | (TA)14 | TGGTTTGTGGGCTGTTGTAA | ACGACGTTTCTTTCGCATTC | 103 |
| chr11:53959426-53961438 | p2 | (TA)17 | AGTTGTGCATCCACCAAACA | GCAAGGAGTTTTCCACGGTA | 221 |
| chr11:53959426-53961438 | p2 | (GT)6 | AGTTGTGCATCCACCAAACA | GCAAGGAGTTTTCCACGGTA | 221 |
| chr11:54029943-54031958 | p3 | (CAT)5 | TAGAGGGATATGACGGGCAC | ATGGTAGCCTTGTTGGCTTG | 125 |
| chr11:54056633-54058649 | p2 | (AT)8 | CACCCCTAAACTCACTCCCA | TCGTCAGGTGCTTTCAGTTG | 119 |
| chr11:54120644-54122658 | p2 | (TA)7 | GGGGCTCAACTTACCCTTTC | GATGTGATCCAGGAGGCTGT | 244 |
| chr11:54120644-54122658 | p5 | (TTCAT)3 | GGGGCTCAACTTACCCTTTC | GATGTGATCCAGGAGGCTGT | 244 |

| Marker ID | SSR type | SSR motif | FORWARD PRIMER (5'-3') | REVERSE PRIMER (5'-3') | size (bp) |
|---|---|---|---|---|---|
| chr11:54187789-54189801 | p2 | (GC)6 | ACCCTGCTGAGACTAGCCAA | CTTGTGCTGGTGTGCTTGTT | 221 |
| chr11:54297701-54299725 | p6 | (ATATAC)4 | GCATCGCTGAAAGCATAACA | AGTTCGTCCTTGGGTGTGTC | 158 |
| chr11:54387116-54389128 | p3 | (CAA)5 | CCGTTTGTTCCGTCAAAAAT | GTTGATTGTTTTGGCCGAGT | 214 |
| chr11:54387116-54389128 | p2 | (AT)6 | CCGTTTGTTCCGTCAAAAAT | GTTGATTGTTTTGGCCGAGT | 214 |
| chr11:54449670-54451714 | p2 | (AT)29 | TTAGCACACCACGACGCTAC | CATGCTGCACTGCGATATTT | 109 |
| chr11:54449670-54451714 | p2 | (TA)22 | TTAGCACACCACGACGCTAC | CATGCTGCACTGCGATATTT | 109 |
| chr11:54621842-54623866 | p6 | (CCCCAC)4 | ATTGCAAAATCCACCAAAGC | TCACCCTTCTCTTTCTCCCA | 140 |
| chr11:54649074-54651092 | p6 | (TGTATA)3 | CCTCACAAATTTGGGCCTTA | CGTTTTCGGGTGATCTGACT | 229 |
| chr11:54676873-54678885 | p2 | (TG)6 | ATGCACCCCATATGACGAGT | TCATACCTGGCCCTACCAAG | 153 |
| chr11:54676873-54678885 | p2 | (AT)9 | ATGCACCCCATATGACGAGT | TCATACCTGGCCCTACCAAG | 153 |
| chr11:54677475-54679493 | p2 | (TG)6 | TTTTATTTGTGGATGCTGCG | CCAAAAGATATGGCTCCGAA | 190 |
| chr11:54677475-54679493 | p2 | (AT)9 | TTTTATTTGTGGATGCTGCG | CCAAAAGATATGGCTCCGAA | 190 |
| chr11:54792407-54794439 | p2 | (AT)16 | CAACACCAGAGAGTCCGTCA | TTGGCTAACCCACTCCAAAC | 186 |
| chr11:54801426-54803440 | p2 | (AT)7 | CGGGCACCTCTTATCACACT | TTAGCCTCGTCGGCTTATGT | 175 |
| chr11:54969050-54971065 | p4 | (GAAT)4 | ACTCATGGCCGTAAGGTGTC | AACAATCCCGCACTTGTTTC | 288 |
| chr11:54969050-54971065 | p3 | (TAT)5 | ACTCATGGCCGTAAGGTGTC | AACAATCCCGCACTTGTTTC | 288 |
| chr11:54974794-54976810 | p2 | (TA)8 | TGCTCAAATTTCCGTAACCC | CATGCCCTAGTTGGATTCGT | 275 |
| chr11:54974794-54976810 | p2 | (AT)25 | TGCTCAAATTTCCGTAACCC | CATGCCCTAGTTGGATTCGT | 275 |
| chr11:55029260-55031275 | p5 | (TGCAA)3 | GGTGATGCGATACCATTCCT | GTGAGCTTTGGCTAGATCCG | 217 |
| chr11:55087435-55089460 | p5 | (GATAT)3 | GTGGATGTCCCAATGTTTCC | GAGGAATCAGAAAAAGGGGC | 181 |
| chr11:55087435-55089460 | p5 | (TATGA)5 | GTGGATGTCCCAATGTTTCC | GAGGAATCAGAAAAAGGGGC | 181 |
| chr11:55164990-55167012 | p2 | (AG)11 | CACATATTCGCCACTCATGG | CTTGCCCTTATTGTGGAGGA | 165 |
| chr11:55187158-55189170 | p2 | (AT)6 | ACTCATGAAATCACGGCCTC | AGAACAACAAGAGGGGGCTT | 194 |
| chr11:55263521-55265536 | p3 | (ACA)5 | TCTCTCTTGGGTTTGGTTGG | GGTGAACTTCTGGTGGCATT | 220 |
| chr11:55278900-55280914 | p2 | (AT)7 | AAAGCTTTGGCGGTCCTAAT | TCTCGGTCCTCTCACTCGTT | 234 |
| chr11:55368648-55370668 | p2 | (AT)10 | GGAGAGCTTCTGTTGTTCCG | AGAACCCATACACAACCCCA | 190 |
| chr11:55368648-55370668 | p2 | (AT)10 | GGAGAGCTTCTGTTGTTCCG | AGAACCCATACACAACCCCA | 190 |
| chr11:55368648-55370668 | p2 | (AC)6 | GGAGAGCTTCTGTTGTTCCG | AGAACCCATACACAACCCCA | 190 |
| chr11:55393155-55395167 | p2 | (AT)6 | TGGTAATACGTCGATGGCAA | TCGGGCCTTTCTGTGTTATC | 293 |
| chr11:55393155-55395167 | p3 | (CAT)5 | TGGTAATACGTCGATGGCAA | TCGGGCCTTTCTGTGTTATC | 293 |
| chr11:55393155-55395167 | p2 | (AT)11 | TGGTAATACGTCGATGGCAA | TCGGGCCTTTCTGTGTTATC | 293 |
| chr11:55403027-55405045 | p2 | (TA)8 | CAGCGTCCTCAATATCCCAT | GAGATGCGAGAGGCTATTCG | 237 |
| chr11:55403043-55405087 | p2 | (TA)8 | CAGCGTCCTCAATATCCCAT | GAGATGCGAGAGGCTATTCG | 221 |
| chr11:55427271-55429293 | p2 | (AT)11 | AAGAGGGAGAACTTCCCCAA | TCCCTTGCTTCCTCTTTGAA | 150 |
| chr11:55462291-55464335 | p2 | (TA)22 | GATGAGCACCGAGCCTAGAG | AATCAAGTAGTCCCACCCCC | 157 |
| chr11:55474906-55476920 | p2 | (AT)6 | CAAACCCCGCTACAGACAAT | GCAGTACAGACACCGCGTAA | 163 |

| Marker ID | SSR type | SSR motif | FORWARD PRIMER (5'-3') | REVERSE PRIMER (5'-3') | size (bp) |
|---|---|---|---|---|---|
| chr11:55474906-55476920 | p2 | (AT)7 | CAAACCCCGCTACAGACAAT | GCAGTACAGACACCGCGTAA | 163 |
| chr11:55494390-55496405 | p5 | (TTCAT)3 | GGTCGTGTTTGAATTTGGCT | TTTTTACCGTTTTCGGCATC | 275 |
| chr11:55592525-55594541 | p2 | (TA)8 | TCGCAAGTAGGCTAAAGGGA | CGGTCCATCCTATTTTGGAA | 215 |
| chr11:55638210-55640266 | p2 | (AT)28 | AGCCGACAAGGCTATCAAAA | GGAGACAGGCTTGACCCATA | 208 |
| chr11:55678047-55680065 | p6 | (CCGAAC)3 | ACATGTGAACCTAGCCCGTC | AAGTTTCATTGGGGCAGATG | 297 |
| chr11:55691513-55693531 | p3 | (CAT)6 | AGACAACCACTGGGACGAAC | TGACGGAAGCGTCCTTTTAT | 280 |
| chr11:55691513-55693531 | p3 | (ACC)5 | AGACAACCACTGGGACGAAC | TGACGGAAGCGTCCTTTTAT | 280 |
| chr11:55740336-55742350 | p2 | (AT)7 | CATTTCGACGGTCCTGATTT | AAGTGAACAGGCCCACATTC | 134 |
| chr11:55768706-55770722 | p2 | (AT)14 | ATGGTCGGGCAGTTTACTTG | AAGAGGGGTCAGTACGAGCA | 175 |
| chr11:55768706-55770722 | p2 | (AT)8 | ATGGTCGGGCAGTTTACTTG | AAGAGGGGTCAGTACGAGCA | 175 |
| chr11:55796077-55798095 | p6 | (TATCGT)3 | ACAAGCCGGACCACATCTAC | GGAGCCATATCTTTTGGGGT | 161 |
| chr11:55808402-55810422 | p2 | (AT)10 | AAGTGTATGGCCTCGGTCAC | GGGAAAAGCTATAGTCGCCC | 289 |
| chr11:55817207-55819221 | p2 | (TA)7 | CAGATAAAAGGGCGCAGAAG | TTGGTTGGGAATGTTGGTTT | 208 |
| chr11:55861143-55863158 | p3 | (CAA)5 | ATTGGTGCATAGCACGATGA | GACCGAAGGTGAATTTGGAA | 221 |
| chr11:55889542-55891557 | p3 | (GAT)5 | GCAACTAGGCAACTCTTCGC | TTGAATCCAAGCCAAGATCC | 158 |
| chr11:55889542-55891557 | p2 | (TA)6 | GCAACTAGGCAACTCTTCGC | TTGAATCCAAGCCAAGATCC | 158 |
| chr11:55935039-55937067 | p2 | (AT)14 | TACCAAACCCAACAGCCTTC | CCAAGACCCAACTCTCCTTG | 214 |
| chr11:55986197-55988213 | p4 | (TTAA)4 | GGCAAATTGCTCAGAAAAGC | TCCACCCTACAACATGGGAT | 125 |
| chr11:56129449-56131464 | p3 | (AAG)5 | TGAGGAGGCTACTTGGGAGA | TGGGGAAACAAGTCCAAAAG | 203 |
| chr11:56141800-56143824 | p2 | (AT)9 | TCTCCCGAGGAGTCACAACT | GTAGTGGCAGCCTTTTCTGC | 269 |
| chr11:56141800-56143824 | p2 | (AT)12 | TCTCCCGAGGAGTCACAACT | GTAGTGGCAGCCTTTTCTGC | 269 |
| chr11:56185245-56187257 | p2 | (AT)6 | CCACCAGGGTTGAGACACTT | TTGGCTCATATCCGGTTTTC | 162 |
| chr11:56204808-56206862 | p2 | (AT)27 | GTATCAGCCAAAATCGGCAT | CCATCATTTGGCTCCTCAGT | 261 |
| chr11:56288225-56290267 | p2 | (AT)21 | TCCCTATGGCAGGTATGAGC | TGACCATCTCCTCTTGGGTC | 289 |
| chr11:56344266-56346278 | p6 | (CTGAAA)3 | CCGATATGATGTGTTTCCCC | AAGAGAGGCCATCCTTCCTC | 152 |
| chr11:56344266-56346278 | p2 | (TG)6 | CCGATATGATGTGTTTCCCC | AAGAGAGGCCATCCTTCCTC | 152 |
| chr11:56348809-56350824 | p3 | (GTT)5 | ATTTGGGAATTTGGGAAACC | TAGCAAGGGAAATGGGTCTG | 219 |
| chr11:56356432-56358447 | p3 | (GTT)5 | ATTTGGGAATTTGGGAAAGC | TAGCAAGGGAAATGGGTCTG | 220 |
| chr11:56367632-56369654 | p2 | (TA)11 | TGATTCCTCACCGCATGATA | GTCGTAGTGGTGTCTTCGCA | 274 |
| chr11:56367632-56369654 | p2 | (AT)50 | TGATTCCTCACCGCATGATA | GTCGTAGTGGTGTCTTCGCA | 274 |
| chr11:56367780-56369880 | p2 | (TA)11 | TGATTCCTCACCGCATGATA | GTCGTAGTGGTGTCTTCGCA | 126 |
| chr11:56367780-56369880 | p2 | (AT)50 | TGATTCCTCACCGCATGATA | GTCGTAGTGGTGTCTTCGCA | 126 |
| chr11:56385461-56387476 | p3 | (CAT)5 | CCATATGGGCAAGTGAAGGT | TTCATGGACCAAATGCTTGA | 129 |
| chr11:56385461-56387476 | p2 | (TA)32 | CCATATGGGCAAGTGAAGGT | TTCATGGACCAAATGCTTGA | 129 |
| chr11:56410037-56412057 | p2 | (TA)10 | CAAAAGATGAAGAGGCCAGC | TTACCGACGACCAGGATAGG | 286 |
| chr11:56439911-56441929 | p3 | (TTG)6 | GTCGGTCCAAACCGACTCTA | AATTTCGTGGCTGGAAGTTG | 237 |

| Marker ID | SSR type | SSR motif | FORWARD PRIMER (5'-3') | REVERSE PRIMER (5'-3') | size (bp) |
|---|---|---|---|---|---|
| chr11:56487425-56489443 | p3 | (CAA)6 | CCCCTTCAGGCATAATAGCA | ATTATGGGGCATAGGAAGGG | 223 |
| chr11:56575605-56577657 | p2 | (AT)26 | AAAAGGCTGCCACTACGAAA | CCCTTTCTGCTGTCTTGGAG | 279 |
| chr11:56582327-56584347 | p2 | (AT)10 | TACCTTACCCGACATGCCTC | CCCAGCCATAGGAAATGAAA | 144 |
| chr11:56582327-56584347 | p2 | (AT)9 | TACCTTACCCGACATGCCTC | CCCAGCCATAGGAAATGAAA | 144 |
| chr11:56618810-56620826 | p2 | (AT)8 | TCTAACACCCCTCGATGGAC | CACCCTTGCCCTGATGTACT | 169 |
| chr11:56700484-56702498 | p2 | (GA)7 | TTTCATGTGCCATGTGTGTG | AGGCTCAAAAGGGGTTGACT | 276 |
| chr11:56736634-56738678 | p2 | (AT)22 | TTACCCGACACGCAACATAA | ATTGCCAGGAATACCTGCTG | 237 |
| chr11:56783035-56785053 | p6 | (TCAGTT)3 | TGCCCACCAAGTGTTTGATA | AATTCCAGCCTTTGAGCGTA | 284 |
| chr11:56809418-56811446 | p2 | (AT)6 | CGTGCATGCCTCAATATGTC | GGCCCTTTAGTGAGGGACTC | 146 |
| chr11:56809418-56811446 | p2 | (AT)16 | CGTGCATGCCTCAATATGTC | GGCCCTTTAGTGAGGGACTC | 146 |
| chr11:56838768-56840780 | p2 | (TA)6 | CGTGCATGCCTCAATATGTC | ACCGTACCTGGCCCTCTAGT | 175 |
| chr11:56907761-56909841 | p2 | (TA)40 | CCATGCTACCCACACATACG | TTTGTTCATGCCCACAGGTA | 126 |
| chr11:56927842-56929900 | p2 | (AT)10 | GTTGCGGGACTAAACTTGGA | CCGGACTCGGAACTTCACTA | 246 |
| chr11:56927842-56929900 | p2 | (AT)6 | GTTGCGGGACTAAACTTGGA | CCGGACTCGGAACTTCACTA | 246 |
| chr11:56927842-56929900 | p2 | (TA)7 | GTTGCGGGACTAAACTTGGA | CCGGACTCGGAACTTCACTA | 246 |
| chr11:57030071-57032111 | p2 | (TA)20 | CGGATATCGACATGCCTCTT | AGCCCCAAAACAAGTAGGGT | 154 |
| chr11:57030071-57032111 | p2 | (AT)8 | CGGATATCGACATGCCTCTT | AGCCCCAAAACAAGTAGGGT | 154 |
| chr11:57040274-57042294 | p2 | (AT)10 | CATCCCGCATATCTTGACCT | AGAGGAGCTTAGAGGGGTGC | 207 |
| chr11:57065410-57067425 | p5 | (AGAAG)3 | CCCATCCCTGCAAGTTAAAA | TTCCCATGGAAAGCTAATGC | 250 |
| chr11:57208940-57210970 | p2 | (AT)10 | GTTACCCCTCGTCGATCAAA | GAAGTCGGTGATCGATTCGT | 198 |
| chr11:57208940-57210970 | p2 | (AT)15 | GTTACCCCTCGTCGATCAAA | GAAGTCGGTGATCGATTCGT | 198 |
| chr11:57235542-57237557 | p3 | (ATC)5 | ATGTGCATTGATTATCGGCA | CACTTGATGATACCCGGACC | 262 |
| chr11:57414084-57416128 | p2 | (TA)22 | GAGTGACGTGAGATGGCTGA | TGACTCGCGAAGTCTAAGCA | 177 |
| chr11:57665223-57667241 | p2 | (AT)9 | GAGTCCTCAAATCAGCTGCC | CCCAACGAAAAATCAAGCAT | 115 |
| chr11:57751002-57753014 | p6 | (TCGTTA)3 | TGGCCTTACCAAGGATTCAG | TATGGAACCCCGAGCTTATG | 178 |
| chr11:57751002-57753014 | p2 | (CA)6 | TGGCCTTACCAAGGATTCAG | TATGGAACCCCGAGCTTATG | 178 |
| chr11:57815186-57817204 | p3 | (TAT)6 | CCTCTTAAAGGGTTGGCTCC | GGTGGTCTGTCATGCACTTG | 211 |
| chr11:57882037-57884053 | p4 | (TACA)4 | ATACATGTCGGGACAGAGCC | ATGCCCGCAGGTGTAGATAG | 287 |
| chr11:57884790-57886805 | p5 | (ATCAT)6 | CATTCGGGATTTGAAATTGG | CTGTTCACCGAGTCCCGTAT | 272 |
| chr11:57884790-57886805 | p5 | (ATCAT)3 | CATTCGGGATTTGAAATTGG | CTGTTCACCGAGTCCCGTAT | 272 |
| chr11:57960332-57962347 | p5 | (ATGAA)3 | GCTTGATTGTTCCCCTTCAA | TTGTCGTGTGCAATGTTGTG | 296 |
| chr11:57966311-57968331 | p5 | (ATGAA)4 | CTATTCCACCGCTTGATCGT | ATTGTCGTGTGCGATGTTGT | 286 |
| chr11:58203800-58205830 | p5 | (TGAAT)6 | TTCATGGGAAGACAAAAGGG | CCATGGCCGAGCACTACTAT | 121 |
| chr11:58203800-58205830 | p6 | (TATATG)4 | TTCATGGGAAGACAAAAGGG | CCATGGCCGAGCACTACTAT | 121 |
| chr11:58203800-58205830 | p6 | (TATATG)3 | TTCATGGGAAGACAAAAGGG | CCATGGCCGAGCACTACTAT | 121 |
| chr11:58317699-58319717 | p2 | (AT)9 | GGCCTCACGGTACACTCATT | ATGCTCGAAAATACGGATGC | 127 |

| Marker ID | SSR type | SSR motif | FORWARD PRIMER (5'-3') | REVERSE PRIMER (5'-3') | size (bp) |
|---|---|---|---|---|---|
| chr11:58488143-58490181 | p2 | (TA)19 | CCTTCTTGCTCCCATGGTTA | AGGAGGAAGAAACCCAAGGA | 190 |
| chr11:58506599-58508637 | p2 | (TA)19 | CCTTCTTGCTCCCATGGTTA | AGGAGGAAGAAACCCAAGGA | 190 |
| chr11:58534502-58536520 | p3 | (TTG)6 | GACAGAAGAAGCCTTCACCG | CACCCGCCAAAGTTTTTCTA | 124 |
| chr11:58622952-58624972 | p2 | (TA)6 | TCCATAACCGCCTTGTTTTC | TCATGGAATGGCTCAAATCA | 295 |
| chr11:58622952-58624972 | p2 | (TA)10 | TCCATAACCGCCTTGTTTTC | TCATGGAATGGCTCAAATCA | 295 |
| chr11:58622952-58624972 | p2 | (TA)6 | TCCATAACCGCCTTGTTTTC | TCATGGAATGGCTCAAATCA | 295 |
| chr11:58636698-58638714 | p4 | (TATG)5 | ACAAACAAGGGAAATGCTGC | GAGCTTTTGAAAACAAGGCG | 233 |
| chr11:58636754-58638770 | p4 | (TATG)5 | ACAAACAAGGGAAATGCTGC | GAGCTTTTGAAAACAAGGCG | 177 |
| chr11:58648583-58650595 | p2 | (AT)6 | AGCACCCCTTGGCTTAATCT | GGCCATGTCAGCCTATGTTT | 148 |
| chr11:58693481-58695521 | p2 | (AT)20 | TGACTATCACACCGATCCGA | GGAAATAATGCATGCTGGCT | 121 |
| chr11:58720385-58722411 | p2 | (AT)24 | GAGAGCATACTGCCCACACA | CGTTGTATTCATGCCCACAG | 283 |
| chr11:58720385-58722411 | p2 | (TA)13 | GAGAGCATACTGCCCACACA | CGTTGTATTCATGCCCACAG | 283 |
| chr11:58791247-58793259 | p2 | (AT)6 | TTATGGGACGCAACGTGTTA | CATGTGGTGACGCTCATTTC | 227 |
| chr11:58791247-58793259 | p3 | (GTT)5 | TTATGGGACGCAACGTGTTA | CATGTGGTGACGCTCATTTC | 227 |
| chr11:58903478-58905490 | p2 | (AT)6 | GGCTCGTAACTCCAGCATTC | ATTGGCAAGTTGACAGGTCC | 108 |
| chr11:58912825-58914841 | p2 | (TA)8 | CCTCAGAAAAGCTCGGACTG | ACGAGCGGAGTAGTGAGGAA | 228 |
| chr11:59007223-59009238 | p3 | (AAC)5 | GGTGTCATCAGCCCTATCGT | GCTATCCCCGACTCCTCTTC | 264 |
| chr11:59217225-59219239 | p2 | (GT)7 | GGCCACATAGGACGAGACAT | TTGCCCCATCTCTTGCTAGT | 279 |
| chr11:59288123-59290138 | p5 | (ACACA)3 | GGGATCATGCCAAAGAAAAA | TTCCGGATTGTTGTTGTTGA | 151 |
| chr11:59309034-59311049 | p5 | (ATTCA)3 | CTTCGACGACACCCTTCATT | GTGTGGTCTAAGTTGGCCGT | 261 |
| chr11:59309088-59311103 | p5 | (ATTCA)3 | CTTCGACGACACCCTTCATT | GTGTGGTCTAAGTTGGCCGT | 207 |
| chr11:59439551-59441571 | p4 | (GAAA)5 | ATACCTCCACCACGTCTTCG | GATGCGATACCCGTTCTTGT | 281 |
| chr11:59439551-59441571 | p2 | (AT)18 | ATACCTCCACCACGTCTTCG | GATGCGATACCCGTTCTTGT | 281 |
| chr11:59439835-59441871 | p4 | (GAAA)5 | CAGCAGCAGTCATAAGAGCG | ATGGCTCCAGCTCAGACACT | 185 |
| chr11:59439835-59441871 | p2 | (AT)18 | CAGCAGCAGTCATAAGAGCG | ATGGCTCCAGCTCAGACACT | 185 |
| chr11:59483736-59485756 | p4 | (GAAA)5 | ATACCTCCACCACGTCTTCG | GATGCGATACCCGTTCTTGT | 281 |
| chr11:59483736-59485756 | p2 | (AT)18 | ATACCTCCACCACGTCTTCG | GATGCGATACCCGTTCTTGT | 281 |
| chr11:59484020-59486056 | p4 | (GAAA)5 | CAGCAGCAGTCATAAGAGCG | ATGGCTCCAGCTCAGACACT | 185 |
| chr11:59484020-59486056 | p2 | (AT)18 | CAGCAGCAGTCATAAGAGCG | ATGGCTCCAGCTCAGACACT | 185 |
| chr11:59507518-59509530 | p2 | (TG)6 | ATCCGCGTAATTTTTGAACG | CCAAGGGTCCTTCATGTGTT | 140 |
| chr11:59514955-59517019 | p2 | (AT)32 | GGCATGTCATTATCGGGTTC | CCAAGGACCACCTTTCATGT | 131 |
| chr11:59525650-59527672 | p2 | (AT)15 | TGGTCATAACTTTGCGCTTG | CCCCGAGATTTTTACGTTGA | 241 |
| chr11:59525650-59527672 | p2 | (AT)6 | TGGTCATAACTTTGCGCTTG | CCCCGAGATTTTTACGTTGA | 241 |
| chr11:59525674-59527686 | p2 | (AT)15 | TGGTCATAACTTTGCGCTTG | CCCCGAGATTTTTACGTTGA | 217 |
| chr11:59525674-59527686 | p2 | (AT)6 | TGGTCATAACTTTGCGCTTG | CCCCGAGATTTTTACGTTGA | 217 |
| chr11:59525788-59527820 | p2 | (AT)15 | TGGTCATAACTTTGCGCTTG | CCCCGAGATTTTTACGTTGA | 103 |

| Marker ID | SSR type | SSR motif | FORWARD PRIMER (5'-3') | REVERSE PRIMER (5'-3') | size (bp) |
|---|---|---|---|---|---|
| chr11:59525788-59527820 | p2 | (AT)6 | TGGTCATAACTTTGCGCTTG | CCCCGAGATTTTTACGTTGA | 103 |
| chr11:59686832-59688847 | p2 | (AT)9 | CTCACTTGGAGGGTCCAAAA | AAGAAACGGGATTTATGGGG | 245 |
| chr11:59686832-59688847 | p3 | (TTG)5 | CTCACTTGGAGGGTCCAAAA | AAGAAACGGGATTTATGGGG | 245 |
| chr11:59686832-59688847 | p4 | (TGTA)4 | CTCACTTGGAGGGTCCAAAA | AAGAAACGGGATTTATGGGG | 245 |
| chr11:59686832-59688847 | p2 | (TA)6 | CTCACTTGGAGGGTCCAAAA | AAGAAACGGGATTTATGGGG | 245 |
| chr11:59690378-59692418 | p2 | (TA)6 | TCCTTCTTTTGGCATGATCC | GTCGGCTTGCGTATCATTTT | 116 |
| chr11:59696292-59698336 | p2 | (TA)9 | CTTTTAGCCGCAAACTCGTC | GGGCTTGTGACACTAAGGGA | 197 |
| chr11:59696338-59698352 | p2 | (TA)9 | CTTTTAGCCGCAAACTCGTC | GGGCTTGTGACACTAAGGGA | 151 |
| chr11:59696350-59698370 | p2 | (TA)9 | CTTTTAGCCGCAAACTCGTC | GGGCTTGTGACACTAAGGGA | 139 |
| chr11:59781380-59783402 | p2 | (TA)11 | CCGAAAGTTGTTTCTGAGCC | ACTCGGGGAACAATGAAGTG | 148 |
| chr11:59781380-59783402 | p2 | (AT)16 | CCGAAAGTTGTTTCTGAGCC | ACTCGGGGAACAATGAAGTG | 148 |
| chr11:59820291-59822305 | p2 | (AT)7 | CTGCCCGATCTCTTGAACTC | CCCCTAGTTCTTCTACCGGG | 167 |
| chr11:59856080-59858108 | p2 | (AT)14 | GGACACGACTTTGCAGACAA | GGCCCTAATTAAGGGTCGAG | 206 |
| chr11:59994476-59996488 | p2 | (AC)6 | CAAACGAAGAAGTGGGAAGC | CGAGCTAAGTCTCTGGGGTG | 247 |
| chr11:60023008-60025034 | p2 | (AT)10 | TACGACCCAAACCGATAAGC | TAGCCTCGTCGGCACTTATT | 253 |
| chr11:60023008-60025034 | p2 | (AT)13 | TACGACCCAAACCGATAAGC | TAGCCTCGTCGGCACTTATT | 253 |
| chr11:60023008-60025034 | p2 | (AT)24 | TACGACCCAAACCGATAAGC | TAGCCTCGTCGGCACTTATT | 253 |
| chr11:60091035-60093049 | p2 | (TA)15 | CTACTCGTGCATGCCTCAAA | CGTACCCGACCCTCTAGTGA | 291 |
| chr11:60091035-60093049 | p2 | (AT)7 | CTACTCGTGCATGCCTCAAA | CGTACCCGACCCTCTAGTGA | 291 |
| chr11:60126188-60128202 | p2 | (TA)7 | CTCGATGTAGCCACTCGTGA | GAACTTGGGAGCATACGAGC | 118 |
| chr11:60127916-60129942 | p4 | (ATAC)4 | ATGAGTGCAGCGTCAACAAG | GCACCTTCCGCAACACTATT | 282 |
| chr11:60127916-60129942 | p3 | (ATC)6 | ATGAGTGCAGCGTCAACAAG | GCACCTTCCGCAACACTATT | 282 |
| chr11:60190791-60192803 | p2 | (TA)6 | CGTGCATCATTCTCTGCATT | AAATCCATACAGAGGCGTGG | 140 |
| chr11:60306346-60308372 | p2 | (AT)13 | GGGCATCTAGGGACTGTTGA | ATCTGTACGCCGCAATTACC | 292 |
| chr11:60374767-60376781 | p2 | (TA)7 | CATTTATCATCCTTCGCGGT | AATCGCAACACTTTCAGGCT | 213 |
| chr11:60376876-60378894 | p5 | (CATAT)3 | CGAACCCGCTGACTCATATT | GGCTATACCAGCCGATGTGT | 178 |
| chr11:60376876-60378894 | p6 | (CATATA)3 | CGAACCCGCTGACTCATATT | GGCTATACCAGCCGATGTGT | 178 |
| chr11:60502801-60504825 | p2 | (AT)12 | TCGGACGTTTGATTTCCTTC | AGCCGCGAGAAAGGTATGTA | 127 |
| chr11:60502801-60504825 | p4 | (TTTC)4 | TCGGACGTTTGATTTCCTTC | AGCCGCGAGAAAGGTATGTA | 127 |
| chr11:60634798-60636826 | p4 | (AATA)4 | TCTTTACCGGCTACCGTTTG | AAAAATGGATGAGACGGGTG | 153 |
| chr11:60634798-60636826 | p2 | (TA)14 | TCTTTACCGGCTACCGTTTG | AAAAATGGATGAGACGGGTG | 153 |
| chr11:60661401-60663419 | p2 | (TA)9 | GCCACTATTGGGAATCTGGA | ACTCAGATGAAATCCGGTGC | 291 |
| chr11:60787668-60789692 | p4 | (TATT)6 | TGCAAAAACACGGGACATAA | GTAGCGAACCTGAAAGTGGC | 273 |
| chr11:60787668-60789692 | p4 | (ATAC)4 | TGCAAAAACACGGGACATAA | GTAGCGAACCTGAAAGTGGC | 273 |
| chr11:60790866-60792881 | p5 | (TTCAT)3 | AACAATCCCACCAACACCAT | TGGTTCAAGTTGGCATGAAA | 218 |
| chr11:61098696-61100742 | p2 | (TA)23 | GTTTGATGCAGTTGGTGTGG | AGGAGTGAACGACATGGACC | 201 |

| Marker ID | SSR type | SSR motif | FORWARD PRIMER (5'-3') | REVERSE PRIMER (5'-3') | size (bp) |
|---|---|---|---|---|---|
| chr11:61166442-61168454 | p2 | (TA)6 | ATAAGATTTCGGGGCCAAAC | AATTTTACGGCACTTCACGG | 280 |
| chr11:61197035-61199047 | p2 | (GT)6 | CCGACGTCCACAGGTAATCT | CTTATGATCATGGCGGGAGT | 176 |
| chr11:61270679-61272694 | p3 | (TTC)5 | ACGTTGCGGGTGACTAAATC | GGCGTAAAATGACCCAGAAA | 158 |
| chr11:61690490-61692502 | p2 | (TA)6 | AGAGACGCGATATCGAAGGA | TGATCCCCCAATTGCATTAT | 164 |
| chr11:61696365-61698380 | p5 | (ATATG)3 | AAGAGAAATGGTGGAATGCG | ATCGGACAACACTCCCTTTG | 254 |
| chr11:61696365-61698380 | p2 | (TA)6 | AAGAGAAATGGTGGAATGCG | ATCGGACAACACTCCCTTTG | 254 |
| chr11:61696365-61698380 | p2 | (AT)6 | AAGAGAAATGGTGGAATGCG | ATCGGACAACACTCCCTTTG | 254 |
| chr11:61761334-61763350 | p4 | (ATAA)4 | CGGGAATTGGTTTCTGATGT | GTTCGAGCCATCTTCTTTGC | 292 |
| chr11:61761334-61763350 | p3 | (AAT)7 | CGGGAATTGGTTTCTGATGT | GTTCGAGCCATCTTCTTTGC | 292 |
| chr11:61777136-61779151 | p5 | (AAAAT)3 | TTGAGACCCCTTTTGTCCAG | GCCCAATTCATAAGCTCCAA | 226 |
| chr11:61809384-61811400 | p4 | (ATAA)4 | CGGGAATTGGTTTCTGATGT | GTTCGAGCCATCTTCTTTGC | 292 |
| chr11:61809384-61811400 | p3 | (AAT)7 | CGGGAATTGGTTTCTGATGT | GTTCGAGCCATCTTCTTTGC | 292 |
| chr11:61825186-61827201 | p5 | (AAAAT)3 | TTGAGACCCCTTTTGTCCAG | GCCCAATTCATAAGCTCCAA | 226 |
| chr11:62001260-62003275 | p3 | (ATC)5 | AGGAATCCATAAAGACGGGG | AGGTAGACGGACCAGACCCT | 216 |
| chr11:62078660-62080675 | p5 | (TATGA)3 | GATGCTGCCCAAATTTCTGT | ACAATTTCAAACGCGGATTC | 121 |
| chr11:62186625-62188643 | p2 | (AG)9 | TAAGGCATGATCCCATGGTT | ATCATACTCGGCCACAAAGG | 282 |
| chr11:62186625-62188643 | p3 | (TGT)5 | TAAGGCATGATCCCATGGTT | ATCATACTCGGCCACAAAGG | 282 |
| chr11:62188037-62190053 | p2 | (TA)8 | CGTATGCGAAACGTATGTGG | CATCGCTCAATTTCCTCACA | 276 |
| chr11:62219690-62221764 | p2 | (TA)37 | CTAGGGGCACTGCATTTCAT | TCTCGACGCACTATCCACAG | 261 |
| chr11:62257877-62259889 | p2 | (AT)6 | AAGTGTATGGCCTCGGTCAC | AGGGCGAATCACAAAATGAG | 287 |
| chr11:62266469-62268485 | p2 | (AT)19 | AAATAGACCGCAGCTCTGGA | CTAACTCCCGTATTGCTCGG | 250 |
| chr11:62266469-62268485 | p4 | (TTCG)4 | AAATAGACCGCAGCTCTGGA | CTAACTCCCGTATTGCTCGG | 250 |
| chr11:62266469-62268485 | p3 | (CAA)9 | AAATAGACCGCAGCTCTGGA | CTAACTCCCGTATTGCTCGG | 250 |
| chr11:62277111-62279131 | p5 | (AATGC)4 | GCACCGACCTTGTGAAATCT | ATCGGTTGTGGATGGTTCAT | 171 |
| chr11:62297966-62299982 | p4 | (TTCG)4 | AAATAGACCGCAGCTCTGGA | CTAACTCCCGTATTGCTCGG | 250 |
| chr11:62297966-62299982 | p3 | (CAA)9 | AAATAGACCGCAGCTCTGGA | CTAACTCCCGTATTGCTCGG | 250 |
| chr11:62308608-62310628 | p5 | (AATGC)4 | GCACCGACCTTGTGAAATCT | ATCGGTTGTGGATGGTTCAT | 171 |
| chr11:62377524-62379536 | p2 | (CA)6 | CTCCACCGGTTTTGACCTAA | CAGGCTCAAAGGCAAAGTTC | 151 |
| chr11:62417568-62419586 | p2 | (TA)9 | TTGGTCGTTAGTAGGGGCTG | ATTTCCCCTAAACATTCGCC | 117 |
| chr11:62471840-62473852 | p2 | (AT)23 | TTGGAAAGCTCCATTGTTCC | ACTTGCCTTCAAAATGCCAC | 273 |
| chr11:62471840-62473852 | p2 | (CT)6 | TTGGAAAGCTCCATTGTTCC | ACTTGCCTTCAAAATGCCAC | 273 |
| chr11:62488422-62490440 | p6 | (ATACAT)3 | TAATCCACTGGGCACCTCTC | AGTGCCTTAGCTCGATGGAA | 202 |
| chr11:62488422-62490440 | p6 | (ATACAT)3 | TAATCCACTGGGCACCTCTC | AGTGCCTTAGCTCGATGGAA | 202 |
| chr11:62488422-62490440 | p2 | (TA)19 | TAATCCACTGGGCACCTCTC | AGTGCCTTAGCTCGATGGAA | 202 |
| chr11:62552569-62554581 | p2 | (AG)6 | AGTTTGGCCATAGAACCACG | AACGTGAGCATCGAAGGAGT | 265 |
| chr11:62552569-62554581 | p5 | (GCATT)3 | AGTTTGGCCATAGAACCACG | AACGTGAGCATCGAAGGAGT | 265 |

| Marker ID | SSR type | SSR motif | FORWARD PRIMER (5'-3') | REVERSE PRIMER (5'-3') | size (bp) |
|---|---|---|---|---|---|
| chr11:62560617-62562633 | p2 | (TC)8 | AAAGGAAAACAAATGCACCG | TTTGCTATTAGGCCGTTGCT | 283 |
| chr11:62707002-62709017 | p2 | (TA)14 | ACATCACAACAAAGGCCACA | AAAAGGCACAGTTGACAGGG | 112 |
| chr11:62707002-62709017 | p2 | (AT)20 | ACATCACAACAAAGGCCACA | AAAAGGCACAGTTGACAGGG | 112 |
| chr11:62707002-62709017 | p3 | (CAA)5 | ACATCACAACAAAGGCCACA | AAAAGGCACAGTTGACAGGG | 112 |
| chr11:62707002-62709017 | p2 | (TA)8 | ACATCACAACAAAGGCCACA | AAAAGGCACAGTTGACAGGG | 112 |
| chr11:62730026-62732046 | p5 | (CATAT)4 | ACCCCTAGGACAAACCTGCT | GGAAAGCATCGAACAAGCAT | 259 |
| chr11:62730026-62732046 | p2 | (AT)7 | ACCCCTAGGACAAACCTGCT | GGAAAGCATCGAACAAGCAT | 259 |
| chr11:62765211-62767249 | p2 | (AT)19 | CTTTTTCCCGTGGGATTTTT | TCCTTGGTGGCCAAGTATTC | 202 |
| chr11:62773923-62775938 | p3 | (GTT)5 | ATTCGAGTGGCATCAAGACC | CAAAACCATCCTCCAATGCT | 138 |
| chr11:62816254-62818286 | p2 | (AT)16 | AAATCCCAACACCAATCCAA | AACGACCCGCTAGGTTGTTA | 246 |
| chr11:62816254-62818286 | p2 | (TA)37 | AAATCCCAACACCAATCCAA | AACGACCCGCTAGGTTGTTA | 246 |
| chr11:62853800-62855824 | p6 | (CATATA)4 | GCTAGAGGCCATGATGGGTA | CCTTGTCCATCCTTGCTGTT | 168 |
| chr11:62896048-62898072 | p6 | (CATATA)4 | GCTAGAGGCCATGATGGGTA | CCTTGTCCATCCTTGCTGTT | 168 |
| chr11:62951049-62953064 | p2 | (AT)13 | ATGTAACGTGTCCCGACCAT | ACTGCATTGTTCACTGCGTC | 182 |
| chr11:62951049-62953064 | p3 | (CAA)5 | ATGTAACGTGTCCCGACCAT | ACTGCATTGTTCACTGCGTC | 182 |
| chr11:63087097-63089115 | p2 | (AT)29 | GGAGATGCTGCTCGATTTTC | CGACCTCCTGTTAGCTACGC | 213 |
| chr11:63087097-63089115 | p2 | (TA)9 | GGAGATGCTGCTCGATTTTC | CGACCTCCTGTTAGCTACGC | 213 |
| chr11:63135899-63137911 | p2 | (AT)6 | AAGAAGTCACGAGCCCTCAA | AGAGATGCGGTTCAGTTGCT | 135 |
| chr11:63149702-63151788 | p2 | (AT)43 | GTGGAGGAACTGGGGGTATT | CGAGCCAGGGTGAGTCTAAG | 151 |
| chr11:63319682-63321698 | p4 | (GAAT)4 | TTCATTTGGTTGAGCATGGA | ATCCGGTCAGATGGTCGTAG | 172 |
| chr11:63420771-63422786 | p3 | (TCA)5 | GCAGGAAAAACAAGCTCCTG | AGCTGGGTATTCTGGGGAGT | 138 |
| chr11:63420771-63422786 | p2 | (AC)14 | GCAGGAAAAACAAGCTCCTG | AGCTGGGTATTCTGGGGAGT | 138 |
| chr11:63421428-63423456 | p3 | (TCA)5 | TCTGAATCCCTGACCAGTCC | GATGATGATGATGACAGGCG | 208 |
| chr11:63421428-63423456 | p2 | (AC)14 | TCTGAATCCCTGACCAGTCC | GATGATGATGATGACAGGCG | 208 |
| chr11:63462929-63464944 | p5 | (ATTGC)3 | AGCGGAAGGGAAAGGATAAA | CACAATACAAAGGGCTGGCT | 174 |
| chr11:63472440-63474452 | p2 | (AG)6 | TGTATGTGTTGGCCGTGAGT | GCATTCCTGTGCCTGTACCT | 157 |
| chr11:63491396-63493412 | p2 | (AT)8 | GGCGAAGGTCAGACGAGTAG | ACCTTGATCGCCGTTACTTG | 240 |
| chr11:63503831-63505864 | p3 | (AAT)11 | ATCATTCTCCCGTCTCATGC | TTTTATCCGCGGTTGAACTC | 219 |
| chr11:63587312-63589330 | p3 | (TTG)6 | GGGCCAAAACGCTATTACAA | GGAAGCTATTTGGCAGCTTG | 244 |
| chr11:63596273-63598301 | p4 | (GAAA)7 | TGAACCCTAGTCAACGGAGG | ATGGGGGATGTGGACAAGTA | 278 |
| chr11:63610344-63612362 | p6 | (GTTATG)3 | GCATGTCGATGAGTGAGGAA | TCTCGTTTGATCTCCGATCC | 163 |
| chr11:63619730-63621794 | p2 | (AT)32 | TTTAGTTGGCAGGGAACAGG | CTCAATTGCCAACAGCTCAA | 275 |
| chr11:63738084-63740096 | p2 | (TA)6 | AGCTTGCTAGCTGGAACTGG | ACGTCTTTTCTTGTGGACGC | 142 |
| chr11:63738084-63740096 | p2 | (AT)6 | AGCTTGCTAGCTGGAACTGG | ACGTCTTTTCTTGTGGACGC | 142 |
| chr11:63738084-63740096 | p3 | (CAA)6 | AGCTTGCTAGCTGGAACTGG | ACGTCTTTTCTTGTGGACGC | 142 |
| chr11:63787028-63789070 | p2 | (GT)8 | ATTTCTCCTGCAGCCATACG | TGAGGCATTTAAGGGTGTCC | 118 |

| Marker ID | SSR type | SSR motif | FORWARD PRIMER (5'-3') | REVERSE PRIMER (5'-3') | size (bp) |
|---|---|---|---|---|---|
| chr11:63787343-63789357 | p2 | (GT)8 | GGCCACCATGGAACTTGTAT | CAATAGGTTTCCCCAGCTCA | 269 |
| chr11:63787343-63789357 | p4 | (ATAC)6 | GGCCACCATGGAACTTGTAT | CAATAGGTTTCCCCAGCTCA | 269 |
| chr11:63787358-63789416 | p2 | (GT)8 | GGCCACCATGGAACTTGTAT | CAATAGGTTTCCCCAGCTCA | 254 |
| chr11:63787358-63789416 | p4 | (ATAC)6 | GGCCACCATGGAACTTGTAT | CAATAGGTTTCCCCAGCTCA | 254 |
| chr11:63797104-63799122 | p6 | (CATATA)3 | ATCTCGCTAGGGTGTGATGG | TCATCAGCCGATGTGTCATT | 173 |
| chr11:63948855-63950879 | p2 | (TA)12 | CAAGGTAGACAGGGAGACGG | TCATAAGGACGGGACAGGAC | 245 |
| chr11:63961753-63963781 | p2 | (AT)14 | TAACATGGCTCTATCGCACG | ATATCTACGAGCCGTGTCCG | 190 |
| chr11:63961753-63963781 | p3 | (ATC)6 | TAACATGGCTCTATCGCACG | ATATCTACGAGCCGTGTCCG | 190 |
| chr11:63978227-63980253 | p2 | (TG)13 | CGAGCCTATATGGACGGGTA | GTAGGTTCGGTGTCATCCGT | 165 |
| chr11:64034903-64036917 | p2 | (AG)7 | AAACATGTGCTGAGGGAAGG | TTGGCATTTGGCTTCTATCC | 141 |
| chr11:64093337-64095353 | p4 | (ACGA)4 | TTTGAGGAGGATCTGGCTGT | CCAAAACAGTCCACCGAACT | 276 |
| chr11:64093337-64095353 | p2 | (AT)8 | TTTGAGGAGGATCTGGCTGT | CCAAAACAGTCCACCGAACT | 276 |
| chr11:64100761-64102809 | p2 | (AT)24 | CATCGCTGAAATCAAAGCAA | TGGGTGTCTAGGCATTAGGG | 291 |
| chr11:64173248-64175268 | p2 | (TA)10 | CAACTCCTTTCCCCAATGAA | GAGAGAGGTTTCCCTTTCGG | 297 |
| chr11:64182287-64184301 | p2 | (AT)7 | TTTGTGGTTTCTATGCAGCG | TTGAGCACCATGAGAGTTGG | 213 |
| chr11:64308199-64310223 | p2 | (AT)25 | TTCCCGAGTTTTGGGTACTG | ACCTCCTTTGCCTTTTCCAT | 159 |
| chr11:64308199-64310223 | p3 | (AAT)8 | TTCCCGAGTTTTGGGTACTG | ACCTCCTTTGCCTTTTCCAT | 159 |
| chr11:64308199-64310223 | p6 | (CACAAG)4 | TTCCCGAGTTTTGGGTACTG | ACCTCCTTTGCCTTTTCCAT | 159 |
| chr11:64315563-64317583 | p2 | (AT)10 | TCAATAAAACCTTGCGGGTC | AATACTTCCTCCACCCCACC | 264 |
| chr11:64315871-64317895 | p2 | (AT)10 | GGTGGGGTGGAGGAAGTATT | AAACAGGAGGCTACAAGGCA | 190 |
| chr11:64315896-64317926 | p2 | (AT)10 | GGTGGGGTGGAGGAAGTATT | AAACAGGAGGCTACAAGGCA | 165 |
| chr11:64379416-64381440 | p2 | (AT)12 | CCTCTCCAGTCAAGCTTTGG | TTTAGGCGGGATTGAAGATG | 255 |
| chr11:64403087-64405117 | p2 | (AT)8 | ATGTTTCCGCAACTTCCTTG | ATGGTATGAGCGGTACTCGG | 224 |
| chr11:64403087-64405117 | p2 | (AT)15 | ATGTTTCCGCAACTTCCTTG | ATGGTATGAGCGGTACTCGG | 224 |
| chr11:64600901-64602919 | p2 | (AT)9 | TGAAAGGGAAACAGGACAGG | CCATAGAATGGGGGAAAGGT | 203 |
| chr11:64668045-64670065 | p2 | (AT)16 | CCCTCTTTAAACGCTTGGTG | GGCCATATCTGCAAACTGGT | 296 |
| chr11:64668045-64670065 | p2 | (AT)10 | CCCTCTTTAAACGCTTGGTG | GGCCATATCTGCAAACTGGT | 296 |
| chr11:64722229-64724251 | p5 | (AAAAT)4 | TGAGCACATCAAATGCCAGT | ACCATACCATGAACACCCGT | 118 |
| chr11:64726491-64728503 | p2 | (AT)6 | TAAGGTTCAAATTCGGGTCG | CGTTACAAGTTCGGCTGACA | 286 |
| chr11:64736437-64738449 | p2 | (AT)6 | TAAGGTTCAAATTCGGGTCG | CGTTACAAGTTCGGCTGACA | 286 |
| chr11:64736437-64738449 | p2 | (AT)12 | TAAGGTTCAAATTCGGGTCG | CGTTACAAGTTCGGCTGACA | 286 |
| chr11:64848418-64850433 | p5 | (TTGCA)3 | CCTCGGGAGATGGATGACTA | ACTAAGTGACCAAAACGCGG | 199 |
| chr11:64863161-64865176 | p3 | (CAA)5 | TGGTTTACGGTCCGTCTTTC | TGAAAATGCGAAAGTTGCAG | 144 |
| chr11:64867714-64869729 | p5 | (TCATA)3 | ATAAGAGGGCGCTCTAAGCC | TATGATGTGTGGTGCTCGGT | 254 |
| chr11:64867714-64869729 | p2 | (TA)9 | ATAAGAGGGCGCTCTAAGCC | TATGATGTGTGGTGCTCGGT | 254 |
| chr11:64867865-64869883 | p5 | (TCATA)3 | ATAAGAGGGCGCTCTAAGCC | TATGATGTGTGGTGCTCGGT | 103 |

| Marker ID | SSR type | SSR motif | FORWARD PRIMER (5'-3') | REVERSE PRIMER (5'-3') | size (bp) |
|---|---|---|---|---|---|
| chr11:64867865-64869883 | p2 | (TA)9 | ATAAGAGGGCGCTCTAAGCC | TATGATGTGTGGTGCTCGGT | 103 |
| chr11:64922191-64924203 | p2 | (AT)6 | TTACGGCTCATAGGTTTGGG | GAACGCTTAACACAGGGCTC | 158 |
| chr11:64933346-64935364 | p2 | (AT)9 | GGGCAGCTGTAGCTTTTCAC | AAACAGGAGGCTACAAGGCA | 148 |
| chr11:64956868-64958883 | p5 | (ATTGC)3 | TGAATACGCGGTTCATCAAA | TGGTGCCAGATGTTTTTCAA | 207 |
| chr11:65122025-65124040 | p2 | (AT)7 | CATATGGACGGGACAAGACC | TCCACAGGTAGACAGGGAGG | 136 |
| chr11:65122025-65124040 | p2 | (TA)8 | CATATGGACGGGACAAGACC | TCCACAGGTAGACAGGGAGG | 136 |
| chr11:65122025-65124040 | p3 | (TCA)5 | CATATGGACGGGACAAGACC | TCCACAGGTAGACAGGGAGG | 136 |
| chr11:65166451-65168469 | p2 | (AT)9 | CCCGTCGTACCCTTAAACAA | TTTCATGCTCGCAGGTACAG | 262 |
| chr11:65177007-65179022 | p3 | (TGT)5 | GGAGAAAACATGGAGGGGTT | GCCTCTCTTGCTCCAACAAC | 102 |
| chr11:65177007-65179022 | p5 | (TGAAT)3 | GGAGAAAACATGGAGGGGTT | GCCTCTCTTGCTCCAACAAC | 102 |
| chr11:65302906-65304918 | p2 | (TA)6 | TAGCAAAAGGAGCGGAAAGA | TCATCGCGTTACTTGTGAGG | 154 |
| chr11:65308041-65310056 | p5 | (ATTGC)3 | AAGGCTCAAAGGAAGGGAAA | GGCTTACATCCATTCTCCGA | 254 |
| chr11:65455146-65457158 | p2 | (TA)6 | TAGGGGAAACCGGTTGTGTA | CTGTCCGACCAGTTTTGGAT | 276 |
| chr11:65457569-65459617 | p2 | (AT)24 | GTCAACACGGTTCGTCAATG | TCCGACAGAATTTTTGGTCC | 227 |
| chr11:65490929-65492991 | p2 | (AT)31 | AGCCGATGAGGCTATCAGAA | TACCTATGGGCATGGAGAGG | 191 |
| chr11:65507155-65509175 | p5 | (TTCAT)4 | TTCCAAGCTAGGGCAAGAAA | ATTCACTGCACATCAACGGA | 264 |
| chr11:65674714-65676732 | p6 | (ATATGT)3 | TTGGATATGATCCTCGGGAA | ACAATAAGGAAAATCCGGGG | 170 |
| chr11:65674714-65676732 | p2 | (AT)8 | TTGGATATGATCCTCGGGAA | ACAATAAGGAAAATCCGGGG | 170 |
| chr11:65697361-65699376 | p2 | (TA)11 | CGCGGGCTAATGATGTTACT | CAACCTGCTGAAAAAGCTCC | 261 |
| chr11:65697361-65699376 | p3 | (ATC)5 | CGCGGGCTAATGATGTTACT | CAACCTGCTGAAAAAGCTCC | 261 |
| chr11:65782661-65784679 | p2 | (TA)9 | CTGAGTGTGCTATGGCCTCA | CGTTTATTTGCTGGGTTCGT | 298 |
| chr11:65853103-65855117 | p2 | (TA)7 | TTTCAAAACCACTCCGAAGG | CTGAGGCCATAGCACACTCA | 192 |
| chr11:65859129-65861147 | p2 | (AT)9 | CTTTCGTCCCAAATCAAGGA | TTTGAACATGTTGTTGCCGT | 114 |
| chr11:65862090-65864105 | p3 | (CAA)5 | CCTTTTAGCCGCACTTATCG | CATAAGTGAGCGAAGCCTCC | 119 |
| chr11:65862090-65864105 | p2 | (AT)10 | CCTTTTAGCCGCACTTATCG | CATAAGTGAGCGAAGCCTCC | 119 |
| chr11:65883439-65885451 | p2 | (AT)6 | AGTGGGCTCCTCTTGGATTT | AGTTGTGAGATGGCTTCGCT | 280 |
| chr11:65893013-65895028 | p5 | (GTGGG)3 | TCCGTATAATCTCATCGGGC | CCCTAGCAAGTCTCCCAACA | 193 |
| chr11:65893013-65895028 | p2 | (AT)7 | TCCGTATAATCTCATCGGGC | CCCTAGCAAGTCTCCCAACA | 193 |
| chr11:65893103-65895133 | p5 | (GTGGG)3 | TCCGTATAATCTCATCGGGC | CCCTAGCAAGTCTCCCAACA | 103 |
| chr11:65893103-65895133 | p2 | (AT)7 | TCCGTATAATCTCATCGGGC | CCCTAGCAAGTCTCCCAACA | 103 |
| chr11:65913958-65915974 | p2 | (AT)6 | ACTTTTGTCACGACCCAACC | CGGCTTGCGTTTTATTTCAT | 155 |
| chr11:65913958-65915974 | p2 | (AT)8 | ACTTTTGTCACGACCCAACC | CGGCTTGCGTTTTATTTCAT | 155 |
| chr11:65913958-65915974 | p2 | (AT)16 | ACTTTTGTCACGACCCAACC | CGGCTTGCGTTTTATTTCAT | 155 |
| chr11:66016676-66018692 | p4 | (ACAT)4 | ATAAGCCGACGAGGCTAACA | TCCCGCTTATGTGACATTGA | 185 |
| chr11:66109973-66111994 | p3 | (TTG)7 | ACCCTTAAGCCAAAAATGGG | TCTGGGCAGTTTCCAAATTC | 291 |
| chr11:66134026-66136090 | p2 | (AT)32 | ATACACGGGATATGGGCAAA | CGTGCAGCCTGAGTCATTTA | 250 |

| Marker ID | SSR type | SSR motif | FORWARD PRIMER (5'-3') | REVERSE PRIMER (5'-3') | size (bp) |
|---|---|---|---|---|---|
| chr11:66172556-66174571 | p5 | (TATAA)3 | CCGTACGACGATAAAGCGAT | CCCCTTGTGCCTCATACACT | 117 |
| chr11:66185809-66187837 | p5 | (ATTTC)3 | CGGGAATGAGTTCGGAGTAA | TTTCGCTGACCGTATGTCTG | 267 |
| chr11:66185809-66187837 | p2 | (TA)11 | CGGGAATGAGTTCGGAGTAA | TTTCGCTGACCGTATGTCTG | 267 |
| chr11:66185809-66187837 | p2 | (AT)14 | CGGGAATGAGTTCGGAGTAA | TTTCGCTGACCGTATGTCTG | 267 |
| chr11:66229611-66231627 | p2 | (AC)8 | TGTGCGTTCGATCAAATCAT | TGTTCAAAAGTGAGGGGGAG | 158 |
| chr11:66229611-66231627 | p2 | (AT)6 | TGTGCGTTCGATCAAATCAT | TGTTCAAAAGTGAGGGGGAG | 158 |
| chr11:66282646-66284662 | p2 | (AT)11 | TTGTCACACCCCACCCTATT | GTCGGCATCATGGGTCTATT | 151 |
| chr11:66282646-66284662 | p4 | (CATA)4 | TTGTCACACCCCACCCTATT | GTCGGCATCATGGGTCTATT | 151 |
| chr11:66330270-66332285 | p5 | (TTGGT)3 | ACCTATGTTGTGGGCCATGT | TACTGCGCAGGTGTATGCTC | 205 |
| chr11:66370898-66372916 | p6 | (TGGGTG)3 | TTCGTCTCATTTTTCAGGGG | AAGAGTGCAAACCCGTGACT | 114 |
| chr11:66405966-66407981 | p3 | (GGT)5 | GGAAATTTGGTTAAACGCGA | TACTGAGCTCTCCCATGCCT | 176 |
| chr11:66412951-66414973 | p2 | (AT)11 | ATCCCCCATTCTCCTTTGAC | CTATGTCGAACAGGGAGGGA | 229 |
| chr11:66412951-66414973 | p3 | (TCA)6 | ATCCCCCATTCTCCTTTGAC | CTATGTCGAACAGGGAGGGA | 229 |
| chr11:66444193-66446208 | p2 | (AT)11 | ATCATCCCATAGACCGGACA | TCTTTTGTTTGTGGACGCTG | 212 |
| chr11:66444193-66446208 | p3 | (TCA)5 | ATCATCCCATAGACCGGACA | TCTTTTGTTTGTGGACGCTG | 212 |
| chr11:66452744-66454762 | p5 | (TTCAT)3 | GGGGACAGGTTTGGCTTATT | GGAACAAGCCTTGGTTGGTA | 216 |
| chr11:66452744-66454762 | p2 | (AT)9 | GGGGACAGGTTTGGCTTATT | GGAACAAGCCTTGGTTGGTA | 216 |
| chr11:66452744-66454762 | p2 | (AT)35 | GGGGACAGGTTTGGCTTATT | GGAACAAGCCTTGGTTGGTA | 216 |
| chr11:66463760-66465800 | p2 | (AT)20 | CCAACCAAGGCTTGTTCCTA | TCCCATGTTTTTCAACGTCA | 111 |
| chr11:66472744-66474756 | p2 | (TA)6 | AACACTTTAACCGACGCCAC | CTTCCAGGGTGTTCCCTACA | 109 |
| chr11:66551336-66553354 | p6 | (CATAAT)3 | AATCATATGCAGCTCGCCTT | CACCAAGGTGAGACATGACG | 191 |
| chr11:66594995-66597009 | p2 | (TA)7 | TCTATAGCCCGCAACAATCC | ATTTCGGGAGACGGAGTTCT | 116 |
| chr11:66616203-66618215 | p2 | (CA)6 | GCTCAAATTCGATTCCGTGT | CGACTAGAGTTCGGGAATGC | 247 |
| chr11:66667204-66669224 | p2 | (AT)10 | GTAACCTGCCAATCCCAGAA | TGCAAAATTTTCAAAGTGCG | 269 |
| chr11:66802679-66804727 | p2 | (AT)24 | TCAACTTCCAACACCCACAA | GATAGCGAACCCACAACCAT | 186 |
| chr11:66847917-66849945 | p2 | (TA)14 | TTGGGGATTTGGTGTTGATT | TTCTAACCGAATTTCGACCG | 104 |
| chr11:67004988-67007000 | p2 | (CA)6 | ACCTTACATACCTGTGCCGC | AGGGGAGATGCTACCGAAAT | 225 |
| chr11:67185605-67187617 | p2 | (TA)6 | GAAAAGGCGAATGACGATGT | CTTCACAGAACATGCGGCTA | 113 |
| chr11:67317910-67319932 | p2 | (AT)10 | ACAACTGGACTGCATCCCTC | AGCTGCCCAGAAGGGTAAAT | 229 |
| chr11:67317910-67319932 | p2 | (AT)11 | ACAACTGGACTGCATCCCTC | AGCTGCCCAGAAGGGTAAAT | 229 |
| chr11:67317910-67319932 | p2 | (AT)6 | ACAACTGGACTGCATCCCTC | AGCTGCCCAGAAGGGTAAAT | 229 |
| chr11:67371874-67373886 | p2 | (TA)6 | TGGGAAGGTTATGAAGGTGG | CGAAAATCCAATTCGGCTAA | 100 |
| chr11:67479880-67481898 | p3 | (TTG)6 | AAGCCCTCCAAAAGGATGAT | TTTGGCAGCTTGGAGTCTTT | 192 |
| chr11:67491603-67493618 | p5 | (GATAT)3 | GATGCTGCCCAAATTTCTGT | CTAGGAATAGAATCGCCCCC | 191 |
| chr11:67491603-67493618 | p2 | (TA)7 | GATGCTGCCCAAATTTCTGT | CTAGGAATAGAATCGCCCCC | 191 |
| chr11:67573220-67575235 | p5 | (ATTTC)3 | AACATGGCAAATCAATGCAA | CGAACGGAGATTTTCTGAGG | 286 |

| Marker ID | SSR type | SSR motif | FORWARD PRIMER (5'-3') | REVERSE PRIMER (5'-3') | size (bp) |
|---|---|---|---|---|---|
| chr11:67599993-67602011 | p2 | (AC)9 | TGCCAAAGCTTGAGGTTCTT | CCATATTTCTTCCCAGCCAA | 296 |
| chr11:67701227-67703242 | p3 | (CAA)5 | AAATGTTTCGCATAGACCCG | TCCTGGTTTTGCCCTTATTG | 228 |
| chr11:67707392-67709418 | p2 | (TA)7 | CATTCGACCCCTCGTACCTA | GTCCCACCTCGACCACTAGA | 131 |
| chr11:67707392-67709418 | p5 | (TGTGT)3 | CATTCGACCCCTCGTACCTA | GTCCCACCTCGACCACTAGA | 131 |
| chr11:67758486-67760501 | p3 | (ATG)5 | ATGGACGTTACAAGTTCGGC | TAGGGTCGGGTCACTGTAGC | 144 |
| chr11:67880253-67882268 | p5 | (ATCAT)3 | TAAAGCACAAGGAAGCCGAT | TGCTCTATTGTTCGGAGCCT | 297 |
| chr11:67880253-67882268 | p5 | (ATCAT)3 | TAAAGCACAAGGAAGCCGAT | TGCTCTATTGTTCGGAGCCT | 297 |
| chr11:67881905-67883923 | p2 | (AT)9 | GACAATTTCAAACGCGGATT | GATGCTGCCCAAATTTCTGT | 196 |
| chr11:67934958-67937015 | p3 | (TTC)19 | GTTTGGGAAAATTGGGGTTT | TTGATTCGGAACCCAAAAAG | 167 |
| chr11:67956298-67958310 | p2 | (TA)6 | GGGCCGTAATTCAAGTCAAA | TCCGTTTGACCCCCATAATA | 115 |
| chr11:67971565-67973587 | p2 | (AT)11 | CGTGACAAAGGATGAACGTG | GAATCCCTTTTGACGCTGAA | 112 |
| chr11:67971565-67973587 | p2 | (AT)6 | CGTGACAAAGGATGAACGTG | GAATCCCTTTTGACGCTGAA | 112 |
| chr11:67987992-67990026 | p2 | (TA)17 | TTTCATTCCCTACGACTCCG | ATTGAGTAAGGCATCGGGTG | 171 |
| chr11:68048288-68050303 | p5 | (ATCAT)3 | GAAGCCGATGAGGCTGTTAC | GCTATTTGGAGCGCTCTTTG | 287 |
| chr11:68077072-68079090 | p2 | (TA)9 | GTAAATCGGGCCGTATCAGA | CACACATTTACGGAGCATGG | 178 |
| chr11:68077072-68079090 | p2 | (GT)6 | GTAAATCGGGCCGTATCAGA | CACACATTTACGGAGCATGG | 178 |
| chr11:68077420-68079432 | p2 | (TA)9 | AAAAAGAAACTCCCCCGCTA | CCGATGTCTCTGCGTCAATA | 235 |
| chr11:68077420-68079432 | p2 | (GT)6 | AAAAAGAAACTCCCCCGCTA | CCGATGTCTCTGCGTCAATA | 235 |
| chr11:68243258-68245273 | p3 | (TGA)5 | GCGCTGAACGATTTTAGAGG | AAAAGTTGGCTTGGTGGTTG | 204 |
| chr11:68243258-68245273 | p4 | (TATG)4 | GCGCTGAACGATTTTAGAGG | AAAAGTTGGCTTGGTGGTTG | 204 |
| chr11:68244264-68246286 | p4 | (TATG)4 | TGATTTCATGTTGCGATGGT | TTTCGGGACTCGGTGTTATC | 210 |
| chr11:68244264-68246286 | p2 | (TA)11 | TGATTTCATGTTGCGATGGT | TTTCGGGACTCGGTGTTATC | 210 |
| chr11:68274869-68276897 | p2 | (AT)14 | GCCGGGAGGTAGTACATCAA | CATCGCCAAAGGTAGTCCAT | 234 |
| chr11:68320300-68322315 | p5 | (ATATA)3 | CATGGTCGCTTTACTCGACA | GGTTCATGCCACACAGTCAC | 259 |
| chr11:68331557-68333579 | p2 | (TA)11 | TAAGGTAAACCCCGCTCCTT | GTAGCACCCACAACGAGGAT | 153 |
| chr11:68470231-68472249 | p6 | (AACTGA)3 | TCCTCCCTTTTCATCATTGC | CTAGGGTTGTCTACAGGCCG | 228 |
| chr11:68522977-68525017 | p2 | (TA)20 | CATGTCATATGCAAGCCGAC | TCTTGGAGAGCTCCGTTGTT | 292 |
| chr11:68535014-68537029 | p3 | (TTG)5 | CTACCTTTTGGTGATGCCGT | GTCTCCCCAACTTCAAACCA | 247 |
| chr11:68602090-68604108 | p6 | (AGTTGA)3 | TATACGGGCCAACATACGGT | CAAACGAGTCGACGAAAACA | 110 |
| chr11:68621900-68623915 | p5 | (ATATG)3 | TTGTGGGTGTTGGAAGTTGA | CGTTCAATGTCACGCCTCTA | 294 |
| chr11:68646455-68648470 | p3 | (CAT)5 | AGAGGAATTGTGAAATGCGG | TTGATAGCCGAGAGGCTTGT | 296 |
| chr11:68656121-68658136 | p5 | (ATCAT)3 | TCCAGATGAACTCCCAGGTC | GAACCGTCCTTCTTTCTCCC | 158 |
| chr11:68941016-68943031 | p2 | (AT)6 | GCGATTCAAAATTGATGCCT | AGTCTTGCTCGGCATTTGTT | 198 |
| chr11:68941016-68943031 | p5 | (CCCAC)3 | GCGATTCAAAATTGATGCCT | AGTCTTGCTCGGCATTTGTT | 198 |
| chr11:68941016-68943031 | p2 | (AT)7 | GCGATTCAAAATTGATGCCT | AGTCTTGCTCGGCATTTGTT | 198 |
| chr11:68964777-68966792 | p3 | (TGA)5 | TTGGGCCTTTGTTTTGTTTC | ACTTCTGTCACGACCCAACC | 230 |

| Marker ID | SSR type | SSR motif | FORWARD PRIMER (5'-3') | REVERSE PRIMER (5'-3') | size (bp) |
|---|---|---|---|---|---|
| chr11:68986864-68988876 | p2 | (CT)6 | ATCCCAGAATTACGCACGAC | GGCCGGTAATGATGAAAGAA | 161 |
| chr11:69187765-69189780 | p5 | (TTTTG)3 | TGCCAAGAGAAAGAAAGGGA | TGGAATATGGAGTTCTCGGG | 189 |
| chr11:69250368-69252386 | p6 | (AAAAGA)3 | TACACCGGTTGGGGATTTTA | CCCTTGAAAACTGGAAACGA | 166 |
| chr11:69252640-69254660 | p2 | (TA)10 | AGCGTCTTGCGTCGTAAAAT | GGATCAAGTCGTGGTGGAGT | 127 |
| chr11:69252640-69254660 | p2 | (AT)6 | AGCGTCTTGCGTCGTAAAAT | GGATCAAGTCGTGGTGGAGT | 127 |
| chr11:69318520-69320532 | p2 | (CA)6 | CATTCAAGAAGATCGACGCA | TTTACAGGGGAGATGCTGCT | 296 |
| chr11:69349370-69351384 | p2 | (AG)7 | CCATGTTTTCGCAACTTCCT | TGCTTGTGAACGCAGTTTTC | 153 |
| chr11:69349370-69351384 | p3 | (TTG)5 | CCATGTTTTCGCAACTTCCT | TGCTTGTGAACGCAGTTTTC | 153 |
| chr11:69372284-69374316 | p3 | (GTT)5 | CGGGCTAGGCATATAAGCAG | CCTTTGAAAGAATGGGACGA | 138 |
| chr11:69372284-69374316 | p5 | (ATTGC)3 | CGGGCTAGGCATATAAGCAG | CCTTTGAAAGAATGGGACGA | 138 |
| chr11:69425876-69427914 | p2 | (AT)19 | GCAACAAACATGGAAGCTGA | GCATGATCCAAAGCCTGATT | 204 |
| chr11:69452402-69454440 | p2 | (AT)19 | GCAACAAACATGGAAGCTGA | GCATGATCCAAAGCCTGATT | 204 |
| chr11:69543558-69545570 | p2 | (AT)6 | TTGGAGAAGAGACACGGCTT | TGCCCTATTGGGCTTAGTTG | 207 |
| chr11:69543558-69545570 | p2 | (TA)8 | TTGGAGAAGAGACACGGCTT | TGCCCTATTGGGCTTAGTTG | 207 |
| chr11:69543558-69545570 | p2 | (AT)10 | TTGGAGAAGAGACACGGCTT | TGCCCTATTGGGCTTAGTTG | 207 |
| chr11:69544390-69546410 | p2 | (AT)6 | CCGTTTGGTAGGCAATCTGT | GCCACTTGGTCAGTGAGACA | 289 |
| chr11:69544390-69546410 | p2 | (TA)8 | CCGTTTGGTAGGCAATCTGT | GCCACTTGGTCAGTGAGACA | 289 |
| chr11:69544390-69546410 | p2 | (AT)10 | CCGTTTGGTAGGCAATCTGT | GCCACTTGGTCAGTGAGACA | 289 |
| chr11:69562146-69564164 | p6 | (CATATA)3 | AACCCTTAGACCGACCTGCT | GTCAGTCTGTAAGCGGCTCC | 142 |
| chr11:69564233-69566257 | p2 | (AT)12 | TGTACCATGGAACCCGAAAT | GTAATTGTTGCGGGCTGTTT | 137 |
| chr11:69673791-69675809 | p6 | (TTTTTA)3 | CGACTAATCGCAAACCCATT | AATTGGGAATTCGGGATAGG | 210 |
| chr11:69789237-69791253 | p2 | (AT)8 | GGTTGACAATTTGATGCGTG | ATATACAGTGCGGAGGACCG | 180 |
| chr11:69879012-69881030 | p2 | (AT)9 | ATCCACACCGGAGAGTCAAC | TTCTGGGTCATGGGTTTTTC | 252 |
| chr11:70002097-70004135 | p2 | (TA)19 | ATAATGAAAGGGGGTTTGGG | AGGCTTGACTTTAGGCCACA | 270 |
| chr11:70028038-70030053 | p3 | (ATT)5 | AATTTCCGGATGTGTTCTCG | GGACAAACAATACGGGTGCT | 225 |
| chr11:70053932-70055950 | p2 | (TA)9 | ATAAGCCGACGAGGCTAACA | CAGGGACTACATAGCGGAGC | 278 |
| chr11:70110632-70112647 | p3 | (CAA)5 | AGTGAGGGACTCGGTGAATG | TGAGCGAACGACGAACATAC | 287 |
| chr11:70214721-70216755 | p2 | (AT)17 | CTCTAACCACCTTTGCTGCC | GGCCTTGGTTGTGTTCTGAT | 122 |
| chr11:70276135-70278150 | p5 | (TTTCA)3 | GCCGGCACTACAAGGTATGT | GTATGGCTCTGCCATCCTGT | 244 |
| chr11:70306698-70308713 | p5 | (ATATA)3 | CGATGACGATAACGATGACG | CATGATAGGCTCGGGTTCAT | 185 |
| chr11:70306698-70308713 | p5 | (ATATA)3 | CGATGACGATAACGATGACG | CATGATAGGCTCGGGTTCAT | 185 |
| chr11:70468594-70470612 | p6 | (TATGAA)3 | CATGGATGTGGATGATTGGA | TTGCACGTTCTCCTTCAATG | 283 |
| chr11:70507196-70509214 | p6 | (TTTTGT)3 | ACCCTTATGCTTGGTAGGGG | AGCATACCCGACAACGTAGG | 281 |
| chr11:70527317-70529343 | p2 | (AT)13 | CTCGTGGTATCTGTCCGGTT | TGCATATGGCACAGGAACAT | 203 |
| chr11:70593972-70595988 | p2 | (TA)8 | TCGAGGGTTCACGACCTTAC | TATCGTATAGCTGCCCGCTT | 207 |
| chr11:70593972-70595988 | p2 | (TA)8 | TCGAGGGTTCACGACCTTAC | TATCGTATAGCTGCCCGCTT | 207 |

| Marker ID | SSR type | SSR motif | FORWARD PRIMER (5'-3') | REVERSE PRIMER (5'-3') | size (bp) |
|---|---|---|---|---|---|
| chr11:70612096-70614108 | p2 | (AG)6 | AGGTACCATGCAATGCAACA | TTTCTTACCGGGAACGATTG | 238 |
| chr11:70767748-70769760 | p2 | (GT)6 | TCTAATACTTCCCCCACCCC | AGATGAATGGCCTCGTTCAC | 100 |
| chr11:70879653-70881669 | p4 | (AGAA)4 | GAACATTTTGTACCACCCCG | CTAATCCCTGATCACCCGAA | 300 |
| chr11:70955323-70957338 | p2 | (AC)8 | GATTGGGCCGAGATCTGTTA | CATTCTGCACTTGCCTGAAA | 171 |
| chr11:70955323-70957338 | p5 | (TTAGT)3 | GATTGGGCCGAGATCTGTTA | CATTCTGCACTTGCCTGAAA | 171 |
| chr11:70970851-70972908 | p6 | (AAAGAG)3 | AAAAATCCCCCAGGTTCTTG | TGGAGTGGGAAATCGAAAAG | 208 |
| chr11:70982319-70984331 | p2 | (AT)6 | TTCCAGATTCCCGATGGTAG | TGGATTAAGCATGGCATGAA | 212 |
| chr11:71033651-71035669 | p6 | (TTTTCC)3 | ACCCTCTGTGGCTATCATGG | AAATGCCTGATTTTGAAGCG | 279 |
| chr11:71057740-71059760 | p2 | (AT)10 | CTCGGGTGGCAAGTAATGAT | GTCCATGCCTCCGAAACTAA | 104 |
| chr11:71206231-71208263 | p2 | (TA)16 | GGCCCAATACCACTTTCTGA | CCAAAGTGCAACGATTTGAA | 119 |
| chr11:71222594-71224606 | p4 | (CAAA)4 | AGACGTCTTACGCTCTCCCA | TTGTTTGATGTGTGGGCTGT | 288 |
| chr11:71222594-71224606 | p2 | (AC)6 | AGACGTCTTACGCTCTCCCA | TTGTTTGATGTGTGGGCTGT | 288 |
| chr11:71246886-71248898 | p2 | (AT)6 | ATCACCGTGAGTAAATGCCC | GCCCTTATCTTTTCCGAAGG | 242 |
| chr11:71340854-71342866 | p2 | (AT)6 | ATAGCAATGAACCCGTTTGC | TCTTATGGTAGGGACCGTGC | 205 |
| chr11:71430107-71432129 | p2 | (AT)11 | AAAGCTGTAGCTCCGGATGA | GAAATGGAAGGTCCCTGTGA | 254 |
| chr11:71606437-71608459 | p2 | (AT)6 | GGGCCCATAAGTGACTCAAA | TGCACTATTCCTTCTGCACG | 107 |
| chr11:71631038-71633053 | p5 | (AATGA)3 | ACCCCATTTATTTCCCTTCG | TTAAAGAACATTCGTCCCCG | 204 |
| chr11:71694082-71696100 | p3 | (TCA)5 | GCCACAAACTCATCCCAAGT | GACTGGTCAGGGCTTCAGAG | 156 |
| chr11:71694082-71696100 | p2 | (AT)9 | GCCACAAACTCATCCCAAGT | GACTGGTCAGGGCTTCAGAG | 156 |
| chr11:71749881-71751899 | p6 | (ATTCAT)3 | TCCTAAAATGTCCGCAGCTT | CCAACGGGACTAGCATGAAT | 153 |
| chr11:71749881-71751899 | p3 | (ATC)5 | TCCTAAAATGTCCGCAGCTT | CCAACGGGACTAGCATGAAT | 153 |
| chr11:71814659-71816675 | p2 | (AC)8 | CAAGCTACTCACGCCTTTCC | TGGGCTGTTGTGGTTTGTTA | 281 |
| chr11:71862822-71864837 | p5 | (CATTT)3 | AGGTTAATGATGAAACGCCG | AACATTCTTCCCCCTTTTGG | 299 |
| chr11:71878134-71880148 | p5 | (TGATA)3 | GATTTGTTGCGTGGGTTCTT | TTGATAACAAATCCTGGCCC | 123 |
| chr11:71878134-71880148 | p2 | (AT)7 | GATTTGTTGCGTGGGTTCTT | TTGATAACAAATCCTGGCCC | 123 |
| chr11:71878134-71880148 | p2 | (TA)10 | GATTTGTTGCGTGGGTTCTT | TTGATAACAAATCCTGGCCC | 123 |
| chr11:71890751-71892763 | p2 | (AT)6 | CTGGCCTAACACAATGCTGA | TTCCATTAAAACGTGGCTCC | 127 |
| chr11:71965024-71967036 | p2 | (AC)6 | CCACAGGTAGACAGGGAGGA | TTGTGAGACCATCTGACCCA | 153 |
| chr11:72005985-72008001 | p2 | (GT)8 | CATCCGGAAATTCGTTCACT | TAGAATGGAAAGGGAATGCG | 117 |
| chr11:72025499-72027517 | p2 | (TA)9 | CCCGTCAGCTCATATACCGT | CTTTTGGGTTAAGCGTGCTC | 114 |
| chr11:72089696-72091732 | p2 | (TA)18 | TTTGAATGAACGAGTGAGCG | GCTACCCGACCATATAGGCA | 261 |
| chr11:72095177-72097213 | p2 | (AT)18 | TCCACTCCAAAATCCCAGAC | CAGCACTTCAGTGATCGGAA | 192 |
| chr11:72101457-72103472 | p3 | (TTG)5 | GGAGGGGCCAAAACACTATT | GGTTCCATGGACCTTCTCAA | 240 |
| chr11:72170481-72172495 | p2 | (AT)7 | CAGGCGGAGATATGGTGATT | AAAAATGTGGAGGGTTGCAG | 279 |
| chr11:72170481-72172495 | p3 | (ATC)6 | CAGGCGGAGATATGGTGATT | AAAAATGTGGAGGGTTGCAG | 279 |
| chr11:72170481-72172495 | p3 | (ACA)6 | CAGGCGGAGATATGGTGATT | AAAAATGTGGAGGGTTGCAG | 279 |

| Marker ID | SSR type | SSR motif | FORWARD PRIMER (5'-3') | REVERSE PRIMER (5'-3') | size (bp) |
|---|---|---|---|---|---|
| chr11:72226242-72228257 | p2 | (AT)9 | CAGTGGAGTACGCACGAGAA | AAAAAGGGTTCTGCAAGGGT | 234 |
| chr11:72226242-72228257 | p3 | (AAC)5 | CAGTGGAGTACGCACGAGAA | AAAAAGGGTTCTGCAAGGGT | 234 |
| chr11:72226242-72228257 | p2 | (TA)10 | CAGTGGAGTACGCACGAGAA | AAAAAGGGTTCTGCAAGGGT | 234 |
| chr11:72255099-72257115 | p4 | (AATA)4 | TCCGCAGGTCTTCCAGTATC | CTCGAGCGTGTTACGCATTA | 256 |
| chr11:72389442-72391457 | p3 | (AAC)5 | GCCTTCAGTGTGGGAAACAT | TTATGATCCCGTCCGACTTC | 198 |
| chr11:72447918-72449936 | p3 | (ACA)6 | TTGGATCCCTTGATTCTTGC | ATTAGTCGGGCCAGCTTTTT | 189 |
| chr11:72463712-72465727 | p5 | (AAACA)3 | CCGCGTCTGCATCCTATTAT | CAAGAAGTGAAAGGCGAAGG | 107 |
| chr11:72476623-72478638 | p5 | (ATCAT)3 | TACATGACCCACCTTGCGTA | AAGTGTCGCCAATGAAAAGG | 109 |
| chr11:72563403-72565423 | p2 | (AC)10 | GACCATGCCAAAAGAAAGGA | TTTCGGCAGACAACATGAAG | 147 |
| chr11:72629655-72631671 | p2 | (AG)8 | AAGTCCCTCCGGAAAAGAAA | CGATGTCACCTCGGGATACT | 168 |
| chr11:72656985-72659000 | p5 | (ATTTC)3 | TGGCCTCACATGACCAGTAA | TTGGAGGAATCGGAATATCG | 113 |
| chr11:72677037-72679052 | p3 | (ACA)5 | AAGGCACACGAATCAAAACC | CGTTTCGCAAAATGAAACCT | 189 |
| chr11:72682190-72684202 | p2 | (AT)6 | CATCATTGTCACGACCCAAC | TGCTGTGGTATGATGTCCGT | 218 |
| chr11:72762380-72764394 | p2 | (AT)7 | TGTATTCCTCTTATGCCCGC | TATTCCGGATCAAGTGGAGC | 107 |
| chr11:72787705-72789720 | p3 | (TTG)5 | CATCGAGGACGGAATGTTCT | GGTTGCCAATCACCCTCTTA | 151 |
| chr11:72788745-72790761 | p2 | (AT)8 | TCCTTTCTTTTGCCATGTCC | GGCAATCCGATTTCATTTGT | 188 |
| chr11:72819609-72821629 | p2 | (TA)10 | TCGTGTCATTCGGGTCAATA | TTTCGGATTTAGGCTCCCTT | 148 |
| chr11:72835066-72837088 | p2 | (TA)11 | CCACTTGCCATTCTACAGCA | TGGAACCATAATAGCCGAGG | 131 |
| chr11:72847067-72849079 | p2 | (TA)6 | CTTCGTTGGCTTTCCATTGT | CCTTGTCGGCTTGCATATTT | 131 |
| chr11:72877059-72879107 | p2 | (TA)24 | GGAACACAGCGTCTGTGAAA | TCGACTAATCTTATGGGGCG | 243 |
| chr11:72877059-72879107 | p3 | (TTG)5 | GGAACACAGCGTCTGTGAAA | TCGACTAATCTTATGGGGCG | 243 |
| chr11:72913254-72915278 | p2 | (AT)12 | AGGAAGCGTGGAAAGGAAAT | TCCTCTTGGCTGCTAATCGT | 232 |
| chr11:73029815-73031833 | p6 | (ATATAC)3 | TCACTGACAATCCTAGGCCC | TATGCGTTGGCTGTGAGTTC | 225 |
| chr11:73129391-73131403 | p2 | (TA)6 | TACGCTTTCACTTGGGCTCT | CATCCAATTAGCCCACGTCT | 152 |
| chr11:73129391-73131403 | p2 | (TA)6 | TACGCTTTCACTTGGGCTCT | CATCCAATTAGCCCACGTCT | 152 |
| chr11:73181602-73183614 | p2 | (GT)6 | AGCCCGCATTTTGTACATTC | CTCGGTAATTCCGCATCAAT | 217 |
| chr11:73214067-73216097 | p2 | (AT)15 | TCCACTGCATTATTCACCGA | ACGTTTCCCGACCCTCTAGT | 194 |
| chr11:73293050-73295062 | p2 | (CA)6 | TCCTCACTTTGAACCCAAGG | GTCAGCCTCAGCTTCTCCAC | 202 |
| chr11:73362086-73364101 | p2 | (AT)6 | GTTTTGGAGAGCCAAATGGA | ATCCACAACACAACAACCCA | 291 |
| chr11:73362086-73364101 | p5 | (TGTGT)3 | GTTTTGGAGAGCCAAATGGA | ATCCACAACACAACAACCCA | 291 |
| chr11:73362086-73364101 | p5 | (TGAAT)3 | GTTTTGGAGAGCCAAATGGA | ATCCACAACACAACAACCCA | 291 |
| chr11:73457803-73459815 | p2 | (TA)6 | ATATAGGGGAAATGCTGCCC | GTTCGTAAATCCGTAGCCCA | 251 |
| chr11:73457803-73459815 | p2 | (AT)9 | ATATAGGGGAAATGCTGCCC | GTTCGTAAATCCGTAGCCCA | 251 |
| chr11:73494309-73496333 | p6 | (TTCATA)4 | TGGGATTGGCAGGCTATTAC | GTGTCAGGACTGGGGCTAAA | 126 |
| chr11:73499192-73501207 | p3 | (TAT)5 | CATGATGTAATCGGGGGTTC | CATTTGGGTACGGTGGAATC | 190 |
| chr11:73562404-73564472 | p2 | (TA)34 | GTCCACCGAGGTAGATTGGA | TTGATCGCCAAGTTCTTTCC | 140 |

| Marker ID | SSR type | SSR motif | FORWARD PRIMER (5'-3') | REVERSE PRIMER (5'-3') | size (bp) |
|---|---|---|---|---|---|
| chr11:73562404-73564472 | p2 | (TA)9 | GTCCACCGAGGTAGATTGGA | TTGATCGCCAAGTTCTTTCC | 140 |
| chr11:73638732-73640790 | p2 | (AT)29 | TGCCTCAGACTGAACCTCCT | AGACAGGTTCGTCGGTTACG | 157 |
| chr11:73644468-73646480 | p2 | (AT)15 | TATCAGAAGCCGACAAGGCT | TCCCGTCATATGATTCGGTT | 147 |
| chr11:73644468-73646480 | p2 | (TA)6 | TATCAGAAGCCGACAAGGCT | TCCCGTCATATGATTCGGTT | 147 |
| chr11:73778772-73780790 | p3 | (CAA)6 | CGACCCTCTTTTGCGAGTAG | CTGTTATTCCTTTCGCCGAG | 183 |
| chr11:73803964-73805976 | p2 | (GT)6 | TATCCTCCTATGTGCCCGAC | TTACCGGCCTTACAAACGTC | 112 |
| chr11:73928506-73930536 | p2 | (AT)15 | TGTTCTCGTCGACTCGTTTG | ACGGTGATTTGGATGGGTAA | 216 |
| chr11:73934366-73936382 | p2 | (GA)8 | CTAGAGGTGGACAGCAAGCC | CACGAAATTTTCTGGGGTGT | 266 |
| chr11:73934366-73936382 | p3 | (GTT)5 | CTAGAGGTGGACAGCAAGCC | CACGAAATTTTCTGGGGTGT | 266 |
| chr11:73978475-73980489 | p2 | (TA)7 | TTCGGGCAAGGCATATAAAC | ATAGGGTAACCCCAAGGTGC | 112 |
| chr11:74007574-74009586 | p2 | (AG)6 | CTAACCCTTTTTCTTCGGGG | CCATTATTCACGGGGACATC | 172 |
| chr11:74007574-74009586 | p3 | (TGT)5 | CTAACCCTTTTTCTTCGGGG | CCATTATTCACGGGGACATC | 172 |
| chr11:74091043-74093058 | p5 | (ATAAA)3 | TATTGACATGGTGGGCTTCA | ACTTCGCTAACGGTAGGGGT | 296 |
| chr11:74161593-74163611 | p2 | (AT)9 | TTCTGCTGTGCCATCAAGTC | CTTAGGTGAGACGCGGTAGG | 248 |
| chr11:74234850-74236866 | p2 | (TA)7 | TCGTGCATGCCTCAATATGT | GGGACTCGGTGAAAAAGTCA | 147 |
| chr11:74234867-74236903 | p2 | (TA)7 | TCGTGCATGCCTCAATATGT | GGGACTCGGTGAAAAAGTCA | 130 |
| chr11:74316074-74318089 | p5 | (AATTC)3 | CGTACATGGCACAGGCATAC | TGTTTCCCTACACTGAGCCC | 201 |
| chr11:74321533-74323551 | p2 | (AT)9 | AGCTCCCCCAAATTCGTACT | GTCATCCATTGCACGTATGC | 172 |
| chr11:74354562-74356577 | p3 | (CAA)5 | AAATCAATGAAGTCGGACGG | CCCGCTTTGCTCTCTTATTG | 245 |
| chr11:74382144-74384159 | p5 | (CATAT)3 | TGGACACTCAAACGTACCCA | ATTGCGTACAAGGATCTGCC | 196 |
| chr11:74418129-74420141 | p2 | (GT)6 | TGATTTCGGAAGGCCATAAC | GATGTCACATGCAGACCCAC | 111 |
| chr11:74418129-74420141 | p3 | (TTG)5 | TGATTTCGGAAGGCCATAAC | GATGTCACATGCAGACCCAC | 111 |
| chr11:74449599-74451614 | p2 | (AG)6 | GGAATTTGGGAAAACCCCTA | TCCCTTAAATGGCATGGGTA | 222 |
| chr11:74449599-74451614 | p3 | (GTT)5 | GGAATTTGGGAAAACCCCTA | TCCCTTAAATGGCATGGGTA | 222 |
| chr11:74449599-74451614 | p2 | (TA)15 | GGAATTTGGGAAAACCCCTA | TCCCTTAAATGGCATGGGTA | 222 |
| chr11:74452781-74454799 | p3 | (CAA)6 | TTTTAACGCAAAACGGGAAC | CGAACCCAGCAAATAAAGGA | 157 |
| chr11:74464319-74466334 | p5 | (GAAGT)3 | AAGTGAAAATCCGTTGACCG | TGGCAAAGAACCAACTTTCC | 118 |
| chr11:74523688-74525700 | p2 | (AC)6 | ATATACGCCTCGCCCTTTTT | ATGGACTCGCTTACTCGCAT | 246 |
| chr11:74523688-74525700 | p2 | (TA)6 | ATATACGCCTCGCCCTTTTT | ATGGACTCGCTTACTCGCAT | 246 |
| chr11:74523688-74525700 | p3 | (ATC)5 | ATATACGCCTCGCCCTTTTT | ATGGACTCGCTTACTCGCAT | 246 |
| chr11:74524512-74526527 | p2 | (TA)6 | TTATCCGTACTGACTCCCCG | GAAGCCGACAAGGCTACATC | 222 |
| chr11:74524512-74526527 | p3 | (ATC)5 | TTATCCGTACTGACTCCCCG | GAAGCCGACAAGGCTACATC | 222 |
| chr11:74547707-74549722 | p3 | (ATC)5 | CCGTTTTGATCCGTTCAACT | CACCTGAGGAGGGAGATGAG | 128 |
| chr11:74619681-74621695 | p2 | (AT)9 | TGCGAAAACATGGGACATAA | TGCGTATGGCATTGATCTGT | 149 |
| chr11:74619681-74621695 | p2 | (AT)7 | TGCGAAAACATGGGACATAA | TGCGTATGGCATTGATCTGT | 149 |
| chr11:74619681-74621695 | p2 | (AT)7 | TGCGAAAACATGGGACATAA | TGCGTATGGCATTGATCTGT | 149 |

| Marker ID | SSR type | SSR motif | FORWARD PRIMER (5'-3') | REVERSE PRIMER (5'-3') | size (bp) |
|---|---|---|---|---|---|
| chr11:74678273-74680297 | p2 | (TA)12 | GTTTACCCATGCTTCGTCGT | AAATGACATGGCGGAATAGC | 246 |
| chr11:74729897-74731921 | p2 | (TA)12 | GTTTACCCATGCTTCGTCGT | AAATGACATGGCGGAATAGC | 246 |
| chr11:74824814-74826826 | p2 | (AT)6 | CAGCCAACTCCACAGACTCA | GTGGTCGTTCGGACAGGTAT | 142 |
| chr11:74838356-74840376 | p2 | (TC)10 | GTTTCGCCCAAAACCTACAA | TCGTTAGAATTTCGGGCATC | 177 |
| chr11:74887462-74889494 | p2 | (AT)16 | GGAGCATTTATCAGCCCAAA | GTCGGCTTGCGTATCATTTT | 112 |
| chr11:74981715-74983729 | p2 | (TG)7 | AAAGGTGCCTACGGGCTATT | CAAGGCTTCAGGGTTATCCA | 219 |
| chr11:74996425-74998439 | p2 | (TG)7 | AAAGGTGCCTACGGGCTATT | CCAACCAAGGCTCAAGGTTA | 221 |
| chr11:75013124-75015136 | p2 | (CA)6 | AATGCCATTCGGAAACAGAC | ATAGGAAGCGGGCAAGGTAT | 150 |
| chr11:75104869-75106884 | p5 | (GTTTT)3 | ATCCCAATTTCGGTTTACCC | ACAAAATTGTCATCCCCCAA | 171 |
| chr11:75190109-75192143 | p2 | (TA)17 | CGAAACCTTTCCTAAACCCC | TCGTCGCGTAGTGAATGAAC | 138 |
| chr11:75324805-75326821 | p2 | (AT)8 | ATTTCTGGGAAAATTTGGGC | AATTGTTAGCCCTGTGTGGC | 197 |
| chr11:75416648-75418663 | p5 | (TATAT)3 | TCGTGTTGTGGTCGTGTTTT | CCCTTCTTAACCCCTTTCCA | 294 |
| chr11:75416648-75418663 | p2 | (AT)11 | TCGTGTTGTGGTCGTGTTTT | CCCTTCTTAACCCCTTTCCA | 294 |
| chr11:75431996-75434008 | p2 | (AT)6 | ATTCGATTTCAGAGGCATGG | ACCGGGCATCAAGTACAAAG | 183 |
| chr11:75578960-75580975 | p3 | (TTG)5 | CTTGTTGTTTTCCCGCATTT | TGGGGATCTACACTTTTGCC | 229 |
| chr11:75662785-75664797 | p2 | (AT)6 | TGACCCAAAGAATTTCGGAG | GTGTCTGAGGCCATCCATTT | 171 |
| chr11:75683297-75685319 | p2 | (AT)11 | CGTCTCCCTATCTACCTGCG | CTCCCCCTTAAGAGGCATTC | 109 |
| chr11:75683297-75685319 | p2 | (TA)7 | CGTCTCCCTATCTACCTGCG | CTCCCCCTTAAGAGGCATTC | 109 |
| chr11:75683449-75685463 | p2 | (AT)11 | GCCTCTTAAGGGGGAGTGTC | CGGATGACACCGATCCTATT | 110 |
| chr11:75683449-75685463 | p2 | (TA)7 | GCCTCTTAAGGGGGAGTGTC | CGGATGACACCGATCCTATT | 110 |
| chr11:75868590-75870605 | p5 | (ATAAA)3 | ATTGCATTTGTCGTTGGTGA | AAGTACGCGCGCTTAAGAAG | 133 |
| chr11:75870766-75872788 | p2 | (AT)11 | AGCCCTTCGGCTATCAAAAT | CCTGTCCCGTCCATATGTCT | 202 |
| chr11:75870766-75872788 | p2 | (AT)40 | AGCCCTTCGGCTATCAAAAT | CCTGTCCCGTCCATATGTCT | 202 |
| chr11:75917901-75919958 | p3 | (TTA)19 | CGGTGCAACCATCTTTAGGT | TCCCTTGAGCAAGGAGTGTT | 150 |
| chr11:75920359-75922401 | p2 | (AT)8 | GCAGTGTTAGGACTCGCCTC | GTGTTTAGGGCGTACGGTTG | 287 |
| chr11:75920359-75922401 | p2 | (GC)7 | GCAGTGTTAGGACTCGCCTC | GTGTTTAGGGCGTACGGTTG | 287 |
| chr11:75920507-75922521 | p2 | (GC)7 | GCAGTGTTAGGACTCGCCTC | GTGTTTAGGGCGTACGGTTG | 139 |
| chr11:76034433-76036477 | p4 | (GAAA)11 | GATAAAGGAAGCGGGTCACA | GAAGGTATTTCGTGGCGTGT | 168 |
| chr11:76034433-76036477 | p2 | (AT)10 | GATAAAGGAAGCGGGTCACA | GAAGGTATTTCGTGGCGTGT | 168 |
| chr11:76034433-76036477 | p5 | (AATGC)3 | GATAAAGGAAGCGGGTCACA | GAAGGTATTTCGTGGCGTGT | 168 |
| chr11:76262707-76264725 | p3 | (TCA)6 | CCTGCTCCTCGTTCTTTCAC | AGGCAAAGAGATGGAAGGGT | 165 |
| chr11:76277309-76279327 | p6 | (AACCGG)3 | TCCTGGATCTTGAACGCTCT | GTTTTTCCCGTCTTGGGTTT | 188 |
| chr11:76279748-76281762 | p2 | (TA)7 | AAAACTCAGAAGCGGAGCAA | TTTCGTTAATTTCCCCCACA | 179 |
| chr11:76279748-76281762 | p2 | (AT)6 | AAAACTCAGAAGCGGAGCAA | TTTCGTTAATTTCCCCCACA | 179 |
| chr11:76282994-76285008 | p2 | (GT)7 | AATTCGCAGCGGAGATAAGA | TTTAAGCCGATTCTATGCGG | 240 |
| chr11:76301267-76303282 | p3 | (TTG)5 | AAGTACATGCGCTTGACACG | ACTCAGGTGCAGATCTCCGT | 110 |

| Marker ID | SSR type | SSR motif | FORWARD PRIMER (5'-3') | REVERSE PRIMER (5'-3') | size (bp) |
|---|---|---|---|---|---|
| chr11:76323724-76325739 | p5 | (AAACC)3 | GTAGTGTGGTGGTGGAGCCT | TTCTCCCATTTTCATAGCCG | 247 |
| chr11:76329948-76331969 | p3 | (TTC)7 | CCAACTTATGCCTTGCGAAT | TGGCCTTTAATTTTTGTCCG | 144 |
| chr11:76329948-76331969 | p3 | (TTG)7 | CCAACTTATGCCTTGCGAAT | TGGCCTTTAATTTTTGTCCG | 144 |
| chr11:76329948-76331969 | p5 | (TTTGG)5 | CCAACTTATGCCTTGCGAAT | TGGCCTTTAATTTTTGTCCG | 144 |
| chr11:76398044-76400068 | p2 | (AT)12 | AAGAAGACGGACAACGAGGA | AATTCGGTTTGGGCATGTAG | 151 |
| chr11:76431092-76433110 | p6 | (TTTTCC)3 | AGAAGCATACACAATGCCCC | GTTTTGTCTGCCTGGTCCAT | 181 |
| chr11:76431835-76433889 | p6 | (TTTTCC)3 | TTCCTTTTCCTTTTTGGGCT | GGGCGATTATGATAGCAGGA | 265 |
| chr11:76431835-76433889 | p2 | (GA)29 | TTCCTTTTCCTTTTTGGGCT | GGGCGATTATGATAGCAGGA | 265 |
| chr11:76455417-76457447 | p6 | (CAATAT)5 | CCCATGGCCTCTCGTAGATA | GTGCAGTTCCTTGGGGATTA | 142 |
| chr11:76505129-76507144 | p3 | (ATA)5 | AGCTTTGAAAAGGCCTCACA | TTCATGCTCAAGCAGACTGG | 181 |
| chr11:76511196-76513211 | p5 | (CCACA)3 | AATGCTTCACCTTCCAATGC | GGGTATGGTCCATCAACCTG | 255 |
| chr11:76511196-76513211 | p2 | (AT)6 | AATGCTTCACCTTCCAATGC | GGGTATGGTCCATCAACCTG | 255 |
| chr11:76511541-76513553 | p5 | (CCACA)3 | ACCTTCGAACGCACTTCATC | CAATGTGCAGGGGAGTAGGT | 234 |
| chr11:76511541-76513553 | p2 | (AT)6 | ACCTTCGAACGCACTTCATC | CAATGTGCAGGGGAGTAGGT | 234 |
| chr11:76569716-76571728 | p2 | (AT)6 | AGTCCCAATTGGACGACAAC | GGAGCTTCATTTCATTCGGA | 289 |
| chr11:76720229-76722257 | p2 | (AT)14 | ATTTTTCCGTCCAGGGTTCT | TGTGAGGCATTGGATGTGTT | 294 |
| chr11:76720229-76722257 | p2 | (TA)9 | ATTTTTCCGTCCAGGGTTCT | TGTGAGGCATTGGATGTGTT | 294 |
| chr11:76720397-76722415 | p2 | (AT)14 | ATTTTTCCGTCCAGGGTTCT | TGTGAGGCATTGGATGTGTT | 126 |
| chr11:76720397-76722415 | p2 | (TA)9 | ATTTTTCCGTCCAGGGTTCT | TGTGAGGCATTGGATGTGTT | 126 |
| chr11:76826524-76828552 | p2 | (TA)14 | GCCTCAAAATCTTCAAAGCG | CATTGCTTATGCTTCCCGAT | 142 |
| chr11:76931438-76933450 | p2 | (TA)6 | CGAAAACGTTAAGGCTACGG | TACAACCGATCCAACGTCAA | 135 |
| chr11:77049676-77051696 | p2 | (TA)10 | TAATGCGGACTCCATGTTGA | ATACGCCTGCGATTTTTCAG | 186 |
| chr11:77247158-77249173 | p5 | (TGGCT)3 | GAACCAGGGACGGAGTACAA | AAATGCCAACGTGAACATGA | 254 |
| chr11:77293195-77295225 | p2 | (TG)15 | AATTGCTGAAAATCGCCAAC | TTGGGCAAGGTTTTCATCTC | 281 |
| chr11:77336667-77338693 | p2 | (TA)13 | TAATTTGGGGGTTTTGCTTG | CTCCTTTGCCTTTCCTTTCC | 115 |
| chr11:77391283-77393298 | p5 | (ATTTC)3 | ACGTTCATGCTGAGTTGCAG | TGGCCCTACTTGAGGAAGAA | 135 |
| chr11:77396157-77398172 | p5 | (ATTTC)3 | GGCCTGATAACAACCGAGAA | TTGTCACAACAGCCCACAAT | 205 |
| chr11:77400821-77402836 | p5 | (AATGC)3 | CTCCTCAAAGCCTTCACGTC | TGACTTTGCAATTGTTGGGA | 165 |
| chr11:77400821-77402836 | p5 | (CCTTT)4 | CTCCTCAAAGCCTTCACGTC | TGACTTTGCAATTGTTGGGA | 165 |
| chr11:77447217-77449233 | p2 | (TA)8 | CCACATTCTGTCATTGTGCC | CAAGACTACGGCAAAGAGGC | 220 |
| chr11:77482301-77484313 | p2 | (CA)6 | CCTCCTACAGTTTCGACCCA | GCCAAGACGTGAGAGAGGTC | 172 |
| chr11:77482301-77484313 | p6 | (ATTCAT)3 | CCTCCTACAGTTTCGACCCA | GCCAAGACGTGAGAGAGGTC | 172 |
| chr11:77514677-77516691 | p2 | (TA)7 | TCCGTGTTTCCCTTTCTTTG | CCGAGCCCATTAGTGTCATT | 106 |
| chr11:77587233-77589251 | p5 | (ATATA)3 | TACGGCCCGATCCATAAATA | ATGTCACCAGAGGGGTTCAG | 289 |
| chr11:77587233-77589251 | p3 | (ACA)6 | TACGGCCCGATCCATAAATA | ATGTCACCAGAGGGGTTCAG | 289 |
| chr11:77587233-77589251 | p2 | (CT)7 | TACGGCCCGATCCATAAATA | ATGTCACCAGAGGGGTTCAG | 289 |

| Marker ID | SSR type | SSR motif | FORWARD PRIMER (5'-3') | REVERSE PRIMER (5'-3') | size (bp) |
|---|---|---|---|---|---|
| chr11:77623175-77625201 | p5 | (AAATT)3 | GTTACGCCTTCGGGTATGAA | TTCACAATCACGGTAAGGCA | 176 |
| chr11:77787927-77789941 | p2 | (TA)7 | CTTTGTAGGGATCGCGAGAG | ACTGGCCAATGTAATCTCCG | 254 |
| chr11:77811440-77813455 | p3 | (ACA)5 | AAGGGATTCCTTGCTTCCAT | GGTCGGGCCAGATTTTATTT | 276 |
| chr11:77902626-77904638 | p2 | (AT)6 | AAAGGAAGGCGTGGAAAAAT | AGATGGTTCGCCTACCAATG | 282 |
| chr11:77941343-77943357 | p2 | (AT)7 | TGTGGCCACTCTTTTAAGCA | TTCTGCACTCCGCAGAATAA | 258 |
| chr11:77948922-77950940 | p3 | (TAA)6 | GGCAACCCTGAAAGCTACTG | GACAAGATTGCCCAAAGCAT | 161 |
| chr11:78156399-78158429 | p2 | (TA)15 | CTCCATGGCCAACTCAGACT | CTCGGAGAGCATAAAGGTGC | 130 |
| chr11:78200120-78202135 | p5 | (GTGTC)3 | GGGTACTTGGTGGCTTTTGA | TGGAAAATGTTGAGCAGCAG | 268 |
| chr11:78210705-78212720 | p3 | (TTG)6 | TTGCATTTGATTGCTGCTTC | TGGGGAGGGTAGAGTGTACG | 277 |
| chr11:78210705-78212720 | p5 | (TTATT)3 | TTGCATTTGATTGCTGCTTC | TGGGGAGGGTAGAGTGTACG | 277 |
| chr11:78233491-78235557 | p2 | (AT)33 | ATTGCGGACTAAGGATGTGG | TAATGCCACACAGGGCTACA | 212 |
| chr11:78263393-78265414 | p3 | (AAC)7 | GGTAGACGGGGAAGAAGACC | TCCCTTAGCCCGTAGATCCT | 163 |
| chr11:78515573-78517588 | p3 | (TTC)6 | AGCTCCTTCATCCCCTTCAT | AGGAAACCCACGAAACCTCT | 300 |
| chr11:78515573-78517588 | p3 | (ACA)5 | AGCTCCTTCATCCCCTTCAT | AGGAAACCCACGAAACCTCT | 300 |
| chr11:78616360-78618380 | p5 | (AAAAG)4 | GCAAAGGTGGCGGTAGTAGA | TTTTGCTGACATGGCAAAAG | 203 |
| chr11:78694363-78696378 | p5 | (AAGAA)3 | CGAGCTAGACTTACCGTCCG | ACCTACAGGCTGCTCTGCAT | 154 |
| chr11:78770476-78772488 | p2 | (AG)6 | TCTACGCGTTCCGTCCTAGT | ACTTTCCGACAGGCTTCTCA | 277 |
| chr11:78770476-78772488 | p2 | (GT)7 | TCTACGCGTTCCGTCCTAGT | ACTTTCCGACAGGCTTCTCA | 277 |
| chr11:78770802-78772816 | p2 | (AG)6 | TGAGAAGCCTGTCGGAAAGT | GGCCACCTTTCATAATTCCA | 194 |
| chr11:78770802-78772816 | p2 | (GT)7 | TGAGAAGCCTGTCGGAAAGT | GGCCACCTTTCATAATTCCA | 194 |
| chr11:78813734-78815746 | p2 | (AC)6 | TCGGGTTTCTAGGAGCAGAA | TGCAGATTTTTGACGAAACG | 191 |
| chr11:78993228-78995242 | p2 | (TA)7 | TTGTAATTTTGGGCTAGGCG | AAGGGGACGAGCCTTAACAT | 139 |
| chr11:78993228-78995242 | p2 | (TA)16 | TTGTAATTTTGGGCTAGGCG | AAGGGGACGAGCCTTAACAT | 139 |
| chr11:79047948-79049968 | p5 | (ATCAT)4 | GAGCGCTCCCAATATAGCAG | GGGACCAGGCTAACGACATA | 267 |
| chr11:79110254-79112276 | p5 | (TTTTC)3 | CAAATCTCCTGCCCACAAAT | GAGAGTGTCCTGACGGCTTC | 244 |
| chr11:79275733-79277748 | p3 | (TTC)5 | ACGTTGCGGGTGACTAAATC | GGCAGAAAATGACCCAGAAA | 158 |
| chr11:79311162-79313177 | p3 | (TTC)5 | ACGTTGCGGGTGACTAAATC | GGCAGAAAATGACCCAGAAA | 158 |
| chr11:79327416-79329442 | p2 | (AT)13 | GTGTATGGCCTCGGTCACTT | CAGAAGGGCAAAGCTACGAC | 299 |
| chr11:79405641-79407656 | p3 | (CAA)5 | CCGGTGAGGCTGTCATAGAT | CCCGCTTTGCTCTCTTATTG | 100 |
| chr11:79440963-79443055 | p2 | (TA)8 | CACGCTTTGAGACGTTTTGA | TTGCCCAATGATCCAATGTA | 208 |
| chr11:79440963-79443055 | p2 | (AT)46 | CACGCTTTGAGACGTTTTGA | TTGCCCAATGATCCAATGTA | 208 |
| chr11:79844260-79846288 | p2 | (AT)14 | CTGAGCGGCTGTAGCTTTCT | GGTTGTCTGCACACCATGTC | 222 |
| chr11:79844260-79846288 | p2 | (AT)6 | CTGAGCGGCTGTAGCTTTCT | GGTTGTCTGCACACCATGTC | 222 |
| chr11:80002032-80004048 | p3 | (ACA)5 | CTTTACGCTCCTCAAATCGC | GGGCTGTTGTTGTTGTTGTG | 253 |
| chr11:80002032-80004048 | p2 | (AC)8 | CTTTACGCTCCTCAAATCGC | GGGCTGTTGTTGTTGTTGTG | 253 |
| chr11:80002032-80004048 | p6 | (TATATG)3 | CTTTACGCTCCTCAAATCGC | GGGCTGTTGTTGTTGTTGTG | 253 |

| Marker ID | SSR type | SSR motif | FORWARD PRIMER (5'-3') | REVERSE PRIMER (5'-3') | size (bp) |
|---|---|---|---|---|---|
| chr11:80031925-80033940 | p5 | (ATCAT)3 | TACATGGAAGCCGATAAGGC | AACGACAGGTTTGCTGATCC | 297 |
| chr11:80100180-80102192 | p2 | (TA)6 | CAGCTATGCAAGCCGATGTA | TACCACAGCCAGCAGAACAC | 111 |
| chr11:80241577-80243592 | p5 | (ATTTC)3 | AAGGGGACAAGAACATGCAC | ACAGCCACGAATAAACGGAC | 129 |
| chr11:80287103-80289121 | p6 | (GTTTTT)3 | AGTGAGACATGGCCGAAAAC | CCCCGAATTCAGTGGTTCTA | 166 |
| chr11:80328199-80330214 | p3 | (ATC)5 | GGATTCTGTCATGCCCAAGT | TTCTTGTGGACGCTGTGTTC | 100 |
| chr11:80328199-80330214 | p3 | (CAA)7 | GGATTCTGTCATGCCCAAGT | TTCTTGTGGACGCTGTGTTC | 100 |
| chr11:80357423-80359438 | p5 | (GAATT)3 | CCTGAGGGGTCATTTTCAGA | TCATCCTCCGCCTGTATTTC | 226 |
| chr11:80440335-80442350 | p5 | (ATGCA)3 | AAAATTTAGCCCGTGAACCC | GAAAAGAAGACCCGAAAGGG | 202 |
| chr11:80440335-80442350 | p2 | (TA)10 | AAAATTTAGCCCGTGAACCC | GAAAAGAAGACCCGAAAGGG | 202 |
| chr11:80464143-80466161 | p6 | (CTTCAA)3 | TCAAGCCCAAATATCGTTCC | GCAGATGGCCTTGGAAATTA | 137 |
| chr11:80488206-80490228 | p2 | (TA)11 | ACAATCAATCCGAGGTCCTG | TAATTCTGACCTCCGTTGGG | 198 |
| chr11:80488206-80490228 | p5 | (AATTT)3 | ACAATCAATCCGAGGTCCTG | TAATTCTGACCTCCGTTGGG | 198 |
| chr11:80516934-80518946 | p2 | (AC)6 | CCGGTGTTTGTCAGAGTCCT | ATGGAATTAGTCGAGGTGCG | 166 |
| chr11:80516934-80518946 | p6 | (GATGAA)3 | CCGGTGTTTGTCAGAGTCCT | ATGGAATTAGTCGAGGTGCG | 166 |
| chr11:80517318-80519336 | p2 | (AC)6 | AACAATGCAATCTCCCAAGG | TTGCCTTTTTGGAGCTTTTG | 229 |
| chr11:80517318-80519336 | p6 | (GATGAA)3 | AACAATGCAATCTCCCAAGG | TTGCCTTTTTGGAGCTTTTG | 229 |
| chr11:80563256-80565271 | p3 | (AAG)5 | TTTCCATTAGCCTTTTTGGC | CCTCCGTAAAACATGGCTGT | 277 |
| chr11:80614270-80616334 | p2 | (TA)32 | GTGATCGTCGGTGTTTGTTG | GGGGGAGGGTCAACCTAATA | 156 |
| chr11:80618142-80620160 | p6 | (ATCCGT)3 | ACGAGGAGGAACTTTCGGAT | ATTCGGGTCGTGACAAAAAG | 218 |
| chr11:80624354-80626369 | p5 | (TGATA)3 | TACCCCGCTAGGGCTATTTT | TCGTCACGAACGTTTCAGAG | 274 |
| chr11:80638655-80640667 | p2 | (TA)6 | CCCCTGTGAAGAAGGTGGTA | GGATCATGCCTTTGCTCAAT | 102 |
| chr11:80688596-80690611 | p5 | (ATTTA)3 | GATGCACCGTTATGTTGTGG | AACTTCGTCACGACCAAACC | 285 |
| chr11:80772555-80774570 | p5 | (TATGA)3 | ATTGACCACTCTGGTAGCGG | AGACCCGAAAAACTCTGGGT | 151 |
| chr11:80788472-80790486 | p2 | (AT)7 | ACTTTTGTCACGACCCAACC | CCTTTACGGCTGGTGATGAT | 157 |
| chr11:80788472-80790486 | p2 | (AT)12 | ACTTTTGTCACGACCCAACC | CCTTTACGGCTGGTGATGAT | 157 |
| chr11:80854558-80856570 | p2 | (AT)6 | TGTACCGAGCACCCCTTATC | ATTTTGGCATCTTAGCGTCG | 170 |
| chr11:80854558-80856570 | p2 | (AT)6 | TGTACCGAGCACCCCTTATC | ATTTTGGCATCTTAGCGTCG | 170 |
| chr11:80861420-80863432 | p2 | (AG)6 | CCCGATTCTCTCGTTATGGA | CCAAATTGCCAGACATCCTT | 123 |
| chr11:80976090-80978112 | p2 | (AT)11 | TCAATGCTTCTCCTCCCATC | GTCGGCTTGCGTATCATTTT | 166 |
| chr11:80981482-80983510 | p2 | (AT)14 | ACTTTTGTCACGACCCAACC | TAGTGGTAGCCTTGTCGGCT | 176 |
| chr11:80981482-80983510 | p3 | (ACA)5 | ACTTTTGTCACGACCCAACC | TAGTGGTAGCCTTGTCGGCT | 176 |
| chr11:81022000-81024030 | p3 | (ATT)10 | CTCTTGGCTTGTTTGCATGA | CAAGCCACAAAGGCCATAAT | 275 |
| chr11:81066140-81068160 | p4 | (TATG)5 | CGACTTCTGGGTTAGCTTCG | TGCAAAAATTTCAAAGTGCG | 295 |
| chr11:81121788-81123806 | p2 | (AT)9 | ATGAATGGTGTGATGGGGTT | CACTGGCTCCTCCTCAAAAG | 189 |
| chr11:81224075-81226093 | p3 | (CAA)6 | GGAAATAAAGCCTTTTCGGC | TCCGAGCATGTTATGATCCA | 172 |
| chr11:81224075-81226093 | p3 | (CAA)6 | GGAAATAAAGCCTTTTCGGC | TCCGAGCATGTTATGATCCA | 172 |

| Marker ID | SSR type | SSR motif | FORWARD PRIMER (5'-3') | REVERSE PRIMER (5'-3') | size (bp) |
|---|---|---|---|---|---|
| chr11:81224075-81226093 | p5 | (CAACA)3 | GGAAATAAAGCCTTTTCGGC | TCCGAGCATGTTATGATCCA | 172 |
| chr11:81233847-81235867 | p2 | (AT)10 | TACTGTTGGGGCATTTGTCA | TGCGGATGTGCATTGATTAT | 131 |
| chr11:81251611-81253623 | p2 | (GA)6 | GACTGGTGTCCACCTCTGGT | GATGGTCGCCGTACCTTCTA | 158 |
| chr11:81275137-81277161 | p6 | (CATATA)4 | GATGGAGCCGCTTACAGAAC | ACATTGCTCGTACTGACCCC | 257 |
| chr11:81275137-81277161 | p5 | (ATTCA)4 | GATGGAGCCGCTTACAGAAC | ACATTGCTCGTACTGACCCC | 257 |
| chr11:81288461-81290495 | p2 | (AT)17 | CTTCCATGTCCCGACCTAAA | GTATGGCCTTGTCGACCTGT | 235 |
| chr11:81343403-81345415 | p2 | (AT)6 | AAAGCTTGTCGGGGAAAGTT | AGATTGCGTCAATTCGATCC | 237 |
| chr11:81349935-81351950 | p5 | (AGGAC)3 | GCCCATTAAGTGGAGACCAA | AACAAGATCGAAGCGGAGAA | 200 |
| chr11:81394359-81396380 | p3 | (ACA)7 | CCGACAAAAGGGATGTCAGT | CTGGAGAGGTATTCAGGGCA | 193 |
| chr11:81407017-81409032 | p5 | (TATAT)3 | CGCTTCGATCTCGTTATTCC | CACTTCCAAAACAGCCCAAT | 228 |
| chr11:81469764-81471780 | p4 | (TGTA)5 | TACTAAGAGGGCCGGGTTCT | CAGGCGAACAGACAGAATCA | 200 |
| chr11:81469764-81471780 | p2 | (AT)8 | TACTAAGAGGGCCGGGTTCT | CAGGCGAACAGACAGAATCA | 200 |
| chr11:81536316-81538328 | p2 | (CT)6 | GGGAAGAGCTAAAACTGGGG | ATTATCGCAGGTTCGTGGAG | 221 |
| chr11:81575962-81577977 | p5 | (ATTTC)3 | CGCATTTATTTCAGCCCCTA | TCGAATGTTAGGTTCTCGGG | 206 |
| chr11:81592371-81594386 | p5 | (TTTCA)3 | CTCGAATCACGCAAGAACAA | CCTTCAACTGACCCGGAGTA | 276 |
| chr11:81609094-81611110 | p4 | (TTTA)4 | GCATACATACGGGGCTGAGT | ACTTGATGCGAGATTGTCCC | 126 |
| chr11:81618265-81620277 | p2 | (AT)6 | CCTTTCAAAAACGGGCAATA | CAAAGAGAGATGCGAGGACC | 181 |
| chr11:81618265-81620277 | p2 | (AT)6 | CCTTTCAAAAACGGGCAATA | CAAAGAGAGATGCGAGGACC | 181 |
| chr11:81625667-81627679 | p2 | (AT)6 | TTTGACCCGAAAAACTCCAG | TTCACCTTCGATTCCTCCAC | 292 |
| chr11:81625667-81627679 | p5 | (TCTTA)3 | TTTGACCCGAAAAACTCCAG | TTCACCTTCGATTCCTCCAC | 292 |
| chr11:81690925-81692940 | p5 | (TTTTC)3 | AACAGCCCACACATCAACAA | ATTTCGTGTATGGCATGGGT | 232 |
| chr11:81957195-81959207 | p2 | (TA)6 | ATGGAAGTTGAGGTCGTTGC | TGCACCCAATAAGCCACATA | 148 |
| chr11:81978554-81980566 | p2 | (AT)6 | CCCTATGTCACCACCCAATC | TATTTTGGCAGCCTTATCGG | 151 |
| chr11:81978554-81980566 | p2 | (AT)6 | CCCTATGTCACCACCCAATC | TATTTTGGCAGCCTTATCGG | 151 |
| chr11:82025477-82027492 | p5 | (TATGA)3 | CCTTACAAGCAACGCTGTGA | TCATTCGGTAAGGAGATCGG | 170 |
| chr11:82027519-82029531 | p2 | (TA)6 | GATGCGGTTCGTTTTGGTAT | AATGAGAGCCGACAAGGCTA | 110 |
| chr11:82281460-82283472 | p2 | (CT)6 | CAATGCAACAGAGAGGGTCA | GATCTCCGATCCTCATGGAA | 277 |
| chr11:82303830-82305850 | p2 | (AT)10 | TTTGAGGGTTCTCATTTGGC | CAGTATGCGGCTCAATCTCA | 263 |
| chr11:82303830-82305850 | p2 | (AT)17 | TTTGAGGGTTCTCATTTGGC | CAGTATGCGGCTCAATCTCA | 263 |
| chr11:82306663-82308679 | p2 | (AT)8 | ATATAGGGGAAATGCTGCCC | CATGCCAAAAGAAAGAAGGG | 284 |
| chr11:82306663-82308679 | p2 | (AT)25 | ATATAGGGGAAATGCTGCCC | CATGCCAAAAGAAAGAAGGG | 284 |
| chr11:82326708-82328726 | p6 | (AGCACC)3 | CAGAATTCTCCGGGACAAAA | GCTCTTCGTAGCTTTGGTGG | 120 |
| chr11:82643282-82645328 | p5 | (AAAAG)3 | GGTTTGTGTTCAGCCGACTT | GGTGTTACATAGGCCGCACT | 225 |
| chr11:82643282-82645328 | p2 | (TC)23 | GGTTTGTGTTCAGCCGACTT | GGTGTTACATAGGCCGCACT | 225 |
| chr11:82707009-82709024 | p5 | (ATTTC)3 | GCCGTATGTTTCCTCCAAAA | AATTGCGGGTTATACGGTCA | 179 |
| chr11:82734642-82736682 | p2 | (AT)20 | TCCATAATGAAATCGGGGAA | GTGTATAGCTGCCCGCCTTA | 284 |

| Marker ID | SSR type | SSR motif | FORWARD PRIMER (5'-3') | REVERSE PRIMER (5'-3') | size (bp) |
|---|---|---|---|---|---|
| chr11:82761619-82763635 | p4 | (AAAT)4 | GACCCTTTCACGCACAAAAT | TGAAAGGCAAAATGGGAAAC | 275 |
| chr11:82761778-82763793 | p4 | (AAAT)4 | GACCCTTTCACGCACAAAAT | TGAAAGGCAAAATGGGAAAC | 116 |
| chr11:82761778-82763793 | p5 | (TGTGT)3 | GACCCTTTCACGCACAAAAT | TGAAAGGCAAAATGGGAAAC | 116 |
| chr11:82762678-82764693 | p5 | (TGTGT)3 | ATGTGACCATAATTGGGGGA | AGTAACTTGGCCGAAACCCT | 161 |
| chr11:82912560-82914580 | p5 | (AATAT)4 | GGAGTATTGGGCCAAACAGA | CGAAGACGTGGTGGAGGTAT | 221 |
| chr11:82938053-82940069 | p2 | (AC)8 | ACCCAGATCTTCCCAGTGTG | CCCCTCCAGAGTCATGGTTA | 187 |
| chr11:82979488-82981506 | p3 | (TGT)6 | TTATTCCCGTTTTTGTTCGG | CTTCACCCATCCTTTCTCCA | 163 |
| chr11:82987189-82989204 | p3 | (ATC)5 | TCCAGAACAACTGGAGCCTT | AAAGATGCATGGTATTCGCC | 239 |
| chr11:83008387-83010403 | p2 | (AT)20 | AGTCTTAAGCTGGCGGTTCA | CACGCTTTACATGCTCGGTA | 239 |
| chr11:83008387-83010403 | p2 | (TA)8 | AGTCTTAAGCTGGCGGTTCA | CACGCTTTACATGCTCGGTA | 239 |
| chr11:83065005-83067038 | p3 | (CAA)5 | GAGGAGCTAGCCCCAGAGTT | TCCAATCAATGGTAGGGAGC | 169 |
| chr11:83223407-83225419 | p2 | (AT)6 | GGAGGAGGTGTATGCGATGT | TGGTGTCATTCTTTCCCTCC | 119 |
| chr11:83243304-83245358 | p3 | (ATA)18 | ATTACAACACGCTCTTGGGG | CAATTTCCAGTCCTTGCCAT | 145 |
| chr11:83270210-83272226 | p6 | (TGTAAG)3 | CTTTTGTTAGCCTTGTCGGC | TCCTTGCCAGGGTTATTGAC | 171 |
| chr11:83270210-83272226 | p4 | (GAAA)4 | CTTTTGTTAGCCTTGTCGGC | TCCTTGCCAGGGTTATTGAC | 171 |
| chr11:83358060-83360076 | p4 | (TTGT)4 | AGGTCAATCGAACATCCGAG | TTTGTTTTGGCTCAAGGGTC | 101 |
| chr11:83420316-83422332 | p2 | (AT)8 | TTAGGTCGGGGACTGAAATG | TAGGAACAAGCCTTGGTTGG | 183 |
| chr11:83492564-83494579 | p2 | (AT)9 | ATACTCCCCCGAAGGACACT | GCCTTACCGGCTTTATGACA | 172 |
| chr11:83492574-83494590 | p2 | (AT)9 | ATACTCCCCCGAAGGACACT | GCCTTACCGGCTTTATGACA | 162 |
| chr11:83492603-83494615 | p2 | (AT)9 | ATACTCCCCCGAAGGACACT | GCCTTACCGGCTTTATGACA | 133 |
| chr11:83514368-83516382 | p2 | (TA)7 | GGAAGCACCTCCTCAATGAA | TCATCGTAACGTGGACCAAA | 229 |
| chr11:83648713-83650729 | p4 | (CCCA)4 | CCTGGAGGAAGATCAACCAA | CATCGGGAAGAGACAAAGGA | 210 |
| chr11:83686609-83688633 | p6 | (TCAGTT)4 | TGCGTGATGGAGAATGATGT | TTCCAGCCTTTGAGCGTAAT | 173 |
| chr11:83693717-83695741 | p2 | (AT)12 | AGGCAACCCACATTTACTGC | ACGCTCCATTATACGAACCG | 263 |
| chr11:83695631-83697649 | p2 | (TA)9 | TTCGTAAATCCATTCGCCTC | ATATAGGGGAATTGCTGCCC | 198 |
| chr11:83742087-83744105 | p3 | (ATC)6 | GGACTTGGCAAAGTTGTGGT | ATCACTCGCGAACCTGTAGC | 214 |
| chr11:83742087-83744105 | p3 | (ATT)6 | GGACTTGGCAAAGTTGTGGT | ATCACTCGCGAACCTGTAGC | 214 |
| chr11:83764695-83766710 | p3 | (GCA)5 | GTTACGGGCCGAGTGTTAGA | CCTGCCATTGGTCTAGGAAA | 166 |
| chr11:83868654-83870672 | p2 | (AT)9 | CACTGCGTGTATGAATTGGC | TTCGCCTAGCTATAACCGGA | 164 |
| chr11:83868654-83870672 | p5 | (AATAA)3 | CACTGCGTGTATGAATTGGC | TTCGCCTAGCTATAACCGGA | 164 |
| chr11:83884475-83886490 | p3 | (ATG)5 | ACATACCCAACCAAACCCAA | CCTTAATCACACGCAAGCAA | 166 |
| chr11:83951637-83953652 | p5 | (AATCT)3 | GTACCCAAAATTTGCAACCG | ATGGCAAAATAGGCAGCAAG | 131 |
| chr11:84034456-84036471 | p3 | (TTG)5 | TGGAAAGGCGTAAAGAGCAT | AGAATACTTTTCGAGGCGCA | 239 |
| chr11:84034456-84036471 | p5 | (TATGA)3 | TGGAAAGGCGTAAAGAGCAT | AGAATACTTTTCGAGGCGCA | 239 |
| chr11:84078633-84080648 | p5 | (AAATG)3 | TATCGCTCGCTTGTTCTCCT | TTGTGGGTGTTGGAAGTTGA | 290 |
| chr11:84155065-84157095 | p3 | (CAA)10 | TGCTTACCCTCGCACTACCT | TTTGTACCACCTCGGAAAGG | 220 |

| Marker ID | SSR type | SSR motif | FORWARD PRIMER (5'-3') | REVERSE PRIMER (5'-3') | size (bp) |
|---|---|---|---|---|---|
| chr11:84155065-84157095 | p5 | (AAGAC)4 | TGCTTACCCTCGCACTACCT | TTTGTACCACCTCGGAAAGG | 220 |
| chr11:84181363-84183375 | p2 | (TA)6 | CATTCTCCTTCCCTGGATGA | TTGTGATGCTTCGAGAGTGG | 130 |
| chr11:84198465-84200480 | p5 | (AAAGA)3 | AGATGCAGATGGAGTGGGTC | GGGAGACAACAGGGAGATGA | 128 |
| chr11:84287549-84289563 | p2 | (AT)6 | CCCGATAACCAGTCTCCTGA | CCTTGTCAGCTTGCGTATCA | 281 |
| chr11:84287549-84289563 | p2 | (AT)7 | CCCGATAACCAGTCTCCTGA | CCTTGTCAGCTTGCGTATCA | 281 |
| chr11:84308940-84310980 | p2 | (AT)20 | CTTTGACCCGAAAAACTCCA | TAACACATCCTCTACCCCGC | 138 |
| chr11:84322337-84324361 | p4 | (AATT)4 | AAAAAGGCAACCCAATTTCC | GGAGGTAGCACGTGTCCAAT | 235 |
| chr11:84322337-84324361 | p2 | (AT)12 | AAAAAGGCAACCCAATTTCC | GGAGGTAGCACGTGTCCAAT | 235 |
| chr11:84350587-84352599 | p2 | (TG)6 | CGCTGTTATATCCCGCATTT | CGCCCCTAGCCTTTCTTAAT | 150 |
| chr11:84397700-84399712 | p2 | (AT)6 | ATCCTGTCGTCTGGGTAACG | CGGTAGGGCAGACTTCTCAG | 177 |
| chr11:84397700-84399712 | p2 | (TA)8 | ATCCTGTCGTCTGGGTAACG | CGGTAGGGCAGACTTCTCAG | 177 |
| chr11:84397700-84399712 | p5 | (ATCAT)3 | ATCCTGTCGTCTGGGTAACG | CGGTAGGGCAGACTTCTCAG | 177 |
| chr11:84421569-84423583 | p2 | (CT)7 | AAGCCATTCTGCTGGGATTA | ATATCTCCATCCATGCTGCC | 221 |
| chr11:84442163-84444181 | p2 | (AT)9 | TTTATGTTGCGGGGGTTTTA | GCCTTCAGCTCTTGGAGAAA | 239 |
| chr11:84485492-84487504 | p2 | (CA)6 | AAATAGGCGTTCTCGGGATT | GGTAGCAATTCTGCTCCTCG | 146 |
| chr11:84511097-84513131 | p2 | (AT)17 | AAACTCATGAAATCACGGGC | GAAAAACAAGAGGGGGCTTC | 169 |
| chr11:84754206-84756226 | p2 | (CA)10 | GATCTGGCCGTCGATATTGT | AAATGAAACTTCACCGCAGG | 202 |
| chr11:84877850-84879865 | p5 | (ATAAA)4 | CAAATAGTGAGGAGGGGCAA | AAAACGAAAAAGGGCCAAAT | 296 |
| chr11:84877850-84879865 | p5 | (TCGGT)3 | CAAATAGTGAGGAGGGGCAA | AAAACGAAAAAGGGCCAAAT | 296 |
| chr11:84897723-84899737 | p2 | (AT)6 | ACGTTACGGCTCCACGTTAC | AGGTCTGCCTTCTGGGAGAT | 102 |
| chr11:84897723-84899737 | p2 | (TA)7 | ACGTTACGGCTCCACGTTAC | AGGTCTGCCTTCTGGGAGAT | 102 |
| chr11:84952271-84954283 | p2 | (GT)6 | ACTACGACGCTGAACCAACC | TGTTGGGTGAAGAAACACCA | 137 |
| chr11:84986672-84988712 | p5 | (TCATA)4 | CGTTTAGGATGTTGCAGGGT | TGATGAGTGGTGCTCGGTAG | 176 |
| chr11:84986672-84988712 | p2 | (AT)20 | CGTTTAGGATGTTGCAGGGT | TGATGAGTGGTGCTCGGTAG | 176 |
| chr11:85000527-85002545 | p3 | (TGT)5 | GCATGCAGAAAAGTCCCAAT | TCGCCAGCTTGCATACATAC | 286 |
| chr11:85000527-85002545 | p2 | (AT)9 | GCATGCAGAAAAGTCCCAAT | TCGCCAGCTTGCATACATAC | 286 |
| chr11:85000527-85002545 | p2 | (AT)29 | GCATGCAGAAAAGTCCCAAT | TCGCCAGCTTGCATACATAC | 286 |
| chr11:85080964-85083008 | p2 | (AT)22 | CAGCTAACACTTGGTTGGCA | AATGTCGGAATTTCACAGCC | 203 |
| chr11:85084574-85086600 | p2 | (TA)13 | TCCCTCCAGCTAACACTTGG | AATGTCGGAATTTCACAGCC | 181 |
| chr11:85128932-85130947 | p5 | (TGCAT)3 | AAACGGAGAAAATACGGCCT | TGTCCGTTTTACTGAAGGGC | 227 |
| chr11:85242778-85244790 | p2 | (CA)6 | GTTCGGCTTAGCACTTGAGG | CTTCCTCCCTCCCCATAGAG | 116 |
| chr11:85242778-85244790 | p2 | (CA)6 | GTTCGGCTTAGCACTTGAGG | CTTCCTCCCTCCCCATAGAG | 116 |
| chr11:85242967-85244981 | p2 | (CA)6 | TAGATCCCGACAGACCAAGG | CTTCCAAACCCTCATCCTCA | 187 |
| chr11:85242967-85244981 | p2 | (CA)6 | TAGATCCCGACAGACCAAGG | CTTCCAAACCCTCATCCTCA | 187 |
| chr11:85242983-85245001 | p2 | (CA)6 | TAGATCCCGACAGACCAAGG | CTTCCAAACCCTCATCCTCA | 171 |
| chr11:85242983-85245001 | p2 | (CA)6 | TAGATCCCGACAGACCAAGG | CTTCCAAACCCTCATCCTCA | 171 |

| Marker ID | SSR type | SSR motif | FORWARD PRIMER (5'-3') | REVERSE PRIMER (5'-3') | size (bp) |
|---|---|---|---|---|---|
| chr11:85274182-85276246 | p2 | (TA)6 | TGAGCGAACCACCCTAAATC | TAGCCTTGTCGGCTCTCATT | 251 |
| chr11:85274182-85276246 | p2 | (AT)32 | TGAGCGAACCACCCTAAATC | TAGCCTTGTCGGCTCTCATT | 251 |
| chr11:85386922-85388934 | p2 | (TG)9 | AACCCTTTGAAACTTGTGCG | ACTTACCCCTTTTGCCTGCT | 182 |
| chr11:85386922-85388934 | p2 | (AT)6 | AACCCTTTGAAACTTGTGCG | ACTTACCCCTTTTGCCTGCT | 182 |
| chr11:85386922-85388934 | p2 | (AT)7 | AACCCTTTGAAACTTGTGCG | ACTTACCCCTTTTGCCTGCT | 182 |
| chr11:85446556-85448571 | p5 | (TTTTG)3 | AAGAAAGCGACTTCCTGCAA | TTGGCCAGTATCCACAACAA | 229 |
| chr11:85446556-85448571 | p5 | (TTTAT)3 | AAGAAAGCGACTTCCTGCAA | TTGGCCAGTATCCACAACAA | 229 |
| chr11:85446919-85448934 | p5 | (TTTTG)3 | GGCATGTTGGTCCTCATTCT | TGTCTTTTTGGTGTGGCAAA | 250 |
| chr11:85446919-85448934 | p5 | (TTTAT)3 | GGCATGTTGGTCCTCATTCT | TGTCTTTTTGGTGTGGCAAA | 250 |
| chr11:85449587-85451602 | p5 | (TTTAT)3 | ACCTGTGACGCCAGAAAAAC | TGGTACCAAAAAGTCTGGGC | 154 |
| chr11:85455781-85457813 | p2 | (TA)16 | TCTCTGATTTGGATACGGGC | GGCATGATCCCTATGGTACG | 157 |
| chr11:85573326-85575341 | p5 | (TTTAT)3 | GTTGAGGTGATTTGGAGGGA | GTTTGGAACCCATACCATGC | 275 |
| chr11:85629339-85631367 | p2 | (TA)14 | TTAACCGCTAGGCATTTTGG | TTATTTGGGGGACGTTGTGT | 218 |
| chr11:85731969-85733984 | p5 | (GATCA)3 | CCCGATAAAACGGAATGCTA | TTTACCCGGTAGCGTTTCAC | 207 |
| chr11:85774229-85776249 | p3 | (TTG)6 | GGAAAACATCTTGCGGGATA | GGCTCAAAGTTATACGCCCA | 156 |
| chr11:85774229-85776249 | p3 | (ATG)6 | GGAAAACATCTTGCGGGATA | GGCTCAAAGTTATACGCCCA | 156 |
| chr11:85774255-85776267 | p3 | (TTG)6 | GGAAAACATCTTGCGGGATA | GGCTCAAAGTTATACGCCCA | 130 |
| chr11:85774255-85776267 | p3 | (ATG)6 | GGAAAACATCTTGCGGGATA | GGCTCAAAGTTATACGCCCA | 130 |
| chr11:85774273-85776285 | p3 | (TTG)6 | GGAAAACATCTTGCGGGATA | GGCTCAAAGTTATACGCCCA | 112 |
| chr11:85774273-85776285 | p3 | (ATG)6 | GGAAAACATCTTGCGGGATA | GGCTCAAAGTTATACGCCCA | 112 |
| chr11:85808892-85810910 | p2 | (TA)9 | GAATGCCCTAACAGACGGAA | GAGGCATTCGGATGTGATTT | 149 |
| chr11:85910390-85912405 | p3 | (TTG)5 | TGGAGACTAAACCGACCCAC | CTCTAGAGGGTTGGGGAAGG | 123 |
| chr11:85910390-85912405 | p3 | (ATG)6 | TGGAGACTAAACCGACCCAC | CTCTAGAGGGTTGGGGAAGG | 123 |
| chr11:85910390-85912405 | p2 | (AT)6 | TGGAGACTAAACCGACCCAC | CTCTAGAGGGTTGGGGAAGG | 123 |
| chr11:85910390-85912405 | p2 | (AT)11 | TGGAGACTAAACCGACCCAC | CTCTAGAGGGTTGGGGAAGG | 123 |
| chr11:85948243-85950258 | p5 | (TTTTA)3 | GCTCTCTTCTTCAGAGCCCA | GCCGATAAGGCCATCATAGA | 113 |
| chr11:85951021-85953036 | p5 | (TTTTA)3 | GCTCTCTTCTTCAGAGCCCA | GCCGATAAGGCCATCATAGA | 113 |
| chr11:86097240-86099256 | p2 | (TA)8 | TGGTGCCCTTTACGGACTAC | CGTGCAGGTTCTGGGATAGT | 258 |
| chr11:86145584-86147608 | p2 | (TA)12 | TTGACGGGCTGTTTTAGGAC | AACATAGCACCAACCCAAGC | 156 |
| chr11:86152608-86154622 | p2 | (TA)7 | GCGTATAAGGGCTGAAGCTG | TACGCCAACCATACAACCAA | 197 |
| chr11:86228256-86230268 | p2 | (CA)6 | TTGAAAGAGGAGGAGCGAAA | TAGCAACAGGCGTGGTGTAG | 241 |
| chr11:86261983-86264017 | p2 | (AT)17 | TGGAAATGCAAATCACGAAA | GAGGTAGGGGTAGGCCAAAG | 118 |
| chr11:86312101-86314185 | p2 | (TA)42 | TGTCACCCTGACATAGCCAC | GCACGGTGGAATCGTATCTT | 190 |
| chr11:86349757-86351772 | p3 | (GTT)5 | CCAAAAGGGAGCAAAAATGA | TGCTAGGGACCTTCCATTTG | 253 |
| chr11:86369022-86371034 | p5 | (TTTAA)3 | CCAATCACGCCTTACCAAGT | CCTTCCTAAATGCCTTGCAG | 168 |
| chr11:86369022-86371034 | p2 | (AT)6 | CCAATCACGCCTTACCAAGT | CCTTCCTAAATGCCTTGCAG | 168 |

| Marker ID | SSR type | SSR motif | FORWARD PRIMER (5'-3') | REVERSE PRIMER (5'-3') | size (bp) |
|---|---|---|---|---|---|
| chr11:86423301-86425319 | p3 | (CAA)6 | GGGGTGGAATTACGAAAGGT | GCCTCTTAAAATTGGGCCTC | 136 |
| chr11:86451107-86453122 | p2 | (TG)6 | ATGGAGGAAGGCTAAGGGAA | TTTTGGACCAAGGATGGAAG | 293 |
| chr11:86451107-86453122 | p3 | (TTG)5 | ATGGAGGAAGGCTAAGGGAA | TTTTGGACCAAGGATGGAAG | 293 |
| chr11:86481209-86483224 | p5 | (GGAGG)3 | AGAGGAGTTGGAGGAGAGCC | CACCCCTTCTACCCCTTCTC | 127 |
| chr11:86481209-86483224 | p2 | (TA)6 | AGAGGAGTTGGAGGAGAGCC | CACCCCTTCTACCCCTTCTC | 127 |
| chr11:86572331-86574349 | p2 | (AT)14 | ATGGCTCGTAGACAGCCCTA | TTTGGGCTTGTGTCCTTTTC | 289 |
| chr11:86572331-86574349 | p2 | (AT)9 | ATGGCTCGTAGACAGCCCTA | TTTGGGCTTGTGTCCTTTTC | 289 |
| chr11:86598245-86600259 | p2 | (TA)7 | TGTACGATAAGGGGGTGCTC | TCCTAAAATGTCCGCAGCTT | 294 |
| chr11:86650494-86652514 | p3 | (TCA)7 | AATACCGCCACATCTTGGAG | GCCCGTGATTTCATGAGTTT | 132 |
| chr11:86650516-86652532 | p3 | (TCA)7 | AATACCGCCACATCTTGGAG | GCCCGTGATTTCATGAGTTT | 110 |
| chr11:86651475-86653496 | p3 | (TCA)7 | GACATATGGACGGAACAGGG | TCATGCCCACAGGTAGACAG | 126 |
| chr11:86699434-86701446 | p2 | (TA)21 | ACAACCGAGTTCCACCTGAC | TATCCTTGTGTTGGGCCTTC | 123 |
| chr11:86699434-86701446 | p2 | (AT)6 | ACAACCGAGTTCCACCTGAC | TATCCTTGTGTTGGGCCTTC | 123 |
| chr11:86737770-86739782 | p2 | (AT)7 | GAACAACGGAGCTCTCCAAG | ATTCCGTACTGACCCCCTTT | 299 |
| chr11:86737770-86739782 | p2 | (TA)6 | GAACAACGGAGCTCTCCAAG | ATTCCGTACTGACCCCCTTT | 299 |
| chr11:86792236-86794248 | p2 | (AT)6 | GAGTCATCATCCGAGGTGGT | GGCTTCTCAATGCCTCAGAC | 198 |
| chr11:86826935-86828947 | p6 | (ATATGT)3 | TGCATATGGTTTCCGCACTA | GGTGCCACAACACAGCATAC | 132 |
| chr11:86826935-86828947 | p2 | (TA)6 | TGCATATGGTTTCCGCACTA | GGTGCCACAACACAGCATAC | 132 |
| chr11:86834949-86836961 | p2 | (TG)6 | TGCATATGGTTTCCGCACTA | ATATCCGTACACCCCGAACA | 292 |
| chr11:86834949-86836961 | p2 | (AT)9 | TGCATATGGTTTCCGCACTA | ATATCCGTACACCCCGAACA | 292 |
| chr11:86835114-86837132 | p2 | (TG)6 | TGCATATGGTTTCCGCACTA | ATATCCGTACACCCCGAACA | 127 |
| chr11:86835114-86837132 | p2 | (AT)9 | TGCATATGGTTTCCGCACTA | ATATCCGTACACCCCGAACA | 127 |
| chr11:86852589-86854601 | p2 | (AG)6 | GCTGATGAATTACGAGGGGA | TCCTTCCTTTTCGTTCCTCA | 101 |
| chr11:86903871-86905886 | p5 | (TTTTC)3 | ACGAGACCTAACTCGGCTCA | TCGTGTATGGCATGGTTTGT | 187 |
| chr11:86941010-86943032 | p2 | (AT)11 | TAGCAAAAGGAGCGGAAAGA | TGAGGGTTCTTGGGTAATCG | 183 |
| chr11:86941010-86943032 | p2 | (TA)8 | TAGCAAAAGGAGCGGAAAGA | TGAGGGTTCTTGGGTAATCG | 183 |
| chr11:86943715-86945730 | p3 | (TTG)5 | TAGAAGGACCCTCAGCCAAA | AAGCCATTTGGCAGCTTAGA | 154 |
| chr11:87009490-87011524 | p2 | (TA)17 | GTTGGGCTGTTTTGGAGTGT | CACACAACCAAGCCAACATC | 262 |
| chr11:87011111-87013127 | p2 | (TA)8 | CGTGTGGATTGTTGGCATAG | ACAACACTCCAAAACAGCCC | 164 |
| chr11:87011111-87013127 | p2 | (AT)17 | CGTGTGGATTGTTGGCATAG | ACAACACTCCAAAACAGCCC | 164 |
| chr11:87073142-87075157 | p3 | (ACA)5 | CAGACCTTGAATCGGTCCAT | TCCCAGGTCATTTCTTTTCG | 261 |
| chr11:87142761-87144785 | p2 | (AT)12 | CTCGTCATCCGCATATTGTG | GATAAGGGGTGCTCGGTACA | 243 |
| chr11:87181756-87183772 | p2 | (TA)8 | TGCTTTTCCGTTTCACACTG | GATGGATTACGAACGGGATG | 227 |
| chr11:87216350-87218374 | p2 | (AT)6 | AATGATTCCCAGATGCCTTG | GGGTGGGGTGTGATAAATTG | 204 |
| chr11:87216350-87218374 | p2 | (TA)12 | AATGATTCCCAGATGCCTTG | GGGTGGGGTGTGATAAATTG | 204 |
| chr11:87260544-87262556 | p2 | (AT)6 | AGCTACATTCTCTTGCCCGA | GGGACCCGATTACCCTAAGA | 199 |

| Marker ID | SSR type | SSR motif | FORWARD PRIMER (5'-3') | REVERSE PRIMER (5'-3') | size (bp) |
|---|---|---|---|---|---|
| chr11:87304285-87306297 | p2 | (CA)6 | AGGTTTCGGACATTTCGTTG | GGAAGGAACGGGATTTGATT | 100 |
| chr11:87304285-87306297 | p2 | (AT)6 | AGGTTTCGGACATTTCGTTG | GGAAGGAACGGGATTTGATT | 100 |
| chr11:87328816-87330831 | p5 | (TTACT)3 | TTCTTGGACAGAAACCCCAG | TCATCCTAGGAAGGCCAATG | 256 |
| chr11:87376045-87378063 | p3 | (CAT)6 | TCATATATGTGGGCGAGGGT | TCACCGGTCATAAGTGCAAA | 163 |
| chr11:87438955-87440970 | p5 | (TATCA)3 | ATTGTTTGACGGCCTCTTTG | CCTAGTCGGGTCGTGACAAT | 213 |
| chr11:87447990-87450011 | p3 | (CAA)7 | ATAAGTGGAACCCGACCCTC | TCCCGCGATTTTTCTTTATG | 170 |
| chr11:87506670-87508682 | p2 | (CA)6 | GGGACATGCCAAAGAAGAAA | GCTTGGTTGTTGGTTGGTTT | 162 |
| chr11:87506670-87508682 | p2 | (TA)6 | GGGACATGCCAAAGAAGAAA | GCTTGGTTGTTGGTTGGTTT | 162 |
| chr11:87681314-87683329 | p5 | (CATAT)3 | TGCAGTGGAATACGCTTGAC | CGTTGTTTGGGTTCTTCGAT | 142 |
| chr11:87681314-87683329 | p3 | (CAA)5 | TGCAGTGGAATACGCTTGAC | CGTTGTTTGGGTTCTTCGAT | 142 |
| chr11:87756497-87758511 | p2 | (TA)7 | CTTCTTTGTCCACGTCCCAT | TGTTCGTCTGTAGTGGCAGC | 151 |
| chr11:87786964-87788979 | p3 | (TTG)5 | CAGGGTGCTGGACCATAGAT | TTGGGCGACGAAAAATAAAG | 192 |
| chr11:87824960-87826974 | p2 | (AT)7 | GCCAAGGACCCATAGGAAAT | GCCAAAAATGGCAAAGTTGT | 190 |
| chr11:87824960-87826974 | p2 | (CT)30 | GCCAAGGACCCATAGGAAAT | GCCAAAAATGGCAAAGTTGT | 190 |
| chr11:87909898-87911913 | p5 | (TGTTT)3 | ATCCGATGAGAAGGCATTTG | TACGCCCAAGCATAATTTCC | 266 |
| chr11:87919399-87921453 | p5 | (CGATA)3 | ACGACCTGCACTTTTTGGTC | CCAAGTCGTGGGAAAACCTA | 106 |
| chr11:87919399-87921453 | p2 | (AT)15 | ACGACCTGCACTTTTTGGTC | CCAAGTCGTGGGAAAACCTA | 106 |
| chr11:88005847-88007901 | p2 | (TA)27 | CTGTACCTTTCTTGCGAGGC | AAGCAATTCCGTCCTAGGGT | 187 |
| chr11:88068733-88070749 | p2 | (AG)8 | GGTCGTGACAATGCCTTTTT | TTTCTTGGTGAACGGCTTCT | 254 |
| chr11:88150959-88152977 | p3 | (TTG)6 | GCATCCTTCAATGCTTCCAT | CTTGATGTTTCCCTCCCTGA | 114 |
| chr11:88229825-88231840 | p3 | (TTG)5 | GCCTTGTCGCAATTACACCT | CCCGAAACATCTTTAAGCCA | 248 |
| chr11:88229825-88231840 | p2 | (AT)6 | GCCTTGTCGCAATTACACCT | CCCGAAACATCTTTAAGCCA | 248 |
| chr11:88303524-88305539 | p5 | (ATATG)3 | AATGGTGGATCGTGGTCATT | AGAGACAATTGGGCAGCACT | 233 |
| chr11:88306807-88308827 | p2 | (AG)10 | GCACAGAACCCAACAAGACA | TCGTATGTGGCTGAGTTTGC | 210 |
| chr11:88306807-88308827 | p2 | (TA)10 | GCACAGAACCCAACAAGACA | TCGTATGTGGCTGAGTTTGC | 210 |
| chr11:88319232-88321254 | p2 | (AT)6 | ATGAGAGCCCACCAATGAAC | TGCCATATGGTGTGGGTATG | 300 |
| chr11:88319232-88321254 | p2 | (AT)12 | ATGAGAGCCCACCAATGAAC | TGCCATATGGTGTGGGTATG | 300 |
| chr11:88319232-88321254 | p2 | (TA)31 | ATGAGAGCCCACCAATGAAC | TGCCATATGGTGTGGGTATG | 300 |
| chr11:88354684-88356699 | p5 | (TATTA)3 | GTCCCAAATGTTTCCCACAC | AGGAATCAGAAAAAGGGCGT | 229 |
| chr11:88354684-88356699 | p6 | (TATGTA)3 | GTCCCAAATGTTTCCCACAC | AGGAATCAGAAAAAGGGCGT | 229 |
| chr11:88354684-88356699 | p2 | (AT)20 | GTCCCAAATGTTTCCCACAC | AGGAATCAGAAAAAGGGCGT | 229 |
| chr11:88366712-88368727 | p6 | (TATACA)3 | ATCGGAGTGACGTAAGGTCG | CCCATCGGGTCCTAAAAGTT | 105 |
| chr11:88379943-88382025 | p3 | (CTT)6 | GACGAGCAGCAAAACAACAA | TTCTTCAAAGGAAGCGTGGT | 257 |
| chr11:88391485-88393567 | p3 | (CTT)6 | GACGAGCAGCAAAACAACAA | TTCTTCAAAGGAAGCGTGGT | 257 |
| chr11:88398939-88400955 | p2 | (GT)8 | TTCGTCACTTCAACCTTCCC | AACCGATTTCCAAAATTCCC | 220 |
| chr11:88423771-88425785 | p2 | (AT)13 | GAAATGCAAGGTAAGGCTGC | GAGGGGTTATCGTTCCCATT | 275 |

| Marker ID | SSR type | SSR motif | FORWARD PRIMER (5'-3') | REVERSE PRIMER (5'-3') | size (bp) |
|---|---|---|---|---|---|
| chr11:88423787-88425799 | p2 | (AT)13 | GAAATGCAAGGTAAGGCTGC | GAGGGGTTATCGTTCCCATT | 259 |
| chr11:88478506-88480518 | p2 | (TA)6 | CGCACCTCAATTTTTCCACT | GGTAACGAACTCGGGATCAA | 253 |
| chr11:88478506-88480518 | p5 | (GCTAT)3 | CGCACCTCAATTTTTCCACT | GGTAACGAACTCGGGATCAA | 253 |
| chr11:88575296-88577311 | p5 | (TTATA)3 | ATATAGGGGAAATGCTGCCC | TTCGTAAACCGTAGCCCATC | 156 |
| chr11:88681803-88683833 | p2 | (TA)15 | ACGTCCGTTTTCTTTGGATG | AATCGTCATAAGCACCTCCG | 187 |
| chr11:88681803-88683833 | p5 | (AATGC)3 | ACGTCCGTTTTCTTTGGATG | AATCGTCATAAGCACCTCCG | 187 |
| chr11:88721558-88723570 | p3 | (CAA)5 | TTGGACACGACTTTGCAGAC | GCGGGTTCGATCGACTATAA | 247 |
| chr11:88721558-88723570 | p2 | (CT)6 | TTGGACACGACTTTGCAGAC | GCGGGTTCGATCGACTATAA | 247 |
| chr11:88737009-88739027 | p6 | (AGAGAA)3 | GGCTTGAAGAACGAATCCTG | TATACGATAGGAGGTGCCCG | 114 |
| chr11:88787778-88789793 | p3 | (AAC)5 | TGATGGGACGTGACGAGATA | CCAGACCCAACCCATAATTG | 135 |
| chr11:88791355-88793403 | p3 | (GAA)16 | AGGTTCATCCAAATCCTCCC | TAAGGTGGTTGTTGAAGGGC | 184 |
| chr11:88791355-88793403 | p6 | (AACAAT)3 | AGGTTCATCCAAATCCTCCC | TAAGGTGGTTGTTGAAGGGC | 184 |
| chr11:88837618-88839630 | p2 | (TG)6 | TTACGATCTTAAGTGGCCCG | CATCGCCAAATTTCACACAC | 286 |
| chr11:88837618-88839630 | p2 | (TG)6 | TTACGATCTTAAGTGGCCCG | CATCGCCAAATTTCACACAC | 286 |
| chr11:88852165-88854179 | p6 | (ATATGG)3 | CATGGCCGGGTATATGAAAC | CCATTTCTAATGCGTGCTCA | 206 |
| chr11:88852165-88854179 | p2 | (TA)7 | CATGGCCGGGTATATGAAAC | CCATTTCTAATGCGTGCTCA | 206 |
| chr11:88883106-88885121 | p3 | (TCA)5 | GGGACCAAGAAGGTCACAGA | TGACGACTTTTTGCTTGTGG | 156 |
| chr11:88907354-88909387 | p3 | (CTT)6 | GCCATGGTGATCCACTTCTT | CTGCTGAGCTGCTGAAATTG | 286 |
| chr11:88907389-88909419 | p3 | (CTT)6 | GCCATGGTGATCCACTTCTT | CTGCTGAGCTGCTGAAATTG | 251 |
| chr11:88907423-88909438 | p3 | (CTT)6 | GCCATGGTGATCCACTTCTT | CTGCTGAGCTGCTGAAATTG | 217 |
| chr11:88907451-88909493 | p3 | (CTT)6 | GCCATGGTGATCCACTTCTT | CTGCTGAGCTGCTGAAATTG | 189 |
| chr11:88907517-88909553 | p3 | (CTT)6 | GCCATGGTGATCCACTTCTT | CTGCTGAGCTGCTGAAATTG | 123 |
| chr11:88907739-88909796 | p3 | (CTT)6 | CAAACTCCCTCAACCCAAAA | TTCCCTGGCAAGATCTCAGT | 166 |
| chr11:88931551-88933566 | p5 | (TTGGT)3 | AAGTCCCGACTTGTGGTTTG | TGGTTTGATTCGGGTTGATT | 134 |
| chr11:88965080-88967142 | p2 | (TA)40 | GGCCGGGCATGTAACTACTA | AATCATCATCGTTAACCCGC | 240 |
| chr11:88965080-88967142 | p2 | (TA)31 | GGCCGGGCATGTAACTACTA | AATCATCATCGTTAACCCGC | 240 |
| chr11:89021715-89023730 | p5 | (TCATT)3 | TCCGAGCTATGGATGGAATC | CATCCCCATACTCGTGGAAC | 143 |
| chr11:89033116-89035128 | p2 | (CT)6 | GCGGGTTCAGGTAGACAGAG | ACTTGCCTTCAAAATGCCAC | 252 |
| chr11:89134898-89136910 | p2 | (TA)6 | ACGAACTGGTTCCGTGGTAG | TCCACAAGAAGGACAGGGAC | 216 |
| chr11:89297343-89299361 | p6 | (TATATG)3 | CCGAGTCTAGTTTCCAACGC | ATAAGGTTCAAATTCGGGGG | 170 |
| chr11:89360645-89362707 | p2 | (TG)10 | ACCGTCCACAACTTCCAAAG | CTTGATCAGAAGAGCTCGGG | 274 |
| chr11:89360709-89362741 | p2 | (TG)10 | ACCGTCCACAACTTCCAAAG | CTTGATCAGAAGAGCTCGGG | 210 |
| chr11:89441748-89443766 | p6 | (AAAATA)3 | AAACGACCCGAGACAATCAG | CCTTGCTTTACTCCGCTACG | 179 |
| chr11:89568896-89570934 | p2 | (AT)19 | TAAGCCATGTGGCACACCTA | ATCGAGTCGAACAAATTCGG | 285 |
| chr11:89779291-89781313 | p2 | (AT)11 | CCGCTCATTTTATGCCTCAT | ATACCAAGCTTTGTCACGCC | 201 |
| chr11:89861394-89863458 | p2 | (TA)32 | GAAGCTCGTTTCGTCGATTC | TTGACCAAGAGAAGGCAAGG | 193 |

| Marker ID | SSR type | SSR motif | FORWARD PRIMER (5'-3') | REVERSE PRIMER (5'-3') | size (bp) |
|---|---|---|---|---|---|
| chr11:89861394-89863458 | p5 | (ACTCA)3 | GAAGCTCGTTTCGTCGATTC | TTGACCAAGAGAAGGCAAGG | 193 |
| chr11:89918723-89920737 | p2 | (TG)7 | CAACGACTCATCCCCGTAGT | AACATTGTCGAAATCCTCCG | 182 |
| chr11:89951916-89953931 | p3 | (TGT)5 | AGACCGACCTTGTTGTACGG | GTGTGCCCACAGATTCATTG | 280 |
| chr11:89962399-89964413 | p2 | (TA)8 | CGTGCATGCCTCAATACATC | GGGACTCGGTGAATAAGGGT | 214 |
| chr11:89962399-89964413 | p2 | (AT)7 | CGTGCATGCCTCAATACATC | GGGACTCGGTGAATAAGGGT | 214 |
| chr11:90108868-90110889 | p3 | (GTT)7 | CCAAGTGTAAGCCCTCCAAA | CTCTTTATCCCGATGTCCCA | 154 |
| chr11:90108868-90110889 | p2 | (TA)6 | CCAAGTGTAAGCCCTCCAAA | CTCTTTATCCCGATGTCCCA | 154 |
| chr11:90589000-90591016 | p2 | (AT)7 | ATTTTGGGACTTGGGTAGGG | ACATACACTGACCCGCATGA | 172 |
| chr11:90589000-90591016 | p2 | (AT)8 | ATTTTGGGACTTGGGTAGGG | ACATACACTGACCCGCATGA | 172 |
| chr11:90607078-90609092 | p2 | (GT)7 | ATCGGGTTCAGGGGATTAAG | CCCTTACTGCACCTCATGGT | 230 |
| chr11:90618186-90620202 | p2 | (AT)8 | AACTTGCGTCGCCAATTATC | CTCAGCCGAGTAGGGTCTTG | 291 |
| chr11:90618186-90620202 | p3 | (CAA)5 | AACTTGCGTCGCCAATTATC | CTCAGCCGAGTAGGGTCTTG | 291 |
| chr11:90788909-90790965 | p2 | (AT)31 | ATCGTGAGACCATGCTACCC | GAGAGCACCCCATTATTCCA | 219 |
| chr11:90788974-90790996 | p2 | (AT)31 | ATCGTGAGACCATGCTACCC | GAGAGCACCCCATTATTCCA | 154 |
| chr11:90829455-90831471 | p4 | (TGAA)4 | TTCCAGGGTAAGCCATGTTC | CCCAAGGAATCTGGTCTGAA | 171 |
| chr11:90869195-90871211 | p4 | (AATT)4 | GGATTGGGTTTTGCTTTCAA | CACGCCTCCTGGGATTATTA | 294 |
| chr11:90893204-90895228 | p6 | (TTGTTT)4 | TGAATCCGAACTGGTTCTCC | TGTTATTTTTAGGACGGCGG | 180 |
| chr11:90942845-90944863 | p3 | (GTT)6 | CCACCAAAACTTAAGGGCAA | TTCACATGCCCAACCATCTA | 266 |
| chr11:90983389-90985440 | p3 | (TAA)17 | CTTGTTTCCGAGCGAGTAGC | TGACCGATGACTCCAAATCA | 124 |
| chr11:91127840-91129860 | p5 | (TTCAT)4 | TGGACTGTGCTACCGAGATG | CTTCGTGCCCTTCGTTAGTC | 287 |
| chr11:91197772-91199786 | p2 | (TC)7 | CGGATGTTTTGGAATTTTGG | AAATAGGGATTTCCCGCAAC | 144 |
| chr11:91197772-91199786 | p2 | (AT)9 | CGGATGTTTTGGAATTTTGG | AAATAGGGATTTCCCGCAAC | 144 |
| chr11:91216646-91218658 | p2 | (AT)6 | CTTTGAATCCGTCCCTTTCA | AAGATGTCACCGAACCAAGG | 238 |
| chr11:91441410-91443425 | p5 | (AATGC)3 | TCGTATGCCTTGCCTTCTCT | GTGCCATCATGACTCGCTTA | 127 |
| chr11:91509461-91511473 | p2 | (AT)6 | AGTGCCTCCTCCAAGACTCA | CCCCATTCACGAAAGAAAAA | 294 |
| chr11:91641329-91643363 | p3 | (CAT)5 | AACTATCACGACCCAAACCG | TGTCTATGGTGGCCTTGTCA | 145 |
| chr11:91804198-91806213 | p5 | (TATAT)3 | GAATGGCCATGACCTTGTCT | TAGGGGCAACGTATCAAAGG | 134 |
| chr11:91882427-91884442 | p5 | (TTTAT)3 | CCTGGAGGAATGAAGCATGT | GATGGTGCAACCAGTTTGTG | 129 |
| chr11:91970528-91972543 | p5 | (ATTTC)3 | TACCCGCATTTATTTCAGCC | ATAACAATGCCTCACAGGGC | 203 |
| chr11:91996678-91998693 | p5 | (AAATA)3 | TTGCAACCCCATAGTTTTCC | ACAATTTGGGGTTTGGGATT | 149 |
| chr11:91996678-91998693 | p5 | (TTATT)3 | TTGCAACCCCATAGTTTTCC | ACAATTTGGGGTTTGGGATT | 149 |
| chr11:91996678-91998693 | p5 | (TTTTA)4 | TTGCAACCCCATAGTTTTCC | ACAATTTGGGGTTTGGGATT | 149 |
| chr11:91996678-91998693 | p5 | (TTTTA)4 | TTGCAACCCCATAGTTTTCC | ACAATTTGGGGTTTGGGATT | 149 |
| chr11:91996837-91998857 | p5 | (AAATA)3 | ATGAAACCCAATCCCAAACC | ATCACGTGTTCAATGGATGG | 209 |
| chr11:91996837-91998857 | p5 | (TTATT)3 | ATGAAACCCAATCCCAAACC | ATCACGTGTTCAATGGATGG | 209 |
| chr11:91996837-91998857 | p5 | (TTTTA)4 | ATGAAACCCAATCCCAAACC | ATCACGTGTTCAATGGATGG | 209 |

| Marker ID | SSR type | SSR motif | FORWARD PRIMER (5'-3') | REVERSE PRIMER (5'-3') | size (bp) |
|---|---|---|---|---|---|
| chr11:91996837-91998857 | p5 | (TTTTA)4 | ATGAAACCCAATCCCAAACC | ATCACGTGTTCAATGGATGG | 209 |
| chr11:92050940-92052952 | p2 | (TA)6 | CGTTGCACCAGTCTCTGAAA | TCAAGAGTGGGAAAAATCGG | 203 |
| chr11:92259724-92261739 | p5 | (ATGAT)3 | TCGTGTTGTTGTTGGGTTGT | ACAATTTCAAACGCGGATTC | 199 |
| chr11:92292280-92294322 | p6 | (TATATG)7 | CCACGTGCACACCATTTAAG | GGGATCATGCCTTGAAAGAA | 211 |
| chr11:92292280-92294322 | p5 | (GATAT)3 | CCACGTGCACACCATTTAAG | GGGATCATGCCTTGAAAGAA | 211 |
| chr11:92299335-92301365 | p5 | (TGAAT)4 | TGAATTCCCGGATTGTTGTT | GGGATCATGCCTTGAAAGAA | 225 |
| chr11:92299335-92301365 | p6 | (TATATG)5 | TGAATTCCCGGATTGTTGTT | GGGATCATGCCTTGAAAGAA | 225 |
| chr11:92299335-92301365 | p5 | (TATGA)7 | TGAATTCCCGGATTGTTGTT | GGGATCATGCCTTGAAAGAA | 225 |
| chr11:92359349-92361367 | p2 | (TA)9 | ACCGTTCAAGCAAATTCCTG | CGGCAATACCTCGAGAGAAG | 223 |
| chr11:92377126-92379138 | p2 | (AT)6 | TATATCTTTCCCCGGTTCCC | CCGTATCCCCGTCACATATC | 146 |
| chr11:92459238-92461262 | p2 | (AG)12 | AGAGGAACCTCTGCCCAAAT | CCCTCCTGAAATGATCTCCA | 296 |
| chr11:92470096-92472108 | p2 | (AT)6 | ATCATATAGGCGGGACACGA | TGGACGCTCCTGAGTAAGGT | 184 |
| chr11:92513864-92515879 | p5 | (TGAAT)3 | AGATAGTTGTGGTCGGGTGC | TTACAGGTTATCCTTCGCCG | 291 |
| chr11:92576702-92578734 | p2 | (TA)16 | CTCAGGGTGAACCGTTGTCT | CACTCTGTCACTCCTGGCAA | 151 |
| chr11:92603765-92605780 | p3 | (ATC)5 | AAGCCGACAAGGCTACAAAA | GTCCTGTCCCGCCTATATGA | 111 |
| chr11:92668929-92670944 | p2 | (AG)6 | ACTCTTGGGTGAACGTTTGG | GAAGAAAGAATTGGGGAGGG | 279 |
| chr11:92747974-92749986 | p2 | (TA)6 | GTATATATGGGTGTGGCGGG | ACAAGCAGAGCATGACAACG | 287 |
| chr11:92747974-92749986 | p2 | (TA)6 | GTATATATGGGTGTGGCGGG | ACAAGCAGAGCATGACAACG | 287 |
| chr11:92768021-92770037 | p3 | (ATG)6 | CGGTGGAGAAATCGGATAAA | ATATCCTGCCGAAAATCGTG | 125 |
| chr11:92768021-92770037 | p4 | (TATG)4 | CGGTGGAGAAATCGGATAAA | ATATCCTGCCGAAAATCGTG | 125 |
| chr11:92768021-92770037 | p2 | (TA)9 | CGGTGGAGAAATCGGATAAA | ATATCCTGCCGAAAATCGTG | 125 |
| chr11:92790443-92792461 | p2 | (TA)9 | CAGCACAAGTTCCCCTTTGT | TCTGGGAGCTATGGAGCACT | 207 |
| chr11:92901331-92903345 | p2 | (AT)6 | ACGAGTCCCGACCCTCTAAT | TATTCTGCTCGTCCATGTCG | 109 |
| chr11:93125088-93127100 | p2 | (CA)6 | TTCAGGCCACTCTTGGACTT | GGCCAAGTCTTCACAGCTTC | 255 |
| chr11:93167790-93169828 | p2 | (AT)14 | ACCACTAGCGACCAAACACC | CCTGTCCCGTCCTTTCTACA | 116 |
| chr11:93167790-93169828 | p2 | (AT)19 | ACCACTAGCGACCAAACACC | CCTGTCCCGTCCTTTCTACA | 116 |
| chr11:93239992-93242006 | p2 | (AC)7 | GGAAAATTCAATAAGCGGCA | GCCGAGCGAACTATTCAGAC | 125 |
| chr11:93266742-93268756 | p2 | (AG)13 | ACCCTTCGAGACGTGTATGG | TGCGTTGTACACTAGGCTCG | 246 |
| chr11:93266742-93268756 | p2 | (AG)7 | ACCCTTCGAGACGTGTATGG | TGCGTTGTACACTAGGCTCG | 246 |
| chr11:93285945-93287969 | p2 | (AT)12 | AACTCTTCTTCCAGCCACCA | TGCAGGGCATTTAGGAAATC | 200 |
| chr11:93302718-93304733 | p3 | (CAA)5 | GAAAAGGGACGTCAGTTGGA | TGTTATTCTCTGCCTTCGGG | 135 |
| chr11:93302718-93304733 | p3 | (ACA)5 | GAAAAGGGACGTCAGTTGGA | TGTTATTCTCTGCCTTCGGG | 135 |
| chr11:93520424-93522439 | p5 | (ATTTC)3 | GAAACAAAAAGGTGGCCTCA | ACACACGACCAAAAGCATGA | 106 |
| chr11:93541885-93543897 | p2 | (CA)6 | ACATGCCAAAGGAAGGAATG | GGTTGTGTGATAATTGGGGG | 244 |
| chr11:93575892-93577969 | p2 | (TA)33 | CGCTCGTCGTGAAATGTTTA | CCGTTCTTCTCATTTTCCCA | 155 |
| chr11:93697439-93699457 | p2 | (AT)9 | GTGTTCTGGATTTCGGAGGA | GCTGAATCAATTCATGGCCT | 120 |

| Marker ID | SSR type | SSR motif | FORWARD PRIMER (5'-3') | REVERSE PRIMER (5'-3') | size (bp) |
|---|---|---|---|---|---|
| chr11:93728258-93730276 | p6 | (TATTCG)3 | TTAAAAAGGGACGCGAAAAA | GGGCAGTGACAAAACATCCT | 100 |
| chr11:93732647-93734661 | p2 | (AT)7 | GAAATGGGGGAGAGAAGAGG | AAGAAAGGAGGAGTCGCACA | 202 |
| chr11:93808169-93810184 | p5 | (TTCAT)3 | TCCGTATATTGTGGGTCGGT | CGACGGTCACAAATCACAAC | 282 |
| chr11:93891073-93893109 | p2 | (TA)18 | TGGTCCACCCAAAAATGACT | CCCTAGGACGAACTGCTCTG | 100 |
| chr11:93944114-93946129 | p2 | (TA)9 | GGTTTGTGGGTTCTTCCAGA | GTTAGCTCTGGGTGCTCGTC | 147 |
| chr11:93944114-93946129 | p5 | (TATCA)3 | GGTTTGTGGGTTCTTCCAGA | GTTAGCTCTGGGTGCTCGTC | 147 |
| chr11:94109693-94111709 | p4 | (TTGA)4 | ACTTGGAAGCCAAAAAGGCT | AGTCATCGTAACGGGTCGTC | 257 |
| chr11:94110472-94112512 | p4 | (TTGA)4 | CCGAGAGTAAAACCGTTGGA | ATAACCTTCGGAAAATCGGG | 103 |
| chr11:94113061-94115111 | p2 | (AC)25 | ACTTTGGATTGCCAAACAGG | CGGCATTACACACACACACA | 103 |
| chr11:94113061-94115111 | p2 | (AT)12 | ACTTTGGATTGCCAAACAGG | CGGCATTACACACACACACA | 103 |
| chr11:94118956-94121012 | p2 | (TA)6 | GACGCTGTATTTCCCTTCCA | TCTATGTCCAAAACTCGGGC | 118 |
| chr11:94161541-94163555 | p2 | (GT)7 | TCAGTTACTGGCTCATCCCC | GGTTCGAAGGACCGTGTAAA | 236 |
| chr11:94199610-94201625 | p5 | (AATAC)3 | CTCTCCTCTTTCTCAGGCGA | GAGCAGCATCCCCAAAATAA | 214 |
| chr11:94263353-94265377 | p2 | (AT)12 | TATGCCTTGGGAATCGAGAC | CGATTCGGTTCGTTGAACTT | 286 |
| chr11:94263353-94265377 | p2 | (TA)7 | TATGCCTTGGGAATCGAGAC | CGATTCGGTTCGTTGAACTT | 286 |
| chr11:94283006-94285022 | p2 | (AG)8 | GATGGCACTAGTCCAACCGT | TGGAACGTCTCCGATACTCC | 294 |
| chr11:94283006-94285022 | p3 | (TTG)6 | GATGGCACTAGTCCAACCGT | TGGAACGTCTCCGATACTCC | 294 |
| chr11:94285786-94287798 | p2 | (TA)6 | AGCAGGTTCGTATACGGGTG | GTGTATCGGGGTGTTTGGTC | 124 |
| chr11:94308345-94310361 | p2 | (TA)8 | TATATGTCCGTCCCCGGTTA | CTTCGAACATCTCCGTCACA | 217 |
| chr11:94308345-94310361 | p3 | (GAT)5 | TATATGTCCGTCCCCGGTTA | CTTCGAACATCTCCGTCACA | 217 |
| chr11:94308583-94310598 | p2 | (TA)8 | AATTGTGAGTATGCCCGTCC | ACCAAAATTACACGCTTCGG | 277 |
| chr11:94308583-94310598 | p3 | (GAT)5 | AATTGTGAGTATGCCCGTCC | ACCAAAATTACACGCTTCGG | 277 |
| chr11:94382081-94384096 | p5 | (TTCTT)3 | TCATAGAACTCGCAACGACG | GTGAAGTTTTGTTTGGCCGT | 183 |
| chr11:94404822-94406840 | p6 | (CATATA)3 | GGTTCATCCAAGCCAAGAAA | TGTGTTGATTGTTCGGGAGA | 103 |
| chr11:94601589-94603601 | p2 | (TA)6 | AATGTTATGCGGGCTATTCG | TGATACGTATACGCTGCCCA | 192 |
| chr11:94717861-94719923 | p2 | (AT)31 | AGCCAAATTCTTGTTGTGGC | TATGTGAGGCATCGGATTGA | 156 |
| chr11:94752248-94754278 | p2 | (AT)15 | TCCTCTCCCGTCCATATGTC | CCCATTGTACACATCAAGCG | 167 |
| chr11:94779815-94781837 | p2 | (TA)11 | CAGACAACCCCTAGGACGAA | GCAAGCCGACGTACCATTAT | 114 |
| chr11:94867800-94869850 | p2 | (TA)8 | TGCACAGGGAGACAGACTTG | AAGCCGACAAGGCTATCAAA | 232 |
| chr11:94877821-94879847 | p2 | (AT)13 | TAGTTCCCGAACTGGACACC | GTCGGCTTGCGTATCATTTT | 160 |
| chr11:94884313-94886325 | p2 | (AT)6 | ATCATAGCCACACAGGGCTC | GCATGCCGGATAAGGTAAGA | 159 |
| chr11:94904765-94906781 | p2 | (AT)8 | TCATTAATAAAGCCCTGCGG | ACGTGGAAGAGCAAAGGAGA | 215 |
| chr11:94939842-94941856 | p2 | (AT)7 | TCATGGAGAGGGTTTTGGAG | ACGTCCCGTACCTCATTCTG | 234 |
| chr11:94998731-95000743 | p2 | (CA)6 | AATGAAGTCACGACCCCTTG | TATTACTGAGGAGCCCGTGG | 188 |
| chr11:95104921-95106939 | p3 | (TTG)6 | ACGTTTTCGGGTGATCTGAC | CTAGCCCACTGACCTTCCTG | 210 |
| chr11:95126678-95128693 | p5 | (TGCAT)3 | GTCGGCTTAATGCCATGAGT | TACCAACAGTGGATACGCCA | 295 |

| Marker ID | SSR type | SSR motif | FORWARD PRIMER (5'-3') | REVERSE PRIMER (5'-3') | size (bp) |
|---|---|---|---|---|---|
| chr11:95268859-95270897 | p2 | (TA)19 | ATAAGATGCCAACTCCACGG | CAAGTACCAAAGCCCGTGAT | 151 |
| chr11:95362462-95364474 | p2 | (TA)6 | TAAACAGGGGAAATGATGCC | CTTACTTCCCGTCTTGCAGC | 216 |
| chr11:95504589-95506605 | p2 | (TA)8 | AATTGTTAGCCCTGTGTGGC | AAAATTTGGGCAGAGGTTCC | 119 |
| chr11:95504589-95506605 | p2 | (AG)10 | AATTGTTAGCCCTGTGTGGC | AAAATTTGGGCAGAGGTTCC | 119 |
| chr11:95531400-95533416 | p2 | (AT)8 | TTTTTCTTGGAACCCGTCAC | TACGAGGAGGGTGTTGATCC | 262 |
| chr11:95614983-95617001 | p3 | (TCA)6 | GGATCACCAACCGGAGTAGA | GATCCCAACTGGTCGAGGTA | 197 |
| chr11:95651359-95653371 | p2 | (GT)6 | TTATATCCCGTTTTCGCGTC | ATAGTTTTCGCCCTTGGCTT | 259 |
| chr11:95661425-95663443 | p3 | (TTG)6 | AAATGGGGGATTCCATAAGC | TTGATCCCGATTTTGTGTGA | 235 |
| chr11:95698571-95700589 | p3 | (TTC)6 | TTTTCGCAAGGCAGAGTTTT | TTCTCATGGCCAAACGTGTA | 161 |
| chr11:95775514-95777528 | p2 | (AT)7 | TGTACCGAGCACCCCTTATC | ACTTTTGATGGTCCTCGTCG | 284 |
| chr11:95803532-95805548 | p2 | (TA)8 | CGTCCTCAGCATCTCATCAA | GTTCAGCCAGTACGAGAGCC | 222 |
| chr11:95817779-95819791 | p2 | (GA)6 | GCAAATTATGGTCCGGCTTA | ACCCTGGGTCATTTACCCTC | 206 |
| chr11:95819615-95821630 | p5 | (ATGAT)3 | AACGATGTTTGAGGTTTGGC | TGGGACAAGGGCAAGTTTAG | 198 |
| chr11:95856947-95858962 | p3 | (AAT)5 | CCCTACAGTCCTGCAAAAGC | CATTTGTTGGTGGCAGTTTG | 155 |
| chr11:95871948-95873963 | p5 | (ATACA)3 | CTTCCCTATCACTCGGACCA | CCACCCCGCTTAATGTAGAA | 170 |
| chr11:95871948-95873963 | p2 | (AT)11 | CTTCCCTATCACTCGGACCA | CCACCCCGCTTAATGTAGAA | 170 |
| chr11:95894166-95896184 | p6 | (CATATA)3 | CCTGAATTGGGCACTCGTAT | TGGGTCGTATTCAGACGTGA | 135 |
| chr11:95894166-95896184 | p2 | (AT)7 | CCTGAATTGGGCACTCGTAT | TGGGTCGTATTCAGACGTGA | 135 |
| chr11:95894460-95896474 | p6 | (CATATA)3 | CCCGTCGTACCCTTAAACAA | ATGCCCGCAGGTACAGATAC | 139 |
| chr11:95894460-95896474 | p2 | (AT)7 | CCCGTCGTACCCTTAAACAA | ATGCCCGCAGGTACAGATAC | 139 |
| chr11:95971248-95973310 | p2 | (TA)6 | GATATGGACGGGCAGTTTGT | ACAACGTCCAAAGAAAACGG | 282 |
| chr11:95971248-95973310 | p2 | (AT)31 | GATATGGACGGGCAGTTTGT | ACAACGTCCAAAGAAAACGG | 282 |
| chr11:95971248-95973310 | p3 | (CAA)6 | GATATGGACGGGCAGTTTGT | ACAACGTCCAAAGAAAACGG | 282 |
| chr11:96020043-96022071 | p2 | (AT)14 | TAGAGGTGATTCGGGTCAGG | GGGCTGGGTCCTACCTCTAC | 140 |
| chr11:96117530-96119545 | p5 | (CAATC)3 | TTTATTATGATCGACCGGGG | TAGTCTCGCACGGTATTCCC | 131 |
| chr11:96125619-96127635 | p2 | (TA)8 | AAACTTTGCCAACGATCTCA | GGTTCCCTAGAGCCGTTACC | 216 |
| chr11:96125619-96127635 | p2 | (TA)7 | AAACTTTGCCAACGATCTCA | GGTTCCCTAGAGCCGTTACC | 216 |
| chr11:96129509-96131533 | p6 | (CCGGTT)4 | GGGGGTTGTGTTTCATCATC | AAACGTGGTGCCGGTAGTAG | 111 |
| chr11:96140974-96142986 | p2 | (CT)6 | GAAAGCGAGATGTTGAAGGC | TTTCTGCAAGTTGTTGGTGG | 202 |
| chr11:96140974-96142986 | p2 | (TA)9 | GAAAGCGAGATGTTGAAGGC | TTTCTGCAAGTTGTTGGTGG | 202 |
| chr11:96172120-96174140 | p3 | (TTG)5 | GGAGAGACATAGATGCCCCA | ACTGCGATAATGCGCCTACT | 207 |
| chr11:96172120-96174140 | p5 | (TTGGT)4 | GGAGAGACATAGATGCCCCA | ACTGCGATAATGCGCCTACT | 207 |
| chr11:96261542-96263557 | p5 | (TATCA)3 | CGAGTTTGGAGAAAGTGGGA | TAAGGGGTGCTCGGTACAAG | 272 |
| chr11:96355655-96357673 | p6 | (CCAATG)3 | CGTGATTCTTTTGGGTCGTT | ACAACATGTACCGAGGGAGC | 186 |
| chr11:96475393-96477417 | p2 | (TA)12 | GTCCGCTTCGTTTTCTCATC | TGAACTCAAGGGAACAACCC | 224 |
| chr11:96499028-96501043 | p5 | (TATTT)3 | CATGGAGAAAGGGCAAATTC | CAACCACCCTTCGTCATCTT | 218 |

| Marker ID | SSR type | SSR motif | FORWARD PRIMER (5'-3') | REVERSE PRIMER (5'-3') | size (bp) |
|---|---|---|---|---|---|
| chr11:96499028-96501043 | p5 | (TAAAA)3 | CATGGAGAAAGGGCAAATTC | CAACCACCCTTCGTCATCTT | 218 |
| chr11:96584364-96586386 | p2 | (TA)11 | GTCAGTCTGTAAGCGGCTCC | GAACCCGCAGACCTTGATTA | 227 |
| chr11:96584364-96586386 | p2 | (TA)7 | GTCAGTCTGTAAGCGGCTCC | GAACCCGCAGACCTTGATTA | 227 |
| chr11:96648595-96650627 | p2 | (AT)16 | GTTGCCTTGTCTTTCGAAGC | AAAATTTCCGTTTTGCCCTT | 143 |
| chr11:96722158-96724174 | p2 | (AC)8 | CATTTAGACGCGATTCCGAT | GTGAATTCTCGGGTGTCGAT | 223 |
| chr11:96725676-96727688 | p2 | (AT)6 | ACGGCTCGTAGACACTTCGT | TGTTGTTTGGGGTACTGGGT | 203 |
| chr11:96736816-96738828 | p2 | (GT)6 | TACAACTGGTTGGCCCTTTC | CCTTCCTCAATATTTCCGCA | 298 |
| chr11:96834123-96836135 | p2 | (CA)6 | TTCCAATTCGGATTTCCAAG | TTTGAGTTTTGACGAGCGTG | 173 |
| chr11:96856061-96858076 | p5 | (ATACG)3 | AGGATAGGGGGCTGGAGTAA | GCCCCTAGAGTAGATTCGGG | 265 |
| chr11:96856061-96858076 | p2 | (AT)6 | AGGATAGGGGGCTGGAGTAA | GCCCCTAGAGTAGATTCGGG | 265 |
| chr11:96881626-96883640 | p2 | (AG)7 | TCCTGTGTGCAGGTTTTGAG | TTCTCGAGGTCAGCCGTACT | 178 |
| chr11:96904512-96906524 | p2 | (AT)6 | ATCAGATAGCCAATGTGCCC | GGAGTCGTCGTGGGAAGTTA | 133 |
| chr11:97067204-97069224 | p5 | (ATGCA)4 | GCGAGTTGAGGCTGAGATTC | TAAGGGAAGGGTTTTCCCAC | 132 |
| chr11:97248724-97250760 | p3 | (GAA)12 | TCCTTCACCATCACCACTCA | GTTTGGTCATTGGTTTCGCT | 297 |
| chr11:97248724-97250760 | p3 | (ACA)5 | TCCTTCACCATCACCACTCA | GTTTGGTCATTGGTTTCGCT | 297 |
| chr11:97269547-97271565 | p6 | (ATCACA)3 | CACTTCCGGGGCATTAGTTA | ATGGCTCCTTGAAAATGTGC | 191 |
| chr11:97281134-97283149 | p3 | (TTC)5 | GTTCTATGGGGTCAAAGCCA | TAACTGCCACATCCCACAAA | 101 |
| chr11:97282818-97284833 | p5 | (TGGGT)3 | TTTTCTCGGGCCAGTACATC | GTGATACCCTGACCGCAAGT | 165 |
| chr11:97449996-97452018 | p2 | (AT)11 | TAAGACAACATATGCGGCCA | AAGGTTCGGGGTTATATCGG | 138 |
| chr11:97528079-97530097 | p3 | (TTG)6 | GGGGGAGGAATGTTACACCT | CCTCCAAAGTTGACCCAGAA | 187 |
| chr11:97528079-97530097 | p2 | (AT)7 | GGGGGAGGAATGTTACACCT | CCTCCAAAGTTGACCCAGAA | 187 |
| chr11:97528079-97530097 | p2 | (TA)7 | GGGGGAGGAATGTTACACCT | CCTCCAAAGTTGACCCAGAA | 187 |
| chr11:97534254-97536266 | p2 | (TA)6 | TAATCACATCAACTGCCCCA | AACCCGTGAGCTGTCTAGGA | 177 |
| chr11:97562919-97564934 | p5 | (TGATA)3 | GCCCAAATTTTTGTAGGCAA | TTTGCCAAAACTCAAATCCC | 124 |
| chr11:97602869-97604895 | p2 | (AT)13 | GTGCTTCGATGTCATGCACT | TTAGTTCAAATTTTCGGGCG | 247 |
| chr11:97602869-97604895 | p5 | (ATGAA)3 | GTGCTTCGATGTCATGCACT | TTAGTTCAAATTTTCGGGCG | 247 |
| chr11:97741091-97743109 | p2 | (TA)9 | GCAAACTCCATAAAGGGCAA | TCGATTGCATGGTGTTCCTA | 295 |
| chr11:97741091-97743109 | p2 | (AT)29 | GCAAACTCCATAAAGGGCAA | TCGATTGCATGGTGTTCCTA | 295 |
| chr11:97741575-97743633 | p2 | (TA)9 | TAATTCGCTCCCATTTCCAC | AGTGGCGCAATATGAAAAGG | 151 |
| chr11:97741575-97743633 | p2 | (AT)29 | TAATTCGCTCCCATTTCCAC | AGTGGCGCAATATGAAAAGG | 151 |
| chr11:97786633-97788651 | p3 | (TTA)6 | GATCTCGAATTGGCTGTGGT | AAGCGCATCGGCTACTACAT | 154 |
| chr11:97851605-97853620 | p5 | (TATCA)3 | GGGGCTCACCAACTAACTGA | ATTGTGAACGGGCTCATAGG | 237 |
| chr11:97851605-97853620 | p5 | (TCATA)3 | GGGGCTCACCAACTAACTGA | ATTGTGAACGGGCTCATAGG | 237 |
| chr11:97989052-97991080 | p2 | (AT)25 | CAGCAGATTGGTAAGCTCCC | ATATCTGAGCAAGCCGAGGA | 224 |
| chr11:97989052-97991080 | p2 | (TA)14 | CAGCAGATTGGTAAGCTCCC | ATATCTGAGCAAGCCGAGGA | 224 |
| chr11:98132335-98134353 | p6 | (CATATA)3 | ACACCGTACCGGAAACTCAC | CATATATGGGTACGGCAGGG | 294 |

| Marker ID | SSR type | SSR motif | FORWARD PRIMER (5'-3') | REVERSE PRIMER (5'-3') | size (bp) |
|---|---|---|---|---|---|
| chr11:98234275-98236287 | p2 | (AT)6 | AACGTAGCCTCAACGAATGG | AACGGAAGGCTATAGGGCAT | 129 |
| chr11:98250833-98252845 | p2 | (AC)6 | AGGGAAGAATTGGGTTTGCT | GTTGAATTCTCGGGGTGCTA | 103 |
| chr11:98277965-98279980 | p3 | (TTG)5 | GTCGTGGGTACAAAAATGGG | TTATTTCTGGGGGTTTTCCC | 227 |
| chr11:98317136-98319151 | p3 | (ATG)5 | CGAGGTTCAAAAATACGGGA | TCTGTTCGAACCTAAACCCG | 201 |
| chr11:98325532-98327544 | p2 | (AT)12 | TTTTCCTTAAATGGGCGAGA | TGGGGGCATGAAAGTAAAAG | 119 |
| chr11:98325532-98327544 | p2 | (TA)6 | TTTTCCTTAAATGGGCGAGA | TGGGGGCATGAAAGTAAAAG | 119 |
| chr11:98370684-98372699 | p5 | (TGCAA)3 | GGAGACACAGAATGGGAGGA | CTTTGTGTGACCCTTTCGGT | 196 |
| chr11:98373293-98375313 | p5 | (ATTTC)4 | GGTCCAGTCGAAACACGAAT | TTCGAGTGTGATTTGGGTCA | 137 |
| chr11:98380579-98382594 | p5 | (ATTTC)3 | CAGGATGGTAGCCCTGTGAT | GCCATCAATTGCCATTTCTT | 296 |
| chr11:98408446-98410461 | p5 | (AAAAC)3 | TTCAAAACATCAACACGGGA | TTTCATTAGGTCGGAGGGTG | 176 |
| chr11:98410880-98412895 | p5 | (GAAAG)3 | TGCATATGCTGCCTTCAGAC | AAATGATCCTGGCCTTCCTT | 265 |
| chr11:98461532-98463550 | p2 | (AT)9 | TCGTGCATGCCTCAATATGT | TATCATATCCGGCCCCATTA | 226 |
| chr11:98541239-98543257 | p6 | (ATTATG)3 | ATCAAGTGCCCAGTCTGTCC | CATCCTCAAGGCCTTACCAA | 101 |
| chr11:98541239-98543257 | p2 | (AT)6 | ATCAAGTGCCCAGTCTGTCC | CATCCTCAAGGCCTTACCAA | 101 |
| chr11:98622404-98624424 | p2 | (AG)7 | GTTTCCTACGCCCACAGGTA | TTCCAAATTTTTCCGAATGC | 211 |
| chr11:98622404-98624424 | p2 | (TG)10 | GTTTCCTACGCCCACAGGTA | TTCCAAATTTTTCCGAATGC | 211 |
| chr11:98641993-98644008 | p2 | (TA)9 | TCGAGGCATGGAGGTATTTC | AAGAAATCAATGCGGAAACG | 143 |
| chr11:98641993-98644008 | p5 | (CCCAC)3 | TCGAGGCATGGAGGTATTTC | AAGAAATCAATGCGGAAACG | 143 |
| chr11:98724750-98726765 | p2 | (AT)13 | GCTCTCTTTCTACCCGCCTT | AAACAAGAGGCTACAGGGCA | 101 |
| chr11:98724750-98726765 | p3 | (CAT)5 | GCTCTCTTTCTACCCGCCTT | AAACAAGAGGCTACAGGGCA | 101 |
| chr11:98827516-98829562 | p2 | (TA)23 | AATGTGGGATTACTTTGGCG | ACCCACCACTTGCTTATTGG | 278 |
| chr11:98839409-98841429 | p2 | (TA)10 | AAGAATGGAGGAGAGGGCAT | ACAACCTTCCATTTGGTGGA | 199 |
| chr11:98839409-98841429 | p5 | (TGAAT)4 | AAGAATGGAGGAGAGGGCAT | ACAACCTTCCATTTGGTGGA | 199 |
| chr11:99021093-99023113 | p2 | (TA)10 | ATGTCACCCCGGTAGAGTTG | GTTCGGTGTCATCCATACCC | 239 |
| chr11:99021093-99023113 | p2 | (TA)17 | ATGTCACCCCGGTAGAGTTG | GTTCGGTGTCATCCATACCC | 239 |
| chr11:99260288-99262306 | p2 | (AT)9 | CGGCTTACATAGCCGCTTAC | GCATTATGGGTATGTTGGGG | 278 |
| chr11:99260288-99262306 | p2 | (AT)14 | CGGCTTACATAGCCGCTTAC | GCATTATGGGTATGTTGGGG | 278 |
| chr11:99317365-99319383 | p3 | (ATG)6 | CTCCCCAAACCTTCCCTAAC | TGATTCGTCGAGTTTCGTTG | 265 |
| chr11:99324315-99326330 | p3 | (ATG)5 | GCAGAAACGCACAGCTACAG | GAAAGTTCTAGGGGGTTCGG | 118 |
| chr11:99334542-99336557 | p3 | (TGT)5 | ATTCCGGGACCAAGAAAATC | GGCTTATCTTGAAATGCCCA | 159 |
| chr11:99334542-99336557 | p6 | (TGATAA)3 | ATTCCGGGACCAAGAAAATC | GGCTTATCTTGAAATGCCCA | 159 |
| chr11:99340626-99342642 | p2 | (AT)8 | ACAACTCAGCACGAATGCAC | ATTTGGGGTGTTTGGAATGA | 228 |
| chr11:99411933-99413948 | p5 | (ATTGC)3 | CTTCGGGTATGAAGTTGGGA | CGCCACCAATATAGCCAAGT | 105 |
| chr11:99440954-99442976 | p2 | (TA)11 | GGGGTCGTTTGGTAGATTGA | AGGGGTCGTTTGGTTTAAGG | 242 |
| chr11:99527869-99529885 | p2 | (TA)8 | CAAGGAGGGCACATGAAGAT | TCTCCCTTGGTTTTCATTGG | 258 |
| chr11:99527869-99529885 | p2 | (TA)7 | CAAGGAGGGCACATGAAGAT | TCTCCCTTGGTTTTCATTGG | 258 |

| Marker ID | SSR type | SSR motif | FORWARD PRIMER (5'-3') | REVERSE PRIMER (5'-3') | size (bp) |
|---|---|---|---|---|---|
| chr11:99527997-99530011 | p2 | (TA)8 | CAAGGAGGGCACATGAAGAT | TCTCCCTTGGTTTTCATTGG | 130 |
| chr11:99527997-99530011 | p2 | (TA)7 | CAAGGAGGGCACATGAAGAT | TCTCCCTTGGTTTTCATTGG | 130 |
| chr11:99555668-99557683 | p5 | (TACAA)3 | GACTTGCCAAGGATATCGGA | AGTTGGCGACAGTCCTCAGT | 134 |
| chr11:99721239-99723266 | p3 | (TTG)9 | AGAAGATGAAAGTCGCGGAA | TCCCGTATCTTGTCCTCCAC | 289 |
| chr11:99761519-99763555 | p2 | (AT)25 | ACTCACGGCCAACACATACA | AGGGCCTTGTCCTGTCTCTT | 203 |
| chr11:99761519-99763555 | p3 | (ATC)5 | ACTCACGGCCAACACATACA | AGGGCCTTGTCCTGTCTCTT | 203 |
| chr11:99761551-99763566 | p2 | (AT)25 | ACTCACGGCCAACACATACA | AGGGCCTTGTCCTGTCTCTT | 171 |
| chr11:99761551-99763566 | p3 | (ATC)5 | ACTCACGGCCAACACATACA | AGGGCCTTGTCCTGTCTCTT | 171 |
| chr11:99803634-99805646 | p2 | (AG)6 | AGATTGTCGGTTCGTCATCC | CATCCCTCAAAATGGTGGTC | 300 |
| chr11:99824398-99826412 | p2 | (TA)7 | TCACTTTCTGCAAGCACCAC | CGCTTTGTTTCTTCCACTCC | 210 |
| chr11:99825855-99827870 | p2 | (AG)6 | AAAGAAGCTCGCAGGAGTTG | TCTTGTCACTGCTTCGTTGG | 224 |
| chr11:99825855-99827870 | p5 | (TATTT)3 | AAAGAAGCTCGCAGGAGTTG | TCTTGTCACTGCTTCGTTGG | 224 |
| chr11:99843517-99845555 | p2 | (TA)19 | TCGCAATTCCTATGCATCAA | AGGCTCGTAGGCAGTTGTGT | 242 |
| chr11:99852087-99854102 | p3 | (ACA)5 | GTTCCTCTTATTCCGGAGCC | AGAAGCCGACAAGGCTATCA | 190 |
| chr11:99857458-99859476 | p6 | (ATACAT)3 | ATGGCTATCGCAAACGAATC | TGATGAGGGGTTCCCAGTAG | 172 |
| chr11:99857458-99859476 | p2 | (TA)6 | ATGGCTATCGCAAACGAATC | TGATGAGGGGTTCCCAGTAG | 172 |
| chr11:99857677-99859689 | p6 | (ATACAT)3 | CTACTGGGAACCCCTCATCA | TTTTAGCCTTGTCGGCTTGT | 183 |
| chr11:99857677-99859689 | p2 | (TA)6 | CTACTGGGAACCCCTCATCA | TTTTAGCCTTGTCGGCTTGT | 183 |
| chr11:99874623-99876637 | p2 | (AT)7 | ATGTCGGAAAGTCCACGAAG | AATTTTGCCCTCGACCTCTT | 198 |
| chr11:99927857-99929875 | p6 | (TATGTA)3 | TTCGTGGTCGCATTTAATCA | GCCATGTCGTATTTTGGCTT | 149 |
| chr11:99944407-99946423 | p2 | (AT)8 | GGAATGCCTTGGTCACTGTT | ACCTTCTAGATCGCCGGATT | 125 |
| chr11:99945053-99947079 | p2 | (AT)8 | ACTTTTGTCACGACCCAACC | GGGAGACAGGTTGCTGGATA | 206 |
| chr11:99945156-99947172 | p2 | (AT)8 | ACTTTTGTCACGACCCAACC | GGGAGACAGGTTGCTGGATA | 103 |
| chr11:99995237-99997252 | p3 | (TAT)5 | AATGGCGGACCTCATCATTA | CCAGGATAAGGCTCATTCCA | 190 |
| chr11:100106677-10010869 | p5 | (TTAAA)3 | CCACTTTCGATTCCGGTTTA | GACGGGTATTGGTGAGAGGA | 207 |
| chr11:100201991-10020400 | p6 | (TGTAAG)3 | GGGCAGGTTCCATAACTGAA | ACATGCCCTTATTCGCATTC | 164 |
| chr11:100201991-10020400 | p5 | (TTGCA)3 | GGGCAGGTTCCATAACTGAA | ACATGCCCTTATTCGCATTC | 164 |
| chr11:100202393-10020440 | p6 | (TGTAAG)3 | GAGCTCCATCTCATTCGGAG | AGCCGACAAGACCAGACAAT | 283 |
| chr11:100202393-10020440 | p5 | (TTGCA)3 | GAGCTCCATCTCATTCGGAG | AGCCGACAAGACCAGACAAT | 283 |
| chr11:100236912-10023900 | p2 | (TA)44 | GGTAACGGGAATGATGTTGG | GCCCTCTAGTTGGGTTGTGA | 293 |
| chr11:100236912-10023900 | p3 | (ATC)8 | GGTAACGGGAATGATGTTGG | GCCCTCTAGTTGGGTTGTGA | 293 |
| chr11:100387119-10038913 | p2 | (TA)7 | CAATTTTGCTGTGGAACGTG | CGACCTTACGTCACTCCGAT | 282 |
| chr11:100387119-10038913 | p3 | (CAA)6 | CAATTTTGCTGTGGAACGTG | CGACCTTACGTCACTCCGAT | 282 |
| chr11:100652538-10065456 | p3 | (TCA)9 | TTTTCATTGGGCTAAGGCAC | TCTTAGCCTTGTCGGCTCAT | 261 |
| chr11:100662112-10066413 | p3 | (ATC)6 | GCGCACTTCATACCCCTAAA | GCCTTGTCGGCTAGCTTATG | 135 |
| chr11:100662112-10066413 | p6 | (ATACAT)3 | GCGCACTTCATACCCCTAAA | GCCTTGTCGGCTAGCTTATG | 135 |

| Marker ID | SSR type | SSR motif | FORWARD PRIMER (5'-3') | REVERSE PRIMER (5'-3') | size (bp) |
|---|---|---|---|---|---|
| chr11:100666204-10066822 | p2 | (AT)7 | CCAGCTACACAAGCCAACAA | TGTGTAGTGGCAGCCTTGTC | 180 |
| chr11:100666204-10066822 | p6 | (CATATA)3 | CCAGCTACACAAGCCAACAA | TGTGTAGTGGCAGCCTTGTC | 180 |
| chr11:100716789-10071880 | p6 | (TTTGCT)3 | GCCACTATCCCTGAACCAAA | AATGACTACATGCCGAAGGG | 158 |
| chr11:100806429-10080844 | p5 | (TGTGT)3 | AATTGCGGGAAGATGACAAG | TTGCGAATTTCCCTTTATGC | 292 |
| chr11:100957398-10095941 | p2 | (TA)6 | TATATTCCCCGTGTTTTCGC | TCTTCAATTTTTCATGGCCC | 298 |
| chr11:100963223-10096524 | p2 | (TA)9 | TGGGCTTGGATTCAAGGTAG | ACCAACAACCAACAACAGCA | 279 |
| chr11:100967212-10096922 | p2 | (CA)6 | TCTTGTTGAGTCCGCTCCTT | AGCAGGATGAAACAGGTTCG | 243 |
| chr11:101049788-10105183 | p2 | (AT)7 | GAATAATTTTTGGGCCGGTT | CTTAATTGTCCAAGCACGCA | 194 |
| chr11:101100126-10110214 | p2 | (AG)7 | GTGGCCGAATACGGATATTG | CGTCCAAAGAAAACGGACAT | 166 |
| chr11:101152320-10115433 | p5 | (AGGAC)3 | CTGGCACTAGAGGCTTGGTC | CAAGACAGCGATAGGAAGGC | 210 |
| chr11:101163073-10116512 | p2 | (TA)24 | TTTTCGCAAGCTTCCAGATT | AGGGTCTTGGGTGTCAGTTG | 260 |
| chr11:101195675-10119768 | p2 | (CA)6 | GGACACCCCAAAGAAACTCA | GTGTGTGAAATTTGGCGATG | 175 |
| chr11:101195675-10119768 | p2 | (CA)6 | GGACACCCCAAAGAAACTCA | GTGTGTGAAATTTGGCGATG | 175 |
| chr11:101275408-10127742 | p5 | (ATCAG)3 | AGAAGCCGACAAGGCTAACA | CGGAGCGTACCTTTTTGGTA | 240 |
| chr11:101323686-10132571 | p2 | (TA)14 | ATTCCTCAGTGGGCAATCAC | CACGGTGTACTCGGTGTGAC | 144 |
| chr11:101438522-10144055 | p2 | (AT)16 | GGAGAACGGTGGCTTACAAA | CGGTTGTTTTCCCGATTAGA | 299 |
| chr11:101467616-10146963 | p5 | (AAAAT)3 | AAAGGAGAACAAGCGAACGA | CAATCAGCACCTCTGGATCA | 247 |
| chr11:101469907-10147192 | p3 | (ATG)6 | TTGACAAGCGGCATAAATGA | AAGCCACCGACTTCTGCTTA | 279 |
| chr11:101469907-10147192 | p2 | (TA)9 | TTGACAAGCGGCATAAATGA | AAGCCACCGACTTCTGCTTA | 279 |
| chr11:101469907-10147192 | p2 | (AT)6 | TTGACAAGCGGCATAAATGA | AAGCCACCGACTTCTGCTTA | 279 |
| chr11:101544084-10154609 | p2 | (TA)6 | CCCTTTCCATGTCTGTTCGT | GCAATGCAGACTCGATAGCA | 196 |
| chr11:101572994-10157500 | p6 | (AAAAGA)3 | CGAAAAATTAGCGGCAAGAC | CCGCATTATTTTCATGACCC | 111 |
| chr11:101573004-10157501 | p6 | (AAAAGA)3 | CGAAAAATTAGCGGCAAGAC | CCGCATTATTTTCATGACCC | 101 |
| chr11:101580447-10158246 | p6 | (CATGTC)3 | ATCTGGGCATCCAAACACAT | TGCTTTGTTAGCCCTGTGTG | 163 |
| chr11:101580447-10158246 | p5 | (CATAG)3 | ATCTGGGCATCCAAACACAT | TGCTTTGTTAGCCCTGTGTG | 163 |
| chr11:101696416-10169843 | p6 | (CATATA)3 | CCCAGACGGCGTATAGTTGT | CGGGAACGATGAAAAGGATA | 114 |
| chr11:101735115-10173713 | p2 | (TA)8 | CCAAGGCAGGCTATGTGTTT | TGCGCTACAAATTCGCTATG | 174 |
| chr11:101770572-10177258 | p5 | (AATAA)3 | AAGTCTGCCAACAATCACCC | AGCTCAATGTCCAAATTGCC | 238 |
| chr11:101791843-10179385 | p2 | (AT)6 | CCAACTCCACGGATTCAGTT | GTTGCTTCGGGTCAACAAAT | 178 |
| chr11:101855082-10185711 | p2 | (TA)16 | CATGGGCTAACTCGTAGGGA | CACCAGATGGCGTATCAATG | 100 |
| chr11:101855082-10185711 | p3 | (CAA)6 | CATGGGCTAACTCGTAGGGA | CACCAGATGGCGTATCAATG | 100 |
| chr11:101932438-10193445 | p6 | (CCAGTT)3 | AAGCGCAAAAAGACCTTCAA | CATTTGAAGAGGGGTTGGAA | 180 |
| chr11:101981778-10198379 | p5 | (AATGC)3 | TTTGTCCTCTTCCCCTTCCT | GTGGTTCGCCTACTCTGAGG | 221 |
| chr11:102162693-10216470 | p5 | (TTTGG)3 | TTTGTGGTATGTTGCCGCTA | CAGCAGGAGGTACACAAGCA | 293 |
| chr11:102190727-10219274 | p5 | (TTTGG)3 | TTTGTGGTATGTTGCCGCTA | CAGCAGGAGGTACACAAGCA | 293 |
| chr11:102231090-10223310 | p5 | (TATAT)3 | TGGCTAGATACGCTCCGACT | CAATATCACGCTCACCCCTT | 222 |

| Marker ID | SSR type | SSR motif | FORWARD PRIMER (5'-3') | REVERSE PRIMER (5'-3') | size (bp) |
|---|---|---|---|---|---|
| chr11:102363046-10236506 | p3 | (ACA)6 | CATCATAATCATACGCCCCC | TGGCATGACCCATGATAGAA | 270 |
| chr11:102363046-10236506 | p5 | (ACTTT)3 | CATCATAATCATACGCCCCC | TGGCATGACCCATGATAGAA | 270 |
| chr11:102363046-10236506 | p2 | (AT)11 | CATCATAATCATACGCCCCC | TGGCATGACCCATGATAGAA | 270 |
| chr11:102363193-10236520 | p3 | (ACA)6 | TTCGGGACTTGGAATCAATC | TGGCATGACCCATGATAGAA | 143 |
| chr11:102363193-10236520 | p5 | (ACTTT)3 | TTCGGGACTTGGAATCAATC | TGGCATGACCCATGATAGAA | 143 |
| chr11:102363193-10236520 | p2 | (AT)11 | TTCGGGACTTGGAATCAATC | TGGCATGACCCATGATAGAA | 143 |
| chr11:102404680-10240669 | p5 | (ATTTC)3 | GAGAGGAAATGACGTGGGAA | AGGTGTAACGTTCATTCGCC | 221 |
| chr11:102488305-10249031 | p2 | (CT)6 | TTCAATCGTCCTTTTGGAGG | GGGTTCGGGATGAAGGTTAT | 251 |
| chr11:102534687-10253672 | p2 | (TA)18 | TCGGACATTATTTGCCATCA | ACGTGGTCTCCCATATCTGC | 103 |
| chr11:102593245-10259526 | p2 | (AT)8 | CGTTATTCACCGAGTCCCAT | TACCTGCGAGCATGAAACAC | 203 |
| chr11:102593245-10259526 | p2 | (AT)39 | CGTTATTCACCGAGTCCCAT | TACCTGCGAGCATGAAACAC | 203 |
| chr11:102593245-10259526 | p2 | (AT)9 | CGTTATTCACCGAGTCCCAT | TACCTGCGAGCATGAAACAC | 203 |
| chr11:102626623-10262863 | p3 | (ACA)5 | GAACCGGTGAACGAAGTCAT | TTTGGAACATCCTCCTACGG | 218 |
| chr11:102626623-10262863 | p5 | (TTTTC)3 | GAACCGGTGAACGAAGTCAT | TTTGGAACATCCTCCTACGG | 218 |
| chr11:102765513-10276753 | p2 | (AT)10 | CAGCATGGAAACTTGCAAAA | GTAGTAGCAGCCTTGTCGGC | 280 |
| chr11:102772633-10277465 | p3 | (TTG)6 | AAAGTTTGGGCACTCAGGTG | ACACACATGCCCGTAGCATA | 116 |
| chr11:102880846-10288285 | p2 | (TG)6 | CACATGGTTGACATTCCGAG | TCTTACCCGGCATCATAAGC | 156 |
| chr11:102994801-10299681 | p2 | (TA)6 | CCAACCCTTTGCACATTTTT | GCTTGCTCCAAAGCTCATTC | 227 |
| chr11:103080744-10308276 | p6 | (ATCACA)3 | CACTTCCGGGGCATTAGTTA | ATGGCTCCTTGAAAATGTGC | 191 |
| chr11:103136907-10313891 | p2 | (GA)6 | TACACCTCGCAGTTCAGACG | TTTCCTTCATTCACTTGCCC | 171 |
| chr11:103215015-10321704 | p6 | (TAGAGG)3 | TAATGAGGCTGCAGCAGATG | CCTCAAAACCCGAAAATTCA | 232 |
| chr11:103215052-10321706 | p6 | (TAGAGG)3 | TAATGAGGCTGCAGCAGATG | CCTCAAAACCCGAAAATTCA | 195 |
| chr11:103215148-10321716 | p6 | (TAGAGG)3 | CCAGGCATACCAGGTACGAT | CCTCAAAACCCGAAAATTCA | 175 |
| chr11:103270724-10327273 | p3 | (AAC)5 | CTAGCTTGAGCACACACCCA | TCAACTAGCGTCCCTCCAGT | 202 |
| chr11:103270724-10327273 | p2 | (AT)7 | CTAGCTTGAGCACACACCCA | TCAACTAGCGTCCCTCCAGT | 202 |
| chr11:103460990-10346300 | p3 | (CAA)5 | GTGCAACCCGATCCACTTAT | TTTGATTTTCCAAAAACGGC | 109 |
| chr11:103708457-10371047 | p5 | (TTGCA)3 | CTGTCAGCTGATGGAAAGCA | CCCACACTGCGCATCTACTA | 199 |
| chr11:103726448-10372846 | p5 | (ATATG)3 | AAGAGAAATGGTGGAATGCG | TCCTTCGATATCGCGTCTCT | 263 |
| chr11:103726448-10372846 | p2 | (TA)9 | AAGAGAAATGGTGGAATGCG | TCCTTCGATATCGCGTCTCT | 263 |
| chr11:103771281-10377332 | p2 | (CT)20 | GAGTTTGCTACAGGATCCGC | TTGTCTCGGAACAATGGACA | 197 |
| chr11:103805142-10380715 | p2 | (CT)6 | GTTGGGCCTGATCCAACTTA | AGCTCTGGCTGTGTGATCCT | 273 |
| chr11:103825774-10382778 | p2 | (AT)6 | TTGCTCCACGCCCTACTACT | TTTGGCTGCATCAAATACGA | 173 |
| chr11:103825774-10382778 | p2 | (AT)26 | TTGCTCCACGCCCTACTACT | TTTGGCTGCATCAAATACGA | 173 |
| chr11:103825774-10382778 | p2 | (AT)11 | TTGCTCCACGCCCTACTACT | TTTGGCTGCATCAAATACGA | 173 |
| chr11:103825774-10382778 | p5 | (CCGGT)3 | TTGCTCCACGCCCTACTACT | TTTGGCTGCATCAAATACGA | 173 |
| chr11:103825774-10382778 | p2 | (TA)7 | TTGCTCCACGCCCTACTACT | TTTGGCTGCATCAAATACGA | 173 |

| Marker ID | SSR type | SSR motif | FORWARD PRIMER (5'-3') | REVERSE PRIMER (5'-3') | size (bp) |
|---|---|---|---|---|---|
| chr11:103825774-10382778 | p4 | (TTTC)4 | TTGCTCCACGCCCTACTACT | TTTGGCTGCATCAAATACGA | 173 |
| chr11:103949642-10395166 | p3 | (TAA)7 | GGAAACAAACGGGTTAGGGT | TGTTTTCTTCCTTCGCCATT | 291 |
| chr11:103949642-10395166 | p3 | (TTG)5 | GGAAACAAACGGGTTAGGGT | TGTTTTCTTCCTTCGCCATT | 291 |
| chr11:103950229-10395224 | p3 | (TAA)7 | GTCGTTTTGGGTTGCAATTT | ACCGGCAATATAACCAAACG | 189 |
| chr11:103950229-10395224 | p3 | (TTG)5 | GTCGTTTTGGGTTGCAATTT | ACCGGCAATATAACCAAACG | 189 |
| chr11:104016839-10401886 | p6 | (TTTTTG)4 | GTCCTAAAACGGACCAACGA | TATCGGAGTAACGGGGAGTG | 126 |
| chr11:104022442-10402445 | p6 | (ATACAT)5 | TAACTCACCGAACACCCCTC | GGGCCTTGTCCCATCTTTAT | 117 |
| chr11:104077642-10407966 | p2 | (TG)6 | CCAAACAATGAAGGGCATTT | TTAAGTGCTTTGTTGCGGTG | 265 |
| chr11:104077673-10407968 | p2 | (TG)6 | CCAAACAATGAAGGGCATTT | TTAAGTGCTTTGTTGCGGTG | 234 |
| chr11:104113509-10411552 | p2 | (TA)9 | CTTCGGTGGGAGAAAATTCA | AACAAGGCAGCGTCTGAGAT | 191 |
| chr11:104182026-10418404 | p5 | (TGTTT)3 | TGCCACTATTTGCCTCATTTC | TTGTCAATTCCACCCCAAAT | 199 |
| chr11:104213126-10421514 | p3 | (TTA)20 | CTGTCTACCTGTGGGCATGA | ACCCCTAAAAGTTCGCAGGT | 222 |
| chr11:104213126-10421514 | p3 | (TCA)5 | CTGTCTACCTGTGGGCATGA | ACCCCTAAAAGTTCGCAGGT | 222 |
| chr11:104385476-10438748 | p2 | (TA)6 | CTGTTGATTGCCCTTTTGGT | AAAATGGCACAGTTGCATGA | 271 |
| chr11:104401820-10440383 | p2 | (AG)7 | ACAAACAACCCTCGTAACCG | TTGGCTTAAGTTTCCCCCTT | 186 |
| chr11:104401820-10440383 | p3 | (TTG)5 | ACAAACAACCCTCGTAACCG | TTGGCTTAAGTTTCCCCCTT | 186 |
| chr11:104401820-10440383 | p2 | (TA)10 | ACAAACAACCCTCGTAACCG | TTGGCTTAAGTTTCCCCCTT | 186 |
| chr11:104415906-10441791 | p2 | (CA)6 | AACCTCTTGACCCCTTCGTT | CCTATTGTGAGGGTCTGGGA | 258 |
| chr11:104445448-10444746 | p2 | (CA)6 | AACCTCTTGACCCCTTCGTT | CCTATTGTGAGGGTCTGGGA | 258 |
| chr11:104527208-10452922 | p3 | (ATG)5 | AAGGCTCGTCCCTTTCTTTC | TATCGGGAAAGGAATCATGC | 187 |
| chr11:104527208-10452922 | p2 | (TA)7 | AAGGCTCGTCCCTTTCTTTC | TATCGGGAAAGGAATCATGC | 187 |
| chr11:104596565-10459859 | p5 | (GCATT)5 | GGTTTACCCCTGTGGTGAGA | TAGCCCGATCAAGCAATACC | 263 |
| chr11:104599946-10460196 | p4 | (TATT)4 | TGTTCCTTGGCATTAAAGGG | GCACAAAAATGCGTGAAAAA | 212 |
| chr11:104607066-10460908 | p3 | (TTG)6 | TATTGGTGCAGGGTTGATGA | TCGCCCCGTACAATAAAGTC | 143 |
| chr11:104684571-10468658 | p2 | (AT)7 | TGCCAAAATTCAGGAAAAGG | ACCAAAAGAACCATGTTCCG | 270 |
| chr11:104695208-10469722 | p2 | (AT)6 | CACCCCAAGGGGAGATACTT | CTTACTTGGCCCATGTTCGT | 216 |
| chr11:104738543-10474055 | p5 | (AAAAT)3 | AAAGCCCGTGGTCTAACCTT | ACAGAGCAAGAAGAAGCGGA | 251 |
| chr11:104738543-10474055 | p2 | (TC)6 | AAAGCCCGTGGTCTAACCTT | ACAGAGCAAGAAGAAGCGGA | 251 |
| chr11:104789777-10479179 | p6 | (ATATGT)3 | CGCATAAGGAGCAGAAAAGG | GGCTTTAGGGTAGTCGGGTC | 201 |
| chr11:104843170-10484518 | p2 | (AT)11 | TCGTGAGACCATACAACCCA | CATTTTTCTGGAGGTGCCAT | 184 |
| chr11:104843170-10484518 | p3 | (TCA)5 | TCGTGAGACCATACAACCCA | CATTTTTCTGGAGGTGCCAT | 184 |
| chr11:104879791-10488180 | p2 | (TA)6 | ACCCGATGTTGATCTTGGAG | AAATGCACAACACAGTCGGA | 118 |
| chr11:104879791-10488180 | p6 | (TTGTTA)3 | ACCCGATGTTGATCTTGGAG | AAATGCACAACACAGTCGGA | 118 |
| chr11:104894145-10489619 | p2 | (TA)25 | TTGAGACTAGGCCTGCTGGT | GCGGGACCAATAATCTGAAA | 269 |
| chr11:104903222-10490524 | p2 | (AT)10 | AAACTCATGAAATCACGGGC | AATTCCTCCACCGGCTACTT | 280 |
| chr11:105030523-10503258 | p2 | (AT)30 | TCCAGCCCTCTATGTTGTCC | CCGGTGGGCTATAAAGTGAA | 127 |

| Marker ID | SSR type | SSR motif | FORWARD PRIMER (5'-3') | REVERSE PRIMER (5'-3') | size (bp) |
|---|---|---|---|---|---|
| chr11:105065023-10506703 | p3 | (TTC)5 | TTGGGACGATTTTTCTCAGG | ATATAGTCCCGGACACTGCG | 271 |
| chr11:105107989-10511000 | p2 | (TA)7 | TTGGGTTCAGTTGGATGACA | TCCGACAACAACATCGGATA | 173 |
| chr11:105118262-10512030 | p2 | (AT)20 | CCTCACAAATTTGGGCCTTA | TGACTTCAAGGGGCAAAAAC | 259 |
| chr11:105214666-10521669 | p2 | (TA)16 | ATCACGACCTTACGTCACCC | TGGTGTGTGTATCTTCCCGA | 270 |
| chr11:105343519-10534553 | p2 | (TG)6 | AAAGGTGCCTACGGGCTATT | CGACCTCATAGGATCGGTGT | 224 |
| chr11:105343519-10534553 | p2 | (TA)8 | AAAGGTGCCTACGGGCTATT | CGACCTCATAGGATCGGTGT | 224 |
| chr11:105343519-10534553 | p2 | (AT)11 | AAAGGTGCCTACGGGCTATT | CGACCTCATAGGATCGGTGT | 224 |
| chr11:105344242-10534625 | p2 | (TG)6 | TTCGGAGCCATATCTTTTGG | AAACATAAGCCGACAATGCC | 168 |
| chr11:105344242-10534625 | p2 | (TA)8 | TTCGGAGCCATATCTTTTGG | AAACATAAGCCGACAATGCC | 168 |
| chr11:105344242-10534625 | p2 | (AT)11 | TTCGGAGCCATATCTTTTGG | AAACATAAGCCGACAATGCC | 168 |
| chr11:105378673-10538069 | p3 | (ATG)5 | ATGGTTATGCCAGGTTTGGA | TCCACATCACTACTCACGCC | 242 |
| chr11:105387213-10538922 | p5 | (ACACA)3 | ACACACGGCCAACTTAGACC | TCATGGAGTGGGTGTGTTGT | 201 |
| chr11:105405258-10540727 | p5 | (TGAAT)4 | TCGAGGATGGTGCATTTACA | GTGGGGATCATGCCTTTAGA | 239 |
| chr11:105405258-10540727 | p6 | (TATATG)3 | TCGAGGATGGTGCATTTACA | GTGGGGATCATGCCTTTAGA | 239 |
| chr11:105436256-10543828 | p4 | (TGAA)4 | GACGATGGGGCAATTTAGAA | TATGCTTCGGTCACTTCACG | 106 |
| chr11:105436256-10543828 | p2 | (TA)16 | GACGATGGGGCAATTTAGAA | TATGCTTCGGTCACTTCACG | 106 |
| chr11:105560921-10556293 | p3 | (TAA)5 | ATCCTTTTTGGTGCCTTTCC | CGATGACGTGGTCAATGAAG | 195 |
| chr11:105560921-10556293 | p6 | (CTTTCT)3 | ATCCTTTTTGGTGCCTTTCC | CGATGACGTGGTCAATGAAG | 195 |
| chr11:105599210-10560124 | p6 | (AGGAGA)6 | TCGTACGCAGATGAAAGGTG | AATTTTGGAAGCACGTGAGG | 139 |
| chr11:105727964-10572999 | p2 | (TA)17 | CGACAGATTCGAGTGTGAGC | AACCCCAAAATCCGAAAGTC | 190 |
| chr11:105738323-10574034 | p2 | (AT)9 | TCCATGGGAACTGAGGAATC | AGTCTGCCAAACTGTTGCCT | 160 |
| chr11:105738323-10574034 | p2 | (TA)7 | TCCATGGGAACTGAGGAATC | AGTCTGCCAAACTGTTGCCT | 160 |
| chr11:105790531-10579254 | p2 | (TA)6 | TCACTACCCCTACCCCCTCT | GTTGATAGGTTGGGGTGGGT | 101 |
| chr11:105790531-10579254 | p3 | (TTA)15 | TCACTACCCCTACCCCCTCT | GTTGATAGGTTGGGGTGGGT | 101 |
| chr11:105790909-10579295 | p2 | (TA)6 | GTTGAGCTTTGCGAATGTTG | AGCTTCGGATGAGTGAAGTGA | 152 |
| chr11:105790909-10579295 | p3 | (TTA)15 | GTTGAGCTTTGCGAATGTTG | AGCTTCGGATGAGTGAAGTGA | 152 |
| chr11:105796383-10579845 | p2 | (AT)36 | CCTTACGCTCTCGCCTTTTA | GGTTTGAGCCAATGCGTTTA | 290 |
| chr11:105808986-10581100 | p4 | (ATCG)4 | TACCTGGTTTAGGCGTCACC | CTTTGTATGGACCCCCAATG | 209 |
| chr11:105841707-10584371 | p2 | (TA)11 | TCCAACTCATCCTTCCATCC | GATGTGACCGAGTCCTGGTT | 213 |
| chr11:105940865-10594288 | p3 | (ACA)7 | ACATGGGCAGAGATGGTAGC | TAGGATCCAGCCATCGGTAG | 279 |
| chr11:105949744-10595176 | p2 | (TA)7 | GCCCTTGTGACCTTTCCATA | GGGCATCTAGGGATTGTTGA | 203 |
| chr11:105962724-10596473 | p2 | (CA)7 | GACATGCCAAAGGAAAGGAA | ATTCCGGCAGGATACAAGTG | 115 |
| chr11:105962724-10596473 | p2 | (TA)6 | GACATGCCAAAGGAAAGGAA | ATTCCGGCAGGATACAAGTG | 115 |
| chr11:105968518-10597053 | p2 | (CA)8 | AAGCCTACATACCTGTGCCG | TTTTTACAGGGGAGGTGCTG | 145 |
| chr11:106043760-10604577 | p5 | (TATTT)3 | CTCGGGTGGCAAATAGTGAT | CCGTGGAGTTGGCATCTTAT | 208 |
| chr11:106060398-10606241 | p2 | (GT)6 | CGTGCTCACTCTATGCGTGT | ATTTCCTAAGCTCGGGGTGT | 254 |

| Marker ID | SSR type | SSR motif | FORWARD PRIMER (5'-3') | REVERSE PRIMER (5'-3') | size (bp) |
|---|---|---|---|---|---|
| chr11:106133185-10613520 | p5 | (ATGAT)3 | GATTTTGGAGGAAACGGGAT | ATGACGCAAGACCCTTTTTG | 170 |
| chr11:106221668-10622368 | p2 | (GT)7 | GGAGCTCCATTCATTTTGGA | GCGCCAAAAACACGAAGTAT | 271 |
| chr11:106336804-10633884 | p2 | (TA)21 | GCATGACGTATCGGGAACTT | TAGCGGAACGCTTGAGAAAT | 198 |
| chr11:106353557-10635557 | p6 | (CATATA)3 | TCTGGAGCGAGTCGAGGTAT | AAGCGAAAGACGAAGGTTGA | 276 |
| chr11:106476550-10647856 | p2 | (AT)9 | GACAGGTGGACGTGATCCTT | GAGTACTGTCGTGCCCGAAT | 283 |
| chr11:106714646-10671666 | p2 | (AT)9 | AAAGTCACCATCAACCCGAC | AAAGCCACATCCTTGGTACG | 200 |
| chr11:106762734-10676474 | p5 | (ATTGC)3 | TGCCCACTGTTGGTACTTGA | TCCAGAATGGATCTTCGGAC | 287 |
| chr11:106911343-10691336 | p2 | (TA)11 | ATGTCATATGCAAGCCGACA | TCTTGGAGAGCTCCGTTGTT | 265 |
| chr11:106968859-10697088 | p2 | (AT)15 | GACCCCACTGGACACAGTTT | CATGTGGTTCACTCGGTCAC | 282 |
| chr11:107031308-10703332 | p2 | (TA)7 | GCTACAATGGGACTCGGTGT | TGTGATGATTGGCATGGACT | 213 |
| chr11:107041148-10704317 | p2 | (AT)7 | GGACCTCGTATTGCCTGAAA | TGTTGATGTGGCGTGATTTT | 208 |
| chr11:107121271-10712328 | p2 | (AT)9 | GACACACCCTAGGACGGAAA | GCCATGTGGCTCACCTAAAT | 104 |
| chr11:107121271-10712328 | p3 | (CAT)5 | GACACACCCTAGGACGGAAA | GCCATGTGGCTCACCTAAAT | 104 |
| chr11:107194474-10719649 | p2 | (TA)11 | AATCTAGGCTCGGGAAGCTC | TGATCCTTGATGGCTTCACA | 202 |
| chr11:107194474-10719649 | p2 | (TG)10 | AATCTAGGCTCGGGAAGCTC | TGATCCTTGATGGCTTCACA | 202 |
| chr11:107194474-10719649 | p2 | (TA)16 | AATCTAGGCTCGGGAAGCTC | TGATCCTTGATGGCTTCACA | 202 |
| chr11:107210347-10721236 | p2 | (AC)7 | CGTTTTCGCCCAAAAGTCTA | GCTGGCCGAGATCTTTGTAG | 193 |
| chr11:107346753-10734876 | p2 | (AT)6 | ACGATTTCCTTTCTGCCCTT | TGTTCAGTGGTGCTCGGTAG | 179 |
| chr11:107346753-10734876 | p2 | (AT)6 | ACGATTTCCTTTCTGCCCTT | TGTTCAGTGGTGCTCGGTAG | 179 |
| chr11:107358060-10736007 | p5 | (TTTTC)3 | TCACGCTCACTCTTTTGTGG | CAGATATAGGGGACCGCGTA | 280 |
| chr11:107378294-10738030 | p5 | (ATTTC)3 | GCCCTCTTGGGAGTTTTTGT | TTCAAGATGCGCCATTCATA | 158 |
| chr11:107389833-10739185 | p6 | (ATTATG)3 | GGTCCCATGAGGATTTGAGA | GTGAAATCACCCCCATGAAC | 106 |
| chr11:107433329-10743537 | p2 | (TA)25 | TCCAAAGAAAGGGTGTTTCG | TAAACCTCCGTCAGTCCTGG | 213 |
| chr11:107573341-10757535 | p6 | (AATTAA)3 | TTCGTAAACCGTAGCCCATC | TGCCCGATTTTCGTTAAGTC | 257 |
| chr11:107583345-10758535 | p2 | (CA)6 | CTCCTCTTCATGGGCATTGT | TTGGGGTCCGAACACTTTAG | 177 |
| chr11:107597831-10759986 | p2 | (TA)16 | ATGTCCTGGAAAATAGCCCC | TGCATGGTCTGGATTGAAAA | 176 |
| chr11:107654975-10765698 | p2 | (AT)6 | TGAATAGTGTGCTCGGCTTG | AGCTCATTTTCTTCCGGTCA | 203 |
| chr11:107654975-10765698 | p2 | (AT)6 | TGAATAGTGTGCTCGGCTTG | AGCTCATTTTCTTCCGGTCA | 203 |
| chr11:107654975-10765698 | p2 | (TA)13 | TGAATAGTGTGCTCGGCTTG | AGCTCATTTTCTTCCGGTCA | 203 |
| chr11:107654975-10765698 | p2 | (AT)6 | TGAATAGTGTGCTCGGCTTG | AGCTCATTTTCTTCCGGTCA | 203 |
| chr11:107692417-10769443 | p2 | (AT)12 | ATCATAGCCACACAGGGCTC | GCTGAAGAACGACATCGACA | 105 |
| chr11:107692417-10769443 | p2 | (AT)7 | ATCATAGCCACACAGGGCTC | GCTGAAGAACGACATCGACA | 105 |
| chr11:107726875-10772890 | p2 | (AT)15 | TACAAGTGTGCTGACGAGGC | GAAGCACCACTACTCCGAGC | 273 |
| chr11:107754514-10775652 | p5 | (AAATG)3 | CTATCGCTCGCTTGTTCTCC | TTGTGGGTGTTGGAAGTTGA | 289 |
| chr11:107958298-10796031 | p2 | (TA)6 | TTGTTCTGATGATGTGGGGA | ACAGACATACAGATCCGCCC | 136 |
| chr11:108258374-10826039 | p4 | (AAGA)4 | AAGCTGAGTGAGGAAGCTCG | TAAAATGAAAGCGGCAGGTC | 229 |

| Marker ID | SSR type | SSR motif | FORWARD PRIMER (5'-3') | REVERSE PRIMER (5'-3') | size (bp) |
|---|---|---|---|---|---|
| chr11:108368795-10837080 | p2 | (GA)6 | CTAGCCAAGTGCTAGGACCG | TTGTATGGTTCAAATGCCGA | 237 |
| chr11:108460285-10846230 | p2 | (AT)11 | TTAATGGGTGTTGGAGCACA | ACCACCCTCACATGAACCAT | 152 |
| chr11:108460285-10846230 | p3 | (GAA)6 | TTAATGGGTGTTGGAGCACA | ACCACCCTCACATGAACCAT | 152 |
| chr11:108473280-10847530 | p2 | (AT)11 | TTAATGGGTGTTGGAGCACA | ACCACCCTCACATGAACCAT | 152 |
| chr11:108473280-10847530 | p3 | (GAA)6 | TTAATGGGTGTTGGAGCACA | ACCACCCTCACATGAACCAT | 152 |
| chr11:108675013-10867702 | p5 | (TGGTT)3 | AAGGTTGCGCAAAGAAAAGA | ACCAAATCAAACCAAACCGA | 159 |
| chr11:108709272-10871128 | p5 | (AAATA)3 | CGGGTGTGAAGTCCGTTACT | CAAGTGCCCTATTCTTCCCA | 146 |
| chr11:108732188-10873420 | p2 | (TG)6 | AGCACCACAAAGCAAAAACC | CGGAAAATGCAAAACTCCAT | 281 |
| chr11:108938656-10894069 | p3 | (AAT)14 | ACTCGCGTATGTTCGGATTC | TAATCGCTCAAGCCGTTCTT | 142 |
| chr11:108953276-10895530 | p3 | (AAT)9 | ACTCGCGTATGTTCGGATTC | GAGCATTTTGAATACCCCGA | 129 |
| chr11:108990353-10899236 | p5 | (TTTTA)3 | GGGGTTGGATAAAGGGTTGT | TCTCATATGCGACCAACCAA | 146 |
| chr11:109001396-10900341 | p5 | (ATTTT)3 | CGACGGACTCAAGACAATCA | CTAGACTCGGGGTCTCGATG | 267 |
| chr11:109065037-10906705 | p2 | (AT)7 | CCATTGAGACGAGGCGTATT | CGTGTCCAGTGGGGTTACTT | 166 |
| chr11:109081877-10908391 | p3 | (TAT)11 | ATGGCAATTGGATGCCTAAC | GGCTCTTGCTTGAGCTTTTG | 285 |
| chr11:109081877-10908391 | p2 | (GA)6 | ATGGCAATTGGATGCCTAAC | GGCTCTTGCTTGAGCTTTTG | 285 |
| chr11:109082060-10908407 | p3 | (TAT)11 | ATGGCAATTGGATGCCTAAC | GGCTCTTGCTTGAGCTTTTG | 102 |
| chr11:109082060-10908407 | p2 | (GA)6 | ATGGCAATTGGATGCCTAAC | GGCTCTTGCTTGAGCTTTTG | 102 |
| chr11:109087339-10908935 | p2 | (GT)6 | CAAACCCTAACCCATTGTGG | ACCCATCACACAACCCACTT | 209 |
| chr11:109087339-10908935 | p3 | (TTG)5 | CAAACCCTAACCCATTGTGG | ACCCATCACACAACCCACTT | 209 |
| chr11:109186707-10918872 | p3 | (TTG)6 | AGCCCAGCACACATATAGGG | CAATTTCTCAACCCGCAACT | 176 |
| chr11:109186758-10918882 | p3 | (TTG)6 | AGCCCAGCACACATATAGGG | CAATTTCTCAACCCGCAACT | 125 |
| chr11:109336404-10933843 | p2 | (AT)9 | CGATGGGCTATTTGCAGTTT | TTATTTTATGCCTGAGCGGG | 129 |
| chr11:109336404-10933843 | p2 | (AT)13 | CGATGGGCTATTTGCAGTTT | TTATTTTATGCCTGAGCGGG | 129 |
| chr11:109466154-10946816 | p5 | (CAATT)3 | CCCAGACAATAGGGGAGTCA | CCTATAGGGTCGGGTACGGT | 264 |
| chr11:109544116-10954612 | p2 | (CT)6 | GATACCTCCGAGGGACCATT | AAACGCATTCTCGACTTGCT | 122 |
| chr11:109575308-10957732 | p2 | (AT)6 | TCAACTTGTCACGACCCAAA | CCGTGTATATTGGCAGCCTT | 146 |
| chr11:109743506-10974552 | p4 | (CATT)4 | ACGCCCAACCAAAGTGTAAG | CATGACCCTTCTGGTCGAAT | 298 |
| chr11:109743608-10974562 | p4 | (CATT)4 | ACGCCCAACCAAAGTGTAAG | CATGACCCTTCTGGTCGAAT | 196 |
| chr11:109825152-10982716 | p2 | (TA)6 | TACCACATGCGTATCAACCC | AATGAAATGGCGGAAAAGC | 204 |
| chr11:109825152-10982716 | p2 | (GT)6 | TACCACATGCGTATCAACCC | AATGAAATGGCGGAAAAGC | 204 |
| chr11:109894910-10989692 | p3 | (AAC)5 | GGAAGGAATTGATTGGCAGA | TCTTTGGCTTGCCTGATTTT | 221 |
| chr11:109920253-10992227 | p2 | (AC)13 | AATTTCTTCGGCGCTAGTGA | AGAAGGGGAGCAGACCAAAT | 104 |
| chr11:110164145-11016615 | p2 | (TC)6 | TTTTCCCATTACTCCCCACA | TCTGTCCTCTCCCCCTTTTT | 200 |
| chr11:110201196-11020321 | p4 | (AGTT)4 | GCAGTACAGGTGTTGAGCCA | ATTCAGCTTCCGCTGACTGT | 265 |
| chr11:110209628-11021166 | p2 | (TA)19 | CTTAAGGCTGAGGTCGATGC | AGGACACGAGTTGTTCACCC | 238 |
| chr11:110224419-11022648 | p5 | (ATCAT)3 | GGAATTGAGATATCGGGGGT | AAGCCAGAATTGCACAGGAC | 159 |

| Marker ID | SSR type | SSR motif | FORWARD PRIMER (5'-3') | REVERSE PRIMER (5'-3') | size (bp) |
|---|---|---|---|---|---|
| chr11:110253562-11025557 | p2 | (AG)6 | CCTATTCCAAGAGCCAGCAC | ATCCCACATTTCTTTGGCAG | 158 |
| chr11:110253562-11025557 | p2 | (AG)6 | CCTATTCCAAGAGCCAGCAC | ATCCCACATTTCTTTGGCAG | 158 |
| chr11:110562416-11056443 | p5 | (ATGAT)3 | CTTATCCTCCCACCCCACTT | AAAACGTTGATGGCTGGAAG | 141 |
| chr11:110592488-11059450 | p2 | (AT)6 | CTTGCCTGTCTCCCTTTCTG | GCAGCCTTACCTTGCACTTC | 195 |
| chr11:110596806-11059882 | p3 | (GGC)5 | AAAAATTGATGCCCAGTTGC | TGGAGGAAGAAAATGGGTTG | 272 |
| chr11:110607071-11060908 | p5 | (CAATG)3 | TTGGAGCCCAACTATATCCG | TGGAGTTAAGGGTTCGCCTA | 158 |
| chr11:110719530-11072154 | p5 | (TTTAT)3 | CATATTCAGGGGGCAGGATA | TGATGTGGTCCTTGGAACCT | 282 |
| chr11:110843730-11084575 | p6 | (GGAAGG)4 | TGCAGCAAGGTAGCATCAAC | CGATCGTGACCCTCAAAAAT | 242 |
| chr11:110871114-11087312 | p2 | (TA)6 | AAACCAGGATGTGTCTTCCG | AAAATAATGCCACACAGGGC | 277 |
| chr11:110882770-11088479 | p2 | (GA)10 | CGGGGTTACCTTTTTGGATT | CCATGGATACCCATACTGCC | 300 |
| chr11:110886632-11088865 | p5 | (GGGGT)3 | TATCGGTTTCACCGCCTTAG | CCTTCCCCTTGGGGTAAATA | 174 |
| chr11:110886632-11088865 | p3 | (CAA)6 | TATCGGTTTCACCGCCTTAG | CCTTCCCCTTGGGGTAAATA | 174 |
| chr11:110927764-11092979 | p2 | (CT)13 | GACCGAGTCAACTCCCGATA | ATTTGGGTGATTTTGTGGGA | 141 |
| chr11:110970686-11097269 | p2 | (TA)6 | GAGAAAACGCGATAACGAGC | CGAGAGAATTTGGGCTAACG | 148 |
| chr11:110995905-11099794 | p2 | (AT)22 | AACGTTCTCAGCATCCCATC | GCCCTCTGGTTGGTTTATGA | 253 |
| chr11:111044331-11104636 | p2 | (AT)15 | TCCAAACAGGGATGGATGAT | TGCGATCTGAAAATCTCACG | 236 |
| chr11:111086422-11108846 | p3 | (TAA)15 | TTTCCGGATTTGTTCTCGTC | AAAGGGTATTCGGCAACCTT | 144 |
| chr11:111159361-11116137 | p3 | (TTG)6 | GTGGAAGATTCCAGGACCAA | GCCGAACTTGTAACGTCCAT | 274 |
| chr11:111159361-11116137 | p2 | (TA)14 | GTGGAAGATTCCAGGACCAA | GCCGAACTTGTAACGTCCAT | 274 |
| chr11:111227060-11122907 | p2 | (TC)13 | GCCATTATTGAGTCTTCCGC | GGAAGAGTGGGAGTGTGGAG | 165 |
| chr11:111227060-11122907 | p2 | (AT)7 | GCCATTATTGAGTCTTCCGC | GGAAGAGTGGGAGTGTGGAG | 165 |
| chr11:111334821-11133683 | p5 | (ATTCA)3 | ATTCCGCAACATAACTGGGA | CATTTCCACCACTTTGCCTT | 299 |
| chr11:111370048-11137206 | p2 | (TA)8 | TTTCCTTCTTCGTAATGGCG | TTTAGCCCAATCCTTCATCG | 167 |
| chr11:111370048-11137206 | p2 | (TA)26 | TTTCCTTCTTCGTAATGGCG | TTTAGCCCAATCCTTCATCG | 167 |
| chr11:111449987-11145201 | p2 | (AT)6 | ATGAAGCAGATCATCCCCAG | TAAAGAAGATGCCCAATGGC | 137 |
| chr11:111449987-11145201 | p3 | (CAA)8 | ATGAAGCAGATCATCCCCAG | TAAAGAAGATGCCCAATGGC | 137 |
| chr11:111620282-11162233 | p2 | (AT)25 | GCTGAAGAGGTCGTTTGGAG | CTGGTCCACCCTTAATCGAA | 244 |
| chr11:111633796-11163581 | p5 | (TGCAA)3 | CAGTTCATGCAAACGCTCAT | ATCGCAGTGGGAATGATAGG | 175 |
| chr11:111633796-11163581 | p2 | (AT)6 | CAGTTCATGCAAACGCTCAT | ATCGCAGTGGGAATGATAGG | 175 |
| chr11:111654109-11165612 | p4 | (ATCT)4 | TGTGGCATTTATTGGAGCAA | AAACGAGTACACACCCTCGC | 176 |
| chr11:111654109-11165612 | p2 | (AT)8 | TGTGGCATTTATTGGAGCAA | AAACGAGTACACACCCTCGC | 176 |
| chr11:111692629-11169467 | p2 | (AT)21 | GGGGTCCAAACAACTTTCCT | GCCGTAAGAGAGCAAACCAG | 278 |
| chr11:111692629-11169467 | p3 | (CAT)6 | GGGGTCCAAACAACTTTCCT | GCCGTAAGAGAGCAAACCAG | 278 |
| chr11:111693279-11169529 | p2 | (AT)21 | TCCCTAGTGACCAAACACCC | TGTACAGCAAATGTAGCGGC | 231 |
| chr11:111693279-11169529 | p3 | (CAT)6 | TCCCTAGTGACCAAACACCC | TGTACAGCAAATGTAGCGGC | 231 |
| chr11:111747826-11174984 | p3 | (TAA)5 | GTGGGTGTGTGTCAGAGGTG | CCTTTACCCCCAAACCCTAA | 133 |

| Marker ID | SSR type | SSR motif | FORWARD PRIMER (5'-3') | REVERSE PRIMER (5'-3') | size (bp) |
|---|---|---|---|---|---|
| chr11:111771964-11177398 | p2 | (AT)11 | GCCAGACCTTTATGCCATGT | CACACAAGATCCAAGGGGTT | 155 |
| chr11:111780417-11178243 | p2 | (AT)10 | AAAGTAGCGCGATGGAAAGA | CTAGGCAAAATGGGTCGTGT | 187 |
| chr11:111780417-11178243 | p5 | (AATGC)3 | AAAGTAGCGCGATGGAAAGA | CTAGGCAAAATGGGTCGTGT | 187 |
| chr11:111794413-11179642 | p5 | (TATGA)3 | TTGGAGACGTGTATGAAGCG | ATAGAGTTGTCCCCGAGGCT | 146 |
| chr11:111824227-11182623 | p2 | (TA)6 | AATGCAACCTCTTCAATGGG | TCCCTAGTCACCCCTGTTTG | 300 |
| chr11:111961944-11196395 | p3 | (GAA)5 | CAAGGTGATGTACTGGGCCT | TGTCCCGGTGGTTAATGAAT | 154 |
| chr11:111970320-11197233 | p3 | (GAA)5 | CAAGGTGATGTACTGGGCCT | TGTCCCGGTGGTTAATGAAT | 154 |
| chr11:111978697-11198071 | p3 | (GAA)5 | CAAGGTGATGTACTGGGCCT | TGTCCCGGTGGTTAATGAAT | 154 |
| chr11:112037537-11203955 | p6 | (ATACAT)3 | CCGAAACTAGCTCACCAAGC | TCTCAGCCTTGTCGTCTCCT | 150 |
| chr11:112037537-11203955 | p3 | (CAT)5 | CCGAAACTAGCTCACCAAGC | TCTCAGCCTTGTCGTCTCCT | 150 |
| chr11:112121186-11212320 | p6 | (ATATAC)3 | TGACTATTCAGGGGTACGGC | CATACAGAAGCCGCTAAGGC | 127 |
| chr11:112121186-11212320 | p5 | (AAAGG)3 | TGACTATTCAGGGGTACGGC | CATACAGAAGCCGCTAAGGC | 127 |
| chr11:112132363-11213439 | p2 | (TA)14 | TCGCCTGGAAAAGAGAATGT | CGATTAGAGCTCACGAAGCC | 275 |
| chr11:112290196-11229221 | p5 | (CATTG)3 | CACCCCATTGTGTTGTGAAC | CAGTACCACACAGGGCACAC | 168 |
| chr11:112357133-11235914 | p2 | (AT)20 | ATCTTCCCACATGGAAATCG | ATTGCTTCAGAGGATGCGTT | 178 |
| chr11:112357133-11235914 | p2 | (TG)8 | ATCTTCCCACATGGAAATCG | ATTGCTTCAGAGGATGCGTT | 178 |
| chr11:112485034-11248706 | p2 | (AG)17 | CCCCACAGCCTATCGATTTA | TTCTGACTCCGCCTCTGAAT | 279 |
| chr11:112485034-11248706 | p6 | (CCAAAT)3 | CCCCACAGCCTATCGATTTA | TTCTGACTCCGCCTCTGAAT | 279 |
| chr11:112577642-11257966 | p6 | (TATATG)3 | AAAGGTTGTCGAAACAACGG | CCTCCAACGTTCGTGATTCT | 282 |
| chr11:112634101-11263611 | p5 | (TGTAG)3 | AAACGGGAAAGCCTTCCTAA | ATCCGAAGCACTCTTCTCCA | 168 |
| chr11:112677706-11267977 | p2 | (AT)32 | CAACCACTAACCCCAGTGCT | GCGAGTGGTACGGTTATGCT | 130 |
| chr11:112953582-11295559 | p3 | (TTG)5 | TTGCGGAAATACCAAGGAAG | ATTTTCCAAGGCCCACTTCT | 207 |
| chr11:113053518-11305553 | p2 | (GA)7 | TCGACTGCAAACCCAATACA | AAGGCCTATGCCCTGATTTT | 248 |
| chr11:113077966-11307998 | p2 | (AG)8 | TCCAGGGTGAGCTTCTTGAT | CTCGTATTCACCCTCTCCGA | 265 |
| chr11:113092143-11309415 | p5 | (ACCAC)3 | AGAAAGGTCGCTTTTTGGGT | TGATAGAAGGCTTCCTGCGT | 126 |
| chr11:113112990-11311500 | p6 | (CATATA)3 | CATGTTGTCACGACCCAAAC | TCACGCGGTTATTCTCAGTG | 138 |
| chr11:113112990-11311500 | p3 | (CAA)6 | CATGTTGTCACGACCCAAAC | TCACGCGGTTATTCTCAGTG | 138 |
| chr11:113152739-11315475 | p3 | (TTG)6 | GAAAACAGCGTGTGGGCTAT | GCAGCATCTCCCCTGTGTAT | 260 |
| chr11:113152739-11315475 | p4 | (TGTA)4 | GAAAACAGCGTGTGGGCTAT | GCAGCATCTCCCCTGTGTAT | 260 |
| chr11:113152739-11315475 | p2 | (TA)9 | GAAAACAGCGTGTGGGCTAT | GCAGCATCTCCCCTGTGTAT | 260 |
| chr11:113152739-11315475 | p2 | (TA)9 | GAAAACAGCGTGTGGGCTAT | GCAGCATCTCCCCTGTGTAT | 260 |
| chr11:113153470-11315548 | p4 | (TGTA)4 | TATTTATGCATTGCGTCCCA | CGGCCATATAGGCTCAGTGT | 283 |
| chr11:113153470-11315548 | p2 | (TA)9 | TATTTATGCATTGCGTCCCA | CGGCCATATAGGCTCAGTGT | 283 |
| chr11:113153470-11315548 | p2 | (TA)9 | TATTTATGCATTGCGTCCCA | CGGCCATATAGGCTCAGTGT | 283 |
| chr11:113169298-11317131 | p5 | (TTATT)3 | CACCCACAAGCTCTCCAAAT | AGAGCCACTTCACGGAGAAA | 126 |
| chr11:113202508-11320457 | p2 | (AT)33 | TTTGGCGAGAGTTCTCGAAT | CTCTGTCCTCCATCTCAGCC | 144 |

| Marker ID | SSR type | SSR motif | FORWARD PRIMER (5'-3') | REVERSE PRIMER (5'-3') | size (bp) |
|---|---|---|---|---|---|
| chr11:113277035-11327905 | p3 | (TGT)6 | GGCATGCATGATTTGTGTTC | CGCTGGATCATCTTCTGACA | 280 |
| chr11:113497125-11349913 | p2 | (AT)6 | ACTCATGAAATCACGGGCTC | AGAAACAGAGCCGCCAACTA | 186 |
| chr11:113595429-11359744 | p5 | (TAATT)3 | CGAAAGGGCCATTGTATGTT | GCAATTCGTCTGGATCTGGT | 207 |
| chr11:113601754-11360376 | p3 | (CAT)5 | TCATAAGGACGGGACAGGAC | TGTTCATGCCCACAGGTAGA | 256 |
| chr11:113634670-11363668 | p2 | (AT)6 | TTGCTCCACGCCCTACTACT | CGAAGCCACGGTTGATACTT | 272 |
| chr11:113634766-11363677 | p2 | (AT)6 | TTGCTCCACGCCCTACTACT | CGAAGCCACGGTTGATACTT | 176 |
| chr11:113635052-11363706 | p2 | (AT)6 | AAAACTCAGAAGCGGAGCAA | TTTCGTTAATTTCCCCCACA | 265 |
| chr11:113666395-11366843 | p2 | (AT)18 | CGACCACTACTGACCAAGCA | GCCTGCTCTTATTTCGTTGC | 279 |
| chr11:113666395-11366843 | p3 | (ACA)7 | CGACCACTACTGACCAAGCA | GCCTGCTCTTATTTCGTTGC | 279 |
| chr11:113770286-11377229 | p2 | (AT)6 | AACTCCACCGATTCAACGTC | CAGCACCTCCATCAAAGGTT | 180 |
| chr11:113804352-11380637 | p6 | (CGTCCT)3 | GTTTTCGGCTAACTTTCCCC | TTGTTTGCTTGGATTTTCCC | 150 |
| chr11:113895235-11389726 | p3 | (TTC)5 | TATATTGCGGTTTCAAGGGG | TAGACGATATTCTTGGCGCAT | 227 |
| chr11:113927141-11392916 | p4 | (TTAT)7 | TTTAAGGCATTTTGGTTCCG | GGGGTAAATCAGGCCCTAAA | 256 |
| chr11:113927141-11392916 | p4 | (AATA)5 | TTTAAGGCATTTTGGTTCCG | GGGGTAAATCAGGCCCTAAA | 256 |
| chr11:113964611-11396662 | p2 | (TA)7 | GTGTGACGGAGATCGGAGAT | CAAAAGCATAATCCGTCGGT | 145 |
| chr11:114000936-11400296 | p2 | (TA)16 | CACCTGGTTCACCACAACAG | CTCCATATTCCCTGTGCGAT | 278 |
| chr11:114108662-11411067 | p2 | (AT)26 | TACGGATCTTATCGGGCAAG | CTCTTATCCTTGTCGGCTCG | 282 |
| chr11:114108662-11411067 | p2 | (AT)6 | TACGGATCTTATCGGGCAAG | CTCTTATCCTTGTCGGCTCG | 282 |
| chr11:114161850-11416386 | p2 | (CA)7 | GGAAAGGGGAAGCAGGATAG | GGAGAGTGAAGGTGAGCCTG | 138 |
| chr11:114203179-11420524 | p2 | (TA)34 | CGTATAAGTCCATCGGGCAT | TCTTGGATTGGGGTGGAATA | 157 |
| chr11:114240514-11424254 | p2 | (TA)6 | TGTTGGCAATTGGATGTGTT | CCTCAAATCGTCCAAGCAAT | 299 |
| chr11:114240546-11424257 | p2 | (TA)6 | TGTTGGCAATTGGATGTGTT | CCTCAAATCGTCCAAGCAAT | 267 |
| chr11:114240578-11424259 | p2 | (TA)6 | TGTTGGCAATTGGATGTGTT | CCTCAAATCGTCCAAGCAAT | 235 |
| chr11:114240595-11424260 | p2 | (TA)6 | TGTTGGCAATTGGATGTGTT | CCTCAAATCGTCCAAGCAAT | 218 |
| chr11:114240608-11424262 | p2 | (TA)6 | TGTTGGCAATTGGATGTGTT | CCTCAAATCGTCCAAGCAAT | 205 |
| chr11:114240629-11424265 | p2 | (TA)6 | TGTTGGCAATTGGATGTGTT | CCTCAAATCGTCCAAGCAAT | 184 |
| chr11:114240660-11424267 | p2 | (TA)6 | TGTTGGCAATTGGATGTGTT | CCTCAAATCGTCCAAGCAAT | 153 |
| chr11:114240677-11424268 | p2 | (TA)6 | TGTTGGCAATTGGATGTGTT | CCTCAAATCGTCCAAGCAAT | 136 |
| chr11:114240690-11424271 | p2 | (TA)6 | TGTTGGCAATTGGATGTGTT | CCTCAAATCGTCCAAGCAAT | 123 |
| chr11:114240711-11424274 | p2 | (TA)6 | TGTTGGCAATTGGATGTGTT | CCTCAAATCGTCCAAGCAAT | 102 |
| chr11:114240841-11424285 | p2 | (TA)6 | GTTGGATTGCTTGGACGATT | TGCATCCTGAATCTGTTGGA | 136 |
| chr11:114240854-11424287 | p2 | (TA)6 | GTTGGATTGCTTGGACGATT | TGCATCCTGAATCTGTTGGA | 123 |
| chr11:114240875-11424290 | p2 | (TA)6 | GTTGGATTGCTTGGACGATT | TGCATCCTGAATCTGTTGGA | 102 |
| chr11:114282906-11428492 | p2 | (TA)8 | CAAGAAATGGCCCTGGACTA | GATACCGGTTTTCAGGAGCA | 258 |
| chr11:114300206-11430221 | p2 | (TG)6 | GCCAAAGGCAAAACAATGAT | ACAACACCAAAACAGCTCCC | 290 |
| chr11:114374593-11437665 | p2 | (AT)33 | ATTTACCGGGAAAACGGTTC | TTCTCGATTTCACCCGATTC | 114 |

| Marker ID | SSR type | SSR motif | FORWARD PRIMER (5'-3') | REVERSE PRIMER (5'-3') | size (bp) |
|---|---|---|---|---|---|
| chr11:114426755-11442876 | p2 | (AT)7 | CTTCTTCCAGTTGGCCCATA | CAAGCATGATTTGGAGCTGA | 204 |
| chr11:114459823-11446184 | p3 | (TGT)5 | GAAGTAGACAGCCGCAAAGG | TGTGAATGTGTGGTGTGGTG | 124 |
| chr11:114459823-11446184 | p2 | (GT)9 | GAAGTAGACAGCCGCAAAGG | TGTGAATGTGTGGTGTGGTG | 124 |
| chr11:114461604-11446361 | p2 | (GT)7 | TTTATTACCGCTTTGCGCTT | ACATCGTGAAGGAATGACCC | 141 |
| chr11:114461604-11446361 | p2 | (GT)7 | TTTATTACCGCTTTGCGCTT | ACATCGTGAAGGAATGACCC | 141 |
| chr11:114461604-11446361 | p2 | (GT)7 | TTTATTACCGCTTTGCGCTT | ACATCGTGAAGGAATGACCC | 141 |
| chr11:114461604-11446361 | p2 | (GT)7 | TTTATTACCGCTTTGCGCTT | ACATCGTGAAGGAATGACCC | 141 |
| chr11:114461604-11446361 | p2 | (TG)9 | TTTATTACCGCTTTGCGCTT | ACATCGTGAAGGAATGACCC | 141 |
| chr11:114493145-11449516 | p2 | (AT)11 | GGGGAACCTGCACAAGTTTA | TGCTTCGGGGAATTGTTAAG | 199 |
| chr11:114493145-11449516 | p3 | (ATA)5 | GGGGAACCTGCACAAGTTTA | TGCTTCGGGGAATTGTTAAG | 199 |
| chr11:114493368-11449538 | p2 | (AT)11 | CTTAACAATTCCCCGAAGCA | AAGGGAATGTTGTTCGCATC | 203 |
| chr11:114493368-11449538 | p3 | (ATA)5 | CTTAACAATTCCCCGAAGCA | AAGGGAATGTTGTTCGCATC | 203 |
| chr11:114589695-11459171 | p4 | (TAAA)4 | CGGCTAAATCGATACGGAAA | TCGAATGAGCCCATCTCTCT | 118 |
| chr11:114639250-11464126 | p2 | (GT)7 | TTGACGTCCCCTTTTATTGC | AAGTGTGATAATGCCCCGAG | 151 |
| chr11:114745659-11474767 | p5 | (CAATG)3 | TCTTCCAAGTTCCCACAACC | CACCCTCTTTGGGGTGTAGA | 290 |
| chr11:114771499-11477351 | p2 | (TA)7 | TCAGAGTTCGGGATGGATTC | CCGAACAACTAACCTCCCAA | 231 |
| chr11:114864058-11486607 | p2 | (TA)9 | TGGAGGATGGGGATAAGTGA | CACCCCTAGCCACAAGTCAT | 170 |
| chr11:114864058-11486607 | p2 | (TA)11 | TGGAGGATGGGGATAAGTGA | CACCCCTAGCCACAAGTCAT | 170 |
| chr11:114989948-11499197 | p2 | (AT)11 | TGAGATTGAGCCGCACTATG | AACAAGAAGCTGCGGACATT | 141 |
| chr11:115021919-11502394 | p2 | (AT)11 | TGAGATTGAGCCGCACTATG | AACAAGAAGCTGCGGACATT | 141 |
| chr11:115086148-11508819 | p3 | (TAT)16 | CCTGGCCCATTTCAAGATTA | ACTTCAACCTAGCAGGCCAA | 262 |
| chr11:115226200-11522821 | p3 | (TCA)6 | GGGGGTTGTGTTTCATCATC | GTTAGGGGTGCGCTTAATGA | 125 |
| chr11:115226200-11522821 | p6 | (CCGGTT)3 | GGGGGTTGTGTTTCATCATC | GTTAGGGGTGCGCTTAATGA | 125 |
| chr11:115264736-11526675 | p5 | (GATTT)3 | ACAATTTCCCAGTCACCGAA | TTTTCGGTTGCAAGATACCC | 149 |
| chr11:115622333-11562435 | p5 | (TGATA)4 | TGGAAGAAACGGGATTTGAG | TGGACAGAATATTCCCCGAG | 209 |
| chr11:115778324-11578038 | p2 | (AT)31 | CAAAGCCTACGACGTGTGAA | TCCATGATCAGCAAATGAACA | 209 |
| chr11:115778324-11578038 | p2 | (TA)7 | CAAAGCCTACGACGTGTGAA | TCCATGATCAGCAAATGAACA | 209 |
| chr11:115796146-11579816 | p5 | (AACCG)3 | GAAGGTGCCAACCATTCAGT | TCTTGGGACAATAACGCTCC | 176 |
| chr11:115903761-11590577 | p3 | (AAG)5 | GCCTCGAGCCAAATCTACAA | ATAACAAGAGCACGGTCGGT | 234 |
| chr11:115903761-11590577 | p3 | (CAA)5 | GCCTCGAGCCAAATCTACAA | ATAACAAGAGCACGGTCGGT | 234 |
| chr11:115943856-11594591 | p3 | (GAA)5 | AGGCCAATTTTTCTCCTGCT | AATCCTCCTAAATGGGTGGG | 129 |
| chr11:115946722-11594875 | p3 | (AAC)6 | GGGAAGCCTAAGATCCCAAC | TTCCCGCTTCAATAATGCTC | 203 |
| chr11:115946722-11594875 | p3 | (GTT)5 | GGGAAGCCTAAGATCCCAAC | TTCCCGCTTCAATAATGCTC | 203 |
| chr11:115977177-11597920 | p2 | (TA)13 | TAACTTGGCACTATCGCACG | TCCTAGGGATGTCCATGAGC | 222 |
| chr11:115983618-11598565 | p2 | (AT)20 | TCTGATCCATCTGTCCTCCC | TGACCTGTGTTTCCTTTCCC | 123 |
| chr11:116009995-11601204 | p2 | (TA)8 | CCCATTAGTGGGAGCAAATG | ATGAGTTTGGAGATGGTGGG | 292 |

| Marker ID | SSR type | SSR motif | FORWARD PRIMER (5'-3') | REVERSE PRIMER (5'-3') | size (bp) |
|---|---|---|---|---|---|
| chr11:116009995-11601204 | p2 | (TA)8 | CCCATTAGTGGGAGCAAATG | ATGAGTTTGGAGATGGTGGG | 292 |
| chr11:116009995-11601204 | p2 | (TA)27 | CCCATTAGTGGGAGCAAATG | ATGAGTTTGGAGATGGTGGG | 292 |
| chr11:116270414-11627242 | p2 | (TC)6 | ACTGTGGGAAAGTCGGACAC | TTTGGGAACCCACACAAGTT | 237 |
| chr11:116275717-11627774 | p4 | (TATT)6 | AGCTTGGGTCTGAGGAGTGA | ATTGGGCATTCAACATCCAT | 170 |
| chr11:116277325-11627934 | p2 | (AT)8 | AGAACCCTTTGCACCACATC | CGAGCGGTTCCATCATATTT | 175 |
| chr11:116504883-11650692 | p2 | (TA)20 | AATGCCGTCTTCTGACCATC | AGGATCCGTGTGGACTATCG | 260 |
| chr11:116530761-11653277 | p5 | (TTGCA)3 | ACATTGTTGGCATACCCCAT | ACATCGGCCTTTTCACAATC | 282 |
| chr11:116552480-11655452 | p2 | (AT)6 | CTTGAGAAGCCACCGCTAAC | TAGACATGTGTGGGTTGCGT | 190 |
| chr11:116552480-11655452 | p2 | (TA)22 | CTTGAGAAGCCACCGCTAAC | TAGACATGTGTGGGTTGCGT | 190 |
| chr11:116552480-11655452 | p3 | (CAA)8 | CTTGAGAAGCCACCGCTAAC | TAGACATGTGTGGGTTGCGT | 190 |
| chr11:116643697-11664571 | p2 | (AT)9 | CCACCATCAAACTACACCCC | GGTGGTGCCTCAGTGTAGGT | 189 |
| chr11:116713394-11671541 | p2 | (AT)8 | TTTTGGCTATCACCCGAGAC | ACTCAGGTGCATTTTGGGAC | 145 |
| chr11:116758781-11676079 | p5 | (AAAGG)3 | GTGTTATTGTTGTCCGGCCT | CCAACTTCATACCCGAAGGA | 288 |
| chr11:116758781-11676079 | p5 | (ATTGC)3 | GTGTTATTGTTGTCCGGCCT | CCAACTTCATACCCGAAGGA | 288 |
| chr11:116789358-11679140 | p2 | (AT)8 | ATGTCAGGGTGCCCAAGTAG | CCGAGGCGCATGTAAGTATT | 290 |
| chr11:116793958-11679597 | p6 | (CCCCCA)3 | TTGTTGGGGTCTTTTGGTTC | GGGTAACACCCCTACACCCT | 186 |
| chr11:116809858-11681187 | p5 | (TTGCA)3 | AAGGTCCGAAGGGAGATGAT | TATGAACGAATCGTGACCCA | 185 |
| chr11:116809858-11681187 | p2 | (TA)8 | AAGGTCCGAAGGGAGATGAT | TATGAACGAATCGTGACCCA | 185 |
| chr11:116809894-11681192 | p5 | (TTGCA)3 | AAGGTCCGAAGGGAGATGAT | TATGAACGAATCGTGACCCA | 149 |
| chr11:116809894-11681192 | p2 | (TA)8 | AAGGTCCGAAGGGAGATGAT | TATGAACGAATCGTGACCCA | 149 |
| chr11:116809920-11681195 | p5 | (TTGCA)3 | AAGGTCCGAAGGGAGATGAT | TATGAACGAATCGTGACCCA | 123 |
| chr11:116809920-11681195 | p2 | (TA)8 | AAGGTCCGAAGGGAGATGAT | TATGAACGAATCGTGACCCA | 123 |
| chr11:116892761-11689478 | p2 | (AT)6 | GCGTTTTCACCATTTTGGTT | CAACTTTCCAAAGGGTCCAA | 254 |
| chr11:116892761-11689478 | p2 | (AT)13 | GCGTTTTCACCATTTTGGTT | CAACTTTCCAAAGGGTCCAA | 254 |
| chr11:116925865-11692788 | p2 | (AT)6 | TTGCTTCGTCGTCAGTCATC | GCTCCCCACTAATGGATTGA | 182 |
| chr11:116930593-11693260 | p2 | (TA)6 | ATACTTAGGCCATTTGGGGC | CAACGGGGCCTATAACTGAA | 215 |
| chr11:116975698-11697772 | p5 | (TGTGT)3 | GAGGGTCCGACATCTGAAAA | AAGGCCGAAGTAAAAGCCAT | 198 |
| chr11:117002697-11700470 | p2 | (TA)6 | ATACTTAGGCCATTTGGGGC | CAACGGGGCCTATAACTGAA | 215 |
| chr11:117047802-11704982 | p5 | (TGTGT)3 | GAGGGTCCGACATCTGAAAA | AAGGCCGAAGTAAAAGCCAT | 198 |
| chr11:117094571-11709658 | p2 | (AT)6 | GGTGTCTTAATCCCTTGGCA | CCCCAGTAATGCAGCAATTT | 103 |
| chr11:117141149-11714316 | p2 | (TA)6 | CCTCAGCCGATATGGTCATT | ACCCATAAAGGGTGGACCTC | 229 |
| chr11:117141579-11714359 | p2 | (TA)6 | GAATACAAAGCCACGCACCT | GTTTTAGACCCCATGCTTCG | 199 |
| chr11:117141645-11714365 | p2 | (TA)6 | GAATACAAAGCCACGCACCT | GTTTTAGACCCCATGCTTCG | 133 |
| chr11:117141658-11714368 | p2 | (TA)6 | GAATACAAAGCCACGCACCT | GTTTTAGACCCCATGCTTCG | 120 |
| chr11:117224477-11722649 | p6 | (TCATGC)3 | GATACGCTCAAAAGCGGAAG | TGGATTAACAGCTGGGGTTC | 170 |
| chr11:117284767-11728678 | p3 | (AGA)5 | GCTAAACCTCCCCAAACACA | TGAACATGAAGCCCATACGA | 237 |

| Marker ID | SSR type | SSR motif | FORWARD PRIMER (5'-3') | REVERSE PRIMER (5'-3') | size (bp) |
|---|---|---|---|---|---|
| chr11:117311862-11731389 | p4 | (AATT)4 | TGGATGACGTTACCACGAAA | ACCCCACCTGATCTCCTTTT | 245 |
| chr11:117311862-11731389 | p2 | (AT)14 | TGGATGACGTTACCACGAAA | ACCCCACCTGATCTCCTTTT | 245 |
| chr11:117389201-11739121 | p5 | (AATTA)3 | ACATCAAATTCGCCCAAAAC | GGCATCAAGAGCCTGCTTAC | 262 |
| chr11:117471117-11747320 | p2 | (AT)44 | TCCGGGGGTAGGTTTATTTC | CTCTGCTGGAGCCATTTCAT | 177 |
| chr11:117540857-11754287 | p2 | (TA)7 | GCTCGTGGGGTTTAATTTGA | GAAAAATCGGGGCATATTCA | 191 |
| chr11:117605618-11760763 | p2 | (GT)6 | TGATTGTGCGAAAAATTGGA | ACTTCGGGGGAGTTTCTGTT | 111 |
| chr11:117605618-11760763 | p3 | (TTG)5 | TGATTGTGCGAAAAATTGGA | ACTTCGGGGGAGTTTCTGTT | 111 |
| chr11:117680471-11768254 | p4 | (AATA)7 | CTTCCTCATGGGGTCCTACA | AACTGAGCAATGAACGGGAC | 138 |
| chr11:117680471-11768254 | p5 | (TGCAA)4 | CTTCCTCATGGGGTCCTACA | AACTGAGCAATGAACGGGAC | 138 |
| chr11:117823402-11782541 | p2 | (AT)8 | CGCTTTAGCTCTTTTCAGGC | AGTGAACGGAGGGAGTAGCA | 131 |
| chr11:117959805-11796181 | p3 | (ATT)5 | CCCGAATAGGCATAGGTTCA | TTGCCCTTCGCTTAAAAAGA | 197 |
| chr11:117959805-11796181 | p2 | (AT)7 | CCCGAATAGGCATAGGTTCA | TTGCCCTTCGCTTAAAAAGA | 197 |
| chr11:117992938-11799495 | p5 | (AATGC)3 | ATGTCTCGGTAGCACAACCC | GCACGCACTCTCATTCTTCA | 134 |
| chr11:117992938-11799495 | p5 | (TTGCC)3 | ATGTCTCGGTAGCACAACCC | GCACGCACTCTCATTCTTCA | 134 |
| chr11:118002815-11800483 | p2 | (TA)12 | GATGCCGCCATAGATCAACT | TTCATCTCAGGCGACCTTTT | 257 |
| chr11:118098991-11810102 | p4 | (ATAG)8 | AAAAGCGAACCGTAACATGG | GATGTGAGGAGGAAATCCCA | 166 |
| chr11:118213884-11821589 | p3 | (GTT)5 | GAGGACAATGCATCTCACCA | GAATGGGCTCAAATGCAGAT | 144 |
| chr11:118213884-11821589 | p3 | (GTT)5 | GAGGACAATGCATCTCACCA | GAATGGGCTCAAATGCAGAT | 144 |
| chr11:118295148-11829718 | p2 | (TA)16 | GGACCTGATGGAAGACCTGA | CCACTCTCACCCATTCCCTA | 159 |
| chr11:118295148-11829718 | p3 | (ACA)5 | GGACCTGATGGAAGACCTGA | CCACTCTCACCCATTCCCTA | 159 |
| chr11:118299331-11830134 | p5 | (ATAAA)3 | TGGGTACATCATTAACGGCA | ACATGAAATTCGCCACCAAT | 228 |
| chr11:118301091-11830310 | p2 | (AT)9 | AAAAAGTGGAACCCACCACA | TTTGCTACGAATACGCATGG | 204 |
| chr11:118301091-11830310 | p2 | (AT)7 | AAAAAGTGGAACCCACCACA | TTTGCTACGAATACGCATGG | 204 |
| chr11:118543771-11854578 | p3 | (ACA)5 | TATTTGATTGGCCGGAACAT | GATCCAATTTCGTTGGGCTA | 219 |
| chr11:118575720-11857773 | p3 | (ATG)6 | GTGGGTGGATTAGAGGCAAA | CTCGGAATGTCAACCATGTG | 203 |
| chr11:118608156-11861021 | p2 | (CA)7 | AACCCTCAACTTAGGCCGAT | ATGTGTGTGTGTGTGCCCTT | 196 |
| chr11:118608156-11861021 | p2 | (AT)31 | AACCCTCAACTTAGGCCGAT | ATGTGTGTGTGTGTGCCCTT | 196 |
| chr11:118626636-11862865 | p5 | (CAACA)3 | TTAGGGGCCAGATTCAAATG | ATTTGTGCCCCAAACTCTTG | 220 |
| chr11:118645181-11864719 | p6 | (ATATAG)3 | AGCCTACCAAGGTGGGAAGT | TTTGTACCTGGGAGGTCTCG | 261 |
| chr11:118667174-11866918 | p3 | (TCT)5 | GGAGAGTCGGAAGCATTGAG | CGCGTCACACTTCACATCTT | 216 |
| chr11:118933202-11893522 | p6 | (TCCTTT)3 | CAATTGGGCGATAAGTTCGT | CAGGCAGACCACCCATACTT | 224 |
| chr11:118972495-11897451 | p2 | (TA)8 | GCAAGAGGCAAAACACCAAT | CCTTCGTCCAACCCTTATGA | 182 |
| chr11:118990607-11899264 | p2 | (GT)17 | AAAGGCCTTGCAAGAGTTGA | GGATATCCGCATCCCCTAAT | 283 |
| chr11:119249470-11925149 | p2 | (AT)10 | ATTTCGGAAAACAGCGATTG | TCTATTTCATCCCCAGGCAC | 229 |
| chr11:119380801-11938282 | p3 | (AAT)19 | ACTGTCTCTAGGAACGCCGA | GGCTTTCAAAGTTCTGTCGC | 180 |
| chr11:119380801-11938282 | p4 | (TCAT)5 | ACTGTCTCTAGGAACGCCGA | GGCTTTCAAAGTTCTGTCGC | 180 |

| Marker ID | SSR type | SSR motif | FORWARD PRIMER (5'-3') | REVERSE PRIMER (5'-3') | size (bp) |
|---|---|---|---|---|---|
| chr11:119477675-11947968 | p2 | (AT)6 | GCCCAGCATAGGAATATCCA | TGTTAGCCTAGGCGGCTTTA | 133 |
| chr11:119477675-11947968 | p3 | (ATC)5 | GCCCAGCATAGGAATATCCA | TGTTAGCCTAGGCGGCTTTA | 133 |
| chr11:119484149-11948616 | p3 | (AAT)19 | ACTGTCTCTAGGAACGCCGA | GGCTTTCAAAGTTCTGTCGC | 180 |
| chr11:119484149-11948616 | p4 | (TCAT)5 | ACTGTCTCTAGGAACGCCGA | GGCTTTCAAAGTTCTGTCGC | 180 |
| chr11:119599998-11960204 | p3 | (ATT)20 | AAGAGGAAAAGCGAGAATGG | TGCATTCGTACCTGCCAATA | 133 |
| chr11:119636818-11963891 | p2 | (AT)13 | CGAGCTGTAGAAGAGACGGG | TTCTTGTGGACACTGCGTTC | 210 |
| chr11:119636818-11963891 | p3 | (ATC)5 | CGAGCTGTAGAAGAGACGGG | TTCTTGTGGACACTGCGTTC | 210 |
| chr11:119662712-11966475 | p2 | (AT)20 | CATTCCGCCATATCGTTTCT | CATCAAACGGGAACCTCCTA | 232 |
| chr11:119668766-11967080 | p5 | (AAAGA)3 | AATGGGAAAGCAATGAATGG | CTTGCCACGTCAGCACTTAG | 196 |
| chr11:119670635-11967265 | p5 | (AATAA)3 | GGCTGCTCCATTGTTCCTAA | CAACAAATGGCGTGTGAAAG | 181 |
| chr11:119891346-11989336 | p2 | (TA)11 | TTGGTGCTTCTCTCCGACTT | TTTGAGGGCAGTCAATTTCC | 147 |
| chr11:119962719-11996473 | p2 | (TA)7 | GCACGGCAAACTTCGATTAT | AAATTTTCGTAGCAAACCCG | 295 |
| chr11:119962719-11996473 | p2 | (TA)6 | GCACGGCAAACTTCGATTAT | AAATTTTCGTAGCAAACCCG | 295 |
| chr11:120059284-12006131 | p2 | (AT)15 | GAAGAACGTGCTACACCCGT | TCTTGCACCATGCTAAGACG | 197 |
| chr11:120059284-12006131 | p2 | (AT)6 | GAAGAACGTGCTACACCCGT | TCTTGCACCATGCTAAGACG | 197 |
| chr11:120059522-12006153 | p2 | (AT)15 | GACGTCTTAGCATGGTGCAA | TCCGCTGCTAAGTTTTGGTT | 136 |
| chr11:120059522-12006153 | p2 | (AT)6 | GACGTCTTAGCATGGTGCAA | TCCGCTGCTAAGTTTTGGTT | 136 |
| chr11:120068316-12007032 | p2 | (TC)6 | CCCAGAACTTACGCTTTTGC | TTTTCATAGGCGTTTTTGGG | 236 |
| chr11:120152325-12015433 | p2 | (GT)6 | GAACCATTCCTGAGGCCATA | GTCCGGATGGTCGTTTAGAA | 251 |
| chr11:120152325-12015433 | p2 | (GT)6 | GAACCATTCCTGAGGCCATA | GTCCGGATGGTCGTTTAGAA | 251 |
| chr11:120197748-12019976 | p2 | (TA)7 | ACCGTAGTTGCTGTGGGAAC | GGGTTGTGCCATCTTTCACT | 138 |
| chr11:120197777-12019978 | p2 | (TA)7 | ACCGTAGTTGCTGTGGGAAC | AGGGTTGTGCCATCTTTCAC | 109 |
| chr11:120205810-12020782 | p5 | (ATTGA)3 | GGGAGTGACCATCTTCCAAA | TTTTTGCCCTTAGCCTCTCA | 272 |
| chr11:120209002-12021110 | p2 | (AT)52 | CAAACAAGCCACTCAAGCAA | GGGTTCATAATCCCGATGTG | 260 |
| chr11:120209002-12021110 | p2 | (TA)16 | CAAACAAGCCACTCAAGCAA | GGGTTCATAATCCCGATGTG | 260 |
| chr11:120228825-12023084 | p5 | (ATGAT)3 | GCTCTTTGAAAGGGAAAGGG | AAGGTGACAAGCCCAATCAC | 248 |
| chr11:120293951-12029596 | p5 | (TTAGT)3 | AGAGGAGGGAGGCAGCTAAG | TCAACGGCTTACAACCCTTC | 286 |
| chr11:120332791-12033480 | p3 | (AAG)5 | GCGTGCTTTCAACAAGCATA | GTGGCCACTCTTGACCATCT | 113 |
| chr11:120596097-12059811 | p2 | (AT)8 | GTTGTTGCCTCTGCACAAAA | AAGCCAAAAGTGATCATGGC | 241 |
| chr11:120612340-12061435 | p6 | (TTTCGT)3 | CTATCCCCACTTCTTTGCGA | CGAGGTCGACCCAGAAAATA | 281 |
| chr11:120622546-12062456 | p5 | (TTCTC)3 | TGGATTGTGACTGAAGCCCT | ATGAAAGATGCAACGGTGGT | 176 |
| chr11:120680958-12068297 | p2 | (TA)7 | ACGTATGTGGACCGAGAAGG | TATTGGTGGCCTTGTTAGCC | 272 |
| chr11:120774453-12077646 | p5 | (TTAGT)3 | TTTGTCCCCTAAACCTGCAC | GGGTCCTGTCACCACACTTT | 113 |
| chr11:120774453-12077646 | p5 | (TGTTG)3 | TTTGTCCCCTAAACCTGCAC | GGGTCCTGTCACCACACTTT | 113 |
| chr11:120860097-12086210 | p2 | (AT)6 | AGGCGCACCAGTCTGTCTAT | GCCTCGTTCAGAGTTTCAGG | 242 |
| chr11:120867282-12086929 | p2 | (TA)7 | ATGATGGATTCATTGCCACA | GACATGGCGGAATAGCAAAT | 120 |

| Marker ID | SSR type | SSR motif | FORWARD PRIMER (5'-3') | REVERSE PRIMER (5'-3') | size (bp) |
|---|---|---|---|---|---|
| chr11:120885302-12088731 | p2 | (AG)6 | AACCAAACGGAAGCAACAAC | ACCTCGACATCCCTGTTCAC | 187 |
| chr11:120974953-12097696 | p5 | (AAAAT)3 | AAGTCGGCATCCCCTCTTAT | ACTCTAACAGGCAGCCGAAA | 285 |
| chr11:121002024-12100403 | p5 | (ACCAA)4 | CCTCTCCCCTTTCAGCTCTT | TTGGTTTGGTTTGGTTTGGT | 136 |
| chr11:121002024-12100403 | p5 | (AACCA)3 | CCTCTCCCCTTTCAGCTCTT | TTGGTTTGGTTTGGTTTGGT | 136 |
| chr11:121020815-12102284 | p2 | (TA)13 | ATATGCCTGCACAATCCACA | CTGCAGTCACTGAGGCCATA | 162 |
| chr11:121066181-12106819 | p5 | (ACCAA)4 | CCTCTCCCCTTTCAGCTCTT | TTGGTTTGGTTTGGTTTGGT | 136 |
| chr11:121066181-12106819 | p5 | (AACCA)3 | CCTCTCCCCTTTCAGCTCTT | TTGGTTTGGTTTGGTTTGGT | 136 |
| chr11:121105546-12110757 | p2 | (TA)14 | TTTTCACATTCTCCAAGCCC | AGGAACGTGGTGTGAGGAAG | 109 |
| chr11:121129107-12113115 | p3 | (AAG)15 | ACAAACCCGGTGATTGTCAT | GCCCTGAAAAAGCCATCATA | 106 |
| chr11:121152658-12115467 | p3 | (CTT)5 | TCTTGCAAGTCCCACATTGA | GCCCAGAAATGCCAAAAATA | 207 |
| chr11:121152658-12115467 | p3 | (GAA)6 | TCTTGCAAGTCCCACATTGA | GCCCAGAAATGCCAAAAATA | 207 |
| chr11:121211368-12121342 | p2 | (TA)23 | TCCGGTCCCTAAGGCTACTT | TGAGACTGTCCCGAAAATCC | 133 |
| chr11:121325136-12132715 | p3 | (TAA)6 | GGGAGAGAGAAGGGAGGAGA | ACCCTAAAAACCATTTGGGC | 243 |
| chr11:121372379-12137441 | p2 | (TG)18 | ATTCGGGACTTGATGACGAC | GAAGATGCTCAAAAGACGGC | 100 |
| chr11:121550176-12155221 | p3 | (ATT)13 | CGACAATATCCATTCCCCAC | GATGCTTAGGTCCCGATGAA | 258 |
| chr11:121639894-12164191 | p6 | (TTTCTA)3 | TGTGCATAAGTTGGCACCTC | TTGACCTCAGCTGGCAAGTA | 279 |
| chr11:121659270-12166129 | p2 | (AC)10 | CGCATCAAGAACTGATTCCA | GCAACCTTTTGCGGTTACAT | 202 |
| chr11:121659270-12166129 | p2 | (TG)9 | CGCATCAAGAACTGATTCCA | GCAACCTTTTGCGGTTACAT | 202 |
| chr11:121659782-12166180 | p2 | (AC)10 | TTTCATCCCAGAATGCATCA | GGGTTAAGGGGGAGAGATTG | 202 |
| chr11:121659782-12166180 | p2 | (TG)9 | TTTCATCCCAGAATGCATCA | GGGTTAAGGGGGAGAGATTG | 202 |
| chr11:121790276-12179229 | p5 | (TTGCA)3 | GATGCCACTTAGGCGATGAT | CATCCCTTAGAGGCCAAACA | 135 |
| chr11:121790276-12179229 | p2 | (AG)9 | GATGCCACTTAGGCGATGAT | CATCCCTTAGAGGCCAAACA | 135 |
| chr11:121874364-12187637 | p2 | (AT)12 | CGTGACCTTACTTGACGGGT | TGGAGGTTGAGTTGGAGGAC | 141 |
| chr11:121874364-12187637 | p2 | (AC)6 | CGTGACCTTACTTGACGGGT | TGGAGGTTGAGTTGGAGGAC | 141 |
| chr11:121882178-12188419 | p6 | (ATCTAT)3 | TGTGATATTGTCCGCTCTGG | GGCTCTCGAAACCAAACAAG | 144 |
| chr11:121940030-12194204 | p2 | (TG)9 | ACAAATGACCACACGCTCTG | TCATGTTGAATGTTAGCGCC | 139 |
| chr11:121940030-12194204 | p5 | (AAATT)3 | ACAAATGACCACACGCTCTG | TCATGTTGAATGTTAGCGCC | 139 |
| chr11:121975552-12197757 | p6 | (TCTTTT)3 | TTCGACCCAGTAACCAAAGC | ACTTCTTCGAGTTGCCTCCA | 244 |
| chr11:122013615-12201563 | p3 | (GAT)5 | CCACAAGCCCATTGAAAGTT | TTGTGAAACGGTGTGTGGAT | 184 |
| chr11:122021248-12202326 | p2 | (TA)9 | TGGGTCTCTCAAGTCCAACC | AATTGCCCAATTGAAAACGA | 127 |
| chr11:122050196-12205221 | p2 | (AG)7 | AGGATTTCAAGGGGCTCAGT | GAGATGAATGTCCCGCTCAT | 208 |
| chr11:122050196-12205221 | p2 | (AT)9 | AGGATTTCAAGGGGCTCAGT | GAGATGAATGTCCCGCTCAT | 208 |
| chr11:122133170-12213520 | p2 | (AT)19 | GAGCCCGTAAGGTTCATTCA | TGTGGTGCGATAGATGAGGA | 220 |
| chr11:122151412-12215342 | p2 | (AT)8 | TTGGTGGTGTTGCTTTTGTC | AAAGTCGATGGCCAAACAAG | 142 |
| chr11:122166069-12216808 | p2 | (TA)6 | AACGAAGAAATTGAAGGGGG | TTGCAGAGAGTACGCAATGG | 221 |
| chr11:122166170-12216818 | p2 | (TA)6 | CCATTGCGTACTCTCTGCAA | TCAATAATGGGCTGCTCACA | 277 |

| Marker ID | SSR type | SSR motif | FORWARD PRIMER (5'-3') | REVERSE PRIMER (5'-3') | size (bp) |
|---|---|---|---|---|---|
| chr11:122166199-12216821 | p2 | (TA)6 | CCATTGCGTACTCTCTGCAA | TCAATAATGGGCTGCTCACA | 248 |
| chr11:122449487-12245150 | p2 | (TA)7 | CCTTTTTCGTCATCCGTTGT | TGTGAGATTTCACTGGGCTG | 194 |
| chr11:122449487-12245150 | p2 | (AT)7 | CCTTTTTCGTCATCCGTTGT | TGTGAGATTTCACTGGGCTG | 194 |
| chr11:122449737-12245175 | p2 | (TA)7 | ACATTGGCTGTTTCCGAGAG | CGTGAGTTTTGGATGTGGTG | 140 |
| chr11:122449737-12245175 | p2 | (AT)7 | ACATTGGCTGTTTCCGAGAG | CGTGAGTTTTGGATGTGGTG | 140 |
| chr11:122590091-12259210 | p3 | (TGT)6 | GGTTGTGAGACCGACCTTGT | CTCGCTTCCCTCATCTTGTC | 270 |
| chr11:122590704-12259271 | p3 | (TGT)6 | TGCATTCTGCTTTGAACTGC | GAGGTCTGGGGAGGGTAAAG | 224 |
| chr11:122590744-12259275 | p3 | (TGT)6 | TGCATTCTGCTTTGAACTGC | GAGGTCTGGGGAGGGTAAAG | 184 |
| chr11:122630999-12263301 | p6 | (TGTAAG)3 | AACTCCGAGCTTATGAGGCA | CCAACCAAGGCTCAAGGTTA | 105 |
| chr11:122672362-12267442 | p3 | (CAA)8 | CACCTTTGGTAAGGCGTGAT | TTTTTGTTCAGGGCATGTCA | 136 |
| chr11:122765422-12276743 | p2 | (TA)7 | AGAAGCCAACAAATCCATGC | GTCGATTTAGAGTCGAGCCG | 217 |
| chr11:122885508-12288752 | p6 | (TGAAAC)3 | CCAAACATTTTGCCTTTGCT | CCCTAGGGAAGGTGTTAGGC | 112 |
| chr11:122885508-12288752 | p2 | (AT)7 | CCAAACATTTTGCCTTTGCT | CCCTAGGGAAGGTGTTAGGC | 112 |
| chr11:122926594-12292861 | p4 | (TTTC)4 | TGCATATCCGAAAATGGTGA | TGAATGAGAGGAGCTTCCGT | 105 |
| chr11:123052365-12305438 | p2 | (TC)8 | TTAGCCATGTGGTCTCCTCC | GTCAAAGGCTTGGCTAAACA | 190 |
| chr11:123084890-12308690 | p2 | (TG)7 | TCTCCTTCTCCTCCTCCACA | TCTCTTCATGGCAGCAACAC | 244 |
| chr11:123092835-12309484 | p2 | (TA)7 | ATTCCCGCCTCATTTCTTTT | CGGGAGAGAGAGACAAGTGG | 199 |
| chr11:123112975-12311499 | p3 | (CAA)6 | TCTTGTGTTTCGGGGCTACT | TTTGGCAAATGCAAAAATCA | 168 |
| chr11:123158867-12316088 | p4 | (TAAA)5 | CCACCAGCACCGTTCTTACT | GCCGAGTCAATTTCTTTCCA | 159 |
| chr11:123188764-12319078 | p2 | (TA)11 | GGATGAGGGTATTCGAGCAA | AAAAAGGTGTCGAATGTGCC | 187 |
| chr11:123213600-12321563 | p3 | (ATA)12 | TGGGCCCTTTGACTATCTTG | TGCTCGATCACTTTTCAACG | 192 |
| chr11:123240388-12324243 | p2 | (AT)14 | CTTAAGACGCGTTGCAAACA | TGGGGTGGTGAAGTTGTGTA | 200 |
| chr11:123271852-12327386 | p5 | (TTGGT)3 | ACCTATGGGAGGTAGGGGTG | TCCAACACCAAACCAAGTCA | 143 |
| chr11:123271892-12327390 | p5 | (TTGGT)3 | ACCTATGGGAGGTAGGGGTG | TCCAACACCAAACCAAGTCA | 103 |
| chr11:123278099-12328012 | p3 | (TTC)9 | CTATGTTCGGGTCTGGTGCT | CAATGTGCTTGCTTCTTCCA | 265 |
| chr11:123278099-12328012 | p6 | (TTTTGA)3 | CTATGTTCGGGTCTGGTGCT | CAATGTGCTTGCTTCTTCCA | 265 |
| chr11:123283524-12328553 | p2 | (TA)6 | GGTGTTCATGGGTCGATTTT | CCGACTACAACCAAACCGAT | 180 |
| chr11:123286544-12328855 | p2 | (AG)20 | GGTCAATTTTGGTGGCATTT | TCTTTGCCCTTCCTTCAAGA | 171 |
| chr11:123286544-12328855 | p2 | (AT)6 | GGTCAATTTTGGTGGCATTT | TCTTTGCCCTTCCTTCAAGA | 171 |
| chr11:123391998-12339404 | p2 | (TA)25 | TTGAAGGGCTAGGATGATGC | AAGGGCCAAAATTACCCTTG | 145 |
| chr11:123440605-12344264 | p2 | (TA)19 | TGGGCGAGCTAAATGTCTCT | TGTGACATGCCGATCAAAGT | 180 |
| chr11:123452031-12345404 | p5 | (CATAT)3 | TGTTCCCCCTGAGAATCAAC | TGAATGGGCTAAAATCGAGG | 200 |
| chr11:123452031-12345404 | p2 | (AC)16 | TGTTCCCCCTGAGAATCAAC | TGAATGGGCTAAAATCGAGG | 200 |
| chr11:123457815-12345983 | p2 | (AT)9 | CTGGCATCCCTGTTCAATTT | ATGGCATTTAGGTTGTTGCC | 160 |
| chr11:123457815-12345983 | p5 | (CATAT)3 | CTGGCATCCCTGTTCAATTT | ATGGCATTTAGGTTGTTGCC | 160 |
| chr11:123458022-12346012 | p2 | (AT)9 | AAGTTTGTCACACCCCAACC | AGATAATGTTGGGTACGGCG | 196 |

| Marker ID | SSR type | SSR motif | FORWARD PRIMER (5'-3') | REVERSE PRIMER (5'-3') | size (bp) |
|---|---|---|---|---|---|
| chr11:123458022-12346012 | p5 | (CATAT)3 | AAGTTTGTCACACCCCAACC | AGATAATGTTGGGTACGGCG | 196 |
| chr11:123558151-12356037 | p2 | (AT)114 | GAGTTGGTCTGGTCCGAAAA | CCCAGAACCTCAGGTATCCA | 183 |
| chr11:123558151-12356037 | p2 | (AT)9 | GAGTTGGTCTGGTCCGAAAA | CCCAGAACCTCAGGTATCCA | 183 |
| chr11:123558151-12356037 | p3 | (ATA)6 | GAGTTGGTCTGGTCCGAAAA | CCCAGAACCTCAGGTATCCA | 183 |
| chr11:123717683-12371969 | p2 | (AT)8 | GGTTAATTGGGTGATCGTGG | CAGCAACCTCCCACGTTATT | 174 |
| chr11:123735899-12373791 | p2 | (AT)7 | TCATGCATGGCTTGTCTTTC | GACCCGATATGCCACGATAG | 184 |
| chr11:123857398-12385942 | p2 | (TA)14 | CGATTTATGAAGTCAGCGCA | CAGGGCATGACATCCTTTCT | 206 |
| chr11:123971385-12397340 | p3 | (TTG)6 | TCCAACTATTCGTTGTGCCA | GGCTAACACACCATGCACAC | 287 |
| chr11:124004404-12400642 | p3 | (ACA)7 | AAACGCATGGTCATCAGACA | TTGTGCCACCGATTTGTTTA | 151 |
| chr11:124026899-12402891 | p2 | (TA)6 | TTAAAGCGGACGAGCCTAAA | ATTGCTTTCAAAATGACCGC | 111 |
| chr11:124026899-12402891 | p2 | (TA)6 | TTAAAGCGGACGAGCCTAAA | ATTGCTTTCAAAATGACCGC | 111 |
| chr11:124059429-12406144 | p2 | (TA)6 | TATTCCGGAGGTGCTAATCG | CCTCTCGGCCATCAAAATAA | 258 |
| chr11:124059429-12406144 | p5 | (AAAAT)3 | TATTCCGGAGGTGCTAATCG | CCTCTCGGCCATCAAAATAA | 258 |
| chr11:124224352-12422639 | p3 | (TCT)14 | GGATCCGTAACGGCTAATCA | GGGATTCGACTTCTATGCCA | 236 |
| chr11:124249866-12425188 | p2 | (AT)6 | GGTTGCTAACCGATCAGCTC | CAAGAGCCTCATCCACCAAT | 257 |
| chr11:124249866-12425188 | p2 | (AT)10 | GGTTGCTAACCGATCAGCTC | CAAGAGCCTCATCCACCAAT | 257 |
| chr11:124249866-12425188 | p2 | (AC)6 | GGTTGCTAACCGATCAGCTC | CAAGAGCCTCATCCACCAAT | 257 |
| chr11:124314684-12431669 | p5 | (ATAAA)3 | CCCAACCTGCTCAAAACAAT | TGATTCCAAGTTAGGTGCCC | 257 |
| chr11:124353300-12435533 | p3 | (CCA)5 | GCTTGGATGAAATGCCCTTA | TGGCCAACAATAACAACCAA | 281 |
| chr11:124394062-12439607 | p5 | (ATTTT)3 | AGGCTTTTGGCCTTGAAAAT | ATTTTTAGGATCCGCTTGGC | 276 |
| chr11:124411024-12441308 | p2 | (AT)30 | CGGGTTATTGACTCGCCTAA | TTCTCCACCGAACCTTTGAC | 215 |
| chr11:124415828-12441784 | p2 | (AT)6 | GGAGTACCCCTAGACATGCG | TCTCAGCGTGTGTATTTCCG | 123 |
| chr11:124668122-12467014 | p2 | (TA)10 | ACTACCAGGATCATTTGCGG | GCAGATACCGGTTTTCAGGA | 209 |
| chr11:124749276-12475129 | p3 | (TCT)12 | CACGTTCCCCAACTAAGCAT | CGGAGTAGGAGTTATGGCGA | 140 |
| chr11:124899922-12490194 | p2 | (AT)11 | TGACACTTTAGGCGAATCCC | ACTGTCACGACCCAAACACA | 130 |
| chr11:125050056-12505207 | p6 | (TCCACA)3 | CTCGAAGGGGTTTGTGGTAA | TCAATTCGACACTGGGACAA | 218 |
| chr11:125050056-12505207 | p6 | (TCCACA)4 | CTCGAAGGGGTTTGTGGTAA | TCAATTCGACACTGGGACAA | 218 |
| chr11:125214878-12521689 | p3 | (CTT)5 | CCGTGCAACAAAATTGCTTA | TTAAACTTGGGCCTGGATCA | 268 |
| chr11:125218976-12522105 | p3 | (CTC)5 | TCTGCCATGGTTGAAAACAA | AATATTACCCGATCCAGCCC | 276 |
| chr11:125223706-12522572 | p3 | (GAT)5 | AACGAATACGCCGAAGCTAA | TTTGGCTGATGAACAAGCTG | 170 |
| chr11:125393747-12539576 | p5 | (TTGGT)3 | CCCTGTAATTGCACGGATCT | CACCCATACCGGGTGAAATA | 198 |
| chr11:125393747-12539576 | p5 | (TTGGT)3 | CCCTGTAATTGCACGGATCT | CACCCATACCGGGTGAAATA | 198 |
| chr11:125425945-12542798 | p2 | (TA)7 | CTAGTTGTGAGCTCGGGTCC | AGCCCAACAACAAGTCAACC | 173 |
| chr11:125425945-12542798 | p2 | (AT)21 | CTAGTTGTGAGCTCGGGTCC | AGCCCAACAACAAGTCAACC | 173 |
| chr11:125465309-12546732 | p2 | (TA)10 | CTTAGAGGTGGAACCGTTGG | CGGTTCCGATTTCCAAAATA | 129 |
| chr11:125465309-12546732 | p2 | (AT)9 | CTTAGAGGTGGAACCGTTGG | CGGTTCCGATTTCCAAAATA | 129 |

| Marker ID | SSR type | SSR motif | FORWARD PRIMER (5'-3') | REVERSE PRIMER (5'-3') | size (bp) |
|---|---|---|---|---|---|
| chr11:125507571-12550958 | p5 | (TTTTG)3 | ATTGTTGGCTCTTCGGATTG | TGTGCATATTCCCATTGCAT | 158 |
| chr11:125509535-12551155 | p5 | (ATTTA)3 | AATCAGACACACGCACAACC | TGGGGTTGCAGTAAGGAAAG | 147 |
| chr11:125615094-12561713 | p5 | (GCATT)3 | ACCCATCGTCTTTGTTTTCG | GGACAGACTCTGCACACCAA | 275 |
| chr11:125615094-12561713 | p4 | (TTTA)4 | ACCCATCGTCTTTGTTTTCG | GGACAGACTCTGCACACCAA | 275 |
| chr11:125615181-12561719 | p4 | (TTTA)4 | CCCATCGTCTTTGTTTTCGT | GGATCACCTAGCAGCTCACC | 189 |
| chr11:125615240-12561725 | p4 | (TTTA)4 | CCCATCGTCTTTGTTTTCGT | GGATCACCTAGCAGCTCACC | 130 |
| chr11:125842130-12584414 | p2 | (TA)6 | GGAGGTTTCGGTGGAAAAAT | ACTCACGCTTACATGGGTCC | 178 |
| chr11:125842130-12584414 | p2 | (AT)8 | GGAGGTTTCGGTGGAAAAAT | ACTCACGCTTACATGGGTCC | 178 |
| chr11:125898939-12590095 | p5 | (TATTT)3 | AGGGTAGCGGGATGAGATTT | AAAGTGCGGCCTAGCAATTA | 241 |
| chr11:125898939-12590095 | p2 | (GT)7 | AGGGTAGCGGGATGAGATTT | AAAGTGCGGCCTAGCAATTA | 241 |
| chr11:125925208-12592722 | p2 | (TC)9 | ATGTCCATCCCCCACTATCA | TCCCATCGTCGGACTTTAAC | 206 |
| chr11:125975534-12597754 | p2 | (TA)6 | CACTAGAGGGTCGGGAAACA | ATCCTTGGCCCATAATCACA | 157 |
| chr11:125975549-12597757 | p2 | (TA)6 | CACTAGAGGGTCGGGAAACA | ATCCTTGGCCCATAATCACA | 142 |
| chr11:126015232-12601724 | p2 | (TA)7 | TGTTGTAATTGATCCCCGGT | TATATGCGCACCTTCGTTTG | 192 |
| chr11:126015232-12601724 | p2 | (TA)6 | TGTTGTAATTGATCCCCGGT | TATATGCGCACCTTCGTTTG | 192 |
| chr11:126015674-12601768 | p2 | (TA)7 | TGGTAGGTTATTGTGCGCTG | TCTTGTCAGACCCCCAAAAG | 171 |
| chr11:126015674-12601768 | p2 | (TA)6 | TGGTAGGTTATTGTGCGCTG | TCTTGTCAGACCCCCAAAAG | 171 |
| chr11:126061025-12606304 | p2 | (TA)8 | TCAATTTCAATTTTCCCCCA | ATTACCCCGGGATTTTTACG | 111 |
| chr11:126098193-12610021 | p5 | (AATGA)3 | TAGGGTGATGTGGGGTTTGT | TTCAGCCATGCCAAAATACA | 265 |
| chr11:126098193-12610021 | p3 | (TTC)6 | TAGGGTGATGTGGGGTTTGT | TTCAGCCATGCCAAAATACA | 265 |
| chr11:126174150-12617616 | p3 | (TGT)5 | GTGTTGGCCGTTCAAGAAAT | ATCTCCTTTGTACATGCCCG | 287 |
| chr11:126184922-12618698 | p2 | (TA)29 | CGGTCCGTTTAACGAAAGAA | TGTTTTGATTTGGCATGGAA | 237 |
| chr11:126249074-12625110 | p2 | (CT)13 | GTGAGACACGTGGTTGATGG | CGGATCGTAGTTTTACCGGA | 300 |
| chr11:126253788-12625580 | p5 | (CAAAT)3 | TCGTGTATGCATGGTCGAAT | CTTAAAACCCGCAAAAGCAG | 167 |
| chr11:126270606-12627262 | p6 | (AAAATA)3 | CTAGCCACTTTTCGACTCGG | TGGTTAACGCTACAAGCCCT | 116 |
| chr11:126320591-12632266 | p3 | (AAG)23 | ATTCACCATGGAAATCAGCC | TCTGTTGGAACAACTGCTGC | 267 |
| chr11:126320591-12632266 | p3 | (GAA)5 | ATTCACCATGGAAATCAGCC | TCTGTTGGAACAACTGCTGC | 267 |
| chr11:126358963-12636098 | p3 | (TGT)6 | AGAAGATGAAAGTCGCGGAA | ATGCCCAAACCATCTCAGTC | 298 |
| chr11:126366250-12636826 | p3 | (AGC)5 | AAGTGAAAATCGAACACCGC | GCCTTTAACACAGGGGCATA | 181 |
| chr11:126412364-12641437 | p5 | (TTGCA)3 | TTTCAGTTTTGCCCTTTTGG | TCCGGAGAGCCTCTAAATGA | 225 |
| chr11:126414078-12641610 | p2 | (TA)13 | GAATCCAAGCCACACTGACA | ATCCATCGCTAAATTGCTCG | 191 |
| chr11:126436717-12643873 | p2 | (TG)7 | GGAGTGGTCTCGCTCTGTTC | AGTGCAGCACAACACTGGAG | 127 |
| chr11:126488572-12649058 | p5 | (ATTTT)3 | CACCCTTCGTCGGAAAATTA | AAACCCAGTGGTGGAGTCAG | 200 |
| chr11:126488572-12649058 | p2 | (AT)6 | CACCCTTCGTCGGAAAATTA | AAACCCAGTGGTGGAGTCAG | 200 |
| chr11:126488572-12649058 | p6 | (AAAGAA)3 | CACCCTTCGTCGGAAAATTA | AAACCCAGTGGTGGAGTCAG | 200 |
| chr11:126547822-12654983 | p5 | (TTGGT)3 | GGAATATTTGCACGGCTGAT | ACCGAACAAATTCTTCACCG | 124 |

| Marker ID | SSR type | SSR motif | FORWARD PRIMER (5'-3') | REVERSE PRIMER (5'-3') | size (bp) |
|---|---|---|---|---|---|
| chr11:126548532-12655054 | p5 | (TTGGT)3 | GTTTGGCTTGGTTTGGTGTT | GTAATGTCGGGTTGGTTTGG | 247 |
| chr11:126548532-12655054 | p2 | (TG)7 | GTTTGGCTTGGTTTGGTGTT | GTAATGTCGGGTTGGTTTGG | 247 |
| chr11:126693679-12669569 | p6 | (TATACA)3 | CAACCCCATACTCAGGGCTA | GCAAGCCGTCGTACCATTAT | 129 |
| chr11:126716869-12671888 | p6 | (TATTTT)3 | CCCTAAAATGACGATCCGAA | AAAAGGTGCCCGAGGTAAGT | 175 |
| chr11:126795353-12679737 | p3 | (AAT)7 | TGCACATCAAAATTCTCCGA | ATCGAAACTTCCGCACTTGT | 202 |
| chr11:126848191-12685020 | p2 | (CT)6 | CTCCATTCGGATGTGGACTT | CAGGGCCAATTTCGATTAAA | 214 |
| chr11:126959495-12696150 | p2 | (TA)7 | AAAGTTGCAATTCGTGGAGG | TTGCCGAAGAGGGTTGTATC | 261 |
| chr11:126959495-12696150 | p2 | (AT)7 | AAAGTTGCAATTCGTGGAGG | TTGCCGAAGAGGGTTGTATC | 261 |
| chr11:126959495-12696150 | p3 | (ATC)5 | AAAGTTGCAATTCGTGGAGG | TTGCCGAAGAGGGTTGTATC | 261 |
| chr11:126987765-12698978 | p5 | (TTGCA)3 | TTTAGGGCTTATTTTGGGGG | ACATGCATGGTAGTGCCAAA | 146 |
| chr11:127094342-12709635 | p5 | (AAAAG)3 | TCATCAAGAAGATGGCGTTG | GCCCTGTTCTAATGGGTGAA | 103 |
| chr11:127124279-12712629 | p3 | (ATG)5 | GTTGTTGCCATTGTTGGTTG | GCCTTTGGAAAGAAAGGGAC | 176 |
| chr11:127124279-12712629 | p2 | (TA)10 | GTTGTTGCCATTGTTGGTTG | GCCTTTGGAAAGAAAGGGAC | 176 |
| chr11:127124279-12712629 | p2 | (AT)6 | GTTGTTGCCATTGTTGGTTG | GCCTTTGGAAAGAAAGGGAC | 176 |
| chr11:127215313-12721732 | p5 | (AAAAT)3 | CGCAAAGCATAAGTTGGGAT | GCTAAAAGCCCCTTGGTTTC | 181 |
| chr11:127295491-12729750 | p4 | (ACTA)4 | TAAGCTCATATGGGAACCCG | TTTCCCTCTGCACACACTTG | 293 |
| chr11:127360885-12736290 | p2 | (AT)9 | CATATCCGGCAAATTATGGG | GTTTCAGGCTGCATTGTTCA | 177 |
| chr11:127360885-12736290 | p3 | (AAC)5 | CATATCCGGCAAATTATGGG | GTTTCAGGCTGCATTGTTCA | 177 |
| chr11:127373652-12737568 | p2 | (AT)13 | TTGGCACCATAACACAGCAT | CGTGCATGCCTCAATATGTC | 195 |
| chr11:127373652-12737568 | p3 | (CAA)12 | TTGGCACCATAACACAGCAT | CGTGCATGCCTCAATATGTC | 195 |
| chr11:127428064-12743007 | p2 | (TA)6 | CAAAAATTAATTCGGGGCAC | AAGAATTTCGTGCGTGAAGG | 214 |
| chr11:127447011-12744902 | p2 | (AT)45 | CCAAATGGCATTACCAACCT | AATGTGTTTGCTTGGATTGCT | 130 |
| chr11:127455461-12745748 | p2 | (TA)6 | CCGTTCTTTTCGAATGGTGT | GTTTGCACTCGGGCTTTCTA | 137 |
| chr11:127455461-12745748 | p5 | (TATTA)4 | CCGTTCTTTTCGAATGGTGT | GTTTGCACTCGGGCTTTCTA | 137 |
| chr11:127463544-12746555 | p5 | (ATTAA)3 | AGGGCATTTTAGGGCTGTCT | CAGTACTTCCACCTCCCCAA | 210 |
| chr11:127476892-12747890 | p5 | (AAAAG)3 | GTCTGCCTGTGCCAACCTAT | CCAAGTTCGGACTTAGACGC | 224 |
| chr11:127505159-12750718 | p3 | (TTG)5 | CTGGGATCGGCATAGTTTGT | TTGGCGTCTCAACTAATCCC | 243 |
| chr11:127530056-12753207 | p5 | (AAATA)3 | AATATTCCCGAGGTTCTGGG | AGGCAGACTTTTGCCTTGAA | 256 |
| chr11:127550738-12755280 | p2 | (TA)31 | AATTTTTGGTGCACCCACTC | ATTAGCCCAAAGGGTCCAAG | 236 |
| chr11:127550738-12755280 | p6 | (TTTAAA)3 | AATTTTTGGTGCACCCACTC | ATTAGCCCAAAGGGTCCAAG | 236 |
| chr11:127598720-12760074 | p2 | (AG)11 | TGCGTTTCCTGGCTATCTTT | GGATCCTATGACCCAAGCAA | 282 |
| chr11:127598720-12760074 | p3 | (TAT)5 | TGCGTTTCCTGGCTATCTTT | GGATCCTATGACCCAAGCAA | 282 |
| chr11:127700607-12770262 | p5 | (AAACC)3 | ATTTTGTCGATTTGGTTCGG | TAGGGGTGTTCATGGTAGGG | 284 |
| chr11:127700631-12770264 | p5 | (AAACC)3 | ATTTTGTCGATTTGGTTCGG | TAGGGGTGTTCATGGTAGGG | 260 |
| chr11:127700655-12770267 | p5 | (AAACC)3 | ATTTTGTCGATTTGGTTCGG | TAGGGGTGTTCATGGTAGGG | 236 |
| chr11:127700679-12770269 | p5 | (AAACC)3 | ATTTTGTCGATTTGGTTCGG | TAGGGGTGTTCATGGTAGGG | 212 |

| Marker ID | SSR type | SSR motif | FORWARD PRIMER (5'-3') | REVERSE PRIMER (5'-3') | size (bp) |
|---|---|---|---|---|---|
| chr11:127700703-12770271 | p5 | (AAACC)3 | ATTTTGTCGATTTGGTTCGG | TAGGGGTGTTCATGGTAGGG | 188 |
| chr11:127700727-12770274 | p5 | (AAACC)3 | ATTTTGTCGATTTGGTTCGG | TAGGGGTGTTCATGGTAGGG | 164 |
| chr11:127700751-12770276 | p5 | (AAACC)3 | ATTTTGTCGATTTGGTTCGG | TAGGGGTGTTCATGGTAGGG | 140 |
| chr11:127700775-12770279 | p5 | (AAACC)3 | ATTTTGTCGATTTGGTTCGG | TAGGGGTGTTCATGGTAGGG | 116 |
| chr11:127807543-12780955 | p3 | (TGT)5 | CCATCATGCCTCGAATTCTT | ATGTGATCTCTGGGAGGCAG | 177 |
| chr11:127844605-12784662 | p5 | (TTTTC)3 | CGACACTCCCCTTTCTTCAG | GCATGTGGTCCCTCTCAGTT | 275 |
| chr11:127844605-12784662 | p4 | (TATG)4 | CGACACTCCCCTTTCTTCAG | GCATGTGGTCCCTCTCAGTT | 275 |
| chr11:127946795-12794881 | p2 | (AT)8 | GTCTGCCCTCTAGCCCTTCT | TCTATGCCGTTGCTACATGC | 160 |
| chr11:128090652-12809266 | p5 | (ATATG)3 | GATTTTGGAGGAAACGGGAT | AATGGAATTTTCCCCAGGAC | 161 |
| chr11:128138094-12814012 | p2 | (AT)8 | CATTAATGTTGGTGAGGGGG | TGGACGGGCCTCTGTTATAG | 120 |
| chr11:128138094-12814012 | p2 | (TA)6 | CATTAATGTTGGTGAGGGGG | TGGACGGGCCTCTGTTATAG | 120 |
| chr11:128138094-12814012 | p2 | (TA)10 | CATTAATGTTGGTGAGGGGG | TGGACGGGCCTCTGTTATAG | 120 |
| chr11:128156479-12815851 | p2 | (TA)17 | AATACAAGGCGAGTCATCCG | ATTTGGTCCACTCCACTTGC | 276 |
| chr11:128182453-12818447 | p6 | (GTTTCA)3 | CATGGCAACATGGAAACAAG | CCCCGTCTTACGAATCAAAA | 190 |
| chr11:128191971-12819398 | p6 | (GTTTCA)3 | CATGGCAACATGGAAACAAG | CCCCGTCTTACGAATCAAAA | 190 |
| chr11:128399120-12840113 | p4 | (AAAC)5 | CCGTGATTTTCTCTCTTCGC | TTTATTCAATTTCCGGTCGC | 244 |
| chr11:128399120-12840113 | p4 | (AAAC)4 | CCGTGATTTTCTCTCTTCGC | TTTATTCAATTTCCGGTCGC | 244 |
| chr11:128399120-12840113 | p4 | (AAAC)4 | CCGTGATTTTCTCTCTTCGC | TTTATTCAATTTCCGGTCGC | 244 |
| chr11:128399952-12840196 | p4 | (AAAC)4 | CCGTGATTTTCTCTCTTCGC | TTCCGGTCGTCGTTTATACC | 247 |
| chr11:128399952-12840196 | p4 | (AAAC)4 | CCGTGATTTTCTCTCTTCGC | TTCCGGTCGTCGTTTATACC | 247 |
| chr11:128399952-12840196 | p4 | (AAAC)4 | CCGTGATTTTCTCTCTTCGC | TTCCGGTCGTCGTTTATACC | 247 |
| chr11:128399952-12840196 | p4 | (AAAC)4 | CCGTGATTTTCTCTCTTCGC | TTCCGGTCGTCGTTTATACC | 247 |
| chr11:128399952-12840196 | p4 | (AAAC)4 | CCGTGATTTTCTCTCTTCGC | TTCCGGTCGTCGTTTATACC | 247 |
| chr11:128412002-12841403 | p2 | (AT)7 | GCTCCAGTTCAGAGGGTGAG | AACCAAAATAAACCCCTCGG | 149 |
| chr11:128477929-12847994 | p3 | (ACA)7 | ATGAATCGGGTTGGTTTGAG | AAAAACCGAGAAAACCCGAG | 161 |
| chr11:128477929-12847994 | p5 | (AAACC)3 | ATGAATCGGGTTGGTTTGAG | AAAAACCGAGAAAACCCGAG | 161 |
| chr11:128477929-12847994 | p5 | (AAACC)3 | ATGAATCGGGTTGGTTTGAG | AAAAACCGAGAAAACCCGAG | 161 |
| chr11:128479179-12848119 | p6 | (AAAAGA)3 | TGAATGCCTTTCAACACGTC | TATATGTCGGAGGCCTGGAG | 229 |
| chr11:128547266-12854928 | p2 | (AC)8 | CAACCCATTTTCTTCCTCCA | ACCCTCCCTTTCTTCCACAT | 235 |
| chr11:128547266-12854928 | p5 | (ATAAA)3 | CAACCCATTTTCTTCCTCCA | ACCCTCCCTTTCTTCCACAT | 235 |
| chr11:128576217-12857823 | p2 | (AT)8 | TAGCGTCTTTGCTACCTCCC | GGCGGACCCATGTATTATTG | 157 |
| chr11:128625546-12862756 | p5 | (GGCTT)3 | GAACCAGGGACGGAGTACAA | CTCATGAAAATGCCAACGTG | 251 |
| chr11:128708503-12871052 | p6 | (CATGAG)3 | GGAGAACGAGCAAGCGATAG | GGCGTGACACATCTTTAGCA | 153 |
| chr11:128785942-12878796 | p6 | (CAACTC)3 | TGTGGAAACCCCTAAAGCTG | AGCACCATGGAGCCATTAAG | 126 |
| chr11:128827556-12882958 | p2 | (AT)11 | TGAATCCTTGCGGCTTATTC | GCTCCGGACTCAGAAAGTTG | 246 |
| chr11:128827629-12882964 | p2 | (AT)11 | TGAATCCTTGCGGCTTATTC | GCTCCGGACTCAGAAAGTTG | 173 |

| Marker ID | SSR type | SSR motif | FORWARD PRIMER (5'-3') | REVERSE PRIMER (5'-3') | size (bp) |
|---|---|---|---|---|---|
| chr11:128830554-12883260 | p2 | (TA)6 | TACAAGAGCCCGTGATTTCC | GCGTCCACAACATATCATCG | 195 |
| chr11:128842349-12884436 | p2 | (TG)7 | CACCCTCCTTTTCTTCCACA | TTTGAAGGACAAACTGTGCG | 121 |
| chr11:128917276-12891929 | p2 | (AT)11 | GAACCCCGAACCTCATGATA | GCAGGAGTGCCCTTATTTTG | 256 |
| chr11:128924305-12892632 | p5 | (CCGTC)4 | CCGTCCACGTGTCAAAGTTA | TCCACACGGAGCTAAAAACC | 231 |
| chr11:129071059-12907312 | p2 | (TA)32 | AAAGATGGGCTAAAACGGGT | GATCGCGAGGTCATGCTAAT | 276 |
| chr11:129071059-12907312 | p2 | (TA)6 | AAAGATGGGCTAAAACGGGT | GATCGCGAGGTCATGCTAAT | 276 |
| chr11:129179539-12918155 | p5 | (TATTT)4 | GCTACGGGGAACTGTGTGAT | GCCTGGAGCAGGATTATTCA | 147 |
| chr11:129206633-12920865 | p3 | (AAC)6 | TCGGTCAAAACAACTTGCAG | AAGTTGGCCCTTTGACACAC | 207 |
| chr11:129337855-12933987 | p3 | (TTC)6 | GATTAATGCTCGGTCACGGT | TGGTCTCATTTTCGCTGATG | 234 |
| chr11:129387531-12938955 | p2 | (AT)13 | ACTTTCTGGGAGAAGCGTGA | TTGAATTGGGTTGAACAGCA | 215 |
| chr11:129407590-12940960 | p3 | (GAA)6 | GCTGAAGTTGAGATCCCTGC | AATTTCCCCAACAACAGTCG | 243 |
| chr11:129407590-12940960 | p5 | (AATTA)3 | GCTGAAGTTGAGATCCCTGC | AATTTCCCCAACAACAGTCG | 243 |
| chr11:129407590-12940960 | p5 | (AATTA)3 | GCTGAAGTTGAGATCCCTGC | AATTTCCCCAACAACAGTCG | 243 |
| chr11:129407590-12940960 | p5 | (TTTAT)3 | GCTGAAGTTGAGATCCCTGC | AATTTCCCCAACAACAGTCG | 243 |
| chr11:129407590-12940960 | p5 | (ATTTA)3 | GCTGAAGTTGAGATCCCTGC | AATTTCCCCAACAACAGTCG | 243 |
| chr11:129447235-12944925 | p5 | (ATTGC)3 | AACACTTTGATGAGCTGGGC | ACATCTCGGTAGCACAACCC | 164 |
| chr11:129447235-12944925 | p2 | (TA)10 | AACACTTTGATGAGCTGGGC | ACATCTCGGTAGCACAACCC | 164 |
| chr11:129464691-12946670 | p2 | (AT)9 | GGTCATGCCAAAAAGGAAGA | TGAATTTACAGGGGAGGTGC | 218 |
| chr11:129464691-12946670 | p2 | (CA)7 | GGTCATGCCAAAAAGGAAGA | TGAATTTACAGGGGAGGTGC | 218 |
| chr11:129464691-12946670 | p4 | (ATTT)5 | GGTCATGCCAAAAAGGAAGA | TGAATTTACAGGGGAGGTGC | 218 |
| chr11:129488236-12949025 | p2 | (AT)9 | GGTCATGCCAAAAAGGAAGA | TGAATTTACAGGGGAGGTGC | 218 |
| chr11:129488236-12949025 | p2 | (CA)7 | GGTCATGCCAAAAAGGAAGA | TGAATTTACAGGGGAGGTGC | 218 |
| chr11:129488236-12949025 | p4 | (ATTT)5 | GGTCATGCCAAAAAGGAAGA | TGAATTTACAGGGGAGGTGC | 218 |
| chr11:129516061-12951807 | p2 | (AT)12 | GTGCAAAAATAATGGTGGGC | ATACACACGGGGCAATGAAT | 265 |
| chr11:129516061-12951807 | p6 | (TTTATT)3 | GTGCAAAAATAATGGTGGGC | ATACACACGGGGCAATGAAT | 265 |
| chr11:129516159-12951819 | p2 | (AT)12 | GTGCAAAAATAATGGTGGGC | ATACACACGGGGCAATGAAT | 167 |
| chr11:129516159-12951819 | p6 | (TTTATT)3 | GTGCAAAAATAATGGTGGGC | ATACACACGGGGCAATGAAT | 167 |
| chr11:129519820-12952184 | p3 | (GTT)9 | TATGGGTTCCCACCGATAAA | AAGGCCGAAGGAAAAACAAT | 110 |
| chr11:129641131-12964314 | p2 | (TA)7 | AAATCCCCGATCATTTCTCC | GGATGAAACGGAAATGGATG | 118 |
| chr11:129644765-12964678 | p3 | (TCA)5 | AGCCCACCCCTAACAATACC | TCTAGCCGCCTTCACTTTGT | 232 |
| chr11:129644765-12964678 | p6 | (CCGGTT)3 | AGCCCACCCCTAACAATACC | TCTAGCCGCCTTCACTTTGT | 232 |
| chr11:129776867-12977888 | p5 | (ACATA)3 | CCACGCGTGAGATCAAAGTA | TACCCCTAAATCCCCCAAAC | 126 |
| chr11:129817423-12981944 | p2 | (TA)9 | TGGTTTTTGGAACGATGACA | TGCTTCGCTATTGCATCAAC | 238 |
| chr11:130003820-13000586 | p3 | (CAC)5 | TTGCATCAACCAGGTACGAA | AGCAAGTGAACTTGATGGGG | 175 |
| chr11:130004515-13000653 | p3 | (CAC)5 | TTGGACTGTTTTCATGCTGC | GCACACATTTCCACCAACAC | 182 |
| chr11:130014270-13001633 | p2 | (TG)8 | TGTGGGAAAAATGCACTTGA | AAACTTATCAGCACCGGCAC | 145 |

| Marker ID | SSR type | SSR motif | FORWARD PRIMER (5'-3') | REVERSE PRIMER (5'-3') | size (bp) |
|---|---|---|---|---|---|
| chr11:130014270-13001633 | p2 | (AT)33 | TGTGGGAAAAATGCACTTGA | AAACTTATCAGCACCGGCAC | 145 |
| chr11:130014270-13001633 | p3 | (TTC)7 | TGTGGGAAAAATGCACTTGA | AAACTTATCAGCACCGGCAC | 145 |
| chr11:130086456-13008847 | p5 | (TTTTA)4 | AACTATCCGCAAGATGCTCG | TTATGTGGCAAGCTGATTCG | 146 |
| chr11:130335665-13033768 | p5 | (TTAGA)3 | CAATTAGGTGGCGACTCGTT | GAACCTAAGTTGCCCAGCAG | 242 |
| chr11:130390795-13039281 | p6 | (GATGAC)3 | CCAGACCCACCCACTAGAAA | CCCCTCAACTTTGCGATTTA | 185 |
| chr11:130444190-13044621 | p3 | (AAT)7 | TAGACGACAATCGCATTCCA | AGAAAATCGGTTCCGGAGAT | 183 |
| chr11:130444190-13044621 | p2 | (AT)9 | TAGACGACAATCGCATTCCA | AGAAAATCGGTTCCGGAGAT | 183 |
| chr11:130487292-13048930 | p2 | (AT)6 | TGATGGCACGTCACCACTAT | ACATGAACAGTCCACCTCCC | 115 |
| chr11:130647834-13064984 | p2 | (TA)7 | AATTTTCCAGGTGAGCATGG | TTTATGTTGGGCCCTTTCTG | 276 |
| chr11:130854894-13085690 | p2 | (AT)6 | CGGTGAACAATGTCGTGAAC | TTTTGTTCAGCCGTGTCTTG | 140 |
| chr11:130854894-13085690 | p2 | (AT)7 | CGGTGAACAATGTCGTGAAC | TTTTGTTCAGCCGTGTCTTG | 140 |
| chr11:130854894-13085690 | p2 | (TA)8 | CGGTGAACAATGTCGTGAAC | TTTTGTTCAGCCGTGTCTTG | 140 |
| chr11:130862639-13086465 | p2 | (AT)19 | ATCCTCGACGCCATAATCAC | AGTTGCATATGGTTGCTCCC | 156 |
| chr11:130862639-13086465 | p2 | (TA)8 | ATCCTCGACGCCATAATCAC | AGTTGCATATGGTTGCTCCC | 156 |
| chr11:130862639-13086465 | p2 | (AT)7 | ATCCTCGACGCCATAATCAC | AGTTGCATATGGTTGCTCCC | 156 |
| chr11:130953678-13095569 | p3 | (TTG)6 | CCAAAAATGTTGTATGGGGC | TGCATCTTATCATCCACCGA | 197 |
| chr11:130979278-13098129 | p5 | (AAACC)3 | ACGGTGTCAACATCTAGGGG | CAAGAAAACCCGAGGTTGAA | 179 |
| chr11:131006770-13100879 | p2 | (AT)10 | AGAAAAATTGATAGGGGGCG | ATGAGGGCATTCATGTGTGA | 190 |
| chr11:131122527-13112455 | p2 | (TA)14 | AAACCGCTCATCGATACGAC | GTGCCAAAGTTGGTGGAACT | 130 |
| chr11:131122527-13112455 | p2 | (AT)7 | AAACCGCTCATCGATACGAC | GTGCCAAAGTTGGTGGAACT | 130 |
| chr11:131130785-13113281 | p5 | (TTTGT)3 | TGCCCACATTTCGTTGATTA | GCTGTCACGAAATGCAAAAA | 269 |
| chr11:131130785-13113281 | p2 | (TA)8 | TGCCCACATTTCGTTGATTA | GCTGTCACGAAATGCAAAAA | 269 |
| chr11:131130785-13113281 | p5 | (AAAAC)3 | TGCCCACATTTCGTTGATTA | GCTGTCACGAAATGCAAAAA | 269 |
| chr11:131130827-13113284 | p5 | (TTTGT)3 | TGCCCACATTTCGTTGATTA | GCTGTCACGAAATGCAAAAA | 227 |
| chr11:131130827-13113284 | p2 | (TA)8 | TGCCCACATTTCGTTGATTA | GCTGTCACGAAATGCAAAAA | 227 |
| chr11:131130827-13113284 | p5 | (AAAAC)3 | TGCCCACATTTCGTTGATTA | GCTGTCACGAAATGCAAAAA | 227 |
| chr11:131130849-13113286 | p5 | (TTTGT)3 | TGCCCACATTTCGTTGATTA | GCTGTCACGAAATGCAAAAA | 205 |
| chr11:131130849-13113286 | p2 | (TA)8 | TGCCCACATTTCGTTGATTA | GCTGTCACGAAATGCAAAAA | 205 |
| chr11:131130849-13113286 | p5 | (AAAAC)3 | TGCCCACATTTCGTTGATTA | GCTGTCACGAAATGCAAAAA | 205 |
| chr11:131130920-13113293 | p5 | (TTTGT)3 | TGCCCACATTTCGTTGATTA | GCTGTCACGAAATGCAAAAA | 134 |
| chr11:131130920-13113293 | p2 | (TA)8 | TGCCCACATTTCGTTGATTA | GCTGTCACGAAATGCAAAAA | 134 |
| chr11:131130920-13113293 | p5 | (AAAAC)3 | TGCCCACATTTCGTTGATTA | GCTGTCACGAAATGCAAAAA | 134 |
| chr11:131130940-13113296 | p5 | (TTTGT)3 | TGCCCACATTTCGTTGATTA | GCTGTCACGAAATGCAAAAA | 114 |
| chr11:131130940-13113296 | p2 | (TA)8 | TGCCCACATTTCGTTGATTA | GCTGTCACGAAATGCAAAAA | 114 |
| chr11:131130940-13113296 | p5 | (AAAAC)3 | TGCCCACATTTCGTTGATTA | GCTGTCACGAAATGCAAAAA | 114 |
| chr11:131159265-13116127 | p2 | (TA)6 | AAGATCCAGATATGTGCCGC | GTTATCGCTGCCCGAAGTAG | 280 |

| Marker ID | SSR type | SSR motif | FORWARD PRIMER (5'-3') | REVERSE PRIMER (5'-3') | size (bp) |
|---|---|---|---|---|---|
| chr11:131159265-13116127 | p2 | (TA)15 | AAGATCCAGATATGTGCCGC | GTTATCGCTGCCCGAAGTAG | 280 |
| chr11:131208510-13121053 | p2 | (AG)9 | CCGGGGAAGAAAACAGTACC | TCTATAAAACAACGCAACCGC | 169 |
| chr11:131208572-13121059 | p2 | (AG)9 | CCGGGGAAGAAAACAGTACC | TCTATAAAACAACGCAACCGC | 107 |
| chr11:131243514-13124557 | p3 | (TTC)24 | CCTTTTCCGGGGTAAACAAT | GCCCAACCAGTCCAACTCTA | 284 |
| chr11:131319670-13132169 | p2 | (AT)10 | GTCAATCCCTCGCATCAAGT | CTTCACATTGCTGCATGCTT | 225 |
| chr11:131353998-13135601 | p2 | (AT)10 | GCTGGGTTTCTATCGTGCTC | GAGCATTCTCAATTGTGCGA | 164 |
| chr11:131353998-13135601 | p2 | (TA)6 | GCTGGGTTTCTATCGTGCTC | GAGCATTCTCAATTGTGCGA | 164 |
| chr11:131376229-13137825 | p2 | (GT)7 | AGATTTCGTTTGATCCGTGG | AATGACCAGATTGGCAAAGG | 248 |
| chr11:131389958-13139199 | p3 | (TAA)5 | AGGAACCAAAATCAAACCCC | GGGTTTCAAGTGTTTTGGGA | 233 |
| chr11:131389958-13139199 | p5 | (GAAGG)3 | AGGAACCAAAATCAAACCCC | GGGTTTCAAGTGTTTTGGGA | 233 |
| chr11:131459761-13146177 | p2 | (AT)7 | AGTTATCGCTGCCCGAAGTA | AGGATCATCAACAGCCCAAC | 195 |
| chr11:131459761-13146177 | p2 | (TA)26 | AGTTATCGCTGCCCGAAGTA | AGGATCATCAACAGCCCAAC | 195 |
| chr11:131515687-13151770 | p2 | (AT)8 | TGGTGTCCATTTCACAGCAT | GTTATGCAGGGGGTTAACGA | 223 |
| chr11:131515707-13151771 | p2 | (AT)8 | TGGTGTCCATTTCACAGCAT | GTTATGCAGGGGGTTAACGA | 203 |
| chr11:131538780-13154079 | p2 | (GA)8 | TGACCTTACCACCAAAAGCC | CTTCAAAGACCAGAGGGTCG | 121 |
| chr11:131538799-13154082 | p2 | (GA)8 | TGACCTTACCACCAAAAGCC | CTTCAAAGACCAGAGGGTCG | 102 |
| chr11:131560270-13156228 | p5 | (TTGGT)3 | CGGGGGTATATTTGACCCTT | GGTTGGTTTGGTTCGATTTG | 106 |
| chr11:131560270-13156228 | p5 | (TTGGT)3 | CGGGGGTATATTTGACCCTT | GGTTGGTTTGGTTCGATTTG | 106 |
| chr11:131577889-13157990 | p3 | (ATG)5 | CTACCCCAATCATTTTCCGA | TGATGGTTTTCTTTGTGGCA | 167 |
| chr11:131583035-13158505 | p3 | (TGT)6 | ATGGATGAAATGGAAGCTCG | TATCCACCACAAAAGCCACA | 222 |
| chr11:131599630-13160164 | p5 | (AAACC)3 | TCGATGAACCAACAAGTGGA | CAAGAAAACCCGAGGTTGAA | 153 |
| chr11:131664700-13166675 | p3 | (AAT)17 | TCGGATTCACAGCAATACCA | TCCCTTACAAAAGTGGTCCG | 151 |
| chr11:131688684-13169073 | p3 | (AAT)17 | CGGACCACTTTTGTAAGGGA | CTCTCAAACGAAAACCCCAA | 270 |
| chr11:131705135-13170717 | p2 | (TA)19 | AAATGGTGGTGGTTGTGGAT | TCTCCCCACCAAGTTTCATC | 172 |
| chr11:131716976-13171900 | p3 | (TTG)8 | CTCAGCCAAGATTTGGAAGG | ATTTTGTCCAGTTCGGCATC | 202 |
| chr11:131981565-13198363 | p2 | (AT)33 | AGCCCAAAACATGGTGAGAC | ATCAAACGTTTATCGTGGGC | 130 |
| chr11:132028243-13203025 | p5 | (AAAAT)3 | TCGAATTTGCGTCAATCTCA | GCCTTAACGTTGGGTTGTGT | 191 |
| chr11:132034121-13203614 | p3 | (TTG)9 | GCCACCAAAACTTAAAGGCA | ATCTTGTCTTCCACCGATGC | 189 |
| chr11:132090855-13209288 | p2 | (AT)16 | TCACAATTTTCATGGCCAAA | AGCCGAGAAGAAATCCGAGT | 251 |
| chr11:132090855-13209288 | p2 | (TA)8 | TCACAATTTTCATGGCCAAA | AGCCGAGAAGAAATCCGAGT | 251 |
| chr11:132112208-13211422 | p6 | (TTTTTA)3 | CTAGGAGGGCCGGAGATTAG | AGCATTTTGTTTTCGGAAGG | 151 |
| chr11:132125440-13212745 | p2 | (CA)6 | TAGCGGGTTGTTACACCACA | ATACTTCGCACCAAACGACC | 280 |
| chr11:132125440-13212745 | p2 | (TA)7 | TAGCGGGTTGTTACACCACA | ATACTTCGCACCAAACGACC | 280 |
| chr11:132187961-13218997 | p2 | (TA)33 | AGTCGTGGGTATTGTTTGCC | TTTACTTAAGGCCGAGCGAA | 209 |
| chr11:132187961-13218997 | p2 | (GA)9 | AGTCGTGGGTATTGTTTGCC | TTTACTTAAGGCCGAGCGAA | 209 |
| chr11:132222227-13222424 | p4 | (AATA)4 | CTGATAGGGCTGGGCATAAA | CGAACTCTAAACCCCGAACA | 145 |

| Marker ID | SSR type | SSR motif | FORWARD PRIMER (5'-3') | REVERSE PRIMER (5'-3') | size (bp) |
|---|---|---|---|---|---|
| chr11:132285153-13228717 | p3 | (AAG)8 | CATCAAAAGCACTTGGGGAT | TTGTGCAATACAACCCCTGA | 159 |
| chr11:132291312-13229332 | p5 | (TTTAT)3 | TATTTATCGGTTCGGTTCGG | TCGGTTTGGTTTGATTTGGT | 192 |
| chr11:132390635-13239265 | p5 | (GGGTT)3 | CTTGTGCCTCCACCTTCAGT | GAAGATTACCGCCAAAACCA | 190 |
| chr11:132390635-13239265 | p4 | (AAAT)5 | CTTGTGCCTCCACCTTCAGT | GAAGATTACCGCCAAAACCA | 190 |
| chr11:132398364-13240038 | p5 | (GGGTT)3 | CATGCTTTCACCCATAGGCT | CCAGACAACTTGCACAATGC | 101 |
| chr11:132398364-13240038 | p4 | (AAAT)5 | CATGCTTTCACCCATAGGCT | CCAGACAACTTGCACAATGC | 101 |
| chr11:132440753-13244276 | p2 | (TC)6 | TTTGGAGGATCTGATACGGG | AGCATTTGTCGTTGCAAGTG | 165 |
| chr11:132440753-13244276 | p4 | (TTAT)4 | TTTGGAGGATCTGATACGGG | AGCATTTGTCGTTGCAAGTG | 165 |
| chr11:132469270-13247128 | p3 | (ACA)6 | CTCCTCTTCCGCACTTTCTG | GGCTCCTTCCTCTGGACTCT | 231 |
| chr11:132495974-13249801 | p5 | (AACCG)3 | GGTTCGGTATTTGGTGTCCA | ATAGAGGGGATTGGGTTTGG | 171 |
| chr11:132495974-13249801 | p5 | (TTTTG)3 | GGTTCGGTATTTGGTGTCCA | ATAGAGGGGATTGGGTTTGG | 171 |
| chr11:132570818-13257283 | p2 | (AT)8 | GACTAGGCGTATTCGGTGGA | GGGCGTGATAAAGCTTGGTA | 169 |
| chr11:132650145-13265216 | p2 | (TA)10 | TTAAAAGGGACAACGGACGA | AAAAGGGTGAGCCCCATACT | 262 |
| chr11:132650145-13265216 | p2 | (AT)8 | TTAAAAGGGACAACGGACGA | AAAAGGGTGAGCCCCATACT | 262 |
| chr11:132650145-13265216 | p5 | (ATTTT)3 | TTAAAAGGGACAACGGACGA | AAAAGGGTGAGCCCCATACT | 262 |
| chr11:132753268-13275529 | p3 | (ATT)9 | GTCTCGTCCTTTGCTTCACC | TTGTTGTTGGTCGTTGGTTC | 114 |
| chr11:132806815-13280884 | p2 | (AT)13 | AAATCAAACGCCCCCTAAGT | TCGACCCTCGTAAATGGAAC | 235 |
| chr11:132822251-13282426 | p2 | (TC)6 | GGCCAGAGCACAAAAATACC | ACCCTGTTGCTAAAACGGTG | 135 |
| chr11:132830227-13283223 | p2 | (TA)6 | TGGGTAAAAGAATCGCATCC | GTCCCTTTCCTTTTGGGTGT | 143 |
| chr11:132830227-13283223 | p6 | (ATTTTT)3 | TGGGTAAAAGAATCGCATCC | GTCCCTTTCCTTTTGGGTGT | 143 |
| chr11:132842084-13284410 | p2 | (AT)12 | GCATGTACGGAGTCAGCAAA | AGGCATGTTCTTGAGCGTTT | 121 |
| chr11:132944793-13294680 | p2 | (TA)7 | TTTGGCAATCAGGTCATTCA | GTGGTACCTTTTATCGCGGA | 231 |
| chr11:133024761-13302677 | p3 | (CTT)5 | TCACAACTCGGATTCCATGA | TAGTTGAGAGCCCTTTTGGC | 279 |
| chr11:133043538-13304555 | p2 | (AT)9 | CGGATTCGACAGAACCAAGT | GCTCTTCCACAAAAAGCTGG | 141 |
| chr11:133043538-13304555 | p2 | (TA)6 | CGGATTCGACAGAACCAAGT | GCTCTTCCACAAAAAGCTGG | 141 |
| chr11:133138494-13314051 | p3 | (AGA)6 | AAGCGCGTACCTCATTTGAC | TTCCACCTGCAGTAGCATTG | 300 |
| chr11:133184033-13318608 | p2 | (AT)25 | TAGAACCAGCTTTGGCGACT | AGTCTGCCCACAAGAAAGGA | 102 |
| chr11:133202592-13320460 | p3 | (TTC)5 | CGGTTCAACTGCTGCATCTA | TTGGCCCTTCATTATCAACC | 143 |
| chr11:133211673-13321368 | p3 | (ATG)5 | CCAACTTATGCCTTGCGAAT | TGGCGGTCTTTAATTTTTGC | 112 |
| chr11:133232474-13323449 | p2 | (TA)6 | CCAACTGAAATCCCGAATTG | AACTGCGATCAAAAGTGCAA | 104 |
| chr11:133232474-13323449 | p4 | (TTTG)4 | CCAACTGAAATCCCGAATTG | AACTGCGATCAAAAGTGCAA | 104 |
| chr11:133398063-13340008 | p6 | (AACTGA)3 | AGAGTGGCACCATCAAGGTC | GTTGATTCAGCCAGCCAGTT | 263 |
| chr11:133401176-13340320 | p2 | (TA)15 | GAGATGCGCCTATCTTACCG | ATGTGCAACCACAAATGAGC | 192 |
| chr11:133437709-13343972 | p3 | (TCA)5 | GTGGATTGCCCAAAATCATC | CGCTGGAAACTCTTAGCCAG | 244 |
| chr11:133437709-13343972 | p2 | (AT)6 | GTGGATTGCCCAAAATCATC | CGCTGGAAACTCTTAGCCAG | 244 |
| chr11:133481081-13348309 | p3 | (TAA)5 | ATGACAGAAGTTTGGACCCG | TTGACACCCAATTTTGTCCA | 110 |

| Marker ID | SSR type | SSR motif | FORWARD PRIMER (5'-3') | REVERSE PRIMER (5'-3') | size (bp) |
|---|---|---|---|---|---|
| chr11:133481081-13348309 | p2 | (TG)6 | ATGACAGAAGTTTGGACCCG | TTGACACCCAATTTTGTCCA | 110 |
| chr11:133481081-13348309 | p2 | (TA)6 | ATGACAGAAGTTTGGACCCG | TTGACACCCAATTTTGTCCA | 110 |
| chr11:133481845-13348385 | p2 | (TG)6 | GTGGGCCCCATACTAAACAA | TTTTTAAGGGACAACGGACG | 286 |
| chr11:133481845-13348385 | p2 | (TA)6 | GTGGGCCCCATACTAAACAA | TTTTTAAGGGACAACGGACG | 286 |
| chr11:133481845-13348385 | p2 | (TA)7 | GTGGGCCCCATACTAAACAA | TTTTTAAGGGACAACGGACG | 286 |
| chr11:133529850-13353186 | p5 | (CAAAC)3 | AACGGCCTCTTGCAGAAATA | AGTCAAACCGGACCATGAAC | 102 |
| chr11:133534233-13353624 | p5 | (AAAAC)3 | TGTCCCCCATCTGTTTCTTC | TCGAGGACGAGCAAGAGTTT | 292 |
| chr11:133782457-13378447 | p5 | (AAAAG)3 | GAAGGGACAGTGGAGAGCAG | ACCACTAAGCCACACCCTTG | 257 |
| chr11:133878631-13388064 | p3 | (TAT)5 | GCAACGGATTTGAGAATGGT | CAATGTCACGTGGTTCAAGG | 146 |
| chr11:133962160-13396417 | p5 | (ATAAA)3 | TTTCTCCACCTCCTGCAACT | AATCTCCGACCACCACTACG | 228 |
| chr11:133987324-13398933 | p2 | (AT)6 | AAATCGGTGGTTTGAACGAA | TTGAAAGGAAATTCATGCCC | 233 |
| chr11:134004393-13400641 | p2 | (TA)9 | TTGTGCCATCAGCAGTGAAT | TGGTTTGTCCCCCAGATTAG | 123 |
| chr11:134004393-13400641 | p2 | (TA)9 | TTGTGCCATCAGCAGTGAAT | TGGTTTGTCCCCCAGATTAG | 123 |
| chr11:134080271-13408229 | p2 | (AT)13 | TAGGAGTTCGGGAGTCGAGA | AATGATCCACCAAAAGCCAG | 229 |
| chr11:134105863-13410788 | p3 | (CAA)7 | ACATCACAACAAAGGCCACA | ACCGTTGATAGGAGGTGACG | 114 |
| chr11:134187971-13418998 | p2 | (AT)8 | CAGTTGATCAGGTGGCAATG | TCTATCATTTGGTCCGCTCC | 105 |
| chr11:134193901-13419591 | p2 | (AT)8 | TGTCATCTTTGGTGTGGCAT | TGAAGGGCAAGTATTCGGAC | 266 |
| chr11:134193901-13419591 | p2 | (AT)8 | TGTCATCTTTGGTGTGGCAT | TGAAGGGCAAGTATTCGGAC | 266 |
| chr11:134193981-13419599 | p2 | (AT)8 | TGTCATCTTTGGTGTGGCAT | TGAAGGGCAAGTATTCGGAC | 186 |
| chr11:134193981-13419599 | p2 | (AT)8 | TGTCATCTTTGGTGTGGCAT | TGAAGGGCAAGTATTCGGAC | 186 |
| chr11:134219283-13422138 | p3 | (ATA)34 | TTTTCGTGTGGAATCGTCAA | GCGAAGTTTGTGGAAAGAGC | 185 |
| chr11:134315917-13431792 | p2 | (TC)6 | TGTAAATTTTCGCCAAAGGG | ATACTGGCTCTGCCACTGCT | 241 |
| chr11:134315917-13431792 | p3 | (GCC)6 | TGTAAATTTTCGCCAAAGGG | ATACTGGCTCTGCCACTGCT | 241 |
| chr11:134328819-13433083 | p2 | (TC)6 | TGTAAATTTTCGCCAAAGGG | ATACTGGCTCTGCCACTGCT | 241 |
| chr11:134328819-13433083 | p3 | (GCC)6 | TGTAAATTTTCGCCAAAGGG | ATACTGGCTCTGCCACTGCT | 241 |
| chr11:134403294-13440530 | p2 | (TG)7 | AAGCCATCTTGGCAGACACT | TAACCCCAAAGAAATGCCAG | 256 |
| chr11:134416593-13441860 | p2 | (AT)7 | AACGTGCACATCTCAAGCAT | ATAGCACCTTTGGCTCCTCA | 225 |
| chr11:134416593-13441860 | p2 | (AT)9 | AACGTGCACATCTCAAGCAT | ATAGCACCTTTGGCTCCTCA | 225 |
| chr11:134416593-13441860 | p2 | (AT)8 | AACGTGCACATCTCAAGCAT | ATAGCACCTTTGGCTCCTCA | 225 |
| chr11:134460856-13446286 | p2 | (TA)6 | AAAAATGCATCGTATTCGGC | AAAACCGGTAAACCGAAACC | 103 |
| chr11:134460856-13446286 | p6 | (CCGGTT)5 | AAAAATGCATCGTATTCGGC | AAAACCGGTAAACCGAAACC | 103 |
| chr11:134470587-13447261 | p6 | (ATAAAA)4 | TCCGTCTGAGGTCAAGGTCT | CAGAAATAGCCAAAGCAGCC | 138 |
| chr11:134523086-13452512 | p3 | (ATT)14 | AGCTTATGCACACGCATCAC | ATGACGCCGTAATAGTTGGG | 162 |
| chr11:134533100-13453517 | p2 | (GT)6 | CCTGATATGCACGGTTTGTG | GCGCTATTTCCCGATCACTA | 200 |
| chr11:134715615-13471763 | p3 | (TAT)6 | ACATCGAGGCTGGAAAATTG | GGACTCGGTAGCAACATCGT | 171 |
| chr11:134800415-13480243 | p6 | (CCGGTT)3 | CTATGGATTTTCGGGGGTTT | GTTAGGGGTGCGCTTAATGA | 115 |

| Marker ID | SSR type | SSR motif | FORWARD PRIMER (5'-3') | REVERSE PRIMER (5'-3') | size (bp) |
|---|---|---|---|---|---|
| chr11:134833612-13483562 | p2 | (TA)6 | AACCCCACAAACTTCACTCG | ATTGAAACGGAGGGAGTCCT | 279 |
| chr11:134848871-13485089 | p2 | (TC)11 | GTGGTCGGGGTAGTTTCTGA | TGTATCCGCGCCATACAATA | 180 |
| chr11:134949509-13495152 | p2 | (TA)7 | AAAATCACGCCACATCAACA | GCATGCCGGTAGTTCAAAGT | 277 |
| chr11:135008467-13501048 | p5 | (TTGAT)3 | AATCAAACTGAACCAAGCCG | CCACCCAAGAAAATTGGCTA | 244 |
| chr11:135008467-13501048 | p6 | (AAAAAT)3 | AATCAAACTGAACCAAGCCG | CCACCCAAGAAAATTGGCTA | 244 |
| chr11:135040094-13504211 | p3 | (TTC)7 | ACCCCACTTGTGGGATTACA | CAAACAAGTCGAGGTCAGCA | 232 |
| chr11:135061142-13506316 | p6 | (TTAAAA)3 | TATTCAATACTCACCGCCCC | TCGGTCCATCATCAATCAAA | 140 |
| chr11:135061142-13506316 | p2 | (AT)14 | TATTCAATACTCACCGCCCC | TCGGTCCATCATCAATCAAA | 140 |
| chr11:135061625-13506365 | p6 | (TTAAAA)3 | CAAATTGAAGGCGTGACTGA | TCCATTTCGTGAATTGCATC | 148 |
| chr11:135061625-13506365 | p2 | (AT)14 | CAAATTGAAGGCGTGACTGA | TCCATTTCGTGAATTGCATC | 148 |
| chr11:135068271-13507028 | p5 | (GGGGT)3 | CTGCAATTAGGCCAAAGGAA | TCAAATGCGAATTGCAAAAG | 173 |
| chr11:135116987-13511899 | p2 | (AG)6 | TTGCACCACATTCTAAGGCA | TGAGCATGACGAAGAAACCA | 119 |
| chr11:135170955-13517297 | p3 | (GAA)5 | AGATGTTCTTTCGCTCCCAA | TTGTTGCAAACACCCCAGTA | 279 |
| chr11:135222085-13522411 | p3 | (CAC)5 | ATTACCACCACCCCAAGTGA | GAATTCACTCCTCCAGCAGC | 193 |
| chr11:135222085-13522411 | p4 | (GTTT)4 | ATTACCACCACCCCAAGTGA | GAATTCACTCCTCCAGCAGC | 193 |
| chr11:135341723-13534373 | p2 | (TA)8 | TCTCATAAACGAGGGGATGC | CCTTCAAGTCCTCTGCTTCG | 113 |
| chr11:135341723-13534373 | p3 | (TTG)5 | TCTCATAAACGAGGGGATGC | CCTTCAAGTCCTCTGCTTCG | 113 |
| chr11:135342000-13534401 | p2 | (TA)8 | CCGTCTAAAGGCTGAACTCG | TTCCAAGCCCACTTCTTTTG | 155 |
| chr11:135342000-13534401 | p3 | (TTG)5 | CCGTCTAAAGGCTGAACTCG | TTCCAAGCCCACTTCTTTTG | 155 |
| chr11:135383231-13538524 | p2 | (CA)6 | ACCCCAAAGCAGCATATCAC | GCGTAGCTAGTAAGGCGTGG | 102 |
| chr11:135567601-13556961 | p3 | (AAC)6 | TCGCGCTCTAATTTCCTCAT | GGCCAATTAAAGGCACAAAA | 281 |
| chr11:135581660-13558367 | p3 | (TGT)5 | AGTGATTTTGGTTGATCGGC | CACTGCACTCCTTCCACAAA | 144 |
| chr11:135633570-13563559 | p4 | (ATAA)4 | CCCATGAGCTGTTCCAAAAT | GTGGGAAGCTTTTTCAACCA | 107 |
| chr11:135633570-13563559 | p2 | (AG)10 | CCCATGAGCTGTTCCAAAAT | GTGGGAAGCTTTTTCAACCA | 107 |
| chr11:135760220-13576225 | p2 | (CT)8 | TTGTGTTTGTGCTGAGGCAT | CTTCTTTGAGGATGGCGAAC | 251 |
| chr11:135760220-13576225 | p2 | (AT)31 | TTGTGTTTGTGCTGAGGCAT | CTTCTTTGAGGATGGCGAAC | 251 |
| chr11:135890842-13589285 | p2 | (TG)10 | GGTTGTGGGTGAATAGGGTG | CATTTATGCCGTCTTCCGTT | 162 |
| chr11:135890842-13589285 | p2 | (TG)7 | GGTTGTGGGTGAATAGGGTG | CATTTATGCCGTCTTCCGTT | 162 |
| chr11:135933237-13593525 | p2 | (TA)7 | AGGTCATTCAAACGATTGGC | CTAAAGACGGCGATTTGAGG | 111 |
| chr11:135985033-13598704 | p5 | (TTTTA)3 | CGAAATTTGGCCCTCTCATA | ATTTTATGCCTGAGCGAGGA | 273 |
| chr11:136016625-13601864 | p3 | (AAC)6 | TACTGTGAACCCTTGTGCCA | CTGCAGCTCAATTGGTGAAA | 253 |
| chr11:136051709-13605372 | p5 | (TGTCT)3 | TTTTGGGAATTGCTGTCTCC | GCTTGAAGAAGCGCTGAGTT | 172 |
| chr11:136066076-13606808 | p2 | (TA)6 | TGAAAGGGAACCAAAGATGG | TTGCTCGACGTCCTCCTAGT | 202 |
| chr11:136066076-13606808 | p2 | (CA)10 | TGAAAGGGAACCAAAGATGG | TTGCTCGACGTCCTCCTAGT | 202 |
| chr11:136073863-13607587 | p3 | (TTG)5 | CCGTTCAACCAAACCTTGAT | CCAAGGTTCAGTTTCCCTGA | 223 |
| chr11:136073863-13607587 | p4 | (AGTG)4 | CCGTTCAACCAAACCTTGAT | CCAAGGTTCAGTTTCCCTGA | 223 |

| Marker ID | SSR type | SSR motif | FORWARD PRIMER (5'-3') | REVERSE PRIMER (5'-3') | size (bp) |
|---|---|---|---|---|---|
| chr11:136073863-13607587 | p2 | (TA)8 | CCGTTCAACCAAACCTTGAT | CCAAGGTTCAGTTTCCCTGA | 223 |
| chr11:136128521-13613053 | p4 | (ATGG)4 | ATGAGTGTCAGCAGTGCAGG | GCCTCTAGAAAACGCACAGG | 294 |
| chr11:136215408-13621742 | p2 | (TA)6 | CGCTTAGCTCCAATTTGTCC | TGTTGATGTGCCGTGAATTT | 177 |
| chr11:136215408-13621742 | p5 | (AACAA)3 | CGCTTAGCTCCAATTTGTCC | TGTTGATGTGCCGTGAATTT | 177 |
| chr11:136359231-13636124 | p2 | (TA)6 | CTGCCACCATAAGCCTGATT | ATGCCCTGGTTGTCTCAGTC | 152 |
| chr11:136359231-13636124 | p2 | (AC)7 | CTGCCACCATAAGCCTGATT | ATGCCCTGGTTGTCTCAGTC | 152 |
| chr11:136509637-13651165 | p5 | (AACCG)3 | CAGTTCGGATATGGTTGGCT | CCAAAAGGGAAGTCAGCTTG | 233 |
| chr11:136587020-13658904 | p2 | (AT)33 | CACACAGAAAAGGCCCAAAT | AATAGCCTTGCCGGCTTTAT | 232 |
| chr11:136587020-13658904 | p2 | (AT)7 | CACACAGAAAAGGCCCAAAT | AATAGCCTTGCCGGCTTTAT | 232 |
| chr11:136587020-13658904 | p2 | (TA)12 | CACACAGAAAAGGCCCAAAT | AATAGCCTTGCCGGCTTTAT | 232 |
| chr11:136689691-13669170 | p3 | (TAT)5 | CACAAGCCGCTCATCAACTA | GGTCTTAATCACAGCCGAGC | 168 |
| chr11:136689691-13669170 | p3 | (TTA)6 | CACAAGCCGCTCATCAACTA | GGTCTTAATCACAGCCGAGC | 168 |
| chr11:136750890-13675290 | p5 | (AAAGA)3 | GGGAAAAGGGCCTGATTTAC | GGGGTTAAAAATAGGGGCAA | 254 |
| chr11:136801116-13680313 | p2 | (TG)6 | CGCAGACCTTACTCCTACCG | GCTCTCCTTCTCGTCTCCAA | 122 |
| chr11:136826894-13682890 | p2 | (AT)7 | GAACGCATCTTTCTCCCTTC | TGCTAGGTTTCACAATGGCA | 241 |
| chr11:136858159-13686018 | p3 | (TCA)6 | GCTCTCCCCAAACATCCATA | GATGATGAAACACAACCCCC | 207 |
| chr11:136858159-13686018 | p6 | (CCGGTT)5 | GCTCTCCCCAAACATCCATA | GATGATGAAACACAACCCCC | 207 |
| chr11:136875761-13687777 | p3 | (AAC)6 | ACGTGACCTTACTTGACGGG | AGAAGTTGCAAAATCCACGG | 261 |
| chr11:136915375-13691739 | p5 | (AATTA)3 | TCTTTGGGCCTTTTTAGCCT | TGCGTATGCCTGTATTCCAA | 228 |
| chr11:137102336-13710435 | p3 | (AAT)5 | AGGGTACAAATGGGGGTAGG | ACATCTCCAACTCACCCCAC | 188 |
| chr11:137102347-13710437 | p3 | (AAT)5 | AGGGTACAAATGGGGGTAGG | ACATCTCCAACTCACCCCAC | 177 |
| chr11:137111873-13711388 | p2 | (CT)6 | AAGGACGAAGTACACCGACG | CTAGTTTGAGTGCATGCCGA | 188 |
| chr11:137158847-13716086 | p2 | (AT)9 | AATCTATGGCCAAACGCCTA | CGGCCAAGGTTTCAAAAGT | 111 |
| chr11:137223882-13722589 | p5 | (AATTT)3 | GTGCTTTGGAGACTTCAGGC | CTGTCCGCTTTGGAGACTTC | 187 |
| chr11:137270219-13727224 | p2 | (AT)12 | AGAGGAATTCACCTCTCGCA | TGCCTAATTTTGTTGGGGTC | 192 |
| chr11:137323630-13732564 | p3 | (AAT)5 | CCTTTTTCGAATCTCTCCCC | CTCTCTCATCTCCACGCTCC | 244 |
| chr11:137326472-13732852 | p2 | (AT)25 | TGGCTTCGGAGAGTGAATTT | TCATTGGTTAATTTCCCCCA | 239 |
| chr11:137338847-13734085 | p2 | (AT)6 | ATTTGTGGGCTGTTTTGAGG | AACGGCTTACTTTCAAGCCA | 285 |
| chr11:137414058-13741607 | p5 | (TTGGT)3 | GTTGGTTGAGCATGGGACTT | CTTGTGCAATAGTCGGCAAA | 288 |
| chr11:137474633-13747664 | p3 | (GGA)5 | CGGATACGTGGGGACTTAAA | TTCCCCCATCCACTACAAAA | 162 |
| chr11:137506088-13750810 | p5 | (TTACT)4 | CCTTGGAAGATCACAACCGT | TATGGAAAGTGGGTTTGGGA | 159 |
| chr11:137506088-13750810 | p2 | (AT)7 | CCTTGGAAGATCACAACCGT | TATGGAAAGTGGGTTTGGGA | 159 |
| chr11:137506088-13750810 | p2 | (TA)8 | CCTTGGAAGATCACAACCGT | TATGGAAAGTGGGTTTGGGA | 159 |
| chr11:137507977-13750998 | p2 | (TA)6 | GTGGGCCCCATACTAAACAA | TTTAAGGGACAACGGACGAC | 298 |
| chr11:137748406-13775044 | p3 | (TAT)14 | AGAGCCACTTTGTGCCTTGT | GCCCTATCTTCCACCAATCA | 298 |
| chr11:137765185-13776720 | p5 | (CGAAC)3 | CTTCACGGCATCAGTCTTGA | TTCGATTTTTCGATTTTCGG | 211 |

| Marker ID | SSR type | SSR motif | FORWARD PRIMER (5'-3') | REVERSE PRIMER (5'-3') | size (bp) |
|---|---|---|---|---|---|
| chr11:137765229-13776724 | p5 | (CGAAC)3 | CTTCACGGCATCAGTCTTGA | TTCGATTTTTCGATTTTCGG | 167 |
| chr11:137848857-13785088 | p4 | (AACA)6 | CGCAGCTCCACCTTCTCTAC | AGAATCTGGGCGACTCAAGA | 164 |
| chr11:137881720-13788373 | p2 | (AT)7 | AGGGAAGAATGGAAAGGGAA | GCGCAGTATTTTAACGCCTC | 234 |
| chr11:137963418-13796543 | p2 | (TA)10 | TGAATGTGAAAGTTGCTGGC | GCGTGTTTAGGGCGTACAAT | 212 |
| chr11:137978512-13798055 | p2 | (AC)9 | AAATGTGTGCTTTTCGTCCC | AGCCCTTTTGGTTCCTCTGT | 117 |
| chr11:138070194-13807221 | p2 | (TA)10 | TGAATGTGAAAGTTGCTGGC | GCGTGTTTAGGGCGTACAAT | 212 |
| chr11:138085288-13808732 | p2 | (AC)9 | AAATGTGTGCTTTTCGTCCC | AGCCCTTTTGGTTCCTCTGT | 117 |
| chr12:74034-76049 | p3 | (TTG)5 | ATGCTGGTGCCCAATAGAAC | GGAAGCTGTCAAGGAGATGC | 190 |
| chr12:240523-242538 | p3 | (AAC)5 | TGCACGAAAAACAGAACCTG | CTTTGATTTGACCCACCGTT | 152 |
| chr12:325516-327532 | p4 | (ATGT)4 | AACCTCGATAATCAGCCACG | TATTTTGATGGCCGAAAAGC | 190 |
| chr12:353579-355595 | p4 | (ATGT)4 | AACCTCGATAATCAGCCACG | TATTTTGATGGCCGAAAAGC | 190 |
| chr12:388990-391002 | p2 | (AG)6 | TCAACAAGGCCCAACTAAGG | TGTTCGGCTTTCCATTTAGG | 271 |
| chr12:388990-391002 | p3 | (GTT)5 | TCAACAAGGCCCAACTAAGG | TGTTCGGCTTTCCATTTAGG | 271 |
| chr12:396848-398863 | p2 | (AG)6 | CTGAATTCGGGAAGCCATAA | GATGTCACATGCAGACCCAC | 137 |
| chr12:396848-398863 | p3 | (GTT)5 | CTGAATTCGGGAAGCCATAA | GATGTCACATGCAGACCCAC | 137 |
| chr12:492997-495015 | p2 | (TA)9 | GAAATGGTGGGTTGAAATGG | AATTGTCCTCCCGTCAACAC | 123 |
| chr12:503790-505856 | p2 | (TA)33 | TATGGCAGGAAATCACACCA | GGCAGGAAATCACACCAGTT | 119 |
| chr12:586451-588517 | p2 | (TA)33 | TATGGCAGGAAATCACACCA | GGCAGGAAATCACACCAGTT | 119 |
| chr12:692519-694534 | p3 | (GTG)5 | CAGCGTGGCTGTTGAATAAA | AAAAACGCGAAATACAACCG | 199 |
| chr12:704900-706920 | p5 | (AAATA)3 | CCTAGGCAACCAACTTTCCA | CTCGAGCCTCTCCCTATCCT | 154 |
| chr12:704900-706920 | p5 | (AAATA)4 | CCTAGGCAACCAACTTTCCA | CTCGAGCCTCTCCCTATCCT | 154 |
| chr12:704900-706920 | p5 | (AAATA)3 | CCTAGGCAACCAACTTTCCA | CTCGAGCCTCTCCCTATCCT | 154 |
| chr12:704900-706920 | p5 | (TTTAT)3 | CCTAGGCAACCAACTTTCCA | CTCGAGCCTCTCCCTATCCT | 154 |
| chr12:707994-710009 | p3 | (TGT)5 | CACATCGTCATACCTGAGCG | CTGGTTTATTTCGCCTCCAA | 201 |
| chr12:741608-743622 | p2 | (AT)7 | TCATCCGAAAACGTCTCCTT | AGCCCTCGAAACCGTAAGTT | 146 |
| chr12:746904-748920 | p4 | (CTGT)4 | TCCTTACCCGATACAGTGGG | TTCGAATTGGTGCAATCTCA | 262 |
| chr12:751330-753345 | p5 | (AAGGG)3 | CTTTGTGGATCATGTGCTGC | TTAGAACAGCGGGAGAACCT | 226 |
| chr12:775663-777681 | p6 | (TTCCAC)3 | CATGATGATGTCCGCTTTTG | TGAGTTTTATGACGTGGGCA | 195 |
| chr12:775924-777942 | p6 | (TTCCAC)3 | CACGGAATTCAATGTGTTGG | TACCGTGGAAGTGGATGTGA | 261 |
| chr12:775991-778009 | p6 | (TTCCAC)3 | CACGGAATTCAATGTGTTGG | TACCGTGGAAGTGGATGTGA | 194 |
| chr12:837526-839541 | p5 | (AAAGG)3 | GGGAGAAGTCCCTGACATGA | CCGCAACTAACACCCAAGAT | 202 |
| chr12:900975-902993 | p6 | (AAGCCG)3 | CAAATTTGTTTTGGGTTGGG | ACTGGTTTTGAACACGGGAG | 138 |
| chr12:905632-907650 | p6 | (CAGTAG)3 | TCTCCGCTGCCTTGTTATCT | TCAAGTTGCGGCAGTGATAG | 281 |
| chr12:1032565-1034586 | p3 | (TTA)12 | TCTCTCATGTCAAGATGCGG | CTTCGGAGAATGGTGGAAGA | 129 |
| chr12:1032565-1034586 | p3 | (TTA)7 | TCTCTCATGTCAAGATGCGG | CTTCGGAGAATGGTGGAAGA | 129 |
| chr12:1117972-1120022 | p2 | (TA)25 | GCCCACTTTGCTAAATCCAA | GGCTTCTTTCGACCAATCAG | 168 |

| Marker ID | SSR type | SSR motif | FORWARD PRIMER (5'-3') | REVERSE PRIMER (5'-3') | size (bp) |
|---|---|---|---|---|---|
| chr12:1173709-1175724 | p5 | (TGTGT)3 | CCCATAGGCTGCTTTCTCAG | TCCATGAACCACCTTGTGAA | 280 |
| chr12:1290917-1292935 | p6 | (ACATGA)3 | AGTGACCCCATTGGAAACAA | CTCGGTCATATGCAGCAAGA | 208 |
| chr12:1367377-1369392 | p5 | (ATTTT)3 | ATGTGGCAATGAGTCCATCA | TGGAACTCACCACATCCAAA | 111 |
| chr12:1383770-1385782 | p2 | (TA)6 | ACCGATCATGTCCTCTACGG | TGGCCCCTGTGGTAGATAAG | 201 |
| chr12:1383770-1385782 | p2 | (GA)10 | ACCGATCATGTCCTCTACGG | TGGCCCCTGTGGTAGATAAG | 201 |
| chr12:1387160-1389180 | p2 | (AT)6 | TGCACACGGAAAAGGTGTAG | TAGGCAAGAACAACCTTCGC | 140 |
| chr12:1387160-1389180 | p3 | (TTA)5 | TGCACACGGAAAAGGTGTAG | TAGGCAAGAACAACCTTCGC | 140 |
| chr12:1387175-1389190 | p2 | (AT)6 | TGCACACGGAAAAGGTGTAG | TAGGCAAGAACAACCTTCGC | 125 |
| chr12:1387175-1389190 | p3 | (TTA)5 | TGCACACGGAAAAGGTGTAG | TAGGCAAGAACAACCTTCGC | 125 |
| chr12:1414363-1416389 | p5 | (ATTTT)3 | GAGTTTGGATGTGGTGGGTT | TGGGCCCCATATTAAACAAA | 110 |
| chr12:1414363-1416389 | p2 | (TA)13 | GAGTTTGGATGTGGTGGGTT | TGGGCCCCATATTAAACAAA | 110 |
| chr12:1475449-1477463 | p2 | (TA)14 | GTTTTGCAAATCTGTCCCGT | TTTGTAGGAAAAGATGCCCG | 231 |
| chr12:1475468-1477488 | p2 | (TA)14 | GTTTTGCAAATCTGTCCCGT | TTTGTAGGAAAAGATGCCCG | 212 |
| chr12:1536390-1538408 | p2 | (AT)9 | GGAGGGGATGGTTAAGTGGT | CAGTTCATTGTCTCCTCCCC | 233 |
| chr12:1536390-1538408 | p2 | (AT)9 | GGAGGGGATGGTTAAGTGGT | CAGTTCATTGTCTCCTCCCC | 233 |
| chr12:1569402-1571417 | p5 | (TATAT)3 | AGACTCCCCCTTTGAGCAAT | ACGGAAATATCGACGTCTCG | 252 |
| chr12:1572424-1574439 | p3 | (AAT)5 | GCAACTCCAAAAGATCCCAA | TTGCTTCGGAGACTGTTCCT | 135 |
| chr12:1620100-1622120 | p6 | (TTCAGT)4 | ATACGTATTGCTTGGGCCTG | ACAGGGAAGCAAGAAGCTGA | 184 |
| chr12:1620100-1622120 | p5 | (CAAAC)4 | ATACGTATTGCTTGGGCCTG | ACAGGGAAGCAAGAAGCTGA | 184 |
| chr12:1714547-1716592 | p3 | (AAG)15 | CCGAACGGTAGAGGAATGAA | TCCCCGACTACGTTATTTGC | 240 |
| chr12:1729326-1731350 | p2 | (AT)12 | ACAACAATTTGCGGTTGACA | AGCCATTGCGTCCATATACC | 139 |
| chr12:1761474-1763489 | p5 | (TTTAT)3 | TATGACAAGTAGCGGGGGAG | TATGTGGGAAGAAGCCAAGG | 294 |
| chr12:1761474-1763489 | p3 | (TAA)10 | TATGACAAGTAGCGGGGGAG | TATGTGGGAAGAAGCCAAGG | 294 |
| chr12:2031061-2033085 | p2 | (AG)12 | CCACCCCTTTCTCTGTTGAA | ATGAGGGATGAAACACGGAG | 126 |
| chr12:2043545-2045569 | p3 | (ATC)8 | ACCAATAGACTGCAGCTCCG | TATAGACGCTGCGTTCATGC | 243 |
| chr12:2054255-2056271 | p2 | (AT)8 | AGGGACCCAAAATTTAACGG | GCAATCGAAAGACGTTTGGT | 168 |
| chr12:2054255-2056271 | p2 | (AT)8 | AGGGACCCAAAATTTAACGG | GCAATCGAAAGACGTTTGGT | 168 |
| chr12:2054255-2056271 | p2 | (AT)8 | AGGGACCCAAAATTTAACGG | GCAATCGAAAGACGTTTGGT | 168 |
| chr12:2054255-2056271 | p2 | (AT)8 | AGGGACCCAAAATTTAACGG | GCAATCGAAAGACGTTTGGT | 168 |
| chr12:2054769-2056785 | p2 | (AT)8 | TCCCTCAAGCTCACCAAGTT | TTGGCTCCGGACTCATTTAC | 126 |
| chr12:2054769-2056785 | p2 | (AT)8 | TCCCTCAAGCTCACCAAGTT | TTGGCTCCGGACTCATTTAC | 126 |
| chr12:2054769-2056785 | p2 | (AT)8 | TCCCTCAAGCTCACCAAGTT | TTGGCTCCGGACTCATTTAC | 126 |
| chr12:2054769-2056785 | p2 | (AT)8 | TCCCTCAAGCTCACCAAGTT | TTGGCTCCGGACTCATTTAC | 126 |
| chr12:2054769-2056785 | p2 | (AT)8 | TCCCTCAAGCTCACCAAGTT | TTGGCTCCGGACTCATTTAC | 126 |
| chr12:2054769-2056785 | p2 | (AT)8 | TCCCTCAAGCTCACCAAGTT | TTGGCTCCGGACTCATTTAC | 126 |
| chr12:2064551-2066565 | p2 | (AT)7 | GTTGCCTTTCGAATCACGTT | ATGGATCGAAGCCATGAAAG | 264 |

| Marker ID | SSR type | SSR motif | FORWARD PRIMER (5'-3') | REVERSE PRIMER (5'-3') | size (bp) |
|---|---|---|---|---|---|
| chr12:2069327-2071342 | p5 | (TTTTG)3 | GCCGAGTAATAGATCGCCAA | TAGTTGACCCAAAGCATCCC | 227 |
| chr12:2127614-2129672 | p2 | (AT)29 | TGACGTGCATGATTTTGGAT | TTGTACTTGGTCTTGCGTGC | 246 |
| chr12:2312703-2314755 | p2 | (TG)26 | CACGATCAGGAAAAAGGGAA | GGTAGCTGAGCAGTTCGTCC | 172 |
| chr12:2312703-2314755 | p2 | (TA)8 | CACGATCAGGAAAAAGGGAA | GGTAGCTGAGCAGTTCGTCC | 172 |
| chr12:2361763-2363775 | p2 | (AC)7 | TTTAAGGGACAACGGACGAC | CCAAAAGGGTGGGTCTCATA | 150 |
| chr12:2361763-2363775 | p2 | (AT)6 | TTTAAGGGACAACGGACGAC | CCAAAAGGGTGGGTCTCATA | 150 |
| chr12:2361763-2363775 | p2 | (AT)6 | TTTAAGGGACAACGGACGAC | CCAAAAGGGTGGGTCTCATA | 150 |
| chr12:2429815-2431869 | p2 | (TA)27 | ATGTGCCCCTTCCAGTTATG | TTTGTCACGACCCAATCAGA | 161 |
| chr12:2564914-2566926 | p2 | (AT)6 | TTCTAAATTCTGGGCATGGG | CGAGGTCCGTCGAACTAGAG | 155 |
| chr12:2600100-2602156 | p2 | (AT)28 | CCTTATTCGGGGTGGTCTTT | ATTGGGTGCTCCATTTTCTG | 122 |
| chr12:2619466-2621481 | p5 | (CCGAA)3 | TATTAGAAGCCCCCATTCCC | CAACGTCATCGTCCACATTC | 282 |
| chr12:2734418-2736458 | p4 | (TTAT)5 | AAGCTATGATCCCCAGCAGA | CTTCCCCTCGGGGTAAATAA | 242 |
| chr12:2734418-2736458 | p2 | (AT)20 | AAGCTATGATCCCCAGCAGA | CTTCCCCTCGGGGTAAATAA | 242 |
| chr12:2736552-2738576 | p6 | (CATCAC)4 | GACGACTTGAAATCCTCCCA | TTTTCAAGCGTGTAAACCCC | 298 |
| chr12:2758845-2760884 | p3 | (CAT)6 | GGAACGTACAGCCCGATCTA | TAGGGTATGGGGAAAGGGAC | 298 |
| chr12:2758845-2760884 | p3 | (AAC)13 | GGAACGTACAGCCCGATCTA | TAGGGTATGGGGAAAGGGAC | 298 |
| chr12:2758845-2760884 | p5 | (TCATT)3 | GGAACGTACAGCCCGATCTA | TAGGGTATGGGGAAAGGGAC | 298 |
| chr12:2794380-2796395 | p6 | (AAACCG)3 | GGAAATGAGCTGGAATCGAA | TGTGCCGTCTCTTCCTCTTT | 210 |
| chr12:2794380-2796395 | p5 | (TAAAA)3 | GGAAATGAGCTGGAATCGAA | TGTGCCGTCTCTTCCTCTTT | 210 |
| chr12:2797653-2799671 | p6 | (AGTTTA)3 | CGACATCAAACGAAACAACG | GGTTGAGGGATTGTTGGAGA | 150 |
| chr12:2882060-2884076 | p4 | (ATTT)4 | TTTGAATGTTGTGCGGGTTA | ACAGAAATTTCGGCAGCATC | 253 |
| chr12:2924881-2926896 | p5 | (CTATT)4 | ACTGATTCCGTAATGGCTGG | TGGGGAGGGTAGAGTGTACG | 248 |
| chr12:2924881-2926896 | p3 | (ACA)5 | ACTGATTCCGTAATGGCTGG | TGGGGAGGGTAGAGTGTACG | 248 |
| chr12:2946838-2948856 | p3 | (TGT)6 | TGATCATTCTTTGCTGACGC | TTTTCATACCACTGCCTCCC | 182 |
| chr12:2957569-2959597 | p2 | (AT)14 | ATACCGACGAAAAAGCGAAA | TTTGGGGATCTCAATGGTTC | 198 |
| chr12:2979868-2981886 | p2 | (TC)9 | GAATGAAGGGGAAAAGAGCC | TCACGTTCCCTCTTTATCGG | 185 |
| chr12:2991199-2993217 | p5 | (AATGG)3 | TATGCACTGGGCATCTGTGT | CAAAAAGTTGAGCTTTCCGC | 138 |
| chr12:2991199-2993217 | p6 | (CCCCCA)3 | TATGCACTGGGCATCTGTGT | CAAAAAGTTGAGCTTTCCGC | 138 |
| chr12:3015661-3017683 | p2 | (TA)7 | TGACCGGAAGTAACAAACCC | TAGGCACCCTTTCTAGGGCT | 109 |
| chr12:3015661-3017683 | p2 | (AT)11 | TGACCGGAAGTAACAAACCC | TAGGCACCCTTTCTAGGGCT | 109 |
| chr12:3044944-3046956 | p2 | (CT)6 | GGAAAGGGGAATTTGTGGTT | GAACCAAAATGGCTCCAAAA | 223 |
| chr12:3069691-3071709 | p3 | (ACA)6 | TTGTGCCCAACACCTTATCA | TTGTCTTCTTGTGCAGACCG | 177 |
| chr12:3097083-3099111 | p4 | (GAAA)7 | GTGGGCAGCAACTAAAGAGC | CCGTTGACTTGGGTTCAGTT | 134 |
| chr12:3162489-3164507 | p3 | (TGT)6 | GCTCCATTACCCACCTCGTA | GATGATTCGACACACATCCG | 154 |
| chr12:3162489-3164507 | p3 | (TGT)6 | GCTCCATTACCCACCTCGTA | GATGATTCGACACACATCCG | 154 |
| chr12:3162489-3164507 | p3 | (TGT)6 | GCTCCATTACCCACCTCGTA | GATGATTCGACACACATCCG | 154 |

| Marker ID | SSR type | SSR motif | FORWARD PRIMER (5'-3') | REVERSE PRIMER (5'-3') | size (bp) |
|---|---|---|---|---|---|
| chr12:3162489-3164507 | p3 | (TGT)6 | GCTCCATTACCCACCTCGTA | GATGATTCGACACACATCCG | 154 |
| chr12:3162489-3164507 | p3 | (TGT)7 | GCTCCATTACCCACCTCGTA | GATGATTCGACACACATCCG | 154 |
| chr12:3292083-3294097 | p2 | (AT)7 | TCCAATTCGGACTGCCTTAC | GCGGTTCTCGCTAAGTATGC | 222 |
| chr12:3292083-3294097 | p2 | (AT)9 | TCCAATTCGGACTGCCTTAC | GCGGTTCTCGCTAAGTATGC | 222 |
| chr12:3355567-3357583 | p4 | (AATA)4 | GTGCTGAACCAATCATGGTG | TGTTCGTGTAGCTCATTGCC | 200 |
| chr12:3355567-3357583 | p4 | (ATAG)4 | GTGCTGAACCAATCATGGTG | TGTTCGTGTAGCTCATTGCC | 200 |
| chr12:3356595-3358610 | p4 | (ATAG)4 | GTTTGCTCCACCAAAGGAAA | GCCCCAAAATAAAATGCTGA | 267 |
| chr12:3356607-3358625 | p4 | (ATAG)4 | GTTTGCTCCACCAAAGGAAA | GCCCCAAAATAAAATGCTGA | 255 |
| chr12:3356633-3358699 | p4 | (ATAG)4 | GTTTGCTCCACCAAAGGAAA | GCCCCAAAATAAAATGCTGA | 229 |
| chr12:3356694-3358760 | p4 | (ATAG)4 | GTTTGCTCCACCAAAGGAAA | GCCCCAAAATAAAATGCTGA | 168 |
| chr12:3398238-3400253 | p5 | (TAAAA)3 | GGTTGGCACTCCTTGGTAGA | GTCAATGGTCTCAGGCTGGT | 294 |
| chr12:3413958-3415976 | p2 | (TA)9 | GACGCCATGAAGAAAGAAGC | TGAATTATCCGATTGCACGA | 173 |
| chr12:3459708-3461723 | p3 | (TCA)5 | CACCTTCGGAACAATGGAGT | ACTGACGTCCGTTTTCCTTG | 127 |
| chr12:3534840-3536855 | p3 | (TCT)5 | AATTCCTGCAAACCGTTGTC | TAGGGCCCAACAAAAACAAG | 125 |
| chr12:3557876-3559894 | p6 | (AACTGA)3 | TCCGTACCGGCCATAGTAAG | GCCGGGTACAGATACCACTG | 103 |
| chr12:3595669-3597719 | p2 | (AT)25 | CTGACTGAGGAGCTGGAACC | ATGTTTACAGCGATGCCTCC | 208 |
| chr12:3685696-3687720 | p6 | (TTTCAG)4 | GAATGGGAATTGGGTGAAGA | CAAGCATGAAGAATGAGGCA | 139 |
| chr12:3788528-3790546 | p2 | (AT)9 | CCTAGGACGGAACTGCTCTG | GGCTTGCACTGTTTTTCCTC | 209 |
| chr12:3788528-3790546 | p2 | (TA)9 | CCTAGGACGGAACTGCTCTG | GGCTTGCACTGTTTTTCCTC | 209 |
| chr12:3801101-3803115 | p2 | (AT)7 | TTTGAGTGCTGCAAATCTGG | CAGAGGATGACGTGTCCCTT | 109 |
| chr12:3801101-3803115 | p2 | (AT)6 | TTTGAGTGCTGCAAATCTGG | CAGAGGATGACGTGTCCCTT | 109 |
| chr12:3873846-3875916 | p2 | (AG)10 | GATTTTGGCGAGAAATGGAA | CAAAACAAACGCAACCCTCT | 185 |
| chr12:3873941-3875955 | p2 | (AG)10 | AGAGGGTTGCGTTTGTTTTG | CCCATTACCTGACTAGCCCA | 170 |
| chr12:3873957-3875971 | p2 | (AG)10 | AGAGGGTTGCGTTTGTTTTG | CCCATTACCTGACTAGCCCA | 154 |
| chr12:3873973-3875987 | p2 | (AG)10 | AGAGGGTTGCGTTTGTTTTG | CCCATTACCTGACTAGCCCA | 138 |
| chr12:3873989-3876003 | p2 | (AG)10 | AGAGGGTTGCGTTTGTTTTG | CCCATTACCTGACTAGCCCA | 122 |
| chr12:3874005-3876019 | p2 | (AG)10 | AGAGGGTTGCGTTTGTTTTG | CCCATTACCTGACTAGCCCA | 106 |
| chr12:3986246-3988264 | p6 | (ACACCC)3 | GGATCAGATTCTCAAGCCCA | TAATTGCTTGGCAAAGACCC | 115 |
| chr12:4087696-4089711 | p5 | (AAACC)3 | CAATCTCCTCAGCCAGCTTC | CATGGGTGACAAATCGACTG | 129 |
| chr12:4106121-4108136 | p5 | (AAACC)3 | CAATCTCCTCAGCCAGCTTC | CATGGGTGACAAATCGACTG | 129 |
| chr12:4210378-4212390 | p2 | (AT)6 | AAGGGAGGAAGGGACAAGAA | GGCCACGAGTAACCGAATTA | 179 |
| chr12:4210378-4212390 | p5 | (TAAAA)3 | AAGGGAGGAAGGGACAAGAA | GGCCACGAGTAACCGAATTA | 179 |
| chr12:4261183-4263195 | p2 | (AT)6 | AAGGGAGGAAGGGACAAGAA | GGCCACGAGTAACCGAATTA | 179 |
| chr12:4261183-4263195 | p5 | (TAAAA)3 | AAGGGAGGAAGGGACAAGAA | GGCCACGAGTAACCGAATTA | 179 |
| chr12:4330051-4332069 | p3 | (AAT)6 | TTGGTGAGAGTTCGATTCCC | GTTGATGTTGCCGAATGATG | 275 |
| chr12:4465136-4467151 | p3 | (TCA)5 | GGGTCCCATCTCAGCAAATA | CCGCTGATTTGGGTGTTAGT | 275 |

| Marker ID | SSR type | SSR motif | FORWARD PRIMER (5'-3') | REVERSE PRIMER (5'-3') | size (bp) |
|---|---|---|---|---|---|
| chr12:4481351-4483366 | p3 | (CAT)5 | CAAGATGTGATCGGCAAGTG | GAGCAGCCTTTCAAAACGTC | 154 |
| chr12:4494053-4496068 | p5 | (AAATA)3 | GGTCGTCTTGACCCTGTGTT | ACATGGTTCCAGAGGCAATC | 279 |
| chr12:4526728-4528756 | p2 | (GA)6 | TCAATATTGGGCCCTTGTTT | AGGATGGTCCACATGGTAGC | 189 |
| chr12:4526766-4528782 | p2 | (GA)6 | TCAATATTGGGCCCTTGTTT | AGGATGGTCCACATGGTAGC | 151 |
| chr12:4669679-4671694 | p2 | (AT)14 | GACACACCCTAGGACGGAAA | GACGACATATGTAGCGGCCT | 280 |
| chr12:4669719-4671777 | p2 | (AT)14 | GACACACCCTAGGACGGAAA | GACGACATATGTAGCGGCCT | 240 |
| chr12:4825336-4827405 | p3 | (TAT)23 | TCGTTTGCCTTGTTTGAATG | GGCTGCAAAGGAGTACAAGC | 259 |
| chr12:5097665-5099677 | p2 | (TC)6 | GGTGGCTTCCATATTGTGCT | AGGAGAGACCACGGTTCCTT | 240 |
| chr12:5158666-5160678 | p2 | (AT)6 | ACATGCACATTGCGTCTCAT | GTGACTATTGGTTGGCCTGC | 234 |
| chr12:5158666-5160678 | p2 | (AT)7 | ACATGCACATTGCGTCTCAT | GTGACTATTGGTTGGCCTGC | 234 |
| chr12:5242854-5244870 | p2 | (TG)6 | TACGAAAGCCAGGTAAGCGT | GATATGTTGCTCCGAAGGGA | 134 |
| chr12:5242872-5244888 | p2 | (TG)6 | TACGAAAGCCAGGTAAGCGT | GATATGTTGCTCCGAAGGGA | 116 |
| chr12:5264583-5266603 | p4 | (TTCT)5 | ACAAGCTTTCAGCAAGGCAT | CCACTTTAATCGGGTGTGGT | 242 |
| chr12:5297777-5299789 | p2 | (AT)6 | TTACATACCTTTCGCGCTCC | CGTTTGGGTTGTTGTTGATG | 287 |
| chr12:5365999-5368019 | p5 | (AAACC)4 | ATTTTCATTCCAGGTCGTGC | TTCACATTAAGCTTCGGGCT | 226 |
| chr12:5419738-5421756 | p6 | (TTCAGT)3 | CTTTTTGCCCACCAAGTGTT | TAACTTCGGCCTTTGAGCAT | 292 |
| chr12:5479246-5481262 | p4 | (TTAA)4 | AGGACGGTTTTCAGGGAGTT | ATGTGGTATGCTCGGTCACA | 104 |
| chr12:5638113-5640129 | p4 | (TGAT)4 | CTCGGAAGGATCTGGAATCA | GGATAGGGTCACGGAGGATT | 180 |
| chr12:5659864-5661882 | p2 | (TA)9 | ATTATTCCGAGTGACGTGCC | GTTCAAGTCGGATCATCGGT | 129 |
| chr12:5703739-5705755 | p2 | (TA)8 | GAACGCATACACGACTTCCA | TTGGAATCTCCAGAAATGGG | 134 |
| chr12:5714070-5716085 | p5 | (ATATA)3 | TTCAATTGGTGGTGTAGCGA | TGAAATCTTGCAGCCAGTTG | 143 |
| chr12:5778191-5780203 | p2 | (GT)6 | GTGAAAACCCTCCAAACGAA | GGGAACCACGACAATAATGG | 255 |
| chr12:5795945-5798001 | p2 | (AT)28 | TCTCCAAGCCCTAGAACGAA | CACACGTGGATAGAAGCCCT | 116 |
| chr12:5813893-5815908 | p5 | (CATTG)3 | CCCCTTGCTCCATAGTTGAA | TCCCAACTTCATACCCGAAG | 231 |
| chr12:5813893-5815908 | p2 | (TA)17 | CCCCTTGCTCCATAGTTGAA | TCCCAACTTCATACCCGAAG | 231 |
| chr12:5824492-5826506 | p2 | (AT)7 | GTAGCTCCGGATGAAATGGA | TGCTGACGTCCTTTTGTTTG | 260 |
| chr12:5870338-5872350 | p2 | (TC)6 | TCTGAAATGACTTGGAGGGG | GGAGCCTCGATCTTAGCCTT | 126 |
| chr12:5872343-5874365 | p2 | (AT)11 | AGCACAATTCCCAAACCAAA | TTGGATTGTGCTTTCTTCCC | 102 |
| chr12:5874796-5876814 | p5 | (AACCG)4 | GATTTGGGTTTGGGATGATG | TTTTGACGATGGACACCAAA | 283 |
| chr12:5874796-5876814 | p6 | (AAACCG)3 | GATTTGGGTTTGGGATGATG | TTTTGACGATGGACACCAAA | 283 |
| chr12:5926214-5928229 | p3 | (TGT)5 | TCCAAAGGGATGATTCAAGG | CAGGGTTAATTCTCTGCCCA | 268 |
| chr12:5926214-5928229 | p6 | (TATATG)3 | TCCAAAGGGATGATTCAAGG | CAGGGTTAATTCTCTGCCCA | 268 |
| chr12:5981930-5983942 | p2 | (TA)6 | TATGAAGAAGTCCCCATCGC | TGGAAAATGGTCTTGCCTTC | 300 |
| chr12:5981930-5983942 | p2 | (GT)6 | TATGAAGAAGTCCCCATCGC | TGGAAAATGGTCTTGCCTTC | 300 |
| chr12:6065683-6067697 | p2 | (TA)7 | GAATGGTAGGGATGGGGACT | TTTTGCCCTTCAAATGGAAC | 244 |
| chr12:6079725-6081741 | p4 | (ATGT)9 | TGCTCATTAGGATCTGCACG | CGGCTAACCAACTCTGGAAG | 227 |

| Marker ID | SSR type | SSR motif | FORWARD PRIMER (5'-3') | REVERSE PRIMER (5'-3') | size (bp) |
|---|---|---|---|---|---|
| chr12:6079725-6081741 | p2 | (TA)8 | TGCTCATTAGGATCTGCACG | CGGCTAACCAACTCTGGAAG | 227 |
| chr12:6281921-6283939 | p2 | (AT)9 | GTAACGGTAGGATGGGGGTT | TTGACCCAGAAGGATGTTCC | 141 |
| chr12:6318845-6320863 | p3 | (CAA)6 | ATACTGCCCAGCCATGTAGG | AATCGGAGGTTACCGACCTT | 227 |
| chr12:6325205-6327223 | p3 | (CAA)6 | TCCATGTAGGCACGGTGTTA | AGCTTTCACTTGGGCTCTCA | 221 |
| chr12:6374548-6376592 | p2 | (TA)22 | TTTTAAAAGGGGAACGACCC | CAACTTGTGCCAAAAACCCT | 181 |
| chr12:6572239-6574257 | p6 | (AAACTG)3 | TGGGCCATTAGAGCTGTCTT | TGTTTCGGAGTTGCTCTGTG | 288 |
| chr12:6572239-6574257 | p2 | (AT)13 | TGGGCCATTAGAGCTGTCTT | TGTTTCGGAGTTGCTCTGTG | 288 |
| chr12:6572239-6574257 | p2 | (AT)9 | TGGGCCATTAGAGCTGTCTT | TGTTTCGGAGTTGCTCTGTG | 288 |
| chr12:6665238-6667256 | p2 | (AT)7 | TGTGTGCAGGCAGTCCTTAG | CTCCCTAGGGCAGACCTTCT | 260 |
| chr12:6665238-6667256 | p2 | (AT)9 | TGTGTGCAGGCAGTCCTTAG | CTCCCTAGGGCAGACCTTCT | 260 |
| chr12:6665238-6667256 | p2 | (AT)8 | TGTGTGCAGGCAGTCCTTAG | CTCCCTAGGGCAGACCTTCT | 260 |
| chr12:6665647-6667663 | p2 | (AT)7 | GTGCCTGAACTGGACACTCA | TGCGTATGGCATTGATCTGT | 178 |
| chr12:6665647-6667663 | p2 | (AT)9 | GTGCCTGAACTGGACACTCA | TGCGTATGGCATTGATCTGT | 178 |
| chr12:6665647-6667663 | p2 | (AT)8 | GTGCCTGAACTGGACACTCA | TGCGTATGGCATTGATCTGT | 178 |
| chr12:6679962-6681978 | p2 | (TA)8 | AAAATGATACGCAAGCCGAC | CTGTCCCGTCATATTTCGCT | 273 |
| chr12:6679962-6681978 | p2 | (AT)8 | AAAATGATACGCAAGCCGAC | CTGTCCCGTCATATTTCGCT | 273 |
| chr12:6691236-6693250 | p2 | (AT)7 | CCCGAACTGGACACTCGTAT | CCGTGTATAGTGGCAGCCTT | 267 |
| chr12:6701728-6703743 | p5 | (TGTGG)3 | CCTATGAGCCACGGAGTTGT | GGTTCATAGGCCAGGCAATA | 224 |
| chr12:6749346-6751364 | p2 | (AT)31 | TGTTTTACGAGCCGATAGGG | CACTCCACTTGTTCCGGAGT | 190 |
| chr12:6749346-6751364 | p2 | (AT)9 | TGTTTTACGAGCCGATAGGG | CACTCCACTTGTTCCGGAGT | 190 |
| chr12:6749346-6751364 | p2 | (AT)9 | TGTTTTACGAGCCGATAGGG | CACTCCACTTGTTCCGGAGT | 190 |
| chr12:6749346-6751364 | p2 | (AT)9 | TGTTTTACGAGCCGATAGGG | CACTCCACTTGTTCCGGAGT | 190 |
| chr12:6749346-6751364 | p2 | (AT)9 | TGTTTTACGAGCCGATAGGG | CACTCCACTTGTTCCGGAGT | 190 |
| chr12:6749346-6751364 | p2 | (AT)9 | TGTTTTACGAGCCGATAGGG | CACTCCACTTGTTCCGGAGT | 190 |
| chr12:6749346-6751364 | p2 | (AT)9 | TGTTTTACGAGCCGATAGGG | CACTCCACTTGTTCCGGAGT | 190 |
| chr12:6749346-6751364 | p2 | (AT)9 | TGTTTTACGAGCCGATAGGG | CACTCCACTTGTTCCGGAGT | 190 |
| chr12:6749514-6751532 | p2 | (AT)31 | ACTCCGGAACAAGTGGAGTG | TGCATGCAGGCATAACATTT | 169 |
| chr12:6749514-6751532 | p2 | (AT)9 | ACTCCGGAACAAGTGGAGTG | TGCATGCAGGCATAACATTT | 169 |
| chr12:6749514-6751532 | p2 | (AT)9 | ACTCCGGAACAAGTGGAGTG | TGCATGCAGGCATAACATTT | 169 |
| chr12:6749514-6751532 | p2 | (AT)9 | ACTCCGGAACAAGTGGAGTG | TGCATGCAGGCATAACATTT | 169 |
| chr12:6749514-6751532 | p2 | (AT)9 | ACTCCGGAACAAGTGGAGTG | TGCATGCAGGCATAACATTT | 169 |
| chr12:6749514-6751532 | p2 | (AT)9 | ACTCCGGAACAAGTGGAGTG | TGCATGCAGGCATAACATTT | 169 |
| chr12:6749514-6751532 | p2 | (AT)9 | ACTCCGGAACAAGTGGAGTG | TGCATGCAGGCATAACATTT | 169 |
| chr12:6749514-6751532 | p2 | (AT)9 | ACTCCGGAACAAGTGGAGTG | TGCATGCAGGCATAACATTT | 169 |
| chr12:6749514-6751532 | p2 | (AT)9 | ACTCCGGAACAAGTGGAGTG | TGCATGCAGGCATAACATTT | 169 |
| chr12:6749682-6751700 | p2 | (AT)31 | CGCTGTGAGCCAATGATATG | AGGGAAAGTGGCTCCGTTAT | 226 |

| Marker ID | SSR type | SSR motif | FORWARD PRIMER (5'-3') | REVERSE PRIMER (5'-3') | size (bp) |
|---|---|---|---|---|---|
| chr12:6749682-6751700 | p2 | (AT)9 | CGCTGTGAGCCAATGATATG | AGGGAAAGTGGCTCCGTTAT | 226 |
| chr12:6749682-6751700 | p2 | (AT)9 | CGCTGTGAGCCAATGATATG | AGGGAAAGTGGCTCCGTTAT | 226 |
| chr12:6749682-6751700 | p2 | (AT)9 | CGCTGTGAGCCAATGATATG | AGGGAAAGTGGCTCCGTTAT | 226 |
| chr12:6749682-6751700 | p2 | (AT)9 | CGCTGTGAGCCAATGATATG | AGGGAAAGTGGCTCCGTTAT | 226 |
| chr12:6749682-6751700 | p2 | (AT)9 | CGCTGTGAGCCAATGATATG | AGGGAAAGTGGCTCCGTTAT | 226 |
| chr12:6749682-6751700 | p2 | (AT)9 | CGCTGTGAGCCAATGATATG | AGGGAAAGTGGCTCCGTTAT | 226 |
| chr12:6749682-6751700 | p2 | (AT)9 | CGCTGTGAGCCAATGATATG | AGGGAAAGTGGCTCCGTTAT | 226 |
| chr12:6749682-6751700 | p2 | (AT)9 | CGCTGTGAGCCAATGATATG | AGGGAAAGTGGCTCCGTTAT | 226 |
| chr12:6749682-6751700 | p2 | (AT)9 | CGCTGTGAGCCAATGATATG | AGGGAAAGTGGCTCCGTTAT | 226 |
| chr12:6771299-6773327 | p2 | (AT)14 | TAAATACCCTTAACCGCCCC | AACCAAAAACCCAAAAACCC | 108 |
| chr12:6822663-6824703 | p2 | (AT)20 | TTGGCATGGCATGAAGAATA | TGTCGACTTTCCCTGAATCC | 276 |
| chr12:6833331-6835345 | p2 | (AT)7 | TTGGCATGGCATGAAGAATA | TGTCGACTTTCCCAGAATCC | 281 |
| chr12:6847783-6849799 | p4 | (TTCT)4 | ATTCATGGACTGTTTTGGGC | CTGTCCACGGTTTTTGGACT | 238 |
| chr12:6874292-6876307 | p3 | (ATG)5 | AGAGCAAAATTGCACCGACT | TTGAATCCAAGCCAAGATCC | 209 |
| chr12:6874292-6876307 | p2 | (AT)8 | AGAGCAAAATTGCACCGACT | TTGAATCCAAGCCAAGATCC | 209 |
| chr12:6883612-6885633 | p3 | (TGT)7 | CCTCTGGAACATTCTCCCAA | TCCACCGCAACACGATACTA | 299 |
| chr12:6883612-6885633 | p3 | (ATG)7 | CCTCTGGAACATTCTCCCAA | TCCACCGCAACACGATACTA | 299 |
| chr12:6883612-6885633 | p2 | (AT)30 | CCTCTGGAACATTCTCCCAA | TCCACCGCAACACGATACTA | 299 |
| chr12:6911425-6913443 | p6 | (AAAAGA)3 | TCCATCCAGATTATGGCCTC | ACAGGGGCAATGGTAGACAG | 118 |
| chr12:6917144-6919156 | p2 | (AT)6 | GGCACTTGAAGCTAACTGGG | GCGAGCTTGCCTTTAGAAGA | 230 |
| chr12:6959113-6961141 | p2 | (AT)14 | AATGGTGCCTCTTACGGATG | GCCCAAATTTTCCCAGAAAT | 144 |
| chr12:7079136-7081148 | p2 | (CT)6 | CCAGCAGTCACACGACAACT | ATTTTGTTGAATTGAGGCCG | 101 |
| chr12:7130808-7132853 | p2 | (TA)9 | GAAAGTAGGGTAATGCCCCC | TTGGTCCGGTTTCTCCATAG | 236 |
| chr12:7130808-7132853 | p2 | (TA)9 | GAAAGTAGGGTAATGCCCCC | TTGGTCCGGTTTCTCCATAG | 236 |
| chr12:7198383-7200401 | p2 | (TA)14 | ACAAGAATGTGCCACCACAA | TCCTAATCCTGTCACCCCTG | 250 |
| chr12:7198383-7200401 | p5 | (ATATA)3 | ACAAGAATGTGCCACCACAA | TCCTAATCCTGTCACCCCTG | 250 |
| chr12:7198456-7200468 | p2 | (TA)14 | ACAAGAATGTGCCACCACAA | TCCTAATCCTGTCACCCCTG | 177 |
| chr12:7198456-7200468 | p5 | (ATATA)3 | ACAAGAATGTGCCACCACAA | TCCTAATCCTGTCACCCCTG | 177 |
| chr12:7324224-7326236 | p2 | (TA)6 | TCCAAACCAAACCTTTACGG | TAATCTCCCACTCCCTCACG | 222 |
| chr12:7510754-7512814 | p2 | (AT)30 | ATTGTTGCCTGATTTCCGAG | AAGGGGCCAGGCTAAATAAA | 144 |
| chr12:7539305-7541323 | p3 | (ATT)6 | TATGTTGATTCAGCCAGCCA | ATACCAAGCTTTGTCACGCC | 239 |
| chr12:7572058-7574078 | p2 | (TA)10 | TGGAATCGAGGGTTTTCAAC | GTGACTACCCGGCCAACTAA | 203 |
| chr12:7572058-7574078 | p2 | (TA)7 | TGGAATCGAGGGTTTTCAAC | GTGACTACCCGGCCAACTAA | 203 |
| chr12:7628962-7630977 | p3 | (AGA)5 | GCAAAAGTGGGGGTGAACTA | TCGTCATCCGTGAATGTGAT | 144 |
| chr12:7693496-7695543 | p2 | (AT)6 | AAATGAGGTGTTATTGGGCG | GAAGGGTTGCTCAGAAATGG | 202 |
| chr12:7693496-7695543 | p2 | (TA)7 | AAATGAGGTGTTATTGGGCG | GAAGGGTTGCTCAGAAATGG | 202 |

| Marker ID | SSR type | SSR motif | FORWARD PRIMER (5'-3') | REVERSE PRIMER (5'-3') | size (bp) |
|---|---|---|---|---|---|
| chr12:7693542-7695556 | p2 | (AT)6 | AAATGAGGTGTTATTGGGCG | GAAGGGTTGCTCAGAAATGG | 156 |
| chr12:7693542-7695556 | p2 | (TA)7 | AAATGAGGTGTTATTGGGCG | GAAGGGTTGCTCAGAAATGG | 156 |
| chr12:7924905-7926937 | p2 | (AG)16 | CCCGAAGAATACCCACATTG | TGAGCTCACATTAAGCCGTG | 174 |
| chr12:7942811-7944839 | p2 | (TA)14 | GCTTCTCATAGGGGCTTTCC | ACCTTCGTGGTCAGGTATGC | 289 |
| chr12:7969489-7971537 | p2 | (TC)24 | TTGGATCGACAGGATCAACA | TAAGGGTAAGCATTTTGGCG | 274 |
| chr12:7981749-7983761 | p2 | (AT)6 | CGTCGTCCGTTATCCCTTTA | AAACGATACAGAGCCCAGGA | 267 |
| chr12:7981749-7983761 | p2 | (AT)6 | CGTCGTCCGTTATCCCTTTA | AAACGATACAGAGCCCAGGA | 267 |
| chr12:7981749-7983761 | p2 | (AT)6 | CGTCGTCCGTTATCCCTTTA | AAACGATACAGAGCCCAGGA | 267 |
| chr12:8124439-8126451 | p3 | (TTG)5 | GTTGATTGTTTTGGCCGAGT | CCCGTTTCACTCCGAATCTA | 261 |
| chr12:8124439-8126451 | p2 | (AT)6 | GTTGATTGTTTTGGCCGAGT | CCCGTTTCACTCCGAATCTA | 261 |
| chr12:8170275-8172296 | p3 | (AAC)7 | ATTGGGCAACACTTCCTTTG | CCTTGAGAGGGCATTTTGAG | 198 |
| chr12:8213549-8215561 | p2 | (AC)6 | AGAACGAAGGGGTAAGCCAT | TCTCGGGGGTGGTGTATTTA | 230 |
| chr12:8262488-8264502 | p2 | (TG)7 | GAATAGGGGGATCACGACCT | GTGTATACGGTGCCTGCCTT | 128 |
| chr12:8262488-8264502 | p2 | (AT)9 | GAATAGGGGGATCACGACCT | GTGTATACGGTGCCTGCCTT | 128 |
| chr12:8373791-8375806 | p3 | (TTC)5 | AGATTTGGCCAGCCCTTATT | CAAGTGGCATGAGCTTCTGA | 208 |
| chr12:8373791-8375806 | p3 | (CTT)14 | AGATTTGGCCAGCCCTTATT | CAAGTGGCATGAGCTTCTGA | 208 |
| chr12:8436009-8438024 | p5 | (TGCAT)3 | CTTGGTAGGGCCAGGTATGA | AACTTTCCGTTTCAAATGCG | 101 |
| chr12:8558908-8560932 | p6 | (CATATA)4 | GATGGAGCCGCTTACAGAAC | GTGTTGCCGGGTCAAAGTAT | 236 |
| chr12:8766292-8768313 | p3 | (TGT)7 | CAGTGAAATCCCACAACGTG | CGCTTTGAATCTCTTAGCCG | 277 |
| chr12:8794954-8796968 | p2 | (CA)7 | CGGTCGATTTTTCCACCTAA | ATGAGAATTCAGGCCGTTTG | 103 |
| chr12:8894208-8896247 | p6 | (AGAAGG)3 | GAACCCTAATTCAGGAGGAGG | CCTTCTCCCCCTTCTTCTTC | 262 |
| chr12:8896700-8898718 | p3 | (AGG)6 | GGGGAGAAGGAGAAGGAGAA | CCTCCTCCTGAATTAGGGTTC | 276 |
| chr12:8896721-8898739 | p3 | (AGG)6 | GGGGAGAAGGAGAAGGAGAA | CCTCCTCCTGAATTAGGGTTC | 255 |
| chr12:8896760-8898778 | p3 | (AGG)6 | GGGGAGAAGGAGAAGGAGAA | CCTCCTCCTGAATTAGGGTTC | 216 |
| chr12:8896787-8898805 | p3 | (AGG)6 | GGGGAGAAGGAGAAGGAGAA | CCTCCTCCTGAATTAGGGTTC | 189 |
| chr12:8896845-8898863 | p3 | (AGG)6 | GGGGAGAAGGAGAAGGAGAA | CCTCCTCCTGAATTAGGGTTC | 131 |
| chr12:8925144-8927190 | p2 | (AT)23 | ACAGCTTCTTCCTGAGCTGC | GCGTCTATGGAACCGTGTCT | 118 |
| chr12:8975855-8977867 | p2 | (TA)12 | CCATTTGGTCAGGTGATCCT | ACGGTCCGTACTTGAAATCG | 148 |
| chr12:8975880-8977900 | p2 | (TA)12 | CCATTTGGTCAGGTGATCCT | ACGGTCCGTACTTGAAATCG | 123 |
| chr12:9064642-9066656 | p6 | (TCCACC)3 | TCCAAACTTCCAGCTCTCGT | CAATGATGGTGAATTGCCAG | 113 |
| chr12:9064642-9066656 | p6 | (TCTTCC)4 | TCCAAACTTCCAGCTCTCGT | CAATGATGGTGAATTGCCAG | 113 |
| chr12:9077527-9079543 | p5 | (TATTT)3 | CACATCCCATCTGACAATCG | GTTGGCTTAACCGAGCAGAG | 161 |
| chr12:9077527-9079543 | p4 | (TAAT)4 | CACATCCCATCTGACAATCG | GTTGGCTTAACCGAGCAGAG | 161 |
| chr12:9159790-9161822 | p2 | (AT)13 | CGTTTTCGCTACTAGGTGGG | CCGTCCTCGTGGATTTATGT | 228 |
| chr12:9159887-9161937 | p2 | (AT)13 | CGTTTTCGCTACTAGGTGGG | CCGTCCTCGTGGATTTATGT | 131 |
| chr12:9241480-9243495 | p3 | (CAA)5 | TCCCTTTTTCCCATATTCCC | AATCGGAGGTTACCGACCTT | 180 |

| Marker ID | SSR type | SSR motif | FORWARD PRIMER (5'-3') | REVERSE PRIMER (5'-3') | size (bp) |
|---|---|---|---|---|---|
| chr12:9298532-9300559 | p3 | (CTT)9 | TGGGCACAGTTGTTGATGTT | TTGGTTCCTTTGCCTGAAAC | 169 |
| chr12:9391596-9393614 | p2 | (AT)9 | TTTACGCTCACAGATGCCAG | GTCGGCTTGCGTATCATTTT | 139 |
| chr12:9477922-9479934 | p2 | (AT)6 | CAGTTGTTGTTTGCAATCGG | GAGGAGGACACCAAGCTGAG | 206 |
| chr12:9487114-9489129 | p5 | (ATCAT)3 | CGAGCCCGGACTAGATGATA | CGTTGTGAACCTTTGGGTCT | 111 |
| chr12:9523932-9525950 | p2 | (AT)16 | AAGGCTGCCACTACACATCC | CTGTCCCGTCATATTTCGCT | 136 |
| chr12:9523932-9525950 | p2 | (AT)6 | AAGGCTGCCACTACACATCC | CTGTCCCGTCATATTTCGCT | 136 |
| chr12:9523932-9525950 | p2 | (AT)9 | AAGGCTGCCACTACACATCC | CTGTCCCGTCATATTTCGCT | 136 |
| chr12:9697310-9699332 | p2 | (TA)11 | TGATGCTTTTCCTTGTGTGC | GTAACCATGGCTATGTCGGG | 178 |
| chr12:9697310-9699332 | p2 | (TA)8 | TGATGCTTTTCCTTGTGTGC | GTAACCATGGCTATGTCGGG | 178 |
| chr12:9704860-9706878 | p2 | (AT)9 | AAGGTGTCATGAGATTCGGC | TGAGGATAACCCCGTCTGTC | 211 |
| chr12:9793066-9795098 | p2 | (AT)16 | CTCCATTTGATCCGGAGTGT | AACCCCGTAACCCACATACA | 202 |
| chr12:9815943-9817993 | p2 | (AT)25 | CAGGTTGATATACGGGTGCC | CGACGACAAGGTGGTAGGAT | 188 |
| chr12:9815943-9817993 | p2 | (AT)6 | CAGGTTGATATACGGGTGCC | CGACGACAAGGTGGTAGGAT | 188 |
| chr12:9849376-9851392 | p4 | (TGTA)4 | AGCCTCGGGGATTATTCTGT | TTTGGAAGCATACCGACTCC | 222 |
| chr12:9859861-9861877 | p4 | (TATG)4 | CACACCCAAATTTCAAGCCT | GTTTATCCCACATTGCCACC | 178 |
| chr12:9891916-9893934 | p3 | (TTG)6 | AAGCCCTCCAAAAGGATGAT | TTTGGCAGCTTGGAGTCTTT | 191 |
| chr12:9951594-9953609 | p3 | (AAC)5 | AACCTCGGTGACCACTCATC | GTGCTTACGCCTAGCCATTC | 246 |
| chr12:9979069-9981083 | p2 | (AT)7 | TCCAAATAACCGGTCTCCTG | CTGTCCCGTCATATTTCGCT | 299 |
| chr12:10039500-10041514 | p2 | (AT)7 | ATTTCAGCAGGGAAGTGGTG | AGAAAAGTTGCCGTCCAATG | 117 |
| chr12:10047406-10049421 | p5 | (TATGA)3 | CAGCTGTCTTGGAAAGCTCC | GCTGCCACTACAGACGAACA | 119 |
| chr12:10123613-10125625 | p2 | (AG)6 | CAATAAAGAATCGGCGGTGT | CAGGGCTATGATGTTGCCTT | 215 |
| chr12:10123613-10125625 | p5 | (TTGTA)3 | CAATAAAGAATCGGCGGTGT | CAGGGCTATGATGTTGCCTT | 215 |
| chr12:10131910-10133925 | p6 | (GGTGGG)3 | TAGCCCTGTGGGCAACTATC | AGTTCTCCCTTGTCGGGAAT | 278 |
| chr12:10169368-10171383 | p5 | (AAAGG)3 | TGGCCTCGGGTATTAGTCTG | TACCGTTTTTACGGTAGCGG | 169 |
| chr12:10207291-10209311 | p2 | (AT)6 | CCTTAAAGGCCGGACAAGAT | TCCTGCATGCTTAGCCTTTT | 235 |
| chr12:10207291-10209311 | p2 | (AT)10 | CCTTAAAGGCCGGACAAGAT | TCCTGCATGCTTAGCCTTTT | 235 |
| chr12:10207291-10209311 | p2 | (AT)6 | CCTTAAAGGCCGGACAAGAT | TCCTGCATGCTTAGCCTTTT | 235 |
| chr12:10253181-10255197 | p3 | (AAC)5 | CCTTTTCGCTCCTTGAACTG | CCCCTTCTTCATCCAAAACA | 281 |
| chr12:10253199-10255215 | p3 | (AAC)5 | CCTTTTCGCTCCTTGAACTG | CCCCTTCTTCATCCAAAACA | 263 |
| chr12:10363488-10365503 | p3 | (GCC)5 | GCTGAAACCCGTGTCATTTT | TGGTGAGGATGACGAACTGA | 270 |
| chr12:10363560-10365575 | p3 | (GCC)5 | GCTGAAACCCGTGTCATTTT | TGGTGAGGATGACGAACTGA | 198 |
| chr12:10379037-10381051 | p2 | (TA)7 | AGTTCCCGAGGTTAGCCATT | AACATGCTCGTGCTCAGTTG | 266 |
| chr12:10381272-10383288 | p2 | (TA)8 | GCGACATCCTGAGTGCTACA | AATGTTGGGACTGGTTTTGC | 119 |
| chr12:10409264-10411282 | p6 | (AAAGAG)3 | CGAATCTGTCCGCCATAGTT | CTTTTCGCTCTTTCACGGAC | 150 |
| chr12:10415217-10417235 | p6 | (AAAGAG)3 | CGAATCTGTCCGCCATAGTT | CTTTTCGCTCTTTCACGGAC | 150 |
| chr12:10447946-10449960 | p2 | (AT)7 | GGGTGGAATATGCTTGGAGA | GTCCATCGGGCAAATTTTTA | 220 |

| Marker ID | SSR type | SSR motif | FORWARD PRIMER (5'-3') | REVERSE PRIMER (5'-3') | size (bp) |
|---|---|---|---|---|---|
| chr12:10483422-10485473 | p5 | (AAGAG)3 | GATCCTGTCATCCGGCTAAA | GCGATGTAGTCAGTGCGGTA | 139 |
| chr12:10483422-10485473 | p3 | (AAT)20 | GATCCTGTCATCCGGCTAAA | GCGATGTAGTCAGTGCGGTA | 139 |
| chr12:10484234-10486294 | p5 | (AAGAG)3 | TTCGGGTCCAAACTTCTGTC | TCCCTGAGTCTCGTCGAAAT | 269 |
| chr12:10484234-10486294 | p3 | (AAT)20 | TTCGGGTCCAAACTTCTGTC | TCCCTGAGTCTCGTCGAAAT | 269 |
| chr12:10530211-10532223 | p2 | (AT)6 | ACTCACGGCCAACACATACA | TCGTGTTGTTATGCAGCCTC | 251 |
| chr12:10530211-10532223 | p2 | (AT)6 | ACTCACGGCCAACACATACA | TCGTGTTGTTATGCAGCCTC | 251 |
| chr12:10557077-10559101 | p2 | (TA)12 | AATCGGAGGTTACCGACCTT | GACCAAATAGGCACGGTGTT | 246 |
| chr12:10643144-10645156 | p2 | (TA)6 | CACTGACCTCATAGCAGCCA | GCCCACTAGGGCAATAGTCA | 104 |
| chr12:10643144-10645156 | p2 | (AT)6 | CACTGACCTCATAGCAGCCA | GCCCACTAGGGCAATAGTCA | 104 |
| chr12:10710939-10712953 | p2 | (AT)6 | GCACGCAGTAAATACCAGCA | GTAAGCGACGACAGGAAAGC | 116 |
| chr12:10710939-10712953 | p2 | (AT)13 | GCACGCAGTAAATACCAGCA | GTAAGCGACGACAGGAAAGC | 116 |
| chr12:10711423-10713435 | p2 | (AT)6 | GCACGCAACACACCATAAAC | TTTGATAGCCGAAGGGCTTA | 246 |
| chr12:10711423-10713435 | p2 | (AT)13 | GCACGCAACACACCATAAAC | TTTGATAGCCGAAGGGCTTA | 246 |
| chr12:10750114-10752154 | p2 | (AT)20 | CCCATAAGAAGCCAACTCCA | ACCAGGATGTGTGGGATGAT | 187 |
| chr12:10839208-10841238 | p6 | (CATATA)5 | AAGGGAGGATAAACGCGACT | TCAGTGGAAGACAAGATGCG | 169 |
| chr12:11083256-11085271 | p3 | (GTT)5 | TATTGGGGTAGGGTGGATGA | CTCCCACCTTGTCTCGGATA | 163 |
| chr12:11083256-11085271 | p4 | (GATG)4 | TATTGGGGTAGGGTGGATGA | CTCCCACCTTGTCTCGGATA | 163 |
| chr12:11083256-11085271 | p2 | (TA)13 | TATTGGGGTAGGGTGGATGA | CTCCCACCTTGTCTCGGATA | 163 |
| chr12:11083407-11085423 | p3 | (GTT)5 | CTATCCGAGACAAGGTGGGA | CTGCGCCATGTCTTGTCTAA | 245 |
| chr12:11083407-11085423 | p4 | (GATG)4 | CTATCCGAGACAAGGTGGGA | CTGCGCCATGTCTTGTCTAA | 245 |
| chr12:11083407-11085423 | p2 | (TA)13 | CTATCCGAGACAAGGTGGGA | CTGCGCCATGTCTTGTCTAA | 245 |
| chr12:11143067-11145085 | p2 | (TA)9 | GGAAATCTGCTACACCCCAG | ATTTTCAACAGCCACCTTGG | 104 |
| chr12:11163603-11165624 | p3 | (GAG)7 | GTTTTGCAAATCTGTCCCGT | TGCGCACACAGGTGTATTTT | 270 |
| chr12:11266830-11268854 | p2 | (AT)12 | CGACGAGAGGAATCAAGGAG | GAAAATGATCGCCTGAGAGC | 120 |
| chr12:11266830-11268854 | p2 | (AT)8 | CGACGAGAGGAATCAAGGAG | GAAAATGATCGCCTGAGAGC | 120 |
| chr12:11279313-11281331 | p6 | (ACTGAA)3 | TGTGTCCAGGTTTTGTTCCA | ACAGCCTCAGACTACCGCAT | 190 |
| chr12:11368390-11370410 | p2 | (AT)10 | GGGGTATGAGTAACGAGCCA | CGCTTACTTGGGAGCAGTTC | 265 |
| chr12:11368390-11370410 | p5 | (AATGC)3 | GGGGTATGAGTAACGAGCCA | CGCTTACTTGGGAGCAGTTC | 265 |
| chr12:11466919-11468931 | p3 | (CAA)6 | CAAACAAGTGTTGAGCGGAA | ACTTCCCTATTGTTTGGGGG | 296 |
| chr12:11473091-11475103 | p2 | (TA)6 | ACGACGGAGAAACGAGTTGT | ACCTTTCCACGATATGCCAA | 137 |
| chr12:11478169-11480183 | p2 | (AT)7 | TGTGTTGGTTTGCCGTGTAT | ACATGGCTGGTCGTTCTTTC | 171 |
| chr12:11520178-11522193 | p5 | (AGAAA)3 | CACACGGACAACAAGCTCAT | GGCCATTGTGAGGTGAGAGT | 290 |
| chr12:11707584-11709599 | p5 | (CTTTT)3 | GAACAGCACTTTTGGTGGGT | CCCAGTTTCAAGGCAGTGAT | 159 |
| chr12:11732124-11734154 | p5 | (TTATG)3 | AGCAGCAGAATGTTCCCACT | AATATGGATTTCGCCGACTG | 105 |
| chr12:11732124-11734154 | p2 | (TA)15 | AGCAGCAGAATGTTCCCACT | AATATGGATTTCGCCGACTG | 105 |
| chr12:11800812-11802824 | p2 | (AT)6 | TACGCCACCTCGACTTTACC | TGCGAATCACAGCAACTAGG | 223 |

| Marker ID | SSR type | SSR motif | FORWARD PRIMER (5'-3') | REVERSE PRIMER (5'-3') | size (bp) |
|---|---|---|---|---|---|
| chr12:11806139-11808154 | p5 | (GATAT)3 | GCTGGCATTAAAACAAGGGA | GGTCGAATTCAAAATCGGAA | 164 |
| chr12:11806139-11808154 | p2 | (TA)8 | GCTGGCATTAAAACAAGGGA | GGTCGAATTCAAAATCGGAA | 164 |
| chr12:11845236-11847248 | p2 | (AT)6 | GCCCTCACGAAGACCAAATA | TAGTAGCCTTGTCGGCTGGT | 204 |
| chr12:11895249-11897293 | p2 | (AT)22 | TACTTCGAGGTTGTGGGCTC | TTCCGAGGAATCACCAGAAC | 112 |
| chr12:11895249-11897293 | p2 | (AC)20 | TACTTCGAGGTTGTGGGCTC | TTCCGAGGAATCACCAGAAC | 112 |
| chr12:11895249-11897293 | p2 | (TA)10 | TACTTCGAGGTTGTGGGCTC | TTCCGAGGAATCACCAGAAC | 112 |
| chr12:12109367-12111379 | p2 | (TA)6 | TTGGCATGATCCTTGTGAAA | TGCCATTCGGAAACATATCA | 100 |
| chr12:12168800-12170818 | p3 | (TTG)6 | TGGTGTTGGAAGGTTGTTGA | ACATGCACGACATAACCCAA | 255 |
| chr12:12168800-12170818 | p2 | (TA)8 | TGGTGTTGGAAGGTTGTTGA | ACATGCACGACATAACCCAA | 255 |
| chr12:12277413-12279431 | p6 | (GTGCGG)3 | AACCATGTTGGCGATAGAGG | AACAGCCCAAATCAAAATGC | 300 |
| chr12:12400081-12402105 | p5 | (TTTTC)3 | AGACCCACAAGCTCTCTCCA | GGATTGTTCATTTGCGTCCT | 212 |
| chr12:12400081-12402105 | p3 | (CTT)8 | AGACCCACAAGCTCTCTCCA | GGATTGTTCATTTGCGTCCT | 212 |
| chr12:12411947-12413963 | p4 | (ATAA)4 | CAAACACACAAACCCCAACA | CAACCTCTTCAAAAGCCGAG | 137 |
| chr12:12468253-12470291 | p2 | (TA)19 | CTACGAGTATTCCTCGCCCA | GGGCGCACAACTGATTTTAT | 273 |
| chr12:12488855-12490894 | p6 | (ATTTTT)3 | CACACTGGGGCAGAATTTTT | TTCTGTCTCAGTTCGGGCTT | 197 |
| chr12:12488924-12491077 | p6 | (ATTTTT)3 | TGGGGCAGAATTTTTGAAAG | GATCCCCCAGGAGGTATGAT | 133 |
| chr12:12519254-12521269 | p3 | (GAA)12 | GGAACAATCCCAGGACAAGA | TCGAGCAATGTGCTCAAAAG | 174 |
| chr12:12519254-12521269 | p5 | (ATTTT)3 | GGAACAATCCCAGGACAAGA | TCGAGCAATGTGCTCAAAAG | 174 |
| chr12:12588566-12590592 | p2 | (CT)13 | CAAGGTTGAAGCTCTTTCCG | TCCCATCGTCGGACTTTAAC | 274 |
| chr12:12615079-12617091 | p2 | (CA)6 | AAGCATGTATCAACCGGAGC | GATAGGGCTGTTAGCGCAAG | 119 |
| chr12:12628162-12630177 | p2 | (AT)6 | TATAGTGGACGGGACAAGCC | TGTTCATGCCCACAGGTAGA | 104 |
| chr12:12628162-12630177 | p3 | (TCA)5 | TATAGTGGACGGGACAAGCC | TGTTCATGCCCACAGGTAGA | 104 |
| chr12:12649144-12651194 | p2 | (TA)25 | CCTTTTCGTCGTAGCTTTGC | AAACAAGAGGCTACAGGGCA | 105 |
| chr12:12657756-12659771 | p5 | (AAAAT)3 | ACTTCCTCGCACGATTCTGA | TTCCAATTTGGTTGGAGAGTG | 257 |
| chr12:12692097-12694113 | p4 | (TCTT)4 | CGCCTGAAGCATCACAATAA | GGGCTTGGCAGGTTATTACA | 228 |
| chr12:12697747-12699762 | p5 | (TTTTA)3 | TCAACCCTAACAGATTCGCC | TGTTATCTGACGGGGAAAGG | 182 |
| chr12:12724512-12726524 | p2 | (CT)6 | ATCCCAGAACTACACACGGC | TGATGAAAGAATTTTCGGGC | 236 |
| chr12:12724512-12726524 | p5 | (TTTTA)3 | ATCCCAGAACTACACACGGC | TGATGAAAGAATTTTCGGGC | 236 |
| chr12:12830267-12832282 | p5 | (TGCAA)3 | CCACAACCCGCTAGAACAAT | ATTTTTCGGATGTATGCCCA | 224 |
| chr12:12862356-12864371 | p2 | (AT)7 | AACACCCCGTCATACCCATA | GCGATCCAGCTCAGTAGGTC | 211 |
| chr12:12862356-12864371 | p2 | (AT)6 | AACACCCCGTCATACCCATA | GCGATCCAGCTCAGTAGGTC | 211 |
| chr12:12862356-12864371 | p3 | (ATC)5 | AACACCCCGTCATACCCATA | GCGATCCAGCTCAGTAGGTC | 211 |
| chr12:13099861-13101876 | p5 | (ATAAA)3 | TTTTCCTGCACCTCGAACTT | TCGAGTCTTCCATGCAGATG | 120 |
| chr12:13099861-13101876 | p5 | (AAAAC)3 | TTTTCCTGCACCTCGAACTT | TCGAGTCTTCCATGCAGATG | 120 |
| chr12:13162509-13164527 | p6 | (TGTAAG)3 | GTAACCCTGACGCCTTGGTA | CATACCCGACCATGAAGGAT | 148 |
| chr12:13315407-13317422 | p5 | (AATAA)3 | GTATGCAGCACGGATGATTG | CCGGCGTATACAGAGGTGTT | 265 |

| Marker ID | SSR type | SSR motif | FORWARD PRIMER (5'-3') | REVERSE PRIMER (5'-3') | size (bp) |
|---|---|---|---|---|---|
| chr12:13315407-13317422 | p3 | (TGG)5 | GTATGCAGCACGGATGATTG | CCGGCGTATACAGAGGTGTT | 265 |
| chr12:13419624-13421639 | p5 | (TTCAT)3 | CGTTCTCCTTCCAAGCTCAC | GCTGCCCAACTTTCATTAGC | 186 |
| chr12:13424762-13426774 | p2 | (AT)10 | TGTCTACCTGTGGGCATGAA | CCCGTAAAAGTTGCACAGGT | 144 |
| chr12:13424762-13426774 | p2 | (AT)6 | TGTCTACCTGTGGGCATGAA | CCCGTAAAAGTTGCACAGGT | 144 |
| chr12:13432338-13434361 | p5 | (GGGTG)3 | GCAGACGAACTCACAGGTCA | CCTTTTCCCAGAAAATGCAA | 136 |
| chr12:13432801-13434816 | p5 | (GGGTG)3 | GAAGGGTGTGTTCGGTAGGA | AAGTTTTTAGGGTGGGGGTG | 172 |
| chr12:13496715-13498730 | p3 | (GAA)5 | GATGTAGCCTTGTCGGCTTC | CTAGCGACCAAACACCCCTA | 252 |
| chr12:13535710-13537728 | p3 | (CAA)6 | ATCAGTGGTAACTGCCCGTC | ATCAGCTAGACATCGTCCCG | 143 |
| chr12:13535710-13537728 | p2 | (CT)6 | ATCAGTGGTAACTGCCCGTC | ATCAGCTAGACATCGTCCCG | 143 |
| chr12:13576938-13578956 | p3 | (AAT)6 | TAAATGCGGAACAATGACGA | TCCGGTGGAAAATTGCTAAG | 224 |
| chr12:13618038-13620053 | p3 | (ATC)5 | GTGCAGCGTCCACAAATAAA | ATATGGTGGGGCAGCATAAC | 207 |
| chr12:13618038-13620053 | p3 | (AAC)5 | GTGCAGCGTCCACAAATAAA | ATATGGTGGGGCAGCATAAC | 207 |
| chr12:13637711-13639726 | p5 | (ATATA)3 | CCCTATGAAAGTCGGTTCCA | AAAACGACATCGGAGTACGG | 192 |
| chr12:13658021-13660045 | p2 | (TA)12 | GAATGTGCAAGGTGGCTTTT | AGGCACCATAACCCACTCAC | 212 |
| chr12:13670009-13672069 | p2 | (TA)30 | CGAAGAGCCGTAGGTGTAGC | CCGTGCATGACACTATGACC | 229 |
| chr12:13676591-13678606 | p5 | (CATAT)3 | CATAGGCAGCTATGCCACAA | GAGCTCCATCTCATTCGGAG | 266 |
| chr12:13687551-13689563 | p2 | (GT)6 | CAGCGTCCACAAACAAAAGA | CCCCTATGGAAAGCAAGTCA | 127 |
| chr12:13878833-13880851 | p6 | (GAATAC)3 | CCTGTTTGGATGGCTATGCT | TTGGCTGGCCAGTTTAAGTC | 263 |
| chr12:14112648-14114705 | p3 | (ATT)14 | GATAGGTTTGCCCGAATTGA | CTTTGACTTAGGGCGTCGAG | 219 |
| chr12:14155830-14157882 | p4 | (AGAA)13 | GTGGGAAGCAACTGAAGAGC | CAACTATGGTGGAGGGCCTA | 104 |
| chr12:14157441-14159459 | p6 | (AAATTC)3 | TTACCGCTGAGCAGGAGAAT | TTCTTTCATTCGTGTCCGTG | 273 |
| chr12:14206402-14208468 | p2 | (TA)33 | AATAACCCCTTCACCATCCC | CTTTCAGCTGCTCGTAACCC | 195 |
| chr12:14224357-14226375 | p3 | (TCC)5 | TTCGGTCTCTCCACATCCTC | GACGTGGCACAAGAAGTTGA | 125 |
| chr12:14224357-14226375 | p2 | (AC)9 | TTCGGTCTCTCCACATCCTC | GACGTGGCACAAGAAGTTGA | 125 |
| chr12:14234142-14236182 | p2 | (CT)20 | TTGGGTCTTTATTTGCGGTC | TGGATTCCGGTCAAATTTTC | 146 |
| chr12:14234142-14236182 | p6 | (TATTTT)3 | TTGGGTCTTTATTTGCGGTC | TGGATTCCGGTCAAATTTTC | 146 |
| chr12:14291543-14293555 | p2 | (AT)6 | GCCAATCAGAACAGCCATTT | GTTCTGTACCCCAGCAGAGC | 174 |
| chr12:14334851-14336985 | p2 | (TA)40 | TCATTCGAAGGTTACCGACC | CATCCCACATCCAGGGTATC | 227 |
| chr12:14406636-14408648 | p2 | (CA)6 | CTCGGTGTCTGAAGCGTGTA | CTTCGGGCCAAGATGTAAAA | 107 |
| chr12:14410799-14412811 | p2 | (CA)6 | CTCGGTGTCTGAAGCGTGTA | TTCGGGCCAAGATGTAAAAG | 107 |
| chr12:14410799-14412811 | p3 | (TAT)5 | CTCGGTGTCTGAAGCGTGTA | TTCGGGCCAAGATGTAAAAG | 107 |
| chr12:14542020-14544035 | p3 | (TGT)5 | TCTTGGGAGGTAAGCCCTTT | ACGTCATCTAGGACGCTGCT | 167 |
| chr12:14542020-14544035 | p3 | (TAT)12 | TCTTGGGAGGTAAGCCCTTT | ACGTCATCTAGGACGCTGCT | 167 |
| chr12:14564363-14566378 | p3 | (TCA)5 | GACATGAGGACGGGATAGGA | ACGGTGCAGATCCATAGGAG | 272 |
| chr12:14657778-14659802 | p2 | (AT)12 | CGGTCAAAATGAGCCGTAAT | TCAAAGTTCCCGAGTCCAAG | 207 |
| chr12:14657997-14660013 | p2 | (AT)12 | TTACTCCGTCTTGGACTCGG | CGTCATGACTCATCGGTTTG | 195 |

| Marker ID | SSR type | SSR motif | FORWARD PRIMER (5'-3') | REVERSE PRIMER (5'-3') | size (bp) |
|---|---|---|---|---|---|
| chr12:14708474-14710492 | p6 | (CATATA)3 | GTGACGAGTACCCGACCACT | TGATGTTATGCAGCCTCGTC | 178 |
| chr12:14708474-14710492 | p3 | (CAA)6 | GTGACGAGTACCCGACCACT | TGATGTTATGCAGCCTCGTC | 178 |
| chr12:14749326-14751338 | p2 | (TA)6 | TGGAATCCGGTATCCGTAAA | TGTTGATGTGGCGTGATTTT | 117 |
| chr12:14763088-14765103 | p3 | (ATC)5 | CTGAAATCAAAAGGACGGGA | AGACAGGGAGACGTTCCAGA | 229 |
| chr12:14842709-14844724 | p5 | (TTCAT)3 | TATGCATTTGGGTCGTGTGT | TTGAGGCGTAACATTTTCCC | 280 |
| chr12:14882070-14884086 | p4 | (CATT)4 | TACACCACAAGTGAACGGGA | TCATTTTGACCCCCTCAGTC | 286 |
| chr12:14882070-14884086 | p4 | (ATGA)5 | TACACCACAAGTGAACGGGA | TCATTTTGACCCCCTCAGTC | 286 |
| chr12:14884384-14886399 | p2 | (TG)7 | TCCAGCCGTTATCCAAAAAC | CCCTCACCCCCTTAAACATT | 270 |
| chr12:14884407-14886425 | p2 | (TG)7 | TCCAGCCGTTATCCAAAAAC | CCCTCACCCCCTTAAACATT | 247 |
| chr12:14884448-14886466 | p2 | (TG)7 | TCCAGCCGTTATCCAAAAAC | CCCTCACCCCCTTAAACATT | 206 |
| chr12:14889508-14891544 | p3 | (TTC)10 | TTGGAATCCTGCTCCTTGAC | TTGAAGGTCAAAAACCCGTC | 158 |
| chr12:14889508-14891544 | p3 | (TTC)12 | TTGGAATCCTGCTCCTTGAC | TTGAAGGTCAAAAACCCGTC | 158 |
| chr12:15018798-15020813 | p3 | (ACA)5 | GCTGTTTAGGTGGTTTCCCA | GGCTGATTTGGAACGTGAAT | 268 |
| chr12:15018798-15020813 | p2 | (TA)11 | GCTGTTTAGGTGGTTTCCCA | GGCTGATTTGGAACGTGAAT | 268 |
| chr12:15018983-15021005 | p3 | (ACA)5 | CTGAATGAAAAAGGGCCAAA | ATTTTGGCTGATTTGGAACG | 133 |
| chr12:15018983-15021005 | p2 | (TA)11 | CTGAATGAAAAAGGGCCAAA | ATTTTGGCTGATTTGGAACG | 133 |
| chr12:15077511-15079526 | p3 | (ATA)5 | AAAAGGAATTGGGAACCACC | TTATTTTGATGGCCGAAAGG | 109 |
| chr12:15100172-15102208 | p4 | (ACAT)5 | AGCCCTTCGGCTATCAAAAT | TCCATTATTCCAGAGGTGCC | 157 |
| chr12:15124780-15126795 | p3 | (TCA)5 | GCGCGATCTTTAAGGATGAA | GGCCGAATACGGATATTGTG | 230 |
| chr12:15201933-15203945 | p2 | (GT)6 | AGACGAATGGACGAAGATGG | TCTTACCCGGCATCATAAGC | 121 |
| chr12:15242478-15244493 | p5 | (CCACA)3 | GTCAAACGGAGCAATAGGGA | TTGCATAAATAAGCCCCGAC | 139 |
| chr12:15278992-15281006 | p2 | (AG)7 | GGTCAAGGTGTTGGTGGACT | ACTTCCCCTCCTCCACTTGT | 217 |
| chr12:15313490-15315514 | p4 | (TTAT)5 | TGCACGCGTTAACTCCATTA | CCCTTCAAAGACAGGCACTC | 145 |
| chr12:15313490-15315514 | p4 | (TCTT)6 | TGCACGCGTTAACTCCATTA | CCCTTCAAAGACAGGCACTC | 145 |
| chr12:15313490-15315514 | p2 | (AT)7 | TGCACGCGTTAACTCCATTA | CCCTTCAAAGACAGGCACTC | 145 |
| chr12:15324408-15326423 | p5 | (CATTG)3 | TTCCCTTTTTAAGGGTCGCT | TGCATACGCTTCACTAACCG | 236 |
| chr12:15349041-15351055 | p2 | (AT)7 | CAGAACCAGATTCCCCAAAA | GGGAAGTTGAAATTGAGGCA | 111 |
| chr12:15386562-15388576 | p2 | (AG)7 | CGTTGATTGAGCGAAACTGA | ACATTGGCCTCCTCACAATC | 209 |
| chr12:15386562-15388576 | p3 | (GTT)5 | CGTTGATTGAGCGAAACTGA | ACATTGGCCTCCTCACAATC | 209 |
| chr12:15386865-15388880 | p2 | (AG)7 | ATTCGGGCATCTGTTACACC | ACCCATCCTTTCTCCGAAGT | 149 |
| chr12:15386865-15388880 | p3 | (GTT)5 | ATTCGGGCATCTGTTACACC | ACCCATCCTTTCTCCGAAGT | 149 |
| chr12:15386865-15388880 | p2 | (TA)7 | ATTCGGGCATCTGTTACACC | ACCCATCCTTTCTCCGAAGT | 149 |
| chr12:15398681-15400699 | p2 | (TA)12 | CGTCCGTTGTCCCTTAAAAA | GTTTTGTGCTCGCGTATCCT | 175 |
| chr12:15398681-15400699 | p2 | (TA)9 | CGTCCGTTGTCCCTTAAAAA | GTTTTGTGCTCGCGTATCCT | 175 |
| chr12:15495856-15497871 | p3 | (TTG)5 | GAGGAGCTCCATTTGATTCG | ACAAGCCGACAAGGCTAAAA | 169 |
| chr12:15510907-15512922 | p3 | (TTG)5 | GAGGAGCTCCATTTGATTCG | ACAAGCCGACAAGGCTAAAA | 169 |

| Marker ID | SSR type | SSR motif | FORWARD PRIMER (5'-3') | REVERSE PRIMER (5'-3') | size (bp) |
|---|---|---|---|---|---|
| chr12:15518532-15520550 | p6 | (ATTCCA)3 | TACGGGGAAGTTTCGAAGTG | ATGGGAATCGAGTTTGAACG | 110 |
| chr12:15536443-15538461 | p6 | (CATATT)3 | AGGGTTCCATGACCTCTCCT | ATGGGCCTAGTTGTTGATGG | 204 |
| chr12:15695266-15697281 | p3 | (AAC)5 | TTGCCTCATAAGGCGGATAC | ATATATGGATCGGGCTGCAC | 128 |
| chr12:15832050-15834066 | p2 | (AG)8 | GAACCGGAAGCATTACAGGA | GCAAAAAGCATAGGGACCAA | 148 |
| chr12:15851205-15853221 | p2 | (AG)8 | GAACCGGAAGCATTACAGGA | GCAAAAAGCATAGGGACCAA | 148 |
| chr12:15902639-15904655 | p2 | (AT)12 | CGCAGTCAGTGGACATCAGT | AGCGACCCAAAGACATCATC | 238 |
| chr12:15902713-15904729 | p2 | (AT)12 | CGCAGTCAGTGGACATCAGT | AACGTCCTCCACATAGCGAC | 164 |
| chr12:15902935-15904951 | p2 | (AT)12 | TGTCTTTGGGTCGCTATGTG | GGCGCCACAAAGATATATTGA | 149 |
| chr12:15903157-15905173 | p2 | (AT)12 | TGTCCGGCGACATATAGGG | TGGCTCCACGGAGATATATTG | 131 |
| chr12:15916857-15918881 | p3 | (ACA)8 | GTTGGTTGCTGGTTGGAGTT | GGTTGCTGGTTAGGAGTGGA | 158 |
| chr12:15959959-15961974 | p3 | (TTG)5 | TCACCTCAGCCTCGAGTTTT | TATTCCACAGCCCCTGATTC | 265 |
| chr12:15989431-15991443 | p2 | (CT)6 | CCCCTTTGCGTAATTCTTGA | GCTTTTTCCGATTCAGTCCA | 140 |
| chr12:16017020-16019044 | p2 | (TA)12 | GACGAGAACGAAAACGGAAC | ATGAATTAAAGCCTGGCCCT | 288 |
| chr12:16051188-16053204 | p2 | (AT)8 | ACACCAACCCCGAGAATATG | TGGATGGTTGTGAGCATTGT | 278 |
| chr12:16051188-16053204 | p2 | (CA)8 | ACACCAACCCCGAGAATATG | TGGATGGTTGTGAGCATTGT | 278 |
| chr12:16061365-16063377 | p2 | (AT)6 | TATCGGGGAAGCAAGCATAC | CAGGCTTGTTAAGGCTCGTC | 161 |
| chr12:16095897-16097912 | p5 | (ATCAG)3 | ACGAAACCCGTATACCCACA | TCTAGGGCACAGGTTCTGCT | 197 |
| chr12:16233218-16235234 | p2 | (CA)8 | AAAGAATGATCCCCGAGCTT | CTGGCAAAATTTCGGTAGGA | 204 |
| chr12:16252980-16254995 | p3 | (TTG)5 | GGGCATAGGCCCTCTAAAAG | GCCTCTAAAATCGTTCAGCG | 300 |
| chr12:16294269-16296281 | p5 | (TATAT)3 | TGCACCATTTGGGAGTTGTA | AATGCTCGATGCGTACTTCC | 286 |
| chr12:16294269-16296281 | p2 | (TA)6 | TGCACCATTTGGGAGTTGTA | AATGCTCGATGCGTACTTCC | 286 |
| chr12:16383023-16385045 | p2 | (TA)11 | GGGCTTACAAGGAATGACGA | CTTCTTCCCGTGTCATGGTT | 278 |
| chr12:16539012-16541030 | p6 | (CATATA)3 | TAACCGAACAATAGCGACCC | TGGATTACTCCCTTGTTCCG | 201 |
| chr12:16540384-16542396 | p2 | (AC)6 | TACATACCTTGCCCGCTTTC | TGTATGTTTGGCCGAGTTGA | 289 |
| chr12:16551090-16553105 | p5 | (ATTTC)3 | CTGGGGGTAAGTCCAGTTCA | ATCTTATGCGGACAACCGTC | 115 |
| chr12:16660232-16662247 | p2 | (TA)6 | CTTGAGCACAAGTTGTCCGA | TGGTGGCAGTATGCCATAGA | 139 |
| chr12:16660232-16662247 | p5 | (CAATA)3 | CTTGAGCACAAGTTGTCCGA | TGGTGGCAGTATGCCATAGA | 139 |
| chr12:16660232-16662247 | p3 | (CAT)8 | CTTGAGCACAAGTTGTCCGA | TGGTGGCAGTATGCCATAGA | 139 |
| chr12:16742823-16744849 | p2 | (TG)13 | TGGGACTCGCTACGAGAACT | GTTACCCCTGTTTTCCCCAT | 133 |
| chr12:16775187-16777199 | p6 | (CATATA)3 | TGTGTGTACCTTCGTGGCAT | CATCGGTATGCATGATTTGC | 297 |
| chr12:16775187-16777199 | p2 | (AT)6 | TGTGTGTACCTTCGTGGCAT | CATCGGTATGCATGATTTGC | 297 |
| chr12:16783899-16785917 | p6 | (TGAACC)3 | ATGGAAAATTGGTGGCATGT | TGAGGTGGAGATTGGAAAGG | 259 |
| chr12:16791241-16793255 | p2 | (GT)7 | CCAATCAAATCAGCCCTTGT | TCCGAGATTCCAATTTTTGC | 276 |
| chr12:16856161-16858175 | p2 | (AG)7 | CGCAATACAACTGACAACCG | TCCCTTCACATGCCTTTTTC | 238 |
| chr12:16856161-16858175 | p3 | (TTG)5 | CGCAATACAACTGACAACCG | TCCCTTCACATGCCTTTTTC | 238 |
| chr12:16859979-16862009 | p2 | (TA)15 | CACGCGTTTCTGCTTTATCA | ACGACGTTGGAGAAAGGATG | 147 |

| Marker ID | SSR type | SSR motif | FORWARD PRIMER (5'-3') | REVERSE PRIMER (5'-3') | size (bp) |
|---|---|---|---|---|---|
| chr12:16859979-16862009 | p3 | (ATA)5 | CACGCGTTTCTGCTTTATCA | ACGACGTTGGAGAAAGGATG | 147 |
| chr12:16911001-16913015 | p2 | (GA)6 | CCCTAACCTAATGCACGGAA | GTGAATCTTGCAGAGTGCCA | 299 |
| chr12:16936515-16938530 | p3 | (TTG)5 | GAATGCCTCCACCGAAGTAA | GGGCATTCTGAACTGGTTGT | 221 |
| chr12:16982705-16984723 | p2 | (AT)12 | CCCGAACTGGACACTCGTAT | ATTGATATGTCGTAGCCGCC | 124 |
| chr12:16982705-16984723 | p2 | (AT)9 | CCCGAACTGGACACTCGTAT | ATTGATATGTCGTAGCCGCC | 124 |
| chr12:17038660-17040728 | p2 | (AT)34 | TTTGACTAATTGCTGCCACG | ATTATTCGCAAGCAAATCGG | 138 |
| chr12:17038660-17040728 | p2 | (AT)34 | TTTGACTAATTGCTGCCACG | ATTATTCGCAAGCAAATCGG | 138 |
| chr12:17059675-17061743 | p2 | (AT)34 | TCCTTGCACCTCACCCTAAC | AGCCGTGACTCCTCAGGATA | 213 |
| chr12:17124868-17126880 | p2 | (AT)6 | GGGGGTCAGAATCGTCTGTA | CCTGGGTCCATAAGAGCAAA | 273 |
| chr12:17124868-17126880 | p2 | (AT)25 | GGGGGTCAGAATCGTCTGTA | CCTGGGTCCATAAGAGCAAA | 273 |
| chr12:17509707-17511725 | p6 | (TTTTTA)3 | CGAAACACCCTTTCTTTGGA | TTTCGAATTGACCTTGGACC | 164 |
| chr12:17523767-17525787 | p5 | (AATAT)4 | CAGGTAGGAGAAGGACAGCG | CTACGGGGTGTTTTTGGAGA | 291 |
| chr12:17613659-17615671 | p2 | (AG)6 | TCAGGTTCAATCCCAAGACC | CAACATACCCAGAGGCGAAT | 260 |
| chr12:17682281-17684293 | p2 | (TC)6 | TCGGAACCCCCTTAATTACC | TTGTCGGACTTCTATTCCCG | 247 |
| chr12:17693907-17695922 | p3 | (TTG)5 | CTGGGAAGGACAACCTTGAA | TCTTTGATTTGCTTGGTCCC | 191 |
| chr12:17865462-17867474 | p2 | (CT)6 | GTCCCACCATAACATGGACC | TCGGAATATGAGGCAAGGAC | 225 |
| chr12:17865540-17867624 | p2 | (CT)6 | GTCCCACCATAACATGGACC | TCGGAATATGAGGCAAGGAC | 147 |
| chr12:17869595-17871635 | p2 | (TA)20 | CAGAGGTAACACACCGAGCA | AAGCCTTGTCAGCTGGTGAT | 257 |
| chr12:17886250-17888268 | p3 | (TTG)6 | TCTGATCCGTCTCTCTGCCT | GTTTCATGGCTTCTTCGAGC | 191 |
| chr12:18005008-18007020 | p2 | (AT)6 | GAGAATTCAACTCGGCCAAA | TCCTTGTTGGAATTGGAAGG | 145 |
| chr12:18073283-18075298 | p2 | (AT)20 | GCGTACCCACAAGTGGTTTT | AACAACGGGTATGCCAACAT | 198 |
| chr12:18073309-18075324 | p2 | (AT)20 | GCGTACCCACAAGTGGTTTT | AACAACGGGTATGCCAACAT | 172 |
| chr12:18073309-18075324 | p2 | (AT)7 | GCGTACCCACAAGTGGTTTT | AACAACGGGTATGCCAACAT | 172 |
| chr12:18261049-18263085 | p4 | (CATA)9 | ACCCCAATTCCCAGATTACC | TTGTGACCCGAACGATGTAA | 218 |
| chr12:18318486-18320504 | p6 | (GAAGAT)3 | TGTCCTGCTCAGTCACCAAG | CACAGTGGCAACAGATTTGG | 299 |
| chr12:18427260-18429278 | p2 | (TA)28 | CTCCCAATCTACTCCACCCC | GCTTTGGACAACACTGCAAA | 200 |
| chr12:18427260-18429278 | p2 | (TA)9 | CTCCCAATCTACTCCACCCC | GCTTTGGACAACACTGCAAA | 200 |
| chr12:18457958-18460004 | p3 | (CAA)6 | CATAGCTCGTAGGCACTCCC | TTTGGTAGCTTTGTCGGCTT | 120 |
| chr12:18457958-18460004 | p2 | (AT)31 | CATAGCTCGTAGGCACTCCC | TTTGGTAGCTTTGTCGGCTT | 120 |
| chr12:18526212-18528240 | p2 | (AT)14 | GACGATCTCGTGGATGGTTT | TTCTGATAGCCTTGTCGGCT | 285 |
| chr12:18538581-18540606 | p5 | (AATAA)5 | TGATGTGGAGGACGTGATGT | TCTTACTTGTCCCGCCAAAC | 183 |
| chr12:18538581-18540606 | p6 | (TAATTT)3 | TGATGTGGAGGACGTGATGT | TCTTACTTGTCCCGCCAAAC | 183 |
| chr12:18538581-18540606 | p5 | (TTTAT)3 | TGATGTGGAGGACGTGATGT | TCTTACTTGTCCCGCCAAAC | 183 |
| chr12:18539175-18541193 | p5 | (AATAA)5 | CGTTTTAGCCCAACACATGA | TATTGTGTGACCCACGGCTA | 156 |
| chr12:18539175-18541193 | p6 | (TAATTT)3 | CGTTTTAGCCCAACACATGA | TATTGTGTGACCCACGGCTA | 156 |
| chr12:18539175-18541193 | p5 | (TTTAT)3 | CGTTTTAGCCCAACACATGA | TATTGTGTGACCCACGGCTA | 156 |

| Marker ID | SSR type | SSR motif | FORWARD PRIMER (5'-3') | REVERSE PRIMER (5'-3') | size (bp) |
|---|---|---|---|---|---|
| chr12:18679849-18681867 | p6 | (AATGCA)3 | TCCAACCTCGGTAGGTGAAC | GATCCACAGCCCTGATGTCT | 295 |
| chr12:18681926-18683948 | p2 | (TA)8 | GCTGTTGGATTACCCGTTGT | GCTATAGCGTGTGGGATGGT | 256 |
| chr12:18681974-18683988 | p2 | (TA)8 | GCTGTTGGATTACCCGTTGT | GCTATAGCGTGTGGGATGGT | 208 |
| chr12:18754379-18756403 | p6 | (ATCTAT)4 | TTGGTCAGCAGAGCAGAAGA | ACTCTGTTCGAGGGGGAGAT | 118 |
| chr12:18836464-18838480 | p2 | (AT)22 | TCTGGTTGGGTCGTGACATA | TAACCAGCTGTCAAACGTGG | 165 |
| chr12:18836464-18838480 | p4 | (TTAT)4 | TCTGGTTGGGTCGTGACATA | TAACCAGCTGTCAAACGTGG | 165 |
| chr12:18874071-18876083 | p2 | (TA)6 | CAATATGCGCCACAACACTC | CAATTTTCAAACGCGCTGTA | 130 |
| chr12:18874071-18876083 | p2 | (TA)10 | CAATATGCGCCACAACACTC | CAATTTTCAAACGCGCTGTA | 130 |
| chr12:18949638-18951650 | p2 | (AT)8 | TTGGATTGGCCTTAGGTGTC | ACAGCGTTAAGGCAACGTCT | 164 |
| chr12:18949638-18951650 | p2 | (AT)6 | TTGGATTGGCCTTAGGTGTC | ACAGCGTTAAGGCAACGTCT | 164 |
| chr12:18949638-18951650 | p2 | (AT)11 | TTGGATTGGCCTTAGGTGTC | ACAGCGTTAAGGCAACGTCT | 164 |
| chr12:18949812-18951834 | p2 | (AT)8 | AGACGTTGCCTTAACGCTGT | TTATGTTTTTGGGGCGTACC | 206 |
| chr12:18949812-18951834 | p2 | (AT)6 | AGACGTTGCCTTAACGCTGT | TTATGTTTTTGGGGCGTACC | 206 |
| chr12:18949812-18951834 | p2 | (AT)11 | AGACGTTGCCTTAACGCTGT | TTATGTTTTTGGGGCGTACC | 206 |
| chr12:18998626-19000644 | p6 | (ATATAC)3 | CGCTGAGGAGAACCAAAATC | TCCAGCACAAATGCAATGAT | 106 |
| chr12:18999934-19001952 | p3 | (ACA)6 | TTCCCCACCCCTTTATTTTC | ATTTACCATTTTCGGCAACG | 238 |
| chr12:19001099-19003151 | p2 | (AT)26 | TACTTCTTCGGCCTACCCCT | GATTGAGATGGTTTGGGCAT | 270 |
| chr12:19032526-19034538 | p2 | (AT)8 | TTGGATTGGCCTTAGGTGTC | ACAGCGTTAAGGCAACGTCT | 164 |
| chr12:19032526-19034538 | p2 | (AT)6 | TTGGATTGGCCTTAGGTGTC | ACAGCGTTAAGGCAACGTCT | 164 |
| chr12:19032526-19034538 | p2 | (AT)11 | TTGGATTGGCCTTAGGTGTC | ACAGCGTTAAGGCAACGTCT | 164 |
| chr12:19032700-19034722 | p2 | (AT)8 | AGACGTTGCCTTAACGCTGT | TTATGTTTTTGGGGCGTACC | 206 |
| chr12:19032700-19034722 | p2 | (AT)6 | AGACGTTGCCTTAACGCTGT | TTATGTTTTTGGGGCGTACC | 206 |
| chr12:19032700-19034722 | p2 | (AT)11 | AGACGTTGCCTTAACGCTGT | TTATGTTTTTGGGGCGTACC | 206 |
| chr12:19081514-19083532 | p6 | (ATATAC)3 | CGCTGAGGAGAACCAAAATC | TCCAGCACAAATGCAATGAT | 106 |
| chr12:19082822-19084840 | p3 | (ACA)6 | TTCCCCACCCCTTTATTTTC | ATTTACCATTTTCGGCAACG | 238 |
| chr12:19083987-19086039 | p2 | (AT)26 | TACTTCTTCGGCCTACCCCT | GATTGAGATGGTTTGGGCAT | 270 |
| chr12:19083987-19086039 | p2 | (AT)17 | TACTTCTTCGGCCTACCCCT | GATTGAGATGGTTTGGGCAT | 270 |
| chr12:19186776-19188791 | p5 | (AAAGG)3 | AGTCAGGTCATCCGAAAACG | GGCCCATATGAAGGGATTTT | 114 |
| chr12:19186776-19188791 | p5 | (GCATT)3 | AGTCAGGTCATCCGAAAACG | GGCCCATATGAAGGGATTTT | 114 |
| chr12:19230115-19232135 | p2 | (CT)10 | TTGAGAAACAGGCAAACACG | AGCTTAGGCATACCGGTGAA | 207 |
| chr12:19265149-19267163 | p2 | (TA)7 | ATGCCCGACATTTTCTTCAC | GACTGGGAACAAAACCCTCA | 193 |
| chr12:19303818-19305832 | p2 | (CT)7 | GGATCTTTTAGGGGCTCGAC | AATGGGCGTCCATCTACTTG | 220 |
| chr12:19348122-19350152 | p2 | (AT)6 | ACCCTTTTCGCGAATTTTCT | CTCGGCGAACAATACAGTGA | 257 |
| chr12:19348122-19350152 | p2 | (AT)15 | ACCCTTTTCGCGAATTTTCT | CTCGGCGAACAATACAGTGA | 257 |
| chr12:19393349-19395365 | p2 | (TA)8 | TATATGTCCGTCCCCGGTTA | TCACACATAGGGTGCAGGAA | 226 |
| chr12:19550597-19552617 | p2 | (AT)10 | CCCGTCGTACCCTTAGTCAA | ATACTCGCTTTGGTGATCCG | 137 |

| Marker ID | SSR type | SSR motif | FORWARD PRIMER (5'-3') | REVERSE PRIMER (5'-3') | size (bp) |
|---|---|---|---|---|---|
| chr12:19550597-19552617 | p2 | (TA)10 | CCCGTCGTACCCTTAGTCAA | ATACTCGCTTTGGTGATCCG | 137 |
| chr12:19560095-19562107 | p2 | (TA)6 | TCGTTCCTAGGGTTCCATTG | TCACAACACCATCTCCTCCA | 230 |
| chr12:19560095-19562107 | p2 | (TA)11 | TCGTTCCTAGGGTTCCATTG | TCACAACACCATCTCCTCCA | 230 |
| chr12:19578786-19580800 | p2 | (GT)7 | CCCGTTATGTTTGTGAGCCT | TCTTAAAATGCCCCGTTTTG | 272 |
| chr12:19647546-19649560 | p2 | (AT)7 | AGAATTGACGGGAGAGAGCA | AATAACGACGACAAGCCCAC | 263 |
| chr12:19710398-19712410 | p2 | (TG)6 | TGTGCAATCGGAAAATTCAA | ATGCAAATTTCGTCCCTTTG | 246 |
| chr12:19777198-19779213 | p5 | (ATCCA)3 | CCTCGGTTCCACTAGTTCCA | TCCTTGTAAAATGGGCCTTG | 222 |
| chr12:19783967-19785983 | p2 | (AT)6 | TGCAATTGGTGCAAGTGATT | TTCCAATTTCGCCTAACACC | 236 |
| chr12:19783967-19785983 | p2 | (AT)8 | TGCAATTGGTGCAAGTGATT | TTCCAATTTCGCCTAACACC | 236 |
| chr12:19941739-19943754 | p5 | (TGAAT)4 | TGATTGATGATTTGAGCCGA | TGGTAGGAACATATCCCCGA | 270 |
| chr12:19941739-19943754 | p6 | (TATATG)4 | TGATTGATGATTTGAGCCGA | TGGTAGGAACATATCCCCGA | 270 |
| chr12:19947141-19949156 | p5 | (ATTCA)3 | TTACCCCAAGGCAGCATATC | TTTGGAGCCGAACCATATTC | 292 |
| chr12:19947141-19949156 | p5 | (ACACA)3 | TTACCCCAAGGCAGCATATC | TTTGGAGCCGAACCATATTC | 292 |
| chr12:19947141-19949156 | p5 | (TTTTC)3 | TTACCCCAAGGCAGCATATC | TTTGGAGCCGAACCATATTC | 292 |
| chr12:19974969-19976985 | p2 | (GT)8 | CCCTTGTTCTGTGGCATTCT | CATAGCCGACACATTCATGC | 295 |
| chr12:19974969-19976985 | p2 | (AT)6 | CCCTTGTTCTGTGGCATTCT | CATAGCCGACACATTCATGC | 295 |
| chr12:20000072-20002084 | p2 | (AT)6 | CCATTGGGGCAACAATACAT | ACTAGCGGAGCTCCATTTCA | 255 |
| chr12:20041930-20043944 | p2 | (TA)6 | AATTCCATCGGGCAACAATA | TTTTGTTAGCTTCGTCGGCT | 124 |
| chr12:20041930-20043944 | p2 | (AT)7 | AATTCCATCGGGCAACAATA | TTTTGTTAGCTTCGTCGGCT | 124 |
| chr12:20083584-20085712 | p2 | (AT)64 | CGGGTAGGGGATTCTACCAT | TTGACGGCCGTTAGAAAAAC | 101 |
| chr12:20224205-20226219 | p2 | (TA)7 | GAACAACGGAGCTCTCCAAG | CCGTATTGACCCCCTTTCTT | 291 |
| chr12:20237279-20239291 | p2 | (AG)6 | CTTGTGCGGCTATGATACGA | TTGGCAGAGTTTCCCTCTGT | 248 |
| chr12:20253859-20255893 | p2 | (CT)17 | TTTTCCAGCAGATTGAACCC | GGAAGATCGGGTCGTTGATA | 208 |
| chr12:20355977-20357992 | p3 | (TGA)5 | TCGTTTGCGTCGTGAGTAAG | GCTTTTGTCCCACCTCAAAA | 187 |
| chr12:20412037-20414059 | p2 | (AC)11 | AACGGAAGTCGAACAACACC | AGATTCGTGGATTCACAGCC | 116 |
| chr12:20448139-20450151 | p2 | (TA)6 | CTTCGTCGTCCGTTAACCAT | GTGAGTTTGGAGATGGTGGG | 248 |
| chr12:20466692-20468712 | p2 | (TA)10 | GCGAACCATTCGGAATAAAA | CCTATCAGCAGCATTCGACA | 269 |
| chr12:20482003-20484021 | p2 | (GA)9 | CAAAGACGATATGCCCACCT | GGCATCTGCCGAGTAGTCTC | 111 |
| chr12:20495676-20497691 | p2 | (AC)9 | TGGATGACCCACAAAATCCT | ACGGGCTGTACTTTCATTGG | 192 |
| chr12:20495676-20497691 | p5 | (AAGAA)3 | TGGATGACCCACAAAATCCT | ACGGGCTGTACTTTCATTGG | 192 |
| chr12:20539271-20541286 | p5 | (TGTGT)3 | GCGGGAAGATAACAAGCAAG | AGTCATATTGCGCATTTCCC | 173 |
| chr12:20545172-20547184 | p6 | (ATGTAT)3 | GCGGGAAGATAACAAGCAAG | GTCATATTGCGCATTTCCCT | 178 |
| chr12:20551946-20553958 | p2 | (AT)6 | ACCGACCTTACATCACTCCG | CCGTGCCACTTATCTCCAAT | 264 |
| chr12:20588453-20590468 | p5 | (ATTGC)3 | CCCCAAATTCTAGGGTTGGT | CTCCTTAGCCTTGCCCTTCT | 105 |
| chr12:20640490-20642510 | p5 | (AATAT)4 | GTGGCTCCAACATCCTGAAT | CCTGAGCAGAGGCATACACA | 130 |
| chr12:20640490-20642510 | p5 | (ATCCA)3 | GTGGCTCCAACATCCTGAAT | CCTGAGCAGAGGCATACACA | 130 |

| Marker ID | SSR type | SSR motif | FORWARD PRIMER (5'-3') | REVERSE PRIMER (5'-3') | size (bp) |
|---|---|---|---|---|---|
| chr12:20947151-20949166 | p3 | (CAA)5 | CCTTCATCCCAGCCTTCATA | TTGGGGTTGAGGTGAGTAGG | 265 |
| chr12:20986676-20988691 | p5 | (GAAAT)3 | GATGATTGGGCAAAAGGAAA | ATTGCGGTGAATGTTGTGAA | 204 |
| chr12:21068536-21070551 | p3 | (CAT)5 | TGAATGGACGCTTGATGTGT | ACATGTCTGGCTAACGGGTC | 135 |
| chr12:21079062-21081104 | p2 | (TA)21 | ACATTGCCTTGCATGTACCA | CATCAATGTGTTGACGGTCC | 103 |
| chr12:21124272-21126287 | p5 | (TCATT)3 | TGTTCCAATCCCCAGAAGAG | CCACCATATCTTCGCCTCAT | 270 |
| chr12:21148070-21150084 | p2 | (AT)7 | TCGCTTTCATCCTCCAACTT | GCTGAAGAACGACATCGACA | 273 |
| chr12:21266569-21268584 | p3 | (ATG)5 | TGATGAAGGGTAAGATGGGG | GCGAACAACATTTCCCTTGT | 252 |
| chr12:21285482-21287497 | p3 | (ACA)5 | AGTGCGCACATCACAAAGTC | ATCGATTCCTACTTCGGCCT | 111 |
| chr12:21285482-21287497 | p2 | (AC)6 | AGTGCGCACATCACAAAGTC | ATCGATTCCTACTTCGGCCT | 111 |
| chr12:21344919-21346935 | p2 | (AT)8 | GCGTACAAGGCTGCATAACA | ATGCCCACAAATAGACAGGG | 143 |
| chr12:21414024-21416062 | p2 | (AT)19 | CTTGTTGAGCCGAAGTAGCC | GAAAGGGAGCCACAAATGAA | 286 |
| chr12:21414024-21416062 | p3 | (TCA)5 | CTTGTTGAGCCGAAGTAGCC | GAAAGGGAGCCACAAATGAA | 286 |
| chr12:21509415-21511453 | p2 | (AT)19 | CTTGTTGAGCCGAAGTAGCC | GAAAGGGAGCCACAAATGAA | 286 |
| chr12:21509415-21511453 | p3 | (TCA)5 | CTTGTTGAGCCGAAGTAGCC | GAAAGGGAGCCACAAATGAA | 286 |
| chr12:21529897-21531912 | p3 | (AAT)5 | GGGCGTTCTGTTGCTAATGT | GGAAAGCCACTGGTTATGGA | 280 |
| chr12:21604101-21606113 | p2 | (GA)6 | ATGAAGGAGGCATGTGAAGG | CCTCGAGCTCACATTAAGCC | 267 |
| chr12:21604101-21606113 | p3 | (GTT)6 | ATGAAGGAGGCATGTGAAGG | CCTCGAGCTCACATTAAGCC | 267 |
| chr12:21604629-21606647 | p2 | (GA)6 | CCAAGTGTAAGCCCTCCAAA | TGTTACATACAGGGCCACGA | 141 |
| chr12:21604629-21606647 | p3 | (GTT)6 | CCAAGTGTAAGCCCTCCAAA | TGTTACATACAGGGCCACGA | 141 |
| chr12:21668444-21670460 | p2 | (TA)8 | TGGCATGAATTTCCTTCCTC | GCCTCACCTTGAAGGAACAA | 128 |
| chr12:21696125-21698140 | p5 | (GTTTT)3 | TACTGACCTGCACTTGCTGC | CGACTCACCCTGGAAAACTC | 255 |
| chr12:21911652-21913667 | p6 | (AAAAAG)3 | TCCCAAGCATAATTTCCAGC | AGCAAGAAGGAGGTTTGGGT | 178 |
| chr12:21911652-21913667 | p3 | (AAT)5 | TCCCAAGCATAATTTCCAGC | AGCAAGAAGGAGGTTTGGGT | 178 |
| chr12:22263605-22265620 | p5 | (GATAT)3 | ACTGCCAAGTCAGGGTATGG | CTCGAATCTTCCAAGCGAAC | 298 |
| chr12:22288270-22290282 | p2 | (CA)6 | CGTCTGTACGGGTTATGGCT | TTTGGACGAAAAGGGAATTG | 130 |
| chr12:22336740-22338788 | p2 | (GA)7 | GATTTTTGCGACTCGAAAGC | TCTTTTCCATCGTCATTCCC | 211 |
| chr12:22336740-22338788 | p3 | (TAA)16 | GATTTTTGCGACTCGAAAGC | TCTTTTCCATCGTCATTCCC | 211 |
| chr12:22467820-22469832 | p2 | (GT)6 | TGTTGGACCTCCCTTGAGAC | TCCGATATTCCCACACCAAT | 219 |
| chr12:22516432-22518450 | p2 | (AT)9 | CCATGGCTTGCTAATTGGTT | GAGATCAAGAAGTGGCGCTC | 249 |
| chr12:22568091-22570103 | p2 | (TA)6 | TTTCCTATATGCCCCACAGC | AAAGTCGGGGTGTGACAAAG | 121 |
| chr12:22584229-22586253 | p6 | (ATACAT)4 | CGGGCCTTTAAATCAAACAA | AGAGTCGTGGGTATTGGTCG | 163 |
| chr12:22584229-22586253 | p3 | (CAT)8 | CGGGCCTTTAAATCAAACAA | AGAGTCGTGGGTATTGGTCG | 163 |
| chr12:22584229-22586253 | p5 | (ATTCA)3 | CGGGCCTTTAAATCAAACAA | AGAGTCGTGGGTATTGGTCG | 163 |
| chr12:22640982-22642997 | p3 | (TTC)5 | CTACTCGGGGTGAACCGTTA | TACGGTCCATCTTTTGCTCC | 240 |
| chr12:22737367-22739382 | p5 | (CACAA)3 | TTCAAGGAACAGAATTGCCC | CATTTGCAGATTCTGAGCGA | 197 |
| chr12:22804880-22806892 | p2 | (AT)6 | CCTTACGTCACCCCGATAGA | TGCCAAAAGGTCATGGTGTA | 278 |

| Marker ID | SSR type | SSR motif | FORWARD PRIMER (5'-3') | REVERSE PRIMER (5'-3') | size (bp) |
|---|---|---|---|---|---|
| chr12:22922121-22924137 | p2 | (AT)8 | CGGAAGGAATGTGCAAAGAT | GAGAGTACACCCCAGGTCCA | 279 |
| chr12:23008075-23010091 | p2 | (AC)8 | GAAATGCTACGGAGGTTCCA | TGCAGATTCTGAACGAAACG | 217 |
| chr12:23022900-23024912 | p4 | (CTTT)4 | TGTGGACTGCTTTGAAGGTG | GGCCAAATCCTCAACAAAAA | 157 |
| chr12:23022900-23024912 | p2 | (TA)6 | TGTGGACTGCTTTGAAGGTG | GGCCAAATCCTCAACAAAAA | 157 |
| chr12:23138000-23140018 | p5 | (ACAAC)3 | CTTCGACGACACCCTTCATT | TGTGTTGTATTGCAAGGGGA | 162 |
| chr12:23138000-23140018 | p2 | (TA)9 | CTTCGACGACACCCTTCATT | TGTGTTGTATTGCAAGGGGA | 162 |
| chr12:23167168-23169180 | p2 | (TG)6 | AACCATATCCGTGTTTGGGA | TCTCCTTACCCCGACATCAC | 286 |
| chr12:23169831-23171855 | p6 | (TGTATA)4 | AACGTGAATTTGGAGAAGCG | TTATTCCCGAGGCTCACATC | 147 |
| chr12:23252041-23254067 | p2 | (AT)13 | TTCTGGGTTTGGCAGGTTAC | GTGTTCACACGCATCATTCC | 267 |
| chr12:23515869-23517883 | p2 | (TA)7 | CATCCCGTTTTTCGTCTGTT | ATCGTGTTTAAACCGCCAAC | 166 |
| chr12:23515869-23517883 | p3 | (TTG)6 | CATCCCGTTTTTCGTCTGTT | ATCGTGTTTAAACCGCCAAC | 166 |
| chr12:23521544-23523562 | p3 | (TTG)6 | CGCTCGTCGTGAGATATTGA | AATCTCCTCGTCTCCCCAAT | 260 |
| chr12:23521544-23523562 | p6 | (TATGTA)3 | CGCTCGTCGTGAGATATTGA | AATCTCCTCGTCTCCCCAAT | 260 |
| chr12:23572329-23574345 | p2 | (TA)8 | ATCGCGGAGCTGAGAATCTA | GGCGTGAAAATTCCATCACT | 169 |
| chr12:23572329-23574345 | p2 | (AG)6 | ATCGCGGAGCTGAGAATCTA | GGCGTGAAAATTCCATCACT | 169 |
| chr12:23604961-23606973 | p2 | (AT)6 | ACATCCTTCACTGAGTCCCG | ACCGTACTCGGCCCTTTAGT | 213 |
| chr12:23711292-23713308 | p2 | (AT)39 | TCTAAGCTTGCTGCTCGTCA | AGTTCAACAATCCCCTGTCG | 149 |
| chr12:23711292-23713308 | p2 | (TG)8 | TCTAAGCTTGCTGCTCGTCA | AGTTCAACAATCCCCTGTCG | 149 |
| chr12:23725486-23727498 | p2 | (GT)6 | GCATTGGAATCAGTGACGTG | GGGACATACATTCTCCCCCT | 263 |
| chr12:23788685-23790697 | p2 | (CA)6 | TTTTGACCCAAAGAATTCCG | AGGAAAGTGAGTCCCAGGGT | 168 |
| chr12:23788685-23790697 | p2 | (AT)7 | TTTTGACCCAAAGAATTCCG | AGGAAAGTGAGTCCCAGGGT | 168 |
| chr12:23826728-23828740 | p2 | (AT)6 | GGGTCCCATCTCAGCAAATA | AGCAGGATTGCGAATGAGAT | 237 |
| chr12:23873921-23875937 | p2 | (AT)9 | ATCGTGCACAGTCTCACTGC | TTGGTGCCATAACACAGCAT | 285 |
| chr12:23873921-23875937 | p2 | (AT)8 | ATCGTGCACAGTCTCACTGC | TTGGTGCCATAACACAGCAT | 285 |
| chr12:23937346-23939358 | p2 | (AT)6 | CGTAAATACAACCATCCCCG | ATTGTTGGTGTGGTTCGTGA | 106 |
| chr12:23960237-23962252 | p5 | (AATGC)3 | GTAGGCAAACCCTCCATTCA | CCTAGACCAGGTCATGGGAA | 145 |
| chr12:23994869-23996881 | p2 | (AT)6 | GCCATTCCCGACTTAACAAA | CTTAGGGTTGTCCGTACCCA | 120 |
| chr12:23996497-23998513 | p2 | (AG)8 | GATGCGGAAATATGGAGGAA | TTTTGATTTTCCAAGCCCAC | 297 |
| chr12:24014183-24016219 | p2 | (TA)11 | CGTGCCAGTAAATGATGTGC | CCTAGTCGGGTCGTGACAAT | 228 |
| chr12:24014183-24016219 | p2 | (TA)10 | CGTGCCAGTAAATGATGTGC | CCTAGTCGGGTCGTGACAAT | 228 |
| chr12:24014255-24016305 | p2 | (TA)11 | CGTGCCAGTAAATGATGTGC | CCTAGTCGGGTCGTGACAAT | 156 |
| chr12:24014255-24016305 | p2 | (TA)10 | CGTGCCAGTAAATGATGTGC | CCTAGTCGGGTCGTGACAAT | 156 |
| chr12:24014660-24016682 | p2 | (TA)11 | TATGGACGAGACAGGAACCC | TGTTCATGCCCACAGGTAGA | 172 |
| chr12:24014660-24016682 | p2 | (TA)10 | TATGGACGAGACAGGAACCC | TGTTCATGCCCACAGGTAGA | 172 |
| chr12:24029753-24031791 | p2 | (TA)19 | TATGAGTCGAGACAAGGCCC | GTGTTCATGCCACACAGTCC | 296 |
| chr12:24117430-24119448 | p6 | (AGAAAA)3 | GACGAGTGGAATGAGGGAAA | TCTTTACGGGAATGAATCGC | 122 |

| Marker ID | SSR type | SSR motif | FORWARD PRIMER (5'-3') | REVERSE PRIMER (5'-3') | size (bp) |
|---|---|---|---|---|---|
| chr12:24140000-24142014 | p3 | (ACA)5 | CCCAAAACAGTCCCTCAAAA | TTAAAGGGCTGGAAACATGG | 267 |
| chr12:24162468-24164482 | p3 | (ACA)5 | CCCAAAACAGTCCCTCAAAA | AGTGGGCAGATTTGGAATTG | 269 |
| chr12:24203074-24205092 | p2 | (AT)9 | CATAGAGTCGTTCCGGTCGT | CCCGAAGACCCTCACAAGTA | 178 |
| chr12:24211307-24213322 | p5 | (ATCAT)3 | TACATGGAAGCCGATAAGGC | ACGACAGATTTGCTTACCCG | 293 |
| chr12:24222539-24224557 | p2 | (TC)9 | CCTTGGTCCCAAGACTTCAA | TTTCTGTGTCAACAGCGGAG | 138 |
| chr12:24241184-24243198 | p2 | (AT)7 | TCTTCCGCTCCTTTTTGCTA | TTCGGTAGCCAAGTGTTTCC | 285 |
| chr12:24251386-24253401 | p3 | (ATA)5 | GAAGACGATGAAGGCCAGAC | ACTTAGTTGTCGGCCCCTTT | 130 |
| chr12:24268222-24270236 | p2 | (AT)7 | TCTTCCGCTCCTTTTTGCTA | TTCGGTAGCCAAGTGTTTCC | 285 |
| chr12:24422542-24424554 | p2 | (AT)7 | TCATGATGACATTCGAGGGA | TTTTCGGATGTCACTGTGGA | 103 |
| chr12:24422542-24424554 | p2 | (AT)6 | TCATGATGACATTCGAGGGA | TTTTCGGATGTCACTGTGGA | 103 |
| chr12:24472611-24474623 | p2 | (AT)7 | TCATGATGACATTCGAGGGA | TTTTCGGATGTCACTGTGGA | 103 |
| chr12:24472611-24474623 | p2 | (AT)6 | TCATGATGACATTCGAGGGA | TTTTCGGATGTCACTGTGGA | 103 |
| chr12:24557271-24559295 | p2 | (TA)12 | ATAATTAGGGGCATAGCGGG | AGCCGACAAGGCTGTAAAAA | 243 |
| chr12:24573540-24575554 | p2 | (TA)7 | AACACATCCAATGCCTCACA | AAGCAGTTGATTATGCCGCT | 201 |
| chr12:24592158-24594186 | p2 | (AT)14 | AATAAAATTCGGACCCGACC | CGAATCCGGAAAAGTATGGA | 217 |
| chr12:24620155-24622167 | p2 | (TA)6 | TACGAGCCTCTGGGACAACT | GGTAATCGGGTCCCACTTTT | 236 |
| chr12:24655816-24657878 | p2 | (TA)31 | CCTATAACAAGCCAGCTCCG | ATTGACCAGATTGGTAGCCG | 197 |
| chr12:24655816-24657878 | p2 | (AT)7 | CCTATAACAAGCCAGCTCCG | ATTGACCAGATTGGTAGCCG | 197 |
| chr12:24655816-24657878 | p2 | (AT)9 | CCTATAACAAGCCAGCTCCG | ATTGACCAGATTGGTAGCCG | 197 |
| chr12:24700456-24702518 | p2 | (TA)31 | CCTATAACAAGCCAGCTCCG | ATTGACCAGATTGGTAGCCG | 197 |
| chr12:24700456-24702518 | p2 | (AT)7 | CCTATAACAAGCCAGCTCCG | ATTGACCAGATTGGTAGCCG | 197 |
| chr12:24700456-24702518 | p2 | (AT)9 | CCTATAACAAGCCAGCTCCG | ATTGACCAGATTGGTAGCCG | 197 |
| chr12:24807264-24809294 | p5 | (ATTCA)3 | CACCATCCCCGAGAATCTAA | AATTTGGAAAGTGTCGGGTG | 156 |
| chr12:24807264-24809294 | p2 | (AT)15 | CACCATCCCCGAGAATCTAA | AATTTGGAAAGTGTCGGGTG | 156 |
| chr12:24807264-24809294 | p2 | (AT)17 | CACCATCCCCGAGAATCTAA | AATTTGGAAAGTGTCGGGTG | 156 |
| chr12:24828438-24830460 | p2 | (AT)11 | ATGTCATTGAGGAACGAGGG | CTTGGACGTCGGAAAAGTGT | 197 |
| chr12:24848214-24850226 | p2 | (TA)6 | GATCGGGAGTTGTCGGTAAA | CGCTTCTGGCCTCTTAACTG | 239 |
| chr12:24850857-24852869 | p2 | (TA)6 | GATCGGGAGTTGTCGGTAAA | CGCTTCTGGCCTCTTAACTG | 239 |
| chr12:24871470-24873485 | p5 | (GAAAG)3 | CAGGGCTGCCAAAGTGTTAT | GAAGCAGGGAGACGAATCTG | 148 |
| chr12:24906381-24908413 | p3 | (CAA)5 | TGGCGTGCTTCATACCAATA | GGTTAGCTCAGCTGGCATTC | 280 |
| chr12:24906446-24908461 | p3 | (CAA)5 | TGGCGTGCTTCATACCAATA | GGTTAGCTCAGCTGGCATTC | 215 |
| chr12:24906503-24908518 | p3 | (CAA)5 | TGGCGTGCTTCATACCAATA | GGTTAGCTCAGCTGGCATTC | 158 |
| chr12:24929208-24931224 | p2 | (GT)8 | GATCCCCAGCTGGTTCACTA | TTCCTCAATATTTCCGCGTC | 243 |
| chr12:24977847-24979863 | p2 | (TA)8 | ACAACAACTCCGGAACAAGG | TATCATTGGCCCTCATAGGC | 176 |
| chr12:24977847-24979863 | p2 | (AT)49 | ACAACAACTCCGGAACAAGG | TATCATTGGCCCTCATAGGC | 176 |
| chr12:25020889-25022901 | p2 | (AG)6 | GCCTGATGCTTGTGAAGTGA | GCACGTTGGTGTTCCTTCTT | 173 |

| Marker ID | SSR type | SSR motif | FORWARD PRIMER (5'-3') | REVERSE PRIMER (5'-3') | size (bp) |
|---|---|---|---|---|---|
| chr12:25054541-25056557 | p2 | (AT)8 | CGTGCATGCCTCAATATGTC | TTGGTGCCATAACACAGCAT | 118 |
| chr12:25081456-25083471 | p5 | (CATAG)3 | ACAAACCCCTAGGACCGAAC | CATACGGCCTTGCTAAGAGG | 166 |
| chr12:25083739-25085759 | p5 | (AAATG)4 | CCAATGGGACTAGCGTGATT | AGCTAACGGTCATGCGACTT | 136 |
| chr12:25113618-25115638 | p5 | (AAATG)4 | CCAATGGGACTAGCGTGATT | AGCTAACGGTCATGCGACTT | 136 |
| chr12:25222050-25224080 | p2 | (AT)15 | TCCTCTTGCTCCTCACACCT | CCCCATATTCCACAAGTTCG | 262 |
| chr12:25285854-25287874 | p2 | (AC)7 | CCGAGAGTACAACTCGGCTC | GAGGTTGGGTTGGAAATGTG | 214 |
| chr12:25285962-25287974 | p2 | (AC)7 | CCGAGAGTACAACTCGGCTC | GAGGTTGGGTTGGAAATGTG | 106 |
| chr12:25443352-25445364 | p2 | (TG)6 | ACGACTGTACTACGCACCCC | CTCATACCTGGCCCTACCAA | 148 |
| chr12:25522801-25524817 | p2 | (AT)6 | CCACCCTTAAAAGTTTCGCA | ACAACGTCCAAAGAACAGGG | 127 |
| chr12:25522801-25524817 | p2 | (CT)8 | CCACCCTTAAAAGTTTCGCA | ACAACGTCCAAAGAACAGGG | 127 |
| chr12:25568869-25570884 | p5 | (TTTCA)3 | AAGAAATGACATGGGAAGCG | ATAACAATGCCTCACAGGGC | 170 |
| chr12:25625192-25627204 | p2 | (AG)6 | CCGTCATGGCCTATCAGTTT | GGTGTTGCTTGATCCGAACT | 171 |
| chr12:25711538-25713550 | p2 | (TA)6 | CAAAGTGGAAACCCTCCAAA | TAGCCCACTGACCTTTCGAC | 278 |
| chr12:25737255-25739282 | p3 | (CAA)9 | AATGGGAGCTCACCAATCAG | GCCACTTGGTCAGTGAGACA | 161 |
| chr12:25770543-25772561 | p6 | (CATATA)3 | ATGAGCCGTGACAAGTACCC | GAAATGATGTCACCCAGCCT | 154 |
| chr12:25770543-25772561 | p3 | (CAA)5 | ATGAGCCGTGACAAGTACCC | GAAATGATGTCACCCAGCCT | 154 |
| chr12:25783460-25785475 | p5 | (ATCAT)3 | GATAAAGCACAAGGAAGCCG | TGCTCTATTGTTCGGAGCCT | 292 |
| chr12:25794847-25796859 | p2 | (GT)6 | CGTATTTTCGAACGTCGGAT | GAAGTCAGGTCACCCGAAAA | 119 |
| chr12:25807299-25809311 | p2 | (GT)6 | CGTATTTTCGAACGTCGGAT | GAAGTCAGGTCACCCGAAAA | 119 |
| chr12:25825205-25827225 | p2 | (TC)10 | AGCTGAGGCCATAGCACACT | GAAGCGAGCTTCCAGGTATG | 250 |
| chr12:25920402-25922422 | p2 | (AT)10 | ATACATTTTCATCGCTCCCG | TCGGGCAGTATGGGAGTTAC | 131 |
| chr12:25955398-25957410 | p2 | (TA)6 | AGCTAAAGGAGCGGAAGGAG | GATTCCGGGATTAGGGTTGT | 186 |
| chr12:26011513-26013531 | p3 | (TGT)6 | GGCTGGTCATCTCAAGAAGC | GTTTCGGACCATAACCGAGA | 108 |
| chr12:26054032-26056050 | p6 | (TATATG)3 | GTTGATGATTTGGCCGAGTT | TTACATACCTTGCCGGCTTC | 274 |
| chr12:26096819-26098879 | p2 | (AT)30 | ACTACCTGTTGAGCACCGCT | ACGTATGTCCTTTGGGCTTG | 285 |
| chr12:26260781-26262819 | p2 | (TA)19 | GCTCTGTGAGGCATTGTTGA | AATCGCCTTTCTCGCTATCA | 116 |
| chr12:26345788-26347800 | p2 | (TA)6 | CTTGTACCGAGCACCCCTTA | TCCGGAGCCTTATATCATCG | 165 |
| chr12:26379723-26381738 | p3 | (ACA)5 | TAAACTGACGCCAGAAGGCT | ATTCGGTTTGTTTGATTCGC | 268 |
| chr12:26379723-26381738 | p6 | (ACACAT)4 | TAAACTGACGCCAGAAGGCT | ATTCGGTTTGTTTGATTCGC | 268 |
| chr12:26465986-26468004 | p6 | (TATATG)3 | GGAGTCGCATAAGAAGCAGG | GGATTTTAGGGTCATCGGGT | 166 |
| chr12:26566049-26568077 | p2 | (TA)14 | GGACACGACTTTGCAGACAA | CTCAGCCGAGTAGGGTCTTG | 179 |
| chr12:26574089-26576105 | p2 | (GA)8 | AGATATCCATGTGCGGGAAG | ACCCAAGTCATTCAGCAACC | 111 |
| chr12:26766580-26768592 | p2 | (AT)6 | GTCCCAAATCATGAGCGAAT | CATGCAGAAGATTGAAGCGA | 179 |
| chr12:26864762-26866777 | p5 | (ATGTT)3 | GGCGAGTTTGTTGTTGGTTT | ACGTTCGTTCGTTTCGTTCT | 154 |
| chr12:26880873-26882888 | p3 | (CAT)5 | CACCAGCCGACAAGGTTACT | GATCAGATTCGTCGGTGCTT | 296 |
| chr12:26885652-26887667 | p3 | (CAT)5 | TACATCCCTCCAAATCCACC | CAACGGATTAGAACGGGAAA | 277 |

| Marker ID | SSR type | SSR motif | FORWARD PRIMER (5'-3') | REVERSE PRIMER (5'-3') | size (bp) |
|---|---|---|---|---|---|
| chr12:27034861-27036876 | p4 | (TATG)4 | TGTGCATGACCAGGTTTCAT | GTACCTGTCGGGGCAATAGA | 215 |
| chr12:27034861-27036876 | p5 | (AATGC)3 | TGTGCATGACCAGGTTTCAT | GTACCTGTCGGGGCAATAGA | 215 |
| chr12:27119654-27121669 | p5 | (TTTTC)3 | GAGGTTGGGTTTTTCGCATA | TGGAACAACTGGGTTGAACA | 190 |
| chr12:27215860-27217876 | p2 | (CA)8 | TGCCAAGAGAAGAAGAGGGA | TGTATAGGGGAAATGCTGCC | 261 |
| chr12:27234519-27236537 | p6 | (ATTATA)3 | GGAGTTTGTGGAATGGGAGA | TTCTTACAAAGGCGACCACC | 196 |
| chr12:27359888-27361903 | p5 | (ACACA)3 | TTCACAAGGATCATGCCAAA | GTTGATGATTTGGCCGAGTT | 281 |
| chr12:27359888-27361903 | p2 | (AT)24 | TTCACAAGGATCATGCCAAA | GTTGATGATTTGGCCGAGTT | 281 |
| chr12:27359888-27361903 | p2 | (AT)9 | TTCACAAGGATCATGCCAAA | GTTGATGATTTGGCCGAGTT | 281 |
| chr12:27360286-27362334 | p5 | (ACACA)3 | AACTCGGCCAAATCATCAAC | CATTTGGAAGGTTTCGGGTA | 111 |
| chr12:27360286-27362334 | p2 | (AT)24 | AACTCGGCCAAATCATCAAC | CATTTGGAAGGTTTCGGGTA | 111 |
| chr12:27360286-27362334 | p2 | (AT)9 | AACTCGGCCAAATCATCAAC | CATTTGGAAGGTTTCGGGTA | 111 |
| chr12:27361961-27363982 | p2 | (GT)8 | CAAGAAGGTGCCACCAAAAT | CTATCGCTCCTAGTGTGCCC | 274 |
| chr12:27361961-27363982 | p3 | (TGT)7 | CAAGAAGGTGCCACCAAAAT | CTATCGCTCCTAGTGTGCCC | 274 |
| chr12:27372171-27374189 | p3 | (GTT)6 | AGGGTTCCACGCTATCTCCT | CCGCCCAAAACTATGGTCTA | 120 |
| chr12:27382331-27384377 | p2 | (TA)23 | AGACTTTGAGCGTTTCGAGC | TCTAACCAGACTCCATGCCC | 137 |
| chr12:27550328-27552346 | p6 | (ATATTG)3 | CGCGTGAAACATCGTCTAAA | TCATATCATTGCGTGAGCGT | 100 |
| chr12:27550328-27552346 | p2 | (TA)7 | CGCGTGAAACATCGTCTAAA | TCATATCATTGCGTGAGCGT | 100 |
| chr12:27550818-27552832 | p6 | (ATATTG)3 | CGTTTTCGTGCGCATATTTA | GGCCCTTTAGTGAGGGACTC | 221 |
| chr12:27550818-27552832 | p2 | (TA)7 | CGTTTTCGTGCGCATATTTA | GGCCCTTTAGTGAGGGACTC | 221 |
| chr12:27574972-27576986 | p2 | (AT)7 | GCCATTACGAATGGGTATGC | TTGTTCCGGAGCTAGCTTGT | 281 |
| chr12:27584860-27586875 | p2 | (AT)11 | TTCGCCCTTAACCTCTCTCA | GGAAAATTCAAGGCAATGGA | 299 |
| chr12:27584860-27586875 | p5 | (TTGAG)3 | TTCGCCCTTAACCTCTCTCA | GGAAAATTCAAGGCAATGGA | 299 |
| chr12:27584860-27586875 | p3 | (ATG)7 | TTCGCCCTTAACCTCTCTCA | GGAAAATTCAAGGCAATGGA | 299 |
| chr12:27602168-27604183 | p5 | (AAATT)3 | TGTGCCTTATACTGCGTTGC | CCTCATCCCTGCATCAATCT | 102 |
| chr12:27602168-27604183 | p2 | (AT)6 | TGTGCCTTATACTGCGTTGC | CCTCATCCCTGCATCAATCT | 102 |
| chr12:27620986-27623001 | p5 | (TTTCA)3 | TGGAGTTTTTCGGGTCAAAG | CCCGGGCACTTATATTTCCT | 180 |
| chr12:27724731-27726747 | p4 | (TGTA)4 | TGCCTCCGAATCATGTTGTA | ATCAACACAGCAACCCAACA | 206 |
| chr12:27923483-27925498 | p3 | (CAA)5 | TTTCGGGACTCGGTGTTATC | GATTTCATCTCGCGATGGTT | 249 |
| chr12:27923483-27925498 | p2 | (AT)6 | TTTCGGGACTCGGTGTTATC | GATTTCATCTCGCGATGGTT | 249 |
| chr12:28094686-28096722 | p2 | (AT)18 | GCAAGTTTTGGCATTCCATT | CCATACCTTTGCAGCCAAAT | 205 |
| chr12:28149109-28151130 | p3 | (TTG)7 | AGAAGATGAAAGTCGCGGAA | CCTAAAACCGTCGATAGCCA | 297 |
| chr12:28165675-28167729 | p2 | (TA)27 | TGGCTCTTCAGGAGACAGGT | ATGACTAGTGCCCGAGTTGG | 200 |
| chr12:28220252-28222264 | p2 | (TA)6 | TGGAGGGATGATTTTCGTTC | CCAAAGTACGCCCAGAATGT | 248 |
| chr12:28361759-28363797 | p2 | (AT)19 | TCCCGAAATCGATAAGATGC | GTTGCTTCGGGTCAACAAAT | 157 |
| chr12:28388297-28390317 | p5 | (CTTTT)4 | GCAAACCGGTAGTTGTTCGT | CTTTGATGCCTTCGCATTTT | 250 |
| chr12:28428960-28430975 | p5 | (AAACG)3 | AAGAAAGGAGAAGAACGGGC | GTGGATTCCAATCGCGTAGT | 200 |

| Marker ID | SSR type | SSR motif | FORWARD PRIMER (5'-3') | REVERSE PRIMER (5'-3') | size (bp) |
|---|---|---|---|---|---|
| chr12:28450390-28452404 | p2 | (TA)7 | CGTCATCTTGCTCGTTTTCA | TTTTGCCCATGGAAATTGAT | 104 |
| chr12:28450390-28452404 | p2 | (TA)12 | CGTCATCTTGCTCGTTTTCA | TTTTGCCCATGGAAATTGAT | 104 |
| chr12:28450390-28452404 | p2 | (TA)7 | CGTCATCTTGCTCGTTTTCA | TTTTGCCCATGGAAATTGAT | 104 |
| chr12:28533060-28535120 | p6 | (AAACTG)3 | GGCTTGCTGAATCATTGTGA | GCCATCTTTGGCTCATTCAT | 258 |
| chr12:28539208-28541234 | p2 | (TA)13 | TTGAAGGGTGGGAGAGAAGA | ATCACCTGCATAAAGTGGGC | 293 |
| chr12:28542527-28544539 | p2 | (GT)6 | AATTTTTCCGGTTCAATCCC | TGCTCTTAATCTCCCGCAAT | 103 |
| chr12:28553164-28555179 | p5 | (CAATG)3 | TTGTACCCCTGAAGACTGGG | CCCGCATGACCTAGGATAAA | 126 |
| chr12:28607929-28609944 | p5 | (AATGC)4 | CCACGAAGCATTATCAAGCA | ATTGGGTCCTTTGAGCTGTG | 202 |
| chr12:28607929-28609944 | p5 | (CCTTT)3 | CCACGAAGCATTATCAAGCA | ATTGGGTCCTTTGAGCTGTG | 202 |
| chr12:28630912-28632924 | p2 | (TA)6 | CCCCGGATTTTCCTTATTGT | AGGCATTTTCTCGTCTTGGA | 271 |
| chr12:28646453-28648468 | p3 | (TGT)5 | ATTCGCAAGCAAATCGACTC | CCCCCTAATGACTCACCAAA | 259 |
| chr12:28646453-28648468 | p2 | (AT)9 | ATTCGCAAGCAAATCGACTC | CCCCCTAATGACTCACCAAA | 259 |
| chr12:28662605-28664620 | p3 | (TTG)7 | TGAAATTGGGAAATGTGGGT | GCTACCCTATCGCTTGAACG | 107 |
| chr12:28677347-28679365 | p2 | (AT)22 | AAGGACCTGGCAACTCTGAA | GGCATCTTTTGTCCTCCAAA | 123 |
| chr12:28677347-28679365 | p3 | (CAA)6 | AAGGACCTGGCAACTCTGAA | GGCATCTTTTGTCCTCCAAA | 123 |
| chr12:28720864-28722886 | p2 | (AT)11 | GTCTAGAAGTGTGGCACGCA | CGGGTCGTGATAAAAGTGGT | 186 |
| chr12:28737398-28739410 | p2 | (AT)6 | TAATAGCTGGCCAAACCCAC | GTAGCATTCTTGGGCCATGT | 188 |
| chr12:28778096-28780108 | p2 | (AT)6 | ACTCGGCCAATTTCAACAAC | TTGAAGTTGGAAAGGTTCGG | 118 |
| chr12:28778096-28780108 | p3 | (TTA)5 | ACTCGGCCAATTTCAACAAC | TTGAAGTTGGAAAGGTTCGG | 118 |
| chr12:28778223-28780238 | p2 | (AT)6 | CCGAACCTTTCCAACTTCAA | GGCCGAATAGGGTATGGTTT | 112 |
| chr12:28778223-28780238 | p3 | (TTA)5 | CCGAACCTTTCCAACTTCAA | GGCCGAATAGGGTATGGTTT | 112 |
| chr12:28841588-28843604 | p2 | (TA)12 | CAAACCCCAACCATTTTCAC | CAGCCACCCTATTAGGCAAA | 211 |
| chr12:28841600-28843615 | p2 | (TA)12 | CAAACCCCAACCATTTTCAC | CAGCCACCCTATTAGGCAAA | 199 |
| chr12:28860134-28862152 | p2 | (AT)9 | TCAACGATTCCTAGATGCCC | CAAGGGGTTGTCTGCAAAGT | 172 |
| chr12:28995575-28997590 | p2 | (AT)11 | TTTGGCCTAAGGGTCCTTCT | GAGCAATTTCAAGGCTCAGG | 275 |
| chr12:28995622-28997637 | p2 | (AT)11 | TTTGGCCTAAGGGTCCTTCT | GAGCAATTTCAAGGCTCAGG | 228 |
| chr12:29015893-29017905 | p2 | (TA)6 | TCCATATTGAACCCCGAGAG | CTGCACTTCACGATGCAGAT | 275 |
| chr12:29032013-29034029 | p2 | (AT)8 | CCAGAGCTTGGAAACCTGAC | TAGCTGTGTTCAAATGCCCA | 201 |
| chr12:29134632-29136650 | p6 | (CATATA)3 | AACAAGTGCCCGACCACTAC | TCGTGATGTTATGCAGCCTC | 112 |
| chr12:29166529-29168541 | p2 | (AT)6 | ACATCCATAGCCAAAGGTCG | TCTGGTTTGGATCGTGACAA | 268 |
| chr12:29274399-29276417 | p6 | (AGCACC)3 | AGAGGACCCCCAGGAGTTTA | TGTCTCAGGTGCAAGAATCG | 121 |
| chr12:29484983-29487005 | p2 | (AT)11 | TGCCAACTCCACAGATTCAG | AGTTCCTCCACCGGCTACTT | 156 |
| chr12:29598610-29600622 | p2 | (TA)6 | GCCGGGATCTGTATGTCTGT | AGAACCCGGCCCTCTTAGTA | 151 |
| chr12:29611685-29613703 | p6 | (ATCCGT)3 | TTTGCGTGTATCATCACGGT | CTCCCTGCAACAATCCAAAT | 179 |
| chr12:29639891-29641906 | p2 | (AT)43 | CCATAAGATGGGGGAGGATT | ATGTCCTTTCTTGTGGACGC | 135 |
| chr12:29639891-29641906 | p3 | (ATC)5 | CCATAAGATGGGGGAGGATT | ATGTCCTTTCTTGTGGACGC | 135 |

| Marker ID | SSR type | SSR motif | FORWARD PRIMER (5'-3') | REVERSE PRIMER (5'-3') | size (bp) |
|---|---|---|---|---|---|
| chr12:29690685-29692703 | p6 | (GGTAGA)3 | CCGGTATGGCAAGAGTTTGT | TTAGGCCTTTCACAAATCGG | 241 |
| chr12:29729811-29731859 | p3 | (GTT)5 | TATTTGATACATGCCGGGGT | ATTAAACCGCTCGTCCCTTT | 170 |
| chr12:29738232-29740247 | p5 | (AAGTG)3 | CGTAAGAGTTGGAGCTTCCG | TCAGCTTCGCATGTCATCTC | 173 |
| chr12:29747268-29749282 | p2 | (AG)7 | TACCGCTGAACAAGCAACTG | TTGTGGTTTAACTGGCCACA | 130 |
| chr12:29747268-29749282 | p4 | (TGTA)4 | TACCGCTGAACAAGCAACTG | TTGTGGTTTAACTGGCCACA | 130 |
| chr12:29781812-29783824 | p2 | (AT)6 | ATGAGAGCGTGGCAAGAACT | GTAGTGGCAGCCTTGTTGGT | 106 |
| chr12:29853313-29855331 | p2 | (AT)9 | TTCCTTGTAAGACGGGATGG | TCGAACTCTAGCCGTGTCCT | 271 |
| chr12:29854512-29856524 | p2 | (AT)6 | GGAAAATAAGTGCAAGCCCA | TTTGAAACCTTCCCTTGGTG | 278 |
| chr12:29854512-29856524 | p3 | (CAA)6 | GGAAAATAAGTGCAAGCCCA | TTTGAAACCTTCCCTTGGTG | 278 |
| chr12:29869296-29871311 | p2 | (TA)21 | TGTAGTGGCAGCCTTGTCAG | TGCGAAAGAACTCCAAAGGT | 275 |
| chr12:29869296-29871311 | p5 | (TATGA)3 | TGTAGTGGCAGCCTTGTCAG | TGCGAAAGAACTCCAAAGGT | 275 |
| chr12:29884210-29886222 | p2 | (AC)6 | TGCCAAATGAAAAGAAAGGG | TTAGATTCTCGGGGATGGTG | 214 |
| chr12:29906962-29908982 | p5 | (TGATA)4 | GAAGATGCTTGCCCGATAAG | GGGTCAAATTCAAACTCGGA | 109 |
| chr12:30000791-30002823 | p2 | (TG)11 | TCCAAAAATCCCCATCAAAA | TGCAACAAAGAGGGGATTTC | 264 |
| chr12:30000791-30002823 | p2 | (AT)16 | TCCAAAAATCCCCATCAAAA | TGCAACAAAGAGGGGATTTC | 264 |
| chr12:30008045-30010085 | p2 | (AT)20 | TCCAAAAATCCCCATCAAAA | TACATACACACACCCACCCG | 236 |
| chr12:30125722-30127734 | p2 | (AT)6 | TAATGTCGGAGGAAGGCATC | TGATAAGGGGTGCTCGGTAG | 173 |
| chr12:30150692-30152708 | p2 | (TA)8 | TTGTACCGAGCACCCCTTAG | TCCGGAGCCTTATATCATCG | 162 |
| chr12:30156481-30158493 | p2 | (TA)6 | TCCAATATCGCGTTACACCA | ATGCCCACAGATAGACAGGG | 227 |
| chr12:30189716-30191731 | p3 | (CAT)5 | GGTTAACCCCAAATTTCCGT | GACCCCGATCAACCCTAAAT | 115 |
| chr12:30269155-30271193 | p2 | (TA)19 | TGTCAGAGAGCCTTTGCTCA | GGAAAGCAAAGACCAAGCAG | 252 |
| chr12:30391190-30393210 | p2 | (TA)7 | AAAATGATACGCAAGCCGAC | AATTAATCCAGGGGGTACGG | 133 |
| chr12:30391219-30393251 | p2 | (TA)7 | AAAATGATACGCAAGCCGAC | AATTAATCCAGGGGGTACGG | 104 |
| chr12:30400403-30402429 | p2 | (AT)13 | CCACCAAATAAAAGCCGAAA | CACACCACATTTCATCCGAG | 122 |
| chr12:30407750-30409768 | p2 | (AT)9 | CCCGAACTGGACACTCGTAT | TGTTCGTCTGTAGTGCCAGC | 261 |
| chr12:30445462-30447482 | p5 | (TCATA)4 | CCATATCGTCACTCGCCTTT | TATAAACAGGAGGTTGCGGG | 164 |
| chr12:30482875-30484905 | p2 | (AT)15 | TCTGTTGGTGATGCTTCTGC | CCGAGTCCGAAACGAAAATA | 247 |
| chr12:30527424-30529439 | p5 | (ATATA)3 | ACAACCCACGCACATGTCTA | TCTTGGAGAGCTCCGTTGTT | 289 |
| chr12:30558988-30561000 | p2 | (CT)6 | TGTAGCAGAATTGCACGGAC | TCACTTCCCATTTCCAGGTC | 137 |
| chr12:30583909-30585921 | p2 | (TA)6 | TTCGAGTCGAGCGTATGATG | TATGCCTTGGAATCTGGGAC | 268 |
| chr12:30594150-30596168 | p6 | (ATATGT)3 | ATTTTGACGGATTTGGAACG | GGGTCCTATTTATGCGGGTT | 211 |
| chr12:30709322-30711344 | p2 | (AT)11 | GTTGTCATAATCCAACCCCG | GGACGTTATGAACCTTCGGA | 119 |
| chr12:30824917-30826932 | p2 | (GT)6 | GATGCGGAAATATCGAGGAA | CCCTTTTTCCCTTCTATGGC | 267 |
| chr12:30824917-30826932 | p3 | (TTG)5 | GATGCGGAAATATCGAGGAA | CCCTTTTTCCCTTCTATGGC | 267 |
| chr12:30871952-30873964 | p2 | (GA)6 | TGTAGGGACTGCCCAAGTTC | CGTTCGCATTTAACCCAACT | 263 |
| chr12:30932679-30934697 | p3 | (TTG)6 | GGAAGTGTTGTTGGAGCGAT | CACACGGGTAACACACCAAG | 178 |

| Marker ID | SSR type | SSR motif | FORWARD PRIMER (5'-3') | REVERSE PRIMER (5'-3') | size (bp) |
|---|---|---|---|---|---|
| chr12:30958501-30960516 | p5 | (ATGCA)3 | GGTCGGGTTTCTTCTTCCTC | TGGTGAGGTTGGTTTGATGA | 295 |
| chr12:30976328-30978343 | p5 | (ATTGC)3 | GAGTGACGTGAGATGGCAGA | GATGTCTAAGTGCCACCCGT | 234 |
| chr12:30976354-30978387 | p5 | (ATTGC)3 | GAGTGACGTGAGATGGCAGA | GATGTCTAAGTGCCACCCGT | 208 |
| chr12:30976405-30978462 | p5 | (ATTGC)3 | GAGTGACGTGAGATGGCAGA | GATGTCTAAGTGCCACCCGT | 157 |
| chr12:30986820-30988832 | p2 | (TA)8 | TAAGCCCGAACGACATAACC | CCAACAATGATAGGCTCGGT | 145 |
| chr12:30986820-30988832 | p2 | (GT)6 | TAAGCCCGAACGACATAACC | CCAACAATGATAGGCTCGGT | 145 |
| chr12:31009186-31011206 | p2 | (TA)13 | TTCCAAATTCCAAGTTTGCC | AAATTCCCAAAGTCGTCGTG | 157 |
| chr12:31009186-31011206 | p2 | (AT)10 | TTCCAAATTCCAAGTTTGCC | AAATTCCCAAAGTCGTCGTG | 157 |
| chr12:31179119-31181139 | p2 | (TA)10 | TGGTCACTTGGGCCTTTTAC | CGGGTTTGAATAGGCAAAGA | 293 |
| chr12:31481297-31483312 | p3 | (CAT)5 | ATACGCTTTTGTCACGACCC | AAAGTGCTCTTGCCCGATAA | 136 |
| chr12:31494281-31496303 | p2 | (AT)11 | ACAAGCCACTTGGCAATTTT | CAGCCGTGGGTTTGTAGTTT | 200 |
| chr12:31494281-31496303 | p2 | (TA)11 | ACAAGCCACTTGGCAATTTT | CAGCCGTGGGTTTGTAGTTT | 200 |
| chr12:31509159-31511174 | p3 | (TGC)5 | TCAGTTCAAGTTTTTGCCCC | AAGGTCACGTCATAGGTCCG | 184 |
| chr12:31532882-31534897 | p5 | (AACAA)3 | CCCCTTATCGACCTAAAGCC | CTGCTGGTATGACTGCCTCA | 161 |
| chr12:31532882-31534897 | p5 | (TTTTA)4 | CCCCTTATCGACCTAAAGCC | CTGCTGGTATGACTGCCTCA | 161 |
| chr12:31533072-31535092 | p5 | (AACAA)3 | AGCAGAGAGGAGGATGACGA | AGTTGGCCACCTTCAAAATG | 209 |
| chr12:31533072-31535092 | p5 | (TTTTA)4 | AGCAGAGAGGAGGATGACGA | AGTTGGCCACCTTCAAAATG | 209 |
| chr12:31650138-31652240 | p2 | (AT)51 | TTGAGAAAAGGCAATGCACA | CCCACCTTGTAGCTGTTGGT | 251 |
| chr12:31696046-31698061 | p3 | (GTT)5 | TGGAGCCAAATCCTTGATTC | ATGGCATACACAACGACCAA | 106 |
| chr12:31706858-31708873 | p3 | (ATC)5 | TTGGTACAGGACCCCAACAT | ATGCCACACAACCCCATATT | 217 |
| chr12:31733602-31735617 | p5 | (AGGAC)3 | GGTTGTCAAACGTCATTCCC | AACATCATTCATAAGCCGCC | 177 |
| chr12:31816050-31818065 | p3 | (ACA)5 | ATTTCGCGTAATCGTGCTCT | GCTTAGCATCACTTTGGGGA | 202 |
| chr12:31852983-31855017 | p2 | (TA)17 | AGGGGAGATGCTGCCTAAAT | CGAATCATGCCTTAGGGTGT | 222 |
| chr12:31889842-31891857 | p3 | (TGT)5 | ACTAAAGTGGGCACGACACC | TCTCGATTCGTTTGACCTCC | 102 |
| chr12:31903380-31905420 | p2 | (TA)20 | AGCGAGGGACTGACTGACAT | ACCGCTAATGGGACTCCTTT | 158 |
| chr12:31969107-31971125 | p3 | (CAA)6 | AATATACGTACCCGGCCCTC | GAGCGAACGACGAACATACA | 183 |
| chr12:32027822-32029846 | p2 | (AT)12 | GAAACCTCCAACCTTCGACA | TTGTGGCCTTGTGGTGTAAA | 156 |
| chr12:32059859-32061874 | p3 | (CAA)5 | GGCATCACTTGTGAGGGTTT | GGACCACTCCATTCCTGAAA | 270 |
| chr12:32205890-32207902 | p2 | (AC)6 | CGTCCAAACTCGATTCCTGT | ATCTCGGGGAGGCTATTTGT | 174 |
| chr12:32248038-32250050 | p2 | (TA)6 | GTTCCTTTGCCATATTCGGA | AAATGATACGCCACCCCATA | 267 |
| chr12:32251446-32253464 | p6 | (TTTTAT)3 | GCACTTTGTGGAGGACCCTA | GAGCTCGACGAACCAATAGC | 125 |
| chr12:32273297-32275315 | p6 | (TATATG)3 | AAGTGTTGTACGGGCAGTCC | CTACGGTTACGCTACGGGAA | 192 |
| chr12:32273297-32275315 | p2 | (TA)6 | AAGTGTTGTACGGGCAGTCC | CTACGGTTACGCTACGGGAA | 192 |
| chr12:32282956-32284982 | p2 | (AT)13 | GCGCTGTAGAGGTGAGGTTC | TATACGGTTTTGAGGACCGC | 251 |
| chr12:32315769-32317781 | p6 | (TATATG)3 | TTACCGAGCCCCTCACTAGA | TAGAACCCAATACGGCAACC | 207 |
| chr12:32315769-32317781 | p2 | (TA)6 | TTACCGAGCCCCTCACTAGA | TAGAACCCAATACGGCAACC | 207 |

| Marker ID | SSR type | SSR motif | FORWARD PRIMER (5'-3') | REVERSE PRIMER (5'-3') | size (bp) |
|---|---|---|---|---|---|
| chr12:32342356-32344380 | p4 | (TATG)6 | CCTTACGTCACCCCGATAGA | GCCCATGCCCTCTAGACATA | 300 |
| chr12:32777271-32779285 | p2 | (AG)7 | TTGCCTTGCTTGTATGTTGC | AGAACCGAACCTTGACATGC | 238 |
| chr12:32893857-32895872 | p5 | (AACCA)3 | GGCAATCGGAGTAAAAACGA | GCTCTCCAACACTCTTTCGG | 257 |
| chr12:33031043-33033065 | p2 | (AT)11 | AAGCCGAGTAAACAGAGCCA | GTGTCTCGGCATGATTGTTG | 146 |
| chr12:33032798-33034813 | p5 | (AACAC)3 | TCTTTCCGCTCCTTTTGCTA | TTCGGTAGCCAAGTGTTTCC | 294 |
| chr12:33035382-33037397 | p5 | (AATGC)3 | TTTTGTGTGGCTGAGTTTCG | AGGTAGGTTCCCGAACGACT | 202 |
| chr12:33120112-33122128 | p2 | (AT)8 | CAACTTGGGAAGCAGAGGAG | CGCAAACGATTCGTGTAAGA | 244 |
| chr12:33127635-33129659 | p2 | (AT)12 | CGGAGGATTCAGGCTCATAA | GTTGCTTCGGGTCAACAAAT | 276 |
| chr12:33159251-33161266 | p3 | (ATT)5 | TTGCTTTCACATGGGTTTCA | TCAGCAGCAACAACAAGTCC | 102 |
| chr12:33313948-33316008 | p2 | (TA)6 | GAAGTGGGATGCGTTGTTTT | TCGGGTATTAGCCTGTTTGG | 240 |
| chr12:33313948-33316008 | p2 | (AT)30 | GAAGTGGGATGCGTTGTTTT | TCGGGTATTAGCCTGTTTGG | 240 |
| chr12:33333946-33335970 | p2 | (TA)12 | GGGTCGGGACACGTTATATG | GTGAGGGACTCGGTGGATAA | 183 |
| chr12:33337801-33339819 | p3 | (ATA)6 | GACCGAGTCAACCCTCGATA | GAATTTCGTCGATTCCGAGA | 185 |
| chr12:33375648-33377663 | p3 | (TTG)5 | GCGGGAAGACAACAAGAAAG | CCTTGGCCCAAAGTGATCTA | 107 |
| chr12:33412001-33414019 | p3 | (CAA)6 | TAATAACTGCCCAACCAGCC | CCACGGGAAGGTCATAGAAA | 162 |
| chr12:33537263-33539287 | p2 | (TC)13 | GGCCAAAATCTGGTGAAAAA | CAACGCAACAAACTGCTCAT | 148 |
| chr12:33537263-33539287 | p4 | (CATA)6 | GGCCAAAATCTGGTGAAAAA | CAACGCAACAAACTGCTCAT | 148 |
| chr12:33537263-33539287 | p2 | (TC)13 | GGCCAAAATCTGGTGAAAAA | CAACGCAACAAACTGCTCAT | 148 |
| chr12:33566652-33568670 | p2 | (TA)31 | AATTAAGCGAGCGAGTTCCA | GCGCGGTGTAAGAAAATAGC | 145 |
| chr12:33566652-33568670 | p2 | (TA)9 | AATTAAGCGAGCGAGTTCCA | GCGCGGTGTAAGAAAATAGC | 145 |
| chr12:33571162-33573212 | p2 | (TA)25 | CCAAGTAATGTGCAGCCTGA | GGCCTACAAATATGCCCAAA | 108 |
| chr12:33571162-33573212 | p3 | (TTG)5 | CCAAGTAATGTGCAGCCTGA | GGCCTACAAATATGCCCAAA | 108 |
| chr12:33578570-33580584 | p2 | (AT)7 | TCTGCGAAGGGTCTACAGGT | GCGAGCTCTTCTGCTGAAAT | 250 |
| chr12:33628657-33630672 | p5 | (TATTT)3 | GCGATTCCTGGTTCACTGAT | GACGATGACTCCTCCAGAGC | 184 |
| chr12:33696104-33698116 | p2 | (AT)6 | GGGGCACGTTACTTGCATAC | CGAGGGACACGGTGAGTAAT | 107 |
| chr12:33709765-33711777 | p2 | (AT)6 | TGGACCGGTAGGGTTATGAA | CAGGTACACGGATTTGACCC | 100 |
| chr12:33709765-33711777 | p3 | (AAT)5 | TGGACCGGTAGGGTTATGAA | CAGGTACACGGATTTGACCC | 100 |
| chr12:33722133-33724148 | p5 | (ATTTC)3 | AAGCCCACGAGGCTAAAAAT | GAGACGGCAAAAACGCTTAG | 124 |
| chr12:33743872-33745892 | p2 | (AT)10 | TCCTGAAATTCCACCTTTCG | CTGTGCAGAAAAGGGGTCAT | 118 |
| chr12:33811858-33813880 | p2 | (AT)6 | TTGTCGGCTTATGTACTGCG | ACTTAATGCCACGACCCAAC | 245 |
| chr12:33811858-33813880 | p2 | (AT)11 | TTGTCGGCTTATGTACTGCG | ACTTAATGCCACGACCCAAC | 245 |
| chr12:33811858-33813880 | p2 | (TA)30 | TTGTCGGCTTATGTACTGCG | ACTTAATGCCACGACCCAAC | 245 |
| chr12:33815540-33817567 | p3 | (TCT)11 | CCTGGGCGAGTATACAAGGA | ATTGACTGCCATCTTCGCTT | 221 |
| chr12:33815540-33817567 | p2 | (TA)7 | CCTGGGCGAGTATACAAGGA | ATTGACTGCCATCTTCGCTT | 221 |
| chr12:33918786-33920828 | p2 | (AT)21 | AATGAGCAAGAGCAAGGGAA | TTTAACGACCGTTTTACCCG | 200 |
| chr12:33918786-33920828 | p6 | (GTCAAC)6 | AATGAGCAAGAGCAAGGGAA | TTTAACGACCGTTTTACCCG | 200 |

| Marker ID | SSR type | SSR motif | FORWARD PRIMER (5'-3') | REVERSE PRIMER (5'-3') | size (bp) |
|---|---|---|---|---|---|
| chr12:33975552-33977567 | p5 | (AAAGG)3 | TGGGAAAAATCTGGTTCTGG | CAACTCCGAATGTCGTGATG | 213 |
| chr12:34003444-34005456 | p2 | (GT)6 | GCAAGACCTGCTTCGATACC | GAGTCTTTGCAGGGAAGTCG | 155 |
| chr12:34003444-34005456 | p3 | (GTT)5 | GCAAGACCTGCTTCGATACC | GAGTCTTTGCAGGGAAGTCG | 155 |
| chr12:34031415-34033433 | p2 | (CT)9 | CTTGTGGGGAAGGGGTATTT | GCGTCCTCAAATGTAGCCTC | 207 |
| chr12:34031415-34033433 | p6 | (TTTATT)3 | CTTGTGGGGAAGGGGTATTT | GCGTCCTCAAATGTAGCCTC | 207 |
| chr12:34040818-34042833 | p5 | (ATATA)3 | CAAAAAGTGTGTGTGTGCCC | ATAATTGACACCCCATCCCA | 228 |
| chr12:34063983-34065998 | p3 | (ACA)5 | TAGATTGGGCATAGGTTCGC | ACGGAGAGTGGTTTCCCTTT | 175 |
| chr12:34130313-34132337 | p2 | (TA)12 | GCCGGACCACACTGTTTACT | AGGCAGTGACGGATTTCATC | 291 |
| chr12:34200615-34202627 | p2 | (TA)6 | CCTAGCTGGGTCGTGAAAAA | TGGCCATTCTTCCTAAATGC | 107 |
| chr12:34200615-34202627 | p6 | (CATATA)3 | CCTAGCTGGGTCGTGAAAAA | TGGCCATTCTTCCTAAATGC | 107 |
| chr12:34311670-34313784 | p3 | (TCT)38 | AACATGAATTTGGGGCGTTA | TTGATCAATGGCCACAAGAA | 118 |
| chr12:34544167-34546181 | p2 | (AT)7 | CGAACACAATCACAAATCCG | CAAAGGAAAACTTCTTGCCG | 104 |
| chr12:34564009-34566021 | p2 | (TA)6 | TTTTCAGGACCCAACCAGAG | ATCTTTTGTCGGCCATTTTG | 131 |
| chr12:34589733-34591754 | p2 | (TA)7 | CATTTTAACCACCCACGTCC | AATTTCCGTTGGTGCGTAAG | 111 |
| chr12:34589733-34591754 | p3 | (ATC)5 | CATTTTAACCACCCACGTCC | AATTTCCGTTGGTGCGTAAG | 111 |
| chr12:34589733-34591754 | p3 | (AAC)7 | CATTTTAACCACCCACGTCC | AATTTCCGTTGGTGCGTAAG | 111 |
| chr12:34592814-34594828 | p2 | (TA)7 | GATTGCATGTTGGGTAAGGG | TAGGGATTCTCGCATGTTCC | 130 |
| chr12:34592814-34594828 | p2 | (TA)23 | GATTGCATGTTGGGTAAGGG | TAGGGATTCTCGCATGTTCC | 130 |
| chr12:34610935-34612967 | p2 | (AT)11 | TCTCCAACCTCAAACCATCC | AGTCGGCTTGCACTGTTCTT | 286 |
| chr12:34610935-34612967 | p3 | (AAC)5 | TCTCCAACCTCAAACCATCC | AGTCGGCTTGCACTGTTCTT | 286 |
| chr12:34702664-34704679 | p3 | (CAA)5 | CGGACGGAATCATAGCAAGT | TTGCTGACTCCCTCGAGTTT | 129 |
| chr12:34743787-34745799 | p2 | (AT)6 | CCCCAACAGGTGTAGCAAGT | GTGAGTATAGGCGGATCGGA | 155 |
| chr12:34764695-34766710 | p5 | (CATTG)3 | TCGTTGCCACGTTGATACAT | AACAACCAATTCAACCCCAA | 278 |
| chr12:34764695-34766710 | p5 | (TCATA)3 | TCGTTGCCACGTTGATACAT | AACAACCAATTCAACCCCAA | 278 |
| chr12:34783672-34785687 | p3 | (ATG)5 | GAGAAGCAACTGGGCAACTC | GTTTGAGGGGTGAATGGATG | 152 |
| chr12:34913658-34915678 | p4 | (AAAT)5 | CACAGGCCCTTTCTTTTTCA | TAAATCGCAAAGTTGAGGGG | 198 |
| chr12:35016269-35018284 | p3 | (TTA)5 | TGAAGAAAGCCCATACCCAC | GCTAATGACAGGTGGAGGGA | 154 |
| chr12:35016269-35018284 | p3 | (TTA)5 | TGAAGAAAGCCCATACCCAC | GCTAATGACAGGTGGAGGGA | 154 |
| chr12:35024881-35026896 | p5 | (GAAAA)3 | ACCGACTTCAAGGAAGGGTT | ACCGAATGAAATGAGCCAAC | 219 |
| chr12:35080256-35082271 | p5 | (TAAAA)3 | TTAAAAACATTTCCACCGCC | ACGGCCCGTACCTTATTTTC | 250 |
| chr12:35104804-35106818 | p2 | (TA)7 | ACTCGGCTGAGTGCCATACT | CTGTGGATAGTGCGTCGAGA | 226 |
| chr12:35106436-35108464 | p2 | (AT)14 | TATGCCTAGCCCGAAATCAC | TGTGGCTGGGATGATTTACA | 209 |
| chr12:35149976-35152004 | p2 | (AG)14 | CAAGTCCCAGCATCTTCTCC | AGACCGTATGGTGGAACAGC | 223 |
| chr12:35187347-35189359 | p2 | (TA)6 | ATCAGAGATCGGAGCAAGGA | TGTATTTGCACGAAATGGGA | 100 |
| chr12:35229787-35231801 | p2 | (GA)7 | CCTGCAATGGATGAGAACCT | CAACCGGATATTTCAGCGAT | 132 |
| chr12:35229787-35231801 | p6 | (GGGGGA)3 | CCTGCAATGGATGAGAACCT | CAACCGGATATTTCAGCGAT | 132 |

| Marker ID | SSR type | SSR motif | FORWARD PRIMER (5'-3') | REVERSE PRIMER (5'-3') | size (bp) |
|---|---|---|---|---|---|
| chr12:35230577-35232595 | p2 | (GA)7 | TGCTGGGACACAAGACTGAC | TCCCAGGTAACCTCCAACTG | 242 |
| chr12:35230577-35232595 | p6 | (GGGGGA)3 | TGCTGGGACACAAGACTGAC | TCCCAGGTAACCTCCAACTG | 242 |
| chr12:35230577-35232595 | p5 | (CACCC)3 | TGCTGGGACACAAGACTGAC | TCCCAGGTAACCTCCAACTG | 242 |
| chr12:35341759-35343771 | p2 | (AT)6 | AGTATAGCGATGCTCGGGAA | TTTTTCATGGACAGCATGGA | 246 |
| chr12:35445323-35447338 | p3 | (TAA)5 | TTGGAGTGTTTTCAAAGGGG | GCGGGATTATGACAGGAAAA | 182 |
| chr12:35453347-35455362 | p5 | (AAGAA)3 | CGTCAGCCCCTTCATATTGT | TGACTGATTTGCTTGCTTGG | 109 |
| chr12:35465501-35467515 | p2 | (AT)7 | ATCAGAAGCCGACAAGGCTA | GATAAGGGGTGCTCGGTACA | 206 |
| chr12:35489084-35491099 | p3 | (GGC)5 | TAAAACAGCCCCAAAACAGC | CTGCCGGTTGTATTCTTGGT | 157 |
| chr12:35492796-35494808 | p2 | (TG)6 | TCTGAGGGCCCTTAAATGTG | CAAATCGTTGCGGGTCTTAT | 275 |
| chr12:35492796-35494808 | p2 | (AT)13 | TCTGAGGGCCCTTAAATGTG | CAAATCGTTGCGGGTCTTAT | 275 |
| chr12:35501774-35503789 | p5 | (AAAGG)3 | GATAGTGCTGTTCGGCCTTC | TATAGGGCCCAACCTGTGAC | 114 |
| chr12:35501774-35503789 | p5 | (GCATT)3 | GATAGTGCTGTTCGGCCTTC | TATAGGGCCCAACCTGTGAC | 114 |
| chr12:35519268-35521283 | p3 | (GGC)5 | TAAAACAGCCCCAAAACAGC | CTGCCGGTTGTATTCTTGGT | 157 |
| chr12:35522980-35524992 | p2 | (TG)6 | TCTGAGGGCCCTTAAATGTG | CAAATCGTTGCGGGTCTTAT | 275 |
| chr12:35522980-35524992 | p2 | (AT)13 | TCTGAGGGCCCTTAAATGTG | CAAATCGTTGCGGGTCTTAT | 275 |
| chr12:35531958-35533973 | p5 | (AAAGG)3 | GATAGTGCTGTTCGGCCTTC | TATAGGGCCCAACCTGTGAC | 114 |
| chr12:35531958-35533973 | p5 | (GCATT)3 | GATAGTGCTGTTCGGCCTTC | TATAGGGCCCAACCTGTGAC | 114 |
| chr12:35772806-35774875 | p3 | (ATA)23 | CGATGAGCCTCTACACTCCC | GACAAAATGTGCCTTCGGTT | 180 |
| chr12:35779277-35781347 | p2 | (AT)35 | TTCTCCGTTGAATCTACCCG | GAAAACACCGGATACTCGGA | 282 |
| chr12:35779277-35781347 | p2 | (AG)25 | TTCTCCGTTGAATCTACCCG | GAAAACACCGGATACTCGGA | 282 |
| chr12:35831101-35833117 | p2 | (AT)8 | AAGCCGACAAGGCTAACAAA | GAGCTCCATCTCATTCGGAG | 221 |
| chr12:36033930-36035942 | p2 | (AT)6 | GGGTCTTCATCCGTCTTTGA | TTAGTTGCTGCTCAGGCTCA | 110 |
| chr12:36159122-36161224 | p2 | (AT)18 | ACAACCCTACGACGGAACTG | TAGCCTTGTCAGCTCCCATT | 249 |
| chr12:36159122-36161224 | p2 | (AT)8 | ACAACCCTACGACGGAACTG | TAGCCTTGTCAGCTCCCATT | 249 |
| chr12:36159236-36161248 | p2 | (AT)18 | ACAACCCTACGACGGAACTG | TAGCCTTGTCAGCTCCCATT | 135 |
| chr12:36159236-36161248 | p2 | (AT)8 | ACAACCCTACGACGGAACTG | TAGCCTTGTCAGCTCCCATT | 135 |
| chr12:36159606-36161622 | p2 | (AT)18 | CTCCGTATCAAATGGAGCGT | ATTATCCGTACTGCCTCCCC | 175 |
| chr12:36159606-36161622 | p2 | (AT)8 | CTCCGTATCAAATGGAGCGT | ATTATCCGTACTGCCTCCCC | 175 |
| chr12:36206211-36208237 | p2 | (TA)13 | GTTTCAAGGATGGAAAGGCA | TGTCGAATGCTGCTGATAGG | 146 |
| chr12:36214508-36216524 | p4 | (TTAC)4 | ATGCACGGGAATATCCAAAA | TGCTTGGAGGGCTCTAGTGT | 238 |
| chr12:36214508-36216524 | p2 | (AT)24 | ATGCACGGGAATATCCAAAA | TGCTTGGAGGGCTCTAGTGT | 238 |
| chr12:36292831-36294843 | p2 | (GA)6 | TTTCGAAAGTTGCTGCCTTT | TGGACCTGGTCTTCCCTATG | 130 |
| chr12:36296679-36298715 | p2 | (AT)18 | GGAAATTTGTCGTGTGCCTT | CTTCCCAATTTCTTGGGTCA | 174 |
| chr12:36307308-36309330 | p2 | (GT)11 | TGGACCGGTACTAAAGCCAC | TTCGCAACCCAGAGACTTTT | 170 |
| chr12:36307867-36309883 | p2 | (GT)11 | ATCTGATCACGTCGAGGACC | GATTTGGTGTAACGACCCGT | 278 |
| chr12:36307894-36309910 | p2 | (GT)11 | ATCTGATCACGTCGAGGACC | ACGACCCGTTTGGTCATTTA | 251 |

| Marker ID | SSR type | SSR motif | FORWARD PRIMER (5'-3') | REVERSE PRIMER (5'-3') | size (bp) |
|---|---|---|---|---|---|
| chr12:36307912-36309928 | p2 | (GT)11 | ATCTGATCACGTCGAGGACC | ACGACCCGTTTGGTCATTTA | 233 |
| chr12:36371696-36373714 | p6 | (GCATTG)3 | CTCGGGAAAAGCTTCAGTTG | TTGCTTCCTGATCCTTCACC | 244 |
| chr12:36553754-36555782 | p2 | (AT)14 | ATAAGATGCCAACTCCACGG | CAAGTACCAAAGCCCGTGAT | 168 |
| chr12:36603138-36605160 | p2 | (AT)11 | CGACTCTTCTTTCAGCCACC | TCGGCGCATTTAGGAATAAC | 145 |
| chr12:36660285-36662313 | p2 | (AT)14 | GGGGTACACAGCCACAGTCT | GGACCCGTAGCTCGATAACA | 275 |
| chr12:36660285-36662313 | p2 | (AT)10 | GGGGTACACAGCCACAGTCT | GGACCCGTAGCTCGATAACA | 275 |
| chr12:36660285-36662313 | p2 | (TA)14 | GGGGTACACAGCCACAGTCT | GGACCCGTAGCTCGATAACA | 275 |
| chr12:36661165-36663193 | p2 | (AT)14 | AGAACAACGGAGCTCTCCAA | ATAATGGGGCCGGATATGAT | 180 |
| chr12:36661165-36663193 | p2 | (AT)10 | AGAACAACGGAGCTCTCCAA | ATAATGGGGCCGGATATGAT | 180 |
| chr12:36661165-36663193 | p2 | (TA)14 | AGAACAACGGAGCTCTCCAA | ATAATGGGGCCGGATATGAT | 180 |
| chr12:36669608-36671622 | p5 | (TATAT)3 | CGGATCACCAAAGCGAGTAT | TAATGGGCCGGATTTGATTA | 185 |
| chr12:36669608-36671622 | p2 | (TA)8 | CGGATCACCAAAGCGAGTAT | TAATGGGCCGGATTTGATTA | 185 |
| chr12:36669640-36671672 | p5 | (TATAT)3 | CGGATCACCAAAGCGAGTAT | TAATGGGCCGGATTTGATTA | 153 |
| chr12:36669640-36671672 | p2 | (TA)8 | CGGATCACCAAAGCGAGTAT | TAATGGGCCGGATTTGATTA | 153 |
| chr12:36748143-36750155 | p5 | (ATGCA)3 | AACCTGAAAGCGTGGTATGG | CAGTTCCTGTCGTCAAGGGT | 215 |
| chr12:36748163-36750189 | p5 | (ATGCA)3 | AACCTGAAAGCGTGGTATGG | CAGTTCCTGTCGTCAAGGGT | 195 |
| chr12:36748224-36750238 | p5 | (ATGCA)3 | AACCTGAAAGCGTGGTATGG | CAGTTCCTGTCGTCAAGGGT | 134 |
| chr12:36748239-36750255 | p5 | (ATGCA)3 | AACCTGAAAGCGTGGTATGG | CAGTTCCTGTCGTCAAGGGT | 119 |
| chr12:36809937-36811961 | p3 | (TGA)5 | TTTGTATAGCAGCGCATTGG | TCCATATGGTGTTTGTCCCA | 154 |
| chr12:36825159-36827177 | p6 | (TGTATA)3 | TAATGTTGAAGGGGCTCGTC | TGCGGAAAAGAAAATCATCC | 296 |
| chr12:36825159-36827177 | p2 | (TA)6 | TAATGTTGAAGGGGCTCGTC | TGCGGAAAAGAAAATCATCC | 296 |
| chr12:36825346-36827358 | p6 | (TGTATA)3 | CAAAGGGCATGGATTCCTAA | TGCGGAAAAGAAAATCATCC | 140 |
| chr12:36825346-36827358 | p2 | (TA)6 | CAAAGGGCATGGATTCCTAA | TGCGGAAAAGAAAATCATCC | 140 |
| chr12:36999319-37001334 | p5 | (ATTTC)3 | AAGGGAAACTCTGGCGAAAT | TTCGTGCCCTTCGTTAGTCT | 265 |
| chr12:37009359-37011375 | p2 | (AT)8 | CATAAGCCCTTCGGCTATCA | TCCATTACTCTGGAGGTGCC | 230 |
| chr12:37070176-37072188 | p2 | (AT)14 | TTTGGAGGGCTTACACTTGG | ACGTTTTCGGGTGATCTGAC | 179 |
| chr12:37070176-37072188 | p2 | (AT)6 | TTTGGAGGGCTTACACTTGG | ACGTTTTCGGGTGATCTGAC | 179 |
| chr12:37091800-37093822 | p2 | (TA)8 | TGGATGTTGGTTGGGATTTT | GCCAAAGGAAAAAGGGAAAC | 280 |
| chr12:37091800-37093822 | p2 | (AC)11 | TGGATGTTGGTTGGGATTTT | GCCAAAGGAAAAAGGGAAAC | 280 |
| chr12:37091800-37093822 | p5 | (TTATA)3 | TGGATGTTGGTTGGGATTTT | GCCAAAGGAAAAAGGGAAAC | 280 |
| chr12:37158981-37161023 | p2 | (TA)21 | GCACCTAGGGTGAATTTCCA | TCACCATTGTAGACCCACGA | 143 |
| chr12:37173022-37175037 | p2 | (CA)9 | GCAGGTCAACAGACAGACGA | CGACGAGGACAATCCAAAGT | 243 |
| chr12:37173022-37175037 | p5 | (TCGGT)3 | GCAGGTCAACAGACAGACGA | CGACGAGGACAATCCAAAGT | 243 |
| chr12:37173022-37175037 | p5 | (AAACC)3 | GCAGGTCAACAGACAGACGA | CGACGAGGACAATCCAAAGT | 243 |
| chr12:37173514-37175529 | p2 | (CA)9 | CCCTAGTTGGGTCGTGACAT | GGCATGGTCTCGAGGTAAAA | 141 |
| chr12:37173514-37175529 | p5 | (TCGGT)3 | CCCTAGTTGGGTCGTGACAT | GGCATGGTCTCGAGGTAAAA | 141 |

| Marker ID | SSR type | SSR motif | FORWARD PRIMER (5'-3') | REVERSE PRIMER (5'-3') | size (bp) |
|---|---|---|---|---|---|
| chr12:37173514-37175529 | p5 | (AAACC)3 | CCCTAGTTGGGTCGTGACAT | GGCATGGTCTCGAGGTAAAA | 141 |
| chr12:37173514-37175529 | p2 | (TA)6 | CCCTAGTTGGGTCGTGACAT | GGCATGGTCTCGAGGTAAAA | 141 |
| chr12:37330954-37332980 | p2 | (TA)13 | TCTTTCGCCACTTCCTCACT | ATTGTGGCGGATAATGAAGG | 297 |
| chr12:37339387-37341437 | p2 | (TA)11 | CTTACGTCACTCGGAAAGGC | TGGTGTGCATATCTACGGGA | 264 |
| chr12:37339387-37341437 | p2 | (TA)25 | CTTACGTCACTCGGAAAGGC | TGGTGTGCATATCTACGGGA | 264 |
| chr12:37483908-37485974 | p2 | (AT)33 | AACGTGTAGCCCGATCCATA | AGAAATCGCAGGTTTACCGA | 195 |
| chr12:37503384-37505396 | p2 | (AT)6 | AGCCCCTATCACACAACAGG | GGAAGGGCAAGGCTAAAAAG | 281 |
| chr12:37503384-37505396 | p2 | (AT)8 | AGCCCCTATCACACAACAGG | GGAAGGGCAAGGCTAAAAAG | 281 |
| chr12:37524867-37526891 | p2 | (GT)12 | AGGGAAATGCGCAATATGAC | ATGATCCCGCTCTCTCCTTT | 294 |
| chr12:37653055-37655077 | p2 | (TA)11 | ATCAGAGGATAACAACGGCG | TGGGATAACTGCATGGCATA | 120 |
| chr12:37664512-37666527 | p5 | (AGAAC)3 | ACCAAGTCAAAGCAACACCC | GGCTTGGAAATTGGAATGAA | 109 |
| chr12:37690074-37692089 | p5 | (CTTTT)3 | TGAAAGAGGGGCTTTAGCAA | GCTTTTGGAAATTGGAAGCA | 138 |
| chr12:37714894-37716909 | p5 | (TATTT)3 | TGCGCATTGCTAACTAGCTG | TTAAAGGGGCAAGATTGGTG | 172 |
| chr12:37747335-37749349 | p2 | (TA)7 | TGCCCGGTCACTTAAGATTC | GGAATGATACCGGTGGATTG | 285 |
| chr12:37859403-37861424 | p3 | (GTT)7 | CTCGTCGACAGGACATCAGA | GTACTGCAGCACGTCGGTTA | 256 |
| chr12:37867513-37869533 | p6 | (AAAAGG)3 | ATTCGAGCCATGAAAAGTGG | CGGACCCTCCTCATAAGTCA | 273 |
| chr12:37867513-37869533 | p5 | (AAAAG)4 | ATTCGAGCCATGAAAAGTGG | CGGACCCTCCTCATAAGTCA | 273 |
| chr12:37888390-37890402 | p2 | (CT)6 | GGGCTAAGATGAAATGCCAA | GTTTGGATGTGGTGGGTTTC | 167 |
| chr12:37890624-37892636 | p2 | (AT)6 | GGAAACAAAAGTGGACCCAA | ATGTGGCAATGAATCCATCA | 292 |
| chr12:37890624-37892636 | p2 | (AT)10 | GGAAACAAAAGTGGACCCAA | ATGTGGCAATGAATCCATCA | 292 |
| chr12:37978230-37980246 | p6 | (TATATG)3 | TCATTAACTCTGCTCGTGCG | ATCTTCATCTGCCATCCTGG | 142 |
| chr12:37978253-37980277 | p6 | (TATATG)3 | TCATTAACTCTGCTCGTGCG | ATCTTCATCTGCCATCCTGG | 119 |
| chr12:38013664-38015680 | p4 | (AAAT)4 | TCCATGTCGCTAACCAAACA | AAGGTTGAGGCGTATGATGC | 199 |
| chr12:38038983-38041029 | p2 | (TA)23 | ATGGCTCTATCGCACGATCT | ATGCGTAGGCTATGTGACCC | 176 |
| chr12:38153458-38155473 | p3 | (TGA)5 | TTCGGAGGTGATACGAAAGG | ATCTCCATTGCACCCTCTTG | 112 |
| chr12:38153458-38155473 | p5 | (TTGCA)3 | TTCGGAGGTGATACGAAAGG | ATCTCCATTGCACCCTCTTG | 112 |
| chr12:38161917-38163935 | p2 | (AT)9 | CAAAAGAATAGAACCCGCCA | GGTATGCCACCCTTTTGAGA | 254 |
| chr12:38240298-38242326 | p2 | (TA)14 | GAAGGAGTGCAACTGCAACA | GAATCAATTCCTTCCAGCCA | 118 |
| chr12:38309064-38311076 | p2 | (TG)6 | TTCCAAGGTTTTCCTTCACG | TCAAATACGTCGTCCAGCAG | 256 |
| chr12:38309064-38311076 | p3 | (GAA)5 | TTCCAAGGTTTTCCTTCACG | TCAAATACGTCGTCCAGCAG | 256 |
| chr12:38309192-38311207 | p2 | (TG)6 | TTCCAAGGTTTTCCTTCACG | TCAAATACGTCGTCCAGCAG | 128 |
| chr12:38309192-38311207 | p3 | (GAA)5 | TTCCAAGGTTTTCCTTCACG | TCAAATACGTCGTCCAGCAG | 128 |
| chr12:38356926-38358944 | p6 | (AGAGAA)3 | TTTCGCATGCCACTGTTTAG | ACCGGTGACTGAATTGCTTC | 261 |
| chr12:38356926-38358944 | p2 | (GT)6 | TTTCGCATGCCACTGTTTAG | ACCGGTGACTGAATTGCTTC | 261 |
| chr12:38359825-38361837 | p2 | (TA)6 | GGATCATCTGACCGACTGGT | GGAAGATAATCCGGACGACA | 209 |
| chr12:38367438-38369466 | p2 | (TA)6 | TGACTTGGGAGGTACCTTGG | CAGGAACTGACGTGTCCTCA | 214 |

| Marker ID | SSR type | SSR motif | FORWARD PRIMER (5'-3') | REVERSE PRIMER (5'-3') | size (bp) |
|---|---|---|---|---|---|
| chr12:38427328-38429342 | p2 | (TA)7 | TCATCCGTTACTGACTCCCC | TATACACACCCGGCATCAGA | 115 |
| chr12:38427328-38429342 | p2 | (AT)6 | TCATCCGTTACTGACTCCCC | TATACACACCCGGCATCAGA | 115 |
| chr12:38427993-38430005 | p2 | (TA)7 | AGCCATTCAATTATGCCCTG | GCCTTGGCTTTAGTGATCCA | 184 |
| chr12:38427993-38430005 | p2 | (AT)6 | AGCCATTCAATTATGCCCTG | GCCTTGGCTTTAGTGATCCA | 184 |
| chr12:38434708-38436736 | p2 | (TA)14 | CCCTCTGGTACAAGCCTCAG | TATGGCACTATGGACCAGCA | 147 |
| chr12:38485473-38487485 | p2 | (TA)6 | CTTCTGGGCTGCTGTAGCTT | GGTTGGGTCGTGACAAAAGT | 115 |
| chr12:38540596-38542608 | p2 | (TA)6 | TCCCCACGGGTATCATAAAA | GCTATGGCACGCTGTATTGA | 171 |
| chr12:38541014-38543034 | p2 | (TA)6 | TATTGGCACCTCGTCTCTCC | TGGGTCGTGACAAGTTGGTA | 240 |
| chr12:38541095-38543110 | p2 | (TA)6 | TATTGGCACCTCGTCTCTCC | TGGGTCGTGACAAGTTGGTA | 159 |
| chr12:38598069-38600109 | p2 | (AT)20 | GAGGTTGGTTTAGCCTTCCC | GAAGAGAGGGACCCGGTTAG | 229 |
| chr12:38665863-38667878 | p5 | (AATGC)3 | TTTTGCCCCTAGCCTTTCTT | TTGTGCGTTCCGATAAATCA | 257 |
| chr12:38665863-38667878 | p5 | (CCTTT)3 | TTTTGCCCCTAGCCTTTCTT | TTGTGCGTTCCGATAAATCA | 257 |
| chr12:38697229-38699247 | p2 | (AT)9 | AACCTCCGGAACAAAGGAGT | TGTTTTCTTTGGACGCTGTG | 103 |
| chr12:38730994-38733078 | p2 | (AT)8 | GACCGTGGTCAACTCCAAAT | TCGTCATTGCATTAGGTCCA | 143 |
| chr12:38812934-38814946 | p2 | (AT)32 | GAAAGATAGGGTGGCCCATT | TAGCCTTGTCGGCTCTCATT | 145 |
| chr12:38812934-38814946 | p2 | (AT)6 | GAAAGATAGGGTGGCCCATT | TAGCCTTGTCGGCTCTCATT | 145 |
| chr12:38857811-38859831 | p5 | (TGATA)4 | GATGCTGCCCAAATTTCTGT | ATGCCAGGAGAAGGAAAGGT | 117 |
| chr12:38895719-38897777 | p2 | (AT)29 | TTGGCTTTTTGGCTGACTTT | CACATGGCCTTGTCCTTCTT | 114 |
| chr12:38899367-38901382 | p5 | (TATAT)3 | CGTATGTGGTTTGCTTGGTG | TGGCAACATGGTGCATAAAT | 229 |
| chr12:38931412-38933427 | p5 | (AATTT)3 | TCCTTTCCCGAAACATGAGT | GGGATTGCCTAGGTTGTTCA | 172 |
| chr12:38977671-38979689 | p6 | (ATTCAT)3 | GGTGAGGCAGAATGCTTAGG | GTGTCTGCCCGACCATATTT | 120 |
| chr12:38991482-38993497 | p3 | (AGG)5 | TATGAAGCGAAACCTGAGGG | GCGGATCGAGGATTGATTTA | 108 |
| chr12:39028050-39030065 | p5 | (TTCCT)3 | CATCTGCTGCATTTTGAGGA | TACAGCATTGGCAGCAACTC | 141 |
| chr12:39060831-39062845 | p2 | (TA)7 | CTGAGCCCAAAATAAACCCA | GGCTTGCTACAGATTTTGGC | 258 |
| chr12:39060831-39062845 | p3 | (TAT)9 | CTGAGCCCAAAATAAACCCA | GGCTTGCTACAGATTTTGGC | 258 |
| chr12:39089318-39091330 | p2 | (AT)11 | TTTTGAAGGGAGGGCTTTTT | GGGGTGACATAAGGTCGAGA | 199 |
| chr12:39089318-39091330 | p2 | (AT)6 | TTTTGAAGGGAGGGCTTTTT | GGGGTGACATAAGGTCGAGA | 199 |
| chr12:39089318-39091330 | p2 | (AT)10 | TTTTGAAGGGAGGGCTTTTT | GGGGTGACATAAGGTCGAGA | 199 |
| chr12:39089593-39091613 | p2 | (AT)11 | TCTCGACCTTATGTCACCCC | ATATATGCAGCATTCGCGTG | 103 |
| chr12:39089593-39091613 | p2 | (AT)6 | TCTCGACCTTATGTCACCCC | ATATATGCAGCATTCGCGTG | 103 |
| chr12:39089593-39091613 | p2 | (AT)10 | TCTCGACCTTATGTCACCCC | ATATATGCAGCATTCGCGTG | 103 |
| chr12:39126208-39128272 | p2 | (TA)32 | GGGCTAGAGGCCCAACTATC | ACCTAAGCCCACCCGTTACT | 186 |
| chr12:39126208-39128272 | p5 | (TCATA)3 | GGGCTAGAGGCCCAACTATC | ACCTAAGCCCACCCGTTACT | 186 |
| chr12:39177393-39179408 | p5 | (TTCAT)3 | ACCTGTCATTTCCAGGCAAG | ATCGGCGTAGCAAGAAGCTA | 170 |
| chr12:39302130-39304176 | p2 | (TA)23 | AGCCGGTCAGAAAAGAGTCA | GTCTGCTTGGGTGTCCAAAT | 137 |
| chr12:39303727-39305741 | p2 | (CA)7 | TTTAGGGGTCATGCCAAAAG | TGAATTTACAGGGGAGGTGC | 219 |

| Marker ID | SSR type | SSR motif | FORWARD PRIMER (5'-3') | REVERSE PRIMER (5'-3') | size (bp) |
|---|---|---|---|---|---|
| chr12:39366018-39368064 | p2 | (TA)23 | AGCCGGTCAGAAAAGAGTCA | GTCTGCTTGGGTGTCCAAAT | 137 |
| chr12:39367615-39369629 | p2 | (CA)7 | TTTAGGGGTCATGCCAAAAG | TGAATTTACAGGGGAGGTGC | 219 |
| chr12:39549041-39551061 | p2 | (AT)10 | CAGCGCACCACTCTTGAATA | TGTGATTTTGCCTTGCCATA | 119 |
| chr12:39562297-39564315 | p6 | (TATATG)3 | TGGTGGTGGTTGAATTGAGA | AGCCTCTTCATCGCACACTT | 163 |
| chr12:39647623-39649643 | p2 | (AT)10 | CAGCGCACCACTCTTGAATA | TGTGATTTTGCCTTGCCATA | 119 |
| chr12:39660879-39662897 | p6 | (TATATG)3 | TGGTGGTGGTTGAATTGAGA | AGCCTCTTCATCGCACACTT | 163 |
| chr12:39685485-39687500 | p5 | (AAAGG)3 | TGTATGCGGCTTCATCTGTC | GAATTGTCACACCCCAATCC | 144 |
| chr12:39824423-39826438 | p5 | (ATTTC)3 | AAGAAATGACATGGGAAGCG | GCTACACGCGACCTAAAAGC | 154 |
| chr12:39840044-39842082 | p2 | (AT)19 | ATACCTCGATCGCTCGCTAA | TGGGCCGAGTTAAATTCTTG | 221 |
| chr12:39881966-39884012 | p2 | (TA)23 | GCTGAGTGACGTGAGATGGA | CCGGATCTTTACCCAACAGA | 230 |
| chr12:39921776-39923812 | p2 | (GA)6 | TGAGGGTAGACCTCAATGGG | GATGCTGCCCAATTTTTGTT | 190 |
| chr12:39921776-39923812 | p2 | (AT)18 | TGAGGGTAGACCTCAATGGG | GATGCTGCCCAATTTTTGTT | 190 |
| chr12:39929162-39931174 | p2 | (GT)7 | AACATGTAATGCGTGCGTGT | TGGTGTATTTCGCCATGAGA | 159 |
| chr12:39929162-39931174 | p2 | (TG)6 | AACATGTAATGCGTGCGTGT | TGGTGTATTTCGCCATGAGA | 159 |
| chr12:40038051-40040072 | p3 | (TGT)7 | TTTGGACAGAAAAGTTGGGC | GAAGCCATTTGGCAGCTTAG | 216 |
| chr12:40038051-40040072 | p3 | (GAT)8 | TTTGGACAGAAAAGTTGGGC | GAAGCCATTTGGCAGCTTAG | 216 |
| chr12:40038051-40040072 | p2 | (AT)23 | TTTGGACAGAAAAGTTGGGC | GAAGCCATTTGGCAGCTTAG | 216 |
| chr12:40048535-40050553 | p3 | (TTG)6 | CGTTTTCGGGTGATCTGACT | TGGGTGTTGTGTCCTGAAAA | 216 |
| chr12:40103257-40105275 | p6 | (AAAAAT)3 | AAACACTCCCGAATGTCTCG | AGAGTGTTTTTGACCCGTGG | 256 |
| chr12:40186070-40188091 | p3 | (AAT)7 | CCAATGGAGGTCTCGTTGAT | TGGGCTTTAAGCGAAAAAGA | 105 |
| chr12:40191431-40193443 | p2 | (CA)6 | CCTAAGTTCCGTTTCGTCCA | TGTTGATGTTGCTGGGTTGT | 206 |
| chr12:40341008-40343026 | p2 | (AT)9 | CTATGGGACAGATGCCGACT | GTCGGTTTGCGTAGCATTTT | 101 |
| chr12:40353903-40355929 | p2 | (AT)13 | ATCAGGGGTGCTGAATAACG | GGAGACGCGAAGACCTAGTG | 171 |
| chr12:40493776-40495794 | p2 | (AT)9 | GCGGCAACTAACTGTGTCAA | AAGGCGGGTAGAAAGAGAGC | 242 |
| chr12:40493776-40495794 | p2 | (AT)23 | GCGGCAACTAACTGTGTCAA | AAGGCGGGTAGAAAGAGAGC | 242 |
| chr12:40494095-40496141 | p2 | (AT)9 | TTTTCACCGTAGTCTTGCCC | GGAAGTGTCTACGAGCCGAG | 113 |
| chr12:40494095-40496141 | p2 | (AT)23 | TTTTCACCGTAGTCTTGCCC | GGAAGTGTCTACGAGCCGAG | 113 |
| chr12:40565052-40567064 | p2 | (AT)6 | AAATCATTGGAGCCTTGTCG | ACCCTGCCCATTCTAGGTTT | 187 |
| chr12:40565052-40567064 | p3 | (TTC)6 | AAATCATTGGAGCCTTGTCG | ACCCTGCCCATTCTAGGTTT | 187 |
| chr12:40613755-40615770 | p2 | (TA)20 | GCTGCAGTCCCATTTGGTAT | AGAACTCACAAGCAGCGGAT | 250 |
| chr12:40613825-40615837 | p2 | (TA)20 | GCTGCAGTCCCATTTGGTAT | AGAACTCACAAGCAGCGGAT | 180 |
| chr12:40613847-40615863 | p2 | (TA)20 | GCTGCAGTCCCATTTGGTAT | AGAACTCACAAGCAGCGGAT | 158 |
| chr12:40644473-40646501 | p6 | (ATACAT)4 | CCGAAACTAACTCACCGAGC | TCTCAGCCTTATCGGCTCAT | 244 |
| chr12:40644473-40646501 | p2 | (TA)14 | CCGAAACTAACTCACCGAGC | TCTCAGCCTTATCGGCTCAT | 244 |
| chr12:40892856-40894871 | p5 | (ATTTC)3 | GAAGGGGTGAAGTGTGGAAA | ATCATTTTGTAGCCTTGCCG | 171 |
| chr12:40998917-41000929 | p2 | (GT)6 | ATACCCGCACTTGTTTCCTG | ACGGGACCTAACATTCTCCC | 190 |

| Marker ID | SSR type | SSR motif | FORWARD PRIMER (5'-3') | REVERSE PRIMER (5'-3') | size (bp) |
|---|---|---|---|---|---|
| chr12:41055330-41057346 | p4 | (AAAG)4 | AGGACGCGGTGAACATAATC | TGTGCATATGGTTTCGCACT | 280 |
| chr12:41071201-41073216 | p3 | (CAA)5 | TAAATGCCCTGCAGCTTCTT | TTGTTACGACCCATCGGTTT | 115 |
| chr12:41071201-41073216 | p6 | (ATATAC)3 | TAAATGCCCTGCAGCTTCTT | TTGTTACGACCCATCGGTTT | 115 |
| chr12:41089950-41091972 | p2 | (AT)9 | CGGATCACCAAAGCGAGTAT | ACTGTTGCGCTTTTCCTTGT | 109 |
| chr12:41111602-41113617 | p3 | (GTT)5 | TGTGTTGTTGTTGTTGGGCT | AACAAAATGGCAGCATCTCC | 200 |
| chr12:41111602-41113617 | p3 | (GAT)5 | TGTGTTGTTGTTGTTGGGCT | AACAAAATGGCAGCATCTCC | 200 |
| chr12:41153223-41155239 | p4 | (GAAA)4 | TCCCCTGACCAATAGTAGCG | CCGTTGACTTGGGTTCAGTT | 202 |
| chr12:41223370-41225406 | p2 | (TA)13 | CCCAATTTTTGGGATTTCCT | AATGGATCGACCACTTGAGG | 254 |
| chr12:41223408-41225430 | p2 | (TA)13 | CCCAATTTTTGGGATTTCCT | AATGGATCGACCACTTGAGG | 216 |
| chr12:41223432-41225454 | p2 | (TA)13 | CCCAATTTTTGGGATTTCCT | AATGGATCGACCACTTGAGG | 192 |
| chr12:41223432-41225454 | p5 | (TGAAT)3 | CCCAATTTTTGGGATTTCCT | AATGGATCGACCACTTGAGG | 192 |
| chr12:41223456-41225478 | p2 | (TA)13 | CCCAATTTTTGGGATTTCCT | AATGGATCGACCACTTGAGG | 168 |
| chr12:41223456-41225478 | p5 | (TGAAT)3 | CCCAATTTTTGGGATTTCCT | AATGGATCGACCACTTGAGG | 168 |
| chr12:41223480-41225502 | p2 | (TA)13 | CCCAATTTTTGGGATTTCCT | AATGGATCGACCACTTGAGG | 144 |
| chr12:41223480-41225502 | p5 | (TGAAT)3 | CCCAATTTTTGGGATTTCCT | AATGGATCGACCACTTGAGG | 144 |
| chr12:41223504-41225526 | p2 | (TA)13 | CCCAATTTTTGGGATTTCCT | AATGGATCGACCACTTGAGG | 120 |
| chr12:41223504-41225526 | p5 | (TGAAT)3 | CCCAATTTTTGGGATTTCCT | AATGGATCGACCACTTGAGG | 120 |
| chr12:41223677-41225703 | p2 | (TA)13 | AGCATTGATTGCTTTACCCG | AAACCTTATCCGCCACTCCT | 205 |
| chr12:41223677-41225703 | p5 | (TGAAT)3 | AGCATTGATTGCTTTACCCG | AAACCTTATCCGCCACTCCT | 205 |
| chr12:41254683-41256705 | p2 | (TA)36 | GAAAGCGGGCAAGGTATGTA | CTCTCGAATGCTATTTCGGC | 260 |
| chr12:41254707-41256729 | p2 | (TA)36 | GAAAGCGGGCAAGGTATGTA | CTCTCGAATGCTATTTCGGC | 236 |
| chr12:41254731-41256753 | p2 | (TA)36 | GAAAGCGGGCAAGGTATGTA | CTCTCGAATGCTATTTCGGC | 212 |
| chr12:41254755-41256777 | p2 | (TA)36 | GAAAGCGGGCAAGGTATGTA | CTCTCGAATGCTATTTCGGC | 188 |
| chr12:41254779-41256801 | p2 | (TA)36 | GAAAGCGGGCAAGGTATGTA | CTCTCGAATGCTATTTCGGC | 164 |
| chr12:41312759-41314775 | p2 | (TA)8 | ACCACCATTATCCTAGCCCC | TTGGTCCGGAATTCATTTGT | 136 |
| chr12:41444181-41446261 | p2 | (TA)6 | GATGTTGCCCAAACTTCGTT | TTGGAACTCGCTCGCTTACT | 270 |
| chr12:41444181-41446261 | p2 | (AT)8 | GATGTTGCCCAAACTTCGTT | TTGGAACTCGCTCGCTTACT | 270 |
| chr12:41444289-41446363 | p2 | (TA)6 | GATGTTGCCCAAACTTCGTT | TTGGAACTCGCTCGCTTACT | 162 |
| chr12:41444289-41446363 | p2 | (AT)8 | GATGTTGCCCAAACTTCGTT | TTGGAACTCGCTCGCTTACT | 162 |
| chr12:41475586-41477601 | p5 | (ATTAA)3 | AACCATTGCCACATCAGACA | TATGCTGCAAGGTGAATTGC | 286 |
| chr12:41525547-41527573 | p2 | (TA)13 | CAACAACATTGGCTTTCCCT | TTTGAAGAGCTGGAGGCATT | 190 |
| chr12:41525547-41527573 | p2 | (AT)13 | CAACAACATTGGCTTTCCCT | TTTGAAGAGCTGGAGGCATT | 190 |
| chr12:41563210-41565230 | p5 | (AAACC)4 | GAGAACCTTTTATGCGCGAG | TATGGCGAGGTTGTGTGAAA | 154 |
| chr12:41563210-41565230 | p4 | (TTAA)4 | GAGAACCTTTTATGCGCGAG | TATGGCGAGGTTGTGTGAAA | 154 |
| chr12:41603686-41605710 | p2 | (AT)12 | GTCCGGATTTCAACCGTCTA | ACACTCTTGGACCCATTTCG | 199 |
| chr12:41603686-41605710 | p2 | (AT)41 | GTCCGGATTTCAACCGTCTA | ACACTCTTGGACCCATTTCG | 199 |

| Marker ID | SSR type | SSR motif | FORWARD PRIMER (5'-3') | REVERSE PRIMER (5'-3') | size (bp) |
|---|---|---|---|---|---|
| chr12:41802240-41804255 | p5 | (GCATT)3 | TGGTGAGACATTTGAGAGCG | CCTTTCGACTTTACTCCCCC | 298 |
| chr12:41862493-41864508 | p5 | (TGAAT)3 | GATCCTGTCGTCCGGTTAAA | TCGCTAGGGTTCAGATGCTT | 300 |
| chr12:41892377-41894395 | p6 | (CATATA)3 | TGGGCCCATATACAACCTGT | CCTGCCTTGTCCTGTCATTT | 258 |
| chr12:42009172-42011188 | p2 | (AT)8 | ATAAGATGCCAACTCCACGG | ACACAATTGGTTTTCGCACA | 168 |
| chr12:42098988-42101000 | p2 | (TA)6 | CAAAGGAGCTTGCATGAACA | TGCTCTTCCTCTCATCCGTT | 115 |
| chr12:42150522-42152552 | p2 | (AT)15 | ACTTTTGTCACAGCCCAACC | GTCGGCTTGCGTATCGTATT | 177 |
| chr12:42181182-42183222 | p2 | (TA)20 | AATCGAGAATCAACATCGGC | AGTTCGGGCACTAGTCATGG | 108 |
| chr12:42403441-42405453 | p2 | (AT)6 | GGGGTATCACCAGAACCCTT | GCGTGGATGGTGCTCTAAAT | 300 |
| chr12:42403441-42405453 | p5 | (TTTGG)3 | GGGGTATCACCAGAACCCTT | GCGTGGATGGTGCTCTAAAT | 300 |
| chr12:42546702-42548720 | p2 | (TA)9 | TCGAGTCCGAAAGAGGAAAA | GTTCTGCATCCATGAGGGTT | 227 |
| chr12:42610758-42612773 | p5 | (GGTTT)3 | CCACATCTTTAGCCGTGGAT | TCCTGCCTGGTCGAGTAGTT | 277 |
| chr12:42664053-42666071 | p6 | (ATATAA)3 | GCTCAGAGACGGCAAAACAT | GGAAAATCTCAAACGGACCA | 125 |
| chr12:42664053-42666071 | p3 | (ATA)5 | GCTCAGAGACGGCAAAACAT | GGAAAATCTCAAACGGACCA | 125 |
| chr12:42684555-42686573 | p3 | (CAA)6 | ATGTCCCCTTCGGGAATTAG | ACGAACCCAGCAAATAAACG | 255 |
| chr12:42695846-42697870 | p2 | (TA)10 | AGCTAACACACCGAGCACCT | TAGTAGCCTTGTCGGCTGGT | 242 |
| chr12:42695846-42697870 | p2 | (TA)12 | AGCTAACACACCGAGCACCT | TAGTAGCCTTGTCGGCTGGT | 242 |
| chr12:42695846-42697870 | p3 | (CAA)6 | AGCTAACACACCGAGCACCT | TAGTAGCCTTGTCGGCTGGT | 242 |
| chr12:42707163-42709217 | p2 | (AT)6 | TCACAGGGCGATCACAAATA | CCCCCATGATTTCTAACCCT | 288 |
| chr12:42734310-42736324 | p2 | (TA)7 | AAGCCGACGTACCATTATGC | GACCCCATACTCAGAGCCAA | 200 |
| chr12:42862286-42864301 | p3 | (TTG)5 | TTAAGGAAGGCTTACGGCAA | GAAGACTAGTGGCCCCTTCC | 243 |
| chr12:42958936-42960952 | p2 | (AT)8 | TTACGAAACTTGGGGAGTGG | GGGTCAAGTCTACCGGACAA | 231 |
| chr12:42958936-42960952 | p2 | (AT)7 | TTACGAAACTTGGGGAGTGG | GGGTCAAGTCTACCGGACAA | 231 |
| chr12:43001070-43003082 | p2 | (CA)6 | GAGTAACCTTCCGGAGGACC | GATGAGGCTATGGCCGTAAA | 176 |
| chr12:43071440-43073464 | p3 | (TTG)10 | ATTGAAGCGTCCGAATCATC | AGGAACGGTTTGGGAAGTTT | 155 |
| chr12:43071483-43073501 | p3 | (TTG)10 | ATTGAAGCGTCCGAATCATC | AGGAACGGTTTGGGAAGTTT | 112 |
| chr12:43121454-43123472 | p6 | (TACATA)3 | ATAAAGCCTTAGCGGCACCT | GTCCCGACCTTTTATGAGCA | 294 |
| chr12:43205466-43207482 | p2 | (AT)8 | TCTTTGTGCTCATCCACTGC | AAACAGGAAGGCGGATAGGT | 297 |
| chr12:43211926-43213948 | p2 | (AT)11 | GCATGGATAATCCTCAGCGT | TTTATGAGTTTGAACCCGCC | 226 |
| chr12:43237959-43239975 | p2 | (AT)8 | TCTTTGTGCTCATCCACTGC | AAACAGGAAGGCGGATAGGT | 297 |
| chr12:43244419-43246441 | p2 | (AT)11 | GCATGGATAATCCTCAGCGT | TTTATGAGTTTGAACCCGCC | 226 |
| chr12:43348232-43350250 | p2 | (AT)9 | CTCACTACTCTGCTCGTGCG | ACAAAACGCCATACACCACA | 223 |
| chr12:43348232-43350250 | p2 | (AT)6 | CTCACTACTCTGCTCGTGCG | ACAAAACGCCATACACCACA | 223 |
| chr12:43353638-43355653 | p5 | (TATCA)3 | TTTTGGTAAGAATCCGCAGG | GATAGGGGTGCTCGGTACAA | 132 |
| chr12:43366955-43368969 | p2 | (AT)7 | TATTTGGTTACGGCGAGTCC | ATGCACAAACATATTGCGGA | 287 |
| chr12:43366955-43368969 | p5 | (ATTCA)3 | TATTTGGTTACGGCGAGTCC | ATGCACAAACATATTGCGGA | 287 |
| chr12:43379017-43381029 | p2 | (AT)6 | TAGTTGAAGGGGCATCTTGG | GCAGCACCAGTTGTTTCAGA | 144 |

| Marker ID | SSR type | SSR motif | FORWARD PRIMER (5'-3') | REVERSE PRIMER (5'-3') | size (bp) |
|---|---|---|---|---|---|
| chr12:43425045-43427059 | p2 | (AT)7 | AACGGATGTCCTTGACTTCG | ACATATTGAGGCATGCACGA | 147 |
| chr12:43425045-43427059 | p5 | (TGAAT)3 | AACGGATGTCCTTGACTTCG | ACATATTGAGGCATGCACGA | 147 |
| chr12:43447208-43449223 | p5 | (CTTTG)3 | ATCTTCGGCACACACATTCA | AAGAGTATTTGGCCGTGTGG | 299 |
| chr12:43454366-43456378 | p2 | (TA)6 | CGAGGACCTGTGAACGGTAT | CTTCGACCACTGGGTCAACT | 111 |
| chr12:43454366-43456378 | p2 | (TA)6 | CGAGGACCTGTGAACGGTAT | CTTCGACCACTGGGTCAACT | 111 |
| chr12:43455060-43457072 | p2 | (TA)6 | AATTTTGGAAGCATTTCCCC | CGAGCGAAGGTTATGTCGTT | 286 |
| chr12:43455060-43457072 | p2 | (TA)6 | AATTTTGGAAGCATTTCCCC | CGAGCGAAGGTTATGTCGTT | 286 |
| chr12:43524987-43527035 | p2 | (TA)12 | TGACGAAATAGGGGAAATGC | TATAGCAAGGGAACCATGCC | 282 |
| chr12:43524987-43527035 | p2 | (AT)24 | TGACGAAATAGGGGAAATGC | TATAGCAAGGGAACCATGCC | 282 |
| chr12:43671046-43673064 | p3 | (CAA)6 | ACCGGCGAACGATATCATAG | GAAAAGCTCGGATTGCTTTG | 259 |
| chr12:43685222-43687236 | p2 | (AT)7 | CTATCACGACCCAAACCACC | TAGTGGTAGCCTTGTCGCCT | 149 |
| chr12:43698645-43700663 | p6 | (TGTGTA)3 | AGGAAACCTCTGCCCAAAGT | CACGAAATCCGTCTCCAAAT | 298 |
| chr12:43811041-43813067 | p2 | (AT)13 | CCTGACTGAGGAGCTGGAAC | AGGGCAAGTCAGTAGCTGGA | 283 |
| chr12:43818850-43820908 | p2 | (AT)29 | CTGACTGAGGAGCTGGAACC | AGGGCAAGTCAGTAGCTGGA | 275 |
| chr12:43860968-43862982 | p6 | (TATATG)3 | TGTTAAGGCTCGTCCCATTC | CGGCATTTTGGAAAACAACT | 164 |
| chr12:43860968-43862982 | p2 | (TA)7 | TGTTAAGGCTCGTCCCATTC | CGGCATTTTGGAAAACAACT | 164 |
| chr12:43887470-43889488 | p2 | (AT)9 | CTCGGTGAAGAGGCAAAAAG | TTTCGGTGTCCCTTTACTCG | 169 |
| chr12:43887470-43889488 | p2 | (AT)16 | CTCGGTGAAGAGGCAAAAAG | TTTCGGTGTCCCTTTACTCG | 169 |
| chr12:43887871-43889903 | p2 | (AT)9 | TGGGGCGAGAATAAAGTACG | TTGCTCACCTTTTCATTCCC | 215 |
| chr12:43887871-43889903 | p2 | (AT)16 | TGGGGCGAGAATAAAGTACG | TTGCTCACCTTTTCATTCCC | 215 |
| chr12:43914336-43916351 | p3 | (ATG)5 | GGACGAATGGGAGAAGTTGA | TACATACCTTTTCGGCGGAC | 100 |
| chr12:43993479-43995494 | p3 | (CAT)5 | CGACCCAAAGTAGAGGGTCA | TGTCGGCATGTCCAGTATGT | 265 |
| chr12:44022370-44024382 | p2 | (AT)6 | ATGTCTGGCCGTAGATTTGG | AACGAACTTCGGAACACTGG | 191 |
| chr12:44022370-44024382 | p2 | (AT)10 | ATGTCTGGCCGTAGATTTGG | AACGAACTTCGGAACACTGG | 191 |
| chr12:44022370-44024382 | p2 | (AT)10 | ATGTCTGGCCGTAGATTTGG | AACGAACTTCGGAACACTGG | 191 |
| chr12:44022941-44024961 | p2 | (AT)6 | GACAAGGCTGCCACTACACA | TCCGTTGTTCCGGAGTTTAG | 223 |
| chr12:44022941-44024961 | p2 | (AT)10 | GACAAGGCTGCCACTACACA | TCCGTTGTTCCGGAGTTTAG | 223 |
| chr12:44022941-44024961 | p2 | (AT)10 | GACAAGGCTGCCACTACACA | TCCGTTGTTCCGGAGTTTAG | 223 |
| chr12:44093061-44095076 | p3 | (ATG)5 | GATGTCACATTGACGAACGG | GTTTGTCGGAACTTTGGGAA | 180 |
| chr12:44171573-44173591 | p3 | (CAA)6 | TCCTTGAAGAAAGGGCAGAA | TTGATTCAAACCCTGCATGA | 108 |
| chr12:44171573-44173591 | p3 | (CAA)7 | TCCTTGAAGAAAGGGCAGAA | TTGATTCAAACCCTGCATGA | 108 |
| chr12:44171573-44173591 | p3 | (TGT)6 | TCCTTGAAGAAAGGGCAGAA | TTGATTCAAACCCTGCATGA | 108 |
| chr12:44171887-44173908 | p3 | (CAA)6 | TGGCTCTTGTTACTGCATCG | TTTGCAGAGTTAGGGCCTGT | 242 |
| chr12:44171887-44173908 | p3 | (CAA)7 | TGGCTCTTGTTACTGCATCG | TTTGCAGAGTTAGGGCCTGT | 242 |
| chr12:44171887-44173908 | p3 | (TGT)6 | TGGCTCTTGTTACTGCATCG | TTTGCAGAGTTAGGGCCTGT | 242 |
| chr12:44191564-44193578 | p2 | (TG)7 | AAAGGTGCCTACGGGCTATT | TCATACCTGGCCCTACCAAG | 224 |

| Marker ID | SSR type | SSR motif | FORWARD PRIMER (5'-3') | REVERSE PRIMER (5'-3') | size (bp) |
|---|---|---|---|---|---|
| chr12:44191564-44193578 | p2 | (CA)6 | AAAGGTGCCTACGGGCTATT | TCATACCTGGCCCTACCAAG | 224 |
| chr12:44235865-44237880 | p5 | (TTCAT)3 | TACCCGCACTTATTTCAGCC | ATAACAATGCCTCACAGGGC | 205 |
| chr12:44455551-44457563 | p2 | (AG)6 | CTTCCAGGGTGTTCCCTACA | ACTTTAACCGACGCCACATC | 250 |
| chr12:44457217-44459232 | p5 | (AAATG)3 | GCCTTGTCGACTTGCTTAGG | ACTCCGAGGATTTAACCGCT | 197 |
| chr12:44504768-44506816 | p2 | (AT)7 | AGCTAACACACCGAGCACCT | TTCATAACCCTACCGGTCCA | 240 |
| chr12:44504768-44506816 | p4 | (ATAC)12 | AGCTAACACACCGAGCACCT | TTCATAACCCTACCGGTCCA | 240 |
| chr12:44536832-44538876 | p2 | (TA)12 | GGGCAAATCATCCTTGCTAA | CATCCTGATATGGATGCGTG | 223 |
| chr12:44536832-44538876 | p2 | (TA)22 | GGGCAAATCATCCTTGCTAA | CATCCTGATATGGATGCGTG | 223 |
| chr12:44570206-44572218 | p2 | (AT)6 | ACTTAGGGTGTGATGGGCAC | TGAGCACGATGAAAAGAACG | 275 |
| chr12:44664945-44666957 | p2 | (TA)6 | AGCCTCGGTCATTTGTTGAC | AGGCCTTGTCTAACGCTTCA | 280 |
| chr12:44683003-44685018 | p3 | (TTG)5 | ATCCACATGTCGGGGTTTTA | TTGGTGTGTTTCCACTTCCA | 123 |
| chr12:44741508-44743523 | p5 | (CATAT)3 | TGGACACTCAAACGTACCCA | ATGCGATTCAGGGGTATGAC | 216 |
| chr12:44746252-44748267 | p5 | (TTACT)3 | TCAGAAATTCCATTCCCTGG | AACAACGTCAGTTTGAGGGG | 239 |
| chr12:44746252-44748267 | p2 | (TA)7 | TCAGAAATTCCATTCCCTGG | AACAACGTCAGTTTGAGGGG | 239 |
| chr12:44746656-44748679 | p5 | (TTACT)3 | GTTTCTGAATGTACGCCGGT | GCATCACTTACATGGGGTCC | 175 |
| chr12:44746656-44748679 | p2 | (TA)7 | GTTTCTGAATGTACGCCGGT | GCATCACTTACATGGGGTCC | 175 |
| chr12:44796826-44798854 | p2 | (AT)14 | CATTTAGAGCCGAGCGAATC | GGCATTCCCCTACACTACGA | 136 |
| chr12:44843279-44845297 | p3 | (TTG)6 | CAAATCAAAGTGGCAACCCT | GGTTGGAAATCTAACGGCAA | 169 |
| chr12:44939339-44941357 | p2 | (AT)9 | CCCTAGGACGAACTGCTCTG | CGTTCGCGAATTTCTATGGT | 189 |
| chr12:44939339-44941357 | p3 | (AAC)8 | CCCTAGGACGAACTGCTCTG | CGTTCGCGAATTTCTATGGT | 189 |
| chr12:44940047-44942071 | p2 | (AT)9 | AATGTCGAGGAATGTACGGC | ACTAGAGGTCCGGGTACGGT | 209 |
| chr12:44940047-44942071 | p3 | (AAC)8 | AATGTCGAGGAATGTACGGC | ACTAGAGGTCCGGGTACGGT | 209 |
| chr12:44944871-44946893 | p4 | (AATT)4 | CCATAGCTGCTCCTTTGTCC | CAAGGAATGAAGTCCCGAAA | 276 |
| chr12:44944931-44946997 | p4 | (AATT)4 | CCATAGCTGCTCCTTTGTCC | CAAGGAATGAAGTCCCGAAA | 216 |
| chr12:45044932-45046952 | p2 | (CT)10 | CTACCGAAATTTGGCAGCAT | GCATACGGGCTACTTGGTGT | 272 |
| chr12:45044932-45046952 | p2 | (AT)6 | CTACCGAAATTTGGCAGCAT | GCATACGGGCTACTTGGTGT | 272 |
| chr12:45069469-45071484 | p3 | (GAT)5 | AAGCAAAAATTGATCGGTGC | TTGGGAGAATGTTCCAGAGG | 218 |
| chr12:45085716-45087728 | p2 | (GT)6 | AAAAAGAAACTCCCCCGAAA | CCTCAATATTTCCGCCTCAA | 239 |
| chr12:45095723-45097738 | p5 | (CAAAG)3 | GCAACAGAGACGGGAAAAAG | GGCATCTGTTTGGTTTGCTT | 133 |
| chr12:45109977-45111992 | p3 | (AAC)5 | AAGGCTCGGTGTCTGACTGT | CCCACTTCTTTCATTGGCAT | 255 |
| chr12:45124201-45126213 | p2 | (TA)6 | TCGTGCATGCCTCAATATGT | ACCGTACCCGACCCTCTAGT | 290 |
| chr12:45124201-45126213 | p2 | (TA)6 | TCGTGCATGCCTCAATATGT | ACCGTACCCGACCCTCTAGT | 290 |
| chr12:45124201-45126213 | p2 | (AT)6 | TCGTGCATGCCTCAATATGT | ACCGTACCCGACCCTCTAGT | 290 |
| chr12:45150748-45152764 | p2 | (AG)8 | TATCTGGATCAAGCTCGCCT | CGATGTCACCTCGGGATACT | 142 |
| chr12:45343172-45345198 | p2 | (AG)13 | CAGCCCAGTTGGTCGTAAAT | CGGGATACTTAAGGCAACCA | 218 |
| chr12:45431750-45433768 | p2 | (CA)9 | CCAAAATAATTCCGGAGGGT | AGTCCTAGAGTGCGGCTCAA | 179 |

| Marker ID | SSR type | SSR motif | FORWARD PRIMER (5'-3') | REVERSE PRIMER (5'-3') | size (bp) |
|---|---|---|---|---|---|
| chr12:45526035-45528053 | p2 | (AT)13 | GCAGGGTGATAGGGCTGTTA | CATCAATAATGTCCTGGGGG | 209 |
| chr12:45526035-45528053 | p2 | (AT)9 | GCAGGGTGATAGGGCTGTTA | CATCAATAATGTCCTGGGGG | 209 |
| chr12:45586340-45588364 | p2 | (AT)12 | ATGCTCCTCTGAAATGGGTG | ATGGCTCATCAGTTTGGGAC | 105 |
| chr12:45615110-45617122 | p5 | (TATTT)3 | TGGAACCTCTATGCAGACCC | CTCCGTATTTCCGCATCAAT | 199 |
| chr12:45615110-45617122 | p2 | (GT)6 | TGGAACCTCTATGCAGACCC | CTCCGTATTTCCGCATCAAT | 199 |
| chr12:45628779-45630794 | p5 | (TTATT)3 | CAGATCTCCGATTGGTTGGT | TTCCTGCCGGATCTCATTAC | 181 |
| chr12:45664497-45666509 | p2 | (CA)6 | TGCACTGAAATTTGCCTCTG | GGTCAATCTCAGTCCTCCCA | 245 |
| chr12:45713459-45715474 | p2 | (TA)7 | TTTCGGTCAACACAGTCCAA | TTTAAGGATGCGTAATGGGC | 217 |
| chr12:45713459-45715474 | p5 | (AGAAA)3 | TTTCGGTCAACACAGTCCAA | TTTAAGGATGCGTAATGGGC | 217 |
| chr12:45733912-45735930 | p2 | (GA)6 | ATTTGCGGTGGAAATAGTCG | ACGCCATTTTTGGCATAATC | 266 |
| chr12:45734016-45736034 | p2 | (GA)6 | GCGGTGGAAATAGTCGAAAA | CCACCCCTCTCTCTCTCTCC | 166 |
| chr12:45792661-45794677 | p2 | (AT)8 | TCTCTGAAGCCCTGACCAGT | GATGATGATGATGACAGGCG | 188 |
| chr12:45811775-45813791 | p2 | (AT)8 | TCTCTGAAGCCCTGACCAGT | GATGATGATGATGACAGGCG | 188 |
| chr12:45845795-45847813 | p2 | (AT)9 | ACTCGAGCGAATAAACCCCT | CATATCCGTGACACCAGTGC | 133 |
| chr12:45896887-45898901 | p2 | (AT)7 | CCCGAACTGGACACTCGTAT | GTCGGCTTGCGTATCGTATT | 258 |
| chr12:45905780-45907840 | p2 | (AT)30 | AGACCATGCGATGAAGGAAC | CCTGTTATGCCAATACGCCT | 214 |
| chr12:45953700-45955712 | p2 | (TA)6 | TGGGGTGTGATAGGTTTGGT | AAGAGAGAAAAAGGACCCCG | 138 |
| chr12:45964414-45966444 | p2 | (TA)15 | TGGGGTGTGATAGGTTTGGT | AAGGACCCCGGTCAGTTTAT | 139 |
| chr12:45978808-45980823 | p5 | (ATCAT)3 | TGAATGCCTCTCATCATCCA | AGATCGACATGTTCCCTTGG | 288 |
| chr12:46045539-46047563 | p2 | (TA)31 | TGACCGTGAAACACTTCCAG | GTCAAACGGATCGGAAAGAA | 299 |
| chr12:46045539-46047563 | p2 | (AT)12 | TGACCGTGAAACACTTCCAG | GTCAAACGGATCGGAAAGAA | 299 |
| chr12:46057971-46059983 | p2 | (CA)6 | GTACCGGGGACTAAGCTTCC | TACGGATAATCGTCGAAGGG | 262 |
| chr12:46098345-46100369 | p2 | (CA)12 | TAAGGTGAGGCAGAACGCTT | TATATGTTGCCCGCCTTAGC | 247 |
| chr12:46130098-46132110 | p2 | (AT)6 | TGATTGCTGCAGAGGTTTTG | ACTACGCAAACCAGCGACTT | 290 |
| chr12:46201779-46203794 | p3 | (AAC)5 | ATTCAAATCTCCGAATCCCC | GGCCGTTTAATGTTTGTCGT | 163 |
| chr12:46272851-46274863 | p2 | (GT)6 | CCCGCCATGATCATAAGACT | GCTCGTTTTGTAATTCCCGA | 264 |
| chr12:46301886-46303898 | p2 | (CA)6 | CTGAACTCTCAGGCTGTCCC | GACTCCAACTCCTCAGCCAG | 224 |
| chr12:46358504-46360519 | p3 | (ACA)5 | TTTATGGGAGGCGTTGTAGG | CAATTCCCTACGCCTCATGT | 165 |
| chr12:46530723-46532741 | p6 | (TTCATA)3 | AATCACTTCCCACTTCCACG | TGTGTCCCTAATGTGGCAGA | 245 |
| chr12:46547683-46549698 | p3 | (CAA)5 | ATGTACGCGTATACCTGCCC | CGACTTCATTGGATCGGTTT | 133 |
| chr12:46565695-46567711 | p2 | (AT)8 | TCGACGTTGTAGTTTCGGTG | AACTCGCCTAACCCGAACTT | 129 |
| chr12:46593270-46595282 | p2 | (AT)6 | TGTGCTCTGAAACCGTTCTG | GTGCCCATCACACCCTAAGT | 176 |
| chr12:46633535-46635555 | p2 | (TA)10 | CACCATCCCCGAGAATCTAA | TTGAATTTGGAAGGTTTCGG | 124 |
| chr12:46689571-46691586 | p2 | (AT)7 | CAAGAAAAAGGGAGAAGGGG | TTCCAAGGAGCCAATTTTTG | 190 |
| chr12:46689571-46691586 | p5 | (TGAAT)3 | CAAGAAAAAGGGAGAAGGGG | TTCCAAGGAGCCAATTTTTG | 190 |
| chr12:46689571-46691586 | p6 | (TGTATA)3 | CAAGAAAAAGGGAGAAGGGG | TTCCAAGGAGCCAATTTTTG | 190 |

| Marker ID | SSR type | SSR motif | FORWARD PRIMER (5'-3') | REVERSE PRIMER (5'-3') | size (bp) |
|---|---|---|---|---|---|
| chr12:46742124-46744144 | p5 | (ACACA)4 | TTACATACCTTGCCCGCTTC | GGTAATTTGGCCGACTTGAA | 287 |
| chr12:46807492-46809504 | p2 | (TA)6 | GAAATGCCGTCGCAATTTAT | ACTCGGCCAAAATCATCAAC | 190 |
| chr12:46903184-46905199 | p5 | (AAAAT)3 | GGCATCAGACACCTCAACCT | AAGAGAGCGAGTTCCCACAA | 239 |
| chr12:46908279-46910307 | p2 | (AT)14 | GCGCATTCTCTCCTCTTGAC | GCAGCAAGTATTAGAGCCCG | 105 |
| chr12:46919361-46921373 | p2 | (TA)6 | AGAATTTGCATGGAAGACCG | TCCATATTTTTCGAGCGTCC | 200 |
| chr12:46919361-46921373 | p2 | (TA)7 | AGAATTTGCATGGAAGACCG | TCCATATTTTTCGAGCGTCC | 200 |
| chr12:46919361-46921373 | p5 | (TGAAT)3 | AGAATTTGCATGGAAGACCG | TCCATATTTTTCGAGCGTCC | 200 |
| chr12:46940955-46942970 | p3 | (CAA)5 | CGGACGGAATCATAGCAAGT | TTTGCTGACTCCCTCGAGTT | 136 |
| chr12:46986685-46988749 | p2 | (TA)32 | CCTGATGCAACACTTCCTGA | GCAAATCTCCCAGTGTTCGT | 223 |
| chr12:47004750-47006764 | p3 | (CAT)8 | CACCAAGGCATGAGAAACCT | AAAATGCTGCCCAATTTTTG | 278 |
| chr12:47004750-47006764 | p2 | (AT)7 | CACCAAGGCATGAGAAACCT | AAAATGCTGCCCAATTTTTG | 278 |
| chr12:47164291-47166307 | p2 | (TA)8 | GTCCTGATCAGACGCAGTCA | CTTCATCTTGATGCCAAGCA | 144 |
| chr12:47183951-47185967 | p4 | (TGTA)7 | CATTCCTTCGAGGTGGGATA | GTGTTATCTTACCCGGCCCT | 190 |
| chr12:47208086-47210101 | p3 | (ATC)5 | TCACGACCCAAACTAGAGGG | ATAGCTCTATCGGCCCACCT | 158 |
| chr12:47329530-47331548 | p3 | (AAT)6 | GAACCTCAACCAAAACCGAA | TAGACCAGGTCATGGGAAGG | 263 |
| chr12:47464402-47466420 | p2 | (AT)12 | CTTCGTGGTCGGGTAGGATA | TTCGATCGTGTGATTGGGTA | 261 |
| chr12:47464402-47466420 | p6 | (TGTAAG)3 | CTTCGTGGTCGGGTAGGATA | TTCGATCGTGTGATTGGGTA | 261 |
| chr12:47492148-47494160 | p2 | (AT)6 | TTTCATCATCCTCCCACCAT | GCATTACATCAGAGCAGCCA | 152 |
| chr12:47492148-47494160 | p3 | (ATC)5 | TTTCATCATCCTCCCACCAT | GCATTACATCAGAGCAGCCA | 152 |
| chr12:47703328-47705343 | p5 | (CATTG)3 | CCTGTCCCGTCCTTTCTACA | TCTACCAACCATTTGTGCCA | 160 |
| chr12:47722467-47724503 | p2 | (AG)18 | TCAATGGTAGGGTTTGGCAT | TTTATGGACCAAGCTACGCC | 104 |
| chr12:47722467-47724503 | p2 | (AG)22 | TCAATGGTAGGGTTTGGCAT | TTTATGGACCAAGCTACGCC | 104 |
| chr12:47781141-47783166 | p5 | (ATGAT)5 | GACCCTTGTCCCGATTATCA | GGGGATAATGCCTTTGGAAT | 181 |
| chr12:47851433-47853451 | p2 | (TA)9 | TATACGCTACCTCATCCGGC | TTCGGTCGTAGGATCATTCC | 234 |
| chr12:47941398-47943458 | p2 | (AT)30 | GCTGTCATGATGGAGTCGTG | TCAGTCATATGCAGCAAGGC | 277 |
| chr12:48046954-48048969 | p5 | (GCATT)3 | GCTAGCCACCCGTTATGTGT | GGATCATCACCAGGTATCGG | 159 |
| chr12:48059071-48061099 | p2 | (TA)14 | GCCCACACGTCAAGATTTTT | GAACCCACGACCTATGGATG | 185 |
| chr12:48103212-48105226 | p2 | (TA)7 | CTTTCTGTCGTAGCCTTGCC | GAGTCTCGCTTACGGGTCTG | 173 |
| chr12:48167493-48169511 | p6 | (CATATA)3 | TCGCCGAACATTCAACATAA | TCGTGTTGTTATGCAGCCTC | 273 |
| chr12:48187382-48189397 | p3 | (ATG)5 | GGGCGTTAAATGGAGCATAA | TTATCGTTCTCCACCCAAGC | 209 |
| chr12:48187382-48189397 | p5 | (CAGAC)3 | GGGCGTTAAATGGAGCATAA | TTATCGTTCTCCACCCAAGC | 209 |
| chr12:48221784-48223796 | p2 | (TA)6 | GAAATTTCACGGCCTTCTGA | CACGAAGATTGAAGCGATCA | 121 |
| chr12:48291029-48293044 | p2 | (TA)16 | TGGGCGGTGTGTGATAGTTA | GGGTCTGCACCATCTTCCTA | 147 |
| chr12:48291029-48293044 | p5 | (CATTG)3 | TGGGCGGTGTGTGATAGTTA | GGGTCTGCACCATCTTCCTA | 147 |
| chr12:48596675-48598703 | p2 | (AT)6 | GACTTGTGCCCGAGGTAGAC | GTCGGCTTGCGTATGGTATT | 205 |
| chr12:48596675-48598703 | p2 | (AT)10 | GACTTGTGCCCGAGGTAGAC | GTCGGCTTGCGTATGGTATT | 205 |

| Marker ID | SSR type | SSR motif | FORWARD PRIMER (5'-3') | REVERSE PRIMER (5'-3') | size (bp) |
|---|---|---|---|---|---|
| chr12:48596675-48598703 | p2 | (AT)14 | GACTTGTGCCCGAGGTAGAC | GTCGGCTTGCGTATGGTATT | 205 |
| chr12:48596675-48598703 | p3 | (ACA)5 | GACTTGTGCCCGAGGTAGAC | GTCGGCTTGCGTATGGTATT | 205 |
| chr12:48624144-48626172 | p2 | (AT)6 | GACTTGTGCCCGAGGTAGAC | GTCGGCTTGCGTATGGTATT | 205 |
| chr12:48624144-48626172 | p2 | (AT)10 | GACTTGTGCCCGAGGTAGAC | GTCGGCTTGCGTATGGTATT | 205 |
| chr12:48624144-48626172 | p2 | (AT)14 | GACTTGTGCCCGAGGTAGAC | GTCGGCTTGCGTATGGTATT | 205 |
| chr12:48624144-48626172 | p3 | (ACA)5 | GACTTGTGCCCGAGGTAGAC | GTCGGCTTGCGTATGGTATT | 205 |
| chr12:48677582-48679598 | p2 | (TA)8 | GAGCGCTGTAGAGTACCGCT | CGCGATATGCTACGACAAGA | 114 |
| chr12:48702211-48704223 | p2 | (TA)6 | GGCCCTCGTACCCAACTTAT | GGAGATTTGCGACGTGAACT | 232 |
| chr12:48799971-48801993 | p2 | (TA)11 | AGGTCGATTCGCTGAAGTGT | CACCTCCCCTATATCGCTCA | 239 |
| chr12:48842046-48844068 | p2 | (TA)11 | AGGTCGATTCGCTGAAGTGT | CACCTCCCCTATATCGCTCA | 239 |
| chr12:48931976-48933988 | p2 | (AT)6 | ACCATGACTAGTGCCCGAAC | TGTCGTGCTTTGGTTAGCTG | 253 |
| chr12:48935048-48937076 | p2 | (TA)14 | TTGTACATGGGTGGTGGTTG | CTTAACGGAAATCGAGCAGC | 129 |
| chr12:48990797-48992817 | p4 | (CATA)5 | CACCCAACCCCTCACTTAGA | CTCTGCAAGCCGATGTATCA | 150 |
| chr12:49064151-49066185 | p2 | (AT)17 | ATAACCCCACTCCCCAAAAC | GGATCATTTGGATGTGGGTC | 233 |
| chr12:49067710-49069730 | p2 | (TA)10 | TGGCATGGCATGAAGAATAA | TGTCGATTTTCCCTGAATCC | 286 |
| chr12:49078222-49080237 | p5 | (ATTGC)3 | ATTCCAATCTTGCCCTTGTG | ACTTCATGCCCGAAGACCTA | 278 |
| chr12:49078222-49080237 | p2 | (TA)6 | ATTCCAATCTTGCCCTTGTG | ACTTCATGCCCGAAGACCTA | 278 |
| chr12:49102140-49104154 | p2 | (TA)7 | TGAAATAGGGGAAATGCTGC | GACAAATGGGAACATGCCTT | 133 |
| chr12:49209162-49211188 | p3 | (CAT)6 | GCCGCTATGTGACAAACTGA | ATCGAACTCCACTTGTTCCG | 276 |
| chr12:49209162-49211188 | p2 | (CA)13 | GCCGCTATGTGACAAACTGA | ATCGAACTCCACTTGTTCCG | 276 |
| chr12:49271690-49273706 | p2 | (TA)8 | AGACATAGGGCTGCTGGAGA | TTCTCGGAGGGATGAATGTC | 179 |
| chr12:49271690-49273706 | p5 | (AATGC)3 | AGACATAGGGCTGCTGGAGA | TTCTCGGAGGGATGAATGTC | 179 |
| chr12:49345776-49347792 | p4 | (CATT)4 | ATGATGAAATTGCGAAAGCC | CCCTCTGGTTGTGATGTCCT | 143 |
| chr12:49415253-49417271 | p6 | (ATATGT)3 | ATGTTGGATATGAGCCTCGG | GTCGGGTCCTATTTATGCGA | 226 |
| chr12:49434248-49436268 | p2 | (TA)10 | CATCCATCCCCGAGAGTCTA | TTGAATTTGGAAGGTTTCGG | 155 |
| chr12:49484302-49486320 | p6 | (ATATGT)3 | ATGTTGGATATGAGCCTCGG | GTCGGGTCCTATTTATGCGA | 226 |
| chr12:49503297-49505317 | p2 | (TA)10 | CATCCATCCCCGAGAGTCTA | TTGAATTTGGAAGGTTTCGG | 155 |
| chr12:49584219-49586234 | p5 | (ACACA)3 | TTCACAAGGATCATGCCAAA | TTTGGCCGAGTTGAATTCTC | 262 |
| chr12:49607426-49609444 | p2 | (TA)9 | TAAACGGGGTCGACTACAGG | TTAATTGATGTGCCGCAAGA | 196 |
| chr12:49608480-49610494 | p2 | (AT)7 | TGCGTACACTTTACCCTCCC | CAATTGATGTTCCCCTTGGT | 253 |
| chr12:49608480-49610494 | p2 | (CT)6 | TGCGTACACTTTACCCTCCC | CAATTGATGTTCCCCTTGGT | 253 |
| chr12:49629567-49631582 | p5 | (TGTGT)3 | CCAAGGGAAGTTGATGGAGA | ACTCGGCTTGCTTGCTAAAA | 151 |
| chr12:49629567-49631582 | p5 | (AATTG)3 | CCAAGGGAAGTTGATGGAGA | ACTCGGCTTGCTTGCTAAAA | 151 |
| chr12:49712973-49714991 | p6 | (TTTCAA)3 | ACGGAATAATCAAACGTGCC | TTGGCATGATCCTTGTGAAA | 212 |
| chr12:49720808-49722823 | p5 | (ATGCA)3 | CACCTACCTTTCCCACCTCA | GGAAGCGGATCGAGTTTGTA | 153 |
| chr12:49742754-49744772 | p3 | (CAA)6 | GGGGAGGACGTATGGAAAAT | ACGTTCCTCGGCACTATGAC | 148 |

| Marker ID | SSR type | SSR motif | FORWARD PRIMER (5'-3') | REVERSE PRIMER (5'-3') | size (bp) |
|---|---|---|---|---|---|
| chr12:49790395-49792409 | p2 | (TA)7 | AGGGGAAGTGCTACCGAAAT | CACGTCGCAATCTCGTTAAA | 162 |
| chr12:49790395-49792409 | p5 | (GCTAT)3 | AGGGGAAGTGCTACCGAAAT | CACGTCGCAATCTCGTTAAA | 162 |
| chr12:49790395-49792409 | p5 | (GCTAT)3 | AGGGGAAGTGCTACCGAAAT | CACGTCGCAATCTCGTTAAA | 162 |
| chr12:49857881-49859896 | p3 | (TTG)5 | TATGCTACGGGCATGTGTGT | GTTTCTCCCACTTCGAACCA | 194 |
| chr12:49884838-49886853 | p5 | (AAAAT)3 | ACTTTCTTCGCAGTGGTCGT | TCCACCAGGTCGTATTAGCC | 195 |
| chr12:49943213-49945227 | p2 | (AT)7 | ATGTCCCTCTTCATGCCATC | GGCTCTAAGTAAGGGTCGGG | 149 |
| chr12:49986944-49988960 | p2 | (TA)8 | AGCTAACACACCGAGCACCT | TCGGCTGGTGATGATAGACA | 287 |
| chr12:50092018-50094030 | p2 | (AT)6 | AGTGTGTGGTTCCAAAAGCC | AAGGTCGGTAACCTCCGAAT | 212 |
| chr12:50201888-50203906 | p2 | (TA)9 | ATAACGCAATAAACCGCTCG | ATTTGGCTGAAATTTCGTGG | 274 |
| chr12:50232095-50234110 | p5 | (GCATT)3 | GGATGGAATTGACATACCCG | TCAACGATACCGGATCATCA | 263 |
| chr12:50232095-50234110 | p4 | (CTTT)4 | GGATGGAATTGACATACCCG | TCAACGATACCGGATCATCA | 263 |
| chr12:50256374-50258386 | p2 | (AT)6 | ATCGCCCAGACATACACACA | TCCGGAGCTCATATTTTTGG | 155 |
| chr12:50268987-50270999 | p2 | (GT)6 | TACACAAGGTGGGCCATGTA | CCGGCACAACTCTAGAAGGA | 114 |
| chr12:50272585-50274597 | p2 | (TA)6 | GGAAGAAGCATGGAATGGAA | CCTTGTCCACTAAAGCCTCG | 196 |
| chr12:50307884-50309899 | p5 | (GAAAG)3 | ATCTGTCCGCCATAGTTTCG | CTTTTCGCTCTTTCACGGAC | 161 |
| chr12:50330974-50333018 | p2 | (TA)22 | TGCCTCTTAGGGTGGATGAC | CTTTCTTTGGGCTCTCATGC | 145 |
| chr12:50465157-50467199 | p3 | (AAT)14 | ACAGCGGGGTTAGAAGGATT | AAACACGAAAGTGGTGGAGG | 104 |
| chr12:50589984-50592002 | p2 | (TA)9 | CCCTCTTGGGTTTCTTCCTC | GTGACTATGGCGAGGGATGT | 281 |
| chr12:50620973-50622985 | p2 | (TA)6 | AGTGCGAGCTGTTTCAAGGT | GCGGCATTTCCCCTATATTT | 289 |
| chr12:50800929-50802944 | p5 | (AAATT)3 | ACTGCAAAATTCCCAACCAG | CTGAGAGGTGATGTCCCGAT | 194 |
| chr12:50848180-50850192 | p4 | (AGAA)5 | CATGCGTTGCCATGCTATAC | GCCATAGGCATCATACGCTT | 274 |
| chr12:50848180-50850192 | p2 | (TG)6 | CATGCGTTGCCATGCTATAC | GCCATAGGCATCATACGCTT | 274 |
| chr12:50910374-50912389 | p3 | (GTT)5 | TGGCCCTGTAGCACTTTCTT | CACGCCTTGGCTCTAATCTC | 131 |
| chr12:50928239-50930251 | p2 | (AT)7 | CCACACATCCAAACTCAACG | CACGGTGTTTAATATGGGGC | 147 |
| chr12:50928239-50930251 | p2 | (TA)6 | CCACACATCCAAACTCAACG | CACGGTGTTTAATATGGGGC | 147 |
| chr12:50942042-50944054 | p2 | (GA)6 | GAGCATTCTCAGAACCTCGG | GTTGCAGGAATAGAGAGCGG | 223 |
| chr12:50959081-50961093 | p2 | (GA)6 | GAGCATTCTCAGAACCTCGG | GTTGCAGGAATAGAGAGCGG | 223 |
| chr12:50961084-50963096 | p2 | (CT)6 | GCGTAGCTCTTTGTGTGCAG | TTCAGGGGCACCATCTTTAC | 275 |
| chr12:51024928-51026949 | p3 | (TTG)7 | TTTTGGGTCAACTTTGGAGG | GATGAGCGGTCCATTGAGTT | 175 |
| chr12:51024928-51026949 | p4 | (TATG)4 | TTTTGGGTCAACTTTGGAGG | GATGAGCGGTCCATTGAGTT | 175 |
| chr12:51029908-51031935 | p3 | (TTG)9 | GGGACCAAGCAAATTGAAGA | GATGAGCGGTCCATTGAGTT | 125 |
| chr12:51029908-51031935 | p4 | (TATG)4 | GGGACCAAGCAAATTGAAGA | GATGAGCGGTCCATTGAGTT | 125 |
| chr12:51193068-51195083 | p5 | (AAAGG)3 | GGAAAATCTGGTGCTGGTGT | GTAACTTTTCGGAAGCAGCG | 198 |
| chr12:51193068-51195083 | p5 | (ATTGC)3 | GGAAAATCTGGTGCTGGTGT | GTAACTTTTCGGAAGCAGCG | 198 |
| chr12:51193831-51195846 | p5 | (AAAGG)3 | GGGTCAAAGGAAAGGAAAGG | GGTGACTAGCCCCAATCAAA | 232 |
| chr12:51193831-51195846 | p5 | (ATTGC)3 | GGGTCAAAGGAAAGGAAAGG | GGTGACTAGCCCCAATCAAA | 232 |

| Marker ID | SSR type | SSR motif | FORWARD PRIMER (5'-3') | REVERSE PRIMER (5'-3') | size (bp) |
|---|---|---|---|---|---|
| chr12:51231348-51233382 | p2 | (TA)17 | ATCACGACCTTACGTCACCC | TGCCAAAAGGTCATGGTGTA | 285 |
| chr12:51274597-51276609 | p2 | (AT)6 | GAAGCCAGCTCCACTGACTC | ATGCACCGGGCTAGAATATG | 203 |
| chr12:51501368-51503394 | p2 | (AT)13 | TCAAGGGATGTTGTGTTCCA | CTTTTCCAAATTCCCAACGA | 236 |
| chr12:51579403-51581418 | p4 | (ACAT)7 | ATGTCACGACCCGAACTAGG | TCGGTCGTCATTTTGTCGTA | 164 |
| chr12:51579403-51581418 | p5 | (ATAAT)3 | ATGTCACGACCCGAACTAGG | TCGGTCGTCATTTTGTCGTA | 164 |
| chr12:51610018-51612033 | p5 | (TGGGG)3 | ACATTTGAACCCGTTTGCTC | GACAAACCGCCTAGAACTCG | 173 |
| chr12:51610018-51612033 | p6 | (CCGGTT)3 | ACATTTGAACCCGTTTGCTC | GACAAACCGCCTAGAACTCG | 173 |
| chr12:51638053-51640081 | p2 | (TA)9 | TGCTCCACGCCCTACTACTT | CGAAGCCACGGTTGATACTT | 182 |
| chr12:51638480-51640498 | p2 | (TA)9 | ACTTAGAAGCGGAGGAAGGC | TTTCGTTAATTTCCCCCACA | 132 |
| chr12:51647963-51649989 | p2 | (TA)6 | CTCGGGCACCTTTTATCGTA | CGGCTTGCGTATATGGGTAT | 159 |
| chr12:51647963-51649989 | p2 | (TA)13 | CTCGGGCACCTTTTATCGTA | CGGCTTGCGTATATGGGTAT | 159 |
| chr12:51677331-51679346 | p3 | (TTG)5 | AAAAGCTGCAGCATCCCTAA | TGCGTGAATGAGTTCGTAGC | 268 |
| chr12:51677331-51679346 | p2 | (TA)31 | AAAAGCTGCAGCATCCCTAA | TGCGTGAATGAGTTCGTAGC | 268 |
| chr12:51756531-51758546 | p5 | (AATGC)3 | CGTCCTTCGTGGTCGTATTT | TAAGCTGCCACGTCGTATTG | 174 |
| chr12:51793551-51795566 | p2 | (GT)6 | CCTTGCCAGGTTGAAGTTGT | AAAAACCATCACCCCTTTCC | 263 |
| chr12:51793551-51795566 | p5 | (TGAAT)3 | CCTTGCCAGGTTGAAGTTGT | AAAAACCATCACCCCTTTCC | 263 |
| chr12:51888225-51890237 | p2 | (TG)6 | TATGTCAGTGCCGAGAAACG | TTTCAAATTTGCCCTTCGTC | 275 |
| chr12:51921450-51923468 | p6 | (TATATG)3 | GATAATTTGGCCGAGTTGGA | TCACAAGGATCATGCCAAAA | 181 |
| chr12:51958818-51960842 | p3 | (TTG)8 | TTTTGGGTCAACTTTGGAGG | AGGTTCGGGTTCCTCTGACT | 170 |
| chr12:51986921-51988957 | p3 | (ATC)6 | TCGTACTGACCGCCTTTCTT | ACGCCCTATCCACAAGACAC | 264 |
| chr12:52014374-52016390 | p2 | (TG)8 | TTCCCCACTTGTTTCCTACG | CTTACAGGCATAACGGGGAA | 157 |
| chr12:52115218-52117230 | p2 | (CA)6 | TGTTGTTCATGTCGTCGGAT | CAAGGATCGTTGGACAGGTT | 197 |
| chr12:52115218-52117230 | p2 | (AT)14 | TGTTGTTCATGTCGTCGGAT | CAAGGATCGTTGGACAGGTT | 197 |
| chr12:52133851-52135877 | p2 | (TA)14 | TGCAAACTATCACGACCCAA | ACTCCAGGGGAGACAGGTTT | 257 |
| chr12:52133851-52135877 | p2 | (AT)13 | TGCAAACTATCACGACCCAA | ACTCCAGGGGAGACAGGTTT | 257 |
| chr12:52168607-52170621 | p2 | (TA)6 | ATCGAGGGTCCACGACATTA | CCGACATATAGGTGCGTGTG | 294 |
| chr12:52168622-52170634 | p2 | (TA)6 | ATCGAGGGTCCACGACATTA | CCGACATATAGGTGCGTGTG | 279 |
| chr12:52168741-52170767 | p2 | (TA)6 | ATCGAGGGTCCACGACATTA | CCGACATATAGGTGCGTGTG | 160 |
| chr12:52184172-52186194 | p2 | (AT)11 | TGCCGGTCAAAATTCACATA | CATCAACTCAAGGGCCAAGT | 107 |
| chr12:52184172-52186194 | p2 | (TA)17 | TGCCGGTCAAAATTCACATA | CATCAACTCAAGGGCCAAGT | 107 |
| chr12:52186312-52188327 | p5 | (AATAA)3 | CCCTACTCGGTCCTGTTCAA | AGACATTGGACAAGCCATCC | 241 |
| chr12:52193361-52195377 | p4 | (TATG)4 | CCCCACTCCCCATAAAACTT | GTTCCACTTTCGACCCAAAA | 184 |
| chr12:52292983-52294998 | p3 | (AAC)5 | GGTTACTCCGACTTCACCCA | AGCTTTTGTGATTCATGGGG | 101 |
| chr12:52319962-52322001 | p6 | (AACATA)3 | AGCTGGAGCAGTCCACAACT | CCTCATTTTGTCGCCACTTT | 278 |
| chr12:52319962-52322001 | p2 | (AT)13 | AGCTGGAGCAGTCCACAACT | CCTCATTTTGTCGCCACTTT | 278 |
| chr12:52545977-52547992 | p3 | (AAC)5 | TCAAGGCCAAAATCATGTCA | CCATCGAGCCTTGGATATGT | 135 |

| Marker ID | SSR type | SSR motif | FORWARD PRIMER (5'-3') | REVERSE PRIMER (5'-3') | size (bp) |
|---|---|---|---|---|---|
| chr12:52545977-52547992 | p3 | (AAC)5 | TCAAGGCCAAAATCATGTCA | CCATCGAGCCTTGGATATGT | 135 |
| chr12:52546163-52548178 | p3 | (AAC)5 | GTTAGCCATCCTAGGGGCTC | GAAAGCGGGCAAGGTATGTA | 227 |
| chr12:52546163-52548178 | p3 | (AAC)5 | GTTAGCCATCCTAGGGGCTC | GAAAGCGGGCAAGGTATGTA | 227 |
| chr12:52624189-52626209 | p2 | (AT)10 | TCCGAGGCCTTAAAGACAAA | ATACGAGTGTCCAGTTCGGG | 140 |
| chr12:52624189-52626209 | p2 | (AT)9 | TCCGAGGCCTTAAAGACAAA | ATACGAGTGTCCAGTTCGGG | 140 |
| chr12:52625518-52627538 | p2 | (AT)9 | TTCGCCCCTTTAGCCTATTT | ATTGGAAGTTCGGGTTAGGC | 142 |
| chr12:52625518-52627538 | p2 | (AT)10 | TTCGCCCCTTTAGCCTATTT | ATTGGAAGTTCGGGTTAGGC | 142 |
| chr12:52625518-52627538 | p2 | (TA)6 | TTCGCCCCTTTAGCCTATTT | ATTGGAAGTTCGGGTTAGGC | 142 |
| chr12:52738042-52740060 | p3 | (TCT)6 | GATACGTCGTGTGGGTGTTG | GCTAAACTACCGAGCATCGC | 261 |
| chr12:52738042-52740060 | p5 | (AACAT)3 | GATACGTCGTGTGGGTGTTG | GCTAAACTACCGAGCATCGC | 261 |
| chr12:52831974-52833988 | p2 | (GA)7 | GGGGATCGACGAGTCAAGTA | TCTCGCTTGCAATATTCCCT | 165 |
| chr12:52833437-52835479 | p2 | (TA)21 | ATCGTAGTTTTGGTGGGCTG | CCCATAACCACAACCACCTC | 168 |
| chr12:52835861-52837879 | p3 | (TTG)6 | GACCCTTGAGCCAAATCAAA | CCATTGACCTTCCTGCCTTA | 161 |
| chr12:52911776-52913798 | p2 | (AT)11 | TGTTGGGAGTTGTTGTGGAA | GAGGCGTCATGGAAGACATT | 110 |
| chr12:52915156-52917170 | p2 | (TA)7 | AAAAGGCTTCGAATTGGGAT | GTGCGAGGGTATGAGGTGTT | 269 |
| chr12:52977522-52979540 | p6 | (CATATA)3 | TGTGAATTTCCCTTTTTGCC | CGGCTTTCCCCTACACTACA | 237 |
| chr12:52994084-52996102 | p2 | (AT)15 | GGAGCCGATTACAATACCGA | ACATTGCTCGTACTGACCCC | 256 |
| chr12:52994084-52996102 | p6 | (CATATA)3 | GGAGCCGATTACAATACCGA | ACATTGCTCGTACTGACCCC | 256 |
| chr12:52994084-52996102 | p5 | (ATTCA)3 | GGAGCCGATTACAATACCGA | ACATTGCTCGTACTGACCCC | 256 |
| chr12:53051341-53053353 | p2 | (AT)6 | GAAATCGGATAAATCGGGGT | ACGAAACATTCACAATGCCA | 281 |
| chr12:53118557-53120581 | p2 | (AT)12 | TTCCAAAACACCAATCGACA | CAGACTTGTCGGCTTGCATA | 216 |
| chr12:53160430-53162448 | p6 | (TATATG)3 | GTCAGTCTGTAAGCGGCTCC | GACCCTTACTTAGGGCCGAG | 163 |
| chr12:53164351-53166371 | p2 | (TA)7 | AAAGGGAGTCCTAAGCTGGC | GTCACCAAGTCCCATAACGG | 271 |
| chr12:53234981-53236997 | p2 | (AT)9 | GTGTTTTCCCCTCCAAGTCA | TGAGCAAGCCCTAAGAAGGA | 195 |
| chr12:53234981-53236997 | p2 | (AT)8 | GTGTTTTCCCCTCCAAGTCA | TGAGCAAGCCCTAAGAAGGA | 195 |
| chr12:53242312-53244330 | p3 | (GTT)6 | GTAACCATCTTGGGCAGCAT | CGTTGCCAATCCTCTTGAAT | 167 |
| chr12:53242312-53244330 | p2 | (TA)8 | GTAACCATCTTGGGCAGCAT | CGTTGCCAATCCTCTTGAAT | 167 |
| chr12:53249607-53251619 | p2 | (AG)6 | TATTCACCCAAGTGCAACCA | CTATTTTTCCCTTGGCCTCC | 277 |
| chr12:53249607-53251619 | p2 | (GT)6 | TATTCACCCAAGTGCAACCA | CTATTTTTCCCTTGGCCTCC | 277 |
| chr12:53256257-53258271 | p2 | (TA)7 | CCTGAACGCAGGGTAATTGT | ATTTGGCCCAAGGGTACTTC | 130 |
| chr12:53268834-53270849 | p3 | (GTT)5 | CCAAGTGTAAGCCCTCCAAA | ATACAGGCCCACGACCTATG | 137 |
| chr12:53302722-53304734 | p2 | (TA)6 | AGACAGCAGGGCAAGAGTGT | CCCTCTATGTCGATCCTCCA | 103 |
| chr12:53630910-53632928 | p2 | (AT)9 | GGCTTGATTTGGAGTTGGAA | CATACCCCGTTGTCTCCCTA | 272 |
| chr12:53636076-53638094 | p6 | (CATATA)3 | AGTGATTCCTGTCACGACCC | ACCTTGTTTGTCCTTGACGC | 194 |
| chr12:53638953-53640969 | p4 | (AAAG)4 | GAACGTACAAAATGCGGGAT | AATTAACCCCGGAATAACCG | 201 |
| chr12:53795530-53797546 | p2 | (TG)9 | CTACTTCCATCGTTCCCAGG | AAGCCGCACGAGCTATAAGA | 124 |

| Marker ID | SSR type | SSR motif | FORWARD PRIMER (5'-3') | REVERSE PRIMER (5'-3') | size (bp) |
|---|---|---|---|---|---|
| chr12:53795548-53797562 | p2 | (TG)9 | CTACTTCCATCGTTCCCAGG | AAGCCGCACGAGCTATAAGA | 106 |
| chr12:53853915-53855933 | p2 | (GA)9 | TGGTGATTCGGTTATGGGAT | CTCGGATCACTTTCGACCAT | 285 |
| chr12:54079346-54081360 | p2 | (AT)7 | AAGAGGAATCACATCACGGG | CATTGTTTGCCACGAATGAG | 190 |
| chr12:54138214-54140230 | p3 | (TAC)5 | AGAGAAGAGCGGACCATTGA | GGTCGCCGACTGCTACTAAG | 222 |
| chr12:54138214-54140230 | p4 | (TGGG)4 | AGAGAAGAGCGGACCATTGA | GGTCGCCGACTGCTACTAAG | 222 |
| chr12:54138214-54140230 | p4 | (AAAT)4 | AGAGAAGAGCGGACCATTGA | GGTCGCCGACTGCTACTAAG | 222 |
| chr12:54141501-54143515 | p2 | (CT)7 | TGGCAGAGCACAATAGCATC | GATTGCAACAGCTCACAGGA | 234 |
| chr12:54164348-54166375 | p2 | (GA)6 | AAACAAAAGCTCACCCCCTT | TGAACAGTTTGACCTTCCCC | 266 |
| chr12:54164348-54166375 | p2 | (GA)6 | AAACAAAAGCTCACCCCCTT | TGAACAGTTTGACCTTCCCC | 266 |
| chr12:54164384-54166399 | p2 | (GA)6 | AAACAAAAGCTCACCCCCTT | TGAACAGTTTGACCTTCCCC | 230 |
| chr12:54164384-54166399 | p2 | (GA)6 | AAACAAAAGCTCACCCCCTT | TGAACAGTTTGACCTTCCCC | 230 |
| chr12:54258659-54260703 | p2 | (TA)22 | GATGGCCACATAAGCGTTTT | TTTCGTCCCCAAACTATTGC | 250 |
| chr12:54301736-54303748 | p2 | (TA)6 | CTGGCCCTGGACTACCATAA | TTCAGGAGCATACATCTGCG | 231 |
| chr12:54356194-54358248 | p3 | (TAT)18 | ACCCATTAAATGACCCGTGA | TCACCCTTGAACTACCCTGC | 252 |
| chr12:54481244-54483267 | p6 | (TTGGAG)3 | CCGAGGAAGAGCTCAAAATG | TGAACCAAACACCCAACAAA | 267 |
| chr12:54514372-54516402 | p3 | (AAG)6 | GCAAGAGTCTATGGGCAAGG | ATTTCGTTCAGTCCAAACGG | 107 |
| chr12:54519139-54521154 | p3 | (AAC)5 | ATATATACCGTAGCCGGCCC | TCTTCACGGCATCATCTCAC | 278 |
| chr12:54545167-54547199 | p2 | (AT)16 | TAACACTTAAACCCGCCAGG | GTGGGTCACCCCTTAGCATA | 118 |
| chr12:54604033-54606048 | p5 | (TGCAT)3 | GAGTCAATCGGAAGGACGAA | CAAGTTCAAGCTGTGCCAAA | 156 |
| chr12:54675813-54677839 | p2 | (TA)13 | TGTTCAGCAAAAGGGGTAGG | ATTTCTACGCCGTGATGGAC | 228 |
| chr12:54722496-54724524 | p2 | (TA)7 | ATTTGGCGAGATAACATGGC | TCATCGGGCATAAAAGTCGT | 198 |
| chr12:54722496-54724524 | p2 | (AT)14 | ATTTGGCGAGATAACATGGC | TCATCGGGCATAAAAGTCGT | 198 |
| chr12:54809474-54811507 | p2 | (TG)9 | CCAAATGGCGGTATTTGACT | GTGCATCACCATTCACCAAG | 190 |
| chr12:54814154-54816166 | p2 | (TA)6 | AACGCTGGAATGGTAGTTGG | ACTCGAGGACATGTTACGGC | 155 |
| chr12:54827311-54829323 | p2 | (TA)6 | GGTCAAAGAATCCCCCAGTT | GGCAAGCGGACTATCTTCAG | 150 |
| chr12:55014582-55016596 | p2 | (TA)7 | ACCTTCTGGGTGCTTTGAGA | ATGAGAGTTCGATTGCCCAC | 276 |
| chr12:55015583-55017601 | p2 | (AT)7 | ATAGGGTTTGGGGAGGTGAC | GACAAGGGCCAAAAATGAAA | 251 |
| chr12:55015583-55017601 | p6 | (TTTCTG)3 | ATAGGGTTTGGGGAGGTGAC | GACAAGGGCCAAAAATGAAA | 251 |
| chr12:55051368-55053386 | p2 | (AT)6 | AAACCCCACAACAAAAACCA | CTCCAGTTGGGGAATCTTGA | 263 |
| chr12:55051368-55053386 | p6 | (TTTTTC)3 | AAACCCCACAACAAAAACCA | CTCCAGTTGGGGAATCTTGA | 263 |
| chr12:55093264-55095279 | p3 | (TTG)5 | TAAAATTCTCCCCAACGCAG | GTCTTCCGCAATCACCATTT | 192 |
| chr12:55136072-55138090 | p2 | (AT)6 | AAACCCCACAACAAAAACCA | CTCCAGTTGGGGAATCTTGA | 263 |
| chr12:55136072-55138090 | p6 | (TTTTTC)3 | AAACCCCACAACAAAAACCA | CTCCAGTTGGGGAATCTTGA | 263 |
| chr12:55177968-55179983 | p3 | (TTG)5 | TAAAATTCTCCCCAACGCAG | GTCTTCCGCAATCACCATTT | 192 |
| chr12:55215899-55217919 | p5 | (GGGGT)4 | AGTCCAAATTCGGGAGTCCT | GCATTTGACAAAATTGCACG | 263 |
| chr12:55272201-55274216 | p5 | (TAGTT)3 | TTAATGATGTGGGGAGGAGC | AACACCCTTTTCTTGACCCC | 299 |

| Marker ID | SSR type | SSR motif | FORWARD PRIMER (5'-3') | REVERSE PRIMER (5'-3') | size (bp) |
|---|---|---|---|---|---|
| chr12:55285226-55287241 | p3 | (CTC)5 | CTTATCCCGACGTCACCTGT | CAAAGTGGAAACCCTCCAAA | 258 |
| chr12:55299890-55301905 | p5 | (TATTT)3 | GGGCCAGGCTCACTATTTTT | AAAGGATCCCCGAATCAACT | 125 |
| chr12:55319419-55321481 | p2 | (AT)6 | GTCACAACACAACAGGGTGG | ATTGTTGTTTGGCCTTCCAG | 146 |
| chr12:55398527-55400543 | p4 | (TTAA)4 | AAATAAGGGAAATGCTGCCC | GGATCATGCCAAAAGAAGGA | 197 |
| chr12:55415519-55417534 | p3 | (ATA)5 | ACCGTTCCAATGATCGAAAG | CGAGGCCTCTTCCTAGACCT | 289 |
| chr12:55433358-55435376 | p6 | (AAAATA)3 | CCAGGGCACCTCTTATCGTA | TCCTGTCCCGCCTATATGAC | 192 |
| chr12:55433358-55435376 | p5 | (AAACC)4 | CCAGGGCACCTCTTATCGTA | TCCTGTCCCGCCTATATGAC | 192 |
| chr12:55447463-55449475 | p2 | (TA)6 | AAAAAGTGGAACCCACCACA | CACGGTGTTTAATATGGGGC | 249 |
| chr12:55638288-55640303 | p3 | (TAT)5 | CCCACAAATGCGAAAAGATT | CCCCAACTCTCCCTTTCTTC | 138 |
| chr12:55645814-55647826 | p2 | (TG)6 | TGGTCCATTCGTTGTGGTAA | CCCCTGCCGTTATACTTTGA | 294 |
| chr12:55676578-55678593 | p5 | (CAAAC)3 | ATTTTGCCCTCAATTCTCCC | CTGCTGCAATTCAAGCACAT | 222 |
| chr12:55694171-55696183 | p2 | (CA)6 | CTACCGCTTGCTTGTTCTCC | GAGAAATGGCGAATTGTGGT | 276 |
| chr12:55704331-55706343 | p2 | (CA)6 | CTACCGCTTGCTTGTTCTCC | GAGAAATGGCGAATTGTGGT | 276 |
| chr12:55768507-55770523 | p2 | (AT)8 | CAACCTTCAGCAGAGAAGGG | ATGAGAAAAGGGTATCGGCA | 247 |
| chr12:55769859-55771871 | p2 | (TA)6 | TCATGCGCTTTAGTGTTGGA | AATAGAGGCGCAAAGCTTCA | 258 |
| chr12:55814594-55816624 | p2 | (GA)15 | TTCACTCCGAAAGCGTTCTT | TGCGGCTACTGTCAGAGCTA | 158 |
| chr12:55854285-55856360 | p3 | (TTC)25 | GTGGGGCTAGGAACCTTCTC | TCAAGCACAGCATTTTCAGG | 196 |
| chr12:55854285-55856360 | p3 | (TTC)20 | GTGGGGCTAGGAACCTTCTC | TCAAGCACAGCATTTTCAGG | 196 |
| chr12:55876397-55878409 | p2 | (TA)6 | TTGGAGTTGGACGATTGTGA | AAACAATTTGACGGCGTTTC | 143 |
| chr12:55958106-55960130 | p2 | (AT)12 | ACCTACGCTTGCTGCATTCT | GTGCTCACCCTGGAATGACT | 291 |
| chr12:55958106-55960130 | p2 | (AT)34 | ACCTACGCTTGCTGCATTCT | GTGCTCACCCTGGAATGACT | 291 |
| chr12:55958106-55960130 | p2 | (TA)13 | ACCTACGCTTGCTGCATTCT | GTGCTCACCCTGGAATGACT | 291 |
| chr12:55958874-55960942 | p2 | (AT)12 | GCCTTTCGGAGTGACGTAAG | TCATTCGGAAGTTACCGACC | 139 |
| chr12:55958874-55960942 | p2 | (AT)34 | GCCTTTCGGAGTGACGTAAG | TCATTCGGAAGTTACCGACC | 139 |
| chr12:55958874-55960942 | p2 | (TA)13 | GCCTTTCGGAGTGACGTAAG | TCATTCGGAAGTTACCGACC | 139 |
| chr12:56087198-56089213 | p3 | (CTG)5 | TGAGGAGTTTGGGGATTGAG | TGCAGTGAAACATTGGGAAA | 288 |
| chr12:56124183-56126203 | p2 | (AT)10 | AAATTAAAAGGCTGGCCCAT | ATTAAATGGGCTAGGACGGG | 100 |
| chr12:56140400-56142418 | p6 | (AAAAAG)3 | ACCAAATGCACCATTTAGCC | TCCATGACCCAACCTAGCTC | 238 |
| chr12:56199091-56201113 | p2 | (TA)11 | TCCTCTCCTTGTTCTTCCCA | AGAAACACGGCTACCCATCA | 217 |
| chr12:56217193-56219208 | p5 | (GGGTG)3 | TGAAGCTTTCCGCTTTCAGT | ATTCTGCCACAATCCCTTTG | 270 |
| chr12:56309424-56311436 | p2 | (AT)6 | AACCCATTAGCAGGAGCAAA | TTGACCGGAGATACTTTGGG | 117 |
| chr12:56393445-56395457 | p2 | (AT)6 | ATAAGCCTTGATGTCGTCGG | GTGAAAACCAAGTTGCGGAT | 202 |
| chr12:56478720-56480738 | p6 | (TCAGTT)3 | ACTTTCACCGTTTTTGGCAG | TGGGATTGCTTACCTTAGCG | 124 |
| chr12:56576652-56578678 | p2 | (GA)13 | CTACCTCCTTGCCCTAGCCT | TTGATGGACGTGTCGGATTA | 251 |
| chr12:56626136-56628150 | p2 | (TA)6 | GACCACGTAACCGACGAAGT | TTAATGAAACCGACGGGAAC | 104 |
| chr12:56658934-56660958 | p4 | (ATGA)6 | CCATCCTTGAGCAACGAAAT | TATGTAGGAACCCCAGCAGC | 206 |

| Marker ID | SSR type | SSR motif | FORWARD PRIMER (5'-3') | REVERSE PRIMER (5'-3') | size (bp) |
|---|---|---|---|---|---|
| chr12:56713323-56715335 | p2 | (TA)8 | GGTTGCTTTGACAGGATGGT | ATAGCAGCCTTGTCGGCTTA | 143 |
| chr12:56713323-56715335 | p2 | (AT)6 | GGTTGCTTTGACAGGATGGT | ATAGCAGCCTTGTCGGCTTA | 143 |
| chr12:56774965-56776980 | p5 | (TTGCA)3 | AGGTCACTTTTTGGGGTGTG | ATTCACCTCCACTTCATGCC | 256 |
| chr12:56774965-56776980 | p5 | (ATTTC)3 | AGGTCACTTTTTGGGGTGTG | ATTCACCTCCACTTCATGCC | 256 |
| chr12:56813518-56815536 | p3 | (TTG)6 | GTGATGTGGCAATGAAGGTG | CTTGCATGCTCCAAAACTCA | 278 |
| chr12:56872391-56874455 | p2 | (AT)8 | CTCGATGCACAGCGAACTTA | TGGACTGATGTTTTGTCCGA | 250 |
| chr12:56872391-56874455 | p2 | (AT)32 | CTCGATGCACAGCGAACTTA | TGGACTGATGTTTTGTCCGA | 250 |
| chr12:56899547-56901568 | p3 | (TAT)7 | AAATGCATCAGGAAGATGCC | GCGTCATGTATGGCACAATC | 212 |
| chr12:57016327-57018347 | p5 | (TCGGT)4 | CGAAAGGATGGAGAAACCAA | CTTCAGGGTAGCCGTAACCA | 123 |
| chr12:57072208-57074220 | p2 | (TA)6 | TGCGCATTAGCGTGAATAAG | ATTTTTAAACCCCCTTCCCC | 153 |
| chr12:57135394-57137409 | p3 | (AAC)5 | AAGCTTGTCACGCCCTAAAA | GGCGTTCGTCACGTTTTAAT | 300 |
| chr12:57148105-57150131 | p2 | (AT)13 | TCCGGGAACTACGAGTCAAC | GGGGTGACATAAGGTCGAGA | 255 |
| chr12:57183766-57185788 | p2 | (AT)11 | TTCTGCAAAGTCGTGTCCAG | TTGGCTCTATCGCTCGATCT | 161 |
| chr12:57220251-57222269 | p3 | (AAT)6 | GGGCATTGCATACATAAGGG | CATGCAGACAGGCACAGAGT | 295 |
| chr12:57388661-57390673 | p2 | (AT)11 | AGCGAAATAGGGGAAATGCT | GTTCGTAAATCCGTAGCCCA | 256 |
| chr12:57388661-57390673 | p2 | (TA)6 | AGCGAAATAGGGGAAATGCT | GTTCGTAAATCCGTAGCCCA | 256 |
| chr12:57388661-57390673 | p2 | (AT)7 | AGCGAAATAGGGGAAATGCT | GTTCGTAAATCCGTAGCCCA | 256 |
| chr12:57423826-57425840 | p2 | (TA)7 | TGTTTCGGTTTTGGAGGAAG | CTTTACTCTGATGCTCGCCC | 120 |
| chr12:57423826-57425840 | p2 | (TA)7 | TGTTTCGGTTTTGGAGGAAG | CTTTACTCTGATGCTCGCCC | 120 |
| chr12:57546155-57548173 | p6 | (ATAGTA)3 | CGTACACAATTTGGGCCTCT | GTTCGGAGCATCACTTCGAT | 156 |
| chr12:57548235-57550250 | p5 | (GATAT)3 | GATGTCCCAATGTTTCCCAC | GAGGAATCAGAAAAAGGGGC | 226 |
| chr12:57548235-57550250 | p3 | (TAT)5 | GATGTCCCAATGTTTCCCAC | GAGGAATCAGAAAAAGGGGC | 226 |
| chr12:57583966-57586004 | p2 | (TA)19 | GTATCGACACATGGCAATCG | GGACCCATGAACAGAAGGAA | 283 |
| chr12:57614465-57616480 | p5 | (GCATT)3 | TAGCGCTTGATGTCGATACG | CGCCAACAGCATAATCAATG | 191 |
| chr12:57619421-57621433 | p2 | (TA)6 | TCAATATCCTCTCGGCCATC | CGGCTTGAATTTGAGATCGT | 293 |
| chr12:57671154-57673170 | p2 | (AT)7 | TAGGCCTCATGAAGAATCGG | GTTGGGCTTGTGTCATGTTG | 275 |
| chr12:57694231-57696255 | p3 | (TTC)8 | GGGTCTAATTGTGGGGGTTT | GGAGCATTGACGGTTTTGTT | 181 |
| chr12:57704241-57706259 | p3 | (TTC)5 | AGCTCCTTCATCCCCTTCAT | AGTGAAACTGGGGCTTCAGA | 144 |
| chr12:57704241-57706259 | p3 | (CAA)5 | AGCTCCTTCATCCCCTTCAT | AGTGAAACTGGGGCTTCAGA | 144 |
| chr12:57704241-57706259 | p3 | (CAA)6 | AGCTCCTTCATCCCCTTCAT | AGTGAAACTGGGGCTTCAGA | 144 |
| chr12:57864327-57866342 | p5 | (TTGGT)3 | CAAACCTTCAGTTTTTCCGC | TTAAGTCCCCACAGGTCTGC | 276 |
| chr12:57864327-57866342 | p5 | (TTGGT)3 | CAAACCTTCAGTTTTTCCGC | TTAAGTCCCCACAGGTCTGC | 276 |
| chr12:57865061-57867076 | p5 | (TTGGT)3 | AACCGACAAACCGCACTAAA | GTAATGTCGGGTTGGTTTGG | 168 |
| chr12:57865061-57867076 | p5 | (TTGGT)3 | AACCGACAAACCGCACTAAA | GTAATGTCGGGTTGGTTTGG | 168 |
| chr12:57965180-57967195 | p3 | (TCA)5 | CCGGTAGCTCGATAGCACTC | TCAGGTTCACAGACAGCCAG | 156 |
| chr12:58031738-58033753 | p5 | (AGAAA)3 | TGCTTGCAATCGGAGTATTG | GAAGGCGAGATTTTAGTGCG | 229 |

| Marker ID | SSR type | SSR motif | FORWARD PRIMER (5'-3') | REVERSE PRIMER (5'-3') | size (bp) |
|---|---|---|---|---|---|
| chr12:58031738-58033753 | p3 | (ACA)6 | TGCTTGCAATCGGAGTATTG | GAAGGCGAGATTTTAGTGCG | 229 |
| chr12:58081469-58083504 | p3 | (CAA)5 | AGACCACCAATGCCACTCTC | GCGTGGACGGATTACTCAAT | 168 |
| chr12:58103442-58105457 | p3 | (TTA)5 | TCTCCTGGTTCGTTCTGCTT | AGAATGGAAACGATGTTGGC | 100 |
| chr12:58175280-58177292 | p2 | (TA)6 | CGAATCCATGACACCAACAG | CACTGGTGAAGAAAACGCAA | 198 |
| chr12:58196642-58198684 | p2 | (AT)21 | TTTTGGACCGTCACCTTTTC | AGGTTGCAGGCCATAAATTG | 243 |
| chr12:58324707-58326723 | p4 | (TGTT)4 | CTAGATCCCCTTTCCCCAAG | TTGGCACATCATGCAAAAAT | 248 |
| chr12:58329812-58331848 | p3 | (TTG)12 | GTAACAAACCGCTCGTCGAT | AATTTCGTGGGTGGAAGTTG | 132 |
| chr12:58329812-58331848 | p2 | (AT)14 | GTAACAAACCGCTCGTCGAT | AATTTCGTGGGTGGAAGTTG | 132 |
| chr12:58329812-58331848 | p2 | (TA)11 | GTAACAAACCGCTCGTCGAT | AATTTCGTGGGTGGAAGTTG | 132 |
| chr12:58332352-58334406 | p2 | (TA)27 | TGGGTCGTGACAAAGTGGTA | AATGACCTGCAGCCTCTTGT | 191 |
| chr12:58355689-58357707 | p3 | (ATA)18 | GCTGATCCGAAGTGTTGGTT | CGGCTATTACAACGCCATTT | 255 |
| chr12:58355689-58357707 | p3 | (ATA)7 | GCTGATCCGAAGTGTTGGTT | CGGCTATTACAACGCCATTT | 255 |
| chr12:58355689-58357707 | p2 | (AT)9 | GCTGATCCGAAGTGTTGGTT | CGGCTATTACAACGCCATTT | 255 |
| chr12:58355689-58357707 | p2 | (AT)8 | GCTGATCCGAAGTGTTGGTT | CGGCTATTACAACGCCATTT | 255 |
| chr12:58356690-58358714 | p2 | (TA)8 | TGTGTCAAGACTCCGAAACG | CGACCACCTTTTCAACACCT | 179 |
| chr12:58356690-58358714 | p2 | (AT)12 | TGTGTCAAGACTCCGAAACG | CGACCACCTTTTCAACACCT | 179 |
| chr12:58356690-58358714 | p2 | (AT)9 | TGTGTCAAGACTCCGAAACG | CGACCACCTTTTCAACACCT | 179 |
| chr12:58360003-58362045 | p2 | (AT)8 | CGTTTTGAATCTCCCCGTAA | AGCGCTTATTCGTCACACAA | 132 |
| chr12:58360003-58362045 | p2 | (TA)15 | CGTTTTGAATCTCCCCGTAA | AGCGCTTATTCGTCACACAA | 132 |
| chr12:58369860-58371899 | p2 | (TA)6 | GAAAGGAGTGGGGTGTCAGA | ACTTCCCTCCACATTACCCC | 243 |
| chr12:58369902-58371929 | p2 | (TA)6 | GAAAGGAGTGGGGTGTCAGA | ACTTCCCTCCACATTACCCC | 201 |
| chr12:58397719-58399734 | p5 | (TATGA)3 | CGTTTTGGGTTAGCCATGTT | GACTGTATGTGACCGAGCGA | 100 |
| chr12:58429559-58431571 | p2 | (AT)6 | TGCTCCACGCCCTACTACTT | CCCTGGCTAAATGCACTCTC | 171 |
| chr12:58502114-58504130 | p2 | (AT)8 | AACACCGGAAGCACCTTATG | TGACTTTTTGGGTGTTGCTG | 195 |
| chr12:58502114-58504130 | p5 | (AATGC)3 | AACACCGGAAGCACCTTATG | TGACTTTTTGGGTGTTGCTG | 195 |
| chr12:58511567-58513587 | p4 | (GAAA)5 | CCCGGTAAAACTGAGTCCAA | ATCCGATCAGGACCTTGTTG | 224 |
| chr12:58511567-58513587 | p2 | (AT)30 | CCCGGTAAAACTGAGTCCAA | ATCCGATCAGGACCTTGTTG | 224 |
| chr12:58540258-58542273 | p5 | (TGGTT)3 | GAACCGGTAAACCGAGTCAA | ACCAAACCAAACCAAACCAA | 174 |
| chr12:58540258-58542273 | p4 | (ATAG)5 | GAACCGGTAAACCGAGTCAA | ACCAAACCAAACCAAACCAA | 174 |
| chr12:58587022-58589042 | p2 | (AT)10 | TTTTACTTTTGGGCGTTTGG | GCCGAGTGTAAGTAGGCGTT | 239 |
| chr12:58593969-58595984 | p5 | (TGGCT)3 | AGCTTCCAAAAGATGAGGCA | CTACTCGCAACACCGAGACA | 147 |
| chr12:58599861-58601876 | p5 | (TATTT)3 | AATGACGACATTGAATCGCA | CCATCAGTCGCCTCATTTCT | 298 |
| chr12:58599861-58601876 | p5 | (TATTT)3 | AATGACGACATTGAATCGCA | CCATCAGTCGCCTCATTTCT | 298 |
| chr12:58599861-58601876 | p3 | (ATA)10 | AATGACGACATTGAATCGCA | CCATCAGTCGCCTCATTTCT | 298 |
| chr12:58658726-58660742 | p2 | (TC)8 | CAGCAGATGTTTGCGACTGT | TTGGGGTTTGTGCTGTTGTA | 228 |
| chr12:58678922-58680938 | p2 | (TA)8 | CTGAAGTTGCCATTGCTGAA | GGCTCATTGTGGGGTTAGAA | 191 |

| Marker ID | SSR type | SSR motif | FORWARD PRIMER (5'-3') | REVERSE PRIMER (5'-3') | size (bp) |
|---|---|---|---|---|---|
| chr12:58678922-58680938 | p3 | (ACA)5 | CTGAAGTTGCCATTGCTGAA | GGCTCATTGTGGGGTTAGAA | 191 |
| chr12:59034119-59036137 | p6 | (AAAAAT)3 | TCGAGAAACTCGAACTGGCT | GGAGGATTTGGACGAGATGA | 200 |
| chr12:59034119-59036137 | p2 | (AG)6 | TCGAGAAACTCGAACTGGCT | GGAGGATTTGGACGAGATGA | 200 |
| chr12:59092200-59094215 | p5 | (TATAT)3 | CTTAGCCCAGACCTGAGTGC | CATCGTCCACCTCCTCGTAT | 109 |
| chr12:59092200-59094215 | p3 | (TCT)5 | CTTAGCCCAGACCTGAGTGC | CATCGTCCACCTCCTCGTAT | 109 |
| chr12:59092356-59094377 | p5 | (TATAT)3 | TGATGGACGATGCGATAAAA | ATGGGACAGTTCGTTTCAGG | 173 |
| chr12:59092356-59094377 | p3 | (TCT)5 | TGATGGACGATGCGATAAAA | ATGGGACAGTTCGTTTCAGG | 173 |
| chr12:59092388-59094403 | p5 | (TATAT)3 | TGATGGACGATGCGATAAAA | ATGGGACAGTTCGTTTCAGG | 141 |
| chr12:59092388-59094403 | p3 | (TCT)5 | TGATGGACGATGCGATAAAA | ATGGGACAGTTCGTTTCAGG | 141 |
| chr12:59205091-59207103 | p2 | (GA)6 | CATCGTTAAGGACGGTTCGT | GAGCCAATGTTGGGTGAAGT | 271 |
| chr12:59205091-59207103 | p3 | (TTA)20 | CATCGTTAAGGACGGTTCGT | GAGCCAATGTTGGGTGAAGT | 271 |
| chr12:59205205-59207265 | p2 | (GA)6 | CATCGTTAAGGACGGTTCGT | GAGCCAATGTTGGGTGAAGT | 157 |
| chr12:59205205-59207265 | p3 | (TTA)20 | CATCGTTAAGGACGGTTCGT | GAGCCAATGTTGGGTGAAGT | 157 |
| chr12:59288233-59290248 | p4 | (ATTT)4 | TTAAGGTTCAGCCATCCAGG | CCCGAAGGTTCACATGCTAT | 232 |
| chr12:59288259-59290274 | p4 | (ATTT)4 | TTAAGGTTCAGCCATCCAGG | CCCGAAGGTTCACATGCTAT | 206 |
| chr12:59401256-59403271 | p5 | (AAAAT)3 | TAGAGCAAGCAGTCTCGCAA | ATCAATGCGACAAGATGCAA | 137 |
| chr12:59446480-59448492 | p2 | (GA)6 | GACCACCAGGTCAGCTTCAT | CAATGTACCAAAACGCCCTT | 267 |
| chr12:59461880-59463898 | p3 | (CTT)6 | CCAGGAATCTCGTCAAGCTC | CAAGGACGAGGATGGAAAAA | 292 |
| chr12:59532226-59534250 | p6 | (AGGTAG)4 | CTTGGACCAGGCAATGAAGT | TCCCCTTTAATGCACACTCC | 215 |
| chr12:59757078-59759098 | p5 | (GGGGT)4 | TCTTTCTCCACCGATCATCC | GCAAAGTTTAGGGGGTTGGT | 134 |
| chr12:59842120-59844138 | p6 | (CAAAAA)3 | TTCGCCCCAATTTATGTGAT | CCTCGGGCAAGTTATGTGAT | 226 |
| chr12:59842120-59844138 | p5 | (TTTTA)3 | TTCGCCCCAATTTATGTGAT | CCTCGGGCAAGTTATGTGAT | 226 |
| chr12:60012620-60014635 | p5 | (TCATT)3 | AAGGTGTGATGAGGTTTGGG | ATCTAGCATCGCAACAGGCT | 291 |
| chr12:60017364-60019434 | p2 | (TA)35 | AGACCTTTTGCTCCCCAAAT | AATTCGGGATTTATCCCACC | 269 |
| chr12:60038090-60040144 | p3 | (AAG)14 | TGAACATAGCCCAGTTCGTG | ATTGCCATTGACACCCATTT | 282 |
| chr12:60070648-60072663 | p2 | (AT)12 | CAAGAGGGAATATCGACCCA | AGTGTTGTCGAGTGGAGGCT | 117 |
| chr12:60070648-60072663 | p5 | (CAATG)3 | CAAGAGGGAATATCGACCCA | AGTGTTGTCGAGTGGAGGCT | 117 |
| chr12:60211106-60213118 | p2 | (CA)9 | AAGGGTCAATGAAGGTGGTG | AGAAATTGCCACGTACCCTG | 282 |
| chr12:60211106-60213118 | p2 | (AG)6 | AAGGGTCAATGAAGGTGGTG | AGAAATTGCCACGTACCCTG | 282 |
| chr12:60211106-60213118 | p2 | (AG)6 | AAGGGTCAATGAAGGTGGTG | AGAAATTGCCACGTACCCTG | 282 |
| chr12:60211106-60213118 | p2 | (AG)9 | AAGGGTCAATGAAGGTGGTG | AGAAATTGCCACGTACCCTG | 282 |
| chr12:60211106-60213118 | p5 | (TTTTG)6 | AAGGGTCAATGAAGGTGGTG | AGAAATTGCCACGTACCCTG | 282 |
| chr12:60261586-60263632 | p2 | (AT)23 | ATGTTCGCTACAGTGGACCC | TCGTATTCGATGTTTTCGCA | 109 |
| chr12:60322742-60324757 | p5 | (AAACC)3 | CCAACCGCTTTCTGCTCTAC | CCCGACGCAATTAGTTTGAT | 187 |
| chr12:60366664-60368682 | p3 | (ACA)6 | CTCCCGTTCTGCGATATCAT | ACTCGTTCATTCAAGGTGGG | 178 |
| chr12:60371724-60373757 | p3 | (TAT)11 | CGCTCCTCCTTGTGTGTGTA | AGGAAAAAGCACAAGCCAGA | 214 |

| Marker ID | SSR type | SSR motif | FORWARD PRIMER (5'-3') | REVERSE PRIMER (5'-3') | size (bp) |
|---|---|---|---|---|---|
| chr12:60371724-60373757 | p2 | (CT)7 | CGCTCCTCCTTGTGTGTGTA | AGGAAAAAGCACAAGCCAGA | 214 |
| chr12:60525517-60527532 | p5 | (TGATA)3 | TGCCACAAACATCTAGCAGC | GGAGGGGATCCTTTCCACTA | 110 |
| chr12:60533838-60535850 | p2 | (GA)6 | AGCACCAAACTGGACCAAAC | AAGCCTGCCTAAACTGAGCA | 102 |
| chr12:60548150-60550168 | p3 | (TAA)5 | TCCGGTAAGTCCAAATACGG | TCCGCATACACATTGCATTT | 209 |
| chr12:60548150-60550168 | p3 | (ATA)6 | TCCGGTAAGTCCAAATACGG | TCCGCATACACATTGCATTT | 209 |
| chr12:60548150-60550168 | p3 | (TAA)6 | TCCGGTAAGTCCAAATACGG | TCCGCATACACATTGCATTT | 209 |
| chr12:60579569-60581585 | p2 | (AT)8 | GCAAACAGATGAGCCCAAAT | AATCAGTCAGCCATCCCAAC | 218 |
| chr12:60607632-60609647 | p5 | (AAATT)3 | AATGGGTCAAAAAGATCCCC | ATGGAGGTCATTTATGGGCA | 121 |
| chr12:60622128-60624143 | p3 | (TAT)5 | TTGGCTTTAGAGGCATGTCA | CAAGATGCCATATTCAAAGGG | 293 |
| chr12:60654388-60656402 | p2 | (AT)7 | TCCCATCCCTCGTGTAACTC | TGCAATTGGAGTTTCGTCAA | 185 |
| chr12:60744091-60746106 | p5 | (ATTGC)3 | TGGGGATTTGGGTCTTGATA | TCCAACTTCATACCCGAAGG | 140 |
| chr12:60783698-60785776 | p2 | (AT)39 | CCCACCCCCTTAAACTGATT | TGCATATTTTGCTCCGTCTG | 262 |
| chr12:60783698-60785776 | p2 | (AT)29 | CCCACCCCCTTAAACTGATT | TGCATATTTTGCTCCGTCTG | 262 |
| chr12:60855700-60857720 | p5 | (TTTTC)4 | CCACTGAAACGCAAGGAAAT | TCGGTTTTTCTTTTTGCCTG | 214 |
| chr12:60873399-60875414 | p2 | (CA)8 | GCTTGGACAAGGTTCCAAAA | TCTGGAGAGCCAAGTGTGTG | 289 |
| chr12:60873399-60875414 | p3 | (TTG)5 | GCTTGGACAAGGTTCCAAAA | TCTGGAGAGCCAAGTGTGTG | 289 |
| chr12:60885153-60887211 | p2 | (AT)29 | TATGGCCAAACACAACTCCA | GATATTTAGGGCCCGTTTGG | 268 |
| chr12:60885153-60887211 | p2 | (TA)12 | TATGGCCAAACACAACTCCA | GATATTTAGGGCCCGTTTGG | 268 |
| chr12:60885153-60887211 | p2 | (TA)11 | TATGGCCAAACACAACTCCA | GATATTTAGGGCCCGTTTGG | 268 |
| chr12:60937943-60940000 | p2 | (AG)6 | CTAAGCCCATCATTGCCATT | GCCTCTCTCTTTGCTTGCAT | 200 |
| chr12:60938003-60940027 | p2 | (AG)6 | CTAAGCCCATCATTGCCATT | GCCTCTCTCTTTGCTTGCAT | 140 |
| chr12:60954136-60956151 | p5 | (ATATA)3 | TTGGGCATCTATGGGTAAGC | TGGTCCCACCATACCCTTTA | 278 |
| chr12:60954136-60956151 | p5 | (CTTTT)3 | TTGGGCATCTATGGGTAAGC | TGGTCCCACCATACCCTTTA | 278 |
| chr12:60954287-60956302 | p5 | (ATATA)3 | TTGGGCATCTATGGGTAAGC | TGGTCCCACCATACCCTTTA | 127 |
| chr12:60954287-60956302 | p5 | (CTTTT)3 | TTGGGCATCTATGGGTAAGC | TGGTCCCACCATACCCTTTA | 127 |
| chr12:61055587-61057655 | p2 | (AT)34 | GCAAGCAGCTCCATGGTTA | ATTTTGGCTGATTGAATTGGA | 104 |
| chr12:61060450-61062466 | p4 | (TTAT)4 | ATGGGTTTGTTGCACTCTCA | CGCATCGGTAATGTAAACCA | 133 |
| chr12:61080407-61082435 | p2 | (TA)6 | GAACGCCCATATCTTTTGGA | GAGTTTGGATGTGGTGGGTT | 120 |
| chr12:61080407-61082435 | p2 | (TA)14 | GAACGCCCATATCTTTTGGA | GAGTTTGGATGTGGTGGGTT | 120 |
| chr12:61090862-61092876 | p2 | (TA)7 | GTCGGGACTTTTGGTTGAGA | ACCCTCAGGACCACACGTAG | 184 |
| chr12:61115658-61117673 | p5 | (TGTGT)3 | CGCATTTGGAGGGGTTAGTA | TCCATGTTCTCCACGAATCA | 229 |
| chr12:61145533-61147553 | p2 | (AT)10 | TATATCGAAGGATCCGCACC | GCTGCAACAGTCCCAAAACT | 144 |
| chr12:61223126-61225172 | p2 | (AT)23 | ATTTGGAGTTTTTGGGGGAA | AGATCAAATCTCGGCTCCAA | 207 |
| chr12:61223126-61225172 | p2 | (AT)23 | ATTTGGAGTTTTTGGGGGAA | AGATCAAATCTCGGCTCCAA | 207 |
| chr12:61224285-61226299 | p2 | (AT)7 | TTTCTATATTTGGCGGGGAG | TTTGGGACCTAAAGCAATGG | 143 |
| chr12:61226390-61228406 | p2 | (TA)6 | CCCATGAGTTTGGAGTTGGT | GTCGTCCGTTGTCCCTTAAA | 296 |

| Marker ID | SSR type | SSR motif | FORWARD PRIMER (5'-3') | REVERSE PRIMER (5'-3') | size (bp) |
|---|---|---|---|---|---|
| chr12:61226390-61228406 | p2 | (TA)8 | CCCATGAGTTTGGAGTTGGT | GTCGTCCGTTGTCCCTTAAA | 296 |
| chr12:61226390-61228406 | p3 | (GTA)5 | CCCATGAGTTTGGAGTTGGT | GTCGTCCGTTGTCCCTTAAA | 296 |
| chr12:61238212-61240226 | p2 | (AT)7 | TTTCTATATTTGGCGGGGAG | TTTGGGACCTAAAGCAATGG | 143 |
| chr12:61240317-61242333 | p2 | (TA)6 | CCCATGAGTTTGGAGTTGGT | GTCGTCCGTTGTCCCTTAAA | 296 |
| chr12:61240317-61242333 | p2 | (TA)8 | CCCATGAGTTTGGAGTTGGT | GTCGTCCGTTGTCCCTTAAA | 296 |
| chr12:61240317-61242333 | p3 | (GTA)5 | CCCATGAGTTTGGAGTTGGT | GTCGTCCGTTGTCCCTTAAA | 296 |
| chr12:61304611-61306623 | p2 | (GT)6 | TTGTTGTTTGGGAGGAGAGG | AACATTGATCAGCTTGTGCG | 275 |
| chr12:61315691-61317719 | p2 | (TA)14 | GCTTAAGGGCACAGTTCCAG | GACCCCAGACAGTAGTGGGA | 218 |
| chr12:61362110-61364125 | p5 | (ATAAA)3 | TCTTTATGGCCGAGTTGGTC | CCCTTAAAAGTGGGCAATGA | 123 |
| chr12:61436638-61438659 | p3 | (AAG)7 | AGAAGCTGGTGCTGGTGACT | CTCCACCCTGAACTACTCGC | 215 |
| chr12:61458211-61460226 | p3 | (AAC)5 | TTGGATTGACGTCCATGTGT | GCCATTGCCGAGTTTGTTAT | 278 |
| chr12:61458211-61460226 | p5 | (AAACC)3 | TTGGATTGACGTCCATGTGT | GCCATTGCCGAGTTTGTTAT | 278 |
| chr12:61514023-61516041 | p6 | (ATTAAT)3 | GACTACAATTCCAGGCGCTC | TAGCGCCTAGGAGCAAAGAG | 293 |
| chr12:61527949-61529977 | p4 | (TATG)7 | TGCCATCCCCAAATTGTTAT | CCTCGAAGCCAAACTGAAAC | 182 |
| chr12:61923755-61925782 | p3 | (GAA)9 | GAGGAGGGATGTTGCCACTA | ATAACAACAAAAATCGCGGC | 116 |
| chr12:61951297-61953313 | p2 | (AG)8 | GTCATCACCAAAGCAGCTCA | TGGCAATTGGATGGGATAAT | 270 |
| chr12:61951580-61953600 | p2 | (AG)8 | TATGTGCAGGCATTATCCCA | CCCTTAAACCCACCAATGTG | 190 |
| chr12:61961914-61963938 | p3 | (TTC)8 | TATTGGTTCGGGTCATGGTT | CTTCTTGCGAGTTGGAGGAG | 124 |
| chr12:62001906-62003920 | p2 | (AT)7 | TAATGAATCATGTCCGTGCG | TGACATGGCGGAATAACAAA | 228 |
| chr12:62001906-62003920 | p2 | (TA)6 | TAATGAATCATGTCCGTGCG | TGACATGGCGGAATAACAAA | 228 |
| chr12:62083623-62085653 | p2 | (TA)15 | TTGAGGCCCCAATTAAACAG | CTTTTTCCCGTGAGTTTGGA | 121 |
| chr12:62087657-62089669 | p2 | (TA)10 | CCACTTTTTCGTCGTTCGTT | TTTTCCCGTAGAGTTGGTGG | 293 |
| chr12:62087657-62089669 | p2 | (TA)6 | CCACTTTTTCGTCGTTCGTT | TTTTCCCGTAGAGTTGGTGG | 293 |
| chr12:62116581-62118596 | p5 | (AGGAC)3 | TTTCGACCCGAAAAATTCTG | AGCGAGGCTCAGAGTGCTAC | 174 |
| chr12:62161089-62163144 | p5 | (TGGGG)3 | AGATATCTCAACATGGGCGG | AAGCGCAAAGAAAAACAGGA | 166 |
| chr12:62223416-62225432 | p2 | (TA)8 | AGCAAATGCGGAGTCAAAAT | ATTGTGTAGCGCCTATTCGG | 219 |
| chr12:62254734-62256748 | p2 | (TA)7 | TGCTACGCAACACGTAAAGG | GCAACCAGCTGAGCCTAAAC | 161 |
| chr12:62329505-62331520 | p5 | (TTTTA)3 | TATCTTCAGCCAGGGGACAC | ACAACTTGGACCGCTTGAAC | 147 |
| chr12:62330599-62332614 | p5 | (TAAAT)3 | CCTATGTTCTTGGGTTTCGC | AATCGCTGGAAATGACTTGC | 223 |
| chr12:62330599-62332614 | p2 | (AT)6 | CCTATGTTCTTGGGTTTCGC | AATCGCTGGAAATGACTTGC | 223 |
| chr12:62425314-62427332 | p6 | (CATGCT)3 | GTAGCAGCCCTATTTCTGCG | GTCAAATTTGGAGGGGCATA | 122 |
| chr12:62430406-62432421 | p3 | (GAA)5 | TGCTCGAGACAACATTGAGG | TTCTCGAGGGCATAAACCAC | 230 |
| chr12:62437213-62439229 | p2 | (TA)8 | ATCCCTTTTCGCCGTAAGAT | GCACAGGGATGGATACTCGT | 179 |
| chr12:62442996-62445037 | p2 | (TC)13 | GCTCTGCTGTTCCTTGAACC | TCCATTTCCTGTTCCTTTGC | 261 |
| chr12:62502963-62504987 | p2 | (TA)6 | TGGGACACAAGTCCAAAACA | ATTCTTAGGCTGCCACTCCA | 218 |
| chr12:62502963-62504987 | p2 | (AT)12 | TGGGACACAAGTCCAAAACA | ATTCTTAGGCTGCCACTCCA | 218 |

| Marker ID | SSR type | SSR motif | FORWARD PRIMER (5'-3') | REVERSE PRIMER (5'-3') | size (bp) |
|---|---|---|---|---|---|
| chr12:62545474-62547492 | p5 | (TGGTT)3 | TTCCGTGGTTACGTGTTTCA | TGATTAATCCAGGGATTCCG | 275 |
| chr12:62546130-62548145 | p5 | (TGGTT)3 | AGCTTTGCACAAAATGGGAT | TTCGGTTCGGTTTTAATTCG | 187 |
| chr12:62551841-62553859 | p6 | (TTTTAC)3 | GGATCCATTATGGTCAACCG | GCGGGGTATAACCCACTTCT | 177 |
| chr12:62592717-62594732 | p5 | (TTTTG)3 | GCACTCGCGACAAACAATAA | TTTTGCCCAACCTATTTTGC | 180 |
| chr12:62637722-62639740 | p3 | (AAG)6 | TGAATGTTTCTCTTGCTGCG | ATGACATCAATGTGCCGAGA | 103 |
| chr12:62647709-62649724 | p2 | (TA)6 | TGCAACAGACGCAAAAGAAG | TTCAAAAACCCAAAAGCCAC | 129 |
| chr12:62647709-62649724 | p5 | (TTGGT)3 | TGCAACAGACGCAAAAGAAG | TTCAAAAACCCAAAAGCCAC | 129 |
| chr12:62673994-62676016 | p2 | (TA)11 | GTCATATCGACGAGCGGTTT | TCAGAGAGGTCATGGAACCC | 208 |
| chr12:62675153-62677177 | p4 | (TTTA)6 | TCCCACCTGTCTTTTGGAAG | AACGATTACCCTCTACCCGC | 241 |
| chr12:62679359-62681377 | p6 | (TTTTCT)3 | TGATCTTTGTTCTCCCACCC | CGAACCCTCCAAGGATGTTA | 160 |
| chr12:62679359-62681377 | p6 | (TTTTCT)3 | TGATCTTTGTTCTCCCACCC | CGAACCCTCCAAGGATGTTA | 160 |
| chr12:62685812-62687874 | p2 | (AT)31 | ACGTCTTACGCCTCTCCAAA | GATGCTGCCCAAATTTCTGT | 106 |
| chr12:62702460-62704475 | p3 | (AAT)5 | TGTCCCCGAGCTCTTTAGAA | ATGCTGCCGAAGTCCTCTTA | 216 |
| chr12:62783953-62785974 | p3 | (CAT)7 | TTTTGGAGGGCTTTCATTTG | GCCCACATGTCGAGATTTTT | 167 |
| chr12:62960715-62962727 | p2 | (AG)8 | GCATGTGCTTGAGCCTTGTA | ATTGTTGCAAGGGAGGAGTG | 191 |
| chr12:62960715-62962727 | p2 | (TA)6 | GCATGTGCTTGAGCCTTGTA | ATTGTTGCAAGGGAGGAGTG | 191 |
| chr12:62979225-62981243 | p5 | (CATAT)3 | CTCCTCTTTTTCCGCCTTCT | TTTTGCATCAACCTCACTCG | 270 |
| chr12:62979225-62981243 | p6 | (TGGGGG)3 | CTCCTCTTTTTCCGCCTTCT | TTTTGCATCAACCTCACTCG | 270 |
| chr12:62979225-62981243 | p2 | (AT)10 | CTCCTCTTTTTCCGCCTTCT | TTTTGCATCAACCTCACTCG | 270 |
| chr12:63005838-63007852 | p2 | (TA)7 | CGCAACCTTGCAATAACTCA | TGCGTAGTCAACCGCATTAG | 291 |
| chr12:63005838-63007852 | p2 | (AT)7 | CGCAACCTTGCAATAACTCA | TGCGTAGTCAACCGCATTAG | 291 |
| chr12:63033004-63035020 | p2 | (TA)8 | TTTCCAATGGCCAAGTCTTC | TCCCAATAATACGCCAAGGA | 216 |
| chr12:63043815-63045851 | p2 | (TA)18 | AAATGGGAACCCATGAAACA | CCATACCCGATCCTTGATTG | 238 |
| chr12:63043815-63045851 | p5 | (AGTTT)4 | AAATGGGAACCCATGAAACA | CCATACCCGATCCTTGATTG | 238 |
| chr12:63072282-63074297 | p5 | (AAACA)3 | ATCTCACTAAGCCGCAAGGA | TCATCATCCTGCACTGAAGC | 183 |
| chr12:63096354-63098376 | p2 | (CA)11 | TTGGGGTTCTAGACACTGGG | AGTTTCCAGCAATCGTGCTT | 170 |
| chr12:63096354-63098376 | p2 | (AT)7 | TTGGGGTTCTAGACACTGGG | AGTTTCCAGCAATCGTGCTT | 170 |
| chr12:63140403-63142415 | p2 | (TA)6 | AATGCCCAAGTTATGACCCA | CGCACAACCATTCTTTCCTT | 192 |
| chr12:63206137-63208149 | p2 | (TA)6 | AGATAGGGCGCCCAGTTAAT | ACTTCCACCATTTTGACGGA | 150 |
| chr12:63262680-63264696 | p2 | (AG)8 | AATCAAATCCCTTGCACCAG | GCTAGCGATCAAGAAAACGG | 106 |
| chr12:63432637-63434663 | p2 | (TA)13 | AACACGCAGTGCTAGGAGGT | CCCCCTCAGGAAAAGAACTC | 217 |
| chr12:63449862-63451900 | p2 | (AT)19 | CATCCCAAGAGAATAGGGCA | TACACCATGCTGGTCCTGAA | 114 |
| chr12:63453163-63455187 | p2 | (TC)9 | AGCAGCCGTGTCTCTCTCTC | GCCACAACTCACTTCCCAAT | 221 |
| chr12:63453163-63455187 | p2 | (AT)12 | AGCAGCCGTGTCTCTCTCTC | GCCACAACTCACTTCCCAAT | 221 |
| chr12:63488166-63490187 | p3 | (AAT)7 | CTGAAACCGAAACAAGCACA | GGGACTCGACTCGCAGTAAG | 122 |
| chr12:63547985-63549999 | p2 | (AT)15 | CGCAAATTTAGGCGATGATT | AAAGTGAGCGTGAACTGCCT | 171 |

| Marker ID | SSR type | SSR motif | FORWARD PRIMER (5'-3') | REVERSE PRIMER (5'-3') | size (bp) |
|---|---|---|---|---|---|
| chr12:63547985-63549999 | p2 | (AT)7 | CGCAAATTTAGGCGATGATT | AAAGTGAGCGTGAACTGCCT | 171 |
| chr12:63547985-63549999 | p2 | (AT)7 | CGCAAATTTAGGCGATGATT | AAAGTGAGCGTGAACTGCCT | 171 |
| chr12:63673977-63675992 | p2 | (TA)20 | TCAGCTGACTTTCCCCTGTT | GGTTTGGTTGGTTGTTTGCT | 297 |
| chr12:63673977-63675992 | p5 | (AGTAA)3 | TCAGCTGACTTTCCCCTGTT | GGTTTGGTTGGTTGTTTGCT | 297 |
| chr12:63751099-63753114 | p3 | (ACA)5 | CGGAAAAACAGAAACCCAAA | GGGAAGGCCATTACATCTGA | 192 |
| chr12:63958590-63960620 | p3 | (TAA)9 | AATGGGAAATGAGGGAAACC | GATCCCAAATCAAAGTCCGA | 215 |
| chr12:64054130-64056168 | p2 | (TA)7 | ACCAACCTTACGTCACTCCG | ACATCACAACAAAGGCCACA | 279 |
| chr12:64054274-64056288 | p2 | (TA)7 | ACCAACCTTACGTCACTCCG | ACATCACAACAAAGGCCACA | 135 |
| chr12:64068381-64070393 | p2 | (AT)6 | GCACGGTTTGTCCTTCAAAT | TGCAAATGGCTAAGTGGACA | 261 |
| chr12:64158798-64160820 | p2 | (TA)11 | TTGGACCCGTTTGACTTAGC | TGGCTCCGAGCTCAGATTAT | 158 |
| chr12:64158798-64160820 | p2 | (TA)8 | TTGGACCCGTTTGACTTAGC | TGGCTCCGAGCTCAGATTAT | 158 |
| chr12:64158798-64160820 | p2 | (AT)8 | TTGGACCCGTTTGACTTAGC | TGGCTCCGAGCTCAGATTAT | 158 |
| chr12:64216191-64218235 | p2 | (AT)22 | TAGGGTGTGACAGTTCCGGT | GCGAACCAAATGACTTGTTG | 153 |
| chr12:64376137-64378152 | p5 | (TCAAT)3 | CTCATCAATCTCTCCCTCGC | CCTATTGGTCACAGAGGGGA | 181 |
| chr12:64376137-64378152 | p5 | (TATGG)3 | CTCATCAATCTCTCCCTCGC | CCTATTGGTCACAGAGGGGA | 181 |
| chr12:64415373-64417388 | p2 | (GA)6 | TTGACCGTTGCTACACGAAG | TTCATCCGACCATTTGAGGT | 300 |
| chr12:64415373-64417388 | p3 | (ATA)5 | TTGACCGTTGCTACACGAAG | TTCATCCGACCATTTGAGGT | 300 |
| chr12:64469106-64471122 | p2 | (AT)8 | GGCACCAATAACAAGCACCT | CAATGGTTTCGTTGGCTTTT | 282 |
| chr12:64475882-64477896 | p2 | (AT)6 | AAAACTCAGAAGCGGAGCAA | TTTCGTTAATTTCCCCCACA | 260 |
| chr12:64477281-64479297 | p5 | (GGGTG)3 | ATCTTCTTCCTGCGCCATTA | GGTGAGGTGAAATGCTCGAT | 206 |
| chr12:64477281-64479297 | p4 | (TTAT)4 | ATCTTCTTCCTGCGCCATTA | GGTGAGGTGAAATGCTCGAT | 206 |
| chr12:64507897-64509909 | p2 | (GA)6 | GCATTTTGAATCTTTCGGGA | TTGGCAAAACGGGTAGAATC | 102 |
| chr12:64610127-64612175 | p3 | (TTA)16 | TAAATTCCAGCAGCTCCACC | CAATCCCTAGCCATGGAAGA | 171 |
| chr12:64610127-64612175 | p2 | (CT)8 | TAAATTCCAGCAGCTCCACC | CAATCCCTAGCCATGGAAGA | 171 |
| chr12:64700259-64702299 | p2 | (TA)20 | AGCCCATCAAACACTATGGC | TTTGGGATGCTTATTTCGCT | 212 |
| chr12:64700259-64702299 | p2 | (TA)30 | AGCCCATCAAACACTATGGC | TTTGGGATGCTTATTTCGCT | 212 |
| chr12:64892554-64894578 | p6 | (AAACCC)4 | AGTCGTAAACCAATCCGACG | TGGTGTGCTTTTGAGTGAGC | 190 |
| chr12:64907014-64909029 | p5 | (TTAAT)3 | CCGAACAAGAGTTGGGATGT | GACCTCCGATTTTCTGTGGA | 291 |
| chr12:65089076-65091127 | p3 | (TCT)17 | AATTCAACCCTTCTGCATGG | TCTGAGCGCAAAACTGAAGA | 183 |
| chr12:65146815-65148829 | p2 | (TC)7 | TGACGAAATTCCGAAAAAGG | TAGCTCAATTTGACCCGAGG | 105 |
| chr12:65187629-65189641 | p2 | (TA)14 | AATGACTCATCATCTCCCGC | TGGGTTTGGCCCATTTATTA | 294 |
| chr12:65187651-65189673 | p2 | (TA)14 | AATGACTCATCATCTCCCGC | TGGGTTTGGCCCATTTATTA | 272 |
| chr12:65327759-65329775 | p2 | (CT)8 | TTGAGATGTCACAACACGCA | ACATGCCACACAGGGTCATA | 243 |
| chr12:65500025-65502079 | p3 | (TTC)18 | TTGGCGAGAGTTCACAGATG | CTCTGTCCTCCATCTCAGCC | 295 |
| chr12:65543631-65545649 | p3 | (TTC)6 | TTTGAAAACCCCAGGCATAG | GGCAGACAAGAAAGCAAAGG | 219 |
| chr12:65557391-65559409 | p3 | (TTC)6 | TTTGAAAACCCCAGGCATAG | GGCAGACAAGAAAGCAAAGG | 219 |

| Marker ID | SSR type | SSR motif | FORWARD PRIMER (5'-3') | REVERSE PRIMER (5'-3') | size (bp) |
|---|---|---|---|---|---|
| chr12:65619536-65621551 | p3 | (ATA)5 | GAAGGAGTGAAAGGGGGAAG | ACCGGGTCCGATACATTTTT | 130 |
| chr12:65632250-65634262 | p2 | (AT)6 | AGGCTGAAGACATGGCAGAT | AGGTTTGCACTTTGACCCAG | 296 |
| chr12:65653069-65655091 | p2 | (AT)6 | AACCCTGAAGAAATGGGGAC | TTGGCACATAGTCTTGCTCG | 255 |
| chr12:65653187-65655199 | p2 | (AT)6 | AACCCTGAAGAAATGGGGAC | TTGGCACATAGTCTTGCTCG | 137 |
| chr12:65655288-65657354 | p2 | (AT)7 | CAAAAGGGTGGCCTCTCATA | CCGCTTTTAAGGGACAACAG | 266 |
| chr12:65691233-65693254 | p3 | (TTA)5 | GGAATTAAACTCGCGATGGA | CCCGGCAAGTTGTTAGAAAA | 181 |
| chr12:65813538-65815550 | p2 | (TA)6 | GTGGGGGTATTCAAATGACG | CCATTCTTGAGGGTATCGGA | 144 |
| chr12:65832663-65834681 | p3 | (AAC)6 | AGCTTCCTAAACCCTAGCCG | GAATAGTGTCGCCCATTCGT | 140 |
| chr12:65875546-65877561 | p3 | (TAA)5 | TTGTTCTCCTCCCTTGTTGC | CATTTGTCTCTCTCTCCCGC | 129 |
| chr12:65890372-65892392 | p4 | (TTTA)5 | ATGTTCGCACTCCATTCTCC | TGCATTAACTGATCGGGACA | 240 |
| chr12:65890372-65892392 | p2 | (AG)7 | ATGTTCGCACTCCATTCTCC | TGCATTAACTGATCGGGACA | 240 |
| chr12:65929848-65931870 | p2 | (AT)11 | GGGAGGGTAAAGTGTACGCA | GGTTGGGAATGAGAAAAATGG | 220 |
| chr12:65946930-65948945 | p5 | (ATACG)3 | GTGTATGTTGTTCGCGGATG | TGGGTACCAAGCCTCACTTC | 112 |
| chr12:65946930-65948945 | p2 | (AT)7 | GTGTATGTTGTTCGCGGATG | TGGGTACCAAGCCTCACTTC | 112 |
| chr12:66002708-66004726 | p6 | (AAAAAT)3 | TTCATTCAGGATGGCATTGA | AGGTCAAACGCCATTGAAAC | 285 |
| chr12:66004475-66006490 | p5 | (TTTTA)3 | TTCAGTCCGGTTCTCTTGCT | GGAGCTGCCCTACAGTATCG | 282 |
| chr12:66108300-66110315 | p3 | (TTG)5 | TTCCTCCCCACAAGAATTTG | CACAATAGGATGAGGGTGGG | 208 |
| chr12:66140951-66142965 | p2 | (TA)7 | CGCCACTTTGTTTCACTTCA | GTAACATCCCACCTCAGCGT | 288 |
| chr12:66167936-66169954 | p3 | (CAA)6 | ACGAGCGGAGTAGTGAGGAA | GAAAAGCTCGGATTGCTTTG | 280 |
| chr12:66176962-66178983 | p3 | (TGT)7 | TATGGGGTCAGGCTTTTGAG | CAGGGCCAGAGTCATGGTAT | 217 |
| chr12:66184499-66186554 | p3 | (AAC)5 | ATCCAAACTCACGGGAAAAA | ACGCATGTGCAATAAATCCA | 232 |
| chr12:66184499-66186554 | p2 | (TA)6 | ATCCAAACTCACGGGAAAAA | ACGCATGTGCAATAAATCCA | 232 |
| chr12:66186498-66188514 | p2 | (AT)8 | GAGCTGCGGTATCCATTCAT | AAATAATGCCGCCCCAATAG | 170 |
| chr12:66192721-66194733 | p2 | (TA)6 | CACGTTTGTGAAAATGGTGG | TTGACATGCCGTCTCATTGT | 115 |
| chr12:66332914-66334929 | p5 | (TTTGG)3 | TTAATGAACCGAAAATCGGC | TAAACGTAAATGGGCTTGGG | 103 |
| chr12:66332914-66334929 | p5 | (TTGGT)3 | TTAATGAACCGAAAATCGGC | TAAACGTAAATGGGCTTGGG | 103 |
| chr12:66473620-66475680 | p3 | (AAT)20 | AAATCCGAGCTTGTGATGCT | TGAATCGTTGAAAGGGGAAG | 103 |
| chr12:66481805-66483820 | p5 | (TTTGA)3 | ACCAATCAAAAATCCCCTCC | ACTTCGTTTTATTCCCCGCT | 285 |
| chr12:66481805-66483820 | p3 | (AAT)5 | ACCAATCAAAAATCCCCTCC | ACTTCGTTTTATTCCCCGCT | 285 |
| chr12:66482091-66484106 | p5 | (TTTGA)3 | AGCGGGGAATAAAACGAAGT | CCGTCACCTCCACTACACCT | 243 |
| chr12:66482091-66484106 | p3 | (AAT)5 | AGCGGGGAATAAAACGAAGT | CCGTCACCTCCACTACACCT | 243 |
| chr12:66509345-66511363 | p3 | (AGA)6 | TGATTGGGAGAGAAAGGTGG | TTCGCATAACCCTCCTCATC | 253 |
| chr12:66515408-66517420 | p2 | (TC)11 | TGAGTGGGGGTGGTTGTAAT | AACCAATGAGATCGGGTCAG | 192 |
| chr12:66515421-66517437 | p2 | (TC)11 | TGAGTGGGGGTGGTTGTAAT | AACCAATGAGATCGGGTCAG | 179 |
| chr12:66640973-66643002 | p5 | (GGTGG)3 | TCCACGGATTAGATCTTGGC | GGCCATAGATTTTGGAAGCA | 126 |
| chr12:66788136-66790154 | p2 | (CT)9 | CGGGAGTTTCCGAATGTAGA | GTTTGGTGCAAATTGCCTTT | 228 |

| Marker ID | SSR type | SSR motif | FORWARD PRIMER (5'-3') | REVERSE PRIMER (5'-3') | size (bp) |
|---|---|---|---|---|---|
| chr12:66788136-66790154 | p5 | (TGAGC)3 | CGGGAGTTTCCGAATGTAGA | GTTTGGTGCAAATTGCCTTT | 228 |
| chr12:66792153-66794168 | p3 | (TAT)5 | CTTTAGAAAAAGTCGGGGGC | GTGCTGAAAAAGTGTTGGCA | 183 |
| chr12:66792179-66794194 | p3 | (TAT)5 | CTTTAGAAAAAGTCGGGGGC | GTGCTGAAAAAGTGTTGGCA | 157 |
| chr12:66808909-66810921 | p2 | (AT)6 | AGGCATTGCTACCCAATGAG | ATGCAAACAGTTTCCCGTTC | 156 |
| chr12:66846263-66848279 | p2 | (TA)8 | TTCGGTAGCCAAATGTTTCC | TCTTTCCGCTCCTTTTGCTA | 136 |
| chr12:66917658-66919670 | p2 | (AG)6 | TCCGAATCATCTCAACCTCC | GGGTGTTCGCCAGTTAAGAA | 271 |
| chr12:66991915-66993936 | p3 | (ACA)7 | TAGTCGAAGCATGATGCAGG | TGAAGGCCTCTCTGTTCCAT | 197 |
| chr12:66991915-66993936 | p3 | (ACA)7 | TAGTCGAAGCATGATGCAGG | TGAAGGCCTCTCTGTTCCAT | 197 |
| chr12:67003377-67005393 | p4 | (CTTA)4 | CAGAAAGGCAATGTCCCCTA | CCGTGTACATTTTTGGAGGG | 148 |
| chr12:67003377-67005393 | p2 | (AT)7 | CAGAAAGGCAATGTCCCCTA | CCGTGTACATTTTTGGAGGG | 148 |
| chr12:67098652-67100672 | p2 | (AC)10 | TCGCTCCAAATTCTCCAACT | ATGCTTTCAACGGGCTAAGA | 173 |
| chr12:67098652-67100672 | p2 | (TA)10 | TCGCTCCAAATTCTCCAACT | ATGCTTTCAACGGGCTAAGA | 173 |
| chr12:67177098-67179114 | p6 | (ATTTGA)3 | ATTCACGCTCATAACCGGAG | CAAAAAGCGTCAATTTGGGT | 157 |
| chr12:67177098-67179114 | p2 | (TC)8 | ATTCACGCTCATAACCGGAG | CAAAAAGCGTCAATTTGGGT | 157 |
| chr12:67317601-67319617 | p2 | (GT)8 | CAAGGCCATCGTTATCTGGT | AGGCTCTTGCACTCTTCCAA | 227 |
| chr12:67317601-67319617 | p3 | (TTG)5 | CAAGGCCATCGTTATCTGGT | AGGCTCTTGCACTCTTCCAA | 227 |
| chr12:67317601-67319617 | p2 | (GT)7 | CAAGGCCATCGTTATCTGGT | AGGCTCTTGCACTCTTCCAA | 227 |
| chr12:67341192-67343216 | p6 | (ATCTAT)4 | ATCGGCTTTTGCTTCTTCAA | GTCCATGGATTATGTTGGGG | 149 |
| chr12:67345030-67347045 | p5 | (ACCAA)3 | CCCCAGACCCTACTTGTGAA | GCTGGATAAGGAAGTGCGTC | 174 |
| chr12:67345030-67347045 | p5 | (AAACC)3 | CCCCAGACCCTACTTGTGAA | GCTGGATAAGGAAGTGCGTC | 174 |
| chr12:67359234-67361249 | p5 | (TTGGT)3 | GGCACGTGGGAATTATCAAC | ACGGTTCAAGCTTCCAAGAA | 165 |
| chr12:67359234-67361249 | p5 | (TAATT)3 | GGCACGTGGGAATTATCAAC | ACGGTTCAAGCTTCCAAGAA | 165 |
| chr12:67457245-67459260 | p3 | (TTA)5 | CGTTGGCAACAGATTCGTAA | TTTAGAGCGTTTTGCAGGCT | 191 |
| chr12:67523811-67525833 | p2 | (AT)11 | TCTCCTTCCTGCCACTGACT | CAGAGTAGTTCGTCCTCGGG | 155 |
| chr12:67523811-67525833 | p3 | (ATC)7 | TCTCCTTCCTGCCACTGACT | CAGAGTAGTTCGTCCTCGGG | 155 |
| chr12:67524052-67526073 | p2 | (AT)11 | TCGTAGACAAACCCGAGGAC | TCTTTCGAGACTCATGCCCT | 142 |
| chr12:67524052-67526073 | p3 | (ATC)7 | TCGTAGACAAACCCGAGGAC | TCTTTCGAGACTCATGCCCT | 142 |
| chr12:67553334-67555352 | p2 | (AT)9 | TGGATGGGTGAAATGAGACA | TCAAAGCATCCAAAATGCTG | 173 |
| chr12:67642387-67644405 | p3 | (ACA)6 | CTCAAACTCAAGCACAGCCA | TTGGAAAAAGGAGCATTTGG | 110 |
| chr12:67855531-67857551 | p2 | (TA)10 | ACCACTGTGGGCAACTTTTC | TATGCCAAAATTCTCGGGAC | 113 |
| chr12:67869424-67871439 | p3 | (TGT)5 | AAACTTGTTCCACCAGCCAG | TGCAGACCTCTCCACAACTG | 294 |
| chr12:68008505-68010523 | p2 | (AT)8 | GGCCCTGTAATTTTGAAGCA | ACAAAGTTGGTGATGGAGGC | 108 |
| chr12:68008505-68010523 | p6 | (TTTATT)3 | GGCCCTGTAATTTTGAAGCA | ACAAAGTTGGTGATGGAGGC | 108 |
| chr12:68022474-68024492 | p2 | (AT)9 | TCAGCGCGAGTAGGATTTTT | ACTCGTCCTGAATTCGTTGG | 223 |
| chr12:68042017-68044035 | p3 | (TAT)6 | TTATTTCGGAGAGAGGCGAA | GGCTTCCCGACAAGGTCTAT | 299 |
| chr12:68042017-68044035 | p3 | (TAT)6 | TTATTTCGGAGAGAGGCGAA | GGCTTCCCGACAAGGTCTAT | 299 |

| Marker ID | SSR type | SSR motif | FORWARD PRIMER (5'-3') | REVERSE PRIMER (5'-3') | size (bp) |
|---|---|---|---|---|---|
| chr12:68042017-68044035 | p5 | (TACTA)3 | TTATTTCGGAGAGAGGCGAA | GGCTTCCCGACAAGGTCTAT | 299 |
| chr12:68055303-68057323 | p5 | (TCTAG)4 | CACCGGAGGAAGAGTTGAAA | CCCGAAACTGCTTTTCCATA | 237 |
| chr12:68059376-68061391 | p3 | (TTA)5 | ATGGGAGGCAAATCAACAAG | GAGTAGGAGGGTGGGAGAGG | 171 |
| chr12:68082564-68084582 | p6 | (CGGTGG)3 | CAACCCACACACTAGCCCTT | GCACCAAAGGACCAAAGAGA | 122 |
| chr12:68092787-68094805 | p3 | (TTC)6 | TTTCGGGCAAAAGCAATAAC | CAACCACCACACCATTTTCA | 286 |
| chr12:68100188-68102212 | p2 | (TA)12 | TTCCCACAAAAGAACCCTTG | CTCCTCACACCTGCAACGTA | 242 |
| chr12:68401501-68403530 | p6 | (TGGGGG)3 | TGCGTACACTCTACCCTCCC | CAAAAACTTGTGGCACCCTT | 169 |
| chr12:68401789-68403818 | p6 | (TGGGGG)3 | CGAGACAATTTGACTTGGCA | GGCGTAAGAGTCAAACCCAA | 273 |
| chr12:68404901-68406923 | p2 | (AT)11 | CTCATTTGTCGCAAAGCAAA | ACGACGGGACATTTCTTCAG | 126 |
| chr12:68404901-68406923 | p2 | (TA)27 | CTCATTTGTCGCAAAGCAAA | ACGACGGGACATTTCTTCAG | 126 |
| chr12:68463377-68465393 | p2 | (AT)8 | TATCGGCCATGAATGCAATA | AAGAAAAACGTGTCGGTTGG | 143 |
| chr12:68490201-68492253 | p2 | (GA)10 | TTTTGCCCGTAACTTTGTCC | GCGGCAGTATGGATTTCATT | 128 |
| chr12:68503597-68505625 | p2 | (TA)14 | ATGGTTGTGGGAGGATGAAG | CACCCAAAGTTGGAGAGCAT | 221 |
| chr12:68565290-68567308 | p3 | (TAA)6 | GAACAGATCCGAACCTCGAC | ACGTTTGAATTTTGAACGGC | 124 |
| chr12:68609672-68611687 | p5 | (CATTT)3 | TGCAGTGGGCATGTTATGAT | TATGCAATGCCTGAGTTTCG | 274 |
| chr12:68628519-68630535 | p4 | (TTAT)4 | TGCTGTGGGTCTGGATATGA | AAATCGCACACGAAGGGTAG | 231 |
| chr12:68718568-68720600 | p2 | (AG)16 | AAACGAAATCCATACTGCCG | TGATGATTTTGTGGCTGCAT | 295 |
| chr12:68881772-68883816 | p2 | (TA)22 | AACAGTGTAATTCTCCGCCG | TAATCGCTGTACGACCACCA | 180 |
| chr12:68887994-68890012 | p2 | (AT)12 | ATTGCTGGACCATCATCACA | GGTGGATGGAGGGGTATTTT | 259 |
| chr12:68887994-68890012 | p2 | (AT)9 | ATTGCTGGACCATCATCACA | GGTGGATGGAGGGGTATTTT | 259 |
| chr12:68999057-69001097 | p2 | (AT)20 | GGGCATTCCTATTGCGTAAA | GCCATTTTCACTCATTGGCT | 114 |
| chr12:69097155-69099180 | p5 | (TAAAA)5 | TGCATTTTGCTTTGAACTGC | GCAAACCTTCCCCCTATCTC | 140 |
| chr12:69103796-69105834 | p2 | (TA)19 | CCCATTAAAAGGAGAAGCCC | AAAGGAGGGAATTTTGGTGG | 170 |
| chr12:69106662-69108690 | p2 | (TA)7 | ATGAGGGCATAACGTAGCCA | CCAGACCATACACATGCTGC | 113 |
| chr12:69141464-69143476 | p5 | (AAAAT)3 | TGAGTCATTCCCACGCATAA | CGCATTAGTTTTGGGCTTGT | 109 |
| chr12:69141464-69143476 | p2 | (GA)6 | TGAGTCATTCCCACGCATAA | CGCATTAGTTTTGGGCTTGT | 109 |
| chr12:69141464-69143476 | p3 | (ATA)18 | TGAGTCATTCCCACGCATAA | CGCATTAGTTTTGGGCTTGT | 109 |
| chr12:69168564-69170578 | p2 | (CA)7 | TGTTCCGCACAATATGAAGC | AGAAAATTCAGGGCAAACCC | 204 |
| chr12:69170712-69172726 | p2 | (CT)7 | AACTGAATTGCCTTTGTGGG | GCAGAGCTGGGACTTTTGAC | 278 |
| chr12:69178921-69180971 | p2 | (TA)25 | TTGTCTGAAGCTCGATGTGG | TATTTTGGGCTAATGGCTGC | 223 |
| chr12:69187137-69189149 | p2 | (AC)6 | AATGGAGGGTGACCTCATGT | TTTGAAGGAAGACTCGCTGG | 110 |
| chr12:69194727-69196745 | p6 | (GGAAGA)3 | ATAGGGCAGCCTGTTTGATG | ATCTCGGTGATTCGGTTCAG | 108 |
| chr12:69220744-69222756 | p2 | (AT)6 | CACAAATACCCGTCCAATCC | TTCACGAAATGCCAAAAACA | 103 |
| chr12:69220744-69222756 | p2 | (TA)8 | CACAAATACCCGTCCAATCC | TTCACGAAATGCCAAAAACA | 103 |
| chr12:69244643-69246655 | p2 | (AT)6 | GCCGATCTGAAGTTACGCTC | GAACAGATTGGGCAGGTCAT | 105 |
| chr12:69307246-69309262 | p4 | (TTTG)4 | AGTTGCTATCAGGCAAGGGA | TGTTTGTTTTTGTGGGCAAG | 111 |

| Marker ID | SSR type | SSR motif | FORWARD PRIMER (5'-3') | REVERSE PRIMER (5'-3') | size (bp) |
|---|---|---|---|---|---|
| chr12:69315635-69317651 | p2 | (CT)8 | CTTATGGGCAGACTTGGCAT | GTAAGGTTTGGATCGCGTGT | 285 |
| chr12:69374870-69376894 | p3 | (ATG)8 | AGATCGTTTGGCTCTTTCCA | CCGTCACATCAGGAGGAACT | 175 |
| chr12:69374870-69376894 | p6 | (TCAGTT)3 | AGATCGTTTGGCTCTTTCCA | CCGTCACATCAGGAGGAACT | 175 |
| chr12:69411693-69413711 | p3 | (TTA)6 | CCTGGAACGAGGTGAACAAT | AGATTTCTTGGGTTGGCCTT | 297 |
| chr12:69416903-69418921 | p6 | (TCAAGT)3 | GCCTTGTGGACCTTTTTGAA | TGATCTTCAACCCTTCCAGG | 136 |
| chr12:69431924-69433939 | p5 | (AAAAG)3 | CAGTGGCAGTGCAAGAAAAA | CAAACCCTGCTACCCCAATA | 208 |
| chr12:69431924-69433939 | p2 | (GA)7 | CAGTGGCAGTGCAAGAAAAA | CAAACCCTGCTACCCCAATA | 208 |
| chr12:69431924-69433939 | p3 | (TCT)5 | CAGTGGCAGTGCAAGAAAAA | CAAACCCTGCTACCCCAATA | 208 |
| chr12:69455707-69457719 | p2 | (AT)6 | AAGTATCAACCGTGGCTTCG | CCCCATATCGTATTCGGCTA | 190 |
| chr12:69455721-69457733 | p2 | (AT)6 | AAGTATCAACCGTGGCTTCG | CCCCATATCGTATTCGGCTA | 176 |
| chr12:69455735-69457747 | p2 | (AT)6 | AAGTATCAACCGTGGCTTCG | CCCCATATCGTATTCGGCTA | 162 |
| chr12:69455749-69457761 | p2 | (AT)6 | AAGTATCAACCGTGGCTTCG | CCCCATATCGTATTCGGCTA | 148 |
| chr12:69455763-69457775 | p2 | (AT)6 | AAGTATCAACCGTGGCTTCG | CCCCATATCGTATTCGGCTA | 134 |
| chr12:69455777-69457789 | p2 | (AT)6 | AAGTATCAACCGTGGCTTCG | CCCCATATCGTATTCGGCTA | 120 |
| chr12:69455791-69457803 | p2 | (AT)6 | AAGTATCAACCGTGGCTTCG | CCCCATATCGTATTCGGCTA | 106 |
| chr12:69473932-69475948 | p2 | (GA)6 | TTGCCAAGCCTTACCAATTC | AAATTGGGGCAAATAGAGGG | 278 |
| chr12:69473932-69475948 | p2 | (TA)8 | TTGCCAAGCCTTACCAATTC | AAATTGGGGCAAATAGAGGG | 278 |
| chr12:69513109-69515124 | p5 | (TTGGT)3 | CCCGACTAGTGGTTTGGTGT | GTAATGTCGGGTTGGTTTGG | 235 |
| chr12:69513149-69515164 | p5 | (TTGGT)3 | CCCGACTAGTGGTTTGGTGT | GTAATGTCGGGTTGGTTTGG | 195 |
| chr12:69811273-69813289 | p2 | (TA)8 | TTCGTTGGTGTGTTGTTCGT | AAGGGGACGATCTTGTGTTG | 274 |
| chr12:69825359-69827377 | p2 | (AC)9 | GCCTTTTGTTTTCTCGATGC | CCCTTGGATCTCCTTGTTGA | 177 |
| chr12:69850208-69852232 | p3 | (TTA)5 | ATCCACTCCCCGGTAAATTC | TCTCAACGTTCCATCTTCCC | 187 |
| chr12:69850208-69852232 | p5 | (ATTTT)3 | ATCCACTCCCCGGTAAATTC | TCTCAACGTTCCATCTTCCC | 187 |
| chr12:69850234-69852262 | p3 | (TTA)5 | ATCCACTCCCCGGTAAATTC | TCTCAACGTTCCATCTTCCC | 161 |
| chr12:69850234-69852262 | p5 | (ATTTT)3 | ATCCACTCCCCGGTAAATTC | TCTCAACGTTCCATCTTCCC | 161 |
| chr12:69850268-69852280 | p3 | (TTA)5 | TGGCACGTGTCGGAATAATA | CGCTGATTTGGTCCAGGTAT | 199 |
| chr12:69850268-69852280 | p5 | (ATTTT)3 | TGGCACGTGTCGGAATAATA | CGCTGATTTGGTCCAGGTAT | 199 |
| chr12:69895268-69897283 | p5 | (ATACC)3 | CAGGGAAACCATGAAAGGAA | GTTGAGGAGGCTATGTGGGA | 228 |
| chr12:69921968-69923983 | p5 | (TTGTA)3 | TGGCTACAAAAGGAAATGGG | CTCTTGCAAAGCAACTTCCC | 277 |
| chr12:69961545-69963559 | p2 | (CT)7 | TTTCTTTGCGTGGATTACCC | GCCTTGAGGTAGAATTTGCG | 167 |
| chr12:69975943-69977961 | p3 | (CAG)6 | AGGATGGGGTCTTACCCTTG | AGCCGGAAAACATTTCACTG | 159 |
| chr12:70430360-70432420 | p2 | (AT)30 | AACGATGTACCCTCACTCCG | GTACCAAAGCACATCGCCTT | 182 |
| chr12:70442388-70444400 | p2 | (AG)6 | CGAAAGTGAAAGCGAACACA | CGAGCTCTTCCGTTGAAATC | 219 |
| chr12:70501225-70503241 | p4 | (TACA)4 | AAAGGCTTAAGGTCACGGGT | CTCTCCCCTTTTGTTCTCCC | 198 |
| chr12:70501225-70503241 | p4 | (ATGT)4 | AAAGGCTTAAGGTCACGGGT | CTCTCCCCTTTTGTTCTCCC | 198 |
| chr12:70501225-70503241 | p4 | (ATGT)4 | AAAGGCTTAAGGTCACGGGT | CTCTCCCCTTTTGTTCTCCC | 198 |

| Marker ID | SSR type | SSR motif | FORWARD PRIMER (5'-3') | REVERSE PRIMER (5'-3') | size (bp) |
|---|---|---|---|---|---|
| chr12:70506090-70508105 | p5 | (TAAAA)3 | TGCCAACGGATGAGTGATAA | ACATAAATTGGGACGAAGCG | 253 |
| chr12:70506090-70508105 | p6 | (TAAAAA)3 | TGCCAACGGATGAGTGATAA | ACATAAATTGGGACGAAGCG | 253 |
| chr12:70506090-70508105 | p5 | (ATTTT)3 | TGCCAACGGATGAGTGATAA | ACATAAATTGGGACGAAGCG | 253 |
| chr12:70524988-70527021 | p5 | (TTTTC)3 | AAGGCCAACAACTAGCTCCA | AAGAGCGTCCCTATGGGTCT | 227 |
| chr12:70550888-70552906 | p6 | (CCCCCA)3 | GAAAGATTCGGTTTGGTTGC | CCACTGCCCTAACAATCACA | 260 |
| chr12:70550888-70552906 | p2 | (AT)6 | GAAAGATTCGGTTTGGTTGC | CCACTGCCCTAACAATCACA | 260 |
| chr12:70626472-70628484 | p2 | (AC)6 | CAAATGCGACAAAAAGGGTT | CGGTAGCCAAATATAGCCCA | 129 |
| chr12:70671012-70673024 | p2 | (TA)6 | CGCAACAATTTGCTCAAAAA | GACGAAAAAGTGAGCCCAAC | 105 |
| chr12:70671012-70673024 | p2 | (TA)8 | CGCAACAATTTGCTCAAAAA | GACGAAAAAGTGAGCCCAAC | 105 |
| chr12:70671012-70673024 | p2 | (TA)7 | CGCAACAATTTGCTCAAAAA | GACGAAAAAGTGAGCCCAAC | 105 |
| chr12:70671012-70673024 | p2 | (CT)7 | CGCAACAATTTGCTCAAAAA | GACGAAAAAGTGAGCCCAAC | 105 |
| chr12:70710406-70712424 | p3 | (CTT)6 | TTTTGCTTAACGGATTTGGC | GTGCACATGCACAACCTTCT | 226 |
| chr12:70738848-70740886 | p2 | (AT)11 | AAGGAAACCTGCGCAGTAAA | AGTTACCCGTGGAGCAGATG | 226 |
| chr12:70738890-70740916 | p2 | (AT)11 | AAGGAAACCTGCGCAGTAAA | AGTTACCCGTGGAGCAGATG | 184 |
| chr12:70770720-70772738 | p6 | (TCACCA)3 | ACAATTTCACCACCCACCAT | TACCTTTCTCTTCCGCTCCA | 102 |
| chr12:70803380-70805395 | p5 | (TTTTA)3 | TGCGTTGCAAACTAGACTCG | TCGTAATCGGGAAGGACAAC | 287 |
| chr12:70803380-70805395 | p2 | (TA)16 | TGCGTTGCAAACTAGACTCG | TCGTAATCGGGAAGGACAAC | 287 |
| chr12:70835521-70837536 | p3 | (TTA)5 | CCTCCAAATGCGGGTACTTA | CCAGCGATTTGATGTTGATG | 202 |
| chr12:70835521-70837536 | p2 | (AT)13 | CCTCCAAATGCGGGTACTTA | CCAGCGATTTGATGTTGATG | 202 |
| chr12:70962545-70964611 | p2 | (AT)33 | TTGAGATGCCTTTCAGCCTT | CTGGAGCTTTCGACTCGTTT | 153 |
| chr12:71021059-71023074 | p2 | (TC)9 | GATGGACAATCGTGCAAATG | ACACACATTTTGCCACCTCA | 243 |
| chr12:71021059-71023074 | p5 | (AAAGA)3 | GATGGACAATCGTGCAAATG | ACACACATTTTGCCACCTCA | 243 |
| chr12:71150242-71152257 | p3 | (AAT)5 | TCTTCATTCGATACCAGCCG | GCCACTTGGCTAAACCACAT | 145 |
| chr12:71151810-71153832 | p2 | (TA)11 | TGTAGCCATGCAGCAACTTC | TTGCTGCCTTTTGTTGTGTC | 255 |
| chr12:71173951-71175973 | p2 | (TA)11 | TAATCGTGAGGTTGCTGCAC | CGAGTCATCTTCTTGTGGCA | 138 |
| chr12:71184199-71186214 | p3 | (TTC)5 | TGATGATTCAGCAAGCCAAG | TTTGGAACATACCGAGAGGC | 242 |
| chr12:71202250-71204274 | p6 | (CCTTTT)4 | TTTGAAAGTTGCGATTTCCC | TTGAAAGTGCTCTTGCCTCA | 292 |
| chr12:71238773-71240788 | p5 | (GTTTG)3 | ACCTGAATTAACCGTGGTGC | TTATTAAGCACCGGGTGTCC | 160 |
| chr12:71253086-71255104 | p3 | (TTC)6 | CAAGCAAGGCAGAATGAACA | TCTGGTTGGATTTTTGGAGC | 210 |
| chr12:71253124-71255155 | p3 | (TTC)6 | CAAGCAAGGCAGAATGAACA | TCTGGTTGGATTTTTGGAGC | 172 |
| chr12:71368019-71370039 | p2 | (CT)10 | TAAGCGTACGACAACCTCCC | AAATCCCGCGTAGATGTCAC | 122 |
| chr12:71450520-71452535 | p3 | (AAC)5 | CCAAGGCGATCAAATTGTTT | GATGTTGAAACGTGTGACGG | 236 |
| chr12:71455275-71457311 | p3 | (ACA)12 | CCTTACCACGAAGCATCCAT | GCGAAGTTTGTGGAAAGAGC | 101 |
| chr12:71471484-71473496 | p2 | (GT)6 | GAAGGATGCGAATTTAGGCA | CTTACCTCCGCGTTAGACCA | 186 |
| chr12:71631580-71633598 | p6 | (ACTGAA)3 | TCGACAATTAGTTGGGAGCC | TATCGAATGATGCGACGAAC | 273 |
| chr12:71648252-71650276 | p2 | (AT)12 | CTTGCACCGAAAAGCATGTA | TGCGTCTGTACGAATCTGGA | 103 |

| Marker ID | SSR type | SSR motif | FORWARD PRIMER (5'-3') | REVERSE PRIMER (5'-3') | size (bp) |
|---|---|---|---|---|---|
| chr12:71684357-71686373 | p2 | (TA)8 | TTATCCCGTCGTAGCTTCATT | TCCAAAAGGGAGCTAAACCA | 270 |
| chr12:71684357-71686373 | p2 | (AT)18 | TTATCCCGTCGTAGCTTCATT | TCCAAAAGGGAGCTAAACCA | 270 |
| chr12:71684357-71686373 | p2 | (AG)7 | TTATCCCGTCGTAGCTTCATT | TCCAAAAGGGAGCTAAACCA | 270 |
| chr12:71686826-71688841 | p3 | (AAT)5 | TGCACATCAAAATTCTCCGA | GTGGGATTACGCTGGGTATG | 203 |
| chr12:71710041-71712053 | p2 | (TA)6 | TTACACCCAAACTCTTCCCG | GTCAGACATCGGGTGTTGTG | 235 |
| chr12:71807748-71809760 | p2 | (TA)6 | AGGATGTGGTAAAGCGGATG | TGCAGATTTGCACTGGGTAA | 200 |
| chr12:71836444-71838464 | p5 | (TCATT)4 | CCTAGTGGAATTGGGGTGTG | ATTGCGGTGACAGGTAGTCC | 174 |
| chr12:71836444-71838464 | p3 | (ATG)6 | CCTAGTGGAATTGGGGTGTG | ATTGCGGTGACAGGTAGTCC | 174 |
| chr12:71872821-71874845 | p3 | (TTC)8 | CATGTTGGTTTGGACTGTCG | CGGACATTTGTAAGGGTCGT | 132 |
| chr12:71883355-71885373 | p3 | (CAT)6 | GGCTAGCCAACGTTTTACCA | TGTTTCAAACGCCCTCTCTT | 143 |
| chr12:71932111-71934126 | p3 | (ACA)5 | GTTTAATTAGCGCGCACACA | TTACTGCTGCTTTTGGCCTT | 232 |
| chr12:71961338-71963350 | p2 | (AT)6 | AGCTGGCCAGTCAAAACACT | GCATGGAAGTAGAGGCAAGG | 126 |
| chr12:72029493-72031545 | p3 | (ATC)8 | TCTTCTTCACTTCGGCCACT | GTCGCTCCATGGGAAGTTTA | 119 |
| chr12:72041906-72043922 | p5 | (AACCG)3 | CCGAATAAATCGAACCGAAA | CGCAATTTCTGTGGGAAAAT | 204 |
| chr12:72041906-72043922 | p2 | (TA)8 | CCGAATAAATCGAACCGAAA | CGCAATTTCTGTGGGAAAAT | 204 |
| chr12:72120708-72122736 | p2 | (TA)14 | AAGACTTGCAAGCCATCGTT | TTTCGCCACAACCCTAGAAT | 101 |
| chr12:72120708-72122736 | p2 | (TA)15 | AAGACTTGCAAGCCATCGTT | TTTCGCCACAACCCTAGAAT | 101 |
| chr12:72244335-72246353 | p6 | (TTTCTC)3 | CCCTTCGAAGCAAAAATGAG | TCCAGCCGAAAAGAAGAAAA | 200 |
| chr12:72407748-72409764 | p4 | (AAAT)4 | TTTTCGAGCCCTGAGTATGG | CAACTTTCAAGCCCACGATT | 240 |
| chr12:72428188-72430208 | p5 | (TGTTG)3 | CGGCTTACCTTTCGGTTACA | CCTTTCCCCCGATGATAAAT | 245 |
| chr12:72428188-72430208 | p5 | (TATGA)4 | CGGCTTACCTTTCGGTTACA | CCTTTCCCCCGATGATAAAT | 245 |
| chr12:72428188-72430208 | p5 | (GCATT)3 | CGGCTTACCTTTCGGTTACA | CCTTTCCCCCGATGATAAAT | 245 |
| chr12:72495219-72497234 | p2 | (TA)17 | TCCCTCCAACCATTTACCAA | TGTGGAGTCCCTTCTCCCTA | 150 |
| chr12:72495219-72497234 | p5 | (AATTA)3 | TCCCTCCAACCATTTACCAA | TGTGGAGTCCCTTCTCCCTA | 150 |
| chr12:72495259-72497274 | p2 | (TA)17 | TCCCTCCAACCATTTACCAA | TGTGGAGTCCCTTCTCCCTA | 110 |
| chr12:72495259-72497274 | p5 | (AATTA)3 | TCCCTCCAACCATTTACCAA | TGTGGAGTCCCTTCTCCCTA | 110 |
| chr12:72532480-72534507 | p3 | (AGA)9 | TCAAAAGCGCATCTTCTCCT | TGGGGTGGGCATTTTAGTAG | 141 |
| chr12:72555535-72557550 | p5 | (TTGGT)3 | GAGGAAAAGGGAAATGGAGG | GGCAGCCAGTCCATAAACAT | 239 |
| chr12:72555535-72557550 | p5 | (TTGGT)3 | GAGGAAAAGGGAAATGGAGG | GGCAGCCAGTCCATAAACAT | 239 |
| chr12:72599264-72601294 | p2 | (TA)6 | CCAAAGTGGTCCCTTCAAGA | CGCGATAAGTTAAGGGCAAG | 225 |
| chr12:72599264-72601294 | p2 | (AT)15 | CCAAAGTGGTCCCTTCAAGA | CGCGATAAGTTAAGGGCAAG | 225 |
| chr12:72599264-72601294 | p2 | (AT)6 | CCAAAGTGGTCCCTTCAAGA | CGCGATAAGTTAAGGGCAAG | 225 |
| chr12:72672484-72674496 | p2 | (GT)6 | TTGCAAACATGAACTTCCCA | GCCTTGTCCCACACTTTCAT | 216 |
| chr12:72804080-72806092 | p5 | (ACCGG)3 | AAGTCGAGCCAGACCTCCTT | TGCCAGGCTTAGTGGAGAGT | 180 |
| chr12:72804080-72806092 | p2 | (TG)16 | AAGTCGAGCCAGACCTCCTT | TGCCAGGCTTAGTGGAGAGT | 180 |
| chr12:72868372-72870387 | p3 | (TGT)5 | TTTGTTGAGTCCGCTTTGTG | CCATCAGCCAATGCCTCTAT | 216 |

| Marker ID | SSR type | SSR motif | FORWARD PRIMER (5'-3') | REVERSE PRIMER (5'-3') | size (bp) |
|---|---|---|---|---|---|
| chr12:72918216-72920236 | p4 | (CTTC)5 | CACGGTGATTTTCAGTGGTG | CATTTTCTCCGATTGCACCT | 145 |
| chr12:72919831-72921845 | p2 | (TG)7 | CAACCATATTCCCATCCCAG | TGAGAATGTCTGCTCATGCC | 117 |
| chr12:72919831-72921845 | p2 | (AC)7 | CAACCATATTCCCATCCCAG | TGAGAATGTCTGCTCATGCC | 117 |
| chr12:73021084-73023096 | p2 | (TA)6 | CGACGCCCTATTTACACTCC | GGCGAGAGCGAATATTGAAG | 218 |
| chr12:73021084-73023096 | p2 | (AT)6 | CGACGCCCTATTTACACTCC | GGCGAGAGCGAATATTGAAG | 218 |
| chr12:73057771-73059786 | p3 | (CAA)5 | AATCAAATCGATGAGGTCGG | TAGTCTACCACCATTCCCGC | 116 |
| chr12:73138508-73140544 | p2 | (AC)7 | TGATGGAATGCCACCAGTTA | AGCTGGGTTGGAATCACTTG | 286 |
| chr12:73142636-73144654 | p2 | (AG)13 | ATTCTAGCATGGTTTTGCGG | CATCCCATGTGGGGTTTATC | 115 |
| chr12:73154027-73156042 | p3 | (TAA)5 | TGACAGAGATGTCGGAGGTG | ATTTCCACTTCATTTCCCCC | 136 |
| chr12:73154027-73156042 | p3 | (AAT)11 | TGACAGAGATGTCGGAGGTG | ATTTCCACTTCATTTCCCCC | 136 |
| chr12:73154164-73156185 | p3 | (TAA)5 | AGTGGGGGTGTTGATGATGT | AAACCCTAAGGATCCCCTCA | 106 |
| chr12:73154164-73156185 | p3 | (AAT)11 | AGTGGGGGTGTTGATGATGT | AAACCCTAAGGATCCCCTCA | 106 |
| chr12:73154368-73156401 | p3 | (TAA)5 | TACGGATCGAGTGCAAGAGA | GCATTTATTTTCGCGCATTT | 234 |
| chr12:73154368-73156401 | p3 | (AAT)11 | TACGGATCGAGTGCAAGAGA | GCATTTATTTTCGCGCATTT | 234 |
| chr12:73167056-73169080 | p3 | (TCT)8 | TGGCTATTTCAGCATCGAGA | TTAGACCCCCAACTGAGACG | 153 |
| chr12:73167056-73169080 | p5 | (ATTTT)3 | TGGCTATTTCAGCATCGAGA | TTAGACCCCCAACTGAGACG | 153 |
| chr12:73181412-73183430 | p6 | (TTATAA)3 | TTACAAGGATTCTCGGGTCG | TAAAACTGTTGGAATCCCGC | 192 |
| chr12:73275505-73277517 | p2 | (CT)6 | TGCCCTTCTCCCAATATACG | TTTCCCAAAACCCCTTTTTC | 263 |
| chr12:73275505-73277517 | p2 | (TC)6 | TGCCCTTCTCCCAATATACG | TTTCCCAAAACCCCTTTTTC | 263 |
| chr12:73290605-73292620 | p2 | (TG)6 | AAATTTCTTATCCACCCCGC | CACCCAACACACACACACAA | 156 |
| chr12:73290605-73292620 | p5 | (TTTTA)3 | AAATTTCTTATCCACCCCGC | CACCCAACACACACACACAA | 156 |
| chr12:73290605-73292620 | p3 | (TGT)5 | AAATTTCTTATCCACCCCGC | CACCCAACACACACACACAA | 156 |
| chr12:73330046-73332064 | p2 | (TA)9 | TCTTGAATTCACCTCCGACC | TTTTGTAGGGTTCAATCCCG | 266 |
| chr12:73330046-73332064 | p5 | (CAACA)3 | TCTTGAATTCACCTCCGACC | TTTTGTAGGGTTCAATCCCG | 266 |
| chr12:73334237-73336251 | p2 | (AT)7 | GCAAAGGGTTAAGGCTAGGG | TCAGCACGACAATGTGAGGT | 250 |
| chr12:73372122-73374134 | p6 | (TATATG)3 | AAATGAGGCAAAAGTCGTGG | TTGCCCAGAATTTCCAGAAC | 278 |
| chr12:73421434-73423449 | p5 | (GGTAT)3 | GTGGATTTTGAAGGAGCGAA | TTGAGGCATTGTTGATCTGG | 239 |
| chr12:73458738-73460753 | p5 | (GGTAT)3 | CTGCCCAGCAAAATATGGTT | GACCATTTGACGGTCCGTAT | 123 |
| chr12:73507178-73509193 | p5 | (TTTTA)3 | GAGTTGCCTTGCCATGTCTT | ATTTCATGAATCCCACGCAT | 169 |
| chr12:73528214-73530229 | p5 | (TTTTA)3 | GAGTTGCCTTGCCATGTCTT | ATTTCATGAATCCCACGCAT | 169 |
| chr12:73549118-73551148 | p2 | (AT)6 | GATTAAGGAGGAGGGTTGCC | AAACCAGTGTATCCCCCACA | 183 |
| chr12:73549118-73551148 | p2 | (AT)10 | GATTAAGGAGGAGGGTTGCC | AAACCAGTGTATCCCCCACA | 183 |
| chr12:73549118-73551148 | p2 | (AT)13 | GATTAAGGAGGAGGGTTGCC | AAACCAGTGTATCCCCCACA | 183 |
| chr12:73549153-73551177 | p2 | (AT)6 | GATTAAGGAGGAGGGTTGCC | AAACCAGTGTATCCCCCACA | 148 |
| chr12:73549153-73551177 | p2 | (AT)10 | GATTAAGGAGGAGGGTTGCC | AAACCAGTGTATCCCCCACA | 148 |
| chr12:73549153-73551177 | p2 | (AT)13 | GATTAAGGAGGAGGGTTGCC | AAACCAGTGTATCCCCCACA | 148 |

| Marker ID | SSR type | SSR motif | FORWARD PRIMER (5'-3') | REVERSE PRIMER (5'-3') | size (bp) |
|---|---|---|---|---|---|
| chr12:73688144-73690168 | p4 | (AAAT)6 | TGTGTTAGCAGTCAGGCCAC | TGCTGGAAGAAGCACATCAC | 159 |
| chr12:73872278-73874300 | p2 | (AT)11 | CCATGACTAGTGCCCGATCT | TGTAGTGGCAGCCTTGTCAG | 160 |
| chr12:73876250-73878265 | p5 | (GTCTT)3 | GGCGCTGTCTCAAATACCAT | GGAAGCGGATGAGAGAGATG | 229 |
| chr12:73912365-73914387 | p2 | (AT)11 | CCATGACTAGTGCCCGATCT | TGTAGTGGCAGCCTTGTCAG | 160 |
| chr12:73916337-73918352 | p5 | (GTCTT)3 | GGCGCTGTCTCAAATACCAT | GGAAGCGGATGAGAGAGATG | 229 |
| chr12:73939770-73941785 | p5 | (TTTAA)3 | ACAACATCCATGGCTTCTCC | TCCATACCAATCCACACCCT | 132 |
| chr12:73967928-73969946 | p3 | (GGA)6 | GCCTATTCCACCCCTTCTTC | TGAAATCTAACAAAGCGGGG | 165 |
| chr12:73970688-73972736 | p2 | (AC)6 | ATGAACCGATGGGGTTATGA | AAGTAGGCAGGGAGACGGAT | 259 |
| chr12:73970765-73972831 | p2 | (AC)6 | ATGAACCGATGGGGTTATGA | AAGTAGGCAGGGAGACGGAT | 182 |
| chr12:73982153-73984169 | p4 | (TTTA)4 | GAACCCAGAACCTCACGGTA | ATACTTCTCTGCCAGCCCAA | 187 |
| chr12:73985013-73987037 | p6 | (TCTTTT)4 | AATGCCGAATACCATACCGA | CTTCGCCGACTTAAGAAACG | 246 |
| chr12:74031921-74033933 | p2 | (TA)9 | TTGTGCTAGGTTTTGTCCCC | GGAGCCCACATAAGCAAAAA | 201 |
| chr12:74031921-74033933 | p2 | (AG)6 | TTGTGCTAGGTTTTGTCCCC | GGAGCCCACATAAGCAAAAA | 201 |
| chr12:74031921-74033933 | p6 | (TATTTG)3 | TTGTGCTAGGTTTTGTCCCC | GGAGCCCACATAAGCAAAAA | 201 |
| chr12:74031921-74033933 | p6 | (TTTTCT)3 | TTGTGCTAGGTTTTGTCCCC | GGAGCCCACATAAGCAAAAA | 201 |
| chr12:74146485-74148523 | p2 | (TA)6 | CCCATCCCAAAACATTCAAC | TCCGTGTTCAATTTGGGTTT | 167 |
| chr12:74146485-74148523 | p2 | (AT)19 | CCCATCCCAAAACATTCAAC | TCCGTGTTCAATTTGGGTTT | 167 |
| chr12:74146485-74148523 | p2 | (AT)8 | CCCATCCCAAAACATTCAAC | TCCGTGTTCAATTTGGGTTT | 167 |
| chr12:74198311-74200326 | p3 | (ATT)5 | GCACCAATTTTATCGCGTTT | GTAGAGCCAAGCGGAGTTTG | 130 |
| chr12:74229031-74231046 | p3 | (TTA)5 | ACCATTTCAGCCTCCACAAC | GACCCAATTACCACCACGAC | 264 |
| chr12:74236453-74238473 | p2 | (TA)10 | TGTTTCCTATGGGAAGCCAC | CCAAAGGCAAGGTATCCAAA | 229 |
| chr12:74289563-74291595 | p2 | (AT)16 | CAGCGACCTTGACAACTTCA | GAAATGGGGTTGCTTTCTCA | 231 |
| chr12:74298200-74300212 | p2 | (AC)6 | ACTGTGCTTCACATTGCTGC | ACAGAACTCTGCCCATCCAC | 138 |
| chr12:74298200-74300212 | p2 | (TA)6 | ACTGTGCTTCACATTGCTGC | ACAGAACTCTGCCCATCCAC | 138 |
| chr12:74302238-74304254 | p4 | (AACA)4 | CGCTTGTCTGGATCATCCTT | TGAGAATGATTTCATTGCGG | 300 |
| chr12:74316706-74318722 | p5 | (GGGTT)3 | GAAAAGGGCCAAAACTACCC | TAAGGGCATAATTGACCCCA | 156 |
| chr12:74316706-74318722 | p2 | (AT)8 | GAAAAGGGCCAAAACTACCC | TAAGGGCATAATTGACCCCA | 156 |
| chr12:74365113-74367129 | p4 | (GACT)4 | GCCTTGGTCGTACCTTTTGA | ACCTTGATTGGCTTTCCCTT | 142 |
| chr12:74427933-74429979 | p2 | (AT)23 | GTGCTTGAAGCTCCTGAACC | GCAGGCCAAGAGAGATACCA | 123 |
| chr12:74521446-74523482 | p2 | (TA)16 | ACTCGGTGCATACCAAAAGC | AGCTCATCATGCACTTCACG | 281 |
| chr12:74521446-74523482 | p2 | (AT)18 | ACTCGGTGCATACCAAAAGC | AGCTCATCATGCACTTCACG | 281 |
| chr12:74548051-74550063 | p2 | (AT)6 | TCTTAAGGGTTGGCTGGATG | AGGGTAAGGTAATGTGCCCC | 177 |
| chr12:74548051-74550063 | p2 | (AT)6 | TCTTAAGGGTTGGCTGGATG | AGGGTAAGGTAATGTGCCCC | 177 |
| chr12:74657066-74659078 | p2 | (TA)6 | TTTGATCTTCCTTGATGGGG | TTGACCACGGAGAGAAAACC | 156 |
| chr12:74716329-74718347 | p2 | (GA)6 | GATCCGAAACAAAACCCAGA | TGGAATCCTCCAAGAACAGG | 214 |
| chr12:74716329-74718347 | p5 | (TCGGG)3 | GATCCGAAACAAAACCCAGA | TGGAATCCTCCAAGAACAGG | 214 |

| Marker ID | SSR type | SSR motif | FORWARD PRIMER (5'-3') | REVERSE PRIMER (5'-3') | size (bp) |
|---|---|---|---|---|---|
| chr12:74716329-74718347 | p3 | (TAA)6 | GATCCGAAACAAAACCCAGA | TGGAATCCTCCAAGAACAGG | 214 |
| chr12:74728037-74730052 | p5 | (TTCTT)3 | TCAAAAACAACTGCGAGTGC | AATGGGAAGGCTGTTTGTTG | 133 |
| chr12:74860955-74862971 | p3 | (TAT)10 | GGTGATAAGCGCTCGTTAGG | CGATTCTAAGCGGTTTCTGG | 140 |
| chr12:74860955-74862971 | p2 | (AT)8 | GGTGATAAGCGCTCGTTAGG | CGATTCTAAGCGGTTTCTGG | 140 |
| chr12:74962769-74964789 | p2 | (TA)6 | CGGGTTGGTCTTGAACGTAT | CGCTAATTCGCCACACAATA | 282 |
| chr12:74962769-74964789 | p2 | (TA)6 | CGGGTTGGTCTTGAACGTAT | CGCTAATTCGCCACACAATA | 282 |
| chr12:75021364-75023378 | p2 | (TA)7 | TACGAAGCTGCTTGCTCAAA | TATGGAGACATCCAATGCCA | 189 |
| chr12:75037855-75039870 | p5 | (ATTTT)3 | TCGGCTACCTTTTCAACACC | CGTGTGTCAAGACTCCGAAA | 152 |
| chr12:75037874-75039908 | p5 | (ATTTT)3 | TCGGCTACCTTTTCAACACC | CGTGTGTCAAGACTCCGAAA | 133 |
| chr12:75062174-75064188 | p2 | (AT)6 | TGGCATTGAATGAAGCAAAG | GTTTGGGGTATGATCGTCGT | 121 |
| chr12:75154402-75156417 | p5 | (TATTT)3 | CATGTGCTGTGCTGGTCTCT | GAACTTCCTCGCAGCACAAT | 160 |
| chr12:75326233-75328248 | p3 | (ATT)5 | TCTTTGGCCTACCCCTACCT | TGGGTGTGTTTGCTTTTGAA | 182 |
| chr12:75426428-75428443 | p3 | (TGT)5 | CAATTGCAAGCTCTGGATGA | CTCCAAAACAACGAGGGGTA | 246 |
| chr12:75473526-75475540 | p2 | (TA)7 | ATATTTGGGCATGCGTGATT | GCAGGTTTTTGTTGAGGGAA | 235 |
| chr12:75541875-75543891 | p4 | (ATTT)4 | TTTCAGAAAAATGCGAAGCA | TTTCCAAAGAAGCCAAATCG | 207 |
| chr12:75635344-75637364 | p2 | (TA)6 | TTTTTCCATTTTGGCACCTC | CAATTCCTGGTTGGCTGAAT | 119 |
| chr12:75635362-75637416 | p2 | (TA)6 | TTTTTCCATTTTGGCACCTC | CAATTCCTGGTTGGCTGAAT | 101 |
| chr12:75677728-75679743 | p5 | (TCTTT)3 | CGTGCTGCAACAGTTGTTCT | GCCCCTATGCAAATTTTCTG | 191 |
| chr12:75719164-75721178 | p2 | (TA)7 | ACACGGGCCCAACATAATAA | CCAATGATGGCGTTCTTGTA | 172 |
| chr12:75729556-75731568 | p2 | (TA)6 | AAAAGAGATCCTCGCCCATT | ATGGCATGATGTCACGAAAA | 213 |
| chr12:75855076-75857092 | p4 | (TTAA)4 | TCCAACGGAGTACTTTGGCT | GGAAACTCAAAGCCTCAACG | 204 |
| chr12:75855076-75857092 | p2 | (AT)16 | TCCAACGGAGTACTTTGGCT | GGAAACTCAAAGCCTCAACG | 204 |
| chr12:75865723-75867738 | p3 | (GTT)5 | CCAGGCAACTGTAGCAGTCA | CCCTGGGAGCTATCACAAAA | 193 |
| chr12:75892827-75894845 | p6 | (TAGCAG)3 | AAAGTCCTCCGCATTAGGGT | ACCACCCCGAATGACAATAA | 103 |
| chr12:75951765-75953780 | p5 | (AAAAT)3 | CTAAATCAGCAGCAAAGGGC | GAGGAAGGCACCTGAATCTG | 154 |
| chr12:75987754-75989766 | p5 | (ATTTT)3 | AAGTGTATGCTGGTGGGGAG | CCACGTTATACCCGTTTTGG | 251 |
| chr12:75987776-75989790 | p5 | (ATTTT)3 | AAGTGTATGCTGGTGGGGAG | CCACGTTATACCCGTTTTGG | 229 |
| chr12:75999490-76001508 | p6 | (GTATGC)3 | CGCTCAACGCAAGATAATCA | TAGGGAGACACATGTTGCGA | 243 |
| chr12:76016403-76018421 | p6 | (GTATGC)3 | CGCTCAACGCAAGATAATCA | TAGGGAGACACATGTTGCGA | 243 |
| chr12:76036071-76038086 | p5 | (TGGCT)3 | GAACCAGGGACGGAGTACAA | ACGCCAACGTGAACATGATA | 280 |
| chr12:76076865-76078880 | p5 | (TATTT)3 | GGGTGTGGGGTTAAAGGAAT | ATTTACCGGAAAGGTCCAGC | 263 |
| chr12:76107717-76109729 | p2 | (TA)6 | GCAAAGAAAAACGGGAGTCA | ACCTTATAGGTTGGGGGTGG | 170 |
| chr12:76234154-76236169 | p5 | (AAACC)3 | TGGCTGCACCATTTTTGTTA | ATGCAAGTATTGCCCCAAAG | 137 |
| chr12:76234154-76236169 | p5 | (TTTGG)3 | TGGCTGCACCATTTTTGTTA | ATGCAAGTATTGCCCCAAAG | 137 |
| chr12:76234154-76236169 | p2 | (TA)6 | TGGCTGCACCATTTTTGTTA | ATGCAAGTATTGCCCCAAAG | 137 |
| chr12:76291198-76293213 | p3 | (TTG)5 | CACTTTCAGCTCACACCGAA | TCTTCTGCCCATTTGACCTC | 214 |

| Marker ID | SSR type | SSR motif | FORWARD PRIMER (5'-3') | REVERSE PRIMER (5'-3') | size (bp) |
|---|---|---|---|---|---|
| chr12:76319136-76321162 | p2 | (TC)13 | CGAATCACCCCTACAGCAAT | CGAAAGGGGGTGATGAAGTA | 150 |
| chr12:76320447-76322465 | p3 | (GAA)6 | TGGGTAGTCCAGGGAATGAG | AATTGGGGATTAAAGGGCAG | 110 |
| chr12:76451875-76453889 | p2 | (AT)7 | CGGTCATGTTTTTGAGCCTT | GGACTACCTGTCACCGCAAT | 136 |
| chr12:76451875-76453889 | p4 | (CATA)4 | CGGTCATGTTTTTGAGCCTT | GGACTACCTGTCACCGCAAT | 136 |
| chr12:76462675-76464690 | p5 | (AAAAG)3 | GGGTTGGCGTAGAGGACATA | TGGGACACATGGAAAGTGAA | 133 |
| chr12:76469351-76471365 | p5 | (ATTTT)3 | GCATGGGAGGTGTCAGAGAT | TCACTCTTTCCCCCTTTCCT | 277 |
| chr12:76487294-76489312 | p6 | (CACCTT)3 | ATCTGGACACGGACATAGGC | GGGTTTGTAGGGTGCATGAT | 234 |
| chr12:76487294-76489312 | p2 | (TG)6 | ATCTGGACACGGACATAGGC | GGGTTTGTAGGGTGCATGAT | 234 |
| chr12:76585108-76587129 | p3 | (TGT)7 | GGTTGTGAGACCGACCTTGT | TCGTTTCCCTCATCTTGTCC | 269 |
| chr12:76641550-76643565 | p3 | (TGT)5 | AGGCAATCACAAGGTCCAAC | TCCATGGTTGTGACTGGAAA | 292 |
| chr12:76819637-76821649 | p2 | (AT)6 | AAGTATCAACCGTGGCTTCG | TCCGCTTCTGAGTTTTGGTT | 206 |
| chr12:76819637-76821649 | p2 | (AT)11 | AAGTATCAACCGTGGCTTCG | TCCGCTTCTGAGTTTTGGTT | 206 |
| chr12:76819863-76821885 | p2 | (AT)6 | AACCAAAACTCAGAAGCGGA | TTTCGTTAATTTCCCCCACA | 114 |
| chr12:76819863-76821885 | p2 | (AT)11 | AACCAAAACTCAGAAGCGGA | TTTCGTTAATTTCCCCCACA | 114 |
| chr12:76859597-76861621 | p2 | (TA)12 | AGAAAACCCAGCCCAAAAAT | TGTCCATGTTTAAGGTGCCA | 144 |
| chr12:76889949-76891964 | p5 | (TTTTA)3 | ATCACCAACACCGGAGAATC | CCATTGAATTTTATCGGGGA | 205 |
| chr12:76973631-76975645 | p2 | (AT)7 | CTTACCCAGTGAGCCGAGAG | CCTCATTCTAGCTGAACCCG | 150 |
| chr12:77214182-77216194 | p2 | (AC)6 | CAACGTGGACGTAGCTCTCA | TTTGTAAGCAGGTTCCCAGG | 229 |
| chr12:77224614-77226634 | p2 | (TG)10 | CATTTCCGACGATCTCCACT | GGTTCCCCTTCATCACAAGA | 137 |
| chr12:77273278-77275330 | p4 | (ATAA)4 | TTACCTTGTCTCGAGGGGTG | ACGCACTAGATAACCGGTGG | 253 |
| chr12:77273278-77275330 | p2 | (TA)7 | TTACCTTGTCTCGAGGGGTG | ACGCACTAGATAACCGGTGG | 253 |
| chr12:77273278-77275330 | p2 | (AT)8 | TTACCTTGTCTCGAGGGGTG | ACGCACTAGATAACCGGTGG | 253 |
| chr12:77273595-77275609 | p4 | (ATAA)4 | ACCACCGGTTATCTAGTGCG | GGGAATAGGGATGCTGATGA | 161 |
| chr12:77273595-77275609 | p2 | (TA)7 | ACCACCGGTTATCTAGTGCG | GGGAATAGGGATGCTGATGA | 161 |
| chr12:77273595-77275609 | p2 | (AT)8 | ACCACCGGTTATCTAGTGCG | GGGAATAGGGATGCTGATGA | 161 |
| chr12:77273864-77275880 | p2 | (TA)7 | CCTATTCCCGCATCAGGTTA | ATGGGGTCGTTGAACTTGAC | 125 |
| chr12:77273864-77275880 | p2 | (AT)8 | CCTATTCCCGCATCAGGTTA | ATGGGGTCGTTGAACTTGAC | 125 |
| chr12:77291106-77293124 | p5 | (ATTTT)3 | TCTTTGTACGATAACGGCGA | ATGGATTTTCAACGGAAACG | 244 |
| chr12:77291122-77293160 | p5 | (ATTTT)3 | TCTTTGTACGATAACGGCGA | ATGGATTTTCAACGGAAACG | 228 |
| chr12:77291183-77293273 | p5 | (ATTTT)3 | TCTTTGTACGATAACGGCGA | ATGGATTTTCAACGGAAACG | 167 |
| chr12:77312028-77314043 | p3 | (TTA)5 | GCACGAACGTAAGATCACCA | AACTGTGGGCAGAGTTGGAC | 184 |
| chr12:77348180-77350195 | p3 | (TAT)7 | TTCAAAATGGGTTTGGGGTA | AACTGCTCATCCCTCTTCCA | 177 |
| chr12:77348180-77350195 | p5 | (TCCTT)3 | TTCAAAATGGGTTTGGGGTA | AACTGCTCATCCCTCTTCCA | 177 |
| chr12:77390883-77392904 | p3 | (TAA)7 | CAGCAAAAGGCAATTCAACA | TCGATGAAGAGCCTCAACCT | 165 |
| chr12:77442966-77444978 | p2 | (AG)6 | AGGACGAAGCCAATCAGAGA | GAATTTCCAGCGAGACAAGC | 154 |
| chr12:77442966-77444978 | p2 | (AG)6 | AGGACGAAGCCAATCAGAGA | GAATTTCCAGCGAGACAAGC | 154 |

| Marker ID | SSR type | SSR motif | FORWARD PRIMER (5'-3') | REVERSE PRIMER (5'-3') | size (bp) |
|---|---|---|---|---|---|
| chr12:77534630-77536645 | p5 | (CATTA)3 | GTTGCACCTTTTTCCCGTAA | AGACAGAGACAGGCCACACC | 235 |
| chr12:77561522-77563567 | p3 | (TAG)6 | AAAGATAACCCAATCTAGGGGG | TCAAATCGACAAGTTTTATGGC | 286 |
| chr12:77561522-77563567 | p2 | (TA)10 | AAAGATAACCCAATCTAGGGGG | TCAAATCGACAAGTTTTATGGC | 286 |
| chr12:77561702-77563768 | p3 | (TAG)6 | AAAGATAACCCAATCTAGGGGG | TCAAATCGACAAGTTTTATGGC | 106 |
| chr12:77561702-77563768 | p2 | (TA)10 | AAAGATAACCCAATCTAGGGGG | TCAAATCGACAAGTTTTATGGC | 106 |
| chr12:77563523-77565537 | p2 | (TA)10 | TTTGTTTCGAAGTTTCGGGT | CGAATTTGCGCTGTGTTAAA | 188 |
| chr12:77563523-77565537 | p2 | (TA)7 | TTTGTTTCGAAGTTTCGGGT | CGAATTTGCGCTGTGTTAAA | 188 |
| chr12:77575618-77577636 | p6 | (ATCTAT)3 | TGCCCATAAATGGACAACAA | CGAATTAAATGGGCTCCAAA | 122 |
| chr12:77637462-77639487 | p3 | (TTA)10 | AAGGCTGGATCCGTCTTTTT | GTGGGACAAAATTGGGTGTC | 219 |
| chr12:77637548-77639560 | p3 | (TTA)10 | AAGGCTGGATCCGTCTTTTT | GTGGGACAAAATTGGGTGTC | 133 |
| chr12:77659614-77661632 | p3 | (TAC)6 | ACTCAATGCACAACCACGAG | TGTTGTTGGAATTGACGGTG | 256 |
| chr12:77659614-77661632 | p3 | (TCT)6 | ACTCAATGCACAACCACGAG | TGTTGTTGGAATTGACGGTG | 256 |
| chr12:77692637-77694658 | p2 | (GA)7 | TATCTTGAAACGGAATGGGC | GTTCCTTGCATTCAGTCCGT | 108 |
| chr12:77692637-77694658 | p3 | (AAT)5 | TATCTTGAAACGGAATGGGC | GTTCCTTGCATTCAGTCCGT | 108 |
| chr12:77692637-77694658 | p3 | (ACA)7 | TATCTTGAAACGGAATGGGC | GTTCCTTGCATTCAGTCCGT | 108 |
| chr12:77831686-77833702 | p2 | (TA)9 | ACATGTCCCAAAAGGTGGAC | CATGAGTTTGGAGACGGTGA | 249 |
| chr12:77831686-77833702 | p2 | (TA)8 | ACATGTCCCAAAAGGTGGAC | CATGAGTTTGGAGACGGTGA | 249 |
| chr12:77831686-77833702 | p2 | (AT)6 | ACATGTCCCAAAAGGTGGAC | CATGAGTTTGGAGACGGTGA | 249 |
| chr12:77928669-77930689 | p4 | (TGGG)5 | GTCGTGTTTTCGATCCGTTT | AACCATATGGGGTTGGGATT | 230 |
| chr12:77935214-77937229 | p3 | (AAG)5 | AACAAGGTGGACAACAAGGC | ACCAACAAACCTCCCACTTG | 187 |
| chr12:78072193-78074213 | p2 | (AT)10 | GGCCCCAACTGGAATAAAAT | AAGCAATTTTGAGAGCCGAA | 243 |
| chr12:78123643-78125658 | p3 | (CAA)5 | ACGAGCGGAGTAGTGAGGAA | TTCGGGTCTATACTCGTCCG | 296 |
| chr12:78151095-78153141 | p2 | (AT)23 | CAGCACCAAAACCTGCACTA | GGAAGCGGTTAGGGTTCATT | 208 |
| chr12:78225031-78227049 | p2 | (AT)13 | GGGTTGGTTCACCGAACTTA | AGTGCTATGTCACACTGGCG | 260 |
| chr12:78225031-78227049 | p3 | (AGT)6 | GGGTTGGTTCACCGAACTTA | AGTGCTATGTCACACTGGCG | 260 |
| chr12:78339096-78341123 | p3 | (TAA)9 | TGTCCAAAATGCACGAAAGA | GCCGATGACACCAGAACTTT | 101 |
| chr12:78360073-78362089 | p2 | (AT)8 | AGGGTTTCCACATCCAACAA | TCGATTTCGGGTCAAAAGTC | 259 |
| chr12:78377474-78379486 | p2 | (TA)6 | TGAACGGAGGGAGTACTTGG | GGCAGATGAGGAAGACCTTG | 165 |
| chr12:78450465-78452479 | p2 | (AG)7 | AAGGAGCACTTCCACGAAAA | TTGTCGATCCCAAGATCCTC | 175 |
| chr12:78524645-78526660 | p5 | (ACCCG)3 | ACTCCGTTCCAATGTCTTCG | GTGGGACAAATTTGGGTGTC | 232 |
| chr12:78534007-78536027 | p4 | (CAAA)5 | ATCCCTTTTAGGCCACCACT | AGGAGTTTTTAGCGAAGGGC | 151 |
| chr12:78549079-78551099 | p4 | (CAAA)5 | ATCCCTTTTAGGCCACCACT | AGGAGTTTTTAGCGAAGGGC | 151 |
| chr12:78572231-78574246 | p5 | (CCGGT)3 | ACTTAGAAGCGGAGGAAGGC | TCATATTCGGCCATGGAGAT | 112 |
| chr12:78779600-78781612 | p2 | (TG)6 | CCTACACCACCAGCTCCAAT | GTGTGGTGAGCGGTTAGTGA | 213 |
| chr12:78870568-78872588 | p2 | (TA)10 | CCATATCTGCATGGTGTTGC | TAAAACCGCATTCGGAACTC | 210 |
| chr12:78881471-78883502 | p3 | (GTT)7 | TCCACTTTGTTGGAGCATGA | ACAAAGAAGTCGAGGGGGTT | 101 |

| Marker ID | SSR type | SSR motif | FORWARD PRIMER (5'-3') | REVERSE PRIMER (5'-3') | size (bp) |
|---|---|---|---|---|---|
| chr12:78881528-78883549 | p3 | (GTT)7 | AACCCCCTCGACTTCTTTGT | AAACCTTTCACCTCTTGGGG | 194 |
| chr12:78883296-78885326 | p2 | (AT)15 | CCCAGAAGGCAGTTGCTAAA | CACCTTGGCTCCAAATACCA | 204 |
| chr12:78994751-78996763 | p2 | (AT)6 | TGAACCAGTCAGGACCACAG | AAACCCGATAATCATGCGTC | 276 |
| chr12:79089146-79091158 | p2 | (AT)6 | ACCAAATTTCGATCACTCGC | GCCCAAATTTCGGTAGACAA | 232 |
| chr12:79102547-79104562 | p5 | (TTGGT)3 | ACCTTGCTGTGCCTGTCTTT | CTGCCAAAAACCAGCAAAAT | 279 |
| chr12:79105143-79107158 | p3 | (AAG)5 | TTGATTGTCCAGTGTCCCAA | TCTGGCGAGCAGGTTAAAGT | 245 |
| chr12:79114268-79116289 | p3 | (TAA)7 | TTGATAGACCCCACACACCA | TTTTTGTTCCACTGCCATCA | 236 |
| chr12:79166820-79168870 | p3 | (TTC)6 | ACCTGGCATGCATAAGGAAC | ACAAGCCCATCATCTTGCTT | 289 |
| chr12:79229720-79231732 | p2 | (TA)6 | TCCCAAAGTATCCCCATCAA | GACATGGCGGAATAGCAAAT | 278 |
| chr12:79229720-79231732 | p3 | (ACA)11 | TCCCAAAGTATCCCCATCAA | GACATGGCGGAATAGCAAAT | 278 |
| chr12:79232701-79234716 | p5 | (ACAAT)3 | ACCCCACTAAAAACCCCATC | GTTGTACGCCGCTTTTTCTC | 252 |
| chr12:79382199-79384214 | p5 | (ATTTT)3 | TCCAAGGCCAGAACAAATTC | TGCATCACCAACTTGTCCAT | 130 |
| chr12:79437952-79439970 | p2 | (TA)6 | TTGTGATCCCACCCATTTTT | TTTCCGTGAGTTTGGAGAGG | 239 |
| chr12:79437952-79439970 | p6 | (AAAATA)3 | TTGTGATCCCACCCATTTTT | TTTCCGTGAGTTTGGAGAGG | 239 |
| chr12:79437952-79439970 | p2 | (TA)12 | TTGTGATCCCACCCATTTTT | TTTCCGTGAGTTTGGAGAGG | 239 |
| chr12:79514501-79516519 | p6 | (TATAAA)3 | AGGTGGCACTGGTTTTATCG | CATTTATCTCGGGGCAAAGA | 131 |
| chr12:79591312-79593334 | p3 | (TGT)5 | TGCAATGATTCAGGTCGAAA | TTGTCCCATACACACGGCTA | 183 |
| chr12:79591312-79593334 | p2 | (TA)11 | TGCAATGATTCAGGTCGAAA | TTGTCCCATACACACGGCTA | 183 |
| chr12:79702730-79704751 | p3 | (TAT)7 | GTGCAAAGCGCGTACACTAA | GAAGTTGACGGCAACCCTAA | 112 |
| chr12:79752446-79754458 | p2 | (AT)6 | GACACCCGTACGTCACACAG | CAGAAAAACGTGATCTGCGA | 248 |
| chr12:79888761-79890779 | p2 | (TA)9 | CAGTCCTAGGAGGGACCACA | TGTTTTGGTGAACCAGGTGA | 127 |
| chr12:80021290-80023305 | p3 | (ACA)5 | TTAAGCGAAGCAACACATGG | GAGCCAACCCAGAAATCGTA | 229 |
| chr12:80021290-80023305 | p2 | (AT)8 | TTAAGCGAAGCAACACATGG | GAGCCAACCCAGAAATCGTA | 229 |
| chr12:80021870-80023886 | p3 | (ACA)5 | ACAACGCTTCTTGTCGATCC | CAGACCCCACTTGTTGGATT | 216 |
| chr12:80021870-80023886 | p2 | (AT)8 | ACAACGCTTCTTGTCGATCC | CAGACCCCACTTGTTGGATT | 216 |
| chr12:80021870-80023886 | p2 | (TA)18 | ACAACGCTTCTTGTCGATCC | CAGACCCCACTTGTTGGATT | 216 |
| chr12:80059814-80061828 | p2 | (AT)7 | CAGCATGCAGGTTCTAAGCA | GAGGATGGGGAAAAGAAAGC | 251 |
| chr12:80122176-80124190 | p2 | (AT)7 | CAGCATGCAGGTTCTAAGCA | GAGGATGGGGAAAAGAAAGC | 251 |
| chr12:80248280-80250295 | p3 | (TCA)5 | TCGACCCGAAAAATGATCTC | AAAAATGTCCTCGCGATCTG | 166 |
| chr12:80287354-80289369 | p3 | (TCA)5 | TCGACCCGAAAAATGATCTC | AAAAATGTCCTCGCGATCTG | 166 |
| chr12:80360392-80362417 | p5 | (CATAT)5 | GGTGGTAATTGTCACACCCC | TCAGCCGATACGTCATTCTG | 247 |
| chr12:80379820-80381836 | p2 | (AT)13 | CTCAAAAGGGGCATACCAAA | TACTGACTCACCGTTGCTCG | 294 |
| chr12:80379842-80381856 | p2 | (AT)13 | CTCAAAAGGGGCATACCAAA | TACTGACTCACCGTTGCTCG | 272 |
| chr12:80379869-80381884 | p2 | (AT)13 | CTCAAAAGGGGCATACCAAA | TACTGACTCACCGTTGCTCG | 245 |
| chr12:80379879-80381891 | p2 | (AT)13 | CTCAAAAGGGGCATACCAAA | TACTGACTCACCGTTGCTCG | 235 |
| chr12:80603381-80605397 | p4 | (GAAA)4 | CCACTGTCTGGCCATATCCT | GCATGCCACCGTTTTCTAAT | 183 |

| Marker ID | SSR type | SSR motif | FORWARD PRIMER (5'-3') | REVERSE PRIMER (5'-3') | size (bp) |
|---|---|---|---|---|---|
| chr12:80603381-80605397 | p5 | (TTTTA)3 | CCACTGTCTGGCCATATCCT | GCATGCCACCGTTTTCTAAT | 183 |
| chr12:80660486-80662498 | p2 | (AT)15 | ATCAAAGGCCCACAAGTGAC | GGACCTTGTCCCTTCCTTTC | 137 |
| chr12:80660486-80662498 | p2 | (AT)6 | ATCAAAGGCCCACAAGTGAC | GGACCTTGTCCCTTCCTTTC | 137 |
| chr12:80660486-80662498 | p6 | (CAACTC)3 | ATCAAAGGCCCACAAGTGAC | GGACCTTGTCCCTTCCTTTC | 137 |
| chr12:80693626-80695650 | p2 | (TA)7 | AAGAAGATTGAGGCCCACCT | CCCGGTAGGGAGAAAGAGAC | 191 |
| chr12:80739291-80741309 | p6 | (GGTGAC)3 | TACAGGTGGCTGAGCAGTTG | CTTCACCAGCAAAGCAATCA | 288 |
| chr12:80739291-80741309 | p2 | (AT)6 | TACAGGTGGCTGAGCAGTTG | CTTCACCAGCAAAGCAATCA | 288 |
| chr12:80739449-80741467 | p6 | (GGTGAC)3 | TACAGGTGGCTGAGCAGTTG | CTTCACCAGCAAAGCAATCA | 130 |
| chr12:80739449-80741467 | p2 | (AT)6 | TACAGGTGGCTGAGCAGTTG | CTTCACCAGCAAAGCAATCA | 130 |
| chr12:80739992-80742004 | p6 | (GGTGAC)3 | GAGGTGGTGACTGTCGAGGT | TACTTGCCTCATTCTTGGGG | 261 |
| chr12:80739992-80742004 | p2 | (AT)6 | GAGGTGGTGACTGTCGAGGT | TACTTGCCTCATTCTTGGGG | 261 |
| chr12:80795045-80797072 | p2 | (TA)6 | CACGAAAATCATGGGCTTCT | ACGTGGAGTCTGGTACCTCG | 195 |
| chr12:80833081-80835096 | p5 | (AAATA)3 | TGTCCAGATCCCAGAGGTTC | TGTCACGTCAGCATTTAGGG | 107 |
| chr12:80890201-80892221 | p5 | (AACCG)4 | GGTGTTGATGGAGAGGAGGA | AGATGGCAGCTTGTTGGTCT | 155 |
| chr12:80912200-80914212 | p2 | (AT)6 | ACTTGGCTTGCCACGTAAAT | CTCGGAAAGCGAAAAGTGTC | 194 |
| chr12:80912200-80914212 | p2 | (AT)8 | ACTTGGCTTGCCACGTAAAT | CTCGGAAAGCGAAAAGTGTC | 194 |
| chr12:80941738-80943762 | p2 | (TA)12 | TGACTGGAGAGAAAGGGGAA | TTGTCATCCAAGCAATCCAA | 288 |
| chr12:80964438-80966555 | p2 | (AT)15 | GTTCTTGGGGAATTTCGTGA | GTCTGTCACCCGCTCTTCTC | 207 |
| chr12:81051240-81053254 | p2 | (AT)7 | CACATCTGTCACGACCCAAC | ATCTGTGATGGCCTTATCGG | 231 |
| chr12:81059801-81061816 | p3 | (TCC)5 | AATGGATTTGGGCCACACTA | TAATGCTAAGACCTCGGGGA | 298 |
| chr12:81118601-81120667 | p2 | (TA)6 | AGTATCAACCGTGGCTTTGG | TCTTCTCTACAGCCTTCGGC | 163 |
| chr12:81185620-81187636 | p2 | (AT)8 | GTGTGTTGTGGGCTTGTTTG | CACAGTTGAACCAGCCATTG | 211 |
| chr12:81199241-81201271 | p2 | (TA)7 | TGAGAAGAAACAGCAGCGAA | TGGAAGTTGAGACATCTGCG | 261 |
| chr12:81199241-81201271 | p2 | (TC)7 | TGAGAAGAAACAGCAGCGAA | TGGAAGTTGAGACATCTGCG | 261 |
| chr12:81353810-81355846 | p3 | (TTA)12 | GCCAAAGGCATCATTCTCAT | GTTCGCTGCCGAAGAAATAG | 182 |
| chr12:81540636-81542651 | p5 | (CAAGC)3 | GCAGATGGAGAAGCTAACGG | GTTTCCAGACTCCCCCTTTC | 148 |
| chr12:81542153-81544168 | p5 | (AAAAT)3 | TCGAAAATAACTGATCGGGC | TCTACGCTGCCAAAAGGAAT | 120 |
| chr12:81542153-81544168 | p5 | (TAAAA)3 | TCGAAAATAACTGATCGGGC | TCTACGCTGCCAAAAGGAAT | 120 |
| chr12:81551752-81553770 | p3 | (ATA)6 | AACCGGACAATGTCTTCCTG | ACATTCTTGAAGGTGCCCAG | 196 |
| chr12:81598667-81600697 | p2 | (AT)15 | CGTCAGTTTGACTTGTCCGA | TCCGAGCTCGTAGGTTGACT | 171 |
| chr12:81628084-81630100 | p2 | (GA)8 | AGCGAAGTGGAAACAGCCTA | GCAAATTTTGATTTTCCCGA | 107 |
| chr12:81726036-81728051 | p5 | (CATGG)3 | CATCAAATGCCACTACCACG | GGGCCAAATATACCCCTTGT | 212 |
| chr12:81726036-81728051 | p5 | (CAAGG)3 | CATCAAATGCCACTACCACG | GGGCCAAATATACCCCTTGT | 212 |
| chr12:81740831-81742845 | p2 | (AG)7 | TGATATGAAGTTGGCCACGA | GGAAAACATGGCTTGGAAGA | 257 |
| chr12:81801246-81803276 | p5 | (TCGGT)3 | CTTGCACTTAGGGTTAGGCG | TTTATTTCGATTCGGTTCGG | 270 |
| chr12:81833458-81835472 | p2 | (TG)7 | AGATCCAGGGTGACTTCGTG | TGGGAAGGGACGATAACAAA | 216 |

| Marker ID | SSR type | SSR motif | FORWARD PRIMER (5'-3') | REVERSE PRIMER (5'-3') | size (bp) |
|---|---|---|---|---|---|
| chr12:82005261-82007276 | p6 | (GGTGGA)3 | TTGGTGGTATGCTGACCTGA | GGGGAGAAAGGGAAAAGTTG | 176 |
| chr12:82005261-82007276 | p3 | (GAG)5 | TTGGTGGTATGCTGACCTGA | GGGGAGAAAGGGAAAAGTTG | 176 |
| chr12:82057804-82059822 | p6 | (ATTTTT)3 | GGCCTCATTTGTTTGCACTT | GTTGGAGTGGAGAGTGGGTG | 263 |
| chr12:82113130-82115142 | p2 | (AT)6 | ACTCGGTCTGTCCCATAACG | TTAAATGTCGTGCCCAGTCA | 192 |
| chr12:82201734-82203749 | p5 | (AGGAG)3 | CGAAATCTGGCTAAGCCTTG | CACTCGGTGACACCAACAAC | 262 |
| chr12:82266408-82268423 | p5 | (AAAAT)3 | GAAATTCGCTGGAAAAACGA | TTTCTTTCGATCCACCATCC | 271 |
| chr12:82294687-82296707 | p2 | (AT)10 | AGTCCAAAGTGAACGGATGG | CATGACCAACGTAACATGCC | 149 |
| chr12:82340111-82342129 | p3 | (TTG)6 | CGTACACTCTACCCTCCCCA | ATTTGCAGGATTGTCCTTCG | 240 |
| chr12:82340111-82342129 | p6 | (TAAAAT)3 | CGTACACTCTACCCTCCCCA | ATTTGCAGGATTGTCCTTCG | 240 |
| chr12:82379827-82381839 | p2 | (AT)6 | GAAGCATGGGGAAGAACGTA | ATGCCCACCAAGTGTTTGAT | 294 |
| chr12:82379827-82381839 | p2 | (AT)6 | GAAGCATGGGGAAGAACGTA | ATGCCCACCAAGTGTTTGAT | 294 |
| chr12:82392763-82394785 | p3 | (CCT)5 | CAGTTTCCGGGCGATACTTA | AAAATTGACTGGCACCGAAG | 282 |
| chr12:82392993-82395008 | p3 | (CCT)5 | TTGCAAATTCGTTCTCATCG | TCCTGGGAGAAGCAGAGAAA | 200 |
| chr12:82392993-82395008 | p2 | (TA)8 | TTGCAAATTCGTTCTCATCG | TCCTGGGAGAAGCAGAGAAA | 200 |
| chr12:82471386-82473400 | p2 | (AT)6 | ATTCTCTGGGATTCCTGCAA | TACAAGACGGGAATCGAACC | 130 |
| chr12:82471386-82473400 | p2 | (TA)7 | ATTCTCTGGGATTCCTGCAA | TACAAGACGGGAATCGAACC | 130 |
| chr12:82490620-82492635 | p5 | (GGGTT)3 | CCTCCATGTCCACCATCTCT | TTGGCATAAATTCTGGCTCC | 140 |
| chr12:82496349-82498364 | p5 | (TATTG)3 | TTGGCAAATAAAGGACCCAG | CCATCCTTGTCTGCTTGGAT | 286 |
| chr12:82706214-82708234 | p2 | (CT)10 | GCCATGTCAAAGTAGCCCAT | TACTTTCCATGGGACTTGGC | 295 |
| chr12:82738890-82740906 | p2 | (AT)8 | TTGTGGAGCCCAAAAAGAAC | TGAAATCCCGACTTTTGACC | 102 |
| chr12:82784812-82786824 | p2 | (TA)6 | AAATGACATGGCGGAATAGC | AACAAAAAGTGCACCCCCTA | 113 |
| chr12:82784812-82786824 | p2 | (AT)13 | AAATGACATGGCGGAATAGC | AACAAAAAGTGCACCCCCTA | 113 |
| chr12:82830008-82832022 | p2 | (TA)7 | CAACCCATTAGTCGAAGCAAA | GACATGGCGGAATACCAAAT | 173 |
| chr12:82869225-82871241 | p5 | (TTTTA)3 | CTCCCAAAGACTTCAGCAGC | GACGATGAAATAGGTCCCGA | 152 |
| chr12:82869248-82871260 | p5 | (TTTTA)3 | CTCCCAAAGACTTCAGCAGC | GACGATGAAATAGGTCCCGA | 129 |
| chr12:82870203-82872218 | p5 | (TTTTA)3 | GGCGGAGAAGATGACGTAAA | ACCACCACCAAAAATCCAAA | 124 |
| chr12:82870203-82872218 | p6 | (ACTGAA)3 | GGCGGAGAAGATGACGTAAA | ACCACCACCAAAAATCCAAA | 124 |
| chr12:82905216-82907232 | p5 | (TTTTA)3 | CTCCCAAAGACTTCAGCAGC | GACGATGAAATAGGTCCCGA | 152 |
| chr12:82905239-82907251 | p5 | (TTTTA)3 | CTCCCAAAGACTTCAGCAGC | GACGATGAAATAGGTCCCGA | 129 |
| chr12:82906194-82908209 | p5 | (TTTTA)3 | GGCGGAGAAGATGACGTAAA | ACCACCACCAAAAATCCAAA | 124 |
| chr12:82906194-82908209 | p6 | (ACTGAA)3 | GGCGGAGAAGATGACGTAAA | ACCACCACCAAAAATCCAAA | 124 |
| chr12:82994010-82996028 | p6 | (CATGAG)3 | GCTCCACCCACCTTCTCATA | CACATTGATGCCGAAGAATG | 182 |
| chr12:82994010-82996028 | p4 | (ATAC)4 | GCTCCACCCACCTTCTCATA | CACATTGATGCCGAAGAATG | 182 |
| chr12:83036624-83038640 | p4 | (TTTA)4 | CACGCAAGGAGTAGCTAGGG | CGGAGTAAGGTTTCGGTCAA | 104 |
| chr12:83071219-83073239 | p4 | (TATT)5 | GCCACCCTTATGTCGGAGTA | AGGACGGATTGAGCTCCTCT | 125 |
| chr12:83093837-83095852 | p5 | (ATTTT)3 | CGTGAAATTGTCACAGGTGG | ACATGCTCTTTCACGCACAG | 205 |

| Marker ID | SSR type | SSR motif | FORWARD PRIMER (5'-3') | REVERSE PRIMER (5'-3') | size (bp) |
|---|---|---|---|---|---|
| chr12:83113580-83115592 | p2 | (AT)6 | CAAGTCCTTCTTGATCCCCA | CGTGAAACACAGGCACATCT | 218 |
| chr12:83225958-83227974 | p2 | (TA)6 | CGACCAGAGATCCATCGAAT | TCATTCTTTTGTTCGTTGCG | 170 |
| chr12:83225958-83227974 | p3 | (GTT)8 | CGACCAGAGATCCATCGAAT | TCATTCTTTTGTTCGTTGCG | 170 |
| chr12:83225958-83227974 | p2 | (AT)8 | CGACCAGAGATCCATCGAAT | TCATTCTTTTGTTCGTTGCG | 170 |
| chr12:83294675-83296708 | p2 | (GT)6 | CCAATAGTCGCTGGACAACA | GCTCATGAACAAAGACGCAA | 191 |
| chr12:83302456-83304471 | p3 | (TCT)7 | TTTTCCAATGGCTTAGGTGG | AAATGGCTGGTCCAAAACTG | 131 |
| chr12:83302456-83304471 | p5 | (AAATA)3 | TTTTCCAATGGCTTAGGTGG | AAATGGCTGGTCCAAAACTG | 131 |
| chr12:83303505-83305520 | p5 | (TTTGA)3 | TTTTTGTCCTTCAAATGGGC | GATTGGTCTTTGCCCTTCAA | 254 |
| chr12:83401098-83403114 | p2 | (AT)9 | GTCGTCTCAAAACTGTCGCA | CGTTCTCATTCGTAGTGGCA | 220 |
| chr12:83401098-83403114 | p2 | (TA)8 | GTCGTCTCAAAACTGTCGCA | CGTTCTCATTCGTAGTGGCA | 220 |
| chr12:83545320-83547356 | p3 | (TTA)12 | TTTGGTAATGGTTCCCCTCA | ATCCGGTGACCCATCAAATA | 214 |
| chr12:83613947-83615963 | p2 | (CT)8 | CAGGTGACCCGAAACGTACT | TATTATGGGGGTCAAACGGA | 120 |
| chr12:83659698-83661736 | p2 | (AG)9 | CCAAATAGGAACCGGTCTGA | GTTCCTCCCTCTCCCTCATC | 109 |
| chr12:83659698-83661736 | p2 | (AT)9 | CCAAATAGGAACCGGTCTGA | GTTCCTCCCTCTCCCTCATC | 109 |
| chr12:83659698-83661736 | p2 | (CT)19 | CCAAATAGGAACCGGTCTGA | GTTCCTCCCTCTCCCTCATC | 109 |
| chr12:83773942-83775954 | p2 | (TA)6 | AATTCACGAAAGTCATGGGC | GTATCGTCCTAACTGGCGGA | 131 |
| chr12:83788722-83790744 | p2 | (TA)11 | TTCCTGTAGCCGGATCATTC | AGATACGTGGCTTCCTGGTG | 283 |
| chr12:83788722-83790744 | p2 | (TA)7 | TTCCTGTAGCCGGATCATTC | AGATACGTGGCTTCCTGGTG | 283 |
| chr12:83864165-83866177 | p2 | (TA)11 | CCGTCACGACCATGTTACAG | ACAATACCAACAAGGCCGAC | 100 |
| chr12:83884897-83886911 | p2 | (TG)7 | TATATCCCCCGCATTTGGTA | TACCGCGAATTCCCTCTATG | 291 |
| chr12:83898828-83900840 | p2 | (TA)6 | AACAATGACGTGCCAGACAA | CCAAACGGGCTCATAGAAAA | 139 |
| chr12:83945379-83947391 | p5 | (ATTTA)3 | CAAATGCACATTCACGTTCC | ATGAACATCTTCCCAGCCAC | 147 |
| chr12:83945379-83947391 | p2 | (TA)6 | CAAATGCACATTCACGTTCC | ATGAACATCTTCCCAGCCAC | 147 |
| chr12:83978907-83980925 | p3 | (AAC)6 | AGAATTAAAACGGCGCATTG | AAACGCCAACCATTCAGAAG | 195 |
| chr12:83978907-83980925 | p5 | (AACAA)3 | AGAATTAAAACGGCGCATTG | AAACGCCAACCATTCAGAAG | 195 |
| chr12:84032288-84034303 | p3 | (ACA)5 | ATATCCCATGCTTGCTCCAC | TTGGTGTGCGGTCACATACT | 154 |
| chr12:84045836-84047851 | p5 | (TAGTT)3 | CCTCCAGAGTCAGAATGGGA | GCGATGCCAGGAATACACTT | 118 |
| chr12:84045836-84047851 | p5 | (AATGA)3 | CCTCCAGAGTCAGAATGGGA | GCGATGCCAGGAATACACTT | 118 |
| chr12:84158098-84160124 | p2 | (TA)13 | GTATGATGTGCCATGTTCGC | GCTGGTGCCACAATACAGAG | 179 |
| chr12:84212715-84214729 | p2 | (AT)7 | ATGGGTGGAAGTGGATGAAG | CGGTCTAACCGGAAAAGTCA | 290 |
| chr12:84212715-84214729 | p2 | (TA)7 | ATGGGTGGAAGTGGATGAAG | CGGTCTAACCGGAAAAGTCA | 290 |
| chr12:84212715-84214729 | p2 | (GA)6 | ATGGGTGGAAGTGGATGAAG | CGGTCTAACCGGAAAAGTCA | 290 |
| chr12:84219718-84221730 | p2 | (TA)12 | GGAACCACATGGCTGAAGTT | ATTCACCAAATTCTTGCCCA | 279 |
| chr12:84219718-84221730 | p2 | (TA)13 | GGAACCACATGGCTGAAGTT | ATTCACCAAATTCTTGCCCA | 279 |
| chr12:84219756-84221768 | p2 | (TA)12 | GGAACCACATGGCTGAAGTT | ATTCACCAAATTCTTGCCCA | 241 |
| chr12:84219756-84221768 | p2 | (TA)13 | GGAACCACATGGCTGAAGTT | ATTCACCAAATTCTTGCCCA | 241 |

| Marker ID | SSR type | SSR motif | FORWARD PRIMER (5'-3') | REVERSE PRIMER (5'-3') | size (bp) |
|---|---|---|---|---|---|
| chr12:84219862-84221880 | p2 | (TA)12 | GGAACCACATGGCTGAAGTT | ATTCACCAAATTCTTGCCCA | 135 |
| chr12:84219862-84221880 | p2 | (TA)13 | GGAACCACATGGCTGAAGTT | ATTCACCAAATTCTTGCCCA | 135 |
| chr12:84269019-84271037 | p6 | (TTTGAA)3 | GAAATGGCCAAAGATGGAAA | CAAATCTGAAAACTTGGGGC | 271 |
| chr12:84374231-84376257 | p2 | (TA)13 | AAATATCAACCGTGGCTTCG | GTTTTGGTTAAGTCGCTCCG | 285 |
| chr12:84374231-84376257 | p2 | (AT)6 | AAATATCAACCGTGGCTTCG | GTTTTGGTTAAGTCGCTCCG | 285 |
| chr12:84374231-84376257 | p5 | (CCGGT)3 | AAATATCAACCGTGGCTTCG | GTTTTGGTTAAGTCGCTCCG | 285 |
| chr12:84508765-84510779 | p2 | (CT)7 | GTTAATGATTCGTGCGCCTT | CGAGCCAGATCACAGCATAA | 216 |
| chr12:84509552-84511577 | p2 | (CT)7 | CGCACATATCGCCTCCTTAT | AGGTCCGGTGAGAAAGGATT | 281 |
| chr12:84577797-84579811 | p2 | (TA)7 | CACGGTGTTTAATATGGGGC | CCACCACATCCATACTCACG | 235 |
| chr12:84577797-84579811 | p2 | (TA)10 | CACGGTGTTTAATATGGGGC | CCACCACATCCATACTCACG | 235 |
| chr12:84577797-84579811 | p5 | (ATTTT)3 | CACGGTGTTTAATATGGGGC | CCACCACATCCATACTCACG | 235 |
| chr12:84623858-84625872 | p2 | (TA)7 | CAAGCTTGGAACCCGTATGT | CATTACTAGCAGCTTCCCGC | 231 |
| chr12:84637821-84639857 | p6 | (CATATA)6 | GGCTGCCACTACGAAATGAT | TCTTGGAGAGCTCCGTTGTT | 279 |
| chr12:84695531-84697549 | p6 | (ATTTTA)3 | AACACAAAGACTGACGGCCT | TTTATCCATCGGTTTCCGTC | 115 |
| chr12:84706606-84708648 | p2 | (TA)6 | GATCGCATTTAGCCAAGGAA | GCCTTCCTCCGCTTCTAAGT | 211 |
| chr12:84706681-84708737 | p2 | (TA)6 | GATCGCATTTAGCCAAGGAA | GCCTTCCTCCGCTTCTAAGT | 136 |
| chr12:84812423-84814441 | p2 | (CT)6 | AACCATCGGTTTCGTGACTC | TTCATGCTCGTGCTTGTTTC | 266 |
| chr12:84812423-84814441 | p2 | (AT)9 | AACCATCGGTTTCGTGACTC | TTCATGCTCGTGCTTGTTTC | 266 |
| chr12:84812423-84814441 | p2 | (TA)31 | AACCATCGGTTTCGTGACTC | TTCATGCTCGTGCTTGTTTC | 266 |
| chr12:84863147-84865173 | p2 | (AT)13 | AAACTCAGAAGCGGAGCAAG | TTTTGGTTATTTCCCCCACA | 109 |
| chr12:84893143-84895157 | p2 | (AT)12 | TTGAAGATGTCACGACCCAA | CCTCAGGAGACAGGTGCCTA | 102 |
| chr12:84893143-84895157 | p2 | (AT)7 | TTGAAGATGTCACGACCCAA | CCTCAGGAGACAGGTGCCTA | 102 |
| chr12:84953889-84955901 | p2 | (CT)6 | CCCACTATGTCCATGAGCCT | GAGTCCAAACCCGAAAATGA | 253 |
| chr12:85129551-85131566 | p3 | (TGT)5 | TGAGCGAACGTGCTAAGAGA | CATAAAACGCCTCGCCTTAC | 131 |
| chr12:85152485-85154500 | p3 | (CCA)5 | TTGACAGTTGCATCAGAGCC | ATTGAAAGTATGCGCAACCC | 107 |
| chr12:85198057-85200072 | p3 | (GTT)7 | GAGAAATCGGCAAAATGGAA | TATTATGCTTCCTCCACGGC | 176 |
| chr12:85198057-85200072 | p5 | (ATAAA)3 | GAGAAATCGGCAAAATGGAA | TATTATGCTTCCTCCACGGC | 176 |
| chr12:85212629-85214644 | p5 | (AGAGG)3 | TTCCTTTGAGAGCGAGCATT | TTTACCCCGCATCACCTAAG | 108 |
| chr12:85218852-85220867 | p2 | (AT)10 | TGAGGATTGGAAATCGGAAG | CCTCAGGCATCCAAGCTAAG | 230 |
| chr12:85218852-85220867 | p2 | (AT)11 | TGAGGATTGGAAATCGGAAG | CCTCAGGCATCCAAGCTAAG | 230 |
| chr12:85218852-85220867 | p5 | (TTTTA)3 | TGAGGATTGGAAATCGGAAG | CCTCAGGCATCCAAGCTAAG | 230 |
| chr12:85297461-85299479 | p3 | (CAA)6 | TCATTTTGTGAAGCATGGGA | CTTCCACCAGGTGATTTTCG | 175 |
| chr12:85309935-85311950 | p5 | (CCCCA)3 | AAGCCTGACATGTTCCCAAC | GAGCTTTGAAATCGGTCGTC | 213 |
| chr12:85443178-85445199 | p3 | (TTC)7 | GCATAATAAAGTGGCACCCG | GCCGGATTGGAGAACTTACA | 129 |
| chr12:85443178-85445199 | p3 | (AAT)5 | GCATAATAAAGTGGCACCCG | GCCGGATTGGAGAACTTACA | 129 |
| chr12:85538770-85540785 | p5 | (AAACC)3 | CGTCATGAGCTCCGACACTA | TCCTACCTGTTCTCCCAACG | 249 |

| Marker ID | SSR type | SSR motif | FORWARD PRIMER (5'-3') | REVERSE PRIMER (5'-3') | size (bp) |
|---|---|---|---|---|---|
| chr12:85538810-85540825 | p5 | (AAACC)3 | CGTCATGAGCTCCGACACTA | TCCTACCTGTTCTCCCAACG | 209 |
| chr12:85538810-85540825 | p2 | (AT)10 | CGTCATGAGCTCCGACACTA | TCCTACCTGTTCTCCCAACG | 209 |
| chr12:85569193-85571208 | p5 | (TTTTC)3 | CCCTATATGAATCCGCCGTA | ATGCCCTTATTTTGTGCTGG | 225 |
| chr12:85591121-85593136 | p5 | (TTTTC)3 | CCCTATATGAATCCGCCGTA | ATGCCCTTATTTTGTGCTGG | 225 |
| chr12:85619835-85621853 | p6 | (TTTTTC)3 | AGAACTTAGCCGCCACTTGA | GTGAACTAGGACGTCAGGGC | 160 |
| chr12:85620842-85622860 | p3 | (TAT)6 | TTGCAGAATATCGCTACCCC | AAAAAGAAGAAACGGGCCAT | 111 |
| chr12:85635032-85637048 | p4 | (TTAT)4 | CACCACCACGATTCTTCCTT | TTCATTGAACCTGGACGTGA | 127 |
| chr12:85675298-85677331 | p3 | (ATA)11 | TTTTCCAAAAGCATGCACAA | AGAGGTTGGAAGATGGAGGG | 143 |
| chr12:85690889-85692910 | p3 | (CAA)7 | CTCGGCGGAAGAAGTAACAG | CATAAAGGCTCGGATTGCTC | 258 |
| chr12:85705200-85707228 | p2 | (TA)14 | CACTGCTTTATCGCATTCCA | AGCATAGAGGGGGAGGTCAT | 207 |
| chr12:85749363-85751377 | p2 | (TA)8 | TGTGCACAAATCGGTTCAAT | AATTTGGTTTGTTCGGATCG | 275 |
| chr12:85749363-85751377 | p2 | (AT)7 | TGTGCACAAATCGGTTCAAT | AATTTGGTTTGTTCGGATCG | 275 |
| chr12:85756659-85758701 | p2 | (TA)21 | ATGTGAAACCCCAAAAACCA | TTATCGAGGCCACCTTATCG | 191 |
| chr12:85782240-85784255 | p5 | (GTATT)3 | CTTTCCGTGTTCTATGGGGA | GCATTCTTCTTTCAGCCCAC | 274 |
| chr12:85789916-85791934 | p2 | (TA)9 | CAGGCCAAGAAGTTGGGTAA | GGCCAAAATAAAGACCAGCA | 209 |
| chr12:85789916-85791934 | p5 | (TTTAT)4 | CAGGCCAAGAAGTTGGGTAA | GGCCAAAATAAAGACCAGCA | 209 |
| chr12:85857649-85859667 | p2 | (TA)9 | CAGGCCAAGAAGTTGGGTAA | GGCCAAAATAAAGACCAGCA | 209 |
| chr12:85857649-85859667 | p5 | (TTTAT)4 | CAGGCCAAGAAGTTGGGTAA | GGCCAAAATAAAGACCAGCA | 209 |
| chr12:85976436-85978451 | p5 | (TATTT)3 | TGGTGTCACTGGGCTGAATA | CTCGAAACAAGGGACTGCTC | 148 |
| chr12:86013749-86015764 | p3 | (AAG)5 | TTTGACGTCTCCCATAGGCT | TGCAGAATTGTTTCAGACGC | 214 |
| chr12:86060173-86062189 | p2 | (AG)8 | GGCACCTATGTTCCTTCCAA | GACCTGTATGCTCCCCAAAA | 209 |
| chr12:86078484-86080499 | p3 | (ATT)5 | AGAGTAGGTCATTGGCGGTG | TTACAATCGAAAAGGTGGGC | 286 |
| chr12:86111982-86113997 | p5 | (TTGGT)3 | TTTCTTCGCCGCTAATGACT | ATCATTACCAGCAACCGGAC | 195 |
| chr12:86130140-86132158 | p3 | (CAA)6 | ACGAGCGGAGTAGTGAGGAA | GAAAAGCTCGGATTGCTTTG | 283 |
| chr12:86245900-86247915 | p3 | (TTC)5 | TGTTCATCCCAAACAGGGTT | CGAAAACCTCAACCCAAAAA | 257 |
| chr12:86283979-86285991 | p2 | (TA)6 | CTTTTCCGCACAATGGAAAT | GAGGTGCCAAATGGTGAGAT | 234 |
| chr12:86283979-86285991 | p2 | (AT)6 | CTTTTCCGCACAATGGAAAT | GAGGTGCCAAATGGTGAGAT | 234 |
| chr12:86283979-86285991 | p6 | (CCGGTT)3 | CTTTTCCGCACAATGGAAAT | GAGGTGCCAAATGGTGAGAT | 234 |
| chr12:86332132-86334147 | p6 | (TTTTTG)3 | TTGCAGGACAGTCCATTTCA | CTAACGTGAACGCACGAGAA | 124 |
| chr12:86332132-86334147 | p5 | (TGTCA)3 | TTGCAGGACAGTCCATTTCA | CTAACGTGAACGCACGAGAA | 124 |
| chr12:86444426-86446460 | p2 | (TA)17 | GAACCACCTGACTCATCCGT | GCTCCACCGGTTCATATTGT | 229 |
| chr12:86517429-86519447 | p6 | (TATATG)3 | AGATTGAGGAAGAGCGGACA | ATGGAATTACCCCGAAGTCC | 286 |
| chr12:86517429-86519447 | p2 | (TA)8 | AGATTGAGGAAGAGCGGACA | ATGGAATTACCCCGAAGTCC | 286 |
| chr12:86517614-86519630 | p6 | (TATATG)3 | AGATTGAGGAAGAGCGGACA | ATGGAATTACCCCGAAGTCC | 101 |
| chr12:86517614-86519630 | p2 | (TA)8 | AGATTGAGGAAGAGCGGACA | ATGGAATTACCCCGAAGTCC | 101 |
| chr12:86530755-86532770 | p3 | (CTA)5 | CCCATCCAAACAGGCTCTAA | ACCTGTTGCTGCTGCTTCTT | 265 |

| Marker ID | SSR type | SSR motif | FORWARD PRIMER (5'-3') | REVERSE PRIMER (5'-3') | size (bp) |
|---|---|---|---|---|---|
| chr12:86546915-86548930 | p5 | (GAAAA)4 | AATCATCAACACAGCCTCCC | AAGGAAGCTTCGGAAAAAGC | 141 |
| chr12:86546915-86548930 | p5 | (TAAAC)3 | AATCATCAACACAGCCTCCC | AAGGAAGCTTCGGAAAAAGC | 141 |
| chr12:86566372-86568384 | p2 | (TA)6 | AAAGCCATGCATGATATCCC | AAAATTGCCAAGTGGCTTGT | 224 |
| chr12:86566372-86568384 | p2 | (AC)6 | AAAGCCATGCATGATATCCC | AAAATTGCCAAGTGGCTTGT | 224 |
| chr12:86612592-86614610 | p3 | (CAA)6 | CGAGTGGCTCGGTAAATCAT | CATGACTCTGTTCGTGCGTT | 197 |
| chr12:86805603-86807631 | p2 | (AT)7 | GAATCCCCACTCAGGCATAA | TGCCCCATAAGGCATAGTTC | 123 |
| chr12:86805603-86807631 | p2 | (AT)14 | GAATCCCCACTCAGGCATAA | TGCCCCATAAGGCATAGTTC | 123 |
| chr12:86805603-86807631 | p2 | (TA)32 | GAATCCCCACTCAGGCATAA | TGCCCCATAAGGCATAGTTC | 123 |
| chr12:86851104-86853122 | p2 | (TA)9 | CTTTCATCCCCGTGTCCTAA | TTGGTGCAACCAATGGAGTA | 251 |
| chr12:86877829-86879847 | p3 | (TTA)6 | CTGTTTGGGGTCACGCTATT | GACAATCTGCAAATCGAGCA | 264 |
| chr12:86877950-86879965 | p3 | (TTA)6 | CTGTTTGGGGTCACGCTATT | TGACAATCTGCAAATCGAGC | 143 |
| chr12:86877987-86880002 | p3 | (TTA)6 | CTGTTTGGGGTCACGCTATT | TGACAATCTGCAAATCGAGC | 106 |
| chr12:86937296-86939311 | p3 | (CCA)5 | AAAATGGCGTGCAATACACA | GAGCAAACTTTCACGCACAA | 213 |
| chr12:86937296-86939311 | p5 | (TTATT)3 | AAAATGGCGTGCAATACACA | GAGCAAACTTTCACGCACAA | 213 |
| chr12:86937296-86939311 | p5 | (TTATT)3 | AAAATGGCGTGCAATACACA | GAGCAAACTTTCACGCACAA | 213 |
| chr12:86937296-86939311 | p5 | (TTATT)3 | AAAATGGCGTGCAATACACA | GAGCAAACTTTCACGCACAA | 213 |
| chr12:86974465-86976495 | p2 | (AT)7 | TTTCGTTTCCGCACTTTTCT | GCACCTGTCGAGCTTCTACC | 191 |
| chr12:86974465-86976495 | p2 | (CT)15 | TTTCGTTTCCGCACTTTTCT | GCACCTGTCGAGCTTCTACC | 191 |
| chr12:87022123-87024141 | p5 | (CCCCA)3 | GAGAGAGAAATGGGGAAGGG | GGATGGTTTTGAGGCGTTTA | 145 |
| chr12:87022123-87024141 | p2 | (TA)9 | GAGAGAGAAATGGGGAAGGG | GGATGGTTTTGAGGCGTTTA | 145 |
| chr12:87024852-87026867 | p2 | (TA)11 | TGTTGCTCCCTCTAATGGGT | CACCACATTCAAACTCACGG | 121 |
| chr12:87024852-87026867 | p5 | (AAATT)3 | TGTTGCTCCCTCTAATGGGT | CACCACATTCAAACTCACGG | 121 |
| chr12:87024852-87026867 | p2 | (AT)6 | TGTTGCTCCCTCTAATGGGT | CACCACATTCAAACTCACGG | 121 |
| chr12:87338226-87340241 | p3 | (TCA)6 | GGGGGTTGTGTTTCATCATC | GTTAGGGGTGCGCTTAATGA | 202 |
| chr12:87338226-87340241 | p5 | (TTTTA)3 | GGGGGTTGTGTTTCATCATC | GTTAGGGGTGCGCTTAATGA | 202 |
| chr12:87343229-87345244 | p3 | (ATA)5 | TGAATTTTGGTGCAAAGCAG | TTGCTCCTACCCATTTGTCC | 134 |
| chr12:87444534-87446549 | p5 | (TTTTA)3 | TCCTTCGTTCTAGTGTGGGG | TTGCAACTTTTGGAACAACG | 299 |
| chr12:87485980-87487996 | p4 | (AAAG)4 | CAAATCCTGCTGATGTGGAA | TCTCTTGAAAACGACCCTGG | 232 |
| chr12:87485980-87487996 | p5 | (TTGGT)3 | CAAATCCTGCTGATGTGGAA | TCTCTTGAAAACGACCCTGG | 232 |
| chr12:87533913-87535928 | p5 | (ATTTT)3 | TAGCGTTCGATTTGTTGCTG | ACTCAAGCCACGTCGATTCT | 209 |
| chr12:87577057-87579081 | p4 | (TATT)6 | ATGAGCGATTCTCAGCTGGT | TTCATCTTCTTCTCGCGGTT | 208 |
| chr12:87609132-87611144 | p2 | (AT)6 | GGGGTCGTTTGGTAAAAGGT | TGGTGCAAGGGATGTTTGTA | 250 |
| chr12:87610721-87612751 | p2 | (TA)15 | CCAAAAGGATTGGCCTCATA | GGATGTGGGTGAGTTCCACT | 189 |
| chr12:87628753-87630771 | p3 | (TCA)6 | ACCGCGCTTTTGTTGTTAGT | TCTTACCGCTGATTTGGGTC | 202 |
| chr12:87717084-87719102 | p3 | (GGT)6 | TTGCAGCCTTTGGAAGAACT | GATTGCGTGGGGATACATTC | 159 |
| chr12:87722246-87724264 | p6 | (TTGATT)3 | CACCCTCAACTTCTTCCTGC | CCAAGTGGAAAAGAAGCAGC | 247 |

| Marker ID | SSR type | SSR motif | FORWARD PRIMER (5'-3') | REVERSE PRIMER (5'-3') | size (bp) |
|---|---|---|---|---|---|
| chr12:87722246-87724264 | p3 | (TGG)6 | CACCCTCAACTTCTTCCTGC | CCAAGTGGAAAAGAAGCAGC | 247 |
| chr12:87745407-87747423 | p2 | (AT)8 | ATAACAATGCGCCAGGGATA | GGACGAAGCATGGGTTTCTA | 280 |
| chr12:87785233-87787257 | p2 | (AT)12 | CACAAGTTTTGGTTAGGCGG | GGCAGGATGCTGATGAAAAT | 187 |
| chr12:87785233-87787257 | p2 | (AT)6 | CACAAGTTTTGGTTAGGCGG | GGCAGGATGCTGATGAAAAT | 187 |
| chr12:87785233-87787257 | p2 | (AT)13 | CACAAGTTTTGGTTAGGCGG | GGCAGGATGCTGATGAAAAT | 187 |
| chr12:87785233-87787257 | p2 | (AT)12 | CACAAGTTTTGGTTAGGCGG | GGCAGGATGCTGATGAAAAT | 187 |
| chr12:87785393-87787405 | p2 | (AT)12 | ATTTTCATCAGCATCCTGCC | TTGGCTAGGCAAACATGAGA | 221 |
| chr12:87785393-87787405 | p2 | (AT)6 | ATTTTCATCAGCATCCTGCC | TTGGCTAGGCAAACATGAGA | 221 |
| chr12:87785393-87787405 | p2 | (AT)13 | ATTTTCATCAGCATCCTGCC | TTGGCTAGGCAAACATGAGA | 221 |
| chr12:87785393-87787405 | p2 | (AT)12 | ATTTTCATCAGCATCCTGCC | TTGGCTAGGCAAACATGAGA | 221 |
| chr12:87785599-87787625 | p2 | (AT)12 | TCCGGAGCCAAAATGACTTA | TCGTGAATTGGACAACGAAA | 285 |
| chr12:87785599-87787625 | p2 | (AT)6 | TCCGGAGCCAAAATGACTTA | TCGTGAATTGGACAACGAAA | 285 |
| chr12:87785599-87787625 | p2 | (AT)13 | TCCGGAGCCAAAATGACTTA | TCGTGAATTGGACAACGAAA | 285 |
| chr12:87785599-87787625 | p2 | (AT)12 | TCCGGAGCCAAAATGACTTA | TCGTGAATTGGACAACGAAA | 285 |
| chr12:87809527-87811542 | p3 | (TTA)5 | AGGCCCCGTCATACCTTTAT | CAAACAGACGATCCAGCTCA | 122 |
| chr12:87809527-87811542 | p3 | (TAT)7 | AGGCCCCGTCATACCTTTAT | CAAACAGACGATCCAGCTCA | 122 |
| chr12:87942572-87944590 | p2 | (AT)9 | AGCGATCATAGGATGTTGGG | TACGGTTCAGTGTCCCCTTC | 217 |
| chr12:87970667-87972695 | p2 | (GT)6 | GCATAATGCAAACAAACCCC | ACACCTCGACTAATCCACCG | 144 |
| chr12:87978747-87980762 | p5 | (TTTTA)3 | TCCCTTCTTAACCACCATGC | TTACCCAATTTCACCCCAAA | 161 |
| chr12:88109772-88111784 | p2 | (AT)6 | ATCGTCGTCGCTTTGAATTT | TTTCTCTCCAGCTACCCGAA | 229 |
| chr12:88129514-88131529 | p5 | (ATAAT)3 | TGGAGGGTGCACAATTTCTT | CGCCCCATATTAAAACCATC | 118 |
| chr12:88383998-88386012 | p2 | (AT)7 | CAAGGATGAGCAGCAACAGA | AGTGGACATGGGTTTGGAAG | 193 |
| chr12:88479908-88481920 | p2 | (TA)6 | CCACCCGCAGAAAATAAGAA | AGAGTTGTTCGCTGTTGGCT | 276 |
| chr12:88559805-88561823 | p3 | (ACA)6 | CCCCTTTCAAATGTGCCTTA | ACCGACCTTACATCACTCCG | 275 |
| chr12:88606998-88609012 | p2 | (AT)7 | AATGCGGAACCAGCATAAAC | TTAAATTTTCCCCTTCGGCT | 121 |
| chr12:88669585-88671603 | p2 | (TA)7 | AGGCCTAAGCCTTGCTTTTC | AAAAGCACATTGAGGATCGG | 190 |
| chr12:88669585-88671603 | p2 | (TA)7 | AGGCCTAAGCCTTGCTTTTC | AAAAGCACATTGAGGATCGG | 190 |
| chr12:88669585-88671603 | p6 | (TACGAC)3 | AGGCCTAAGCCTTGCTTTTC | AAAAGCACATTGAGGATCGG | 190 |
| chr12:88673170-88675182 | p2 | (GT)6 | CTTCCACCCTCCCTTTCTTC | ATGGTTGTCGTTTTCGAAGC | 289 |
| chr12:88711292-88713306 | p2 | (AT)7 | TATTCTGCTTGGTGCCTCCT | CCTTCGCCTTCTTCTTTGTG | 165 |
| chr12:88754106-88756132 | p2 | (AT)13 | CCACGATTTATGCACGTCAC | ATGGAGAGAAGCGAAATGGA | 215 |
| chr12:88796851-88798871 | p5 | (TACAA)4 | TGACCTAGGCTCCTTCCTGA | AAAGGGAGATTGCATTCGTG | 258 |
| chr12:88855134-88857154 | p2 | (CT)10 | AGGTGAGAGTACCGGTGGTG | ATCAACGCCACAATCTCCTC | 235 |
| chr12:88859605-88861693 | p2 | (AT)44 | GTTGACGGAAAACCCAAGAA | CCGGGTAAGCCTGTGATATG | 274 |
| chr12:88875401-88877416 | p5 | (TTCAT)3 | GGAGGAACTGGACCAGATGA | AATCCCGACGCATGTATTTC | 100 |
| chr12:88903930-88905944 | p2 | (AG)7 | TGGATGGGAAAGTGAAGGAG | AATGCTTTCCGTTCAACGAC | 251 |

| Marker ID | SSR type | SSR motif | FORWARD PRIMER (5'-3') | REVERSE PRIMER (5'-3') | size (bp) |
|---|---|---|---|---|---|
| chr12:88937864-88939880 | p4 | (TGTT)4 | TTCTTGTACTCGGGGAGTGG | TGCGTACAAGGCATAAAGCA | 115 |
| chr12:88947210-88949228 | p6 | (TTGGAG)3 | CGAACCCCAAACCTATGATG | AAGTTGCACGTGAGCTTCCT | 115 |
| chr12:88962385-88964397 | p2 | (TA)7 | GATAAGCCATCCAAACAGGC | TTCCGATTGGAACCTCAGTT | 280 |
| chr12:88962385-88964397 | p2 | (TA)6 | GATAAGCCATCCAAACAGGC | TTCCGATTGGAACCTCAGTT | 280 |
| chr12:88962385-88964397 | p2 | (TA)9 | GATAAGCCATCCAAACAGGC | TTCCGATTGGAACCTCAGTT | 280 |
| chr12:88962385-88964397 | p2 | (TA)7 | GATAAGCCATCCAAACAGGC | TTCCGATTGGAACCTCAGTT | 280 |
| chr12:88962385-88964397 | p2 | (TA)7 | GATAAGCCATCCAAACAGGC | TTCCGATTGGAACCTCAGTT | 280 |
| chr12:89045514-89047528 | p2 | (AT)7 | TTTGACTGAAAAGGGGTTCG | CCGTGTCTATGGGCTGAAAT | 252 |
| chr12:89124018-89126033 | p5 | (CAAAC)3 | GATCTCAAACTTTGGGGCAG | CTGCAACTCAAGCACATGGT | 266 |
| chr12:89133245-89135263 | p3 | (TGT)6 | CAAGGAAATGGGGAGATTGA | CAGCATTCTCATTTCTGCCA | 182 |
| chr12:89133245-89135263 | p3 | (TTG)5 | CAAGGAAATGGGGAGATTGA | CAGCATTCTCATTTCTGCCA | 182 |
| chr12:89172602-89174617 | p5 | (AAAAT)3 | TGAGCTCTCAGACCAGCAAA | TAAAGAATCGGAATGACGGC | 262 |
| chr12:89172602-89174617 | p3 | (AAC)6 | TGAGCTCTCAGACCAGCAAA | TAAAGAATCGGAATGACGGC | 262 |
| chr12:89206242-89208257 | p5 | (CAAAC)3 | GATCTCAAACTTTGGGGCAG | CTGCAACTCAAGCACATGGT | 266 |
| chr12:89215469-89217487 | p3 | (TGT)6 | CAAGGAAATGGGGAGATTGA | CAGCATTCTCATTTCTGCCA | 182 |
| chr12:89215469-89217487 | p3 | (TTG)5 | CAAGGAAATGGGGAGATTGA | CAGCATTCTCATTTCTGCCA | 182 |
| chr12:89291762-89293782 | p2 | (TA)10 | GGACCGTCATTTAGCTCCAA | TGTTGATGTGCCGTGAATTT | 170 |
| chr12:89315603-89317615 | p2 | (TA)6 | TCTAATGAATGGGGAGTCCG | GCCAAAGTCACTGAATGGGT | 289 |
| chr12:89320447-89322462 | p5 | (GTTTG)3 | ACAAGTTCGAACCCACAACC | AACCCACCCAATTCCCTTAC | 159 |
| chr12:89460677-89462692 | p5 | (TTGGT)3 | CCGACGCAATTGGTTTAGTT | AAACCACCATCCAAACAAGG | 152 |
| chr12:89460677-89462692 | p5 | (TACAA)3 | CCGACGCAATTGGTTTAGTT | AAACCACCATCCAAACAAGG | 152 |
| chr12:89460677-89462692 | p2 | (TA)20 | CCGACGCAATTGGTTTAGTT | AAACCACCATCCAAACAAGG | 152 |
| chr12:89470524-89472536 | p2 | (TA)6 | CCTCAATTCTGCGGAAAAAG | TAGCACCATCACACCACCAT | 102 |
| chr12:89602140-89604158 | p6 | (AAAAAT)3 | TAATAGGCATGGGCTTGGAC | TACTGAAACCCGAAAAACCG | 270 |
| chr12:89688117-89690132 | p5 | (CTTTT)3 | TGGCAACGACCATTTTTGTA | GTTCAAACCGATAACCGGAA | 229 |
| chr12:89744921-89746942 | p3 | (ATT)7 | CACCAAATTCTCCACCTGCT | AGGGCCATTGTTTTCATCAG | 100 |
| chr12:89777069-89779083 | p2 | (AT)7 | TGTGGACATCCGTGAAAAGA | TGGTGATCGGGTAAGGAAAG | 269 |
| chr12:89805897-89807927 | p5 | (CCGGT)3 | GGGCTTTTCGGGTTTAAAAG | CATTCTTGAAAGTGCCGGAT | 140 |
| chr12:89805897-89807927 | p2 | (TA)25 | GGGCTTTTCGGGTTTAAAAG | CATTCTTGAAAGTGCCGGAT | 140 |
| chr12:89888938-89890953 | p5 | (AAAAG)3 | TTAGTGGCGACGTTTTTGTG | CATTGTGTGTCCACTCAGCC | 171 |
| chr12:89900823-89902838 | p5 | (CCCGA)3 | TGTGCGTGAAGCCTACTTTG | TCCCATCTGTAAGTTTGGGC | 152 |
| chr12:90066605-90068623 | p2 | (AT)9 | GGCAAACCTCTGCCTGAATA | CCGCCACGTACTTCATTTTT | 135 |
| chr12:90066605-90068623 | p2 | (AT)15 | GGCAAACCTCTGCCTGAATA | CCGCCACGTACTTCATTTTT | 135 |
| chr12:90079130-90081148 | p2 | (AT)9 | GGCAAACCTCTGCCTGAATA | CCGCCACGTACTTCATTTTT | 135 |
| chr12:90079130-90081148 | p2 | (AT)15 | GGCAAACCTCTGCCTGAATA | CCGCCACGTACTTCATTTTT | 135 |
| chr12:90160519-90162534 | p5 | (TAATT)3 | CTTTGCTGCTGCTTCTTTCC | TCGTTTGACTTAATTGCCCC | 173 |

| Marker ID | SSR type | SSR motif | FORWARD PRIMER (5'-3') | REVERSE PRIMER (5'-3') | size (bp) |
|---|---|---|---|---|---|
| chr12:90216610-90218634 | p2 | (AT)12 | GCTCCACGCCGTACTACTTC | CTAAATGCGCTCTCCGAAAC | 111 |
| chr12:90367803-90369851 | p5 | (AAAAT)3 | ATTTTCTTGCGGTGAGGATG | AGGTTCCGGACTCTGGTTTT | 277 |
| chr12:90367803-90369851 | p2 | (TC)9 | ATTTTCTTGCGGTGAGGATG | AGGTTCCGGACTCTGGTTTT | 277 |
| chr12:90374123-90376138 | p2 | (AT)6 | AATTCCACCGTGCCTAAATG | ATCATCGTTAACCTGCGTCC | 149 |
| chr12:90374123-90376138 | p2 | (TA)15 | AATTCCACCGTGCCTAAATG | ATCATCGTTAACCTGCGTCC | 149 |
| chr12:90374123-90376138 | p3 | (CTT)5 | AATTCCACCGTGCCTAAATG | ATCATCGTTAACCTGCGTCC | 149 |
| chr12:90374123-90376138 | p3 | (TAA)10 | AATTCCACCGTGCCTAAATG | ATCATCGTTAACCTGCGTCC | 149 |
| chr12:90374123-90376138 | p3 | (GGC)5 | AATTCCACCGTGCCTAAATG | ATCATCGTTAACCTGCGTCC | 149 |
| chr12:90374649-90376664 | p2 | (TA)15 | GGGTAGCATGGTCTCACGAT | GAAGGAGGAGGAGGAAGGTG | 221 |
| chr12:90374649-90376664 | p3 | (CTT)5 | GGGTAGCATGGTCTCACGAT | GAAGGAGGAGGAGGAAGGTG | 221 |
| chr12:90374649-90376664 | p3 | (TAA)10 | GGGTAGCATGGTCTCACGAT | GAAGGAGGAGGAGGAAGGTG | 221 |
| chr12:90374649-90376664 | p3 | (GGC)5 | GGGTAGCATGGTCTCACGAT | GAAGGAGGAGGAGGAAGGTG | 221 |
| chr12:90438829-90440844 | p3 | (TCT)5 | AACGAATCCCCATATGACCA | TCGGTTTTGCTAATTCACCC | 295 |
| chr12:90461202-90463217 | p5 | (TTTTC)3 | AGTGACAACAAAGATGGGGC | GCGGCGGAGTTAACATAAAG | 112 |
| chr12:90511300-90513318 | p6 | (TTTTAT)3 | GTGCAAAGGATCGGCTAGAG | TCTTGGCAGCTTCATCTTCA | 177 |
| chr12:90540568-90542586 | p6 | (CAGTAG)3 | TAGTTCCTTGTCCCGATTGC | AGGTTTGGTGATTAGTGGCG | 165 |
| chr12:90560942-90562960 | p2 | (GT)9 | TGCCCATGCCACATAAGATA | TGCTCGAGTTTCGAAGGTTT | 121 |
| chr12:90576280-90578295 | p5 | (TATTT)3 | GCGTACTTCCGCCAAATAAA | AATGTATCCGTGTGCACCAA | 162 |
| chr12:90606929-90608963 | p5 | (CTTTT)3 | ATGGTCGTTAGTCGAGGGTG | GCCTATTCCTTCCACGTCTG | 248 |
| chr12:90606929-90608963 | p2 | (AT)17 | ATGGTCGTTAGTCGAGGGTG | GCCTATTCCTTCCACGTCTG | 248 |
| chr12:90736766-90738778 | p2 | (AT)6 | TGCAAGACCATTTTCGTCAA | CGCACATAGGCCAAGTTTTT | 238 |
| chr12:90788622-90790646 | p3 | (AAT)8 | ATTTTCAGGGGGATTTTTGG | CTTTTCTCTCTCCCGCTCCT | 291 |
| chr12:90860939-90862957 | p3 | (ACA)6 | GCGCCAGATATTATGGAGGA | GCACGTTTCTGGCATAAGGT | 113 |
| chr12:91006283-91008305 | p2 | (AT)11 | AGCTACGAAAAGGTGCCTGA | TCGGGGTCAGATTAAGTTGC | 120 |
| chr12:91038545-91040565 | p3 | (AAC)5 | GGGTTCGAACATGGATATGG | GGTTTTCTGGTAGGTGCTCG | 288 |
| chr12:91038545-91040565 | p2 | (AT)10 | GGGTTCGAACATGGATATGG | GGTTTTCTGGTAGGTGCTCG | 288 |
| chr12:91038545-91040565 | p2 | (AC)12 | GGGTTCGAACATGGATATGG | GGTTTTCTGGTAGGTGCTCG | 288 |
| chr12:91038688-91040712 | p2 | (AT)10 | ATTAATTGGGCTCAAATGCG | CGAACCTGGGAGCTTTTGTA | 226 |
| chr12:91038688-91040712 | p2 | (AC)12 | ATTAATTGGGCTCAAATGCG | CGAACCTGGGAGCTTTTGTA | 226 |
| chr12:91038688-91040712 | p2 | (AT)8 | ATTAATTGGGCTCAAATGCG | CGAACCTGGGAGCTTTTGTA | 226 |
| chr12:91048825-91050839 | p2 | (AT)8 | CGAGCTTCCTGTTCTGAGGT | CCATCCTCGAAGGCATAAAA | 203 |
| chr12:91048825-91050839 | p2 | (TA)7 | CGAGCTTCCTGTTCTGAGGT | CCATCCTCGAAGGCATAAAA | 203 |
| chr12:91114258-91116274 | p2 | (AC)8 | CTGATGACACCAAGCCTTCA | CTGCGCTATAGTGCCTCCTC | 199 |
| chr12:91209699-91211717 | p3 | (AAC)6 | TATAGGCATAAGCCCGTCCA | AAATCATGTCCAAACGGCTC | 205 |
| chr12:91213883-91215904 | p5 | (AAACC)4 | AGAGCGGTTGCTGTTCATCT | GTTTTGTGAGTGCACCATGC | 215 |
| chr12:91213883-91215904 | p3 | (AAC)7 | AGAGCGGTTGCTGTTCATCT | GTTTTGTGAGTGCACCATGC | 215 |

| Marker ID | SSR type | SSR motif | FORWARD PRIMER (5'-3') | REVERSE PRIMER (5'-3') | size (bp) |
|---|---|---|---|---|---|
| chr12:91237073-91239085 | p2 | (AT)6 | CCTAAGTTCATGCCCCTTGA | TAAGGAGAAGCGAGAGGCAA | 236 |
| chr12:91429785-91431801 | p4 | (TATT)4 | GGGACCCTTTCAGATGGTTT | TCAACTCAGTCAGGCTCACG | 300 |
| chr12:91452368-91454382 | p2 | (TA)7 | CTGGTTCACAAAGATCGGCT | AAAGGCCGCAACATAAGCTA | 218 |
| chr12:91462332-91464396 | p3 | (TTG)5 | CATTGTTCCGAAGCCATCTT | TTTAATCGGAGGCATTCCAG | 112 |
| chr12:91545125-91547164 | p3 | (TTG)13 | CATCGTATTGGTGCAGGATG | CTCATTTCCGCCACTTTCAT | 134 |
| chr12:91568338-91570353 | p3 | (TTA)5 | CATTCTAGGCGTGGAAGAGG | GCTCAACTAGGTGCTTTGGC | 188 |
| chr12:91568354-91570372 | p3 | (TTA)5 | CATTCTAGGCGTGGAAGAGG | GCTCAACTAGGTGCTTTGGC | 172 |
| chr12:91595159-91597175 | p4 | (AAAT)4 | AGCGCCCAATCATGCTATAC | GGCCCGACGATAGATCAGTA | 283 |
| chr12:91619638-91621677 | p3 | (TTG)13 | CATCGTATTGGTGCAGGATG | CTCATTTCCGCCACTTTCAT | 134 |
| chr12:91642851-91644866 | p3 | (TTA)5 | CATTCTAGGCGTGGAAGAGG | GCTCAACTAGGTGCTTTGGC | 188 |
| chr12:91642867-91644885 | p3 | (TTA)5 | CATTCTAGGCGTGGAAGAGG | GCTCAACTAGGTGCTTTGGC | 172 |
| chr12:91669672-91671688 | p4 | (AAAT)4 | AGCGCCCAATCATGCTATAC | GGCCCGACGATAGATCAGTA | 283 |
| chr12:91692800-91694852 | p2 | (TA)6 | CCTCTTCAGTCCACTGAGCC | CTCCAGCAAAAAGAGGATGC | 136 |
| chr12:91731797-91733812 | p5 | (AACAA)3 | CTTGCGTGCACCTCAACTAA | TGTGCAAGAAGCCACAAGAC | 270 |
| chr12:91830592-91832606 | p2 | (AT)6 | GTGAGACCCAGGAGATCGAA | GAACCCTTGGAAAGAGTCCC | 134 |
| chr12:91830592-91832606 | p2 | (TA)7 | GTGAGACCCAGGAGATCGAA | GAACCCTTGGAAAGAGTCCC | 134 |
| chr12:91932840-91934855 | p3 | (AAG)5 | CCCCGCTTCTTCTTGTTCTT | CGGCAGATTTGGAAAACAGT | 252 |
| chr12:91932840-91934855 | p5 | (AACTA)3 | CCCCGCTTCTTCTTGTTCTT | CGGCAGATTTGGAAAACAGT | 252 |
| chr12:91965249-91967271 | p2 | (TA)11 | CCACCAGCTATATCCTCCGA | TGGGACACGATTCGTTATGA | 297 |
| chr12:92020308-92022328 | p2 | (TA)10 | GGGCTGTTGTTGCTCAGAAT | CTTGAAAAGGTGGGCGAATA | 198 |
| chr12:92025475-92027495 | p2 | (TA)10 | GGGCTGTTGTTGCTCAGAAT | CTTGAAAAGGTGGGCGAATA | 198 |
| chr12:92141219-92143270 | p5 | (CGGGT)3 | CTGAGACGATTCAACCCGAT | CCACTGCTTCTCTCTCCCAC | 279 |
| chr12:92141219-92143270 | p3 | (ATA)17 | CTGAGACGATTCAACCCGAT | CCACTGCTTCTCTCTCCCAC | 279 |
| chr12:92288006-92290060 | p6 | (AGAAAA)3 | CAACACTTGACGGGTGTGAG | GGGAGAAGTTGCAAAATCCA | 106 |
| chr12:92318368-92320383 | p5 | (TTTTA)3 | TGTTTTGGTGGATCCTGTCA | TTGACTTTTCACCTTTGGGG | 216 |
| chr12:92338276-92340291 | p3 | (TAA)7 | CAGGAGTCAATCGGGCTTTA | GCATTTATTTTCGCGCATTT | 156 |
| chr12:92338276-92340291 | p3 | (TAA)5 | CAGGAGTCAATCGGGCTTTA | GCATTTATTTTCGCGCATTT | 156 |
| chr12:92441944-92443959 | p5 | (GGATC)3 | TTTTCGGTGAACGTGTGGTA | TCCATGACACATCCGTGTTT | 108 |
| chr12:92503851-92505866 | p5 | (AAACA)3 | ATTCGAGCCACGGTTATCAG | CACAACATCCAAAGTGGTCG | 149 |
| chr12:92527240-92529255 | p5 | (TATTT)3 | TCGAAAATTCACGCCCTAAC | AACGAATGAGGAGGTTCAACA | 210 |
| chr12:92579817-92581831 | p2 | (AT)19 | GACGACGAAGCTGGGTTCTA | AATGTCCAAAAGGGAGAGCC | 295 |
| chr12:92579817-92581831 | p5 | (AGTAA)3 | GACGACGAAGCTGGGTTCTA | AATGTCCAAAAGGGAGAGCC | 295 |
| chr12:92615216-92617230 | p2 | (AT)7 | TGGAAGAAGAGGAGCTTGGA | CAAACTGTGACCACATCCCTT | 233 |
| chr12:92638501-92640522 | p3 | (TCT)7 | ACACGGAGTTGTCCTCCAAC | GTTTCATGTGGGGAAGTGCT | 216 |
| chr12:92650930-92652950 | p5 | (GTTTA)4 | ACCACACACGCGAGATATGA | TCTGGTGGAACACGGTATGA | 221 |
| chr12:92658343-92660365 | p6 | (CCCCCA)3 | AAGAAAAGGGGCCGTTAAAA | TAAAAAGGGGTTTGGGTGTG | 273 |

| Marker ID | SSR type | SSR motif | FORWARD PRIMER (5'-3') | REVERSE PRIMER (5'-3') | size (bp) |
|---|---|---|---|---|---|
| chr12:92658437-92660451 | p6 | (CCCCCA)3 | AAGAAAAGGGGCCGTTAAAA | TAAAAAGGGGTTTGGGTGTG | 179 |
| chr12:92659786-92661812 | p2 | (AT)13 | TGACATGGCGGAATAACAAA | GCGTGCGTATCAATCCATTA | 291 |
| chr12:92659786-92661812 | p2 | (TA)9 | TGACATGGCGGAATAACAAA | GCGTGCGTATCAATCCATTA | 291 |
| chr12:92678031-92680046 | p5 | (TACAT)3 | TATTGCGTGTTTTGAGCCAC | GATGAGTTGCTGAGTGGCAA | 203 |
| chr12:92689376-92691394 | p6 | (TTTTTA)3 | CCCTTAGGGGTAAGGTCTGC | CGGTATATGGGGGCTGTATG | 103 |
| chr12:92714056-92716068 | p2 | (TA)6 | GAGTACAAATACCGGGCGAA | TATGAATTCCCTGGTCCCAA | 286 |
| chr12:92757495-92759511 | p2 | (AT)8 | CGGAGTTTGCCTCAGTTAGC | AAGATCCGGTTCTCGGTTTT | 214 |
| chr12:92761176-92763196 | p2 | (GT)10 | GAAATGACCTGGAAAGCCAA | CATTCCGGGGAGTCAGAGTA | 149 |
| chr12:92803813-92805837 | p2 | (AT)12 | AATTGCTACACCCGTTGCTC | AGCTCGAGCCATTCTTGAAA | 186 |
| chr12:93000383-93002398 | p5 | (GAAAA)3 | GAGGTGACGATGAAGAGGGA | AACATTTTCAAATGCCAGCC | 263 |
| chr12:93000383-93002398 | p5 | (AATAA)3 | GAGGTGACGATGAAGAGGGA | AACATTTTCAAATGCCAGCC | 263 |
| chr12:93000383-93002398 | p3 | (GAA)5 | GAGGTGACGATGAAGAGGGA | AACATTTTCAAATGCCAGCC | 263 |
| chr12:93070602-93072618 | p2 | (TA)8 | TTCCGGTCTTCATCATCCTC | TGAGCAGCCACGACAGTAAC | 272 |
| chr12:93118058-93120073 | p5 | (CCAAA)3 | GTTGAACCCGCCAAAATAAA | TCCCAGTGATAAACATGCCA | 108 |
| chr12:93118058-93120073 | p5 | (ATAAA)3 | GTTGAACCCGCCAAAATAAA | TCCCAGTGATAAACATGCCA | 108 |
| chr12:93129368-93131386 | p2 | (TA)9 | TGTAGATTGTGGTGCGGAGA | CTTGCTGGGTGAGTGAAACA | 206 |
| chr12:93233155-93235171 | p2 | (AT)8 | AATTGTTGGCCCCATATTCA | GTTTGCTTCGAATACCGCAT | 147 |
| chr12:93233155-93235171 | p2 | (TA)10 | AATTGTTGGCCCCATATTCA | GTTTGCTTCGAATACCGCAT | 147 |
| chr12:93293090-93295108 | p3 | (ATC)6 | CAATGAGATCCTGCGATTGA | TTAGATGCCATCAGCGTGTC | 222 |
| chr12:93368373-93370388 | p3 | (TTG)5 | AGAAGATGAAAGTCGCGGAA | ATGCCCAAACCATCTCAGTC | 277 |
| chr12:93379182-93381196 | p2 | (CT)7 | GAGCGGGAACACTGATGAAT | TTTCCGACCTGATTTTGAGG | 177 |
| chr12:93468721-93470733 | p2 | (TA)6 | GACAAAATTTCAGCGAGGGA | ACATCAAACGGGCTGGATAG | 151 |
| chr12:93511696-93513712 | p4 | (ATTT)4 | CGATTCGTTCAACCTCCAAT | TCATACGAATCGGTGGAACA | 232 |
| chr12:93619919-93621943 | p3 | (AAT)8 | GGAAAAGAGGAAGTGGGAGG | CCCTTAGGCCATTCCATTTT | 251 |
| chr12:93679104-93681140 | p3 | (TAA)12 | ATTGGTCCTTCCTCCGCTAT | CGTCATTTCACTTTCGGGTT | 241 |
| chr12:93706422-93708437 | p3 | (TAT)7 | GTCTGCTGGAGGAGGCATAG | AATAACTGCTGGGATTTGCG | 228 |
| chr12:93706422-93708437 | p5 | (CATAA)3 | GTCTGCTGGAGGAGGCATAG | AATAACTGCTGGGATTTGCG | 228 |
| chr12:93735696-93737711 | p3 | (TTA)5 | GATCGTCGTCAAGGGTGTTT | AAAAGGTAAAAAGGCGCCAC | 230 |
| chr12:93820369-93822387 | p3 | (TGT)6 | TATGTTGTATGGGGCGGAAT | CCATGTCGTCCTTGTGTGAC | 236 |
| chr12:93827323-93829339 | p4 | (TATT)4 | TTTGCTTTCATCTCCTGGCT | TGTCTTTTTCCTGTTTCGGG | 157 |
| chr12:93835294-93837309 | p5 | (ACCAA)3 | CCATTGCTAGCTGCTTGTGA | AGGAGTTGGTCAAGTGGCTG | 172 |
| chr12:93864343-93866376 | p3 | (AAT)11 | CCATCAACATAGCCTGCCTT | GTCGGTAAGACAGGAGCCAA | 100 |
| chr12:93937568-93939590 | p2 | (AG)11 | GTTCAAGTTGTGGGAGGGAA | CCAATAATTCAATGGACCGC | 273 |
| chr12:93937568-93939590 | p2 | (AG)11 | GTTCAAGTTGTGGGAGGGAA | CCAATAATTCAATGGACCGC | 273 |
| chr12:94009340-94011374 | p2 | (AT)17 | AGCTTGTGGAACCGTAGTGG | CAGCATTTCTAAAGGCTCCG | 206 |
| chr12:94015386-94017402 | p2 | (AT)6 | GTACCCCTTTTTCTCCGCTC | AAATGCTGGGAATTTCGTTG | 290 |

| Marker ID | SSR type | SSR motif | FORWARD PRIMER (5'-3') | REVERSE PRIMER (5'-3') | size (bp) |
|---|---|---|---|---|---|
| chr12:94015386-94017402 | p2 | (TA)8 | GTACCCCTTTTTCTCCGCTC | AAATGCTGGGAATTTCGTTG | 290 |
| chr12:94015386-94017402 | p5 | (TTTTA)3 | GTACCCCTTTTTCTCCGCTC | AAATGCTGGGAATTTCGTTG | 290 |
| chr12:94040638-94042650 | p2 | (GA)6 | GCGGTGCTTATTTATGCGAT | AACCCAGGTCGTCTGAACAC | 137 |
| chr12:94094480-94096492 | p2 | (AT)6 | GGTCCGGCATAACTTTGGTA | TTTTTAAATTTTGCCCGTCG | 206 |
| chr12:94174729-94176769 | p5 | (AAATA)3 | ACCTGCAAGGAATGACCATC | CAGCATACAGGCAGAATGGA | 269 |
| chr12:94193390-94195405 | p5 | (AAAAT)3 | TCGGTAGTGTGCTCTGCATT | GCTCGTATGAGTGGGGTGTT | 168 |
| chr12:94204763-94206781 | p3 | (TGT)6 | GAAATGAATTGCCTTGCGAT | GCCCAAAATGCAACACCTAT | 246 |
| chr12:94218807-94220821 | p2 | (TC)7 | CTGCAGCCAAAAACTCATCA | GCAAACAAAAAGCGTCCTTC | 104 |
| chr12:94254933-94256957 | p2 | (TA)10 | ATTTAAGTGGACGGGCATCA | CTTGCTCATTTGTGCCTTCA | 275 |
| chr12:94255037-94257061 | p2 | (TA)10 | ATTTAAGTGGACGGGCATCA | CTTGCTCATTTGTGCCTTCA | 171 |
| chr12:94313565-94315577 | p2 | (AT)6 | CAAACCTCATCACACCCCTT | TGATTCAAGCCAAGCTGTTG | 121 |
| chr12:94313565-94315577 | p2 | (AT)9 | CAAACCTCATCACACCCCTT | TGATTCAAGCCAAGCTGTTG | 121 |
| chr12:94379254-94381278 | p6 | (GTGGGG)4 | GACACCTCGCAACTCTCTCC | CCATTACTCGGTCCTCCAAA | 286 |
| chr12:94453696-94455726 | p3 | (TCT)10 | AGTCAGCCACTCCAAGCCTA | TTGAACAAAGGAGGGGACAG | 234 |
| chr12:94487984-94490008 | p2 | (AT)13 | TAAGAATGGTGCCCCTTACG | ACAGAGGAAACTCTGCCCAA | 122 |
| chr12:94571244-94573256 | p2 | (TG)6 | AAGGAAACCTCTGCCGAAAT | TTGCCCGGTATCATAAGCTC | 170 |
| chr12:94571244-94573256 | p2 | (TA)14 | AAGGAAACCTCTGCCGAAAT | TTGCCCGGTATCATAAGCTC | 170 |
| chr12:94571731-94573759 | p2 | (TG)6 | CGGTGTATGCAAGTAAGCGA | CCAAAAGATATGGCTCCGAA | 169 |
| chr12:94571731-94573759 | p2 | (TA)14 | CGGTGTATGCAAGTAAGCGA | CCAAAAGATATGGCTCCGAA | 169 |
| chr12:94574793-94576821 | p2 | (TG)6 | CGGTGTATGCAAGTAAGCGA | CCAAAAGATATGGCTCCGAA | 169 |
| chr12:94574793-94576821 | p2 | (TA)14 | CGGTGTATGCAAGTAAGCGA | CCAAAAGATATGGCTCCGAA | 169 |
| chr12:94636532-94638544 | p2 | (TA)6 | CCAACGGTATATTATGGGCG | TTCTTGATGATGGGAGAGGG | 250 |
| chr12:94689719-94691739 | p2 | (AT)10 | TCCTATGATATTGGGAGGCG | ACTTTCACCACCGTCCAGTC | 152 |
| chr12:94713227-94715242 | p3 | (GAC)5 | AGCTCTAGTGCTTCACCCGA | AGGAGCAATTTTGGTTGTGG | 265 |
| chr12:94732817-94734829 | p2 | (AC)6 | ATGGCCGCCTTTATTTTCTT | GCTGCCACTGAACTTTCACA | 298 |
| chr12:94732863-94734885 | p2 | (AC)6 | ATGGCCGCCTTTATTTTCTT | GCTGCCACTGAACTTTCACA | 252 |
| chr12:94749574-94751592 | p2 | (AT)9 | ACGGACCAAGATACGGTCTG | ACGAGTCGACGAAAACGAGT | 298 |
| chr12:94771945-94773957 | p2 | (TA)6 | CAACCCATTAGAGGGAGCAA | TTTGGATGTGGTGAGTTCCA | 176 |
| chr12:94834860-94836872 | p2 | (TA)6 | TATTATGGGGGTCAAACGGA | GTGCTGGCCGTATTTTCAGT | 268 |
| chr12:94935821-94937839 | p5 | (TAAAT)4 | TTGCTCCCTCTAATGGGTTG | TGATGGATTTATTGCCCCAT | 168 |
| chr12:94935821-94937839 | p6 | (AGGGGA)3 | TTGCTCCCTCTAATGGGTTG | TGATGGATTTATTGCCCCAT | 168 |
| chr12:94966707-94968722 | p5 | (AAACC)3 | ATTGTCATGAGGGGCAAAAG | CCGACTACAACCAAACCGAT | 238 |
| chr12:94966707-94968722 | p5 | (GATTT)3 | ATTGTCATGAGGGGCAAAAG | CCGACTACAACCAAACCGAT | 238 |
| chr12:95012495-95014513 | p6 | (AAAAAT)3 | TCTCAATGGGCTCTGAATCC | CGTCCAGTTATTCCCTGCAT | 234 |
| chr12:95024666-95026690 | p3 | (TTG)8 | TCGTATTGGTGCAGAATGGA | CTCATTTCCGCCACTTTCAT | 120 |
| chr12:95146136-95148148 | p2 | (AT)6 | CCTTTGCTCCATTGGTGACT | GACGCAATCTTAATCGAGGC | 156 |

| Marker ID | SSR type | SSR motif | FORWARD PRIMER (5'-3') | REVERSE PRIMER (5'-3') | size (bp) |
|---|---|---|---|---|---|
| chr12:95169891-95171906 | p5 | (ATAGA)3 | GGTAGGTGTTGAACCCCCTT | AGGTTTGATTTCCAGGGACC | 273 |
| chr12:95476474-95478489 | p3 | (TAT)5 | CCAAGAAAGAGACTGGACCG | TGCAGTTGCACTCACAAATG | 102 |
| chr12:95476474-95478489 | p3 | (TTG)5 | CCAAGAAAGAGACTGGACCG | TGCAGTTGCACTCACAAATG | 102 |
| chr12:95521256-95523271 | p3 | (CAA)5 | TCCTCTCCCACGTCCATAAC | CAGATGTCTGTGTCGAGCGT | 234 |
| chr12:95658250-95660264 | p2 | (AC)7 | AGCACGATTCAAGCCAATTC | TTACAAGTTTGGGGCTTTGC | 155 |
| chr12:95658250-95660264 | p2 | (CA)7 | AGCACGATTCAAGCCAATTC | TTACAAGTTTGGGGCTTTGC | 155 |
| chr12:95658250-95660264 | p5 | (TATAA)3 | AGCACGATTCAAGCCAATTC | TTACAAGTTTGGGGCTTTGC | 155 |
| chr12:95842984-95845036 | p2 | (TA)26 | GTAGGGTTCGCTAAAAGGGG | TTTTGGAGCCCAACAGAAAG | 287 |
| chr12:95851831-95853845 | p2 | (CT)7 | CACACGGAAAAGGCTCAAAT | TTTGGTAACGGACGATGACA | 118 |
| chr12:95864487-95866505 | p5 | (AGAAA)3 | ATCTGCACCCGAAATTGAAC | GCGAAATTTGGCAGAGAGAG | 281 |
| chr12:95864487-95866505 | p6 | (ATTTTT)3 | ATCTGCACCCGAAATTGAAC | GCGAAATTTGGCAGAGAGAG | 281 |
| chr12:95866316-95868348 | p2 | (TA)16 | TATTCGTCCCTCCCAGATTG | CCCACTTCCTCTTCATCGAG | 154 |
| chr12:95866316-95868348 | p2 | (AT)16 | TATTCGTCCCTCCCAGATTG | CCCACTTCCTCTTCATCGAG | 154 |
| chr12:96097171-96099186 | p5 | (GAACC)3 | GTTTTACGTTGGCTCCCTGA | CATTTTCCCCGTATTGATGC | 122 |
| chr12:96097171-96099186 | p5 | (AACCG)3 | GTTTTACGTTGGCTCCCTGA | CATTTTCCCCGTATTGATGC | 122 |
| chr12:96142492-96144507 | p3 | (GAA)5 | GTCCGTATCAGGCCTACCAA | TGATTTTACGGTCCCCAAAG | 244 |
| chr12:96166779-96168794 | p5 | (TTGGT)4 | CAAAAGCAAACCGACAAACC | AACCAAACCAAACCAAACCA | 136 |
| chr12:96166779-96168794 | p5 | (TTGGT)3 | CAAAAGCAAACCGACAAACC | AACCAAACCAAACCAAACCA | 136 |
| chr12:96241874-96243922 | p3 | (TAA)16 | AGCCAAGCACCAATTTCATC | ACCCATGGCTGCTACAAATC | 133 |
| chr12:96271678-96273694 | p4 | (TATT)4 | GTGGAAATGGAGAGGGGTTT | TAATAATAGCCGTGGGCCAG | 120 |
| chr12:96271678-96273694 | p2 | (AT)7 | GTGGAAATGGAGAGGGGTTT | TAATAATAGCCGTGGGCCAG | 120 |
| chr12:96393851-96395865 | p2 | (TA)7 | GCACCGCCACCTATTAAAAA | CAAATAAATGACATGCCGGA | 142 |
| chr12:96393851-96395865 | p2 | (TA)7 | GCACCGCCACCTATTAAAAA | CAAATAAATGACATGCCGGA | 142 |
| chr12:96462219-96464239 | p5 | (GGCAT)4 | CGAGAATGTAAGCCATGCAA | CCAATGCAATCTTCTCGGAT | 299 |
| chr12:96496960-96498975 | p5 | (ACCAA)3 | CGGCAATATCCCTCATCTGT | AAAACCCGAGAAAACCGAAT | 198 |
| chr12:96496960-96498975 | p5 | (AAACC)3 | CGGCAATATCCCTCATCTGT | AAAACCCGAGAAAACCGAAT | 198 |
| chr12:96500864-96502878 | p2 | (AT)7 | CTCCCAAAAGCTTGGTCAAA | CTAGGCACTGATGCAGACGA | 208 |
| chr12:96728797-96730909 | p2 | (AG)6 | TGAGGATCGAACCCATATCC | GAGTGTTTTGGAGGGTGCAT | 152 |
| chr12:96782942-96784956 | p2 | (GA)7 | GCCACAACAATGCATAGGAA | AATAATGACGGCCCACAAAG | 154 |
| chr12:96825311-96827326 | p5 | (CTTTT)3 | AATTAGAACCCCGAACCCAC | ACCACCATTCACCAAGCCTA | 297 |
| chr12:96825311-96827326 | p2 | (TA)7 | AATTAGAACCCCGAACCCAC | ACCACCATTCACCAAGCCTA | 297 |
| chr12:96848996-96851008 | p2 | (TA)6 | CCCTTCTCCTACCCCCTATG | AGTCGAGTTGGCCCATTTTA | 230 |
| chr12:96889949-96891967 | p2 | (AG)9 | CGCGGGGTTAAAGTATACCA | TGAATTGAACTTGCCGTTTG | 293 |
| chr12:96898215-96900233 | p2 | (GA)9 | AGATTCAGGGCCACTAGGGT | TTTGTGTCAGCCCCCATTAT | 229 |
| chr12:96916443-96918459 | p6 | (CAAAAA)3 | CTCGAGCTTCGGGTTCTATG | CCCATGACGGCTTTAAACAT | 245 |
| chr12:96916443-96918459 | p2 | (TC)8 | CTCGAGCTTCGGGTTCTATG | CCCATGACGGCTTTAAACAT | 245 |

| Marker ID | SSR type | SSR motif | FORWARD PRIMER (5'-3') | REVERSE PRIMER (5'-3') | size (bp) |
|---|---|---|---|---|---|
| chr12:96941757-96943771 | p2 | (TA)7 | TTGTTCCCATTTGCACTTGA | ATTCATGAGTGGAGCCAACC | 189 |
| chr12:96941757-96943771 | p5 | (TTTTG)3 | TTGTTCCCATTTGCACTTGA | ATTCATGAGTGGAGCCAACC | 189 |
| chr12:96963449-96965464 | p5 | (ATAAA)3 | ATAGGCGTTTGGACGACATC | GGTGATTGATGATTTCCGCT | 225 |
| chr12:97095906-97097924 | p6 | (TTTCAT)3 | TGATCACGAACCTTGTCGAA | GGATACAACCATACCGGGTG | 147 |
| chr12:97161165-97163180 | p5 | (TTGGT)3 | CCACTTTTTATGGCCTTGGA | CAGATCGGTTTTGTTCGGTT | 187 |
| chr12:97161165-97163180 | p5 | (TTGGT)3 | CCACTTTTTATGGCCTTGGA | CAGATCGGTTTTGTTCGGTT | 187 |
| chr12:97216864-97218882 | p3 | (TGT)6 | GTGGGCAACGAAAACAAGAT | TTCAGGTGCATCAGGAAACA | 241 |
| chr12:97236694-97238712 | p3 | (TGT)6 | GTGGGCAACGAAAACAAGAT | TTCAGGTGCATCAGGAAACA | 241 |
| chr12:97418930-97420946 | p6 | (GGGGGC)3 | GAGGACTCGTTGGGTTTGAA | TAACGGCGAGGGTATTTTTG | 143 |
| chr12:97418930-97420946 | p4 | (CCAC)4 | GAGGACTCGTTGGGTTTGAA | TAACGGCGAGGGTATTTTTG | 143 |
| chr12:97476846-97478868 | p2 | (AT)10 | CAAGGCCTTATTCCACCAGA | AGTCCTGCCTTGCGAAATTA | 144 |
| chr12:97476846-97478868 | p2 | (AT)11 | CAAGGCCTTATTCCACCAGA | AGTCCTGCCTTGCGAAATTA | 144 |
| chr12:97542970-97544990 | p5 | (CTTTT)3 | TCGTCGTCAACAAACCAAAA | GATTGATACGCATGTGGCAA | 173 |
| chr12:97542970-97544990 | p2 | (TA)10 | TCGTCGTCAACAAACCAAAA | GATTGATACGCATGTGGCAA | 173 |
| chr12:97542970-97544990 | p3 | (AAG)5 | TCGTCGTCAACAAACCAAAA | GATTGATACGCATGTGGCAA | 173 |
| chr12:97557435-97559450 | p5 | (AACCG)3 | AGCTCGGTGTCACCATAACC | CTGGGCAGATGCAGTCAATA | 208 |
| chr12:97602873-97604888 | p3 | (CAG)5 | AAGAATGGGTTCAGGCCTTT | CCGAGGTGCATATAGGCTGT | 233 |
| chr12:97675989-97678001 | p2 | (TA)6 | CCATTAGTGGGAGCAAATGAA | AAATGACATGGCGGAATAGC | 272 |
| chr12:97699404-97701416 | p2 | (TA)6 | GGTAGGGCACAATTTGAGGA | GGTACCTGAGCGTTTTTCCA | 160 |
| chr12:97736615-97738655 | p2 | (AT)20 | TTGCATCTATGCCTGAGCAC | GTCCTGTCAAGCAATGCAAA | 167 |
| chr12:97798542-97800560 | p3 | (GTT)6 | ATGTTCGAACCCCTAATCCC | TCAAATCAAATGTGGAGGCA | 299 |
| chr12:97798542-97800560 | p2 | (TA)8 | ATGTTCGAACCCCTAATCCC | TCAAATCAAATGTGGAGGCA | 299 |
| chr12:97798542-97800560 | p2 | (TA)9 | ATGTTCGAACCCCTAATCCC | TCAAATCAAATGTGGAGGCA | 299 |
| chr12:97798675-97800693 | p3 | (GTT)6 | ATGTTCGAACCCCTAATCCC | TCAAATCAAATGTGGAGGCA | 166 |
| chr12:97798675-97800693 | p2 | (TA)8 | ATGTTCGAACCCCTAATCCC | TCAAATCAAATGTGGAGGCA | 166 |
| chr12:97798675-97800693 | p2 | (TA)9 | ATGTTCGAACCCCTAATCCC | TCAAATCAAATGTGGAGGCA | 166 |
| chr12:97924143-97926155 | p2 | (TA)7 | TATTCGACCAGCACACAAGC | GAAACAAGCCGGCAAAAATA | 178 |
| chr12:97924240-97926254 | p2 | (TA)7 | GAGGGGATTCAAAGAGAGGC | GAAACAAGCCGGCAAAAATA | 145 |
| chr12:97941084-97943099 | p3 | (AAC)5 | TGCCAAAAGGTCATGGTGTA | ATCACGACCTTACGTCACCC | 232 |
| chr12:98080042-98082057 | p5 | (AAAGC)3 | AAGGAGAAGAAGAGGGTCGC | CCCAGGACATCTCTAGCTGC | 111 |
| chr12:98080042-98082057 | p5 | (TTTCA)3 | AAGGAGAAGAAGAGGGTCGC | CCCAGGACATCTCTAGCTGC | 111 |
| chr12:98148528-98150546 | p2 | (GT)9 | GCTCCTTACCAGTGCTCCAC | CACATGGTCGCTCGACTAAA | 159 |
| chr12:98158120-98160142 | p2 | (AT)11 | TTTGGAAAACCCCAGTGAAC | TGGATAAGGGATTCCCCATT | 194 |
| chr12:98198671-98200692 | p3 | (GAA)7 | ATTCAGCTTTTGGGGCTTTT | TTTAGGGGCCTGTTTACGTG | 160 |
| chr12:98202093-98204111 | p3 | (CTT)10 | GTTGCACACTCTCCGCTACA | TGGAGGAAGAAAATGGGTTG | 291 |
| chr12:98202093-98204111 | p2 | (AC)9 | GTTGCACACTCTCCGCTACA | TGGAGGAAGAAAATGGGTTG | 291 |

| Marker ID | SSR type | SSR motif | FORWARD PRIMER (5'-3') | REVERSE PRIMER (5'-3') | size (bp) |
|---|---|---|---|---|---|
| chr12:98202093-98204111 | p3 | (AGA)6 | GTTGCACACTCTCCGCTACA | TGGAGGAAGAAAATGGGTTG | 291 |
| chr12:98372340-98374372 | p5 | (CAAAC)3 | TAACCCGAACCATTAACCCA | ACCATGAACACCCCTAGCTG | 157 |
| chr12:98382276-98384306 | p3 | (AAT)10 | GGGTTTCAAACTTGGGGAAT | CATATCACTGTCACGCCCAC | 101 |
| chr12:98399460-98401484 | p2 | (AT)12 | AGGACACTTTGATTGCCCAG | GATGATTTATTCTCCGGGGG | 259 |
| chr12:98417072-98419084 | p2 | (AT)6 | TTTAATAGCAATGCCCCGAC | CCGAAAAATGCCTTATCCCT | 217 |
| chr12:98419321-98421339 | p2 | (AT)6 | CCCACATGAGAGGTGCTTTT | CATAACTGTGCCGCTTACGA | 165 |
| chr12:98419321-98421339 | p6 | (TGTTTT)3 | CCCACATGAGAGGTGCTTTT | CATAACTGTGCCGCTTACGA | 165 |
| chr12:98421882-98423897 | p5 | (AACCG)3 | GTTGTCCTTGCAAGCCTCTC | ATGTTTGCACCAGCATTTGA | 268 |
| chr12:98441690-98443720 | p2 | (AT)19 | GAATTACACCGGTTGGGCTA | TAGGGGGTGTCAAATGGGTA | 202 |
| chr12:98441737-98443749 | p2 | (AT)19 | TACCCATTTGACACCCCCTA | CGTCTTATTTTCTTCGGGCA | 276 |
| chr12:98475507-98477523 | p2 | (TA)8 | CCGCAGTCTGTTTCTTGTCA | TTTCCGACGACTTTTGCTCT | 204 |
| chr12:98561697-98563739 | p3 | (TTA)14 | TTGAGGTCGGCATATTAGGG | CGAGGTTAGCCGAAGAAGTG | 157 |
| chr12:98745871-98747885 | p2 | (AT)7 | AGCCCTAGATCCCCAAAGAA | TTTGACTGTGAATTTGGGCA | 166 |
| chr12:98789702-98791718 | p2 | (AT)8 | TCCTCCCTGTACTTGCCATC | TGCCACCTTGCAATGAATTA | 252 |
| chr12:98808998-98811018 | p2 | (AT)12 | CTACGCAGCTCCTAAGGTCG | TGCGGAGATTCCTTTTTCAG | 271 |
| chr12:98929540-98931560 | p4 | (TTTA)5 | ATTGGAACCTCGTGTCTTGC | GGCCAAGGTACTGGAAATCA | 248 |
| chr12:99064860-99066875 | p3 | (AAC)5 | AATAAGGGAGCCAAGTGCAA | TCAACAATCGGAGCTGTCTG | 106 |
| chr12:99067837-99069849 | p2 | (TC)6 | CAGGTGCCCAGGTTGATACT | ACGTTTGGTGGGGGAATAAT | 165 |
| chr12:99130856-99132912 | p2 | (CT)28 | CAGTTCGTTGGTGTTGAACG | AAGTATGCATGGGGCACATT | 231 |
| chr12:99145552-99147566 | p2 | (AT)7 | GGTGAACGCGTAATGGAAGT | CTGCAAGCGGGTCATAAAAT | 112 |
| chr12:99165523-99167579 | p2 | (CT)28 | CAGTTCGTTGGTGTTGAACG | AAGTATGCATGGGGCACATT | 231 |
| chr12:99180219-99182233 | p2 | (AT)7 | GGTGAACGCGTAATGGAAGT | CTGCAAGCGGGTCATAAAAT | 112 |
| chr12:99266295-99268313 | p2 | (AT)15 | AATCAACTTGCTTGGGGTTG | CGGCCTTTTCTTTCTCATTG | 222 |
| chr12:99266295-99268313 | p2 | (AT)9 | AATCAACTTGCTTGGGGTTG | CGGCCTTTTCTTTCTCATTG | 222 |
| chr12:99519795-99521813 | p3 | (TTC)6 | TTTCTTTCGCCGTTCTCTGT | AGGGTTAAGGCGGCTAAAAA | 196 |
| chr12:99524926-99526938 | p2 | (AT)6 | GGGAATCCACGACTTCTTGA | AGCTCCACTTGCAGTTCGAT | 145 |
| chr12:99594862-99596892 | p2 | (AT)15 | GCAGTATGGCACAACCAAGA | TTCCCTATCGTCCTATCCCC | 154 |
| chr12:99598730-99600758 | p2 | (AT)14 | GCAGAAGGCAGAAATGAAGG | TTCTGTACAAAACCCAGCCC | 280 |
| chr12:99610472-99612484 | p2 | (AT)6 | GCCATGAACAATGGGAGTTT | TGGGATCTTGGGAGATGGTA | 209 |
| chr12:99680430-99682444 | p2 | (TA)8 | CATGGGCATAGATGCGGTAT | TTCGGAGCACCATAATAGCC | 287 |
| chr12:99680459-99682473 | p2 | (TA)8 | CATGGGCATAGATGCGGTAT | TTCGGAGCACCATAATAGCC | 258 |
| chr12:99775200-99777215 | p5 | (AAAAT)3 | TTAGGGCTCTTACCACCACG | TGAGTGCCCTCTCAAGTCCT | 180 |
| chr12:99775200-99777215 | p2 | (AT)34 | TTAGGGCTCTTACCACCACG | TGAGTGCCCTCTCAAGTCCT | 180 |
| chr12:99775200-99777215 | p3 | (GTT)6 | TTAGGGCTCTTACCACCACG | TGAGTGCCCTCTCAAGTCCT | 180 |
| chr12:99775668-99777736 | p5 | (AAAAT)3 | TTTGTAACCCGAAAAGGACG | CTGAAGCCGCCACTAATAGG | 193 |
| chr12:99775668-99777736 | p2 | (AT)34 | TTTGTAACCCGAAAAGGACG | CTGAAGCCGCCACTAATAGG | 193 |

| Marker ID | SSR type | SSR motif | FORWARD PRIMER (5'-3') | REVERSE PRIMER (5'-3') | size (bp) |
|---|---|---|---|---|---|
| chr12:99775668-99777736 | p3 | (GTT)6 | TTTGTAACCCGAAAAGGACG | CTGAAGCCGCCACTAATAGG | 193 |
| chr12:99775668-99777736 | p2 | (TA)7 | TTTGTAACCCGAAAAGGACG | CTGAAGCCGCCACTAATAGG | 193 |
| chr12:99776082-99778100 | p5 | (AAAAT)3 | TGCTGATCAACAGGCATCAT | TTGGTAGAACACCTACCGGC | 272 |
| chr12:99776082-99778100 | p2 | (AT)34 | TGCTGATCAACAGGCATCAT | TTGGTAGAACACCTACCGGC | 272 |
| chr12:99776082-99778100 | p3 | (GTT)6 | TGCTGATCAACAGGCATCAT | TTGGTAGAACACCTACCGGC | 272 |
| chr12:99776082-99778100 | p2 | (TA)7 | TGCTGATCAACAGGCATCAT | TTGGTAGAACACCTACCGGC | 272 |
| chr12:99813839-99815863 | p3 | (TTA)5 | ATGATGATGTTCGGGGATGT | AAGGGGGAAGCGAACAGTAT | 255 |
| chr12:99813839-99815863 | p3 | (TTA)6 | ATGATGATGTTCGGGGATGT | AAGGGGGAAGCGAACAGTAT | 255 |
| chr12:99813945-99815966 | p3 | (TTA)5 | ATGATGATGTTCGGGGATGT | AAGGGGGAAGCGAACAGTAT | 149 |
| chr12:99813945-99815966 | p3 | (TTA)6 | ATGATGATGTTCGGGGATGT | AAGGGGGAAGCGAACAGTAT | 149 |
| chr12:99814224-99816242 | p3 | (TTA)5 | CCCGTCATATGTCTTCGCTT | GCAAACATATGGGCCAGTTT | 295 |
| chr12:99814224-99816242 | p3 | (TTA)6 | CCCGTCATATGTCTTCGCTT | GCAAACATATGGGCCAGTTT | 295 |
| chr12:99832041-99834056 | p5 | (ATAAT)3 | GAGGAAGTGGGATGTCCAGA | CTCCCCTTCATCCTAAAGCC | 201 |
| chr12:99849432-99851448 | p4 | (AAAT)4 | TTGGAGAGAGAAACGTGGGT | GGAAATACAAAAATCCGGCA | 156 |
| chr12:99950521-99952536 | p5 | (TGTTG)3 | TGTTCTGTCTCGTCCCATGA | GATGGACCCACAGCTCAAAT | 160 |
| chr12:99950521-99952536 | p3 | (GTT)6 | TGTTCTGTCTCGTCCCATGA | GATGGACCCACAGCTCAAAT | 160 |
| chr12:99961210-99963226 | p4 | (ATAA)4 | GCGGGATCTTTAAACCATGA | CAAATTTCGCCGCTAGTCTC | 275 |
| chr12:99961210-99963226 | p5 | (GGTTT)3 | GCGGGATCTTTAAACCATGA | CAAATTTCGCCGCTAGTCTC | 275 |
| chr12:99961333-99963348 | p4 | (ATAA)4 | GCGGGATCTTTAAACCATGA | CAAATTTCGCCGCTAGTCTC | 152 |
| chr12:99961333-99963348 | p5 | (GGTTT)3 | GCGGGATCTTTAAACCATGA | CAAATTTCGCCGCTAGTCTC | 152 |
| chr12:100046772-10004878 | p2 | (TG)8 | CAAGGCGGAGTTTTCTATCG | TTTCCCTTTGCAGACCAAAC | 242 |
| chr12:100067125-10006916 | p5 | (GGTTC)3 | GCCTTGCCTGGTAAATGGTA | TAGCTGGTTAGCAGCCCACT | 158 |
| chr12:100067125-10006916 | p3 | (TTA)14 | GCCTTGCCTGGTAAATGGTA | TAGCTGGTTAGCAGCCCACT | 158 |
| chr12:100067125-10006916 | p2 | (CT)6 | GCCTTGCCTGGTAAATGGTA | TAGCTGGTTAGCAGCCCACT | 158 |
| chr12:100119004-10012102 | p6 | (CCGGTT)3 | TGGCTATTGCAACGATGTGT | AGCTATCCATCTCGTGTGCC | 282 |
| chr12:100188548-10019056 | p2 | (GA)10 | TCCTCTGCAGTATCCCCATC | TGGATAACGGTGCTCCTACC | 251 |
| chr12:100205092-10020710 | p2 | (TA)7 | GTATAAACCGTGGCTTCGGA | TTTTGGTCCGGTCCACTTAC | 227 |
| chr12:100205092-10020710 | p2 | (AT)7 | GTATAAACCGTGGCTTCGGA | TTTTGGTCCGGTCCACTTAC | 227 |
| chr12:100216167-10021818 | p3 | (TGG)5 | TGACTACTTTTTCCGGTGCC | CAATCCGCACACCCTTCTAT | 150 |
| chr12:100416575-10041858 | p2 | (AT)7 | CTCCACAATGGGGTATTTGG | TTGAACGGGATTGAAGAAGG | 272 |
| chr12:100428282-10043029 | p5 | (AAAAG)3 | TGAGAAGGTCCAATTCCCAG | CGATGTCCAAAGACAGACGA | 162 |
| chr12:100518445-10052046 | p4 | (TATT)4 | GAATTTCAAGGAAGAGGGGG | TCCCCCTAAAGAGGAAAGGA | 175 |
| chr12:100518445-10052046 | p3 | (TAA)5 | GAATTTCAAGGAAGAGGGGG | TCCCCCTAAAGAGGAAAGGA | 175 |
| chr12:100536095-10053811 | p4 | (ATTT)4 | GCCTAACTTTGACCCGAACA | AATTTTTGTCCCTTTCGCCT | 238 |
| chr12:100536095-10053811 | p2 | (TA)6 | GCCTAACTTTGACCCGAACA | AATTTTTGTCCCTTTCGCCT | 238 |
| chr12:100674377-10067638 | p2 | (AT)6 | TATTGAGGGAAAATGCCTGG | AATCCGGCTATGAACAGTGC | 278 |

| Marker ID | SSR type | SSR motif | FORWARD PRIMER (5'-3') | REVERSE PRIMER (5'-3') | size (bp) |
|---|---|---|---|---|---|
| chr12:100674377-10067638 | p2 | (AT)8 | TATTGAGGGAAAATGCCTGG | AATCCGGCTATGAACAGTGC | 278 |
| chr12:100709644-10071166 | p4 | (ATTA)4 | AGCGGAAGGGAAAAATTCAT | TAATGACTTCGGCCTGGTTC | 283 |
| chr12:100709644-10071166 | p2 | (AT)8 | AGCGGAAGGGAAAAATTCAT | TAATGACTTCGGCCTGGTTC | 283 |
| chr12:100750557-10075256 | p2 | (AT)6 | AACCCTCGGGTAAATGATCC | GCCAGTAGAACCCCAGCATA | 169 |
| chr12:100750557-10075256 | p2 | (AT)9 | AACCCTCGGGTAAATGATCC | GCCAGTAGAACCCCAGCATA | 169 |
| chr12:100805699-10080771 | p2 | (TA)10 | AACTCGTAAGCCAACCGCTA | TACCCGTTCCACCATCCTTA | 233 |
| chr12:100808078-10081009 | p2 | (AT)10 | GGTCCTTACTGGGATCGTGA | ATGCCCGTACACACATGCTA | 207 |
| chr12:100808078-10081009 | p2 | (AT)9 | GGTCCTTACTGGGATCGTGA | ATGCCCGTACACACATGCTA | 207 |
| chr12:100859252-10086127 | p6 | (TTAAAA)3 | TCTCTGAAAGGGGTGCAGTT | TATCATGAAGCAAAAGCCCC | 230 |
| chr12:100874303-10087631 | p5 | (AAAAT)3 | TCAAAGAGAATGGGATTGGG | TCATAATTCGTCCTCCTGCC | 144 |
| chr12:100918504-10092051 | p3 | (GTG)5 | AGAAGGCAACGAGCAGAAAA | AGCAGCCACCTAGCGATAAA | 198 |
| chr12:101001336-10100334 | p2 | (TA)6 | GTGGAAGTCCTCCAAACGAA | GAACCCACGACCTATGGATG | 279 |
| chr12:101003754-10100576 | p5 | (TGATA)3 | ACGTTCTGTTTGCCGAGTCT | GAGTGCGTGTAACATCCGAA | 145 |
| chr12:101003754-10100576 | p5 | (TTGGT)3 | ACGTTCTGTTTGCCGAGTCT | GAGTGCGTGTAACATCCGAA | 145 |
| chr12:101081305-10108331 | p2 | (TA)6 | GTGGAAGTCCTCCAAACGAA | GAACCCACGACCTATGGATG | 279 |
| chr12:101083723-10108573 | p5 | (TGATA)3 | ACGTTCTGTTTGCCGAGTCT | GAGTGCGTGTAACATCCGAA | 145 |
| chr12:101083723-10108573 | p5 | (TTGGT)3 | ACGTTCTGTTTGCCGAGTCT | GAGTGCGTGTAACATCCGAA | 145 |
| chr12:101225786-10122781 | p3 | (TTG)8 | TCGGAGGATAACGAATCGAC | AGGCTCAGATGCAGGTGTTT | 191 |
| chr12:101243620-10124563 | p3 | (TGA)5 | CAAGTTTTCCCACCTCCAAA | TTGGGACCTTTGTGAAAAGC | 270 |
| chr12:101277884-10127989 | p5 | (AAACC)3 | GTAGGGGTGTACATGGACCG | ACCCGACTTTTGTTGGTTTG | 149 |
| chr12:101282162-10128418 | p3 | (ATA)9 | CAATTGCAGGATGATTGCAC | CTGTCCTGTTGCCAGTCAGA | 212 |
| chr12:101311296-10131331 | p5 | (AAACC)3 | GTAGGGGTGTACATGGACCG | ACCCGACTTTTGTTGGTTTG | 149 |
| chr12:101315574-10131760 | p3 | (ATA)9 | CAATTGCAGGATGATTGCAC | CTGTCCTGTTGCCAGTCAGA | 212 |
| chr12:101344801-10134681 | p2 | (TA)6 | ACCCACCATCTCCAAACTCA | TGTGGGCCCTCAATTAACAT | 295 |
| chr12:101604574-10160659 | p4 | (TTTA)4 | GTTCGAACCCCAGCTCAATA | TTTGACCCGCCTAAATTCTG | 122 |
| chr12:101661183-10166319 | p2 | (TA)6 | GATAGCGGTGAAGAGCCAAG | AGAGCAAGTTGTGGGAGGAA | 166 |
| chr12:101712150-10171416 | p5 | (TATTA)4 | CAGAAGTGGGGCAATGAGAT | CTGCTTGGCCATTTTCATTT | 161 |
| chr12:101747159-10174918 | p2 | (AT)13 | GAGATCGAGCGTCCACATTT | AACCAAACCGAACCATTGAC | 108 |
| chr12:101765706-10176773 | p2 | (AT)13 | GAGATCGAGCGTCCACATTT | AACCAAACCGAACCATTGAC | 108 |
| chr12:101878174-10188020 | p2 | (AT)8 | ACATGCGGTATCAACCCATT | ATGGTGGTTTATATGGGGGC | 129 |
| chr12:101878174-10188020 | p2 | (TA)14 | ACATGCGGTATCAACCCATT | ATGGTGGTTTATATGGGGGC | 129 |
| chr12:101878174-10188020 | p2 | (AT)10 | ACATGCGGTATCAACCCATT | ATGGTGGTTTATATGGGGGC | 129 |
| chr12:101941044-10194305 | p2 | (TA)8 | GGACCGGTAACGTGTTAGGA | TCAAACCGGTTAAACGGAAG | 270 |
| chr12:101941068-10194308 | p2 | (TA)8 | GGACCGGTAACGTGTTAGGA | TCAAACCGGTTAAACGGAAG | 246 |
| chr12:102062211-10206422 | p2 | (TA)7 | TTGCTCCACGCCCTACTACT | CGGCATTCTTGAGTACCGAT | 293 |
| chr12:102116259-10211827 | p2 | (AT)8 | AAATGACATGGCGGAATAGC | TTCCCAAAATATCCCCATCA | 287 |

| Marker ID | SSR type | SSR motif | FORWARD PRIMER (5'-3') | REVERSE PRIMER (5'-3') | size (bp) |
|---|---|---|---|---|---|
| chr12:102116259-10211827 | p5 | (ATATA)3 | AAATGACATGGCGGAATAGC | TTCCCAAAATATCCCCATCA | 287 |
| chr12:102154703-10215672 | p2 | (AT)6 | TCACAGATGGGGAGAGAAGG | CAAGCTGCCTTGTCACAAAA | 286 |
| chr12:102154703-10215672 | p6 | (ATCTGA)4 | TCACAGATGGGGAGAGAAGG | CAAGCTGCCTTGTCACAAAA | 286 |
| chr12:102221010-10222303 | p2 | (AG)10 | GTTTGCCGCTCTTTTCAGTC | ATACATCAGGCAAAGTGGGG | 125 |
| chr12:102223450-10222548 | p4 | (AAAC)4 | TCCACTAACCATTTGGGCTC | TGCTTGCAGGTGAACAACTC | 168 |
| chr12:102223450-10222548 | p2 | (AT)16 | TCCACTAACCATTTGGGCTC | TGCTTGCAGGTGAACAACTC | 168 |
| chr12:102279800-10228186 | p4 | (ATCA)4 | GGGCAACCATGTTGCTAACT | CTGTACCGGGAGAAAATCCA | 207 |
| chr12:102310595-10231262 | p5 | (TGTTG)6 | TTCCTTGGCCAATCAAACTC | TGCAGTTGGAAAAACTGCTG | 147 |
| chr12:102310595-10231262 | p5 | (GTTTT)4 | TTCCTTGGCCAATCAAACTC | TGCAGTTGGAAAAACTGCTG | 147 |
| chr12:102310817-10231283 | p5 | (TGTTG)6 | CAATCACCACTAATGCCACG | CCAAAAGTAGTCGCCTTTGC | 207 |
| chr12:102310817-10231283 | p5 | (GTTTT)4 | CAATCACCACTAATGCCACG | CCAAAAGTAGTCGCCTTTGC | 207 |
| chr12:102458339-10246036 | p2 | (AT)11 | GAGGAGGTATCACGACCCAA | GTTAGCCTCGTCGGCTTATG | 201 |
| chr12:102458339-10246036 | p5 | (CATAT)3 | GAGGAGGTATCACGACCCAA | GTTAGCCTCGTCGGCTTATG | 201 |
| chr12:102560325-10256233 | p2 | (TA)7 | GGCCCACCTGAAGTGTCTAA | ACCCTAAACCCATGGAAAGG | 119 |
| chr12:102560325-10256233 | p2 | (AT)10 | GGCCCACCTGAAGTGTCTAA | ACCCTAAACCCATGGAAAGG | 119 |
| chr12:102560524-10256254 | p2 | (TA)7 | CCTTTCCATGGGTTTAGGGT | GACCCCGTCATACTCGTTGT | 130 |
| chr12:102560524-10256254 | p2 | (AT)10 | CCTTTCCATGGGTTTAGGGT | GACCCCGTCATACTCGTTGT | 130 |
| chr12:102581642-10258365 | p2 | (TA)6 | TTTCCCTTTGTTCCTTGTGC | TTTTCGCTGTGTTAGGACCC | 257 |
| chr12:102591966-10259398 | p2 | (AT)15 | GGAATGTGGTCAGTCCAACC | CGACACGCGAAGCAATATAA | 176 |
| chr12:102591966-10259398 | p2 | (AT)7 | GGAATGTGGTCAGTCCAACC | CGACACGCGAAGCAATATAA | 176 |
| chr12:102591966-10259398 | p2 | (AT)11 | GGAATGTGGTCAGTCCAACC | CGACACGCGAAGCAATATAA | 176 |
| chr12:102608157-10261016 | p2 | (AT)6 | TTGCAGATGGTTGGTGGATA | GCCAGAATATTCCCCAGGTT | 288 |
| chr12:102608207-10261021 | p2 | (AT)6 | TTGCAGATGGTTGGTGGATA | GCCAGAATATTCCCCAGGTT | 238 |
| chr12:102716313-10271832 | p2 | (TG)8 | GTTGTTGGTTCCCTCTCCAA | ATGGGAGTGCATTAGGCAAC | 137 |
| chr12:102739726-10274174 | p5 | (TTTTC)3 | CCCAGGAGCATCAGCATATT | ATCCCCTCCCTTCTCTCAAA | 153 |
| chr12:102803674-10280569 | p3 | (TTA)7 | TCGATCAAAGCTGCAGAAGA | AAGCATAACCACCAGGAACG | 254 |
| chr12:102834222-10283623 | p5 | (ATATA)3 | CCTAACGCCAAAGTACGAGG | GTGAGCCCCATCTCATTCAT | 207 |
| chr12:102851869-10285390 | p2 | (TA)19 | TTTATTTCTGGGGGTTTCCC | TAATTGACGAAAAGCCCTCG | 200 |
| chr12:102858900-10286091 | p6 | (TTCATG)3 | CGCCTTGATTTGGTACGAGT | GAACGTGGCGTAGGGAATAA | 255 |
| chr12:102945536-10294755 | p3 | (GTT)6 | TTCGGAAACAACCGTCCTAC | GAGCAGATGGCATGAACAGA | 139 |
| chr12:102945536-10294755 | p3 | (TGT)7 | TTCGGAAACAACCGTCCTAC | GAGCAGATGGCATGAACAGA | 139 |
| chr12:102974191-10297622 | p2 | (AT)15 | TTATTTGCCTTGGCTTGGAC | TAGCCGCATAGGTAGCTCGT | 105 |
| chr12:103143823-10314583 | p3 | (TGA)5 | TCATGGTTGCACGATTGTCT | CTACCTTGCATGTGCTTCCA | 299 |
| chr12:103224059-10322609 | p3 | (AAT)12 | AGCAATATCAAGGCCAAACG | CGCCATAAAATCCCTTGAAA | 196 |
| chr12:103297654-10329967 | p4 | (TATT)4 | CCGATATCGTGTGATAGCCC | GACTTGGAAGGGCAACAAAG | 142 |
| chr12:103571203-10357323 | p3 | (AAC)9 | AGATCAGGGCAGCAAAAAGA | CAATCTGTTCCTGAAGGGGA | 194 |

| Marker ID | SSR type | SSR motif | FORWARD PRIMER (5'-3') | REVERSE PRIMER (5'-3') | size (bp) |
|---|---|---|---|---|---|
| chr12:103689768-10369178 | p4 | (TAAA)4 | GGTGGTGCAGGAACAGAGTT | TGAGAGGCACAGAAGGTGTG | 174 |
| chr12:103798729-10380074 | p2 | (TA)7 | TTCGGACTTCAACGATGTGA | TACTGGTCAATCTACGGGGC | 195 |
| chr12:103827897-10382991 | p6 | (AATAAA)3 | ACATTGAAAGCCATTGAGGC | AACTTCTTCTCAACAGCGGC | 208 |
| chr12:103827897-10382991 | p5 | (TTTCT)3 | ACATTGAAAGCCATTGAGGC | AACTTCTTCTCAACAGCGGC | 208 |
| chr12:103900825-10390284 | p3 | (CAA)6 | CTTCCCCACATATGTCCCAC | AATCGGAGGTTACCGACCTT | 179 |
| chr12:104016523-10401855 | p3 | (TTA)11 | GTGGGTGCCTGAACTGAAAT | TGGATGTGGACGTGGAAGTA | 269 |
| chr12:104054974-10405703 | p2 | (TA)10 | AAAAGGAGGAAGAAGCGTCC | GCTGCTGACTTGGCATCATA | 281 |
| chr12:104055080-10405714 | p2 | (TA)10 | AAAAGGAGGAAGAAGCGTCC | GCTGCTGACTTGGCATCATA | 175 |
| chr12:104055292-10405733 | p2 | (TA)10 | GACACGTAGCTCCACCCCTA | GGTTTTTGGGTTGGTTGAGA | 206 |
| chr12:104079937-10408195 | p5 | (GGATT)3 | TAACCTGATGCAATGGGACA | GGTTCTAGGGTGGGGCTTAG | 147 |
| chr12:104096603-10409861 | p2 | (TA)11 | ATAGGAAGCCCAAAGCCACT | TGACAGCACCAACAATCCAT | 139 |
| chr12:104096603-10409861 | p2 | (AT)8 | ATAGGAAGCCCAAAGCCACT | TGACAGCACCAACAATCCAT | 139 |
| chr12:104132153-10413417 | p2 | (AT)10 | AAATATCAACGGTGGCTTCG | ATTTGGTCCATTCCACTTGC | 184 |
| chr12:104287709-10428972 | p5 | (TTGGT)3 | AGAGGAGGACCCGTTTTGTT | TACAACAAAGTGCGAAACGC | 186 |
| chr2:16499-18514 | p5 | (GTTGG)3 | GATGAATGAGGAAAGCGAGC | GAATGCAGGCTTTGGAATGT | 183 |
| chr2:221310-223325 | p3 | (TGT)5 | CTTGCCCGGTTAGTCTTTCA | ACACCCCCAACTTAGGCTTT | 279 |
| chr2:261024-263039 | p3 | (GTT)5 | GCGGGAAGTTAACAAGCAAG | TTTCAAATTTGCCCTTCGTC | 281 |
| chr2:347763-349778 | p3 | (TGT)5 | CTTGCCCGGTTAGTCTTTCA | ACACCCCCAACTTAGGCTTT | 279 |
| chr2:387477-389492 | p3 | (GTT)5 | GCGGGAAGTTAACAAGCAAG | TTTCAAATTTGCCCTTCGTC | 281 |
| chr2:480375-482447 | p6 | (AAAAAT)4 | TCCACCCATTAGGGTTACCA | AAGCGGAAGCGTTGTTTCTA | 137 |
| chr2:485260-487278 | p6 | (CAGTAG)3 | TGTGTCCGATTTTTCCACAA | CTCATCGTCCGGTTGAGAAT | 126 |
| chr2:519403-521429 | p2 | (AT)13 | TCAGAAGGACGTGACGTGAG | TTTCCGGACCAAAGTTTCAC | 245 |
| chr2:523776-525788 | p2 | (AG)6 | TAACTAAAACCTGGCGTGGC | TTTAAGCCTGTCAAGTGGGC | 254 |
| chr2:523776-525788 | p5 | (GGCCC)3 | TAACTAAAACCTGGCGTGGC | TTTAAGCCTGTCAAGTGGGC | 254 |
| chr2:523790-525806 | p2 | (AG)6 | TAACTAAAACCTGGCGTGGC | TTTAAGCCTGTCAAGTGGGC | 240 |
| chr2:523790-525806 | p5 | (GGCCC)3 | TAACTAAAACCTGGCGTGGC | TTTAAGCCTGTCAAGTGGGC | 240 |
| chr2:523811-525865 | p5 | (GGCCC)3 | TAACTAAAACCTGGCGTGGC | TTTAAGCCTGTCAAGTGGGC | 219 |
| chr2:523893-525905 | p5 | (GGCCC)3 | TAACTAAAACCTGGCGTGGC | TTTAAGCCTGTCAAGTGGGC | 137 |
| chr2:523916-525928 | p5 | (GGCCC)3 | TAACTAAAACCTGGCGTGGC | TTTAAGCCTGTCAAGTGGGC | 114 |
| chr2:584458-586484 | p2 | (AT)13 | TCAGAAGGACGTGACGTGAG | TTTCCGGACCAAAGTTTCAC | 245 |
| chr2:588831-590843 | p2 | (AG)6 | TAACTAAAACCTGGCGTGGC | TTTAAGCCTGTCAAGTGGGC | 254 |
| chr2:588831-590843 | p5 | (GGCCC)3 | TAACTAAAACCTGGCGTGGC | TTTAAGCCTGTCAAGTGGGC | 254 |
| chr2:588845-590861 | p2 | (AG)6 | TAACTAAAACCTGGCGTGGC | TTTAAGCCTGTCAAGTGGGC | 240 |
| chr2:588845-590861 | p5 | (GGCCC)3 | TAACTAAAACCTGGCGTGGC | TTTAAGCCTGTCAAGTGGGC | 240 |
| chr2:588866-590920 | p5 | (GGCCC)3 | TAACTAAAACCTGGCGTGGC | TTTAAGCCTGTCAAGTGGGC | 219 |
| chr2:588948-590960 | p5 | (GGCCC)3 | TAACTAAAACCTGGCGTGGC | TTTAAGCCTGTCAAGTGGGC | 137 |

| Marker ID | SSR type | SSR motif | FORWARD PRIMER (5'-3') | REVERSE PRIMER (5'-3') | size (bp) |
|---|---|---|---|---|---|
| chr2:588971-590983 | p5 | (GGCCC)3 | TAACTAAAACCTGGCGTGGC | TTTAAGCCTGTCAAGTGGGC | 114 |
| chr2:713424-715439 | p5 | (AAAAT)3 | AACGACCGTGGAGACTTGAC | CAATCCAAACGGGCTCTTAG | 256 |
| chr2:794225-796243 | p2 | (AT)9 | TATGACACCGAGTCCCATGA | ATACCGTACACGGCCCATTA | 110 |
| chr2:837485-839497 | p2 | (AG)6 | TAGGAAAGTCCGCAGGCTAA | TGGCCTTCATTTCCTCTTTG | 210 |
| chr2:839438-841456 | p2 | (TA)9 | GGCTGTTTGGAATATTGCGT | TTTCCCCACGTTAGCAACTC | 277 |
| chr2:970126-972138 | p2 | (CA)6 | AAGCCTTTGGCTTCACAAGA | CATTTCAGGGGAGATGCTGT | 249 |
| chr2:983565-985591 | p2 | (TA)6 | AAATGCATGGGTCTGTTGCT | GGCATAGACAAAGTAATGCGG | 282 |
| chr2:1006618-1008630 | p2 | (AG)6 | GAGCCAAAGGATGCATTGAT | TTGCATGATTTTTGGGATGA | 156 |
| chr2:1021569-1023593 | p6 | (GGTGCA)4 | TAGCCTAAGCGGAAGGTCAA | GGTAGGTCGATTGCGGAGTA | 107 |
| chr2:1021569-1023593 | p5 | (AATGC)3 | TAGCCTAAGCGGAAGGTCAA | GGTAGGTCGATTGCGGAGTA | 107 |
| chr2:1023335-1025347 | p5 | (AATGC)3 | AATGCAACAGTCGCCTCTCT | TGATTGAAAGCCTCCGAGAT | 165 |
| chr2:1023356-1025368 | p5 | (AATGC)3 | AATGCAACAGTCGCCTCTCT | TGATTGAAAGCCTCCGAGAT | 144 |
| chr2:1025124-1027138 | p2 | (AT)6 | AAAACCATGCTTCGTCATCC | GATAGTATAGCGGGCGTCCA | 268 |
| chr2:1025124-1027138 | p2 | (TC)6 | AAAACCATGCTTCGTCATCC | GATAGTATAGCGGGCGTCCA | 268 |
| chr2:1025159-1027173 | p2 | (AT)6 | AAAACCATGCTTCGTCATCC | GATAGTATAGCGGGCGTCCA | 233 |
| chr2:1025159-1027173 | p2 | (TC)6 | AAAACCATGCTTCGTCATCC | GATAGTATAGCGGGCGTCCA | 233 |
| chr2:1027935-1029947 | p2 | (TA)14 | TTAATGTCCAAAAGGGTGGG | TTTTAGGGACAACGGACGAC | 219 |
| chr2:1027935-1029947 | p2 | (AT)6 | TTAATGTCCAAAAGGGTGGG | TTTTAGGGACAACGGACGAC | 219 |
| chr2:1147923-1149938 | p5 | (AGTAT)3 | ATCGACGAGCGGTTTGTTAC | CATGGAATCTATTGGGCGTT | 125 |
| chr2:1157933-1159948 | p2 | (AC)8 | CTTTTGCCCTTCATACGGAA | TGACAGCGAAAACGACACTC | 173 |
| chr2:1157933-1159948 | p5 | (AAAAT)3 | CTTTTGCCCTTCATACGGAA | TGACAGCGAAAACGACACTC | 173 |
| chr2:1157945-1159963 | p2 | (AC)8 | CTTTTGCCCTTCATACGGAA | TGACAGCGAAAACGACACTC | 161 |
| chr2:1157945-1159963 | p5 | (AAAAT)3 | CTTTTGCCCTTCATACGGAA | TGACAGCGAAAACGACACTC | 161 |
| chr2:1221685-1223705 | p2 | (TA)10 | ATCAAGGTACGGCTTGGATG | ATACTTGCTGAGGCGGAGAA | 237 |
| chr2:1342274-1344290 | p2 | (AC)8 | TAGACCCTCGGCTCAAAAGA | TGTTTATGTTTCTCGGCGTG | 265 |
| chr2:1408385-1410403 | p3 | (GTT)6 | GCCAGCTTAAAGCCATTTTG | CCTGTTTGGATGGGCTTAAA | 164 |
| chr2:1456742-1458756 | p4 | (TTTA)4 | GGGCATTTTCGTCCATACAC | GTCGACGGATCAGATTGGTT | 239 |
| chr2:1456742-1458756 | p2 | (TG)6 | GGGCATTTTCGTCCATACAC | GTCGACGGATCAGATTGGTT | 239 |
| chr2:1456742-1458756 | p2 | (TG)7 | GGGCATTTTCGTCCATACAC | GTCGACGGATCAGATTGGTT | 239 |
| chr2:1477496-1479511 | p5 | (GTTAT)3 | TCAGAGTCGAACGCTTCCTT | AGCCCTTCTCTTTGACCCTC | 150 |
| chr2:1495071-1497086 | p5 | (TACAA)3 | TTGTCCCGTGGTGTTCATTA | TGAGTGTAGAATGGGGAGGG | 202 |
| chr2:1524203-1526236 | p3 | (AAT)11 | TCAGACCAATTTGCTTGCAC | CTTAGGGGAGAGCAGCAATG | 172 |
| chr2:1835534-1837546 | p2 | (TA)6 | TTTTGCCAGACTTTTGAGGG | ACCCCACCTTGAATCTCCTT | 294 |
| chr2:1888749-1890809 | p2 | (AT)6 | GAGAGTAAAAGCGCAAACCG | GACCAGTAGGGCCTCTTTCC | 268 |
| chr2:1888749-1890809 | p2 | (TA)30 | GAGAGTAAAAGCGCAAACCG | GACCAGTAGGGCCTCTTTCC | 268 |
| chr2:1888749-1890809 | p3 | (CAA)5 | GAGAGTAAAAGCGCAAACCG | GACCAGTAGGGCCTCTTTCC | 268 |

| Marker ID | SSR type | SSR motif | FORWARD PRIMER (5'-3') | REVERSE PRIMER (5'-3') | size (bp) |
|---|---|---|---|---|---|
| chr2:1893796-1895812 | p2 | (TA)8 | GCCCATCATCCTCGTTCTTA | CGATTCAGGGTCGATTTTGT | 127 |
| chr2:1910356-1912374 | p2 | (AT)9 | ATTGGACGTTTTCGCATTTC | CTTTTTCAAAACTCGGGTGG | 106 |
| chr2:1988753-1990771 | p2 | (AT)9 | TATGACACCGAGTCCCATGA | ATACCGTACACGGCCCATTA | 110 |
| chr2:2032013-2034025 | p2 | (AG)6 | TAGGAAAGTCCGCAGGCTAA | TGGCCTTCATTTCCTCTTTG | 210 |
| chr2:2033966-2035984 | p2 | (TA)9 | GGCTGTTTGGAATATTGCGT | TTTCCCCACGTTAGCAACTC | 277 |
| chr2:2071208-2073223 | p3 | (ATT)5 | TATTTTGGCAGAGAGACGGG | TGTGTACATCTTGCTCAAGGC | 101 |
| chr2:2071208-2073223 | p2 | (AT)23 | TATTTTGGCAGAGAGACGGG | TGTGTACATCTTGCTCAAGGC | 101 |
| chr2:2226052-2228064 | p2 | (AC)6 | TTGTCTGATCCGTTCCCTTC | TGGTTCCCCAGATCTTCATC | 297 |
| chr2:2420978-2422993 | p3 | (CAA)5 | GGGAACAATGCAGTGGAAGT | ATTGTTTTTGAGGCACTCGG | 109 |
| chr2:2584823-2586835 | p2 | (TA)6 | TTCCGGTATTCAAATGGCTC | ATCGGTCTCGGATGATCTTG | 102 |
| chr2:2632579-2634597 | p6 | (TATATG)3 | TTAGATTCTCGGGGATGGTG | TCCACGCCCTATTAGCTACG | 168 |
| chr2:2692718-2694733 | p3 | (TAA)5 | AGTGGTGTAGTGGAGGGTGC | ATTTCCCCTCTTTCGTCGTT | 133 |
| chr2:2692993-2695008 | p3 | (TAA)5 | TTTTGGATAAAGACGGGTCG | ACTCGGTTCGCATTTTTGAC | 111 |
| chr2:2765605-2767650 | p2 | (AG)6 | CGGGAATTTCTTGAATCGAA | CGCTCCTCTCCTTCATCATC | 232 |
| chr2:2765605-2767650 | p3 | (AAT)15 | CGGGAATTTCTTGAATCGAA | CGCTCCTCTCCTTCATCATC | 232 |
| chr2:2765605-2767650 | p2 | (AT)19 | CGGGAATTTCTTGAATCGAA | CGCTCCTCTCCTTCATCATC | 232 |
| chr2:2765842-2767880 | p2 | (AG)6 | GAAGAAAATAAGGGAGCGGG | GGACTAGCCCATACCCGTTT | 235 |
| chr2:2765842-2767880 | p3 | (AAT)15 | GAAGAAAATAAGGGAGCGGG | GGACTAGCCCATACCCGTTT | 235 |
| chr2:2765842-2767880 | p2 | (AT)19 | GAAGAAAATAAGGGAGCGGG | GGACTAGCCCATACCCGTTT | 235 |
| chr2:2814224-2816239 | p5 | (TACCA)3 | ACCCCCGACATACTTGACTG | AAGAGGTGCCTAGGTGGGTT | 138 |
| chr2:2912130-2914152 | p2 | (TA)7 | CAAGCCGATTTTTGGTTCAT | CGTACCTAACTCCGCCTCTG | 191 |
| chr2:2912130-2914152 | p2 | (TA)11 | CAAGCCGATTTTTGGTTCAT | CGTACCTAACTCCGCCTCTG | 191 |
| chr2:2912130-2914152 | p2 | (TC)21 | CAAGCCGATTTTTGGTTCAT | CGTACCTAACTCCGCCTCTG | 191 |
| chr2:3082110-3084125 | p5 | (TTAAT)3 | CCAGATTCTTGTTCCCTCCA | AAAACGCATCATGCAACAAA | 102 |
| chr2:3082110-3084125 | p3 | (CTT)5 | CCAGATTCTTGTTCCCTCCA | AAAACGCATCATGCAACAAA | 102 |
| chr2:3117699-3119717 | p3 | (AAT)6 | TTATGGAGATTTTGAGGCGG | AGAATCGCAAGAATCCCCTT | 197 |
| chr2:3117699-3119717 | p2 | (AT)6 | TTATGGAGATTTTGAGGCGG | AGAATCGCAAGAATCCCCTT | 197 |
| chr2:3174004-3176032 | p2 | (TA)14 | ATATGGGGCCCACAATTTTT | CGTCGTCCATTGATGTTGAC | 196 |
| chr2:3174004-3176032 | p2 | (TA)8 | ATATGGGGCCCACAATTTTT | CGTCGTCCATTGATGTTGAC | 196 |
| chr2:3179413-3181425 | p2 | (TA)6 | CCATGACTTTGGATGTGGTG | CAACCCATAAGAGGGAGCAA | 183 |
| chr2:3179413-3181425 | p2 | (TA)7 | CCATGACTTTGGATGTGGTG | CAACCCATAAGAGGGAGCAA | 183 |
| chr2:3179413-3181425 | p2 | (AT)8 | CCATGACTTTGGATGTGGTG | CAACCCATAAGAGGGAGCAA | 183 |
| chr2:3232996-3235014 | p6 | (ATCTAT)3 | CATAACCCTGCCCATTTCAC | TCGGGATCAAGTATTTTCCG | 147 |
| chr2:3309074-3311089 | p5 | (TTTCT)3 | TGGCATTTTTGAAGCTGTTG | ACATGTCGTTGTCCGAAACA | 225 |
| chr2:3447784-3449800 | p2 | (AT)8 | AAAGTCCAAACGGCTTCAAA | ACCAATGGACCCATCTTTCA | 186 |
| chr2:3453605-3455617 | p2 | (TA)6 | TGATGCGCTTTGTCTTTGTC | TCTCGGACCAACAGTAACCC | 239 |

| Marker ID | SSR type | SSR motif | FORWARD PRIMER (5'-3') | REVERSE PRIMER (5'-3') | size (bp) |
|---|---|---|---|---|---|
| chr2:3453605-3455617 | p2 | (AT)8 | TGATGCGCTTTGTCTTTGTC | TCTCGGACCAACAGTAACCC | 239 |
| chr2:3454177-3456193 | p2 | (TA)6 | CTTGACTGCAGCATCAGGAA | GGCCCCATATTAAACAGGCT | 106 |
| chr2:3454177-3456193 | p2 | (AT)8 | CTTGACTGCAGCATCAGGAA | GGCCCCATATTAAACAGGCT | 106 |
| chr2:3486423-3488438 | p5 | (TAAAA)3 | ACCCAATTGTAACAAGCCCA | CGGTTCGGTTCAATTCAGTT | 175 |
| chr2:3490327-3492342 | p5 | (TAAAA)3 | CGGGGTGAATGCTAAGTGAT | TGTGATAAGGAGGTCCCGAG | 217 |
| chr2:3507597-3509612 | p3 | (ATA)5 | ACACACTTCCAGGATGAGGC | GACACGGACTTGGAGGAAAA | 120 |
| chr2:3625398-3627414 | p2 | (AT)11 | TCCCAAAGTATCCCGATCAA | GACATGGCGGAATAGCAAAT | 223 |
| chr2:3625398-3627414 | p2 | (TA)8 | TCCCAAAGTATCCCGATCAA | GACATGGCGGAATAGCAAAT | 223 |
| chr2:3678482-3680500 | p3 | (TGT)6 | ATGGATACTGTGCCCCTGAG | AAGCCTTTCGCCCTGTTATT | 267 |
| chr2:3706845-3708892 | p2 | (TG)7 | ATGCAATCGTGCAGCAAATA | TCACGTGAACACATGGTCCT | 131 |
| chr2:3726239-3728255 | p4 | (ATTT)4 | AAATGCCTACGAGCAGCAAT | TAATGACAGGCAAAATCGCA | 164 |
| chr2:3726239-3728255 | p5 | (TTATT)3 | AAATGCCTACGAGCAGCAAT | TAATGACAGGCAAAATCGCA | 164 |
| chr2:3914069-3916084 | p5 | (GAAAT)3 | CATCTACCTCAACGGGCCTA | CTAGCATTGGAGGAGAACGC | 111 |
| chr2:3922638-3924653 | p3 | (TTG)5 | CAAAAATTTGCCCTCCAAAA | CTAATGCTCCCCACTCCAAA | 234 |
| chr2:3979227-3981242 | p5 | (GGGGT)3 | ACCAAATCGGGTTGTTTTCA | AAACCCCTTACCATTTCGCT | 198 |
| chr2:3988031-3990046 | p3 | (AAG)5 | GGGGCCCAAAATTAGAACTC | AGGCGTTCATTGTAGGTTGG | 237 |
| chr2:3994684-3996699 | p6 | (AGGCAA)3 | GCTGATAGGCAAAGGCAAAG | AACCGGTGTATGATTTGGGA | 149 |
| chr2:3994684-3996699 | p5 | (ACAAC)3 | GCTGATAGGCAAAGGCAAAG | AACCGGTGTATGATTTGGGA | 149 |
| chr2:4044507-4046525 | p5 | (CGGGT)3 | GGGGAACAGAAGTGAGTGGA | TACCTAACGAACCAGCCCAC | 176 |
| chr2:4044507-4046525 | p3 | (TAA)9 | GGGGAACAGAAGTGAGTGGA | TACCTAACGAACCAGCCCAC | 176 |
| chr2:4044507-4046525 | p2 | (TA)9 | GGGGAACAGAAGTGAGTGGA | TACCTAACGAACCAGCCCAC | 176 |
| chr2:4077655-4079670 | p5 | (TTTGG)3 | CGTTAGCGACGACATTTTCA | GGCCCAAAATCATCACTCAC | 107 |
| chr2:4120528-4122576 | p3 | (AAT)9 | CATTAAAATGGGCTGGGATG | TGCACGGCTATTACAACGTC | 169 |
| chr2:4120528-4122576 | p3 | (ATA)16 | CATTAAAATGGGCTGGGATG | TGCACGGCTATTACAACGTC | 169 |
| chr2:4280770-4282785 | p3 | (TTG)5 | CCACCGAAACTTAAAGGCAA | CGCATCTTGTCTTCCACTGA | 200 |
| chr2:4287939-4289965 | p3 | (GAT)6 | TGTTGTTTCCATTGTTGCGT | CCCATCGCAGCTTCATTTAT | 233 |
| chr2:4307621-4309636 | p5 | (AAACC)3 | CTGGGATGGAGGGAGTATCA | CCCATGAACACCCCTAGTTG | 231 |
| chr2:4319409-4321425 | p4 | (AAAT)4 | TCGGGCAAGTTTTAATGAGC | TTGTGACGTGGAGCAGTCTC | 294 |
| chr2:4389485-4391517 | p2 | (TA)16 | TGGTCCAGGAGGTTACTTGC | TTAGGTTATCATGGGCCTGG | 198 |
| chr2:4448136-4450151 | p3 | (TGT)5 | GAGTTCATTTCATTCGCGGT | TTTATGCGGTACTTCTGCCC | 243 |
| chr2:4503003-4505021 | p6 | (TAAAAT)3 | GATCCGCGAGTGACTTTAGC | GCTCTGGAAACGGCTAGATG | 161 |
| chr2:4503003-4505021 | p5 | (CAAAC)3 | GATCCGCGAGTGACTTTAGC | GCTCTGGAAACGGCTAGATG | 161 |
| chr2:4535086-4537132 | p2 | (TA)8 | TTCATTACCACGTGCGTGTC | AAATAAATGACATGGCGGGA | 234 |
| chr2:4535086-4537132 | p2 | (AT)6 | TTCATTACCACGTGCGTGTC | AAATAAATGACATGGCGGGA | 234 |
| chr2:4535086-4537132 | p2 | (AT)23 | TTCATTACCACGTGCGTGTC | AAATAAATGACATGGCGGGA | 234 |
| chr2:4632026-4634044 | p5 | (ACCAA)3 | CGGATTAGGGGTGTACATGG | TCGGTCGTGACACCTGAATA | 262 |

| Marker ID | SSR type | SSR motif | FORWARD PRIMER (5'-3') | REVERSE PRIMER (5'-3') | size (bp) |
|---|---|---|---|---|---|
| chr2:4632026-4634044 | p2 | (AT)7 | CGGATTAGGGGTGTACATGG | TCGGTCGTGACACCTGAATA | 262 |
| chr2:4632026-4634044 | p6 | (AAAAAT)3 | CGGATTAGGGGTGTACATGG | TCGGTCGTGACACCTGAATA | 262 |
| chr2:4689108-4691126 | p2 | (AT)12 | CACCCCTTATCTCGCATCAT | TGGTTGTCCCTGTCGATGTA | 183 |
| chr2:4689108-4691126 | p6 | (CATATA)3 | CACCCCTTATCTCGCATCAT | TGGTTGTCCCTGTCGATGTA | 183 |
| chr2:4733499-4735533 | p3 | (GAA)5 | GCAAGTGGTGTTGCTTGCTA | TTTTCATCTCCCTCCACCAG | 285 |
| chr2:5007057-5009073 | p2 | (AT)8 | TCCTGCCAAGAAAATTGACC | CAGCCTTACATGGTGGGTTT | 165 |
| chr2:5022719-5024735 | p2 | (AT)8 | TCCTGCCAAGAAAATTGACC | CAGCCTTACATGGTGGGTTT | 165 |
| chr2:5085555-5087585 | p2 | (TA)15 | TTTGCACTTGAAGCATCCAC | TATTGAGCCACCCATGATGA | 235 |
| chr2:5210544-5212560 | p4 | (CTTT)4 | ATTTACACAGCAGCAGTGCG | GTCTTCCAAGGGAAACCTCC | 148 |
| chr2:5222317-5224329 | p2 | (TC)6 | CGTGAACAAGTACCGACCCT | CCAGTTGACACCCCTTGATT | 190 |
| chr2:5294083-5296099 | p3 | (AAG)5 | AAATCCTTCACAAGGCGATG | GCTCCGAAGATTCAAACTGG | 207 |
| chr2:5294083-5296099 | p4 | (TAAA)4 | AAATCCTTCACAAGGCGATG | GCTCCGAAGATTCAAACTGG | 207 |
| chr2:5294083-5296099 | p3 | (AAG)5 | AAATCCTTCACAAGGCGATG | GCTCCGAAGATTCAAACTGG | 207 |
| chr2:5706981-5709001 | p6 | (ATTTTA)4 | TTGTCGTTTGTCGCCATTAG | GTATGCCGACAAGGGTCTGT | 130 |
| chr2:5706981-5709001 | p5 | (AAATA)4 | TTGTCGTTTGTCGCCATTAG | GTATGCCGACAAGGGTCTGT | 130 |
| chr2:5715495-5717519 | p6 | (TTCCCC)4 | GAGAGTAGCCCCAGAGGACC | ACCTCTGCCAAAAGAAGCAA | 168 |
| chr2:5715495-5717519 | p6 | (TTCCCC)4 | GAGAGTAGCCCCAGAGGACC | ACCTCTGCCAAAAGAAGCAA | 168 |
| chr2:5820241-5822274 | p3 | (TAA)11 | TTGCATACGCATTGCTCTTC | TGTGTCTGGTAGGCTGTCCA | 265 |
| chr2:5831952-5833993 | p3 | (AAC)5 | AGCCATGGTTGTCCTATTGC | AACTGCTGGTCTGCCTCAGT | 266 |
| chr2:5835026-5837046 | p2 | (TA)9 | GCTGCATTTCCACCAAAAGT | GGAAAGGCCATGATAAAGCA | 125 |
| chr2:5835026-5837046 | p5 | (TTTGT)4 | GCTGCATTTCCACCAAAAGT | GGAAAGGCCATGATAAAGCA | 125 |
| chr2:5845817-5847832 | p3 | (TTG)5 | GATGCAGGTGTTGCTGAAGA | GGAACTACGGGAGGAAGGAC | 171 |
| chr2:5845817-5847832 | p2 | (TC)10 | GATGCAGGTGTTGCTGAAGA | GGAACTACGGGAGGAAGGAC | 171 |
| chr2:5985194-5987216 | p2 | (AG)11 | CATGACGGAATTGGAGCTTT | AACTCATCCCATTTGCTTGG | 207 |
| chr2:5985194-5987216 | p3 | (TTG)5 | CATGACGGAATTGGAGCTTT | AACTCATCCCATTTGCTTGG | 207 |
| chr2:6029598-6031613 | p3 | (CAT)5 | GTGGAAGGCATCGTCAAAAT | TACGCGCATTGTTAATTCCA | 135 |
| chr2:6050583-6052598 | p5 | (TTTTC)3 | GACACAAGTGCGTCAGCATT | TTTCAGTTTTGGGGTTGAGG | 205 |
| chr2:6063397-6065413 | p4 | (CTTT)4 | CATCTTTTTGCAACTGGGGT | TTTGGTTGGACTTAGGGTGC | 286 |
| chr2:6110263-6112279 | p4 | (CTTT)4 | CATCTTTTTGCAACTGGGGT | TTTGGTTGGACTTAGGGTGC | 286 |
| chr2:6192511-6194558 | p2 | (CT)11 | TTTTCCTTTTCCGTGAATGC | ATCCCACTAGCCAGTGATGG | 183 |
| chr2:6244397-6246409 | p2 | (GA)6 | CATGTTGAGATCGCTCCTGA | GATGCTTCTTCTCTCCACCG | 224 |
| chr2:6251904-6253918 | p2 | (TC)7 | TTTCATCATCACCTGCCAAA | GAACGAAAACGCCGGTATTA | 242 |
| chr2:6267888-6269903 | p5 | (TTGAT)3 | GAAGGGTTTGTGGATCTGGA | TGTACCCTTTTTCGGCTGAC | 263 |
| chr2:6316024-6318042 | p2 | (AT)6 | CCAATAGTGGAAGCGCAAGT | CTCGCTCGAATTGGTTGATT | 228 |
| chr2:6316059-6318083 | p2 | (AT)6 | CCAATAGTGGAAGCGCAAGT | CTCGCTCGAATTGGTTGATT | 193 |
| chr2:6317747-6319787 | p2 | (TA)20 | GATGCAATTCATTACCCCGT | AAATGACATGGCGGAATAGC | 239 |

| Marker ID | SSR type | SSR motif | FORWARD PRIMER (5'-3') | REVERSE PRIMER (5'-3') | size (bp) |
|---|---|---|---|---|---|
| chr2:6545639-6547654 | p5 | (ATAGT)3 | CGGGCTTATCGAAGAATCAA | TTTCCTCAATCATGGGCTTC | 260 |
| chr2:6565845-6567871 | p2 | (AG)13 | TAGGTAGTGAGTGGGGGTGG | CGATGTCACCTCGGGATACT | 186 |
| chr2:6676267-6678281 | p2 | (TG)7 | TAACCCGGGACAATACCAAA | TGATGAGAGAGGGAGGGAAA | 290 |
| chr2:6705973-6707989 | p4 | (TAAA)4 | TTGGGCATTCGTGTTTTGTA | AGGACAAACGATAGGCCCTT | 158 |
| chr2:6732469-6734483 | p2 | (AT)7 | CCCCTGCATCAACATTCTTT | GCGGGTACAGAAGGTACCAA | 189 |
| chr2:6783171-6785183 | p2 | (TA)6 | ACATGCGCATTTGCTGAATA | ATAATCCCACAGCAAGGCAC | 157 |
| chr2:6806585-6808600 | p5 | (GAAAA)3 | CCTCAACTTCAACCGTTGGT | CGACAGATCGTACCGGATTT | 256 |
| chr2:6812515-6814536 | p3 | (AAT)7 | TTCGACCGCCAATAGAGAAC | TTCCTCCTCCATTCTCAGGA | 163 |
| chr2:6902368-6904383 | p3 | (TTG)5 | TGTAAAATCCCGTTTTGGGA | ATTTGGAGCATGGAACTTGG | 172 |
| chr2:6955155-6957170 | p3 | (TAA)5 | CCCTTTTGTGATGGAGCATT | AAATTCACACTTTGGCTGGC | 214 |
| chr2:7012118-7014130 | p2 | (TA)6 | TGCTTTTTCGGCGTAGATTT | GACCTTTCACCACCGTCACT | 128 |
| chr2:7012118-7014130 | p2 | (TA)6 | TGCTTTTTCGGCGTAGATTT | GACCTTTCACCACCGTCACT | 128 |
| chr2:7123320-7125332 | p2 | (AT)6 | TGATTACCGTTGCGTTTCAA | CATATGCCGAAGGAGAGAGC | 175 |
| chr2:7130585-7132603 | p3 | (ACA)5 | CAAGCGGTATGCCCTTGTAT | AGCATTTTCATTGCTCACCC | 128 |
| chr2:7130585-7132603 | p3 | (CAA)6 | CAAGCGGTATGCCCTTGTAT | AGCATTTTCATTGCTCACCC | 128 |
| chr2:7153795-7155816 | p3 | (TTC)7 | TGACACAACCATGGCAAAGT | TTTTTGATCCGCCTGTTTTC | 268 |
| chr2:7153795-7155816 | p3 | (ACA)5 | TGACACAACCATGGCAAAGT | TTTTTGATCCGCCTGTTTTC | 268 |
| chr2:7154401-7156416 | p3 | (TTC)7 | AATGTGTCGGATTGGATGGT | GGTTCAAAGGCCACAGACAT | 172 |
| chr2:7154401-7156416 | p3 | (ACA)5 | AATGTGTCGGATTGGATGGT | GGTTCAAAGGCCACAGACAT | 172 |
| chr2:7310781-7312801 | p2 | (AT)10 | TCATCAAGCTCCGGGTAAAT | GGGGTGCAATTAACGACATT | 153 |
| chr2:7444836-7446851 | p3 | (ACA)5 | GAAGCAGGAATCCTTTGTGC | TTTGGGCCACCCATTTATTA | 243 |
| chr2:7488222-7490234 | p2 | (CT)6 | CGGTGGTAGCTGGTAATGCT | TCTGGACAATGCAGAGCAAC | 156 |
| chr2:7534408-7536424 | p4 | (GTTT)4 | TCAAACATTGGTGCTTGCAT | CCTTGCCGTCTGGGTAAATA | 144 |
| chr2:7577778-7579799 | p3 | (TAA)7 | GTCGAGGTTTTGGGTTCAGA | ATCGTAATCGTGGTTGGAGC | 233 |
| chr2:7577778-7579799 | p3 | (TTA)5 | GTCGAGGTTTTGGGTTCAGA | ATCGTAATCGTGGTTGGAGC | 233 |
| chr2:7723938-7725953 | p3 | (AAC)5 | TATAGCCTCTGGCCCATTTG | AATTGGAGACGGAACAGACG | 208 |
| chr2:7878224-7880239 | p5 | (TTCTT)3 | TGAAACTCATAACCTCCGGG | AACCTTGGTTGGGCCTTACT | 112 |
| chr2:7927727-7929742 | p5 | (TAAAA)3 | GATTGATTTCCGGTTCATGC | GTAATTTTCCCCGCAATGTG | 121 |
| chr2:7966247-7968263 | p4 | (TTTC)4 | CGCGAGATTGGAGGTTAGAG | ATCTAGCTGCACGTCCTCGT | 187 |
| chr2:8310785-8312797 | p2 | (TA)6 | TGGCCTACCAAAGGTTCTTG | TAGTTCGCTGTGCAGTTTGG | 143 |
| chr2:8446743-8448761 | p6 | (AATTCA)3 | GCATATGCCCCCATTAACAT | TAGAAACCCATCTGCTGCCT | 131 |
| chr2:8498301-8500361 | p3 | (TTA)5 | ACACGACAAACCTCCACCTC | TTTCTACAATGAGGGCCCAG | 227 |
| chr2:8498301-8500361 | p3 | (TTA)8 | ACACGACAAACCTCCACCTC | TTTCTACAATGAGGGCCCAG | 227 |
| chr2:8498619-8500643 | p3 | (TTA)5 | CTGGGCCCTCATTGTAGAAA | ATTCAATTTGGCCGCATAAG | 124 |
| chr2:8498619-8500643 | p3 | (TTA)8 | CTGGGCCCTCATTGTAGAAA | ATTCAATTTGGCCGCATAAG | 124 |
| chr2:8503767-8505782 | p5 | (CTTTT)3 | CGCTGAGGATGAGATTGACA | TCTGATCAACAGGGAGGACC | 129 |

| Marker ID | SSR type | SSR motif | FORWARD PRIMER (5'-3') | REVERSE PRIMER (5'-3') | size (bp) |
|---|---|---|---|---|---|
| chr2:8508504-8510522 | p3 | (GTT)6 | CGCCCCAATTAGTAGAACCA | TGCAAGCATGTTTTTCCATC | 272 |
| chr2:8596254-8598270 | p4 | (TTAA)4 | CTCTCCCGATCCTGTGTGTT | CAAGGGAAAGAGGGGTTAGG | 149 |
| chr2:8623845-8625860 | p6 | (TTTTTA)3 | CTTGTGGGATTTCACCGAGT | AAAATGGCAGCATGAAGGAC | 265 |
| chr2:8623845-8625860 | p5 | (AAAAG)3 | CTTGTGGGATTTCACCGAGT | AAAATGGCAGCATGAAGGAC | 265 |
| chr2:8625164-8627180 | p4 | (TTTA)4 | GATCACTCCGTCGTGACCTT | ACAGTTGCTGGAAATGGGAC | 223 |
| chr2:8693101-8695127 | p2 | (TA)13 | GATTGGGCATGCAAAAGACT | ACGGTTCCTGGCAATTAGTG | 173 |
| chr2:8721150-8723162 | p2 | (AG)6 | AAGAATGGTCCCGACAACAC | ACTCCATATCAGGGTGGCAG | 280 |
| chr2:8870390-8872402 | p5 | (ATCAT)3 | GGAAATTGATGTCAACCGCT | TCCGAGTGGGTTGTTTCTTC | 221 |
| chr2:8870390-8872402 | p2 | (TA)6 | GGAAATTGATGTCAACCGCT | TCCGAGTGGGTTGTTTCTTC | 221 |
| chr2:8872940-8874955 | p3 | (AAG)5 | CCTGGATTCTTAAATCGCCA | CACCACCAGAGAAAGGCAAT | 225 |
| chr2:8888178-8890193 | p2 | (TG)8 | GTTCCCCCTTTTTGCTGAAT | TCATCCTTGCTTGCTGTCAC | 190 |
| chr2:8888178-8890193 | p3 | (AAT)5 | GTTCCCCCTTTTTGCTGAAT | TCATCCTTGCTTGCTGTCAC | 190 |
| chr2:8915228-8917243 | p3 | (TAA)5 | TTTGGAGGGTTTCCACTTTG | TCCCGTATTTTTGAACCTCG | 114 |
| chr2:8970270-8972306 | p5 | (TTTTA)3 | TGGGTGTGCGTGAGATAAAA | AACGCATAACAAGGACACCC | 204 |
| chr2:9038570-9040594 | p3 | (TCT)8 | CGTCACACGGCAGCTAATAA | TTGCCCTGATTTTCGTTAGG | 180 |
| chr2:9331269-9333287 | p3 | (ACA)6 | GCTTGGCCAAACATGCTATT | CTGAAGCCCATCAAACCAGT | 207 |
| chr2:9369254-9371272 | p3 | (ACA)6 | GCTTGGCCAAACATGCTATT | CTGAAGCCCATCAAACCAGT | 207 |
| chr2:9417126-9419141 | p5 | (AATTT)3 | GGATTAGCTGAAGATTGCCG | TATGCTCCACGCTGAAAGTG | 204 |
| chr2:9434208-9436226 | p6 | (AATTGT)3 | CAGTTCTGAAGCTCTTGGGC | CAAATTCACTTTCCCCCTCA | 249 |
| chr2:9515070-9517082 | p2 | (AT)6 | AAGGGTGTCATGAGATTCGG | ATCTGCCGATCAAGTATGGC | 123 |
| chr2:9543708-9545723 | p5 | (AAAAT)3 | ACCTACAGATGAGATGGCGG | ACTTGGACGCTCTGCTCATT | 169 |
| chr2:9549117-9551135 | p6 | (TATATG)3 | AGGTAAGCGGGTCAGGGTAT | TTGATCAAGGATACCGGCTC | 160 |
| chr2:9689918-9691952 | p3 | (ACT)5 | AGAGCCTGGTTGGAGAGTGA | CGTATGCAACAGCATCCACT | 244 |
| chr2:9689918-9691952 | p2 | (TA)17 | AGAGCCTGGTTGGAGAGTGA | CGTATGCAACAGCATCCACT | 244 |
| chr2:9689918-9691952 | p2 | (AT)11 | AGAGCCTGGTTGGAGAGTGA | CGTATGCAACAGCATCCACT | 244 |
| chr2:9741519-9743537 | p6 | (TTTTTA)3 | AGAATCAAATCAGGTTGCGG | ATCAGGGAACTATACGGGGG | 269 |
| chr2:9781717-9783757 | p2 | (CA)7 | GCCACTTTCACAGGCATTTT | GGCTCACGTGGGTAAAACAT | 149 |
| chr2:9781717-9783757 | p2 | (TA)16 | GCCACTTTCACAGGCATTTT | GGCTCACGTGGGTAAAACAT | 149 |
| chr2:9781717-9783757 | p5 | (AAAAT)3 | GCCACTTTCACAGGCATTTT | GGCTCACGTGGGTAAAACAT | 149 |
| chr2:9807040-9809061 | p3 | (GAT)7 | AAATTTGAAACACCGCCTTG | ACTTCTGTCACGACCCAACC | 285 |
| chr2:9845095-9847141 | p2 | (TA)6 | AAGATGAAACACACGGGGAG | TCCAAACTCATGGGGAAAAA | 122 |
| chr2:9852040-9854052 | p2 | (TA)6 | TCAAACCCACTGATTGTGGA | AGGAGCCATCCCAAGATACC | 200 |
| chr2:9852040-9854052 | p2 | (TG)6 | TCAAACCCACTGATTGTGGA | AGGAGCCATCCCAAGATACC | 200 |
| chr2:9927035-9929050 | p3 | (AAC)5 | GCTGGTGGTGGTACAGAGGT | GGAATCAAAGAGCAATGGGA | 267 |
| chr2:10018484-10020496 | p2 | (AT)6 | CCCATTTAGGCCGAACCTAT | CGATGAACGCGTAGCTAACA | 279 |
| chr2:10103545-10105593 | p2 | (TA)24 | GGGAACCCCAAACTTCAAAT | GGAGTACGTGTTTCCTCCCA | 185 |

| Marker ID | SSR type | SSR motif | FORWARD PRIMER (5'-3') | REVERSE PRIMER (5'-3') | size (bp) |
|---|---|---|---|---|---|
| chr2:10107890-10109905 | p5 | (TCATA)3 | AATGGATCTGATCAGGGCAG | ATTGCGGTGACAGGTAGTCC | 148 |
| chr2:10115946-10117962 | p2 | (TA)8 | TGGAGGGAATCTGTTCATCC | GGGAGCAAACAGAGCTTCAC | 156 |
| chr2:10115946-10117962 | p6 | (TAATCC)3 | TGGAGGGAATCTGTTCATCC | GGGAGCAAACAGAGCTTCAC | 156 |
| chr2:10122315-10124331 | p2 | (TA)6 | GGACGACAAAACCCATTTGA | GCCACCACTAGTTTGCATGA | 235 |
| chr2:10122315-10124331 | p2 | (TA)7 | GGACGACAAAACCCATTTGA | GCCACCACTAGTTTGCATGA | 235 |
| chr2:10122315-10124331 | p2 | (TA)6 | GGACGACAAAACCCATTTGA | GCCACCACTAGTTTGCATGA | 235 |
| chr2:10122391-10124403 | p2 | (TA)6 | GGACGACAAAACCCATTTGA | GCCACCACTAGTTTGCATGA | 159 |
| chr2:10122391-10124403 | p2 | (TA)7 | GGACGACAAAACCCATTTGA | GCCACCACTAGTTTGCATGA | 159 |
| chr2:10122391-10124403 | p2 | (TA)6 | GGACGACAAAACCCATTTGA | GCCACCACTAGTTTGCATGA | 159 |
| chr2:10230547-10232563 | p2 | (TA)8 | GAAAAATCGAACCAAACCGA | GTTTGACTTTAGGCCCCCAT | 235 |
| chr2:10232833-10234865 | p2 | (AT)16 | CCAAAGAAAGGAAACCACCA | TACAGAGTTGCACTCCGCAC | 106 |
| chr2:10252753-10254768 | p5 | (ACCAA)3 | GGGCCATGACTGATGAAACT | ATTTCTACCCCTCCACCACC | 282 |
| chr2:10270746-10272758 | p2 | (AT)6 | AAGTATCAACCGTGGCTTCG | TCTTCCATTTGGTCCAGTCC | 236 |
| chr2:10300344-10302362 | p3 | (ACC)6 | TATTTCTATGGCCAAACGCC | CAAGTTTAGGGGGCTGCATA | 289 |
| chr2:10300344-10302362 | p3 | (GGC)5 | TATTTCTATGGCCAAACGCC | CAAGTTTAGGGGGCTGCATA | 289 |
| chr2:10393293-10395305 | p3 | (GTT)5 | AGCTTTGTCAAGGCGACTGT | GATTCGGCTGGGTCTTGTTA | 166 |
| chr2:10393329-10395345 | p3 | (GTT)5 | AGCTTTGTCAAGGCGACTGT | GATTCGGCTGGGTCTTGTTA | 130 |
| chr2:10406982-10408997 | p3 | (TTC)5 | TTTGTTTGAATCGTCGTGGA | GGCAAGCCCCTCCTAGATAC | 280 |
| chr2:10455093-10457109 | p4 | (TATT)4 | TGCACTTCTCTGCAATTTGG | GTTCATGCCTCTTGACCGTT | 101 |
| chr2:10527161-10529176 | p2 | (AT)12 | CAAGAAGCAGCCTTGGTAGG | GGGACCCACAAAATAAAGGG | 259 |
| chr2:10527161-10529176 | p5 | (ATAAA)3 | CAAGAAGCAGCCTTGGTAGG | GGGACCCACAAAATAAAGGG | 259 |
| chr2:10607572-10609587 | p3 | (TCA)5 | TGGACAAATTTCCCGACTTC | TTGCTGTTGCAACTTCCTTG | 234 |
| chr2:10615120-10617134 | p2 | (TA)7 | TTGGGGTATTTCGGTTCAAA | ATGACGCCCTGATTATTTCG | 153 |
| chr2:10615120-10617134 | p4 | (TAAA)4 | TTGGGGTATTTCGGTTCAAA | ATGACGCCCTGATTATTTCG | 153 |
| chr2:10656759-10658774 | p5 | (GACCC)3 | GAGTAATTCAGCCCAACCCA | GGTTACTGTGCTGAGGGAGC | 289 |
| chr2:10668594-10670612 | p2 | (AT)9 | CACAAGTGTATGTCGGTGGG | CGAATTGCCCGAAAAAGATA | 109 |
| chr2:10791138-10793154 | p5 | (TGGTT)3 | TTGGGACAAGAGGGTTGAAG | ATGAATATCGGACACCGAGC | 251 |
| chr2:10791240-10793255 | p5 | (TGGTT)3 | TTGGGACAAGAGGGTTGAAG | ATGAATATCGGACACCGAGC | 149 |
| chr2:10923162-10925177 | p2 | (AT)11 | GCCTGCAGGTAGAGTTTTGC | ATTGCACGTCAAAAGCACTG | 297 |
| chr2:10923172-10925218 | p2 | (AT)11 | GCCTGCAGGTAGAGTTTTGC | ATTGCACGTCAAAAGCACTG | 287 |
| chr2:11033435-11035453 | p3 | (TGT)6 | GAGAGAATCCAAGGGGAAGG | GAAGCAGTTGAAAAGAGCCG | 256 |
| chr2:11052332-11054358 | p2 | (AT)13 | GATCCAAACGGGCTCTTACA | GGACCAAAGGGAACAGTTGA | 146 |
| chr2:11091516-11093530 | p4 | (TTTA)4 | GGATTCAATGGACTTGGGTG | ATGTTTGGCGCTCGAGTATC | 225 |
| chr2:11091516-11093530 | p2 | (AT)7 | GGATTCAATGGACTTGGGTG | ATGTTTGGCGCTCGAGTATC | 225 |
| chr2:11091516-11093530 | p2 | (AT)7 | GGATTCAATGGACTTGGGTG | ATGTTTGGCGCTCGAGTATC | 225 |
| chr2:11193855-11195889 | p2 | (AT)17 | CTTCAAATCAAATGCGGAGC | AAGGGAGGACCAACTCCATC | 223 |

| Marker ID | SSR type | SSR motif | FORWARD PRIMER (5'-3') | REVERSE PRIMER (5'-3') | size (bp) |
|---|---|---|---|---|---|
| chr2:11194178-11196192 | p2 | (AT)17 | ATTAATTGGGCTCAAATGCG | GGGCAAACTTAGGGACTTTTG | 153 |
| chr2:11215411-11217427 | p2 | (AT)8 | TCATCCAACAAAATGCCAAA | CCACAATTCACGAAATTCCA | 234 |
| chr2:11215411-11217427 | p2 | (TA)7 | TCATCCAACAAAATGCCAAA | CCACAATTCACGAAATTCCA | 234 |
| chr2:11215411-11217427 | p2 | (TA)7 | TCATCCAACAAAATGCCAAA | CCACAATTCACGAAATTCCA | 234 |
| chr2:11259165-11261180 | p5 | (AAATC)3 | ATTACTCGTTTGGCCACCAG | AGGCGTTTGGTCATGAAAAT | 127 |
| chr2:11259165-11261180 | p3 | (AAG)5 | ATTACTCGTTTGGCCACCAG | AGGCGTTTGGTCATGAAAAT | 127 |
| chr2:11303939-11305954 | p5 | (TTGGG)3 | GGCAGCCAGAAAGAGACAAC | TCAAGGCACACTTATGCTGC | 249 |
| chr2:11303939-11305954 | p3 | (ACA)5 | GGCAGCCAGAAAGAGACAAC | TCAAGGCACACTTATGCTGC | 249 |
| chr2:11315086-11317098 | p5 | (AGTAA)3 | TTGGGTTTAATCCCTTGACG | TTCAAGTCTCATTTTGCCCC | 213 |
| chr2:11315086-11317098 | p2 | (TA)6 | TTGGGTTTAATCCCTTGACG | TTCAAGTCTCATTTTGCCCC | 213 |
| chr2:11315086-11317098 | p5 | (AAATA)3 | TTGGGTTTAATCCCTTGACG | TTCAAGTCTCATTTTGCCCC | 213 |
| chr2:11316043-11318058 | p2 | (TA)6 | GAGTGAAAGCGAAGGCTCAC | GGGTGGAGACTGGAGATTGA | 111 |
| chr2:11316043-11318058 | p5 | (AAATA)3 | GAGTGAAAGCGAAGGCTCAC | GGGTGGAGACTGGAGATTGA | 111 |
| chr2:11353019-11355053 | p2 | (TA)17 | CCTGCTTGCATGATTTCAGA | TACCTTGCCAGCAACAACTG | 171 |
| chr2:11498605-11500626 | p3 | (TGT)7 | GTGCACCATCTTAGCGTTGA | GTGTGTTGTTTGCGCACTCT | 274 |
| chr2:11526487-11528499 | p2 | (GA)6 | AGGCAGTACCCTTTGCCTTT | TACTTTGTCCCGAAACCAGG | 269 |
| chr2:11549616-11551628 | p2 | (TA)6 | GCTTAGCCATGACCGAAGAG | ACAAAAGCCCTTAGCAACCA | 200 |
| chr2:11620597-11622615 | p3 | (TTA)5 | CCCTTATTCAGGGGTGGTCT | AGGCAAACTCTACCTTGCGA | 260 |
| chr2:11620597-11622615 | p3 | (TTA)6 | CCCTTATTCAGGGGTGGTCT | AGGCAAACTCTACCTTGCGA | 260 |
| chr2:11620719-11622734 | p3 | (TTA)5 | CCCTTATTCAGGGGTGGTCT | AGGCAAACTCTACCTTGCGA | 138 |
| chr2:11620719-11622734 | p3 | (TTA)6 | CCCTTATTCAGGGGTGGTCT | AGGCAAACTCTACCTTGCGA | 138 |
| chr2:11693721-11695736 | p5 | (AAATA)3 | ATCGCATGGGGATCAAATAC | GAAAATAATGGGTTTGGGGG | 251 |
| chr2:11699848-11701869 | p3 | (GGT)7 | AATTCGAATGAACCCCCTTC | GTGTGGAAGAAAGTTGGGGA | 193 |
| chr2:11863294-11865316 | p2 | (TA)11 | ACCAGTGAACGGACTGCTCT | TTTCGGGTATTTGCGTTAGG | 245 |
| chr2:11888920-11890932 | p2 | (AT)6 | GCACATTTAGGTGCGGATTT | TAGATTTTGGACGAAACGGG | 116 |
| chr2:11915620-11917634 | p3 | (ACA)5 | ATGCGGAGCAAGAATTTGAC | TCGATATCCGCATTTAAGCC | 257 |
| chr2:11929109-11931125 | p2 | (AT)8 | GCAGAAGCTTCCGAGAACAC | CGTTCCTTTCGCTCTTCATC | 280 |
| chr2:11946210-11948226 | p2 | (CA)8 | AAAGAGGTCCTCGGATGGTT | ACTCCTTACCTTACGCCGGT | 290 |
| chr2:11954704-11956719 | p3 | (ACA)5 | AGGGTTAGGATGGGAACCAC | GATCCAATGCACCCTCATCT | 240 |
| chr2:12006630-12008648 | p6 | (TTTTTA)3 | TTGATGCTCCAAGAAAACCC | GTCACGGAATTCTCTCGCA | 167 |
| chr2:12006649-12008709 | p6 | (TTTTTA)3 | TTGATGCTCCAAGAAAACCC | GTCACGGAATTCTCTCGCA | 148 |
| chr2:12256891-12258906 | p3 | (ACA)5 | CGAAACGAAAGAAGACAGCC | CAGCCAAAGCATTGATGAGA | 105 |
| chr2:12398915-12400930 | p5 | (TAAAA)3 | CTCGGACGAGCAATACATGA | CGGGTAAGATGGGGGTAAAT | 145 |
| chr2:12439272-12441287 | p3 | (TAA)5 | TGTTAAGGGCCTGTTTGGAA | GAAACAAGGCCAGTTTGGAA | 125 |
| chr2:12453222-12455234 | p2 | (AT)6 | ACGTCAACACCTCGGATCTC | GTCCTCAGTTTTGTGCCCAT | 286 |
| chr2:12504585-12506600 | p5 | (TTATT)3 | TTCAGCCCCACTCTCATCTC | CAAAAATCCCCTTGTCGTTG | 137 |

| Marker ID | SSR type | SSR motif | FORWARD PRIMER (5'-3') | REVERSE PRIMER (5'-3') | size (bp) |
|---|---|---|---|---|---|
| chr2:12539713-12541725 | p2 | (CT)6 | GGCTAATTATGCCGGAGGAT | TTACACATTTGACCAGCCGA | 244 |
| chr2:12539713-12541725 | p2 | (AT)9 | GGCTAATTATGCCGGAGGAT | TTACACATTTGACCAGCCGA | 244 |
| chr2:12689657-12691683 | p2 | (AT)13 | GGTGTGCGTAAGCATTTCCT | GGCAATGAGCCATCTCAACT | 227 |
| chr2:12740887-12742905 | p6 | (TAACTG)3 | CTGCCTACACCGGCATACTT | TCTCTCGGTCACATGCAGTC | 275 |
| chr2:12813356-12815371 | p3 | (TGT)5 | TTTATGGCATGCATGGTTGT | CCACGGATCACCAAAAGTCT | 212 |
| chr2:12828557-12830575 | p6 | (GGGGGT)3 | TGGGAAAAGTTCCACAGTCC | GATTGCAAGGCTGAAGAAGC | 151 |
| chr2:12828557-12830575 | p2 | (CT)7 | TGGGAAAAGTTCCACAGTCC | GATTGCAAGGCTGAAGAAGC | 151 |
| chr2:12828557-12830575 | p5 | (AAAAG)3 | TGGGAAAAGTTCCACAGTCC | GATTGCAAGGCTGAAGAAGC | 151 |
| chr2:12828557-12830575 | p4 | (AGAA)8 | TGGGAAAAGTTCCACAGTCC | GATTGCAAGGCTGAAGAAGC | 151 |
| chr2:13040539-13042555 | p6 | (TTTTTA)3 | ATGGCTCACAAGGAACCAAG | CAGAGTGCGAATTTGGAACA | 129 |
| chr2:13040539-13042555 | p4 | (AAGA)4 | ATGGCTCACAAGGAACCAAG | CAGAGTGCGAATTTGGAACA | 129 |
| chr2:13149113-13151128 | p5 | (CATAT)3 | TAGGCTCGGGAATTTTTCCT | GCAAATATGCTCGTTTGGGT | 128 |
| chr2:13193757-13195789 | p2 | (CT)16 | GACACCCAATTTTGTCCCAC | AACAGGATAATGAAACGCCG | 200 |
| chr2:13266521-13268571 | p5 | (GGGGT)3 | TCTGGCATGTTCTCAGTTGC | ACCGCCTCCACAAATAAAAA | 207 |
| chr2:13342536-13344551 | p5 | (CCACC)3 | GAACCCCCACTCAGTCAAAA | ATGCCAGACCCTGCATAAAC | 209 |
| chr2:13382491-13384506 | p5 | (AAATA)3 | AGGGTTTCCACATCCAACAA | TCGATTTCGGGTCAAAAGTC | 245 |
| chr2:13399170-13401182 | p2 | (AT)6 | AGTAGGGTCTCGCAAAGGGT | TGTGATCTTGGCTTTTGTCG | 188 |
| chr2:13399170-13401182 | p2 | (GA)7 | AGTAGGGTCTCGCAAAGGGT | TGTGATCTTGGCTTTTGTCG | 188 |
| chr2:13415883-13417915 | p2 | (TA)16 | TTTGAACCAAGTCATGCGAA | TTGCATTCCTATCCCACCAT | 100 |
| chr2:13525037-13527052 | p4 | (ATAA)4 | AAATGTGGAACTTTCGCCAG | GGCCCTTACAAGAAGCAATG | 101 |
| chr2:13525037-13527052 | p5 | (AAAAG)3 | AAATGTGGAACTTTCGCCAG | GGCCCTTACAAGAAGCAATG | 101 |
| chr2:13525037-13527052 | p3 | (TGA)5 | AAATGTGGAACTTTCGCCAG | GGCCCTTACAAGAAGCAATG | 101 |
| chr2:13644102-13646124 | p2 | (TC)11 | CCCAAGTGTCTTAAGCCCAA | TGAAACTGCCCCTAGACCAG | 153 |
| chr2:13667944-13669959 | p5 | (TAAAA)3 | TTTCTGCACCAACCAATTCA | TCCATTTCAAGAATCCTGGG | 273 |
| chr2:13741486-13743501 | p5 | (CATGT)3 | TGCTGATTAAGAACCCCGTC | TTGTGCTGAAACCTGGATCA | 119 |
| chr2:13756418-13758436 | p6 | (CCCCAC)3 | TTACTGCTGCCGAAACACAG | TTCCAAAAGCTCGGTCAATC | 117 |
| chr2:13761670-13763700 | p2 | (AT)15 | ACTGCCTTCCGTAAGATGGA | GAAAATCCCTGCCTTTCCTT | 238 |
| chr2:13883636-13885654 | p3 | (TCT)5 | CAGTGGCGGATCCAGTATTT | TTTGTCAAATTCATCGCAGC | 171 |
| chr2:13883636-13885654 | p6 | (AATTGG)3 | CAGTGGCGGATCCAGTATTT | TTTGTCAAATTCATCGCAGC | 171 |
| chr2:13901425-13903440 | p5 | (TATTA)3 | AATGTTTCCCACACTGAGCC | TGGTGCCACAATACAGAGGA | 220 |
| chr2:14011110-14013152 | p2 | (TA)6 | AAGGTCGCAGACATTGATCC | CTCATTTTGCACTTGCCTCA | 138 |
| chr2:14011110-14013152 | p5 | (TATTT)3 | AAGGTCGCAGACATTGATCC | CTCATTTTGCACTTGCCTCA | 138 |
| chr2:14012231-14014279 | p2 | (TA)6 | TTCCCCTCTTCATTTTGCTC | AAGGCAGCCCTATTTTTGGT | 153 |
| chr2:14012231-14014279 | p5 | (TATTT)3 | TTCCCCTCTTCATTTTGCTC | AAGGCAGCCCTATTTTTGGT | 153 |
| chr2:14030317-14032331 | p2 | (TA)11 | GGTGGGCCCAGACTCTAAAT | CCCGTAAGTTTGGATGTGGT | 169 |
| chr2:14030317-14032331 | p2 | (AT)7 | GGTGGGCCCAGACTCTAAAT | CCCGTAAGTTTGGATGTGGT | 169 |

| Marker ID | SSR type | SSR motif | FORWARD PRIMER (5'-3') | REVERSE PRIMER (5'-3') | size (bp) |
|---|---|---|---|---|---|
| chr2:14121027-14123045 | p3 | (ACC)6 | CAGAAAATTCGGCGAGAAAC | GCAAAATGAACCTTCGGAAA | 161 |
| chr2:14158294-14160306 | p2 | (AT)6 | TATTAGCCGCCGCAATTAAC | AATCCCCAATTTCCCTCAAC | 158 |
| chr2:14229297-14231309 | p2 | (AT)6 | ATGCCGCTTGATTTTGTCA | CATTGCCACGTGCGTATTAG | 281 |
| chr2:14229297-14231309 | p2 | (AT)6 | ATGCCGCTTGATTTTGTCA | CATTGCCACGTGCGTATTAG | 281 |
| chr2:14229297-14231309 | p6 | (TACTAA)3 | ATGCCGCTTGATTTTGTCA | CATTGCCACGTGCGTATTAG | 281 |
| chr2:14255415-14257429 | p2 | (AG)7 | TTTGTGCCAAGTCAAAACCA | CGTTTGATTTCCATGAACCC | 271 |
| chr2:14257108-14259122 | p2 | (AG)7 | CACGACAAGTATGGAAGCGA | TCGACGTTTGATTTCCATGA | 253 |
| chr2:14296386-14298401 | p5 | (ATTTT)3 | AGCTAGTCCCATTCCACACG | TGTTCGTCCATCTTTCATGC | 210 |
| chr2:14296386-14298401 | p5 | (TATTT)3 | AGCTAGTCCCATTCCACACG | TGTTCGTCCATCTTTCATGC | 210 |
| chr2:14360827-14362839 | p2 | (AC)6 | TATCACACGCTGCCACAAAT | ATTTTGATGCGGCCTAAATG | 146 |
| chr2:14472458-14474476 | p2 | (TA)9 | TTGGGTCATGACAGCAAAAA | CAAACAACAGCTTGACGTGG | 234 |
| chr2:14472458-14474476 | p5 | (TTTAA)3 | TTGGGTCATGACAGCAAAAA | CAAACAACAGCTTGACGTGG | 234 |
| chr2:14488774-14490786 | p2 | (TA)6 | TTGCACTTGACAACCCAAAA | CATCAAAATAGACCGGCGTT | 266 |
| chr2:14490764-14492776 | p2 | (CA)6 | AATACCAAAGCCCGTGAATG | CTCGAACGCAACAAATCAGA | 196 |
| chr2:14566850-14568862 | p2 | (AT)6 | ACCCTAGTTTGTGAACCCCC | GACCAGGCTGCAGAGGATAG | 286 |
| chr2:14566850-14568862 | p5 | (AATGC)3 | ACCCTAGTTTGTGAACCCCC | GACCAGGCTGCAGAGGATAG | 286 |
| chr2:14604805-14606871 | p2 | (TA)33 | CTCTTTATGCGAGTGCGTGA | TCCCATTTGACAGGCTTAGG | 175 |
| chr2:14622526-14624538 | p2 | (TA)6 | GCCTATAAAAGCGTGCAAGG | ATTTCTTGGAAGCCAACCCT | 213 |
| chr2:14622526-14624538 | p6 | (GGCTTT)3 | GCCTATAAAAGCGTGCAAGG | ATTTCTTGGAAGCCAACCCT | 213 |
| chr2:14695317-14697332 | p5 | (ATCAT)3 | CCAAGTTTGACGGAGAAGGA | CAAAACCCCGGGAATAAAAT | 199 |
| chr2:14746749-14748765 | p2 | (CT)8 | TATTTTCGGGCAAATGAAGC | AATTTGACTGGCGTGGAGAG | 199 |
| chr2:14775327-14777343 | p2 | (CT)8 | TATTTTCGGGCAAATGAAGC | AATTTGACTGGCGTGGAGAG | 202 |
| chr2:14812336-14814354 | p3 | (CTC)6 | TTCTCCTCCTCCTCCTCCTC | TCACGCAAGCCAAAATACAA | 183 |
| chr2:14812336-14814354 | p2 | (AT)9 | TTCTCCTCCTCCTCCTCCTC | TCACGCAAGCCAAAATACAA | 183 |
| chr2:14841309-14843324 | p3 | (GAT)5 | AAAGACAACAAGGTCCACGG | GGACCGACATGGGATGTAAG | 126 |
| chr2:14916267-14918287 | p5 | (ATAAA)4 | TTGGCAGAAAATTGACGTTG | ATGCGTGTTTTTGGCTAAGG | 171 |
| chr2:14931253-14933269 | p2 | (AT)8 | AGAATCCAACCCTTCTGGCT | AGAGTCGCTTTTGTCAGGGA | 228 |
| chr2:14968515-14970539 | p3 | (TTG)8 | CATGCTGGATCTTTTCCCTC | AGCTACAGAGCTCCCGTGAA | 215 |
| chr2:14978133-14980149 | p2 | (AT)8 | ATTTTCTTTGCTCCTTCGCA | TCCCAAAATTCTGGCAGTTC | 155 |
| chr2:15202334-15204358 | p2 | (TG)7 | GAATTGAAACGAAACCGGAA | TGTGCGGTTACGAGACAAAG | 101 |
| chr2:15259749-15261773 | p3 | (CAA)6 | ACTGATCCCCATTCCATCAA | TGCCTTTCTGCTTCTCCAAT | 210 |
| chr2:15259749-15261773 | p2 | (AT)12 | ACTGATCCCCATTCCATCAA | TGCCTTTCTGCTTCTCCAAT | 210 |
| chr2:15259749-15261773 | p2 | (AT)6 | ACTGATCCCCATTCCATCAA | TGCCTTTCTGCTTCTCCAAT | 210 |
| chr2:15421650-15423664 | p5 | (CGGGT)3 | GGAGCGTGAGAGAGGAGAGA | TTTTAATTCACCCCAACCCA | 298 |
| chr2:15421650-15423664 | p3 | (TAA)7 | GGAGCGTGAGAGAGGAGAGA | TTTTAATTCACCCCAACCCA | 298 |
| chr2:15421650-15423664 | p2 | (CT)7 | GGAGCGTGAGAGAGGAGAGA | TTTTAATTCACCCCAACCCA | 298 |

| Marker ID | SSR type | SSR motif | FORWARD PRIMER (5'-3') | REVERSE PRIMER (5'-3') | size (bp) |
|---|---|---|---|---|---|
| chr2:15431007-15433022 | p5 | (AATGC)3 | TGTTTTCTACGAGCAGCGTG | CCCTCCCTTAGAAGGTTTGC | 161 |
| chr2:15431007-15433022 | p2 | (AT)11 | TGTTTTCTACGAGCAGCGTG | CCCTCCCTTAGAAGGTTTGC | 161 |
| chr2:15444825-15446840 | p5 | (AAGGC)3 | TCGAGAAAAGGCAAAGAGGA | TCAGTAAGGGTTTGCCGTCT | 231 |
| chr2:15465595-15467623 | p2 | (AT)14 | CACAACGTACTGTTCCGGTG | TTGATTCGCGAAATGCATAG | 206 |
| chr2:15552160-15554175 | p5 | (TATAA)3 | CTCTGCCACATCAGCAAAAA | CAACGTTTGACCAAAATCCC | 184 |
| chr2:15719521-15721539 | p3 | (GAT)6 | TTTTAAGCCCATGTTCCAGC | TATCCCAGATGCCGAAAAAG | 226 |
| chr2:15788629-15790644 | p3 | (CTA)5 | GACGTTAAGCCTTGGGATGA | CTTAATCCCGTTGCCGATTA | 187 |
| chr2:15806959-15808975 | p5 | (TTCTC)3 | AGATGCGGGGTGTTACATTC | TTTTCCACAAGATTCGGGTC | 172 |
| chr2:15806959-15808975 | p6 | (TGTTTT)4 | AGATGCGGGGTGTTACATTC | TTTTCCACAAGATTCGGGTC | 172 |
| chr2:15815020-15817038 | p2 | (AT)9 | GATTTTTGTGATGGCAGGCT | CCACCATCTGAGAGGCATTT | 272 |
| chr2:15840720-15842740 | p2 | (TA)10 | ACAGCCCTTCACAAAGAGGA | GAACCAGTCACCAAATGCCT | 186 |
| chr2:15940217-15942241 | p3 | (CAT)8 | AGAGGGTCATGACAGGCATC | ATATCCCTCAGTGTCGTCGG | 142 |
| chr2:16044630-16046650 | p2 | (TA)10 | AGGCATTTGGTGACTGGTTC | CCACGTATACTTGCGTGTGC | 195 |
| chr2:16054568-16056582 | p2 | (TA)7 | TTGGTTCGATCCTCTCCTTG | AACACCAAATGCGGAAGAAG | 161 |
| chr2:16063212-16065227 | p3 | (AAT)5 | GCCCTTATCGACCTAAAGCC | GACCGCCTCAATTCCATAAA | 247 |
| chr2:16063212-16065227 | p2 | (AT)21 | GCCCTTATCGACCTAAAGCC | GACCGCCTCAATTCCATAAA | 247 |
| chr2:16077464-16079484 | p4 | (AAAT)5 | TAAATGACCCGGCCTACTTG | CTCAGTGCCTTGTTGCATGT | 228 |
| chr2:16079831-16081843 | p2 | (CA)6 | TGCAAAAAGAAGGAGTGGGT | TTGTGGGCTATTGTGGTTGA | 262 |
| chr2:16125791-16127809 | p2 | (AG)9 | ATCCAACGAGTATCTCCCCC | GTACCATGAGAAATGGCGGT | 223 |
| chr2:16125791-16127809 | p2 | (TC)8 | ATCCAACGAGTATCTCCCCC | GTACCATGAGAAATGGCGGT | 223 |
| chr2:16125791-16127809 | p2 | (TC)8 | ATCCAACGAGTATCTCCCCC | GTACCATGAGAAATGGCGGT | 223 |
| chr2:16126680-16128696 | p2 | (AG)9 | GGGGTTTTGCGTGAATAAAA | GGGAAAACAAAGCGCAGTAA | 288 |
| chr2:16126680-16128696 | p2 | (TC)8 | GGGGTTTTGCGTGAATAAAA | GGGAAAACAAAGCGCAGTAA | 288 |
| chr2:16126680-16128696 | p2 | (TC)8 | GGGGTTTTGCGTGAATAAAA | GGGAAAACAAAGCGCAGTAA | 288 |
| chr2:16213455-16215483 | p2 | (TC)6 | ATGTCCTGCATCTTGGGTTC | GCATCTCCTGCTCCAGAAAC | 194 |
| chr2:16226512-16228527 | p5 | (GTTCG)3 | GCCTCCAAACAAACCAAAAA | CCGAAACCGAAATAGCCATA | 234 |
| chr2:16228224-16230238 | p2 | (AT)7 | TTTTTCCGTGTTTGCAGTTG | TTTCCCACTTGACCTTTTGG | 218 |
| chr2:16255983-16257998 | p3 | (TGT)5 | TGCCTCCTCTCTGCTTCTTC | CTCTTTCCATAAATGGGGCA | 297 |
| chr2:16272106-16274121 | p5 | (CTTTT)3 | TACCCCCGACATTTGAAGAG | GACAAACGCGCCTAGAACTC | 226 |
| chr2:16355995-16358010 | p3 | (ACA)5 | TGGCATGTTGTCTCCTTCAA | ACACTCCGGTTTTTGGAATG | 128 |
| chr2:16391563-16393578 | p5 | (GGATG)3 | ACCTCGTGCCGTTTTATCAC | AAATGAACGAATAGGGGGCT | 155 |
| chr2:16476220-16478238 | p2 | (AT)8 | GCGATCGGGAACATCTCTAC | TGCGACTTTGGGAAAACTTC | 127 |
| chr2:16476220-16478238 | p2 | (AT)9 | GCGATCGGGAACATCTCTAC | TGCGACTTTGGGAAAACTTC | 127 |
| chr2:16476220-16478238 | p2 | (AT)7 | GCGATCGGGAACATCTCTAC | TGCGACTTTGGGAAAACTTC | 127 |
| chr2:16485426-16487444 | p6 | (CCCACC)3 | TTGGTTAGGCGCTACGATTT | GTGGCGTACGCAGAATTTTT | 250 |
| chr2:16495727-16497743 | p4 | (TATT)4 | ACCGTTTGAGGCCTTGTATG | CTTCCGATCCGCATACATCT | 263 |

| Marker ID | SSR type | SSR motif | FORWARD PRIMER (5'-3') | REVERSE PRIMER (5'-3') | size (bp) |
|---|---|---|---|---|---|
| chr2:16495727-16497743 | p2 | (AT)6 | ACCGTTTGAGGCCTTGTATG | CTTCCGATCCGCATACATCT | 263 |
| chr2:16622783-16624801 | p3 | (TCA)5 | TCATTAAGCGCACCCCTAAC | TTTGCAATTCTAGCCGCTTT | 295 |
| chr2:16622783-16624801 | p6 | (CCGGTT)3 | TCATTAAGCGCACCCCTAAC | TTTGCAATTCTAGCCGCTTT | 295 |
| chr2:16711146-16713158 | p2 | (TA)7 | TTGTTCCACGAAATGCAAAA | TTGTTCCACGAAATGCAAAA | 230 |
| chr2:16711204-16713224 | p2 | (TA)7 | TTGTTCCACGAAATGCAAAA | TTGTTCCACGAAATGCAAAA | 172 |
| chr2:16711431-16713447 | p2 | (TA)7 | CCGCAGAAGTGAAATTGACA | CCGATTTACGCGATTTGTTT | 265 |
| chr2:16711453-16713471 | p2 | (TA)7 | CCGCAGAAGTGAAATTGACA | CCGATTTACGCGATTTGTTT | 243 |
| chr2:16730617-16732631 | p2 | (TA)7 | TGTCTCTGTGTGTGTGGGGT | TCACCCCTTCAATCTTGCTT | 115 |
| chr2:16730617-16732631 | p5 | (AATTT)3 | TGTCTCTGTGTGTGTGGGGT | TCACCCCTTCAATCTTGCTT | 115 |
| chr2:16749668-16751683 | p5 | (GTTCG)3 | TCCAATCGAACCACACTTCA | ACCCCTATTTGTCACCACCA | 259 |
| chr2:16749668-16751683 | p5 | (GTTCG)3 | TCCAATCGAACCACACTTCA | ACCCCTATTTGTCACCACCA | 259 |
| chr2:16830433-16832465 | p2 | (AT)16 | CTACTCGTACCGCTCGCTTT | TGCGCTCTTAGTCCGTTTTT | 209 |
| chr2:16857338-16859352 | p2 | (AT)7 | GGAACCCATCACATCCAAAC | GATACGCATGTGGCAATGAA | 176 |
| chr2:16857338-16859352 | p2 | (TA)6 | GGAACCCATCACATCCAAAC | GATACGCATGTGGCAATGAA | 176 |
| chr2:16927862-16929874 | p2 | (AT)6 | GAAGCGCTTTACCAGTGAGC | ACCAAGGGGTGTCATCAGTC | 296 |
| chr2:16968692-16970754 | p2 | (AG)31 | AAAATCATGCTCCGGTTTTG | GAAACACCAGTTGGGAGGAA | 295 |
| chr2:16998313-17000327 | p2 | (TA)7 | CGCACTTATGGCCAATTTTT | GGCACCGATGACATGTAGTG | 275 |
| chr2:17008679-17010697 | p6 | (GAATCC)3 | AAGATCTGCTTGCCGACACT | ATGGAAGCCCTTCCTGATTT | 219 |
| chr2:17018834-17020846 | p2 | (AT)6 | TTGAAAAAGGGAGGAGCAAA | TCATTTGTATCGGTTCGCAA | 237 |
| chr2:17066337-17068355 | p3 | (TTA)6 | TCCTCACGCTCACACTCATC | TTGCTTTAGCCTTTGGGAAA | 232 |
| chr2:17090298-17092316 | p2 | (AG)9 | GAGTTTGGGCTGAAGCTTTG | TTGGGATTGCTAAGTCTGGG | 170 |
| chr2:17130193-17132205 | p5 | (GGAAA)3 | ATTTAGGGGTAATGGGCTGG | ATAGCGGAGGAAGGACCAAT | 172 |
| chr2:17130193-17132205 | p2 | (AT)6 | ATTTAGGGGTAATGGGCTGG | ATAGCGGAGGAAGGACCAAT | 172 |
| chr2:17145334-17147349 | p2 | (AT)7 | ATGATTAACGCGCTACCACC | CTAGTTTAAGCAGGCCCGTG | 127 |
| chr2:17145334-17147349 | p3 | (CAA)5 | ATGATTAACGCGCTACCACC | CTAGTTTAAGCAGGCCCGTG | 127 |
| chr2:17149069-17151085 | p2 | (TC)8 | CACCACATCCAAACTCATGG | GCACGGTGTTTAATATGGGG | 174 |
| chr2:17244279-17246297 | p3 | (ACA)6 | GTCCGGAAATATGATGGGTG | GCTTCAACCCATTTGGAGAA | 161 |
| chr2:17323265-17325285 | p5 | (AAACC)4 | CTGAGTTCGCTCTCCCTCAC | GAGAAAACCCATGGTCGAAA | 116 |
| chr2:17323265-17325285 | p5 | (TAAAA)3 | CTGAGTTCGCTCTCCCTCAC | GAGAAAACCCATGGTCGAAA | 116 |
| chr2:17451577-17453592 | p5 | (AAATA)3 | GAGCCGTCGATGATATGGAT | GCGCCCAATGTAAAAGAGAA | 268 |
| chr2:17451577-17453592 | p2 | (AG)8 | GAGCCGTCGATGATATGGAT | GCGCCCAATGTAAAAGAGAA | 268 |
| chr2:17559348-17561366 | p2 | (GA)7 | GAGAGTATCGAAGCATCGCC | TTTGTCCACAGCCATTTTGA | 123 |
| chr2:17559348-17561366 | p5 | (AAATA)5 | GAGAGTATCGAAGCATCGCC | TTTGTCCACAGCCATTTTGA | 123 |
| chr2:17559348-17561366 | p3 | (AAT)11 | GAGAGTATCGAAGCATCGCC | TTTGTCCACAGCCATTTTGA | 123 |
| chr2:17559348-17561366 | p3 | (ATA)6 | GAGAGTATCGAAGCATCGCC | TTTGTCCACAGCCATTTTGA | 123 |
| chr2:17690687-17692705 | p6 | (GAAAAA)3 | CAACCGACACATCAGCTACG | GAAAAACTCACATGCAGCCA | 221 |

| Marker ID | SSR type | SSR motif | FORWARD PRIMER (5'-3') | REVERSE PRIMER (5'-3') | size (bp) |
|---|---|---|---|---|---|
| chr2:17694465-17696483 | p2 | (AT)21 | GTTTGGATTGTGGCGAGTTT | CCATGCTTCATCGTCTGTTG | 185 |
| chr2:17694465-17696483 | p6 | (ATTTTT)3 | GTTTGGATTGTGGCGAGTTT | CCATGCTTCATCGTCTGTTG | 185 |
| chr2:17724398-17726418 | p2 | (AG)10 | ACTCGAGTGTTGGCCAGAGT | AACACCGAGGTTGCTAATGG | 206 |
| chr2:17729674-17731696 | p2 | (TA)11 | AGTGTTGGGCAACCAAGAAC | CTAGCTTCCCGCATCTTGTC | 278 |
| chr2:18012079-18014097 | p6 | (TTTTTA)3 | GATTTCCCTTCATTTGGGGT | AGTTTTGTGCCTTGCGAATC | 247 |
| chr2:18015228-18017240 | p2 | (AT)6 | ACCTCTTCCTCGTTGGGAGT | TCAAACAGCATCTCGTCCAG | 187 |
| chr2:18107189-18109201 | p5 | (GCAGG)3 | TTGATCACATGACGGACGAT | GTCCTCAACGATGTCCAGGT | 185 |
| chr2:18107189-18109201 | p2 | (TA)6 | TTGATCACATGACGGACGAT | GTCCTCAACGATGTCCAGGT | 185 |
| chr2:18122088-18124103 | p5 | (GCAGG)3 | TTTTTGTCCAATTTGGCCTC | CCAGCTGTAGTCTTCCAGGC | 276 |
| chr2:18122088-18124103 | p2 | (TA)6 | TTTTTGTCCAATTTGGCCTC | CCAGCTGTAGTCTTCCAGGC | 276 |
| chr2:18122742-18124754 | p5 | (GCAGG)3 | TTGATCACATGACGGACGAT | GTCCTCAACGATGTCCAGGT | 185 |
| chr2:18122742-18124754 | p2 | (TA)6 | TTGATCACATGACGGACGAT | GTCCTCAACGATGTCCAGGT | 185 |
| chr2:18163412-18165444 | p2 | (TA)10 | TCCAATCACAACTTGCCAAA | GTGAGTTCAAAGGTGGAGCC | 264 |
| chr2:18194838-18196856 | p6 | (ACCCAA)3 | ATGCCTTATGGGCAGACTTG | ACTTGGCATAAGTATGCGGG | 250 |
| chr2:18194838-18196856 | p2 | (AT)11 | ATGCCTTATGGGCAGACTTG | ACTTGGCATAAGTATGCGGG | 250 |
| chr2:18195558-18197580 | p6 | (ACCCAA)3 | CAAAAGAAGGGCAATTCTGC | AAAAGGAGAAGAGGGAAGCG | 172 |
| chr2:18195558-18197580 | p2 | (AT)11 | CAAAAGAAGGGCAATTCTGC | AAAAGGAGAAGAGGGAAGCG | 172 |
| chr2:18227246-18229258 | p2 | (TA)6 | CACTCCCCGTTACTCCGATA | GAACGGTAGCTCCACCACAT | 196 |
| chr2:18241955-18243970 | p2 | (AT)15 | GGCTCATTATGCTTAGCCCA | GTTTGCTTCTTGGTGGAGGA | 211 |
| chr2:18242047-18244062 | p2 | (AT)15 | GGCTCATTATGCTTAGCCCA | GTTTGCTTCTTGGTGGAGGA | 119 |
| chr2:18252528-18254544 | p4 | (AATA)4 | GCACTCAGAAGAGCCGAGTT | CATTTGGTCATGTCCATTGC | 176 |
| chr2:18418812-18420827 | p5 | (TATTA)3 | GAGGGGGAAAGAGAAGGAA | AGCTCGCTCACCAAAAACAT | 286 |
| chr2:18489743-18491755 | p2 | (TA)9 | CCGGTTAAAATAGGGTGCAA | TGTGGGACAATCCACTTTGA | 296 |
| chr2:18489743-18491755 | p2 | (TA)6 | CCGGTTAAAATAGGGTGCAA | TGTGGGACAATCCACTTTGA | 296 |
| chr2:18507079-18509099 | p4 | (ATTT)5 | AAGTGATGCCAGAGCCAAGT | TGTTGCTTGCCATGGTTAAA | 175 |
| chr2:18507079-18509099 | p2 | (AC)11 | AAGTGATGCCAGAGCCAAGT | TGTTGCTTGCCATGGTTAAA | 175 |
| chr2:18604805-18606817 | p2 | (AT)6 | TTGGCCGTCTTTAGTTGACC | TATGGCCAAACACAACTCCA | 192 |
| chr2:18604805-18606817 | p5 | (AAAAT)3 | TTGGCCGTCTTTAGTTGACC | TATGGCCAAACACAACTCCA | 192 |
| chr2:18604805-18606817 | p3 | (TGT)8 | TTGGCCGTCTTTAGTTGACC | TATGGCCAAACACAACTCCA | 192 |
| chr2:18604805-18606817 | p3 | (AAT)5 | TTGGCCGTCTTTAGTTGACC | TATGGCCAAACACAACTCCA | 192 |
| chr2:18653468-18655488 | p2 | (AT)10 | TGCCACATGGCAGAATTAGT | ACTTAGCCATTTGCACACCC | 205 |
| chr2:18725350-18727390 | p2 | (TA)20 | CTGCCCCTTTGTGTCATTTT | AACTCCCCACACACAGCTTC | 286 |
| chr2:18739973-18741991 | p2 | (AT)9 | TGCTCTTTCGGCTCGTTACT | GAATGCAAACAGCAAGCAGA | 236 |
| chr2:18755944-18757959 | p3 | (TTG)5 | ACCCCCAAAAATCTTGTTCC | CCAAATGCTCCTTCTTCCAA | 117 |
| chr2:18781883-18783905 | p2 | (TA)11 | TCAGCAATAAAGTTGCCACG | GAACCGTACCATTGACACCC | 235 |
| chr2:18822814-18824832 | p3 | (TCG)5 | TCTTCCATCACTGATTGCCA | ATCACTGGAAGGCATTGGAC | 223 |

| Marker ID | SSR type | SSR motif | FORWARD PRIMER (5'-3') | REVERSE PRIMER (5'-3') | size (bp) |
|---|---|---|---|---|---|
| chr2:18822814-18824832 | p3 | (ATG)6 | TCTTCCATCACTGATTGCCA | ATCACTGGAAGGCATTGGAC | 223 |
| chr2:18847003-18849045 | p3 | (ATT)14 | GAAAAGGACAACGAGACCCA | CCCAAAGTTGAGGGTTTGAA | 214 |
| chr2:18847003-18849045 | p5 | (ACCCG)3 | GAAAAGGACAACGAGACCCA | CCCAAAGTTGAGGGTTTGAA | 214 |
| chr2:18847378-18849393 | p3 | (ATT)14 | CCACCCAACCCATTCACTAC | TCTTAGTCCCCGCAAATCAC | 198 |
| chr2:18847378-18849393 | p5 | (ACCCG)3 | CCACCCAACCCATTCACTAC | TCTTAGTCCCCGCAAATCAC | 198 |
| chr2:18847378-18849393 | p5 | (TAGTT)3 | CCACCCAACCCATTCACTAC | TCTTAGTCCCCGCAAATCAC | 198 |
| chr2:19057820-19059832 | p2 | (AT)6 | TATTGGTGTCGTTGGGTTCA | AACAGTGCGTTGTGTGGGTA | 289 |
| chr2:19057820-19059832 | p2 | (TA)13 | TATTGGTGTCGTTGGGTTCA | AACAGTGCGTTGTGTGGGTA | 289 |
| chr2:19058931-19060943 | p2 | (AT)7 | TTTTTCGAGATCTATGCGGG | CCGCAGAAGTGAAATTGACA | 235 |
| chr2:19058968-19060984 | p2 | (AT)7 | TCAATTTCAATTTTCCCCCA | TGTTTAAAAGGTGGCCGAAG | 278 |
| chr2:19313807-19315822 | p3 | (CTT)5 | CGGGCAATTTCAAGCTAAGA | TTCATGCACGACCAAAACAT | 206 |
| chr2:19373438-19375450 | p2 | (TG)6 | ATAGGCAGAATTTGCATGGC | AGCGAGAGTTCGAAAAACCA | 176 |
| chr2:19398098-19400128 | p2 | (TA)6 | TCATAAAATGGCGCGATACA | TTTGTGCTGGCATTGATTGT | 153 |
| chr2:19398098-19400128 | p3 | (AAC)10 | TCATAAAATGGCGCGATACA | TTTGTGCTGGCATTGATTGT | 153 |
| chr2:19549002-19551017 | p5 | (TCGGT)3 | AAAAGAGATCACTTCGGGCA | ACTGAGAAAACCGAAACCGA | 180 |
| chr2:19600024-19602046 | p2 | (AT)11 | CACAAGACACTGTGCAGCCT | AGGGAAAACCACAGCATGAC | 223 |
| chr2:19614300-19616315 | p5 | (ACCAA)3 | TTCCGATCATGACACCTTCA | AACCCGACGCAATTAGTTTG | 128 |
| chr2:19614300-19616315 | p5 | (AAACC)3 | TTCCGATCATGACACCTTCA | AACCCGACGCAATTAGTTTG | 128 |
| chr2:19626931-19628945 | p2 | (AT)7 | GTGTACGCAGACCATACCCC | CCTATCCATTTGGCGCTTTA | 168 |
| chr2:19626931-19628945 | p4 | (AATA)4 | GTGTACGCAGACCATACCCC | CCTATCCATTTGGCGCTTTA | 168 |
| chr2:19626931-19628945 | p2 | (AG)7 | GTGTACGCAGACCATACCCC | CCTATCCATTTGGCGCTTTA | 168 |
| chr2:19626931-19628945 | p2 | (AT)7 | GTGTACGCAGACCATACCCC | CCTATCCATTTGGCGCTTTA | 168 |
| chr2:19655313-19657333 | p2 | (TA)10 | ATTCCCGAAAAGAGACCGTT | CAGCACCTGCTGTACCAAGA | 228 |
| chr2:19665019-19667059 | p2 | (AG)20 | TGACCATGAACTCGGAAACA | TCTTTATGTTGCGCTCCCTT | 296 |
| chr2:19860014-19862029 | p5 | (CAAAC)3 | CTTTGGGGCAGTACTTGCAT | CTGCAACTCAAGCACATGGT | 296 |
| chr2:19909150-19911165 | p5 | (CAAAC)3 | CTTTGGGGCAGTACTTGCAT | CTGCAACTCAAGCACATGGT | 296 |
| chr2:19985606-19987618 | p2 | (AT)6 | AATTCGCAACGGATTTTCAC | GTTGGGAATTTGGAAAAGCA | 218 |
| chr2:20004764-20006779 | p5 | (TTAGT)3 | TACCTATATGAGCGGGCAGC | CGAATCTGGGTCAAGCTCTC | 284 |
| chr2:20103520-20105538 | p3 | (AGA)31 | ACCATAGGTGACTTGTGGGG | GAAGAAGTGCGAGGGAGTTG | 176 |
| chr2:20103520-20105538 | p6 | (TGTTGG)3 | ACCATAGGTGACTTGTGGGG | GAAGAAGTGCGAGGGAGTTG | 176 |
| chr2:20138093-20140109 | p2 | (AT)8 | CTCCCGGAATTGAATTGAGA | GCTTGAAGGCCCATTCTTTA | 198 |
| chr2:20144234-20146268 | p2 | (CA)7 | TTTTTGAGGGCCATATCAGC | TTGTCCATGAAGCACTCGTC | 131 |
| chr2:20145471-20147483 | p2 | (AC)6 | GACAATTGAGACACTGCGGA | GACGAACGCAGCAACAATAA | 212 |
| chr2:20366724-20368740 | p2 | (TA)8 | AAGTGGATGTCCTCCAAACG | ACTCGGTTGGTCCAGAGTTG | 285 |
| chr2:20491308-20493320 | p2 | (AT)8 | CCCATCAAAACAACTCCACC | AGAAAATTTGAGCGGGGATT | 244 |
| chr2:20491418-20493470 | p2 | (AT)8 | CCCATCAAAACAACTCCACC | AGAAAATTTGAGCGGGGATT | 134 |

| Marker ID | SSR type | SSR motif | FORWARD PRIMER (5'-3') | REVERSE PRIMER (5'-3') | size (bp) |
|---|---|---|---|---|---|
| chr2:20491619-20493647 | p2 | (AT)8 | CCCCGCTCAAATTTTCTTTT | GGCGGCTGGTTTATCTGTTA | 121 |
| chr2:20491619-20493647 | p2 | (AT)15 | CCCCGCTCAAATTTTCTTTT | GGCGGCTGGTTTATCTGTTA | 121 |
| chr2:20532881-20534897 | p2 | (TA)8 | ACACATTTCAAACGCAACCA | TATTTGTTCAGGGGTACGGC | 140 |
| chr2:20604340-20606352 | p2 | (TA)6 | TTTGTCCAAATTCAAGCCCT | ACAATGGGTCAATGCTCACA | 100 |
| chr2:20612146-20614164 | p5 | (ATTTT)3 | TTGAGTTGTCCAAAACGCTG | GCTAAACGTGAAGGAGCTGG | 221 |
| chr2:20612146-20614164 | p6 | (AAAAAT)3 | TTGAGTTGTCCAAAACGCTG | GCTAAACGTGAAGGAGCTGG | 221 |
| chr2:20654793-20656805 | p2 | (AT)6 | CAGGTGAGCAGACATTTGGA | AATTTTGCCCCAAAAAGACA | 151 |
| chr2:20660207-20662243 | p2 | (AG)18 | TTGATGTGCAACAAATCCGT | GCTTTGAGAGCTGCCTCATT | 267 |
| chr2:20696523-20698538 | p5 | (TGCCC)3 | AGCAAATGGAGAAGCGAAAA | CCACATGCCATCCATATCAA | 220 |
| chr2:20717616-20719640 | p6 | (TTTTCC)4 | AGGGAAGCATACACAATGCC | ATGTTTTGTCTGCCTGGTCC | 166 |
| chr2:20803519-20805533 | p2 | (TA)7 | GTGCGTGTCAATCCATGAAA | GACATGGCGGAATAGCAAAT | 212 |
| chr2:20844168-20846192 | p3 | (TCT)9 | TTTGTTGGTGTAGCACTGGC | CCCAGCTGTTGCAGGTTATT | 297 |
| chr2:20844193-20846209 | p3 | (TCT)9 | TTTGTTGGTGTAGCACTGGC | CCCAGCTGTTGCAGGTTATT | 272 |
| chr2:20844210-20846222 | p3 | (TTC)5 | TTTGTTGGTGTAGCACTGGC | CCCAGCTGTTGCAGGTTATT | 255 |
| chr2:20844210-20846222 | p3 | (TCT)9 | TTTGTTGGTGTAGCACTGGC | CCCAGCTGTTGCAGGTTATT | 255 |
| chr2:20844234-20846264 | p3 | (TTC)5 | TTTGTTGGTGTAGCACTGGC | CCCAGCTGTTGCAGGTTATT | 231 |
| chr2:20844234-20846264 | p3 | (TCT)9 | TTTGTTGGTGTAGCACTGGC | CCCAGCTGTTGCAGGTTATT | 231 |
| chr2:20923662-20925676 | p2 | (AT)7 | AAGGCTCCTCATGCATTCAC | AAGTGACCCCACTGGATACG | 177 |
| chr2:21078730-21080742 | p2 | (TA)6 | ATGAACCCTGCATCGAGAAC | TGTAGCCTTGTCGGCTCTTT | 244 |
| chr2:21098104-21100122 | p6 | (TCGGCT)3 | TCCGATATTCTCAGGGCATC | CAGGGTTGAGGACGATCAAT | 219 |
| chr2:21101689-21103710 | p3 | (TGT)5 | ATTCCCCTCAACCTGTCCTT | CAGGTGTCGGTTTGTGATTG | 275 |
| chr2:21101689-21103710 | p3 | (TGT)7 | ATTCCCCTCAACCTGTCCTT | CAGGTGTCGGTTTGTGATTG | 275 |
| chr2:21101689-21103710 | p4 | (AAAT)8 | ATTCCCCTCAACCTGTCCTT | CAGGTGTCGGTTTGTGATTG | 275 |
| chr2:21113106-21115121 | p5 | (AATAG)3 | GTCCCTTCCACCAAGTCTGA | TTTGATTCGAGCCACAACTG | 122 |
| chr2:21113106-21115121 | p5 | (AGAAG)3 | GTCCCTTCCACCAAGTCTGA | TTTGATTCGAGCCACAACTG | 122 |
| chr2:21135542-21137562 | p5 | (CTTTT)4 | AAATTTTGTATGCCCCCTCC | GACTGGGGGAATGTATGTCG | 236 |
| chr2:21161045-21163099 | p2 | (AT)27 | GTGTACGTGCACCAGACCAC | GCAGCAAACTCGCTAAAACC | 125 |
| chr2:21176067-21178082 | p5 | (TAAAA)3 | TCACATTTCAAAACGTCGGA | CCCTAAAACGACGATCCAAA | 131 |
| chr2:21234220-21236235 | p5 | (ACAAA)3 | CATGAACGTATGATGGCGAC | TTGCTGAGGACTTTGTGTGC | 147 |
| chr2:21235330-21237345 | p5 | (AAACA)3 | AATGAGGACCTGGCAAACTG | GGTAGCAACAATGCCAGGAT | 189 |
| chr2:21245289-21247304 | p5 | (TTTGT)3 | GCTGAAGCTGCAAAGCTACC | GTTTTCTGATGTGGCGTTGA | 260 |
| chr2:21245289-21247304 | p3 | (TCT)10 | GCTGAAGCTGCAAAGCTACC | GTTTTCTGATGTGGCGTTGA | 260 |
| chr2:21245289-21247304 | p4 | (TCTT)4 | GCTGAAGCTGCAAAGCTACC | GTTTTCTGATGTGGCGTTGA | 260 |
| chr2:21245431-21247461 | p5 | (TTTGT)3 | GCTGAAGCTGCAAAGCTACC | GTTTTCTGATGTGGCGTTGA | 118 |
| chr2:21245431-21247461 | p3 | (TCT)10 | GCTGAAGCTGCAAAGCTACC | GTTTTCTGATGTGGCGTTGA | 118 |
| chr2:21245431-21247461 | p4 | (TCTT)4 | GCTGAAGCTGCAAAGCTACC | GTTTTCTGATGTGGCGTTGA | 118 |

| Marker ID | SSR type | SSR motif | FORWARD PRIMER (5'-3') | REVERSE PRIMER (5'-3') | size (bp) |
|---|---|---|---|---|---|
| chr2:21311616-21313628 | p2 | (AG)6 | ATGGAGGTTTCTTGGTCGTG | CGGAACTTCATGTGGGTCTT | 136 |
| chr2:21365982-21367997 | p3 | (TTA)5 | GTAGAAAGTTGCTCGCGCTT | GGTGTTGCTGTCTGCACAGT | 177 |
| chr2:21373398-21375410 | p4 | (AACA)5 | GCGGGTATCAGCTCAGATGT | TTTGGATAACGTGGCAAACA | 182 |
| chr2:21373408-21375424 | p4 | (AACA)5 | GCGGGTATCAGCTCAGATGT | TTTGGATAACGTGGCAAACA | 172 |
| chr2:21378733-21380745 | p2 | (CA)6 | CTTGAGGCAACAAGAGGGAG | TCCACTATGGAAGGACCCTG | 119 |
| chr2:21474013-21476049 | p2 | (TA)18 | CCCAATCTTTCGTGAGCAAT | ATCTGAAAGTTCGTCCACGC | 182 |
| chr2:21583854-21585869 | p5 | (CGGGT)3 | CTCGGAACCGAAACAGTAGC | GTTTGGGACAAGTTTTCGGA | 172 |
| chr2:21610368-21612386 | p6 | (TTTTTA)3 | GGACGACGATCTTGCCTCTA | CCCATTAGTGGGAGCAAATG | 139 |
| chr2:21610368-21612386 | p2 | (AT)6 | GGACGACGATCTTGCCTCTA | CCCATTAGTGGGAGCAAATG | 139 |
| chr2:21653613-21655649 | p2 | (TA)18 | AGATGGTGATTTTCCAAGCG | GCCACCTTCTTCTCCAAACA | 223 |
| chr2:21830836-21832851 | p3 | (TTA)5 | ACATGCTTGCCATGTGTTGT | AGAAGGTGGTTGGTTTGTGG | 229 |
| chr2:21830836-21832851 | p4 | (AAAT)4 | ACATGCTTGCCATGTGTTGT | AGAAGGTGGTTGGTTTGTGG | 229 |
| chr2:21875734-21877749 | p5 | (AAAGC)3 | GGCCAACGGATAGATCTTGA | CCGAACCGAAGTTCGATAAA | 216 |
| chr2:21875802-21877817 | p5 | (AAAGC)3 | GGCCAACGGATAGATCTTGA | CCGAACCGAAGTTCGATAAA | 148 |
| chr2:21888368-21890402 | p2 | (CT)12 | CCGAGGGACGTGACATAACT | CCAGAAGGAAAAACCGATCA | 201 |
| chr2:21888368-21890402 | p2 | (AT)17 | CCGAGGGACGTGACATAACT | CCAGAAGGAAAAACCGATCA | 201 |
| chr2:21888368-21890402 | p5 | (TATTT)3 | CCGAGGGACGTGACATAACT | CCAGAAGGAAAAACCGATCA | 201 |
| chr2:21888605-21890620 | p2 | (CT)12 | TGATCGGTTTTTCCTTCTGG | TGGAGAAACCGGACTAATGG | 173 |
| chr2:21888605-21890620 | p2 | (AT)17 | TGATCGGTTTTTCCTTCTGG | TGGAGAAACCGGACTAATGG | 173 |
| chr2:21888605-21890620 | p5 | (TATTT)3 | TGATCGGTTTTTCCTTCTGG | TGGAGAAACCGGACTAATGG | 173 |
| chr2:21890134-21892149 | p3 | (TCT)5 | ATGCTTGGGATTGATAGCCA | TCGAGCGACCTTTTAAGGAA | 127 |
| chr2:21897925-21899943 | p6 | (AACTGA)3 | AAATGTGGATGCATTCGTGA | GATGAAAAATTGGCATGGCT | 255 |
| chr2:22211766-22213781 | p5 | (TATTA)3 | TTGCACAAACAAAAGCAAGG | ACCATTTTGCTCATTCCCAG | 186 |
| chr2:22213229-22215247 | p3 | (GTT)6 | ACGTGGTCTTCAAATGAGGG | TGTGTTTACGAGGTTTCCCC | 235 |
| chr2:22213269-22215291 | p3 | (GTT)6 | ACGTGGTCTTCAAATGAGGG | TGTGTTTACGAGGTTTCCCC | 195 |
| chr2:22288796-22290812 | p4 | (AACT)4 | TTCATGCCACGCAGATGTAT | TCGGCTTAAATAGCCGCTTA | 113 |
| chr2:22309501-22311589 | p2 | (AT)44 | AATTCGTGCTGGAGGACCTA | AATGGGAGGATATTTGCGTG | 231 |
| chr2:22309501-22311589 | p3 | (TAT)12 | AATTCGTGCTGGAGGACCTA | AATGGGAGGATATTTGCGTG | 231 |
| chr2:22376111-22378123 | p2 | (AT)6 | TTTTTGATCCGGAGGGAGTA | TGGTCTCTACAGGTGCATGG | 235 |
| chr2:22415594-22417612 | p6 | (AAAATA)3 | CAGCTCAGGCGGATTACATT | AGCGATATGCTCTTGGAGGA | 168 |
| chr2:22509167-22511182 | p5 | (TTTGG)3 | TGGATGACATACCGTTGGAA | TCACCAAACCGTGAATAGCA | 216 |
| chr2:22509167-22511182 | p5 | (TTTGG)3 | TGGATGACATACCGTTGGAA | TCACCAAACCGTGAATAGCA | 216 |
| chr2:22539577-22541592 | p3 | (CAT)5 | GAATCTGGGTAAAGGCCACA | GGAAGGAAATAGAGCTCGCC | 180 |
| chr2:22633322-22635337 | p5 | (ATCAT)3 | TGAGGTTTTGAGTGTGCCAG | TCCCTAGGGCTAAGGCTCAT | 116 |
| chr2:22633322-22635337 | p3 | (TCT)5 | TGAGGTTTTGAGTGTGCCAG | TCCCTAGGGCTAAGGCTCAT | 116 |
| chr2:22641336-22643351 | p5 | (TAAGA)3 | TTTAGGGGACCCTCGCTAGT | CCTAAAGGATTGGACACCCA | 266 |

| Marker ID | SSR type | SSR motif | FORWARD PRIMER (5'-3') | REVERSE PRIMER (5'-3') | size (bp) |
|---|---|---|---|---|---|
| chr2:22659129-22661151 | p2 | (TA)11 | TGCCACATGCATATTGAACC | AAATGACATGGCGGAATAGC | 261 |
| chr2:22659129-22661151 | p2 | (AT)8 | TGCCACATGCATATTGAACC | AAATGACATGGCGGAATAGC | 261 |
| chr2:22683709-22685724 | p5 | (CCCCA)3 | ACAACCCCCAAAAATGACAA | GTAGGCCAGTGCAAAACCAT | 175 |
| chr2:22683709-22685724 | p5 | (TTTCT)3 | ACAACCCCCAAAAATGACAA | GTAGGCCAGTGCAAAACCAT | 175 |
| chr2:22706411-22708438 | p2 | (AT)32 | AGGTAATTTTTACCGGCGCT | GGTTGGTAGCAGTGGTGGTT | 203 |
| chr2:22707243-22709307 | p2 | (AT)32 | AAAACAATTGAGGCAGCGTT | CGGGGAAGAAATTGTTGAAA | 110 |
| chr2:22777842-22779875 | p3 | (AAT)7 | TGGGTCGTGACAAGTTGGTA | CATCATCCGGGAGAAGGTAA | 188 |
| chr2:22777842-22779875 | p2 | (AT)6 | TGGGTCGTGACAAGTTGGTA | CATCATCCGGGAGAAGGTAA | 188 |
| chr2:22778785-22780797 | p3 | (AAT)7 | TTGGCAAAGGGTACTTCGAC | CAAATTTTGCAAGCAGGTCA | 108 |
| chr2:22778785-22780797 | p2 | (AT)6 | TTGGCAAAGGGTACTTCGAC | CAAATTTTGCAAGCAGGTCA | 108 |
| chr2:22815060-22817075 | p5 | (TATTT)3 | AGGTGGTTTATGAGTTGGCG | GGATTTCTACCGGCTCTTCC | 293 |
| chr2:22835932-22837950 | p6 | (AAAAGG)3 | TTGCCTTTGGCAGAGTTTTT | TTTCCAAGGATTGCTGAACC | 128 |
| chr2:22896095-22898110 | p5 | (TTGGT)3 | AGTAAACCGACAAACCGCAC | AAACCAATTATGGTCGGGTG | 190 |
| chr2:22896095-22898110 | p5 | (TTGGT)3 | AGTAAACCGACAAACCGCAC | AAACCAATTATGGTCGGGTG | 190 |
| chr2:22991585-22993600 | p3 | (ACA)5 | CGCTTGAACCGGAGTAAAAG | GGAGGTCAATGCCCCTTATT | 235 |
| chr2:23260813-23262828 | p5 | (TGATA)3 | GAATTGAGTTTGGAGAGGCG | CTGAATGAAAAAGGGGCAAA | 178 |
| chr2:23424692-23426707 | p3 | (TGT)5 | TATCAATGGAGCACAGGCAG | TGCAGGCAGATGAAACTGAG | 284 |
| chr2:23454010-23456025 | p3 | (TGT)5 | TATCAATGGAGCACAGGCAG | TGCAGGCAGATGAAACTGAG | 284 |
| chr2:23493409-23495429 | p5 | (GGCCC)3 | CCCACTAGTCCCGCTAATGA | CGGGCAAAATAATTCCACAC | 289 |
| chr2:23493409-23495429 | p5 | (TTGGT)3 | CCCACTAGTCCCGCTAATGA | CGGGCAAAATAATTCCACAC | 289 |
| chr2:23493426-23495441 | p5 | (GGCCC)3 | CCCACTAGTCCCGCTAATGA | CGGGCAAAATAATTCCACAC | 272 |
| chr2:23493426-23495441 | p5 | (TTGGT)3 | CCCACTAGTCCCGCTAATGA | CGGGCAAAATAATTCCACAC | 272 |
| chr2:23511279-23513294 | p5 | (TTATA)3 | ACGGGGTTGAGATGCTATTG | CGTAGCCGAGCTCGTAAATC | 142 |
| chr2:23614183-23616208 | p3 | (TGT)5 | ATGGTGGTGGAGTGAAGGAG | ATCTGACTTCGTCGTGAGGG | 260 |
| chr2:23614183-23616208 | p2 | (TA)6 | ATGGTGGTGGAGTGAAGGAG | ATCTGACTTCGTCGTGAGGG | 260 |
| chr2:23629862-23631877 | p5 | (ATTTT)3 | GAGACATTAGTTCAGGCGGG | TTTTATTGAGCTGGGGTTGG | 268 |
| chr2:23846162-23848186 | p4 | (GTTT)6 | AGGGATGAGTGGTTGTCGTC | CTATATCCGGCCGAGTGTGT | 131 |
| chr2:23873332-23875344 | p3 | (ACA)8 | GTTAGTGCACGACCAGCAGA | GGGCATGTTGAATCGTTGAT | 109 |
| chr2:23883100-23885118 | p2 | (TA)6 | ATACTTCACGTGCCAGGGAG | TGTGCACCCGACAATTATGT | 248 |
| chr2:23883100-23885118 | p6 | (CCGGTT)4 | ATACTTCACGTGCCAGGGAG | TGTGCACCCGACAATTATGT | 248 |
| chr2:23883124-23885150 | p2 | (TA)6 | ATACTTCACGTGCCAGGGAG | TGTGCACCCGACAATTATGT | 224 |
| chr2:23883124-23885150 | p6 | (CCGGTT)4 | ATACTTCACGTGCCAGGGAG | TGTGCACCCGACAATTATGT | 224 |
| chr2:24002154-24004172 | p2 | (AG)9 | TCTTTCGAGCTTAACCGCAT | TGGACAGGCTGAAAATTGTG | 127 |
| chr2:24002154-24004172 | p2 | (TA)6 | TCTTTCGAGCTTAACCGCAT | TGGACAGGCTGAAAATTGTG | 127 |
| chr2:24002154-24004172 | p3 | (CTA)11 | TCTTTCGAGCTTAACCGCAT | TGGACAGGCTGAAAATTGTG | 127 |
| chr2:24003128-24005188 | p2 | (AG)9 | TTGACGGGTCGGGTACTAAA | GGGTCAAATATGCCCTTGAG | 102 |

| Marker ID | SSR type | SSR motif | FORWARD PRIMER (5'-3') | REVERSE PRIMER (5'-3') | size (bp) |
|---|---|---|---|---|---|
| chr2:24003128-24005188 | p2 | (TA)6 | TTGACGGGTCGGGTACTAAA | GGGTCAAATATGCCCTTGAG | 102 |
| chr2:24003128-24005188 | p3 | (ATA)15 | TTGACGGGTCGGGTACTAAA | GGGTCAAATATGCCCTTGAG | 102 |
| chr2:24003527-24005583 | p2 | (TA)6 | CAAGGACAATATGAATCCCAAAA | CAGCCGCCTAGCTAAAAATG | 264 |
| chr2:24003527-24005583 | p3 | (ATA)15 | CAAGGACAATATGAATCCCAAAA | CAGCCGCCTAGCTAAAAATG | 264 |
| chr2:24004173-24006205 | p3 | (ATA)15 | GCCACTTGGCAACAAATTCT | AGCTTGTTCATATTACGCCACTT | 209 |
| chr2:24004173-24006205 | p3 | (AAT)12 | GCCACTTGGCAACAAATTCT | AGCTTGTTCATATTACGCCACTT | 209 |
| chr2:24004173-24006205 | p5 | (ATAAA)3 | GCCACTTGGCAACAAATTCT | AGCTTGTTCATATTACGCCACTT | 209 |
| chr2:24004204-24006246 | p3 | (ATA)15 | GCCACTTGGCAACAAATTCT | AGCTTGTTCATATTACGCCACTT | 178 |
| chr2:24004204-24006246 | p3 | (AAT)12 | GCCACTTGGCAACAAATTCT | AGCTTGTTCATATTACGCCACTT | 178 |
| chr2:24004204-24006246 | p5 | (ATAAA)3 | GCCACTTGGCAACAAATTCT | AGCTTGTTCATATTACGCCACTT | 178 |
| chr2:24004247-24006275 | p3 | (ATA)15 | GCCACTTGGCAACAAATTCT | AGCTTGTTCATATTACGCCACTT | 135 |
| chr2:24004247-24006275 | p3 | (AAT)12 | GCCACTTGGCAACAAATTCT | AGCTTGTTCATATTACGCCACTT | 135 |
| chr2:24004247-24006275 | p5 | (ATAAA)3 | GCCACTTGGCAACAAATTCT | AGCTTGTTCATATTACGCCACTT | 135 |
| chr2:24004277-24006319 | p2 | (TA)22 | GCCACTTGGCAACAAATTCT | AGCTTGTTCATATTACGCCACTT | 105 |
| chr2:24004277-24006319 | p3 | (ATA)15 | GCCACTTGGCAACAAATTCT | AGCTTGTTCATATTACGCCACTT | 105 |
| chr2:24004277-24006319 | p3 | (AAT)12 | GCCACTTGGCAACAAATTCT | AGCTTGTTCATATTACGCCACTT | 105 |
| chr2:24004277-24006319 | p5 | (ATAAA)3 | GCCACTTGGCAACAAATTCT | AGCTTGTTCATATTACGCCACTT | 105 |
| chr2:24007341-24009356 | p5 | (TTTGG)3 | CAACAATCCATCCCCATCTC | CCGAAGTCTGCATTCCTTTC | 157 |
| chr2:24341965-24343979 | p2 | (TA)7 | TGGTACATCTCCCTTTTGCC | TCTCGTGGAAACGACTTGAA | 243 |
| chr2:24341965-24343979 | p2 | (AT)7 | TGGTACATCTCCCTTTTGCC | TCTCGTGGAAACGACTTGAA | 243 |
| chr2:24378427-24380454 | p3 | (TTC)7 | TCACCTCATTCATCCCCTTC | AGAGGAAACCCACGAAACCT | 202 |
| chr2:24378427-24380454 | p3 | (CAA)9 | TCACCTCATTCATCCCCTTC | AGAGGAAACCCACGAAACCT | 202 |
| chr2:24378427-24380454 | p3 | (CAA)5 | TCACCTCATTCATCCCCTTC | AGAGGAAACCCACGAAACCT | 202 |
| chr2:24378427-24380454 | p2 | (GA)8 | TCACCTCATTCATCCCCTTC | AGAGGAAACCCACGAAACCT | 202 |
| chr2:24379137-24381153 | p3 | (CAA)9 | CAATTGCCAAACAAGTGCTG | CATCTGTTGAGGTTGCATGG | 252 |
| chr2:24379137-24381153 | p3 | (CAA)5 | CAATTGCCAAACAAGTGCTG | CATCTGTTGAGGTTGCATGG | 252 |
| chr2:24379137-24381153 | p2 | (GA)8 | CAATTGCCAAACAAGTGCTG | CATCTGTTGAGGTTGCATGG | 252 |
| chr2:24379137-24381153 | p3 | (AAC)7 | CAATTGCCAAACAAGTGCTG | CATCTGTTGAGGTTGCATGG | 252 |
| chr2:24436356-24438428 | p2 | (AT)8 | ACAGTGACATCCACAGGCAA | ATAATGTGGCCAACAACGGT | 229 |
| chr2:24436356-24438428 | p3 | (ATT)24 | ACAGTGACATCCACAGGCAA | ATAATGTGGCCAACAACGGT | 229 |
| chr2:24543634-24545648 | p2 | (TA)8 | GCGACCAACTTCGTGTCTTT | GACCAAAAAGTCATCACGCA | 258 |
| chr2:24543650-24545664 | p2 | (TA)8 | GCGACCAACTTCGTGTCTTT | GACCAAAAAGTCATCACGCA | 242 |
| chr2:24543666-24545680 | p2 | (TA)8 | GCGACCAACTTCGTGTCTTT | GACCAAAAAGTCATCACGCA | 226 |
| chr2:24543682-24545696 | p2 | (TA)8 | GCGACCAACTTCGTGTCTTT | GACCAAAAAGTCATCACGCA | 210 |
| chr2:24546809-24548831 | p2 | (TA)18 | ATGTGGCAATGAGTCCATCA | GGGTTTAGACCCATGCTTCA | 287 |
| chr2:24546809-24548831 | p2 | (TA)11 | ATGTGGCAATGAGTCCATCA | GGGTTTAGACCCATGCTTCA | 287 |

| Marker ID | SSR type | SSR motif | FORWARD PRIMER (5'-3') | REVERSE PRIMER (5'-3') | size (bp) |
|---|---|---|---|---|---|
| chr2:24582577-24584589 | p2 | (AT)6 | GCCAAGCTTCTCTGGACAAC | TTTGTCTGCCCCTAACTTGG | 188 |
| chr2:24659919-24661941 | p2 | (TA)11 | GCTCTGGGTATTTGCTTTGC | CAATTGCTGCTTTGCTTTCA | 181 |
| chr2:24659919-24661941 | p2 | (AT)52 | GCTCTGGGTATTTGCTTTGC | CAATTGCTGCTTTGCTTTCA | 181 |
| chr2:24687677-24689689 | p2 | (GT)6 | TCTGCTTACCGTGCAAGATG | CTCGGTAATTCCGCATCAAT | 275 |
| chr2:24707277-24709289 | p2 | (TA)9 | GGAAAATTAAGGGCGAGACC | ATTTTTCCCCTAGCCTTCCA | 245 |
| chr2:24707277-24709289 | p2 | (GT)6 | GGAAAATTAAGGGCGAGACC | ATTTTTCCCCTAGCCTTCCA | 245 |
| chr2:24707277-24709289 | p5 | (AATTG)3 | GGAAAATTAAGGGCGAGACC | ATTTTTCCCCTAGCCTTCCA | 245 |
| chr2:24873876-24875888 | p2 | (AT)7 | TTACACTGAAGTCCGGAGCC | TGTCTGCGGCTATGAACAGT | 189 |
| chr2:24873876-24875888 | p2 | (TA)6 | TTACACTGAAGTCCGGAGCC | TGTCTGCGGCTATGAACAGT | 189 |
| chr2:24873937-24875953 | p2 | (AT)7 | TTACACTGAAGTCCGGAGCC | ATTGTCTGCGGCTATGAACA | 128 |
| chr2:24873937-24875953 | p2 | (TA)6 | TTACACTGAAGTCCGGAGCC | ATTGTCTGCGGCTATGAACA | 128 |
| chr2:24890678-24892693 | p5 | (TTCTT)3 | CAAGTGACCAGAGGCAGACA | AAAAATGCTGCTGTGCAGTG | 263 |
| chr2:24892855-24894870 | p5 | (TTCTT)3 | CAAGTGACCAGAGGCAGACA | AAAAATGCTGCTGTGCAGTG | 263 |
| chr2:24958796-24960814 | p3 | (TTG)6 | TTCTGATGCGATTTAAGCCC | CAGACGAAGAACATGAGGCA | 288 |
| chr2:24996625-24998650 | p6 | (CAAAAA)3 | TCAACATGGATGCAAAAGGA | TTTCTTTCCTCGTTCTCCGA | 131 |
| chr2:25004299-25006314 | p2 | (CT)6 | TTCATGTTTCACCGTACCGA | TTCTGCATCATCACAGCACA | 209 |
| chr2:25004299-25006314 | p5 | (TTTTG)3 | TTCATGTTTCACCGTACCGA | TTCTGCATCATCACAGCACA | 209 |
| chr2:25034989-25037017 | p2 | (TA)14 | TTGGTTTTCTGGTTGGTGCT | GGGGTTCGAACATGGATATG | 109 |
| chr2:25034989-25037017 | p2 | (AT)7 | TTGGTTTTCTGGTTGGTGCT | GGGGTTCGAACATGGATATG | 109 |
| chr2:25034989-25037017 | p2 | (TA)15 | TTGGTTTTCTGGTTGGTGCT | GGGGTTCGAACATGGATATG | 109 |
| chr2:25137885-25139900 | p3 | (TGT)5 | CCACCAAAACTTAAGGGCAA | TTCCGCGACTTTCATCTTCT | 247 |
| chr2:25152118-25154130 | p2 | (AT)6 | AAAAGCAGCCTACGGTCTCA | TATGCTTCGTGTGCTCATCC | 185 |
| chr2:25217538-25219550 | p3 | (TTG)5 | AAAACTCAGAAGCGGAGCAA | TCCCCCACATCATATTCGTT | 133 |
| chr2:25272636-25274650 | p4 | (GGGT)4 | AACAAAAGGGCCCTAAATGG | TTAGGAAGCAGCGTTGAGGT | 211 |
| chr2:25272636-25274650 | p2 | (GA)7 | AACAAAAGGGCCCTAAATGG | TTAGGAAGCAGCGTTGAGGT | 211 |
| chr2:25272636-25274650 | p4 | (GATA)4 | AACAAAAGGGCCCTAAATGG | TTAGGAAGCAGCGTTGAGGT | 211 |
| chr2:25273388-25275404 | p2 | (GA)7 | CGCTTTCAGGGTTGTGGTAT | ATTTCAGTTGCCTGCCTTTG | 263 |
| chr2:25273388-25275404 | p4 | (GATA)4 | CGCTTTCAGGGTTGTGGTAT | ATTTCAGTTGCCTGCCTTTG | 263 |
| chr2:25293495-25295510 | p5 | (CCGGC)3 | AAGTGGATCAGCAACCAAGG | ATACCTGCCACTTGGGATGA | 257 |
| chr2:25294269-25296297 | p5 | (CCGGC)3 | TGTAAGGGGCTGGGTTAGGT | TTTTAATTTGAAGTGGGCCG | 121 |
| chr2:25306863-25308875 | p2 | (TA)6 | GCAGATCATTCAGGTGCAGA | TCTGGCCAATTTTTCCTTTG | 159 |
| chr2:25358422-25360437 | p5 | (AAAAT)3 | ATTTGGTCAAAGCACTTGCC | GCCAATTAAATCACCATGCC | 273 |
| chr2:25406017-25408057 | p2 | (AT)20 | AGCGATTTTCACTCGTGGAC | AAAGCTGGTTAGCTTTGCCA | 253 |
| chr2:25406017-25408057 | p2 | (AT)21 | AGCGATTTTCACTCGTGGAC | AAAGCTGGTTAGCTTTGCCA | 253 |
| chr2:25413194-25415206 | p5 | (ACAAC)3 | AACACATGCACGCAGAAAAG | CCTTGCCGGCTTCATAATAA | 212 |
| chr2:25413212-25415224 | p5 | (ACAAC)3 | AACACATGCACGCAGAAAAG | CCTTGCCGGCTTCATAATAA | 194 |

| Marker ID | SSR type | SSR motif | FORWARD PRIMER (5'-3') | REVERSE PRIMER (5'-3') | size (bp) |
|---|---|---|---|---|---|
| chr2:25413226-25415248 | p5 | (ACAAC)3 | AACACATGCACGCAGAAAAG | CCTTGCCGGCTTCATAATAA | 180 |
| chr2:25424272-25426298 | p2 | (AT)13 | ATCATAGCCACACAGGGCTC | GCTGAAGAACGACATCGACA | 230 |
| chr2:25424272-25426298 | p2 | (TA)6 | ATCATAGCCACACAGGGCTC | GCTGAAGAACGACATCGACA | 230 |
| chr2:25424272-25426298 | p2 | (AT)12 | ATCATAGCCACACAGGGCTC | GCTGAAGAACGACATCGACA | 230 |
| chr2:25430133-25432151 | p3 | (AAT)7 | ACGCATCCTCATGGTTTTTC | ATCCATGGCACCACATCTTT | 197 |
| chr2:25430133-25432151 | p2 | (AG)9 | ACGCATCCTCATGGTTTTTC | ATCCATGGCACCACATCTTT | 197 |
| chr2:25449746-25451764 | p6 | (TTCAGT)3 | TGCCCACCAAGTGTTTGATA | AATTCCAGCCTTTGAGCGTA | 271 |
| chr2:25475116-25477128 | p2 | (GA)6 | TCAAGAACGGGGTTGATCTC | ATACCCACGACTTGAATCCG | 247 |
| chr2:25559756-25561774 | p3 | (AAG)6 | AAACGCAAGTTGTTAACGGG | CTTTTGTTGGGTGTGCCTCT | 229 |
| chr2:25559756-25561774 | p5 | (TAAAT)3 | AAACGCAAGTTGTTAACGGG | CTTTTGTTGGGTGTGCCTCT | 229 |
| chr2:25561484-25563498 | p5 | (TAAAT)3 | TTGTACGAAGGGAGTCGTCC | GATTCTTGATTCGAACCCCA | 212 |
| chr2:25561484-25563498 | p2 | (AT)7 | TTGTACGAAGGGAGTCGTCC | GATTCTTGATTCGAACCCCA | 212 |
| chr2:25561484-25563498 | p2 | (TG)14 | TTGTACGAAGGGAGTCGTCC | GATTCTTGATTCGAACCCCA | 212 |
| chr2:25583864-25585879 | p5 | (TCTTT)3 | TAATCTCGCCCTTCGACAGT | GTTGCCGAATCTTATGGGAA | 244 |
| chr2:25583864-25585879 | p2 | (TC)6 | TAATCTCGCCCTTCGACAGT | GTTGCCGAATCTTATGGGAA | 244 |
| chr2:25665506-25667524 | p6 | (TTTTTA)3 | AATTCACAGGGCAAAATTCG | CATGGGTTCGAGTTCAGGAT | 273 |
| chr2:25730463-25732495 | p2 | (TA)16 | TTGGAGTCCACCTTGGTTTC | GCGTATCAAATTTTGGCGTT | 227 |
| chr2:25798962-25800977 | p3 | (TGT)5 | ATGCTTATGGACGAATTGGC | AGCGGCAACAATAAAAATGC | 150 |
| chr2:25821790-25823802 | p2 | (AT)6 | AATGGTGTGTTTAGGGCAGG | GGGGAACAAGTTCCAGATCA | 141 |
| chr2:25832519-25834571 | p2 | (AT)26 | ACCTTACGTCACCCGATGAG | CGACCATATAGCCGGATGTT | 269 |
| chr2:25895380-25897400 | p2 | (TA)10 | CCGCATGTCGAAACTAACAA | ATTGGTTTTCATGGCCAAAC | 165 |
| chr2:25991709-25993725 | p2 | (AG)8 | CAACCATCCCCAAGCTATGT | TTGCTCTGTTTTGTGGCAAG | 164 |
| chr2:26049740-26051755 | p3 | (CTT)5 | CTTTACATCCCTTGCCCAAA | ATTCATCCAAGGCAACCAAG | 245 |
| chr2:26065942-26067960 | p6 | (ATAAAA)3 | CGTTGCCTGCATATCTTTGA | ACTTGAGGCCAATCCGTACA | 127 |
| chr2:26068981-26071027 | p2 | (AG)23 | CGCGTGCCTTCTCAATAAAT | GGGGGAACAAGGAAGAGAAG | 183 |
| chr2:26118146-26120166 | p5 | (TGGTT)4 | GATCCGAACCAGTAAACCGA | CGGTTTGGTTCAGTTTTGGT | 153 |
| chr2:26182493-26184508 | p4 | (TAAA)4 | TTGTAAAATAAATGCGCCCC | TCTCAATCCCCTCAATCTGG | 263 |
| chr2:26182493-26184508 | p5 | (TTTTC)3 | TTGTAAAATAAATGCGCCCC | TCTCAATCCCCTCAATCTGG | 263 |
| chr2:26254191-26256207 | p4 | (TACA)4 | TCAGGCAGGAAACTCACAAA | CCTTAAGCAGTGGCTGAACC | 170 |
| chr2:26254191-26256207 | p4 | (TACA)4 | TCAGGCAGGAAACTCACAAA | CCTTAAGCAGTGGCTGAACC | 170 |
| chr2:26348206-26350221 | p5 | (TTGGT)3 | CATTTCGGGTGAAACCACTT | TTTACAAAACACGGATGGCA | 144 |
| chr2:26358757-26360773 | p2 | (AT)8 | GTTCGGTTGGGTAGTGGAAA | ATGGCCCACTCAGGTATGAC | 290 |
| chr2:26358757-26360773 | p2 | (TA)6 | GTTCGGTTGGGTAGTGGAAA | ATGGCCCACTCAGGTATGAC | 290 |
| chr2:26559247-26561277 | p3 | (TAT)10 | TCTATGTGGCAAAGTGCAGG | AAAACCCTGTAACTCCGGCT | 279 |
| chr2:26559247-26561277 | p2 | (CT)6 | TCTATGTGGCAAAGTGCAGG | AAAACCCTGTAACTCCGGCT | 279 |
| chr2:26559247-26561277 | p2 | (AT)18 | TCTATGTGGCAAAGTGCAGG | AAAACCCTGTAACTCCGGCT | 279 |

| Marker ID | SSR type | SSR motif | FORWARD PRIMER (5'-3') | REVERSE PRIMER (5'-3') | size (bp) |
|---|---|---|---|---|---|
| chr2:26658451-26660469 | p6 | (TTTTTA)3 | AAAGGGAAGGGAAACAGGAA | GACGAAAAAGTGAGCCAACC | 284 |
| chr2:26658451-26660469 | p2 | (AT)6 | AAAGGGAAGGGAAACAGGAA | GACGAAAAAGTGAGCCAACC | 284 |
| chr2:26661016-26663038 | p2 | (TA)11 | TTCCCAAAGTATCCCCATCA | AAATGACATGGCGGAATAGC | 171 |
| chr2:26688924-26690939 | p5 | (ACCAA)3 | TATAAGCCAATCCAAACGGG | CAAACCGAATAACCCGAGAA | 117 |
| chr2:26688924-26690939 | p5 | (AAACC)3 | TATAAGCCAATCCAAACGGG | CAAACCGAATAACCCGAGAA | 117 |
| chr2:26721701-26723758 | p2 | (AG)8 | TTGAAGCAGAGGGGAAAGAA | TCACCCCTCTTTATTCGTCG | 292 |
| chr2:26721701-26723758 | p3 | (AAT)19 | TTGAAGCAGAGGGGAAAGAA | TCACCCCTCTTTATTCGTCG | 292 |
| chr2:26749798-26751812 | p2 | (TA)7 | CAAATCATGTCTCGGTGCAA | GGAGCGTCAGCCATATTTTC | 216 |
| chr2:26749798-26751812 | p2 | (AT)8 | CAAATCATGTCTCGGTGCAA | GGAGCGTCAGCCATATTTTC | 216 |
| chr2:26750027-26752042 | p2 | (TA)7 | GAAAATATGGCTGACGCTCC | TTACTTGGGTTGCCACATCC | 187 |
| chr2:26750027-26752042 | p2 | (AT)8 | GAAAATATGGCTGACGCTCC | TTACTTGGGTTGCCACATCC | 187 |
| chr2:26750027-26752042 | p2 | (TG)7 | GAAAATATGGCTGACGCTCC | TTACTTGGGTTGCCACATCC | 187 |
| chr2:26756294-26758309 | p5 | (CTTTT)3 | GGAAACTCTGCAACTGAGCC | CCTTATATGGACTCGGGCAA | 187 |
| chr2:26758213-26760247 | p2 | (AT)17 | TCCACATTGAGCATGCATTT | TAGGCTTGCACTTTCCGTTT | 173 |
| chr2:26839388-26841410 | p2 | (TC)11 | AACATGAATTGGGACGAAGC | TAAAAGGGCTGGACCACTTG | 114 |
| chr2:26871970-26873982 | p2 | (AT)6 | TTTGCAAAGCCTTACCTTGC | GTTGGGGGTGGTCTTTAGGT | 273 |
| chr2:26871970-26873982 | p6 | (CAAATC)3 | TTTGCAAAGCCTTACCTTGC | GTTGGGGGTGGTCTTTAGGT | 273 |
| chr2:26901648-26903663 | p5 | (ATAAT)3 | GGAAAAGAGGAAGTGGGAGG | TAAGGGAAACACTTACGGCG | 182 |
| chr2:26901668-26903683 | p5 | (ATAAT)3 | GGAAAAGAGGAAGTGGGAGG | TAAGGGAAACACTTACGGCG | 162 |
| chr2:26939602-26941620 | p6 | (CTCTTT)3 | CTACAACCCCCGCAAGATTA | GGGCGAAGAAGTTGACGTAG | 222 |
| chr2:27030535-27032551 | p4 | (TTAT)4 | TTTTAATGGAGGAAGGGGCT | GTTGCTACATGGCAAAGCAA | 126 |
| chr2:27050826-27052844 | p6 | (TGGGGT)3 | CCCTGTCGGAAAAGTACCAA | GTGGGTGTTCGTTTTCGAGT | 208 |
| chr2:27123112-27125139 | p3 | (CAT)9 | AGCTTGTACCCGTCCACATC | GGGTTGAGGGAGCTTCTAGG | 219 |
| chr2:27123112-27125139 | p3 | (TAA)19 | AGCTTGTACCCGTCCACATC | GGGTTGAGGGAGCTTCTAGG | 219 |
| chr2:27123613-27125670 | p3 | (CAT)9 | GACCGGAAACCTACTGGTGA | CGGCCAGATCTCTCTCTCTG | 281 |
| chr2:27123613-27125670 | p3 | (TAA)19 | GACCGGAAACCTACTGGTGA | CGGCCAGATCTCTCTCTCTG | 281 |
| chr2:27220796-27222810 | p2 | (AC)7 | CCTATCATGGACCCTGGATG | ATTTCCAATGTTTGGTGGGA | 203 |
| chr2:27249617-27251632 | p5 | (AGTTA)3 | TTTCCTTTCTCCGTCCCTTT | CCTTTTGGGTTGAATGTGCT | 183 |
| chr2:27446824-27448839 | p3 | (AAG)5 | GGTGGGGGCAAACTTTTAAT | CTTATGGGCAGACTTGGCAT | 232 |
| chr2:27446824-27448839 | p5 | (TTTTA)3 | GGTGGGGGCAAACTTTTAAT | CTTATGGGCAGACTTGGCAT | 232 |
| chr2:27534130-27536148 | p6 | (AAAAAC)3 | CCAAACGCCAATTAAGGAAA | TCTCTTTGAGCCCCGACTAA | 249 |
| chr2:27588944-27590988 | p2 | (AT)22 | TCTTAAACTCCGTGCCCAGT | GGGTCGTTTTGGCAAAGTTA | 180 |
| chr2:27588944-27590988 | p5 | (AAAAT)3 | TCTTAAACTCCGTGCCCAGT | GGGTCGTTTTGGCAAAGTTA | 180 |
| chr2:27608497-27610521 | p6 | (AACTCA)4 | CTCTCCCGCGATATTGTGTT | TCATGCATAATAAGCGCAGC | 248 |
| chr2:27621529-27623549 | p2 | (TC)10 | GTTGGAACGATGGAACGATT | CCTTCAAATGCGTTGGTCTT | 215 |
| chr2:27694215-27696227 | p2 | (TA)6 | TTTTGGGATTGGCTGAACAT | TGATGCTGGTGGTGTCATTT | 238 |

| Marker ID | SSR type | SSR motif | FORWARD PRIMER (5'-3') | REVERSE PRIMER (5'-3') | size (bp) |
|---|---|---|---|---|---|
| chr2:27749105-27751121 | p2 | (AT)8 | ATGCTTTATGCGGGCAATAC | CCCGGCATACTATTGCCTTA | 279 |
| chr2:27759227-27761242 | p3 | (ATA)5 | TTTTGTTGGACAAGCTGACG | AATCGCTGAGACGAGATGCT | 292 |
| chr2:27860510-27862534 | p2 | (AT)12 | TGGGGTGGGTAGTAGCGTAG | GATGGTTGTTTCCGTGGTTC | 216 |
| chr2:27920945-27922960 | p3 | (CAA)5 | TCATGCCTCTGCTCTGTTTG | ACCGAAAGTCACAAAGCGTC | 237 |
| chr2:27940365-27942385 | p2 | (TA)10 | TATACCTGATGTGTCGCGGA | GCTTATCGACGCTTCAAAGG | 212 |
| chr2:27978833-27980848 | p3 | (TAT)5 | TGTGTTTGCCTTGCTATTCG | TACAGGCCCAAGAACCAATC | 125 |
| chr2:28023987-28026003 | p4 | (AATA)4 | GGGAGGGCCCAATAATTAAA | TGCTTGCATTCGTATTGGAC | 218 |
| chr2:28023987-28026003 | p5 | (ATTTT)3 | GGGAGGGCCCAATAATTAAA | TGCTTGCATTCGTATTGGAC | 218 |
| chr2:28028545-28030563 | p2 | (TA)9 | AAATGGGGACGGAAGGAATA | TGAAGTCAAAAACTTCGGGG | 276 |
| chr2:28044822-28046836 | p3 | (TTG)8 | AAAACAGCCGCCTTACCTTT | CCCAAGTCTGGCTGTCTCTC | 259 |
| chr2:28044822-28046836 | p2 | (TA)7 | AAAACAGCCGCCTTACCTTT | CCCAAGTCTGGCTGTCTCTC | 259 |
| chr2:28179695-28181707 | p2 | (TA)6 | CCTGGAGAACTCTTCCGTTG | GTGAACGACCAGCTGTGAGA | 196 |
| chr2:28179695-28181707 | p2 | (AG)7 | CCTGGAGAACTCTTCCGTTG | GTGAACGACCAGCTGTGAGA | 196 |
| chr2:28238661-28240676 | p5 | (ACCAA)3 | GTGGGGTGGACATGAGAAGT | AGAGGGTAAAGTTGCAGCGA | 145 |
| chr2:28238661-28240676 | p5 | (AAACC)3 | GTGGGGTGGACATGAGAAGT | AGAGGGTAAAGTTGCAGCGA | 145 |
| chr2:28270097-28272115 | p3 | (CAT)6 | TTACCACGAAGCAAAAAGGG | TCATTTTTGGTAGCCTTGCC | 188 |
| chr2:28332022-28334037 | p5 | (ACCAA)3 | GTGGGGTGGACATGAGAAGT | AGAGGGTAAAGTTGCAGCGA | 145 |
| chr2:28332022-28334037 | p5 | (AAACC)3 | GTGGGGTGGACATGAGAAGT | AGAGGGTAAAGTTGCAGCGA | 145 |
| chr2:28363458-28365476 | p3 | (CAT)6 | TTACCACGAAGCAAAAAGGG | TCATTTTTGGTAGCCTTGCC | 188 |
| chr2:28409070-28411086 | p2 | (TC)8 | CTTTCCCTAAACTAGCCGCC | AATTGAAGCGGAACAACCAC | 244 |
| chr2:28409070-28411086 | p6 | (GGGCCG)3 | CTTTCCCTAAACTAGCCGCC | AATTGAAGCGGAACAACCAC | 244 |
| chr2:28409766-28411784 | p2 | (TC)8 | GCCATGGCTCCAATTCTTTA | ACTCTTCACCCGAAATTCCC | 270 |
| chr2:28409766-28411784 | p6 | (GGGCCG)3 | GCCATGGCTCCAATTCTTTA | ACTCTTCACCCGAAATTCCC | 270 |
| chr2:28421610-28423625 | p5 | (TACAA)3 | ATACCTGGTAACGAACCCCC | TGAACCACGGGTTTACAACA | 284 |
| chr2:28441245-28443260 | p2 | (AG)15 | GCCTTTTGTCCAAGTGCCTA | ACCTCCTCCAGTCCGAAGAT | 143 |
| chr2:28441245-28443260 | p5 | (CCGGC)3 | GCCTTTTGTCCAAGTGCCTA | ACCTCCTCCAGTCCGAAGAT | 143 |
| chr2:28441245-28443260 | p5 | (GGCCC)3 | GCCTTTTGTCCAAGTGCCTA | ACCTCCTCCAGTCCGAAGAT | 143 |
| chr2:28441245-28443260 | p5 | (ACCAA)3 | GCCTTTTGTCCAAGTGCCTA | ACCTCCTCCAGTCCGAAGAT | 143 |
| chr2:28441245-28443260 | p5 | (TTTAG)3 | GCCTTTTGTCCAAGTGCCTA | ACCTCCTCCAGTCCGAAGAT | 143 |
| chr2:28512260-28514272 | p2 | (AT)6 | TTTTGCCCAAGTCTTGCTTT | GGGATCAAGCCAAGTGAAAA | 276 |
| chr2:28514615-28516630 | p5 | (AATGC)3 | ATATACCTCACGGTCCAGCG | ATACATGGCACGCATTCTCA | 239 |
| chr2:28514615-28516630 | p5 | (AAATT)3 | ATATACCTCACGGTCCAGCG | ATACATGGCACGCATTCTCA | 239 |
| chr2:28551874-28553896 | p2 | (TG)11 | CTTCGGTTCGGAATTTGTGT | GTCCAGCCGAACTTCTTGTC | 223 |
| chr2:28600500-28602524 | p2 | (TA)12 | TGATGAGCCCCATCTTCTTC | AGCCAAAGATGGCTAAACGA | 114 |
| chr2:28600500-28602524 | p3 | (TCT)6 | TGATGAGCCCCATCTTCTTC | AGCCAAAGATGGCTAAACGA | 114 |
| chr2:28600500-28602524 | p2 | (AG)30 | TGATGAGCCCCATCTTCTTC | AGCCAAAGATGGCTAAACGA | 114 |

| Marker ID | SSR type | SSR motif | FORWARD PRIMER (5'-3') | REVERSE PRIMER (5'-3') | size (bp) |
|---|---|---|---|---|---|
| chr2:28718462-28720477 | p5 | (TTATT)3 | TTGATTCTGATTTGCCATGC | AGTGCGAAATACAACCTGGG | 298 |
| chr2:28754040-28756056 | p4 | (AATC)4 | GAGGCACCGAAAGTGAAGAG | TCCCCTCTGAAATGCTCCTA | 173 |
| chr2:28754040-28756056 | p3 | (ACA)6 | GAGGCACCGAAAGTGAAGAG | TCCCCTCTGAAATGCTCCTA | 173 |
| chr2:28810357-28812372 | p5 | (ATGCA)3 | GTGACTTCAGTCCTTGGGGA | ATCGCCAACTAGGAGGGTTT | 176 |
| chr2:28893750-28895770 | p2 | (TA)10 | CTGGATGGCAGTGACTCAGA | TGCACTCTACTCCGCCTTCT | 180 |
| chr2:28940285-28942300 | p5 | (ATCAT)3 | ATCGGAATCATATGACGGGA | CTCTGGAGTGCTCCCTTGTC | 245 |
| chr2:28962170-28964184 | p2 | (TA)7 | ATGGGTAACGCCCAAACATA | TTGTTCCGGAGCTAGCTTGT | 285 |
| chr2:28974678-28976692 | p2 | (TA)7 | ACGTCTTATCATGGCGGAAG | AGCTAAATGCGTTCTCCGAA | 211 |
| chr2:28974678-28976692 | p2 | (AT)6 | ACGTCTTATCATGGCGGAAG | AGCTAAATGCGTTCTCCGAA | 211 |
| chr2:28974823-28976835 | p2 | (TA)7 | GCTTCGGAGAACGCATTTAG | GTCGCTCCGATCTAATCCAA | 161 |
| chr2:28974823-28976835 | p2 | (AT)6 | GCTTCGGAGAACGCATTTAG | GTCGCTCCGATCTAATCCAA | 161 |
| chr2:28982102-28984118 | p4 | (TCTT)4 | GTTTGGTGCCCAACATATCC | TGGAATCGAACACTGGGAAT | 211 |
| chr2:29179325-29181343 | p6 | (TTAACA)3 | GGTCCATTAAGGGGTCGAAT | AATCCTAAAACCCAGCCACC | 226 |
| chr2:29188878-29190893 | p5 | (ACCGA)3 | CCTCGAGTGGGGACTTGATA | TCCGGGGTAGCATCTAACAG | 127 |
| chr2:29241456-29243468 | p2 | (AT)6 | CCACCTGTCTCCTGAGGTGT | GATGTTGTGTGTAGTGGCGG | 150 |
| chr2:29256784-29258802 | p2 | (TA)6 | ACGTGCGTATTCGTAGCAAA | AAACGGACGACGAAAATGAG | 136 |
| chr2:29256805-29258819 | p2 | (TA)6 | ACGTGCGTATTCGTAGCAAA | AAACGGACGACGAAAATGAG | 115 |
| chr2:29275726-29277741 | p3 | (TGG)5 | CCTGATTCTGCTGTGCCATA | TATAAGTGGCCAACCCAAGC | 255 |
| chr2:29275726-29277741 | p2 | (AT)6 | CCTGATTCTGCTGTGCCATA | TATAAGTGGCCAACCCAAGC | 255 |
| chr2:29276432-29278444 | p3 | (TGG)5 | CATTGTTGGTGATGACGGAG | ACCACAACCATCAACCACCT | 221 |
| chr2:29276432-29278444 | p2 | (AT)6 | CATTGTTGGTGATGACGGAG | ACCACAACCATCAACCACCT | 221 |
| chr2:29276432-29278444 | p2 | (TA)7 | CATTGTTGGTGATGACGGAG | ACCACAACCATCAACCACCT | 221 |
| chr2:29278205-29280223 | p6 | (TCCTAT)3 | CTTAGGTGCCGTTTGGACAT | TGTTTGCACAAAAGGTGGAA | 256 |
| chr2:29291814-29293832 | p6 | (TCCTAT)3 | ATGTTTGGAAGCAACGGAAC | TCGGAACTTCAAGCCCTATG | 120 |
| chr2:29300531-29302555 | p2 | (AT)12 | TGCATAGCACCTTCCTTGTG | ACTCGCTCCAAGTCCAAGAA | 124 |
| chr2:29374598-29376610 | p2 | (CT)6 | GGCATGGTAACTGTAGCGGT | TTTAGTTTTGGCGGGAACAG | 222 |
| chr2:29435570-29437604 | p2 | (TC)7 | GGACCGAAGGTATCGCACTA | ACTGATCACCCGGACTAACG | 232 |
| chr2:29435570-29437604 | p2 | (TA)17 | GGACCGAAGGTATCGCACTA | ACTGATCACCCGGACTAACG | 232 |
| chr2:29623422-29625434 | p2 | (TA)6 | CCAACATAGATACCGGGTGG | GCAAGATCGAAGCCAAAGTC | 268 |
| chr2:29629139-29631154 | p3 | (TAT)5 | GCCAACGTGGAAAAGAAAAA | GGGAAAATTCTGATGCCCTT | 212 |
| chr2:29633976-29636021 | p3 | (TTA)15 | GATATGTTTGGCGTTGGCTT | ATTTTTATCCGGGGGTTCAA | 157 |
| chr2:29734179-29736194 | p3 | (AAT)5 | TTTTATGGGGTTTATGGGCA | GGCACCTGACGAGCTTCTAC | 200 |
| chr2:29734179-29736194 | p5 | (TTTCT)3 | TTTTATGGGGTTTATGGGCA | GGCACCTGACGAGCTTCTAC | 200 |
| chr2:29734773-29736788 | p3 | (AAT)5 | TCGTCGTCAGACACCAAAAA | CATGCGGACCTCACCTTAAT | 170 |
| chr2:29734773-29736788 | p5 | (TTTCT)3 | TCGTCGTCAGACACCAAAAA | CATGCGGACCTCACCTTAAT | 170 |
| chr2:29774378-29776393 | p3 | (ATC)5 | ATTCCGGATAACTCGACCCT | ACCCACATGGCTCTCAAAAC | 276 |

| Marker ID | SSR type | SSR motif | FORWARD PRIMER (5'-3') | REVERSE PRIMER (5'-3') | size (bp) |
|---|---|---|---|---|---|
| chr2:29796327-29798339 | p2 | (AC)6 | GGTCAACTCCCAGCAGAGAC | CCAGAATTAGGGGTGCAAAA | 296 |
| chr2:29796327-29798339 | p2 | (AT)9 | GGTCAACTCCCAGCAGAGAC | CCAGAATTAGGGGTGCAAAA | 296 |
| chr2:29810461-29812476 | p3 | (CAA)5 | CACAGCATAACACCGGAATG | TATGTTTTTCATAGGCGGGC | 216 |
| chr2:29825323-29827338 | p5 | (AAATA)3 | GAATTGTTGGTGACGACGTG | CCTTTCCTTTCCATTCTCCC | 287 |
| chr2:29909892-29911910 | p6 | (TATACA)3 | CTCCTTCAATTCCTCCACCA | TCAATGGCTGAGAAAGACCC | 145 |
| chr2:30093993-30096008 | p3 | (GTT)5 | AGGTGGTGGTTAATTGCAGC | TACTTCTTCGGCCTACCCCT | 269 |
| chr2:30093993-30096008 | p6 | (ATTTTC)4 | AGGTGGTGGTTAATTGCAGC | TACTTCTTCGGCCTACCCCT | 269 |
| chr2:30164017-30166029 | p2 | (TA)13 | TACATTTCCTTGGGTGTGGG | TTAACCACTCCAAACCGGAC | 204 |
| chr2:30164017-30166029 | p2 | (TA)6 | TACATTTCCTTGGGTGTGGG | TTAACCACTCCAAACCGGAC | 204 |
| chr2:30220412-30222427 | p5 | (TTTTC)3 | GATGCAGTCCGTCGATTTTT | AACACGGTCCGTCAGTTTTC | 161 |
| chr2:30253819-30255841 | p2 | (AT)11 | CGCATTTAGCCAAGGAAGAC | GTCGCTCCGATCTAATCCAA | 222 |
| chr2:30253819-30255841 | p2 | (AT)11 | CGCATTTAGCCAAGGAAGAC | GTCGCTCCGATCTAATCCAA | 222 |
| chr2:30253819-30255841 | p2 | (AT)11 | CGCATTTAGCCAAGGAAGAC | GTCGCTCCGATCTAATCCAA | 222 |
| chr2:30298010-30300036 | p2 | (AT)13 | CGCTTTCTTTGGAGGCATAG | CCAGTTCGGGCACTAGTCAT | 106 |
| chr2:30303614-30305632 | p2 | (AT)8 | AGCCACACACACATATCCGA | CTTGGAGTGCTCCCTTGTTC | 218 |
| chr2:30303614-30305632 | p6 | (CATATA)3 | AGCCACACACACATATCCGA | CTTGGAGTGCTCCCTTGTTC | 218 |
| chr2:30307601-30309621 | p2 | (AT)10 | CCCTAGGACGAACTGCTCTG | AAATCGGTTTGCGCTGTTAC | 208 |
| chr2:30326833-30328845 | p2 | (AC)6 | TACATGCCTCGTACGCACTC | AATTCTCGGGTTGGCCTATT | 168 |
| chr2:30430904-30432918 | p5 | (ATAAA)3 | CAAAACAACCGGCTCAAAAT | CCCCCATCAATCATTTACCA | 258 |
| chr2:30430904-30432918 | p2 | (TA)7 | CAAAACAACCGGCTCAAAAT | CCCCCATCAATCATTTACCA | 258 |
| chr2:30430904-30432918 | p3 | (TTC)6 | CAAAACAACCGGCTCAAAAT | CCCCCATCAATCATTTACCA | 258 |
| chr2:30512726-30514741 | p5 | (AAACC)3 | TTATGATGATCTGGGGTGGG | AGCGTTTGATATTCGGTTCG | 266 |
| chr2:30523432-30525444 | p2 | (TA)6 | AATTGGATCCACCGTTACCA | TGCAAACTTTTCAATGTGGC | 129 |
| chr2:30551192-30553222 | p3 | (TAT)10 | GGTGGTACATTTCCCTGTGG | TTGGGAAAAACTGTGGAAGC | 194 |
| chr2:30609386-30611406 | p2 | (TA)10 | GATTGGGAGGTGGCTACTGA | ACAAGCAAACTGTGTTCCCC | 203 |
| chr2:30615006-30617030 | p2 | (TC)6 | TGCAAAGACAGATTCAACGC | GGCCTGTGTTGTCCTCTGAT | 132 |
| chr2:30661161-30663179 | p3 | (TAT)6 | CCAATAAAAGGCGGTGAGAA | CAAAAGCAAGGCCTCAACTC | 120 |
| chr2:30669856-30671868 | p2 | (TG)6 | GCCCAGGAATATCAGGAACA | ACAAGGAGCAAACGCCTAGA | 191 |
| chr2:30785134-30787152 | p6 | (ATATCC)3 | TATTGGGTTTGTGTGGGGTT | CCCTAGTTGGGTCGTGACAT | 175 |
| chr2:30792733-30794771 | p2 | (GA)19 | CTTGCCCCTTATTCCTCTCC | TTCCGACAATTTCTCAACCC | 105 |
| chr2:30865215-30867249 | p2 | (AT)6 | CAAAATCCGAAAGTCACCGT | CATTTAGCGTGGAGGACGTT | 242 |
| chr2:30865215-30867249 | p2 | (TA)32 | CAAAATCCGAAAGTCACCGT | CATTTAGCGTGGAGGACGTT | 242 |
| chr2:30944861-30946876 | p3 | (TGT)5 | CGACTCTGTTGTACGGAGCA | TGGGGCATCTATGTCTCTCC | 297 |
| chr2:31110186-31112201 | p5 | (TTGTA)3 | TGGATTAGGGGAGAGAACGA | GGTTGAATCATGCCACAATG | 179 |
| chr2:31126518-31128533 | p5 | (GTTCG)3 | CTGTCCACCCCAGACTTCAT | CATCGTTGAAGTTGTCCCCT | 101 |
| chr2:31126518-31128533 | p5 | (GTTCG)3 | CTGTCCACCCCAGACTTCAT | CATCGTTGAAGTTGTCCCCT | 101 |

| Marker ID | SSR type | SSR motif | FORWARD PRIMER (5'-3') | REVERSE PRIMER (5'-3') | size (bp) |
|---|---|---|---|---|---|
| chr2:31127032-31129047 | p5 | (GTTCG)3 | CATGCTCAGGGACCGACTAT | TGTTTGGTGTCCACCTTCAA | 259 |
| chr2:31127032-31129047 | p5 | (GTTCG)3 | CATGCTCAGGGACCGACTAT | TGTTTGGTGTCCACCTTCAA | 259 |
| chr2:31205191-31207209 | p6 | (CTTACA)3 | CTAGAAAGGGCCATTCCACA | GTGTCATGGCAAAACTGGTG | 184 |
| chr2:31288594-31290646 | p2 | (TA)26 | AGCTGAATCCGAACTCGCTA | CGACAAGCCCTTTTTGCTAC | 117 |
| chr2:31366715-31368760 | p3 | (TTC)15 | ACGAACGCAAACCAAAAATC | TCCCCAGTCATAATGCTTCC | 197 |
| chr2:31403496-31405510 | p2 | (TA)7 | CGCTTAGCTCCAATTTGTCC | TGTTGATGTGCCGTGAATTT | 160 |
| chr2:31502882-31504900 | p3 | (TAG)6 | ACGGTTGGCAGTATTCTTGG | GCAGAAGTAGTGTCCGGTCC | 246 |
| chr2:31520610-31522636 | p2 | (AT)13 | CGTCAGATCAAGGTTCACGA | GAACGCCGACTAAGAGCATC | 165 |
| chr2:31520610-31522636 | p2 | (AT)8 | CGTCAGATCAAGGTTCACGA | GAACGCCGACTAAGAGCATC | 165 |
| chr2:31587968-31589980 | p2 | (TA)6 | TTGTCACTAAATGTCGGGCA | ACTTAGCGACAGGGTAGCGA | 102 |
| chr2:31622525-31624543 | p2 | (TA)6 | CATCTGCAATACAACGTGGG | TGGTTGAACAAAAAGACCCC | 284 |
| chr2:31622525-31624543 | p2 | (TA)6 | CATCTGCAATACAACGTGGG | TGGTTGAACAAAAAGACCCC | 284 |
| chr2:31622525-31624543 | p2 | (AT)6 | CATCTGCAATACAACGTGGG | TGGTTGAACAAAAAGACCCC | 284 |
| chr2:31622525-31624543 | p3 | (AAG)7 | CATCTGCAATACAACGTGGG | TGGTTGAACAAAAAGACCCC | 284 |
| chr2:31622555-31624567 | p2 | (TA)6 | CATCTGCAATACAACGTGGG | TGGTTGAACAAAAAGACCCC | 254 |
| chr2:31622555-31624567 | p2 | (AT)6 | CATCTGCAATACAACGTGGG | TGGTTGAACAAAAAGACCCC | 254 |
| chr2:31622555-31624567 | p3 | (AAG)7 | CATCTGCAATACAACGTGGG | TGGTTGAACAAAAAGACCCC | 254 |
| chr2:31664321-31666336 | p5 | (TAATA)3 | CCAGCCCAATTAAACCAAAA | CGAATTTTGAGAAAAAGCGG | 233 |
| chr2:31674560-31676641 | p3 | (TTA)27 | ACTCTGTTGCTGCTGTCCCT | GGCCTACTCAGAGGCAGTTG | 131 |
| chr2:31727533-31729548 | p5 | (ATTTC)3 | AACGCTTGTATCCCGTGTTC | ATACTTCTTGTCCGCCCCTT | 294 |
| chr2:31758044-31760062 | p3 | (GTT)7 | CCGAATTTTCCGTAGAACGA | CCTCTGCTGTCTCGCTAACC | 274 |
| chr2:31758148-31760160 | p3 | (GTT)7 | CCGAATTTTCCGTAGAACGA | CCTCTGCTGTCTCGCTAACC | 170 |
| chr2:31758148-31760160 | p2 | (TA)30 | CCGAATTTTCCGTAGAACGA | CCTCTGCTGTCTCGCTAACC | 170 |
| chr2:31794129-31796144 | p3 | (TCA)5 | TTGCACCAAATGAAAATGGA | GGCAACCCTCATAGAATGGA | 204 |
| chr2:31809924-31811939 | p5 | (ATTTT)3 | CGCAACACATACTGCTTCGT | GCGAGTCAGTCCCTCTTTTG | 258 |
| chr2:31836472-31838487 | p3 | (TCA)5 | TTGCACCAAATGAAAATGGA | GGCAACCCTCATAGAATGGA | 204 |
| chr2:31852267-31854282 | p5 | (ATTTT)3 | CGCAACACATACTGCTTCGT | GCGAGTCAGTCCCTCTTTTG | 258 |
| chr2:31863286-31865298 | p2 | (TC)6 | TCACCCTCACGATGTTCTCA | TCACGTACGGGGCTTAATTC | 181 |
| chr2:31938378-31940394 | p2 | (AT)8 | CTCGGTTACATGCAGCAAGA | GCTTCCGAAAGCAGCATAAG | 126 |
| chr2:32032039-32034097 | p2 | (AT)29 | TTCATTAACGGGCCATTAGC | TTACTCGGGGTTCAAATCCA | 250 |
| chr2:32096169-32098184 | p5 | (TTTTA)3 | GGACATCCAAGAGGGAGTGT | GAATGTGGGGAAGATCATGG | 122 |
| chr2:32096169-32098184 | p2 | (AT)34 | GGACATCCAAGAGGGAGTGT | GAATGTGGGGAAGATCATGG | 122 |
| chr2:32214248-32216264 | p2 | (AT)8 | TGCATGTCTGAGAGAGGGTG | TCGGCTAGCGAATTCAGTTT | 231 |
| chr2:32232413-32234467 | p3 | (GAA)7 | CCTCGCGAAAACCTACTTTG | TCCCCAGTAAAGGTGCAAAC | 210 |
| chr2:32232413-32234467 | p2 | (TA)17 | CCTCGCGAAAACCTACTTTG | TCCCCAGTAAAGGTGCAAAC | 210 |
| chr2:32340690-32342702 | p2 | (AT)6 | GAAGTGAAACAAATGCGGGT | CCTTCCAGAATTGTGCCTGT | 233 |

| Marker ID | SSR type | SSR motif | FORWARD PRIMER (5'-3') | REVERSE PRIMER (5'-3') | size (bp) |
|---|---|---|---|---|---|
| chr2:32368973-32370988 | p5 | (AACAA)3 | TACGGATCCCGTCATTTAGC | GCTTAAGTTGGGGGTGTTGA | 152 |
| chr2:32515142-32517154 | p2 | (AT)6 | TCAGAATTTCCAATGGAGGC | AGACCGGATTAGGGCAACTT | 117 |
| chr2:32652991-32655006 | p3 | (GTT)5 | CCACCAAAACTTAAGGGCAA | CTGGGGCTTCTGTGTCTCTC | 269 |
| chr2:32656080-32658095 | p3 | (GAT)5 | CCGAGTTTTCTTGGGAAACA | ATCCTGATTGGGAGAAGGCT | 258 |
| chr2:32749163-32751178 | p3 | (TTA)5 | TGGTTTGCCTTGTAGCATTG | AGGCTCTCCAGTGTGTTGCT | 286 |
| chr2:32762125-32764140 | p3 | (TTA)5 | TGGTTTGCCTTGTAGCATTG | AGGCTCTCCAGTGTGTTGCT | 286 |
| chr2:32808755-32810779 | p2 | (TA)12 | CCCACCATCTCCAAACTCAC | CGCAAACAAAGTCTGCACAT | 242 |
| chr2:32815291-32817305 | p2 | (AG)7 | AGGGTGCGTAGGCTGATAGA | TCAAGATGGTGCGGAATACA | 165 |
| chr2:32823595-32825609 | p2 | (AG)7 | AGGGTGCGTAGGCTGATAGA | TCAAGATGGTGCGGAATACA | 165 |
| chr2:32870864-32872888 | p2 | (TA)6 | CAGTCCTCGATTTGACACGA | TTTCCTGAGTTTGGGATTGG | 220 |
| chr2:32870864-32872888 | p2 | (TA)7 | CAGTCCTCGATTTGACACGA | TTTCCTGAGTTTGGGATTGG | 220 |
| chr2:32870864-32872888 | p2 | (AT)19 | CAGTCCTCGATTTGACACGA | TTTCCTGAGTTTGGGATTGG | 220 |
| chr2:32878140-32880154 | p2 | (CA)7 | TTGGTGGGATTGAGACATCA | AAAACAAGGAAGGGCCTGTT | 181 |
| chr2:32886920-32888935 | p3 | (GTT)5 | CCACCAAAACTTAAGGGCAA | GGCTACCCCCACCTTATCTC | 259 |
| chr2:32955946-32957976 | p2 | (AT)15 | CACGTGTGGGCTAAGCAGTA | TTTTATTTGGGACGGACGAC | 202 |
| chr2:32955946-32957976 | p2 | (AT)8 | CACGTGTGGGCTAAGCAGTA | TTTTATTTGGGACGGACGAC | 202 |
| chr2:33058666-33060678 | p2 | (TA)6 | TCCACAAACCCTTCCATAGC | AGGGGCATATTTGAGCCTTT | 116 |
| chr2:33117887-33119945 | p2 | (AT)29 | AAAGAAGGGCAGTTATGCGA | TCCCCTACAATCAGTCCTGC | 118 |
| chr2:33117887-33119945 | p2 | (TA)6 | AAAGAAGGGCAGTTATGCGA | TCCCCTACAATCAGTCCTGC | 118 |
| chr2:33138342-33140356 | p2 | (CT)7 | TGAATCCTTGGAGCAAATCC | AACAATCCTTGAAGAGGGCA | 231 |
| chr2:33198506-33200526 | p2 | (TA)10 | TAGCTTCCCATTCCCCTTTT | AGCATTGGTATATTTGCGCC | 236 |
| chr2:33261063-33263078 | p3 | (TGT)5 | ATACAAACCCGAAGCCCTTT | TGGCCAAACGAGCTATAACC | 295 |
| chr2:33296467-33298479 | p2 | (GA)6 | GGCTGACGGCGTTATTATGT | GCGAGTCAATTCTTTGGGAC | 287 |
| chr2:33314162-33316174 | p2 | (AT)6 | TCGCGACTGCGTTATATCAG | TCTTGTGCAATGCCCAGTAG | 139 |
| chr2:33395972-33397987 | p3 | (TGT)5 | ACACCATATTGGTGCAGGGT | ACCGAGGCTATTCCAACCTT | 216 |
| chr2:33395972-33397987 | p2 | (TA)25 | ACACCATATTGGTGCAGGGT | ACCGAGGCTATTCCAACCTT | 216 |
| chr2:33473724-33475736 | p2 | (TA)6 | TGATGTTAACGCCTTTTCCC | GCCAGCTCCAAGTTGATTTC | 127 |
| chr2:33491875-33493901 | p6 | (ATTTTT)3 | AATCAGCCCAGCTCTCACAT | CAATGAGCCCGCTTATGAAT | 168 |
| chr2:33491934-33493986 | p6 | (ATTTTT)3 | AATCAGCCCAGCTCTCACAT | CAATGAGCCCGCTTATGAAT | 109 |
| chr2:33550309-33552325 | p2 | (TA)8 | GACAGGTGGCTGGAGTTGAT | CCAAGGCTCTCGATCTTGTC | 194 |
| chr2:33583307-33585325 | p3 | (CAC)6 | TGTTCATGCCCACAGGTAGA | TCGTGAAACCACATAACCCA | 137 |
| chr2:33583307-33585325 | p3 | (ACC)6 | TGTTCATGCCCACAGGTAGA | TCGTGAAACCACATAACCCA | 137 |
| chr2:33590948-33592960 | p3 | (ACA)5 | TGGGGAGGGTAGAGTGTACG | GTGTCGCTTGCGTTACTTCA | 119 |
| chr2:33590948-33592960 | p2 | (AC)6 | TGGGGAGGGTAGAGTGTACG | GTGTCGCTTGCGTTACTTCA | 119 |
| chr2:33603783-33605801 | p3 | (TTG)6 | GCCACCAAAACTTAAAGGCA | CTCCCACCTTGTCTCGGATA | 203 |
| chr2:33603783-33605801 | p6 | (AGGGGA)5 | GCCACCAAAACTTAAAGGCA | CTCCCACCTTGTCTCGGATA | 203 |

| Marker ID | SSR type | SSR motif | FORWARD PRIMER (5'-3') | REVERSE PRIMER (5'-3') | size (bp) |
|---|---|---|---|---|---|
| chr2:33650315-33652330 | p6 | (TTTATT)3 | CACAAACATCACGGATGGAG | CCAAGCCCGACTATCTTGAC | 123 |
| chr2:33768592-33770607 | p5 | (TTTCA)3 | ATGGCGGAATAACGATCAAG | TCCTGCACAACAACCCATAA | 177 |
| chr2:33912087-33914099 | p2 | (AT)6 | TTCGGTTCGGTACGGTATTC | AGCACAGAGGGTTGTCGTTT | 110 |
| chr2:33912087-33914099 | p2 | (AG)9 | TTCGGTTCGGTACGGTATTC | AGCACAGAGGGTTGTCGTTT | 110 |
| chr2:33912617-33914635 | p2 | (AT)6 | TTATCAGGCCTTAGCCCGTA | TTGAGCAAGTCTTTTGGGGT | 195 |
| chr2:33912617-33914635 | p2 | (AG)9 | TTATCAGGCCTTAGCCCGTA | TTGAGCAAGTCTTTTGGGGT | 195 |
| chr2:33990534-33992549 | p5 | (AAAAT)3 | TCATAATGGAGCCCACACAA | TTTGCCGCTACTGAACTCCT | 151 |
| chr2:34012462-34014477 | p3 | (TCA)5 | ATTGGATATCATCCGCTTCG | CAACAAGAAAACGCACAAGC | 101 |
| chr2:34052648-34054663 | p5 | (GGGGT)3 | TATGGCGACAAGCCACATAA | ACACAAAAATGGCCAAGGAG | 275 |
| chr2:34055889-34057907 | p6 | (AATAAA)3 | TATTGATTTCTTGGGACCCG | AATCAAAAGTGTTGAGCCCG | 292 |
| chr2:34055889-34057907 | p6 | (AAAAAT)3 | TATTGATTTCTTGGGACCCG | AATCAAAAGTGTTGAGCCCG | 292 |
| chr2:34115578-34117593 | p5 | (TTTAA)3 | TTCGCACTTGGAGTGTCTTG | TAGCTGCGGAGCAAGAATTT | 258 |
| chr2:34115578-34117593 | p2 | (AT)10 | TTCGCACTTGGAGTGTCTTG | TAGCTGCGGAGCAAGAATTT | 258 |
| chr2:34243938-34245974 | p2 | (AT)18 | AAAAGGCCAATCTGCTTTCA | TGTCAATGCACCATCGATCT | 213 |
| chr2:34243938-34245974 | p2 | (AT)9 | AAAAGGCCAATCTGCTTTCA | TGTCAATGCACCATCGATCT | 213 |
| chr2:34424702-34426717 | p5 | (AATGC)3 | GTAGAGTTGCGGGTGAGGAG | TTAGATCGGACTCTGGGTGG | 207 |
| chr2:34460622-34462637 | p2 | (AT)36 | GTGGGCTCCACACGTTATTT | ATACCGTACTGTGCCCACCT | 105 |
| chr2:34460622-34462637 | p5 | (TTTGG)3 | GTGGGCTCCACACGTTATTT | ATACCGTACTGTGCCCACCT | 105 |
| chr2:34481531-34483549 | p3 | (TTC)6 | TGGTTAGGCACCACCTTTTC | AGAACAGCAACGGGAGAGAA | 110 |
| chr2:34481531-34483549 | p3 | (CTT)7 | TGGTTAGGCACCACCTTTTC | AGAACAGCAACGGGAGAGAA | 110 |
| chr2:34533801-34535817 | p5 | (ATATG)3 | TACGAACGTAACTTGCGACG | GGCAATCGTAAAAGTGGGAA | 259 |
| chr2:34533818-34535830 | p5 | (ATATG)3 | TACGAACGTAACTTGCGACG | GGCAATCGTAAAAGTGGGAA | 242 |
| chr2:34533831-34535875 | p5 | (ATATG)3 | TACGAACGTAACTTGCGACG | GGCAATCGTAAAAGTGGGAA | 229 |
| chr2:34536968-34538988 | p2 | (AG)10 | GGGCTGCAATGTTCATCTTT | CCGGAGACATTTTGTGGAGT | 226 |
| chr2:34544210-34546225 | p3 | (TTA)5 | TCATGAACTGATCGAACCCA | TTCCTTCCCCTTCTCTCCAT | 211 |
| chr2:34607560-34609576 | p2 | (TG)8 | TTCTTGGACGTGCTGAAGTG | GCAATTTTACAAGGACGGGA | 222 |
| chr2:34609511-34611526 | p5 | (AAAGA)3 | TTCAGCGAACTCAACGAATG | CGAAAAGGCATTTCACACCT | 210 |
| chr2:34613064-34615076 | p2 | (CA)6 | TGGTTGACTGCTGTCTTTGG | TCTCATCAGTGGGGCTTTTC | 217 |
| chr2:34640047-34642071 | p2 | (AT)9 | AAAATCCACACAGGCCAAAG | CTTCCATGGCTACCCCTACA | 102 |
| chr2:34640047-34642071 | p6 | (AGAAGG)4 | AAAATCCACACAGGCCAAAG | CTTCCATGGCTACCCCTACA | 102 |
| chr2:34683471-34685575 | p3 | (ACA)5 | ATGCAGATTCGTTTTCCACC | TGTGCAACTTCATTTGGCAT | 249 |
| chr2:34717323-34719338 | p5 | (TGGTT)3 | CAATGTTAGGCTCGCAGTCA | TGCAACTTCACTTCACGGAC | 135 |
| chr2:34828079-34830115 | p2 | (GA)7 | AATATTTTCTCCCCCGGTTG | CCGCTCCTCTCTTCCTTTCT | 161 |
| chr2:34828079-34830115 | p3 | (ATA)12 | AATATTTTCTCCCCCGGTTG | CCGCTCCTCTCTTCCTTTCT | 161 |
| chr2:34853624-34855682 | p2 | (CT)29 | GTGTTAGGCACCTCGGAAAA | AGTTTGACCCCACCACTCAC | 139 |
| chr2:34862648-34864666 | p5 | (TCATA)3 | AAGCCGACAGGCTAACAAAA | ATTCGGAAGCCATAGCCTTT | 208 |

| Marker ID | SSR type | SSR motif | FORWARD PRIMER (5'-3') | REVERSE PRIMER (5'-3') | size (bp) |
|---|---|---|---|---|---|
| chr2:34862648-34864666 | p2 | (TA)9 | AAGCCGACAGGCTAACAAAA | ATTCGGAAGCCATAGCCTTT | 208 |
| chr2:34895503-34897515 | p2 | (TG)6 | ATGATGCTAGGTGCCTACGG | TCATACCTGGCCCTACAAGG | 198 |
| chr2:35010058-35012082 | p2 | (TA)12 | CAACTCCCCAACGGTCTAAA | CCCGTTTGACTCACAGTTGA | 162 |
| chr2:35055946-35058006 | p3 | (AAT)20 | CCAAGGCAAATAAAAACGGA | TGTATGAGGGAGCCAAAACC | 158 |
| chr2:35210639-35212651 | p2 | (GT)6 | GAGATGGTTGCTGCAGATGA | GATGCCTTACAACCGAGGAA | 198 |
| chr2:35220068-35222086 | p6 | (AACAAT)3 | CGTCCACACTCCACAGTACG | TCTATGCCACCTCAGCATTG | 213 |
| chr2:35282009-35284021 | p6 | (TGGGGA)3 | AATCCATGGCATTGGTTCAT | CCCACTACCACTGCCAAAAT | 240 |
| chr2:35282009-35284021 | p2 | (AT)6 | AATCCATGGCATTGGTTCAT | CCCACTACCACTGCCAAAAT | 240 |
| chr2:35297059-35299071 | p2 | (GA)6 | ACTGCCAGCTAGGCAAACAT | CGGTCTTAGCTAGGTTCCCC | 123 |
| chr2:35340400-35342415 | p3 | (AAG)5 | ACAATAAGGCGGTTGTGTCC | ACTGCTGGAGCGATGAAGAT | 274 |
| chr2:35340530-35342580 | p3 | (AAG)5 | ACAATAAGGCGGTTGTGTCC | ACTGCTGGAGCGATGAAGAT | 144 |
| chr2:35487884-35489899 | p5 | (CGACC)3 | CTCGTGGTGGAAGTGGAGTT | ACAAAGGATTGGCCACTGTC | 224 |
| chr2:35592721-35594736 | p5 | (TTGGT)3 | TCCGTGAGCAAAGGACTCTT | GTAATGTCGGGTTGGTTTGG | 103 |
| chr2:35651411-35653426 | p5 | (TTTTA)3 | ATGGTCGGAGTTGAATGGTC | TAACACAGCACCCAAAACGA | 118 |
| chr2:35802564-35804576 | p2 | (AT)6 | TAATATTCAAAGCGGCGGAC | TAAGTCAGGTGGTTCCCGAC | 277 |
| chr2:35829026-35831041 | p5 | (CAAAC)3 | TAGTGGTTGCACCGTTTTTG | ATGCAAGTATTGCCCCAAAG | 113 |
| chr2:36083169-36085187 | p3 | (AAC)6 | ACGAGCGGAGTAGTGAGGAA | GAAAAGCTCGGATTGCTTTG | 276 |
| chr2:36127896-36129941 | p3 | (TCT)15 | AAAATGGAATCGGATGCAAG | ACTCGTTACATTTGGCCAGG | 227 |
| chr2:36127896-36129941 | p2 | (GT)11 | AAAATGGAATCGGATGCAAG | ACTCGTTACATTTGGCCAGG | 227 |
| chr2:36139276-36141292 | p2 | (TA)6 | TAGAGCCTCTTTGGATTGGC | TAAGAGCCTGTTTGGATGGG | 175 |
| chr2:36139276-36141292 | p2 | (TA)23 | TAGAGCCTCTTTGGATTGGC | TAAGAGCCTGTTTGGATGGG | 175 |
| chr2:36139296-36141308 | p2 | (TA)6 | TAGAGCCTCTTTGGATTGGC | TAAGAGCCTGTTTGGATGGG | 155 |
| chr2:36139296-36141308 | p2 | (TA)23 | TAGAGCCTCTTTGGATTGGC | TAAGAGCCTGTTTGGATGGG | 155 |
| chr2:36239327-36241342 | p5 | (CTACC)3 | TTCCAAAATTCTTTGACGCC | CATTGAGGAGGAAGAGCCAG | 102 |
| chr2:36239327-36241342 | p5 | (TACAA)3 | TTCCAAAATTCTTTGACGCC | CATTGAGGAGGAAGAGCCAG | 102 |
| chr2:36284990-36287006 | p2 | (AT)8 | ACGGACAGTTTCAAACAGGG | TATTCTGGCGGGAACAAGTC | 136 |
| chr2:36284990-36287006 | p2 | (AT)12 | ACGGACAGTTTCAAACAGGG | TATTCTGGCGGGAACAAGTC | 136 |
| chr2:36290389-36292401 | p2 | (AG)6 | ATTGCATTCCGGAAAGTGTC | CCTCTACCTCGACTTGCTGG | 186 |
| chr2:36316193-36318213 | p2 | (TA)8 | AAGCAACGGCTAGCAGACAT | AAAGGTGGTTGCATGCTCTT | 181 |
| chr2:36316220-36318235 | p2 | (TA)8 | AAGCAACGGCTAGCAGACAT | AAAGGTGGTTGCATGCTCTT | 154 |
| chr2:36316436-36318452 | p2 | (TA)8 | AGTTGAAGTCCACCAGACGC | AAGGGATTGAGCATTTGACG | 132 |
| chr2:36384885-36386903 | p6 | (CCACCC)3 | GTCTTAACACGGCACGGATT | TTCCCCTTTATGTGCTTTGG | 225 |
| chr2:36407894-36409916 | p2 | (TA)11 | TGGAAGAACTTTGAGCAGCA | CCCCAAGCTCTAGCTCACAC | 125 |
| chr2:36469602-36471623 | p3 | (ACA)7 | AACGAATCACCTCCAGCATC | TGGTCTAGGTGTTGATGGCA | 224 |
| chr2:36469602-36471623 | p3 | (ATT)22 | AACGAATCACCTCCAGCATC | TGGTCTAGGTGTTGATGGCA | 224 |
| chr2:36500120-36502135 | p5 | (ACCAA)3 | ACAATTGATTGCTGAAGGGG | CAAGAAAACCCGAGGTTGAA | 200 |

| Marker ID | SSR type | SSR motif | FORWARD PRIMER (5'-3') | REVERSE PRIMER (5'-3') | size (bp) |
|---|---|---|---|---|---|
| chr2:36500120-36502135 | p5 | (AAACC)3 | ACAATTGATTGCTGAAGGGG | CAAGAAAACCCGAGGTTGAA | 200 |
| chr2:36520072-36522087 | p3 | (ACA)5 | GCAGCAGAAATCGAAAAACC | ACCGAAGAACCATTGACCTG | 191 |
| chr2:36541976-36543991 | p5 | (TTGGT)3 | ACGGAGCTCTCCAAGACAAA | ACCGAATAACCCGAAAAACC | 115 |
| chr2:36601372-36603394 | p2 | (TA)11 | ATGTCGACTAGCCGTTGACC | TCAAACTCCAACTCCATCCC | 107 |
| chr2:36601372-36603394 | p2 | (TA)13 | ATGTCGACTAGCCGTTGACC | TCAAACTCCAACTCCATCCC | 107 |
| chr2:36602096-36604122 | p2 | (TA)11 | CGAGCGAGTTCCAAAGACTC | TCATCATCCATAAGCCTCCC | 134 |
| chr2:36602096-36604122 | p2 | (TA)13 | CGAGCGAGTTCCAAAGACTC | TCATCATCCATAAGCCTCCC | 134 |
| chr2:36602096-36604122 | p2 | (TA)13 | CGAGCGAGTTCCAAAGACTC | TCATCATCCATAAGCCTCCC | 134 |
| chr2:36759139-36761157 | p6 | (TGAGTA)3 | GTGGTGGTTGGAGAATGCTT | TAGTACCACAAGCCCTTCCG | 265 |
| chr2:36761448-36763464 | p2 | (TA)8 | GCAAATTTGAGTAGAGCGGC | ACCCGCTTGTTCTGATTGTC | 184 |
| chr2:36775173-36777207 | p2 | (AT)17 | GAGAACGACGATTAGCAGCC | TTCCCATATTCCCGATGGTA | 298 |
| chr2:36792845-36794905 | p3 | (AAT)20 | GCTCCTGCTATTCCTGTTGC | ACCTCTCGTGAGCCAAAGAA | 193 |
| chr2:36807749-36809761 | p2 | (TA)6 | TCTGCGAAGGGTCTACAGGT | GCGAGCTCTTCTGCTGAAAT | 299 |
| chr2:36807749-36809761 | p2 | (AT)10 | TCTGCGAAGGGTCTACAGGT | GCGAGCTCTTCTGCTGAAAT | 299 |
| chr2:36892560-36894572 | p2 | (AT)6 | CCTTGGGTTTCTTCCTCCTC | GCCTTGTACGATAGGGGGAT | 296 |
| chr2:36908278-36910290 | p2 | (TA)6 | TTGGCCAATTTCAATCAACA | TTTAATTTGGAAGGTTGCGG | 118 |
| chr2:36952284-36954300 | p4 | (ATCT)4 | TCTCAGCTAATGGTGGTCCC | CACTTTTTGCCCAGCTCTTC | 155 |
| chr2:36957865-36959877 | p2 | (AG)6 | AATCCGTGCCATTGTTCTTC | TCCCTAGCCACCTCAGAAAA | 230 |
| chr2:36957865-36959877 | p2 | (AG)7 | AATCCGTGCCATTGTTCTTC | TCCCTAGCCACCTCAGAAAA | 230 |
| chr2:37024752-37026767 | p3 | (AAT)5 | TTATTTTGCTTTTGGTCCCG | TTGATGCGTCTTATGCAAGC | 130 |
| chr2:37080157-37082175 | p5 | (AAGAG)3 | CTCACACCTTACTGGCCCAT | AAGGTGGGAGTGGCCTCTAT | 129 |
| chr2:37080157-37082175 | p3 | (ACA)6 | CTCACACCTTACTGGCCCAT | AAGGTGGGAGTGGCCTCTAT | 129 |
| chr2:37128587-37130602 | p5 | (TTTTA)3 | ATATGCCAAACGAGCCAATC | TATGTGAAGGGGTGGGAGAG | 156 |
| chr2:37274661-37276675 | p2 | (AT)7 | TTTCTTATTTGCCGTGAGGG | CATGGCGACAGAAAGGAAAT | 122 |
| chr2:37367491-37369583 | p2 | (AT)33 | CTGCCCGAATTTTGACAGAT | CGCTTATTCTCCCGAACTTG | 169 |
| chr2:37371757-37373772 | p5 | (ATGCA)3 | TACGGCCCATCAGTTAGGTC | AGACATTGTGGGTGTCACGA | 149 |
| chr2:37434640-37436656 | p2 | (TA)8 | CCCACCATCTCCAAACTCAC | AAATGACATGGCGGAGTAGC | 282 |
| chr2:37455253-37457293 | p2 | (CT)20 | TTTACGAAAGTTGGCAACCC | AACGGCATTGAGGATGAGTC | 227 |
| chr2:37465757-37467771 | p3 | (TGT)5 | AGGTTGAATTCTCGGGGTTT | TTACATCCCTTGTCCGCTTC | 240 |
| chr2:37465757-37467771 | p2 | (AT)8 | AGGTTGAATTCTCGGGGTTT | TTACATCCCTTGTCCGCTTC | 240 |
| chr2:37567486-37569516 | p2 | (AT)7 | CCAAGCTGCAGTTGTTTCAA | AAAGTACGCATCCCATCGTC | 220 |
| chr2:37567486-37569516 | p2 | (TC)15 | CCAAGCTGCAGTTGTTTCAA | AAAGTACGCATCCCATCGTC | 220 |
| chr2:37608463-37610478 | p3 | (ATA)5 | ACCACTGCCCATCAAATTGT | GAAGTCACTCAAGCGGAACC | 217 |
| chr2:37608463-37610478 | p5 | (TTCAT)3 | ACCACTGCCCATCAAATTGT | GAAGTCACTCAAGCGGAACC | 217 |
| chr2:37614492-37616512 | p2 | (AT)14 | AAGGAAAAGTTTTGCCCCAT | GCAGAATTGCCCTTCTTTTG | 174 |
| chr2:37614492-37616512 | p5 | (AACCC)4 | AAGGAAAAGTTTTGCCCCAT | GCAGAATTGCCCTTCTTTTG | 174 |

| Marker ID | SSR type | SSR motif | FORWARD PRIMER (5'-3') | REVERSE PRIMER (5'-3') | size (bp) |
|---|---|---|---|---|---|
| chr2:37665905-37667920 | p5 | (TTGGT)3 | TGCAAAAACCATGAAAACCA | GATTGCTTTGAGCAACCACA | 130 |
| chr2:37666609-37668624 | p5 | (TTGGT)3 | GGTTTGACTTGATTTGGCGT | ATGTCGGATTGGTTTGGTTC | 253 |
| chr2:37666649-37668664 | p5 | (TTGGT)3 | GGTTTGACTTGATTTGGCGT | ATGTCGGATTGGTTTGGTTC | 213 |
| chr2:37666689-37668704 | p5 | (TTGGT)3 | GGTTTGACTTGATTTGGCGT | ATGTCGGATTGGTTTGGTTC | 173 |
| chr2:37733710-37735726 | p2 | (TA)8 | CAGCGAAAAGACAAAGGAGG | CCCCCTTTTAACCCATGTTT | 261 |
| chr2:37733710-37735726 | p3 | (TTA)5 | CAGCGAAAAGACAAAGGAGG | CCCCCTTTTAACCCATGTTT | 261 |
| chr2:37733833-37735870 | p2 | (TA)8 | CAGCGAAAAGACAAAGGAGG | CCCCCTTTTAACCCATGTTT | 138 |
| chr2:37733833-37735870 | p3 | (TTA)5 | CAGCGAAAAGACAAAGGAGG | CCCCCTTTTAACCCATGTTT | 138 |
| chr2:37849325-37851343 | p6 | (TTTTCT)3 | GCAGAGCCCCTCTATCTCCT | AAACAAGTATGGTGCCGGTC | 175 |
| chr2:37924840-37926862 | p2 | (AT)11 | AGCGGTAGGCTGGTTACTGA | ACCGCTTATGTGCTTGACCT | 231 |
| chr2:37939851-37941866 | p3 | (CTT)5 | GGGTCGAGACCTGATCAAAA | GTTATGCGCAAAGCAGTGAA | 150 |
| chr2:37966346-37968364 | p6 | (CATGAG)3 | TTGTGTTTTGGGCATTCAAA | AGCTCCATTTGCATCCATTC | 206 |
| chr2:37995208-37997240 | p2 | (AT)6 | TTTACACAGCATCCCAACCA | TTGGAGTGCTCCTTTGTTCC | 232 |
| chr2:37995244-37997258 | p2 | (AT)6 | TTTACACAGCATCCCAACCA | TTGGAGTGCTCCTTTGTTCC | 196 |
| chr2:37995271-37997299 | p2 | (AT)6 | TTTACACAGCATCCCAACCA | TTGGAGTGCTCCTTTGTTCC | 169 |
| chr2:38103011-38105044 | p3 | (TAA)11 | TAATCCGAACGTTGAAAGCC | GTCTTGCCTTTCCATTTCCA | 226 |
| chr2:38103011-38105044 | p5 | (ATAAA)3 | TAATCCGAACGTTGAAAGCC | GTCTTGCCTTTCCATTTCCA | 226 |
| chr2:38109286-38111301 | p3 | (ATA)5 | TTCAATCAATTTGTGCCGAG | CCAATGGAAATCCCACAAAG | 127 |
| chr2:38186521-38188577 | p2 | (AT)28 | GAAGATCCGTGCTAAGTCCG | AACCACATGTGTCCCTTTGA | 240 |
| chr2:38254110-38256122 | p2 | (TA)6 | CCGGACTGAGCTTCTTTCTG | ACCTTCACGTCTTCCACCAC | 232 |
| chr2:38286714-38288777 | p2 | (GA)7 | TAGGGTGACGAGATTTTGGG | ATGGATTTTTGTCCCCTTCC | 168 |
| chr2:38286714-38288777 | p3 | (AAT)21 | TAGGGTGACGAGATTTTGGG | ATGGATTTTTGTCCCCTTCC | 168 |
| chr2:38289928-38291952 | p3 | (TGT)8 | CACACATCATATTGGAGCGG | TCCATCTCAGCATCTGCATC | 210 |
| chr2:38468264-38470280 | p2 | (AT)8 | ATTTGCATTGAAGGCTCACC | ATTTGGGCGACTTCCTAACA | 239 |
| chr2:38483583-38485601 | p2 | (AT)19 | GGAGGACAATGCCCCTTATT | CCTGTTTTATCGTGGCAGGT | 137 |
| chr2:38483583-38485601 | p3 | (TAT)6 | GGAGGACAATGCCCCTTATT | CCTGTTTTATCGTGGCAGGT | 137 |
| chr2:38513665-38515686 | p6 | (AAAGAG)3 | CGGCCATGGAGAAGAAAATA | ACACACACATTGAAGGGCAA | 190 |
| chr2:38513665-38515686 | p2 | (TG)7 | CGGCCATGGAGAAGAAAATA | ACACACACATTGAAGGGCAA | 190 |
| chr2:38513665-38515686 | p3 | (GAA)6 | CGGCCATGGAGAAGAAAATA | ACACACACATTGAAGGGCAA | 190 |
| chr2:38513665-38515686 | p3 | (AAC)7 | CGGCCATGGAGAAGAAAATA | ACACACACATTGAAGGGCAA | 190 |
| chr2:38513665-38515686 | p2 | (TA)8 | CGGCCATGGAGAAGAAAATA | ACACACACATTGAAGGGCAA | 190 |
| chr2:38513665-38515686 | p2 | (TG)10 | CGGCCATGGAGAAGAAAATA | ACACACACATTGAAGGGCAA | 190 |
| chr2:38513900-38515916 | p2 | (TG)7 | TGGAGGAAGAAAATGGGTTG | AAAATTTGATGCCCAGTTGC | 128 |
| chr2:38513900-38515916 | p3 | (GAA)6 | TGGAGGAAGAAAATGGGTTG | AAAATTTGATGCCCAGTTGC | 128 |
| chr2:38513900-38515916 | p3 | (AAC)7 | TGGAGGAAGAAAATGGGTTG | AAAATTTGATGCCCAGTTGC | 128 |
| chr2:38513900-38515916 | p2 | (TA)8 | TGGAGGAAGAAAATGGGTTG | AAAATTTGATGCCCAGTTGC | 128 |

| Marker ID | SSR type | SSR motif | FORWARD PRIMER (5'-3') | REVERSE PRIMER (5'-3') | size (bp) |
|---|---|---|---|---|---|
| chr2:38513900-38515916 | p2 | (TG)10 | TGGAGGAAGAAAATGGGTTG | AAAATTTGATGCCCAGTTGC | 128 |
| chr2:38662800-38664812 | p2 | (AC)6 | TAGGAGCAGAAAGGATCCGA | AAGCCTCGGGGTGTTAGATT | 218 |
| chr2:38749714-38751729 | p3 | (ATA)5 | CCCTCTCCATACCCCACTTT | GCATGTCCAAACGCCTATTT | 157 |
| chr2:38986388-38988408 | p5 | (GAATG)4 | CCCACCCTCTCCCTTAAAAA | CATTTGCCTTGATTGGTGTG | 109 |
| chr2:39044073-39046088 | p5 | (TAAAA)3 | CAAGCAACCTTGACTGGACA | GAATACTGTCGGCACACACG | 193 |
| chr2:39147759-39149779 | p2 | (AG)10 | TTGGGCATGTTTGTTTTGAA | GATGTTTTGGGGAGAACGAA | 114 |
| chr2:39147759-39149779 | p6 | (ATTAAA)3 | TTGGGCATGTTTGTTTTGAA | GATGTTTTGGGGAGAACGAA | 114 |
| chr2:39427685-39429700 | p5 | (TTTAT)3 | TGCAAGATGAATTAGCCACG | AAAAACGAGCCGCCTAAAAT | 112 |
| chr2:39427685-39429700 | p2 | (TA)8 | TGCAAGATGAATTAGCCACG | AAAAACGAGCCGCCTAAAAT | 112 |
| chr2:39427685-39429700 | p3 | (TGT)8 | TGCAAGATGAATTAGCCACG | AAAAACGAGCCGCCTAAAAT | 112 |
| chr2:39456315-39458327 | p2 | (TA)6 | TGAAGTCAGATGAGGATGCG | TGACACTGACTGCCTTACGC | 193 |
| chr2:39456315-39458327 | p2 | (AT)6 | TGAAGTCAGATGAGGATGCG | TGACACTGACTGCCTTACGC | 193 |
| chr2:39488285-39490313 | p2 | (TA)14 | TGCCCACTTGTGACCACTTA | ACAAACGGACGACGAAAAAG | 108 |
| chr2:39488285-39490313 | p2 | (TA)6 | TGCCCACTTGTGACCACTTA | ACAAACGGACGACGAAAAAG | 108 |
| chr2:39489107-39491119 | p2 | (TA)14 | CAACCCATTAGAGGGAGCAA | TGGGTTGATACGCATGTGAC | 278 |
| chr2:39489107-39491119 | p2 | (TA)6 | CAACCCATTAGAGGGAGCAA | TGGGTTGATACGCATGTGAC | 278 |
| chr2:39489107-39491119 | p6 | (ATAAAA)3 | CAACCCATTAGAGGGAGCAA | TGGGTTGATACGCATGTGAC | 278 |
| chr2:39489107-39491119 | p2 | (TA)22 | CAACCCATTAGAGGGAGCAA | TGGGTTGATACGCATGTGAC | 278 |
| chr2:39577067-39579082 | p3 | (AAC)5 | TAACGTCCTGCCATGTTCAA | ACACAACAAATGTGGACCGA | 114 |
| chr2:39710492-39712524 | p2 | (TA)16 | CGAGGTTTGCAAAGTGAACA | TGGGAAGATATGAGCGAAGC | 125 |
| chr2:39731001-39733016 | p2 | (TA)7 | AAATTCCAAAGTCGTCGTGG | CTTGCCTCAGTTCCCACATT | 156 |
| chr2:39731001-39733016 | p3 | (TTG)5 | AAATTCCAAAGTCGTCGTGG | CTTGCCTCAGTTCCCACATT | 156 |
| chr2:39943128-39945142 | p2 | (AT)15 | ACATATGGAAGTTGAGGCCG | AAATATGCACCCAACAAGCC | 149 |
| chr2:39943128-39945142 | p2 | (AT)9 | ACATATGGAAGTTGAGGCCG | AAATATGCACCCAACAAGCC | 149 |
| chr2:39943137-39945152 | p2 | (AT)15 | ACATATGGAAGTTGAGGCCG | AAATATGCACCCAACAAGCC | 140 |
| chr2:39943137-39945152 | p2 | (AT)9 | ACATATGGAAGTTGAGGCCG | AAATATGCACCCAACAAGCC | 140 |
| chr2:39943147-39945161 | p2 | (AT)15 | ACATATGGAAGTTGAGGCCG | AAATATGCACCCAACAAGCC | 130 |
| chr2:39943147-39945161 | p2 | (AT)9 | ACATATGGAAGTTGAGGCCG | AAATATGCACCCAACAAGCC | 130 |
| chr2:40016167-40018182 | p3 | (AAC)5 | TCACCTTCCCTCCTAACCCT | CCAGAGCGGAAGATTAGCTG | 172 |
| chr2:40020031-40022043 | p5 | (CGTCC)3 | ATACAGATGCAGATTGGGGC | CCAAAAGAAAGTTGCGAAGC | 176 |
| chr2:40020031-40022043 | p2 | (CA)6 | ATACAGATGCAGATTGGGGC | CCAAAAGAAAGTTGCGAAGC | 176 |
| chr2:40020031-40022043 | p5 | (CCACC)3 | ATACAGATGCAGATTGGGGC | CCAAAAGAAAGTTGCGAAGC | 176 |
| chr2:40034551-40036566 | p5 | (TTTTA)3 | TCGGCCTATGATGCCTACTC | AGGAAATCGAACCAGTCACG | 189 |
| chr2:40182493-40184508 | p3 | (CAA)5 | AAACACAGCATAACTCCGGC | TATGTTTTCCATAAGCGGGC | 150 |
| chr2:40198925-40200940 | p5 | (AATAA)3 | TGGGGAGTAGCAGAAGCAGT | GCCAAATACATAGAGCCCCA | 296 |
| chr2:40251311-40253357 | p4 | (TAAT)4 | CGATTTCTCGTCCTCTTTGC | GGTGGCTGAAAAGCCAGTAG | 235 |

| Marker ID | SSR type | SSR motif | FORWARD PRIMER (5'-3') | REVERSE PRIMER (5'-3') | size (bp) |
|---|---|---|---|---|---|
| chr2:40251311-40253357 | p2 | (TA)23 | CGATTTCTCGTCCTCTTTGC | GGTGGCTGAAAAGCCAGTAG | 235 |
| chr2:40360645-40362666 | p2 | (AT)6 | TTGGCTAATTGGAGGTGGAG | CGGGAAAAGGCAAGTTTACA | 111 |
| chr2:40360645-40362666 | p3 | (ATT)7 | TTGGCTAATTGGAGGTGGAG | CGGGAAAAGGCAAGTTTACA | 111 |
| chr2:40410613-40412649 | p2 | (TA)18 | CCGTCGACAATTTGGTAGGT | GTAGCCTTTTCACTCGGCTG | 212 |
| chr2:40455205-40457223 | p3 | (CAA)6 | ACCGGCGAACGATATCATAG | GAAAAGCTCGGATTGCTTTG | 259 |
| chr2:40487026-40489076 | p2 | (AT)25 | AGGCGTATGCGGAGTATGAC | CATTCGCAAGATTTGCTTGA | 283 |
| chr2:40487026-40489076 | p5 | (AGAAA)3 | AGGCGTATGCGGAGTATGAC | CATTCGCAAGATTTGCTTGA | 283 |
| chr2:40494628-40496679 | p3 | (TTG)17 | GTTGCTGGAGAGATCAAGCC | ACACACACCTCACACGGCTA | 183 |
| chr2:40688519-40690583 | p2 | (TA)32 | TAAAAACCTTTTGGCAACGG | TAGGCCCAATACATCAAGGG | 121 |
| chr2:40706249-40708267 | p2 | (TA)9 | CCACCTGTGGGGAATTAAAA | CCCTTTCGCCCCATATTACT | 134 |
| chr2:40740960-40742999 | p5 | (GGAAG)3 | CATCCTCAGTTTTTGGGGAA | CATGTTCTTTGTCCCCGTCT | 189 |
| chr2:40741008-40743023 | p5 | (GGAAG)3 | CATCCTCAGTTTTTGGGGAA | CATGTTCTTTGTCCCCGTCT | 141 |
| chr2:40741039-40743054 | p5 | (GGAAG)3 | CATCCTCAGTTTTTGGGGAA | CATGTTCTTTGTCCCCGTCT | 110 |
| chr2:41086987-41089001 | p2 | (TA)7 | GACCAATTGGGCTCAAAAGA | CTCTTTCTCTCTTGGCCGTG | 128 |
| chr2:41086987-41089001 | p2 | (TA)7 | GACCAATTGGGCTCAAAAGA | CTCTTTCTCTCTTGGCCGTG | 128 |
| chr2:41087014-41089029 | p2 | (TA)7 | GACCAATTGGGCTCAAAAGA | CTCTTTCTCTCTTGGCCGTG | 101 |
| chr2:41087014-41089029 | p2 | (TA)7 | GACCAATTGGGCTCAAAAGA | CTCTTTCTCTCTTGGCCGTG | 101 |
| chr2:41087014-41089029 | p2 | (AT)7 | GACCAATTGGGCTCAAAAGA | CTCTTTCTCTCTTGGCCGTG | 101 |
| chr2:41103681-41105696 | p5 | (TGCAT)3 | ACGGGGAAAGCATGTTAGTG | GCAGCAAGCCAAACATATCA | 105 |
| chr2:41108112-41110126 | p3 | (ACA)6 | TCCTCAAGAACAGCCCCTAA | ATGGACTGTTTTGGGTTTGC | 286 |
| chr2:41108196-41110210 | p3 | (ACA)6 | TTCCTCAAGAACAGCCCCTA | ATGGACTGTTTTGGGTTTGC | 201 |
| chr2:41274648-41276672 | p4 | (AACA)6 | AATTAAGGGGTCCGACCAAG | ACCTGTTGCCATTAAGGTGG | 220 |
| chr2:41353808-41355856 | p4 | (ATAA)5 | CACCTTGCACCAATACGATG | CTTGATTGACGAAACTCGCA | 125 |
| chr2:41353808-41355856 | p3 | (ATA)16 | CACCTTGCACCAATACGATG | CTTGATTGACGAAACTCGCA | 125 |
| chr2:41589849-41591865 | p2 | (CT)8 | GTAACTCGATCGTGGGGAGA | CCGCAAAACTCAAACAACAA | 226 |
| chr2:41589849-41591865 | p2 | (GT)6 | GTAACTCGATCGTGGGGAGA | CCGCAAAACTCAAACAACAA | 226 |
| chr2:41589849-41591865 | p2 | (GT)6 | GTAACTCGATCGTGGGGAGA | CCGCAAAACTCAAACAACAA | 226 |
| chr2:41648388-41650406 | p3 | (CAT)6 | CCGCAAGGCTACAAAATGAT | CCGTCCTCATGACTTCCAGT | 109 |
| chr2:41658109-41660121 | p2 | (TA)6 | TCCGTCTAGGCCAGGAAAAT | TGACATAGTTCCGGGGGATA | 109 |
| chr2:41684700-41686714 | p2 | (TA)7 | TGAGGCTCTCCAAAGAGGAA | GGTGACGATCCCACAAAATC | 116 |
| chr2:41731650-41733670 | p5 | (TCTTT)4 | TCACATCATATGCGGTGCTT | CCTGTTGAAATCCTTTGGGA | 212 |
| chr2:41748362-41750392 | p2 | (AG)15 | AAGGGGGAAGGTATGCTTGT | CAGCTGGACGAATACCATGA | 216 |
| chr2:41796875-41798887 | p2 | (AT)6 | ATGAAACCCACGAAAAGTGC | CCAAACGAGGGCATTACATT | 296 |
| chr2:41799287-41801325 | p2 | (TA)19 | TCCTCGCGGCTCTTATTTTA | ACGGTGCAAATCACACTTGA | 219 |
| chr2:41810763-41812778 | p5 | (ACCAA)3 | GACTAGCAATTTGAGGGGCA | CGAGAAAACCGAATAACCCA | 100 |
| chr2:41856690-41858705 | p5 | (CAATT)3 | GCATGAGAGCCCAAAGAAAG | GATTGGCTCGAAACGTGATT | 225 |

| Marker ID | SSR type | SSR motif | FORWARD PRIMER (5'-3') | REVERSE PRIMER (5'-3') | size (bp) |
|---|---|---|---|---|---|
| chr2:41909935-41911951 | p4 | (TATT)6 | GCGTTGGTGTTTCATTGTTG | CCGACCAAGATTAGCATCGT | 240 |
| chr2:41909935-41911951 | p4 | (TTTC)4 | GCGTTGGTGTTTCATTGTTG | CCGACCAAGATTAGCATCGT | 240 |
| chr2:41918816-41920834 | p6 | (ATACAT)3 | CTCTGTGGCTTTCCTTGGTC | TTCTGTTAGGGTCGTGGGTC | 177 |
| chr2:42049671-42051692 | p3 | (TGT)7 | AAAGCTTGCGGAAACACACT | AGAACCCGATCCAAACACAG | 187 |
| chr2:42108206-42110218 | p2 | (TA)6 | GATTTGTGGCACATTCAACG | CTTTGCTTTTTGCTTTTGGG | 230 |
| chr2:42117880-42119895 | p3 | (TGA)5 | TAGAGCAGTGGTTGTGGCTG | TAGCTGCAAATGCATCAAGG | 103 |
| chr2:42331331-42333349 | p6 | (CATGCC)3 | TCCCAACGGAGGTCTATTTG | TTTCGGAAATTCGTCGATTC | 281 |
| chr2:42375260-42377275 | p3 | (TAA)5 | TCGCCCCTCTTTATAGGGTT | CCTCTCCTCTTCAATTCCCC | 116 |
| chr2:42375260-42377275 | p3 | (ATA)5 | TCGCCCCTCTTTATAGGGTT | CCTCTCCTCTTCAATTCCCC | 116 |
| chr2:42425364-42427376 | p2 | (CT)6 | ACCATCATTGTTGTGGCAGA | TTGCATCAGGGTTGCAATTA | 254 |
| chr2:42448162-42450176 | p2 | (TA)7 | CAGAGTCACCGTCAGGGATT | TCACGGAACCAATTTCAACA | 109 |
| chr2:42538771-42540785 | p2 | (AT)6 | CCGGAGCCAAAATGGTTATT | CGAAAAACAACGAACTGCAA | 160 |
| chr2:42538771-42540785 | p2 | (TA)10 | CCGGAGCCAAAATGGTTATT | CGAAAAACAACGAACTGCAA | 160 |
| chr2:42538771-42540785 | p2 | (AT)8 | CCGGAGCCAAAATGGTTATT | CGAAAAACAACGAACTGCAA | 160 |
| chr2:42538771-42540785 | p2 | (TA)6 | CCGGAGCCAAAATGGTTATT | CGAAAAACAACGAACTGCAA | 160 |
| chr2:42538813-42540829 | p2 | (AT)6 | CCGGAGCCAAAATGGTTATT | CGAAAAACAACGAACTGCAA | 118 |
| chr2:42538813-42540829 | p2 | (TA)10 | CCGGAGCCAAAATGGTTATT | CGAAAAACAACGAACTGCAA | 118 |
| chr2:42538813-42540829 | p2 | (AT)8 | CCGGAGCCAAAATGGTTATT | CGAAAAACAACGAACTGCAA | 118 |
| chr2:42538813-42540829 | p2 | (TA)6 | CCGGAGCCAAAATGGTTATT | CGAAAAACAACGAACTGCAA | 118 |
| chr2:42612552-42614582 | p3 | (TAA)18 | TAATCCGGACCATCCATAGC | GGAGGAATCAGGTTGGAACA | 286 |
| chr2:42612602-42614620 | p3 | (TAA)18 | TCACTTTCCTCCGCATTAGG | TCCAACAACTCAAAGCGATG | 173 |
| chr2:42612721-42614775 | p3 | (TAA)18 | TAATCCGGACCATCCATAGC | GATGGAGGAATCAGGTTGGA | 117 |
| chr2:42662973-42664988 | p3 | (AAG)5 | CGTAGAACACCATGACCGTG | TAAGGTGGTTCTGCTTTGGG | 167 |
| chr2:42669008-42671038 | p2 | (AT)15 | AGGGTCATCAGATGGCAAAC | TGAGTTGCTGTCGTTCTTGG | 235 |
| chr2:42674202-42676214 | p2 | (AC)6 | TGAGGCTCAGCAAAGGAAGT | GTCCTGCTGGTTTCCCATAA | 284 |
| chr2:42710495-42712525 | p2 | (AT)15 | AAGCTGTAGCTCCGGATGAA | CTGACGTCCGTTTGTTTGTG | 288 |
| chr2:42735547-42737562 | p5 | (AATGC)3 | GTGCCTCAAAGCCATGAACT | CCGAGGTCTTTTGTCGGATA | 173 |
| chr2:42735547-42737562 | p5 | (AAAAC)3 | GTGCCTCAAAGCCATGAACT | CCGAGGTCTTTTGTCGGATA | 173 |
| chr2:42797490-42799556 | p2 | (AT)33 | TGATTGCTACATCAGCAGGC | CCAGATTCCCCAGAAACAAA | 109 |
| chr2:42904742-42906769 | p2 | (TA)7 | GGGTTTAGCTGTGCTTCTCG | CGCTCGTGATTTTGTTGAAA | 265 |
| chr2:42935848-42937882 | p2 | (AT)17 | GCGTCTTCTCAAGGATTTGC | GCAATTCACAGAAATGCCCT | 119 |
| chr2:42935848-42937882 | p2 | (AT)14 | GCGTCTTCTCAAGGATTTGC | GCAATTCACAGAAATGCCCT | 119 |
| chr2:42949405-42951419 | p2 | (AT)7 | CCTTTCTGTCGTAGCCTTGC | GTTTATGGTGTGTTGCGTGC | 136 |
| chr2:42976643-42978667 | p6 | (CCAAAT)3 | TCCAATCCAAAGTCCAAAGC | TGAAAGTGTTACGTGCCTGC | 244 |
| chr2:42976643-42978667 | p6 | (CCAAAT)3 | TCCAATCCAAAGTCCAAAGC | TGAAAGTGTTACGTGCCTGC | 244 |
| chr2:42976643-42978667 | p6 | (AAGTCC)4 | TCCAATCCAAAGTCCAAAGC | TGAAAGTGTTACGTGCCTGC | 244 |

| Marker ID | SSR type | SSR motif | FORWARD PRIMER (5'-3') | REVERSE PRIMER (5'-3') | size (bp) |
|---|---|---|---|---|---|
| chr2:43078102-43080122 | p2 | (AT)10 | TTGGACGTCTTAGCATGGTG | AATCCAATCACGAATGCACA | 266 |
| chr2:43113167-43115185 | p3 | (TTG)6 | TGGACCGATTTTCTCTGGTC | CCAAGCCCCTCTCTCTTCTT | 116 |
| chr2:43138877-43140889 | p2 | (TA)6 | CTTCATGTATTCGCCCACCT | CCGTGTCAATTCCCCAATAG | 153 |
| chr2:43169164-43171180 | p2 | (TA)30 | GGTCGTGCCGTTCTAATTGT | TTCCCATTGCAGTTTTGTACC | 199 |
| chr2:43169164-43171180 | p2 | (TA)8 | GGTCGTGCCGTTCTAATTGT | TTCCCATTGCAGTTTTGTACC | 199 |
| chr2:43169164-43171180 | p2 | (AG)7 | GGTCGTGCCGTTCTAATTGT | TTCCCATTGCAGTTTTGTACC | 199 |
| chr2:43180914-43182930 | p2 | (TA)30 | GGTCGTGCCGTTCTAATTGT | TTCCCATTGCAGTTTTGTACC | 199 |
| chr2:43180914-43182930 | p2 | (TA)8 | GGTCGTGCCGTTCTAATTGT | TTCCCATTGCAGTTTTGTACC | 199 |
| chr2:43180914-43182930 | p2 | (AG)7 | GGTCGTGCCGTTCTAATTGT | TTCCCATTGCAGTTTTGTACC | 199 |
| chr2:43222802-43224816 | p2 | (AT)7 | TGCTCCACGCCCTACTTACT | CTTCTTCCACCCATGGCTAA | 217 |
| chr2:43244232-43246253 | p3 | (AAC)7 | GATGGCTTCTCGCCTTACTG | CGCGTTCATTTTCTCAACAA | 173 |
| chr2:43311291-43313311 | p5 | (CTTTT)4 | AGTTTGCGGGAGAGGGTATT | GAGACAAGAATCCCGCGTAG | 144 |
| chr2:43525637-43527652 | p3 | (GAA)5 | TTCGTTTGACTTCCAATCCC | TTGGTAATGTCTTGAGGGGC | 192 |
| chr2:43545807-43547825 | p5 | (TTTTC)3 | ACCAACTTCGCTGGAACATC | ACATTGCTCGTACTGACCCC | 243 |
| chr2:43545807-43547825 | p6 | (TTCATA)3 | ACCAACTTCGCTGGAACATC | ACATTGCTCGTACTGACCCC | 243 |
| chr2:43814338-43816390 | p2 | (AT)7 | TCGGTTAGAGAAGATGGCGT | ATGGGAAGAAAGGCCAAGAT | 110 |
| chr2:43814338-43816390 | p5 | (AGGAC)3 | TCGGTTAGAGAAGATGGCGT | ATGGGAAGAAAGGCCAAGAT | 110 |
| chr2:43814338-43816390 | p2 | (AT)26 | TCGGTTAGAGAAGATGGCGT | ATGGGAAGAAAGGCCAAGAT | 110 |
| chr2:43814338-43816390 | p3 | (ATC)5 | TCGGTTAGAGAAGATGGCGT | ATGGGAAGAAAGGCCAAGAT | 110 |
| chr2:43814808-43816823 | p5 | (AGGAC)3 | AGTGCTCCTGCGTATCTGCT | TCCTTTCTTTTGTGGACGCT | 272 |
| chr2:43814808-43816823 | p2 | (AT)26 | AGTGCTCCTGCGTATCTGCT | TCCTTTCTTTTGTGGACGCT | 272 |
| chr2:43814808-43816823 | p3 | (ATC)5 | AGTGCTCCTGCGTATCTGCT | TCCTTTCTTTTGTGGACGCT | 272 |
| chr2:44061019-44063034 | p5 | (AAAGG)3 | GGTTTGTGAACAGATGGCCT | GCTATGGTGGTACCATTGGG | 199 |
| chr2:44141863-44143879 | p2 | (AC)8 | GGCATTCCAGCCAAAATAGA | CTTTCGTTTTGGGCATGTCT | 139 |
| chr2:44246816-44248838 | p2 | (AT)11 | GGCTAGTGCTTTTGCGCTAC | GATGGAAATTCTGAGGCCAA | 153 |
| chr2:44359204-44361219 | p4 | (CATA)5 | GGGACTCGATGCTATCCGTA | CCCTGACATCGCCATAGTTT | 182 |
| chr2:44359204-44361219 | p5 | (AAGAA)3 | GGGACTCGATGCTATCCGTA | CCCTGACATCGCCATAGTTT | 182 |
| chr2:44359204-44361219 | p2 | (TG)6 | GGGACTCGATGCTATCCGTA | CCCTGACATCGCCATAGTTT | 182 |
| chr2:44493883-44495895 | p2 | (AT)6 | CTCGATACGGGGAATTCAAA | ATGAGTATACCGTGAGGCCG | 125 |
| chr2:44533516-44535531 | p3 | (TCA)5 | CGCGTTCTACGAGCCTCTAC | GATCCGTAGTAGCTGTCCGC | 120 |
| chr2:44569183-44571198 | p5 | (GCATT)3 | AGCGGAAGAGGAAGGCTAAG | AGTGGCATCAGGCCAATAAC | 205 |
| chr2:44569183-44571198 | p5 | (ATATT)3 | AGCGGAAGAGGAAGGCTAAG | AGTGGCATCAGGCCAATAAC | 205 |
| chr2:44589145-44591163 | p6 | (GTATCG)3 | TGAGTCTGACATGCGTAGCC | AACCAAGCTCAGGAAAGGGT | 260 |
| chr2:44607428-44609461 | p2 | (AT)7 | TTGCGCCATAATCACATCAT | TGCTCGTGCATGCCTTAATA | 165 |
| chr2:44607428-44609461 | p3 | (ACA)11 | TTGCGCCATAATCACATCAT | TGCTCGTGCATGCCTTAATA | 165 |
| chr2:44616375-44618393 | p2 | (CT)9 | AGATAAGACGAACCGCGAAA | CATTGACAATCATTTCCCCC | 237 |

| Marker ID | SSR type | SSR motif | FORWARD PRIMER (5'-3') | REVERSE PRIMER (5'-3') | size (bp) |
|---|---|---|---|---|---|
| chr2:44662602-44664617 | p3 | (TTG)5 | TGGTTTATGGGCAGCAATTT | AGCCTTTCTTGGAGGGTTGT | 245 |
| chr2:44815052-44817070 | p6 | (TAAAAA)3 | CATGGCTGTTCATTTTGCAG | TCCAGCCTGACTGTGATCTG | 239 |
| chr2:44839424-44841448 | p2 | (AT)25 | GCCGACAAGGCTAAACACAT | CAGGGACTACATAGCGGAGC | 168 |
| chr2:44839424-44841448 | p2 | (AT)12 | GCCGACAAGGCTAAACACAT | CAGGGACTACATAGCGGAGC | 168 |
| chr2:44844172-44846198 | p2 | (AT)13 | TGTCAGTGGTTCTTGCCTTG | ATATGAGTGTCCAGCTCGGG | 106 |
| chr2:44869083-44871095 | p2 | (CA)6 | GGTGGTGCAACAAATGACAC | GCAATTGGGAAATGGTATGG | 159 |
| chr2:44977115-44979131 | p4 | (GAAA)4 | GGTCCCTCAAAATACGACGA | TGATGCCAAGTTATGGTGGA | 130 |
| chr2:45027796-45029812 | p2 | (AT)8 | CATGCATTTCCATTCATTCG | GTGGGGGTAGATGTGTTTGG | 148 |
| chr2:45139913-45141935 | p2 | (TA)11 | GATTTCATGCAAGTGTCGGA | CGAAAACAAAGGGAGATCCA | 263 |
| chr2:45190768-45192824 | p2 | (TA)28 | GGGTAGAGCCACACCCAATA | TGCAAGATAGTTTGGGCCTC | 280 |
| chr2:45190768-45192824 | p2 | (TA)28 | GGGTAGAGCCACACCCAATA | TGCAAGATAGTTTGGGCCTC | 280 |
| chr2:45190768-45192824 | p3 | (TAA)5 | GGGTAGAGCCACACCCAATA | TGCAAGATAGTTTGGGCCTC | 280 |
| chr2:45212839-45214854 | p3 | (AGA)5 | TGCTCTCGACTGTTGATTGG | CACCCCAACTGGTCTTCCTA | 123 |
| chr2:45254528-45256561 | p3 | (ATA)11 | CCATTCGCAGCCTTGATATT | ATATTCGGCGAAGTTTGTGG | 141 |
| chr2:45264253-45266269 | p2 | (TA)8 | AGGCCTCGTTATGTGATTCG | AAAAGCTACAGCCGACCAGA | 233 |
| chr2:45326260-45328275 | p3 | (CAT)5 | TTCTGGGCTGTCGTAGCTTT | TACCCGTCACGACCCTAGTC | 129 |
| chr2:45432804-45434870 | p3 | (CAA)5 | CTTGCCCTCTGCCTAACTTG | TTGCTTGATTTGGCAATGAG | 157 |
| chr2:45484880-45486892 | p2 | (GT)6 | GCATTTGTTCGAAAAGGGAG | CAAGCCAACAGAAAACACCA | 234 |
| chr2:45609817-45611835 | p3 | (CAT)6 | ACTTTTGTCACGACCCAACC | TAGCCTCGTCGGCTTAGGTA | 110 |
| chr2:45760164-45762180 | p4 | (GAAA)4 | CCTCCTCAGTAGGCAGCAAC | TGATGCCAAGTTATGGTGGA | 102 |
| chr2:45857673-45859691 | p4 | (CTTT)4 | TTGGCCGAGTTAAATTCTCG | GCTTATTTGTCCAAGCTCGC | 144 |
| chr2:45857673-45859691 | p6 | (TATATG)3 | TTGGCCGAGTTAAATTCTCG | GCTTATTTGTCCAAGCTCGC | 144 |
| chr2:45912311-45914349 | p2 | (AT)10 | GGCATGATCGTTGTGAAATG | ACCTTTCGGATGTCCTTGTG | 111 |
| chr2:45912311-45914349 | p2 | (AT)6 | GGCATGATCGTTGTGAAATG | ACCTTTCGGATGTCCTTGTG | 111 |
| chr2:45912311-45914349 | p2 | (AT)19 | GGCATGATCGTTGTGAAATG | ACCTTTCGGATGTCCTTGTG | 111 |
| chr2:45924885-45926911 | p2 | (AT)13 | AAAACACCATACGACTCGCC | CTGCATCAAGCCCAACTGTA | 271 |
| chr2:45947330-45949352 | p2 | (AT)6 | GTATTCCTGTCTCGACCCCA | GGGCTTGTGTCCGTTACAGT | 161 |
| chr2:45947330-45949352 | p2 | (TA)11 | GTATTCCTGTCTCGACCCCA | GGGCTTGTGTCCGTTACAGT | 161 |
| chr2:46029997-46032013 | p2 | (TA)8 | ACTGTCGTGCCCGAAAATAC | TTCGTACTGACGTCCCTTCC | 154 |
| chr2:46041917-46043935 | p6 | (ATACAT)3 | TGCCACTGACTTTCCCTTCT | GGTGTGTTTACGAGCCGTTT | 119 |
| chr2:46066033-46068045 | p2 | (TG)6 | TGACGAAGATAGCGATGACG | ACTTAGCCAACCAAGGCTCA | 215 |
| chr2:46092061-46094079 | p6 | (TATTCA)3 | ATTCCAAATTTACCCTCGGC | ATTGTTGCCCATTGGGATTA | 159 |
| chr2:46171931-46173945 | p2 | (AT)7 | GGAACAAGCCTTGGTTGAAA | TCCACATTTCACCCCTTCTC | 109 |
| chr2:46313140-46315170 | p2 | (TC)15 | CTAGAACAACACCGCCCCTA | GGCTCCACTTTCTCTGTTGC | 229 |
| chr2:46337977-46339992 | p5 | (GTTTT)3 | GTTCCGCCCAATTTCAACTA | TGCAGATTTGCTGAACAAGG | 265 |
| chr2:46350563-46352613 | p2 | (TA)8 | TTGCCCGAATTCTTGTAACC | TATTCCGATGCTCGTCCTTC | 289 |

| Marker ID | SSR type | SSR motif | FORWARD PRIMER (5'-3') | REVERSE PRIMER (5'-3') | size (bp) |
|---|---|---|---|---|---|
| chr2:46506286-46508298 | p2 | (TA)7 | ATGAGATGGATTTTGGCTCG | TTGTGGAACAATCCCGAAAT | 234 |
| chr2:46506286-46508298 | p2 | (AT)10 | ATGAGATGGATTTTGGCTCG | TTGTGGAACAATCCCGAAAT | 234 |
| chr2:46547383-46549417 | p2 | (AT)17 | GACGGGCGGCTTATACATTA | TGTCTACCTGTGGGCATGAA | 231 |
| chr2:46547383-46549417 | p2 | (TA)6 | GACGGGCGGCTTATACATTA | TGTCTACCTGTGGGCATGAA | 231 |
| chr2:46547383-46549417 | p3 | (TCA)5 | GACGGGCGGCTTATACATTA | TGTCTACCTGTGGGCATGAA | 231 |
| chr2:46654078-46656090 | p2 | (TA)6 | ACTAACGGCCCTGTCATTTG | ATAGTATTTGCTGCCGGACG | 228 |
| chr2:46739664-46741682 | p6 | (AAAATA)3 | TGAAGACCACCCCAAATGAT | AAGCTCCGTGCCTATTCAAA | 234 |
| chr2:46773913-46775933 | p2 | (AT)10 | TCAAGGCCAACGCATACATA | ATGACAGGGGCCTTGTACTG | 264 |
| chr2:46817208-46819222 | p2 | (TA)7 | TTAGCCAGGCAAGCAACTTT | CCGTGTGTTATGCAATGGAG | 172 |
| chr2:46822670-46824778 | p2 | (AT)54 | CTACCTGGAAAGCCGAACAG | CCTAGTCGGGTCGTGACATT | 156 |
| chr2:46838613-46840635 | p2 | (TA)11 | AACACCCAGTCCACTATGGC | TGTGTTCGCATGGTTGTTTT | 130 |
| chr2:46840887-46842902 | p5 | (TGATA)3 | TCAGGTGAAACTCCCCTTTG | CTCGATTCAACCCCTTTTCA | 193 |
| chr2:46866106-46868118 | p2 | (AC)6 | AAATAGGACAGGGTGGGGTC | CCTGCCTATGACCTTTAGCG | 227 |
| chr2:46891868-46893910 | p3 | (TAT)14 | GCATTCATGGCCCATTTTAC | CCAAGGTCAAGAAAACCCAA | 263 |
| chr2:46961679-46963694 | p5 | (TGCAA)3 | TGGGAAAAATTGGGTTTTGA | CCTGCAAGTGCAAACTGAAA | 114 |
| chr2:46980114-46982132 | p2 | (AC)8 | TTGTTGGCTGTTAGGTGTGC | GGATCATGCCAAAAGAAGGA | 269 |
| chr2:46980114-46982132 | p2 | (AC)9 | TTGTTGGCTGTTAGGTGTGC | GGATCATGCCAAAAGAAGGA | 269 |
| chr2:47032720-47034738 | p6 | (GGGGGT)3 | CTTTTCACCTCCTCACCCAA | CCACGGCTAAAGATGTGGAT | 184 |
| chr2:47065849-47067869 | p5 | (TTCAT)4 | GAGGTCGTTCTATACGCCCA | ACTCCGTATCACCCAAGTGC | 238 |
| chr2:47256318-47258340 | p2 | (AT)11 | CCATAGCGGCTAGAGGACAG | GTTAAGTCCCGAAATGGGGT | 164 |
| chr2:47272804-47274818 | p2 | (TA)7 | GAGATGCGTTACGAGCATGA | AGTCATGGAAGAGGCAGGAA | 120 |
| chr2:47338116-47340160 | p2 | (TA)22 | GGTTTGTTGCGTTGGAAACT | TATGAGGGGCAATTTTGGTC | 118 |
| chr2:47407647-47409671 | p2 | (AG)12 | AACCATGTTCTAAGGCGTGG | CGAGGGAGTCCAAATACCAA | 132 |
| chr2:47421958-47423976 | p2 | (AG)9 | AATTGGTGGTTTTGGTGGAA | AGCCACGAACTAATCCGCTA | 279 |
| chr2:47566503-47568521 | p2 | (AT)9 | GACAGAGGGCAGTTGGAGAG | CCGGTTGCTACCTATGCACT | 177 |
| chr2:47603702-47605720 | p2 | (GA)9 | ACCGCTCTAAGCCTCACGTA | CATATTCCCTGTGGGATTCG | 134 |
| chr2:47619262-47621276 | p2 | (AT)25 | TAAATTCTGCACATTGCCCA | GCCACATTCAGACCCTCATT | 229 |
| chr2:47619262-47621276 | p2 | (GT)7 | TAAATTCTGCACATTGCCCA | GCCACATTCAGACCCTCATT | 229 |
| chr2:47662576-47664672 | p2 | (TA)48 | TAGCCGAATCAATCCTGGAG | ACAAGGATGCTTGAAATGGG | 125 |
| chr2:47668350-47670406 | p5 | (ATTGC)4 | GTCTCGTCTGTCTCTTGGGC | TACAATGCCATGCAAATGCT | 196 |
| chr2:47668350-47670406 | p2 | (AT)28 | GTCTCGTCTGTCTCTTGGGC | TACAATGCCATGCAAATGCT | 196 |
| chr2:47695929-47697947 | p2 | (TA)6 | CCCTAGGGAAGGTGTTAGGC | ACCTTTTGCGGGAAATAACC | 163 |
| chr2:47695929-47697947 | p6 | (AAAAAT)3 | CCCTAGGGAAGGTGTTAGGC | ACCTTTTGCGGGAAATAACC | 163 |
| chr2:47807094-47809116 | p2 | (TA)11 | CATTATCTGGCGGAGCATTT | ACAGTGGATACCAAGCTGCC | 118 |
| chr2:47885482-47887502 | p2 | (TA)6 | GAGGTGCTCGACAGGTTAGC | TAAGTGTGGCCCATTTCACA | 299 |
| chr2:47885519-47887531 | p2 | (AT)14 | GAGGTGCTCGACAGGTTAGC | TAAGTGTGGCCCATTTCACA | 262 |

| Marker ID | SSR type | SSR motif | FORWARD PRIMER (5'-3') | REVERSE PRIMER (5'-3') | size (bp) |
|---|---|---|---|---|---|
| chr2:47963438-47965466 | p2 | (AT)14 | GATTTCACCTTGCGATGGTT | TTTCGGACTCGGTGTTATCC | 127 |
| chr2:47963438-47965466 | p3 | (ATT)5 | GATTTCACCTTGCGATGGTT | TTTCGGACTCGGTGTTATCC | 127 |
| chr2:48108237-48110274 | p5 | (ACAAA)3 | CACGATCAGGTAAAGCAGCA | TTTTGCTCAGGAGGCTCTGT | 185 |
| chr2:48111500-48113512 | p2 | (AT)6 | AAAGCTGTAGCTCCGGATGA | GACTGCTCGACTATCTCCCG | 243 |
| chr2:48174316-48176334 | p6 | (ACACTC)3 | ACGGGCTACTTCTTCAGCAA | TCCCATACATGTCTCGCGTA | 259 |
| chr2:48220539-48222557 | p2 | (AT)9 | GAAGACCAATGCACCCTCAT | TGGCTTGAGGATGTTGTGAG | 296 |
| chr2:48220539-48222557 | p5 | (AATGC)3 | GAAGACCAATGCACCCTCAT | TGGCTTGAGGATGTTGTGAG | 296 |
| chr2:48246307-48248323 | p2 | (TA)8 | GCCTTAAAAGAGCAGCAGGA | CAAAACAAACCCATTTTGGC | 127 |
| chr2:48246307-48248323 | p6 | (GATGAG)3 | GCCTTAAAAGAGCAGCAGGA | CAAAACAAACCCATTTTGGC | 127 |
| chr2:48370325-48372359 | p2 | (AG)17 | CAGGAGAGAGGCACATAGCC | TCCAAATCCTGGTTGGAGAG | 127 |
| chr2:48475415-48477435 | p4 | (TAAT)5 | CCGGATTATTTTCGGTCCTT | TGCAGTTGAGTCCTATTGCG | 249 |
| chr2:48538834-48540849 | p5 | (AAATA)3 | CGCAGGGAGGAAATTACTGA | ACCTGCACATCCTGGAAAAG | 197 |
| chr2:48714716-48716734 | p2 | (TA)9 | CAACCCGGCCAAACTATAAA | GTGTTGATGGCTTGTTGTGG | 152 |
| chr2:48729611-48731623 | p2 | (TA)6 | TCCCTCCAAGTTCACGATTC | CTTGAAAGCGGTGTTTGTGA | 195 |
| chr2:48826930-48828945 | p5 | (ATCAA)3 | AGGGATGAAAATGCCACAAG | GGTCAAGTATGCCGTTCGTT | 142 |
| chr2:48860589-48862604 | p5 | (AAAAT)3 | GTTTTCGCTGGACCTGGTAA | AACTTTCCGTCCCTATCCGT | 298 |
| chr2:48885736-48887766 | p6 | (CATATA)5 | TCAACAATCCCTAGATGCCC | AAGATCGGGGTGTGACAAAG | 120 |
| chr2:49012655-49014670 | p3 | (GTT)5 | ACCATAAGTCGTGGGTCTGC | TCCCTTAAATGGCATGGGTA | 238 |
| chr2:49301613-49303625 | p2 | (TA)14 | ATGAAATCGGGGATCACAAA | GCATATAGGCGCGGTGTTAT | 262 |
| chr2:49371524-49373539 | p3 | (TCA)5 | ATAACACTCTACCGGGTGCG | CCCACACGACCAGTACACAC | 109 |
| chr2:49398551-49400583 | p2 | (TA)13 | CGACGTGAAGCCATACTCAA | TGTCCTTCCCAGGTCATTTC | 264 |
| chr2:49398551-49400583 | p2 | (TA)16 | CGACGTGAAGCCATACTCAA | TGTCCTTCCCAGGTCATTTC | 264 |
| chr2:49435527-49437539 | p2 | (TA)6 | GACCAAACGGAACTCCAAGA | AGTCGTGAACCTATGGACGG | 112 |
| chr2:49435527-49437539 | p5 | (CATAT)3 | GACCAAACGGAACTCCAAGA | AGTCGTGAACCTATGGACGG | 112 |
| chr2:49470207-49472221 | p2 | (AT)7 | ATACGTCCCAACCACATCGT | TAAGGCATTCTCCAACCACC | 180 |
| chr2:49479057-49481075 | p6 | (ACATAT)3 | CAAACAAGAGGGCCATGACT | TAGTGACCATTTCACCGCAA | 243 |
| chr2:49479057-49481075 | p3 | (CAA)6 | CAAACAAGAGGGCCATGACT | TAGTGACCATTTCACCGCAA | 243 |
| chr2:49554594-49556612 | p6 | (ATACAT)3 | TGCATAACCATCCGAACAAA | CCTTAATGACCTCGGCGATA | 222 |
| chr2:49624977-49626989 | p2 | (AT)6 | TACCAATTTGCCTCCACCTC | CAGAATTCTCCGGGACAGAA | 101 |
| chr2:49637961-49639979 | p6 | (CATTAT)3 | ATAATGCCCAATTCATCCGA | AAAGCAAGGGATATGTTGCG | 166 |
| chr2:49672265-49674277 | p2 | (AT)6 | GGCCCTGAGTTGATTCAAAA | ATAGGCTACTTGGCCCACCT | 276 |
| chr2:49734880-49736895 | p3 | (ATC)5 | CAGGGACTACATAGCGGAGC | ATAAGCCGAAAAGGCTGACA | 270 |
| chr2:49734880-49736895 | p2 | (AT)24 | CAGGGACTACATAGCGGAGC | ATAAGCCGAAAAGGCTGACA | 270 |
| chr2:49753647-49755659 | p2 | (AT)8 | ATAAGCCGACGAGGCTAACA | CAGGGACTACATAGCGGAGC | 226 |
| chr2:49753647-49755659 | p2 | (AT)6 | ATAAGCCGACGAGGCTAACA | CAGGGACTACATAGCGGAGC | 226 |
| chr2:49859700-49861712 | p2 | (TA)6 | TCCAAATTGCCAACAAGTGA | TCGCATAATCTTCTGTTGCG | 153 |

| Marker ID | SSR type | SSR motif | FORWARD PRIMER (5'-3') | REVERSE PRIMER (5'-3') | size (bp) |
|---|---|---|---|---|---|
| chr2:49953123-49955138 | p2 | (AT)15 | TGTCACCGCTAAGGTAGGCT | GTCGTGACACTGCGTGACTT | 212 |
| chr2:49953123-49955138 | p5 | (TCATA)3 | TGTCACCGCTAAGGTAGGCT | GTCGTGACACTGCGTGACTT | 212 |
| chr2:50055480-50057502 | p2 | (AT)11 | TATCAGTGTGAAAGCGGCTG | CAGTGCCGTCTCGGATTATT | 105 |
| chr2:50187837-50189849 | p2 | (AT)6 | GCGAACCATTCGGAATAAAA | CCTATCAGCAGCATTCGACA | 215 |
| chr2:50265219-50267231 | p2 | (TA)6 | TCCTCCAGCATTAGCGACTT | ATGCTGGCCACTATCTACGG | 191 |
| chr2:50311546-50313560 | p2 | (CA)7 | TTGTGGGGCATCTAGGAAAC | CCAACAGGATTGCACAACAC | 125 |
| chr2:50311546-50313560 | p5 | (AAAAC)3 | TTGTGGGGCATCTAGGAAAC | CCAACAGGATTGCACAACAC | 125 |
| chr2:50317924-50319936 | p2 | (CA)6 | CTCTGGAGGGCCACAAAATA | CGGGCTCGTGTTTATGATTT | 210 |
| chr2:50340544-50342558 | p2 | (TA)7 | AACCCGAGAGCTCCTCTTTC | GATGAGAAATGAGGCCCAAA | 176 |
| chr2:50358869-50360893 | p2 | (AT)12 | TTTATCGACGGCCAGGATAG | ACCGAGTCGGAAGAGTCTCA | 138 |
| chr2:50367165-50369189 | p2 | (TA)12 | ATCAAATCTTGCAACCTCCG | CCCAGTCGGGTAATTTCGTA | 151 |
| chr2:50382337-50384352 | p5 | (ATGAT)3 | CTTGGGGAAAACTCCATCCT | TACGGACGTTTTATGGCACA | 173 |
| chr2:50388857-50390872 | p5 | (ATTTC)3 | GAAATCCATGTACCCGCACT | ATTTCGCCAAAGTTTTCCCT | 206 |
| chr2:50388857-50390872 | p2 | (AT)9 | GAAATCCATGTACCCGCACT | ATTTCGCCAAAGTTTTCCCT | 206 |
| chr2:50441216-50443248 | p2 | (AT)16 | GCTTCATGTCCACTTTCGGT | GACTCATCGGTTTTGGTCGT | 243 |
| chr2:50460090-50462108 | p6 | (CAGCTC)3 | CAGGTGCACAAGGAGACAGA | GCCAACATAGCCGAACATCT | 178 |
| chr2:50480790-50482836 | p2 | (AT)6 | CATATCACAGCCACACAGGG | GGTAAGCAAAGGCACGAAAG | 257 |
| chr2:50492757-50494791 | p2 | (TA)17 | TTGAGTTGTTTATGCGCTGG | AACTCGCCTAACCCGAACTT | 255 |
| chr2:50537159-50539177 | p3 | (TTG)5 | CCGATTTTCGGCAGAATTTA | TGCCTTTGAAAGAAAGGGAA | 236 |
| chr2:50537159-50539177 | p6 | (TATATG)3 | CCGATTTTCGGCAGAATTTA | TGCCTTTGAAAGAAAGGGAA | 236 |
| chr2:50632829-50634853 | p3 | (CAA)8 | GAAAATGCCTCTGTGGGTGT | GCTATTATGCCCGAAGTGGA | 297 |
| chr2:50644590-50646605 | p5 | (ATATA)3 | TCGATAACATATGCAAGCCG | TCTTGGAGAGCTCCGTTGTT | 258 |
| chr2:50725354-50727372 | p6 | (CATATT)3 | CCCCGCCTACCTACTGGTAT | TATTGTTGCCCAATGGGATT | 152 |
| chr2:50725354-50727372 | p2 | (TA)7 | CCCCGCCTACCTACTGGTAT | TATTGTTGCCCAATGGGATT | 152 |
| chr2:50732271-50734295 | p6 | (TATATG)4 | GTTGTATGGCATGTTGGTCG | CGCCTCCAATAGTATCCCAA | 163 |
| chr2:50738697-50740715 | p3 | (ATA)6 | GATAATCGGCGTCGACACTT | AACACTGGGTGTACCGCTTC | 136 |
| chr2:50789975-50791987 | p2 | (TA)6 | TCCTGCCACTTCTTGTTCCT | GGATTGCACTTTTGAAGGGA | 145 |
| chr2:50821406-50823418 | p2 | (TG)6 | TAGGGTTATATCCCGCCCTT | CCAGATGGCCCCTTAATTTT | 286 |
| chr2:50863329-50865345 | p2 | (AT)8 | CCGTATTTTTGAACCTCGGA | CAAAGTAGGGTCACCCGAAA | 134 |
| chr2:51021732-51023746 | p2 | (TA)7 | GACTTACGCTTGCCCAAAAC | TTGCTTGGGTTGTTGATTGA | 177 |
| chr2:51055739-51057751 | p2 | (TA)6 | CCAGAATCCGCACCTACAGT | TCCTTAGGGTAATCGGGTCC | 253 |
| chr2:51072590-51074608 | p2 | (TA)9 | CCAAATCTGTTCCGTCACCT | GAAAATTGGGGTGGTCCTTT | 174 |
| chr2:51253249-51255261 | p2 | (TA)6 | AGTGGGCCTCAGGGAGTATT | ACCCAAGAGAGTGGATGGTG | 263 |
| chr2:51376585-51378597 | p2 | (TA)14 | GAATACAATCGGCTCTCCCA | TGTATGTGGGGAAAAAGGGA | 105 |
| chr2:51376585-51378597 | p2 | (AT)6 | GAATACAATCGGCTCTCCCA | TGTATGTGGGGAAAAAGGGA | 105 |
| chr2:51383830-51385850 | p2 | (TA)10 | TTGTACCGAGCACCCCTTAG | TCCGGAGCCTTATTACATCG | 161 |

| Marker ID | SSR type | SSR motif | FORWARD PRIMER (5'-3') | REVERSE PRIMER (5'-3') | size (bp) |
|---|---|---|---|---|---|
| chr2:51610372-51612384 | p2 | (AT)6 | GTCCTGTCCCGCCTATATGA | TCACCCGGACACCTCTTAAC | 197 |
| chr2:51610372-51612384 | p3 | (CAT)5 | GTCCTGTCCCGCCTATATGA | TCACCCGGACACCTCTTAAC | 197 |
| chr2:51610695-51612710 | p2 | (AT)6 | GTTAAGAGGTGTCCGGGTGA | TTGTCGCCATAGCTTCAATG | 129 |
| chr2:51610695-51612710 | p3 | (CAT)5 | GTTAAGAGGTGTCCGGGTGA | TTGTCGCCATAGCTTCAATG | 129 |
| chr2:51665216-51667231 | p5 | (ATCAT)3 | ATTTGGACACCCCAAACAAA | GCTTAAATGGCCTGCATGTT | 157 |
| chr2:51665216-51667231 | p2 | (TA)7 | ATTTGGACACCCCAAACAAA | GCTTAAATGGCCTGCATGTT | 157 |
| chr2:51713486-51715500 | p2 | (TA)7 | GATAATCGACGCCGACACTT | ACTGCTCCCAAATCAGATGG | 105 |
| chr2:51751262-51753280 | p3 | (CAA)6 | GTCCGGGACGACATCATAGT | TGGAGGTTACCGACCTTACG | 290 |
| chr2:51774067-51776079 | p2 | (TA)6 | GGTTCAAGATGAAACGCCAT | TCGAATGTTAAGCTCTCGGG | 223 |
| chr2:51780886-51782902 | p2 | (TA)8 | TAATGGTGGATTCATTGCCA | AAATGACATGGCGGAATAGC | 117 |
| chr2:51780886-51782902 | p3 | (AAT)14 | TAATGGTGGATTCATTGCCA | AAATGACATGGCGGAATAGC | 117 |
| chr2:51806925-51808975 | p2 | (AT)25 | TCACAAGGATTACGAAGCCC | GGAAATTTGCATTTGCTGCT | 113 |
| chr2:51865978-51867994 | p2 | (TG)8 | AGGTGCCTACGGGCTATTTT | TCATACCTGGCCCTACCAAG | 221 |
| chr2:51916037-51918089 | p5 | (AGAAA)3 | CTCTGCTGCGTCTCCTTTTC | ACAAGCCTAATCGCTTGCAT | 205 |
| chr2:51916037-51918089 | p2 | (AT)26 | CTCTGCTGCGTCTCCTTTTC | ACAAGCCTAATCGCTTGCAT | 205 |
| chr2:52066236-52068251 | p3 | (TTC)5 | CAAACACACAAACCCCAACA | ACGAACTGCTGCTCTCCATT | 249 |
| chr2:52077332-52079347 | p5 | (ATGCA)3 | CTAACCCAAACCCAACCCTT | GTTCCACGACGCTAGACTCC | 200 |
| chr2:52189041-52191061 | p2 | (AT)10 | ATTCTGCTCGTGCTTACCGT | GGCCCTTTAATGAGGGACTC | 103 |
| chr2:52224502-52226520 | p2 | (TA)33 | TATGATACGCAAGCCGACAA | TGTTCGTCTGTAATGGCAGC | 250 |
| chr2:52224502-52226520 | p2 | (AT)9 | TATGATACGCAAGCCGACAA | TGTTCGTCTGTAATGGCAGC | 250 |
| chr2:52229305-52231357 | p2 | (TA)8 | AGCTCGGCGAATGAATAGAA | ATGCTAGGTGGCGACTCTGT | 199 |
| chr2:52411186-52413202 | p2 | (AT)8 | CTTCCTGCTCAAGGTATCCG | GCTCATGGCTTAGAGTTGGC | 160 |
| chr2:52413272-52415300 | p3 | (TTG)5 | ATTGTTCTGAACGGTTTCGG | ATGGCTCGGCTCTAAATCAA | 169 |
| chr2:52553345-52555399 | p3 | (ATT)18 | TTTCATCAGCCGATTCTTCC | TAAGGCTGATTTCTGGGTGG | 254 |
| chr2:52816401-52818416 | p5 | (AATGC)3 | CCTCACGTGGCTATGGAAGT | CGTGACAAAGGGTTGTCCTT | 172 |
| chr2:52849573-52851588 | p5 | (AATGC)3 | CGACCTCGTGAGTGATCTGA | TGAGGTGGGAAAGGTAGGTG | 221 |
| chr2:52870346-52872442 | p2 | (TA)17 | TTATTTTGGTGGCCAAGAGG | CCACAATGCCCCATACTTCT | 271 |
| chr2:52903785-52905821 | p2 | (AT)18 | ATTGAGATGGAGGGAAGGGT | TTAGCTCGGATTGCCAGTTT | 161 |
| chr2:52968653-52970669 | p2 | (TG)8 | GGAAATTCGTGGCTGAATGT | CGGGGTAAGCCTGTATTGAA | 129 |
| chr2:52968653-52970669 | p2 | (TA)6 | GGAAATTCGTGGCTGAATGT | CGGGGTAAGCCTGTATTGAA | 129 |
| chr2:52986627-52988645 | p2 | (AT)21 | TCTGGTACTAGCTGCCCGAT | CCCTGCCCAATGTAGAAGAA | 159 |
| chr2:52986627-52988645 | p2 | (AT)9 | TCTGGTACTAGCTGCCCGAT | CCCTGCCCAATGTAGAAGAA | 159 |
| chr2:53006006-53008018 | p2 | (AT)6 | GAGGGGTACTCCTAGTCCCG | GCAGCAATTTTTGAATCGGT | 280 |
| chr2:53076887-53078911 | p3 | (AAT)8 | ATCATGAAGCGACGAGAACC | TTTGGACACGTATCATCGGA | 162 |
| chr2:53137523-53139538 | p5 | (GAAAT)3 | TAGCCCCTCACATCCTTACG | TGACATTGTGAATTGCGGAT | 271 |
| chr2:53213335-53215353 | p3 | (TTG)6 | ATCGCCGACGAGACTTAGAA | GGATTCACTCCAAGTCCGAA | 247 |

| Marker ID | SSR type | SSR motif | FORWARD PRIMER (5'-3') | REVERSE PRIMER (5'-3') | size (bp) |
|---|---|---|---|---|---|
| chr2:53221735-53223791 | p2 | (TC)28 | CGAACGACGAAATGTGCTAA | AGACCCTGTGTTACCATGCC | 259 |
| chr2:53262245-53264259 | p2 | (GA)7 | GCGTTTGGGAAATGTTTTGT | TCTCTCCATTCCCGAACATC | 115 |
| chr2:53414579-53416591 | p2 | (TG)6 | TAGTGCGTGCACAAAAGACC | AAGGTGGCATGTGATTGTGA | 283 |
| chr2:53519687-53521713 | p2 | (TA)13 | CCTGAAACTGGAAGCTCTGG | ATTTCCTTCCCACGCTTCTT | 266 |
| chr2:53525397-53527411 | p2 | (TA)13 | AAGAAGCGTGGGAAGGAAAT | TTCGACCCTCAACTTTCACC | 178 |
| chr2:53525397-53527411 | p2 | (AG)7 | AAGAAGCGTGGGAAGGAAAT | TTCGACCCTCAACTTTCACC | 178 |
| chr2:53525397-53527411 | p2 | (AT)7 | AAGAAGCGTGGGAAGGAAAT | TTCGACCCTCAACTTTCACC | 178 |
| chr2:53526898-53528913 | p2 | (AT)7 | GGACATGTGTCCACCTTGTG | TGGCCCTCCTAGTTTTCCTT | 238 |
| chr2:53526898-53528913 | p5 | (TGAAT)3 | GGACATGTGTCCACCTTGTG | TGGCCCTCCTAGTTTTCCTT | 238 |
| chr2:53537848-53539866 | p6 | (TCATGC)3 | GAACGTCACCCCACTGAACT | TTCTGCACCATTCCAAAACA | 186 |
| chr2:53745370-53747384 | p3 | (CAA)5 | CGTTATCCATCCTTGTGGCT | AGTCGGGATATGTTGGCAAG | 127 |
| chr2:53745370-53747384 | p2 | (CA)7 | CGTTATCCATCCTTGTGGCT | AGTCGGGATATGTTGGCAAG | 127 |
| chr2:53789516-53791570 | p2 | (AT)27 | AAACTCATGAAATCACGGGC | TATGCCTCCAGTTCCTCCAC | 283 |
| chr2:53852407-53854423 | p2 | (TA)8 | AACAGCATGCGGCTAACTTT | TTTGGGCTTGTCTCCCATAC | 284 |
| chr2:53852407-53854423 | p2 | (TA)7 | AACAGCATGCGGCTAACTTT | TTTGGGCTTGTCTCCCATAC | 284 |
| chr2:53887680-53889694 | p2 | (AT)7 | AGAGGGCAGGGTTGAAGATT | CATCTGAGCAGTTTGTCCGA | 180 |
| chr2:53903522-53905538 | p2 | (TA)8 | CTTAGAGTTCCAGGTTGGCG | GTAGGATGGCTCCTCCATCA | 147 |
| chr2:53937593-53939605 | p2 | (TA)6 | TGTCCGGATCTAGAGACGCT | CAAACGCGATGGTACAAGTG | 285 |
| chr2:53937593-53939605 | p2 | (AT)7 | TGTCCGGATCTAGAGACGCT | CAAACGCGATGGTACAAGTG | 285 |
| chr2:53937718-53939732 | p2 | (TA)6 | TGTCCGGATCTAGAGACGCT | CAAACGCGATGGTACAAGTG | 160 |
| chr2:53937718-53939732 | p2 | (AT)7 | TGTCCGGATCTAGAGACGCT | CAAACGCGATGGTACAAGTG | 160 |
| chr2:53950720-53952736 | p2 | (AT)7 | GAACTGGTGGAGGAATTGGA | AAGAGATCCGCCAATGAATG | 275 |
| chr2:53950720-53952736 | p2 | (AT)8 | GAACTGGTGGAGGAATTGGA | AAGAGATCCGCCAATGAATG | 275 |
| chr2:53989910-53991922 | p2 | (TA)6 | TGTCCGGATCTAGAGACGCT | CAAACGCGATGGTACAAGTG | 285 |
| chr2:53989910-53991922 | p2 | (AT)7 | TGTCCGGATCTAGAGACGCT | CAAACGCGATGGTACAAGTG | 285 |
| chr2:53990035-53992049 | p2 | (TA)6 | TGTCCGGATCTAGAGACGCT | CAAACGCGATGGTACAAGTG | 160 |
| chr2:53990035-53992049 | p2 | (AT)7 | TGTCCGGATCTAGAGACGCT | CAAACGCGATGGTACAAGTG | 160 |
| chr2:54003037-54005053 | p2 | (AT)7 | GAACTGGTGGAGGAATTGGA | AAGAGATCCGCCAATGAATG | 275 |
| chr2:54003037-54005053 | p2 | (AT)8 | GAACTGGTGGAGGAATTGGA | AAGAGATCCGCCAATGAATG | 275 |
| chr2:54234778-54236790 | p2 | (TA)8 | CGTGCATGCCTCAATATGTC | ACCGTACTCGGCCCTCTAGT | 131 |
| chr2:54234778-54236790 | p2 | (TA)6 | CGTGCATGCCTCAATATGTC | ACCGTACTCGGCCCTCTAGT | 131 |
| chr2:54296903-54298917 | p2 | (AT)7 | TTTGATTGGAGCATGGATGA | GGCGTTGTGAATCTCTCGTT | 142 |
| chr2:54354388-54356404 | p2 | (AT)8 | TGACGGACGTACAACCAGAA | CCAGTTCGGGCACTAGTCAT | 100 |
| chr2:54421936-54423964 | p2 | (TA)14 | CCTTAGACCGACCTGCTCTG | GTCAGTCTGTAAGCGGCTCC | 256 |
| chr2:54430359-54432375 | p2 | (TA)8 | GAGAGGGGTATTTTGTGCGA | TTCTGCAGGTTGTGCGTTAG | 118 |
| chr2:54484961-54486973 | p2 | (TG)6 | ATGATGCCAAAGGTGCCTAC | TACATACATAGCGTGCCCGA | 246 |

| Marker ID | SSR type | SSR motif | FORWARD PRIMER (5'-3') | REVERSE PRIMER (5'-3') | size (bp) |
|---|---|---|---|---|---|
| chr2:54491239-54493255 | p2 | (TA)8 | ATTGCTCCATAACGGAATCG | GGTGACTTCCTGGCCAAATA | 220 |
| chr2:54495842-54497870 | p2 | (AT)14 | ACTTTTGTCACGACCCAACC | TGTAGTGGCAGTCTTGTCGG | 272 |
| chr2:54558016-54560032 | p5 | (ATTCA)3 | TACCAAAAATCGTGCCAACA | GAAGGTTTCGGGGGAAGTAG | 170 |
| chr2:54558016-54560032 | p2 | (AG)6 | TACCAAAAATCGTGCCAACA | GAAGGTTTCGGGGGAAGTAG | 170 |
| chr2:54561263-54563277 | p2 | (AT)7 | GGCCCTGTCCTGACTCATAA | TCCACAAATGACCCAACTGA | 112 |
| chr2:54617757-54619787 | p2 | (AT)15 | CCGTGAAATGATCCTGTCCT | AAGTTCAGTGGACTGGTGGG | 262 |
| chr2:54659552-54661596 | p2 | (AT)22 | CTGTCCCGTCATATTTCGCT | GACAAGGCTGCCACTACACA | 126 |
| chr2:54686739-54688771 | p2 | (AT)16 | AGGATTCTCGAATACGGGCT | GCATGCCTCTGGTAACCTGT | 296 |
| chr2:54686739-54688771 | p3 | (AAT)5 | AGGATTCTCGAATACGGGCT | GCATGCCTCTGGTAACCTGT | 296 |
| chr2:54686878-54688893 | p2 | (AT)16 | AGGATTCTCGAATACGGGCT | GCATGCCTCTGGTAACCTGT | 157 |
| chr2:54686878-54688893 | p3 | (AAT)5 | AGGATTCTCGAATACGGGCT | GCATGCCTCTGGTAACCTGT | 157 |
| chr2:54707824-54709898 | p3 | (TGT)5 | GAAGCTCCTTTGTGCAGTCC | TATAGCTGGGACCGCATTTC | 131 |
| chr2:54725292-54727312 | p2 | (AT)10 | ACACTTCGAGGATGAACGCT | ACTCCAGGGGAGACAGGTTT | 230 |
| chr2:54725292-54727312 | p2 | (CT)7 | ACACTTCGAGGATGAACGCT | ACTCCAGGGGAGACAGGTTT | 230 |
| chr2:54763100-54765120 | p2 | (AT)10 | AACTGCCCAGTCGAATATGG | AGTTTCCAACGCAACAAACC | 111 |
| chr2:54860179-54862193 | p2 | (TA)7 | GTTGTTGGCTTATGGGCTGT | TGCCATAATCCCCTTTTCAA | 156 |
| chr2:54942055-54944087 | p2 | (AT)16 | CGGTTCGGTAGAATGACGTT | TGAGGGCATGAGAAGAATCC | 102 |
| chr2:55005820-55007844 | p6 | (CATAAG)4 | ACAAGGTAATTGGGCAGCAC | GAATGAAGGTGGAGAAGCCA | 152 |
| chr2:55093751-55095765 | p2 | (CA)7 | GCCAAATGAAGAAGTGGGAA | TGTGGGCTGTTGTTGTTGAT | 264 |
| chr2:55100287-55102301 | p2 | (CA)7 | TTTTCTGTGGTTCGGGTTTC | ACCATTTGCAGATTTCGGAC | 191 |
| chr2:55154344-55156358 | p2 | (AT)6 | TTCATTTCTGTCACGACCCA | AGGCGACAGTTCCGAGACTA | 231 |
| chr2:55154344-55156358 | p2 | (AT)7 | TTCATTTCTGTCACGACCCA | AGGCGACAGTTCCGAGACTA | 231 |
| chr2:55154344-55156358 | p2 | (AT)6 | TTCATTTCTGTCACGACCCA | AGGCGACAGTTCCGAGACTA | 231 |
| chr2:55154344-55156358 | p2 | (AT)6 | TTCATTTCTGTCACGACCCA | AGGCGACAGTTCCGAGACTA | 231 |
| chr2:55154467-55156479 | p2 | (AT)6 | TTCATTTCTGTCACGACCCA | AGGCGACAGTTCCGAGACTA | 108 |
| chr2:55154467-55156479 | p2 | (AT)7 | TTCATTTCTGTCACGACCCA | AGGCGACAGTTCCGAGACTA | 108 |
| chr2:55154467-55156479 | p2 | (AT)6 | TTCATTTCTGTCACGACCCA | AGGCGACAGTTCCGAGACTA | 108 |
| chr2:55154467-55156479 | p2 | (AT)6 | TTCATTTCTGTCACGACCCA | AGGCGACAGTTCCGAGACTA | 108 |
| chr2:55161363-55163383 | p2 | (TA)10 | ACTTTTGTCACGACCCAACC | TAGGCGACTGTTCCGAGACT | 212 |
| chr2:55161363-55163383 | p2 | (AT)9 | ACTTTTGTCACGACCCAACC | TAGGCGACTGTTCCGAGACT | 212 |
| chr2:55298017-55300029 | p2 | (TG)6 | TTCAGAGATTCCGACAGCCT | TGTTGGCACGGTGTGATTAT | 136 |
| chr2:55389613-55391629 | p2 | (AT)8 | CGCAGCATCTGTCGAATAAA | ACACGACAGTTTGAGGGTCC | 151 |
| chr2:55495751-55497765 | p2 | (AG)7 | AAGGAAAGGTGCCATGTTTG | TCCTCGAATGTTGTTGCTTG | 178 |
| chr2:55495751-55497765 | p3 | (TTG)5 | AAGGAAAGGTGCCATGTTTG | TCCTCGAATGTTGTTGCTTG | 178 |
| chr2:55495751-55497765 | p4 | (TATG)4 | AAGGAAAGGTGCCATGTTTG | TCCTCGAATGTTGTTGCTTG | 178 |
| chr2:55504192-55506206 | p2 | (AT)7 | CTACCGAGCACCACTCATCA | TGTGTGGGTAGCGGATGTTA | 222 |

| Marker ID | SSR type | SSR motif | FORWARD PRIMER (5'-3') | REVERSE PRIMER (5'-3') | size (bp) |
|---|---|---|---|---|---|
| chr2:55537422-55539437 | p5 | (TTCAT)3 | GCCTGAATCAATTAGGCGAA | GACAAAGGCGGTGGTTAGAA | 239 |
| chr2:55702641-55704659 | p6 | (AACATG)3 | AGGGAATGACGTTTGACGAC | GACTGGACCGAAGTGCAAAT | 154 |
| chr2:55702641-55704659 | p2 | (TA)6 | AGGGAATGACGTTTGACGAC | GACTGGACCGAAGTGCAAAT | 154 |
| chr2:55708171-55710187 | p2 | (AT)8 | CTTGTAACGTGGGGAGTGCT | TTGCCATCGTAGGATAAGGG | 101 |
| chr2:55852078-55854096 | p2 | (AT)9 | CGACCAAGGTATGTGAGGCT | AACATTTGTGCTCTACCGGG | 139 |
| chr2:55855755-55857781 | p3 | (ACA)5 | GAATCTTTGGAACTCGCTCG | GGAACGGGAATTATAGGGGA | 295 |
| chr2:55855755-55857781 | p2 | (AT)13 | GAATCTTTGGAACTCGCTCG | GGAACGGGAATTATAGGGGA | 295 |
| chr2:55862237-55864252 | p5 | (ATTTC)3 | TCACTGAACAACAGGGTGGA | TACGTAGCCTTGTCGGCTCT | 206 |
| chr2:56037010-56039025 | p5 | (GAAAG)3 | GAGCGGGAATGAAATGAAAA | GGCCGTATAAAATTCACGGA | 107 |
| chr2:56119761-56121788 | p3 | (TTG)5 | ATACACATTTTGCCTTCCCG | TGGCCAGATAGAAAGGCTGT | 112 |
| chr2:56119761-56121788 | p3 | (ATA)9 | ATACACATTTTGCCTTCCCG | TGGCCAGATAGAAAGGCTGT | 112 |
| chr2:56191199-56193217 | p3 | (ACA)6 | AGGTCAATGGGCGTAAAGTG | GAAGCACCGTTAAGTCGAGC | 173 |
| chr2:56197345-56199360 | p5 | (ATGTT)3 | AGGAGGCGGAAAAGGTATGT | AAGTTTCAAGGGGATTCGGT | 246 |
| chr2:56207743-56209758 | p5 | (ATGTT)3 | AGGAGGCGGAAAAGGTATGT | AAGTTTCAAGGGGATTCGGT | 246 |
| chr2:56222739-56224754 | p3 | (ATA)5 | GAAACAGCTATCTCGGGCAG | CAATAAACCTGGGGCTCAAA | 162 |
| chr2:56225675-56227689 | p2 | (TA)7 | TGTTTCGACTTCTTCATGCG | TGGAGCCTGGATCGTACTCT | 129 |
| chr2:56225675-56227689 | p2 | (TA)11 | TGTTTCGACTTCTTCATGCG | TGGAGCCTGGATCGTACTCT | 129 |
| chr2:56226618-56228640 | p2 | (TA)7 | ATGATGGATTCATTGCCACA | GACATGGCGGAATAGCAAAT | 170 |
| chr2:56226618-56228640 | p2 | (TA)11 | ATGATGGATTCATTGCCACA | GACATGGCGGAATAGCAAAT | 170 |
| chr2:56229170-56231212 | p2 | (AT)21 | GGGTCACAGTAAGGCGGTAA | CCACAAGTCTGCCAAATCCT | 293 |
| chr2:56330106-56332122 | p2 | (AG)8 | GACCTCTCGGAATCGATGAA | CCCTTTCCGAGCATTGATTA | 177 |
| chr2:56393519-56395541 | p3 | (TGA)5 | GAAACAAAGGAGATGCTGCC | ATGCCTTAGGGAAGAAGGGA | 266 |
| chr2:56393519-56395541 | p2 | (TA)11 | GAAACAAAGGAGATGCTGCC | ATGCCTTAGGGAAGAAGGGA | 266 |
| chr2:56428951-56430966 | p5 | (CAAAA)3 | TGACCGATCTATGGACACCA | AACCAACAACCAAGGCTCAC | 283 |
| chr2:56430784-56432796 | p2 | (AT)6 | TTACAGGGGAAATGCTACCG | ATCAAGAACGTTTCGCCATC | 203 |
| chr2:56432572-56434593 | p3 | (TTG)7 | AAATTCGTGGTTATGAGCGG | ATATGCGGTTCGTACTTGGG | 282 |
| chr2:56520155-56522170 | p5 | (CAGGA)3 | CAGAATGCGAGTCCAAGTGA | TTTGTGGGTCTTTATTCGGC | 264 |
| chr2:56570544-56572562 | p6 | (ACATTC)3 | AGGCATCCACAAATTCTTGC | TGGTTAATGCATGTTTCGGA | 185 |
| chr2:56709018-56711036 | p2 | (AT)9 | ATAACCCTAGGCCCATGTCC | TCATGGATTACGGACGGATT | 289 |
| chr2:56750607-56752621 | p3 | (ACA)5 | ACAGCCCACAACAACAACAA | ATGGACTGTTTTGGGTTTGC | 277 |
| chr2:56750607-56752621 | p2 | (TC)7 | ACAGCCCACAACAACAACAA | ATGGACTGTTTTGGGTTTGC | 277 |
| chr2:56760641-56762653 | p2 | (AT)6 | AGCTTTGACAGTCTTGGGGA | GAAGGCGTGTTAGAACCCAA | 259 |
| chr2:56814545-56816560 | p3 | (ATG)5 | TTTCGCTGAACCTACTGCCT | TGAAAAATGGGGTCTTTTGG | 119 |
| chr2:56951885-56953905 | p2 | (AT)10 | AGACAGGGAGGTGAGCTTGA | CTTTTCGGCCGTCAAAATAA | 140 |
| chr2:56993420-56995435 | p5 | (ATTTC)3 | CAACCCTTGGAAGAGATCCA | AACAGGGCACTACCATCCTG | 202 |
| chr2:56993420-56995435 | p6 | (ATGAAT)3 | CAACCCTTGGAAGAGATCCA | AACAGGGCACTACCATCCTG | 202 |

| Marker ID | SSR type | SSR motif | FORWARD PRIMER (5'-3') | REVERSE PRIMER (5'-3') | size (bp) |
|---|---|---|---|---|---|
| chr2:57092844-57094876 | p2 | (AT)6 | GAGTTTCCCCATCACAAGGA | TCGGAATTGGGGTGTAACAT | 186 |
| chr2:57092921-57094947 | p2 | (AT)6 | GAGTTTCCCCATCACAAGGA | TCGGAATTGGGGTGTAACAT | 109 |
| chr2:57203009-57205027 | p6 | (CATGAG)3 | AAATGAATTCAGCTGGTGCC | CTCAGGCCCATGCAAGTAAT | 257 |
| chr2:57216086-57218101 | p3 | (TAT)5 | AGCAAGGCCTGAGTTTGTGT | AGGGCCATGATACGCATAAG | 272 |
| chr2:57216086-57218101 | p2 | (AT)33 | AGCAAGGCCTGAGTTTGTGT | AGGGCCATGATACGCATAAG | 272 |
| chr2:57227474-57229490 | p2 | (TA)8 | CTTTCACCACTTGCTGACGA | ATATGGGCTCAGAGAGCGAA | 268 |
| chr2:57234225-57236243 | p6 | (AACTGA)3 | CTTGCCATGCTGCATTAAGA | TTGACTTCTGCTCCGGTTTT | 171 |
| chr2:57245327-57247345 | p2 | (AT)9 | TACGACTTGAATTACGGCCC | TCTAAATACACCGAAGGCGG | 179 |
| chr2:57245327-57247345 | p3 | (TAT)5 | TACGACTTGAATTACGGCCC | TCTAAATACACCGAAGGCGG | 179 |
| chr2:57335865-57337877 | p2 | (AT)6 | TTGCTCCACGCCCTACTACT | CGAAGCCACGGTTGATACTT | 118 |
| chr2:57335865-57337877 | p2 | (AT)7 | TTGCTCCACGCCCTACTACT | CGAAGCCACGGTTGATACTT | 118 |
| chr2:57336000-57338012 | p2 | (AT)6 | AAGTATCAACCGTGGCTTCG | TTTCGTTAATTTCCCCCACA | 220 |
| chr2:57336000-57338012 | p2 | (AT)7 | AAGTATCAACCGTGGCTTCG | TTTCGTTAATTTCCCCCACA | 220 |
| chr2:57336033-57338048 | p2 | (AT)6 | AAGTATCAACCGTGGCTTCG | TTTCGTTAATTTCCCCCACA | 187 |
| chr2:57336033-57338048 | p2 | (AT)7 | AAGTATCAACCGTGGCTTCG | TTTCGTTAATTTCCCCCACA | 187 |
| chr2:57336043-57338055 | p2 | (AT)6 | AAGTATCAACCGTGGCTTCG | TTTCGTTAATTTCCCCCACA | 177 |
| chr2:57336043-57338055 | p2 | (AT)7 | AAGTATCAACCGTGGCTTCG | TTTCGTTAATTTCCCCCACA | 177 |
| chr2:57346566-57348582 | p4 | (GAAA)4 | CCAAGAAATAAGCCCGACAA | GGAATTAAACTCGCGATGGA | 258 |
| chr2:57557678-57559693 | p2 | (TA)32 | TGGAATTCGTGGATTGTTGA | TCCAACCGAACCTACAAAGG | 126 |
| chr2:57569058-57571072 | p2 | (AT)7 | ACCGCCACATCTCTGAGTCT | AAGACCCGCAAGGCTTTATT | 153 |
| chr2:57623187-57625223 | p3 | (TTA)5 | GAACCTTGATCAGATCCCCA | CCCTGATCTTGGGTTGCTTA | 241 |
| chr2:57652351-57654363 | p2 | (TG)6 | ATGATGCCAAAGGTGCCTAC | GCGAACCAAGGTTCAATGTT | 223 |
| chr2:57687963-57689981 | p6 | (ATACAT)3 | TCGATCCTGGCTATCTGCTT | TTTTCCTTGTCGGGCATAAG | 288 |
| chr2:57730343-57732355 | p2 | (AT)13 | GGCACAAATATCACACGCTG | GATAGCCTTGTCGGCTTCTG | 101 |
| chr2:57730343-57732355 | p2 | (TC)6 | GGCACAAATATCACACGCTG | GATAGCCTTGTCGGCTTCTG | 101 |
| chr2:57730343-57732355 | p5 | (ATTAT)6 | GGCACAAATATCACACGCTG | GATAGCCTTGTCGGCTTCTG | 101 |
| chr2:57830092-57832108 | p2 | (TA)7 | GAACGGTCCAAACTTCCAGA | GGGTCTGCAACTGGGTTTTA | 177 |
| chr2:57830092-57832108 | p2 | (AT)8 | GAACGGTCCAAACTTCCAGA | GGGTCTGCAACTGGGTTTTA | 177 |
| chr2:57886535-57888547 | p2 | (AT)6 | GACTCTGAATTGGCCCAAAA | AGGGTCAGGTCTTCAGGGTT | 179 |
| chr2:57904471-57906489 | p3 | (CAC)6 | TGCCATCAGCAAAGTTCTTG | GTTGAAGCCGGTTCTCGTAG | 166 |
| chr2:57912767-57914785 | p2 | (AG)9 | TCGGGTCCTCCTATTTCCTT | CTGATGGTTCCCGTCAAAAT | 108 |
| chr2:57917425-57919437 | p2 | (AT)6 | AACCTGCCATGTCAACATCA | CAGGAACCTTGGCAAAAAGA | 231 |
| chr2:57919414-57921429 | p3 | (CTT)5 | CACAATTCCCACAAAAAGGG | GTTGCTTAAATTGGGCCAGA | 113 |
| chr2:58002064-58004082 | p3 | (ACC)6 | ATTGAGCATCAAAAGCCAGG | TTAAATTGTTGGTGGACGCA | 287 |
| chr2:58002064-58004082 | p2 | (AT)13 | ATTGAGCATCAAAAGCCAGG | TTAAATTGTTGGTGGACGCA | 287 |
| chr2:58002308-58004334 | p3 | (ACC)6 | GAGGCATGCATCTTTCGTTT | AGGAGGCTGAGATTCCCATT | 297 |

| Marker ID | SSR type | SSR motif | FORWARD PRIMER (5'-3') | REVERSE PRIMER (5'-3') | size (bp) |
|---|---|---|---|---|---|
| chr2:58002308-58004334 | p2 | (AT)13 | GAGGCATGCATCTTTCGTTT | AGGAGGCTGAGATTCCCATT | 297 |
| chr2:58071482-58073526 | p2 | (TA)22 | GGCAAACCGGTATGAAGAGA | GAGGCACTTCTTCGATGGTC | 265 |
| chr2:58101014-58103030 | p3 | (GAA)5 | TCAACTACTATGCCCCGACC | GTTCAGAGGGCCTACCCTTC | 280 |
| chr2:58101014-58103030 | p2 | (AG)8 | TCAACTACTATGCCCCGACC | GTTCAGAGGGCCTACCCTTC | 280 |
| chr2:58214611-58216623 | p2 | (GC)6 | TCATGGAGGAAAGGTTTTGG | GCGCTCAAACAATTTAGGGA | 109 |
| chr2:58232671-58234686 | p2 | (TA)10 | CCTCGTCAATCTCCCCATTA | TCGACATCAAAGTGCTCGTC | 240 |
| chr2:58232671-58234686 | p3 | (ACC)5 | CCTCGTCAATCTCCCCATTA | TCGACATCAAAGTGCTCGTC | 240 |
| chr2:58392278-58394344 | p5 | (TCATT)3 | TGGGAATGTCCTGGAAGAAG | CCATGGGTCTGCTCTAGCTC | 294 |
| chr2:58392278-58394344 | p2 | (TA)33 | TGGGAATGTCCTGGAAGAAG | CCATGGGTCTGCTCTAGCTC | 294 |
| chr2:58442835-58444905 | p3 | (ATC)5 | TACATCGCAAGGTAATGGCA | CGTTCCAGTTGTTTCGGAGT | 257 |
| chr2:58442835-58444905 | p3 | (CAA)5 | TACATCGCAAGGTAATGGCA | CGTTCCAGTTGTTTCGGAGT | 257 |
| chr2:58498475-58500489 | p2 | (AT)7 | GCCAAAATATTGCCCAAAGA | AACCGTCACAATGGCTAAGG | 289 |
| chr2:58498475-58500489 | p6 | (TACAAA)3 | GCCAAAATATTGCCCAAAGA | AACCGTCACAATGGCTAAGG | 289 |
| chr2:58551461-58553515 | p5 | (TTCTC)3 | CAACCAGTCGTGATGACACC | TTTAATGAGGTCCGCCGTAG | 215 |
| chr2:58551461-58553515 | p3 | (TAA)19 | CAACCAGTCGTGATGACACC | TTTAATGAGGTCCGCCGTAG | 215 |
| chr2:58551530-58553587 | p5 | (TTCTC)3 | CAACCAGTCGTGATGACACC | TTTAATGAGGTCCGCCGTAG | 146 |
| chr2:58551530-58553587 | p3 | (TAA)19 | CAACCAGTCGTGATGACACC | TTTAATGAGGTCCGCCGTAG | 146 |
| chr2:58578545-58580561 | p2 | (AG)8 | TCGAGGACGAGCAAGAGTTT | TTATCCAGGGAACAAGTCGG | 269 |
| chr2:58660888-58662954 | p2 | (AT)33 | CCTTCCTTCCAAACACGGTA | CTAAGAACGCATGGCCAGAT | 130 |
| chr2:58685013-58687028 | p5 | (ATTGC)3 | AAGGCAAGGCAAAAGAGTGA | CAAGTCACAACACAATGGGG | 185 |
| chr2:58689244-58691260 | p2 | (TA)8 | GTGGTGGTGTGTCCTCCTCT | AGTCCGTGTTGCATTTTTCC | 166 |
| chr2:58741956-58743968 | p5 | (TTTAA)3 | AGCAGTTCGGCCTAGGGTAT | TGAGAAGGAAGGGTCAATGG | 213 |
| chr2:58741956-58743968 | p2 | (TA)6 | AGCAGTTCGGCCTAGGGTAT | TGAGAAGGAAGGGTCAATGG | 213 |
| chr2:58824283-58826298 | p5 | (AAAAC)3 | GTTCCTACGTCACTCCCCAA | GTTGGCTAGTTTGAGCCTGG | 260 |
| chr2:59003348-59005364 | p2 | (AT)8 | ATAAGCCCGAGCCAAAAGAT | CATCTGAAGGGCAAAATCGT | 130 |
| chr2:59102546-59104588 | p3 | (ATT)14 | CCGAAAGGCATCATTCTCAT | GTTCGCTGCCGAAGAAATAG | 218 |
| chr2:59123269-59125311 | p3 | (ATT)14 | CCGAAAGGCATCATTCTCAT | GTTCGCTGCCGAAGAAATAG | 218 |
| chr2:59163127-59165142 | p5 | (TTGGT)3 | TCTCCTCATTTCGGGTATGC | TGACCAATCTAACCTCCCCA | 252 |
| chr2:59202101-59204116 | p5 | (TGAAT)3 | GAAGTACCACTTGTGGCCGT | CACATGGCCTCTTCTCCATT | 109 |
| chr2:59202101-59204116 | p6 | (TATATG)3 | GAAGTACCACTTGTGGCCGT | CACATGGCCTCTTCTCCATT | 109 |
| chr2:59205249-59207263 | p2 | (AT)7 | GTTTGCAGTCACATCTCGGA | TACTGAAACCTGTTTGGGGC | 105 |
| chr2:59210371-59212383 | p2 | (TC)6 | ACATTCTTTGGCCGATATGC | GGCATTACGACCTTCGTGTT | 225 |
| chr2:59381737-59383752 | p5 | (TTAGT)3 | TCAAAGCCCAATCAACAACA | ACGAACGTGGGAGAGAGAAA | 274 |
| chr2:59389570-59391585 | p5 | (TGATA)3 | CCATGCCATTGTCTTTGATG | CTGCGGTAGGGAAGTTCTCA | 101 |
| chr2:59440543-59442558 | p5 | (TTTTA)3 | GAAAGGTTTTGCCAACCAAA | ATCACCTTGACCATGAGGGA | 233 |
| chr2:59499141-59501167 | p2 | (AT)13 | AGTTCTTGCCACGCTCTCAT | CATGACTGAGGCAGGGGTAT | 243 |

| Marker ID | SSR type | SSR motif | FORWARD PRIMER (5'-3') | REVERSE PRIMER (5'-3') | size (bp) |
|---|---|---|---|---|---|
| chr2:59509119-59511134 | p5 | (TTTCT)3 | ATTACACGAGGGCGACAATC | TCGACTTTCCGGTTCAATTC | 225 |
| chr2:59509119-59511134 | p3 | (TGT)5 | ATTACACGAGGGCGACAATC | TCGACTTTCCGGTTCAATTC | 225 |
| chr2:59839881-59841901 | p2 | (AG)10 | GCCCATCTGTGGAAGACCTA | CAATTCAACATTCCCCGACT | 182 |
| chr2:59850478-59852508 | p2 | (AT)15 | AGACCCATGCTTCATCATCC | GATGTGGCGAGTTCCACTTT | 300 |
| chr2:59850478-59852508 | p2 | (TA)24 | AGACCCATGCTTCATCATCC | GATGTGGCGAGTTCCACTTT | 300 |
| chr2:59900478-59902496 | p6 | (ATCTAT)3 | TCCGAGTGGAAAACGAAATC | CATGGGGTTTTTGTTCAAGG | 275 |
| chr2:59900478-59902496 | p4 | (ATTT)4 | TCCGAGTGGAAAACGAAATC | CATGGGGTTTTTGTTCAAGG | 275 |
| chr2:59952988-59955006 | p6 | (CAACTC)3 | GCATTTGAAATAGCCCAAGC | TTCTGTAGCGCCATCCTTTT | 213 |
| chr2:59957180-59959195 | p5 | (TCTTC)3 | GGACGGAGGGGGTAATAAGA | CAGGGGATGGTCAGTTTGAT | 139 |
| chr2:60084409-60086425 | p2 | (TA)8 | GCCGGTGTATCAATATGCCT | CCCGACCCTTACTTAGAGCC | 125 |
| chr2:60091028-60093070 | p2 | (AT)21 | AAGCTGTAGCTCCGGATGAA | CGTACCGACGTCCTTTTGTT | 280 |
| chr2:60280384-60282399 | p5 | (TTTTG)3 | AGTTGATAGGGTGGACTGCG | TTGATTGGTGCCAAACACAT | 142 |
| chr2:60280384-60282399 | p3 | (GTT)5 | AGTTGATAGGGTGGACTGCG | TTGATTGGTGCCAAACACAT | 142 |
| chr2:60293732-60295748 | p3 | (AAT)6 | CACAATAACCGGCAGACCTT | CCGCTTCTACGGTCAAACAT | 241 |
| chr2:60293732-60295748 | p4 | (TTAA)4 | CACAATAACCGGCAGACCTT | CCGCTTCTACGGTCAAACAT | 241 |
| chr2:60293732-60295748 | p2 | (CT)6 | CACAATAACCGGCAGACCTT | CCGCTTCTACGGTCAAACAT | 241 |
| chr2:60302411-60304453 | p2 | (TA)21 | ATGGTCTCACGATCTCCCAC | GGCTAACCACCGAGCACTAC | 123 |
| chr2:60378163-60380178 | p5 | (ATTTC)3 | TACCCGCACTTATTTCAGCC | ATAACAATGCCTCACAGGGC | 202 |
| chr2:60399442-60401460 | p2 | (AT)9 | CAGTCTTTTCCCCCTCAACA | AAACCGGGTTCTTGTCCTTT | 235 |
| chr2:60461478-60463522 | p2 | (GA)22 | AACCATTTTGACCCTGGTTG | GAATTTGTAGCCGGTGTCGT | 131 |
| chr2:60480617-60482631 | p2 | (AG)7 | ATCTTGGCTGAATTTGTGGC | TAACGCGAAATTTTCTTGGG | 220 |
| chr2:60649055-60651067 | p2 | (AT)6 | GGAAGATGATACGATCCCGA | GAATAGCCTCTTTGCATGGC | 216 |
| chr2:60662409-60664427 | p6 | (TCAAGA)3 | CATGGCACGAACCACTAAGA | GTTATGCGATTTTGGGCATT | 236 |
| chr2:60707208-60709223 | p5 | (TCATA)3 | CCATCCACAACACGTGAAAC | TCTTGGAGTTGGAGCGTTCT | 277 |
| chr2:60707208-60709223 | p2 | (TA)9 | CCATCCACAACACGTGAAAC | TCTTGGAGTTGGAGCGTTCT | 277 |
| chr2:60708506-60710526 | p2 | (TA)9 | ACTTTTGTCACGACCCAACC | TGTAATGGCAGCCTTGTCAG | 181 |
| chr2:60708506-60710526 | p2 | (AT)10 | ACTTTTGTCACGACCCAACC | TGTAATGGCAGCCTTGTCAG | 181 |
| chr2:60737834-60739846 | p2 | (TA)6 | GCATGCATATCCCTCAAGGT | AGGAGATGGTGTTGACCAGG | 237 |
| chr2:60750989-60753021 | p2 | (AT)16 | ACGAAGTGGACTTTGGTTGG | TCGGGCTTGGTTCACTATTC | 105 |
| chr2:60750989-60753021 | p2 | (AT)7 | ACGAAGTGGACTTTGGTTGG | TCGGGCTTGGTTCACTATTC | 105 |
| chr2:60759596-60761658 | p2 | (AT)31 | AAACAGCAGACAAGGGGAAA | CGAATGGCTCCGAATTGTAT | 120 |
| chr2:60759596-60761658 | p3 | (CAA)5 | AAACAGCAGACAAGGGGAAA | CGAATGGCTCCGAATTGTAT | 120 |
| chr2:60767603-60769619 | p2 | (CA)8 | AGCGGCATAATCAACAGCTT | TGGTGTCAGAGCAGTTCGTC | 217 |
| chr2:60792496-60794528 | p2 | (AT)16 | ACGAAGTGGACTTTGGTTGG | TCGGGCTTGGTTCACTATTC | 105 |
| chr2:60792496-60794528 | p2 | (AT)7 | ACGAAGTGGACTTTGGTTGG | TCGGGCTTGGTTCACTATTC | 105 |
| chr2:60801103-60803165 | p2 | (AT)31 | AAACAGCAGACAAGGGGAAA | CGAATGGCTCCGAATTGTAT | 120 |

| Marker ID | SSR type | SSR motif | FORWARD PRIMER (5'-3') | REVERSE PRIMER (5'-3') | size (bp) |
|---|---|---|---|---|---|
| chr2:60801103-60803165 | p3 | (CAA)5 | AAACAGCAGACAAGGGGAAA | CGAATGGCTCCGAATTGTAT | 120 |
| chr2:60809110-60811126 | p2 | (CA)8 | AGCGGCATAATCAACAGCTT | TGGTGTCAGAGCAGTTCGTC | 217 |
| chr2:60814023-60816051 | p2 | (TA)11 | AATAGAATGAGCGGTGCTCG | CCCGCAGGTAAACAGGTAGA | 271 |
| chr2:60814023-60816051 | p5 | (AAAGA)3 | AATAGAATGAGCGGTGCTCG | CCCGCAGGTAAACAGGTAGA | 271 |
| chr2:60818546-60820561 | p5 | (GATAT)3 | TGCGATGGCATCTGTATGAT | GACGCTTGCTGCACATAAAA | 280 |
| chr2:60896269-60898299 | p2 | (AT)15 | ACAGATGATCCAGAGGCACC | TAGCCTATCTCCCATGCACC | 211 |
| chr2:60896269-60898299 | p3 | (TGT)5 | ACAGATGATCCAGAGGCACC | TAGCCTATCTCCCATGCACC | 211 |
| chr2:60911663-60913681 | p6 | (CGTCCC)3 | GCCCCCATATCCTTCAATTT | CATTCCGAGGGACAACCTTA | 128 |
| chr2:61024470-61026485 | p5 | (TTCAT)3 | CACGAGAAACACCGAACCTT | CAGCTGACCCGGAGTATCAT | 114 |
| chr2:61081487-61083514 | p2 | (AT)11 | CATCATCACAACCCGACAAG | CCTTAGGAGCTTCATCAGCG | 114 |
| chr2:61081487-61083514 | p3 | (ATC)9 | CATCATCACAACCCGACAAG | CCTTAGGAGCTTCATCAGCG | 114 |
| chr2:61127665-61129713 | p2 | (TA)24 | CAACCTTCACTTCCGAGTCC | ATGGCTACTGTTTCGTTGGG | 162 |
| chr2:61145792-61147810 | p3 | (TTG)6 | TGGCTTATTGCCATGTGTGT | CCCCTCCTTTCTCCAAAGTC | 127 |
| chr2:61190970-61192990 | p3 | (CAA)5 | CCAACCTAATCCGAGAAGCA | GAGGCAGTCAGAGATTTGGC | 172 |
| chr2:61190999-61193014 | p3 | (CAA)5 | CCAACCTAATCCGAGAAGCA | GAGGCAGTCAGAGATTTGGC | 143 |
| chr2:61196342-61198354 | p2 | (TA)6 | AGCTTGCCAGAGAATTCCAA | ATTGCCGGGATTCACAATTA | 267 |
| chr2:61228665-61230679 | p2 | (AT)7 | ACCGAGCACCGCTTATTCTA | TATGTGGATAGCCAAAGGGC | 111 |
| chr2:61228665-61230679 | p2 | (TA)7 | ACCGAGCACCGCTTATTCTA | TATGTGGATAGCCAAAGGGC | 111 |
| chr2:61291152-61293190 | p2 | (AT)19 | TATGGACGGGCAGCCTATAC | CCCTGTCTACCTGTTGGCAT | 128 |
| chr2:61379627-61381647 | p2 | (AT)10 | TATGCCGGAGACTATCCCTG | GGATAGGGGAAGAGGAGACG | 141 |
| chr2:61400773-61402788 | p5 | (TTTTC)3 | GCGGGTCAAACAGAGAAATC | TGCCATTGGGTTAATTTGGT | 174 |
| chr2:61413405-61415428 | p2 | (AG)6 | GAAGCGATTCACTGGAGGAG | GTTCTCGTGGTGACAGCAAA | 167 |
| chr2:61685420-61687434 | p2 | (TA)7 | CCTTACGTCACCCCGATAGA | GAAGGTCGTGGTGTATCCGT | 277 |
| chr2:61775780-61777795 | p3 | (GAT)5 | CATCCCAACCCAAGAGAAAA | AGGAATGAGCTTTGGAGCAA | 149 |
| chr2:61847473-61849491 | p6 | (TGTAAG)3 | CGATAACGATGACGACGATG | TGCAGGATCGGTGTTTCATA | 154 |
| chr2:61887489-61889505 | p2 | (TA)14 | CAGCAGACAGGTGGACTTGA | GAGTACTGTCGTGCCCGAAT | 159 |
| chr2:61887489-61889505 | p2 | (AT)8 | CAGCAGACAGGTGGACTTGA | GAGTACTGTCGTGCCCGAAT | 159 |
| chr2:61928081-61930093 | p2 | (TA)6 | ACCCGGCCATAATATCAACA | GAGAGTGGGCTGATTTGGAG | 127 |
| chr2:61935634-61937652 | p3 | (ATC)6 | GCCCACGAGGCTACAAGATA | GTAGACAGGGAGACGGTCCA | 202 |
| chr2:61948804-61950818 | p2 | (TA)7 | TCACACAATTCAACGGCATT | GAATTTGGAAGGTTTCGGGT | 218 |
| chr2:61972146-61974164 | p2 | (TA)11 | CAAGCCGACAAGGATGCTAT | TCCGATGTGGTCAGTTGTGT | 271 |
| chr2:61972146-61974164 | p6 | (CATATA)3 | CAAGCCGACAAGGATGCTAT | TCCGATGTGGTCAGTTGTGT | 271 |
| chr2:62040474-62042499 | p5 | (CAATT)5 | TTACGAACATTTCGGAAGCC | TATGGTCCCAAGGCATTGAT | 251 |
| chr2:62054426-62056438 | p2 | (AT)6 | TCATATAGGCGGGACAGGAC | ACAGGTGGACAGGGAGACAG | 212 |
| chr2:62090100-62092112 | p2 | (AG)6 | ATACGTCGAGTTATTCGCGG | CCTTGGACGTTGATGCTCTT | 228 |
| chr2:62187866-62189878 | p2 | (GT)6 | CTTTGTGGGTGCGTAAACCT | GAAGTCGATCCAAAAGGCAA | 236 |

| Marker ID | SSR type | SSR motif | FORWARD PRIMER (5'-3') | REVERSE PRIMER (5'-3') | size (bp) |
|---|---|---|---|---|---|
| chr2:62218204-62220216 | p2 | (AT)6 | TTTGTACATATGACCGGCGA | TTTGTACCCGTTTGTCACGA | 126 |
| chr2:62248474-62250489 | p3 | (ATC)5 | TTCGCTGCAGTTATTCATGC | TGCCGCAGATTTAACTCCTT | 206 |
| chr2:62338286-62340314 | p2 | (AT)14 | TCTCGGGGATAACTCCCTCT | CCCGTCACTTGCAAAGAAAT | 239 |
| chr2:62346225-62348303 | p2 | (AT)39 | GACGCAGTATTCCCTTTCCA | TGGATTGGTTGGCTTCTTTC | 124 |
| chr2:62346225-62348303 | p2 | (AT)14 | GACGCAGTATTCCCTTTCCA | TGGATTGGTTGGCTTCTTTC | 124 |
| chr2:62410904-62412916 | p2 | (CA)6 | TGCCAAAAGGAATGGTTAGC | TTTGTGATGATTGCTGGCAT | 245 |
| chr2:62532930-62534945 | p3 | (GAA)5 | CAAGGTGATGTACTGGGCCT | ATGAACTTCCAGGCCTTCCT | 154 |
| chr2:62535335-62537349 | p2 | (AT)7 | GACTAGTGCCCGAGTTGGAC | CCCATTCGTAATGACAGCCT | 131 |
| chr2:62762741-62764755 | p2 | (AT)7 | GGGTTATGACTAGTGCCCGA | CGGCTTGCGTAGTATGTTGA | 155 |
| chr2:62772030-62774045 | p3 | (ACA)5 | TGAGCAAGCAAATGCAAATC | TTTGGAACATCCTCCTACGG | 256 |
| chr2:62780606-62782660 | p2 | (AT)27 | CGTCCTTTCGCTTGTTTAGC | TTAGGCTTGCGATGATGTTG | 292 |
| chr2:62780606-62782660 | p5 | (ATTCA)3 | CGTCCTTTCGCTTGTTTAGC | TTAGGCTTGCGATGATGTTG | 292 |
| chr2:62781309-62783324 | p2 | (AT)27 | TGAACTTCGGTACCCTTTGC | AGGCTTGAATAGCGTTTGGA | 190 |
| chr2:62781309-62783324 | p5 | (ATTCA)3 | TGAACTTCGGTACCCTTTGC | AGGCTTGAATAGCGTTTGGA | 190 |
| chr2:62805174-62807189 | p3 | (ATC)5 | TTATCCGTACTGACTCCCCG | TGTAGAAAGGACGGGACAGG | 260 |
| chr2:62814061-62816073 | p2 | (AT)6 | ACTCTGCTCGTGCTTACCGT | GTGTCCCGACCCTTTAGTGA | 169 |
| chr2:62887786-62889801 | p5 | (CATTT)3 | TTGGGAAGCGGAAGAGAGTA | ATATCATTGCCTCACAGGGC | 175 |
| chr2:63010330-63012342 | p2 | (AT)6 | GGTGAACCGCCAGTTTAAGA | GCTGCCACTGTAGACGAACA | 204 |
| chr2:63105936-63107950 | p2 | (TA)7 | TGGGAGCGAATTGAGAAATC | CGTCCTCCTCTGAAGCTCAC | 257 |
| chr2:63138146-63140158 | p2 | (GT)6 | CCGGAGGTGCTCATGTTTAT | ACGAAAGCACGGGACATAAC | 220 |
| chr2:63140185-63142287 | p2 | (AT)51 | ACTCCGCTCGTGCCTTATTA | GCCACACACAGCATAACACC | 250 |
| chr2:63140185-63142287 | p2 | (AT)6 | ACTCCGCTCGTGCCTTATTA | GCCACACACAGCATAACACC | 250 |
| chr2:63174717-63176733 | p2 | (AT)18 | CAGGAAGAAATGGGGACAGA | AAGGATCACACCTTCATCGC | 181 |
| chr2:63174717-63176733 | p2 | (TA)8 | CAGGAAGAAATGGGGACAGA | AAGGATCACACCTTCATCGC | 181 |
| chr2:63190781-63192797 | p2 | (AT)18 | CAGGAAGAAATGGGGACAGA | AAGGATCACACCTTCATCGC | 181 |
| chr2:63190781-63192797 | p2 | (TA)8 | CAGGAAGAAATGGGGACAGA | AAGGATCACACCTTCATCGC | 181 |
| chr2:63200563-63202577 | p2 | (AG)7 | ACATTCGAGGACGAATGCTC | CCTTGCACGTTGATTTCCTT | 164 |
| chr2:63337355-63339385 | p4 | (TGTA)4 | GATTTGTGTATGGGGATGGC | AGTATGATCCAAACGGACGG | 227 |
| chr2:63337355-63339385 | p2 | (TA)15 | GATTTGTGTATGGGGATGGC | AGTATGATCCAAACGGACGG | 227 |
| chr2:63341460-63343472 | p2 | (GT)6 | ACAGGTTGCTACAATTCGGG | TGCAGCGCGCTATAACATAC | 281 |
| chr2:63341460-63343472 | p6 | (TCATTA)3 | ACAGGTTGCTACAATTCGGG | TGCAGCGCGCTATAACATAC | 281 |
| chr2:63447883-63449897 | p2 | (GT)7 | ACCCCCATTTATTCCCTACG | TAACACGAAAACACGGGACA | 162 |
| chr2:63508581-63510593 | p2 | (AT)6 | GACGAGGCCAACAAAATGAT | CTTGACCCATAGCTGCCAAT | 251 |
| chr2:63508581-63510593 | p5 | (GAAAT)3 | GACGAGGCCAACAAAATGAT | CTTGACCCATAGCTGCCAAT | 251 |
| chr2:63510770-63512782 | p2 | (TC)6 | ACCCCACCTTGCCTTATCTT | ATTGAGGCCGTGTAGTGGAC | 110 |
| chr2:63584311-63586326 | p5 | (GTTGG)3 | GATGAATGAGGAAAGCGAGC | GAATGCAGGCTTTGGAATGT | 183 |

| Marker ID | SSR type | SSR motif | FORWARD PRIMER (5'-3') | REVERSE PRIMER (5'-3') | size (bp) |
|---|---|---|---|---|---|
| chr2:63618094-63620106 | p2 | (AT)6 | TTTGGTTTGGAAATGGGAAG | GTAAATACAACGCCCCGAGA | 166 |
| chr2:63730707-63732721 | p2 | (AT)7 | GCTGATTATTGCCCTCGGTA | CTATTAGGGTCAGGTGCCCA | 154 |
| chr2:63730747-63732763 | p2 | (AT)7 | GCTGATTATTGCCCTCGGTA | CTATTAGGGTCAGGTGCCCA | 114 |
| chr2:63778064-63780124 | p2 | (AT)30 | AACATATTGCTGGACAGGGC | TTCATGCTTTACATGCTCGG | 110 |
| chr2:63843409-63845427 | p3 | (TTG)6 | GACCCTTGAGCCAAATCAAA | ATAGGTCATGGGCCTAACCC | 158 |
| chr2:63846422-63848438 | p4 | (AGAA)4 | TCTGCAAAGTTGTGTCCAGC | CGCACGATCTGAAACAGAAA | 206 |
| chr2:63846422-63848438 | p2 | (TG)6 | TCTGCAAAGTTGTGTCCAGC | CGCACGATCTGAAACAGAAA | 206 |
| chr2:63846422-63848438 | p3 | (TTG)6 | TCTGCAAAGTTGTGTCCAGC | CGCACGATCTGAAACAGAAA | 206 |
| chr2:63874856-63876876 | p2 | (AT)10 | CGCTTCCGGCTCAATAATAA | TTTCAAAACCACTCCGAAGG | 146 |
| chr2:63884893-63886909 | p2 | (TA)8 | ATCGTGAGACCATGCTACCC | TGTTCATGCCCACAGGTAGA | 241 |
| chr2:63884893-63886909 | p6 | (ATTATC)3 | ATCGTGAGACCATGCTACCC | TGTTCATGCCCACAGGTAGA | 241 |
| chr2:63887518-63889548 | p2 | (AT)6 | GTCGAATCACCAAAGCGAAT | TTCATGCTTTACATGCTCGG | 232 |
| chr2:63887518-63889548 | p2 | (TA)15 | GTCGAATCACCAAAGCGAAT | TTCATGCTTTACATGCTCGG | 232 |
| chr2:63934877-63936889 | p2 | (AG)6 | GATTGACGAAATCAGCCCAT | GGGCACCTCGGTCAGTAATA | 160 |
| chr2:63987617-63989635 | p2 | (AT)9 | ACTTTTGTCACGACCCAACC | TCGTGGGTCATGTTTGCTTA | 163 |
| chr2:64082116-64084131 | p5 | (AAATG)3 | GTCGTTCCGCTAGGCTATTG | GACGATGCGTTTGTAGACGA | 270 |
| chr2:64174682-64176700 | p6 | (GAATAT)3 | ACACGGCTAGAACACCCATC | TGTACCCATTTGCAAACCCT | 139 |
| chr2:64176488-64178503 | p5 | (AAATA)3 | TTATCGTAGCGCCGGTTACT | CCATAAAAGGCCACGTAGGA | 209 |
| chr2:64176488-64178503 | p5 | (CATAT)3 | TTATCGTAGCGCCGGTTACT | CCATAAAAGGCCACGTAGGA | 209 |
| chr2:64177122-64179137 | p5 | (AAATA)3 | CCCAATTCCACTAGGGTGTG | GGCTTCATCAGCCGATATGT | 265 |
| chr2:64177122-64179137 | p5 | (CATAT)3 | CCCAATTCCACTAGGGTGTG | GGCTTCATCAGCCGATATGT | 265 |
| chr2:64180947-64182959 | p2 | (CT)6 | CTTCGCACTTCCAAACCAAT | ACCTTGAGGAGTTTTCGGGT | 294 |
| chr2:64184420-64186435 | p5 | (AAATA)3 | TTGCAGACAACCCCTAGGAC | GGCTTCATCAGCCGATATGT | 233 |
| chr2:64184420-64186435 | p5 | (CATAT)3 | TTGCAGACAACCCCTAGGAC | GGCTTCATCAGCCGATATGT | 233 |
| chr2:64256711-64258723 | p2 | (TA)6 | TCTTTCTGACGGGCCATATC | TGTCTCACTGACCAAGTGGC | 160 |
| chr2:64338845-64340860 | p3 | (TTG)6 | TTAAGAATTCGGGCTAGGCA | ATAGAATCGTCCCCAAGGCT | 152 |
| chr2:64338845-64340860 | p5 | (GATAT)3 | TTAAGAATTCGGGCTAGGCA | ATAGAATCGTCCCCAAGGCT | 152 |
| chr2:64393890-64395904 | p2 | (AC)7 | ATGCCAAATGAAAGAGTGGG | GTGAAGCCGAGCTAAGTTGG | 257 |
| chr2:64467207-64469222 | p5 | (CATAT)3 | GGAAAAATGTCACGCCCTAA | CTTCCGTGCGCTATATCCAT | 287 |
| chr2:64545195-64547211 | p2 | (TA)8 | AATCCAAACCAAGTTTCCCC | CCCAACTTCCGTCCACTAGA | 243 |
| chr2:64566404-64568416 | p2 | (TA)6 | CTTAACAAAGGGTCGGGACA | GGTGAATGCCTTATCCGAGA | 278 |
| chr2:64587554-64589570 | p4 | (CATA)4 | TATACAAGTCGGGACAGGGC | ATGCCCGCAGGTATAGATTG | 216 |
| chr2:64590976-64593034 | p2 | (AT)29 | TGGTGAACCACGACCACTTA | TGACTCCTCGGGAGACAGTT | 125 |
| chr2:64600288-64602392 | p2 | (TA)52 | TATAGGGGAAGTGCTGTCCG | TATTCGTGAATCCTTTCGCC | 105 |
| chr2:64640088-64642120 | p6 | (ATATGA)4 | AGTTCGGTACGGATAACCCC | GACTATGGCTAGGCTCGGTG | 286 |
| chr2:64681719-64683735 | p4 | (TCTG)4 | TTTTGTTGTTGTGTCCGCAT | ACGATTGCCTTTCATATCGG | 119 |

| Marker ID | SSR type | SSR motif | FORWARD PRIMER (5'-3') | REVERSE PRIMER (5'-3') | size (bp) |
|---|---|---|---|---|---|
| chr2:64683172-64685190 | p6 | (TGTATG)3 | GGTTGGGTCGTGACAGAAGT | AGCTGGCTCTATCGCATGAT | 231 |
| chr2:64692641-64694673 | p6 | (ATATGA)4 | AGTTCGGTACGGATAACCCC | GACTATGGCTAGGCTCGGTG | 286 |
| chr2:64734272-64736288 | p4 | (TCTG)4 | TTTTGTTGTTGTGTCCGCAT | ACGATTGCCTTTCATATCGG | 119 |
| chr2:64766032-64768056 | p2 | (TA)12 | ACAAATCCAAGAGTGGCCTG | CCTCTCCCAAGGATTGATGA | 276 |
| chr2:64819177-64821201 | p2 | (TA)12 | ACAAATCCAAGAGTGGCCTG | CCTCTCCCAAGGATTGATGA | 276 |
| chr2:64847627-64849651 | p2 | (TA)12 | GTTGATGTTTTGGCCGAGTT | TTACATACCTTGCCCGCTTC | 275 |
| chr2:65001422-65003434 | p3 | (ATG)5 | TCGGCAGATGCTAAGAGGAT | CTTCCGGTCTCGCATTCTAC | 273 |
| chr2:65001422-65003434 | p2 | (TA)9 | TCGGCAGATGCTAAGAGGAT | CTTCCGGTCTCGCATTCTAC | 273 |
| chr2:65001422-65003434 | p2 | (TA)6 | TCGGCAGATGCTAAGAGGAT | CTTCCGGTCTCGCATTCTAC | 273 |
| chr2:65004367-65006379 | p2 | (AC)6 | ATTCAAAAGGAGAGCCCGAT | GAAACCCGATCCACAGAAAA | 137 |
| chr2:65142283-65144307 | p2 | (AT)12 | AGCCGACAAGGCTATCAAAA | TGCACAGGGAGACAGACTTG | 232 |
| chr2:65169874-65171898 | p2 | (AT)12 | AGCCGACAAGGCTATCAAAA | TGCACAGGGAGACAGACTTG | 232 |
| chr2:65207862-65209877 | p5 | (TATTT)3 | GACCGCGGTATCTACGACAT | TCCCGGAAGACAGATACGAC | 256 |
| chr2:65236616-65238631 | p5 | (ACACA)3 | ACGGCCAACTTAGACCACAC | TTGGACGAATGAGGTGTTGA | 259 |
| chr2:65483671-65485707 | p2 | (AT)18 | TGCCTGCTTTAGAGGAGCAT | ACTATGACCGTACCTTGCCG | 211 |
| chr2:65483671-65485707 | p2 | (TA)6 | TGCCTGCTTTAGAGGAGCAT | ACTATGACCGTACCTTGCCG | 211 |
| chr2:65534641-65536653 | p2 | (AC)6 | CGAAACCATATGAAACGGCT | TGGTGCGCATTAATGTTGTT | 187 |
| chr2:65564238-65566253 | p3 | (CAA)5 | CACAAATGTATTTGCTGCGG | CACAAGGAGTCGGATACCGT | 105 |
| chr2:65589121-65591133 | p2 | (AT)6 | GCTGCAAATCAATGGGAAGT | GTGCCCAGTTCGGGTACTTA | 259 |
| chr2:65592039-65594054 | p5 | (ATTTC)3 | ATCTACCCGAATGCCATCAG | TGCAGATCACTTGAGAACGG | 201 |
| chr2:65603916-65605932 | p2 | (AT)7 | GAGCAACACGCTGTAGCAAG | GCCTGTAGGCCGTTGATTTA | 133 |
| chr2:65603916-65605932 | p2 | (TA)8 | GAGCAACACGCTGTAGCAAG | GCCTGTAGGCCGTTGATTTA | 133 |
| chr2:65603916-65605932 | p2 | (TA)8 | GAGCAACACGCTGTAGCAAG | GCCTGTAGGCCGTTGATTTA | 133 |
| chr2:65603916-65605932 | p2 | (TA)8 | GAGCAACACGCTGTAGCAAG | GCCTGTAGGCCGTTGATTTA | 133 |
| chr2:65603916-65605932 | p2 | (TA)8 | GAGCAACACGCTGTAGCAAG | GCCTGTAGGCCGTTGATTTA | 133 |
| chr2:65855300-65857320 | p2 | (TA)10 | CGATTAGCTACGCTCCAAGG | TTTCACAACTACGTGCCTGC | 152 |
| chr2:65880894-65882914 | p2 | (AT)10 | TGTGGCCCTCATAAAGTTCC | CCCGTCACTTGCAAAGAAAT | 292 |
| chr2:66050594-66052610 | p2 | (AT)8 | TTTCGGCCATATCTCCTTTG | ATTGGGTTCGATGTTTCCAA | 210 |
| chr2:66074917-66076932 | p6 | (CATATA)3 | TGTATGTACCTGCGTGGCAT | CACGCATGATTTCATTCACC | 172 |
| chr2:66074917-66076932 | p5 | (ATCAA)3 | TGTATGTACCTGCGTGGCAT | CACGCATGATTTCATTCACC | 172 |
| chr2:66105084-66107112 | p2 | (TA)14 | ATCCAAGCCAAAATCCACAG | GAGCTCAGGAAACATAGCGG | 219 |
| chr2:66177705-66179721 | p5 | (AAAGA)3 | TTAAGTTTCCGCAAAATGCC | CTAGAGAAATCCCGGGTGTG | 275 |
| chr2:66177723-66179739 | p5 | (AAAGA)3 | TTAAGTTTCCGCAAAATGCC | CTAGAGAAATCCCGGGTGTG | 257 |
| chr2:66177741-66179757 | p5 | (AAAGA)3 | TTAAGTTTCCGCAAAATGCC | CTAGAGAAATCCCGGGTGTG | 239 |
| chr2:66177759-66179775 | p5 | (AAAGA)3 | TTAAGTTTCCGCAAAATGCC | CTAGAGAAATCCCGGGTGTG | 221 |
| chr2:66177777-66179793 | p5 | (AAAGA)3 | TTAAGTTTCCGCAAAATGCC | CTAGAGAAATCCCGGGTGTG | 203 |

| Marker ID | SSR type | SSR motif | FORWARD PRIMER (5'-3') | REVERSE PRIMER (5'-3') | size (bp) |
|---|---|---|---|---|---|
| chr2:66177795-66179811 | p5 | (AAAGA)3 | TTAAGTTTCCGCAAAATGCC | CTAGAGAAATCCCGGGTGTG | 185 |
| chr2:66177813-66179829 | p5 | (AAAGA)3 | TTAAGTTTCCGCAAAATGCC | CTAGAGAAATCCCGGGTGTG | 167 |
| chr2:66177831-66179847 | p5 | (AAAGA)3 | TTAAGTTTCCGCAAAATGCC | CTAGAGAAATCCCGGGTGTG | 149 |
| chr2:66177849-66179865 | p5 | (AAAGA)3 | TTAAGTTTCCGCAAAATGCC | CTAGAGAAATCCCGGGTGTG | 131 |
| chr2:66177867-66179883 | p5 | (AAAGA)3 | TTAAGTTTCCGCAAAATGCC | CTAGAGAAATCCCGGGTGTG | 113 |
| chr2:66188567-66190589 | p2 | (TA)11 | CACCACGTTCCTGTGCTCTA | ATTATGGTTCAGGCGGACAG | 208 |
| chr2:66188567-66190589 | p5 | (AAACA)3 | CACCACGTTCCTGTGCTCTA | ATTATGGTTCAGGCGGACAG | 208 |
| chr2:66222416-66224430 | p2 | (AT)7 | TCATGCCTTTGAAAGAAGGG | GAAACAAAGGAAATGCTGCC | 210 |
| chr2:66336315-66338327 | p2 | (TG)6 | ACCCAAGCTCCTAAAAGGGA | TCTCGAATTATCCGATTGCC | 167 |
| chr2:66353207-66355247 | p2 | (AT)20 | GTACTGGCGGAACGATCACT | AGAAGGGCAAAGCTACGACA | 228 |
| chr2:66364088-66366103 | p6 | (CCCCCT)3 | GGCCCAATTGGCATAGTTTA | CATTGAAATGACACCAACGC | 181 |
| chr2:66364088-66366103 | p5 | (TATGA)3 | GGCCCAATTGGCATAGTTTA | CATTGAAATGACACCAACGC | 181 |
| chr2:66370409-66372421 | p2 | (TG)6 | CCCGTCTCTATTTGCCATGT | TTTTTCCCCTAGCCTTCGTT | 271 |
| chr2:66416973-66418991 | p2 | (AT)6 | ACTTCTGTCACGACCCAACC | CAGGTTAATGATGTGCGACG | 240 |
| chr2:66416973-66418991 | p2 | (AT)9 | ACTTCTGTCACGACCCAACC | CAGGTTAATGATGTGCGACG | 240 |
| chr2:66502741-66504767 | p2 | (AT)13 | TGGCGGGTTCAAATATTGTT | TATAAATGCGTTGTTGCCCA | 246 |
| chr2:66507326-66509340 | p2 | (AT)7 | TTTCATCTCCTTTGGCCATC | AATTCCCGGATTGTGATTGA | 291 |
| chr2:66521419-66523455 | p2 | (AT)18 | TCGCACGATCTCGTAAGATG | GCTGGATTTGTACGATGGGT | 246 |
| chr2:66532187-66534199 | p6 | (CAACAT)3 | ACATGCCAAAAGAAGGATCG | GTGATGGTTTTGAGCCGAGT | 169 |
| chr2:66532187-66534199 | p2 | (AC)6 | ACATGCCAAAAGAAGGATCG | GTGATGGTTTTGAGCCGAGT | 169 |
| chr2:66602031-66604045 | p2 | (TA)7 | CTCGCTTCGCCTAACAAATC | TTTACAGGGGAGATGCTGCT | 174 |
| chr2:66602031-66604045 | p2 | (CT)8 | CTCGCTTCGCCTAACAAATC | TTTACAGGGGAGATGCTGCT | 174 |
| chr2:66672556-66674568 | p2 | (AT)6 | GACGAGGCCAACAAAATGAT | GGTTGTATAGCAGAGCCCCA | 256 |
| chr2:66719728-66721740 | p2 | (AT)10 | TTGCAAACAATGGTTCGTGT | GGCCCTTTAATGAGGGACTC | 178 |
| chr2:66719728-66721740 | p2 | (TG)6 | TTGCAAACAATGGTTCGTGT | GGCCCTTTAATGAGGGACTC | 178 |
| chr2:66754087-66756101 | p2 | (TA)7 | CTGTGGCCTTATTGGATCGT | ACCTTTTTCTTCCTGCCCAT | 275 |
| chr2:66762376-66764390 | p2 | (AT)7 | CCGACCTCTTCTGATCAAGC | AGTTATTGGCACCGAGTTGG | 150 |
| chr2:66762376-66764390 | p3 | (CAA)5 | CCGACCTCTTCTGATCAAGC | AGTTATTGGCACCGAGTTGG | 150 |
| chr2:66763014-66765029 | p2 | (AT)7 | TAACGTACCCAGCCTTTTCG | CGGAGGTTACCGACCTTACA | 249 |
| chr2:66763014-66765029 | p3 | (CAA)5 | TAACGTACCCAGCCTTTTCG | CGGAGGTTACCGACCTTACA | 249 |
| chr2:66823730-66825750 | p2 | (TA)10 | ACGACCAAGGTATGTGAGGC | TGCCTCACCTTGAAAGGACT | 126 |
| chr2:66823730-66825750 | p4 | (TATT)5 | ACGACCAAGGTATGTGAGGC | TGCCTCACCTTGAAAGGACT | 126 |
| chr2:66854261-66856275 | p2 | (AT)7 | CCGACCTCTTCTGATCAAGC | AGTTATTGGCACCGAGTTGG | 150 |
| chr2:66854261-66856275 | p3 | (CAA)5 | CCGACCTCTTCTGATCAAGC | AGTTATTGGCACCGAGTTGG | 150 |
| chr2:66854899-66856914 | p2 | (AT)7 | TAACGTACCCAGCCTTTTCG | CGGAGGTTACCGACCTTACA | 249 |
| chr2:66854899-66856914 | p3 | (CAA)5 | TAACGTACCCAGCCTTTTCG | CGGAGGTTACCGACCTTACA | 249 |

| Marker ID | SSR type | SSR motif | FORWARD PRIMER (5'-3') | REVERSE PRIMER (5'-3') | size (bp) |
|---|---|---|---|---|---|
| chr2:66915615-66917635 | p2 | (TA)10 | ACGACCAAGGTATGTGAGGC | TGCCTCACCTTGAAAGGACT | 126 |
| chr2:66915615-66917635 | p4 | (TATT)5 | ACGACCAAGGTATGTGAGGC | TGCCTCACCTTGAAAGGACT | 126 |
| chr2:67008407-67010419 | p4 | (GAAA)4 | GAAACGATGGCGTTTTGTTT | AACCCCACTTCCCTTTGTTC | 246 |
| chr2:67008407-67010419 | p2 | (TA)6 | GAAACGATGGCGTTTTGTTT | AACCCCACTTCCCTTTGTTC | 246 |
| chr2:67097289-67099307 | p6 | (AAAAAG)3 | AAGGAGATTCCAGCTCAGCA | TCATGAACTGAGTCGCCAAC | 291 |
| chr2:67131749-67133764 | p5 | (ATATA)3 | TGACAAGGCTGCCACTACAG | GGTGACCCTTCAGCATAGGA | 248 |
| chr2:67235616-67237628 | p2 | (GT)6 | CAAGCGAAAGACGTGGTACA | TCGAATTTTCCGAGTATGCC | 103 |
| chr2:67305658-67307673 | p5 | (ATATC)3 | CCCTGCCAAACCTCTGTCTA | ATATGGGATAGGGGGTGCTC | 154 |
| chr2:67318561-67320579 | p2 | (TA)9 | TGGACTTGGTATGGCATGAA | TTCCCTGAATCTCCCAACAG | 272 |
| chr2:67318561-67320579 | p2 | (AT)9 | TGGACTTGGTATGGCATGAA | TTCCCTGAATCTCCCAACAG | 272 |
| chr2:67337154-67339174 | p2 | (AT)10 | AGGTGATGACGGATACTCGG | GGAAACTCGAGTGGAAGCTG | 263 |
| chr2:67349287-67351305 | p2 | (TA)9 | AGATTGTTATCGGTGGACGC | GGCCCTTTTAGAACTGAGGC | 245 |
| chr2:67429834-67431849 | p5 | (AAAAT)3 | TTGGTGATAAGCAAATGGCA | CCGTGCTTATAGAAAGGGCA | 234 |
| chr2:67510245-67512281 | p2 | (AT)13 | CTATAGGTCTGCGCCGTCTC | AAGGTCACACAGGTTTTGCC | 204 |
| chr2:67510245-67512281 | p2 | (TA)18 | CTATAGGTCTGCGCCGTCTC | AAGGTCACACAGGTTTTGCC | 204 |
| chr2:67549073-67551088 | p3 | (ATC)5 | AGCCGACAAGGCTAAACAAA | CAGGGACTACATAGCGGAGC | 101 |
| chr2:67610021-67612039 | p2 | (TA)16 | AGCCCTTCGGCTATCAAAAT | TCCTGTCCCGTCCATATGTT | 283 |
| chr2:67610021-67612039 | p2 | (AT)9 | AGCCCTTCGGCTATCAAAAT | TCCTGTCCCGTCCATATGTT | 283 |
| chr2:67632650-67634668 | p2 | (AT)9 | ATGTCGAAGGGGATGATGAG | ACCCATTCCTCGTTAGCCTT | 236 |
| chr2:67700260-67702296 | p2 | (AT)18 | TGCGTAATCGGGTTCCTATC | AGAATGGACCTGCGAGCTTA | 147 |
| chr2:67790827-67792841 | p2 | (AT)7 | TCGGGATTTGATTAGCGAAC | ATGGAGATCAAAGACGTCGG | 124 |
| chr2:67810759-67812774 | p5 | (TGATA)3 | ATGGAACATCCCTAAGCCCT | CAATAAGAGAGCAAAGCGGG | 267 |
| chr2:67898319-67900337 | p3 | (TGT)6 | AAGCCCTCCAAAAGGATGAT | TTTGGCAGCTTGGAGTCTTT | 190 |
| chr2:67898319-67900337 | p2 | (TA)15 | AAGCCCTCCAAAAGGATGAT | TTTGGCAGCTTGGAGTCTTT | 190 |
| chr2:67901751-67903767 | p2 | (TA)8 | TCTGAGAGCACAATGGTTCG | GGCGAACATACCTTCATCGT | 250 |
| chr2:67923245-67925260 | p5 | (ATTCA)3 | CGACACATGAAGGCTCAGAA | GATGATGCCAAGGAGGAGAA | 224 |
| chr2:67923245-67925260 | p2 | (AT)6 | CGACACATGAAGGCTCAGAA | GATGATGCCAAGGAGGAGAA | 224 |
| chr2:67932421-67934436 | p5 | (ATTCA)3 | CGACACATGAAGGCTCAGAA | GATGATGCCAAGGAGGAGAA | 224 |
| chr2:67932421-67934436 | p2 | (AT)6 | CGACACATGAAGGCTCAGAA | GATGATGCCAAGGAGGAGAA | 224 |
| chr2:68026905-68028917 | p2 | (AT)6 | ACATTCAGCCCCTTTCTCCT | AAGCGATAAAGGATTGGCCT | 114 |
| chr2:68028424-68030448 | p2 | (AT)12 | TGTTTCTGAGTCGAGAGCGA | ACGTGAAGGGCAAAAATGTC | 181 |
| chr2:68120816-68122834 | p6 | (CATATA)3 | GAGGCTGCCACTACACATCA | CGCTATAGAAGCCTTGTCGG | 191 |
| chr2:68171094-68173106 | p2 | (TG)6 | AAATCTGTCCCGACACGAAC | CAAACGAGTCGACGAGAACA | 271 |
| chr2:68171094-68173106 | p2 | (TA)6 | AAATCTGTCCCGACACGAAC | CAAACGAGTCGACGAGAACA | 271 |
| chr2:68171255-68173267 | p2 | (TG)6 | AAATCTGTCCCGACACGAAC | CAAACGAGTCGACGAGAACA | 110 |
| chr2:68171255-68173267 | p2 | (TA)6 | AAATCTGTCCCGACACGAAC | CAAACGAGTCGACGAGAACA | 110 |

| Marker ID | SSR type | SSR motif | FORWARD PRIMER (5'-3') | REVERSE PRIMER (5'-3') | size (bp) |
|---|---|---|---|---|---|
| chr2:68216984-68218999 | p5 | (TTTAA)3 | CATGTGTGCCCAGTTCAGAC | AATAAGGCCAACCTGCACAC | 234 |
| chr2:68304177-68306192 | p3 | (GAT)5 | TTTCCCTTTTCACTCCCCTT | GATTCCTTGGCTTGGATGAA | 270 |
| chr2:68440044-68442062 | p6 | (ATATGT)3 | GTGTTGTATCGCAGCCCTTT | AACCTTGATGGAAGCAATGG | 299 |
| chr2:68463149-68465207 | p2 | (TG)6 | TCCGCTGGTTTCTAATGTCC | GCTCTTCAACTTCTCCCACG | 281 |
| chr2:68463209-68465223 | p2 | (TG)6 | TCCGCTGGTTTCTAATGTCC | GCTCTTCAACTTCTCCCACG | 221 |
| chr2:68463251-68465285 | p2 | (TG)6 | TCCGCTGGTTTCTAATGTCC | GCTCTTCAACTTCTCCCACG | 179 |
| chr2:68635620-68637690 | p2 | (AT)35 | CGATGGCCAGGAAAATTCTA | CGCCTAGAATCCAAACCAAA | 289 |
| chr2:68635620-68637690 | p2 | (TA)10 | CGATGGCCAGGAAAATTCTA | CGCCTAGAATCCAAACCAAA | 289 |
| chr2:68674267-68676279 | p2 | (AG)6 | GTGATTTTATGTTGCCCGCT | CTCATCCCTTTGCTAGCCTG | 187 |
| chr2:68675898-68677914 | p2 | (AT)6 | CGTGTTGTTGTTTTGGTTGG | ATTTAAACATGACGCGAGGG | 125 |
| chr2:68805708-68807720 | p2 | (TC)6 | TACGCACACGCATCGTATTT | AAATGGCGTTGTAATAGCCG | 232 |
| chr2:68805708-68807720 | p2 | (TA)6 | TACGCACACGCATCGTATTT | AAATGGCGTTGTAATAGCCG | 232 |
| chr2:68874961-68876976 | p5 | (AATAA)3 | CGCCATAGACCAACCAAAGT | GAAACTGAGCGACAGTGCAG | 201 |
| chr2:68949770-68951812 | p2 | (AT)21 | CGTTGTTCCATTTTGTCGTG | ACAACATATCCGATTTCCGC | 247 |
| chr2:68949770-68951812 | p5 | (TGCAT)3 | CGTTGTTCCATTTTGTCGTG | ACAACATATCCGATTTCCGC | 247 |
| chr2:68949770-68951812 | p2 | (TA)6 | CGTTGTTCCATTTTGTCGTG | ACAACATATCCGATTTCCGC | 247 |
| chr2:68956378-68958392 | p2 | (AT)7 | CTCCGGGTATATCAGGGGTT | GGTCACCACATAAGTGGGCT | 127 |
| chr2:68992095-68994123 | p3 | (ATC)5 | GAAGCCGACCTCTTCATCTG | GGTTGGGTCGTGACAAAAGT | 149 |
| chr2:68992624-68994639 | p3 | (ATC)5 | ACAGGAATTCGTGAAGACGG | CTTAGGAGCTTCATCAGCGG | 150 |
| chr2:68992624-68994639 | p2 | (TA)6 | ACAGGAATTCGTGAAGACGG | CTTAGGAGCTTCATCAGCGG | 150 |
| chr2:69004141-69006153 | p2 | (TA)6 | ACCATCTGATTTGGGAGCAG | ACCTGACGATCCAGTATGGC | 266 |
| chr2:69058921-69060936 | p4 | (ATAA)4 | ACATCCGGAAACGTATCGAC | TTAGCGAGCGATCGAGGTAT | 300 |
| chr2:69058921-69060936 | p3 | (CAA)5 | ACATCCGGAAACGTATCGAC | TTAGCGAGCGATCGAGGTAT | 300 |
| chr2:69058921-69060936 | p2 | (CT)7 | ACATCCGGAAACGTATCGAC | TTAGCGAGCGATCGAGGTAT | 300 |
| chr2:69117742-69119757 | p5 | (ATTTC)3 | AAGAATTGACATGGGAAGCG | AACAGGGCACTACCATCCTG | 156 |
| chr2:69122089-69124109 | p2 | (AT)10 | CAACCTGTGCTGTCACGACT | GTTTTGGCAATTTCCTGCAT | 287 |
| chr2:69130327-69132347 | p2 | (TA)10 | AGACGCTAGCATTGGAAGGA | CGTCATATGAGGGGCGTAGT | 220 |
| chr2:69206717-69208785 | p2 | (AT)34 | ACCGACCTTACATCACTCCG | GGGGTAATCCGTACCCAACT | 283 |
| chr2:69223222-69225237 | p3 | (ATC)5 | AGCCGACAAGGCTAACAAAA | TCCATTTCATTCGGAACCAT | 141 |
| chr2:69284234-69286248 | p2 | (TA)7 | AACGAGTGTAAGGCGCAAGT | AAGTTTCAAGGGGATTCGGT | 130 |
| chr2:69290427-69292455 | p2 | (AT)14 | TTCCATATTTCACAACGCCA | TTTGACCAAAATCTGAGCCC | 153 |
| chr2:69314045-69316115 | p2 | (AT)35 | ATAAGATGCCAACTCCACCG | ATGTGGGCCCAAGTGATTAG | 171 |
| chr2:69341309-69343379 | p2 | (AT)35 | ATAAGATGCCAACTCCACCG | ATGTGGGCCCAAGTGATTAG | 171 |
| chr2:69368011-69370027 | p2 | (AT)17 | CTGGAATTGTGCGTAGGGTT | TTTTGTCAAGTCTTTGGGGC | 214 |
| chr2:69368011-69370027 | p2 | (TA)50 | CTGGAATTGTGCGTAGGGTT | TTTTGTCAAGTCTTTGGGGC | 214 |
| chr2:69368011-69370027 | p2 | (TA)8 | CTGGAATTGTGCGTAGGGTT | TTTTGTCAAGTCTTTGGGGC | 214 |

| Marker ID | SSR type | SSR motif | FORWARD PRIMER (5'-3') | REVERSE PRIMER (5'-3') | size (bp) |
|---|---|---|---|---|---|
| chr2:69368011-69370027 | p2 | (TA)9 | CTGGAATTGTGCGTAGGGTT | TTTTGTCAAGTCTTTGGGGC | 214 |
| chr2:69565684-69567708 | p3 | (TAT)20 | AATGGGGACACCGATTCATA | GGAGTTGCGTATGACGCTTT | 104 |
| chr2:69575074-69577086 | p2 | (CA)6 | ATTCCCCCAAAGGTTGATTC | CGATGAGTTCGTCATTCCCT | 165 |
| chr2:69692575-69694590 | p5 | (CATAT)3 | GTCCCTGATGTGGCAAGTTT | TGTCAGTTGTGTAAAGCGGC | 262 |
| chr2:69770091-69772106 | p3 | (AAC)5 | ATTCCCAGCATTTCTTGACG | TCTCGCTGTGCATTTACCAG | 292 |
| chr2:69783109-69785125 | p4 | (TATT)4 | GACCAGCCGTGTATCCCTTA | GCCCCAAAAGTATTTCACGA | 226 |
| chr2:69783109-69785125 | p3 | (ATA)17 | GACCAGCCGTGTATCCCTTA | GCCCCAAAAGTATTTCACGA | 226 |
| chr2:69873971-69875986 | p5 | (TCATT)3 | TCGGAGATATGGAAGGCAAG | CCCCTCACAGATTTAGGCAA | 108 |
| chr2:69991193-69993208 | p5 | (TGTGG)3 | TGTACGATAAGGGGGTGCTC | AAGCTGAGGAGGCAATGAGA | 130 |
| chr2:70070633-70072687 | p2 | (GA)27 | CAGCAGATACGCCAACAAGA | CTCTCTCACCTGATGAGCCC | 180 |
| chr2:70072528-70074543 | p3 | (ATA)5 | GATATACGGCCAAGATGCGT | TCTTGACCTTGCATTTTCCC | 154 |
| chr2:70104323-70106335 | p2 | (AT)7 | GATTTTGAGGAGCGGAAATG | GTTTGGGGAGCGTAATCGTA | 104 |
| chr2:70104323-70106335 | p2 | (CA)6 | GATTTTGAGGAGCGGAAATG | GTTTGGGGAGCGTAATCGTA | 104 |
| chr2:70134198-70136218 | p5 | (ATATC)4 | CATCGAATAGCTGAGGCTCC | GACGATAAGCTTGCGACCTC | 180 |
| chr2:70352894-70354906 | p3 | (CAA)5 | CGAAACCATATGAAACGGCT | TGGTGCGCATTAATGTTGTT | 144 |
| chr2:70352894-70354906 | p2 | (CA)6 | CGAAACCATATGAAACGGCT | TGGTGCGCATTAATGTTGTT | 144 |
| chr2:70358750-70360762 | p3 | (CAA)5 | CGAAACCATATGAAACGGCT | TGGTGCGCATTAATGTTGTT | 105 |
| chr2:70358750-70360762 | p2 | (CA)6 | CGAAACCATATGAAACGGCT | TGGTGCGCATTAATGTTGTT | 105 |
| chr2:70389935-70391971 | p2 | (AT)18 | GTTCTTCCAAATTCTGGGCA | TTGGATAGTTATGCTCCGGC | 131 |
| chr2:70417121-70419153 | p2 | (AT)16 | ATGTGGCTGGGCTGATTTAC | CCCGAAATCACAATACCGAC | 173 |
| chr2:70484136-70486160 | p2 | (TA)12 | TAAACATTGCGGGATCATCA | TCGATACGTTGAGCGACAAG | 226 |
| chr2:70528247-70530259 | p2 | (CA)6 | ACATGCCAAAGGAAGGAATG | TGTTGATGTTGCTGGGTTGT | 142 |
| chr2:70539867-70541882 | p5 | (TGATA)3 | CGATACAAGTCTCGGGGAAA | TATAGGGTCACCCACGAAGG | 233 |
| chr2:70544531-70546637 | p2 | (TA)53 | CCAATAATCTCCAGCCTCCA | CGCACAGCCAGAGTACAAAA | 225 |
| chr2:70598195-70600207 | p2 | (TA)6 | TAAGAGTCTGTGGGTCCGCT | TTTCACAAGGATCATGCCAA | 194 |
| chr2:70664801-70666823 | p2 | (AT)11 | CCAAAGGGGGTTAAGGATGT | CGGGATACTTAAGGCAACCA | 173 |
| chr2:70671895-70673917 | p2 | (AT)6 | TCGAGGGTTCTCGACCATAC | GTTGTGACACATGTCCGTCC | 272 |
| chr2:70671895-70673917 | p2 | (AT)11 | TCGAGGGTTCTCGACCATAC | GTTGTGACACATGTCCGTCC | 272 |
| chr2:70671895-70673917 | p3 | (ATC)6 | TCGAGGGTTCTCGACCATAC | GTTGTGACACATGTCCGTCC | 272 |
| chr2:70672340-70674358 | p2 | (AT)6 | AGGAAATCGTGAGACCATGC | GATTCGTAGGAGCCATCAGC | 159 |
| chr2:70672340-70674358 | p2 | (AT)11 | AGGAAATCGTGAGACCATGC | GATTCGTAGGAGCCATCAGC | 159 |
| chr2:70672340-70674358 | p3 | (ATC)6 | AGGAAATCGTGAGACCATGC | GATTCGTAGGAGCCATCAGC | 159 |
| chr2:70834901-70836921 | p4 | (AAAT)5 | ATACGGCCAGTATGTTTGGC | AGAGAAGGCAAGGCATACGA | 100 |
| chr2:70834901-70836921 | p5 | (TTTTA)3 | ATACGGCCAGTATGTTTGGC | AGAGAAGGCAAGGCATACGA | 100 |
| chr2:70854778-70856796 | p6 | (AGAGGC)3 | CTCCTCCTGTTTGGCAAGAG | AATCTATGGACACGATCGCC | 236 |
| chr2:70925603-70927618 | p5 | (TGAAT)3 | TGTGGCACATGCCATATTTT | TTTGGGGGAAATTGTCCATA | 106 |

| Marker ID | SSR type | SSR motif | FORWARD PRIMER (5'-3') | REVERSE PRIMER (5'-3') | size (bp) |
|---|---|---|---|---|---|
| chr2:70951918-70953936 | p2 | (AG)9 | CAACAATCCACCAGGCTTTT | CCCTGGAGCAACAAAAATGT | 274 |
| chr2:70951918-70953936 | p3 | (TGT)5 | CAACAATCCACCAGGCTTTT | CCCTGGAGCAACAAAAATGT | 274 |
| chr2:70954013-70956033 | p2 | (AG)10 | GTTTGGTGACCCTTCAGCAT | GACAAGGCTCCCACCTCATA | 141 |
| chr2:70954013-70956033 | p3 | (TGT)5 | GTTTGGTGACCCTTCAGCAT | GACAAGGCTCCCACCTCATA | 141 |
| chr2:71012855-71014870 | p5 | (TTGCA)3 | AGCAATATGGGCATCTACGG | GGACCCCTTTTAGTACCCCA | 166 |
| chr2:71175387-71177399 | p2 | (CA)6 | GAACTCTGGAGGGCCACATA | GACTCCAGTTCCTCAGCCAG | 188 |
| chr2:71280335-71282431 | p2 | (TG)22 | AACCAATCGGAGATCTGCAC | GATATGCTACGGGCCTGTGT | 244 |
| chr2:71280441-71282456 | p2 | (TG)22 | AACCAATCGGAGATCTGCAC | GATATGCTACGGGCCTGTGT | 138 |
| chr2:71407774-71409792 | p2 | (AT)9 | CTGACAAGCTCTGTGGGTGA | CGCCGTACTATCTCCTCGTC | 192 |
| chr2:71548773-71550788 | p5 | (GAAAA)3 | TTTGGTTGGTGACATGGAAA | CAAGGCCATAAGAAAACCCA | 122 |
| chr2:71598679-71600693 | p2 | (AG)7 | AGGACCCGAGCTTGATAGGT | ATACGGAAATGTCCATCCCA | 134 |
| chr2:71702892-71704904 | p2 | (AC)6 | GTTTCGAGGAGCAGAAATGC | ACCATTTGCAGATTTCGGAG | 190 |
| chr2:71766080-71768096 | p2 | (TA)8 | GGCAGCATTTCCCGTATAAA | GGAAGATTGCGGATTGTGTT | 167 |
| chr2:71819975-71821990 | p3 | (CAA)5 | CATACTGAGGCTTGCACGAA | CCGATCGATTCGAAACTTGT | 122 |
| chr2:71819975-71821990 | p2 | (CA)6 | CATACTGAGGCTTGCACGAA | CCGATCGATTCGAAACTTGT | 122 |
| chr2:71823956-71825974 | p6 | (TATATG)3 | TTTGGCCGAGTTGAATTCTC | GCTTAATTGTCCAAGCTCGC | 171 |
| chr2:71884121-71886139 | p3 | (ACA)6 | TCGGAAACGGTATCACATCA | TCTGCACTACTCCGTTCGTG | 106 |
| chr2:71899109-71901121 | p2 | (TA)6 | GGACCGTAAACCAGGTCGTA | AATTCTCGTCGGGTGAACAG | 121 |
| chr2:71993495-71995507 | p2 | (TA)9 | CCCGACATAGCCATAGTGGT | GTAGCCCGTATTAGCGGTGA | 109 |
| chr2:71993495-71995507 | p2 | (AT)6 | CCCGACATAGCCATAGTGGT | GTAGCCCGTATTAGCGGTGA | 109 |
| chr2:72019471-72021489 | p6 | (ATTTTC)3 | AAAATGCCCTCCCAAATAGC | TCACAAGGCCACAACACAAT | 121 |
| chr2:72025456-72027500 | p2 | (TA)22 | CATAAGCCGACGAGGCTAAC | GGAGATCCATTTCATTCGGA | 279 |
| chr2:72041870-72043918 | p2 | (AT)24 | TTCCCTGAACTGGGAATCTG | GGGATGGATATCTCCCGAAT | 262 |
| chr2:72142753-72144771 | p6 | (ATACAT)3 | GACAAGGCCGACAAAGTGAT | CAGAGCTCCATTTCTTTCGG | 300 |
| chr2:72163275-72165290 | p5 | (CATAG)3 | GTGCTCTCAGCTCGGCTAGT | TACGGCCCGCTAGAGAAGTA | 249 |
| chr2:72163275-72165290 | p2 | (TA)7 | GTGCTCTCAGCTCGGCTAGT | TACGGCCCGCTAGAGAAGTA | 249 |
| chr2:72176337-72178352 | p5 | (ATTGC)3 | CCTCTTCGATCTTGTGGCTC | GCGCCCAACCTAAGACAATA | 180 |
| chr2:72210399-72212419 | p5 | (TCATT)4 | AAAAAGGGGCATCCCTAAGA | AAACAACAACACTGCCCTCC | 129 |
| chr2:72261808-72263820 | p2 | (TG)6 | ATGTTATGAGCTTCGCCCAC | TCGTCAGCTAGGCTTGGAGT | 229 |
| chr2:72357147-72359162 | p3 | (CAT)5 | TCATAAGGACGGGACAGGAC | TGTGTTCATGCCCTCAGGTA | 212 |
| chr2:72357147-72359162 | p2 | (TA)18 | TCATAAGGACGGGACAGGAC | TGTGTTCATGCCCTCAGGTA | 212 |
| chr2:72400149-72402161 | p2 | (GA)15 | TGCGGAAATATTGAGGAAGG | CCACCTTGGCCATTTTCTTA | 290 |
| chr2:72400149-72402161 | p2 | (GT)6 | TGCGGAAATATTGAGGAAGG | CCACCTTGGCCATTTTCTTA | 290 |
| chr2:72515583-72517595 | p2 | (TG)6 | GATCTTGTAACGCGCAGACA | CTCCATATTTCCGCACCAAT | 274 |
| chr2:72515583-72517595 | p3 | (TTG)5 | GATCTTGTAACGCGCAGACA | CTCCATATTTCCGCACCAAT | 274 |
| chr2:72577784-72579796 | p2 | (AT)6 | TTCTCGCTTTCATTTCCGTT | TTCGAAGTCCTGTGTGCTTG | 136 |

| Marker ID | SSR type | SSR motif | FORWARD PRIMER (5'-3') | REVERSE PRIMER (5'-3') | size (bp) |
|---|---|---|---|---|---|
| chr2:72671307-72673327 | p5 | (ATCAT)4 | GCTGGACACCCAAACGTACT | TTAGTGATGGCCTTATCGGC | 198 |
| chr2:72748849-72750861 | p2 | (CT)6 | ACTTCACTCACGACTCCGCT | ACTCTTTCCCAAAATGCCCT | 295 |
| chr2:72797913-72799925 | p2 | (TA)6 | GGTATCGTTGTTGGTTTGGC | CTCGCTACGCTTATTCGTCC | 247 |
| chr2:72880850-72882871 | p2 | (AT)20 | TGCGACAAATTTGACACCAT | CTAAGTACCCGAACTGGGCA | 102 |
| chr2:72880850-72882871 | p3 | (ATC)7 | TGCGACAAATTTGACACCAT | CTAAGTACCCGAACTGGGCA | 102 |
| chr2:72921870-72923888 | p2 | (AT)9 | GGTCTGTGGTGTTGACGTTG | CTCATTCGAAAAAGCCAAGC | 162 |
| chr2:73003721-73005737 | p2 | (AT)8 | TTTAAAAGGCTCAACGTGGG | CCCCTCAAAGAGAGGAAACC | 118 |
| chr2:73024149-73026203 | p2 | (AT)27 | TTTCTCCGTTCCTTTGATGG | AGCCAACCCAGAGTCTCAAA | 120 |
| chr2:73048511-73050551 | p2 | (TA)20 | TATAGGGGAAGTGCTGTCCG | TATTCGTGAATCCTTTCGCC | 226 |
| chr2:73064341-73066353 | p2 | (TC)6 | GAATTTCCAATTGCCCTCCT | TGGGGGCTGTTTTGATATGT | 154 |
| chr2:73103766-73105781 | p5 | (GCATT)3 | GTTGGGTTGGACATTGCTTT | CGTGGCATCAAGCTAACAGA | 298 |
| chr2:73130741-73132756 | p5 | (GTAAT)3 | AGTAAAACGAGTCGGCGAGA | ACCAGGGATCACCCTTCTCT | 263 |
| chr2:73130741-73132756 | p2 | (AT)6 | AGTAAAACGAGTCGGCGAGA | ACCAGGGATCACCCTTCTCT | 263 |
| chr2:73130741-73132756 | p2 | (AT)8 | AGTAAAACGAGTCGGCGAGA | ACCAGGGATCACCCTTCTCT | 263 |
| chr2:73142049-73144083 | p2 | (TA)17 | GGTCGGGTCGGATTTTATTT | GCTTTGGAGCAGCTTTCAAC | 161 |
| chr2:73248264-73250279 | p3 | (TGT)5 | GACTTGAAGAGGCCAAAACG | GTGCATAACGTGTGATTCCG | 282 |
| chr2:73253521-73255539 | p3 | (CAA)6 | ACGAGCGGAGTAGTGAGGAA | GAAAAGCTCGGATTGCTTTG | 282 |
| chr2:73286632-73288647 | p5 | (GCTAT)3 | GGGAACTTTTGATCCCGATT | GTCGTTCCGGTAGCACGTAT | 178 |
| chr2:73296863-73298875 | p2 | (AT)6 | ATCATAGCCACACAGGGCTC | GCTGAAGAACGACATCGACA | 109 |
| chr2:73296863-73298875 | p5 | (ATATA)3 | ATCATAGCCACACAGGGCTC | GCTGAAGAACGACATCGACA | 109 |
| chr2:73311343-73313358 | p2 | (AT)9 | TACCAATAAGCCGCAACTCC | CCGCTTTAAGAGGCAATCAG | 284 |
| chr2:73311343-73313358 | p3 | (ATC)5 | TACCAATAAGCCGCAACTCC | CCGCTTTAAGAGGCAATCAG | 284 |
| chr2:73311343-73313358 | p2 | (AT)8 | TACCAATAAGCCGCAACTCC | CCGCTTTAAGAGGCAATCAG | 284 |
| chr2:73533245-73535260 | p5 | (AATAT)3 | CCCCAACCCTAGTCTGTGAA | CCAGCAAGGAGAGAGGTGTC | 295 |
| chr2:73631668-73633686 | p6 | (CATATA)3 | AGCTTTATTGTCACACCCCG | ACAGAGTCACGGGTACTGGG | 183 |
| chr2:73700605-73702617 | p2 | (AT)6 | CCTGTAATACCACCATCCCG | TGCTGATTGGGTCATTTTGA | 200 |
| chr2:73700605-73702617 | p2 | (AC)7 | CCTGTAATACCACCATCCCG | TGCTGATTGGGTCATTTTGA | 200 |
| chr2:73722882-73724934 | p5 | (TTTAA)3 | CTGTCCCGTCATATTTCGCT | GTCTCCTGCGGAGTCTCAAC | 200 |
| chr2:73722882-73724934 | p2 | (TA)26 | CTGTCCCGTCATATTTCGCT | GTCTCCTGCGGAGTCTCAAC | 200 |
| chr2:73781923-73783938 | p5 | (AATAA)3 | TTACCCGGCATCATAAGCTC | CACATGGTTGACATTCCGAG | 104 |
| chr2:73806950-73808962 | p2 | (AT)6 | GACCCTTACTTAGGGCCGAG | AAGCCGATGTACCATTTTGC | 249 |
| chr2:73919924-73921936 | p2 | (TA)6 | ATTGGAAATTCGGGATAGGC | GCCAAAAGAATGAAGGGACA | 232 |
| chr2:73955969-73957984 | p5 | (AACAC)3 | TCCTCACATCCTATGGCCTC | AGGTTGGGCATAGTTTCACG | 133 |
| chr2:74018035-74020047 | p2 | (CT)6 | GAAGCGGAAGAGAATATGCG | TCGAATGTTAAAGGCTTGGG | 105 |
| chr2:74018035-74020047 | p2 | (TA)6 | GAAGCGGAAGAGAATATGCG | TCGAATGTTAAAGGCTTGGG | 105 |
| chr2:74109084-74111100 | p4 | (TGTA)4 | ACCCAGAAGCATCACCAGAC | TGATTCGGGAGTTCGTTACC | 278 |

| Marker ID | SSR type | SSR motif | FORWARD PRIMER (5'-3') | REVERSE PRIMER (5'-3') | size (bp) |
|---|---|---|---|---|---|
| chr2:74197370-74199384 | p2 | (TC)7 | CGGCATTAGCATTTCCCTTA | TCGTTAGATTTTCGGGCATC | 251 |
| chr2:74224115-74226127 | p2 | (AT)6 | ACATTGTGGGAAATTTGGGA | AACCCCCGATACCTCAAAAC | 285 |
| chr2:74252283-74254317 | p2 | (AT)17 | GGTGCGTAATCGGGTTTCTA | TGGTCGAAGGAAGAGAGGAA | 130 |
| chr2:74264356-74266374 | p6 | (TCATAT)3 | GATAAGGGGTGCTCGGTACA | GCCCACATGGGAGCTTAATA | 230 |
| chr2:74264356-74266374 | p4 | (TATT)4 | GATAAGGGGTGCTCGGTACA | GCCCACATGGGAGCTTAATA | 230 |
| chr2:74290903-74292915 | p2 | (AT)6 | CTGCTCGAGGAAGAAAATGG | TATCCTCTACCTTCGGGGCT | 272 |
| chr2:74290903-74292915 | p2 | (GT)7 | CTGCTCGAGGAAGAAAATGG | TATCCTCTACCTTCGGGGCT | 272 |
| chr2:74291249-74293263 | p2 | (AT)6 | CCCCAAAGGCATTGACTAGA | TCACGATTTCCAATTACGCA | 185 |
| chr2:74291249-74293263 | p2 | (GT)7 | CCCCAAAGGCATTGACTAGA | TCACGATTTCCAATTACGCA | 185 |
| chr2:74314766-74316778 | p3 | (CAA)5 | TATGGAGGGAAAGGACATCG | GGGATTCGTTTACCGAGTGA | 144 |
| chr2:74314766-74316778 | p2 | (CA)6 | TATGGAGGGAAAGGACATCG | GGGATTCGTTTACCGAGTGA | 144 |
| chr2:74333492-74335507 | p5 | (TGATA)3 | CACGCCTCGGATCATATTTT | GGCGGTCTCAAATTACGTGT | 132 |
| chr2:74402314-74404329 | p3 | (CAG)5 | GACTCCAGTTCCTCAGCCAG | CAGTCTCTGAAGCCCTGACC | 249 |
| chr2:74413838-74415866 | p5 | (CATAG)3 | ATCAAGTCCACCTGTCTGCC | CGCGATATGCTACGACAAGA | 201 |
| chr2:74413838-74415866 | p2 | (TA)14 | ATCAAGTCCACCTGTCTGCC | CGCGATATGCTACGACAAGA | 201 |
| chr2:74523395-74525435 | p2 | (TA)20 | ACTAGTGCGCCTTCACGATT | ATACGGGCTATAACACCCCC | 281 |
| chr2:74523395-74525435 | p5 | (AATTG)5 | ACTAGTGCGCCTTCACGATT | ATACGGGCTATAACACCCCC | 281 |
| chr2:74584808-74586868 | p3 | (ATG)7 | TGCCCAAATTCCATTAGCTC | CCTCCCAAGTGCACAAGTCT | 282 |
| chr2:74617940-74619952 | p2 | (AT)6 | ATGAGCCATGACTAGTGCCC | ATGGGTTTTGGTCAGCTACG | 267 |
| chr2:74678790-74680806 | p2 | (GT)8 | ACAACAACCGGGAAGAGATG | TTATGAGCATAACGGGGGAG | 109 |
| chr2:74722467-74724497 | p2 | (TA)15 | ACCAAACAACGCACAAATCA | AGGGTAGTAATGCTCCGGCT | 255 |
| chr2:74725326-74727360 | p2 | (TA)17 | CTTCCTCCACCACCAAGTGT | CACTACCGCCAGTACCACCT | 123 |
| chr2:74725326-74727360 | p2 | (TA)29 | CTTCCTCCACCACCAAGTGT | CACTACCGCCAGTACCACCT | 123 |
| chr2:74752378-74754408 | p2 | (TA)15 | ACCAAACAACGCACAAATCA | AGGGTAGTAATGCTCCGGCT | 255 |
| chr2:74755237-74757271 | p2 | (TA)17 | CTTCCTCCACCACCAAGTGT | CACTACCGCCAGTACCACCT | 123 |
| chr2:74755237-74757271 | p2 | (TA)29 | CTTCCTCCACCACCAAGTGT | CACTACCGCCAGTACCACCT | 123 |
| chr2:74786501-74788519 | p6 | (ATACAT)3 | AAAGCTTGACATCACCCAGG | TGCTTTATGGTGCTGAGTCG | 283 |
| chr2:74828951-74830963 | p2 | (AT)6 | GTACTGGCGGAACGATCACT | CAGAAGGGCAAAGCTACGAC | 226 |
| chr2:74864710-74866722 | p2 | (AT)20 | TGCGAAAGAACTCCAAAGGT | CCTTGTCAGCTTGCGTATCA | 247 |
| chr2:74864710-74866722 | p2 | (TA)6 | TGCGAAAGAACTCCAAAGGT | CCTTGTCAGCTTGCGTATCA | 247 |
| chr2:74864710-74866722 | p3 | (ACA)6 | TGCGAAAGAACTCCAAAGGT | CCTTGTCAGCTTGCGTATCA | 247 |
| chr2:74899115-74901133 | p6 | (ATATGT)3 | ATGTTGGATATGAGCCTCGG | GTCGGGTCCTATTTATGCGA | 226 |
| chr2:75051658-75053670 | p2 | (TA)6 | AAAAGGGCAAGGAGTTCGAT | CCTCGGAATGGTCAACTGAT | 272 |
| chr2:75073341-75075353 | p2 | (AT)6 | AAAATGTGGAGGAGGAGGCT | ACGTTTTCGGATGACCTGAC | 222 |
| chr2:75261855-75263933 | p2 | (TA)7 | CGCAACGACATCGACTAGAA | TGTGGTGTGTGGGAGTGAAT | 228 |
| chr2:75261855-75263933 | p2 | (AT)39 | CGCAACGACATCGACTAGAA | TGTGGTGTGTGGGAGTGAAT | 228 |

| Marker ID | SSR type | SSR motif | FORWARD PRIMER (5'-3') | REVERSE PRIMER (5'-3') | size (bp) |
|---|---|---|---|---|---|
| chr2:75335412-75337430 | p2 | (AT)9 | CCAGTTGTTCCGTTTTCGTT | TGTCATGTGTCGGGTAGGAA | 212 |
| chr2:75367613-75369625 | p2 | (GT)6 | TTGAGAACGGGGTAATCCTG | GCCTACCCCGAGGTCTTAAC | 256 |
| chr2:75374041-75376055 | p2 | (CT)7 | AGCATACAATTGGTGCTCCC | GGTTAGTGATGCCGTTTGGT | 139 |
| chr2:75420109-75422124 | p3 | (ATC)5 | CCGACGAGGATACAAAAGGA | GCTCCATTTCATTCAGAGCC | 180 |
| chr2:75423842-75425857 | p5 | (GAAAT)3 | AGCACATCAAGCATCGTGAG | ACGTGAGCTTCAGAGGAGGA | 154 |
| chr2:75501380-75503395 | p5 | (TTTTG)3 | GAGCTACACATTCCGCAACA | TAATGAAACGCAAGCACGTC | 299 |
| chr2:75559615-75561630 | p5 | (TTTCA)3 | TCGGTCAAAGTGTTGTAGCG | GAAGGTTCCCGAAAGTAGGG | 291 |
| chr2:75603581-75605637 | p2 | (AT)28 | TTGGTGTTGGGGTCAATTTT | ATGTGGAAACACGGAGGAAG | 266 |
| chr2:75707253-75709269 | p5 | (TCATA)3 | GCTGGGGATTTCCTAGGTTC | TTCCAAAGAAGGGTGTTTCG | 259 |
| chr2:75707253-75709269 | p2 | (AT)8 | GCTGGGGATTTCCTAGGTTC | TTCCAAAGAAGGGTGTTTCG | 259 |
| chr2:75707253-75709269 | p5 | (CCCTA)3 | GCTGGGGATTTCCTAGGTTC | TTCCAAAGAAGGGTGTTTCG | 259 |
| chr2:75743446-75745471 | p5 | (ATATC)5 | AGGGAAATTTGCTCCCAACT | TAAGGGGTGCTCGGTACAAG | 247 |
| chr2:75819902-75821914 | p2 | (CA)6 | GAGTTCTTTTCCTGAGGGGG | TTTGTCAGTTTCGAGGGGTC | 161 |
| chr2:75821956-75823968 | p3 | (ACA)5 | TTACCCAAATGGGACCAAAA | TTTTGGCATGTCCTAGACCC | 246 |
| chr2:75821956-75823968 | p2 | (AC)6 | TTACCCAAATGGGACCAAAA | TTTTGGCATGTCCTAGACCC | 246 |
| chr2:75875418-75877430 | p2 | (AG)6 | GTCTGCCAACACATCAATGG | ATTTCCGGGAGAACCAGTCT | 298 |
| chr2:76046806-76048870 | p2 | (TA)32 | CTCTAGCCATCGCCTCTGAC | GTGTCGTTGTTTGGTGATGC | 192 |
| chr2:76046806-76048870 | p3 | (ATC)5 | CTCTAGCCATCGCCTCTGAC | GTGTCGTTGTTTGGTGATGC | 192 |
| chr2:76084789-76086804 | p3 | (CAA)5 | CAACTGTCGCTATGGAGCAA | CTATGATGGCCGACGTCTTT | 205 |
| chr2:76194825-76196840 | p5 | (ATTTC)3 | ACGAGAGGAAATGACATGGG | ATAACAATGCCTCACAGGGC | 153 |
| chr2:76419709-76421721 | p2 | (TA)6 | TTTGCGTACGAATTTGGTGA | CCAAAACAGTCCCTAACCGA | 146 |
| chr2:76474296-76476314 | p2 | (AT)9 | GGACCCTCAAACTGTCGTGT | TATCAGAGGAGCCTAGGGCA | 237 |
| chr2:76508484-76510499 | p5 | (TATGA)3 | AGCCTCTGGGACAACCCTAT | TAGGAAAATCCGGGGGTAAC | 123 |
| chr2:76514486-76516504 | p6 | (CATATA)3 | GTACACGTTCCAGCACCCTT | AGTAGACTTGAGCCACGGGA | 209 |
| chr2:76547088-76549103 | p5 | (ATGCA)3 | CCCAAAATCGCTTAGCTTTG | GACCAGATGCCTTGGGTAGA | 101 |
| chr2:76564720-76566742 | p3 | (CAT)5 | GTTTTGGCTTGTGTGGAGGT | TATGGGCCTCAGTCAAAACC | 178 |
| chr2:76564737-76566752 | p3 | (CAT)5 | GTTTTGGCTTGTGTGGAGGT | TATGGGCCTCAGTCAAAACC | 161 |
| chr2:76564747-76566759 | p3 | (CAT)5 | GTTTTGGCTTGTGTGGAGGT | TATGGGCCTCAGTCAAAACC | 151 |
| chr2:76564762-76566778 | p3 | (CAT)5 | GTTTTGGCTTGTGTGGAGGT | TATGGGCCTCAGTCAAAACC | 136 |
| chr2:76587070-76589085 | p3 | (TTG)5 | ATTTTGGCAATTTGTTTCGG | AAGACTTGGTGGCCCTTTTT | 283 |
| chr2:76769865-76771879 | p2 | (TA)7 | ACGTGGAAAATCGTGAGACC | CGTCCGTTTTCTTTGGATGT | 264 |
| chr2:76899452-76901472 | p2 | (TA)10 | TGCCCGATTTTCGTTAAGTC | TTTTCGCCTCGTCACTCTTT | 116 |
| chr2:76899452-76901472 | p2 | (TA)7 | TGCCCGATTTTCGTTAAGTC | TTTTCGCCTCGTCACTCTTT | 116 |
| chr2:76899573-76901587 | p2 | (TA)10 | AAAAGAGTGACGAGGCGAAA | AAACTGCATAGGGAGGCTCA | 144 |
| chr2:76899573-76901587 | p2 | (TA)7 | AAAAGAGTGACGAGGCGAAA | AAACTGCATAGGGAGGCTCA | 144 |
| chr2:77022361-77024376 | p2 | (TA)6 | TTGGGAGAAGATTCAAACGG | AGTCGATACGTTTCCGGATG | 185 |

| Marker ID | SSR type | SSR motif | FORWARD PRIMER (5'-3') | REVERSE PRIMER (5'-3') | size (bp) |
|---|---|---|---|---|---|
| chr2:77022361-77024376 | p3 | (CAA)5 | TTGGGAGAAGATTCAAACGG | AGTCGATACGTTTCCGGATG | 185 |
| chr2:77022361-77024376 | p2 | (CA)6 | TTGGGAGAAGATTCAAACGG | AGTCGATACGTTTCCGGATG | 185 |
| chr2:77156579-77158594 | p3 | (ACA)5 | GGCCCTTTAATGAGGGACTC | CTCACTAGAGGGTCGGGACA | 136 |
| chr2:77156579-77158594 | p2 | (CT)7 | GGCCCTTTAATGAGGGACTC | CTCACTAGAGGGTCGGGACA | 136 |
| chr2:77159743-77161771 | p2 | (TA)14 | TTGGTCTCACCCACATCAAA | TTGTGGGCCTGTGTTATTGA | 293 |
| chr2:77226174-77228189 | p3 | (CAA)5 | AGGTCTCGAGAGCCGTAACA | CCCTCGTGCCCTTATTATGA | 140 |
| chr2:77226174-77228189 | p2 | (CA)6 | AGGTCTCGAGAGCCGTAACA | CCCTCGTGCCCTTATTATGA | 140 |
| chr2:77230447-77232471 | p6 | (CATATA)4 | ACCCCTAGGACAAACCTGCT | GTCGACGTGTCATTTTGTGC | 242 |
| chr2:77237407-77239427 | p2 | (AT)10 | GTGTCCTGTGGATAAGGCGT | CGACTCCCTTCAGAGTCGAG | 144 |
| chr2:77305894-77307906 | p2 | (TA)6 | TTTATTGGTGTGGTTCGGGT | TGGAGCAGGGAGTTATGGAC | 298 |
| chr2:77322767-77324782 | p3 | (GTT)5 | GGCTATGGTTCGAACTGCTC | TATTATTCCCACTCACCGCC | 154 |
| chr2:77361348-77363360 | p2 | (TA)6 | AACCGGATTGACTTGAAACG | ACATATTGAGGCATGCACGA | 106 |
| chr2:77445522-77447537 | p5 | (GATAT)3 | AAGCAACCCAATTCTTCCCT | GGGTCAAATTGTGCCCTCTA | 145 |
| chr2:77455005-77457029 | p2 | (AT)12 | GGAAATCACGGGCTCTTGTA | GGCAAGGCTACGACAAAAAG | 204 |
| chr2:77622850-77624868 | p2 | (TA)6 | AGCACCGAGTCCCATGATAG | ACGTACCCAGGCCCTCTAGT | 107 |
| chr2:77648569-77650625 | p2 | (AT)28 | TTTCGTGCACAGTTTTCAGC | CACGTATACCATCCGGCTCT | 124 |
| chr2:77691523-77693539 | p2 | (AT)8 | CGGCAAGGCTAACAGAAAAG | GGCCCTGACAGGATTATTGA | 253 |
| chr2:77750311-77752326 | p3 | (GAT)5 | GTTTCTGCCAGTTCCCTGAG | TTAAGTTCCTCACCTCGCGT | 119 |
| chr2:77872796-77874820 | p2 | (AT)12 | GAGGCGCATTCTCTTGTCTC | GATGAAGACCCGCAATGTTT | 152 |
| chr2:77952125-77954140 | p5 | (AAAAG)3 | TCAGTGTCATCCGTACCCAA | CCAAGGATAGACGTAGCGGA | 152 |
| chr2:78095939-78097964 | p5 | (ATGAT)5 | AACGCGATTTGAGTTTGGAC | AAAGGATGCAAGACAATCCG | 211 |
| chr2:78110468-78112488 | p5 | (ATGAT)4 | GATTTTGGAGGAAACGGGAT | TTGCTCACAAAGGATGCAAG | 199 |
| chr2:78157453-78159469 | p4 | (CATA)4 | GGAACTGTCTGCAAAGCCTC | TCCACTTCTTCCTGGAGCAT | 298 |
| chr2:78157453-78159469 | p3 | (ATC)5 | GGAACTGTCTGCAAAGCCTC | TCCACTTCTTCCTGGAGCAT | 298 |
| chr2:78157453-78159469 | p2 | (TC)9 | GGAACTGTCTGCAAAGCCTC | TCCACTTCTTCCTGGAGCAT | 298 |
| chr2:78160073-78162099 | p2 | (AT)9 | GCCTCTTTCCTTTTCGTTCC | TCCCCACACCATGATAACCT | 200 |
| chr2:78180794-78182818 | p2 | (AT)32 | CGACGTACCCTTAACGCAAT | ATTGCTCGTATTGACGTCCC | 184 |
| chr2:78180794-78182818 | p4 | (CATA)6 | CGACGTACCCTTAACGCAAT | ATTGCTCGTATTGACGTCCC | 184 |
| chr2:78180794-78182818 | p3 | (ATC)5 | CGACGTACCCTTAACGCAAT | ATTGCTCGTATTGACGTCCC | 184 |
| chr2:78180794-78182818 | p2 | (CT)7 | CGACGTACCCTTAACGCAAT | ATTGCTCGTATTGACGTCCC | 184 |
| chr2:78218313-78220347 | p2 | (AT)17 | CCGTCATGGCCCTCTAGTTA | ATATGGAGGGGTCCTGTTCC | 184 |
| chr2:78256145-78258167 | p3 | (TTG)6 | TGTTATGAAGGGTGCGGATT | TAAGGGTCATCAAACAGCCC | 162 |
| chr2:78256145-78258167 | p2 | (TA)27 | TGTTATGAAGGGTGCGGATT | TAAGGGTCATCAAACAGCCC | 162 |
| chr2:78256177-78258205 | p3 | (TTG)6 | TGTTATGAAGGGTGCGGATT | TAAGGGTCATCAAACAGCCC | 130 |
| chr2:78256177-78258205 | p2 | (TA)27 | TGTTATGAAGGGTGCGGATT | TAAGGGTCATCAAACAGCCC | 130 |
| chr2:78269666-78271684 | p2 | (AT)9 | AAGGCCGATGGACACTTATG | CATGTATTTGGCACGCAATC | 271 |

| Marker ID | SSR type | SSR motif | FORWARD PRIMER (5'-3') | REVERSE PRIMER (5'-3') | size (bp) |
|---|---|---|---|---|---|
| chr2:78275162-78277180 | p6 | (CATATA)3 | ATGTCCCACCTTGAAGCAAC | ATGTTCAAATGTTCCGAGCC | 271 |
| chr2:78275162-78277180 | p3 | (CAA)6 | ATGTCCCACCTTGAAGCAAC | ATGTTCAAATGTTCCGAGCC | 271 |
| chr2:78290420-78292435 | p5 | (GATAT)3 | CGATACGGGCATTACGAGTT | ACGAACAGGTCAAGGAATGG | 255 |
| chr2:78306555-78308573 | p3 | (CAA)6 | ACATCACAACAAAGGCCACA | AGGCATAGTTGACAGGGGTG | 125 |
| chr2:78315467-78317481 | p2 | (AC)7 | ATGCCAAAATGAAGGAGTGG | TGTTGTTGGGCTGTTGTTGT | 230 |
| chr2:78475357-78477373 | p2 | (AT)6 | ATGGCCTCTTGTGTTGCTCT | GAAAAAGGCCCAGTGTGTGT | 232 |
| chr2:78475357-78477373 | p2 | (AT)8 | ATGGCCTCTTGTGTTGCTCT | GAAAAAGGCCCAGTGTGTGT | 232 |
| chr2:78475357-78477373 | p2 | (AT)8 | ATGGCCTCTTGTGTTGCTCT | GAAAAAGGCCCAGTGTGTGT | 232 |
| chr2:78683879-78685904 | p5 | (GATAT)3 | GCCCCTTTTTCTGATTCCTC | TCCAAAACAGACAGAATGCG | 184 |
| chr2:78683914-78685944 | p5 | (GATAT)3 | GCCCCTTTTTCTGATTCCTC | TCCAAAACAGACAGAATGCG | 149 |
| chr2:78697345-78699357 | p2 | (AT)6 | CACATCACAACAAAGGCCAC | TAAGGCACAGTTGAAAGGGG | 115 |
| chr2:78697345-78699357 | p2 | (TA)6 | CACATCACAACAAAGGCCAC | TAAGGCACAGTTGAAAGGGG | 115 |
| chr2:78697345-78699357 | p2 | (TA)11 | CACATCACAACAAAGGCCAC | TAAGGCACAGTTGAAAGGGG | 115 |
| chr2:78725947-78727962 | p5 | (TATAT)3 | GTTGTTCCGTTTTATCGCGT | AATAGCCCCCAACTCCAACT | 166 |
| chr2:78725947-78727962 | p5 | (ATATT)3 | GTTGTTCCGTTTTATCGCGT | AATAGCCCCCAACTCCAACT | 166 |
| chr2:78737605-78739617 | p2 | (CA)6 | GAATTCAACTCGGCCAAAAA | TCGAGACGCTGTTGAAGTTG | 162 |
| chr2:78739845-78741857 | p2 | (AC)6 | GAGAAGGGTTATTCCCGAGG | CGGTAGCCAAATATAGCCCA | 178 |
| chr2:78746375-78748391 | p4 | (CATT)4 | CTGGCATAAGCGGAGAGAAC | CGGGAATAACTCTGCTCTCG | 156 |
| chr2:78754853-78756871 | p2 | (AT)7 | TGTAGACACACCCTGGGACA | TGTTAGCCTCGTCGGCTTAT | 237 |
| chr2:78754853-78756871 | p6 | (CATATT)3 | TGTAGACACACCCTGGGACA | TGTTAGCCTCGTCGGCTTAT | 237 |
| chr2:78754853-78756871 | p2 | (AT)6 | TGTAGACACACCCTGGGACA | TGTTAGCCTCGTCGGCTTAT | 237 |
| chr2:78761309-78763321 | p2 | (AT)6 | TGAGCGGAAACTCCTCACTT | CCCTTTTCACTCCCCTTCTC | 256 |
| chr2:78821341-78823356 | p5 | (ATTTC)3 | ACAGGCAGCAGTTCCCTAGA | ACTGACCCCAATGCAACTTC | 113 |
| chr2:78848283-78850298 | p5 | (ATTTC)3 | CCGGAAGAGATCCAAGATGA | TTAAGAACATCCGTCCTCGC | 241 |
| chr2:78952469-78954484 | p6 | (TTATTG)3 | GGCCATGTGGCTTATGAGTT | AATTGCTCGGTGATCTGGAC | 180 |
| chr2:78952469-78954484 | p5 | (TGTGT)3 | GGCCATGTGGCTTATGAGTT | AATTGCTCGGTGATCTGGAC | 180 |
| chr2:78952524-78954539 | p6 | (TTATTG)3 | GGCCATGTGGCTTATGAGTT | AATTGCTCGGTGATCTGGAC | 125 |
| chr2:78952524-78954539 | p5 | (TGTGT)3 | GGCCATGTGGCTTATGAGTT | AATTGCTCGGTGATCTGGAC | 125 |
| chr2:78960580-78962625 | p3 | (ATG)15 | TTACATCTTGGGGCCTTGTC | TGGGGAAATAAGTGTGGGAG | 136 |
| chr2:78960580-78962625 | p2 | (TA)20 | TTACATCTTGGGGCCTTGTC | TGGGGAAATAAGTGTGGGAG | 136 |
| chr2:79054768-79056814 | p2 | (TA)23 | CTCAAGGGCAGTGAGCTAGG | GTCTGCCCGTGCCATATAGT | 279 |
| chr2:79148350-79150362 | p3 | (CAA)5 | CAAAAATTTGGGCAGCATCT | GATTGTTGTTGTTGTTGGCG | 299 |
| chr2:79148350-79150362 | p2 | (AC)6 | CAAAAATTTGGGCAGCATCT | GATTGTTGTTGTTGTTGGCG | 299 |
| chr2:79227430-79229442 | p2 | (AT)6 | GCCAGGAATGTTGTCAAGGT | TCAGAACAAGCGTACATGGC | 190 |
| chr2:79285680-79287695 | p5 | (GCAAG)3 | GAGTGAAGCCCAACATCCAT | TCATTCCTTAAAGGCCGATG | 140 |
| chr2:79449359-79451377 | p2 | (TA)9 | GGGTGGAAAATAAGGGGGTA | TTTACTGAATGAACGCGCAG | 158 |

| Marker ID | SSR type | SSR motif | FORWARD PRIMER (5'-3') | REVERSE PRIMER (5'-3') | size (bp) |
|---|---|---|---|---|---|
| chr2:79449515-79451527 | p2 | (TA)9 | CTGCGCGTTCATTCAGTAAA | TGGAAACACACCGAAAGTGA | 125 |
| chr2:79449577-79451591 | p2 | (TA)9 | TCACTTTCGGTGTGTTTCCA | CGTAGGTTACGGTGAGGCAT | 163 |
| chr2:79449612-79451626 | p2 | (TA)9 | TCACTTTCGGTGTGTTTCCA | CGTAGGTTACGGTGAGGCAT | 128 |
| chr2:79449637-79451651 | p2 | (TA)9 | TCACTTTCGGTGTGTTTCCA | CGTAGGTTACGGTGAGGCAT | 103 |
| chr2:79466356-79468382 | p5 | (ATACG)3 | CGGGTATTCGGTACTAGCCA | TGCCTCGTCCTTATGGATTC | 274 |
| chr2:79469422-79471440 | p3 | (ATC)6 | CCAAGGATCAATTCCACGTT | GTAAGGTTTGTGGGCAGCAT | 246 |
| chr2:79493326-79495352 | p5 | (ATACG)3 | CGGGTATTCGGTACTAGCCA | TGCCTCGTCCTTATGGATTC | 274 |
| chr2:79496392-79498410 | p3 | (ATC)6 | CCAAGGATCAATTCCACGTT | GTAAGGTTTGTGGGCAGCAT | 246 |
| chr2:79533832-79535844 | p2 | (AT)6 | ATTCCGCCACTTTGTTTGAG | CGAACGTGCTAGCATTTGAA | 110 |
| chr2:79544512-79546526 | p2 | (AC)7 | CAGACGTCTTACGCTTTCCC | TGGGCTGTTGTTGTTGATGT | 291 |
| chr2:79705403-79707415 | p3 | (ACA)5 | CAAATGCGACAAAAAGGGTT | CGGTAGCCAAATATAGCCCA | 129 |
| chr2:79705403-79707415 | p2 | (AC)6 | CAAATGCGACAAAAAGGGTT | CGGTAGCCAAATATAGCCCA | 129 |
| chr2:79732135-79734149 | p2 | (GT)7 | GCCCACACGTCAAGATTTTT | GATTTGTCCCCTCCAAGTCA | 182 |
| chr2:79732135-79734149 | p3 | (GTT)5 | GCCCACACGTCAAGATTTTT | GATTTGTCCCCTCCAAGTCA | 182 |
| chr2:79745433-79747445 | p2 | (AT)7 | AATGTCAGCTTTGTCGGCTT | GTTGGCCATGACTAATGCCT | 140 |
| chr2:79745433-79747445 | p2 | (AT)6 | AATGTCAGCTTTGTCGGCTT | GTTGGCCATGACTAATGCCT | 140 |
| chr2:79745433-79747445 | p2 | (GT)6 | AATGTCAGCTTTGTCGGCTT | GTTGGCCATGACTAATGCCT | 140 |
| chr2:79774055-79776073 | p6 | (CATATA)3 | TGCAGACAACCCCTTAGACC | GTCAGTCTGTAAGCGGCTCC | 132 |
| chr2:79799261-79801273 | p2 | (GT)6 | TAGGTTGGAGAGGGCTGCTA | CTGTTTACCGACGACCCAAT | 120 |
| chr2:79874646-79876662 | p6 | (TGTATA)3 | AGGGTTATACCATGACCCCC | AAGACATCGCCGGTCATAAG | 157 |
| chr2:79927052-79929067 | p3 | (TAT)5 | ACATTCTTCGTATGGCCAGG | ATAAGCCGACGAGGCTAACA | 210 |
| chr2:79958596-79960620 | p6 | (CATACA)4 | ATACCCCACGTAATCACCGA | ACGTCTGGCCCTAGTTCTGA | 114 |
| chr2:79964804-79966822 | p6 | (TTCATA)3 | ATGAACTGAGGGGCAATCAC | GAGATGATTCGGGTCAGGAA | 138 |
| chr2:79991373-79993388 | p2 | (TA)8 | CACGATCCGTCCTTTCCTTA | TGCATACGAGCAAGCTATGG | 232 |
| chr2:79991373-79993388 | p5 | (TTAAA)3 | CACGATCCGTCCTTTCCTTA | TGCATACGAGCAAGCTATGG | 232 |
| chr2:80081710-80083728 | p2 | (AT)9 | GGGCACCCCTTTAACATACA | ATTTTGTTGGCCTTGTCGTC | 300 |
| chr2:80125168-80127182 | p2 | (AT)7 | ACAATGGCTGTCTTCCCAAC | AATTTGAGGCCACGAATGAG | 201 |
| chr2:80156482-80158497 | p3 | (AAC)5 | CCAAAGAATGCCCTGAACAT | GGCCGCTTAGTCAGTGAGAC | 136 |
| chr2:80209140-80211155 | p5 | (TAAAA)3 | AAATCTTGCATGGGCAAATC | TTGGAGTCAACACCAACCAA | 190 |
| chr2:80252945-80254960 | p3 | (AAC)5 | TTTCTTGGATTTTGGCTTGG | TTTGAAAGCCCGTAAAATGG | 179 |
| chr2:80477511-80479525 | p2 | (TA)7 | CGGGGAAATTACGCTACAAA | GACTCCTCAAACGCCTTACG | 171 |
| chr2:80477511-80479525 | p2 | (TA)24 | CGGGGAAATTACGCTACAAA | GACTCCTCAAACGCCTTACG | 171 |
| chr2:80540604-80542622 | p2 | (AT)9 | TCGGCAGAATTTACAAAGGG | GTGGACAGGCTCGGATCTTA | 224 |
| chr2:80583843-80585858 | p3 | (ATG)5 | ACACTTGGATTGTGGAAGGG | TCTTCTCAAGGTTCGGTGCT | 199 |
| chr2:80590592-80592610 | p2 | (TA)9 | TGTGACAGGACAGTGGTGGT | AGGTGTAACATTCCTCCCCC | 182 |
| chr2:80590592-80592610 | p3 | (TTG)6 | TGTGACAGGACAGTGGTGGT | AGGTGTAACATTCCTCCCCC | 182 |

| Marker ID | SSR type | SSR motif | FORWARD PRIMER (5'-3') | REVERSE PRIMER (5'-3') | size (bp) |
|---|---|---|---|---|---|
| chr2:80619519-80621531 | p2 | (TA)6 | TCGGGATTTGATTAGCGAAC | ACTCGAACATCATAAGGGCG | 149 |
| chr2:80691612-80693624 | p2 | (GT)6 | GCAGAGCTCCATTTCTTTCG | CCATTATTTTGGCCCAATG | 287 |
| chr2:80774148-80776162 | p2 | (TA)7 | ATCGTGCGATAGAGCCAAGT | TCAACAATGCCTCACAGAGC | 117 |
| chr2:80774148-80776162 | p2 | (CT)6 | ATCGTGCGATAGAGCCAAGT | TCAACAATGCCTCACAGAGC | 117 |
| chr2:80826077-80828089 | p2 | (TA)6 | ATCGATGCTTCTCTTTGCGT | TATGGCTATGTCGGGGCTAC | 207 |
| chr2:80826077-80828089 | p3 | (TTG)7 | ATCGATGCTTCTCTTTGCGT | TATGGCTATGTCGGGGCTAC | 207 |
| chr2:80878683-80880701 | p6 | (TGTGTA)3 | CTGCTTAAGGGCAGTTCAGG | AGAAGCACACCACGTCCTCT | 188 |
| chr2:80972441-80974457 | p2 | (AT)8 | GCAAAACTCTGCAGGTCCTC | GACTCAGTTATTCGCCGCTC | 249 |
| chr2:81014676-81016688 | p2 | (AT)6 | CTTCCGGATTGTGTTGGAAT | GAATCAAACTCGGCCAAAAA | 140 |
| chr2:81038609-81040627 | p6 | (AGAGAA)3 | TTCATTACAGGCTTACCCCG | ACCATGGAGTCGCACTATCC | 131 |
| chr2:81085742-81087754 | p2 | (TA)6 | TGTTAAGGTCATGTGGCGAA | CTTACAAGCATTTTGGGGGA | 191 |
| chr2:81085742-81087754 | p3 | (TTG)5 | TGTTAAGGTCATGTGGCGAA | CTTACAAGCATTTTGGGGGA | 191 |
| chr2:81095146-81097164 | p6 | (CATAAT)3 | ATTTAGGCATGGTGGAATCG | TCCAAGGTGAGACATGACGA | 231 |
| chr2:81108471-81110486 | p3 | (TTG)5 | CCAAGGGAAAAATTGGGAAT | TTGGGCCCTTTTATTTTCCT | 267 |
| chr2:81133209-81135227 | p6 | (AAAAAT)3 | TGTGGTAGCTGTCCCATTGA | TCCAATTTTGTGTTTTGGCA | 182 |
| chr2:81147463-81149475 | p2 | (CA)6 | GAATAGAGTTTCCCCGAGGC | ACCATTGTAGATTCTGGGCG | 202 |
| chr2:81220832-81222858 | p2 | (TA)13 | TCACCAAAGCAGCAAACAAG | CATATTCCCTGTGGGATTCG | 285 |
| chr2:81220832-81222858 | p2 | (AT)8 | TCACCAAAGCAGCAAACAAG | CATATTCCCTGTGGGATTCG | 285 |
| chr2:81276635-81278647 | p2 | (AT)6 | CCCCGTAAGAGTCCGTGATA | CGGGATACTTAAGGCAACCA | 282 |
| chr2:81279461-81281476 | p5 | (TATGA)3 | TGTTCGTCTGTAGTGGCAGC | TCCAAATAACCGGTCTCCTG | 216 |
| chr2:81302919-81304933 | p2 | (TG)7 | ACCCCAAAATAACCAAACCC | GTTCGGGGCGTGATAACTAA | 187 |
| chr2:81347961-81349976 | p5 | (ATACA)3 | AAATGCCATTCACCCTCTTG | TATCTTTATGGTCCGCCCAG | 269 |
| chr2:81347961-81349976 | p3 | (CAT)5 | AAATGCCATTCACCCTCTTG | TATCTTTATGGTCCGCCCAG | 269 |
| chr2:81387416-81389428 | p2 | (TA)6 | TGCTCTTGTTGTTGCCTCAC | ATCGGAGTGACGTAAGGTCG | 229 |
| chr2:81391334-81393362 | p2 | (AT)14 | ATGAACTCACTCGCCCATTC | AGTGCCGTATTTCACCCAAG | 270 |
| chr2:81479704-81481742 | p2 | (AT)19 | CAGCATTTGTGGAGAAGCAA | GGAGAGAGGTTTCCCTTTCG | 248 |
| chr2:81479704-81481742 | p2 | (AT)19 | CAGCATTTGTGGAGAAGCAA | GGAGAGAGGTTTCCCTTTCG | 248 |
| chr2:81479852-81481890 | p2 | (AT)19 | TCACGAACAAGAGCACCAAC | GGAGAGAGGTTTCCCTTTCG | 125 |
| chr2:81479852-81481890 | p2 | (AT)19 | TCACGAACAAGAGCACCAAC | GGAGAGAGGTTTCCCTTTCG | 125 |
| chr2:81502882-81504960 | p2 | (AT)39 | TAACAGGCTCCAGAGGAGGA | TTCATGTCGACCTCACGGTA | 131 |
| chr2:81560515-81562533 | p3 | (TTG)6 | AGAACGCTCGCCAAGTCTAA | TCATCTTCACCCTTTCGACC | 248 |
| chr2:81560515-81562533 | p2 | (TA)13 | AGAACGCTCGCCAAGTCTAA | TCATCTTCACCCTTTCGACC | 248 |
| chr2:81573105-81575117 | p2 | (AT)10 | TTAGGGCTTTTTGCCATGAA | GGTCCACAAGGACGAGAAAA | 298 |
| chr2:81573105-81575117 | p2 | (TA)6 | TTAGGGCTTTTTGCCATGAA | GGTCCACAAGGACGAGAAAA | 298 |
| chr2:81664254-81666269 | p2 | (TA)21 | CGCGGGCTAATGTTATTGAT | CGTCCAAAGAAAACGGATGT | 140 |
| chr2:81664254-81666269 | p3 | (ATA)5 | CGCGGGCTAATGTTATTGAT | CGTCCAAAGAAAACGGATGT | 140 |

| Marker ID | SSR type | SSR motif | FORWARD PRIMER (5'-3') | REVERSE PRIMER (5'-3') | size (bp) |
|---|---|---|---|---|---|
| chr2:81677734-81679748 | p2 | (TA)6 | CCCTTCTTTGTGCTCTCCAC | CCTCCCTTTTCCCCAATAAA | 284 |
| chr2:81677734-81679748 | p2 | (AT)7 | CCCTTCTTTGTGCTCTCCAC | CCTCCCTTTTCCCCAATAAA | 284 |
| chr2:81697634-81699652 | p3 | (CAA)6 | ACAGTAGGCACAAGCGGAGT | TTCGGGTCTATATTCGTCCG | 270 |
| chr2:81697634-81699652 | p2 | (CA)6 | ACAGTAGGCACAAGCGGAGT | TTCGGGTCTATATTCGTCCG | 270 |
| chr2:81745169-81747184 | p3 | (TTG)5 | GGGATGATCACTTGCCACTT | GACCAAGTCTGGCCCTATCA | 230 |
| chr2:81754363-81756397 | p2 | (TA)6 | TGGGCTAGATCCATGAGGAC | CAAATCCACGACACCAACAG | 212 |
| chr2:81754442-81756476 | p2 | (TA)6 | TGGGCTAGATCCATGAGGAC | CAAATCCACGACACCAACAG | 133 |
| chr2:81799825-81801849 | p2 | (TA)6 | CCGACACTAACCTGGTCGAT | TACCAGAGAAGCCTAGGGCA | 211 |
| chr2:81799894-81801924 | p2 | (TA)6 | CCGACACTAACCTGGTCGAT | TACCAGAGAAGCCTAGGGCA | 142 |
| chr2:81826583-81828639 | p2 | (TA)28 | GACGCCATTTGGTAACACCT | GTGCCCGTCATACTCCTCAT | 201 |
| chr2:81863409-81865421 | p2 | (TG)6 | TGATGTGTTACGGTTTCCGA | TGATAAACAAAGCACCTGCG | 252 |
| chr2:82028179-82030213 | p2 | (TA)17 | TGCCCAGGGAGATAGACTTG | TGGCATCTATGCCCATATCA | 117 |
| chr2:82104271-82106309 | p2 | (TA)6 | GGTGCATTGACTCCCCTAGA | CATTAAAAGCCGGATGAGGA | 270 |
| chr2:82104322-82106392 | p2 | (TA)6 | GGTGCATTGACTCCCCTAGA | CATTAAAAGCCGGATGAGGA | 219 |
| chr2:82162927-82164942 | p3 | (CAT)5 | ACATATCTCCCAAAGGCGTG | TCTGGATTCCTCGGAGCTTA | 136 |
| chr2:82248450-82250465 | p3 | (TTG)5 | TTCGGAGGGAGAAGCACTTA | CATTGGCATATCATGCAAGC | 185 |
| chr2:82253359-82255373 | p2 | (AG)7 | AAACACAGGTGTTGGTGCAA | CCCATCCCTCTTCAGACAAA | 290 |
| chr2:82253359-82255373 | p2 | (AT)7 | AAACACAGGTGTTGGTGCAA | CCCATCCCTCTTCAGACAAA | 290 |
| chr2:82289338-82291353 | p5 | (CATAT)3 | AGACACGCCCCTAGACTGAA | CTTGTTGTCCGATGGGAAGT | 272 |
| chr2:82306644-82308660 | p2 | (AT)8 | TATCTCAGATGGCCCCTCAC | TTATGCGTTCGCTTCTGATG | 136 |
| chr2:82393665-82395677 | p2 | (TG)6 | AAACCGTCGAACCTTCCTTT | CCTCAATATTTCCGCCTCAA | 198 |
| chr2:82393665-82395677 | p3 | (TTG)5 | AAACCGTCGAACCTTCCTTT | CCTCAATATTTCCGCCTCAA | 198 |
| chr2:82452490-82454505 | p5 | (ATTTC)3 | GACAGGGTGGATGAGGAAAA | CCTCTGCATAGGGGAATCAA | 111 |
| chr2:82452490-82454505 | p2 | (TA)20 | GACAGGGTGGATGAGGAAAA | CCTCTGCATAGGGGAATCAA | 111 |
| chr2:82524992-82527004 | p2 | (TC)6 | CCAGCAGTCACACGACAACT | ATTTTGTTGAATTGAGGCCG | 101 |
| chr2:82595216-82597228 | p2 | (AC)6 | CTGTGTCGGCTGGGATAGAT | GAGTCAGGATCGGAGACAGC | 279 |
| chr2:82647115-82649135 | p2 | (TA)7 | GGAACTGTCTGCAAAGCCTC | GAATCTTCCGCAGTAGGCTG | 285 |
| chr2:82647115-82649135 | p4 | (ACAT)4 | GGAACTGTCTGCAAAGCCTC | GAATCTTCCGCAGTAGGCTG | 285 |
| chr2:82647115-82649135 | p4 | (CATA)5 | GGAACTGTCTGCAAAGCCTC | GAATCTTCCGCAGTAGGCTG | 285 |
| chr2:82647115-82649135 | p3 | (ATC)5 | GGAACTGTCTGCAAAGCCTC | GAATCTTCCGCAGTAGGCTG | 285 |
| chr2:82649710-82651752 | p2 | (AT)21 | TTCTGTACCAAACCTTGCCC | TATCTTTGGGCTTGTGTCCC | 270 |
| chr2:82811336-82813351 | p3 | (TTG)5 | GGAAGGCTAAGGGCAAAATC | TACATGGGCTTGGGACATTT | 213 |
| chr2:82905700-82907718 | p6 | (CATATA)3 | ACTGAGTGGGCCGTATGAAC | ATTGAGAGCGCTCCAGTTGT | 287 |
| chr2:82917899-82919925 | p2 | (AT)13 | GGTTCTGAAAGCGGTCTTTG | GCCCTATAGTTGGGTCGTGA | 179 |
| chr2:82950040-82952142 | p3 | (AAG)34 | GGTATGGCTATCACGGCCTA | GCGTTGCTAAAGCGTCTTCT | 165 |
| chr2:82984375-82986402 | p3 | (ACA)9 | GTTGCAGGGATGCTTCATTT | ACGGGAACAAAGGGAGAACT | 110 |

| Marker ID | SSR type | SSR motif | FORWARD PRIMER (5'-3') | REVERSE PRIMER (5'-3') | size (bp) |
|---|---|---|---|---|---|
| chr2:83029890-83031914 | p2 | (AT)12 | TCAAACCCAAGTAAGGTCGG | GCGTGCTTTTCAGCACATAA | 133 |
| chr2:83101537-83103549 | p2 | (CA)6 | CTTGTGGAGAAGCCAGAAGG | CCTGACATGGGCTATGGAAT | 227 |
| chr2:83108346-83110366 | p2 | (AC)6 | TTTCTTGAAACCCCGACTTG | ACACCACAAAAGAGGGTTCG | 147 |
| chr2:83108346-83110366 | p2 | (AT)10 | TTTCTTGAAACCCCGACTTG | ACACCACAAAAGAGGGTTCG | 147 |
| chr2:83144247-83146262 | p5 | (AATGA)3 | TGCCATTGGAAAGAAAGGAC | AAGGAGATGCCGCCTAATTT | 234 |
| chr2:83369990-83372006 | p4 | (TCTG)4 | CAAGTGACGTGTAGGTGCGT | CACGATATACATACCGGCCC | 254 |
| chr2:83441247-83443267 | p2 | (AT)10 | CCCGATTTGGACACTCGTAT | GTCGGCTTGCGTATGGTATT | 154 |
| chr2:83441247-83443267 | p3 | (AAC)5 | CCCGATTTGGACACTCGTAT | GTCGGCTTGCGTATGGTATT | 154 |
| chr2:83441247-83443267 | p2 | (TA)13 | CCCGATTTGGACACTCGTAT | GTCGGCTTGCGTATGGTATT | 154 |
| chr2:83496008-83498024 | p2 | (AT)8 | TGCAACGTGGTCAAATCCTA | ACAGAGGAAACTCTGCCGAA | 173 |
| chr2:83496008-83498024 | p2 | (AT)7 | TGCAACGTGGTCAAATCCTA | ACAGAGGAAACTCTGCCGAA | 173 |
| chr2:83496707-83498721 | p2 | (AT)8 | TAGGCCATGACTAAGTGCCC | ACCTTGGAGTTCTTTTGCGA | 181 |
| chr2:83496707-83498721 | p2 | (AT)7 | TAGGCCATGACTAAGTGCCC | ACCTTGGAGTTCTTTTGCGA | 181 |
| chr2:83525983-83528013 | p2 | (TA)9 | ACACCTACCGAGAGCTGGAA | CCAAGAAGGACGATGGGTAA | 196 |
| chr2:83539677-83541692 | p5 | (AGCTT)3 | AAACATCGGGATCAACTTCG | CCGTTGGGTGTTTTATGGAC | 137 |
| chr2:83596111-83598126 | p3 | (CAA)5 | CCGAGGAACGTATAGTCCGA | TGATGATGATGGCCGTAGAA | 257 |
| chr2:83609043-83611058 | p3 | (CAA)5 | CCGAGGAACGTATAGTCCGA | TGATGATGATGGCCGTAGAA | 257 |
| chr2:83623053-83625075 | p2 | (AT)11 | GTGAAGTGTGCTCCTCGTCA | ACAGATTAATCCAGCCCCCT | 153 |
| chr2:83623053-83625075 | p2 | (AT)6 | GTGAAGTGTGCTCCTCGTCA | ACAGATTAATCCAGCCCCCT | 153 |
| chr2:83735593-83737607 | p2 | (CA)7 | TCACCCCGACAATTTTTCAT | CCCTTTGTGGGATTTCTTGA | 104 |
| chr2:83758114-83760142 | p2 | (AT)14 | TCGGCCTATGATGCCTACTC | GCCATGTGCTCACCCTAAAT | 145 |
| chr2:83827759-83829774 | p5 | (TTTTA)3 | CCAACCAAGCCCAATAGAAA | CCCAATTTGTTGGTGGATCT | 166 |
| chr2:83851634-83853660 | p2 | (CA)6 | CATGTTCGTGACACTACCCG | GGAGAAATAGCCCTGGGAAG | 215 |
| chr2:83851953-83853965 | p2 | (CA)6 | TGGAGACCTTTCGTCTGCTT | GGCATTGAGGATTGTCACCT | 294 |
| chr2:83889465-83891480 | p5 | (GATAT)3 | TGCTCTATTGTTCGGAGCCT | CGGATAAAGCACAAGGAAGC | 214 |
| chr2:83967823-83969841 | p2 | (AC)9 | ATACCTCGATCGCTCGCTAA | GATGAATTTGGCCGAGTTGT | 264 |
| chr2:84028269-84030293 | p6 | (ATACAC)4 | CAGGGTCAGACGTACGGAAT | TGCGAACCTGTCTGATATGC | 215 |
| chr2:84036859-84038881 | p2 | (TA)11 | ACTCGTTTTCGTCGACTCGT | GGCAAGGCATACGGTAGGTA | 200 |
| chr2:84109278-84111292 | p2 | (AT)7 | ACAGTAGCGCAAAGCAAGGT | GATCTTGTAACGCGCAGACA | 125 |
| chr2:84125902-84127914 | p2 | (TA)6 | CTTGTACCGAGCACCCCTTA | TCCGGAGCCTTATTTCATTG | 255 |
| chr2:84129830-84131842 | p2 | (TA)6 | GGACGCGAACAATTCAAAAT | GCGACAGTACTCAGCCCTTC | 261 |
| chr2:84164112-84166127 | p3 | (ATG)5 | GGGGGTTCCATGACATTCTA | CTCAACTCCTTCCAGGACCA | 267 |
| chr2:84187807-84189825 | p6 | (ATACAT)3 | TACCTGCCACTATCTTGCCC | GGTTGGGTCGTGACAAAAGT | 116 |
| chr2:84218939-84220957 | p6 | (ATACAT)3 | TACCTGCCACTATCTTGCCC | GGTTGGGTCGTGACAAAAGT | 116 |
| chr2:84246507-84248523 | p4 | (CCTT)4 | TTGCGTGTTTCTGATTCTCG | TTTCGATCAAAGGAGCAAGC | 125 |
| chr2:84302201-84304216 | p3 | (GTT)6 | GGCGAAAAGAGCATACAAGG | TGGGGTCACCTACAGAGGTC | 243 |

| Marker ID | SSR type | SSR motif | FORWARD PRIMER (5'-3') | REVERSE PRIMER (5'-3') | size (bp) |
|---|---|---|---|---|---|
| chr2:84302201-84304216 | p5 | (TGATA)3 | GGCGAAAAGAGCATACAAGG | TGGGGTCACCTACAGAGGTC | 243 |
| chr2:84315076-84317092 | p3 | (CAT)6 | ACTGGACTATGCCAACCTGC | CATAGAGGCCGGAAATGAAA | 225 |
| chr2:84315157-84317173 | p3 | (CAT)6 | ACTGGACTATGCCAACCTGC | CATAGAGGCCGGAAATGAAA | 144 |
| chr2:84315183-84317199 | p3 | (CAT)6 | ACTGGACTATGCCAACCTGC | CATAGAGGCCGGAAATGAAA | 118 |
| chr2:84388511-84390529 | p6 | (CATATA)3 | TGGTAGAGTTGGGGTTCAGC | GTCTTGACATACACGGGGCT | 150 |
| chr2:84451907-84453921 | p2 | (TA)7 | TGAATCCCAAGCAACCTTTC | CCCAAAAGAAGACACCCAAA | 114 |
| chr2:84465876-84467888 | p3 | (CAA)7 | ATGGAGATCAAAGACGTCGG | ATTAGCGAGCGATCGAGGTA | 189 |
| chr2:84465876-84467888 | p4 | (TTTC)4 | ATGGAGATCAAAGACGTCGG | ATTAGCGAGCGATCGAGGTA | 189 |
| chr2:84471423-84473439 | p2 | (TA)8 | AATCCCAAGCAACCTTTCCT | CAATTTCGAGTTCTTCCCCA | 142 |
| chr2:84480731-84482745 | p2 | (TA)7 | TGAATCCCAAGCAACCTTTC | CCCAAAAGAAGACACCCAAA | 142 |
| chr2:84730162-84732180 | p3 | (GAT)6 | TAAACAAAGGAATGCTGCCC | ATTTTCCTTCGATGTCCACG | 163 |
| chr2:84735935-84737950 | p5 | (AAGAA)3 | GCAAGATTTGCCTCCAAGAG | AGAGCGGTCTGAGCCATTTA | 180 |
| chr2:84779144-84781159 | p5 | (AAGAA)3 | GCAAGATTTGCCTCCAAGAG | AGAGCGGTCTGAGCCATTTA | 180 |
| chr2:84832735-84834759 | p2 | (AT)12 | AAATGCCATGTAGCCTCCTG | AGTGGTGTTCGGTGGTTAGC | 288 |
| chr2:84832735-84834759 | p2 | (AT)44 | AAATGCCATGTAGCCTCCTG | AGTGGTGTTCGGTGGTTAGC | 288 |
| chr2:84832735-84834759 | p2 | (AT)34 | AAATGCCATGTAGCCTCCTG | AGTGGTGTTCGGTGGTTAGC | 288 |
| chr2:84833187-84835275 | p2 | (AT)12 | AGCCCTTCGGCTATCAAAAT | AGACAGGGAGGTGATCCAGA | 114 |
| chr2:84833187-84835275 | p2 | (AT)44 | AGCCCTTCGGCTATCAAAAT | AGACAGGGAGGTGATCCAGA | 114 |
| chr2:84833187-84835275 | p2 | (AT)34 | AGCCCTTCGGCTATCAAAAT | AGACAGGGAGGTGATCCAGA | 114 |
| chr2:84833187-84835275 | p2 | (TA)34 | AGCCCTTCGGCTATCAAAAT | AGACAGGGAGGTGATCCAGA | 114 |
| chr2:84972831-84974845 | p2 | (GT)7 | ACAAGCCCATGCTCATAACC | CCTGAGTACCTGGAATGCGT | 181 |
| chr2:84993624-84995636 | p2 | (GT)6 | GTTTCCTACGCCCACAGGTA | AGCATAACGGGGGAGTCTCT | 182 |
| chr2:85045146-85047158 | p2 | (AT)6 | ACTCGGTACGTTGCAAATCC | TTCATGTCGTTGCACAGTGA | 145 |
| chr2:85114776-85116791 | p5 | (TGAAT)3 | CGTGTGACATAAGGTGTGGG | TTGCTTCAACAACCATTCCA | 196 |
| chr2:85246743-85248755 | p2 | (AT)6 | ACCGCATTAAGACGCAAATC | CCTTTTGATTCGGAGCACAT | 128 |
| chr2:85248050-85250066 | p2 | (CA)8 | AAACTCAATCCCCATTTCCC | GGCCTAGTAAATTCTCGGGG | 286 |
| chr2:85394532-85396552 | p2 | (TA)10 | TGGGGCTGTTTGGTATGATT | ACTCCGTCCAACTCACCAAC | 290 |
| chr2:85560049-85562079 | p2 | (AT)15 | GAGTGGCACACTCAAGGTCA | TGCCCTGTAGCTTCCTGTTT | 222 |
| chr2:85560049-85562079 | p2 | (AT)6 | GAGTGGCACACTCAAGGTCA | TGCCCTGTAGCTTCCTGTTT | 222 |
| chr2:85660887-85662899 | p2 | (AC)6 | TTGTAGATCGGGTTTCGAGG | ACCATTTGCAGATTTCGGAC | 189 |
| chr2:85668645-85670657 | p2 | (AC)6 | ACATGCCAAACGAAAGAAGG | TTTATAGGGGAAATGCTGCG | 243 |
| chr2:85770123-85772175 | p2 | (TA)26 | TACGGGGCATAACATCCAAT | TTGCACGCTCAGACATAAGC | 111 |
| chr2:85770123-85772175 | p2 | (AT)11 | TACGGGGCATAACATCCAAT | TTGCACGCTCAGACATAAGC | 111 |
| chr2:85820375-85822387 | p2 | (TG)6 | GGAGACTCAGGCAGAGATGC | CCTTCCTCAATATTTCCGCA | 178 |
| chr2:85833223-85835238 | p3 | (GAA)5 | AGCATGGAGATAACGGATGG | ATGCCATTAATGAGGCAAGG | 236 |
| chr2:85844010-85846024 | p2 | (AC)7 | TCCCAATATTCCGCGTAGAG | TGCACGCAGAAAATAGCAAC | 166 |

| Marker ID | SSR type | SSR motif | FORWARD PRIMER (5'-3') | REVERSE PRIMER (5'-3') | size (bp) |
|---|---|---|---|---|---|
| chr2:85903214-85905228 | p2 | (GT)7 | AATATCCTCGCCTGTTCCCT | CAAATGTGTGAAAACACGGC | 127 |
| chr2:85903214-85905228 | p3 | (TTG)5 | AATATCCTCGCCTGTTCCCT | CAAATGTGTGAAAACACGGC | 127 |
| chr2:85904789-85906804 | p3 | (TTG)5 | TTTGAAGGAAACGGGAATTG | TTCCATTGAACGAAAAAGGG | 211 |
| chr2:85904789-85906804 | p5 | (TGATA)3 | TTTGAAGGAAACGGGAATTG | TTCCATTGAACGAAAAAGGG | 211 |
| chr2:86020578-86022596 | p3 | (AAT)6 | CTGCAACTTTTTCGGTGACA | ATTCTTTCGGGGTTTTGCTT | 206 |
| chr2:86119743-86121757 | p2 | (TA)7 | CATCGCTAAGATGGGCTGAT | AGGCACGGTGGAATCATAAG | 291 |
| chr2:86231645-86233660 | p5 | (ACCCC)3 | GTATGAGCCAACTTGGGGAA | GTATCGACGAGCGGTTTGTT | 175 |
| chr2:86231645-86233660 | p3 | (ATG)5 | GTATGAGCCAACTTGGGGAA | GTATCGACGAGCGGTTTGTT | 175 |
| chr2:86231645-86233660 | p2 | (TA)6 | GTATGAGCCAACTTGGGGAA | GTATCGACGAGCGGTTTGTT | 175 |
| chr2:86285165-86287183 | p3 | (TGT)6 | ATGGATGAAATGGAAGCTCG | ATATCCATCTCAGCATCCGC | 221 |
| chr2:86403060-86405072 | p2 | (CA)6 | TTCAAGGAACAGAATTGCCC | AGCTAAGTCTCGGGGGTGTT | 203 |
| chr2:86416279-86418299 | p2 | (AT)10 | GATCCTCTGCTGGCTTTGAC | GACAGGGCTATGAGAGACGC | 128 |
| chr2:86432552-86434564 | p2 | (TG)6 | CCTGCGGGTTCAAGTGTTAT | TTGGCCCAAATTCGTAAAAG | 263 |
| chr2:86475684-86477698 | p2 | (TA)7 | TTCACCTTAAATTCCAGCCG | CAACCATAGGTTGTGGTCCC | 199 |
| chr2:86538774-86540789 | p5 | (ATTTC)3 | TTGCTCGTACTGACCCCTCT | GGAGCCGCTTACAGACTGAC | 285 |
| chr2:86613713-86615733 | p5 | (TGTGT)3 | AAATGAGCCATGGGAAGTTG | TGCTAAAACGCAACAACGAG | 135 |
| chr2:86613713-86615733 | p5 | (AATTG)4 | AAATGAGCCATGGGAAGTTG | TGCTAAAACGCAACAACGAG | 135 |
| chr2:86641644-86643658 | p2 | (AT)7 | TGACACTGACTGCCTTACGC | ATGCTAGGTGGCGACTCTGT | 149 |
| chr2:86657240-86659298 | p3 | (CAA)5 | GTTCGCTAATCAAATCCCGA | TTGGCCGAGTTAAATTCTCG | 236 |
| chr2:86657240-86659298 | p2 | (AT)29 | GTTCGCTAATCAAATCCCGA | TTGGCCGAGTTAAATTCTCG | 236 |
| chr2:86698612-86700624 | p2 | (AT)6 | ATTAGCGAGCGATCGAGGTA | ATGGAGATCAAAGACGTCGG | 155 |
| chr2:86729636-86731654 | p6 | (TGAACC)3 | TCCTTTCCTTCATTGGCAAC | TATGGAACCCTCAGGACGAG | 118 |
| chr2:86729636-86731654 | p2 | (AT)43 | TCCTTTCCTTCATTGGCAAC | TATGGAACCCTCAGGACGAG | 118 |
| chr2:86775252-86777270 | p6 | (TGAACC)3 | TCCTTTCCTTCATTGGCAAC | TATGGAACCCTCAGGACGAG | 118 |
| chr2:86775252-86777270 | p2 | (AT)43 | TCCTTTCCTTCATTGGCAAC | TATGGAACCCTCAGGACGAG | 118 |
| chr2:86890558-86892573 | p5 | (TGTGG)3 | GACGAAAGTTCCTAAGGGGG | TTTCCAAATTTGCCTTCGTC | 187 |
| chr2:86912610-86914624 | p2 | (AC)7 | GTTTCGAGGAGCAGAAATGC | AAGCCTCGGGGATGTTAGAT | 146 |
| chr2:86922582-86924594 | p2 | (AT)6 | CTACCGGAACGGATTGAAGA | AGGTCGGGGTGTGACAGTAG | 266 |
| chr2:86934180-86936196 | p4 | (TCAA)4 | TTTGCATTTGCCGATTTGTA | CGGCAAAGAGTGAAGCCTAC | 167 |
| chr2:86992335-86994347 | p2 | (AT)6 | AATCATATGACGGACAGGGC | CGGGTACAGAGGTTCGTGAT | 244 |
| chr2:87001094-87003116 | p2 | (AT)11 | CACAACCTAAACTCCCGCAT | TGTTGTGACAGGATGGCAGT | 202 |
| chr2:87092133-87094148 | p2 | (CA)6 | TACACCAATTTCAATGCCGA | GTGGTGAGGGGAAGAGATGA | 274 |
| chr2:87092133-87094148 | p5 | (AAATA)3 | TACACCAATTTCAATGCCGA | GTGGTGAGGGGAAGAGATGA | 274 |
| chr2:87116393-87118429 | p2 | (AT)6 | ACACAACACCCAAAGCAACA | ATTGGGTTGGAGGTGTGAAG | 277 |
| chr2:87134423-87136435 | p2 | (TA)6 | GGGACCCGAATACCCTAAAA | CCCTTTGGACTCACTTCGAG | 158 |
| chr2:87166029-87168044 | p5 | (AATTG)3 | AGAAAGGGGGCCAGTAAGAA | CTAAGCGTTGCGATGTTTCA | 119 |

| Marker ID | SSR type | SSR motif | FORWARD PRIMER (5'-3') | REVERSE PRIMER (5'-3') | size (bp) |
|---|---|---|---|---|---|
| chr2:87286725-87288741 | p2 | (AG)8 | TTTGGGCAAAAATTCAAAGG | CGGGATACTTAAGGCAACCA | 161 |
| chr2:87364375-87366390 | p5 | (TGCAT)3 | GAGGCGCTACAGAAGGATTG | CGACAAGCCACAATCAACAG | 105 |
| chr2:87384070-87386094 | p3 | (AAC)8 | AGGGTAATCGAAATGGTCCC | GAACGAGATTGCAACGTGAA | 285 |
| chr2:87420287-87422302 | p3 | (TAT)5 | CGTCCGACTTCATTGGATTT | GGCTCAGTGTGGGAAACATT | 131 |
| chr2:87487114-87489126 | p2 | (TG)6 | ATAGGTCGTGGGCCTGTATG | AATGTGCCTAACCCGACTTG | 198 |
| chr2:87489613-87491635 | p2 | (GA)11 | TCGGAAAATAAGCACCTTGG | TACCAAAAGTATGGGCTCCG | 283 |
| chr2:87489613-87491635 | p3 | (GTT)6 | TCGGAAAATAAGCACCTTGG | TACCAAAAGTATGGGCTCCG | 283 |
| chr2:87563855-87565873 | p2 | (AT)9 | CGCACCTTGGGGTAATTCTA | GGCCTCGGAAACATATCAAA | 104 |
| chr2:87590279-87592297 | p6 | (TTCATA)3 | AGTGTCACGACCCGACTAGG | TGATAAAGGCCTTGTCGTCC | 257 |
| chr2:87621859-87623874 | p5 | (GAAAT)3 | CAATCGGGCGTATCAAGAAT | GAAACGCTAGCATTGGAAGG | 153 |
| chr2:87675059-87677071 | p2 | (AT)6 | TTGTATAGGCCCAAACAGCC | ATATGAGTGTCCTGCTCGGG | 155 |
| chr2:87675059-87677071 | p2 | (AT)6 | TTGTATAGGCCCAAACAGCC | ATATGAGTGTCCTGCTCGGG | 155 |
| chr2:87675059-87677071 | p6 | (TAGAAT)3 | TTGTATAGGCCCAAACAGCC | ATATGAGTGTCCTGCTCGGG | 155 |
| chr2:87738458-87740473 | p3 | (GTT)5 | ACCCTATCACAACAGCCCAC | CATCTATTTCAGCCCCCAGA | 240 |
| chr2:87738458-87740473 | p2 | (AT)23 | ACCCTATCACAACAGCCCAC | CATCTATTTCAGCCCCCAGA | 240 |
| chr2:87932969-87934981 | p2 | (AT)6 | TCACATGGTTCCTCATTCCA | TGGCTTCCATTAAGGTACGG | 112 |
| chr2:87948376-87950394 | p3 | (ATA)6 | CCACTGTTCACCCAACACAG | TGCTCTTTGAGCAAAACCCT | 157 |
| chr2:88233197-88235254 | p3 | (AAT)19 | CGCTCCTGCTATTCCTTTTG | TTATCCACCTCTCGTGACCC | 195 |
| chr2:88317986-88320001 | p5 | (ATGAT)3 | TAGGGGAGATGCTGTCCAAC | TTGCCAAAAGAAGAATTGGG | 128 |
| chr2:88335631-88337653 | p2 | (AT)11 | AAGCCCATTTTGTAAACCCC | CTTGCTGGGAATGATGGAGT | 121 |
| chr2:88457394-88459410 | p2 | (TG)8 | AAATTTAGCCAGCACGTTCG | ACTCGGACTGGTCCTGAAGA | 122 |
| chr2:88464682-88466697 | p5 | (TTTTA)3 | AGCTTTTTGCTCGTCGTGTT | GCCTCCAAAGTACAACCCAA | 118 |
| chr2:88555916-88557931 | p3 | (TGT)5 | GGAAAGAAAGGCATTGGTGA | CCCTCATCCAAGCAATCAAT | 283 |
| chr2:88562843-88564858 | p3 | (ATC)6 | AGAGCCTATAAGGCCGGGTA | CCAGCCATCTATGTTCGGTT | 106 |
| chr2:88675454-88677518 | p2 | (TA)32 | GACCCAAATCGATGAGTCGT | TCAGATGATGCAGCCTTGTC | 298 |
| chr2:88705373-88707393 | p2 | (TG)6 | GCTTATGTTTTGGGCCATGT | CTTGTTGTGCACCCCCTACT | 197 |
| chr2:88705373-88707393 | p2 | (AT)10 | GCTTATGTTTTGGGCCATGT | CTTGTTGTGCACCCCCTACT | 197 |
| chr2:88823280-88825294 | p2 | (AT)7 | TGTGACAATCCTACGGTCCA | GTGTGTGTTGGGCAAGAATG | 126 |
| chr2:88941214-88943238 | p2 | (AT)12 | AATAACCGAACTCGCCATTG | GCCTTAGCCTGATCATCTGC | 277 |
| chr2:89168225-89170300 | p3 | (AAT)25 | GACTTAACAGCGCTTACCCG | AGAAGGTCCGGTTCTTGGTT | 126 |
| chr2:89242200-89244216 | p4 | (TCTT)4 | TCCTGAATTCGCCTTTGAAC | CCTAATTGAATGCCCTCGAA | 230 |
| chr2:89264983-89266995 | p2 | (GA)10 | TACCACTCAAAGGTTTCGGG | CAATCCGAAAATTGCTCCAT | 153 |
| chr2:89264983-89266995 | p2 | (TA)11 | TACCACTCAAAGGTTTCGGG | CAATCCGAAAATTGCTCCAT | 153 |
| chr2:89264983-89266995 | p3 | (CAT)5 | TACCACTCAAAGGTTTCGGG | CAATCCGAAAATTGCTCCAT | 153 |
| chr2:89264983-89266995 | p2 | (TA)32 | TACCACTCAAAGGTTTCGGG | CAATCCGAAAATTGCTCCAT | 153 |
| chr2:89265016-89267038 | p2 | (GA)10 | TACCACTCAAAGGTTTCGGG | CAATCCGAAAATTGCTCCAT | 120 |

| Marker ID | SSR type | SSR motif | FORWARD PRIMER (5'-3') | REVERSE PRIMER (5'-3') | size (bp) |
|---|---|---|---|---|---|
| chr2:89265016-89267038 | p2 | (TA)11 | TACCACTCAAAGGTTTCGGG | CAATCCGAAAATTGCTCCAT | 120 |
| chr2:89265016-89267038 | p3 | (CAT)5 | TACCACTCAAAGGTTTCGGG | CAATCCGAAAATTGCTCCAT | 120 |
| chr2:89265016-89267038 | p2 | (TA)32 | TACCACTCAAAGGTTTCGGG | CAATCCGAAAATTGCTCCAT | 120 |
| chr2:89265146-89267168 | p2 | (GA)10 | CTTCGAGTTGCTGAGGCTCT | GTGACAAAGGCCGATTGATT | 159 |
| chr2:89265146-89267168 | p2 | (TA)11 | CTTCGAGTTGCTGAGGCTCT | GTGACAAAGGCCGATTGATT | 159 |
| chr2:89265146-89267168 | p3 | (CAT)5 | CTTCGAGTTGCTGAGGCTCT | GTGACAAAGGCCGATTGATT | 159 |
| chr2:89265146-89267168 | p2 | (TA)32 | CTTCGAGTTGCTGAGGCTCT | GTGACAAAGGCCGATTGATT | 159 |
| chr2:89307517-89309557 | p2 | (TG)20 | CAGTTCCCACGGGATAGAGA | ACAACTCTATCGGGGTGACG | 128 |
| chr2:89338123-89340150 | p3 | (ATG)9 | TCGAATGTGCCGTAATTTCA | ATATTCGGCGAAGTTTGTGG | 123 |
| chr2:89395631-89397646 | p3 | (ACA)5 | AACCCCAGCTTCCAAAACTT | TTTGCAAAAGCTGTCCAGTG | 175 |
| chr2:89485911-89487929 | p6 | (ATGTAT)3 | GAGCAGGTCTGATTTCCTGC | TCAAGGACAAAAACCAAGGC | 230 |
| chr2:89605950-89607976 | p2 | (AT)14 | GGGAACGTCCCAAAACATAA | ACAGATAGCCGGTTGGTGAC | 214 |
| chr2:89605950-89607976 | p2 | (AT)13 | GGGAACGTCCCAAAACATAA | ACAGATAGCCGGTTGGTGAC | 214 |
| chr2:89640985-89642997 | p2 | (AT)8 | CCTACCTGTGGAGCTGGAAG | ACCAGTAGGACCCGGAGAGT | 190 |
| chr2:89641004-89643030 | p2 | (AT)8 | CCTACCTGTGGAGCTGGAAG | ACCAGTAGGACCCGGAGAGT | 171 |
| chr2:89652353-89654365 | p2 | (AT)6 | AGTTGGGAAACGAGTTGTGG | CGAGTCTAGTTTGCAACGCA | 298 |
| chr2:89840029-89842059 | p2 | (AT)15 | CGTCAAATGTGCCACAAATC | TTGAAAGGACCTTGTTTGCC | 296 |
| chr2:89968094-89970109 | p5 | (TAAAA)3 | TACTTTCCGCATAACACCCC | GCTTTAGACAGTCCCCCTCC | 187 |
| chr2:89981439-89983467 | p2 | (GA)14 | AGCAGCTTATGTTTGTGGGG | GCAACCCTAATGAGAACCCA | 120 |
| chr2:90045436-90047451 | p5 | (ATTGC)3 | GTGATTGTTGGTGTTTTGCG | TCACGTGGCATCAAGTCAAT | 119 |
| chr2:90061125-90063140 | p5 | (ATTGC)3 | GTGATTGTTGGTGTTTTGCG | TCACGTGGCATCAAGTCAAT | 119 |
| chr2:90100591-90102606 | p5 | (AAAAT)3 | TTAGTTCATCTCAACGGGGG | GATCCATGTTAGGTTGCGCT | 135 |
| chr2:90100591-90102606 | p2 | (AT)10 | TTAGTTCATCTCAACGGGGG | GATCCATGTTAGGTTGCGCT | 135 |
| chr2:90219280-90221295 | p5 | (CATTG)3 | CCTGAAACTCAGCAGAAGGG | GCTAGCCCCAATCACACAAT | 200 |
| chr2:90264277-90266292 | p5 | (TCATT)3 | GAAATGGTTTCAATGGTGGG | ATGGACAAAGCTATGCCTGG | 199 |
| chr2:90318021-90320037 | p2 | (TA)8 | TGGGTCGTGACAAAGTGGTA | TACATGCCCTTCAGCCTCTT | 182 |
| chr2:90322577-90324595 | p3 | (TGT)6 | AAGCCTAGTTTCCAACGCAA | AACAATGAGGGAAAAACCCC | 117 |
| chr2:90322577-90324595 | p2 | (TA)7 | AAGCCTAGTTTCCAACGCAA | AACAATGAGGGAAAAACCCC | 117 |
| chr2:90353797-90355812 | p5 | (ATTCA)3 | TAAGGGGGAGACTGTCAACG | CGGATTGACCTGGAACTTGT | 127 |
| chr2:90353797-90355812 | p2 | (TA)10 | TAAGGGGGAGACTGTCAACG | CGGATTGACCTGGAACTTGT | 127 |
| chr2:90421335-90423353 | p6 | (ATTATG)3 | CCTGAGTGAGTGAAGCCTCC | CCCCACACATTCCTCTTTTG | 158 |
| chr2:90440079-90442094 | p5 | (AAAAT)3 | TACGCACTTTTGCAATCAGC | GAGATGTGCCAATCGGAGTT | 244 |
| chr2:90486727-90488741 | p2 | (AT)7 | CCTTGTTGTTGGGCTTGAGT | ATTAGGGGAGCAAGAAGGGA | 229 |
| chr2:90530163-90532188 | p5 | (ATTTT)5 | TGATCGTTGCTTGTCTTTGC | GCACCAAAGGAGCTCAAATC | 273 |
| chr2:90548508-90550523 | p5 | (ATTTC)3 | AGCGCTAAGTGGTTCTCCAA | GAGATCCAAGAATGGCCTGA | 128 |
| chr2:90612455-90614471 | p5 | (TATTT)3 | TGATGTGGAAGGCGTCATTA | TTTGGAGGGTGGTTGAAAAG | 169 |

| Marker ID | SSR type | SSR motif | FORWARD PRIMER (5'-3') | REVERSE PRIMER (5'-3') | size (bp) |
|---|---|---|---|---|---|
| chr2:90612455-90614471 | p4 | (ATTT)4 | TGATGTGGAAGGCGTCATTA | TTTGGAGGGTGGTTGAAAAG | 169 |
| chr2:90619291-90621305 | p2 | (AT)7 | TCTAGGCCGGATGTTAGTGG | GCGAACCCTCAACCAAAATA | 220 |
| chr2:90638262-90640304 | p2 | (AT)21 | CCCAAAAGGACGATTCTTCA | AGTGGACAGACACACGCATC | 142 |
| chr2:90676587-90678603 | p2 | (AT)8 | ACGCTGCGTTCTTTCTTGTT | AAACATGTCGGCACATCAAA | 189 |
| chr2:90714230-90716246 | p2 | (AT)8 | ACGCTGCGTTCTTTCTTGTT | AAACATGTCGGCACATCAAA | 189 |
| chr2:90857467-90859482 | p3 | (TTC)5 | CCTTCCTATGGTGCGTGAAT | CTTTGTCGTTATCGCGGTTT | 198 |
| chr2:90894427-90896442 | p2 | (TA)7 | CGTTAAATGCACACACGTCC | TAAGGCTGCATGGTTTCCTC | 211 |
| chr2:90894427-90896442 | p5 | (TTTCT)3 | CGTTAAATGCACACACGTCC | TAAGGCTGCATGGTTTCCTC | 211 |
| chr2:90894427-90896442 | p5 | (TTTTA)3 | CGTTAAATGCACACACGTCC | TAAGGCTGCATGGTTTCCTC | 211 |
| chr2:90894587-90896602 | p5 | (TTTCT)3 | GAGGAAACCATGCAGCCTTA | ATCGTGAAAATGAAGGCGAC | 274 |
| chr2:90894587-90896602 | p5 | (TTTTA)3 | GAGGAAACCATGCAGCCTTA | ATCGTGAAAATGAAGGCGAC | 274 |
| chr2:90900051-90902079 | p3 | (TTG)5 | ACTCTCGTTCCTTGCGTGTT | GGCCATCACGCCATAGTAGT | 255 |
| chr2:90935586-90937601 | p2 | (TA)11 | CAGACAACCCCCTAGGACAA | TCCCAAAGGTCTGTTGGTTC | 165 |
| chr2:90935635-90937650 | p2 | (TA)11 | CAGACAACCCCCTAGGACAA | TCCCAAAGGTCTGTTGGTTC | 116 |
| chr2:90936041-90938063 | p2 | (TA)11 | GAGGGGTCAGTACGCAAAAA | CATAATGGGACCGGATATGG | 206 |
| chr2:90969865-90971883 | p2 | (TA)6 | TAAATGGCCGAGCAGGATAC | CTCCCTGTCTACTTGCAGGC | 287 |
| chr2:90969865-90971883 | p2 | (TA)9 | TAAATGGCCGAGCAGGATAC | CTCCCTGTCTACTTGCAGGC | 287 |
| chr2:91149559-91151574 | p5 | (AATGC)3 | GTCTCGGTAGCACAACCCAT | CTCGATGTTTCTGTCGGGAT | 249 |
| chr2:91149559-91151574 | p5 | (CCTTT)3 | GTCTCGGTAGCACAACCCAT | CTCGATGTTTCTGTCGGGAT | 249 |
| chr2:91201442-91203457 | p3 | (ATC)5 | GGAGTGCTCTTGAGAATCCG | GATAGGGGGTGATGATGTGG | 140 |
| chr2:91208223-91210238 | p2 | (AT)10 | TCATCATTACCAGCCGACAA | GCAAGTAGGCAGGGAGACTG | 259 |
| chr2:91208223-91210238 | p3 | (ATC)5 | TCATCATTACCAGCCGACAA | GCAAGTAGGCAGGGAGACTG | 259 |
| chr2:91398749-91400767 | p2 | (AT)9 | ATCGTGGAACTGGTGGTCTC | TCAAGGCAGATTTATTCCCG | 146 |
| chr2:91561321-91563387 | p2 | (AT)33 | CTTCAAATCGGGGATAAGCA | GAAAACCGATAGTGGGACGA | 182 |
| chr2:91561321-91563387 | p5 | (AATGC)3 | CTTCAAATCGGGGATAAGCA | GAAAACCGATAGTGGGACGA | 182 |
| chr2:91574650-91576710 | p2 | (TA)30 | CTTTTTGTGCTTCCAAAGCC | TCGAGTTCAGGCCTCTCAGT | 181 |
| chr2:91592632-91594644 | p2 | (TG)6 | CCTTGACTGAAGGAAACCCA | CCTTCCTCCGTATTTCCACA | 277 |
| chr2:91640798-91642813 | p3 | (TTG)5 | AGTGAGTATAGAGGCGGGCA | CAACTTGGCCTTTTGTGGTT | 131 |
| chr2:91660572-91662587 | p5 | (TTCAT)3 | TACCCGCACTTATTTCAGCC | ACGGGGCCACAACATAATAG | 206 |
| chr2:91713412-91715427 | p5 | (TGATA)3 | TTGAAGATTTTGGGCGAAAC | TCACCTACGGAGGTACTGGG | 204 |
| chr2:91866729-91868747 | p3 | (CAA)6 | ACCGTACCCGGCTCTTTAGT | CGGGCTGGATACATAATCGT | 122 |
| chr2:91866729-91868747 | p2 | (TC)6 | ACCGTACCCGGCTCTTTAGT | CGGGCTGGATACATAATCGT | 122 |
| chr2:91888458-91890518 | p3 | (AAG)20 | GAGCAGTGGTACGTGCTGAA | AGCGACCAAACATGGAAAAG | 180 |
| chr2:91954889-91956905 | p2 | (AT)8 | GAGATTTAGCCGCACTCTGG | AAACAAGAAGCTGCGGACAT | 264 |
| chr2:92004536-92006550 | p2 | (TA)7 | CATCCCTATCACCATGACCC | AGTATCGCCGACGGTTATTG | 165 |
| chr2:92005060-92007074 | p2 | (TA)7 | CACCCTATGACGGAACTGCT | GTGCCTTAGCCCAATGAAAA | 115 |

| Marker ID | SSR type | SSR motif | FORWARD PRIMER (5'-3') | REVERSE PRIMER (5'-3') | size (bp) |
|---|---|---|---|---|---|
| chr2:92024615-92026666 | p5 | (TTGCA)3 | CAGGTCATGGCTACTGAGCA | CATTCACCTTGCTACTGCGA | 222 |
| chr2:92024615-92026666 | p3 | (TTA)17 | CAGGTCATGGCTACTGAGCA | CATTCACCTTGCTACTGCGA | 222 |
| chr2:92123289-92125303 | p2 | (AT)7 | TACAAATAGGGCCGACAAGG | GATGACCTCGTCAGTCAGCA | 239 |
| chr2:92225324-92227354 | p2 | (AT)15 | CTCCACGGACTTGACCTCAT | CAGATACACCGAGCCAGGAT | 200 |
| chr2:92263258-92265276 | p6 | (TATGTA)3 | TATTCCACCGAGGCTTTGAC | TCCAAAACAAGTTTCGGGTC | 257 |
| chr2:92372918-92374982 | p2 | (TA)32 | TTGTTGCTGGGTGTGTTGAT | GTTGCTCGCTTGCTTTTCTT | 296 |
| chr2:92419249-92421294 | p3 | (ACA)11 | CAAGGTTGGTGCATTCATTG | CATTTGGGGATTACACTGGG | 116 |
| chr2:92419249-92421294 | p3 | (ATT)15 | CAAGGTTGGTGCATTCATTG | CATTTGGGGATTACACTGGG | 116 |
| chr2:92494436-92496451 | p3 | (CTT)5 | AAATATTGAAGGTTCGGGGG | GCGCTAAAAAGCCAGTAACG | 263 |
| chr2:92494436-92496451 | p5 | (TTTTC)3 | AAATATTGAAGGTTCGGGGG | GCGCTAAAAAGCCAGTAACG | 263 |
| chr2:92685226-92687240 | p2 | (TA)7 | GATAGAGACCGCTCATTCGC | ACCTGCTCTCTGTGCACCTT | 111 |
| chr2:92795353-92797371 | p6 | (TGTAAG)3 | TTGGTAGGGCCAGGTAGATG | TACCCGACTACGAAGGATCG | 140 |
| chr2:92867556-92869574 | p2 | (AT)8 | ATTCTATCGATGTGCCAGGG | AGAAGAGGGCGAGTCACAAA | 220 |
| chr2:92867556-92869574 | p6 | (ATACAT)3 | ATTCTATCGATGTGCCAGGG | AGAAGAGGGCGAGTCACAAA | 220 |
| chr2:92900197-92902247 | p2 | (AT)15 | GCATGAGCAGCATTTTTCAA | TTGGGTCGATGCAGAATACA | 254 |
| chr2:92900283-92902295 | p2 | (AT)15 | GCATGAGCAGCATTTTTCAA | TTGGGTCGATGCAGAATACA | 168 |
| chr2:92911954-92913968 | p2 | (AT)7 | CTCACGGGCAACCAAAGTAT | ATTCATGCGCAAGCCTAGTT | 211 |
| chr2:92970372-92972387 | p3 | (GTT)5 | TATTTGGGCATTTGGGAAAA | CCGCCTTAAAACGTCCATAA | 263 |
| chr2:93070850-93073015 | p3 | (TTC)55 | GCCCAAGGGAAGCCTTATAG | AAAGCACAAGGAGGCACTGT | 119 |
| chr2:93070850-93073015 | p3 | (CAA)5 | GCCCAAGGGAAGCCTTATAG | AAAGCACAAGGAGGCACTGT | 119 |
| chr2:93096078-93098243 | p3 | (TTC)55 | GCCCAAGGGAAGCCTTATAG | AAAGCACAAGGAGGCACTGT | 119 |
| chr2:93096078-93098243 | p3 | (CAA)5 | GCCCAAGGGAAGCCTTATAG | AAAGCACAAGGAGGCACTGT | 119 |
| chr2:93106663-93108678 | p5 | (TTTAA)3 | ACAAGAGGGTATATCGGGGG | TTACGAAAACCAAGCGAACC | 150 |
| chr2:93164545-93166559 | p2 | (TG)6 | GTTGTTGTGGCCTTGTGATG | AACCTCCAATCTTCGACACG | 114 |
| chr2:93164545-93166559 | p2 | (TA)7 | GTTGTTGTGGCCTTGTGATG | AACCTCCAATCTTCGACACG | 114 |
| chr2:93164545-93166559 | p2 | (TA)16 | GTTGTTGTGGCCTTGTGATG | AACCTCCAATCTTCGACACG | 114 |
| chr2:93164545-93166559 | p2 | (AT)12 | GTTGTTGTGGCCTTGTGATG | AACCTCCAATCTTCGACACG | 114 |
| chr2:93165167-93167191 | p2 | (TA)7 | ATTAGCGAGCGATCGAGGTA | ACATCCGGAAACGTATCGAC | 125 |
| chr2:93165167-93167191 | p2 | (TA)16 | ATTAGCGAGCGATCGAGGTA | ACATCCGGAAACGTATCGAC | 125 |
| chr2:93165167-93167191 | p2 | (AT)12 | ATTAGCGAGCGATCGAGGTA | ACATCCGGAAACGTATCGAC | 125 |
| chr2:93265402-93267417 | p3 | (TTC)17 | CTTGCAATCTGCATTCTCCA | AGACCTAGTTGCGCTCGTGT | 266 |
| chr2:93265402-93267417 | p5 | (AATAA)3 | CTTGCAATCTGCATTCTCCA | AGACCTAGTTGCGCTCGTGT | 266 |
| chr2:93265402-93267417 | p5 | (TTTAT)3 | CTTGCAATCTGCATTCTCCA | AGACCTAGTTGCGCTCGTGT | 266 |
| chr2:93284158-93286182 | p3 | (TGT)8 | CTTTCGTCGCAAGAAAGTCC | TTTCCTGAGCAGCTCCAACT | 224 |
| chr2:93295099-93297117 | p6 | (AAAGTG)3 | CTATTCTATTCACCCGGGCA | ACCTCCGGGCTTCTAATGTT | 276 |
| chr2:93298787-93300805 | p6 | (AAAGTG)3 | TCATTATCGAGATACGGCCC | CCGGGTGAATAGAATAGCGA | 132 |

| Marker ID | SSR type | SSR motif | FORWARD PRIMER (5'-3') | REVERSE PRIMER (5'-3') | size (bp) |
|---|---|---|---|---|---|
| chr2:93341927-93343939 | p2 | (AG)6 | CAGGTCTTGGCAAGGTGTTT | TCACAACCCAAAATCCCATT | 275 |
| chr2:93367535-93369550 | p5 | (ATTCA)3 | AGCCCAAGGAAAGTCATGTG | GTTCCTTCAAGGCGTGACAT | 181 |
| chr2:93425719-93427744 | p2 | (TC)6 | GTCCCAAAAGTCACGCAAAT | TGTTGGGGGATGGAGTAGAG | 224 |
| chr2:93472205-93474219 | p2 | (AT)9 | AACAACTCGGGTGGTTCTTG | TTACCATCCGAGCCATTCTC | 165 |
| chr2:93472205-93474219 | p2 | (CT)7 | AACAACTCGGGTGGTTCTTG | TTACCATCCGAGCCATTCTC | 165 |
| chr2:93497636-93499654 | p3 | (CAA)6 | ACGAGCGGAGTAGTGAGGAA | AAAAGCTCGGATTGCTTTGA | 284 |
| chr2:93563607-93565619 | p3 | (CAA)6 | CTGCAAACTGCGATGGTCTA | GAGTAGGATCTTGGGTGCCA | 154 |
| chr2:93599457-93601475 | p4 | (CATA)4 | AGGCACGGTGTTATATTGCC | GAGCCTACATGGCTGGGTAA | 177 |
| chr2:93599457-93601475 | p3 | (CAA)6 | AGGCACGGTGTTATATTGCC | GAGCCTACATGGCTGGGTAA | 177 |
| chr2:93666911-93668923 | p2 | (GT)6 | AATGTAATCGGAGGGGGAAC | CTGGTGCTATCTGCCCAACT | 281 |
| chr2:93768758-93770773 | p5 | (CACCA)3 | AAGGCAGTCGGGTCCTATTT | TATTTTCCGGAATCTGCACC | 249 |
| chr2:93789629-93791647 | p6 | (ATGTAT)3 | ACAGGATAGCCGTATGTCGG | ATGCCGGTCCCAACATAATA | 148 |
| chr2:93830894-93832960 | p3 | (AAT)22 | TTTGCTTGGGGTTCAAAAAC | CTTCCTCTTCTGTCCGCATC | 175 |
| chr2:93861319-93863335 | p2 | (AT)14 | AAACCTGTCCCCTGAGGAGT | CTTGTCGGCTTGCGTACATA | 252 |
| chr2:93861343-93863361 | p2 | (AT)14 | AAACCTGTCCCCTGAGGAGT | CTTGTCGGCTTGCGTACATA | 228 |
| chr2:94039819-94041843 | p3 | (TCA)8 | ACAAAGGAAATCCGTGCATC | GGCCTGTTTTCATCCTCAAA | 162 |
| chr2:94042010-94044025 | p3 | (TTA)5 | AGTTTGACGCGCTTTTGAAT | GACAAGTGGGATTGGCTGAT | 207 |
| chr2:94092379-94094395 | p3 | (CAA)5 | ACAACTCGGCCAAATTCATC | GGTTTCGCGTGAAGTATGGT | 216 |
| chr2:94092379-94094395 | p2 | (CA)6 | ACAACTCGGCCAAATTCATC | GGTTTCGCGTGAAGTATGGT | 216 |
| chr2:94092379-94094395 | p4 | (TTTC)4 | ACAACTCGGCCAAATTCATC | GGTTTCGCGTGAAGTATGGT | 216 |
| chr2:94199039-94201051 | p2 | (GT)6 | GAATGGACGAAGTTGGGTGT | TACCTCGATATTTCCGCGTC | 216 |
| chr2:94199039-94201051 | p3 | (TTG)5 | GAATGGACGAAGTTGGGTGT | TACCTCGATATTTCCGCGTC | 216 |
| chr2:94234008-94236023 | p5 | (TCTCC)3 | ATCGAGATGAGTTTGGGTGG | AGCTGCCCATATATCGTTGG | 204 |
| chr2:94234008-94236023 | p5 | (TCTCC)3 | ATCGAGATGAGTTTGGGTGG | AGCTGCCCATATATCGTTGG | 204 |
| chr2:94256073-94258091 | p6 | (TATATG)3 | TGGATTCTCGGGGATAGTTG | CCCGCTTCCTAACCTAATCC | 202 |
| chr2:94295847-94297867 | p5 | (TATCA)4 | CACCCAACGTACCCAATACC | ATCTGTGATGGCCTTATCGG | 196 |
| chr2:94444182-94446202 | p5 | (ACCCC)4 | GGAGTGGGCAACGAATAGAA | TTCTCCAAAATCGGAACACC | 121 |
| chr2:94444182-94446202 | p6 | (CCCACC)3 | GGAGTGGGCAACGAATAGAA | TTCTCCAAAATCGGAACACC | 121 |
| chr2:94447786-94449801 | p5 | (AAACC)3 | ACTTCCTAGCACATCCCCCT | CCAAACCATAAACACCCCTG | 173 |
| chr2:94479633-94481645 | p2 | (GT)6 | TTATACCCCGCGTTTTATGC | ATTTTGCCCTTCGTCTTCCT | 144 |
| chr2:94520551-94522563 | p2 | (AT)6 | TCCCTCGTTTCTTTCACCTG | TTCGAATGAAAATAAGCCCG | 223 |
| chr2:94677759-94679779 | p2 | (TA)10 | ACGCTTGACAGGGGTTAATG | AAGCACCCATTGTCACATCA | 281 |
| chr2:94711574-94713586 | p2 | (CT)6 | TGACGGCTCTTGTTTTGTTG | CAGATGACCAAGCACACACC | 260 |
| chr2:94735477-94737492 | p3 | (ACA)5 | CAAACGCTAATGTCTGGCAA | ATGTTGATGCGATGACCAAA | 245 |
| chr2:94758744-94760820 | p2 | (AT)38 | AAAAATCTCGATGTGTGGGC | TATACCCCGCGTTTTCAGAG | 174 |
| chr2:94758744-94760820 | p2 | (AT)9 | AAAAATCTCGATGTGTGGGC | TATACCCCGCGTTTTCAGAG | 174 |

| Marker ID | SSR type | SSR motif | FORWARD PRIMER (5'-3') | REVERSE PRIMER (5'-3') | size (bp) |
|---|---|---|---|---|---|
| chr2:94889756-94891771 | p3 | (TTC)5 | TGCAATTTTTATGAGCAGCG | ATTAAACACCAGCCCATTCG | 124 |
| chr2:95010403-95012415 | p2 | (TA)6 | TGATGGGATGGGAATATCGT | AATCCAAGTAACGTTCCCCC | 225 |
| chr2:95010403-95012415 | p2 | (TA)12 | TGATGGGATGGGAATATCGT | AATCCAAGTAACGTTCCCCC | 225 |
| chr2:95012867-95014882 | p3 | (ATC)5 | TTCCCGACGAGATAAAATCG | GACGAGTAAGGGAATGAGCG | 300 |
| chr2:95047654-95049668 | p2 | (TA)7 | CGGGCCCTCAAACTGTAATA | AGGTTTGGTTTCAGACCACG | 284 |
| chr2:95199249-95201281 | p2 | (AT)16 | CCGGATAACCTGTCTCCTGA | TTATGTTCGTCTGCTGTGGC | 275 |
| chr2:95264024-95266040 | p2 | (AT)8 | ACTTTTCCTCCCGAATTGCT | ACTAGTCATGGCTCATCGGG | 108 |
| chr2:95287826-95289838 | p2 | (GT)6 | GGCGTCCAACCAAAGAATAA | TGCAAAAACACGGGACATAA | 217 |
| chr2:95290586-95292601 | p3 | (GAT)5 | AAAGCGGTCCGTTTGTTATG | TCATATCGAGTCCGAAAGGG | 223 |
| chr2:95295719-95297731 | p2 | (GT)6 | CCGCCATGATCATAAGACCT | TTCCCCAATGACTTAGCACC | 261 |
| chr2:95308482-95310588 | p2 | (TA)7 | CTCAATGGCCGGGTAAGATA | AGGTACGGTGTTATGCTGCC | 213 |
| chr2:95308482-95310588 | p2 | (TA)53 | CTCAATGGCCGGGTAAGATA | AGGTACGGTGTTATGCTGCC | 213 |
| chr2:95308482-95310588 | p2 | (TA)7 | CTCAATGGCCGGGTAAGATA | AGGTACGGTGTTATGCTGCC | 213 |
| chr2:95324018-95326042 | p2 | (AT)12 | CCCGAGATGGACACTCGTAT | CAGGAGACAGGTTATCGGGA | 209 |
| chr2:95395493-95397519 | p2 | (AT)13 | ACTCGGTCAACGAGACCTTG | TTAGCCTCGTCGGCTTATGT | 112 |
| chr2:95418982-95421003 | p3 | (GTT)7 | GTTGGAGCCGTACATCAGGT | TGGCATGGGATAAGAGTTCC | 236 |
| chr2:95433426-95435444 | p6 | (CATAAT)3 | CCAACTAGTGTGTCTGCCCA | CTATAACGGAATCGGGGGTT | 221 |
| chr2:95493198-95495210 | p2 | (TA)6 | AAACAACGAGATCAAAGCGG | GGCCCTACGTGTAAGGACAA | 247 |
| chr2:95520553-95522573 | p2 | (TA)10 | TTGATCCGTGCAGTCAGAAG | ATATAGCGTCGTGTGCCTCC | 232 |
| chr2:95585134-95587149 | p4 | (CATA)5 | CTGATCAGTGGGATGCAATG | TTTCATGAATCCCTTAGCCG | 300 |
| chr2:95585134-95587149 | p3 | (CAA)5 | CTGATCAGTGGGATGCAATG | TTTCATGAATCCCTTAGCCG | 300 |
| chr2:95605590-95607616 | p2 | (TA)13 | TCTCCGATGCCACTATACCC | CGCTTTATCCTCATACCCCA | 169 |
| chr2:95605590-95607616 | p2 | (TA)13 | TCTCCGATGCCACTATACCC | CGCTTTATCCTCATACCCCA | 169 |
| chr2:95655483-95657499 | p4 | (AATT)4 | ATGTCTGGCCGTAGATCTGG | ACGAGAACGAGTTTCGGAGA | 161 |
| chr2:95733861-95735877 | p2 | (TA)8 | GGTCTTGGAAAGGGGTGAAT | ACCTCATGGCCATCAACTTC | 181 |
| chr2:95737828-95739843 | p3 | (GAT)5 | CCCCAAGACATCTCCAAAAA | AATTTTAGGCCACAACACGC | 282 |
| chr2:95737828-95739843 | p4 | (TATG)4 | CCCCAAGACATCTCCAAAAA | AATTTTAGGCCACAACACGC | 282 |
| chr2:95753382-95755410 | p2 | (AT)14 | CAATCACGGAGCCTTTTCTC | CAAACATTCACTGCGGGTAA | 244 |
| chr2:95753382-95755410 | p2 | (AT)42 | CAATCACGGAGCCTTTTCTC | CAAACATTCACTGCGGGTAA | 244 |
| chr2:95765046-95767062 | p2 | (TA)8 | ATGCATTGAAAGGTTGAGGC | TCGGGATCACAGAATGGAAT | 222 |
| chr2:95798647-95800662 | p6 | (GGAGTT)3 | CCCTTGTTCTTACACCCCAA | CTCCAACTCCAACTCCGAAA | 164 |
| chr2:95798647-95800662 | p5 | (AAACC)3 | CCCTTGTTCTTACACCCCAA | CTCCAACTCCAACTCCGAAA | 164 |
| chr2:95798687-95800702 | p6 | (GGAGTT)3 | CCCTTGTTCTTACACCCCAA | CTCCAACTCCAACTCCGAAA | 124 |
| chr2:95798687-95800702 | p5 | (AAACC)3 | CCCTTGTTCTTACACCCCAA | CTCCAACTCCAACTCCGAAA | 124 |
| chr2:95907402-95909426 | p2 | (AT)12 | TGAACCTCGAGGACACCTCT | GTCGGGTCCTATTTATGCGA | 265 |
| chr2:95946554-95948569 | p2 | (CA)6 | ACCCGAAATCTCCACAAATG | TGTAAGTGGGTGGAGGAAGG | 289 |

| Marker ID | SSR type | SSR motif | FORWARD PRIMER (5'-3') | REVERSE PRIMER (5'-3') | size (bp) |
|---|---|---|---|---|---|
| chr2:95946554-95948569 | p5 | (TTTTA)3 | ACCCGAAATCTCCACAAATG | TGTAAGTGGGTGGAGGAAGG | 289 |
| chr2:96154250-96156264 | p2 | (AT)7 | GGCATGATCGTTGTGAAATG | TCGGATGTACCTGTGGATCA | 120 |
| chr2:96182468-96184504 | p2 | (TA)18 | ATCCAATGCTCTTGGTGGTC | GGGATTCTCCCTTATGCTCC | 265 |
| chr2:96182468-96184504 | p2 | (AT)8 | ATCCAATGCTCTTGGTGGTC | GGGATTCTCCCTTATGCTCC | 265 |
| chr2:96182468-96184504 | p3 | (ATC)5 | ATCCAATGCTCTTGGTGGTC | GGGATTCTCCCTTATGCTCC | 265 |
| chr2:96199315-96201371 | p3 | (ATC)5 | AAAATCATGGGCTCTTGCAC | ATACCTCAGGCCAGCAAATG | 135 |
| chr2:96200476-96202494 | p3 | (ATC)5 | GTGCAGCCCGATTAACAAAT | AAATCGGAGGTTACCGACCT | 211 |
| chr2:96200476-96202494 | p3 | (ACA)6 | GTGCAGCCCGATTAACAAAT | AAATCGGAGGTTACCGACCT | 211 |
| chr2:96202555-96204570 | p5 | (TATGA)3 | CAGTCGGCACAAGTCTTGAA | GAGACAGGATGACTTGGGGA | 139 |
| chr2:96202555-96204570 | p2 | (AT)11 | CAGTCGGCACAAGTCTTGAA | GAGACAGGATGACTTGGGGA | 139 |
| chr2:96268064-96270096 | p2 | (TA)16 | TCTACAAAACCTTGCGGGTC | ACAAGTGACCGAGGCCATAC | 227 |
| chr2:96283284-96285340 | p2 | (TA)28 | ATCCGAGTGCATTGAAAACC | CGCTTCCCGAGATATTTGAA | 142 |
| chr2:96345524-96347608 | p2 | (TA)42 | TATGGTAGTGCCCCGATGAT | TGTTGAGGTTGTTGTTGGGA | 202 |
| chr2:96377653-96379668 | p3 | (AAG)5 | ATCGATTGGGTCGTTACCCT | CCTCTGTTTGGATGGTCGTT | 267 |
| chr2:96497377-96499395 | p6 | (AGGAAG)3 | GGAATCACGGGGATTAACCT | CCACCCAGGACCTTTCTGTA | 243 |
| chr2:96665892-96667910 | p6 | (AGCACC)3 | AAACCAACAGACCAACCTCG | GTTGGAGCAATCGTTCACCT | 156 |
| chr2:96709332-96711348 | p2 | (TA)10 | CCTTACGTCACCCCGATAGA | CACGGTGGAATTGTATGCAG | 198 |
| chr2:96709372-96711384 | p2 | (TA)10 | CCTTACGTCACCCCGATAGA | CACGGTGGAATTGTATGCAG | 158 |
| chr2:96709409-96711421 | p2 | (TA)10 | CCTTACGTCACCCCGATAGA | CACGGTGGAATTGTATGCAG | 121 |
| chr2:96865068-96867098 | p2 | (TC)15 | GGTCAAGTGCTGGGATTGTT | ACTTGGGTCCTTTTCGAGGT | 152 |
| chr2:96904051-96906099 | p2 | (AT)24 | AGGAAGGGCAATCCTCCTTA | TGGTCCTTAGGTTCCACCAG | 232 |
| chr2:96904051-96906099 | p2 | (AT)10 | AGGAAGGGCAATCCTCCTTA | TGGTCCTTAGGTTCCACCAG | 232 |
| chr2:96949551-96951573 | p2 | (AT)11 | GCTGCAAAATTGGAAAGGAA | CCGTTAAAAAGAGGGCATGA | 262 |
| chr2:96949551-96951573 | p3 | (ATT)6 | GCTGCAAAATTGGAAAGGAA | CCGTTAAAAAGAGGGCATGA | 262 |
| chr2:96958151-96960166 | p3 | (AAT)5 | CCCATAAAGCTTTGGCTGAT | CCATCAACGAAAAGAATGCC | 178 |
| chr2:96979380-96981392 | p2 | (GA)6 | GGCAGCCAAACAACAGATTT | CTTGAAGCATTGCCCGTATT | 134 |
| chr2:96991010-96993036 | p2 | (AT)13 | AAGTATCAACCGTGGCTTCG | TCCGCTTCTGAGTTTTGGTT | 202 |
| chr2:96993936-96995964 | p2 | (TA)14 | AGCAAGGCACGAGAGTGAAT | TTTTCAATCGAACCCTCACC | 118 |
| chr2:96999432-97001460 | p2 | (TA)14 | AGCAAGGCACGAGAGTGAAT | TTTTCAATCGAACCCTCACC | 118 |
| chr2:97021680-97023714 | p2 | (TA)17 | CCCCTAGAATGCTCCTACCC | CCGACAGTTGAAGGGTTCAT | 119 |
| chr2:97059797-97061849 | p2 | (AT)26 | TCCGTTTGACCCCCATAATA | TGGGAGTTTACGCACATGAA | 300 |
| chr2:97077616-97079628 | p2 | (TA)6 | GGATCACCAAAGCCGAGTTA | CTGTTCACCGAGTCGCATAA | 169 |
| chr2:97091552-97093564 | p2 | (AC)6 | ACGAAATCTGCGAATTGGTC | TGTACAGAGGAAATGCTGCG | 271 |
| chr2:97190794-97192809 | p5 | (ACCCG)3 | GCAAGGGAACGGTATGAAAA | AACCCGTGAGTTGGAGATTG | 210 |
| chr2:97292454-97294469 | p5 | (ATTTA)3 | GAAGGTCCCCATGAAAGACA | GTAGCTCCGGATGAAATGGA | 165 |
| chr2:97331302-97333322 | p5 | (GAAAT)4 | AAAGCCATAGCTCCGAATGA | TAAGGCCCTTTTGTTTGTGG | 160 |

| Marker ID | SSR type | SSR motif | FORWARD PRIMER (5'-3') | REVERSE PRIMER (5'-3') | size (bp) |
|---|---|---|---|---|---|
| chr2:97473976-97475991 | p3 | (TGT)5 | GGGCAAGGCTTGTACGATAA | ATTCACTTCATGCCACCTCC | 256 |
| chr2:97473976-97475991 | p3 | (GAT)5 | GGGCAAGGCTTGTACGATAA | ATTCACTTCATGCCACCTCC | 256 |
| chr2:97474347-97476362 | p3 | (TGT)5 | GAAGCAACCGAGGAACTCTG | TTGGGATGTTTTGAGGGGTA | 148 |
| chr2:97474347-97476362 | p3 | (GAT)5 | GAAGCAACCGAGGAACTCTG | TTGGGATGTTTTGAGGGGTA | 148 |
| chr2:97483249-97485453 | p2 | (GA)7 | AAGAATGGATTTGGGAGCCT | CAAGTTTTGGAAGGGTGGAA | 111 |
| chr2:97483249-97485453 | p2 | (TA)102 | AAGAATGGATTTGGGAGCCT | CAAGTTTTGGAAGGGTGGAA | 111 |
| chr2:97598457-97600485 | p2 | (TA)33 | TTGTGCCAAAGCAACTTCTG | GCAGCCTGGTATATGAAGCG | 290 |
| chr2:97598457-97600485 | p2 | (AT)14 | TTGTGCCAAAGCAACTTCTG | GCAGCCTGGTATATGAAGCG | 290 |
| chr2:97629072-97631120 | p2 | (TA)24 | TTTTGGGGAACACTTTGACC | CTCGAGACCCGAACCAAATA | 259 |
| chr2:97652023-97654050 | p3 | (TAA)9 | AAGCAACGGCTAGCAGACAT | GTCTGGCGGACTTCAACTTC | 135 |
| chr2:97652023-97654050 | p2 | (AT)14 | AAGCAACGGCTAGCAGACAT | GTCTGGCGGACTTCAACTTC | 135 |
| chr2:97673850-97675877 | p3 | (ATC)9 | TGAGAGCACCCCATTATTCC | AGCCCTTCGGCTATCAAAAT | 241 |
| chr2:97739123-97741135 | p2 | (CT)6 | CCTCCATAGTGGTCATGCCT | TCAGCGTCAAATGCTAGTGG | 139 |
| chr2:97823812-97825830 | p2 | (TG)9 | TAGACAGGGAGGTGGTCCAG | TGTAGAAAGGACGGGACAGG | 136 |
| chr2:97924800-97926818 | p2 | (TA)9 | ATATAGGGGAAATGCTGCCC | TCCGTTAACTTCGTCGCTCT | 230 |
| chr2:97924800-97926818 | p2 | (TA)9 | ATATAGGGGAAATGCTGCCC | TCCGTTAACTTCGTCGCTCT | 230 |
| chr2:97973670-97975688 | p2 | (TA)9 | ATATAGGGGAAATGCTGCCC | TCCGTTAACTTCGTCGCTCT | 230 |
| chr2:97973670-97975688 | p2 | (TA)9 | ATATAGGGGAAATGCTGCCC | TCCGTTAACTTCGTCGCTCT | 230 |
| chr2:98002726-98004741 | p5 | (CATAT)3 | GCCAAGACCCACAACTCCTA | ATTTTGTGACGGCCTTATCG | 229 |
| chr2:98002726-98004741 | p3 | (CAA)5 | GCCAAGACCCACAACTCCTA | ATTTTGTGACGGCCTTATCG | 229 |
| chr2:98019764-98021779 | p5 | (CATAT)3 | GCCAAGACCCACAACTCCTA | ATTTTGTGACGGCCTTATCG | 229 |
| chr2:98019764-98021779 | p3 | (CAA)5 | GCCAAGACCCACAACTCCTA | ATTTTGTGACGGCCTTATCG | 229 |
| chr2:98129598-98131610 | p2 | (TA)6 | CATCATCTTGTCATGCGTCC | ACTGGCTTGAAAGAGCCAAA | 146 |
| chr2:98139261-98141303 | p2 | (TA)21 | TGCACTGACCGCTCTATCAC | GACCTCTCCAACTCCATCCA | 150 |
| chr2:98215587-98217607 | p3 | (CAA)5 | CGGCCATCCTAGTGGTTCTA | TTAGCGAGCGATCGAGGTAT | 102 |
| chr2:98252391-98254425 | p2 | (AT)17 | CCATAACGCAATCGGAAGTT | GCTGTGCTATACTCCTGCCC | 218 |
| chr2:98332050-98334096 | p2 | (TA)23 | CGCTGCAACTGAAGGTGTTA | TGCTTGGTCTTCGAGTGATG | 182 |
| chr2:98366732-98368744 | p2 | (TA)6 | ATTTTGGTTGCTGTGTGCAG | GGGGTGTAACATTCTCCCCT | 112 |
| chr2:98366732-98368744 | p2 | (TA)7 | ATTTTGGTTGCTGTGTGCAG | GGGGTGTAACATTCTCCCCT | 112 |
| chr2:98395389-98397401 | p2 | (GA)6 | GTTTCAGATTTCGGGTGGAA | CAACTTCGCATCTGTGGAGA | 139 |
| chr2:98395389-98397401 | p5 | (TTGCA)3 | GTTTCAGATTTCGGGTGGAA | CAACTTCGCATCTGTGGAGA | 139 |
| chr2:98399335-98401361 | p2 | (TG)13 | AGTACATCGGGGATCCTTCC | AACCTACAAACCTGGCGATG | 215 |
| chr2:98399335-98401361 | p2 | (TG)14 | AGTACATCGGGGATCCTTCC | AACCTACAAACCTGGCGATG | 215 |
| chr2:98424439-98426454 | p5 | (GCATT)3 | AAGGCAAGGCAAAAGAGTGA | AAGGTCCACATGCTGTCTCC | 197 |
| chr2:98424439-98426454 | p2 | (TA)28 | AAGGCAAGGCAAAAGAGTGA | AAGGTCCACATGCTGTCTCC | 197 |
| chr2:98451162-98453178 | p3 | (CAA)5 | ATGGAGGTCAAAGACATCGG | GGGATTTGGTTAGTGAGCGA | 179 |

| Marker ID | SSR type | SSR motif | FORWARD PRIMER (5'-3') | REVERSE PRIMER (5'-3') | size (bp) |
|---|---|---|---|---|---|
| chr2:98451162-98453178 | p2 | (AT)8 | ATGGAGGTCAAAGACATCGG | GGGATTTGGTTAGTGAGCGA | 179 |
| chr2:98472325-98474337 | p2 | (GT)6 | TTTTGCCGTAGCAAGTGTTG | GACCTTGCAGCTCGGATAAG | 195 |
| chr2:98656567-98658582 | p5 | (ACCAA)3 | TGTCATGTCCTTCGGATTGA | TGCTCTGCATGATCAACCTC | 219 |
| chr2:98656567-98658582 | p5 | (AAACC)3 | TGTCATGTCCTTCGGATTGA | TGCTCTGCATGATCAACCTC | 219 |
| chr2:98672700-98674718 | p5 | (GCATT)3 | TGTTCGATGATCAATTCCGA | CCATCCTGAATCTTTTCCCA | 191 |
| chr2:98672700-98674718 | p2 | (TA)9 | TGTTCGATGATCAATTCCGA | CCATCCTGAATCTTTTCCCA | 191 |
| chr2:98683811-98685863 | p2 | (AT)26 | GGCCTTAGCCCTCAAAATTC | GACCAAAAACCAATCGGAAA | 157 |
| chr2:98717511-98719526 | p5 | (AATGC)3 | TCGAGTCCTCACCATCTGTG | CCGTACTTTTCTGCGAGAGG | 105 |
| chr2:98763416-98765431 | p5 | (ATAAA)3 | AGTACCGTGATGGCCAAAAG | TTGGAGTTTCCATTCTTCCG | 273 |
| chr2:98904228-98906244 | p2 | (TA)8 | CGGAATAGCAAATGCAACAG | CGTGGACCCCATATTATTGC | 148 |
| chr2:98904228-98906244 | p6 | (TTTTTG)3 | CGGAATAGCAAATGCAACAG | CGTGGACCCCATATTATTGC | 148 |
| chr2:98956355-98958371 | p2 | (TA)8 | TGGGCCCCATACTAAACAAC | TTTAAGGGACAACGGACGAC | 282 |
| chr2:98956355-98958371 | p2 | (TA)8 | TGGGCCCCATACTAAACAAC | TTTAAGGGACAACGGACGAC | 282 |
| chr2:98956355-98958371 | p2 | (TA)11 | TGGGCCCCATACTAAACAAC | TTTAAGGGACAACGGACGAC | 282 |
| chr2:99027881-99029893 | p2 | (AG)6 | CACACCTGGACTCTCTGCAA | AAGGCTGTGTCTTTGCCACT | 155 |
| chr2:99031994-99034006 | p2 | (AG)6 | CACACCTGGACTCTCTGCAA | AAGGCTGTGTCTTTGCCACT | 155 |
| chr2:99157000-99159029 | p2 | (AT)10 | CCGTGAGTTGTTGGTTGTTG | GCCATGGACACTTGGAGAAT | 285 |
| chr2:99157000-99159029 | p2 | (AT)6 | CCGTGAGTTGTTGGTTGTTG | GCCATGGACACTTGGAGAAT | 285 |
| chr2:99179573-99181585 | p2 | (TA)27 | ACCGCTCGTGCTATCATTTC | CCATTATTCCGGAGGTGCTA | 266 |
| chr2:99179864-99181918 | p2 | (TA)27 | TCTCCCTGTCTACCTGTGGG | ATATATGGATCGGGCTGCAC | 273 |
| chr2:99244064-99246080 | p4 | (TTTA)4 | AAACCCATGCTTTGCATTTC | GCACTTGAGCCATGGTTTTT | 263 |
| chr2:99260361-99262379 | p6 | (TCCACT)3 | ACCTCATGACGAACGAATCC | GAACTATTTCTTCGGCAGCG | 260 |
| chr2:99260361-99262379 | p6 | (TCCACA)3 | ACCTCATGACGAACGAATCC | GAACTATTTCTTCGGCAGCG | 260 |
| chr2:99338593-99340617 | p2 | (AT)12 | CCCAACCGCTATGTTACGTT | TCACACCCTAGGGGCATTAG | 216 |
| chr2:99349676-99351690 | p2 | (TA)7 | GCCATTAAGGCTCGGTGTTA | GCCGTATTTTCCAAACCAGA | 148 |
| chr2:99349676-99351690 | p2 | (TA)7 | GCCATTAAGGCTCGGTGTTA | GCCGTATTTTCCAAACCAGA | 148 |
| chr2:99408125-99410155 | p3 | (TTA)5 | TACCTCCCAAAGCCAGCTAA | CTTTGTGATGCTCTGGGGAT | 291 |
| chr2:99408125-99410155 | p3 | (TTA)5 | TACCTCCCAAAGCCAGCTAA | CTTTGTGATGCTCTGGGGAT | 291 |
| chr2:99408156-99410171 | p3 | (TTA)5 | TACCTCCCAAAGCCAGCTAA | CTTTGTGATGCTCTGGGGAT | 260 |
| chr2:99408156-99410171 | p3 | (TTA)5 | TACCTCCCAAAGCCAGCTAA | CTTTGTGATGCTCTGGGGAT | 260 |
| chr2:99437427-99439460 | p2 | (GA)7 | CCTAGATCCCATTCCCGTTT | AACCGTCGTGAGGTGATTTC | 271 |
| chr2:99437467-99439482 | p2 | (GA)7 | CCTAGATCCCATTCCCGTTT | AACCGTCGTGAGGTGATTTC | 231 |
| chr2:99437526-99439544 | p2 | (GA)7 | CCTAGATCCCATTCCCGTTT | AACCGTCGTGAGGTGATTTC | 172 |
| chr2:99437570-99439588 | p2 | (GA)7 | CCTAGATCCCATTCCCGTTT | AACCGTCGTGAGGTGATTTC | 128 |
| chr2:99498912-99500924 | p2 | (AG)6 | TAGACAGGAGGGTGACCCAG | TGGGACACAACTTACCCACA | 164 |
| chr2:99498912-99500924 | p4 | (TATG)4 | TAGACAGGAGGGTGACCCAG | TGGGACACAACTTACCCACA | 164 |

| Marker ID | SSR type | SSR motif | FORWARD PRIMER (5'-3') | REVERSE PRIMER (5'-3') | size (bp) |
|---|---|---|---|---|---|
| chr2:99580177-99582192 | p5 | (TATTT)3 | TTTAGGTGGGTGTGCAGTGA | ATTTACACACGAATTCGCCC | 268 |
| chr2:99688823-99690838 | p5 | (AATAT)3 | GAAATCTTCCCGCAATTTGA | TAATCCACCTCCACATCCGT | 197 |
| chr2:99822156-99824174 | p5 | (ATGAT)3 | GGAGTGTTTCGGGCAAAGTA | GCCAAACCAACAACGTTACC | 115 |
| chr2:99849056-99851074 | p6 | (ATTTGC)3 | GCATTGATCGATGTGGTGAC | GCTCTGTCATGTCTGCTCCA | 288 |
| chr2:99849056-99851074 | p5 | (AAAAC)3 | GCATTGATCGATGTGGTGAC | GCTCTGTCATGTCTGCTCCA | 288 |
| chr2:99887788-99889803 | p3 | (ACA)5 | CCAACACATGTTTCGTGAGG | AGGTCCTTCCAAGATTGCCT | 209 |
| chr2:99904867-99906887 | p2 | (TA)10 | ACGCTGCCAGCTATGCTATT | GCCCCTCTTTCTGTTTTCCT | 257 |
| chr2:99962025-99964037 | p2 | (AT)6 | CAATGATGTATGCGGTCCTG | TTGGCATATTCAGCATCACC | 235 |
| chr2:99994717-99996731 | p2 | (TG)7 | AGGCAAAGCGAATGATATGG | GGGGAGGTCTCTGTCTTTCC | 203 |
| chr2:99994717-99996731 | p3 | (TTG)5 | AGGCAAAGCGAATGATATGG | GGGGAGGTCTCTGTCTTTCC | 203 |
| chr2:100028805-100030869 | p2 | (TA)11 | ATCTCCCCCGCAATAATACC | GGGCTCCAACACCTTATCAA | 235 |
| chr2:100028925-100030989 | p2 | (TA)11 | ATCTCCCCCGCAATAATACC | GGGCTCCAACACCTTATCAA | 115 |
| chr2:100039827-100041843 | p2 | (TA)8 | ATAAAATGAGGGGAATGGGG | TAACTTGAGGGAGGTGTGGC | 233 |
| chr2:100039827-100041843 | p2 | (AT)13 | ATAAAATGAGGGGAATGGGG | TAACTTGAGGGAGGTGTGGC | 233 |
| chr2:100040426-100042452 | p2 | (TA)8 | CTCCAAGTTGCCCTCATCTC | TATTGGACGAAATTCACGCA | 184 |
| chr2:100040426-100042452 | p2 | (AT)13 | CTCCAAGTTGCCCTCATCTC | TATTGGACGAAATTCACGCA | 184 |
| chr2:100040426-100042452 | p6 | (TCATCG)3 | CTCCAAGTTGCCCTCATCTC | TATTGGACGAAATTCACGCA | 184 |
| chr2:100057309-100059321 | p2 | (TA)11 | TTGGCCTAAAACCTCTGGTG | AACAGCTCAAAAGGGCAAGA | 188 |
| chr2:100057309-100059321 | p2 | (TA)6 | TTGGCCTAAAACCTCTGGTG | AACAGCTCAAAAGGGCAAGA | 188 |
| chr2:100077933-100079966 | p3 | (TAT)11 | GCCAAAGGCATCATTCTCAT | AATTGCCCCAGTTCACTGTC | 140 |
| chr2:100081694-100083714 | p2 | (TA)10 | ACGCTGCCTGCTATGCTATT | GCAGGCCTTGAGTTTAGCAG | 269 |
| chr2:100094337-100096365 | p2 | (AT)14 | CCTCAGATTCCGGACGATAA | TTGACCGGCATTTGATTGTA | 184 |
| chr2:100116582-100118598 | p2 | (AT)8 | TGTTGTTGTGGTTGGTTGCT | CTTATCTATCCGATCCCGCA | 192 |
| chr2:100116639-100118651 | p2 | (AT)8 | TGTTGTTGTGGTTGGTTGCT | CTTATCTATCCGATCCCGCA | 135 |
| chr2:100133970-100135984 | p2 | (TA)7 | TGTTGTTGTGGTTGGTTGCT | GCCTTTGAAAGAAAGGGGAC | 139 |
| chr2:100133970-100135984 | p2 | (TA)10 | TGTTGTTGTGGTTGGTTGCT | GCCTTTGAAAGAAAGGGGAC | 139 |
| chr2:100265276-100267291 | p5 | (TATCG)3 | TGATATTGAGCCGCACTCTG | TTCTTCATAAGAATCGGGCG | 147 |
| chr2:100265276-100267291 | p2 | (TA)11 | TGATATTGAGCCGCACTCTG | TTCTTCATAAGAATCGGGCG | 147 |
| chr2:100270416-100272428 | p2 | (TA)6 | TAACCAAATGTGGCCCTAGC | GATATCATCGAGGGGAGCAA | 216 |
| chr2:100270416-100272428 | p5 | (TATCG)3 | TAACCAAATGTGGCCCTAGC | GATATCATCGAGGGGAGCAA | 216 |
| chr2:100368876-100370891 | p3 | (ATC)5 | GTGTTGCCAAAATTGTCACG | CGGCTTGCATTTTTCTTTTC | 171 |
| chr2:100614844-100616872 | p2 | (AT)7 | TTACATCGCATTTGCTCTGC | GGGCAAAGACATGTTCCATT | 119 |
| chr2:100614844-100616872 | p2 | (AT)14 | TTACATCGCATTTGCTCTGC | GGGCAAAGACATGTTCCATT | 119 |
| chr2:100614844-100616872 | p2 | (AT)7 | TTACATCGCATTTGCTCTGC | GGGCAAAGACATGTTCCATT | 119 |
| chr2:100761113-100763131 | p2 | (AT)9 | AATATGGCGCTACCATCCTG | TCAGTCAAAGTCTTGTGGCG | 169 |
| chr2:100778201-100780222 | p3 | (GAA)7 | CAAAGCAAGGCACAAATTCA | TCAAAGACACTGCCATCAGC | 245 |

| Marker ID | SSR type | SSR motif | FORWARD PRIMER (5'-3') | REVERSE PRIMER (5'-3') | size (bp) |
|---|---|---|---|---|---|
| chr2:100810229-100812259 | p2 | (TA)15 | GTCAAACTTAGGCCTTCCCC | TGTGGTTTGCTTGGTCACAT | 128 |
| chr2:100885220-100887238 | p6 | (ATAAAA)3 | TCCCGTAGTTTCATTTTGGC | GCGGTTCACATGAAATCCTT | 162 |
| chr2:100945908-100947922 | p4 | (ATAA)4 | CTCCCCATTGCTCCTAATCA | CATCCATTTCAATCCAACCC | 286 |
| chr2:100945928-100947940 | p4 | (ATAA)4 | CTCCCCATTGCTCCTAATCA | CATCCATTTCAATCCAACCC | 266 |
| chr2:101039839-101041867 | p2 | (TA)14 | TCAAATTCGTGATTCCAGCA | TCACCGACGTGGATAATGAA | 299 |
| chr2:101058551-101060571 | p5 | (TCATT)4 | TCAGTCAAAGTTTTGTGGCG | TATTGCCTCACAGGGTCACA | 145 |
| chr2:101066416-101068432 | p5 | (TTGCA)3 | GTTGTTCTTTTGCCGGTGAT | GCTACTGGCAATTTTCGCTC | 293 |
| chr2:101066416-101068432 | p4 | (TTAT)4 | GTTGTTCTTTTGCCGGTGAT | GCTACTGGCAATTTTCGCTC | 293 |
| chr2:101081516-101083542 | p2 | (AT)13 | GACGAGGCCAACAAAATGAT | CGTAGCTGCTTATTCCGAGG | 245 |
| chr2:101081516-101083542 | p5 | (GAAAT)3 | GACGAGGCCAACAAAATGAT | CGTAGCTGCTTATTCCGAGG | 245 |
| chr2:101121832-101123844 | p2 | (GA)6 | GACGACATGTTGATCGTTGG | TGTTGAGCCGGTCCTAAGTC | 289 |
| chr2:101151974-101154018 | p2 | (AT)22 | AGTAACCATGAGCAATGGGG | TTGGTTACGGATGGTTGGTT | 230 |
| chr2:101151974-101154018 | p2 | (AT)20 | AGTAACCATGAGCAATGGGG | TTGGTTACGGATGGTTGGTT | 230 |
| chr2:101191948-101193994 | p4 | (TATG)12 | TCTTGTCACGACCCAAATCA | CGTTCTATCATGGTGGCCTT | 180 |
| chr2:101191948-101193994 | p3 | (AAC)5 | TCTTGTCACGACCCAAATCA | CGTTCTATCATGGTGGCCTT | 180 |
| chr2:101196338-101198380 | p5 | (TACAT)3 | CGGAGATTCAAGAGCACTCC | AATCACAAAGGACGGGACAG | 277 |
| chr2:101196455-101198481 | p5 | (TACAT)3 | CGGAGATTCAAGAGCACTCC | AATCACAAAGGACGGGACAG | 160 |
| chr2:101196489-101198501 | p5 | (TACAT)3 | CGGAGATTCAAGAGCACTCC | AATCACAAAGGACGGGACAG | 126 |
| chr2:101254524-101256539 | p2 | (AT)7 | AGTGCACCCACACCCATATT | AAATTGATACGTGCAAGGGC | 101 |
| chr2:101254524-101256539 | p5 | (TTTCC)3 | AGTGCACCCACACCCATATT | AAATTGATACGTGCAAGGGC | 101 |
| chr2:101281764-101283776 | p2 | (TC)6 | CATAGCGTGACCCTGGAGAT | CCTCGAACGACAGGGAATTA | 170 |
| chr2:101329988-101332008 | p3 | (TAA)8 | TCCTTGTTAAGTGGAAGGCG | AAATTTCTTCCATCCAGCCC | 143 |
| chr2:101330009-101332025 | p3 | (TAA)8 | TCCTTGTTAAGTGGAAGGCG | AAATTTCTTCCATCCAGCCC | 122 |
| chr2:101341624-101343674 | p2 | (TA)25 | TTCCATGACCTCCCTTTCTG | TGCAAATCATCTGGGCATTA | 138 |
| chr2:101358959-101360973 | p2 | (CA)7 | CGCTCATTTGTGCAACTTGT | AGCCATATTCTCGGTGTTGG | 182 |
| chr2:101358959-101360973 | p2 | (TA)7 | CGCTCATTTGTGCAACTTGT | AGCCATATTCTCGGTGTTGG | 182 |
| chr2:101397134-101399149 | p3 | (TGT)5 | AGGAAGAGGCATTAAACGCA | GCGGGATTTGTTTGTTGAGT | 261 |
| chr2:101421794-101423822 | p2 | (GA)14 | TTCGCTCGAAGATACGAGGT | ACTGCATCGCATAAGGGAAC | 261 |
| chr2:101458033-101460048 | p2 | (AG)8 | CGGGTTTAGGGTTCTGATGA | CTCCTGCTCCCTCCTCTTTT | 143 |
| chr2:101458049-101460121 | p2 | (AG)8 | CGGGTTTAGGGTTCTGATGA | CTCCTGCTCCCTCCTCTTTT | 127 |
| chr2:101771414-101773428 | p2 | (CT)7 | CCGTCGAATGAGTTACGGTT | TGTACGGGGGATTTTCTGAG | 278 |
| chr2:101784698-101786712 | p2 | (CT)7 | CCGTCGAATGAGTTACGGTT | TGTACGGGGGATTTTCTGAG | 278 |
| chr2:101842725-101844743 | p6 | (ATATTC)3 | CACTTCCTTCGGAGCCATAG | ACATTGCCATTCATTCGTCA | 266 |
| chr2:101862355-101864373 | p6 | (CTTACA)3 | CGATGCTAACTCCACCGATT | GGCAATGTGATAGGGCTGTT | 273 |
| chr2:101956147-101958161 | p2 | (TG)7 | AAAAGGATGCGAAGAGGGAT | TTGGGATGGGTATCGAACAT | 224 |
| chr2:102038578-102040594 | p2 | (TG)7 | TCCCAAAGGGAAGAATGTTG | TTCCCATTTTTCCCCTATCC | 125 |

| Marker ID | SSR type | SSR motif | FORWARD PRIMER (5'-3') | REVERSE PRIMER (5'-3') | size (bp) |
|---|---|---|---|---|---|
| chr2:102038578-102040594 | p2 | (GT)8 | TCCCAAAGGGAAGAATGTTG | TTCCCATTTTTCCCCTATCC | 125 |
| chr2:102105171-102107189 | p3 | (CAA)6 | GTAAACATTGGGGCATCCAC | CCGTCCGACTTCATTTGATT | 172 |
| chr2:102205610-102207625 | p5 | (TGAAA)3 | ACGATCCGTCTTTCATGACC | TGAAAATGCGAAAGTTGCAG | 280 |
| chr2:102205610-102207625 | p5 | (ATTTC)3 | ACGATCCGTCTTTCATGACC | TGAAAATGCGAAAGTTGCAG | 280 |
| chr2:102215812-102217830 | p3 | (ATC)6 | AGTCACTCGGAGCCAACCTA | CCCGCCCTTACAGATTATGA | 124 |
| chr2:102293331-102295433 | p2 | (CA)6 | GATCTAAGCCTGTCGAACCG | CCTACCCCCAACCAAAAATC | 102 |
| chr2:102293331-102295433 | p2 | (AT)15 | GATCTAAGCCTGTCGAACCG | CCTACCCCCAACCAAAAATC | 102 |
| chr2:102293331-102295433 | p2 | (TG)6 | GATCTAAGCCTGTCGAACCG | CCTACCCCCAACCAAAAATC | 102 |
| chr2:102293331-102295433 | p2 | (TA)51 | GATCTAAGCCTGTCGAACCG | CCTACCCCCAACCAAAAATC | 102 |
| chr2:102333537-102335561 | p6 | (TTCCTC)3 | AATGCATTGGGAACTGAACC | GCATTGGTTTGCTCGTTTTT | 101 |
| chr2:102348970-102350982 | p2 | (GT)6 | TTCCGTTAAGGTCTTGTGGC | CCTGGAGGCATAGGAAACAA | 101 |
| chr2:102372149-102374161 | p2 | (TA)6 | CGACCACGTTCTTGTACCCT | ATGCGGACATAGAGAATCGG | 225 |
| chr2:102472026-102474041 | p6 | (CATATA)3 | GTGCCCTTTTTCCCCATATT | ATCGACCTTACGTCACTCCG | 137 |
| chr2:102472026-102474041 | p3 | (ACA)5 | GTGCCCTTTTTCCCCATATT | ATCGACCTTACGTCACTCCG | 137 |
| chr2:102479020-102481032 | p3 | (ACA)5 | CAAAAACATGTCCGTCGTTG | AAGGCTCGTCCCTTTCTTTC | 258 |
| chr2:102479020-102481032 | p2 | (CT)6 | CAAAAACATGTCCGTCGTTG | AAGGCTCGTCCCTTTCTTTC | 258 |
| chr2:102542814-102544836 | p2 | (AT)11 | AGCTGGGAGGAAAGGGATTA | GTTAGTGGCCGAGGAACGTA | 265 |
| chr2:102563855-102565867 | p2 | (CA)6 | ACATGCCAAAGGAAGGAATG | TGATGTTGCTGGGTTGTTGT | 141 |
| chr2:102571583-102573595 | p2 | (AT)6 | TGATATGTTTCCGAGTGGCA | GTGCACAAAAACGTGTCCTG | 291 |
| chr2:102628896-102630911 | p3 | (CAG)5 | GGTCTGCTACACCCCCACTA | GAACTCTGGAGGGCCACATA | 173 |
| chr2:102709927-102711943 | p2 | (AC)8 | GGTTTCGAGGAGCAGACTTG | TTTGGGGAAAGCATAAGACG | 191 |
| chr2:102709927-102711943 | p5 | (TTTTG)3 | GGTTTCGAGGAGCAGACTTG | TTTGGGGAAAGCATAAGACG | 191 |
| chr2:102765306-102767321 | p3 | (TGT)5 | GTTAGCCTCGTCGGCTTATG | CACTTAGTGGACTGTGCGGA | 197 |
| chr2:102765306-102767321 | p3 | (TGA)6 | GTTAGCCTCGTCGGCTTATG | CACTTAGTGGACTGTGCGGA | 197 |
| chr2:102927162-102929186 | p2 | (TA)12 | CCGTGAGTTTGGGACTGATT | ATTTTGACGCTCACAATCCC | 203 |
| chr2:102938878-102940922 | p3 | (TGT)7 | ACAGTTCATTCCCGACGAAG | CATGCTTGTTAGGACGGGTT | 272 |
| chr2:102938930-102940942 | p3 | (TGT)7 | ACAGTTCATTCCCGACGAAG | CATGCTTGTTAGGACGGGTT | 220 |
| chr2:102980866-102982926 | p2 | (AT)30 | TTTGGGTTCGTCATCCTCTC | CAAGAGAGCACATTACGCCA | 110 |
| chr2:103006838-103008856 | p2 | (TA)9 | TTTGGGTTCGTCATCCTCTC | TGGGCAAAAATTGTGTCGTA | 127 |
| chr2:103038938-103040953 | p5 | (ATTTC)3 | GAAGCGGAAGAAAGTATGCG | CAACCCAAAATATCATCGCC | 143 |
| chr2:103067599-103069614 | p2 | (TA)6 | AGAGAAACGGATTCAAGCGA | TCTTTTCCTTGTCATGTGCG | 294 |
| chr2:103067609-103069625 | p2 | (TA)6 | AGAGAAACGGATTCAAGCGA | TCTTTTCCTTGTCATGTGCG | 284 |
| chr2:103157404-103159416 | p2 | (CA)6 | CAATCCCCTCATCGAACTGT | ACGTGATTTGGAGTTGAGGG | 258 |
| chr2:103221992-103224007 | p2 | (AC)13 | TAAACGATCATCCGGACCTC | TTAGATTTGATTCGGGCGAC | 115 |
| chr2:103221992-103224007 | p3 | (CAA)5 | TAAACGATCATCCGGACCTC | TTAGATTTGATTCGGGCGAC | 115 |
| chr2:103222077-103224089 | p2 | (AC)13 | GTCGCCCGAATCAAATCTAA | TATGTAACGACCCGCTTGGT | 143 |

| Marker ID | SSR type | SSR motif | FORWARD PRIMER (5'-3') | REVERSE PRIMER (5'-3') | size (bp) |
|---|---|---|---|---|---|
| chr2:103222077-103224089 | p3 | (CAA)5 | GTCGCCCGAATCAAATCTAA | TATGTAACGACCCGCTTGGT | 143 |
| chr2:103235098-103237116 | p6 | (TATATG)3 | TTAGATTCTCGGGGATGGTG | ACTCTTGGCCTTCCCAAAAT | 183 |
| chr2:103339674-103341690 | p4 | (AATA)4 | TGGTTGTTGTACCAGCCAAA | GGACAGGATGGGTTTCTTGA | 274 |
| chr2:103367797-103369815 | p6 | (AAAACA)3 | AAAAGGTGCCATTCGTGAAC | AGGAGTCCCGAAAGAAGAGG | 144 |
| chr2:103367797-103369815 | p6 | (TTTTTG)3 | AAAAGGTGCCATTCGTGAAC | AGGAGTCCCGAAAGAAGAGG | 144 |
| chr2:103493368-103495386 | p6 | (CATATA)3 | ATAGGCACCCGACCCTTACT | AAGTCTTGTAAGCGGCTCCA | 190 |
| chr2:103529627-103531641 | p5 | (TCAAT)3 | GAGAATTCAACTCGGCCAAA | GCATGAAATTGGAAGGGAGA | 154 |
| chr2:103529667-103531693 | p5 | (TCAAT)3 | GAGAATTCAACTCGGCCAAA | GCATGAAATTGGAAGGGAGA | 114 |
| chr2:103543974-103545992 | p6 | (CATATA)3 | ATCCATCTCCTTGGGCTTCT | CATGGCATGAAGAACGATTG | 275 |
| chr2:103750269-103752284 | p2 | (AT)6 | AGCCCAAGGAAAGTCATGTG | AATTGTTCCTTCAAGGCGTG | 189 |
| chr2:103750269-103752284 | p5 | (AATTC)3 | AGCCCAAGGAAAGTCATGTG | AATTGTTCCTTCAAGGCGTG | 189 |
| chr2:103752444-103754458 | p2 | (AT)7 | TAATGTCGTTTTTGCCCCTC | GCCCTCGTAACAATAAGCCA | 275 |
| chr2:103766113-103768147 | p2 | (GA)17 | GACCAATGAAGGAGGCATGT | CTTTCCCCCTTTGGAACATT | 263 |
| chr2:103766113-103768147 | p3 | (TGT)5 | GACCAATGAAGGAGGCATGT | CTTTCCCCCTTTGGAACATT | 263 |
| chr2:103778269-103780281 | p2 | (AT)6 | AGCTACATTCTCTTGCCCGA | GGGACCCGATTACCCTAAGA | 199 |
| chr2:103941107-103943143 | p2 | (AT)18 | GCTTTCGACCTCAACTTTCG | GGAGTTCATCCGGTCTTTCA | 209 |
| chr2:103949609-103951653 | p2 | (TA)22 | TTTGGGTACATGCCCTTCTC | CGACAGGAGTTGCAAACAAA | 132 |
| chr2:103949609-103951653 | p2 | (TC)15 | TTTGGGTACATGCCCTTCTC | CGACAGGAGTTGCAAACAAA | 132 |
| chr2:104205688-104207710 | p2 | (AT)11 | CAGGTTTCCTATATCGCGGA | CCAAAATCGGGAAGTGAAGA | 205 |
| chr2:104290900-104292912 | p2 | (AT)6 | GAATTTGAGCGAGCGTAAGG | AGGCATTCCGACTTCATTTG | 134 |
| chr2:104306661-104308673 | p2 | (CA)6 | GAGTGAGGCGACGTACACAA | CTCCGGAATACGCACCTAAA | 134 |
| chr2:104316579-104318593 | p2 | (AT)7 | CAGTTTCGGAAGCCTGAGTC | CAGTTTGAGGGTCCGTGATT | 134 |
| chr2:104316579-104318593 | p2 | (AT)6 | CAGTTTCGGAAGCCTGAGTC | CAGTTTGAGGGTCCGTGATT | 134 |
| chr2:104328773-104330785 | p2 | (AT)7 | TTCTTTACCGTAGCGCCAGT | CTGGAGACCAAGAGAGGTCG | 162 |
| chr2:104328773-104330785 | p2 | (TA)6 | TTCTTTACCGTAGCGCCAGT | CTGGAGACCAAGAGAGGTCG | 162 |
| chr2:104338287-104340305 | p3 | (CAA)6 | TCACTCCCTGTCAACTGTGC | ACTGACCTTACACCGTTCCG | 117 |
| chr2:104516485-104518553 | p2 | (TA)34 | TGCCCGATTTTCGTTAAGTC | TTGTGAATCCTTTTCGCCTT | 267 |
| chr2:104516485-104518553 | p2 | (TA)11 | TGCCCGATTTTCGTTAAGTC | TTGTGAATCCTTTTCGCCTT | 267 |
| chr2:104516485-104518553 | p2 | (TA)13 | TGCCCGATTTTCGTTAAGTC | TTGTGAATCCTTTTCGCCTT | 267 |
| chr2:104541669-104543687 | p2 | (AT)9 | ATCGATTCTGGGAAGAGGGT | TATTACCAGACACCGAGCCC | 256 |
| chr2:104541669-104543687 | p2 | (AT)6 | ATCGATTCTGGGAAGAGGGT | TATTACCAGACACCGAGCCC | 256 |
| chr2:104548197-104550215 | p6 | (ATACAT)3 | CGCTGCATCTGTCGAATAAA | GGGTCGTGACACAGTTGTTG | 197 |
| chr2:104550459-104552471 | p2 | (AT)7 | CAACAACATTAATGCGCACC | GCGCATTTACAGGGGAAATA | 292 |
| chr2:104550510-104552526 | p2 | (AT)7 | CAACAACATTAATGCGCACC | GCGCATTTACAGGGGAAATA | 241 |
| chr2:104573531-104575546 | p3 | (CAA)5 | ATCTCTCGAACACCATTCGG | GAAGCCCGAGACTTGAGTTG | 297 |
| chr2:104573531-104575546 | p2 | (CA)6 | ATCTCTCGAACACCATTCGG | GAAGCCCGAGACTTGAGTTG | 297 |

| Marker ID | SSR type | SSR motif | FORWARD PRIMER (5'-3') | REVERSE PRIMER (5'-3') | size (bp) |
|---|---|---|---|---|---|
| chr2:104664360-104666378 | p2 | (TA)7 | TTCCTTCTGTTCATGGGTCC | ATATAACGCCTCGCCCTTTT | 229 |
| chr2:104664360-104666378 | p2 | (TA)9 | TTCCTTCTGTTCATGGGTCC | ATATAACGCCTCGCCCTTTT | 229 |
| chr2:104808700-104810722 | p2 | (TA)11 | ATTGTACGGGCTGTCCAGAG | ACACGCTGCCCAATAGTACC | 190 |
| chr2:104825453-104827468 | p3 | (GAT)5 | GTGGGTAAGGGGGATGTTTT | TGGGTTGTGTTCTTCATCCA | 282 |
| chr2:104863228-104865258 | p2 | (TA)6 | TGCAACGAGACTGCATAAGG | TTGATCCGGAGCTCATTTTT | 229 |
| chr2:104863228-104865258 | p2 | (AT)15 | TGCAACGAGACTGCATAAGG | TTGATCCGGAGCTCATTTTT | 229 |
| chr2:104863228-104865258 | p3 | (CAA)5 | TGCAACGAGACTGCATAAGG | TTGATCCGGAGCTCATTTTT | 229 |
| chr2:104883523-104885541 | p2 | (AG)20 | CAGATAGGGGAACCGTGAAA | CCCTAAGGACCAGATCCCTC | 236 |
| chr2:104883551-104885583 | p2 | (AG)20 | CAGATAGGGGAACCGTGAAA | CCCTAAGGACCAGATCCCTC | 208 |
| chr2:104898911-104900929 | p3 | (TGT)5 | AGAAGCCAACAACCACAAGG | TAAATGATCACCCGCAACAA | 113 |
| chr2:104898911-104900929 | p6 | (TTTTTA)3 | AGAAGCCAACAACCACAAGG | TAAATGATCACCCGCAACAA | 113 |
| chr2:104907193-104909211 | p2 | (CT)9 | CTTGACATGGCATCAAATGG | CTTTGTGCAGCTGATAGGCA | 229 |
| chr2:104951613-104953627 | p2 | (AT)7 | ACAATGTGCCCCAAAAATGT | GCACCAGCAGTGTTTATGGA | 155 |
| chr2:105075190-105077202 | p2 | (GT)7 | TTGTTGTGGCCTTGTGATGT | ATATTCAACAATCCCGCCAA | 120 |
| chr2:105075190-105077202 | p2 | (AT)6 | TTGTTGTGGCCTTGTGATGT | ATATTCAACAATCCCGCCAA | 120 |
| chr2:105075190-105077202 | p5 | (ATATG)3 | TTGTTGTGGCCTTGTGATGT | ATATTCAACAATCCCGCCAA | 120 |
| chr2:105075653-105077668 | p2 | (AT)6 | TGGAGAGCCCGATATTTGAG | CATCCGGAAACGTATCGACT | 243 |
| chr2:105075653-105077668 | p5 | (ATATG)3 | TGGAGAGCCCGATATTTGAG | CATCCGGAAACGTATCGACT | 243 |
| chr2:105100228-105102243 | p5 | (CATAT)3 | CAAACTCTGACTCGGCAACA | TACTGACCCCTGTTCTTCGG | 290 |
| chr2:105100228-105102243 | p2 | (AT)8 | CAAACTCTGACTCGGCAACA | TACTGACCCCTGTTCTTCGG | 290 |
| chr2:105258710-105260728 | p6 | (ATCAAT)3 | GGACTGAGATTGAACCGCAT | AGACCAAAAGGGCATGACAG | 144 |
| chr2:105258710-105260728 | p6 | (ATCCGT)3 | GGACTGAGATTGAACCGCAT | AGACCAAAAGGGCATGACAG | 144 |
| chr2:105328741-105330756 | p5 | (ATCAT)3 | TGTCGGAAAAAGGTGAAAGG | ATCTGTGATGGCCTTATCGG | 175 |
| chr2:105328741-105330756 | p3 | (CAA)5 | TGTCGGAAAAAGGTGAAAGG | ATCTGTGATGGCCTTATCGG | 175 |
| chr2:105337826-105339853 | p5 | (ATCAT)3 | GCATCCACGAACGGAGTAGT | TTTGAGCGGCTTTAAAGGAA | 197 |
| chr2:105337826-105339853 | p3 | (CAA)9 | GCATCCACGAACGGAGTAGT | TTTGAGCGGCTTTAAAGGAA | 197 |
| chr2:105346583-105348598 | p3 | (ACA)5 | TCCGGCTTAATAATGATGGC | GATTTCCAAAACCACTCCGA | 123 |
| chr2:105418401-105420417 | p2 | (TA)8 | GTCGGAACTGAAGCCCATTA | AGAGGGAGCCAATAACCGAT | 229 |
| chr2:105418401-105420417 | p3 | (AAT)9 | GTCGGAACTGAAGCCCATTA | AGAGGGAGCCAATAACCGAT | 229 |
| chr2:105479280-105481294 | p2 | (TA)7 | CCCCAAAGACCGGTTATTCT | ACAATAAATCGGGCAGCATC | 217 |
| chr2:105501510-105503525 | p5 | (TTCAT)3 | ATACAAAATGAAGCGCCGAC | CCTCACAGGGCCATATCACT | 291 |
| chr2:105503828-105505843 | p2 | (TA)10 | GGGCATGACGAAGTCTTGTT | CCTCTCGGCCATCAAAATAA | 275 |
| chr2:105503828-105505843 | p5 | (TATGA)3 | GGGCATGACGAAGTCTTGTT | CCTCTCGGCCATCAAAATAA | 275 |
| chr2:105570166-105572190 | p2 | (AC)7 | CCCACTTTGCAAACTTCCAT | TTTGGCTTTGGAAAATCACC | 142 |
| chr2:105570199-105572211 | p2 | (AC)7 | CCCACTTTGCAAACTTCCAT | TTTGGCTTTGGAAAATCACC | 109 |
| chr2:105625128-105627140 | p2 | (GT)6 | AAGTTGAGGCTCCCCTTGTT | AATTTGCCGTTAGCCTTCCT | 265 |

| Marker ID | SSR type | SSR motif | FORWARD PRIMER (5'-3') | REVERSE PRIMER (5'-3') | size (bp) |
|---|---|---|---|---|---|
| chr2:105644297-105646312 | p5 | (TTACT)3 | TAAGTGGTTCTGACGGGGAG | CCGTTGACCAGTGCCTAAAT | 102 |
| chr2:105654395-105656410 | p5 | (CATAT)3 | CCCAAATGTACCCAACAACC | TCTATGATGGCCTTATCGGC | 226 |
| chr2:105758814-105760832 | p6 | (ATCCAT)3 | AACCCATTAGGGTAAACCGC | CCTCTACGGGTCGTGACATT | 109 |
| chr2:105758814-105760832 | p2 | (TA)12 | AACCCATTAGGGTAAACCGC | CCTCTACGGGTCGTGACATT | 109 |
| chr2:105961579-105963597 | p3 | (ACA)6 | AAAAGACACAACGTCACCCC | GTGGCCTTTGTTGTGATGTG | 138 |
| chr2:106059957-106061971 | p2 | (TG)7 | AACCACGAGGATCGATCAAG | CGCCTACCTCCATATTTCCA | 237 |
| chr2:106228546-106230561 | p5 | (GGGAC)3 | GACTCGTGCTAGGGAAGTCG | CAAAACGACCCAAACTTGCT | 293 |
| chr2:106228546-106230561 | p3 | (TCA)5 | GACTCGTGCTAGGGAAGTCG | CAAAACGACCCAAACTTGCT | 293 |
| chr2:106229448-106231463 | p5 | (GGGAC)3 | ACCCCTTCGTGCCTAAAAAT | TCATGCCCACAGGTAGACAG | 111 |
| chr2:106229448-106231463 | p3 | (TCA)5 | ACCCCTTCGTGCCTAAAAAT | TCATGCCCACAGGTAGACAG | 111 |
| chr2:106247358-106249384 | p2 | (TG)13 | AGCCTCACATACCTTGGTCG | TTTCGAATTTTTCCGAATGC | 151 |
| chr2:106264878-106266896 | p6 | (ATATAC)3 | ACTGACTCCCAATACCACGC | AAGATGTTCCCACAAAAGCG | 297 |
| chr2:106293754-106295768 | p2 | (AT)7 | GTCGTGATTTGTGACCATCG | TTCCTCGCCTTGAGATAGGA | 244 |
| chr2:106362546-106364576 | p2 | (AT)15 | GGGGAAGTCTCCCTTATTGC | GATGTGGCTTCCGTCAAAGT | 260 |
| chr2:106362546-106364576 | p2 | (AT)7 | GGGGAAGTCTCCCTTATTGC | GATGTGGCTTCCGTCAAAGT | 260 |
| chr2:106365436-106367458 | p2 | (TA)11 | AGGGTAATTGGAAATTCGGG | CGCCTCGTCACTCCTTTAAC | 218 |
| chr2:106622984-106624999 | p5 | (CCACA)3 | TGACTATTTGGGTAAGGCGG | ATAGACATCGGCGTCCAAAG | 281 |
| chr2:106726785-106728811 | p2 | (AT)6 | ATAGGATCCTCGTACCGGCT | GCTAGCCTAGGAGTTCGGGT | 101 |
| chr2:106743528-106745540 | p2 | (AT)6 | TTAGGTACTGGAACTGCGGG | TTGCTCCTGATATTTTGCCC | 195 |
| chr2:106800775-106802787 | p2 | (AT)6 | CTCGGAGGTAGATTGGGTGA | CCCCATGTTCACTGGATACC | 153 |
| chr2:106834241-106836263 | p2 | (TA)11 | ATGCCCGCAGGTACAGATAC | ACAACCCACACATGCCTACA | 238 |
| chr2:107003023-107005037 | p2 | (AT)7 | TACGAAATCGGGATCAAAGG | GCTACCTCATTCAGGCCAAG | 283 |
| chr2:107034219-107036233 | p2 | (AT)7 | TACGAAATCGGGATCAAAGG | GCTACCTCATTCAGGCCAAG | 283 |
| chr2:107063299-107065314 | p5 | (TTTTC)3 | CGGTCTCAACACAACCACAC | GAGCTTAGGTGAAGCCATGC | 141 |
| chr2:107270009-107272021 | p2 | (TA)6 | CAAGCAAAGGGAGCTGAATC | TCCAAAGGTCGAATTCCAAG | 165 |
| chr2:107270009-107272021 | p5 | (TGAAT)5 | CAAGCAAAGGGAGCTGAATC | TCCAAAGGTCGAATTCCAAG | 165 |
| chr2:107270830-107272855 | p2 | (TA)6 | CAAGAAGGAGAAGCATTCGG | TGCACCAATAAATGGCTTGA | 243 |
| chr2:107270830-107272855 | p5 | (TGAAT)5 | CAAGAAGGAGAAGCATTCGG | TGCACCAATAAATGGCTTGA | 243 |
| chr2:107270830-107272855 | p6 | (TATATG)6 | CAAGAAGGAGAAGCATTCGG | TGCACCAATAAATGGCTTGA | 243 |
| chr2:107319510-107321528 | p4 | (ATAA)4 | GGTCCAAGCCCATGTAGAAA | TGATTTTCCACTTGCAACCA | 168 |
| chr2:107319510-107321528 | p2 | (AG)9 | GGTCCAAGCCCATGTAGAAA | TGATTTTCCACTTGCAACCA | 168 |
| chr2:107360840-107362855 | p5 | (AAAGG)3 | GGGTGAAAGAGCATAAGGCA | TGGTATCTTTATCCCAGCGG | 145 |
| chr2:107364742-107366794 | p2 | (TA)9 | CTGCTCGTGCATGCATAAAT | ACTGTACCCGGCCCTCTAGT | 171 |
| chr2:107364742-107366794 | p2 | (AT)26 | CTGCTCGTGCATGCATAAAT | ACTGTACCCGGCCCTCTAGT | 171 |
| chr2:107392702-107394718 | p4 | (TAAA)4 | CCGAAGTACATTGGGCCTTA | AACCTGCCGATCAAGTATGG | 259 |
| chr2:107703755-107705818 | p3 | (AAT)21 | AACAAGTGACGGGTAGGTCG | ATCACCTCAACCAAAGCAGG | 268 |

| Marker ID | SSR type | SSR motif | FORWARD PRIMER (5'-3') | REVERSE PRIMER (5'-3') | size (bp) |
|---|---|---|---|---|---|
| chr2:107716229-107718253 | p6 | (GGAGAA)4 | AAGAGCCACCTTTCTGGGAT | AGAGAAAGGGGAGAGCAAGG | 250 |
| chr2:107716229-107718253 | p6 | (CTCTAC)3 | AAGAGCCACCTTTCTGGGAT | AGAGAAAGGGGAGAGCAAGG | 250 |
| chr2:107876502-107878520 | p2 | (AT)25 | TCCTGAGCAACAACAACAGG | AGCTATGTGGCTCGCCTAGA | 173 |
| chr2:107876502-107878520 | p6 | (CATATT)3 | TCCTGAGCAACAACAACAGG | AGCTATGTGGCTCGCCTAGA | 173 |
| chr2:107876502-107878520 | p2 | (AT)6 | TCCTGAGCAACAACAACAGG | AGCTATGTGGCTCGCCTAGA | 173 |
| chr2:107985842-107987857 | p3 | (TTG)5 | ACGTTTTCGGATGACCTGAC | TATTGTGTGAGGGACCACGA | 230 |
| chr2:107985842-107987857 | p4 | (TGTA)6 | ACGTTTTCGGATGACCTGAC | TATTGTGTGAGGGACCACGA | 230 |
| chr2:108024220-108026235 | p5 | (TTTTA)3 | ACATGTCCGTCCATTTAGGC | CCCCTCCAGGTGAGATATGA | 151 |
| chr2:108034652-108036668 | p2 | (CA)8 | ACAGAATATTCCCCGAGGCT | GAGGAAACGGGGATTGAGTT | 211 |
| chr2:108034652-108036668 | p2 | (AT)6 | ACAGAATATTCCCCGAGGCT | GAGGAAACGGGGATTGAGTT | 211 |
| chr2:108184156-108186168 | p2 | (AT)6 | GCCAGGATGGCATATGATCT | GATGGCACACGCATACATTC | 191 |
| chr2:108196517-108198541 | p2 | (TA)9 | CCACAGGTAGACAGGGTGGT | ATGCGCATCTATGAGCCTCT | 255 |
| chr2:108196517-108198541 | p2 | (TA)12 | CCACAGGTAGACAGGGTGGT | ATGCGCATCTATGAGCCTCT | 255 |
| chr2:108253998-108256014 | p2 | (CA)8 | GACCATGCCAAAAGAAAGGA | GCAATTACACGGGAAATGCT | 260 |
| chr2:108317506-108319520 | p2 | (AG)7 | CCCTATATGTGTGCTGGGCT | CGTTACAAGTTCGGCTGACA | 134 |
| chr2:108317506-108319520 | p3 | (TGT)5 | CCCTATATGTGTGCTGGGCT | CGTTACAAGTTCGGCTGACA | 134 |
| chr2:108348118-108350134 | p2 | (AG)8 | TGCTGGGCTTTAGGATCTGT | ATGGACGTTACAAGTTCGGC | 145 |
| chr2:108410459-108412474 | p3 | (TTG)5 | ATACTAACCCCCGCAAGCTC | TTCCCTGTTTGATCCCTCTG | 182 |
| chr2:108410459-108412474 | p2 | (AT)12 | ATACTAACCCCCGCAAGCTC | TTCCCTGTTTGATCCCTCTG | 182 |
| chr2:108434623-108436637 | p2 | (TA)7 | ACATATCGCACCCTCGAGAC | GTGAAAGAGGCATGTCGGAT | 147 |
| chr2:108435424-108437438 | p2 | (TA)7 | ACATCCTTCACCGAGTGTCC | ATGCCTCTAGCGCCATAAAA | 106 |
| chr2:108435424-108437438 | p2 | (AT)8 | ACATCCTTCACCGAGTGTCC | ATGCCTCTAGCGCCATAAAA | 106 |
| chr2:108457966-108459981 | p3 | (ATT)8 | TGGACAAAAACTGTGCAAGC | AGCACTCCTCTTGTTTCGGA | 268 |
| chr2:108458071-108460101 | p3 | (ATT)8 | TGGACAAAAACTGTGCAAGC | AGCACTCCTCTTGTTTCGGA | 163 |
| chr2:108458332-108460356 | p3 | (ATT)8 | CTCCGAAACAAGAGGAGTGC | GGCTTTCAGAGGCAGTTCAG | 101 |
| chr2:108466313-108468365 | p2 | (AG)26 | TGTTATGCGTTCTCCCCTTC | GTTTGGTGGATGCATAGCCT | 153 |
| chr2:108492971-108494995 | p2 | (CT)12 | TGAGGGGAACGGTGATAAAG | ATATTTGTTCCCCCTCAGGC | 239 |
| chr2:108495216-108497231 | p2 | (AC)6 | CTGTCGAGCACTTCCAAACA | TGTCTTTTTGGGCTCATTCC | 261 |
| chr2:108495216-108497231 | p5 | (TATCA)3 | CTGTCGAGCACTTCCAAACA | TGTCTTTTTGGGCTCATTCC | 261 |
| chr2:108558626-108560672 | p5 | (TATGA)3 | ACCGCCCAGTCTACTTCTCA | TAACGGTTCACCCCGAGTAG | 108 |
| chr2:108558626-108560672 | p2 | (TA)23 | ACCGCCCAGTCTACTTCTCA | TAACGGTTCACCCCGAGTAG | 108 |
| chr2:108632290-108634326 | p4 | (TTTA)9 | ACTCCTCGATAAGGCCCATT | TTTTCTTGAGGCGCAAAACT | 196 |
| chr2:108637690-108639712 | p2 | (AT)11 | AAGTATCAACCGTGGCTTCG | CTCTTCCTCTTCGCCTTCCT | 288 |
| chr2:108637690-108639712 | p2 | (AT)17 | AAGTATCAACCGTGGCTTCG | CTCTTCCTCTTCGCCTTCCT | 288 |
| chr2:108637690-108639712 | p2 | (AT)13 | AAGTATCAACCGTGGCTTCG | CTCTTCCTCTTCGCCTTCCT | 288 |
| chr2:108653202-108655220 | p6 | (TCTGAT)3 | GTGATCGGCGAAAGGTTTTA | TTCGGCTTATTCGCACTTCT | 287 |

| Marker ID | SSR type | SSR motif | FORWARD PRIMER (5'-3') | REVERSE PRIMER (5'-3') | size (bp) |
|---|---|---|---|---|---|
| chr2:108723314-108725348 | p2 | (AT)17 | TGAACCTCTTCGGGATCAAC | ACCAAAAGGGGTTACCGAAC | 179 |
| chr2:108838447-108840477 | p2 | (AT)15 | TTGGGAAGTTGGATTTGAGG | CGCTGCTGCAGTAATGATGT | 139 |
| chr2:108919200-108921218 | p2 | (AT)9 | CCTCAGGCCAGAGATGAGAC | TTTCCCCCAACTCATGCTAC | 160 |
| chr2:108975911-108977927 | p2 | (TA)8 | GTTGGTTGCTGAGTTGGGAT | CAGCCTCCAACTACCTACGC | 193 |
| chr2:109065387-109067399 | p2 | (AT)6 | CATATGGAGCCCTCACGTTT | TTGTGGGGTCTAAGATTCCG | 113 |
| chr2:109065387-109067399 | p2 | (TA)9 | CATATGGAGCCCTCACGTTT | TTGTGGGGTCTAAGATTCCG | 113 |
| chr2:109191877-109193905 | p2 | (AT)14 | CGAGCCTCCGTAATCAAATC | CGCTCGTCTGAAGGGAATAG | 258 |
| chr2:109414378-109416394 | p2 | (AT)8 | CCCGTCGTACCCTTAAACAA | TGCAGATACTCGCTTTGGTG | 235 |
| chr2:109451056-109453098 | p2 | (AT)13 | CTTGTTGTTTTCCCGCATTT | CGGGTCAGGTTCTAAAACGA | 156 |
| chr2:109551936-109553950 | p2 | (GT)7 | GTTTCCTACGCCCACAGGTA | CTTATGAGCATAACGGGGGA | 168 |
| chr2:109830076-109832102 | p6 | (TCATGT)3 | GTCGGTTTATGAAATGCGCT | GCAACCGGAAGAAACAACAT | 126 |
| chr2:109830076-109832102 | p2 | (AT)13 | GTCGGTTTATGAAATGCGCT | GCAACCGGAAGAAACAACAT | 126 |
| chr2:109832639-109834660 | p3 | (ATG)7 | CCACAGGTAGACAGGAAGGC | ACGAGCCCTTCGACTATCAA | 104 |
| chr2:109832639-109834660 | p2 | (AT)7 | CCACAGGTAGACAGGAAGGC | ACGAGCCCTTCGACTATCAA | 104 |
| chr2:109832639-109834660 | p2 | (TA)22 | CCACAGGTAGACAGGAAGGC | ACGAGCCCTTCGACTATCAA | 104 |
| chr2:109855309-109857324 | p5 | (AAAAT)3 | CTGCCCTAAACCCAACAAGA | AGCTTCCTTTTCCTCAAGGC | 193 |
| chr2:109881456-109883471 | p5 | (CGAAC)3 | GGGTTGGATCGTGACAGAGT | CGGTTTGTTCAAGTTTCGGT | 230 |
| chr2:110086108-110088178 | p2 | (TA)35 | CAGGACCCTGCCATACTCAT | GTCGGTCGTTATATTGGCGT | 144 |
| chr2:110086108-110088178 | p3 | (CAA)6 | CAGGACCCTGCCATACTCAT | GTCGGTCGTTATATTGGCGT | 144 |
| chr2:110086329-110088389 | p2 | (TA)35 | TATACAAGTCGGGACAGGGC | GATCTTCCATCGTAGGCTGC | 189 |
| chr2:110086329-110088389 | p3 | (CAA)6 | TATACAAGTCGGGACAGGGC | GATCTTCCATCGTAGGCTGC | 189 |
| chr2:110086401-110088425 | p2 | (TA)35 | TATACAAGTCGGGACAGGGC | GATCTTCCATCGTAGGCTGC | 117 |
| chr2:110086401-110088425 | p3 | (CAA)6 | TATACAAGTCGGGACAGGGC | GATCTTCCATCGTAGGCTGC | 117 |
| chr2:110205436-110207502 | p2 | (TA)33 | AACTATGGCGGACAGATTCG | ATACGTCTGCGCCTCTGACT | 206 |
| chr2:110227473-110229539 | p2 | (TA)33 | GCTCGCTTACACAAACCACA | TCGAACTGTGCTGAGAATGG | 120 |
| chr2:110270996-110273011 | p3 | (TTG)5 | TTCTTGAAATGCGAATGCTG | GCATCTTGTCTTCACCCGAT | 265 |
| chr2:110304022-110306034 | p2 | (TA)6 | TGAGCTATGCTGACAGTGGG | TAACACCTAGCCCCAAATCG | 198 |
| chr2:110304022-110306034 | p2 | (AT)9 | TGAGCTATGCTGACAGTGGG | TAACACCTAGCCCCAAATCG | 198 |
| chr2:110314379-110316397 | p3 | (TCA)6 | GGGGGTTGTGTTTCATCATC | GTTAGGGGTGCGCTTAATGA | 251 |
| chr2:110314475-110316493 | p3 | (TCA)6 | GGGGGTTGTGTTTCATCATC | GTTAGGGGTGCGCTTAATGA | 155 |
| chr2:110344488-110346532 | p2 | (AT)22 | ACGTGCATATCTCCCTACCG | GTAAAAACAAAAGCCGGCAA | 121 |
| chr2:110388254-110390270 | p4 | (AAAT)4 | TCTCGTCGGAAACGAGAAAT | TCCCTTCCATCGTTGATTGT | 104 |
| chr2:110496162-110498177 | p5 | (GCATT)3 | CGAGGATCAAGGACAGGAAA | ACTAACCACACGTGGCATCA | 199 |
| chr2:110529580-110531604 | p3 | (TGT)8 | GCGACACACAAACAACCATC | ATGTATGCCAAAGGGCAAAG | 132 |
| chr2:110563742-110565754 | p2 | (TA)6 | CTTGCAAGATGGAAAGCTCC | TCTGGCCAATTTTTCCTTTG | 175 |
| chr2:110570485-110572497 | p2 | (TC)6 | ATGAATTGGGACGAAGCAAC | ATCACGCGGTTGAAAATCTC | 108 |

| Marker ID | SSR type | SSR motif | FORWARD PRIMER (5'-3') | REVERSE PRIMER (5'-3') | size (bp) |
|---|---|---|---|---|---|
| chr2:110604106-110606121 | p5 | (TTGGT)3 | GTTGAGAACCCCCAGTTCAA | GAGAAAGCTGATTTCCGCAC | 182 |
| chr2:110607216-110609231 | p3 | (TGT)5 | CCTCGTTATCGCCGATCTTA | AGACTTCTGGTGGGGCCTAT | 139 |
| chr2:110621469-110623481 | p2 | (TG)6 | TATGATTTCGTGCGTCGGTA | GAAAAATCGCATCACACACG | 182 |
| chr2:110662976-110664991 | p3 | (TTG)5 | GAGGCTTGTTTCTGGAGTGC | CTTCTCGCATCTGGTCTTCC | 208 |
| chr2:110703426-110705440 | p2 | (TA)7 | AAAAAGTGGAACCCACCACA | GTGCGGTGTTTAATATGGGG | 141 |
| chr2:110721337-110723349 | p2 | (AT)6 | ACTTTCTCCACGAAGGCGTA | CCAGGGTTCGGTTTGAGTAA | 152 |
| chr2:110753590-110755608 | p3 | (TCT)6 | ACAACGGAAGTCAATCGTCC | TGTTTGTGTTTGGGCCTGTA | 267 |
| chr2:110774720-110776738 | p6 | (ATTTTT)3 | GTCAATTTTTGGCTCCGAGA | TTTCGGGATTTATTCGATGG | 118 |
| chr2:110807669-110809684 | p3 | (GAA)5 | CGCCAAAACCTTTAAGGACA | AAGGGTTTTGAAAAGGGGAA | 176 |
| chr2:110822281-110824299 | p3 | (GTT)6 | CGGAGTGTTGGCCAGTTAAT | ACTTCTTCGGCCTACCCCTA | 239 |
| chr2:110822281-110824299 | p3 | (CTC)5 | CGGAGTGTTGGCCAGTTAAT | ACTTCTTCGGCCTACCCCTA | 239 |
| chr2:110853923-110855938 | p3 | (ATA)5 | CACGCCATAAACAAGACCCT | TTATTTTGATGGCCGAAAGG | 103 |
| chr2:110868249-110870261 | p6 | (ATATTT)3 | GAACCATTCTTAAACGGGCA | CCCAGGAGACAGGTTGTTGT | 148 |
| chr2:110868249-110870261 | p2 | (TA)6 | GAACCATTCTTAAACGGGCA | CCCAGGAGACAGGTTGTTGT | 148 |
| chr2:110868249-110870261 | p2 | (AT)6 | GAACCATTCTTAAACGGGCA | CCCAGGAGACAGGTTGTTGT | 148 |
| chr2:110915255-110917273 | p5 | (TGAAT)3 | TTAGATTCTCGGGGATGGTG | AACATACCTGCTCCACGTCC | 279 |
| chr2:110915255-110917273 | p6 | (TATATG)3 | TTAGATTCTCGGGGATGGTG | AACATACCTGCTCCACGTCC | 279 |
| chr2:111002157-111004177 | p5 | (GATTT)4 | CCGGGATCTGTTATCTGCAT | GAAGGTACCGAATGCCAAAA | 199 |
| chr2:111020298-111022318 | p5 | (ATGAC)4 | ACCCTTCCATGACGAGAATG | AAAATGAAATTGGGGCATCA | 263 |
| chr2:111082065-111084083 | p6 | (CATATA)3 | CGTTCTTTTCATCGTGCTCA | AGATGTTCCAGCGAAATTGG | 299 |
| chr2:111182837-111184855 | p6 | (CATATA)3 | TTGCAGACAACCCCTAGGAC | GGCTTCATCAGCCGATATGT | 279 |
| chr2:111182837-111184855 | p5 | (CATAT)3 | TTGCAGACAACCCCTAGGAC | GGCTTCATCAGCCGATATGT | 279 |
| chr2:111245158-111247170 | p2 | (GT)6 | ATTGCGGGAGATTAAGAGCA | CAGTTGGGCCTCAATTTTGT | 238 |
| chr2:111272375-111274387 | p2 | (AT)6 | AACAACAAACCCGAGAATGC | GAACCTGAATGAAATGCCGT | 135 |
| chr2:111298910-111300922 | p2 | (AT)6 | CCAAAGCATAGTCTCGGAGC | ATAGCAGCCTTGTCGGCTTA | 176 |
| chr2:111298910-111300922 | p2 | (AT)6 | CCAAAGCATAGTCTCGGAGC | ATAGCAGCCTTGTCGGCTTA | 176 |
| chr2:111319306-111321324 | p2 | (TA)9 | TCCCGACTTGTCCCTTATTG | TGAGGTCGCTTCCATATTCC | 124 |
| chr2:111355357-111357372 | p5 | (GAAAT)3 | TTTGGCTGATGGACATGGTA | TCCAGCTTGTCAAAAATCCC | 272 |
| chr2:111469613-111471659 | p2 | (AT)23 | ATCATACCGAAGCCACCAAG | GTCTAAGCGGTTCCACTTGC | 217 |
| chr2:111483222-111485237 | p5 | (ATTCA)3 | TGCACGAAAACATACTTGGC | TGTTTCTGAGCCATCAGACG | 161 |
| chr2:111499782-111501794 | p2 | (TA)6 | CTTAGAGTTCCAGGTTGGCG | TTCCTCGTAGGATAGCCCCT | 134 |
| chr2:111505234-111507254 | p2 | (AT)10 | AGAAATTGGCACTGAATGGG | GCTGCCCAATGGGATTAGTA | 103 |
| chr2:111535018-111537033 | p3 | (TCA)5 | CGAGCCTCTACTGGACTGCT | CACCCACAGGTAGAAAGGGA | 200 |
| chr2:111582318-111584336 | p2 | (AT)9 | TAGCCTGTCCCGATACCAAC | AGCTCCGCTACACCATCACT | 280 |
| chr2:111600336-111602358 | p2 | (TA)11 | GTGGAAGTCCTCCAAACGAA | GAACCACGACCTATGGATGG | 280 |
| chr2:111600336-111602358 | p5 | (GAAAT)3 | GTGGAAGTCCTCCAAACGAA | GAACCACGACCTATGGATGG | 280 |

| Marker ID | SSR type | SSR motif | FORWARD PRIMER (5'-3') | REVERSE PRIMER (5'-3') | size (bp) |
|---|---|---|---|---|---|
| chr2:111677371-111679385 | p2 | (TA)7 | ACGACCTGGTTTACAGACCG | GATGTTTTGTTAAGGCGGGA | 106 |
| chr2:111725110-111727125 | p5 | (ACACA)3 | GCTTAATTGTCCAAGCTCGC | GTTGCATTCTCGGGTTTGTT | 269 |
| chr2:111768914-111770929 | p5 | (TTTCA)3 | TATGCATTTGGGTCGTGTGT | TCGAATGTTAAGTCCTCGGG | 286 |
| chr2:111785307-111787322 | p3 | (CAA)5 | CATGGCACAGCATACTCCAC | CGACTTCATTGGATCGGTTT | 106 |
| chr2:111835701-111837717 | p2 | (TA)8 | GTTTTGGCACGTTGTTGTTG | TCAAACAGCCCACATTCAAA | 111 |
| chr2:111838941-111840957 | p2 | (AT)8 | TCCGGCTTAATAATGATGGC | TTGGGTAGGGCAAGAAGATG | 274 |
| chr2:111857821-111859836 | p5 | (GATAT)3 | GGTCAGCAGACAGGTGGATT | TGCCCGAATATACATACGCA | 261 |
| chr2:111875712-111877724 | p2 | (AC)6 | ATGTCCCACGTCCAACTAGC | AGATTCTCGGGGATCATGTG | 253 |
| chr2:112011790-112013808 | p6 | (ATATGT)3 | TTAGGAGGAACGTGGATTGG | TTATTCCCGAGGCTCACATC | 138 |
| chr2:112044923-112046938 | p2 | (AG)6 | GGCCAAGGTGAAAAACAAGA | CACACTCACGGCTCACTCTC | 249 |
| chr2:112044923-112046938 | p3 | (TGT)5 | GGCCAAGGTGAAAAACAAGA | CACACTCACGGCTCACTCTC | 249 |
| chr2:112044923-112046938 | p2 | (TA)6 | GGCCAAGGTGAAAAACAAGA | CACACTCACGGCTCACTCTC | 249 |
| chr2:112102432-112104452 | p5 | (ATTCA)4 | AAAAGACCGTATCCATCCCC | CCGAGGCTTGTTTTTGAGTC | 158 |
| chr2:112102432-112104452 | p2 | (CA)6 | AAAAGACCGTATCCATCCCC | CCGAGGCTTGTTTTTGAGTC | 158 |
| chr2:112102432-112104452 | p2 | (AT)16 | AAAAGACCGTATCCATCCCC | CCGAGGCTTGTTTTTGAGTC | 158 |
| chr2:112207308-112209324 | p2 | (GT)8 | CTCTGAAGGGAGTCAGGTGC | GGAGGGATTAGGGACTCAGG | 223 |
| chr2:112325197-112327211 | p2 | (AT)7 | GGGACGAGACATTTGGAAGA | CGAGGAGCAGAAATACTCCG | 103 |
| chr2:112406154-112408184 | p2 | (TA)34 | CGAAGCATCGAGACATGAAA | GTCGGTTATTAGCAGCCCAA | 284 |
| chr2:112406154-112408184 | p2 | (TA)12 | CGAAGCATCGAGACATGAAA | GTCGGTTATTAGCAGCCCAA | 284 |
| chr2:112406154-112408184 | p6 | (TATGAA)5 | CGAAGCATCGAGACATGAAA | GTCGGTTATTAGCAGCCCAA | 284 |
| chr2:112406154-112408184 | p2 | (AT)28 | CGAAGCATCGAGACATGAAA | GTCGGTTATTAGCAGCCCAA | 284 |
| chr2:112431254-112433318 | p6 | (AATATG)3 | GAAGCCTTGGTAGGGCTAGG | ACGAAGGTTCGGTGTTTCAC | 241 |
| chr2:112623255-112625269 | p2 | (AT)7 | GCGGGATATAACAAGGACGA | GCAAGCCGATGTACCATTTT | 253 |
| chr2:112728747-112730759 | p2 | (TA)6 | CCTTATTCTGCGTCATGGGT | CTTCCTCCGTGTTTCCACAT | 222 |
| chr2:112733383-112735395 | p2 | (AT)6 | ACCTCATGTGTTGGTGCTCA | GCACAAGCTGGAAAAGAAGG | 107 |
| chr2:112784631-112786643 | p2 | (AC)6 | TGCCATTCGGAAACATATCA | GCGAGCGAGGTATGTAAAGC | 109 |
| chr2:112799574-112801590 | p5 | (TGAAT)3 | TCGCCACAAGACTTTGACTG | CCTCGATCCATTGGACCTTA | 276 |
| chr2:112799604-112801632 | p5 | (TGAAT)3 | TCGCCACAAGACTTTGACTG | CCTCGATCCATTGGACCTTA | 246 |
| chr2:112819564-112821579 | p2 | (TA)17 | AAGCTTGGAGAAGGGGACAT | CCAACTTGCACAACAAATGC | 229 |
| chr2:112819564-112821579 | p5 | (TGAAT)3 | AAGCTTGGAGAAGGGGACAT | CCAACTTGCACAACAAATGC | 229 |
| chr2:112916573-112918588 | p5 | (ATCAG)3 | GAGTCACAAAATCGTTCGCA | CCGGAGCATACACTTTTGGT | 140 |
| chr2:112980215-112982230 | p5 | (ACATA)3 | AACTTCCGGGATGCATGTAG | TTCTTTGGGTCGAAATCTGG | 132 |
| chr2:113015352-113017378 | p2 | (AT)13 | TTAAGCGCTTTCCTCCAATG | TTTGTGATTGATGCTTGGGA | 206 |
| chr2:113020805-113022827 | p2 | (AT)13 | AGCTGCTCCATTAGCCGATA | GCCAAGTGACGAAAGTGTCA | 244 |
| chr2:113020805-113022827 | p2 | (AT)11 | AGCTGCTCCATTAGCCGATA | GCCAAGTGACGAAAGTGTCA | 244 |
| chr2:113075699-113077711 | p2 | (TA)6 | GCTTACTCAAGAGGTTGGCG | ATTTGCGCCACAACAATGTA | 291 |

| Marker ID | SSR type | SSR motif | FORWARD PRIMER (5'-3') | REVERSE PRIMER (5'-3') | size (bp) |
|---|---|---|---|---|---|
| chr2:113106330-113108354 | p6 | (TATATG)4 | AATGCCGTTGAATTGTGTGA | AATTTCGGTAGCATTTCCCC | 166 |
| chr2:113106330-113108354 | p2 | (AT)6 | AATGCCGTTGAATTGTGTGA | AATTTCGGTAGCATTTCCCC | 166 |
| chr2:113132089-113134104 | p5 | (AAGTT)3 | TGGGAAGGTATGAAAGGCTG | CGGCAACCAACTAACGAAAT | 170 |
| chr2:113194205-113196223 | p6 | (AAGAAA)3 | ATGTCTTGCAGCCTGACCTT | CCAACCATTGGTCGTAAACC | 214 |
| chr2:113250510-113252534 | p5 | (ATTCA)3 | TCGACCCAAATCATCAATCA | TGGGTTGGGTTGTGGTTATT | 100 |
| chr2:113250510-113252534 | p2 | (TA)12 | TCGACCCAAATCATCAATCA | TGGGTTGGGTTGTGGTTATT | 100 |
| chr2:113291663-113293681 | p2 | (TA)9 | GCTCGGACGAATAAGCGTAG | CGAGCTTTTGAAAACAAGGC | 254 |
| chr2:113295658-113297670 | p2 | (CA)6 | TGAGATAAATGGGCGAAAGG | CGTCTCTCTTCCCGACTTTG | 232 |
| chr2:113299495-113301507 | p2 | (TA)37 | GCTCGGACGAATAAGCGTAG | CGAGCTTTTGAAAACAAGGC | 103 |
| chr2:113337500-113339542 | p2 | (TA)21 | AGTCCTATGGCCGAACCTCT | ACCAGCGTGCAATACACAAA | 280 |
| chr2:113353617-113355632 | p5 | (AAAAG)3 | CTGTTCACCTCGGATGGTTT | TGAACGTCGATGTCACCATT | 131 |
| chr2:113364047-113366059 | p3 | (TGA)6 | GCATATTCCAGACAAACGCA | TGGAATCATGCCTTGTGAAA | 258 |
| chr2:113364069-113366083 | p3 | (TGA)6 | GCATATTCCAGACAAACGCA | TGGAATCATGCCTTGTGAAA | 236 |
| chr2:113444226-113446244 | p6 | (CATATA)3 | GAAAGCGAAGGTCAACAAGC | CGCATCACACCCTAGGAGAT | 135 |
| chr2:113455055-113457073 | p5 | (CATAT)4 | TTGATAACAAATCCTGGCCC | AATTCGATATAATCCCCCGC | 157 |
| chr2:113455055-113457073 | p3 | (CAA)6 | TTGATAACAAATCCTGGCCC | AATTCGATATAATCCCCCGC | 157 |
| chr2:113455055-113457073 | p2 | (AC)6 | TTGATAACAAATCCTGGCCC | AATTCGATATAATCCCCCGC | 157 |
| chr2:113461302-113463314 | p2 | (AT)6 | TAGAGGTCGTGGTGGAGCTT | TCGATTCAGGTTCTATCCCG | 200 |
| chr2:113465998-113468013 | p5 | (ATTCA)3 | TACTTACCGACCCGTGAAGG | TTTCCCAAAAATGGCTTCTG | 138 |
| chr2:113465998-113468013 | p2 | (TA)8 | TACTTACCGACCCGTGAAGG | TTTCCCAAAAATGGCTTCTG | 138 |
| chr2:113465998-113468013 | p2 | (AT)6 | TACTTACCGACCCGTGAAGG | TTTCCCAAAAATGGCTTCTG | 138 |
| chr2:113514929-113516944 | p2 | (TA)16 | TGGAGAAGGGGAAATGTGTC | CCTTGCACAAAGGAATGGTT | 222 |
| chr2:113514929-113516944 | p5 | (TGAAT)3 | TGGAGAAGGGGAAATGTGTC | CCTTGCACAAAGGAATGGTT | 222 |
| chr2:113549538-113551553 | p5 | (TGAAT)3 | AAGCTTGGAGAAGGGGACAT | CCTTGCACAAAGGAATGGTT | 218 |
| chr2:113640421-113642439 | p6 | (AGAAAA)3 | CGTTCGAAAGATAAGCGAGG | ACGGAGGGCACTCCTTTAGT | 130 |
| chr2:113656793-113658808 | p5 | (TTTTG)3 | GACGGCGTTCCATTTATGTT | TTGGAAGGTTTCGGATGAAG | 210 |
| chr2:113666543-113668559 | p2 | (TA)8 | GTGGCATCACACTATGTCCG | CCGTAGAAATCTGTGCCCAT | 294 |
| chr2:113738261-113740276 | p5 | (TCATA)3 | GAAAGTTTCAGGGCAATCCA | AGTCGGGGTGTGACAGATTC | 270 |
| chr2:113790123-113792141 | p2 | (TA)6 | CGTGTGAGCGTAGGTTTTCA | TAATCGCATCATGTTTCCCA | 239 |
| chr2:113790142-113792160 | p2 | (TA)6 | CGTGTGAGCGTAGGTTTTCA | TAATCGCATCATGTTTCCCA | 220 |
| chr2:113849202-113851217 | p2 | (GT)6 | CCAACTATCTTGCCCCAAAA | CCCACTTTGCAAACTTCCAT | 243 |
| chr2:113849202-113851217 | p3 | (TTG)5 | CCAACTATCTTGCCCCAAAA | CCCACTTTGCAAACTTCCAT | 243 |
| chr2:113849202-113851217 | p5 | (TATGA)3 | CCAACTATCTTGCCCCAAAA | CCCACTTTGCAAACTTCCAT | 243 |
| chr2:113878509-113880524 | p5 | (AAATG)3 | TCACGCTCATAGGTGACTCG | AGGGCGTCCTTATGGCTAAT | 172 |
| chr2:113890453-113892471 | p2 | (AT)34 | ATGCCCACAAGTAGACAGGG | CCTCTACTGGAGCGCTGAAC | 176 |
| chr2:113890453-113892471 | p2 | (TA)9 | ATGCCCACAAGTAGACAGGG | CCTCTACTGGAGCGCTGAAC | 176 |

| Marker ID | SSR type | SSR motif | FORWARD PRIMER (5'-3') | REVERSE PRIMER (5'-3') | size (bp) |
|---|---|---|---|---|---|
| chr2:113904576-113906591 | p2 | (GT)6 | CCAACTATCTTGCCCCAAAA | CCCACTTTGCAAACTTCCAT | 243 |
| chr2:113904576-113906591 | p3 | (TTG)5 | CCAACTATCTTGCCCCAAAA | CCCACTTTGCAAACTTCCAT | 243 |
| chr2:113904576-113906591 | p5 | (TATGA)3 | CCAACTATCTTGCCCCAAAA | CCCACTTTGCAAACTTCCAT | 243 |
| chr2:113933883-113935898 | p5 | (AAATG)3 | TCACGCTCATAGGTGACTCG | AGGGCGTCCTTATGGCTAAT | 172 |
| chr2:113946007-113948019 | p2 | (GT)6 | GTCCTTATGTTCAGCGCTCC | CCCGCGATTACCTCAAACTA | 152 |
| chr2:113971041-113973056 | p3 | (AAC)5 | TGTCAGTTTTGTGAGCAGGC | CGATCGAACCCACTGAATCT | 209 |
| chr2:113988892-113990908 | p2 | (TA)8 | AACAACAAACCCGAGAATGC | GTTGTGCGGGTTTTTGAAGT | 197 |
| chr2:114050396-114052438 | p2 | (AT)21 | ACTCGCCTTCTTCCGTCATA | GTGGTATCCGAGCAGGTTGT | 115 |
| chr2:114067127-114069139 | p2 | (CA)6 | CACGAAAGGACTATCGGCAT | GGCCTCACGGTACACTCATT | 185 |
| chr2:114077711-114079727 | p2 | (AG)8 | GGTAAAGAAGCCATGTCGGA | CGGGATACTTAAGGCAACCA | 112 |
| chr2:114086618-114088638 | p2 | (AT)10 | GAAATTCCCATTTTGCCCTT | TGCGTCTGGACATTTCAAAG | 135 |
| chr2:114140859-114142897 | p2 | (AT)19 | AGAAACCCGAGCCAGATTTT | TGGCGAACATAAATCACGAA | 261 |
| chr2:114172444-114174462 | p3 | (CAT)6 | CAATACCATTTCCAACGGCT | CCCTGTCGGCTCATATTGTT | 248 |
| chr2:114239318-114241346 | p2 | (AT)14 | GACCATTTGATGCGTGATTG | ACCTGTAGTCACGTCTGGGG | 184 |
| chr2:114346312-114348324 | p2 | (AT)6 | TCATCTTAAGCGGTCACACG | ATATGTGGGTCATGGGTCGT | 166 |
| chr2:114346312-114348324 | p2 | (AT)6 | TCATCTTAAGCGGTCACACG | ATATGTGGGTCATGGGTCGT | 166 |
| chr2:114356045-114358063 | p3 | (CAA)5 | TGGTAGAGGGCTTTTGCACT | GATGCTGCCCAAATTTCTGT | 259 |
| chr2:114356045-114358063 | p6 | (CATTTC)3 | TGGTAGAGGGCTTTTGCACT | GATGCTGCCCAAATTTCTGT | 259 |
| chr2:114366163-114368181 | p3 | (CAA)5 | TAGAGGGCTTTGCACTTGGT | GCCCAAATTTCTGTAGGCAA | 263 |
| chr2:114366163-114368181 | p6 | (CATTTC)3 | TAGAGGGCTTTGCACTTGGT | GCCCAAATTTCTGTAGGCAA | 263 |
| chr2:114388954-114390969 | p5 | (CATAG)3 | TTTGGCATGCTTCACACCTA | GTAAGGGGTGCTCGGTACAA | 167 |
| chr2:114388954-114390969 | p2 | (AT)7 | TTTGGCATGCTTCACACCTA | GTAAGGGGTGCTCGGTACAA | 167 |
| chr2:114413356-114415372 | p2 | (AT)8 | TACGAAAATGCTCGGTGTCA | TGATTCCATTGCATCCTTCA | 216 |
| chr2:114442636-114444651 | p5 | (TCATA)3 | CGTTGACTGGTGCCCTATTT | TTTATTCGCCAGTCCTACCG | 169 |
| chr2:114607029-114609063 | p2 | (AT)17 | CTCGATAATTGCTCCGGGTA | TAACACTTACTCCACCCCGC | 284 |
| chr2:114622122-114624136 | p2 | (TA)7 | TATCGGTAGGCCAGAGGTTG | TTTGTTTGGGGTGTCCAAAT | 264 |
| chr2:114662160-114664182 | p2 | (AT)11 | CCTCCATATCTTTCCAGCCA | TCCTAAACCAGAATCGGCAC | 165 |
| chr2:114662160-114664182 | p2 | (AT)6 | CCTCCATATCTTTCCAGCCA | TCCTAAACCAGAATCGGCAC | 165 |
| chr2:114735598-114737610 | p2 | (GT)6 | TCTGTCGGGGAAAGAATGTC | TAATCCCCTTTCCCATTTCC | 151 |
| chr2:114746054-114748322 | p2 | (AT)134 | TCCTCGTTGTAGTATGGGGC | CGTCCGTTGTCCCTTAAAAA | 152 |
| chr2:114746054-114748322 | p2 | (AT)9 | TCCTCGTTGTAGTATGGGGC | CGTCCGTTGTCCCTTAAAAA | 152 |
| chr2:114746054-114748322 | p2 | (AT)20 | TCCTCGTTGTAGTATGGGGC | CGTCCGTTGTCCCTTAAAAA | 152 |
| chr2:114758831-114760873 | p2 | (TC)21 | AATCGAACGAAACAAATGCC | AAACCAACACCAAAACCCAA | 114 |
| chr2:114767157-114769205 | p3 | (AAT)16 | TTGCGGAGTGCTCCTAAAAT | AATACGGCGAGTAACCGAAA | 214 |
| chr2:114767157-114769205 | p2 | (TA)33 | TTGCGGAGTGCTCCTAAAAT | AATACGGCGAGTAACCGAAA | 214 |
| chr2:114808600-114810642 | p2 | (TC)21 | AATCGAACGAAACAAATGCC | AAACCAACACCAAAACCCAA | 114 |

| Marker ID | SSR type | SSR motif | FORWARD PRIMER (5'-3') | REVERSE PRIMER (5'-3') | size (bp) |
|---|---|---|---|---|---|
| chr2:114816926-114818974 | p3 | (AAT)16 | TTGCGGAGTGCTCCTAAAAT | AATACGGCGAGTAACCGAAA | 214 |
| chr2:114816926-114818974 | p2 | (TA)33 | TTGCGGAGTGCTCCTAAAAT | AATACGGCGAGTAACCGAAA | 214 |
| chr2:114972982-114975030 | p2 | (TA)24 | ACACCTTGCACTCTTGCCTT | CAATTGGTATCAAAGCGGGT | 238 |
| chr2:115025462-115027474 | p2 | (TA)9 | CCGGTTAAAATAGGGTGCAA | TGTGGGACAATCCACTTTGA | 296 |
| chr2:115025462-115027474 | p2 | (TA)6 | CCGGTTAAAATAGGGTGCAA | TGTGGGACAATCCACTTTGA | 296 |
| chr2:115042798-115044818 | p4 | (ATTT)5 | AAGTGATGCCAGAGCCAAGT | TGTTGCTTGCCATGGTTAAA | 175 |
| chr2:115042798-115044818 | p2 | (AC)11 | AAGTGATGCCAGAGCCAAGT | TGTTGCTTGCCATGGTTAAA | 175 |
| chr2:115061232-115063280 | p2 | (TA)24 | ACACCTTGCACTCTTGCCTT | CAATTGGTATCAAAGCGGGT | 238 |
| chr2:115183056-115185071 | p5 | (TTGGT)3 | TCGACTTGGTTTGGTGTTGA | TTCGGTTTGGGCTTTATTTG | 281 |
| chr2:115205445-115207460 | p5 | (TTGGT)3 | AGTGGTTTGGTTCGACTTGG | GGTTTCGGTTTGGGCTTTAT | 270 |
| chr2:115215002-115217018 | p4 | (AAAC)4 | CAAGACAGACACGGCACCTA | ACTCTCGAGCCGTCCATCTA | 134 |
| chr2:115232384-115234402 | p6 | (TTGGAT)3 | TGGATGCTGGGGATAGAGTC | GCAGAAAGAGCGGATGATTC | 263 |
| chr2:115242074-115244119 | p3 | (TAT)15 | CAAAAGTGCAAATGACCCCT | AGATGGACCTCCGACAGCTA | 199 |
| chr2:115276658-115278688 | p2 | (AT)15 | GGTGCTGCGTTCTTTCTTTC | AGAAAATGGCCACGAGCTAA | 127 |
| chr2:115317580-115319598 | p3 | (GTT)6 | AGGCTTCGGAAGGATATGGT | TCTCATCTCGGCTAAGCGAT | 207 |
| chr2:115400317-115402341 | p3 | (ACA)8 | AGGGGAAAGTTCCATTGTCC | ACTTCTCATTTCAATGGCCG | 186 |
| chr2:115421364-115423379 | p5 | (CTTCT)3 | AAACCGGGTGCTTGTAAGTG | AATCGGCGTTAAGACTCGAA | 124 |
| chr2:115426518-115428530 | p2 | (GA)6 | GGACAACGGAAAAGGCAATA | AGTTGGAAGGACAATCGGTG | 281 |
| chr2:115460623-115462638 | p2 | (AT)19 | AGCGAAATATGACGGGACAG | TGCAGATACTCGCTTTGGTG | 272 |
| chr2:115460623-115462638 | p5 | (AAATG)3 | AGCGAAATATGACGGGACAG | TGCAGATACTCGCTTTGGTG | 272 |
| chr2:115477847-115479859 | p2 | (TA)6 | CGTGGTGGGAGTAACGACTT | CAATAGGCCCCACAAACACT | 199 |
| chr2:115540109-115542127 | p2 | (AT)9 | GGAGAGTGCATTTAGCCGAG | ATTCTTCCTCTTCGCCTTCC | 137 |
| chr2:115540109-115542127 | p2 | (TA)8 | GGAGAGTGCATTTAGCCGAG | ATTCTTCCTCTTCGCCTTCC | 137 |
| chr2:115714766-115716784 | p3 | (TTG)6 | AGTTTCCAACGCAACAAACC | TGTGGGATGAGAAATGGTCA | 245 |
| chr2:115802951-115804973 | p2 | (AT)11 | CACTGGAGCACCCTCTTAGC | GGCATAGAGGCCTATGGTGA | 169 |
| chr2:115999970-116001984 | p2 | (AT)7 | CAGCAACCCACAACAACAAC | ATTTCCGAATTCCAGTGCAG | 209 |
| chr2:116011292-116013304 | p2 | (TA)6 | TTGGGGATTGAGAACCAAAG | GGTCCCACACGACCAATAAC | 266 |
| chr2:116050616-116052630 | p2 | (AC)7 | AACGCCAACGAATACCAAAC | GAGCTGTGATGTGGCTTGAA | 249 |
| chr2:116185533-116187565 | p2 | (AG)16 | CCCACTTAGGCAGACTCAGG | TATGTGCGATGCTCCTCTTG | 162 |
| chr2:116256695-116258710 | p6 | (ATATAC)3 | TGTTTTCTTCAATTTCCGCC | AGCAGCTCACCCTGACAACT | 156 |
| chr2:116256695-116258710 | p5 | (TTATT)3 | TGTTTTCTTCAATTTCCGCC | AGCAGCTCACCCTGACAACT | 156 |
| chr2:116422411-116424425 | p2 | (TA)7 | AATTGCTCCTCTAGTCGCCA | GGGAGTTACAAAATGGGGGT | 164 |
| chr2:116422411-116424425 | p5 | (GAAAA)3 | AATTGCTCCTCTAGTCGCCA | GGGAGTTACAAAATGGGGGT | 164 |
| chr2:116423087-116425102 | p2 | (TA)7 | TCTTCGGAGTTGCCTCTTGT | CTCCATTACGAAGGACCAGC | 135 |
| chr2:116423087-116425102 | p5 | (GAAAA)3 | TCTTCGGAGTTGCCTCTTGT | CTCCATTACGAAGGACCAGC | 135 |
| chr2:116448132-116450147 | p5 | (ATGCA)3 | TGCCGATGACAGACATGATT | CAGATACGGTCTTTCGGCAT | 132 |

| Marker ID | SSR type | SSR motif | FORWARD PRIMER (5'-3') | REVERSE PRIMER (5'-3') | size (bp) |
|---|---|---|---|---|---|
| chr2:116498509-116500521 | p2 | (TA)6 | GGCAACCAAAGTCTAGCGAC | CATTGTTTCAACGCATTTGG | 154 |
| chr2:116498509-116500521 | p2 | (TA)6 | GGCAACCAAAGTCTAGCGAC | CATTGTTTCAACGCATTTGG | 154 |
| chr2:116498509-116500521 | p3 | (TAT)6 | GGCAACCAAAGTCTAGCGAC | CATTGTTTCAACGCATTTGG | 154 |
| chr2:116604089-116606101 | p2 | (AT)6 | AAAGCGAACCTGTCTCCTGA | GTTACCCTCCCGCTTAGGAC | 282 |
| chr2:116611419-116613437 | p3 | (CAA)6 | CAACACACCCTTGTCCACTG | TGGCCTTTGTTGTGATGTGT | 192 |
| chr2:116816348-116818362 | p2 | (CA)7 | TGAATAGAATTGCCTTCGGG | GATATTGAGCCGAGCAAAGC | 203 |
| chr2:116857228-116859240 | p2 | (TA)6 | TATGGCTTGTTGGGTGCATA | AATTAGGCAGCACTTCCCCT | 239 |
| chr2:116857228-116859240 | p2 | (TA)7 | TATGGCTTGTTGGGTGCATA | AATTAGGCAGCACTTCCCCT | 239 |
| chr2:116857228-116859240 | p2 | (AT)13 | TATGGCTTGTTGGGTGCATA | AATTAGGCAGCACTTCCCCT | 239 |
| chr2:116857955-116859981 | p2 | (TA)6 | CGTGCATGCCTCAATGTATC | GTCATCACATGCCATCCTTG | 167 |
| chr2:116857955-116859981 | p2 | (TA)7 | CGTGCATGCCTCAATGTATC | GTCATCACATGCCATCCTTG | 167 |
| chr2:116857955-116859981 | p2 | (AT)13 | CGTGCATGCCTCAATGTATC | GTCATCACATGCCATCCTTG | 167 |
| chr2:116973008-116975022 | p2 | (AT)7 | ATAGGTTGGGGATTTGGGTC | TGTGTTAGCTCTGTGTGCCC | 220 |
| chr2:116973008-116975022 | p2 | (TA)8 | ATAGGTTGGGGATTTGGGTC | TGTGTTAGCTCTGTGTGCCC | 220 |
| chr2:116994662-116996680 | p2 | (AT)9 | AAGGTTTGATTGTTCGTGGC | ACAAGAGGGAGAGGGGAAAA | 206 |
| chr2:116994662-116996680 | p5 | (CATTG)3 | AAGGTTTGATTGTTCGTGGC | ACAAGAGGGAGAGGGGAAAA | 206 |
| chr2:117065985-117067997 | p2 | (TA)6 | CACCCCGACATAGCCTTAAA | ATGCCTGTCCATCTAGGCAC | 252 |
| chr2:117078922-117080937 | p5 | (ATGAT)3 | GATGCTGCCCAAATTTCTGT | GACAATTTCAAACGCGGATT | 114 |
| chr2:117136611-117138626 | p3 | (GAA)5 | AAGGTGATGTACTGGGCCTG | ATGAACTTCCAGGCCTTCCT | 155 |
| chr2:117206424-117208439 | p2 | (AG)10 | GAAGGGATTCGACCAATGAA | TACAAGCAAGTCGTGCCAAG | 253 |
| chr2:117206424-117208439 | p3 | (TTG)5 | GAAGGGATTCGACCAATGAA | TACAAGCAAGTCGTGCCAAG | 253 |
| chr2:117214699-117216715 | p2 | (TA)8 | TTTGTCGGATTTACGGGAAG | GTCAAGCCCGGATACGATTA | 112 |
| chr2:117225026-117227044 | p2 | (AT)9 | GCGTGCATAACCCTCAAAGT | GCATAGACCGCCTCTAGCAC | 254 |
| chr2:117251408-117253420 | p3 | (CAA)5 | GGTCATGCCAAAGAAAGGAA | GGTGAATTTTGCCGAGTTGT | 232 |
| chr2:117251408-117253420 | p2 | (CA)6 | GGTCATGCCAAAGAAAGGAA | GGTGAATTTTGCCGAGTTGT | 232 |
| chr2:117253257-117255269 | p2 | (AT)6 | CCCTTTGGGCGATATTACCT | GCTCCATCTCATTCAGAGCC | 164 |
| chr2:117473971-117475986 | p2 | (AT)15 | ACGCGTTGCAATACTGACTG | TACACATATGCTTCGAGCGG | 129 |
| chr2:117494099-117496159 | p2 | (AT)30 | TCAGAAGCCGACAAGGCTAT | CCGGAGCCTACATTTTGGTA | 159 |
| chr2:117521549-117523565 | p2 | (AT)8 | CCAAACCGTGGTCTCAGATT | TTGTTTAAGGGTACGACGGG | 245 |
| chr2:117658907-117660925 | p3 | (GAT)6 | TCATTCCCATTTCTCCTCCA | CTTGCACCAAAACCCTTGTT | 124 |
| chr2:117660604-117662620 | p2 | (TA)8 | TAACCCTGAAGCCTTGGTTG | CACATACATATCCTCCGGGC | 294 |
| chr2:117674200-117676218 | p3 | (GAT)6 | TCATTCCCATTTCTCCTCCA | CTTGCACCAAAACCCTTGTT | 124 |
| chr2:117675897-117677913 | p2 | (TA)8 | TAACCCTGAAGCCTTGGTTG | CACATACATATCCTCCGGGC | 294 |
| chr2:117731562-117733577 | p5 | (TTCAT)3 | AGTTCCGGTTGATGACGTTC | CGCTTCCCAAGTCATTTGTT | 159 |
| chr2:117739176-117741194 | p2 | (AT)16 | TTTTGTCACGACCTAACCCC | TGTTCGTCTGTAGTGGCAGC | 139 |
| chr2:117739176-117741194 | p2 | (AT)9 | TTTTGTCACGACCTAACCCC | TGTTCGTCTGTAGTGGCAGC | 139 |

| Marker ID | SSR type | SSR motif | FORWARD PRIMER (5'-3') | REVERSE PRIMER (5'-3') | size (bp) |
|---|---|---|---|---|---|
| chr2:117746677-117748722 | p2 | (AG)17 | TTTTTGACCACCGGAAACTC | CCCACCAGTTGTCAACCTTT | 138 |
| chr2:117746677-117748722 | p2 | (AT)10 | TTTTTGACCACCGGAAACTC | CCCACCAGTTGTCAACCTTT | 138 |
| chr2:117773632-117775646 | p2 | (TA)7 | CTCCTGTTCAGAGGGTGAGC | CCCCCTCAGGAAAAGAAATC | 144 |
| chr2:117778094-117780109 | p5 | (TATTT)3 | CCTGGTATGACCATCTGCCT | CTCATCATCCCTTTGGGCTA | 261 |
| chr2:117908123-117910138 | p3 | (CAA)5 | TGTGGAGCAAATCAAATGGA | CACTAGGATGGCCGATGTCT | 242 |
| chr2:117908123-117910138 | p2 | (CA)6 | TGTGGAGCAAATCAAATGGA | CACTAGGATGGCCGATGTCT | 242 |
| chr2:118005725-118007746 | p3 | (GAA)7 | TCTGGCAAATGCAGACAACT | ACGAGGATGCACATGCACTA | 134 |
| chr2:118005725-118007746 | p2 | (TC)18 | TCTGGCAAATGCAGACAACT | ACGAGGATGCACATGCACTA | 134 |
| chr2:118046448-118048463 | p3 | (AAG)5 | TTCTCCATCATCCGTTGTGA | AATTGGGATTGGGTTTGTGA | 275 |
| chr2:118132661-118134673 | p2 | (TA)6 | GGAGGCCACGTGTCATATCT | CGCGGTTACAAGATATTGGG | 178 |
| chr2:118333717-118335732 | p5 | (TTTTC)3 | TTCATGCTGCTTTGAACTGC | AACAACAACATGCCCAGTGA | 252 |
| chr2:118363972-118365990 | p3 | (TTG)6 | TTTCGTCGCAAGAAAGTCCT | TACAACGATGGCTCGTCAAC | 246 |
| chr2:118504502-118506520 | p6 | (TTCGGT)3 | ACCGGTGAGTGAGTTTACCG | TCTTTCGATGTGGTGGATGA | 145 |
| chr2:118505028-118507043 | p6 | (TTCGGT)3 | CCCGTTATCACACAAAGCCT | CCCACCCCTATCCAAGAGAT | 116 |
| chr2:118522637-118524665 | p2 | (AT)8 | CCTCACAACTCAGCCACAGA | GTCCCACATCGTTTGGACTT | 214 |
| chr2:118522680-118524694 | p2 | (AT)8 | CCTCACAACTCAGCCACAGA | GTCCCACATCGTTTGGACTT | 171 |
| chr2:118652967-118654987 | p5 | (TATCG)4 | GATGGGTCCCCAAATAAGGT | CGACTGGCAGTTGACTTGAA | 243 |
| chr2:118652967-118654987 | p2 | (AT)15 | GATGGGTCCCCAAATAAGGT | CGACTGGCAGTTGACTTGAA | 243 |
| chr2:118708132-118710150 | p2 | (TA)9 | GCAAGGAGTGCACAGATCAA | TTCTCCTCAAGATGGTCGCT | 273 |
| chr2:118709165-118711187 | p2 | (AT)11 | CTTAGCTGCACCCTTCGTTC | TTATGAGGTGGACGCACAAA | 122 |
| chr2:118732060-118734084 | p2 | (AT)12 | ACCGACCTTATGTCACTCCG | TACGGCCCGATCCATAAATA | 165 |
| chr2:118774823-118776835 | p2 | (AT)6 | GCGCTAAGCAATTCCAAAAG | AGGCTGCCAAGATTGTTTGT | 101 |
| chr2:118957050-118959086 | p2 | (TG)18 | TTTCCAAGCTAGGGCAAGAA | AATTCACCATTGGAAACCCA | 150 |
| chr2:119065489-119067501 | p2 | (TA)6 | AGTGCTCAAGGAAGCAAGGA | CCTGCATGTTATTGAAGCGA | 290 |
| chr2:119217641-119219662 | p3 | (ATG)7 | GCCCGCAGATAGACAAAGAG | AAAGACGGGGCAAGACTTTT | 298 |
| chr2:119278095-119280125 | p2 | (GT)15 | AGTGTTGGATTCAAGGCTGG | TCGGTCCATACAGTTGTCCA | 293 |
| chr2:119278095-119280125 | p3 | (CAA)7 | AGTGTTGGATTCAAGGCTGG | TCGGTCCATACAGTTGTCCA | 293 |
| chr2:119350286-119352310 | p3 | (GTT)8 | CCACAAAATTCTGCAAGGGT | GGTTGAACAATCAGGCGAAT | 247 |
| chr2:119414918-119416933 | p3 | (GAG)5 | AAGGTACAATGCCATCTGCC | CGAAGTAGGTGGAGCTGGAG | 195 |
| chr2:119414918-119416933 | p2 | (TA)7 | AAGGTACAATGCCATCTGCC | CGAAGTAGGTGGAGCTGGAG | 195 |
| chr2:119414918-119416933 | p5 | (TGTGT)3 | AAGGTACAATGCCATCTGCC | CGAAGTAGGTGGAGCTGGAG | 195 |
| chr2:119415870-119417885 | p3 | (GAG)5 | GCCTGATACCACCATTGCTT | AACCTACCACGACCACCTTG | 206 |
| chr2:119415870-119417885 | p2 | (TA)7 | GCCTGATACCACCATTGCTT | AACCTACCACGACCACCTTG | 206 |
| chr2:119415870-119417885 | p5 | (TGTGT)3 | GCCTGATACCACCATTGCTT | AACCTACCACGACCACCTTG | 206 |
| chr2:119415870-119417885 | p5 | (TGAAT)3 | GCCTGATACCACCATTGCTT | AACCTACCACGACCACCTTG | 206 |
| chr2:119434830-119436850 | p2 | (CT)10 | CCAATGGCGTTTTTGATTTT | GGAGCCGTTAAGTGCTTCAG | 102 |

| Marker ID | SSR type | SSR motif | FORWARD PRIMER (5'-3') | REVERSE PRIMER (5'-3') | size (bp) |
|---|---|---|---|---|---|
| chr2:119434830-119436850 | p4 | (TATC)4 | CCAATGGCGTTTTTGATTTT | GGAGCCGTTAAGTGCTTCAG | 102 |
| chr2:119471263-119473283 | p5 | (AAAAC)4 | GAACACTCACCACACATGGC | ATGAGAGCCCTTGAAAGCAA | 237 |
| chr2:119519999-119522017 | p3 | (TCT)6 | GTCCAACAAACGCCCTAAAA | ATGCATCATGATCAAAGCCA | 260 |
| chr2:119583260-119585276 | p2 | (AT)8 | TTCTGTTGGGTTGAGGGTTC | CTGAGGTTGTCCATCGAGGT | 122 |
| chr2:119583260-119585276 | p5 | (AATAA)3 | TTCTGTTGGGTTGAGGGTTC | CTGAGGTTGTCCATCGAGGT | 122 |
| chr2:119595544-119597574 | p2 | (TA)15 | GGAATGTCTACGAGCCGTGT | GGTTAAAATCAAGGGCGTCA | 227 |
| chr2:119615557-119617581 | p2 | (TA)12 | CTACTTCCCCCAAAGCCTTC | GGACGGTGAGACTTGTGGTT | 167 |
| chr2:119698793-119700805 | p3 | (TTC)10 | CAGACCACGCGCAGTAGATA | GAGAATGGGAACGAGTTGGA | 117 |
| chr2:119766201-119768215 | p2 | (TC)7 | TTTGCCTGGATTGCTCTTTT | CCCCCAACTTCACTTCGTTA | 266 |
| chr2:119783419-119785434 | p5 | (TGATA)3 | CGCGAAAAAGGTTGTAAAGC | TTGAAGCCTATGAGCCATCC | 237 |
| chr2:119795521-119797536 | p5 | (ATTTC)3 | CATGTCATCCCACACCTCAG | TCGCTCGCCTATTCACTTTT | 275 |
| chr2:119797256-119799314 | p2 | (TA)29 | ATTTCCTTCAAGGTGAGGCA | CCATATAGGCGCGGTGTAAT | 252 |
| chr2:119906131-119908143 | p2 | (AT)6 | AGCTCGAGGAGTAGACGCTG | ATATCAAAATGGACGGCTGG | 114 |
| chr2:119906131-119908143 | p2 | (TA)6 | AGCTCGAGGAGTAGACGCTG | ATATCAAAATGGACGGCTGG | 114 |
| chr2:119961001-119963022 | p3 | (TGT)7 | TTGGAGGGTATTCGCATTTC | GTTTTGTTCGCATGTTGTGG | 233 |
| chr2:120062446-120064461 | p5 | (TACAA)3 | ACCCATGTTTGGATGGTTGT | GGTATGGGTAGGGTGGAGGT | 291 |
| chr2:120069294-120071306 | p2 | (TC)6 | ATTTTTGCATTCAATTCGGC | CAAATTGAAGCCTATGGGGA | 132 |
| chr2:120069294-120071306 | p6 | (AAAAAT)3 | ATTTTTGCATTCAATTCGGC | CAAATTGAAGCCTATGGGGA | 132 |
| chr2:120091590-120093606 | p4 | (TATG)4 | AGGTTGTGGGTTCACAGGAG | TCCTTTACGAGCTTTGGCAT | 162 |
| chr2:120152380-120154395 | p5 | (ATAAC)3 | CTGCGCTAGAGCTGTTTCCT | CAGAGGGTTGCACCAGGTAT | 272 |
| chr2:120173810-120175825 | p3 | (ATA)5 | GGAGTAAGACAAACGCCCAA | ATCCAAATGCATGGGTCAAT | 206 |
| chr2:120179019-120181034 | p5 | (ATTGC)3 | AGCGAAGGAATGATTGTTGG | AACACAACGGGGTGACTAGC | 246 |
| chr2:120198364-120200398 | p2 | (TA)17 | TAGTATTGCCAAGCAAGGGG | CCTTCTTGTTGCTTAAGGCG | 293 |
| chr2:120199913-120201928 | p5 | (TTTAT)3 | CGTGGGAAGGTTACCTCAGA | TCTCCTCCTTTCACCACCAC | 175 |
| chr2:120409910-120411925 | p3 | (TTG)5 | ATGGATGAAATGGAGGCTTG | CGCATCTTGTCTTCCACTGA | 135 |
| chr2:120409910-120411925 | p2 | (TC)6 | ATGGATGAAATGGAGGCTTG | CGCATCTTGTCTTCCACTGA | 135 |
| chr2:120438442-120440457 | p5 | (TACAA)3 | TCTTAAATCGCACAAACCCC | GAATGCAGCGAACTGGGTAT | 294 |
| chr2:120463676-120465706 | p3 | (CTT)10 | TGAAATCTGCGAGCATTCAG | TGATAAGTGGGGCATGTGAA | 181 |
| chr2:120487567-120489630 | p3 | (ATT)21 | GCCAAAGGCATCATTCTCAT | TAGTTCGCTGCCGAAGAAAT | 279 |
| chr2:120487567-120489630 | p2 | (TC)6 | GCCAAAGGCATCATTCTCAT | TAGTTCGCTGCCGAAGAAAT | 279 |
| chr2:120492566-120494578 | p2 | (AT)6 | GCGATAATCAGATGGGCAGT | CCGAGTATGTAAGGCGGAAA | 295 |
| chr2:120616303-120618318 | p3 | (TAT)5 | CTGTCCGTCGAACATGCTTA | ATAGTTTATGGTGCGGGCTG | 193 |
| chr2:120631304-120633374 | p6 | (TAAAAA)3 | CTCCTTCACCAAACACAGCA | AATTTTCTACCATTGCCCCC | 240 |
| chr2:120631384-120633420 | p6 | (TAAAAA)3 | TGCTCCTTCACCAAACACAG | AATTTTCTACCATTGCCCCC | 158 |
| chr2:120631799-120633817 | p6 | (TAAAAA)3 | TAAGTGGGAGGGAGGATGAA | AAAGGTGGAGTTGTGCCAAG | 100 |
| chr2:120689849-120691867 | p6 | (AGGAAA)3 | CTCTCACGGTTGCATTTTCA | TGGGCATGAAATTGTTCTCA | 136 |

| Marker ID | SSR type | SSR motif | FORWARD PRIMER (5'-3') | REVERSE PRIMER (5'-3') | size (bp) |
|---|---|---|---|---|---|
| chr2:120697477-120699491 | p2 | (CA)10 | CACACACGACATGCACACAT | AACCCTCTCCGTTGCTTTTT | 272 |
| chr2:120697477-120699491 | p2 | (AT)7 | CACACACGACATGCACACAT | AACCCTCTCCGTTGCTTTTT | 272 |
| chr2:120779376-120781390 | p2 | (TA)7 | TTGTGTTGGGTTTTTGGGTT | TGTCGCAATTCACAAAGGA | 221 |
| chr2:120814912-120816936 | p5 | (ATTTT)3 | GCAAATTACTATGCGAGCCC | GAGTCCTCACATCGGTGGAT | 186 |
| chr2:120821964-120823979 | p5 | (GAAAT)3 | CTCGTCACAAGAGGTGCGTA | CGCTAGCATTGGAGAGGAAC | 128 |
| chr2:120862569-120864584 | p5 | (AATTA)3 | TTTTTCTTCACCACCGGAAC | CGTTAGCTGCAATCCTGTCA | 151 |
| chr2:120933921-120935939 | p2 | (AT)7 | AGCCCTTCGGCTATCAAAAT | TTGCAGATCCATATGAGCCA | 264 |
| chr2:120933921-120935939 | p6 | (GGCATA)3 | AGCCCTTCGGCTATCAAAAT | TTGCAGATCCATATGAGCCA | 264 |
| chr2:121010224-121012240 | p2 | (TA)8 | GAAGAAATGACTGGGAGCCA | TTACAAGCATTTTGGGGGAG | 123 |
| chr2:121010224-121012240 | p3 | (TTG)5 | GAAGAAATGACTGGGAGCCA | TTACAAGCATTTTGGGGGAG | 123 |
| chr2:121022148-121024163 | p3 | (GTT)5 | CCCCCAAACTCAGTTCAAAA | GATCTGGAAAACCCCTCTCC | 251 |
| chr2:121163513-121165531 | p2 | (CT)6 | CGACATGTCAGTCCACCAAC | GGATCGTGCTCATTTGGACT | 229 |
| chr2:121180368-121182386 | p6 | (CATATA)3 | ACCACTTTGTCACACACCGA | GTCTTGAAAGCGGCTTCATC | 117 |
| chr2:121199930-121201960 | p2 | (AT)15 | ACATGTGGACCCCACTTAGC | CCCAGATTCGGGATAAAGGT | 253 |
| chr2:121235069-121237084 | p5 | (AACCA)3 | ACAAGGCAACAAATTCTGGG | CCCTTTGGTTGTCGAATCAT | 212 |
| chr2:121407122-121409137 | p3 | (TGT)9 | GTGGATCTCGCTCAGCTACC | TCCCAACTTCATACCCGAAG | 273 |
| chr2:121408620-121410644 | p3 | (TGT)9 | AGGGGGCTTACCCCTTATTT | ACCGCAACTATACGCTGTCC | 109 |
| chr2:121408620-121410644 | p3 | (GAT)8 | AGGGGGCTTACCCCTTATTT | ACCGCAACTATACGCTGTCC | 109 |
| chr2:121408620-121410644 | p2 | (TA)14 | AGGGGGCTTACCCCTTATTT | ACCGCAACTATACGCTGTCC | 109 |
| chr2:121410845-121412861 | p4 | (AATT)4 | GGGGTGTTGTTATTGTTCCG | CCTCGGTAAGTGAACCCTCA | 183 |
| chr2:121591739-121593755 | p4 | (CATA)4 | ATGTCTGAGCGATACCGTCC | CCCCTGTTGTCGTCTCTTGT | 202 |
| chr2:121601888-121603904 | p4 | (AATT)4 | TCTTTTTCCAGTGTTTCCCG | TATGTCTTTCCGTGCCACAA | 171 |
| chr2:121601888-121603904 | p2 | (TG)7 | TCTTTTTCCAGTGTTTCCCG | TATGTCTTTCCGTGCCACAA | 171 |
| chr2:121696507-121698521 | p2 | (AT)13 | CAGATCGACCCGAAACTTGT | TCGGAAACCGTAACACATCA | 288 |
| chr2:121696507-121698521 | p2 | (TA)7 | CAGATCGACCCGAAACTTGT | TCGGAAACCGTAACACATCA | 288 |
| chr2:121702780-121704792 | p2 | (GT)6 | TGAATTCTATGTGCCGTCCA | CCTCACTATTTGCGCATCAA | 229 |
| chr2:121710109-121712124 | p5 | (CAATG)3 | GGTAGGCGGACCCACTTATT | CATGGGAAGTTGGATCGAGT | 175 |
| chr2:121756382-121758446 | p2 | (AT)32 | GGGCAGCTGTAGCTTTCTTG | ACACTCGTCACGACTCATCG | 222 |
| chr2:121807551-121809575 | p4 | (GAAA)6 | GAACCCTAGCGAGCTTCCTT | GTCCCGACTGGCTGAGTAAG | 203 |
| chr2:121971998-121974024 | p2 | (TA)13 | TTGGTAAACCAGGAGTTGCC | ATATTTTTGCACGCGAGACC | 260 |
| chr2:122232351-122234366 | p5 | (AAACC)3 | TATGGACCGGGTTGGTTAGA | CTGAGGTTGAAAAACCCGAA | 203 |
| chr2:122310041-122312056 | p5 | (CAAAC)3 | GGCAGTAGAATAAGCCTGCG | GGGTGTTTGGCCTGAAAATA | 186 |
| chr2:122409125-122411140 | p5 | (TTCAT)3 | GCCTCTCTTACCCATTCACG | GCAAGCAATGCAACAAGAAA | 156 |
| chr2:122409125-122411140 | p5 | (TTTGT)3 | GCCTCTCTTACCCATTCACG | GCAAGCAATGCAACAAGAAA | 156 |
| chr2:122437059-122439077 | p6 | (CATATT)3 | CCTGACGCTGACTATCACGA | GTTAGCCTCGTCGGCTTATG | 286 |
| chr2:122502805-122504859 | p2 | (TA)27 | CCCTTTGAGCCACAAACAGT | TATGAAGCGTGGGGAAGAAC | 222 |

| Marker ID | SSR type | SSR motif | FORWARD PRIMER (5'-3') | REVERSE PRIMER (5'-3') | size (bp) |
|---|---|---|---|---|---|
| chr2:122514614-122516628 | p2 | (AC)6 | GTCTTGGAAATCGACGGAAA | GATTTGCGGCTATCAGCTTC | 163 |
| chr2:122514614-122516628 | p2 | (CA)7 | GTCTTGGAAATCGACGGAAA | GATTTGCGGCTATCAGCTTC | 163 |
| chr2:122552827-122554842 | p5 | (TAGTT)3 | CCTCCAAAGTCAGAATGGGA | ATGCCAGGAATACGCTTCAC | 104 |
| chr2:122552827-122554842 | p5 | (AAATA)3 | CCTCCAAAGTCAGAATGGGA | ATGCCAGGAATACGCTTCAC | 104 |
| chr2:122552827-122554842 | p5 | (TTTAT)4 | CCTCCAAAGTCAGAATGGGA | ATGCCAGGAATACGCTTCAC | 104 |
| chr2:122553450-122555465 | p5 | (TAGTT)3 | ATGCCTAGAAGATTCCGCCT | CCTGCAAGAGGACACACAAA | 146 |
| chr2:122553450-122555465 | p5 | (AAATA)3 | ATGCCTAGAAGATTCCGCCT | CCTGCAAGAGGACACACAAA | 146 |
| chr2:122553450-122555465 | p5 | (TTTAT)4 | ATGCCTAGAAGATTCCGCCT | CCTGCAAGAGGACACACAAA | 146 |
| chr2:122553450-122555465 | p5 | (TTTAT)3 | ATGCCTAGAAGATTCCGCCT | CCTGCAAGAGGACACACAAA | 146 |
| chr2:122665429-122667444 | p3 | (AGA)5 | GGTGGATGCCATCTTTGACT | TTCGCATAACCCTCCTCATC | 269 |
| chr2:122730728-122732746 | p3 | (TTG)6 | ATGGAAGTTGGAAAACAGCG | TATGCCTAGCCCCAAATCAC | 102 |
| chr2:122730728-122732746 | p6 | (TATATG)3 | ATGGAAGTTGGAAAACAGCG | TATGCCTAGCCCCAAATCAC | 102 |
| chr2:122762877-122764919 | p2 | (TA)21 | CGATAAGGCGAGCAAACTTC | ATGTTGCTTGTCATCGGTCA | 102 |
| chr2:122828081-122830117 | p3 | (TTC)12 | GTCCACCATTGAAGGCTGAT | TCCTCTTGATGTTTCCCGTC | 215 |
| chr2:122841870-122843882 | p2 | (TG)7 | TGTCTAAGGGTTCGCTCGAC | CCCCTTCAAAGTTCAAACCA | 144 |
| chr2:122852806-122854842 | p3 | (GAA)11 | AAGTACCGTGGCATCCAAAG | ACGGATCATTATTCGGCTTG | 174 |
| chr2:122852806-122854842 | p4 | (AGTG)9 | AAGTACCGTGGCATCCAAAG | ACGGATCATTATTCGGCTTG | 174 |
| chr2:122852806-122854842 | p5 | (ATTTT)3 | AAGTACCGTGGCATCCAAAG | ACGGATCATTATTCGGCTTG | 174 |
| chr2:122861011-122863026 | p5 | (TTTTA)3 | CCAAAACACCACCAGACCTT | AATTTCAGCCGCCTCCTTAT | 137 |
| chr2:122896874-122898886 | p2 | (TA)33 | TAACACGTCGGAACCAGACA | GTTTCGCCACTGGTCATTCT | 132 |
| chr2:122896874-122898886 | p2 | (TC)6 | TAACACGTCGGAACCAGACA | GTTTCGCCACTGGTCATTCT | 132 |
| chr2:122924388-122926403 | p5 | (ATTTC)3 | AGAAGTGCCCATTGCCATAC | CGCTTCCCATGTCATTTCTT | 120 |
| chr2:122935137-122937173 | p2 | (AT)18 | CTGAGGTTTGTCCCACCACT | AAATTCCGCTCATGTTTTGC | 241 |
| chr2:122964872-122966887 | p5 | (ATTTC)3 | AGAAGTGCCCATTGCCATAC | CGCTTCCCATGTCATTTCTT | 120 |
| chr2:122975621-122977657 | p2 | (AT)18 | CTGAGGTTTGTCCCACCACT | AAATTCCGCTCATGTTTTGC | 241 |
| chr2:123007597-123009612 | p5 | (ATCAT)3 | CTGCAACTTTCGCATTTTCA | ACTCACGGCCAACACATACA | 224 |
| chr2:123213252-123215267 | p3 | (AAT)5 | CGGAAATAGTTGGAATGCGT | GTTGAAAAAGTCAACCCCGA | 170 |
| chr2:123230432-123232450 | p6 | (CATGTT)3 | AGAAAATGCCTCGAAGACGA | GAATCAACGTAGCTTGGGGA | 294 |
| chr2:123231896-123233911 | p3 | (TTG)5 | GCAGGTTAGGATTCACGCAT | CCACAAGGCCGAACATAACT | 108 |
| chr2:123302366-123304380 | p2 | (GA)7 | TCGTGCTAAACAAACCCACA | GCATTGGTTTGAATCCGAGT | 197 |
| chr2:123302964-123305042 | p2 | (GA)7 | CGATCAGAATACCCCGAAGA | ATGAGCTTTTGTCCGCTTGT | 138 |
| chr2:123310346-123312364 | p2 | (AT)9 | TTTCCAAAATGCCCTGTAGC | GATACCCGTCATGGCCTCTA | 289 |
| chr2:123324137-123326155 | p6 | (TCATAT)3 | CATTTCCGAACATTGTCACG | TGTTAGCCTCGTCGGCTTAT | 276 |
| chr2:123355589-123357604 | p5 | (AGAAA)3 | ATGCGGTTGGGTTGTGTATT | CTGTTTCCTCGGCATGATTT | 197 |
| chr2:123360005-123362020 | p2 | (TA)15 | AATCGGAGGTTTACCGACCT | CTATAGCGCCTTCCCTTTCC | 239 |
| chr2:123360005-123362020 | p5 | (GTCCT)3 | AATCGGAGGTTTACCGACCT | CTATAGCGCCTTCCCTTTCC | 239 |

| Marker ID | SSR type | SSR motif | FORWARD PRIMER (5'-3') | REVERSE PRIMER (5'-3') | size (bp) |
|---|---|---|---|---|---|
| chr2:123505533-123507545 | p2 | (AT)6 | AAACCATGATCTCAGCCTGC | GCGAGTTTGAGATGGAGGAG | 218 |
| chr2:123514709-123516724 | p5 | (TATTT)3 | CAACAGCGTCTTCTTGACCA | TGAGGAAGCAGTGGAGGAAT | 266 |
| chr2:123525028-123527044 | p4 | (TTTC)4 | AATGATGAAAAATCGGCTGG | AAGAGAGGCTTCAAAAGGGG | 103 |
| chr2:123525028-123527044 | p2 | (AT)13 | AATGATGAAAAATCGGCTGG | AAGAGAGGCTTCAAAAGGGG | 103 |
| chr2:123539382-123541416 | p2 | (CA)10 | AATTTTGGAAGCTTGGCCTT | GTCTGGGACAGGTATCCGAA | 243 |
| chr2:123539382-123541416 | p3 | (TTC)5 | AATTTTGGAAGCTTGGCCTT | GTCTGGGACAGGTATCCGAA | 243 |
| chr2:123551141-123553183 | p3 | (CAA)5 | ACCTACCGAGCACCTCTGAA | ATATTTTGGCAGCCTTGTCG | 272 |
| chr2:123551199-123553241 | p3 | (CAA)5 | ACCTACCGAGCACCTCTGAA | ATATTTTGGCAGCCTTGTCG | 214 |
| chr2:123582364-123584384 | p5 | (ATAAA)3 | TGTCGGAGAATGGAGAACAA | GGTTTGTTTGGTACGGAGGA | 106 |
| chr2:123582364-123584384 | p2 | (TA)10 | TGTCGGAGAATGGAGAACAA | GGTTTGTTTGGTACGGAGGA | 106 |
| chr2:123648868-123650888 | p2 | (AT)10 | TGGCCATTAGGGTTGTTCAT | GGAGCGCCAGCTAAAATATG | 164 |
| chr2:123648868-123650888 | p2 | (AT)10 | TGGCCATTAGGGTTGTTCAT | GGAGCGCCAGCTAAAATATG | 164 |
| chr2:123690217-123692233 | p2 | (AT)8 | AGGATCAAAGGGGAGGCTTA | TTCTCCAACCCCTGTTATCG | 170 |
| chr2:123710832-123712847 | p2 | (TG)11 | TGGGTACGTGAGACATCGAA | TATTCTTTTTCCACCTGCGG | 164 |
| chr2:123710832-123712847 | p3 | (TTC)5 | TGGGTACGTGAGACATCGAA | TATTCTTTTTCCACCTGCGG | 164 |
| chr2:123726332-123728350 | p2 | (TA)12 | TCCGCAACACGAACATACAT | CACTTGCTGACACCGCTAAA | 256 |
| chr2:123726332-123728350 | p2 | (AT)16 | TCCGCAACACGAACATACAT | CACTTGCTGACACCGCTAAA | 256 |
| chr2:123726332-123728350 | p2 | (AT)9 | TCCGCAACACGAACATACAT | CACTTGCTGACACCGCTAAA | 256 |
| chr2:123726332-123728350 | p2 | (AT)15 | TCCGCAACACGAACATACAT | CACTTGCTGACACCGCTAAA | 256 |
| chr2:123726467-123728497 | p2 | (AT)16 | TCCGCAACACGAACATACAT | CACTTGCTGACACCGCTAAA | 121 |
| chr2:123726467-123728497 | p2 | (AT)9 | TCCGCAACACGAACATACAT | CACTTGCTGACACCGCTAAA | 121 |
| chr2:123726467-123728497 | p2 | (AT)15 | TCCGCAACACGAACATACAT | CACTTGCTGACACCGCTAAA | 121 |
| chr2:123739404-123741428 | p6 | (AAGAAA)4 | ACTGCCAATGATCGAGGTTC | ACCCCAACTGTTTCCATCAA | 165 |
| chr2:123911626-123913644 | p4 | (TATG)4 | CATATCCATGTTCGAACCCC | CAAACGTGGAGCCAGAATTT | 154 |
| chr2:123911626-123913644 | p2 | (AT)9 | CATATCCATGTTCGAACCCC | CAAACGTGGAGCCAGAATTT | 154 |
| chr2:123913729-123915741 | p2 | (TC)6 | TGAAAAATGAGGACTTGGGG | AAATTTCACAGCTTCGACGG | 223 |
| chr2:123913729-123915741 | p3 | (AAC)5 | TGAAAAATGAGGACTTGGGG | AAATTTCACAGCTTCGACGG | 223 |
| chr2:123964874-123966908 | p2 | (AT)17 | CTGTAGGTCCCTGCTTGAGG | AGTCTTGTCGGCTTGCGTAT | 249 |
| chr2:124076610-124078628 | p3 | (CAA)6 | CTGACGAACCACAGCCACTA | CCGTCATCATCACTGTCACC | 104 |
| chr2:124076610-124078628 | p3 | (AAG)6 | CTGACGAACCACAGCCACTA | CCGTCATCATCACTGTCACC | 104 |
| chr2:124076610-124078628 | p4 | (AAAT)5 | CTGACGAACCACAGCCACTA | CCGTCATCATCACTGTCACC | 104 |
| chr2:124170519-124172539 | p4 | (TTTA)5 | TTCCTAAAACTTCTGCCCCA | CCTTTTTCTTCTTTTGCGGA | 232 |
| chr2:124296583-124298598 | p5 | (TTTCT)3 | CGCCGGGAAATCAATATCTA | GTATGGGGCCCACTCTTTTT | 106 |
| chr2:124296583-124298598 | p2 | (TA)17 | CGCCGGGAAATCAATATCTA | GTATGGGGCCCACTCTTTTT | 106 |
| chr2:124306616-124308636 | p5 | (CATTG)4 | AAGCATGTTACACCTTCGGG | TCCTTCCCCACAACATCAAT | 292 |
| chr2:124389518-124391538 | p4 | (ATTT)5 | ATCACTTTGCCCCTTACGTG | TGCTAAGTCCGAACCCAAAC | 197 |

| Marker ID | SSR type | SSR motif | FORWARD PRIMER (5'-3') | REVERSE PRIMER (5'-3') | size (bp) |
|---|---|---|---|---|---|
| chr2:124409000-124411015 | p3 | (AAG)5 | GAACCCAACTACCTGCCAAA | TAACATCCATGTCCGAAGCA | 242 |
| chr2:124511966-124513982 | p2 | (TA)8 | CAAGGCAGCCCAAATACAAT | TTGAACGGAGGAAGTCAAGG | 152 |
| chr2:124638134-124640154 | p2 | (AT)10 | ATATCATGCTCGGATTTCGG | AGATTGGCCATCTTGGAGTG | 163 |
| chr2:124678001-124680023 | p3 | (AAT)7 | CGAGAAATGGCGTTCAACTT | TGTAGCAAACTCTTTGGCCC | 218 |
| chr2:124678001-124680023 | p2 | (GA)6 | CGAGAAATGGCGTTCAACTT | TGTAGCAAACTCTTTGGCCC | 218 |
| chr2:124678001-124680023 | p2 | (AT)11 | CGAGAAATGGCGTTCAACTT | TGTAGCAAACTCTTTGGCCC | 218 |
| chr2:124678001-124680023 | p4 | (AATT)4 | CGAGAAATGGCGTTCAACTT | TGTAGCAAACTCTTTGGCCC | 218 |
| chr2:124742011-124744045 | p2 | (AT)17 | CACATCCCGACGAGGAGTAT | CTTTTTCTGCGTGCATGTGT | 121 |
| chr2:124807538-124809576 | p3 | (CAA)8 | CCGACTTTTCACACCCTGAT | TGTTGTAGCAGCTTTGTCGG | 131 |
| chr2:124807577-124809593 | p3 | (CAA)8 | CCGACAAAGCTGCTACAACA | AGCTCCACCGGTTCATATTG | 264 |
| chr2:124807605-124809621 | p3 | (CAA)8 | CCGACAAAGCTGCTACAACA | AGCTCCACCGGTTCATATTG | 236 |
| chr2:124807637-124809649 | p3 | (CAA)8 | CCGACAAAGCTGCTACAACA | AGCTCCACCGGTTCATATTG | 204 |
| chr2:124953948-124956000 | p2 | (TA)26 | ACACGGGAAGCAGAAGAAGA | CATCAATAACCATGGCCACA | 164 |
| chr2:125014106-125016118 | p2 | (GT)6 | GGCAATAACAATCGGGAAGA | AGCCTCACAAACATAACGGG | 120 |
| chr2:125077609-125079624 | p6 | (TATTGG)3 | AGACAAGCGTAAAAGGCGAA | CTGAGTGAAAAAGGGGCAAA | 226 |
| chr2:125086180-125088192 | p2 | (AT)6 | AGTATTTGGGGGCTCTTCGT | CACACACATGGAAACAAGCC | 274 |
| chr2:125139580-125141595 | p3 | (ACA)5 | ATGAAACAATTCCACCGAGC | TGCATAACAGGCAAATCCAA | 198 |
| chr2:125184228-125186240 | p2 | (AT)6 | TTCCGGAATCAACATACGGT | TCACGGTCCGTAAAATCCAT | 236 |
| chr2:125184228-125186240 | p2 | (TA)7 | TTCCGGAATCAACATACGGT | TCACGGTCCGTAAAATCCAT | 236 |
| chr2:125305668-125307680 | p2 | (AT)6 | TCGAGGCGTACCTGATCTTT | GCTGAAGAACGACATCGACA | 186 |
| chr2:125305668-125307680 | p2 | (AT)7 | TCGAGGCGTACCTGATCTTT | GCTGAAGAACGACATCGACA | 186 |
| chr2:125306089-125308103 | p2 | (AT)6 | ACTTTTGTCACGACCCAACC | CAGCCTTGTCAGCTTGCATA | 162 |
| chr2:125306089-125308103 | p2 | (AT)7 | ACTTTTGTCACGACCCAACC | CAGCCTTGTCAGCTTGCATA | 162 |
| chr2:125376676-125378694 | p6 | (TGATTC)3 | AACAAGAAATGGTGAACGCC | GAAGCTGTCAAGGAGATGCC | 106 |
| chr2:125421735-125423787 | p2 | (AT)26 | AGCAAAATTTCAATGGACCG | GGTATCTGTTGGCCGTGAGT | 219 |
| chr2:125421735-125423787 | p2 | (TA)10 | AGCAAAATTTCAATGGACCG | GGTATCTGTTGGCCGTGAGT | 219 |
| chr2:125485092-125487122 | p5 | (TTTTG)6 | CTGCTTCAACCCAGTCCATT | CCAAGGGAAACTCTTCCTCC | 216 |
| chr2:125566434-125568449 | p5 | (TGATA)3 | GGAGTTTGGGAATTGGGTTT | AGTGATGGACGTTTTACGGC | 132 |
| chr2:125674663-125676675 | p2 | (AC)6 | TCCATCAAGATCATGCCAAA | GGATGTTGCATTTACAGGGG | 256 |
| chr2:125692037-125694052 | p2 | (AT)8 | CGTCCAGCCAAAGCATAAAT | CCTAGTCGGGTCGTGACAAT | 143 |
| chr2:125692037-125694052 | p3 | (CAT)5 | CGTCCAGCCAAAGCATAAAT | CCTAGTCGGGTCGTGACAAT | 143 |
| chr2:125888740-125890776 | p2 | (AT)18 | GCTAAAACTTTACCGCGCAC | ATACGAGTGTCCAGTTCGGG | 105 |
| chr2:125908002-125910014 | p2 | (TA)6 | GCCTAACCCGAACTTCCAAT | CGTGTTGTCGTTGATATGGC | 287 |
| chr2:125944051-125946069 | p2 | (AT)9 | AACAAAGGGAGAGGGCAGAT | ATCAAAGAGGAGGGGTTGGT | 193 |
| chr2:125968869-125970887 | p2 | (AC)6 | GAATTTGGGGTCCATGGTTT | GGGTTTCATTCCTTGCTTGA | 269 |
| chr2:125968869-125970887 | p6 | (ATATAG)3 | GAATTTGGGGTCCATGGTTT | GGGTTTCATTCCTTGCTTGA | 269 |

| Marker ID | SSR type | SSR motif | FORWARD PRIMER (5'-3') | REVERSE PRIMER (5'-3') | size (bp) |
|---|---|---|---|---|---|
| chr2:126023572-126025584 | p2 | (TC)6 | CTTACGCTTCAGAAGGACGG | TTGCCTTAAATGCTCCAACC | 181 |
| chr2:126178243-126180263 | p2 | (AT)10 | TTTGAATTGCTCCTCCTTGG | CAGCCCGCATTTATGAAAGT | 244 |
| chr2:126232711-126234745 | p2 | (GA)6 | GGGTTAATCGAATCGTCGAA | TTTTCAACTTCAAATCCCGC | 141 |
| chr2:126288284-126290378 | p2 | (TA)13 | AGGCAAAGGTGTGGTTGTTC | TCCCAACTTCATACCCGAAG | 246 |
| chr2:126288284-126290378 | p2 | (TA)17 | AGGCAAAGGTGTGGTTGTTC | TCCCAACTTCATACCCGAAG | 246 |
| chr2:126354064-126356079 | p5 | (GCATT)3 | GACTGCTTTGGTTCTGAGGC | TCGTGGCATCAAGCTAACAC | 201 |
| chr2:126355504-126357519 | p5 | (AATAA)3 | GGCTGGCAAGAAACTCTGAC | CTACGGACTTGGCTTTACCG | 266 |
| chr2:126358592-126360616 | p6 | (CCGTGA)4 | TTGGGGTGATAAGCGAAAAC | ACCGTAGGAGTGGACGTTTG | 186 |
| chr2:126358592-126360616 | p3 | (GGT)5 | TTGGGGTGATAAGCGAAAAC | ACCGTAGGAGTGGACGTTTG | 186 |
| chr2:126552735-126554750 | p3 | (AGA)5 | CCGTTCGACGTACAACAATG | TTGCGCGAAGGACTCTCTAT | 117 |
| chr2:126710778-126712807 | p2 | (TA)15 | ATCGAGGATTACCTTGCCCT | ACTGTTGTTGCACACCCAAA | 170 |
| chr2:126727873-126729895 | p2 | (TA)6 | AAAAATGCACGAGTCCGAAG | GGAGGGGTATTTCGTTGGTT | 140 |
| chr2:126727873-126729895 | p2 | (CA)11 | AAAAATGCACGAGTCCGAAG | GGAGGGGTATTTCGTTGGTT | 140 |
| chr2:126727873-126729895 | p2 | (AT)12 | AAAAATGCACGAGTCCGAAG | GGAGGGGTATTTCGTTGGTT | 140 |
| chr2:126728995-126731015 | p2 | (AT)12 | ATCTATGCGGACAAACTGCC | TCGTGTGTCAAGAATCCGAA | 247 |
| chr2:126840524-126842548 | p4 | (TATT)6 | TCTGAACTGGAGGAACGCTT | AAGTTCCACAATCCATTGCC | 176 |
| chr2:126854963-126856978 | p5 | (TTGGT)3 | TCTGCGTACGCCACTGATAC | TCCAACACCAAACCAAGTCA | 108 |
| chr2:126854963-126856978 | p5 | (ACCAA)3 | TCTGCGTACGCCACTGATAC | TCCAACACCAAACCAAGTCA | 108 |
| chr2:126953699-126955714 | p5 | (AAAAT)3 | GGGTATTTGGTTGGATGTGG | AAACAAACACCGCCAAGTTC | 291 |
| chr2:127064023-127066059 | p2 | (AT)11 | GGAAGTATCGCTTGGGTTCA | GCCACGCCTCTCTAGATTTG | 170 |
| chr2:127064023-127066059 | p2 | (AT)18 | GGAAGTATCGCTTGGGTTCA | GCCACGCCTCTCTAGATTTG | 170 |
| chr2:127369715-127371730 | p3 | (ATC)5 | CCTAGTCGGGTCGTGACACT | CGGAGTTGCAGGTTATTGGT | 156 |
| chr2:127512847-127514863 | p2 | (TG)8 | CATAACGGTTGGAGATGGCT | ACTCCCCAATTTTACCCCAC | 136 |
| chr2:127556098-127558122 | p2 | (CA)6 | GGCCGCGTTTGATTTAGTAG | ATTGGCCCACAGAAGAAGTG | 164 |
| chr2:127558672-127560734 | p2 | (TA)31 | CAACCAAAGACACAACACGG | CTCAAATTGGGACGGAGAGA | 207 |
| chr2:127676432-127678458 | p2 | (CT)8 | AAACTTCTGACGGCCATTTG | TTTGTACTCCAGGGGCAAAG | 225 |
| chr2:127676432-127678458 | p2 | (TC)6 | AAACTTCTGACGGCCATTTG | TTTGTACTCCAGGGGCAAAG | 225 |
| chr2:127731646-127733664 | p3 | (GAA)6 | AACTTTGCGGGGTTCCTACT | TTTGGGTTCCTCCCTCTCTT | 187 |
| chr2:127731646-127733664 | p6 | (ATAAAA)3 | AACTTTGCGGGGTTCCTACT | TTTGGGTTCCTCCCTCTCTT | 187 |
| chr2:127750758-127752772 | p2 | (TG)7 | AGCCCACATTTGAGATACCG | GGTGTTGCCCAAAATGAATC | 162 |
| chr2:127767098-127769113 | p5 | (TTATT)3 | GGAGCCTTTAGAGTTCGGCT | TTCATTCCTCTGCGTCTGTG | 282 |
| chr2:127767098-127769113 | p5 | (TTATT)3 | GGAGCCTTTAGAGTTCGGCT | TTCATTCCTCTGCGTCTGTG | 282 |
| chr2:127767098-127769113 | p5 | (TTATT)3 | GGAGCCTTTAGAGTTCGGCT | TTCATTCCTCTGCGTCTGTG | 282 |
| chr2:127767098-127769113 | p5 | (TTATT)3 | GGAGCCTTTAGAGTTCGGCT | TTCATTCCTCTGCGTCTGTG | 282 |
| chr2:127835231-127837243 | p2 | (AC)6 | TTGTCTGATCCGTTCCCTTC | TGGTTCCCCAGATCTTCATC | 297 |
| chr2:127893065-127895079 | p2 | (TG)7 | AGCCCACATTTGAGATACCG | GGTGTTGCCCAAAATGAATC | 162 |

| Marker ID | SSR type | SSR motif | FORWARD PRIMER (5'-3') | REVERSE PRIMER (5'-3') | size (bp) |
|---|---|---|---|---|---|
| chr2:127909405-127911420 | p5 | (TTATT)3 | GGAGCCTTTAGAGTTCGGCT | TTCATTCCTCTGCGTCTGTG | 282 |
| chr2:127909405-127911420 | p5 | (TTATT)3 | GGAGCCTTTAGAGTTCGGCT | TTCATTCCTCTGCGTCTGTG | 282 |
| chr2:127909405-127911420 | p5 | (TTATT)3 | GGAGCCTTTAGAGTTCGGCT | TTCATTCCTCTGCGTCTGTG | 282 |
| chr2:127909405-127911420 | p5 | (TTATT)3 | GGAGCCTTTAGAGTTCGGCT | TTCATTCCTCTGCGTCTGTG | 282 |
| chr2:127953968-127955983 | p3 | (TAT)5 | TCGTCATACCTGATCGTGGA | TCCTCTTTTCATCTCCCCCT | 252 |
| chr2:127953968-127955983 | p3 | (TTA)6 | TCGTCATACCTGATCGTGGA | TCCTCTTTTCATCTCCCCCT | 252 |
| chr2:128000959-128003004 | p2 | (AT)9 | CAGAGGGGTTAGTGTGGGTG | AAGCAAGCAGCTACGTGACA | 125 |
| chr2:128182499-128184511 | p2 | (AC)6 | TCGAGAACGTTAGCCATCCT | GCGAGCGAGGTATGTAAAGC | 297 |
| chr2:128223569-128225584 | p5 | (ATCAT)3 | TAGCGTAGGGACAAACTGGG | ACTTACAGTGGTGATCCGCC | 208 |
| chr2:128232027-128234039 | p2 | (TA)6 | GTTGTTTGGCCGTGTGTATG | ATTTTGACGCTCACAATCCC | 192 |
| chr2:128247225-128249239 | p2 | (TA)7 | TCGCTTCACACCCATAAACA | TTTATGGTCCACCCAGGAAA | 213 |
| chr2:128247225-128249239 | p3 | (ATC)7 | TCGCTTCACACCCATAAACA | TTTATGGTCCACCCAGGAAA | 213 |
| chr2:128247560-128249581 | p2 | (TA)7 | CATCATCACCAGCCAACAAG | CGGAGCTTCAGTCTATTGGG | 140 |
| chr2:128247560-128249581 | p3 | (ATC)7 | CATCATCACCAGCCAACAAG | CGGAGCTTCAGTCTATTGGG | 140 |
| chr2:128293692-128295713 | p3 | (GTT)7 | AAGCAGTGTGGTTGAGGGTC | CGGAGGTAGTCAGTGTGCAA | 122 |
| chr2:128293692-128295713 | p6 | (CTTGCT)4 | AAGCAGTGTGGTTGAGGGTC | CGGAGGTAGTCAGTGTGCAA | 122 |
| chr2:128305249-128307264 | p5 | (ATATA)3 | GGCTGCCATTACAGAAGAGC | TCTTGGAGAGCTCCGTTGTT | 143 |
| chr2:128305249-128307264 | p3 | (ACA)7 | GGCTGCCATTACAGAAGAGC | TCTTGGAGAGCTCCGTTGTT | 143 |
| chr2:128467061-128469077 | p2 | (TA)8 | GGGGAAAAGGATAGGCGTTA | AGTGCCGAGGAACATACGAC | 293 |
| chr2:128502095-128504164 | p3 | (TTA)23 | GGATGCAACTTCCGAAACAT | GTTCGCTGCCGAAGAAATAG | 282 |
| chr2:128504809-128506821 | p2 | (TA)6 | TCATCAGGCCAGCCTAAGTT | TCTCGGATGAAGCAACAGTG | 234 |
| chr2:128513789-128515804 | p5 | (TGTAT)3 | GAAAAATGGCACGGATGAGT | TATGCGTCATAGCCACCAAC | 111 |
| chr2:128537993-128540005 | p2 | (AG)6 | ATCTTGCCTTGTGGGAAATG | TCGAAATTTCAATTTTGCCC | 245 |
| chr2:128537993-128540005 | p2 | (GT)6 | ATCTTGCCTTGTGGGAAATG | TCGAAATTTCAATTTTGCCC | 245 |
| chr2:128572493-128574547 | p2 | (AT)27 | GAGGGTTCACAACGTCCACT | AGGTGGGACTCTCATCATGC | 203 |
| chr2:128572493-128574547 | p2 | (AT)7 | GAGGGTTCACAACGTCCACT | AGGTGGGACTCTCATCATGC | 203 |
| chr2:128612001-128614027 | p2 | (TA)8 | AGGCAAACCGTAAGGTGACA | TCCTCACTGTTGACCCACAA | 159 |
| chr2:128612029-128614047 | p2 | (TA)8 | AGGCAAACCGTAAGGTGACA | TCCTCACTGTTGACCCACAA | 131 |
| chr2:128612058-128614074 | p2 | (TA)8 | AGGCAAACCGTAAGGTGACA | TCCTCACTGTTGACCCACAA | 102 |
| chr2:128626524-128628596 | p3 | (AAT)24 | CCATTCGCAGCCTTGATATT | GCGAAGTTTGTGGAAAGAGC | 130 |
| chr2:128776122-128778140 | p3 | (TTG)6 | TTTCGATTCGCAATTGTCTG | CGGCCACTTCTACAACAACA | 139 |
| chr2:128776122-128778140 | p6 | (AATTCA)3 | TTTCGATTCGCAATTGTCTG | CGGCCACTTCTACAACAACA | 139 |
| chr2:128776122-128778140 | p2 | (AT)11 | TTTCGATTCGCAATTGTCTG | CGGCCACTTCTACAACAACA | 139 |
| chr2:128821426-128823438 | p2 | (AT)6 | TTCTTGCCACACTCAGCATC | TCGGCAAGCCCTCTATTAAA | 237 |
| chr2:128821538-128823550 | p2 | (AT)6 | TTCTTGCCACACTCAGCATC | TCGGCAAGCCCTCTATTAAA | 125 |
| chr2:128821701-128823725 | p2 | (AT)6 | TTTAATAGAGGGCTTGCCGA | CGTCTCAAATCCGAACACCT | 193 |

| Marker ID | SSR type | SSR motif | FORWARD PRIMER (5'-3') | REVERSE PRIMER (5'-3') | size (bp) |
|---|---|---|---|---|---|
| chr2:128821771-128823783 | p2 | (AT)6 | TTTAATAGAGGGCTTGCCGA | CGTCTCAAATCCGAACACCT | 123 |
| chr2:128821771-128823783 | p2 | (AT)15 | TTTAATAGAGGGCTTGCCGA | CGTCTCAAATCCGAACACCT | 123 |
| chr2:128821812-128823834 | p2 | (AT)6 | AAACTCCAGAGGTGTTCGGA | TGAAGGATCTGCTTGACGTG | 194 |
| chr2:128821812-128823834 | p2 | (AT)15 | AAACTCCAGAGGTGTTCGGA | TGAAGGATCTGCTTGACGTG | 194 |
| chr2:128924933-128926949 | p2 | (AC)13 | ACCGTTGGTTTTATTGCTCG | TGATTTTCAGCCAAACCACA | 212 |
| chr2:128924995-128927009 | p2 | (AC)13 | ACCGTTGGTTTTATTGCTCG | TGATTTTCAGCCAAACCACA | 150 |
| chr2:129030477-129032495 | p3 | (TTA)6 | TGTGTTACCGGCATTTTTGA | CTGATAGACAACTGCGCCAA | 156 |
| chr2:129032907-129034922 | p5 | (TTTCA)3 | TGACCGCTTAGCATCCTCTT | GGATAACGAAGCCTGTCCAA | 159 |
| chr2:129100999-129103013 | p2 | (GA)7 | TGCCATGTGTGTTGAGGTTT | TGGATCAATTAAGAACGGGG | 279 |
| chr2:129110754-129112769 | p5 | (CATAT)3 | ATATCGGCTGATGAAGCCAC | GTTGGCAAGCTCCATTTGTT | 170 |
| chr2:129201054-129203100 | p2 | (TC)23 | CAAAAGGGATTTTCAACCGA | CATTTCCAGTTCAGGACCGT | 249 |
| chr2:129206213-129208279 | p4 | (ATTA)7 | ACCAGGTTTTGGAGGGAAAT | GCTTCGGATTTGTCAACGAT | 232 |
| chr2:129206213-129208279 | p2 | (TA)22 | ACCAGGTTTTGGAGGGAAAT | GCTTCGGATTTGTCAACGAT | 232 |
| chr2:129206289-129208337 | p4 | (ATTA)7 | ACCAGGTTTTGGAGGGAAAT | GCTTCGGATTTGTCAACGAT | 156 |
| chr2:129206289-129208337 | p2 | (TA)22 | ACCAGGTTTTGGAGGGAAAT | GCTTCGGATTTGTCAACGAT | 156 |
| chr2:129206289-129208337 | p5 | (TATTT)3 | ACCAGGTTTTGGAGGGAAAT | GCTTCGGATTTGTCAACGAT | 156 |
| chr2:129233823-129235850 | p3 | (GTA)9 | GTGGGTGAAGGTAGGGGTTT | TTGCAAAATTCTGCCTTGTG | 269 |
| chr2:129271632-129273676 | p2 | (AT)22 | CGTTACAACGGAAGGACACA | GCAGTAGGCATGCATGATTTT | 149 |
| chr2:129277885-129280229 | p2 | (AT)7 | ATCCATGGGGTTACAGTCCA | TTTTGTGAAACTTGCAGCCA | 269 |
| chr2:129277885-129280229 | p2 | (AT)6 | ATCCATGGGGTTACAGTCCA | TTTTGTGAAACTTGCAGCCA | 269 |
| chr2:129299396-129301454 | p2 | (AT)29 | ACCCGCAACACTTCTACACC | CGTGGGAATAAGAGAACCGA | 186 |
| chr2:129299396-129301454 | p3 | (TTG)5 | ACCCGCAACACTTCTACACC | CGTGGGAATAAGAGAACCGA | 186 |
| chr2:129380866-129382918 | p2 | (AT)26 | GGTTCACCTCTGCTCGCTAC | CGACTCATCGGTTTGTGTTG | 294 |
| chr2:129423977-129425995 | p2 | (TA)16 | ATTGGAAATTCGGGTTAGGC | CGTCGTAATCTCGTTCGCTT | 232 |
| chr2:129424001-129426093 | p2 | (TA)16 | ATTGGAAATTCGGGTTAGGC | CGTCGTAATCTCGTTCGCTT | 208 |
| chr2:129459683-129461721 | p2 | (AT)19 | ATATAGGCGGGACAGGAACC | CGTACGGACGTCCTTTTGTT | 258 |
| chr2:129516519-129518535 | p2 | (AC)8 | AGGGGTAGAGGTCCTGCCTA | TTTCGCAAGTCCAGGTAAGG | 107 |
| chr2:129523480-129525495 | p2 | (GT)6 | TCCTACCAATTTGAGTGCCC | CCCACTTTGCAAACTTCCAT | 158 |
| chr2:129523480-129525495 | p3 | (TTG)5 | TCCTACCAATTTGAGTGCCC | CCCACTTTGCAAACTTCCAT | 158 |
| chr2:129528331-129530346 | p3 | (TTG)5 | ATGGAAGTTTGCAAAGTGGG | CACAACTCATACCACACGGC | 279 |
| chr2:129534386-129536402 | p4 | (TTAT)4 | CCTCCGCATGGTGAACTACT | TTTTGAGCCAGATTCATCCC | 232 |
| chr2:129547998-129550064 | p2 | (TA)33 | TCCTATCGGGATGAATCTGG | TGGCAACAACTCCTAGCACA | 194 |
| chr2:129567147-129569159 | p2 | (TA)6 | CGATTGAGGATCCGTTTGTT | AGCGTATCTGTTACACCCCG | 164 |
| chr2:129583749-129585761 | p2 | (AT)6 | TCACCTGGTTCACCAAAACA | AGGAATAGTGGCCGGAGAAT | 282 |
| chr2:129583749-129585761 | p2 | (TA)27 | TCACCTGGTTCACCAAAACA | AGGAATAGTGGCCGGAGAAT | 282 |
| chr2:129585814-129587830 | p2 | (GA)7 | ATTGCAGAATCTTTGGTGGG | AGCATATGCATGAGGAAGGG | 193 |

| Marker ID | SSR type | SSR motif | FORWARD PRIMER (5'-3') | REVERSE PRIMER (5'-3') | size (bp) |
|---|---|---|---|---|---|
| chr2:129585814-129587830 | p2 | (AT)8 | ATTGCAGAATCTTTGGTGGG | AGCATATGCATGAGGAAGGG | 193 |
| chr2:129591463-129593479 | p4 | (CATA)4 | ACAAGGCCCCTGATATACCC | ACTGACGTCCCTTCTTGTGG | 264 |
| chr2:129604193-129606215 | p2 | (AT)11 | AGTTTCAAGTGACGATGGGG | AAAAGGAGTTGGGGCATTTT | 107 |
| chr2:129650713-129652725 | p2 | (AT)6 | CGTGAAATCTATTGGGCGTT | CTTGGTCCCAAAGTTTTCCA | 235 |
| chr2:129650713-129652725 | p2 | (TA)11 | CGTGAAATCTATTGGGCGTT | CTTGGTCCCAAAGTTTTCCA | 235 |
| chr2:129676811-129678827 | p2 | (AT)8 | TACGGATCGTATTGGCTTCC | CCAATCCAAGATCACAGCCT | 141 |
| chr2:129767247-129769262 | p5 | (GTTCG)3 | GGGAAATTGTCGGGTATGTG | GGGTTAAATGGCCCAAAGAT | 151 |
| chr2:129767247-129769262 | p2 | (AT)15 | GGGAAATTGTCGGGTATGTG | GGGTTAAATGGCCCAAAGAT | 151 |
| chr2:129824652-129826670 | p6 | (CTGCTT)3 | TTGCAGAAATGCAAGGTCAG | AAAGGGCGTGTTTGATATGG | 188 |
| chr2:129889485-129891539 | p2 | (TA)29 | TACCACAGCCATTCCAGATG | TCATTCTTCTTGTTCCGCCT | 131 |
| chr2:129889485-129891539 | p2 | (AT)27 | TACCACAGCCATTCCAGATG | TCATTCTTCTTGTTCCGCCT | 131 |
| chr2:129889485-129891539 | p4 | (TTAT)4 | TACCACAGCCATTCCAGATG | TCATTCTTCTTGTTCCGCCT | 131 |
| chr2:129943544-129945559 | p5 | (ATATT)3 | CGGAGGCAGGATTTTTGTTA | ATCGTGTCCTTTCAACCAGG | 157 |
| chr2:129977173-129979187 | p2 | (CA)7 | CGAAGAACAGAATTGCTCCC | CTGAGCTAAGTCTCGGGGTG | 200 |
| chr2:130068542-130070558 | p4 | (GAGC)4 | AGGCGTGATTCATCCACTTC | ACGATCAAACGGACTAACGG | 179 |
| chr2:130123176-130125224 | p6 | (TAAAAA)4 | AAATTTCTTCGCCCCTGTTT | GTGGGATGGGCTCTAAGACA | 145 |
| chr2:130123176-130125224 | p3 | (TTA)16 | AAATTTCTTCGCCCCTGTTT | GTGGGATGGGCTCTAAGACA | 145 |
| chr2:130285884-130287896 | p2 | (TA)6 | TCGCCACATATCGTTCACAT | TGGAGGACACAGATCCACAA | 195 |
| chr2:130374513-130376529 | p5 | (AAATA)3 | CCCAGCTATGCCCTCAAGTA | GTACCTGCTACCTCCCACCA | 257 |
| chr2:130374513-130376529 | p2 | (TA)8 | CCCAGCTATGCCCTCAAGTA | GTACCTGCTACCTCCCACCA | 257 |
| chr2:130442477-130444492 | p3 | (TGG)6 | AACACGCAGTGAAGCAACTG | GGATGTTTTTGAGCGGTTGT | 126 |
| chr2:130442477-130444492 | p3 | (ATG)5 | AACACGCAGTGAAGCAACTG | GGATGTTTTTGAGCGGTTGT | 126 |
| chr2:130572620-130574632 | p2 | (CT)6 | TTTTGACACATACCAGGCCA | GTTGCTTCTTGGCTTTCCAG | 194 |
| chr2:130572620-130574632 | p2 | (TG)6 | TTTTGACACATACCAGGCCA | GTTGCTTCTTGGCTTTCCAG | 194 |
| chr2:130572828-130574840 | p2 | (CT)6 | TCTGTCCCTCACTGCCTTCT | AGTGAGGTCAGCAACAGCCT | 143 |
| chr2:130572828-130574840 | p2 | (TG)6 | TCTGTCCCTCACTGCCTTCT | AGTGAGGTCAGCAACAGCCT | 143 |
| chr2:130608799-130610815 | p2 | (TG)8 | TGGTTTCAGACGAGGTAGGG | TTCCTCCAAAGCCCTCCTAT | 152 |
| chr2:130632832-130634847 | p2 | (AT)6 | TTTCGGGACTCGGTGTTATC | GCGTTATAGACGCACAACCA | 115 |
| chr2:130632832-130634847 | p3 | (CAA)5 | TTTCGGGACTCGGTGTTATC | GCGTTATAGACGCACAACCA | 115 |
| chr2:130634507-130636529 | p2 | (AT)11 | CGTTCCTGGTTGGTGCTATT | AAAGAGCGTCACCTGCTCAT | 162 |
| chr2:130634507-130636529 | p2 | (TA)10 | CGTTCCTGGTTGGTGCTATT | AAAGAGCGTCACCTGCTCAT | 162 |
| chr2:130673262-130675274 | p2 | (AT)6 | ATTTGCCGGGTGTTCATTAG | ATGTAAATTGGGGCAGCAGT | 188 |
| chr2:130686580-130688620 | p2 | (AT)20 | TCCGTGACAACATCGTCAAT | AAAGAAAGCAGCCTTTTCCC | 113 |
| chr2:130742589-130744609 | p2 | (AT)10 | AGTCCATTTGAAGGGTGTGC | AACCTCGGAGTATTTCCGGT | 239 |
| chr2:130761213-130763228 | p2 | (TA)8 | GAACCTGTTTGGCATGGACT | TCTTGCGCAACTAAGAGCCT | 101 |
| chr2:130761213-130763228 | p3 | (TTG)5 | GAACCTGTTTGGCATGGACT | TCTTGCGCAACTAAGAGCCT | 101 |

| Marker ID | SSR type | SSR motif | FORWARD PRIMER (5'-3') | REVERSE PRIMER (5'-3') | size (bp) |
|---|---|---|---|---|---|
| chr2:130921002-130923092 | p3 | (TAA)30 | AGCTAGCGTTTGGCCATAGA | TGCTCTGCCGACTGTAGAAA | 151 |
| chr2:130964880-130966900 | p2 | (TA)10 | CACCATCTTGCTTGCATTTG | TTTCGTTGGGTTTTAGCCAC | 141 |
| chr2:131060935-131062953 | p3 | (TCT)6 | TACTCGCGCACTGAAATCAC | AGTTGAGTCATTGGTTCGGG | 269 |
| chr2:131172373-131174417 | p2 | (AT)22 | ATAAAATGTAAGGTGCGCCG | TTGTGGCAGGACAAGTTCAG | 266 |
| chr2:131198936-131200951 | p5 | (GATTT)3 | AGACAATTTCCCAGTCACCG | TCAACTTTTCGGTTGCAAGA | 167 |
| chr2:131319112-131321132 | p5 | (ATTGC)4 | CTTTGATGAGCTAGGCTCGG | ACATCTCGGTAGCACAACCC | 186 |
| chr2:131319112-131321132 | p2 | (TA)10 | CTTTGATGAGCTAGGCTCGG | ACATCTCGGTAGCACAACCC | 186 |
| chr2:131374371-131376386 | p5 | (TTCAT)3 | GGATGACCACCTGCCTCTTA | GACCAAATCTGGCCCTAACA | 198 |
| chr2:131420989-131423004 | p5 | (CGGGT)3 | TTAGGGGATTTTTGCCACTG | CCTCTCCCCCTTTCTTCAAC | 226 |
| chr2:131421006-131423024 | p5 | (CGGGT)3 | TTAGGGGATTTTTGCCACTG | CCTCTCCCCCTTTCTTCAAC | 209 |
| chr2:131421033-131423051 | p5 | (CGGGT)3 | TTAGGGGATTTTTGCCACTG | CCTCTCCCCCTTTCTTCAAC | 182 |
| chr2:131456139-131458155 | p4 | (TTTA)4 | AGTTCCATCCACCAGTTTCG | CTTGAGGCATTGGTTCAGGT | 120 |
| chr2:131524742-131526778 | p2 | (TA)18 | GCAAATGCCAGTCAAGGATT | ACATACCCCACGATCTCAGC | 204 |
| chr2:131524742-131526778 | p2 | (TA)6 | GCAAATGCCAGTCAAGGATT | ACATACCCCACGATCTCAGC | 204 |
| chr2:131525686-131527698 | p2 | (TA)18 | GCCCACACGTCAAGATTTTT | ACTTTGATTTGGACCAAGCG | 186 |
| chr2:131525686-131527698 | p2 | (TA)6 | GCCCACACGTCAAGATTTTT | ACTTTGATTTGGACCAAGCG | 186 |
| chr2:131525686-131527698 | p3 | (TTG)5 | GCCCACACGTCAAGATTTTT | ACTTTGATTTGGACCAAGCG | 186 |
| chr2:131579166-131581182 | p2 | (AT)8 | ATCTCATCACCAGGAGTGGG | GAGAGAGATCTCCATCCCCC | 221 |
| chr2:131583463-131585483 | p2 | (AT)10 | AATAGTGTCACGACCCAGCC | TGTGTAGTGGCAGCCTTGTC | 108 |
| chr2:131599485-131601509 | p4 | (TTAT)6 | GGAACGATTGAGTGTCCGAT | CATGTCCCGGATGGAATAAC | 259 |
| chr2:131616151-131618175 | p6 | (CCATCT)3 | CACCACCTCCACCAAAGTCT | GGTGATGGCGATTTATTGCT | 175 |
| chr2:131616151-131618175 | p6 | (CCATCG)4 | CACCACCTCCACCAAAGTCT | GGTGATGGCGATTTATTGCT | 175 |
| chr2:131651762-131653777 | p3 | (ATA)19 | TCACTGCGCCTTTGTGTATC | GATCTTTAACGGCTTGCTGC | 229 |
| chr2:131651762-131653777 | p3 | (TGT)5 | TCACTGCGCCTTTGTGTATC | GATCTTTAACGGCTTGCTGC | 229 |
| chr2:131652056-131654080 | p3 | (TGT)5 | AGGGGTACTCTTTTCGCCAT | ACTGAATGTCCATCGCAACA | 234 |
| chr2:131652088-131654124 | p3 | (TGT)5 | AGGGGTACTCTTTTCGCCAT | ACTGAATGTCCATCGCAACA | 202 |
| chr2:131668335-131670350 | p3 | (ATA)19 | TCACTGCGCCTTTGTGTATC | GATCTTTAACGGCTTGCTGC | 229 |
| chr2:131668335-131670350 | p3 | (TGT)5 | TCACTGCGCCTTTGTGTATC | GATCTTTAACGGCTTGCTGC | 229 |
| chr2:131668629-131670653 | p3 | (TGT)5 | AGGGGTACTCTTTTCGCCAT | ACTGAATGTCCATCGCAACA | 234 |
| chr2:131668661-131670697 | p3 | (TGT)5 | AGGGGTACTCTTTTCGCCAT | ACTGAATGTCCATCGCAACA | 202 |
| chr2:131674142-131676156 | p2 | (AT)7 | TGCTACACCAGTTGCTCCAG | AAGACGTCCACCTCATCCAG | 102 |
| chr2:131674142-131676156 | p2 | (TA)6 | TGCTACACCAGTTGCTCCAG | AAGACGTCCACCTCATCCAG | 102 |
| chr2:131674142-131676156 | p5 | (ATATA)3 | TGCTACACCAGTTGCTCCAG | AAGACGTCCACCTCATCCAG | 102 |
| chr2:131674290-131676302 | p2 | (AT)7 | CATTATCCGGCTTAGCTTGC | TTCGTTAATTTCCCCCACAA | 272 |
| chr2:131674290-131676302 | p2 | (TA)6 | CATTATCCGGCTTAGCTTGC | TTCGTTAATTTCCCCCACAA | 272 |
| chr2:131674290-131676302 | p5 | (ATATA)3 | CATTATCCGGCTTAGCTTGC | TTCGTTAATTTCCCCCACAA | 272 |

| Marker ID | SSR type | SSR motif | FORWARD PRIMER (5'-3') | REVERSE PRIMER (5'-3') | size (bp) |
|---|---|---|---|---|---|
| chr2:131800900-131802924 | p3 | (GTT)8 | ACCATAGGTCGTGAACCTGC | TCTTCATGAGCTTCGGGATT | 219 |
| chr2:131800900-131802924 | p2 | (TA)10 | ACCATAGGTCGTGAACCTGC | TCTTCATGAGCTTCGGGATT | 219 |
| chr2:131816294-131818312 | p6 | (ACATGA)3 | TTGAAGCCTCGTGACAACTG | CAACCAGATTTGTCCTCGGT | 100 |
| chr2:131842528-131844554 | p2 | (AT)13 | GGGTCCCATTTAGGCAAGTT | TTTTGACGAGATTGTGACGC | 214 |
| chr2:131941129-131943144 | p5 | (AAAAT)3 | ATCCAGTGAATCCCTATGCG | TTCTTGTGCAGTTGCAGACC | 153 |
| chr2:131941129-131943144 | p2 | (TG)7 | ATCCAGTGAATCCCTATGCG | TTCTTGTGCAGTTGCAGACC | 153 |
| chr2:132051229-132053263 | p5 | (TTTGT)5 | GTATGAAGGGGAGTGGGGTT | ATAATATCCGGTCGTTCCCC | 231 |
| chr2:132051229-132053263 | p2 | (TA)17 | GTATGAAGGGGAGTGGGGTT | ATAATATCCGGTCGTTCCCC | 231 |
| chr2:132085363-132087384 | p3 | (TAA)7 | AGTGGTCATTCGGAAAAACG | TTGAATGATCGCTTGTCTGC | 281 |
| chr2:132085363-132087384 | p2 | (TA)6 | AGTGGTCATTCGGAAAAACG | TTGAATGATCGCTTGTCTGC | 281 |
| chr2:132124689-132126713 | p2 | (TA)12 | TGTGGTCCTCTAACCCTTGG | GTTCCACGGACCATGAAAGT | 172 |
| chr2:132271141-132273159 | p3 | (ACA)6 | GAATCAGCGGATATGGAGGA | AGGGATGAACCCTCGACTCT | 228 |
| chr2:132271141-132273159 | p2 | (GA)7 | GAATCAGCGGATATGGAGGA | AGGGATGAACCCTCGACTCT | 228 |
| chr2:132271141-132273159 | p5 | (CAACA)3 | GAATCAGCGGATATGGAGGA | AGGGATGAACCCTCGACTCT | 228 |
| chr2:132538091-132540125 | p2 | (AT)17 | CGACCTGTTTTGGACCTGTT | GAGGGACATCATTGGCTTGT | 199 |
| chr2:132567178-132569193 | p5 | (AAAAT)3 | GAACCGCTTGTTGGACACTT | CGCGTCACTTTTGCTGATAA | 296 |
| chr2:132590789-132592805 | p4 | (GAAA)4 | CTCAGGTCAGCACTCCACAA | CCTGGTTGGGAGATTCAAGA | 260 |
| chr2:132591710-132593725 | p4 | (GAAA)4 | ATTTTGCCTAAGTCATGCGG | TACCCTGGGGGTGTTTATCA | 233 |
| chr2:132591737-132593752 | p4 | (GAAA)4 | ATTTTGCCTAAGTCATGCGG | TACCCTGGGGGTGTTTATCA | 206 |
| chr2:132676673-132678701 | p2 | (TA)14 | GGGCATCTAGGGATTGTTGA | CAGAATCTGTACGCAGCCAA | 105 |
| chr2:132679378-132681396 | p2 | (AG)7 | ACGTTTTCGGGTGATCTGAC | GACCCATGACCTATGGTTGG | 208 |
| chr2:132679378-132681396 | p3 | (TTG)6 | ACGTTTTCGGGTGATCTGAC | GACCCATGACCTATGGTTGG | 208 |
| chr2:132756595-132758621 | p2 | (AG)35 | GCCAGTCGAACTACGTGTGA | TTGGAGGAAACAATTCGGAG | 212 |
| chr2:132756595-132758621 | p2 | (TA)13 | GCCAGTCGAACTACGTGTGA | TTGGAGGAAACAATTCGGAG | 212 |
| chr2:132756595-132758621 | p2 | (TG)13 | GCCAGTCGAACTACGTGTGA | TTGGAGGAAACAATTCGGAG | 212 |
| chr2:132820789-132822805 | p4 | (GAAA)4 | CTCAGGTCTGCACTCCACAA | CATGTGAGGATGCCAATGAG | 260 |
| chr2:132820789-132822805 | p2 | (AT)7 | CTCAGGTCTGCACTCCACAA | CATGTGAGGATGCCAATGAG | 260 |
| chr2:132858774-132860798 | p4 | (GTAT)6 | AAACGTTGACCATCTGGACC | AGTGGGCTATCGGAGAGGAT | 275 |
| chr2:132858774-132860798 | p6 | (CCACCC)3 | AAACGTTGACCATCTGGACC | AGTGGGCTATCGGAGAGGAT | 275 |
| chr2:132957793-132959808 | p6 | (TTCGGT)3 | TATAAGCCAATCCAAACGGG | TGTTCGGTGTCCACTTTCAA | 135 |
| chr2:132957793-132959808 | p6 | (TAGAAC)4 | TATAAGCCAATCCAAACGGG | TGTTCGGTGTCCACTTTCAA | 135 |
| chr2:132996238-132998253 | p5 | (AATAA)3 | AGGTCCAGAGTTGGGATGTG | TATCCCAGGCTATCTGGACG | 152 |
| chr2:132996238-132998253 | p3 | (TGG)5 | AGGTCCAGAGTTGGGATGTG | TATCCCAGGCTATCTGGACG | 152 |
| chr2:133048989-133051004 | p5 | (TTTTA)3 | CACCCACTATCCCTCACCAC | TTTTTACACCACCACCCCAT | 266 |
| chr2:133062584-133064599 | p5 | (AATGC)3 | TGAACAATCTCGCACGAAAG | ACCCATCCTCGAGAGGCTAT | 210 |
| chr2:133133380-133135404 | p6 | (TCTACC)4 | AAAATGGGAACACAGCCTTG | GTCGAATTTTCATCGTGCCT | 271 |

| Marker ID | SSR type | SSR motif | FORWARD PRIMER (5'-3') | REVERSE PRIMER (5'-3') | size (bp) |
|---|---|---|---|---|---|
| chr2:133147369-133149381 | p2 | (TA)6 | CCAGTAGACGTGGACGGAAT | GACGGATTTCTTCCCCACTT | 243 |
| chr2:133147369-133149381 | p2 | (TA)6 | CCAGTAGACGTGGACGGAAT | GACGGATTTCTTCCCCACTT | 243 |
| chr2:133147936-133149954 | p2 | (TA)6 | CCGGTAACCGTTTAGGAACC | TCCGTATTGCTGACTTGCTG | 111 |
| chr2:133147936-133149954 | p2 | (TA)6 | CCGGTAACCGTTTAGGAACC | TCCGTATTGCTGACTTGCTG | 111 |
| chr2:133147936-133149954 | p2 | (TA)6 | CCGGTAACCGTTTAGGAACC | TCCGTATTGCTGACTTGCTG | 111 |
| chr2:133147936-133149954 | p6 | (GTTCCG)3 | CCGGTAACCGTTTAGGAACC | TCCGTATTGCTGACTTGCTG | 111 |
| chr2:133150409-133152424 | p5 | (TTTTA)3 | ATCCCTCACCACCACTCAAC | ACCCCTCACTCCCCTCTAAA | 273 |
| chr2:133166687-133168703 | p2 | (TA)7 | AGAAGCGGAGGAAGGAGAAG | TTCCCCACATCATATTCGGT | 219 |
| chr2:133166687-133168703 | p2 | (AT)8 | AGAAGCGGAGGAAGGAGAAG | TTCCCCACATCATATTCGGT | 219 |
| chr2:133166687-133168703 | p2 | (TA)9 | AGAAGCGGAGGAAGGAGAAG | TTCCCCACATCATATTCGGT | 219 |
| chr2:133166687-133168703 | p2 | (TA)12 | AGAAGCGGAGGAAGGAGAAG | TTCCCCACATCATATTCGGT | 219 |
| chr2:133252654-133254680 | p2 | (AT)11 | CGGGCACCTCTGTCATACTT | GTTAGCCTCGTCGGCTTATG | 160 |
| chr2:133252654-133254680 | p2 | (TA)13 | CGGGCACCTCTGTCATACTT | GTTAGCCTCGTCGGCTTATG | 160 |
| chr2:133276902-133278929 | p3 | (AAT)9 | AAGGACATGCAGAAAATGGG | AATCAAAATCAAATCCCGCA | 145 |
| chr2:133313760-133315781 | p2 | (TA)7 | CGGGAAAGGCTAATGTGGTA | AAACTGCTGGGCCCTAATCT | 223 |
| chr2:133313794-133315806 | p2 | (TA)7 | CGGGAAAGGCTAATGTGGTA | AAACTGCTGGGCCCTAATCT | 189 |
| chr2:133315556-133317574 | p2 | (TA)7 | GGCTGAATCTGGCAACAAAT | ATTTCCAAGTGGCTTGTTCA | 185 |
| chr2:133315556-133317574 | p2 | (TA)6 | GGCTGAATCTGGCAACAAAT | ATTTCCAAGTGGCTTGTTCA | 185 |
| chr2:133315556-133317574 | p2 | (AT)9 | GGCTGAATCTGGCAACAAAT | ATTTCCAAGTGGCTTGTTCA | 185 |
| chr2:133315556-133317574 | p2 | (AT)8 | GGCTGAATCTGGCAACAAAT | ATTTCCAAGTGGCTTGTTCA | 185 |
| chr2:133315556-133317574 | p3 | (AAT)5 | GGCTGAATCTGGCAACAAAT | ATTTCCAAGTGGCTTGTTCA | 185 |
| chr2:133315556-133317574 | p5 | (AATAA)3 | GGCTGAATCTGGCAACAAAT | ATTTCCAAGTGGCTTGTTCA | 185 |
| chr2:133315575-133317587 | p2 | (TA)7 | GGCTGAATCTGGCAACAAAT | ATTTCCAAGTGGCTTGTTCA | 166 |
| chr2:133315575-133317587 | p2 | (TA)6 | GGCTGAATCTGGCAACAAAT | ATTTCCAAGTGGCTTGTTCA | 166 |
| chr2:133315575-133317587 | p2 | (AT)9 | GGCTGAATCTGGCAACAAAT | ATTTCCAAGTGGCTTGTTCA | 166 |
| chr2:133315575-133317587 | p2 | (AT)8 | GGCTGAATCTGGCAACAAAT | ATTTCCAAGTGGCTTGTTCA | 166 |
| chr2:133315575-133317587 | p3 | (AAT)5 | GGCTGAATCTGGCAACAAAT | ATTTCCAAGTGGCTTGTTCA | 166 |
| chr2:133315575-133317587 | p5 | (AATAA)3 | GGCTGAATCTGGCAACAAAT | ATTTCCAAGTGGCTTGTTCA | 166 |
| chr2:133315779-133317794 | p2 | (TA)6 | AATGATAAGACCAGAGTGGCG | AATTTCCCGCTAAGTGGCTT | 262 |
| chr2:133315779-133317794 | p2 | (AT)9 | AATGATAAGACCAGAGTGGCG | AATTTCCCGCTAAGTGGCTT | 262 |
| chr2:133315779-133317794 | p2 | (AT)8 | AATGATAAGACCAGAGTGGCG | AATTTCCCGCTAAGTGGCTT | 262 |
| chr2:133315779-133317794 | p3 | (AAT)5 | AATGATAAGACCAGAGTGGCG | AATTTCCCGCTAAGTGGCTT | 262 |
| chr2:133315779-133317794 | p5 | (AATAA)3 | AATGATAAGACCAGAGTGGCG | AATTTCCCGCTAAGTGGCTT | 262 |
| chr2:133315827-133317853 | p2 | (TA)6 | AATGATAAGACCAGAGTGGCG | AATTTCCCGCTAAGTGGCTT | 214 |
| chr2:133315827-133317853 | p2 | (AT)9 | AATGATAAGACCAGAGTGGCG | AATTTCCCGCTAAGTGGCTT | 214 |
| chr2:133315827-133317853 | p2 | (AT)8 | AATGATAAGACCAGAGTGGCG | AATTTCCCGCTAAGTGGCTT | 214 |

| Marker ID | SSR type | SSR motif | FORWARD PRIMER (5'-3') | REVERSE PRIMER (5'-3') | size (bp) |
|---|---|---|---|---|---|
| chr2:133315827-133317853 | p3 | (AAT)5 | AATGATAAGACCAGAGTGGCG | AATTTCCCGCTAAGTGGCTT | 214 |
| chr2:133315827-133317853 | p5 | (AATAA)3 | AATGATAAGACCAGAGTGGCG | AATTTCCCGCTAAGTGGCTT | 214 |
| chr2:133357843-133359858 | p3 | (CAA)5 | AGAAGGCAAACCAGCAAGAA | CACGGTACCAGAAAGGAGGA | 102 |
| chr2:133361351-133363367 | p2 | (TA)8 | TGGCAAAATCTATATCCGCC | CATACCGAGCTGTAGGGCAT | 231 |
| chr2:133374143-133376158 | p3 | (CAA)5 | AGAAGGCAAACCAGCAAGAA | CACGGTACCAGAAAGGAGGA | 102 |
| chr2:133382522-133384538 | p2 | (TC)8 | GCCTGCAGTAATATGCCCTC | CAGATTCGGACATGGTGTTG | 215 |
| chr2:133382522-133384538 | p2 | (CT)9 | GCCTGCAGTAATATGCCCTC | CAGATTCGGACATGGTGTTG | 215 |
| chr2:133686888-133688904 | p2 | (TA)8 | ATGATTGGCAAGGGAAATTG | TACGGCATACAGCCATGGTA | 284 |
| chr2:133686997-133689029 | p2 | (TA)8 | ATGATTGGCAAGGGAAATTG | TACGGCATACAGCCATGGTA | 175 |
| chr2:133841160-133843176 | p4 | (TAAT)4 | ATGGCAAAAAGGTTCTGTGG | AGGTTTGGATTGAGCGTTTG | 268 |
| chr2:133894196-133896211 | p3 | (TGT)5 | TTTTCCCACTCACTGCCTTC | TTAAACAGCAAACCGAGGCT | 138 |
| chr2:133965675-133967690 | p5 | (AAACC)3 | AGCTGGTAGGGGTGTACGTG | CAGAGGTTGAAAAACCCGAA | 193 |
| chr2:134030688-134032706 | p2 | (AC)9 | GCAATTATGACAGCAGCCAA | CATTCCCCATTTCATTCCAC | 142 |
| chr2:134039266-134041287 | p3 | (AGA)7 | CGACATTTGTTCGGAAACCT | GAGAATCCTTACACGCAGCC | 279 |
| chr2:134178761-134180783 | p2 | (AT)11 | GGACGCCGTATGATCAGATT | ATAATGGTCGTGCTCCCAAG | 288 |
| chr2:134254495-134256519 | p5 | (CTTTC)3 | GGCAAGAATTGGAAGGTTCA | AGAAGACTTTCCGCCAGTCA | 173 |
| chr2:134254515-134256530 | p5 | (CTTTC)3 | GGCAAGAATTGGAAGGTTCA | AGAAGACTTTCCGCCAGTCA | 153 |
| chr2:134376365-134378377 | p2 | (AG)6 | GAAAATTCTGCCCATCTGGA | CCCCCTGACTTCTTCGTGTA | 132 |
| chr2:134376365-134378377 | p5 | (TACCA)4 | GAAAATTCTGCCCATCTGGA | CCCCCTGACTTCTTCGTGTA | 132 |
| chr2:134454265-134456283 | p3 | (AAT)6 | GAGCATTCTCAGAACCTCGG | GGACACAAGTCCGGAAAGAA | 166 |
| chr2:134461320-134463340 | p5 | (ATCAT)4 | AGGCTCAATCATGTTCAGGAG | GGGTGTGCAAATGGCTAAGT | 202 |
| chr2:134491975-134493990 | p5 | (TTTTA)3 | CCAAGCATGATGAAGAGGGT | GCTCTATGGCAACATGAGCA | 136 |
| chr2:134493459-134495483 | p3 | (TTG)8 | AGGATTTGGTAGATGCGACG | TGTCAGCAGTGAAATGCCTC | 153 |
| chr2:134574141-134576156 | p3 | (AGT)5 | ATAATAACCCATGCGGCAAA | TTGTGATTGTGTTTGGGGTG | 182 |
| chr2:134574141-134576156 | p6 | (TTGGAG)3 | ATAATAACCCATGCGGCAAA | TTGTGATTGTGTTTGGGGTG | 182 |
| chr2:134600400-134602424 | p6 | (CATGAG)4 | ACAAGTCCAACACACCCCAT | GGGGCGTGACACCTACTTTA | 163 |
| chr2:134695022-134697040 | p6 | (TTCGGT)3 | CACCAATGCGGGAAAATTAC | AACACTTGCGCATCTCTCCT | 248 |
| chr2:134695022-134697040 | p3 | (AGC)6 | CACCAATGCGGGAAAATTAC | AACACTTGCGCATCTCTCCT | 248 |
| chr2:134705649-134707679 | p2 | (AT)15 | GAATGCTGACTCCCTACCCA | GGGGTACAGTCTGGGTTTGA | 263 |
| chr2:134737841-134739856 | p5 | (TATGA)3 | TATCCGGCATTTTGAACCTC | CTCGAAACTTTGGAAAAGCG | 153 |
| chr2:134804308-134806324 | p2 | (AT)8 | TCAGGGCCCTCACATTTATC | CCATTAGCCCACTCGGATTA | 180 |
| chr2:134866200-134868212 | p2 | (CA)6 | TTTGAAGGATCGGAGGAAGA | GCACTCTGCATGAGTCCAAA | 192 |
| chr2:134898887-134900947 | p2 | (CT)10 | GCACAAAGGCCTAACACCAT | ATGGACTGGATTGGCCATAA | 118 |
| chr2:134898887-134900947 | p5 | (TTTTA)3 | GCACAAAGGCCTAACACCAT | ATGGACTGGATTGGCCATAA | 118 |
| chr2:134898887-134900947 | p3 | (ATA)20 | GCACAAAGGCCTAACACCAT | ATGGACTGGATTGGCCATAA | 118 |
| chr2:134901153-134903213 | p2 | (CT)10 | GCACAAAGGCCTAACACCAT | GAAGACGCAAACACACGAAA | 118 |

| Marker ID | SSR type | SSR motif | FORWARD PRIMER (5'-3') | REVERSE PRIMER (5'-3') | size (bp) |
|---|---|---|---|---|---|
| chr2:134901153-134903213 | p5 | (TTTTA)3 | GCACAAAGGCCTAACACCAT | GAAGACGCAAACACACGAAA | 118 |
| chr2:134901153-134903213 | p3 | (ATA)20 | GCACAAAGGCCTAACACCAT | GAAGACGCAAACACACGAAA | 118 |
| chr2:134901153-134903213 | p2 | (AT)16 | GCACAAAGGCCTAACACCAT | GAAGACGCAAACACACGAAA | 118 |
| chr2:134953777-134955789 | p2 | (TG)6 | GCCCTTCTTGTTATCACCGA | TCAAAATTTTCCATTTCGCC | 283 |
| chr2:135071547-135073559 | p2 | (AT)6 | TGTATGAAAACCTCGTTGCG | TCGAGTTGAGGAAGATGGCT | 276 |
| chr2:135080012-135082027 | p3 | (TGT)5 | CACCATGCCTTCAAGGTTTT | ATGCAAAATTTGAGCTTGCC | 290 |
| chr2:135104422-135106437 | p5 | (ATCAT)3 | TACATGGAAGCCGATAAGGC | AACGACAGGTTTGCTGATCC | 298 |
| chr2:135155428-135157485 | p3 | (TAT)19 | AAGAGAGAGGTGAGGGAGGC | TATCCTAAATTACCCGGCCC | 291 |
| chr2:135233845-135235865 | p5 | (GGGGT)4 | CAAAGTCCTGTGCCTGTCAA | GCATACAAGCTGGCCAAAAT | 124 |
| chr2:135238891-135240936 | p3 | (AAT)15 | AAGAGGCTCCTTCCACTTCC | GGGAGAGAATCTGGTGGTGA | 140 |
| chr2:135338023-135340043 | p2 | (AT)23 | GGATGACCTGGCACAAACTT | CTCAAATGATCGGTGGCTTT | 215 |
| chr2:135338023-135340043 | p5 | (CTTTT)4 | GGATGACCTGGCACAAACTT | CTCAAATGATCGGTGGCTTT | 215 |
| chr2:135422638-135424650 | p2 | (TA)6 | CCACATTGCTCCTCCAAAGT | AAGGCCTAGACCCTTGTGGT | 188 |
| chr2:135538154-135540169 | p5 | (CCACC)3 | GGGAAGGAGAAATCGGAAAG | TGTTGGGCAAGCTTAAAACC | 247 |
| chr2:135614771-135616786 | p5 | (ATTTT)3 | GGCTTTGAGATTCCCTGTGA | TCTGTCTTCACAAACCTGCG | 192 |
| chr2:135614811-135616826 | p5 | (ATTTT)3 | GGCTTTGAGATTCCCTGTGA | TCTGTCTTCACAAACCTGCG | 152 |
| chr2:135652640-135654655 | p5 | (ATTTT)3 | GGCTTTGAGATTCCCTGTGA | TCTGTCTTCACAAACCTGCG | 192 |
| chr2:135652680-135654695 | p5 | (ATTTT)3 | GGCTTTGAGATTCCCTGTGA | TCTGTCTTCACAAACCTGCG | 152 |
| chr2:135675326-135677346 | p2 | (TG)10 | GCCGCTTTTGGTGTTATTGT | GAATCCAGGCTTCACGTTGT | 183 |
| chr2:135813289-135815301 | p2 | (AT)6 | ACACGTGCCGAGAAGCTAGT | AGAACAGAGGTGGCCGAATA | 251 |
| chr2:135827774-135829792 | p2 | (AG)6 | TGAGAGGGAGGGCACATTAC | TTCGTTGTGAACTTCGTTCG | 275 |
| chr2:135827799-135829811 | p2 | (AG)6 | TGAGAGGGAGGGCACATTAC | TTCGTTGTGAACTTCGTTCG | 250 |
| chr2:135877192-135879207 | p2 | (TA)9 | GGTTTAAGGAAGAAAGGCGG | TCAATCTTTTTGGGACGGAG | 270 |
| chr2:135877192-135879207 | p3 | (TTG)9 | GGTTTAAGGAAGAAAGGCGG | TCAATCTTTTTGGGACGGAG | 270 |
| chr2:135877267-135879282 | p2 | (TA)9 | GGTTTAAGGAAGAAAGGCGG | TCAATCTTTTTGGGACGGAG | 195 |
| chr2:135877267-135879282 | p3 | (TTG)9 | GGTTTAAGGAAGAAAGGCGG | TCAATCTTTTTGGGACGGAG | 195 |
| chr2:135877510-135879562 | p2 | (TA)9 | TCATGTTACTCCCTCCGTCC | TCTGTCCCAAAAAGATTGGC | 188 |
| chr2:135877510-135879562 | p3 | (TTG)9 | TCATGTTACTCCCTCCGTCC | TCTGTCCCAAAAAGATTGGC | 188 |
| chr2:135877596-135879614 | p2 | (TA)9 | TCATGTTACTCCCTCCGTCC | TCTGTCCCAAAAAGATTGGC | 102 |
| chr2:135877596-135879614 | p3 | (TTG)9 | TCATGTTACTCCCTCCGTCC | TCTGTCCCAAAAAGATTGGC | 102 |
| chr2:135916952-135918967 | p2 | (CT)6 | TAACGTATCGATGATGCCGA | CCGCCTTTCTTCCTTAAACC | 299 |
| chr2:135963663-135965679 | p3 | (CAA)5 | TACCAAACAACCCCTAACGG | AGGGTAAGGCCTCCACTTGT | 252 |
| chr2:135963663-135965679 | p4 | (ATCG)4 | TACCAAACAACCCCTAACGG | AGGGTAAGGCCTCCACTTGT | 252 |
| chr2:135963663-135965679 | p6 | (GAAAAT)3 | TACCAAACAACCCCTAACGG | AGGGTAAGGCCTCCACTTGT | 252 |
| chr2:135981573-135983589 | p3 | (CAA)5 | TACCAAACAACCCCTAACGG | AGGGTAAGGCCTCCACTTGT | 252 |
| chr2:135981573-135983589 | p4 | (ATCG)4 | TACCAAACAACCCCTAACGG | AGGGTAAGGCCTCCACTTGT | 252 |

| Marker ID | SSR type | SSR motif | FORWARD PRIMER (5'-3') | REVERSE PRIMER (5'-3') | size (bp) |
|---|---|---|---|---|---|
| chr2:135981573-135983589 | p6 | (GAAAAT)3 | TACCAAACAACCCCTAACGG | AGGGTAAGGCCTCCACTTGT | 252 |
| chr2:136038419-136040439 | p5 | (AAACC)4 | CTTAGGGGTCGTTTGGTTCA | TATCTGCCACAGGGTGGAAT | 219 |
| chr2:136059942-136061956 | p2 | (TA)7 | GAGGAAAATCCTCAGTGCCA | AATTGGTAGACATCGTCGCC | 185 |
| chr2:136078101-136080117 | p2 | (AT)8 | AGTGTCGCCTTAGCCAAAAA | GGATTTCACTGGGTTGCTGT | 287 |
| chr2:136202305-136204317 | p2 | (AT)6 | CAGCCGGTCAGAAAAGAGTC | TTAAGTGTGGGATGCGATCA | 139 |
| chr2:136206642-136208676 | p5 | (CCTTG)3 | ATTGCTCCAAGGTGACGAAC | TGCAATGCAAGACTGACCTC | 147 |
| chr2:136206674-136208694 | p5 | (CCTTG)3 | ATTGCTCCAAGGTGACGAAC | TGCAATGCAAGACTGACCTC | 115 |
| chr2:136259311-136261329 | p3 | (GTT)6 | TCTTGAAGTGGCAGACATGG | TTGCAAATGATGGAGGACAA | 119 |
| chr2:136286784-136288799 | p3 | (AAT)5 | AGTGCTGTAACTGCAGGGGT | AAAATCTTGGATCCGCCTCT | 238 |
| chr2:136288522-136290536 | p2 | (AT)7 | TTTTTGGCCAACGAGATACC | AATTCCATCCGTTCATCTGG | 234 |
| chr2:136288522-136290536 | p5 | (TTTGG)3 | TTTTTGGCCAACGAGATACC | AATTCCATCCGTTCATCTGG | 234 |
| chr2:136402319-136404334 | p3 | (AAC)5 | ACATGCGTTAGGGTAGGCTG | TCCCCTCTTCCTGATCATTG | 174 |
| chr2:136402319-136404334 | p3 | (ACA)5 | ACATGCGTTAGGGTAGGCTG | TCCCCTCTTCCTGATCATTG | 174 |
| chr2:136421939-136423989 | p2 | (TA)6 | GCAGTAACCAAACGGTGGAT | GCCCCCTAATCCTTGTAACC | 144 |
| chr2:136429751-136431766 | p5 | (AATAC)3 | GCTTGTTTGGCCATGAAAAT | TGGGAATGGGCTTCATAAAA | 142 |
| chr2:136471467-136473482 | p5 | (AAAGA)3 | AAAAGCGAATCTCCAATCCA | TTTATGAACCAGGAAACCCG | 173 |
| chr2:136471467-136473482 | p5 | (AAAAT)3 | AAAAGCGAATCTCCAATCCA | TTTATGAACCAGGAAACCCG | 173 |
| chr2:136476205-136478220 | p5 | (AACCA)3 | AGATATCATCGCTGCAACCC | CCAATTTTTCACGTGGTGTG | 119 |
| chr2:136476205-136478220 | p5 | (TTGAT)3 | AGATATCATCGCTGCAACCC | CCAATTTTTCACGTGGTGTG | 119 |
| chr2:136491697-136493709 | p2 | (AG)6 | CTGCATAAGTGAGATGCGGA | TAGGTAGGGTGAATTTGCGG | 265 |
| chr2:136550050-136552065 | p5 | (ATTGC)3 | TTTGATGAGCTAGGCTCGGT | ACATCTCGGTAGCACAACCC | 201 |
| chr2:136550050-136552065 | p3 | (TTA)5 | TTTGATGAGCTAGGCTCGGT | ACATCTCGGTAGCACAACCC | 201 |
| chr2:136603193-136605208 | p5 | (TTTAA)3 | AGCTCCACCGGTTCATATTG | GAACATCTGCACGCAGAAAA | 146 |
| chr2:136603193-136605208 | p2 | (TG)8 | AGCTCCACCGGTTCATATTG | GAACATCTGCACGCAGAAAA | 146 |
| chr2:136715138-136717212 | p2 | (TA)10 | CAAAATCCCGAATGAGATGG | GTCCCCTTTTAATTCCTCGG | 139 |
| chr2:136715138-136717212 | p2 | (AT)37 | CAAAATCCCGAATGAGATGG | GTCCCCTTTTAATTCCTCGG | 139 |
| chr2:136715138-136717212 | p2 | (TA)10 | CAAAATCCCGAATGAGATGG | GTCCCCTTTTAATTCCTCGG | 139 |
| chr2:136772147-136774163 | p2 | (AT)8 | TTGAACTGGGGGTTCTCAAC | AAATCCAAGCATGGAGCAAC | 129 |
| chr2:136879276-136881294 | p2 | (AT)9 | TGAATCCTTCTGGCCATCTC | GTCATTTTCACCGATTGGCT | 140 |
| chr2:136879276-136881294 | p2 | (AT)9 | TGAATCCTTCTGGCCATCTC | GTCATTTTCACCGATTGGCT | 140 |
| chr2:137089260-137091275 | p5 | (AAAAC)3 | TTCTTAGCGTGCCACATCAG | TGCGTGGTGCTTCAAATAAG | 121 |
| chr2:137099052-137101066 | p2 | (AT)7 | CAAGTGGCATACCAAGGGTT | TCAAGCCTTTTTCACATCCC | 126 |
| chr2:137099052-137101066 | p3 | (AAT)12 | CAAGTGGCATACCAAGGGTT | TCAAGCCTTTTTCACATCCC | 126 |
| chr2:137175586-137177622 | p2 | (TA)18 | TTGGTGAGTGCATTGTGGAT | AGGAATTTCAGGGCAGGATT | 251 |
| chr2:137183647-137185669 | p2 | (TA)11 | CTAGGAATGGGGCCAGAAAT | CTTGTCGGCGAACAATCTTT | 122 |
| chr2:137184268-137186334 | p2 | (TA)11 | AAAGGGTGTTTCCATTGTGG | ATGGCCCACATGTCCATTAG | 180 |

| Marker ID | SSR type | SSR motif | FORWARD PRIMER (5'-3') | REVERSE PRIMER (5'-3') | size (bp) |
|---|---|---|---|---|---|
| chr2:137184268-137186334 | p2 | (TA)8 | AAAGGGTGTTTCCATTGTGG | ATGGCCCACATGTCCATTAG | 180 |
| chr2:137232640-137234680 | p2 | (TA)20 | AGGGTAATTGGAATTTCGGG | GGAGGGACAGCTTACAATGC | 153 |
| chr2:137267761-137269776 | p3 | (AAG)5 | TGTACCAAGATGGGCAACAA | TGGAATGTAGGGGTTTGAGC | 134 |
| chr2:137329930-137331942 | p2 | (AT)6 | AAAGAGCCGCCCTAACTTTC | GCTATTCCGCCATGTCATTT | 204 |
| chr2:137329930-137331942 | p5 | (TTTGG)3 | AAAGAGCCGCCCTAACTTTC | GCTATTCCGCCATGTCATTT | 204 |
| chr2:137334527-137336575 | p2 | (AT)9 | ATATGGGGCCCACAATTTTT | TTGCTTGATTTAGTTGGGGG | 135 |
| chr2:137412294-137414309 | p5 | (ACAAT)3 | TCATTGAGGCTTTTTCCCAC | TTAGTTGCCAAGCCACATGA | 238 |
| chr2:137513091-137515111 | p2 | (TA)10 | CACGAATTCGAGCTTTGAGA | GCTTTTTGCGGTCAAGTTTT | 156 |
| chr2:137537385-137539403 | p3 | (GTT)5 | TCGGACGGATAAGCGTAACT | GGCGTGACAAACGTAATCCT | 255 |
| chr2:137537385-137539403 | p6 | (AACTGA)3 | TCGGACGGATAAGCGTAACT | GGCGTGACAAACGTAATCCT | 255 |
| chr2:137593427-137595484 | p3 | (TTC)19 | GGCCGGAAAAGCTTTAATTC | CGACCGGAGAAAATCAATGT | 161 |
| chr2:137602858-137604900 | p3 | (TAA)14 | GGAGATTGCATTACGTGGCT | CTGGAGCGACAAAAACAACA | 239 |
| chr2:137622264-137624276 | p2 | (GA)6 | GGATCCGTATCCCGAATCTT | GGTATCCTGCACGAACACCT | 240 |
| chr2:137682750-137684764 | p2 | (AT)7 | GGGGGTGGAAGTTAGGAAAA | TGGAAAGTAATGCGGGTAGG | 298 |
| chr2:137691000-137693016 | p2 | (AT)9 | CCCATGAGTTTGGAGATGGT | ACCCATGCTTCGTCGAATAC | 279 |
| chr2:137691000-137693016 | p2 | (AT)6 | CCCATGAGTTTGGAGATGGT | ACCCATGCTTCGTCGAATAC | 279 |
| chr2:137693254-137695266 | p2 | (AT)6 | ATGCTTTACGGACGACGAAG | TCCAAAGAGTGAGCCCCATA | 133 |
| chr2:137693254-137695266 | p2 | (AT)7 | ATGCTTTACGGACGACGAAG | TCCAAAGAGTGAGCCCCATA | 133 |
| chr2:137704809-137706821 | p2 | (AT)10 | ACGACGAAGCATGGGTCTAA | CCAAAAGAGTGGGCCTCATA | 127 |
| chr2:137704809-137706821 | p2 | (AT)12 | ACGACGAAGCATGGGTCTAA | CCAAAAGAGTGGGCCTCATA | 127 |
| chr2:137704809-137706821 | p2 | (AT)6 | ACGACGAAGCATGGGTCTAA | CCAAAAGAGTGGGCCTCATA | 127 |
| chr2:137704809-137706821 | p2 | (AT)7 | ACGACGAAGCATGGGTCTAA | CCAAAAGAGTGGGCCTCATA | 127 |
| chr2:137714188-137716204 | p4 | (ATTA)4 | TGTGTTGCTTCTTTTTCCCC | TACCACCCCATCACAACCTT | 260 |
| chr2:137756323-137758338 | p5 | (TTTTA)3 | CAAATTCGTGCTGCGTAAGA | ATGGATATGATCCGTCCACC | 280 |
| chr2:137756323-137758338 | p5 | (TTTTA)3 | CAAATTCGTGCTGCGTAAGA | ATGGATATGATCCGTCCACC | 280 |
| chr2:137773135-137775150 | p5 | (TACAA)3 | AGGCGGAACATTTCATTCAC | GTCGGTTACATAAATCCCCG | 168 |
| chr2:137791567-137793582 | p3 | (AAC)5 | GATATCTTACGTAGGCGGCG | GCTGGTTTGAGGTCATGGTT | 192 |
| chr2:137795386-137797401 | p5 | (AACCG)3 | AATATGTCCAAGCCCGTGTC | AGTTGCAGCAAATGCTGATG | 155 |
| chr2:137845469-137847481 | p2 | (TA)6 | CACAGCTAAAGGCAGGCTCT | TACGCCTGTGACTCCATCAA | 172 |
| chr2:137848152-137850170 | p6 | (GGGTGG)3 | AAAGATGCAGCAGCAGGAAT | CGCCAATGGTTGAAAGAGAT | 242 |
| chr2:137848152-137850170 | p5 | (GGGGT)3 | AAAGATGCAGCAGCAGGAAT | CGCCAATGGTTGAAAGAGAT | 242 |
| chr2:137910754-137912769 | p5 | (TTTTC)3 | GAGAAAGCGCGAATCTTTTG | GAAAATAGCCGAAAACGCAA | 112 |
| chr2:137922245-137924293 | p3 | (CTT)16 | CACCACGGCTATTTGATCCT | GTGGAATCAAAGCCACATGA | 176 |
| chr2:137934970-137936998 | p5 | (AAAAT)3 | GCATGTGTCATGTGTGGTGA | AAGCACAAAAACGGTCCAAG | 270 |
| chr2:137935001-137937013 | p5 | (AAAAT)3 | TCTAGAACTTGCCCGGAATG | ATCACCCCCAAACTTAGGCT | 291 |
| chr2:137984072-137986087 | p3 | (AAT)7 | TGTACCAAGGGTGCCAGAAT | CATTATTGCACGGTCATTGC | 267 |

| Marker ID | SSR type | SSR motif | FORWARD PRIMER (5'-3') | REVERSE PRIMER (5'-3') | size (bp) |
|---|---|---|---|---|---|
| chr2:137984083-137986099 | p3 | (AAT)7 | TGTACCAAGGGTGCCAGAAT | CATTATTGCACGGTCATTGC | 256 |
| chr2:137984101-137986117 | p3 | (AAT)7 | TGTACCAAGGGTGCCAGAAT | CATTATTGCACGGTCATTGC | 238 |
| chr2:137990079-137992127 | p2 | (AT)24 | ATCGTGAGACCATGCTACCC | TGTTCATGCCCACAGGTAGA | 281 |
| chr2:138041022-138043040 | p6 | (TTTTTG)3 | CACCGTCCTCCTGTCATTCT | AGATGCAACAACAACCACCA | 123 |
| chr2:138041022-138043040 | p3 | (TTG)7 | CACCGTCCTCCTGTCATTCT | AGATGCAACAACAACCACCA | 123 |
| chr2:138124330-138126345 | p3 | (TGT)5 | ATTTGAAGCTTGTGGGTTCG | AGGTTATAAGTCAGGGGCGG | 191 |
| chr2:138124330-138126345 | p2 | (AT)6 | ATTTGAAGCTTGTGGGTTCG | AGGTTATAAGTCAGGGGCGG | 191 |
| chr2:138313046-138315067 | p3 | (AAT)8 | AACGATGTACCAAGGTTGCC | CGTTTTACCAACTTCCGCAT | 179 |
| chr2:138321647-138323662 | p3 | (TTA)5 | AATGGTCGTCACCTCTTTGC | CATTTGCATTACCTGGCCTT | 256 |
| chr2:138374909-138376924 | p2 | (CT)6 | GAAACAGGGGTTGCAAGTGT | TGTTGTTCAAAGCCAACAGC | 220 |
| chr2:138385309-138387333 | p3 | (ATG)8 | TCTTCATCATGGGGACACAA | TTGCCTAAAACTTCCCATGC | 196 |
| chr2:138385309-138387333 | p2 | (AT)6 | TCTTCATCATGGGGACACAA | TTGCCTAAAACTTCCCATGC | 196 |
| chr2:138415770-138417786 | p2 | (AT)8 | GAGCCGGCCACATAATAAGA | AGGAGAGGCTGTGCAGGTTA | 100 |
| chr2:138556784-138558842 | p2 | (AT)29 | AAAATTAAAGACCACCCCCG | CTTTTTGCTGTCGGGACATT | 201 |
| chr2:138556784-138558842 | p2 | (TA)7 | AAAATTAAAGACCACCCCCG | CTTTTTGCTGTCGGGACATT | 201 |
| chr2:138801603-138803641 | p2 | (TA)19 | GTTCAGTGGGCTGAATTTCG | CCGTCGTGTACCAGGAAGAT | 155 |
| chr2:138807354-138809384 | p2 | (AT)15 | ACCCTCGCGACTCAGAATAA | CCCCATGGCAAAGTTAATGT | 139 |
| chr2:138815558-138817574 | p4 | (TTTG)4 | TCAGGGCAAAACCCTAACAC | CACATTATCTCCTCCGCCAT | 256 |
| chr2:138847787-138849805 | p3 | (TCT)6 | GACATTTTTGGGGCGAACTA | TCAAACAAGCCTTACGGACC | 123 |
| chr2:138971013-138973031 | p6 | (TATGTG)3 | TTACTTCATGTGCATGGGGA | CACGTAGTTGATTTTGGCCC | 158 |
| chr2:138979950-138981980 | p2 | (AT)15 | GAGTATCCCAATTTTCGCCA | ATTTGGATGAAGCCACCTTG | 249 |
| chr2:139061329-139063343 | p2 | (AT)7 | TGTGTCGGGCAAGAATTACA | GATATGGCTGGACATGGCTT | 217 |
| chr2:139061329-139063343 | p2 | (TA)10 | TGTGTCGGGCAAGAATTACA | GATATGGCTGGACATGGCTT | 217 |
| chr2:139083796-139085816 | p5 | (AAAAG)4 | CAACAACAACACCACCTTGC | AAGAGCCCTCTCCTTCAAGC | 272 |
| chr2:139168373-139170388 | p5 | (ATTGT)3 | CATCACCTACAAGGGTGGCT | TCGGGTGCACACAAGATAAA | 160 |
| chr2:139195954-139197974 | p5 | (TCGGT)3 | GTAATGTCGGGTTGGTTTGG | TTGCGAGAATGCTTCATTTG | 263 |
| chr2:139195954-139197974 | p4 | (ATTT)5 | GTAATGTCGGGTTGGTTTGG | TTGCGAGAATGCTTCATTTG | 263 |
| chr2:139297689-139299735 | p2 | (AT)23 | TTTTCATGGCAAAACGTTGA | GCAGCATATTCTGGGACCAT | 294 |
| chr2:139349604-139351619 | p5 | (TATTT)3 | TGAGGCAATTGATGGGATTT | AAAAGTTCATCCCTCCACCC | 188 |
| chr2:139349604-139351619 | p5 | (TATTT)3 | TGAGGCAATTGATGGGATTT | AAAAGTTCATCCCTCCACCC | 188 |
| chr2:139349604-139351619 | p5 | (TATTT)3 | TGAGGCAATTGATGGGATTT | AAAAGTTCATCCCTCCACCC | 188 |
| chr2:139359204-139361222 | p2 | (CA)9 | TCGCTTTACTCGTCGTCCTT | AACCGAACCTGTAATGCTGG | 231 |
| chr2:139359204-139361222 | p2 | (CA)9 | TCGCTTTACTCGTCGTCCTT | AACCGAACCTGTAATGCTGG | 231 |
| chr2:139366882-139368898 | p3 | (TGT)5 | CTTTGCCCCTGGAGTACAAA | AGGAGCAGGCATGACAAGAC | 271 |
| chr2:139366882-139368898 | p2 | (TC)8 | CTTTGCCCCTGGAGTACAAA | AGGAGCAGGCATGACAAGAC | 271 |
| chr2:139374084-139376098 | p2 | (AT)7 | GGCTTTGACTTTGGTGCCTA | CGTACCGTGTGGTGTCTTTG | 110 |

| Marker ID | SSR type | SSR motif | FORWARD PRIMER (5'-3') | REVERSE PRIMER (5'-3') | size (bp) |
|---|---|---|---|---|---|
| chr2:139375100-139377116 | p2 | (TA)8 | AGATGGCATCATGGAGGACT | GCATGCCCCTCAAATAAAAA | 174 |
| chr2:139447318-139449333 | p5 | (ATTGT)3 | GCTCAGTTGGTTGAGCATGA | AACCCTCAGTCGCTACCCTT | 222 |
| chr2:139450626-139452641 | p5 | (CCACC)3 | GCGTTCTTGTTACCTCTGGC | AAACAGGGGGTAAATCGTCC | 153 |
| chr2:139450626-139452641 | p2 | (AT)6 | GCGTTCTTGTTACCTCTGGC | AAACAGGGGGTAAATCGTCC | 153 |
| chr2:139450626-139452641 | p5 | (TACAA)3 | GCGTTCTTGTTACCTCTGGC | AAACAGGGGGTAAATCGTCC | 153 |
| chr2:139461526-139463542 | p4 | (ATTT)4 | TTTTCAAGGGGCATACGTTC | TCTGGAGATTGGGAACAACC | 228 |
| chr2:139517130-139519145 | p5 | (GGTGG)3 | ATTTTTCAATCATGGAGGCG | CTAGCTACACCCTCAAGCCG | 180 |
| chr2:139545820-139547838 | p5 | (TGGTT)3 | AGCCGACAAGGCTATCAAAA | GCGGTTCAGCTTGGTATGAT | 174 |
| chr2:139545820-139547838 | p6 | (ATAACC)3 | AGCCGACAAGGCTATCAAAA | GCGGTTCAGCTTGGTATGAT | 174 |
| chr2:139545820-139547838 | p2 | (AT)24 | AGCCGACAAGGCTATCAAAA | GCGGTTCAGCTTGGTATGAT | 174 |
| chr2:139579667-139581683 | p4 | (TGCA)4 | TGGGCAAAAATATCCGCTAA | ACATCATGCAAAATGCTGGA | 269 |
| chr2:139652351-139654374 | p2 | (TA)6 | CCTCCATTGTCTCCCAGGTA | TATGGGAGAAACAAATCCGC | 100 |
| chr2:139656010-139658025 | p5 | (TTTCT)3 | AGAAGCATACACAATGCCCC | ATGTTTTGTCTGCCTGGTCC | 246 |
| chr2:139660532-139662547 | p5 | (TTTCT)3 | GGGAAGCATACACAATGCCT | GGCCAGCCGCATATACTTTA | 239 |
| chr2:139670103-139672118 | p3 | (CAA)5 | CCGCTTCCGGTTAATAATGA | TTCCAAAACCACTCCGAAAG | 206 |
| chr2:139700697-139702712 | p5 | (AAAAT)3 | TTACGTTAAGCAGATCCCCG | CCCAACTTATGCTTTGCGAT | 235 |
| chr2:139700697-139702712 | p3 | (AAT)5 | TTACGTTAAGCAGATCCCCG | CCCAACTTATGCTTTGCGAT | 235 |
| chr2:139704214-139706230 | p4 | (AATA)4 | AAACTTGAAGCTCCTGGCAA | GCACATATAGGGGCGAATGT | 211 |
| chr2:139704214-139706230 | p4 | (TTTA)4 | AAACTTGAAGCTCCTGGCAA | GCACATATAGGGGCGAATGT | 211 |
| chr2:139710070-139712082 | p2 | (TA)6 | TAAAGAGCCCGTTTGGATTG | TTGAGCATGTTTGGATGAGC | 287 |
| chr2:139710070-139712082 | p2 | (TG)6 | TAAAGAGCCCGTTTGGATTG | TTGAGCATGTTTGGATGAGC | 287 |
| chr2:139778640-139780655 | p3 | (TTC)5 | CAAGCAAATCAATTCGGGTT | TCGGGATGAGGAATCTTGTC | 283 |
| chr2:139786782-139788797 | p3 | (ATC)5 | ACGACCAAAACCGATGAGTC | GAATGGGCCACCCTATCTTT | 237 |
| chr2:139870848-139872860 | p2 | (AT)6 | CAATGCGAGCATGCATAAGT | ATCATGCTGGCCAAAGAGAG | 232 |
| chr2:139870848-139872860 | p2 | (AT)6 | CAATGCGAGCATGCATAAGT | ATCATGCTGGCCAAAGAGAG | 232 |
| chr2:139870848-139872860 | p2 | (AT)10 | CAATGCGAGCATGCATAAGT | ATCATGCTGGCCAAAGAGAG | 232 |
| chr2:139887021-139889036 | p3 | (ATT)5 | ATCAAAAGGGAAAAATGGGG | ATTTTTCCACCTGTCCCTCC | 113 |
| chr2:139900773-139902788 | p5 | (AAAAT)3 | CCATTTGAACCCTCCAAGAA | CCCATCCAAACAGGCTCTAA | 280 |
| chr2:139973691-139975707 | p2 | (TA)8 | GGATTGACTCGAAACGTGGT | GGACGGTGCGGTGTTATACT | 174 |
| chr2:139976245-139978308 | p2 | (TA)7 | TCCTAGGCTCGTTCGATTGT | CATCGCAGTCGAAAACTTGA | 284 |
| chr2:139976245-139978308 | p3 | (AAT)21 | TCCTAGGCTCGTTCGATTGT | CATCGCAGTCGAAAACTTGA | 284 |
| chr2:139976245-139978308 | p3 | (CTC)7 | TCCTAGGCTCGTTCGATTGT | CATCGCAGTCGAAAACTTGA | 284 |
| chr2:140001148-140003169 | p3 | (ACA)7 | GGATGTGCCATTGGATTTTC | TCCCAGCTTCTCTACCTCCA | 169 |
| chr2:140060668-140062689 | p2 | (TA)6 | ATCTCACAAGTGGGGTCTGG | ATTTCTTTTGAGCCGAGGGT | 198 |
| chr2:140060668-140062689 | p3 | (ACA)7 | ATCTCACAAGTGGGGTCTGG | ATTTCTTTTGAGCCGAGGGT | 198 |
| chr2:140098453-140100465 | p2 | (TA)6 | GTTGAAGGGGTTGAAGGACA | GAAACGAATCATGGGGAAGA | 189 |

| Marker ID | SSR type | SSR motif | FORWARD PRIMER (5'-3') | REVERSE PRIMER (5'-3') | size (bp) |
|---|---|---|---|---|---|
| chr2:140120833-140122845 | p2 | (TA)6 | GTTGAAGGGGTTGAAGGACA | GAAACGAATCATGGGGAAGA | 189 |
| chr2:140286591-140288651 | p2 | (AT)30 | TTCCCCATTGAACCAATCAT | ATTGATGAGCAAGAAGCCGT | 198 |
| chr2:140394486-140396504 | p2 | (TA)9 | ACTGGAAAGGTCACACCAGC | GTGCCAGCTTCTTCCATGTT | 235 |
| chr2:140421721-140423736 | p2 | (AT)33 | GGCCAAGTGTGTGTACGTTG | CAACAAGCCCAAGCAAAAAT | 300 |
| chr2:140421834-140423846 | p2 | (AT)33 | GGCCAAGTGTGTGTACGTTG | CAACAAGCCCAAGCAAAAAT | 187 |
| chr2:140450010-140452028 | p2 | (TA)9 | GAACCCACGTTCTGTCGTTT | AATCCGCGCGATATTAAATG | 219 |
| chr2:140452726-140454740 | p3 | (AAT)5 | ACCCGATCCTTTATGGAACC | AGCGAATGTGCAAAATAGGC | 262 |
| chr2:140452726-140454740 | p2 | (AC)7 | ACCCGATCCTTTATGGAACC | AGCGAATGTGCAAAATAGGC | 262 |
| chr2:140458030-140460042 | p2 | (AT)6 | GCCAAGTGTGTGTACGTTGG | AATTGGCAAGGTCAAAATGG | 149 |
| chr2:140460428-140462444 | p4 | (TTAT)4 | CGTGAACTGATATGGGAGGG | GCTGAAGTTGGTCGGGTTTA | 283 |
| chr2:140460428-140462444 | p2 | (AT)28 | CGTGAACTGATATGGGAGGG | GCTGAAGTTGGTCGGGTTTA | 283 |
| chr2:140460428-140462444 | p4 | (ATTA)4 | CGTGAACTGATATGGGAGGG | GCTGAAGTTGGTCGGGTTTA | 283 |
| chr2:140468817-140470833 | p4 | (TTAT)4 | CGTGAACTGATATGGGAGGG | GCTGAAGTTGGTCGGGTTTA | 283 |
| chr2:140468817-140470833 | p2 | (AT)28 | CGTGAACTGATATGGGAGGG | GCTGAAGTTGGTCGGGTTTA | 283 |
| chr2:140468817-140470833 | p4 | (ATTA)4 | CGTGAACTGATATGGGAGGG | GCTGAAGTTGGTCGGGTTTA | 283 |
| chr2:140548156-140550170 | p2 | (AT)7 | ATCTGGTAACAAGGTTGCGG | CAGTTTCCTTCCAATTCCCA | 111 |
| chr2:140604221-140606320 | p3 | (TAA)33 | ATTTCCTGCGATTCCTCCTT | CTCCTTCATCTCGCTTCACC | 244 |
| chr2:140604221-140606320 | p2 | (AT)7 | ATTTCCTGCGATTCCTCCTT | CTCCTTCATCTCGCTTCACC | 244 |
| chr2:140656608-140658629 | p3 | (ACA)7 | GAAAAATGTCGGTGCCACTT | GACACCGAAGGCACATTTTT | 261 |
| chr2:140658578-140660596 | p6 | (CGTCCC)3 | CCCTCGGAATGACCTTACAA | CTTTTGGATTTCGCAAGAGC | 286 |
| chr2:140685970-140687985 | p2 | (GT)13 | GATGCGGAAATATCGAAGGA | TTGGGCCTTTTGTTTGTTTC | 287 |
| chr2:140685970-140687985 | p3 | (TTG)5 | GATGCGGAAATATCGAAGGA | TTGGGCCTTTTGTTTGTTTC | 287 |
| chr2:140788163-140790181 | p6 | (AATAAA)3 | TGGGATGACACGTAGGATGA | CCTCAAGGTATTTTCGGCAA | 265 |
| chr2:140788163-140790181 | p5 | (CAAAC)3 | TGGGATGACACGTAGGATGA | CCTCAAGGTATTTTCGGCAA | 265 |
| chr2:140815304-140817318 | p2 | (TA)7 | CACGAATGCTACGCGTCTTA | CCTATTAGAAAGCAGCCCCA | 125 |
| chr2:140904519-140906561 | p2 | (AT)21 | TCGGCCGAACTCAGTAGTTT | GACTTAGCGAGATGGTTCGG | 219 |
| chr2:140931386-140933401 | p5 | (AAAAT)3 | GCAATGGAAGTCACCAACCT | CCACATCAGCTGCACAAAAT | 245 |
| chr2:140931397-140933425 | p5 | (AAAAT)3 | GCAATGGAAGTCACCAACCT | CCACATCAGCTGCACAAAAT | 234 |
| chr2:141049303-141051323 | p2 | (TA)10 | CTGAATTGCCTGGGATCACT | CTCAACAAATACCGCAGGGT | 107 |
| chr2:141076927-141078945 | p6 | (GAGTTT)3 | AATCGTTAGCGAATCCGTTG | TCAAATTTCATGGCCAAACA | 213 |
| chr2:141084100-141086120 | p2 | (AT)10 | TTGTAGTAAATGGGCTGGGC | GGTTGGGGTCCTGGTAAAAT | 160 |
| chr2:141084100-141086120 | p3 | (AAT)5 | TTGTAGTAAATGGGCTGGGC | GGTTGGGGTCCTGGTAAAAT | 160 |
| chr2:141154629-141156647 | p3 | (GTT)6 | CAACACAAAATTCATTGCCG | TGGGCCTGAGTCTCAACTCT | 119 |
| chr2:141167249-141169305 | p2 | (TA)28 | TGGGCCCAATAACTATTCCA | GCCGGTCTAATTTTGTCCCT | 277 |
| chr2:141228112-141230130 | p2 | (TC)9 | ACATAAATCTGGGCAGCACC | GTAAAGCTCCGTTTTGAGCG | 213 |
| chr2:141284799-141286817 | p6 | (ATTTAT)3 | TGAACGTGCCACCAAGATAA | CGTAAAAGGGGTCAAGCAGA | 161 |

| Marker ID | SSR type | SSR motif | FORWARD PRIMER (5'-3') | REVERSE PRIMER (5'-3') | size (bp) |
|---|---|---|---|---|---|
| chr2:141289769-141291789 | p2 | (AT)10 | AATTTCATGTCCCGAGGTTG | CCTCTTAGTCTTCGCCAACG | 125 |
| chr2:141289769-141291789 | p2 | (TA)6 | AATTTCATGTCCCGAGGTTG | CCTCTTAGTCTTCGCCAACG | 125 |
| chr2:141332896-141334910 | p2 | (AT)7 | GATCCAGGTGTAAACCCACG | GCCCTCAAGATTACAGAGCG | 186 |
| chr2:141337680-141339712 | p5 | (ATCAA)3 | ATTCGAACCTTGCTTGCATC | ATTTGTGCACCTTTTCAGGG | 226 |
| chr2:141337680-141339712 | p2 | (TA)16 | ATTCGAACCTTGCTTGCATC | ATTTGTGCACCTTTTCAGGG | 226 |
| chr2:141393033-141395054 | p3 | (TGT)5 | TGAGGGGAAGAAGGCATAGA | TAAAGGCACCAAACGAAAGG | 149 |
| chr2:141683679-141685697 | p3 | (TGT)6 | ATGGATGAAATGGAGGCTTG | CGCATCTTGTCTTCCACTGA | 226 |
| chr2:141698435-141700447 | p4 | (TAAA)4 | ATTCCCAGAACTCGAACACG | TATTAAACATGGCATCGGCA | 152 |
| chr2:141698435-141700447 | p2 | (TA)6 | ATTCCCAGAACTCGAACACG | TATTAAACATGGCATCGGCA | 152 |
| chr2:141831128-141833143 | p5 | (TTTAA)3 | ACCCCCAATATAAAGGACCG | CAAAGGGGAGGGGATTAAAA | 219 |
| chr2:141839511-141841526 | p5 | (TTTAA)3 | ACCCCCAATATAAAGGACCG | CAAAGGGGAGGGGATTAAAA | 219 |
| chr2:141918925-141920943 | p2 | (CT)9 | CTCACCGTTGGAATTCCTGT | AGTCAACCAAAGGAGGGGAT | 111 |
| chr2:141970669-141972684 | p3 | (TTC)5 | ACATTGTAATCCCCTTCCCC | AATCCATTGCAACCGATCAT | 115 |
| chr2:141970669-141972684 | p4 | (ATTT)4 | ACATTGTAATCCCCTTCCCC | AATCCATTGCAACCGATCAT | 115 |
| chr2:142050047-142052067 | p2 | (TA)10 | TGAGAGCACTCCATTGTTCG | TTGTGAGACCATCTGACCCA | 250 |
| chr2:142051303-142053319 | p2 | (AT)8 | TCATAAGCTGAGGTGGGGTC | GACCTCCCGAAATCAAATCA | 147 |
| chr2:142051303-142053319 | p2 | (TA)8 | TCATAAGCTGAGGTGGGGTC | GACCTCCCGAAATCAAATCA | 147 |
| chr2:142187756-142189771 | p5 | (TAACA)3 | TGTCAATCTTGCGCTCTCAC | TGATCGGGCTAAACATTTCC | 149 |
| chr2:142265801-142267825 | p2 | (TA)12 | CATCAAACAACAACAACGGC | GTCTGAGGGTTTTCGTAGCG | 193 |
| chr2:142299585-142301600 | p5 | (AAATA)3 | AACTTTGGTCCCATGCGTTA | TTAGGCCCGAAAGCCTAAAT | 267 |
| chr2:142312640-142314658 | p3 | (ATT)6 | CCTTTTGGCAAACAGATGGT | CCTCTGGTATGAGCGGGTAA | 255 |
| chr2:142324492-142326507 | p5 | (TTGTA)3 | GTGCCTTTCCCGTTGTATGT | AATGGAGTTAATGCGGCAAC | 221 |
| chr2:142478096-142480108 | p2 | (AT)6 | GATCCCAAGTGGAAGGAACA | TCAGTGCCTGGGAAAATACC | 204 |
| chr2:142479153-142481199 | p2 | (AT)23 | TCAAAAGCTCCAAAACCAGG | CCAAACCCTTCCTTCAAAAA | 287 |
| chr2:142479153-142481199 | p2 | (AT)23 | TCAAAAGCTCCAAAACCAGG | CCAAACCCTTCCTTCAAAAA | 287 |
| chr2:142528697-142530712 | p5 | (TATTG)3 | TGTGGCTGCTTAGATGGATG | ATGCAGCATCAACGACTGAG | 293 |
| chr2:142535755-142537776 | p3 | (AAG)7 | ACCTTTGCGGGAGACCTATT | GCGAACTGGACTGTGTTCAA | 182 |
| chr2:142537236-142539256 | p2 | (TG)10 | AAAAGGTGGCCGAAAATACC | AGTCTTCGCGAAAAGTCTGC | 272 |
| chr2:142537236-142539256 | p2 | (AT)23 | AAAAGGTGGCCGAAAATACC | AGTCTTCGCGAAAAGTCTGC | 272 |
| chr2:142540908-142542926 | p6 | (TAAAAA)3 | AGGAAAATGTGGGGAAAAGG | ACCTGATTTAAACGGTGGCA | 224 |
| chr2:142543924-142545962 | p2 | (AT)19 | TTCGAACCCAAAACCTCAAG | AATTTGCACGATTGTCCCTC | 170 |
| chr2:142552221-142554539 | p2 | (TA)159 | TGCTCCACGCCCTACTACTT | CGAAGCCACGGTTGATACTT | 187 |
| chr2:142602766-142604788 | p2 | (AG)11 | TCGAAACTCGGGATGATAGG | TCCCACCAAAGGAAAACAAA | 180 |
| chr2:142604460-142606480 | p4 | (TTAT)5 | GAGGTGAAGGTGAGTGGGTG | GGCTGAAAGGCCAGTAGTTG | 198 |
| chr2:142645573-142647623 | p5 | (TATGA)3 | ATGTGTTGTGGCACCAGTGT | CGGAATATCCGGAATAGCAA | 188 |
| chr2:142728571-142730583 | p2 | (AT)6 | ATCAGCACCAACGTCTACCC | AAAACAAAGGCTTCTTGGCA | 276 |

| Marker ID | SSR type | SSR motif | FORWARD PRIMER (5'-3') | REVERSE PRIMER (5'-3') | size (bp) |
|---|---|---|---|---|---|
| chr2:142852695-142854713 | p2 | (AT)29 | GATCACTCAAACACCCCACC | TATGGACTTGGGCAAACCTC | 266 |
| chr2:142852750-142854765 | p2 | (AT)29 | GATCACTCAAACACCCCACC | TATGGACTTGGGCAAACCTC | 211 |
| chr2:142856136-142858152 | p2 | (AC)8 | CCCCCTACCTTTGGTTTCTC | AAAGCTGTGATGGGATGAGG | 188 |
| chr2:142927855-142929869 | p2 | (TC)7 | GCACATGTTGGAGAAAAGCA | CAATTCGTGTGTGAAATGGC | 268 |
| chr2:142942939-142944955 | p4 | (ATGT)4 | GTCAATCCAAGGGAGGCATA | CTTGGTCGTGAGCTTCCTTC | 236 |
| chr2:142974628-142976642 | p2 | (AT)7 | GAACCCACCACATCCAAACT | TTGCTAGCGAATACACACGC | 208 |
| chr2:142974628-142976642 | p2 | (TA)6 | GAACCCACCACATCCAAACT | TTGCTAGCGAATACACACGC | 208 |
| chr2:142974628-142976642 | p2 | (AT)7 | GAACCCACCACATCCAAACT | TTGCTAGCGAATACACACGC | 208 |
| chr2:142978839-142980854 | p5 | (AAAAT)3 | GTTCACTCCGGCTTAGTTCG | GCCTTTTAGCCCTAGCCATC | 290 |
| chr2:142978839-142980854 | p5 | (AAAAG)3 | GTTCACTCCGGCTTAGTTCG | GCCTTTTAGCCCTAGCCATC | 290 |
| chr2:142992808-142994860 | p2 | (AT)26 | GATGCGCTAATGGCTGAAAT | CATGTGGGCTCAATGATTTG | 156 |
| chr2:143065222-143067260 | p2 | (TA)19 | TGGCTGAAAAATCATGGTCA | TTTTGTTGGATTCACAGGCA | 114 |
| chr2:143110641-143112665 | p3 | (AAT)8 | ATTGCAACTTTCAACCTCCG | CCTTTGCTAAAGCACTTGCC | 300 |
| chr2:143150659-143152683 | p3 | (AAT)8 | ATTGCAACTTTCAACCTCCG | CCTTTGCTAAAGCACTTGCC | 300 |
| chr2:143185134-143187158 | p3 | (AAT)8 | ATTGACAGACCGGAAGTTGG | TTGAGCAATTCCACCAAATG | 182 |
| chr2:143185134-143187158 | p3 | (AAT)8 | ATTGACAGACCGGAAGTTGG | TTGAGCAATTCCACCAAATG | 182 |
| chr2:143185134-143187158 | p3 | (AAT)9 | ATTGACAGACCGGAAGTTGG | TTGAGCAATTCCACCAAATG | 182 |
| chr2:143185134-143187158 | p3 | (AAT)12 | ATTGACAGACCGGAAGTTGG | TTGAGCAATTCCACCAAATG | 182 |
| chr2:143201246-143203264 | p3 | (TGT)6 | GTGGGCTCAGGGCATAAATA | ACCCTAATGGCGACAAACAG | 175 |
| chr2:143222970-143224991 | p3 | (ATA)7 | CTGTCGATGAGTTTGGCCTT | TGCTTTGTGCTTGTCTCACA | 165 |
| chr2:143250471-143252486 | p5 | (AAATA)3 | ATTTAGGCTCGACTTCACGC | TCATGTGTTCAATGGATGGG | 129 |
| chr2:143250471-143252486 | p5 | (AAAAT)3 | ATTTAGGCTCGACTTCACGC | TCATGTGTTCAATGGATGGG | 129 |
| chr2:143250471-143252486 | p5 | (AATAA)4 | ATTTAGGCTCGACTTCACGC | TCATGTGTTCAATGGATGGG | 129 |
| chr2:143250471-143252486 | p5 | (TTTAT)3 | ATTTAGGCTCGACTTCACGC | TCATGTGTTCAATGGATGGG | 129 |
| chr2:143250471-143252486 | p4 | (TTTA)4 | ATTTAGGCTCGACTTCACGC | TCATGTGTTCAATGGATGGG | 129 |
| chr2:143299835-143301853 | p3 | (TAA)6 | CCCTCTTTCGAATCTCTCCC | CTGACACCCCACACTTTCCT | 211 |
| chr2:143330410-143332425 | p5 | (CCTTT)3 | TTAGAGGCACAGGGAGAGGA | CCTTTTGCAACAATTCAGCA | 174 |
| chr2:143410649-143412667 | p2 | (TC)9 | ATTTTGAAAGGATTTGCCCC | CTGGGGTGGAGTTCCAATTA | 147 |
| chr2:143452085-143454100 | p5 | (AATGC)3 | TTTAACACGCGTAACGCAAC | ACGACTCGTAAATTGGGTCG | 220 |
| chr2:143524462-143526477 | p3 | (AAC)5 | TGTGGTGCAACTACCTCTGC | CGGATCCTCGGTAACCACTA | 207 |
| chr2:143524462-143526477 | p3 | (TTC)6 | TGTGGTGCAACTACCTCTGC | CGGATCCTCGGTAACCACTA | 207 |
| chr2:143626392-143628428 | p3 | (TCT)12 | AACCACTCCAAGCCTACCCT | ACGAAGGAGGGGACAGAGAT | 242 |
| chr2:143713686-143715698 | p2 | (AT)6 | CTTATCAGCAACCTCGGAGC | ATCGCGTTTTAGGGGAAGAT | 129 |
| chr2:143715538-143717552 | p2 | (AT)6 | CCGGTAACCGTAGGAAAACC | TTAACGAGTCAAACGGGACC | 117 |
| chr2:143715538-143717552 | p2 | (TA)7 | CCGGTAACCGTAGGAAAACC | TTAACGAGTCAAACGGGACC | 117 |
| chr2:143715538-143717552 | p2 | (TA)7 | CCGGTAACCGTAGGAAAACC | TTAACGAGTCAAACGGGACC | 117 |

| Marker ID | SSR type | SSR motif | FORWARD PRIMER (5'-3') | REVERSE PRIMER (5'-3') | size (bp) |
|---|---|---|---|---|---|
| chr2:143765306-143767370 | p2 | (TA)32 | ACCCACCCATTCATGTTGAT | CATATGCGCATTCTCTGGTG | 246 |
| chr2:143827324-143829360 | p2 | (AT)18 | ACCGTTTGTAAATGGGGTCA | TCACAAAGTCGCCACAAAAG | 160 |
| chr2:143886390-143888410 | p2 | (AT)10 | TTAACTTGGCATGTTGCGTC | ACATGAGCCCGCTTCATTAG | 120 |
| chr2:144080202-144082217 | p3 | (ACA)5 | AACTTGCAAGCAGACAAGCA | GTTGGCCTCAAGCTGACTTC | 269 |
| chr2:144080202-144082217 | p5 | (TGACA)4 | AACTTGCAAGCAGACAAGCA | GTTGGCCTCAAGCTGACTTC | 269 |
| chr2:144154847-144156865 | p6 | (TAACCC)3 | ATCGATTGTCCATCCCGATA | TGACCAAAGTGGACAGGTGA | 260 |
| chr2:144175432-144177446 | p2 | (AC)7 | CAGATGCGTATCGGATCCTT | TTTTTCTTAAAGCAGCGGGA | 231 |
| chr2:144175432-144177446 | p4 | (AATA)4 | CAGATGCGTATCGGATCCTT | TTTTTCTTAAAGCAGCGGGA | 231 |
| chr2:144185921-144187961 | p2 | (TA)20 | TCCCAAAGTATCCCCATCAA | AAATGACATGGCGGAATAGC | 227 |
| chr2:144196191-144198259 | p2 | (TA)34 | ATTTTGATGGTCAGGCTTGG | CAAGAGTCAACAAGGACGCA | 122 |
| chr2:144250412-144252427 | p5 | (AAAAT)3 | TCTTGAAAAGTACCGCCCAC | TTTCGGCCATCCTAAATGTC | 185 |
| chr2:144251667-144253685 | p2 | (TA)9 | TTTGTCATGCAAACTCCCAT | TGCCTGTATGCATCGCTTAG | 158 |
| chr2:144354021-144356039 | p2 | (TG)8 | ACACACACTTCGGTCGTTCA | TTGCATTTTGGAGTTAGGGC | 285 |
| chr2:144354021-144356039 | p2 | (TG)9 | ACACACACTTCGGTCGTTCA | TTGCATTTTGGAGTTAGGGC | 285 |
| chr2:144354021-144356039 | p2 | (TG)8 | ACACACACTTCGGTCGTTCA | TTGCATTTTGGAGTTAGGGC | 285 |
| chr2:144354146-144356162 | p2 | (TG)8 | ACACACACTTCGGTCGTTCA | TTGCATTTTGGAGTTAGGGC | 160 |
| chr2:144354146-144356162 | p2 | (TG)9 | ACACACACTTCGGTCGTTCA | TTGCATTTTGGAGTTAGGGC | 160 |
| chr2:144354146-144356162 | p2 | (TG)8 | ACACACACTTCGGTCGTTCA | TTGCATTTTGGAGTTAGGGC | 160 |
| chr2:144490736-144492751 | p3 | (GTG)5 | TTTCGAAAACCAAAAGGTCG | ATCCCAGTCATTTGCCACTC | 130 |
| chr2:144511927-144513965 | p2 | (AT)19 | TCCATTCCTTTTGGTGCTTC | GCAGCCTTACCTTGCATTTC | 118 |
| chr2:144521408-144523423 | p5 | (TTGGT)3 | GGGTTTTGTTTTTCGGGTCT | TTCAAACCATGTCCAAACGA | 167 |
| chr2:144540030-144542048 | p6 | (TTTTTA)3 | CAAAAGAGGAGGTGGCAAAA | AATCAGGCCAATGACTTTCG | 219 |
| chr2:144540030-144542048 | p3 | (CAC)5 | CAAAAGAGGAGGTGGCAAAA | AATCAGGCCAATGACTTTCG | 219 |
| chr2:144622360-144624384 | p2 | (AT)12 | CCTCTCAGACGCAAATCACA | CTCGCAGGTACAGACGTTCA | 188 |
| chr2:144640066-144642081 | p3 | (TGT)5 | AGTGTTTGTACGCTGCGTTG | CCCGTTCTATTCCGTTGAAA | 212 |
| chr2:144669506-144671521 | p3 | (TAT)5 | TGAGCCGAGAGTCTTTTGGT | TTTTGTCGACGATAGCCCAT | 198 |
| chr2:144669506-144671521 | p2 | (TA)11 | TGAGCCGAGAGTCTTTTGGT | TTTTGTCGACGATAGCCCAT | 198 |
| chr2:144722470-144724488 | p6 | (CTCCGA)3 | TTTGTCTCCTCTGCCGAACT | TCCCGAGTTCAAATGAAAGG | 131 |
| chr2:144757981-144759996 | p5 | (TTATT)3 | TGGACCAACCAGACATCAAA | TGCATGGTCACCAAGTTCAT | 161 |
| chr2:144813490-144815510 | p5 | (CATAG)4 | TCCGATTCACAAAGTCCACA | GCTTCACATCCTCTCGTTCC | 176 |
| chr2:144824087-144826102 | p5 | (ATAGC)3 | CAAGGTCCGATCCACAAAGT | CAAAGTTCACTCCTTTCCCG | 166 |
| chr2:144856560-144858575 | p3 | (TGA)5 | GGGTTGGTTGCTCGTGTAGT | TCTCTCCCAGCGAAACTGAT | 132 |
| chr2:144947465-144949481 | p2 | (TA)8 | CTTTCCCCTTGTCTGCTGTC | AGCAGTAGAGGACGATGGCT | 104 |
| chr2:144989885-144991897 | p2 | (TA)6 | ATTGTCACGACCCGACTAGG | TGGCACTTTAATGTTGCTGC | 126 |
| chr2:144989885-144991897 | p2 | (TA)6 | ATTGTCACGACCCGACTAGG | TGGCACTTTAATGTTGCTGC | 126 |
| chr2:144996565-144998580 | p6 | (CATATA)3 | AAGGTCGGTAACCTCCGATT | TCCGAAATGTTCGAAAAAGG | 298 |

| Marker ID | SSR type | SSR motif | FORWARD PRIMER (5'-3') | REVERSE PRIMER (5'-3') | size (bp) |
|---|---|---|---|---|---|
| chr2:144996565-144998580 | p3 | (CAA)5 | AAGGTCGGTAACCTCCGATT | TCCGAAATGTTCGAAAAAGG | 298 |
| chr2:145400531-145402546 | p5 | (AATTA)3 | CGAACCCTCACCAACAAAGT | ATTAGTGATGGCAAATGGGC | 142 |
| chr2:145416607-145418711 | p3 | (ACA)5 | ATGCAGATTCGTTTTCCACC | TGTGCAACTTCATTTGGCAT | 249 |
| chr2:145442189-145444205 | p2 | (CT)8 | ATTTTACATGCCTTCGGACG | CTCAATTCTGCCCAATGCTT | 185 |
| chr2:145449640-145451655 | p5 | (AATTA)3 | CGAACCCTCACCAACAAAGT | ATTAGTGATGGCAAATGGGC | 142 |
| chr2:145471185-145473197 | p2 | (TG)6 | TCGAGATAGTGGCGAAAGGT | CCTTGGCCCAAAGTGATCTA | 173 |
| chr2:145518523-145520535 | p2 | (TG)6 | TCGAGATAGTGGCGAAAGGT | CCTTGGCCCAAAGTGATCTA | 173 |
| chr2:145570158-145572172 | p2 | (AC)7 | TTTTGAGACGGAAATGCTCC | TATAGGGGAAATGCTGCTGG | 152 |
| chr2:145758386-145760401 | p5 | (AACCG)3 | CCGAATCGACTCGAACAAAT | GCTTGAGGCCAACAAGAAAG | 259 |
| chr2:145762198-145764218 | p2 | (AG)10 | TGGTCACGGAAAAATTGTCA | CGCGGTTGATATGAAAAGAA | 279 |
| chr2:145797860-145799872 | p2 | (CA)6 | CGATCGCTGCATACACAAGT | TCTAAACGCGAACCTTGACC | 165 |
| chr2:145812360-145814375 | p3 | (TAC)5 | GGTGGTTCCCATTTGAACAT | GGACTGGCCGGATTAAGATT | 150 |
| chr2:145816771-145818783 | p2 | (TA)6 | GGAAGGGAGTGCTAGCTGTG | TGCTTGCATTTGGACTTTTG | 280 |
| chr2:145847451-145849469 | p3 | (TCT)6 | AGTCAGCCACTCCAAGCCTA | ATAGGCGTCGCGACTTTAGA | 245 |
| chr2:145892549-145894564 | p5 | (AACCG)3 | CCGAATCGACTCGAACAAAT | GCTTGAGGCCAACAAGAAAG | 259 |
| chr2:145896361-145898381 | p2 | (AG)10 | TGGTCACGGAAAAATTGTCA | CGCGGTTGATATGAAAAGAA | 279 |
| chr2:145932023-145934035 | p2 | (CA)6 | CGATCGCTGCATACACAAGT | TCTAAACGCGAACCTTGACC | 165 |
| chr2:145946523-145948538 | p3 | (TAC)5 | GGTGGTTCCCATTTGAACAT | GGACTGGCCGGATTAAGATT | 150 |
| chr2:145950934-145952946 | p2 | (TA)6 | GGAAGGGAGTGCTAGCTGTG | TGCTTGCATTTGGACTTTTG | 280 |
| chr2:145981614-145983632 | p3 | (TCT)6 | AGTCAGCCACTCCAAGCCTA | ATAGGCGTCGCGACTTTAGA | 245 |
| chr2:145987728-145989756 | p2 | (TC)6 | ATGTCCTGCATCTTGGGTTC | GCATCTCCTGCTCCAGAAAC | 194 |
| chr2:146000785-146002800 | p5 | (GTTCG)3 | GCCTCCAAACAAACCAAAAA | CCGAAACCGAAATAGCCATA | 234 |
| chr2:146002497-146004511 | p2 | (AT)7 | TTTTTCCGTGTTTGCAGTTG | TTTCCCACTTGACCTTTTGG | 218 |
| chr2:146030256-146032271 | p3 | (TGT)5 | TGCCTCCTCTCTGCTTCTTC | CTCTTTCCATAAATGGGGCA | 297 |
| chr2:146033391-146034416 | p2 | (AT)8 | GGACAGCACACCGCTATGTA | ATTGCACAATTGCACAGGAA | 290 |
| chr3:1704-3719 | p5 | (GAAAG)3 | GCTGCCACTACACGTCAGAA | TCCGATGTGGTCAGTTGTGT | 258 |
| chr3:17063-19078 | p5 | (GAAAG)3 | AGGCTTCCACTACGAAACGA | TCCGATGTGGTCAGTTGTGT | 294 |
| chr3:52044-54060 | p2 | (AT)9 | GCAAGTTAACGGAGCAGACC | TCATCGACTGTCACGAGAGG | 271 |
| chr3:52044-54060 | p4 | (GAAA)4 | GCAAGTTAACGGAGCAGACC | TCATCGACTGTCACGAGAGG | 271 |
| chr3:54134-56150 | p2 | (AT)8 | TTCATCCCATGAGCAACAAA | CCATTCCAAGGGTAGAACGA | 237 |
| chr3:54134-56150 | p2 | (TA)6 | TTCATCCCATGAGCAACAAA | CCATTCCAAGGGTAGAACGA | 237 |
| chr3:214396-216444 | p2 | (TA)24 | CAGCATCCATGTCCCTTTTT | ATCTCTGTGCATCTCCGCTT | 216 |
| chr3:258984-260999 | p5 | (ATTGC)3 | ATTGCCTTAGGTTGGGTCCT | GTACGCCAACAATGCATCAC | 113 |
| chr3:258984-260999 | p5 | (TATAT)3 | ATTGCCTTAGGTTGGGTCCT | GTACGCCAACAATGCATCAC | 113 |
| chr3:259561-261576 | p5 | (ATTGC)3 | GGCTGATGTCCGATGGTTAT | ATGCCATGCTATGCAAATGA | 126 |
| chr3:259561-261576 | p5 | (TATAT)3 | GGCTGATGTCCGATGGTTAT | ATGCCATGCTATGCAAATGA | 126 |

| Marker ID | SSR type | SSR motif | FORWARD PRIMER (5'-3') | REVERSE PRIMER (5'-3') | size (bp) |
|---|---|---|---|---|---|
| chr3:270883-272898 | p5 | (TTTTA)3 | TGGTCTTCTCGACCAACCTC | TCACATGGGCACACATTCTT | 212 |
| chr3:270883-272898 | p2 | (TA)6 | TGGTCTTCTCGACCAACCTC | TCACATGGGCACACATTCTT | 212 |
| chr3:480595-482613 | p6 | (CATGAG)3 | CACATCTAAGGGGTGACCGT | AAGTCATGTGGTTCCTTCGG | 222 |
| chr3:531147-533195 | p2 | (AT)24 | CTGATACCGAACCAACCGAT | GCCTTGTTAGGGTTAGGGCT | 123 |
| chr3:531147-533195 | p2 | (AT)6 | CTGATACCGAACCAACCGAT | GCCTTGTTAGGGTTAGGGCT | 123 |
| chr3:698474-700486 | p5 | (CCGGT)4 | TTGCTCCACGCCCTACTACT | CCATGCTAAAACGTCCCACT | 231 |
| chr3:698533-700547 | p5 | (CCGGT)4 | TTGCTCCACGCCCTACTACT | CCATGCTAAAACGTCCCACT | 172 |
| chr3:698553-700595 | p5 | (CCGGT)4 | TTGCTCCACGCCCTACTACT | CCATGCTAAAACGTCCCACT | 152 |
| chr3:698596-700610 | p5 | (CCGGT)4 | TTGCTCCACGCCCTACTACT | CCATGCTAAAACGTCCCACT | 109 |
| chr3:703763-705778 | p5 | (AAAAG)3 | GTTAGGTGGGGATCTGGTGA | TGATGTTACCGGCAGAGATG | 290 |
| chr3:741974-744006 | p2 | (AT)16 | ATTTTCGAACAAATGCTGCC | CGTCAAGCCAAATAAGGGAA | 257 |
| chr3:856839-858860 | p3 | (ATT)7 | CTGCAACAAAATCCCCAAGT | AAATTTGGCGTCATTTAGCG | 146 |
| chr3:869045-871103 | p2 | (AT)29 | CGGAATTCGAAAAACTCCAA | GACCAGTAATTTGAGGGGCA | 155 |
| chr3:917650-919670 | p4 | (TATG)5 | ACTGAGACACGTCATCGCAC | GCCTCTCCTCCGAAATACCT | 252 |
| chr3:944741-946799 | p2 | (AT)29 | CAAGGATTGATCTCCACCGT | TTTTCCCACATCGTTGATGA | 169 |
| chr3:959269-961303 | p2 | (TA)17 | TGTCGAGATTCGAACCAGTG | TACTATCACACGCAGAGGCG | 181 |
| chr3:959269-961303 | p2 | (TA)6 | TGTCGAGATTCGAACCAGTG | TACTATCACACGCAGAGGCG | 181 |
| chr3:961386-963398 | p2 | (TA)6 | ATGAGCGTCACCCTTTATGG | AGTGGCTGGCTTTGTATTCG | 200 |
| chr3:961386-963398 | p2 | (TA)6 | ATGAGCGTCACCCTTTATGG | AGTGGCTGGCTTTGTATTCG | 200 |
| chr3:961386-963398 | p2 | (AT)7 | ATGAGCGTCACCCTTTATGG | AGTGGCTGGCTTTGTATTCG | 200 |
| chr3:1226203-1228223 | p2 | (AC)10 | CCCCAATTGCAAATAACCAC | GAAGCGAGAAGAAGCAATGG | 105 |
| chr3:1233100-1235114 | p2 | (AT)7 | TCGAACTCAGATGCTTGGTG | CATCAGGGGTGGACCTACAT | 144 |
| chr3:1275383-1277398 | p2 | (TA)18 | AATGGCAGGCTACCAGTGAC | TCTATCGATTCGGTTCGGTT | 101 |
| chr3:1275383-1277398 | p5 | (TGGTT)3 | AATGGCAGGCTACCAGTGAC | TCTATCGATTCGGTTCGGTT | 101 |
| chr3:1423009-1425041 | p2 | (AT)16 | ACTCCTGAAACGCGACAAGT | CGAGGAGGGGTTGATAGTGA | 276 |
| chr3:1571257-1573369 | p3 | (TGG)5 | TAATGATGGTGGTGGTGGTG | CAAACCCAAAAATTCCCAAC | 138 |
| chr3:1571257-1573369 | p3 | (TAA)5 | TAATGATGGTGGTGGTGGTG | CAAACCCAAAAATTCCCAAC | 138 |
| chr3:1627932-1629948 | p2 | (AT)8 | GGTGAAACCGGAGATCAAGA | CAAATGACCTGCCAATGATG | 162 |
| chr3:1676025-1678041 | p5 | (GGGCA)7 | TGTAGGAGCCGATAGGGATG | TGTTTAAGATCGTCCTGCCC | 151 |
| chr3:1678321-1680347 | p2 | (AT)13 | CACATGCGTATCAACCCATT | GACATGGCGGAATAGCAAAT | 156 |
| chr3:1678321-1680347 | p2 | (TA)7 | CACATGCGTATCAACCCATT | GACATGGCGGAATAGCAAAT | 156 |
| chr3:1696759-1698771 | p2 | (AT)6 | TTGCTGTCTTTCAACGCATC | CAGCCCAATTTTAAATCCCA | 160 |
| chr3:1721635-1723649 | p2 | (AT)7 | TTGGGTCAAAAAGGATCAGC | GCGCCAAAACTTTGATTGTT | 262 |
| chr3:1729090-1731105 | p5 | (TTTTC)3 | AAGCCCGGAGGTTACTTTGT | AGCCCGAACATAGTGAGACG | 172 |
| chr3:1767369-1769387 | p2 | (TA)9 | AAATGGGGTTTGGTGCAATA | TTTTGGGAGTGATTTGGAGC | 299 |
| chr3:1767369-1769387 | p4 | (TCTT)4 | AAATGGGGTTTGGTGCAATA | TTTTGGGAGTGATTTGGAGC | 299 |

| Marker ID | SSR type | SSR motif | FORWARD PRIMER (5'-3') | REVERSE PRIMER (5'-3') | size (bp) |
|---|---|---|---|---|---|
| chr3:1767485-1769500 | p2 | (TA)9 | AAATGGGGTTTGGTGCAATA | TTTTGGGAGTGATTTGGAGC | 183 |
| chr3:1767485-1769500 | p4 | (TCTT)4 | AAATGGGGTTTGGTGCAATA | TTTTGGGAGTGATTTGGAGC | 183 |
| chr3:1779167-1781182 | p3 | (TAA)5 | AGCGCCATCCTTTTCAACTA | GGACTATTGGCTTTCCGTGA | 142 |
| chr3:1894310-1896342 | p2 | (TA)16 | GTGTCGTTCTGCATGCTTGT | TCCAAACCGAATAATCCGAA | 154 |
| chr3:2008585-2010605 | p5 | (AAGAA)4 | TTTAGTCGGTTGGCCAAAAA | AGCTTTGCTTCACCTTGTGC | 195 |
| chr3:2120624-2122651 | p3 | (AAC)9 | TGCTATAAACCTTTTGGCGG | TCCATGCGGTGTGTACAGTT | 235 |
| chr3:2123594-2125609 | p3 | (ACC)5 | AAATGCATCTCGACCAAACC | TTCCATTGGCTTATTCCAGC | 144 |
| chr3:2166623-2168638 | p3 | (AAT)5 | TGGAGTGGAGAGTAGGTGGG | AAGGCCTCATTTGTTTGCAC | 114 |
| chr3:2166623-2168638 | p3 | (ATT)5 | TGGAGTGGAGAGTAGGTGGG | AAGGCCTCATTTGTTTGCAC | 114 |
| chr3:2166623-2168638 | p2 | (TG)6 | TGGAGTGGAGAGTAGGTGGG | AAGGCCTCATTTGTTTGCAC | 114 |
| chr3:2166623-2168638 | p2 | (TG)6 | TGGAGTGGAGAGTAGGTGGG | AAGGCCTCATTTGTTTGCAC | 114 |
| chr3:2212610-2214660 | p2 | (AT)25 | ATCATTGCCTGTTTCAAGCC | CCATTTATGAGGGAGCCAAA | 136 |
| chr3:2293736-2295778 | p2 | (TA)14 | AGGGAGCGTAGGTGATGATG | CTCTCTTCGCTCCCCTCTTT | 280 |
| chr3:2293736-2295778 | p5 | (TCGGT)3 | AGGGAGCGTAGGTGATGATG | CTCTCTTCGCTCCCCTCTTT | 280 |
| chr3:2293847-2295859 | p2 | (TA)14 | AGGGAGCGTAGGTGATGATG | CTCTCTTCGCTCCCCTCTTT | 169 |
| chr3:2293847-2295859 | p5 | (TCGGT)3 | AGGGAGCGTAGGTGATGATG | CTCTCTTCGCTCCCCTCTTT | 169 |
| chr3:2332179-2334197 | p6 | (GTTTTT)3 | AACTGAGCCGAGTCATTGCT | ACAGCCTCGCCTTTAACAGA | 262 |
| chr3:2379855-2381881 | p2 | (AT)13 | CAACTCGATTCCAAGCCTTC | GCTCTGGAAACGGCTAGATG | 196 |
| chr3:2412323-2414335 | p3 | (CTT)6 | AAACCCTTTCCTAGGGAGCA | ACCTCCAGCTTAGGCGTTTT | 194 |
| chr3:2412323-2414335 | p2 | (TA)6 | AAACCCTTTCCTAGGGAGCA | ACCTCCAGCTTAGGCGTTTT | 194 |
| chr3:2412323-2414335 | p5 | (GAACC)3 | AAACCCTTTCCTAGGGAGCA | ACCTCCAGCTTAGGCGTTTT | 194 |
| chr3:2448486-2450500 | p2 | (TA)7 | CAGATCTCGGTGAGGATGGT | CCAAGTCCATGAATTCGCTT | 201 |
| chr3:2461933-2463945 | p2 | (AG)6 | CCCATCCAAACAGGCTCTAA | CCATTTGCCCAACGACTATT | 240 |
| chr3:2473521-2475536 | p5 | (AAAAT)3 | TCATTTCGTCAAAATTGCCA | ACTACTCGCTCATTGCGGAT | 198 |
| chr3:2625259-2627275 | p2 | (TA)8 | CCGGCTTTAGCTTGCAATAC | TACGGCCTGCTAAGAGGAAA | 265 |
| chr3:2628665-2630679 | p2 | (TA)7 | TAGCCCACTGACCTTTCACC | ACCTTGGCCAAATCAAAGTG | 249 |
| chr3:2628665-2630679 | p2 | (AT)10 | TAGCCCACTGACCTTTCACC | ACCTTGGCCAAATCAAAGTG | 249 |
| chr3:2804162-2806177 | p5 | (ATTTT)3 | TTTGTGAATTGGGTCAGCAA | CATATCGTTGGTCGGAGGTC | 134 |
| chr3:2880753-2882769 | p2 | (AT)8 | CACTGCCCACTTCAAAAACA | GCCTGTGGTTCTGAGGGTTA | 281 |
| chr3:2886749-2888767 | p2 | (AT)9 | ACCACACAACAAAACTGGCA | TAGCCAAGGCATCAAAACCT | 252 |
| chr3:2886749-2888767 | p2 | (AT)9 | ACCACACAACAAAACTGGCA | TAGCCAAGGCATCAAAACCT | 252 |
| chr3:2981063-2983083 | p5 | (ATTGT)4 | GTGCTCAAAACGAGAAAGGG | CGATGATGCTCCTCTCCATT | 234 |
| chr3:3042144-3044164 | p3 | (AAT)10 | TTGGTTCGGTATCGGGTTAG | TGAAGTGATCACCATCGGAA | 111 |
| chr3:3042144-3044164 | p5 | (AAACC)4 | TTGGTTCGGTATCGGGTTAG | TGAAGTGATCACCATCGGAA | 111 |
| chr3:3107340-3109356 | p2 | (GT)8 | CCGACGTCCACAGGTAATCT | TATGGGAAAACACGGACAT | 187 |
| chr3:3159246-3161290 | p2 | (AT)22 | TCCAATTTCTTTGCCGATTC | AATTTCAACGTGCTTCGTCC | 103 |

| Marker ID | SSR type | SSR motif | FORWARD PRIMER (5'-3') | REVERSE PRIMER (5'-3') | size (bp) |
|---|---|---|---|---|---|
| chr3:3162021-3164041 | p2 | (TA)10 | CTTGGGGTCATTTGTGCTTT | TTGCATGGGTTGTCTTTCAA | 218 |
| chr3:3233354-3235370 | p2 | (TA)8 | CACCTTCTTATTGGCGCATT | AGTCACATGCAGTCAGGCAC | 233 |
| chr3:3266579-3268597 | p2 | (AT)6 | AGTCGTTGGGCAGATAATGC | GCCGAATTATCCCATCCTTT | 140 |
| chr3:3266579-3268597 | p2 | (GT)9 | AGTCGTTGGGCAGATAATGC | GCCGAATTATCCCATCCTTT | 140 |
| chr3:3314387-3316405 | p2 | (TA)9 | TACCAGCAACACTGCTGAGG | TGTGATGCAATTACGGAGGA | 298 |
| chr3:3335811-3337829 | p2 | (TG)9 | TCCGACCCATGAAAGGTAAG | TTCTGATATTGCCGAGGCTT | 230 |
| chr3:3393421-3395436 | p5 | (AATTT)3 | TTATCCCAGGTTTGGTTGGA | GCTCGTTTGGTTGGGAAATA | 117 |
| chr3:3420402-3422417 | p3 | (GTA)5 | TGCTGAATGTGGTGAAGGAG | CCATGGTGAAATGCACTGAC | 100 |
| chr3:3420402-3422417 | p3 | (GTA)5 | TGCTGAATGTGGTGAAGGAG | CCATGGTGAAATGCACTGAC | 100 |
| chr3:3510526-3512541 | p5 | (TTTTA)3 | TGGAAAAGCCCTTCTCATTG | TGCAACTTGTATTGACGGGA | 142 |
| chr3:3518251-3520269 | p2 | (TA)6 | GAGGAAAAGATGCACCCGTA | AAAGGGTTTGTTTTTGCGTG | 192 |
| chr3:3518251-3520269 | p6 | (TTCATC)3 | GAGGAAAAGATGCACCCGTA | AAAGGGTTTGTTTTTGCGTG | 192 |
| chr3:3543163-3545175 | p2 | (TA)6 | CAATTCGCAGGATTGTCCTT | TGGTTCAAACCCAGAACCTC | 252 |
| chr3:3543163-3545175 | p3 | (ATA)5 | CAATTCGCAGGATTGTCCTT | TGGTTCAAACCCAGAACCTC | 252 |
| chr3:3543856-3545871 | p2 | (TA)6 | TATCGGGTCTGGCATACACA | TAGTGGCAGAGCCAGGATTT | 194 |
| chr3:3543856-3545871 | p3 | (ATA)5 | TATCGGGTCTGGCATACACA | TAGTGGCAGAGCCAGGATTT | 194 |
| chr3:3559935-3561995 | p2 | (AT)7 | ATCCAAATCCGGGGTAAAAG | TTATGGGTGCCTTGCCTTAC | 108 |
| chr3:3585634-3587646 | p2 | (AT)6 | TCGGAGCTCACAATTCAAAA | CATGGCATCATGTTGTGTGA | 263 |
| chr3:3682991-3685003 | p2 | (AT)6 | TTCTGTGTGCCAAGTCTGCT | ATTACGTACCGTCACGACCC | 174 |
| chr3:3692140-3694204 | p2 | (TA)32 | ACCCACTCACCACCTAACCA | GCCAGCTGCCAACCTACTAC | 154 |
| chr3:3731730-3733814 | p2 | (CT)10 | ACCATCTGGTAGATGAGGCG | CCTGGCTCCCTCTAAATTCC | 128 |
| chr3:3740550-3742565 | p5 | (CATAT)3 | AGTTCCGCTAGGGTGTGATG | GGCTTCATCAGCCGATATGT | 268 |
| chr3:3744762-3746777 | p2 | (AG)6 | CCAAGTGTAAGCCCTCCAAA | CACATACAGGCCCAAGACCT | 139 |
| chr3:3744762-3746777 | p3 | (GTT)5 | CCAAGTGTAAGCCCTCCAAA | CACATACAGGCCCAAGACCT | 139 |
| chr3:3811348-3813364 | p2 | (AT)8 | CAACAATGGAAGGTGTGTCG | GAAGCGCGTTCGCTATACTC | 198 |
| chr3:3822209-3824224 | p3 | (ACA)5 | ATTCCTGACACTGGGAATGC | ACACCTCAGCGTTGCTCTTT | 118 |
| chr3:3942535-3944547 | p2 | (TA)12 | AAATTTCCGTTGCTTTTGGA | GTGAGTTTGGGAAATGGTGG | 289 |
| chr3:4002080-4004095 | p5 | (TTGTC)3 | CCACCGAAACTTAAAGGCAA | CGCATCTTGTCTTCCACTGA | 197 |
| chr3:4002144-4004159 | p5 | (TTGTC)3 | CCACCGAAACTTAAAGGCAA | CGCATCTTGTCTTCCACTGA | 133 |
| chr3:4080496-4082508 | p2 | (TA)6 | CATGCTTCGTGGTCAGTCAC | TTACGCATGTGGCAATGAAT | 290 |
| chr3:4080496-4082508 | p2 | (AT)6 | CATGCTTCGTGGTCAGTCAC | TTACGCATGTGGCAATGAAT | 290 |
| chr3:4080496-4082508 | p2 | (AT)7 | CATGCTTCGTGGTCAGTCAC | TTACGCATGTGGCAATGAAT | 290 |
| chr3:4102455-4104479 | p2 | (AT)8 | CCTGGGGCTGAGATTATCAA | CCGTGTTTGCATGGAGTAGA | 124 |
| chr3:4123829-4125843 | p2 | (TA)7 | GAGTGAGCAGTGATCCCACA | AAATTGGGAAGAAACATGCG | 165 |
| chr3:4136573-4138585 | p2 | (CA)6 | TCGCATGTGTGACATCTTCA | TCTGCTGACAGAAAAGGGCT | 152 |
| chr3:4158237-4160257 | p5 | (TAATT)4 | CTTCAAATGGGCTGGTGTTT | TTTGCATGGCTTGGTATTCA | 238 |

| Marker ID | SSR type | SSR motif | FORWARD PRIMER (5'-3') | REVERSE PRIMER (5'-3') | size (bp) |
|---|---|---|---|---|---|
| chr3:4183760-4185772 | p2 | (CA)6 | GGGAAAGGAAGGATGGGTTA | AGAAATTGACACAGCGACCC | 210 |
| chr3:4315770-4317828 | p3 | (TTG)6 | TTGCTTTGAACTGCTTGTGC | ACAATGTGGAGTCTGGGGAG | 299 |
| chr3:4315770-4317828 | p2 | (AT)29 | TTGCTTTGAACTGCTTGTGC | ACAATGTGGAGTCTGGGGAG | 299 |
| chr3:4315770-4317828 | p2 | (AT)12 | TTGCTTTGAACTGCTTGTGC | ACAATGTGGAGTCTGGGGAG | 299 |
| chr3:4401102-4403126 | p2 | (GT)6 | ATATGCTCGCTGTAGGCGTT | AGAAAATCTGCTCGCGTGTT | 217 |
| chr3:4424553-4426579 | p2 | (TA)13 | AGCATATTGTTCTCCGTGGG | GCGGCAATGAATCCACTATT | 127 |
| chr3:4523571-4525633 | p2 | (TA)31 | AAATGGGCATGGAATGTTGT | TCCACCTATCCAAACAAGCC | 225 |
| chr3:4543537-4545599 | p2 | (TA)31 | AAATGGGCATGGAATGTTGT | TCCACCTATCCAAACAAGCC | 225 |
| chr3:4584063-4586081 | p3 | (AAT)6 | ACCTGGCGATGGTTGATTAG | AAAGACGGAAAAGTGCATGG | 223 |
| chr3:4711367-4713415 | p6 | (CATGTT)3 | ACAGGACGACGCTCTCTGAT | TGAGTGTGCCAAAATGGCTA | 261 |
| chr3:4711417-4713429 | p6 | (CATGTT)3 | ACAGGACGACGCTCTCTGAT | TGAGTGTGCCAAAATGGCTA | 211 |
| chr3:5057856-5059870 | p2 | (TA)7 | CCATCTCCGCAAAGAAACAT | GCATGGACTTCAACCAACCT | 296 |
| chr3:5057856-5059870 | p5 | (CATAT)3 | CCATCTCCGCAAAGAAACAT | GCATGGACTTCAACCAACCT | 296 |
| chr3:5057856-5059870 | p2 | (AC)7 | CCATCTCCGCAAAGAAACAT | GCATGGACTTCAACCAACCT | 296 |
| chr3:5057856-5059870 | p2 | (TA)8 | CCATCTCCGCAAAGAAACAT | GCATGGACTTCAACCAACCT | 296 |
| chr3:5061096-5063153 | p3 | (AAT)19 | AGGCCTTAGATTTGGTGGGT | CAGCGAGTTGGGAATACGAT | 242 |
| chr3:5189091-5191106 | p3 | (AAC)5 | TGCAAGCTCTTTGGACACAC | GCAGAGTGTGCATTATGCGT | 237 |
| chr3:5205547-5207562 | p3 | (CTT)5 | AATTAATGCAAAACGCGGAG | TCTTCAATGAGCTGAGGGGT | 261 |
| chr3:5275223-5277241 | p3 | (GAA)6 | GGCATTCTTGGACTTGGAAA | CCCTTCAAATAGCGCGTAAG | 219 |
| chr3:5281221-5283235 | p2 | (TA)7 | CAACAGCTTCTTTGGCAACA | TAAACTTTTCCCAACGGCTG | 160 |
| chr3:5301776-5303802 | p2 | (TA)8 | AACGTCCTGTCCTGGTCAAC | CAGGGAGCCAACCTTTACAA | 148 |
| chr3:5345574-5347592 | p6 | (CAACTC)3 | TAAACTTCGGGCATTTTTGC | TTACTGCATTGCCCACACAT | 110 |
| chr3:5472717-5474735 | p6 | (ACCCCC)3 | GGCTACCCTCGAATTCTTCC | AATTCGTCGAACCACCATTC | 147 |
| chr3:5504852-5506870 | p6 | (ACCCCC)3 | GGCTACCCTCGAATTCTTCC | AATTCGTCGAACCACCATTC | 147 |
| chr3:5546274-5548298 | p3 | (AAT)8 | TATGAGGGGGTTTATGCTGC | TATCCCAAGGGAATTGTCCA | 128 |
| chr3:5566581-5568596 | p3 | (TGT)5 | ACTTTGCTGCTTTGATGGCT | TCCACCTCAGCATCCCTAAC | 100 |
| chr3:5593670-5595690 | p5 | (ATAAA)4 | CAACCGATTTGAAGGGAAAA | CAAGCAATTTGAGGGGAAAA | 267 |
| chr3:5593670-5595690 | p5 | (TAAAA)3 | CAACCGATTTGAAGGGAAAA | CAAGCAATTTGAGGGGAAAA | 267 |
| chr3:5681325-5683341 | p2 | (TA)6 | TTGGGATGGGAGGTAAGAAA | GTGGAGGTGGCAATTTGAGT | 219 |
| chr3:5681350-5683402 | p2 | (TA)6 | TTGGGATGGGAGGTAAGAAA | GTGGAGGTGGCAATTTGAGT | 194 |
| chr3:5681410-5683422 | p2 | (TA)6 | TTGGGATGGGAGGTAAGAAA | GTGGAGGTGGCAATTTGAGT | 134 |
| chr3:5681423-5683443 | p2 | (TA)6 | TTGGGATGGGAGGTAAGAAA | GTGGAGGTGGCAATTTGAGT | 121 |
| chr3:5887084-5889096 | p2 | (TC)6 | GGGGGCATTTTATGAACCTT | TGGAGTCAGTGTGGATCGAG | 101 |
| chr3:5887084-5889096 | p2 | (AG)10 | GGGGGCATTTTATGAACCTT | TGGAGTCAGTGTGGATCGAG | 101 |
| chr3:5897960-5899975 | p5 | (ATTAA)3 | TTCCAATGCTCTCCACCTTC | GGAAGACTGGGTCAATTCCA | 155 |
| chr3:5996351-5998366 | p5 | (TTCTC)3 | ACAGCCCGAGATATGTACGC | TAGTGGCCTATTTTGTCCCG | 195 |

| Marker ID | SSR type | SSR motif | FORWARD PRIMER (5'-3') | REVERSE PRIMER (5'-3') | size (bp) |
|---|---|---|---|---|---|
| chr3:6015514-6017535 | p3 | (TAT)7 | CGCTCCTATGAGATTTTGGC | AATGCAGTAGGATTGGTGCC | 168 |
| chr3:6033347-6035365 | p6 | (ATATGT)3 | AAGCGGAAAAGGTATGGGTT | TGACGTCACTTGTGTGGGTT | 162 |
| chr3:6072319-6074346 | p3 | (AAC)9 | GGTTAAAGGCTCAAAAGGGG | TGTGCCATGTGTGGTAAGGT | 283 |
| chr3:6100301-6102325 | p2 | (AT)6 | TGGTTCCTTCCGATCAGTTC | ATGATGCAACACATGCGACT | 123 |
| chr3:6104367-6106389 | p2 | (TA)12 | CCGCCTAATTCTTTGCTCAC | AGCGGGATTGAAAGGAATTT | 235 |
| chr3:6104367-6106389 | p2 | (AT)11 | CCGCCTAATTCTTTGCTCAC | AGCGGGATTGAAAGGAATTT | 235 |
| chr3:6136296-6138324 | p2 | (AT)32 | TTCGGCCACCTTTTTAAGTG | TAATCCGAAGTGTTAGCCCG | 116 |
| chr3:6136296-6138324 | p2 | (TA)14 | TTCGGCCACCTTTTTAAGTG | TAATCCGAAGTGTTAGCCCG | 116 |
| chr3:6151637-6153655 | p3 | (GTT)6 | TGTGGAATTGGGGAAGAAAG | CCATACACGACCCAACTTCC | 203 |
| chr3:6163087-6165103 | p4 | (AAGA)4 | CCACAAAGCTGCACAAAGAA | CACCAGCACTGGAAGTGAGA | 217 |
| chr3:6168406-6170426 | p4 | (TTTG)5 | TTGGCAGCCATATAGCACAA | CTGCTCATAAAGTAGCCGCC | 284 |
| chr3:6186036-6188051 | p5 | (AAGAG)3 | TTTGATGACTAACGGGCACA | TTTCTTCTCGCCCGTCTAAA | 188 |
| chr3:6225282-6227300 | p6 | (CTTTTT)3 | AGGTTTGGTGGAAGAGAGCA | CCTTTAAGCGCACCTTTGAC | 204 |
| chr3:6234803-6236823 | p2 | (AT)10 | TAACCACCAAACGATGCAAA | GCATTCTGCGGGTCATATTT | 259 |
| chr3:6334540-6336558 | p3 | (TTG)6 | CAATGAAAAGTTGCTTGGCA | TCCTGAAGATAAGCGCCACT | 138 |
| chr3:6373511-6375523 | p2 | (AT)6 | GCTTCCCCATAGAAACTCCC | AAGAAGTGGCGTAAGCTGGA | 264 |
| chr3:6407714-6409726 | p2 | (AT)6 | GCTTCCCCATAGAAACTCCC | AAGAAGTGGCGTAAGCTGGA | 264 |
| chr3:6496588-6498600 | p2 | (TA)6 | GTAAGCCTCGAGACACAGGG | TTGAGGTCGGGAAGGACTAA | 200 |
| chr3:6534533-6536565 | p5 | (TTGGT)3 | ACCCGACTTTTGTTGGTTTG | GTAGGGGTGTACATGGACCG | 169 |
| chr3:6610696-6612711 | p5 | (TTGTC)3 | TCACGCAATTAGTGGGTCAA | TCGGGGATCAAAAATGGATA | 274 |
| chr3:6610696-6612711 | p3 | (TTG)5 | TCACGCAATTAGTGGGTCAA | TCGGGGATCAAAAATGGATA | 274 |
| chr3:6610799-6612811 | p5 | (TTGTC)3 | TCACGCAATTAGTGGGTCAA | TCGGGGATCAAAAATGGATA | 171 |
| chr3:6610799-6612811 | p3 | (TTG)5 | TCACGCAATTAGTGGGTCAA | TCGGGGATCAAAAATGGATA | 171 |
| chr3:6610808-6612829 | p5 | (TTGTC)3 | TCACGCAATTAGTGGGTCAA | TCGGGGATCAAAAATGGATA | 162 |
| chr3:6610808-6612829 | p3 | (TTG)5 | TCACGCAATTAGTGGGTCAA | TCGGGGATCAAAAATGGATA | 162 |
| chr3:6622125-6624182 | p3 | (ATA)19 | TGTCAGCATCCCCTATCTCC | TGAGATTATTAGGCGGACGG | 172 |
| chr3:6628170-6630185 | p3 | (CTT)6 | AAACTTCCTGCCTGGTCTTG | CCCAAATCCATGCAACTCTC | 176 |
| chr3:6628170-6630185 | p5 | (AAAAT)3 | AAACTTCCTGCCTGGTCTTG | CCCAAATCCATGCAACTCTC | 176 |
| chr3:6628170-6630185 | p5 | (TCTCA)3 | AAACTTCCTGCCTGGTCTTG | CCCAAATCCATGCAACTCTC | 176 |
| chr3:6657410-6659443 | p3 | (GAA)6 | TTCGAGCTGAAAATCATCCC | AGAGTTGACGCCGATTCAGT | 258 |
| chr3:6657473-6659491 | p3 | (GAA)6 | TTCGAGCTGAAAATCATCCC | AGAGTTGACGCCGATTCAGT | 195 |
| chr3:6657492-6659510 | p3 | (GAA)6 | TTCGAGCTGAAAATCATCCC | AGAGTTGACGCCGATTCAGT | 176 |
| chr3:6820865-6822877 | p2 | (TA)7 | TTGCTGCTGTTGTTGAGGTC | TCGTGATTATCGTCATCCCA | 204 |
| chr3:6820865-6822877 | p2 | (TA)6 | TTGCTGCTGTTGTTGAGGTC | TCGTGATTATCGTCATCCCA | 204 |
| chr3:6914194-6916209 | p5 | (AAAAT)3 | TTGGGAGGGTAGGATGTACG | CGACTTTGTTGGGAGGAGAG | 172 |
| chr3:6914194-6916209 | p2 | (AT)10 | TTGGGAGGGTAGGATGTACG | CGACTTTGTTGGGAGGAGAG | 172 |

| Marker ID | SSR type | SSR motif | FORWARD PRIMER (5'-3') | REVERSE PRIMER (5'-3') | size (bp) |
|---|---|---|---|---|---|
| chr3:6936885-6939064 | p2 | (TA)82 | ATGGCATCTCTTCCAGCAGT | GTTTGTCGGCCCATTAAAGA | 241 |
| chr3:7057494-7059521 | p3 | (TTC)9 | GGCAAACTTGGCCAATAAGA | GTGGACATCCTCAGCAGGTT | 174 |
| chr3:7150009-7152029 | p4 | (TTTC)5 | AGAACGACGACTCGAGCATT | ATTTGGTGGGATTGGACTGA | 220 |
| chr3:7217907-7219925 | p3 | (GTG)6 | AAAATTAAGCAACCCCCACC | TTTTTCGTGGGACCAGAGTC | 264 |
| chr3:7266520-7268538 | p3 | (TCT)6 | CCGTCAATCCATAAAACCGT | AGAGAAAGTGCTGCAAGGGA | 289 |
| chr3:7354139-7356159 | p2 | (TA)10 | GATATGCTTTCTTTGAGCCGA | AGAAGTGGGGTTTGGAGAGG | 187 |
| chr3:7375389-7377407 | p2 | (TA)9 | CACGGATTTGGTGGGATTAG | CCGACTGATCTGCCTTCTTC | 267 |
| chr3:7383719-7385784 | p2 | (AT)6 | TTTCTTCGACCTCAATTGGC | TGATGCCCTTATAGGGGTTG | 126 |
| chr3:7398529-7400557 | p2 | (TA)6 | CCCACGTCTTTAAAACCCAA | TGAGCAATCAAATCACGAGC | 184 |
| chr3:7398529-7400557 | p2 | (TA)14 | CCCACGTCTTTAAAACCCAA | TGAGCAATCAAATCACGAGC | 184 |
| chr3:7398529-7400557 | p2 | (TA)8 | CCCACGTCTTTAAAACCCAA | TGAGCAATCAAATCACGAGC | 184 |
| chr3:7398529-7400557 | p2 | (TA)7 | CCCACGTCTTTAAAACCCAA | TGAGCAATCAAATCACGAGC | 184 |
| chr3:7442546-7444561 | p5 | (AAAAT)3 | AGAGGATCTGCTTTCTCCCC | CCTGGCATTGTTTCCTGTCT | 118 |
| chr3:7449340-7451355 | p5 | (TTGTA)3 | TCATGGCATTTTCCTCACAA | CAAAAGTGTCAGCAACGGAA | 141 |
| chr3:7449340-7451355 | p2 | (TA)15 | TCATGGCATTTTCCTCACAA | CAAAAGTGTCAGCAACGGAA | 141 |
| chr3:7490312-7492330 | p2 | (TA)9 | GGCCGGTCAAACACTTAAAA | CTGCTCCTGGTGGAGACATT | 161 |
| chr3:7490312-7492330 | p2 | (TC)7 | GGCCGGTCAAACACTTAAAA | CTGCTCCTGGTGGAGACATT | 161 |
| chr3:7546373-7548419 | p6 | (CTCATG)3 | TGACGTCCCCTTTTTATTGC | TAAAAGTTGACCAAACCCCG | 169 |
| chr3:7546373-7548419 | p3 | (CTT)8 | TGACGTCCCCTTTTTATTGC | TAAAAGTTGACCAAACCCCG | 169 |
| chr3:7548350-7550364 | p2 | (TA)6 | TCCTCCTCCTCCTTCTCCTC | CTGGGGTATCTTTCCGTTGA | 166 |
| chr3:7548350-7550364 | p2 | (AT)7 | TCCTCCTCCTCCTTCTCCTC | CTGGGGTATCTTTCCGTTGA | 166 |
| chr3:7548350-7550364 | p2 | (AT)8 | TCCTCCTCCTCCTTCTCCTC | CTGGGGTATCTTTCCGTTGA | 166 |
| chr3:7549317-7551333 | p2 | (AT)7 | TTCCACGAAAATGCCAAAAT | TTGTTCCACGAAATGCAAAA | 174 |
| chr3:7549317-7551333 | p3 | (CTC)5 | TTCCACGAAAATGCCAAAAT | TTGTTCCACGAAATGCAAAA | 174 |
| chr3:7549386-7551420 | p3 | (CTC)5 | TTCCACGAAAATGCCAAAAT | TTGTTCCACGAAATGCAAAA | 105 |
| chr3:7549492-7551504 | p3 | (CTC)5 | TTGCATTTCGTGGAACAATC | TGTCAATTTCACTTCTGCGG | 157 |
| chr3:7792033-7794048 | p5 | (AAAAC)3 | GGCCTTTGATCTGGTGATGT | GAGCTGCTAACCGGAAAATG | 257 |
| chr3:7935302-7937316 | p2 | (TA)7 | ACTTGGGTAAATTCCCGTCC | CGACCCTCTTTTGCGAGTAG | 148 |
| chr3:7939522-7941554 | p2 | (AT)16 | GAAACCCACCACATCCAAAC | TACGCATGTGGCAATGAATC | 165 |
| chr3:7939522-7941554 | p2 | (TA)19 | GAAACCCACCACATCCAAAC | TACGCATGTGGCAATGAATC | 165 |
| chr3:7998097-8000109 | p2 | (TA)6 | CCGACCCTAGTGATCCTTCA | GGAGCATGGTATAGTCGGGA | 113 |
| chr3:7998097-8000109 | p5 | (TTAAA)4 | CCGACCCTAGTGATCCTTCA | GGAGCATGGTATAGTCGGGA | 113 |
| chr3:8033915-8035927 | p2 | (AC)6 | TTTTTATCAAAACCCTGCCG | GCGCTCTTTAACGAATCAGG | 175 |
| chr3:8039817-8041841 | p3 | (TTC)8 | GAGGCGCATAAGGAAGACAC | TCTTCGCTGATGGTGCATAG | 180 |
| chr3:8049847-8051862 | p5 | (TTCTT)3 | TACCGAAGCAATTCAGGCTT | AAAAGATGAGCAGCCACGTT | 241 |
| chr3:8049847-8051862 | p6 | (AAAAAT)4 | TACCGAAGCAATTCAGGCTT | AAAAGATGAGCAGCCACGTT | 241 |

| Marker ID | SSR type | SSR motif | FORWARD PRIMER (5'-3') | REVERSE PRIMER (5'-3') | size (bp) |
|---|---|---|---|---|---|
| chr3:8097700-8099715 | p3 | (TTG)5 | AAGTGGCGGAAATGAGAATG | CGCATCTTGTCTTCCACTGA | 290 |
| chr3:8126449-8128465 | p2 | (CT)8 | ATGAAACTTTGGCAGCCACT | CCACCCCTATGTCTCCTTCA | 235 |
| chr3:8132184-8134199 | p5 | (TTCGG)3 | CGAGTGGTACAACCACGATG | TTAGCCTTGTGGGATTTTGC | 260 |
| chr3:8133410-8135422 | p2 | (AT)6 | ACAAATTAGGGGTGGGCATT | CCGAAAATCGAAAAACCAGA | 202 |
| chr3:8151572-8153590 | p2 | (AT)9 | TGATCCCCAAATCCAATCAT | TCTGTGTTTTTGTGGCGTGT | 108 |
| chr3:8151572-8153590 | p4 | (AATA)5 | TGATCCCCAAATCCAATCAT | TCTGTGTTTTTGTGGCGTGT | 108 |
| chr3:8187492-8189510 | p2 | (AT)11 | CCCCCTATGGGTAAACATCC | CTTGTCCAGTGGGGTCACTT | 143 |
| chr3:8187513-8189533 | p2 | (AT)11 | CCCCCTATGGGTAAACATCC | CTTGTCCAGTGGGGTCACTT | 122 |
| chr3:8188048-8190070 | p2 | (AT)11 | AGGCCCCCAACATACCTTAG | TGACGTCCCCTTTTTATTGC | 160 |
| chr3:8208899-8210914 | p3 | (AAT)5 | TTGGCTGCTCGATTTTCTTT | GTAGCAGCACCCCAAACAAT | 106 |
| chr3:8260803-8262819 | p4 | (ATGT)4 | TGACGTCCCCTTTTTATTGC | GAGATGGGGCTCACCAGTAA | 157 |
| chr3:8317907-8319922 | p3 | (TAA)5 | ATCGAAATGGGCAGAGATTG | CGTTCCCAAGCCTAATTCAA | 144 |
| chr3:8350107-8352122 | p3 | (TAA)5 | ATCGAAATGGGCAGAGATTG | CGTTCCCAAGCCTAATTCAA | 144 |
| chr3:8409465-8411483 | p3 | (ATA)6 | GTATCCCCAAGCCAAACTGA | TCAGGCAAATAGCAGCAATG | 114 |
| chr3:8445582-8447597 | p3 | (ATA)5 | TTTTCGTGTGGAATCGTCAA | GCGAAGTTTGTGGAAAGAGC | 207 |
| chr3:8558634-8560654 | p2 | (AT)10 | AGAGCTCAAAATATGGCCGA | GATAAGGGGTGCTCGGTACA | 145 |
| chr3:8615084-8617160 | p2 | (AT)38 | AAGAACAGAGCCGAGAAGCA | TGTTCAACAGCAATTTTGGC | 102 |
| chr3:8617987-8620002 | p5 | (AATGC)3 | ATCAAAACCGAACCGAACTG | CGCTCTTGTACCCTTTGACC | 298 |
| chr3:8724970-8727008 | p2 | (AT)19 | TGCAGGTGTACGCCATAAAA | AACATGTGCAATGGTGGCTA | 147 |
| chr3:8724970-8727008 | p2 | (AT)8 | TGCAGGTGTACGCCATAAAA | AACATGTGCAATGGTGGCTA | 147 |
| chr3:8768017-8770029 | p6 | (AACCGG)4 | AGCCCAACTCATTTATCCCC | TGGGCATCCCTAAAATCAAG | 281 |
| chr3:8768017-8770029 | p2 | (TA)6 | AGCCCAACTCATTTATCCCC | TGGGCATCCCTAAAATCAAG | 281 |
| chr3:8768017-8770029 | p2 | (TA)7 | AGCCCAACTCATTTATCCCC | TGGGCATCCCTAAAATCAAG | 281 |
| chr3:8768017-8770029 | p2 | (AT)7 | AGCCCAACTCATTTATCCCC | TGGGCATCCCTAAAATCAAG | 281 |
| chr3:8768322-8770336 | p6 | (AACCGG)4 | GTCAGGACGCGCTTCTACAT | TGGGCATCCCTAAAATCAAG | 106 |
| chr3:8768322-8770336 | p2 | (TA)6 | GTCAGGACGCGCTTCTACAT | TGGGCATCCCTAAAATCAAG | 106 |
| chr3:8768322-8770336 | p2 | (TA)7 | GTCAGGACGCGCTTCTACAT | TGGGCATCCCTAAAATCAAG | 106 |
| chr3:8768322-8770336 | p2 | (AT)7 | GTCAGGACGCGCTTCTACAT | TGGGCATCCCTAAAATCAAG | 106 |
| chr3:8794015-8796030 | p3 | (AAC)5 | GTCTGTCCATGCCAAAAGGT | CAAGGTGAGGCAGAATGTCA | 222 |
| chr3:8976739-8978759 | p4 | (TTAA)5 | CCCTAAAACGACGATCCAAA | TGCGTGAAAAGACCAAACTG | 297 |
| chr3:8996877-8998889 | p2 | (AC)6 | GCCCATTCAAATGGATTCAC | CCGATTCAAATCGAAAATGG | 126 |
| chr3:9018052-9020067 | p5 | (AAATA)3 | ATGAAAACCGTCACCTCCAG | TCTAGAAGCACACGCAGCAT | 101 |
| chr3:9073474-9075489 | p2 | (TA)6 | CCGGGTTGACGGTAGTCTTA | TAATCACTCCAAACCGGACC | 154 |
| chr3:9073474-9075489 | p3 | (ACA)5 | CCGGGTTGACGGTAGTCTTA | TAATCACTCCAAACCGGACC | 154 |
| chr3:9213922-9215946 | p3 | (TGT)8 | CGGAAAGCACAGGGTAATGT | TCGTCTGAAACCATCCACAA | 247 |
| chr3:9224526-9226550 | p3 | (TGT)8 | CGGAAAGCACAGGGTAATGT | TCGTCTGAAACCATCCACAA | 247 |

| Marker ID | SSR type | SSR motif | FORWARD PRIMER (5'-3') | REVERSE PRIMER (5'-3') | size (bp) |
|---|---|---|---|---|---|
| chr3:9288643-9290658 | p5 | (AAAAT)3 | ATCTGATTGCAGTGGGCTCT | GTCCGAGGAACGACAACATT | 125 |
| chr3:9335804-9337819 | p3 | (TAA)5 | AGCTCCATCGAAACTGTCGT | GCCCATTCTGGTGCTAACAT | 170 |
| chr3:9471298-9473319 | p3 | (CTT)5 | ATCGAAAACACTGGAGCCAC | GAGGAAGAGGAGGAGGAGGA | 241 |
| chr3:9471298-9473319 | p3 | (CTT)16 | ATCGAAAACACTGGAGCCAC | GAGGAAGAGGAGGAGGAGGA | 241 |
| chr3:9471322-9473340 | p3 | (CTT)5 | ATCGAAAACACTGGAGCCAC | GAGGAAGAGGAGGAGGAGGA | 217 |
| chr3:9471322-9473340 | p3 | (CTT)16 | ATCGAAAACACTGGAGCCAC | GAGGAAGAGGAGGAGGAGGA | 217 |
| chr3:9519930-9521942 | p2 | (AT)6 | GAGGATGCTCAGGTGGATGT | TGGAGCTGTTCATGTCTTGC | 283 |
| chr3:9546626-9548647 | p3 | (TAA)5 | AAAAGAGGGAAAGGGATGGA | ATTTTGACTTACCGGACCCC | 176 |
| chr3:9546626-9548647 | p3 | (ATA)7 | AAAAGAGGGAAAGGGATGGA | ATTTTGACTTACCGGACCCC | 176 |
| chr3:9546753-9548777 | p3 | (TAA)5 | GGGGTCCGGTAAGTCAAAAT | TCTACCAACTTCCGCACACA | 287 |
| chr3:9546753-9548777 | p3 | (ATA)7 | GGGGTCCGGTAAGTCAAAAT | TCTACCAACTTCCGCACACA | 287 |
| chr3:9546832-9548850 | p3 | (TAA)5 | GGGGTCCGGTAAGTCAAAAT | CGCACACACATCGCATTTAT | 208 |
| chr3:9546832-9548850 | p3 | (ATA)7 | GGGGTCCGGTAAGTCAAAAT | CGCACACACATCGCATTTAT | 208 |
| chr3:9565187-9567215 | p2 | (TA)13 | TGGAACTCACCACATCCAAA | ATGTGGCAATGAGTCCATCA | 271 |
| chr3:9565187-9567215 | p2 | (TA)6 | TGGAACTCACCACATCCAAA | ATGTGGCAATGAGTCCATCA | 271 |
| chr3:9569621-9571648 | p2 | (TA)7 | CAACCCATTAGAGGGAGCAA | GCATGTGGCCATAAATCCAT | 165 |
| chr3:9625930-9627960 | p2 | (CT)15 | GAATCGACTTTGGGCTGGTA | TTAGGTTTGCTGCTGTGTGC | 155 |
| chr3:9746900-9748932 | p2 | (TA)16 | TTCCATAGAAATTCGCCCAC | GGCTCTTGATGCCAATTGTT | 296 |
| chr3:10037059-10039074 | p5 | (AAACC)3 | GTAGGGGTGTACATGGACCG | AAAACCCGAGAAAACCCAAG | 229 |
| chr3:10153437-10155452 | p3 | (GCA)5 | AATATTTCTCCGCTCAGCCA | GGCGCATCTTAAAGTTTTCG | 282 |
| chr3:10361445-10363460 | p5 | (AATAA)3 | GCCCGACTTGAAAACGTAAA | TAATTTCCCAACCCCAACAA | 235 |
| chr3:10361445-10363460 | p5 | (TTTAT)3 | GCCCGACTTGAAAACGTAAA | TAATTTCCCAACCCCAACAA | 235 |
| chr3:10398921-10400936 | p5 | (ATTTT)3 | CCCTTGGCGAAAAATTACCT | CAGAATTTAAGCGGAGCCAC | 214 |
| chr3:10448461-10450476 | p5 | (ATATG)3 | GGCGTGAAATTAAATCCGAA | CCACCGACATTCAGTGAAAA | 161 |
| chr3:10469758-10471782 | p6 | (GTGGAC)4 | GTTCCCCAACAAGCACAAGT | AACAACGTCCACCGTTTAGC | 231 |
| chr3:10469758-10471782 | p6 | (GTGGAC)3 | GTTCCCCAACAAGCACAAGT | AACAACGTCCACCGTTTAGC | 231 |
| chr3:10731610-10733625 | p5 | (ATCAT)3 | AATCCAAACGGGCTCTAGGT | TTCGAAGTGTGTCAGCCAAA | 284 |
| chr3:10737314-10739329 | p3 | (TGC)5 | GAGGACGATGGTTACCCTGA | TTCCCATTGCCCACATTATT | 296 |
| chr3:10761048-10763063 | p5 | (ATAAA)3 | GGGCGGATCTAGTCATTGTC | AACCAAACAACGCTTATGGC | 280 |
| chr3:10761048-10763063 | p2 | (AT)8 | GGGCGGATCTAGTCATTGTC | AACCAAACAACGCTTATGGC | 280 |
| chr3:10798631-10800649 | p6 | (TTTTTA)3 | ACCCTCGACTTAAGGGGAAA | ATTGGCTCCTCCATCATCTG | 139 |
| chr3:10844613-10846628 | p5 | (AAACC)4 | ATGCATCCAACCGTTCTTTC | GCAATCTCAACCTGACGGAT | 133 |
| chr3:10845428-10847448 | p5 | (AAACC)4 | GTAGGGGTGTACATGGACCG | CGACTTTCGTTGGTTTGGTT | 143 |
| chr3:10883562-10885578 | p5 | (ACCCC)3 | GCCGAGCTCTTATTGCATTC | TTCCAGGGTCATGGTTTGTT | 230 |
| chr3:10883562-10885578 | p2 | (AT)8 | GCCGAGCTCTTATTGCATTC | TTCCAGGGTCATGGTTTGTT | 230 |
| chr3:10956759-10958787 | p2 | (AT)14 | CTAGTGCACTTTGGCCCATT | TGGGACTTAAACCCAACGTC | 297 |

| Marker ID | SSR type | SSR motif | FORWARD PRIMER (5'-3') | REVERSE PRIMER (5'-3') | size (bp) |
|---|---|---|---|---|---|
| chr3:10983674-10985690 | p2 | (TA)8 | AAAAAGGAGTGCCGATCTCA | TCGGTAGCTTAGCATTCGCT | 155 |
| chr3:11024249-11026285 | p3 | (AAT)12 | AAGGAGTGGGGTGTCAGAGA | TCATCATCGTTCCCTTCCTC | 267 |
| chr3:11129631-11131651 | p5 | (AAAAT)4 | GGTGCACAGTTGCATTGAAG | GCTGCTTTCAGATGCCTTTC | 227 |
| chr3:11203177-11205203 | p2 | (TA)7 | CACCCCATAAAGGGTGGAC | CCCCCACCTTCAATCTTTTT | 230 |
| chr3:11203177-11205203 | p2 | (TA)13 | CACCCCATAAAGGGTGGAC | CCCCCACCTTCAATCTTTTT | 230 |
| chr3:11203177-11205203 | p2 | (TA)27 | CACCCCATAAAGGGTGGAC | CCCCCACCTTCAATCTTTTT | 230 |
| chr3:11212532-11214544 | p2 | (AC)6 | CTGCCAGTGTGGACAGAAGA | TGTTTAATAGCCCGCAAACC | 152 |
| chr3:11261479-11263505 | p2 | (TA)7 | CACCCCATAAAGGGTGGAC | CCCCCACCTTCAATCTTTTT | 230 |
| chr3:11261479-11263505 | p2 | (TA)13 | CACCCCATAAAGGGTGGAC | CCCCCACCTTCAATCTTTTT | 230 |
| chr3:11261479-11263505 | p2 | (TA)27 | CACCCCATAAAGGGTGGAC | CCCCCACCTTCAATCTTTTT | 230 |
| chr3:11270834-11272846 | p2 | (AC)6 | CTGCCAGTGTGGACAGAAGA | TGTTTAATAGCCCGCAAACC | 152 |
| chr3:11320044-11322065 | p3 | (AAT)7 | TGGAGGTGTCAGAGCAAGTG | CTTTCGTCGTCATCTCCCAT | 236 |
| chr3:11340861-11342873 | p2 | (AT)6 | TCAAGGGATGTCTCCAGTCC | GATTCGTGACGGGACAAAGT | 190 |
| chr3:11340861-11342873 | p2 | (TA)13 | TCAAGGGATGTCTCCAGTCC | GATTCGTGACGGGACAAAGT | 190 |
| chr3:11350537-11352558 | p3 | (TAA)7 | CACCTCAGCATCTAGGGAGC | TCCTGCCTGGATATGGATTC | 277 |
| chr3:11354022-11356037 | p3 | (TTG)5 | ACTCCCAAAAATCACCCCTC | AACAAGTGCCCTTTCTTCCA | 104 |
| chr3:11397504-11399522 | p3 | (TTG)6 | TTTTGGGTCAATTTTGGAGG | GGTTTGTTGCGTTGGAAACT | 218 |
| chr3:11397504-11399522 | p5 | (TTGTA)3 | TTTTGGGTCAATTTTGGAGG | GGTTTGTTGCGTTGGAAACT | 218 |
| chr3:11601843-11603863 | p3 | (AAC)6 | TGCATTAATGGAACCCGATT | TTTGAGCCTTGGGTATGGAG | 178 |
| chr3:11601843-11603863 | p5 | (AACCA)4 | TGCATTAATGGAACCCGATT | TTTGAGCCTTGGGTATGGAG | 178 |
| chr3:11613695-11615710 | p5 | (AAGAA)3 | TACCAGGTGGCTTTCCAAAC | TTTTTGGGCATTCATTGGTT | 287 |
| chr3:11656513-11658525 | p2 | (GT)6 | TTCATTCCCCAATCGAGAAG | GTGAAGTGGGAAAAACCGAA | 233 |
| chr3:11672570-11674582 | p2 | (TA)6 | GAGAGCGCATATAGCCAAGG | AAAGTCTTCCTCCGGTCCAT | 297 |
| chr3:11672570-11674582 | p2 | (TA)7 | GAGAGCGCATATAGCCAAGG | AAAGTCTTCCTCCGGTCCAT | 297 |
| chr3:11672752-11674766 | p2 | (TA)6 | GAGAGCGCATATAGCCAAGG | AAAGTCTTCCTCCGGTCCAT | 115 |
| chr3:11672752-11674766 | p2 | (TA)7 | GAGAGCGCATATAGCCAAGG | AAAGTCTTCCTCCGGTCCAT | 115 |
| chr3:11944719-11946735 | p2 | (TA)8 | CAGCCCATTTTAGAACTGCC | AGAACGCTCTGAAAAGCAGC | 201 |
| chr3:12034725-12036740 | p3 | (CTG)5 | GTTTTCCCATACCGGGATCT | GGGATGAGTTCACTTTCGGA | 142 |
| chr3:12045546-12047561 | p5 | (TTTTA)3 | AGCTAGTAGGCGCAAGGGAT | TGAGGAAGTCAACGGGATTC | 259 |
| chr3:12063027-12065057 | p2 | (AT)7 | CTCTCATTTTTGGGTTGGGA | TCATGCAAAATTAAAGCCCC | 292 |
| chr3:12063027-12065057 | p3 | (TAA)10 | CTCTCATTTTTGGGTTGGGA | TCATGCAAAATTAAAGCCCC | 292 |
| chr3:12065179-12067251 | p5 | (AAACC)3 | TACACCCCTAGGCACTGGTC | AGAGATTTGCCTTCAAGCGA | 273 |
| chr3:12065179-12067251 | p2 | (CA)6 | TACACCCCTAGGCACTGGTC | AGAGATTTGCCTTCAAGCGA | 273 |
| chr3:12083160-12085178 | p6 | (TTTAGT)3 | TATAAGCCAATCCAAACGGG | CGTACTAGTTGCACAGGCGA | 278 |
| chr3:12125587-12127599 | p2 | (AT)6 | GGTGAGAGACCCCTTTTTCC | CGGGTTTAAATGAAGGAGCA | 294 |
| chr3:12134689-12136713 | p6 | (TATATG)4 | TTGGCCGAATTAGATTTTCG | AACATACCTGCTCCACGTCC | 157 |

| Marker ID | SSR type | SSR motif | FORWARD PRIMER (5'-3') | REVERSE PRIMER (5'-3') | size (bp) |
|---|---|---|---|---|---|
| chr3:12251316-12253328 | p5 | (TTGGT)4 | CGTGTGTCAAGACTCCGAAA | GGGCACCTTTTCAACACCTA | 125 |
| chr3:12254223-12256239 | p5 | (TTGGT)4 | ACCGCATCAAACCGATAAAC | AACCAAACCAAACCAAACCA | 170 |
| chr3:12254223-12256239 | p2 | (AT)6 | ACCGCATCAAACCGATAAAC | AACCAAACCAAACCAAACCA | 170 |
| chr3:12265693-12267708 | p3 | (AAC)5 | CGAACCAAAATATGGAGCGT | CCAAATGTTACTGCTTCGCA | 271 |
| chr3:12287530-12289545 | p5 | (ATTTT)3 | ATCCAGCCAGCCTCTCACTA | GTGCACTACGCACCAGAAAA | 195 |
| chr3:12287530-12289545 | p5 | (TATTT)4 | ATCCAGCCAGCCTCTCACTA | GTGCACTACGCACCAGAAAA | 195 |
| chr3:12384052-12386070 | p6 | (AAAAAT)3 | CCTCCATGGCTATGAGCTTC | TGGCTGATTAGGATTGGTCC | 165 |
| chr3:12504666-12506684 | p2 | (AT)7 | TTCACTCATTCAATCGCGTC | GAGACTGTCCCTCGACTTGC | 295 |
| chr3:12504666-12506684 | p2 | (AC)9 | TTCACTCATTCAATCGCGTC | GAGACTGTCCCTCGACTTGC | 295 |
| chr3:12523185-12525197 | p2 | (CT)6 | TAAAAAGGGGACGTCAGCAC | ATTGGTTGGCAACACTCTCC | 173 |
| chr3:12581483-12583537 | p3 | (TTA)18 | CTGCCATGTAACCACACGTC | TTATGCTGCTGTGAAGGCTG | 145 |
| chr3:12628976-12631006 | p2 | (GA)15 | TTGGAGAACAGTCCCTGAGC | CACTTCCCACAACTCCCAGT | 138 |
| chr3:12773455-12775470 | p3 | (AAG)5 | AGAGCTGGAGTTGGTGGAGA | TTTGTTGGAGCAAATTGCAG | 298 |
| chr3:12773477-12775516 | p3 | (AAG)5 | AGAGCTGGAGTTGGTGGAGA | TTTGTTGGAGCAAATTGCAG | 276 |
| chr3:12773525-12775552 | p3 | (AAG)5 | AGAGCTGGAGTTGGTGGAGA | TTTGTTGGAGCAAATTGCAG | 228 |
| chr3:13017831-13019847 | p2 | (AT)8 | CAACGCTAGCCAAAGGAAAG | ACTTTCGATGTGTTCGGAGG | 204 |
| chr3:13169807-13171828 | p3 | (TGT)7 | CCACAACTGAAGGGCAAGTT | CACTCTCCCACCTTATCCGA | 101 |
| chr3:13243349-13245367 | p6 | (CATGAG)3 | GAAAACACGTAGACGCACGA | TGAGTGCCCCATGCATATAA | 215 |
| chr3:13265230-13267245 | p5 | (CATAT)3 | CTGTGGCAATGCTCAGTGTT | TGGGTTGATATGTTCTGCGA | 186 |
| chr3:13269371-13271398 | p3 | (TGT)9 | AATCTGGTGTTGCCTTTTGG | ACACCATTAGCCCCATCAAA | 225 |
| chr3:13271646-13273661 | p3 | (TGT)5 | ACAATTCAAACCCCATGGAA | GGCTCATGTGAAACATTGGA | 197 |
| chr3:13419278-13421306 | p2 | (AT)6 | TCTGTTGGGGTTGTTTCACA | CGTGGCCTTGTTAGCCTTTA | 175 |
| chr3:13419278-13421306 | p2 | (AT)14 | TCTGTTGGGGTTGTTTCACA | CGTGGCCTTGTTAGCCTTTA | 175 |
| chr3:13419278-13421306 | p2 | (TA)14 | TCTGTTGGGGTTGTTTCACA | CGTGGCCTTGTTAGCCTTTA | 175 |
| chr3:13425811-13427825 | p2 | (AT)14 | GAGGCAATGGACAGAAATGG | TCCCGACTTAGCACAGACCT | 156 |
| chr3:13425811-13427825 | p2 | (TA)14 | GAGGCAATGGACAGAAATGG | TCCCGACTTAGCACAGACCT | 156 |
| chr3:13425811-13427825 | p2 | (AT)6 | GAGGCAATGGACAGAAATGG | TCCCGACTTAGCACAGACCT | 156 |
| chr3:13425827-13427869 | p2 | (AT)14 | GAGGCAATGGACAGAAATGG | TCCCGACTTAGCACAGACCT | 140 |
| chr3:13425827-13427869 | p2 | (TA)14 | GAGGCAATGGACAGAAATGG | TCCCGACTTAGCACAGACCT | 140 |
| chr3:13425827-13427869 | p2 | (AT)28 | GAGGCAATGGACAGAAATGG | TCCCGACTTAGCACAGACCT | 140 |
| chr3:13440733-13442783 | p2 | (AT)11 | GTCACGATGCCATCAACATC | TGTCCTATTAAAGGCGGACG | 110 |
| chr3:13440733-13442783 | p2 | (TA)25 | GTCACGATGCCATCAACATC | TGTCCTATTAAAGGCGGACG | 110 |
| chr3:13446160-13448178 | p6 | (ATAAAA)3 | CCTTCTTTCCGGTCAACAAA | GTGACGGTTTTGAAAAGGGA | 232 |
| chr3:13525820-13527832 | p2 | (AT)6 | CAACGCTAGCCAAAGGAAAG | CTTTCGATGTGTTCGGAGGT | 168 |
| chr3:13591674-13593688 | p2 | (AT)6 | GCATGCATGTTTGCACTTTT | TAATATTGGACCCCCGCATA | 287 |
| chr3:13591674-13593688 | p2 | (GA)7 | GCATGCATGTTTGCACTTTT | TAATATTGGACCCCCGCATA | 287 |

| Marker ID | SSR type | SSR motif | FORWARD PRIMER (5'-3') | REVERSE PRIMER (5'-3') | size (bp) |
|---|---|---|---|---|---|
| chr3:13661031-13663049 | p6 | (TTTCTT)3 | TTACAAAATGGCGCTGACTG | GTCCATCTATCTGCCACGGT | 268 |
| chr3:13671030-13673045 | p3 | (ATG)5 | CAACGTTTGACCAAAATCCC | GCTCTTGTCCACATCAGCAA | 250 |
| chr3:13746093-13748105 | p2 | (GA)6 | ACCGAACCTGCAATCAAAAG | TGTTCCATGTTCAAAGGCAA | 201 |
| chr3:13815562-13817582 | p4 | (AATA)5 | AAGAAGTGGACGACCACAGG | CGTCTTCTCCCCCACTATCA | 106 |
| chr3:13829725-13831803 | p2 | (AT)39 | GAGCTGTGGTGCAGTTGAAA | GGTTTCTCCCGACCCTAGAC | 174 |
| chr3:13957343-13959363 | p4 | (TTTC)5 | CCATGCTGGACTTCAATCCT | CGAGGAATTGGTGCTTCATT | 138 |
| chr3:13964401-13966428 | p3 | (TGT)9 | AGTGGCTCCGACAACTCTGT | TTTCACTTGGCAGCTAAGCC | 210 |
| chr3:13974732-13976744 | p2 | (TG)6 | GCAAGACCCATACCGGAGTA | TTCTAAGGGCGAGACCTCAA | 262 |
| chr3:13974732-13976744 | p5 | (AAGTC)3 | GCAAGACCCATACCGGAGTA | TTCTAAGGGCGAGACCTCAA | 262 |
| chr3:13990752-13992772 | p4 | (ATAC)5 | AGGAGAGCTTAGTGCGGGTA | TGTTAGAGGGGTGAAGTGGG | 173 |
| chr3:13990752-13992772 | p2 | (TA)27 | AGGAGAGCTTAGTGCGGGTA | TGTTAGAGGGGTGAAGTGGG | 173 |
| chr3:14035746-14037758 | p2 | (TC)6 | AATGACGAAATGAAGGACGG | TTGCCTTAAATGCTCCAACC | 255 |
| chr3:14057797-14059831 | p2 | (AT)6 | TTGATGGAAATGCGATGAAA | GGATTCGTGCCAAAGAAAAA | 272 |
| chr3:14057797-14059831 | p2 | (AT)17 | TTGATGGAAATGCGATGAAA | GGATTCGTGCCAAAGAAAAA | 272 |
| chr3:14083011-14085025 | p3 | (GAA)7 | TGCAAAAATCCCAAATCTCC | TCATTCCAGGCAGCTTATCC | 282 |
| chr3:14083011-14085025 | p2 | (GT)7 | TGCAAAAATCCCAAATCTCC | TCATTCCAGGCAGCTTATCC | 282 |
| chr3:14122416-14124431 | p5 | (GGCCC)3 | CGGAACAAACGATGCTGTTA | CAAAAGGGACCAAATAGCCA | 195 |
| chr3:14122416-14124431 | p2 | (AG)21 | CGGAACAAACGATGCTGTTA | CAAAAGGGACCAAATAGCCA | 195 |
| chr3:14131323-14133341 | p3 | (GAA)6 | TTTCCATCAGCAAAGCACAG | TGATCTGAATCAATGAGCGG | 226 |
| chr3:14131323-14133341 | p2 | (GT)7 | TTTCCATCAGCAAAGCACAG | TGATCTGAATCAATGAGCGG | 226 |
| chr3:14131806-14133841 | p3 | (GAA)6 | TTACGCAAATTCACCGAACA | TCATTCCAGGCAGCTTATCC | 129 |
| chr3:14131806-14133841 | p2 | (GT)7 | TTACGCAAATTCACCGAACA | TCATTCCAGGCAGCTTATCC | 129 |
| chr3:14173412-14175427 | p2 | (AG)7 | GGAATGAAGACTCCCAACGA | CTCCATCGATGCTAGCGTTT | 207 |
| chr3:14173412-14175427 | p5 | (GGCCC)3 | GGAATGAAGACTCCCAACGA | CTCCATCGATGCTAGCGTTT | 207 |
| chr3:14247292-14249307 | p3 | (GAA)5 | ATCACCAAAGCACAGAACGA | TCCCCCTAAATAATCCCTGC | 232 |
| chr3:14261447-14263462 | p3 | (GAA)5 | ATCACCAAAGCACAGAACGA | TCCCCCTAAATAATCCCTGC | 232 |
| chr3:14298646-14300688 | p2 | (AT)21 | AGACATGGGCCATCCTAGTG | TCGGGATCTGGTTAGTGAGC | 107 |
| chr3:14311996-14314016 | p5 | (GTGCA)4 | GGCCGGTAATGATGAAAGAA | GCTCATTCGCAAGTGTCGTA | 141 |
| chr3:14429539-14431571 | p2 | (TA)9 | CAATCAACAAGACGTGCAGC | GCGACGAAGAAGCATAGGTC | 100 |
| chr3:14429539-14431571 | p2 | (AT)16 | CAATCAACAAGACGTGCAGC | GCGACGAAGAAGCATAGGTC | 100 |
| chr3:14533188-14535203 | p3 | (TGG)5 | GGAACTCGAATGGGGGTATT | GGCGAGGTGCCAATTTAGTA | 237 |
| chr3:14565887-14567902 | p5 | (CCACC)3 | TTTGCTTGAAGGGCAAGAGT | CTCGGTTTCATTTTAGGCCA | 237 |
| chr3:14565887-14567902 | p5 | (TACAA)3 | TTTGCTTGAAGGGCAAGAGT | CTCGGTTTCATTTTAGGCCA | 237 |
| chr3:14630163-14632178 | p5 | (CTTTT)3 | TAGCAACCAAGGTCCAAACC | TTGTGTATGTTGCCGTGGAT | 154 |
| chr3:14654890-14656908 | p2 | (TA)12 | AAGGGGGAAAGAGACGTGAT | TCGTGGCCCAATTTATTGTT | 258 |
| chr3:14654890-14656908 | p6 | (TTCAGT)3 | AAGGGGGAAAGAGACGTGAT | TCGTGGCCCAATTTATTGTT | 258 |

| Marker ID | SSR type | SSR motif | FORWARD PRIMER (5'-3') | REVERSE PRIMER (5'-3') | size (bp) |
|---|---|---|---|---|---|
| chr3:14686212-14688236 | p2 | (AT)12 | GCCAGTGGCCTTGATATGTT | GGTGGTCACAATTCATGCAG | 247 |
| chr3:14720814-14722838 | p3 | (ACA)8 | CACCAGCAAAAGCAGAATCA | GATTCTTGCTTCCAAGTCGG | 128 |
| chr3:14724036-14726048 | p2 | (AT)6 | TACACTAGCGACAGCCCCTT | CTCCAGCTTTGTCCACATGA | 229 |
| chr3:14729494-14731510 | p4 | (AAAT)4 | ATATGGGCAATGCATTAGGG | TTCCTAAATGCCACACATGC | 233 |
| chr3:14766601-14768653 | p4 | (AGAA)13 | ATGTGTCATTGTCCTGGGGT | GGGTTCCGGAAAGGATAGAG | 214 |
| chr3:14774603-14776621 | p3 | (TGT)6 | CAGGCAAAGCATATCAAGCA | TGATAGTTGGGGGATGAAGC | 206 |
| chr3:14774603-14776621 | p3 | (ACA)5 | CAGGCAAAGCATATCAAGCA | TGATAGTTGGGGGATGAAGC | 206 |
| chr3:14774603-14776621 | p3 | (TGT)6 | CAGGCAAAGCATATCAAGCA | TGATAGTTGGGGGATGAAGC | 206 |
| chr3:14896308-14898326 | p2 | (AT)9 | AGTGTTGAGACACGTGGCAG | TATACAGATAGCCCCGACGC | 147 |
| chr3:15157567-15159588 | p3 | (CAA)7 | ACCAACAGGAATTGCTCCAC | CTTCGGGCTTTGGTGTTAAA | 244 |
| chr3:15184340-15186396 | p2 | (TA)28 | TGTCGTGTTACGCTCAGACC | GGATTAGTGTGGGTGCCATC | 124 |
| chr3:15327533-15329545 | p2 | (AG)6 | CTGAAGCACCCTGAGTGACA | TGCTTGGCTCAGTAATTCCC | 298 |
| chr3:15327533-15329545 | p4 | (AAAT)4 | CTGAAGCACCCTGAGTGACA | TGCTTGGCTCAGTAATTCCC | 298 |
| chr3:15338715-15340730 | p3 | (CAA)5 | AACAAAAGACCACACAGGGC | ATTCTCGTTCATGGGTACGG | 171 |
| chr3:15340620-15342636 | p2 | (AT)8 | GGGCCTCAATTTTTCCATTC | TCGAGGAAGACTAAGGGCAA | 179 |
| chr3:15340620-15342636 | p4 | (AAAT)5 | GGGCCTCAATTTTTCCATTC | TCGAGGAAGACTAAGGGCAA | 179 |
| chr3:15419953-15421973 | p5 | (TTGTA)4 | GCCGGAGTTCAAATTCAAAA | ATCCTTCTTTTGGGGTGGTC | 159 |
| chr3:15452517-15454537 | p5 | (TTGTA)4 | GCCGGAGTTCAAATTCAAAA | ATCCTTCTTTTGGGGTGGTC | 159 |
| chr3:15455166-15457181 | p2 | (AT)7 | GAAAGAGTATGGAACGGGCA | ACGGGCTTTAATATGGGGTC | 285 |
| chr3:15455166-15457181 | p2 | (TA)7 | GAAAGAGTATGGAACGGGCA | ACGGGCTTTAATATGGGGTC | 285 |
| chr3:15455182-15457196 | p2 | (AT)7 | GAAAGAGTATGGAACGGGCA | ACGGGCTTTAATATGGGGTC | 269 |
| chr3:15455182-15457196 | p2 | (TA)7 | GAAAGAGTATGGAACGGGCA | ACGGGCTTTAATATGGGGTC | 269 |
| chr3:15618072-15620087 | p5 | (TTTTC)3 | CTTCGTACCAAATGCACCCT | TAAATTGGTGGCTCGTCACA | 136 |
| chr3:15618072-15620087 | p5 | (GAAAA)3 | CTTCGTACCAAATGCACCCT | TAAATTGGTGGCTCGTCACA | 136 |
| chr3:15627595-15629613 | p3 | (TGT)6 | AGGAGTGGATGTGGAACCTG | ATCATGCAACTCTGAACCCC | 218 |
| chr3:15763662-15765682 | p5 | (AATAC)4 | CCCTTTCTGAACTTTTCCCC | GCAAAAATCACCAAATCCGT | 245 |
| chr3:15780348-15782363 | p3 | (TTG)5 | ACGTTTTCGGATGACCTGAC | CGGAGTTGATGGGGTCTTTA | 242 |
| chr3:15879765-15881781 | p2 | (TA)8 | TGTCATGCCAAGAAGAAACG | AATCAAGAAAAGCACGGTGG | 275 |
| chr3:15879765-15881781 | p3 | (TGT)5 | TGTCATGCCAAGAAGAAACG | AATCAAGAAAAGCACGGTGG | 275 |
| chr3:15970538-15972550 | p2 | (AT)6 | ATAGGACAACCCAAAAGGGG | GTCAATCCAAACGGGCTCTA | 181 |
| chr3:15993412-15995426 | p2 | (TA)7 | TTCGTCCTAGGGGTTGTCTG | GAAGCAATTTCCTCGTAGCG | 220 |
| chr3:16044051-16046069 | p6 | (AAAAAT)3 | AATCGTCACCAACCTATCGC | GGGACTCTAACGGGGAAGTC | 214 |
| chr3:16044121-16046149 | p6 | (AAAAAT)3 | AATCGTCACCAACCTATCGC | GGGACTCTAACGGGGAAGTC | 144 |
| chr3:16061603-16063630 | p3 | (AGA)9 | CAGCCGTGCAAAAACTTGTA | TCTGGAAGGAAAAATGGCAC | 170 |
| chr3:16108596-16110644 | p2 | (AT)24 | CGCTCCCATAATCGTTGAAT | GTGGCTGCTGTGATTGAGAA | 194 |
| chr3:16108596-16110644 | p4 | (TAAT)5 | CGCTCCCATAATCGTTGAAT | GTGGCTGCTGTGATTGAGAA | 194 |

| Marker ID | SSR type | SSR motif | FORWARD PRIMER (5'-3') | REVERSE PRIMER (5'-3') | size (bp) |
|---|---|---|---|---|---|
| chr3:16129136-16131148 | p2 | (CT)6 | TCGAGACGGTTACAGATCCC | TTTGCGGACCGTAGTTTACC | 262 |
| chr3:16260356-16262390 | p5 | (ATTTT)3 | TTGGCATCAGATGGAACAAA | TTATGCTTCGATCAGCAACG | 300 |
| chr3:16260356-16262390 | p2 | (AT)17 | TTGGCATCAGATGGAACAAA | TTATGCTTCGATCAGCAACG | 300 |
| chr3:16275375-16277390 | p5 | (TCGGG)3 | AAAGCGGAAACAACATCAGG | CCTTTGGATCGCTCTTTCAA | 166 |
| chr3:16309059-16311081 | p2 | (AT)8 | TGGTATTGTTATGGGTGGGC | TTGCATTGCCTCATGACCTA | 186 |
| chr3:16319661-16321676 | p3 | (TTG)5 | AGAAGATGAAAGTCGCGGAA | CCCGCATATTGTCTTCCACT | 296 |
| chr3:16353921-16355965 | p4 | (AAAT)7 | TTAGGCAAATCGTGTACCCC | TTCCCGTCTGTTCCATCTTC | 299 |
| chr3:16353921-16355965 | p2 | (AT)9 | TTAGGCAAATCGTGTACCCC | TTCCCGTCTGTTCCATCTTC | 299 |
| chr3:16353921-16355965 | p2 | (TA)22 | TTAGGCAAATCGTGTACCCC | TTCCCGTCTGTTCCATCTTC | 299 |
| chr3:16367127-16369145 | p3 | (CAA)6 | TCACAGTAGGCACTGAACGC | AAAAGCTCGGATTGCTTTGA | 259 |
| chr3:16380188-16382212 | p2 | (AG)8 | TCCAGCCACTCAAGTCCTCT | GCATTTTGGCATTTCCAGAT | 261 |
| chr3:16380188-16382212 | p6 | (CAACCC)4 | TCCAGCCACTCAAGTCCTCT | GCATTTTGGCATTTCCAGAT | 261 |
| chr3:16388242-16390310 | p2 | (TA)6 | GTCTTGTTTTGCGGTGGATT | ACACGCTGTTTTCCAACTCC | 109 |
| chr3:16418072-16420090 | p2 | (AT)6 | TTGGTAAGGTGTGATTGGCA | GAGGCTTTGCAGACAGTTCC | 138 |
| chr3:16418072-16420090 | p6 | (CATTAT)3 | TTGGTAAGGTGTGATTGGCA | GAGGCTTTGCAGACAGTTCC | 138 |
| chr3:16429832-16431852 | p2 | (AT)10 | AGGCAACAACTCCAAACACC | GCATTTTGGCATTTCCAGAT | 282 |
| chr3:16465819-16467857 | p2 | (TA)33 | TTCCCAAAGTATCCCCATCA | AAATGACATGGCGGAATAGC | 148 |
| chr3:16530528-16532543 | p5 | (TTCAT)3 | AGCCATGGATCAGGAAGTTG | ACAAGTGCCATTTGCTCCTT | 168 |
| chr3:16590176-16592196 | p2 | (TA)10 | CGAACATGAAAGCACCACTG | TCAACTTATTGGTGAGCCCC | 164 |
| chr3:16670139-16672154 | p5 | (TAAAA)3 | AGCCACCGTCTTCTTCTTCA | ACAGAGAACGCTCTTCGCAT | 159 |
| chr3:16671843-16673857 | p2 | (AT)7 | TGCCCTTATTTTGGATGGTC | AGGAGATTCGAACCCGAAAT | 231 |
| chr3:16676668-16678680 | p2 | (AG)6 | TTGAACGTGACATGAAGGGA | GCAGCTAGCGGAATGCTAGT | 276 |
| chr3:16676668-16678680 | p2 | (TA)6 | TTGAACGTGACATGAAGGGA | GCAGCTAGCGGAATGCTAGT | 276 |
| chr3:16715013-16717025 | p2 | (CT)6 | GCTCGTTTGGTGGTTCAGTT | TACTCGTTCCCATTTCTGCC | 207 |
| chr3:16749696-16751711 | p5 | (AAAAC)3 | GATGTATTGGAGGGCTGGAA | TCGAGGAAGGAAGTTTTGGA | 223 |
| chr3:16749696-16751711 | p2 | (TA)9 | GATGTATTGGAGGGCTGGAA | TCGAGGAAGGAAGTTTTGGA | 223 |
| chr3:16786559-16788579 | p5 | (TAATA)4 | TTTCCTTGAACCTCCTCCTGT | TTTCTGATTCGTCAAATTGGG | 113 |
| chr3:16844272-16846287 | p5 | (CAAAC)3 | TGTGCAAGGCAAGACAAAAG | ATCCCAATGCCTTGTGTAGC | 247 |
| chr3:16865728-16867743 | p5 | (ACCAA)3 | GGAACTTGAGAACCATGGGA | AACCAAACAACGCGAAAAAC | 168 |
| chr3:16865728-16867743 | p5 | (AAACC)3 | GGAACTTGAGAACCATGGGA | AACCAAACAACGCGAAAAAC | 168 |
| chr3:16865728-16867743 | p2 | (TA)7 | GGAACTTGAGAACCATGGGA | AACCAAACAACGCGAAAAAC | 168 |
| chr3:16996347-16998381 | p2 | (TA)17 | ACTTCCCCCTCCAGCTACAT | TAAGGGATGCAACACAAGCA | 141 |
| chr3:17022079-17024113 | p2 | (TA)17 | ACTTCCCCCTCCAGCTACAT | TAAGGGATGCAACACAAGCA | 141 |
| chr3:17059266-17061296 | p2 | (AT)15 | TCCGACTGTTGGAATGTGAA | TGTGTGGGTATTTCGTTGGA | 191 |
| chr3:17059266-17061296 | p2 | (AT)9 | TCCGACTGTTGGAATGTGAA | TGTGTGGGTATTTCGTTGGA | 191 |
| chr3:17094034-17096082 | p2 | (CT)24 | TTTAGCTCCAGGCACGACTT | AAGAAAATGAAGCGAGCCAA | 169 |

| Marker ID | SSR type | SSR motif | FORWARD PRIMER (5'-3') | REVERSE PRIMER (5'-3') | size (bp) |
|---|---|---|---|---|---|
| chr3:17120875-17122893 | p3 | (TGT)6 | AGACCGACCTTGTTGTACGG | ACTGGAGCATCTGTGCCTCT | 278 |
| chr3:17176933-17178957 | p6 | (TAGCAC)4 | TCCATCAAACCTGCATCAAA | CCCACCACCTATTCATCCTG | 241 |
| chr3:17196465-17198480 | p5 | (AAAAC)3 | CTTTTTCCATATCCACCCGA | TTTATGGTCAAACGCCCACT | 187 |
| chr3:17263718-17265739 | p3 | (AAG)7 | CCCAAATGAGGGAAAACCTT | TATGCGCCCTTTTCTCAGAT | 245 |
| chr3:17392284-17394300 | p4 | (TCTT)4 | ATCGCGAGAAGTCTCCTTCA | AGGCAAAAGAAGAAAACGCA | 189 |
| chr3:17418609-17420635 | p2 | (AT)13 | AGGGCCCAACCTAAGACAAT | ATATTGAGGCTCAGCGGAAA | 206 |
| chr3:17462564-17464580 | p4 | (TATT)4 | AAACGCCGGTAGTGGTGTAG | ACCCACACAACACCCAAAAT | 202 |
| chr3:17462564-17464580 | p6 | (CTTTTT)3 | AAACGCCGGTAGTGGTGTAG | ACCCACACAACACCCAAAAT | 202 |
| chr3:17557494-17559508 | p2 | (AT)7 | AGCTTTGTCAGTCTTGGGGA | GAAGGCGTGTTAGAACCCAA | 256 |
| chr3:17585276-17587291 | p5 | (ATTGC)3 | ATTCCAATCTTGCCCTTGTG | CTATCCTGCTATCCCCGTCA | 222 |
| chr3:17651213-17653238 | p3 | (TCA)8 | GGGGGTTGTGTTTCATCATC | GACGTGGTGCCGGTAGTAGT | 248 |
| chr3:17706440-17708455 | p3 | (TTA)5 | ATACACACCAGCACATGGGA | GTAGCAGGCATGGAAAAGGA | 221 |
| chr3:17726426-17728478 | p2 | (TA)26 | TTCAAAAATCCTCGCATTCC | TTGATCCAGTGTGTGGGCTA | 168 |
| chr3:17778900-17780912 | p2 | (AT)6 | GACAAAAGGTACGGGACCAA | TCCAATGTGGAAAATGCTGA | 162 |
| chr3:17856610-17858624 | p2 | (TA)7 | TGTAACTGTATGCCCGCTGA | CCAAGTGAGGTCTGGAGAGG | 239 |
| chr3:17882750-17884766 | p4 | (TTTC)4 | GGATCAGCTTGGCTGCTATT | TTGTGCAATGGGAGTTTGAA | 221 |
| chr3:18101477-18103503 | p2 | (TA)6 | AGTATCAATCGTGGCTTCGG | TCCGCTTCTGAGTTTTGGTT | 164 |
| chr3:18101477-18103503 | p2 | (AT)13 | AGTATCAATCGTGGCTTCGG | TCCGCTTCTGAGTTTTGGTT | 164 |
| chr3:18124111-18126133 | p2 | (TA)8 | CAAGCCAACAAGGCTACCAT | TAGGGGCGTACCATTCGTAG | 185 |
| chr3:18124134-18126180 | p2 | (TA)8 | CAAGCCAACAAGGCTACCAT | TAGGGGCGTACCATTCGTAG | 162 |
| chr3:18191568-18193610 | p3 | (CTT)14 | TACCATTCGACACACTCCCA | TGAAAGCAAACCAAGACGTG | 188 |
| chr3:18316765-18318781 | p4 | (AGAA)4 | TGAGGGAAGGATTGTCCAAG | AAACTGTCCACGCTCCAGAT | 171 |
| chr3:18437086-18439104 | p6 | (CATATA)3 | CAATTCCATGTCACACCTCG | TGGCCACAGATTTTGTTTGA | 208 |
| chr3:18503329-18505355 | p2 | (AT)13 | TATGCAAGTCGACAAGGCTG | CCGTTGTTCCGGAGTTTAGA | 281 |
| chr3:18596924-18598939 | p5 | (TGCAT)3 | TGGACCCATCTATGACAGCA | TCCCCCACTCAAAAATCAAG | 189 |
| chr3:18621274-18623300 | p2 | (AC)13 | CAGAGAAATCCTGCGGAGAC | TTACAACCCCAAAAATCCGA | 258 |
| chr3:18627024-18629040 | p4 | (TTTG)4 | ACGAGCTAGATCCAGAGCCA | CGGAACAAAACTCTACCCCA | 287 |
| chr3:18639217-18641235 | p3 | (CAA)6 | CCTTTAACCCGGAATCATCA | TCCAAGGCATAATTCCGTTC | 200 |
| chr3:18639217-18641235 | p2 | (CT)9 | CCTTTAACCCGGAATCATCA | TCCAAGGCATAATTCCGTTC | 200 |
| chr3:18681265-18683280 | p3 | (AAC)5 | TGTGTGGTTGAATGATGCCT | CCAGTAGAACCCCAGCATGT | 196 |
| chr3:18802041-18804059 | p2 | (AT)9 | GACCGTCTTTTTGCTCCAAG | TGTTGATGTGCCGTGAATTT | 165 |
| chr3:18843747-18845817 | p2 | (AT)35 | AACCAAAACTCAGAAGCGGA | TTTCGTTAATTTCCCCCACA | 267 |
| chr3:18843747-18845817 | p2 | (TA)8 | AACCAAAACTCAGAAGCGGA | TTTCGTTAATTTCCCCCACA | 267 |
| chr3:18859391-18861403 | p2 | (AT)6 | TATAGGCTGCTTCCGTCCAT | ACTGGTTCAGGCCTTCAATG | 280 |
| chr3:18962300-18964315 | p5 | (AGAAA)3 | GTACCTGCTACCTCCCACCA | CTTTCCCATTCCGTTGAAGA | 208 |
| chr3:19009968-19011983 | p3 | (CAA)5 | ATGTACGCCGACCTTGCTAC | TGTAATGCTTTTCCCCCAAG | 108 |

| Marker ID | SSR type | SSR motif | FORWARD PRIMER (5'-3') | REVERSE PRIMER (5'-3') | size (bp) |
|---|---|---|---|---|---|
| chr3:19009968-19011983 | p5 | (GGGGT)3 | ATGTACGCCGACCTTGCTAC | TGTAATGCTTTTCCCCCAAG | 108 |
| chr3:19035906-19037918 | p2 | (GA)6 | CGCTTCGTCACCGAAATACT | TTTGTGACGTCATCACCGTT | 200 |
| chr3:19088828-19090846 | p6 | (CTAATT)3 | ATTCCGGAGATCATGTTTGG | TTTGTGTTTGGCCGTGAATA | 121 |
| chr3:19136999-19139014 | p3 | (TGT)5 | TATTTCTGCATCCAGAGGGG | CGGGGTTTTGAGCATAAAGA | 119 |
| chr3:19136999-19139014 | p2 | (AT)31 | TATTTCTGCATCCAGAGGGG | CGGGGTTTTGAGCATAAAGA | 119 |
| chr3:19163911-19165927 | p4 | (AAAC)4 | TGAATTGCTTGTTCTGTGGC | GGTCTGGAGAGGGTAGAGGG | 156 |
| chr3:19201148-19203168 | p2 | (AT)10 | TTGCTCCACGCCCTACTACT | ATTCACATTTGGCTGCATCA | 185 |
| chr3:19201148-19203168 | p2 | (AT)16 | TTGCTCCACGCCCTACTACT | ATTCACATTTGGCTGCATCA | 185 |
| chr3:19201148-19203168 | p2 | (AT)6 | TTGCTCCACGCCCTACTACT | ATTCACATTTGGCTGCATCA | 185 |
| chr3:19247137-19249152 | p5 | (TTTAT)3 | TGCCCCTCAAATTAGTGGTC | AGTTTTGCCCTTCGCCTAAT | 239 |
| chr3:19253117-19255143 | p2 | (AT)13 | TTGTCGCATTGTGTGTGAGA | TGGGGTCTAAGGAGGGTAGG | 198 |
| chr3:19336417-19338431 | p2 | (CA)7 | CGGACACATGTTCAGAGGTTT | CTTGTTGGCTCAACACGAAA | 107 |
| chr3:19364029-19366047 | p6 | (AAAGAG)3 | GTATCCACGCACTTCCCCTA | TTACTCTCGGTTCGTGCCTT | 145 |
| chr3:19364029-19366047 | p2 | (TA)33 | GTATCCACGCACTTCCCCTA | TTACTCTCGGTTCGTGCCTT | 145 |
| chr3:19364379-19366445 | p6 | (AAAGAG)3 | TCCATTAAGAAACCCAACGG | CTCTTCCTACCGCAACAAGC | 162 |
| chr3:19364379-19366445 | p2 | (TA)33 | TCCATTAAGAAACCCAACGG | CTCTTCCTACCGCAACAAGC | 162 |
| chr3:19512592-19514646 | p2 | (TC)7 | CTGGTGTCCCATGTTGTCTG | CAGCAGTGTATGGGTTGCAC | 241 |
| chr3:19532702-19534714 | p2 | (GA)6 | TGGGTTCTCAAGAAATTCGC | GGTGGAAGGTCCAAGCATAA | 124 |
| chr3:19585223-19587287 | p2 | (AG)8 | GCCATTCGTTATCCGACAAG | ATGCCGATTATCGTTTCACC | 123 |
| chr3:19585223-19587287 | p2 | (AT)32 | GCCATTCGTTATCCGACAAG | ATGCCGATTATCGTTTCACC | 123 |
| chr3:19593951-19596031 | p2 | (AT)40 | CCTTACTTAGAGCCGAGCGA | ATGCCAGTATTGTAAGCGGC | 202 |
| chr3:19604202-19606266 | p2 | (AG)8 | GCCATTCGTTATCCGACAAG | ATGCCGATTATCGTTTCACC | 123 |
| chr3:19604202-19606266 | p2 | (AT)32 | GCCATTCGTTATCCGACAAG | ATGCCGATTATCGTTTCACC | 123 |
| chr3:19612931-19614943 | p5 | (TTTTA)3 | GACGAACCGCTGTTTTTGTT | ATGCCAGTATTGTAAGCGGC | 227 |
| chr3:19612945-19614961 | p5 | (TTTTA)3 | GACGAACCGCTGTTTTTGTT | ATGCCAGTATTGTAAGCGGC | 213 |
| chr3:19612965-19614997 | p5 | (TTTTA)3 | GACGAACCGCTGTTTTTGTT | ATGCCAGTATTGTAAGCGGC | 193 |
| chr3:19705098-19707110 | p2 | (AG)6 | AGGGTCGAAGATTGTTGGTG | TCCATTAAATGCTGCCCAAT | 241 |
| chr3:19705098-19707110 | p2 | (TA)28 | AGGGTCGAAGATTGTTGGTG | TCCATTAAATGCTGCCCAAT | 241 |
| chr3:19778591-19780657 | p2 | (AT)33 | GAACATCTGCACGCAGAAAA | GCCTCGTCAGCCTTATTTTG | 213 |
| chr3:19778799-19780811 | p2 | (AT)33 | CAAAATAAGGCTGACGAGGC | TAACGTCCCTTCCTGTGGAC | 144 |
| chr3:19778813-19780833 | p2 | (AT)33 | ACAATATGAACCGGTGGAGC | CTAACGTCCCTTCCTGTGGA | 167 |
| chr3:19778855-19780871 | p2 | (AT)33 | ACAATATGAACCGGTGGAGC | CTAACGTCCCTTCCTGTGGA | 125 |
| chr3:19778869-19780903 | p2 | (AT)33 | ACAATATGAACCGGTGGAGC | CTAACGTCCCTTCCTGTGGA | 111 |
| chr3:19782761-19784773 | p2 | (GA)6 | ACCAAGGAAATGTGGAGACG | TAAACCAGGAGAGCTGCCAT | 272 |
| chr3:19820065-19822081 | p2 | (TG)8 | CCGAACAGCAACAACTAGCA | CCGCATTCATGCACATTTAC | 134 |
| chr3:19820065-19822081 | p3 | (TGT)5 | CCGAACAGCAACAACTAGCA | CCGCATTCATGCACATTTAC | 134 |

| Marker ID | SSR type | SSR motif | FORWARD PRIMER (5'-3') | REVERSE PRIMER (5'-3') | size (bp) |
|---|---|---|---|---|---|
| chr3:19828160-19830175 | p5 | (TTAAA)3 | TGCTCCTTCAGCCGTAAGTT | GAGAACAAGAATCGGTGGGA | 113 |
| chr3:19854516-19856531 | p5 | (GTGTC)3 | CCCTGAAAATCTCCAAGCTG | CAAATGGATGTGCCTCATTG | 254 |
| chr3:19901022-19903036 | p2 | (CT)7 | TCCTTGCGCTTACCACTTCT | TGATCACTCTGAATCCGCTG | 210 |
| chr3:19977812-19979824 | p2 | (AT)6 | ACAAATTCAACAAAAGGCGG | TCTTGCCAACGTACCATTCA | 281 |
| chr3:19992334-19994361 | p3 | (TGT)9 | TCTGCGCAATGACAGAAAAC | CTTTGGCCTACCCCTACCTC | 264 |
| chr3:19992334-19994361 | p6 | (CAACTC)3 | TCTGCGCAATGACAGAAAAC | CTTTGGCCTACCCCTACCTC | 264 |
| chr3:19993050-19995068 | p3 | (TGT)9 | GTCTCTCGGAAACAGCCATC | TAAAGACCTGGCCCCACATA | 192 |
| chr3:19993050-19995068 | p6 | (CAACTC)3 | GTCTCTCGGAAACAGCCATC | TAAAGACCTGGCCCCACATA | 192 |
| chr3:20002734-20004749 | p3 | (TTA)5 | CAATGCCAATGAATCAGGTG | AGAAATGGCCTTTGTCATGG | 176 |
| chr3:20019800-20021814 | p2 | (AG)7 | TTTTATGCTCTTCGGGCATC | GCAGCGTTATTGACAAGCAA | 150 |
| chr3:20055111-20057141 | p2 | (TA)6 | TCCCTCCGTTCCAATTTATG | GTGACGCACTACTGTCCCAA | 210 |
| chr3:20055111-20057141 | p2 | (TA)15 | TCCCTCCGTTCCAATTTATG | GTGACGCACTACTGTCCCAA | 210 |
| chr3:20078548-20080560 | p2 | (TG)6 | TACACACTTCATCTTCGCGG | GGTGTTAGGGAAGAGGGAGG | 276 |
| chr3:20078689-20080743 | p2 | (TG)6 | TACACACTTCATCTTCGCGG | GGTGTTAGGGAAGAGGGAGG | 135 |
| chr3:20105573-20107603 | p2 | (TA)6 | TCCCTCCGTTCCAATTTATG | GTGACGCACTACTGTCCCAA | 210 |
| chr3:20105573-20107603 | p2 | (TA)15 | TCCCTCCGTTCCAATTTATG | GTGACGCACTACTGTCCCAA | 210 |
| chr3:20231093-20233159 | p2 | (TA)33 | CATAAATTCAACCAACCCCG | GATTGCGGATTGTGTTGTTG | 229 |
| chr3:20234740-20236768 | p2 | (TA)8 | AGGCACCTGAGCAAACAACT | AGGTTTCCAAAACAAGCCAA | 183 |
| chr3:20238085-20240100 | p5 | (TATTT)3 | AGCAGCTTCGCGATACAAAT | CAAGTGGTTCTAGCGAAGGG | 237 |
| chr3:20245268-20247294 | p2 | (AT)13 | GAAGTGGATCGTCGACCAAG | ATTTTTATGAGCAGGGGCAA | 203 |
| chr3:20354135-20356191 | p2 | (TA)28 | AATCACTTTCGACTTCGCGT | GGCCTTGCCACATAGAAAAA | 261 |
| chr3:20379792-20381806 | p2 | (AG)7 | TTTGATCATGTGGGGAACCT | TTGCAGATCAAAACGCACTC | 130 |
| chr3:20424557-20426573 | p4 | (AAAT)4 | TAATCTTGAGGGGCTGTTGC | ACAAGTTGAACGACCGAAGC | 100 |
| chr3:20424856-20426895 | p4 | (AAAT)4 | GGGACTATTTCGCCATGAAC | AAAGTTAGCCAAGCGACGAA | 124 |
| chr3:20490915-20492936 | p2 | (TA)9 | CAAAAACCCCAACTTCTCCA | GCACAAAGTGCACTCCTGAA | 160 |
| chr3:20490915-20492936 | p3 | (ATT)7 | CAAAAACCCCAACTTCTCCA | GCACAAAGTGCACTCCTGAA | 160 |
| chr3:20494794-20496812 | p2 | (TA)9 | GGGGATGAAATGTGGAAATG | TACGGACGGTAAACCGTGAT | 119 |
| chr3:20536160-20538174 | p2 | (TA)7 | AATTCATCGATGGGCTTGAG | CCCCACCCTAGTTGGGTACT | 245 |
| chr3:20537194-20539210 | p2 | (TA)8 | TATTCAATGCTCACCCACCA | TTGATCTTGCTATCAGGGGC | 170 |
| chr3:20725803-20727818 | p3 | (AAC)5 | CGGTAAGTTTACAACCCCCA | CTATACCCTCCGGTGTTGGA | 139 |
| chr3:20725803-20727818 | p2 | (GA)7 | CGGTAAGTTTACAACCCCCA | CTATACCCTCCGGTGTTGGA | 139 |
| chr3:20744504-20746519 | p5 | (AAACC)3 | GTGTTCATAGACCGGGTTGG | AAAACTGAAAAACCCGAGCA | 124 |
| chr3:20744504-20746519 | p5 | (TTTGA)3 | GTGTTCATAGACCGGGTTGG | AAAACTGAAAAACCCGAGCA | 124 |
| chr3:20744504-20746519 | p3 | (ATT)15 | GTGTTCATAGACCGGGTTGG | AAAACTGAAAAACCCGAGCA | 124 |
| chr3:20757249-20759264 | p2 | (TA)10 | TTGCTCCCTCTAATGGGTTG | AAGTGAAACCCACCACATCC | 113 |
| chr3:20757249-20759264 | p3 | (ATT)5 | TTGCTCCCTCTAATGGGTTG | AAGTGAAACCCACCACATCC | 113 |

| Marker ID | SSR type | SSR motif | FORWARD PRIMER (5'-3') | REVERSE PRIMER (5'-3') | size (bp) |
|---|---|---|---|---|---|
| chr3:20767939-20769951 | p2 | (GT)6 | CGTTGAAAGGGATTTTGGAA | GGTCTCTCCGTATTTTCGCA | 220 |
| chr3:20774604-20776619 | p5 | (AAAAT)3 | GGGAGGGGGAGAATTGTTAG | GAATCGGGAAACGCAATTTA | 105 |
| chr3:20780697-20782709 | p2 | (TA)15 | AATATCAACCGTGGCTTTGG | TCTTCCATTTGGTCCACTCC | 185 |
| chr3:20794987-20796999 | p2 | (AT)6 | CAGTTTGGCCACATTGAGTG | GGTCCTCTGATGTGGTTGCT | 141 |
| chr3:20794987-20796999 | p5 | (GAGTT)3 | CAGTTTGGCCACATTGAGTG | GGTCCTCTGATGTGGTTGCT | 141 |
| chr3:20825398-20827422 | p3 | (CAC)8 | GAAGAAATCGTGGGCTACCA | TTCTGGGGAATTTTCACGAC | 201 |
| chr3:20862358-20864376 | p2 | (AT)17 | CGCATTTAGCCAAGGAAGAC | GCCTTCCTCCGCTTCTAAGT | 174 |
| chr3:20903454-20905468 | p2 | (TA)7 | GTGGGCCCCATACTAAACAA | ATGAGTTTGGAGATGGCGAG | 210 |
| chr3:20903454-20905468 | p2 | (TA)7 | GTGGGCCCCATACTAAACAA | ATGAGTTTGGAGATGGCGAG | 210 |
| chr3:20903454-20905468 | p2 | (TA)6 | GTGGGCCCCATACTAAACAA | ATGAGTTTGGAGATGGCGAG | 210 |
| chr3:20913416-20915434 | p3 | (ATC)6 | CCCAGAGCTAGCCTATCGTG | ATCCTTCGATAGCCATGTCG | 295 |
| chr3:20913578-20915608 | p3 | (ATC)6 | CCCAGAGCTAGCCTATCGTG | ATCCTTCGATAGCCATGTCG | 133 |
| chr3:20944870-20946926 | p2 | (AT)28 | AGGAGCATGCCAAAAGAAGA | TAAAACAGGGGAAATGCTGC | 296 |
| chr3:20944870-20946926 | p2 | (AT)9 | AGGAGCATGCCAAAAGAAGA | TAAAACAGGGGAAATGCTGC | 296 |
| chr3:20993578-20995593 | p3 | (CTT)5 | CAAGGACCTAATCCTTGGCA | TTGAGGGTTTAATTCGCAGG | 148 |
| chr3:21082474-21084514 | p2 | (AT)20 | TTTTTCGAGGCCACTCCTTA | GGTTCGAATCCAGAACCTCA | 169 |
| chr3:21110143-21112158 | p5 | (AAACC)3 | GCCTCTAGTGGGCAAAACAG | CTAAAAATACACCGGCGGAA | 141 |
| chr3:21110143-21112158 | p5 | (AAACC)3 | GCCTCTAGTGGGCAAAACAG | CTAAAAATACACCGGCGGAA | 141 |
| chr3:21118134-21120158 | p3 | (CTT)8 | TTTACCACTGGAGGGACCAG | TGTGCTAAAGCACTTCACGG | 221 |
| chr3:21191791-21193806 | p5 | (GACCC)3 | TGTGGCGTACATTAGGGTGA | AACCCCATCACAACTGGAAA | 275 |
| chr3:21251742-21253757 | p5 | (GTTCG)3 | AAATTGCAAGACTTGTGCCC | TCGGTCAATGATGGATGAGA | 108 |
| chr3:21258565-21260577 | p4 | (AACA)4 | TTTCCGTCCCAAAAAGATTG | TCCGTCCCAAAAAGATTGTC | 266 |
| chr3:21258613-21260625 | p4 | (AACA)4 | TTTCCGTCCCAAAAAGATTG | TCCGTCCCAAAAAGATTGTC | 218 |
| chr3:21258951-21260967 | p4 | (AACA)4 | GAACGCATCTTTGCCTCATT | GTGCCCACGAAAGACAAAAT | 135 |
| chr3:21413634-21415652 | p3 | (CAA)6 | CTTTCCTTTCCCCATCTTCC | TGGAGGTTACCGACCTTACG | 182 |
| chr3:21498540-21500556 | p4 | (TGTA)4 | TTCCTTCTGTTCATGGGTCC | ATAACGCCTTCTGGTCATGG | 163 |
| chr3:21505842-21507856 | p3 | (TTA)5 | GCAGAGTTTCGCATGTTTCA | AGAGATTTCGCAAGGCAGAA | 193 |
| chr3:21505842-21507856 | p2 | (TA)7 | GCAGAGTTTCGCATGTTTCA | AGAGATTTCGCAAGGCAGAA | 193 |
| chr3:21505842-21507856 | p2 | (AT)7 | GCAGAGTTTCGCATGTTTCA | AGAGATTTCGCAAGGCAGAA | 193 |
| chr3:21521536-21523548 | p2 | (TC)6 | AGAGACGGGTGTCGATTCAG | CCAGCATAAGAGATCGGAGC | 243 |
| chr3:21531011-21533026 | p5 | (ATTTG)3 | GAAGTTGCGGAAATGTGGAT | TACTTTTTCGGCCTCCCTTT | 227 |
| chr3:21549518-21551533 | p3 | (CAG)5 | TTCCTTGCAAATAGGCCATC | GGAGTTGCACGGTGGATAGT | 237 |
| chr3:21593172-21595208 | p2 | (TA)18 | AGTTTCGGTTCGAGCTTTTG | TGTGTTCAAGCATTTCCTCG | 102 |
| chr3:21593172-21595208 | p6 | (ATTTTT)3 | AGTTTCGGTTCGAGCTTTTG | TGTGTTCAAGCATTTCCTCG | 102 |
| chr3:21719175-21721187 | p2 | (TC)6 | CGCTGGATCGTATTCCGTAT | GAACCAAAATGGCTCCAAAA | 132 |
| chr3:21734321-21736339 | p6 | (GTGGAC)3 | GGTGGACGTTGTTCCACTCT | AACTCCACGATTATGTGCCC | 123 |

| Marker ID | SSR type | SSR motif | FORWARD PRIMER (5'-3') | REVERSE PRIMER (5'-3') | size (bp) |
|---|---|---|---|---|---|
| chr3:21734321-21736339 | p6 | (GTGGAC)3 | GGTGGACGTTGTTCCACTCT | AACTCCACGATTATGTGCCC | 123 |
| chr3:21905889-21907937 | p2 | (TA)31 | GCAGCGGACCTATAGCAGAG | TGCAGTTTGAATGTTGGGAA | 202 |
| chr3:21905889-21907937 | p2 | (AT)24 | GCAGCGGACCTATAGCAGAG | TGCAGTTTGAATGTTGGGAA | 202 |
| chr3:21905889-21907937 | p2 | (AT)18 | GCAGCGGACCTATAGCAGAG | TGCAGTTTGAATGTTGGGAA | 202 |
| chr3:21905889-21907937 | p3 | (CTA)9 | GCAGCGGACCTATAGCAGAG | TGCAGTTTGAATGTTGGGAA | 202 |
| chr3:21909975-21911996 | p3 | (CTA)7 | ACATACTTAAGGGCATGGCG | CATAAGCCGACGAGGCTAAC | 248 |
| chr3:21909975-21911996 | p2 | (AT)52 | ACATACTTAAGGGCATGGCG | CATAAGCCGACGAGGCTAAC | 248 |
| chr3:21909975-21911996 | p3 | (CTA)6 | ACATACTTAAGGGCATGGCG | CATAAGCCGACGAGGCTAAC | 248 |
| chr3:21946485-21948503 | p2 | (TC)9 | AACATGAATTGGGACGAAGC | ATCACGCGGTTGAAAATCTC | 101 |
| chr3:22038422-22040443 | p3 | (TTC)7 | AATAAAATTAGCCCACCCCG | ATTCCACATACTTCTGGCCG | 157 |
| chr3:22063196-22065210 | p2 | (AT)7 | TGTTAATGGCCGAGTGGAAT | AAAACCTCCGCAAATTACCC | 138 |
| chr3:22065239-22067259 | p2 | (AT)10 | TGCTACGAATACAAAGGCAGC | TGATGGATTCATTGCCAAAA | 245 |
| chr3:22111283-22113298 | p5 | (TTATT)3 | GCTTCATAATCACCCCCAGA | ATAAGGGTGGCGGAAGCTAT | 277 |
| chr3:22240720-22242744 | p6 | (GGTGCA)4 | GAAGGTCCAAACCCTTCACA | CCTGGACCCAGTTCTACCAA | 113 |
| chr3:22284379-22286401 | p2 | (AT)11 | GAGCCACAGACCCACAATCT | TGCAAGCTCATCTTTGGTTG | 249 |
| chr3:22334866-22336890 | p3 | (TAT)8 | CCTGTTTGGCCAAGCTTTAT | CTGCTTCTGCTACTCCCCAA | 148 |
| chr3:22383341-22385365 | p2 | (AT)12 | ATCGTATTCGAATGGCCAAG | AGTACGTCTCGAGGTGCGAT | 124 |
| chr3:22417166-22419181 | p3 | (ATA)5 | GAAATGGGGAAAGGGAGAAG | AGTCTAAAATTTGCCCCGGT | 150 |
| chr3:22417375-22419390 | p3 | (ATA)5 | TAATCGGATTTTGATTGGGG | CAACTTTCGCATACACACCG | 175 |
| chr3:22468285-22470300 | p2 | (TA)19 | TTTGTTGGCAAGTGGTTTGT | TGGCGTAATATGAACGAGCC | 159 |
| chr3:22468285-22470300 | p2 | (TA)15 | TTTGTTGGCAAGTGGTTTGT | TGGCGTAATATGAACGAGCC | 159 |
| chr3:22468322-22470340 | p2 | (TA)19 | TTTGTTGGCAAGTGGTTTGT | TGGCGTAATATGAACGAGCC | 122 |
| chr3:22468322-22470340 | p2 | (TA)15 | TTTGTTGGCAAGTGGTTTGT | TGGCGTAATATGAACGAGCC | 122 |
| chr3:22533709-22535727 | p3 | (GTT)6 | ACTGGAGAAACGTGGGTGAC | AATCCTACAAGGGGGTCTGG | 213 |
| chr3:22573014-22575029 | p5 | (ATTGC)3 | TGTTGGCGTACCCCATTATT | AATTGAAACTCGGACATCGG | 295 |
| chr3:22705417-22707441 | p2 | (TA)12 | CTGCACTCTTTTTCCCTTCG | GATCGGTCTGCTCTATTCGC | 225 |
| chr3:22725471-22727483 | p2 | (TA)6 | TGTTTGGTCATGCAAGTTGG | GAATATCCTAGGGCGCAACA | 229 |
| chr3:22780371-22782386 | p5 | (CCCCA)3 | AATGATGATAGGCAGGTCGG | CCCTCCGTTAGCGATAGTGA | 219 |
| chr3:22810040-22812055 | p3 | (TTC)9 | CCACCCCTTCTCTCTTCTCC | AATCGTTCTTTCCGGTGATG | 168 |
| chr3:22810040-22812055 | p5 | (AAAAT)3 | CCACCCCTTCTCTCTTCTCC | AATCGTTCTTTCCGGTGATG | 168 |
| chr3:22849918-22851938 | p2 | (TA)10 | TGAGTACCCACGGCACATTA | GCTGTCACCCCGTCTATCAT | 214 |
| chr3:22849918-22851938 | p3 | (TTG)5 | TGAGTACCCACGGCACATTA | GCTGTCACCCCGTCTATCAT | 214 |
| chr3:22926270-22928285 | p5 | (TATTT)3 | TTTGTGGTCAAATCCTTCCC | GTTTTGCGAAATGCACTGAA | 254 |
| chr3:23002065-23004122 | p3 | (ATA)19 | AATTCAAACAGGTGCCAAGG | TGAACCCAAAAATCCACCAT | 142 |
| chr3:23113943-23115965 | p2 | (TA)11 | ATGATAGCCACACAGGGCTC | GCTGAAGAACGACATCGACA | 116 |
| chr3:23207598-23209620 | p2 | (AC)11 | AGGGGCATTGCTTTAGTCCT | TCCTTTGCACTGTACGACCA | 275 |

| Marker ID | SSR type | SSR motif | FORWARD PRIMER (5'-3') | REVERSE PRIMER (5'-3') | size (bp) |
|---|---|---|---|---|---|
| chr3:23222104-23224118 | p2 | (CT)7 | AGATGTGATACGAGGGGACG | TCCGGATCGTAGTTTTACCG | 285 |
| chr3:23225416-23227430 | p2 | (TA)8 | TGCTCCACGCCCTACTACTT | CGAAGCCACGGTTGATACTT | 198 |
| chr3:23225416-23227430 | p6 | (CCGGTT)3 | TGCTCCACGCCCTACTACTT | CGAAGCCACGGTTGATACTT | 198 |
| chr3:23225437-23227457 | p2 | (TA)8 | TGCTCCACGCCCTACTACTT | CGAAGCCACGGTTGATACTT | 177 |
| chr3:23225437-23227457 | p6 | (CCGGTT)3 | TGCTCCACGCCCTACTACTT | CGAAGCCACGGTTGATACTT | 177 |
| chr3:23225486-23227500 | p2 | (TA)8 | TGCTCCACGCCCTACTACTT | CGAAGCCACGGTTGATACTT | 128 |
| chr3:23225486-23227500 | p6 | (CCGGTT)3 | TGCTCCACGCCCTACTACTT | CGAAGCCACGGTTGATACTT | 128 |
| chr3:23227623-23229639 | p2 | (TA)8 | TCGCTTCTCCCGACTTCTAA | CAGAGCTTTACGGATCCTGG | 241 |
| chr3:23240807-23242825 | p6 | (TATTTT)3 | TGAAGCAATGTTGCCAGAAG | GAAGCCTCCAAACAAGCAAC | 222 |
| chr3:23345224-23347236 | p2 | (AT)6 | ATGGTCAAACGGCACCTAAG | GATTTTGCCGAAACAGAGGA | 252 |
| chr3:23365863-23367875 | p2 | (CT)6 | CAATGCAATCTTTTCAGCCA | TTGAAGCCGTAGAGGCTGTT | 220 |
| chr3:23378349-23380361 | p2 | (TA)6 | CAGCTCAACAAACGGACAAA | GTTCCTCCACAAATGCTGGT | 225 |
| chr3:23406022-23408036 | p2 | (AT)7 | TAGGGGAGCAAAACTCCTCA | CGTTTTGAGAACGGAAGCTC | 262 |
| chr3:23412213-23414228 | p5 | (CTTTT)3 | TACCCCTTTCGGGATTTCTC | TTCCAAAGGAATGGTGAAGC | 259 |
| chr3:23421529-23423561 | p2 | (AC)16 | TCGTAGATACGCAGTGTGGG | TCCGTCCTAGGGTTGTTGTC | 296 |
| chr3:23456006-23458021 | p5 | (TGTGT)3 | TGAATCGTACCGGGAAAGTC | ACGGATCTTTAATACCGGGG | 285 |
| chr3:23456116-23458130 | p5 | (TGTGT)3 | TGAATCGTACCGGGAAAGTC | ACGGATCTTTAATACCGGGG | 175 |
| chr3:23496479-23498535 | p2 | (TA)28 | TAGGGTAGTCACCAGCACCC | GGTCACACCTGAGTGAAGCA | 137 |
| chr3:23585574-23587594 | p5 | (ACCTT)3 | CTCTGTCCCGACTCAGAAGG | GAAAAGGTCAAGGTGGGACA | 234 |
| chr3:23585610-23587624 | p5 | (ACCTT)3 | CTCTGTCCCGACTCAGAAGG | GAAAAGGTCAAGGTGGGACA | 198 |
| chr3:23609620-23611638 | p2 | (TA)9 | AGTTCCAATTCCTGTCGTCG | GACCCAAAATCTGGTCTGGA | 148 |
| chr3:23651141-23653155 | p2 | (TA)7 | TGACCACCAGTGAAACTTGC | TTGCGCATGCTCTAATCATC | 199 |
| chr3:23669137-23671151 | p2 | (TA)7 | TGACCACCAGTGAAACTTGC | TTGCGCATGCTCTAATCATC | 199 |
| chr3:23682543-23684557 | p2 | (AT)7 | CACCACATTCAAACTCACGG | GCTCCCCCTAATGGGTTAAT | 211 |
| chr3:23767979-23769995 | p2 | (AC)8 | GTTCACTTCTCGCCTTGACC | TCACTTAATTGGGCCTCTGG | 187 |
| chr3:23772372-23774384 | p2 | (GT)6 | ATTGCGGATTTGATAGCTGC | CTCCCAAAGGAGACAATGGA | 184 |
| chr3:23784194-23786227 | p3 | (AAT)11 | GGCTTACAATTCAACCCCCT | CCATTCTCTGCAACCCAAAT | 198 |
| chr3:23784194-23786227 | p6 | (TCCTTC)3 | GGCTTACAATTCAACCCCCT | CCATTCTCTGCAACCCAAAT | 198 |
| chr3:23836824-23838839 | p5 | (GTTCG)3 | GGCTATACTTTGAAGCCGCA | CCGAAATAGCCAAATCGAAA | 202 |
| chr3:23850792-23852806 | p2 | (TA)7 | GAGAATGGTGGGGAGGAAAT | GACCGCATTTTTGGTCACTT | 151 |
| chr3:23855062-23857077 | p5 | (AAACC)3 | GGGACACCATGAGTGGAAAG | AACCGACTACAACCAAACCG | 251 |
| chr3:23870509-23872527 | p2 | (AT)9 | GCCGTTGAAGGCTACGTAAA | TTGAAGAATGCTGGTAGGGG | 203 |
| chr3:23870509-23872527 | p2 | (AT)20 | GCCGTTGAAGGCTACGTAAA | TTGAAGAATGCTGGTAGGGG | 203 |
| chr3:23874776-23876794 | p3 | (AAT)6 | GGTGCAGTTTCAAGGAGAGC | AGGGCAGCAAAGTAGCAAGA | 116 |
| chr3:23877318-23879336 | p3 | (AAT)6 | TAATGCGCACTTCAGAGTCG | GCACAAATAAAGCATGGGCT | 109 |
| chr3:23940091-23942106 | p3 | (ACA)5 | GAAGGCACCTTCCGATAACA | TTGCCACCACTACGGTTACA | 238 |

| Marker ID | SSR type | SSR motif | FORWARD PRIMER (5'-3') | REVERSE PRIMER (5'-3') | size (bp) |
|---|---|---|---|---|---|
| chr3:23940091-23942106 | p4 | (AAAG)5 | GAAGGCACCTTCCGATAACA | TTGCCACCACTACGGTTACA | 238 |
| chr3:23957528-23959546 | p3 | (GAA)6 | GTGTTTCCAAATTCGCCATT | GGAACTTCGTAAGTGCAGGC | 201 |
| chr3:24340094-24342124 | p3 | (TAG)10 | CCGACCTCCCTCACAAATAA | CTTCCCCCTTTAAACTTCCG | 112 |
| chr3:24355387-24357405 | p6 | (TATTTT)3 | CCAAGCCTTTTCGAAGTCAA | TTGTGGTTTGTGAGGGACAA | 182 |
| chr3:24511304-24513328 | p3 | (TTC)8 | GTCCCGGACGATCTCTCATA | CATGGTCTGGTGCTTGTGTC | 110 |
| chr3:24533164-24535179 | p5 | (CTTTT)3 | TTTTTGGTTGAGGAAACGCT | CACTCTTTTTGAACCGGAGG | 267 |
| chr3:24635827-24637842 | p5 | (AATTA)3 | AACAGAGCGGAAAGCAAAAA | ATTCGATTCTTGGTCGCAGT | 235 |
| chr3:24807165-24809195 | p2 | (TA)15 | GCTACAAACGATTCGCTTCC | TTAAGACCATGAATTCGCCC | 161 |
| chr3:24862487-24864507 | p5 | (AAAAT)4 | GATCGCTGTTTTCAGGGGTA | CAAATGGCGAGAAAAGTGGT | 254 |
| chr3:24868978-24871000 | p2 | (AT)11 | TTTGGTGATGTGTCACGGAT | TCCAGGACGAGCAAAAATTC | 137 |
| chr3:24878974-24881022 | p3 | (TAA)16 | TTCACATTTGAAGCGGTCTG | TGACAAAATGGAGCACCAAA | 197 |
| chr3:24893083-24895097 | p2 | (AT)7 | AAAGCTGTAGCTCCGGATGA | GAAGCCCTTTTGTTTGTGGA | 256 |
| chr3:24979497-24981513 | p2 | (CT)8 | TCCTACGATTTCGAGGGATG | AGACCTCAAAGCCTCGTTCA | 278 |
| chr3:24979497-24981513 | p2 | (TA)11 | TCCTACGATTTCGAGGGATG | AGACCTCAAAGCCTCGTTCA | 278 |
| chr3:24994999-24997011 | p2 | (AT)6 | GAAAAAGGAGTCCCATGCAA | AGTTGCAGCCTTGTCGACTT | 132 |
| chr3:25016767-25018783 | p2 | (TA)8 | TTCAAGAAAACCCAACCCTG | GGAAGAAGAATCCCAAAGCC | 195 |
| chr3:25031713-25033731 | p6 | (TAATGA)3 | TGGTTTTGAAAGGAGTTGGG | TCCAAAGTCCCCATTTTCAG | 266 |
| chr3:25036183-25038195 | p2 | (AT)6 | GTCTTGTCTCTCTCCGTGCC | TGGTAGGGCACACACACACT | 272 |
| chr3:25153553-25155569 | p4 | (AATA)4 | GCGCCGAGAGCTATAATTTG | CTTCTCGGGAAGTGACCAAG | 228 |
| chr3:25201289-25203304 | p5 | (CATAG)3 | ATTGTCACGACCCGTCTAGG | AGAATGACGCCGCTAATGTT | 237 |
| chr3:25201289-25203304 | p2 | (TA)15 | ATTGTCACGACCCGTCTAGG | AGAATGACGCCGCTAATGTT | 237 |
| chr3:25237282-25239344 | p4 | (ATCT)7 | CATGACAAGGCTGTTCGAGA | CTATCATGTTGAGGCGGGTT | 161 |
| chr3:25237282-25239344 | p2 | (AT)31 | CATGACAAGGCTGTTCGAGA | CTATCATGTTGAGGCGGGTT | 161 |
| chr3:25307398-25309416 | p6 | (TTGGTA)3 | TGGACTCTGGAGTTGCCTCT | TTGCCCCTCAAACATTCTCT | 178 |
| chr3:25307398-25309416 | p2 | (TA)6 | TGGACTCTGGAGTTGCCTCT | TTGCCCCTCAAACATTCTCT | 178 |
| chr3:25582929-25584949 | p2 | (AT)10 | TGTATGTGTGTGTGGAGGGG | AAAATTGCCAAGTGGCTTGT | 199 |
| chr3:25666091-25668109 | p2 | (AG)9 | AACTATTTCTTCGGCAGCGA | AACAAATCCCGATTAGTGCG | 100 |
| chr3:25681985-25684000 | p3 | (TAT)5 | TTTTGAGGCATTTTGGTTCC | CATGTCTCGACCAGCATGAC | 253 |
| chr3:25778860-25780887 | p3 | (AGA)9 | TGATGGACTTCTTCGTTCCC | GACATATAACGGGAAGGGCA | 129 |
| chr3:25800070-25802082 | p2 | (GT)6 | CCCGAAACTGATGCAAAAAT | TTATGCTGCCCAGAATTTCC | 274 |
| chr3:25902486-25904508 | p2 | (TA)11 | CCTGTCAGAACACTCAGCCA | TTTCTTTGTGGAGATGGCCT | 179 |
| chr3:25910028-25912040 | p2 | (AT)6 | GAGGGGATTTCTACTTCGGC | ATGCATTTCGATTAGGGTCG | 139 |
| chr3:26105581-26107595 | p2 | (TG)7 | GGCTATAATGACACCCCACG | AGGCTTCAGGGTTTATCCGT | 236 |
| chr3:26304120-26306144 | p3 | (CAT)8 | CAACCTACCGAGCACCTCTC | GCCCTGCCCTGTATTTATGA | 163 |
| chr3:26360882-26362897 | p5 | (AAGAA)3 | CATGACCGAGTTGATTGTGG | AGTTTTGCCTTATGGGCCTT | 144 |
| chr3:26496111-26498123 | p2 | (TG)6 | CACATGGTTGACATTCCGAG | TCTTACCCGGCATCATAAGC | 104 |

| Marker ID | SSR type | SSR motif | FORWARD PRIMER (5'-3') | REVERSE PRIMER (5'-3') | size (bp) |
|---|---|---|---|---|---|
| chr3:26624810-26626825 | p3 | (ATT)5 | TCCTTCCCCTTCTCTCCATT | TTACCACCATGCCAATCTCA | 225 |
| chr3:26669511-26671525 | p2 | (AT)7 | TCAAGGGAGGTAAGCCATTG | AAGAAAAGGAGAAGGGCGAG | 155 |
| chr3:26669511-26671525 | p2 | (AT)13 | TCAAGGGAGGTAAGCCATTG | AAGAAAAGGAGAAGGGCGAG | 155 |
| chr3:26932965-26934980 | p5 | (ATAAA)3 | CCATCTCCCAAACATCCTTC | TCTGTTAACCGAGGGTGGAC | 101 |
| chr3:26958326-26960386 | p2 | (TA)30 | ATTTTTCAAATGGACGCAGG | GTCAATCCAAACGGGCTCTA | 114 |
| chr3:26958326-26960386 | p2 | (TA)16 | ATTTTTCAAATGGACGCAGG | GTCAATCCAAACGGGCTCTA | 114 |
| chr3:26970068-26972086 | p6 | (TCTTTT)3 | CGAGGCCCACTACTAAGTGC | ACTTCCCAATGACATGGAGC | 190 |
| chr3:27137459-27139475 | p2 | (TA)8 | AACGTCCGAATCTTGAATGC | ATTTGATGAGGTGGTGCCAT | 266 |
| chr3:27270214-27272234 | p6 | (CTATTA)4 | AAACGGAAAAGGGCCTAAAA | GGACGAAAGGATGGACAAAA | 152 |
| chr3:27270214-27272234 | p5 | (TTCTC)4 | AAACGGAAAAGGGCCTAAAA | GGACGAAAGGATGGACAAAA | 152 |
| chr3:27368699-27370714 | p5 | (ACTTG)3 | CGAATATGTGGGTGCCTTCT | GTAATGTCGGGTTGGTTTGG | 115 |
| chr3:27373766-27375781 | p3 | (TTG)5 | ATGGCAACCTCTTTCCCTTT | ACCAGAACCCGAAACAACAG | 184 |
| chr3:27462944-27464956 | p2 | (TC)6 | TGCAAGACTTGTTGACCAGC | AGTGTGGGTTGAGAAATGGC | 219 |
| chr3:27542198-27544226 | p2 | (AT)14 | GGTCTGCAGCTGATTTTTCC | GTAGGTCTGTCTTCTGGGCG | 131 |
| chr3:27560260-27562306 | p2 | (TA)23 | TCGAGACTCAAAACATCCCC | GGAGAGAGGTTTCCCTTTCG | 134 |
| chr3:27617021-27619033 | p2 | (AT)6 | ATCGTAGCCCTCACATCCTG | AAGCCGACGTACCATTATGC | 127 |
| chr3:27658604-27660616 | p2 | (TA)6 | CAGGTGGGTTCACTTGGAGT | CATATTCCCTGTGGGATTCG | 215 |
| chr3:27683198-27685212 | p2 | (CT)7 | TCTGCTGGCACAAGAAGCTA | CCCTTTCTTTCATTTTGGCA | 268 |
| chr3:27715783-27717798 | p5 | (CACCC)3 | AACCACAACAAACGACACCA | TGGTGAAAAGAGGGTTGGAG | 203 |
| chr3:27801760-27803784 | p4 | (AAAG)6 | CCGACATGGCCTTATCTTGT | GCTGCGGTCTGTTTTGGTAT | 103 |
| chr3:27924647-27926671 | p3 | (AAT)8 | ACTTCGGGCATACAATCCAG | TTGCACTATGTCCGTGCTTC | 296 |
| chr3:28000215-28002233 | p2 | (AG)7 | CATTCCGGAAGACGATGAGT | CCTCGAAGATACGGGGTGTA | 113 |
| chr3:28000215-28002233 | p3 | (TTG)6 | CATTCCGGAAGACGATGAGT | CCTCGAAGATACGGGGTGTA | 113 |
| chr3:28083167-28085201 | p2 | (AT)10 | GGCTGCCACTATGAAACGAT | TTTTGTCCAGGGGTACGAAG | 178 |
| chr3:28083167-28085201 | p2 | (TA)17 | GGCTGCCACTATGAAACGAT | TTTTGTCCAGGGGTACGAAG | 178 |
| chr3:28103607-28105625 | p3 | (TTG)6 | CAGCGGCGTATCCTAAAGAG | GAGATTGCCACACCGCTTAT | 125 |
| chr3:28159358-28161388 | p2 | (GA)11 | GGGGAAGGGAGAGAGAGAGA | CCTCTAAATACCTGCCCGTG | 164 |
| chr3:28159358-28161388 | p2 | (AG)15 | GGGGAAGGGAGAGAGAGAGA | CCTCTAAATACCTGCCCGTG | 164 |
| chr3:28215753-28217765 | p2 | (AT)6 | TAGGAGCAACTGGTACGGCT | AGGTGCAGATTGCGAGAGAT | 191 |
| chr3:28243115-28245130 | p5 | (TTTTA)3 | TTCATTGGCTAGGTATGCCC | TGTCTGTGGAAGTCCTGACG | 103 |
| chr3:28261039-28263054 | p5 | (TTTTA)3 | TTGACAAAAACCCTGGTTCC | GGAAGTCCTGACAGGCTGAG | 105 |
| chr3:28339326-28341352 | p5 | (GATTA)3 | TGTTCATGCCCACAGGTAGA | TCGTGCCCAAAACAAATACA | 141 |
| chr3:28339354-28341380 | p5 | (GATTA)3 | TGTTCATGCCCACAGGTAGA | TCGTGCCCAAAACAAATACA | 113 |
| chr3:28345417-28347445 | p2 | (TA)33 | TTATTGCCCTCTTTTGGGTG | ACGCAAAAGAAACACCAAGG | 124 |
| chr3:28345417-28347445 | p2 | (TA)14 | TTATTGCCCTCTTTTGGGTG | ACGCAAAAGAAACACCAAGG | 124 |
| chr3:28354390-28356412 | p2 | (TA)11 | ATCTGGGGTTAATCAGTGCG | GCAAGGTTTACAGACCCCAA | 124 |

| Marker ID | SSR type | SSR motif | FORWARD PRIMER (5'-3') | REVERSE PRIMER (5'-3') | size (bp) |
|---|---|---|---|---|---|
| chr3:28372292-28374343 | p3 | (CTT)20 | GAGGGTTGGCGAAGATGATA | GGACCGTCCTTCTCACCATA | 115 |
| chr3:28398471-28400489 | p3 | (TTG)6 | ACGTGAAGGACAAACCTTGG | TGCGTTGGAAAGTAGACACG | 254 |
| chr3:28410561-28412593 | p2 | (TA)16 | CCCCGAGAGTCAGTCACATT | CGGGACTATCCATGAGCAAT | 178 |
| chr3:28410561-28412593 | p2 | (TC)8 | CCCCGAGAGTCAGTCACATT | CGGGACTATCCATGAGCAAT | 178 |
| chr3:28410561-28412593 | p2 | (AT)7 | CCCCGAGAGTCAGTCACATT | CGGGACTATCCATGAGCAAT | 178 |
| chr3:28450564-28452580 | p2 | (GT)8 | CCTCAGCGAGATTTCAAAGG | CAAACACTCGTTGCTCGAAA | 212 |
| chr3:28457935-28459950 | p5 | (CATTT)3 | GCCACTGTAGGCCACACTTT | AGCGAACACACATAGGACCC | 185 |
| chr3:28578078-28580136 | p2 | (AT)29 | TCAATAAAACCTTGCGGGTC | TACGGGTCTAGGTGACAGGG | 283 |
| chr3:28629155-28631170 | p5 | (ATTTC)3 | AGAAGTGCCCATTGCCATAC | TGAATCTCTTTTCGGGGTTG | 112 |
| chr3:28683517-28685565 | p2 | (AT)10 | CCGAGCTTGACACTCGTACA | AGCCTTGTCGGCTTGTGTAT | 213 |
| chr3:28683577-28685635 | p2 | (AT)10 | CCGAGCTTGACACTCGTACA | AGCCTTGTCGGCTTGTGTAT | 153 |
| chr3:28691483-28693501 | p2 | (AT)9 | ACTTTTGTCACGACCCAACC | GTTGGCCTGCGTATCGTATT | 132 |
| chr3:28807308-28809324 | p2 | (TA)8 | GCACCAACGGAAGAGGAGTA | GGTGGACGACACTTGGATTC | 124 |
| chr3:28844206-28846224 | p6 | (TAAAAA)3 | AAATCCTCTTGCCTGCTGAA | TTTATACCCCGCATCGAAAG | 225 |
| chr3:28865072-28867092 | p2 | (AT)10 | CCTGATCAATTTGGGCTGAT | ACCTCAGTGAAATCGGCAAC | 248 |
| chr3:28865072-28867092 | p5 | (TTTTA)3 | CCTGATCAATTTGGGCTGAT | ACCTCAGTGAAATCGGCAAC | 248 |
| chr3:28865072-28867092 | p2 | (AT)7 | CCTGATCAATTTGGGCTGAT | ACCTCAGTGAAATCGGCAAC | 248 |
| chr3:28871163-28873178 | p5 | (TTAAA)3 | GCATTGGCTTCAGTTCCATT | GTAAGCCCTTGCTTCACACC | 242 |
| chr3:28894118-28896130 | p2 | (TC)6 | TACAACCAACCCCGAGAATC | TCGTCGAATCGTGTAGCAAG | 205 |
| chr3:28905645-28907659 | p2 | (TC)7 | TCTGGAAGGGTCAAAACTGG | AGTGGATGCCCAGAAGATTG | 101 |
| chr3:28921819-28923837 | p6 | (CATATA)3 | ATGACCCAAACTGATGAGCC | CCGTAAGTTCCCGTCACAGT | 181 |
| chr3:29088573-29090588 | p3 | (CTT)5 | TGAAGGCAAATCAAACACCA | GAAGACACCTTCCCTCCCTC | 288 |
| chr3:29088573-29090588 | p3 | (CTT)5 | TGAAGGCAAATCAAACACCA | GAAGACACCTTCCCTCCCTC | 288 |
| chr3:29088573-29090588 | p3 | (TGC)7 | TGAAGGCAAATCAAACACCA | GAAGACACCTTCCCTCCCTC | 288 |
| chr3:29187000-29189018 | p6 | (AAAAAT)3 | GTTTGCCCGTGAAAGAAAAA | CTTTGAATGCCCATGTGATG | 196 |
| chr3:29187000-29189018 | p6 | (AAAAAT)3 | GTTTGCCCGTGAAAGAAAAA | CTTTGAATGCCCATGTGATG | 196 |
| chr3:29233829-29235847 | p5 | (ATTGC)3 | GTTGTTTGGCCTTCCAGGTA | GGATGGCTAGCCCCTAAATC | 109 |
| chr3:29233829-29235847 | p2 | (AT)9 | GTTGTTTGGCCTTCCAGGTA | GGATGGCTAGCCCCTAAATC | 109 |
| chr3:29233829-29235847 | p5 | (ATTGC)3 | GTTGTTTGGCCTTCCAGGTA | GGATGGCTAGCCCCTAAATC | 109 |
| chr3:29233829-29235847 | p2 | (AT)9 | GTTGTTTGGCCTTCCAGGTA | GGATGGCTAGCCCCTAAATC | 109 |
| chr3:29234679-29236697 | p2 | (AT)9 | ATTGATGGATTCGGATTGGA | CAGCCACAAGGGGTAGACAT | 219 |
| chr3:29234679-29236697 | p5 | (ATTGC)3 | ATTGATGGATTCGGATTGGA | CAGCCACAAGGGGTAGACAT | 219 |
| chr3:29234679-29236697 | p2 | (AT)9 | ATTGATGGATTCGGATTGGA | CAGCCACAAGGGGTAGACAT | 219 |
| chr3:29408321-29410347 | p2 | (AC)13 | CTGGAAAATGATCCGCCTTA | CCTCGGATTTGAGGACTGAG | 122 |
| chr3:29408321-29410347 | p2 | (TG)13 | CTGGAAAATGATCCGCCTTA | CCTCGGATTTGAGGACTGAG | 122 |
| chr3:29462294-29464312 | p6 | (AATTGT)3 | TGGGGCAAAAGACTTGAAAC | GGTATCGACCTCCGATTTGA | 181 |

| Marker ID | SSR type | SSR motif | FORWARD PRIMER (5'-3') | REVERSE PRIMER (5'-3') | size (bp) |
|---|---|---|---|---|---|
| chr3:29532959-29534971 | p2 | (AT)6 | TGGTGACTGATGGTGGTGAT | CGATAAAGAGGGCAGCTCAC | 116 |
| chr3:29693559-29695579 | p2 | (TG)10 | AGACCTCCGTTGAGCTTGAA | CCAGCGCTTAGTTAAGGTCG | 223 |
| chr3:29763933-29765945 | p2 | (CT)6 | CTCTCGATGGTCCACCAAAT | CAGAATGCAAGTGTAGCCGA | 238 |
| chr3:29855842-29857857 | p3 | (TTC)5 | TCACTTGCTGCAACAACCTC | GAAGAACATGGGCAAGGTGT | 218 |
| chr3:29979725-29981753 | p4 | (TATG)7 | TATTACTATGCCGGAAGCGG | ATAGCATTTTGAACCGTGCC | 282 |
| chr3:30190262-30192289 | p2 | (GA)7 | AAGAGCTCCAAACCGAGTCA | TCAGTTAACCCCTCCCCTCT | 290 |
| chr3:30190292-30192325 | p2 | (GA)7 | AAGAGCTCCAAACCGAGTCA | AAATGCATTCTCACTTCGGG | 260 |
| chr3:30190328-30192349 | p2 | (GA)7 | AAGAGCTCCAAACCGAGTCA | AAATGCATTCTCACTTCGGG | 224 |
| chr3:30199955-30201970 | p5 | (AAATT)3 | TAGTGGCTGCACCATTTTTG | AGACCCTAGCCTTGGGGATA | 144 |
| chr3:30242647-30244662 | p3 | (GTT)5 | GCGCGTTCCCTAATCATAAA | TCACAAATCCATCGCAACAT | 214 |
| chr3:30303978-30305994 | p2 | (GA)8 | CTTCATCCAAGTCCTCCCAA | TTTGGTTCTCGGGTAAGTCG | 101 |
| chr3:30323639-30325705 | p3 | (ATA)22 | TTTAAACTCCCTCATCCCCC | GCGAAGTTTGTGGAAAGAGC | 265 |
| chr3:30387557-30389572 | p3 | (CAT)5 | AGCGCGAAGAATGAAAAGAA | TTGTTGGGTTGAGGGTTAGC | 114 |
| chr3:30399061-30401079 | p6 | (ATAAAA)3 | GGTCCGGCATAACTTTGGTA | TTCACGCAGAAATCATGAGG | 163 |
| chr3:30411265-30413280 | p3 | (GTT)5 | TGCAAAGATCCATCAGTTCG | CGGGTAGACTTGCTGGAGAC | 206 |
| chr3:30470091-30472115 | p2 | (GA)12 | TGTGATATTGGGCCTTAGCC | TTCGGGGTTCGTAGTGTCTC | 261 |
| chr3:30521693-30523711 | p2 | (AT)9 | TCTCTCCTACTGGGCCTGAA | GGGAGGAAGTGTTTGACGAA | 100 |
| chr3:30664967-30666985 | p2 | (TA)9 | TTACGTCACCCCGATATGGT | AGGCGCGGTATAATCACAAG | 275 |
| chr3:30686424-30688439 | p5 | (ATATG)3 | ATTAACAGCGAACACTGGGG | CATGGCAAAAGAAAGAAGGG | 164 |
| chr3:30686424-30688439 | p2 | (TA)6 | ATTAACAGCGAACACTGGGG | CATGGCAAAAGAAAGAAGGG | 164 |
| chr3:30744767-30746779 | p2 | (AT)6 | ATTTCTGGGCAGGTTGTTTG | ACCCCCGAATTACCAAACTC | 162 |
| chr3:30825427-30827439 | p2 | (AT)6 | ATTTCTGGGCAGGTTGTTTG | ACCCCCGAATTACCAAACTC | 162 |
| chr3:30895492-30897507 | p5 | (TATTT)3 | CAAAACGCACACACCAAATC | TGGGTTTATGGAGGTAAGCG | 151 |
| chr3:30905011-30907103 | p2 | (AT)16 | GAGAGGTCCCGCAATAACAA | TGGGTAAGATTTTTCAGCGG | 147 |
| chr3:30905011-30907103 | p2 | (AT)46 | GAGAGGTCCCGCAATAACAA | TGGGTAAGATTTTTCAGCGG | 147 |
| chr3:30913764-30915776 | p2 | (CA)6 | GCACTGCCGATAGAAGGAAG | GGTTGAATTCCCGGATTCTT | 247 |
| chr3:30949590-30951614 | p5 | (ATTGC)3 | CACAGCATTGTATGGGTTGC | TCGGAACCAAGTACACGTCA | 121 |
| chr3:30949590-30951614 | p3 | (TAT)8 | CACAGCATTGTATGGGTTGC | TCGGAACCAAGTACACGTCA | 121 |
| chr3:30949590-30951614 | p2 | (CT)6 | CACAGCATTGTATGGGTTGC | TCGGAACCAAGTACACGTCA | 121 |
| chr3:30969533-30971548 | p3 | (AAT)5 | AACGTTGGAGTCCGTTTCAC | AATACAGCCGCACCTACGTC | 292 |
| chr3:31099957-31101975 | p3 | (TTC)6 | ACAATGCTTTTGGCTTTGCT | GCTGGCTGTCCATGGTTATT | 266 |
| chr3:31115124-31117136 | p2 | (TA)6 | AGATGAATGCCATGGTGACA | ATTCCATGAATTATGCCCCA | 132 |
| chr3:31115124-31117136 | p2 | (AT)6 | AGATGAATGCCATGGTGACA | ATTCCATGAATTATGCCCCA | 132 |
| chr3:31236010-31238044 | p2 | (AT)17 | TTTCCATGGCTTCGAGTACC | CTTCGTTTTCCTCTTTCCCC | 120 |
| chr3:31313237-31315252 | p5 | (AATGC)3 | GCTGTGTGCGGAGTGACTAA | GTTGAGCTTCGCTAACCGAG | 290 |
| chr3:31508881-31510903 | p2 | (TA)11 | GCCAAACTGGAGGAGGTACA | CCAGATCCTATGCCCACAGT | 171 |

| Marker ID | SSR type | SSR motif | FORWARD PRIMER (5'-3') | REVERSE PRIMER (5'-3') | size (bp) |
|---|---|---|---|---|---|
| chr3:31508881-31510903 | p2 | (AT)14 | GCCAAACTGGAGGAGGTACA | CCAGATCCTATGCCCACAGT | 171 |
| chr3:31508881-31510903 | p3 | (TCA)5 | GCCAAACTGGAGGAGGTACA | CCAGATCCTATGCCCACAGT | 171 |
| chr3:31509718-31511733 | p2 | (TA)11 | TCGTCGTGCCCAAATATGTA | TCCTTTTCTTTGGACGCTGT | 236 |
| chr3:31509718-31511733 | p2 | (AT)14 | TCGTCGTGCCCAAATATGTA | TCCTTTTCTTTGGACGCTGT | 236 |
| chr3:31509718-31511733 | p3 | (TCA)5 | TCGTCGTGCCCAAATATGTA | TCCTTTTCTTTGGACGCTGT | 236 |
| chr3:31523840-31525858 | p3 | (CAA)6 | CCGTATCCCCGTCACATATC | TGTTGTCGTGCACACTTTGA | 112 |
| chr3:31530475-31532487 | p2 | (AT)6 | AGCACGAACATCCCAAAAAC | TTTGGGTTGCTCCTTTGTTC | 259 |
| chr3:31541503-31543517 | p2 | (TC)7 | TTGGCTCCCTATGAACCTTG | TCTGGAAACCAACGGAAAAC | 256 |
| chr3:31587964-31589978 | p2 | (TA)7 | GACGTGTTGTTGTTGGTTGG | TGACTTAAGGGAGCTTCGGA | 155 |
| chr3:31587964-31589978 | p2 | (AT)6 | GACGTGTTGTTGTTGGTTGG | TGACTTAAGGGAGCTTCGGA | 155 |
| chr3:31671201-31673219 | p3 | (TCT)6 | ACGGGCGACATTGTTTAGTC | TTTATGAATCGAACCCGGAG | 119 |
| chr3:31695432-31697472 | p2 | (TA)20 | TTACATACCTTGCCCGCTTC | TTGGGGTTGGTTGAATTTGT | 100 |
| chr3:31720533-31722551 | p3 | (TCT)6 | ACGGGCGACATTGTTTAGTC | TTTATGAATCGAACCCGGAG | 119 |
| chr3:31754183-31756195 | p2 | (TA)6 | GCCATCGAAAAACTACGGAA | TAGTCGGGTCGTGACATGAA | 142 |
| chr3:31790343-31792355 | p2 | (TA)6 | GCCATCGAAAAACTACGGAA | TAGTCGGGTCGTGACATGAA | 142 |
| chr3:31855981-31857995 | p2 | (AT)7 | ACAAAGTGAAGGCGGCTAGA | TCCCCCACCCCTATTAAAAC | 237 |
| chr3:31981083-31983095 | p2 | (GA)6 | AATCAAAGGCCATATGCTGC | TTAGGAAGAGGTCGCCAAGA | 224 |
| chr3:31997257-31999272 | p5 | (ATGCA)3 | TGGCTGGCACTATTTACACG | CTAGTGGGTTGTGGATGGCT | 164 |
| chr3:32133862-32135874 | p2 | (TA)7 | GGTTCAGGTATCAACCGTGG | GTCGCTCCGATCTAATCCAA | 182 |
| chr3:32133862-32135874 | p2 | (AT)6 | GGTTCAGGTATCAACCGTGG | GTCGCTCCGATCTAATCCAA | 182 |
| chr3:32133862-32135874 | p2 | (TA)12 | GGTTCAGGTATCAACCGTGG | GTCGCTCCGATCTAATCCAA | 182 |
| chr3:32138960-32140974 | p2 | (GA)7 | CGCCTTTAAAAGCAGCAAAG | TATTTGCTTGCGAAAATCCC | 215 |
| chr3:32138960-32140974 | p2 | (TA)7 | CGCCTTTAAAAGCAGCAAAG | TATTTGCTTGCGAAAATCCC | 215 |
| chr3:32182085-32184097 | p2 | (GT)6 | CTCCGCAAAGTCTCTTTTGG | ACCGTAACAGATGGCGACTC | 148 |
| chr3:32322413-32324428 | p5 | (ATTTC)3 | GAAGCGGAAGAAAGTATGCG | AATATCATCGCCTCACAGGG | 166 |
| chr3:32362342-32364396 | p2 | (AT)27 | TTCCAATGGCATAGGTCACA | AGTTTGGCAAAGGATTGGTG | 246 |
| chr3:32362342-32364396 | p2 | (TG)6 | TTCCAATGGCATAGGTCACA | AGTTTGGCAAAGGATTGGTG | 246 |
| chr3:32439330-32441344 | p2 | (AT)7 | AGGGGATTGTAGCTCCTGCT | GGGGGAAAACTGAAAATGGT | 160 |
| chr3:32439330-32441344 | p2 | (AT)9 | AGGGGATTGTAGCTCCTGCT | GGGGGAAAACTGAAAATGGT | 160 |
| chr3:32532812-32534830 | p6 | (AATGAA)3 | TCTTCTTTAACCACACCCCG | GGAAACCTCCCTCCATCTTC | 265 |
| chr3:32532812-32534830 | p2 | (TA)6 | TCTTCTTTAACCACACCCCG | GGAAACCTCCCTCCATCTTC | 265 |
| chr3:32580730-32582750 | p2 | (TA)18 | TCCAAACGCTTGTCCACATA | CCAAAGCTGAATTGCACGTA | 250 |
| chr3:32580730-32582750 | p4 | (AAAG)5 | TCCAAACGCTTGTCCACATA | CCAAAGCTGAATTGCACGTA | 250 |
| chr3:32642747-32644761 | p2 | (TA)7 | TATGCAAGTCGACAAGGCTG | TGGAGATCCACCAGCTTAGG | 287 |
| chr3:32689885-32691913 | p6 | (TATATG)3 | TATGTGTTGCGGCACGTAAT | TTATTTTGATGGCCGAGAGG | 280 |
| chr3:32689924-32691942 | p6 | (TATATG)3 | TATGTGTTGCGGCACGTAAT | TTATTTTGATGGCCGAGAGG | 241 |

| Marker ID | SSR type | SSR motif | FORWARD PRIMER (5'-3') | REVERSE PRIMER (5'-3') | size (bp) |
|---|---|---|---|---|---|
| chr3:32689943-32691957 | p6 | (TATATG)3 | TATGTGTTGCGGCACGTAAT | TTATTTTGATGGCCGAGAGG | 222 |
| chr3:32793157-32795178 | p5 | (ATTGC)3 | CGTCGTTTGCGCTGTAAGTA | TTCTTCGATTTGCTTGGTCC | 189 |
| chr3:32793157-32795178 | p3 | (TTG)7 | CGTCGTTTGCGCTGTAAGTA | TTCTTCGATTTGCTTGGTCC | 189 |
| chr3:32811163-32813175 | p3 | (AAC)5 | CAAAAACATGTCCGTCGTTG | TGATTCCCTTCCTGACAACC | 267 |
| chr3:32811163-32813175 | p2 | (CT)6 | CAAAAACATGTCCGTCGTTG | TGATTCCCTTCCTGACAACC | 267 |
| chr3:32877550-32879574 | p2 | (TA)14 | TTCCTTCTGTTCATGGGTCC | CCTCCGGGCATAATAGCATA | 151 |
| chr3:32877550-32879574 | p2 | (TA)12 | TTCCTTCTGTTCATGGGTCC | CCTCCGGGCATAATAGCATA | 151 |
| chr3:32877550-32879574 | p2 | (AG)7 | TTCCTTCTGTTCATGGGTCC | CCTCCGGGCATAATAGCATA | 151 |
| chr3:32887426-32889441 | p5 | (AATGC)3 | CTCCGATTTCGTAGACCAGC | CATGGTATTAACGCCAGGCT | 120 |
| chr3:32887426-32889441 | p2 | (TA)27 | CTCCGATTTCGTAGACCAGC | CATGGTATTAACGCCAGGCT | 120 |
| chr3:32902087-32904102 | p5 | (TGATC)3 | GCTCCAATTCCTTCCATTCA | CCCGAGGCCAAACTATGTAA | 294 |
| chr3:32927242-32929257 | p5 | (TTTGG)3 | TTTACAATCATCACCGCCAA | CACCAATTGGGGACTCTGTT | 224 |
| chr3:32980844-32982910 | p2 | (TA)33 | AACCCACTTTTCCCCAAAAC | GGTTTCGCAACAAATGACCT | 167 |
| chr3:33007399-33009417 | p6 | (GGAATA)3 | CACATGGACATCGTTGAAGG | CATGACCACCCTTCGCTAAT | 102 |
| chr3:33222261-33224277 | p2 | (TA)8 | TATTTCTGCTATTTGGGGCG | GAGCAGACTTGGCACACAGA | 214 |
| chr3:33289612-33291627 | p2 | (AT)7 | AGCATTTATCCGTCATTCGG | TGTGGGTTTAATAGGGCCAG | 273 |
| chr3:33289612-33291627 | p3 | (TTA)5 | AGCATTTATCCGTCATTCGG | TGTGGGTTTAATAGGGCCAG | 273 |
| chr3:33318758-33320810 | p2 | (AT)26 | TGAAGAACAGTGCAAGCCAG | GGTATGCCACCCTTTTGAGA | 277 |
| chr3:33372903-33374923 | p3 | (AAT)6 | GGTCTTGACAGGAGAGCGAC | CCGTTGTGTTGGTTGTTGAC | 234 |
| chr3:33470634-33472646 | p2 | (AC)6 | ACTCGGGTTTCGATGAACAG | GGAACTGAATTTGAGCGAGC | 234 |
| chr3:33530953-33532981 | p2 | (AT)41 | GATGAAAACCCTTCCACGAA | TGATGGCATTCGAGTGGATA | 149 |
| chr3:33530953-33532981 | p2 | (TA)14 | GATGAAAACCCTTCCACGAA | TGATGGCATTCGAGTGGATA | 149 |
| chr3:33541335-33543405 | p2 | (AT)35 | TATCCGGCATCTTGAACCTC | GATGGTTGAGAGCAATGGGT | 106 |
| chr3:33696825-33698843 | p6 | (AATATG)3 | GCCACAATGTGTTTTTGCAC | CCTCCCGCAATTGTCTTAAA | 294 |
| chr3:33709728-33711740 | p2 | (GT)6 | GTCCCACGTTCTGAGTGGAT | TCCACGAGTCATATTGCGAA | 126 |
| chr3:33764496-33766511 | p5 | (TTTTG)3 | AGCACGGGGGAGACTTTATT | GTGCCATCATTTGTGTGAGG | 161 |
| chr3:33954803-33956817 | p2 | (TA)7 | GGTCAAGTATCAACCGTGGC | TTCCCCACATCATATTCGGT | 163 |
| chr3:33954803-33956817 | p2 | (TA)7 | GGTCAAGTATCAACCGTGGC | TTCCCCACATCATATTCGGT | 163 |
| chr3:33966801-33968816 | p2 | (AG)7 | TTCACGGCTAAACAGGCTCT | CTCTTCTCCTCCACCTTCCC | 197 |
| chr3:33966830-33968857 | p2 | (AG)7 | TTCACGGCTAAACAGGCTCT | CTCTTCTCCTCCACCTTCCC | 168 |
| chr3:33995194-33997209 | p2 | (AG)7 | TTCACGGCTAAACAGGCTCT | CTCTTCTCCTCCACCTTCCC | 197 |
| chr3:33995223-33997250 | p2 | (AG)7 | TTCACGGCTAAACAGGCTCT | CTCTTCTCCTCCACCTTCCC | 168 |
| chr3:34023137-34025165 | p2 | (TA)9 | CCTCCTCCCCATTACCATTT | CGATTTGGTTTGCTACGACA | 155 |
| chr3:34036771-34038786 | p5 | (ACCAA)3 | ATATCTCCCCATCGAGTCCC | AATGAATCCACCGACTTTCG | 169 |
| chr3:34036771-34038786 | p5 | (AAACC)3 | ATATCTCCCCATCGAGTCCC | AATGAATCCACCGACTTTCG | 169 |
| chr3:34136135-34138156 | p5 | (AATGC)3 | CCGGGACAAAGAAAATCAAA | GATTCCACCGACCTTCTCAA | 235 |

| Marker ID | SSR type | SSR motif | FORWARD PRIMER (5'-3') | REVERSE PRIMER (5'-3') | size (bp) |
|---|---|---|---|---|---|
| chr3:34136135-34138156 | p3 | (TTG)7 | CCGGGACAAAGAAAATCAAA | GATTCCACCGACCTTCTCAA | 235 |
| chr3:34189064-34191085 | p5 | (AATGC)3 | CCGGGACAAAGAAAATCAAA | GATTCCACCGACCTTCTCAA | 235 |
| chr3:34189064-34191085 | p3 | (TTG)7 | CCGGGACAAAGAAAATCAAA | GATTCCACCGACCTTCTCAA | 235 |
| chr3:34506161-34508173 | p2 | (AT)9 | TTGATAAATGCACTGCGGAA | CTTTGTGGCCCACCTAAAAA | 153 |
| chr3:34506161-34508173 | p2 | (AT)6 | TTGATAAATGCACTGCGGAA | CTTTGTGGCCCACCTAAAAA | 153 |
| chr3:34651425-34653455 | p3 | (TAT)10 | TTCATAGGTCCCCCAGAGTG | ACATCTCGGTAGCACAACCC | 298 |
| chr3:34651425-34653455 | p2 | (AG)6 | TTCATAGGTCCCCCAGAGTG | ACATCTCGGTAGCACAACCC | 298 |
| chr3:34751788-34753804 | p2 | (GA)8 | TGCCTCACAGAGCATACGAC | TGTGAAGCCGACCACATTTA | 105 |
| chr3:34835333-34837351 | p2 | (CT)6 | CCCTCTCTCTGCTCCCTTCT | GTTGGAAGGTTTTTGGACGA | 242 |
| chr3:34835333-34837351 | p6 | (GTTGTA)3 | CCCTCTCTCTGCTCCCTTCT | GTTGGAAGGTTTTTGGACGA | 242 |
| chr3:34835333-34837351 | p5 | (TTAGT)3 | CCCTCTCTCTGCTCCCTTCT | GTTGGAAGGTTTTTGGACGA | 242 |
| chr3:34848012-34850042 | p2 | (AT)15 | CAGCGGGCTTTAGTTCGTAG | CATTCAGCAATTCACGTTGG | 180 |
| chr3:34882838-34884854 | p4 | (AAAT)4 | AAAGGTTTCAAGAGGCAGCA | GCTTACATTTGGGCCCTTTT | 139 |
| chr3:34890370-34892388 | p6 | (TATATG)3 | GGGTCGGGACACGTTATATG | GTGAGGGACTCGGTGGATAA | 108 |
| chr3:34890370-34892388 | p6 | (ATTATG)3 | GGGTCGGGACACGTTATATG | GTGAGGGACTCGGTGGATAA | 108 |
| chr3:34939584-34941598 | p2 | (TA)7 | TGGTTATACGGTCCCGACAT | ATTGGAGACCAAACGAAACG | 209 |
| chr3:35004008-35006056 | p2 | (AT)24 | CTTTGGCCCCCTCATTTTAT | CCCTTGGAATCCCCAATACT | 189 |
| chr3:35004008-35006056 | p2 | (AT)23 | CTTTGGCCCCCTCATTTTAT | CCCTTGGAATCCCCAATACT | 189 |
| chr3:35021000-35023015 | p5 | (TCATT)3 | GAAGCGTCAAAGAATAGCGG | CCGCTTTTGGTATCGTGATT | 253 |
| chr3:35038122-35040188 | p3 | (ATA)22 | CCTCAAATCGATGTGGGACT | CAGATTCACCAAACCCGACT | 286 |
| chr3:35158242-35160258 | p5 | (ATATC)3 | GTCCCGATACCAACATGGTC | TGTCAGCGACTGCTACAACC | 296 |
| chr3:35181985-35184006 | p3 | (TAA)7 | CCCGAGAATACAAATCGCAT | TCCGTTTGATATTGTGGCAA | 208 |
| chr3:35285092-35287118 | p2 | (AT)6 | GGAACTGTCTGCAAAGCCTC | TTCGTACTGACGTCCCCTTC | 151 |
| chr3:35285092-35287118 | p2 | (TA)13 | GGAACTGTCTGCAAAGCCTC | TTCGTACTGACGTCCCCTTC | 151 |
| chr3:35399910-35401930 | p2 | (TA)10 | GCTCGTGTCTAGCGTCCTTC | CAGCCATTTTTCCACAGGTT | 201 |
| chr3:35655296-35657314 | p2 | (AT)27 | GGCAATTTGACCATTTTTCG | ATCACCAAGCCTTTGACCAC | 253 |
| chr3:35655296-35657314 | p2 | (AT)9 | GGCAATTTGACCATTTTTCG | ATCACCAAGCCTTTGACCAC | 253 |
| chr3:35655296-35657314 | p3 | (TTC)18 | GGCAATTTGACCATTTTTCG | ATCACCAAGCCTTTGACCAC | 253 |
| chr3:35719975-35721993 | p3 | (TAT)6 | CCATTTTCTTTGGACGCTGT | CCCAATGGGATTCGTATGAC | 239 |
| chr3:35759297-35761309 | p2 | (CA)6 | CTTATCCGCTCCAAATCTGC | GTAAACCACCCCATTCGTTG | 201 |
| chr3:35953595-35955613 | p6 | (TATATG)3 | TCTGAGATACCCAAGGCTCG | TGCATATCATAACTCCGCCA | 167 |
| chr3:35953595-35955613 | p6 | (TATATG)3 | TCTGAGATACCCAAGGCTCG | TGCATATCATAACTCCGCCA | 167 |
| chr3:36066480-36068496 | p4 | (AATT)4 | CAACTGCAGATGCCAAGGTA | CCAAGGGTGCTTTTTGGTTA | 243 |
| chr3:36066480-36068496 | p5 | (ATAAA)3 | CAACTGCAGATGCCAAGGTA | CCAAGGGTGCTTTTTGGTTA | 243 |
| chr3:36275221-36277233 | p2 | (AG)6 | GAGGCACAAAGTCTGAAGGC | AAACTGCCAGGTGACAGACC | 250 |
| chr3:36301475-36303490 | p5 | (TTTAA)3 | GATAACACCGAGTCCCGAAA | GAAGGGACGTCAGTACGAGC | 165 |

| Marker ID | SSR type | SSR motif | FORWARD PRIMER (5'-3') | REVERSE PRIMER (5'-3') | size (bp) |
|---|---|---|---|---|---|
| chr3:36356370-36358382 | p2 | (AT)6 | TTCCCTTTGGCTTTGTTTTG | GATTTGGGACTTCTCCGTCA | 179 |
| chr3:36412500-36414516 | p2 | (AT)8 | CTCAGCCACACAAAGCGTAA | CACCATTTCACACAGCCTTG | 146 |
| chr3:36475240-36477264 | p3 | (AAG)8 | TTGCTGAAATGGATGGTCAA | ACAAGCAAAGGGAATGCAAC | 192 |
| chr3:36492885-36494903 | p6 | (GGGTTG)3 | GGCCAGGTTAAATGGTTTGA | CATGTTAAGCTTCGTTGGCA | 226 |
| chr3:36560814-36562826 | p2 | (AT)6 | ACCCTTACCCGTTTCTCTGG | CTTGGGGGTAACCTCGTCTT | 286 |
| chr3:36713339-36715353 | p3 | (AAG)12 | CGGTGGTATCAACCAAAACC | ATGCCATTGAAATTGGGAAA | 207 |
| chr3:36713339-36715353 | p2 | (TA)7 | CGGTGGTATCAACCAAAACC | ATGCCATTGAAATTGGGAAA | 207 |
| chr3:36761606-36763624 | p2 | (AT)9 | GTCCATTCTTCGCTTCCAAC | GGCGTTGTGAATCTTTCGTT | 156 |
| chr3:36854816-36856836 | p5 | (AATGC)4 | AAGAACGTGTTCCCACCATC | CCGTACCCATCCTTGAGAGA | 171 |
| chr3:36985051-36987063 | p2 | (TA)6 | ACGGTGGAAGAAAAGGCTCT | ATGTTTGGATGGTCCCATGT | 139 |
| chr3:37066453-37068477 | p2 | (TA)12 | CCTTTTTGGATGATTTGGGA | GGTTTCTTGCGGTTGATGAT | 225 |
| chr3:37126859-37128874 | p3 | (ATC)5 | ACAGGAATTCACAAGGACGG | CCTAAGAGCTTCATCAGCGG | 248 |
| chr3:37146167-37148182 | p5 | (ATACG)3 | TCAGCAACTGAACTGCATCC | CCCAGAAGGGCAAATCAGTA | 180 |
| chr3:37146167-37148182 | p2 | (AT)9 | TCAGCAACTGAACTGCATCC | CCCAGAAGGGCAAATCAGTA | 180 |
| chr3:37186054-37188114 | p2 | (AT)30 | TTTTTCAAGTCCCCACCAAG | AACACGTGGCATGATCTGAA | 275 |
| chr3:37197290-37199308 | p3 | (GAT)6 | TTTCAAACCCCATCATCCTC | TTGTTCCGTTTCATGGTTGA | 224 |
| chr3:37197290-37199308 | p6 | (AAAGTC)3 | TTTCAAACCCCATCATCCTC | TTGTTCCGTTTCATGGTTGA | 224 |
| chr3:37293267-37295282 | p5 | (ATTTC)3 | CCACCTTCTAATGCGGGTAA | GTGGGTGCCTGAACTGAAAT | 247 |
| chr3:37355392-37357407 | p5 | (TTTTC)3 | ATCATCATAAAACTCGCCCG | TGTCCTCTCTTGGCCATTTC | 251 |
| chr3:37368399-37370414 | p3 | (ATT)5 | ACCCAGTGGTTATGGAGCTG | AGTCTGGAATCGGGAACCTT | 141 |
| chr3:37406966-37408986 | p2 | (TA)10 | ATGTAACTGCCCACTTTCGC | TGTTTTTGCGTTGACATGGT | 126 |
| chr3:37419317-37421335 | p6 | (CATATA)3 | TGCTTAACTGTCACACCCCA | ATTGTAAACGGCTCCACCAG | 222 |
| chr3:37419317-37421335 | p5 | (ATTCA)3 | TGCTTAACTGTCACACCCCA | ATTGTAAACGGCTCCACCAG | 222 |
| chr3:37427402-37429420 | p6 | (CATATA)3 | ATTGGCCAAGCACTCCTATG | GTTGACCGTGAGTTTCCGAT | 199 |
| chr3:37465095-37467123 | p2 | (TA)14 | CCAAAAGCCTTCTCCAAACA | CAACTTGTGCCAAAAACCCT | 250 |
| chr3:37474546-37476561 | p5 | (TTCTG)3 | TGGCATGCCCTAGTGTGTTA | CATTAAGGCAAAACCGGAAA | 299 |
| chr3:37506304-37508318 | p2 | (AG)7 | TTGATTACTCCCTCCGTCCA | GCCCTGTTTGGCAGTGTAAT | 266 |
| chr3:37506304-37508318 | p2 | (AG)7 | TTGATTACTCCCTCCGTCCA | GCCCTGTTTGGCAGTGTAAT | 266 |
| chr3:37570908-37572923 | p3 | (ACA)5 | GTTTTCAGCCAGCATTGACA | CCTGGAAACTAAAGCACCCA | 205 |
| chr3:37617496-37619514 | p4 | (TTAT)4 | TTCACTTTCCCGCCATTAAG | GCTTTTGATAGGTGACCGGA | 199 |
| chr3:37617496-37619514 | p6 | (TTTTTA)3 | TTCACTTTCCCGCCATTAAG | GCTTTTGATAGGTGACCGGA | 199 |
| chr3:37617496-37619514 | p2 | (GT)10 | TTCACTTTCCCGCCATTAAG | GCTTTTGATAGGTGACCGGA | 199 |
| chr3:37645769-37647793 | p2 | (TA)12 | CATAAGCCCTTCGGCTATCA | CCGTTTTATTTGGACGCTGT | 165 |
| chr3:37736158-37738172 | p5 | (TGTTT)3 | AGCTCCACCTCTTCAGTCCA | GTACAATCACCCCAAGGTGG | 122 |
| chr3:37736176-37738212 | p5 | (TGTTT)3 | AGCTCCACCTCTTCAGTCCA | GTACAATCACCCCAAGGTGG | 104 |
| chr3:37742066-37744080 | p2 | (AC)7 | GCCGTCGATCTCTTGTTAGC | GTCTTTCCCACCGTCTTTCA | 230 |

| Marker ID | SSR type | SSR motif | FORWARD PRIMER (5'-3') | REVERSE PRIMER (5'-3') | size (bp) |
|---|---|---|---|---|---|
| chr3:37742066-37744080 | p2 | (AG)12 | GCCGTCGATCTCTTGTTAGC | GTCTTTCCCACCGTCTTTCA | 230 |
| chr3:37742183-37744207 | p2 | (AC)7 | GCCGTCGATCTCTTGTTAGC | GTCTTTCCCACCGTCTTTCA | 113 |
| chr3:37742183-37744207 | p2 | (AG)12 | GCCGTCGATCTCTTGTTAGC | GTCTTTCCCACCGTCTTTCA | 113 |
| chr3:37854226-37856241 | p5 | (ATTTC)3 | CCGGAAGAGATCCAAGATGA | ATAACAATGCCTCACAGGGC | 224 |
| chr3:38053598-38055628 | p2 | (TA)15 | GGTGTGCAAGGTTTTGGAGT | ACCTCTCCTCCATGTTGTGG | 182 |
| chr3:38128196-38130208 | p3 | (TGT)5 | ATTTTTGCGTGGATTCAAGG | CGTGACAACCAACATCCAAC | 195 |
| chr3:38128261-38130279 | p3 | (TGT)5 | ATTTTTGCGTGGATTCAAGG | CGTGACAACCAACATCCAAC | 130 |
| chr3:38138979-38141031 | p2 | (TA)6 | TTTTATTTGGGCTCCAATGC | CTTTCGAGTGTCGGTGTGAA | 214 |
| chr3:38139032-38141088 | p2 | (TA)6 | TTTTATTTGGGCTCCAATGC | CTTTCGAGTGTCGGTGTGAA | 161 |
| chr3:38291638-38293656 | p6 | (CATAAT)3 | GTATACGCTGCCCGCTTTAG | ACGACCTTACCCAATCATGC | 270 |
| chr3:38323190-38325202 | p2 | (TA)6 | ATACGACCCATGACCCACAT | CGTCATCTTAGGCCATCCAT | 264 |
| chr3:38347720-38349734 | p2 | (AT)7 | AAGACGATAAGGGCTTGGGT | AACGGGTTTGAATGGACAAA | 161 |
| chr3:38382063-38384089 | p3 | (TTG)6 | CCTATAAAAGGGGCAAAGCC | AGCCAAAATCCACCACAAAG | 119 |
| chr3:38382063-38384089 | p2 | (TA)13 | CCTATAAAAGGGGCAAAGCC | AGCCAAAATCCACCACAAAG | 119 |
| chr3:38434123-38436135 | p2 | (TG)6 | AAGGGAGACCTCCACGAAGT | CGCCTTTCTCCATATTTCCA | 239 |
| chr3:38468227-38470257 | p2 | (TA)15 | ACTGCGGAGAAGATAGCCAA | GTCGTGACACAATGACCACC | 102 |
| chr3:38471237-38473255 | p3 | (CAA)6 | CCGTCACACATCACAACACA | TTCCTCACTACTCCGCTCGT | 109 |
| chr3:38476724-38478739 | p3 | (CTG)5 | ACTTACCCCTCGGCGATACT | AATCCGAACCGATAGAGCCT | 147 |
| chr3:38479665-38481680 | p3 | (CAA)5 | ACGAGCGGAGTAGTGAGGAA | CCTCAGAAAAGCTCGGATTG | 282 |
| chr3:38546863-38548881 | p6 | (ATATCT)3 | CGGTGCATTAAGCTCCAGTT | GAGCTGTGGAAACATTGCCT | 283 |
| chr3:38601304-38603318 | p2 | (TC)7 | GCGCTCGTACAACATACCCT | TCAGTTCGTCTCAACATCGC | 170 |
| chr3:38649377-38651392 | p5 | (TTGCA)3 | ATTGGGATCGTTAGGAGGCT | CCCATTTCATTACCCCATTG | 231 |
| chr3:38683905-38685921 | p2 | (AT)8 | TGAGTTGCCACAAGAGTTGG | ACCTGACGATCAAGAATGGC | 283 |
| chr3:38686322-38688340 | p2 | (AT)9 | AATCCTCTTGCCCCATCTCT | ATTCGGGCAGCATCTGTTAC | 181 |
| chr3:38692917-38694933 | p2 | (TA)8 | CCTTGGGTTTCTTCCTCCTC | AATGAATAAGCCCTGCGATG | 293 |
| chr3:38795535-38797550 | p5 | (CATTG)3 | TCTGCCGTAGTGGACAAGTG | TGGCTTACAAAAGGTTTGGG | 122 |
| chr3:38898412-38900424 | p6 | (TATTAC)3 | ACACCAAGAGGAGCCTCAAA | GGAGACGAATTTGAGGTGGA | 205 |
| chr3:38898412-38900424 | p2 | (AT)6 | ACACCAAGAGGAGCCTCAAA | GGAGACGAATTTGAGGTGGA | 205 |
| chr3:38954287-38956305 | p5 | (TTTTG)3 | CATAAAACTCTGGCGGCTTC | GCAGGAAAGTGTTCCAGCTC | 263 |
| chr3:38954287-38956305 | p6 | (TTAACA)3 | CATAAAACTCTGGCGGCTTC | GCAGGAAAGTGTTCCAGCTC | 263 |
| chr3:39007711-39009729 | p2 | (TA)9 | CCTGTTTTCCTCCCTTTTCC | TTGGTGCAACCATGTTCTGT | 298 |
| chr3:39045376-39047396 | p4 | (ATAC)5 | AGCCCTTCGGCTATCAAAAT | CCCACAGGTAGACAGGGAGA | 239 |
| chr3:39119440-39121455 | p5 | (TAACA)3 | GCCACTCAAACGTACCCAAT | GATAGCCTCATCGGTTTCCA | 219 |
| chr3:39492095-39494110 | p5 | (CAAAT)3 | GCTTGATTCTTGGGCCTGTA | GGTTCAATAGTCGCCTCCAA | 113 |
| chr3:39501435-39503453 | p3 | (CAA)6 | TCCGAGTTAGAGCCTGCATT | TGACAGCGTTGTTTGTTTCC | 196 |
| chr3:39517950-39519966 | p4 | (TTAG)4 | AAACCATTTGAATGCAAGGC | TGGGAGATAGCAGGTATCCG | 236 |

| Marker ID | SSR type | SSR motif | FORWARD PRIMER (5'-3') | REVERSE PRIMER (5'-3') | size (bp) |
|---|---|---|---|---|---|
| chr3:39517950-39519966 | p2 | (TA)6 | AAACCATTTGAATGCAAGGC | TGGGAGATAGCAGGTATCCG | 236 |
| chr3:39518319-39520331 | p4 | (TTAG)4 | CCCACCATCTCTCCTGTGTT | CTTGAAGAAGTGGTGCCCAT | 169 |
| chr3:39518319-39520331 | p2 | (TA)6 | CCCACCATCTCTCCTGTGTT | CTTGAAGAAGTGGTGCCCAT | 169 |
| chr3:39683251-39685265 | p2 | (TA)7 | GGGTGTAAAGAGATTCGGCA | CTGGAAGGGTCGCCTATACA | 212 |
| chr3:39692848-39694864 | p4 | (ATGA)4 | CGTTCTTAAGCGTTGCATGA | CCCAAGAAGGGACGTCAGTA | 231 |
| chr3:39704134-39706162 | p2 | (TA)14 | TTGTACATGGATGGTGGTCG | AAGCCCGCAATAACAACAAC | 216 |
| chr3:39704134-39706162 | p2 | (AT)18 | TTGTACATGGATGGTGGTCG | AAGCCCGCAATAACAACAAC | 216 |
| chr3:39707608-39709623 | p5 | (GAAAT)3 | CGGAGTCCATGGCCTAATAA | GGCGGTTGATATGTGATGTG | 211 |
| chr3:39714281-39716296 | p5 | (GAAAT)3 | TCTATCGCTTGATCGTTCCC | ACGGGGATTGTTGGTTATGA | 286 |
| chr3:39758968-39760996 | p2 | (TA)8 | TCGAGGGTTCACGACCTTAC | TATCGTATAGCTGCCCGCTT | 199 |
| chr3:39758968-39760996 | p2 | (TA)14 | TCGAGGGTTCACGACCTTAC | TATCGTATAGCTGCCCGCTT | 199 |
| chr3:39792584-39794598 | p2 | (TA)7 | AAGTATGTCGGCTGCGAACT | CACACGACCGACCACAATAG | 240 |
| chr3:39839112-39841148 | p2 | (AT)18 | ATCACTCGCCAAACAAATCC | TGAATTTACAGGGGAGGTGC | 255 |
| chr3:39888765-39890783 | p6 | (TCATAT)3 | TTTCTAGGGAGTGCCCAATG | TGGTCGTCATATCCTGCTCA | 155 |
| chr3:39898392-39900412 | p2 | (AT)10 | AAGCCGACAAGGACAAACAC | TAGCGGAGCTCCATTTCATT | 274 |
| chr3:39911988-39914003 | p2 | (AT)9 | AGACAAAGCAAAAGCCCAGA | TATTTTCCTGGCTTGCCATC | 144 |
| chr3:39911988-39914003 | p3 | (TTG)5 | AGACAAAGCAAAAGCCCAGA | TATTTTCCTGGCTTGCCATC | 144 |
| chr3:39932587-39934602 | p3 | (GAA)5 | CAAGGTGATGTACTGGGCCT | TGTCCCGGTGGTTAATGAAT | 154 |
| chr3:40008792-40010807 | p2 | (TA)9 | GCAGGTACACCGACAGGTTT | ATAATAACGGGGATCAGGGC | 117 |
| chr3:40008792-40010807 | p3 | (ATC)5 | GCAGGTACACCGACAGGTTT | ATAATAACGGGGATCAGGGC | 117 |
| chr3:40063111-40065123 | p2 | (AC)6 | GCGAAGTGTGAAAGAGGGAA | TGGGAGAAAATCAATCTGCC | 162 |
| chr3:40113594-40115612 | p6 | (ATATAC)3 | CTTGTGGCAATTCAAGCTCA | AAGGGTGTCATGAGGTTTGG | 141 |
| chr3:40166523-40168535 | p2 | (AT)6 | TGATAGGCAACGATCAACCA | ACGAATTGTGTCCCAGCTTC | 100 |
| chr3:40207018-40209033 | p2 | (AG)6 | ACCATATGTGATGGGCCTGT | CGTGATTCGGGACTCAGATT | 277 |
| chr3:40207018-40209033 | p3 | (GTT)5 | ACCATATGTGATGGGCCTGT | CGTGATTCGGGACTCAGATT | 277 |
| chr3:40207018-40209033 | p2 | (TA)8 | ACCATATGTGATGGGCCTGT | CGTGATTCGGGACTCAGATT | 277 |
| chr3:40209068-40211083 | p5 | (TATGA)3 | AGACTTGTCGGCTTGCGTAT | CCTGGGGAGTCACAACTGAT | 273 |
| chr3:40214087-40216102 | p2 | (AG)6 | CTTAAGCACGACCCTTGAGC | AGGAGTGTTCCCAAATTCCC | 113 |
| chr3:40214087-40216102 | p3 | (GTT)5 | CTTAAGCACGACCCTTGAGC | AGGAGTGTTCCCAAATTCCC | 113 |
| chr3:40214087-40216102 | p3 | (TGT)5 | CTTAAGCACGACCCTTGAGC | AGGAGTGTTCCCAAATTCCC | 113 |
| chr3:40214087-40216102 | p2 | (TA)6 | CTTAAGCACGACCCTTGAGC | AGGAGTGTTCCCAAATTCCC | 113 |
| chr3:40224246-40226261 | p3 | (ATA)5 | TGCAAGTTCATGGAAGCAAG | ACCCTTATCCTACGGTGGCT | 198 |
| chr3:40460677-40462731 | p2 | (TA)27 | GCTCTCTTTCTACCCGCCTT | CGCGCCACACTAATAAACAA | 192 |
| chr3:40566596-40568618 | p2 | (TA)11 | ATAAGCCGACGAGGCTAACA | TAGCGGAGCTCCATTTCATT | 298 |
| chr3:40602590-40604612 | p2 | (AT)8 | CGATGTCCATGAGGTTGATG | ACTATAGCGATGTCGGGGTG | 156 |
| chr3:40602590-40604612 | p2 | (TA)11 | CGATGTCCATGAGGTTGATG | ACTATAGCGATGTCGGGGTG | 156 |

| Marker ID | SSR type | SSR motif | FORWARD PRIMER (5'-3') | REVERSE PRIMER (5'-3') | size (bp) |
|---|---|---|---|---|---|
| chr3:40647037-40649052 | p5 | (AAAGG)3 | GGATTTCGGCATGAGATTGT | CCCAATACCAATACGGTTGC | 175 |
| chr3:40660174-40662186 | p2 | (AT)6 | CGATGAAGCGAACAAAATGA | CAGGGACTACATAGCGGAGC | 238 |
| chr3:40686732-40688744 | p2 | (CA)6 | TCCGGAAGGTCTCATATTCG | AAGCCTCGGGGATGTTAGAT | 220 |
| chr3:40693397-40695409 | p2 | (CA)6 | TCCGGAAGGTCTCATATTCG | AAGCCTCGGGGATGTTAGAT | 220 |
| chr3:40782286-40784318 | p2 | (AT)16 | ACAATAAGGAAAATCCGGGG | GAATTGGGCCCTTTTTCATT | 125 |
| chr3:40782286-40784318 | p2 | (AT)8 | ACAATAAGGAAAATCCGGGG | GAATTGGGCCCTTTTTCATT | 125 |
| chr3:40793836-40795854 | p6 | (TAAAAT)3 | ACTGCATGTGTTCACCGTGT | ATATATACCGTACCCGGCCC | 160 |
| chr3:40793836-40795854 | p5 | (CAAAC)4 | ACTGCATGTGTTCACCGTGT | ATATATACCGTACCCGGCCC | 160 |
| chr3:40794588-40796608 | p6 | (TAAAAT)3 | CTTTGGGGCAATACTTGCAT | GTTGCAACTCAAGCACATGG | 300 |
| chr3:40794588-40796608 | p5 | (CAAAC)4 | CTTTGGGGCAATACTTGCAT | GTTGCAACTCAAGCACATGG | 300 |
| chr3:40827384-40829410 | p2 | (AT)13 | CAGGGTCATGGTAAGAGGGA | TACCACACTGGACACACCGT | 107 |
| chr3:40827384-40829410 | p5 | (AGGAC)3 | CAGGGTCATGGTAAGAGGGA | TACCACACTGGACACACCGT | 107 |
| chr3:41050382-41052406 | p3 | (GTT)6 | CGCGAAAATATGTGTGTTGG | AAAACAGCCCACACGGTTAC | 242 |
| chr3:41050382-41052406 | p4 | (CTTT)6 | CGCGAAAATATGTGTGTTGG | AAAACAGCCCACACGGTTAC | 242 |
| chr3:41164370-41166384 | p2 | (AT)7 | TCTTCTTCCCCAAATTGTCG | TGCTTTCTCCCATCAATTCC | 237 |
| chr3:41170641-41172653 | p2 | (TA)6 | GGGTCTTCATCCGTCTTTGA | AGATGAACAAGAGGGGGCTT | 249 |
| chr3:41171115-41173135 | p2 | (TA)6 | TGGTATTCTTGCCCTTCTGG | TGATAAGGGGTGCTCGGTAG | 100 |
| chr3:41171235-41173249 | p2 | (TA)6 | ACACAAACCGGTCAGGCTAC | GTGACCTGCCAGCTTAGGAC | 240 |
| chr3:41183022-41185048 | p2 | (AT)13 | CTGCCATATCCCGGAGTCTA | CTAGTTGCCACTCAGGCACA | 200 |
| chr3:41211516-41213528 | p2 | (AT)6 | ATCAAGCCATGAAGTTCGCT | CAGCTCCATTGCACGATTTA | 218 |
| chr3:41227676-41229688 | p2 | (TA)6 | AAAACGGGGTGTGAAGTTTG | AGTCGGGTGGAGTGAAAATG | 167 |
| chr3:41231899-41233914 | p5 | (TAAAA)3 | TCTTTTGTTCTGTGCGTTCG | GCCCAATTCATAAGCTCCAA | 235 |
| chr3:41276821-41278871 | p3 | (TTG)6 | TTGAAAAACATCGTGTGGGA | CGCTTATTCTCCCGAACTTG | 211 |
| chr3:41276821-41278871 | p2 | (TA)25 | TTGAAAAACATCGTGTGGGA | CGCTTATTCTCCCGAACTTG | 211 |
| chr3:41345974-41348004 | p2 | (AT)15 | TCGAACCCCTGGTCTGTAAC | GTAAGAGGGCCAGATCCACA | 152 |
| chr3:41386457-41388475 | p2 | (AT)9 | CGATTGTCACATTTCCCCTT | CTGAGGCAAATGTTACGGGT | 275 |
| chr3:41389057-41391072 | p5 | (TGATT)3 | ATTTGGAGTGACGACCAAGC | AAGGAAATTGGAGCGGTTTT | 168 |
| chr3:41470011-41472051 | p2 | (AT)20 | GCAAGAACCTATCTGCTCGG | TGCAGAGGACTTTGAGGGTT | 157 |
| chr3:41518075-41520095 | p2 | (AT)10 | CCCCTGGGAAGTGTACAAAA | TACCCGTTGATACAGCCACA | 251 |
| chr3:41600682-41602694 | p2 | (TG)6 | CCAAGATGCCTCACCAAGAT | TAGCGCTATATCGGGACCAC | 140 |
| chr3:41659889-41661901 | p2 | (AC)6 | TAGGGCCCTTAGGCTTCAAT | AGTTCCCCACTTCTTCGGTT | 151 |
| chr3:41681678-41683690 | p2 | (TA)6 | AATTCGATCTTGTCGCTCGT | ACGTCGTTCAAGAGCTTGGT | 187 |
| chr3:41681678-41683690 | p2 | (AT)24 | AATTCGATCTTGTCGCTCGT | ACGTCGTTCAAGAGCTTGGT | 187 |
| chr3:41684061-41686083 | p2 | (TA)11 | GCCTTGGATGTGCCATAACT | TGTGTGCCGGTAGTTGAGAG | 270 |
| chr3:41684061-41686083 | p2 | (AT)7 | GCCTTGGATGTGCCATAACT | TGTGTGCCGGTAGTTGAGAG | 270 |
| chr3:41715275-41717301 | p2 | (TA)10 | CCATGACTAGTGCCCGATCT | TGTTCGTCTGTAGTGGCAGC | 165 |

| Marker ID | SSR type | SSR motif | FORWARD PRIMER (5'-3') | REVERSE PRIMER (5'-3') | size (bp) |
|---|---|---|---|---|---|
| chr3:41715275-41717301 | p2 | (AT)13 | CCATGACTAGTGCCCGATCT | TGTTCGTCTGTAGTGGCAGC | 165 |
| chr3:41806991-41809023 | p2 | (TA)6 | GCACAGTGTCGCTGCATTAT | ACCGTACCTGGCCCTTTAGT | 126 |
| chr3:41806991-41809023 | p2 | (TA)16 | GCACAGTGTCGCTGCATTAT | ACCGTACCTGGCCCTTTAGT | 126 |
| chr3:41878511-41880526 | p3 | (TTG)5 | GACCGGATTACCATTTGACG | TCTCCCTGCCTCATTTTGAC | 246 |
| chr3:41878511-41880526 | p2 | (TA)9 | GACCGGATTACCATTTGACG | TCTCCCTGCCTCATTTTGAC | 246 |
| chr3:41878511-41880526 | p2 | (TA)9 | GACCGGATTACCATTTGACG | TCTCCCTGCCTCATTTTGAC | 246 |
| chr3:41955845-41957860 | p5 | (AGGAC)3 | TTGCCATGAAAGAGTGCAAG | TTCTCATGATTTTCCCCCAG | 126 |
| chr3:41994104-41996120 | p4 | (TTAC)4 | TTCCTTAATGCCCTGCAATC | TGTGGACTCATGCCCTTACA | 111 |
| chr3:42065832-42067938 | p2 | (AT)6 | TGAAGGTATCCCTGGTCGAG | ACAGTCCCAAAGGCAAACTG | 185 |
| chr3:42133357-42135372 | p5 | (ATTTT)3 | TCCTTGATTCACTTGCCCTC | TGGACTCTAGTGGCTGCCTT | 226 |
| chr3:42144696-42146711 | p5 | (ATTTT)3 | TCCTTGATTCACTTGCCCTC | TGGACTCTAGTGGCTGCCTT | 226 |
| chr3:42192481-42194495 | p2 | (GA)6 | TGACACGTGCTTCCTCAGAC | TTTGGAGGGTTTCCACTTTG | 154 |
| chr3:42192481-42194495 | p2 | (AT)7 | TGACACGTGCTTCCTCAGAC | TTTGGAGGGTTTCCACTTTG | 154 |
| chr3:42199088-42201100 | p2 | (GA)6 | GCCGTAGAAAATAAACGCCA | TCCCTTTTGGAACATTCGTC | 288 |
| chr3:42204596-42206608 | p2 | (TA)6 | TAGATTCTCGGGGTTGGTTG | TGCCATTCGGAAACATATCA | 101 |
| chr3:42204596-42206608 | p3 | (ATG)5 | TAGATTCTCGGGGTTGGTTG | TGCCATTCGGAAACATATCA | 101 |
| chr3:42205191-42207206 | p2 | (TA)6 | TATCGTGCGCACTCCACTAC | ACCGTACCTGGCCCTCTAGT | 131 |
| chr3:42205191-42207206 | p3 | (ATG)5 | TATCGTGCGCACTCCACTAC | ACCGTACCTGGCCCTCTAGT | 131 |
| chr3:42240801-42242819 | p6 | (TATATG)3 | TAAACAAGGGAAATGCTGCC | TTTCACAAGGATCATGCCAA | 154 |
| chr3:42240801-42242819 | p2 | (TA)21 | TAAACAAGGGAAATGCTGCC | TTTCACAAGGATCATGCCAA | 154 |
| chr3:42279593-42281608 | p2 | (TG)7 | CACGTATTTTGCACATTCGG | TTTTGCCCTTAGCCTTCCTT | 110 |
| chr3:42279593-42281608 | p3 | (TTG)5 | CACGTATTTTGCACATTCGG | TTTTGCCCTTAGCCTTCCTT | 110 |
| chr3:42384261-42386273 | p2 | (TA)6 | TGACGACGAGAAACTTGGTG | ATCATTTTGTTGGCCTCGTC | 277 |
| chr3:42403072-42405087 | p2 | (GT)6 | TTCAACAAAAACCCCTTTCG | GCTTCACAACCATGCAAAAA | 207 |
| chr3:42403072-42405087 | p3 | (GTT)5 | TTCAACAAAAACCCCTTTCG | GCTTCACAACCATGCAAAAA | 207 |
| chr3:42542172-42544193 | p3 | (CAT)7 | TCGTCACCAGTCAGCAAGAC | ACTGACGTCCCTTCTTGTGG | 292 |
| chr3:42547112-42549126 | p2 | (TA)7 | CCACTTGCCATTCTACAGCA | AGTTGCAATTAGGACGTGGG | 180 |
| chr3:42841160-42843172 | p2 | (AT)6 | TTCCGAGGATGAACTGCTCT | CGGCTTGCGTTTCATTTTAT | 173 |
| chr3:42857929-42859973 | p2 | (AT)9 | TAAAGGCCCGTGAATAGCAC | TTTGGTATGCCCCTTTTGAG | 122 |
| chr3:42857929-42859973 | p2 | (AT)17 | TAAAGGCCCGTGAATAGCAC | TTTGGTATGCCCCTTTTGAG | 122 |
| chr3:42902272-42904287 | p5 | (ACCAA)3 | ACGGGAAAATGTTCGAGTTG | TTTTCCTCCTTCGTGGTACG | 244 |
| chr3:42902272-42904287 | p5 | (AAACC)3 | ACGGGAAAATGTTCGAGTTG | TTTTCCTCCTTCGTGGTACG | 244 |
| chr3:42961984-42964010 | p2 | (AT)13 | TGCACTGGTACCACGTTTGT | CTGGGGTATGTGGATATGGG | 144 |
| chr3:42989734-42991764 | p3 | (ATA)10 | TTCTTGCGATTCTCAGCCTT | CACAAGAAACCCTAGCTCGC | 259 |
| chr3:43066872-43068887 | p5 | (AAATG)3 | ATCGCTCACTTGTTCTCCGT | GAAGCGTTGTCCGACTTCTC | 289 |
| chr3:43144126-43146146 | p4 | (TATG)5 | CCTTCAAGGTGAGGCAGAAC | CATATAGCTGCCCGCCTTAG | 255 |

| Marker ID | SSR type | SSR motif | FORWARD PRIMER (5'-3') | REVERSE PRIMER (5'-3') | size (bp) |
|---|---|---|---|---|---|
| chr3:43193205-43195217 | p2 | (AT)6 | ACGACAAGCAGTGGAGGACT | CCCCCTTTTTAAACCCTCAA | 251 |
| chr3:43193218-43195232 | p2 | (AT)6 | ACGACAAGCAGTGGAGGACT | CCCCCTTTTTAAACCCTCAA | 238 |
| chr3:43193233-43195249 | p2 | (AT)6 | ACGACAAGCAGTGGAGGACT | CCCCCTTTTTAAACCCTCAA | 223 |
| chr3:43240037-43242075 | p2 | (AT)19 | GTTGAACCAACGGAGAGGAA | GCCGTTTGGTTATCCCTACA | 212 |
| chr3:43240037-43242075 | p2 | (AT)7 | GTTGAACCAACGGAGAGGAA | GCCGTTTGGTTATCCCTACA | 212 |
| chr3:43240428-43242458 | p2 | (AT)19 | GAATGGAGGTCATAGAGCGG | GCCTTGTGTGGGTATTTCGT | 275 |
| chr3:43240428-43242458 | p2 | (AT)7 | GAATGGAGGTCATAGAGCGG | GCCTTGTGTGGGTATTTCGT | 275 |
| chr3:43240593-43242605 | p2 | (AT)19 | GAATGGAGGTCATAGAGCGG | GCCTTGTGTGGGTATTTCGT | 110 |
| chr3:43240593-43242605 | p2 | (AT)7 | GAATGGAGGTCATAGAGCGG | GCCTTGTGTGGGTATTTCGT | 110 |
| chr3:43307369-43309407 | p6 | (AAAACA)3 | CGGTTAAGGGTCCAATAGCA | TTTAGCGAAAAGGGTCCAAA | 170 |
| chr3:43337281-43339296 | p5 | (GTTCG)3 | CCGCTGGGACTTTTTCATAA | TACCCCTATTTTGCACTGGC | 119 |
| chr3:43337281-43339296 | p5 | (ATTTG)3 | CCGCTGGGACTTTTTCATAA | TACCCCTATTTTGCACTGGC | 119 |
| chr3:43350898-43352916 | p2 | (TA)9 | AACACTCCCCTCAACCCTTT | TACCTTATGCAGCCCCAAAC | 235 |
| chr3:43358636-43360672 | p3 | (TAT)12 | AGGGTGATCCTTCATTTCCC | AGGCAAGCTCACCCTAGACA | 189 |
| chr3:43362544-43364562 | p3 | (TAT)6 | TGGTCACAAGATGACATCCC | CTCAAATTTGGGGGAAATGA | 195 |
| chr3:43458966-43460984 | p3 | (GTT)6 | TGTGTCGCTTGCGTTACTTC | GTGAAATCCCATAACGTGGG | 274 |
| chr3:43458983-43461001 | p3 | (GTT)6 | TGTGTCGCTTGCGTTACTTC | GTGAAATCCCATAACGTGGG | 257 |
| chr3:43525251-43527265 | p2 | (CA)7 | GCGAGCCTTACATACCTTGC | GGCCGAGTTGTAATTCTGGA | 221 |
| chr3:43556249-43558263 | p2 | (TA)7 | ATATATGCAAGCCGACAGGG | TTGTTCCGGAGCTAGCTTGT | 247 |
| chr3:43716691-43718709 | p3 | (TTC)6 | AGCCTCAGTTTCAGCTTCCA | GCAATACAGATGGGGCAAGT | 206 |
| chr3:43778066-43780081 | p5 | (TGCAT)3 | GACAATTCTTTTACGGCCCA | GGAAGGAAACCGTTTTGTGA | 222 |
| chr3:43784385-43786417 | p2 | (TA)16 | GAATTCAACTCGGCCACATT | GCATGTTGATTGGGTTGTTG | 119 |
| chr3:43784771-43786791 | p2 | (TA)16 | CATTTCAAGCCACCAAACCT | TAAGTATAGGGGTGTGGCCG | 137 |
| chr3:43784793-43786877 | p2 | (TA)16 | CATTTCAAGCCACCAAACCT | TAAGTATAGGGGTGTGGCCG | 115 |
| chr3:43789783-43791798 | p3 | (TCA)5 | TTGCAGACAACCCCTAGGAC | GAGTCAGTAGGTTCGCTCCG | 101 |
| chr3:43795960-43797975 | p3 | (TCA)5 | TTGCAGACAACCCCTAGGAC | GAGTCAGTAGGTTCGCTCCG | 101 |
| chr3:43812284-43814334 | p2 | (TA)6 | GTAAGGGTTTTTGTGGCGAA | TAACCCGTCATGGTCCTAGC | 300 |
| chr3:43812465-43814477 | p2 | (TA)6 | AGAAATGACAATGGGAAGCG | TAACCCGTCATGGTCCTAGC | 151 |
| chr3:43815466-43817480 | p2 | (AT)7 | GGAGAGTTGTCGGGTCAGAG | CACAATAACACATCGGCTGG | 224 |
| chr3:43815466-43817480 | p2 | (AT)8 | GGAGAGTTGTCGGGTCAGAG | CACAATAACACATCGGCTGG | 224 |
| chr3:43818823-43820838 | p5 | (ATTTC)3 | TTGCTTCCTCTTGGCAACTT | TCCTGTCACAACAGCCCATA | 131 |
| chr3:43818823-43820838 | p2 | (AT)11 | TTGCTTCCTCTTGGCAACTT | TCCTGTCACAACAGCCCATA | 131 |
| chr3:43861089-43863128 | p3 | (ATA)13 | CCAATTGCAACATGACCTTG | CAATTTCCTGTCCTTGCCAT | 142 |
| chr3:43872402-43874430 | p6 | (ATACAT)3 | ATGCCCGCTGGTATAGACAC | ATGCACGGGAATATCCAAAA | 260 |
| chr3:43872402-43874430 | p2 | (TA)14 | ATGCCCGCTGGTATAGACAC | ATGCACGGGAATATCCAAAA | 260 |
| chr3:43872402-43874430 | p2 | (AT)6 | ATGCCCGCTGGTATAGACAC | ATGCACGGGAATATCCAAAA | 260 |

| Marker ID | SSR type | SSR motif | FORWARD PRIMER (5'-3') | REVERSE PRIMER (5'-3') | size (bp) |
|---|---|---|---|---|---|
| chr3:43873289-43875301 | p2 | (TA)14 | TGATTTTCGGAAGGAAGTCG | GGGGAAATGCTACCCAATTT | 270 |
| chr3:43873289-43875301 | p2 | (AT)6 | TGATTTTCGGAAGGAAGTCG | GGGGAAATGCTACCCAATTT | 270 |
| chr3:43881602-43883616 | p2 | (AT)7 | AGGAGGCCAACTTTACCGAT | TGGTGATAGGGCTGTTAGGG | 159 |
| chr3:43922221-43924233 | p2 | (CT)6 | CAACAACCAAGGCACGTCTA | GAAGCGTATTGGGCATTGTT | 242 |
| chr3:43960949-43962967 | p2 | (AT)20 | GACTCAACCAGATGGGCACT | CCCGCCATATGACACTCTTT | 264 |
| chr3:44000379-44002397 | p3 | (ATG)6 | GAAAATTCTGGGCCAGTGAA | TGTCCGAACCTAAATCCGAG | 224 |
| chr3:44012224-44014250 | p2 | (AT)13 | GCGGCTATTGTGAGGGATAA | ACCATGGCATCTTGGAGAAC | 166 |
| chr3:44029404-44031418 | p2 | (TA)7 | AAGAGAGCGGCCATGAAATA | GGCGACAAATTCGATGAGAT | 203 |
| chr3:44029404-44031418 | p2 | (GT)6 | AAGAGAGCGGCCATGAAATA | GGCGACAAATTCGATGAGAT | 203 |
| chr3:44097009-44099023 | p2 | (AT)7 | CATTGGACGGTTCATAGCCT | ATCCTGTCGAATGCCTTTTG | 153 |
| chr3:44106484-44108502 | p3 | (GTT)6 | GTGTCGCTTGCGTTACTTCA | TGGGGAGGGTAGAGTGTACG | 124 |
| chr3:44106484-44108502 | p6 | (ATACAT)3 | GTGTCGCTTGCGTTACTTCA | TGGGGAGGGTAGAGTGTACG | 124 |
| chr3:44106484-44108502 | p2 | (TA)6 | GTGTCGCTTGCGTTACTTCA | TGGGGAGGGTAGAGTGTACG | 124 |
| chr3:44133817-44135833 | p4 | (ATTC)4 | GTGCAAAACATTCCATGACG | CGCAATTGAACCCTATTCGT | 274 |
| chr3:44133817-44135833 | p2 | (CA)7 | GTGCAAAACATTCCATGACG | CGCAATTGAACCCTATTCGT | 274 |
| chr3:44292242-44294262 | p2 | (AT)6 | GACTATAGTGCCCGAGCTGC | GCGTATGTGGATATGATGCG | 229 |
| chr3:44292242-44294262 | p2 | (TA)10 | GACTATAGTGCCCGAGCTGC | GCGTATGTGGATATGATGCG | 229 |
| chr3:44302494-44304528 | p2 | (AT)17 | TGACTCCTTTGATCTCCGCT | TGCCCGGTTATCTACAGTCC | 164 |
| chr3:44302494-44304528 | p2 | (AT)18 | TGACTCCTTTGATCTCCGCT | TGCCCGGTTATCTACAGTCC | 164 |
| chr3:44304035-44306047 | p2 | (TA)6 | CCATTATGGGACTCGGTGAC | GCCGGGTACGGTATATGATG | 159 |
| chr3:44355237-44357261 | p4 | (TATT)6 | CGTGCTTCCTCAGACATTGA | ATTCAGATCACCCGAAAACG | 156 |
| chr3:44387095-44389110 | p3 | (TTG)5 | AGGGGTTATGGGCTAGTGCT | CAACGTGTCACTTCCTCCCT | 285 |
| chr3:44406689-44408701 | p2 | (AT)6 | GGTCCGAGATTTGGAGTTCA | TCAAACAATACCGTGTCCCA | 158 |
| chr3:44406689-44408701 | p2 | (TA)29 | GGTCCGAGATTTGGAGTTCA | TCAAACAATACCGTGTCCCA | 158 |
| chr3:44412963-44414979 | p2 | (AT)8 | GCTCAATTTTCTGGGGTTCA | CAAGACGGGGAGTGTTAGGA | 119 |
| chr3:44576287-44578299 | p2 | (GT)6 | TGCAATAGACCTCCAGGCTT | TAGCCTTGTCCCACACTTCC | 289 |
| chr3:44604060-44606074 | p2 | (GA)7 | CAGATTTGAAGGACGGGAAA | GACAAAAACCGTGGAAGGAA | 167 |
| chr3:44719588-44721603 | p3 | (ATC)5 | TATATAATTTCAGCCCGGCG | TCGATGCTCCGTCATTTGTA | 265 |
| chr3:44769822-44771834 | p2 | (TA)6 | TCAACTCGGCCAATGTATCA | GTTTGAAAGGTTTCGGGTGA | 115 |
| chr3:44769822-44771834 | p2 | (AT)16 | TCAACTCGGCCAATGTATCA | GTTTGAAAGGTTTCGGGTGA | 115 |
| chr3:44772493-44774505 | p6 | (TATATG)3 | CGACCTTACGTCACTCCGAT | TCCCTTTTTCCCCATATTCC | 218 |
| chr3:44772493-44774505 | p2 | (AT)6 | CGACCTTACGTCACTCCGAT | TCCCTTTTTCCCCATATTCC | 218 |
| chr3:44772493-44774505 | p2 | (AT)8 | CGACCTTACGTCACTCCGAT | TCCCTTTTTCCCCATATTCC | 218 |
| chr3:44787202-44789236 | p3 | (AAT)16 | GCTGGAGATCATGGCAATTT | GTTGCAGGTTTGCACTTTGA | 187 |
| chr3:44787202-44789236 | p2 | (AT)17 | GCTGGAGATCATGGCAATTT | GTTGCAGGTTTGCACTTTGA | 187 |
| chr3:44894722-44896734 | p2 | (AT)8 | TACTTGTGCCACACGACCAT | TGGGACATTTTAAGCCAACC | 204 |

| Marker ID | SSR type | SSR motif | FORWARD PRIMER (5'-3') | REVERSE PRIMER (5'-3') | size (bp) |
|---|---|---|---|---|---|
| chr3:44894722-44896734 | p2 | (AT)6 | TACTTGTGCCACACGACCAT | TGGGACATTTTAAGCCAACC | 204 |
| chr3:44908517-44910553 | p2 | (TA)18 | TTAAGACCCCAACTTTCCCC | TAAGGCTTCTAACCGCTCCA | 108 |
| chr3:44908517-44910553 | p2 | (AC)6 | TTAAGACCCCAACTTTCCCC | TAAGGCTTCTAACCGCTCCA | 108 |
| chr3:44917288-44919312 | p2 | (AT)12 | ACGTCGTTCCCTTCTCATTG | AGCGAAACTCTGGCGAAATA | 178 |
| chr3:44950041-44952056 | p5 | (AAAGA)3 | GCCAAAACATACTGGCCCTA | AATTGTCGGTCCGTTTCAAG | 253 |
| chr3:44950041-44952056 | p2 | (GT)6 | GCCAAAACATACTGGCCCTA | AATTGTCGGTCCGTTTCAAG | 253 |
| chr3:45088113-45090131 | p2 | (AT)9 | ATGAGCATGACACGTGCTTC | AAAATCTCGACGTGTGGGAC | 147 |
| chr3:45157604-45159625 | p6 | (CATATA)3 | AATAACACATCGGCTGAGGG | ATGCCACGCAGGTACATACA | 196 |
| chr3:45205543-45207555 | p2 | (TG)6 | ATGTTCCTCGCTTGACGAGT | CCGGTGTTATAGCTCGGGTA | 165 |
| chr3:45282767-45284779 | p2 | (AG)6 | AAGTGTGACGACGGTGATTG | ACGTCGATTCTTTCGCAAAT | 128 |
| chr3:45397283-45399315 | p2 | (AT)16 | CTAGCTCCTCCACCAGATGC | TCACCTTGACCTGCACTTTG | 176 |
| chr3:45445847-45447868 | p3 | (ATG)7 | TTCTTTTGGCATGATCCCTC | TCCAGTCAGGCCATAACCTC | 119 |
| chr3:45483587-45485605 | p6 | (TCAGTT)3 | TGGGAATTGGGTGAAGAAAC | CTCAATTCCAGCCTTTGAGC | 273 |
| chr3:45489324-45491339 | p5 | (ATATC)3 | TCACAAGCCCTTTCCAAGTT | TTGTGGTGGTTTTGGAGGAT | 165 |
| chr3:45493077-45495115 | p2 | (AT)19 | CAAATACGAACGGTGCCTCT | CGTGCAGGTTTTGGGATAGT | 134 |
| chr3:45493077-45495115 | p2 | (AT)8 | CAAATACGAACGGTGCCTCT | CGTGCAGGTTTTGGGATAGT | 134 |
| chr3:45567379-45569393 | p2 | (AT)7 | TGTTGTGATGAATGCCGTTT | CACCGTACTCGACATGGTTG | 186 |
| chr3:45595528-45597543 | p5 | (ATTTC)3 | GCCCCTAAAAGAGATCCCAG | ATAACAATGCCTCACAGGGC | 232 |
| chr3:45996591-45998613 | p2 | (AT)11 | ATTAGCGAGCGATCGAGGTA | ATGGAGATCAAAGACGTCGG | 157 |
| chr3:46028105-46030117 | p2 | (CT)6 | ACAAATAGGCTTTTGGGGCT | CGGATTTGTGTTCCCAGAGT | 221 |
| chr3:46115064-46117079 | p5 | (AAAAT)3 | GGAGGATGATGGAGGATCAA | TGGTGCTTTTTCTTGAAGGG | 103 |
| chr3:46152888-46154902 | p2 | (AT)7 | AAAGAAGAATCGGGATGGCT | GCCGGCATATTTCTGTAGGA | 255 |
| chr3:46379542-46381572 | p3 | (AAT)10 | GCATGCAATACTCCAGGGAT | TATTTCGCAGTCGGTTGTCA | 228 |
| chr3:46476556-46478576 | p4 | (TATG)5 | TACCTCGGAGAAGCACTGGT | CTTGATCCCGAAATCTTCCA | 123 |
| chr3:46476556-46478576 | p2 | (AT)6 | TACCTCGGAGAAGCACTGGT | CTTGATCCCGAAATCTTCCA | 123 |
| chr3:46535874-46537920 | p2 | (TA)23 | TGAAGCTACAGGGGAAATGC | TCACAAGGATCATGCCAAAA | 176 |
| chr3:46572501-46574516 | p3 | (ATG)5 | TGGGTTGTTGTCGTTGTTGT | TGCTCCCTTATTCGACTGCT | 193 |
| chr3:46585023-46587041 | p3 | (ATG)6 | CGTTGATTGGGAAGAAAGGA | GGGTAGCATCTCCCCTGTTT | 249 |
| chr3:46585023-46587041 | p5 | (TTGTA)4 | CGTTGATTGGGAAGAAAGGA | GGGTAGCATCTCCCCTGTTT | 249 |
| chr3:46586686-46588702 | p2 | (TA)8 | GGTTGGGTCGTGACAAAAGT | TACACTCCCTGCAGCCTCTT | 185 |
| chr3:46718910-46720925 | p5 | (ATTTC)3 | GTCGTTACAGGATCGCGTTT | CCGTTGTCAGTTGTATTGCG | 178 |
| chr3:46747306-46749378 | p2 | (AT)36 | AGGGGGTGCGAGATTTTACT | TCCGGCTTAATAATGATGGC | 291 |
| chr3:46762283-46764359 | p2 | (AT)38 | TGCCCGATTTTCGTTAAGTC | TTCGCTTTGTCGCTCTTTTT | 233 |
| chr3:46796101-46798113 | p2 | (CA)6 | TGATCAGAATAACCCCCGAG | AGGAAACGGGGATTGAGTTT | 207 |
| chr3:46812454-46814466 | p2 | (AT)6 | TCGAGATGGGCACTCATACA | TGATTATTCAGGGGTACGGC | 214 |
| chr3:46829527-46831539 | p2 | (AG)6 | GTACTTGTTGCCCGGTCACT | AAAGCTACAGTCGCCCAGAA | 221 |

| Marker ID | SSR type | SSR motif | FORWARD PRIMER (5'-3') | REVERSE PRIMER (5'-3') | size (bp) |
|---|---|---|---|---|---|
| chr3:46838809-46840825 | p2 | (TA)8 | TGTGTATGGGGCAGTTTTGA | TGACCCTCACAATCCCTTTC | 206 |
| chr3:46844724-46846736 | p2 | (AG)6 | TCCGGTCTTTCATGTGTCAA | AACACTTTAACCGACGCCAC | 200 |
| chr3:46866418-46868434 | p2 | (TA)8 | GAGAATGGGAAATTCGGGTT | ATTACGTTCGCTTTGTCGCT | 207 |
| chr3:46925691-46927711 | p2 | (TA)14 | CGTATCGTCGTTGGAATCCT | GTAGCTCCGAGTACCCGTCA | 293 |
| chr3:46925712-46927760 | p2 | (TA)14 | CGTATCGTCGTTGGAATCCT | GTAGCTCCGAGTACCCGTCA | 272 |
| chr3:47052274-47054289 | p5 | (ATTTC)3 | AAGCATGCGATCCAGATACC | ATAACAATGCCTCACAGGGC | 183 |
| chr3:47075974-47077989 | p5 | (CATAG)3 | AGCCATCTGCTCCTTTTGAA | GTAAGGGGTGCTCGGTACAA | 113 |
| chr3:47075974-47077989 | p2 | (TA)9 | AGCCATCTGCTCCTTTTGAA | GTAAGGGGTGCTCGGTACAA | 113 |
| chr3:47225574-47227592 | p2 | (AT)9 | TAACCTTTGATTTCAGCGGG | CGGGTCGTGACACTTACCTT | 224 |
| chr3:47244302-47246324 | p2 | (AT)11 | TCCATTGCGCAACAATACAT | TTTATGGGCATACTGGGGTC | 237 |
| chr3:47247965-47249979 | p2 | (AT)7 | CATAAGCCGACGAGGCTAAC | TAGCGGAGCTCCATTTCATT | 293 |
| chr3:47345356-47347371 | p5 | (GATAT)3 | ATAAGGGACCCCAAGCTCAT | TTGGAACAAGCCTTGGGTAG | 120 |
| chr3:47358611-47360623 | p2 | (CA)6 | TATTCTCCTTCGATGCCCAC | AGGAAATGCTTCCCGATTTT | 290 |
| chr3:47361893-47363905 | p2 | (CA)6 | TAACAATCGTGGTCAGGCAA | TGAGTTTGTGCCTCGTTCTG | 242 |
| chr3:47445582-47447594 | p2 | (TA)6 | CGAGATGGCTATGGGACATT | CCATTCAGGGGAGCATAAGA | 197 |
| chr3:47594345-47596359 | p2 | (TA)7 | CTGTATAGCGCTCTGGACCC | CGACCTTACGTCACTCCGAT | 295 |
| chr3:47673947-47675961 | p2 | (AT)7 | ACTGCCCAGTCATATAGGCG | AAACCACTGCAATGGGTAGC | 189 |
| chr3:47718093-47720105 | p2 | (TA)15 | GTGCCCTCTTAAGGTTTCCC | ACTTGTTCCAGGTTATGCGG | 200 |
| chr3:47718093-47720105 | p2 | (TA)6 | GTGCCCTCTTAAGGTTTCCC | ACTTGTTCCAGGTTATGCGG | 200 |
| chr3:47737765-47739777 | p2 | (TA)15 | GTGCCCTCTTAAGGTTTCCC | ACTTGTTCCAGGTTATGCGG | 200 |
| chr3:47737765-47739777 | p2 | (TA)6 | GTGCCCTCTTAAGGTTTCCC | ACTTGTTCCAGGTTATGCGG | 200 |
| chr3:47757459-47759473 | p2 | (TA)7 | ACTAACGGGTCACGGAGTTG | CTACGTGGGTTGCCTCTCAT | 111 |
| chr3:47819290-47821308 | p2 | (TA)6 | TCCTTCTTTTGGCATGATCC | TACACCTTTCCACACCCCAT | 265 |
| chr3:47819290-47821308 | p2 | (TA)9 | TCCTTCTTTTGGCATGATCC | TACACCTTTCCACACCCCAT | 265 |
| chr3:47819290-47821308 | p2 | (AG)7 | TCCTTCTTTTGGCATGATCC | TACACCTTTCCACACCCCAT | 265 |
| chr3:47862575-47864595 | p2 | (AT)10 | AACCTCGTAAGCCGTGATTG | TGCTGTGGTATGGTGTTCGT | 160 |
| chr3:48023342-48025357 | p3 | (GTT)5 | GGGAAAATGGTCCAGACTGA | CACAACCCACATTGAGCATC | 204 |
| chr3:48066014-48068036 | p2 | (TA)11 | GGTCTCGGAGGCATGAATAA | GCCCGATGTAGAAGAGCAAG | 247 |
| chr3:48076993-48079011 | p3 | (TTG)6 | CACCTCGTATTTTCAGGGGA | TCCAATCCTTCCGACACTTC | 248 |
| chr3:48184502-48186514 | p2 | (TG)6 | TGAGTGCTGGACACATGACA | CCTTCCTCAATATTTCCGCA | 261 |
| chr3:48184502-48186514 | p3 | (GTT)5 | TGAGTGCTGGACACATGACA | CCTTCCTCAATATTTCCGCA | 261 |
| chr3:48301866-48303904 | p2 | (TA)19 | CGGGAACACACAACCTACCT | CCGCTTCCTACACCAACAAT | 181 |
| chr3:48308409-48310424 | p5 | (ACAAC)3 | CCGAGGGTGAGGCATAGATA | ACCTACGGGTCATGGAGTTG | 296 |
| chr3:48342633-48344657 | p3 | (TTG)8 | TTTGGTCCAACTTGAGGGAG | ACCGAACTGTTTTGTGGTCC | 113 |
| chr3:48342633-48344657 | p2 | (TA)10 | TTTGGTCCAACTTGAGGGAG | ACCGAACTGTTTTGTGGTCC | 113 |
| chr3:48342633-48344657 | p3 | (AAC)8 | TTTGGTCCAACTTGAGGGAG | ACCGAACTGTTTTGTGGTCC | 113 |

| Marker ID | SSR type | SSR motif | FORWARD PRIMER (5'-3') | REVERSE PRIMER (5'-3') | size (bp) |
|---|---|---|---|---|---|
| chr3:48343289-48345313 | p3 | (TTG)8 | CACTGATGGGAAGAAAGGGA | TCGATCTTAAAGCGAAACGG | 236 |
| chr3:48343289-48345313 | p2 | (TA)10 | CACTGATGGGAAGAAAGGGA | TCGATCTTAAAGCGAAACGG | 236 |
| chr3:48343289-48345313 | p3 | (AAC)8 | CACTGATGGGAAGAAAGGGA | TCGATCTTAAAGCGAAACGG | 236 |
| chr3:48370728-48372794 | p2 | (AT)33 | TCCTCATCTGGTTTCGATCC | GCATAGAAGAGCTCATGCCC | 273 |
| chr3:48379758-48381776 | p2 | (AT)9 | GTAGGATGGCCAGATCCTGA | TCGGGTTTCACCCAATACAT | 299 |
| chr3:48392428-48394446 | p6 | (CATATA)3 | GTATTGTCACGACCCAACCC | AGCCTTGTCGGCTTCTGTTA | 295 |
| chr3:48423204-48425222 | p6 | (CATATA)3 | AGATCGGGCACTAGTCATGG | AGCCTTGTCGGCTTCTGTTA | 207 |
| chr3:48515312-48517334 | p2 | (AT)11 | CCCCCTTCCTTTTGTTTTTC | CGAGGAAACGCCTAGAGATG | 173 |
| chr3:48515312-48517334 | p2 | (AT)7 | CCCCCTTCCTTTTGTTTTTC | CGAGGAAACGCCTAGAGATG | 173 |
| chr3:48517060-48519074 | p2 | (AT)7 | ACATATGGACGGGACAGGAC | CCGTTTTCTTTGGACGTTGT | 200 |
| chr3:48517060-48519074 | p2 | (TA)7 | ACATATGGACGGGACAGGAC | CCGTTTTCTTTGGACGTTGT | 200 |
| chr3:48517060-48519074 | p2 | (AT)7 | ACATATGGACGGGACAGGAC | CCGTTTTCTTTGGACGTTGT | 200 |
| chr3:48557556-48559572 | p2 | (TA)8 | TTTCCCAAATAAGAGTGGCG | GCCCAAATTTTCCCAGAAAT | 130 |
| chr3:48557556-48559572 | p2 | (AT)8 | TTTCCCAAATAAGAGTGGCG | GCCCAAATTTTCCCAGAAAT | 130 |
| chr3:48721200-48723214 | p2 | (AT)7 | ATTCAGTTTCGGAAGCATGG | CAAGTACCAAAGCCCGTGAT | 186 |
| chr3:48737624-48739656 | p2 | (AT)8 | GTAACGTAGTCGGGGGTTCA | ACGACGTTACGAACCTTTGG | 110 |
| chr3:48737624-48739656 | p2 | (AT)16 | GTAACGTAGTCGGGGGTTCA | ACGACGTTACGAACCTTTGG | 110 |
| chr3:48819826-48821838 | p2 | (AT)8 | ACCACTTTTTGTCACGACCC | TGACGACGTTATGAACCCAA | 102 |
| chr3:48843770-48845800 | p2 | (AT)15 | CTTCCTTCCAACTCTGTGCC | AAACGCGCAGTTAAGGAAGA | 177 |
| chr3:48931963-48933975 | p3 | (ACA)5 | CTAAGGAACGTACGACCCGA | CGACCTTACGTCACTCCGAT | 107 |
| chr3:48956348-48958363 | p3 | (TTG)5 | CACTAGCCGCCTAGATTTGG | GACGGCTCGTCAATCTTCTC | 257 |
| chr3:49024709-49026721 | p2 | (GT)6 | CATTGCCACCTGATCCTTCT | TTTTCATCTGTTTCAGCCCC | 130 |
| chr3:49073154-49075174 | p2 | (AT)10 | GACTAGTGCCCGAGTTGGAC | TGCAACACCTCAGGAGACAG | 206 |
| chr3:49073154-49075174 | p2 | (AT)6 | GACTAGTGCCCGAGTTGGAC | TGCAACACCTCAGGAGACAG | 206 |
| chr3:49289856-49291871 | p5 | (TTTCA)3 | AATTATGCGATCCAAGCACC | TGTTAAGTCGTCGGGGATTC | 197 |
| chr3:49318111-49320126 | p5 | (TATTG)3 | CAAATTGCTGACTGCCTCAA | AAGCTGACCAGCCAAACACT | 203 |
| chr3:49472785-49474797 | p5 | (TATAT)3 | GTCACCATGAAGCGTCCTCT | AGCTTATGTCGAAGGGCAAA | 209 |
| chr3:49472785-49474797 | p2 | (AT)6 | GTCACCATGAAGCGTCCTCT | AGCTTATGTCGAAGGGCAAA | 209 |
| chr3:49489401-49491415 | p2 | (TA)7 | AGGCCAAGGATTTGGATTCT | TTTATGGAAGAAGGGCATGG | 237 |
| chr3:49499897-49501911 | p2 | (TA)7 | TTACTCCAGCCCCCTATCCT | GCCGATCGGTCTCTAGTCTG | 155 |
| chr3:49502191-49504211 | p2 | (TG)10 | TCAGACCATACGATGACGGA | TATGTTTCGAACTCTGGCCC | 215 |
| chr3:49508166-49510210 | p2 | (AT)15 | AAGATCACCGCCTATGGTTG | GATGTGATGTTCCATCGTCG | 281 |
| chr3:49508166-49510210 | p2 | (AT)22 | AAGATCACCGCCTATGGTTG | GATGTGATGTTCCATCGTCG | 281 |
| chr3:49508166-49510210 | p2 | (AT)9 | AAGATCACCGCCTATGGTTG | GATGTGATGTTCCATCGTCG | 281 |
| chr3:49559178-49561204 | p2 | (TA)13 | AAAGTGGCCAAAGTCACCAC | GGACAGTCATCCAGGCCTTA | 199 |
| chr3:49618511-49620527 | p2 | (TC)8 | GACAGCGATCAAAACAAGCA | ACCGCTACAGTTACCATGCC | 112 |

| Marker ID | SSR type | SSR motif | FORWARD PRIMER (5'-3') | REVERSE PRIMER (5'-3') | size (bp) |
|---|---|---|---|---|---|
| chr3:49640578-49642608 | p2 | (AT)15 | TTAATTGGCTCGGGTTTCAC | TCAGCCCCGTATTTATGCTC | 229 |
| chr3:49640578-49642608 | p3 | (AAT)5 | TTAATTGGCTCGGGTTTCAC | TCAGCCCCGTATTTATGCTC | 229 |
| chr3:49668074-49670089 | p5 | (CAGAT)3 | GCACGTTCCCGTACATACCT | GTCCCGTCATCTGATTTCGT | 112 |
| chr3:49829199-49831214 | p5 | (CATTG)3 | CGCCATGATTGTTGAGATTG | CCGTACACTTCCTCAGCCTC | 262 |
| chr3:49951495-49953523 | p2 | (TA)14 | CGATTCCATCGTGCCTAGAT | CATATTCCCTGTGGGATTCG | 211 |
| chr3:50030200-50032212 | p2 | (TA)6 | TTCCAAACACATTTGAGGCA | CCAAAGTTGATCCCGTTGTT | 261 |
| chr3:50139665-50141681 | p3 | (ATC)5 | GACTCGCCTTCCTCCCTATC | TATAAACAGGAGGTTGCGGG | 121 |
| chr3:50156755-50158770 | p2 | (AT)7 | AGTCGGGGCCTTAACAACTT | ATTTACAAGGAGCGACACGG | 210 |
| chr3:50156755-50158770 | p2 | (TA)6 | AGTCGGGGCCTTAACAACTT | ATTTACAAGGAGCGACACGG | 210 |
| chr3:50156755-50158770 | p5 | (CATAT)3 | AGTCGGGGCCTTAACAACTT | ATTTACAAGGAGCGACACGG | 210 |
| chr3:50192968-50194988 | p2 | (AC)10 | CAAATGGCGCAAATTTTTCT | CCTTGTTTGCTGGAATAGGC | 151 |
| chr3:50206106-50208121 | p5 | (ATTAT)3 | TTGTTAGTGCATGAGCTCCG | TCACCACGAATCGAAATCAA | 241 |
| chr3:50237140-50239158 | p6 | (TATATG)3 | TTAGATTCTCGGGGATGGTG | ATATCTGCTGCGCGTCCTAT | 166 |
| chr3:50487472-50489487 | p5 | (ATGCA)3 | CGTCATGTGTCCTCACCATC | GTGTGGGTGCCACTAAAGGT | 150 |
| chr3:50498654-50500666 | p2 | (AT)6 | TCACCTACGCCTAAAATGGC | GTACCCTGTTGCCCGTAGAA | 289 |
| chr3:50514156-50516171 | p3 | (GAT)5 | TTGGAAAGGTTTTCCCAATG | TGGACCCTATAATCCCACCA | 160 |
| chr3:50668473-50670529 | p2 | (AT)28 | CATTCTAGACCGTGCCGATT | ACCTGGACGCTTCTCTTTGA | 156 |
| chr3:50724821-50726883 | p2 | (AT)31 | ATCATCATTAGCCTGCGTCC | TGATCACACCGTGCCTACAT | 179 |
| chr3:50732432-50734446 | p2 | (AT)7 | TTTGTTGACCCGAAGTAGCC | CGGAAGAGGGTGAATCACAT | 286 |
| chr3:50796905-50798933 | p2 | (AT)14 | TCTATGAAACCTTGCGGGTC | CAACAAGTAACCGAGGCCAT | 233 |
| chr3:50853931-50855949 | p3 | (TGT)6 | CCAACACAACAAACCACTCG | TGCACCGACCTAGGATTTTC | 135 |
| chr3:50929634-50931648 | p2 | (AT)7 | TGTTCATGCCCACAGGTAGA | ACAAAACGCATAGACGGGAC | 260 |
| chr3:51085464-51087482 | p2 | (AT)6 | GTCCGCTATTGGACCAGTGT | AGCATTGACATCCACGAACA | 205 |
| chr3:51085464-51087482 | p6 | (GATTCG)3 | GTCCGCTATTGGACCAGTGT | AGCATTGACATCCACGAACA | 205 |
| chr3:51152264-51154278 | p2 | (AT)7 | AATCAGACACCCTGGTGGAG | GGAAAGAGAGCATACCGCAG | 138 |
| chr3:51193866-51195932 | p5 | (ATTCA)3 | TTCACACGACACCTTCCAAA | AATGAAGGTTTGGTGGCTTG | 124 |
| chr3:51193866-51195932 | p2 | (TA)33 | TTCACACGACACCTTCCAAA | AATGAAGGTTTGGTGGCTTG | 124 |
| chr3:51199422-51201440 | p5 | (ATTCA)3 | TACTTCACCCGACACCTTCC | AATGAAGGTTTGGTGGCTTG | 181 |
| chr3:51199422-51201440 | p2 | (AT)10 | TACTTCACCCGACACCTTCC | AATGAAGGTTTGGTGGCTTG | 181 |
| chr3:51199422-51201440 | p2 | (AT)9 | TACTTCACCCGACACCTTCC | AATGAAGGTTTGGTGGCTTG | 181 |
| chr3:51261044-51263059 | p5 | (ATTTC)3 | TATGAGCAAATCGGAGTCCC | TTACCCGATGGTCGTAAAGC | 227 |
| chr3:51273197-51275209 | p2 | (TA)6 | TTTCACAAAAGACCGGGAAC | CTTCGCGGATCATACATTCA | 198 |
| chr3:51320670-51322720 | p2 | (AG)25 | TGGATGAAGAGTGGAGAGGG | GCAATTTCACGGTCGCTTAT | 179 |
| chr3:51383463-51385500 | p4 | (ATAA)4 | ATCCCCGTGTATAGTGGCAG | CCCGAACTGGACACTCGTAT | 202 |
| chr3:51433911-51435925 | p2 | (AT)7 | TGATACGCAAGTCGACAAGG | GATTGACATGTTGTGGCTGC | 286 |
| chr3:51454288-51456302 | p2 | (AT)7 | TGATACGCAAGTCGACAAGG | GATTGACATGTTGTGGCTGC | 286 |

| Marker ID | SSR type | SSR motif | FORWARD PRIMER (5'-3') | REVERSE PRIMER (5'-3') | size (bp) |
|---|---|---|---|---|---|
| chr3:51483543-51485563 | p2 | (AT)10 | TTTCATTTCCTATGGCTGGG | GGTTGGGTCGTGACAAAAGT | 113 |
| chr3:51521838-51523852 | p2 | (TA)7 | GAGTTGTTTGTGGGCTGGAT | AACTCGCCTAACCCGAACTT | 285 |
| chr3:51521838-51523852 | p2 | (TA)14 | GAGTTGTTTGTGGGCTGGAT | AACTCGCCTAACCCGAACTT | 285 |
| chr3:51522225-51524253 | p2 | (TA)7 | CGACACAGCGAACTTGGTTA | ACGGCCGAAATCATAGACAC | 182 |
| chr3:51522225-51524253 | p2 | (TA)14 | CGACACAGCGAACTTGGTTA | ACGGCCGAAATCATAGACAC | 182 |
| chr3:51573857-51575872 | p3 | (TTG)6 | GCCCGATTTTCGACAAGATA | ACGGAACTCAACGTCCAAAC | 127 |
| chr3:51573857-51575872 | p3 | (TGT)5 | GCCCGATTTTCGACAAGATA | ACGGAACTCAACGTCCAAAC | 127 |
| chr3:51591943-51594023 | p3 | (CAA)6 | TTATCCCATATTTCCGCCAA | CCAACTCCCAACACCAACTT | 234 |
| chr3:51591943-51594023 | p2 | (TA)72 | TTATCCCATATTTCCGCCAA | CCAACTCCCAACACCAACTT | 234 |
| chr3:51682115-51684133 | p2 | (AT)6 | ACGCTGTACCTATATGGCCG | TGTCTACCTGTGGGCATGAA | 218 |
| chr3:51682115-51684133 | p3 | (ATG)6 | ACGCTGTACCTATATGGCCG | TGTCTACCTGTGGGCATGAA | 218 |
| chr3:51684939-51686957 | p6 | (CTTACA)3 | TCTCGTATTTAGGCCCATGC | AATGTTGTTGCCTCTGACCC | 260 |
| chr3:51693733-51695745 | p2 | (TA)11 | TGTGGTTGGGCTGATTTACA | AGCTTTTGGGAACAAGGACA | 162 |
| chr3:51693733-51695745 | p2 | (TA)6 | TGTGGTTGGGCTGATTTACA | AGCTTTTGGGAACAAGGACA | 162 |
| chr3:51809379-51811400 | p3 | (TGT)7 | GGTTGTGAGACCGACCTTGT | ATCCTGTCCTCCACTGATGC | 265 |
| chr3:51863673-51865727 | p2 | (AT)27 | AAACACCATGTCCGAAGACC | TGGAGTAGTTCGTGGCTCCT | 119 |
| chr3:51879985-51882007 | p2 | (AT)8 | GTGCACAGTTCACTCGCATT | GGCGCCTATTAATCACTCCA | 255 |
| chr3:51879985-51882007 | p2 | (AT)11 | GTGCACAGTTCACTCGCATT | GGCGCCTATTAATCACTCCA | 255 |
| chr3:51879985-51882007 | p3 | (TTG)5 | GTGCACAGTTCACTCGCATT | GGCGCCTATTAATCACTCCA | 255 |
| chr3:51887543-51889558 | p3 | (TTG)5 | AGGGGCCAAAACGCTATTAT | TTTGGCAGCTTGGAGTCTTT | 242 |
| chr3:52086638-52088658 | p4 | (TGTA)5 | TGGAAATCTGCTCCAATTCC | GGACCAACTCGAGACGACAT | 271 |
| chr3:52205130-52207142 | p2 | (AT)6 | TTCACAAGGATCATGCCAAA | TTAGATTCTCCGGGATGGTG | 219 |
| chr3:52213019-52215034 | p5 | (ATTTC)3 | GAAAAGGGTGAGTCCCACAA | ACACCCAGTGGAGGAACTTG | 139 |
| chr3:52254095-52256110 | p5 | (ATGAT)3 | ATAGGAAGCGGGCAAGGTAT | TTACGGATGTTTACGACGCA | 144 |
| chr3:52282760-52284782 | p2 | (TA)18 | TAACGATGCCGACGATGATA | AGCCAAGGCTCAAGGTTACA | 251 |
| chr3:52282760-52284782 | p2 | (AT)11 | TAACGATGCCGACGATGATA | AGCCAAGGCTCAAGGTTACA | 251 |
| chr3:52345988-52348020 | p2 | (AT)16 | ATATAGGGGGAACGTCCCAG | GATGTTGTGTGTAGTGGCGG | 277 |
| chr3:52379173-52381185 | p2 | (CA)6 | ATCTTCTCCACGCCTCTCAA | GGAATGAGCTTTGGATGGAA | 245 |
| chr3:52401323-52403349 | p2 | (AC)13 | TGCCGGTTCTAGTAAGTGGG | ATCTCATGGCAGCCCTAATG | 209 |
| chr3:52401323-52403349 | p2 | (AT)7 | TGCCGGTTCTAGTAAGTGGG | ATCTCATGGCAGCCCTAATG | 209 |
| chr3:52442187-52444199 | p2 | (CT)6 | TGGTGGAACAACAATCGAGA | TCTTGCCACAACAACCCATA | 129 |
| chr3:52473182-52475258 | p2 | (TA)38 | AATGAGGCAAGTGCAAATCC | CATAATCCTCCACCAGCGTT | 245 |
| chr3:52522291-52524315 | p6 | (ATTCAT)4 | TCGCACTATTCTGTTCGTGC | GACACAGATCATGCGTACCG | 202 |
| chr3:52536009-52538027 | p2 | (TA)9 | GTGATTTGGCCGAGTTGAAT | CATATCTTGCCCGCTTCCTA | 279 |
| chr3:52536009-52538027 | p2 | (TA)17 | GTGATTTGGCCGAGTTGAAT | CATATCTTGCCCGCTTCCTA | 279 |
| chr3:52649819-52651837 | p2 | (TA)6 | GGCCTTCCTCCATATTCACA | GAATGTTGCCATTGATGACG | 192 |

| Marker ID | SSR type | SSR motif | FORWARD PRIMER (5'-3') | REVERSE PRIMER (5'-3') | size (bp) |
|---|---|---|---|---|---|
| chr3:52649819-52651837 | p2 | (TA)9 | GGCCTTCCTCCATATTCACA | GAATGTTGCCATTGATGACG | 192 |
| chr3:52661181-52663196 | p5 | (ATATC)3 | CATGAAGGCCTCCTCCATAA | GAAACGCGACCAAGAATCAT | 271 |
| chr3:52675555-52677567 | p2 | (TG)6 | TTCTCCCATAGACCTGCACC | TTCCCTTAGCCTTCCTCCAT | 248 |
| chr3:52748755-52750769 | p2 | (CT)7 | CTGTCCACGGTTTTGGACTT | TCATGAAATGGTGGAAGGGT | 268 |
| chr3:52817470-52819485 | p5 | (CACAA)3 | TTCAAGGAACAGAATTGCCC | CATTTGCAGATTCTGAGCGA | 185 |
| chr3:52843611-52845625 | p2 | (GT)7 | AAATACCCCCACTTGTTCCC | AAGTCTTATGATCGTGGCGG | 146 |
| chr3:52902563-52904575 | p2 | (TG)6 | TGATAGCCAATTTCCCCTTG | TGAGAATTGAAATGAGGCCC | 209 |
| chr3:52902563-52904575 | p2 | (TA)21 | TGATAGCCAATTTCCCCTTG | TGAGAATTGAAATGAGGCCC | 209 |
| chr3:52929752-52931766 | p2 | (AC)7 | CATCCAGGTTCAGTTTCCGT | TGTATAGGGGAAATGCTGCC | 293 |
| chr3:53045463-53047481 | p6 | (TCAGTT)3 | TTGGGAGAATCGTAAATGGG | ACACTTGGTGGGCATAAAGC | 143 |
| chr3:53061261-53063276 | p3 | (TGT)5 | ATACTCCCACACATGCCACA | CCGTGGTAAGTTAAGGGCAA | 119 |
| chr3:53061261-53063276 | p3 | (GAT)5 | ATACTCCCACACATGCCACA | CCGTGGTAAGTTAAGGGCAA | 119 |
| chr3:53061261-53063276 | p2 | (TA)13 | ATACTCCCACACATGCCACA | CCGTGGTAAGTTAAGGGCAA | 119 |
| chr3:53327616-53329636 | p2 | (TA)7 | TCGGGTGGTTGGTTTGTAAT | GCAAAGGAACCATGCGTTAT | 291 |
| chr3:53327616-53329636 | p2 | (AT)10 | TCGGGTGGTTGGTTTGTAAT | GCAAAGGAACCATGCGTTAT | 291 |
| chr3:53409865-53411880 | p5 | (ATATC)3 | TCGCTTTCATTTCCGTTACC | GACGTGGTCTATGCGACTGA | 166 |
| chr3:53411170-53413214 | p2 | (AT)22 | ACATTCCGAGGATGAACTGC | TATGTAGTGGCGGCCTTTTC | 147 |
| chr3:53424423-53426505 | p2 | (AT)28 | TGCCAGGGATGGTCTTTAAC | AGGGTATTAGGCGAAGGGAA | 237 |
| chr3:53424423-53426505 | p2 | (AT)20 | TGCCAGGGATGGTCTTTAAC | AGGGTATTAGGCGAAGGGAA | 237 |
| chr3:53424510-53426540 | p2 | (AT)28 | TGCCAGGGATGGTCTTTAAC | AGGGTATTAGGCGAAGGGAA | 150 |
| chr3:53424510-53426540 | p2 | (AT)20 | TGCCAGGGATGGTCTTTAAC | AGGGTATTAGGCGAAGGGAA | 150 |
| chr3:53440338-53442362 | p3 | (TTG)6 | ATGGAAATTGGAAAACAGCG | CTGTTTATACGCCTAGCCCG | 103 |
| chr3:53440338-53442362 | p6 | (TATATG)4 | ATGGAAATTGGAAAACAGCG | CTGTTTATACGCCTAGCCCG | 103 |
| chr3:53456834-53458852 | p2 | (TA)9 | CCGCTAGGTTGAAAATCTCG | TGGTTTGGGGTTACTGGGTA | 218 |
| chr3:53462259-53464274 | p5 | (ATATC)3 | TCGCTTTCATTTCCGTTACC | GACGTGGTCTATGCGACTGA | 166 |
| chr3:53463564-53465608 | p2 | (AT)22 | ACATTCCGAGGATGAACTGC | TATGTAGTGGCGGCCTTTTC | 147 |
| chr3:53476817-53478899 | p2 | (AT)28 | TGCCAGGGATGGTCTTTAAC | AGGGTATTAGGCGAAGGGAA | 237 |
| chr3:53476817-53478899 | p2 | (AT)20 | TGCCAGGGATGGTCTTTAAC | AGGGTATTAGGCGAAGGGAA | 237 |
| chr3:53476904-53478934 | p2 | (AT)28 | TGCCAGGGATGGTCTTTAAC | AGGGTATTAGGCGAAGGGAA | 150 |
| chr3:53476904-53478934 | p2 | (AT)20 | TGCCAGGGATGGTCTTTAAC | AGGGTATTAGGCGAAGGGAA | 150 |
| chr3:53492732-53494756 | p3 | (TTG)6 | ATGGAAATTGGAAAACAGCG | CTGTTTATACGCCTAGCCCG | 103 |
| chr3:53492732-53494756 | p6 | (TATATG)4 | ATGGAAATTGGAAAACAGCG | CTGTTTATACGCCTAGCCCG | 103 |
| chr3:53535877-53537889 | p2 | (AT)6 | GAATTGGAGGAATTGCCAGA | AAGGGATGCCATATGAACCA | 108 |
| chr3:53623577-53625591 | p2 | (TA)18 | CCACTAGGATGGCCAATGTT | TAACATACCCGAGCAGGGAC | 264 |
| chr3:53623577-53625591 | p2 | (AT)7 | CCACTAGGATGGCCAATGTT | TAACATACCCGAGCAGGGAC | 264 |
| chr3:53623577-53625591 | p2 | (AT)7 | CCACTAGGATGGCCAATGTT | TAACATACCCGAGCAGGGAC | 264 |

| Marker ID | SSR type | SSR motif | FORWARD PRIMER (5'-3') | REVERSE PRIMER (5'-3') | size (bp) |
|---|---|---|---|---|---|
| chr3:53623577-53625591 | p5 | (TGATA)3 | CCACTAGGATGGCCAATGTT | TAACATACCCGAGCAGGGAC | 264 |
| chr3:53639882-53641897 | p3 | (CAA)5 | GTCATCACTATGCCCGTCCT | TTTTCGTGCACAAGTTTCCA | 291 |
| chr3:53653910-53655924 | p2 | (AT)7 | TGCCCTTTCTGCATTTATCC | TCTTGACAGCGACAAGGATG | 200 |
| chr3:53687223-53689238 | p5 | (ATATC)3 | TCCCTTAGTGTCACAAGCCC | TTGTGGTGGTTTTGGAGGAT | 140 |
| chr3:53689087-53691099 | p2 | (AT)6 | GTATCTGTACCTGCGGGCAT | CCCTAGTTGGGTCGTGACAT | 169 |
| chr3:53692074-53694089 | p3 | (CAA)5 | TAGGCGCCATAATCACATCA | ACTCTTCTCGTGCATGCCTT | 290 |
| chr3:53778330-53780390 | p2 | (TA)30 | TGATCCATAACGACATCCGA | TTCGGGTCAATCCGACTTAG | 177 |
| chr3:53778330-53780390 | p2 | (AT)7 | TGATCCATAACGACATCCGA | TTCGGGTCAATCCGACTTAG | 177 |
| chr3:53790096-53792114 | p2 | (AT)9 | GAGTGCGTCTCTCATAGCCC | CAGTGAGACCCGGAGGATTA | 274 |
| chr3:53802683-53804705 | p2 | (AT)11 | GCAACCGATATGAAGCCAAT | GAGGGTCCGTGATTTCTTGA | 114 |
| chr3:53824220-53826235 | p2 | (TA)34 | CGCAATTGATGAAAATCCCT | GAGGATTGAGGCAGTTCGAG | 242 |
| chr3:53824220-53826235 | p3 | (AAT)5 | CGCAATTGATGAAAATCCCT | GAGGATTGAGGCAGTTCGAG | 242 |
| chr3:53849449-53851461 | p2 | (AT)6 | TGCCAACTCCACAGATTCAG | TTCGGGTTAAAAAGTGACCG | 227 |
| chr3:53912818-53914840 | p2 | (TA)11 | CAGCCACCAGAATCAAAGGT | TGTTCATGACGTCGGATGTT | 274 |
| chr3:53912818-53914840 | p2 | (AT)8 | CAGCCACCAGAATCAAAGGT | TGTTCATGACGTCGGATGTT | 274 |
| chr3:53961935-53963963 | p2 | (AT)14 | AGGCGCACTAGCTGTTGAAT | TCAGAGCAAGAGGTCCAGGT | 101 |
| chr3:53961935-53963963 | p2 | (AT)8 | AGGCGCACTAGCTGTTGAAT | TCAGAGCAAGAGGTCCAGGT | 101 |
| chr3:54011107-54013121 | p2 | (AT)7 | TTCTGCAATTCCCCTCAATC | TGGTTATTTCAATCGCGACA | 260 |
| chr3:54028995-54031039 | p3 | (ATG)5 | TTTGGAAAGCATCGTGTGAG | TATGCCTAGCCCGAAATCAC | 181 |
| chr3:54028995-54031039 | p2 | (TA)22 | TTTGGAAAGCATCGTGTGAG | TATGCCTAGCCCGAAATCAC | 181 |
| chr3:54028995-54031039 | p2 | (AT)14 | TTTGGAAAGCATCGTGTGAG | TATGCCTAGCCCGAAATCAC | 181 |
| chr3:54028995-54031039 | p2 | (AT)14 | TTTGGAAAGCATCGTGTGAG | TATGCCTAGCCCGAAATCAC | 181 |
| chr3:54036327-54038339 | p2 | (TA)6 | ATTGATTGGGGTAGTGCAGG | ATGGATGTTTCAAATGGGGA | 230 |
| chr3:54174760-54176792 | p2 | (TA)16 | GTACCCGTCATGGCCTTCTA | TACGGCACAATCGAAGATGA | 157 |
| chr3:54198932-54201010 | p2 | (AT)21 | AAATCGGAGGTTACCGACCT | AGGGTAAACATCTCATGCCG | 221 |
| chr3:54487557-54489569 | p2 | (CT)6 | AGAAACACAACCCCTACCCC | AGGTGTAACAGCATCCCAGG | 226 |
| chr3:54511205-54513223 | p5 | (TGAAT)3 | GGCCCGAGTTAAGATTCTCC | GCCTTTTGAAGGAAAGGGAC | 186 |
| chr3:54511205-54513223 | p6 | (TATATG)3 | GGCCCGAGTTAAGATTCTCC | GCCTTTTGAAGGAAAGGGAC | 186 |
| chr3:54550315-54552345 | p2 | (TA)8 | CCGTGAGTTTGGGACTGATT | ATTTTGACGCTCACAATCCC | 189 |
| chr3:54550354-54552376 | p2 | (TA)8 | CCGTGAGTTTGGGACTGATT | ATTTTGACGCTCACAATCCC | 150 |
| chr3:54555638-54557660 | p2 | (TA)11 | GCCGTGAGTTTTGGACTGAT | ATTTTGACGCTCACAATCCC | 198 |
| chr3:54555663-54557677 | p2 | (TA)11 | GCCGTGAGTTTTGGACTGAT | ATTTTGACGCTCACAATCCC | 173 |
| chr3:54555698-54557712 | p2 | (TA)11 | GCCGTGAGTTTTGGACTGAT | ATTTTGACGCTCACAATCCC | 138 |
| chr3:54679173-54681188 | p5 | (CTTTT)3 | AGGGAAGCATACACATTGCC | GGCCGACCGCATATACTTTA | 299 |
| chr3:54683170-54685192 | p2 | (TA)11 | CTCGGAAGGGCAATAATCAA | GCCCGTTTGATCGCTATAAG | 283 |
| chr3:54683320-54685336 | p2 | (TA)11 | CTCGGAAGGGCAATAATCAA | GCCCGTTTGATCGCTATAAG | 133 |

| Marker ID | SSR type | SSR motif | FORWARD PRIMER (5'-3') | REVERSE PRIMER (5'-3') | size (bp) |
|---|---|---|---|---|---|
| chr3:54683338-54685378 | p2 | (TA)11 | CTCGGAAGGGCAATAATCAA | GCCCGTTTGATCGCTATAAG | 115 |
| chr3:54785993-54788023 | p6 | (GGAAGT)3 | GTGGAAGTGGAAGTGCCTGT | TCGGCCTATGATGCCTATTC | 249 |
| chr3:55001452-55003464 | p2 | (AT)6 | TCCTGAGGAGCTGCACTTTT | AGTGTTTATTGGGTACGCGG | 201 |
| chr3:55135694-55137709 | p5 | (TGCAA)3 | CTGCAAGTGGGATTGGCTAT | CCACTTGCGGGTAAACAAGT | 200 |
| chr3:55183539-55185551 | p5 | (TATGA)3 | GAGCCCTGTGTGGCTATGAT | ATTTTGGCAGAGATTCCCCT | 287 |
| chr3:55195872-55197887 | p5 | (TTCAT)3 | GTTCTATTGTCCCGGCTTGA | CCAGCTCATCGAGTGTTTCA | 158 |
| chr3:55210282-55212294 | p2 | (TA)6 | AGAGACCTCCCCGACGTATT | TTCCGCATCAACATTTTTCA | 103 |
| chr3:55243664-55245676 | p2 | (GT)6 | GAAAGCTGCTATCTGTGCCC | TTATGCGAAAACACGGAACA | 163 |
| chr3:55270120-55272136 | p2 | (AT)8 | TGGACGACCGTCATCATTTA | GCAGAGCTCCAATTCTTTCG | 190 |
| chr3:55343875-55345893 | p6 | (GTATAT)3 | GAATTCCCGGATTGTTGTTG | CCTCGGAATCATGCCTTAAA | 185 |
| chr3:55343875-55345893 | p2 | (TA)6 | GAATTCCCGGATTGTTGTTG | CCTCGGAATCATGCCTTAAA | 185 |
| chr3:55367964-55369979 | p5 | (ATTGA)3 | CCAATGCTTACAAGTGCGAA | TTTCACTCGCTAGGGCTTGT | 285 |
| chr3:55373439-55375454 | p5 | (ATTGC)3 | TGATGAATGATGGCGAGTGT | TGGCTAGTCCCAAACACACA | 121 |
| chr3:55539219-55541245 | p2 | (TA)13 | GCGATATTCCCTCAGACCAA | CCGCACCTAGATCCATGATT | 187 |
| chr3:55542219-55544249 | p2 | (TA)15 | GGCGAATATGATTCTGGCAT | CACCTTTCGAACCACAACCT | 109 |
| chr3:55588475-55590493 | p2 | (TA)9 | TGATTAGGGGAAAATGCTGC | CCTCCCAAGCTTGAAAATCA | 297 |
| chr3:55623125-55625139 | p2 | (AG)9 | CGGAGACGTTTTGAACGAAT | AAATAAGTGGAACCCGACCC | 130 |
| chr3:55623125-55625139 | p3 | (TTG)6 | CGGAGACGTTTTGAACGAAT | AAATAAGTGGAACCCGACCC | 130 |
| chr3:55623148-55625170 | p2 | (AG)9 | CGGAGACGTTTTGAACGAAT | AAATAAGTGGAACCCGACCC | 107 |
| chr3:55623148-55625170 | p3 | (TTG)6 | CGGAGACGTTTTGAACGAAT | AAATAAGTGGAACCCGACCC | 107 |
| chr3:55774173-55776188 | p5 | (GGTGC)3 | GAGGCGAGTGTTCGAGGTAG | TCGGCCTAAAACCTCATTTG | 253 |
| chr3:55801674-55803702 | p2 | (AT)14 | GGGATCAAGAACGTTAGCCA | CGGACTAGCATAGAAAGCGG | 169 |
| chr3:55813380-55815395 | p5 | (TGAAA)3 | TTTAGTGCAATCGTGGCGTA | GCTAGCATTGGAGGAGAACG | 143 |
| chr3:55813380-55815395 | p5 | (CATGC)3 | TTTAGTGCAATCGTGGCGTA | GCTAGCATTGGAGGAGAACG | 143 |
| chr3:55830373-55832393 | p5 | (CATGC)4 | GACTTCCCCGAAACTAAGCC | GAAGGTTGCAAGACTGGAGG | 165 |
| chr3:55940262-55942274 | p2 | (CA)6 | ATACCTCGAACGCTCGCTAA | TGAATTTACAGGGGAGGTGC | 285 |
| chr3:55988755-55990769 | p2 | (TA)24 | AAATAGGGGAGATGCTGCCT | AAGTTTGTCCATCCACGCTC | 158 |
| chr3:55988772-55990864 | p2 | (TA)24 | AAATAGGGGAGATGCTGCCT | AAGTTTGTCCATCCACGCTC | 141 |
| chr3:55996777-55998792 | p5 | (TTTTA)3 | AACTGGCCATTGGTATTGGA | AAGGACTCCAAACGAACCCT | 133 |
| chr3:56152796-56154811 | p3 | (ATG)5 | AGGAGGTTTCGAACGGATTT | TTCCTCCAAATTCCAAACCA | 299 |
| chr3:56220213-56222228 | p3 | (ATG)5 | AATGGTTGCAAATGGTGGAT | AACAAGCGTCATTCAAACCC | 278 |
| chr3:56270136-56272163 | p3 | (TTG)9 | GAGGGCAAGTGAATCAAGGA | TCTGCTCTGGTGGCCTAACT | 241 |
| chr3:56358534-56360549 | p3 | (GAA)5 | GGTTCGCAAGTAAGCAGAGG | GAACCATTCTTAAACGGGCA | 128 |
| chr3:56382160-56384174 | p2 | (TA)7 | ACTGACGTCCCTTTGTGGAC | GCAAAATGATACGTCGGCTT | 270 |
| chr3:56382178-56384208 | p2 | (TA)7 | ACTGACGTCCCTTTGTGGAC | GCAAAATGATACGTCGGCTT | 252 |
| chr3:56495918-56497948 | p2 | (AT)15 | CTCTATCGCACGACCTCACA | ATGGCCCTTAGTTGGCTCTT | 270 |

| Marker ID | SSR type | SSR motif | FORWARD PRIMER (5'-3') | REVERSE PRIMER (5'-3') | size (bp) |
|---|---|---|---|---|---|
| chr3:56514040-56516052 | p2 | (CT)6 | ATGTACCCCCATTTGTTCCA | CATCTATGGCCACACACGAC | 224 |
| chr3:56598147-56600163 | p2 | (AT)8 | TTTATCTTAAGGGCCGGCTT | GGGTCGTGACAGTTAAGGGA | 132 |
| chr3:56800968-56803000 | p2 | (AT)16 | GGCGAGACACCAGTAACCAT | TTTTGGGCTATTGGTTGGAG | 165 |
| chr3:56823034-56825062 | p2 | (AT)14 | TTACATTCACAGACGCCGAG | AAACCCCTTGCCTCATATCC | 166 |
| chr3:56847803-56849828 | p2 | (AT)15 | CCCTAGGGAAGGTTTTACGC | AAGGGAATTTTTGCCGAACT | 209 |
| chr3:56849652-56851670 | p6 | (ATATAC)3 | AACTGATGGGCCGTAACAAG | TGTGTTGATTGTTCGGGAGA | 177 |
| chr3:56932850-56934868 | p6 | (TTTTAA)3 | GCTCCTCGCACACATCACTA | GCTTCATTGTTGGCTTCTCC | 210 |
| chr3:57095870-57097885 | p3 | (CAT)5 | AGGAACATCGACCACGATTC | CTATAGACGCATGGCCACCT | 113 |
| chr3:57095870-57097885 | p3 | (CAA)5 | AGGAACATCGACCACGATTC | CTATAGACGCATGGCCACCT | 113 |
| chr3:57095870-57097885 | p2 | (AT)8 | AGGAACATCGACCACGATTC | CTATAGACGCATGGCCACCT | 113 |
| chr3:57181890-57183905 | p5 | (ACATG)3 | TAAACGACCATTCGGACCTC | AGATCGGATTCGGGAAACTT | 257 |
| chr3:57201783-57203815 | p2 | (AT)13 | CCCTCAAAGGTTGGTGAAGA | ATAACCGCAACCAATTCGAG | 242 |
| chr3:57201817-57203849 | p2 | (AT)13 | CCCTCAAAGGTTGGTGAAGA | ATAACCGCAACCAATTCGAG | 208 |
| chr3:57201851-57203883 | p2 | (AT)13 | CCCTCAAAGGTTGGTGAAGA | ATAACCGCAACCAATTCGAG | 174 |
| chr3:57201885-57203917 | p2 | (AT)13 | CCCTCAAAGGTTGGTGAAGA | ATAACCGCAACCAATTCGAG | 140 |
| chr3:57201919-57203951 | p2 | (AT)13 | CCCTCAAAGGTTGGTGAAGA | ATAACCGCAACCAATTCGAG | 106 |
| chr3:57202055-57204089 | p2 | (AT)13 | GAAAGAAACGGCCAATCAAA | TCTACCGCAGGTTCAATTCC | 113 |
| chr3:57210086-57212101 | p5 | (ACATG)3 | TAAACGACCATTCGGACCTC | AGATCGGATTCGGGAAACTT | 257 |
| chr3:57229979-57232011 | p2 | (AT)13 | CCCTCAAAGGTTGGTGAAGA | ATAACCGCAACCAATTCGAG | 242 |
| chr3:57230013-57232045 | p2 | (AT)13 | CCCTCAAAGGTTGGTGAAGA | ATAACCGCAACCAATTCGAG | 208 |
| chr3:57230047-57232079 | p2 | (AT)13 | CCCTCAAAGGTTGGTGAAGA | ATAACCGCAACCAATTCGAG | 174 |
| chr3:57230081-57232113 | p2 | (AT)13 | CCCTCAAAGGTTGGTGAAGA | ATAACCGCAACCAATTCGAG | 140 |
| chr3:57230115-57232147 | p2 | (AT)13 | CCCTCAAAGGTTGGTGAAGA | ATAACCGCAACCAATTCGAG | 106 |
| chr3:57230251-57232285 | p2 | (AT)13 | GAAAGAAACGGCCAATCAAA | TCTACCGCAGGTTCAATTCC | 113 |
| chr3:57237272-57239284 | p2 | (AT)6 | TCTCGCTAGGGTATGATGGG | GACTTGAGAACGCCGAAAAG | 292 |
| chr3:57237272-57239284 | p3 | (AAC)5 | TCTCGCTAGGGTATGATGGG | GACTTGAGAACGCCGAAAAG | 292 |
| chr3:57293610-57295630 | p4 | (ATTA)5 | GGCCATCCAGGTAGGTTACA | TCAATAACACAACGGGGTGA | 217 |
| chr3:57326301-57328329 | p2 | (AT)14 | TTTGGTTTTGGAAAATGGGA | GAGTGCATTAAGGGAGAGCG | 134 |
| chr3:57326301-57328329 | p2 | (TA)6 | TTTGGTTTTGGAAAATGGGA | GAGTGCATTAAGGGAGAGCG | 134 |
| chr3:57331743-57333755 | p2 | (TA)6 | AGCCCAAATCAAGAAGGGTT | TCTTGATCATTGGGTGACGA | 214 |
| chr3:57406624-57408686 | p3 | (GAT)5 | CAACATACACATGGCAAGGC | CCGTAGGGTGACGTTTAGGA | 259 |
| chr3:57552034-57554049 | p5 | (ATTTC)3 | TCACGAGGAACCCTTGAAAC | AGTTCTCACCGAGGATGTGG | 159 |
| chr3:57594269-57596284 | p6 | (CATATA)3 | TTCGGGTCAATCCGACTTAG | CCGTCAATGCTTTCCAAAAT | 246 |
| chr3:57594269-57596284 | p5 | (ATTCA)3 | TTCGGGTCAATCCGACTTAG | CCGTCAATGCTTTCCAAAAT | 246 |
| chr3:57648408-57650420 | p2 | (AT)6 | GTCCTCCGCAATCGTTTTTA | ACAGTGGTCGTAAAGGGCAC | 217 |
| chr3:57653777-57655791 | p2 | (TA)7 | GTCTCCCGAACAATCGTCAT | CTTGTCCTGTCCCTTCCAAA | 184 |

| Marker ID | SSR type | SSR motif | FORWARD PRIMER (5'-3') | REVERSE PRIMER (5'-3') | size (bp) |
|---|---|---|---|---|---|
| chr3:57653777-57655791 | p3 | (ATC)6 | GTCTCCCGAACAATCGTCAT | CTTGTCCTGTCCCTTCCAAA | 184 |
| chr3:57688669-57690745 | p2 | (AT)38 | GCCGAGCCTATTACATCCAG | TGGGTTGCTGTGTTGATGAT | 120 |
| chr3:57713373-57715385 | p2 | (AT)6 | GTCCTCCGCAATCGTTTTTA | ACAGTGGTCGTAAAGGGCAC | 217 |
| chr3:57718742-57720756 | p2 | (TA)7 | GTCTCCCGAACAATCGTCAT | CTTGTCCTGTCCCTTCCAAA | 184 |
| chr3:57718742-57720756 | p3 | (ATC)6 | GTCTCCCGAACAATCGTCAT | CTTGTCCTGTCCCTTCCAAA | 184 |
| chr3:57753634-57755710 | p2 | (AT)38 | GCCGAGCCTATTACATCCAG | TGGGTTGCTGTGTTGATGAT | 120 |
| chr3:57753634-57755710 | p2 | (TA)9 | GCCGAGCCTATTACATCCAG | TGGGTTGCTGTGTTGATGAT | 120 |
| chr3:57753634-57755710 | p2 | (CT)7 | GCCGAGCCTATTACATCCAG | TGGGTTGCTGTGTTGATGAT | 120 |
| chr3:57798098-57800116 | p2 | (AG)8 | TGGGCATATGCGATGAGTTA | TCATGTCGATTGGCTTACGA | 189 |
| chr3:57798098-57800116 | p3 | (TTG)6 | TGGGCATATGCGATGAGTTA | TCATGTCGATTGGCTTACGA | 189 |
| chr3:57799198-57801216 | p2 | (TA)9 | GAAGCGTATTGGGCATTGTT | CTAGACGATTGGGCTGCATT | 267 |
| chr3:57801268-57803283 | p5 | (ATATA)3 | TTACTCCAGCCCCCTATCCT | GTGAAGTGTGCTCCTCGTCA | 294 |
| chr3:57821171-57823183 | p2 | (TA)6 | CAATTCCATTTTCCACACCC | CCCATTTGTGGAATTGAAGG | 116 |
| chr3:57958426-57960441 | p5 | (TTCAT)3 | TGGGCAGCATCTCCTTAGTT | TGGTTGACGAAGATGGTTGA | 140 |
| chr3:57967626-57969640 | p2 | (AT)7 | GGCGGAGTGTGTGAGGTATT | AGAAAAGCTACAGTCGCCCA | 113 |
| chr3:57976337-57978352 | p3 | (CAT)5 | AAAGTTGGTGATGGTGGAGG | TGATAGCCGAAGGGCTTATG | 286 |
| chr3:58003769-58005781 | p2 | (AT)6 | TAGTCATCCATCTCCCGAGG | TTTGACTTAAGTTTGGGGCG | 171 |
| chr3:58009700-58011715 | p3 | (GAA)5 | CAAGGTGATGTACTGGGCCT | TGTCCCGGTGGTTAATGAAT | 154 |
| chr3:58262697-58264712 | p3 | (TTG)5 | AAGGCGGGTGGCTTATAGAT | GAGGCATCAAATGAACAGCA | 164 |
| chr3:58301100-58303125 | p5 | (AAAAC)3 | AAGGCAGACGTATCGCAAGT | CGGGTTAGGCTTGAATGTGT | 120 |
| chr3:58383393-58385408 | p5 | (AACCG)3 | CGAATACGGTTGGCTTCAAT | GTGAGAAGAACGCCGAAAAG | 235 |
| chr3:58401345-58403387 | p2 | (TA)7 | GGGTTCCATGACCTCTCTGA | AAAGTGAGAAAAGGCGGTTG | 133 |
| chr3:58401345-58403387 | p2 | (AT)21 | GGGTTCCATGACCTCTCTGA | AAAGTGAGAAAAGGCGGTTG | 133 |
| chr3:58401345-58403387 | p2 | (AT)24 | GGGTTCCATGACCTCTCTGA | AAAGTGAGAAAAGGCGGTTG | 133 |
| chr3:58456718-58458816 | p2 | (TA)49 | TAAACGCCCTCTTTTGTCCA | TGGGTTCTTGATCCATTGGT | 299 |
| chr3:58549763-58551775 | p5 | (ACACA)3 | ACACAGCCTCTGCTTGGAAT | GATTCTTGAGGGTCCGTTGA | 145 |
| chr3:58549763-58551775 | p2 | (AT)6 | ACACAGCCTCTGCTTGGAAT | GATTCTTGAGGGTCCGTTGA | 145 |
| chr3:58620898-58622913 | p5 | (TCATT)3 | CGCTATACCTTTGGATGGGA | TCCTGGGGAATACACCCATA | 238 |
| chr3:58635975-58638011 | p2 | (GA)10 | TCAGGTGCCCAATCATACAA | AGAGCTGGACTCCCCAATTT | 219 |
| chr3:58811599-58813623 | p2 | (TA)12 | ATTGTTGCTGTTGCTGTTCG | CAAGGCCCAACCTAAGACAA | 119 |
| chr3:58811599-58813623 | p5 | (CATTG)3 | ATTGTTGCTGTTGCTGTTCG | CAAGGCCCAACCTAAGACAA | 119 |
| chr3:58811599-58813623 | p4 | (ATTT)4 | ATTGTTGCTGTTGCTGTTCG | CAAGGCCCAACCTAAGACAA | 119 |
| chr3:58812509-58814525 | p2 | (TA)12 | TTGACAGGGTGTGACGATGT | GCACATGTACTCTGGACGGA | 241 |
| chr3:58812509-58814525 | p5 | (CATTG)3 | TTGACAGGGTGTGACGATGT | GCACATGTACTCTGGACGGA | 241 |
| chr3:58812509-58814525 | p4 | (ATTT)4 | TTGACAGGGTGTGACGATGT | GCACATGTACTCTGGACGGA | 241 |
| chr3:58861234-58863250 | p5 | (TTTTG)3 | ATGTGGGCCAGAATTGAAAG | TATCAGAGCGGGTCAGTCCT | 127 |

| Marker ID | SSR type | SSR motif | FORWARD PRIMER (5'-3') | REVERSE PRIMER (5'-3') | size (bp) |
|---|---|---|---|---|---|
| chr3:58899852-58901864 | p2 | (TG)6 | AAGAAAATTCCCCCGAAATG | CCTTCCTCAATATTTCCGCA | 243 |
| chr3:58899852-58901864 | p3 | (TTG)5 | AAGAAAATTCCCCCGAAATG | CCTTCCTCAATATTTCCGCA | 243 |
| chr3:58951912-58953932 | p2 | (AT)10 | TTGATATTTTTCCGAACGGC | TTTGGAAAATCGAAGCAACC | 263 |
| chr3:59020027-59022045 | p3 | (TGG)6 | CCAGAATCACCACACGTCAC | TTCGGGTATTGGATCGAGAG | 224 |
| chr3:59121483-59123501 | p6 | (AATGGT)3 | GTGCTCCTTGATTGGGAAGA | ACGAAAAACCACAATCCAGC | 246 |
| chr3:59139833-59141845 | p2 | (TA)6 | CACTCCGATGGACACATGAC | ACTAGGATGTGATGGGCACC | 171 |
| chr3:59139833-59141845 | p2 | (TA)28 | CACTCCGATGGACACATGAC | ACTAGGATGTGATGGGCACC | 171 |
| chr3:59145494-59147506 | p2 | (TA)6 | CTTCCGTCCGGTAACAAAAA | TCTGTGGATAGTGCGTCGAG | 298 |
| chr3:59158472-59160487 | p5 | (ATCAT)3 | CGTTGAGGCTGTCATAGCAA | GCTCGCAGGTATAGACGCTC | 128 |
| chr3:59175272-59177287 | p2 | (AT)11 | CGATTAGCGATGGGAAATGT | TCGGTATCGGTCTTACCCTG | 218 |
| chr3:59175272-59177287 | p5 | (CAAAA)3 | CGATTAGCGATGGGAAATGT | TCGGTATCGGTCTTACCCTG | 218 |
| chr3:59195546-59197572 | p2 | (AT)13 | TGAGTTGTTTTGTGGGCTTG | AGTCAGGAAGCATTTCCCCT | 193 |
| chr3:59195546-59197572 | p2 | (CA)6 | TGAGTTGTTTTGTGGGCTTG | AGTCAGGAAGCATTTCCCCT | 193 |
| chr3:59220858-59222874 | p2 | (AT)6 | TTCCTTCTCGCGACACTTTT | AACCGTCGAGGGAGGTTAGT | 241 |
| chr3:59220881-59222893 | p2 | (AT)6 | TTCCTTCTCGCGACACTTTT | AACCGTCGAGGGAGGTTAGT | 218 |
| chr3:59256863-59258881 | p2 | (AC)9 | CTACCCAAGCTCACTCTCCG | TGTAAAAAGGGGAGGTGCTG | 179 |
| chr3:59290217-59292265 | p6 | (CATATT)8 | AAGATTCTGAGCGTGTGCCT | TTTGTGAGAGGTTTGGGACC | 165 |
| chr3:59290217-59292265 | p2 | (TA)11 | AAGATTCTGAGCGTGTGCCT | TTTGTGAGAGGTTTGGGACC | 165 |
| chr3:59318819-59320877 | p2 | (AT)29 | CGGAAGTACCTGAAAGGCTG | TTGCCAATTCTCACACGAAG | 171 |
| chr3:59392433-59394447 | p2 | (AC)8 | CCAAATCATGCAACACCTTG | CTGACCTAGAGGTCGGTTGC | 102 |
| chr3:59463704-59465720 | p2 | (AT)8 | TACCCCATGTGGTCTCCAAT | ATCGGGGTAAGGAGGTATGG | 138 |
| chr3:59491926-59493942 | p4 | (AACA)4 | TGAGTGGCAACATCTCGAAG | CGATGATGATGATGACAGGC | 106 |
| chr3:59689504-59691524 | p2 | (AT)10 | CCTACATGAGGGCCATGACT | CACTTGGTCGGGTTCAGTTT | 223 |
| chr3:59689504-59691524 | p2 | (CA)6 | CCTACATGAGGGCCATGACT | CACTTGGTCGGGTTCAGTTT | 223 |
| chr3:59976355-59978385 | p2 | (AT)15 | GATCTTCAGGTCGCATGACA | GCACCCTTTTGAGTGCAAAT | 269 |
| chr3:60025840-60027855 | p5 | (AATTG)3 | GAAAAAGGGAAATGCCAACA | CTCGTATGTGTGTGTTCGGG | 225 |
| chr3:60025840-60027855 | p2 | (AC)6 | GAAAAAGGGAAATGCCAACA | CTCGTATGTGTGTGTTCGGG | 225 |
| chr3:60111876-60113888 | p2 | (AG)6 | GGTCATGTAGGAAAAGGGCA | GGGCCCTTATGTGTCTTGAA | 200 |
| chr3:60178195-60180215 | p2 | (AT)10 | GTACGAGCCTCGGAGACAAC | AGGGTCCTATGGTAATCGGG | 217 |
| chr3:60178195-60180215 | p5 | (GCTAT)3 | GTACGAGCCTCGGAGACAAC | AGGGTCCTATGGTAATCGGG | 217 |
| chr3:60190414-60192430 | p4 | (AAAT)4 | GCTCCAGCAAGAAAATGGAG | TCGTACTGACGTCCTGTTGC | 158 |
| chr3:60306731-60308746 | p2 | (AT)6 | CGTACGTTCCTCGGCACTAT | TGAGCATGTAAGGCATGGAA | 262 |
| chr3:60306731-60308746 | p2 | (AT)8 | CGTACGTTCCTCGGCACTAT | TGAGCATGTAAGGCATGGAA | 262 |
| chr3:60306731-60308746 | p6 | (TATATG)3 | CGTACGTTCCTCGGCACTAT | TGAGCATGTAAGGCATGGAA | 262 |
| chr3:60306814-60308888 | p2 | (AT)8 | GACATATGATGTTGGCACGC | CCCAAAGAAAGGGGTTCAGT | 239 |
| chr3:60306814-60308888 | p6 | (TATATG)3 | GACATATGATGTTGGCACGC | CCCAAAGAAAGGGGTTCAGT | 239 |

| Marker ID | SSR type | SSR motif | FORWARD PRIMER (5'-3') | REVERSE PRIMER (5'-3') | size (bp) |
|---|---|---|---|---|---|
| chr3:60349905-60351932 | p3 | (TCT)9 | GGGGCTAACGAGACCTTTTC | CCCATTACACTTTCGGGCTA | 293 |
| chr3:60394465-60396477 | p2 | (TC)6 | TCACCGTAACATCATCCGAA | CCGTCCTTCATTTCGTCATT | 130 |
| chr3:60408363-60410375 | p2 | (TC)6 | TCACCGTAACATCATCCGAA | CCGTCCTTCATTTCGTCATT | 130 |
| chr3:60454088-60456106 | p5 | (CGGGT)4 | AAACTCGTGCAAATCCGAAC | TCCCTCTCCCTCTCCATTTT | 207 |
| chr3:60518187-60520207 | p4 | (ACAT)5 | TCGAGAATGCATTTGGTTGA | GGCAATGCCTTAGTGGACAT | 285 |
| chr3:60577679-60579694 | p3 | (TCA)5 | TCAACCCCTTGACGAATCTC | AGCTGAGTTTCCTCCGTTGA | 296 |
| chr3:60993419-60995447 | p2 | (TA)13 | ACAAAGCTTTTTGGGCGTTA | AGGCTCCCAACCTCATTTCT | 159 |
| chr3:60993627-60995653 | p2 | (TA)13 | TGGCGGAAACAAGTTTAAGG | CTTCATGAGATGCTCCGGTT | 262 |
| chr3:61006673-61008688 | p3 | (AAT)5 | TTAGCTTCGCCTTCTCCAAA | GAGTTGGGACGAAAAATCCA | 161 |
| chr3:61037806-61039824 | p6 | (CCTTTT)3 | GCCGCCCTACAAAAACACTA | AGGCTTTGGGAAAAACCACT | 215 |
| chr3:61147222-61149246 | p2 | (AT)18 | GATAACCACCGAGCACCACT | CGCATGTGTGGGTAGTATGC | 288 |
| chr3:61147222-61149246 | p6 | (TATACA)4 | GATAACCACCGAGCACCACT | CGCATGTGTGGGTAGTATGC | 288 |
| chr3:61149496-61151520 | p2 | (AT)17 | TCGGTTCCACCGACTTACTC | CCTTAAGCCAAATATGGGCA | 119 |
| chr3:61149496-61151520 | p6 | (TATACA)4 | TCGGTTCCACCGACTTACTC | CCTTAAGCCAAATATGGGCA | 119 |
| chr3:61154746-61156758 | p2 | (AT)6 | TCCCTCCTTGGGTTTCTTCT | TTAACGCTCCGGATTTCATC | 133 |
| chr3:61155750-61157786 | p2 | (AT)18 | GGACACGACTTTGCAGACAA | GCAAGCCGTCGTACCATTAT | 281 |
| chr3:61155750-61157786 | p6 | (TATACA)4 | GGACACGACTTTGCAGACAA | GCAAGCCGTCGTACCATTAT | 281 |
| chr3:61163528-61165542 | p2 | (AT)7 | TCTGGTACTAGCTGCCCGAT | CCTCGCCTAATGTGGAAAAA | 166 |
| chr3:61177677-61179695 | p3 | (TTC)6 | GGCATGCCATTTTGTCTTCT | AGCGAATTTGAAAATGGTGC | 198 |
| chr3:61181804-61183850 | p2 | (AT)23 | ACCCACTCGAGCACCTCTTA | GTCCTGTCCCGCCTATATGA | 151 |
| chr3:61210229-61212279 | p2 | (TA)25 | CGGATGTTTATAGGAGGGCA | ACAGCAGCGGAGATTAGGAA | 231 |
| chr3:61292228-61294336 | p2 | (TA)14 | ATTTCGTGGTGAGGTTCCAC | AAAGGAAAATCCAACCCGTC | 177 |
| chr3:61361014-61363030 | p2 | (TC)8 | AATGGCTCGACCACTTATGG | TGGTACCAAATTCGCCTTTC | 129 |
| chr3:61364390-61366408 | p3 | (CAA)6 | ACCGGCGAACGATATCATAG | GAAAAGCTCGGATTGCTTTG | 259 |
| chr3:61433530-61435545 | p3 | (TGT)5 | TATGACCCCCTGAACGAGAC | ATGTACGCAGACCTTACCCG | 268 |
| chr3:61433530-61435545 | p2 | (AT)6 | TATGACCCCCTGAACGAGAC | ATGTACGCAGACCTTACCCG | 268 |
| chr3:61461379-61463391 | p2 | (TA)6 | GGACTTTGAAAACGGTTGGA | TTTCCGGACTACGAAACACC | 103 |
| chr3:61461379-61463391 | p2 | (TA)6 | GGACTTTGAAAACGGTTGGA | TTTCCGGACTACGAAACACC | 103 |
| chr3:61508520-61510536 | p2 | (AT)8 | TCCCTGATAAAGCACCAACC | GGGGAGAAAGGGATCAGAAG | 152 |
| chr3:61524259-61526291 | p2 | (AT)17 | TCCACCATTGAAACCCAAAT | AATGTGTTTTGGTCCTCCCA | 151 |
| chr3:61524259-61526291 | p2 | (TA)16 | TCCACCATTGAAACCCAAAT | AATGTGTTTTGGTCCTCCCA | 151 |
| chr3:61542617-61544641 | p2 | (AT)12 | TTTTTCATTAAAGCCCACGC | GCCTGAACTTGTGGAATGGT | 266 |
| chr3:61587459-61589474 | p6 | (TAAAAA)3 | ACAATCCACCTGGGTCAGTC | TGCTGAGCACAACCTTCATC | 299 |
| chr3:61587459-61589474 | p3 | (TCT)5 | ACAATCCACCTGGGTCAGTC | TGCTGAGCACAACCTTCATC | 299 |
| chr3:61589873-61591900 | p3 | (TTG)9 | TTCCGCTTTCAAGAAACTCC | AAACATGGCAGGTAGGTGGA | 204 |
| chr3:61608446-61610464 | p6 | (ACTTTT)3 | GATCCATTATGGTCCAACCG | ACAAGGCATATTTTGGGACG | 189 |

| Marker ID | SSR type | SSR motif | FORWARD PRIMER (5'-3') | REVERSE PRIMER (5'-3') | size (bp) |
|---|---|---|---|---|---|
| chr3:61681233-61683261 | p2 | (TA)14 | TACCCTTAGCCTCCCTCGAT | GCACCTTCGGAAAGATTTGA | 188 |
| chr3:61844800-61846814 | p2 | (TA)7 | AATGAAATCGAGGGTTCACG | CGTCCGGAGTACCATCTCAT | 203 |
| chr3:61844800-61846814 | p2 | (AT)18 | AATGAAATCGAGGGTTCACG | CGTCCGGAGTACCATCTCAT | 203 |
| chr3:61844800-61846814 | p2 | (AT)16 | AATGAAATCGAGGGTTCACG | CGTCCGGAGTACCATCTCAT | 203 |
| chr3:61844980-61847008 | p2 | (TA)7 | GCGGGCAAATGTTACTGATT | TGTCTACCTGTGGGCATGAA | 296 |
| chr3:61844980-61847008 | p2 | (AT)18 | GCGGGCAAATGTTACTGATT | TGTCTACCTGTGGGCATGAA | 296 |
| chr3:61844980-61847008 | p2 | (AT)16 | GCGGGCAAATGTTACTGATT | TGTCTACCTGTGGGCATGAA | 296 |
| chr3:61895498-61897519 | p3 | (ACA)7 | GAGAATTGGCATCCCAAAAA | CGTCATTTAGGCCCATCTGT | 145 |
| chr3:61899127-61901139 | p2 | (TA)6 | GCATTCTGGAATTTGGCAAT | TCCGTTTGACCCCCATAATA | 141 |
| chr3:61978261-61980273 | p2 | (AT)6 | CATCAGGAAGATTAGCCCCA | ATAGGATGAGTGGTGCTCGG | 271 |
| chr3:62060201-62062216 | p5 | (ATGAT)3 | GGGTTTGGACAAGCGTAAAA | ATGACGCAAGACCCTTTTTG | 192 |
| chr3:62074472-62076487 | p5 | (AAACA)3 | TCCATGCAGCAGTTCAAAAG | TGCCCTGGAGCTAGGTTCTA | 129 |
| chr3:62234084-62236102 | p2 | (TA)14 | GAATTCGAACCCTCACCAAA | CAAACCCAAGAGCGGTTTTA | 103 |
| chr3:62234084-62236102 | p6 | (TTTTTA)3 | GAATTCGAACCCTCACCAAA | CAAACCCAAGAGCGGTTTTA | 103 |
| chr3:62342902-62344914 | p2 | (CA)6 | ATCGGAGCGTCCTTAGCATA | CAAGAGCCCGTGATTTCATT | 129 |
| chr3:62352780-62354816 | p2 | (AC)24 | AGACAAGTCAGGCGCAAACT | ACACCCCGTACTGTTTGACC | 203 |
| chr3:62352819-62354841 | p2 | (AC)24 | AGACAAGTCAGGCGCAAACT | ACACCCCGTACTGTTTGACC | 164 |
| chr3:62433048-62435096 | p3 | (TAA)16 | AAGCCTCCCTTCAGTGGATT | TCTCTCTTCCACCTCCCTCA | 139 |
| chr3:62460100-62462118 | p6 | (TGTTTG)3 | CACCTGTAGACCCACGACCT | ACCCATGAGCCAAATCAAAG | 147 |
| chr3:62462283-62464299 | p4 | (ACAT)4 | ATTTGCGGAACTCATATCGG | TGGCCCAAAAGTCTCATTTC | 196 |
| chr3:62648740-62650833 | p3 | (TTC)31 | ATTACCAATTGTCACCCCCA | TTTCTGAGTTGGTTTGCACG | 275 |
| chr3:62698561-62700595 | p2 | (GA)6 | CTTTGCTCACATGCGTCAGT | GGCGTATTGTTGGAGCTGAT | 234 |
| chr3:62744993-62747011 | p6 | (CCCACC)3 | GCCTGCATTTTCAAAACGAT | GAATGAGATGGGGCTAACCA | 289 |
| chr3:62847167-62849183 | p4 | (GAGT)4 | TGTAGGGATGCAGCTCTGTG | TGAGCCAACCATGAACAAAA | 207 |
| chr3:62847167-62849183 | p3 | (AAC)5 | TGTAGGGATGCAGCTCTGTG | TGAGCCAACCATGAACAAAA | 207 |
| chr3:62912109-62914172 | p2 | (GA)7 | TCACAAATCGACAAAACCCA | CTTTCGTGGTAACAGGCCAT | 122 |
| chr3:62912109-62914172 | p3 | (AAT)21 | TCACAAATCGACAAAACCCA | CTTTCGTGGTAACAGGCCAT | 122 |
| chr3:62938968-62940982 | p2 | (TG)6 | TATTCCCGTCAACAAGCACA | AGCTTTATGAGGCTGCAGGA | 116 |
| chr3:62938968-62940982 | p2 | (TG)7 | TATTCCCGTCAACAAGCACA | AGCTTTATGAGGCTGCAGGA | 116 |
| chr3:62942247-62944269 | p2 | (TA)11 | TCGGTGAATCTCCCAACAAT | GCGTAAGGTGGGGTGTTAAA | 116 |
| chr3:63005987-63008002 | p3 | (CAA)5 | TGCGCACAGATAACAGAACC | CGCCCGAATTCCTTTATGTA | 194 |
| chr3:63010161-63012176 | p5 | (AAGCT)3 | AGCCTCGGGGACAACTCTAT | GTCAAACGGAGCAATAGGGA | 246 |
| chr3:63087248-63089262 | p2 | (TA)7 | TGCATGTGTCTAAGCGAACC | GGGCCATGTCCTGGTAGTTA | 177 |
| chr3:63125921-63127933 | p2 | (CA)6 | GTGGGACCACCCTACTGCTA | GCCCTTAACCCCTAGTTTCG | 133 |
| chr3:63240698-63242718 | p2 | (TA)10 | CTCATGACGACATCCAAACG | CTTCGACCTCCAACAGCTTC | 241 |
| chr3:63444944-63446958 | p2 | (AT)7 | TTTTGAACCTCGGAGCATCT | TTTGGAGGGTTTCCACTTTG | 148 |

| Marker ID | SSR type | SSR motif | FORWARD PRIMER (5'-3') | REVERSE PRIMER (5'-3') | size (bp) |
|---|---|---|---|---|---|
| chr3:63678332-63680347 | p5 | (TTTTA)4 | CTCCAAGCCCCAAACATCTA | ACCCAACGAACTTGACTTGG | 232 |
| chr3:63678370-63680385 | p5 | (TTTTA)4 | CTCCAAGCCCCAAACATCTA | ACCCAACGAACTTGACTTGG | 194 |
| chr3:63924466-63926504 | p2 | (AT)9 | CCTCAGGCAGTGTCCGTATT | TCACAGATGGTGAGGACTCG | 294 |
| chr3:63924628-63926646 | p2 | (AT)9 | CCTCAGGCAGTGTCCGTATT | TCACAGATGGTGAGGACTCG | 132 |
| chr3:63954008-63956030 | p2 | (AT)11 | AAGAGAGGCCCGGTAAGGTA | TCGGAAAATTCTCAAGGGTG | 300 |
| chr3:63954008-63956030 | p4 | (AACT)5 | AAGAGAGGCCCGGTAAGGTA | TCGGAAAATTCTCAAGGGTG | 300 |
| chr3:64102506-64104521 | p5 | (ATTTC)3 | GGTTGGCCTCATGTTATGCT | AGGAAGGCCTGGAAGTTCAT | 141 |
| chr3:64121794-64123812 | p2 | (TC)9 | TACCGAAAATTTCCAGGCAG | GATGCGTCCGATGTATGTTG | 150 |
| chr3:64156088-64158100 | p2 | (TG)6 | CATTCCGAGCTCCTATGCTC | AATAGCCCGTAGGCACCTTT | 101 |
| chr3:64156088-64158100 | p2 | (TA)31 | CATTCCGAGCTCCTATGCTC | AATAGCCCGTAGGCACCTTT | 101 |
| chr3:64157828-64159876 | p2 | (TA)15 | AAGTCCAGTAGCCGAAAGCA | TACTTCGCTATCGGTCACCC | 184 |
| chr3:64157828-64159876 | p2 | (TA)24 | AAGTCCAGTAGCCGAAAGCA | TACTTCGCTATCGGTCACCC | 184 |
| chr3:64200753-64202765 | p2 | (AT)6 | TTCTCCAAACTCAAATCCCG | TTGTTGTGGGTGCTGTTGTT | 266 |
| chr3:64248342-64250356 | p2 | (TA)7 | TTCATGATTCTCCCTACGCC | GGTATGCTCCTCTACGTGCC | 299 |
| chr3:64301548-64303563 | p5 | (ATTCA)3 | TGTTCATGCCCACAGGTAGA | GCCTCTTGGCCATCAAAATA | 161 |
| chr3:64301548-64303563 | p2 | (TA)10 | TGTTCATGCCCACAGGTAGA | GCCTCTTGGCCATCAAAATA | 161 |
| chr3:64497040-64499082 | p5 | (CATAT)4 | TCCGTAGGTATCATAGGCCG | TCGTGAGTCCCCCATAAGTC | 261 |
| chr3:64497040-64499082 | p2 | (TA)21 | TCCGTAGGTATCATAGGCCG | TCGTGAGTCCCCCATAAGTC | 261 |
| chr3:64609922-64611946 | p2 | (AT)12 | TTCGGGTGGCAAATAGTCTC | GTAAGGGGTGCTCGGTACAA | 157 |
| chr3:64712056-64714070 | p2 | (AT)12 | TAGACGGTGGGAAGACAAGG | CCCCTTTAGGATTTTCGCTC | 247 |
| chr3:64712071-64714141 | p2 | (AT)12 | TAGACGGTGGGAAGACAAGG | CCCCTTTAGGATTTTCGCTC | 232 |
| chr3:64735363-64737379 | p3 | (CAA)6 | CACCAATGATCCCACATTCA | TGCTGATTGGGTTGTTTTGA | 175 |
| chr3:64735382-64737394 | p3 | (CAA)6 | CACCAATGATCCCACATTCA | TGCTGATTGGGTTGTTTTGA | 156 |
| chr3:64752918-64754933 | p5 | (TCTTT)3 | AAAACAAGGCGATTTGATGG | TCCTCGGATTTGCTGACTCT | 244 |
| chr3:64754119-64756134 | p5 | (TCTTT)3 | AAAACAAGGCGATTTGATGG | TCCTCGGATTTGCTGACTCT | 190 |
| chr3:64758591-64760606 | p5 | (TCTTT)3 | TCTACATGTGAAGGGGAGCC | AAATTTTGGGCCAATTCCTC | 151 |
| chr3:64764862-64766877 | p5 | (TCTTT)3 | AAAACAAGGCGATTTGATGG | TCCTCGGATTTGCTGACTCT | 244 |
| chr3:64766063-64768078 | p5 | (TCTTT)3 | AAAACAAGGCGATTTGATGG | TCCTCGGATTTGCTGACTCT | 190 |
| chr3:64772248-64774263 | p5 | (AAAAG)3 | GCTCCCCTGCACTTGTAGAC | TGGGATTTGCTTCAACAACA | 101 |
| chr3:64772248-64774263 | p5 | (AAAAG)3 | GCTCCCCTGCACTTGTAGAC | TGGGATTTGCTTCAACAACA | 101 |
| chr3:64779508-64781523 | p5 | (AAAAG)3 | ATGTGGACCTCCCATGATGT | GGGGCCATAAGAGTCAACAA | 291 |
| chr3:64781947-64783962 | p5 | (AAAAG)3 | TTCAGTAAGGGCCTTGGTTG | AGCTTGGATGGAAAGCTTCA | 118 |
| chr3:64783113-64785128 | p5 | (AAAAG)3 | GCCTCCTTCTCCAAATAGCC | AGAAGGACACGTACGCGAAT | 254 |
| chr3:64788762-64790777 | p5 | (AAAAG)3 | CCTTCTCCAAGTAGCCTCCC | TGGCCCAAAATTTGAAGAAG | 269 |
| chr3:64808803-64810818 | p5 | (AAAGA)3 | TCAGTAAGGGCCTTGTTTGG | CGAGCTTCATGAAAAGGGAG | 124 |
| chr3:64812447-64814462 | p5 | (AAAAG)3 | CTCCCATATCTTGAAGGCCA | GGGGCCATAAGAGTCAACAA | 283 |

| Marker ID | SSR type | SSR motif | FORWARD PRIMER (5'-3') | REVERSE PRIMER (5'-3') | size (bp) |
|---|---|---|---|---|---|
| chr3:64816834-64818849 | p5 | (AAAAG)3 | CTCCCATATCTTGAAGGCCA | GGGGCCATAAGAGTCAACAA | 285 |
| chr3:64816834-64818849 | p5 | (AAAAG)3 | CTCCCATATCTTGAAGGCCA | GGGGCCATAAGAGTCAACAA | 285 |
| chr3:64826763-64828778 | p5 | (AAAAG)3 | CCTTCTCCAAATAGCCTCCC | GGGGCCATAAGAGTCAACAA | 219 |
| chr3:64829653-64831668 | p5 | (AAAAG)3 | CAGTAAGGGCCTTGTTTGGA | TTTGAAGATCGTGGGAGGTC | 126 |
| chr3:64846698-64848713 | p5 | (AAAAG)3 | CAGTAAGGGCCTTGTTTGGA | GGGAGGCTATTTGGAGAAGG | 122 |
| chr3:64865473-64867488 | p5 | (AAAAG)3 | CAGTAAGGGCCTTGTTTGGA | TTCGAAGATCATGGGAGGTC | 122 |
| chr3:64886721-64888736 | p5 | (AAAGA)3 | CTCCCATATCTTGAAGCCCA | AAGTTGTGTGGGGCCATAAG | 288 |
| chr3:64886721-64888736 | p5 | (AAAGA)3 | CTCCCATATCTTGAAGCCCA | AAGTTGTGTGGGGCCATAAG | 288 |
| chr3:64908157-64910172 | p5 | (AAAGA)3 | CAGTAAGGGCCTTGTTTGGA | GGGAGGCTATTTGGAGAAGG | 127 |
| chr3:64909352-64911372 | p5 | (AAAGA)4 | CAGTAAGGGCCTTGTTTGGA | GCTTGGATGGAAGGTTTCAA | 128 |
| chr3:64998809-65000839 | p2 | (TA)15 | GTTCATCATCCGCTCGTTTT | TGGAGTAGGTAACATCGGGG | 196 |
| chr3:65007331-65009349 | p6 | (GGCGGT)3 | GAACCCAAAGAAATGCCAGA | CAGCTTCCATTCGTGACTGA | 227 |
| chr3:65039041-65041059 | p6 | (TGTATA)3 | AGTTGTTGTAGCCGAATGGG | TCATCGGGTCCTATTTACGC | 136 |
| chr3:65039041-65041059 | p2 | (TA)10 | AGTTGTTGTAGCCGAATGGG | TCATCGGGTCCTATTTACGC | 136 |
| chr3:65218916-65220931 | p5 | (ATACG)3 | CCCTCCTCATGAACAATGCT | TAGATCCTCAGCTAGGGGCA | 222 |
| chr3:65340516-65342530 | p2 | (TG)7 | CGTTCTGTATTCCGTCCGAT | TCGCACATACCGATACCAGA | 177 |
| chr3:65470032-65472052 | p6 | (TATATG)3 | CGTACGTTCCTCGGCACTAT | CGGAACAACGGAACAAAAGT | 244 |
| chr3:65470032-65472052 | p5 | (GAAAT)4 | CGTACGTTCCTCGGCACTAT | CGGAACAACGGAACAAAAGT | 244 |
| chr3:65561417-65563432 | p5 | (TGAAT)3 | CAAAGGAAACTCTGCCGAAG | GGGACATTCCTCTTGGTTGA | 159 |
| chr3:65579581-65581596 | p3 | (TTG)5 | TTTTGGTGCATTCGGTATGA | TCGCATTTAAATCCCGAGTC | 127 |
| chr3:65593730-65595744 | p2 | (GT)7 | CTGCTCAGCTGTGTGGAGAG | CCTTCCTCAATATTTCCGCA | 217 |
| chr3:65595157-65597172 | p5 | (TATGA)3 | AGTTAAATTTCGGGGGCTGT | ATAGAGTTGTCCCCGAGGCT | 105 |
| chr3:65751004-65753016 | p2 | (AT)6 | TGTCTACCTGTGGGCATGAA | GCCTATGACATGTGTGCAGG | 289 |
| chr3:65751004-65753016 | p2 | (TC)6 | TGTCTACCTGTGGGCATGAA | GCCTATGACATGTGTGCAGG | 289 |
| chr3:65880055-65882070 | p3 | (GAA)5 | GAATCAGACATTCCCGTGCT | AAGCTGCTGAAATGATGCCT | 225 |
| chr3:65880055-65882070 | p5 | (GCCTC)3 | GAATCAGACATTCCCGTGCT | AAGCTGCTGAAATGATGCCT | 225 |
| chr3:65915720-65917735 | p2 | (AT)18 | AATCATCCTTTTGGAGGGCT | ACCATAAGGTAAGGTGGGGC | 278 |
| chr3:65915720-65917735 | p3 | (CTT)5 | AATCATCCTTTTGGAGGGCT | ACCATAAGGTAAGGTGGGGC | 278 |
| chr3:65928213-65930225 | p2 | (CT)6 | TGGGATGCTGTGACAGGTTA | TAACGGAAAACATTCCGAGG | 279 |
| chr3:65962184-65964202 | p3 | (ATT)6 | GACGAGGCCAACAAAATGAT | GGGAGACAGACCTGATCCAA | 203 |
| chr3:65968238-65970256 | p3 | (ATT)6 | CACAACAATGCTACATGGGC | GAGCATACGGCTTTTGGGTA | 176 |
| chr3:66054543-66056559 | p2 | (AT)8 | AAAATTATTGCCATGGCTGC | GTTGCTATCAACCCCTTGGA | 150 |
| chr3:66054543-66056559 | p2 | (TA)11 | AAAATTATTGCCATGGCTGC | GTTGCTATCAACCCCTTGGA | 150 |
| chr3:66076496-66078512 | p6 | (CCTCCC)4 | CTTTCCTACCCATCCCCATT | GTCCGGATTACGCTCATGTT | 162 |
| chr3:66076496-66078512 | p2 | (AT)8 | CTTTCCTACCCATCCCCATT | GTCCGGATTACGCTCATGTT | 162 |
| chr3:66095268-66097280 | p2 | (AT)13 | AACCAGAGGGTCATGACAGG | GTGGGTCATGGGTGGTTATC | 246 |

| Marker ID | SSR type | SSR motif | FORWARD PRIMER (5'-3') | REVERSE PRIMER (5'-3') | size (bp) |
|---|---|---|---|---|---|
| chr3:66095268-66097280 | p2 | (TA)6 | AACCAGAGGGTCATGACAGG | GTGGGTCATGGGTGGTTATC | 246 |
| chr3:66095268-66097280 | p2 | (AT)8 | AACCAGAGGGTCATGACAGG | GTGGGTCATGGGTGGTTATC | 246 |
| chr3:66095394-66097410 | p2 | (AT)13 | AACCAGAGGGTCATGACAGG | TAGCCTGACCGACTTGTGTG | 120 |
| chr3:66095394-66097410 | p2 | (TA)6 | AACCAGAGGGTCATGACAGG | TAGCCTGACCGACTTGTGTG | 120 |
| chr3:66095394-66097410 | p2 | (AT)8 | AACCAGAGGGTCATGACAGG | TAGCCTGACCGACTTGTGTG | 120 |
| chr3:66126553-66128568 | p5 | (GGTGT)3 | TTCCTACCCCTCCCTGTTCT | GAACGTACAAAATGCGGGAT | 149 |
| chr3:66213844-66215864 | p2 | (TA)13 | TTTCTAGCCATACGCCAACC | TATATGATGATGGCCGCAGA | 192 |
| chr3:66213844-66215864 | p2 | (AT)10 | TTTCTAGCCATACGCCAACC | TATATGATGATGGCCGCAGA | 192 |
| chr3:66432872-66434886 | p2 | (TA)7 | AAGCATCGGATCAACAATCC | GCATTGGAAAGAAGACTCGC | 179 |
| chr3:66500697-66502715 | p2 | (AT)15 | GGTGTGATGGGTACCTGACC | CCAGCCGATGTGTTATTGTG | 189 |
| chr3:66500697-66502715 | p6 | (ATATAC)3 | GGTGTGATGGGTACCTGACC | CCAGCCGATGTGTTATTGTG | 189 |
| chr3:66528656-66530671 | p5 | (TTCAT)3 | GGTTCCAGATGAAACGCTGT | TCAGGGCTACAACACCACAA | 247 |
| chr3:66572004-66574019 | p5 | (GACAT)3 | CATTGGGCGAAATTTTCAGT | ACGGAGATCAAACGGACATC | 241 |
| chr3:66615290-66617302 | p2 | (GT)6 | ACAAGCATCTGAACCCAAGC | TTCCCTTAGCCTTCCTCCAT | 277 |
| chr3:66740104-66742119 | p5 | (ATTTC)3 | CGCGGATCATTTGTGTACTG | TAGCCTGCTAAGCCCAGAAA | 187 |
| chr3:66777062-66779088 | p2 | (AT)13 | TTTGGGCCTCATTTCATCTC | GCAAGACAAGCTTAATGGGC | 292 |
| chr3:66802481-66804496 | p3 | (ATC)5 | TTCGTATTTCTCCTCCCACG | GTGTTGTTGAATTCGGGCTT | 257 |
| chr3:66819026-66821040 | p2 | (AT)7 | CCCAAGCTGTACACCCGTAT | AGCCTTGTCAGCTTGCGTAT | 217 |
| chr3:66894538-66896558 | p2 | (TA)10 | TCCACGAATCATACTGCGAA | GCGGGAAGACTACAAGCAAG | 295 |
| chr3:67031862-67033886 | p2 | (AT)12 | TGAGTCAGCAGTTGAATGGC | GGGAAAAACAATGGCAGAAA | 244 |
| chr3:67085573-67087585 | p2 | (AT)6 | ACGTACCTGATCCTTGGACG | GTGTGATAGAGCCCAGCCAT | 193 |
| chr3:67085573-67087585 | p2 | (AT)6 | ACGTACCTGATCCTTGGACG | GTGTGATAGAGCCCAGCCAT | 193 |
| chr3:67157042-67159060 | p3 | (TCA)6 | CGCCTATCATCACAAGCTGA | TAAGAGCGCTCCATTTGCTT | 162 |
| chr3:67168917-67170932 | p3 | (TGA)5 | ATCATAGCCGCCCACTACAC | CGACCTTACGTCACTCCGAT | 277 |
| chr3:67197458-67199473 | p3 | (GTT)5 | GCATGATTGAGCCGGACTAT | CCGAGTCCGAAACGAAAATA | 167 |
| chr3:67363873-67365897 | p5 | (GAAAT)3 | TGTTTGGCCGTATAACTCCC | AGAGAAGGCAAGGCATACGA | 278 |
| chr3:67363873-67365897 | p2 | (TA)15 | TGTTTGGCCGTATAACTCCC | AGAGAAGGCAAGGCATACGA | 278 |
| chr3:67363873-67365897 | p2 | (AT)12 | TGTTTGGCCGTATAACTCCC | AGAGAAGGCAAGGCATACGA | 278 |
| chr3:67363873-67365897 | p2 | (AT)17 | TGTTTGGCCGTATAACTCCC | AGAGAAGGCAAGGCATACGA | 278 |
| chr3:67364774-67366792 | p2 | (AT)12 | TTGGGCCTCACTTTATCGTC | ATATGGGGCCCACAATTTTT | 238 |
| chr3:67364774-67366792 | p2 | (AT)17 | TTGGGCCTCACTTTATCGTC | ATATGGGGCCCACAATTTTT | 238 |
| chr3:67364774-67366792 | p2 | (TA)13 | TTGGGCCTCACTTTATCGTC | ATATGGGGCCCACAATTTTT | 238 |
| chr3:67364774-67366792 | p2 | (AT)6 | TTGGGCCTCACTTTATCGTC | ATATGGGGCCCACAATTTTT | 238 |
| chr3:67382755-67384770 | p5 | (TGTGG)3 | GCATAATTTCCCTTGGGGTT | GCAACCTCCCATATGCCTTA | 103 |
| chr3:67470965-67472980 | p5 | (AATAT)3 | CGGGTCCGAGATTTTATTGA | GTTTCGTCGAAGCTGTGTGA | 279 |
| chr3:67555765-67557783 | p3 | (TTG)6 | GACCCTTGAGCCAAATCAAA | CACCTGTAGACCCACGACCT | 158 |

| Marker ID | SSR type | SSR motif | FORWARD PRIMER (5'-3') | REVERSE PRIMER (5'-3') | size (bp) |
|---|---|---|---|---|---|
| chr3:67575419-67577434 | p3 | (TGT)5 | CGTTTTCGGGTGATCTGACT | ATAGCAAGGAAAGTGGGCCT | 205 |
| chr3:67660963-67662983 | p2 | (TG)10 | ATGTGAATGCCAGAGGGTTC | TAAATGCCCTGTAGCAGCCT | 294 |
| chr3:67684166-67686208 | p2 | (TG)21 | TGGCGGAACCTTCTTAACAC | CGTTGGTACGCAAGTGAATG | 247 |
| chr3:67718462-67720504 | p2 | (TG)21 | TGGCGGAACCTTCTTAACAC | CGTTGGTACGCAAGTGAATG | 247 |
| chr3:67847574-67849588 | p2 | (TC)7 | CCGAATTCAGTGGTTGCTTT | AAAATGGGTAACATTCGGCA | 184 |
| chr3:67893447-67895469 | p2 | (AT)11 | ATGAGCCATGACTAGTGCCC | GGTTAGCCGTGACTCCTCAG | 266 |
| chr3:67946693-67948709 | p2 | (TA)8 | GACTCCGGTTCGGACTATGA | TGACAGCTAGGCTGGAGGAT | 223 |
| chr3:67946693-67948709 | p2 | (TA)8 | GACTCCGGTTCGGACTATGA | TGACAGCTAGGCTGGAGGAT | 223 |
| chr3:68110618-68112634 | p2 | (AT)8 | GATTGGCCGATGTTGATTCT | TTTAATTGAGTCGCCCGTTC | 152 |
| chr3:68171154-68173170 | p2 | (AG)8 | TCCAGTGAATGTTGGTGCAT | TCTGCAGTCTCGGTAACACG | 276 |
| chr3:68171154-68173170 | p3 | (TTG)5 | TCCAGTGAATGTTGGTGCAT | TCTGCAGTCTCGGTAACACG | 276 |
| chr3:68243683-68245787 | p2 | (AT)52 | TAACTGACGGTTACGAGGGC | TCTTAGGCTCGAATCAGGGA | 257 |
| chr3:68522697-68524709 | p2 | (CA)6 | GCCCATGACTACTACCCCAA | TGACCCTTACATGGAGCACA | 108 |
| chr3:68598233-68600251 | p6 | (ATATGT)3 | ATGTTGGATATGAGCCTCGG | GTCGGGTCCTATTTATGCGA | 257 |
| chr3:68619802-68621814 | p2 | (TC)6 | ATGAGTTCAAGTGATCGGGC | GCCAACTCCATGGATTCACT | 134 |
| chr3:68672924-68674950 | p2 | (TA)17 | AGGAGAACAAGCAGGCGATA | AGACTCTTAGCTTTCCGGGG | 267 |
| chr3:68672959-68675003 | p2 | (TA)17 | AGGAGAACAAGCAGGCGATA | AGACTCTTAGCTTTCCGGGG | 232 |
| chr3:68690291-68692303 | p2 | (CA)6 | GACACCGAAATCAGCCAAGT | CCATGTTGCATCAACACACA | 106 |
| chr3:68721080-68723100 | p2 | (AT)10 | ATGCACATTCTCAAGGAGCC | ACGGGCTATAACATGTTGCC | 287 |
| chr3:68721080-68723100 | p5 | (AATTG)4 | ATGCACATTCTCAAGGAGCC | ACGGGCTATAACATGTTGCC | 287 |
| chr3:68731829-68733879 | p5 | (TTGCA)3 | ATACGGTCCCAACATACGGA | AAGTTGGCCAAGAGAAAGCA | 220 |
| chr3:68731829-68733879 | p2 | (TC)14 | ATACGGTCCCAACATACGGA | AAGTTGGCCAAGAGAAAGCA | 220 |
| chr3:68731829-68733879 | p2 | (TA)10 | ATACGGTCCCAACATACGGA | AAGTTGGCCAAGAGAAAGCA | 220 |
| chr3:68751084-68753099 | p3 | (TAA)5 | GGGGTATATTTGGCGCTTTT | TCTGGCTATTAGGCGAAGGA | 195 |
| chr3:68751084-68753099 | p3 | (TTA)5 | GGGGTATATTTGGCGCTTTT | TCTGGCTATTAGGCGAAGGA | 195 |
| chr3:68759072-68761084 | p2 | (AG)6 | TACGACTGGGTTTACGGTCC | TTTGGAAATAAGAGGTGGCG | 226 |
| chr3:68854845-68856860 | p5 | (CATAT)3 | TTGCAGACAACCCCTAGGAC | GGCTTCGTCAACCGATATGT | 244 |
| chr3:68979464-68981490 | p2 | (TA)13 | ACTCTTTCGACGATGTCGCT | GTGGAGCTCGATAAACCCAA | 260 |
| chr3:68993681-68995705 | p2 | (AT)12 | TGTCGTGTGTGTTATGGCCT | ACTCCTTCGATGCTCACGTT | 193 |
| chr3:69023166-69025186 | p2 | (AC)10 | ATCCAAAAAGAGAGCCCGAT | TCTGCTCCTCGAAACTCGAT | 268 |
| chr3:69065991-69068017 | p2 | (TA)13 | ACTCTTTCGACGATGTCGCT | GTGGAGCTCGATAAACCCAA | 260 |
| chr3:69080208-69082232 | p2 | (AT)12 | TGTCGTGTGTGTTATGGCCT | ACTCCTTCGATGCTCACGTT | 193 |
| chr3:69146271-69148293 | p2 | (TA)11 | ACCACGGCAGAAGCATAATC | ACCAAGGTGGACTCCAAGTG | 140 |
| chr3:69216986-69218998 | p2 | (AT)6 | AGCCCTCTATCCTGGTCGTT | CAGCCGATCGGTCTCTAGTC | 164 |
| chr3:69350225-69352237 | p2 | (AT)6 | AGCATAGAGGGGGAGGTCAT | AACCGCACTCAAAGGTATGG | 209 |
| chr3:69357356-69359424 | p2 | (AT)6 | GGTGTTTCCTTGTTCTGGGA | CGCAAAAAGGGTATGTTGCT | 187 |

| Marker ID | SSR type | SSR motif | FORWARD PRIMER (5'-3') | REVERSE PRIMER (5'-3') | size (bp) |
|---|---|---|---|---|---|
| chr3:69357356-69359424 | p2 | (TA)34 | GGTGTTTCCTTGTTCTGGGA | CGCAAAAAGGGTATGTTGCT | 187 |
| chr3:69446481-69448493 | p5 | (TGAAT)3 | GTTGTGTGTTGATGGGCAAG | CTAACCACCTCCTCTGTCGC | 232 |
| chr3:69446499-69448513 | p5 | (TGAAT)3 | GTTGTGTGTTGATGGGCAAG | CTAACCACCTCCTCTGTCGC | 214 |
| chr3:69446514-69448530 | p5 | (TGAAT)3 | GTTGTGTGTTGATGGGCAAG | CTAACCACCTCCTCTGTCGC | 199 |
| chr3:69458963-69460978 | p2 | (TA)6 | TTGAGGTTGATTCCAAAGGG | CATGGAGTCGTTCCGGTAGT | 169 |
| chr3:69458963-69460978 | p5 | (GCTAT)3 | TTGAGGTTGATTCCAAAGGG | CATGGAGTCGTTCCGGTAGT | 169 |
| chr3:69496050-69498062 | p2 | (TA)6 | TGACTCGCCCTCCTCTACAT | GATTTTAGAAATGCCGCCAA | 105 |
| chr3:69496050-69498062 | p2 | (AT)6 | TGACTCGCCCTCCTCTACAT | GATTTTAGAAATGCCGCCAA | 105 |
| chr3:69545457-69547482 | p2 | (TA)8 | TGTTTAGTGCGCTCCACTTG | GGAACGGTGTAAGCCGACTA | 138 |
| chr3:69545457-69547482 | p5 | (TCATA)5 | TGTTTAGTGCGCTCCACTTG | GGAACGGTGTAAGCCGACTA | 138 |
| chr3:69576864-69578879 | p3 | (TAA)5 | TGTAGCTGGCACTGGTTCTG | GACCCACAAAAAGAGTCGGA | 125 |
| chr3:69649727-69651739 | p2 | (AT)6 | TTGACTTACGGTCCGTCCTC | TGCAATTGTCCTTCAGCTTG | 284 |
| chr3:69666452-69668480 | p2 | (TA)27 | AAATGTGGGCTATTTGCAGG | CATGCCAAAAGAAGGAAAGC | 182 |
| chr3:69666483-69668501 | p2 | (TA)27 | AAATGTGGGCTATTTGCAGG | CATGCCAAAAGAAGGAAAGC | 151 |
| chr3:69666504-69668522 | p2 | (TA)27 | AAATGTGGGCTATTTGCAGG | CATGCCAAAAGAAGGAAAGC | 130 |
| chr3:69666525-69668543 | p2 | (TA)27 | AAATGTGGGCTATTTGCAGG | CATGCCAAAAGAAGGAAAGC | 109 |
| chr3:69893166-69895194 | p2 | (AT)6 | ATTGGAGTTCGCCTTCTTGA | ATGAACACCGAGCTTATGGC | 223 |
| chr3:69914617-69916638 | p3 | (CAT)7 | CCAACACCAATTACCGTTCC | AGACAGTTGTGTTTTGGCCC | 152 |
| chr3:69945987-69948043 | p2 | (AT)28 | GAGGAAACGCCCTCAACATA | CGTAAGGTGTCGGGTCCTAA | 211 |
| chr3:69945987-69948043 | p3 | (ATC)5 | GAGGAAACGCCCTCAACATA | CGTAAGGTGTCGGGTCCTAA | 211 |
| chr3:70008126-70010192 | p3 | (TTC)22 | TTAAGAGGCAGTCGCTTGGT | CTTGGCGAAATGGGTAAGAA | 265 |
| chr3:70039205-70041220 | p2 | (TA)9 | ATGGGGTCCAGACTACAACG | CCAATGGTATCCAACCGAAC | 158 |
| chr3:70039205-70041220 | p3 | (TAT)5 | ATGGGGTCCAGACTACAACG | CCAATGGTATCCAACCGAAC | 158 |
| chr3:70061319-70063353 | p2 | (TA)17 | AACTCGGCCAAATCATCAAC | GGAAGCGAGGAATGTGTTGT | 239 |
| chr3:70096459-70098474 | p3 | (AAC)6 | GATCCTTATGTTCCGCCAAA | TATCGGGTACGTTTGGGTGT | 143 |
| chr3:70096459-70098474 | p5 | (CATAT)3 | GATCCTTATGTTCCGCCAAA | TATCGGGTACGTTTGGGTGT | 143 |
| chr3:70108492-70110510 | p3 | (CAT)6 | AAGTTCCATCAAACATCGGC | AAAGTGCTCCATTTGATCCG | 244 |
| chr3:70136402-70138436 | p2 | (TA)17 | AACTCGGCCAAATCATCAAC | GGAAGCGAGGAATGTGTTGT | 239 |
| chr3:70171542-70173557 | p3 | (AAC)6 | GATCCTTATGTTCCGCCAAA | TATCGGGTACGTTTGGGTGT | 143 |
| chr3:70171542-70173557 | p5 | (CATAT)3 | GATCCTTATGTTCCGCCAAA | TATCGGGTACGTTTGGGTGT | 143 |
| chr3:70183575-70185593 | p3 | (CAT)6 | AAGTTCCATCAAACATCGGC | AAAGTGCTCCATTTGATCCG | 244 |
| chr3:70340397-70342411 | p3 | (ATC)5 | TCCTGTCCCGTCCTCATAAC | CTACCGAGCACCACTCATCA | 272 |
| chr3:70340397-70342411 | p2 | (AT)7 | TCCTGTCCCGTCCTCATAAC | CTACCGAGCACCACTCATCA | 272 |
| chr3:70467291-70469305 | p2 | (TA)7 | GACGAGACCTCAGGACAAGC | CACCCCCAACTAAAGAAGCA | 154 |
| chr3:70491953-70493965 | p2 | (TA)6 | TAAATAGAACCACGCCGACC | GGCAAATCAGGTCCATCTGT | 161 |
| chr3:70491953-70493965 | p2 | (AT)11 | TAAATAGAACCACGCCGACC | GGCAAATCAGGTCCATCTGT | 161 |

| Marker ID | SSR type | SSR motif | FORWARD PRIMER (5'-3') | REVERSE PRIMER (5'-3') | size (bp) |
|---|---|---|---|---|---|
| chr3:70491953-70493965 | p2 | (AT)24 | TAAATAGAACCACGCCGACC | GGCAAATCAGGTCCATCTGT | 161 |
| chr3:70511605-70513620 | p5 | (ATCAT)3 | TGTAGCGTAGGGGTAAACCG | CCGAGCCCATACTTTTGGTA | 125 |
| chr3:70538553-70540583 | p2 | (AT)7 | GATAGCCTTGTCGGCTTCTG | TGGTCACCCGAACATACTCA | 145 |
| chr3:70538553-70540583 | p2 | (AT)15 | GATAGCCTTGTCGGCTTCTG | TGGTCACCCGAACATACTCA | 145 |
| chr3:70698627-70700639 | p5 | (AAGAA)3 | CTGTTAAAGCACTGTGGCGA | CTTACAAGCATTTTGGGGGA | 107 |
| chr3:70698627-70700639 | p2 | (TG)6 | CTGTTAAAGCACTGTGGCGA | CTTACAAGCATTTTGGGGGA | 107 |
| chr3:70800066-70802080 | p2 | (AG)7 | GAAGCAGTGCATCCGGTATT | CGCTTCCCATGTCATTTCTT | 205 |
| chr3:70867353-70869381 | p2 | (AT)14 | ATCTTGAGTCTTCCCCCGAT | TGGAATATAGCCTTGTCGGC | 260 |
| chr3:70925122-70927138 | p2 | (TA)10 | AGCGTGCTAAGGTGGGAGTA | TTTAGTCTCCAACCATCCCG | 225 |
| chr3:70925122-70927138 | p2 | (AG)8 | AGCGTGCTAAGGTGGGAGTA | TTTAGTCTCCAACCATCCCG | 225 |
| chr3:70932230-70934248 | p2 | (AT)9 | GTACTAAGAGGGCCGGGTTC | AATCGGAATCCATACGGACA | 108 |
| chr3:71064127-71066145 | p2 | (AC)9 | GGGGGTTTTGACTTGAATGA | GAAGAAGGGGAAAATGAGCC | 127 |
| chr3:71079066-71081078 | p2 | (AT)6 | GTGGAGCCCGAATCTATCAA | ATATCATGCGTAACTGCCCC | 123 |
| chr3:71108969-71110984 | p5 | (CATAT)3 | ATTAGCATAAGGAGGGGCGT | AGCCTTATCGGCTTCCTTGT | 160 |
| chr3:71173806-71175830 | p4 | (ATAC)6 | AATTGTTGTCGGGTCCTCTG | AGATGAAAAGGGCAGCTTCA | 274 |
| chr3:71236637-71238652 | p3 | (TCA)5 | TCGTGAGACCATGTAACCCA | TGTTCATGCCCACAGGTAGA | 191 |
| chr3:71248827-71250959 | p2 | (TA)66 | GAACTTGGCTGCCTTCTGAG | ACTTCACACCAACCCTTTGC | 199 |
| chr3:71264941-71266961 | p2 | (AT)10 | TGATATGCTTATGGCACCGA | ATATATCGTACCCGGCCCTC | 262 |
| chr3:71264941-71266961 | p2 | (AT)9 | TGATATGCTTATGGCACCGA | ATATATCGTACCCGGCCCTC | 262 |
| chr3:71310248-71312300 | p2 | (TA)26 | GCTTGCTCCAAAGCTCATTC | CCAACCCTTTGCACATTTTT | 170 |
| chr3:71310248-71312300 | p2 | (TA)32 | GCTTGCTCCAAAGCTCATTC | CCAACCCTTTGCACATTTTT | 170 |
| chr3:71310248-71312300 | p3 | (ACA)5 | GCTTGCTCCAAAGCTCATTC | CCAACCCTTTGCACATTTTT | 170 |
| chr3:71356416-71358430 | p2 | (TA)7 | TTGAGGGCTGGAATTTATGG | CTAAGCTTGCTTCGTCGCTT | 126 |
| chr3:71356416-71358430 | p2 | (AT)11 | TTGAGGGCTGGAATTTATGG | CTAAGCTTGCTTCGTCGCTT | 126 |
| chr3:71452789-71454801 | p2 | (AT)6 | TGAATCACCATGTTCTGGGA | GCAGGTTGTAGGCCATGATT | 261 |
| chr3:71465092-71467107 | p5 | (AAACA)3 | AAAATGTGGCACCCAAAGAG | GATTTCTCCGGACCTCCTTC | 217 |
| chr3:71576991-71579016 | p5 | (TCATA)5 | TCGCATATTCTTCTTCGGCT | CCGAGTTGGGTCGTGATACT | 130 |
| chr3:71702751-71704767 | p2 | (TA)8 | GGAACGATGCTACGGCTAAG | AATCATGGTCGCCTTGTCTC | 166 |
| chr3:71731827-71733842 | p6 | (CATATA)4 | TAAGTTGAGACGAGCAGCGA | CGACATTGATGATGTCGAGG | 182 |
| chr3:71731827-71733842 | p5 | (ATTCA)3 | TAAGTTGAGACGAGCAGCGA | CGACATTGATGATGTCGAGG | 182 |
| chr3:71908900-71910914 | p5 | (ATCAT)3 | ATGAAACGCAGTGTCCACAA | CGCGATATGCTACGACAAGA | 259 |
| chr3:71908917-71910971 | p5 | (ATCAT)3 | ATGAAACGCAGTGTCCACAA | CGCGATATGCTACGACAAGA | 242 |
| chr3:71910486-71912504 | p2 | (AT)9 | TGCAAGTTCTAGGGGATTGG | CAGATTTTGACGAGATGGCA | 100 |
| chr3:71956151-71958169 | p6 | (TATACA)3 | AGCGAACCCCTTTAGCTTTC | GCAAGCCGACGTACCATTAT | 205 |
| chr3:72163374-72165389 | p2 | (AT)7 | GACCTGCTGGTATCTGGGAA | TTTTGTTCTCAGTTTGGGGG | 137 |
| chr3:72163374-72165389 | p3 | (CAT)5 | GACCTGCTGGTATCTGGGAA | TTTTGTTCTCAGTTTGGGGG | 137 |

| Marker ID | SSR type | SSR motif | FORWARD PRIMER (5'-3') | REVERSE PRIMER (5'-3') | size (bp) |
|---|---|---|---|---|---|
| chr3:72250742-72252760 | p6 | (TATATG)3 | CTTGACCTCCATGAGCCATT | TCCAATCACGCAACAAACTC | 293 |
| chr3:72304992-72307010 | p2 | (TA)8 | TCGTGCATGCCGTAATGTAT | TACCCGACCGTCTAGTGAGG | 109 |
| chr3:72304992-72307010 | p2 | (TA)43 | TCGTGCATGCCGTAATGTAT | TACCCGACCGTCTAGTGAGG | 109 |
| chr3:72304992-72307010 | p2 | (AT)9 | TCGTGCATGCCGTAATGTAT | TACCCGACCGTCTAGTGAGG | 109 |
| chr3:72359373-72361385 | p2 | (AT)6 | TAGCCATGTGAGGCTTTGTG | TTCCTGGTCCAAAAGTACGG | 219 |
| chr3:72359373-72361385 | p3 | (GTT)5 | TAGCCATGTGAGGCTTTGTG | TTCCTGGTCCAAAAGTACGG | 219 |
| chr3:72359373-72361385 | p2 | (AT)8 | TAGCCATGTGAGGCTTTGTG | TTCCTGGTCCAAAAGTACGG | 219 |
| chr3:72372866-72374881 | p5 | (CATTG)3 | AAGACGAAGCTCGAGATGGA | TAACATGGTGACAAGCCCAA | 116 |
| chr3:72393086-72395102 | p2 | (AT)8 | AAATCTAACCGGACCTGCCT | CTGGCACTTGGTCAGTGAGA | 165 |
| chr3:72425373-72427385 | p2 | (TC)6 | CAAAAACATGTCCGTCGTTG | ATTCCCTTCCTGATAACCCG | 264 |
| chr3:72465051-72467101 | p2 | (AT)25 | TCCCAAGTAAGAATGGTGCC | CGTGCAGGTTTTGGGATAGT | 132 |
| chr3:72465051-72467101 | p2 | (AT)11 | TCCCAAGTAAGAATGGTGCC | CGTGCAGGTTTTGGGATAGT | 132 |
| chr3:72465051-72467101 | p5 | (ATCAT)3 | TCCCAAGTAAGAATGGTGCC | CGTGCAGGTTTTGGGATAGT | 132 |
| chr3:72480482-72482494 | p2 | (AC)6 | GGTCATGCCAAAAGAAAGGA | TTACAGGGGAGATGCTACCG | 217 |
| chr3:72490788-72492802 | p5 | (ATATC)3 | CGAAGTCAGTCGTACTCGCA | TCAAAATTGGGAGCCTTTTG | 173 |
| chr3:72490788-72492802 | p2 | (AT)7 | CGAAGTCAGTCGTACTCGCA | TCAAAATTGGGAGCCTTTTG | 173 |
| chr3:72490788-72492802 | p2 | (TA)15 | CGAAGTCAGTCGTACTCGCA | TCAAAATTGGGAGCCTTTTG | 173 |
| chr3:72491390-72493420 | p2 | (AT)7 | ACTTTTGTCACGACCCAACC | CCTTGTCAGCTTGCGTATCA | 256 |
| chr3:72491390-72493420 | p2 | (TA)15 | ACTTTTGTCACGACCCAACC | CCTTGTCAGCTTGCGTATCA | 256 |
| chr3:72521160-72523172 | p2 | (TG)6 | AAAGGTGCCAAAGGGCTATT | TCATACCTGGCCCTACCAAG | 237 |
| chr3:72521160-72523172 | p2 | (TA)20 | AAAGGTGCCAAAGGGCTATT | TCATACCTGGCCCTACCAAG | 237 |
| chr3:72807722-72809736 | p2 | (TA)7 | GTCCAGTAAGGCCCATGAGA | GCCTTTCACTGTACCCCTGA | 171 |
| chr3:72807722-72809736 | p3 | (ACC)5 | GTCCAGTAAGGCCCATGAGA | GCCTTTCACTGTACCCCTGA | 171 |
| chr3:72817157-72819172 | p5 | (ATATC)3 | AGCTGCTTCTTCATTGGCAT | GAAGCCGGCCACACTTATAG | 120 |
| chr3:73008205-73010220 | p3 | (AAT)5 | TTTGTTGCTGGCAAGTTTTG | GTAGGAAGCCAACCAATCCA | 104 |
| chr3:73115521-73117545 | p6 | (TATATG)4 | GTTGCATTCTCGGGTTTGTT | TGCCATTGAAAGAAAGGGAC | 190 |
| chr3:73190936-73192958 | p6 | (TATATG)3 | GAAGCGGGCAAGGTATGTAA | ACAGCTTTCGGGAGTCAAGA | 289 |
| chr3:73190936-73192958 | p2 | (AT)11 | GAAGCGGGCAAGGTATGTAA | ACAGCTTTCGGGAGTCAAGA | 289 |
| chr3:73190936-73192958 | p2 | (AT)9 | GAAGCGGGCAAGGTATGTAA | ACAGCTTTCGGGAGTCAAGA | 289 |
| chr3:73234836-73236851 | p5 | (AAGAA)3 | TTGCAGCAGCAATACTGACC | CAGAAAGGCCACGGTAATGT | 130 |
| chr3:73266484-73268506 | p2 | (AT)11 | GCTGTTGTTGTGTCGTTGCT | ACAATTGGGCAGGACTTCAC | 210 |
| chr3:73267708-73269723 | p5 | (TATGA)3 | CTGTCCCGTCATATTTCGCT | CTGACAAGCCTGCCACTACA | 207 |
| chr3:73302663-73304683 | p2 | (AT)10 | TTGTACCGAGCACCCTTACC | ATTTTGAGATGTTCGCGTCC | 206 |
| chr3:73353293-73355329 | p2 | (AT)18 | GTGAGCATCTGCTGAGTCCA | AACCCGTCCCTTAACCATTC | 106 |
| chr3:73383146-73385168 | p2 | (AT)11 | GCCACCCTGTTTGATAGGAA | CGTGGCAATGAAGACCTTTT | 133 |
| chr3:73389255-73391270 | p2 | (GT)7 | GGTTTCGGCCAACATAAGAA | TCGGAGAGCGTCTTGAACTT | 223 |

| Marker ID | SSR type | SSR motif | FORWARD PRIMER (5'-3') | REVERSE PRIMER (5'-3') | size (bp) |
|---|---|---|---|---|---|
| chr3:73389255-73391270 | p3 | (GTT)5 | GGTTTCGGCCAACATAAGAA | TCGGAGAGCGTCTTGAACTT | 223 |
| chr3:73401685-73403727 | p3 | (CAA)5 | TTGGGAAAAGGTTTGGATTG | GCTCGCTATCGCATTTTCTC | 283 |
| chr3:73439870-73441884 | p2 | (AT)7 | TCACACTCAAACCTTCTGCG | GTTCCTCCACATGATGCCTT | 165 |
| chr3:73489019-73491035 | p2 | (TA)8 | TCCGATGGGAATTCTGAGAC | GGCGCACAGAAACCAGTAAT | 213 |
| chr3:73537010-73539025 | p5 | (GCATT)3 | TCGAGGCTCAAGGAAAGAAA | CGCTACACGAAGTCTCACCA | 225 |
| chr3:73581914-73583928 | p2 | (TA)7 | CCCTAAGGCTCTCCAAAACC | TTTTTCCTCCGTCAACCAAG | 129 |
| chr3:73814429-73816441 | p2 | (AG)13 | TGGTGAATAAGGGAACTGGC | GCAGTATGTTGGGGCTGATT | 285 |
| chr3:73814443-73816461 | p2 | (AG)13 | TGGTGAATAAGGGAACTGGC | GCAGTATGTTGGGGCTGATT | 271 |
| chr3:73814596-73816622 | p2 | (AG)13 | TGGTGAATAAGGGAACTGGC | GCAGTATGTTGGGGCTGATT | 118 |
| chr3:73891275-73893290 | p5 | (AACAC)3 | GGAATTACCTGTGGGGGTTT | GAGGCATGTAAGGCTTCACC | 267 |
| chr3:73926027-73928042 | p5 | (TCATA)3 | CAAGAGCCTCCTGAAACTGG | TTGGGTCGTGACATGTTTGT | 246 |
| chr3:73948165-73950177 | p2 | (AG)6 | AGAAACCCAGAGTTGAGCGA | GGTAATGGAGGGACCAGGTT | 292 |
| chr3:73981668-73983686 | p6 | (TTATCA)3 | CATAGTGACGGGACAGGACC | CCCACAGGTAGACAGGGAGA | 282 |
| chr3:74060018-74062030 | p2 | (CT)6 | TCAGTCAAAGTCTTGTGGCG | ATGGCACTGTCATCCTGTCA | 136 |
| chr3:74107010-74109074 | p2 | (TA)32 | TTCGATACCTACCGTACGCC | TCCCGTTGATGCATAGTGAA | 100 |
| chr3:74109198-74111213 | p5 | (ATTCA)3 | AAGGTCGGTAACCTCCGATT | AACGTTCGTGGAAGGTTCTG | 115 |
| chr3:74119774-74121788 | p2 | (TA)7 | TTCCACTCGGCCAAATTATC | TGGAAGCGAGAGATGTGTTG | 163 |
| chr3:74119774-74121788 | p2 | (TA)6 | TTCCACTCGGCCAAATTATC | TGGAAGCGAGAGATGTGTTG | 163 |
| chr3:74119774-74121788 | p2 | (TA)14 | TTCCACTCGGCCAAATTATC | TGGAAGCGAGAGATGTGTTG | 163 |
| chr3:74135570-74137594 | p2 | (AT)12 | AACGTGTCTTTGCTGCTCCT | AACAACGAGGTCGAAGCAGT | 110 |
| chr3:74186666-74188700 | p2 | (AT)17 | ACACCAACATTTTGATGCCA | CCGCCCTCTCAAATAACAAA | 197 |
| chr3:74215191-74217249 | p6 | (TATATG)3 | GTTCTGACCCATGCTACCGT | CCATGAGAGCCCAAAGAAAG | 162 |
| chr3:74215191-74217249 | p2 | (AT)29 | GTTCTGACCCATGCTACCGT | CCATGAGAGCCCAAAGAAAG | 162 |
| chr3:74290679-74292695 | p2 | (AC)8 | AGAGGAATGTGTGGGGTCAG | CGAGGCAACCGTGTTAATTT | 149 |
| chr3:74309767-74311801 | p2 | (TA)17 | TTTCTTGGGTCATGTGGTGA | GGACGAAAAACCTTCCACAA | 218 |
| chr3:74315134-74317149 | p3 | (TGT)5 | TAGCTGGCCGAGAATGAACT | CTCAATCCCTCAATCGGCTA | 272 |
| chr3:74315134-74317149 | p2 | (TA)11 | TAGCTGGCCGAGAATGAACT | CTCAATCCCTCAATCGGCTA | 272 |
| chr3:74536495-74538510 | p5 | (TGTGG)3 | CGTGGAAATATGGATGGAGG | TGGGCCCTTTTTAATTTTCC | 287 |
| chr3:74556821-74558949 | p2 | (TA)6 | GCCGGGTACGGTATATGATG | GCAAGGGACTCGGTGAATAA | 104 |
| chr3:74556821-74558949 | p2 | (AT)64 | GCCGGGTACGGTATATGATG | GCAAGGGACTCGGTGAATAA | 104 |
| chr3:74594511-74596523 | p2 | (AC)8 | ATCCTCTTCCATCCAATCCC | TTGTTGACGTGCCAATTCAT | 184 |
| chr3:74594511-74596523 | p2 | (AC)6 | ATCCTCTTCCATCCAATCCC | TTGTTGACGTGCCAATTCAT | 184 |
| chr3:74594511-74596523 | p2 | (AC)6 | ATCCTCTTCCATCCAATCCC | TTGTTGACGTGCCAATTCAT | 184 |
| chr3:74594511-74596523 | p2 | (GA)7 | ATCCTCTTCCATCCAATCCC | TTGTTGACGTGCCAATTCAT | 184 |
| chr3:74624172-74626190 | p6 | (CATATA)3 | CAATAGCTCTAGCCATCGCC | GAATTAGCGGCGGTTATTGA | 105 |
| chr3:74643037-74645051 | p2 | (AT)7 | TGGCCTTTACACGAATCCTC | AATGCGAACAAAAGGGTGAC | 180 |

| Marker ID | SSR type | SSR motif | FORWARD PRIMER (5'-3') | REVERSE PRIMER (5'-3') | size (bp) |
|---|---|---|---|---|---|
| chr3:74744887-74746907 | p2 | (AT)10 | CTCATGCTGGTCTTCGATCA | GCCAGTGGAGCATAGAGGAG | 208 |
| chr3:74744887-74746907 | p5 | (TTCAT)3 | CTCATGCTGGTCTTCGATCA | GCCAGTGGAGCATAGAGGAG | 208 |
| chr3:74769307-74771323 | p2 | (AT)8 | GGTGAAGGCCCAATTATGTG | GTTTTGAGGTGGCTTGCCTA | 198 |
| chr3:74846238-74848253 | p5 | (ATTTC)3 | ACAGAGGAAACTCTGGCGAA | ACATCGTATGCCTTGCCTTC | 275 |
| chr3:74886174-74888186 | p3 | (ATG)5 | AGCTCCACTTCATTCGGAGA | ATAAGCCGACGAGGCTAACA | 170 |
| chr3:74886174-74888186 | p2 | (TA)6 | AGCTCCACTTCATTCGGAGA | ATAAGCCGACGAGGCTAACA | 170 |
| chr3:74890499-74892515 | p2 | (TA)15 | GGACATGTGTCCACCTTGTG | GCTTAATTGTCCAAGCTCGC | 175 |
| chr3:74890499-74892515 | p2 | (TA)15 | GGACATGTGTCCACCTTGTG | GCTTAATTGTCCAAGCTCGC | 175 |
| chr3:74890499-74892515 | p2 | (AT)8 | GGACATGTGTCCACCTTGTG | GCTTAATTGTCCAAGCTCGC | 175 |
| chr3:74894773-74896791 | p2 | (TA)9 | CGTCAGAAGGATACCCGAAA | TTAGCTCTCCCGTGTGGACT | 151 |
| chr3:74894773-74896791 | p2 | (AT)11 | CGTCAGAAGGATACCCGAAA | TTAGCTCTCCCGTGTGGACT | 151 |
| chr3:74919777-74921843 | p2 | (TA)6 | TTGCTGTTTTGAGCTTGTGG | CTTGTCAGTCCAATCTCGCA | 258 |
| chr3:74920793-74922805 | p2 | (TA)25 | CTTGCATAGTTCAGCGGACA | GTAACATAGCGGTCGGGAAA | 233 |
| chr3:74920793-74922805 | p2 | (TA)6 | CTTGCATAGTTCAGCGGACA | GTAACATAGCGGTCGGGAAA | 233 |
| chr3:75031876-75033938 | p2 | (TA)31 | CAAGCTTCAGTTCAGCAACG | AACATGATTTCCCCCATCAA | 201 |
| chr3:75043308-75045326 | p5 | (ATTGC)3 | ATGGGTGGCACTCAGACTTC | GCAATGCAATGCAATAATGG | 157 |
| chr3:75043308-75045326 | p3 | (ATG)6 | ATGGGTGGCACTCAGACTTC | GCAATGCAATGCAATAATGG | 157 |
| chr3:75043308-75045326 | p2 | (AT)25 | ATGGGTGGCACTCAGACTTC | GCAATGCAATGCAATAATGG | 157 |
| chr3:75101078-75103108 | p2 | (AT)12 | GCCCCTCTTGTTATTCAAACG | ACTCGTGAATAGGCAGTCGC | 211 |
| chr3:75101078-75103108 | p2 | (TA)15 | GCCCCTCTTGTTATTCAAACG | ACTCGTGAATAGGCAGTCGC | 211 |
| chr3:75128109-75130129 | p2 | (CT)18 | CAACGAAGAAGTGCCAAACA | ACCAGCTAAGCCAGTCCTGA | 232 |
| chr3:75128109-75130129 | p5 | (GTTTA)4 | CAACGAAGAAGTGCCAAACA | ACCAGCTAAGCCAGTCCTGA | 232 |
| chr3:75140232-75142260 | p2 | (TA)6 | CGTCACCCCGATAGAGTTGT | GTGTTATGTCCGGCCCTCTA | 289 |
| chr3:75173305-75175320 | p5 | (TGCAA)3 | TGAAGTCGCACGAGAGAATG | ACGGTTCGACTACCAAGGTG | 190 |
| chr3:75245095-75247147 | p2 | (TA)26 | TCCTTCCGAATAGCTAGGCA | CTTTGTTGTTGTGTTTGGCG | 213 |
| chr3:75245095-75247147 | p2 | (TA)8 | TCCTTCCGAATAGCTAGGCA | CTTTGTTGTTGTGTTTGGCG | 213 |
| chr3:75245095-75247147 | p2 | (TA)8 | TCCTTCCGAATAGCTAGGCA | CTTTGTTGTTGTGTTTGGCG | 213 |
| chr3:75245095-75247147 | p2 | (TA)24 | TCCTTCCGAATAGCTAGGCA | CTTTGTTGTTGTGTTTGGCG | 213 |
| chr3:75327111-75329125 | p2 | (TA)7 | TGATGTGGCACCTCTCCATA | TGAGGTTTGTCAAGGAAGGC | 230 |
| chr3:75332384-75334396 | p2 | (AT)6 | CCCGTTTCGTCCAGAATCTA | GATTCTCGGCGATGTGATTT | 272 |
| chr3:75359758-75361806 | p5 | (ATTGC)3 | AGGTGTCGATGGATACTCGG | GTACACGGGGTAACGCACTT | 149 |
| chr3:75382334-75384382 | p5 | (ATTGC)3 | AGGTGTCGATGGATACTCGG | GTACACGGGGTAACGCACTT | 149 |
| chr3:75426811-75428829 | p6 | (ACAAAA)3 | ACAAACCCCCAAAATTCACA | TTGCTACATGGTGGCAAAAG | 256 |
| chr3:75473884-75475948 | p2 | (AT)32 | ACTTCTGTCACGACCCAACC | TGATGTTGCCTTGTCAGCTC | 295 |
| chr3:75568856-75570872 | p2 | (GT)8 | TGGAGTAGTTCGTGGCTCCT | ATCCAACCCAAACACCATGT | 188 |
| chr3:75648505-75650523 | p2 | (TC)9 | TGTTAAACTTAGTGGCCCCG | TCCAGCCCAATAGCCAATAG | 196 |

| Marker ID | SSR type | SSR motif | FORWARD PRIMER (5'-3') | REVERSE PRIMER (5'-3') | size (bp) |
|---|---|---|---|---|---|
| chr3:75811930-75813942 | p2 | (AG)6 | CCGATGAGAGAGAGAGTGGG | GCGGTGCTAAGGACTACTCG | 253 |
| chr3:75830728-75832818 | p2 | (AT)45 | TCTGAAAACCCATTCTTGGC | GGGATCAGTCCCATCGTTTA | 156 |
| chr3:75868586-75870607 | p3 | (ATC)7 | GCTGGCTACCTGATCCTACG | GGCGTCCAGATGTACAGGTT | 220 |
| chr3:75868586-75870607 | p3 | (CAA)6 | GCTGGCTACCTGATCCTACG | GGCGTCCAGATGTACAGGTT | 220 |
| chr3:76061514-76063529 | p3 | (CAA)5 | TCCTGGTGTGGCCTTTTTAG | GATTATCGCGAAGGGAATCA | 126 |
| chr3:76220846-76222928 | p2 | (AT)41 | TCCTGTAGCGACCATCCTTT | CTGGTAGGGCATAGCCTTCA | 268 |
| chr3:76235795-76237810 | p5 | (TGTGT)3 | TTATTGCTCTTGGCCACTCC | TACCATGAACCACTTTGCGA | 223 |
| chr3:76329987-76332007 | p2 | (AT)10 | GCGTGATCGGGTTCCTATTA | GTGTTCTCCGGAAAGGATGA | 171 |
| chr3:76356578-76358596 | p2 | (TA)7 | GTGAACTCAAAGTTCGGGGA | TCAAACCGGTAAACCGAAAC | 292 |
| chr3:76356578-76358596 | p2 | (TA)9 | GTGAACTCAAAGTTCGGGGA | TCAAACCGGTAAACCGAAAC | 292 |
| chr3:76364122-76366137 | p5 | (TTTTA)3 | AAAGCCTAGCGACTGTCGAA | CTTTCAAATAGGCTCAGGCG | 136 |
| chr3:76364122-76366137 | p6 | (TTTTTA)3 | AAAGCCTAGCGACTGTCGAA | CTTTCAAATAGGCTCAGGCG | 136 |
| chr3:76371603-76373645 | p5 | (AACCA)3 | AATAATCGGACGAATGCAGG | ATCTGCAATCCATGCACAAA | 212 |
| chr3:76371603-76373645 | p2 | (TA)21 | AATAATCGGACGAATGCAGG | ATCTGCAATCCATGCACAAA | 212 |
| chr3:76390446-76392462 | p2 | (AT)8 | ATCCAATGACTCGGTTGAGG | GCCATCCACTTTTCTTGGAA | 259 |
| chr3:76390446-76392462 | p2 | (AT)30 | ATCCAATGACTCGGTTGAGG | GCCATCCACTTTTCTTGGAA | 259 |
| chr3:76396820-76398865 | p3 | (TTC)15 | AAACGCATCTTCCATTTTCG | ATGAGCAGATTACGGATGGC | 277 |
| chr3:76404139-76406185 | p2 | (TA)11 | AAAAACCCATGCTTTGCATC | TGCAACTCCCAAAAACATGA | 231 |
| chr3:76470650-76472666 | p2 | (TA)8 | CCAACAGCTGCACAACAACT | ACCCACTTCGGGATAAAACC | 135 |
| chr3:76470650-76472666 | p2 | (AT)6 | CCAACAGCTGCACAACAACT | ACCCACTTCGGGATAAAACC | 135 |
| chr3:76505710-76507728 | p3 | (ACA)6 | ACTTAGGCCACATCGTCACC | CGACCTTACGTCACTCCGAT | 201 |
| chr3:76514513-76516525 | p2 | (TA)6 | CTGACGGATCCTCCTCCATA | ATGAAGAGGTCGGCTTCTGA | 267 |
| chr3:76730074-76732089 | p5 | (TTTTA)3 | ACCTTACCCACCCACAATCA | CGGAAAAGGCTCAAATATGC | 178 |
| chr3:76852299-76854355 | p2 | (AT)28 | CCCTTGCATCAAAATCTCGT | CATGGGATTGTTCGACCTCT | 136 |
| chr3:76867877-76869897 | p5 | (TGCAA)3 | GTCCATAATCCGATCCATGC | CGTGACAGTGACGATATGGC | 193 |
| chr3:76867877-76869897 | p2 | (AT)10 | GTCCATAATCCGATCCATGC | CGTGACAGTGACGATATGGC | 193 |
| chr3:76867877-76869897 | p2 | (TA)21 | GTCCATAATCCGATCCATGC | CGTGACAGTGACGATATGGC | 193 |
| chr3:76881891-76883906 | p3 | (TTC)5 | GAGTTTTGAACTTGCCCCAA | AAACGACACTGAAGGGGATG | 190 |
| chr3:76881891-76883906 | p5 | (TCGGT)3 | GAGTTTTGAACTTGCCCCAA | AAACGACACTGAAGGGGATG | 190 |
| chr3:76881891-76883906 | p6 | (ATTGGG)3 | GAGTTTTGAACTTGCCCCAA | AAACGACACTGAAGGGGATG | 190 |
| chr3:76889909-76891924 | p5 | (TTTTG)3 | TCAACCCTATCCAAAGCCAC | CTCAACTCACCGTCGACTCA | 167 |
| chr3:76931870-76933894 | p4 | (TTAT)6 | TATGAGTGTAGCGATTGCCG | TGTACTCCGGTGATGGAACA | 232 |
| chr3:76961113-76963125 | p2 | (CA)6 | ACGCCAAAGAAAGAAAGGGT | TTAGATTCTCGGGGATGGTG | 150 |
| chr3:76983251-76985266 | p3 | (ACA)5 | CTCCTCTACCCTTCTCCGCT | CAACCGAATTTGTACCCCAC | 276 |
| chr3:76983251-76985266 | p2 | (AT)10 | CTCCTCTACCCTTCTCCGCT | CAACCGAATTTGTACCCCAC | 276 |
| chr3:77251474-77253486 | p2 | (TA)6 | GTGCACCACAGGAAGCTACA | ACCTTCCCCCTCAACTGTCT | 270 |

| Marker ID | SSR type | SSR motif | FORWARD PRIMER (5'-3') | REVERSE PRIMER (5'-3') | size (bp) |
|---|---|---|---|---|---|
| chr3:77326552-77328564 | p5 | (AATGC)3 | CTCGCCAAATGATCAGTCAA | CTCCGAGGTCTTTGCTGAAG | 206 |
| chr3:77326552-77328564 | p2 | (TA)6 | CTCGCCAAATGATCAGTCAA | CTCCGAGGTCTTTGCTGAAG | 206 |
| chr3:77556915-77558927 | p2 | (CA)6 | TGGATGGCTGGTAACTCCTC | ATGCAGTGCCATCCACATAA | 163 |
| chr3:77564298-77566310 | p2 | (AT)6 | TTCATACCCGAAGGCGTAAC | ATTCGAAAGATGGCCTAGCA | 161 |
| chr3:77587578-77589592 | p2 | (CA)7 | CAGGCAATTCCTGGTTCACT | AGCATTCTTCTGGGACATCG | 185 |
| chr3:77587578-77589592 | p2 | (AC)20 | CAGGCAATTCCTGGTTCACT | AGCATTCTTCTGGGACATCG | 185 |
| chr3:77592814-77594832 | p2 | (AT)9 | TACTGGCAATTAGGCGAACC | GGCAGAGCTTGTGTCATTCA | 104 |
| chr3:77592814-77594832 | p2 | (TA)8 | TACTGGCAATTAGGCGAACC | GGCAGAGCTTGTGTCATTCA | 104 |
| chr3:77638970-77640985 | p3 | (TTG)5 | CTTTCCATCCGCAGTCAAAT | GGGATCTACATCTCCGTCCA | 128 |
| chr3:77656704-77658719 | p6 | (TATATG)3 | ACCTTTTGTCACTCCGATGG | TCCGGGATGATAAGTTACGC | 117 |
| chr3:77656704-77658719 | p5 | (CATTG)3 | ACCTTTTGTCACTCCGATGG | TCCGGGATGATAAGTTACGC | 117 |
| chr3:77706150-77708168 | p6 | (AGAAGG)3 | ATGGAATGAGGAAATGTCGC | ACGAGCTGAATGAACGGTTT | 128 |
| chr3:77706150-77708168 | p3 | (AGA)5 | ATGGAATGAGGAAATGTCGC | ACGAGCTGAATGAACGGTTT | 128 |
| chr3:77706885-77708912 | p6 | (AGAAGG)3 | GGAGGAGCAGGAGAATGAGA | TCCTCCTCCTTTCGCTTCTT | 245 |
| chr3:77706885-77708912 | p3 | (AGA)5 | GGAGGAGCAGGAGAATGAGA | TCCTCCTCCTTTCGCTTCTT | 245 |
| chr3:77706885-77708912 | p3 | (AGA)14 | GGAGGAGCAGGAGAATGAGA | TCCTCCTCCTTTCGCTTCTT | 245 |
| chr3:77712634-77714661 | p6 | (AGAAGG)3 | GGACTGAGAAGGGGAAGGAG | TTCTCCTTTTACCGTCCTGC | 145 |
| chr3:77788767-77790797 | p2 | (AT)15 | GGGAAGGTGTAGCCTTGTCA | TTGGGCTCCAAATGATGAGT | 128 |
| chr3:77808448-77810466 | p2 | (TA)6 | ATACCAACATCGTGCCAACA | TTGAATTTGGAAGGTCTCGG | 141 |
| chr3:77808468-77810488 | p2 | (TA)6 | ATACCAACATCGTGCCAACA | TTGAATTTGGAAGGTCTCGG | 121 |
| chr3:77815746-77817758 | p2 | (AT)6 | CCATGAGCCATGTTTTTGTG | GCGAAAGTCAAAGGCAAGAC | 103 |
| chr3:77815746-77817758 | p2 | (TA)6 | CCATGAGCCATGTTTTTGTG | GCGAAAGTCAAAGGCAAGAC | 103 |
| chr3:77892982-77895056 | p2 | (AT)37 | GCCGTCTTTTGAGCATCTTC | CATAGCCCCGATACCAAAGA | 253 |
| chr3:77892982-77895056 | p3 | (TTC)7 | GCCGTCTTTTGAGCATCTTC | CATAGCCCCGATACCAAAGA | 253 |
| chr3:77918242-77920266 | p6 | (CACCCC)4 | ACTACTACCGGCACCACGTC | TTGTTGGAAATTAGTCCGGC | 201 |
| chr3:77921945-77923960 | p5 | (TTATT)3 | AAAAACCCGAATAACCCGAG | ACATGGATCGGGTTGGTTTA | 230 |
| chr3:77974803-77976817 | p2 | (TA)7 | ATCCAACACTGGGCTATTCG | AGCTGGGCCTTTACCATTTT | 103 |
| chr3:78013527-78015542 | p5 | (ATTCA)3 | GGAAACATTGGGACATCCAC | GACCCCATTGGATTTGTTTG | 110 |
| chr3:78137170-78139202 | p2 | (TA)16 | GCCCAAATTTTCCCAGAAAT | ATTCGCTGCTCACATAGGCT | 132 |
| chr3:78137170-78139202 | p2 | (TC)7 | GCCCAAATTTTCCCAGAAAT | ATTCGCTGCTCACATAGGCT | 132 |
| chr3:78192554-78194566 | p2 | (AT)15 | AAAATCTTGGAAGCATTGCG | AATTAAGGTGCCACACCCAA | 296 |
| chr3:78192554-78194566 | p2 | (AT)26 | AAAATCTTGGAAGCATTGCG | AATTAAGGTGCCACACCCAA | 296 |
| chr3:78192554-78194566 | p2 | (TA)6 | AAAATCTTGGAAGCATTGCG | AATTAAGGTGCCACACCCAA | 296 |
| chr3:78247799-78249811 | p2 | (TA)6 | TGGTGGCCCATTTCTAACTC | CCTAATTTCCGTACAGGGCA | 275 |
| chr3:78262413-78264428 | p3 | (ATT)5 | AACAAACAAGACTGGGGCAC | ATGTTATTCCCGACACCCAA | 160 |
| chr3:78342170-78344185 | p3 | (TTG)5 | ATTGGGCAACCCATTAACAA | ATGCTCCCTACCGTTGATTG | 116 |

| Marker ID | SSR type | SSR motif | FORWARD PRIMER (5'-3') | REVERSE PRIMER (5'-3') | size (bp) |
|---|---|---|---|---|---|
| chr3:78388823-78390843 | p5 | (CATTG)4 | TAGCAGAAGGGCAAGGCTAA | CACACAACAGGGCCTAACCT | 213 |
| chr3:78401799-78403814 | p5 | (TATAT)3 | CCTCAAGTGGCTGTGCAATA | ACCAATTTTCCACCTCTCCC | 269 |
| chr3:78438887-78440915 | p4 | (TTAT)7 | GCCGAGAATGAATGGAGTGT | TACAACATGTCCTGGACGGA | 213 |
| chr3:78465487-78467509 | p2 | (AT)11 | AGGGCAACAATCAAGACCTG | GTGACTGTGCTGGTAGGGGT | 212 |
| chr3:78465487-78467509 | p2 | (TA)14 | AGGGCAACAATCAAGACCTG | GTGACTGTGCTGGTAGGGGT | 212 |
| chr3:78477834-78479850 | p2 | (AT)8 | CAAGCCGAACGCTAAAATTC | TTGGAGGCAAATCACAACAA | 268 |
| chr3:78490792-78492807 | p3 | (TTC)5 | ATCGCAGCAGGACTCAGATT | AGCGGGGTTACGAACTCTTT | 267 |
| chr3:78490792-78492807 | p5 | (ATTTA)3 | ATCGCAGCAGGACTCAGATT | AGCGGGGTTACGAACTCTTT | 267 |
| chr3:78515032-78517047 | p2 | (AT)10 | CATCTGTGTCACGACCCAAC | CCCGTGTATAGTGGCAGCTT | 204 |
| chr3:78515032-78517047 | p3 | (CAT)5 | CATCTGTGTCACGACCCAAC | CCCGTGTATAGTGGCAGCTT | 204 |
| chr3:78526707-78528727 | p4 | (AATA)5 | CTTCTCAACCCAAGTTCCCA | TTCAGAGCAGATGATGCAGG | 223 |
| chr3:78529641-78531656 | p5 | (ATTAA)3 | TCTATCTGCCCGATACCCAC | GCAAATAGCCTCGGGTGATA | 166 |
| chr3:78548079-78550111 | p2 | (AT)16 | CAGCATTTGTGGAGAAGCAA | GTTTTTGACTGGGCAAAGGA | 237 |
| chr3:78555988-78558003 | p5 | (AAAAT)3 | AGGTCGGGAAAAAGAAGGAA | ACTCCGCCACACGAAAATAA | 131 |
| chr3:78845741-78847757 | p2 | (AG)8 | TCTTCAGGATTGCTTGGGTC | AGTCGCTGATCACGAAACCT | 259 |
| chr3:78847630-78849645 | p5 | (CTGAT)3 | CAAAAATGCTGCCCAAATCT | GAACAAAGGGAAATGCTCCA | 113 |
| chr3:78916091-78918106 | p3 | (TTC)5 | TGTGTCGCAAAGTTCTGACC | AATCCATTTCCACGACTTGC | 273 |
| chr3:78942649-78944683 | p2 | (TA)17 | TATACGCTACCTCATCCGGC | CGAAATCAAGGGATCAGAGG | 227 |
| chr3:79114683-79116711 | p2 | (TA)14 | GTCCCAATTTTCGAACCAGA | AATGGAAGCCAGTGGTTGAC | 170 |
| chr3:79279082-79281102 | p3 | (ACA)5 | TCAGGCATGTTGAGACCTTG | GCCAAAATGAAAAGGCAAAA | 254 |
| chr3:79279106-79281122 | p3 | (ACA)5 | TCAGGCATGTTGAGACCTTG | GCCAAAATGAAAAGGCAAAA | 230 |
| chr3:79432034-79434058 | p6 | (TGTAAG)4 | CCACTGCAGTTGGGTACAGA | TCACATAGCATACCCGACCA | 223 |
| chr3:79443388-79445418 | p2 | (AT)6 | ACTTTTGTCACGACCCAACC | GGGGTTGTGTATGTATGGGC | 197 |
| chr3:79443428-79445526 | p2 | (AT)6 | ACTTTTGTCACGACCCAACC | GGGGTTGTGTATGTATGGGC | 157 |
| chr3:79454102-79456116 | p2 | (CA)7 | TTTTGACCGAAAGAATTCCG | CGGGCCAAGATGTAAAAGAA | 181 |
| chr3:79466812-79468830 | p6 | (TATATG)3 | GTTGATGATTTGGCCGAGTT | CTTGGCTTCGCAAGACCTAC | 204 |
| chr3:79466812-79468830 | p2 | (TA)16 | GTTGATGATTTGGCCGAGTT | CTTGGCTTCGCAAGACCTAC | 204 |
| chr3:79475157-79477173 | p4 | (TAAT)4 | CACGGTCTGGCCGTATATTT | TGGTGAGACCATCCCTAAGC | 149 |
| chr3:79475157-79477173 | p2 | (TC)8 | CACGGTCTGGCCGTATATTT | TGGTGAGACCATCCCTAAGC | 149 |
| chr3:79482545-79484614 | p3 | (CTT)23 | ATCCCATGGCTTCTGACTTG | AAAATCTTGGATGTCGACGC | 109 |
| chr3:79491862-79493878 | p4 | (TATG)4 | TTGGCATGGCATGAAGAATA | TGTCGACTTTCCCTGAATCC | 273 |
| chr3:79569603-79571647 | p2 | (AT)22 | GTATCCTTGGACGTCGGAAA | AGGCAAAAGTATTCGAGCCA | 268 |
| chr3:79783526-79785580 | p2 | (AG)6 | ATTTCATCGATTCAAAGCCG | GCTCCCTCCACTCACTTTTG | 208 |
| chr3:79783526-79785580 | p3 | (AAT)18 | ATTTCATCGATTCAAAGCCG | GCTCCCTCCACTCACTTTTG | 208 |
| chr3:79862529-79864544 | p3 | (TTC)5 | TCATGACTGACCTTGCGAAC | GAGCGTTTCAAATGCCAAAT | 236 |
| chr3:79862529-79864544 | p5 | (TGGTT)3 | TCATGACTGACCTTGCGAAC | GAGCGTTTCAAATGCCAAAT | 236 |

| Marker ID | SSR type | SSR motif | FORWARD PRIMER (5'-3') | REVERSE PRIMER (5'-3') | size (bp) |
|---|---|---|---|---|---|
| chr3:79862529-79864544 | p3 | (CAG)5 | TCATGACTGACCTTGCGAAC | GAGCGTTTCAAATGCCAAAT | 236 |
| chr3:79892069-79894087 | p6 | (ATACAT)3 | CGGTCGCACATATCAAACAC | GGGTCGCTATTGTTCGGTAA | 299 |
| chr3:79892069-79894087 | p3 | (CAT)7 | CGGTCGCACATATCAAACAC | GGGTCGCTATTGTTCGGTAA | 299 |
| chr3:79892276-79894297 | p6 | (ATACAT)3 | CCAAGGCCGTCATAGAACAT | GGGTCGCTATTGTTCGGTAA | 127 |
| chr3:79892276-79894297 | p3 | (CAT)7 | CCAAGGCCGTCATAGAACAT | GGGTCGCTATTGTTCGGTAA | 127 |
| chr3:79893686-79895700 | p2 | (CA)7 | TACCTTGCCCGCTTTTTATG | GGGGATGTTGCATTTACAGG | 273 |
| chr3:79893686-79895700 | p2 | (AT)9 | TACCTTGCCCGCTTTTTATG | GGGGATGTTGCATTTACAGG | 273 |
| chr3:79932054-79934120 | p3 | (TTA)22 | TTCCTTCTCTCCATTCCCCT | AGAGCTGGGTTGGGAGTTTT | 246 |
| chr3:79932054-79934120 | p5 | (CATAT)3 | TTCCTTCTCTCCATTCCCCT | AGAGCTGGGTTGGGAGTTTT | 246 |
| chr3:80038983-80040998 | p5 | (TGGGG)3 | GCTCCTGTCACGTTACGGTT | TGGGGAAAATGGAGAGAGTG | 183 |
| chr3:80128032-80130047 | p5 | (GGGAC)3 | TCTTCATCCGACTTTGACCC | TATGCCTCCAGTTCCTCCAC | 184 |
| chr3:80140857-80142875 | p3 | (TTG)6 | GCGTTTTGTGGCCACTATTT | TTCTTTATCCCCTCCGTCCT | 299 |
| chr3:80160039-80162054 | p5 | (AGAAA)3 | TTTTTGAGTTGCGAAACGTG | AATTCGAAGTGGGTGACTGG | 140 |
| chr3:80186761-80188777 | p2 | (AT)8 | GGCCGGACATATCTAATCCA | TGCACAGGGAGACAGACTTG | 226 |
| chr3:80209737-80211769 | p2 | (AT)6 | TCTCCTCTTCCTTCGACCAA | CTCGGACATGGAGGATGTTT | 256 |
| chr3:80209775-80211833 | p2 | (AT)6 | TCTCCTCTTCCTTCGACCAA | CTCGGACATGGAGGATGTTT | 218 |
| chr3:80228610-80230642 | p2 | (AT)6 | TCTCCTCTTCCTTCGACCAA | CTCGGACATGGAGGATGTTT | 256 |
| chr3:80228648-80230706 | p2 | (AT)6 | TCTCCTCTTCCTTCGACCAA | CTCGGACATGGAGGATGTTT | 218 |
| chr3:80256250-80258268 | p2 | (GT)6 | GAACGATGTTTGACGACCCT | CGGTCCGTATTTCAAGTCGT | 196 |
| chr3:80335126-80337141 | p5 | (ATGCA)3 | GTAATCTCTGACCCGCCAAA | AAAATGACAGCAGCAACACG | 163 |
| chr3:80360628-80362726 | p2 | (AT)8 | GACCTATCGGAATCGACGAA | CTCACTTTTGACTCGAGGGG | 226 |
| chr3:80433488-80435504 | p2 | (AT)8 | CATCCGCCAACAGCTTCTAT | GGCACTTAAATTTGGACGGA | 168 |
| chr3:80477553-80479568 | p5 | (TGATA)3 | TCGGCGTTTGTGAATGTAAA | GGGGTCAGTACGAGCAATGT | 160 |
| chr3:80500645-80502659 | p3 | (AAT)6 | AGGGCTGGCCTACGATTACT | TCAAGAGGTGCTCGGTAGGT | 240 |
| chr3:80500645-80502659 | p2 | (AT)7 | AGGGCTGGCCTACGATTACT | TCAAGAGGTGCTCGGTAGGT | 240 |
| chr3:80532982-80534997 | p5 | (CCTTT)3 | CACGTGTACCCAGCAACATC | CTGGTGCATATTTGCCATTG | 179 |
| chr3:80699631-80701653 | p2 | (TA)11 | TGGGAGAGAAAAACGCAGAT | CTTCACACTCAACCTTGGCA | 117 |
| chr3:80699631-80701653 | p6 | (ATGTGA)3 | TGGGAGAGAAAAACGCAGAT | CTTCACACTCAACCTTGGCA | 117 |
| chr3:80996700-80998715 | p5 | (TGCAT)3 | CCCTTATGGTCCCGATTTTT | TGGGTGCCACGAACTATACA | 179 |
| chr3:80996700-80998715 | p5 | (TCATG)3 | CCCTTATGGTCCCGATTTTT | TGGGTGCCACGAACTATACA | 179 |
| chr3:81095946-81097964 | p2 | (TA)9 | TCCCGAGGTACAGACGTTTC | GCCGTACCGAACAAACACTT | 115 |
| chr3:81143163-81145178 | p3 | (GAA)5 | AAGGTGATGTACTGGGCCTG | TGTCCCGGTGGTTAATGAAT | 167 |
| chr3:81152908-81154920 | p2 | (TA)6 | CAACTCCATGGATTCGGTCT | TGCTTTGGGTCAACAAATGA | 241 |
| chr3:81152908-81154920 | p2 | (AT)8 | CAACTCCATGGATTCGGTCT | TGCTTTGGGTCAACAAATGA | 241 |
| chr3:81159928-81161943 | p3 | (TGG)5 | TGACAGATTGCAAATCCGAA | AAAAAGAGCTGCACTGCCAT | 297 |
| chr3:81159928-81161943 | p2 | (TA)21 | TGACAGATTGCAAATCCGAA | AAAAAGAGCTGCACTGCCAT | 297 |

| Marker ID | SSR type | SSR motif | FORWARD PRIMER (5'-3') | REVERSE PRIMER (5'-3') | size (bp) |
|---|---|---|---|---|---|
| chr3:81159928-81161943 | p2 | (TA)7 | TGACAGATTGCAAATCCGAA | AAAAAGAGCTGCACTGCCAT | 297 |
| chr3:81197983-81199999 | p2 | (CT)8 | TTCCAATTTCGTCCTTCCAG | AGGCGAGGTTTATAGGGGAA | 145 |
| chr3:81202145-81204185 | p2 | (TA)6 | CCATGCCCTCTAGACATGGT | CCTTACGTCACCCCGATAGA | 107 |
| chr3:81202145-81204185 | p2 | (TA)20 | CCATGCCCTCTAGACATGGT | CCTTACGTCACCCCGATAGA | 107 |
| chr3:81254532-81256547 | p2 | (AT)6 | TTGGCATGCATGATTTGTTT | CAATCGGAGTCAGAAGCACA | 205 |
| chr3:81254532-81256547 | p5 | (TTCAT)3 | TTGGCATGCATGATTTGTTT | CAATCGGAGTCAGAAGCACA | 205 |
| chr3:81298215-81300227 | p2 | (CA)6 | AATCATCCTTTTGGAGGGCT | CATTTGGAGATGGTATGGGG | 206 |
| chr3:81298215-81300227 | p2 | (AT)9 | AATCATCCTTTTGGAGGGCT | CATTTGGAGATGGTATGGGG | 206 |
| chr3:81298215-81300227 | p2 | (GT)6 | AATCATCCTTTTGGAGGGCT | CATTTGGAGATGGTATGGGG | 206 |
| chr3:81407500-81409528 | p4 | (TTAT)7 | CTTGAGGACCGATGTTCGAT | TGGAACATCATATCTCGGCA | 183 |
| chr3:81543136-81545150 | p2 | (TC)7 | CACTTGTCCTGACCCCACTT | CTGGACCAGTGCCAATTTTT | 126 |
| chr3:81543136-81545150 | p5 | (TTTTA)3 | CACTTGTCCTGACCCCACTT | CTGGACCAGTGCCAATTTTT | 126 |
| chr3:81616784-81618799 | p5 | (TTCAT)3 | TCCTACTGGGGTAGGTCGTG | ATTCACTGTGCATCAACGGA | 245 |
| chr3:81832980-81835050 | p2 | (TA)12 | ACTTCTGTCACGACCCAACC | AGCCCTGTTGGCTTACAATG | 143 |
| chr3:81832980-81835050 | p3 | (CAA)9 | ACTTCTGTCACGACCCAACC | AGCCCTGTTGGCTTACAATG | 143 |
| chr3:81855733-81857748 | p3 | (CAA)5 | TCGTAAACGGACTCGGAAAC | GGGATTTGATTAGCGAACGA | 300 |
| chr3:81855733-81857748 | p2 | (AC)6 | TCGTAAACGGACTCGGAAAC | GGGATTTGATTAGCGAACGA | 300 |
| chr3:81908244-81910264 | p2 | (TA)10 | GCGTACCGCTGTTGGTACTT | GAAAATGAGAGTGCGTGCAA | 228 |
| chr3:82186449-82188467 | p3 | (TTG)6 | TGAGCTTTGCATGATTTTCG | TCTCCCAAACGAGCTTCTGT | 278 |
| chr3:82272783-82274809 | p2 | (AT)13 | ATTTTGTGCTGCAGGCTCTT | AGTCGGCAAGGCTATCAGAA | 171 |
| chr3:82339887-82341901 | p2 | (AC)7 | CAGACGTCGTAAGCTCTCCC | TTCGGTAGGCAAAATTGGAG | 285 |
| chr3:82347363-82349377 | p2 | (AC)7 | CAGACGTCGTAAGCTCTCCC | TTCGGTAGGCAAAATTGGAG | 285 |
| chr3:82452776-82454788 | p2 | (AT)6 | CCGCCACTACACACAACATC | TTGGAGTGCTCCTTTGTTCC | 252 |
| chr3:82564514-82566526 | p2 | (TA)6 | TGTACCGAGCACCCCTTATC | GATGGTCCTAGTCGAGTCGC | 154 |
| chr3:82761744-82763766 | p2 | (TA)11 | CTCCAGTTCAGAGGGTGAGC | AAGAAAAGCCGAGGAGGTTC | 156 |
| chr3:82844164-82846194 | p2 | (AT)15 | ATGTGGGTTACGGGTCATGT | CATCATTGTCACGACCCAAC | 101 |
| chr3:82946912-82948927 | p5 | (TCATA)3 | TTCCAAAGGTATGATCCCCA | TGGCTTTCGGAAAATACTGG | 133 |
| chr3:82946933-82948948 | p5 | (TCATA)3 | TTCCAAAGGTATGATCCCCA | TGGCTTTCGGAAAATACTGG | 112 |
| chr3:82967479-82969497 | p6 | (CATATA)3 | TATTGACTACCGAGCACCCC | CCTTTTGGCAGTTCCCTACA | 194 |
| chr3:83005199-83007213 | p2 | (TA)7 | GCGACCTCTTATGGTCTCCA | GGTGGGAGAAATTTTGGGAT | 107 |
| chr3:83060976-83062992 | p2 | (AT)8 | CTCGCAGGCATAGACTAGGG | TCCCATACTGTTGGAGGAGC | 271 |
| chr3:83252349-83254364 | p5 | (TATAT)3 | AGAACGACATACGGGCTTTG | TCTCGTCCCATGATTTAGCC | 111 |
| chr3:83252349-83254364 | p2 | (AT)18 | AGAACGACATACGGGCTTTG | TCTCGTCCCATGATTTAGCC | 111 |
| chr3:83260305-83262383 | p5 | (TATAT)3 | TCGTTGGCCTCCTATGTTTT | TACGGCCCGATCCATAAATA | 267 |
| chr3:83260305-83262383 | p2 | (AT)39 | TCGTTGGCCTCCTATGTTTT | TACGGCCCGATCCATAAATA | 267 |
| chr3:83279130-83281142 | p5 | (ATATT)4 | TTTCCTTTTTGGCATGATCC | ATGTTAGTGCCAAGGGAACG | 119 |

| Marker ID | SSR type | SSR motif | FORWARD PRIMER (5'-3') | REVERSE PRIMER (5'-3') | size (bp) |
|---|---|---|---|---|---|
| chr3:83279147-83281159 | p5 | (ATATT)4 | TTTCCTTTTTGGCATGATCC | ATGTTAGTGCCAAGGGAACG | 102 |
| chr3:83294177-83296215 | p2 | (TA)19 | GCAGGATAGGGGTCATAGCA | GCATAGACCCCAATGCAACT | 239 |
| chr3:83301537-83303549 | p2 | (AT)6 | TTGAGTGCAGATCGTGTTCC | CCCCGAAAATCTGAAAGTCA | 126 |
| chr3:83312752-83314767 | p3 | (GTT)5 | GGGTGTGGAAAATGGAATTG | CCATATATACGGCCACACCC | 181 |
| chr3:83361570-83363588 | p2 | (AT)8 | GTATATGTCACGGCCCAACC | TGGAGTCCTTTCGCAAGTTT | 128 |
| chr3:83361570-83363588 | p2 | (TA)9 | GTATATGTCACGGCCCAACC | TGGAGTCCTTTCGCAAGTTT | 128 |
| chr3:83383286-83385304 | p6 | (GGAGAA)3 | ATGTGCAAGGTTGGATGTGA | GTTTGTCTTGAGGGGCAAAA | 268 |
| chr3:83402785-83404803 | p2 | (AT)9 | GGTTCAGCAGCTGGTTCTTC | AAAGCATATACACGGGGCTG | 229 |
| chr3:83441096-83443110 | p2 | (TA)7 | ATATAGGGGAAATGCTGCCC | ATAGCAAAACCGTTCGCTTG | 224 |
| chr3:83474407-83476423 | p2 | (TA)8 | CTAAGGGTGGATGCCTTTCA | CACGGATGTATGCGCTCTAA | 189 |
| chr3:83503544-83505559 | p5 | (ATCAT)3 | ATTTGGACACCCCAAACAAA | GCGATGTATATTCCGCCAAT | 288 |
| chr3:83594300-83596315 | p5 | (TGTGT)3 | AACGCCTGTAAAGCCCCTAT | CTGATCAGCTGGTGCACATT | 159 |
| chr3:83596811-83598826 | p5 | (TGTGT)3 | GATAAGGAATGGGCGTGAAA | GGCGAATTTCCCTTTTTACC | 199 |
| chr3:83629952-83631964 | p2 | (TC)6 | TCAGAAGGACGGACCGTAAC | TTGCCTTAAATGCTCCAACC | 189 |
| chr3:83679325-83681337 | p2 | (AC)6 | TCCACAAGATCATGCCAAAA | GTTGCATTCTCGGGTTTGTT | 228 |
| chr3:83767823-83769838 | p3 | (CAA)5 | AGAGTGTCGCTGCTTTTGGT | ATTTCAGTTCCTCCCCGTTT | 226 |
| chr3:83790062-83792076 | p2 | (TA)7 | ATAAGATGCCAACTCCACCG | ACAGTTAGTTGCCACCCAGG | 215 |
| chr3:83870073-83872103 | p2 | (AT)15 | ACAACTCCATTTGCTCCTGG | TGGAGCCGCTTACAAAACTT | 285 |
| chr3:83876391-83878409 | p2 | (CA)9 | CCCCAAGGTTCAAACTCAAA | TAGATTTTGGACGAAACGGG | 222 |
| chr3:84007498-84009516 | p6 | (ATATGT)3 | TGAGCCTCGCGAATAACTCT | GTCGGGTCCTATTTATGCGA | 240 |
| chr3:84017761-84019777 | p4 | (TATG)4 | CATATAGCGGCGTGTTGTTG | CGCTGTTTTCCATCTTCCAT | 116 |
| chr3:84183106-84185120 | p2 | (AT)7 | CTTCTCTTCCTGTGCTCGCT | GGAGCCTCGTGATGCTAGTC | 169 |
| chr3:84204546-84206558 | p2 | (AC)6 | CTAGGAAAAATTGGGCAGCA | CATTGATGTGGCGTAGATGC | 219 |
| chr3:84482984-84485008 | p3 | (ATC)8 | AAAGTTGTCACGACCCAACC | TAAATGTGTCGTTGCCCAAA | 108 |
| chr3:84482984-84485008 | p2 | (TG)6 | AAAGTTGTCACGACCCAACC | TAAATGTGTCGTTGCCCAAA | 108 |
| chr3:84491991-84494005 | p2 | (TG)7 | CCCGCCATGATCATAAGACT | ATTCCAAATTTACCCTCGGC | 255 |
| chr3:84500661-84502676 | p5 | (ATTTC)3 | CCCCAGGAGATTCAAGATGA | TTTCCTCATAGGGCCACAAC | 234 |
| chr3:84523397-84525417 | p2 | (TA)11 | CGCACGATCTCACAAGAAGA | GGGTCGTGAAAGGAGTGGTA | 212 |
| chr3:84523397-84525417 | p2 | (TA)10 | CGCACGATCTCACAAGAAGA | GGGTCGTGAAAGGAGTGGTA | 212 |
| chr3:84568255-84570269 | p2 | (AT)7 | CCGCTACATCCCTCAACCTA | TCAAAGTCGGATGAAGACCC | 126 |
| chr3:84597836-84599878 | p2 | (AT)21 | GGTCATGCCAAAAAGGAAGA | AGAGCTCGAGACCTGAGTCG | 174 |
| chr3:84597836-84599878 | p2 | (TC)8 | GGTCATGCCAAAAAGGAAGA | AGAGCTCGAGACCTGAGTCG | 174 |
| chr3:84689337-84691355 | p6 | (ATATGT)3 | AGTTGCATAAGGAGCGGAAA | ACTTGCGAGGGTTTTAGGGT | 159 |
| chr3:84791831-84793849 | p3 | (ATC)6 | ACCAATAAACTGCAGCTCCG | CCAGAGGCACAGGTTGATCT | 153 |
| chr3:84871852-84873872 | p5 | (TTACT)3 | CTTTGGATGCTGTGTTCACG | ACAACCAATTAGCACCTCCG | 180 |
| chr3:84871852-84873872 | p2 | (AT)10 | CTTTGGATGCTGTGTTCACG | ACAACCAATTAGCACCTCCG | 180 |

| Marker ID | SSR type | SSR motif | FORWARD PRIMER (5'-3') | REVERSE PRIMER (5'-3') | size (bp) |
|---|---|---|---|---|---|
| chr3:84999336-85001351 | p5 | (ATATA)3 | CTTAGAGATGAGCGAACCCG | GGGGACCTGTCCCAAATTAT | 241 |
| chr3:85015010-85017028 | p2 | (TC)9 | CGGATTACGATGAGCAGGAT | TGAGTTTGGACAAGCGTGAG | 112 |
| chr3:85046321-85048355 | p2 | (TA)7 | CAAAACAAAGGCCACAGGAT | CCTCCTTCCGTAATTCCACA | 253 |
| chr3:85046321-85048355 | p2 | (AT)17 | CAAAACAAAGGCCACAGGAT | CCTCCTTCCGTAATTCCACA | 253 |
| chr3:85117845-85119860 | p5 | (TGATA)3 | TGAGCTTAACGGGATTTTGG | GACAATTTCAAACGCGGATT | 201 |
| chr3:85177420-85179456 | p2 | (AT)18 | TCTAGAAACGCGAGGAAGGA | GCCATTTATTACCCTTCGCA | 177 |
| chr3:85185351-85187375 | p3 | (AAG)8 | GCTTTCAATTCATGGCGATT | CACAAGATGGCTAAGGGCTC | 284 |
| chr3:85189987-85191999 | p2 | (AT)6 | CCCAAATCGAGTGGGACTAA | TGGTCGGAAAATTAAAACGC | 189 |
| chr3:85202500-85204512 | p2 | (AT)6 | AAGGGAGAATTGGGTTTGCT | TGAATTCTCGTGGTGCCATA | 128 |
| chr3:85317078-85319096 | p6 | (TATATG)3 | TGTATGTTTGGCCGAGTTGA | TACATACCTTGCCCGCTTTC | 156 |
| chr3:85317078-85319096 | p2 | (AT)20 | TGTATGTTTGGCCGAGTTGA | TACATACCTTGCCCGCTTTC | 156 |
| chr3:85356315-85358329 | p2 | (AT)7 | ATCCTGCTAGGGTGTGATGG | AGTCTTGTGAGCAGCTCCGT | 280 |
| chr3:85378273-85380291 | p6 | (GTATAT)3 | TGGCGTATTTACAGGGGAAG | ACTCTTGGCCTTCCCAAAAT | 247 |
| chr3:85439591-85441629 | p2 | (TA)8 | TGGATCAACTTTAGGCCACC | TTGAGTGGGACACATTTCCA | 225 |
| chr3:85483896-85485908 | p2 | (AT)6 | GGGCGAGATCTACCATTTGA | AGGCGAGGACACCTCTTTTT | 231 |
| chr3:85483896-85485908 | p2 | (GT)6 | GGGCGAGATCTACCATTTGA | AGGCGAGGACACCTCTTTTT | 231 |
| chr3:85484301-85486313 | p2 | (AT)6 | AAAGAAGCTCCCCCGAAGTA | TCAAATTTTCCCGAATACGC | 177 |
| chr3:85484301-85486313 | p2 | (GT)6 | AAAGAAGCTCCCCCGAAGTA | TCAAATTTTCCCGAATACGC | 177 |
| chr3:85495114-85497134 | p2 | (AT)45 | ATTGCGGTAAATCTTGGCTG | ATACCAACACCGCAATCCAT | 207 |
| chr3:85495135-85497147 | p2 | (AT)45 | ATTGCGGTAAATCTTGGCTG | ATACCAACACCGCAATCCAT | 186 |
| chr3:85520505-85522535 | p3 | (ACA)5 | CACCCGGAGCTAAACTACCA | CTTGCCCTGTCGTTATGGAT | 199 |
| chr3:85540780-85542795 | p5 | (AAAGT)3 | TAGGGATTTCCATTGCTTGC | GCTGTGTGTTCCGAATGTTG | 266 |
| chr3:85553252-85555270 | p6 | (TTCCCG)3 | ACGACGAGTTGGGTCGTAAC | CTTTCTTGGGGATGACCTGA | 231 |
| chr3:85553252-85555270 | p3 | (TGT)10 | ACGACGAGTTGGGTCGTAAC | CTTTCTTGGGGATGACCTGA | 231 |
| chr3:85558598-85560613 | p5 | (TTGTA)3 | CATATGCGGCCCACTAAACT | ACCGACACATCCATGAACCT | 233 |
| chr3:85558598-85560613 | p5 | (GGGGT)3 | CATATGCGGCCCACTAAACT | ACCGACACATCCATGAACCT | 233 |
| chr3:85578452-85580464 | p2 | (AT)6 | TCCTTCAAGGCCATTCACTC | AAGGCCCAAAGTGACCTTTT | 282 |
| chr3:85654021-85656033 | p2 | (AG)6 | AGCTCAATGTCCAAATTGCC | GAGTCATACTTGCCTCGGGA | 295 |
| chr3:85712234-85714246 | p2 | (AT)6 | ATTCCTAAGCAAGCGGACAA | TTGACTTAAGTTTTGGGGCG | 220 |
| chr3:85789809-85791821 | p2 | (TA)6 | TAATTGGAAGTTCGGGTTCG | TCGTAAATCCGTAGCCCATC | 236 |
| chr3:85789809-85791821 | p2 | (TA)28 | TAATTGGAAGTTCGGGTTCG | TCGTAAATCCGTAGCCCATC | 236 |
| chr3:85834420-85836435 | p5 | (ATATA)3 | ATGTCGCTGAGGGATGGTAG | ATTGACGTCCCCTTCTTGTG | 193 |
| chr3:85929576-85931590 | p2 | (AT)7 | AGGTACCACAATTCTCCCCC | TTGTGATGCAAATGGCTTGT | 172 |
| chr3:85948987-85950999 | p2 | (AC)6 | ATGAAGGAGTGGGAGGCTTT | TGTTGATTGTGTGGGCTGTT | 238 |
| chr3:85972230-85974242 | p2 | (GT)6 | TCTGTCAAAGTTCTGTGGCG | ACATGGTCCATCCGTTCATT | 101 |
| chr3:85999883-86001895 | p2 | (AT)6 | ACGAGGAGCACACTTCACCT | ATTCCCATTGGTGGTGAAAA | 276 |

| Marker ID | SSR type | SSR motif | FORWARD PRIMER (5'-3') | REVERSE PRIMER (5'-3') | size (bp) |
|---|---|---|---|---|---|
| chr3:85999883-86001895 | p2 | (AT)6 | ACGAGGAGCACACTTCACCT | ATTCCCATTGGTGGTGAAAA | 276 |
| chr3:86065955-86067971 | p2 | (AT)8 | CACCCTCCTCATTCGTGTTT | GGAAAGCCGTAACGATGGTA | 224 |
| chr3:86100330-86102346 | p4 | (TTAA)4 | GAGTGGCCTCAGTGGAAGAG | CTTTGGCCTACCCCTACCTC | 116 |
| chr3:86100330-86102346 | p5 | (TTTAA)3 | GAGTGGCCTCAGTGGAAGAG | CTTTGGCCTACCCCTACCTC | 116 |
| chr3:86100330-86102346 | p2 | (TC)10 | GAGTGGCCTCAGTGGAAGAG | CTTTGGCCTACCCCTACCTC | 116 |
| chr3:86100712-86102728 | p4 | (TTAA)4 | TCGCTTGCATTATTTCGTTG | ACAACAGCCCAGTGAAATCC | 167 |
| chr3:86100712-86102728 | p5 | (TTTAA)3 | TCGCTTGCATTATTTCGTTG | ACAACAGCCCAGTGAAATCC | 167 |
| chr3:86100712-86102728 | p2 | (TC)10 | TCGCTTGCATTATTTCGTTG | ACAACAGCCCAGTGAAATCC | 167 |
| chr3:86100725-86102767 | p4 | (TTAA)4 | TCGCTTGCATTATTTCGTTG | ACAACAGCCCAGTGAAATCC | 154 |
| chr3:86100725-86102767 | p5 | (TTTAA)3 | TCGCTTGCATTATTTCGTTG | ACAACAGCCCAGTGAAATCC | 154 |
| chr3:86100725-86102767 | p2 | (TC)10 | TCGCTTGCATTATTTCGTTG | ACAACAGCCCAGTGAAATCC | 154 |
| chr3:86109236-86111250 | p2 | (AG)7 | GCTCTCCGTACCAACCAGAG | AAGGGCGCACTACAGAAGAA | 111 |
| chr3:86175474-86177486 | p4 | (TTAA)4 | TTGTGCCTTCCAAAACATCA | TGTCAACAACTGCTCCTTCG | 113 |
| chr3:86175474-86177486 | p2 | (AT)6 | TTGTGCCTTCCAAAACATCA | TGTCAACAACTGCTCCTTCG | 113 |
| chr3:86175474-86177486 | p2 | (TC)25 | TTGTGCCTTCCAAAACATCA | TGTCAACAACTGCTCCTTCG | 113 |
| chr3:86234880-86236894 | p2 | (TG)7 | ATGATCCTCATGCTCTGCCT | GGGAGAAGTCCCTGACATGA | 252 |
| chr3:86268645-86270665 | p5 | (TGCAA)4 | AAGTGCAAAGGAAATGTGCC | GGTAGGAAAGGTAGGTGCCC | 194 |
| chr3:86318553-86320571 | p3 | (ATT)6 | TTGGTTGGCTGCAAACATTA | GGTCCAATTGGTGCAAAAGT | 246 |
| chr3:86432079-86434091 | p2 | (AC)6 | GAGCAGAAATGCTCTGGAGG | TGCAGATTTCAGACGAAACG | 197 |
| chr3:86531074-86533106 | p5 | (AAGAA)3 | ATAAGGTGGGGAATCGAACC | ACTGGGGTGATGCACTAAGC | 149 |
| chr3:86531074-86533106 | p2 | (GT)16 | ATAAGGTGGGGAATCGAACC | ACTGGGGTGATGCACTAAGC | 149 |
| chr3:86531074-86533106 | p3 | (TTG)5 | ATAAGGTGGGGAATCGAACC | ACTGGGGTGATGCACTAAGC | 149 |
| chr3:86531309-86533324 | p5 | (AAGAA)3 | GAAGCATGGAGGAAGGTCAA | TGGTGACTAAACAAAGGCCC | 247 |
| chr3:86531309-86533324 | p2 | (GT)16 | GAAGCATGGAGGAAGGTCAA | TGGTGACTAAACAAAGGCCC | 247 |
| chr3:86531309-86533324 | p3 | (TTG)5 | GAAGCATGGAGGAAGGTCAA | TGGTGACTAAACAAAGGCCC | 247 |
| chr3:86537232-86539280 | p3 | (AGA)16 | ATGTTCACAGGTCCCGAGAG | TCCACCAAAGAAATCTTGCC | 197 |
| chr3:86621053-86623071 | p2 | (TA)9 | CTCTCAATGCGTAATCGGGT | AGGAGGAAGAAACCCAAGGA | 182 |
| chr3:86650506-86652528 | p2 | (TA)11 | AAATTTGGGCAGCATTTCAC | GGAAGCGAGGAATGTGTTGT | 183 |
| chr3:86767395-86769415 | p2 | (AG)10 | CTCCCGAAGTAGTTTGACGC | ACAAGTTTCCATTCGCCAAC | 209 |
| chr3:86767419-86769433 | p2 | (AG)10 | CTCCCGAAGTAGTTTGACGC | ACAAGTTTCCATTCGCCAAC | 185 |
| chr3:86767435-86769451 | p2 | (AG)10 | CTCCCGAAGTAGTTTGACGC | ACAAGTTTCCATTCGCCAAC | 169 |
| chr3:86842450-86844471 | p3 | (TTG)7 | TTGGGTCAATTTTGGAGAGG | CATTGACGAACGGTTTGTTG | 175 |
| chr3:86842450-86844471 | p2 | (AT)7 | TTGGGTCAATTTTGGAGAGG | CATTGACGAACGGTTTGTTG | 175 |
| chr3:86913892-86915907 | p5 | (TGATC)3 | AGGCGCTATCGAGGTATGTG | TGGATTGAAGGAATTGGAGC | 231 |
| chr3:86913892-86915907 | p2 | (TA)11 | AGGCGCTATCGAGGTATGTG | TGGATTGAAGGAATTGGAGC | 231 |
| chr3:86952752-86954796 | p2 | (TA)22 | CAGGTACCGACGACAGGTTT | ATCATTATGACGGGACAGGC | 232 |

| Marker ID | SSR type | SSR motif | FORWARD PRIMER (5'-3') | REVERSE PRIMER (5'-3') | size (bp) |
|---|---|---|---|---|---|
| chr3:87242567-87244582 | p5 | (TTTAA)3 | GGCAAGATAACATCGCACCT | GATCATGACGAGCACTCGAA | 292 |
| chr3:87286333-87288351 | p3 | (TTC)6 | AGGCCAACCCTCATCTTCTT | AAAATCTGCACGCAAAAAGG | 115 |
| chr3:87463025-87465043 | p2 | (AT)9 | GTATTTTCTGTGCGCAGGGT | TACACCTGCAGATGCAAAGC | 267 |
| chr3:87463025-87465043 | p2 | (TA)24 | GTATTTTCTGTGCGCAGGGT | TACACCTGCAGATGCAAAGC | 267 |
| chr3:87524740-87526758 | p2 | (AT)9 | TTCAAACAACCCAACCAACA | TTCATGAATTCGGAAGGGAG | 165 |
| chr3:87585708-87587780 | p2 | (AT)6 | CACCCCAAAAGAAGGACAAA | GTTTCGAACCCAGAACCTCA | 298 |
| chr3:87685895-87687923 | p2 | (AT)14 | CGACCCATCATCCTTGTTCT | ATGGATTTGCGCATTTAAGG | 187 |
| chr3:87685895-87687923 | p2 | (TG)6 | CGACCCATCATCCTTGTTCT | ATGGATTTGCGCATTTAAGG | 187 |
| chr3:87691965-87693983 | p2 | (AT)6 | TCAACTATGTTCGGCCACAA | GGAGTGGTTGTTGGAGCAAT | 300 |
| chr3:87691992-87694008 | p2 | (AT)6 | TCAACTATGTTCGGCCACAA | GGAGTGGTTGTTGGAGCAAT | 273 |
| chr3:87781009-87783023 | p2 | (CA)7 | TTCTTGGAACAGAATTGCCC | GATATTGAGCCGAGCAAAGC | 209 |
| chr3:87781009-87783023 | p2 | (CT)8 | TTCTTGGAACAGAATTGCCC | GATATTGAGCCGAGCAAAGC | 209 |
| chr3:87811224-87813244 | p5 | (AATAA)4 | ACGCCATAGACCAACCAAAG | TACCCCTTTGAAACTGAGCG | 201 |
| chr3:87812960-87814978 | p2 | (TA)9 | TCGGCTGATATGACCATGAA | TCAACTCCCAAACGTCAACA | 195 |
| chr3:87852200-87854212 | p2 | (CA)6 | TATGAGCAGAATAACCCCCG | TGGGTTGCTGTGTTGATGTT | 223 |
| chr3:87936873-87938888 | p3 | (ATA)5 | AAGGGTGTCATGAGATTCGG | AACCTGCCGATCAAGTATGG | 193 |
| chr3:88043367-88045439 | p2 | (AT)36 | AATGCAAACTCGGATTGACC | ATTGAGGCATCCACGAGAAG | 291 |
| chr3:88163237-88165337 | p3 | (ACA)5 | CACGAACGGAGTAGTGCAGA | TGAACGTTTTGCACTTCGAG | 192 |
| chr3:88263466-88265480 | p2 | (AT)7 | ACGCAAACTCGATACAACCC | TTCCGGAGGGACCTTACTCT | 103 |
| chr3:88286736-88288754 | p6 | (TATATG)3 | GAATTTCGGCAGGACCATAA | ACTAGGGATCATGCCGTTTG | 225 |
| chr3:88327915-88329929 | p2 | (TA)7 | TCCTCCCAGTCTACCTGTGG | AAGGTGGAGTAGCCGATGTG | 144 |
| chr3:88332071-88334089 | p2 | (AT)9 | TCATCCCTCGCCAAATAAAG | TTGCCACATGAAAGGTCAAA | 289 |
| chr3:88410164-88412179 | p5 | (TTCAT)3 | TCATTTTGATAGCCGAAGGG | GCTCGTAGACACTTCCTCGG | 146 |
| chr3:88443135-88445150 | p5 | (CAAAA)3 | GCAATGGCTTGGCTCATATT | GCCCCAAAGGACACTTTACA | 262 |
| chr3:88447593-88449605 | p2 | (AT)6 | AATCCCATTGGGCAACAATA | ATACTTATGAGCATGGCGGG | 165 |
| chr3:88474219-88476234 | p3 | (CAA)5 | AACTCGAGGGAGTCAGCAAA | GGCCCACTATTTCCGATTTT | 232 |
| chr3:88494064-88496088 | p2 | (AT)12 | TATGGATGACGTGGGGGTAT | GAGCCCCTGTATGGCTATGA | 154 |
| chr3:88562761-88564776 | p3 | (CAT)5 | CCATCGGGCTATGACACTTT | TGCAGCTGTTGGCACTATTC | 294 |
| chr3:88620478-88622493 | p5 | (ATTCA)3 | GGGACGCAGTGAAAAGACAT | GGATAGCTTGAAAATGCCCA | 128 |
| chr3:88620478-88622493 | p5 | (ACACA)3 | GGGACGCAGTGAAAAGACAT | GGATAGCTTGAAAATGCCCA | 128 |
| chr3:88620478-88622493 | p2 | (TA)17 | GGGACGCAGTGAAAAGACAT | GGATAGCTTGAAAATGCCCA | 128 |
| chr3:88647264-88649282 | p2 | (AT)9 | TTCCAGAGGTTTGGGATGTC | GGCTAGGAAAACACTCTGCG | 162 |
| chr3:88706124-88708139 | p5 | (TCATA)3 | CCACTAGCATGCATCTTCCA | CCCCGTATTTTTGTACGTCG | 140 |
| chr3:88722009-88724024 | p5 | (TCATA)3 | CCACTAGCATGCATCTTCCA | CCCCGTATTTTTGTACGTCG | 140 |
| chr3:88802914-88804940 | p2 | (TA)13 | GCGCATTACGGGTTACATCT | ATACAAGAGTCGGCTCGCAT | 186 |
| chr3:88818066-88820084 | p2 | (AT)9 | CACAAACAGGCTTGGGAAAT | TGCTATGCACATGACCCCTA | 227 |

| Marker ID | SSR type | SSR motif | FORWARD PRIMER (5'-3') | REVERSE PRIMER (5'-3') | size (bp) |
|---|---|---|---|---|---|
| chr3:88836911-88838927 | p2 | (AT)6 | TGATTATGGCTTGGCTTTCC | AGTACACTCACCGACCCCTG | 149 |
| chr3:88836911-88838927 | p2 | (TA)8 | TGATTATGGCTTGGCTTTCC | AGTACACTCACCGACCCCTG | 149 |
| chr3:88849304-88851370 | p2 | (AT)10 | AATTTCGATGGACTCGATGC | CCGAACACATCATATGCCAC | 141 |
| chr3:88889998-88892014 | p2 | (AT)8 | ATTCCTGGTTCACTGATGGC | CCAGGACCGAGTAAGCAGAG | 242 |
| chr3:88923385-88925401 | p2 | (TG)8 | AGACCTCCCCGAAGCATTAT | AATTTGCCCTTCGTCTTCCT | 221 |
| chr3:88945770-88947786 | p4 | (TTAA)4 | ACGTGCATAGCACACAGCTC | GGAGAATACCCAATGCTCCA | 170 |
| chr3:88945770-88947786 | p4 | (TTAT)5 | ACGTGCATAGCACACAGCTC | GGAGAATACCCAATGCTCCA | 170 |
| chr3:88961231-88963243 | p2 | (TA)6 | TTGGGTCCTTCAATTCTTGC | TTTGATGAAGACCCACCACA | 228 |
| chr3:89046763-89048777 | p2 | (AG)7 | AACCAATTTGTTCTGCGACC | GGTCAGTCTCAGCTTCAGCC | 139 |
| chr3:89070066-89072120 | p3 | (TAA)18 | AGGAAGAGAGAGACGGAGCC | GGAAACCCTCATTTTCCCAT | 281 |
| chr3:89336225-89338249 | p2 | (TA)12 | TGTCTCGGAGCTCTAGGGAA | TACAGCCCCTTTTGGTATGC | 115 |
| chr3:89436466-89438518 | p2 | (AT)26 | AGGCGATTTTAGGTGTCCCT | TCCAATTCGAACCCAAATGT | 300 |
| chr3:89515659-89517687 | p2 | (AT)14 | AGATCCTGAACCAACCCTCC | CCCACCAGTTAGGTCGTGAT | 154 |
| chr3:89530442-89532457 | p2 | (AG)9 | GAAGGGATCGACCAATGAAA | GTCCTTGCCTCATATCCCAA | 102 |
| chr3:89530442-89532457 | p3 | (TGT)5 | GAAGGGATCGACCAATGAAA | GTCCTTGCCTCATATCCCAA | 102 |
| chr3:89651987-89654002 | p5 | (TCATT)3 | TGGAATGAACAGTCGTCGAA | CGTAAACGGCGATCGTAAAT | 119 |
| chr3:89747600-89749669 | p3 | (ATA)23 | GGTGCGGTACCATCAAGTCT | GGCCGCAAAGTATTCCACTA | 172 |
| chr3:89764661-89766677 | p4 | (CCTT)4 | TCAGATTGACCCAAGAAGGG | GAACCCCGAAAGTTCAACAA | 256 |
| chr3:89778216-89780231 | p5 | (ATTCA)3 | TCCAGAAACAAGTTCCGGTC | AAATAGAAGCCCGTCCGATT | 206 |
| chr3:89778216-89780231 | p2 | (TA)6 | TCCAGAAACAAGTTCCGGTC | AAATAGAAGCCCGTCCGATT | 206 |
| chr3:89827130-89829151 | p6 | (CTTCTC)3 | CTTATGCTCCTTCTGCTCCTG | GAGGGGGAAGAAGAGGAAGA | 142 |
| chr3:89827166-89829205 | p6 | (CTTCTC)3 | CTTATGCTCCTTCTGCTCCTG | GAGGGGGAAGAAGAGGAAGA | 106 |
| chr3:89868770-89870786 | p5 | (CCTTT)3 | GATTGACATCGGCTCCAGTT | CCACACCGAGGTTGAAAGAT | 161 |
| chr3:89868770-89870786 | p2 | (AT)8 | GATTGACATCGGCTCCAGTT | CCACACCGAGGTTGAAAGAT | 161 |
| chr3:89918429-89920441 | p2 | (GT)6 | TGGGCAACTTGTGACTTGAA | GCCATGGTTGTTGTTCTCCT | 240 |
| chr3:90002624-90004638 | p2 | (AG)7 | ATCCGGCTTGAAAGGGTATT | ACCGCTGATGAAGCTCCTAA | 197 |
| chr3:90008878-90010892 | p2 | (TG)7 | TTGACCTACGGCATAAAGGC | TGTTAAACCGGGCCTCATAG | 102 |
| chr3:90132219-90134239 | p2 | (AT)10 | CGGCTGAGTTGAGAGAGCTT | ACCCTGGCTTGGGTCTCTAT | 166 |
| chr3:90132219-90134239 | p2 | (TA)10 | CGGCTGAGTTGAGAGAGCTT | ACCCTGGCTTGGGTCTCTAT | 166 |
| chr3:90143367-90145382 | p2 | (CA)6 | ACTCGGCCAAATTCATCAAC | TGTCGAAGGTTGGAGGTTTC | 243 |
| chr3:90143367-90145382 | p5 | (TTTTC)3 | ACTCGGCCAAATTCATCAAC | TGTCGAAGGTTGGAGGTTTC | 243 |
| chr3:90181187-90183199 | p2 | (GA)6 | TCATCAATCCTTGGGAGAGG | GGCTCCAGCAGAATTGAAAG | 236 |
| chr3:90186518-90188538 | p2 | (TA)10 | GAAGCAGCTGGTGAAGGAAC | AGAAAGAGAGCCGCAATGAA | 107 |
| chr3:90191119-90193135 | p4 | (TATG)4 | TAGGTCCGGGTAACGCATAG | CCCATTTCTCCCTTCACAAA | 225 |
| chr3:90191119-90193135 | p2 | (TA)6 | TAGGTCCGGGTAACGCATAG | CCCATTTCTCCCTTCACAAA | 225 |
| chr3:90191119-90193135 | p2 | (AT)21 | TAGGTCCGGGTAACGCATAG | CCCATTTCTCCCTTCACAAA | 225 |

| Marker ID | SSR type | SSR motif | FORWARD PRIMER (5'-3') | REVERSE PRIMER (5'-3') | size (bp) |
|---|---|---|---|---|---|
| chr3:90201336-90203351 | p5 | (AAAAT)3 | CCCTTCTCTCCATCTTTCCC | AATCACCATGCCACCAATTT | 268 |
| chr3:90258109-90260124 | p5 | (TGGAA)3 | AGGTGGCTAAGGGCAATTTT | TAAAAATTTGTGGGCTTGGG | 263 |
| chr3:90279501-90281517 | p2 | (AT)8 | AGGATGGCTAGCGTTCTTGA | ATGTATCAAGGGCTCGATGG | 299 |
| chr3:90298814-90300832 | p6 | (GATCGG)3 | ACATTCCAGCCATCCTGAAC | GGCTACAGGACATCTAGCGG | 145 |
| chr3:90301231-90303261 | p3 | (CAT)10 | CAACGCGACGTACAACACTT | TTCAGACCAAAACACACCCA | 270 |
| chr3:90307746-90309782 | p3 | (ACA)12 | ACCTTTACCCATTCGCTCCT | GGGAAGGGCTATGGCATTAT | 258 |
| chr3:90427629-90429644 | p5 | (TTCTG)3 | CTTGGCACCTTATGGGTGTT | TGGCATGATCCTTGTCGTTA | 183 |
| chr3:90442389-90444455 | p4 | (ATAC)6 | AGAGCTCCACTTCATTCGGA | AAGCCGACAAGGCTAACAAA | 192 |
| chr3:90442389-90444455 | p2 | (AT)33 | AGAGCTCCACTTCATTCGGA | AAGCCGACAAGGCTAACAAA | 192 |
| chr3:90536196-90538212 | p4 | (TTTC)4 | ATTCAATTACCACCACCCGA | GGCTTAATGGCTCGAAATGA | 245 |
| chr3:90543065-90545080 | p2 | (AT)7 | CACACCGGACAGCATAACAC | CGCGTGCCTTATTTTGATTT | 299 |
| chr3:90543065-90545080 | p5 | (GAATT)3 | CACACCGGACAGCATAACAC | CGCGTGCCTTATTTTGATTT | 299 |
| chr3:90567382-90569402 | p2 | (TA)10 | AAATGGTGGGTTGAAATGGA | GGGCAGCATTTCCCTACATA | 118 |
| chr3:90567382-90569402 | p6 | (TATATG)3 | AAATGGTGGGTTGAAATGGA | GGGCAGCATTTCCCTACATA | 118 |
| chr3:90765101-90767119 | p2 | (AT)9 | TGCACTGTTATCTCGCTTGC | TTCGGGTTCCTCTTCTTGTG | 189 |
| chr3:90791878-90793910 | p2 | (TA)11 | GCTTACGTCACCCCGTTAAA | TCATAACATGAGCCTGCGTC | 269 |
| chr3:90791878-90793910 | p2 | (AT)16 | GCTTACGTCACCCCGTTAAA | TCATAACATGAGCCTGCGTC | 269 |
| chr3:90899629-90901641 | p2 | (CT)7 | AGTTGGTTTCGTGCGGATAC | CTTGCTGCCCAATGGTACTT | 273 |
| chr3:90899629-90901641 | p2 | (TA)6 | AGTTGGTTTCGTGCGGATAC | CTTGCTGCCCAATGGTACTT | 273 |
| chr3:90912188-90914248 | p2 | (AT)30 | AACTCGGCCAAATCATCAAC | GAAATGAAATGCCGTCGAAT | 218 |
| chr3:90963740-90965762 | p5 | (AAGAA)3 | GCACATAAGGAACCCCTCAA | TTATAATCTGCTGGGCCGAC | 198 |
| chr3:90963740-90965762 | p2 | (AT)11 | GCACATAAGGAACCCCTCAA | TTATAATCTGCTGGGCCGAC | 198 |
| chr3:91021223-91023238 | p3 | (ACA)5 | TATCTGGGAACACCTCTGGG | GGAAACAGCTTTGGAATGGA | 224 |
| chr3:91087476-91089491 | p3 | (TGA)5 | TACTTGTTCAGGGGTACGGC | TCAGAAGCCGACAAGGCTAT | 119 |
| chr3:91277603-91279618 | p5 | (ATATG)3 | GAGAAACGGTGGATTGTGGT | TTCGCTTGTTCTCCTTCGAT | 228 |
| chr3:91277603-91279618 | p2 | (AT)8 | GAGAAACGGTGGATTGTGGT | TTCGCTTGTTCTCCTTCGAT | 228 |
| chr3:91282319-91284337 | p3 | (ATC)6 | TGCCCCAGATATGAACACAA | TTAGCATGCATGACTCCGTC | 106 |
| chr3:91362470-91364484 | p2 | (AT)7 | GGACAACTCCATTCTTCGGA | ACAATAAGGAAAATCCGGGG | 130 |
| chr3:91389408-91391420 | p2 | (TA)19 | TCCACCATGTTCAGCCTACA | AAATCGTACGCCAACACACA | 288 |
| chr3:91389453-91391471 | p2 | (TA)19 | TCCACCATGTTCAGCCTACA | AAATCGTACGCCAACACACA | 243 |
| chr3:91389574-91391586 | p2 | (TA)19 | TCCACCATGTTCAGCCTACA | AAATCGTACGCCAACACACA | 122 |
| chr3:91389740-91391752 | p2 | (TA)19 | ACGCCTTGGATGAGGTTTTA | ATTCACCACCTCGTCACACA | 121 |
| chr3:91447991-91450003 | p2 | (TA)6 | TGAAAAACAATGCAAGCCAG | GGTATGCTAGCGCCTTTGAG | 209 |
| chr3:91486080-91488096 | p2 | (AT)8 | TACGAAGAGAATCGGGGTTG | CTGCTTTCCTGCTTTCATCC | 281 |
| chr3:91489129-91491144 | p3 | (TGT)5 | TGGAGGGCAAGTGGATTAAG | GCGGTTTGTTAGGTTGGAAA | 241 |
| chr3:91490244-91492259 | p5 | (ATTTC)3 | TAGCTGCCTTTCAGTGAGCA | CTTTGCCTCAAAACTCCTCG | 103 |

| Marker ID | SSR type | SSR motif | FORWARD PRIMER (5'-3') | REVERSE PRIMER (5'-3') | size (bp) |
|---|---|---|---|---|---|
| chr3:91494133-91496148 | p3 | (ATA)5 | AGCTTTTGATTGGGCTCTCA | CAAGTAAACTGCCCGACCAT | 260 |
| chr3:91603759-91605777 | p6 | (TATGTA)3 | GAGCCCCTAGGATGGCTAAC | CAAAGGCTCGATGGAACAAT | 296 |
| chr3:91643177-91645193 | p2 | (AT)8 | CGGGTCCGAGATTTTATTGA | TTTCGTCGAAGCTGTGTGAG | 109 |
| chr3:91650431-91652446 | p3 | (CAA)5 | CGTCACCCCCTGTAAACTGT | GTGGCCTTTGTTGTGATGTG | 122 |
| chr3:91664548-91666563 | p3 | (CAA)5 | CCCGATGTCTTAGCACCAAT | TGTTATGTCCCGTGTTTTCG | 238 |
| chr3:91667547-91669562 | p3 | (CAA)5 | TGGGCCTCAATTCTTGTTTC | ATTGGTGCTAAGACATCGGG | 132 |
| chr3:91667547-91669562 | p2 | (TA)14 | TGGGCCTCAATTCTTGTTTC | ATTGGTGCTAAGACATCGGG | 132 |
| chr3:91732497-91734509 | p2 | (TA)6 | GTTTATTGATGTGGCTCGCA | AATTAAGTGGTGCTCGGTGG | 285 |
| chr3:91735931-91737946 | p5 | (AATAA)3 | AATGCCATTTTGAGACCGAG | GTGGAAGAAAAGGTGGCAAA | 169 |
| chr3:91887817-91889835 | p6 | (TACAAC)3 | GCCAGTATAATGCATCCGCT | CCGAAAAGGACAAAACGGTA | 284 |
| chr3:91887817-91889835 | p4 | (ATAC)4 | GCCAGTATAATGCATCCGCT | CCGAAAAGGACAAAACGGTA | 284 |
| chr3:91914436-91916448 | p2 | (GT)6 | CCAGAGTATCGGAGACCCAA | AAGTAGGGCCTTGCTTGTCA | 154 |
| chr3:91920332-91922344 | p2 | (GT)6 | AAGGAGACCTCCCCGAAGTA | TCCATGTTCTCCACGAATCA | 219 |
| chr3:91924654-91926690 | p2 | (TA)18 | TTGAGGTAGGGTTTTGGTGC | TTCACCCATTCAAGACCCAT | 292 |
| chr3:91924654-91926690 | p2 | (AT)31 | TTGAGGTAGGGTTTTGGTGC | TTCACCCATTCAAGACCCAT | 292 |
| chr3:91984906-91986918 | p2 | (TA)6 | ACAAAATATGCAAGCCGGAC | TTGTTCCGGAGCTTGAGTTT | 242 |
| chr3:91984906-91986918 | p3 | (CAA)5 | ACAAAATATGCAAGCCGGAC | TTGTTCCGGAGCTTGAGTTT | 242 |
| chr3:92026960-92028972 | p2 | (AG)6 | TGACATTCTGCTTCACCTCG | AGCGACCAAGGTATGTGAGG | 143 |
| chr3:92138138-92140168 | p2 | (AT)15 | GCGATAGAGCCAGCTGTAGG | CCTGCTACACCATTCCGTTT | 224 |
| chr3:92181353-92183365 | p2 | (AT)6 | TCCGCAGACTGATGAACAAG | CATTGATCTTTTCCACCGCT | 279 |
| chr3:92225365-92227381 | p2 | (AT)8 | TAGTAAGCCGGCCAACAATC | GGTACCGAGGCAGATGATGT | 177 |
| chr3:92225365-92227381 | p2 | (AT)11 | TAGTAAGCCGGCCAACAATC | GGTACCGAGGCAGATGATGT | 177 |
| chr3:92233096-92235108 | p2 | (AT)17 | AATAACATACGCCAGCCGAC | TTTGTCTAGGGGTACGACGG | 283 |
| chr3:92233096-92235108 | p2 | (AT)6 | AATAACATACGCCAGCCGAC | TTTGTCTAGGGGTACGACGG | 283 |
| chr3:92244726-92246742 | p2 | (AT)8 | CAAGTCAGCGGTAGGTAGGC | GCCTAGAGCATCTGGTGGAG | 113 |
| chr3:92249768-92251780 | p2 | (AT)6 | TTACCAACTTGCCCTTCTGG | GGTTGGGTCGTGACAAAAGT | 209 |
| chr3:92311070-92313082 | p2 | (TA)10 | GATGGGCTACGGATTTACGA | TCGAGGGGAAACTCAATCAC | 243 |
| chr3:92311070-92313082 | p2 | (TA)6 | GATGGGCTACGGATTTACGA | TCGAGGGGAAACTCAATCAC | 243 |
| chr3:92335612-92337627 | p5 | (TGAAA)3 | GGCGTATCAAGAATTGGGAA | TTAAGGCATGGTTCTTTGGC | 167 |
| chr3:92446228-92448243 | p5 | (AATGC)3 | CGTTCCATTCCACGAGATTT | ACGATAGGTGCCCATACGAC | 265 |
| chr3:92463691-92465705 | p2 | (AT)7 | GTTGTCGCTCTGCCTCTACC | TCGGTTTTGCTATGCTTGTG | 103 |
| chr3:92463691-92465705 | p6 | (ATACAT)3 | GTTGTCGCTCTGCCTCTACC | TCGGTTTTGCTATGCTTGTG | 103 |
| chr3:92478164-92480176 | p2 | (TA)6 | CCATTACACGCAACATTCCA | TGAAGTGCTCATTTGTTCCG | 257 |
| chr3:92545006-92547018 | p2 | (TA)6 | TGCATGGCATGACGAATAAT | CTCGCCACTCTTGCATAACA | 249 |
| chr3:92617472-92619496 | p2 | (AT)12 | GTCCCTTTCTTCAAATGGCA | ATCCCCCGATCTCTTTATGG | 244 |
| chr3:92641798-92643816 | p2 | (AT)13 | CCCCGTCGTACCTTTAGTCA | ATATTCGCTTTGGTGATCCG | 135 |

| Marker ID | SSR type | SSR motif | FORWARD PRIMER (5'-3') | REVERSE PRIMER (5'-3') | size (bp) |
|---|---|---|---|---|---|
| chr3:92641798-92643816 | p2 | (TA)9 | CCCCGTCGTACCTTTAGTCA | ATATTCGCTTTGGTGATCCG | 135 |
| chr3:92717003-92719019 | p2 | (AT)8 | CCCGATAACCTGTCTCCTGA | GGCCTGCGTATGGTATTGAT | 297 |
| chr3:92893016-92895031 | p5 | (TTCAT)3 | TGATAGGGCCAGACTTGGTC | CCTTCGACAATTATCGGCAT | 130 |
| chr3:92893016-92895031 | p2 | (AT)17 | TGATAGGGCCAGACTTGGTC | CCTTCGACAATTATCGGCAT | 130 |
| chr3:92927394-92929409 | p5 | (ATGCC)3 | CAATTTTCCCACGACTTGGT | GTATGCGGAAGTTATGCGGT | 254 |
| chr3:92948385-92950419 | p2 | (TA)17 | GGCAAATTGAGGAATCCAGA | GAGCTCGGAATGTCAACCAT | 159 |
| chr3:92966887-92968902 | p3 | (CAT)5 | GGGCACCCTGAGCTAACATA | CTCTCAGTGTCGTCGGTTCA | 137 |
| chr3:93144832-93146856 | p2 | (AT)12 | TTGAAGCCATGCAAGATGAG | ACCAAGAACTGTCCGAATGG | 116 |
| chr3:93144832-93146856 | p2 | (AT)6 | TTGAAGCCATGCAAGATGAG | ACCAAGAACTGTCCGAATGG | 116 |
| chr3:93160589-93162613 | p2 | (AT)12 | TTGAAGCCATGCAAGATGAG | ACCAAGAACTGTCCGAATGG | 116 |
| chr3:93160589-93162613 | p2 | (AT)6 | TTGAAGCCATGCAAGATGAG | ACCAAGAACTGTCCGAATGG | 116 |
| chr3:93168831-93170899 | p2 | (TA)34 | GAACCACTCCAAATCCGAGA | TCTAGGCACCCAATTCGTTC | 118 |
| chr3:93251122-93253140 | p2 | (TA)24 | ACTGAGCGGCACTGTTTTCT | GGATATATGAAGCGGCCAGA | 131 |
| chr3:93251122-93253140 | p6 | (CATATA)3 | ACTGAGCGGCACTGTTTTCT | GGATATATGAAGCGGCCAGA | 131 |
| chr3:93294597-93296627 | p2 | (AT)15 | GTTCGGTTTTGACCCAAAGA | CTCAGGCCAAGAGGTGAAAG | 102 |
| chr3:93298552-93300564 | p2 | (AT)49 | ATCCCCAAAATCCGAAAGTC | TATCTAGCGTTGAGGGCGTT | 129 |
| chr3:93298552-93300564 | p2 | (TA)6 | ATCCCCAAAATCCGAAAGTC | TATCTAGCGTTGAGGGCGTT | 129 |
| chr3:93463458-93465476 | p3 | (ATG)6 | GAGGTTATGGCCTGATTGGA | TATTGCTGCGGAACGTGTAG | 177 |
| chr3:93463458-93465476 | p2 | (TA)9 | GAGGTTATGGCCTGATTGGA | TATTGCTGCGGAACGTGTAG | 177 |
| chr3:93468117-93470135 | p2 | (TA)16 | ATGCGCCTTCTTTCTCTTCA | TAGCATTAACTCCCCGGATG | 207 |
| chr3:93468117-93470135 | p2 | (AT)9 | ATGCGCCTTCTTTCTCTTCA | TAGCATTAACTCCCCGGATG | 207 |
| chr3:93547435-93549453 | p2 | (AT)12 | CATGGTGGCCGAATACTCTT | GAAGGTTGGAGAAGCATGGA | 212 |
| chr3:93547435-93549453 | p2 | (TC)9 | CATGGTGGCCGAATACTCTT | GAAGGTTGGAGAAGCATGGA | 212 |
| chr3:93561710-93563740 | p2 | (AT)15 | TTTATGGAAACCGAGCGAAC | GAGGCTTTGCAAACAGTTCC | 239 |
| chr3:93691142-93693158 | p2 | (AT)8 | ACATCCTTCACTGAGTCCCG | GTGTCCCGACCCTTTAGTGA | 222 |
| chr3:93815375-93817390 | p5 | (AATGC)3 | GTATTGTCACGACCCAACCC | CGGGGTTTTATTTCGGATTT | 184 |
| chr3:93932948-93934963 | p3 | (ACA)5 | ACCTCACCTTGGATTTGTCG | ATTTGGGGCACATTTATGGA | 261 |
| chr3:94062961-94064973 | p2 | (AT)6 | AGGCTCCATTTTCGGCTATT | AAGAACCCAACCCACAAACA | 180 |
| chr3:94176144-94178159 | p5 | (TTGCA)3 | TGGTGTTGCTGTTCAACCAT | CCCAACTTCATACCCGAAGA | 203 |
| chr3:94296938-94298962 | p2 | (AG)12 | AATGATTGGCATCTTCCAGC | CGACATCGAGGAATCGAAAT | 277 |
| chr3:94322738-94324750 | p2 | (AG)6 | TGGCCAAGACAGTGTTTCAC | TGTCGGGCTTACTTGTTTCC | 265 |
| chr3:94322738-94324750 | p5 | (TAAAA)3 | TGGCCAAGACAGTGTTTCAC | TGTCGGGCTTACTTGTTTCC | 265 |
| chr3:94350505-94352521 | p2 | (TA)6 | GGAGTGGATTCGAGGATTGA | CACTAGTTGCCGATCACGAA | 145 |
| chr3:94350505-94352521 | p2 | (TA)8 | GGAGTGGATTCGAGGATTGA | CACTAGTTGCCGATCACGAA | 145 |
| chr3:94350505-94352521 | p5 | (TGAAT)3 | GGAGTGGATTCGAGGATTGA | CACTAGTTGCCGATCACGAA | 145 |
| chr3:94464110-94466128 | p3 | (ATT)6 | GCTTCAGTCTTCACAAGCCC | GCAAAGGACGAAGCAGAAAG | 111 |

| Marker ID | SSR type | SSR motif | FORWARD PRIMER (5'-3') | REVERSE PRIMER (5'-3') | size (bp) |
|---|---|---|---|---|---|
| chr3:94464110-94466128 | p6 | (TTATTT)3 | GCTTCAGTCTTCACAAGCCC | GCAAAGGACGAAGCAGAAAG | 111 |
| chr3:94477324-94479380 | p2 | (TA)28 | ATGAAACTATGGACGTGCCC | ATATACCATACCCGGCCCAT | 193 |
| chr3:94488620-94490638 | p3 | (AAC)6 | GTGTGACTTACACCATGCCG | GCGACTACCTTTGGGGTGTA | 162 |
| chr3:94488620-94490638 | p2 | (CT)6 | GTGTGACTTACACCATGCCG | GCGACTACCTTTGGGGTGTA | 162 |
| chr3:94506015-94508030 | p4 | (TATT)4 | ACTCTCGGGCACGTTGTTAC | GGTAAGGGTTGCTCGGTACA | 149 |
| chr3:94506015-94508030 | p5 | (CATAT)3 | ACTCTCGGGCACGTTGTTAC | GGTAAGGGTTGCTCGGTACA | 149 |
| chr3:94642207-94644219 | p2 | (AT)6 | AAAGCGGGACAAATTGTGAG | TCCACCTCCACTTGGATTTC | 181 |
| chr3:94685171-94687183 | p2 | (AT)6 | CAGACGGTCTCGGATTGTTT | TAAGTTGAGGCATGCACGAG | 171 |
| chr3:94919791-94921809 | p2 | (AT)9 | CCTAGGACCGAACTGCTCTG | TGATGGTCCTCGTCGTTGTA | 214 |
| chr3:94919791-94921809 | p2 | (TA)11 | CCTAGGACCGAACTGCTCTG | TGATGGTCCTCGTCGTTGTA | 214 |
| chr3:94934819-94936831 | p2 | (AT)6 | CCCATCGTACCCCTAGACAA | TGCAGATATTCGCTTTGGTG | 167 |
| chr3:94936673-94938699 | p2 | (TA)13 | TTATACGGACCCCAAGGTCA | AGGTTGCTGCTCAACGTTTT | 262 |
| chr3:94938960-94940972 | p2 | (TA)6 | TATTTTATGGCCTTGTCGGC | ACCAATACCAAGGCAAGACG | 168 |
| chr3:95037671-95039689 | p2 | (TA)9 | GGATAATTTTGAGCGACGGA | AGCCAACAACACCAACACAA | 226 |
| chr3:95051582-95053600 | p3 | (CAA)6 | ATGCCCGCAGGTGTAGATAG | GGAACTGTCTGCAAAGCCTC | 216 |
| chr3:95072995-95075010 | p3 | (ATG)5 | TGACGTAAGGTCGGTAGCCT | TTCGTTGGTAGTCTCGCCTT | 101 |
| chr3:95144837-95146852 | p5 | (CATTT)3 | TGTATAGGCGACCCTTCCAG | ACGTTGACGGAAGCCATATC | 194 |
| chr3:95175329-95177341 | p2 | (AG)6 | ATGCTGTAGACCCGAACCAC | CCTACCAAAACGTAGGCGAA | 132 |
| chr3:95175329-95177341 | p3 | (TTG)5 | ATGCTGTAGACCCGAACCAC | CCTACCAAAACGTAGGCGAA | 132 |
| chr3:95183837-95185849 | p2 | (AT)6 | GCCGTTGTAATCTTTCGCAT | GCATTATGGCCAACTCAGGT | 124 |
| chr3:95316224-95318236 | p2 | (TG)6 | AAAGGTGCCTACGGGCTATT | CTCATACCTGGCCCTACCAA | 224 |
| chr3:95331303-95333315 | p2 | (AT)6 | TTGAACCTGAACCCTATCGC | AAAGCTACAGTCGCCCAGAA | 206 |
| chr3:95372219-95374240 | p3 | (AGA)9 | ACCCGGCCATAATATCAACA | GAGAGTGGGCTGATTTGGAG | 218 |
| chr3:95382851-95384863 | p2 | (AT)6 | GATATGCTTCGGGCTTGTGT | GCTGCCAAAAGCTTACCTTG | 271 |
| chr3:95395802-95397817 | p5 | (TGCAA)3 | GTTGTGCGGACACCTACCTT | GGGGCGGATTTCATTTACTT | 260 |
| chr3:95395802-95397817 | p2 | (AT)18 | GTTGTGCGGACACCTACCTT | GGGGCGGATTTCATTTACTT | 260 |
| chr3:95415547-95417562 | p3 | (AAC)5 | GTGACCCGTCGGTCAAGTAT | TCGCCTAGCCTACGAGTCAT | 217 |
| chr3:95531610-95533628 | p6 | (ATTATG)3 | CGCGTTAAGGATTGTTTGGT | GTTGTCCTCGTCGTTTCGTT | 199 |
| chr3:95542662-95544678 | p2 | (AT)8 | ATGCTCTTTACGCCTTTCCA | GTTGATTTGTTTGGCCGAGT | 298 |
| chr3:95665343-95667358 | p5 | (TGATC)3 | CTTTTGCTGTTGTGTCCGAA | CATGCCTAGGAACAAAGGGA | 157 |
| chr3:95667599-95669611 | p2 | (TG)6 | CCTCCGATGTTGTTCGAGTT | GTATCGCGGAACCTGTTTGT | 102 |
| chr3:95672657-95674672 | p5 | (ATGCA)3 | TCGGTCAAATTGTCCTTTCC | GACTTGAGGAGGAGCTGTGG | 247 |
| chr3:95678617-95680632 | p5 | (TATGA)3 | TTCGGCTGTGTTAGCATGAC | TGCCACTACATACGGGAACA | 162 |
| chr3:95775715-95777730 | p5 | (TTCAT)3 | CAACCAATGCAATCCCTTCT | ATCGCGGTACGAGAAATGAC | 200 |
| chr3:95779051-95781066 | p2 | (CA)6 | CCAAAAGATATGGCTCCGAA | TCTTCGTACTGGCGTCCTTT | 295 |
| chr3:95779051-95781066 | p3 | (ATC)5 | CCAAAAGATATGGCTCCGAA | TCTTCGTACTGGCGTCCTTT | 295 |

| Marker ID | SSR type | SSR motif | FORWARD PRIMER (5'-3') | REVERSE PRIMER (5'-3') | size (bp) |
|---|---|---|---|---|---|
| chr3:95809504-95811522 | p6 | (TATATG)3 | TTAGATTCTCGGGGATGGTG | GCTTAATTGTCCAAGCTCGC | 161 |
| chr3:95818709-95820724 | p5 | (TTTCA)3 | GTTGTTCCATGGCCAATTTC | TTGCCAAACAGACAGACGAG | 165 |
| chr3:95837106-95839130 | p2 | (TA)12 | TCGTGTTGTTGTCGGATTGT | CGCCTAACCAGAACTTCCAA | 124 |
| chr3:95837106-95839130 | p2 | (AT)16 | TCGTGTTGTTGTCGGATTGT | CGCCTAACCAGAACTTCCAA | 124 |
| chr3:95865296-95867320 | p2 | (AT)11 | ATGACTAGTGCCCGACTTGG | CAGACTTGTCGGCTTGCATA | 172 |
| chr3:95865296-95867320 | p2 | (AT)12 | ATGACTAGTGCCCGACTTGG | CAGACTTGTCGGCTTGCATA | 172 |
| chr3:95872570-95874586 | p2 | (AT)7 | CCCGACTTGGACACTCGTAT | AGCCTTGTCGACTTGCGTAT | 188 |
| chr3:95872570-95874586 | p2 | (AT)8 | CCCGACTTGGACACTCGTAT | AGCCTTGTCGACTTGCGTAT | 188 |
| chr3:95888886-95890901 | p5 | (ATCAT)3 | CGAAATTCGGGTCGATTCTA | TGAGCCGACACACATCAAAT | 118 |
| chr3:95898900-95900921 | p3 | (ATT)7 | GTTTCCTCCGGATCTTAGCC | ACTCACGGCTTTCACTCACC | 216 |
| chr3:95906847-95908871 | p2 | (AT)11 | ATGACTAGTGCCCGACTTGG | CAGACTTGTCGGCTTGCATA | 172 |
| chr3:95906847-95908871 | p2 | (AT)12 | ATGACTAGTGCCCGACTTGG | CAGACTTGTCGGCTTGCATA | 172 |
| chr3:95914121-95916137 | p2 | (AT)7 | CCCGACTTGGACACTCGTAT | AGCCTTGTCGACTTGCGTAT | 188 |
| chr3:95914121-95916137 | p2 | (AT)8 | CCCGACTTGGACACTCGTAT | AGCCTTGTCGACTTGCGTAT | 188 |
| chr3:95978366-95980406 | p2 | (TA)20 | CATTGATCGCTTTGTTGTGG | ACAAGGGCACCTCTAACCCT | 202 |
| chr3:96022900-96024922 | p2 | (TA)11 | ATGGCAATGTCATGCTCAAA | TGTCCGTTTAACAGCCATCA | 133 |
| chr3:96051628-96053643 | p5 | (CGACA)3 | CTTTTGGGAGGACTTAGGGC | CTCCGCCATGACCCTAGATA | 182 |
| chr3:96091573-96093589 | p2 | (AT)8 | AGGTGAAGTGTGCTCCTCGT | AACCCCCTATCCTGGTCATC | 265 |
| chr3:96091573-96093589 | p2 | (AT)9 | AGGTGAAGTGTGCTCCTCGT | AACCCCCTATCCTGGTCATC | 265 |
| chr3:96109170-96111182 | p2 | (AC)6 | GTTTCGAGGAGCAGAAATGC | TTACGCCTCGGGATGTTTAG | 153 |
| chr3:96160318-96162333 | p3 | (ATG)5 | CGTGCGAGATGTCTTATGGA | CTACTGAACGCCTGTCCTCC | 279 |
| chr3:96172539-96174551 | p2 | (TA)6 | TGAGGACATGATCAGCTTCG | CTCGCCCTCTACCTTATCCC | 129 |
| chr3:96333395-96335435 | p2 | (TA)20 | TTGCTAGGGCTTGTGGAACT | TTTTCCTTTACCCACATGGC | 149 |
| chr3:96353683-96355695 | p2 | (AT)6 | GGATAGCTTAGGAAACGGGC | ATTCGGAAACATATCAGCCG | 238 |
| chr3:96353683-96355695 | p2 | (AT)12 | GGATAGCTTAGGAAACGGGC | ATTCGGAAACATATCAGCCG | 238 |
| chr3:96405451-96407465 | p2 | (TA)7 | GTTGATGATTTGGCCGAGTT | TTCACAAGGATCATGCCAAA | 227 |
| chr3:96473091-96475111 | p5 | (AAATG)4 | GGGTCAAGCCAAGTGAAAAA | ATGGCCGAAGAGATAACGTG | 281 |
| chr3:96611456-96613554 | p2 | (GA)14 | GCTGAGCTCCCAATGTGATT | GTCCAACTTGGTGGACGAAT | 232 |
| chr3:96611456-96613554 | p4 | (TTAA)4 | GCTGAGCTCCCAATGTGATT | GTCCAACTTGGTGGACGAAT | 232 |
| chr3:96616657-96618669 | p3 | (CAA)6 | GATCGGGGGTCCTTTTGTAT | TTTGTGGTCCACCCAAAAAT | 147 |
| chr3:96616683-96618701 | p3 | (CAA)6 | GATCGGGGGTCCTTTTGTAT | TTTGTGGTCCACCCAAAAAT | 121 |
| chr3:96680179-96682191 | p2 | (AT)6 | CCCGGCCCTATCAAAATAAT | TGTTCATGCCCATCAGGTAA | 132 |
| chr3:96701625-96703641 | p2 | (TG)31 | GCATGTCAGCCCAGGTTTAT | ACACACACACACACCCTGCT | 278 |
| chr3:96701625-96703641 | p4 | (TTAA)4 | GCATGTCAGCCCAGGTTTAT | ACACACACACACACCCTGCT | 278 |
| chr3:96705396-96707424 | p2 | (AT)14 | AGAGTGTCGATCTGCTCCGT | TCACGGAAAGGAGGTTTGAC | 142 |
| chr3:96754930-96756974 | p2 | (AT)22 | TGCCATATATGTTTTCCGCA | ATGGCGAGGTCTAGGGAACT | 178 |

| Marker ID | SSR type | SSR motif | FORWARD PRIMER (5'-3') | REVERSE PRIMER (5'-3') | size (bp) |
|---|---|---|---|---|---|
| chr3:96765495-96767513 | p6 | (AGGTAG)3 | TTATTCATGCCTCCGAGACC | AGCATCCACAAATTCTTGCC | 132 |
| chr3:96804297-96806309 | p2 | (GT)6 | AATGAGGCTTTGGTCCCTTT | TAGCCTTGTCCCACACTTCC | 190 |
| chr3:96900163-96902175 | p2 | (CT)6 | TATATGCTTCCGGTTCAGGC | CTAAACCAGCGGCGTATCTC | 288 |
| chr3:96900163-96902175 | p2 | (TA)6 | TATATGCTTCCGGTTCAGGC | CTAAACCAGCGGCGTATCTC | 288 |
| chr3:96924572-96926590 | p2 | (AT)9 | AAGAAATGACATGGGAAGCG | TTACAAGCATTTTGGGGGAG | 201 |
| chr3:96924640-96926652 | p2 | (AT)9 | AAGAAATGACATGGGAAGCG | TTACAAGCATTTTGGGGGAG | 133 |
| chr3:96924640-96926652 | p5 | (TCATA)3 | AAGAAATGACATGGGAAGCG | TTACAAGCATTTTGGGGGAG | 133 |
| chr3:96947130-96949145 | p3 | (GAA)5 | GTGCGTTGGACAACTGCTTA | TCAACCAAAAAGCACCCTTC | 261 |
| chr3:97064659-97066673 | p2 | (TA)7 | CTCGAGGCCAGAGTTCAAAG | TTTTTCTAATCCATTGCGCC | 274 |
| chr3:97098393-97100407 | p2 | (AT)7 | AGCGAATTGCAGCTACTGGT | CATCCTCGGAATTTCCTTCA | 107 |
| chr3:97129701-97131717 | p3 | (ACA)6 | GGTTGATTGATGCTCTCGGT | TGTGCTTCCTCCTGGAGACT | 157 |
| chr3:97129701-97131717 | p2 | (AT)8 | GGTTGATTGATGCTCTCGGT | TGTGCTTCCTCCTGGAGACT | 157 |
| chr3:97170387-97172415 | p2 | (AT)14 | AGACGTTGCCTTAACGCTGT | ATTCGTAGTGGCAGCCTTGT | 228 |
| chr3:97329551-97331566 | p3 | (TTG)5 | GCCACCAAAACTTAAAGGCA | CCAAACTATCTCAGCCTCGC | 203 |
| chr3:97401143-97403157 | p2 | (AT)8 | TGCTAACACAGCCGAACAAC | AGCCTTGTCGACTTGCGTAT | 223 |
| chr3:97401143-97403157 | p2 | (TA)7 | TGCTAACACAGCCGAACAAC | AGCCTTGTCGACTTGCGTAT | 223 |
| chr3:97401143-97403157 | p2 | (AT)7 | TGCTAACACAGCCGAACAAC | AGCCTTGTCGACTTGCGTAT | 223 |
| chr3:97435093-97437120 | p2 | (TG)12 | TTGGGAGGCCTATCAATCTG | CTTTGGCCTCCAAATTGTGT | 122 |
| chr3:97435093-97437120 | p3 | (ACA)9 | TTGGGAGGCCTATCAATCTG | CTTTGGCCTCCAAATTGTGT | 122 |
| chr3:97532338-97534354 | p2 | (AT)6 | AGCGAAACTAGACGAACCGA | GAGTAGTGGCAGCCTTTTCG | 265 |
| chr3:97532338-97534354 | p2 | (AT)8 | AGCGAAACTAGACGAACCGA | GAGTAGTGGCAGCCTTTTCG | 265 |
| chr3:97532338-97534354 | p2 | (AT)15 | AGCGAAACTAGACGAACCGA | GAGTAGTGGCAGCCTTTTCG | 265 |
| chr3:97594736-97596760 | p2 | (AG)12 | CCTCCCATGCATCTGACTTT | CTCTTCGGTCCCTTACCACA | 152 |
| chr3:97594736-97596760 | p2 | (AT)10 | CCTCCCATGCATCTGACTTT | CTCTTCGGTCCCTTACCACA | 152 |
| chr3:97692215-97694227 | p2 | (AG)6 | AGTCAGATCACCCGAAAACG | TTAATTGCTGAGGTGGGACC | 121 |
| chr3:97700142-97702156 | p2 | (AG)7 | TGGTTGCCTTAAGTATCCCG | GAAGGGTTCGGAGAATCACA | 242 |
| chr3:97717792-97719804 | p2 | (AT)6 | TATCTACGCACTTACCGGGC | AGAGCCTGGGTCAATCAATG | 300 |
| chr3:97774288-97776304 | p4 | (ATAC)4 | CAGGAGCAAATGGAGCTTTC | TACTGACCCCCTGTTCTTCG | 259 |
| chr3:97790441-97792456 | p5 | (AGGAC)3 | GCGGCTTATGATTACACCGT | AGAAAAGGACGCCAGTACGA | 112 |
| chr3:97797386-97799401 | p6 | (AAGAGC)3 | GCCCTCTTACTCTGCCCTCT | TTGGTATTTCCATAAGCCGC | 207 |
| chr3:97797386-97799401 | p5 | (TGTGT)3 | GCCCTCTTACTCTGCCCTCT | TTGGTATTTCCATAAGCCGC | 207 |
| chr3:97797386-97799401 | p2 | (GT)6 | GCCCTCTTACTCTGCCCTCT | TTGGTATTTCCATAAGCCGC | 207 |
| chr3:97848699-97850777 | p2 | (AT)39 | AGTCAGGTCATCCGAAAACG | CCCTCGTAACCATAAGCCAA | 116 |
| chr3:97962953-97964971 | p2 | (AT)8 | GACAAGCGCTGCCATACATA | GGTACGATGACCCATATGCC | 253 |
| chr3:97962953-97964971 | p2 | (AT)9 | GACAAGCGCTGCCATACATA | GGTACGATGACCCATATGCC | 253 |
| chr3:97971799-97973817 | p3 | (CAA)6 | TTGCCTCATAAGGCGGATAC | TGAAGCGTTTAATTGGGCTC | 155 |

| Marker ID | SSR type | SSR motif | FORWARD PRIMER (5'-3') | REVERSE PRIMER (5'-3') | size (bp) |
|---|---|---|---|---|---|
| chr3:97984646-97986658 | p2 | (GT)6 | TGACACCTACTCCGAGCCTT | CATTATGAATGGCACGAACG | 122 |
| chr3:98003290-98005308 | p2 | (TA)9 | GTAACCTGCCAATCCCAGAA | TGCAAAATTTTCAAAGTGCG | 194 |
| chr3:98076281-98078296 | p5 | (GATAC)3 | GATGCCCACGATTTCATTCT | CATATAGCTATCGGAGCGGC | 121 |
| chr3:98099281-98101293 | p2 | (TC)6 | CGTAAGTGGTTATTCGCGGT | GGCTTCTCGTGCATGGTTAT | 298 |
| chr3:98177173-98179185 | p2 | (TG)6 | AAAGGTGCCTACGGGCTATT | CCAACCAAGGCTCAAGGTTA | 254 |
| chr3:98194188-98196206 | p6 | (GTGGAC)3 | ATTCAAGGGACAAGTTCCCC | AGAGTGGAACAACGTCCACC | 233 |
| chr3:98202611-98204626 | p3 | (ATT)16 | GGTGGCGACTCCTTAAAACA | TGATGAAAGAATTTTCGGGC | 154 |
| chr3:98202611-98204626 | p5 | (ACCCG)3 | GGTGGCGACTCCTTAAAACA | TGATGAAAGAATTTTCGGGC | 154 |
| chr3:98202611-98204626 | p2 | (CT)6 | GGTGGCGACTCCTTAAAACA | TGATGAAAGAATTTTCGGGC | 154 |
| chr3:98281487-98283505 | p2 | (TA)9 | AATTGGAATTCGGGTTAGGC | TCGTAATTCCGTAGCCCATC | 242 |
| chr3:98315760-98317772 | p2 | (AT)6 | TACCCGACCCCATAATCAGA | TGCAAGCCGATGTACCATTA | 237 |
| chr3:98324081-98326096 | p5 | (GAAAT)3 | GGCCCTTTTCTTTTTCTTGC | CAAGGCCCAAGTGGAATTTA | 252 |
| chr3:98338989-98341007 | p6 | (TATATG)3 | TAACGATGGCACTTCCATGA | TAGAATGAAATCGGGGCATC | 166 |
| chr3:98417872-98419890 | p6 | (AGTTGA)3 | CAGCAACAAAAGGCGTAACA | ACTACAACGAACACGAGGGG | 242 |
| chr3:98438975-98441035 | p2 | (TA)30 | GCACTGACCACTGCTGGATA | ATATTGTTCGCCCACTGAGG | 241 |
| chr3:98453287-98455309 | p2 | (AG)11 | CCGAGGTTAGGAATGAGCAG | TGTAGTGAGACCCCTCCTCG | 169 |
| chr3:98609686-98611701 | p5 | (GAAGA)3 | TACCCGTCCACATCCTTAGC | TTTGACGGGGAAGTCCATAG | 226 |
| chr3:98609686-98611701 | p5 | (GGGGT)3 | TACCCGTCCACATCCTTAGC | TTTGACGGGGAAGTCCATAG | 226 |
| chr3:98627456-98629468 | p2 | (AT)6 | GAAGAAGGCACCAAGACGAG | ACCTCCGAGGGGAGATCTTA | 196 |
| chr3:98651755-98653783 | p4 | (AAAT)4 | TGAGTTTCGTCATGCTTTCG | CAATGACTCATGCGTTGCTT | 197 |
| chr3:98651755-98653783 | p4 | (TTTC)5 | TGAGTTTCGTCATGCTTTCG | CAATGACTCATGCGTTGCTT | 197 |
| chr3:98708262-98710280 | p2 | (TA)9 | ACATCACGCAACCGATATGA | GGAGGATAGGGCTATGAGGG | 182 |
| chr3:98708262-98710280 | p6 | (CATATA)3 | ACATCACGCAACCGATATGA | GGAGGATAGGGCTATGAGGG | 182 |
| chr3:98775675-98777693 | p2 | (AT)6 | ACGAAATAATACGCAAGCCG | GGAGAGCTCTGTTGTTTCGG | 291 |
| chr3:98775675-98777693 | p6 | (CATATA)3 | ACGAAATAATACGCAAGCCG | GGAGAGCTCTGTTGTTTCGG | 291 |
| chr3:98923257-98925272 | p5 | (TATGA)3 | ATGTGGGTCTTGGGTCATGT | TGACAGATTGGCGGAATGTA | 152 |
| chr3:98995301-98997337 | p4 | (TCTT)9 | AGGGAAGCATACACAATGCC | ATGTTTTGTCTGCCTGGTCC | 197 |
| chr3:99006577-99008613 | p4 | (TCTT)9 | AGGGAAGCATACACAATGCC | ATGTTTTGTCTGCCTGGTCC | 197 |
| chr3:99179068-99181095 | p3 | (TGT)9 | GGATGTCACGTCCCGTACTT | AAACCTTGCCCACCTTTTCT | 114 |
| chr3:99179068-99181095 | p2 | (TA)6 | GGATGTCACGTCCCGTACTT | AAACCTTGCCCACCTTTTCT | 114 |
| chr3:99236495-99238522 | p3 | (TGT)9 | GGATGTCACGTCCCGTACTT | AAACCTTGCCCACCTTTTCT | 114 |
| chr3:99236495-99238522 | p2 | (TA)6 | GGATGTCACGTCCCGTACTT | AAACCTTGCCCACCTTTTCT | 114 |
| chr3:99252710-99254722 | p2 | (AT)6 | GAGCTTACGTCACCCCGATA | CGTGCAATATAGGCATGGTG | 249 |
| chr3:99252710-99254722 | p2 | (TG)6 | GAGCTTACGTCACCCCGATA | CGTGCAATATAGGCATGGTG | 249 |
| chr3:99257025-99259079 | p2 | (AT)27 | CCTAGGACGGAACTGCTCTG | GATGTAGCCTTGTCGGCTTC | 251 |
| chr3:99257025-99259079 | p2 | (AT)8 | CCTAGGACGGAACTGCTCTG | GATGTAGCCTTGTCGGCTTC | 251 |

| Marker ID | SSR type | SSR motif | FORWARD PRIMER (5'-3') | REVERSE PRIMER (5'-3') | size (bp) |
|---|---|---|---|---|---|
| chr3:99272657-99274675 | p2 | (CT)9 | GGTACGTTTGGGTGTCCAAC | GAGAAGTCTGCGGAAACCAG | 188 |
| chr3:99309246-99311288 | p2 | (AT)21 | CTTTGGCAGCATCACTGAAA | CTGTGGGGCATTTAGGAAAA | 241 |
| chr3:99510959-99513015 | p2 | (AT)28 | TGGATGACGTGGGGATATTT | CTACGTGCAGGTTCTGGGAT | 153 |
| chr3:99510959-99513015 | p2 | (AT)13 | TGGATGACGTGGGGATATTT | CTACGTGCAGGTTCTGGGAT | 153 |
| chr3:99588914-99590934 | p4 | (TGTA)5 | GTTAGCCTCGTCGGCTTATG | CAAACACACCTGTCACGACC | 277 |
| chr3:99674040-99676078 | p2 | (AT)7 | TGGCATCTATGCCCATATCA | GTCCTGTCCCGCCTATATGA | 191 |
| chr3:99674122-99676140 | p2 | (AT)7 | TGGCATCTATGCCCATATCA | GTCCTGTCCCGCCTATATGA | 109 |
| chr3:99682304-99684318 | p2 | (AT)7 | TCAAGAAATCACGGACCCTC | AGGAACCTAGAGCAGCACCA | 130 |
| chr3:99682304-99684318 | p2 | (AT)8 | TCAAGAAATCACGGACCCTC | AGGAACCTAGAGCAGCACCA | 130 |
| chr3:99739919-99741935 | p2 | (AT)8 | TCAATAAAACCTTGCGGGTC | ACAATTAGTTGCTGCCCAGG | 112 |
| chr3:99801595-99803610 | p2 | (AT)7 | CATATTCCCTGTGGGATTCG | CTTGTCGGCTTGCGTATTTT | 133 |
| chr3:99801595-99803610 | p5 | (ATCAT)3 | CATATTCCCTGTGGGATTCG | CTTGTCGGCTTGCGTATTTT | 133 |
| chr3:99887835-99889847 | p2 | (TG)6 | AGTGCCGTTTTGACACTTCC | ATGCACAAATCCGATGTCAA | 198 |
| chr3:99935171-99937201 | p2 | (GA)9 | TGTGACGGCAAAACCAACTA | GGCCAAATGACTGCTTGATT | 138 |
| chr3:99935171-99937201 | p5 | (TTACT)5 | TGTGACGGCAAAACCAACTA | GGCCAAATGACTGCTTGATT | 138 |
| chr3:99960138-99962262 | p2 | (GT)62 | ATGAAGCAACCATCACCCAT | GTACACGGGGCAAGACACTT | 234 |
| chr3:100022934-100024952 | p2 | (TA)6 | CATAAGCACCTCCGGACAAT | CGTCCGTTTTCTTTAGACGC | 184 |
| chr3:100022934-100024952 | p2 | (TA)7 | CATAAGCACCTCCGGACAAT | CGTCCGTTTTCTTTAGACGC | 184 |
| chr3:100022934-100024952 | p6 | (TTATTT)3 | CATAAGCACCTCCGGACAAT | CGTCCGTTTTCTTTAGACGC | 184 |
| chr3:100025256-100027276 | p4 | (GTTT)5 | TCCGTCCCATATTACTTGGC | TTACTTGCCCCTTTCCCTTT | 210 |
| chr3:100065030-100067048 | p2 | (AT)32 | TGGGAGGATTGAAGATCAGG | GTGCTAAGCAATTTCCAGGC | 220 |
| chr3:100065030-100067048 | p3 | (TTG)6 | TGGGAGGATTGAAGATCAGG | GTGCTAAGCAATTTCCAGGC | 220 |
| chr3:100088669-100090684 | p5 | (CAATG)3 | ATGATTTCAAAGGGGACACG | ACTCGGGGTGAACTGTTGTC | 133 |
| chr3:100135231-100137243 | p2 | (AG)6 | CACCTGGGCAGCTTCTATTC | TCTGCAGTCTCGGTAACACG | 263 |
| chr3:100135231-100137243 | p3 | (TTG)5 | CACCTGGGCAGCTTCTATTC | TCTGCAGTCTCGGTAACACG | 263 |
| chr3:100258096-100260111 | p3 | (ATC)5 | AAGGTGTTCCAGCGAGATTG | TGATTCACACTTCTCGACGC | 262 |
| chr3:100258096-100260111 | p5 | (TTTTA)4 | AAGGTGTTCCAGCGAGATTG | TGATTCACACTTCTCGACGC | 262 |
| chr3:100314664-100316676 | p3 | (AAC)5 | TAAGTTCAACCAACCCCGAG | CTTGGAAGATTGCGGATTGT | 200 |
| chr3:100314664-100316676 | p2 | (TA)6 | TAAGTTCAACCAACCCCGAG | CTTGGAAGATTGCGGATTGT | 200 |
| chr3:100368775-100370787 | p2 | (TG)6 | CCCCATTTGGCGAGTAAGTA | GCCCTTCAAAGGCAAACATA | 127 |
| chr3:100465066-100467078 | p2 | (CA)6 | GTTTTCAATGCTCCCTTCCA | AGCCGGAGATTGATTTGCTA | 204 |
| chr3:100533388-100535454 | p3 | (GAA)22 | TCGCGTCCACTTCTTCTTCT | TGTCCAATTCACAACAGGGA | 136 |
| chr3:100549601-100551616 | p5 | (TCTTT)3 | TACACGATGCCCCAGTATCA | GGCAGAATTGGTGGTCCTTA | 263 |
| chr3:100613863-100615878 | p3 | (TTG)5 | ACGCTTTAGGCGGAATACCT | GGACACTGTAGCACCCGATT | 120 |
| chr3:100613863-100615878 | p2 | (TA)6 | ACGCTTTAGGCGGAATACCT | GGACACTGTAGCACCCGATT | 120 |
| chr3:100642911-100644925 | p2 | (AT)20 | TTGCAAAAGGACTCCAAAGG | TGTTCGTCTGTAGTGGCAGC | 215 |

| Marker ID | SSR type | SSR motif | FORWARD PRIMER (5'-3') | REVERSE PRIMER (5'-3') | size (bp) |
|---|---|---|---|---|---|
| chr3:100642911-100644925 | p2 | (TA)7 | TTGCAAAAGGACTCCAAAGG | TGTTCGTCTGTAGTGGCAGC | 215 |
| chr3:100661071-100663083 | p2 | (AT)7 | GACAAGGCTGCCACTACACA | TGTTCCGGAGCCTAGCTTTA | 271 |
| chr3:100661071-100663083 | p2 | (AT)6 | GACAAGGCTGCCACTACACA | TGTTCCGGAGCCTAGCTTTA | 271 |
| chr3:100669512-100671532 | p2 | (AT)7 | ACATAAGCTGGCAAGGCTGT | ATGATTTATGTGTGGGGCGT | 145 |
| chr3:100669512-100671532 | p2 | (AT)9 | ACATAAGCTGGCAAGGCTGT | ATGATTTATGTGTGGGGCGT | 145 |
| chr3:100669512-100671532 | p2 | (AT)10 | ACATAAGCTGGCAAGGCTGT | ATGATTTATGTGTGGGGCGT | 145 |
| chr3:100686740-100688758 | p3 | (TTG)6 | ACGTTTTCGGGTGATCTGAC | CTAGCCCACTGACCTTCCTG | 230 |
| chr3:100927470-100929485 | p3 | (CAT)5 | TTCTGGGTGGACCATAAAGG | GTAGTCCTGCCCCATCTTCA | 241 |
| chr3:100940767-100942779 | p2 | (AT)6 | CCATCAGTCGCCTCATTTCT | GGAAAGGTCCGGTCAAAACT | 160 |
| chr3:100967170-100969185 | p5 | (AAGTA)3 | TTGTTAAGTGCATGCGGAAG | GCAGAACGGGAAAAAGATCA | 224 |
| chr3:101042350-101044364 | p2 | (AT)7 | CATGAATCTCTTCTGGGGGA | AAAAGGTACCCATTGCCACA | 100 |
| chr3:101052373-101054399 | p2 | (TA)16 | TATGCAAGACGACAAGGCTG | CTGTCCCGTCATATTTCGCT | 128 |
| chr3:101052373-101054399 | p2 | (TA)13 | TATGCAAGACGACAAGGCTG | CTGTCCCGTCATATTTCGCT | 128 |
| chr3:101133565-101135583 | p2 | (TA)9 | CGTTTACGAGCACGAACAAA | CCCGGAATGTAAGGAAGTCA | 132 |
| chr3:101169387-101171402 | p3 | (TGT)7 | GGGTGTAACTAGCGATCGGA | ACGATTTGTGGAAGTTTGCC | 292 |
| chr3:101169387-101171402 | p5 | (ATGAT)3 | GGGTGTAACTAGCGATCGGA | ACGATTTGTGGAAGTTTGCC | 292 |
| chr3:101181177-101183192 | p5 | (CATTG)3 | TGGCTTACCCTTGGTGAAAC | TAAGACAACAGGGCCCAATC | 170 |
| chr3:101299489-101301501 | p2 | (TG)6 | AGATAATGTCTTGCCCCGTG | CTGGTGTCCTGCAGGTTTTT | 293 |
| chr3:101299489-101301501 | p2 | (TG)6 | AGATAATGTCTTGCCCCGTG | CTGGTGTCCTGCAGGTTTTT | 293 |
| chr3:101303778-101305794 | p2 | (AT)40 | TATTGTGCGCATTGTGAGGT | TCCATAACCAAAGGTCGCTC | 151 |
| chr3:101303778-101305794 | p2 | (AT)8 | TATTGTGCGCATTGTGAGGT | TCCATAACCAAAGGTCGCTC | 151 |
| chr3:101466815-101468863 | p2 | (AT)24 | ATCATCGGACCACTGGAAAG | GGATCATGGTTGATCCCAAG | 216 |
| chr3:101514707-101516719 | p2 | (AT)6 | CCCAACATATTCCACAACCC | AGCCTTGTCAGCTTGCGTAT | 127 |
| chr3:101548280-101550302 | p2 | (AT)11 | CCCGATAACCAGTCTCCTGA | TCGGCTTGCGTATCATATTG | 274 |
| chr3:101573411-101575431 | p2 | (AT)10 | AGGATAATGGGAAATTCGGG | TCGTAAATCCGTAGCCCATC | 224 |
| chr3:101618094-101620115 | p3 | (ACA)7 | AAGTCTCGGGAACCAAGGAT | TGTGCCCTTCGTATGATTGA | 180 |
| chr3:101638772-101640787 | p3 | (AGC)5 | TCTTTTCTGCCATTTGGACC | CCTCTAAGCAGTACGGCGAC | 125 |
| chr3:101660960-101662975 | p3 | (GAT)5 | TCAATCACCGTGGTCACACT | GCCCGAAAAATTCGATCATA | 296 |
| chr3:101781761-101783788 | p3 | (TGA)9 | CTCCCACACTTATTTCCCGA | TCAACACCCTTCATGACCAA | 255 |
| chr3:101887894-101889906 | p2 | (TC)6 | TTACTACTGCGTGCCACCAC | TGAGGGGAAGATTGTCCAAG | 295 |
| chr3:102184764-102186782 | p2 | (AT)9 | GGCTCAAGCGTGAGGTAGAC | CCCCATTTTAGCCCAAATCT | 144 |
| chr3:102325277-102327295 | p2 | (AT)9 | GGCTCAAGCGTGAGGTAGAC | CCCCATTTTAGCCCAAATCT | 144 |
| chr3:102335135-102337155 | p3 | (TTG)5 | TTGAAAAACATCGTGTCCGA | GTCCAAGCTCGCAGGTCTAC | 150 |
| chr3:102335135-102337155 | p4 | (TATG)5 | TTGAAAAACATCGTGTCCGA | GTCCAAGCTCGCAGGTCTAC | 150 |
| chr3:102342395-102344411 | p3 | (TTG)5 | TTGAAAAACATCGTGTGGGA | GTCCAAGCTCGCAGGTCTAC | 144 |
| chr3:102342395-102344411 | p4 | (TATG)4 | TTGAAAAACATCGTGTGGGA | GTCCAAGCTCGCAGGTCTAC | 144 |

| Marker ID | SSR type | SSR motif | FORWARD PRIMER (5'-3') | REVERSE PRIMER (5'-3') | size (bp) |
|---|---|---|---|---|---|
| chr3:102342395-102344411 | p6 | (GTAAGT)3 | TTGAAAAACATCGTGTGGGA | GTCCAAGCTCGCAGGTCTAC | 144 |
| chr3:102371833-102373851 | p3 | (CAA)6 | CGGAGTGACATAAGGTCGGT | GTCCCAAGGCTACGTTTTCA | 241 |
| chr3:102465843-102467858 | p5 | (ATCAT)3 | TACATGGAAGCCGATAAGGC | AACGACAGGTTTGCTGATCC | 298 |
| chr3:102468635-102470655 | p2 | (CT)6 | GAAGACAACGCAGCGTACAA | GGCCCATTACCCAAGTTTTT | 133 |
| chr3:102468635-102470655 | p2 | (TA)10 | GAAGACAACGCAGCGTACAA | GGCCCATTACCCAAGTTTTT | 133 |
| chr3:102528477-102530492 | p5 | (AAAAG)3 | CCTTTGAACTCGCCCATTTA | GGAGCATTAAGAACACCCGA | 131 |
| chr3:102550191-102552203 | p2 | (AG)6 | ATGTGAGGATTGGGGTCAAA | TTGTTTGCTGGCTTGTGTTC | 201 |
| chr3:102550191-102552203 | p5 | (AATAA)3 | ATGTGAGGATTGGGGTCAAA | TTGTTTGCTGGCTTGTGTTC | 201 |
| chr3:102556701-102558717 | p2 | (AT)8 | AGAGGAAAGCAAGCCAATCA | TATGTTAAAGGGGTGCTCGG | 197 |
| chr3:102574662-102576674 | p5 | (TATTT)3 | GCAGTAGTCTGAGGGTTCGC | TGTATAGGGGAAATGCTGCC | 148 |
| chr3:102574662-102576674 | p2 | (AC)6 | GCAGTAGTCTGAGGGTTCGC | TGTATAGGGGAAATGCTGCC | 148 |
| chr3:102584987-102587015 | p2 | (AT)14 | ATGATTATGGGAGTGCGGAG | AGGGGAGCGGAATTCAGTAT | 121 |
| chr3:102621853-102623883 | p2 | (TA)15 | TAAACAAGGGAAATGCTGCC | AGGTGCGTGCCATACCTTAC | 285 |
| chr3:102623768-102625790 | p2 | (TA)11 | TTGGCATGGCATGAAGAATA | TGTCGACTTTCCCTGAATCC | 268 |
| chr3:102645016-102647040 | p6 | (ATAATG)3 | TCGTTGTTGTTGTTGCTGGT | TCCTTAGCCACTCAACGCTT | 132 |
| chr3:102645016-102647040 | p2 | (TA)9 | TCGTTGTTGTTGTTGCTGGT | TCCTTAGCCACTCAACGCTT | 132 |
| chr3:102645016-102647040 | p2 | (TA)12 | TCGTTGTTGTTGTTGCTGGT | TCCTTAGCCACTCAACGCTT | 132 |
| chr3:102755825-102757843 | p2 | (AG)9 | TTCCTTCGTTGGGATTCTTG | CCAGAAACAGAGGGTGGAAA | 101 |
| chr3:102778576-102780596 | p2 | (TA)10 | AGTAAAATCTCGCACCCCCT | GGTCTCGGATTGTTTTGGAA | 254 |
| chr3:102802245-102804257 | p2 | (TC)6 | TCAGAAGGACGGACCGTAAC | TTGCCTTAAATGCTCCAACC | 189 |
| chr3:102802245-102804257 | p3 | (ATT)6 | TCAGAAGGACGGACCGTAAC | TTGCCTTAAATGCTCCAACC | 189 |
| chr3:102930348-102932362 | p2 | (AT)7 | GGAAACCGAACCAAATGATG | CGTAGTTTGCCTCTTGCTCC | 106 |
| chr3:102959380-102961395 | p5 | (CCACA)3 | GTCCGTCTGAGGCTCGTATC | GATGCTGCCCAAATTTCTGT | 252 |
| chr3:103026412-103028427 | p5 | (GAGAG)3 | CGGGTAGCCATTTCCTTGTA | CTTTTTCCTTCACGAGCCTG | 139 |
| chr3:103026412-103028427 | p3 | (TTG)5 | CGGGTAGCCATTTCCTTGTA | CTTTTTCCTTCACGAGCCTG | 139 |
| chr3:103070874-103072889 | p5 | (GAAAT)3 | CATCTACCTCAACGGGCCTA | CTAGCATTGGAGGAGAACGC | 148 |
| chr3:103125479-103127497 | p2 | (AT)9 | GACATATGGATGGGACAGGG | TCATGCCCACAGGTAGACAA | 126 |
| chr3:103125479-103127497 | p3 | (TCA)6 | GACATATGGATGGGACAGGG | TCATGCCCACAGGTAGACAA | 126 |
| chr3:103141970-103143988 | p2 | (AT)15 | CAAATTCATGAACACTGCCG | GTAGAAGGAGTGGTGCTCGG | 186 |
| chr3:103141970-103143988 | p2 | (AT)9 | CAAATTCATGAACACTGCCG | GTAGAAGGAGTGGTGCTCGG | 186 |
| chr3:103187194-103189209 | p5 | (TTCAT)3 | TCATGCTTTTGGTCGTGTGT | AAACATCCTTCCCCCTTTTG | 260 |
| chr3:103194110-103196128 | p3 | (TCA)6 | AATGAGTCAATGGCACCTCC | TGTTCATGCCCACAGGTAGA | 238 |
| chr3:103227761-103229791 | p2 | (AT)15 | TCAACAATGCCTCACAGAGC | GGTTGGGTCGTGACAAAAGT | 177 |
| chr3:103228834-103230854 | p2 | (CA)10 | CCGGAACTCGGTGAAAGATA | GTGCAAACGACCTCAGGAAT | 147 |
| chr3:103241987-103244005 | p3 | (TGT)6 | GGGCTACTTTCGGACATGAA | TGGCACCTACAAAGTGGACA | 203 |
| chr3:103255480-103257498 | p3 | (AAG)6 | GGGACGAAATGACGAAAAGA | AAATGTCACGTTTTAGGGCG | 233 |

| Marker ID | SSR type | SSR motif | FORWARD PRIMER (5'-3') | REVERSE PRIMER (5'-3') | size (bp) |
|---|---|---|---|---|---|
| chr3:103353208-103355229 | p3 | (AAG)7 | CCAGCAAGGAAAACCATCAT | AGGAAGAGCGTCCAGCATTA | 253 |
| chr3:103353208-103355229 | p5 | (AAAAT)3 | CCAGCAAGGAAAACCATCAT | AGGAAGAGCGTCCAGCATTA | 253 |
| chr3:103355448-103357466 | p6 | (CATATA)4 | CCCGACCCTTACTTAGAGCC | AATGTCAGTTTTGGAAGCGG | 283 |
| chr3:103355467-103357479 | p6 | (CATATA)4 | CCCGACCCTTACTTAGAGCC | AATGTCAGTTTTGGAAGCGG | 264 |
| chr3:103355481-103357497 | p6 | (CATATA)4 | CCCGACCCTTACTTAGAGCC | AATGTCAGTTTTGGAAGCGG | 250 |
| chr3:103439839-103441851 | p2 | (TA)6 | TCCCATAGCAATTTTGGGAA | TTTCACCAGCATCACTGCTC | 177 |
| chr3:103655516-103657528 | p2 | (AT)6 | ATGTCATAGCCACAGAGGGC | GCTGAAGAACGACATCGACA | 136 |
| chr3:103655516-103657528 | p2 | (AT)18 | ATGTCATAGCCACAGAGGGC | GCTGAAGAACGACATCGACA | 136 |
| chr3:103717926-103719944 | p2 | (AT)9 | CATAAGCCGACGAGGCTAAC | GAGCTCCATCTCATTCGGAG | 236 |
| chr3:103875411-103877423 | p2 | (TA)6 | ACGACCAAGGTATGTGAGGC | GTAAGGTCGTGATCCCCTGA | 135 |
| chr3:103922230-103924245 | p3 | (TTC)5 | AGACCATTTTCCACACCTGC | CGGACATCGTATGAGGGAGT | 182 |
| chr3:103949367-103951403 | p2 | (AT)18 | GCTGCCGAAGACTCCTCTTA | ATTTCACTAAGTCGTGCGGG | 131 |
| chr3:104048297-104050312 | p5 | (TTTTG)3 | AATTCGCCTTGGAACCTCTT | CATGAGCTTGCGGAGTATCA | 249 |
| chr3:104114241-104116256 | p3 | (CAA)5 | CCACCTTTCTCTCATCCCAA | AAGGTGCAATTCACCCTGTC | 240 |
| chr3:104128638-104130652 | p2 | (TG)7 | TGGAACCTAGCACTGCCTTT | AAGTAAAATGGACCGGAGGG | 293 |
| chr3:104178150-104180162 | p2 | (TA)6 | TTTTGAACCTCGGAGCATCT | TTTGGAGGGTTTCCACTTTG | 147 |
| chr3:104284027-104286059 | p4 | (CATT)4 | TCCTGCTGTAGGGAAGGCTA | CCTAGTTTGGCTGGTTGCAT | 132 |
| chr3:104312721-104314742 | p2 | (TA)6 | GAACACCACTACCCCCAATG | GGCTTTTGTGTGTCACATGG | 217 |
| chr3:104312721-104314742 | p3 | (GAT)7 | GAACACCACTACCCCCAATG | GGCTTTTGTGTGTCACATGG | 217 |
| chr3:104376385-104378397 | p2 | (TA)6 | TATGAGCCTCTGGGACAACC | TAGTCGGGTCCCACTTATGC | 279 |
| chr3:104376404-104378440 | p2 | (TA)6 | TATGAGCCTCTGGGACAACC | TAGTCGGGTCCCACTTATGC | 260 |
| chr3:104404376-104406392 | p4 | (TTAA)4 | AGGGTATCCTTGCTCCGTCT | TGGACCTTGTTGCAAATTGA | 232 |
| chr3:104404376-104406392 | p4 | (TTAA)4 | AGGGTATCCTTGCTCCGTCT | TGGACCTTGTTGCAAATTGA | 232 |
| chr3:104448857-104450875 | p2 | (AT)22 | AACCACTGAGCCATCCTCAC | TAATGCATCCCCAACACTCA | 258 |
| chr3:104448871-104450886 | p2 | (AT)22 | AACCACTGAGCCATCCTCAC | TAATGCATCCCCAACACTCA | 244 |
| chr3:104612236-104614264 | p2 | (TA)14 | CTCCACCACCAGGTCCTCTA | GGAGGCTGAGTATCAGGCAG | 133 |
| chr3:104757278-104759290 | p2 | (TA)6 | GCTCGGCCTAATTAAGGGTC | GGACACGACTTTGCAGACAA | 266 |
| chr3:104865947-104867971 | p3 | (TTG)8 | TTGGGGTGTCACTGTTTTGA | TTGGGGGTTGTAGCACTTTC | 262 |
| chr3:104868182-104870197 | p5 | (TAATA)3 | CCGAGACCTCCCGTACATAA | TTAAAGTTCCGTCAAACCCG | 193 |
| chr3:104906667-104908691 | p3 | (TTG)8 | TTGGGGTGTCACTGTTTTGA | TTGGGGGTTGTAGCACTTTC | 262 |
| chr3:104908902-104910917 | p5 | (TAATA)3 | CCGAGACCTCCCGTACATAA | TTAAAGTTCCGTCAAACCCG | 193 |
| chr3:104997269-104999281 | p2 | (AT)9 | TGCTCATTAGGATCTGCACG | CGGCTAACCAACTCTGGAAG | 209 |
| chr3:104997269-104999281 | p2 | (TA)6 | TGCTCATTAGGATCTGCACG | CGGCTAACCAACTCTGGAAG | 209 |
| chr3:104997269-104999281 | p2 | (AT)6 | TGCTCATTAGGATCTGCACG | CGGCTAACCAACTCTGGAAG | 209 |
| chr3:105014070-105016096 | p2 | (TA)13 | TCTCGATCAGAAGCAGAGCA | TGTAATAGCCTGCCAAACCC | 190 |
| chr3:105114522-105116540 | p6 | (AATAAA)3 | ATGCATTTCGTTGAAGGACC | ATCAGTGGAGATGGAGGTGG | 150 |

| Marker ID | SSR type | SSR motif | FORWARD PRIMER (5'-3') | REVERSE PRIMER (5'-3') | size (bp) |
|---|---|---|---|---|---|
| chr3:105133538-105135553 | p5 | (TTCTG)3 | TATATTGGTCAACGCCAGCA | GGAGGCTGAAACCATGTGAT | 190 |
| chr3:105138115-105140130 | p5 | (ATATA)3 | GCACGACATATCAAAAGCCA | CGAATTCCCCTCAATTCTCA | 124 |
| chr3:105171107-105173121 | p2 | (AT)7 | TTGAGCTTCCTGGACTTCCT | AGGTATAGCCAATCCCCACC | 253 |
| chr3:105171107-105173121 | p2 | (AT)7 | TTGAGCTTCCTGGACTTCCT | AGGTATAGCCAATCCCCACC | 253 |
| chr3:105171429-105173441 | p2 | (AT)7 | GGATTGGCTATACCTGGCAA | ACATGTTTGGCACTTGGGA | 141 |
| chr3:105171429-105173441 | p2 | (AT)7 | GGATTGGCTATACCTGGCAA | ACATGTTTGGCACTTGGGA | 141 |
| chr3:105171444-105173470 | p2 | (AT)7 | GGATTGGCTATACCTGGCAA | ACATGTTTGGCACTTGGGA | 126 |
| chr3:105171444-105173470 | p2 | (AT)7 | GGATTGGCTATACCTGGCAA | ACATGTTTGGCACTTGGGA | 126 |
| chr3:105374427-105376441 | p2 | (TC)7 | GGAGTGGAGGAATCTGTGGA | AGGTCGACTCTTGCCTTGAA | 162 |
| chr3:105380746-105382761 | p3 | (CAC)5 | AAACCATCAGGACCACAAGC | ATAAGGAGCAGCTTCCAGCA | 187 |
| chr3:105428090-105430130 | p2 | (TC)20 | TTTAGCCCGATTTTTCATCG | GCGAGCAAAGTCTAAGTGGG | 168 |
| chr3:105430556-105432576 | p5 | (AAAAT)4 | TTGGGCATCAGACGTATCAA | ACTTCATGGGTTTCTCCGTG | 235 |
| chr3:105467151-105469169 | p2 | (CT)9 | TCATGAGGTCGGAAATAGCC | CCCAAACTAATCCCACTGGA | 120 |
| chr3:105495386-105497398 | p2 | (AT)6 | ATCGTGAGACCATGCTACCC | TGTTCATGCCCACAGGTAGA | 221 |
| chr3:105566652-105568667 | p5 | (GCATT)3 | GTTGCGGTTCGACTATCCAT | CCCGAAGATTCACATGCTTT | 253 |
| chr3:105957354-105959368 | p2 | (AT)7 | TCGGTGGTAGAGGCATTTTC | CTTGGAAACTTTGGGGAACA | 220 |
| chr3:106076317-106078353 | p2 | (AT)18 | GGTTTGTTGCGTTGGAATCT | TTCTGAGGCTTGGAAGGAAA | 198 |
| chr3:106081105-106083120 | p3 | (GAT)5 | TTTCTTTGGCCCATACCAAG | TTCCTCCCTTCCTCCAAGTT | 270 |
| chr3:106081105-106083120 | p2 | (TA)15 | TTTCTTTGGCCCATACCAAG | TTCCTCCCTTCCTCCAAGTT | 270 |
| chr3:106097688-106099712 | p6 | (CATATA)4 | CCCAAAGCACTGCCTATGTT | GAATGCGAGAAAAGAGGTGC | 175 |
| chr3:106108978-106110993 | p6 | (CATATA)6 | ACGCGTATATAATGCTCGGG | TAAATCGCAGGTTACCGACC | 149 |
| chr3:106108978-106110993 | p3 | (CAA)5 | ACGCGTATATAATGCTCGGG | TAAATCGCAGGTTACCGACC | 149 |
| chr3:106199737-106201755 | p2 | (TA)9 | TGAAAATGTTGAAGTGCCCA | CCTCTTTCAATCCCCCATTT | 176 |
| chr3:106201109-106203137 | p2 | (AT)14 | ACGTTCTCCTTCCAAGCTCA | TGCCCAATTTTCGTTAGCTC | 270 |
| chr3:106316747-106318765 | p3 | (CAA)6 | CGGAATTCATACATGGGACC | GTGCTTTCCCAAGCAGGATA | 102 |
| chr3:106402273-106404339 | p3 | (TTA)22 | GCACGCACTCTCATTTTTCA | AGACGGGACCTCGAATCTTT | 167 |
| chr3:106402273-106404339 | p4 | (CTTT)4 | GCACGCACTCTCATTTTTCA | AGACGGGACCTCGAATCTTT | 167 |
| chr3:106439420-106441438 | p2 | (AT)7 | GAACGTAGCGTCAGCATGAA | ATGCATGGTATCCGCCTTAC | 292 |
| chr3:106439420-106441438 | p3 | (CAT)6 | GAACGTAGCGTCAGCATGAA | ATGCATGGTATCCGCCTTAC | 292 |
| chr3:106460228-106462243 | p5 | (CATGT)3 | TTTCTGATCGTGGTGCTCTG | TCTGTCTGGGGATGAAAAGC | 136 |
| chr3:106493610-106495672 | p2 | (AT)31 | ACGTTCTCCTTCCAAGCTCA | CAAAGGAAATGTTGCCCAAT | 244 |
| chr3:106499952-106501986 | p2 | (TA)17 | TTCTGGGAAGCTGTTGCTTT | GTGTCTACGAGCCGTGTCAA | 161 |
| chr3:106614057-106616077 | p2 | (TA)10 | AAGAGGCCAACTCCAAGGAT | CACAGGCATAGCGGAAAAAT | 167 |
| chr3:106621461-106623481 | p2 | (TA)21 | AACCTCTCGTGGGGAAAGAT | ACATCTCGGTAGCACAACCC | 126 |
| chr3:106621461-106623481 | p4 | (CTTT)5 | AACCTCTCGTGGGGAAAGAT | ACATCTCGGTAGCACAACCC | 126 |
| chr3:106660579-106662617 | p6 | (CATATA)3 | GGATGGATTTTGTCACACCC | GTCTTGAGAGCGGCTACACC | 299 |

| Marker ID | SSR type | SSR motif | FORWARD PRIMER (5'-3') | REVERSE PRIMER (5'-3') | size (bp) |
|---|---|---|---|---|---|
| chr3:106660651-106662673 | p6 | (CATATA)3 | GGGATGGATTTTGTCACACC | GTCTTGAGAGCGGCTACACC | 226 |
| chr3:106660969-106662987 | p6 | (CATATA)3 | TGACACTTCGGCTGGTGTAG | ACATTGCTCGTACTGACCCC | 134 |
| chr3:106674641-106676657 | p3 | (TTA)18 | GGAGATGGAGATCTCTTGCG | GCACCAGAGGGAGAGAACAG | 227 |
| chr3:106674641-106676657 | p4 | (CTTT)4 | GGAGATGGAGATCTCTTGCG | GCACCAGAGGGAGAGAACAG | 227 |
| chr3:106719129-106721169 | p4 | (CTTT)4 | ACACTTTGATGAGCTGGGCT | TTCTCACCAGCACCACTCTG | 217 |
| chr3:106767745-106769763 | p6 | (ATGATA)3 | ATATGAGTGTCCAGCTCGGG | ACCCAAATGTGCCAAGTCTC | 179 |
| chr3:106996971-106998992 | p3 | (TTG)7 | GTTGCTTGTTTGAATCCGCT | ACTGGGAGAAGCAAAGCAAA | 167 |
| chr3:107023762-107025864 | p3 | (TAT)5 | ACAATTTCCATTCCCAACGA | CTTTTGGAGGCATGCTCATT | 124 |
| chr3:107045387-107047423 | p2 | (TA)18 | CGGTTCCAGCAGAAGTAAGG | GGCTATGTAGGCACGGTGTT | 229 |
| chr3:107056454-107058514 | p3 | (TTA)20 | AGGTCGAGCCTAGAACCCAT | CTCGAGGACATGTGAAAGCA | 218 |
| chr3:107070193-107072205 | p2 | (AG)6 | CAGAGGACAGAGCAAGAGGG | AGTCTCGAACTCTGACCCGA | 132 |
| chr3:107070193-107072205 | p2 | (GT)6 | CAGAGGACAGAGCAAGAGGG | AGTCTCGAACTCTGACCCGA | 132 |
| chr3:107078678-107080690 | p2 | (AG)6 | CAGAGGACAGAGCAAGAGGG | GAGTCTCGAACTCTGACCCG | 118 |
| chr3:107078678-107080690 | p2 | (GT)6 | CAGAGGACAGAGCAAGAGGG | GAGTCTCGAACTCTGACCCG | 118 |
| chr3:107084340-107086352 | p2 | (AG)6 | TCTACGCCCCTGATCTTGAC | GATTCGTAATCGGAGGCAAA | 164 |
| chr3:107084340-107086352 | p2 | (GT)6 | TCTACGCCCCTGATCTTGAC | GATTCGTAATCGGAGGCAAA | 164 |
| chr3:107141212-107143228 | p2 | (TA)6 | AAAAGCAACCGTGATTCCAC | GCGGAAACAACCTCTCTCAC | 133 |
| chr3:107195538-107197556 | p5 | (AATGC)3 | TACTCCGGACGGAAGTATCG | GTGGTAGCGGTTGTTGAGGT | 233 |
| chr3:107195538-107197556 | p6 | (CCTTAG)3 | TACTCCGGACGGAAGTATCG | GTGGTAGCGGTTGTTGAGGT | 233 |
| chr3:107251225-107253259 | p2 | (TA)17 | AGGTCGTCAAACGTCATTCC | CGTGTTACCGATCGTCCTTT | 291 |
| chr3:107406936-107408954 | p2 | (AT)9 | TGGACCTCTTGGAAATTGGA | GTGGGCATAGTTTGGGCTAA | 206 |
| chr3:107487470-107489486 | p2 | (AT)8 | AGTAAGGTGCGCGATGATTC | ATGCCAAAAGATTGACCTCG | 136 |
| chr3:107500293-107502311 | p6 | (GAATCT)3 | TACACGCGCTGCAAGTTTAG | CACTACTCCGGCACTCCATT | 205 |
| chr3:107500293-107502311 | p5 | (CATTT)3 | TACACGCGCTGCAAGTTTAG | CACTACTCCGGCACTCCATT | 205 |
| chr3:107525668-107527684 | p2 | (TA)8 | TATAACAAAAGGGCAAGCCG | TGCTCTCGCCACTCTCACTA | 266 |
| chr3:107525668-107527684 | p2 | (TA)7 | TATAACAAAAGGGCAAGCCG | TGCTCTCGCCACTCTCACTA | 266 |
| chr3:107545223-107547241 | p2 | (AT)6 | ACTCCCAAATGGGGATAAGG | AATTTGTCCTCTATGGCCCC | 148 |
| chr3:107565782-107567804 | p2 | (AT)11 | GGTGTTGTAAAGTGCGCTGA | CCAACGAAACGTGGGATAAG | 219 |
| chr3:107586729-107588753 | p2 | (TA)12 | GCCTTTAAACGAAAATGGCA | TAATCATGCTTCGAGACCCC | 106 |
| chr3:107775995-107778007 | p2 | (TA)6 | TGTGCTTGTGGCACTTTTTC | ATCTCTTCCTCTCCGCCTTC | 278 |
| chr3:107794667-107796687 | p5 | (AAACC)4 | TCCTCCTTCGAATCTCCAAA | ACACCTTTGAACCAGGATGG | 212 |
| chr3:107892412-107894442 | p2 | (AT)11 | TTTGATGGTGTTTTGGGTCA | ACGCGGTTGAATCAAGAAGT | 203 |
| chr3:107892412-107894442 | p2 | (AT)15 | TTTGATGGTGTTTTGGGTCA | ACGCGGTTGAATCAAGAAGT | 203 |
| chr3:107949886-107951958 | p3 | (ATT)24 | GATGTTGTTCATGACGTCGG | ATCTGGACGTGTGCATACCA | 127 |
| chr3:107953154-107955175 | p3 | (ATT)7 | CTCCGAATGCGGGTATCTTA | TCCACGAGTTGGTGTGTGAT | 138 |
| chr3:107985410-107987425 | p3 | (ATA)5 | TAGTGGTTTGTGGAGGGGAG | TGTTATTATTTCCGCCGTCC | 149 |

| Marker ID | SSR type | SSR motif | FORWARD PRIMER (5'-3') | REVERSE PRIMER (5'-3') | size (bp) |
|---|---|---|---|---|---|
| chr3:108023033-108025047 | p2 | (AG)7 | TCCGAATCCAATCTTGAAGG | AAATGGAGCAGCGGATTCTA | 169 |
| chr3:108023033-108025047 | p3 | (TTG)6 | TCCGAATCCAATCTTGAAGG | AAATGGAGCAGCGGATTCTA | 169 |
| chr3:108023033-108025047 | p4 | (TGTA)5 | TCCGAATCCAATCTTGAAGG | AAATGGAGCAGCGGATTCTA | 169 |
| chr3:108060077-108062095 | p3 | (ACA)5 | CATCAGGTTCCTTCCCTTGA | GCCAGGAAAAACAAAAGCAA | 259 |
| chr3:108060077-108062095 | p3 | (ACA)6 | CATCAGGTTCCTTCCCTTGA | GCCAGGAAAAACAAAAGCAA | 259 |
| chr3:108060077-108062095 | p3 | (AAC)5 | CATCAGGTTCCTTCCCTTGA | GCCAGGAAAAACAAAAGCAA | 259 |
| chr3:108060077-108062095 | p3 | (AAC)5 | CATCAGGTTCCTTCCCTTGA | GCCAGGAAAAACAAAAGCAA | 259 |
| chr3:108178158-108180179 | p3 | (ATA)8 | TAGTCTACGCGTTTTGTGCG | CAAGCAATTTAGCGAGGGAT | 109 |
| chr3:108179165-108181181 | p3 | (ATA)8 | GAGCACTTTCAGGCATGTGA | GCTGCAGGAAAAATGTGAAA | 282 |
| chr3:108179165-108181181 | p4 | (TTTA)4 | GAGCACTTTCAGGCATGTGA | GCTGCAGGAAAAATGTGAAA | 282 |
| chr3:108179165-108181181 | p3 | (TTA)14 | GAGCACTTTCAGGCATGTGA | GCTGCAGGAAAAATGTGAAA | 282 |
| chr3:108179291-108181336 | p3 | (ATA)8 | GAGCACTTTCAGGCATGTGA | GCTGCAGGAAAAATGTGAAA | 156 |
| chr3:108179291-108181336 | p4 | (TTTA)4 | GAGCACTTTCAGGCATGTGA | GCTGCAGGAAAAATGTGAAA | 156 |
| chr3:108179291-108181336 | p3 | (TTA)14 | GAGCACTTTCAGGCATGTGA | GCTGCAGGAAAAATGTGAAA | 156 |
| chr3:108179337-108181370 | p3 | (ATA)8 | GAGCACTTTCAGGCATGTGA | GCTGCAGGAAAAATGTGAAA | 110 |
| chr3:108179337-108181370 | p4 | (TTTA)4 | GAGCACTTTCAGGCATGTGA | GCTGCAGGAAAAATGTGAAA | 110 |
| chr3:108179337-108181370 | p3 | (TTA)14 | GAGCACTTTCAGGCATGTGA | GCTGCAGGAAAAATGTGAAA | 110 |
| chr3:108264241-108266292 | p3 | (TAT)17 | AGAAATTGCACAAACTGCCC | AAACAATGAAGGGCAACCAG | 112 |
| chr3:108386252-108388264 | p2 | (AT)6 | CGTTCAAAGAGAGCCACCTC | GTGGAGGCATCAGTTTTGGT | 273 |
| chr3:108386252-108388264 | p2 | (AT)6 | CGTTCAAAGAGAGCCACCTC | GTGGAGGCATCAGTTTTGGT | 273 |
| chr3:108386252-108388264 | p2 | (TA)9 | CGTTCAAAGAGAGCCACCTC | GTGGAGGCATCAGTTTTGGT | 273 |
| chr3:108386365-108388377 | p2 | (AT)6 | CGTTCAAAGAGAGCCACCTC | GTGGAGGCATCAGTTTTGGT | 160 |
| chr3:108386365-108388377 | p2 | (AT)6 | CGTTCAAAGAGAGCCACCTC | GTGGAGGCATCAGTTTTGGT | 160 |
| chr3:108386365-108388377 | p2 | (TA)9 | CGTTCAAAGAGAGCCACCTC | GTGGAGGCATCAGTTTTGGT | 160 |
| chr3:108457765-108459790 | p5 | (GCATT)5 | AAGGCAAGGCAAAAGAGTGA | AAATCACAACACAACGGGGT | 281 |
| chr3:108475746-108477762 | p4 | (CAAA)4 | GGTTATTTGGGGTGTGTTCG | TATTTTCCGCTACCCAAACG | 217 |
| chr3:108569599-108571650 | p3 | (TTC)17 | CATCATGTCAGGTGAATCCG | GTCACTTCATCCGCCTCTGT | 254 |
| chr3:108573480-108575495 | p5 | (AAACC)3 | CCTAGCTGGGTCGTGACAGT | CGAACCATGAACACCCCTAC | 215 |
| chr3:108628324-108630374 | p2 | (TA)25 | TGATATTGACTATGGCCGCA | TATGCATGCTCACGCCTATC | 284 |
| chr3:108678126-108680156 | p2 | (AG)7 | TGCAAATGGACAAAACCTCA | CTTCTCTTCAGAACACCCGC | 267 |
| chr3:108678126-108680156 | p3 | (ATA)10 | TGCAAATGGACAAAACCTCA | CTTCTCTTCAGAACACCCGC | 267 |
| chr3:108695432-108697444 | p2 | (AT)6 | GGCCGTATAATCCCCAGTTT | GTTTGCCAAGAGAAGGCAAG | 112 |
| chr3:108695432-108697444 | p5 | (AATGC)4 | GGCCGTATAATCCCCAGTTT | GTTTGCCAAGAGAAGGCAAG | 112 |
| chr3:108850810-108852830 | p5 | (CTAAG)4 | TTCAGGTTTTACCCAGTCCG | ACTTGCCAGGAGTTGAATGG | 226 |
| chr3:108917394-108919466 | p2 | (TA)10 | ATGGCCTGAGCTGTGCTTAT | ATTTGCCCAACTGGTCGTAG | 264 |
| chr3:108924177-108926189 | p2 | (AT)6 | GGCACCGCTCGTACTATCAT | GAAGAGCACTCCATCTTCCG | 251 |

| Marker ID | SSR type | SSR motif | FORWARD PRIMER (5'-3') | REVERSE PRIMER (5'-3') | size (bp) |
|---|---|---|---|---|---|
| chr3:108970229-108972259 | p2 | (AT)15 | ATCCTAGGTCATTGCAACCG | TCTCACGTGTTTTTGGGTGA | 147 |
| chr3:108970229-108972259 | p2 | (TA)25 | ATCCTAGGTCATTGCAACCG | TCTCACGTGTTTTTGGGTGA | 147 |
| chr3:109079439-109081454 | p5 | (CCCAC)3 | TCCTGCAGATTTGTTCACCA | CGCTTAGAGAGCTGCCTCAT | 182 |
| chr3:109079439-109081454 | p5 | (AAAAT)3 | TCCTGCAGATTTGTTCACCA | CGCTTAGAGAGCTGCCTCAT | 182 |
| chr3:109093921-109095948 | p3 | (TTA)9 | CCCATTACATCTTCTGCGGT | TGTGAGCAGATGGTCTCAGG | 297 |
| chr3:109148400-109150415 | p5 | (AATGC)3 | GACAAGATGAGGGAAGCGAG | CATGGGAAGTTGGATCGAGT | 111 |
| chr3:109193044-109195059 | p3 | (TGT)5 | GCGGAGATGAGGATGCTTAG | CTCCGCAACTCTTTACTGGC | 297 |
| chr3:109260017-109262032 | p3 | (TAA)5 | AGAGGAGAGGGGACAAAAGC | GTTTCCCCTTCATTTCGTCA | 189 |
| chr3:109260017-109262032 | p3 | (ATA)5 | AGAGGAGAGGGGACAAAAGC | GTTTCCCCTTCATTTCGTCA | 189 |
| chr3:109291918-109293930 | p2 | (AT)6 | GTGTATGGACACCCCAGCTT | GGGCGTAACATTTGCCTAAA | 138 |
| chr3:109291918-109293930 | p2 | (AT)9 | GTGTATGGACACCCCAGCTT | GGGCGTAACATTTGCCTAAA | 138 |
| chr3:109297780-109299792 | p2 | (AT)6 | AGATGGGGCTACCCAAAATC | CGATGTCCCAGTTTCCAGTT | 260 |
| chr3:109394048-109396064 | p2 | (AG)8 | TTTCTATGGGGGAAGCCTCT | TGGATTGTTTGATTGCAGGA | 247 |
| chr3:109399424-109401439 | p5 | (ATTGC)3 | CGAAAGGCAAGACAAAGGAG | TGGCTAGTCCCAAACACACA | 206 |
| chr3:109417601-109419616 | p3 | (ATC)5 | GATCCGCATGGCTTACTCAT | CTGTCCCGTTCTTGGATGAT | 188 |
| chr3:109723679-109725693 | p2 | (TA)7 | TGACTTGCATGAGGACAAGC | CTTTGCTTTTTGCTTTTGGG | 182 |
| chr3:109731485-109733501 | p2 | (TA)8 | ATGCTCATATCCCATTCCCA | TGTTTGATGTGGGCTGTTGT | 297 |
| chr3:109733226-109735242 | p4 | (GAAA)4 | CGGGGTCAACAATGACTCTT | CAAATTCGGGAAAAACGAAA | 261 |
| chr3:109741718-109743736 | p2 | (AT)9 | CGGCAACCCTAGCTTAACTG | GTTGTGAGATTTGTGCGTGG | 156 |
| chr3:109744366-109746381 | p5 | (TGAAA)3 | CCTTGGATCATTTTGTGCCT | TCCACATCACGACTTTTCCA | 131 |
| chr3:109749721-109751737 | p4 | (TCTT)4 | CGCTTACAGACTCCGCTCTT | GGTGGAGTCAACGAAGGTGT | 147 |
| chr3:109775491-109777507 | p2 | (TG)8 | GCAAAGATAAGCACCCAAGC | CCGGTTCGTGTTGGTAAGTT | 190 |
| chr3:109781329-109783345 | p2 | (TG)8 | GCAAAGATAAGCACCCAAGC | CCGGTTCGTGTTGGTAAGTT | 190 |
| chr3:109824100-109826124 | p4 | (TTTC)6 | CGGTAGCATCTCCCCTGTAA | TTTGAATGTTGTGCGGGTTA | 275 |
| chr3:109861460-109863478 | p6 | (TTTTTG)3 | TCACTGATTTCCAACCCACA | AATTAATCCAGCGACCATGC | 124 |
| chr3:109877366-109879384 | p2 | (TG)9 | GGTTAGGGCAACCAGCTGTA | TTGTTGCTGCTTTCCTTGTG | 136 |
| chr3:109879211-109881226 | p5 | (TTTGT)3 | CTCAAAATCAGGGGGTGAGA | TCAGGGAATGAAGAAGGTGG | 192 |
| chr3:109929428-109931446 | p2 | (TG)9 | ATTTAGACAAAGCATGCCCG | CATCCGGATTTGGTTGTACC | 147 |
| chr3:109982476-109984491 | p5 | (AAGAA)3 | CAAAGACGGTGGCAAAAGTT | CCTTTGCCTTTGACATTGGT | 291 |
| chr3:109998656-110000707 | p3 | (ATT)17 | TTTCCTGGGAACTGAAATCG | TTTCGCCCTTTGTTGCTAGT | 196 |
| chr3:109998656-110000707 | p2 | (CT)16 | TTTCCTGGGAACTGAAATCG | TTTCGCCCTTTGTTGCTAGT | 196 |
| chr3:109999182-110001214 | p3 | (ATT)17 | AAGATGAACCAGCAACCGAC | ACATCCTCGATGTGGTAGGC | 157 |
| chr3:109999182-110001214 | p2 | (CT)16 | AAGATGAACCAGCAACCGAC | ACATCCTCGATGTGGTAGGC | 157 |
| chr3:110002375-110004393 | p2 | (AC)9 | AGTGGCGGTGTAGCAGAACT | CTTCCAAAGAAAGCGACAGC | 229 |
| chr3:110002375-110004393 | p2 | (AT)7 | AGTGGCGGTGTAGCAGAACT | CTTCCAAAGAAAGCGACAGC | 229 |
| chr3:110002375-110004393 | p2 | (AT)11 | AGTGGCGGTGTAGCAGAACT | CTTCCAAAGAAAGCGACAGC | 229 |

| Marker ID | SSR type | SSR motif | FORWARD PRIMER (5'-3') | REVERSE PRIMER (5'-3') | size (bp) |
|---|---|---|---|---|---|
| chr3:110002528-110004542 | p2 | (AC)9 | AGCTGTCGCTTTCTTTGGAA | GGATGGGTCGTGACAAAAGT | 195 |
| chr3:110002528-110004542 | p2 | (AT)7 | AGCTGTCGCTTTCTTTGGAA | GGATGGGTCGTGACAAAAGT | 195 |
| chr3:110002528-110004542 | p2 | (AT)11 | AGCTGTCGCTTTCTTTGGAA | GGATGGGTCGTGACAAAAGT | 195 |
| chr3:110054538-110056566 | p2 | (AT)14 | ATAACCACGACTGCCACCTC | AAGCGAGTTCAAAGTTCCCA | 228 |
| chr3:110078681-110080697 | p2 | (TA)8 | CCATTAGTGGGAGCCAATGA | AAATGACATGGCGGAATAGC | 182 |
| chr3:110135814-110137829 | p3 | (TAC)5 | AGCCCTTTGGATTGGCTTAT | TTAGAGCCTGTTTGGATGGG | 173 |
| chr3:110157900-110159915 | p2 | (TG)7 | GAATTCCTCGTTGCCACAAT | AAAGCTAAGGAGGCCACACA | 280 |
| chr3:110157900-110159915 | p3 | (TGT)5 | GAATTCCTCGTTGCCACAAT | AAAGCTAAGGAGGCCACACA | 280 |
| chr3:110193770-110195854 | p3 | (AAT)28 | CCCATTTTCGTATTGCTGCT | CGACCCAATTTCCTTCAAAA | 230 |
| chr3:110259611-110261653 | p2 | (AG)6 | CGAATTGTCACAACCACTGC | AGAGCCTGTTTAGCCGTGAA | 257 |
| chr3:110259611-110261653 | p3 | (ATA)14 | CGAATTGTCACAACCACTGC | AGAGCCTGTTTAGCCGTGAA | 257 |
| chr3:110305429-110307495 | p2 | (TA)8 | TCTTGTTTGTGCAGTGAGCC | AAGTTGTTGCACGTTGTTGC | 227 |
| chr3:110305429-110307495 | p2 | (AT)33 | TCTTGTTTGTGCAGTGAGCC | AAGTTGTTGCACGTTGTTGC | 227 |
| chr3:110333217-110335229 | p2 | (TG)6 | GGTTTCGTATAGGTTGCGGA | CGAGCTTAGCATTTTGGCTC | 227 |
| chr3:110445333-110447363 | p2 | (AT)6 | TGAAACCCTCGAATCTCCTG | AACTCGGGCACTAGTCATGG | 110 |
| chr3:110445333-110447363 | p2 | (AT)15 | TGAAACCCTCGAATCTCCTG | AACTCGGGCACTAGTCATGG | 110 |
| chr3:110566424-110568439 | p5 | (AAACC)3 | TAGCTTCCGATACACCCACC | GCAAAGGAAATACGTCGGAA | 196 |
| chr3:110566464-110568479 | p5 | (AAACC)3 | TAGCTTCCGATACACCCACC | GCAAAGGAAATACGTCGGAA | 156 |
| chr3:110583880-110585892 | p2 | (AT)6 | TTCCCGATTTTGCAATTTGT | AGGGTTAGGTGCCATCACAG | 162 |
| chr3:110632222-110634237 | p3 | (AAG)5 | GCAGACAATGAGGTGGGTTT | TCGGTCAGTCTGAATGTTGG | 193 |
| chr3:110718588-110720606 | p3 | (TTC)6 | AACGAGCTTGAAAAGAGGCA | ACCCCTTATTGTTCGGTTCC | 106 |
| chr3:110718588-110720606 | p3 | (TAT)5 | AACGAGCTTGAAAAGAGGCA | ACCCCTTATTGTTCGGTTCC | 106 |
| chr3:110718588-110720606 | p3 | (TTC)5 | AACGAGCTTGAAAAGAGGCA | ACCCCTTATTGTTCGGTTCC | 106 |
| chr3:110719051-110721066 | p3 | (TTC)6 | TTTTGTGTGGGAAGGAAAGC | GCCGAATGGTAGGAAGTTTG | 135 |
| chr3:110719051-110721066 | p3 | (TAT)5 | TTTTGTGTGGGAAGGAAAGC | GCCGAATGGTAGGAAGTTTG | 135 |
| chr3:110719051-110721066 | p3 | (TTC)5 | TTTTGTGTGGGAAGGAAAGC | GCCGAATGGTAGGAAGTTTG | 135 |
| chr3:110757572-110759590 | p6 | (ATTTTT)3 | CAGTGGGGACAAGTTGAGGT | GGCAGATTCCAAGTCCAAAA | 258 |
| chr3:110762842-110764858 | p2 | (TA)8 | GTCGTCCGTTGTCCCTTAAA | TTTTCCCATGAGTTTGGAGG | 262 |
| chr3:110762842-110764858 | p2 | (TA)8 | GTCGTCCGTTGTCCCTTAAA | TTTTCCCATGAGTTTGGAGG | 262 |
| chr3:110813890-110815908 | p3 | (GAA)6 | TGTGCACAAATCGGTTCAAT | ACAAATCCACCAACACCACA | 240 |
| chr3:110822517-110824537 | p2 | (AT)10 | TCACCGATCATAACAACCCA | CAGGTACCGACGACAGGTTT | 247 |
| chr3:110908627-110910699 | p3 | (TCA)5 | TTTTGGGTGAAAAATGAGGC | TACGTCCCGTACTTTCTGGG | 148 |
| chr3:111007198-111009242 | p5 | (TTTAA)3 | AATGTCTACAAGCCGTGCCT | GAAGCAATTTCCTCGTAGCG | 230 |
| chr3:111007198-111009242 | p2 | (TA)22 | AATGTCTACAAGCCGTGCCT | GAAGCAATTTCCTCGTAGCG | 230 |
| chr3:111166723-111168737 | p2 | (AG)7 | TTCAACCGTGGACATTACGA | GTCTCCTCTTTCCCTTCGCT | 205 |
| chr3:111230707-111232721 | p2 | (AT)7 | GCAGCCTCCACACTCCTAAG | AGTTCGGGCACTAGTCATGG | 194 |

| Marker ID | SSR type | SSR motif | FORWARD PRIMER (5'-3') | REVERSE PRIMER (5'-3') | size (bp) |
|---|---|---|---|---|---|
| chr3:111270125-111272141 | p4 | (AATT)4 | GTGGGAAATACCGTCCACAC | TGCCGCTTCATTGATTTGTA | 283 |
| chr3:111273680-111275692 | p2 | (AC)6 | GCGTAGATCCATAGGGCGTA | ATCAAATTTAACGGCCGGAG | 231 |
| chr3:111279939-111281955 | p4 | (AAGA)4 | CAAGGGTGATCTTTGCGACT | GGGATGTTCCGAAAAACTCA | 173 |
| chr3:111666309-111668324 | p5 | (TTTTA)3 | TTAAACTGGGACGGATGGAG | CCCCTTTTACCCTGCTTTTC | 231 |
| chr3:111694375-111696389 | p2 | (AT)7 | GAAAAGGGAAAAGGACGCTC | AAACCAACAAGGCAATCCAA | 231 |
| chr3:111694375-111696389 | p2 | (TA)6 | GAAAAGGGAAAAGGACGCTC | AAACCAACAAGGCAATCCAA | 231 |
| chr3:111694375-111696389 | p2 | (AT)6 | GAAAAGGGAAAAGGACGCTC | AAACCAACAAGGCAATCCAA | 231 |
| chr3:111694375-111696389 | p2 | (AT)7 | GAAAAGGGAAAAGGACGCTC | AAACCAACAAGGCAATCCAA | 231 |
| chr3:111822049-111824079 | p2 | (TA)6 | CCATTAAGCCTCGGTGTGAT | TGCAGTGGGCATGTTATGAT | 200 |
| chr3:111822049-111824079 | p3 | (CAA)10 | CCATTAAGCCTCGGTGTGAT | TGCAGTGGGCATGTTATGAT | 200 |
| chr3:111845617-111847635 | p6 | (TTTTAT)3 | TACCCTGTCGCAATCATCAA | GAGCTGGAATGACTTGGCTC | 206 |
| chr3:111893325-111895345 | p2 | (AT)10 | GGCCATTAGGACTTCTTCCC | ACCGACCTTACATCACTCCG | 204 |
| chr3:112001844-112003866 | p2 | (AT)7 | AAAACTCAGAAGCGGAGCAA | TTTCGTTAATTTCCCCCACA | 147 |
| chr3:112001844-112003866 | p2 | (AT)6 | AAAACTCAGAAGCGGAGCAA | TTTCGTTAATTTCCCCCACA | 147 |
| chr3:112001884-112003902 | p2 | (AT)7 | AAAACTCAGAAGCGGAGCAA | TTTCGTTAATTTCCCCCACA | 107 |
| chr3:112001884-112003902 | p2 | (AT)6 | AAAACTCAGAAGCGGAGCAA | TTTCGTTAATTTCCCCCACA | 107 |
| chr3:112047171-112049185 | p2 | (AG)7 | GCAGTGCATCCAGTCTTTCA | CGTCCTTAGTTCGCAACCTC | 193 |
| chr3:112053261-112055275 | p3 | (ATC)5 | CCAGGTCTCCCTAGTGTGGA | CCCCGAAAGCTTATGGTACA | 102 |
| chr3:112061254-112063268 | p2 | (TA)7 | GTGATGCAGGTCGAGTGAGA | TTTCATGACAACTTTGCCGA | 206 |
| chr3:112278613-112280628 | p2 | (TC)7 | TTCTTGGAGGGAATCACCTG | ACTCAGAGGAGCAGCGAGAG | 194 |
| chr3:112278670-112280685 | p2 | (TC)7 | TTCTTGGAGGGAATCACCTG | ACTCAGAGGAGCAGCGAGAG | 137 |
| chr3:112278699-112280738 | p2 | (TC)7 | TTCTTGGAGGGAATCACCTG | ACTCAGAGGAGCAGCGAGAG | 108 |
| chr3:112345664-112347679 | p5 | (TTAAA)3 | CATGAGACTTCATGCCCCTT | CAGGGCAAGAAAATGGAAAA | 240 |
| chr3:112412349-112414361 | p2 | (CT)6 | GTCGGGTAGACACCGCTAAG | AATGAGGAGGCATCAAATGG | 171 |
| chr3:112542955-112544971 | p2 | (AT)8 | AGGAAATCCTGGCATCAGTG | AAATCCCTTCGTCACACACC | 267 |
| chr3:112542955-112544971 | p5 | (TGGTT)3 | AGGAAATCCTGGCATCAGTG | AAATCCCTTCGTCACACACC | 267 |
| chr3:112570521-112572535 | p2 | (GT)7 | GCTTAATACCTTCCCCTCGG | TTACGCCCTATTTTGAACGG | 110 |
| chr3:112570521-112572535 | p3 | (TTG)5 | GCTTAATACCTTCCCCTCGG | TTACGCCCTATTTTGAACGG | 110 |
| chr3:112604075-112606090 | p5 | (AATGC)3 | TTTGATCTTCGAGGGTTTGG | TGGGTTGAAGGTTCGCTTAC | 219 |
| chr3:112675537-112677552 | p5 | (GTTCG)3 | GGCTGAAAGGTCCCAATGTA | AAAACCTCCAAAATCACCCC | 100 |
| chr3:112726584-112728623 | p3 | (AAT)13 | TCTTTGCGATTTGAAAGCCT | CCCCATCCCTTCTCTCTCTC | 256 |
| chr3:112733349-112735367 | p6 | (ATCCAC)3 | TGGATATGCTGGATTGCAGA | TGTTGCGGATTCAACTTTCA | 134 |
| chr3:112733349-112735367 | p6 | (TCCACG)3 | TGGATATGCTGGATTGCAGA | TGTTGCGGATTCAACTTTCA | 134 |
| chr3:112745750-112747789 | p3 | (AAT)13 | TCTTTGCGATTTGAAAGCCT | CCCCATCCCTTCTCTCTCTC | 256 |
| chr3:112778180-112780195 | p3 | (CAA)5 | CCGTCACACATCACAACACA | TTACTCCGCTCGTGCCTACT | 115 |
| chr3:112811389-112813404 | p3 | (CAA)5 | CCGTCACACATCACAACACA | TTACTCCGCTCGTGCCTACT | 115 |

| Marker ID | SSR type | SSR motif | FORWARD PRIMER (5'-3') | REVERSE PRIMER (5'-3') | size (bp) |
|---|---|---|---|---|---|
| chr3:112868921-112870933 | p2 | (AT)6 | TGACGTACGTGTCATGCTCA | GTATGCGAGTGTCCAGCTCA | 233 |
| chr3:113123919-113125955 | p2 | (AG)8 | TGGTTCAAAATAAGAGCCCG | TTAGAGCCTGTTTGGATGGG | 132 |
| chr3:113123919-113125955 | p6 | (TTTTTG)6 | TGGTTCAAAATAAGAGCCCG | TTAGAGCCTGTTTGGATGGG | 132 |
| chr3:113174480-113176542 | p2 | (TG)6 | TGCGGAATATAACAAAGCCC | TTTGGTCCGGTTTTTCCATA | 200 |
| chr3:113174480-113176542 | p2 | (AT)31 | TGCGGAATATAACAAAGCCC | TTTGGTCCGGTTTTTCCATA | 200 |
| chr3:113213302-113215324 | p4 | (AAAC)4 | CCAGAGCTCATGCAAAAACA | AAGGAAATGGCATTCACAGG | 204 |
| chr3:113213302-113215324 | p2 | (AT)11 | CCAGAGCTCATGCAAAAACA | AAGGAAATGGCATTCACAGG | 204 |
| chr3:113263472-113265487 | p5 | (CTCGC)3 | TCGGAAAGTCACCAAAGGAG | GAGCCTGTCTCGCCATAGAC | 259 |
| chr3:113263472-113265487 | p5 | (TTAAT)3 | TCGGAAAGTCACCAAAGGAG | GAGCCTGTCTCGCCATAGAC | 259 |
| chr3:113310596-113312611 | p3 | (TCA)5 | ACCGTAGCATGTGGTGACAA | TTTGTAGGAAAAGATGCCCG | 177 |
| chr3:113311373-113313409 | p3 | (TCA)5 | TTCCAAATTTTAGCGGTTGG | TGAGGCAATGCAACAACATT | 121 |
| chr3:113341178-113343193 | p3 | (TCA)5 | ACCGTAGCATGTGGTGACAA | TTTGTAGGAAAAGATGCCCG | 177 |
| chr3:113341955-113343991 | p3 | (TCA)5 | TTCCAAATTTTAGCGGTTGG | TGAGGCAATGCAACAACATT | 121 |
| chr3:113356618-113358630 | p2 | (AT)6 | TTGGTGGACAACAACCTTGA | TCGGGTACCAAACACTAGCC | 119 |
| chr3:113356618-113358630 | p2 | (TA)6 | TTGGTGGACAACAACCTTGA | TCGGGTACCAAACACTAGCC | 119 |
| chr3:113367126-113369151 | p5 | (TTTGG)5 | ACATGTGTGCATTTGGGCTA | TGAGGGGGTTCATGTGAAAT | 205 |
| chr3:113367126-113369151 | p5 | (TTATA)3 | ACATGTGTGCATTTGGGCTA | TGAGGGGGTTCATGTGAAAT | 205 |
| chr3:113391696-113393778 | p2 | (GA)8 | TATCCCTCCTTAGTTCGGGG | TATTGCAGGAACAGGGAACC | 264 |
| chr3:113391902-113393918 | p2 | (GA)8 | GGTTCCCTGTTCCTGCAATA | CAACGAAATTGTGGCTGAAA | 158 |
| chr3:113429429-113431445 | p2 | (AT)8 | AATCTTTGCACTGAGTCCCG | TTGTTCCACGAAATGCAAAA | 221 |
| chr3:113429459-113431487 | p2 | (AT)8 | AATCTTTGCACTGAGTCCCG | TTGTTCCACGAAATGCAAAA | 191 |
| chr3:113429540-113431554 | p2 | (AT)8 | AATCTTTGCACTGAGTCCCG | TTGTTCCACGAAATGCAAAA | 110 |
| chr3:113464604-113466618 | p2 | (TC)7 | TAACAATACGGCGTAAGGGC | TTGCCTTAAATGCTCCAACC | 298 |
| chr3:113489224-113491239 | p5 | (ATGAT)3 | GCCCAAATTTCTGTAGGCAA | GGACATGCCAAGAGAAGGAA | 116 |
| chr3:113608747-113610759 | p2 | (AG)6 | AAATCGTTGGTGCTTTTTCG | AGGATTTTTCATGGCGAATG | 107 |
| chr3:113614831-113616867 | p2 | (AT)18 | TGCGTAACTGACAATGCCTC | ATGCCATAGAGGCTCACAGG | 240 |
| chr3:113620488-113622503 | p3 | (ATC)5 | GAATAACTGGGTGGCCCATA | GGCCTAGTCGGCTTACACTG | 243 |
| chr3:113637543-113639609 | p2 | (AT)7 | GGGTGGCCTCTCATAAAACA | AAGGGACAACAGACGACCAC | 271 |
| chr3:113637543-113639609 | p2 | (TA)11 | GGGTGGCCTCTCATAAAACA | AAGGGACAACAGACGACCAC | 271 |
| chr3:113653485-113655507 | p2 | (TA)11 | TTGTGCACATGGTTGCTTTT | TTTGGTTGGGAGTTTCGTTC | 119 |
| chr3:113653485-113655507 | p3 | (GGT)5 | TTGTGCACATGGTTGCTTTT | TTTGGTTGGGAGTTTCGTTC | 119 |
| chr3:113747351-113749366 | p3 | (CTC)5 | TGTTGAATGGCCAAAGATGA | AAATGGAGCAAAAATGGCTG | 278 |
| chr3:113750573-113752588 | p5 | (TATTT)3 | GAACCAAATGGTGCCAGTCT | ACATGGTTTATCTGCCAGCC | 272 |
| chr3:113855033-113857045 | p2 | (TA)6 | ACATCAGGGCTTTCATTTGG | GGTGACTCCGTACCCTTGTC | 146 |
| chr3:113857620-113859636 | p2 | (TC)8 | CCAACCTGATTCAAATGCCT | CATTTTCACACAATGGGCAC | 222 |
| chr3:113869300-113871324 | p6 | (ACTGAA)4 | AGCGTGGAAGAATCTGAGGA | TGCATAGCAACGAAGGTGTC | 119 |

| Marker ID | SSR type | SSR motif | FORWARD PRIMER (5'-3') | REVERSE PRIMER (5'-3') | size (bp) |
|---|---|---|---|---|---|
| chr3:114006746-114008761 | p5 | (GGTGC)3 | GATGAAAGAATTTTCGGGCA | ATCCCAGAACTACACACGGC | 167 |
| chr3:114024675-114026690 | p3 | (CAA)5 | TCCCAATTATCAGCCGAGAC | AGGAGACTTCCGTTGGGAAT | 296 |
| chr3:114058046-114060060 | p2 | (TA)7 | ATGCAAGATTCGCTCAGTCA | TTGGCACGGTTTGTCAGTTA | 164 |
| chr3:114063824-114065842 | p2 | (TA)9 | TTTTTCCCGTGAGTTTGGAG | TGGGCCCCATACTAAACAAC | 112 |
| chr3:114063824-114065842 | p2 | (TA)9 | TTTTTCCCGTGAGTTTGGAG | TGGGCCCCATACTAAACAAC | 112 |
| chr3:114123973-114125989 | p3 | (TGT)6 | TGTTGTTGTTGTTGGCCATT | TGATAAAACGACGATGACCG | 103 |
| chr3:114123973-114125989 | p2 | (TA)8 | TGTTGTTGTTGTTGGCCATT | TGATAAAACGACGATGACCG | 103 |
| chr3:114178024-114180040 | p2 | (TA)8 | GGGCTGATTTGATGCTTGTT | ACAATTTGCTCCAAAGGTGG | 288 |
| chr3:114240179-114242194 | p2 | (AT)6 | TTTAAGCCCATCCAAACAGG | TGGACCATCCAAACAACTCA | 143 |
| chr3:114240179-114242194 | p3 | (ATA)5 | TTTAAGCCCATCCAAACAGG | TGGACCATCCAAACAACTCA | 143 |
| chr3:114357743-114359761 | p2 | (AT)9 | GCGAACGTAAGAGCAAGGTC | GTCGGATCGAAAGTTGGAGA | 174 |
| chr3:114388823-114390857 | p2 | (AT)17 | TGCACAAATAGGCACCTCAA | TGGATTTTGAAAGGCAAAGG | 246 |
| chr3:114396829-114398881 | p6 | (ATTTTT)3 | TCTCGTTTCTCATCGAACCC | GCCAAGTTTGAAGGCATGTT | 203 |
| chr3:114403962-114405978 | p2 | (TA)8 | TAGCACCTCCAAAACCGAAC | CAATTACGGGCGATGAATCT | 101 |
| chr3:114403962-114405978 | p2 | (TA)6 | TAGCACCTCCAAAACCGAAC | CAATTACGGGCGATGAATCT | 101 |
| chr3:114424489-114426507 | p3 | (CAG)6 | GTGATGATTTTGGCATGACG | TAGGTCTGCTCCAGCAAGGT | 125 |
| chr3:114440588-114442608 | p2 | (AG)10 | GGAAGCTGATGTTCTGAGGG | ACTAGGTGCGCTTTCCAAAA | 257 |
| chr3:114492726-114494740 | p2 | (TC)7 | AAGCTGACCAGCCAAACACT | AGTGGCTACTCCATCCATGC | 128 |
| chr3:114519269-114521284 | p3 | (TAC)9 | CAAATGCAACCAAATTGCAG | TGAGGAAATCGAAGACCCAC | 113 |
| chr3:114519269-114521284 | p5 | (GAGAA)3 | CAAATGCAACCAAATTGCAG | TGAGGAAATCGAAGACCCAC | 113 |
| chr3:114519821-114521866 | p3 | (TAC)9 | CGGGTCCAAACTTCTGTCAT | TCAAACATTCCCCGATTCTC | 294 |
| chr3:114519821-114521866 | p5 | (GAGAA)3 | CGGGTCCAAACTTCTGTCAT | TCAAACATTCCCCGATTCTC | 294 |
| chr3:114519868-114521889 | p3 | (TAC)9 | CGGGTCCAAACTTCTGTCAT | TCAAACATTCCCCGATTCTC | 247 |
| chr3:114519868-114521889 | p5 | (GAGAA)3 | CGGGTCCAAACTTCTGTCAT | TCAAACATTCCCCGATTCTC | 247 |
| chr3:114519958-114521991 | p3 | (TAC)9 | CGGGTCCAAACTTCTGTCAT | TCAAACATTCCCCGATTCTC | 157 |
| chr3:114519958-114521991 | p5 | (GAGAA)3 | CGGGTCCAAACTTCTGTCAT | TCAAACATTCCCCGATTCTC | 157 |
| chr3:114610407-114612475 | p2 | (AT)34 | AAAAAGGTGTCGAATGTGCC | AAGACGAAATTTCCCTGGGT | 108 |
| chr3:114643394-114645412 | p2 | (GA)9 | CGATCTCGTCGTCAATCTCA | CCTCGGCCAATGAAGTGTAT | 200 |
| chr3:114742077-114744092 | p3 | (TCT)5 | TTTTATCCCACTCGGGTCAG | TGGACTACGCTGTTGTCTGC | 187 |
| chr3:114742077-114744092 | p2 | (TG)7 | TTTTATCCCACTCGGGTCAG | TGGACTACGCTGTTGTCTGC | 187 |
| chr3:114827718-114829733 | p2 | (GA)6 | CGGACCACTTTTGTAAGGGA | CCTCTGCACCTTCACTCTCC | 293 |
| chr3:114827741-114829756 | p2 | (GA)6 | CGGACCACTTTTGTAAGGGA | CCTCTGCACCTTCACTCTCC | 270 |
| chr3:114852921-114854941 | p2 | (TA)10 | GACACATTCGACGCCTTTTT | CGGACCGTTGATTCTCATTT | 242 |
| chr3:114971707-114973725 | p2 | (TA)9 | GGTTGAGAAGCTTGGTCTGC | CTTCTTGTGCGGTGGAGATT | 172 |
| chr3:114990375-114992406 | p6 | (TTACTT)4 | CAAATAAATTGAGCCGGGAA | TCCAGGAGCAGGTGACTTTT | 239 |
| chr3:114990375-114992406 | p2 | (AT)11 | CAAATAAATTGAGCCGGGAA | TCCAGGAGCAGGTGACTTTT | 239 |

| Marker ID | SSR type | SSR motif | FORWARD PRIMER (5'-3') | REVERSE PRIMER (5'-3') | size (bp) |
|---|---|---|---|---|---|
| chr3:114990375-114992406 | p2 | (AT)8 | CAAATAAATTGAGCCGGGAA | TCCAGGAGCAGGTGACTTTT | 239 |
| chr3:114990375-114992406 | p2 | (TA)10 | CAAATAAATTGAGCCGGGAA | TCCAGGAGCAGGTGACTTTT | 239 |
| chr3:114990375-114992406 | p2 | (AT)12 | CAAATAAATTGAGCCGGGAA | TCCAGGAGCAGGTGACTTTT | 239 |
| chr3:115028132-115030160 | p3 | (ATT)5 | CATCTCCCTCCTGCAAAAAG | CCCTAATTCATGGAGACCCA | 285 |
| chr3:115264339-115266354 | p2 | (AG)7 | ACAAAATGCATAGGGCGTTC | GCAAAGGGAAAATCAGCAAA | 236 |
| chr3:115264417-115266435 | p2 | (AG)7 | ACAAAATGCATAGGGCGTTC | GCAAAGGGAAAATCAGCAAA | 158 |
| chr3:115265111-115267126 | p2 | (AG)7 | GGGGTGTTCAAATGAGATCG | GGACTTGGAATGAGCCAAAA | 105 |
| chr3:115340643-115342661 | p6 | (TCTTTT)3 | ATGACTTAGCGAGCCGAAAA | TGCGGAATTATCAACATCCA | 173 |
| chr3:115385507-115387525 | p2 | (AT)9 | TATGAATGTGCCGTTGGAAA | GAGTCAAAGGAGCAGCAACC | 213 |
| chr3:115562955-115564971 | p4 | (AAAT)4 | TTGAAGTGAGGTGCGTTGAA | CAGCATGGCCGTTATTCTTT | 219 |
| chr3:115562955-115564971 | p2 | (TA)11 | TTGAAGTGAGGTGCGTTGAA | CAGCATGGCCGTTATTCTTT | 219 |
| chr3:115583968-115585980 | p2 | (AT)17 | TACACTCGTTGCTCCACGTC | CGAAGCCACGGTTGATACTT | 188 |
| chr3:115583981-115585995 | p2 | (AT)17 | TACACTCGTTGCTCCACGTC | CGAAGCCACGGTTGATACTT | 175 |
| chr3:115584002-115586028 | p2 | (AT)17 | TACACTCGTTGCTCCACGTC | CGAAGCCACGGTTGATACTT | 154 |
| chr3:115584175-115586195 | p2 | (AT)17 | CGGCTCGAGATATCCTTACG | ACTTGCAAGCTAAAACCGGA | 211 |
| chr3:115584210-115586230 | p2 | (AT)17 | CGGCTCGAGATATCCTTACG | ACTTGCAAGCTAAAACCGGA | 176 |
| chr3:115584245-115586265 | p2 | (AT)17 | CGGCTCGAGATATCCTTACG | ACTTGCAAGCTAAAACCGGA | 141 |
| chr3:115584280-115586300 | p2 | (AT)17 | CGGCTCGAGATATCCTTACG | ACTTGCAAGCTAAAACCGGA | 106 |
| chr3:115629799-115631811 | p2 | (AT)6 | TATAGGCAAGGGACTGTGGC | GCTGGTGACTCCGTTAGCTC | 270 |
| chr3:115839749-115841764 | p5 | (TTAAG)3 | TTGCATGAACACTTCGGAGA | GCCAAAATGAAGACTCTCGC | 117 |
| chr3:115914896-115916950 | p3 | (AAT)18 | TGTGCCAATGGACTACCAAA | TGCCTCAAGGGTCTATCACC | 264 |
| chr3:115983271-115985287 | p2 | (TA)6 | TTCGTGCAAAGTTTCACTGC | CTTTAATGAGGGACCGGTGA | 199 |
| chr3:115983341-115985363 | p2 | (TA)6 | TTCGTGCAAAGTTTCACTGC | CTTTAATGAGGGACCGGTGA | 129 |
| chr3:116008354-116010366 | p2 | (AT)6 | GGGCCGTAATTCAAGTCGTA | ATGTTCGTTTGACCCCCATA | 249 |
| chr3:116154377-116156395 | p6 | (TTTTTA)3 | AGCTCAAATGATCCAATCCG | GATGCAGTGAAGGGACACAA | 188 |
| chr3:116154377-116156395 | p2 | (AT)15 | AGCTCAAATGATCCAATCCG | GATGCAGTGAAGGGACACAA | 188 |
| chr3:116406822-116408867 | p2 | (TA)7 | GCACATACATCGGCAATGAC | AATCGCGAGCACAATCTCTT | 209 |
| chr3:116406822-116408867 | p2 | (TC)6 | GCACATACATCGGCAATGAC | AATCGCGAGCACAATCTCTT | 209 |
| chr3:116502788-116504806 | p6 | (CTTTTT)3 | TGAAGAGGTGATCATGGTGG | CCTTAGCACCCCTAACCCAT | 198 |
| chr3:116514462-116516477 | p2 | (AG)9 | CGACCAGCTGAAACAACAAA | TCTTTCTTTAGCCGCCTTGA | 127 |
| chr3:116514462-116516477 | p3 | (TAA)5 | CGACCAGCTGAAACAACAAA | TCTTTCTTTAGCCGCCTTGA | 127 |
| chr3:116650442-116652475 | p3 | (TGT)11 | GGTTGTGAGACCGACCTTGT | CTCGCTTCCCTCATCTTGTC | 269 |
| chr3:116650442-116652475 | p2 | (AG)7 | GGTTGTGAGACCGACCTTGT | CTCGCTTCCCTCATCTTGTC | 269 |
| chr3:116650596-116652610 | p3 | (TGT)11 | GGTTGTGAGACCGACCTTGT | CTCGCTTCCCTCATCTTGTC | 115 |
| chr3:116650596-116652610 | p2 | (AG)7 | GGTTGTGAGACCGACCTTGT | CTCGCTTCCCTCATCTTGTC | 115 |
| chr3:116736298-116738386 | p2 | (TC)7 | CAGCGAAACGAGTCACAAAA | GCTTTACAAGCGACCTCGAC | 156 |

| Marker ID | SSR type | SSR motif | FORWARD PRIMER (5'-3') | REVERSE PRIMER (5'-3') | size (bp) |
|---|---|---|---|---|---|
| chr3:116736298-116738386 | p2 | (TA)44 | CAGCGAAACGAGTCACAAAA | GCTTTACAAGCGACCTCGAC | 156 |
| chr3:116786854-116788905 | p3 | (TAT)17 | GAATTGGTTGATGGCAGAAAA | AGAAAATCTTCCCGGCATTT | 111 |
| chr3:116786854-116788905 | p3 | (ATT)18 | GAATTGGTTGATGGCAGAAAA | AGAAAATCTTCCCGGCATTT | 111 |
| chr3:116786854-116788905 | p2 | (TA)22 | GAATTGGTTGATGGCAGAAAA | AGAAAATCTTCCCGGCATTT | 111 |
| chr3:116797944-116799966 | p2 | (AT)11 | GGCATGTCCTCTTTTCTCCA | ACAATGCTGACAGTGAACGC | 189 |
| chr3:116814827-116816878 | p3 | (TAT)17 | GAATTGGTTGATGGCAGAAAA | AGAAAATCTTCCCGGCATTT | 111 |
| chr3:116814827-116816878 | p3 | (ATT)18 | GAATTGGTTGATGGCAGAAAA | AGAAAATCTTCCCGGCATTT | 111 |
| chr3:116814827-116816878 | p2 | (TA)22 | GAATTGGTTGATGGCAGAAAA | AGAAAATCTTCCCGGCATTT | 111 |
| chr3:116856698-116858710 | p2 | (AT)6 | ACCAAAGTGTGGTTCCAAGC | TTCTTCTCCGATGAACCACC | 236 |
| chr3:116931553-116933568 | p5 | (ACCAA)3 | TCGGGTTGGTGTGAATTTTT | CGAGAAAATCCGAGGTTGAA | 231 |
| chr3:116931553-116933568 | p5 | (AAACC)3 | TCGGGTTGGTGTGAATTTTT | CGAGAAAATCCGAGGTTGAA | 231 |
| chr3:116994640-116996655 | p2 | (TA)6 | TATGCACTCCGGTAGAGGCT | GTAAGGGTCACAAGCCCAAA | 153 |
| chr3:116994640-116996655 | p5 | (CATTT)3 | TATGCACTCCGGTAGAGGCT | GTAAGGGTCACAAGCCCAAA | 153 |
| chr3:116994640-116996655 | p5 | (CATAT)3 | TATGCACTCCGGTAGAGGCT | GTAAGGGTCACAAGCCCAAA | 153 |
| chr3:117063397-117065415 | p3 | (ATT)5 | CCAATGGTATATTGTGGGGG | TTGTCCTTAGGGTTTGGGTG | 156 |
| chr3:117063450-117065465 | p3 | (ATT)5 | CCAATGGTATATTGTGGGGG | TTGTCCTTAGGGTTTGGGTG | 103 |
| chr3:117173932-117175946 | p2 | (TC)7 | TCATTTTGAGAGGGATTGCC | AGTCCCACTCGTCTGCTCAT | 203 |
| chr3:117190952-117192968 | p2 | (TA)8 | TGGACCCGTCTCCTATTGAG | GAACCTTACCACCCTCACCA | 147 |
| chr3:117405397-117407409 | p2 | (AC)6 | TTTTAAGCGTCTGGGCTGTT | AACGGTGGCTGTATTTCGAC | 180 |
| chr3:117413234-117415249 | p5 | (AAGGA)3 | AGTCGAGCTTACTGCCCAAA | ATAGCGGTACAATGCTCGCT | 157 |
| chr3:117484746-117486768 | p2 | (TA)11 | CTGGAATTTGGCAGTTGGAT | TCGGCCCAATTAAATACTGG | 138 |
| chr3:117505627-117507645 | p5 | (AATAT)3 | ACCTCTCTTGAAACGCCAAA | GGACTTTGAATCCGAGGTGA | 207 |
| chr3:117505648-117507660 | p5 | (AATAT)3 | ACCTCTCTTGAAACGCCAAA | GGACTTTGAATCCGAGGTGA | 186 |
| chr3:117505704-117507719 | p5 | (AATAT)3 | ACCTCTCTTGAAACGCCAAA | GGACTTTGAATCCGAGGTGA | 130 |
| chr3:117514168-117516183 | p5 | (AAAGA)3 | ATGTATTGATGGGGCGTGTT | CTTGGCGAGATGCCTAAGAC | 190 |
| chr3:117558797-117560813 | p2 | (TA)7 | TTGACACATTCTGAACCCCA | CAAGAAAACGAATGCCCAAT | 229 |
| chr3:117558797-117560813 | p2 | (TA)8 | TTGACACATTCTGAACCCCA | CAAGAAAACGAATGCCCAAT | 229 |
| chr3:117558797-117560813 | p2 | (TA)6 | TTGACACATTCTGAACCCCA | CAAGAAAACGAATGCCCAAT | 229 |
| chr3:117558797-117560813 | p2 | (AT)8 | TTGACACATTCTGAACCCCA | CAAGAAAACGAATGCCCAAT | 229 |
| chr3:117701553-117703568 | p5 | (TTGGT)3 | AAAGTGGCCGAAAATATCCC | TTCGAACTCAGAATTTCGGG | 290 |
| chr3:117701553-117703568 | p5 | (TTTCT)3 | AAAGTGGCCGAAAATATCCC | TTCGAACTCAGAATTTCGGG | 290 |
| chr3:117708881-117710905 | p2 | (AT)6 | AAGTATCAACCGTGGCTTCG | TCCGCTTCTGAGTTTTGGTT | 190 |
| chr3:117722450-117724470 | p5 | (ATTTT)4 | GCTTCTTCGACATGCTGACA | CGGGCTGTTATCCAGTTCAT | 229 |
| chr3:117786200-117788215 | p5 | (AAAAT)3 | ATCCCACAATGCCTATGCTC | GCCAAAGTGGCCAGTAATGT | 264 |
| chr3:117788022-117790050 | p2 | (AT)14 | ACTAAGCGGGAATTCGAACC | ATCTGAGATGGTGGCAAAGG | 257 |
| chr3:117788022-117790050 | p2 | (AT)14 | ACTAAGCGGGAATTCGAACC | ATCTGAGATGGTGGCAAAGG | 257 |

| Marker ID | SSR type | SSR motif | FORWARD PRIMER (5'-3') | REVERSE PRIMER (5'-3') | size (bp) |
|---|---|---|---|---|---|
| chr3:117893621-117895633 | p2 | (TA)6 | TGAACGTTCTTATGAGCCCC | GTGAGTTTAGCGATGGTGGG | 102 |
| chr3:117893621-117895633 | p2 | (AT)6 | TGAACGTTCTTATGAGCCCC | GTGAGTTTAGCGATGGTGGG | 102 |
| chr3:117893621-117895633 | p2 | (TA)6 | TGAACGTTCTTATGAGCCCC | GTGAGTTTAGCGATGGTGGG | 102 |
| chr3:117893863-117895875 | p2 | (TA)6 | GGGAAAGAATGCCCCATATT | TCGTTGCCCTACGAATACAA | 176 |
| chr3:117893863-117895875 | p2 | (AT)6 | GGGAAAGAATGCCCCATATT | TCGTTGCCCTACGAATACAA | 176 |
| chr3:117893863-117895875 | p2 | (TA)6 | GGGAAAGAATGCCCCATATT | TCGTTGCCCTACGAATACAA | 176 |
| chr3:117906420-117908435 | p6 | (AAGCCG)3 | CATCCCATCATTTCCCATTC | AGGGCACAATGGTCAAACTC | 182 |
| chr3:117906420-117908435 | p5 | (AGAAA)3 | CATCCCATCATTTCCCATTC | AGGGCACAATGGTCAAACTC | 182 |
| chr3:117919209-117921235 | p2 | (AT)13 | GCAGGCTCTCTCCTAGGGTT | GAATCACTGTTCACGCAGGA | 124 |
| chr3:117925505-117927517 | p2 | (AT)6 | GACCGATCGAGAATTTGCAT | ATAAGAGCTACACGCGAGGC | 231 |
| chr3:118002215-118004233 | p3 | (TTG)6 | CCACCGAAACTTAAAGGCAA | CGCATCTTGTCTTCCACTGA | 278 |
| chr3:118091846-118093862 | p4 | (ATTT)4 | TATACGGATCGGAAGGTTCG | ACACCTGCCAAACCTAGTGG | 153 |
| chr3:118186756-118188776 | p3 | (GGA)5 | AGCCTTCATCTCTTCTGGCA | AGCCATGGAGAAACATCACC | 215 |
| chr3:118186756-118188776 | p5 | (CTATG)3 | AGCCTTCATCTCTTCTGGCA | AGCCATGGAGAAACATCACC | 215 |
| chr3:118186869-118188885 | p3 | (GGA)5 | AGCCTTCATCTCTTCTGGCA | AGCCATGGAGAAACATCACC | 102 |
| chr3:118186869-118188885 | p5 | (CTATG)3 | AGCCTTCATCTCTTCTGGCA | AGCCATGGAGAAACATCACC | 102 |
| chr3:118212171-118214195 | p4 | (ATTT)6 | TGAAGTTTGAGGGAATTGGG | TACCCCTGTCTTGACGGAAA | 262 |
| chr3:118222926-118224988 | p2 | (AT)31 | GGTACCTTAGGGCTTAGGCG | TTTTGATCGGAAACCTACCG | 293 |
| chr3:118306854-118308866 | p2 | (TC)6 | TCCCTCATCTTGTCCCTCAC | TGAGGCTTTCTTCCGTGTCT | 153 |
| chr3:118363672-118365696 | p2 | (AT)12 | ATGAATTCCCCCGAGATAGC | CGTTGCCTGATATGGGATCT | 115 |
| chr3:118363672-118365696 | p3 | (ACA)5 | ATGAATTCCCCCGAGATAGC | CGTTGCCTGATATGGGATCT | 115 |
| chr3:118373164-118375188 | p2 | (AT)12 | ATGAATTCCCCCGAGATAGC | CGTTGCCTGATATGGGATCT | 115 |
| chr3:118373164-118375188 | p3 | (ACA)5 | ATGAATTCCCCCGAGATAGC | CGTTGCCTGATATGGGATCT | 115 |
| chr3:118442289-118444313 | p6 | (AACCTA)4 | CAAGGACACATGCATTTTGG | GGCCGTCAAAAACACTTTTC | 157 |
| chr3:118442289-118444313 | p2 | (AT)25 | CAAGGACACATGCATTTTGG | GGCCGTCAAAAACACTTTTC | 157 |
| chr3:118499446-118501491 | p3 | (AAT)15 | AATTGGATCGGCTTTAGGCT | TGTGAAAAAGTGGCTATGCG | 202 |
| chr3:118529371-118531386 | p5 | (ATAAA)3 | GATGAAACCAGCGTTTGGAC | GGTTGGTAAATGGTTGGTGG | 138 |
| chr3:118548524-118550542 | p3 | (TAA)6 | TTGACAATCCAAATGGCAAA | TGGAGGAAAATTTTGGGATG | 204 |
| chr3:118572949-118574964 | p5 | (TAAAT)3 | TTTAGGCCCCTAAGGTGGAT | TGGATTGTCAGGCAATGAAA | 242 |
| chr3:118574716-118576736 | p5 | (AAACC)4 | TTTTTGAAGAACCCCGTCAG | AATCCGAGAAAATCCGAGGT | 180 |
| chr3:118585547-118587562 | p5 | (CCCGC)3 | ATGGCATTCAGTTGCCTTTC | GGTCAGCTCATCCTAACCCA | 108 |
| chr3:118601969-118603985 | p2 | (AT)8 | ATGGGAGGCGGAAGTAGATT | CTTTTGCCCCTTTACAACCA | 144 |
| chr3:118717458-118719473 | p5 | (TGGGT)3 | CACCAATTTAACCCAATCCG | ATGTTTTCCAGGACCGTTTG | 258 |
| chr3:118717458-118719473 | p3 | (TTA)9 | CACCAATTTAACCCAATCCG | ATGTTTTCCAGGACCGTTTG | 258 |
| chr3:118794510-118796524 | p2 | (AT)7 | CAACCCATTTGAAGGGAAAA | AAACGCCTGGTGTGGATAAG | 252 |
| chr3:118794510-118796524 | p2 | (AT)11 | CAACCCATTTGAAGGGAAAA | AAACGCCTGGTGTGGATAAG | 252 |

| Marker ID | SSR type | SSR motif | FORWARD PRIMER (5'-3') | REVERSE PRIMER (5'-3') | size (bp) |
|---|---|---|---|---|---|
| chr3:118809152-118811167 | p3 | (AAT)14 | ACGAACGGAAGAAGGGTTTT | CCTGATCCGACCCATCTTTA | 189 |
| chr3:118809550-118811580 | p3 | (AAT)14 | GCAATTGTAAAAGACCCGGA | CTCGGCATTTTATCCACCAC | 212 |
| chr3:118809577-118811592 | p3 | (AAT)14 | GCAATTGTAAAAGACCCGGA | CTCGGCATTTTATCCACCAC | 185 |
| chr3:118833226-118835240 | p2 | (TA)7 | ATCTTATCCCGCTTTTGGCT | AAGGACAACGGACGATGAAG | 190 |
| chr3:118851981-118853996 | p5 | (ATTTT)3 | GAGATCTAGCCGTGCAAACC | AAAACTGAAAATTTCCCGCC | 288 |
| chr3:118851981-118853996 | p6 | (ATTTTT)3 | GAGATCTAGCCGTGCAAACC | AAAACTGAAAATTTCCCGCC | 288 |
| chr3:118874084-118876099 | p5 | (ATTTT)3 | GAGATCTAGCCGTGCAAACC | AAAACTGAAAATTTCCCGCC | 288 |
| chr3:118874084-118876099 | p6 | (ATTTTT)3 | GAGATCTAGCCGTGCAAACC | AAAACTGAAAATTTCCCGCC | 288 |
| chr3:118915703-118917745 | p2 | (AT)6 | AAGTATCAACCGTGGCTTCG | TCGCCTTCCTCCACTTCTAA | 268 |
| chr3:118915746-118917762 | p2 | (AT)6 | AAGTATCAACCGTGGCTTCG | TCGCCTTCCTCCACTTCTAA | 225 |
| chr3:118962880-118964892 | p2 | (GA)6 | TGTGGCGTGTGAATTGAACT | CCTATGGTCGCCAGAGGTAA | 121 |
| chr3:119080774-119082790 | p4 | (TTAT)4 | TTCCAAAAACGGGAACAAAA | TGTCACCACATGCTACGGTT | 114 |
| chr3:119107361-119109376 | p5 | (ATTGC)3 | TCATTCGTGATCGGATTTCA | GCCTCGAAAAGTGACCTCAG | 141 |
| chr3:119159629-119161644 | p5 | (TTTAG)3 | GACCTATGCACGGTCGATTT | AACAGATTTACCCGCACGTC | 124 |
| chr3:119266983-119268998 | p5 | (AATGC)3 | GCGATGAATCAAAGGGGATA | CCACCACGACTTGTGAATTG | 240 |
| chr3:119352081-119354096 | p5 | (TTTTA)3 | TCACAAGAATTTTCGGGCTC | ATCATGGAATTGCTCCAAGC | 251 |
| chr3:119366461-119368485 | p4 | (ATAA)6 | CAAACTTCTCAAATGGGGGA | AGTGAGCACGAATTGGCTTT | 160 |
| chr3:119388025-119390043 | p6 | (TACAAT)3 | CAACCCCATAGGAAAGGGAT | TCTCTTGGACCTTTCCCTCA | 111 |
| chr3:119390944-119392962 | p3 | (AGG)6 | GGGTTGGAATCCAGAACCTT | CTGTCAAACTTCGGTCCCAT | 245 |
| chr3:119393305-119395320 | p5 | (ATATG)3 | GAATGAAGAAATCGAGCGGA | CTGCTTAGCCCTTCGTTTTG | 224 |
| chr3:119421954-119423966 | p2 | (TA)6 | AACTTGCATCCGGAGTTTTG | CCCAAGCATCTTGAATTGGT | 104 |
| chr3:119430629-119432647 | p3 | (TTG)14 | GAACCACCCCACATCAAAAC | GGAGTTGTGGGAAGGGAACT | 272 |
| chr3:119430652-119432676 | p3 | (TTG)14 | GAACCACCCCACATCAAAAC | GGAGTTGTGGGAAGGGAACT | 249 |
| chr3:119486901-119488916 | p5 | (AAACC)3 | GCATCAGATTGGTTCGGATT | CGAGAAAACCCGAAGTTGAA | 223 |
| chr3:119486901-119488916 | p2 | (TA)6 | GCATCAGATTGGTTCGGATT | CGAGAAAACCCGAAGTTGAA | 223 |
| chr3:119569756-119571774 | p6 | (AACCGG)3 | TTGTCGTAGAACTTTCGCCC | TTTGCGGCTAGTAGTGACGA | 114 |
| chr3:119587888-119589912 | p2 | (AT)12 | CCACCTTCTTCCCCTTTTTC | GTAGAAGGGTGGCCAAACAA | 189 |
| chr3:119590439-119592454 | p5 | (AAAAT)3 | GAAGCTGCAAAAACTTGGCT | CCTTACGTTTGAGGCTGGAG | 172 |
| chr3:119590439-119592454 | p2 | (TA)6 | GAAGCTGCAAAAACTTGGCT | CCTTACGTTTGAGGCTGGAG | 172 |
| chr3:119603003-119605019 | p2 | (TG)8 | GCCATGGACCCTTATGAAAA | ATTTTCCAAACCATGTCCGA | 168 |
| chr3:119621894-119623906 | p2 | (AT)6 | TCAACTCAAGCAAATGGCAC | TGGTGTGTACGCAGCCTTAC | 175 |
| chr3:119676476-119678494 | p3 | (ACA)7 | GGCAACTTTTCTTGAGCAGG | TCCCATAAGGTGAGGTCTGG | 150 |
| chr3:119676476-119678494 | p6 | (TATCTA)3 | GGCAACTTTTCTTGAGCAGG | TCCCATAAGGTGAGGTCTGG | 150 |
| chr3:119737858-119739870 | p2 | (AC)6 | TTGTGATGATGTGGCACCTT | TCCTACTCCTTCCGATCCCT | 108 |
| chr3:119737858-119739870 | p2 | (TA)17 | TTGTGATGATGTGGCACCTT | TCCTACTCCTTCCGATCCCT | 108 |
| chr3:119802054-119804072 | p6 | (TTTATT)3 | CAGCTTCAGGATTAGCAGCC | CGGGGAATTAATAGCCCAAT | 282 |

| Marker ID | SSR type | SSR motif | FORWARD PRIMER (5'-3') | REVERSE PRIMER (5'-3') | size (bp) |
|---|---|---|---|---|---|
| chr3:119802054-119804072 | p2 | (AT)8 | CAGCTTCAGGATTAGCAGCC | CGGGGAATTAATAGCCCAAT | 282 |
| chr3:119847913-119849925 | p3 | (GTT)6 | GAAGCTTGTTGAGTCGAGGG | CAAACATGCCTCAACCCTTT | 225 |
| chr3:119847913-119849925 | p2 | (AT)6 | GAAGCTTGTTGAGTCGAGGG | CAAACATGCCTCAACCCTTT | 225 |
| chr3:119847913-119849925 | p3 | (TCT)5 | GAAGCTTGTTGAGTCGAGGG | CAAACATGCCTCAACCCTTT | 225 |
| chr3:119847913-119849925 | p3 | (CAA)5 | GAAGCTTGTTGAGTCGAGGG | CAAACATGCCTCAACCCTTT | 225 |
| chr3:119848750-119850765 | p2 | (AT)6 | TATTGGGAGCCCCTTTTCTT | TTTTTGGAGGTCAACGGTTC | 281 |
| chr3:119848750-119850765 | p3 | (TCT)5 | TATTGGGAGCCCCTTTTCTT | TTTTTGGAGGTCAACGGTTC | 281 |
| chr3:119848750-119850765 | p3 | (CAA)5 | TATTGGGAGCCCCTTTTCTT | TTTTTGGAGGTCAACGGTTC | 281 |
| chr3:120097007-120099022 | p5 | (AATTA)3 | GTTATGTTCCAAGCCGCCTA | TCGATCCTGGCTATTTGCTT | 200 |
| chr3:120174656-120176671 | p5 | (ATTTT)3 | CTTCCCCTCACTTACCCCTC | TTTCACTCCATCATCGACCA | 111 |
| chr3:120210662-120212704 | p3 | (ATT)14 | TTGGGGAAGGAAGGAGTTTT | CCCCCTTTGTTGGTAGATCA | 132 |
| chr3:120224909-120226923 | p2 | (TC)7 | ATGATGAAAAACGGCCAAAG | GAGGCCAAATTGGACAAAAA | 203 |
| chr3:120287659-120289689 | p3 | (TTA)10 | CCGAAAGGCATCATTCTCAT | GACTCTCGTCGGGAAGTCTG | 185 |
| chr3:120368874-120370907 | p3 | (AAT)11 | GGATGGAAAAAGTTTCGCAG | CCCTCGTGCACATTTAGCTT | 250 |
| chr3:120570448-120572464 | p2 | (AT)8 | GGCCTGAGGTTCATATTCCA | TATTCCACGGGATACTTGCC | 172 |
| chr3:120609259-120611323 | p2 | (TA)32 | TAAGCTAGTGGGGGTGATGG | TGTTCCACGAAATGCAAAAA | 100 |
| chr3:120627714-120629726 | p2 | (AT)6 | TAAGCTAGTGGGGGTGATGG | CGTGGAACAATCCACGAAAT | 239 |
| chr3:120706127-120708151 | p2 | (TC)12 | TGTTGAACCATGCAGACCAT | TTCCTGTTGTTTGCGTTGAG | 104 |
| chr3:120713946-120715960 | p2 | (AT)7 | AGCGGAATATTCTCCGAGGT | GATGATTGTTTTGGCCGAGT | 204 |
| chr3:120715392-120717407 | p5 | (TCGGG)3 | TAGGAACCCACGACCTATCG | AAACGTTTTCGGGTGATCTG | 115 |
| chr3:120716727-120718745 | p3 | (TTG)6 | TCCGAGGAAGCACTACCATT | ACCAACAAAATGACATGCGA | 124 |
| chr3:120764988-120767006 | p2 | (AC)9 | CATTAGATTGCCCCAAGGAA | TCGTTGACGTGGTCATTTGT | 185 |
| chr3:120767696-120769750 | p6 | (CAGTTT)3 | AACATCATTGTTCTTCCCCG | AGGGGACGTCAGTACATTCG | 132 |
| chr3:120767696-120769750 | p6 | (TTAGGG)3 | AACATCATTGTTCTTCCCCG | AGGGGACGTCAGTACATTCG | 132 |
| chr3:120768856-120770880 | p6 | (TTAGGG)3 | ACCCCCACAACATTCTCAAA | TGCGGGGCTTAATTTTTATG | 291 |
| chr3:120768856-120770880 | p6 | (ACCACA)4 | ACCCCCACAACATTCTCAAA | TGCGGGGCTTAATTTTTATG | 291 |
| chr3:120768856-120770880 | p3 | (GAT)7 | ACCCCCACAACATTCTCAAA | TGCGGGGCTTAATTTTTATG | 291 |
| chr3:120814344-120816364 | p5 | (TAAAA)4 | TGTACCACCTGGGATCAACA | ATCTCTTTGGCAATGGCATC | 264 |
| chr3:120847530-120849578 | p2 | (TA)7 | TGCATTGCTATTGTTCGTCC | TAGCGTGGGATCCTCATCA | 175 |
| chr3:120849265-120851279 | p2 | (TA)7 | TGGCCAAGTAATATGGACGA | TGAAGTACTCCTTCCGTTCCA | 223 |
| chr3:120849265-120851279 | p3 | (ATT)5 | TGGCCAAGTAATATGGACGA | TGAAGTACTCCTTCCGTTCCA | 223 |
| chr3:120860725-120862737 | p2 | (AT)6 | ATGACATGTTCGATGGGGAT | GGTAGGTGCCTTGGAACAAA | 101 |
| chr3:120932097-120934118 | p3 | (CAC)7 | TTTCCTGGGGCATGAAAATA | CGTGCACAGTCAAATTGGTT | 292 |
| chr3:120995458-120997473 | p5 | (AAAAT)3 | AGAACCCTCCTGGAAAATGG | GGCCCTGCTAACTAAATCCC | 182 |
| chr3:121015903-121017924 | p3 | (GAA)7 | TTAAACTGGCGCATTTCCTT | GGAAATCATGGCTCCAGAAA | 261 |
| chr3:121076981-121078995 | p2 | (TA)7 | ACCTCGCTGCAACTCTGATT | TTGGTTGCTGTCAAGCAGTC | 163 |

| Marker ID | SSR type | SSR motif | FORWARD PRIMER (5'-3') | REVERSE PRIMER (5'-3') | size (bp) |
|---|---|---|---|---|---|
| chr3:121106392-121108404 | p2 | (AT)6 | CCCGCAAGTATAGGCAACAT | TGATAGGGGTGTGACAAGCA | 112 |
| chr3:121114967-121116985 | p2 | (AG)6 | GAAATTCGTGCGATTAGGGA | TCCCCACTTCACTTGTCTCC | 238 |
| chr3:121114967-121116985 | p3 | (AAT)6 | GAAATTCGTGCGATTAGGGA | TCCCCACTTCACTTGTCTCC | 238 |
| chr3:121232762-121234780 | p6 | (AAAAAT)3 | TTCATGAGGCAATCCTAGCC | CAATCAACAGGGCCATTTCT | 288 |
| chr3:121300102-121302116 | p2 | (GA)7 | CGGACCAAGATACGGTCACT | AACGGTATCTTACGGGGGTC | 123 |
| chr3:121300102-121302116 | p2 | (GA)7 | CGGACCAAGATACGGTCACT | AACGGTATCTTACGGGGGTC | 123 |
| chr3:121300969-121302983 | p2 | (GA)7 | GGTCAAAAGGCATCACCAGT | GTCGAAAGTTTGGGGTTCAA | 271 |
| chr3:121300969-121302983 | p2 | (GA)7 | GGTCAAAAGGCATCACCAGT | GTCGAAAGTTTGGGGTTCAA | 271 |
| chr3:121329640-121331655 | p5 | (ATTTT)3 | AACGCCAACCTACCCTCTCT | CCATTTCAATGTGCTTGGTG | 180 |
| chr3:121356324-121358336 | p5 | (ACCGG)3 | TTGGTGTTTCTCTTTTGGGC | CTTAGGGCAGATGGACCAGA | 156 |
| chr3:121376125-121378145 | p5 | (AATAA)4 | CAATCCAAACGGGCTCTAAG | GCGTGCCTCTGTATCCAAAT | 276 |
| chr3:121468596-121470611 | p5 | (AACCG)3 | CCAATAGCGGCTGGTTAAAA | CGGTTCGGTTTGTTGAAGTT | 299 |
| chr3:121582787-121584803 | p4 | (CAAT)4 | TTTGTCCTCTCCGCATAACC | CTTTTCCTTTTCCCCTTTGC | 225 |
| chr3:121636981-121639001 | p4 | (GGGT)5 | GGTCCAATTGGTGCAAAAGT | TTGGTTGGCTGCAAACATTA | 203 |
| chr3:121724860-121726898 | p2 | (TA)19 | TCTATGAAACCTTGCGGGTC | GGTTGCTTCGGGTCAATAAA | 207 |
| chr3:121812742-121814757 | p5 | (TTTTG)3 | TACGCGAAAATGCAAATGAA | AACCCACAGCCAAACAAGTC | 211 |
| chr3:121838770-121840782 | p2 | (AT)6 | CAGTGGTCACATCAAGGTCG | AGATGTTGGCCTCACAATCC | 295 |
| chr3:121843658-121845676 | p2 | (TA)10 | AGAGTGCATTTAGCCAGGGA | CTTCCTCTTCGCCTTCCTCT | 218 |
| chr3:121843658-121845676 | p2 | (AT)9 | AGAGTGCATTTAGCCAGGGA | CTTCCTCTTCGCCTTCCTCT | 218 |
| chr3:121857790-121859835 | p3 | (TAA)15 | CCAACCTGTCCTCAACGATT | TTTTCCCTTCGTTCCCTTTT | 267 |
| chr3:121925708-121927732 | p2 | (TA)7 | TTTTGGATGGAAAAACGGAC | GAACCTCGACCCACTTTTGA | 211 |
| chr3:121925708-121927732 | p2 | (TA)12 | TTTTGGATGGAAAAACGGAC | GAACCTCGACCCACTTTTGA | 211 |
| chr3:121925708-121927732 | p2 | (TA)12 | TTTTGGATGGAAAAACGGAC | GAACCTCGACCCACTTTTGA | 211 |
| chr3:121925708-121927732 | p3 | (GTT)10 | TTTTGGATGGAAAAACGGAC | GAACCTCGACCCACTTTTGA | 211 |
| chr3:121930840-121932897 | p3 | (TTA)19 | ATAAGGGAGCGAGATGGGTT | TTAACCAAGGAACGGGTCTG | 107 |
| chr3:122147733-122149755 | p2 | (TC)11 | AAGCCAGTAACGCAAGTGCT | ATTGGGACGGTTGTTTCTTG | 133 |
| chr3:122147733-122149755 | p5 | (ACCGG)3 | AAGCCAGTAACGCAAGTGCT | ATTGGGACGGTTGTTTCTTG | 133 |
| chr3:122165170-122167184 | p2 | (AT)13 | GCAGCCATTAGCCCAAAATA | TCGCATTTGCTTACCACTTG | 246 |
| chr3:122165170-122167184 | p2 | (AT)9 | GCAGCCATTAGCCCAAAATA | TCGCATTTGCTTACCACTTG | 246 |
| chr3:122689655-122691667 | p2 | (AG)6 | CGTGGAAGCTGGAAAATCAT | GAACCACCTCGAAGACTCCA | 166 |
| chr3:122701966-122703990 | p3 | (AAT)8 | CTTGAAACTTGTGTCGGCAA | ACCTGCTCTCTGTGCACCTT | 157 |
| chr3:122701966-122703990 | p2 | (TA)30 | CTTGAAACTTGTGTCGGCAA | ACCTGCTCTCTGTGCACCTT | 157 |
| chr3:122702902-122704920 | p3 | (AAT)8 | TGTGCAAATTCAGTGCTGC | AACTGCACAATGGGAGAGGT | 171 |
| chr3:122702902-122704920 | p2 | (TA)30 | TGTGCAAATTCAGTGCTGC | AACTGCACAATGGGAGAGGT | 171 |
| chr3:122702923-122704941 | p3 | (AAT)8 | TGTGCAAATTCAGTGCTGC | AACTGCACAATGGGAGAGGT | 150 |
| chr3:122702923-122704941 | p2 | (TA)30 | TGTGCAAATTCAGTGCTGC | AACTGCACAATGGGAGAGGT | 150 |

| Marker ID | SSR type | SSR motif | FORWARD PRIMER (5'-3') | REVERSE PRIMER (5'-3') | size (bp) |
|---|---|---|---|---|---|
| chr3:122702944-122704962 | p3 | (AAT)8 | TGTGCAAATTCAGTGCTGC | AACTGCACAATGGGAGAGGT | 129 |
| chr3:122702944-122704962 | p2 | (TA)30 | TGTGCAAATTCAGTGCTGC | AACTGCACAATGGGAGAGGT | 129 |
| chr3:122702965-122704983 | p3 | (AAT)8 | TGTGCAAATTCAGTGCTGC | AACTGCACAATGGGAGAGGT | 108 |
| chr3:122702965-122704983 | p2 | (TA)30 | TGTGCAAATTCAGTGCTGC | AACTGCACAATGGGAGAGGT | 108 |
| chr3:122798446-122800470 | p5 | (TATTT)3 | GTTCGGTATCGGGTTAGCAA | GTGATCACCATCGAAAGGGT | 225 |
| chr3:122798446-122800470 | p3 | (ATA)8 | GTTCGGTATCGGGTTAGCAA | GTGATCACCATCGAAAGGGT | 225 |
| chr3:122798446-122800470 | p4 | (ATAG)6 | GTTCGGTATCGGGTTAGCAA | GTGATCACCATCGAAAGGGT | 225 |
| chr3:122957254-122959269 | p5 | (AAGAA)3 | CAATAAACAAGGCGCCAGAT | ATCAATGCTGAAACCATCCC | 154 |
| chr3:122966196-122968208 | p2 | (TA)6 | TCGCAATCAAAACAAATGGA | TTAAAAGCCCACCCTTCCTT | 129 |
| chr3:122992540-122994562 | p3 | (AAT)7 | GGAGAAGAGGCGATGATGAG | TGTTTCTTGCACTCAGTCCG | 274 |
| chr3:123010195-123012213 | p6 | (TTCTTT)3 | TCGTGGGGAAAAATGTCACT | ATGAAATTCAAACCCCCTCC | 185 |
| chr3:123019690-123021702 | p2 | (TG)6 | GGCTCAGTGTGGGAAACATT | CCCCATTGGATTTGTTTGTC | 268 |
| chr3:123043946-123045985 | p3 | (TGT)9 | CGCTTTCGTCTCTCGTTTTC | GCATTGCATTGTGGCTTATG | 290 |
| chr3:123043946-123045985 | p3 | (GTT)13 | CGCTTTCGTCTCTCGTTTTC | GCATTGCATTGTGGCTTATG | 290 |
| chr3:123113046-123115061 | p3 | (CAA)5 | GGGTATTTTGGTTCCGGACT | ATACAGTGGGATTTCGCTGG | 298 |
| chr3:123113046-123115061 | p5 | (TTTTA)3 | GGGTATTTTGGTTCCGGACT | ATACAGTGGGATTTCGCTGG | 298 |
| chr3:123250893-123252913 | p2 | (TA)10 | ATACCACACCCCGAAAATCA | CGTGCTGGAACTTTAGGGAG | 283 |
| chr3:123254905-123256923 | p4 | (ATAA)4 | GGGAGTTTCCTTTGCAACAA | CCATTGAAGCTCAGCTCCTC | 170 |
| chr3:123254905-123256923 | p6 | (ATTTTT)3 | GGGAGTTTCCTTTGCAACAA | CCATTGAAGCTCAGCTCCTC | 170 |
| chr3:123254905-123256923 | p3 | (AGA)5 | GGGAGTTTCCTTTGCAACAA | CCATTGAAGCTCAGCTCCTC | 170 |
| chr3:123254905-123256923 | p2 | (AT)6 | GGGAGTTTCCTTTGCAACAA | CCATTGAAGCTCAGCTCCTC | 170 |
| chr3:123255247-123257262 | p4 | (ATAA)4 | AATGGGATTGGGTCAGAACA | GGCCAGCCAAACACTTAAAA | 244 |
| chr3:123255247-123257262 | p6 | (ATTTTT)3 | AATGGGATTGGGTCAGAACA | GGCCAGCCAAACACTTAAAA | 244 |
| chr3:123255247-123257262 | p3 | (AGA)5 | AATGGGATTGGGTCAGAACA | GGCCAGCCAAACACTTAAAA | 244 |
| chr3:123255247-123257262 | p2 | (AT)6 | AATGGGATTGGGTCAGAACA | GGCCAGCCAAACACTTAAAA | 244 |
| chr3:123330266-123332292 | p2 | (AT)13 | GGATGCGTTTGTCAGTGCTA | TCAAATCCAAGAACCCAAGG | 223 |
| chr3:123352598-123354614 | p4 | (CATA)4 | ACATTAAGTGGCCGGAACAG | CGTACCAACGTCCTTTTGCT | 299 |
| chr3:123519412-123521436 | p3 | (AAT)8 | CGAACCCTGGACCTTTATCA | AAAGTTGCAGGCATTTCAGG | 178 |
| chr3:123535492-123537516 | p3 | (ACA)8 | GCCAATCCAAACACCCTCTA | CCATGGCTTTGTTGGACTCT | 128 |
| chr3:123579584-123581599 | p5 | (ATAAC)3 | TCAATCCAAAGCCCAAAAAG | GAGAGGATGGTGGAAAACGA | 213 |
| chr3:123610579-123612594 | p4 | (AAGA)4 | TGATGAAGCAGTTTGCAAGG | GGCGCAGTTATAGCTTACCG | 164 |
| chr3:123610579-123612594 | p3 | (CCG)5 | TGATGAAGCAGTTTGCAAGG | GGCGCAGTTATAGCTTACCG | 164 |
| chr3:123792504-123794519 | p5 | (AAGAT)3 | CTAGAGACCACGCCAAAAGC | GTGAGTGAATGAGGCCAACA | 138 |
| chr3:123794960-123796978 | p6 | (TTTTCC)3 | CCGTGTTCAAAACTTGGTCA | AACACCAACCCCAACACATT | 258 |
| chr3:123817128-123819146 | p6 | (ATTAAG)3 | GGGGTTTAGGAGAAATTGGC | TTTGGAGAGTCCGAGCAACT | 185 |
| chr3:123817128-123819146 | p2 | (TA)10 | GGGGTTTAGGAGAAATTGGC | TTTGGAGAGTCCGAGCAACT | 185 |

| Marker ID | SSR type | SSR motif | FORWARD PRIMER (5'-3') | REVERSE PRIMER (5'-3') | size (bp) |
|---|---|---|---|---|---|
| chr3:123867738-123869758 | p5 | (TTTAT)4 | AGGGGGTTCCTTTCCCTATT | GCTCCAACTCACGCACTGTA | 213 |
| chr3:123896266-123898278 | p2 | (CA)6 | ATCACTCGCCAAACAAATCC | GATGCTGCCGAAATTTTTGT | 299 |
| chr3:123920109-123922123 | p2 | (AG)7 | AATCGTAGGTACGGGGCTTT | TCGGCCGCTGTTAGAGTATT | 147 |
| chr3:124059999-124062014 | p3 | (AAC)6 | AGCTGATTCGGTGAAATGCT | CTTTGAGTCGTCAAGGGAGC | 243 |
| chr3:124059999-124062014 | p2 | (TC)6 | AGCTGATTCGGTGAAATGCT | CTTTGAGTCGTCAAGGGAGC | 243 |
| chr3:124124003-124126018 | p3 | (AAG)5 | TGTTTCCAAAAGATCCTCGG | AAAATTACCCGATGAGCACG | 184 |
| chr3:124172261-124174279 | p6 | (TTTTTA)3 | TCAACGTGTGCTGCTTTTTC | AGCATTCACGCGCTAAACTT | 156 |
| chr3:124179729-124181744 | p5 | (TTGTA)3 | TTCCTCTCCACTTCAATGCC | GTGCAGGAACAAAGGGTGTT | 216 |
| chr3:124179729-124181744 | p5 | (GGGTG)3 | TTCCTCTCCACTTCAATGCC | GTGCAGGAACAAAGGGTGTT | 216 |
| chr3:124187301-124189325 | p6 | (CCGAAG)4 | GGGAGGACAATGACAAGGAA | CAAAGGGGTAACGGAAGACA | 200 |
| chr3:124329230-124331242 | p2 | (AT)6 | GGACGGTTTGGGTAAAGTCA | TAAAAAGGGGTGTGACAGGG | 198 |
| chr3:124329230-124331242 | p2 | (AT)8 | GGACGGTTTGGGTAAAGTCA | TAAAAAGGGGTGTGACAGGG | 198 |
| chr3:124459098-124461118 | p5 | (GGCAT)4 | ATGCAAGGGACCTTTTTGTG | TGGGCCTTCAATTAATAGCG | 280 |
| chr3:124467717-124469732 | p5 | (AGATA)3 | CGCCCAGAAGACAGACCTAC | CCCACAACTCATACACACGG | 204 |
| chr3:124467717-124469732 | p2 | (TG)7 | CGCCCAGAAGACAGACCTAC | CCCACAACTCATACACACGG | 204 |
| chr3:124485900-124487915 | p5 | (TATTT)3 | TGCAAGATACGTGGAGGTCA | AACTTCCCTGAGAGGCCATT | 131 |
| chr3:124494782-124496797 | p3 | (ACA)5 | CCATGACCATCAAACACAGC | AGGGTGTGAGATGCTCGACT | 181 |
| chr3:124520028-124522043 | p5 | (ATACC)3 | TTGGGGGAAATTAGAAACCC | TGATCCCTTTTGTTTTTGCC | 176 |
| chr3:124520028-124522043 | p3 | (TTC)5 | TTGGGGGAAATTAGAAACCC | TGATCCCTTTTGTTTTTGCC | 176 |
| chr3:124520242-124522267 | p5 | (ATACC)3 | AATATGCACCTTACACGCCC | GCAACGATGCACCTAACCTT | 231 |
| chr3:124520242-124522267 | p3 | (TTC)5 | AATATGCACCTTACACGCCC | GCAACGATGCACCTAACCTT | 231 |
| chr3:124598638-124600653 | p3 | (TTA)5 | GTGGATCGGTGCTGATTTTT | TTTGTCTCGGGATTTGAACC | 199 |
| chr3:124607844-124609859 | p5 | (TATTT)3 | CGATTGTCCCCTCTGGAATA | TTTGAGGGTAGTGGGTGGAG | 240 |
| chr3:124640490-124642510 | p3 | (TTA)9 | TCGACCTTGTTTGGTTTTCC | GCGGTCCCTAATTTGTTTGA | 223 |
| chr3:124640490-124642510 | p2 | (CA)6 | TCGACCTTGTTTGGTTTTCC | GCGGTCCCTAATTTGTTTGA | 223 |
| chr3:124640549-124642605 | p3 | (TTA)9 | TCGACCTTGTTTGGTTTTCC | GCGGTCCCTAATTTGTTTGA | 164 |
| chr3:124640549-124642605 | p2 | (CA)6 | TCGACCTTGTTTGGTTTTCC | GCGGTCCCTAATTTGTTTGA | 164 |
| chr3:124649988-124652002 | p2 | (TA)7 | TCCACCAGCAACCTACCTTC | AAAGCGCAATTTGTGGAAAG | 222 |
| chr3:124660434-124662452 | p6 | (AAAAAT)3 | TGATTCGGAGTACGAGGGTC | GGAGCACCCAAAATTTGCTA | 154 |
| chr3:124756237-124758255 | p2 | (AG)9 | AAGAAGGAACAGTGCGGCTA | AAGTCGTCTGCAGTCTCGGT | 216 |
| chr3:124756237-124758255 | p3 | (TTG)5 | AAGAAGGAACAGTGCGGCTA | AAGTCGTCTGCAGTCTCGGT | 216 |
| chr3:124837472-124839490 | p3 | (TTG)6 | GATAAAAAGGCCACCGACAA | ATGGGACAGCAGGGTGTTAC | 104 |
| chr3:124850106-124852127 | p3 | (TTC)7 | ATTGCATCTCATCAAAGGGG | TTTCTTTGCCCAGAGCAACT | 142 |
| chr3:124864089-124866109 | p5 | (AAAAT)4 | TAAACGGGCCTTACATTTGC | CACCAATTTTCCCAATACGG | 133 |
| chr3:124982182-124984197 | p5 | (ACCAA)3 | ATTATGGGATGGACCCAACA | GAGAAAATCCAACAACCCGA | 199 |
| chr3:124982182-124984197 | p5 | (AAACC)3 | ATTATGGGATGGACCCAACA | GAGAAAATCCAACAACCCGA | 199 |

| Marker ID | SSR type | SSR motif | FORWARD PRIMER (5'-3') | REVERSE PRIMER (5'-3') | size (bp) |
|---|---|---|---|---|---|
| chr3:124982182-124984197 | p6 | (CTTCAA)4 | ATTATGGGATGGACCCAACA | GAGAAAATCCAACAACCCGA | 199 |
| chr3:124993800-124995812 | p2 | (AT)6 | TATTTCAAACCCCAAATGGC | TGGGACAGAACCAGTGTACG | 225 |
| chr3:125085414-125087432 | p3 | (TTC)5 | GAACTGCCACATCAGCAAAA | CACTCATTCAGTCACGCGAT | 147 |
| chr3:125085414-125087432 | p6 | (TTTTAT)3 | GAACTGCCACATCAGCAAAA | CACTCATTCAGTCACGCGAT | 147 |
| chr3:125116773-125118788 | p2 | (GA)7 | TTGATTATTCAAATCCGGGG | CAGTGGATGGAATTCTTGCT | 243 |
| chr3:125116773-125118788 | p5 | (AATAA)3 | TTGATTATTCAAATCCGGGG | CAGTGGATGGAATTCTTGCT | 243 |
| chr3:125191383-125193398 | p2 | (AT)12 | AAAGCCCAAAACAATAGGGG | AGCTAATCCCACAGCGAGAC | 155 |
| chr3:125191383-125193398 | p5 | (AACAA)3 | AAAGCCCAAAACAATAGGGG | AGCTAATCCCACAGCGAGAC | 155 |
| chr3:125228664-125230676 | p2 | (AT)6 | ATCCCGTGGGTACGGATACT | GGGTGCCATGTTATTGCTTT | 100 |
| chr3:125239997-125242012 | p5 | (AACCG)3 | TTGGAGGCAGGATTACAACC | CGGTTCGGTTTGTTGAAGTT | 261 |
| chr3:125239997-125242012 | p5 | (AACCG)3 | TTGGAGGCAGGATTACAACC | CGGTTCGGTTTGTTGAAGTT | 261 |
| chr3:125284786-125286798 | p2 | (AT)6 | AAAATCGCCCAGAAGACAGA | CGCCATGTCATTGTAACACC | 120 |
| chr3:125291875-125293890 | p3 | (CAA)5 | ATATGCTAAGAGCCGCGAAA | ATCTGCTTCCAAGGGATGTG | 143 |
| chr3:125305754-125307766 | p2 | (TA)6 | ACGCTTGAACCTGTGAAACC | TGGAAAAACACCATAGGGGA | 286 |
| chr3:125305754-125307766 | p4 | (AAAT)8 | ACGCTTGAACCTGTGAAACC | TGGAAAAACACCATAGGGGA | 286 |
| chr3:125305932-125307964 | p2 | (TA)6 | CGCTTGAACCTGTGAAACCT | TGGAAAAACACCATAGGGGA | 109 |
| chr3:125305932-125307964 | p4 | (AAAT)8 | CGCTTGAACCTGTGAAACCT | TGGAAAAACACCATAGGGGA | 109 |
| chr3:125408240-125410255 | p2 | (AT)10 | TTTTTAAGGGACAACGGACG | GGGCCCCATACTAAACAACA | 223 |
| chr3:125408240-125410255 | p2 | (TA)6 | TTTTTAAGGGACAACGGACG | GGGCCCCATACTAAACAACA | 223 |
| chr3:125408240-125410255 | p2 | (AT)7 | TTTTTAAGGGACAACGGACG | GGGCCCCATACTAAACAACA | 223 |
| chr3:125408240-125410255 | p5 | (ATTTT)3 | TTTTTAAGGGACAACGGACG | GGGCCCCATACTAAACAACA | 223 |
| chr3:125408240-125410255 | p2 | (AT)12 | TTTTTAAGGGACAACGGACG | GGGCCCCATACTAAACAACA | 223 |
| chr3:125408240-125410255 | p2 | (AT)17 | TTTTTAAGGGACAACGGACG | GGGCCCCATACTAAACAACA | 223 |
| chr3:125408908-125410942 | p2 | (AT)7 | TGGGGCTCACAATTCTTTTT | GAACCCACCACATCCAAACT | 127 |
| chr3:125408908-125410942 | p5 | (ATTTT)3 | TGGGGCTCACAATTCTTTTT | GAACCCACCACATCCAAACT | 127 |
| chr3:125408908-125410942 | p2 | (AT)12 | TGGGGCTCACAATTCTTTTT | GAACCCACCACATCCAAACT | 127 |
| chr3:125408908-125410942 | p2 | (AT)17 | TGGGGCTCACAATTCTTTTT | GAACCCACCACATCCAAACT | 127 |
| chr3:125408908-125410942 | p2 | (AT)15 | TGGGGCTCACAATTCTTTTT | GAACCCACCACATCCAAACT | 127 |
| chr3:125408908-125410942 | p2 | (AT)6 | TGGGGCTCACAATTCTTTTT | GAACCCACCACATCCAAACT | 127 |
| chr3:125408908-125410942 | p2 | (TA)7 | TGGGGCTCACAATTCTTTTT | GAACCCACCACATCCAAACT | 127 |
| chr3:125455376-125457391 | p5 | (AAACC)3 | ACCAAACCAAACCAAACCAA | ATAACCCGAGAAAACCCGAG | 264 |
| chr3:125459027-125461042 | p3 | (ATG)5 | CACTTTCTTGCCAGTCTCCC | AGATCTTCGTGGAATGTCGG | 166 |
| chr3:125496632-125498644 | p2 | (TA)6 | GGGTTCGGTTGCATTTAGAA | ATGTGGGCTGCCTACGTATC | 231 |
| chr3:125500318-125502330 | p2 | (CT)6 | CTACGTCAACTTCTTCGCCC | GACTCCAAACCCGAAAATGA | 161 |
| chr3:125581349-125583377 | p2 | (AT)14 | GGACGAGAGGCAACAAGAAG | CGGGTAGGGTTGTGACAGAT | 153 |
| chr3:125581349-125583377 | p2 | (AT)38 | GGACGAGAGGCAACAAGAAG | CGGGTAGGGTTGTGACAGAT | 153 |

| Marker ID | SSR type | SSR motif | FORWARD PRIMER (5'-3') | REVERSE PRIMER (5'-3') | size (bp) |
|---|---|---|---|---|---|
| chr3:125790794-125792864 | p2 | (AT)35 | TACAGCAGATGAGGTGTGGC | TACAGCTTGTCATTGGCGTC | 173 |
| chr3:125869190-125871205 | p3 | (TTA)6 | TCGGAAATCAATTTGAAGGG | ACTTCAAAACAAACACCGCC | 235 |
| chr3:125869190-125871205 | p5 | (TATTG)3 | TCGGAAATCAATTTGAAGGG | ACTTCAAAACAAACACCGCC | 235 |
| chr3:125997682-125999697 | p5 | (GTTTG)3 | TCGATCCGAACCAGTAAACC | CCAAACCAAATGTTATCGGG | 136 |
| chr3:126030218-126032233 | p5 | (TTGGT)3 | TATTGCAAGGACTCACGACG | TGGATCGATATTCAGCCTCC | 181 |
| chr3:126030218-126032233 | p5 | (TTGGT)3 | TATTGCAAGGACTCACGACG | TGGATCGATATTCAGCCTCC | 181 |
| chr3:126032961-126034976 | p5 | (GGTTT)3 | TTCGTGATGCAATCCGTTTA | TTCCCAGTGGACAACACTCA | 296 |
| chr3:126033001-126035016 | p5 | (GGTTT)3 | TTCGTGATGCAATCCGTTTA | TTCCCAGTGGACAACACTCA | 256 |
| chr3:126055976-126057991 | p3 | (AAC)5 | CACATAAGTTGCGCGAAAAA | CACCAAGAACGCAGTTTCAA | 277 |
| chr3:126126018-126128056 | p2 | (TA)19 | ATGATGCCAAGACCAAAAGG | ACCCTAACCCCATCCTCAAT | 220 |
| chr3:126129334-126131354 | p2 | (AC)9 | ACTTTAGTCCGCAGCCTTGA | CCCAAGGCCTAGACATGAAA | 254 |
| chr3:126129334-126131354 | p2 | (TA)6 | ACTTTAGTCCGCAGCCTTGA | CCCAAGGCCTAGACATGAAA | 254 |
| chr3:126129356-126131368 | p2 | (TA)6 | ACTTTAGTCCGCAGCCTTGA | CCCAAGGCCTAGACATGAAA | 232 |
| chr3:126209755-126211770 | p3 | (ATT)5 | TGCCGTTGGTTACTGAAACA | CCCGACAATCTCTTGGAAAA | 108 |
| chr3:126209755-126211770 | p2 | (AT)7 | TGCCGTTGGTTACTGAAACA | CCCGACAATCTCTTGGAAAA | 108 |
| chr3:126209755-126211770 | p2 | (AT)13 | TGCCGTTGGTTACTGAAACA | CCCGACAATCTCTTGGAAAA | 108 |
| chr3:126324171-126326191 | p2 | (AT)10 | CCAGCTGGTAGCCAAATGAT | TCGGAATTTACGATCCCATC | 242 |
| chr3:126387017-126389037 | p2 | (AT)10 | CCAGCTGGTAGCCAAATGAT | TCGGAATTTACGATCCCATC | 242 |
| chr3:126423059-126425073 | p2 | (TA)7 | TAAGCTGGCTTGACCGACTT | GCGGGTATGCATCTTTTGAT | 184 |
| chr3:126528177-126530193 | p4 | (ATAC)4 | AGTGTGGGCTTCTTGCCTAA | TATGCAACTGGAAACACCGA | 194 |
| chr3:126567982-126569996 | p2 | (AC)7 | GAATCATTCTTTCCCCAGCA | AGAGGGCAGACGTTGGTAGA | 139 |
| chr3:126567982-126569996 | p5 | (TTATT)3 | GAATCATTCTTTCCCCAGCA | AGAGGGCAGACGTTGGTAGA | 139 |
| chr3:126748854-126750869 | p3 | (AAC)5 | ATTATTTGAAGCTGCGGTGG | GATGTGCGGCTATTCAAGGT | 141 |
| chr3:126768025-126770037 | p2 | (GC)6 | TTCTGCCACCCATTTTCTTC | TGCCACATAAGATGCCACAT | 151 |
| chr3:126790601-126792616 | p5 | (AAATG)3 | TTTTATGGGGTTTATGGGCA | GGCACCTGACGAGCTTCTAC | 163 |
| chr3:126799793-126801808 | p5 | (AAATG)3 | TTTTATGGGGTTTATGGGCA | GGCACCTGACGAGCTTCTAC | 163 |
| chr3:126807119-126809157 | p2 | (AT)30 | CCACCTTATCCCGAGTACCA | ACTTGGGGGAATGGAAAAAC | 222 |
| chr3:126807119-126809157 | p2 | (TA)19 | CCACCTTATCCCGAGTACCA | ACTTGGGGGAATGGAAAAAC | 222 |
| chr3:126807119-126809157 | p2 | (AT)8 | CCACCTTATCCCGAGTACCA | ACTTGGGGGAATGGAAAAAC | 222 |
| chr3:126863599-126865683 | p2 | (AT)70 | TTCCTTTTGGACTTTGTGCC | CTGCTGCTTCTGACCATTGA | 145 |
| chr3:126902083-126904099 | p4 | (TTTA)4 | ATTCATGCGCCACTAACCAT | CCCGATCAAGCTACCAAAAA | 249 |
| chr3:126913686-126915706 | p5 | (ATTTT)3 | TGTGCCAATGGCTAAATGAA | TTTCGGTCCAAGTTTATCCG | 300 |
| chr3:126960538-126962550 | p2 | (AT)6 | GTGGTGGTGGAGCTACTGGT | AGAAGCCTAAAGAGCCCGAG | 256 |
| chr3:126993315-126995330 | p5 | (TTCTT)3 | AGGGAACAAGAGCATAGCCA | GCATTCAGGTTCTCTTTCGC | 197 |
| chr3:127059716-127061740 | p3 | (AAT)8 | TCCTTGCGTGAGTGTCTTTG | TCAAACGGTAGCCGGTAAAG | 146 |
| chr3:127075736-127077752 | p2 | (TA)8 | AAAAGCTGTTCATTCCCACG | GCAGACTGAAAGGGGTATGC | 294 |

| Marker ID | SSR type | SSR motif | FORWARD PRIMER (5'-3') | REVERSE PRIMER (5'-3') | size (bp) |
|---|---|---|---|---|---|
| chr3:127075736-127077752 | p6 | (GAGACA)3 | AAAAGCTGTTCATTCCCACG | GCAGACTGAAAGGGGTATGC | 294 |
| chr3:127092722-127094738 | p4 | (CCCA)4 | GAAAGGGACCAAAAAGCCTC | AAGCCTTGCGTCTCACACTT | 245 |
| chr3:127105607-127107619 | p2 | (AG)6 | TAAAAAGGCTGCCCAGAAAA | TTCCCATGGCCAAGATAGAG | 229 |
| chr3:127243356-127245372 | p2 | (TC)8 | TTCACATGGTGTTTGCCTGT | CAATGCGTTCAATGACAACC | 170 |
| chr3:127308496-127310518 | p2 | (TA)17 | TGTGTAGATCTTCGCGTTGC | TGTTGGTCCTCCTTAGCCAC | 242 |
| chr3:127308496-127310518 | p2 | (TA)11 | TGTGTAGATCTTCGCGTTGC | TGTTGGTCCTCCTTAGCCAC | 242 |
| chr3:127384489-127386503 | p2 | (AT)7 | AGTGGAGGAAACATGAACGG | CTCTCGGGTTAAAATGGGGT | 247 |
| chr3:127466548-127468563 | p5 | (ATTTT)3 | ATGCCCCTATTTTAGGGTGG | GCCTTATCCGGCATAATTCA | 291 |
| chr3:127469823-127471837 | p2 | (TA)7 | AAATTTTGATTCCGCCACTG | GATGTGGCTCATCTTCCCAT | 229 |
| chr3:127542071-127544086 | p5 | (ATATT)3 | GATGTTGTTCATGACGTCGG | CAAAATCGCCAATCTTCCAT | 117 |
| chr3:127542071-127544086 | p4 | (AAAT)4 | GATGTTGTTCATGACGTCGG | CAAAATCGCCAATCTTCCAT | 117 |
| chr3:127665980-127667995 | p6 | (AAGAAA)3 | CTCCATTTGAGGAGAGACGC | TTTGTGCTGATGAGAGGCTG | 228 |
| chr3:127666085-127668109 | p6 | (AAGAAA)3 | CTCCATTTGAGGAGAGACGC | TTTGTGCTGATGAGAGGCTG | 123 |
| chr3:127812895-127814911 | p4 | (ATTA)4 | CTTAACTAGGGGTCTCGGGC | ATTCAGTACCCGCATTGGAG | 277 |
| chr3:127864590-127866606 | p4 | (ATTA)4 | CTTAACTAGGGGTCTCGGGC | ATTCAGTACCCGCATTGGAG | 277 |
| chr3:127979365-127981380 | p3 | (TTA)11 | TCGCCTAGCATATTGTTCCC | CGCCATTTTTGATAACGTCC | 300 |
| chr3:128125156-128127201 | p3 | (TTA)15 | TACGCGGGTATGGGATATGT | GGTTGGCTCCTTTACCTTCC | 293 |
| chr3:128125156-128127201 | p2 | (CT)7 | TACGCGGGTATGGGATATGT | GGTTGGCTCCTTTACCTTCC | 293 |
| chr3:128173762-128175774 | p2 | (TA)6 | TGCCCTCTTGTTATATCCCG | TTGGAGGGTTTTCACTTTGG | 132 |
| chr3:128242789-128244810 | p3 | (GTT)7 | ATGTTTGTATGGGGATCGGA | ATCTTGTCATTCCACCGAGG | 268 |
| chr3:128281068-128283083 | p5 | (AAACC)3 | ACACCTCGGAGCAAAAACAG | AGTCAAACCGAACCGTGAAC | 132 |
| chr3:128283794-128285890 | p2 | (AT)48 | GGTGGCCTAATGGGACTGTA | CATAAAAACTAAGCCCCGCA | 199 |
| chr3:128393151-128395163 | p2 | (TA)6 | TGCAGCACCACACATAAACA | GGTGGGTTGAGTTCTCCAAA | 284 |
| chr3:128423525-128425543 | p3 | (AAG)6 | ATACACTTTGGGCAGGGTTG | AAACGTAAGGGTGGCAAATG | 202 |
| chr3:128500010-128502064 | p3 | (AAT)29 | CCACGCCCAACAATATATCC | TAAAATGGGAAAGGGTCACG | 192 |
| chr3:128500010-128502064 | p3 | (ATA)18 | CCACGCCCAACAATATATCC | TAAAATGGGAAAGGGTCACG | 192 |
| chr3:128587481-128589493 | p2 | (AT)6 | TCTTTTCTCGGATCAGCCAC | GGGCTTGTCACAACCAACTT | 295 |
| chr3:128596007-128598023 | p2 | (AG)8 | AAACATGGTTCTTTTTGCCG | TGTCTGCATGGAGAAAGCTG | 128 |
| chr3:128598829-128600845 | p2 | (AG)8 | AAACATGGTTCTTTTTGCCG | TGTCTGCATGGAGAAAGCTG | 128 |
| chr3:128616879-128618894 | p3 | (ATA)5 | TGAGGCTCGACGTGACATAG | TTAAAGTAGCTCGGGTGCGT | 174 |
| chr3:128667123-128669147 | p2 | (TC)7 | AACGTAGCCACTTTCCCCTT | ACCCCCGCTGATTCTTATTT | 244 |
| chr3:128667123-128669147 | p3 | (ACA)8 | AACGTAGCCACTTTCCCCTT | ACCCCCGCTGATTCTTATTT | 244 |
| chr3:128716323-128718339 | p5 | (AATAA)3 | CTCAAAAGCCCTCTCACCAG | TCCACAGACGCAGAAGAATG | 106 |
| chr3:128742889-128744904 | p3 | (ACA)5 | CCCCATCTGTGTCCTCACTT | CGTTCAAATCGGCCTACTGT | 100 |
| chr3:128742889-128744904 | p3 | (TTC)12 | CCCCATCTGTGTCCTCACTT | CGTTCAAATCGGCCTACTGT | 100 |
| chr3:128766871-128768883 | p2 | (GT)8 | CCAAGGCTTCACATTAGGGA | AGATCCGATTCGTTCGAAAA | 285 |

| Marker ID | SSR type | SSR motif | FORWARD PRIMER (5'-3') | REVERSE PRIMER (5'-3') | size (bp) |
|---|---|---|---|---|---|
| chr3:128766871-128768883 | p2 | (GT)7 | CCAAGGCTTCACATTAGGGA | AGATCCGATTCGTTCGAAAA | 285 |
| chr3:128766871-128768883 | p2 | (GT)6 | CCAAGGCTTCACATTAGGGA | AGATCCGATTCGTTCGAAAA | 285 |
| chr3:128766871-128768883 | p2 | (CA)6 | CCAAGGCTTCACATTAGGGA | AGATCCGATTCGTTCGAAAA | 285 |
| chr3:128798649-128800665 | p4 | (TTGC)4 | TGAGTGAATGAGCGTCAACC | ATAAAACGACGCGTCAATCC | 300 |
| chr3:128895090-128897105 | p5 | (TAATT)3 | GATGTTGAACCTCGGCTTGT | AAGGCCCTATTGTGTGATGC | 102 |
| chr3:128952106-128954124 | p6 | (TATAAA)3 | ATACATTGCCGAAACACCGT | AGGGGAATTGCCTCCTTAAA | 165 |
| chr3:129085004-129087016 | p2 | (AG)6 | GAGGGATGGCTCAGTGGTTA | TTTTCCCCTCCCATTTTAGG | 175 |
| chr3:129100073-129102089 | p5 | (TGATA)4 | ACCAGACTCCACGTACCCAC | TCCATACCTGCATCGTGAAA | 298 |
| chr3:129100091-129102105 | p5 | (TGATA)4 | ACCAGACTCCACGTACCCAC | TCCATACCTGCATCGTGAAA | 280 |
| chr3:129118093-129120105 | p2 | (GA)6 | AGCACCTTTCCATGCAGTCT | TGACCCTCTGGAGGAGAAGA | 272 |
| chr3:129142849-129144881 | p2 | (AT)16 | TCCACCTCGATAGACGAACC | TAGTGGCTCTTGTACGGCCT | 113 |
| chr3:129171952-129173968 | p2 | (CA)9 | ACTATCAAGCAACCATGCCC | GAGAGCGATTCCTGGTGGTA | 162 |
| chr3:129171952-129173968 | p2 | (AC)8 | ACTATCAAGCAACCATGCCC | GAGAGCGATTCCTGGTGGTA | 162 |
| chr3:129175025-129177041 | p4 | (AATA)4 | ATGCATGCAGCCTTACATCA | CATTGAAACGCTGGGAAACT | 255 |
| chr3:129183220-129185234 | p2 | (CA)7 | TCGGGATCACTCAAAACCTC | CATGAGGTTTTCAATTCGGG | 284 |
| chr3:129183220-129185234 | p2 | (AC)7 | TCGGGATCACTCAAAACCTC | CATGAGGTTTTCAATTCGGG | 284 |
| chr3:129183353-129185367 | p2 | (CA)7 | TCGGGATCACTCAAAACCTC | CATGAGGTTTTCAATTCGGG | 151 |
| chr3:129183353-129185367 | p2 | (AC)7 | TCGGGATCACTCAAAACCTC | CATGAGGTTTTCAATTCGGG | 151 |
| chr3:129209854-129211866 | p2 | (GA)6 | TTATTTCGGGCAGCGATAAC | TTCTGGAAATTTCGTCTGGG | 252 |
| chr3:129253983-129256008 | p5 | (TTTCC)3 | GTAATGGAAGGGAATGGGCT | GCCGGTTTGAAGAGTGAATC | 127 |
| chr3:129294974-129296998 | p6 | (TAGATG)4 | GTGCTTCAATTGTCACCACG | GGCATGGTGGTGAGTCTTTT | 195 |
| chr3:129342358-129344382 | p5 | (TTCTC)3 | TGAGCGACGTGAAACTTTTG | TGCATGGACTTCTATGCTGC | 293 |
| chr3:129342710-129344725 | p5 | (TTCTC)3 | GGGCCTAAAATACCCTCCAA | TTGACCCCATAGGTGGATGT | 271 |
| chr3:129472890-129474905 | p5 | (TATTG)3 | AAGATGTTGCGAGGGAAATG | CGCAGTCACGAAACCTCTTT | 122 |
| chr3:129472890-129474905 | p5 | (CAATA)3 | AAGATGTTGCGAGGGAAATG | CGCAGTCACGAAACCTCTTT | 122 |
| chr3:129527327-129529363 | p2 | (TA)18 | GAACTGGAATGCGACCAAAT | GTAACCCAGGGCGGATCTAT | 182 |
| chr3:129549521-129551549 | p2 | (AT)14 | TTGTTTGCACGACAAGAAGG | TAACGAACGAACTCGGGAAC | 240 |
| chr3:129583280-129585295 | p5 | (AAATA)3 | CCCCTACCTTGTGACGGTAA | CCCACAAAGTGGGGTACTGT | 154 |
| chr3:129583280-129585295 | p2 | (TA)16 | CCCCTACCTTGTGACGGTAA | CCCACAAAGTGGGGTACTGT | 154 |
| chr3:129583280-129585295 | p5 | (TTTCT)3 | CCCCTACCTTGTGACGGTAA | CCCACAAAGTGGGGTACTGT | 154 |
| chr3:129602450-129604465 | p3 | (ACA)5 | ACGTAGTTCAAATGGACGCC | TAGCTGGGTTGGTTCTGCTT | 298 |
| chr3:129626695-129628710 | p2 | (AT)13 | TCCGTTTGACCCCCATAATA | GCATTCTGGAATTTGGCATT | 275 |
| chr3:129753686-129755722 | p3 | (TAA)5 | ACCGGGCTTTAATTTGGAGT | TACCAACTTCCGCACAAACA | 204 |
| chr3:129753686-129755722 | p3 | (ATA)12 | ACCGGGCTTTAATTTGGAGT | TACCAACTTCCGCACAAACA | 204 |
| chr3:129766790-129768808 | p3 | (ACA)6 | TTCACACTTGACGGGTTTCA | AGCAGAGGTCAGTGCAGGTT | 117 |
| chr3:129798040-129800052 | p2 | (AT)6 | AGCCCGTTTGGATTGACTTA | TAAGAGCCTGTTTGGATGGG | 193 |

| Marker ID | SSR type | SSR motif | FORWARD PRIMER (5'-3') | REVERSE PRIMER (5'-3') | size (bp) |
|---|---|---|---|---|---|
| chr3:129915133-129917185 | p2 | (AT)26 | TATTGTGGTCAGCAAGCAGC | TTGTGGTACGTGAACATCGG | 143 |
| chr3:130000295-130002313 | p2 | (AT)9 | GTTCGTTTGGTTCCGTTGAT | GCCAGAAAATGGTGGAAAAA | 113 |
| chr3:130038274-130040314 | p2 | (AT)20 | CCCATTGAAGGTGTTGGACT | GTTGGGTGCGAAAAGTTCAT | 123 |
| chr3:130041881-130043899 | p2 | (TA)9 | CTGCCTGAGGCCTAACTTTG | CCCTTATTTTGGGGTGGTCT | 194 |
| chr3:130048426-130050444 | p6 | (TTTTAT)3 | TACTAGATGGGTATGGCCCG | TGAATTCTTTTCCCCGTGTC | 142 |
| chr3:130060490-130062517 | p5 | (AAAAT)3 | GGGGTGGATTTTGAGGATTT | TTTCTGCTTCACGCTTTCCT | 255 |
| chr3:130060616-130062640 | p5 | (AAAAT)3 | GGGGTGGATTTTGAGGATTT | TTTCTGCTTCACGCTTTCCT | 129 |
| chr3:130070767-130072835 | p2 | (AT)6 | CACCGGTACCACTCTCCCTA | TTTCGACAATTGGATCGTGA | 263 |
| chr3:130070767-130072835 | p6 | (TATTTT)3 | CACCGGTACCACTCTCCCTA | TTTCGACAATTGGATCGTGA | 263 |
| chr3:130184565-130186589 | p2 | (AC)12 | TGATGGCACTTCATGGTTGT | CCCGAACCCCAGGTATCTAT | 192 |
| chr3:130252019-130254079 | p2 | (TA)6 | GAGAAGCTAACAAGGTGGCG | CCTTTGACCTTTAGGGGGAG | 180 |
| chr3:130252019-130254079 | p2 | (AT)30 | GAGAAGCTAACAAGGTGGCG | CCTTTGACCTTTAGGGGGAG | 180 |
| chr3:130264484-130266502 | p6 | (GTCCAT)3 | ACCCAATCAAGACATGGGAA | GCCCGTAGATTTCACCGTAA | 196 |
| chr3:130276291-130278305 | p2 | (AT)7 | GACAATAGCCAGCGAAGAGG | CAATGGCAGAGCCACATAGA | 194 |
| chr3:130276291-130278305 | p2 | (AT)7 | GACAATAGCCAGCGAAGAGG | CAATGGCAGAGCCACATAGA | 194 |
| chr3:130284538-130286560 | p2 | (TA)11 | CTCACTAAGGAGAATCGCCG | GTGCGTGTGTAGGTACCGTG | 176 |
| chr3:130303983-130305995 | p6 | (TTTTTA)3 | ATGATGGATTCATTGCCACA | AAATGACATGGCGGATTAGC | 136 |
| chr3:130304001-130306015 | p6 | (TTTTTA)3 | ATGATGGATTCATTGCCACA | AAATGACATGGCGGATTAGC | 118 |
| chr3:130313767-130315783 | p2 | (AT)8 | TGGCTTATGGGTCCTTCTTG | ATATGTTGTGTGTCGGGCAA | 175 |
| chr3:130313767-130315783 | p2 | (AT)7 | TGGCTTATGGGTCCTTCTTG | ATATGTTGTGTGTCGGGCAA | 175 |
| chr3:130333213-130335245 | p2 | (TA)7 | TTCTCCACCGGATGAAAAAG | AAGGACATCCCACAACTTGC | 244 |
| chr3:130333213-130335245 | p2 | (TA)16 | TTCTCCACCGGATGAAAAAG | AAGGACATCCCACAACTTGC | 244 |
| chr3:130383011-130385025 | p2 | (GA)7 | TGTGAAATTTGCCTCGTCTG | AACAATCCAAGCCAACCAAG | 137 |
| chr3:130506780-130508798 | p2 | (TA)6 | AGCGATCTTGTTGTATGGGG | TCCCTCATCTTTTCTGCCAC | 235 |
| chr3:130509170-130511188 | p6 | (CTCTTG)3 | CAAAAGGGAGAAAGAAGGGG | ACACAAGAGGCCCAAAAGTG | 169 |
| chr3:130518086-130520101 | p5 | (AGGAC)3 | TGACCCGAAAAATTCTGGAG | GAGGCTCCATCTCATGCTTC | 163 |
| chr3:130591649-130593664 | p3 | (ACA)5 | GGGTCGTCTGGTTACAAGGA | AAACGACTCCCCTGAGTGTG | 112 |
| chr3:130600909-130602924 | p5 | (AAGCC)3 | CATACACTAGGGGCGCATTT | ACCCTTGATTTTGCCTCCTT | 112 |
| chr3:130649356-130651368 | p2 | (TA)6 | AATTGGTCTCGCCGTTACTG | TAGTCGTATGACCGTTTGCG | 234 |
| chr3:130670250-130672268 | p3 | (ATT)6 | TGAACACCATCTGAACCCAA | AATTTGACAACAACCGGAGC | 153 |
| chr3:130707187-130709202 | p3 | (TAA)6 | AGCGTATGAGAGAGACGGGA | CATTTTCCTCCATACCCACG | 213 |
| chr3:130707187-130709202 | p3 | (ATA)6 | AGCGTATGAGAGAGACGGGA | CATTTTCCTCCATACCCACG | 213 |
| chr3:130707212-130709227 | p3 | (TAA)6 | AGCGTATGAGAGAGACGGGA | CATTTTCCTCCATACCCACG | 188 |
| chr3:130707212-130709227 | p3 | (ATA)6 | AGCGTATGAGAGAGACGGGA | CATTTTCCTCCATACCCACG | 188 |
| chr3:130756828-130758844 | p4 | (CAAT)4 | GGTTTATGGGCTTGAAGCAA | GGCGGTTCTTTGAATCTCAG | 160 |
| chr3:130762067-130764093 | p2 | (TA)13 | CCCCATCCTCTTCAAACTCA | TTGATGGGGATACTTTGGGA | 174 |

| Marker ID | SSR type | SSR motif | FORWARD PRIMER (5'-3') | REVERSE PRIMER (5'-3') | size (bp) |
|---|---|---|---|---|---|
| chr3:130763146-130765166 | p2 | (TA)10 | CCATTCGATATTGGACCCTG | TCGAAGGAGGTTTGAGTTGC | 126 |
| chr3:130805788-130807808 | p2 | (TA)10 | CCATTCGATATTGGACCCTG | TCGAAGGAGGTTTGAGTTGC | 126 |
| chr3:130964576-130966591 | p6 | (GTGGGG)4 | AAAATATCTCACCCACCCCC | TAATATCTCACCCACCCCCA | 123 |
| chr3:130964576-130966591 | p5 | (CCCCA)3 | AAAATATCTCACCCACCCCC | TAATATCTCACCCACCCCCA | 123 |
| chr3:130964576-130966591 | p5 | (TACAA)3 | AAAATATCTCACCCACCCCC | TAATATCTCACCCACCCCCA | 123 |
| chr3:131026082-131028110 | p2 | (AT)14 | GTTGCCTGTTTTTGCCACTT | CTGGATTTGGCATAGCGATT | 189 |
| chr3:131072016-131074034 | p3 | (AAC)6 | GAAGGATAGAGGGGCAGACC | TGGGTAGGGAGCTGAGAAGA | 122 |
| chr3:131093780-131095795 | p3 | (TGT)5 | GACGTCACACGTCGTATTGG | AAGTTGCCACACATACGCAG | 150 |
| chr3:131121483-131123498 | p3 | (TAA)5 | TAATCGGACCTTGATTTGGG | CCGCATTTATTTCGCACTTT | 278 |
| chr3:131121501-131123516 | p3 | (TAA)5 | TAATCGGACCTTGATTTGGG | CCGCATTTATTTCGCACTTT | 260 |
| chr3:131180111-131182133 | p2 | (AG)11 | AATACCCAAAATCGTGACGC | TGAAGTGCATCAAGGACGAG | 170 |
| chr3:131272528-131274540 | p2 | (TA)6 | GATGTCCGATACCCGCTTTA | TAACATGGGGCACTACACGA | 193 |
| chr3:131272528-131274540 | p2 | (TA)6 | GATGTCCGATACCCGCTTTA | TAACATGGGGCACTACACGA | 193 |
| chr3:131329941-131332013 | p2 | (GA)7 | AAGATCCACCAACAGCAACC | GGAAGAGCCCAATTGAATGA | 162 |
| chr3:131368644-131370665 | p3 | (TTA)7 | ATGTGAGGCTTGGGAAAAGA | ATTGCCGTAGGAGTTGATCG | 134 |
| chr3:131368644-131370665 | p5 | (AAACT)3 | ATGTGAGGCTTGGGAAAAGA | ATTGCCGTAGGAGTTGATCG | 134 |
| chr3:131404179-131406241 | p2 | (TA)7 | ATGCATCATGTCCTCGATCA | ACTGCACACAGTCAAAACGC | 296 |
| chr3:131404179-131406241 | p2 | (TA)12 | ATGCATCATGTCCTCGATCA | ACTGCACACAGTCAAAACGC | 296 |
| chr3:131404179-131406241 | p2 | (TA)8 | ATGCATCATGTCCTCGATCA | ACTGCACACAGTCAAAACGC | 296 |
| chr3:131404179-131406241 | p2 | (AT)31 | ATGCATCATGTCCTCGATCA | ACTGCACACAGTCAAAACGC | 296 |
| chr3:131404179-131406241 | p2 | (GA)20 | ATGCATCATGTCCTCGATCA | ACTGCACACAGTCAAAACGC | 296 |
| chr3:131449726-131451746 | p5 | (GGGGT)4 | AAGTGCGTTCGTCGAAAAAT | AAGTGAGTCAGCGACGGAGT | 243 |
| chr3:131458993-131461008 | p5 | (GAAAA)3 | AAACCATGGGCTGAGACTTG | ACTCCCAAAAATGTTGTCGC | 149 |
| chr3:131504946-131506962 | p2 | (AT)8 | ATAAACTGGACCCCACCCTC | TGATGGGGATACTTTGGGAA | 122 |
| chr3:131514699-131516714 | p5 | (TCGGG)3 | CCACTAGCAACTAGCGGAGG | TTGACCAGGTTGCATTTTGA | 202 |
| chr3:131516008-131518026 | p2 | (AT)9 | ACTTTGGACCGCTTGAGCTA | AAGGGTCTGGCAACACTTTG | 190 |
| chr3:131529400-131531416 | p2 | (TA)8 | ACGGCCATCTTTAGTGCTTG | GGTGCACGATATACGCATTG | 104 |
| chr3:131588109-131590125 | p2 | (TA)6 | TCTTAATTTTGCCACCCCAG | CGGCAAACCCATTTATCCTA | 164 |
| chr3:131588109-131590125 | p4 | (GCTA)4 | TCTTAATTTTGCCACCCCAG | CGGCAAACCCATTTATCCTA | 164 |
| chr3:131588109-131590125 | p2 | (TA)14 | TCTTAATTTTGCCACCCCAG | CGGCAAACCCATTTATCCTA | 164 |
| chr3:131627422-131629438 | p4 | (ATAG)4 | TACAAACGACACTCACCGGA | TCTGTGTTTCGCTTGGTCAG | 271 |
| chr3:131719007-131721037 | p2 | (TC)15 | AGCCTTCGCTTTCTTTGACA | AGCCCCTTTGGATGTCTTTT | 206 |
| chr3:131721530-131723548 | p2 | (AT)9 | AATAAAGTCCGTCGCCAATG | GTGAACACTCTCACCCCGTT | 206 |
| chr3:131817995-131820055 | p5 | (TTTAA)3 | AAGCCGGCTCTTTAATCTGG | AGGACTCTCATCATCGCACAT | 201 |
| chr3:131817995-131820055 | p2 | (AT)9 | AAGCCGGCTCTTTAATCTGG | AGGACTCTCATCATCGCACAT | 201 |
| chr3:131833963-131835978 | p5 | (TATTT)3 | CCATGGCATTATTTGAACCC | ATGACTCCTTCGGTCCTCCT | 104 |

| Marker ID | SSR type | SSR motif | FORWARD PRIMER (5'-3') | REVERSE PRIMER (5'-3') | size (bp) |
|---|---|---|---|---|---|
| chr3:131872339-131874366 | p3 | (AAT)9 | CGGCTGATGGCAGATATTTT | TTGTGATGGTCTTTGCTGGA | 234 |
| chr3:131882325-131884385 | p5 | (TTTAA)3 | AAGCCGGCTCTTTAATCTGG | AGGACTCTCATCATCGCACAT | 201 |
| chr3:131882325-131884385 | p2 | (AT)9 | AAGCCGGCTCTTTAATCTGG | AGGACTCTCATCATCGCACAT | 201 |
| chr3:131898293-131900308 | p5 | (TATTT)3 | CCATGGCATTATTTGAACCC | ATGACTCCTTCGGTCCTCCT | 104 |
| chr3:132247155-132249173 | p3 | (TCA)6 | GGGGGTTGTGTTTCATCATC | GTTAGGGGTGCGCTTAATGA | 275 |
| chr3:132247155-132249173 | p6 | (CCGGTT)3 | GGGGGTTGTGTTTCATCATC | GTTAGGGGTGCGCTTAATGA | 275 |
| chr3:132247155-132249173 | p2 | (TA)16 | GGGGGTTGTGTTTCATCATC | GTTAGGGGTGCGCTTAATGA | 275 |
| chr4:103715-105729 | p2 | (AT)7 | AGTGAGGTGAGTGGAATGGC | GCTGGTGAAATGCACAAAAA | 269 |
| chr4:211719-213767 | p2 | (TA)24 | ATAGCACTTCGCAAAAGGGA | TTGAATCGTGTTTGTGGAGC | 259 |
| chr4:240214-242236 | p3 | (GAA)8 | TTTATGGTCCGAAACAAGCC | ATTTATGCGGCACACTTTCC | 215 |
| chr4:240214-242236 | p2 | (AG)11 | TTTATGGTCCGAAACAAGCC | ATTTATGCGGCACACTTTCC | 215 |
| chr4:260585-262600 | p5 | (TTTTA)3 | CCGGTCACGACTTACCAGAT | ATTGTTACGCCTCGCACTTT | 133 |
| chr4:341249-343265 | p2 | (AT)8 | CAAATGAACAGCTTCAGCCA | GGTAACGGAGATGGGTCAAA | 205 |
| chr4:358910-360925 | p5 | (TTTGT)3 | CAGGGGATAGATTGCTCCAA | AAGACGTGGCAGATGATTCC | 107 |
| chr4:368536-370551 | p3 | (TGT)5 | TGGTTGGGTTTTTGGTTTGT | GGTTTAAGGACCGCACACAT | 259 |
| chr4:443640-445655 | p5 | (ATAAA)3 | GGGTGTGTTTGGTATGGAGG | TTAGAGGAGTTGGGTGGGTG | 260 |
| chr4:487267-489291 | p2 | (AT)8 | AGCGGATTGTCCTTCATTTG | ATTCTGCCCATGCAAAACTC | 131 |
| chr4:602545-604563 | p6 | (TTTATA)3 | CCGGGTGAAAGCTATGGTAA | GCGATTAGGCCTTCATTGAG | 272 |
| chr4:610159-612174 | p5 | (TCCAA)3 | AACCACGGAGTAGCTCGAAA | CAAACTGTGCCCACCCTAGT | 106 |
| chr4:648975-650987 | p2 | (TA)6 | CCAACAGCAAATGAAAAGCA | AGGCAGAGTTTTACCTGCGA | 147 |
| chr4:666653-668671 | p3 | (ATT)6 | GAATCCGAGCAACCAGGATA | TGGTGTGGGATCAACTCAAA | 184 |
| chr4:683589-685604 | p3 | (CTT)5 | AAGGAATGCTACGAAAGCGA | TAAAAATGGGCAGAGGGATG | 281 |
| chr4:686568-688583 | p3 | (CTT)5 | AGGAATGCTACGAAAGCGAA | TAAAAATGGGCAGAGGGATG | 282 |
| chr4:754530-756548 | p2 | (GA)8 | CCTGTTCACCACCGAGAAAT | TTTCCGGGTAATTTCTGCAC | 264 |
| chr4:754530-756548 | p6 | (CAAAAG)3 | CCTGTTCACCACCGAGAAAT | TTTCCGGGTAATTTCTGCAC | 264 |
| chr4:833124-835180 | p2 | (CA)7 | GCAACGTGACAACAAAATGG | TCACTCCGAGTTGGATCCTC | 276 |
| chr4:866693-868705 | p2 | (TA)6 | ACCGGCCATGTCTATCAAAG | CATATGCACGGCTAAGCAAA | 130 |
| chr4:875292-877304 | p3 | (TTG)11 | AACCCGACGCAATTAGTTTG | TCAGCCTCAAACGTCTTTCA | 279 |
| chr4:875292-877304 | p2 | (TA)6 | AACCCGACGCAATTAGTTTG | TCAGCCTCAAACGTCTTTCA | 279 |
| chr4:875292-877304 | p2 | (TA)6 | AACCCGACGCAATTAGTTTG | TCAGCCTCAAACGTCTTTCA | 279 |
| chr4:933835-935850 | p3 | (ACA)5 | ACCAGTGGAGAGCGGTTAGA | TCACAGTGTGCAAAGGCTTC | 268 |
| chr4:1130960-1132972 | p2 | (TA)6 | GATTTGCCATAAAGCGTTGC | GATAATATGGTCCGCCGTGT | 109 |
| chr4:1130960-1132972 | p2 | (AT)7 | GATTTGCCATAAAGCGTTGC | GATAATATGGTCCGCCGTGT | 109 |
| chr4:1130960-1132972 | p4 | (GGGT)7 | GATTTGCCATAAAGCGTTGC | GATAATATGGTCCGCCGTGT | 109 |
| chr4:1139670-1141682 | p2 | (TA)6 | TGAATCACACGGAGAAGTGC | ATCGGGTCTCCTACACCCTT | 200 |
| chr4:1149396-1151411 | p5 | (ACCGA)3 | CCCGTGAGTTTGAGTGATGA | CCCCCGATATTAAACAGGCT | 264 |

| Marker ID | SSR type | SSR motif | FORWARD PRIMER (5'-3') | REVERSE PRIMER (5'-3') | size (bp) |
|---|---|---|---|---|---|
| chr4:1149396-1151411 | p5 | (AACCG)3 | CCCGTGAGTTTGAGTGATGA | CCCCCGATATTAAACAGGCT | 264 |
| chr4:1157905-1159927 | p2 | (AT)9 | TTCCGAATACGTTAAACGCC | GAAAATTCTGACTCCGCCAC | 177 |
| chr4:1157905-1159927 | p2 | (AT)11 | TTCCGAATACGTTAAACGCC | GAAAATTCTGACTCCGCCAC | 177 |
| chr4:1262848-1264864 | p2 | (AT)8 | TGATTATACTCGGGCGTTCC | CTGACATTGAGAGATGGGCA | 162 |
| chr4:1436114-1438132 | p3 | (TGT)5 | CACGCAGAAGGATGTGAGAA | TACCAAGGAGGAGATCGTGG | 232 |
| chr4:1436114-1438132 | p3 | (ACC)6 | CACGCAGAAGGATGTGAGAA | TACCAAGGAGGAGATCGTGG | 232 |
| chr4:1477013-1479028 | p5 | (TAAAA)3 | TTTTCAAGGTGCCCCTTATG | TGTATCTCCCACGTCGTTCA | 143 |
| chr4:1591940-1593958 | p6 | (TTAATT)3 | AAAAAGTGGCGTTGACTCGT | GCCAAAAATGTGATGCATTG | 114 |
| chr4:1601064-1603100 | p3 | (TAT)12 | CATGTCTGCACCAAATGAGG | CCCACACGAGGAAGAACAAT | 101 |
| chr4:1640816-1642831 | p5 | (CATCC)3 | TTAAGCCTTGGCTCAGCAAT | TTAGCCCGAAGGCTAGTTGA | 186 |
| chr4:1648454-1650520 | p2 | (AT)33 | ATACGCTCCTTCGTTTGTGG | AATTTGTTGGCACCCAGTGT | 113 |
| chr4:1657775-1659790 | p3 | (TCT)5 | GTTCAATTGGGTCATCCCAC | CCAGTGGGGTCTTACAGGAA | 150 |
| chr4:1682347-1684371 | p3 | (AGA)8 | CCCATTGATCCCTCCCTAAT | TGGCCAAACACTGATTTTCA | 299 |
| chr4:1691291-1693315 | p6 | (AATTTA)4 | CCGCAACTTGACATGAGAGA | CCGGAAAAGAGAAAGCTCCT | 281 |
| chr4:1700285-1702300 | p5 | (TAAAA)3 | AAACCGAAAATTCCCTCTGC | CAGCCTGTTGAAAGTGGGTT | 188 |
| chr4:1789953-1791975 | p2 | (TA)8 | AGATACGGAGCACGAACCAT | GACGAAGCATGGCTCTAACC | 191 |
| chr4:1789953-1791975 | p2 | (AT)15 | AGATACGGAGCACGAACCAT | GACGAAGCATGGCTCTAACC | 191 |
| chr4:1789979-1791995 | p2 | (TA)8 | AGATACGGAGCACGAACCAT | GACGAAGCATGGCTCTAACC | 165 |
| chr4:1789979-1791995 | p2 | (AT)15 | AGATACGGAGCACGAACCAT | GACGAAGCATGGCTCTAACC | 165 |
| chr4:1789999-1792035 | p2 | (TA)8 | AGATACGGAGCACGAACCAT | GACGAAGCATGGCTCTAACC | 145 |
| chr4:1789999-1792035 | p2 | (AT)15 | AGATACGGAGCACGAACCAT | GACGAAGCATGGCTCTAACC | 145 |
| chr4:1795199-1797217 | p6 | (TAAAAA)3 | GAACTCTCGCCAAAGATTGC | AAAAAGCAGGCCAAATTCCT | 188 |
| chr4:1853682-1855694 | p2 | (GT)6 | ACTTCCAGTGGAGGACCCTT | TTCCTCAACGCACACAAGTC | 287 |
| chr4:2101462-2103480 | p6 | (CACCAG)3 | TGAAACCCAGGATGTTCCTC | CGTTGGCGTTATTTGGAACT | 108 |
| chr4:2121909-2123927 | p3 | (ACA)6 | GTTAAACCGGTGAGACCGAA | ACGATTGGCAGTTGAACACA | 131 |
| chr4:2139572-2141592 | p5 | (AAAAG)3 | CCGAGCGCCATAGATTAAAG | GTCTGGTTCTGCAGCTAGGG | 252 |
| chr4:2139572-2141592 | p2 | (CT)10 | CCGAGCGCCATAGATTAAAG | GTCTGGTTCTGCAGCTAGGG | 252 |
| chr4:2140579-2142594 | p2 | (TC)6 | AGGGTCTCTTGCTTGGGATT | TTTCGGCAAGCTACCAAGTT | 254 |
| chr4:2140579-2142594 | p3 | (TTG)5 | AGGGTCTCTTGCTTGGGATT | TTTCGGCAAGCTACCAAGTT | 254 |
| chr4:2211011-2213026 | p3 | (ATC)5 | GGACGCAAGTACCAGACCAT | ATGAAGAACACGGCACAACA | 104 |
| chr4:2268475-2270493 | p2 | (GA)9 | AGCGTTCTCAAATGCCTTGT | ACTTGGGTTTTCCCCCATAC | 178 |
| chr4:2282917-2284931 | p2 | (TA)7 | TCATCTCCTGGTCCTCAACC | CGGAGCCTCACAAAAGTAGG | 219 |
| chr4:2298465-2300477 | p2 | (AT)6 | CGAGTATGTTTGGATGGGCT | CACTTTGAATTTCGGGTCGT | 215 |
| chr4:2307766-2309814 | p2 | (TA)24 | CCTTGTCACACAATTCCACG | TTGAATTTGGAAGGTTTCGG | 202 |
| chr4:2321401-2323416 | p3 | (GTT)5 | TCTATTGTGCAAGACAGCGG | TCGTTCCTAATCCTGTCGCT | 137 |
| chr4:2357655-2359673 | p3 | (ACA)6 | TGCAGCAACCAATTCATGTT | TTTGTCTGGATTCTCCCTGG | 132 |

| Marker ID | SSR type | SSR motif | FORWARD PRIMER (5'-3') | REVERSE PRIMER (5'-3') | size (bp) |
|---|---|---|---|---|---|
| chr4:2359527-2361539 | p2 | (AT)6 | AAACGCCTACGAACAACACC | TGCGACATGAAGGAAGTTTG | 221 |
| chr4:2468745-2470791 | p5 | (CCGGT)3 | ACTTAGAAGCGGAGGAAGGC | CGCCGCAAGTTGTAGTTTTT | 240 |
| chr4:2684114-2686144 | p2 | (TA)15 | ACGACTGCCAGTTTACGGAC | TGGGGCCCATATTTTTGTTA | 187 |
| chr4:2684114-2686144 | p2 | (AT)9 | ACGACTGCCAGTTTACGGAC | TGGGGCCCATATTTTTGTTA | 187 |
| chr4:2684114-2686144 | p2 | (AT)52 | ACGACTGCCAGTTTACGGAC | TGGGGCCCATATTTTTGTTA | 187 |
| chr4:2702400-2704415 | p5 | (AATAA)3 | TCCCATCGACTTCTCATTCC | GTCCTCGAACATGGAGTCGT | 211 |
| chr4:2702400-2704415 | p3 | (TGG)5 | TCCCATCGACTTCTCATTCC | GTCCTCGAACATGGAGTCGT | 211 |
| chr4:2850815-2852830 | p5 | (AAAAG)3 | GAGCTAGCGTTTGAGGATGG | TGCCAAGCTTTGAGGAATCT | 142 |
| chr4:2905238-2907250 | p2 | (AC)6 | CCACCCCAAAAGCTAGAACA | TGTGGGGTTTCACTGGGTAT | 218 |
| chr4:2947872-2949910 | p2 | (TA)19 | TCCCAAAGTATCCCCATCAA | GACATGGCGGAATAGCAAAT | 238 |
| chr4:2998746-3000766 | p2 | (TA)10 | CTCCTCCTCCTTTTTCGCTT | GGCAACCCTTCAAAAATCAA | 147 |
| chr4:3009606-3011630 | p6 | (GAAAAA)3 | AATTAAAGACCACCCCCAGC | GGTAGCCCAGTTTCTGGTGA | 256 |
| chr4:3009606-3011630 | p2 | (TC)12 | AATTAAAGACCACCCCCAGC | GGTAGCCCAGTTTCTGGTGA | 256 |
| chr4:3016195-3018211 | p2 | (TA)7 | AAAGCAGGCAGATGGAGCTA | GGCCAACACCAACAAAAATC | 195 |
| chr4:3016195-3018211 | p2 | (TA)13 | AAAGCAGGCAGATGGAGCTA | GGCCAACACCAACAAAAATC | 195 |
| chr4:3050512-3052536 | p2 | (AT)12 | TACTTCTGGGCATCGGAAAC | ATTTTGCGTCTTGATTTGGC | 276 |
| chr4:3206086-3208101 | p3 | (TTG)5 | CATCGTATTGGTGCAGGATG | CTCCCACCTTGTCTCGGATA | 148 |
| chr4:3302291-3304305 | p2 | (TA)7 | TGCAGAACCGACACTCGTAG | ACCCTACCTGTTACGCGATG | 177 |
| chr4:3310120-3312138 | p2 | (TA)9 | CGCCATTACTATCCGTCGAT | CCTTTCCACGACCTGACATT | 209 |
| chr4:3346660-3348675 | p3 | (ATC)5 | ACACCTTGGCGTTCATTAGG | CCAAGAACTTTGCCACTGGT | 151 |
| chr4:3492581-3494593 | p2 | (GA)6 | TTTTTCGTTTGGTTATGGGG | TTGTCGCGAAATACATGAGG | 142 |
| chr4:3535263-3537277 | p2 | (TA)7 | TATTGCTCCGCATTTCATCA | TAGCACTGAATTGGCCATGA | 134 |
| chr4:3576751-3578766 | p5 | (AAATA)3 | GGTTAGTCGGTTTGGCTCAC | CACTCAGTGCCTGACTGCAT | 140 |
| chr4:3576751-3578766 | p5 | (AAATA)3 | GGTTAGTCGGTTTGGCTCAC | CACTCAGTGCCTGACTGCAT | 140 |
| chr4:3576751-3578766 | p5 | (AATAA)3 | GGTTAGTCGGTTTGGCTCAC | CACTCAGTGCCTGACTGCAT | 140 |
| chr4:3636216-3638234 | p2 | (AT)7 | CGCATTTAGCCAAGGAAGAC | TTCCCCACATCATATTCGGT | 282 |
| chr4:3636354-3638368 | p2 | (AT)7 | CGCATTTAGCCAAGGAAGAC | TTCCCCACATCATATTCGGT | 144 |
| chr4:3803416-3805434 | p3 | (AAT)6 | GACACTTGCTCCCACCAGAT | ACTCTGGATGCTGGACTGCT | 183 |
| chr4:3831193-3833213 | p2 | (TA)6 | GCGGCATATCCTTTCACAGT | TTGACGGGCTTATTGGAGAC | 209 |
| chr4:3831193-3833213 | p2 | (TA)9 | GCGGCATATCCTTTCACAGT | TTGACGGGCTTATTGGAGAC | 209 |
| chr4:3831270-3833282 | p2 | (TA)6 | GCGGCATATCCTTTCACAGT | TTGACGGGCTTATTGGAGAC | 132 |
| chr4:3831270-3833282 | p2 | (TA)9 | GCGGCATATCCTTTCACAGT | TTGACGGGCTTATTGGAGAC | 132 |
| chr4:3843882-3845902 | p2 | (AT)10 | TACACGAGGATACGTGGCAG | TCGAACCCGAGAGTCTGATT | 128 |
| chr4:3843882-3845902 | p4 | (ATGT)5 | TACACGAGGATACGTGGCAG | TCGAACCCGAGAGTCTGATT | 128 |
| chr4:3862582-3864600 | p2 | (AT)9 | TATCAACCCATTAGCGGGAG | CAAATAAATGACATGGCGGA | 142 |
| chr4:3920483-3922499 | p2 | (AT)8 | ACCTTGCACATCACTCACCA | GTGGCGCAGTAAGAACCATT | 143 |

| Marker ID | SSR type | SSR motif | FORWARD PRIMER (5'-3') | REVERSE PRIMER (5'-3') | size (bp) |
|---|---|---|---|---|---|
| chr4:4005742-4007757 | p3 | (TTA)5 | AGTCCCCAATGATGAAGCTG | TCCAAATTCACAAGGCAACA | 296 |
| chr4:4005742-4007757 | p5 | (TTTAT)3 | AGTCCCCAATGATGAAGCTG | TCCAAATTCACAAGGCAACA | 296 |
| chr4:4005742-4007757 | p5 | (TTTAT)3 | AGTCCCCAATGATGAAGCTG | TCCAAATTCACAAGGCAACA | 296 |
| chr4:4005847-4007867 | p3 | (TTA)5 | AGTCCCCAATGATGAAGCTG | TGTCCAAATTCACAAGGCAA | 191 |
| chr4:4005847-4007867 | p5 | (TTTAT)3 | AGTCCCCAATGATGAAGCTG | TGTCCAAATTCACAAGGCAA | 191 |
| chr4:4005847-4007867 | p5 | (TTTAT)3 | AGTCCCCAATGATGAAGCTG | TGTCCAAATTCACAAGGCAA | 191 |
| chr4:4048315-4050330 | p3 | (TTA)5 | AGTCCCCAATGATGAAGCTG | TCCAAATTCACAAGGCAACA | 296 |
| chr4:4048315-4050330 | p5 | (TTTAT)3 | AGTCCCCAATGATGAAGCTG | TCCAAATTCACAAGGCAACA | 296 |
| chr4:4048315-4050330 | p5 | (TTTAT)3 | AGTCCCCAATGATGAAGCTG | TCCAAATTCACAAGGCAACA | 296 |
| chr4:4048420-4050440 | p3 | (TTA)5 | AGTCCCCAATGATGAAGCTG | TGTCCAAATTCACAAGGCAA | 191 |
| chr4:4048420-4050440 | p5 | (TTTAT)3 | AGTCCCCAATGATGAAGCTG | TGTCCAAATTCACAAGGCAA | 191 |
| chr4:4048420-4050440 | p5 | (TTTAT)3 | AGTCCCCAATGATGAAGCTG | TGTCCAAATTCACAAGGCAA | 191 |
| chr4:4135687-4137702 | p5 | (ACAAA)3 | AGATCTATGCGGGCAAACTG | GCTACATTACCCCGGGATTT | 281 |
| chr4:4222467-4224489 | p2 | (TA)11 | TTCCATAACAGCATGGCAAA | ATGAGCAATCTCATACCGGC | 126 |
| chr4:4222467-4224489 | p2 | (TA)15 | TTCCATAACAGCATGGCAAA | ATGAGCAATCTCATACCGGC | 126 |
| chr4:4272494-4274506 | p2 | (AT)8 | GTGCCAACGCTTGGTAGTTT | TATTGTTGGGCATTGCTTCA | 141 |
| chr4:4272515-4274531 | p2 | (AT)8 | GTGCCAACGCTTGGTAGTTT | TATTGTTGGGCATTGCTTCA | 120 |
| chr4:4272532-4274552 | p2 | (AT)8 | GTGCCAACGCTTGGTAGTTT | TATTGTTGGGCATTGCTTCA | 103 |
| chr4:4272954-4274970 | p2 | (AT)8 | AAAACCCATTGACGTGGAAG | TTGCCTTATGACATCCTCCC | 118 |
| chr4:4356268-4358286 | p3 | (TTG)6 | TGCCACCCAAAACTTAAAGG | CTCGCATCTTGTCATCTCCA | 145 |
| chr4:4384901-4386915 | p2 | (AT)7 | TTTTGCGGCTACCTGTCTTT | TCATCGAGCAATTGGAATGA | 138 |
| chr4:4390750-4392818 | p2 | (TA)34 | ATGCCATGTCACTAAAGCCC | TTGATTCTTTTGCCGCTTTT | 276 |
| chr4:4443952-4445967 | p5 | (AAACC)5 | ACCAAACCAAACCAAACCAA | GGATCACGTCAAAGCCGTAT | 231 |
| chr4:4443952-4445967 | p5 | (ATTTT)3 | ACCAAACCAAACCAAACCAA | GGATCACGTCAAAGCCGTAT | 231 |
| chr4:4443952-4445967 | p5 | (CAAAC)3 | ACCAAACCAAACCAAACCAA | GGATCACGTCAAAGCCGTAT | 231 |
| chr4:4443990-4446015 | p5 | (AAACC)5 | ACCAAACCAAACCAAACCAA | GGATCACGTCAAAGCCGTAT | 193 |
| chr4:4443990-4446015 | p5 | (ATTTT)3 | ACCAAACCAAACCAAACCAA | GGATCACGTCAAAGCCGTAT | 193 |
| chr4:4443990-4446015 | p5 | (CAAAC)3 | ACCAAACCAAACCAAACCAA | GGATCACGTCAAAGCCGTAT | 193 |
| chr4:4567322-4569334 | p2 | (TA)6 | TGGCCCGTATTTCAAAATGT | TCCGTTTGACCCCCATAATA | 193 |
| chr4:4575192-4577216 | p5 | (TATTT)3 | GTGAATTTAAGGGCGGATCA | CTTGACCGAAAAGCGACAAT | 246 |
| chr4:4575192-4577216 | p2 | (TA)12 | GTGAATTTAAGGGCGGATCA | CTTGACCGAAAAGCGACAAT | 246 |
| chr4:4575192-4577216 | p2 | (TA)16 | GTGAATTTAAGGGCGGATCA | CTTGACCGAAAAGCGACAAT | 246 |
| chr4:4575974-4578006 | p2 | (TA)12 | CGTGGCAGAGCTACCCTTAT | CTGCACCTCTTTCCTCCAAC | 139 |
| chr4:4575974-4578006 | p2 | (TA)16 | CGTGGCAGAGCTACCCTTAT | CTGCACCTCTTTCCTCCAAC | 139 |
| chr4:4656546-4658558 | p2 | (TA)6 | ATATTTTTGGCCAAGGGGAG | CTTACGAAGCTTGCCTGGTC | 230 |
| chr4:4656546-4658558 | p2 | (AT)8 | ATATTTTTGGCCAAGGGGAG | CTTACGAAGCTTGCCTGGTC | 230 |

| Marker ID | SSR type | SSR motif | FORWARD PRIMER (5'-3') | REVERSE PRIMER (5'-3') | size (bp) |
|---|---|---|---|---|---|
| chr4:4656546-4658558 | p2 | (AG)6 | ATATTTTTGGCCAAGGGGAG | CTTACGAAGCTTGCCTGGTC | 230 |
| chr4:4656546-4658558 | p6 | (TTGCAT)3 | ATATTTTTGGCCAAGGGGAG | CTTACGAAGCTTGCCTGGTC | 230 |
| chr4:4724867-4726924 | p2 | (AG)6 | GCTCGAATTCACAGCAACAA | CCCAAAATTTCCTTGCAAAA | 117 |
| chr4:4724867-4726924 | p3 | (AAT)19 | GCTCGAATTCACAGCAACAA | CCCAAAATTTCCTTGCAAAA | 117 |
| chr4:4770199-4772214 | p3 | (TTA)5 | CATGCACCGCCTTTCTAAAT | TTTTCCTTCCCGACGTATTG | 264 |
| chr4:4770215-4772230 | p3 | (TTA)5 | CATGCACCGCCTTTCTAAAT | TTTTCCTTCCCGACGTATTG | 248 |
| chr4:4848113-4850143 | p2 | (AT)15 | GTATCCTTGTGCTTCGCTCC | ATGTGATCAGGAGGTCAGGG | 230 |
| chr4:4887496-4889511 | p5 | (AATAA)3 | TTCGTGAACACAAAAACCCA | TTCCAAACAGGCCCTTAGTG | 110 |
| chr4:4887496-4889511 | p5 | (TTTTA)3 | TTCGTGAACACAAAAACCCA | TTCCAAACAGGCCCTTAGTG | 110 |
| chr4:4921946-4923960 | p2 | (AG)15 | CAAACAGGGCTAGTTCGCTC | ATTTATCTGGGCCTCTGGCT | 166 |
| chr4:4921977-4923991 | p2 | (AG)16 | CAAACAGGGCTAGTTCGCTC | ATTTATCTGGGCCTCTGGCT | 135 |
| chr4:5003274-5005289 | p5 | (ATTGT)3 | GTTGTTGTTATGGTTCCGGG | GGACTGGGTTCCATGCTAAA | 242 |
| chr4:5058063-5060087 | p2 | (AC)8 | TGCCATGCCATTATACAGGA | CACAACCGTTTCAAATCCCT | 203 |
| chr4:5058063-5060087 | p2 | (AT)6 | TGCCATGCCATTATACAGGA | CACAACCGTTTCAAATCCCT | 203 |
| chr4:5058063-5060087 | p2 | (AT)12 | TGCCATGCCATTATACAGGA | CACAACCGTTTCAAATCCCT | 203 |
| chr4:5139439-5141451 | p2 | (TA)6 | TGGCCATCAATTTTCAAACC | CCAAACCATGCTTTTTGGAT | 132 |
| chr4:5253587-5255605 | p6 | (CATATA)3 | CTAACCCGAGCACCACTCAT | TGTATGTTTCGAGCAGGACG | 246 |
| chr4:5264788-5266815 | p3 | (AAT)9 | AAAAGGAAGAGGAGCGTGGT | TCGCCTCTCCTTCATCATCT | 287 |
| chr4:5264788-5266815 | p3 | (AAT)9 | AAAAGGAAGAGGAGCGTGGT | TCGCCTCTCCTTCATCATCT | 287 |
| chr4:5300962-5302990 | p4 | (AATA)7 | CTTTCGTTCCAAATCCTCCA | AGATGGGGTTGCAAGATGAC | 118 |
| chr4:5300962-5302990 | p5 | (GTGAC)3 | CTTTCGTTCCAAATCCTCCA | AGATGGGGTTGCAAGATGAC | 118 |
| chr4:5344414-5346434 | p4 | (AAAT)5 | GACGAAGGATGACAAGAGGC | AGCCTCAGGTGAAGCAGAAA | 148 |
| chr4:5360886-5362916 | p2 | (TA)15 | CGAACCGCAACTCACATTTA | ATTTAGAGGCTCTGGGTGCC | 273 |
| chr4:5384089-5386104 | p6 | (GACGGA)3 | CGGACTGCGTCGATTATTTT | TGTTCCTTGGCATTAAAGGG | 208 |
| chr4:5384089-5386104 | p3 | (AAG)5 | CGGACTGCGTCGATTATTTT | TGTTCCTTGGCATTAAAGGG | 208 |
| chr4:5438705-5440723 | p6 | (CCCCCA)3 | AGCTCTTGGACCCTGTGAGA | CCTTGACTGACCTTCCCTGA | 176 |
| chr4:5438705-5440723 | p6 | (CCCCCA)3 | AGCTCTTGGACCCTGTGAGA | CCTTGACTGACCTTCCCTGA | 176 |
| chr4:5487629-5489644 | p5 | (CACTT)3 | TGCTTGATTCGCATTGTCTC | GCCTTGAAGATATCCTCCCC | 192 |
| chr4:5516223-5518239 | p2 | (AG)8 | AACTTCGGTGGATGAACCAG | AAGGCTGGAAAGTACCGGAT | 217 |
| chr4:5563350-5565362 | p2 | (AT)6 | TTTGGGGCTACAAGATTTCG | ATACATACCGCCACCACCAT | 192 |
| chr4:5567568-5569588 | p2 | (AT)7 | AATTTCAACCATGGCAGACC | TTGGGTGAGCTGCAATATGA | 239 |
| chr4:5567568-5569588 | p2 | (CT)10 | AATTTCAACCATGGCAGACC | TTGGGTGAGCTGCAATATGA | 239 |
| chr4:5567568-5569588 | p3 | (ACA)6 | AATTTCAACCATGGCAGACC | TTGGGTGAGCTGCAATATGA | 239 |
| chr4:5715594-5717612 | p3 | (AAT)6 | TGTTTTGTTGGCCTCATTCA | TTGTTTTCCTCCGATTTTGC | 262 |
| chr4:5715594-5717612 | p4 | (TATG)4 | TGTTTTGTTGGCCTCATTCA | TTGTTTTCCTCCGATTTTGC | 262 |
| chr4:5793457-5795475 | p2 | (GT)7 | AATCATTTCATCCCTTCCCC | ACACACACACACACGGGTTC | 290 |

| Marker ID | SSR type | SSR motif | FORWARD PRIMER (5'-3') | REVERSE PRIMER (5'-3') | size (bp) |
|---|---|---|---|---|---|
| chr4:5793457-5795475 | p6 | (TTTTTG)3 | AATCATTTCATCCCTTCCCC | ACACACACACACACGGGTTC | 290 |
| chr4:5793457-5795475 | p5 | (TATTT)3 | AATCATTTCATCCCTTCCCC | ACACACACACACACGGGTTC | 290 |
| chr4:5802625-5804655 | p3 | (TGT)10 | GAAACAAACAAGCACGAGCA | AAGGTGGCAGAACGGTATTG | 269 |
| chr4:5852848-5854868 | p4 | (ATTA)5 | CAGCCACCACCTTTGAATCT | ATTCGAGCGGAGTTGCTAAA | 123 |
| chr4:5866931-5868961 | p2 | (AT)9 | CGAGATATCCTTCCGGTTCA | GTCGCTCCGATCTAATCCAA | 255 |
| chr4:5866931-5868961 | p2 | (TA)15 | CGAGATATCCTTCCGGTTCA | GTCGCTCCGATCTAATCCAA | 255 |
| chr4:5866931-5868961 | p2 | (AT)6 | CGAGATATCCTTCCGGTTCA | GTCGCTCCGATCTAATCCAA | 255 |
| chr4:5910325-5912355 | p5 | (GCGGG)6 | CTTCAACTCGGGTTGCATTT | TTCTTGGACGATCAAGACCC | 291 |
| chr4:5988467-5990491 | p3 | (TAT)8 | ATGACGTAATTCGGGAGACG | CGATATCTTCCCTCCTTCCC | 293 |
| chr4:6011812-6013826 | p2 | (TA)7 | CAATGCTTTTGAAGGCCAAT | CAGGCCCATGTTCTTCATTT | 298 |
| chr4:6093844-6095868 | p5 | (AAAAG)3 | AGTTTCGGGTTCGCCTATTT | ACCCCGAATGCTAGGTTTTT | 285 |
| chr4:6093844-6095868 | p2 | (TA)12 | AGTTTCGGGTTCGCCTATTT | ACCCCGAATGCTAGGTTTTT | 285 |
| chr4:6149848-6151860 | p2 | (TC)8 | TGCTAAGGAGAAGGGGGAAT | CGTTCTTTCTTTGTGCCCAT | 234 |
| chr4:6149848-6151860 | p2 | (AG)6 | TGCTAAGGAGAAGGGGGAAT | CGTTCTTTCTTTGTGCCCAT | 234 |
| chr4:6237931-6239946 | p5 | (ACCAA)3 | GACGGTTGAGGGATCAAAGA | TCGCCAATAGTATGCCCTTT | 149 |
| chr4:6251055-6253067 | p2 | (AT)6 | CAGATTGGGATGGACAAGGT | ATACTGCTGCCCTACTGCGT | 109 |
| chr4:6251055-6253067 | p2 | (CT)7 | CAGATTGGGATGGACAAGGT | ATACTGCTGCCCTACTGCGT | 109 |
| chr4:6282363-6284377 | p2 | (CA)7 | GTGGATGTGTGCCAAGTGAG | AATGTCGAAATGAGGCGAAC | 167 |
| chr4:6320601-6322627 | p2 | (TA)11 | TTGCCCATTTTTCTTTCTGC | TGACCCCTGCGGTAAATTAG | 290 |
| chr4:6447490-6449511 | p3 | (AAT)6 | CCAAAAGACCCCGATGAGTA | GCGACCCAAAGTCAATCAGT | 219 |
| chr4:6483869-6485883 | p2 | (AT)7 | ATTGCTTCGAAACACCAACC | TTTTCATCGATTCCCCTCAC | 286 |
| chr4:6494317-6496341 | p3 | (TGT)8 | ACAGAAGAATTCACCGCCAC | CGGAAGAGCGAGATTCTTTG | 104 |
| chr4:6499596-6501611 | p5 | (CAAAA)3 | TTGTTCCAACAATGGCTTCA | TCACATGAGGTGGAGACTCG | 285 |
| chr4:6530278-6532293 | p3 | (ACA)5 | GACATGCCACCTTGGATCTT | TCAAACTCGCTTTTCGGAGT | 199 |
| chr4:6629695-6631713 | p3 | (AAG)6 | TCTCATCTGCTCAGCATTGG | CAAAGCCGTCAACAAAGACA | 136 |
| chr4:6681491-6683506 | p3 | (GAT)5 | AGCCATATGAAGTCCCATCG | AATGGGGGCCAAAAATTAAG | 160 |
| chr4:6831018-6833033 | p5 | (AAATA)3 | CTGCAAATTGCCCCATAAGT | TAAATTTATGCTCAGCGGGG | 210 |
| chr4:6878238-6880253 | p3 | (AAC)11 | ACAAGTCCCCAAACAGATGC | ACATGCAATGGTCGTTTTGA | 236 |
| chr4:6878238-6880253 | p3 | (AGT)5 | ACAAGTCCCCAAACAGATGC | ACATGCAATGGTCGTTTTGA | 236 |
| chr4:6969572-6971590 | p6 | (CTAATT)3 | ACACATATTAGCGGGCTTGG | CGTGAAGGGTTGTGTTGTTG | 126 |
| chr4:7060075-7062089 | p2 | (AT)7 | GAAGAGCGCATTTAGCCAAG | CCATTTCGTTAATTTCCCCC | 175 |
| chr4:7060075-7062089 | p2 | (TA)6 | GAAGAGCGCATTTAGCCAAG | CCATTTCGTTAATTTCCCCC | 175 |
| chr4:7248631-7250647 | p4 | (AAGG)4 | AACACAAGACCCTGACTGGC | CGGAAGGAACTAAATTGGCA | 100 |
| chr4:7361283-7363298 | p3 | (ACA)5 | GTTCATTCTATCGGGGCAGA | CACATGGCACTAACCCAATG | 254 |
| chr4:7400624-7402648 | p3 | (TTG)8 | TCCGGAATGCTGTGTGATAA | CTCCCACCTTGTCTCGGATA | 163 |
| chr4:7443220-7445238 | p2 | (TA)9 | AGGGGAAAATGGGAGAAAGA | TCCCTTATCCACACAGCTCC | 287 |

| Marker ID | SSR type | SSR motif | FORWARD PRIMER (5'-3') | REVERSE PRIMER (5'-3') | size (bp) |
|---|---|---|---|---|---|
| chr4:7443285-7445323 | p2 | (TA)9 | AGGGGAAAATGGAGAAAGA | TCCCTTATCCACACAGCTCC | 222 |
| chr4:7453287-7455301 | p2 | (CA)7 | GTGCATACGAAGCAACATGG | TTGCCTTCAAACTCTGCCTT | 124 |
| chr4:7477751-7479766 | p5 | (TTTTC)3 | TGCCGTACTAGCAATGATGC | AAGCCTGCCAAATGAAGCTA | 291 |
| chr4:7489322-7491337 | p5 | (GTATT)3 | TGTATACCCCGGAATGGAAA | GGAAAACAAGTCCTCCCCAT | 231 |
| chr4:7489322-7491337 | p5 | (GGTGG)3 | TGTATACCCCGGAATGGAAA | GGAAAACAAGTCCTCCCCAT | 231 |
| chr4:7489872-7491887 | p5 | (GTATT)3 | GGTGAAAGGTGGCAAACAAT | CAACCATCCAAACAGAGGGT | 283 |
| chr4:7489872-7491887 | p5 | (GGTGG)3 | GGTGAAAGGTGGCAAACAAT | CAACCATCCAAACAGAGGGT | 283 |
| chr4:7489872-7491887 | p5 | (CCCCA)3 | GGTGAAAGGTGGCAAACAAT | CAACCATCCAAACAGAGGGT | 283 |
| chr4:7512333-7514351 | p2 | (AT)9 | CCATCGATGTGCTAGGAGTG | CATATAGGCTGCCTTGTGGC | 171 |
| chr4:7537057-7539075 | p2 | (TA)9 | AATCACTCCATGCCTTTTCG | GTCTGTTCGAGGTGGAGAGC | 175 |
| chr4:7575842-7577862 | p2 | (TA)10 | CCCCATATTAAACTGGCCCT | ATATGGGGCCCACAATTTTT | 207 |
| chr4:7607931-7609947 | p2 | (TA)7 | TGGACGTGATCAGTTCAGGA | CTTAAAGCACTAGCACCGCA | 218 |
| chr4:7607931-7609947 | p2 | (AT)8 | TGGACGTGATCAGTTCAGGA | CTTAAAGCACTAGCACCGCA | 218 |
| chr4:7671638-7673656 | p6 | (TCTTTC)3 | CCCGAAGCTACGAGGTGTTA | TTGACATGGGAATGATGTGG | 257 |
| chr4:7671638-7673656 | p3 | (CCA)5 | CCCGAAGCTACGAGGTGTTA | TTGACATGGGAATGATGTGG | 257 |
| chr4:7671638-7673656 | p6 | (AAAATC)3 | CCCGAAGCTACGAGGTGTTA | TTGACATGGGAATGATGTGG | 257 |
| chr4:7671638-7673656 | p3 | (GAA)13 | CCCGAAGCTACGAGGTGTTA | TTGACATGGGAATGATGTGG | 257 |
| chr4:7671638-7673656 | p2 | (AT)15 | CCCGAAGCTACGAGGTGTTA | TTGACATGGGAATGATGTGG | 257 |
| chr4:7675303-7677319 | p4 | (AAAT)4 | AACATGGTTCCAATGAAGGC | TGCAGCCGAAACAGAGTATG | 223 |
| chr4:7683680-7685692 | p2 | (AT)6 | CAAAGTCGTTCCTCAGCCTC | GCACTACCCTTCTCAACCCA | 105 |
| chr4:7916632-7918652 | p2 | (AT)9 | TGATGCAAGCCAAATGTGAT | CCTAAATGCAGACTCTCGCC | 267 |
| chr4:7916740-7918752 | p2 | (AT)9 | TGATGCAAGCCAAATGTGAT | CCTAAATGCAGACTCTCGCC | 159 |
| chr4:7950288-7952303 | p5 | (AATTA)3 | CGTCAATTGGCAGAGCTGTA | TTGCTTCTGCTGCGTATGTC | 158 |
| chr4:7975391-7977406 | p5 | (ATTTA)3 | AGGATTGTTGCGTGTTTTCC | TTTTCAGCAAGCTGACATGC | 102 |
| chr4:8005088-8007118 | p3 | (ATA)10 | AGAGAGAGAGAAGCGGGGAC | CCTTCCCTCATCACCTTCAA | 264 |
| chr4:8070833-8072855 | p5 | (AAAAT)5 | ACATTCCCCACTGCTTCATC | TTTTGCCTAGGGTGTTCAGG | 235 |
| chr4:8070833-8072855 | p2 | (TA)11 | ACATTCCCCACTGCTTCATC | TTTTGCCTAGGGTGTTCAGG | 235 |
| chr4:8105626-8107648 | p5 | (ACAAC)3 | TGGCAATAAACGAATGAGCA | GCAGTTTCAGCTTACCGAGG | 293 |
| chr4:8105626-8107648 | p2 | (TA)11 | TGGCAATAAACGAATGAGCA | GCAGTTTCAGCTTACCGAGG | 293 |
| chr4:8115670-8117688 | p6 | (AATAAA)3 | CATTTATGGGTCCCACTCCA | TCTAAAGTATTGGGCCCGTG | 149 |
| chr4:8115670-8117688 | p2 | (AT)21 | CATTTATGGGTCCCACTCCA | TCTAAAGTATTGGGCCCGTG | 149 |
| chr4:8117806-8119818 | p2 | (TA)7 | TCCCATGAGTTGGAGAAGGT | CATCGTCGTCCGTTATCCTT | 179 |
| chr4:8117806-8119818 | p2 | (AT)6 | TCCCATGAGTTGGAGAAGGT | CATCGTCGTCCGTTATCCTT | 179 |
| chr4:8123168-8125200 | p2 | (AG)16 | AATCTGCCTCGCTACAATGG | AAATCATCTTGGACCCACCA | 158 |
| chr4:8140328-8142386 | p2 | (TA)9 | TCGCGATAGCATTCAAAGTG | TCCTGCTATGCGGAAAAAGT | 157 |
| chr4:8176860-8178894 | p2 | (TC)6 | GCTACAATCAGAAGGCGGAG | CTTTAGTCGGGTGGGCTACA | 187 |

| Marker ID | SSR type | SSR motif | FORWARD PRIMER (5'-3') | REVERSE PRIMER (5'-3') | size (bp) |
|---|---|---|---|---|---|
| chr4:8176860-8178894 | p2 | (TA)17 | GCTACAATCAGAAGGCGGAG | CTTTAGTCGGGTGGGCTACA | 187 |
| chr4:8192810-8194832 | p2 | (TA)7 | TGCATCCAAAATTAGCTCACC | CAGTACTGGGCCATTCGTTT | 224 |
| chr4:8192810-8194832 | p2 | (AT)11 | TGCATCCAAAATTAGCTCACC | CAGTACTGGGCCATTCGTTT | 224 |
| chr4:8192810-8194832 | p2 | (TA)9 | TGCATCCAAAATTAGCTCACC | CAGTACTGGGCCATTCGTTT | 224 |
| chr4:8195937-8197961 | p2 | (AT)12 | ATCACAAGCCACACATGGAA | TGCCTATTAATTTGGAGCCG | 177 |
| chr4:8253837-8255852 | p5 | (AAGGA)3 | CAGCCTACCCTAATGCAAGC | ACCGAGCAACATCACTTTCC | 271 |
| chr4:8266732-8268747 | p5 | (AATTT)3 | AAGAACAAAAATGCCCCTGA | CAAGGGGAATAAATGAGCCA | 206 |
| chr4:8327194-8329206 | p2 | (TC)6 | GTCCGTAACCCTTCACCGTA | CCGTCCTTCATTTCGTCATT | 118 |
| chr4:8390746-8392758 | p2 | (GT)6 | AAGAGAACCCTCCCGGAGTA | ACGAAAGCACGGGACATAAC | 260 |
| chr4:8435009-8437041 | p2 | (TA)12 | AAAGGGAGGCCGAAAAAGTA | GAGAAGCCACCTAGCCCTCT | 291 |
| chr4:8435060-8437072 | p2 | (TA)12 | AAAGGGAGGCCGAAAAAGTA | GAGAAGCCACCTAGCCCTCT | 240 |
| chr4:8489590-8491648 | p2 | (AT)13 | GCAACTCAAGGGTGTGACCT | TGCACCTCGACCATTTATCA | 136 |
| chr4:8489590-8491648 | p4 | (TTTA)4 | GCAACTCAAGGGTGTGACCT | TGCACCTCGACCATTTATCA | 136 |
| chr4:8490141-8492167 | p2 | (AT)13 | GATACGTGGACTTTGCACGA | CAACTATGGCAAGGGGAAAA | 227 |
| chr4:8490141-8492167 | p4 | (TTTA)4 | GATACGTGGACTTTGCACGA | CAACTATGGCAAGGGGAAAA | 227 |
| chr4:8559356-8561380 | p2 | (TA)12 | AGGACTGTACTGGAAGGGCA | TGCTTTTTGATTGCTCATGC | 164 |
| chr4:8575636-8577652 | p2 | (TC)8 | AAGGGTATTCATCCAAGCCA | GGGCCACCAATATTCAGATG | 117 |
| chr4:8575636-8577652 | p3 | (AAT)12 | AAGGGTATTCATCCAAGCCA | GGGCCACCAATATTCAGATG | 117 |
| chr4:8579477-8581492 | p3 | (TTA)5 | CTGGACAGAGCCACAGAACA | GGTTCATGACCACTTCCACC | 204 |
| chr4:8579477-8581492 | p2 | (GA)16 | CTGGACAGAGCCACAGAACA | GGTTCATGACCACTTCCACC | 204 |
| chr4:8648824-8650838 | p2 | (AC)7 | TTTGGTGGGAAATTTGTGGT | CAACGCGAAGTAAGCAGTCA | 151 |
| chr4:8679521-8681533 | p2 | (AG)6 | CGCTGTTACAAAATGGCTCA | GCACGTTGGTGTTCCTTCTT | 181 |
| chr4:8745736-8747751 | p3 | (ATG)5 | TAGACGGTCTCGGATTGCTT | GAGGAGTGTGCACGAGATCA | 300 |
| chr4:8785267-8787297 | p5 | (ATTTT)3 | TGCTACGAATACGCACGGTA | ACCCACCGTATCCAAATTCA | 113 |
| chr4:8785267-8787297 | p2 | (TA)15 | TGCTACGAATACGCACGGTA | ACCCACCGTATCCAAATTCA | 113 |
| chr4:8802417-8804432 | p3 | (CAA)5 | GAATGGCTTTCCTTGAGCTG | AAGGCTGGACAGACGAGAAA | 186 |
| chr4:8850938-8852952 | p2 | (TA)7 | GGGGAGACAATGTGAGCTGT | TTACCCGCCATTATGGTGTT | 187 |
| chr4:8895399-8897413 | p2 | (AT)7 | CCCTTCGCCTAATACCATGA | ATTGCCCTTCAAAAGCACTG | 204 |
| chr4:9024442-9026464 | p2 | (AC)11 | ACATGAAATCGCATGTCCAA | AACCGGTCAAAAGTTTGGTG | 137 |
| chr4:9065853-9067868 | p5 | (AAAAT)3 | GCCCATTTACCGCTTTGATA | CAAAGAAAGGTTCAGGAGCG | 105 |
| chr4:9065853-9067868 | p2 | (AT)6 | GCCCATTTACCGCTTTGATA | CAAAGAAAGGTTCAGGAGCG | 105 |
| chr4:9065853-9067868 | p2 | (CT)32 | GCCCATTTACCGCTTTGATA | CAAAGAAAGGTTCAGGAGCG | 105 |
| chr4:9075249-9077283 | p3 | (TAT)7 | GAAATTTGCATGGGCTGATT | CAAAGCCAAAGAAAAGTCGC | 141 |
| chr4:9075249-9077283 | p2 | (CA)8 | GAAATTTGCATGGGCTGATT | CAAAGCCAAAGAAAAGTCGC | 141 |
| chr4:9106125-9108157 | p2 | (AT)16 | ATCCCAAATCAGCAAGATCG | GCAACCCTCCTCCTTTATCC | 182 |
| chr4:9107248-9109263 | p5 | (GATCT)3 | ACTCAACTCCGTCCAGCACT | GGGATCTCGAGACTTGCTTG | 258 |

| Marker ID | SSR type | SSR motif | FORWARD PRIMER (5'-3') | REVERSE PRIMER (5'-3') | size (bp) |
|---|---|---|---|---|---|
| chr4:9197999-9200019 | p5 | (TATTG)4 | GCAAACACCAACGAAAAGGT | GAAGGCGAGAGTAGCCATTG | 287 |
| chr4:9345194-9347212 | p2 | (AT)9 | TTCAATGCCCAGTGTCAAAA | TAACCTTCTCTGGCTTGCGT | 297 |
| chr4:9426048-9428074 | p2 | (TA)13 | AGGAACAAATGCCCTCAGAA | AGTGGTCCTTTCCCGAAATC | 159 |
| chr4:9432151-9434166 | p3 | (GTT)5 | GTCTATCATGCAAGGCAGCA | TCCGCCCCATACAACATAAT | 157 |
| chr4:9538261-9540276 | p3 | (CAA)5 | AGGGCACAATGTTCTTACGG | TTTCGGGAGATGTAGTTGCC | 168 |
| chr4:9556295-9558310 | p3 | (TCT)5 | CACGTCATTGCAAGAAGCAT | GCACCTTATTTCCCACCTGA | 131 |
| chr4:9562448-9564500 | p2 | (TA)26 | GCAGATGCAAGAAGAGGGTC | TTGAGAAGCCCACACATGAC | 192 |
| chr4:9725954-9727972 | p3 | (TGA)6 | TTCTTACCGGTCCGATTTTG | CGTGCAACGGTCAATAACAC | 199 |
| chr4:9728589-9730601 | p2 | (TA)6 | AAGTATCAACCGTGGCTTCG | CTCTTCGCCTCTCGCTTCTA | 221 |
| chr4:9728589-9730601 | p2 | (AT)8 | AAGTATCAACCGTGGCTTCG | CTCTTCGCCTCTCGCTTCTA | 221 |
| chr4:9728589-9730601 | p2 | (AT)7 | AAGTATCAACCGTGGCTTCG | CTCTTCGCCTCTCGCTTCTA | 221 |
| chr4:9728589-9730601 | p5 | (CCGGT)3 | AAGTATCAACCGTGGCTTCG | CTCTTCGCCTCTCGCTTCTA | 221 |
| chr4:9728824-9730840 | p2 | (TA)6 | TAGAAGCGAGAGGCGAAGAG | TCATTCGTTTAATTCCCCCA | 130 |
| chr4:9728824-9730840 | p2 | (AT)8 | TAGAAGCGAGAGGCGAAGAG | TCATTCGTTTAATTCCCCCA | 130 |
| chr4:9728824-9730840 | p2 | (AT)7 | TAGAAGCGAGAGGCGAAGAG | TCATTCGTTTAATTCCCCCA | 130 |
| chr4:9728824-9730840 | p5 | (CCGGT)3 | TAGAAGCGAGAGGCGAAGAG | TCATTCGTTTAATTCCCCCA | 130 |
| chr4:9759493-9761521 | p2 | (AT)14 | TGCATCAGGCATTTTTCTCA | GGTCACGCTGATAAGTGGCT | 262 |
| chr4:9777708-9779728 | p2 | (TA)10 | ACAAAGGGAAAGGGAAAGGA | TAGACCGTCCACCCAGTACC | 214 |
| chr4:9794639-9796654 | p5 | (AAGAA)3 | CAGCAGGACCAATTTCATCA | ACGATCGATGCTGGATTTTC | 162 |
| chr4:9800348-9802360 | p2 | (AT)6 | ATTGGTCAGGTTCGGATGAG | AACGCAGAAACCACCAAATC | 174 |
| chr4:9801442-9803457 | p3 | (ATT)5 | ATGCGGGTGCCATTATAGAG | GTTCTTCCTTAGCTTCCGGG | 200 |
| chr4:9801442-9803457 | p3 | (ATT)5 | ATGCGGGTGCCATTATAGAG | GTTCTTCCTTAGCTTCCGGG | 200 |
| chr4:9801442-9803457 | p3 | (TTA)6 | ATGCGGGTGCCATTATAGAG | GTTCTTCCTTAGCTTCCGGG | 200 |
| chr4:9801721-9803739 | p3 | (ATT)5 | TCCAAGAGAACCATGTGCTG | CCTCGTTAAAAACCTTGCCA | 126 |
| chr4:9801721-9803739 | p3 | (ATT)5 | TCCAAGAGAACCATGTGCTG | CCTCGTTAAAAACCTTGCCA | 126 |
| chr4:9801721-9803739 | p3 | (TTA)6 | TCCAAGAGAACCATGTGCTG | CCTCGTTAAAAACCTTGCCA | 126 |
| chr4:9816990-9819005 | p5 | (CTTTT)3 | GAAGTTGGAAGGGGCTCAAT | ATTTTCGCCTGAGTAGGGGT | 163 |
| chr4:9845017-9847032 | p5 | (GGTTC)3 | GAAAAGGCGAGAGAGAGGGT | TTTGAGGAGCTGACAAGCCT | 126 |
| chr4:9888797-9890821 | p2 | (TA)18 | TGCTACGAATACAAAATCGGC | GCTATTCCGCCATGTGATTT | 277 |
| chr4:9888797-9890821 | p2 | (AT)8 | TGCTACGAATACAAAATCGGC | GCTATTCCGCCATGTGATTT | 277 |
| chr4:9888797-9890821 | p2 | (AT)12 | TGCTACGAATACAAAATCGGC | GCTATTCCGCCATGTGATTT | 277 |
| chr4:9888797-9890821 | p2 | (AT)9 | TGCTACGAATACAAAATCGGC | GCTATTCCGCCATGTGATTT | 277 |
| chr4:9888797-9890821 | p2 | (TA)9 | TGCTACGAATACAAAATCGGC | GCTATTCCGCCATGTGATTT | 277 |
| chr4:9907206-9909218 | p2 | (TA)6 | CACCACTTTCCTTCCACCAT | CATCACTGCCCTTCTCCATT | 236 |
| chr4:9911528-9913544 | p2 | (TA)8 | GGTCGAAACAAATGAGGGAA | GCCGTAGTGGTGGAAGGTAA | 118 |
| chr4:9949025-9951037 | p2 | (AG)6 | GCTCACAAGAAACACGACCA | TTCACTCTGGCCCCTTTAGA | 300 |

| Marker ID | SSR type | SSR motif | FORWARD PRIMER (5'-3') | REVERSE PRIMER (5'-3') | size (bp) |
|---|---|---|---|---|---|
| chr4:10028545-10030563 | p6 | (TTTTTC)3 | AATCCAAACGGGCTCTAGGT | ATTTGCTGATCCAACAAGGG | 268 |
| chr4:10161520-10163534 | p6 | (CATCAC)4 | CCCTTCATCCTCACCTGAAA | TGGGTCCATCTCCAAGAATC | 137 |
| chr4:10161520-10163534 | p2 | (AG)7 | CCCTTCATCCTCACCTGAAA | TGGGTCCATCTCCAAGAATC | 137 |
| chr4:10340515-10342539 | p5 | (CCGGT)4 | TTCGGAGAGTGCATTTAGCC | GCCTTCCTCCGCTTCTAAGT | 289 |
| chr4:10340630-10342642 | p5 | (CCGGT)4 | TTCGGAGAGTGCATTTAGCC | GCCTTCCTCCGCTTCTAAGT | 174 |
| chr4:10366940-10368956 | p4 | (TATT)4 | GGGACCCTTTCAGATGGTTT | TCAACTCAGTCAGGCTCACG | 300 |
| chr4:10369889-10371909 | p2 | (AT)9 | TTTCTTACGCCCCTTTTGTG | ATGTTGGGGTGTGAGTGGAT | 191 |
| chr4:10369889-10371909 | p2 | (AT)7 | TTTCTTACGCCCCTTTTGTG | ATGTTGGGGTGTGAGTGGAT | 191 |
| chr4:10369889-10371909 | p5 | (CCGGT)4 | TTTCTTACGCCCCTTTTGTG | ATGTTGGGGTGTGAGTGGAT | 191 |
| chr4:10421195-10423210 | p2 | (TA)7 | AAGAGCATTGTTTTGGACGG | CCTCACTTCAAAAAGGCGAG | 262 |
| chr4:10421195-10423210 | p5 | (TCGGG)3 | AAGAGCATTGTTTTGGACGG | CCTCACTTCAAAAAGGCGAG | 262 |
| chr4:10421226-10423244 | p2 | (TA)7 | AAGAGCATTGTTTTGGACGG | CCTCACTTCAAAAAGGCGAG | 231 |
| chr4:10421226-10423244 | p5 | (TCGGG)3 | AAGAGCATTGTTTTGGACGG | CCTCACTTCAAAAAGGCGAG | 231 |
| chr4:10421246-10423261 | p5 | (TCGGG)3 | AAGAGCATTGTTTTGGACGG | CCTCACTTCAAAAAGGCGAG | 211 |
| chr4:10421743-10423758 | p5 | (TCGGG)3 | GCTACGGGCGTAATGTAGGA | ATCTGGCGACGACTCCTCTA | 222 |
| chr4:10628008-10630023 | p2 | (CT)11 | TCCCGAAGAAGGAAAAGGAT | GTTGAAACCCAGCTTCCGTA | 157 |
| chr4:10628008-10630023 | p3 | (TGT)5 | TCCCGAAGAAGGAAAAGGAT | GTTGAAACCCAGCTTCCGTA | 157 |
| chr4:10681922-10683937 | p5 | (TTTGG)3 | TTAATTATCCGCTCATGCCC | ACGGAGTCATGAAGGACTGG | 295 |
| chr4:10686831-10688849 | p3 | (ACA)6 | TCTAAGCACGGTCCCTGTTC | CCAGCTGCATTGCATAAAAA | 187 |
| chr4:10702346-10704362 | p2 | (AT)8 | CATCGTCTCCATCGACCTTT | GTGCCAGCGAGACTAGATCC | 175 |
| chr4:10724342-10726354 | p2 | (CT)6 | TCCAATCAAAAACAGCCCTC | CTTGCGTACCAACGGAAGAT | 254 |
| chr4:10728165-10730177 | p2 | (TC)6 | ATGCTATGGCTCGAAGCTGT | ACAACTTAGCCACACAGGGC | 286 |
| chr4:10774834-10776852 | p2 | (AT)7 | CTTCGGAGAGCGCATTTATC | TTGCTCCGCTTTTGAGTTTT | 101 |
| chr4:10774834-10776852 | p2 | (TA)9 | CTTCGGAGAGCGCATTTATC | TTGCTCCGCTTTTGAGTTTT | 101 |
| chr4:10774834-10776852 | p2 | (AT)9 | CTTCGGAGAGCGCATTTATC | TTGCTCCGCTTTTGAGTTTT | 101 |
| chr4:10943841-10945861 | p5 | (AGAAA)4 | CGATGCAGCCACTGAACTTA | CCTAAGCAAGAGGCTTGGTG | 206 |
| chr4:10964016-10966028 | p4 | (AACT)4 | TGAATGCATTGGGAACTGAA | AGCATTGATTTGCTCGCTTT | 156 |
| chr4:10964016-10966028 | p2 | (AT)6 | TGAATGCATTGGGAACTGAA | AGCATTGATTTGCTCGCTTT | 156 |
| chr4:10972223-10974238 | p3 | (AAC)5 | ACAACAACAACAACAGCCCA | CTCACTTCAAAAAGGCGAGG | 214 |
| chr4:10979398-10981413 | p3 | (AAC)5 | CTGCTGCAAGTTTTGTTCCA | TCCGCAAGCCGTAATTTATC | 100 |
| chr4:10988500-10990514 | p2 | (AT)7 | TGGAAACTTTGGATTTTGCC | CAGATTAGCGCGTTCAACAA | 181 |
| chr4:11055012-11057068 | p4 | (TTAT)14 | AAGGATTGGGAAAGTTGCCT | GGCTACTGGATGGACAAGGA | 122 |
| chr4:11055012-11057068 | p2 | (AT)12 | AAGGATTGGGAAAGTTGCCT | GGCTACTGGATGGACAAGGA | 122 |
| chr4:11056965-11058981 | p2 | (AT)8 | GGGAGCAACCTCAAAAATGA | GAGCCTCCAATCATCCACAT | 191 |
| chr4:11071349-11073361 | p2 | (CT)6 | GTTGCTTCGAGGAAAACTCG | ATTGGGTGTCAACAATGGGT | 262 |
| chr4:11202523-11204538 | p5 | (CGACC)3 | ATTCAACACTGGGGCAAAAG | TAGCCAATTTTGTTCCGACC | 209 |

| Marker ID | SSR type | SSR motif | FORWARD PRIMER (5'-3') | REVERSE PRIMER (5'-3') | size (bp) |
|---|---|---|---|---|---|
| chr4:11235925-11237943 | p2 | (AT)9 | TCCGCAAACTTTCGTCTTCT | CCTTAAGCTGGGTTCGTGAC | 209 |
| chr4:11264313-11266329 | p4 | (TCTT)4 | AGCATTCTTACCGTAACCGC | CCATTCCATTTCAGCCAAGT | 228 |
| chr4:11299661-11301677 | p4 | (TCTT)4 | AGCATTCTTACCGTAACCGC | CCATTCCATTTCAGCCAAGT | 228 |
| chr4:11387199-11389217 | p6 | (CCACTC)3 | TTTTAACACCAGCTCCACCC | AGGGTTGGGGTAGTGTAGGG | 285 |
| chr4:11402486-11404530 | p2 | (TA)22 | CCGTTCTTTCAACAATGGCT | GAACATTCATTGCGTTGCTG | 154 |
| chr4:11425513-11427525 | p2 | (TA)6 | CCAGGAAGAGGCCATAATGA | TTTTCCTCCTCCTTGCTTGA | 148 |
| chr4:11425513-11427525 | p2 | (AC)6 | CCAGGAAGAGGCCATAATGA | TTTTCCTCCTCCTTGCTTGA | 148 |
| chr4:11443292-11445314 | p2 | (TA)11 | AATTGGCCATTTAACACCCA | ATGTGGCAATGAGTCCATCA | 169 |
| chr4:11517460-11519475 | p2 | (GA)8 | ACGTTTTCGGGTGATCTGAC | CAGGCTCACGACCTATGGTT | 169 |
| chr4:11517460-11519475 | p3 | (GTT)5 | ACGTTTTCGGGTGATCTGAC | CAGGCTCACGACCTATGGTT | 169 |
| chr4:11540119-11542133 | p2 | (TA)7 | CCCCGTAAGAACATAGCATGA | TTGTTGAATCCTGGGGGATA | 188 |
| chr4:11540119-11542133 | p2 | (AT)6 | CCCCGTAAGAACATAGCATGA | TTGTTGAATCCTGGGGGATA | 188 |
| chr4:11546163-11548178 | p3 | (CTT)5 | GCTTGTGGCTGCTTCCTATC | CCAATCAGGCCAAGCTTTAG | 167 |
| chr4:11546163-11548178 | p2 | (TC)9 | GCTTGTGGCTGCTTCCTATC | CCAATCAGGCCAAGCTTTAG | 167 |
| chr4:11586485-11588503 | p2 | (AT)7 | ATGCTTTGGTTCACTCCCAC | AGAAAACCCTGAAGGCCAAT | 254 |
| chr4:11586485-11588503 | p3 | (AAC)6 | ATGCTTTGGTTCACTCCCAC | AGAAAACCCTGAAGGCCAAT | 254 |
| chr4:11707481-11709513 | p2 | (AT)16 | CTAGGGTTCTTGCAATCCCA | CTCGGGGACTTCATCAGTGT | 173 |
| chr4:11818607-11820625 | p3 | (AAT)6 | GCTACTGCTTTTGAGACCCG | AATTTCGAACCCCCAAAAAC | 228 |
| chr4:11859232-11861283 | p5 | (AGTAA)3 | AATGTAACCCGCCCTCTTTT | CGAAGGACAATCCTGCAAAT | 207 |
| chr4:11859232-11861283 | p6 | (AAAAAG)3 | AATGTAACCCGCCCTCTTTT | CGAAGGACAATCCTGCAAAT | 207 |
| chr4:11859232-11861283 | p3 | (AAT)17 | AATGTAACCCGCCCTCTTTT | CGAAGGACAATCCTGCAAAT | 207 |
| chr4:11873772-11875788 | p2 | (AT)8 | GAGGGAAAGAAGTTGGAGGG | AAACGCTTCCCATACGACAC | 250 |
| chr4:11892169-11894189 | p2 | (AT)10 | CAAGTCTGCCCATAAGCCAT | TTTTGGCTATAAGATGGCCG | 113 |
| chr4:11904101-11906116 | p3 | (CTT)6 | TCGTCTTTCTCTCCGACGAT | TCCATGGATGAGCAATTGAA | 194 |
| chr4:11904101-11906116 | p3 | (TTG)5 | TCGTCTTTCTCTCCGACGAT | TCCATGGATGAGCAATTGAA | 194 |
| chr4:11906512-11908527 | p3 | (TAC)5 | AGCAACCGTTAGCAGGAAGA | ATTTCAATTCATCCTTGCCG | 294 |
| chr4:11962179-11964194 | p3 | (CTT)6 | TCGTCTTTCTCTCCGACGAT | TCCATGGATGAGCAATTGAA | 194 |
| chr4:11962179-11964194 | p3 | (TTG)5 | TCGTCTTTCTCTCCGACGAT | TCCATGGATGAGCAATTGAA | 194 |
| chr4:11964590-11966605 | p3 | (TAC)5 | AGCAACCGTTAGCAGGAAGA | ATTTCAATTCATCCTTGCCG | 294 |
| chr4:12037790-12039808 | p6 | (TCAGTT)3 | TGTTGGGTGATGGAGAATGA | TGGGCATAAAGCTCCACAAT | 168 |
| chr4:12037790-12039808 | p2 | (AT)8 | TGTTGGGTGATGGAGAATGA | TGGGCATAAAGCTCCACAAT | 168 |
| chr4:12075662-12077677 | p5 | (TTTTC)3 | TCCAGGCAATGGATAGGTTC | GATTCTGCACCAACACGCTA | 140 |
| chr4:12156645-12158660 | p5 | (TATAT)3 | AGACGGTGGTGTAGTGGAGG | TCCCCTCATTTTCCTTCTCA | 148 |
| chr4:12177522-12179550 | p2 | (AT)14 | TCTCACCAAGGGCGTTTATC | AGGATTCCCCGAATCTCAAT | 106 |
| chr4:12336783-12338798 | p5 | (CCCCT)3 | CGTCCACAGTAGGGGAAAGA | CGGAATTGTTCGGTTCTGTT | 270 |
| chr4:12378271-12380297 | p3 | (TTG)5 | TAGGCACCTGCATAAAAGGG | ACCACACGTGCTCATCAAAA | 188 |

| Marker ID | SSR type | SSR motif | FORWARD PRIMER (5'-3') | REVERSE PRIMER (5'-3') | size (bp) |
|---|---|---|---|---|---|
| chr4:12378271-12380297 | p2 | (AC)6 | TAGGCACCTGCATAAAAGGG | ACCACACGTGCTCATCAAAA | 188 |
| chr4:12378271-12380297 | p2 | (AT)13 | TAGGCACCTGCATAAAAGGG | ACCACACGTGCTCATCAAAA | 188 |
| chr4:12378271-12380297 | p2 | (AT)17 | TAGGCACCTGCATAAAAGGG | ACCACACGTGCTCATCAAAA | 188 |
| chr4:12402115-12404139 | p3 | (TTC)8 | ATCGGTCAGATCAGATTGGG | GGGCGAACAGAAAATTTGAA | 137 |
| chr4:12402115-12404139 | p6 | (AAAAAT)4 | ATCGGTCAGATCAGATTGGG | GGGCGAACAGAAAATTTGAA | 137 |
| chr4:12654148-12656163 | p5 | (AAAAT)3 | TGGCTCTTGATGCTGTATGC | CCGGAGTGCAGTTGTGTAGA | 236 |
| chr4:12749398-12751413 | p5 | (ATATG)3 | AAATGGACCGCAGATGGTAG | GGGTTCCCTCTCGTCTTTTC | 184 |
| chr4:12754800-12756812 | p2 | (TA)6 | CACAACGGCACCTCTAACAA | AGGTCGTGTGGTTGTGTGAA | 237 |
| chr4:12785942-12787962 | p2 | (TA)7 | GTGATCGGATTGCAATGATG | ACGGAGATCGTATTTTCCCC | 197 |
| chr4:12785942-12787962 | p2 | (AG)10 | GTGATCGGATTGCAATGATG | ACGGAGATCGTATTTTCCCC | 197 |
| chr4:12823443-12825461 | p2 | (AG)9 | TACTGCGATAATGCGACTGC | AGGGGTAGGCCGAAGAAGTA | 275 |
| chr4:12906910-12908955 | p3 | (ATA)15 | GTGATGACGGAGAGATGGGT | ACCGGGTCCAATTCATTTTT | 189 |
| chr4:12920770-12922788 | p2 | (AT)9 | AGCGTTTCATCTCATTTCCG | TGTTGATGTGCCGTGAATTT | 194 |
| chr4:13202184-13204199 | p5 | (AAGAA)3 | ATCGATGATCCTGAGGCAAC | TCAAAATTGGGCCTATCGAG | 139 |
| chr4:13271169-13273225 | p2 | (TA)8 | AATGCAACGAAATAGGTGGG | ATAGTGTTCACGAATGGCGA | 240 |
| chr4:13271169-13273225 | p2 | (TA)15 | AATGCAACGAAATAGGTGGG | ATAGTGTTCACGAATGGCGA | 240 |
| chr4:13313787-13315805 | p6 | (TTTGAA)3 | ATCGGTTCGATCCATTTTGA | ATGCGACGTTAAGAATTGCC | 115 |
| chr4:13333131-13335149 | p2 | (AT)9 | TTGCAATGTGTTTGGCATTT | TCTCCAGATTGGTTTCTGGG | 154 |
| chr4:13333131-13335149 | p2 | (TA)22 | TTGCAATGTGTTTGGCATTT | TCTCCAGATTGGTTTCTGGG | 154 |
| chr4:13426522-13428536 | p2 | (AT)7 | TGGCCATCATTCTGTCACTG | TTGGCTACAATTGCAAGGTTC | 175 |
| chr4:13445438-13447477 | p3 | (TAT)7 | AACTCGTATGCACCTCGGTC | CGAGTATCGAACACCGGATT | 198 |
| chr4:13445529-13447550 | p3 | (TAT)7 | AACTCGTATGCACCTCGGTC | CGAGTATCGAACACCGGATT | 107 |
| chr4:13468681-13470713 | p2 | (TA)15 | ATGATTGATTCATTGCCACG | AAATGACATGGCGGAATAGC | 207 |
| chr4:13468681-13470713 | p2 | (AT)16 | ATGATTGATTCATTGCCACG | AAATGACATGGCGGAATAGC | 207 |
| chr4:13499966-13501978 | p2 | (TA)6 | GCCGTTTCACTTGGATTTGT | CCAAGCTTTCCCGTTCATTA | 288 |
| chr4:13520032-13522098 | p2 | (AT)10 | GTGGAAAATTGTCACACCCC | ATATTTGGGCACTTTCTGCG | 140 |
| chr4:13520185-13522201 | p2 | (AT)10 | AGTGTCATAAACCGGCAAGG | TCAGCGACGTCAGCAACTAC | 224 |
| chr4:13520246-13522261 | p2 | (AT)10 | AGTGTCATAAACCGGCAAGG | TCAGCGACGTCAGCAACTAC | 163 |
| chr4:13520289-13522305 | p2 | (AT)10 | AGTGTCATAAACCGGCAAGG | TCAGCGACGTCAGCAACTAC | 120 |
| chr4:13546993-13549059 | p2 | (AT)10 | GTGGAAAATTGTCACACCCC | ATATTTGGGCACTTTCTGCG | 140 |
| chr4:13547146-13549162 | p2 | (AT)10 | AGTGTCATAAACCGGCAAGG | TCAGCGACGTCAGCAACTAC | 224 |
| chr4:13547207-13549222 | p2 | (AT)10 | AGTGTCATAAACCGGCAAGG | TCAGCGACGTCAGCAACTAC | 163 |
| chr4:13547250-13549266 | p2 | (AT)10 | AGTGTCATAAACCGGCAAGG | TCAGCGACGTCAGCAACTAC | 120 |
| chr4:13805145-13807160 | p5 | (TTGGT)3 | AATGGCATGTGTGGTGAAGA | ATGGCCTAATTCATTGCAGG | 131 |
| chr4:13805145-13807160 | p5 | (TTGGT)3 | AATGGCATGTGTGGTGAAGA | ATGGCCTAATTCATTGCAGG | 131 |
| chr4:13810248-13812266 | p3 | (TTC)6 | TGGGTGGATGCATGCTATTA | AGAACTGGTTGGCGCTAGAA | 218 |

| Marker ID | SSR type | SSR motif | FORWARD PRIMER (5'-3') | REVERSE PRIMER (5'-3') | size (bp) |
|---|---|---|---|---|---|
| chr4:13810248-13812266 | p2 | (AT)6 | TGGGTGGATGCATGCTATTA | AGAACTGGTTGGCGCTAGAA | 218 |
| chr4:13811152-13813164 | p3 | (TTC)6 | TTAGCTGCTTCCTGTGCGTA | TATTGTTGATCGGATGGCAA | 142 |
| chr4:13811152-13813164 | p2 | (AT)6 | TTAGCTGCTTCCTGTGCGTA | TATTGTTGATCGGATGGCAA | 142 |
| chr4:13811152-13813164 | p3 | (GTG)6 | TTAGCTGCTTCCTGTGCGTA | TATTGTTGATCGGATGGCAA | 142 |
| chr4:13813364-13815382 | p3 | (TTC)6 | TGGGTGGATGCATGCTATTA | AGAACTGGTTGGCGCTAGAA | 219 |
| chr4:13813364-13815382 | p2 | (AT)7 | TGGGTGGATGCATGCTATTA | AGAACTGGTTGGCGCTAGAA | 219 |
| chr4:13924431-13926451 | p5 | (TTTTC)4 | CTCCCAGTCCTCCCTTCTTC | TGATCCATGAGCCACAAAAA | 202 |
| chr4:13973949-13975964 | p5 | (TAAAA)3 | CACATCCAAACAACTCACGG | ATATGGGGCCCACAATTTTT | 149 |
| chr4:13975401-13977437 | p2 | (AT)18 | CACCACATCCAAACTTCACG | AGTTGATACGCATGTGGCAA | 227 |
| chr4:14108796-14110810 | p2 | (AT)6 | GGATGCTGTCCTCCATGAAT | TTGGAGTTGCACCTGTTGTC | 146 |
| chr4:14108796-14110810 | p2 | (AT)7 | GGATGCTGTCCTCCATGAAT | TTGGAGTTGCACCTGTTGTC | 146 |
| chr4:14108796-14110810 | p6 | (ATTTTG)3 | GGATGCTGTCCTCCATGAAT | TTGGAGTTGCACCTGTTGTC | 146 |
| chr4:14141247-14143263 | p4 | (TTTA)4 | ACGGATTTAACACGCAGGTC | AGGGCGGGCTATATGCTACT | 277 |
| chr4:14163089-14165104 | p5 | (AAAAG)3 | GGAAAATTCAGCTTTGCAGG | TCTTTCAAGCCAAACCATCC | 195 |
| chr4:14266992-14269006 | p2 | (AG)7 | AAAAAGCACTTTTGGCGAGA | TGGCCAAACTACTTCAAGGC | 146 |
| chr4:14301923-14303947 | p3 | (AAT)8 | TTCGAATCTTGACCCGACTC | ACGTTCAACCGGCTTTTATG | 175 |
| chr4:14381387-14383411 | p3 | (TTA)5 | CACCAGTTGAGCAGTGGAGA | TTGATCCGTTTTATCCGAGG | 136 |
| chr4:14381387-14383411 | p3 | (TTA)6 | CACCAGTTGAGCAGTGGAGA | TTGATCCGTTTTATCCGAGG | 136 |
| chr4:14387949-14389965 | p2 | (TG)8 | TGATGAAAATCGGCAAGACA | CGACTACATGTTGGTCCACG | 234 |
| chr4:14454563-14456585 | p2 | (TA)11 | TCCTGTCAATAAGGGCGAAG | ATACCGAACACGCCAAAGAG | 112 |
| chr4:14464135-14466159 | p3 | (TTG)8 | CGCTGTGCCTTATGTCTGAA | ATCACAATCTTCCGAGTGCC | 254 |
| chr4:14464135-14466159 | p3 | (TTG)8 | CGCTGTGCCTTATGTCTGAA | ATCACAATCTTCCGAGTGCC | 254 |
| chr4:14481123-14483138 | p5 | (TTTGG)3 | GCCTGAAAAACCGCAAATTA | TTGCCGTGTTTCTACCTTCC | 220 |
| chr4:14482595-14484611 | p4 | (TGGG)4 | TCCTTGTAGTCGAGGCCCTA | CTTGCATTTTGCCCTCATTT | 116 |
| chr4:14681721-14683763 | p2 | (TA)7 | ATTCATCGTCCGTAATTGGC | GTGGTCCTGAAATAGGGCAA | 245 |
| chr4:14681721-14683763 | p2 | (TA)12 | ATTCATCGTCCGTAATTGGC | GTGGTCCTGAAATAGGGCAA | 245 |
| chr4:14827279-14829294 | p3 | (TGT)5 | GAAATGGAGGCTTGCTTCAG | GGTGTGCTCACACATCCATC | 195 |
| chr4:14834540-14836552 | p2 | (TA)6 | TCATTGTTTGATACACCCGAA | TCGAGCTACGATCAGCAAGA | 210 |
| chr4:14893409-14895430 | p3 | (AGA)7 | TTTCTTGTAGTGACGTGGCG | ATTGCAGGAGTGGATTTTCG | 126 |
| chr4:14967365-14969383 | p5 | (ATTTT)3 | ATTTGTGGATCCAAAGCTGG | CGACCCCTACCCATTTTCTT | 190 |
| chr4:14967365-14969383 | p2 | (AT)6 | ATTTGTGGATCCAAAGCTGG | CGACCCCTACCCATTTTCTT | 190 |
| chr4:14967365-14969383 | p6 | (CCCCTG)3 | ATTTGTGGATCCAAAGCTGG | CGACCCCTACCCATTTTCTT | 190 |
| chr4:14976834-14978849 | p5 | (ACCAA)3 | TCCAACAACACACACCAACC | AATCGGAGGTTGAAAAACCC | 135 |
| chr4:14976834-14978849 | p5 | (AAACC)3 | TCCAACAACACACACCAACC | AATCGGAGGTTGAAAAACCC | 135 |
| chr4:15007689-15009703 | p2 | (AT)7 | TTCCGGCTAAGACTTGCCTA | CTTTCGCTTTGGCGTTACAT | 169 |
| chr4:15028413-15030437 | p3 | (ATA)8 | AGGAGTGGTGGTGACAGAGG | CTTCCTCCCCTACGCTCTCT | 290 |

| Marker ID | SSR type | SSR motif | FORWARD PRIMER (5'-3') | REVERSE PRIMER (5'-3') | size (bp) |
|---|---|---|---|---|---|
| chr4:15057351-15059371 | p2 | (GA)10 | TTTCTTGCAAAAATTTCCCG | ATTGGGAGCTGTGAATCCTG | 166 |
| chr4:15111545-15113559 | p5 | (CCGGT)3 | ACGCCCTGCTACTTCTCAAA | TGCCTAGGTCCAACTCATCC | 191 |
| chr4:15111585-15113599 | p5 | (CCGGT)3 | ACGCCCTGCTACTTCTCAAA | TGCCTAGGTCCAACTCATCC | 151 |
| chr4:15270928-15272943 | p5 | (AAACC)3 | TGTATATGGACCGGGTTGGT | CGAACAACCCGAGAAAACTC | 106 |
| chr4:15320966-15322984 | p3 | (AGA)6 | GGTAGCTGGAATATTGGGCA | GGGTCCACCCATTAGTCCTT | 300 |
| chr4:15389380-15391406 | p2 | (TA)6 | AACCAAAACTCAGAAGCGGA | TTTCGTTAATTTCCCCCACA | 163 |
| chr4:15389380-15391406 | p2 | (AT)13 | AACCAAAACTCAGAAGCGGA | TTTCGTTAATTTCCCCCACA | 163 |
| chr4:15405603-15407617 | p2 | (AG)7 | GTAGGCGTTTGGCCATAGAA | AAAAGAATGGAGGGGTTTGG | 132 |
| chr4:15405603-15407617 | p6 | (TCTATA)3 | GTAGGCGTTTGGCCATAGAA | AAAAGAATGGAGGGGTTTGG | 132 |
| chr4:15578873-15580919 | p3 | (TGT)6 | TCTCCCCTTTCTCCGAAAAT | AAAATCGTACACCGCACGTT | 208 |
| chr4:15578873-15580919 | p2 | (AT)23 | TCTCCCCTTTCTCCGAAAAT | AAAATCGTACACCGCACGTT | 208 |
| chr4:15640800-15642815 | p3 | (TTC)5 | TCGGCAAGTTTAATCGATCC | AAAGCACAAGGAGGCACTGT | 260 |
| chr4:15640800-15642815 | p3 | (CAA)6 | TCGGCAAGTTTAATCGATCC | AAAGCACAAGGAGGCACTGT | 260 |
| chr4:15658822-15660840 | p3 | (AAG)6 | TGGCCAAGTGTGTGAAATACA | TCTTGGACGGATTTTCTTGG | 128 |
| chr4:15658822-15660840 | p3 | (ATT)14 | TGGCCAAGTGTGTGAAATACA | TCTTGGACGGATTTTCTTGG | 128 |
| chr4:15668219-15670234 | p5 | (GCTTG)3 | GAACCAGGGACGGAGTACAA | AGGGATTCGAAAAGAGCCAT | 260 |
| chr4:15711051-15713091 | p4 | (ATTC)4 | CAACCAACAACTCCCATCCT | TTTCTGGCTTCTTGCTGCTT | 226 |
| chr4:15711051-15713091 | p2 | (AG)20 | CAACCAACAACTCCCATCCT | TTTCTGGCTTCTTGCTGCTT | 226 |
| chr4:15816391-15818409 | p2 | (CA)9 | CCTCAGCAGATTTTGCATCA | TTCCCAAGAAGCTGATTGCT | 142 |
| chr4:15816391-15818409 | p3 | (TGT)6 | CCTCAGCAGATTTTGCATCA | TTCCCAAGAAGCTGATTGCT | 142 |
| chr4:15885945-15888105 | p2 | (AT)80 | TTGTGGCACGAATTGATGAT | CTTGCAGGGCTTATCAGAGG | 140 |
| chr4:15885945-15888105 | p2 | (TA)8 | TTGTGGCACGAATTGATGAT | CTTGCAGGGCTTATCAGAGG | 140 |
| chr4:15886360-15888424 | p2 | (AT)80 | GTACTCCCTCCGGATCAAAA | CGGTTGCCAGGCTTACTTAG | 137 |
| chr4:15886360-15888424 | p2 | (TA)8 | GTACTCCCTCCGGATCAAAA | CGGTTGCCAGGCTTACTTAG | 137 |
| chr4:15993654-15995675 | p3 | (TGT)7 | GGGGATCAACCACAATCATC | TGCTTGTAAAGACAATGGCG | 162 |
| chr4:16144114-16146126 | p3 | (TGC)5 | ACCTCATGTTGCTGCTGTTG | CCTTCCTCCCTTTCTTCCAC | 263 |
| chr4:16144114-16146126 | p5 | (AAGAA)3 | ACCTCATGTTGCTGCTGTTG | CCTTCCTCCCTTTCTTCCAC | 263 |
| chr4:16144114-16146126 | p2 | (CT)7 | ACCTCATGTTGCTGCTGTTG | CCTTCCTCCCTTTCTTCCAC | 263 |
| chr4:16144140-16146156 | p3 | (TGC)5 | ACCTCATGTTGCTGCTGTTG | CCTTCCTCCCTTTCTTCCAC | 237 |
| chr4:16144140-16146156 | p5 | (AAGAA)3 | ACCTCATGTTGCTGCTGTTG | CCTTCCTCCCTTTCTTCCAC | 237 |
| chr4:16144140-16146156 | p2 | (CT)7 | ACCTCATGTTGCTGCTGTTG | CCTTCCTCCCTTTCTTCCAC | 237 |
| chr4:16197650-16199665 | p2 | (GA)6 | AAGAGGAGGAAGCGGAGAAG | TCCCCTTCATTTCGTCATTC | 145 |
| chr4:16197650-16199665 | p3 | (ATA)5 | AAGAGGAGGAAGCGGAGAAG | TCCCCTTCATTTCGTCATTC | 145 |
| chr4:16263236-16265251 | p5 | (AACCG)3 | CACTCCCCCTTGGATGTCTA | TACCGAAGAACCATTGACCC | 258 |
| chr4:16314802-16316820 | p3 | (GTT)5 | CGCTTTGAATCTCTTAGCCG | TGGGGAGGGTAGAGTGTACG | 158 |
| chr4:16314802-16316820 | p2 | (AT)9 | CGCTTTGAATCTCTTAGCCG | TGGGGAGGGTAGAGTGTACG | 158 |

| Marker ID | SSR type | SSR motif | FORWARD PRIMER (5'-3') | REVERSE PRIMER (5'-3') | size (bp) |
|---|---|---|---|---|---|
| chr4:16415587-16417607 | p2 | (TA)10 | AGCAGGTTCCCAGGAAAGAT | CAACGTGGACGTAGCTCTCA | 177 |
| chr4:16415587-16417607 | p2 | (GT)6 | AGCAGGTTCCCAGGAAAGAT | CAACGTGGACGTAGCTCTCA | 177 |
| chr4:16564032-16566074 | p2 | (TA)12 | GAGCGTTAGGCGTGTTTAGG | AATCGGTAACCGTCCACAAC | 106 |
| chr4:16566375-16568395 | p2 | (GT)10 | TTGCACCAAATTCCTCACCT | GTTGCAACATCTGCAACAGC | 196 |
| chr4:16566375-16568395 | p2 | (TA)6 | TTGCACCAAATTCCTCACCT | GTTGCAACATCTGCAACAGC | 196 |
| chr4:16566375-16568395 | p2 | (TA)31 | TTGCACCAAATTCCTCACCT | GTTGCAACATCTGCAACAGC | 196 |
| chr4:16566375-16568395 | p2 | (GT)8 | TTGCACCAAATTCCTCACCT | GTTGCAACATCTGCAACAGC | 196 |
| chr4:16602092-16604106 | p2 | (AT)7 | CCGAAGGATGAACCTGAAAA | ATCACACCCATCTCAGGAGG | 178 |
| chr4:16626004-16628022 | p3 | (TGG)6 | TGAAAACCATTGACACCCCT | TGCACTTTCTTAACGGGAGG | 144 |
| chr4:16687535-16689550 | p3 | (ATT)5 | CATGGTTATTCCGGGATTTG | ACTGGCTCCCAAATTTTCCT | 148 |
| chr4:16846418-16848436 | p3 | (TCT)6 | GGGAGGTTGAAGGGCTAAAG | ACGAAGACGAGGAAGACGAA | 112 |
| chr4:16846418-16848436 | p6 | (GGGGGT)3 | GGGAGGTTGAAGGGCTAAAG | ACGAAGACGAGGAAGACGAA | 112 |
| chr4:16895093-16897105 | p2 | (AG)6 | GATACTCAAGGGATGGGGGT | GTTAGCTCAAACGGTCCAGG | 168 |
| chr4:16898510-16900540 | p3 | (TGT)5 | TTTGTGTGTTTCCATCGCAT | TGGCCTTTGTTGGGATAAAC | 273 |
| chr4:16898538-16900570 | p3 | (TGT)5 | TTTGTGTGTTTCCATCGCAT | TGGCCTTTGTTGGGATAAAC | 245 |
| chr4:16898573-16900593 | p3 | (TGT)5 | TTTGTGTGTTTCCATCGCAT | TGGCCTTTGTTGGGATAAAC | 210 |
| chr4:16898573-16900593 | p4 | (TAAC)4 | TTTGTGTGTTTCCATCGCAT | TGGCCTTTGTTGGGATAAAC | 210 |
| chr4:16948550-16950586 | p2 | (TA)18 | TGAAGGTTGTGGTGACGGTA | GCCGTTATATTTTGAGGCCA | 261 |
| chr4:16948550-16950586 | p2 | (AT)7 | TGAAGGTTGTGGTGACGGTA | GCCGTTATATTTTGAGGCCA | 261 |
| chr4:16948693-16950707 | p2 | (TA)18 | TGAAGGTTGTGGTGACGGTA | GCCGTTATATTTTGAGGCCA | 118 |
| chr4:16948693-16950707 | p2 | (AT)7 | TGAAGGTTGTGGTGACGGTA | GCCGTTATATTTTGAGGCCA | 118 |
| chr4:16986481-16988493 | p2 | (AT)6 | GGCCAAAAAGCAGTTGGTAA | ACGACATCCCGAAACAATTC | 101 |
| chr4:17007425-17009445 | p5 | (TTGTA)4 | TACACTCCCCCAGTCCTGTC | TGAGTAAATCAACCCAGGGC | 219 |
| chr4:17009862-17011880 | p2 | (TA)6 | TGGTGCCGAATGATTACTTG | GCACAACGAAAATGCAAAAA | 105 |
| chr4:17009862-17011880 | p3 | (TAT)6 | TGGTGCCGAATGATTACTTG | GCACAACGAAAATGCAAAAA | 105 |
| chr4:17053569-17055621 | p2 | (AT)6 | CGCTTGGGTATTTCGTTCAT | CGGAGGAAGTATCGCTTACG | 267 |
| chr4:17053672-17055738 | p2 | (AT)6 | CGCTTGGGTATTTCGTTCAT | CGGAGGAAGTATCGCTTACG | 164 |
| chr4:17094920-17096972 | p2 | (AT)6 | CGCTTGGGTATTTCGTTCAT | CGGAGGAAGTATCGCTTACG | 267 |
| chr4:17095023-17097089 | p2 | (AT)6 | CGCTTGGGTATTTCGTTCAT | CGGAGGAAGTATCGCTTACG | 164 |
| chr4:17223174-17225189 | p3 | (TGA)5 | TTAGGAGCCTGTTTGGATGG | TCTAGAGCCCGTTTGGATTG | 243 |
| chr4:17378977-17380993 | p2 | (AT)8 | CATTTTGAACCTCTTCGGGA | GAAGAGATGAGGCACGAAGG | 245 |
| chr4:17388131-17390147 | p3 | (TCT)5 | CCTTTTTCTGGATCGCCATA | GTCTGCACTTCCAGACCCAT | 252 |
| chr4:17388131-17390147 | p2 | (TC)8 | CCTTTTTCTGGATCGCCATA | GTCTGCACTTCCAGACCCAT | 252 |
| chr4:17412067-17414083 | p4 | (TTTA)4 | TGCTGGCAGATCCATACTTG | CATGTGCACGGTTTTTGAAC | 152 |
| chr4:17424076-17426100 | p2 | (AT)12 | GTTGCAGTGTCGACTGAGGA | TTTCCGTTTCCATTCTCACC | 290 |
| chr4:17426329-17428341 | p2 | (TA)6 | CAGTACCCGCATTGGAGTTT | TTCCTCCTTTAACCAGGGGT | 107 |

| Marker ID | SSR type | SSR motif | FORWARD PRIMER (5'-3') | REVERSE PRIMER (5'-3') | size (bp) |
|---|---|---|---|---|---|
| chr4:17436810-17438876 | p2 | (TA)33 | ATATCGGCAACAATTGCCAC | ATCAACTGCCAAGAACGGAT | 146 |
| chr4:17592181-17594199 | p6 | (GAACAG)3 | CCCCTCCCCAACAATTTACT | TTCGACCTTTGGGCTAGATG | 249 |
| chr4:17709044-17711058 | p2 | (AG)7 | AGCTTCTATTCCAGTGGGCA | GCAACTTGGCTTCTTTTTCG | 156 |
| chr4:17723558-17725573 | p3 | (CAT)5 | CCTCTCGGCCATCAAAATAA | TGTTCATGCCCACAGGTAGA | 147 |
| chr4:17723558-17725573 | p4 | (TTCT)4 | CCTCTCGGCCATCAAAATAA | TGTTCATGCCCACAGGTAGA | 147 |
| chr4:17727302-17729320 | p6 | (TTTGGG)3 | TTTCGTGCAGAATTCCATCA | TAAACACAAGTCCAAGGCCC | 131 |
| chr4:17765989-17768004 | p4 | (AATA)6 | CGGATTGGTTTGGTTTAGGA | TACATCAATAAAGCCGTGCG | 173 |
| chr4:17766002-17768046 | p4 | (AATA)6 | CGGATTGGTTTGGTTTAGGA | TACATCAATAAAGCCGTGCG | 160 |
| chr4:17819265-17821280 | p5 | (TTTGT)3 | GCCTTGTCCTAAGCGTGAAG | CCTAAATCTTTCATGGGGCA | 108 |
| chr4:17829537-17831553 | p2 | (GT)6 | TAGCAAATTTTGGTGACCCC | CCTCGCTTGTAGTTGCCTTC | 225 |
| chr4:17829537-17831553 | p2 | (TA)8 | TAGCAAATTTTGGTGACCCC | CCTCGCTTGTAGTTGCCTTC | 225 |
| chr4:17841722-17843794 | p3 | (TTA)7 | CGAGTCGCAGACATAGACCA | GTCTCCTCCACATTTGGCAT | 267 |
| chr4:17841722-17843794 | p3 | (TTA)5 | CGAGTCGCAGACATAGACCA | GTCTCCTCCACATTTGGCAT | 267 |
| chr4:17841871-17843931 | p3 | (TTA)7 | CGAGTCGCAGACATAGACCA | GTCTCCTCCACATTTGGCAT | 118 |
| chr4:17841871-17843931 | p3 | (TTA)5 | CGAGTCGCAGACATAGACCA | GTCTCCTCCACATTTGGCAT | 118 |
| chr4:17925114-17927130 | p2 | (TA)8 | CAGCTCTAGTTCCGAGGGTG | TGTCCTCACTGCATCCCATA | 107 |
| chr4:17974113-17976128 | p5 | (TTGTA)3 | AAGTTGCATGGAAGAATGCC | TGTCGAGTTGTTCATTGGGA | 110 |
| chr4:17974113-17976128 | p5 | (GGGGT)5 | AAGTTGCATGGAAGAATGCC | TGTCGAGTTGTTCATTGGGA | 110 |
| chr4:17974653-17976678 | p5 | (TTGTA)3 | CTGAAACATGAAGCACCGAA | CGCAGACTCTTCCTTTTTGC | 151 |
| chr4:17974653-17976678 | p5 | (GGGGT)5 | CTGAAACATGAAGCACCGAA | CGCAGACTCTTCCTTTTTGC | 151 |
| chr4:18058735-18060813 | p3 | (CTT)5 | TTCGTCGTCCCCGTAGATAG | AAGGCAGAGTTGAAGTCGGA | 231 |
| chr4:18058735-18060813 | p3 | (TAA)8 | TTCGTCGTCCCCGTAGATAG | AAGGCAGAGTTGAAGTCGGA | 231 |
| chr4:18075601-18077633 | p2 | (AG)7 | GTGGGGATTGGGATTCTTTT | ACGGGTCCGGGTATATTTGT | 122 |
| chr4:18075601-18077633 | p2 | (AC)6 | GTGGGGATTGGGATTCTTTT | ACGGGTCCGGGTATATTTGT | 122 |
| chr4:18122244-18124262 | p5 | (AAAAT)3 | GAATTTTGCTTGCTTCAGGC | AGGTCCCTTCAAATGCACTG | 118 |
| chr4:18122244-18124262 | p3 | (TCA)6 | GAATTTTGCTTGCTTCAGGC | AGGTCCCTTCAAATGCACTG | 118 |
| chr4:18154743-18156755 | p2 | (AT)6 | AGTTTGAAGTCGGGTTTCCA | ATCCAAGCACCAAATCCATC | 298 |
| chr4:18205981-18208011 | p3 | (AAT)10 | GAGATGGAGGAGAGAAGGGG | GTCCACAAATGGAATGGGTC | 112 |
| chr4:18254646-18256664 | p3 | (CAA)6 | ACCGGCGAACGATATCATAG | GAAAAGCTCGGATTGCTTTG | 259 |
| chr4:18350459-18352475 | p4 | (AAAT)4 | AAGTCGACCAACCTCAATGC | GTATGGGATGCACCGGTATC | 273 |
| chr4:18400173-18402185 | p2 | (CA)6 | CTTTTAGACCAGCACGAGCC | AGCGCTAGGTTTTGGTTCAA | 289 |
| chr4:18428798-18430810 | p2 | (AT)6 | TAGCGGTTGGCTTACGAGTT | AAACGTGGTGCCGGTAGTAG | 193 |
| chr4:18495892-18497922 | p3 | (CTT)10 | GTCCAACAAAACGCCCTAAA | ATGCATCATGATCAAAGCCA | 258 |
| chr4:18534748-18536760 | p2 | (AT)7 | CCAAACGACCCCTAATGGTA | GAGGATGGAGAGAAGGGTCC | 262 |
| chr4:18534764-18536804 | p2 | (AT)7 | CCAAACGACCCCTAATGGTA | GAGGATGGAGAGAAGGGTCC | 246 |
| chr4:18537623-18539641 | p3 | (TTG)6 | GCCGGTAAGTCCCTAAAAGC | GAAGGAGGAGAGGAGGAGGA | 114 |

| Marker ID | SSR type | SSR motif | FORWARD PRIMER (5'-3') | REVERSE PRIMER (5'-3') | size (bp) |
|---|---|---|---|---|---|
| chr4:18593706-18595718 | p2 | (CT)6 | TGATCCACCAGCACCTTGTA | TCCCTCTTGCCTTAGCTTGA | 259 |
| chr4:18653383-18655398 | p3 | (GAA)5 | AAATTAGGGCAGTGTGTGGC | TCACTTTCTCTCTCGCCCAT | 105 |
| chr4:18665322-18667340 | p6 | (ATGGGT)3 | TCCTCACCAACCCTATCACC | GCACCACATGGTGAATGAAC | 105 |
| chr4:18764238-18766250 | p2 | (CT)6 | AATACCGTTGCAATGCATGA | GCCATGTTTGCCCTCATAGT | 208 |
| chr4:18786232-18788250 | p6 | (ACTAGT)3 | TGCATGTGCTCTCTGCTCTT | TGGGATTGAAATTCCCAAAG | 145 |
| chr4:18786232-18788250 | p2 | (TA)9 | TGCATGTGCTCTCTGCTCTT | TGGGATTGAAATTCCCAAAG | 145 |
| chr4:18816567-18818582 | p5 | (AATTT)3 | TTTCCACCACCAAACCAAAT | CCTGAGGTAAGGCGGTACAA | 147 |
| chr4:18873050-18875070 | p2 | (AT)10 | CAGGCCGGTTCTGTTATTGT | CTCCGAATCCCCATCTCATA | 229 |
| chr4:18958460-18960478 | p6 | (TTTTTC)3 | ATAGGATTCTTTGGGTGGGC | GATATCAACCTCGGCATGGT | 240 |
| chr4:18976976-18978994 | p6 | (TTTTTC)3 | ATAGGATTCTTTGGGTGGGC | GATATCAACCTCGGCATGGT | 240 |
| chr4:19069629-19071653 | p3 | (ATA)8 | GAGTGGGTTGACGACCAAAC | GGCCAAACGTAGATTTCCAA | 209 |
| chr4:19122704-19124724 | p5 | (TTCAT)4 | CCCCGGAAGATATCCAAGAT | GAATGTTAAGTGCTCGGGGA | 240 |
| chr4:19151110-19153128 | p2 | (AG)19 | TGGATTAGGCCATTTTGGAA | ACCGGCAATTACACTTTTGC | 285 |
| chr4:19151110-19153128 | p6 | (TGTATA)3 | TGGATTAGGCCATTTTGGAA | ACCGGCAATTACACTTTTGC | 285 |
| chr4:19151110-19153128 | p4 | (TTAA)4 | TGGATTAGGCCATTTTGGAA | ACCGGCAATTACACTTTTGC | 285 |
| chr4:19211078-19213098 | p5 | (TTCAT)4 | CCCCGGAAGATATCCAAGAT | GAATGTTAAGTGCTCGGGGA | 240 |
| chr4:19220748-19222772 | p3 | (TTA)8 | GTCTCAGGAAGGTGCCATGT | CCAGGCCATCTAACCTTTCA | 106 |
| chr4:19220748-19222772 | p3 | (TCT)5 | GTCTCAGGAAGGTGCCATGT | CCAGGCCATCTAACCTTTCA | 106 |
| chr4:19228627-19230642 | p2 | (AT)8 | CCTTCCTTTTACACACCCCA | CTTCCCCTCGCTTATTCACA | 190 |
| chr4:19228627-19230642 | p5 | (CCCCA)3 | CCTTCCTTTTACACACCCCA | CTTCCCCTCGCTTATTCACA | 190 |
| chr4:19228627-19230642 | p3 | (TTC)5 | CCTTCCTTTTACACACCCCA | CTTCCCCTCGCTTATTCACA | 190 |
| chr4:19230416-19232431 | p3 | (AAG)5 | CGTGGCTGGGGTGTATTACT | GGCAACCCAAAAGAAGGATT | 214 |
| chr4:19354827-19356845 | p6 | (AAGAAA)3 | TCCACAGCAATAGGTTTCCC | CGGCAGTATGGATTTCGTTT | 113 |
| chr4:19366822-19368837 | p5 | (GACCC)3 | TGAGCTGCTGACGCAACTAT | CTGCAAGAAGTGCGAACAAA | 295 |
| chr4:19366822-19368837 | p2 | (AT)6 | TGAGCTGCTGACGCAACTAT | CTGCAAGAAGTGCGAACAAA | 295 |
| chr4:19399957-19401972 | p3 | (AAG)5 | CGTGGCTGGGGTGTATTACT | AAAGGCAACCCAAAAGAAGG | 223 |
| chr4:19403187-19405201 | p5 | (AAAAT)3 | TTTTCCCTTGTTGCCCTAGA | TTGGGGATTTCCATCCATTA | 119 |
| chr4:19431733-19433803 | p3 | (ACA)5 | TGTTAGCGAGGGGCACTAAT | GGATTTCACTGGGCTGTTGT | 213 |
| chr4:19431733-19433803 | p2 | (GA)20 | TGTTAGCGAGGGGCACTAAT | GGATTTCACTGGGCTGTTGT | 213 |
| chr4:19447306-19449321 | p5 | (ACAAT)3 | TGCGGACGAATAATACCACA | TCGCATCTTCTCTTCTCCGT | 159 |
| chr4:19447306-19449321 | p2 | (AT)6 | TGCGGACGAATAATACCACA | TCGCATCTTCTCTTCTCCGT | 159 |
| chr4:19456651-19458677 | p2 | (AT)13 | TTTGAAGGCATTTTGGTTCC | TTGCATCGCAACTAGTGCTT | 175 |
| chr4:19488644-19490662 | p6 | (CATATA)3 | GGAGCCGCTTACAGACTGAC | CGCAGGTACTCCCAGATGAT | 168 |
| chr4:19604508-19606523 | p5 | (TTGGT)3 | TAAATCGACAAATCGCACCA | GTAATGTCGGGTTGGTTTGG | 192 |
| chr4:19604508-19606523 | p5 | (TAAAA)3 | TAAATCGACAAATCGCACCA | GTAATGTCGGGTTGGTTTGG | 192 |
| chr4:19672902-19674922 | p5 | (TTCTT)4 | TCCCTGCACTCAGACACTTG | ACTTGGAGACGGTCACATCC | 180 |

| Marker ID | SSR type | SSR motif | FORWARD PRIMER (5'-3') | REVERSE PRIMER (5'-3') | size (bp) |
|---|---|---|---|---|---|
| chr4:19684768-19686783 | p5 | (GAAAA)3 | TGTTTCTCCCTACCCCACAG | GAACCCTTTAGCCTTCCACC | 246 |
| chr4:19702718-19704732 | p2 | (AG)7 | TGGTCAATGAAGTGGGTTGA | GAACCTGCTACCTCCCATCA | 214 |
| chr4:19728955-19730969 | p2 | (AT)7 | TTGCTTCCCATTTCCAAGTC | TTACCCCAGCAGATTATGGC | 161 |
| chr4:19728955-19730969 | p2 | (AT)7 | TTGCTTCCCATTTCCAAGTC | TTACCCCAGCAGATTATGGC | 161 |
| chr4:19728955-19730969 | p6 | (CATATA)3 | TTGCTTCCCATTTCCAAGTC | TTACCCCAGCAGATTATGGC | 161 |
| chr4:19755292-19757310 | p6 | (CTGAAA)3 | AAAAGGGACCAAGGCTTCAT | TTTACCAGCTTACGCACACG | 162 |
| chr4:19755292-19757310 | p5 | (ATAAT)3 | AAAAGGGACCAAGGCTTCAT | TTTACCAGCTTACGCACACG | 162 |
| chr4:19767737-19769813 | p2 | (TA)6 | AATGTACGGGGACTCTGCTG | GCAATAAAAGGGGACGTCAA | 114 |
| chr4:19767737-19769813 | p6 | (CTGAAA)3 | AATGTACGGGGACTCTGCTG | GCAATAAAAGGGGACGTCAA | 114 |
| chr4:19842216-19844230 | p2 | (GA)6 | CAAGCCTGCTACTTGCATCA | GAGGTCTCTGCGTTTTCCTG | 211 |
| chr4:19842216-19844230 | p2 | (AG)7 | CAAGCCTGCTACTTGCATCA | GAGGTCTCTGCGTTTTCCTG | 211 |
| chr4:19933910-19935925 | p5 | (CAAAA)3 | TGCCTTGAGCCAAGGTCTAT | CGCCCGATTAGTTTGAGTGT | 138 |
| chr4:19974798-19976819 | p3 | (TTC)7 | TGGTGGTAATGGTGAGGGTT | ATTAGCGGTGTTCGAGGTTG | 177 |
| chr4:19980367-19982387 | p5 | (AATCA)4 | GGGTGGTTGCAGCTGTCTAT | AATTTGCCATCATAGCTCGG | 183 |
| chr4:20118878-20120896 | p2 | (CT)9 | TCAGAACTCAGCCCTCCTGT | TACTTTTTGGCACTCCCGAC | 202 |
| chr4:20132244-20134258 | p2 | (AT)7 | AAATAACCCGAACCGTAGCC | AAGGAGAAGAAGGCCACCAT | 111 |
| chr4:20139212-20141227 | p3 | (AAT)5 | TTGCCCATAAACCCCATAAA | GGAGGTGAGCCACCATTAGA | 255 |
| chr4:20172573-20174621 | p3 | (TTA)16 | CTCCGAATGCGGGTATCTTA | TCGCTGCCGAAGAAAATAGT | 201 |
| chr4:20183002-20185014 | p2 | (AT)7 | GCTACTGGGCTTGTTTTGCT | TTAAATTGCAACGGACGACA | 267 |
| chr4:20183002-20185014 | p2 | (AT)23 | GCTACTGGGCTTGTTTTGCT | TTAAATTGCAACGGACGACA | 267 |
| chr4:20229294-20231309 | p5 | (TCATA)3 | TCACCCGTATTTTGGGTGTT | TTCGCTAATCTGGGTATCGG | 287 |
| chr4:20417539-20419566 | p3 | (TGT)9 | GGAATACTGGGTTTGGAGCA | AAAGTGTGGTGGCAAGATCC | 254 |
| chr4:20427137-20429153 | p4 | (TACA)4 | TCACAGGGTGACTTGTTCCA | ACCAGGACTGGAGACACCTG | 105 |
| chr4:20471765-20473791 | p2 | (AT)13 | TTTGGCTTCGCTCTACCCTA | TTCCTTAAACTGGCATTCCG | 192 |
| chr4:20504123-20506191 | p2 | (AT)34 | TTTGTGGAATTGGCTGATGA | ATTGGACCCCAACCCTTTAG | 211 |
| chr4:20523997-20526015 | p3 | (GAA)6 | AACCTCACGTTCACGCTCTT | GCAGTGTTCTCCAAAAAGGG | 189 |
| chr4:20523997-20526015 | p2 | (AC)7 | AACCTCACGTTCACGCTCTT | GCAGTGTTCTCCAAAAAGGG | 189 |
| chr4:20625128-20627196 | p2 | (AT)34 | TTTGTGGAATTGGCTGATGA | ATTGGACCCCAACCCTTTAG | 211 |
| chr4:20686081-20688096 | p5 | (ACCCG)3 | CCGACAGGCCAGAATAAAAA | TTCGAGGCACCCATTTCTAC | 139 |
| chr4:20686081-20688096 | p2 | (CT)6 | CCGACAGGCCAGAATAAAAA | TTCGAGGCACCCATTTCTAC | 139 |
| chr4:20699189-20701213 | p4 | (TATT)6 | AAAACCAACCCATGAACACC | CGAACTACATGGCTTGCTGA | 276 |
| chr4:20717522-20719540 | p6 | (CAACTC)3 | TGGGCCATGCCTAAGACTAC | TCAAATCCCCTATATCCCCC | 100 |
| chr4:20738681-20740695 | p2 | (AG)7 | GATGTATGGAGCGCTGGAGT | GGCCAAGGTTCAGATTCAAA | 240 |
| chr4:20777672-20779684 | p2 | (CT)6 | ATCTGCAACGGTAGCCAAGT | GTAAACCCCGCTTCGATGTA | 235 |
| chr4:20777672-20779684 | p3 | (AAT)7 | ATCTGCAACGGTAGCCAAGT | GTAAACCCCGCTTCGATGTA | 235 |
| chr4:20829596-20831632 | p2 | (AT)9 | TTCCCAAGGAAGTGAGAGGA | GGAATGGTGGCCATAGAAAA | 223 |

| Marker ID | SSR type | SSR motif | FORWARD PRIMER (5'-3') | REVERSE PRIMER (5'-3') | size (bp) |
|---|---|---|---|---|---|
| chr4:20829596-20831632 | p2 | (AT)18 | TTCCCAAGGAAGTGAGAGGA | GGAATGGTGGCCATAGAAAA | 223 |
| chr4:20834601-20836621 | p2 | (TC)10 | GTTTTCCATCGTGCCTTCTC | TATGCAGAGGCACATTCTGG | 133 |
| chr4:20838058-20840070 | p2 | (AT)6 | CCTTTGTGGTGCTACCCACT | TGGAGCAGTAATGATCCGGT | 186 |
| chr4:20838058-20840070 | p2 | (AT)6 | CCTTTGTGGTGCTACCCACT | TGGAGCAGTAATGATCCGGT | 186 |
| chr4:20906438-20908450 | p2 | (TA)6 | CTCGGGGAGAGTGCATTTAG | CTTCTTGGTTTTGGTTGCGT | 281 |
| chr4:20926108-20928123 | p5 | (GATTG)3 | AAGAAAAGTGGGATTGGCCT | GGCAGAAAACTCGTCCACAT | 194 |
| chr4:20985098-20987116 | p2 | (AG)8 | ACACAACGATTCTCTTCGGG | ATCTTAGCCGGAGATCAGCA | 211 |
| chr4:20985098-20987116 | p6 | (CACCTC)3 | ACACAACGATTCTCTTCGGG | ATCTTAGCCGGAGATCAGCA | 211 |
| chr4:20985098-20987116 | p5 | (GGGGT)3 | ACACAACGATTCTCTTCGGG | ATCTTAGCCGGAGATCAGCA | 211 |
| chr4:20992197-20994209 | p2 | (AG)6 | AACAGAGGGCCAAAGGAACT | AAATTTGCTCAATATGGGCG | 286 |
| chr4:21031987-21034011 | p6 | (TTTTTC)4 | ACGTCCACCACCAGAAGAAG | GAAGTGAAAGCAGGAGCAGG | 282 |
| chr4:21052970-21054985 | p3 | (CAC)5 | CCCAAGAAGATGTGGCAAGT | TAGTGGTGGCCATGACAAAA | 231 |
| chr4:21081372-21083390 | p2 | (TA)9 | TTGTCTACCGCAAACTCGTC | TTTACAGCCAGAAACCCACC | 105 |
| chr4:21132494-21134509 | p3 | (AGA)5 | GAGCCGAACTGGCTCTTCTA | TGTCGAGTGAGTGGAAATCG | 265 |
| chr4:21182472-21184490 | p6 | (CCTCTG)3 | TTGTTCTTTATTGTCGCCCC | GGCTCCTGCTGAATTGAAAG | 125 |
| chr4:21373983-21375997 | p2 | (TG)7 | GGGCACCAGCTAATTATCCA | CCTTTGGCTTTGAGAAAACG | 226 |
| chr4:21430735-21432789 | p2 | (TA)27 | CCGTACCGATACCGAAGAAA | CATGCTTTGCGAAAAATCAA | 119 |
| chr4:21434593-21436608 | p5 | (TTATT)3 | GAGGTTTGCCTCACCAAAAA | AGAAAGCCCGTGACCCTATT | 254 |
| chr4:21508841-21510859 | p3 | (ACA)6 | AGTAATTAGGGGTGGTGGGG | TAAAGCTATGAAACGCCGCT | 289 |
| chr4:21523059-21525074 | p5 | (AACCG)3 | GATGTCACCGACATTTTCCC | ATTCGGATTGAACCGATTTG | 270 |
| chr4:21523059-21525074 | p2 | (TA)16 | GATGTCACCGACATTTTCCC | ATTCGGATTGAACCGATTTG | 270 |
| chr4:21544653-21546668 | p6 | (TTTCGG)3 | GCTCTCCAAGCAGCCTCTAA | GTCCACCTTCAAAAACCGAA | 180 |
| chr4:21544653-21546668 | p5 | (GTTCG)3 | GCTCTCCAAGCAGCCTCTAA | GTCCACCTTCAAAAACCGAA | 180 |
| chr4:21584740-21586755 | p5 | (CTTTT)3 | TACGGACCAGTGCCAATACA | GGCCTGGTTATCGTTTTGAA | 224 |
| chr4:21632796-21634816 | p2 | (AC)10 | CTCACACCTGAGTTGCCTGA | TTTTGGGCTTGAGAACTGCT | 101 |
| chr4:21684819-21686867 | p2 | (CT)24 | TGTCAAGAGTGCGCAAATTC | TGAAACTCCTCTGACCACCC | 185 |
| chr4:21844934-21846952 | p2 | (TG)9 | GGGATTATGATCTGCGGAAA | TTCCAGGACAATTTTGGAGC | 214 |
| chr4:22106122-22108137 | p5 | (CCCGC)3 | ATGGCGTGAGCAAGTTCTTT | ATAGAAAATGCGGGACATCG | 122 |
| chr4:22128742-22130802 | p3 | (GTT)6 | GGGAAGTTTCGGCAATATCA | TTCTGGGAATTCTCGGATTG | 282 |
| chr4:22145914-22147934 | p2 | (TA)10 | GTTCAAGGGGGTGCAAATTA | GGCGAGCTACTACGATGAGC | 233 |
| chr4:22238370-22240386 | p2 | (TG)8 | GCACTTAACGGCTTCGTCTC | CCCAATTGATTCGTGAGGTT | 100 |
| chr4:22287662-22289677 | p2 | (AT)8 | ACAGGAATCCGTGAAGATGG | CAGCGGAGTTTCAAAAGAGC | 161 |
| chr4:22287662-22289677 | p3 | (ATC)5 | ACAGGAATCCGTGAAGATGG | CAGCGGAGTTTCAAAAGAGC | 161 |
| chr4:22345330-22347345 | p5 | (GTTTT)3 | ATTCTTGGGTGAGTGGATGC | GGCGGTGGTAGAAAAACAAA | 251 |
| chr4:22349163-22351212 | p2 | (AT)7 | TTTTGCAATTCGTGGTGAAA | GCATTTCGTGAATTGTGGTG | 224 |
| chr4:22349163-22351212 | p2 | (TA)17 | TTTTGCAATTCGTGGTGAAA | GCATTTCGTGAATTGTGGTG | 224 |

| Marker ID | SSR type | SSR motif | FORWARD PRIMER (5'-3') | REVERSE PRIMER (5'-3') | size (bp) |
|---|---|---|---|---|---|
| chr4:22349163-22351212 | p6 | (GAAGAT)3 | TTTTGCAATTCGTGGTGAAA | GCATTTCGTGAATTGTGGTG | 224 |
| chr4:22371572-22373602 | p6 | (CCACCC)5 | ATTTCGTCTCCAAATGCCAC | CTTGGGCTAGGCATCTTGTC | 180 |
| chr4:22386569-22388599 | p2 | (AT)15 | CTACCCACCAACGCTAGGAG | ACACCTGCTGGAAAATTTGG | 209 |
| chr4:22420490-22422510 | p4 | (TATC)5 | CTCCCAGCCTCTTCCTCTTT | CATGTGCTTAATCCGGGAAA | 276 |
| chr4:22508209-22510224 | p5 | (AATTT)3 | CAGTTGCTTCCGAGTGTTGA | AATTCTGCAAGGGGAAATGA | 206 |
| chr4:22542996-22545011 | p5 | (AATTT)3 | CAGTTGCTTCCGAGTGTTGA | AATTCTGCAAGGGGAAATGA | 206 |
| chr4:22552162-22554186 | p2 | (AT)12 | AACTGAAATTTGATTGGCCG | TTCGCCTTCGTTCTTGATCT | 202 |
| chr4:22556302-22558328 | p2 | (AT)13 | AGAGTTTTATGCGAAGGGCA | CTCCTATACAAGCCCCACCA | 288 |
| chr4:22599914-22601932 | p6 | (AAGGAG)3 | GCAAAGAGCCAACTTTCTGG | AGGAAGGGGAGTGCAAAGTT | 261 |
| chr4:22636908-22638920 | p2 | (TC)6 | TAAATGATTCGCACTGCTCG | ATTCCGTACTGACCCCCTTT | 265 |
| chr4:22672802-22674814 | p2 | (TC)6 | TAAATGATTCGCACTGCTCG | ATTCCGTACTGACCCCCTTT | 265 |
| chr4:22693468-22695558 | p2 | (AT)45 | AAGAGTTTTAGGCGAAGGGC | CTTACACAAGCCCCACCACT | 289 |
| chr4:22693468-22695558 | p2 | (AT)8 | AAGAGTTTTAGGCGAAGGGC | CTTACACAAGCCCCACCACT | 289 |
| chr4:22693831-22695847 | p2 | (AT)45 | AGCTAGTTACGGGTTTGCGA | GGCTACATACGCCTCTCAGG | 119 |
| chr4:22693831-22695847 | p2 | (AT)8 | AGCTAGTTACGGGTTTGCGA | GGCTACATACGCCTCTCAGG | 119 |
| chr4:22722676-22724691 | p3 | (ATA)7 | CCCATGTTGAGCCTTTGACT | GCTTTAGGCTCTTGGCCTTT | 160 |
| chr4:22722676-22724691 | p3 | (ATT)5 | CCCATGTTGAGCCTTTGACT | GCTTTAGGCTCTTGGCCTTT | 160 |
| chr4:22726420-22728432 | p2 | (AT)6 | AATGAAGAATCGACGTTCCG | TCTCCGTCCTTTGTCGTCTT | 274 |
| chr4:22726420-22728432 | p2 | (AT)7 | AATGAAGAATCGACGTTCCG | TCTCCGTCCTTTGTCGTCTT | 274 |
| chr4:22828474-22830498 | p2 | (AG)34 | TGTCCCGAAAAATCAAAAGG | TTTATCCGCTTCTTTCCCCT | 175 |
| chr4:22886665-22888680 | p3 | (TAA)5 | ATGTTGAGGCAAGCTCGATT | TTTGTTGGGGCTCTGATACC | 111 |
| chr4:23108850-23110874 | p4 | (AAAG)5 | TCCTCGAGAAAAGGACCAAA | CAAGGGGAATAAATGAGCCA | 284 |
| chr4:23108850-23110874 | p2 | (TA)12 | TCCTCGAGAAAAGGACCAAA | CAAGGGGAATAAATGAGCCA | 284 |
| chr4:23108850-23110874 | p2 | (AG)8 | TCCTCGAGAAAAGGACCAAA | CAAGGGGAATAAATGAGCCA | 284 |
| chr4:23108850-23110874 | p2 | (AT)13 | TCCTCGAGAAAAGGACCAAA | CAAGGGGAATAAATGAGCCA | 284 |
| chr4:23124063-23126075 | p2 | (TA)6 | TCATTGCCATCCATTGTCAT | CAATAGCCTGAGCCTTGGAG | 293 |
| chr4:23144509-23146523 | p2 | (TA)7 | CGTAGATTTGGGGCGTAAAA | TTAAAGTTGCTGTCGGACCC | 283 |
| chr4:23288655-23290670 | p5 | (ATTTT)3 | ATGAAGGGTTTTTCCCCATC | AAAGAAAAGTGCACCCCGTA | 268 |
| chr4:23301583-23303607 | p5 | (TAAAA)3 | CAAAAGGGTGGCCCATACTA | CCCATGAGTTTGGAGATGGT | 299 |
| chr4:23301583-23303607 | p2 | (TA)12 | CAAAAGGGTGGCCCATACTA | CCCATGAGTTTGGAGATGGT | 299 |
| chr4:23301583-23303607 | p2 | (TA)6 | CAAAAGGGTGGCCCATACTA | CCCATGAGTTTGGAGATGGT | 299 |
| chr4:23315206-23317226 | p2 | (AT)10 | CACAAAAGCGAGTTATCGCA | TAGGCGTCAATAGGTGGAGG | 209 |
| chr4:23346007-23348022 | p3 | (TGT)5 | GTTGTACGGGGCAGAGTGTT | TTCTGAAACCGTCCATAGCC | 258 |
| chr4:23419236-23421251 | p5 | (AATTT)3 | CTTCGATGCCAATCTTGGTT | CGAAGCATCTAAGACCCGAG | 253 |
| chr4:23449488-23451503 | p5 | (AATAA)3 | AATATCCCTGGAAAGGGTGG | TCCGATTAACAGCAACACCA | 158 |
| chr4:23449488-23451503 | p5 | (ACCAA)3 | AATATCCCTGGAAAGGGTGG | TCCGATTAACAGCAACACCA | 158 |

| Marker ID | SSR type | SSR motif | FORWARD PRIMER (5'-3') | REVERSE PRIMER (5'-3') | size (bp) |
|---|---|---|---|---|---|
| chr4:23481701-23483716 | p3 | (AAT)5 | GTGGGTGGGTAGGGTGTATG | CAATCGCCCGTTATCAAAAT | 145 |
| chr4:23666731-23668745 | p2 | (AG)7 | CTGATGCTGTGGCTGTAGGA | GGATTCAGTACCTCAGGGCA | 261 |
| chr4:23746579-23748615 | p2 | (AT)18 | GACTAGTGCCTGGAAGCTGG | TGTCTGTAGTGGCAGCCTTG | 182 |
| chr4:23757765-23759780 | p3 | (TCA)5 | GGGGGTTGTGTTTCATCATC | GTGGTGCCGGTAGTAGTGGT | 263 |
| chr4:23757765-23759780 | p3 | (GTT)5 | GGGGGTTGTGTTTCATCATC | GTGGTGCCGGTAGTAGTGGT | 263 |
| chr4:23757791-23759809 | p3 | (TCA)5 | GGGGGTTGTGTTTCATCATC | GTGGTGCCGGTAGTAGTGGT | 237 |
| chr4:23757791-23759809 | p3 | (GTT)5 | GGGGGTTGTGTTTCATCATC | GTGGTGCCGGTAGTAGTGGT | 237 |
| chr4:23758651-23760666 | p3 | (GTT)5 | GTTCCGGTTCCGGTTTTACT | AGGCCCTAAGTTTGCCCTTA | 143 |
| chr4:23791033-23793051 | p3 | (ATA)6 | ATTCAACTCCGCCCCTTTAT | CCCTTGTTCCCTCCTTCTTC | 158 |
| chr4:23808018-23810033 | p5 | (ATCAT)3 | CGGGATTTCACTAGGCATGT | TGCTCTGCAATCTTGACACC | 296 |
| chr4:23841794-23843822 | p3 | (GTG)5 | TGGGAATTCGCCTATTTCTG | GACACCTTCCACCATCATCC | 139 |
| chr4:23841794-23843822 | p2 | (AT)14 | TGGGAATTCGCCTATTTCTG | GACACCTTCCACCATCATCC | 139 |
| chr4:23883887-23885953 | p2 | (AT)33 | CTTCCTCCGTTCCAAAACAA | CGAGAAACGAAACGTGACAA | 284 |
| chr4:23912064-23914084 | p2 | (AT)10 | AAAACTGACCCACATTTGCC | GTACGGTACGGAGGATTGGA | 196 |
| chr4:23918134-23920152 | p5 | (AAACC)3 | CCACCGGTTTGGTTATTGTC | GCAAGTTTGGTCCACCTGAT | 270 |
| chr4:23918134-23920152 | p6 | (ATTTTT)3 | CCACCGGTTTGGTTATTGTC | GCAAGTTTGGTCCACCTGAT | 270 |
| chr4:23963398-23965416 | p6 | (TATATG)3 | TTGTATGGTGTGTTGGTCGG | CGTATACGCTGCCCAAAAGT | 164 |
| chr4:23978428-23980458 | p5 | (ATTCA)6 | CGGGGATGGACATATACCAC | CGGTATGTATCGTGCGAATG | 171 |
| chr4:24065405-24067417 | p6 | (TAAAAA)3 | ATTCAAGACAATTGCGGGAC | ATTTTGCCCTTAGCGTTCCT | 238 |
| chr4:24065405-24067417 | p2 | (GT)6 | ATTCAAGACAATTGCGGGAC | ATTTTGCCCTTAGCGTTCCT | 238 |
| chr4:24184686-24186716 | p5 | (TGAAT)6 | CCCCTTGAAATCTTGTTCCA | CACATACACCATGCACACGA | 243 |
| chr4:24239523-24241541 | p6 | (TATATG)3 | GCTAGCGTGAATGACGATGA | CCTTGGAAAACGCTTATGGA | 237 |
| chr4:24245249-24247279 | p6 | (TATATG)5 | CCAAAATTTCGGCAGATCAT | GAAGGAAATCATGCTTGGGA | 192 |
| chr4:24329754-24331780 | p2 | (GT)11 | ATGGATTGGTTGGCTTCTTG | GCACCCTTTCGTGATCATTT | 139 |
| chr4:24329754-24331780 | p2 | (TA)13 | ATGGATTGGTTGGCTTCTTG | GCACCCTTTCGTGATCATTT | 139 |
| chr4:24349883-24351895 | p2 | (TA)6 | TAGTCGGTTGGTGACCCTTC | CAAAGCTTCCATTACACGCA | 160 |
| chr4:24368504-24370522 | p2 | (AT)9 | CAACCGAGAGCTACACCACA | GGCCGAATGGAATAGGGTAT | 228 |
| chr4:24466411-24468435 | p3 | (TTG)8 | GCGATCAAGCCATCAATTTT | CCAATCACACCCCCATTTAC | 201 |
| chr4:24478556-24480574 | p3 | (GAT)10 | TTACGGGATATAACAGCCGC | CCAATCTCCAAATCTCCCAA | 191 |
| chr4:24478556-24480574 | p2 | (TA)13 | TTACGGGATATAACAGCCGC | CCAATCTCCAAATCTCCCAA | 191 |
| chr4:24478625-24480637 | p3 | (GAT)10 | TTACGGGATATAACAGCCGC | CCAATCTCCAAATCTCCCAA | 122 |
| chr4:24478625-24480637 | p2 | (TA)13 | TTACGGGATATAACAGCCGC | CCAATCTCCAAATCTCCCAA | 122 |
| chr4:24604533-24606553 | p5 | (ACAAT)4 | CTTTGAGAGGCGATTTCAGG | CAAGATGGCAAAGGGCTAAG | 237 |
| chr4:24646337-24648355 | p6 | (AACAAA)3 | GGTGGACATCAACCAAATCC | AGGAGGAACATCCCTTGGTT | 245 |
| chr4:24646337-24648355 | p5 | (AAAAG)3 | GGTGGACATCAACCAAATCC | AGGAGGAACATCCCTTGGTT | 245 |
| chr4:24684602-24686617 | p5 | (TATTT)3 | ATTCACGTCATTTGCCTTCC | TGATAGGGGTCAAGCCAAAC | 246 |

| Marker ID | SSR type | SSR motif | FORWARD PRIMER (5'-3') | REVERSE PRIMER (5'-3') | size (bp) |
|---|---|---|---|---|---|
| chr4:24687904-24689920 | p4 | (ATAC)4 | AACAGATTCAACAGGCCCAG | TGGGTCGTGACAGTCAACAT | 127 |
| chr4:24787748-24789768 | p5 | (TCTTT)4 | CGAAAATCACCTCTTCCCAA | TGTCAAATTTGGGGGCTATC | 263 |
| chr4:24968573-24970587 | p2 | (TA)7 | GTCGCGACAGATTGAAAACA | ATGAAGTGGGATTGGCTCAC | 185 |
| chr4:24968573-24970587 | p2 | (AT)6 | GTCGCGACAGATTGAAAACA | ATGAAGTGGGATTGGCTCAC | 185 |
| chr4:25025834-25027849 | p5 | (TGCAT)3 | GCATAGCAACGAAAAGCACA | AATTCCTGCGCTCAAAGAAA | 256 |
| chr4:25221885-25223900 | p3 | (AGA)5 | GGACTGGCAACACAAGGAAT | TACGCTATGGCTGTTTGCTG | 172 |
| chr4:25263682-25265697 | p3 | (AGA)5 | GGACTGGCAACACAAGGAAT | TACGCTATGGCTGTTTGCTG | 172 |
| chr4:25302557-25304569 | p2 | (TA)6 | TCAGTTCGGGCTTCACATAA | GAGTTTGCGGTAGACGAAGC | 125 |
| chr4:25398750-25400768 | p2 | (TA)6 | TAGGTTGTGAATGCACCCAA | GCCTTCGAATTTGTGCCTTA | 233 |
| chr4:25398750-25400768 | p6 | (GGGTTG)3 | TAGGTTGTGAATGCACCCAA | GCCTTCGAATTTGTGCCTTA | 233 |
| chr4:25496899-25498913 | p3 | (TTC)6 | ATGCTGGTGCCACATATCAA | CAGCCCTATCGAACACCATT | 123 |
| chr4:25496899-25498913 | p2 | (AT)7 | ATGCTGGTGCCACATATCAA | CAGCCCTATCGAACACCATT | 123 |
| chr4:25504005-25506017 | p2 | (AC)6 | GGGACGAGTGTCCATCACTT | TTCTGGTTAGCGCTTCGTTT | 203 |
| chr4:25552167-25554182 | p6 | (GGGTGG)3 | TGAATGGACGGATGGATTTT | GCTGCTTCTGGATTTGAAGG | 143 |
| chr4:25552205-25554220 | p6 | (GGGTGG)3 | TGAATGGACGGATGGATTTT | GCTGCTTCTGGATTTGAAGG | 105 |
| chr4:25565767-25567783 | p2 | (TA)8 | AAACCGAAACCAAAGGAACC | TTATCAAAATTCGGTTCGGC | 282 |
| chr4:25744727-25746739 | p2 | (TC)6 | GACGGACGGTAACACAGGTT | GGCCTTAAATGCTCCAACAA | 190 |
| chr4:25777554-25779566 | p2 | (CT)6 | TTTAGAGCCCGTTTGGATTG | GCATGTTTGGATGGGCTTAT | 293 |
| chr4:25794293-25796330 | p3 | (ATA)5 | ACGGGACAAATTGCTAATGG | AGTTTGGTTTACCCTGTGCG | 217 |
| chr4:25795428-25797446 | p3 | (ATA)5 | CCCACCCTGTAACCCTACAA | CGGGGAAAAGTTGAAAAACA | 272 |
| chr4:25795428-25797446 | p3 | (TAA)6 | CCCACCCTGTAACCCTACAA | CGGGGAAAAGTTGAAAAACA | 272 |
| chr4:25860503-25862521 | p6 | (CCTTGG)3 | ACGAGCTAAATCATGGTGCC | GCTGACCAAGAAGAAGACGG | 104 |
| chr4:25880852-25882870 | p2 | (AG)9 | TTTGGTCGGTCAGGCTAAAC | TTTTCATTTACGCGTCCCTT | 153 |
| chr4:25880852-25882870 | p2 | (TG)9 | TTTGGTCGGTCAGGCTAAAC | TTTTCATTTACGCGTCCCTT | 153 |
| chr4:25921977-25923993 | p2 | (CT)8 | CAAGCCAACATCAACGCTTA | TCGTTAGATTTTCGGGCATC | 170 |
| chr4:25976708-25978720 | p2 | (AT)6 | TTTTCATGGGCCAAATTCAT | ACCCTTAATGTGTCCTTGCG | 229 |
| chr4:25976708-25978720 | p2 | (TC)6 | TTTTCATGGGCCAAATTCAT | ACCCTTAATGTGTCCTTGCG | 229 |
| chr4:26011294-26013328 | p2 | (AG)17 | ATGAGAGCTGCGGTGACTTT | CGCTCCAGCTCCATTTTAAC | 133 |
| chr4:26120371-26122383 | p2 | (TA)6 | TAGCGGCACGTATCCTCTCT | AGTTGGGGGTGTTGATGTGT | 101 |
| chr4:26194470-26196485 | p2 | (AT)14 | TTTGTGGGTGTGTTTGCCTA | GGCACAAATATCACACGCTG | 222 |
| chr4:26194491-26196509 | p2 | (AT)14 | TTTGTGGGTGTGTTTGCCTA | GGCACAAATATCACACGCTG | 201 |
| chr4:26236664-26238680 | p2 | (AT)8 | GAAATTTCACGGCCTTCTGA | ACATAGACCGAAATACGCGG | 142 |
| chr4:26267299-26269314 | p2 | (AT)6 | CCTCGATAAGGGGTGCATAA | AGGTGCTTCATCAGCGACTT | 245 |
| chr4:26267299-26269314 | p2 | (AT)7 | CCTCGATAAGGGGTGCATAA | AGGTGCTTCATCAGCGACTT | 245 |
| chr4:26267299-26269314 | p3 | (ATC)5 | CCTCGATAAGGGGTGCATAA | AGGTGCTTCATCAGCGACTT | 245 |
| chr4:26547298-26549344 | p2 | (AT)9 | CAAGGTTGAAGCTCTTTCCG | GAAAGTACGCGTCCCATCAT | 297 |

| Marker ID | SSR type | SSR motif | FORWARD PRIMER (5'-3') | REVERSE PRIMER (5'-3') | size (bp) |
|---|---|---|---|---|---|
| chr4:26547298-26549344 | p2 | (TC)23 | CAAGGTTGAAGCTCTTTCCG | GAAAGTACGCGTCCCATCAT | 297 |
| chr4:26600141-26602173 | p2 | (AT)16 | CTTCCATCTTCCTTCCACCA | TGCGGGTGGAAAATTAAAAG | 149 |
| chr4:26629940-26632046 | p2 | (TA)53 | GGGCCTAATTCTAGTTGGGG | GAGCACTGGATGGGACTGTT | 146 |
| chr4:26629940-26632046 | p2 | (AT)22 | GGGCCTAATTCTAGTTGGGG | GAGCACTGGATGGGACTGTT | 146 |
| chr4:26673873-26675888 | p5 | (AAACA)3 | CCCGGATCTGTTCTCATGTT | CGTAAACCGGTCACCAGAAT | 258 |
| chr4:26673873-26675888 | p5 | (AACCG)3 | CCCGGATCTGTTCTCATGTT | CGTAAACCGGTCACCAGAAT | 258 |
| chr4:26689601-26691613 | p2 | (AC)6 | CACATAAGCAGCACACCCAC | GCTTTGCGATTATTTGGAGG | 206 |
| chr4:26734575-26736587 | p2 | (AT)6 | CAACGGAAACTGCATACGAA | CGTTGTGTGTTTGTAACGGC | 239 |
| chr4:26739795-26741810 | p5 | (TTTTA)3 | ATCCCATAGGTTTCGGGTTC | CTTATGCCAAGTCTGCCCAT | 280 |
| chr4:26828787-26830802 | p5 | (TTGTT)3 | AAGAACAAATGGTTGCGGAC | ACTTCCTGGGCCTTCAATTT | 220 |
| chr4:26942811-26944831 | p2 | (TC)10 | TATGTAGGCAACCCGGAGAC | TGATGAAAGAATTTTCGGGC | 239 |
| chr4:26986316-26988334 | p6 | (CCCACC)3 | AACGTGGTGTGGTCCCTAAG | ACAAGTGGGACGTATCCAGC | 216 |
| chr4:26986316-26988334 | p6 | (TTATTT)3 | AACGTGGTGTGGTCCCTAAG | ACAAGTGGGACGTATCCAGC | 216 |
| chr4:27017534-27019549 | p3 | (ACA)5 | CACCAGCCCCAGAATAAAAA | GGGACTTGTCCTTGTGGAGA | 159 |
| chr4:27042648-27044684 | p2 | (TA)18 | GCAGGAACTTCAGGCAAAAA | ACCGTTTCAAACAACTTCGG | 228 |
| chr4:27187950-27189965 | p5 | (ATATC)3 | TAGCTTTCCTGTGATCGCCT | GACCCAAAGGGTTAGCATGA | 131 |
| chr4:27194698-27196752 | p3 | (ATT)5 | TCCACCAGTTTAGGACACCC | CATATACAAAATGCACGCCG | 300 |
| chr4:27303898-27305913 | p2 | (AT)11 | ATAAGGCCGATAAGGCCACT | ACTGACGTCCCTTCTTGTGG | 130 |
| chr4:27303898-27305913 | p3 | (ATC)5 | ATAAGGCCGATAAGGCCACT | ACTGACGTCCCTTCTTGTGG | 130 |
| chr4:27303898-27305913 | p3 | (CAA)7 | ATAAGGCCGATAAGGCCACT | ACTGACGTCCCTTCTTGTGG | 130 |
| chr4:27308475-27310487 | p2 | (AT)6 | GACGCGGGAGATGTTTTCTA | GAAGGCACACAAGTGACCAA | 231 |
| chr4:27308475-27310487 | p5 | (TGAAT)3 | GACGCGGGAGATGTTTTCTA | GAAGGCACACAAGTGACCAA | 231 |
| chr4:27314293-27316308 | p2 | (AT)16 | TCAAAATAAGGCCGACAAGG | AGGTAGGCAGGGAGATGGAT | 173 |
| chr4:27314293-27316308 | p3 | (ATC)5 | TCAAAATAAGGCCGACAAGG | AGGTAGGCAGGGAGATGGAT | 173 |
| chr4:27314293-27316308 | p3 | (CAA)8 | TCAAAATAAGGCCGACAAGG | AGGTAGGCAGGGAGATGGAT | 173 |
| chr4:27345632-27347650 | p6 | (AACTGA)3 | GAATACAAGCCTCGGACGAC | GTAGCGACCAAATACCCGAA | 135 |
| chr4:27455105-27457129 | p5 | (ACCCG)3 | ATACCCTACCAGGATTGGGG | ATTTCTGGATGAGGCGTGAC | 158 |
| chr4:27455132-27457147 | p5 | (ACCCG)3 | ATACCCTACCAGGATTGGGG | ATTTCTGGATGAGGCGTGAC | 131 |
| chr4:27455150-27457171 | p5 | (ACCCG)3 | ATACCCTACCAGGATTGGGG | ATTTCTGGATGAGGCGTGAC | 113 |
| chr4:27482427-27484453 | p2 | (TA)6 | TTACAGGGTGAGGTTCCAGC | GCGTGTTTAGGGCGTACAAT | 249 |
| chr4:27735516-27737534 | p2 | (TA)6 | TGGTGGGATTTGGTTAGTCC | AACAAGCCATTTTGATTCCG | 289 |
| chr4:27735570-27737586 | p2 | (TA)6 | TGGTGGGATTTGGTTAGTCC | AACAAGCCATTTTGATTCCG | 235 |
| chr4:27775515-27777533 | p3 | (AAC)6 | CACATTGATCATATGGGGCA | CGTCATGCTCTTGCGAGATA | 293 |
| chr4:27789984-27792002 | p2 | (TA)9 | TTTGCAAATTGACCCTTTCC | GGGGGAGTTGTGACATATCG | 275 |
| chr4:27966303-27968318 | p3 | (AAC)5 | TCAACAACTTGCGGAGACTG | TCAAAGCTGGCATCACAAAG | 124 |
| chr4:28025880-28027895 | p3 | (AAC)5 | TCAACAACTTGCGGAGACTG | TCAAAGCTGGCATCACAAAG | 124 |

| Marker ID | SSR type | SSR motif | FORWARD PRIMER (5'-3') | REVERSE PRIMER (5'-3') | size (bp) |
|---|---|---|---|---|---|
| chr4:28147355-28149367 | p2 | (AT)6 | GCCTTGTCAGGACGAGTAGG | ACGTGTCGGAGTTATGGAGG | 177 |
| chr4:28210591-28212623 | p2 | (TA)16 | GCCATTCCACGTCATTCTTT | ATATGTGGTAGCGGCGAGTT | 195 |
| chr4:28272760-28274784 | p4 | (TATT)6 | TAATCTTGGGAGCACTTGGG | ACACATGTCCCATCAGACGA | 177 |
| chr4:28275641-28277665 | p5 | (TTTGT)3 | ATAAATGTTGGATCGTCGGC | TCCGGAAATTATGAACCGTC | 250 |
| chr4:28275683-28277707 | p5 | (TTTGT)3 | ATAAATGTTGGATCGTCGGC | TCCGGAAATTATGAACCGTC | 208 |
| chr4:28275767-28277779 | p5 | (TTTGT)3 | ATAAATGTTGGATCGTCGGC | TCCGGAAATTATGAACCGTC | 124 |
| chr4:28276031-28278045 | p5 | (TTTGT)3 | TGTTTGCGTTCGTGAACTGT | TAATTGGTACGAACATGCGG | 183 |
| chr4:28276049-28278061 | p5 | (TTTGT)3 | TGTTTGCGTTCGTGAACTGT | TAATTGGTACGAACATGCGG | 165 |
| chr4:28300932-28302944 | p2 | (AC)6 | TCGCTCAAAAACTGCAAATG | TTTGGAGCCGAGTTGAATCT | 204 |
| chr4:28335946-28337964 | p3 | (TAT)6 | TGGACTTGTCGCTTTGTTTG | GGGTGTCTGGCAGATGTGTA | 186 |
| chr4:28407046-28409064 | p6 | (TCTATA)3 | CCAAAGAATTCGAGGGTTCA | TTCGGGCCAAGATGTAAAAG | 153 |
| chr4:28468614-28470629 | p6 | (TCATCC)3 | TTTTCGTGTGGAATCGTCAA | TGATGATCATAATCGGCGAA | 131 |
| chr4:28468644-28470662 | p6 | (TCATCC)3 | TTTTCGTGTGGAATCGTCAA | TGATGATCATAATCGGCGAA | 101 |
| chr4:28477259-28479287 | p2 | (AT)14 | CAGAGTGTGATTGGTGGGTG | TTGAGCTCTTTGGCTGGATT | 209 |
| chr4:28520709-28522725 | p2 | (TA)8 | GGCGGACAAAACAAAGAAAA | TCAAATCGTGGGCACTATGA | 151 |
| chr4:28785041-28787101 | p2 | (TA)30 | ATATTCCACCATTCCCGACA | GCAACGGCCATAGCAATAAT | 153 |
| chr4:28878571-28880607 | p2 | (AT)6 | CATGTTATGCCTCGTGATGG | ATACCAAGCTTTGTCACGCC | 205 |
| chr4:28878571-28880607 | p2 | (AT)18 | CATGTTATGCCTCGTGATGG | ATACCAAGCTTTGTCACGCC | 205 |
| chr4:28898883-28900901 | p2 | (TA)9 | ATACAAGCGAGTCATCCGGT | AGTTTTGATTGCGTCGCTCT | 226 |
| chr4:28898883-28900901 | p2 | (AT)6 | ATACAAGCGAGTCATCCGGT | AGTTTTGATTGCGTCGCTCT | 226 |
| chr4:28898883-28900901 | p2 | (AT)16 | ATACAAGCGAGTCATCCGGT | AGTTTTGATTGCGTCGCTCT | 226 |
| chr4:28907505-28909526 | p3 | (CAA)7 | TTTCCCGTTTTCCAACACTC | GTTTTCGGGATTACCCGTTT | 135 |
| chr4:28913111-28915125 | p2 | (AT)7 | AATGAACCCTAACCGCTTCC | CGTGTGGGTGTACACGATG | 178 |
| chr4:28913111-28915125 | p2 | (AT)7 | AATGAACCCTAACCGCTTCC | CGTGTGGGTGTACACGATG | 178 |
| chr4:28921817-28923833 | p4 | (TTAT)4 | TCCTTTTCCTCTAACGCGAA | TACGCATAATTTGGGAAGGG | 221 |
| chr4:29014987-29017011 | p2 | (AT)12 | GCTCACAATCCCTTTTGGAA | TCCATAATCCGAACAAAGCC | 281 |
| chr4:29028840-29030856 | p2 | (TA)8 | AACAGCCCACACATCATCAA | TTTTTGGAGCCAATTCCTTG | 138 |
| chr4:29028871-29030885 | p2 | (TA)8 | AACAGCCCACACATCATCAA | TTTTTGGAGCCAATTCCTTG | 107 |
| chr4:29074923-29076935 | p2 | (CA)6 | CTCAAACACAAGAGGACGCA | AGTGCTTCCCATGCTCAGAT | 172 |
| chr4:29081634-29083654 | p2 | (CT)10 | CAGAGACCAGAGAGTTCGGC | GATGAAAGAATTTTCGGGCA | 146 |
| chr4:29155535-29157559 | p4 | (ATTT)6 | GCCCTTGCAACTGTACCAAT | TATCAACAACCCCAACAGCA | 242 |
| chr4:29167736-29169763 | p3 | (TAA)9 | TGGAGATTTTGAGGCGATTC | AAGAACCGCCCCTATTTTGT | 186 |
| chr4:29326189-29328204 | p3 | (ATT)5 | TAGAACGACCGTCGAACTCC | TTACACCTCGGAACTTTGGG | 132 |
| chr4:29376210-29378222 | p2 | (TA)6 | TAAATGCTCCCTCCATTCCA | AAATGGAAAACCCCAAAAGG | 259 |
| chr4:29376210-29378222 | p2 | (TA)8 | TAAATGCTCCCTCCATTCCA | AAATGGAAAACCCCAAAAGG | 259 |
| chr4:29376210-29378222 | p2 | (AT)6 | TAAATGCTCCCTCCATTCCA | AAATGGAAAACCCCAAAAGG | 259 |

| Marker ID | SSR type | SSR motif | FORWARD PRIMER (5'-3') | REVERSE PRIMER (5'-3') | size (bp) |
|---|---|---|---|---|---|
| chr4:29434688-29436733 | p3 | (AAC)5 | CTTTGTGCACTTGGTCGAGA | AAGGTTGCGTTTCCATCTTG | 100 |
| chr4:29434688-29436733 | p3 | (AAG)15 | CTTTGTGCACTTGGTCGAGA | AAGGTTGCGTTTCCATCTTG | 100 |
| chr4:29486256-29488272 | p4 | (TTTG)4 | TCCCGCAAGATAGTAGGTGC | AATCCGCGCAAAAATTGTAA | 166 |
| chr4:29500417-29502448 | p2 | (AT)6 | AGTCGCGAAATCTGGCTCTA | TAAATTCCCCGGTGCATAAG | 204 |
| chr4:29500511-29502545 | p2 | (AT)6 | AGTCGCGAAATCTGGCTCTA | TAAATTCCCCGGTGCATAAG | 110 |
| chr4:29589919-29591935 | p4 | (AATG)4 | TACGAAGCGCGGACTAGTTT | TTTCCGATGTGGTTCAGTCA | 256 |
| chr4:29648140-29650212 | p5 | (TTGCA)3 | CAATTCCTCCCGAAAATCAA | CCAAACGGGTCGTTACATTC | 287 |
| chr4:29670763-29672778 | p5 | (AATTA)3 | AGATGAAACCAGCGTTGGAC | CAATCTCTTCACGTCGAGCA | 244 |
| chr4:29764951-29766966 | p3 | (TGT)5 | GGACCAAGCAAATCGAAGAA | GGTTTGTTGCGTTGGAAACT | 162 |
| chr4:29785453-29787471 | p3 | (TTC)6 | TTCATGTTTCCGGTAAAGCC | GGAGACAATAAGCCCGTCAA | 229 |
| chr4:29785453-29787471 | p5 | (TTATT)3 | TTCATGTTTCCGGTAAAGCC | GGAGACAATAAGCCCGTCAA | 229 |
| chr4:29785579-29787594 | p3 | (TTC)6 | TTCATGTTTCCGGTAAAGCC | GGAGACAATAAGCCCGTCAA | 103 |
| chr4:29785579-29787594 | p5 | (TTATT)3 | TTCATGTTTCCGGTAAAGCC | GGAGACAATAAGCCCGTCAA | 103 |
| chr4:29796857-29798885 | p2 | (AT)13 | CCCCATTATTGGCATCTTTG | CTTGCTCATGCACCGTCTAA | 174 |
| chr4:29796888-29798900 | p2 | (AT)13 | CCCCATTATTGGCATCTTTG | CTTGCTCATGCACCGTCTAA | 143 |
| chr4:29796905-29798981 | p2 | (AT)13 | CCCCATTATTGGCATCTTTG | CTTGCTCATGCACCGTCTAA | 126 |
| chr4:29824776-29826788 | p2 | (TA)6 | GCATTCTGGAATTTGGCATT | TGCACAGATGTCCGTTTGAT | 139 |
| chr4:29838604-29840626 | p2 | (TA)8 | ATTCGGATGCACCAACTTTC | TGCTCTCCCAAATTTTCACC | 213 |
| chr4:29838604-29840626 | p2 | (AT)11 | ATTCGGATGCACCAACTTTC | TGCTCTCCCAAATTTTCACC | 213 |
| chr4:29860185-29862200 | p3 | (CAA)5 | TCCAACAATGAACCCGAAAT | GTCAAAAGTGCCAGGGGTTA | 146 |
| chr4:29881029-29883051 | p2 | (TA)11 | ATTCTCGGGCTACCCAACTT | CCGACCACCAATTCATCTCT | 248 |
| chr4:29896150-29898170 | p2 | (TC)10 | AGTGACTGGTTTCCCGTGAG | GGCAATTAACTTCCGGCATA | 275 |
| chr4:29961335-29963349 | p2 | (AT)7 | CGAAGGGAACTGAACCTGAA | ATTTTCAGATGATGGAGGCG | 238 |
| chr4:30133182-30135232 | p2 | (AT)6 | AAGCTTCCAAGAAATGGGGT | GCAACAAGAAGAGGGCTTTG | 167 |
| chr4:30133182-30135232 | p2 | (TA)17 | AAGCTTCCAAGAAATGGGGT | GCAACAAGAAGAGGGCTTTG | 167 |
| chr4:30143886-30145919 | p3 | (AAC)11 | TGATCACACGCATTTCCACT | GCGGAAGACCCAAATAGTGA | 156 |
| chr4:30201507-30203531 | p3 | (GTT)8 | GTATGAGGCGGAGTGTTGGT | GCACTGGGGCATCTGTATCT | 251 |
| chr4:30282376-30284394 | p6 | (CTCCGT)3 | TGACGAGGAAGAGGTCGAGT | GCCCTCACCATCGTCTACAT | 209 |
| chr4:30426371-30428386 | p5 | (TTGTA)3 | TTGGAAACGGATACGGAAAG | GGATTCTGGAAAAATGCGAA | 162 |
| chr4:30511167-30513179 | p2 | (TA)6 | GGGGACTTGATGCTGTGTCT | CTCCGAAGACAATCGGAGAG | 270 |
| chr4:30534338-30536358 | p2 | (TA)10 | ACATGAACCCGTCAAAAAGG | TCTAAATCCACCGGCTCAAC | 100 |
| chr4:30540908-30542923 | p5 | (TTATT)3 | TAAAGCAATGATCCTTCGGC | AGAAATCATGTGGGCTTTGC | 280 |
| chr4:30580816-30582828 | p2 | (TG)6 | AGCTGGAGAATGCGACAAAT | TGTGAAAGCTGAAGCAGGTG | 113 |
| chr4:30586179-30588194 | p3 | (TGT)5 | CCACCAAAACTTAAGGGCAA | CCCGGATGTCTTCATTCCTA | 294 |
| chr4:30597548-30599566 | p6 | (TAAAAA)3 | TTTCAGATCATTCGACAGCG | CACCTTAATGCGATGGGTCT | 225 |
| chr4:30640829-30642844 | p5 | (AAAAT)3 | AAAATGGCGAAAAGTGTTGG | TGAGGGATATCTTTGCGGTC | 255 |

| Marker ID | SSR type | SSR motif | FORWARD PRIMER (5'-3') | REVERSE PRIMER (5'-3') | size (bp) |
|---|---|---|---|---|---|
| chr4:30677631-30679651 | p2 | (TA)10 | TTCCAGTTAGCAGCCTTCGT | TTCACAAAATGCGCTTTGAC | 197 |
| chr4:30696389-30698433 | p2 | (TC)22 | ATTTGATCGCTCCAATTGCT | GGGGTGTTTAGATTGCCTGA | 150 |
| chr4:30710325-30712346 | p3 | (AAT)7 | TGGTGAAACATGAGGTGCAT | GTCCTCAATGGCGGAAATAA | 114 |
| chr4:30710325-30712346 | p5 | (TTGGT)3 | TGGTGAAACATGAGGTGCAT | GTCCTCAATGGCGGAAATAA | 114 |
| chr4:30711533-30713553 | p5 | (TTGGT)3 | CCTCTCGAGGGGTGTACAAA | GTAATGTCGGGTTGGTTTGG | 168 |
| chr4:30711578-30713593 | p5 | (TTGGT)3 | CCTCTCGAGGGGTGTACAAA | GTAATGTCGGGTTGGTTTGG | 123 |
| chr4:30735407-30737422 | p5 | (GGGGT)3 | TTGGAGAGGAAAACATGGCT | GTGTAGTATGGGGTGGTGGG | 217 |
| chr4:30740170-30742188 | p3 | (AGA)6 | CAAATTTTTGGGCTTGGGTA | CGATCCGTCAGGATTTGTCT | 287 |
| chr4:30747365-30749380 | p5 | (GATTA)3 | TCAATGACGTGGTGGTTGTT | CACTCGCCCTCTAAAGATGC | 252 |
| chr4:30803157-30805181 | p3 | (TAA)8 | TCGTCCGAAAAAGGAAGAGA | CCTTTACCCCCAAACCCTAA | 138 |
| chr4:30803157-30805181 | p2 | (TA)9 | TCGTCCGAAAAAGGAAGAGA | CCTTTACCCCCAAACCCTAA | 138 |
| chr4:30803157-30805181 | p2 | (TA)7 | TCGTCCGAAAAAGGAAGAGA | CCTTTACCCCCAAACCCTAA | 138 |
| chr4:30829629-30831647 | p6 | (TTTTTA)3 | ACTTCGGCTATAGAACGGCA | GTGAGGCAAATGGGATGAGT | 242 |
| chr4:30829629-30831647 | p2 | (AT)34 | ACTTCGGCTATAGAACGGCA | GTGAGGCAAATGGGATGAGT | 242 |
| chr4:30829881-30831949 | p6 | (TTTTTA)3 | TGCACATTAGAGATCGCCAG | TGTACCGGAACCCCTGAATA | 289 |
| chr4:30829881-30831949 | p2 | (AT)34 | TGCACATTAGAGATCGCCAG | TGTACCGGAACCCCTGAATA | 289 |
| chr4:30867269-30869289 | p2 | (AT)10 | TTGAGACGGAGGGAGTAGGA | CCAAATCGTCCCTCTTGAAA | 211 |
| chr4:30940521-30942542 | p3 | (ACA)7 | AGTTATTCCCGAGGCCATCT | GTCCGGATGGTCGTTTAGAA | 173 |
| chr4:30942878-30944893 | p5 | (TTTTA)3 | CAACCAAACACCATCAGCAC | CTTCCACACACAGCCAAAGA | 143 |
| chr4:30979085-30981097 | p2 | (CT)6 | GCTACCGTCCAAAGAGCAAG | TGAGTCCACACGAAACCAAA | 187 |
| chr4:30998752-31000772 | p2 | (AT)10 | CTGGGGCACGATCCTACTTA | AAAATCAGTGCCAACAAGGC | 105 |
| chr4:31209556-31211568 | p2 | (TA)6 | ACACTATGCCGAGGGTGTTC | ACGAAGAAGTCTGAGGGGGT | 246 |
| chr4:31209556-31211568 | p2 | (GT)6 | ACACTATGCCGAGGGTGTTC | ACGAAGAAGTCTGAGGGGGT | 246 |
| chr4:31290982-31293040 | p2 | (TA)29 | CCGCTTTTACTTGCCCTCTA | TTGGCAAATGCATTGAGAAG | 271 |
| chr4:31326527-31328543 | p2 | (TG)8 | TTTTCAAATCAAACTCCCCC | AAGGAAAGTTGCGAAGGTGA | 293 |
| chr4:31326527-31328543 | p2 | (TG)9 | TTTTCAAATCAAACTCCCCC | AAGGAAAGTTGCGAAGGTGA | 293 |
| chr4:31326527-31328543 | p5 | (TTTTA)3 | TTTTCAAATCAAACTCCCCC | AAGGAAAGTTGCGAAGGTGA | 293 |
| chr4:31330985-31333001 | p2 | (TA)8 | GACAAACCGCCTAGAACTCG | TTAGTGAGCCGTTCCAAACC | 201 |
| chr4:31330985-31333001 | p2 | (TA)6 | GACAAACCGCCTAGAACTCG | TTAGTGAGCCGTTCCAAACC | 201 |
| chr4:31330985-31333001 | p2 | (AT)6 | GACAAACCGCCTAGAACTCG | TTAGTGAGCCGTTCCAAACC | 201 |
| chr4:31338623-31340635 | p2 | (TA)6 | AGCGTTAGCTCGATGGTGTT | ACTTTTCCAAAGGCCGATCT | 174 |
| chr4:31408026-31410044 | p6 | (TCTTTT)3 | AAACCGAGTTATTTTGGGGG | TGGCCCTCTACTTGGAGAAA | 291 |
| chr4:31464205-31466217 | p2 | (AT)6 | GACATGGCGGAATAGCAAAT | TGATGGATTCATTGCCACAT | 202 |
| chr4:31530925-31532940 | p5 | (AAAAT)3 | GCCAATGCCAAGAAGAAGAG | TCATGATTTCTGTGCGAAGC | 114 |
| chr4:31530925-31532940 | p2 | (TA)11 | GCCAATGCCAAGAAGAAGAG | TCATGATTTCTGTGCGAAGC | 114 |
| chr4:31757123-31759153 | p3 | (AAT)10 | ATCCAAACGCCGTGACTAAC | ATGATGCATTGGAGGAGAGC | 133 |

| Marker ID | SSR type | SSR motif | FORWARD PRIMER (5'-3') | REVERSE PRIMER (5'-3') | size (bp) |
|---|---|---|---|---|---|
| chr4:31763327-31765345 | p3 | (AAC)6 | TACCCGGTCTTTTGTTTTCG | GTTAGTCCCCAGCAGAGTCG | 154 |
| chr4:31810737-31812752 | p2 | (TA)11 | TAGATTTTGGACGAAACGGG | ACGCGGGTTTCTATGAACAG | 198 |
| chr4:31810787-31812832 | p2 | (TA)11 | TAGATTTTGGACGAAACGGG | ACGCGGGTTTCTATGAACAG | 148 |
| chr4:31823255-31825273 | p4 | (AAAG)7 | ATTAGTTGGGGGTTGAAGGG | ACTTCGGATTGTCGTTTTGG | 136 |
| chr4:31823284-31825302 | p4 | (AAAG)7 | ATTAGTTGGGGGTTGAAGGG | ACTTCGGATTGTCGTTTTGG | 107 |
| chr4:31823492-31825515 | p4 | (AAAG)7 | ACCAAAACGACAATCCGAAG | GATTTTCATGCGTCTCCGTT | 122 |
| chr4:31885214-31887230 | p4 | (TTAT)4 | TTAGCCGACAAAACTGCCTT | TACATCCCTTCTCCGGTGTC | 280 |
| chr4:32093492-32095508 | p2 | (TA)8 | TGGGTATAGCCGGACACTTC | CATCCTTTGCACCATCCTCT | 287 |
| chr4:32096061-32098117 | p2 | (TA)28 | ACATCGTGTGGGATGTTTCA | TTTGGGCAGTATCTTCCCTG | 166 |
| chr4:32113359-32115374 | p5 | (ACCAA)3 | TCTGAATTCGTCCCCTTGTC | ATTGTGTGTACAATGCCCGA | 255 |
| chr4:32113359-32115374 | p5 | (AAACC)3 | TCTGAATTCGTCCCCTTGTC | ATTGTGTGTACAATGCCCGA | 255 |
| chr4:32166267-32168283 | p2 | (AT)8 | TGCAGGCTATTTCGCTTTTT | ACGATTCGATTACCGCAAAG | 254 |
| chr4:32395014-32397026 | p2 | (TA)6 | TGGATTGATCTGCCTGTGAG | CTCAGCGAGTGTCTAAGGGG | 116 |
| chr4:32411655-32413667 | p2 | (AT)6 | TAGTGTGCATTCAGAAGCGG | TTTCGTTAATTTCCCCCACA | 159 |
| chr4:32467990-32470022 | p2 | (AT)20 | TCCTCCTGCTCTGCAATTTT | ATGGATCAGAAAGACCACCG | 135 |
| chr4:32482991-32485006 | p5 | (CATTT)3 | TTTCTCCGAGGAAATTGTGG | ACCCGAATTTCCAATTACCC | 118 |
| chr4:32489709-32491725 | p4 | (TTTG)4 | TGCTCCTTTTGTGTGGTTTG | CTGGGCTATCTAGGGTGCTG | 141 |
| chr4:32489709-32491725 | p2 | (CT)8 | TGCTCCTTTTGTGTGGTTTG | CTGGGCTATCTAGGGTGCTG | 141 |
| chr4:32499976-32502000 | p2 | (AT)12 | AACCGACACTTGTGCTTGTG | CGACGAATGGGAAGATTGAT | 200 |
| chr4:32503285-32505299 | p6 | (ATATAG)4 | AGTGAATTAGCTCGCGCTTC | TTCACTCAAAGAGATGGGGG | 242 |
| chr4:32503285-32505299 | p2 | (TA)7 | AGTGAATTAGCTCGCGCTTC | TTCACTCAAAGAGATGGGGG | 242 |
| chr4:32503285-32505299 | p2 | (AC)6 | AGTGAATTAGCTCGCGCTTC | TTCACTCAAAGAGATGGGGG | 242 |
| chr4:32513696-32515714 | p2 | (TA)9 | TTGTCATGACCCAAACAGGA | TTGATGGAAGCGTCATTTCA | 300 |
| chr4:32513696-32515714 | p2 | (TA)6 | TTGTCATGACCCAAACAGGA | TTGATGGAAGCGTCATTTCA | 300 |
| chr4:32513696-32515714 | p3 | (ACA)5 | TTGTCATGACCCAAACAGGA | TTGATGGAAGCGTCATTTCA | 300 |
| chr4:32513833-32515845 | p2 | (TA)9 | TTGTCATGACCCAAACAGGA | TTGATGGAAGCGTCATTTCA | 163 |
| chr4:32513833-32515845 | p2 | (TA)6 | TTGTCATGACCCAAACAGGA | TTGATGGAAGCGTCATTTCA | 163 |
| chr4:32513833-32515845 | p3 | (ACA)5 | TTGTCATGACCCAAACAGGA | TTGATGGAAGCGTCATTTCA | 163 |
| chr4:32519487-32521511 | p6 | (TGCTTA)4 | AGAGCTGCCACCCTTATTCA | TGGAGACGATGTGCTTGTTC | 248 |
| chr4:32529527-32531545 | p2 | (AT)9 | TTCGGATCGTCATTTTAGGG | CCGCATAGATCTCGGAAAAA | 168 |
| chr4:32591938-32593956 | p6 | (TTGGGG)3 | AAAGGGAAGAAAGGGCTGAA | CCAAGGATCCTGAACTTCCA | 150 |
| chr4:32591938-32593956 | p6 | (AAAAAG)3 | AAAGGGAAGAAAGGGCTGAA | CCAAGGATCCTGAACTTCCA | 150 |
| chr4:32672585-32674599 | p2 | (TA)7 | AGCTCAAGGCAATCCTCGTA | CATCAATGTGACGCTCCAAC | 246 |
| chr4:32767805-32769838 | p2 | (TG)6 | CCTTTCTTTCTTTCGGGGTC | GCAGCCTTACCTTGCATTTC | 206 |
| chr4:32792299-32794317 | p6 | (ATGAGT)3 | ATATCAGGCAACTGCCAACC | TAGACCCTCGGCTCAGAAAA | 267 |
| chr4:32798804-32800843 | p2 | (AT)11 | CATGCAGTTTGAATGTTGGG | GAGTTAACGGAGGTTGAGCG | 266 |

| Marker ID | SSR type | SSR motif | FORWARD PRIMER (5'-3') | REVERSE PRIMER (5'-3') | size (bp) |
|---|---|---|---|---|---|
| chr4:32805586-32807616 | p2 | (TA)15 | GAATTGGGAGCGAATTGAGA | CCGAAGCTCACGTTTCTACC | 222 |
| chr4:32806526-32808547 | p2 | (TA)15 | CAACAGAATTGGGCTGTCCT | TATGGATCGAGCCTGTCTCC | 271 |
| chr4:32806526-32808547 | p2 | (TA)16 | CAACAGAATTGGGCTGTCCT | TATGGATCGAGCCTGTCTCC | 271 |
| chr4:32806562-32808577 | p2 | (TA)15 | CAACAGAATTGGGCTGTCCT | TATGGATCGAGCCTGTCTCC | 235 |
| chr4:32806562-32808577 | p2 | (TA)16 | CAACAGAATTGGGCTGTCCT | TATGGATCGAGCCTGTCTCC | 235 |
| chr4:32929753-32931767 | p2 | (GA)7 | TTGCTGGACTTCCTTCACCT | TTTCCGGAAGAAAACGTGTC | 101 |
| chr4:32981648-32983723 | p3 | (TTC)25 | TCCTGGAAAAGAACACTGGG | TCATCTCCTTGCCAGCTTTT | 279 |
| chr4:33001627-33003647 | p2 | (TA)10 | CTGGGGCTCGATATCCTGTA | GTAAATGCGAAACCGGCTAA | 107 |
| chr4:33030702-33032720 | p6 | (TTCATG)3 | GCGTGCCAAAAATCATAGGT | TGCTGGTAGGAGGAGAGGAA | 157 |
| chr4:33131370-33133421 | p3 | (ATT)17 | TGAATCATCGGCCTAGAACC | TTGATCCTTGTCGGGAAGTC | 229 |
| chr4:33131370-33133421 | p5 | (TTATT)3 | TGAATCATCGGCCTAGAACC | TTGATCCTTGTCGGGAAGTC | 229 |
| chr4:33167439-33169455 | p2 | (AT)8 | AACCCATTAGTGGGAGCAAA | AAATGACACGGCGGAATAGT | 123 |
| chr4:33222810-33224828 | p6 | (ATACAT)3 | GGGTGACGTAAGGTCGTGAT | TCACACCCTAGGGGAATCAG | 160 |
| chr4:33227856-33229878 | p2 | (AT)33 | GGGTTATGACCTCCCGAAAT | CAACCCAAGAAGGACCTTGA | 201 |
| chr4:33227856-33229878 | p2 | (AC)11 | GGGTTATGACCTCCCGAAAT | CAACCCAAGAAGGACCTTGA | 201 |
| chr4:33227856-33229878 | p2 | (TC)6 | GGGTTATGACCTCCCGAAAT | CAACCCAAGAAGGACCTTGA | 201 |
| chr4:33230047-33232062 | p3 | (ACA)5 | CGCTCAAAAATCCCCTTACA | ATACCTCGAAACACCGCAAG | 291 |
| chr4:33275475-33277493 | p3 | (ATG)6 | GAAATGCCGTCGAATTTTGT | GGCAGCATATTCCCCTGTAA | 272 |
| chr4:33286840-33288878 | p2 | (TA)6 | GTAGGCCATGACTAGTGCCC | ATATGCGACCGTGTTAAGGC | 135 |
| chr4:33286840-33288878 | p2 | (AT)6 | GTAGGCCATGACTAGTGCCC | ATATGCGACCGTGTTAAGGC | 135 |
| chr4:33286840-33288878 | p2 | (AT)19 | GTAGGCCATGACTAGTGCCC | ATATGCGACCGTGTTAAGGC | 135 |
| chr4:33299353-33301379 | p3 | (AAC)8 | AAGACGTTGCCTTAACACGG | TTTTTGGGATGTTGTGCGTA | 233 |
| chr4:33299380-33301400 | p3 | (AAC)8 | AAGACGTTGCCTTAACACGG | TTTTTGGGATGTTGTGCGTA | 206 |
| chr4:33351551-33353569 | p6 | (TTTAAT)3 | ATGTTGCTTTTGCTCGGAAG | CCCATCCAATCAGGCTCTAA | 101 |
| chr4:33368413-33370439 | p6 | (CCGGTT)3 | TTAACGGCATCAATGGAACA | GTGTCGCTGTATGTGGTTGG | 254 |
| chr4:33368413-33370439 | p2 | (TA)13 | TTAACGGCATCAATGGAACA | GTGTCGCTGTATGTGGTTGG | 254 |
| chr4:33662431-33664446 | p3 | (GTT)5 | TGGGAGGCTTTGAAACAATC | GGCAAGTGGTGTTTGGAAGT | 282 |
| chr4:33693219-33695237 | p2 | (TA)9 | ATCGCATCGTTAAATCCGTC | ATCGTTTCCCATCGTCACAT | 300 |
| chr4:33693219-33695237 | p3 | (AAT)6 | ATCGCATCGTTAAATCCGTC | ATCGTTTCCCATCGTCACAT | 300 |
| chr4:33709295-33711331 | p3 | (TTG)12 | GCCACCAAAACTTAAAGGCA | TTCCGCGACTTTCATCTTCT | 203 |
| chr4:33709295-33711331 | p2 | (AT)14 | GCCACCAAAACTTAAAGGCA | TTCCGCGACTTTCATCTTCT | 203 |
| chr4:33718653-33720665 | p2 | (TG)6 | GTTTGGTCCTTTCACGCACT | AGTCCGCGACTTTTGTAGGA | 257 |
| chr4:33762010-33764025 | p5 | (ATTTA)3 | GACGAGTACGATCAGCACGA | CTATGGAGAAACCGGACCAA | 121 |
| chr4:33762380-33764395 | p5 | (ATTTA)3 | CAAATAAGGGGAGGCCAAAT | CGTGAGCAAATGAATCGAAA | 153 |
| chr4:33762402-33764420 | p5 | (ATTTA)3 | CAAATAAGGGGAGGCCAAAT | CGTGAGCAAATGAATCGAAA | 131 |
| chr4:33762426-33764438 | p5 | (ATTTA)3 | CAAATAAGGGGAGGCCAAAT | CGTGAGCAAATGAATCGAAA | 107 |

| Marker ID | SSR type | SSR motif | FORWARD PRIMER (5'-3') | REVERSE PRIMER (5'-3') | size (bp) |
|---|---|---|---|---|---|
| chr4:33874618-33876663 | p3 | (ATA)15 | GATACTCAGCGATGTTGCGA | CCCTTAGCGTGAAAATGGAA | 217 |
| chr4:33884794-33886809 | p5 | (CCTTT)3 | TAAAGCCATAGTGCAATGCG | AGTGATTCCCGAGGTTTGTG | 257 |
| chr4:33978477-33980489 | p2 | (AC)6 | GGAATCTGGAAAACAGCCAA | GATTTGAAGATGGGGTCGAA | 221 |
| chr4:33982958-33984972 | p2 | (TA)7 | GGGGGAGTATTTAGCCAACC | TTGAAGTCCCTGCTGCTTTT | 158 |
| chr4:34059773-34061809 | p3 | (AAG)12 | AAATGGAGAAGGGAGGGAAA | AAATCCCATTGTTTGGGACA | 151 |
| chr4:34059808-34061840 | p3 | (AAG)12 | AAATGGAGAAGGGAGGGAAA | AAATCCCATTGTTTGGGACA | 116 |
| chr4:34075796-34077811 | p5 | (AAGAG)3 | TCCTCTTTTGGTTTGCAAGG | GGAATTGGAGGTGTTCCAGA | 138 |
| chr4:34075796-34077811 | p3 | (ATC)6 | TCCTCTTTTGGTTTGCAAGG | GGAATTGGAGGTGTTCCAGA | 138 |
| chr4:34102501-34104513 | p2 | (GT)6 | AATACCGATGACCCATGGAA | TCAAAAATGTAGCCTTGCCC | 289 |
| chr4:34126176-34128191 | p5 | (GTTTG)3 | ATTGCTGCCCATAGGAAATG | AACCTCCGTACCAACACAGG | 188 |
| chr4:34126176-34128191 | p5 | (AATAC)3 | ATTGCTGCCCATAGGAAATG | AACCTCCGTACCAACACAGG | 188 |
| chr4:34168579-34170594 | p5 | (CCTTT)3 | CTCCTCGCTAGATGTGAGGG | CCTTGGTTGACTTGCTGTGA | 155 |
| chr4:34311936-34313966 | p6 | (AAAGAG)3 | AGGAATGATTTTCGGGGAAG | CAAACACATGGAGGCCTTTT | 100 |
| chr4:34311936-34313966 | p3 | (AAC)10 | AGGAATGATTTTCGGGGAAG | CAAACACATGGAGGCCTTTT | 100 |
| chr4:34311936-34313966 | p2 | (TA)22 | AGGAATGATTTTCGGGGAAG | CAAACACATGGAGGCCTTTT | 100 |
| chr4:34361430-34363445 | p5 | (TGCAT)3 | GCTTCATTTCCAGGCATGTT | TGAGCATCAGAACCCAATCA | 277 |
| chr4:34390026-34392042 | p4 | (TCTT)4 | ATTGGCATGGCTTTCAAGAG | CCCACTTGCCATTCAAACTT | 216 |
| chr4:34505093-34507108 | p5 | (TTTGG)3 | TGCGTGAGTTATGGATCACC | TTGCATTTGCCAATGTGTTT | 280 |
| chr4:34661018-34663033 | p3 | (ATT)8 | CCAAGTTTGCCCTTGTGATT | TTATGCCCGTTAGGCTATGG | 229 |
| chr4:34661018-34663033 | p5 | (ATATA)3 | CCAAGTTTGCCCTTGTGATT | TTATGCCCGTTAGGCTATGG | 229 |
| chr4:34725317-34727331 | p2 | (TA)8 | AACACGCGCAATTTTTCTCT | AAAAACCGGTTGACACCCTT | 297 |
| chr4:34725317-34727331 | p2 | (TA)6 | AACACGCGCAATTTTTCTCT | AAAAACCGGTTGACACCCTT | 297 |
| chr4:34725317-34727331 | p2 | (AT)7 | AACACGCGCAATTTTTCTCT | AAAAACCGGTTGACACCCTT | 297 |
| chr4:34742316-34744328 | p2 | (AT)10 | GGGATGTGACACTTATGGGG | CGGTAAGAAGTTTGGCGAAG | 246 |
| chr4:34742316-34744328 | p2 | (CT)6 | GGGATGTGACACTTATGGGG | CGGTAAGAAGTTTGGCGAAG | 246 |
| chr4:34775996-34778011 | p5 | (AATGC)3 | TGGCTTCTGAAAATGTCACG | CAGACCAGGTCATGGGAGTT | 132 |
| chr4:34803581-34805593 | p2 | (TA)6 | AGCTCCAGTTCAGAGGGTGA | CCCCCTCAGGAAAAGAACTC | 117 |
| chr4:34825519-34827534 | p3 | (TGT)5 | CCCAACGGAGGTCTATTTGA | ACTGTTTCGAGATGATCGGG | 157 |
| chr4:35001158-35003176 | p3 | (ATC)6 | TCATAAGGACGGGACAGGAC | GAGACGGCACAGAGGCTTAG | 188 |
| chr4:35049642-35051660 | p6 | (AAACTG)3 | CAGGTTCGAGGTAGTGGCAT | TTTCTACTCCTCGCGTCCAT | 147 |
| chr4:35049642-35051660 | p6 | (ACATGA)3 | CAGGTTCGAGGTAGTGGCAT | TTTCTACTCCTCGCGTCCAT | 147 |
| chr4:35051361-35053376 | p2 | (GA)19 | ATTCCTAGGGCTTGAGGGAA | TACTTCCCCCATCGAAATCA | 252 |
| chr4:35051379-35053430 | p2 | (GA)19 | ATTCCTAGGGCTTGAGGGAA | TACTTCCCCCATCGAAATCA | 234 |
| chr4:35051433-35053493 | p2 | (GA)19 | ATTCCTAGGGCTTGAGGGAA | TACTTCCCCCATCGAAATCA | 180 |
| chr4:35127078-35129096 | p6 | (AAACTG)3 | CAGGTTCGAGGTAGTGGCAT | TTTCTACTCCTCGCGTCCAT | 147 |
| chr4:35127078-35129096 | p6 | (ACATGA)3 | CAGGTTCGAGGTAGTGGCAT | TTTCTACTCCTCGCGTCCAT | 147 |

| Marker ID | SSR type | SSR motif | FORWARD PRIMER (5'-3') | REVERSE PRIMER (5'-3') | size (bp) |
|---|---|---|---|---|---|
| chr4:35128797-35130812 | p2 | (GA)19 | ATTCCTAGGGCTTGAGGGAA | TACTTCCCCCATCGAAATCA | 252 |
| chr4:35128815-35130866 | p2 | (GA)19 | ATTCCTAGGGCTTGAGGGAA | TACTTCCCCCATCGAAATCA | 234 |
| chr4:35128869-35130929 | p2 | (GA)19 | ATTCCTAGGGCTTGAGGGAA | TACTTCCCCCATCGAAATCA | 180 |
| chr4:35215150-35217214 | p2 | (AT)32 | TTTGATGCAGTTGCCTTCAG | AGGATGAAATGGTGCTTTGG | 150 |
| chr4:35333697-35335717 | p3 | (ATG)7 | CCCAGTTAAGAGGTGTGGGA | GCCTCCACCCTGATACTTGA | 279 |
| chr4:35333697-35335717 | p4 | (CTAT)5 | CCCAGTTAAGAGGTGTGGGA | GCCTCCACCCTGATACTTGA | 279 |
| chr4:35393224-35395239 | p5 | (GTTCG)3 | TGACGTCCCCTTTTATTTGC | AGATGGGTCGCACCAATAAG | 125 |
| chr4:35393224-35395239 | p5 | (TTCGG)3 | TGACGTCCCCTTTTATTTGC | AGATGGGTCGCACCAATAAG | 125 |
| chr4:35500877-35502937 | p3 | (TAT)20 | TTTGCAGTCAAACCAACTGC | TTCATCGGGTGTCCTCCTTA | 233 |
| chr4:35521420-35523434 | p2 | (TA)7 | GATATGCTTCGGGCTTGTGT | TTTCCTAAAATTTGGTGGCG | 194 |
| chr4:35669702-35671723 | p2 | (TA)11 | TTTTCTTTTCCATTTTGGCG | TGAAGTCACATGCTATCCGC | 288 |
| chr4:35669702-35671723 | p3 | (AAT)7 | TTTTCTTTTCCATTTTGGCG | TGAAGTCACATGCTATCCGC | 288 |
| chr4:35703007-35705022 | p5 | (TTCAT)3 | TACCTCGGAGAAGCACTGGT | TTTCAAGATTGTCCTTCCCG | 129 |
| chr4:35841823-35843837 | p2 | (TC)7 | GGTGGGGGTGGCTTTATAGT | CCAAATCCCGAGGAAAGAAT | 256 |
| chr4:35911353-35913369 | p2 | (AT)7 | GGTAGGGCCTCCGTACAGTT | AATGCGATGTTCGCTTTCTT | 165 |
| chr4:35911353-35913369 | p2 | (AT)7 | GGTAGGGCCTCCGTACAGTT | AATGCGATGTTCGCTTTCTT | 165 |
| chr4:35911405-35913417 | p2 | (AT)7 | GGTAGGGCCTCCGTACAGTT | AATGCGATGTTCGCTTTCTT | 113 |
| chr4:35911405-35913417 | p2 | (AT)7 | GGTAGGGCCTCCGTACAGTT | AATGCGATGTTCGCTTTCTT | 113 |
| chr4:36000470-36002500 | p3 | (CAA)10 | CATCCCCTTCACCTCTTTCA | AACCTCTGTTGGGCATGAAG | 124 |
| chr4:36000470-36002500 | p2 | (GA)7 | CATCCCCTTCACCTCTTTCA | AACCTCTGTTGGGCATGAAG | 124 |
| chr4:36039159-36041171 | p6 | (AACTCC)3 | AACGGGGCCTTAGAAAGAAA | GGGGACAAACGTATGCCTTA | 239 |
| chr4:36039172-36041200 | p6 | (AACTCC)3 | AACGGGGCCTTAGAAAGAAA | GGGGACAAACGTATGCCTTA | 226 |
| chr4:36088229-36090250 | p2 | (TA)9 | TGTGGTGGGTTCCACTTTTT | TTTCGGGTTGTTACAGGAGC | 163 |
| chr4:36088229-36090250 | p2 | (TA)7 | TGTGGTGGGTTCCACTTTTT | TTTCGGGTTGTTACAGGAGC | 163 |
| chr4:36098240-36100254 | p2 | (AT)7 | CCCTCCCCACCATTTTACTT | ATGAGTCGGAAGCTCGTCAT | 235 |
| chr4:36307600-36309615 | p5 | (GAACC)3 | ACAACACTTGGGGATGAAGC | CGGTTCGGTTTGTTGAAGTT | 280 |
| chr4:36380337-36382349 | p2 | (AT)6 | ATCCAGATAAAGGCCCGAGT | AATCAGCGGCAAATAGTTCG | 131 |
| chr4:36474654-36476674 | p2 | (AT)10 | TGTCGTAAATCGGCAGAACA | AAACGGTGTTTGAAACCCAA | 201 |
| chr4:36483339-36485365 | p2 | (TA)13 | TGCAATTTAGCAGCTGATGG | CATCCAAAGTTTGCTTGGGT | 129 |
| chr4:36557960-36559975 | p5 | (TTTTA)3 | ACCCCAACTCTCCACCCTAC | CGCGGTGGAAGATAAGTGAT | 290 |
| chr4:36557960-36559975 | p2 | (TA)10 | ACCCCAACTCTCCACCCTAC | CGCGGTGGAAGATAAGTGAT | 290 |
| chr4:36557960-36559975 | p2 | (AT)18 | ACCCCAACTCTCCACCCTAC | CGCGGTGGAAGATAAGTGAT | 290 |
| chr4:36558939-36561021 | p5 | (TTTTA)3 | AATTTGTTGTTTGGCTCGCT | TAATTGCCACCCAGCATTTT | 194 |
| chr4:36558939-36561021 | p2 | (TA)10 | AATTTGTTGTTTGGCTCGCT | TAATTGCCACCCAGCATTTT | 194 |
| chr4:36585726-36587741 | p5 | (GGGGT)3 | ATGGATGACCAGGCCAATAA | CTCCCATCGTCATGCCTAAT | 211 |
| chr4:36612052-36614066 | p2 | (AT)7 | TTTCACTCTCCCCACAGACC | CGACTTCTACGGGCTCTACG | 177 |

| Marker ID | SSR type | SSR motif | FORWARD PRIMER (5'-3') | REVERSE PRIMER (5'-3') | size (bp) |
|---|---|---|---|---|---|
| chr4:36621633-36623651 | p3 | (CTG)5 | CTGCATTTGCTGATTGCTGT | TTTGAAGTTCGGTTCGGTTC | 250 |
| chr4:36621633-36623651 | p5 | (AACCG)3 | CTGCATTTGCTGATTGCTGT | TTTGAAGTTCGGTTCGGTTC | 250 |
| chr4:36621633-36623651 | p6 | (AAACCG)3 | CTGCATTTGCTGATTGCTGT | TTTGAAGTTCGGTTCGGTTC | 250 |
| chr4:36694498-36696510 | p2 | (TC)6 | AATGACGAAATGAAGGACGG | TTGCCTTAAATGCTCCAACC | 255 |
| chr4:36722392-36724410 | p2 | (AT)10 | GGGCTAAAACGGGCTAAAAG | TTCGGCCATTATTTTAAGCG | 238 |
| chr4:36722392-36724410 | p6 | (TTTACC)3 | GGGCTAAAACGGGCTAAAAG | TTCGGCCATTATTTTAAGCG | 238 |
| chr4:36810482-36812494 | p2 | (TA)6 | TTGTTTTGATAGCCGAAGGG | GACGGCTCGTAGACACTTCC | 170 |
| chr4:36976973-36978988 | p5 | (TTGTA)3 | TCTGAATTGCCTTGGAGCTT | ATTTTGGGAAAGGAGACGCT | 278 |
| chr4:36976973-36978988 | p5 | (GGGTG)4 | TCTGAATTGCCTTGGAGCTT | ATTTTGGGAAAGGAGACGCT | 278 |
| chr4:37026932-37028950 | p3 | (TAA)6 | TTAAACGATCCAAACTCGCC | CTGTCTCTCTCCACTTGCCC | 177 |
| chr4:37052957-37054993 | p2 | (TA)18 | TACATATGCAAGGCCGACAA | TGTCGCACTCCACTTCTCTG | 286 |
| chr4:37078528-37080543 | p5 | (TCGGT)3 | CAGGCCTACTGTGCTTGACA | AACCGAAATAGCCGAACAGA | 206 |
| chr4:37078528-37080543 | p5 | (GTTCG)3 | CAGGCCTACTGTGCTTGACA | AACCGAAATAGCCGAACAGA | 206 |
| chr4:37117264-37119279 | p3 | (TAT)5 | TGTGGAACTGAAGGATGACG | TGGCTGCACATAGAGACACC | 145 |
| chr4:37260607-37262627 | p5 | (AAAGA)4 | CGTCTTCGTCCCATTCTTGT | AAAGGACCACCCCAATTTTC | 253 |
| chr4:37298251-37300263 | p2 | (AT)6 | AGGTGGACGTCTTAGCATGG | TTTGCTGTCTTCCTTGGCTT | 259 |
| chr4:37319181-37321254 | p4 | (TTTC)4 | ATGCCAAGTCTGCCCATAAG | ATCATGAGGTTCTGGGTTCG | 269 |
| chr4:37344765-37346777 | p2 | (TG)6 | AAGTACAAGGCGTGGAGGAA | ATTGGAATCACCGCTTCTTG | 177 |
| chr4:37382750-37384770 | p5 | (AAACC)4 | ACCCAACTGTGCCAAGTCTC | GGCTGTGTGTGGAATGACTG | 142 |
| chr4:37407673-37409693 | p5 | (AAACC)4 | ACCCAACTGTGCCAAGTCTC | GGCTGTGTGTGGAATGACTG | 142 |
| chr4:37492989-37495015 | p2 | (AT)13 | TAGCAAAAGGAGCGGAAAGA | GGGTCATAGGGTAATCGGGT | 183 |
| chr4:37492989-37495015 | p5 | (TATGA)3 | TAGCAAAAGGAGCGGAAAGA | GGGTCATAGGGTAATCGGGT | 183 |
| chr4:37492989-37495015 | p5 | (GATAC)3 | TAGCAAAAGGAGCGGAAAGA | GGGTCATAGGGTAATCGGGT | 183 |
| chr4:37508988-37511004 | p4 | (AGTA)4 | CGCATTTAGGCAAGGAAGAC | CGTCTTCCATTTGGTCCACT | 292 |
| chr4:37513729-37515756 | p6 | (TGGGGG)3 | TGAATTGCTCGTTCTTGTGC | ACCCAGTGAAATCCCACAAA | 256 |
| chr4:37586036-37588051 | p5 | (TTTTG)3 | AGAAGGAGGGAAAGGAGCAG | GTTCGGCCCAATCTTGTTTA | 199 |
| chr4:37586036-37588051 | p3 | (AAT)7 | AGAAGGAGGGAAAGGAGCAG | GTTCGGCCCAATCTTGTTTA | 199 |
| chr4:37586067-37588082 | p5 | (TTTTG)3 | AGGGAAAAATGAGAAGGGGA | ATTTTGACTTACCGGACCCC | 211 |
| chr4:37586067-37588082 | p3 | (AAT)7 | AGGGAAAAATGAGAAGGGGA | ATTTTGACTTACCGGACCCC | 211 |
| chr4:37683403-37685418 | p5 | (TGACA)3 | GACTGCGTTGCTGTGAAAAA | CTGAAGTTGTCCGTCGGTTT | 195 |
| chr4:37768613-37770628 | p5 | (TAATT)3 | CCAGTCAGATGGCTGCAGTA | TCTCATGGGCACCCTATCTC | 178 |
| chr4:37834684-37836698 | p2 | (AT)7 | TTCGTACCGTCGCTTACCTT | TTGATTGCAGCAGAAGAACG | 109 |
| chr4:37977035-37979067 | p5 | (CAATG)3 | ACCTAGGATCACGAGCCCTT | CGACCAAATCGAGAGAGAGG | 146 |
| chr4:37977035-37979067 | p2 | (TA)16 | ACCTAGGATCACGAGCCCTT | CGACCAAATCGAGAGAGAGG | 146 |
| chr4:37985881-37987897 | p4 | (ATTT)4 | ATTAAGTAATCCGTCCCCCG | TAGGGCTGTTCAAAACCGAC | 237 |
| chr4:37987069-37989084 | p5 | (ACCAA)3 | TGTGTTGCACTTAGGAACGC | CCTATAGGGGCCACCAATTT | 246 |

| Marker ID | SSR type | SSR motif | FORWARD PRIMER (5'-3') | REVERSE PRIMER (5'-3') | size (bp) |
|---|---|---|---|---|---|
| chr4:37987069-37989084 | p3 | (TGT)5 | TGTGTTGCACTTAGGAACGC | CCTATAGGGGCCACCAATTT | 246 |
| chr4:38019634-38021654 | p2 | (TA)8 | TTTGCATTTCGGTGAATCAA | TTTGCTCGATATGCCACAAC | 263 |
| chr4:38019634-38021654 | p2 | (AT)10 | TTTGCATTTCGGTGAATCAA | TTTGCTCGATATGCCACAAC | 263 |
| chr4:38019634-38021654 | p2 | (AT)9 | TTTGCATTTCGGTGAATCAA | TTTGCTCGATATGCCACAAC | 263 |
| chr4:38023706-38025722 | p5 | (TTTAT)3 | CTTGAGATGGCCCATGATCT | TGAACGGAGGGAGTAAATGC | 206 |
| chr4:38023706-38025722 | p4 | (AAAT)4 | CTTGAGATGGCCCATGATCT | TGAACGGAGGGAGTAAATGC | 206 |
| chr4:38075411-38077435 | p3 | (TTG)8 | AGAAGATGAAAGTCGCGGAA | CTCTTCACATGCCCAAACCT | 298 |
| chr4:38099197-38101209 | p3 | (CAA)5 | GTTTCGGACATTTGGTTGCT | AAAGCTCCCCTTCATTCGAT | 105 |
| chr4:38099197-38101209 | p2 | (CA)6 | GTTTCGGACATTTGGTTGCT | AAAGCTCCCCTTCATTCGAT | 105 |
| chr4:38180486-38182501 | p5 | (AATAT)3 | CAACTTCCTGAATCTCCCCA | ACTGATAGTACATCCCGCCG | 134 |
| chr4:38199633-38201648 | p3 | (TGT)5 | CTTGCCCGGTTAGTCTTTCA | GGTTTTTGGCCTCAAATTCA | 253 |
| chr4:38246038-38248092 | p2 | (TA)27 | CGCCAGGAGCAGATTAAAAA | CAGTTTACTCGCAACGACCA | 244 |
| chr4:38246038-38248092 | p3 | (CAA)7 | CGCCAGGAGCAGATTAAAAA | CAGTTTACTCGCAACGACCA | 244 |
| chr4:38376038-38378056 | p2 | (AG)19 | TGGATTAGGCCATTTTGGAA | ACCGGCAATTACACTTTTGC | 285 |
| chr4:38376038-38378056 | p6 | (TGTATA)3 | TGGATTAGGCCATTTTGGAA | ACCGGCAATTACACTTTTGC | 285 |
| chr4:38376038-38378056 | p4 | (TTAA)4 | TGGATTAGGCCATTTTGGAA | ACCGGCAATTACACTTTTGC | 285 |
| chr4:38519337-38521352 | p5 | (AGAAA)3 | ACTCTCTCATCCAAAGGCGA | TTCTTTGGCTATAAGGGGCA | 243 |
| chr4:38619158-38621192 | p2 | (AT)7 | CCATGCTACCCACACATACG | ATGCCCACAGGTAGACAAGG | 208 |
| chr4:38619158-38621192 | p2 | (AT)17 | CCATGCTACCCACACATACG | ATGCCCACAGGTAGACAAGG | 208 |
| chr4:38622955-38624971 | p2 | (AT)8 | TTGCTCCACGCCCTACTACT | CGAAGCCACGGTTGATACTT | 246 |
| chr4:38622955-38624971 | p2 | (AT)8 | TTGCTCCACGCCCTACTACT | CGAAGCCACGGTTGATACTT | 246 |
| chr4:38622955-38624971 | p2 | (AT)6 | TTGCTCCACGCCCTACTACT | CGAAGCCACGGTTGATACTT | 246 |
| chr4:38622955-38624971 | p5 | (CCGGT)3 | TTGCTCCACGCCCTACTACT | CGAAGCCACGGTTGATACTT | 246 |
| chr4:38812296-38814316 | p3 | (TGT)5 | ATTGCTTTTCCATGTCGCTT | TGGGGAGGGTAGAGTGTACG | 129 |
| chr4:38812296-38814316 | p2 | (TA)10 | ATTGCTTTTCCATGTCGCTT | TGGGGAGGGTAGAGTGTACG | 129 |
| chr4:38812296-38814316 | p3 | (ACA)5 | ATTGCTTTTCCATGTCGCTT | TGGGGAGGGTAGAGTGTACG | 129 |
| chr4:38857593-38859623 | p2 | (AT)15 | TTTGTACTGACCTGCGCTTG | CCCTGAAGGACTCGAATCAG | 126 |
| chr4:38857593-38859623 | p2 | (TA)8 | TTTGTACTGACCTGCGCTTG | CCCTGAAGGACTCGAATCAG | 126 |
| chr4:38858438-38860454 | p2 | (AT)15 | TAATGTCCAAAAGGATGGGC | TGAGTTTGGAGATGGTGGGT | 212 |
| chr4:38858438-38860454 | p2 | (TA)8 | TAATGTCCAAAAGGATGGGC | TGAGTTTGGAGATGGTGGGT | 212 |
| chr4:38858438-38860454 | p2 | (AT)27 | TAATGTCCAAAAGGATGGGC | TGAGTTTGGAGATGGTGGGT | 212 |
| chr4:38954100-38956130 | p2 | (AT)15 | TTGTGACCCATTTCTCCACA | AATGGGTTGCGTAAGTCCAC | 149 |
| chr4:39015861-39017876 | p5 | (TGATA)3 | ATTCCATATGGCGGGATACA | CCATATAGGAACCTTGCGGA | 127 |
| chr4:39086612-39088627 | p3 | (TTG)5 | GAGAGCCAAGGGAGGAGACT | GCACCAACACACCACATTTC | 219 |
| chr4:39112730-39114746 | p2 | (AT)8 | ACCTCTTTGCGAATTTCCCT | GTGATAAGTTGGGGGCAAGA | 216 |
| chr4:39127046-39129068 | p2 | (AT)11 | GAACCTGGGAAAATCAGCAA | CTTGGTCCGCCATTATTGTT | 223 |

| Marker ID | SSR type | SSR motif | FORWARD PRIMER (5'-3') | REVERSE PRIMER (5'-3') | size (bp) |
|---|---|---|---|---|---|
| chr4:39127046-39129068 | p2 | (AT)8 | GAACCTGGGAAAATCAGCAA | CTTGGTCCGCCATTATTGTT | 223 |
| chr4:39138926-39140941 | p5 | (AAGAG)3 | GGATGGTTCTGGTTTGATGG | TGTATCTGGTGGCGTGTCAT | 232 |
| chr4:39214847-39216887 | p3 | (ATG)5 | AGCTTGGGAGGATAAGCGTT | AGCTTTTCGGGAACAAGGAT | 266 |
| chr4:39214847-39216887 | p2 | (AT)20 | AGCTTGGGAGGATAAGCGTT | AGCTTTTCGGGAACAAGGAT | 266 |
| chr4:39363343-39365361 | p2 | (TA)9 | GAACCATTAGGGCTGTTGGA | TTTGTTCCAAAGGCGATTTC | 189 |
| chr4:39365183-39367198 | p5 | (TTTTA)3 | CGAAGCATGGGTAAAACGAT | CTTTGTGGTCCCATCCACTT | 256 |
| chr4:39432621-39434641 | p2 | (TA)10 | AGTTGCTCCACGCCCTACTA | TCCGAAGCCACGGTTAATAC | 284 |
| chr4:39541068-39543083 | p5 | (AATTA)3 | CAATCACACCACACACACCA | CATCCTCACTCGCTGTCAAA | 125 |
| chr4:39670259-39672277 | p3 | (AAT)6 | CCCTTTGAAGGTCAGTCCAA | CCTTCCCATTTTTGCTTCAA | 228 |
| chr4:39728437-39730453 | p2 | (CT)8 | ATAACGTAATGGCGGTGAGC | GACTCGGGTGGAGTAGTTCG | 274 |
| chr4:39742211-39744226 | p5 | (AAAGA)3 | GCTGGTGTGGCTGTATCAGA | TGCATGATCCCAACTTGCTA | 208 |
| chr4:39758026-39760044 | p6 | (AACTGA)3 | GTGTCACCTTGGCCTTTCAT | ACGTCCATGCCTAGGTTCAG | 226 |
| chr4:39921957-39923972 | p3 | (CAA)5 | TGTATACACCTGCGTGGCAT | TCACTAGAGGGTCGGGTACG | 168 |
| chr4:39921957-39923972 | p2 | (TC)20 | TGTATACACCTGCGTGGCAT | TCACTAGAGGGTCGGGTACG | 168 |
| chr4:39929986-39932074 | p2 | (TA)44 | TCGACGAGAGGTTTGTTGTG | ACTTCGGAGAAAGGATGGGT | 134 |
| chr4:39972615-39974630 | p5 | (AACAA)3 | AACCCGCATCAGGAAGTATG | GTCCGAGTTGTAGGAGACGC | 245 |
| chr4:39998392-40000408 | p2 | (AT)8 | TGTACTTGCGATCTTCGTCG | ACAACAGGATAAAAGGCCCC | 242 |
| chr4:40012545-40014575 | p6 | (CATATA)5 | AGCTCCCAAATTGTCACACC | GTCTGTCAGCGACTCCATCA | 165 |
| chr4:40012545-40014575 | p5 | (ATTCA)3 | AGCTCCCAAATTGTCACACC | GTCTGTCAGCGACTCCATCA | 165 |
| chr4:40031202-40033217 | p5 | (ATTCA)4 | TACTAGGCGTCTGCGGAAAT | AAAGTCGGGGTGTGACAAAG | 176 |
| chr4:40031232-40033247 | p5 | (ATTCA)4 | TACTAGGCGTCTGCGGAAAT | AAAGTCGGGGTGTGACAAAG | 146 |
| chr4:40032110-40034130 | p5 | (ATTCA)4 | CCATCGGAGTGACGTAAGGT | TTGGGGATATGCTCCTACCA | 182 |
| chr4:40032110-40034130 | p2 | (TA)30 | CCATCGGAGTGACGTAAGGT | TTGGGGATATGCTCCTACCA | 182 |
| chr4:40032767-40034827 | p5 | (ATTCA)4 | CAACAATCCGGGAATTCAAC | GGAAGGTGTTGGACGGAGTA | 271 |
| chr4:40032767-40034827 | p2 | (TA)30 | CAACAATCCGGGAATTCAAC | GGAAGGTGTTGGACGGAGTA | 271 |
| chr4:40217388-40219406 | p6 | (ACCTCT)3 | GAGCATATCGGGACAAGGAA | CCGTTGACTTGGGTTCAGTT | 249 |
| chr4:40240704-40242722 | p6 | (ATATGT)3 | TGGTATTAGGGCCTGTTTGC | TTTAAAGCAATGGCACACCA | 199 |
| chr4:40297450-40299468 | p6 | (GAAAGT)3 | TGAGCTTAGCCACGACAATG | TTCCCCCACATTTGTGATCT | 156 |
| chr4:40297450-40299468 | p6 | (CTTTCC)3 | TGAGCTTAGCCACGACAATG | TTCCCCCACATTTGTGATCT | 156 |
| chr4:40313455-40315489 | p2 | (AT)17 | TTTGGACCGTCCTTTAGTGG | CTAGCCCAAGCAAAGTTTCG | 195 |
| chr4:40441868-40443880 | p2 | (TA)6 | TAGCAAAAGGAGCGGAAAGA | TGAGGGTTCTAGGGTGATCG | 159 |
| chr4:40462830-40464842 | p2 | (TA)6 | TAGCAAAAGGAGCGGAAAGA | TGAGGGTTCTAGGGTGATCG | 159 |
| chr4:40501012-40503027 | p5 | (TCCTT)3 | TCCTTGCTTTACTCCGCTGT | AAGCGGGTTACTGGTTGCTA | 219 |
| chr4:40514119-40516134 | p3 | (ATC)5 | TGGCCTGCTCCTGAGTATCT | AGCTGGGTATTCTGGGGAGT | 272 |
| chr4:40515429-40517443 | p2 | (AT)7 | CATGCATTTCCATTGATTCG | TTCGGAGCCATATCTTTTGG | 186 |
| chr4:40515429-40517443 | p5 | (GAAAT)3 | CATGCATTTCCATTGATTCG | TTCGGAGCCATATCTTTTGG | 186 |

| Marker ID | SSR type | SSR motif | FORWARD PRIMER (5'-3') | REVERSE PRIMER (5'-3') | size (bp) |
|---|---|---|---|---|---|
| chr4:40533786-40535800 | p2 | (AT)7 | AAGTGGACCCTACCCTCTCC | ATATGGGGTGCACGTGTTTT | 132 |
| chr4:40544213-40546228 | p5 | (AATTT)3 | CTTCGATGCCAATCTTGGTT | CGAAGCATCTAAGACCCGAG | 235 |
| chr4:40555287-40557301 | p2 | (TA)7 | CAAAGCGCGTATCTCATTTG | CGGATACGACATCTTGGGTT | 247 |
| chr4:40555287-40557301 | p5 | (AAAAT)3 | CAAAGCGCGTATCTCATTTG | CGGATACGACATCTTGGGTT | 247 |
| chr4:40601931-40603943 | p2 | (AT)6 | TGGCCGAACTCTCTGTCTCT | CAAAAGAGGATTCCGAACCA | 231 |
| chr4:40601931-40603943 | p2 | (AT)6 | TGGCCGAACTCTCTGTCTCT | CAAAAGAGGATTCCGAACCA | 231 |
| chr4:40701100-40703136 | p2 | (TA)18 | GAACCTCCTCGGCTTTTCTT | CAAATTTTCTGAGGGGTCCA | 288 |
| chr4:40754891-40756906 | p5 | (TGCAT)3 | GGTTTTGAGGTTCAACGGAA | ATACGACGTCTCACCAAGGG | 224 |
| chr4:40808229-40810244 | p5 | (AATGC)3 | GGCGAACCCTCTCATTCATA | AGGCCTCCTCCAACTTCATT | 205 |
| chr4:40876015-40878029 | p2 | (AT)7 | CCCTGGGTCTATTCCTCCTC | GAATGGTTAGACCGTGCGAT | 300 |
| chr4:41014832-41016846 | p2 | (AT)7 | GCTCAAACCCACAACACCTT | ATGTGACCTTGGACCAGAGC | 127 |
| chr4:41056432-41058452 | p4 | (TTGT)5 | AGCGCATCTAACCAAGAGGA | AATCGGATTTACATCGCCTG | 124 |
| chr4:41221850-41223862 | p6 | (GGAGCT)3 | ACCAACAACAAAGTCCTGGG | CTTCGGGCCAAGATGTAAAA | 143 |
| chr4:41221850-41223862 | p2 | (CA)6 | ACCAACAACAAAGTCCTGGG | CTTCGGGCCAAGATGTAAAA | 143 |
| chr4:41369163-41371179 | p4 | (TTCT)4 | AGTTATGGGCTCGTGATTCG | ACTCAGCCCGTGTCATTACC | 119 |
| chr4:41461616-41463644 | p2 | (AT)14 | AAAGCATGTTATGCCTTCGG | ATCAAATCAATCGTGCCACA | 230 |
| chr4:41507574-41509594 | p2 | (TA)10 | CTGCCCTATGCATTCACCTT | CCTCTGAGAAGCACCACCTC | 132 |
| chr4:41507574-41509594 | p2 | (TA)10 | CTGCCCTATGCATTCACCTT | CCTCTGAGAAGCACCACCTC | 132 |
| chr4:41507574-41509594 | p2 | (TA)10 | CTGCCCTATGCATTCACCTT | CCTCTGAGAAGCACCACCTC | 132 |
| chr4:41507574-41509594 | p2 | (TA)10 | CTGCCCTATGCATTCACCTT | CCTCTGAGAAGCACCACCTC | 132 |
| chr4:41507574-41509594 | p2 | (TA)10 | CTGCCCTATGCATTCACCTT | CCTCTGAGAAGCACCACCTC | 132 |
| chr4:41507574-41509594 | p2 | (TA)10 | CTGCCCTATGCATTCACCTT | CCTCTGAGAAGCACCACCTC | 132 |
| chr4:41666862-41668876 | p2 | (TA)7 | AATCAACGGTGTTGACCCAT | TCGGGAATGGAAGTGTTTTC | 164 |
| chr4:41704309-41706323 | p2 | (TA)7 | AATCAACGGTGTTGACCCAT | TCGGGAATGGAAGTGTTTTC | 164 |
| chr4:41778377-41780421 | p2 | (TA)21 | CTTGTTTTGCGGTGGATTTT | CGCTGTTTTCCAACTTCCAT | 138 |
| chr4:41849236-41851254 | p3 | (TCA)6 | TGTCTACCTGTGGGCATGAA | CTTCCTCCCATGGCTTATGA | 146 |
| chr4:41865737-41867752 | p4 | (ACAT)4 | CTAGGGCCATGACAAGCAAT | TTGGTGTAAGGCTTTTTGGC | 258 |
| chr4:41865830-41867845 | p4 | (ACAT)4 | AATTCTGTCACGACCCGAAC | TTGGTGTAAGGCTTTTTGGC | 146 |
| chr4:41870933-41872961 | p2 | (TA)11 | GGGTAATTGGGATCGTGATG | GCCGAGGTTGAGTACGAAAG | 232 |
| chr4:41870971-41872985 | p2 | (TA)11 | GGGTAATTGGGATCGTGATG | GCCGAGGTTGAGTACGAAAG | 194 |
| chr4:41885140-41887162 | p2 | (CT)11 | GTAACTCGATCGTGGGGAGA | AACCGCAAACCTCAAACAAC | 226 |
| chr4:41885140-41887162 | p2 | (GT)6 | GTAACTCGATCGTGGGGAGA | AACCGCAAACCTCAAACAAC | 226 |
| chr4:41885140-41887162 | p2 | (GT)7 | GTAACTCGATCGTGGGGAGA | AACCGCAAACCTCAAACAAC | 226 |
| chr4:41929329-41931341 | p2 | (TG)6 | GGAGGAGGAGACCTGGAAAC | TTACAAGCATTTCGGGGAAG | 102 |
| chr4:41984221-41986251 | p2 | (TA)15 | AAAAGGTCCTTATGGCCCAG | GTAACATCTGCCCGACTGGT | 139 |
| chr4:42007261-42009275 | p2 | (TC)7 | AGTGTCCGGCCATACTTTTG | CTTTGAAGGCCATCCTTGAG | 273 |

| Marker ID | SSR type | SSR motif | FORWARD PRIMER (5'-3') | REVERSE PRIMER (5'-3') | size (bp) |
|---|---|---|---|---|---|
| chr4:42027167-42029191 | p2 | (TA)12 | ACATGAGGCGTAGGGAATTG | CTCAGCGGAGTAGGGTCTTG | 161 |
| chr4:42027167-42029191 | p3 | (CAA)5 | ACATGAGGCGTAGGGAATTG | CTCAGCGGAGTAGGGTCTTG | 161 |
| chr4:42067095-42069111 | p4 | (AATA)4 | GAGCGTGCAAGTGCTAGATG | CCCGATTTTACTTATGGGCA | 141 |
| chr4:42067095-42069111 | p3 | (ACA)5 | GAGCGTGCAAGTGCTAGATG | CCCGATTTTACTTATGGGCA | 141 |
| chr4:42089643-42091655 | p2 | (TA)9 | TCACATTCGGAATCCTGTCA | TATGTCGGAACGCATGGATA | 252 |
| chr4:42089643-42091655 | p2 | (TA)6 | TCACATTCGGAATCCTGTCA | TATGTCGGAACGCATGGATA | 252 |
| chr4:42134385-42136400 | p5 | (ATTTC)3 | TCCTTGGGGTGATCTCGTAG | ATTTGTCCCTTGCGAACAAC | 236 |
| chr4:42245581-42247605 | p2 | (TA)12 | CTCAGCCGAGTAGGGTCTTG | CCTGTTTATAACTGCGGCGT | 119 |
| chr4:42289259-42291271 | p2 | (AT)6 | AACCTGCACGCAGGAAATAC | TGCGGTTAAAGGTTGAAAGG | 206 |
| chr4:42289259-42291271 | p2 | (AT)6 | AACCTGCACGCAGGAAATAC | TGCGGTTAAAGGTTGAAAGG | 206 |
| chr4:42303913-42305963 | p2 | (AT)28 | GAATGAATTTGCGAGCCATT | GGGTGTGACAGAATCACCCT | 279 |
| chr4:42522676-42524696 | p2 | (AT)10 | GCACTGACCACTGCTGGATA | ATGAGCCCACAGAGGAATTG | 109 |
| chr4:42522676-42524696 | p6 | (CATATA)3 | GCACTGACCACTGCTGGATA | ATGAGCCCACAGAGGAATTG | 109 |
| chr4:42563273-42565313 | p2 | (TA)20 | TTATAATGCCGTTTTTGCCC | CCTGATAATCGCCCTCGTAA | 198 |
| chr4:42571900-42573915 | p5 | (TGTTG)3 | TTAATTGCTGAGGTGGGACC | AGTCAGATCACCCGAAAACG | 180 |
| chr4:42711382-42713394 | p2 | (AC)6 | TACATACCTTGCCCGCTTTC | GGTAAATTGGCCGAGTTGAA | 248 |
| chr4:42729778-42731793 | p5 | (AATGC)3 | TCTACCCCAGAAGAGGCTCA | ATGGTAGGTGCCCACATGAT | 210 |
| chr4:42805626-42807642 | p2 | (TA)6 | AGGGAAAGCCAAGTCAGGAT | TGAAGCCTCGGGTAATATGG | 277 |
| chr4:42808156-42810174 | p2 | (TA)9 | AGTGTATGGGACACCCAAGC | TAGATGTCAGGCACTCGTCG | 130 |
| chr4:42940972-42943040 | p2 | (AT)34 | TATTCCATGGATTTTGGGGA | GGTTCGAACTCAGAACCTCG | 285 |
| chr4:43121312-43123327 | p3 | (GAA)7 | ATGCCCACCCCTACACAATA | TACCCCAGAGGAAAATGTGG | 264 |
| chr4:43121312-43123327 | p3 | (GAA)5 | ATGCCCACCCCTACACAATA | TACCCCAGAGGAAAATGTGG | 264 |
| chr4:43203110-43205125 | p3 | (GAT)5 | TACCAACCAAGGCTTGTTCC | TGCTCATCCCATGTTCTCAA | 225 |
| chr4:43305501-43307515 | p2 | (TA)7 | TTTGTCACGACCCAATCTGA | CGGCTTGTATGTTTTTGGGT | 188 |
| chr4:43390657-43392681 | p6 | (TTGGTG)4 | GTCGTGTGTGGCCATAATTG | GCCAGGGTTTTCCCTGTAAT | 130 |
| chr4:43416227-43418242 | p5 | (AAATA)4 | ATGACCAATCATGCCACTCA | ATTACACGCAGAGAATGCCC | 169 |
| chr4:43416227-43418242 | p5 | (AAATA)3 | ATGACCAATCATGCCACTCA | ATTACACGCAGAGAATGCCC | 169 |
| chr4:43416227-43418242 | p5 | (TTTAT)3 | ATGACCAATCATGCCACTCA | ATTACACGCAGAGAATGCCC | 169 |
| chr4:43416227-43418242 | p4 | (TTAT)4 | ATGACCAATCATGCCACTCA | ATTACACGCAGAGAATGCCC | 169 |
| chr4:43416227-43418242 | p5 | (TATTT)3 | ATGACCAATCATGCCACTCA | ATTACACGCAGAGAATGCCC | 169 |
| chr4:43443459-43445483 | p6 | (AACCGG)4 | CTTATCTCGTTTGACCCCGA | GGTGAACGGACCTTTTTCAA | 200 |
| chr4:43443459-43445483 | p3 | (GAT)7 | CTTATCTCGTTTGACCCCGA | GGTGAACGGACCTTTTTCAA | 200 |
| chr4:43446036-43448070 | p2 | (AT)17 | ATCTCAGAAGCGGAGGATGA | ATCATATTCGGCCATGGAAA | 147 |
| chr4:43446036-43448070 | p2 | (AT)9 | ATCTCAGAAGCGGAGGATGA | ATCATATTCGGCCATGGAAA | 147 |
| chr4:43463894-43465912 | p2 | (AT)6 | TGCAGAAAAGGCCCAAATAC | CGTCAGCCTTGTTTGGTTTT | 278 |
| chr4:43463894-43465912 | p2 | (AT)18 | TGCAGAAAAGGCCCAAATAC | CGTCAGCCTTGTTTGGTTTT | 278 |

| Marker ID | SSR type | SSR motif | FORWARD PRIMER (5'-3') | REVERSE PRIMER (5'-3') | size (bp) |
|---|---|---|---|---|---|
| chr4:43463894-43465912 | p2 | (AT)9 | TGCAGAAAAGGCCCAAATAC | CGTCAGCCTTGTTTGGTTTT | 278 |
| chr4:43794929-43796941 | p2 | (AT)6 | ATCTCGAAGAACGCCAAGAA | CCCTTAACTTTCCACGACGA | 279 |
| chr4:44013123-44015138 | p2 | (AG)9 | ATGGCGTGAGCAAGTTCTCT | AAGCGTTAATGGTGTTCGCT | 192 |
| chr4:44013123-44015138 | p5 | (CCCGC)3 | ATGGCGTGAGCAAGTTCTCT | AAGCGTTAATGGTGTTCGCT | 192 |
| chr4:44020564-44022579 | p5 | (TTATT)3 | GCGGATTTATGTAGTGGCGT | TAAGCAACAGGGCTCGAACT | 224 |
| chr4:44020564-44022579 | p3 | (TTG)5 | GCGGATTTATGTAGTGGCGT | TAAGCAACAGGGCTCGAACT | 224 |
| chr4:44070798-44072812 | p5 | (AAACC)4 | ACCAAACCAAACCAAACCAA | CAAGGTCACTGGCCAAGTTT | 226 |
| chr4:44070798-44072812 | p2 | (AT)7 | ACCAAACCAAACCAAACCAA | CAAGGTCACTGGCCAAGTTT | 226 |
| chr4:44070798-44072812 | p2 | (AT)11 | ACCAAACCAAACCAAACCAA | CAAGGTCACTGGCCAAGTTT | 226 |
| chr4:44080781-44082795 | p2 | (TA)7 | TTAAGCTGGCAGGAGAGGAA | ATTTCAGCCCTTCCCAAGAT | 173 |
| chr4:44119035-44121050 | p5 | (ATATA)3 | AATACAAGGCGAGTCATCCG | ATTTGGTCCACTCCACTTGC | 261 |
| chr4:44119035-44121050 | p2 | (TA)11 | AATACAAGGCGAGTCATCCG | ATTTGGTCCACTCCACTTGC | 261 |
| chr4:44119035-44121050 | p2 | (AC)6 | AATACAAGGCGAGTCATCCG | ATTTGGTCCACTCCACTTGC | 261 |
| chr4:44168096-44170111 | p5 | (AAGAA)3 | AAACGAAAACATAAGCCCCA | CATGCAATGACACTTCACCC | 272 |
| chr4:44168096-44170111 | p2 | (AT)179 | AAACGAAAACATAAGCCCCA | CATGCAATGACACTTCACCC | 272 |
| chr4:44336509-44338521 | p2 | (TG)6 | TAGTAACACCGGGAGAACGG | TTTTTCTTCCGTGGTGGTTC | 155 |
| chr4:44359536-44361551 | p5 | (ATGTG)3 | AACAAAGGAGGTGCTGCCTA | ATCGCTTGCGTGTTCTTCTT | 154 |
| chr4:44359536-44361551 | p2 | (TA)8 | AACAAAGGAGGTGCTGCCTA | ATCGCTTGCGTGTTCTTCTT | 154 |
| chr4:44359536-44361551 | p2 | (AT)36 | AACAAAGGAGGTGCTGCCTA | ATCGCTTGCGTGTTCTTCTT | 154 |
| chr4:44367637-44369651 | p2 | (AT)7 | CCAACTTATGCCTTGCGATT | AATTATGGCCTCAAAGCACG | 202 |
| chr4:44408442-44410457 | p5 | (GAAAA)3 | TCTGCAAAGGTTTTTCCACC | TGCTATACGGTGGAGGGAAG | 153 |
| chr4:44605609-44607624 | p5 | (AAATA)3 | CAAGATGTCCATGTGCCAAG | TTCATTCGCAACTTCAGCAG | 163 |
| chr4:44791723-44793737 | p2 | (AG)7 | TTGAAATCGACCCTCGAAAC | GACTTGGGTTGGGAAGTTGA | 120 |
| chr4:44851318-44853333 | p5 | (GTTTG)3 | TAATAGCCCAAACGCTGTCC | ACCAAAGGGGAAAGAGGAGA | 102 |
| chr4:44973153-44975183 | p2 | (TA)11 | TTGTTCCACGAAATGCAAAA | TTGTTCCACGAAATGCAAAA | 116 |
| chr4:44973153-44975183 | p2 | (AT)10 | TTGTTCCACGAAATGCAAAA | TTGTTCCACGAAATGCAAAA | 116 |
| chr4:45028028-45030046 | p2 | (AT)25 | CCGTCCAAGTTTACTCGTCC | TACACTGCCCAGAGAAAGGC | 126 |
| chr4:45028028-45030046 | p6 | (TAAAAA)3 | CCGTCCAAGTTTACTCGTCC | TACACTGCCCAGAGAAAGGC | 126 |
| chr4:45028028-45030046 | p5 | (AAGAA)3 | CCGTCCAAGTTTACTCGTCC | TACACTGCCCAGAGAAAGGC | 126 |
| chr4:45028028-45030046 | p2 | (AC)6 | CCGTCCAAGTTTACTCGTCC | TACACTGCCCAGAGAAAGGC | 126 |
| chr4:45186842-45188857 | p5 | (GCATT)3 | GTGAGGCCTTCAAGTTCGAG | GGGCTTCCTACGTATCCCTC | 233 |
| chr4:45186842-45188857 | p2 | (AT)33 | GTGAGGCCTTCAAGTTCGAG | GGGCTTCCTACGTATCCCTC | 233 |
| chr4:45226571-45228616 | p2 | (AT)34 | TGTTTTGGGCTAAGGGTTTG | TGTCGAATTGTCGACTCTGG | 128 |
| chr4:45230273-45232288 | p2 | (AT)8 | AGACCGACCTTGTTGTACGG | CTCGCTTCCCTCATCTTGTC | 249 |
| chr4:45255295-45257310 | p5 | (CCACC)4 | TTTATTGGGGCATCGACTTC | CCGTGGTTACAAAAATTGGG | 141 |
| chr4:45255295-45257310 | p5 | (TACAA)3 | TTTATTGGGGCATCGACTTC | CCGTGGTTACAAAAATTGGG | 141 |

| Marker ID | SSR type | SSR motif | FORWARD PRIMER (5'-3') | REVERSE PRIMER (5'-3') | size (bp) |
|---|---|---|---|---|---|
| chr4:45308342-45310386 | p2 | (TA)22 | TGTGGGGGAGGTAGCTTATG | TTATGTACGAACCATCGCCA | 167 |
| chr4:45319289-45321304 | p5 | (TGCAA)3 | TGGTAAGCCGATCACACAAA | TATCGCGGAGGGAATGATAG | 230 |
| chr4:45322240-45324255 | p5 | (TATCA)3 | AATCACCGTCATAACACGCA | TAAGGGGTGCTCGGTACAAG | 225 |
| chr4:45444266-45446288 | p2 | (TA)11 | AGGCTGGAAGGGCATTATCT | GTTATGCACGCGGATAAGGT | 146 |
| chr4:45444266-45446288 | p2 | (AT)20 | AGGCTGGAAGGGCATTATCT | GTTATGCACGCGGATAAGGT | 146 |
| chr4:45535883-45537901 | p6 | (TATAAA)3 | TGGATCGTGACAAACCAAAA | GCCATCATCGGTTTGAAAAT | 214 |
| chr4:45576706-45578726 | p2 | (AT)10 | AGGCATGCTCAGGTAGTGGT | TGGGACAGGTTATGCGTGTA | 195 |
| chr4:45743538-45745550 | p2 | (TA)6 | CTCGAAGAAGAAGCTCGGAA | CGGCCTAAGTCTTGCCTAAA | 222 |
| chr4:45743538-45745550 | p2 | (TA)6 | CTCGAAGAAGAAGCTCGGAA | CGGCCTAAGTCTTGCCTAAA | 222 |
| chr4:45802878-45804893 | p3 | (TTA)5 | ATGCCCAAACCATCTCAGTC | ATGTCGCACATCGTATTGGA | 295 |
| chr4:45841214-45843232 | p6 | (AAATAA)3 | TCGCAAAATGGCAGATGATA | TGATATTTCGCTGCGTCTTG | 292 |
| chr4:45999471-46001507 | p2 | (AT)18 | CCACCAGCTCGATTCTCTTC | CCAGATTGAGGTTTACCCGA | 240 |
| chr4:46071069-46073195 | p3 | (TTG)9 | TAAACGAGAGGGTTCGCCTA | TATGCCTAGCCCGAAATCAC | 202 |
| chr4:46114186-46116244 | p2 | (AT)29 | GGAAGACGTGTGGAAAGGAA | TTCCTTGAATCGCCCAATAG | 207 |
| chr4:46227178-46229206 | p2 | (TA)20 | TCCCTGTCAGTGGTGAGTGA | CCTACTGCAACATGCCAAGA | 268 |
| chr4:46227178-46229206 | p2 | (AT)14 | TCCCTGTCAGTGGTGAGTGA | CCTACTGCAACATGCCAAGA | 268 |
| chr4:46227178-46229206 | p6 | (AAAGTA)4 | TCCCTGTCAGTGGTGAGTGA | CCTACTGCAACATGCCAAGA | 268 |
| chr4:46270878-46272890 | p2 | (TG)6 | TATAATGCACCATGAGGCCA | ATCTCGCGAATTTCCCTTTT | 229 |
| chr4:46348368-46350380 | p2 | (CT)6 | TGCCTTAGTTTATGTCGGGG | CCTTTGAGCCTAGCCAGTTG | 172 |
| chr4:46555399-46557414 | p5 | (TTTTG)3 | ACTTCAACGGCGTTTCACTT | TTGAATTTGGAAGGTTTCGG | 119 |
| chr4:46555399-46557414 | p5 | (CAATG)3 | ACTTCAACGGCGTTTCACTT | TTGAATTTGGAAGGTTTCGG | 119 |
| chr4:46639672-46641706 | p2 | (AT)18 | CGGGCACCTCTCATCATACT | TCTTAGCCTTGTCGGCTCAT | 206 |
| chr4:46639672-46641706 | p2 | (AT)17 | CGGGCACCTCTCATCATACT | TCTTAGCCTTGTCGGCTCAT | 206 |
| chr4:46675156-46677172 | p4 | (AAAG)4 | AATCTTTTCCCTTCGCCCTA | CACGATCCCTTCCAAACCTA | 131 |
| chr4:46721450-46723468 | p6 | (TGTGTA)3 | TAGGGCACAAGCTTTCCTGT | TTCCTCTTTCTCCAGCCGTA | 226 |
| chr4:46746283-46748299 | p2 | (TA)8 | GGCCCAATACCCTTCTTGAT | GCCTACCAAACCCATCAGTG | 162 |
| chr4:46768209-46770247 | p5 | (TTTTG)4 | CCCGATGCTACAGTTGGAAT | CATGGAACGGAGCTTATCGT | 251 |
| chr4:46768209-46770247 | p2 | (AT)19 | CCCGATGCTACAGTTGGAAT | CATGGAACGGAGCTTATCGT | 251 |
| chr4:46833119-46835145 | p2 | (AG)13 | TTCCTTAGGTTGGTGATCGG | GGTGTAACAAATGGGCATCC | 126 |
| chr4:46875383-46877413 | p6 | (ATAATT)5 | TTTCCTTACCACGGAGCATC | AAAATCATTGCAAAGGGCTG | 117 |
| chr4:47035516-47037532 | p2 | (TC)8 | ACAACTTGTCCATGCCAACA | AGCCGGGAGAAGTAATTGGT | 225 |
| chr4:47044471-47046528 | p3 | (AAG)19 | CCCCAACAACTTTTGCAGTT | CCTTAGCAGCTGATCGGAAG | 159 |
| chr4:47114725-47116739 | p2 | (AT)7 | AACCCATGCTTAGTCGTTCG | CTTTTCCGTGAGTTTGGGAG | 109 |
| chr4:47114725-47116739 | p2 | (TA)7 | AACCCATGCTTAGTCGTTCG | CTTTTCCGTGAGTTTGGGAG | 109 |
| chr4:47180003-47182018 | p5 | (CACAA)3 | AGAATTATCCCCGATGGGTC | AACGGGAATTGAGAATGACG | 199 |
| chr4:47320118-47322133 | p4 | (CATT)4 | CATACTGCCGAGAGGGTCAT | TTGCTCTGCAATTTGTCTGG | 161 |

| Marker ID | SSR type | SSR motif | FORWARD PRIMER (5'-3') | REVERSE PRIMER (5'-3') | size (bp) |
|---|---|---|---|---|---|
| chr4:47320118-47322133 | p5 | (TATTT)3 | CATACTGCCGAGAGGGTCAT | TTGCTCTGCAATTTGTCTGG | 161 |
| chr4:47418667-47420679 | p2 | (TA)6 | CAATGTGGCAGTGATATGGC | TTTGAAATCATTTTTCGCCC | 156 |
| chr4:47423841-47425875 | p5 | (ATATC)3 | TCGATTTTTGGGGTTTTGAG | CCATTTTTGGGATTAGGGCT | 228 |
| chr4:47423873-47425909 | p5 | (ATATC)3 | TCGATTTTTGGGGTTTTGAG | CCATTTTTGGGATTAGGGCT | 196 |
| chr4:47463092-47465104 | p2 | (AT)6 | TTAGCGTAGAAAGCGGGAAA | TGAATGCCATCGGAAACATA | 288 |
| chr4:47587403-47589418 | p5 | (TATCA)3 | AATGAGGCTTTGGTCCCTTT | GATAAGAGTCAGCGGGTTCG | 178 |
| chr4:47608498-47610534 | p2 | (TA)18 | TGGATAAACATCGTGTGGGA | CCCATTTAGAATCTGCCGAA | 179 |
| chr4:47623822-47625860 | p2 | (TA)19 | AATGAACCCTAACCGCTTCC | TGCAATGTCACTTCACCCTT | 143 |
| chr4:47656234-47658254 | p5 | (ATGAT)4 | GATGCTGCCGAAATTTTTGT | GGACATGCCAAAAGAAGGAA | 197 |
| chr4:47691019-47693034 | p5 | (TTTAT)3 | GGCTTGGTAAGCCCCTAGTC | AACTGTTGAATTTGCTCGGG | 215 |
| chr4:47740777-47742789 | p2 | (AT)14 | GACGAAGGGACGAGCAGTAG | GTTCGAATGGAGGAGCATGT | 253 |
| chr4:47740777-47742789 | p2 | (TA)6 | GACGAAGGGACGAGCAGTAG | GTTCGAATGGAGGAGCATGT | 253 |
| chr4:47789837-47791849 | p2 | (AT)14 | GACGAAGGGACGAGCAGTAG | GTTCGAATGGAGGAGCATGT | 253 |
| chr4:47789837-47791849 | p2 | (TA)6 | GACGAAGGGACGAGCAGTAG | GTTCGAATGGAGGAGCATGT | 253 |
| chr4:47801558-47803576 | p6 | (ATATCT)3 | CTTCCGTCGTACCCATGAAT | ATCCGCCTGCTTAGGATTTT | 285 |
| chr4:47833330-47835350 | p4 | (TTTA)5 | TGCCTCTTCCTCTTATGCGT | GCCTTTCAACCACAACCAAT | 269 |
| chr4:47902440-47904455 | p5 | (ATTTC)3 | ACTCAGGAGGCTGACTTGGA | TTGCCCTTAAGTTTTGGTGG | 179 |
| chr4:47902440-47904455 | p3 | (GTT)7 | ACTCAGGAGGCTGACTTGGA | TTGCCCTTAAGTTTTGGTGG | 179 |
| chr4:47967871-47969893 | p2 | (AT)11 | TCCAAATTTCTGATTTCGCC | GGTGGCGACTCCTTAAAACA | 174 |
| chr4:48063552-48065608 | p2 | (AT)28 | GATCATGTCTTGCCATCCCT | AATCAACTTCCGAACATGGC | 119 |
| chr4:48084433-48086448 | p5 | (ATATG)3 | AGAGAAATGGTGGATCGTGG | AGAGACAATTGGGCAGCACT | 242 |
| chr4:48091905-48093925 | p5 | (ATATG)4 | GATAGCGAACCCACAACCAT | TCAACTTCCAACACCCACAA | 204 |
| chr4:48109599-48111614 | p5 | (AATGA)3 | CTAAAGTGGCCCTTGTTCCA | CACCCACGTGTTTTTGACTG | 100 |
| chr4:48147736-48149750 | p2 | (AT)7 | GCCAGTCACGATCCAACTTT | TGGGCATATTTAACCCCAAA | 142 |
| chr4:48179559-48181579 | p5 | (CATAT)4 | CAGAAAACCCCAAGGACAAA | AATTTGTGGGTTCGCTCAAC | 139 |
| chr4:48202843-48204859 | p4 | (GTAT)4 | ATCATGCGAAGCGAAAGACT | GTTAGCCATCCTAGGGGCTC | 119 |
| chr4:48202843-48204859 | p6 | (TATGTA)3 | ATCATGCGAAGCGAAAGACT | GTTAGCCATCCTAGGGGCTC | 119 |
| chr4:48244058-48246076 | p6 | (CATATA)3 | GAAAATCCAGATTCGGCGTA | GCTATGACAGCCTAAACGGC | 285 |
| chr4:48304486-48306506 | p2 | (TA)10 | GCATAAGCCATCCAAAAGGA | ACATGGTCGTGGAACTCACA | 171 |
| chr4:48400307-48402322 | p5 | (TTCAT)3 | TGCGAATAACAACAACCGAG | ATGGCCTCACACACAACAAA | 186 |
| chr4:48553494-48555509 | p5 | (ATGCA)3 | ACGAGTTTCCACCCTCTGTG | TGGGTCGTGACAAGTTGGTA | 116 |
| chr4:48591916-48593943 | p3 | (GTT)9 | CTGGGATCAAGCGGATAAAA | GGCTTCTGCCATTTCATTGT | 263 |
| chr4:48609493-48611508 | p3 | (GAT)5 | TTGCTCCAAAGCTCATCCTT | AATCCTTTGCACATTTTCGC | 148 |
| chr4:48629865-48631895 | p2 | (AT)15 | GCCGCAACGAATTACTTCTC | GAATCCATAAAGGGCGCATA | 132 |
| chr4:48673114-48675129 | p3 | (GAT)5 | GCTCCAAACGAATTCAAGGA | CTCTTGTTTTGTGGCCCATT | 160 |
| chr4:48673114-48675129 | p2 | (TG)11 | GCTCCAAACGAATTCAAGGA | CTCTTGTTTTGTGGCCCATT | 160 |

| Marker ID | SSR type | SSR motif | FORWARD PRIMER (5'-3') | REVERSE PRIMER (5'-3') | size (bp) |
|---|---|---|---|---|---|
| chr4:48673690-48675712 | p3 | (GAT)5 | GAAAAACCAAGCGATGGAAA | TTAGGAACCATCATGGGGAG | 242 |
| chr4:48673690-48675712 | p2 | (TG)11 | GAAAAACCAAGCGATGGAAA | TTAGGAACCATCATGGGGAG | 242 |
| chr4:48673690-48675712 | p2 | (TA)6 | GAAAAACCAAGCGATGGAAA | TTAGGAACCATCATGGGGAG | 242 |
| chr4:48674514-48676526 | p2 | (TG)11 | CGCGCGCTTATGTATGACTA | CAAGGCTTCAGGGTTATTGC | 198 |
| chr4:48674514-48676526 | p2 | (TA)6 | CGCGCGCTTATGTATGACTA | CAAGGCTTCAGGGTTATTGC | 198 |
| chr4:48674514-48676526 | p2 | (TA)6 | CGCGCGCTTATGTATGACTA | CAAGGCTTCAGGGTTATTGC | 198 |
| chr4:48675369-48677381 | p2 | (TA)6 | ACAGCCGAAGGGCTTATGTA | CCCTCACTCGCCATAGTCAT | 168 |
| chr4:48675369-48677381 | p2 | (TA)6 | ACAGCCGAAGGGCTTATGTA | CCCTCACTCGCCATAGTCAT | 168 |
| chr4:48687029-48689093 | p2 | (AT)32 | CGAAGCGTATATTGGTCGGT | AGCAGCACCAACAATGACAG | 230 |
| chr4:48699569-48701623 | p2 | (TA)6 | GTTCGTGTTGTGGTCGTGTC | CCGTCTAACCCAAACTTCCA | 191 |
| chr4:48799659-48801671 | p2 | (CA)6 | TATTTCGGAGCTTTCGCACT | GTCTTCCACGTTTCGATGGT | 174 |
| chr4:48807053-48809065 | p2 | (CA)6 | CTGGGGGTTTCAAAGCAATA | TTGAGTCCTAGATTGCGGCT | 186 |
| chr4:48825527-48827541 | p2 | (AT)7 | GGAAGGATTCATGCCTTTGA | AGTGCGAATTCCCTCTTGAA | 141 |
| chr4:48877634-48879648 | p2 | (AT)6 | CATACATGTCTTTCCGGGCT | GGCGTTGTGAATCTTTCGTT | 179 |
| chr4:48877656-48879700 | p2 | (AT)6 | CATACATGTCTTTCCGGGCT | GGCGTTGTGAATCTTTCGTT | 157 |
| chr4:48931520-48933532 | p2 | (AT)6 | AATTTACAGGGGTGGTGCTG | ATACCTCGATCGCTCGCTAA | 226 |
| chr4:49036305-49038335 | p2 | (AG)15 | TCGGTCGTTATATTGGCGAT | AGCCGTCTGCAGCCTTAATA | 183 |
| chr4:49278991-49281006 | p5 | (GCAGT)3 | TCAAGCTTGGCATTCAACTG | TACCCCAGTGCATCCTTTTC | 199 |
| chr4:49291466-49293484 | p3 | (TTG)6 | CAGCAACATGGCAAGCTAAA | ACCAACTGCACCACTTCCTC | 261 |
| chr4:49291466-49293484 | p3 | (GTT)6 | CAGCAACATGGCAAGCTAAA | ACCAACTGCACCACTTCCTC | 261 |
| chr4:49291466-49293484 | p2 | (TA)6 | CAGCAACATGGCAAGCTAAA | ACCAACTGCACCACTTCCTC | 261 |
| chr4:49521851-49523865 | p2 | (GA)7 | CGAAAGTGAAAGCGAACACA | ACGTGCCTTTTTCATTGGTC | 236 |
| chr4:49533141-49535159 | p2 | (AG)9 | TTGAAGGGAATCGACCAAAG | TTCCGTCCTTTTGGAACATC | 244 |
| chr4:49553262-49555277 | p3 | (CAA)5 | ACCAAACACGAGGGAGTCAC | GGGTTGCATTTAGTCTCCCA | 228 |
| chr4:49568378-49570412 | p2 | (TA)14 | TACCCACACTGCGCATCTAC | TGAGAGCACTCCATTGTTCG | 180 |
| chr4:49568378-49570412 | p2 | (AT)17 | TACCCACACTGCGCATCTAC | TGAGAGCACTCCATTGTTCG | 180 |
| chr4:49568378-49570412 | p3 | (CAT)5 | TACCCACACTGCGCATCTAC | TGAGAGCACTCCATTGTTCG | 180 |
| chr4:49575591-49577603 | p2 | (AG)7 | CAGCTGTCACGACCCAACTA | GGCCTTGTCGGCCTATATTT | 228 |
| chr4:49575591-49577603 | p2 | (AT)6 | CAGCTGTCACGACCCAACTA | GGCCTTGTCGGCCTATATTT | 228 |
| chr4:49627829-49629859 | p2 | (AT)15 | TTGCAAAAGAACCTCCAAGG | TCGGCTTGCGTATCACATTA | 267 |
| chr4:49684355-49686369 | p2 | (TA)7 | CGTCATCGTCTACCCACCTT | TCCGAGTATTCGTCTGGGAC | 119 |
| chr4:49744838-49746850 | p2 | (AC)6 | CTGACTTGGATGGGTTTCGT | CCGGAAAATTATCGGTAGCA | 198 |
| chr4:49834220-49836236 | p2 | (AT)8 | AATTAACAAGCCTTGCCCCT | AGGAAGGTCAAGGGCAAAAT | 227 |
| chr4:49834502-49836528 | p2 | (AT)8 | ATTTTGCCCTTGACCTTCCT | CTAGGGTGTTATGCCCCGTA | 144 |
| chr4:49834537-49836587 | p2 | (AT)8 | ATTTTGCCCTTGACCTTCCT | CTAGGGTGTTATGCCCCGTA | 109 |
| chr4:49863929-49865947 | p2 | (AT)9 | AACAACGGAGCTCTCCAAGA | ATTCAAGATGATTGTCGCCC | 187 |

| Marker ID | SSR type | SSR motif | FORWARD PRIMER (5'-3') | REVERSE PRIMER (5'-3') | size (bp) |
|---|---|---|---|---|---|
| chr4:49899138-49901154 | p3 | (ACA)5 | TTCCCCTGAAAGCTTAGCAA | GTGGCAATATTTTGGGGTTG | 282 |
| chr4:49899138-49901154 | p2 | (CT)8 | TTCCCCTGAAAGCTTAGCAA | GTGGCAATATTTTGGGGTTG | 282 |
| chr4:49903872-49905887 | p5 | (ATATC)3 | CATCAACAACGCATGAAACC | AAGTCCCGTGGTGCTTGTAG | 192 |
| chr4:49949157-49951181 | p6 | (TATATG)4 | TGTTGAGGTGCTGTTTGAGG | GCAGCATTTCCCCTATGTGT | 110 |
| chr4:49954499-49956511 | p2 | (AT)6 | GGTACGTCTTGAGGTGCGAT | ATTTCTGGGAAAATTTGGGC | 107 |
| chr4:49962633-49964647 | p2 | (TA)7 | TATACGCTACCTCATCCCGC | GTGTCGTTCCGGTCGTACTT | 235 |
| chr4:50035614-50037626 | p2 | (AT)6 | ATGGTGTCCATGCGGATATT | AGCCACAACATTTGCCTTTC | 148 |
| chr4:50086043-50088057 | p2 | (AT)7 | TTCCACTATACGGCTCGTCC | CCATGCAAGGTTTCCACTTT | 228 |
| chr4:50098070-50100136 | p2 | (AT)33 | ATGCCCACAGGTAGACAAGG | GCCCTTTGGCTATCCACATA | 152 |
| chr4:50098070-50100136 | p2 | (AT)13 | ATGCCCACAGGTAGACAAGG | GCCCTTTGGCTATCCACATA | 152 |
| chr4:50129616-50131631 | p5 | (CAACA)3 | CATGGCGTTACTTGTGATGG | TGAATTTGGAGCAGCATACG | 144 |
| chr4:50130014-50132026 | p5 | (CAACA)3 | GCAGCATTTCCCCTTGTAAA | ATGATGACGTGCATGGTTGT | 112 |
| chr4:50202542-50204557 | p5 | (TTTAA)3 | TAGTAGCCTTGTCGGCTGGT | ACTTTTGTCACGACCCAACC | 225 |
| chr4:50202542-50204557 | p2 | (TA)11 | TAGTAGCCTTGTCGGCTGGT | ACTTTTGTCACGACCCAACC | 225 |
| chr4:50211137-50213169 | p2 | (TA)16 | CCTTTGCCCCTTAGCTTTTC | CATGGGACGTCTCCGATACT | 280 |
| chr4:50211137-50213169 | p3 | (GTT)5 | CCTTTGCCCCTTAGCTTTTC | CATGGGACGTCTCCGATACT | 280 |
| chr4:50281172-50283187 | p5 | (TTTCT)3 | AGTTCCTCTATTGCAGGCCA | TGCAAAACCCTAACCCTACG | 170 |
| chr4:50353197-50355215 | p6 | (TTTCTC)3 | TACAACATACGTCGGAGGCA | GTTGTGGGTGAATTATGCCC | 252 |
| chr4:50462025-50464040 | p5 | (GCGGG)3 | CCATGGCCCTAAGCATAGAA | CCCACCCCCGTACTAATTTT | 147 |
| chr4:50468388-50470402 | p2 | (AT)7 | AAAACACGAAAAGTGCCACC | ACTATCGCTCGCTTGCTCTC | 275 |
| chr4:50468388-50470402 | p2 | (AT)7 | AAAACACGAAAAGTGCCACC | ACTATCGCTCGCTTGCTCTC | 275 |
| chr4:50468388-50470402 | p2 | (TA)7 | AAAACACGAAAAGTGCCACC | ACTATCGCTCGCTTGCTCTC | 275 |
| chr4:50468388-50470402 | p2 | (TA)10 | AAAACACGAAAAGTGCCACC | ACTATCGCTCGCTTGCTCTC | 275 |
| chr4:50468766-50470780 | p2 | (AT)7 | GTTACGACGCTGTACCCCAT | GCCATGGTTGATCGCTTATT | 295 |
| chr4:50468766-50470780 | p2 | (TA)7 | GTTACGACGCTGTACCCCAT | GCCATGGTTGATCGCTTATT | 295 |
| chr4:50468766-50470780 | p2 | (TA)10 | GTTACGACGCTGTACCCCAT | GCCATGGTTGATCGCTTATT | 295 |
| chr4:50551731-50553746 | p2 | (AC)6 | TCGGCCAAATATCAACAACA | GATTTTGGAAGGTTTCGGGT | 122 |
| chr4:50563178-50565196 | p3 | (TGT)6 | AGGAATGAAGTGCCGAAATG | TGCCACCTGCCACATAGATA | 187 |
| chr4:50582775-50584789 | p2 | (TA)7 | CTCCCATATCTTGAAGGCCA | TTCCCACAATTCCCTACCAA | 160 |
| chr4:50785897-50787923 | p2 | (AT)13 | ATAAGATGCCAACTCCACGG | AGTGCCAGAGCCAGTGATTT | 169 |
| chr4:50790980-50792998 | p6 | (GGTAGA)3 | CTCCGAAATTCTTTGGGTCA | GCCATCCATCCGTATACCAC | 172 |
| chr4:50868367-50870381 | p5 | (TATAC)3 | TAAACAAGCACGACCCATGA | GTGCTCCTTTGATCCGTAGC | 202 |
| chr4:50868367-50870381 | p2 | (AT)7 | TAAACAAGCACGACCCATGA | GTGCTCCTTTGATCCGTAGC | 202 |
| chr4:50997877-50999904 | p2 | (TC)7 | TTCCTTCACCGTCATCATCA | TAAAAGGCAAGAGGGCTTGA | 205 |
| chr4:51024149-51026169 | p3 | (ATG)5 | CACGACCTCATGTCACCTTG | ATAATCACAAGCAGTCCGCC | 275 |
| chr4:51024149-51026169 | p2 | (TA)10 | CACGACCTCATGTCACCTTG | ATAATCACAAGCAGTCCGCC | 275 |

| Marker ID | SSR type | SSR motif | FORWARD PRIMER (5'-3') | REVERSE PRIMER (5'-3') | size (bp) |
|---|---|---|---|---|---|
| chr4:51028684-51030699 | p3 | (TCA)5 | GGAACTGTTTGCAAAGCCTC | ATTTGCATGCCCGTAGTAGG | 173 |
| chr4:51090398-51092414 | p2 | (TA)8 | GTTCGGGACTGGAGTTCAAA | TTATTCCTCATGTCTCCGCC | 179 |
| chr4:51110733-51112748 | p5 | (ATGTT)3 | ACCTCTTCGGGCGTTCTATT | TTTAATCTCTGGCCCTCCCT | 251 |
| chr4:51113818-51115834 | p2 | (CT)8 | CCAACGCACAGAAAACTGAA | GGGTTACGAATAGCCACCAA | 174 |
| chr4:51350714-51352735 | p3 | (ATC)7 | GTCAAGCTCACCTCTCGGTC | GGAGGCCGATATCACCACTA | 228 |
| chr4:51371440-51373458 | p2 | (AC)9 | TCTGGTACTAGCTGCCCGAT | AGAAGAGGGCGAGTCACAAA | 189 |
| chr4:51371440-51373458 | p3 | (TCA)6 | TCTGGTACTAGCTGCCCGAT | AGAAGAGGGCGAGTCACAAA | 189 |
| chr4:51372335-51374353 | p2 | (AC)9 | TGTCTACCTGTGGGCATGAA | ACTAGTGGGCGGCATATGAG | 231 |
| chr4:51372335-51374353 | p3 | (TCA)6 | TGTCTACCTGTGGGCATGAA | ACTAGTGGGCGGCATATGAG | 231 |
| chr4:51379913-51381977 | p2 | (TA)7 | CAACGTGAACTTCCAGCAAA | AACCCGTACCGTAGTCGATG | 126 |
| chr4:51379913-51381977 | p2 | (TA)7 | CAACGTGAACTTCCAGCAAA | AACCCGTACCGTAGTCGATG | 126 |
| chr4:51491988-51494000 | p2 | (AT)9 | TACACAACCCACGCATGTCT | CTTGGAGAGCTCCGTTGTTC | 118 |
| chr4:51491988-51494000 | p2 | (AT)14 | TACACAACCCACGCATGTCT | CTTGGAGAGCTCCGTTGTTC | 118 |
| chr4:51491988-51494000 | p2 | (TA)6 | TACACAACCCACGCATGTCT | CTTGGAGAGCTCCGTTGTTC | 118 |
| chr4:51536610-51538622 | p2 | (TA)6 | TTTGGACGCAAGAATTAGGC | GGGGCTAAGCCTTCATTTTC | 174 |
| chr4:51549214-51551228 | p5 | (ATTCA)3 | CCAACAATCATGCCAACAAC | TTGAATTTGGAAGGTTTCGG | 174 |
| chr4:51549214-51551228 | p2 | (AT)7 | CCAACAATCATGCCAACAAC | TTGAATTTGGAAGGTTTCGG | 174 |
| chr4:51630136-51632151 | p3 | (AAC)5 | TATAACGCCACACCTTTCCC | TGATACTTGGTCTTTCGGGG | 201 |
| chr4:51649734-51651746 | p5 | (TTTAT)3 | TATTGTGGATAGCCGAAGGG | CTTTTTGTCACGACCCGACT | 128 |
| chr4:51777013-51779027 | p2 | (AT)7 | ACTGGCCTGAATGCCTAGAA | AGGTTTGTGAGTGGTCTGGG | 278 |
| chr4:51807963-51809978 | p5 | (ATTCA)3 | TCGTGGAATCATTCTTGTCG | ACCTCAATCGGACAAAGGTG | 247 |
| chr4:51807963-51809978 | p3 | (AAC)6 | TCGTGGAATCATTCTTGTCG | ACCTCAATCGGACAAAGGTG | 247 |
| chr4:51853092-51855116 | p2 | (TA)12 | AATCATGCATGTTGGGGTTT | TTAAAGAAACGGCACCGAAG | 223 |
| chr4:51854041-51856056 | p2 | (TA)12 | ATGGGACCAATGGTATCGTG | TACGGACGTTTTATGGCACA | 165 |
| chr4:51854077-51856092 | p2 | (TA)12 | ATGGGACCAATGGTATCGTG | TACGGACGTTTTATGGCACA | 129 |
| chr4:51870451-51872466 | p3 | (CTT)5 | TCCCCACTCAACGTTCTACC | TGTACGCGTCGCTAATGAAG | 204 |
| chr4:51876425-51878441 | p2 | (AT)8 | TACGGAGGATTTCCAAGGTG | GCAATGGAAGGATTTGAGGA | 164 |
| chr4:51876425-51878441 | p2 | (AT)16 | TACGGAGGATTTCCAAGGTG | GCAATGGAAGGATTTGAGGA | 164 |
| chr4:51945299-51947314 | p5 | (CATTT)3 | GGGAAGACATGACTTGGGAA | TTTCCCCTGTAACTTGGCAC | 145 |
| chr4:52017783-52019803 | p2 | (TA)10 | CAGAACCTATTAGCTGCGCC | AGTCCACCACCACCTCAGTC | 211 |
| chr4:52043085-52045105 | p2 | (TA)10 | ACCATCTGATTTGGGAGCAG | ATTCATCCCTTCTTCGGCTT | 282 |
| chr4:52060362-52062374 | p2 | (AC)6 | CTCACAACCCTGAGTGACGA | AGCCTTATTCTTTGCGGGAT | 118 |
| chr4:52169811-52171827 | p2 | (AT)8 | GCATATACACGGGGCTGAGT | GGTTCAGCAGCTGGTTCTTC | 176 |
| chr4:52175158-52177173 | p5 | (ATGAT)3 | TTTGAAGGAAACGGGATTTG | TGTCTCAACAAGGCACGAAG | 205 |
| chr4:52217810-52219826 | p4 | (AAGA)4 | CGTGCCACAGAGCTAAATCA | AGAAAACACATCCAATGCCC | 264 |
| chr4:52269151-52271165 | p2 | (TG)7 | ATCAGGTTCAGACGCTTGCT | AAGATCTTAGCGTCCAGCCA | 156 |

| Marker ID | SSR type | SSR motif | FORWARD PRIMER (5'-3') | REVERSE PRIMER (5'-3') | size (bp) |
|---|---|---|---|---|---|
| chr4:52428584-52430596 | p2 | (AT)6 | GTCGAGGCCAACCAAAGATA | GCAGGATTTTGGGGTACTCA | 205 |
| chr4:52444805-52446821 | p6 | (CATACA)3 | CTCAATTTCCGACGGTTGTT | TTTCGGTGTGCTCAAGACAG | 104 |
| chr4:52444805-52446821 | p4 | (TCTA)4 | CTCAATTTCCGACGGTTGTT | TTTCGGTGTGCTCAAGACAG | 104 |
| chr4:52506714-52508732 | p3 | (CAA)6 | CAAAGGATACTCCCAACGGA | TCCATTTGACTTATTCCCGC | 256 |
| chr4:52514918-52516930 | p2 | (TA)6 | AAAGCGGAACAACCAAACAC | TTTTATTCCGAATGGTTCGC | 154 |
| chr4:52514936-52516948 | p2 | (TA)6 | AAAGCGGAACAACCAAACAC | TTTTATTCCGAATGGTTCGC | 136 |
| chr4:52514949-52516963 | p2 | (TA)6 | AAAGCGGAACAACCAAACAC | TTTTATTCCGAATGGTTCGC | 123 |
| chr4:52514964-52517002 | p2 | (TA)6 | AAAGCGGAACAACCAAACAC | TTTTATTCCGAATGGTTCGC | 108 |
| chr4:52515182-52517194 | p2 | (TA)6 | ACGACTGCTGTACGATGCTG | GCATTGTTTTTCCATGCCTT | 174 |
| chr4:52518574-52520589 | p5 | (ATTTC)3 | TGCTCTTAATCTCCCGCAAT | AGTGGAAACTTGCCTGCTGT | 114 |
| chr4:52567715-52569730 | p3 | (TTG)5 | AAGCGGTTCTGCTCCAGTAA | AATGTCAAAAGGCCACAACC | 254 |
| chr4:52617766-52619781 | p5 | (CATAT)3 | CTCTGGGGAAAGGTCAACAA | TCCGAAAGAAAGATGGATCG | 105 |
| chr4:52624469-52626481 | p2 | (AT)6 | GCCCTGATGGGTTGAGATTA | ATGGTTGTTTTGGGTGTGGT | 154 |
| chr4:52661177-52663189 | p2 | (AT)6 | TGGGTTGTGTTGTGTGTGTG | TATGCACCATACTCCACCGA | 278 |
| chr4:52696973-52699003 | p2 | (CA)15 | TGGAAACAAGCCAAATCACA | GGTGCTTGATCCGTAGTGGT | 167 |
| chr4:52701161-52703173 | p2 | (AC)6 | TTACCCAAATGGGACCAAAA | CATTTGCAGTTTCTGGACGA | 116 |
| chr4:52742667-52744682 | p5 | (ATCAT)3 | GGAACGTACACCCCGATCTA | TTTTCGGGTACGTAAGGTGC | 126 |
| chr4:52785711-52787729 | p4 | (TATG)4 | TAGACTCCCGCAATCTCGTT | CACTTTTGGGACTCGGTGTT | 191 |
| chr4:52785711-52787729 | p2 | (GA)8 | TAGACTCCCGCAATCTCGTT | CACTTTTGGGACTCGGTGTT | 191 |
| chr4:52785731-52787747 | p4 | (TATG)4 | TAGACTCCCGCAATCTCGTT | CACTTTTGGGACTCGGTGTT | 171 |
| chr4:52785731-52787747 | p2 | (GA)8 | TAGACTCCCGCAATCTCGTT | CACTTTTGGGACTCGGTGTT | 171 |
| chr4:52908003-52910049 | p2 | (AT)23 | GATGCGTAATCGGGTTCCTA | GAGGAGGAAGAAACCCAAGG | 112 |
| chr4:52941130-52943145 | p5 | (AAATA)3 | AATTCCAAAGTTGTGGCGTC | TTTAAGGAACCGAAACGCAC | 116 |
| chr4:53165493-53167589 | p5 | (GTTCG)3 | TGTTCATGCCCACAGGTAGA | CCTTGGGATACCACACAACC | 294 |
| chr4:53165493-53167589 | p2 | (AC)7 | TGTTCATGCCCACAGGTAGA | CCTTGGGATACCACACAACC | 294 |
| chr4:53165607-53167699 | p5 | (GTTCG)3 | TGTTCATGCCCACAGGTAGA | CCTTGGGATACCACACAACC | 180 |
| chr4:53165607-53167699 | p2 | (AC)7 | TGTTCATGCCCACAGGTAGA | CCTTGGGATACCACACAACC | 180 |
| chr4:53241227-53243242 | p6 | (CATATA)3 | ATCGGAGTGACGTAAGGTCG | CCGAAACTTGTTTCTGAGCC | 247 |
| chr4:53241227-53243242 | p5 | (ATTCA)3 | ATCGGAGTGACGTAAGGTCG | CCGAAACTTGTTTCTGAGCC | 247 |
| chr4:53300990-53303006 | p2 | (AT)8 | GACTACCCATCAACGCAGGT | AGCCCGATTAAGATCTCGGT | 300 |
| chr4:53384970-53386984 | p2 | (GT)7 | AGGGAGAGTAGAGGGGTTCG | TACCACGTTGAAGAGGACCC | 127 |
| chr4:53388787-53390802 | p5 | (ATGAT)3 | GATCGGGACTTGAGTTTGGA | GGGTCGTAACTATGCTCGGA | 165 |
| chr4:53509963-53511975 | p2 | (AT)6 | CGGCACAGGTATGTAAGGCT | CATGAAGGTTTCAGGGCAAT | 162 |
| chr4:53509963-53511975 | p3 | (TAA)28 | CGGCACAGGTATGTAAGGCT | CATGAAGGTTTCAGGGCAAT | 162 |
| chr4:53556428-53558440 | p2 | (AT)6 | ACATATTGACAGGCGAAGCC | ACATTGCTCGTACTGACCCC | 270 |
| chr4:53557132-53559152 | p2 | (AT)6 | TGCATGCTTATCGGGTCATA | ACACATGACTTTCCTTGGGC | 128 |

| Marker ID | SSR type | SSR motif | FORWARD PRIMER (5'-3') | REVERSE PRIMER (5'-3') | size (bp) |
|---|---|---|---|---|---|
| chr4:53557132-53559152 | p5 | (TCATA)4 | TGCATGCTTATCGGGTCATA | ACACATGACTTTCCTTGGGC | 128 |
| chr4:53733209-53735243 | p2 | (AT)17 | CAACGCAGCGTACAACACTT | CAACCATAGGTTGTGGGTCC | 124 |
| chr4:53754612-53756630 | p2 | (TA)7 | ACTTCAGAGCCGACAAGGAA | ATATGGCCACAGACTGCTCC | 288 |
| chr4:53754632-53756656 | p2 | (TA)7 | ACTTCAGAGCCGACAAGGAA | ATATGGCCACAGACTGCTCC | 268 |
| chr4:53754775-53756789 | p2 | (TA)7 | ACTTCAGAGCCGACAAGGAA | ATATGGCCACAGACTGCTCC | 125 |
| chr4:53758419-53760457 | p2 | (AT)19 | GTGCACTAAGGGTCAGAGGG | TCCAGTCCGCGAGTTCTACT | 188 |
| chr4:53784637-53786667 | p2 | (AT)15 | TGCTTTGTTAGCCCTGTGTG | GCCGACAAGGCTATATTCCA | 121 |
| chr4:53801170-53803194 | p2 | (AT)12 | CCATTGGAAAACGCATACCT | AGGCCTTGTCGGCCTATATT | 253 |
| chr4:53925315-53927327 | p2 | (AG)6 | TGAATAGTTATGGCACGGCA | TTCGTCCTTGAATGTTGCAG | 174 |
| chr4:53934491-53936503 | p6 | (AACAAT)3 | ACGCCGTTATACGATTTTGC | TGGAAACTTCCCCACACTTC | 293 |
| chr4:53934491-53936503 | p2 | (AT)6 | ACGCCGTTATACGATTTTGC | TGGAAACTTCCCCACACTTC | 293 |
| chr4:54029051-54031065 | p2 | (AT)7 | GACATCTCTCATCGCCCTGT | TCAGAGGAGCCTAGAGCAGC | 124 |
| chr4:54080471-54082483 | p2 | (AC)6 | CTTGCAACTAGGACATGCCA | AGGGGAGATGCTACCGAAAT | 264 |
| chr4:54263749-54265767 | p6 | (TATTGT)3 | TCCCTTGATGTTTTGGAAGG | GCATTCCGGCATTTCTTAAA | 117 |
| chr4:54279719-54281734 | p2 | (TA)6 | TGAAAATGTTGATGCGGAAA | AAAATTTTATGGGCTTGGGC | 227 |
| chr4:54279719-54281734 | p3 | (TTG)5 | TGAAAATGTTGATGCGGAAA | AAAATTTTATGGGCTTGGGC | 227 |
| chr4:54434341-54436356 | p5 | (ATCAT)3 | CCTGTCACAGCATAGCCAAA | AGGTCAGGAGCTGGGGTAAT | 247 |
| chr4:54450093-54452105 | p2 | (CA)6 | GACGATCCCATAACGCTGAT | ACGTAAGGTCGTAAACCCCC | 177 |
| chr4:54450093-54452105 | p2 | (AT)7 | GACGATCCCATAACGCTGAT | ACGTAAGGTCGTAAACCCCC | 177 |
| chr4:54484291-54486306 | p2 | (AT)7 | TGATGCCATGAGTTGTTGGT | GTACCACAGGGGGTACGCTA | 267 |
| chr4:54484291-54486306 | p2 | (TA)14 | TGATGCCATGAGTTGTTGGT | GTACCACAGGGGGTACGCTA | 267 |
| chr4:54484375-54486390 | p2 | (AT)7 | TGATGCCATGAGTTGTTGGT | GTACCACAGGGGGTACGCTA | 183 |
| chr4:54484375-54486390 | p2 | (TA)14 | TGATGCCATGAGTTGTTGGT | GTACCACAGGGGGTACGCTA | 183 |
| chr4:54614733-54616773 | p2 | (AT)20 | AATATGGCCACAGGCTGTTC | GTCGACATGAAGGGAAAGGA | 178 |
| chr4:54805501-54807516 | p5 | (TTCAT)3 | CAAGAAAACCCAATCGCCTA | ATGGTTTGATGTTGGGGAAA | 222 |
| chr4:54821677-54823695 | p2 | (AT)9 | CCACATCCTATGACCTGGCT | AGCTTCGGGTTCTCAGCATA | 138 |
| chr4:54846026-54848041 | p5 | (TTCAT)3 | CAAGAAAACCCAATCGCCTA | ATGGTTTGATGTTGGGGAAA | 222 |
| chr4:54862202-54864220 | p2 | (AT)9 | CCACATCCTATGACCTGGCT | AGCTTCGGGTTCTCAGCATA | 138 |
| chr4:54973978-54975992 | p2 | (AG)7 | TGTCACTCTCCTCGGTTCCT | TCCGTCTGACTCTGAACACG | 282 |
| chr4:54973978-54975992 | p2 | (GT)9 | TGTCACTCTCCTCGGTTCCT | TCCGTCTGACTCTGAACACG | 282 |
| chr4:54988957-54990973 | p2 | (AC)8 | GGCCGAGCCTATTATGTTCA | TTTCTGGGTTGCTGTGTTGA | 148 |
| chr4:55053006-55055020 | p2 | (AT)7 | TTTTTCACGACCCGACTAGG | GAGGTTGTTGTCCAAAGGGA | 181 |
| chr4:55077386-55079402 | p2 | (GT)8 | TGCAGACTGCTATGACCCAG | AACATCATCGCCCTTTTCTG | 174 |
| chr4:55175240-55177252 | p2 | (AT)6 | TAGCTTGGGTGAGGGAAGAA | AGCATCGTTCAAACACCTCC | 184 |
| chr4:55217431-55219446 | p2 | (AT)13 | GGCAAAAAGGGAAATTCACA | TTGATCAATTTCCATGGGGT | 107 |
| chr4:55217431-55219446 | p5 | (TGAAT)3 | GGCAAAAAGGGAAATTCACA | TTGATCAATTTCCATGGGGT | 107 |

| Marker ID | SSR type | SSR motif | FORWARD PRIMER (5'-3') | REVERSE PRIMER (5'-3') | size (bp) |
|---|---|---|---|---|---|
| chr4:55217431-55219446 | p6 | (TATATG)4 | GGCAAAAAGGGAAATTCACA | TTGATCAATTTCCATGGGGT | 107 |
| chr4:55218398-55220422 | p5 | (TGAAT)3 | GTTGTTGGCATGATTGTTGG | GCTTAATTGTCCAAGCTCGC | 120 |
| chr4:55218398-55220422 | p6 | (TATATG)4 | GTTGTTGGCATGATTGTTGG | GCTTAATTGTCCAAGCTCGC | 120 |
| chr4:55283787-55285801 | p2 | (CT)7 | AGGGGTCAGTACGAGCAATG | TCTCATGAATGCTCAGGCAG | 103 |
| chr4:55448994-55451036 | p2 | (AT)21 | AAACTCATGAAATCACGGGC | AAGGCGGGTAGAAAGAGAGC | 276 |
| chr4:55466814-55468835 | p3 | (CAA)7 | TACAATTTGGAGGCCAAAGG | CGACGGGTCAGTGAGATTTT | 263 |
| chr4:55518390-55520404 | p2 | (TA)7 | AGGTCGCAGCTTATCCTTCA | AACCTTTGCCTCAGCAGAAA | 216 |
| chr4:55530017-55532053 | p2 | (TA)6 | GCCTCATTAGGCCATCAAAA | GGCATGACGGAGTCTTGTTT | 240 |
| chr4:55530017-55532053 | p2 | (AT)18 | GCCTCATTAGGCCATCAAAA | GGCATGACGGAGTCTTGTTT | 240 |
| chr4:55532396-55534410 | p2 | (TA)6 | AGAGAGATGTTGGGTCGTGG | AAAGGGAAGTCACTGGAGCA | 208 |
| chr4:55532396-55534410 | p2 | (AT)7 | AGAGAGATGTTGGGTCGTGG | AAAGGGAAGTCACTGGAGCA | 208 |
| chr4:55651236-55653258 | p2 | (TA)16 | GACGATGATGATGTTGACGC | AGATATCATACCCGGGCACA | 246 |
| chr4:55651236-55653258 | p2 | (TA)11 | GACGATGATGATGTTGACGC | AGATATCATACCCGGGCACA | 246 |
| chr4:55657475-55659539 | p2 | (AT)6 | CCTAGGACGGAACTGCTCTG | TAGCCTTGTCGGCTCGTATT | 158 |
| chr4:55657475-55659539 | p2 | (AT)32 | CCTAGGACGGAACTGCTCTG | TAGCCTTGTCGGCTCGTATT | 158 |
| chr4:55702016-55704066 | p2 | (TA)25 | GATGAAACTCCCGTCTGCAT | AAAAAGAGAGGCGATGATGG | 234 |
| chr4:55702016-55704066 | p2 | (TA)21 | GATGAAACTCCCGTCTGCAT | AAAAAGAGAGGCGATGATGG | 234 |
| chr4:55732281-55734299 | p3 | (ATA)6 | GAGCGGTGTATGGGAAGAAA | TGCCTGGTTTTAGTCGTTCC | 158 |
| chr4:55799856-55801904 | p3 | (TAT)16 | CGTAACATGGCCAAAGAGGT | CAACGGATCCGATCTTCAGT | 204 |
| chr4:55892636-55894651 | p3 | (AAC)5 | GACGTTGTGGTATGCACAGG | GACGGAACAAAACTCCTCCA | 281 |
| chr4:56034484-56036499 | p2 | (TA)8 | GCCATAAGGAAGCAATAGCG | TGACCCTCTTTTTACGTCGC | 286 |
| chr4:56034484-56036499 | p5 | (TTTTA)3 | GCCATAAGGAAGCAATAGCG | TGACCCTCTTTTTACGTCGC | 286 |
| chr4:56103160-56105174 | p2 | (TG)7 | CACCCTCCTTTTCTTCCACA | ATTTGAAGGACAAACCGTGC | 252 |
| chr4:56344598-56346614 | p2 | (TA)8 | CTGGATTGTGATTTTGGGCT | CGCTTATTCTCCCGAACTTG | 222 |
| chr4:56536978-56539004 | p2 | (AT)13 | ACCGGAGTAAGCGTCAAGAA | TCCAGGAGGTCAAGTAACCG | 254 |
| chr4:56536978-56539004 | p2 | (CT)6 | ACCGGAGTAAGCGTCAAGAA | TCCAGGAGGTCAAGTAACCG | 254 |
| chr4:56552135-56554160 | p5 | (AAGAG)5 | AAATCTTCCCAAAAACGCCT | GCTAAGGATGTGGACGGGTA | 154 |
| chr4:56695480-56697520 | p4 | (CATA)10 | TATACAAGTCGGGACAGGGC | ATCGTTCGTACTGACGTCCC | 129 |
| chr4:56695480-56697520 | p3 | (ATC)7 | TATACAAGTCGGGACAGGGC | ATCGTTCGTACTGACGTCCC | 129 |
| chr4:56695480-56697520 | p3 | (CAA)6 | TATACAAGTCGGGACAGGGC | ATCGTTCGTACTGACGTCCC | 129 |
| chr4:56731681-56733696 | p2 | (AT)36 | AGAGTTTGGGTAGCAAGGCA | GCACGGCTTAGGCAAAATAG | 149 |
| chr4:56731681-56733696 | p5 | (AATGC)3 | AGAGTTTGGGTAGCAAGGCA | GCACGGCTTAGGCAAAATAG | 149 |
| chr4:56750356-56752368 | p2 | (AG)6 | GAGCAGACACCATCCGATCT | AAGGGTACATCTTTGGTGCG | 205 |
| chr4:56894048-56896110 | p2 | (AT)31 | GCGGGGTGTGAGAGGTATTA | AGAAAAGCTACAGTCGCCCA | 121 |
| chr4:56936426-56938441 | p3 | (CAT)5 | ACCGAGACACGAAGAGAGGA | CTAAGCGGTTCCATTTTCCA | 157 |
| chr4:56940510-56942533 | p4 | (TTTA)4 | GCGGGGTTAATTCCCTACTC | TGGAGGGCAAGTGGATTAAG | 231 |

| Marker ID | SSR type | SSR motif | FORWARD PRIMER (5'-3') | REVERSE PRIMER (5'-3') | size (bp) |
|---|---|---|---|---|---|
| chr4:56979320-56981335 | p5 | (TTTCA)3 | GGCACACCCATCCATAGAGT | CAATCCAAAACGCAGACAAA | 189 |
| chr4:56999621-57001637 | p4 | (TTAA)4 | CCGGATGAGTGTCCATCTCT | ATCCCTGCCTTAAGAACGGT | 101 |
| chr4:57140703-57142735 | p2 | (TA)16 | GGGGGTTTTGCTTCAATTTT | ATTTAGGCCAAATCCAACCC | 166 |
| chr4:57140703-57142735 | p5 | (TGAAT)3 | GGGGGTTTTGCTTCAATTTT | ATTTAGGCCAAATCCAACCC | 166 |
| chr4:57153552-57155568 | p2 | (AT)8 | GAGCTGAAACTTCATGCCAAC | TCCCATTCAGCAAAAACAAA | 248 |
| chr4:57247066-57249130 | p2 | (AT)32 | CCCATGTTTGGACCAATTTC | CCACCATCTGGGAAAAAGAA | 175 |
| chr4:57284551-57286566 | p5 | (TTTAT)3 | AAAGGAAGGTTGTTGCAGGA | GTCAAAAGTTGTCCCCGAAA | 160 |
| chr4:57362919-57364933 | p2 | (CT)7 | CACAGTCCAAAACAGCTCCA | CTAGCCACTTGGCATTTGGT | 271 |
| chr4:57362919-57364933 | p2 | (TA)24 | CACAGTCCAAAACAGCTCCA | CTAGCCACTTGGCATTTGGT | 271 |
| chr4:57363805-57365853 | p2 | (CT)7 | AGACAGTTGAGGCAACCACC | ACCCCCACAAAACCATCATA | 239 |
| chr4:57363805-57365853 | p2 | (TA)24 | AGACAGTTGAGGCAACCACC | ACCCCCACAAAACCATCATA | 239 |
| chr4:57378103-57380131 | p2 | (AT)14 | TAAGATTTTGGCGGATTTGC | GTCGGGTCCTATTTATGCGA | 147 |
| chr4:57398360-57400386 | p3 | (CAT)5 | CGAGCACCGCTAATCATACA | TAGACGATGTATGGCCTCCC | 184 |
| chr4:57398360-57400386 | p2 | (AT)13 | CGAGCACCGCTAATCATACA | TAGACGATGTATGGCCTCCC | 184 |
| chr4:57436016-57438044 | p2 | (TA)21 | CATCTGGTAGGGGAACTGGA | AAAAGCTACAGCAGGCCAAA | 228 |
| chr4:57436016-57438044 | p2 | (AT)14 | CATCTGGTAGGGGAACTGGA | AAAAGCTACAGCAGGCCAAA | 228 |
| chr4:57442484-57444502 | p3 | (CAT)6 | ACAGGAGGCTTGCAGAAAAA | TGTCTTTCAGGCTCATCCCT | 203 |
| chr4:57448518-57450530 | p2 | (AT)6 | TTGGAGAGCTCCATTGTTCC | ATATACGCAAGCCGACAAGG | 212 |
| chr4:57448518-57450530 | p2 | (AC)16 | TTGGAGAGCTCCATTGTTCC | ATATACGCAAGCCGACAAGG | 212 |
| chr4:57501624-57503636 | p5 | (AAAGG)3 | GGCTCAAAGGAAAGGAAAGG | AACCCAACATCATACCCGAA | 260 |
| chr4:57501624-57503636 | p2 | (TA)6 | GGCTCAAAGGAAAGGAAAGG | AACCCAACATCATACCCGAA | 260 |
| chr4:57554822-57556843 | p3 | (TGT)7 | TGACTGTAAAGGCGATGTGC | TATACGTAGATCGCCCCTGG | 193 |
| chr4:57590927-57592979 | p2 | (TA)26 | TCATATCTGCCACCACGGTA | CGCATAGACTTCGGGATGTT | 155 |
| chr4:57644077-57646098 | p3 | (TGT)7 | TGACTGTAAAGGCGATGTGC | TATACGTAGATCGCCCCTGG | 193 |
| chr4:57726043-57728061 | p6 | (TGAACA)3 | CGCTTGAGGACCCTATTCTG | TGGATGTCAAGGGAAAGACC | 182 |
| chr4:57726043-57728061 | p2 | (AT)8 | CGCTTGAGGACCCTATTCTG | TGGATGTCAAGGGAAAGACC | 182 |
| chr4:57755443-57757463 | p2 | (TA)10 | AATTAGGCAGCATTTCCCCT | TAGGAAGGGCTGTTGGATTG | 285 |
| chr4:57856713-57858731 | p6 | (TATATG)3 | GGACGTCGTGTGTGTTATGG | TTGCACGTTCTCTTTCAACG | 226 |
| chr4:57881539-57883587 | p2 | (TA)16 | GCCAAAGAATGCCCTGAATA | TAGATATGCCTGGCCTACGG | 225 |
| chr4:57881539-57883587 | p3 | (ACA)16 | GCCAAAGAATGCCCTGAATA | TAGATATGCCTGGCCTACGG | 225 |
| chr4:57891625-57893637 | p2 | (AT)6 | GGCAGCGAAATTGGTTAGAG | ACCATAACAATGCTGGGCTC | 290 |
| chr4:58116992-58119062 | p2 | (TA)35 | GGAGGGCACCTCTAGGATTC | GGGGTCGCCTACAAGTGATA | 300 |
| chr4:58298795-58300811 | p2 | (GA)6 | CCGCATTTTCAGAGCCTAAG | CCCACCTCCGCTTATGACTA | 137 |
| chr4:58298795-58300811 | p4 | (TATT)4 | CCGCATTTTCAGAGCCTAAG | CCCACCTCCGCTTATGACTA | 137 |
| chr4:58305090-58307114 | p6 | (TACATA)4 | ACCCGCTGACTATTATTGCG | AACAGAGTCGTGGGTATCGG | 286 |
| chr4:58305090-58307114 | p5 | (ATTCA)3 | ACCCGCTGACTATTATTGCG | AACAGAGTCGTGGGTATCGG | 286 |

| Marker ID | SSR type | SSR motif | FORWARD PRIMER (5'-3') | REVERSE PRIMER (5'-3') | size (bp) |
|---|---|---|---|---|---|
| chr4:58323343-58325375 | p2 | (TA)16 | GCCCTTTGGCTATCAAAACA | TGTTCATGCCCACAGGTAAA | 269 |
| chr4:58327950-58329965 | p5 | (GAAAT)3 | TTTACACCACAAGGCCACAA | AGTCGCTTGTTTCGTCGATT | 230 |
| chr4:58429667-58431682 | p5 | (GATAT)3 | TGTTCGTTTGTAGTGGCAGC | CCCAAGCTGGACACTCGTAT | 214 |
| chr4:58511547-58513562 | p5 | (TTTCA)3 | CTTATTCCAACCGCCAGAAG | TAAAATTTGGCCACACACGA | 234 |
| chr4:58536631-58538665 | p2 | (AT)17 | TAGAAGTGGGCAGCAATGTG | ACAGCCCATAAGCCAACAAC | 187 |
| chr4:58688296-58690308 | p2 | (TA)6 | TTTTGGGGTGAGCGTAAGAC | GTTTCGAGGAGCAGAATTGC | 114 |
| chr4:58786398-58788414 | p2 | (AT)8 | AAATGCGATAAACGTGGCTC | AAAGAAAACGCCCTTCACCT | 176 |
| chr4:58799399-58801414 | p2 | (AT)26 | CGTACATAACGCCTCACCCT | TTGGAGGTCACCAGCCTTAC | 280 |
| chr4:58799399-58801414 | p3 | (CAA)5 | CGTACATAACGCCTCACCCT | TTGGAGGTCACCAGCCTTAC | 280 |
| chr4:58875402-58877420 | p5 | (TTGTC)3 | AGATGAAAGTGGCGGAAATG | CGCATCTTGTCTTCCACTGA | 283 |
| chr4:58875402-58877420 | p3 | (TTG)6 | AGATGAAAGTGGCGGAAATG | CGCATCTTGTCTTCCACTGA | 283 |
| chr4:58891153-58893168 | p5 | (TAAAA)3 | CATAGCGTTGAGGTCCCATT | ACCCTGATCGTAGCCTTGTG | 173 |
| chr4:58948948-58950981 | p3 | (TAA)11 | ATGAATGTGGGAAAAGTGCC | CGTCATTATCCAATCCCCAC | 118 |
| chr4:58952068-58954080 | p2 | (AT)6 | TTGCCACATCCCACAGTCTA | CCCGTGTTCGTGATTTTCTT | 141 |
| chr4:59100173-59102185 | p2 | (TA)6 | CAGTTCTAGGGGGAACCACA | ACCAAGGTGGACTCCAAGTG | 138 |
| chr4:59231310-59233322 | p2 | (AT)6 | TCAATAAAACCTTGCGGGTC | GGCACAGCGGAAATGTAAGT | 238 |
| chr4:59247805-59249820 | p5 | (TTCAT)3 | GGGTCGTGTGTGGCTAAACT | ATTCACTGTGCATCAACGGA | 292 |
| chr4:59267539-59269557 | p6 | (TCCGTA)3 | TTAATGGCCTTGCTATTCGG | TTAGCCTCGTCGGCTTATGT | 297 |
| chr4:59328573-59330588 | p5 | (ATTAA)3 | AATTAGCCCCGTCAATTTCC | TTGCGTGTCGTCTATTCCAG | 141 |
| chr4:59345511-59347526 | p5 | (GGGCC)3 | TGGAGAGCCCGTAAAATGAC | CTTTCCAAACAGACCCCAAA | 160 |
| chr4:59345511-59347526 | p5 | (CCGGG)3 | TGGAGAGCCCGTAAAATGAC | CTTTCCAAACAGACCCCAAA | 160 |
| chr4:59381564-59383588 | p5 | (ATGAT)3 | AGGTTTAAGGGTTCGTGGCT | CCACAACCTCTAAGGTGGGA | 170 |
| chr4:59381564-59383588 | p4 | (CATT)6 | AGGTTTAAGGGTTCGTGGCT | CCACAACCTCTAAGGTGGGA | 170 |
| chr4:59687794-59689812 | p6 | (TTTCAT)3 | TACATCGCCAATGAGACCAA | TAATTCTGCCTCACAGGGCT | 222 |
| chr4:59705689-59707703 | p2 | (AC)7 | CCCCCTTTTGATTCACTTCA | TAACCCTAATCCCCGGAATC | 113 |
| chr4:59719032-59721050 | p2 | (AT)9 | AAACAAGTATGCAGGCCCAC | TTTGGAAAACAGTCTTGCCC | 130 |
| chr4:59719032-59721050 | p2 | (TG)9 | AAACAAGTATGCAGGCCCAC | TTTGGAAAACAGTCTTGCCC | 130 |
| chr4:59719032-59721050 | p2 | (AT)8 | AAACAAGTATGCAGGCCCAC | TTTGGAAAACAGTCTTGCCC | 130 |
| chr4:59719155-59721171 | p2 | (AT)9 | GATAGGGGCAAAATGGGAAT | GCTTTGGTCCACACACACAC | 233 |
| chr4:59719155-59721171 | p2 | (TG)9 | GATAGGGGCAAAATGGGAAT | GCTTTGGTCCACACACACAC | 233 |
| chr4:59719155-59721171 | p2 | (AT)8 | GATAGGGGCAAAATGGGAAT | GCTTTGGTCCACACACACAC | 233 |
| chr4:59733633-59735651 | p6 | (TATATG)3 | GTTGTTGTTGTGGTTGGCTG | ACGCTTATGTGTCCGAGCTT | 200 |
| chr4:59781292-59783310 | p3 | (CAA)6 | TACCTGGGTTGCCTTTCATC | CCCGACCTATGAGCTATGGA | 288 |
| chr4:59814346-59816372 | p2 | (AT)13 | AAGGTGAGGCAGGATGTTTG | TGGTGTGTATATCTGCCCGA | 263 |
| chr4:59849357-59851372 | p5 | (CAATG)3 | ACAGGTTGGGGGTCATCATA | TTGGGTTAAAGGTTCGGTTG | 178 |
| chr4:59873129-59875153 | p2 | (TA)12 | AGCTTGGGGAAGGGTTAAAA | GTGAAGCAGTCCCACCATTT | 131 |

| Marker ID | SSR type | SSR motif | FORWARD PRIMER (5'-3') | REVERSE PRIMER (5'-3') | size (bp) |
|---|---|---|---|---|---|
| chr4:59897865-59899895 | p5 | (CCCCA)3 | GGAAACCGAACCAAGGTACA | GCAAATCGTGAGCCTAAAGC | 120 |
| chr4:59939022-59941037 | p3 | (TGT)5 | TGGAGGGCAAGTGGATTAAG | GGTTTGTTGCGTTGGAAACT | 241 |
| chr4:59940280-59942296 | p2 | (TA)8 | AGCTTGGCCACTTTGTCAGT | TCGGTCAAGCAACAACTCTG | 240 |
| chr4:59947358-59949370 | p2 | (TA)6 | TTCTGGCAACTTGGTCAGTG | ACAACTCCATGGTCCGAAAG | 242 |
| chr4:59972487-59974502 | p5 | (TTCAT)3 | AAGAAATGACATGGGAAGCG | ATAACAATGCCTCACAGGGC | 204 |
| chr4:60017218-60019233 | p5 | (ATCAT)3 | GCTCTTTCTACCCGCCTCTT | AAACAAGAGGCTACAGGGCA | 116 |
| chr4:60025988-60028002 | p2 | (TA)7 | AATCTTCGACGTGAAGCGAT | TATAGTACCCCGGGAGGGTC | 166 |
| chr4:60151685-60153700 | p5 | (TGATA)3 | AATTTGCCCCTTTTCCATTC | AGCCAAGAGTTGCCTTTGAA | 216 |
| chr4:60190019-60192034 | p3 | (TCA)5 | CACACCCACACACACACAAA | TCCACCCTTAAACGTTACGC | 161 |
| chr4:60212191-60214206 | p5 | (ATCAT)3 | ATCCGTACCCAACTGCAGAG | TGATGCCGACGTTAATGGTA | 255 |
| chr4:60261961-60263976 | p5 | (TTTTA)3 | GTAGGTGATGTTCCGAGGGA | TTTGGGAGGATGCCACTTAC | 104 |
| chr4:60520074-60522086 | p2 | (CT)6 | GTGTCACCTATGAAGGGCGT | ACCCAAATCAGCTGCAAATC | 284 |
| chr4:60538692-60540716 | p4 | (TATT)6 | GATGGTAGGGAGGGAAGAGG | TGCCCATACACCCACTTACA | 214 |
| chr4:60538692-60540716 | p2 | (TA)7 | GATGGTAGGGAGGGAAGAGG | TGCCCATACACCCACTTACA | 214 |
| chr4:60542773-60544788 | p3 | (CAA)5 | CAAGTAAACTGCCCGACCAT | GTCCTCTCATGCATTCCGTT | 207 |
| chr4:60550909-60552924 | p5 | (TTTAT)3 | TTTCATGTATCACCCCCGTT | AGTTGTGCCTGCACACTCAG | 185 |
| chr4:60602938-60604953 | p5 | (ATATT)3 | TGGTTTAGATGCTGCACTCG | ATGTGTAAATGGCCCCAAAA | 197 |
| chr4:60667959-60669974 | p3 | (CAT)5 | ACCTCCCGTCGTACATTCAC | GCCCTGTCTCGTCCTTATGA | 153 |
| chr4:61087500-61089564 | p2 | (AT)21 | TCCGAGGTTCAAAAATACGG | GTTACCTCGGACATGGAGGA | 224 |
| chr4:61087569-61089585 | p2 | (AT)21 | TCCGAGGTTCAAAAATACGG | GTTACCTCGGACATGGAGGA | 155 |
| chr4:61089412-61091476 | p2 | (AT)21 | TCCGAGGTTCAAAAATACGG | GTTACCTCGGACATGGAGGA | 224 |
| chr4:61089412-61091476 | p2 | (AT)20 | TCCGAGGTTCAAAAATACGG | GTTACCTCGGACATGGAGGA | 224 |
| chr4:61089481-61091497 | p2 | (AT)21 | TCCGAGGTTCAAAAATACGG | GTTACCTCGGACATGGAGGA | 155 |
| chr4:61089481-61091497 | p2 | (AT)20 | TCCGAGGTTCAAAAATACGG | GTTACCTCGGACATGGAGGA | 155 |
| chr4:61128792-61130804 | p6 | (CTCCAG)3 | CACCAGTTCCTCCACCTGAT | CTGGACCTCTTGCTCTGACC | 208 |
| chr4:61128792-61130804 | p2 | (TA)6 | CACCAGTTCCTCCACCTGAT | CTGGACCTCTTGCTCTGACC | 208 |
| chr4:61181140-61183152 | p2 | (TA)6 | ATGTGACGGGGTTCCAATAA | TTTACCCTAATTGCCCCTCC | 160 |
| chr4:61248917-61250935 | p6 | (CAAAAA)3 | AGAGGGAGATGAGGCAGACA | GTGTTAAACTGGCTCCCCAA | 138 |
| chr4:61417387-61419402 | p3 | (TAT)5 | TACCCGAAACGAGGAGTTTG | CGCTGCAAAAATTCTCACAA | 146 |
| chr4:61419168-61421186 | p6 | (ATCTAT)3 | GATCAACCCCAACAAGAGGA | GTGAGCCAAAGGGAAAATGA | 119 |
| chr4:61464934-61466949 | p3 | (TTG)5 | AGATGAAAGTGGCGGAAATG | CGCATCTTGTCTTCCACTGA | 286 |
| chr4:61539334-61541352 | p5 | (AGGAC)3 | TCTACGAGCCTCTAACCGGA | TGTTCATGCCCACAGGTAGA | 220 |
| chr4:61539334-61541352 | p6 | (TTATCA)3 | TCTACGAGCCTCTAACCGGA | TGTTCATGCCCACAGGTAGA | 220 |
| chr4:61647274-61649286 | p2 | (TC)6 | AACTGATGGACTTCCGTTGG | GACGACGGACTCTATGCGAT | 233 |
| chr4:61647274-61649286 | p3 | (TAA)9 | AACTGATGGACTTCCGTTGG | GACGACGGACTCTATGCGAT | 233 |
| chr4:61694996-61697032 | p6 | (AAAACA)6 | CCATGGTTGCGGATCTACTT | TAACCGCGGAATTACTGGAA | 271 |

| Marker ID | SSR type | SSR motif | FORWARD PRIMER (5'-3') | REVERSE PRIMER (5'-3') | size (bp) |
|---|---|---|---|---|---|
| chr4:61694996-61697032 | p5 | (AAAAG)3 | CCATGGTTGCGGATCTACTT | TAACCGCGGAATTACTGGAA | 271 |
| chr4:61745235-61747247 | p2 | (AT)6 | TCAATAAAACCTTGCGGGTC | CAGCTCTAGTTCCTCCACCG | 153 |
| chr4:61758294-61760309 | p5 | (AAAAG)3 | TCATGCTCCATTCTTCTCCC | GCAGTGAAGAGAGTTTGGGC | 225 |
| chr4:61758294-61760309 | p2 | (TC)8 | TCATGCTCCATTCTTCTCCC | GCAGTGAAGAGAGTTTGGGC | 225 |
| chr4:61758294-61760309 | p2 | (CA)8 | TCATGCTCCATTCTTCTCCC | GCAGTGAAGAGAGTTTGGGC | 225 |
| chr4:61758294-61760309 | p2 | (TA)6 | TCATGCTCCATTCTTCTCCC | GCAGTGAAGAGAGTTTGGGC | 225 |
| chr4:61817525-61819546 | p3 | (GAA)7 | TGGAGATCATTGCTCGTCTG | CAAAGTCTGACGAACCCCAT | 225 |
| chr4:61872471-61874483 | p3 | (GAG)5 | GAAGAGGAGGAGGAGGAGGA | GGTTCGGACACATAATCCCA | 265 |
| chr4:61902399-61904415 | p2 | (GT)8 | CCCCAACCCTAGTCTGTGAA | CAGTCGGATAGAGACCAGGC | 268 |
| chr4:61907002-61909016 | p2 | (TA)7 | AGTGGTCATGGGTTTGAAGC | ATTCGGGAAGCATTTCCTCT | 274 |
| chr4:61913293-61915311 | p6 | (TTTTTA)3 | TTGTCTCCTTTTTCGTTGGG | AGCAACCGGATAATCTCGTG | 245 |
| chr4:61913293-61915311 | p5 | (ATGCA)3 | TTGTCTCCTTTTTCGTTGGG | AGCAACCGGATAATCTCGTG | 245 |
| chr4:61923142-61925190 | p6 | (TATGAA)8 | TTCCAACCAAAGTCCAATCC | ATCATGACTCCATCCAAGCC | 142 |
| chr4:62000925-62002940 | p3 | (CAA)5 | CAGCCACAACATTCCCTTTT | ATGCTTCAAGGCAATTACGG | 255 |
| chr4:62054425-62056437 | p2 | (AT)6 | GTAATTTCCCCCGATTCGAT | TGAACCTTGCCCATCCTAAG | 239 |
| chr4:62093240-62095272 | p2 | (TA)16 | TTCTCAAACAATTCTGCCCC | AAACTCCACGGAGAGCAAGA | 269 |
| chr4:62143491-62145503 | p2 | (AT)7 | ATCACGACTCAACTGGAGGG | GGAACCCACTCAGGTACGAA | 214 |
| chr4:62143491-62145503 | p2 | (TG)6 | ATCACGACTCAACTGGAGGG | GGAACCCACTCAGGTACGAA | 214 |
| chr4:62304234-62306262 | p2 | (AT)14 | TGTTTCCACCCAAGGTTCAT | GACCCGACCCATTTATCCTT | 172 |
| chr4:62391317-62393361 | p2 | (TA)22 | ATCCCCAAAATCCGAAAGTC | GCCCTCTATTTTGGTCGTGA | 202 |
| chr4:62391317-62393361 | p2 | (AT)14 | ATCCCCAAAATCCGAAAGTC | GCCCTCTATTTTGGTCGTGA | 202 |
| chr4:62431713-62433725 | p2 | (AC)9 | ACTCCCGAAAGGGCACTAAT | CATCCCTAACGCGTCAAAAT | 160 |
| chr4:62431713-62433725 | p2 | (AC)6 | ACTCCCGAAAGGGCACTAAT | CATCCCTAACGCGTCAAAAT | 160 |
| chr4:62651204-62653216 | p6 | (TCATAT)4 | ATAAGCCGACGAGGCTAACA | GTCCTGTCCCGCCTATATGA | 293 |
| chr4:62651204-62653216 | p2 | (TA)6 | ATAAGCCGACGAGGCTAACA | GTCCTGTCCCGCCTATATGA | 293 |
| chr4:62658893-62660908 | p5 | (ATATT)3 | AATCCAGCAAGCGGTAGAGA | GGCCCAATATACCAGGGACT | 147 |
| chr4:62756053-62758068 | p2 | (TA)32 | ATGGAAGGCCAGAACAAGTG | CCAAGTGAGAGCAAGGGTTC | 102 |
| chr4:62756053-62758068 | p2 | (TC)9 | ATGGAAGGCCAGAACAAGTG | CCAAGTGAGAGCAAGGGTTC | 102 |
| chr4:62756053-62758068 | p5 | (TTTTA)3 | ATGGAAGGCCAGAACAAGTG | CCAAGTGAGAGCAAGGGTTC | 102 |
| chr4:62774675-62776690 | p5 | (ATCAT)3 | ATTCATTGGGGGCTCTCTTT | AGGTCGTGCGATAGAGTCGT | 152 |
| chr4:62810643-62812658 | p2 | (AT)48 | ATCTCGCATTCTAGGGCTGA | ATCCGATATCAAGCCAAACG | 251 |
| chr4:62810664-62812703 | p2 | (AT)48 | ATCTCGCATTCTAGGGCTGA | ATCCGATATCAAGCCAAACG | 230 |
| chr4:63041122-63043140 | p6 | (AGCACC)3 | CAGAATTCTCCGGGACAAAA | GCTCTTCGTAGCTTTGGTGG | 120 |
| chr4:63155188-63157203 | p2 | (AT)18 | GTGCCCCGTTGAGGTACTTA | CGGCCTTTTTCACAATCACT | 277 |
| chr4:63155188-63157203 | p5 | (ATTGC)3 | GTGCCCCGTTGAGGTACTTA | CGGCCTTTTTCACAATCACT | 277 |
| chr4:63155188-63157203 | p5 | (TTATA)3 | GTGCCCCGTTGAGGTACTTA | CGGCCTTTTTCACAATCACT | 277 |

| Marker ID | SSR type | SSR motif | FORWARD PRIMER (5'-3') | REVERSE PRIMER (5'-3') | size (bp) |
|---|---|---|---|---|---|
| chr4:63182064-63184084 | p2 | (AT)10 | CATAAGCCGACGAGGCTAAC | CAGGGACTACATAGCGGAGC | 200 |
| chr4:63335992-63338055 | p3 | (TAT)21 | TTTTGTATCAACCGCAACCA | GATGGACTTATTGTCCGCGT | 254 |
| chr4:63449873-63451979 | p2 | (TA)53 | GGCATGGCGTGAAAGATAAT | TTCGGGTATCAGCCTATTCG | 280 |
| chr4:63511974-63513990 | p2 | (TA)8 | TTTGGGAACTGTTGGCTTTC | AAGCAATACCAACAAACCGC | 167 |
| chr4:63619024-63621046 | p2 | (AT)11 | TCATCCCCAAACCTGAGAAC | TTATGGGCTGATTTGGAAGC | 240 |
| chr4:63674136-63676151 | p5 | (CCTTT)3 | GTGGGACTCGTGAAGTCCAT | GCATGCACTCTCATCCTTCA | 239 |
| chr4:63683807-63685837 | p2 | (AT)15 | ACCCGCAAAGTCCATGTAAG | TTATTTTGATGGCCGAGAGG | 171 |
| chr4:63683807-63685837 | p2 | (AT)15 | ACCCGCAAAGTCCATGTAAG | TTATTTTGATGGCCGAGAGG | 171 |
| chr4:63691642-63693660 | p6 | (CATATA)3 | TGACTTGGCTCTATTGCACG | TTGTCCTAGGGGTTGTCTGC | 157 |
| chr4:63713211-63715225 | p2 | (AG)7 | ACAATTCGTTGCTACCCAGG | CGATGTCACCTCGGGATACT | 206 |
| chr4:63717035-63719050 | p5 | (ATATC)3 | TCGTCGTAATTTACCCCTGC | GTGGTTGGAGAATGCCTCAT | 223 |
| chr4:63814014-63816028 | p3 | (AAC)7 | TCCAAAACAGAAAGGTTGGC | AAATCACCATCGTCCTCCAG | 170 |
| chr4:63814029-63816041 | p3 | (AAC)7 | TCCAAAACAGAAAGGTTGGC | AAATCACCATCGTCCTCCAG | 155 |
| chr4:63820536-63822552 | p2 | (AT)8 | GATGGGATCATGGTGGATTC | TGGAAGGGAACATCCTTCTG | 123 |
| chr4:63898103-63900118 | p2 | (AT)7 | CACAATGGATTGGAAAACCC | TTGAAAATGGTCTTGCCCTC | 267 |
| chr4:63898103-63900118 | p5 | (TTGAA)3 | CACAATGGATTGGAAAACCC | TTGAAAATGGTCTTGCCCTC | 267 |
| chr4:63898103-63900118 | p6 | (TATATG)3 | CACAATGGATTGGAAAACCC | TTGAAAATGGTCTTGCCCTC | 267 |
| chr4:64020013-64022029 | p2 | (AT)8 | ATAGTTCGTGGCATCGGTTC | CCTTGGAGTTCTTTTGCGAG | 219 |
| chr4:64020013-64022029 | p3 | (ACA)5 | ATAGTTCGTGGCATCGGTTC | CCTTGGAGTTCTTTTGCGAG | 219 |
| chr4:64046649-64048661 | p2 | (TA)6 | GAAGGAAACGTGCAAGGGTA | GTAGGCACCTTTGCCATCAT | 247 |
| chr4:64054293-64056305 | p2 | (TA)6 | AGACGAATGGACGAAGTTGG | CTTGCACGTTTCCTTCCAAT | 107 |
| chr4:64092629-64094651 | p2 | (AT)11 | GAAAGTGCGCAAGATGATGA | GAGAGTTAAGGGCCCACCTC | 296 |
| chr4:64094439-64096451 | p2 | (AT)6 | GGACATGCCAAAAGAAGGAA | TTATAGGGAAAATGCTGCCG | 190 |
| chr4:64177549-64179561 | p2 | (TA)6 | CCGATGATGTGATGTTCCAG | TAATCGCAATCCACGAAACA | 116 |
| chr4:64204987-64207011 | p2 | (AT)12 | AAATGGCCACCTGCACTTAG | CTTTCCTGCCCAAGACTCAG | 158 |
| chr4:64204987-64207011 | p3 | (CAA)5 | AAATGGCCACCTGCACTTAG | CTTTCCTGCCCAAGACTCAG | 158 |
| chr4:64240957-64242972 | p3 | (GTT)5 | ATTCGGAACACATAGCCTGC | TTTCCGCATCCGTATTTCTC | 122 |
| chr4:64241978-64243996 | p2 | (AT)9 | TTGAATTTTGGGGATGGTGT | CCTTACGCTCGTCCAAACTC | 286 |
| chr4:64257375-64259387 | p2 | (AC)6 | CAGCAGGCGTAGCTTAATCC | TGTTGATGAATTGAGCCGAG | 122 |
| chr4:64257375-64259387 | p5 | (TTTTC)3 | CAGCAGGCGTAGCTTAATCC | TGTTGATGAATTGAGCCGAG | 122 |
| chr4:64269600-64271625 | p5 | (TCATA)3 | ACCAACTTCGCTGGAACATC | ACATTGCTCGTACTGACCCC | 243 |
| chr4:64269653-64271668 | p5 | (TCATA)3 | ACCAACTTCGCTGGAACATC | ACATTGCTCGTACTGACCCC | 190 |
| chr4:64429423-64431451 | p2 | (TA)25 | TCCCACAAGGATTGGAAGTC | TGCCATGGTTACATGGAGAA | 268 |
| chr4:64429423-64431451 | p2 | (AT)14 | TCCCACAAGGATTGGAAGTC | TGCCATGGTTACATGGAGAA | 268 |
| chr4:64473555-64475569 | p2 | (CT)6 | AAATGATGAGCCCTCAAACG | ATCGAGTCTGGTTTCCAACG | 245 |
| chr4:64473555-64475569 | p2 | (AT)7 | AAATGATGAGCCCTCAAACG | ATCGAGTCTGGTTTCCAACG | 245 |

| Marker ID | SSR type | SSR motif | FORWARD PRIMER (5'-3') | REVERSE PRIMER (5'-3') | size (bp) |
|---|---|---|---|---|---|
| chr4:64573206-64575224 | p3 | (TAT)6 | GCTTGCTCCAAGCTCATTCT | CCAACCCTTTGCACATTTTT | 106 |
| chr4:64591424-64593438 | p2 | (TA)7 | TTTGGGTTACACGGACCAAT | ATTGGAGACCAAACGAAACG | 245 |
| chr4:64694888-64696903 | p3 | (TTG)5 | GCTTAAGCCAAATATGGGCA | TTTGGCAGCTTGGAGTCTTT | 171 |
| chr4:64694888-64696903 | p2 | (AT)14 | GCTTAAGCCAAATATGGGCA | TTTGGCAGCTTGGAGTCTTT | 171 |
| chr4:64714791-64716803 | p2 | (AT)13 | CCGTACCGGTCCTCTAGTGA | CTCGTGCATGCTCAAAATGT | 216 |
| chr4:64714791-64716803 | p2 | (AT)6 | CCGTACCGGTCCTCTAGTGA | CTCGTGCATGCTCAAAATGT | 216 |
| chr4:64729725-64731737 | p2 | (TA)6 | GAGACGTGAGCGTTGAATGA | GTCGCCATTTTCATGTCCTT | 147 |
| chr4:64729725-64731737 | p2 | (AT)19 | GAGACGTGAGCGTTGAATGA | GTCGCCATTTTCATGTCCTT | 147 |
| chr4:64729725-64731737 | p2 | (TA)8 | GAGACGTGAGCGTTGAATGA | GTCGCCATTTTCATGTCCTT | 147 |
| chr4:64829655-64831667 | p2 | (AT)6 | CGTCACATCCCGGAGTCTAT | ATGGCAGAGTATGGGTCCAG | 211 |
| chr4:64883857-64885872 | p5 | (ATTTC)3 | TCAGTCAAAGTCTTGTGGCG | TGAATCTCTCTTGGGGGTTG | 148 |
| chr4:64907972-64909990 | p6 | (CATATA)3 | CCTTAGACCGACCTGCTCTG | GCAATGGGTATTGGGTTGTC | 142 |
| chr4:64913287-64915299 | p2 | (TA)6 | ACAACCCTTAGACCGACGTG | TGGGTTGTCAGTCTGTAGCG | 190 |
| chr4:64968084-64970099 | p5 | (TATGA)3 | TGTGGGTAATGGGTCATGTG | CATTTAAGCCGCATCCAAAT | 166 |
| chr4:64968084-64970099 | p5 | (TGCAA)3 | TGTGGGTAATGGGTCATGTG | CATTTAAGCCGCATCCAAAT | 166 |
| chr4:65023900-65025915 | p5 | (TATGA)3 | TGTGGGTAATGGGTCATGTG | CATTTAAGCCGCATCCAAAT | 166 |
| chr4:65023900-65025915 | p5 | (TGCAA)3 | TGTGGGTAATGGGTCATGTG | CATTTAAGCCGCATCCAAAT | 166 |
| chr4:65179672-65181702 | p2 | (TA)15 | CGCGTATGGTTCCAAAAGTT | GCTCAAAAACAAGTCTCGGG | 268 |
| chr4:65210145-65212160 | p3 | (ACA)5 | ACTACATAATTGCCCAGCCG | ATGCATCTTCAGTTACCGCC | 213 |
| chr4:65215993-65218005 | p2 | (AG)8 | TCTGGTCACGGGTCAATGTA | CTATTTTTCCCTTGGCCTCC | 254 |
| chr4:65215993-65218005 | p2 | (GT)6 | TCTGGTCACGGGTCAATGTA | CTATTTTTCCCTTGGCCTCC | 254 |
| chr4:65225060-65227076 | p2 | (AT)8 | AAGAAAGGAACCTCCCGAAA | TCCCATTTTGTAATTCCCGA | 236 |
| chr4:65337987-65339999 | p2 | (AT)6 | CCAACTCCACGGATTCAGTT | CAAGTACCAAAGCCCGTGAT | 173 |
| chr4:65420248-65422260 | p2 | (AC)6 | CGAGACTCGTACCCGACTTC | AATTCCCGGATTGTGTTTGA | 233 |
| chr4:65434869-65436935 | p2 | (AT)33 | ATTTACGGCAAAGACGATGG | CCTCATACTGGCGCTCTTTC | 259 |
| chr4:65454792-65456810 | p5 | (GAAAT)4 | CCGTATAATGCCCAGCTGTT | TTAAAGATGTTTCGGGGACG | 255 |
| chr4:65454792-65456810 | p6 | (CAACTC)3 | CCGTATAATGCCCAGCTGTT | TTAAAGATGTTTCGGGGACG | 255 |
| chr4:65470032-65472044 | p2 | (AC)6 | CGAGACTCGTACCCGACTTC | AATTCCCGGATTGTGTTTGA | 233 |
| chr4:65484653-65486719 | p2 | (AT)33 | ATTTACGGCAAAGACGATGG | CCTCATACTGGCGCTCTTTC | 259 |
| chr4:65504576-65506594 | p5 | (GAAAT)4 | CCGTATAATGCCCAGCTGTT | TTAAAGATGTTTCGGGGACG | 255 |
| chr4:65504576-65506594 | p6 | (CAACTC)3 | CCGTATAATGCCCAGCTGTT | TTAAAGATGTTTCGGGGACG | 255 |
| chr4:65549776-65551798 | p2 | (AT)12 | TGCATGAAGGACACAACGAT | CCAATCCAAACAGGCTCCTA | 146 |
| chr4:65549776-65551798 | p2 | (TC)11 | TGCATGAAGGACACAACGAT | CCAATCCAAACAGGCTCCTA | 146 |
| chr4:65560331-65562346 | p2 | (CA)11 | GGCATATTCGCGTTGGTACT | AGCTCAGCTTACCACGGTGT | 138 |
| chr4:65560331-65562346 | p5 | (ATTGC)3 | GGCATATTCGCGTTGGTACT | AGCTCAGCTTACCACGGTGT | 138 |
| chr4:65560331-65562346 | p4 | (ATTT)4 | GGCATATTCGCGTTGGTACT | AGCTCAGCTTACCACGGTGT | 138 |

| Marker ID | SSR type | SSR motif | FORWARD PRIMER (5'-3') | REVERSE PRIMER (5'-3') | size (bp) |
|---|---|---|---|---|---|
| chr4:65637151-65639169 | p6 | (CATATA)3 | CCTTAGACCGACCTGCTCTG | GCAATGGGTATTGGGTTGTC | 267 |
| chr4:65753684-65755696 | p2 | (AT)6 | AAATCGGGATCAAAGGTTCC | TATACGCTACCTCATCCGGC | 297 |
| chr4:65753684-65755696 | p2 | (CA)6 | AAATCGGGATCAAAGGTTCC | TATACGCTACCTCATCCGGC | 297 |
| chr4:65833829-65835841 | p2 | (GT)6 | ATGAGGCCAGTCCAACATTC | CCTTCCTCAATATTTCCGCA | 295 |
| chr4:65954383-65956411 | p2 | (AT)14 | GGTGGACCCACAACCATATC | TCCCGCTATTTGCTATTTGG | 168 |
| chr4:65960780-65962795 | p5 | (TCATT)3 | ACCTTCGTGGTCAGGTATGC | TCATCGGCGACGTACAAATA | 126 |
| chr4:66024096-66026111 | p5 | (TTCAT)3 | GACCCAAACCAATGAGTCGT | GCATATGTCGTTCCCTTCGT | 253 |
| chr4:66056170-66058182 | p2 | (AC)6 | TTCTCAATTCCCGTTTCGTC | TTTGTTGATGTGTGGGCTGT | 295 |
| chr4:66060313-66062327 | p2 | (TA)7 | CTCGACTCGACTCTTGTCCC | TGCCCTTAAGGAGTCGAGAA | 265 |
| chr4:66094570-66096585 | p2 | (TA)6 | CCTTAAGCCAAATATGGGCA | GGTTTCACCGACTTCCTCAA | 182 |
| chr4:66094570-66096585 | p3 | (TTG)5 | CCTTAAGCCAAATATGGGCA | GGTTTCACCGACTTCCTCAA | 182 |
| chr4:66094570-66096585 | p2 | (TA)7 | CCTTAAGCCAAATATGGGCA | GGTTTCACCGACTTCCTCAA | 182 |
| chr4:66113065-66115109 | p2 | (TA)11 | GAGCGATCACAATTACCCGT | GTTGATTTTGCCGCTAGGAG | 278 |
| chr4:66113065-66115109 | p2 | (AT)22 | GAGCGATCACAATTACCCGT | GTTGATTTTGCCGCTAGGAG | 278 |
| chr4:66143112-66145138 | p2 | (AT)15 | ATTGGAAATTCGGGGTAAGG | TCCGTTCTCTTTGTCGCTCT | 151 |
| chr4:66278817-66280855 | p2 | (TA)19 | GGGGTCCACAACCATATGTC | GGTGTCTGCCTATGCCATCT | 294 |
| chr4:66278817-66280855 | p5 | (TGTGT)3 | GGGGTCCACAACCATATGTC | GGTGTCTGCCTATGCCATCT | 294 |
| chr4:66522614-66524660 | p2 | (AT)23 | CCGGATGTGGTTTGAGAAAT | GGCTCAAGGTAAGCATCGAG | 236 |
| chr4:66542576-66544588 | p2 | (AT)9 | TATGGCTGGACAACGATTCA | CCTAGTCGGGTCGTGACAAT | 166 |
| chr4:66566224-66568236 | p2 | (AT)9 | TATGGCTGGACAACGATTCA | CCTAGTCGGGTCGTGACAAT | 166 |
| chr4:66568648-66570663 | p3 | (TTG)5 | TGGCATGTGCCTTAGTCAAG | TCGGCCGAATAGTGGTTTAG | 116 |
| chr4:66711615-66713633 | p3 | (ACA)6 | ACATTTGCTTTTCCAGGGTG | CAGCTGTGGAAGCATGAAAA | 160 |
| chr4:66743181-66745221 | p2 | (TA)7 | ACATCACAACAAAGGCCACA | AACAGTTGACAGGGGGTGAC | 135 |
| chr4:66743181-66745221 | p2 | (TA)20 | ACATCACAACAAAGGCCACA | AACAGTTGACAGGGGGTGAC | 135 |
| chr4:66760340-66762404 | p2 | (AT)32 | CTTCCTAAATGCCTTGCAGC | TCATAGTTGGCCTTCTTGGC | 201 |
| chr4:66760340-66762404 | p3 | (CAA)5 | CTTCCTAAATGCCTTGCAGC | TCATAGTTGGCCTTCTTGGC | 201 |
| chr4:66865819-66867831 | p2 | (AT)6 | TGTGTGGTTGGGCTGATTTA | CATCACACGCGATGTTTTTC | 109 |
| chr4:66865819-66867831 | p2 | (AT)27 | TGTGTGGTTGGGCTGATTTA | CATCACACGCGATGTTTTTC | 109 |
| chr4:66889666-66891678 | p2 | (AT)6 | TGTGTGGTTGGGCTGATTTA | CATCACACGCGATGTTTTTC | 109 |
| chr4:66992418-66994442 | p2 | (AT)12 | CCTGCCACCTTTGCTCTTAG | AATGCCATAAATGGCGAAAG | 152 |
| chr4:67016113-67018129 | p4 | (AGAA)4 | CCGCACACATCATACTCCAG | TGCTCGTGCAGTTCATTACC | 231 |
| chr4:67029110-67031122 | p2 | (AC)6 | TTCTGGCTCTTCCTCCTCAA | GCATCGTAGGGTTGTGTGTG | 108 |
| chr4:67029110-67031122 | p2 | (TG)6 | TTCTGGCTCTTCCTCCTCAA | GCATCGTAGGGTTGTGTGTG | 108 |
| chr4:67039050-67041062 | p2 | (AG)6 | CGATTACACGCACTCCAAGA | CCTCGAGCTCACATTAAGCC | 219 |
| chr4:67039050-67041062 | p3 | (TTG)5 | CGATTACACGCACTCCAAGA | CCTCGAGCTCACATTAAGCC | 219 |
| chr4:67039050-67041062 | p5 | (TGTTG)3 | CGATTACACGCACTCCAAGA | CCTCGAGCTCACATTAAGCC | 219 |

| Marker ID | SSR type | SSR motif | FORWARD PRIMER (5'-3') | REVERSE PRIMER (5'-3') | size (bp) |
|---|---|---|---|---|---|
| chr4:67039608-67041623 | p2 | (AG)6 | CCAAGTGTAAGCCCTCCAAA | TAACAAGGAAAATGGGCCTG | 115 |
| chr4:67039608-67041623 | p3 | (TTG)5 | CCAAGTGTAAGCCCTCCAAA | TAACAAGGAAAATGGGCCTG | 115 |
| chr4:67039608-67041623 | p5 | (TGTTG)3 | CCAAGTGTAAGCCCTCCAAA | TAACAAGGAAAATGGGCCTG | 115 |
| chr4:67039608-67041623 | p2 | (TA)7 | CCAAGTGTAAGCCCTCCAAA | TAACAAGGAAAATGGGCCTG | 115 |
| chr4:67041288-67043303 | p2 | (TA)7 | GCTGTGTTCTCATCGCATGT | TGGAACCACTTTTGTCACGA | 278 |
| chr4:67041288-67043303 | p5 | (GATAT)3 | GCTGTGTTCTCATCGCATGT | TGGAACCACTTTTGTCACGA | 278 |
| chr4:67052536-67054548 | p2 | (AG)6 | ACTGAAGGGATCGACCAATG | CCTCGAGCTCACATTAAGCC | 249 |
| chr4:67052536-67054548 | p3 | (GTT)6 | ACTGAAGGGATCGACCAATG | CCTCGAGCTCACATTAAGCC | 249 |
| chr4:67053067-67055085 | p2 | (AG)6 | CGTTTTCGGGTGATCTGACT | CTGAAAATGAGGAGGCGAAG | 173 |
| chr4:67053067-67055085 | p3 | (GTT)6 | CGTTTTCGGGTGATCTGACT | CTGAAAATGAGGAGGCGAAG | 173 |
| chr4:67110952-67112966 | p2 | (TC)6 | TATTCGTCCTTTCGCTTGCT | GTGAGCAAAAGTAGCGGGAC | 295 |
| chr4:67110952-67112966 | p2 | (AT)7 | TATTCGTCCTTTCGCTTGCT | GTGAGCAAAAGTAGCGGGAC | 295 |
| chr4:67136882-67138896 | p2 | (TA)6 | AAACCGGAGAGTCATGATGG | ACATATGTCCTTTGGGCTCG | 234 |
| chr4:67136920-67138984 | p2 | (TA)6 | GGAGAGTCATGATGGGCACT | ACATATGTCCTTTGGGCTCG | 201 |
| chr4:67143037-67145059 | p2 | (GA)11 | TTAAGAGGATCTGATGGCGG | GCATCACACACCACAAAAGG | 208 |
| chr4:67244664-67246679 | p5 | (TTGGT)3 | CAAAGGGGGTTCAACGTCTA | AATCCCAACACAATCCCAAA | 236 |
| chr4:67282329-67284363 | p2 | (AT)17 | GGGAAATAGCTTGGAAAGGC | TTTGCAAAATTACCCCCAAA | 260 |
| chr4:67310290-67312305 | p5 | (TTGGT)3 | CAAAGGGGGTTCAACGTCTA | AATCCCAACACAATCCCAAA | 236 |
| chr4:67347955-67349989 | p2 | (AT)17 | GGGAAATAGCTTGGAAAGGC | TTTGCAAAATTACCCCCAAA | 260 |
| chr4:67366300-67368318 | p6 | (TTTGTT)3 | AACTTCGTTGAACCATTGGC | AGGGGGACTTATTCTCCGAA | 294 |
| chr4:67366300-67368318 | p3 | (TGT)6 | AACTTCGTTGAACCATTGGC | AGGGGGACTTATTCTCCGAA | 294 |
| chr4:67374005-67376020 | p5 | (GTTTG)3 | TTATGCGCCAAAAACACAAA | TGGTCGGTCAATTGGAGAAT | 257 |
| chr4:67408440-67410464 | p6 | (GTGGAC)4 | CATGCAATTTTGATCGCAAC | GTCCTTGATCCACCGAGAAA | 171 |
| chr4:67469486-67471501 | p5 | (TTTTA)3 | ATGCTGGCGCATAAGAAACT | CACCGTTCTTACGTGGTGAA | 251 |
| chr4:67492976-67494996 | p4 | (TCCA)5 | CTTGGCTCTGAAACAGGGAC | CAACCTCGGAATTTGCCTAA | 172 |
| chr4:67644865-67646880 | p3 | (ATA)5 | GCACGATCGTATCGGAACTT | TGCCAAAGACACCCCTTATC | 210 |
| chr4:67650951-67652966 | p3 | (AAC)5 | CAATGTCCTCAGCCAACCTT | CCATGGTCGTAAACACCTCC | 171 |
| chr4:67700711-67702726 | p5 | (TATAT)3 | CAAAAATACCCACACGGGTC | AGGATGCCTTGGAGGAGATT | 190 |
| chr4:67709484-67711499 | p5 | (TTCAT)3 | CCGGAAGAGATCCAAGATGA | ACGAAAATGTTTCCGAGGTG | 226 |
| chr4:67720360-67722375 | p3 | (TTG)5 | CGGACAGCATCTCCGTTTAT | TGGCCTTTTCTTTTTGGTTG | 200 |
| chr4:67720360-67722375 | p2 | (AT)16 | CGGACAGCATCTCCGTTTAT | TGGCCTTTTCTTTTTGGTTG | 200 |
| chr4:67743412-67745424 | p2 | (TA)6 | GAACAGTAAAGGCCGATGGA | TGGATCCTCATTTCGTGTCA | 281 |
| chr4:67825158-67827184 | p2 | (AT)13 | GGTACGGTAAAGGTGGGGTT | GGGCTGAGTAGCACCTGAAG | 260 |
| chr4:67825158-67827184 | p2 | (AT)20 | GGTACGGTAAAGGTGGGGTT | GGGCTGAGTAGCACCTGAAG | 260 |
| chr4:67825675-67827715 | p2 | (AT)13 | CTCAAAAGGGGCATACCAAA | CGTACTGACTCCCCATTGCT | 288 |
| chr4:67825675-67827715 | p2 | (AT)20 | CTCAAAAGGGGCATACCAAA | CGTACTGACTCCCCATTGCT | 288 |

| Marker ID | SSR type | SSR motif | FORWARD PRIMER (5'-3') | REVERSE PRIMER (5'-3') | size (bp) |
|---|---|---|---|---|---|
| chr4:67829928-67831940 | p2 | (AT)6 | TATCATACGCTTTTTCGCCC | TGCCCAAAATTTCGGTAGAC | 283 |
| chr4:67860770-67862802 | p2 | (TA)12 | CGATTTGGATTGGAAAAGGA | CTTAACGGAAATCTGGCAGC | 205 |
| chr4:67867958-67869996 | p2 | (AT)19 | GTGTGTTGGTTTGTTCGGTG | CTTAACGGAAATCTGGCAGC | 177 |
| chr4:67867958-67869996 | p2 | (TA)8 | GTGTGTTGGTTTGTTCGGTG | CTTAACGGAAATCTGGCAGC | 177 |
| chr4:68049047-68051075 | p2 | (AT)14 | ACGGACTGTAACAGCGGAGT | TCCCTTGTGCGCTTAGTCTT | 101 |
| chr4:68414295-68416313 | p2 | (TA)7 | ACATCACAACAAAGGCCACA | TAGTTGACAGGGGGTAACGC | 226 |
| chr4:68414295-68416313 | p2 | (TA)9 | ACATCACAACAAAGGCCACA | TAGTTGACAGGGGGTAACGC | 226 |
| chr4:68414295-68416313 | p2 | (TA)9 | ACATCACAACAAAGGCCACA | TAGTTGACAGGGGGTAACGC | 226 |
| chr4:68437148-68439162 | p2 | (TA)7 | AGCCCAGTGGTCGTAAAATG | TGGTTTACGGTCCGTCTTTC | 227 |
| chr4:68437148-68439162 | p2 | (TA)16 | AGCCCAGTGGTCGTAAAATG | TGGTTTACGGTCCGTCTTTC | 227 |
| chr4:68459983-68461995 | p2 | (AG)6 | GGAGAGTAAAGACAACGGCG | GGTGTAACATCCTTCCCCCT | 146 |
| chr4:68505910-68507924 | p2 | (AG)8 | TTAGGCAGAAAGGTCACGCT | CTCGTCATGAGTTTCGGGAT | 206 |
| chr4:68505910-68507924 | p2 | (TA)7 | TTAGGCAGAAAGGTCACGCT | CTCGTCATGAGTTTCGGGAT | 206 |
| chr4:68682361-68684381 | p2 | (TA)8 | CAAGGGAGAAGAGTTGGACG | GGTAATACCATAGCCGGGGT | 165 |
| chr4:68682385-68684417 | p2 | (TA)8 | CCCTTTCTTTCATTTTGGCA | CCTCTCATGGGCCTTACTGA | 193 |
| chr4:68734113-68736125 | p2 | (TA)6 | GTTCGCTCGGTTCCTGTAAG | AGTGCTTCTGTGAATCCGCT | 154 |
| chr4:68734113-68736125 | p2 | (AT)6 | GTTCGCTCGGTTCCTGTAAG | AGTGCTTCTGTGAATCCGCT | 154 |
| chr4:68734113-68736125 | p2 | (AT)6 | GTTCGCTCGGTTCCTGTAAG | AGTGCTTCTGTGAATCCGCT | 154 |
| chr4:68760828-68762858 | p2 | (TA)15 | AGAAGCCGACAAGGCTATCA | TCCCGTCATATGATTCGGTT | 154 |
| chr4:68760828-68762858 | p2 | (CA)13 | AGAAGCCGACAAGGCTATCA | TCCCGTCATATGATTCGGTT | 154 |
| chr4:68768959-68770971 | p2 | (AT)15 | TCCTCAACCTTACGTCACCC | ACGGTGTAATTCATAGCGCC | 274 |
| chr4:68778097-68780109 | p2 | (AT)6 | GAGTGTGCTCCTCGTCATCA | CCTATCCTGGTCGTCGGTAA | 266 |
| chr4:68859305-68861317 | p2 | (AT)6 | GGCGATCAAGAAGCTGAATC | CTAACTGCCCAGGCTCAAAG | 207 |
| chr4:68938162-68940174 | p5 | (ATTCA)3 | TACTTCACCCGACACCTTCC | AATGAAGGTTTGGTGGCTTG | 243 |
| chr4:68938162-68940174 | p2 | (AT)6 | TACTTCACCCGACACCTTCC | AATGAAGGTTTGGTGGCTTG | 243 |
| chr4:68938162-68940174 | p2 | (AT)19 | TACTTCACCCGACACCTTCC | AATGAAGGTTTGGTGGCTTG | 243 |
| chr4:69000610-69002664 | p2 | (AT)27 | TCAGATTCGCATCGCATAAG | CATGCGAAAGTTTGTGATGG | 207 |
| chr4:69050846-69052858 | p2 | (AG)6 | GATATTCAGGTAACGGCGGA | TTTTTACCTGGAGACACGGG | 299 |
| chr4:69053350-69055362 | p2 | (AG)6 | GATATTCAGGTAACGGCGGA | TTTTTACCTGGAGACACGGG | 299 |
| chr4:69071182-69073200 | p6 | (GTAGAG)3 | GGACTCTCCAAAACTGCTGC | AATTGTGCATTGGGGACATT | 276 |
| chr4:69100525-69102541 | p2 | (TG)7 | GGGAGTCCATGAAGCAGTGT | TGCTCGTTTCTTGTGCATTC | 291 |
| chr4:69100525-69102541 | p2 | (AT)8 | GGGAGTCCATGAAGCAGTGT | TGCTCGTTTCTTGTGCATTC | 291 |
| chr4:69168406-69170418 | p2 | (AT)6 | TCATCTGCTCCATCTGTTGC | CCCTACTTCTCCTGACACGC | 251 |
| chr4:69221088-69223106 | p6 | (AAGCAG)3 | CATATGCCAAGGCCAAACTT | ATCAATCCATTTGAAGGGCA | 213 |
| chr4:69275182-69277197 | p5 | (AAAAG)3 | CATGGCAATCCTCTCCAACT | TATTCCATTCATCTGGGGGA | 184 |
| chr4:69339183-69341198 | p5 | (AAAAG)3 | CATGGCAATCCTCTCCAACT | TATTCCATTCATCTGGGGGA | 184 |

| Marker ID | SSR type | SSR motif | FORWARD PRIMER (5'-3') | REVERSE PRIMER (5'-3') | size (bp) |
|---|---|---|---|---|---|
| chr4:69373245-69375261 | p2 | (TA)8 | GTATGCCGGGTAAGGTGAGA | TCCCAATGATAGCGAACACA | 209 |
| chr4:69453826-69455841 | p3 | (ACA)5 | CTGAATGAAAAAGGGCCAAA | GTAAGGCTTCACCCCTCCTC | 241 |
| chr4:69563441-69565456 | p3 | (CCG)5 | ACCAGCCAACTTGGTCAAAC | ATGGGGTCATGAGGTTGTGT | 148 |
| chr4:69691709-69693727 | p2 | (AT)9 | GATGTTCCAGCGGATTTGAT | CATTCTTTTCGTCGACCGTT | 223 |
| chr4:69828831-69830846 | p5 | (AAAAC)3 | GTGGAAGCTGGCTCTTCATC | GATCCACTCAGACTGGGCAT | 114 |
| chr4:69850689-69852703 | p2 | (GT)7 | GTTTGTGGCATTGATTGACG | AAATTCCAAAGTTTCGCCCT | 222 |
| chr4:69922092-69924107 | p5 | (TTCAA)3 | ACGAAGGTTCGGTGTTTCAC | CTGAGACTTGGTAGGGCCAG | 236 |
| chr4:69936400-69938415 | p3 | (ATG)8 | TAGTCATTTGTGACGCCGAG | TCGGTATTATCCGTACCCGA | 125 |
| chr4:69936400-69938415 | p5 | (TTTAA)3 | TAGTCATTTGTGACGCCGAG | TCGGTATTATCCGTACCCGA | 125 |
| chr4:69936400-69938415 | p2 | (TA)7 | TAGTCATTTGTGACGCCGAG | TCGGTATTATCCGTACCCGA | 125 |
| chr4:69995336-69997360 | p3 | (TCT)8 | ATTCGGGGGAAGAATACCTG | GTCACATGGCTTTCCTTGGT | 284 |
| chr4:70030916-70032928 | p2 | (AT)6 | GATATTTTCCCAAGCGGGAT | AAGGCGCGAGATGATAGAGA | 139 |
| chr4:70039244-70041306 | p2 | (AT)31 | CGTTTGGTCTCCGATCCTTA | AGAGAAGGCAAGGCATACGA | 155 |
| chr4:70061520-70063538 | p2 | (AT)9 | AATCCAACATACATGCCCGT | GTCGGCTTGCGTATCATTTT | 168 |
| chr4:70067600-70069612 | p2 | (AT)6 | CCAATATACCGCAACATCCC | TACAGGGCAAGGTGATAGGG | 198 |
| chr4:70067600-70069612 | p2 | (AT)6 | CCAATATACCGCAACATCCC | TACAGGGCAAGGTGATAGGG | 198 |
| chr4:70112810-70114822 | p2 | (TA)8 | CCAACCACATGTTGCTTCAG | AACATTCTCCAACCACCAGC | 161 |
| chr4:70112810-70114822 | p2 | (AT)6 | CCAACCACATGTTGCTTCAG | AACATTCTCCAACCACCAGC | 161 |
| chr4:70164784-70166796 | p2 | (AT)6 | GTGGTGTCTGCCTAGCCAAT | GGCCCCAGAGTCATGAATTA | 164 |
| chr4:70188930-70190944 | p2 | (GT)7 | AATTGTGGAGGTTCACTCCG | TTGAGTCCTGGGAGTTGTCC | 118 |
| chr4:70197124-70199138 | p2 | (GT)7 | ATAGAGCAACAACAACCGGG | CTTATGAGCATAACGGGGGA | 104 |
| chr4:70299210-70301231 | p3 | (ACA)7 | ATTTGGACACCCCAAACAGA | GGGTCCTATGTGTGTTCGCT | 254 |
| chr4:70318238-70320254 | p4 | (AGAA)4 | TCAATGCACACACATCATGG | GGCTGGTTCTTCTTCAGTGC | 258 |
| chr4:70318238-70320254 | p5 | (AATTG)4 | TCAATGCACACACATCATGG | GGCTGGTTCTTCTTCAGTGC | 258 |
| chr4:70320209-70322233 | p6 | (ATATGT)4 | TGTTGGATATAAGCCTCGGG | GGCGTTACTTGTGTGGGTTT | 229 |
| chr4:70553528-70555543 | p5 | (AATTT)3 | ACTCTCGCAAACAGGCCTAA | ATCCTCGTTGCTGAAGGAGA | 110 |
| chr4:70593782-70595794 | p2 | (AT)6 | GATGGCCGATGTCTTTGATT | ATGGAACAAACCAAATGGGA | 272 |
| chr4:70812115-70814139 | p4 | (TGTA)6 | AAGCCTACCCCTTCATTCGT | TTGGAAGTATTCCGACTCCG | 179 |
| chr4:70879164-70881188 | p4 | (TGTA)6 | AAGCCTACCCCTTCATTCGT | TTGGAAGTATTCCGACTCCG | 179 |
| chr4:70898192-70900207 | p5 | (AATTC)3 | TGCTGTTTCTCCCCCTAATG | CGAGCTTAACCATGGGATGT | 146 |
| chr4:70898192-70900207 | p2 | (AC)10 | TGCTGTTTCTCCCCCTAATG | CGAGCTTAACCATGGGATGT | 146 |
| chr4:70906526-70908541 | p3 | (TCA)5 | TAGAACTCCGGCCTGCTTTA | GACATGGACATAGCCCGACT | 265 |
| chr4:70931411-70933426 | p5 | (TTATT)3 | GCCAGCACAAGGAAAGTAGG | AGCTGGTTTCCTTCCCTCAT | 249 |
| chr4:70953450-70955462 | p2 | (AT)6 | TATGGCCGAGTTGGATTCTC | TTACATACCTTGCCCGCTTC | 233 |
| chr4:71013616-71015628 | p2 | (CT)6 | TTGGGAAGCAGAGGAGAAGA | GTTTCCCCTGTAATCTGGCA | 193 |
| chr4:71096096-71098108 | p2 | (TA)6 | TATTCCCTTCATGGTGAGGC | GCCGTCTAGCCATGGTGTAT | 272 |

| Marker ID | SSR type | SSR motif | FORWARD PRIMER (5'-3') | REVERSE PRIMER (5'-3') | size (bp) |
|---|---|---|---|---|---|
| chr4:71096096-71098108 | p5 | (AATGC)3 | TATTCCCTTCATGGTGAGGC | GCCGTCTAGCCATGGTGTAT | 272 |
| chr4:71170500-71172514 | p2 | (TA)7 | GATGTTGACCGCTGGATTCT | ACAATTACCGCGTCCAAAAG | 192 |
| chr4:71254100-71256118 | p6 | (TTTCTC)3 | GCTTCACTTCCGACTTCGTC | AGCTAAGTCTCGGGGGTGTT | 216 |
| chr4:71309203-71311215 | p2 | (TA)6 | GTCCGGAGCATTGGATAAGA | CCATGGTTGATTTGCATGAG | 212 |
| chr4:71309203-71311215 | p2 | (GA)13 | GTCCGGAGCATTGGATAAGA | CCATGGTTGATTTGCATGAG | 212 |
| chr4:71375475-71377493 | p3 | (CAT)6 | ACATTGCTCGTACTGACCCC | CTCATTGCTCTGTCTGCCAA | 119 |
| chr4:71504736-71506748 | p2 | (AT)6 | CCCACCCCCTTTAAAACATT | TGACACGGAAGGGTAAGGAC | 109 |
| chr4:71688003-71690017 | p5 | (CATAG)3 | GAGCGCTGTAGAGTACCGCT | ACTGACGTCCCTTCTTGTGG | 152 |
| chr4:71701467-71703509 | p6 | (CATATA)7 | ACCCGACCCCATATTAGGAG | TGGCCACAGATTTTGTTTGA | 124 |
| chr4:71709156-71711171 | p5 | (TGTGT)3 | GTACACAAATGATCCGCGTG | ACAATGTGTTCAGGCCCTTC | 158 |
| chr4:71757721-71759739 | p6 | (TATATG)3 | TTCGGGCTAGGCACATAAAC | TGGAAACATTTGTGAAGCCA | 213 |
| chr4:71896035-71898053 | p2 | (AT)7 | GGTTTCGGGTTTCGTCTGTA | TACGTACCTACCCGACCGAC | 136 |
| chr4:71896035-71898053 | p6 | (ATAATG)3 | GGTTTCGGGTTTCGTCTGTA | TACGTACCTACCCGACCGAC | 136 |
| chr4:71896035-71898053 | p5 | (ATGAT)3 | GGTTTCGGGTTTCGTCTGTA | TACGTACCTACCCGACCGAC | 136 |
| chr4:71906364-71908376 | p2 | (CA)6 | ACGGATACTGTCAAGCCACC | AGGTGAAGGCAAAACCTGTG | 207 |
| chr4:71953860-71955874 | p2 | (GT)7 | CGGAGGACGAGATGAAGAAG | TATGCGAAACACGGGACATA | 138 |
| chr4:72043407-72045423 | p2 | (AT)9 | ATGACACATCGGTTGGTGAA | GTGCTGCCGAGTCAGATGTA | 149 |
| chr4:72043407-72045423 | p2 | (TA)8 | ATGACACATCGGTTGGTGAA | GTGCTGCCGAGTCAGATGTA | 149 |
| chr4:72143512-72145532 | p2 | (AT)6 | CCCTAGCCTTTGCTCTCCTT | GAAGTGATTGGGCTTGGAAA | 177 |
| chr4:72143512-72145532 | p2 | (AT)10 | CCCTAGCCTTTGCTCTCCTT | GAAGTGATTGGGCTTGGAAA | 177 |
| chr4:72143512-72145532 | p2 | (AT)6 | CCCTAGCCTTTGCTCTCCTT | GAAGTGATTGGGCTTGGAAA | 177 |
| chr4:72157256-72159304 | p2 | (AT)24 | TTTGTCACAACCCAACCAGA | ATGGCTCGCTCAGTACCTGT | 107 |
| chr4:72157256-72159304 | p2 | (TA)6 | TTTGTCACAACCCAACCAGA | ATGGCTCGCTCAGTACCTGT | 107 |
| chr4:72165638-72167654 | p2 | (TA)8 | GTTTGGGTTAGGCGGGTTAT | TGTAGACCAACCGTCACCAA | 113 |
| chr4:72165638-72167654 | p2 | (AT)9 | GTTTGGGTTAGGCGGGTTAT | TGTAGACCAACCGTCACCAA | 113 |
| chr4:72174246-72176260 | p2 | (AT)18 | CCCAGGAGACAGGTTGTTGT | ACTGCGCCATCTCATTATCC | 184 |
| chr4:72174246-72176260 | p2 | (TA)7 | CCCAGGAGACAGGTTGTTGT | ACTGCGCCATCTCATTATCC | 184 |
| chr4:72174246-72176260 | p5 | (TGAAT)3 | CCCAGGAGACAGGTTGTTGT | ACTGCGCCATCTCATTATCC | 184 |
| chr4:72224675-72226693 | p6 | (ACATAT)3 | ACCAGATGATGCTCCAATCC | CTTTCTTGGCCACTTTCTGC | 167 |
| chr4:72362803-72364821 | p3 | (ACA)6 | TGGGTTTTGGCTTGGATTAG | ACGTTTCACTTTGACGGACC | 214 |
| chr4:72362803-72364821 | p2 | (AG)6 | TGGGTTTTGGCTTGGATTAG | ACGTTTCACTTTGACGGACC | 214 |
| chr4:72419352-72421367 | p5 | (TACAA)3 | CTTGCGGATCTGTTGGATTT | TGCTTTGTGTGTTAGCTCCG | 245 |
| chr4:72513394-72515426 | p2 | (AT)16 | CGATGATTGATCGGTTTCCT | ACCAAGCAGGCATAATCACC | 222 |
| chr4:72540971-72542983 | p2 | (AC)6 | CAAATTCAAATCCCGTTTCG | GATGCTGCCCAAATTTCTGT | 244 |
| chr4:72553818-72555830 | p2 | (CA)6 | GGACATGCCAAAAGAGGAAA | AGATGATTCTCGGGGATGTG | 221 |
| chr4:72601820-72603832 | p2 | (TA)6 | CCAAAACTGGGGGAGGTATT | AACTGAATCCGTGGAGTTGG | 151 |

| Marker ID | SSR type | SSR motif | FORWARD PRIMER (5'-3') | REVERSE PRIMER (5'-3') | size (bp) |
|---|---|---|---|---|---|
| chr4:72601820-72603832 | p3 | (ATC)6 | CCAAAACTGGGGGAGGTATT | AACTGAATCCGTGGAGTTGG | 151 |
| chr4:72609877-72611889 | p2 | (TA)6 | TGTACCATTAAAGCAGCCCC | TTGGGGATGAACCCAACTTA | 101 |
| chr4:72609877-72611889 | p2 | (AT)6 | TGTACCATTAAAGCAGCCCC | TTGGGGATGAACCCAACTTA | 101 |
| chr4:72612160-72614174 | p2 | (TA)7 | CGACAACCCTAAAACCCTCA | TAAAATCGGGAAACAGGTGG | 216 |
| chr4:72702915-72704933 | p6 | (CTGAAA)3 | GCCTTGGTGGTCAACTGTTT | ATTTCGCTCACGTTTATGCC | 133 |
| chr4:72702915-72704933 | p6 | (TGAACA)3 | GCCTTGGTGGTCAACTGTTT | ATTTCGCTCACGTTTATGCC | 133 |
| chr4:72771777-72773797 | p4 | (TAAT)5 | CGGAGGTGCTAATTGGTTGT | ACTACCCACGTTGTGGAACC | 184 |
| chr4:72777118-72779133 | p5 | (ATCAT)3 | TCATTGTCACGACCCGTCTA | GGCCTTGTCGGCCTATATTT | 175 |
| chr4:72828898-72830916 | p2 | (AT)6 | AATATAACGCCTCGCCCTTT | TTCCCGACCTTACGTCACTC | 205 |
| chr4:72828898-72830916 | p3 | (ATC)6 | AATATAACGCCTCGCCCTTT | TTCCCGACCTTACGTCACTC | 205 |
| chr4:72828898-72830916 | p2 | (AT)9 | AATATAACGCCTCGCCCTTT | TTCCCGACCTTACGTCACTC | 205 |
| chr4:72828898-72830916 | p2 | (AT)13 | AATATAACGCCTCGCCCTTT | TTCCCGACCTTACGTCACTC | 205 |
| chr4:72829291-72831317 | p3 | (ATC)6 | AGGACATTATTGCCACGACC | CGTCGGCTTGCGTATTTATT | 210 |
| chr4:72829291-72831317 | p2 | (AT)9 | AGGACATTATTGCCACGACC | CGTCGGCTTGCGTATTTATT | 210 |
| chr4:72829291-72831317 | p2 | (AT)13 | AGGACATTATTGCCACGACC | CGTCGGCTTGCGTATTTATT | 210 |
| chr4:72914530-72916542 | p2 | (CA)6 | ATAGAATTGTCCCCGAGGCT | CGAGCGTAAGACGCACATAA | 229 |
| chr4:72930209-72932224 | p3 | (CAA)5 | TCAAAGGCCAATAGGAATCG | GAAAGCGGGCAAGGTATGTA | 255 |
| chr4:72930209-72932224 | p2 | (CA)6 | TCAAAGGCCAATAGGAATCG | GAAAGCGGGCAAGGTATGTA | 255 |
| chr4:73015208-73017220 | p2 | (AG)6 | CTGACGCATGTGAGCAAAGT | ATTCCTTTCGTGCTTCCTGA | 182 |
| chr4:73067292-73069358 | p2 | (TA)33 | AACCCCTGTCATCGTGACTC | GAGCCGTCTACGGATATGGA | 177 |
| chr4:73069633-73071645 | p2 | (CA)6 | CGCCCTCCAAGTCGTATTTA | GATATGGGCGAGGGGTATTT | 278 |
| chr4:73092558-73094570 | p6 | (TATATG)3 | GTGTTCGTGGTTCCCAAAGT | AGGCGTCGCTTTCCTATACA | 250 |
| chr4:73092586-73094600 | p6 | (TATATG)3 | GTGTTCGTGGTTCCCAAAGT | AGGCGTCGCTTTCCTATACA | 222 |
| chr4:73092601-73094615 | p6 | (TATATG)3 | GTGTTCGTGGTTCCCAAAGT | AGGCGTCGCTTTCCTATACA | 207 |
| chr4:73136477-73138493 | p2 | (AT)10 | CGATGTTTTGTGTCACAGCC | GTATGGCCCGCTCAGTACAT | 120 |
| chr4:73136495-73138509 | p2 | (AT)10 | CGATGTTTTGTGTCACAGCC | GTATGGCCCGCTCAGTACAT | 102 |
| chr4:73146295-73148319 | p2 | (TA)12 | AGCCGATAAGGCTACCCAAT | GCACTCCTCTCTTGTCGGAG | 246 |
| chr4:73176514-73178532 | p2 | (TA)9 | AATGGAAATAAGGTGCGACG | CGTGGGTATGTGGTTGTCAG | 265 |
| chr4:73176514-73178532 | p6 | (CATATA)3 | AATGGAAATAAGGTGCGACG | CGTGGGTATGTGGTTGTCAG | 265 |
| chr4:73203821-73205836 | p5 | (TTCAT)3 | ATATTTTCGAGGAATCGGGG | GCCACTCAGGGCTACAACAT | 100 |
| chr4:73235386-73237410 | p6 | (CATATA)4 | CCAGTTTACGGTCCGTCCTA | CTTAGGCGAGATTGCAAAGG | 214 |
| chr4:73235386-73237410 | p2 | (AT)7 | CCAGTTTACGGTCCGTCCTA | CTTAGGCGAGATTGCAAAGG | 214 |
| chr4:73237642-73239654 | p2 | (AT)6 | CACTCACCTTTTTCCGTCGT | AGCAGGTTGTCCTAGGGGTT | 206 |
| chr4:73237642-73239654 | p6 | (CATATA)3 | CACTCACCTTTTTCCGTCGT | AGCAGGTTGTCCTAGGGGTT | 206 |
| chr4:73239691-73241703 | p5 | (ACACA)3 | GTTAACACCAATGATCCCGC | GAATTTGGAAGGTTTCGGGT | 235 |
| chr4:73239691-73241703 | p2 | (AT)6 | GTTAACACCAATGATCCCGC | GAATTTGGAAGGTTTCGGGT | 235 |

| Marker ID | SSR type | SSR motif | FORWARD PRIMER (5'-3') | REVERSE PRIMER (5'-3') | size (bp) |
|---|---|---|---|---|---|
| chr4:73241925-73243943 | p3 | (CAA)6 | CCCCTTCTGGCATAATAGCA | GAACGGCATCATTTCATGTG | 287 |
| chr4:73345714-73347726 | p2 | (TA)6 | CGGAAATTAGTAGCACCCCA | ATTCTGGCCTCAAGATCCCT | 212 |
| chr4:73355848-73357868 | p2 | (TA)10 | CTTGGCACCAAACTTGTCCT | GCCCAATCCTTGAGTTTTGA | 203 |
| chr4:73365677-73367695 | p6 | (ATATAG)3 | CGTGCTTCTCGTACTCACCA | AAACAACCAAAATCCACCCA | 230 |
| chr4:73476804-73478840 | p2 | (AT)18 | ACTTTTGTCACGACCCAACC | TTAGCCTCGTCGGCTTATGT | 103 |
| chr4:73488659-73490677 | p6 | (TTTTTA)3 | ACACCATCCTTCAAAGTCGG | CCCATGCATTTGAGTCATTG | 114 |
| chr4:73547455-73549467 | p2 | (CA)6 | CGTCATCTTGCTCGTTTTCA | GGTAGATTTTGCGAAGCCAG | 215 |
| chr4:73654187-73656202 | p5 | (TATGA)3 | GGTGTGCCTGTTGTTGATTG | AGCCTTACATACCTGTGCCG | 135 |
| chr4:73681411-73683426 | p2 | (AT)16 | CCCAGGAAAAGAGACGTCAG | CCACAGAGGCTTAGACAGGC | 203 |
| chr4:73681411-73683426 | p3 | (CAA)5 | CCCAGGAAAAGAGACGTCAG | CCACAGAGGCTTAGACAGGC | 203 |
| chr4:73681411-73683426 | p2 | (TA)20 | CCCAGGAAAAGAGACGTCAG | CCACAGAGGCTTAGACAGGC | 203 |
| chr4:73693810-73695824 | p2 | (AT)7 | TTTGACCGGAAAAACTCTGG | AGTGACCGAAGCCATTCATC | 132 |
| chr4:73716528-73718540 | p5 | (ATATG)3 | CTAAAGAAGGACCCTTGGGC | CACATGCCCAACTCAAACTG | 300 |
| chr4:73716528-73718540 | p2 | (AT)6 | CTAAAGAAGGACCCTTGGGC | CACATGCCCAACTCAAACTG | 300 |
| chr4:73716528-73718540 | p2 | (AT)6 | CTAAAGAAGGACCCTTGGGC | CACATGCCCAACTCAAACTG | 300 |
| chr4:73716721-73718733 | p5 | (ATATG)3 | CTAAAGAAGGACCCTTGGGC | CACATGCCCAACTCAAACTG | 107 |
| chr4:73716721-73718733 | p2 | (AT)6 | CTAAAGAAGGACCCTTGGGC | CACATGCCCAACTCAAACTG | 107 |
| chr4:73716721-73718733 | p2 | (AT)6 | CTAAAGAAGGACCCTTGGGC | CACATGCCCAACTCAAACTG | 107 |
| chr4:73744434-73746448 | p2 | (AT)7 | TACGTTTGCACTTTCAACGC | ATTTTCCCTAGTCAACCCCG | 118 |
| chr4:73809218-73811236 | p6 | (GACAAA)3 | AGCCATCTACAAGCCGAAGA | AAAGGCACCCCTTGAGATTT | 141 |
| chr4:73809218-73811236 | p2 | (TG)11 | AGCCATCTACAAGCCGAAGA | AAAGGCACCCCTTGAGATTT | 141 |
| chr4:73810026-73812048 | p6 | (GACAAA)3 | TCCCTCAAAGGTTGGTGAAG | GGAAAGCATCAAGGAATGGA | 274 |
| chr4:73810026-73812048 | p2 | (TG)11 | TCCCTCAAAGGTTGGTGAAG | GGAAAGCATCAAGGAATGGA | 274 |
| chr4:73935262-73937274 | p3 | (ATG)5 | GTGAGGCAGAATGCTTGTGA | TGTGTATATCTGCCCGACCA | 261 |
| chr4:73935262-73937274 | p2 | (TA)6 | GTGAGGCAGAATGCTTGTGA | TGTGTATATCTGCCCGACCA | 261 |
| chr4:73935262-73937274 | p2 | (TA)10 | GTGAGGCAGAATGCTTGTGA | TGTGTATATCTGCCCGACCA | 261 |
| chr4:74003613-74005625 | p2 | (TA)6 | CCGAAACTTGTTTCTGAGCC | TTTATCGGGGTGACGTAAGG | 267 |
| chr4:74066233-74068248 | p5 | (ATCAT)3 | TACATGGAAGCCGATAAGGC | AACGACAGGTTTGCTGATCC | 298 |
| chr4:74070158-74072170 | p2 | (AC)6 | GGCCGAGCCTATTATGTTCA | TTGTTGTTGGGTGCTGTGTT | 117 |
| chr4:74113094-74115110 | p4 | (ATTT)4 | GGGGTTGTTGATGAATTTGG | CAAGTCTCGGGTTCTCCAAG | 150 |
| chr4:74134540-74136558 | p3 | (TTG)6 | TCCACTCCAGCACCTCTTCT | GGCCAAAATTGCCTCTCATA | 188 |
| chr4:74284840-74286856 | p2 | (AT)8 | TCTCCTCTTCCTTCGACCAA | AAAGGGACATTTCACGAACG | 242 |
| chr4:74292371-74294386 | p3 | (TGT)5 | AAAATCTCGACGTGTGGACC | TGAGCGTGATACGTGCTTTC | 282 |
| chr4:74303741-74305757 | p2 | (TG)8 | CGTTTGGGAGTTGGATAGGA | CCCAAAAATGAAGGGAAGGT | 146 |
| chr4:74313116-74315164 | p2 | (TA)24 | TCTGTGTTGTGCACCGTTTT | TGTCCGATGTGTACTTCCCA | 188 |
| chr4:74313116-74315164 | p2 | (AT)6 | TCTGTGTTGTGCACCGTTTT | TGTCCGATGTGTACTTCCCA | 188 |

| Marker ID | SSR type | SSR motif | FORWARD PRIMER (5'-3') | REVERSE PRIMER (5'-3') | size (bp) |
|---|---|---|---|---|---|
| chr4:74327250-74329262 | p2 | (TA)6 | ACATTCCGGAAAACGATGAG | ATTCTGCCTCACCATGAAGG | 177 |
| chr4:74358977-74360993 | p6 | (TATTCA)5 | CCATACCGAGAGAGGTGCAT | GACGTTGAATCGGTGGAGTT | 111 |
| chr4:74358977-74360993 | p2 | (AT)6 | CCATACCGAGAGAGGTGCAT | GACGTTGAATCGGTGGAGTT | 111 |
| chr4:74358977-74360993 | p4 | (ATAC)4 | CCATACCGAGAGAGGTGCAT | GACGTTGAATCGGTGGAGTT | 111 |
| chr4:74404505-74406525 | p2 | (TA)10 | AGCCTCGGGGTTTATTCTGT | AGTCTCAAGGGGATTCGGTT | 159 |
| chr4:74432391-74434406 | p5 | (TTCAT)3 | TATGCATTTGGGTCGTGTGT | CCTCGAATGTTAGGCTCTCG | 284 |
| chr4:74437622-74439640 | p6 | (TATATG)3 | CGTGGCACGTGTATGTAAGG | ACGCATCAAAACCACATGAA | 202 |
| chr4:74437622-74439640 | p2 | (TA)7 | CGTGGCACGTGTATGTAAGG | ACGCATCAAAACCACATGAA | 202 |
| chr4:74445342-74447398 | p6 | (TATGAA)4 | GAGAATTCAACTCGGCCAAA | AAATTTGGAAGGTTTCGGGT | 126 |
| chr4:74461212-74463226 | p2 | (TA)7 | TTTGTTGGCTAGGTATGCCC | CCTAGAGTGAGGCCGTCAAG | 209 |
| chr4:74461212-74463226 | p2 | (TA)18 | TTTGTTGGCTAGGTATGCCC | CCTAGAGTGAGGCCGTCAAG | 209 |
| chr4:74535227-74537242 | p5 | (TCATT)3 | TCCCTCCCTCCATTTCTTCT | AACAACACGGTTAGCGGAAG | 133 |
| chr4:74535227-74537242 | p3 | (ATG)5 | TCCCTCCCTCCATTTCTTCT | AACAACACGGTTAGCGGAAG | 133 |
| chr4:74589590-74591612 | p2 | (TA)11 | ATGGGCTGCTTTTGTCAATC | GAACATCAAAAGCCAGGAGG | 279 |
| chr4:74676298-74678318 | p5 | (ATCAT)4 | CCCTAAACCCTCTTGGAAGG | TCTTTCCAGTCAGAATGGGC | 107 |
| chr4:74685623-74687638 | p5 | (CCATA)3 | AAAGGTCTTGATGCCATTCG | GTCCGGAATTTGAGTTTGGA | 229 |
| chr4:74711954-74713969 | p5 | (CCATA)3 | GGAATAGAGTTTCCCCCGAG | TCTCAGGGATGGCCTATTTG | 219 |
| chr4:74731151-74733166 | p3 | (TTC)5 | GGCCAGCATCGTTATCCTTA | CGAATCTTGGTGGTGACCTT | 107 |
| chr4:74793252-74795266 | p2 | (AT)7 | TCGTTTTCCTATGACTGGGC | AGCAGGTTGTCCTAGGGGTT | 159 |
| chr4:74844697-74846729 | p2 | (TA)16 | ATATTCGGCGAAGTTTGTGG | CAACCCTTTGCGATGATTTT | 106 |
| chr4:74844697-74846729 | p2 | (AT)7 | ATATTCGGCGAAGTTTGTGG | CAACCCTTTGCGATGATTTT | 106 |
| chr4:75054857-75056872 | p5 | (AAAGT)3 | ATGAAAATGCCAGCCGTTAG | CGGGCATTAGGAAACTGAAA | 194 |
| chr4:75297370-75299416 | p2 | (CT)23 | AGTAAAATCTCGCACCCCCT | TTCCAAAGGGAACACCCATA | 186 |
| chr4:75346194-75348240 | p2 | (CT)23 | AGTAAAATCTCGCACCCCCT | TTCCAAAGGGAACACCCATA | 186 |
| chr4:75398494-75400506 | p2 | (TC)6 | TCAGAAGGACGGACCGTAAC | TTGCCTTAAATGCTCCAACC | 189 |
| chr4:75408571-75410586 | p5 | (TTGTT)3 | CCTACAAATCCACTCCCGAA | ACTTCCTGGGCCTTCAATTT | 243 |
| chr4:75423898-75425925 | p2 | (GA)6 | AACATGTTTTGAGCCTCGCT | GGGAAGTCATTTGCTTGGAA | 138 |
| chr4:75687037-75689052 | p5 | (AAGAA)3 | CACATGCTTGAAGAAGCCAA | CGATTTGATGGGCAGTTTTT | 111 |
| chr4:75759806-75761833 | p3 | (ACA)9 | CGATCAAGCAAGCAAACTGA | GCCGTTCACCAGCATCTAAT | 193 |
| chr4:75846458-75848474 | p4 | (TGTA)4 | AGCCTCGGGGATTATTCTGT | TTTGGAAGCATACCGACTCC | 222 |
| chr4:75874314-75876330 | p2 | (AT)8 | GACCATAGTGTCATGCACGG | CCAACGGGACTAGCATGATT | 140 |
| chr4:75930154-75932172 | p6 | (TATTCA)3 | CACGCAACACACCCATAAAC | TATAAATGCGTTGTTGCCCA | 222 |
| chr4:75930154-75932172 | p2 | (TA)30 | CACGCAACACACCCATAAAC | TATAAATGCGTTGTTGCCCA | 222 |
| chr4:75930154-75932172 | p2 | (AT)6 | CACGCAACACACCCATAAAC | TATAAATGCGTTGTTGCCCA | 222 |
| chr4:76047517-76049543 | p2 | (TA)13 | GATAGCCTTGTCGGCTTCTG | AACTAGGCACCCGAACACAC | 202 |
| chr4:76047517-76049543 | p2 | (GT)6 | GATAGCCTTGTCGGCTTCTG | AACTAGGCACCCGAACACAC | 202 |

| Marker ID | SSR type | SSR motif | FORWARD PRIMER (5'-3') | REVERSE PRIMER (5'-3') | size (bp) |
|---|---|---|---|---|---|
| chr4:76054307-76056345 | p2 | (AT)7 | CCACACACACACAACCACAA | GTTCGGAGGTGTCCACTGAT | 250 |
| chr4:76159193-76161205 | p5 | (ATTCA)3 | GGCCCAAATCATCAATCAAC | TCCTTGTTGGAATTGGAAGG | 124 |
| chr4:76159193-76161205 | p2 | (AT)6 | GGCCCAAATCATCAATCAAC | TCCTTGTTGGAATTGGAAGG | 124 |
| chr4:76284953-76286968 | p3 | (ACA)5 | TTGATTTTTGTCACGACCCA | AGCCTTGTCGACTTGCGTAT | 180 |
| chr4:76284966-76286978 | p3 | (ACA)5 | TTGATTTTTGTCACGACCCA | AGCCTTGTCGACTTGCGTAT | 167 |
| chr4:76442173-76444185 | p2 | (CA)6 | AAGCATGTATCAACCGGAGC | ATTACTGAGGAGCACGTGGG | 118 |
| chr4:76615668-76617683 | p5 | (ATTCA)3 | ACCACGTTTCGAGTCAATCC | ACTCGGAGGCGTAATTCCTT | 299 |
| chr4:76615668-76617683 | p2 | (TA)7 | ACCACGTTTCGAGTCAATCC | ACTCGGAGGCGTAATTCCTT | 299 |
| chr4:76759893-76761961 | p2 | (TA)34 | GAACTTGGAAGCGAGGAATG | TCGACGGCATTTCATTTACA | 149 |
| chr4:76759893-76761961 | p2 | (TA)7 | GAACTTGGAAGCGAGGAATG | TCGACGGCATTTCATTTACA | 149 |
| chr4:76784975-76786993 | p6 | (ATCCGT)3 | GGCAGGTTTACGCCACTAAA | CCTAGTCGGGTCGTGACATT | 204 |
| chr4:76958756-76960771 | p3 | (ATG)5 | CCACCCTCCAAATATACCCC | TTTGAATCCAAGCCAAGACC | 291 |
| chr4:76960311-76962331 | p2 | (AT)14 | ATGAGATTCATCGGTGGAGG | GACTCATTGGTTTGGGTCGT | 144 |
| chr4:76960311-76962331 | p2 | (TA)7 | ATGAGATTCATCGGTGGAGG | GACTCATTGGTTTGGGTCGT | 144 |
| chr4:76960333-76962349 | p2 | (AT)14 | ATGAGATTCATCGGTGGAGG | GACTCATTGGTTTGGGTCGT | 122 |
| chr4:76960333-76962349 | p2 | (TA)7 | ATGAGATTCATCGGTGGAGG | GACTCATTGGTTTGGGTCGT | 122 |
| chr4:76987710-76989726 | p4 | (AAAG)4 | GAGGGATGCGGTAAACAGAA | GTGTCGTGCAAGCCTCAGTA | 261 |
| chr4:76990471-76992486 | p3 | (AAT)5 | GGGTGACGGTGCATAGAAGT | GGAATTAGAGCCCGTTGACA | 136 |
| chr4:77031414-77033452 | p2 | (AT)19 | AGGGTCGTCAAACGTCATTC | CCCCAGGATGCAAGTTAAGA | 249 |
| chr4:77260381-77262397 | p6 | (CTGGAG)3 | CGCTGAAATCATAACGCAGA | TGTCTACGAGCCGTGTCAAG | 202 |
| chr4:77260381-77262397 | p4 | (CATA)4 | CGCTGAAATCATAACGCAGA | TGTCTACGAGCCGTGTCAAG | 202 |
| chr4:77285622-77287634 | p2 | (AT)6 | TATGCTACCTCATCCGGCTC | TCATAAGAGTCGTCCCGGTC | 108 |
| chr4:77302994-77305022 | p2 | (AT)14 | CATGCTCGCAGGTTCAAGTA | GAAGCCGACAAGGCTACATC | 215 |
| chr4:77302994-77305022 | p2 | (AT)14 | CATGCTCGCAGGTTCAAGTA | GAAGCCGACAAGGCTACATC | 215 |
| chr4:77471030-77473044 | p2 | (AC)7 | CAGACGTCGTATGCTCTCCA | TGTTTGTTGATGTGTGGGCT | 280 |
| chr4:77523584-77525630 | p2 | (TA)23 | TTCCGGTTAAAGTATTGCGG | TCCGAATGTACAAAATGCGA | 269 |
| chr4:77523584-77525630 | p2 | (AT)31 | TTCCGGTTAAAGTATTGCGG | TCCGAATGTACAAAATGCGA | 269 |
| chr4:77531171-77533185 | p2 | (AC)7 | TCGCCCAATCTTGCTAGTCT | TTGGCCGAGTTAAATTCTCG | 292 |
| chr4:77542078-77544102 | p5 | (TGAAT)5 | TTCCCGGATTGTTGTTGATT | GGATCATGCCTTTGGAAGAA | 201 |
| chr4:77542078-77544102 | p6 | (TATATG)4 | TTCCCGGATTGTTGTTGATT | GGATCATGCCTTTGGAAGAA | 201 |
| chr4:77592681-77594693 | p2 | (TG)6 | CCCATATGGCGAGTAAGCAT | TCATACCTGGCCCTACCAAG | 142 |
| chr4:77735654-77737666 | p2 | (GT)6 | GGAGGATGTTATATCCCGCA | ACATGGCCCCTTTAATTTCC | 300 |
| chr4:77739595-77741609 | p2 | (TA)6 | AGGTTTAGCCGGAACAAGGT | TCGGGTATCAGCCTATCCAG | 175 |
| chr4:77739619-77741643 | p2 | (TA)6 | AGGTTTAGCCGGAACAAGGT | TCGGGTATCAGCCTATCCAG | 151 |
| chr4:77838141-77840153 | p2 | (TA)6 | GCTCCTCAGAATAGGCATCG | AAGGCGGGCTGAAAGAGTAT | 212 |
| chr4:77914059-77916077 | p2 | (AT)9 | CGTGTCCCCTTTGAAACAAT | TCTGTTGGGGATTTGCTAGG | 247 |

| Marker ID | SSR type | SSR motif | FORWARD PRIMER (5'-3') | REVERSE PRIMER (5'-3') | size (bp) |
|---|---|---|---|---|---|
| chr4:77943741-77945753 | p2 | (TG)6 | ATTTTTGCGTGTTCGGGTAG | TTTTGCCCCTAGCCTTTCTT | 300 |
| chr4:77970785-77972800 | p5 | (ACACA)3 | CCACGCCTTATTAGCTACGC | GTTGTTGGCACGATTGTTTG | 295 |
| chr4:77970785-77972800 | p2 | (AT)7 | CCACGCCTTATTAGCTACGC | GTTGTTGGCACGATTGTTTG | 295 |
| chr4:77970785-77972800 | p2 | (TA)6 | CCACGCCTTATTAGCTACGC | GTTGTTGGCACGATTGTTTG | 295 |
| chr4:77971182-77973196 | p5 | (ACACA)3 | AAACAATCGTGCCAACAACA | AACTTGCGAAGTGAGGCATT | 147 |
| chr4:77971182-77973196 | p2 | (AT)7 | AAACAATCGTGCCAACAACA | AACTTGCGAAGTGAGGCATT | 147 |
| chr4:77971182-77973196 | p2 | (TA)6 | AAACAATCGTGCCAACAACA | AACTTGCGAAGTGAGGCATT | 147 |
| chr4:78053177-78055197 | p2 | (TA)10 | GCTTTTTCTGTCCTCGATGC | TGTATGTTTCAGCGGCAGAG | 148 |
| chr4:78075014-78077026 | p2 | (TA)6 | TTGAAGTCGAATCCAAAGGG | CTTAGCCGTAGCATCGTTCC | 129 |
| chr4:78075014-78077026 | p2 | (TA)6 | TTGAAGTCGAATCCAAAGGG | CTTAGCCGTAGCATCGTTCC | 129 |
| chr4:78075345-78077357 | p2 | (TA)6 | TCGGAAGTCACCGACCTTAC | GCCATGTACATACACCAGCG | 201 |
| chr4:78075345-78077357 | p2 | (TA)6 | TCGGAAGTCACCGACCTTAC | GCCATGTACATACACCAGCG | 201 |
| chr4:78091688-78093704 | p2 | (AT)8 | GTTGTGCATCCTTCCTCGAT | AAGCAGCAAAGGAGTACCGA | 228 |
| chr4:78152642-78154672 | p2 | (AT)15 | TGTGATAATTCTGCGGACCA | AGAGAAGGCAAGGCATACGA | 176 |
| chr4:78280103-78282121 | p2 | (AT)9 | TGAACTGCATCTCTCATGGC | CAAATCGGTAGCAGGGGATA | 202 |
| chr4:78280103-78282121 | p2 | (AT)21 | TGAACTGCATCTCTCATGGC | CAAATCGGTAGCAGGGGATA | 202 |
| chr4:78409265-78411280 | p3 | (TGT)5 | CTCAGTTGCAGTGGTGGAGA | CTATCGCTCCTAGTGTGCCC | 267 |
| chr4:78465931-78467947 | p4 | (GAAA)4 | GGATCTCCCATGAGCTGAAA | GAGGTGGAGGATGCCTATGA | 247 |
| chr4:78535315-78537331 | p4 | (AAAT)4 | TTAAATGTTTTTCCCCGTCG | TGACGAAAAGGGGTTAATGC | 213 |
| chr4:78752102-78754117 | p5 | (GATAT)3 | CGAGGTTAAGCGTGCTAAGG | TGCTTTCTCCCCTTCTGCTA | 282 |
| chr4:78757763-78759777 | p2 | (TA)7 | TATCGCCCAACTGAAAATCC | TTTTAGTAAGGTGCCGGGTG | 176 |
| chr4:78769174-78771238 | p5 | (GATTG)3 | AGACGCAAGGAAATTCGCTA | TGCTTTGTCGTAATTGACCG | 100 |
| chr4:78788875-78790890 | p3 | (AGA)5 | TGAAACAGTGGCTCAGATCG | CATTTTACCCAGCTCCTCCA | 253 |
| chr4:78793248-78795316 | p2 | (AT)34 | GAATGGATCCGCAAGAGGTA | TTGACCATCCATTTCAGCAA | 268 |
| chr4:78793248-78795316 | p3 | (GAA)15 | GAATGGATCCGCAAGAGGTA | TTGACCATCCATTTCAGCAA | 268 |
| chr4:78793564-78795609 | p2 | (AT)34 | AATGGATGGTCAAGAGTGGC | AAAGCGCCTACATATCCCCT | 214 |
| chr4:78793564-78795609 | p3 | (GAA)15 | AATGGATGGTCAAGAGTGGC | AAAGCGCCTACATATCCCCT | 214 |
| chr4:78806206-78808218 | p2 | (AT)6 | AATCCTGGTACTGCCCTCCT | CCTCTATTGAGACGGACCCA | 290 |
| chr4:78806206-78808218 | p2 | (TA)6 | AATCCTGGTACTGCCCTCCT | CCTCTATTGAGACGGACCCA | 290 |
| chr4:78922326-78924346 | p2 | (AT)10 | CTCCTACTTGCGAATCGAGG | AGCTTCGCTACAAGATGGGA | 166 |
| chr4:79040587-79042613 | p2 | (TA)13 | CTCGATGTTGCCACTCTTGA | TCGAGGGAAGTTACGAGCAT | 149 |
| chr4:79271313-79273325 | p2 | (TA)6 | TTATGGCCCAGTTAGATGCC | CGTTCCCTCGATTGAGATGT | 123 |
| chr4:79354600-79356612 | p2 | (AT)6 | ATGAAAGTCTGTGGGTTCGC | AAGCCGACGTACCATTATGC | 181 |
| chr4:79385233-79387245 | p2 | (AT)6 | ATGAAAGTCTGTGGGTTCGC | AAGCCGACGTACCATTATGC | 181 |
| chr4:79435454-79437469 | p5 | (AGGAC)3 | ATTCAGTATCGGAGGCATGG | AGTTCCCCCACTAGGTGCTT | 138 |
| chr4:79456891-79458907 | p2 | (TA)8 | CACTGGCACGAAGACTGTGT | AATACTCGGCCCCTTTGAGT | 195 |

| Marker ID | SSR type | SSR motif | FORWARD PRIMER (5'-3') | REVERSE PRIMER (5'-3') | size (bp) |
|---|---|---|---|---|---|
| chr4:79509834-79511846 | p2 | (TA)6 | TCAAATTTACCTTGGACCCG | CGAAACACCCTTTCTTTGGA | 186 |
| chr4:79532994-79535006 | p2 | (AT)6 | TCTTCAATTTCCCACTTGCC | GCCCATGTGGGAGCTAAATA | 251 |
| chr4:79642592-79644654 | p3 | (CAT)6 | CGCTACACGAAGTCTCACCA | TCTTGATCTTGGGGCTCATC | 195 |
| chr4:79759286-79761302 | p2 | (AT)8 | GGCTGATGTCCGAGGTTAAA | AGAATCGGCCATGAAACAAC | 105 |
| chr4:79775360-79777382 | p2 | (AT)11 | TCCATGTTGATCACTTCCCA | GATGGCATGAAGAGGGACAT | 170 |
| chr4:79866355-79868379 | p2 | (TG)12 | TCTCCACGACCCAATAGCTC | ACCAAAACCCTCGTTCACTG | 281 |
| chr4:79866484-79868560 | p2 | (TG)12 | TCTCCACGACCCAATAGCTC | ACCAAAACCCTCGTTCACTG | 152 |
| chr4:79913420-79915435 | p5 | (AAATA)3 | AATATTTTTGCGGAGATGCG | ACCAAAAAGCTTCGTGCAGT | 281 |
| chr4:79940253-79942324 | p4 | (AAAT)4 | TTACTTGGTCCACCCGTCTC | CGGCACTTGACATGTATGCT | 107 |
| chr4:79940253-79942324 | p2 | (AT)7 | TTACTTGGTCCACCCGTCTC | CGGCACTTGACATGTATGCT | 107 |
| chr4:79940498-79942512 | p4 | (AAAT)4 | AGCATACATGTCAAGTGCCG | GTGGTCCTGAAATAGGGCAA | 110 |
| chr4:79940498-79942512 | p2 | (AT)7 | AGCATACATGTCAAGTGCCG | GTGGTCCTGAAATAGGGCAA | 110 |
| chr4:79959543-79961586 | p6 | (CATGAA)3 | ACTGAAACCTGGGCATGAAC | GCAAGCCGACGTACCATTAT | 292 |
| chr4:79959543-79961586 | p6 | (TATACA)4 | ACTGAAACCTGGGCATGAAC | GCAAGCCGACGTACCATTAT | 292 |
| chr4:79959597-79961609 | p6 | (CATGAA)3 | ACTGAAACCTGGGCATGAAC | GCAAGCCGACGTACCATTAT | 238 |
| chr4:79959597-79961609 | p6 | (TATACA)4 | ACTGAAACCTGGGCATGAAC | GCAAGCCGACGTACCATTAT | 238 |
| chr4:79959836-79961860 | p6 | (CATGAA)3 | GCATAATGGTACGTCGGCTT | GCAGGTGGATACCGATGAGT | 246 |
| chr4:79959836-79961860 | p6 | (TATACA)4 | GCATAATGGTACGTCGGCTT | GCAGGTGGATACCGATGAGT | 246 |
| chr4:79981338-79983352 | p2 | (AT)7 | GTGACAAACCCCTAGGACGA | GCAAGCCGACGTACCATTAT | 286 |
| chr4:79997251-79999265 | p2 | (AT)7 | CATGACCTAACTAGGGGCCA | CCAGCTCTGACTTCACTCCC | 297 |
| chr4:80023102-80025138 | p2 | (AT)18 | TTTCAGATCGTGGTGCTCAG | CCCACACAGAGCTTCAAACA | 148 |
| chr4:80046878-80048892 | p5 | (TGCAA)3 | TGCGAAGTCAATGTAGCCTG | GGACCGATATCCAATGGTTG | 244 |
| chr4:80046878-80048892 | p2 | (TA)7 | TGCGAAGTCAATGTAGCCTG | GGACCGATATCCAATGGTTG | 244 |
| chr4:80075761-80077779 | p6 | (ACTGAA)3 | AGTCCACTTCTAGGGCAGCA | GATACTCCGCGGGACACTTA | 146 |
| chr4:80203800-80205822 | p2 | (AT)11 | TTGTCGAGTGTGGCTAGTGG | GAGCCATGTGTTAGGCGTTT | 185 |
| chr4:80203800-80205822 | p2 | (TA)15 | TTGTCGAGTGTGGCTAGTGG | GAGCCATGTGTTAGGCGTTT | 185 |
| chr4:80207029-80209047 | p2 | (TA)9 | TTCTCGACCTTACGTCACCC | TAGCCCACGTCAGGGTAATC | 275 |
| chr4:80236342-80238357 | p3 | (AAT)5 | CAGGGGCTGAACTCTCTTTG | GGGGAGGTTATGGGAGAGTC | 140 |
| chr4:80236342-80238357 | p2 | (TA)6 | CAGGGGCTGAACTCTCTTTG | GGGGAGGTTATGGGAGAGTC | 140 |
| chr4:80274306-80276320 | p2 | (AT)7 | TAATAACATACGCACGCCGA | GATGTCCTGTCGCTGTAGCA | 201 |
| chr4:80393304-80395316 | p2 | (AC)6 | GTCCTAGGGTAATCGGGTCC | GCAGATTTTGGAGGATTGGA | 207 |
| chr4:80478578-80480593 | p5 | (TTTTA)3 | GTAGGTGATGTTCCGAGGGA | TTTGGGAGGATGCCACTTAC | 112 |
| chr4:80491732-80493752 | p2 | (AT)6 | CTCGTTACTAGCACCCGGAG | TGATAAGGGGTGCTCGGTAG | 138 |
| chr4:80491732-80493752 | p2 | (AT)10 | CTCGTTACTAGCACCCGGAG | TGATAAGGGGTGCTCGGTAG | 138 |
| chr4:80678361-80680376 | p5 | (TTCCA)3 | CGGTGTAACGTGCACAACTC | GTGAGCTAAACAAGCTCGGG | 156 |
| chr4:80836908-80838923 | p5 | (GCTTT)3 | TCCTCGTGAAACCTTCTTGG | TCGGAAAGGTTTTCGCTCTA | 291 |

| Marker ID | SSR type | SSR motif | FORWARD PRIMER (5'-3') | REVERSE PRIMER (5'-3') | size (bp) |
|---|---|---|---|---|---|
| chr4:80836908-80838923 | p2 | (TA)32 | TCCTCGTGAAACCTTCTTGG | TCGGAAAGGTTTTCGCTCTA | 291 |
| chr4:80859647-80861662 | p3 | (ATC)5 | TCATAAGGACGGGACAGGAC | GACGGCATAGACGCTTAGGA | 276 |
| chr4:80989286-80991310 | p6 | (TATTCA)4 | GCTCGAGAAAGGGTCAACAG | CTTTTCCTCCGTTCCTTTCC | 131 |
| chr4:81032611-81034626 | p5 | (TTCAT)3 | AGGGCCATATCGCCTAGAAT | GCGACTGAAATCTGATGGGT | 281 |
| chr4:81108614-81110626 | p2 | (AT)6 | TATATTCACCCTCCCCCTCC | CACCACACACACCCTTTTTG | 144 |
| chr4:81129679-81131691 | p2 | (AT)6 | TATATTCACCCTCCCCCTCC | CACCACACACACCCTTTTTG | 144 |
| chr4:81232391-81234406 | p3 | (TTG)5 | ATGAAGGACCACTACCGTCG | GAAGGTGAAGGGGTCACAGA | 148 |
| chr4:81247585-81249597 | p2 | (TA)6 | ATCAATGGCGAAAGTCAACC | AATGTAGCATCACCGAAGGG | 283 |
| chr4:81247585-81249597 | p2 | (AT)6 | ATCAATGGCGAAAGTCAACC | AATGTAGCATCACCGAAGGG | 283 |
| chr4:81287895-81289907 | p3 | (TTG)6 | ACATTCTCACCAACCAAGGC | GTTCGAGGAGCCAAGAGATG | 241 |
| chr4:81287895-81289907 | p2 | (CA)6 | ACATTCTCACCAACCAAGGC | GTTCGAGGAGCCAAGAGATG | 241 |
| chr4:81347830-81349844 | p5 | (ATCAT)3 | CACACATGACCCATTACCCA | CTTTTGATCCGAAGCCCATA | 234 |
| chr4:81347857-81349873 | p5 | (ATCAT)3 | CACACATGACCCATTACCCA | CTTTTGATCCGAAGCCCATA | 207 |
| chr4:81347927-81349945 | p5 | (ATCAT)3 | CACACATGACCCATTACCCA | CTTTTGATCCGAAGCCCATA | 137 |
| chr4:81351383-81353398 | p5 | (AATGC)3 | GATTTCCAGTCTGCTACGGC | TCGTGAAAGAAATTCTGGGG | 168 |
| chr4:81391013-81393028 | p2 | (TC)8 | GGATAAGTTGGGCCCTTTTC | GAGAGAGTTTCAAGGCGGTG | 108 |
| chr4:81391013-81393028 | p5 | (TTTAG)3 | GGATAAGTTGGGCCCTTTTC | GAGAGAGTTTCAAGGCGGTG | 108 |
| chr4:81432311-81434326 | p5 | (AATGC)3 | GATTTCCAGTCTGCTACGGC | TCGTGAAAGAAATTCTGGGG | 168 |
| chr4:81471941-81473956 | p2 | (TC)8 | GGATAAGTTGGGCCCTTTTC | GAGAGAGTTTCAAGGCGGTG | 108 |
| chr4:81471941-81473956 | p5 | (TTTAG)3 | GGATAAGTTGGGCCCTTTTC | GAGAGAGTTTCAAGGCGGTG | 108 |
| chr4:81640317-81642331 | p3 | (TGT)5 | GTGTGTGGCCATGTTTTCTG | CATGAAATGTCCGAGGTGTG | 290 |
| chr4:81640317-81642331 | p2 | (TA)7 | GTGTGTGGCCATGTTTTCTG | CATGAAATGTCCGAGGTGTG | 290 |
| chr4:81676657-81678675 | p6 | (CATATA)3 | TAAAGCCCACACCTTTCCTG | GCAAGCCGACGTACCATTAT | 270 |
| chr4:81716478-81718492 | p2 | (AT)7 | AACTGAGCCCCTCTATCCGT | CCTACGCTTGAAATGGGAAA | 221 |
| chr4:81716478-81718492 | p2 | (GT)6 | AACTGAGCCCCTCTATCCGT | CCTACGCTTGAAATGGGAAA | 221 |
| chr4:81734304-81736319 | p3 | (TTC)5 | TTTACCATGAAAGCCCAAGG | CCGTATCGCTTGGATTGATT | 127 |
| chr4:81740706-81742730 | p6 | (CATATA)4 | GTCACACCCCGACTTTCCTA | TCCACCAGACGATGTGTCAT | 216 |
| chr4:81743157-81745196 | p3 | (CAA)13 | ACAATACAAACGGCCCACAT | AGGTGGAAAAATTGGGGAAG | 172 |
| chr4:81743157-81745196 | p2 | (GA)6 | ACAATACAAACGGCCCACAT | AGGTGGAAAAATTGGGGAAG | 172 |
| chr4:81796989-81799004 | p5 | (ATCAT)3 | GATAAAGCACAAGGAAGCCG | ATTCTGCCATCTTGGAGTGG | 272 |
| chr4:81800282-81802298 | p4 | (AATA)4 | TAGCGGAGATCAATCCATCC | ACAACGATTCATCAGTGGCA | 259 |
| chr4:81875017-81877032 | p5 | (TGCAA)3 | TACATTCCATACGGGGCTTC | ATAAGGTAGGTGCCCACACG | 111 |
| chr4:81927549-81929561 | p2 | (CA)6 | CAACACCTTAAGCATCGCAA | CGGAGGTTCGTAGCTTTCTG | 268 |
| chr4:82033548-82035568 | p2 | (TA)6 | TTTGTCATGACCCAACTGGA | TTATCGGCTGGTGATGATGA | 118 |
| chr4:82033548-82035568 | p4 | (CATA)5 | TTTGTCATGACCCAACTGGA | TTATCGGCTGGTGATGATGA | 118 |
| chr4:82061745-82063760 | p5 | (TTTTC)3 | ACAGGGACTAAAAGCGCAAA | CAATGCAGTGAATGATTGGG | 144 |

| Marker ID | SSR type | SSR motif | FORWARD PRIMER (5'-3') | REVERSE PRIMER (5'-3') | size (bp) |
|---|---|---|---|---|---|
| chr4:82093672-82095688 | p2 | (TA)8 | CGAATAGCAGTCCGAAGGAG | TGGTCGCCATTACTCACAAA | 299 |
| chr4:82120012-82122024 | p2 | (AT)6 | AAAACAATTCCCTTGGACCC | CGCAAGAAAATGCTAGGAGG | 197 |
| chr4:82120012-82122024 | p3 | (ATA)5 | AAAACAATTCCCTTGGACCC | CGCAAGAAAATGCTAGGAGG | 197 |
| chr4:82149614-82151626 | p2 | (AT)6 | CCCTTTGGCACATAGTCGTT | CGGCCGATATTCTTCTTCAG | 132 |
| chr4:82149614-82151626 | p2 | (TA)6 | CCCTTTGGCACATAGTCGTT | CGGCCGATATTCTTCTTCAG | 132 |
| chr4:82159090-82161105 | p5 | (TTTAT)3 | GAGCCCTTTAATTTTTCCCG | CCCTGTAACTATGGCCTCCA | 230 |
| chr4:82159090-82161105 | p5 | (TTATT)3 | GAGCCCTTTAATTTTTCCCG | CCCTGTAACTATGGCCTCCA | 230 |
| chr4:82159369-82161384 | p5 | (TTTAT)3 | TGGAGGCCATAGTTACAGGG | AGTTCAATTCACACGCCACA | 112 |
| chr4:82159369-82161384 | p5 | (TTATT)3 | TGGAGGCCATAGTTACAGGG | AGTTCAATTCACACGCCACA | 112 |
| chr4:82260858-82262912 | p2 | (TA)27 | GTTGGTTCGTGACAATGACG | GCCACGTGGGCTTATACATT | 166 |
| chr4:82355418-82357436 | p6 | (AACTGA)3 | TATGGCAACAAGCCCACATA | AGCGGACTAGTTGTGGCACT | 187 |
| chr4:82541942-82543957 | p3 | (TCC)5 | AACCAAGACCTCGTAAGGCA | TCCATCACCCACCTATGGAT | 152 |
| chr4:82541942-82543957 | p5 | (GTTTT)3 | AACCAAGACCTCGTAAGGCA | TCCATCACCCACCTATGGAT | 152 |
| chr4:82625911-82627929 | p3 | (CAA)6 | ACGAGCGGAGTAGTGAGGAA | AAAAGCTCGGATTGCTTTGA | 282 |
| chr4:82652454-82654466 | p2 | (AT)6 | TGCCTAGAGAATCGTTGCCT | CCCACCTCTGTTTTCCCATA | 242 |
| chr4:82680514-82682532 | p2 | (AT)33 | AGGTGTAACATCCTTCCCCC | TTGCCCTGTATGGCATGTAA | 141 |
| chr4:82680514-82682532 | p2 | (AT)9 | AGGTGTAACATCCTTCCCCC | TTGCCCTGTATGGCATGTAA | 141 |
| chr4:82680514-82682532 | p2 | (AT)17 | AGGTGTAACATCCTTCCCCC | TTGCCCTGTATGGCATGTAA | 141 |
| chr4:82684444-82686459 | p3 | (CAA)5 | TGTCACAAGGCCAATATGGA | TTGCAAAATGGTTCACGAAA | 280 |
| chr4:82741638-82743653 | p2 | (AT)8 | GCTCCGCTATGTAGTCCCTG | TACGTACACGGTCACGCATT | 281 |
| chr4:82741638-82743653 | p3 | (CAT)5 | GCTCCGCTATGTAGTCCCTG | TACGTACACGGTCACGCATT | 281 |
| chr4:82795446-82797461 | p5 | (TTTTC)3 | ATACTTGTACCGAGCACCCG | TGTTTGGGCATTGTGATGTT | 162 |
| chr4:82820397-82822415 | p6 | (CATATA)3 | TCCAAAGAAAGGGCAACATC | TAGCCTTGTCGGCTTCTGTT | 164 |
| chr4:82837155-82839169 | p5 | (GATAT)3 | AGAGGAACCTCTGCCCAAAT | GCTGTGCTAAAGCAAAAGGG | 132 |
| chr4:82837155-82839169 | p2 | (AT)7 | AGAGGAACCTCTGCCCAAAT | GCTGTGCTAAAGCAAAAGGG | 132 |
| chr4:82848590-82850620 | p2 | (AT)15 | TTAGCCCCAGTCCAGACATC | GAACTCTGGCAAAGAAACGC | 113 |
| chr4:82875307-82877321 | p2 | (AT)9 | TCTTGGAGAGCTCCGTTGTT | AGCGAAATATGACGGGACAG | 233 |
| chr4:82875307-82877321 | p2 | (AT)7 | TCTTGGAGAGCTCCGTTGTT | AGCGAAATATGACGGGACAG | 233 |
| chr4:82904840-82906855 | p5 | (TCTTT)3 | ATGGGCGATTTTCAGTATGC | ACATGCTCCACCCTCACTTC | 132 |
| chr4:82962670-82964685 | p4 | (AATA)5 | GTTCCATCATTTCACGACCC | CATTCCGCGTATTCCATCTT | 281 |
| chr4:82962670-82964685 | p5 | (TGCAA)3 | GTTCCATCATTTCACGACCC | CATTCCGCGTATTCCATCTT | 281 |
| chr4:82962670-82964685 | p2 | (AT)7 | GTTCCATCATTTCACGACCC | CATTCCGCGTATTCCATCTT | 281 |
| chr4:82969295-82971337 | p6 | (CATATA)7 | ATTGAGTGGGCCGTATGAAC | ATTGAGAGCGCTCCAGTTGT | 236 |
| chr4:82986561-82988576 | p2 | (AT)16 | GAGGCTTTGCAGACAGTTCC | TCACACAGCACCCTTGGTAA | 209 |
| chr4:82986561-82988576 | p5 | (TTTAA)3 | GAGGCTTTGCAGACAGTTCC | TCACACAGCACCCTTGGTAA | 209 |
| chr4:83044827-83046843 | p2 | (AT)6 | GGAACTGTCTGCAAAGCCTC | GCCTTCAGGTACTGGTCTCG | 247 |

| Marker ID | SSR type | SSR motif | FORWARD PRIMER (5'-3') | REVERSE PRIMER (5'-3') | size (bp) |
|---|---|---|---|---|---|
| chr4:83044827-83046843 | p4 | (CATA)4 | GGAACTGTCTGCAAAGCCTC | GCCTTCAGGTACTGGTCTCG | 247 |
| chr4:83065620-83067632 | p2 | (GT)6 | CTTGTTTTCGAGCCGTTTTC | GTTCCGAGTGCCAAAACAAT | 190 |
| chr4:83145050-83147062 | p2 | (GT)6 | TGTGGCGTACATTAGGGTGA | CTTGGCCCAATTCGTAAAAA | 269 |
| chr4:83175965-83177980 | p2 | (AT)22 | CCACCAAAACTTAAGGGCAA | GCCTCTCGCATCTTGTCTTC | 245 |
| chr4:83175981-83177993 | p2 | (AT)22 | CCACCAAAACTTAAGGGCAA | GCCTCTCGCATCTTGTCTTC | 229 |
| chr4:83176135-83178179 | p2 | (AT)22 | GGAAGACAAGATGCGAGAGG | CTTCTTCGGCCTACCCCTAC | 277 |
| chr4:83226987-83229002 | p5 | (TATTT)3 | TGAGACCCTCATATCGGACC | TAGTCGTTCCCGTCCAAAAC | 143 |
| chr4:83284327-83286339 | p2 | (AT)6 | CAGTTCAAAGGACGAACCGT | TCCAACGAACATAACCCCAT | 120 |
| chr4:83312968-83314998 | p5 | (GAAAT)3 | GAAGGGGTGTCAAGGGAAAT | GATTGGCCAAACTTTCTGGA | 169 |
| chr4:83312968-83314998 | p2 | (TA)15 | GAAGGGGTGTCAAGGGAAAT | GATTGGCCAAACTTTCTGGA | 169 |
| chr4:83399344-83401374 | p2 | (AT)6 | ATAAGCCGACGAGGCTAACA | GTCCTGTCCCGCCTATATGA | 264 |
| chr4:83399344-83401374 | p2 | (TA)15 | ATAAGCCGACGAGGCTAACA | GTCCTGTCCCGCCTATATGA | 264 |
| chr4:83429334-83431352 | p2 | (AT)18 | AATCCCATTGGGCAACAATA | GGATCTCCATTTCATTCGGA | 216 |
| chr4:83429334-83431352 | p2 | (AT)9 | AATCCCATTGGGCAACAATA | GGATCTCCATTTCATTCGGA | 216 |
| chr4:83429334-83431352 | p6 | (TCCGTA)3 | AATCCCATTGGGCAACAATA | GGATCTCCATTTCATTCGGA | 216 |
| chr4:83546096-83548114 | p2 | (AT)10 | CAAATAACACGGGTGCTCCT | TCCTTGTAGAATGGATCCCG | 238 |
| chr4:83546096-83548114 | p2 | (AT)6 | CAAATAACACGGGTGCTCCT | TCCTTGTAGAATGGATCCCG | 238 |
| chr4:83546096-83548114 | p2 | (AT)9 | CAAATAACACGGGTGCTCCT | TCCTTGTAGAATGGATCCCG | 238 |
| chr4:83546096-83548114 | p2 | (AT)6 | CAAATAACACGGGTGCTCCT | TCCTTGTAGAATGGATCCCG | 238 |
| chr4:83565573-83567585 | p2 | (TA)39 | TGTGTAGTGGCAGCCTTGTC | GCCATGACTAGTGCCCAAAC | 275 |
| chr4:83565573-83567585 | p6 | (ATCATT)3 | TGTGTAGTGGCAGCCTTGTC | GCCATGACTAGTGCCCAAAC | 275 |
| chr4:83665106-83667118 | p2 | (AT)15 | TTACGCTAATCGGCCTTGTT | GCCCAGGTAAGTCAGCCATA | 217 |
| chr4:83665106-83667118 | p2 | (AT)6 | TTACGCTAATCGGCCTTGTT | GCCCAGGTAAGTCAGCCATA | 217 |
| chr4:83665119-83667167 | p2 | (AT)15 | TTACGCTAATCGGCCTTGTT | GCCCAGGTAAGTCAGCCATA | 204 |
| chr4:83665119-83667167 | p2 | (AT)6 | TTACGCTAATCGGCCTTGTT | GCCCAGGTAAGTCAGCCATA | 204 |
| chr4:83687914-83689926 | p2 | (AT)15 | TTACGCTAATCGGCCTTGTT | GCCCAGGTAAGTCAGCCATA | 217 |
| chr4:83687914-83689926 | p2 | (AT)6 | TTACGCTAATCGGCCTTGTT | GCCCAGGTAAGTCAGCCATA | 217 |
| chr4:83687927-83689975 | p2 | (AT)15 | TTACGCTAATCGGCCTTGTT | GCCCAGGTAAGTCAGCCATA | 204 |
| chr4:83687927-83689975 | p2 | (AT)6 | TTACGCTAATCGGCCTTGTT | GCCCAGGTAAGTCAGCCATA | 204 |
| chr4:83723446-83725458 | p2 | (CA)6 | GAATTGGTTCAGCAAACGGT | AGCTCTGAAGGCAGATCCAA | 270 |
| chr4:83756947-83758971 | p5 | (TGAAT)3 | TTGATGTTGTTGGCACGATT | GCTTAATTGTCCAAGCTCGC | 179 |
| chr4:83756947-83758971 | p6 | (TATATG)4 | TTGATGTTGTTGGCACGATT | GCTTAATTGTCCAAGCTCGC | 179 |
| chr4:83805326-83807341 | p3 | (CAA)5 | CGGACGGAATCATAGCAAGT | TTGCTGACTCCCTCGAGTTT | 135 |
| chr4:83920423-83922479 | p2 | (TA)28 | GCCCTCGAAAGGTGTATGAA | GTGGAGCAATCCTCTCTTGC | 289 |
| chr4:83931752-83933782 | p2 | (AT)15 | CGTGCTGACCTCCTTTCTTC | AGCGAAATATGACGGGACAG | 169 |
| chr4:83988255-83990275 | p4 | (TTAT)5 | CAATATCCGATGGTTGTCCC | TCCATGTACCTAACGGCTCC | 197 |

| Marker ID | SSR type | SSR motif | FORWARD PRIMER (5'-3') | REVERSE PRIMER (5'-3') | size (bp) |
|---|---|---|---|---|---|
| chr4:84025348-84027384 | p4 | (TTAT)9 | GCTTTCTTCTCTCGCCTCAA | ATGGATCCGACAACAACCTC | 156 |
| chr4:84047290-84049305 | p5 | (TATCA)3 | CCTAGGACCGAACTGCTCTG | GATGGTCCTCGTCGCTGTAT | 286 |
| chr4:84102332-84104344 | p2 | (AT)6 | CCACCTGTCTCCTGAGGTGT | ACCCATTCGTAGTGGCAGAC | 134 |
| chr4:84173947-84175965 | p3 | (AAC)6 | GGCACGGTGTTATACTGCCT | CGACCTTACGTCACTCCGAT | 243 |
| chr4:84173947-84175965 | p2 | (CT)10 | GGCACGGTGTTATACTGCCT | CGACCTTACGTCACTCCGAT | 243 |
| chr4:84211801-84213816 | p3 | (ATC)5 | TCCATCATCATTAGCCGACA | ATGCTCGCAGGTAGACAGGT | 133 |
| chr4:84220997-84223009 | p2 | (CT)6 | TGGGGTCAAGGAAGAGAGAA | TCTGAACCCTCCAAAACCAC | 282 |
| chr4:84278778-84280816 | p2 | (TA)19 | TTTCGAATTGACCTTGGACC | ACTCCGCTACTTCTGGCAAA | 124 |
| chr4:84278778-84280816 | p2 | (TA)20 | TTTCGAATTGACCTTGGACC | ACTCCGCTACTTCTGGCAAA | 124 |
| chr4:84289431-84291446 | p2 | (TA)8 | GAGAGCTTGACCCAGATTCG | CCTCTCGGCCATCAAAATAA | 287 |
| chr4:84289431-84291446 | p5 | (AATGC)3 | GAGAGCTTGACCCAGATTCG | CCTCTCGGCCATCAAAATAA | 287 |
| chr4:84294388-84296400 | p5 | (TGAAT)3 | GAGCCACTATGTTGCCGAAT | ACAACGTCCAAAGAAAACGG | 289 |
| chr4:84294497-84296509 | p5 | (TGAAT)3 | GAGCCACTATGTTGCCGAAT | ACAACGTCCAAAGAAAACGG | 180 |
| chr4:84365478-84367493 | p3 | (CAT)5 | CACCATCCGACAAGGCTACT | ACTGACGTCCCTTCTTGTGG | 203 |
| chr4:84522866-84524880 | p2 | (TA)7 | TATTATGGGGGTCAAACGGA | TGATGATGGAGGAGGGTAGG | 295 |
| chr4:84528893-84530905 | p2 | (TA)6 | TTTACGCCCATGAACAGTGA | TCCGTTTGACCCCCATAATA | 122 |
| chr4:84535647-84537659 | p2 | (TA)6 | CCCGGGGTAATGTAGTTGTG | AAAGGTGCCCGAAGAAAGTT | 240 |
| chr4:84583887-84585903 | p2 | (AG)8 | GGTAAAGAAGCCATGTCGGA | CGGGATACTTAAGGCAACCA | 112 |
| chr4:84629989-84632004 | p5 | (CATTG)3 | TGTTCGGCTATCCAGGTAGG | CAAAAGTGAGGGACCACCAT | 225 |
| chr4:84677457-84679472 | p5 | (AACCG)3 | TGGTCCATGTGCACGATACT | ATATGGGCTGCATATCTGGC | 201 |
| chr4:84681794-84683816 | p2 | (TG)11 | GCGGAGGATGAGATGAAAAA | TTATAAGTATGGCGGGGCAG | 146 |
| chr4:84715554-84717572 | p6 | (TAAAAA)3 | AATGGCCAAACTTGCTCTTG | AATTGCAACCGACGAAAAAC | 168 |
| chr4:84750547-84752562 | p5 | (GAAAT)3 | GACGTTAGGCCAAAGAACCA | TGTTGACGTTTGGGAGTTGA | 242 |
| chr4:84779233-84781248 | p5 | (CAATG)3 | ATATAGGGAGTTGGGGCAGG | TGGGTCGTGACAAGTTGGTA | 221 |
| chr4:84850208-84852220 | p2 | (TA)6 | GTGACACAGGATGCTGAGGA | CCCTGGAGTATCGCATGTTT | 158 |
| chr4:84869163-84871177 | p2 | (AT)7 | AGGCCGTACAAGAGCCACTA | GGATCGATTTCTGTGCCCTA | 266 |
| chr4:84894974-84896992 | p3 | (CAT)6 | GCCGACTAGGCATACCAAAA | TCCGGAGCTGCAGTTTAGTT | 281 |
| chr4:85075041-85077061 | p2 | (AT)10 | GCTCTCTTTCTCCAACCCCT | CAACACCTCCAGTCCATCCT | 293 |
| chr4:85075041-85077061 | p2 | (AT)18 | GCTCTCTTTCTCCAACCCCT | CAACACCTCCAGTCCATCCT | 293 |
| chr4:85216088-85218100 | p2 | (CA)6 | AAATCAATGAAGTCACGGGC | ATTTCTGAGGAGCATGTGGG | 202 |
| chr4:85216088-85218100 | p2 | (AT)7 | AAATCAATGAAGTCACGGGC | ATTTCTGAGGAGCATGTGGG | 202 |
| chr4:85307263-85309283 | p6 | (TATATG)4 | ACCTGGTCAGTTGGCGTAAC | ATGCATGCTTGGCTTTCTTT | 137 |
| chr4:85307263-85309283 | p2 | (AC)10 | ACCTGGTCAGTTGGCGTAAC | ATGCATGCTTGGCTTTCTTT | 137 |
| chr4:85322704-85324719 | p6 | (ATACAT)3 | TTAGACGCAACCCACAGACA | GTGCTCCTTTGTTCTGGAGC | 226 |
| chr4:85322704-85324719 | p3 | (CAT)5 | TTAGACGCAACCCACAGACA | GTGCTCCTTTGTTCTGGAGC | 226 |
| chr4:85439383-85441401 | p6 | (TTTTCT)3 | CGCTGGAGTAAGGAAACAGG | GAAACCCCAAATCCAATCCT | 163 |

| Marker ID | SSR type | SSR motif | FORWARD PRIMER (5'-3') | REVERSE PRIMER (5'-3') | size (bp) |
|---|---|---|---|---|---|
| chr4:85535784-85537796 | p2 | (AT)6 | TATTTGCTGCGTGCAGATTC | ATTTCCTTTCCACGCTTCCT | 114 |
| chr4:85674990-85677006 | p2 | (AT)8 | ATCACGACCTTACGTCACCC | CCAAATAGGCGCGATGTTAT | 236 |
| chr4:85674990-85677006 | p2 | (AT)8 | ATCACGACCTTACGTCACCC | CCAAATAGGCGCGATGTTAT | 236 |
| chr4:85774295-85776329 | p2 | (TA)17 | CTTCGGTGGGAGAAAATTCA | ACGGAAGCGTCATTCCATAG | 230 |
| chr4:85811387-85813402 | p5 | (CACTA)3 | CCTTTCCTTAAGGCCCTTTG | TATGTGCATTTGGAACCCCT | 104 |
| chr4:85811387-85813402 | p2 | (AT)9 | CCTTTCCTTAAGGCCCTTTG | TATGTGCATTTGGAACCCCT | 104 |
| chr4:85906407-85908419 | p2 | (AC)6 | CCTTTGCGGGTGACCTATTA | TGGCAACAATGGAGTCAAAA | 188 |
| chr4:85914846-85916861 | p5 | (AAACC)3 | AGTAGTGCCTGCACCGTTTT | AATTGAGGGCAAAATGCAAG | 117 |
| chr4:85914846-85916861 | p5 | (TAAAT)3 | AGTAGTGCCTGCACCGTTTT | AATTGAGGGCAAAATGCAAG | 117 |
| chr4:86008138-86010153 | p3 | (TGT)5 | GAAGATTCGGCCAAAGTGAA | ACCACTACTCACGGCTCACC | 249 |
| chr4:86008138-86010153 | p2 | (TA)15 | GAAGATTCGGCCAAAGTGAA | ACCACTACTCACGGCTCACC | 249 |
| chr4:86054416-86056430 | p2 | (AT)7 | TACGGCCCGATCCATAAATA | GACGTACATTCCTCGGCACT | 178 |
| chr4:86069036-86071054 | p2 | (GT)9 | AAGCTATGGCTCCGAATGAA | TCCTTTTGTTTGAGGACGCT | 167 |
| chr4:86069036-86071054 | p2 | (TA)9 | AAGCTATGGCTCCGAATGAA | TCCTTTTGTTTGAGGACGCT | 167 |
| chr4:86202035-86204071 | p2 | (AT)18 | TCCGGATTGTCCCAATGTAT | CGTCGTGGCTTTTAGTAGGG | 275 |
| chr4:86202035-86204071 | p2 | (AT)6 | TCCGGATTGTCCCAATGTAT | CGTCGTGGCTTTTAGTAGGG | 275 |
| chr4:86217400-86219436 | p6 | (AGAGAA)3 | CGAAACTTGTCCCAAACGAT | CCCTACGCTTCTCTCCCTCT | 283 |
| chr4:86217400-86219436 | p3 | (AAT)12 | CGAAACTTGTCCCAAACGAT | CCCTACGCTTCTCTCCCTCT | 283 |
| chr4:86302953-86304975 | p2 | (TA)12 | GCCGAATAGGCCTTCTCTTT | CGTTTGTATGGAGTGGGCTT | 157 |
| chr4:86302953-86304975 | p2 | (TA)11 | GCCGAATAGGCCTTCTCTTT | CGTTTGTATGGAGTGGGCTT | 157 |
| chr4:86305951-86307973 | p2 | (AT)11 | TCAGGTGCCTGTTGACTCAG | CGTACCGGGAAGAAGATGAG | 138 |
| chr4:86330260-86332275 | p3 | (CAT)5 | TATAAAGAATGGCTTGCGGG | CCCCTTTTTCACGCTCAATA | 223 |
| chr4:86332036-86334105 | p3 | (TAT)23 | GTTTGGGGAAAGTGGGAAAT | TTCCAGAAGGTGTGAATCCC | 132 |
| chr4:86399863-86401877 | p2 | (TA)7 | CCGTTCGATTTCATTGCTTT | ATACCTGCCCTCACATCGAG | 174 |
| chr4:86435163-86437178 | p5 | (ATTCA)3 | ATGGAAGTATGCTCGTTGGG | CTCGCATAAATAGGACCCGA | 259 |
| chr4:86486727-86488742 | p5 | (TAAAT)3 | CCTACCGTGCAAACTTCCAT | TCAGCCCTAGTTGGGTGAAG | 119 |
| chr4:86659054-86661066 | p2 | (AC)6 | TTGCGCTGAGTGAATAGTGC | CGGTAGCCAATATAGCCCAA | 157 |
| chr4:86701828-86703844 | p4 | (TTTC)4 | TACTTCACCCGACACCTTCC | AATGAAGGTTTGGTGGCTTG | 200 |
| chr4:86746092-86748106 | p6 | (TGTATA)3 | CCCTTCCCTTTTTCCTTCAG | ACCTCGGATCAAATGAGGTG | 187 |
| chr4:86746092-86748106 | p2 | (AT)7 | CCCTTCCCTTTTTCCTTCAG | ACCTCGGATCAAATGAGGTG | 187 |
| chr4:86838493-86840523 | p2 | (AT)15 | AACTCGCAACAATGCTGATG | GGTACCCGTCACGACCTCTA | 165 |
| chr4:86838493-86840523 | p6 | (CATATA)3 | AACTCGCAACAATGCTGATG | GGTACCCGTCACGACCTCTA | 165 |
| chr4:86840361-86842376 | p5 | (CATAT)3 | CGCTGTTTTCCAACTTCCAT | GGTTATTGGGTACGTGTGGG | 188 |
| chr4:86952705-86954727 | p2 | (TA)11 | TGTTAAAAGGAGGTCGGGTG | GGATTCATGCCAAAGAAGGA | 139 |
| chr4:86952705-86954727 | p2 | (AT)11 | TGTTAAAAGGAGGTCGGGTG | GGATTCATGCCAAAGAAGGA | 139 |
| chr4:86952705-86954727 | p2 | (TA)10 | TGTTAAAAGGAGGTCGGGTG | GGATTCATGCCAAAGAAGGA | 139 |

| Marker ID | SSR type | SSR motif | FORWARD PRIMER (5'-3') | REVERSE PRIMER (5'-3') | size (bp) |
|---|---|---|---|---|---|
| chr4:86959898-86961913 | p3 | (TGT)5 | AAAAGGGGAATTCGCAAGAT | GGTCCCTTGGCCATTTATTT | 207 |
| chr4:86963251-86965263 | p2 | (TA)6 | AGGAGACCCGGTCAGAAGTT | AACTACGGCAAAACCGATTG | 147 |
| chr4:87012498-87014513 | p5 | (ATGCA)3 | CCTGAAATGATCTCCGTCGT | AGTTTGCCAAGAGAAGGCAA | 236 |
| chr4:87048031-87050052 | p3 | (ATG)7 | GCATTCTCGGGTTTGTTGTT | CCACGCCTTATTAGCTACGC | 133 |
| chr4:87158271-87160295 | p3 | (TCA)8 | TGGGTCAAGCTCACCTCTCT | CCTGGACGCAGAATAACGAT | 256 |
| chr4:87168654-87170722 | p2 | (TA)6 | AAGGCCGCAAATATCAACAG | AATCATTTGATTTGGCTCGG | 258 |
| chr4:87183626-87185646 | p5 | (CATTT)4 | GATTCAAGATGAAGCCCCAA | CATTTCGGAACATTCGTCCT | 232 |
| chr4:87286802-87288826 | p2 | (AT)12 | CGAACTGCACTGACACCACT | TTTGGTAGCCTTGTTGGCTT | 108 |
| chr4:87327362-87329408 | p2 | (AC)9 | ACTACACAAACACACGGCCA | GCGAACCATTCTTTTGGTGT | 125 |
| chr4:87359479-87361494 | p3 | (ATG)5 | AGCCGTCTTTCTTCCTCACA | ATGCAGAAGTAAAGCCACCG | 146 |
| chr4:87365027-87367042 | p5 | (TTTTA)3 | ATATGGGAGACTGCGTCCAC | TACTCACGCTCACTCATCCG | 190 |
| chr4:87367406-87369431 | p5 | (ATTCA)5 | GCCCAAGGAAAGTCATGTGT | AATTGTTCCTTCAAGGCGTG | 181 |
| chr4:87367406-87369431 | p5 | (ATTCA)4 | GCCCAAGGAAAGTCATGTGT | AATTGTTCCTTCAAGGCGTG | 181 |
| chr4:87382259-87384274 | p5 | (CAAAA)3 | AACAACTGAGGGCGCAATAC | ACCTTTGTCGCTATCCCCTT | 135 |
| chr4:87445676-87447691 | p5 | (TATGA)3 | ATTCCATATGGCGGGATACA | TCGGACATCGTTATGGATCA | 265 |
| chr4:87459827-87461841 | p2 | (TA)7 | GGTTCGCATCACACCCTAGT | CTCGGTAAAAACACCGAGGA | 104 |
| chr4:87480233-87482248 | p5 | (GGTTC)3 | ATGGTTTGTGGCCATGTTTT | ACAAATCGCTTTGCTCGACT | 232 |
| chr4:87495578-87497594 | p2 | (AC)8 | CAAACGAAGAAGTGGGAAGC | TTGATGTGTGGGCTGTTGTT | 255 |
| chr4:87495578-87497594 | p2 | (AT)12 | CAAACGAAGAAGTGGGAAGC | TTGATGTGTGGGCTGTTGTT | 255 |
| chr4:87558368-87560392 | p2 | (AT)12 | ACTTTTGTCACGACCCAACC | GCCTTGTCGGCTTGTGTATT | 179 |
| chr4:87675869-87677884 | p3 | (ACA)5 | AGCTTCGCTTATTCGTCCAA | TCTCGGCTTGTTGATTGTTG | 268 |
| chr4:87695626-87697638 | p2 | (TA)6 | ACCTCCTAGGGCCTTCATGT | TGCTCCCATGGCTCCTATAC | 140 |
| chr4:87708937-87710949 | p2 | (TA)6 | AAATCAAAGTGGGAAGCCCT | ACGTCACTTAGGAACCCACG | 230 |
| chr4:87710365-87712380 | p5 | (TGATA)3 | GCCCAAATTTCTGTAGGCAA | AGAATGCTTTTCGAGGCTCA | 120 |
| chr4:87840503-87842517 | p2 | (TA)7 | GGTAATCGTCACAGGCCCTA | CGAAACGACGAAAACCATTT | 218 |
| chr4:87847184-87849196 | p2 | (CA)6 | ACTTGCGAGGGTTCTAGGGT | GGATTTTGCGACGTGAATTT | 226 |
| chr4:87946611-87948639 | p2 | (AT)14 | TGCGTAATCGGGTTTCTACC | CTTGGAAGGATGACTTGGGA | 160 |
| chr4:87961399-87963427 | p2 | (AT)14 | TGCGTAATCGGGTTTCTACC | CTTGGAAGGATGACTTGGGA | 160 |
| chr4:87973706-87975726 | p2 | (AT)10 | ACCCCCTAGGATGAACTGCT | AAGCCGACGTACCATTATGC | 119 |
| chr4:88103479-88105495 | p2 | (TA)8 | GCCGACGTACCATTATGCTT | GACCCTTATTTAGGGCCGAG | 186 |
| chr4:88103479-88105495 | p4 | (AAAG)6 | GCCGACGTACCATTATGCTT | GACCCTTATTTAGGGCCGAG | 186 |
| chr4:88104387-88106411 | p2 | (TA)8 | CTTTCTTCGAGAGCTGCGTT | GTGAAACATGGGGGTACAGG | 298 |
| chr4:88104387-88106411 | p4 | (AAAG)6 | CTTTCTTCGAGAGCTGCGTT | GTGAAACATGGGGGTACAGG | 298 |
| chr4:88232940-88234958 | p3 | (CAT)6 | AATCAGCCCCTTGTTTGTTG | GGATGGCACGAGTCTGAAAT | 285 |
| chr4:88242763-88244779 | p2 | (AT)8 | TTCCACGTGTCTATGCTTCG | TCCTCCCTGTTATTGTTCCG | 266 |
| chr4:88291219-88293231 | p2 | (GT)6 | GCCTCCAAGCTGCAAAATAG | AATTCCCGAAACCAATTTCC | 135 |

| Marker ID | SSR type | SSR motif | FORWARD PRIMER (5'-3') | REVERSE PRIMER (5'-3') | size (bp) |
|---|---|---|---|---|---|
| chr4:88293998-88296016 | p3 | (TGT)6 | CATGTGATTCGGTTGGTCTG | TCAGAAGCCGACAAGGCTAT | 103 |
| chr4:88316929-88318947 | p3 | (CAA)5 | ATTACGAAATGACGCTTCCG | TGCAGACAGTGTGTTGTGGA | 121 |
| chr4:88316949-88318983 | p3 | (CAA)5 | ATTACGAAATGACGCTTCCG | TGCAGACAGTGTGTTGTGGA | 101 |
| chr4:88329146-88331158 | p2 | (TG)6 | TCGGCACTACTCTGTTCGTG | TCGGAAACCGTAACACATCA | 162 |
| chr4:88562121-88564205 | p3 | (AAT)28 | AGTGGGGAACAGAGGGAAGT | GGTTGGACCCGACCTAATTT | 269 |
| chr4:88644227-88646241 | p2 | (AT)7 | TTCAAATCAACCGCATACGA | GCTTGTGTTATTGGTGGGCT | 124 |
| chr4:88647631-88649647 | p2 | (AT)6 | CCTTAGACCGACCTGCTCTG | GTAAGTGGCTTTAGCAGCCG | 235 |
| chr4:88647631-88649647 | p2 | (TA)8 | CCTTAGACCGACCTGCTCTG | GTAAGTGGCTTTAGCAGCCG | 235 |
| chr4:88647631-88649647 | p6 | (CATATA)3 | CCTTAGACCGACCTGCTCTG | GTAAGTGGCTTTAGCAGCCG | 235 |
| chr4:88649778-88651796 | p2 | (AT)9 | TTCGCCGATATTCCACTTTC | GCTTGTGTATTGGATGGGCT | 105 |
| chr4:88649778-88651796 | p5 | (AGGGT)3 | TTCGCCGATATTCCACTTTC | GCTTGTGTATTGGATGGGCT | 105 |
| chr4:88661885-88663899 | p2 | (AG)7 | CTCCTTGCATCGTATGCCTT | TCACCGGTGTTCGTAATTCA | 262 |
| chr4:88677922-88679937 | p3 | (AAT)5 | TCAGAATACGGACCGTCACA | GACGTCCTCCATCGATTCAT | 141 |
| chr4:88677922-88679937 | p2 | (AG)7 | TCAGAATACGGACCGTCACA | GACGTCCTCCATCGATTCAT | 141 |
| chr4:88821060-88823078 | p3 | (TTG)6 | CGTTTTCGGGTGATCTGACT | TGGGTGTTGTGTCCTGAAAA | 212 |
| chr4:88869466-88871480 | p2 | (AT)7 | CTCACGAAGGCCCATTAAAA | ATAGTGGGAAAAACCCCGAC | 227 |
| chr4:88874336-88876352 | p2 | (TA)8 | AAAGGAGACCCGGTCAGAAT | CCTGTCCTTGCTAGACCTCG | 152 |
| chr4:88911297-88913315 | p2 | (AT)8 | GACAAGGCTGCCACTACACA | ATGTTTGTTCAGGGTACGGC | 260 |
| chr4:88911297-88913315 | p2 | (AT)9 | GACAAGGCTGCCACTACACA | ATGTTTGTTCAGGGTACGGC | 260 |
| chr4:88942550-88944568 | p6 | (ATCCGT)3 | TGTTGGTGGTGCTCTTTCAG | ACGAACCGTGTCCCATAGAG | 264 |
| chr4:89135312-89137332 | p2 | (AT)10 | GCTCTGGCGCTAGTCACTCT | TGTAAAGGAGCATGAAGGGG | 109 |
| chr4:89135312-89137332 | p5 | (TTGAA)4 | GCTCTGGCGCTAGTCACTCT | TGTAAAGGAGCATGAAGGGG | 109 |
| chr4:89151160-89153172 | p2 | (TA)6 | TTTACAAGGGAAATGCTGCC | ACAAGGATCATGCCCAAAAG | 149 |
| chr4:89333090-89335108 | p6 | (ATGTAT)3 | TTGCATAAAAAGGACCCGAC | CTCGGTGAACGGAATAAGGA | 287 |
| chr4:89338679-89340697 | p6 | (ATATGT)3 | ATGTTGGATATGAGCCTCGG | GTCGGGTCCTATTTATGCGA | 172 |
| chr4:89338679-89340697 | p2 | (AT)6 | ATGTTGGATATGAGCCTCGG | GTCGGGTCCTATTTATGCGA | 172 |
| chr4:89366185-89368200 | p3 | (ATC)5 | GGGTACCCGCTTAGTTCTCC | CCACCTTGAGAGGCATTCAT | 196 |
| chr4:89418205-89420217 | p2 | (AT)6 | TCTTCTTGATGGTCCAAGCC | GATCAAAGAATTGAGCCCCA | 252 |
| chr4:89483800-89485840 | p2 | (AT)20 | ACCATTGTACGAGCCGAAAC | CGGAGTTGCCAGGTCTATGT | 267 |
| chr4:89483800-89485840 | p3 | (AAC)10 | ACCATTGTACGAGCCGAAAC | CGGAGTTGCCAGGTCTATGT | 267 |
| chr4:89534177-89536195 | p6 | (ACACAT)3 | TCCGACAACGGTTTCTTACC | GCCGGCTTAGTCTGAATCTG | 231 |
| chr4:89550051-89552091 | p2 | (AT)20 | TGTATCTTCCATGAAGGCCC | ACGTTTTCGGGTGATCTGAC | 101 |
| chr4:89562719-89564734 | p2 | (AT)6 | ACCTTGGAGTTCTTTTGCGA | ATGGCTAGTTGTCACGACCC | 142 |
| chr4:89562719-89564734 | p5 | (AAATG)3 | ACCTTGGAGTTCTTTTGCGA | ATGGCTAGTTGTCACGACCC | 142 |
| chr4:89586030-89588054 | p2 | (AT)12 | ACTCGTCCGACGGTACAAAG | TTTGTAGAATCCCCGAGTGC | 283 |
| chr4:89605397-89607412 | p3 | (TGA)5 | ATAGCCTTGTCGGCTTCTGA | ACTTCTGTCACGACCCAACC | 298 |

| Marker ID | SSR type | SSR motif | FORWARD PRIMER (5'-3') | REVERSE PRIMER (5'-3') | size (bp) |
|---|---|---|---|---|---|
| chr4:89637118-89639132 | p4 | (AAAG)4 | GTATTTTCCGCCGTTCGATA | TGCTAATTCCAATTTTGCCC | 177 |
| chr4:89637118-89639132 | p2 | (TG)7 | GTATTTTCCGCCGTTCGATA | TGCTAATTCCAATTTTGCCC | 177 |
| chr4:89695025-89697088 | p3 | (AAT)21 | GGGCGTACCTGAGATTGGTA | AGTGGCTTCAAGTCGAGGAA | 208 |
| chr4:89743904-89745988 | p3 | (ATG)5 | CACCTCGGCATAGCCATAGT | ACATGTTTGGGCATTTAGGC | 230 |
| chr4:89896462-89898582 | p2 | (TA)60 | ACGACGGAAGAAAGCGAGTA | CACGCTTCTTCCTGTTAGGC | 170 |
| chr4:89939062-89941084 | p2 | (AT)11 | AGGCCTTTGGGTGTACCTCT | AGAGGGGATGAAACTCGGAT | 174 |
| chr4:89939062-89941084 | p2 | (AT)20 | AGGCCTTTGGGTGTACCTCT | AGAGGGGATGAAACTCGGAT | 174 |
| chr4:90061157-90063169 | p2 | (CA)6 | TGGAACAACATTCGGAAACA | TATTGTCGATCTTGGGGAGC | 214 |
| chr4:90135677-90137713 | p2 | (AT)18 | GGCTTGGAAGTTGTTGCATT | GGCAGCATCTCCCCTATACA | 157 |
| chr4:90135677-90137713 | p5 | (ATGAT)3 | GGCTTGGAAGTTGTTGCATT | GGCAGCATCTCCCCTATACA | 157 |
| chr4:90200363-90202378 | p3 | (TGA)5 | GATAGCCTTGTCGGCTTCTG | ACTTCTGTCACGACCCAACC | 300 |
| chr4:90321426-90323444 | p6 | (TACATA)3 | GAGATGAAGGGGTGCAAAAA | CAGTTTGAAAGGCTTCCCAG | 151 |
| chr4:90328346-90330361 | p5 | (TTTTA)3 | AATTGCGTGGTTCGAGGTAG | AGGATTTCTGACGGCTCTGA | 252 |
| chr4:90496172-90498187 | p3 | (GTT)5 | GGACTTGCACGTGGCTATTT | TATTTTGCTAATGGGCCGAG | 224 |
| chr4:90504990-90507006 | p6 | (ATCCGT)3 | TCATATAGGCGGGACAGGAC | TTTTGTTTGTGGACTCTGCG | 136 |
| chr4:90504990-90507006 | p4 | (AGAA)4 | TCATATAGGCGGGACAGGAC | TTTTGTTTGTGGACTCTGCG | 136 |
| chr4:90504990-90507006 | p3 | (GTT)5 | TCATATAGGCGGGACAGGAC | TTTTGTTTGTGGACTCTGCG | 136 |
| chr4:90559656-90561668 | p2 | (CA)6 | AAAACTCTGGAGGGCCAAAT | CACCACATTCTGAGGGGTCT | 127 |
| chr4:90593446-90595466 | p2 | (AT)10 | GACCCTTAATTAGGGCCGAG | AAGCCGACGTACCATTATGC | 254 |
| chr4:90636229-90638261 | p3 | (TTG)5 | TTTATGCCTATTTGCGGAGG | AGCCCGAAATCACAATTCAG | 200 |
| chr4:90636229-90638261 | p2 | (AT)16 | TTTATGCCTATTTGCGGAGG | AGCCCGAAATCACAATTCAG | 200 |
| chr4:90636229-90638261 | p3 | (CAA)6 | TTTATGCCTATTTGCGGAGG | AGCCCGAAATCACAATTCAG | 200 |
| chr4:90669720-90671734 | p6 | (AGTTGA)3 | CCCCTCGTGTTTGTAGTGGT | CGACGATGACTGATGTCCTG | 233 |
| chr4:90669720-90671734 | p2 | (TG)7 | CCCCTCGTGTTTGTAGTGGT | CGACGATGACTGATGTCCTG | 233 |
| chr4:90684625-90686637 | p2 | (TA)6 | GTCCCATCCAAGCTGTCAAT | AAAAGTTCGGGGTGTGTCAG | 179 |
| chr4:90773650-90775678 | p2 | (TA)14 | GCCACAATCTTCCAAATGCT | CTGGCAGAGGAAGAAAATGC | 290 |
| chr4:90784022-90786042 | p2 | (TA)10 | CCGATTGAAAGTGGTTCGAT | AATTCATTCACTGCCGGAAC | 171 |
| chr4:90807814-90809829 | p3 | (TGT)5 | GCAAAACGATCAACGGAAAT | GTCTTGCCATGCAGTTTTGA | 182 |
| chr4:90809104-90811116 | p2 | (AT)6 | GGGAATTTGGAGTCGCATAA | TTAGATGCGGATTTTGGAGG | 225 |
| chr4:90876517-90878532 | p5 | (ATCAT)3 | CACACCCCTATGAGGAGCAT | GGCGGCTATGTTTCATTCAT | 227 |
| chr4:90975767-90977782 | p5 | (TCCTT)3 | GACGCCCTTCATAGGTGAAA | AAGGGTTATTTGGCAGCCTT | 166 |
| chr4:90984065-90986080 | p5 | (CATAT)3 | ATGTGCCGAGGAACTACGAC | CGATAGTTTGCGATTCCGTT | 182 |
| chr4:91011012-91013028 | p2 | (AT)8 | CTTCACTGAGTGCCAGGACA | GGCCCTTTAGTGAGGGACTC | 203 |
| chr4:91015105-91017121 | p2 | (AT)8 | CTTCACTGAGTGCCAGGACA | GGCCCTTTAGTGAGGGACTC | 203 |
| chr4:91070057-91072069 | p2 | (TG)6 | GCGGAGGAACAGATTGTTGT | TAGGTGCTTTTGGGGAACTG | 176 |
| chr4:91080702-91082720 | p5 | (ATTCA)6 | CACCATCCCCGAGTTTCTAA | CTTGTTGGATCGGGAAGTGT | 118 |

| Marker ID | SSR type | SSR motif | FORWARD PRIMER (5'-3') | REVERSE PRIMER (5'-3') | size (bp) |
|---|---|---|---|---|---|
| chr4:91080702-91082720 | p2 | (AT)9 | CACCATCCCCGAGTTTCTAA | CTTGTTGGATCGGGAAGTGT | 118 |
| chr4:91146221-91148236 | p3 | (AAC)5 | CGTAACTGCCGGGAAGAATA | CAACCCCAGTCCTGACACTT | 233 |
| chr4:91146221-91148236 | p5 | (AAGAC)3 | CGTAACTGCCGGGAAGAATA | CAACCCCAGTCCTGACACTT | 233 |
| chr4:91157889-91159909 | p2 | (AG)10 | TCCTCTCCCAAGGATTGATG | CGATGTCACCTCGGGATACT | 230 |
| chr4:91229013-91231031 | p6 | (AAAGAG)3 | GATGCTTCTGGTGTTGGGTT | TAATTCGCGGTCATGAGTTG | 132 |
| chr4:91302231-91304243 | p2 | (TA)6 | GAGGGTTCACGACCTGACAT | TATCGTATAGCTGCCCGCTT | 206 |
| chr4:91302231-91304243 | p2 | (AT)9 | GAGGGTTCACGACCTGACAT | TATCGTATAGCTGCCCGCTT | 206 |
| chr4:91412586-91414604 | p6 | (TATATG)3 | TGATATGTTTCCGAATGGCA | TTCGGGTCAATCCGACTTAG | 198 |
| chr4:91518602-91520614 | p2 | (AT)6 | ATCACGGGCCTTTGTACTTG | AGAGGAAGGAGAATCGCACA | 298 |
| chr4:91536426-91538446 | p2 | (AT)10 | CGCACTGCCCGATTTATTAT | GGACAGCCTTTACACACCGT | 137 |
| chr4:91624407-91626449 | p2 | (AT)21 | TCGATGCTTATGTGACTCGC | TGGAACCACTATAACCCCCA | 299 |
| chr4:91624407-91626449 | p2 | (TA)30 | TCGATGCTTATGTGACTCGC | TGGAACCACTATAACCCCCA | 299 |
| chr4:91663820-91665832 | p2 | (GT)6 | GGGGGTGTTGTATTTCATGC | GACAGGTCCCTAGCCTACCC | 292 |
| chr4:91714055-91716070 | p5 | (TCAAT)3 | CGGGAGCTCCGTATTTATGA | TGTTGGATAAGAACCTCGGG | 134 |
| chr4:91714055-91716070 | p5 | (CAACA)3 | CGGGAGCTCCGTATTTATGA | TGTTGGATAAGAACCTCGGG | 134 |
| chr4:91735695-91737710 | p5 | (TTCAT)3 | TGTGTGGGAGATGTTTTGGA | ACGTTTATCCTCCCGAGCTT | 255 |
| chr4:91746337-91748361 | p2 | (CT)6 | AGTGACAAAGGGTGGTGGAG | GGTGATCTGACTTGGGGAGA | 108 |
| chr4:91771199-91773217 | p6 | (ACACAT)3 | GTATCAGACGGGTTCGCAAT | CGTCCGATTCTGGATACGTT | 248 |
| chr4:91826707-91828719 | p2 | (AT)6 | AAACAGCCCGCAACAATTAC | TGTGTGTAGCCGTGTGTGTG | 202 |
| chr4:91854373-91856388 | p3 | (TTG)5 | CGATCGCATCCACTATGCTA | GGAATCTCCCGAAACTCTCC | 139 |
| chr4:91854373-91856388 | p2 | (TA)7 | CGATCGCATCCACTATGCTA | GGAATCTCCCGAAACTCTCC | 139 |
| chr4:91866228-91868244 | p2 | (CT)8 | GGACATGCCAAAAGAAGGAA | GTTCGTTAGAATTTCGGGCA | 170 |
| chr4:91873481-91875501 | p2 | (TA)13 | ACACCAAAACCCGATGGATA | AACAAAAGCCACATCCTTGG | 203 |
| chr4:91873481-91875501 | p2 | (AT)10 | ACACCAAAACCCGATGGATA | AACAAAAGCCACATCCTTGG | 203 |
| chr4:91884376-91886394 | p3 | (ATG)6 | GTGGGCTGTTTTATGGCACT | GTCCCGGATCTTCCACCTAT | 291 |
| chr4:91931992-91934102 | p2 | (AT)55 | AACCCGCTGGTTCTCCTTAT | ATCGGCCTTGTAAGCAGCTA | 189 |
| chr4:91975258-91977274 | p4 | (TATG)4 | GTCCAATAGCGACGGAAAAA | TAACCCTTGTGGAAAAACGC | 184 |
| chr4:92031109-92033121 | p2 | (TA)6 | TGCAGACGGCTTAATGTGAG | AAATCTCGACATGTGGACCC | 271 |
| chr4:92031109-92033121 | p5 | (TGAAT)3 | TGCAGACGGCTTAATGTGAG | AAATCTCGACATGTGGACCC | 271 |
| chr4:92064493-92066505 | p2 | (GT)6 | GAGGATGTTACACCGGCATT | CTCTCCCTTTTGGGCCTTAC | 199 |
| chr4:92085103-92087121 | p3 | (CAA)6 | CCTCTTCCGACCCTCATGTA | TGGTTGCCTTAAGTATCCCG | 132 |
| chr4:92088494-92090506 | p2 | (TG)6 | TGAAACCCCGAGCTTATGAC | CTCATACCTGGCCCTACCAA | 269 |
| chr4:92159246-92161261 | p5 | (ACACA)3 | TTACATACCTTGCCCGCTTC | GTTGATGATTTGGCCGAGTT | 300 |
| chr4:92200817-92202831 | p2 | (TA)7 | GGCCGATGCCTTCTGTATAA | ACGGCGAGTCAAGTCAGAGT | 170 |
| chr4:92211666-92213684 | p6 | (CATATA)3 | TAATGATCCCACGGTTTCGT | CTCTGAAGAGAGTCGGGTGC | 161 |
| chr4:92211666-92213684 | p2 | (AT)11 | TAATGATCCCACGGTTTCGT | CTCTGAAGAGAGTCGGGTGC | 161 |

| Marker ID | SSR type | SSR motif | FORWARD PRIMER (5'-3') | REVERSE PRIMER (5'-3') | size (bp) |
|---|---|---|---|---|---|
| chr4:92247484-92249496 | p2 | (AT)23 | TTGACTAAGGGTACGACGGG | GTAAAAGCAAGCCGACAAGG | 132 |
| chr4:92247497-92249517 | p2 | (AT)23 | TTGACTAAGGGTACGACGGG | GTAAAAGCAAGCCGACAAGG | 119 |
| chr4:92319714-92321734 | p2 | (TA)10 | CTATAACGGAATCGGGGGTT | GGTGTCTGCCCATGCTATCT | 280 |
| chr4:92351427-92353443 | p4 | (CATA)4 | TCCATTTTCTCCTGGAGTGC | TGAAGCCGCTTACACAACTG | 288 |
| chr4:92398720-92400754 | p2 | (AT)17 | TGAGCATCGGGGAATAACTC | GTCGGGTCCTATTTATGCGA | 200 |
| chr4:92580885-92582897 | p2 | (AG)6 | TGGGTGGAATGTAGGGGTTA | CCTGAGGGGATGTACCAAGA | 148 |
| chr4:92667845-92669863 | p6 | (ATAAAA)3 | TCACTGGAGCTTGGTGAGTG | CCAAAAGCAAGGGTTTTGAA | 236 |
| chr4:92791299-92793317 | p2 | (AC)6 | CAACCCATTTTCTTCCTCCA | TCGCTCAGTTCCTTGGAGTT | 300 |
| chr4:92791319-92793337 | p2 | (AC)6 | CAACCCATTTTCTTCCTCCA | TCGCTCAGTTCCTTGGAGTT | 280 |
| chr4:92791338-92793374 | p2 | (AC)6 | CAACCCATTTTCTTCCTCCA | TCGCTCAGTTCCTTGGAGTT | 261 |
| chr4:92791393-92793408 | p2 | (AC)6 | CAACCCATTTTCTTCCTCCA | TCGCTCAGTTCCTTGGAGTT | 206 |
| chr4:93067128-93069142 | p2 | (AT)7 | TGGAGGATACCGACCTTACG | TAATCCGCGTCCAGGATAAC | 295 |
| chr4:93067128-93069142 | p2 | (AT)8 | TGGAGGATACCGACCTTACG | TAATCCGCGTCCAGGATAAC | 295 |
| chr4:93110030-93112046 | p2 | (AT)8 | TCCTCCGGTACTGGAAACTG | ACATAGACCGAAATACGCGG | 103 |
| chr4:93142656-93144674 | p3 | (TTG)6 | GAAGATATCGGGACCAAGCA | TGTCGTATCGACGTGTGGTT | 290 |
| chr4:93157297-93159315 | p3 | (TTG)6 | GAAGATATCGGGACCAAGCA | TGTCGTATCGACGTGTGGTT | 290 |
| chr4:93198112-93200124 | p2 | (TA)6 | TCCCACAGGTAGACAGGGAG | CACGTCGTGGAACCACATAG | 147 |
| chr4:93198112-93200124 | p2 | (AG)6 | TCCCACAGGTAGACAGGGAG | CACGTCGTGGAACCACATAG | 147 |
| chr4:93198781-93200793 | p2 | (TA)6 | CCCCATAAGATTCCGTGCTA | AGGGCACTCACATTACCTCG | 240 |
| chr4:93198781-93200793 | p2 | (AG)6 | CCCCATAAGATTCCGTGCTA | AGGGCACTCACATTACCTCG | 240 |
| chr4:93225322-93227376 | p2 | (AT)27 | CCCCATCAAGGCTGCTATTA | CGAGCAGCTATTCGACCTTC | 250 |
| chr4:93247821-93249836 | p5 | (ATGAT)3 | TGTGTCGCCACTGGATATGT | CGCGAGGACTATAGGGTCAG | 129 |
| chr4:93267107-93269127 | p2 | (CA)10 | CTCTACCGGACATGGATCGT | TTTCCGTAGTCTCGTTGGCT | 172 |
| chr4:93267107-93269127 | p3 | (CAA)5 | CTCTACCGGACATGGATCGT | TTTCCGTAGTCTCGTTGGCT | 172 |
| chr4:93277622-93279638 | p2 | (TA)8 | GCCAGTGTAGGCCACTCATT | CACATGACCCATTACCCACA | 258 |
| chr4:93285440-93287460 | p2 | (AT)10 | CCCGTTTCGTCAAAAACCTA | TGTTGCTTTGGGTGTTGTGT | 118 |
| chr4:93290720-93292754 | p2 | (AT)17 | CCCTCTCTTTTAGCTGGCCT | TCCTTTCAACCATACCTGCC | 271 |
| chr4:93315997-93318012 | p3 | (ATT)5 | CCTTTTGTTTGTGGGATGCT | CGAAAAGGCCAAACACATCT | 201 |
| chr4:93369506-93371518 | p2 | (AT)6 | TAAGAATGGTGCCCCTTACG | GAGCCCTGTGTGGCTATGAT | 140 |
| chr4:93624722-93626737 | p5 | (ACACA)3 | TGGAAGGAGTTACCCAGACG | GGGATTAGCTTAGGAAGCGG | 224 |
| chr4:93635948-93637963 | p5 | (ACACA)3 | AAGGCATGCCAAAAGAAAGA | GTTGATGATTTGGCCGAGTT | 269 |
| chr4:93639041-93641057 | p2 | (AT)17 | TCAACCGAGTTAAACTGCCC | TTAGTGGTGCTCAGCAGGTG | 261 |
| chr4:93649869-93651881 | p2 | (AT)6 | TAGAGTTCGCCAATCCTGCT | AAGCGCTTTGGTAATCTGGA | 153 |
| chr4:93685818-93687833 | p3 | (GAA)5 | AAAATCCAACAGCATCAGGC | GGGTTGGGGGTAGTAGTGGT | 140 |
| chr4:93889134-93891150 | p2 | (AG)8 | TTAGGATGGGTGACCCTCTG | CGATGTCACCTCGGGATACT | 216 |
| chr4:93908827-93910843 | p2 | (AC)8 | ATACCTCGATCGCTCGCTAA | GGTATGGGTTGTTGGTTTGG | 272 |

| Marker ID | SSR type | SSR motif | FORWARD PRIMER (5'-3') | REVERSE PRIMER (5'-3') | size (bp) |
|---|---|---|---|---|---|
| chr4:93925900-93927914 | p2 | (AC)7 | AGACATAGGCCATCCCAGTG | TTAGCGAGCGATCGAGGTAT | 146 |
| chr4:94032996-94035010 | p6 | (TATATG)3 | CACGTTCCGCAGCAGTATAA | GGAACGTACGACCCGATCTA | 216 |
| chr4:94032996-94035010 | p2 | (TA)7 | CACGTTCCGCAGCAGTATAA | GGAACGTACGACCCGATCTA | 216 |
| chr4:94091047-94093065 | p6 | (ATATGT)3 | ATGTTGGATATGAGCCTCGG | TCGGGTCCTATTTATGCCAG | 224 |
| chr4:94102692-94104714 | p2 | (TA)11 | CCGTTTGGTCGTTATGGTCT | TTGGAATCGAACCAAACACA | 250 |
| chr4:94102692-94104714 | p6 | (ATGATA)3 | CCGTTTGGTCGTTATGGTCT | TTGGAATCGAACCAAACACA | 250 |
| chr4:94102692-94104714 | p2 | (TA)10 | CCGTTTGGTCGTTATGGTCT | TTGGAATCGAACCAAACACA | 250 |
| chr4:94169597-94171612 | p3 | (GTT)5 | CCACCAAAACTTAAGGGCAA | CTTCTCGCATCTCTTCCACC | 169 |
| chr4:94387339-94389351 | p2 | (AG)6 | ACTGAAAGCACCCCATTGTC | CATGAGACCACATGACCCAC | 156 |
| chr4:94417770-94419788 | p6 | (CATATA)3 | AACAAGTGCCCGACCACTAC | TGTGTTGATTGTTCGGGAGA | 153 |
| chr4:94417770-94419788 | p3 | (CAA)5 | AACAAGTGCCCGACCACTAC | TGTGTTGATTGTTCGGGAGA | 153 |
| chr4:94490785-94492797 | p2 | (TA)6 | ACTCCATAAGGTAACCGGGG | GACATACCCACGGTGGAATC | 226 |
| chr4:94516266-94518281 | p3 | (ATG)5 | CGGTGTAAACCGACCCTAGA | TATACGCTTCCCGAGCCTAA | 215 |
| chr4:94516266-94518281 | p2 | (TA)6 | CGGTGTAAACCGACCCTAGA | TATACGCTTCCCGAGCCTAA | 215 |
| chr4:94581244-94583260 | p5 | (TTTGA)3 | GTATTTTCTGCGTGCAGGGT | TCACAAGGATCATGCCAAAA | 166 |
| chr4:94581244-94583260 | p2 | (AT)8 | GTATTTTCTGCGTGCAGGGT | TCACAAGGATCATGCCAAAA | 166 |
| chr4:94586216-94588232 | p2 | (AT)6 | GTTACATGAGCCCAGTCCGT | ATACCTTGTCGGTTGCTTGC | 106 |
| chr4:94586216-94588232 | p2 | (TA)8 | GTTACATGAGCCCAGTCCGT | ATACCTTGTCGGTTGCTTGC | 106 |
| chr4:94799112-94801126 | p2 | (GA)7 | TGTTTTGTGCCCCACATCTA | ATCCCCATCATCAGAAGCAC | 217 |
| chr4:94799112-94801126 | p5 | (TGAAA)3 | TGTTTTGTGCCCCACATCTA | ATCCCCATCATCAGAAGCAC | 217 |
| chr4:94813010-94815028 | p2 | (AT)9 | TTCCTCACTACTCCGCTCGT | GTCTCGGGAACCGTACACAT | 231 |
| chr4:94813010-94815028 | p2 | (AC)8 | TTCCTCACTACTCCGCTCGT | GTCTCGGGAACCGTACACAT | 231 |
| chr4:94847486-94849514 | p2 | (AT)14 | TAGACCGACACAACCGAACA | CAGGGACTACATAGCGGAGC | 252 |
| chr4:94852777-94854807 | p2 | (AT)15 | TAGACCGACACAACCGAACA | CAGGGACTACATAGCGGAGC | 252 |
| chr4:94892984-94894998 | p2 | (TG)8 | GAAGGGCCAAAGTGTTCAAA | TGTCCCTTTTCCCCTTTTCT | 275 |
| chr4:94984042-94986066 | p3 | (AAC)8 | GTCCCCGAGTATGTAAGGCA | ACATTATGGCTTGGCCTACG | 246 |
| chr4:94984042-94986066 | p2 | (TA)19 | GTCCCCGAGTATGTAAGGCA | ACATTATGGCTTGGCCTACG | 246 |
| chr4:95005012-95007028 | p2 | (TA)8 | CCAGTTCGGGCACTAGTCAT | TATGCAACACTTCCCCATCA | 194 |
| chr4:95005012-95007028 | p2 | (TA)9 | CCAGTTCGGGCACTAGTCAT | TATGCAACACTTCCCCATCA | 194 |
| chr4:95025833-95027845 | p2 | (CA)6 | CGGGTTTCGATGAACAGAAT | GAATTTGAGCGAGCGTAAGG | 189 |
| chr4:95040241-95042273 | p2 | (TA)16 | ATATCGGAACACCGTGAAGC | CTTACCCCGTTCTCGACGTA | 296 |
| chr4:95057846-95059860 | p2 | (AT)7 | GTACGTTCGGGTCATGGTTC | AGTCGGCTTGCACTGTTCTT | 113 |
| chr4:95069335-95071457 | p2 | (AT)61 | AACTCGGCCAAATCATCAAC | GGGTTGTTGTGATTGCATTG | 212 |
| chr4:95184671-95186683 | p2 | (AT)6 | CAAATCTCTCACGCTCCACA | GGATTTGCGCGTTTAAGGTA | 228 |
| chr4:95244734-95246746 | p2 | (AC)6 | AGACGTCTTACGCTCTCCCA | TGTTTGATGATGTGTGGGCT | 290 |
| chr4:95288881-95290903 | p2 | (TA)11 | TTTTGGCATGATCCTTGTGA | TGCAATGCAAAGCGTAAAAC | 226 |

| Marker ID | SSR type | SSR motif | FORWARD PRIMER (5'-3') | REVERSE PRIMER (5'-3') | size (bp) |
|---|---|---|---|---|---|
| chr4:95302057-95304072 | p5 | (TGCAT)3 | AATTGCATGTTCCATGGGTT | AGCGCCTCCAAGTACTCAAA | 109 |
| chr4:95349990-95352005 | p2 | (AT)11 | TGACGTCCCTTTTTATTGCC | TAATGCCTAAACCCCGAATG | 225 |
| chr4:95349990-95352005 | p5 | (TAGTT)3 | TGACGTCCCTTTTTATTGCC | TAATGCCTAAACCCCGAATG | 225 |
| chr4:95384591-95386603 | p2 | (AC)6 | TATCCCCGAGGTTCGTATCA | GTTGATGTTTTGGCCGAGTT | 242 |
| chr4:95427689-95429704 | p5 | (ATATG)3 | AGAGAAATGGTGGATCGTGG | CCTCACATCCCTTTGACGTT | 265 |
| chr4:95454452-95456464 | p2 | (AT)6 | TCGGACGGATCTGTGTCATA | CGCTTCTGATCCTTTTCCTG | 281 |
| chr4:95463988-95466000 | p2 | (AT)6 | CGTCCTTCGTGGTCGTATTT | GATCTGCATTTTGGCCAGTT | 261 |
| chr4:95463988-95466000 | p2 | (TA)10 | CGTCCTTCGTGGTCGTATTT | GATCTGCATTTTGGCCAGTT | 261 |
| chr4:95516176-95518192 | p2 | (AT)8 | GAACATGAACCTGGTTGGCT | TAATCCATGTATGCGCGTGT | 101 |
| chr4:95594002-95596020 | p6 | (AGGTAG)3 | CGTCACTACTTGCCACCTGA | GTTGTCGTTCCTCGAGCTTC | 256 |
| chr4:95655756-95657774 | p3 | (ATG)6 | CCAACAATCACTCCATGCTG | AGAAAAAGAGGGGCATTCGT | 104 |
| chr4:95687928-95689944 | p2 | (AT)8 | TGGTCAATAACTGAACCGCA | CAGAAGCTGCTAATCGGGAG | 180 |
| chr4:95741402-95743420 | p5 | (GTCCT)3 | GACGGACCGTAAAAGGTTGA | TGAGGAACTTCAACACGCAC | 239 |
| chr4:95741512-95743533 | p5 | (GTCCT)3 | GACGGACCGTAAAAGGTTGA | TGAGGAACTTCAACACGCAC | 129 |
| chr4:95741994-95744009 | p5 | (GTCCT)3 | TTTGGCATGATCCTTGTGAA | TTCACCTTTCCCATATCCCA | 226 |
| chr4:95767896-95769918 | p2 | (TA)11 | AGCCGACAAGGCTAAAGACA | GTGTACATACATGGGCACGG | 162 |
| chr4:95799588-95801610 | p2 | (TA)11 | AGCCGACAAGGCTAAAGACA | GTGTACATACATGGGCACGG | 162 |
| chr4:95872358-95874376 | p2 | (AT)9 | ACTCATTGGAATTCAACCGC | CAGTGCCATAGACCAGCAGA | 137 |
| chr4:95872358-95874376 | p2 | (TA)9 | ACTCATTGGAATTCAACCGC | CAGTGCCATAGACCAGCAGA | 137 |
| chr4:95872681-95874699 | p2 | (AT)9 | ATCGTGAGACCATGCTACCC | TGTTCATGCCCACAGGTAGA | 273 |
| chr4:95872681-95874699 | p2 | (TA)9 | ATCGTGAGACCATGCTACCC | TGTTCATGCCCACAGGTAGA | 273 |
| chr4:95914081-95916099 | p2 | (AT)9 | ACTCATTGGAATTCAACCGC | CAGTGCCATAGACCAGCAGA | 137 |
| chr4:95914081-95916099 | p2 | (TA)9 | ACTCATTGGAATTCAACCGC | CAGTGCCATAGACCAGCAGA | 137 |
| chr4:95914404-95916422 | p2 | (AT)9 | ATCGTGAGACCATGCTACCC | TGTTCATGCCCACAGGTAGA | 273 |
| chr4:95914404-95916422 | p2 | (TA)9 | ATCGTGAGACCATGCTACCC | TGTTCATGCCCACAGGTAGA | 273 |
| chr4:95944206-95946222 | p2 | (AT)32 | AGCCCTTCGGCTATCAAAAT | CGTTGTGTTTATGCCCACAG | 234 |
| chr4:95944206-95946222 | p2 | (AT)8 | AGCCCTTCGGCTATCAAAAT | CGTTGTGTTTATGCCCACAG | 234 |
| chr4:95944206-95946222 | p2 | (TA)12 | AGCCCTTCGGCTATCAAAAT | CGTTGTGTTTATGCCCACAG | 234 |
| chr4:95961145-95963163 | p3 | (CAA)6 | GGCACGGTGTTATACTGCCT | CGACCTTACGTCACTCCGAT | 246 |
| chr4:96005875-96007909 | p2 | (TA)17 | GGACTTCCAGAACTTGGCTG | ATTTTCTCCCCGTCAGGACT | 232 |
| chr4:96005875-96007909 | p2 | (AT)11 | GGACTTCCAGAACTTGGCTG | ATTTTCTCCCCGTCAGGACT | 232 |
| chr4:96072438-96074456 | p2 | (AT)9 | GGAGGCATTGCTGAAATCAT | GGTTGGGTCGTGACAAAAGT | 128 |
| chr4:96223848-96225912 | p2 | (TA)32 | CACCCATCGTTGTCGTAGTG | AGATCGCGTCGATTTGATTC | 216 |
| chr4:96223848-96225912 | p2 | (AT)14 | CACCCATCGTTGTCGTAGTG | AGATCGCGTCGATTTGATTC | 216 |
| chr4:96314373-96316395 | p2 | (AT)11 | GTCACGCCTATATGCTCGGT | TTGCTCCTGATATTTTGCCC | 284 |
| chr4:96314373-96316395 | p6 | (CAACAT)3 | GTCACGCCTATATGCTCGGT | TTGCTCCTGATATTTTGCCC | 284 |

| Marker ID | SSR type | SSR motif | FORWARD PRIMER (5'-3') | REVERSE PRIMER (5'-3') | size (bp) |
|---|---|---|---|---|---|
| chr4:96316256-96318270 | p2 | (TA)7 | AGACGATGGGCAAAGAAATG | TTGTAGGTGACCCCGTAAGC | 247 |
| chr4:96343673-96345709 | p3 | (ATC)12 | ACTTTTGTCACGACCCAACC | TCTTGTCGGCTCCTGTTCTT | 234 |
| chr4:96359665-96361681 | p4 | (TTTC)4 | TTTCCGTTCCGACCTTACAC | TAGAGGGGTTTGTGGCTTTG | 143 |
| chr4:96397641-96399683 | p5 | (GCATT)3 | ATGGCCCATATCCGATGTTA | GAAGTCTAAGTGCCACCCGA | 217 |
| chr4:96397641-96399683 | p2 | (TA)21 | ATGGCCCATATCCGATGTTA | GAAGTCTAAGTGCCACCCGA | 217 |
| chr4:96397756-96399771 | p5 | (GCATT)3 | ATGGCCCATATCCGATGTTA | GAAGTCTAAGTGCCACCCGA | 102 |
| chr4:96397756-96399771 | p2 | (TA)21 | ATGGCCCATATCCGATGTTA | GAAGTCTAAGTGCCACCCGA | 102 |
| chr4:96482373-96484395 | p2 | (AT)18 | TTCTGCACCACTCACAAAGC | ATCAACACACAGCCTAGCCC | 206 |
| chr4:96482877-96484913 | p2 | (AT)18 | TCATTATGCTATGCAAGCCG | TTTTTGTAGCAGCCTTTCCG | 164 |
| chr4:96517751-96519767 | p2 | (AT)7 | CCCGAACTGGACACTCGTAT | ATGTTGTGACAGCCTTGTCG | 248 |
| chr4:96517751-96519767 | p2 | (AT)8 | CCCGAACTGGACACTCGTAT | ATGTTGTGACAGCCTTGTCG | 248 |
| chr4:96517751-96519767 | p3 | (CAA)5 | CCCGAACTGGACACTCGTAT | ATGTTGTGACAGCCTTGTCG | 248 |
| chr4:96725799-96727811 | p2 | (TA)6 | ACGACCAGGACGAATCTACG | GGAACGATCGTGGTCACTTT | 282 |
| chr4:96735821-96737833 | p2 | (AC)6 | ATGCCAAACGAAGTAGTGGG | TTTGATGATGTGTGGGCAGT | 243 |
| chr4:96748444-96750456 | p2 | (CA)6 | AAACCTCCTCCGCTTCTTTC | AAATCTTCGGTACGCAATGG | 281 |
| chr4:96748444-96750456 | p2 | (TA)6 | AAACCTCCTCCGCTTCTTTC | AAATCTTCGGTACGCAATGG | 281 |
| chr4:96905785-96907803 | p6 | (CATATA)3 | CACATCCACAAAACCATTCG | CCTAGACGGGTCGTGACATT | 241 |
| chr4:96917699-96919715 | p4 | (AAAC)4 | GCACATAACACATCGCGTTC | TGTCGTGGAGCATCCAATTA | 185 |
| chr4:96919988-96922003 | p5 | (ATATG)3 | AATGGTGGATCGTGGTCATT | AGAGACAATTGGGCAGCACT | 245 |
| chr4:96928188-96930257 | p3 | (AGA)5 | CTCCCGCTTCTTCTTCAATG | CGACGACAGAACCCTAGCAT | 124 |
| chr4:96928188-96930257 | p5 | (CGGGT)3 | CTCCCGCTTCTTCTTCAATG | CGACGACAGAACCCTAGCAT | 124 |
| chr4:96985639-96987654 | p5 | (ATATG)3 | CCGGATTCGTGTTTCTGTTT | CGCTCCCTTCTCCTCTTCTT | 134 |
| chr4:96987227-96989243 | p5 | (ATATG)3 | ATATTCGTATGGACGCTCCG | TAATCATAAGCACCTCCGGG | 246 |
| chr4:96987227-96989243 | p2 | (AT)8 | ATATTCGTATGGACGCTCCG | TAATCATAAGCACCTCCGGG | 246 |
| chr4:96996320-96998332 | p2 | (TA)6 | TTCGTAAACCGTAGCCCATC | AGGACCGCAATAAACACAGG | 178 |
| chr4:97046987-97049002 | p5 | (ATTGT)3 | GCCATAGTTGGCCGTGTAAT | TTGGGATCAACATCACCTCA | 191 |
| chr4:97098402-97100417 | p3 | (TTG)5 | ATATGAGGCTTGGGTCCCTT | TTCCTCCATATTTTCGCGTC | 179 |
| chr4:97139282-97141294 | p2 | (AT)6 | AATAAAGTCCGTCGCCAATG | AACACTCTCACCCCGTTCAC | 208 |
| chr4:97202617-97204629 | p2 | (AG)10 | GCGGGTAGGGGTGTTATTTT | TTCAAAATTGGGACGGAGTC | 282 |
| chr4:97202617-97204629 | p2 | (CT)6 | GCGGGTAGGGGTGTTATTTT | TTCAAAATTGGGACGGAGTC | 282 |
| chr4:97230491-97232507 | p2 | (AC)8 | CGCACACACCTAATCACCTG | TATTTGCCTCCTGAAGGGTG | 250 |
| chr4:97270099-97272115 | p2 | (AT)8 | CGGCTTGAAAATTCGTGATT | ACTCGGCCAAATTCATCAAC | 207 |
| chr4:97270099-97272115 | p2 | (TA)6 | CGGCTTGAAAATTCGTGATT | ACTCGGCCAAATTCATCAAC | 207 |
| chr4:97272482-97274498 | p2 | (AT)8 | CGGCTTGAAAATTCGTGATT | ACTCGGCCAAATTCATCAAC | 207 |
| chr4:97272482-97274498 | p2 | (TA)6 | CGGCTTGAAAATTCGTGATT | ACTCGGCCAAATTCATCAAC | 207 |
| chr4:97315219-97317235 | p4 | (TGTA)4 | TAGATTCTGGGCGAAATTGG | CCGAGCCTATTACGTTTGGA | 118 |

| Marker ID | SSR type | SSR motif | FORWARD PRIMER (5'-3') | REVERSE PRIMER (5'-3') | size (bp) |
|---|---|---|---|---|---|
| chr4:97337044-97339062 | p2 | (TA)9 | TATGGACTGCCCAAGTTTCC | CCATTCGAACCAACTAGCGT | 219 |
| chr4:97387211-97389235 | p2 | (AT)12 | CGAAAATCCTTTTGTCGGAA | ATTCCTTCACATGATTCGGC | 204 |
| chr4:97455037-97457055 | p3 | (AAC)6 | GTTTAACGCCAAGCGACTTC | AAAAATGGGGTTGATAGGGG | 161 |
| chr4:97455037-97457055 | p3 | (ATG)6 | GTTTAACGCCAAGCGACTTC | AAAAATGGGGTTGATAGGGG | 161 |
| chr4:97457699-97459714 | p5 | (TGTGG)3 | CGCATTTTTGTACAGTCGGA | TCAAAATTCCAAACTTGCCC | 138 |
| chr4:97473701-97475713 | p2 | (AT)9 | AGAAATACCAACAATGCCGC | AAAGGGGAGGTAGATCGGAA | 284 |
| chr4:97473720-97475732 | p2 | (AT)9 | AGAAATACCAACAATGCCGC | AAAGGGGAGGTAGATCGGAA | 265 |
| chr4:97491892-97493946 | p2 | (TA)27 | ACGTCCTTTCTTGTGGATGC | ACCAGCCGACAAGGCTACTA | 135 |
| chr4:97503898-97505914 | p4 | (GAAA)4 | AACTGAACCCAAGTCAACGG | TCATCGACTGTCACGAGAGG | 283 |
| chr4:97551233-97553249 | p2 | (AT)8 | CCTTGTCTAGGTATGCCCCA | ATCAGAATCACGCTCATCCC | 175 |
| chr4:97599847-97601862 | p3 | (TTG)5 | CACGAGAACATGACCATTGC | CTGGAGATATGCCCTTCCAA | 237 |
| chr4:97614656-97616671 | p5 | (TTTAA)3 | GTTGTGGCGACCTTATCGTT | ATTCCCTAGGACGACCTGCT | 252 |
| chr4:97614656-97616671 | p2 | (TA)14 | GTTGTGGCGACCTTATCGTT | ATTCCCTAGGACGACCTGCT | 252 |
| chr4:97716772-97718786 | p2 | (TA)7 | GTCTTGTCGATGGGTGTGTG | GGAACATGGAAGCCTGGTAA | 187 |
| chr4:97746501-97748519 | p2 | (TA)9 | GGTTGATGTTGTTGTCGTGC | GAATCAGGCAGCATTTCCTC | 153 |
| chr4:97746501-97748519 | p2 | (AT)9 | GGTTGATGTTGTTGTCGTGC | GAATCAGGCAGCATTTCCTC | 153 |
| chr4:97780979-97783004 | p5 | (ATATC)5 | TCCATTGGTCTGTGGATGAA | GCCTTATAGGCTTCTGTGCG | 143 |
| chr4:97837029-97839041 | p2 | (TG)6 | AAAGGTGCCTACGGGCTATT | CATAGCGTGCTCGACCATAA | 255 |
| chr4:97905788-97907802 | p2 | (TA)7 | GGACTGCGTCTCTCATAGCC | TAGGGCTTATGCCCAATCTG | 250 |
| chr4:97905788-97907802 | p2 | (AT)6 | GGACTGCGTCTCTCATAGCC | TAGGGCTTATGCCCAATCTG | 250 |
| chr4:97906788-97908806 | p2 | (TA)7 | CTCCGGAACAAATGAGCAAT | CACGCTTTACATGCTCGGTA | 224 |
| chr4:97906788-97908806 | p2 | (AT)6 | CTCCGGAACAAATGAGCAAT | CACGCTTTACATGCTCGGTA | 224 |
| chr4:98048992-98051017 | p5 | (ATCAT)5 | ACTTCTGTCACGACCCAACC | TCTGTGCGCTACATCTGCTT | 279 |
| chr4:98116273-98118287 | p2 | (AT)7 | CAGTACAGGGGGCTGGAATA | TGATATCCAGAGAGGCGGTC | 105 |
| chr4:98116273-98118287 | p2 | (TA)24 | CAGTACAGGGGGCTGGAATA | TGATATCCAGAGAGGCGGTC | 105 |
| chr4:98187622-98189646 | p2 | (TA)6 | TACACGGCCACAACACAACT | TTCCAAGCAATTGGAAGGTC | 205 |
| chr4:98205480-98207494 | p2 | (AT)7 | AGCCAACGAGGCTTACAAAA | AGAGCTCCACTTCATTCGGA | 206 |
| chr4:98272122-98274154 | p2 | (AT)16 | CAGCCGTTGGAAATGAATCT | ACTTACCTCACGTGCCCAAC | 260 |
| chr4:98272122-98274154 | p2 | (TA)15 | CAGCCGTTGGAAATGAATCT | ACTTACCTCACGTGCCCAAC | 260 |
| chr4:98307315-98309353 | p2 | (AT)19 | GTCCGCACCGTTCATTCTAT | CATATGTGTGGGTAGCGTGG | 160 |
| chr4:98405843-98407857 | p2 | (AT)7 | CATGGCATGAAGAACGATTG | TTCCCTGAATCTCCCAACAG | 275 |
| chr4:98451184-98453212 | p2 | (TA)14 | ATAGCTTGCCCATACATGCC | GTCATGGTCCTTTGATTGGG | 100 |
| chr4:98461438-98463450 | p2 | (AT)6 | CACATGACCCATTACCCACA | GAGTGCTCCTCTTTGATCCG | 251 |
| chr4:98732679-98734705 | p2 | (TA)13 | CGTCAACCTTTTCCCACTGT | AAGTCCTATGATTGCGTCGG | 133 |
| chr4:98763965-98765977 | p2 | (CA)6 | TTTTACATCTTGGCCCGAAC | CGGCTCAATCTCAGTTCTCC | 287 |
| chr4:98769352-98771367 | p2 | (AT)7 | TATGATGCCCCGATTTCATT | TCGTGATCGCTTTGTCTCAC | 265 |

| Marker ID | SSR type | SSR motif | FORWARD PRIMER (5'-3') | REVERSE PRIMER (5'-3') | size (bp) |
|---|---|---|---|---|---|
| chr4:98769352-98771367 | p5 | (GCTAT)3 | TATGATGCCCCGATTTCATT | TCGTGATCGCTTTGTCTCAC | 265 |
| chr4:98790480-98792495 | p5 | (TTTTG)3 | TTTTTCTTATGGCCGAATGG | AAAAAGGAAAAGAGGCCCAA | 272 |
| chr4:98824963-98826979 | p2 | (TA)8 | CGTGCTAGCTGAAAGATCCC | AGAAGGGCAAAGCTACGACA | 214 |
| chr4:98863929-98865949 | p2 | (TA)10 | TAACGGTGCACACCATTCAT | TCCGCACTTCGTTAAAATCC | 274 |
| chr4:98867505-98869520 | p4 | (AAAG)4 | AGTTGGAGAGTGATGGCGAC | AAGTGCAATTGTTCTTGCCC | 129 |
| chr4:98867505-98869520 | p5 | (AAAAT)3 | AGTTGGAGAGTGATGGCGAC | AAGTGCAATTGTTCTTGCCC | 129 |
| chr4:98867505-98869520 | p5 | (AAAAT)3 | AGTTGGAGAGTGATGGCGAC | AAGTGCAATTGTTCTTGCCC | 129 |
| chr4:98867505-98869520 | p5 | (AAAAT)3 | AGTTGGAGAGTGATGGCGAC | AAGTGCAATTGTTCTTGCCC | 129 |
| chr4:98867505-98869520 | p5 | (ATTTT)3 | AGTTGGAGAGTGATGGCGAC | AAGTGCAATTGTTCTTGCCC | 129 |
| chr4:98876500-98878515 | p3 | (TTG)5 | ATCGCCTAACCCTTTCCAAT | CCACCAAGCAAACTTCCATT | 117 |
| chr4:98899316-98901334 | p6 | (CATATA)3 | GCCGCTTACAATACCGACAT | ACTGACGTCCCTTCTTGTGG | 261 |
| chr4:98899316-98901334 | p3 | (ACA)9 | GCCGCTTACAATACCGACAT | ACTGACGTCCCTTCTTGTGG | 261 |
| chr4:98927141-98929161 | p4 | (TTCT)5 | AATCGTGCCAACAACATCAA | GAAGGTTTGGTGGCTTGAAA | 279 |
| chr4:99020159-99022177 | p2 | (TA)9 | ATTGGAAATTCGGGTTAGGC | GCCAAATGAAAGAAGGGACA | 210 |
| chr4:99219151-99221169 | p6 | (CATATA)3 | GTGAGGGACTCGGTGGATAA | GCACTCCTCTGCATTGTTCA | 144 |
| chr4:99219151-99221169 | p6 | (CAAAAA)3 | GTGAGGGACTCGGTGGATAA | GCACTCCTCTGCATTGTTCA | 144 |
| chr4:99219151-99221169 | p2 | (GT)6 | GTGAGGGACTCGGTGGATAA | GCACTCCTCTGCATTGTTCA | 144 |
| chr4:99228137-99230152 | p2 | (CA)6 | ACCTCCACATGCACATTTCA | CATGGCTAGCCGAATATGGT | 132 |
| chr4:99259855-99261870 | p5 | (ATATG)3 | AGAGAAATGGTGGATCGTGG | CACGCCTCTTCCTTCAAAAG | 227 |
| chr4:99343897-99345933 | p2 | (AG)18 | AAAGGTCCCATTGTTAGGGG | AAAAGCTGCAATCCAGTGCT | 183 |
| chr4:99410361-99412379 | p6 | (TATATG)3 | TGGCCAAGTTAAATTCTCGG | TACATACCTTGCCCGCTTTC | 127 |
| chr4:99519254-99521276 | p2 | (AT)11 | CGGATACCAGTCATGGCTCT | ATCGCATACGCACCACATTA | 274 |
| chr4:99554112-99556124 | p2 | (AC)6 | TTGGGCTATATTGGCTACCG | TGATGTTGGGCTGTTGTTGT | 158 |
| chr4:99560513-99562525 | p2 | (AC)6 | AACCAAATGCGACAAAAAGG | CATTTGCAGTTTCTGGACGA | 134 |
| chr4:99560513-99562525 | p2 | (AT)19 | AACCAAATGCGACAAAAAGG | CATTTGCAGTTTCTGGACGA | 134 |
| chr4:99598097-99600109 | p2 | (AC)6 | AACCAAATGCGACAAAAAGG | CATTTGCAGTTTCTGGACGA | 134 |
| chr4:99598097-99600109 | p2 | (AT)19 | AACCAAATGCGACAAAAAGG | CATTTGCAGTTTCTGGACGA | 134 |
| chr4:99684774-99686792 | p6 | (CATATA)3 | AAGAATGTCGTGAAATGGGC | ACGGGTCGTGACAGTAAACC | 167 |
| chr4:99703482-99705500 | p3 | (TGT)6 | AAATTGTGGAAGGGTAGGGG | TTAGCCCCCAAAGCAATATG | 122 |
| chr4:99712823-99714838 | p5 | (TCATT)3 | CAATCCCCAGAGGAGATTCA | TTTGAACACTGACGTCCTCG | 273 |
| chr4:99741286-99743304 | p3 | (TGT)6 | GAAGGCCACACATGAAAGGT | AGTAAGGGCCTTGTTTGGCT | 261 |
| chr4:99741286-99743304 | p3 | (TGA)5 | GAAGGCCACACATGAAAGGT | AGTAAGGGCCTTGTTTGGCT | 261 |
| chr4:99741286-99743304 | p2 | (TA)8 | GAAGGCCACACATGAAAGGT | AGTAAGGGCCTTGTTTGGCT | 261 |
| chr4:99741678-99743693 | p3 | (TGT)6 | GCGAACGCGTTGAGTAATTT | AGAGGTTTTGGGGTGCTTTT | 174 |
| chr4:99741678-99743693 | p3 | (TGA)5 | GCGAACGCGTTGAGTAATTT | AGAGGTTTTGGGGTGCTTTT | 174 |
| chr4:99741678-99743693 | p2 | (TA)8 | GCGAACGCGTTGAGTAATTT | AGAGGTTTTGGGGTGCTTTT | 174 |

| Marker ID | SSR type | SSR motif | FORWARD PRIMER (5'-3') | REVERSE PRIMER (5'-3') | size (bp) |
|---|---|---|---|---|---|
| chr4:99785772-99787790 | p6 | (ATACAT)3 | TCGTATCTGTCACGACCCAA | GCCTTAGCGGCTTCTGTATG | 152 |
| chr4:99785772-99787790 | p3 | (CAT)5 | TCGTATCTGTCACGACCCAA | GCCTTAGCGGCTTCTGTATG | 152 |
| chr4:99785979-99787994 | p6 | (ATACAT)3 | CATACAGAAGCCGCTAAGGC | GGGTCGCTAGTGTTCGGTTA | 106 |
| chr4:99785979-99787994 | p3 | (CAT)5 | CATACAGAAGCCGCTAAGGC | GGGTCGCTAGTGTTCGGTTA | 106 |
| chr4:99789750-99791765 | p5 | (GAAAT)3 | CGTCATATGAGGGGCGTAGT | TGGTTTCTTTGCCAAATTCC | 261 |
| chr4:99813680-99815695 | p5 | (ACCCA)3 | AGCAAATGGATTGCGAGACT | TTGGAAAACCAACATGCAAA | 102 |
| chr4:99813680-99815695 | p6 | (ATTTTT)3 | AGCAAATGGATTGCGAGACT | TTGGAAAACCAACATGCAAA | 102 |
| chr4:99813680-99815695 | p2 | (GT)7 | AGCAAATGGATTGCGAGACT | TTGGAAAACCAACATGCAAA | 102 |
| chr4:99852114-99854134 | p2 | (AT)10 | TCTTGTGGTGACGAATTGGA | AGCTGTGACTTCAGGGGAGA | 264 |
| chr4:99899421-99901465 | p2 | (AT)22 | TCGGTGAGTAATCGGTTTCC | AGGAGGAAGAAACCCAAGGA | 136 |
| chr4:99899421-99901465 | p2 | (AT)18 | TCGGTGAGTAATCGGTTTCC | AGGAGGAAGAAACCCAAGGA | 136 |
| chr4:99930534-99932549 | p5 | (ATTTC)3 | CGCTACAAGATGGGATGGAT | CTGTCCTGAACGAAAGGAGC | 145 |
| chr4:99934076-99936100 | p6 | (CATATA)4 | AGCAAACCCGCTGACTCTTA | GCTTCACCAGCCGATATGTT | 215 |
| chr4:99937915-99939931 | p4 | (TTCT)4 | CACCAATGATCCCACATTCA | GAATTTGGAAGATTCCGGGT | 139 |
| chr4:99937915-99939931 | p2 | (CA)8 | CACCAATGATCCCACATTCA | GAATTTGGAAGATTCCGGGT | 139 |
| chr4:99963080-99965095 | p5 | (ATTTC)3 | CAACCCCCAAAAGAGATTCA | ATAACAATGCCTCACAGGGC | 194 |
| chr4:99989022-99991036 | p2 | (AT)7 | CCTAACAGGAAGAAGCGTGG | TGTCGACTTTCCCTGAATCC | 155 |
| chr4:99989022-99991036 | p5 | (GAAAT)3 | CCTAACAGGAAGAAGCGTGG | TGTCGACTTTCCCTGAATCC | 155 |
| chr4:100012197-100014209 | p2 | (TG)6 | ATGCCATAGGTTCGAGGATG | CAATATCCGTATTCGGCCAC | 182 |
| chr4:100012197-100014209 | p2 | (TA)7 | ATGCCATAGGTTCGAGGATG | CAATATCCGTATTCGGCCAC | 182 |
| chr4:100021311-100023326 | p2 | (TA)8 | GGGAACTTTTGATCCCGATT | GTCGTTCCGGTAGCACGTAT | 111 |
| chr4:100021311-100023326 | p5 | (GCTAT)3 | GGGAACTTTTGATCCCGATT | GTCGTTCCGGTAGCACGTAT | 111 |
| chr4:100085661-100087676 | p3 | (ATG)5 | TCTCAGGTTGTTTTCGGAGG | CGGAGACATCATTCGGCTAT | 191 |
| chr4:100085661-100087676 | p6 | (TATGTA)3 | TCTCAGGTTGTTTTCGGAGG | CGGAGACATCATTCGGCTAT | 191 |
| chr4:100108772-100110787 | p5 | (TTTCA)3 | ATGAGTTAGCATGATGGCCC | CGAGTCGCATGCCTATTACA | 230 |
| chr4:100187894-100189960 | p2 | (AT)17 | GATAGCCTTGTCGGCTTCTG | CCTTCCCTAGGGGATTACCA | 142 |
| chr4:100298655-100300673 | p3 | (TCA)6 | GTTCCATTGCCTCATGACCT | GTTAGGGGTGCGCTTAATGA | 106 |
| chr4:100298655-100300673 | p6 | (AACCGG)3 | GTTCCATTGCCTCATGACCT | GTTAGGGGTGCGCTTAATGA | 106 |
| chr4:100308508-100310523 | p5 | (GAAAA)3 | ATCTCCCGATTCTCCCATCT | AGCTGTAGCTCCGGATGAAA | 286 |
| chr4:100317558-100319573 | p5 | (TATCA)3 | GAGATGGGGCTCACCAATAA | TGCTCGGTAAAAGTATCGGG | 207 |
| chr4:100331084-100333099 | p4 | (TTTA)4 | TACACGAAGAAGTCAGGGGG | TCACATCAAGGGGAGCTCTT | 100 |
| chr4:100331084-100333099 | p3 | (ATG)5 | TACACGAAGAAGTCAGGGGG | TCACATCAAGGGGAGCTCTT | 100 |
| chr4:100341432-100343470 | p2 | (TA)19 | CTGCTGAAGCACACACCACT | GGAATGGAACAAACCAATCG | 259 |
| chr4:100367356-100369563 | p3 | (GAA)69 | ACATTGGCCGAGATTGTAGG | CGCCAATTAGGTCACTGGTT | 256 |
| chr4:100423271-100425283 | p5 | (GAAAT)3 | GTTGAATTTCCAGGGAGCAA | GTTGAGTTAGCGCGAAGACC | 176 |
| chr4:100423271-100425283 | p2 | (CA)6 | GTTGAATTTCCAGGGAGCAA | GTTGAGTTAGCGCGAAGACC | 176 |

| Marker ID | SSR type | SSR motif | FORWARD PRIMER (5'-3') | REVERSE PRIMER (5'-3') | size (bp) |
|---|---|---|---|---|---|
| chr4:100437116-100439132 | p2 | (AT)8 | CCGAATTCGAGAATTGGAAA | AGATCATGAGCGTAGGGGTG | 292 |
| chr4:100516169-100518181 | p2 | (AT)6 | ACGACACCACAACGATTTCA | GGCCGTGTATGTGTGACTTG | 272 |
| chr4:100522251-100524319 | p2 | (TA)34 | TCCAAAGAAAGGGTGTTTCG | ATATGGTGACTCTGCTGGGG | 216 |
| chr4:100522251-100524319 | p2 | (TA)10 | TCCAAAGAAAGGGTGTTTCG | ATATGGTGACTCTGCTGGGG | 216 |
| chr4:100592257-100594275 | p2 | (GT)9 | TCCCACTAATTCAAGGGTGC | CATATGCACACCCACTTTGC | 236 |
| chr4:100592257-100594275 | p3 | (TTG)6 | TCCCACTAATTCAAGGGTGC | CATATGCACACCCACTTTGC | 236 |
| chr4:100592257-100594275 | p5 | (TATGA)3 | TCCCACTAATTCAAGGGTGC | CATATGCACACCCACTTTGC | 236 |
| chr4:100593226-100595241 | p3 | (TTG)6 | ATGTGTAGGGGAGATGCTGC | CGGATTTCGATGGACAGAAT | 108 |
| chr4:100593226-100595241 | p5 | (TATGA)3 | ATGTGTAGGGGAGATGCTGC | CGGATTTCGATGGACAGAAT | 108 |
| chr4:100599039-100601059 | p2 | (TA)10 | GATGATCCCTTCGTTCCTCA | AATTAGGGTTTGGGGTTTGG | 219 |
| chr4:100638286-100640300 | p2 | (TA)6 | TAAGCATGCATGGTATCCGA | AACTGCAGCTCCGAACAACT | 284 |
| chr4:100638286-100640300 | p2 | (TA)7 | TAAGCATGCATGGTATCCGA | AACTGCAGCTCCGAACAACT | 284 |
| chr4:100661767-100663785 | p3 | (TGT)6 | ATTGGAGACGTCCCATGAAG | AGGTTTGTTGCGTTGGAAAC | 282 |
| chr4:100661767-100663785 | p2 | (TG)10 | ATTGGAGACGTCCCATGAAG | AGGTTTGTTGCGTTGGAAAC | 282 |
| chr4:100662614-100664634 | p3 | (TGT)6 | CCGTGTAGTTGGGCTGATTT | GGGCAGTATCTCCCCTGTTT | 200 |
| chr4:100662614-100664634 | p2 | (TG)10 | CCGTGTAGTTGGGCTGATTT | GGGCAGTATCTCCCCTGTTT | 200 |
| chr4:100684504-100686525 | p3 | (AGA)7 | TCCCGGTTACGACTTACAGG | GTACCTCGGAGAAGCACTGG | 116 |
| chr4:100698217-100700232 | p3 | (AAT)5 | CCCACGAAAGTTAACCGAAA | TTCAGGCACACTCGCAATAG | 106 |
| chr4:100709773-100711791 | p2 | (AT)9 | TTTTTCGCCTCATTCAAACC | TAGGAAAGTGGCCAAACTGG | 232 |
| chr4:100759261-100761276 | p3 | (TTG)5 | GTGGGGTTTTGAGATGAGGA | TATGGCATGGACCAATCAGA | 179 |
| chr4:100836560-100838575 | p3 | (TGT)5 | AATATGGGCATAAGCCCTCC | GTTCACGCAGGGTGATTTTT | 293 |
| chr4:100858362-100860374 | p2 | (TA)6 | TCACACTCAGCCACACACAA | GGTTGTCCCACATCGAAATC | 231 |
| chr4:100930419-100932431 | p2 | (TA)6 | TGCTTCAGTTGCACAAAACC | CTCTCTCCACGACCCAACAT | 124 |
| chr4:100986732-100988747 | p2 | (TA)29 | AGAAGAACCGTTGAGCCAAA | CCCACTATTGTGTGAGGGCT | 242 |
| chr4:100986749-100988767 | p2 | (TA)29 | AGAAGAACCGTTGAGCCAAA | CCCACTATTGTGTGAGGGCT | 225 |
| chr4:101087330-101089344 | p2 | (TA)7 | TACGTGGCATTCTCATGCTC | GGGCATAAATTTCGGCTACA | 154 |
| chr4:101198905-101200920 | p5 | (TATGA)3 | CTCTTCGGAATCTACGTCCG | TTCAAATGAGGTGGCGTACA | 155 |
| chr4:101223335-101225353 | p2 | (TA)6 | ACAACAAAGGCCATAGCGTC | TCAAGCTGGGATAGAGCGAT | 286 |
| chr4:101223335-101225353 | p3 | (CAA)6 | ACAACAAAGGCCATAGCGTC | TCAAGCTGGGATAGAGCGAT | 286 |
| chr4:101233586-101235598 | p2 | (AG)6 | GGTTACCAGAGGCATGCAAT | TTCGCAGTCGGTTGTAGTTG | 145 |
| chr4:101249642-101251698 | p2 | (AT)28 | TCTTGCATTGGGTAAGAGGG | TCTTCAAGAGGAAGTGGCGT | 294 |
| chr4:101279709-101281771 | p2 | (AT)31 | TCGGGACGTAATGGCATATT | TCACCCTTTCCATCGTTCTC | 299 |
| chr4:101292706-101294730 | p2 | (AT)12 | GCACCGGATTTCATCAGAGT | CCACTCCGCATACTCCAAAT | 111 |
| chr4:101306246-101308282 | p2 | (AT)18 | CTTCCTGAATCTCCCAACCA | GGTGCAGACTCCTTAGCAGG | 156 |
| chr4:101326808-101328823 | p5 | (TTGGT)3 | GTTGCTTTGAGCAAAAAGGG | ATATCTCCCTCATGCGATGG | 239 |
| chr4:101378375-101380393 | p6 | (ATGTAT)3 | TTTCGGGACTTCATTCCTTG | AATGAGGTTATCCCCCAAGC | 300 |

| Marker ID | SSR type | SSR motif | FORWARD PRIMER (5'-3') | REVERSE PRIMER (5'-3') | size (bp) |
|---|---|---|---|---|---|
| chr4:101478205-101480220 | p5 | (AATAA)3 | AGAGCCGTTTGATTTTCCCT | AACTGACACGGATACCGGAC | 151 |
| chr4:101478205-101480220 | p2 | (GT)9 | AGAGCCGTTTGATTTTCCCT | AACTGACACGGATACCGGAC | 151 |
| chr4:101625802-101627814 | p2 | (TG)6 | GGAAACATTGGGACATCCAC | GACCCCATTGGATTTGTTTG | 122 |
| chr4:101640269-101642287 | p6 | (TAAAAA)3 | ACTGGGAGGTGAAAACCTGA | GACACCTCTCTATGGCCTCG | 187 |
| chr4:101680864-101682876 | p6 | (TATATG)3 | CCCACAGGTAGACAGGGAGA | ATATACACAAGCCGGCAAGG | 205 |
| chr4:101680864-101682876 | p2 | (TA)6 | CCCACAGGTAGACAGGGAGA | ATATACACAAGCCGGCAAGG | 205 |
| chr4:101680864-101682876 | p2 | (TC)6 | CCCACAGGTAGACAGGGAGA | ATATACACAAGCCGGCAAGG | 205 |
| chr4:101895727-101897741 | p6 | (ATCCGT)3 | CATAAGCCGACGAGGCTAAC | AGGAGGCTAAGGAGACAGGC | 200 |
| chr4:101895727-101897741 | p2 | (TA)7 | CATAAGCCGACGAGGCTAAC | AGGAGGCTAAGGAGACAGGC | 200 |
| chr4:102002801-102004817 | p4 | (GAAA)5 | TCCTGAATCTCCCAACAAGG | CGGCAACCGGTGTTACTACT | 161 |
| chr4:102002801-102004817 | p4 | (GATA)4 | TCCTGAATCTCCCAACAAGG | CGGCAACCGGTGTTACTACT | 161 |
| chr4:102115718-102117733 | p5 | (ACATA)3 | AAAGACTGACGCTCCCAGAA | TCCATGAGGATGTGCATTGT | 189 |
| chr4:102115718-102117733 | p3 | (CAA)9 | AAAGACTGACGCTCCCAGAA | TCCATGAGGATGTGCATTGT | 189 |
| chr4:102209430-102211468 | p2 | (AT)19 | AACCGATCTGCCAATCTTTG | GTTGAGCTTCAGTTGCACCA | 116 |
| chr4:102228761-102230779 | p6 | (CATATA)3 | CTGAGAATAACAGCGGGAGC | ATCAGGTATCCAGTCGGACG | 257 |
| chr4:102346351-102348375 | p2 | (TA)6 | GCCAAGTATCCAGCGGTAAA | CCGCGGTATTTGTATGTGTG | 266 |
| chr4:102373571-102375589 | p3 | (CAA)6 | TGACTGTGGCTGAGGAGATG | ATTTCACAGCGCCTTATTGG | 116 |
| chr4:102383300-102385318 | p6 | (TTCAGT)3 | CAACCCGCACAAGAACCTAT | CTTGATGGTGGAGGAGGAGA | 203 |
| chr4:102564215-102566233 | p3 | (TGT)6 | TCCAGGCAGAATACCTCCAC | TTAGTATTGGATTTGGCGGC | 245 |
| chr4:102624024-102626042 | p6 | (TTCAGT)3 | CAACCCGCACAAGAACCTAT | CTTGATGGTGGAGGAGGAGA | 203 |
| chr4:102637709-102639745 | p2 | (AT)18 | TTCGGAGGCATGACTAATCC | GAGCCCGTGACTTTATGAGC | 190 |
| chr4:102736998-102739012 | p2 | (AT)7 | CCCCCAAGCTTATAGGTGGT | CTTATGTTGGGCGGTTCAAT | 226 |
| chr4:102828108-102830122 | p2 | (TA)7 | ACGGCATTATGGGTGTTGTT | TTCGTACCATGGTCAGGTCA | 203 |
| chr4:102840924-102842942 | p2 | (AT)9 | CCCTGGGAAATATGCTGAAA | TTCTCCTTCATTCCCAATGC | 145 |
| chr4:102845853-102847865 | p2 | (AT)6 | GTTAAGGCTCTCGGGGATTC | CTTCAGAGGAAACTCCGCAC | 131 |
| chr4:102860007-102862021 | p2 | (AT)6 | CAGTTGAATCCCACAACGTG | TCATTTGAACCTACCCTCGG | 151 |
| chr4:102860017-102862032 | p2 | (AT)6 | CAGTTGAATCCCACAACGTG | TCATTTGAACCTACCCTCGG | 141 |
| chr4:102993608-102995620 | p2 | (TA)6 | TTTGTTTCGAAGGGAACACC | AGTAAAATCTCGCACCCCCT | 123 |
| chr4:103013034-103015052 | p6 | (AACTGA)3 | CACGTACGAAACTACGGGGT | ACATGCTCATGAAGGGGAAC | 293 |
| chr4:103043697-103045721 | p5 | (AATGC)3 | GCTGCAAACCTCGTATGTGA | GAACAGTGGCGAGAGAGGTC | 279 |
| chr4:103055868-103057886 | p2 | (AT)6 | TCATGTCATGTGCAAGCTGA | TCTCGGAGAGCTCGTTTGTT | 295 |
| chr4:103168402-103170416 | p2 | (AT)7 | TTGGTGGAAATCAAGTGCAA | CCAAGTCTTTCAACCCCTCA | 222 |
| chr4:103248630-103250645 | p3 | (TAA)5 | TCAGCCAAAGCTCTATGGGA | TCACTATTTGGGTCTTCCGC | 267 |
| chr4:103253780-103255800 | p3 | (AAT)5 | CATCAAATGCTCCAATCCCT | ATGTTTTCCTTTCGGGGAAT | 295 |
| chr4:103253780-103255800 | p2 | (AG)10 | CATCAAATGCTCCAATCCCT | ATGTTTTCCTTTCGGGGAAT | 295 |
| chr4:103285631-103287649 | p2 | (AC)9 | ACATACCTCTTTCCGCCCTT | AGTTGAACTTGGTGGGATGG | 281 |

| Marker ID | SSR type | SSR motif | FORWARD PRIMER (5'-3') | REVERSE PRIMER (5'-3') | size (bp) |
|---|---|---|---|---|---|
| chr4:103290706-103292724 | p6 | (AAGCTA)3 | GCATCATATCCGGCCTTCTA | CATGTCACTGATAATGCGGG | 277 |
| chr4:103379667-103381683 | p2 | (GT)9 | AAGGCGATTAGCAGTCCAGA | CGACCCTCAATGTCACCTTT | 278 |
| chr4:103379690-103381706 | p2 | (GT)9 | AAGGCGATTAGCAGTCCAGA | CGACCCTCAATGTCACCTTT | 255 |
| chr4:103402113-103404127 | p2 | (AT)7 | TCCAAAAAGCGAGGCTATGT | CCGTTACAAAGGGCAGTCAT | 296 |
| chr4:103402113-103404127 | p2 | (TA)7 | TCCAAAAAGCGAGGCTATGT | CCGTTACAAAGGGCAGTCAT | 296 |
| chr4:103402113-103404127 | p2 | (TA)6 | TCCAAAAAGCGAGGCTATGT | CCGTTACAAAGGGCAGTCAT | 296 |
| chr4:103406795-103408855 | p2 | (CA)7 | CAAAATCAAGCTCAGCCACA | GAGATGCGAAAAGTCCCAAA | 204 |
| chr4:103406795-103408855 | p2 | (AG)7 | CAAAATCAAGCTCAGCCACA | GAGATGCGAAAAGTCCCAAA | 204 |
| chr4:103406856-103408894 | p2 | (CA)7 | CAAAATCAAGCTCAGCCACA | GAGATGCGAAAAGTCCCAAA | 143 |
| chr4:103406856-103408894 | p2 | (AG)7 | CAAAATCAAGCTCAGCCACA | GAGATGCGAAAAGTCCCAAA | 143 |
| chr4:103510115-103512133 | p3 | (GAA)11 | CTGGTACTGTGGTGGGAGGT | CTGCCGGATAAATGTTGCTT | 205 |
| chr4:103510115-103512133 | p6 | (GAAAAA)3 | CTGGTACTGTGGTGGGAGGT | CTGCCGGATAAATGTTGCTT | 205 |
| chr4:103548890-103550929 | p3 | (ATA)13 | ATACACATCGCCTTCCCAAG | AGATGGTAGACCGCTGGATG | 155 |
| chr4:103605314-103607330 | p2 | (AC)8 | GCTTCCACATCCCACAATCT | GACCCTCCTATTTTCTCGGG | 148 |
| chr4:103613527-103615557 | p2 | (AT)15 | GTGTGCGGAACCGATAACTT | ACCTTTCTTTTGGGTGCAGA | 269 |
| chr4:103615488-103617518 | p2 | (AT)15 | AGGCGAACCATCCCTAAATC | GCAGCCCTAGATCTCCTCCT | 179 |
| chr4:103693746-103695764 | p3 | (TTG)6 | AAGCCCTCCAAAAGGACAAT | TTTATGTCTGGGCGTTTTCC | 250 |
| chr4:103743297-103745311 | p3 | (TCA)5 | GCTCCCAGGACTCATACCAA | GATGATGATGATGACAGGCG | 118 |
| chr4:103743297-103745311 | p2 | (AT)7 | GCTCCCAGGACTCATACCAA | GATGATGATGATGACAGGCG | 118 |
| chr4:103793781-103795796 | p5 | (GGTTC)3 | AATACGAGGCTACGGGGAGT | TGACCGCTATCACGAGTCAG | 160 |
| chr4:103793781-103795796 | p3 | (GTT)5 | AATACGAGGCTACGGGGAGT | TGACCGCTATCACGAGTCAG | 160 |
| chr4:103852468-103854483 | p6 | (TCCTCT)3 | CTTCGTATGCCCATGCCTAT | ATGGGGCAAGGAATGTATCA | 111 |
| chr4:103852889-103854907 | p6 | (TCCTCT)3 | ATGGGTCCGGCTTACTTTTT | CTGTTTGGGCCATTTCTTGT | 127 |
| chr4:103910490-103912508 | p6 | (CATATT)3 | CGTGCAACACACCCATAAAC | ATTGTTGCCCAATGGGATTA | 208 |
| chr4:103942417-103944445 | p2 | (TA)14 | GGAGCGCCAATATTGTTTGT | CCTTCTCTTTGACTCGCACC | 275 |
| chr4:103984889-103986904 | p5 | (TTTTA)3 | AATCATGACCAAACCGCTTC | CCCCATCATTTGTCTTGTCC | 103 |
| chr4:104152990-104155002 | p2 | (AT)6 | CCGAGAATATAACTCGGCCA | GCATGTTGATTGGGTTGTTG | 100 |
| chr4:104152990-104155002 | p2 | (TA)16 | CCGAGAATATAACTCGGCCA | GCATGTTGATTGGGTTGTTG | 100 |
| chr4:104170927-104172963 | p2 | (AT)18 | TGGATTGCTTGTTGGTTTCA | GCTAACGGATGGGATCAAGA | 142 |
| chr4:104199089-104201104 | p3 | (TTG)5 | CGTCGTTTGCGCTGTAAGTA | GGTTTGTTGCGTTGGAAACT | 227 |
| chr4:104259139-104261151 | p2 | (AG)6 | ACGTTTGCCCAAAAGTCAAC | CCTCGTTGACCCATTCCTAA | 283 |
| chr4:104414768-104416802 | p2 | (AT)17 | TCCTCCCCTTCTTGGAGAAT | AGGCGAAATGAATAGCATGG | 259 |
| chr4:104417704-104419740 | p5 | (AATGC)3 | TTGGGACCAGCAACACAATA | CCGAGACGTCGATATTCCAT | 176 |
| chr4:104417704-104419740 | p2 | (AT)18 | TTGGGACCAGCAACACAATA | CCGAGACGTCGATATTCCAT | 176 |
| chr4:104461530-104463546 | p2 | (TA)8 | GCGTTGATAATGAGGGAGGA | GCAGTTACAGGTGCGGTTTT | 119 |
| chr4:104602041-104604056 | p2 | (CA)6 | CGGTCAACTGTGCCATACAC | ACAGTTAACAGGGGGTGACG | 202 |

| Marker ID | SSR type | SSR motif | FORWARD PRIMER (5'-3') | REVERSE PRIMER (5'-3') | size (bp) |
|---|---|---|---|---|---|
| chr4:104602041-104604056 | p2 | (CA)6 | CGGTCAACTGTGCCATACAC | ACAGTTAACAGGGGGTGACG | 202 |
| chr4:104602041-104604056 | p3 | (CAA)5 | CGGTCAACTGTGCCATACAC | ACAGTTAACAGGGGGTGACG | 202 |
| chr4:104671498-104673516 | p2 | (AT)9 | GCGAACCCTCAACCAAAATA | AAAGTTGCACGGTGCCTATC | 178 |
| chr4:104834920-104836935 | p3 | (CAT)5 | TTGATGTGGCATAGATGCGT | CCCTCCGGCTAGGACTTATC | 299 |
| chr4:104894328-104896343 | p3 | (TCC)5 | TGGTTGGGGTTGACTACCAT | TGTAAACAAGGCAAAAGGGG | 174 |
| chr4:104980769-104982781 | p2 | (TA)6 | TGGTTGTATGGCAGTGGCTA | GGAGCTCTGACCGACTTTTG | 283 |
| chr4:105015224-105017242 | p6 | (CATATA)3 | AACAAGTGCCCGACCACTAC | CATCGGCGTGTATTTTGTTG | 189 |
| chr4:105062980-105065014 | p2 | (TA)17 | CGGAAGCTTACTTGGGTTTG | CTCTCCCTCCGTTTTTCCTC | 160 |
| chr4:105140831-105142843 | p2 | (AT)6 | TTGCGGTTGTTGATGTTGTT | CTTACTTGCGCCTTACGCTC | 200 |
| chr4:105379314-105381366 | p2 | (TA)26 | ATCGATGCTTCTCTTTGCGT | ATGGCTATATCGGGGTGACA | 254 |
| chr4:105436611-105438641 | p2 | (AT)15 | TCAGCATGCATAACCCTCAA | GGCCAGGATATAGGGGATGT | 247 |
| chr4:105495346-105497358 | p2 | (CA)6 | GGAGGAGTCGTAGAGGGGAC | AAAACAATTTTGGCGTCGAG | 265 |
| chr4:105501436-105503450 | p2 | (AT)7 | AATGGTGGATCGTGGTCATT | TCGTTTATTCCCCTTCAACG | 114 |
| chr4:105501436-105503450 | p2 | (AT)15 | AATGGTGGATCGTGGTCATT | TCGTTTATTCCCCTTCAACG | 114 |
| chr4:105516094-105518110 | p2 | (GT)8 | CGTGGAAAGTTAAAGGCGAG | TGAAATTCTCATTTTGCCCC | 128 |
| chr4:105535859-105537937 | p6 | (ATATGT)4 | CTATGATATCGTTCGCCGGT | CTATGGTGAGGTCTCGGGAA | 170 |
| chr4:105535859-105537937 | p2 | (TA)39 | CTATGATATCGTTCGCCGGT | CTATGGTGAGGTCTCGGGAA | 170 |
| chr4:105541406-105543432 | p2 | (AG)13 | TAAGCTTAGCCGCGGTACAT | GCGCAGTCTCCTCACCTTAC | 137 |
| chr4:105542978-105544996 | p6 | (ATATGT)3 | TTAACTCTCGGGGGATGTTG | TGCCAAGAGAAGAAAAGGGA | 285 |
| chr4:105542978-105544996 | p2 | (AT)9 | TTAACTCTCGGGGGATGTTG | TGCCAAGAGAAGAAAAGGGA | 285 |
| chr4:105557965-105559985 | p2 | (AT)10 | CTCCGGAGGATTCAAACTCA | AAGGCGGGTAGAAAGAGAGC | 143 |
| chr4:105557965-105559985 | p2 | (AT)6 | CTCCGGAGGATTCAAACTCA | AAGGCGGGTAGAAAGAGAGC | 143 |
| chr4:105576658-105578678 | p2 | (AT)10 | CTCCGGAGGATTCAAACTCA | AAGGCGGGTAGAAAGAGAGC | 143 |
| chr4:105576658-105578678 | p2 | (AT)6 | CTCCGGAGGATTCAAACTCA | AAGGCGGGTAGAAAGAGAGC | 143 |
| chr4:105596449-105598470 | p5 | (TTGCA)3 | GCCAGGGTCCGATATGTCTA | ATGCCATGCTATGCAAATGA | 123 |
| chr4:105596449-105598470 | p3 | (AGA)7 | GCCAGGGTCCGATATGTCTA | ATGCCATGCTATGCAAATGA | 123 |
| chr4:105712693-105714707 | p2 | (TA)7 | GCATCGTATGTGAATGGACG | TCAATTCCCAAACGTCAACA | 210 |
| chr4:105834501-105836535 | p2 | (AT)7 | AATGCCCTGAAACCTCCTTT | GCTTGGGTGGTGACAAAAGT | 252 |
| chr4:105917614-105919634 | p2 | (AT)10 | CTTGGCCTAGACCTACGACG | GCATCGGTCCATTGAAACTT | 272 |
| chr4:105941272-105943284 | p2 | (AT)6 | CCCTTTCTGCTGTCTTGGAG | ACCATATAGGCACCACTGGC | 132 |
| chr4:105941272-105943284 | p2 | (TC)6 | CCCTTTCTGCTGTCTTGGAG | ACCATATAGGCACCACTGGC | 132 |
| chr4:105948516-105950534 | p6 | (TTGCTA)3 | CGGAGTTAAGTTTCGGTGGA | GCAAGTCATGCAACATCACC | 203 |
| chr4:105962006-105964021 | p3 | (GTT)5 | CGAACCCTCCAAAACAAAAA | CAAACGTCGCACTCTTGCTA | 216 |
| chr4:106011753-106013771 | p4 | (TCAT)4 | AAAGGGTCGCAATGAATGAG | AAAAATAAACTGCATGGCGG | 121 |
| chr4:106011753-106013771 | p2 | (GA)9 | AAAGGGTCGCAATGAATGAG | AAAAATAAACTGCATGGCGG | 121 |
| chr4:106209567-106211585 | p6 | (TATGGA)3 | ATTTCCACTTTGGCCAGTTG | TAGTTGACGGGCCGTATTTC | 201 |

| Marker ID | SSR type | SSR motif | FORWARD PRIMER (5'-3') | REVERSE PRIMER (5'-3') | size (bp) |
|---|---|---|---|---|---|
| chr4:106279385-106281400 | p3 | (ATC)5 | ACTGCAGCTCCGAAACAAAT | ATCGCTCGTACTGACGTCCT | 290 |
| chr4:106326313-106328328 | p5 | (AAACA)3 | CGAAGAAATTAGCCCAACCA | GAGCCTTAGACAGGTGCCAG | 176 |
| chr4:106341889-106343901 | p2 | (TG)6 | GATGCTAAAGGTGCCTACGG | CGACCTCATAGGATCGGTGT | 208 |
| chr4:106450799-106452817 | p2 | (GA)9 | CAGTCGCGAACAAGATTGAA | CCACACGATGGACTAGGGTT | 158 |
| chr4:106450799-106452817 | p6 | (ATGTAT)3 | CAGTCGCGAACAAGATTGAA | CCACACGATGGACTAGGGTT | 158 |
| chr4:106453726-106455741 | p3 | (TTG)5 | CCCAAACTCCCTCGTTCATA | TAACTGTTCATCCGCGTCAC | 295 |
| chr4:106529664-106531679 | p5 | (AAATA)3 | AATCGATGAGGGATGTCTGC | CGTTGTCTCTGCTCTCCTCC | 266 |
| chr4:106531697-106533715 | p2 | (TA)6 | GCGACCTTATTCACCGAGTC | ATGCTGAAGGGTCACCAAAC | 288 |
| chr4:106531697-106533715 | p2 | (AT)9 | GCGACCTTATTCACCGAGTC | ATGCTGAAGGGTCACCAAAC | 288 |
| chr4:106668698-106670716 | p3 | (CAA)6 | AGGCACGGTGTTATACTGCC | CGACCTTACGTCACTCCGAT | 226 |
| chr4:106668698-106670716 | p2 | (TC)10 | AGGCACGGTGTTATACTGCC | CGACCTTACGTCACTCCGAT | 226 |
| chr4:106858675-106860691 | p4 | (TATG)4 | TGGTGTCTTGGGTGTTGGTA | TGGTCTCTTCAAGCTCGGAT | 187 |
| chr4:106862136-106864164 | p2 | (AT)14 | TCCCATCGGGATGAGTTTAG | TGCCAATCACACCTTACCAA | 141 |
| chr4:106895829-106897841 | p2 | (CT)6 | CATGGTTGGGAAGCTTAGGA | CTTGCGTACCAACGGAAGAT | 217 |
| chr4:106935275-106937287 | p2 | (AG)6 | AACTGCGCTAGTCCAAAGGA | CTTGAGCCCAAAAGGCATTA | 243 |
| chr4:106943989-106946001 | p2 | (AT)6 | CTACCGAGCACCCCTTATCA | GTGGGTCATGGGTGGTTATC | 192 |
| chr4:106964507-106966519 | p2 | (TA)6 | TTCAACTCGGCCAAATAACC | GTTGGAAGCGAGGAATGTGT | 255 |
| chr4:106991492-106993508 | p2 | (TA)17 | GGGGATTAAATGGTTCGCTT | AACATTCTCCAACCACCAGC | 243 |
| chr4:106991492-106993508 | p2 | (AT)8 | GGGGATTAAATGGTTCGCTT | AACATTCTCCAACCACCAGC | 243 |
| chr4:107072161-107074173 | p2 | (GT)6 | CCTGGACATGGATTTCGACT | CAAGGCTCACTGGTCTCACA | 166 |
| chr4:107113169-107115181 | p2 | (CA)6 | ATAGAGTTGTCCCCGAGGCT | GTTGAATTTTGGGGATGGTG | 207 |
| chr4:107215143-107217155 | p2 | (AT)6 | GGATGGCCAAGAGCAATAAA | GCGACTGAGCTTGGAAAAAC | 119 |
| chr4:107258800-107260815 | p5 | (ATTTC)3 | AGAAGTGCCCATTGCCATAC | TGGCCACAACACAATAGCAT | 108 |
| chr4:107258800-107260815 | p2 | (AT)10 | AGAAGTGCCCATTGCCATAC | TGGCCACAACACAATAGCAT | 108 |
| chr4:107259088-107261108 | p5 | (ATTTC)3 | AAGGCATCCGATGTAAGACG | CAGTGATCCCGAAATTCGTT | 288 |
| chr4:107259088-107261108 | p2 | (AT)10 | AAGGCATCCGATGTAAGACG | CAGTGATCCCGAAATTCGTT | 288 |
| chr4:107326468-107328484 | p2 | (GT)8 | AAACGGACCAGTGCATTTTC | ATCAAGCACGGGAGAAACAC | 216 |
| chr4:107410529-107412547 | p3 | (ATC)6 | TTTCACCATGCCCAAGTGTA | GCCCGCAGGTAGACAAGTAG | 176 |
| chr4:107438935-107440965 | p2 | (TA)15 | GAGATGAATAAGGCCGACCA | TTTTTCCGTTTGCTTCTGCT | 207 |
| chr4:107500538-107502562 | p2 | (AT)12 | CGTACGCTGCATCTATCGAA | GAGACAGGGCGATGAGAGAG | 217 |
| chr4:107519258-107521273 | p5 | (TTTCA)3 | GACGGGACACAAAGAGGTGT | TGTAAAGTGGAAGGAACGGC | 108 |
| chr4:107542875-107544905 | p3 | (GTT)6 | ACTCCAAGACTGCTCGGAAA | TTTACGCAAACCCAAACACA | 159 |
| chr4:107542875-107544905 | p2 | (AT)15 | ACTCCAAGACTGCTCGGAAA | TTTACGCAAACCCAAACACA | 159 |
| chr4:107577252-107579292 | p2 | (AT)20 | GTGCTAGAGGCGAGGTATGC | AAGAAAGCTACAGCTGCCCA | 114 |
| chr4:107577252-107579292 | p2 | (AC)22 | GTGCTAGAGGCGAGGTATGC | AAGAAAGCTACAGCTGCCCA | 114 |
| chr4:107629876-107631892 | p2 | (AT)8 | AGATCTTGCGAAGCCAAAGA | CATATGCGTCATCTGCTCGT | 111 |

| Marker ID | SSR type | SSR motif | FORWARD PRIMER (5'-3') | REVERSE PRIMER (5'-3') | size (bp) |
|---|---|---|---|---|---|
| chr4:107795696-107797708 | p5 | (GGGGT)4 | TCAAAATCTGCTTGAATGCG | AATCGATCATCACTCGCTCC | 184 |
| chr4:107795775-107797797 | p5 | (GGGGT)4 | TCAAAATCTGCTTGAATGCG | AATCGATCATCACTCGCTCC | 105 |
| chr4:107986796-107988812 | p4 | (ATTT)4 | CCTGGCCACTCATCTAAGGA | ATCAGCAATGGTGGTTCACA | 197 |
| chr4:108090496-108092508 | p2 | (TA)6 | ATTAATCGCAAACCCGTTCA | TAGGCGAATTGGAGGAAATG | 187 |
| chr4:108090496-108092508 | p3 | (ATC)6 | ATTAATCGCAAACCCGTTCA | TAGGCGAATTGGAGGAAATG | 187 |
| chr4:108328413-108330427 | p2 | (AT)7 | GGACCTCTTAGCATGGTGGA | CGAAGCCACGGTTGATATTT | 274 |
| chr4:108516721-108518741 | p2 | (TA)10 | TACCAACCAAGGCTTGTTCC | TTGCTCATCCCATGTTTTCA | 104 |
| chr4:108519636-108521651 | p5 | (CATTT)3 | CTCTAAACCGACCCGCAATA | CCAGTTCTAGCGGTCCTCTG | 217 |
| chr4:108577394-108579430 | p2 | (AT)18 | TGATCAGCAAGATGGATTGC | CTTTTCAGCTTAGCGATGGG | 202 |
| chr4:108587843-108589857 | p2 | (GA)7 | CCCTCAACAATGTACACCCC | TACTGGTGCTGCTGGTTTTG | 169 |
| chr4:108656300-108658320 | p5 | (GAAAT)3 | CCAAACATACGGACCCAAAC | ACGAGACGACGAAACGAACT | 215 |
| chr4:108656300-108658320 | p5 | (ATGAA)4 | CCAAACATACGGACCCAAAC | ACGAGACGACGAAACGAACT | 215 |
| chr4:108738263-108740279 | p4 | (AGAA)4 | TCGACAGTGATGACTCCAGC | GAAAGGGCCATTCAGTACCA | 240 |
| chr4:108777693-108779709 | p4 | (AGAA)4 | TCGACAGTGATGACTCCAGC | GAAAGGGCCATTCAGTACCA | 240 |
| chr4:108817400-108819415 | p5 | (AAGCC)3 | GGAAATGGAAAAGAATGCGA | ACAGGTAAGTCAACGTCGGG | 237 |
| chr4:108902383-108904431 | p6 | (TGGGTG)4 | CTCCAAGCTTCTCGATCCAC | GGTACTGCTACCTCCCTCCC | 231 |
| chr4:108902383-108904431 | p2 | (TA)24 | CTCCAAGCTTCTCGATCCAC | GGTACTGCTACCTCCCTCCC | 231 |
| chr4:108902383-108904431 | p2 | (AT)8 | CTCCAAGCTTCTCGATCCAC | GGTACTGCTACCTCCCTCCC | 231 |
| chr4:108902383-108904431 | p2 | (AT)8 | CTCCAAGCTTCTCGATCCAC | GGTACTGCTACCTCCCTCCC | 231 |
| chr4:108902383-108904431 | p2 | (TA)8 | CTCCAAGCTTCTCGATCCAC | GGTACTGCTACCTCCCTCCC | 231 |
| chr4:108902383-108904431 | p2 | (TA)22 | CTCCAAGCTTCTCGATCCAC | GGTACTGCTACCTCCCTCCC | 231 |
| chr4:108902383-108904431 | p2 | (TA)11 | CTCCAAGCTTCTCGATCCAC | GGTACTGCTACCTCCCTCCC | 231 |
| chr4:108903011-108905055 | p6 | (TGGGTG)4 | CGCGGACTCTTCAAAATCTC | CCCGAGGTAAGTTGACCAAA | 196 |
| chr4:108903011-108905055 | p2 | (TA)24 | CGCGGACTCTTCAAAATCTC | CCCGAGGTAAGTTGACCAAA | 196 |
| chr4:108903011-108905055 | p2 | (AT)8 | CGCGGACTCTTCAAAATCTC | CCCGAGGTAAGTTGACCAAA | 196 |
| chr4:108903011-108905055 | p2 | (AT)8 | CGCGGACTCTTCAAAATCTC | CCCGAGGTAAGTTGACCAAA | 196 |
| chr4:108903011-108905055 | p2 | (TA)8 | CGCGGACTCTTCAAAATCTC | CCCGAGGTAAGTTGACCAAA | 196 |
| chr4:108903011-108905055 | p2 | (TA)22 | CGCGGACTCTTCAAAATCTC | CCCGAGGTAAGTTGACCAAA | 196 |
| chr4:108903011-108905055 | p2 | (TA)11 | CGCGGACTCTTCAAAATCTC | CCCGAGGTAAGTTGACCAAA | 196 |
| chr4:108993901-108995919 | p6 | (ACACAT)3 | ATGGACAGAATTGTTCCCCA | AAGGCGCAGCTAAGGTATGA | 175 |
| chr4:109071700-109073732 | p2 | (TA)16 | CCCCTTCAGCACGTAGTCAT | AGGTAATGAAACCTCGGCCT | 246 |
| chr4:109210408-109212444 | p2 | (TA)18 | AGCGGTACAAACTGTGGTCC | CGTTGTTTTGGGAAGAGGAA | 232 |
| chr4:109277038-109279053 | p5 | (TGAAA)3 | AACTTTAGCGCGTTGGCTTA | GTTAGGGGTGCGCTTAATGA | 268 |
| chr4:109305351-109307405 | p2 | (AT)27 | AGACCCCTAAAGACCCCAGA | CAATTGATCCTCGATGGCTT | 161 |
| chr4:109305351-109307405 | p2 | (AC)6 | AGACCCCTAAAGACCCCAGA | CAATTGATCCTCGATGGCTT | 161 |
| chr4:109305351-109307405 | p2 | (TA)8 | AGACCCCTAAAGACCCCAGA | CAATTGATCCTCGATGGCTT | 161 |

| Marker ID | SSR type | SSR motif | FORWARD PRIMER (5'-3') | REVERSE PRIMER (5'-3') | size (bp) |
|---|---|---|---|---|---|
| chr4:109305531-109307543 | p2 | (AT)27 | TGGATGAATTGGTGGTTGAA | TTCGGGTTACAAGAATTCGG | 293 |
| chr4:109305531-109307543 | p2 | (AC)6 | TGGATGAATTGGTGGTTGAA | TTCGGGTTACAAGAATTCGG | 293 |
| chr4:109305531-109307543 | p2 | (TA)8 | TGGATGAATTGGTGGTTGAA | TTCGGGTTACAAGAATTCGG | 293 |
| chr4:109521835-109523863 | p4 | (TATT)7 | GCAACTTGAAGTGGGCTCAT | AAGCTAGCGTTTGCGGAATA | 138 |
| chr4:109596829-109598843 | p2 | (TA)8 | TGGGCCCCATACTAAACAAC | CCCATGAGTTTGGAGATGGT | 251 |
| chr4:109596829-109598843 | p2 | (TA)7 | TGGGCCCCATACTAAACAAC | CCCATGAGTTTGGAGATGGT | 251 |
| chr4:109596829-109598843 | p2 | (TA)6 | TGGGCCCCATACTAAACAAC | CCCATGAGTTTGGAGATGGT | 251 |
| chr4:109605058-109607073 | p5 | (TTGGT)3 | AAAGAATGGATGGGCAAGTG | CCAAACCAACCAATTATGGC | 107 |
| chr4:109674267-109676297 | p2 | (TA)15 | ACTTCCCCCAACATTGAACA | AGACGGCAGAAAGCGTTAGA | 290 |
| chr4:109674267-109676297 | p3 | (TAT)6 | ACTTCCCCCAACATTGAACA | AGACGGCAGAAAGCGTTAGA | 290 |
| chr4:109810378-109812392 | p2 | (TA)7 | AACAAATTCTTGCCAAACGG | CAGATGAGGACCCACAAGGT | 141 |
| chr4:109976245-109978261 | p2 | (TA)8 | CTTGTCTGCCCTCAGGACTC | TGTTGTGGATGAAGCAAGGA | 203 |
| chr4:109999373-110001394 | p3 | (TCT)7 | TTACACCGCCATTGAATGAA | TTATAATCGTGCGAAGCGTG | 280 |
| chr4:110043469-110045483 | p2 | (AG)7 | GTGCAGCGAAGACATGGTAA | AAACAGGTCAGCAATGTCCC | 152 |
| chr4:110076984-110079028 | p2 | (AT)22 | TTGAAAAACATCGTGTGGGA | GGGCAGTATTTCCCCTGTTT | 220 |
| chr4:110076984-110079028 | p2 | (TA)15 | TTGAAAAACATCGTGTGGGA | GGGCAGTATTTCCCCTGTTT | 220 |
| chr4:110350979-110352997 | p2 | (AT)9 | ATCACTCGCCAAACAAATCC | TGAATTTACAGGGGAGGTGC | 244 |
| chr4:110350979-110352997 | p2 | (CT)9 | ATCACTCGCCAAACAAATCC | TGAATTTACAGGGGAGGTGC | 244 |
| chr4:110371024-110373036 | p2 | (AT)6 | CCCCTATCATCATGACCCAC | AAAGCTACAGCGACCCAGAA | 173 |
| chr4:110416537-110418552 | p3 | (TTG)5 | CAAGAAGGTTTGGGGACTCA | CGAACCACCCAATGGTTATC | 106 |
| chr4:110487114-110489126 | p2 | (AT)6 | GTTTTTGGGCAAAAGGCATA | AAAGCTGCCCATCAAAACAC | 115 |
| chr4:110487114-110489126 | p5 | (GTGGT)3 | GTTTTTGGGCAAAAGGCATA | AAAGCTGCCCATCAAAACAC | 115 |
| chr4:110494987-110496999 | p2 | (TA)6 | GCGCATTCTCTCCTCTTGAC | CAGTGATGGTCAGCTAGGCA | 105 |
| chr4:110523757-110525773 | p4 | (AAAT)4 | CACGGAGTCTTGCGTCCTAT | GGCTTGGTAGCAAAAATGGA | 171 |
| chr4:110607590-110609606 | p2 | (AT)8 | CGGGTGATAATAAGGGCAAA | GTTGTCACATGACTGGTCGG | 204 |
| chr4:110615988-110618003 | p3 | (AAG)5 | AAATTTTCGACCACGTGAGG | TTGGGCAACTGTGAGCATTA | 168 |
| chr4:110693431-110695446 | p5 | (CATTT)3 | GCATTCATATTTGGATGGGG | TAAGAATCAACCTGGCGCTT | 176 |
| chr4:110744269-110746293 | p2 | (AC)12 | TACTTCCCACACCCTCCAAG | TTGGATTTTTGAGGTCAGGC | 130 |
| chr4:110845260-110847276 | p2 | (TA)12 | CCCCTAACTCGTATTTGCCA | TGGCCATGTCAGACTGTGTT | 158 |
| chr4:110845277-110847301 | p2 | (TA)12 | CCCCTAACTCGTATTTGCCA | TGGCCATGTCAGACTGTGTT | 141 |
| chr4:110850051-110852083 | p2 | (TA)32 | TTCATTCGGAGGTTACCGAC | CGCGGCACAGTGTAGTAAAA | 250 |
| chr4:110850051-110852083 | p2 | (AT)16 | TTCATTCGGAGGTTACCGAC | CGCGGCACAGTGTAGTAAAA | 250 |
| chr4:110910698-110912713 | p3 | (TGT)5 | GGGATGATCATTTGCGACTT | CCGATTGTCCGCATAAGACT | 228 |
| chr4:110948843-110950887 | p2 | (AT)22 | GGTGGAGATACAAAGCCCAA | TTACCCCCATACTTTGCAGC | 228 |
| chr4:110948843-110950887 | p2 | (AT)8 | GGTGGAGATACAAAGCCCAA | TTACCCCCATACTTTGCAGC | 228 |
| chr4:110984157-110986173 | p2 | (AT)6 | ACTCCTCGGGTCAGGTAGGT | TCATCATTAGCCCGTGTCAA | 163 |

| Marker ID | SSR type | SSR motif | FORWARD PRIMER (5'-3') | REVERSE PRIMER (5'-3') | size (bp) |
|---|---|---|---|---|---|
| chr4:110984176-110986188 | p2 | (AT)6 | ACTCCTCGGGTCAGGTAGGT | TCATCATTAGCCCGTGTCAA | 144 |
| chr4:110984193-110986211 | p2 | (AT)6 | ACTCCTCGGGTCAGGTAGGT | TCATCATTAGCCCGTGTCAA | 127 |
| chr4:111014250-111016262 | p6 | (ATCAAT)3 | AGACGGAAGGGCAAAGAAAT | TGCACCAGATCCTACCATCA | 187 |
| chr4:111031943-111033955 | p6 | (ATCAAT)3 | AGACGGAAGGGCAAAGAAAT | TGCACCAGATCCTACCATCA | 186 |
| chr4:111031943-111033955 | p2 | (TA)6 | AGACGGAAGGGCAAAGAAAT | TGCACCAGATCCTACCATCA | 186 |
| chr4:111039433-111041448 | p5 | (TTTTC)3 | GGGGATAGAACACCTGCTGA | GCCCAAGCGTAGTATCGTGT | 267 |
| chr4:111119646-111121661 | p3 | (CAG)5 | GCTCCACCCCCACTAGTACA | GAACTCTGGAGGGCCACATA | 178 |
| chr4:111164525-111166540 | p5 | (CTTTT)3 | TTGCCAATTCATGTGGAGAA | TGATTGCAGGTCCATGTGTT | 105 |
| chr4:111173456-111175468 | p2 | (TC)6 | AACACTAGCTCCCGTCCAGA | TCCCCTTTTGACATCTTTGC | 211 |
| chr4:111173456-111175468 | p2 | (TA)7 | AACACTAGCTCCCGTCCAGA | TCCCCTTTTGACATCTTTGC | 211 |
| chr4:111193945-111195969 | p6 | (AACAAA)4 | TTGGCCTTATTTTTCGCATC | ATGAGCCTAGCCCGTACAGA | 236 |
| chr4:111231982-111234018 | p2 | (AT)18 | TCGAGAAATGTGAGGGAAGG | GTTCTCTTTTCTTTGCCCCC | 159 |
| chr4:111280387-111282415 | p2 | (AT)12 | TGCATATGAATGCTCAAGCC | ACATGCCAGCTTCCAGTTTC | 219 |
| chr4:111280387-111282415 | p2 | (AT)8 | TGCATATGAATGCTCAAGCC | ACATGCCAGCTTCCAGTTTC | 219 |
| chr4:111280476-111282491 | p2 | (AT)12 | TGCATATGAATGCTCAAGCC | ACATGCCAGCTTCCAGTTTC | 130 |
| chr4:111280476-111282491 | p2 | (AT)8 | TGCATATGAATGCTCAAGCC | ACATGCCAGCTTCCAGTTTC | 130 |
| chr4:111281544-111283564 | p2 | (TA)6 | TATGCACACAAACTGCCACA | ATAACCCCGGGATTTTTACG | 296 |
| chr4:111281544-111283564 | p2 | (AT)8 | TATGCACACAAACTGCCACA | ATAACCCCGGGATTTTTACG | 296 |
| chr4:111281544-111283564 | p2 | (TA)10 | TATGCACACAAACTGCCACA | ATAACCCCGGGATTTTTACG | 296 |
| chr4:111281544-111283564 | p2 | (AT)10 | TATGCACACAAACTGCCACA | ATAACCCCGGGATTTTTACG | 296 |
| chr4:111328395-111330410 | p5 | (CCACC)3 | AACGGTGCTAAACCCTTGTG | TGCTGCCAAATTGTCGTTTA | 156 |
| chr4:111328395-111330410 | p5 | (TACAA)3 | AACGGTGCTAAACCCTTGTG | TGCTGCCAAATTGTCGTTTA | 156 |
| chr4:111330204-111332219 | p5 | (TACAA)3 | CCCACCCCAATACAACTCAC | TGGGATGAGGGTGAAGGTAG | 278 |
| chr4:111330204-111332219 | p5 | (CCACC)3 | CCCACCCCAATACAACTCAC | TGGGATGAGGGTGAAGGTAG | 278 |
| chr4:111382515-111384529 | p2 | (AC)7 | ATTTCACTGATGGGCAGTCC | GCACCAGCTCAGATGTTCAA | 206 |
| chr4:111425849-111427863 | p2 | (TA)7 | TCTGCCTGGTGGTGTTACAA | CAACCAACTGAACCACCCTT | 223 |
| chr4:111498415-111500429 | p2 | (TA)7 | TCTGCCTGGTGGTGTTACAA | CAACCAACTGAACCACCCTT | 223 |
| chr4:111674971-111676989 | p5 | (AAGAG)3 | TTTTGTCCATCCTTTCGTCC | GAAGTTTGGACCCGCATCTA | 277 |
| chr4:111674985-111677018 | p5 | (AAGAG)3 | TTTTGTCCATCCTTTCGTCC | GAAGTTTGGACCCGCATCTA | 263 |
| chr4:111675024-111677042 | p5 | (AAGAG)3 | TTTTGTCCATCCTTTCGTCC | GAAGTTTGGACCCGCATCTA | 224 |
| chr4:111675051-111677081 | p5 | (AAGAG)3 | TTTTGTCCATCCTTTCGTCC | GAAGTTTGGACCCGCATCTA | 197 |
| chr4:111837091-111839109 | p2 | (AG)9 | CTTTTCACCTCCTCACCCAA | CCACGGCTAAAGATGTGGAT | 101 |
| chr4:111911108-111913128 | p2 | (AT)28 | CGGTTGACCTACGGGTTCTA | TTGGGCCGTGACATTTCTAT | 188 |
| chr4:111911108-111913128 | p2 | (TA)10 | CGGTTGACCTACGGGTTCTA | TTGGGCCGTGACATTTCTAT | 188 |
| chr4:111921839-111923857 | p6 | (GGGGGT)3 | CTGGGGAGTGTTTTCGTGTT | TATATGCGGTTCGAAGGACC | 150 |
| chr4:111953540-111955552 | p2 | (TA)6 | ATGTGGTTTGAAGGCTGTCC | TAGGTCCCATGACCACCAAT | 229 |

| Marker ID | SSR type | SSR motif | FORWARD PRIMER (5'-3') | REVERSE PRIMER (5'-3') | size (bp) |
|---|---|---|---|---|---|
| chr4:111982236-111984250 | p2 | (AT)7 | TAGCTCCCATGTCCGATACC | GGCAGGGAGATTGTGTTGAT | 144 |
| chr4:112015921-112017935 | p2 | (AT)7 | TAGCTCCCATGTCCGATACC | GGCAGGGAGATTGTGTTGAT | 144 |
| chr4:112172050-112174065 | p5 | (TGCAA)3 | AAGAATTGTTGCACCTTGGG | CTATCGCGATGGGAAAGGTA | 254 |
| chr4:112185635-112187650 | p5 | (GTTCG)3 | ACCATCATAAAGCCCTGACG | GTTTATTGTGGGGGTTCGTG | 185 |
| chr4:112244747-112246762 | p3 | (ATG)5 | GCTGACATAGCAGAAACGCA | ATCCACTTTGAATTCACGCC | 203 |
| chr4:112272548-112274563 | p2 | (AT)16 | CAAACAAAAAGGGCGTCAGT | AACACCGAGCCTTGATATGG | 191 |
| chr4:112272548-112274563 | p3 | (AAC)5 | CAAACAAAAAGGGCGTCAGT | AACACCGAGCCTTGATATGG | 191 |
| chr4:112287146-112289161 | p2 | (AT)22 | CAAACAAAAAGGGCGTCAAT | AAGCATTCATGTTATCCGCC | 143 |
| chr4:112287146-112289161 | p3 | (AAC)5 | CAAACAAAAAGGGCGTCAAT | AAGCATTCATGTTATCCGCC | 143 |
| chr4:112439103-112441121 | p5 | (TTTCA)3 | GCCTTTGGTTGTATCTCGGA | AAGAACTCCAACCCTCAGCA | 102 |
| chr4:112454100-112456114 | p2 | (AT)7 | GCGTCGAGGTACCATCTCAT | ACCGTATGTCACCCCGATAA | 160 |
| chr4:112454100-112456114 | p2 | (AT)6 | GCGTCGAGGTACCATCTCAT | ACCGTATGTCACCCCGATAA | 160 |
| chr4:112476645-112478661 | p4 | (TATC)4 | TCCCGGACTAATGGTGGTTA | TCGTCTGCACCATAGCTCAC | 129 |
| chr4:112487136-112489192 | p2 | (AT)28 | ACTCGACAGATGCTCACCCT | CTTGGGTTGTGTGGTCTGTG | 221 |
| chr4:112487136-112489192 | p6 | (ATACGG)3 | ACTCGACAGATGCTCACCCT | CTTGGGTTGTGTGGTCTGTG | 221 |
| chr4:112557176-112559196 | p4 | (ATAC)5 | GGAGCCGCTTACAGACTGAC | TACTGACCCCCTGTTCTTCG | 277 |
| chr4:112566167-112568201 | p2 | (AT)17 | TTCCACGTTGACGGTTCATA | GGAAAGAATGAACGCTGCTC | 217 |
| chr4:112600862-112602880 | p6 | (CTCATG)3 | GAGTCTAAGGCCATTGCAGC | TCAATTAGACCTCCGTTGGG | 143 |
| chr4:112600862-112602880 | p3 | (ATA)6 | GAGTCTAAGGCCATTGCAGC | TCAATTAGACCTCCGTTGGG | 143 |
| chr4:112611574-112613586 | p2 | (TA)6 | AAAATTGCGAAGCGAAAATG | TTTCGTACGCATTCAAACCA | 156 |
| chr4:112663943-112665963 | p5 | (ATTAT)4 | CTTGTACCGAGCACCCCTTA | CGGAGCCTCATATCATTGGT | 152 |
| chr4:112725028-112727040 | p2 | (AT)14 | CCCGAACTGGACACTCGTAT | TATGATATGCGGCGTTTTGA | 244 |
| chr4:112725028-112727040 | p2 | (AT)6 | CCCGAACTGGACACTCGTAT | TATGATATGCGGCGTTTTGA | 244 |
| chr4:112755367-112757385 | p2 | (AT)9 | CGGGAACGATGAAAAGGATA | TGTCTACCTGTGGGCATGAA | 133 |
| chr4:112766868-112768882 | p2 | (TA)7 | GATTTGTGTATGGGGATGGC | AATAGAACCCGGCCCTCTTA | 137 |
| chr4:112815740-112817755 | p2 | (TA)8 | CGGTTGCAGCTACATCAGAA | GGGTCTTGACATGATGGTCC | 197 |
| chr4:112858759-112860791 | p5 | (GCATT)3 | TTGATGAGCTAGGCTCGGTT | ATCCCGGATCTTTACCCAAC | 163 |
| chr4:112865128-112867148 | p2 | (AT)10 | GAGTAGGGCCAAGGTAAGGG | TCCCTTCACGGAGGTTTATG | 177 |
| chr4:112885947-112887989 | p2 | (TA)21 | AGTTCCGGAATCGTTGATTG | AGATAGACCGGACCCAGGAT | 235 |
| chr4:112894578-112896596 | p3 | (CAA)6 | GGCTTGATCCATAGCCACAT | TTGTTCCTTCAAGGTGGGAC | 281 |
| chr4:112935922-112937934 | p2 | (GT)6 | TGCGGGAACATGTCTAGTGA | GGGGAGGTCTCTTTCGTAGG | 120 |
| chr4:113006505-113008517 | p2 | (TA)6 | CGTAGATACGCATGTGTGGG | TCTAACCACCGAGCACCTCT | 147 |
| chr4:113036708-113038724 | p2 | (TA)8 | GAGAGGTGCTCGGTGGTTAG | CATACCTGTTCGGCCTCCTA | 191 |
| chr4:113061060-113063108 | p2 | (TG)24 | CCGAGTTAGACGAAATGGGA | ATTGCCAGGAATACCTGCTG | 119 |
| chr4:113226545-113228557 | p2 | (AT)7 | GCCAACAGATGACAGAGGGT | AAAGCCCTACGGGTCTTCAT | 284 |
| chr4:113227571-113229591 | p2 | (TA)6 | CAACAAAGGGGGTTCAGAGA | GGGGTATGCTGTGTTATGGC | 160 |

| Marker ID | SSR type | SSR motif | FORWARD PRIMER (5'-3') | REVERSE PRIMER (5'-3') | size (bp) |
|---|---|---|---|---|---|
| chr4:113288187-113290199 | p2 | (AT)6 | CTGTCGTCCTGTCAGTGCAT | TCCGGAGCCTTATATCATCG | 187 |
| chr4:113323002-113325018 | p2 | (TA)8 | GCCCTTTGGCTATCCACATA | CTATTATTCCGGAGGTGCCA | 164 |
| chr4:113329655-113331695 | p2 | (TA)20 | GCCCTTTGGCTATCCACATA | TAGACAGGGAGGTGACCCAG | 164 |
| chr4:113394174-113396188 | p2 | (TA)7 | CTATGTCTCGCTTCATGGCA | ATATCGAAGCCGGCTATGTG | 273 |
| chr4:113490783-113492798 | p3 | (GTT)5 | GGAAGATTGGCCATCAAAAA | CCCACTTTGCAAACTTCCAT | 242 |
| chr4:113511707-113513722 | p3 | (ATG)5 | GCACGCTACAATGCTCAAAA | TATACGCTTCCCGAGCCTAA | 184 |
| chr4:113511707-113513722 | p2 | (TA)8 | GCACGCTACAATGCTCAAAA | TATACGCTTCCCGAGCCTAA | 184 |
| chr4:113538368-113540386 | p3 | (ATG)6 | CGCGAAAAGTTCTAGATCGG | GCTTAGGGAAATTTTGGGGA | 191 |
| chr4:113591180-113593212 | p2 | (TA)16 | CGCGCAGAAGGATTTTATGT | GGCTAAAAAGTGGGTGTGGA | 102 |
| chr4:113603935-113605953 | p3 | (TTG)6 | AGTTTGGACGTTGAGTTCCG | AAGTTGGGCCAAAATTACCC | 148 |
| chr4:113726879-113728899 | p2 | (AG)10 | CGATTTCAAGTACGGGCTGT | CGGGATACTTAAGGCAACCA | 192 |
| chr4:113780772-113782788 | p4 | (ATTA)4 | TTCCAAGCTCAAAAGAGGGA | AGTCCGGAATGCAACGTATC | 152 |
| chr4:113848417-113850438 | p3 | (CAA)7 | TGGGAACTTCTTCCCCTTTT | GAACCCAGCAAATAAACGGA | 201 |
| chr4:113848417-113850438 | p3 | (CAA)6 | TGGGAACTTCTTCCCCTTTT | GAACCCAGCAAATAAACGGA | 201 |
| chr4:113892352-113894367 | p5 | (ATATT)3 | TATTCCTAAAATGCCCGCAG | TGTACGATAAGGGGGTGCTC | 155 |
| chr4:113919817-113921837 | p2 | (AT)10 | CACCATCCCCGAGAATCTAA | GAATTTGGAAGGTTTCGGGT | 126 |
| chr4:113948187-113950202 | p5 | (ATCCT)3 | CGTACATGCGTAGCGTCCTA | AATGGAGTGGTGGAACTTCG | 204 |
| chr4:113993240-113995252 | p2 | (TA)6 | ACAAACTCTTGCCAAATGGG | TACAAGAGCCCGTGATTTCC | 143 |
| chr4:113993240-113995252 | p2 | (TA)7 | ACAAACTCTTGCCAAATGGG | TACAAGAGCCCGTGATTTCC | 143 |
| chr4:114026111-114028123 | p2 | (CA)6 | ATGCCAAAAGAAGGAAGGGT | CGAGTTAAATCTCGGGGATG | 130 |
| chr4:114026111-114028123 | p2 | (AT)10 | ATGCCAAAAGAAGGAAGGGT | CGAGTTAAATCTCGGGGATG | 130 |
| chr4:114055008-114057062 | p2 | (AT)7 | ATCCTTGGAAGCGGGTACTT | TGAGTTGAGGGAATTGGAGG | 161 |
| chr4:114162285-114164303 | p2 | (CA)9 | AGTGCAGCGGGAAAGTAAGA | GTGCATCTCATAGGGCCATT | 286 |
| chr4:114345218-114347284 | p3 | (TTG)6 | TTGAAAAACATCGTGTGGGA | TATGCCTAGCCCGAAATCAC | 292 |
| chr4:114345312-114347376 | p3 | (TTG)6 | TTGAAAAACATCGTGTGGGA | TATGCCTAGCCCGAAATCAC | 198 |
| chr4:114385089-114387103 | p2 | (TA)7 | AAATTCCAGTCGCAATGCTC | TATTCCAAGGATCGATTCGG | 290 |
| chr4:114458015-114460027 | p2 | (TA)6 | AAAGAACGAATGGTGCTTGG | CTGCAATCAGGGGTTCGTAT | 284 |
| chr4:114472958-114474976 | p6 | (ACATAT)3 | AATAAGGCTCCGGAACGACT | TGCCCTGTGACTCCATGATA | 281 |
| chr4:114488467-114490482 | p5 | (GAGAA)4 | GAGAATCGGGGAATGTTTGA | CTACCCAGCCCATAATCCCT | 232 |
| chr4:114488467-114490482 | p3 | (TAA)8 | GAGAATCGGGGAATGTTTGA | CTACCCAGCCCATAATCCCT | 232 |
| chr4:114488467-114490482 | p3 | (TAA)7 | GAGAATCGGGGAATGTTTGA | CTACCCAGCCCATAATCCCT | 232 |
| chr4:114488497-114490524 | p5 | (GAGAA)4 | GAGAATCGGGGAATGTTTGA | CTACCCAGCCCATAATCCCT | 202 |
| chr4:114488497-114490524 | p3 | (TAA)8 | GAGAATCGGGGAATGTTTGA | CTACCCAGCCCATAATCCCT | 202 |
| chr4:114488497-114490524 | p3 | (TAA)7 | GAGAATCGGGGAATGTTTGA | CTACCCAGCCCATAATCCCT | 202 |
| chr4:114488543-114490558 | p5 | (GAGAA)4 | GAGAATCGGGGAATGTTTGA | CTACCCAGCCCATAATCCCT | 156 |
| chr4:114488543-114490558 | p3 | (TAA)8 | GAGAATCGGGGAATGTTTGA | CTACCCAGCCCATAATCCCT | 156 |

| Marker ID | SSR type | SSR motif | FORWARD PRIMER (5'-3') | REVERSE PRIMER (5'-3') | size (bp) |
|---|---|---|---|---|---|
| chr4:114488543-114490558 | p3 | (TAA)7 | GAGAATCGGGGAATGTTTGA | CTACCCAGCCCATAATCCCT | 156 |
| chr4:114544304-114546316 | p2 | (GA)6 | GTAGGTCGTCCGAGCAAGAG | AGTGCAGCAACAACAACCTG | 191 |
| chr4:114612781-114614803 | p2 | (AT)11 | ATCGTGAGACCATGCTACCC | ATGCCACAGGTATAGACGGG | 235 |
| chr4:114612781-114614803 | p2 | (TA)6 | ATCGTGAGACCATGCTACCC | ATGCCACAGGTATAGACGGG | 235 |
| chr4:114652337-114654355 | p3 | (ACA)6 | CGATGAGGTGGCATAATCCT | CTGCTCATCCATCCCTCATT | 227 |
| chr4:114656837-114658879 | p2 | (TA)21 | ATGCCTGCCTCAAGACAGAT | GCACAGTGAAACCAAGCAAA | 257 |
| chr4:114685843-114687861 | p3 | (ACA)6 | CGATGAGGTGGCATAATCCT | CTGCTCATCCATCCCTCATT | 227 |
| chr4:114690343-114692385 | p2 | (TA)21 | ATGCCTGCCTCAAGACAGAT | GCACAGTGAAACCAAGCAAA | 257 |
| chr4:114690343-114692385 | p2 | (TA)23 | ATGCCTGCCTCAAGACAGAT | GCACAGTGAAACCAAGCAAA | 257 |
| chr4:114791684-114793699 | p3 | (TTG)5 | TACGATCGAATTCTCCAGGC | CATACGTTGGAAAGTGTGCG | 199 |
| chr4:114797126-114799144 | p6 | (TTTCAT)3 | ACTGACGTCCCTTCTTGTGG | ATGAATCGGTGGGGTTATGA | 230 |
| chr4:114806054-114808072 | p2 | (TA)9 | GTGTGTTCGTCTAGCCGTGA | GATTGTGGTGTGTCGGTCAG | 122 |
| chr4:114850837-114852857 | p2 | (TA)10 | GAAGCATGCAAAAGGTGACA | TATCACAATGCCAGCCAAAA | 271 |
| chr4:114885028-114887044 | p3 | (GTT)5 | TGTTGTTGTTGGTTGCTGGT | CGCTTCGCACTTGTCATAGA | 246 |
| chr4:114885028-114887044 | p2 | (TA)8 | TGTTGTTGTTGGTTGCTGGT | CGCTTCGCACTTGTCATAGA | 246 |
| chr4:114964090-114966108 | p6 | (TATATG)3 | TTAGATTCTCGGGGATGGTG | GCTACGCTTTTTCGTCCAAG | 160 |
| chr4:114975998-114978016 | p6 | (CATATA)3 | ACTTTTGTCACGACCCAACC | GGGTCGCTATTGTTCGGTTA | 125 |
| chr4:114975998-114978016 | p5 | (ATCAA)3 | ACTTTTGTCACGACCCAACC | GGGTCGCTATTGTTCGGTTA | 125 |
| chr4:114977484-114979502 | p5 | (ATCAA)3 | TCCATCAAGATCATGCCAAA | TGTAGATCTTGCGAAGCGAA | 297 |
| chr4:114977484-114979502 | p3 | (ACA)6 | TCCATCAAGATCATGCCAAA | TGTAGATCTTGCGAAGCGAA | 297 |
| chr4:115024565-115026580 | p5 | (AAAGA)3 | AGGGCCCTATCATACCCATC | GTTTATGCCCGCAGGTTTAG | 177 |
| chr4:115059695-115061710 | p5 | (ATTTC)3 | AATGCATAAATACGGGGCTG | CATCGGTGGGTGAAAGAGTT | 299 |
| chr4:115115869-115117891 | p2 | (TA)11 | CAGACAAACGATCCAGCTCA | AGAGCCGGCAAAGCTACATA | 116 |
| chr4:115202761-115204785 | p6 | (TATATC)4 | TTTCATGGCCCAACTTGAAT | AATTACCCGCCATAGCCTTT | 300 |
| chr4:115202761-115204785 | p2 | (AT)13 | TTTCATGGCCCAACTTGAAT | AATTACCCGCCATAGCCTTT | 300 |
| chr4:115272572-115274592 | p2 | (AT)10 | AGCCGATGAGGCTAACAAAA | GCGGAGCTCTATTTCATTCG | 206 |
| chr4:115297857-115299877 | p2 | (AT)10 | AGCCGATGAGGCTAACAAAA | GCGGAGCTCTATTTCATTCG | 206 |
| chr4:115363246-115365258 | p2 | (TA)6 | AGCAACCCAAAATACATCGC | CTCAGAACCAAAGGCGTAGC | 118 |
| chr4:115363260-115365274 | p2 | (TA)6 | AGCAACCCAAAATACATCGC | CTCAGAACCAAAGGCGTAGC | 104 |
| chr4:115369465-115371480 | p5 | (TATCA)3 | AAACGTCGTCGTATTTTGCC | GGAGATGGAATTCGTGGAAA | 191 |
| chr4:115383164-115385188 | p2 | (TA)8 | CCTCCTCCTCCTCTTTTGCT | CCAAATGAGCGGAAATAACC | 203 |
| chr4:115383187-115385209 | p2 | (TA)8 | CCTCCTCCTCCTCTTTTGCT | CCAAATGAGCGGAAATAACC | 180 |
| chr4:115384302-115386404 | p2 | (TA)8 | ACAGTTGCTGCAGATGTTGC | TTTTTACCGACAAGGTCAGGA | 149 |
| chr4:115384302-115386404 | p2 | (TA)51 | ACAGTTGCTGCAGATGTTGC | TTTTTACCGACAAGGTCAGGA | 149 |
| chr4:115384302-115386404 | p2 | (AT)32 | ACAGTTGCTGCAGATGTTGC | TTTTTACCGACAAGGTCAGGA | 149 |
| chr4:115440112-115442124 | p2 | (AT)6 | ACCGCTCGTGCTATCATTTC | GGTGTGGGTTATGTGGTTCC | 290 |

| Marker ID | SSR type | SSR motif | FORWARD PRIMER (5'-3') | REVERSE PRIMER (5'-3') | size (bp) |
|---|---|---|---|---|---|
| chr4:115443433-115445448 | p3 | (CTT)5 | TTCCTGCCCGGAGTATTATG | GACGGGAAGTCGTTGTGAAT | 237 |
| chr4:115483803-115485823 | p2 | (AT)10 | AATCATTTTCCCCCTCCAAG | TGTGAGGCATTGGATGTGTT | 256 |
| chr4:115506322-115508337 | p5 | (TTTAT)3 | CAAACACAAAACCCCAATCC | ACACCATGGGGATGATTGTT | 208 |
| chr4:115506322-115508337 | p4 | (TTTA)4 | CAAACACAAAACCCCAATCC | ACACCATGGGGATGATTGTT | 208 |
| chr4:115555366-115557381 | p3 | (CAT)5 | CAAGCCGACGAGTGTATCAA | ATGGAGGGTTCCTATCCCAC | 253 |
| chr4:115568271-115570323 | p2 | (AT)26 | CGTACGCATGGGATAGGACT | GCCCATGGTAGTATGGGATG | 199 |
| chr4:115568271-115570323 | p3 | (ATC)6 | CGTACGCATGGGATAGGACT | GCCCATGGTAGTATGGGATG | 199 |
| chr4:115787875-115789893 | p6 | (ATTCTT)3 | CGTGTGTGTATGGTCAAGCC | AAGCAAAACTGTTCGTTGGG | 279 |
| chr4:115800999-115803025 | p2 | (TA)13 | ATCCAAGGTCTCCCGAAGAT | GGAGAGATCGTGTTGCCAAT | 111 |
| chr4:115800999-115803025 | p2 | (TA)8 | ATCCAAGGTCTCCCGAAGAT | GGAGAGATCGTGTTGCCAAT | 111 |
| chr4:115851913-115853925 | p3 | (TTG)5 | GCGTTCCAGCAGGAGTTAAG | ATTCGAGCAGCATCTCCTGT | 132 |
| chr4:115851913-115853925 | p2 | (TA)6 | GCGTTCCAGCAGGAGTTAAG | ATTCGAGCAGCATCTCCTGT | 132 |
| chr4:115921941-115923956 | p5 | (CCCCT)3 | TCCCTTTGTCGAATCCTTTG | TGCTGCTACACTGATTTGGC | 152 |
| chr4:115935621-115937637 | p4 | (TATG)4 | GATGGCCATGAGGTGAGTTT | TCCCACACAATGTTTCTCCA | 166 |
| chr4:115935621-115937637 | p2 | (AT)74 | GATGGCCATGAGGTGAGTTT | TCCCACACAATGTTTCTCCA | 166 |
| chr4:115935621-115937637 | p3 | (TTA)5 | GATGGCCATGAGGTGAGTTT | TCCCACACAATGTTTCTCCA | 166 |
| chr4:116043227-116045248 | p3 | (CAA)7 | CATCATAGCATAATGGCCCC | CCCCTGACCAGAAGGTGTTA | 298 |
| chr4:116209342-116211356 | p2 | (AG)7 | TCCGTACCTCTGATTTTGCC | TCCCTCTTCATACCGGACAC | 197 |
| chr4:116251686-116253702 | p2 | (TA)8 | ATCGATGCTTCTCTTTGCGT | GGGGTGAGATAAGGTCGTGA | 207 |
| chr4:116277958-116279976 | p3 | (TTG)6 | CCTTAAGCCAAATATGGGCA | TGGTGTGACTCCAAATTCCA | 241 |
| chr4:116318031-116320046 | p2 | (TA)6 | AGCTACCCCGTTATGTGTGC | TAGTGGGTACGCCAACAACA | 186 |
| chr4:116318079-116320094 | p2 | (TA)6 | AGCTACCCCGTTATGTGTGC | TAGTGGGTACGCCAACAACA | 138 |
| chr4:116351197-116353211 | p2 | (TA)27 | ACAGATTCGATATGGACGGG | CCTTGTCTACCTGTGGGCAT | 148 |
| chr4:116351197-116353211 | p2 | (TA)7 | ACAGATTCGATATGGACGGG | CCTTGTCTACCTGTGGGCAT | 148 |
| chr4:116390300-116392318 | p6 | (TATACA)3 | AGCGAACCCCTTTAGCTCTC | GCAAGCCGACGTACCATTAT | 180 |
| chr4:116408989-116411009 | p5 | (AAAAT)4 | CCTCGATAAGCTGGAACTGC | TGGTCTCGATGGAGGAAAAG | 129 |
| chr4:116517095-116519107 | p2 | (TA)6 | TGGAATATCGTGGGATTGGT | AACGATATTTGGGCAGCATC | 198 |
| chr4:116658390-116660411 | p3 | (GAT)7 | GTTCGGAGGTGCTTGTGATT | CCTCTCGGCCATCAAAATAA | 219 |
| chr4:116671540-116673555 | p5 | (CCTCC)3 | GCGCCCTCTCTTTCTTCTTT | AGGGAACTGATTTGAATGCG | 137 |
| chr4:116671540-116673555 | p2 | (AT)6 | GCGCCCTCTCTTTCTTCTTT | AGGGAACTGATTTGAATGCG | 137 |
| chr4:116711954-116713972 | p6 | (CTGCTA)3 | CTGCTGAGGCACATCGATTA | AACATAGCCGCATAATTGGC | 212 |
| chr4:116972436-116974454 | p2 | (TA)9 | ATTTTGGCTCCGTCACTGAT | CGGGTAAACAACGGTGATCT | 148 |
| chr4:116972683-116974695 | p2 | (TA)9 | CACCGTTGTTTACCCGAAAT | GACAATGACCAACCTTTGGC | 100 |
| chr4:116976505-116978523 | p3 | (TCA)5 | TTCCTCATTAAGCGACACCC | CGGCAATTTTGTACCTTCGT | 221 |
| chr4:116976505-116978523 | p6 | (CCGGTT)3 | TTCCTCATTAAGCGACACCC | CGGCAATTTTGTACCTTCGT | 221 |
| chr4:117035761-117037777 | p2 | (AT)8 | CTGGATATTGGAGCTTGGGA | TCGTTGTCCATTCACTCCAA | 229 |

| Marker ID | SSR type | SSR motif | FORWARD PRIMER (5'-3') | REVERSE PRIMER (5'-3') | size (bp) |
|---|---|---|---|---|---|
| chr4:117121529-117123543 | p2 | (TA)7 | ACTATTCGCAAACCCATTCG | TTGGAAATTCGGCTTAGGTG | 236 |
| chr4:117174466-117176481 | p3 | (ATG)5 | GTCCCGGAGCCTAGACTTTT | AATACCATACGCAAGCCGAC | 119 |
| chr4:117237196-117239214 | p6 | (CATATA)3 | ACTTCTGTCACGACCCAACC | GCAGTTCCCTGCATTACGAT | 124 |
| chr4:117248775-117250787 | p2 | (TG)6 | CACATGGTTGACATTCCGAG | AATAGCCCGTAGGCACCTTT | 204 |
| chr4:117297302-117299328 | p2 | (AT)13 | TCCTCCGGTACTGGAAACTG | CATAGACCGAAATACGCGGT | 103 |
| chr4:117350148-117352174 | p2 | (AT)13 | GGCGAGACCAACAAAATGAT | GTCCTGTCCCGCCTATATGA | 215 |
| chr4:117535141-117537159 | p3 | (ATG)6 | TATAAAAGGGGTTTTGCCCC | CCTTGTGTCCAAGCCAAGAT | 240 |
| chr4:117595938-117597976 | p2 | (AT)19 | ATATGTAACGGACGGTTCGC | ACGAAAACCAACGGAGTTTG | 278 |
| chr4:117670739-117672754 | p3 | (TCA)5 | TCCATATACACAAGCCGACG | GAGGAGCTCCATTTGATTCG | 291 |
| chr4:117788098-117790186 | p2 | (GA)6 | TTGGTGTGGCTGTGGATAGA | AGCAAGCCAAAAGTTAGCCA | 238 |
| chr4:117788530-117790542 | p2 | (GA)6 | CTCGGAATGAGTTTGAAGGC | TTACCCCCGAACTTAGGCTT | 255 |
| chr4:117824813-117826849 | p2 | (AT)18 | CGTTTTCCCTGACTTTCTCG | GATGGAGGAAGATGCCAAAA | 115 |
| chr4:117862114-117864129 | p5 | (ATATC)3 | TTCGAGCATTCTCGGAACTT | CCCCTAGTTGGGTCGTAACA | 125 |
| chr4:117865229-117867251 | p2 | (AG)11 | CGTCTTCCTCCATTTTTCCA | GCCGAACAAGGAATCACAAT | 183 |
| chr4:117867712-117869738 | p2 | (AT)13 | TAGTCCCATGCCCAGACTTC | TGGCCAATCAAAACAAGTCA | 129 |
| chr4:117878226-117880256 | p2 | (TA)15 | GCTCGGTTTTCACCATTTGT | TTAGCCGAAACCCGAGTAGA | 118 |
| chr4:117902066-117904084 | p6 | (CTGTTT)3 | ATTAACCATCCAAGGGGACC | AACTTTGGGTTGTCGTGAGG | 155 |
| chr4:117909018-117911034 | p4 | (ACTA)4 | TGTACGCAGACCTTAGCCCT | AGAAGCATGTGGCTCGTTTT | 167 |
| chr4:117941879-117943894 | p5 | (ATATA)3 | AGAGTCGCGTCAATTCTGGT | CTTTTTCAGTCCCTCTCCCC | 182 |
| chr4:117951702-117953714 | p2 | (AT)6 | GGTCTCGGAGGCATGAATAA | GCTCAGGCACAATGACAGAA | 214 |
| chr4:118027678-118029702 | p2 | (TA)12 | TCAAGGTGAGGCAGGCTACT | CATAGCTCGGCATTTTCACA | 114 |
| chr4:118088604-118090619 | p3 | (ATG)5 | TTCAGCGATTCTGGTCCTCT | GAGATTGTCGGCAAAACGAT | 267 |
| chr4:118159103-118161118 | p3 | (GTT)5 | CGTTTTCGGGTGATCTGACT | GAGCCCACTACCCAGTTTGA | 210 |
| chr4:118265941-118267957 | p2 | (TA)7 | TCGTGCATGCCTCAATATGT | ACCGTACCCAGCCCTCTAGT | 277 |
| chr4:118265941-118267957 | p2 | (AT)8 | TCGTGCATGCCTCAATATGT | ACCGTACCCAGCCCTCTAGT | 277 |
| chr4:118311045-118313063 | p3 | (GTT)5 | GAATATCGTGGGATTGGCTG | AACTTAACGAAATTCGGGCA | 195 |
| chr4:118311045-118313063 | p2 | (TA)9 | GAATATCGTGGGATTGGCTG | AACTTAACGAAATTCGGGCA | 195 |
| chr4:118325971-118327986 | p3 | (GTT)5 | GATCTGAGTTTGGGAAGCCA | AATAACAAGGGGAAATGGGC | 172 |
| chr4:118325971-118327986 | p2 | (TA)8 | GATCTGAGTTTGGGAAGCCA | AATAACAAGGGGAAATGGGC | 172 |
| chr4:118346260-118348272 | p2 | (AT)7 | TTTGAAGCCTTCGAGCAAAT | CGACAAAGAACTGGAGCACA | 233 |
| chr4:118346277-118348289 | p2 | (AT)7 | TTTGAAGCCTTCGAGCAAAT | CGACAAAGAACTGGAGCACA | 216 |
| chr4:118346384-118348398 | p2 | (AT)7 | TTTGAAGCCTTCGAGCAAAT | CGACAAAGAACTGGAGCACA | 109 |
| chr4:118467460-118469476 | p2 | (TG)8 | GGAGCCTTACACATTCGAGC | TCAGGCTGTGAAAGATGTGC | 179 |
| chr4:118476742-118478780 | p2 | (AT)19 | CTCACAGCTCTATCCTCCGC | CCTCCGAAGAAAGTGACTGC | 170 |
| chr4:118476742-118478780 | p2 | (AT)6 | CTCACAGCTCTATCCTCCGC | CCTCCGAAGAAAGTGACTGC | 170 |
| chr4:118486336-118488398 | p2 | (TA)31 | AACCCTTCACCTCAATCACG | GTGGCTCGAGGTTGTTTCAT | 135 |

| Marker ID | SSR type | SSR motif | FORWARD PRIMER (5'-3') | REVERSE PRIMER (5'-3') | size (bp) |
|---|---|---|---|---|---|
| chr4:118486336-118488398 | p5 | (TTATT)3 | AACCCTTCACCTCAATCACG | GTGGCTCGAGGTTGTTTCAT | 135 |
| chr4:118486692-118488707 | p2 | (TA)31 | TTTGGCAAAGTAAAGGTGCC | AAATTACATGCTCGGATGGC | 225 |
| chr4:118486692-118488707 | p5 | (TTATT)3 | TTTGGCAAAGTAAAGGTGCC | AAATTACATGCTCGGATGGC | 225 |
| chr4:118768921-118770941 | p4 | (ATTT)5 | TGACGATGTGGCCTATGTGT | CATATCACTCGCTCCGGAAT | 142 |
| chr4:118848346-118850364 | p6 | (AAAGGG)3 | CATATGGGTATGGCGGAGTC | GCCCGTATATAAGCCGACAA | 243 |
| chr4:118870651-118872663 | p2 | (TC)6 | ACATGCCAAAGAACAAAGGG | GAGAGCAATTGGAAATTCGG | 118 |
| chr4:118965997-118968015 | p6 | (GCAACA)3 | CACTCAAAGTTGGAGGGCAT | TGTTTTCCGTTGAACTTCCC | 253 |
| chr4:119019682-119021697 | p3 | (TTG)5 | TTGGGACCAAGCAAAGAAAC | GACGTCGTATTGATGAGCGA | 296 |
| chr4:119058555-119060570 | p5 | (CATTT)3 | GCTTCAACTTGCATCCAACA | TTCGCATTTTCACGAGACAG | 134 |
| chr4:119074699-119076714 | p2 | (TG)6 | ATTCCGCCTTCCTTCTGATT | AACGAACTCACACCCAGACC | 146 |
| chr4:119074699-119076714 | p5 | (CATTT)3 | ATTCCGCCTTCCTTCTGATT | AACGAACTCACACCCAGACC | 146 |
| chr4:119132221-119134233 | p2 | (AT)6 | TAAGGACTCCAACCACTCCG | AAGTTAAACGTGCTGGGGTG | 191 |
| chr4:119132221-119134233 | p2 | (AT)11 | TAAGGACTCCAACCACTCCG | AAGTTAAACGTGCTGGGGTG | 191 |
| chr4:119134500-119136515 | p3 | (CAA)5 | CTATTGGAAGGTGCGGTGTT | AGCTTTCACTTGGGCTCTCA | 148 |
| chr4:119140537-119142633 | p2 | (AC)14 | CACTTCTCCTCACCTCGTCC | TGGTGTGGGCAAGAACAATA | 109 |
| chr4:119152810-119154878 | p5 | (TATAT)3 | CGAGGTAGAGGCCTCAGATG | CGGACTCTTACGGGGTGTAA | 140 |
| chr4:119355620-119357642 | p2 | (TA)11 | CCAACTCCACGGATTCAGTT | CAAGTACCAAAGCCCGTGAT | 210 |
| chr4:119387627-119389643 | p2 | (AT)8 | TGTTCCCCCTCACACTTAGC | GTGATGGAAGGCAAGGAAAA | 274 |
| chr4:119387627-119389643 | p2 | (AT)6 | TGTTCCCCCTCACACTTAGC | GTGATGGAAGGCAAGGAAAA | 274 |
| chr4:119403027-119405039 | p2 | (CA)6 | GGAGGACCGCATAACAAGAA | GATAGGAACCTGAGGGAGGC | 183 |
| chr4:119404801-119406816 | p3 | (ACA)5 | CCGCTACGGCCTAAGTAGTG | ATGGTGCGTTTGAAACTTCC | 173 |
| chr4:119543020-119545032 | p2 | (CA)6 | GGCCGAAACCTCTCTTTCTT | GGGGAATTATTTCCTTGGGA | 257 |
| chr4:119630229-119632244 | p5 | (TTTAT)3 | AGCCAATCCGACAATTTGAG | ACACGTCCGATGTCTCAGTG | 226 |
| chr4:119702003-119704019 | p4 | (ATAC)4 | GGCAGCTAGCTCGACAGACT | GATAGGGCTGCTAGTGCCAG | 133 |
| chr4:119777932-119779946 | p2 | (AG)7 | ACCTGAAGACACTGCATCCC | GGTGGACCCACAACCATATC | 127 |
| chr4:119802900-119804915 | p3 | (TGT)5 | AGGAGAATCATTCGTGGCAG | TCAACGCAAGTAGGCATGAG | 193 |
| chr4:119805059-119807071 | p2 | (TC)6 | CTGACACCAGCCGAGGTAAT | CACTCAGCATCTTCCCCATT | 241 |
| chr4:119856510-119858525 | p2 | (TC)9 | AACCAGCTTGGGTCCCTTAT | GCACTCATTGAAGCAGGTCA | 191 |
| chr4:119856510-119858525 | p3 | (TCA)5 | AACCAGCTTGGGTCCCTTAT | GCACTCATTGAAGCAGGTCA | 191 |
| chr4:119873346-119875361 | p5 | (AAAAG)3 | CACCCTCCCACAAGGATAAA | GGCTCATTTGGTATGAGGGA | 222 |
| chr4:119873346-119875361 | p2 | (TA)6 | CACCCTCCCACAAGGATAAA | GGCTCATTTGGTATGAGGGA | 222 |
| chr4:119922584-119924599 | p3 | (TCA)5 | CTCGCCTCCCTGTCTACTTG | AGCCAGCATGCATTATTTCC | 294 |
| chr4:119948934-119950959 | p5 | (TCATA)5 | TGTGATTCCACCTCCTCTCC | AAAGGAGAGGTCAGCCCATT | 166 |
| chr4:119948934-119950959 | p6 | (TACATA)3 | TGTGATTCCACCTCCTCTCC | AAAGGAGAGGTCAGCCCATT | 166 |
| chr4:119954345-119956385 | p5 | (TCATA)8 | CCCTTCTTCCGTCACCACTA | GAGTCGGGGTGTGACAAAGT | 175 |
| chr4:119954345-119956385 | p6 | (TACATA)3 | CCCTTCTTCCGTCACCACTA | GAGTCGGGGTGTGACAAAGT | 175 |

| Marker ID | SSR type | SSR motif | FORWARD PRIMER (5'-3') | REVERSE PRIMER (5'-3') | size (bp) |
|---|---|---|---|---|---|
| chr4:120120281-120122302 | p6 | (CATATA)3 | GCATGAGAGCCCAAAGAAAG | CTTTAAGGTGAGCCAAAGCG | 209 |
| chr4:120120281-120122302 | p3 | (CAA)7 | GCATGAGAGCCCAAAGAAAG | CTTTAAGGTGAGCCAAAGCG | 209 |
| chr4:120125258-120127273 | p2 | (AT)9 | TTGCACGATTATGTACCCGA | CTGAGCGAACGACGAACATA | 253 |
| chr4:120125258-120127273 | p3 | (CAA)5 | TTGCACGATTATGTACCCGA | CTGAGCGAACGACGAACATA | 253 |
| chr4:120127080-120129106 | p2 | (TA)13 | TATGAGAGGGTCCAACGAGG | GAATGCAATGCACTAGCCAA | 286 |
| chr4:120162695-120164707 | p2 | (AT)6 | TCTGCATGCTCATGGCTATC | TGATATAAGGATCGGGCTGC | 198 |
| chr4:120280050-120282068 | p6 | (TATATG)3 | TTAGATTCTCGGGGATGGTG | TTCACAAGGATCATGCCAAA | 160 |
| chr4:120290456-120292468 | p2 | (AT)6 | TGTAGCTGGTGTGCAAGGTC | TGACATGATCCCTCCACGTA | 297 |
| chr4:120290456-120292468 | p2 | (AT)9 | TGTAGCTGGTGTGCAAGGTC | TGACATGATCCCTCCACGTA | 297 |
| chr4:120290937-120292955 | p2 | (AT)6 | TGGGAACTTGCGAAAGAACT | TGATGTCCGTCTGTAGTGGC | 200 |
| chr4:120290937-120292955 | p2 | (AT)9 | TGGGAACTTGCGAAAGAACT | TGATGTCCGTCTGTAGTGGC | 200 |
| chr4:120408631-120410687 | p3 | (TTG)6 | AAATAGGGGAGATGCTGCCT | GCTACGCTTATTCGTCCGAG | 164 |
| chr4:120408631-120410687 | p4 | (TGTA)4 | AAATAGGGGAGATGCTGCCT | GCTACGCTTATTCGTCCGAG | 164 |
| chr4:120408631-120410687 | p2 | (AT)12 | AAATAGGGGAGATGCTGCCT | GCTACGCTTATTCGTCCGAG | 164 |
| chr4:120423453-120425465 | p2 | (AG)6 | TTGCAGACAACCCCTAGGAC | CCTCGAGCTCACATTAAGCC | 196 |
| chr4:120428669-120430709 | p2 | (AT)20 | TTGTGAGTATCGGAGGGGAG | GTGTTGCACTAAGGCGTCAA | 136 |
| chr4:120500833-120502875 | p6 | (TATACA)7 | GTCTCCCGAACAATCGTCAT | CATATATGGGTACGGCAGGG | 197 |
| chr4:120507957-120509981 | p6 | (CATATA)4 | TCACGGCCAACACATACCTA | CTCAGCAGCAGTCAGTACGC | 240 |
| chr4:120509491-120511503 | p2 | (AT)6 | TTCGAGCTCGCAAGTCTACA | AGTTTCCAACGCAACAAACC | 139 |
| chr4:120608307-120610325 | p6 | (TATATG)3 | TCGTGTGGGATGTTTATGGA | GCTTATTTGTCCAAGCTCGC | 130 |
| chr4:120847675-120849695 | p2 | (AT)10 | TGCATATAGACCCGTGACCA | GCCAATAATAGTGCCGAGGA | 156 |
| chr4:120877599-120879649 | p2 | (TA)25 | TCCCTTATTCATCGGACTCG | GCTCGTGTGTCACCTTTTGA | 204 |
| chr4:120910412-120912452 | p3 | (ATT)23 | GGGTTTAGGACGGGGTTTTA | ACCCCTAATTCCCCATTACG | 136 |
| chr4:121048441-121050455 | p2 | (CT)7 | CGTCTAGCCTAACAGCCCAC | GTTGAAGCCATGTAAGGGGA | 300 |
| chr4:121101806-121103818 | p2 | (TA)6 | CTAGAAGATATGCTGCGGGC | GACTCTGGGCTGCCAGTAAC | 186 |
| chr4:121127166-121129186 | p5 | (TCATA)5 | AGCGCAACATATGTCCCTTC | ACCTTACGTCACTCCGATGG | 182 |
| chr4:121127166-121129186 | p6 | (TACATA)4 | AGCGCAACATATGTCCCTTC | ACCTTACGTCACTCCGATGG | 182 |
| chr4:121127166-121129186 | p5 | (ATTCA)4 | AGCGCAACATATGTCCCTTC | ACCTTACGTCACTCCGATGG | 182 |
| chr4:121131985-121134010 | p6 | (TACATA)3 | ACATATGTCCCTTCCCCTCC | CGACCTTACGTCACTCCGAT | 210 |
| chr4:121131985-121134010 | p5 | (ATTCA)5 | ACATATGTCCCTTCCCCTCC | CGACCTTACGTCACTCCGAT | 210 |
| chr4:121169229-121171244 | p5 | (ATTCA)3 | ATGGGCGTTTGAAGTACAGG | TGATATGTTTCCGAATGGCA | 235 |
| chr4:121243266-121245284 | p3 | (GTT)6 | TTGTTTGGGAATTTGGGAAA | AGATGAGGAAGACAAGCGGA | 241 |
| chr4:121243266-121245284 | p2 | (TA)6 | TTGTTTGGGAATTTGGGAAA | AGATGAGGAAGACAAGCGGA | 241 |
| chr4:121409005-121411020 | p5 | (ATTGC)3 | TGCCCACTGTTGGTACTTGA | GGATCATCACCGGATATTGG | 293 |
| chr4:121618098-121620113 | p5 | (TGATA)3 | TTGGAAAGTTCCGTTGTTCC | CAAACCGACAAGGCTACCAT | 133 |
| chr4:121685060-121687080 | p2 | (AT)10 | GATCTACGTGCAGCTCCTCC | TACGCTTGTTGCACCTTTTG | 138 |

| Marker ID | SSR type | SSR motif | FORWARD PRIMER (5'-3') | REVERSE PRIMER (5'-3') | size (bp) |
|---|---|---|---|---|---|
| chr4:121823560-121825578 | p3 | (ATG)6 | GGACCAAGGGTCAACTCAAA | GGGAATTTTGGAGGGATGTT | 184 |
| chr4:121826341-121828359 | p3 | (ATG)6 | GGACCAAGGGTCAACTCAAA | GGGAATTTTGGAGGGATGTT | 184 |
| chr4:121856192-121858210 | p3 | (CAA)6 | TGCAAACAGAGTCGTGGGTA | GTGCACCCGACCCTTACTTA | 282 |
| chr4:121863684-121865696 | p2 | (TA)6 | AAGTGCTATGTCGGACACCC | GTTCGATGGACCTCGTGTCT | 193 |
| chr4:121924358-121926376 | p6 | (TATATG)3 | TTGGACCAAGGCCTAAAGTG | ATGAAAGTGTGGGACAAGGC | 154 |
| chr4:122026366-122028382 | p2 | (AT)8 | TCAACAATCCCTAGATGCCC | GCAGGTTGTCTGCAAAGTCA | 254 |
| chr4:122026366-122028382 | p6 | (CATATA)4 | TCAACAATCCCTAGATGCCC | GCAGGTTGTCTGCAAAGTCA | 254 |
| chr4:122036282-122038312 | p2 | (AT)33 | CGCTGGTTCTCGTTAGGTTC | AAGCCGACGTACCATTATGC | 149 |
| chr4:122036282-122038312 | p2 | (TA)15 | CGCTGGTTCTCGTTAGGTTC | AAGCCGACGTACCATTATGC | 149 |
| chr4:122043465-122045515 | p2 | (AT)25 | GGACACGACTTTGCAGACAA | AAGCCGACGTACCATTATGC | 237 |
| chr4:122097243-122099259 | p2 | (TA)8 | GAGGTGCTCGGTGAGTTAGC | CCCTGCAGCCACTTGTTTAT | 145 |
| chr4:122180034-122182049 | p5 | (TGCAA)3 | AGCTCACCCTGACCTCTCAA | GTTCGACTATCGAGGTGGGA | 131 |
| chr4:122217396-122219418 | p2 | (TA)11 | GAATGCGAGCATGATGAGAA | GAGCAAAGGCAACAATGTCA | 251 |
| chr4:122217396-122219418 | p3 | (AGA)6 | GAATGCGAGCATGATGAGAA | GAGCAAAGGCAACAATGTCA | 251 |
| chr4:122268933-122270947 | p2 | (TA)7 | GGAAGCGAGGAATGTGTTGT | AACTCGGCCAAATCATCAAC | 110 |
| chr4:122276840-122278852 | p3 | (TTG)5 | TTATGGGTCGAGAGATTGGG | TACATGCCTAGCCCGAAATC | 229 |
| chr4:122276840-122278852 | p2 | (TA)9 | TTATGGGTCGAGAGATTGGG | TACATGCCTAGCCCGAAATC | 229 |
| chr4:122276873-122278889 | p3 | (TTG)5 | TTATGGGTCGAGAGATTGGG | TACATGCCTAGCCCGAAATC | 196 |
| chr4:122276873-122278889 | p2 | (TA)9 | TTATGGGTCGAGAGATTGGG | TACATGCCTAGCCCGAAATC | 196 |
| chr4:122303572-122305590 | p2 | (TA)6 | TTGAAAATTCGGATTAGGCG | ACTATTCGCAAATCCGTTCG | 228 |
| chr4:122303572-122305590 | p2 | (TA)9 | TTGAAAATTCGGATTAGGCG | ACTATTCGCAAATCCGTTCG | 228 |
| chr4:122375307-122377319 | p2 | (TA)6 | TGTTGTTTGGGGGATGTTTT | TCCTTCAAAGCCTGCAAAAT | 183 |
| chr4:122456425-122458439 | p2 | (AT)6 | CCGCACTAGACACTCGCATA | CTTATCGGCTTGCGTATGGT | 227 |
| chr4:122456425-122458439 | p2 | (AT)7 | CCGCACTAGACACTCGCATA | CTTATCGGCTTGCGTATGGT | 227 |
| chr4:122601396-122603440 | p2 | (TA)10 | GAGACTTGAGCCGATTGGAG | TGTCGACGATGACTGGTGTT | 252 |
| chr4:122621333-122623348 | p5 | (AATGC)3 | ATCGTACCGATCCACGAGAC | CCGCATGACCTAGGGTAAAA | 225 |
| chr4:122644105-122646129 | p2 | (TA)12 | CAGCGCAAGTCCTACATCAA | CGAGTAGGCCCATGAGATGT | 147 |
| chr4:122644105-122646129 | p2 | (AT)24 | CAGCGCAAGTCCTACATCAA | CGAGTAGGCCCATGAGATGT | 147 |
| chr4:123041192-123043232 | p2 | (AT)14 | AAAGGACTGTCACGACCCAG | GTCGGCTTGCGTATCGTATT | 147 |
| chr4:123041192-123043232 | p2 | (AT)20 | AAAGGACTGTCACGACCCAG | GTCGGCTTGCGTATCGTATT | 147 |
| chr4:123114913-123116963 | p4 | (AATA)4 | TGGGTCGTGACAAAAGATGA | GTGTGGTGTCATTGGAGCAC | 201 |
| chr4:123114992-123117016 | p4 | (AATA)4 | TGGGTCGTGACAAAAGATGA | GTGTGGTGTCATTGGAGCAC | 122 |
| chr4:123115133-123117149 | p4 | (AATA)4 | AGAAACCCCAGAATGCCTCT | GTCCCATCGTTGGAGAGTGT | 275 |
| chr4:123297662-123299692 | p6 | (CATATT)5 | AAATGCTCGTAGCCTCTTGC | GGTTGGGTCGTGACAAAAGT | 164 |
| chr4:123297662-123299692 | p2 | (AT)16 | AAATGCTCGTAGCCTCTTGC | GGTTGGGTCGTGACAAAAGT | 164 |
| chr4:123308829-123310847 | p3 | (ACA)6 | CTTAAAATCCCGTCAGGCAA | GTTCATCAATGTGACGTCGG | 281 |

| Marker ID | SSR type | SSR motif | FORWARD PRIMER (5'-3') | REVERSE PRIMER (5'-3') | size (bp) |
|---|---|---|---|---|---|
| chr4:123314146-123316166 | p5 | (AATGC)4 | TTGGACCTGAAGAATTTCGG | TCAACCCCAGATCTAGCACC | 201 |
| chr4:123323169-123325197 | p2 | (AT)7 | TAGGGACCTCGTATTGCCTG | TGTTGATGTGGCGTGATTTT | 205 |
| chr4:123326131-123328149 | p6 | (TTTTTA)3 | GCGAGCATTCTGTTTGTGAA | CTTGGGAGAATTGGAAACGA | 290 |
| chr4:123375901-123377919 | p3 | (CAT)6 | ACTGTACAACATCCCGCACA | GGCAAGGGAGTATCTGGTCA | 142 |
| chr4:123381480-123383540 | p2 | (AC)30 | ACATTGGACGATTCGTAGCC | CAAGATGGGAGGATTGGAAA | 173 |
| chr4:123424584-123426600 | p3 | (ATG)5 | CTGGCGGGTTCAAGTGTTAT | GGAAAACTAGGAAAACCGGG | 121 |
| chr4:123424584-123426600 | p4 | (TATG)4 | CTGGCGGGTTCAAGTGTTAT | GGAAAACTAGGAAAACCGGG | 121 |
| chr4:123502464-123504493 | p3 | (TTG)5 | TTGGGCCGAGTTAGATTGTC | AACATACCTGCTCCACGTCC | 164 |
| chr4:123502464-123504493 | p3 | (AAC)5 | TTGGGCCGAGTTAGATTGTC | AACATACCTGCTCCACGTCC | 164 |
| chr4:123602496-123604516 | p5 | (CAATG)4 | TCCTACCGATCCACAAGACC | AATGTTCGCCTATCGAGGTG | 268 |
| chr4:123624669-123626681 | p2 | (TG)6 | CCATGATTTGGGAATATCGG | AATCTGCTCACGTTGATCCC | 226 |
| chr4:123637204-123639219 | p3 | (ACA)5 | TACGGACTACCAAACCGAGG | ACCTACGGGTCATGGAGTTG | 201 |
| chr4:123651199-123653214 | p3 | (TCT)5 | TTGTGGACATCAACCTTGGA | TGCCATCGAGGTAAGCTTCT | 170 |
| chr4:123698547-123700563 | p2 | (TA)8 | AGCGAGTCTTTCCTCACCAA | TGCAAAGAAATAGCAATGCG | 102 |
| chr4:123698547-123700563 | p2 | (AT)15 | AGCGAGTCTTTCCTCACCAA | TGCAAAGAAATAGCAATGCG | 102 |
| chr4:123707954-123709978 | p6 | (TCCCTT)4 | GTCGGGCAGTTACCACTGAT | CTATCTACACCTGCGGGCAT | 154 |
| chr4:123876880-123878894 | p2 | (AT)7 | AGAACCCACCTTGGATCCTT | GAACGGAGCTTTGTGCTAGG | 292 |
| chr4:123934748-123936784 | p3 | (AAT)12 | CAACTTTGGCTTGTGCTTCA | ACACTGCATCTCTTTGCCCT | 238 |
| chr4:123934748-123936784 | p2 | (AT)11 | CAACTTTGGCTTGTGCTTCA | ACACTGCATCTCTTTGCCCT | 238 |
| chr4:123944977-123946989 | p5 | (TTCAA)3 | GAAAGAAAGGGACGAGGCTT | ATTTACAGGGGTGATGCTGC | 186 |
| chr4:123944977-123946989 | p2 | (CA)6 | GAAAGAAAGGGACGAGGCTT | ATTTACAGGGGTGATGCTGC | 186 |
| chr4:124234137-124236155 | p6 | (CCTCGG)3 | GGGGGATGGGATACTTGTTT | CGCTGAGTACCTCTCCTTGG | 186 |
| chr4:124235680-124237696 | p4 | (GAAA)4 | CCTCCTCAGTAGGCAGCAAC | GTTTTTGGGCTATGCCATGT | 135 |
| chr4:124317230-124319262 | p2 | (AT)16 | TGAACAAAACTACCCCCGAG | GATTTTGGATGAAACGGGAA | 147 |
| chr4:124325005-124327017 | p2 | (AT)6 | TGAACAAAACTACCCCCGAG | GATTTTGGATGAAACGGGAA | 147 |
| chr4:124393690-124395708 | p2 | (AT)9 | GGGTTTTCATCCAGCTTTGA | GAAAAACAAGAGGGGGCTTC | 121 |
| chr4:124433808-124435820 | p2 | (CA)6 | GGTCATGCCAAAAGAAGGAA | CGGGATATGTGATTTCAGGG | 265 |
| chr4:124433808-124435820 | p2 | (TG)11 | GGTCATGCCAAAAGAAGGAA | CGGGATATGTGATTTCAGGG | 265 |
| chr4:124459652-124461678 | p2 | (TA)13 | CTCGAACCAAAACCCTCAAA | TTTTGTTTTTGCTTTTGGGC | 104 |
| chr4:124470385-124472405 | p2 | (AT)10 | TCCCTCCACCAATCTCTGTC | GCACTTCTCATCAAACGCAA | 286 |
| chr4:124470385-124472405 | p2 | (AT)9 | TCCCTCCACCAATCTCTGTC | GCACTTCTCATCAAACGCAA | 286 |
| chr4:124470507-124472525 | p2 | (AT)10 | TCCCTCCACCAATCTCTGTC | GCACTTCTCATCAAACGCAA | 164 |
| chr4:124470507-124472525 | p2 | (AT)9 | TCCCTCCACCAATCTCTGTC | GCACTTCTCATCAAACGCAA | 164 |
| chr4:124491173-124493188 | p5 | (AAACT)3 | CCCTATTTCCCAACCCAAGT | GTCTAGGCTTGTTTCAGGCG | 278 |
| chr4:124645621-124647707 | p2 | (AG)7 | CAACCTCAACAACTCAGCCA | CCCCACTCTCACAAAAAGGA | 146 |
| chr4:124680196-124682210 | p2 | (TA)7 | TGCCCGATTTTCGTTAAGTC | AAACTGTTCGCTTTGTCGCT | 161 |

| Marker ID | SSR type | SSR motif | FORWARD PRIMER (5'-3') | REVERSE PRIMER (5'-3') | size (bp) |
|---|---|---|---|---|---|
| chr4:124714973-124717030 | p3 | (TAT)19 | GAACAGATGCAAATTCGGGT | TATGGGGATGGACTGGTTGT | 196 |
| chr4:124763561-124765576 | p5 | (ATTTC)3 | AAGCGGAAGAAAGTATGCGA | TCATCGCCTCAAAGGGATAC | 164 |
| chr4:124842220-124844238 | p2 | (AT)30 | GCCTGCACTATCAATCCGTT | CGACCTAGTTGGGCATGTTT | 296 |
| chr4:124842220-124844238 | p6 | (TATATG)3 | GCCTGCACTATCAATCCGTT | CGACCTAGTTGGGCATGTTT | 296 |
| chr4:124903914-124905938 | p2 | (AC)12 | TGTATGGCTTTGTTTTGCCA | TTATCCGGGGATTCTGAGTG | 183 |
| chr4:124909291-124911305 | p2 | (AT)7 | AATGGTTGGGGTTTGGTACA | TGCAGCGTTTTCCTATGATG | 215 |
| chr4:124909291-124911305 | p6 | (ACTTCA)3 | AATGGTTGGGGTTTGGTACA | TGCAGCGTTTTCCTATGATG | 215 |
| chr4:124926516-124928534 | p3 | (AAT)6 | GATGCAATCCCGTCAACTCT | TCTGTAGACTGCTGGCATGG | 196 |
| chr4:124953956-124955968 | p2 | (AT)6 | GCAGATTTTATGCTCCCCAA | TCTCGGTGCTATCATTGCTG | 285 |
| chr4:125009041-125011056 | p5 | (ATTTC)3 | CATGGGAAGCGGAAGAATTA | ATAACAATGCCTCACAGGGC | 170 |
| chr4:125023057-125025072 | p5 | (ATTTC)3 | CATGGGAAGCGGAAGAATTA | ATAACAATGCCTCACAGGGC | 170 |
| chr4:125071240-125073255 | p5 | (TTTAG)3 | CAGCCTTGTCGACTTGCATA | CGACAGACCATCAACACGTC | 166 |
| chr4:125169182-125171197 | p5 | (GGTTC)3 | TTATAAACTACGGTGGCGGC | CACACTTTGGGACTCACCCT | 217 |
| chr4:125187482-125189497 | p3 | (AAC)5 | TCCCTTCGGAGCAACTTTTA | ATGTTTAAATCCCGAAGGGG | 125 |
| chr4:125187482-125189497 | p2 | (TA)10 | TCCCTTCGGAGCAACTTTTA | ATGTTTAAATCCCGAAGGGG | 125 |
| chr4:125209375-125211405 | p2 | (TC)15 | GACAATCTTTCTGGGACGGA | TTTTGTGACTGGCATTGTGG | 220 |
| chr4:125269438-125271454 | p2 | (AG)8 | TTGCAGGTTTGGAATTAGCC | GTACCTTTGCATCCGTTCGT | 207 |
| chr4:125271093-125273117 | p6 | (TATATG)4 | TTTGGCCGAGTTGAATTCTC | AACATACCTGCTCCACGTCC | 213 |
| chr4:125322929-125324943 | p2 | (TA)7 | TTGGGAAACCTTCCTTGATG | CTGGCCGTGATTTTCAGTTT | 186 |
| chr4:125348681-125350705 | p6 | (ATCATT)4 | GCATACAGAGACGTAGGCCC | TAATTATGGGCGTAGCGAGG | 157 |
| chr4:125368052-125370064 | p2 | (TA)6 | GCTCAAGATTTCTTGCCTGG | ATAAGGGACCCACGAAGTCC | 146 |
| chr4:125471048-125473066 | p6 | (AACTCC)3 | CTGTACCACATTGGGCCTTT | GGCCCAAAGCAGAATAATCA | 229 |
| chr4:125540507-125542522 | p5 | (TAACA)3 | TCATGTGCACCTTTGAGAGC | GGCAGGACTTCATCACTGGT | 117 |
| chr4:125542722-125544734 | p2 | (GA)6 | AAGGCGAGAAGCCATAGTCA | GATGCGAATGTTGGTGTCAG | 126 |
| chr4:125607095-125609119 | p4 | (TTGA)6 | CTCCAAGGTTTGCATCCACT | TCTCCCTCTACACCACCCAG | 196 |
| chr4:125619880-125621916 | p3 | (AAC)7 | TGGAGCATTTACCAGTTCCC | CTAGGGAGGGTGGGGTGTAT | 239 |
| chr4:125679037-125681055 | p2 | (AT)10 | TTTACGCACGTGGAAAATGA | GTGGGCCTCATAATTCTTGG | 145 |
| chr4:125679037-125681055 | p2 | (AT)7 | TTTACGCACGTGGAAAATGA | GTGGGCCTCATAATTCTTGG | 145 |
| chr4:125679037-125681055 | p2 | (AT)9 | TTTACGCACGTGGAAAATGA | GTGGGCCTCATAATTCTTGG | 145 |
| chr4:125679037-125681055 | p6 | (TAATTC)3 | TTTACGCACGTGGAAAATGA | GTGGGCCTCATAATTCTTGG | 145 |
| chr4:125707288-125709306 | p2 | (TA)9 | TGGTGTTCCAACCCTGTTTT | TGATTCACCAGCTTCACAGG | 285 |
| chr4:125736461-125738497 | p2 | (TA)18 | AAGCATGCTCGTATCCCATC | TTGGGAGGTTGAGGTTTGTC | 296 |
| chr4:125740356-125742371 | p5 | (CTAAA)3 | ATGCGCATTCAGATGTTCAA | TACATTTTTCTCCCCTCCCC | 182 |
| chr4:125784285-125786305 | p2 | (TA)10 | TCCGGTGTTAACCAACCTTC | GGTGTGGGCTGTCAAAGTCT | 224 |
| chr4:125784285-125786305 | p5 | (GGGCC)3 | TCCGGTGTTAACCAACCTTC | GGTGTGGGCTGTCAAAGTCT | 224 |
| chr4:125921186-125923202 | p2 | (TA)6 | GAAGGCTTCCTCCCGATAAG | TGATCGAACTCTTGGGGATT | 159 |

| Marker ID | SSR type | SSR motif | FORWARD PRIMER (5'-3') | REVERSE PRIMER (5'-3') | size (bp) |
|---|---|---|---|---|---|
| chr4:125983837-125985852 | p5 | (AATGC)3 | ATCGTACCGATCCACGAGAC | GTTGAGGGTTCGCTTACCAA | 227 |
| chr4:126051969-126053984 | p5 | (TGCAA)3 | CCTCGGTTGGTCAATCCTTA | AGCTACTGGGGTGGATTCCT | 267 |
| chr4:126072029-126074059 | p3 | (AAT)10 | GCCAACGGCTATTCAAATGT | ACATCGCAAACCATACGTGA | 241 |
| chr4:126144635-126146651 | p4 | (TTTC)4 | TTTTCAAAAACACGGAAGCC | GGCACTAGCAGAGTTGAGGG | 105 |
| chr4:126266910-126268948 | p2 | (TA)19 | AGGCCGCTCTTGGATTTATT | CTTTTGTAGGCGTCGTAGGC | 174 |
| chr4:126306522-126308534 | p2 | (AC)6 | GGATGGGGGACTGAAGTTTT | TTCTACCCATGGCAATCACA | 178 |
| chr4:126505432-126507447 | p3 | (CAA)5 | CACACCCCTTGTCAACTGTG | TGGCCTTTGTTGTGATGTGT | 124 |
| chr4:126587911-126589926 | p3 | (CAA)5 | CACACCCCTTGTCAACTGTG | TGGCCTTTGTTGTGATGTGT | 124 |
| chr4:126710951-126712963 | p2 | (AT)6 | CGAACGCTGAGAAAGAAACC | GGGTGTGATTTGGATCTTGG | 215 |
| chr4:126710951-126712963 | p6 | (GTATTT)3 | CGAACGCTGAGAAAGAAACC | GGGTGTGATTTGGATCTTGG | 215 |
| chr4:126724774-126726801 | p3 | (TCT)9 | ACCCAGTCAACCACTCCAAG | TCTTGAACGAAGGATGGGAC | 229 |
| chr4:126877561-126879577 | p2 | (AT)7 | CCAAACTCAAGCCCCATAAA | TGGAGATGGTGGTTCCACTT | 297 |
| chr4:126893554-126895618 | p2 | (AT)32 | TTTGATGCAGTTGCCTTCAG | AGGATGAAATGGTGCTTTGG | 150 |
| chr4:127138593-127140609 | p2 | (AT)8 | GGGGAAAATAGCCATTCCAT | TATAAGCCGATCCAAACGGA | 191 |
| chr4:127202038-127204054 | p2 | (AT)8 | GGGGAAAATAGCCATTCCAT | TATAAGCCGATCCAAACGGA | 191 |
| chr4:127412791-127414833 | p2 | (TA)21 | TCATGCTTAATAGTCCCCCG | AAATTACGGCTGAAGGAGCA | 272 |
| chr4:127412791-127414833 | p5 | (TTAAA)3 | TCATGCTTAATAGTCCCCCG | AAATTACGGCTGAAGGAGCA | 272 |
| chr4:127457033-127459075 | p2 | (TA)21 | TCATGCTTAATAGTCCCCCG | AAATTACGGCTGAAGGAGCA | 272 |
| chr4:127457033-127459075 | p5 | (TTAAA)3 | TCATGCTTAATAGTCCCCCG | AAATTACGGCTGAAGGAGCA | 272 |
| chr4:127485970-127487990 | p2 | (AT)10 | CCTCCTTGGGTTTCTTCCTC | AAATTAACACGGTTGCCTCG | 294 |
| chr4:127577851-127579877 | p2 | (AT)13 | AAACTCATGAAATCACGGGC | AAAGGGAGCCTCCAATGAAT | 166 |
| chr4:127577851-127579877 | p2 | (AT)11 | AAACTCATGAAATCACGGGC | AAAGGGAGCCTCCAATGAAT | 166 |
| chr4:127700796-127702808 | p2 | (AG)6 | TCTCACCCCCTAAAATGACG | ACGGATTATCCTGGTGCAAG | 274 |
| chr4:127700796-127702808 | p2 | (TA)15 | TCTCACCCCCTAAAATGACG | ACGGATTATCCTGGTGCAAG | 274 |
| chr4:127716643-127718658 | p6 | (CACGAC)3 | ACTAGACCACGACCCCCTCT | GATATGTAGCGCGGATGGTT | 190 |
| chr4:127716643-127718658 | p3 | (AAT)5 | ACTAGACCACGACCCCCTCT | GATATGTAGCGCGGATGGTT | 190 |
| chr4:127761430-127763451 | p3 | (TTG)7 | ACGTTCAGGTCCGTTAGGTG | CTTAATCCACTTGCCCTCCA | 140 |
| chr4:127787494-127789510 | p2 | (CT)8 | CAAGGTTGAAGCTCTTTCCG | CCCTGAAAGTACGAGTCCCA | 295 |
| chr4:127816762-127818776 | p6 | (ATCCGT)3 | TACCACTCGTTCCGGAAAAC | AAGGCTGAGTATGGGCATTG | 258 |
| chr4:127816780-127818794 | p6 | (ATCCGT)3 | TACCACTCGTTCCGGAAAAC | AAGGCTGAGTATGGGCATTG | 240 |
| chr4:127816812-127818832 | p6 | (ATCCGT)3 | TACCACTCGTTCCGGAAAAC | AAGGCTGAGTATGGGCATTG | 208 |
| chr4:128061345-128063359 | p2 | (AT)7 | TTCGACTATTTGGGTACGGC | GCCATTTTTGCCACCATAAC | 141 |
| chr4:128061345-128063359 | p2 | (AT)7 | TTCGACTATTTGGGTACGGC | GCCATTTTTGCCACCATAAC | 141 |
| chr4:128082482-128084497 | p2 | (AT)6 | ATTTCAACAAGGCACAAGGG | GGTCGTTCAATGATGGTGTG | 185 |
| chr4:128082482-128084497 | p5 | (GTTTT)3 | ATTTCAACAAGGCACAAGGG | GGTCGTTCAATGATGGTGTG | 185 |
| chr4:128152846-128154870 | p6 | (GGAGAA)4 | AAGAGCCACCTTTCTGGGAT | AGAGAAAGGGGAGAGCAAGG | 250 |

| Marker ID | SSR type | SSR motif | FORWARD PRIMER (5'-3') | REVERSE PRIMER (5'-3') | size (bp) |
|---|---|---|---|---|---|
| chr4:128152846-128154870 | p6 | (CTCTAC)3 | AAGAGCCACCTTTCTGGGAT | AGAGAAAGGGGAGAGCAAGG | 250 |
| chr4:128274821-128276836 | p3 | (TTG)5 | ATGGATGAAATGGAGGCTTG | ACTCCGCCCAAACAACATAG | 147 |
| chr4:128277282-128279297 | p3 | (ACA)5 | TGAAAGTCCCTTTCTCCCCT | AAATAGGCTGAGGCCCAAAT | 185 |
| chr4:128290328-128292343 | p5 | (TTCGG)3 | GTTTAGGGCCAGGCATTGTA | ACCGAACCGTACTGATACCG | 158 |
| chr4:128332152-128334166 | p2 | (AT)7 | AGGCCAATCCAGCTTATGTG | TGGAAAGTTCGGAAAATTGG | 256 |
| chr4:128475722-128477737 | p3 | (ATC)5 | GCCCACGAGGCTACAAGATA | GGGTCCTGTCCCATCCTTAT | 186 |
| chr4:128494325-128496357 | p2 | (AT)16 | ATGGGCCATCATTGTTTGTT | CTCCGGTCAAAACGTATGCT | 235 |
| chr4:128513368-128515390 | p2 | (AT)11 | AACAACAACACCGACAACCA | TTTTGGAAGCTCTCGGGATA | 124 |
| chr4:128595236-128597251 | p5 | (TGAAC)3 | CAAAGTGCTTTTGCCAATGA | GCGCAAGGACTTTCGTTAAG | 111 |
| chr4:128613209-128615235 | p2 | (AT)13 | CTGAGGACAGCCTCGGTTAG | ACGCTTGACCGTGCTTACTT | 162 |
| chr4:128672647-128674662 | p5 | (ATTTT)3 | TTCGTCACCGCACAAGTATC | TCGCTTGAGTTTCGGACTTT | 130 |
| chr4:128677156-128679258 | p5 | (GCATT)3 | GGGTTGTGCTACCGAGATGT | AAACAAGCACACGTCATTGC | 150 |
| chr4:128726782-128728797 | p5 | (GGGGT)3 | CCGTAACCCATACACAACCC | TGGGGTGGTTGAAAAACAAT | 235 |
| chr4:128726782-128728797 | p5 | (GGGGT)3 | CCGTAACCCATACACAACCC | TGGGGTGGTTGAAAAACAAT | 235 |
| chr4:128726782-128728797 | p3 | (CAA)7 | CCGTAACCCATACACAACCC | TGGGGTGGTTGAAAAACAAT | 235 |
| chr4:128796627-128798661 | p2 | (AT)17 | GGCAGCATCTCCCTTGTTTA | TGACCAGTGACCAGAAGACG | 196 |
| chr4:128856954-128859000 | p2 | (AT)23 | TGTGGAGAAACCCTAATCGC | ACACGAGGGGCCTCTTTAAT | 116 |
| chr4:128915877-128917892 | p5 | (GAAAA)3 | GGGTTGGAGACCACTGAAAA | GAATCCGCGACTTTTGTAGC | 267 |
| chr4:128915877-128917892 | p2 | (TA)7 | GGGTTGGAGACCACTGAAAA | GAATCCGCGACTTTTGTAGC | 267 |
| chr4:128940086-128942104 | p6 | (CCCCCA)3 | GCAACCATGGCAAGAAAGTT | AACGAGGCATGGTTGAAAAG | 229 |
| chr4:128941169-128943184 | p5 | (ATTTT)3 | TCCAGGATTTTCATTCAGGG | TAGATGAGAGCAGGGGAGGA | 156 |
| chr4:129098525-129100543 | p2 | (AT)9 | TCGACGGATTCCTGAAAAAG | TGACCCTTTCTCGAGCTTGT | 252 |
| chr4:129133269-129135284 | p5 | (TTTAT)3 | TGCCGCTTCTTCAAAAACTT | AAATGAAGATGGGCAATTCG | 191 |
| chr4:129134503-129136518 | p5 | (TTGGT)3 | CCATATTAAGTCGCCAGGGA | GCCAAAAGGCCACAAGTTTA | 138 |
| chr4:129254500-129256515 | p3 | (TAT)5 | AATTACAAGGCGTTATGCGG | GGCCAATTGGTCAAAATCAT | 241 |
| chr4:129294614-129296634 | p4 | (AAGA)5 | GTGATCACATGGACGTTTGC | GGTTGAACCTGATCCATGCT | 218 |
| chr4:129323498-129325518 | p2 | (AT)10 | GAGCCCCACCTAAACATCAA | TACGCATGTGGCAATGAATC | 188 |
| chr4:129325861-129327876 | p5 | (AAATA)3 | AAGATCGTGTCCGTCCTTTG | GAGGGTGGGGTTTCATTTTT | 116 |
| chr4:129360222-129362278 | p2 | (AT)28 | CTGTTACTCCTTTCGCCGAG | CGACCCTCTTTTGCGAGTAG | 185 |
| chr4:129360222-129362278 | p3 | (TGA)5 | CTGTTACTCCTTTCGCCGAG | CGACCCTCTTTTGCGAGTAG | 185 |
| chr4:129396101-129398115 | p5 | (TTATT)3 | GGGATTTTGCTGCGTTATGT | ATCCTTTTAGTTGCTGCGGA | 234 |
| chr4:129396101-129398115 | p2 | (AT)7 | GGGATTTTGCTGCGTTATGT | ATCCTTTTAGTTGCTGCGGA | 234 |
| chr4:129533239-129535254 | p5 | (ATTGC)3 | TCAAAGGCAAGGAAAGGCTA | CTAATCAAAACAGGGCCCAA | 240 |
| chr4:129554617-129556632 | p5 | (GCATT)3 | GATCCAGGACAGTTGGCATT | CCGAATCCAAAACCACAAGT | 257 |
| chr4:129554617-129556632 | p5 | (GGGGT)3 | GATCCAGGACAGTTGGCATT | CCGAATCCAAAACCACAAGT | 257 |
| chr4:129661783-129663798 | p3 | (CCA)5 | TTGCCTTAGCTGGCTGTCTT | AATGAGTTGGTTGTTTCCGC | 286 |

| Marker ID | SSR type | SSR motif | FORWARD PRIMER (5'-3') | REVERSE PRIMER (5'-3') | size (bp) |
|---|---|---|---|---|---|
| chr4:129686534-129688552 | p2 | (TA)15 | AAATGAATCAAGGGGCAACA | GATCCGACACTTTTGGTCGT | 110 |
| chr4:129686534-129688552 | p6 | (TAAAAA)3 | AAATGAATCAAGGGGCAACA | GATCCGACACTTTTGGTCGT | 110 |
| chr4:129686534-129688552 | p2 | (AT)6 | AAATGAATCAAGGGGCAACA | GATCCGACACTTTTGGTCGT | 110 |
| chr4:129686534-129688552 | p3 | (TTA)10 | AAATGAATCAAGGGGCAACA | GATCCGACACTTTTGGTCGT | 110 |
| chr4:129793627-129795653 | p2 | (AT)13 | CCCTCAGACTCTTAGCGCAC | TTCCATCAGCCGATATGTCA | 242 |
| chr4:129846167-129848182 | p5 | (AAAAC)3 | CTGGCTCTTGTTGGTGGAAT | GAATGGGAACAACGGAGAAA | 222 |
| chr4:129865473-129867501 | p2 | (TA)14 | GCGAAAGTGTAGATCCCCAA | GCGTAATCTCATAAGGCCCA | 168 |
| chr4:129997176-129999198 | p4 | (TAAA)6 | TGAGCTTGTCAAAGACACGG | CAAAGGATCACCTTTCGCAT | 184 |
| chr4:129997176-129999198 | p2 | (TA)11 | TGAGCTTGTCAAAGACACGG | CAAAGGATCACCTTTCGCAT | 184 |
| chr4:129999916-130001934 | p3 | (TAA)5 | ACTGAAATTTGTCCCGTTGG | GGACCATTTGTTTGTTTGGG | 167 |
| chr4:129999937-130001951 | p3 | (TAA)5 | ACTGAAATTTGTCCCGTTGG | GGACCATTTGTTTGTTTGGG | 146 |
| chr4:130001708-130003735 | p3 | (CAT)9 | TACAAGCCCACAAGGCTACC | CAGACCCCTAGGCTGTTTCA | 284 |
| chr4:130090708-130092735 | p3 | (ACA)9 | AGGACGAAGAGGAAACGGAT | AAACGAGGAACGAGGGGTAT | 239 |
| chr4:130093442-130095460 | p6 | (TTTTTA)3 | CCACCACCCTGAAAGAAAAA | ATAGGGGGAGGGTTGAGAGA | 165 |
| chr4:130096244-130098256 | p5 | (TTTGA)3 | TTAATGGATGGTCGTGGTCA | TCCGCCTTCACTATCAAACC | 196 |
| chr4:130096244-130098256 | p2 | (CA)6 | TTAATGGATGGTCGTGGTCA | TCCGCCTTCACTATCAAACC | 196 |
| chr4:130191383-130193409 | p2 | (TG)13 | CAAAAATTCGCCCCAGATTA | TTTCTATGGCTTTCCCGTTG | 182 |
| chr4:130240475-130242487 | p2 | (AT)6 | GGGTCGTTATGTGCGTTTTT | ATTCCGCTCATGTATGCCTC | 271 |
| chr4:130270674-130272688 | p2 | (AT)7 | ATTTACCATGTCATCCCCCA | CTGTAACGACCCGCTAGCTC | 100 |
| chr4:130286831-130288847 | p4 | (ATAA)4 | GCGTTATTTTCCCGCATTTA | TTAAACGGATTGATCCGAGC | 110 |
| chr4:130384887-130386902 | p3 | (TGA)5 | ACGGACCATCCTACTCAACG | CTCTGTGGTGGACAATCCCT | 205 |
| chr4:130527135-130529201 | p5 | (AAACC)3 | TTTCGGATGAGTTTCAAGGG | TCACTATGCGTAGGTGCGAC | 271 |
| chr4:130688691-130690709 | p5 | (CATTC)3 | TGAAATTCTGAAGGGGCATT | AATGCTGAACAGGAACGAGG | 235 |
| chr4:130688769-130690784 | p5 | (CATTC)3 | TGAAATTCTGAAGGGGCATT | AATGCTGAACAGGAACGAGG | 157 |
| chr4:130688801-130690816 | p5 | (CATTC)3 | TGAAATTCTGAAGGGGCATT | AATGCTGAACAGGAACGAGG | 125 |
| chr4:130688822-130690837 | p5 | (CATTC)3 | TGAAATTCTGAAGGGGCATT | AATGCTGAACAGGAACGAGG | 104 |
| chr4:130859741-130861756 | p5 | (AAAGG)3 | TTCGGGGAATTCTGGTACTG | GGTCCCTCCTCTACGGTCTC | 185 |
| chr4:130924302-130926362 | p3 | (AAT)20 | GCACAACTGAACTCCGTGAA | TTATTGGTGAGCCCCACTTC | 270 |
| chr4:131059297-131061312 | p5 | (TTTTA)3 | TTTCGCATTTTCACCTTTCC | TGACCCGATTGTGTCTTTCA | 153 |
| chr4:131094536-131096551 | p3 | (TTC)5 | TAGGTTTTGTGTCTTCCCCG | AAAGCACAAGGAGGCACTGT | 155 |
| chr4:131382689-131384723 | p2 | (AT)17 | CAAACCATTGCCCCATAATC | GTTGGAGGCCACTTTGACAT | 216 |
| chr4:131508819-131510834 | p5 | (ATGCA)3 | GGAACCTCTCAAGTCAAGCG | TCGAACTCGCTTGTTGATTG | 133 |
| chr4:131600398-131602413 | p3 | (AGA)5 | ATGACGCTTGCAAATAACCC | TTTGTTGGCCCATTTTTAGG | 112 |
| chr4:131664723-131666756 | p3 | (AAC)11 | CGCACAAGGATTTTTCCATT | TCCCTCGTCAACAAAATTCC | 198 |
| chr4:131664723-131666756 | p5 | (AAAAT)4 | CGCACAAGGATTTTTCCATT | TCCCTCGTCAACAAAATTCC | 198 |
| chr4:131671390-131673405 | p5 | (GCATT)3 | TGGTGAGACTTCGTGTAGCG | GTCACAAAATAACGGGGTGG | 276 |

| Marker ID | SSR type | SSR motif | FORWARD PRIMER (5'-3') | REVERSE PRIMER (5'-3') | size (bp) |
|---|---|---|---|---|---|
| chr4:131703083-131705098 | p3 | (TTC)5 | TGTGAAATTAGCGATTGGCA | ACAATGTGTTTTCGAAGCCC | 213 |
| chr4:131703083-131705098 | p3 | (TTC)6 | TGTGAAATTAGCGATTGGCA | ACAATGTGTTTTCGAAGCCC | 213 |
| chr4:131726646-131728667 | p3 | (TTC)7 | TTGGCTATGTTGAGCTCCCT | TGGTAAATGCGACTGCTCTG | 124 |
| chr4:131766155-131768170 | p5 | (TACCA)3 | GCCAGACCTCAAGAAGCAAC | CATGGCCAGCAATGTATCAG | 179 |
| chr4:131770765-131772783 | p5 | (AAAAT)3 | TCATCATCACCGGAGACAAA | ATCTACACCGTTGTCGGAGG | 251 |
| chr4:131770765-131772783 | p3 | (GTG)6 | TCATCATCACCGGAGACAAA | ATCTACACCGTTGTCGGAGG | 251 |
| chr4:131770765-131772783 | p4 | (AGAA)5 | TCATCATCACCGGAGACAAA | ATCTACACCGTTGTCGGAGG | 251 |
| chr4:131823670-131825684 | p2 | (AG)7 | AGCTTCAGGCTTCCTTTGCT | TTCCACAAGTTTCAATGGCA | 162 |
| chr4:131851018-131853066 | p2 | (AG)24 | TTCAGATGATTTTGGTCCGAG | GTCAAAGCAAAGGAAGCCTG | 105 |
| chr4:131861161-131863185 | p2 | (TA)30 | TAGCCTTTGTTCCTCCTCCA | TGGCACCAAAGTTCATCAAA | 176 |
| chr4:131910267-131912315 | p3 | (AAT)16 | CCATAGTGGAGGTTGCAGGT | AGGAAGAAGAAGAGCCCAGG | 114 |
| chr4:131950844-131952859 | p5 | (TTTTG)3 | CTGCTTCTGCTACTCCCCAG | TTTTGGGAATGTGTGGTTGA | 282 |
| chr4:131950844-131952859 | p3 | (ACA)7 | CTGCTTCTGCTACTCCCCAG | TTTTGGGAATGTGTGGTTGA | 282 |
| chr4:132034090-132036124 | p2 | (TA)17 | ACTAAGCCCTGCACGTTAGC | TCGCTTGCGGAAATAGAGTT | 239 |
| chr4:132036228-132038243 | p5 | (TTTTA)3 | TCAACGACCACGTGTGAAAT | CACCTGCCAGTGATGGTATG | 227 |
| chr4:132036228-132038243 | p3 | (TGT)5 | TCAACGACCACGTGTGAAAT | CACCTGCCAGTGATGGTATG | 227 |
| chr4:132115161-132117177 | p2 | (AT)8 | CGCTGTGCTCTTCTTATCCC | GCACCGAGCTAAAATGCTTC | 157 |
| chr4:132181058-132183073 | p5 | (AATTA)3 | TTTAAATTAGACGTGGGCCG | AATAAGCGGAAAAGGGCCTA | 127 |
| chr4:132301719-132303755 | p3 | (GGA)11 | GAAAATGAGGAGAAAGGGGG | TTCTTCTGCTGCTCCTCCTC | 149 |
| chr4:132301719-132303755 | p3 | (GAA)12 | GAAAATGAGGAGAAAGGGGG | TTCTTCTGCTGCTCCTCCTC | 149 |
| chr4:132302686-132304785 | p3 | (GAA)12 | GGAGGAAAATGGGGGAAATA | CTCCTCCTCCTCCTCCTCAT | 278 |
| chr4:132302686-132304785 | p3 | (AGA)11 | GGAGGAAAATGGGGGAAATA | CTCCTCCTCCTCCTCCTCAT | 278 |
| chr4:132302827-132304866 | p3 | (AGA)38 | GGAGGAAAATGGGGGAAATA | CTCCTCCTCCTCCTCCTCAT | 137 |
| chr4:132304567-132306585 | p3 | (AGA)39 | GAGCGGGATGAGATGAAGAA | ATCCTCCTCCACCTCCTTGT | 137 |
| chr4:132304567-132306585 | p3 | (AGG)5 | GAGCGGGATGAGATGAAGAA | ATCCTCCTCCACCTCCTTGT | 137 |
| chr4:132304567-132306585 | p3 | (AGG)6 | GAGCGGGATGAGATGAAGAA | ATCCTCCTCCACCTCCTTGT | 137 |
| chr4:132304589-132306607 | p3 | (AGA)39 | GAGCGGGATGAGATGAAGAA | ATCCTCCTCCACCTCCTTGT | 115 |
| chr4:132304589-132306607 | p3 | (AGG)5 | GAGCGGGATGAGATGAAGAA | ATCCTCCTCCACCTCCTTGT | 115 |
| chr4:132304589-132306607 | p3 | (AGG)6 | GAGCGGGATGAGATGAAGAA | ATCCTCCTCCACCTCCTTGT | 115 |
| chr4:132316048-132318066 | p6 | (TATTTT)3 | AGCCTCTTGCCGATAACGTA | TGGCCCACTTGTGTATCGTA | 267 |
| chr4:132420057-132422072 | p5 | (TATTT)3 | GTGCAGCGAGGACACTGTAA | ACGAAAGAATTTGGAGGGGT | 279 |
| chr4:132420057-132422072 | p2 | (AT)10 | GTGCAGCGAGGACACTGTAA | ACGAAAGAATTTGGAGGGGT | 279 |
| chr4:132420057-132422072 | p3 | (ACC)5 | GTGCAGCGAGGACACTGTAA | ACGAAAGAATTTGGAGGGGT | 279 |
| chr4:132523660-132525674 | p2 | (TA)7 | GTGAAGACCGAGGGATTTGA | CAGCTGTGCATCCATGTTCT | 134 |
| chr4:132525264-132527276 | p2 | (TA)6 | GTTGATGATTTGGCCGAGTT | TTTCACAAGGATCATGCCAA | 268 |
| chr4:132525401-132527429 | p2 | (TA)6 | GTTGATGATTTGGCCGAGTT | TTTCACAAGGATCATGCCAA | 131 |

| Marker ID | SSR type | SSR motif | FORWARD PRIMER (5'-3') | REVERSE PRIMER (5'-3') | size (bp) |
|---|---|---|---|---|---|
| chr4:132554849-132556865 | p2 | (TA)8 | CCGTACTAGCGTAGGAAGCG | GAAGGCCATTCGGAAACATA | 131 |
| chr4:132577239-132579254 | p5 | (ACCGA)3 | ATGCCAACTCATTCATTCCC | CAAACAGGTGCGTAGCAGAA | 287 |
| chr4:132577239-132579254 | p5 | (AAATA)3 | ATGCCAACTCATTCATTCCC | CAAACAGGTGCGTAGCAGAA | 287 |
| chr4:132610226-132612268 | p2 | (AC)6 | GACATGGCGGAATAGCAAAT | CATGCGTATCAACCCATCAG | 128 |
| chr4:132610226-132612268 | p2 | (AC)8 | GACATGGCGGAATAGCAAAT | CATGCGTATCAACCCATCAG | 128 |
| chr4:132786188-132788202 | p2 | (TA)7 | ATTCCGAACAACGTCAGACC | TCAAAGACGTACCCCGATTC | 120 |
| chr4:132818646-132820660 | p2 | (TA)7 | ATTCCGAACAACGTCAGACC | TCAAAGACGTACCCCGATTC | 120 |
| chr4:132886409-132888455 | p5 | (TTTAA)3 | GCAGTTCGTCCTAGGGAGTG | AAATTCCCTGCAACCACTTG | 209 |
| chr4:132886409-132888455 | p2 | (TA)23 | GCAGTTCGTCCTAGGGAGTG | AAATTCCCTGCAACCACTTG | 209 |
| chr4:132900465-132902491 | p2 | (AT)13 | TCCTCCATTAACAACGCACA | GGCGACCTGAAATGAAGAAG | 254 |
| chr4:132900465-132902491 | p3 | (ATG)5 | TCCTCCATTAACAACGCACA | GGCGACCTGAAATGAAGAAG | 254 |
| chr4:132991590-132993624 | p2 | (TA)6 | CGCTGCGTATCAACCAATTA | AATACCATGCCGGAATACCA | 252 |
| chr4:132991637-132993651 | p2 | (TA)6 | CGCTGCGTATCAACCAATTA | AATACCATGCCGGAATACCA | 205 |
| chr4:133072356-133074382 | p2 | (AG)13 | TGGGATACAGTTAGGGCGAG | ATGGAGCCGAAGAGTCTCAA | 165 |
| chr4:133092575-133094587 | p3 | (ATC)5 | CTCAACATTGTCATGGTGGC | CTAGGGACCCACAACTTCCA | 243 |
| chr4:133092575-133094587 | p2 | (GA)6 | CTCAACATTGTCATGGTGGC | CTAGGGACCCACAACTTCCA | 243 |
| chr4:133147067-133149091 | p2 | (TA)12 | ACCTCGGGTATATTGAGGGG | GGCTTACTATCTGCGCAACC | 243 |
| chr4:133147067-133149091 | p2 | (GA)11 | ACCTCGGGTATATTGAGGGG | GGCTTACTATCTGCGCAACC | 243 |
| chr4:133190653-133192668 | p5 | (TGTGT)3 | TATGCCCACCAAGTGTTTGA | CTCCGGCCATTGAGTGTAAT | 152 |
| chr4:133220965-133223015 | p2 | (AT)17 | GCCGATAAGGCCATCATAAA | TGCAAGTTCAGTTGGACAGG | 273 |
| chr4:133230405-133232420 | p3 | (GTT)5 | CCACCAAAACTTAAGGGCAA | CTTATCTTCCACCAAGGCCA | 269 |
| chr4:133240779-133242803 | p3 | (GTT)8 | TTCACGCTCTTTGTCACGTC | ACTTCAGACCCCCTGAGCTT | 150 |
| chr4:133336174-133338204 | p2 | (TA)15 | AAACGAATGAAATTTTGCGG | CGAGGTAAGGGTTCTGGTGA | 196 |
| chr4:133336174-133338204 | p2 | (AT)8 | AAACGAATGAAATTTTGCGG | CGAGGTAAGGGTTCTGGTGA | 196 |
| chr4:133410209-133412225 | p4 | (TGAT)4 | TCTAGCGGCCTGCTATTTGT | CCTAAAGGAAATTCGCTCCC | 102 |
| chr4:133715381-133717459 | p3 | (AAT)26 | AAAATCCACTCCACGGACAG | GCCTGACGATCAGAGAGGTC | 125 |
| chr4:133763055-133765088 | p3 | (AAT)5 | AATCATGTAGCTCAACCCCG | ATCTCCATTGTCGTTAGGCG | 294 |
| chr4:133763089-133765107 | p3 | (AAT)5 | AATCATGTAGCTCAACCCCG | ATCTCCATTGTCGTTAGGCG | 260 |
| chr4:133865633-133867651 | p6 | (CTTCCG)3 | TGGGCTAATTGTGCCCTTAC | ATGAGCCATGAAGGTTGGAG | 236 |
| chr4:133919373-133921397 | p3 | (TAA)5 | TACCTCCTTTGTTTGCCACC | AAATCAAGGTGGTGGAGCTG | 220 |
| chr4:133919373-133921397 | p4 | (AAGA)4 | TACCTCCTTTGTTTGCCACC | AAATCAAGGTGGTGGAGCTG | 220 |
| chr4:133919373-133921397 | p3 | (AAT)8 | TACCTCCTTTGTTTGCCACC | AAATCAAGGTGGTGGAGCTG | 220 |
| chr4:133931899-133933911 | p2 | (GA)6 | CGACGACAACAACAACAACC | TGGGTTTACGTGACCCATTT | 199 |
| chr4:133941654-133943669 | p5 | (GAAAA)3 | GGCTTCTACCGAAGGGAACT | ATCCAGAGGAATGACGCAAG | 285 |
| chr4:133941654-133943669 | p2 | (TA)18 | GGCTTCTACCGAAGGGAACT | ATCCAGAGGAATGACGCAAG | 285 |
| chr4:133945446-133947462 | p4 | (ATAG)4 | TATCTCTTTGCCCGTTGCTT | TGTCCACCAAAGAATCCACA | 117 |

| Marker ID | SSR type | SSR motif | FORWARD PRIMER (5'-3') | REVERSE PRIMER (5'-3') | size (bp) |
|---|---|---|---|---|---|
| chr4:134019275-134021293 | p6 | (ATTTTT)3 | TCTTCCTGACTCCCGATTTG | AGGGGACGACCTAGGAAAGA | 267 |
| chr4:134040877-134042891 | p2 | (AT)7 | TTGAGATTGGGACTTCGGTC | CCGCGTCGAAAGCTAGATAC | 131 |
| chr4:134102938-134104953 | p6 | (CAGTTT)3 | ATGTTTTGCGAGTTGCACAG | CGTAAGCATGTGGCTTCTGA | 242 |
| chr4:134102938-134104953 | p5 | (ATATT)3 | ATGTTTTGCGAGTTGCACAG | CGTAAGCATGTGGCTTCTGA | 242 |
| chr4:134135139-134137187 | p2 | (AT)24 | GCGGCTACGCTAATCAGTTC | AAGGCCACTAGCCGTACTCA | 184 |
| chr4:134198919-134200934 | p5 | (GTTTG)3 | ATCAAAATTAGGCCGAACCC | CACTTTCACCAATCACACCG | 269 |
| chr4:134249574-134251634 | p2 | (GA)6 | TGAGCGGATCCTTAATACCG | GGGTCGAAAAGGGTTCATTT | 257 |
| chr4:134249636-134251676 | p2 | (GA)6 | TGAGCGGATCCTTAATACCG | GGGTCGAAAAGGGTTCATTT | 195 |
| chr4:134359434-134361448 | p2 | (AT)7 | TTATCACCCGAGTTCATCCC | GCTCACCCTGAAATCTCCAA | 122 |
| chr4:134359607-134361661 | p2 | (AT)7 | GGAGATTTCAGGGTGAGCTG | CCTTCCTCTTACGAGCTCCA | 149 |
| chr4:134397090-134399102 | p2 | (AG)6 | ATTGGCATTACTGCATGTGG | GAGGATCCAACACTCACCGT | 112 |
| chr4:134477583-134479598 | p2 | (CT)6 | CGTTTTCTGTGCTACCCGTT | TATAGCATCAGCGCCATGAG | 295 |
| chr4:134477583-134479598 | p5 | (CATGC)3 | CGTTTTCTGTGCTACCCGTT | TATAGCATCAGCGCCATGAG | 295 |
| chr4:134485360-134487384 | p2 | (TA)12 | TGTAATGCAGCCAAGACGAG | CAGGTCATGGATATTTGGGC | 299 |
| chr4:134485360-134487384 | p2 | (AT)12 | TGTAATGCAGCCAAGACGAG | CAGGTCATGGATATTTGGGC | 299 |
| chr4:134485360-134487384 | p2 | (AT)12 | TGTAATGCAGCCAAGACGAG | CAGGTCATGGATATTTGGGC | 299 |
| chr4:134485360-134487384 | p2 | (AT)12 | TGTAATGCAGCCAAGACGAG | CAGGTCATGGATATTTGGGC | 299 |
| chr4:134485360-134487384 | p2 | (AT)12 | TGTAATGCAGCCAAGACGAG | CAGGTCATGGATATTTGGGC | 299 |
| chr4:134485360-134487384 | p2 | (AT)12 | TGTAATGCAGCCAAGACGAG | CAGGTCATGGATATTTGGGC | 299 |
| chr4:134485360-134487384 | p2 | (AT)12 | TGTAATGCAGCCAAGACGAG | CAGGTCATGGATATTTGGGC | 299 |
| chr4:134485360-134487384 | p2 | (AT)12 | TGTAATGCAGCCAAGACGAG | CAGGTCATGGATATTTGGGC | 299 |
| chr4:134485360-134487384 | p2 | (AT)12 | TGTAATGCAGCCAAGACGAG | CAGGTCATGGATATTTGGGC | 299 |
| chr4:134485360-134487384 | p2 | (AT)12 | TGTAATGCAGCCAAGACGAG | CAGGTCATGGATATTTGGGC | 299 |
| chr4:134485513-134487537 | p2 | (TA)12 | TGTAATGCAGCCAAGACGAG | GCAGGTCATGGATATTTGGG | 146 |
| chr4:134485513-134487537 | p2 | (AT)12 | TGTAATGCAGCCAAGACGAG | GCAGGTCATGGATATTTGGG | 146 |
| chr4:134485513-134487537 | p2 | (AT)12 | TGTAATGCAGCCAAGACGAG | GCAGGTCATGGATATTTGGG | 146 |
| chr4:134485513-134487537 | p2 | (AT)12 | TGTAATGCAGCCAAGACGAG | GCAGGTCATGGATATTTGGG | 146 |
| chr4:134485513-134487537 | p2 | (AT)12 | TGTAATGCAGCCAAGACGAG | GCAGGTCATGGATATTTGGG | 146 |
| chr4:134485513-134487537 | p2 | (AT)12 | TGTAATGCAGCCAAGACGAG | GCAGGTCATGGATATTTGGG | 146 |
| chr4:134485513-134487537 | p2 | (AT)12 | TGTAATGCAGCCAAGACGAG | GCAGGTCATGGATATTTGGG | 146 |
| chr4:134485513-134487537 | p2 | (AT)12 | TGTAATGCAGCCAAGACGAG | GCAGGTCATGGATATTTGGG | 146 |
| chr4:134485513-134487537 | p2 | (AT)12 | TGTAATGCAGCCAAGACGAG | GCAGGTCATGGATATTTGGG | 146 |
| chr4:134485513-134487537 | p2 | (AT)12 | TGTAATGCAGCCAAGACGAG | GCAGGTCATGGATATTTGGG | 146 |
| chr4:134485666-134487690 | p2 | (TA)12 | CCCAAATATCCATGACCTGC | AGTTGTGGCATGATGCTTTG | 101 |
| chr4:134485666-134487690 | p2 | (AT)12 | CCCAAATATCCATGACCTGC | AGTTGTGGCATGATGCTTTG | 101 |
| chr4:134485666-134487690 | p2 | (AT)12 | CCCAAATATCCATGACCTGC | AGTTGTGGCATGATGCTTTG | 101 |

| Marker ID | SSR type | SSR motif | FORWARD PRIMER (5'-3') | REVERSE PRIMER (5'-3') | size (bp) |
|---|---|---|---|---|---|
| chr4:134485666-134487690 | p2 | (AT)12 | CCCAAATATCCATGACCTGC | AGTTGTGGCATGATGCTTTG | 101 |
| chr4:134485666-134487690 | p2 | (AT)12 | CCCAAATATCCATGACCTGC | AGTTGTGGCATGATGCTTTG | 101 |
| chr4:134485666-134487690 | p2 | (AT)12 | CCCAAATATCCATGACCTGC | AGTTGTGGCATGATGCTTTG | 101 |
| chr4:134485666-134487690 | p2 | (AT)12 | CCCAAATATCCATGACCTGC | AGTTGTGGCATGATGCTTTG | 101 |
| chr4:134485666-134487690 | p2 | (AT)12 | CCCAAATATCCATGACCTGC | AGTTGTGGCATGATGCTTTG | 101 |
| chr4:134485666-134487690 | p2 | (AT)12 | CCCAAATATCCATGACCTGC | AGTTGTGGCATGATGCTTTG | 101 |
| chr4:134485666-134487690 | p2 | (AT)12 | CCCAAATATCCATGACCTGC | AGTTGTGGCATGATGCTTTG | 101 |
| chr4:134513010-134515025 | p3 | (TTA)6 | GAACTATTTCTTCGGCAGCG | CTGGAATGTGAACGGGAGAT | 144 |
| chr4:134513010-134515025 | p3 | (TTA)14 | GAACTATTTCTTCGGCAGCG | CTGGAATGTGAACGGGAGAT | 144 |
| chr4:134513010-134515025 | p3 | (TTA)5 | GAACTATTTCTTCGGCAGCG | CTGGAATGTGAACGGGAGAT | 144 |
| chr4:134851254-134853336 | p3 | (CAA)6 | TGTCACGACCCAAACTGAAG | GGTCCTATTGGCCCACCTAT | 109 |
| chr4:134922536-134924605 | p2 | (AT)19 | CATCGACTCTTGCTGGTGAA | AAGACCTGCTTTGGATGGTG | 277 |
| chr4:135072202-135074222 | p4 | (TATG)5 | CCGTTCCTCGGAATACGATA | CACCGACTCCGTTTAGGAAA | 224 |
| chr4:135072202-135074222 | p2 | (TA)7 | CCGTTCCTCGGAATACGATA | CACCGACTCCGTTTAGGAAA | 224 |
| chr4:135090029-135092044 | p5 | (TTTTA)3 | TTCGTTCCTCCACAAGGATT | CTTGGTGTTTCGGCAGTGTA | 284 |
| chr4:135090029-135092044 | p2 | (AT)7 | TTCGTTCCTCCACAAGGATT | CTTGGTGTTTCGGCAGTGTA | 284 |
| chr4:135132751-135134787 | p2 | (TA)18 | CGACGGAAGAAGGTGAGAAG | TTAGGCGTATCCATTCTGGC | 190 |
| chr4:135132751-135134787 | p2 | (AT)23 | CGACGGAAGAAGGTGAGAAG | TTAGGCGTATCCATTCTGGC | 190 |
| chr4:135133232-135135278 | p2 | (TA)18 | CCCCTGGAAAGTATGCAAGA | GCCCACAAAGTACTTCCCAA | 289 |
| chr4:135133232-135135278 | p2 | (AT)23 | CCCCTGGAAAGTATGCAAGA | GCCCACAAAGTACTTCCCAA | 289 |
| chr4:135152487-135154502 | p5 | (ATTTC)3 | CTGCATAGGAGGTCCATCGT | TGCTGATATTGTACCCTGCG | 230 |
| chr4:135170848-135172860 | p2 | (AT)6 | TTGACCATGCGTGCCTATTA | GAAGCCTCAGACAAAGCAGG | 134 |
| chr4:135285442-135287458 | p4 | (AGAA)4 | TGCCTTAGCACGCTTAACCT | GGAGTGGCACCCACTAAGAA | 262 |
| chr4:135292973-135295013 | p2 | (TA)20 | TCCTTCTCCGTGAACTGCTT | TTTGGATTTGATTGTCGGCT | 167 |
| chr4:135292973-135295013 | p5 | (AATTA)3 | TCCTTCTCCGTGAACTGCTT | TTTGGATTTGATTGTCGGCT | 167 |
| chr4:135319695-135321735 | p2 | (TA)20 | TCCTTCTCCGTGAACTGCTT | TTTGGATTTGATTGTCGGCT | 167 |
| chr4:135319695-135321735 | p5 | (AATTA)3 | TCCTTCTCCGTGAACTGCTT | TTTGGATTTGATTGTCGGCT | 167 |
| chr4:135601031-135603053 | p2 | (TA)7 | AGTTTTATTTTCGGGGTGGG | CTAACCGCTTACCAGTTCGG | 251 |
| chr4:135601031-135603053 | p2 | (TA)11 | AGTTTTATTTTCGGGGTGGG | CTAACCGCTTACCAGTTCGG | 251 |
| chr4:135639280-135641294 | p2 | (TC)7 | TCTTGTCTTTGGCCGGTATC | TGGAATTCACTTGGGACACA | 276 |
| chr4:135700302-135702320 | p3 | (CAA)6 | CAATTGGCTCTCACAAGCAA | GGCTTCATTTCCGTCACCTA | 151 |
| chr4:135790374-135792410 | p3 | (TTC)12 | TCTCCTTGCTTCACATCCCT | GGCAAGCCAGCAAATGTTAT | 182 |
| chr4:135819470-135821482 | p2 | (TA)6 | GACTTCTTGACCAAGGCAGC | GGGCTGTTTTGGTTCCTTTT | 166 |
| chr4:135967513-135969528 | p5 | (CGCAC)3 | TATTGCTTTGAATACCCCGC | GGCGTACTAGCCTAAATGCG | 112 |
| chr4:135981773-135983788 | p5 | (TCTTT)3 | AAGTGAAGGGGAGCCAAGAT | ACTCGCGTACGTGTCCTTCT | 286 |
| chr4:135984144-135986159 | p5 | (TCTTT)3 | TGTACAAGTGAAGGGGAGCC | ACTCGCGTACGTGTCCTTCT | 151 |

| Marker ID | SSR type | SSR motif | FORWARD PRIMER (5'-3') | REVERSE PRIMER (5'-3') | size (bp) |
|---|---|---|---|---|---|
| chr4:136108226-136110241 | p5 | (AAAGG)3 | CACGGGTTTATTCGTAGCGT | CCAAATGGTACAGGTCGCTT | 126 |
| chr4:136142165-136144181 | p4 | (TTGT)4 | TAAGATCCCGTTATTCCCCC | GCTCCATCATTTAAACCCGA | 150 |
| chr4:136156220-136158235 | p3 | (TGT)5 | GGTTGTGAGACCGACCTTGT | ATCCTATCTCGCCTGGTGTG | 271 |
| chr4:136180006-136182026 | p5 | (AACCA)4 | TTTTAAAGGGGAATCGGGTC | AGGTGCAAAAATGTCCTTGG | 275 |
| chr4:136357504-136359519 | p5 | (TTTAA)3 | ATTAGCCTTGCCGGCTTTAT | GAACCACCTGACTCATCCGT | 223 |
| chr4:136357504-136359519 | p2 | (TA)8 | ATTAGCCTTGCCGGCTTTAT | GAACCACCTGACTCATCCGT | 223 |
| chr4:136357744-136359760 | p5 | (TTTAA)3 | TGTCTATGAGCCGTGTCCAG | CGAAGCAATTTCCTCGTAGC | 232 |
| chr4:136357744-136359760 | p2 | (TA)8 | TGTCTATGAGCCGTGTCCAG | CGAAGCAATTTCCTCGTAGC | 232 |
| chr4:136418018-136420041 | p2 | (AT)6 | TGCGTGAAATTAAACACGGA | CCATCTGCAGCAATCAGAAA | 264 |
| chr4:136418018-136420041 | p3 | (GGA)5 | TGCGTGAAATTAAACACGGA | CCATCTGCAGCAATCAGAAA | 264 |
| chr4:136447579-136449630 | p3 | (ATT)17 | CAGTAGAGGGTGGGAGGACA | CATTTAGCGTGTGTTCGGTG | 245 |
| chr4:136453225-136455246 | p2 | (TC)8 | GTTCAGCTGCACAGTTCCAA | GTTATCTCTCCCACCCCCAT | 250 |
| chr4:136469848-136471862 | p2 | (TA)7 | CCTTACTGAGAGGGCACGAG | TGGGACTCCGTGAATTAAGC | 170 |
| chr4:136563880-136565895 | p5 | (ACCAA)3 | CTCATGCCCAACAGGTTTTT | CTGAGAAAACCCGAGGTTGA | 106 |
| chr4:136563880-136565895 | p5 | (AAACC)3 | CTCATGCCCAACAGGTTTTT | CTGAGAAAACCCGAGGTTGA | 106 |
| chr4:136563880-136565895 | p2 | (AT)10 | CTCATGCCCAACAGGTTTTT | CTGAGAAAACCCGAGGTTGA | 106 |
| chr4:136632760-136634787 | p2 | (CA)9 | ACATACGGAGCAAAATTGGG | GCTTTGACAAAAGTAGGCGG | 269 |
| chr4:136633263-136635281 | p2 | (CA)9 | ATTTGCTTTTTCATCAGCCC | GGGATGGAGAAGGAGTGTGA | 138 |
| chr4:136635172-136637184 | p2 | (AT)6 | AGGAAATCCTGCTACACCCC | TGGCTCCAAAAACCACAACT | 180 |
| chr4:136643306-136645318 | p2 | (CA)6 | GTTTCGGACGTTTGATTGCT | AGGTTGATATTCCGGAACCC | 102 |
| chr4:136719610-136721662 | p2 | (AT)26 | AGCTGTAGCTCCGGATGAAA | TTTTTGTTTGTGGACGTTGC | 286 |
| chr4:136768491-136770506 | p5 | (ATATC)3 | ATGTGTCACACCCCAACCTT | GTCGGCCTATGATTCACGTT | 261 |
| chr4:136800555-136802571 | p2 | (TA)8 | GCAAATTGAAATTGGGGAGA | TACCTTTTTCCCCTTGGACC | 233 |
| chr4:136800555-136802571 | p2 | (AT)11 | GCAAATTGAAATTGGGGAGA | TACCTTTTTCCCCTTGGACC | 233 |
| chr4:136880697-136882715 | p3 | (CAA)6 | CCGAGCATCGGATTATTGTT | GATCACCCTGAGGATAGCCA | 265 |
| chr4:136946396-136948412 | p2 | (CA)6 | AAACAGCCCGCAACAATTAC | TGTAGCCGTGTGTGTGTGTG | 240 |
| chr4:136946481-136948497 | p2 | (CA)6 | AAACAGCCCGCAACAATTAC | TGTAGCCGTGTGTGTGTGTG | 155 |
| chr4:137099919-137101935 | p2 | (CT)8 | ACCCTCGATCAAAAGTGCAG | TGCTGCTGCTCTGAGTGTTT | 174 |
| chr4:137122649-137124663 | p2 | (AT)9 | GTTGTCCTCAGTCATGGCCT | TGACACCGCACAAAAACAAT | 152 |
| chr4:137122649-137124663 | p2 | (TA)7 | GTTGTCCTCAGTCATGGCCT | TGACACCGCACAAAAACAAT | 152 |
| chr4:137141917-137143929 | p2 | (CA)6 | TTTAATCGCACAAAAGCACG | GGGTGCGTATCCAGTTATGG | 262 |
| chr4:137161820-137163835 | p2 | (TC)7 | AAATGTGGAGGAGGAGGCTT | GGGAAAATGCCAAGAGCATA | 246 |
| chr4:137161820-137163835 | p5 | (TGATA)3 | AAATGTGGAGGAGGAGGCTT | GGGAAAATGCCAAGAGCATA | 246 |
| chr4:137165624-137167640 | p2 | (TA)9 | CTGGGTGGAAAAAGAGACGA | CTCGTGTGCATTTTGCTGAT | 286 |
| chr4:137165624-137167640 | p2 | (AG)8 | CTGGGTGGAAAAAGAGACGA | CTCGTGTGCATTTTGCTGAT | 286 |
| chr4:137165624-137167640 | p5 | (CTAGA)3 | CTGGGTGGAAAAAGAGACGA | CTCGTGTGCATTTTGCTGAT | 286 |

| Marker ID | SSR type | SSR motif | FORWARD PRIMER (5'-3') | REVERSE PRIMER (5'-3') | size (bp) |
|---|---|---|---|---|---|
| chr4:137165746-137167761 | p2 | (TA)9 | CTGGGTGGAAAAAGAGACGA | CTCGTGTGCATTTTGCTGAT | 164 |
| chr4:137165746-137167761 | p2 | (AG)8 | CTGGGTGGAAAAAGAGACGA | CTCGTGTGCATTTTGCTGAT | 164 |
| chr4:137165746-137167761 | p5 | (CTAGA)3 | CTGGGTGGAAAAAGAGACGA | CTCGTGTGCATTTTGCTGAT | 164 |
| chr4:137226421-137228435 | p2 | (AT)7 | TCTGCTGAAGCTTTTGCCTT | TCCATGGAGGCTTTGATTTC | 199 |
| chr4:137254455-137256477 | p2 | (AT)11 | TTGAAAAACATCGTGTGGGA | GAAATTCGAGCAGCATCTCC | 124 |
| chr4:137290833-137292851 | p3 | (CAT)6 | ACTTTTGTCACGACCCAACC | TGTTAGCCTCGCCGACTAAT | 104 |
| chr4:137450392-137452404 | p2 | (TG)6 | TGTCACGATGTGCAAACAAA | CTCGATGTTTTGGCAGATGA | 189 |
| chr4:137450392-137452404 | p4 | (ATTA)4 | TGTCACGATGTGCAAACAAA | CTCGATGTTTTGGCAGATGA | 189 |
| chr4:137453957-137455969 | p2 | (TC)6 | CCGAGCACCAAGTTATGACC | CCAAGGCATGGTTGAAAAGT | 119 |
| chr4:137464953-137466965 | p2 | (AT)6 | ACTTGGCAAGAGGTGGAAGA | GGTTCATAACGGTTCGGTTG | 189 |
| chr4:137464953-137466965 | p2 | (TA)7 | ACTTGGCAAGAGGTGGAAGA | GGTTCATAACGGTTCGGTTG | 189 |
| chr4:137510667-137512682 | p3 | (TAA)5 | GGGGTTTTCCGATTTCTCTC | ACCCGACACAAACACCTCTC | 230 |
| chr4:137510667-137512682 | p2 | (TA)6 | GGGGTTTTCCGATTTCTCTC | ACCCGACACAAACACCTCTC | 230 |
| chr4:137616388-137618403 | p5 | (ATCAA)3 | ACAAGAAGGACAATCCACGC | CTCCAGTCGCGTATGACTGA | 277 |
| chr4:137658266-137660280 | p2 | (GT)7 | GTTGCTGCACAAGAACCAGA | TCCTCAAGCGAACTCAAGGT | 157 |
| chr4:137708878-137710896 | p6 | (TTTTTG)3 | GCCAGGGCATTTGAAAATAA | ATTTGGCCCTGTGTATGCTC | 156 |
| chr4:137766730-137768754 | p4 | (TTTC)6 | AAACGCAAGGATGACGAAAC | CTCTTCACATTGCCTCACCA | 107 |
| chr4:137908353-137910369 | p2 | (TA)8 | GCCACCAAAACTTAAAGGCA | CTCCCACCTTGTCTCGGATA | 178 |
| chr4:137935101-137937116 | p5 | (ATTTA)3 | TATGATCCAAATCCTCCCCA | AGGATGTTGGGTATGCAAGC | 228 |
| chr4:137942169-137944195 | p2 | (AT)13 | AGGGGTAGGCCGAAGAAGTA | GGTCCTCCAAACCCACCTAT | 121 |
| chr4:137951228-137953243 | p3 | (TTG)5 | GCTGCTCTCGAGCCTTCTAA | ACTGAAAGGCAAGGGGATTT | 113 |
| chr4:137954310-137956325 | p5 | (TTGTA)3 | GGAGACGAAGCCGTTAAGTG | CTCCAATGGCGTTTCAACTT | 173 |
| chr4:138068345-138070366 | p3 | (AGA)7 | AACAAATCAGGCTACCACCG | ACATGAAGGCAGCCCTACAC | 255 |
| chr4:138098593-138100608 | p5 | (TTTGG)3 | GAAAAGGGACTTGAAGCGTG | GTGAAATGGGGGAATGTTTG | 109 |
| chr4:138277534-138279558 | p4 | (TCTT)6 | TGATGTTCTGCAGGTTCTCG | ATGAGGTGACGAGTCGAAGG | 150 |
| chr4:138297868-138299884 | p2 | (AT)9 | CATATTCGGGCTTGCCATAG | AATGTGGGAATCTCTTTGCG | 230 |
| chr4:138297868-138299884 | p2 | (AG)8 | CATATTCGGGCTTGCCATAG | AATGTGGGAATCTCTTTGCG | 230 |
| chr4:138297868-138299884 | p2 | (AT)14 | CATATTCGGGCTTGCCATAG | AATGTGGGAATCTCTTTGCG | 230 |
| chr4:138333860-138335876 | p2 | (AG)8 | CGCAGACCACACAGAAGAGA | AGGGCTTTCATAACGGAGGT | 149 |
| chr4:138347773-138349791 | p2 | (AC)9 | ATGGCCCATCCATGAGTAAA | TTTGAAATTCGTTTGGGGAC | 196 |
| chr4:138347773-138349791 | p2 | (AC)6 | ATGGCCCATCCATGAGTAAA | TTTGAAATTCGTTTGGGGAC | 196 |
| chr4:138347929-138349961 | p2 | (AC)9 | GCCGTCTTAGGACAACGGTA | TGCAAGACGCATGATTTGTT | 201 |
| chr4:138347929-138349961 | p2 | (AC)6 | GCCGTCTTAGGACAACGGTA | TGCAAGACGCATGATTTGTT | 201 |
| chr4:138347962-138349976 | p2 | (AC)9 | GCCGTCTTAGGACAACGGTA | TGCAAGACGCATGATTTGTT | 168 |
| chr4:138347962-138349976 | p2 | (AC)6 | GCCGTCTTAGGACAACGGTA | TGCAAGACGCATGATTTGTT | 168 |
| chr4:138374703-138376718 | p3 | (TGT)6 | CATTGTTGGCATACCCACTG | TCCTGAATGGATCTTCGGAC | 249 |

| Marker ID | SSR type | SSR motif | FORWARD PRIMER (5'-3') | REVERSE PRIMER (5'-3') | size (bp) |
|---|---|---|---|---|---|
| chr4:138374728-138376743 | p3 | (TGT)6 | CATTGTTGGCATACCCACTG | TCCTGAATGGATCTTCGGAC | 224 |
| chr4:138385495-138387519 | p6 | (AGGAGA)4 | ATGCGAGGATGAAAAGGATG | ACATCCGGGGCTAGAAGAGT | 114 |
| chr4:138385495-138387519 | p2 | (AT)17 | ATGCGAGGATGAAAAGGATG | ACATCCGGGGCTAGAAGAGT | 114 |
| chr4:138402463-138404501 | p5 | (ATTGC)3 | TGGTGTGGATGTTGAATGCT | CCAAGTCCCAGAATATCCCA | 227 |
| chr4:138406023-138408039 | p2 | (AT)8 | TCACAGGAGCGAGACACAAG | CCCAACTCCGGAAATTGTTA | 143 |
| chr4:138408615-138410711 | p6 | (CACCCC)4 | TTTCCCACCATTACGAGGAG | CCAACTATTTTCCCACCCCT | 268 |
| chr4:138430747-138432763 | p2 | (AT)8 | CTTTGCAAGCCTGCACATTA | TTCGCCTCCTTGTATTCACC | 175 |
| chr4:138509910-138511934 | p2 | (AT)12 | CTCTTTGGCTGTTCTTTGCC | AACATGGACCACCACCACTT | 230 |
| chr4:138549015-138551063 | p2 | (CT)8 | TCAACGCAGAGCAACAATTC | CATACCCTTTTGTCCCCTCA | 199 |
| chr4:138549015-138551063 | p2 | (AT)24 | TCAACGCAGAGCAACAATTC | CATACCCTTTTGTCCCCTCA | 199 |
| chr4:138569057-138571073 | p2 | (CT)8 | GAAAACCCAGAAGTAGCCCC | GTTAAGGCTCGTCCCCTTTC | 273 |
| chr4:138598007-138600075 | p2 | (AT)34 | CCCTCACGACGTTCTCAAAT | GCCCAACTGTACGCCTTAAA | 169 |
| chr4:138731902-138733918 | p2 | (CT)8 | AAAGAGACTCGGCTCCACAA | CAGTGAACTTCGGTTGCTCA | 162 |
| chr4:138731902-138733918 | p2 | (CT)8 | AAAGAGACTCGGCTCCACAA | CAGTGAACTTCGGTTGCTCA | 162 |
| chr4:138732627-138734643 | p2 | (CT)8 | CTCCCTCCTCACTACAACCG | ATGGGTGCCGAGTGAATTAG | 198 |
| chr4:138732627-138734643 | p2 | (CT)8 | CTCCCTCCTCACTACAACCG | ATGGGTGCCGAGTGAATTAG | 198 |
| chr4:138767448-138769469 | p3 | (GAT)7 | ACCAGTTGCGACCAAGATTC | CGCCAGGCATAAGAAGAAAG | 291 |
| chr4:138779735-138781747 | p2 | (AC)6 | TTTTTGCGAATTTTTCCTCG | TCGGAACACAAAGACTGACG | 161 |
| chr4:138805423-138807451 | p6 | (GGGTGG)3 | TTGAGTGTACCCGGCCTTAG | ATGCACTTCGTTTGTGCAAG | 185 |
| chr4:138805423-138807451 | p2 | (TA)14 | TTGAGTGTACCCGGCCTTAG | ATGCACTTCGTTTGTGCAAG | 185 |
| chr4:138805423-138807451 | p2 | (AT)10 | TTGAGTGTACCCGGCCTTAG | ATGCACTTCGTTTGTGCAAG | 185 |
| chr4:138805423-138807451 | p2 | (AT)14 | TTGAGTGTACCCGGCCTTAG | ATGCACTTCGTTTGTGCAAG | 185 |
| chr4:138816347-138818369 | p2 | (GT)11 | GTGTAGGCAGTCTTCAGCCC | GAACCCCAATGAAGGGAAAT | 139 |
| chr4:138858528-138860596 | p6 | (ATTTTT)3 | CGAAATATCCGCCATCAGTT | CCACCCAAAGTAAAAGGGGT | 277 |
| chr4:138867605-138869619 | p2 | (TA)7 | GGCCTTTTTCCTGAGAGTCC | TGGAATGTCGACCACAAAAA | 180 |
| chr4:138867605-138869619 | p2 | (TA)7 | GGCCTTTTTCCTGAGAGTCC | TGGAATGTCGACCACAAAAA | 180 |
| chr4:138873946-138875961 | p5 | (GCATT)3 | TGGTGAGACTTCGTGTAGCG | CAAGGCCCAACCTAAGACAA | 146 |
| chr4:138902162-138904186 | p2 | (AT)8 | CATAAGCCGACCAGGCTAAC | GTCCTGTCCCGCCTATATGA | 295 |
| chr4:138902162-138904186 | p2 | (AT)12 | CATAAGCCGACCAGGCTAAC | GTCCTGTCCCGCCTATATGA | 295 |
| chr4:138956276-138958294 | p3 | (TTG)6 | CCCTTGAAATTCCTGCGTTA | CCAAGTGTTCTTCACCGGAT | 196 |
| chr4:138980831-138982849 | p2 | (TA)9 | CTCTTCCTCCTCCTCGTCCT | CCTCTTTTTATCTGCTGGCG | 252 |
| chr4:138980831-138982849 | p3 | (CTC)6 | CTCTTCCTCCTCCTCGTCCT | CCTCTTTTTATCTGCTGGCG | 252 |
| chr4:138980831-138982849 | p3 | (ACA)6 | CTCTTCCTCCTCCTCGTCCT | CCTCTTTTTATCTGCTGGCG | 252 |
| chr4:139158498-139160548 | p2 | (AT)10 | ATTGGACAAGGGTATGGGCT | GATAAGACCCCACCTCACCA | 105 |
| chr4:139158498-139160548 | p2 | (TA)25 | ATTGGACAAGGGTATGGGCT | GATAAGACCCCACCTCACCA | 105 |
| chr4:139172964-139174980 | p2 | (TA)8 | GGCCATTTTACTGAAGAGCG | CATTTTTCGGCCCAGTTAAA | 227 |

| Marker ID | SSR type | SSR motif | FORWARD PRIMER (5'-3') | REVERSE PRIMER (5'-3') | size (bp) |
|---|---|---|---|---|---|
| chr4:139225852-139227867 | p5 | (CTTTT)3 | ACTGTCCTTCATTTGGGGTG | ATTTGCAGGATTGTCCTTCG | 239 |
| chr4:139233452-139235470 | p6 | (TAACTG)3 | CTGCCTACACCGGCATACTT | TCTCTCGGTCACATGCAGTC | 275 |
| chr4:139278742-139280763 | p3 | (ACA)7 | GATCGATTTGGATTCTCGGA | GGGAAGGACAGGGAGGATTA | 286 |
| chr4:139282437-139284452 | p3 | (AAC)5 | GGAGGCAGAAGTTCACCTTG | TGCCTCCTCCAGCATACTTT | 219 |
| chr4:139282437-139284452 | p5 | (AAAAG)3 | GGAGGCAGAAGTTCACCTTG | TGCCTCCTCCAGCATACTTT | 219 |
| chr4:139302556-139304570 | p2 | (AT)7 | GTGGCATATTCGATGCCTTT | CAACACAACACGAATCGGTC | 273 |
| chr4:139378979-139380993 | p2 | (TA)7 | GAACAAACATTTGCGTCCCT | TACATAAATAGCCGCCCACC | 180 |
| chr4:139427320-139429335 | p5 | (TGCAA)3 | AAGTCCATGTACCTCACGGC | ATATCCGATGGTTGTTCGGA | 227 |
| chr4:139427360-139429375 | p5 | (TGCAA)3 | AAGTCCATGTACCTCACGGC | ATATCCGATGGTTGTTCGGA | 187 |
| chr4:139522587-139524601 | p2 | (TG)7 | ACCATTTTTGGCTTTGTTGG | AAATGTTGCAGGAACAAGGG | 115 |
| chr4:139529052-139531066 | p2 | (GA)7 | TGGTTTTATAATGCGTGGCA | ACCAGGGGTGTTGTATTGGA | 102 |
| chr4:139546703-139548731 | p2 | (TA)14 | CTTCATCTCCATGCCGATCT | TATAAATGGCCTTGCTTGCC | 142 |
| chr4:139572233-139574251 | p6 | (CTTCAA)3 | TCCACTCTGCATCCGTTAAA | TGGAGCTTTCTTCCTCTGGA | 189 |
| chr4:139613206-139615224 | p3 | (ACA)6 | TGAATCAAACCCATTGCTCA | TCAGAATTGACACCCTTCCC | 180 |
| chr4:139622413-139624428 | p3 | (TGT)5 | TAGAATCCCGAACCCACAAG | CCAAAATTGCTGCACAAGAA | 210 |
| chr4:139662613-139664631 | p6 | (TCATCC)3 | CGGGAGCATGAGATGGTATT | TTTGCTCAATTGCAAAGCAC | 192 |
| chr4:139662613-139664631 | p2 | (TA)28 | CGGGAGCATGAGATGGTATT | TTTGCTCAATTGCAAAGCAC | 192 |
| chr4:139676976-139678992 | p2 | (TA)8 | TTGAGAGATGGGGGTAGCTG | TCATCAATGTTTTGCCTCCA | 284 |
| chr4:139685041-139687071 | p2 | (TA)11 | AGGCCGAACTGCTAAAGTGA | GTGCATGCGTGAATTTTGTC | 134 |
| chr4:139719255-139721277 | p2 | (AT)11 | AAGCAATGTCAAAACTGGGC | AAATCAGTCCGCCAAATGAC | 208 |
| chr4:139960291-139962306 | p5 | (ACCGG)3 | TGACCCTCCAATCTTTCCTG | GCATCGATATTGCACGTCAC | 192 |
| chr4:139960291-139962306 | p2 | (TA)6 | TGACCCTCCAATCTTTCCTG | GCATCGATATTGCACGTCAC | 192 |
| chr4:140003373-140005391 | p3 | (ACA)6 | GCATCACACTGTCCCATTTG | AGTGTGCTTGGTAGCATCCC | 214 |
| chr4:140053215-140055227 | p2 | (AT)6 | GTTTGACCCAAAGCGGATAA | TTACCACGATCCACAGGTCA | 148 |
| chr4:140203385-140205397 | p2 | (AT)6 | ATGTCCGCAGCTTCTTGTTT | TGGCTAGAACGCATGATTTG | 153 |
| chr4:140226697-140228712 | p2 | (TA)6 | TGTTGTTTTGGAGCTTGCTG | CCCCAATGCATAGACACAAA | 278 |
| chr4:140226697-140228712 | p2 | (AT)10 | TGTTGTTTTGGAGCTTGCTG | CCCCAATGCATAGACACAAA | 278 |
| chr4:140226784-140228808 | p2 | (TA)6 | TGTTGTTTTGGAGCTTGCTG | CCCCAATGCATAGACACAAA | 191 |
| chr4:140226784-140228808 | p2 | (AT)10 | TGTTGTTTTGGAGCTTGCTG | CCCCAATGCATAGACACAAA | 191 |
| chr4:140226804-140228870 | p2 | (TA)6 | TGTTGTTTTGGAGCTTGCTG | CCCCAATGCATAGACACAAA | 171 |
| chr4:140226804-140228870 | p2 | (AT)10 | TGTTGTTTTGGAGCTTGCTG | CCCCAATGCATAGACACAAA | 171 |
| chr4:140232453-140234485 | p2 | (TA)16 | GAAACCACCATCCAAACAGG | GCAAAAGTGAGCCAGAAAGG | 220 |
| chr4:140245671-140247687 | p2 | (TA)8 | GACCCGTAACCAAAGTGCAT | TTCTTCGGAGGATTGTCCAC | 260 |
| chr4:140251041-140253057 | p4 | (TTTA)5 | TTTTTGCTTTTGGGCTATCG | GTGGTACATTCGACGCCTTT | 256 |
| chr4:140251041-140253057 | p4 | (AAAT)4 | TTTTTGCTTTTGGGCTATCG | GTGGTACATTCGACGCCTTT | 256 |
| chr4:140367874-140369888 | p2 | (AG)7 | CCCTTCAAATGCATTGGTCT | GGCCAAAGTTAGGTCTGCAA | 144 |

| Marker ID | SSR type | SSR motif | FORWARD PRIMER (5'-3') | REVERSE PRIMER (5'-3') | size (bp) |
|---|---|---|---|---|---|
| chr4:140376774-140378789 | p5 | (GGGCA)3 | TGGAAGCAGATGGAAAATCC | GCTTCAGAGCATCCGAGAAC | 266 |
| chr4:140377636-140379648 | p5 | (GGGCA)3 | AAACACACCTAACGCCCAAC | CCCTCACGACGTTCTCAAAT | 235 |
| chr4:140377636-140379648 | p5 | (TAGAA)3 | AAACACACCTAACGCCCAAC | CCCTCACGACGTTCTCAAAT | 235 |
| chr4:140377636-140379648 | p3 | (GAA)5 | AAACACACCTAACGCCCAAC | CCCTCACGACGTTCTCAAAT | 235 |
| chr4:140403180-140405196 | p4 | (TAAT)4 | AATCACAAAAGGCGGCATAC | AGATCCGCTCAATGGAGAAA | 207 |
| chr4:140418377-140420392 | p3 | (TCA)5 | GCAAGGCAGAATTTCGTAGC | CCTCATGGCTAGCTGTGTCA | 201 |
| chr4:140421058-140423074 | p2 | (TA)9 | AATCATGCCCGACCAGTAAG | GCATGTTATGATATCGCCCC | 287 |
| chr4:140421058-140423074 | p2 | (TA)8 | AATCATGCCCGACCAGTAAG | GCATGTTATGATATCGCCCC | 287 |
| chr4:140446861-140448951 | p2 | (AT)45 | GATGCTCAAAAGGGAATCCA | ACCCTCTGCCATGTTATTGC | 184 |
| chr4:140472538-140474552 | p2 | (CA)7 | ATAACAGGAGCCACGGACTG | TTGCAGGCAAAGATGTGAAC | 225 |
| chr4:140514958-140516973 | p5 | (AATAA)3 | ACAGAATGGGGTGGTGTTGT | CCAGAATTTGAGGTTTGGGA | 168 |
| chr4:140558849-140560861 | p2 | (AT)6 | TTCATCATCACAGCAACGGT | CGCCGTAGATCTGGTTTAGC | 147 |
| chr4:140577900-140579918 | p2 | (AT)9 | TTCAGTTGACCCGAGGTAGG | TGAGGTCGCTTCCATATTCC | 201 |
| chr4:140606126-140608138 | p2 | (AT)6 | TAGGTCACTGCTGCAAGGTG | TTTCATCACTTGGATTGGCA | 256 |
| chr4:140606126-140608138 | p4 | (CAGA)4 | TAGGTCACTGCTGCAAGGTG | TTTCATCACTTGGATTGGCA | 256 |
| chr4:140745424-140747444 | p2 | (TA)10 | GTGAAGATTGCGGGTTGTTT | CACGCTGCCCAATAGTACCT | 151 |
| chr4:140824555-140826570 | p5 | (CGAAC)3 | AGAATCTGTGCGTGAAGCCT | GCAGCATATTTGGCCTTGAT | 193 |
| chr4:140824555-140826570 | p5 | (AACCG)3 | AGAATCTGTGCGTGAAGCCT | GCAGCATATTTGGCCTTGAT | 193 |
| chr4:140892101-140894123 | p2 | (TA)11 | GGCAATTTGGTCAAAGCACT | TGACACATGGCCAAAAACAT | 129 |
| chr4:141127658-141129700 | p2 | (AT)21 | GGGTTCGAGCTTGATCCATA | CCCATTATATCACGCCCTTG | 132 |
| chr4:141132032-141134050 | p6 | (AACCGG)4 | CGACGTAGGCTTTCCTCAAG | CCCAACTCGGATGAGACATT | 290 |
| chr4:141132032-141134050 | p6 | (CCGGTT)3 | CGACGTAGGCTTTCCTCAAG | CCCAACTCGGATGAGACATT | 290 |
| chr4:141132032-141134050 | p6 | (CCGGTT)3 | CGACGTAGGCTTTCCTCAAG | CCCAACTCGGATGAGACATT | 290 |
| chr4:141132032-141134050 | p6 | (CCGGTT)3 | CGACGTAGGCTTTCCTCAAG | CCCAACTCGGATGAGACATT | 290 |
| chr4:141132032-141134050 | p6 | (CCGGTT)3 | CGACGTAGGCTTTCCTCAAG | CCCAACTCGGATGAGACATT | 290 |
| chr4:141132032-141134050 | p6 | (CCGGTT)3 | CGACGTAGGCTTTCCTCAAG | CCCAACTCGGATGAGACATT | 290 |
| chr4:141132032-141134050 | p6 | (CCGGTT)3 | CGACGTAGGCTTTCCTCAAG | CCCAACTCGGATGAGACATT | 290 |
| chr4:141132032-141134050 | p6 | (CCGGTT)3 | CGACGTAGGCTTTCCTCAAG | CCCAACTCGGATGAGACATT | 290 |
| chr4:141132189-141134207 | p6 | (AACCGG)4 | CGACGTAGGCTTTCCTCAAG | CCCAACTCGGATGAGACATT | 133 |
| chr4:141132189-141134207 | p6 | (CCGGTT)3 | CGACGTAGGCTTTCCTCAAG | CCCAACTCGGATGAGACATT | 133 |
| chr4:141132189-141134207 | p6 | (CCGGTT)3 | CGACGTAGGCTTTCCTCAAG | CCCAACTCGGATGAGACATT | 133 |
| chr4:141132189-141134207 | p6 | (CCGGTT)3 | CGACGTAGGCTTTCCTCAAG | CCCAACTCGGATGAGACATT | 133 |
| chr4:141132189-141134207 | p6 | (CCGGTT)3 | CGACGTAGGCTTTCCTCAAG | CCCAACTCGGATGAGACATT | 133 |
| chr4:141132189-141134207 | p6 | (CCGGTT)3 | CGACGTAGGCTTTCCTCAAG | CCCAACTCGGATGAGACATT | 133 |
| chr4:141132189-141134207 | p6 | (CCGGTT)3 | CGACGTAGGCTTTCCTCAAG | CCCAACTCGGATGAGACATT | 133 |
| chr4:141132189-141134207 | p6 | (CCGGTT)3 | CGACGTAGGCTTTCCTCAAG | CCCAACTCGGATGAGACATT | 133 |

| Marker ID | SSR type | SSR motif | FORWARD PRIMER (5'-3') | REVERSE PRIMER (5'-3') | size (bp) |
|---|---|---|---|---|---|
| chr4:141132189-141134207 | p6 | (CCGGTT)3 | CGACGTAGGCTTTCCTCAAG | CCCAACTCGGATGAGACATT | 133 |
| chr4:141132346-141134364 | p6 | (AACCGG)4 | GCGCCTGTCATGAATTCTCT | AGCTCGTCTTTGGTAAGGCA | 243 |
| chr4:141132346-141134364 | p6 | (CCGGTT)3 | GCGCCTGTCATGAATTCTCT | AGCTCGTCTTTGGTAAGGCA | 243 |
| chr4:141132346-141134364 | p6 | (CCGGTT)3 | GCGCCTGTCATGAATTCTCT | AGCTCGTCTTTGGTAAGGCA | 243 |
| chr4:141132346-141134364 | p6 | (CCGGTT)3 | GCGCCTGTCATGAATTCTCT | AGCTCGTCTTTGGTAAGGCA | 243 |
| chr4:141132346-141134364 | p6 | (CCGGTT)3 | GCGCCTGTCATGAATTCTCT | AGCTCGTCTTTGGTAAGGCA | 243 |
| chr4:141132346-141134364 | p6 | (CCGGTT)3 | GCGCCTGTCATGAATTCTCT | AGCTCGTCTTTGGTAAGGCA | 243 |
| chr4:141132346-141134364 | p6 | (CCGGTT)3 | GCGCCTGTCATGAATTCTCT | AGCTCGTCTTTGGTAAGGCA | 243 |
| chr4:141132346-141134364 | p6 | (CCGGTT)3 | GCGCCTGTCATGAATTCTCT | AGCTCGTCTTTGGTAAGGCA | 243 |
| chr4:141132346-141134364 | p6 | (CCGGTT)3 | GCGCCTGTCATGAATTCTCT | AGCTCGTCTTTGGTAAGGCA | 243 |
| chr4:141132346-141134364 | p6 | (CCGGTT)3 | GCGCCTGTCATGAATTCTCT | AGCTCGTCTTTGGTAAGGCA | 243 |
| chr4:141274370-141276394 | p6 | (TGGGGG)4 | TCCAAGTTCTAGGCCGTGTC | CAACCGCAAGGAGCACTTAT | 215 |
| chr4:141339508-141341520 | p2 | (TG)6 | AAAGGTGCCTACGGGCTATT | TCATACCTGCCCTACCAAGC | 255 |
| chr4:141385227-141387241 | p2 | (GA)7 | TGAGCACACATGATGCAAGA | TCTATGCGACCAAGCATACG | 259 |
| chr4:141397314-141399328 | p2 | (AC)7 | TAAAAAGGGAACGAGCCAGA | TTGTTCTGGAGTTGCTGTCG | 274 |
| chr4:141397314-141399328 | p3 | (CAA)5 | TAAAAAGGGAACGAGCCAGA | TTGTTCTGGAGTTGCTGTCG | 274 |
| chr4:141397434-141399516 | p2 | (AC)7 | TAAAAAGGGAACGAGCCAGA | TTGTTCTGGAGTTGCTGTCG | 154 |
| chr4:141397434-141399516 | p3 | (CAA)5 | TAAAAAGGGAACGAGCCAGA | TTGTTCTGGAGTTGCTGTCG | 154 |
| chr4:141488810-141490822 | p2 | (AT)6 | GTGAAGTGTGCTCCTCGTCA | ACAGATTATTCCAGCCCCCT | 118 |
| chr4:141510451-141512469 | p6 | (AGGTGC)3 | GGCATGAGATAGTACCCCGA | ACAACGTCCAAAGAAAACGG | 246 |
| chr4:141651975-141653989 | p2 | (AT)7 | TGCGTACACTCTACCTCCCC | AAATAAATGGACACTGGCGG | 262 |
| chr4:141849712-141851724 | p2 | (AG)6 | GACACAATTGCCCCTGAAAT | GACAATGTGCATCTGCGTTT | 243 |
| chr4:141850537-141852621 | p2 | (AG)6 | TTTGATAGCATTGTTGCCCA | CGCTAATCACGTCGGACATT | 242 |
| chr4:141925699-141927714 | p5 | (ACCAA)3 | CATTTTGGGGCAGATGAGTT | CTCGGCTCCCAATAACAAGA | 156 |
| chr4:141925699-141927714 | p5 | (AAACC)3 | CATTTTGGGGCAGATGAGTT | CTCGGCTCCCAATAACAAGA | 156 |
| chr4:141945298-141947314 | p4 | (AAAT)4 | TGGGTGGTTCAATCATTTCA | AAGGTCCGAGAGCAGGTTTT | 120 |
| chr4:141945298-141947314 | p2 | (TA)7 | TGGGTGGTTCAATCATTTCA | AAGGTCCGAGAGCAGGTTTT | 120 |
| chr4:141945298-141947314 | p5 | (TAAAT)4 | TGGGTGGTTCAATCATTTCA | AAGGTCCGAGAGCAGGTTTT | 120 |
| chr4:141948926-141950938 | p5 | (ACCGG)4 | ACCGATTTTGTATTCGCAGC | CCTTCGAAGCATTTGGATGT | 130 |
| chr4:141948926-141950938 | p2 | (TA)12 | ACCGATTTTGTATTCGCAGC | CCTTCGAAGCATTTGGATGT | 130 |
| chr4:141948926-141950938 | p2 | (TA)6 | ACCGATTTTGTATTCGCAGC | CCTTCGAAGCATTTGGATGT | 130 |
| chr4:141948926-141950938 | p2 | (TA)103 | ACCGATTTTGTATTCGCAGC | CCTTCGAAGCATTTGGATGT | 130 |
| chr4:141969332-141971347 | p3 | (ACA)5 | ATACATCAACGGGCTTCTGC | TAATCAGGCTGTTGTCGTCG | 276 |
| chr4:141969332-141971347 | p2 | (TA)7 | ATACATCAACGGGCTTCTGC | TAATCAGGCTGTTGTCGTCG | 276 |
| chr4:142035706-142037720 | p2 | (CA)7 | AGAAAGGAGGCGGATGAGAT | CTAAAATCCCTAAACCCGCC | 114 |
| chr4:142043089-142045101 | p2 | (TC)6 | CGGTGCCTTTTTAAAACCAA | ATCGAATAAGCAATCCTGCG | 284 |

| Marker ID | SSR type | SSR motif | FORWARD PRIMER (5'-3') | REVERSE PRIMER (5'-3') | size (bp) |
|---|---|---|---|---|---|
| chr4:142091901-142093915 | p2 | (AT)7 | CCCAGCATAATCCCACAAGT | CGCAACAAAGATGTTCCTCA | 125 |
| chr4:142152660-142154682 | p2 | (AT)11 | TGTGGAGTTACCTTCAGCCA | ATTGATTCTTATGGCCGTGG | 112 |
| chr4:142223581-142225596 | p5 | (TCATA)3 | TGTCTACCTGTGGGCATGAA | TATGGCGGTCGTGACAGATA | 198 |
| chr4:142246959-142248984 | p6 | (TCGGTT)3 | TCCACCTTTGCTCTTGCTTT | AAAACTGAACGCTCACCCAC | 161 |
| chr4:142246959-142248984 | p5 | (GTTCG)5 | TCCACCTTTGCTCTTGCTTT | AAAACTGAACGCTCACCCAC | 161 |
| chr4:142374195-142376213 | p6 | (TTTTAT)3 | GAGGGAGATGAAGTGTGGGA | ACGGTCCGTATAACACCCAA | 178 |
| chr4:142377795-142379811 | p2 | (TA)28 | GCCAAGTCAACCACCGTAAT | CACATCAATTGAAACGGCAA | 113 |
| chr4:142377795-142379811 | p2 | (GA)8 | GCCAAGTCAACCACCGTAAT | CACATCAATTGAAACGGCAA | 113 |
| chr4:142399997-142402057 | p2 | (GA)9 | ACAGATTCCCATTGAAACGC | GATTTCCCATTGAATGTCGG | 300 |
| chr4:142399997-142402057 | p2 | (TA)29 | ACAGATTCCCATTGAAACGC | GATTTCCCATTGAATGTCGG | 300 |
| chr4:142401845-142403857 | p2 | (TC)6 | TGCACCATTTTATAACGGCA | GATGCATTGAAACAAGGGCT | 295 |
| chr4:142401845-142403857 | p2 | (TA)8 | TGCACCATTTTATAACGGCA | GATGCATTGAAACAAGGGCT | 295 |
| chr4:142453349-142455367 | p6 | (CTCTTT)3 | CTTGCAACAACGCACAGAAT | TGTCAACAATGGGAACTGGA | 227 |
| chr4:142460379-142462430 | p2 | (GA)6 | TCGAAGGAAAAAGCCAAAGA | GTGAAGTCGCAGACCCATTT | 292 |
| chr4:142460379-142462430 | p3 | (ATA)17 | TCGAAGGAAAAAGCCAAAGA | GTGAAGTCGCAGACCCATTT | 292 |
| chr4:142464503-142466521 | p3 | (ACA)6 | GAGTCAGACAGGAGCCGAAG | GTGATGCCTAACGAGGGTGT | 229 |
| chr4:142467940-142469955 | p3 | (TTC)5 | GAGAGAACGGTTTGGTGGAA | CCAATTTTGGACCTGCCTTA | 180 |
| chr4:142467940-142469955 | p2 | (TC)7 | GAGAGAACGGTTTGGTGGAA | CCAATTTTGGACCTGCCTTA | 180 |
| chr4:142524955-142526973 | p2 | (GA)6 | AACAGGATGTGGTGGAGAGG | CCAGCCCATTGCCTAAAATA | 271 |
| chr4:142524955-142526973 | p3 | (AAT)6 | AACAGGATGTGGTGGAGAGG | CCAGCCCATTGCCTAAAATA | 271 |
| chr4:142549234-142551256 | p2 | (AT)11 | GCGGTTCAACAAGTATGCCT | AGTAGGTTGCCGTTTGGATG | 220 |
| chr4:142549234-142551256 | p3 | (AGA)5 | GCGGTTCAACAAGTATGCCT | AGTAGGTTGCCGTTTGGATG | 220 |
| chr4:142586022-142588054 | p2 | (AT)16 | GGCTGGTGGTATGAGTGGTT | CCAAGCCTTTGCCTACTGAG | 136 |
| chr4:142586022-142588054 | p3 | (ATA)14 | GGCTGGTGGTATGAGTGGTT | CCAAGCCTTTGCCTACTGAG | 136 |
| chr4:142680951-142682966 | p5 | (TTTCT)3 | GCCGAAGTTTGCTCAACTCT | GCAGACTAACCGCGACAATA | 184 |
| chr4:142774793-142776805 | p2 | (AT)6 | TAAAAAGAGTCCGGAGCCAA | TGTTGCTTTCGCTTCACTTG | 156 |
| chr4:142774821-142776841 | p2 | (AT)6 | TAAAAAGAGTCCGGAGCCAA | TGTTGCTTTCGCTTCACTTG | 128 |
| chr4:142774837-142776852 | p2 | (AT)6 | TAAAAAGAGTCCGGAGCCAA | TGTTGCTTTCGCTTCACTTG | 112 |
| chr4:142774847-142776859 | p2 | (AT)6 | TAAAAAGAGTCCGGAGCCAA | TGTTGCTTTCGCTTCACTTG | 102 |
| chr4:142864480-142866538 | p2 | (TA)7 | GTCCCGCAAGTGGGTAGTTA | CATCCGCTGCACAAGTTAAA | 145 |
| chr4:142969799-142971879 | p2 | (TA)40 | ATGATGGATTCATTGCCACA | GACATGGCGGAATAGCAAAT | 170 |
| chr4:142978842-142980860 | p3 | (ACA)6 | TTCAACAGCTGTCATGAGCC | TGCCATCTTTTTAATTGGGC | 204 |
| chr4:143069501-143071539 | p2 | (AT)34 | TTTTGCTCCCTTGGTGATTC | TGTGGCCTAAAGCCAAATTC | 108 |
| chr4:143159427-143161439 | p2 | (TA)6 | GCCTAATCGATCCTGCATGT | TATAGAAGTTGTCCGCCGCT | 207 |
| chr4:143186644-143188660 | p2 | (TG)8 | CGATGCATTAAACTCCCGAT | TTATTTTCCCACCGACGTTC | 200 |
| chr4:143186644-143188660 | p2 | (GT)6 | CGATGCATTAAACTCCCGAT | TTATTTTCCCACCGACGTTC | 200 |

| Marker ID | SSR type | SSR motif | FORWARD PRIMER (5'-3') | REVERSE PRIMER (5'-3') | size (bp) |
|---|---|---|---|---|---|
| chr4:143186644-143188660 | p2 | (GT)6 | CGATGCATTAAACTCCCGAT | TTATTTTCCCACCGACGTTC | 200 |
| chr4:143187080-143189092 | p2 | (TG)8 | GCCCAATGACCAAACATTTC | ACGGAATGGAGGGTTAGGAT | 298 |
| chr4:143187080-143189092 | p2 | (GT)6 | GCCCAATGACCAAACATTTC | ACGGAATGGAGGGTTAGGAT | 298 |
| chr4:143187080-143189092 | p2 | (GT)6 | GCCCAATGACCAAACATTTC | ACGGAATGGAGGGTTAGGAT | 298 |
| chr4:143235273-143237297 | p2 | (AT)12 | CCAGAAGCTTGGCCAAATAG | ATGTACCCAATGCCCTTTGA | 236 |
| chr4:143244114-143246128 | p4 | (AATA)4 | TCAATAAGAAGCCGCGAAAT | CACGAAATGCGTTCGTCTAA | 286 |
| chr4:143244114-143246128 | p2 | (AT)7 | TCAATAAGAAGCCGCGAAAT | CACGAAATGCGTTCGTCTAA | 286 |
| chr4:143244210-143246250 | p4 | (AATA)4 | TCAATAAGAAGCCGCGAAAT | CACGAAATGCGTTCGTCTAA | 190 |
| chr4:143244210-143246250 | p2 | (AT)7 | TCAATAAGAAGCCGCGAAAT | CACGAAATGCGTTCGTCTAA | 190 |
| chr4:143253968-143255983 | p3 | (TTA)5 | TGCCTGGCTTTTTCGATTAC | TCATGCATTTTAGCACTCGC | 211 |
| chr4:143293231-143295246 | p3 | (TAT)5 | TCAACTTTGTCATCACCGGA | GCCCAAAATCAAAAAGCAAA | 112 |
| chr4:143305351-143307366 | p3 | (TAT)5 | TGTCATCACCGGAAAATCAA | GCCCAAAATCAAAAAGCAAA | 120 |
| chr4:143329055-143331067 | p2 | (TA)6 | ACGGGTTGGTCTCACTTGAC | TTGAAGAATCAAAGGCCACC | 239 |
| chr4:143329055-143331067 | p4 | (AGAA)4 | ACGGGTTGGTCTCACTTGAC | TTGAAGAATCAAAGGCCACC | 239 |
| chr4:143329178-143331194 | p2 | (TA)6 | ACGGGTTGGTCTCACTTGAC | TTGAAGAATCAAAGGCCACC | 116 |
| chr4:143329178-143331194 | p4 | (AGAA)4 | ACGGGTTGGTCTCACTTGAC | TTGAAGAATCAAAGGCCACC | 116 |
| chr4:143329754-143331806 | p2 | (TA)6 | TGCCAAGAAATGTGATCTCG | TTGGGGTGAAAAACGTTAGC | 247 |
| chr4:143329754-143331806 | p4 | (AGAA)4 | TGCCAAGAAATGTGATCTCG | TTGGGGTGAAAAACGTTAGC | 247 |
| chr4:143422187-143424253 | p3 | (AAT)22 | GCCGGTTTGGTTCATTAAAA | ATACCCTTTTCGCTGCCATA | 237 |
| chr4:143660156-143662170 | p2 | (AT)7 | GAACCACCTGACTCATCCGT | CCTTGTCGGCCTTATTTTGA | 127 |
| chr4:143670032-143672046 | p5 | (GGTTT)3 | AAGTGTAGGGGTGTTCGTGG | GGATCACGTCAAAGCCGTAT | 237 |
| chr4:143670032-143672046 | p2 | (AT)7 | AAGTGTAGGGGTGTTCGTGG | GGATCACGTCAAAGCCGTAT | 237 |
| chr4:143682776-143684791 | p3 | (TGT)5 | TTTGGCACCCAAGTCATACA | TTTTCATTTCCGCCACTTTC | 114 |
| chr4:143698372-143700393 | p3 | (AAC)7 | ACCTGCAACGACATCAAACA | AACGACCCCTAAGGCTAGGA | 288 |
| chr4:143709653-143711668 | p3 | (TGT)5 | TTTGGCACCCAAGTCATACA | TTTTCATTTCCGCCACTTTC | 114 |
| chr4:143738644-143740660 | p4 | (TTTC)4 | TTGACCAAAGAAAGGCTGCT | TGTTTCCCTTTGATCTTCCG | 112 |
| chr4:143750303-143752321 | p2 | (TC)9 | AAACCGAAGAGCAACCCTTT | TGTGTAGTGGGGCCAGTGTA | 232 |
| chr4:143753503-143755517 | p2 | (AG)7 | ACAAACACGATTTTGGAGGC | TTGCAATGGGTTCCTTTCTC | 162 |
| chr4:143755592-143757607 | p3 | (ACA)5 | TTCCAGAACATTGCAAGCTG | GGACCGTGCTATTATGCGAT | 115 |
| chr4:143824906-143826924 | p2 | (TA)19 | TAAACCAGTGTCACGACCCA | TTTCTCATCTATGACCCCCG | 252 |
| chr4:143873068-143875097 | p5 | (CACCC)3 | TGGGGAGTGGTGAAAAGATT | GGGTGGGTGGAGAAAAGATT | 136 |
| chr4:143876541-143878556 | p5 | (AAACC)3 | GTAGGGGTGTACATGGACCG | AAAACCCGAGAAAACCGAAT | 187 |
| chr4:143958016-143960034 | p3 | (ACA)6 | GTGCCATTCCACCAGAAGTT | TGGTTACTCTACGCCATCCC | 164 |
| chr4:143965548-143967564 | p2 | (TA)8 | CCCACCCCAGTTTTTAGGTT | TGCCCGAGGTTATTTTTGTC | 278 |
| chr4:144038769-144040801 | p2 | (CT)13 | ATGAAGTGGTCCAGGAATGC | GCTAAAATGGGGGTTGGTTT | 136 |
| chr4:144038802-144040820 | p2 | (CT)13 | ATGAAGTGGTCCAGGAATGC | GCTAAAATGGGGGTTGGTTT | 103 |

| Marker ID | SSR type | SSR motif | FORWARD PRIMER (5'-3') | REVERSE PRIMER (5'-3') | size (bp) |
|---|---|---|---|---|---|
| chr4:144088773-144090791 | p6 | (CATAAT)3 | CGTCCGGGGTAACTATCTCA | CGACCTTATGTCACCCCAAT | 126 |
| chr4:144091694-144093708 | p2 | (TA)7 | TTTGCCATGATTTTCATCCA | GCTTGTTGGATTCCGTGATT | 130 |
| chr4:144094393-144096405 | p2 | (TA)6 | CGGGTCAAAATGGACTAAGC | GGGATCCCTACTTTTGCTCC | 158 |
| chr4:144113772-144115787 | p5 | (AGGAG)3 | TTCCGTGAACCATACGTGAA | GGTCCAGCACCAAATGTTCT | 296 |
| chr4:144115064-144117082 | p6 | (TGAGTG)3 | TTGATCACGCTAGTTGGTGG | GCTCCATTGAGGCCAATTTA | 275 |
| chr4:144130770-144132785 | p3 | (ACA)5 | TGAAACAGAACCCACCACAA | GATCAAGCAAATCTCTGCCC | 119 |
| chr4:144152139-144154166 | p3 | (CAA)9 | GATCCGGTGAGAAGAACTCG | TCTTTCCCTTTCCCCTCACT | 144 |
| chr4:144152139-144154166 | p5 | (AAGAC)3 | GATCCGGTGAGAAGAACTCG | TCTTTCCCTTTCCCCTCACT | 144 |
| chr4:144154691-144156709 | p3 | (TTG)6 | TGCCACCAAAACTTAAAGGC | GTCTCTCCTCTTCACGTGCC | 184 |
| chr4:144216869-144218887 | p6 | (CTGAAA)3 | GGAGAAGAACGTTGGAGCAG | ACTGATCATGTCCACGGTGA | 227 |
| chr4:144216869-144218887 | p5 | (ATAAC)3 | GGAGAAGAACGTTGGAGCAG | ACTGATCATGTCCACGGTGA | 227 |
| chr4:144288928-144290958 | p2 | (TA)15 | ACGCGCTAGTAACGTCCTGT | GAACACAATGATTCCGGACC | 266 |
| chr4:144332288-144334312 | p6 | (GCAATA)4 | CCTTCATCAAGTGCCCAAAT | TTGCTCTCCTCCTTTTTCCA | 108 |
| chr4:144339173-144341207 | p2 | (AT)17 | AACGGGCCCTTAGAAACAAT | AAACGGGGCCTTAGTTTGAT | 254 |
| chr4:144365191-144367209 | p3 | (TTA)6 | CGCTTGAAAAGAGGTTTTGG | CGGAGACGATAGACTTTGCC | 267 |
| chr4:144391134-144393149 | p2 | (CT)9 | GCAGATGAGGTTGCTCCTTT | AAACAACAAGTGGCTCTGGG | 222 |
| chr4:144391134-144393149 | p5 | (AAAAT)3 | GCAGATGAGGTTGCTCCTTT | AAACAACAAGTGGCTCTGGG | 222 |
| chr4:144391134-144393149 | p5 | (TAATT)3 | GCAGATGAGGTTGCTCCTTT | AAACAACAAGTGGCTCTGGG | 222 |
| chr4:144401948-144403966 | p6 | (TTCTTT)3 | CCTCTGCCCGTTTTCATTTA | GCCCTGAACGGAATATCAAA | 246 |
| chr4:144456152-144458168 | p4 | (ATAC)4 | TTCCAGCGTCAAATAGGAGG | CCTTGACATGATGCTGCACT | 124 |
| chr4:144456152-144458168 | p2 | (TA)9 | TTCCAGCGTCAAATAGGAGG | CCTTGACATGATGCTGCACT | 124 |
| chr4:144522465-144524481 | p4 | (ATAC)4 | TTCCAGCGTCAAATAGGAGG | CCTTGACATGATGCTGCACT | 124 |
| chr4:144522465-144524481 | p2 | (TA)9 | TTCCAGCGTCAAATAGGAGG | CCTTGACATGATGCTGCACT | 124 |
| chr4:144555676-144557688 | p2 | (AT)6 | GGGAGCTAAGAATGGGAAGG | CGACGAATTACAGCGACAGA | 256 |
| chr4:144560203-144562219 | p4 | (CTCC)4 | GGATGTCGATGTAGCGGAAT | GCTTGCCTTCCATTTCAGTC | 226 |
| chr4:144581307-144583331 | p6 | (TAAAAA)3 | CCCTTGGTTTCAGGTTCAAA | AAGGCAGATGTTTGGATTCG | 110 |
| chr4:144581307-144583331 | p6 | (AAAAAG)4 | CCCTTGGTTTCAGGTTCAAA | AAGGCAGATGTTTGGATTCG | 110 |
| chr4:144747215-144749229 | p2 | (CT)7 | CCCGACTTGTTTGGGACTTA | AAAACTATCGTCGCGTTTGG | 146 |
| chr4:144749508-144751526 | p6 | (AACAAA)3 | GGAACCTTGACAGGACCTCA | GTTGGGTGCACGAAGAGTTT | 100 |
| chr4:144878033-144880048 | p5 | (AAAAC)3 | ACTTGTCCTAACGGAGCCAA | ATCCAAGGTCGGTAAAGCCT | 184 |
| chr4:144891478-144893493 | p5 | (AAAAC)3 | ACTTGTCCTAACGGAGCCAA | ATCCAAGGTCGGTAAAGCCT | 184 |
| chr4:144905372-144907392 | p2 | (TA)10 | AGAGCATATCCCCACACCAG | TGATGGCGTCTCTGTGAGTC | 252 |
| chr4:144929255-144931270 | p2 | (AT)9 | TCCGAAAGACCGAAACAATC | CACAACGTGAGGTCTGAGGA | 148 |
| chr4:144929255-144931270 | p3 | (GTT)5 | TCCGAAAGACCGAAACAATC | CACAACGTGAGGTCTGAGGA | 148 |
| chr4:144929255-144931270 | p3 | (AAC)5 | TCCGAAAGACCGAAACAATC | CACAACGTGAGGTCTGAGGA | 148 |
| chr4:144950535-144952549 | p2 | (AT)7 | GTTGTCAAATATCGGGTCGC | CCCCAACATCACACAAGAAA | 292 |

| Marker ID | SSR type | SSR motif | FORWARD PRIMER (5'-3') | REVERSE PRIMER (5'-3') | size (bp) |
|---|---|---|---|---|---|
| chr4:144950535-144952549 | p2 | (TA)7 | GTTGTCAAATATCGGGTCGC | CCCCAACATCACACAAGAAA | 292 |
| chr4:144971218-144973233 | p5 | (ACCAA)3 | CATTGGCCATTCTGTTGTTG | AGGTCCAATGACAGCACCAT | 247 |
| chr4:144971218-144973233 | p5 | (AAACC)3 | CATTGGCCATTCTGTTGTTG | AGGTCCAATGACAGCACCAT | 247 |
| chr4:144984215-144986230 | p5 | (TTCGG)3 | TTCGAAATGGCAGTAAACCC | TTTCTCAACAACCCCTTTGC | 113 |
| chr4:144984215-144986230 | p6 | (AAATAT)3 | TTCGAAATGGCAGTAAACCC | TTTCTCAACAACCCCTTTGC | 113 |
| chr4:145005749-145007764 | p5 | (GACCC)3 | CGATACCTGACCCGTTCACT | CCTTAGGGGGACGGTTAGAC | 206 |
| chr4:145005749-145007764 | p5 | (GACCC)4 | CGATACCTGACCCGTTCACT | CCTTAGGGGGACGGTTAGAC | 206 |
| chr4:145005749-145007764 | p5 | (GACCC)3 | CGATACCTGACCCGTTCACT | CCTTAGGGGGACGGTTAGAC | 206 |
| chr4:145005749-145007764 | p5 | (TATTG)3 | CGATACCTGACCCGTTCACT | CCTTAGGGGGACGGTTAGAC | 206 |
| chr4:145034310-145036328 | p2 | (TA)8 | GGATTCATTGTCACATGCGT | AAATGACATGGCGGAATAGC | 208 |
| chr4:145034310-145036328 | p2 | (AT)9 | GGATTCATTGTCACATGCGT | AAATGACATGGCGGAATAGC | 208 |
| chr4:145034310-145036328 | p2 | (AT)15 | GGATTCATTGTCACATGCGT | AAATGACATGGCGGAATAGC | 208 |
| chr4:145039318-145041336 | p6 | (CACAAT)3 | AATTGGCTCGAGCATTTTGT | AATTGGCATATAGCAACGCC | 180 |
| chr4:145060310-145062335 | p2 | (AT)6 | AAAGTGGAACCCAGCACATC | TTGCTCCCTCTTATGGGTTG | 100 |
| chr4:145060310-145062335 | p2 | (AT)9 | AAAGTGGAACCCAGCACATC | TTGCTCCCTCTTATGGGTTG | 100 |
| chr4:145061248-145063266 | p2 | (AT)9 | AATGGAACTCACCATCTCGC | ATTTCATGGATTGACACGCA | 127 |
| chr4:145135299-145137317 | p3 | (AAT)6 | CTTTTTAGCGCGATTTTTCG | TTGAGGGAGCTTTTTGGAGA | 130 |
| chr4:145135299-145137317 | p3 | (AAT)6 | CTTTTTAGCGCGATTTTTCG | TTGAGGGAGCTTTTTGGAGA | 130 |
| chr4:145154491-145156503 | p2 | (TA)6 | TCCTCTCACGGTCTCCAAGT | TGTCAAAGGAGGGAGGAATG | 166 |
| chr4:145252078-145254092 | p2 | (CT)7 | TTTTCGAACCTCGACCAAAC | TCGTTCTTCTGTCTCCACCC | 202 |
| chr4:145306073-145308088 | p3 | (TGT)5 | AAAAGATGATTGCTGGACGG | GGGTGAGAGGGAAAGGAAAG | 257 |
| chr4:145320727-145322745 | p4 | (TTTA)4 | CCCCAAATCGCTACTGCTAC | CCACCATCGTTGTTCTCCTT | 258 |
| chr4:145320727-145322745 | p3 | (ATT)5 | CCCCAAATCGCTACTGCTAC | CCACCATCGTTGTTCTCCTT | 258 |
| chr4:145320727-145322745 | p3 | (AAC)6 | CCCCAAATCGCTACTGCTAC | CCACCATCGTTGTTCTCCTT | 258 |
| chr4:145353321-145355341 | p2 | (TA)10 | TGAGAGACCCCCTCAAAAGA | ATGTGTCCCTCCCACTTCTG | 170 |
| chr4:145353321-145355341 | p5 | (CAAAA)3 | TGAGAGACCCCCTCAAAAGA | ATGTGTCCCTCCCACTTCTG | 170 |
| chr4:145353321-145355341 | p3 | (ATT)6 | TGAGAGACCCCCTCAAAAGA | ATGTGTCCCTCCCACTTCTG | 170 |
| chr4:145405401-145407437 | p2 | (TA)18 | ATTATGAAGTGCGCCTGTCC | GGTGCCACATCACCTCTCTT | 224 |
| chr4:145457970-145459982 | p2 | (TA)6 | ACGACGAAGCATGGGTAAAC | GCAAGATCATCGTTCGTCCT | 127 |
| chr4:145545126-145547141 | p5 | (TTGGT)3 | TTCCACCAATTTCACGAACA | CAAGAGGGTAACCTGGAGGA | 153 |
| chr4:145579314-145581328 | p2 | (AT)7 | TGGGTATCACCGGGACTTTA | GCCAAAATATAACATGCGGC | 111 |
| chr4:145653165-145655180 | p5 | (TATTG)3 | AGAAGACGAACCCGATCAAA | TCCATCCTCATCGTGTGAAA | 205 |
| chr4:145711628-145713642 | p2 | (AT)7 | CATTTGCTATTTCACCGCCT | GACGAGCATGTGCGAGTTTA | 191 |
| chr4:145721613-145723627 | p2 | (AT)7 | ACCTAAAATCGCCCGCTACT | CATCAAAATCACAAACCCCC | 203 |
| chr4:145721613-145723627 | p2 | (AT)7 | ACCTAAAATCGCCCGCTACT | CATCAAAATCACAAACCCCC | 203 |
| chr4:145721613-145723627 | p2 | (AT)7 | ACCTAAAATCGCCCGCTACT | CATCAAAATCACAAACCCCC | 203 |

| Marker ID | SSR type | SSR motif | FORWARD PRIMER (5'-3') | REVERSE PRIMER (5'-3') | size (bp) |
|---|---|---|---|---|---|
| chr4:145721838-145723853 | p2 | (AT)7 | TGGGGGTTTGTGATTTTGAT | AGCACCCCTCTACACTTGGA | 139 |
| chr4:145721838-145723853 | p2 | (AT)7 | TGGGGGTTTGTGATTTTGAT | AGCACCCCTCTACACTTGGA | 139 |
| chr4:145721838-145723853 | p2 | (AT)7 | TGGGGGTTTGTGATTTTGAT | AGCACCCCTCTACACTTGGA | 139 |
| chr4:145752097-145754111 | p2 | (AT)7 | CATTTGCTATTTCACCGCCT | GACGAGCATGTGCGAGTTTA | 191 |
| chr4:145762082-145764096 | p2 | (AT)7 | ACCTAAAATCGCCCGCTACT | CATCAAAATCACAAACCCCC | 203 |
| chr4:145762082-145764096 | p2 | (AT)7 | ACCTAAAATCGCCCGCTACT | CATCAAAATCACAAACCCCC | 203 |
| chr4:145762082-145764096 | p2 | (AT)7 | ACCTAAAATCGCCCGCTACT | CATCAAAATCACAAACCCCC | 203 |
| chr4:145762307-145764322 | p2 | (AT)7 | TGGGGGTTTGTGATTTTGAT | AGCACCCCTCTACACTTGGA | 139 |
| chr4:145762307-145764322 | p2 | (AT)7 | TGGGGGTTTGTGATTTTGAT | AGCACCCCTCTACACTTGGA | 139 |
| chr4:145762307-145764322 | p2 | (AT)7 | TGGGGGTTTGTGATTTTGAT | AGCACCCCTCTACACTTGGA | 139 |
| chr4:145787171-145789192 | p3 | (TCA)7 | TGAAATCCCTAAAACACCCG | GGGTTCTTCCGAAATCAACA | 101 |
| chr4:145793828-145795848 | p2 | (TA)10 | GGTTCATGCGTGAGAGGAAT | ACACAACCTTTGCCACAACA | 163 |
| chr4:145910301-145912334 | p4 | (TCTT)4 | ATGCCATTTCCCTGTTTCTG | ACCACCAACTCTCCATCAGG | 100 |
| chr4:145910477-145912507 | p4 | (TCTT)4 | GCTGGGACTGACTCACATCA | GCGTGAGGGTACGCTACCTA | 154 |
| chr4:145992938-145994956 | p3 | (TTG)6 | AATTGGGCTGGTCATTCTTG | CGGTACTTACGTGCGTCTGA | 197 |
| chr4:146110970-146112988 | p5 | (TTGGT)3 | TCATTGACCACTAGGCCACA | GTAATGTCGGGTTGGTTTGG | 266 |
| chr4:146110970-146112988 | p5 | (TTGGT)3 | TCATTGACCACTAGGCCACA | GTAATGTCGGGTTGGTTTGG | 266 |
| chr4:146110970-146112988 | p3 | (ATG)6 | TCATTGACCACTAGGCCACA | GTAATGTCGGGTTGGTTTGG | 266 |
| chr4:146142042-146144072 | p2 | (AT)15 | AGCAAACGTTCACAAGCAAA | GATCGGTCTGACCCTTGAAA | 219 |
| chr4:146202917-146204959 | p2 | (TA)21 | AAACCCAGTGCAGGACTTTG | CACATGGCAATGAAACCATC | 127 |
| chr4:146202917-146204959 | p2 | (TA)8 | AAACCCAGTGCAGGACTTTG | CACATGGCAATGAAACCATC | 127 |
| chr4:146214559-146216577 | p6 | (ATATAC)3 | AAAGCCAAAAGGGTCCTCAT | CTCCGCTCACATCTCATCAA | 298 |
| chr4:146223723-146225743 | p5 | (AGATT)4 | ACAGTTACAATCCGCCGAAC | GTGGTCCCTATGCGAGATGT | 238 |
| chr4:146264485-146266497 | p2 | (AT)6 | GACAATTTCAAACGCGGATT | GATGCTGCCCAAATTTCTGT | 181 |
| chr4:146309206-146311221 | p3 | (ATT)5 | AAACCAAAGGGGGTTTATCG | AGTAAAACGTTGGCCACCAC | 239 |
| chr4:146342933-146344948 | p3 | (TTC)5 | CAACCAAACGACCCCTAAGA | ATCCGCTTCTGCTGTTATCG | 245 |
| chr4:146462653-146464671 | p6 | (AAAAAG)3 | CTATGCAAGGATGCAGGGAT | CCGCTAATCAACCAACCACT | 220 |
| chr4:146519414-146521432 | p3 | (ACA)6 | TTTCAAATTGGTTTGCCACA | GTTTGCTGTCCGTATGCTCA | 134 |
| chr4:146519414-146521432 | p3 | (ACA)6 | TTTCAAATTGGTTTGCCACA | GTTTGCTGTCCGTATGCTCA | 134 |
| chr4:146536139-146538160 | p3 | (TCT)7 | TGCCATCTGTTTTGTCTTGC | TTCGAAAACGCCATTTTTCT | 287 |
| chr4:146599502-146601536 | p2 | (AT)6 | AGGACAATCATCCTCATGCC | AATTACACTCGTGATCCGCC | 221 |
| chr4:146599538-146601576 | p2 | (AT)6 | AGGACAATCATCCTCATGCC | AATTACACTCGTGATCCGCC | 185 |
| chr4:146670168-146672183 | p5 | (ATTGT)3 | CCTGTTTGGATGGGCTAAAA | TTTGGCTCCACCTTCGTAAC | 293 |
| chr4:146713999-146716017 | p3 | (TAA)6 | TTCGCTTGGCCAAAATAATC | AAGTCTGGAGAACGTGCCAT | 222 |
| chr4:146713999-146716017 | p5 | (AAACC)3 | TTCGCTTGGCCAAAATAATC | AAGTCTGGAGAACGTGCCAT | 222 |
| chr4:146749455-146751491 | p2 | (TA)18 | CTTTCACAAATGCCCCAACT | ACTGAATTCCATCTTTGCGG | 209 |

| Marker ID | SSR type | SSR motif | FORWARD PRIMER (5'-3') | REVERSE PRIMER (5'-3') | size (bp) |
|---|---|---|---|---|---|
| chr4:146758385-146760397 | p2 | (GT)6 | ACGCTCTCCCAACTGAGCTA | CCGACCAGCACACCTTAAAT | 299 |
| chr4:146771340-146773360 | p2 | (TA)10 | TGCGCTATAAGGCGAAACTT | TGGGCGAATACTTGAGGTTC | 274 |
| chr4:146783849-146785867 | p3 | (CGG)9 | TTGATGATGCCACGGTTTTA | GTTCAGCCGATTGCATTTTT | 243 |
| chr4:146783849-146785867 | p3 | (TGT)6 | TTGATGATGCCACGGTTTTA | GTTCAGCCGATTGCATTTTT | 243 |
| chr4:146836400-146838424 | p2 | (AT)12 | TGGTGACATGTGATCGGTTT | AAGCAAAGTTGGGGAGTGAA | 164 |
| chr4:146861486-146863502 | p4 | (TTGA)4 | AGGTTCGAGCAAACTGGCTA | CCACATCCACCACCCTCTAC | 270 |
| chr4:146911630-146913660 | p5 | (AAACC)3 | GGGACTGCTTCAGCTTCATC | ACAGCGATGTAGGCTTCGTT | 154 |
| chr4:146957082-146959097 | p5 | (TTTTC)3 | ATGAACCCATCATCTGGAGC | GATTTTGACACGGCCATCTT | 124 |
| chr4:146998846-147000876 | p5 | (AAACC)3 | GGGACTGCTTCAGCTTCATC | ACAGCGATGTAGGCTTCGTT | 154 |
| chr4:147044298-147046313 | p5 | (TTTTC)3 | ATGAACCCATCATCTGGAGC | GATTTTGACACGGCCATCTT | 124 |
| chr4:147112769-147114799 | p2 | (TG)11 | CGGACCCGACATACTCTTGT | TTAGGCAATCCTCCTCCCTT | 195 |
| chr4:147112804-147114820 | p2 | (TG)11 | CGGACCCGACATACTCTTGT | TTAGGCAATCCTCCTCCCTT | 160 |
| chr4:147112823-147114839 | p2 | (TG)11 | CGGACCCGACATACTCTTGT | TTAGGCAATCCTCCTCCCTT | 141 |
| chr4:147112861-147114873 | p2 | (TG)11 | CGGACCCGACATACTCTTGT | TTAGGCAATCCTCCTCCCTT | 103 |
| chr4:147146722-147148743 | p3 | (ATT)7 | TTGTCGGATATCGAGCTGTG | AATGATTCCATGCCTCAAGC | 182 |
| chr4:147208116-147210134 | p3 | (TGA)5 | TATCGTCCAACAGCACCAAA | TTCTTGTCCCTGTCCTGTCC | 258 |
| chr4:147208116-147210134 | p3 | (ACA)5 | TATCGTCCAACAGCACCAAA | TTCTTGTCCCTGTCCTGTCC | 258 |
| chr4:147208173-147210188 | p3 | (TGA)5 | TATCGTCCAACAGCACCAAA | TTCTTGTCCCTGTCCTGTCC | 201 |
| chr4:147208173-147210188 | p3 | (ACA)5 | TATCGTCCAACAGCACCAAA | TTCTTGTCCCTGTCCTGTCC | 201 |
| chr4:147211986-147214001 | p5 | (GATTC)3 | TGGGAACTTGGAATCGAAAC | TAAGAAAACCGACGGACTGC | 160 |
| chr4:147211986-147214001 | p3 | (AAC)6 | TGGGAACTTGGAATCGAAAC | TAAGAAAACCGACGGACTGC | 160 |
| chr4:147313449-147315464 | p3 | (AAT)5 | TCTCGAGAGTCGATTTGGCT | AGGTGGGACATATCGGTTGA | 289 |
| chr4:147322424-147324438 | p2 | (AG)7 | CCCTAAAACTGGACAGCCAA | TTGGGAACAGAGGATTCCAG | 177 |
| chr4:147326194-147328212 | p3 | (ACA)6 | ATCCCAAGAAGAGCCGAAGT | CCCACTTGTGGGATTACACC | 192 |
| chr4:147485365-147487380 | p5 | (AACAC)3 | GGGAGCAACTTGAAGGTGAA | ATATGAGAGGCGACCATTGC | 134 |
| chr4:147493756-147495770 | p2 | (TA)7 | TAGCTGTCCATGATGTGGGA | TGGTTTGTGGTAGACTGGCA | 101 |
| chr4:147498107-147500137 | p2 | (AT)15 | CTCAAATGGCACCTGAGGAT | ATCCAGGCAGATTCACCATC | 282 |
| chr4:147505956-147507968 | p6 | (ATTTTT)3 | TCCCAAACAATCATTGCATC | TGGAGTTGAGGAGAAGGGAA | 117 |
| chr4:147505956-147507968 | p2 | (TA)6 | TCCCAAACAATCATTGCATC | TGGAGTTGAGGAGAAGGGAA | 117 |
| chr4:147505956-147507968 | p2 | (AT)10 | TCCCAAACAATCATTGCATC | TGGAGTTGAGGAGAAGGGAA | 117 |
| chr4:147507924-147509938 | p2 | (TA)7 | CGGCCTAACCATTGTGACTT | TTTGAATTTGGTAGGCTCGG | 241 |
| chr4:147522862-147524900 | p2 | (AT)19 | GCTGGGCATAAATACCGAAA | GTGCGTTTGGTTCTTCGATT | 273 |
| chr4:147602120-147604135 | p3 | (TGA)5 | GTATTGCATCGAGGGCATCT | GAAGCAATCAACGAGGAAGC | 117 |
| chr4:147665717-147667757 | p2 | (TA)20 | CATTGATCGCTTTGTTGTGG | ACAAGGGCACCTCTAACCCT | 202 |
| chr4:147710251-147712273 | p2 | (TA)11 | ATGGCAATGTCATGCTCAAA | TGTCCGTTTAACAGCCATCA | 133 |
| chr4:147738979-147740994 | p5 | (CGACA)3 | CTTTTGGGAGGACTTAGGGC | CTCCGCCATGACCCTAGATA | 182 |

| Marker ID | SSR type | SSR motif | FORWARD PRIMER (5'-3') | REVERSE PRIMER (5'-3') | size (bp) |
|---|---|---|---|---|---|
| chr4:147789375-147791397 | p2 | (TA)16 | GACGATGATGATGTTGACGC | AGATATCATACCCGGGCACA | 246 |
| chr4:147789375-147791397 | p2 | (TA)11 | GACGATGATGATGTTGACGC | AGATATCATACCCGGGCACA | 246 |
| chr4:147795614-147797678 | p2 | (AT)6 | CCTAGGACGGAACTGCTCTG | TAGCCTTGTCGGCTCGTATT | 158 |
| chr4:147795614-147797678 | p2 | (AT)32 | CCTAGGACGGAACTGCTCTG | TAGCCTTGTCGGCTCGTATT | 158 |
| chr4:147840155-147842205 | p2 | (TA)25 | GATGAAACTCCCGTCTGCAT | AAAAAGAGAGGCGATGATGG | 234 |
| chr4:147840155-147842205 | p2 | (TA)21 | GATGAAACTCCCGTCTGCAT | AAAAAGAGAGGCGATGATGG | 234 |
| chr4:147886586-147888598 | p2 | (AT)6 | CAGGCTCTAGTTCAGGCACC | CATCTCCAAACTGCCTCCAT | 137 |
| chr4:147911705-147913717 | p2 | (AT)6 | CAGGCTCTAGTTCAGGCACC | CATCTCCAAACTGCCTCCAT | 137 |
| chr4:147973720-147975740 | p2 | (AT)8 | GCCGACAAGGCTAACAGAAG | TTGGAGTGGTCCTTTGTTCC | 257 |
| chr4:147973720-147975740 | p2 | (AT)10 | GCCGACAAGGCTAACAGAAG | TTGGAGTGGTCCTTTGTTCC | 257 |
| chr4:148061648-148063676 | p2 | (TA)29 | AAACCATGGTCTCGGAACTG | TTGACTAAGGGTACGACGGG | 111 |
| chr4:148061648-148063676 | p2 | (TA)14 | AAACCATGGTCTCGGAACTG | TTGACTAAGGGTACGACGGG | 111 |
| chr4:148081937-148083953 | p2 | (TA)7 | TCTTTGGGCCACTTTAATGC | CCGCCTCAAGGATATTACGA | 102 |
| chr4:148081937-148083953 | p2 | (AT)8 | TCTTTGGGCCACTTTAATGC | CCGCCTCAAGGATATTACGA | 102 |
| chr4:148091632-148093646 | p2 | (TA)7 | AATGAATGGACGAAGTTGGG | CGACAATTAGCACACATGGC | 212 |
| chr4:148132382-148134412 | p3 | (ATA)7 | TCACCAACAAGAGCATCAGC | GGAGAGAGGTTTCCCTTTCG | 248 |
| chr4:148132382-148134412 | p2 | (AT)15 | TCACCAACAAGAGCATCAGC | GGAGAGAGGTTTCCCTTTCG | 248 |
| chr4:148156652-148158667 | p5 | (TAATT)3 | TGTTAAAAGTGCCTACGGGC | TACACGTACCTGGCCCTACC | 228 |
| chr4:148213686-148215701 | p5 | (TAATT)3 | TGTTAAAAGTGCCTACGGGC | TACACGTACCTGGCCCTACC | 228 |
| chr5:120334-122346 | p2 | (TA)6 | AATGCCAAATTCCAGAATGC | GGTTTACGCCCATGAACAGT | 287 |
| chr5:142835-144850 | p5 | (TTTTA)3 | GGTGTGAGTTAGGCCCAAGA | GGTTGCAAATCGGTGAAGAT | 184 |
| chr5:191404-193416 | p2 | (CT)6 | TCCCCCAAATTTCTTCTCCT | GATTGTCACCGCCAGTTCTT | 180 |
| chr5:191404-193416 | p2 | (CT)6 | TCCCCCAAATTTCTTCTCCT | GATTGTCACCGCCAGTTCTT | 180 |
| chr5:196918-198946 | p2 | (AT)14 | GAATGCTGACTCCCTACCCA | GGAGAGACTGATGGGGTCAA | 268 |
| chr5:262124-264138 | p5 | (ACCGG)3 | GCAGTTTTGCGGTAGTAGCC | GGCAAAACCCACTGCTCTTA | 238 |
| chr5:262124-264138 | p2 | (AT)11 | GCAGTTTTGCGGTAGTAGCC | GGCAAAACCCACTGCTCTTA | 238 |
| chr5:262188-264206 | p5 | (ACCGG)3 | GCAGTTTTGCGGTAGTAGCC | GGCAAAACCCACTGCTCTTA | 174 |
| chr5:262188-264206 | p2 | (AT)11 | GCAGTTTTGCGGTAGTAGCC | GGCAAAACCCACTGCTCTTA | 174 |
| chr5:285222-287238 | p2 | (AT)8 | CAAGTTACTCCCTCCGACCA | ATACCACCCCAAAAGAAGGG | 206 |
| chr5:434358-436376 | p2 | (AT)9 | TGGGTTGCTTAATCGGGTAG | TCATCGTTTCATTTTCGGGT | 234 |
| chr5:493011-495026 | p5 | (ATTTT)3 | TTGCTTGCATGTGTTTGGAT | CGATGATAACCGGCCTAAAA | 136 |
| chr5:533645-535661 | p2 | (TG)8 | ATGATCTCTTCCCATTTGCG | TAAAAAGGCCCGAGACACAC | 277 |
| chr5:542031-544049 | p3 | (CAA)6 | CGATTTTAATGCCCTGGAAA | TATTTTGGGCGTTTTTGCTC | 197 |
| chr5:546903-548915 | p5 | (AAAAT)3 | CCCGCTCAGGCATAAAATAA | TGCCCCATAAGGCATAGTTC | 141 |
| chr5:546903-548915 | p2 | (TA)6 | CCCGCTCAGGCATAAAATAA | TGCCCCATAAGGCATAGTTC | 141 |
| chr5:734312-736330 | p2 | (TC)6 | CCATATTTCCAGCCCTTCAA | GTGGGCAGATTTGGAGTTGT | 161 |

| Marker ID | SSR type | SSR motif | FORWARD PRIMER (5'-3') | REVERSE PRIMER (5'-3') | size (bp) |
|---|---|---|---|---|---|
| chr5:734312-736330 | p6 | (CGGAAC)3 | CCATATTTCCAGCCCTTCAA | GTGGGCAGATTTGGAGTTGT | 161 |
| chr5:734312-736330 | p3 | (GAT)5 | CCATATTTCCAGCCCTTCAA | GTGGGCAGATTTGGAGTTGT | 161 |
| chr5:813839-815859 | p2 | (TA)10 | GATTAATTGTCGCGTGCGTA | AAATAAATGACATGGCGGGA | 233 |
| chr5:820212-822224 | p2 | (AG)6 | AAACGTGGAAACTCCGTTTG | ATTGTCTGCAGAGTCCAGGC | 126 |
| chr5:887419-889445 | p2 | (TA)13 | TTTATTATGGTCCCGTCCGA | GCGAACAATGCAGAAGATCA | 186 |
| chr5:896561-898575 | p2 | (TA)11 | GCGCATTTAGCCAAGGAATA | GCCTTCCTCCGCTTCTAAGT | 265 |
| chr5:896561-898575 | p2 | (TA)7 | GCGCATTTAGCCAAGGAATA | GCCTTCCTCCGCTTCTAAGT | 265 |
| chr5:896561-898575 | p2 | (AT)9 | GCGCATTTAGCCAAGGAATA | GCCTTCCTCCGCTTCTAAGT | 265 |
| chr5:896561-898575 | p2 | (TA)8 | GCGCATTTAGCCAAGGAATA | GCCTTCCTCCGCTTCTAAGT | 265 |
| chr5:896695-898713 | p2 | (TA)11 | GCGCATTTAGCCAAGGAATA | GCCTTCCTCCGCTTCTAAGT | 131 |
| chr5:896695-898713 | p2 | (TA)7 | GCGCATTTAGCCAAGGAATA | GCCTTCCTCCGCTTCTAAGT | 131 |
| chr5:896695-898713 | p2 | (AT)9 | GCGCATTTAGCCAAGGAATA | GCCTTCCTCCGCTTCTAAGT | 131 |
| chr5:896695-898713 | p2 | (TA)8 | GCGCATTTAGCCAAGGAATA | GCCTTCCTCCGCTTCTAAGT | 131 |
| chr5:994742-996780 | p2 | (TA)19 | TGGAATCGTGTTTGCTGGTA | TCCCAACTTCCCATTTTTGA | 146 |
| chr5:1052413-1054435 | p2 | (TA)11 | AGGAACTTTGCAGGTCGAAA | TTAATGGCGCTTAATGAGGG | 282 |
| chr5:1116394-1118424 | p3 | (TAT)10 | GTGAAGCATTATCGGCCATT | GGCTCATAGGAGTAAGGCCC | 295 |
| chr5:1241641-1243677 | p2 | (AG)18 | CCGAATCTGGGCGTAGTTTA | TCATGTGTCTTGCTTCAGGC | 168 |
| chr5:1305607-1307619 | p2 | (TC)6 | GGGAACAGTTGCTGTTGGAT | ACAATGTGGCAGTGAAAGCA | 181 |
| chr5:1357536-1359554 | p6 | (GGGGTG)3 | CTGCTTTGATGTTGGGAGGT | CATCCGCATTAAAAAGGGAA | 240 |
| chr5:1377326-1379341 | p3 | (GAA)5 | CCCTCTAAATCCGCCCTATC | TGGTGCCAAATAACACTCCA | 153 |
| chr5:1464714-1466732 | p6 | (CAACTC)3 | TGGAAGGCTAATAAAGGCGA | CATGATACAAGGGGAAGCGT | 154 |
| chr5:1478453-1480471 | p6 | (CAACTC)3 | TGGAAGGCTAATAAAGGCGA | CATGATACAAGGGGAAGCGT | 154 |
| chr5:1515738-1517752 | p2 | (AC)7 | GGGTACAATTCGTGCAGCTT | AAATCTTGGATCGCCTCTGA | 168 |
| chr5:1583351-1585372 | p3 | (TGT)7 | TGACCGCTGTTGAATACCAA | GAGGATTTCAACTGCACGGT | 180 |
| chr5:1590317-1592333 | p2 | (AT)15 | TACGAAGCAGCTCACTGGAA | TAGTGAATTTACCCGCGCTT | 118 |
| chr5:1590317-1592333 | p2 | (TA)8 | TACGAAGCAGCTCACTGGAA | TAGTGAATTTACCCGCGCTT | 118 |
| chr5:1591138-1593150 | p2 | (TA)8 | TAATGTCCAAAAGGGTGGCT | TTTTTCCCCATGAGTTTGGA | 223 |
| chr5:1591162-1593180 | p2 | (TA)8 | TAATGTCCAAAAGGGTGGCT | TTTTTCCCCATGAGTTTGGA | 199 |
| chr5:1618956-1620972 | p2 | (AG)8 | TCAGTCGAGTCTACCCGCTT | AAAAGTGTTGCCAAACCCAG | 141 |
| chr5:1655560-1657572 | p2 | (TC)6 | CCGGAATTCTACCTTCCACA | ACCAACATACCCAGCTCCAC | 120 |
| chr5:1722558-1724576 | p3 | (TTG)6 | AGATGAAAGTGGCGGAAATG | CGCATCTTGTCTTCCACTGA | 298 |
| chr5:1722558-1724576 | p5 | (TTGGT)3 | AGATGAAAGTGGCGGAAATG | CGCATCTTGTCTTCCACTGA | 298 |
| chr5:1722677-1724692 | p3 | (TTG)6 | AGATGAAAGTGGCGGAAATG | CGCATCTTGTCTTCCACTGA | 179 |
| chr5:1722677-1724692 | p5 | (TTGGT)3 | AGATGAAAGTGGCGGAAATG | CGCATCTTGTCTTCCACTGA | 179 |
| chr5:1745919-1747937 | p3 | (TTG)6 | AGATGAAAGTGGCGGAAATG | CGCATCTTGTCTTCCACTGA | 298 |
| chr5:1745919-1747937 | p5 | (TTGGT)3 | AGATGAAAGTGGCGGAAATG | CGCATCTTGTCTTCCACTGA | 298 |

| Marker ID | SSR type | SSR motif | FORWARD PRIMER (5'-3') | REVERSE PRIMER (5'-3') | size (bp) |
|---|---|---|---|---|---|
| chr5:1746038-1748053 | p3 | (TTG)6 | AGATGAAAGTGGCGGAAATG | CGCATCTTGTCTTCCACTGA | 179 |
| chr5:1746038-1748053 | p5 | (TTGGT)3 | AGATGAAAGTGGCGGAAATG | CGCATCTTGTCTTCCACTGA | 179 |
| chr5:1909235-1911253 | p5 | (TTTTA)3 | ACCGCCTAGCATTGAGCTAC | AAGCTGGCCAGTCAAACACT | 250 |
| chr5:1992396-1994414 | p6 | (TTTTGC)3 | GCCATTGTTGAGGAACCCTA | ATGGCGTGCCAATTACTCTC | 272 |
| chr5:1992396-1994414 | p6 | (AAAAAG)3 | GCCATTGTTGAGGAACCCTA | ATGGCGTGCCAATTACTCTC | 272 |
| chr5:1992396-1994414 | p5 | (TAAAA)3 | GCCATTGTTGAGGAACCCTA | ATGGCGTGCCAATTACTCTC | 272 |
| chr5:2017219-2019234 | p5 | (CCCCA)3 | ATACCGAACAGTGGCTGGAC | TGGCATGTGAGCTCTTGAAC | 194 |
| chr5:2032361-2034376 | p5 | (TTTTA)3 | ATGTGGCAATGAATCCATCA | AAAAATGGCCCCCATAATTC | 227 |
| chr5:2032361-2034376 | p5 | (TTTAT)3 | ATGTGGCAATGAATCCATCA | AAAAATGGCCCCCATAATTC | 227 |
| chr5:2040250-2042274 | p2 | (TA)12 | ACTGAAAATGCCTCTGTGGG | TACGCACATACATCAGGCGT | 263 |
| chr5:2165981-2168032 | p3 | (TAT)17 | CTCCGAATGCGGGTATTTTA | TTCGCTGCCGAAGAAATAGT | 200 |
| chr5:2202326-2204341 | p5 | (TTTTG)3 | AGAAAGGAAAGCCTGAAGCC | TCTTGGATCGTACATTGGCA | 146 |
| chr5:2203844-2205878 | p2 | (AT)17 | TGCTTTGAACTGCTTGTGCT | GGGAGGGTAAAGTGTACGCA | 141 |
| chr5:2203844-2205878 | p5 | (AAATT)3 | TGCTTTGAACTGCTTGTGCT | GGGAGGGTAAAGTGTACGCA | 141 |
| chr5:2215940-2217964 | p2 | (TA)12 | TGTTCACGTGATTGGGCTTA | TGTGCAATCCATTGGTCTGT | 195 |
| chr5:2215940-2217964 | p5 | (TTTTA)3 | TGTTCACGTGATTGGGCTTA | TGTGCAATCCATTGGTCTGT | 195 |
| chr5:2216205-2218220 | p2 | (TA)12 | ACAGACCAATGGATTGCACA | GCATTGCTATGCCTGATTGA | 157 |
| chr5:2216205-2218220 | p5 | (TTTTA)3 | ACAGACCAATGGATTGCACA | GCATTGCTATGCCTGATTGA | 157 |
| chr5:2223718-2225733 | p3 | (ATT)5 | AGGGACTCGTTTTCTTCCGT | TTGGAAATTGCTGCACTCTG | 159 |
| chr5:2242952-2244970 | p2 | (TA)9 | GGGTAGATGTAAGCCCGTCA | CAACCCTTTCCTCTTAGCCC | 236 |
| chr5:2264564-2266576 | p2 | (TA)6 | TTTTAAAGGGACAACGGACG | GCCCCATACTAAACGACGAC | 206 |
| chr5:2264564-2266576 | p2 | (TA)6 | TTTTAAAGGGACAACGGACG | GCCCCATACTAAACGACGAC | 206 |
| chr5:2269975-2271993 | p3 | (GAG)5 | GACGTTGCCCCAGTATGTCT | ATAAGCACAACCCCGTCAAG | 198 |
| chr5:2269975-2271993 | p2 | (CT)9 | GACGTTGCCCCAGTATGTCT | ATAAGCACAACCCCGTCAAG | 198 |
| chr5:2314981-2316993 | p2 | (CT)6 | ATGCCCGCAGGTACAGATAG | GCTGCCACTACAGACGAACA | 289 |
| chr5:2335646-2337668 | p2 | (AT)11 | GCCCAGCAATACAAGAGCAT | TTCAAATTCCCGCCTATGTC | 201 |
| chr5:2343051-2345066 | p5 | (GTTTA)3 | GGGGTAATTGTGGCCTTCTT | AACCAGCCACGTAATTTTGC | 207 |
| chr5:2343051-2345066 | p5 | (GTTTA)3 | GGGGTAATTGTGGCCTTCTT | AACCAGCCACGTAATTTTGC | 207 |
| chr5:2343051-2345066 | p3 | (ACA)5 | GGGGTAATTGTGGCCTTCTT | AACCAGCCACGTAATTTTGC | 207 |
| chr5:2358876-2360888 | p2 | (TA)6 | CGTGTGACGAAAGTGAAGGA | CGCCTAATCCGAATTTTCAA | 184 |
| chr5:2370945-2373029 | p2 | (TA)10 | GAAATCGGACTGGCTGTGAT | CGGGAGGGAGTAATGAACAA | 277 |
| chr5:2372743-2374758 | p5 | (ATTCC)3 | CATTTGCAAACAATCATGCC | TCGACCTCCTACAAGCGAGT | 281 |
| chr5:2420137-2422152 | p3 | (TAT)5 | CAGCTCTTCTGGTCCACTCC | AAGGATCTAAACAAGCCGCA | 233 |
| chr5:2430774-2432789 | p3 | (AAG)10 | GTCGTAGCCAGGCTTTTCAG | TACAAGCCCCTTCCCTTTTT | 175 |
| chr5:2430774-2432789 | p5 | (AACCG)3 | GTCGTAGCCAGGCTTTTCAG | TACAAGCCCCTTCCCTTTTT | 175 |
| chr5:2533873-2535887 | p2 | (AT)7 | ACGTTCGGTTCATTCTTTCG | CGACAACTGCGGAGAGTACA | 136 |

| Marker ID | SSR type | SSR motif | FORWARD PRIMER (5'-3') | REVERSE PRIMER (5'-3') | size (bp) |
|---|---|---|---|---|---|
| chr5:2536066-2538096 | p3 | (CAA)10 | CTTTCGGCCAACATCGTTAT | CCCATGTGGGAAAGAAGAAA | 282 |
| chr5:2538025-2540041 | p3 | (TTC)8 | ATTCGTGTGCTTTCCATTCC | TTGTCAGGCAATGGTTGGTA | 204 |
| chr5:2538025-2540041 | p2 | (AT)11 | ATTCGTGTGCTTTCCATTCC | TTGTCAGGCAATGGTTGGTA | 204 |
| chr5:2538025-2540041 | p2 | (TA)8 | ATTCGTGTGCTTTCCATTCC | TTGTCAGGCAATGGTTGGTA | 204 |
| chr5:2593737-2595761 | p2 | (CT)12 | GCGTTTTTCCCTTTTTGACA | TGCGCTCCATGATATGGTTA | 159 |
| chr5:2626097-2628109 | p2 | (TA)6 | CGTTGATTACCGGGAGAAAA | TAGGTGGCAACTTCGAATCC | 142 |
| chr5:2639957-2641972 | p3 | (GAG)5 | TCTTGTGTGGCTGTCTCCTG | CAATAGCCGAAGTCAGAGGC | 241 |
| chr5:2639957-2641972 | p3 | (AGA)5 | TCTTGTGTGGCTGTCTCCTG | CAATAGCCGAAGTCAGAGGC | 241 |
| chr5:2679531-2681545 | p2 | (TA)7 | GTCGACCCACTTTTGATGCT | ACTTAAACGGACCAAATGCG | 174 |
| chr5:2748833-2750859 | p2 | (AT)13 | AATGTCTCCGGTTGTCTTCG | ATATGCATAAGCCAGACGGC | 140 |
| chr5:2816364-2818378 | p2 | (AT)7 | AGTTATGTGGTTCGCTCGGT | TGACATGGCGGAATAACAAA | 185 |
| chr5:2871552-2873567 | p2 | (TA)6 | CAGGTACCGACGACAGGTTT | CCGTACCCATGAACGAGAAT | 161 |
| chr5:2871552-2873567 | p5 | (AGAAA)3 | CAGGTACCGACGACAGGTTT | CCGTACCCATGAACGAGAAT | 161 |
| chr5:2918341-2920353 | p2 | (TA)6 | AGGCTCAATGATTCCACCAC | CCGCACTCTTGCTGATTACA | 254 |
| chr5:2946821-2948836 | p3 | (TGA)5 | AAACCGGTAAAACCGGAATC | GGGGGATGGGTTTAAAGTGT | 218 |
| chr5:3025402-3027420 | p2 | (TA)9 | AATTTGCTGCTGGGGTACTG | CTGCACAAGATTTGCCTTGA | 253 |
| chr5:3025402-3027420 | p2 | (AT)7 | AATTTGCTGCTGGGGTACTG | CTGCACAAGATTTGCCTTGA | 253 |
| chr5:3110519-3112533 | p6 | (CCGGTT)4 | GCGAAAACGACGACGTAAAT | TACAACCGGTAAACCGGAAC | 222 |
| chr5:3110633-3112647 | p6 | (CCGGTT)4 | GCGAAAACGACGACGTAAAT | TACAACCGGTAAACCGGAAC | 108 |
| chr5:3168856-3170886 | p2 | (TA)6 | GACTCATGCTTCATCGTCCA | AGTTTGGATGTGGCGAGTTC | 297 |
| chr5:3169810-3171822 | p2 | (TA)6 | GGGTTTAGACCCATGCTTCA | CGTGAGTTTGGATGTGGTGA | 194 |
| chr5:3181508-3183523 | p5 | (AAACC)3 | GAAACTGCTACTTGGCCAGC | CCCGAGAAAACCTGAGATTG | 184 |
| chr5:3194344-3196368 | p2 | (TA)12 | AGTGCATGTAGGGTTGAGGG | GGAGAATGACAAAGCGCAAT | 114 |
| chr5:3194344-3196368 | p2 | (TA)13 | AGTGCATGTAGGGTTGAGGG | GGAGAATGACAAAGCGCAAT | 114 |
| chr5:3229367-3231379 | p2 | (AG)6 | TTTGGTTGATGAAGTGGGGT | CTTACAGCCAAGATGCGTCA | 138 |
| chr5:3229367-3231379 | p2 | (TA)9 | TTTGGTTGATGAAGTGGGGT | CTTACAGCCAAGATGCGTCA | 138 |
| chr5:3237437-3239449 | p6 | (TTATAT)4 | GCGGAAGTTGGAGTTGTGTT | GATAGTGGAGCGCTCTTTCG | 159 |
| chr5:3237437-3239449 | p2 | (TG)6 | GCGGAAGTTGGAGTTGTGTT | GATAGTGGAGCGCTCTTTCG | 159 |
| chr5:3274962-3276980 | p5 | (CCACC)3 | CGCAACCAAAGAGTTGATGA | GGTTAAGCGTTGCATGGAGT | 163 |
| chr5:3274962-3276980 | p6 | (ACCCCC)3 | CGCAACCAAAGAGTTGATGA | GGTTAAGCGTTGCATGGAGT | 163 |
| chr5:3387931-3389946 | p2 | (GA)6 | GCCCACACGTCAAGATTTTT | TCGTTTGGAGGACTTCCACT | 189 |
| chr5:3387931-3389946 | p5 | (TACAA)3 | GCCCACACGTCAAGATTTTT | TCGTTTGGAGGACTTCCACT | 189 |
| chr5:3439557-3441575 | p3 | (TTG)6 | GCCACCAAAACTTAAAGGCA | CATTCTCATTTCCGCCACTT | 200 |
| chr5:3501081-3503099 | p3 | (AAT)6 | AATTCTCATGTCCAAACGGG | ATCGTGCCCAGTCAAATAGG | 100 |
| chr5:3565756-3567771 | p5 | (TTGTA)3 | GCTGACATTCGGCAAGTGTA | CCAGTGGTTTTGTGCATGTC | 282 |
| chr5:3565756-3567771 | p5 | (GGGGT)3 | GCTGACATTCGGCAAGTGTA | CCAGTGGTTTTGTGCATGTC | 282 |

| Marker ID | SSR type | SSR motif | FORWARD PRIMER (5'-3') | REVERSE PRIMER (5'-3') | size (bp) |
|---|---|---|---|---|---|
| chr5:3565756-3567771 | p2 | (AT)7 | GCTGACATTCGGCAAGTGTA | CCAGTGGTTTTGTGCATGTC | 282 |
| chr5:3566497-3568511 | p5 | (GGGGT)3 | GGTGACATGAGTGGTTGTGG | TGGTGGTTTCTCATGGTTCA | 211 |
| chr5:3566497-3568511 | p2 | (AT)7 | GGTGACATGAGTGGTTGTGG | TGGTGGTTTCTCATGGTTCA | 211 |
| chr5:3657982-3660006 | p3 | (TCC)5 | AAATGCAAGAAATCATCGGC | TCCGGAGCCAAAATAGCTTA | 105 |
| chr5:3679769-3681785 | p5 | (TTGGT)3 | GGTTGAAAACCCCGAATCTT | GTGTGGTCCAAAACAAGCCT | 104 |
| chr5:3679769-3681785 | p4 | (TTAA)4 | GGTTGAAAACCCCGAATCTT | GTGTGGTCCAAAACAAGCCT | 104 |
| chr5:3916839-3918867 | p5 | (TTATT)3 | TCAAACAATGCCCACTTCAA | TCCTCCTCCTCCTCCTTCTC | 265 |
| chr5:3916839-3918867 | p3 | (TTG)5 | TCAAACAATGCCCACTTCAA | TCCTCCTCCTCCTCCTTCTC | 265 |
| chr5:3916898-3918912 | p5 | (TTATT)3 | TCAAACAATGCCCACTTCAA | TCCTCCTCCTCCTCCTTCTC | 206 |
| chr5:3916898-3918912 | p3 | (TTG)5 | TCAAACAATGCCCACTTCAA | TCCTCCTCCTCCTCCTTCTC | 206 |
| chr5:3967183-3969217 | p2 | (AT)17 | ATTTCACAGCTTCCCATTGC | TGAGCAAAGAGATGTGTCCG | 286 |
| chr5:3967183-3969217 | p6 | (CCGGTT)4 | ATTTCACAGCTTCCCATTGC | TGAGCAAAGAGATGTGTCCG | 286 |
| chr5:3986669-3988681 | p2 | (AC)6 | AGGTTGTGACAAGTCCCCAG | ATGCCTCCAACAGTCTTGCT | 242 |
| chr5:4030432-4032454 | p2 | (AT)11 | TGAACATCCACCAAAGAGCA | CATAGCCACCATCATCACCA | 176 |
| chr5:4030670-4032688 | p2 | (AT)11 | TTTGGTGATGATGGTGGCTA | TCACAACCACTGCATTTTCC | 179 |
| chr5:4030670-4032688 | p2 | (AT)7 | TTTGGTGATGATGGTGGCTA | TCACAACCACTGCATTTTCC | 179 |
| chr5:4039380-4041392 | p2 | (AC)6 | AGGTTGTGACAAGTCCCCAG | ATGCCTCCAACAGTCTTGCT | 242 |
| chr5:4083143-4085165 | p2 | (AT)11 | TGAACATCCACCAAAGAGCA | CATAGCCACCATCATCACCA | 176 |
| chr5:4083381-4085399 | p2 | (AT)11 | TTTGGTGATGATGGTGGCTA | TCACAACCACTGCATTTTCC | 179 |
| chr5:4083381-4085399 | p2 | (AT)7 | TTTGGTGATGATGGTGGCTA | TCACAACCACTGCATTTTCC | 179 |
| chr5:4129629-4131641 | p5 | (AAACC)3 | AGCTTGAAGATTGGGCAAAA | AGGCTGGAGTTATGGGGATT | 295 |
| chr5:4165565-4167581 | p2 | (GA)8 | ACCCGGTTGTTACCTTCCTC | TTGGGTCCCATTTGATTGAT | 180 |
| chr5:4179098-4181113 | p5 | (CTTTT)3 | TTCATGCCCATTTTGTGAGA | CCTTCAAATTTTGGCCTTCA | 106 |
| chr5:4200502-4202517 | p5 | (ACCAA)3 | GGGGCAATACTTGCATTTTG | AGACCCTAGCCTTGGGGATA | 289 |
| chr5:4204041-4206053 | p2 | (AT)17 | TCTCGTGAATCCCTTTACGG | AAGTTTTGGTTGCGTTGCTC | 243 |
| chr5:4204076-4206090 | p2 | (AT)17 | TCTCGTGAATCCCTTTACGG | TCAATTTCTTCGTCTTCGCC | 208 |
| chr5:4284254-4286266 | p2 | (CT)6 | AGACGGCCAATTGTATCGAC | ACTAAAGCTTCTGCACCGGA | 264 |
| chr5:4286612-4288627 | p5 | (TTACT)4 | TCAGCAAGTGTTCATGGGAA | GCACCAACATTGTGTATCGC | 206 |
| chr5:4286612-4288627 | p5 | (TTTCT)3 | TCAGCAAGTGTTCATGGGAA | GCACCAACATTGTGTATCGC | 206 |
| chr5:4295940-4297960 | p2 | (CT)10 | TTCTTTGTGCAGGACTTCCC | GCACGATTTTGGGATAGGAA | 238 |
| chr5:4315657-4317675 | p2 | (GA)9 | TCTGAGGGTTTTTGTCTGGC | ACGAAGAAAAGGAGTTCGCA | 158 |
| chr5:4326319-4328397 | p2 | (TA)16 | CACAGGCCCAAAGAACAGAT | TAACAGACGACGAAGCATGG | 236 |
| chr5:4393325-4395337 | p2 | (AT)6 | TCGCTTCATCTCCTCTGGTT | TGGGAACCTGCTTACAAAGG | 182 |
| chr5:4428967-4430982 | p3 | (AAC)5 | CATGCAACCATTTTCACAGC | AAAGCGGATTGGATGTTCAC | 274 |
| chr5:4434310-4436325 | p5 | (TTTTA)3 | ACTTCCTTATGTGCTCGCGT | TCTAGCGAGGATTCAATGGC | 144 |
| chr5:4475175-4477193 | p6 | (AAAAAT)3 | AGGGGTGTAATGCATGTGGT | TCCTTCTTGCAACCTCTGCT | 100 |

| Marker ID | SSR type | SSR motif | FORWARD PRIMER (5'-3') | REVERSE PRIMER (5'-3') | size (bp) |
|---|---|---|---|---|---|
| chr5:4475608-4477658 | p6 | (AAAAAT)3 | AGATAGCTGAATGAGCCCCA | TGTAGTTTTGGGCTTTTGGC | 163 |
| chr5:4506168-4508180 | p2 | (AT)6 | GACAATTTCAAACGCGGATT | GATGCTGCCCAAATTTCTGT | 216 |
| chr5:4519500-4521516 | p2 | (AT)7 | GGAATCCACCACATCCAAAC | TGCTCCCTCTAATGGGTTGA | 161 |
| chr5:4519500-4521516 | p2 | (TA)8 | GGAATCCACCACATCCAAAC | TGCTCCCTCTAATGGGTTGA | 161 |
| chr5:4533020-4535032 | p2 | (TA)6 | AGTGGAACCCATCATCTCCA | ATATGGGCGCACTTTTTGTT | 271 |
| chr5:4533020-4535032 | p2 | (TA)6 | AGTGGAACCCATCATCTCCA | ATATGGGCGCACTTTTTGTT | 271 |
| chr5:4552514-4554529 | p3 | (TAA)5 | TGTGGTAGGGGTGTGTCAGA | CACCTACACCCCTTCATCGT | 177 |
| chr5:4554702-4556723 | p5 | (AAATT)3 | CACACATTCATTCGTGGGAC | GTCAAACATGACAACGTGCC | 118 |
| chr5:4554702-4556723 | p3 | (AAT)7 | CACACATTCATTCGTGGGAC | GTCAAACATGACAACGTGCC | 118 |
| chr5:4560875-4562887 | p2 | (TA)6 | TCCAAGAACTTGCGAATGTG | CAGCAATGCAGCATGAAGAT | 117 |
| chr5:4583008-4585024 | p2 | (GA)8 | AGCTCGGGTTAAGCTCACAA | GTCTCTTCGAAGCTCGTTGG | 159 |
| chr5:4592220-4594244 | p2 | (TA)12 | CCAAATGCGGAGAAAATGAT | CATGTGCACACAACATGGAA | 216 |
| chr5:4631683-4633713 | p5 | (TATCA)6 | GACCAAATGAGGGGTGAGAA | AGCAGCAACACCAAAAATCC | 141 |
| chr5:4636773-4638805 | p2 | (AT)15 | AAATGTGTCATCTCCAGGGG | CGTTGGATGTGGTGTTGTTC | 209 |
| chr5:4636808-4638822 | p2 | (AT)15 | AAATGTGTCATCTCCAGGGG | CACGTTGGATGTGGTGTTGT | 174 |
| chr5:4676844-4678859 | p3 | (TTG)5 | GCTTTTCTTGCATTTCCAGC | GTCGAGAAAAGCGTTGAAGC | 248 |
| chr5:4844283-4846301 | p2 | (AT)8 | TGGAGAAACTCGATTGCCTT | TCGATCAGTTGAACCAGCAG | 253 |
| chr5:4844283-4846301 | p3 | (GTT)6 | TGGAGAAACTCGATTGCCTT | TCGATCAGTTGAACCAGCAG | 253 |
| chr5:4887982-4890000 | p2 | (AT)8 | TGGAGAAACTCGATTGCCTT | TCGATCAGTTGAACCAGCAG | 253 |
| chr5:4887982-4890000 | p3 | (GTT)6 | TGGAGAAACTCGATTGCCTT | TCGATCAGTTGAACCAGCAG | 253 |
| chr5:5005325-5007349 | p6 | (AGTTGG)4 | TGAAGCAAGCTCCCAAAACT | CCCGTTGTACTCGAATTTGG | 190 |
| chr5:5097804-5099828 | p2 | (TG)7 | GTACCCGCTCCGATATGAAA | GACCAGTGCATTTGAAGGGT | 159 |
| chr5:5114017-5116038 | p3 | (AAG)7 | TGGGACCGGCATAAGTTTAG | CTTATGGGCAGACTTGGCAT | 149 |
| chr5:5123761-5125776 | p3 | (AAG)5 | TCCGAAAAGGAAGGGTTTTT | TCAAAGAAAATCAAGCCCGT | 299 |
| chr5:5167560-5169575 | p3 | (CAA)5 | AGGCACGACGACCAAAATAC | TTCATTGAATTGGTTTGGCA | 102 |
| chr5:5178951-5180965 | p3 | (TGA)6 | GTTTAGGCCATCCACAGAGC | TGACGTTAGAGGGGTTTTGG | 143 |
| chr5:5178969-5180981 | p3 | (TGA)6 | GTTTAGGCCATCCACAGAGC | TGACGTTAGAGGGGTTTTGG | 125 |
| chr5:5234583-5236599 | p2 | (AT)8 | GGTTTGGTTGGTCTTGTCGT | CTGTTTCTTGGTCCCCACAT | 177 |
| chr5:5261310-5263322 | p3 | (CAA)6 | GATAGGGGCCCAACCTAGAC | ATCCGCAATTGAAAAACTCG | 167 |
| chr5:5261310-5263322 | p2 | (AT)6 | GATAGGGGCCCAACCTAGAC | ATCCGCAATTGAAAAACTCG | 167 |
| chr5:5261310-5263322 | p2 | (AC)8 | GATAGGGGCCCAACCTAGAC | ATCCGCAATTGAAAAACTCG | 167 |
| chr5:5275887-5277901 | p2 | (AT)7 | GCTAATCCGCGAGAGACTTG | CTCTGGAAACGGCTAGATGC | 287 |
| chr5:5275887-5277901 | p2 | (TA)8 | GCTAATCCGCGAGAGACTTG | CTCTGGAAACGGCTAGATGC | 287 |
| chr5:5275887-5277901 | p2 | (AT)8 | GCTAATCCGCGAGAGACTTG | CTCTGGAAACGGCTAGATGC | 287 |
| chr5:5359590-5361606 | p2 | (AT)8 | TCACCCGTAATCCCATTGTT | TCTCCAAGTGCCAAGGATGT | 191 |
| chr5:5359590-5361606 | p2 | (AC)6 | TCACCCGTAATCCCATTGTT | TCTCCAAGTGCCAAGGATGT | 191 |

| Marker ID | SSR type | SSR motif | FORWARD PRIMER (5'-3') | REVERSE PRIMER (5'-3') | size (bp) |
|---|---|---|---|---|---|
| chr5:5390863-5392878 | p5 | (ATGAC)3 | ATGCGCCCAAATTTTTGTAG | CGCCAAGATTATGCCCTAAA | 195 |
| chr5:5397665-5399681 | p2 | (TA)8 | GGTCAGTCGGGCAAAGATAA | CTCCGCCACTAGTGATTGCT | 118 |
| chr5:5397665-5399681 | p2 | (TA)12 | GGTCAGTCGGGCAAAGATAA | CTCCGCCACTAGTGATTGCT | 118 |
| chr5:5397854-5399878 | p2 | (TA)8 | ACGAAGAAGTCAGGGGGTTC | CATCCAAACGGGTTGCTAAT | 141 |
| chr5:5397854-5399878 | p2 | (TA)12 | ACGAAGAAGTCAGGGGGTTC | CATCCAAACGGGTTGCTAAT | 141 |
| chr5:5397854-5399878 | p2 | (AT)9 | ACGAAGAAGTCAGGGGGTTC | CATCCAAACGGGTTGCTAAT | 141 |
| chr5:5465824-5467846 | p2 | (AT)11 | TTCCAACGGTATATTGGGGA | GGTCAAATAAACCCCAGGCT | 119 |
| chr5:5491097-5493112 | p5 | (ATTTT)3 | ACCTTGTCGGACAGCGTTAG | AATTTTGTCGGTTGTCCCAC | 282 |
| chr5:5528103-5530127 | p3 | (TGT)8 | CCACCAAAACTTAAGGGCAA | TTCCGCGACTTTCATCTTCT | 253 |
| chr5:5581465-5583480 | p5 | (ATTAT)3 | CATGATAGTTCAGCGAGCCA | CGTTCGATTTTTGCCATTTT | 127 |
| chr5:5581465-5583480 | p2 | (AT)20 | CATGATAGTTCAGCGAGCCA | CGTTCGATTTTTGCCATTTT | 127 |
| chr5:5581465-5583480 | p5 | (GGTTT)3 | CATGATAGTTCAGCGAGCCA | CGTTCGATTTTTGCCATTTT | 127 |
| chr5:5581827-5583867 | p5 | (ATTAT)3 | CCCTAAAACGACGATCCAAA | GCGCAAAAGACTAAACTGGC | 195 |
| chr5:5581827-5583867 | p2 | (AT)20 | CCCTAAAACGACGATCCAAA | GCGCAAAAGACTAAACTGGC | 195 |
| chr5:5581827-5583867 | p5 | (GGTTT)3 | CCCTAAAACGACGATCCAAA | GCGCAAAAGACTAAACTGGC | 195 |
| chr5:5597494-5599568 | p2 | (TA)37 | CCCTCGGAACCATATCAGAA | TTTAAAGGCTTGTTGCGGAT | 248 |
| chr5:5610238-5612278 | p2 | (AG)20 | TATGAAGGACAATCCGCACA | TACCCTTTCAAATGGCGGTA | 139 |
| chr5:5719975-5721991 | p4 | (ATAA)4 | TATGCCCCTCCAAAGTTGAC | GGAAGATACAAGCCATGGGA | 160 |
| chr5:5762155-5764170 | p3 | (TGT)5 | CTCACAAAGGCGAGGGTAAG | ATTTTCAGAGCTCGAGCCAA | 166 |
| chr5:5762155-5764170 | p3 | (AAG)5 | CTCACAAAGGCGAGGGTAAG | ATTTTCAGAGCTCGAGCCAA | 166 |
| chr5:5762155-5764170 | p3 | (TTC)5 | CTCACAAAGGCGAGGGTAAG | ATTTTCAGAGCTCGAGCCAA | 166 |
| chr5:5772312-5774368 | p2 | (AT)19 | ATGTATTTTCCCACGGCTGA | AACTTTTCGGTTGGCAAGAA | 222 |
| chr5:5772370-5774384 | p2 | (AT)19 | ATGTATTTTCCCACGGCTGA | AACTTTTCGGTTGGCAAGAA | 164 |
| chr5:5805822-5807846 | p2 | (AT)6 | GCATTTATCCAAGGAAGGCA | GTCGCTCCGATCTAATCCAA | 233 |
| chr5:5805927-5807953 | p2 | (AT)6 | GCATTTATCCAAGGAAGGCA | GTCGCTCCGATCTAATCCAA | 128 |
| chr5:5845774-5847996 | p3 | (ATT)7 | GCCCTCGAGTTCCCTTATTC | CAGATGATTGGCGTGTGAAG | 237 |
| chr5:5845774-5847996 | p2 | (TA)111 | GCCCTCGAGTTCCCTTATTC | CAGATGATTGGCGTGTGAAG | 237 |
| chr5:5846822-5848840 | p2 | (TA)87 | GTGGGCCCCATACTAAACAA | TTAAGGGATAACGGACGACG | 216 |
| chr5:5868895-5870919 | p2 | (AT)12 | GCTTTGCTTTTGTGCTGTTG | CCGGTTGCCACCTTTAGATA | 112 |
| chr5:5895637-5897655 | p2 | (TG)9 | CAGTCAGGTGCACAAAGCAT | ATAAAGGGGAGCCTTGGAGA | 184 |
| chr5:5899643-5901659 | p2 | (AT)8 | AAGTTGGACTGGATTGCACC | TTGCGAACTCAAACATCTGC | 121 |
| chr5:5909485-5911500 | p5 | (AAATA)3 | AACCGAACCGAATTGTTGAA | AAAACCGACCGAACCTAACC | 241 |
| chr5:5909485-5911500 | p5 | (TTTAT)3 | AACCGAACCGAATTGTTGAA | AAAACCGACCGAACCTAACC | 241 |
| chr5:5917616-5919630 | p2 | (TA)7 | ATCCCATTGATCCACTTCCA | ACTGGAGCAATTGCCTATGG | 129 |
| chr5:5946191-5948203 | p2 | (TC)6 | GACTTAACGAAAATCGGGCA | TGTTCGTTAGAATTTCGGGC | 103 |
| chr5:5973640-5975652 | p2 | (AC)6 | TTCGAACATCATTCGGAACA | TGAGTCGGGATTTGGTTAGC | 144 |

| Marker ID | SSR type | SSR motif | FORWARD PRIMER (5'-3') | REVERSE PRIMER (5'-3') | size (bp) |
|---|---|---|---|---|---|
| chr5:5985482-5987497 | p3 | (TGT)5 | GTGGTTGTGAGACCGACCTT | CTAAACCATCTCAGCCTCGC | 271 |
| chr5:6129822-6131844 | p2 | (AT)11 | TTGCTTTCGTGAATTGGTCA | CCGTTTCCAATGTCGTTTTT | 214 |
| chr5:6129822-6131844 | p5 | (AAAAT)4 | TTGCTTTCGTGAATTGGTCA | CCGTTTCCAATGTCGTTTTT | 214 |
| chr5:6161847-6163859 | p2 | (GA)6 | TTTTCAGGTTCATCCTTCGG | TCGCCCCTTTTCCTCTTAAT | 219 |
| chr5:6433262-6435277 | p5 | (ACCAA)3 | GAACAAGCCGTGGTTCAAGT | AGATCGTGTCCATCTCAGGG | 183 |
| chr5:6433262-6435277 | p5 | (AAACC)3 | GAACAAGCCGTGGTTCAAGT | AGATCGTGTCCATCTCAGGG | 183 |
| chr5:6434002-6436017 | p5 | (ACCAA)3 | GGGTGTACATGGACCGAGTT | CTGAGGTTGAAAAACCCGAA | 182 |
| chr5:6434002-6436017 | p5 | (AAACC)3 | GGGTGTACATGGACCGAGTT | CTGAGGTTGAAAAACCCGAA | 182 |
| chr5:6501387-6503401 | p2 | (AT)7 | AATTGTGATCAGCCCGAGTC | TCCCTCTTTCTTTTGCTGGA | 209 |
| chr5:6542868-6544883 | p3 | (TGT)5 | CACGTCCGGAAGATGAAAAT | TGCCACTCCCACCTTATCTC | 299 |
| chr5:6724412-6726427 | p3 | (CTT)5 | ATCAGGGGAAAAGGTGGAAC | TACATGTCTCAGTTTGCCGC | 210 |
| chr5:6724412-6726427 | p3 | (ACA)5 | ATCAGGGGAAAAGGTGGAAC | TACATGTCTCAGTTTGCCGC | 210 |
| chr5:6796896-6798911 | p5 | (AAAAT)3 | ATCACCCATCTGACGACCTC | CATGTGCATAACAGGGTGGA | 142 |
| chr5:6796896-6798911 | p3 | (GTT)65 | ATCACCCATCTGACGACCTC | CATGTGCATAACAGGGTGGA | 142 |
| chr5:6892622-6894679 | p3 | (TAT)19 | ATAAGGCGTAAGTGCCGATG | ATTGGCGTTCTTTTCCCTTT | 157 |
| chr5:6978756-6980774 | p6 | (CGGCTA)3 | TACCCCTTTCCCTTGTTCCT | AGACTCAAGCTCGAAGACGC | 196 |
| chr5:7049805-7051820 | p5 | (ATTGC)3 | GCTCGAGGTCACTTTTCGAG | CACGAACGATCAAACCAATG | 293 |
| chr5:7138342-7140357 | p3 | (CAC)5 | GTTCATGACACTCCCGACCT | TATTGACTTCGCGCCTAACC | 294 |
| chr5:7174140-7176206 | p3 | (AAT)22 | ACTTGATTTTCAAGGGGGCT | TCCTCCACTTGTCCCTGTTC | 278 |
| chr5:7192358-7194373 | p5 | (GGCCC)3 | TTGAATGTCTCGCAATGAGC | AGGGGGATACCAGTGTGTTG | 248 |
| chr5:7278950-7280968 | p6 | (AACCGG)3 | AATTACCCCTCCGGATGAAG | GGAGGATCTGGTGAGGTCAA | 299 |
| chr5:7278950-7280968 | p2 | (TA)7 | AATTACCCCTCCGGATGAAG | GGAGGATCTGGTGAGGTCAA | 299 |
| chr5:7304084-7306096 | p2 | (TG)6 | TAGTCGGAAATTGGTGGCTT | TGCAATGCTCCTGCAATAAG | 202 |
| chr5:7335618-7337644 | p2 | (AT)13 | GGGGAACCTGCACAAGTTTA | AAGGCTGTGAGTGGTTGCTT | 199 |
| chr5:7335618-7337644 | p3 | (ATA)19 | GGGGAACCTGCACAAGTTTA | AAGGCTGTGAGTGGTTGCTT | 199 |
| chr5:7335831-7337888 | p2 | (AT)13 | TAACAATTCCCCGAAGCAAC | CGCCTAAAAACCTGGATTGA | 215 |
| chr5:7335831-7337888 | p3 | (ATA)19 | TAACAATTCCCCGAAGCAAC | CGCCTAAAAACCTGGATTGA | 215 |
| chr5:7374574-7376600 | p2 | (AT)13 | GGGGAACCTGCACAAGTTTA | AAGGCTGTGAGTGGTTGCTT | 199 |
| chr5:7374574-7376600 | p3 | (ATA)19 | GGGGAACCTGCACAAGTTTA | AAGGCTGTGAGTGGTTGCTT | 199 |
| chr5:7374787-7376844 | p2 | (AT)13 | TAACAATTCCCCGAAGCAAC | CGCCTAAAAACCTGGATTGA | 215 |
| chr5:7374787-7376844 | p3 | (ATA)19 | TAACAATTCCCCGAAGCAAC | CGCCTAAAAACCTGGATTGA | 215 |
| chr5:7420116-7422132 | p4 | (ATGC)4 | TCAATTTCCAGTGAAAGCCC | AAACGAATCAAAATCAGCCG | 120 |
| chr5:7464318-7466332 | p2 | (AT)13 | TTGGACGTCTTAGCATGGTG | TTCGTCTTCCTCCGCTTCTA | 255 |
| chr5:7464318-7466332 | p2 | (TA)7 | TTGGACGTCTTAGCATGGTG | TTCGTCTTCCTCCGCTTCTA | 255 |
| chr5:7464394-7466414 | p2 | (AT)13 | TTGGACGTCTTAGCATGGTG | ATCCAATCGCGAATGAAAAC | 179 |
| chr5:7464394-7466414 | p2 | (TA)7 | TTGGACGTCTTAGCATGGTG | ATCCAATCGCGAATGAAAAC | 179 |

| Marker ID | SSR type | SSR motif | FORWARD PRIMER (5'-3') | REVERSE PRIMER (5'-3') | size (bp) |
|---|---|---|---|---|---|
| chr5:7596585-7598603 | p3 | (CTT)6 | CCTCAAGTGCAACACAGGAA | CAACGAATTTCCCCTCAAAA | 289 |
| chr5:7605577-7607592 | p5 | (ATTTT)3 | TCTCATGAATCTTCCCCGAC | AATGGGTGGTTGAAGGACTG | 157 |
| chr5:7691256-7693271 | p3 | (TCT)5 | GGGAAAACCCATCAAAGGAT | AGGACTCACTTGGGGGAAAC | 166 |
| chr5:7719943-7721969 | p2 | (CT)13 | CCAGCCCACGGTAAAGACTA | ATGCTTGCTGCTTCGAAAAT | 188 |
| chr5:7731714-7733729 | p5 | (TAAAA)3 | ACTAATAACCCGATGCTGCG | TGTGGTCAACCTTCAACCAA | 241 |
| chr5:7731714-7733729 | p2 | (AC)9 | ACTAATAACCCGATGCTGCG | TGTGGTCAACCTTCAACCAA | 241 |
| chr5:7776845-7778860 | p5 | (CTTTT)3 | TCCAGTTTGACACCTACCCC | ACTCACTCTGAGGCAGCCAT | 258 |
| chr5:7823184-7825229 | p3 | (ATT)15 | GAAGGTGGCAGAGCTGAATC | TTGGGCGTACATTGAGTGAA | 118 |
| chr5:7847731-7849791 | p2 | (AT)8 | TTATCGACAAGGCAAGGGAC | TGTCGCGAACACGTAACACT | 250 |
| chr5:7847801-7849909 | p2 | (AT)8 | TTATCGACAAGGCAAGGGAC | TGTCGCGAACACGTAACACT | 180 |
| chr5:7862590-7864635 | p3 | (ATT)15 | GAAGGTGGCAGAGCTGAATC | TTGGGCGTACATTGAGTGAA | 118 |
| chr5:7887137-7889197 | p2 | (AT)8 | TTATCGACAAGGCAAGGGAC | TGTCGCGAACACGTAACACT | 250 |
| chr5:7887207-7889315 | p2 | (AT)8 | TTATCGACAAGGCAAGGGAC | TGTCGCGAACACGTAACACT | 180 |
| chr5:7976528-7978550 | p2 | (AT)6 | TCGGTTGATTGCATTTATCG | TTGATTTTCCCCCAAATTGA | 279 |
| chr5:7976528-7978550 | p2 | (AT)7 | TCGGTTGATTGCATTTATCG | TTGATTTTCCCCCAAATTGA | 279 |
| chr5:7976528-7978550 | p2 | (TA)17 | TCGGTTGATTGCATTTATCG | TTGATTTTCCCCCAAATTGA | 279 |
| chr5:7976528-7978550 | p2 | (AT)11 | TCGGTTGATTGCATTTATCG | TTGATTTTCCCCCAAATTGA | 279 |
| chr5:7976557-7978573 | p2 | (AT)6 | TCGGTTGATTGCATTTATCG | TTGATTTTCCCCCAAATTGA | 250 |
| chr5:7976557-7978573 | p2 | (AT)7 | TCGGTTGATTGCATTTATCG | TTGATTTTCCCCCAAATTGA | 250 |
| chr5:7976557-7978573 | p2 | (TA)17 | TCGGTTGATTGCATTTATCG | TTGATTTTCCCCCAAATTGA | 250 |
| chr5:7976557-7978573 | p2 | (AT)11 | TCGGTTGATTGCATTTATCG | TTGATTTTCCCCCAAATTGA | 250 |
| chr5:7993897-7995915 | p2 | (TA)9 | AAAATCACGCCACATCAACA | GGTGGTCAGTTGAAGGGAAA | 299 |
| chr5:8012949-8014967 | p2 | (AT)15 | GGGATACGTAGACGTTCGGA | AACCTCGGGGGTAATGTAGC | 299 |
| chr5:8012981-8014995 | p2 | (AT)15 | GGGATACGTAGACGTTCGGA | AACCTCGGGGGTAATGTAGC | 267 |
| chr5:8012996-8015012 | p2 | (AT)15 | GGGATACGTAGACGTTCGGA | AACCTCGGGGGTAATGTAGC | 252 |
| chr5:8184100-8186118 | p2 | (AT)32 | ACCAAATTGCATGCACAAAA | CGTTTATGCCTCAAACGTCA | 293 |
| chr5:8184100-8186118 | p6 | (AATTTT)3 | ACCAAATTGCATGCACAAAA | CGTTTATGCCTCAAACGTCA | 293 |
| chr5:8347737-8349755 | p6 | (AGTTGG)3 | TTGGAAGCACTCAACAACCA | GGCAGTTGACCAGGATCATT | 238 |
| chr5:8364906-8366921 | p5 | (CACCC)3 | CCGTGAGAAAGAAAGAACGC | AAACATGCTACCTTGGCAGG | 107 |
| chr5:8415856-8417871 | p5 | (TAAAA)3 | CAGAAGTCCAGCCCATTGAT | TCCACCAGGACCAATACCAT | 298 |
| chr5:8457744-8459762 | p3 | (GTA)6 | TTGCTGATTTTGTGTGGGAA | CCACCATCAAAAGAATTGGG | 260 |
| chr5:8557159-8559177 | p6 | (TATTTT)3 | TGGTGAAGCCACCCTTATTC | CGCTTTTGTAACGAGGGGTA | 247 |
| chr5:8557159-8559177 | p2 | (TG)6 | TGGTGAAGCCACCCTTATTC | CGCTTTTGTAACGAGGGGTA | 247 |
| chr5:8557159-8559177 | p6 | (TATTTT)3 | TGGTGAAGCCACCCTTATTC | CGCTTTTGTAACGAGGGGTA | 247 |
| chr5:8557159-8559177 | p5 | (ATTTT)3 | TGGTGAAGCCACCCTTATTC | CGCTTTTGTAACGAGGGGTA | 247 |
| chr5:8583101-8585116 | p3 | (TTC)5 | TGTGAGTATCGTCGAGCCAC | TCTTGCCGCCCTCTCTAGTA | 124 |

| Marker ID | SSR type | SSR motif | FORWARD PRIMER (5'-3') | REVERSE PRIMER (5'-3') | size (bp) |
|---|---|---|---|---|---|
| chr5:8608034-8610046 | p2 | (CA)6 | TGCGAGCGAGGGTAACTAAT | GTGTCCACATTCCGCTCTTT | 122 |
| chr5:8640593-8642611 | p6 | (CAACTC)3 | ATTTCCCTTCTTTTGGGGTG | GTGTTTTCGGGCACCTTTTA | 116 |
| chr5:8655536-8657552 | p5 | (TATAC)3 | ATCATTGCTCTTCCCCACAC | GCATCTAAGAACCCACGGAA | 247 |
| chr5:8655536-8657552 | p6 | (CATGTT)3 | ATCATTGCTCTTCCCCACAC | GCATCTAAGAACCCACGGAA | 247 |
| chr5:8659004-8661070 | p2 | (AT)33 | GATATTTGGAAACGGCGAAA | AGTTGGGATTCGGTGTCAAC | 246 |
| chr5:8718948-8720962 | p2 | (TA)6 | GACGATCTTGGCCTTGAAAA | TTGTTCCACGAAATGCAAAA | 282 |
| chr5:8718966-8720980 | p2 | (TA)6 | GACGATCTTGGCCTTGAAAA | TTGTTCCACGAAATGCAAAA | 264 |
| chr5:8718966-8720980 | p2 | (AT)7 | GACGATCTTGGCCTTGAAAA | TTGTTCCACGAAATGCAAAA | 264 |
| chr5:8719057-8721069 | p2 | (TA)6 | GACGATCTTGGCCTTGAAAA | TTGTTCCACGAAATGCAAAA | 173 |
| chr5:8719961-8721975 | p2 | (TA)6 | CGTGAGATTGTTCCACGAAA | TTGTTCCACGAAATGCAAAA | 186 |
| chr5:8719961-8721975 | p2 | (AT)8 | CGTGAGATTGTTCCACGAAA | TTGTTCCACGAAATGCAAAA | 186 |
| chr5:8720016-8722028 | p2 | (AT)6 | CGTGAGATTGTTCCACGAAA | TTGTTCCACGAAATGCAAAA | 131 |
| chr5:8720016-8722028 | p2 | (TA)6 | CGTGAGATTGTTCCACGAAA | TTGTTCCACGAAATGCAAAA | 131 |
| chr5:8720016-8722028 | p2 | (AT)8 | CGTGAGATTGTTCCACGAAA | TTGTTCCACGAAATGCAAAA | 131 |
| chr5:8722532-8724552 | p5 | (GGTGG)4 | TTTCCAAATTGGGACCTTTG | TCCTCCTCCAACCACAAAAC | 164 |
| chr5:8722532-8724552 | p2 | (AT)9 | TTTCCAAATTGGGACCTTTG | TCCTCCTCCAACCACAAAAC | 164 |
| chr5:8724498-8726510 | p2 | (TA)6 | TTAAAGGGACAACGGACGAT | GGGTTTAGACCCATGCTTCA | 282 |
| chr5:8724498-8726510 | p2 | (AT)8 | TTAAAGGGACAACGGACGAT | GGGTTTAGACCCATGCTTCA | 282 |
| chr5:8783943-8785961 | p6 | (AAGTTA)3 | CGGACAAATCTGGAGACGAT | CCATTGGAGAAGGGCAGTTA | 247 |
| chr5:8902617-8904632 | p2 | (TA)42 | TAACCGATTTTCGCGATTTC | TTTCCTCAAAACGAACACCC | 172 |
| chr5:8902633-8904648 | p2 | (TA)42 | TAACCGATTTTCGCGATTTC | TTTCCTCAAAACGAACACCC | 156 |
| chr5:8914325-8916339 | p2 | (AG)7 | AGCCCTCCACAACCTCCTAT | GGCATGTGAAGAGAAGAGGC | 129 |
| chr5:9132988-9135024 | p2 | (AT)8 | TGTGGGCCCCATAATTCTAA | GGGTGCATTTGTTTTTGACA | 193 |
| chr5:9132988-9135024 | p2 | (AT)18 | TGTGGGCCCCATAATTCTAA | GGGTGCATTTGTTTTTGACA | 193 |
| chr5:9132988-9135024 | p2 | (AT)12 | TGTGGGCCCCATAATTCTAA | GGGTGCATTTGTTTTTGACA | 193 |
| chr5:9199687-9201702 | p3 | (ATA)5 | CGGGAGAGAGAGACAAGTGG | TCAGCAAAATGAGGGAAACC | 130 |
| chr5:9199687-9201702 | p6 | (ATAATG)4 | CGGGAGAGAGAGACAAGTGG | TCAGCAAAATGAGGGAAACC | 130 |
| chr5:9210426-9212441 | p3 | (TAA)17 | AAGATGTTCGAAGCGAGGAA | GATTTTGGCCTTGCATCATT | 262 |
| chr5:9210426-9212441 | p3 | (AAT)5 | AAGATGTTCGAAGCGAGGAA | GATTTTGGCCTTGCATCATT | 262 |
| chr5:9225185-9227197 | p2 | (TA)6 | TGGGAGGCATGTCCTACTTC | GAGACCGGACAACCCTATGA | 163 |
| chr5:9225185-9227197 | p2 | (TA)13 | TGGGAGGCATGTCCTACTTC | GAGACCGGACAACCCTATGA | 163 |
| chr5:9225955-9227973 | p2 | (TA)6 | ATCAGCTAAGCCAAACGAGC | CAGGCCCGTCTACCAAAGTA | 204 |
| chr5:9225955-9227973 | p2 | (TA)13 | ATCAGCTAAGCCAAACGAGC | CAGGCCCGTCTACCAAAGTA | 204 |
| chr5:9225955-9227973 | p2 | (AT)9 | ATCAGCTAAGCCAAACGAGC | CAGGCCCGTCTACCAAAGTA | 204 |
| chr5:9226023-9228035 | p2 | (TA)6 | ATCAGCTAAGCCAAACGAGC | CAGGCCCGTCTACCAAAGTA | 136 |
| chr5:9226023-9228035 | p2 | (TA)13 | ATCAGCTAAGCCAAACGAGC | CAGGCCCGTCTACCAAAGTA | 136 |

| Marker ID | SSR type | SSR motif | FORWARD PRIMER (5'-3') | REVERSE PRIMER (5'-3') | size (bp) |
|---|---|---|---|---|---|
| chr5:9226023-9228035 | p2 | (AT)9 | ATCAGCTAAGCCAAACGAGC | CAGGCCCGTCTACCAAAGTA | 136 |
| chr5:9344451-9346471 | p5 | (AAATA)3 | CGGGGTTAACTTTTTACGCA | ACCCAACGCTTTTGATATGG | 117 |
| chr5:9344451-9346471 | p5 | (TTTAT)4 | CGGGGTTAACTTTTTACGCA | ACCCAACGCTTTTGATATGG | 117 |
| chr5:9344451-9346471 | p5 | (TAAAT)3 | CGGGGTTAACTTTTTACGCA | ACCCAACGCTTTTGATATGG | 117 |
| chr5:9344451-9346471 | p3 | (AAT)11 | CGGGGTTAACTTTTTACGCA | ACCCAACGCTTTTGATATGG | 117 |
| chr5:9378701-9380731 | p2 | (AT)15 | TCCTACTGGTGTGAGAGGGG | AGGGCATACACACCCAAGAC | 264 |
| chr5:9386007-9388022 | p6 | (CCCCAA)3 | TCTTAATTTTGCACCCCCAG | GGGTTAGGGTGGGTAGTGGT | 181 |
| chr5:9386007-9388022 | p5 | (GTGGG)3 | TCTTAATTTTGCACCCCCAG | GGGTTAGGGTGGGTAGTGGT | 181 |
| chr5:9458963-9460978 | p4 | (TAAT)5 | GTACGGACCGTTCAGACGAT | CTACGGAGCTCCATCACCTC | 187 |
| chr5:9458963-9460978 | p5 | (AATAA)3 | GTACGGACCGTTCAGACGAT | CTACGGAGCTCCATCACCTC | 187 |
| chr5:9497739-9499757 | p6 | (TTTTTG)3 | CCTTGGTGGAACATCCACTT | GGATCCATTTTCCATGGTTG | 128 |
| chr5:9582363-9584378 | p5 | (AAATA)3 | CGGCAGTTAGAACCCAGAAC | TAATGTGAAGTTGTGGCGGA | 152 |
| chr5:9651119-9653185 | p2 | (TA)33 | CCCTAAACTACCCCCACACA | AAAAGTCTTGGTCGTGGGAA | 268 |
| chr5:9759534-9761546 | p2 | (TA)6 | ACCTTCAAATCATGTTCCGC | AATTAGCGAGCACTCCGAGA | 245 |
| chr5:9777570-9779585 | p5 | (GAAAA)3 | CTTAAAGCATATTCGGGCCA | TGAACGTTTATCATGCCCAA | 112 |
| chr5:9858345-9860366 | p3 | (TAA)7 | TTAAGGGACAGAGCAATGGG | GAATGGGCATCTCTTGGTGT | 117 |
| chr5:9862511-9864541 | p2 | (TA)15 | TTGATCATAACCCCTCCTCG | CAATTTCCTGAGCGTTAGGC | 116 |
| chr5:9862511-9864541 | p2 | (TA)7 | TTGATCATAACCCCTCCTCG | CAATTTCCTGAGCGTTAGGC | 116 |
| chr5:9958478-9960496 | p6 | (TTCATG)3 | CCGCGCAAGTACAGACTACA | AACACATGGAGCCGATAAGG | 137 |
| chr5:9988984-9990999 | p3 | (TTG)5 | AGAAGATGAAAGTCGCGGAA | CGTCTAATCACCTCTCCCCA | 299 |
| chr5:10066901-10068913 | p2 | (TA)6 | GCATTCTGGAATTTGGCAAT | ACGGACAACACTGCCTCTCT | 133 |
| chr5:10126499-10128514 | p3 | (TTG)5 | AGAAGATGAAAGTCGCGGAA | TTCCTGAAACCGTCCATAGC | 268 |
| chr5:10163920-10165932 | p2 | (CG)6 | ACCATCCCGGCTATTTTACC | TTTAGAGGCGTATCGGTTGG | 161 |
| chr5:10177887-10179902 | p3 | (ACA)5 | CTCACGCAGTCTGTTTCCAA | TTGACTACGAGGCGACACTG | 136 |
| chr5:10246985-10249003 | p6 | (TTCTCT)3 | GAGGCTGCTTATCGCTTAGG | AATAATTACCTCCCGCCCAT | 128 |
| chr5:10246985-10249003 | p3 | (ACA)5 | GAGGCTGCTTATCGCTTAGG | AATAATTACCTCCCGCCCAT | 128 |
| chr5:10283740-10285755 | p3 | (TTG)5 | TGCGACCAGGGTGTTATGTA | ACTTGCACCTTCTGGCGTAT | 201 |
| chr5:10283740-10285755 | p3 | (GTT)6 | TGCGACCAGGGTGTTATGTA | ACTTGCACCTTCTGGCGTAT | 201 |
| chr5:10296545-10298563 | p6 | (TTTTAT)3 | TGTATGTGGCTCAATTCCCA | GAGGCCAATCATGAAAAGGA | 255 |
| chr5:10304201-10306231 | p2 | (AT)15 | CTAACTGCGGTAACCTGGGA | CATTTGGATCAGGGAGGAGA | 242 |
| chr5:10319502-10321518 | p2 | (AT)8 | GCGTAGGGTAACTTGTGGGA | TATTCTCAACATGCCCCCTC | 178 |
| chr5:10329511-10331527 | p2 | (CT)8 | CCGTCGAATGAGTTACGGTT | TTGAAACCTCGTTGAGGGTC | 278 |
| chr5:10335556-10337568 | p2 | (CT)6 | GAATGTGAAGTGCGCAAAGA | TTTCACCAGAGTTTCCCCTG | 171 |
| chr5:10397463-10399483 | p2 | (TA)10 | GATCTATGTTGCTCGGGCTC | ACCAAGCAGCCTTTCCTTTT | 235 |
| chr5:10397463-10399483 | p5 | (TTTCT)3 | GATCTATGTTGCTCGGGCTC | ACCAAGCAGCCTTTCCTTTT | 235 |
| chr5:10624032-10626047 | p5 | (GAAAA)3 | TTTCCAGCAAGAAGATGGCT | TGCCGAACTGAACAAAAGTG | 202 |

| Marker ID | SSR type | SSR motif | FORWARD PRIMER (5'-3') | REVERSE PRIMER (5'-3') | size (bp) |
|---|---|---|---|---|---|
| chr5:10640484-10642504 | p5 | (TTGGT)4 | TTTTGGCCTTTCATACGGAC | GTCGAGCGAAACACATCTGA | 123 |
| chr5:10649554-10651566 | p2 | (TA)6 | ATAACGCCCATACGAGTTGC | TGAATGGATTGTTCGGTGAA | 166 |
| chr5:10672241-10674277 | p2 | (AT)18 | TTAGTCCATTTCAGCTGCCC | ATGGCGTGTATTAGGGGGAT | 269 |
| chr5:10695001-10697013 | p2 | (TA)6 | CCCCTCCATGAACGTAGAAA | TACGACCTGATATGCCACGA | 164 |
| chr5:10753200-10755216 | p2 | (TA)8 | CCCCTCCCTCTATCTTCGTC | CGCTGTTAACCGGTGACTTT | 121 |
| chr5:10753200-10755216 | p2 | (TC)6 | CCCCTCCCTCTATCTTCGTC | CGCTGTTAACCGGTGACTTT | 121 |
| chr5:10884071-10886083 | p2 | (TA)6 | TCATTGCCCAATTATGTCCA | TTTTGTATCCAGACCCCACC | 240 |
| chr5:10914518-10916532 | p2 | (TA)7 | AAATAGGGACAAAAGCGCAA | CAGGACTATGAATGCGCTGA | 195 |
| chr5:10971503-10973518 | p3 | (AGC)5 | CAGACCAGACGCAACAAAGA | CCGAATCACTGAAGTGCTGA | 143 |
| chr5:11134382-11136397 | p5 | (ACAAT)4 | ACAATGGGTGACAACGTTCC | CAACGACTTTCGTGCAAATG | 264 |
| chr5:11134382-11136397 | p3 | (ATT)5 | ACAATGGGTGACAACGTTCC | CAACGACTTTCGTGCAAATG | 264 |
| chr5:11164096-11166114 | p2 | (AT)9 | GTGGGTTTTGGGGTTTTCTT | GCGTGTTTAGGGCGTACAAT | 226 |
| chr5:11184158-11186173 | p5 | (TGCAA)3 | GGCAAAAGTTGTCCATTGCT | CAATTGAGGTAGTTGGGGGA | 102 |
| chr5:11186096-11188110 | p2 | (AT)7 | CACCATGCCTGAAAGCCTAT | CAAGACTCCTGTTCGGCTTC | 105 |
| chr5:11241127-11243167 | p2 | (AT)20 | TGCTTAAGCCTGGTGATGTG | GCTTATCGACGCTTCAAAGG | 204 |
| chr5:11241127-11243167 | p2 | (TC)14 | TGCTTAAGCCTGGTGATGTG | GCTTATCGACGCTTCAAAGG | 204 |
| chr5:11431566-11433598 | p2 | (AT)16 | CCTGAGTCCTGGGAATGAAA | CCAATTGTTGGCGTCTTTCT | 162 |
| chr5:11431566-11433598 | p2 | (AT)16 | CCTGAGTCCTGGGAATGAAA | CCAATTGTTGGCGTCTTTCT | 162 |
| chr5:11530890-11532914 | p6 | (CCCCCA)4 | GGCAAAAGTGCAAATGACCT | GGGGGTAGTCGGATGAAAAG | 223 |
| chr5:11556328-11558343 | p3 | (GTT)5 | TCCAAAAGGGTGATTCAAGG | ATATAGGGGTCTTGGGTCGG | 184 |
| chr5:11558784-11560798 | p2 | (CA)6 | GTCTGAGGTTCGCTCGACTC | TTTTGCTTGCATGCAGTTTC | 126 |
| chr5:11558800-11560818 | p2 | (CA)6 | GTCTGAGGTTCGCTCGACTC | TTTTGCTTGCATGCAGTTTC | 110 |
| chr5:11571391-11573406 | p3 | (TGG)5 | CTTTCTCTTCCATTGTGCCC | TAAAATGCGAGAAGGTCGCT | 147 |
| chr5:11601397-11603449 | p5 | (TATTT)3 | CTTGCCGCAACTCTCTCTCT | TAAAATTATTCCGCCAAGCG | 120 |
| chr5:11705809-11707824 | p5 | (TTTTA)3 | CGTCGTTTTAGGGGTTGAAA | AACAAAATCGCGTAAATCGG | 269 |
| chr5:11724991-11727012 | p3 | (GTT)7 | CCCTTTTGGGACGTGAAATA | ACAAATCTCCGGAACCAGTG | 206 |
| chr5:11931575-11933590 | p5 | (TCGGT)3 | GATTGCGCAAGACCCTACAT | CAGGCCCACTCAATCTTGTT | 160 |
| chr5:11931575-11933590 | p2 | (AT)33 | GATTGCGCAAGACCCTACAT | CAGGCCCACTCAATCTTGTT | 160 |
| chr5:11974866-11976880 | p2 | (GA)7 | CGTTGCAAAACTCACCGTAA | TCGTCCGCCTCTAGGAGTAA | 136 |
| chr5:11989828-11991842 | p2 | (TA)9 | GCACCATTGCTGTCTGCTTA | CCTAGACTTACGCCCCAACA | 196 |
| chr5:11989828-11991842 | p2 | (TA)7 | GCACCATTGCTGTCTGCTTA | CCTAGACTTACGCCCCAACA | 196 |
| chr5:11993276-11995291 | p3 | (ATA)5 | CGGGTTCTCGTCAACAATTT | TGGATTCGTTTCGTTTGTGA | 118 |
| chr5:12067003-12069017 | p2 | (CA)7 | TTGTTGTTGTGCAGGTTGGT | TTGTGAGATTCCTGCTGTGC | 126 |
| chr5:12067003-12069017 | p2 | (CA)7 | TTGTTGTTGTGCAGGTTGGT | TTGTGAGATTCCTGCTGTGC | 126 |
| chr5:12213988-12216000 | p2 | (TA)6 | TGGATGAAAATTCTGGCTCC | CTCTGCCCTCCATTGAAAAA | 247 |
| chr5:12234321-12236363 | p2 | (AT)21 | ACCACTGCCCATCAAATTGT | GGAAGCCCGAAATAGACCTC | 151 |

| Marker ID | SSR type | SSR motif | FORWARD PRIMER (5'-3') | REVERSE PRIMER (5'-3') | size (bp) |
|---|---|---|---|---|---|
| chr5:12261183-12263198 | p3 | (TAA)5 | GACAGCTGAAGCACCAAACA | GTCAAGAAACACGGCCAAAT | 272 |
| chr5:12277949-12279969 | p2 | (TA)10 | CATAGATTGGGATGGAGGGA | CACGGAGGATTTACTTCCCA | 295 |
| chr5:12297923-12299935 | p2 | (TA)6 | TGGTGATGGGAACTTTGACA | ATTAGCAACTCCACCTCCCA | 149 |
| chr5:12347424-12349436 | p2 | (TA)10 | GTGGGCCCCATACTAAACAA | TTAAGGGACCACAGACGGAC | 232 |
| chr5:12347424-12349436 | p2 | (TA)6 | GTGGGCCCCATACTAAACAA | TTAAGGGACCACAGACGGAC | 232 |
| chr5:12347453-12349485 | p2 | (TA)10 | GTGGGCCCCATACTAAACAA | TTAAGGGACCACAGACGGAC | 203 |
| chr5:12347453-12349485 | p2 | (TA)6 | GTGGGCCCCATACTAAACAA | TTAAGGGACCACAGACGGAC | 203 |
| chr5:12348259-12350271 | p2 | (TA)6 | AAGTGCACCGCAATTAAAAA | GTTTGCCTACGAATACGCAC | 282 |
| chr5:12348259-12350271 | p2 | (CT)6 | AAGTGCACCGCAATTAAAAA | GTTTGCCTACGAATACGCAC | 282 |
| chr5:12457619-12459658 | p2 | (AT)8 | GTGGATCTTCGGGTTCAAGA | ACAAACCCTAGAATCGCCCT | 164 |
| chr5:12457619-12459658 | p3 | (ATA)13 | GTGGATCTTCGGGTTCAAGA | ACAAACCCTAGAATCGCCCT | 164 |
| chr5:12491770-12493790 | p2 | (TA)10 | GGAGTTCTGACCACCCTTGA | GGCGGTCTTCCCTTATCTTC | 254 |
| chr5:12493329-12495353 | p2 | (AT)12 | TCCCTTCGGCAGAGACTTTA | GTTGAGAGAAGCACCCTTGC | 241 |
| chr5:12503247-12505265 | p6 | (TATCTA)3 | GGTGTGTACGCCTTTCAGGT | CCTCAACTTAGGCCGATGAG | 116 |
| chr5:12547947-12549971 | p2 | (TA)12 | TACATGCATCCAAAAAGGCA | AGCGACCGACCTGAGAACTA | 146 |
| chr5:12547947-12549971 | p3 | (CAC)6 | TACATGCATCCAAAAAGGCA | AGCGACCGACCTGAGAACTA | 146 |
| chr5:12582450-12584466 | p2 | (TA)7 | AAATATCAACCGTGGCTTCG | TCTTTCATTTGGTCCGCTTC | 255 |
| chr5:12636363-12638379 | p2 | (AT)8 | ACATGCGGAGTCGTATAGCC | TTTACACTAACGCCCCGAAC | 208 |
| chr5:12636363-12638379 | p2 | (TA)11 | ACATGCGGAGTCGTATAGCC | TTTACACTAACGCCCCGAAC | 208 |
| chr5:12637246-12639268 | p2 | (AT)8 | GAGACAGCGTCGTTCTTTCC | TTTGTGCTTGGCTCATTACG | 150 |
| chr5:12637246-12639268 | p2 | (TA)11 | GAGACAGCGTCGTTCTTTCC | TTTGTGCTTGGCTCATTACG | 150 |
| chr5:12637246-12639268 | p2 | (AT)6 | GAGACAGCGTCGTTCTTTCC | TTTGTGCTTGGCTCATTACG | 150 |
| chr5:12637246-12639268 | p2 | (TA)10 | GAGACAGCGTCGTTCTTTCC | TTTGTGCTTGGCTCATTACG | 150 |
| chr5:12652879-12654891 | p2 | (TA)6 | TCAGGAACTGACAAAGCACG | AGAAATGGGGACGAAGGTCT | 166 |
| chr5:12661026-12663038 | p2 | (AG)6 | CCCAGGTGAGGACTCGATAA | ACGATGCGTGGAAGTAAAGC | 121 |
| chr5:12661026-12663038 | p2 | (TA)9 | CCCAGGTGAGGACTCGATAA | ACGATGCGTGGAAGTAAAGC | 121 |
| chr5:12668671-12670686 | p5 | (TGTAT)3 | TCCTCTCTGGTGCTTGAGGT | ACAATACGGCTGTCCAATCC | 190 |
| chr5:12745094-12747158 | p2 | (AT)12 | CCATTCGACTGGCCACTAAT | CAATAGCCACAGGCAGGAAT | 258 |
| chr5:12745094-12747158 | p2 | (TA)32 | CCATTCGACTGGCCACTAAT | CAATAGCCACAGGCAGGAAT | 258 |
| chr5:12745094-12747158 | p6 | (TAATGT)3 | CCATTCGACTGGCCACTAAT | CAATAGCCACAGGCAGGAAT | 258 |
| chr5:12784021-12786041 | p3 | (TTA)7 | GTTTCGCAGGAACTTTAGCG | CTGGAATGTGAACGGGAGAT | 295 |
| chr5:12784037-12786052 | p3 | (TTA)7 | GTTTCGCAGGAACTTTAGCG | CTGGAATGTGAACGGGAGAT | 279 |
| chr5:12784060-12786075 | p3 | (TTA)7 | GTTTCGCAGGAACTTTAGCG | CTGGAATGTGAACGGGAGAT | 256 |
| chr5:12784177-12786198 | p3 | (TTA)7 | GTTTCGCAGGAACTTTAGCG | CTGGAATGTGAACGGGAGAT | 139 |
| chr5:12841498-12843516 | p2 | (TA)6 | ATTGCTTCAGCACAAAGGCT | ATACAGATGGGGCAAAGCAC | 164 |
| chr5:12841498-12843516 | p2 | (TA)21 | ATTGCTTCAGCACAAAGGCT | ATACAGATGGGGCAAAGCAC | 164 |

| Marker ID | SSR type | SSR motif | FORWARD PRIMER (5'-3') | REVERSE PRIMER (5'-3') | size (bp) |
|---|---|---|---|---|---|
| chr5:12894877-12896919 | p2 | (AT)21 | TCACAATTTCGAACCCAACA | CGCACGACCAAATTGTTTTA | 161 |
| chr5:12952100-12954115 | p3 | (ATT)5 | GCATCCAACGGCTAGGAATA | TAGTGGAGCAGTGAAATGCG | 209 |
| chr5:13123009-13125021 | p2 | (AT)6 | TAACGGGATGGGACTTGAAC | TGAAAGCTGACTCGGGAAAT | 118 |
| chr5:13185126-13187141 | p5 | (TTTTA)3 | CTCCTCCATTCCCTTTTTCC | GTCCAAACTTGTAACGGGGA | 191 |
| chr5:13416849-13418864 | p6 | (AAAATA)3 | CTTTGGGGCAGTACTTGCAT | GTTGCAACTCAAGCACATGG | 291 |
| chr5:13416849-13418864 | p5 | (CAAAC)3 | CTTTGGGGCAGTACTTGCAT | GTTGCAACTCAAGCACATGG | 291 |
| chr5:13538852-13540867 | p5 | (ATTTT)3 | TTGAATCCTGGATCTGCCTC | ACTATGCGAAGTTGGTTCCG | 200 |
| chr5:13572644-13574660 | p2 | (TA)8 | ACCAAGGCTTGCTCCTACAA | ACATCCGACTTCGTCCATTC | 107 |
| chr5:13586282-13588328 | p2 | (TA)23 | GGGGAACCTGCACAAGTTTA | GTTGCTTCGGGGAATTGTTA | 200 |
| chr5:13672060-13674076 | p2 | (AT)7 | ACACTCCGAGGATGAACCTG | CCCAGGAGACCGGTTATTTT | 106 |
| chr5:13672060-13674076 | p2 | (AT)8 | ACACTCCGAGGATGAACCTG | CCCAGGAGACCGGTTATTTT | 106 |
| chr5:13674359-13676425 | p3 | (ATC)8 | TCAAATTCCGCCATGTGTTA | TCCCCGATTTTAATGAGTGC | 256 |
| chr5:13674784-13676808 | p3 | (ATC)8 | CTGATCGCGATATGGGACTT | GAGTTTCAAAAGCGCTCCAG | 125 |
| chr5:13747664-13749682 | p5 | (TGTTG)3 | AAACATTGGGTGGGACACAT | AGCTCAGTTCTCGGCCACTA | 151 |
| chr5:13747664-13749682 | p3 | (TGT)6 | AAACATTGGGTGGGACACAT | AGCTCAGTTCTCGGCCACTA | 151 |
| chr5:13747664-13749682 | p2 | (TA)9 | AAACATTGGGTGGGACACAT | AGCTCAGTTCTCGGCCACTA | 151 |
| chr5:13751072-13753154 | p2 | (TA)41 | TTCATAGTGCGACAAATGCC | CGGGGAAGTAACCTATGCAA | 217 |
| chr5:13818947-13820962 | p5 | (AACCA)3 | CAAAATGCAAAGAACGCAAA | GTCCTGGGAGAAGTCAGTGC | 119 |
| chr5:13818947-13820962 | p5 | (AAACC)3 | CAAAATGCAAAGAACGCAAA | GTCCTGGGAGAAGTCAGTGC | 119 |
| chr5:13946216-13948231 | p5 | (AACCA)3 | CAAAATGCAAAGAACGCAAA | GTCCTGGGAGAAGTCAGTGC | 119 |
| chr5:13946216-13948231 | p5 | (AAACC)3 | CAAAATGCAAAGAACGCAAA | GTCCTGGGAGAAGTCAGTGC | 119 |
| chr5:14064494-14066510 | p2 | (TA)8 | AAGTGCAAATCGTAAACCGC | GCTTCTGCTCATCCACACAA | 201 |
| chr5:14064871-14066973 | p2 | (TA)8 | TGCTGCTCTCCTCCTTTTGT | ACGCCCTCTCATTCTTGCTA | 291 |
| chr5:14078735-14080755 | p2 | (AT)6 | ACATGTGTCCCTTCCCCATA | GTTCCTTCAAGGCGTGACAT | 107 |
| chr5:14078735-14080755 | p5 | (ATTCA)4 | ACATGTGTCCCTTCCCCATA | GTTCCTTCAAGGCGTGACAT | 107 |
| chr5:14120069-14122083 | p6 | (ATAAAA)3 | GCGTAAACCCGAATTTTTGA | TATGGGTCTGAACAGGGAGC | 276 |
| chr5:14120069-14122083 | p2 | (TA)7 | GCGTAAACCCGAATTTTTGA | TATGGGTCTGAACAGGGAGC | 276 |
| chr5:14120069-14122083 | p2 | (CA)7 | GCGTAAACCCGAATTTTTGA | TATGGGTCTGAACAGGGAGC | 276 |
| chr5:14144370-14146385 | p3 | (TGT)5 | AGCACTTGTTTAGCCAGCGT | AGACGGGGTTTAGGATTCGT | 239 |
| chr5:14159185-14161200 | p3 | (TTG)5 | CTGGGGTTGGCTATGTGAGT | GGAGCAGTCCACTCCAGTTC | 151 |
| chr5:14432512-14434528 | p4 | (TAAA)4 | TTAATCCGCAAGAAATTCGG | GCATCAAATTTGGGCCTCTA | 215 |
| chr5:14432512-14434528 | p6 | (TTAAAA)3 | TTAATCCGCAAGAAATTCGG | GCATCAAATTTGGGCCTCTA | 215 |
| chr5:14433489-14435507 | p4 | (TAAA)4 | TTGTTATGCCCCTTGGAGAG | GATTCTCCGCCTACTCCCTC | 163 |
| chr5:14433489-14435507 | p6 | (TTAAAA)3 | TTGTTATGCCCCTTGGAGAG | GATTCTCCGCCTACTCCCTC | 163 |
| chr5:14438195-14440213 | p6 | (TAAAAA)3 | CAATGTAATCCCCATCCCTG | AGAAGGAGGAAAACGCAACA | 128 |
| chr5:14493164-14495188 | p3 | (TCT)8 | TTCCTTAACTGCAACCTGGG | TGCCCCTGTTATTTAGGTGC | 135 |

| Marker ID | SSR type | SSR motif | FORWARD PRIMER (5'-3') | REVERSE PRIMER (5'-3') | size (bp) |
|---|---|---|---|---|---|
| chr5:14493164-14495188 | p2 | (AC)6 | TTCCTTAACTGCAACCTGGG | TGCCCCTGTTATTTAGGTGC | 135 |
| chr5:14493164-14495188 | p2 | (TG)6 | TTCCTTAACTGCAACCTGGG | TGCCCCTGTTATTTAGGTGC | 135 |
| chr5:14602321-14604363 | p6 | (ATCAGA)7 | GTCCATTGCAGCAAAAAGGT | TGCTGTTCAAAAGGATGCTG | 100 |
| chr5:14647592-14649625 | p3 | (AAT)11 | AAGGGGAAGGCTGATAAAGC | TCGTCTTAGGCCACGAGATT | 173 |
| chr5:14694256-14696274 | p6 | (AAGATA)3 | AGGGCCATGTCCTGATAGTG | ACATGGGCATCAAAACAACA | 136 |
| chr5:14694256-14696274 | p5 | (AACCG)3 | AGGGCCATGTCCTGATAGTG | ACATGGGCATCAAAACAACA | 136 |
| chr5:14696634-14698667 | p3 | (AAT)11 | AAGGGGAAGGCTGATAAAGC | TCGTCTTAGGCCACGAGATT | 173 |
| chr5:14746178-14748193 | p5 | (CCCCT)3 | TGCACAGGGAGACAGACTTG | ACGTGGCTACGAGCCTCTAA | 127 |
| chr5:14746178-14748193 | p5 | (ATTTT)3 | TGCACAGGGAGACAGACTTG | ACGTGGCTACGAGCCTCTAA | 127 |
| chr5:14746178-14748193 | p2 | (GA)7 | TGCACAGGGAGACAGACTTG | ACGTGGCTACGAGCCTCTAA | 127 |
| chr5:14762892-14764922 | p6 | (AAAAAT)5 | ACGCGTTATTGGATACTGCC | GTCACACCTTCAGATGCCCT | 175 |
| chr5:14886055-14888069 | p5 | (AAAAT)3 | GAAACCCACCACATCCAAAC | GACATGGCGGAATAGCAGAT | 276 |
| chr5:14886055-14888069 | p2 | (AT)7 | GAAACCCACCACATCCAAAC | GACATGGCGGAATAGCAGAT | 276 |
| chr5:14970559-14972574 | p5 | (CCACC)3 | GTTGGATGGGGGAGAGGTAT | CTCCATACCGAACACACCCT | 192 |
| chr5:14970559-14972574 | p5 | (CCACC)3 | GTTGGATGGGGGAGAGGTAT | CTCCATACCGAACACACCCT | 192 |
| chr5:14972378-14974393 | p5 | (AAACC)3 | GCTTCACGAATGAAAGCACA | CAAGAAAACCCGAGGTTGAA | 135 |
| chr5:14976409-14978424 | p3 | (CAA)5 | GCTCCAATCATGCCAAAAAT | AATAGCTTTAGGCCGCCAAT | 264 |
| chr5:14976409-14978424 | p3 | (TAA)5 | GCTCCAATCATGCCAAAAAT | AATAGCTTTAGGCCGCCAAT | 264 |
| chr5:14988119-14990134 | p3 | (ATT)5 | GTGAAAATGGGACACCCAAA | TGCATGTGTAGTGCCCAAAT | 217 |
| chr5:14988119-14990134 | p3 | (TTA)5 | GTGAAAATGGGACACCCAAA | TGCATGTGTAGTGCCCAAAT | 217 |
| chr5:15064021-15066039 | p2 | (TA)9 | GGCGTCCATTTCAACAATCT | GGGTCAGTTTGTTGTCCACC | 242 |
| chr5:15277222-15279234 | p2 | (TA)6 | TGGATGCCTCTTGTTTGATG | ATGACATACTAGCCGGCACC | 275 |
| chr5:15277222-15279234 | p2 | (TA)36 | TGGATGCCTCTTGTTTGATG | ATGACATACTAGCCGGCACC | 275 |
| chr5:15277711-15279783 | p2 | (TA)6 | TTGAACCCCTAACTTCCACG | GAACTTCCCAGAGGACGATG | 215 |
| chr5:15277711-15279783 | p2 | (TA)36 | TTGAACCCCTAACTTCCACG | GAACTTCCCAGAGGACGATG | 215 |
| chr5:15281712-15283732 | p2 | (AT)10 | TCCATTGGCAATTTCTTTCC | GGCATGAAGAGAACGTGGTT | 276 |
| chr5:15302159-15304174 | p3 | (CAA)5 | ATGGCCTCAGTTTTCAGTGG | AGAGAGGATACGTGCCGCTA | 284 |
| chr5:15380755-15382767 | p2 | (CT)6 | TCCCGAATATCAGGACGAAG | TTACTTGGATGGTGAAGCCC | 127 |
| chr5:15421182-15423235 | p2 | (TG)7 | ATTCCAGTCCAACCAACTGC | GGGAGGGTAAGCAAAACACA | 297 |
| chr5:15512847-15514862 | p3 | (GTT)5 | CCACCAAAACTTAAGGGCAA | CCATCTCAATCTCGCTCCTC | 270 |
| chr5:15514232-15516244 | p2 | (AT)6 | ACATCTCATTCACTTCGGGG | ACGTTCTCCACCTTTCATGC | 148 |
| chr5:15516020-15518035 | p2 | (TA)7 | ATTGCGGGTAAACATCCAGA | CTTGGTTGCTTTTCAGGCTC | 190 |
| chr5:15548206-15550226 | p4 | (TGTA)5 | CCCACCGTTATGGACTCTGT | CTCCTCCTCTTTGCCTTCCT | 259 |
| chr5:15548360-15550376 | p4 | (TGTA)5 | CCCACCGTTATGGACTCTGT | CTCCTCCTCTTTGCCTTCCT | 105 |
| chr5:15575942-15577958 | p4 | (GATA)4 | GAGAGTGAGCCGAACGAAAG | CATGGGCACTAACTGAAGCA | 182 |
| chr5:15629337-15631349 | p2 | (AC)6 | TGACCGATCAAGGCATGTTA | ATTGTACGTGGAAGGATGGC | 101 |

| Marker ID | SSR type | SSR motif | FORWARD PRIMER (5'-3') | REVERSE PRIMER (5'-3') | size (bp) |
|---|---|---|---|---|---|
| chr5:15655167-15657179 | p2 | (AT)6 | AAACTCAGAAGCGAGGCAAG | CCCCCACATCATATTCGTTC | 190 |
| chr5:15655193-15657221 | p2 | (AT)6 | AAACTCAGAAGCGAGGCAAG | CCCCCACATCATATTCGTTC | 164 |
| chr5:15730743-15732787 | p5 | (TTATG)3 | ATCCCTTTTGAGGTCGGTCT | CGGGTGTCTATGACACGATG | 262 |
| chr5:15730743-15732787 | p2 | (AT)22 | ATCCCTTTTGAGGTCGGTCT | CGGGTGTCTATGACACGATG | 262 |
| chr5:15730743-15732787 | p4 | (TTAT)4 | ATCCCTTTTGAGGTCGGTCT | CGGGTGTCTATGACACGATG | 262 |
| chr5:15751033-15753059 | p2 | (AT)13 | TACAGATGTCCCCTTGACCC | TTTCGGGTACTTGCGTTAGG | 232 |
| chr5:15900157-15902223 | p2 | (GA)10 | AACAGCGGTGTTGCTTTCTT | GTCTCTCACCCCTCTCTCCC | 145 |
| chr5:15900157-15902223 | p2 | (AT)33 | AACAGCGGTGTTGCTTTCTT | GTCTCTCACCCCTCTCTCCC | 145 |
| chr5:16027353-16029368 | p5 | (AAAAT)3 | GATCAGGCACGTCAACAGAA | GCCCAAAAACGTTCATTCAC | 246 |
| chr5:16038937-16040952 | p5 | (AACCG)3 | TGCTAAGCGTGGATGATGAG | TGGGACGGATTTTCTAATCG | 115 |
| chr5:16181248-16183264 | p2 | (TA)8 | TTAATCAAGGGGCTGCAATC | TTGGTCAGTAGCAAAACCCC | 191 |
| chr5:16183374-16185390 | p2 | (TA)8 | TTAATCAAGGGGCTGCAATC | TTGGTCAGTAGCAAAACCCC | 191 |
| chr5:16251568-16253582 | p2 | (AT)7 | TCAACGCTCTTAGCCATGTG | CGAAGCCACGGTTGATACTT | 198 |
| chr5:16439056-16441068 | p2 | (CT)6 | GGAGCTTCAATCAGTTGGGA | AACCGCAAACCTCAATCAAC | 256 |
| chr5:16459087-16461102 | p5 | (TATTT)3 | TCCCCTTGGCTCATTATCTG | GGCTTGGACAACAACCATTT | 103 |
| chr5:16472656-16474671 | p5 | (AATAA)3 | AGGAAGAGAGCCACGACAGA | GAAACGGTTGGCATTGAGTT | 212 |
| chr5:16495700-16497727 | p3 | (AGA)9 | GAGGTTCAATGCAGAGCACA | GCCTTAATGGCCACAAACAT | 176 |
| chr5:16536738-16538754 | p4 | (TATT)4 | CTTACCCCCTGCTTGTGAAA | TGCGCGGTCATTATTAAACA | 132 |
| chr5:16538573-16540585 | p2 | (CT)6 | AAAGCACACCAAGTTGAGGG | GCATTTCTAAGGATGCCCAA | 264 |
| chr5:16592498-16594513 | p2 | (GA)8 | TTTTGGGGTTCAAGGAACTG | GAAATTCCCTAGACCGGAGG | 110 |
| chr5:16852530-16854575 | p5 | (CGGGT)3 | GTTTGCTTGGGGTTCAAAAA | CTTCCTCTTCTGTCCGCATC | 165 |
| chr5:16852530-16854575 | p3 | (AAT)15 | GTTTGCTTGGGGTTCAAAAA | CTTCCTCTTCTGTCCGCATC | 165 |
| chr5:16897856-16899871 | p3 | (TGT)5 | TCTTTTTCGTCGAATCCTCG | TTTCAGCCTTGGTCCCTCTA | 155 |
| chr5:16931206-16933218 | p2 | (AG)6 | GCTGGGTGCAAACAATTTCT | TCTCTTTGAGCCCCGACTAA | 168 |
| chr5:16966793-16968809 | p2 | (AT)8 | GGAAACAAAAGTGAACCCCA | TTGCTACGAATACACACGGC | 187 |
| chr5:16998632-17000644 | p2 | (TA)6 | CTTTCTTCCAACCTCCCCTC | ACCAATGGATCCCAAAAACA | 120 |
| chr5:16998632-17000644 | p2 | (TC)8 | CTTTCTTCCAACCTCCCCTC | ACCAATGGATCCCAAAAACA | 120 |
| chr5:17017265-17019279 | p2 | (TG)7 | AGGTGGTCAAGAATTGCAGC | TGGTTTGTGAGTTCTGGGGT | 182 |
| chr5:17050751-17052763 | p2 | (TA)6 | CTTTCTTCCAACCTCCCCTC | ACCAATGGATCCCAAAAACA | 120 |
| chr5:17050751-17052763 | p2 | (TC)8 | CTTTCTTCCAACCTCCCCTC | ACCAATGGATCCCAAAAACA | 120 |
| chr5:17118674-17120689 | p3 | (AGA)5 | ATGGTTCAAATTTTGCTGCC | TGGGTTCAACCGAACGTAAT | 167 |
| chr5:17136719-17138737 | p6 | (GCACCA)3 | AGGATCATTCCAACACAGGC | CCAATCTGGAAGCGGAATTA | 122 |
| chr5:17207442-17209458 | p5 | (GACCC)3 | TCATTCACATCGGGTAACGA | GCGAAAATGGTGTCGAATTT | 162 |
| chr5:17289714-17291726 | p2 | (TA)6 | AACAAATTCCAAGGGAAGGG | CTTGGGAATCGCCACACTAT | 208 |
| chr5:17289714-17291726 | p2 | (TA)10 | AACAAATTCCAAGGGAAGGG | CTTGGGAATCGCCACACTAT | 208 |
| chr5:17308284-17310296 | p2 | (AT)9 | AATGAGGGATTCATTGCCAC | GTATGGGGCCACTCTTTTGA | 135 |

| Marker ID | SSR type | SSR motif | FORWARD PRIMER (5'-3') | REVERSE PRIMER (5'-3') | size (bp) |
|---|---|---|---|---|---|
| chr5:17308284-17310296 | p2 | (AT)7 | AATGAGGGATTCATTGCCAC | GTATGGGGCCACTCTTTTGA | 135 |
| chr5:17308284-17310296 | p2 | (TA)6 | AATGAGGGATTCATTGCCAC | GTATGGGGCCACTCTTTTGA | 135 |
| chr5:17308793-17310807 | p2 | (AT)7 | AAGTGCATCCGCATAAAAGG | AGTGAGTTTGGAGACGGTGG | 248 |
| chr5:17308793-17310807 | p2 | (TA)34 | AAGTGCATCCGCATAAAAGG | AGTGAGTTTGGAGACGGTGG | 248 |
| chr5:17329563-17331578 | p3 | (AAG)5 | CGGAATAGACCGAATCCTCA | GAAATGGTGGTAGGCGAAAA | 129 |
| chr5:17365511-17367527 | p4 | (TTAA)4 | TCCAAACAAGTGCAGAGTGC | GGATTGGTTTTTCGGTCTCA | 163 |
| chr5:17402120-17404135 | p5 | (ATTTG)3 | GCCTTTGTTGAAGGAATCCA | TTGACGGAAAAAGAGCGAGT | 143 |
| chr5:17587168-17589184 | p3 | (TTA)24 | GACTCTGCTGGGGACTTCAG | ATAACTGCATTCCTGTGCCC | 202 |
| chr5:17587168-17589184 | p4 | (ATTT)4 | GACTCTGCTGGGGACTTCAG | ATAACTGCATTCCTGTGCCC | 202 |
| chr5:17612975-17614991 | p6 | (ATATAG)3 | GAGCCAAAACGACATCACAA | CGAGGTCCGTTCATATGGTT | 296 |
| chr5:17613000-17615014 | p6 | (ATATAG)3 | GAGCCAAAACGACATCACAA | CGAGGTCCGTTCATATGGTT | 271 |
| chr5:17641105-17643120 | p3 | (TTG)5 | AGGTGGGAACGATCAGAATG | TGTTCCCCTCAGTTCCAGTC | 100 |
| chr5:17768284-17770302 | p3 | (AGT)6 | TTCGTTGATCTGTGGGCATA | TTCTCTCCTGTCTCGCCAAT | 106 |
| chr5:17926822-17928837 | p5 | (AAATA)3 | AGCCAATCATGAATTTTCGC | AACCTGCGAAGTCACCAAAT | 141 |
| chr5:17966813-17968838 | p5 | (TTATA)5 | ATGCATGCAAATTGGAACCT | TCGTCCCTTTTCTCCCTCTT | 238 |
| chr5:17987302-17989320 | p6 | (TTTTAT)3 | ACGGCTTGAAGCATATCCAG | ATTGGGTCAACCATGCAAAT | 278 |
| chr5:18073452-18075467 | p5 | (AAACC)3 | GGATCGGGTTGGTTTAGGTT | CCCAACTTTTGTTGGTTTGG | 220 |
| chr5:18079980-18081994 | p2 | (AC)7 | TTAACAGGACCATCTTCGGC | TTTCCTGAGCTCGGATTGAT | 118 |
| chr5:18098783-18100801 | p6 | (CATTAC)3 | GTTACGAATTTGGAAGGCCA | CCAATTTTTGTGTCGGGAGT | 192 |
| chr5:18102188-18104238 | p2 | (AT)25 | GGATGCGAACAACATTCCTT | TGCAATAACAGTCGTTGGGA | 240 |
| chr5:18193433-18195445 | p2 | (GT)6 | CAATCGCAGATTCGTCACTG | TTCAAATGAACCCTCTTGGC | 224 |
| chr5:18228754-18230769 | p3 | (TGA)5 | TGAATCTTGCTGCCTTTCCT | CAGGTGGTCGTCCATTCTTT | 198 |
| chr5:18250965-18252989 | p6 | (TTTCTT)4 | CAAGGAGGTTGGGATGAGAA | CGCATCATTTTGGGAAGACT | 286 |
| chr5:18289563-18291581 | p6 | (CATTCT)3 | GGCAGCCTAACCTTTAACCC | GCTTTAAGTTGTGCAGGGGA | 105 |
| chr5:18314682-18316702 | p5 | (GTTTG)3 | TTCGTGATGCAATCCGTTTA | TTCCCAGTGGACAACTCACA | 258 |
| chr5:18314682-18316702 | p5 | (TTGGT)4 | TTCGTGATGCAATCCGTTTA | TTCCCAGTGGACAACTCACA | 258 |
| chr5:18337082-18339122 | p2 | (AT)20 | GAGCGGCCACTAACAGAGTC | TAGCTAGCCAGAGAGGCAGG | 248 |
| chr5:18337082-18339122 | p5 | (TAAAG)3 | GAGCGGCCACTAACAGAGTC | TAGCTAGCCAGAGAGGCAGG | 248 |
| chr5:18404400-18406418 | p3 | (ACA)6 | TTGACGTCCCCTTTTATTGC | GCTCGCCAAAAGCTAATACG | 188 |
| chr5:18421677-18423704 | p3 | (TGT)9 | TAACCTAACTCGTCGCCCAC | ATGCCACAAGTAGGGTCTGG | 294 |
| chr5:18423586-18425604 | p6 | (ATAAAA)3 | CGACGATGCCCACTTCTACT | CGACTCCAGACAAAACCCAT | 117 |
| chr5:18438744-18440762 | p4 | (AAAG)4 | GATGATTGGGACGCAGATTT | GAGACAACTCTCGTGGAGGC | 278 |
| chr5:18438744-18440762 | p2 | (AT)6 | GATGATTGGGACGCAGATTT | GAGACAACTCTCGTGGAGGC | 278 |
| chr5:18438744-18440762 | p3 | (AAC)6 | GATGATTGGGACGCAGATTT | GAGACAACTCTCGTGGAGGC | 278 |
| chr5:18455483-18457498 | p5 | (TATTT)3 | CGCCATGCAACTCTAACTCA | GGGAAGCCTACTGGTGTTCA | 115 |
| chr5:18470740-18472762 | p2 | (TG)8 | GCCTTTCTCTCCATTCCCTT | TACCACACCGACAAACCAAA | 129 |

| Marker ID | SSR type | SSR motif | FORWARD PRIMER (5'-3') | REVERSE PRIMER (5'-3') | size (bp) |
|---|---|---|---|---|---|
| chr5:18503814-18505832 | p6 | (TGTTGG)3 | ACTAAAGGTGACGTGTGGGG | AACGGTTTGGGATGTTTTGA | 267 |
| chr5:18503814-18505832 | p3 | (TTG)8 | ACTAAAGGTGACGTGTGGGG | AACGGTTTGGGATGTTTTGA | 267 |
| chr5:18523169-18525184 | p2 | (TA)6 | TGCATGCCACAACTCATTTT | ATATACGCAAGCCGACAAGG | 101 |
| chr5:18536458-18538488 | p3 | (TTA)5 | CCCCAAGAGTGATCGAGAAA | TTAAGTGCTCATGTGCTCGG | 223 |
| chr5:18552956-18554976 | p2 | (TA)7 | TGAGATCACGGAGGTGTGAG | AGGATTTGTGTGGCCAATTT | 229 |
| chr5:18553011-18555027 | p2 | (TA)7 | TGAGATCACGGAGGTGTGAG | AGGATTTGTGTGGCCAATTT | 174 |
| chr5:18586399-18588414 | p5 | (TTTCA)3 | TATCCTCCAGCTCCATGACC | TGAAGGAGGAGGAAGTGGTG | 137 |
| chr5:18599727-18601743 | p2 | (AT)8 | TACGTTTGGGATACGTGCAA | TTAGAGCCTGGTGCAGCTTT | 118 |
| chr5:18653290-18655305 | p5 | (TTTCA)3 | TATCCTCCAGCTCCATGACC | TGAAGGAGGAGGAAGTGGTG | 137 |
| chr5:18666618-18668634 | p2 | (AT)8 | TACGTTTGGGATACGTGCAA | TTAGAGCCTGGTGCAGCTTT | 118 |
| chr5:18810295-18812310 | p5 | (TTTTG)3 | AGCTTGGTGCACGAGAGAAT | GATCCTCATTGTTTGGGTGG | 101 |
| chr5:18885990-18888035 | p2 | (GA)6 | CCTCCCTTCACAGATTCCAA | CCTTCTCTCCTCCATCTCCC | 296 |
| chr5:18885990-18888035 | p3 | (ATA)15 | CCTCCCTTCACAGATTCCAA | CCTTCTCTCCTCCATCTCCC | 296 |
| chr5:18941410-18943428 | p2 | (AT)9 | GCCCAAAAACTGTCAACACA | GCCAGTGGTAACGAGTCTCC | 288 |
| chr5:19018462-19020480 | p2 | (AT)9 | GCCCAAAAACTGTCAACACA | GCCAGTGGTAACGAGTCTCC | 288 |
| chr5:19253848-19255924 | p5 | (TTGGT)3 | TGGAATTGAAAAATGCACGA | ATTTGAAAGCCTTACCCCGT | 215 |
| chr5:19255328-19257343 | p5 | (TTGGT)3 | TAGTCAGGAAGGGGCCTACA | TCCAACACCAAACCAAGTCA | 119 |
| chr5:19274869-19276887 | p6 | (CATATA)3 | TAGTTCGAGAGCCCGTGAGT | AAAAATCGCCCCAAGCTACT | 248 |
| chr5:19289314-19291332 | p6 | (CATATA)3 | TAGTTCGAGAGCCCGTGAGT | AAAAATCGCCCCAAGCTACT | 248 |
| chr5:19310362-19312374 | p2 | (AT)6 | CGCAGCATCTGTCGATTAAA | GATAGGGCTATGCGTGAAGC | 138 |
| chr5:19369815-19371830 | p2 | (AT)10 | CGACCTTACGTCACCTCGAT | TATCGTATAGCTGCCCGCTT | 243 |
| chr5:19369815-19371830 | p5 | (TATAT)3 | CGACCTTACGTCACCTCGAT | TATCGTATAGCTGCCCGCTT | 243 |
| chr5:19419145-19421157 | p2 | (TA)11 | TACCCCCGGACAAAAACATA | ATTCCGTACTGACCCCCTTT | 258 |
| chr5:19419145-19421157 | p2 | (AC)6 | TACCCCCGGACAAAAACATA | ATTCCGTACTGACCCCCTTT | 258 |
| chr5:19425972-19428032 | p2 | (AT)30 | ACTTCTGACCGGCTTTGAGA | AGTGGGTATGGGGAGGGTAG | 129 |
| chr5:19425972-19428032 | p2 | (TC)10 | ACTTCTGACCGGCTTTGAGA | AGTGGGTATGGGGAGGGTAG | 129 |
| chr5:19428213-19430228 | p5 | (ATACT)3 | TCTCCAAACGCCACTCTCTT | CTGAGGGTGGATCGTCATCT | 205 |
| chr5:19441445-19443463 | p3 | (TTG)6 | GCCCACACGTCAAGATTTTT | AATTGTCCTTTTGGAGGGCT | 168 |
| chr5:19445040-19447058 | p6 | (ACCAGC)3 | GAGTCCGTTGAGTTGGCTTC | TTTCGGCCATCTCTGCTACT | 177 |
| chr5:19446921-19448936 | p3 | (TGT)5 | ATCCTGCATTTTCGAACGTC | AATTGTCCTTTTGGAGGGCT | 136 |
| chr5:19451908-19453922 | p2 | (TC)7 | AATGCTCGCACACTCAAATG | GTATTGGGAAGGTTTCGGGT | 297 |
| chr5:19466628-19468654 | p2 | (TA)13 | CTGAATTGGAATTCGGGCTA | TTTCACAAAGACCATGCCAA | 255 |
| chr5:19479658-19481672 | p2 | (TA)7 | TGACAACATTAAGCGGTGGA | CCATTTGAAATTGGGTTTGG | 191 |
| chr5:19496851-19498881 | p2 | (AT)11 | AAAAGAGTTGCACCCCCATA | TTTGCTAACGAATACGCACG | 143 |
| chr5:19496888-19498910 | p2 | (AT)11 | AAAAGAGTTGCACCCCCATA | TTTGCTAACGAATACGCACG | 106 |
| chr5:19500191-19502207 | p2 | (CT)6 | GGAGAGAGAGTTTGAGGGGG | TCTTTTCTCTACGAGCCCCA | 170 |

| Marker ID | SSR type | SSR motif | FORWARD PRIMER (5'-3') | REVERSE PRIMER (5'-3') | size (bp) |
|---|---|---|---|---|---|
| chr5:19500191-19502207 | p5 | (ATAAA)3 | GGAGAGAGAGTTTGAGGGGG | TCTTTTCTCTACGAGCCCCA | 170 |
| chr5:19500191-19502207 | p2 | (TA)8 | GGAGAGAGAGTTTGAGGGGG | TCTTTTCTCTACGAGCCCCA | 170 |
| chr5:19563090-19565114 | p2 | (AT)12 | ACGGCTCGTAGACACATCCT | ATCATTTTGTTGGCCTCGTC | 205 |
| chr5:19587637-19589667 | p2 | (TA)7 | TTAGACCCTGTTTGGATGGG | TTGGCTGGCAAGCTTAAAGT | 117 |
| chr5:19587637-19589667 | p2 | (AT)13 | TTAGACCCTGTTTGGATGGG | TTGGCTGGCAAGCTTAAAGT | 117 |
| chr5:19618333-19620348 | p5 | (TTTTA)3 | ACCAGACATCTCCTATGCGG | AGTAGGTTGCCGTTTGGATG | 138 |
| chr5:19684624-19686642 | p2 | (AG)9 | CAAATCACTTTGCTCCCGTT | ACTTGCACCTCACCCAAAAC | 201 |
| chr5:19795608-19797623 | p5 | (AAAAT)3 | CATTGCCAGTGCATAACGAC | CTTTTGCCTTGCTCCTTTTG | 154 |
| chr5:19807600-19809618 | p2 | (TG)9 | AATCACCAAAGCCATTCGAG | CTCCACATGCGATCTCTCAA | 229 |
| chr5:19818243-19820267 | p2 | (TA)7 | TGAAAAATTGTGCATGGTGG | GAAAATATGGCTGACGCTCC | 264 |
| chr5:19818243-19820267 | p2 | (TA)8 | TGAAAAATTGTGCATGGTGG | GAAAATATGGCTGACGCTCC | 264 |
| chr5:19821712-19823730 | p6 | (AGAATT)3 | TCAAGGGATGAAAGAGACCG | ACTCAGGCGTGTAACCTGCT | 112 |
| chr5:19821712-19823730 | p2 | (AT)9 | TCAAGGGATGAAAGAGACCG | ACTCAGGCGTGTAACCTGCT | 112 |
| chr5:19846714-19848729 | p5 | (TTTAT)3 | ACCATGGGGGTAGAAGAAGG | AGTGCAACTCGCTCAAACCT | 168 |
| chr5:19924756-19926774 | p2 | (TA)9 | GTATCAATGCGAGGAGCCAT | TCATTTCATCTTGATGCCCA | 228 |
| chr5:19972296-19974311 | p3 | (AAT)5 | AGGGATTCAATGTGCCAGAC | GCACCCAGGTTCAAAGAGTC | 187 |
| chr5:19982018-19984033 | p5 | (TATTT)3 | AAAGGCATCCAAAGTTGCTG | TTGAAAATCGCTAAAAGCGG | 165 |
| chr5:19988648-19990666 | p6 | (TTTATT)3 | AAAGAGGAACGTCGCAGAAA | GCCTCCACAGTAGCTCCAAG | 178 |
| chr5:20037416-20039462 | p2 | (AT)23 | TCTTTTTCCCTTAGCCCGAT | TTACACATCTCACGGGGTCA | 163 |
| chr5:20062741-20064759 | p3 | (TAA)6 | AAGGTAGGCTTCCTCCAAGC | GGCTGCTGATGAAAAAGAGG | 209 |
| chr5:20089691-20091711 | p2 | (AT)10 | TGCAGCAATCATCTGGAAAC | CCTTAAGGATGCTTCCACCA | 240 |
| chr5:20089691-20091711 | p6 | (ATGATA)3 | TGCAGCAATCATCTGGAAAC | CCTTAAGGATGCTTCCACCA | 240 |
| chr5:20161816-20163844 | p2 | (GA)9 | CCATAGCACCTAGCGCTTCT | ATAACTGCGTTTTTGGACGG | 176 |
| chr5:20161816-20163844 | p2 | (TA)14 | CCATAGCACCTAGCGCTTCT | ATAACTGCGTTTTTGGACGG | 176 |
| chr5:20168141-20170156 | p3 | (TAA)5 | CCCATCCGATCCGTTATATG | TTGGAGTTGGACGTGAACAA | 251 |
| chr5:20186384-20188399 | p5 | (TATTT)3 | CGGTTTTTGGATCGCTTTTA | ATTGACAAGGCTGAATCGCT | 129 |
| chr5:20211462-20213496 | p2 | (AT)17 | GCACGTGTAAAATACCGCCT | AAAAGGGCGAAACCAGAAGT | 241 |
| chr5:20227806-20229866 | p3 | (TAA)5 | GGGGTCCGGTAAGTCAAAAT | CAACTTCCGCATACACATCG | 282 |
| chr5:20227929-20229959 | p3 | (TAA)5 | GGGGTCCGGTAAGTCAAAAT | CAACTTCCGCATACACATCG | 159 |
| chr5:20263304-20265316 | p2 | (AT)6 | AGGATTGATCCCTTGACCCT | ATAACTGCGTTTTTGGACGG | 108 |
| chr5:20313819-20315839 | p5 | (TATTT)4 | GATCGAATCGTCTTCTTCGC | GGTGCAAGGAAAAGCAAAAA | 173 |
| chr5:20313819-20315839 | p2 | (TA)8 | GATCGAATCGTCTTCTTCGC | GGTGCAAGGAAAAGCAAAAA | 173 |
| chr5:20373112-20375127 | p3 | (TTG)5 | TGGAAGATTCCAGGACCAAG | GCAGTCACTGTAGCACGCAT | 281 |
| chr5:20388836-20390851 | p5 | (TTTTA)3 | CGGATTTACTGCCTGCATTT | GCCAGTACTTAGCAGCGGAC | 185 |
| chr5:20388836-20390851 | p5 | (TTTTA)3 | CGGATTTACTGCCTGCATTT | GCCAGTACTTAGCAGCGGAC | 185 |
| chr5:20408246-20410260 | p2 | (TA)7 | TATAAAGCGATGGGCTACGG | ATTAATCGCAAAACCGTTCG | 234 |

| Marker ID | SSR type | SSR motif | FORWARD PRIMER (5'-3') | REVERSE PRIMER (5'-3') | size (bp) |
|---|---|---|---|---|---|
| chr5:20408246-20410260 | p2 | (AT)11 | TATAAAGCGATGGGCTACGG | ATTAATCGCAAAACCGTTCG | 234 |
| chr5:20442326-20444390 | p2 | (TA)11 | TGCAAACATTGAACGACGAT | GGAGGAAAATGTTTTGGACG | 113 |
| chr5:20442326-20444390 | p2 | (TA)32 | TGCAAACATTGAACGACGAT | GGAGGAAAATGTTTTGGACG | 113 |
| chr5:20458425-20460437 | p2 | (TA)6 | AACAAAATTACGGCACGCTT | ACGTTCTCACCATTGGAAGG | 210 |
| chr5:20462524-20464544 | p5 | (ATAAA)4 | CAAGTCTGCCCATAAGGCAT | ACATTCGCAAGGCAGAGTTT | 100 |
| chr5:20463350-20465390 | p5 | (ATAAA)4 | TGCAAAAAGGGAGAGAGGAA | TCAAAGATCCACGCAGTTTG | 271 |
| chr5:20463350-20465390 | p2 | (AT)6 | TGCAAAAAGGGAGAGAGGAA | TCAAAGATCCACGCAGTTTG | 271 |
| chr5:20463350-20465390 | p2 | (AT)8 | TGCAAAAAGGGAGAGAGGAA | TCAAAGATCCACGCAGTTTG | 271 |
| chr5:20463738-20465754 | p2 | (AT)6 | AACGGTGCTTTATTTGCAGG | GCCTTGTGTGGGTATTTCGT | 143 |
| chr5:20463738-20465754 | p2 | (AT)8 | AACGGTGCTTTATTTGCAGG | GCCTTGTGTGGGTATTTCGT | 143 |
| chr5:20463738-20465754 | p3 | (AAT)7 | AACGGTGCTTTATTTGCAGG | GCCTTGTGTGGGTATTTCGT | 143 |
| chr5:20547376-20549391 | p3 | (GGT)5 | GACCCAACAAAACAACCACC | AGGAAAAACGTCAGGGCTTT | 119 |
| chr5:20638859-20640877 | p2 | (AT)9 | TGTCGCACCCCATTTTAATC | AAAGGATGCCTAACCCCTTC | 164 |
| chr5:20677519-20679531 | p6 | (CCTCCC)3 | TTGCTCCCCTCAAACAATTC | AGGAAACGAGGAGGAGGAAA | 238 |
| chr5:20677538-20679558 | p6 | (CCTCCC)3 | TTGCTCCCCTCAAACAATTC | AGGAAACGAGGAGGAGGAAA | 219 |
| chr5:20715075-20717129 | p2 | (AT)7 | TACCAGCCCATTTGAAGGAC | CGATCCACTTCTGCAACTCA | 206 |
| chr5:20715075-20717129 | p3 | (AAT)18 | TACCAGCCCATTTGAAGGAC | CGATCCACTTCTGCAACTCA | 206 |
| chr5:20783120-20785135 | p3 | (ATT)5 | AAGTCAGCCAAACCAGGAAA | AGAATGATGCAAATGGGCTC | 291 |
| chr5:20783120-20785135 | p4 | (TTAA)4 | AAGTCAGCCAAACCAGGAAA | AGAATGATGCAAATGGGCTC | 291 |
| chr5:20910282-20912296 | p2 | (TA)7 | TTTGCATGGATTGTCCTTCA | CGAACCCACAACCTCAGAGT | 233 |
| chr5:20932992-20935006 | p2 | (AT)6 | TCTTCTCGGACCAAAACACC | TTATGTTGGCTCCTTTTCCG | 272 |
| chr5:20933007-20935025 | p2 | (AT)6 | TCTTCTCGGACCAAAACACC | TTATGTTGGCTCCTTTTCCG | 257 |
| chr5:20951100-20953120 | p2 | (TA)10 | TGTGGCTCAGATGGTCTCAC | TACCACTTTGTCACGACCCA | 170 |
| chr5:20985335-20987361 | p2 | (AT)13 | GGAGCAGTGGGGTGAAGTAA | CCGTCTCTCTCTTCCTCCCT | 195 |
| chr5:21035732-21037748 | p2 | (AT)7 | TCCGTTGACGTCATTGTGAT | CAACGCACATGTCGAAAAAC | 212 |
| chr5:21035732-21037748 | p4 | (TCTT)4 | TCCGTTGACGTCATTGTGAT | CAACGCACATGTCGAAAAAC | 212 |
| chr5:21129576-21131592 | p2 | (AT)7 | TCCGTTGACGTCATTGTGAT | CAACGCACATGTCGAAAAAC | 212 |
| chr5:21129576-21131592 | p4 | (TCTT)4 | TCCGTTGACGTCATTGTGAT | CAACGCACATGTCGAAAAAC | 212 |
| chr5:21194049-21196064 | p2 | (AT)15 | GACCGTACGACCCAACCTTA | GAATCCGGACCAAAGTCAGA | 257 |
| chr5:21194049-21196064 | p5 | (ACCAA)3 | GACCGTACGACCCAACCTTA | GAATCCGGACCAAAGTCAGA | 257 |
| chr5:21194049-21196064 | p5 | (AAACC)3 | GACCGTACGACCCAACCTTA | GAATCCGGACCAAAGTCAGA | 257 |
| chr5:21401563-21403578 | p5 | (CAAAC)3 | CTTTGGGGCAGTACTTGCAT | GGGTAATCCTCCCAAAAAGC | 290 |
| chr5:21504164-21506178 | p2 | (GT)7 | CATCCGAAGAAGGTCCGTAA | ACATGAACACGCTTGACTGC | 185 |
| chr5:21508814-21510829 | p5 | (TTTAT)3 | ATTGCGCTGAGGTGAAAAAT | TTGTGCCTGCACACTAGGTC | 185 |
| chr5:21512603-21514627 | p6 | (ATTATG)4 | TTGGTAAGGCCTTGAGGATG | TTTCGTTTGGTCTCCAATCC | 232 |
| chr5:21587329-21589341 | p2 | (AT)6 | GGGAGCACTTCAGCTTCTTG | CCCAAGCAATAAAAGGGACA | 139 |

| Marker ID | SSR type | SSR motif | FORWARD PRIMER (5'-3') | REVERSE PRIMER (5'-3') | size (bp) |
|---|---|---|---|---|---|
| chr5:21597632-21599647 | p5 | (ATAAA)3 | TTTTAATGAGTTTTGGGGCG | TGAGCAATCATCATCGTGGT | 183 |
| chr5:21666942-21668974 | p2 | (AT)9 | GATATCATGCAAACCCGACC | CGTGTGAGGGGTCTTCAGTT | 151 |
| chr5:21666942-21668974 | p2 | (AT)16 | GATATCATGCAAACCCGACC | CGTGTGAGGGGTCTTCAGTT | 151 |
| chr5:21799833-21801848 | p5 | (ACCCG)3 | TCACATGACTGAGACCTCGC | TTGACCATGATGTCGCACTT | 171 |
| chr5:21815001-21817067 | p2 | (AT)33 | TCTTATTGGTTGGCTAGCGG | TGTGGCTAGGGGTGCTTATC | 115 |
| chr5:21830328-21832344 | p4 | (TTTC)4 | ACGTTTTCGTGATGAGTCCC | CCCTGAGTCTTGGTCATGGT | 186 |
| chr5:21864076-21866094 | p6 | (TTTATA)3 | CATCTAAACACGCAGAGGCA | CGTCGGAGTGTGCATTTCTA | 150 |
| chr5:21943897-21945909 | p2 | (AT)6 | TAGTTGTAGTCCGCGCTCCT | TCAATTTCAATTTTCCCCCA | 128 |
| chr5:21943897-21945909 | p5 | (TTCCA)3 | TAGTTGTAGTCCGCGCTCCT | TCAATTTCAATTTTCCCCCA | 128 |
| chr5:22027776-22029791 | p5 | (AAAAG)3 | AACATGTGGCATCAAACCAA | ATCGCCCCATCTAGTAGGCT | 146 |
| chr5:22115803-22117818 | p5 | (AAAAT)3 | CAACAGGATATGGTGCAACG | CTCTCTGTCTTCCACCAGGC | 245 |
| chr5:22115803-22117818 | p5 | (AACCG)3 | CAACAGGATATGGTGCAACG | CTCTCTGTCTTCCACCAGGC | 245 |
| chr5:22148133-22150148 | p5 | (AAAAG)3 | CCTAACATCGCTGCCCTAGA | CTCTCGCCAAATGGACTCTC | 214 |
| chr5:22166535-22168575 | p2 | (AT)20 | ATAGGGGTCAAATTCGGGTC | TTGGAAGAAAACATGAGGGG | 166 |
| chr5:22168051-22170065 | p2 | (AT)7 | TCTTCACGTGGACCCTACCT | TGATGGCATGAAACTCGGTA | 280 |
| chr5:22257944-22259968 | p3 | (ACA)5 | CTTTCCCCATATCCCCTCAT | ACGTAATTACCCGCTTGTGG | 211 |
| chr5:22297240-22299255 | p5 | (AAGTA)3 | GCAGCAGAGTGTCCCAATTT | TGGTACCCCTCTGGAGTCTG | 130 |
| chr5:22494735-22496750 | p3 | (TAT)5 | CACCTGCCCTGTTTTCATCT | CGTTAAAGCTCCTGCGAAAC | 213 |
| chr5:22553098-22555113 | p5 | (TTTAC)3 | CCATCTACCATATGGCCACC | GTAGTGGGGAACGTGGTTGA | 100 |
| chr5:22585543-22587573 | p3 | (ACA)10 | TATGACAATCGCCAGAGTGG | ATCCAAGGTTGTGATAGGCG | 246 |
| chr5:22605754-22607774 | p2 | (TA)10 | GGCACATTCGACGTCTTTTT | CGGAACGTCGATTCTTCATT | 197 |
| chr5:22652689-22654707 | p3 | (GAT)6 | ACGTTTGACCAAAATCCCAA | CTCTGCCACATCAGCAAAAA | 250 |
| chr5:22693639-22695726 | p3 | (ATA)29 | CCTTACCACGAAGCATCCAT | AAAATCATTGCAAAGGGCTG | 150 |
| chr5:22711733-22713748 | p3 | (ACA)5 | GCCCTTGACACCAGAAATGT | TGCTCAGACAATCAACCAGG | 235 |
| chr5:22724806-22726824 | p3 | (TTG)6 | CCACATGTCGGAGTTTTGTG | ATCGTCCTTTTGGAGGGTCT | 131 |
| chr5:22894729-22896749 | p2 | (CA)8 | ATGAACTCGCAGAATACGCC | GATCTGGACGGAGTTGTCGT | 210 |
| chr5:22894729-22896749 | p2 | (AT)10 | ATGAACTCGCAGAATACGCC | GATCTGGACGGAGTTGTCGT | 210 |
| chr5:22900558-22902578 | p2 | (CA)8 | ATGAACTCGCAGAATACGCC | GATCTGGACGGAGTTGTCGT | 210 |
| chr5:22900558-22902578 | p2 | (AT)10 | ATGAACTCGCAGAATACGCC | GATCTGGACGGAGTTGTCGT | 210 |
| chr5:23022245-23024263 | p2 | (TA)9 | TCGTTCTGGGAAAAACCATC | TGTGCACACAATGAAAACGA | 100 |
| chr5:23022245-23024263 | p5 | (TTGTT)3 | TCGTTCTGGGAAAAACCATC | TGTGCACACAATGAAAACGA | 100 |
| chr5:23056429-23058447 | p6 | (ATTCTC)3 | CACCAAATACCGAAGGCAAT | CGATGATATTCGCCCTGTTT | 201 |
| chr5:23059260-23061278 | p6 | (CAAAAA)3 | GAGGGAGACATAACAGCGGA | CCAAAACCGTACAGTGCCTT | 292 |
| chr5:23070899-23072935 | p3 | (TTC)12 | CCTCATGATCCTCCGACAGT | GCAAAGCCAAGGTTGATGAT | 201 |
| chr5:23131706-23133744 | p2 | (TA)19 | GCCATCGATATTTCATCCCA | TGGTGTTGCAGCATCTCTTC | 219 |
| chr5:23251382-23253397 | p5 | (AATTC)3 | TTACTTCCCGTATTGCGACC | TTCGGGCTAGGCACATAAAC | 111 |

| Marker ID | SSR type | SSR motif | FORWARD PRIMER (5'-3') | REVERSE PRIMER (5'-3') | size (bp) |
|---|---|---|---|---|---|
| chr5:23311630-23313645 | p5 | (TTGGT)3 | GCGTTCGAGAATACAGCCTC | TGTTCTTCCAGATTTTGGGC | 163 |
| chr5:23321862-23323886 | p2 | (TA)14 | TTTATTTCATTGATGCCCCC | ATAGGCACCTCCGAACAATG | 234 |
| chr5:23321862-23323886 | p6 | (TCATAT)4 | TTTATTTCATTGATGCCCCC | ATAGGCACCTCCGAACAATG | 234 |
| chr5:23321862-23323886 | p2 | (AT)19 | TTTATTTCATTGATGCCCCC | ATAGGCACCTCCGAACAATG | 234 |
| chr5:23364583-23366598 | p5 | (ATAAA)3 | ATACACGTGTCCGCACGATA | GACCGGTGAGTAGACCCCTT | 104 |
| chr5:23364583-23366598 | p5 | (TTTTA)3 | ATACACGTGTCCGCACGATA | GACCGGTGAGTAGACCCCTT | 104 |
| chr5:23364583-23366598 | p3 | (AAT)5 | ATACACGTGTCCGCACGATA | GACCGGTGAGTAGACCCCTT | 104 |
| chr5:23467678-23469711 | p3 | (AAT)11 | GTCGGTGATGAAGTGGGAGT | TCATTTCTCGTCGTCTCCCT | 112 |
| chr5:23467678-23469711 | p3 | (AAT)7 | GTCGGTGATGAAGTGGGAGT | TCATTTCTCGTCGTCTCCCT | 112 |
| chr5:23467678-23469711 | p3 | (TTA)6 | GTCGGTGATGAAGTGGGAGT | TCATTTCTCGTCGTCTCCCT | 112 |
| chr5:23467852-23469873 | p3 | (AAT)11 | TTAGGGGTTAATCGGGCTTT | ACGGTCATTGCAACAGCATA | 284 |
| chr5:23467852-23469873 | p3 | (AAT)7 | TTAGGGGTTAATCGGGCTTT | ACGGTCATTGCAACAGCATA | 284 |
| chr5:23467852-23469873 | p3 | (TTA)6 | TTAGGGGTTAATCGGGCTTT | ACGGTCATTGCAACAGCATA | 284 |
| chr5:23490187-23492251 | p2 | (AT)8 | TCATGATGATTCAGCAAGCC | ACTTTTGTCACGTCCCGAAC | 131 |
| chr5:23490187-23492251 | p2 | (AT)32 | TCATGATGATTCAGCAAGCC | ACTTTTGTCACGTCCCGAAC | 131 |
| chr5:23490187-23492251 | p3 | (CAA)6 | TCATGATGATTCAGCAAGCC | ACTTTTGTCACGTCCCGAAC | 131 |
| chr5:23512599-23514611 | p2 | (TA)6 | AAGGAAAGAGTGAAGGGGGA | TACCGGATCCCAAATCAAAA | 192 |
| chr5:23512599-23514611 | p3 | (ATA)5 | AAGGAAAGAGTGAAGGGGGA | TACCGGATCCCAAATCAAAA | 192 |
| chr5:23556756-23558774 | p2 | (AG)9 | TTCGGATTCTGGTTCGATTC | AGGGGGTCAGTACGGAAAAT | 277 |
| chr5:23563497-23565521 | p4 | (ATTT)6 | TATGCCTATTCGGGCAAAAG | TGGCCGAATATACCCTTGAG | 284 |
| chr5:23608622-23610637 | p5 | (TAATA)3 | AAGAGGGGCGGAGAGTTTAG | ATGTCGGAATTCCACTTTGC | 215 |
| chr5:23651798-23653813 | p5 | (AAAAT)3 | CGTCTTAAGCGTGTGGACAA | CATTTCGACGCAATCAGCTA | 181 |
| chr5:23674342-23676354 | p2 | (AT)6 | ATTCGAAACTTGTGGGTTCG | AACCCAGCAGCTTTTCTTCA | 132 |
| chr5:23758446-23760460 | p2 | (AT)7 | TCCAAAGGAACGGAGAGAAA | CAAAGGTGAAGCGTGTTCAA | 130 |
| chr5:23761209-23763247 | p2 | (TA)19 | AGTGTACGCTGCATCTGTCG | GACCAGGGGGTGATTATGTG | 190 |
| chr5:23812312-23814332 | p2 | (TA)6 | CGTCTGCAAGGGACACTGTA | TTTTGGCTCCGGACTCTATG | 122 |
| chr5:23877548-23879563 | p5 | (TTTTC)3 | ATGCGATTCAAAGCGAGACT | TTAGAGCTGAGCTTTGGGGA | 156 |
| chr5:23877548-23879563 | p5 | (TCTTT)3 | ATGCGATTCAAAGCGAGACT | TTAGAGCTGAGCTTTGGGGA | 156 |
| chr5:23934784-23936802 | p6 | (GTATAT)3 | TTAGAATCTCGGGGATGGTG | GCCTTTGGAAGAAAAGGGAC | 227 |
| chr5:23934784-23936802 | p5 | (ATGAT)6 | TTAGAATCTCGGGGATGGTG | GCCTTTGGAAGAAAAGGGAC | 227 |
| chr5:23940282-23942312 | p2 | (AT)15 | GCCCCACGACTATGAAATGA | CATGCAATGTCACTTCACCC | 282 |
| chr5:23940282-23942312 | p2 | (AT)11 | GCCCCACGACTATGAAATGA | CATGCAATGTCACTTCACCC | 282 |
| chr5:23948993-23951005 | p2 | (GA)6 | TTCTGGTCCCGTTCCTAGTG | CTCAAGTTGCCAAAGGAAGC | 176 |
| chr5:24070709-24072724 | p5 | (AAGCC)3 | AGGGGCACATTTTGTCACTC | GGCTTCTCATTGCCTCTTTG | 119 |
| chr5:24096513-24098533 | p4 | (TTTG)5 | CCAGTCCCAATTTATGTGGC | CCTCCGTCCCAATTTATGTG | 113 |
| chr5:24136510-24138528 | p2 | (TA)11 | CTGCTGGACCCATTTTCTGT | TCAATATTCTGGGCCAAAGC | 102 |

| Marker ID | SSR type | SSR motif | FORWARD PRIMER (5'-3') | REVERSE PRIMER (5'-3') | size (bp) |
|---|---|---|---|---|---|
| chr5:24136510-24138528 | p3 | (TAT)6 | CTGCTGGACCCATTTTCTGT | TCAATATTCTGGGCCAAAGC | 102 |
| chr5:24163751-24165763 | p2 | (TA)6 | TTATCTCCAAACTGGGCCAC | GCACATTAGCTGGGTTTGGT | 276 |
| chr5:24177961-24179975 | p2 | (TG)7 | TTCAATTTTGGACTTTGGGC | TATGATTGGTGATCGGGGTT | 136 |
| chr5:24258865-24260886 | p3 | (TAA)7 | AACGATGTACTGAGGGTGCC | CAACTTCCGCATACACATCG | 229 |
| chr5:24258865-24260886 | p3 | (ATA)5 | AACGATGTACTGAGGGTGCC | CAACTTCCGCATACACATCG | 229 |
| chr5:24342363-24344377 | p2 | (TG)7 | GGAGTGGGGACACAGAGAAA | GAGGTTGGCAGATTCCCATA | 179 |
| chr5:24396960-24398974 | p2 | (TG)7 | AAGGTTTAAGCACAATGCGG | TTTTTCTACCGCACCTTTGG | 138 |
| chr5:24431255-24433291 | p3 | (ATT)12 | AACAAGGGCCCATTTAACCT | GCCATACAGGTTGCACATTG | 246 |
| chr5:24460784-24462798 | p2 | (TA)33 | CCTTTCAAATGGTCATCCGT | TCCCGAAGAAAGTTCCTCAA | 243 |
| chr5:24460800-24462826 | p2 | (TA)33 | CCTTTCAAATGGTCATCCGT | TCCCGAAGAAAGTTCCTCAA | 227 |
| chr5:24460827-24462893 | p2 | (TA)33 | CCTTTCAAATGGTCATCCGT | TCCCGAAGAAAGTTCCTCAA | 200 |
| chr5:24476484-24478506 | p2 | (TA)11 | TGGACTTGAGGGAAACCATC | TCTGATTTGGCAGCTTACCC | 222 |
| chr5:24480022-24482112 | p3 | (TAT)30 | AACCAGGACACCACTTACCG | TCATTTCCAGATCCTCGGAC | 275 |
| chr5:24485806-24487824 | p6 | (AACTGA)3 | GGATACCTCGCCTTTGATGA | GTACCCCATCGGAAGTGAGA | 146 |
| chr5:24504901-24506917 | p2 | (TA)8 | TTCGAGTCGAGCGTATGATG | GGCGAGCCTTAACATACCTG | 267 |
| chr5:24564003-24566015 | p2 | (AT)6 | CAACTCCAATGTGATGGACG | CCAAGGGAATTAACTCGCAA | 107 |
| chr5:24564003-24566015 | p3 | (CAA)10 | CAACTCCAATGTGATGGACG | CCAAGGGAATTAACTCGCAA | 107 |
| chr5:24616943-24618955 | p2 | (TA)15 | ATATTGGCGTGCAAATAGCC | GTTAAGCGGTTCCGGTTTTT | 151 |
| chr5:24616943-24618955 | p2 | (TA)6 | ATATTGGCGTGCAAATAGCC | GTTAAGCGGTTCCGGTTTTT | 151 |
| chr5:24616943-24618955 | p2 | (CT)6 | ATATTGGCGTGCAAATAGCC | GTTAAGCGGTTCCGGTTTTT | 151 |
| chr5:24684803-24686839 | p2 | (TA)18 | CTGTTTTCGCGGACCATAAT | CATGGCAACCACGTACTCAG | 141 |
| chr5:24752997-24755017 | p5 | (TATAA)4 | TGGCTATTTGATCCTGCACA | TGGAATCAAAGCAACACGAG | 144 |
| chr5:24766190-24768205 | p5 | (ATTGC)3 | CTTCGGGTATGAAGTTGGGA | ACATATCACATTCGGCGACA | 108 |
| chr5:24847801-24849819 | p5 | (AATGC)3 | CTCTCAGCCTTTGTGGGAAG | CACGGGGTGGATTATGTTTC | 193 |
| chr5:24847891-24849909 | p5 | (AATGC)3 | CTCTCAGCCTTTGTGGGAAG | CACGGGGTGGATTATGTTTC | 103 |
| chr5:24909820-24911848 | p2 | (AT)14 | AGACTTCCGGGTCTATGGCT | TATGGCTGGGTCAAAGGAAG | 165 |
| chr5:24918563-24920609 | p6 | (TAATTC)3 | CAACAAAAACGTGGCAGCTA | TGTTTTTCTTGTGTGGGGGT | 195 |
| chr5:24920427-24922445 | p6 | (TTTTAT)3 | ATGTGGCAATGAATCCATCA | TGGCCCCATATTAAAAACCA | 168 |
| chr5:24920427-24922445 | p2 | (AT)13 | ATGTGGCAATGAATCCATCA | TGGCCCCATATTAAAAACCA | 168 |
| chr5:24920427-24922445 | p2 | (AT)33 | ATGTGGCAATGAATCCATCA | TGGCCCCATATTAAAAACCA | 168 |
| chr5:24927007-24929053 | p6 | (TAATTC)3 | CAACAAAAACGTGGCAGCTA | TGTTTTTCTTGTGTGGGGGT | 195 |
| chr5:24928871-24930889 | p6 | (TTTTAT)3 | ATGTGGCAATGAATCCATCA | TGGCCCCATATTAAAAACCA | 168 |
| chr5:24928871-24930889 | p5 | (ATATA)3 | ATGTGGCAATGAATCCATCA | TGGCCCCATATTAAAAACCA | 168 |
| chr5:24968988-24971016 | p2 | (TA)37 | CCTTACGTCACCCCGATAGA | AATAATGTGAGCCTGCGTCC | 184 |
| chr5:24968988-24971016 | p2 | (AT)14 | CCTTACGTCACCCCGATAGA | AATAATGTGAGCCTGCGTCC | 184 |
| chr5:24991702-24993729 | p3 | (TAA)9 | ACCACGAAAATCGAACTTGC | TCTTGGTGACGAAAATTCCC | 130 |

| Marker ID | SSR type | SSR motif | FORWARD PRIMER (5'-3') | REVERSE PRIMER (5'-3') | size (bp) |
|---|---|---|---|---|---|
| chr5:24996566-24998581 | p5 | (ATTGC)3 | TGTCGCCGATTGTGATATGT | TATGTCGTGCATGGTGACCT | 190 |
| chr5:25010717-25012747 | p2 | (AT)15 | GTGGGGACTTGCAATCAACT | ATTCCAGCCAGACCAGAGAA | 280 |
| chr5:25010717-25012747 | p2 | (AT)6 | GTGGGGACTTGCAATCAACT | ATTCCAGCCAGACCAGAGAA | 280 |
| chr5:25168389-25170404 | p5 | (TTTTA)3 | CTCCCCACACCCCTATTTTT | CACTTCCCAGACGGTAAGGA | 208 |
| chr5:25203313-25205327 | p2 | (AT)7 | AAGATTTTGATGTCGCGCTT | TCCTTGCAAATTTTTGGACC | 169 |
| chr5:25275394-25277410 | p2 | (AT)8 | TTCCGAGGTTGAGCAGAGAT | CCCCCTCAGGAAAAGAACTC | 115 |
| chr5:25333686-25335701 | p5 | (TTAAA)3 | AAAACAAAACGAGTGGACGG | TGTCTTGATGGCACCACAGT | 257 |
| chr5:25370649-25372691 | p3 | (AAT)14 | AAGAAAACGATGGGTTCACG | TGCATGGCCAAATTCAGTAA | 231 |
| chr5:25379979-25381991 | p2 | (TA)6 | CCAGTTGACCGAACCTCAAT | CGGTCCGTACCTTTCGTTTA | 190 |
| chr5:25411112-25413130 | p2 | (TA)7 | AAAGACAAGCGACGAAGCAT | AACAATGTCCAAAAGGGTGG | 269 |
| chr5:25444824-25446878 | p3 | (ATT)13 | TCCTATCCCGTCTCCATGTC | CGTGTGCATTGGATGGTAAG | 299 |
| chr5:25444824-25446878 | p2 | (CT)27 | TCCTATCCCGTCTCCATGTC | CGTGTGCATTGGATGGTAAG | 299 |
| chr5:25483378-25485400 | p2 | (TA)11 | AGACTTCCGGGTCTATGGCT | TATGGCTGGGTCAAAGGAAG | 174 |
| chr5:25525392-25527407 | p5 | (TAAAT)3 | AATGTGGAGGCCACTACCAG | ACCGCAGTTCTAGCACAACC | 234 |
| chr5:25545667-25547691 | p2 | (AT)12 | CCGAGCCCAAACTCAAAATA | GGGAAGGTAGGATGTACGCA | 181 |
| chr5:25548354-25550366 | p5 | (ATTTT)3 | TCTAACCCCCAAGGGCTAAC | TAGGTTGGCCCACCGTATAA | 171 |
| chr5:25548354-25550366 | p2 | (AT)6 | TCTAACCCCCAAGGGCTAAC | TAGGTTGGCCCACCGTATAA | 171 |
| chr5:25549988-25552000 | p2 | (CA)6 | ACGATGAGAATCGCTAACGG | TTCGTCATTTGTCGGAACAG | 222 |
| chr5:25549988-25552000 | p2 | (AT)20 | ACGATGAGAATCGCTAACGG | TTCGTCATTTGTCGGAACAG | 222 |
| chr5:25566897-25568912 | p5 | (AAATA)3 | TGGGCTCCTTTGAGTGTTTT | TCTTGAAGACATCACGCACC | 156 |
| chr5:25640757-25642775 | p3 | (CAA)6 | CATATATCCCTTCCCCCACC | ACCTTACGTCACTCCGATGG | 163 |
| chr5:25640757-25642775 | p2 | (AT)6 | CATATATCCCTTCCCCCACC | ACCTTACGTCACTCCGATGG | 163 |
| chr5:25716973-25718993 | p2 | (AT)10 | TTAACGTTCCACGTGTTCCA | GCTGGGGAGGTTCTAGGTTC | 237 |
| chr5:25716973-25718993 | p2 | (TG)6 | TTAACGTTCCACGTGTTCCA | GCTGGGGAGGTTCTAGGTTC | 237 |
| chr5:25744228-25746244 | p2 | (TA)55 | TATGCCCCATAAGGCAGAAC | TACCAAAGTTATGCCGGACC | 123 |
| chr5:25744228-25746244 | p2 | (AT)8 | TATGCCCCATAAGGCAGAAC | TACCAAAGTTATGCCGGACC | 123 |
| chr5:25754542-25756558 | p2 | (TA)8 | TGGATTTGAACGTGGTTTCA | CCTCGTAGCTCTCGCAAAGT | 206 |
| chr5:25813570-25815585 | p2 | (TA)13 | AAGCCCACAAGGCTACAAAA | TATTCGGTGTATGACGGGGT | 174 |
| chr5:25813570-25815585 | p3 | (ATC)5 | AAGCCCACAAGGCTACAAAA | TATTCGGTGTATGACGGGGT | 174 |
| chr5:25913507-25915522 | p3 | (TGA)5 | GTAACCGGACCCGGATAAAT | TAATGGTCCATAAGCCCAGC | 187 |
| chr5:25935685-25937700 | p3 | (CAA)5 | AACGCGGGGTGTAACAATAA | ACCCCGAGATTTCTTTCGTT | 297 |
| chr5:25941117-25943132 | p5 | (TTAGA)3 | AGCCCTAAAACGCTCCTCTC | TGGCCTTTGAATTTCGAATC | 153 |
| chr5:25944960-25946990 | p2 | (AT)15 | TTGCATATAAGGGACCCCAA | AAAACACATTGGGGCTCAAG | 197 |
| chr5:25958305-25960377 | p3 | (TTG)6 | GGCCAAAAATTAGCGTGAAA | GTTTATGTGCCTAGCCCGAA | 234 |
| chr5:25958305-25960377 | p2 | (AT)36 | GGCCAAAAATTAGCGTGAAA | GTTTATGTGCCTAGCCCGAA | 234 |
| chr5:25958305-25960377 | p2 | (TA)10 | GGCCAAAAATTAGCGTGAAA | GTTTATGTGCCTAGCCCGAA | 234 |

| Marker ID | SSR type | SSR motif | FORWARD PRIMER (5'-3') | REVERSE PRIMER (5'-3') | size (bp) |
|---|---|---|---|---|---|
| chr5:25978821-25980836 | p3 | (TGT)5 | GCCCGTCATACTCCTCGTAG | AATGGCTCCGAGCTCACTTA | 142 |
| chr5:25991901-25993916 | p2 | (TA)10 | GATGGGTTTGGGGTTTCTTT | CCGGTCCATTTGAAGGATAA | 172 |
| chr5:25991917-25993947 | p2 | (TA)10 | GATGGGTTTGGGGTTTCTTT | CCGGTCCATTTGAAGGATAA | 156 |
| chr5:26004865-26006925 | p2 | (AT)7 | ACTGCCTATTTGCATTCGCT | TTTCGTTGATTTGGTCACGA | 244 |
| chr5:26004865-26006925 | p2 | (TA)10 | ACTGCCTATTTGCATTCGCT | TTTCGTTGATTTGGTCACGA | 244 |
| chr5:26004865-26006925 | p5 | (AAAAC)3 | ACTGCCTATTTGCATTCGCT | TTTCGTTGATTTGGTCACGA | 244 |
| chr5:26004977-26006995 | p2 | (TA)10 | ACTGCCTATTTGCATTCGCT | TTTCGTTGATTTGGTCACGA | 132 |
| chr5:26004977-26006995 | p5 | (AAAAC)3 | ACTGCCTATTTGCATTCGCT | TTTCGTTGATTTGGTCACGA | 132 |
| chr5:26228907-26230931 | p2 | (AT)12 | AAAAGGCCTCCATTCCAATC | TTCACCAATTTGGGGTTCAT | 207 |
| chr5:26287149-26289164 | p3 | (ATG)5 | ACATGAAGGCAATCCGAATC | ATAGGAAGCATCATGCGGAC | 296 |
| chr5:26301296-26303310 | p2 | (AT)7 | TGGAGGATCAAGAGACGACC | GCGTTCCGGAATACAGAAAA | 188 |
| chr5:26335549-26337576 | p3 | (TGT)9 | CGAGCTATGGCTGGGTAAAA | TGCTTGGTCCCAAAATCTTC | 126 |
| chr5:26335549-26337576 | p3 | (GAT)5 | CGAGCTATGGCTGGGTAAAA | TGCTTGGTCCCAAAATCTTC | 126 |
| chr5:26335549-26337576 | p2 | (TA)15 | CGAGCTATGGCTGGGTAAAA | TGCTTGGTCCCAAAATCTTC | 126 |
| chr5:26363871-26365883 | p2 | (TG)6 | ATCTGGAGGCAGTACATCCG | GCGCAGTCTCCTTACCTGAC | 179 |
| chr5:26409792-26411810 | p2 | (AT)9 | GGGACATGAATGACCCAATC | AATTCTCCGCAGTGACGACT | 252 |
| chr5:26564454-26566469 | p5 | (TGATA)3 | ATAGGCGAAAAGAGCAACGA | GAGCAATCCGAGCCTTTATG | 255 |
| chr5:26569281-26571299 | p6 | (GTGATT)3 | TTTTATGGGGTTTATGGGCA | TAAGGCGAACCCTCTAAGCA | 160 |
| chr5:26572714-26574729 | p6 | (AAAAAT)3 | ATTCATCGCTAATTCGACGG | AATAAATTTTGGGGCTTGGC | 295 |
| chr5:26572714-26574729 | p5 | (ATTTT)3 | ATTCATCGCTAATTCGACGG | AATAAATTTTGGGGCTTGGC | 295 |
| chr5:26630270-26632285 | p3 | (GTT)5 | GTGGGGGCTTTTGTAGTCCT | TCTCCAGGTTCGATGCTTTT | 162 |
| chr5:26666344-26668359 | p5 | (ACCAA)3 | TACATGGATCGGGTTGGTTC | AGGTTGAAAACCCCGAATTT | 249 |
| chr5:26666344-26668359 | p5 | (AAACC)3 | TACATGGATCGGGTTGGTTC | AGGTTGAAAACCCCGAATTT | 249 |
| chr5:26696519-26698552 | p3 | (AAT)11 | TTCAAAAGGGAAATTCGTGC | CACAAGAAACCCTAGCTCGC | 176 |
| chr5:26725640-26727666 | p2 | (TA)8 | CGTGAATTTTGATGTGGTGG | CTGTTTCGTCCTCCGTTGTT | 135 |
| chr5:26807702-26809717 | p5 | (ACCAA)3 | CCAGACACCATGGTTATCCC | AGCAATTGTGGATTTGAGCC | 256 |
| chr5:26884968-26886983 | p5 | (ATTTT)3 | TATTGCGACCATTTGCATGT | GAGGCTCCTACAAGCCACAC | 286 |
| chr5:27029533-27031547 | p2 | (AT)7 | TCCAAAACAAGTGCACTCCA | AAGCAATTTCATTTGCTCCC | 162 |
| chr5:27086004-27088018 | p2 | (TC)7 | TGGGATGTCTTGATTCGACA | ATGTCATGGGTGGAATTGCT | 156 |
| chr5:27106360-27108381 | p3 | (ATT)7 | TTGCCATTTTCTCGGATTTC | AACCACGGAAGGATTCAATG | 227 |
| chr5:27221745-27223760 | p5 | (AAATT)3 | GAGAACCGAGATGATCGGAA | TGGATGACAACTCGTCCAAA | 159 |
| chr5:27221745-27223760 | p5 | (AATAA)3 | GAGAACCGAGATGATCGGAA | TGGATGACAACTCGTCCAAA | 159 |
| chr5:27221745-27223760 | p3 | (AGA)5 | GAGAACCGAGATGATCGGAA | TGGATGACAACTCGTCCAAA | 159 |
| chr5:27221745-27223760 | p5 | (TTTAT)3 | GAGAACCGAGATGATCGGAA | TGGATGACAACTCGTCCAAA | 159 |
| chr5:27350867-27352879 | p2 | (TA)6 | GACTGCACATAATTGGGGCT | TTCAATTCTCGGGTCGTAGG | 212 |
| chr5:27449563-27451583 | p5 | (TTTCT)4 | CATGGGTGAAAGGGAGAGAA | AAAGGAGCCAACTCCAATCA | 186 |

| Marker ID | SSR type | SSR motif | FORWARD PRIMER (5'-3') | REVERSE PRIMER (5'-3') | size (bp) |
|---|---|---|---|---|---|
| chr5:27523769-27525789 | p2 | (TA)10 | CCGCCATGCTAATATAGGGA | AGCGAGATTGGTAAGCTCCA | 145 |
| chr5:27523769-27525789 | p2 | (TA)10 | CCGCCATGCTAATATAGGGA | AGCGAGATTGGTAAGCTCCA | 145 |
| chr5:27567783-27569795 | p2 | (TA)6 | GCTCGTAACCCCCATACTCA | TACTTCACCCCTCGCAAATC | 264 |
| chr5:27576441-27578457 | p2 | (AT)8 | GACATGGCGGAATAGCAAAT | TTCATTACCACGTGCGTGTC | 203 |
| chr5:27576441-27578457 | p2 | (AT)7 | GACATGGCGGAATAGCAAAT | TTCATTACCACGTGCGTGTC | 203 |
| chr5:27609889-27611904 | p3 | (AAC)5 | CACACTGGGGGTAAAGTGCT | CGTTCTTCAATCATCCGACA | 215 |
| chr5:27612704-27614743 | p3 | (TTC)13 | TAGAGAAAGTTTTCCGGCGA | CCAATTAAACCCCAACCCTT | 205 |
| chr5:27612704-27614743 | p3 | (TTC)8 | TAGAGAAAGTTTTCCGGCGA | CCAATTAAACCCCAACCCTT | 205 |
| chr5:27622019-27624037 | p3 | (TTC)6 | TTTGAAAACCCCAGGCATAG | GCAGACAAGAAAGCAAAGGG | 219 |
| chr5:27684755-27686771 | p2 | (AG)6 | ATGGTGCAAGGCGAAAATAC | GGAAGGAGGAACTTTTTGGC | 143 |
| chr5:27684755-27686771 | p4 | (TTTA)4 | ATGGTGCAAGGCGAAAATAC | GGAAGGAGGAACTTTTTGGC | 143 |
| chr5:27690608-27692624 | p2 | (AT)7 | TGAGGGTGAGGTGGAAAATC | CAGCATACCTCGAGCACAAA | 260 |
| chr5:27690654-27692674 | p2 | (AT)7 | TGAGGGTGAGGTGGAAAATC | CAGCATACCTCGAGCACAAA | 214 |
| chr5:27712086-27714101 | p5 | (AAAAT)3 | CGCAAGGCATAAGTTGGATT | CTTCGCTTAACACCCAGAGG | 259 |
| chr5:27715239-27717257 | p3 | (TAA)6 | GTGGAGAAGTGGAAGGTGGA | ACCCTTTTTAGGGTTTCCCC | 208 |
| chr5:27715239-27717257 | p3 | (ATA)6 | GTGGAGAAGTGGAAGGTGGA | ACCCTTTTTAGGGTTTCCCC | 208 |
| chr5:27715239-27717257 | p3 | (ATA)14 | GTGGAGAAGTGGAAGGTGGA | ACCCTTTTTAGGGTTTCCCC | 208 |
| chr5:27715405-27717423 | p3 | (TAA)6 | GGAGCAGTGAGAAAGAGGGA | ATTTTGACTTACCGGACCCC | 146 |
| chr5:27715405-27717423 | p3 | (ATA)6 | GGAGCAGTGAGAAAGAGGGA | ATTTTGACTTACCGGACCCC | 146 |
| chr5:27715405-27717423 | p3 | (ATA)14 | GGAGCAGTGAGAAAGAGGGA | ATTTTGACTTACCGGACCCC | 146 |
| chr5:27797776-27799800 | p2 | (AT)12 | ATGACCCATGACCCACAAAT | AGGTACGGACGATCGTTTTG | 256 |
| chr5:27880415-27882430 | p5 | (GTCTA)3 | AATCGGGACTCTCTGGGTTT | TGTCTGGTTCCATCACCAAA | 152 |
| chr5:27931180-27933196 | p4 | (ATTT)4 | GGCCTCTACAAAGGCACAAG | TGCAAAAGTGAGGTTCAACG | 259 |
| chr5:27949905-27951927 | p2 | (TG)11 | GGTCGCATTTCAAAAGGTGT | ACCACCCCCTAAAAATCACC | 253 |
| chr5:28026463-28028478 | p2 | (AG)18 | CAAAACAAACATGCCGTGTC | CTAAACACTACCCGACCCGA | 285 |
| chr5:28026479-28028524 | p2 | (AG)18 | CAAAACAAACATGCCGTGTC | CTAAACACTACCCGACCCGA | 269 |
| chr5:28217867-28219879 | p2 | (AT)7 | ATGCCCTTTGATCGCTACAT | TCGTGGAACAATCCACAAAA | 253 |
| chr5:28217867-28219879 | p2 | (AT)16 | ATGCCCTTTGATCGCTACAT | TCGTGGAACAATCCACAAAA | 253 |
| chr5:28239589-28241604 | p6 | (TATATG)3 | AAGGCTCGTCCCTTTCTTTC | ATCATTGCCTTGGGACCATA | 211 |
| chr5:28239589-28241604 | p5 | (TATGA)3 | AAGGCTCGTCCCTTTCTTTC | ATCATTGCCTTGGGACCATA | 211 |
| chr5:28258226-28260241 | p5 | (TTTTG)3 | AGCCCATGTTTTTCCAACTG | TTGTTCCAACTCTGCACTGG | 138 |
| chr5:28280012-28282076 | p2 | (TA)32 | TTTTGCAAGATCTGTCGGTG | AGCAGGTATCGGCTCAGCTA | 243 |
| chr5:28380140-28382208 | p2 | (CT)7 | TACGGGTGTCAGACCGTACA | ACCACAAATCCGGACCAATA | 131 |
| chr5:28380140-28382208 | p2 | (TA)34 | TACGGGTGTCAGACCGTACA | ACCACAAATCCGGACCAATA | 131 |
| chr5:28380140-28382208 | p6 | (TGAACA)3 | TACGGGTGTCAGACCGTACA | ACCACAAATCCGGACCAATA | 131 |
| chr5:28483385-28485397 | p2 | (AT)6 | AAAGGGGAAAGGTTAGGGGT | TCCAGCCCATTACTCCTTTG | 117 |

| Marker ID | SSR type | SSR motif | FORWARD PRIMER (5'-3') | REVERSE PRIMER (5'-3') | size (bp) |
|---|---|---|---|---|---|
| chr5:28536426-28538438 | p2 | (TA)6 | CTGTACTACGGGCCAACCAT | TCAACGACGGGGAATTTAAG | 116 |
| chr5:28578122-28580140 | p3 | (TTA)6 | GGAAATGCTTTCCATTGAGC | CAACAATCCGGTGAAAATCC | 286 |
| chr5:28686630-28688646 | p4 | (TCTT)4 | TTGTTGCATTTCGCTTCAAC | CACGTGGCTTGTTTTGATTG | 151 |
| chr5:28768417-28770431 | p2 | (CT)7 | ATAAGGCTTCATACGGGGCT | GTGCATTGGACTTTGGAGGT | 151 |
| chr5:28790060-28792072 | p2 | (AT)6 | GAACCGCTGGAATAACCGTA | AAAATGAATGTAAACCGCCG | 101 |
| chr5:28809167-28811183 | p4 | (TTTC)4 | CAGCCATAACACCCAGAGGT | GAGGGACCATCATTCCTCAA | 126 |
| chr5:28870241-28872256 | p5 | (GTAAT)3 | GCCCACACAAATAAAATCCG | CCCCAACCAGAGCATATTGT | 153 |
| chr5:28935701-28937719 | p3 | (TTG)6 | TCGTAATCGGGAAGGACAAC | TTGATCCCGAAATCTTCCAG | 217 |
| chr5:28975053-28977075 | p2 | (AT)11 | AACCACTTGACTCTTTTGGCA | GGGGTGTTGTGGTCTTCTGT | 170 |
| chr5:28988748-28990760 | p2 | (TA)6 | AGGGGAAGCTGTCCAAAAAT | AGAAAAGAGCGCGGTAGACA | 228 |
| chr5:29029246-29031258 | p2 | (AG)6 | CCGTCGACTTCTTCTTCCAG | TCCAAGTTCTCATCTCCGCT | 209 |
| chr5:29040922-29042944 | p2 | (TA)11 | GGACCCCCATGATTATTGC | TGACATGGCGGAATAACAAA | 210 |
| chr5:29100108-29102134 | p6 | (ATTTTT)3 | ATTGTGGGGTTACCGGTGTA | GACCAGTGCACGAAAAGTCA | 225 |
| chr5:29100108-29102134 | p2 | (TC)13 | ATTGTGGGGTTACCGGTGTA | GACCAGTGCACGAAAAGTCA | 225 |
| chr5:29100108-29102134 | p2 | (GT)10 | ATTGTGGGGTTACCGGTGTA | GACCAGTGCACGAAAAGTCA | 225 |
| chr5:29302518-29304539 | p3 | (TTG)7 | CTGGGCGGATTTTTAGTCAA | TGTTTCTGCGTCGACAGTTC | 206 |
| chr5:29302518-29304539 | p2 | (TA)6 | CTGGGCGGATTTTTAGTCAA | TGTTTCTGCGTCGACAGTTC | 206 |
| chr5:29344047-29346063 | p4 | (TTTA)4 | GGTAATAGCAACCTTGCCGA | AAATTCGATTTGCTGTTCCG | 115 |
| chr5:29351862-29353874 | p2 | (AT)6 | CCGTTCTTTCAACAATGGCT | GCCCAATGAGTGTCGGTTAT | 132 |
| chr5:29351862-29353874 | p5 | (GGCCC)3 | CCGTTCTTTCAACAATGGCT | GCCCAATGAGTGTCGGTTAT | 132 |
| chr5:29518327-29520351 | p3 | (CAA)8 | GTGGTAACATCTGCCCGACT | ATGGCCTGAGCTGTGCTTAT | 171 |
| chr5:29518327-29520351 | p3 | (CAA)7 | GTGGTAACATCTGCCCGACT | ATGGCCTGAGCTGTGCTTAT | 171 |
| chr5:29523056-29525071 | p3 | (ACA)5 | GGAGGAGGAGAGAAGGAGGA | CATTTTCGGACTTTCTCCCA | 196 |
| chr5:29640892-29642908 | p2 | (AT)8 | TTTAAATTCCCGCAGAGTGG | TTTAGGACCCGTTTGCCATA | 217 |
| chr5:29647855-29649876 | p3 | (ACA)7 | TTGGTCCTTTCACCTTTTGC | AAAGCATCTGGCCATTTTTG | 187 |
| chr5:29647855-29649876 | p5 | (TCATA)3 | TTGGTCCTTTCACCTTTTGC | AAAGCATCTGGCCATTTTTG | 187 |
| chr5:29682884-29684899 | p5 | (TGCTT)3 | GATACCTCTTTCCGCCCTTC | GCACTGCATTCTAGGGAAGC | 180 |
| chr5:29713140-29715178 | p2 | (AT)19 | GTCATGACTGCCGAGACAAA | CGAAGCATGCAAATGCTAGA | 255 |
| chr5:29713140-29715178 | p2 | (AT)6 | GTCATGACTGCCGAGACAAA | CGAAGCATGCAAATGCTAGA | 255 |
| chr5:29733075-29735095 | p4 | (ATAA)5 | CCTGGGTCTGATTGTTTGCT | TTGGTTGTGATTCCTGTGGA | 138 |
| chr5:29778959-29780981 | p2 | (AT)11 | TTTTCCCAGTTTGAGATCGG | CCTAATGCAGAAGCTCGGTT | 295 |
| chr5:29778959-29780981 | p2 | (AT)11 | TTTTCCCAGTTTGAGATCGG | CCTAATGCAGAAGCTCGGTT | 295 |
| chr5:29778959-29780981 | p2 | (AT)11 | TTTTCCCAGTTTGAGATCGG | CCTAATGCAGAAGCTCGGTT | 295 |
| chr5:29778959-29780981 | p2 | (AT)11 | TTTTCCCAGTTTGAGATCGG | CCTAATGCAGAAGCTCGGTT | 295 |
| chr5:29778959-29780981 | p2 | (AT)11 | TTTTCCCAGTTTGAGATCGG | CCTAATGCAGAAGCTCGGTT | 295 |
| chr5:29778959-29780981 | p2 | (AT)11 | TTTTCCCAGTTTGAGATCGG | CCTAATGCAGAAGCTCGGTT | 295 |

| Marker ID | SSR type | SSR motif | FORWARD PRIMER (5'-3') | REVERSE PRIMER (5'-3') | size (bp) |
|---|---|---|---|---|---|
| chr5:29778959-29780981 | p2 | (AT)11 | TTTTCCCAGTTTGAGATCGG | CCTAATGCAGAAGCTCGGTT | 295 |
| chr5:29778959-29780981 | p2 | (AT)11 | TTTTCCCAGTTTGAGATCGG | CCTAATGCAGAAGCTCGGTT | 295 |
| chr5:29778959-29780981 | p2 | (AT)11 | TTTTCCCAGTTTGAGATCGG | CCTAATGCAGAAGCTCGGTT | 295 |
| chr5:29779108-29781130 | p2 | (AT)11 | TTTTCCCAGTTTGAGATCGG | CCAAAGCTCGGTTTTGAACT | 146 |
| chr5:29779108-29781130 | p2 | (AT)11 | TTTTCCCAGTTTGAGATCGG | CCAAAGCTCGGTTTTGAACT | 146 |
| chr5:29779108-29781130 | p2 | (AT)11 | TTTTCCCAGTTTGAGATCGG | CCAAAGCTCGGTTTTGAACT | 146 |
| chr5:29779108-29781130 | p2 | (AT)11 | TTTTCCCAGTTTGAGATCGG | CCAAAGCTCGGTTTTGAACT | 146 |
| chr5:29779108-29781130 | p2 | (AT)11 | TTTTCCCAGTTTGAGATCGG | CCAAAGCTCGGTTTTGAACT | 146 |
| chr5:29779108-29781130 | p2 | (AT)11 | TTTTCCCAGTTTGAGATCGG | CCAAAGCTCGGTTTTGAACT | 146 |
| chr5:29779108-29781130 | p2 | (AT)11 | TTTTCCCAGTTTGAGATCGG | CCAAAGCTCGGTTTTGAACT | 146 |
| chr5:29779108-29781130 | p2 | (AT)11 | TTTTCCCAGTTTGAGATCGG | CCAAAGCTCGGTTTTGAACT | 146 |
| chr5:29779108-29781130 | p2 | (AT)11 | TTTTCCCAGTTTGAGATCGG | CCAAAGCTCGGTTTTGAACT | 146 |
| chr5:29779108-29781130 | p2 | (AT)11 | TTTTCCCAGTTTGAGATCGG | CCAAAGCTCGGTTTTGAACT | 146 |
| chr5:29791227-29793241 | p2 | (AT)7 | TCCCAAGCATTCTCGTCTCT | ATGTTAGGAGATCCCGGCTT | 219 |
| chr5:29859374-29861389 | p3 | (CTC)5 | ACAAAATAGGGGCAAACGTG | GTGGTGAATGGTAGGGATGG | 104 |
| chr5:29859374-29861389 | p5 | (TTTTA)3 | ACAAAATAGGGGCAAACGTG | GTGGTGAATGGTAGGGATGG | 104 |
| chr5:29940876-29942891 | p3 | (TTA)5 | GACTGCATGCGTGGCTACTA | GCCACATTTCTTGCCACTTT | 294 |
| chr5:29952407-29954423 | p2 | (TC)8 | ACAAAAGGGGTCATTTGCAG | TAATGGGCTGAAGCTTTTGG | 136 |
| chr5:29952407-29954423 | p4 | (TATT)6 | ACAAAAGGGGTCATTTGCAG | TAATGGGCTGAAGCTTTTGG | 136 |
| chr5:29956273-29958311 | p2 | (AT)19 | CTCGAGGTTTTGGGTTCGTA | CGAACCCCTACTCAGGTGAA | 155 |
| chr5:30007877-30009915 | p2 | (AT)19 | ACGGAGAGGGAGTTTTTGGT | GGTCTCGCCCTTAACTTTCC | 271 |
| chr5:30010934-30012948 | p2 | (AT)7 | AGAGCTGATACCGCGTCTGT | TATGGAATGTCGTTCAGGCA | 136 |
| chr5:30077601-30079623 | p2 | (AG)11 | GAGAAGACCGCGAATGAGTC | GGTCCAGGATAGAGGCATCA | 245 |
| chr5:30098345-30100365 | p5 | (TTTTC)3 | AATGGAGGGGTGGGATTTAG | CCAAAATGCCCAATTTGTTC | 285 |
| chr5:30098345-30100365 | p2 | (AT)10 | AATGGAGGGGTGGGATTTAG | CCAAAATGCCCAATTTGTTC | 285 |
| chr5:30186308-30188322 | p2 | (TA)7 | GTTCGATATTGGGCCGTTTA | AAAGCCAATGCAAACTCACC | 248 |
| chr5:30226520-30228534 | p2 | (TA)7 | TTGGGCCGAAAGTTATCAAA | CGGCAGCTTCAAAATCAAAT | 162 |
| chr5:30240892-30242907 | p5 | (ATTTA)3 | CACATAGAGCCCGTTTGGAT | TCCAAAACCGAGAAGTTGCT | 248 |
| chr5:30261478-30263493 | p5 | (TTTTA)3 | GCGGTCACTTGTGAGTTGAA | ATGTTGAAACGTGTGACGGA | 121 |
| chr5:30261478-30263493 | p5 | (AAAAT)3 | GCGGTCACTTGTGAGTTGAA | ATGTTGAAACGTGTGACGGA | 121 |
| chr5:30261478-30263493 | p2 | (AT)9 | GCGGTCACTTGTGAGTTGAA | ATGTTGAAACGTGTGACGGA | 121 |
| chr5:30261893-30263908 | p5 | (TTTTA)3 | GACAACCCCAAAAGTGGGTA | TTTTGGTCGGAAAGGAATTG | 122 |
| chr5:30261893-30263908 | p5 | (AAAAT)3 | GACAACCCCAAAAGTGGGTA | TTTTGGTCGGAAAGGAATTG | 122 |
| chr5:30261893-30263908 | p2 | (AT)9 | GACAACCCCAAAAGTGGGTA | TTTTGGTCGGAAAGGAATTG | 122 |
| chr5:30297213-30299231 | p6 | (ATAAAA)3 | CTCCGCGAAATATCGTTTGT | CCGTGGTAAGAATGGGAAAA | 114 |
| chr5:30297213-30299231 | p5 | (TAAAA)3 | CTCCGCGAAATATCGTTTGT | CCGTGGTAAGAATGGGAAAA | 114 |

| Marker ID | SSR type | SSR motif | FORWARD PRIMER (5'-3') | REVERSE PRIMER (5'-3') | size (bp) |
|---|---|---|---|---|---|
| chr5:30478016-30480031 | p5 | (TTTTA)3 | TCGACATCAATACAGCCGAG | CCTGCGAGAGGATGGAATAA | 106 |
| chr5:30508400-30510421 | p3 | (ATA)7 | AGATGCTCTTGATGGCGACT | AGGAGCAACACCTCCTCAAA | 182 |
| chr5:30535023-30537038 | p2 | (TC)7 | TGGCAATCGAAAATCTCCTC | AGGGGAGCCAAATCTGAAAT | 108 |
| chr5:30535023-30537038 | p5 | (CTAAG)3 | TGGCAATCGAAAATCTCCTC | AGGGGAGCCAAATCTGAAAT | 108 |
| chr5:30734215-30736241 | p3 | (GAA)5 | AAAATGCCAACTGAACTGCC | GCCTAATCGAACTTCACCCA | 288 |
| chr5:30788129-30790144 | p5 | (AAACA)3 | TTAGTTGCAGGGCAAAACCT | CTGACGAGTTTTTGGGTGGT | 233 |
| chr5:30848138-30850152 | p2 | (TA)7 | AGCAGTTGGAAAAGTGTGGG | AGCAGAAGTCAGTGCCCCTA | 278 |
| chr5:30909648-30911660 | p2 | (AT)6 | TTTGCAGACAGGGTCACGTA | ATCCTGATAGGGCATGATGG | 127 |
| chr5:30918376-30920391 | p5 | (AGGAG)4 | AGCCCGTTCTTGATTTCCTT | CTCTATCTCCGGCCTCTCCT | 126 |
| chr5:30918376-30920391 | p3 | (TTG)5 | AGCCCGTTCTTGATTTCCTT | CTCTATCTCCGGCCTCTCCT | 126 |
| chr5:30956848-30958863 | p2 | (TA)22 | AAAAGCAAAGGGGGAAGTGT | CGGTCCCTACTTTAGTGCCA | 285 |
| chr5:30956848-30958863 | p3 | (ACT)5 | AAAAGCAAAGGGGGAAGTGT | CGGTCCCTACTTTAGTGCCA | 285 |
| chr5:31245131-31247146 | p3 | (AAT)5 | AATGGTTGGCTTTGATTGGA | GAAGAGACACCACTTGTTACAT | 294 |
| chr5:31521432-31523448 | p2 | (AT)8 | TGGTCATGTCGGGATCTTTT | TGCAGTGTCTCGATTGTCCT | 217 |
| chr5:31521432-31523448 | p2 | (TA)9 | TGGTCATGTCGGGATCTTTT | TGCAGTGTCTCGATTGTCCT | 217 |
| chr5:31521432-31523448 | p2 | (TA)6 | TGGTCATGTCGGGATCTTTT | TGCAGTGTCTCGATTGTCCT | 217 |
| chr5:31638123-31640191 | p2 | (TA)10 | CCCTTCAAATGCACTGGTCT | CTCCCTTCAGGCAGAGTTTG | 114 |
| chr5:31638357-31640377 | p2 | (TA)10 | AAACCCGAAATCAGGAGGTT | TGTGTGGTGCCCAAATAAGA | 211 |
| chr5:31643256-31645268 | p2 | (TA)6 | AGAAAAACAGAGGCAAGGCA | ACGCAACCATCCACATTTTT | 147 |
| chr5:31653071-31655083 | p2 | (CT)6 | ATGTACCCCCATTTGTTCCA | TTTTGGAACATTCCTCCTCG | 217 |
| chr5:31762400-31764416 | p2 | (AC)8 | GCAAAAGGGAAAGTGACCAA | CGATTCCTACAGCCAATGGT | 252 |
| chr5:31847464-31849532 | p2 | (TA)34 | GGCCAAGACAAATGGGTAGA | CCTGAGGCTGCACCATATTT | 241 |
| chr5:31882537-31884555 | p6 | (AAAATT)3 | CCGGATTGAATGGATAGGTG | ACTCCGGATGAGCAATTGAG | 256 |
| chr5:32018145-32020163 | p2 | (AT)9 | TTGCGTGCATGTTCTTTCTC | TAAGTGACTTCCGGGTCAGC | 103 |
| chr5:32031146-32033161 | p5 | (GAAAT)3 | TTTGGCTAAAGAACCATGCC | TAAAGGAGCTGCTGTCCGAT | 243 |
| chr5:32151003-32153021 | p2 | (TA)9 | ATGCCAACTCATTTCCCAAC | AGAACCCAAATCTGGTCGTG | 189 |
| chr5:32173168-32175194 | p2 | (TA)13 | ATTACTTGCAACGCGGAATC | AATTGTGCTTCCGACCAAAG | 262 |
| chr5:32283052-32285067 | p3 | (TTG)5 | AAGGGCCTATCGGAAACATC | ACCCATTATGATCCCACGAG | 272 |
| chr5:32283052-32285067 | p5 | (AAATA)3 | AAGGGCCTATCGGAAACATC | ACCCATTATGATCCCACGAG | 272 |
| chr5:32307458-32309473 | p5 | (TTTTC)3 | AGACACTCCTCGGTCTTCCA | TTAAAGCCTAGCCCGTTTGA | 285 |
| chr5:32312798-32314813 | p5 | (CTTTT)3 | TTGGCTTCAAAAAGGATTGG | GTCGGTTTTTCTAAGCGACG | 174 |
| chr5:32312939-32314972 | p5 | (CTTTT)3 | CTGCGTCGCTTAGAAAAACC | AAAACCGACGTTTTCTGTCG | 237 |
| chr5:32312939-32314972 | p2 | (AT)12 | CTGCGTCGCTTAGAAAAACC | AAAACCGACGTTTTCTGTCG | 237 |
| chr5:32331910-32333925 | p5 | (AAACC)3 | AAGGGTGTACATGAACCGGA | CTGAGAAAACCCGAGGTTGA | 218 |
| chr5:32335183-32337198 | p5 | (TACAA)3 | CATAACCCCCTGTTTGGATG | GTGGTAGGGTGGGTTGAATG | 101 |
| chr5:32359086-32361104 | p3 | (TCT)6 | TTGCTTGCTTGGCATCATAG | CCATAACATAAACCGGTGCC | 197 |

| Marker ID | SSR type | SSR motif | FORWARD PRIMER (5'-3') | REVERSE PRIMER (5'-3') | size (bp) |
|---|---|---|---|---|---|
| chr5:32422706-32424726 | p5 | (TTCGG)4 | CCCTGAACCATACGAGGAAA | GTTCGGTCCAGTTTTCGGTA | 250 |
| chr5:32422706-32424726 | p3 | (GGT)5 | CCCTGAACCATACGAGGAAA | GTTCGGTCCAGTTTTCGGTA | 250 |
| chr5:32524432-32526447 | p5 | (ATACT)3 | TCTCCACAACACCAAACGAA | AGGAAGGAAGGAAGGAGCAG | 106 |
| chr5:32533413-32535428 | p3 | (TAA)5 | CTCGCCCCCTCTTTTATAGG | GGTTTCCCCTTCATCTCCTC | 230 |
| chr5:32533413-32535428 | p3 | (ATA)5 | CTCGCCCCCTCTTTTATAGG | GGTTTCCCCTTCATCTCCTC | 230 |
| chr5:32610392-32612438 | p2 | (AT)23 | AGGTAACCGTGTCACCTTGG | AGGGGGTGATAGAATCCCAC | 214 |
| chr5:32614052-32616067 | p6 | (GATTCT)3 | AAGGCGTGGAGTAGTGAGGA | TCATGGTGAGTGCAAGAAGC | 298 |
| chr5:32661288-32663302 | p2 | (TA)7 | TTATTGCATCAGGCCTACCC | AAGGTATTGGCTGAACACGG | 153 |
| chr5:32661288-32663302 | p3 | (TCA)5 | TTATTGCATCAGGCCTACCC | AAGGTATTGGCTGAACACGG | 153 |
| chr5:32794152-32796167 | p6 | (TAAAAA)3 | AACGACATGATGAACAGGCA | TTGAATGTCGGTCCTCAACA | 267 |
| chr5:32794152-32796167 | p3 | (ATA)5 | AACGACATGATGAACAGGCA | TTGAATGTCGGTCCTCAACA | 267 |
| chr5:32794152-32796167 | p5 | (TTTTA)3 | AACGACATGATGAACAGGCA | TTGAATGTCGGTCCTCAACA | 267 |
| chr5:33084259-33086279 | p5 | (CATTT)4 | AAGGCTCGCTGACACATTCT | ACACAGGTCATCCCCAAGAG | 278 |
| chr5:33146302-33148320 | p3 | (TGT)6 | AGGCACTGATTTCATTTGGG | TACCTGGGTAGGGAGGCTTT | 175 |
| chr5:33154088-33156104 | p2 | (CT)8 | ATGAGGTTGAGGATGGCAAG | AAATTATGGTCCACCCCTCC | 255 |
| chr5:33154088-33156104 | p5 | (TTGGT)3 | ATGAGGTTGAGGATGGCAAG | AAATTATGGTCCACCCCTCC | 255 |
| chr5:33301648-33303660 | p2 | (AT)6 | GGGAGAGCTAGGCAAGGACT | CTCTCCCTTTTGAGTGCCAG | 258 |
| chr5:33335624-33337638 | p2 | (TA)7 | TAACCGCACTCTCGTCACAG | AAGTTGCCCTTATTTGTGCG | 206 |
| chr5:33335624-33337638 | p2 | (TA)7 | TAACCGCACTCTCGTCACAG | AAGTTGCCCTTATTTGTGCG | 206 |
| chr5:33335624-33337638 | p2 | (TA)7 | TAACCGCACTCTCGTCACAG | AAGTTGCCCTTATTTGTGCG | 206 |
| chr5:33335624-33337638 | p2 | (AC)6 | TAACCGCACTCTCGTCACAG | AAGTTGCCCTTATTTGTGCG | 206 |
| chr5:33353463-33355478 | p3 | (ATT)5 | AAGACCATCATTCGAGGACG | ACTTTCCCAAATTCCCAACC | 291 |
| chr5:33353516-33355534 | p3 | (ATT)5 | AAGACCATCATTCGAGGACG | ACTTTCCCAAATTCCCAACC | 238 |
| chr5:33422039-33424054 | p5 | (AAAAG)3 | TCTTGCAAAAACTGCACACC | CCTACCCCATCCAAATTTCA | 184 |
| chr5:33422039-33424054 | p2 | (AT)6 | TCTTGCAAAAACTGCACACC | CCTACCCCATCCAAATTTCA | 184 |
| chr5:33446111-33448137 | p2 | (AT)13 | CCTCAGATCCAATCCCTCAA | TTCACCATGCATAAGAAGCG | 286 |
| chr5:33492707-33494722 | p5 | (AAAAG)3 | TCTTGCAAAAACTGCACACC | CCTACCCCATCCAAATTTCA | 184 |
| chr5:33492707-33494722 | p2 | (AT)7 | TCTTGCAAAAACTGCACACC | CCTACCCCATCCAAATTTCA | 184 |
| chr5:33496560-33498572 | p2 | (AT)6 | AAATTAATTGGGGAGGACCG | TGATTCAGGGCGTGACAGTA | 112 |
| chr5:33534573-33536591 | p6 | (CGGTTC)3 | TGCAATCAATCTGCGCTTAC | TTTCCCCCATCAATCAGGTA | 117 |
| chr5:33553230-33555246 | p2 | (TA)8 | ACTGCAATCCCCAAGTTTTG | TGTGTCCGCGTAAAAGTGAA | 161 |
| chr5:33615008-33617023 | p3 | (TAT)5 | GTTTGGACCTACACACCCGT | AAGCACGCAAGAAGGAAAAA | 234 |
| chr5:33632350-33634365 | p5 | (CATAT)3 | GGACTCCAAATAACCCCGAT | GGGTACGTATGGGTGTCCAG | 227 |
| chr5:33658097-33660117 | p2 | (AT)7 | CATGACCCATGACCCACATA | TATTCCGCCACTTGGAGTTC | 248 |
| chr5:33658097-33660117 | p2 | (AT)10 | CATGACCCATGACCCACATA | TATTCCGCCACTTGGAGTTC | 248 |
| chr5:33664984-33666996 | p2 | (AT)6 | GTTGGGCAGATTGTATGCCT | ACCAGGGAGGGTTCAAACTT | 146 |

| Marker ID | SSR type | SSR motif | FORWARD PRIMER (5'-3') | REVERSE PRIMER (5'-3') | size (bp) |
|---|---|---|---|---|---|
| chr5:33706534-33708566 | p2 | (TA)16 | TCTTGCAAGGGACAATGAGA | GCGTGTTTAGGGCGTACAAT | 190 |
| chr5:33722460-33724476 | p4 | (AACA)4 | TTCAACCCAAATCGACACAA | TTCGTCCCTCCACAAAATTC | 255 |
| chr5:33764356-33766368 | p2 | (AG)6 | CCACAAACGAACAGGGTTTT | TACAGCTGGCCAGGAGAAAT | 111 |
| chr5:33803470-33805488 | p6 | (TAAAAA)3 | GGACCCATATTTCCAAACCC | GACGACGAAAAAGTGCCCTA | 175 |
| chr5:33814105-33816165 | p3 | (TAA)20 | GATCTGGATAGGGCGGTACA | CTCTTTCTCACTAACGCCGC | 244 |
| chr5:33834652-33836716 | p2 | (AT)32 | CTTTGGCCTACCCCTACCTC | ATTATGTGCCCCAAGCTGAG | 243 |
| chr5:33962184-33964199 | p3 | (ACA)5 | CGGCAAAGGAGTCAAATCAT | CCATCCTTTTCCCGTCCTAT | 236 |
| chr5:33985955-33987979 | p3 | (ATC)8 | CAAGGGCATTTACTCACGGT | CATGACATGGGACATGGGTA | 267 |
| chr5:34168583-34170622 | p3 | (TAA)8 | CGAATAGGCGCGATAAATGT | AGCTATCGAACCTGGGGACT | 251 |
| chr5:34168583-34170622 | p3 | (ATA)12 | CGAATAGGCGCGATAAATGT | AGCTATCGAACCTGGGGACT | 251 |
| chr5:34200858-34202890 | p2 | (TA)12 | AAGAGTGGGCTCCATGCTAA | TTTAAGGGACAACGGACGAC | 263 |
| chr5:34200858-34202890 | p2 | (TA)16 | AAGAGTGGGCTCCATGCTAA | TTTAAGGGACAACGGACGAC | 263 |
| chr5:34271915-34273930 | p5 | (TTGGT)3 | AAAGCAATGAGGCAACCAAC | GGACAGATGAAAGGGGATGA | 178 |
| chr5:34279654-34281696 | p6 | (TGTTGG)3 | CCCACAACTCCTTTGCACTT | AAGAAGAAGAAGCAGCAGCG | 202 |
| chr5:34279654-34281696 | p3 | (TTG)14 | CCCACAACTCCTTTGCACTT | AAGAAGAAGAAGCAGCAGCG | 202 |
| chr5:34279654-34281696 | p2 | (TA)10 | CCCACAACTCCTTTGCACTT | AAGAAGAAGAAGCAGCAGCG | 202 |
| chr5:34310655-34312675 | p2 | (TA)10 | TCAGTGGCGAGGATGTAGTG | TGGCCTGTTGCAATCATTTA | 277 |
| chr5:34329552-34331564 | p2 | (TA)6 | ATCCCTAGAGCGGGATCATT | CAAACCTTCGCCTGAGCTAC | 144 |
| chr5:34329552-34331564 | p2 | (TG)6 | ATCCCTAGAGCGGGATCATT | CAAACCTTCGCCTGAGCTAC | 144 |
| chr5:34652419-34654441 | p2 | (TG)11 | GAAAAACCCAAAGCTTGCTG | CCAATTTGGGAAAAATGACG | 235 |
| chr5:34691944-34693966 | p2 | (TG)11 | GAAAAACCCAAAGCTTGCTG | CCAATTTGGGAAAAATGACG | 235 |
| chr5:34724098-34726114 | p4 | (TAAT)4 | GCTCCATCATTTAAACCCGA | CCGCTCCATCATTTACATCC | 263 |
| chr5:34752831-34754861 | p2 | (TA)15 | ATTGACCCGTGATCAGAAGG | CCGATCCATATCCCATCATC | 195 |
| chr5:34782803-34784821 | p3 | (CAA)6 | GGGCAAGCATGTTTTCTGTT | AGTTTCCCGATCAAGCATGT | 189 |
| chr5:34782803-34784821 | p2 | (GT)6 | GGGCAAGCATGTTTTCTGTT | AGTTTCCCGATCAAGCATGT | 189 |
| chr5:34856724-34858760 | p2 | (AT)18 | TTCCCAAAGTATCCCCATCA | AAATGACATGGCGGAATAGC | 183 |
| chr5:34904193-34906241 | p2 | (AT)6 | ATTCTACCACTGTGCCTGGG | AAGAGAACCGCCAATGAATG | 193 |
| chr5:34942427-34944493 | p2 | (AT)33 | AGCTGCTGAAGCTGGAAGTC | CGTCCTCAGATGGATCAGGT | 141 |
| chr5:35117985-35120001 | p2 | (TA)8 | TCAAATGGTGTGGCTTGGTA | GCAACGGGTCATAGGACAAT | 188 |
| chr5:35117985-35120001 | p5 | (ATATT)3 | TCAAATGGTGTGGCTTGGTA | GCAACGGGTCATAGGACAAT | 188 |
| chr5:35146592-35148604 | p2 | (TA)6 | AAGTGCATGAAAAGGCCAAC | AAAGGCTGGCACAAAGAAGA | 300 |
| chr5:35201132-35203144 | p2 | (TA)6 | AAGTGCATGAAAAGGCCAAC | AAAGGCTGGCACAAAGAAGA | 300 |
| chr5:35255899-35257917 | p2 | (AT)9 | AAAGGATTGCCATTTGATGC | AAGAGTTTCGTTTGGTGGGA | 133 |
| chr5:35306272-35308286 | p2 | (AT)7 | TTGGAAAAACACCCTCAAGC | GAACAGCTCTGTCCGGAGTC | 130 |
| chr5:35385348-35387360 | p2 | (AT)6 | CTGGCACACTCAGAACCTCA | CACCATCACACCCATCTCAG | 107 |
| chr5:35390188-35392200 | p2 | (TA)6 | TAGGCCAAGGACCAACAGTC | ATCTCCAATTCCAACCCCTT | 170 |

| Marker ID | SSR type | SSR motif | FORWARD PRIMER (5'-3') | REVERSE PRIMER (5'-3') | size (bp) |
|---|---|---|---|---|---|
| chr5:35422284-35424296 | p2 | (GT)6 | TCTGCACTACTCCGTTCGTG | CCTCGGAAACCGTAACACAT | 216 |
| chr5:35439140-35441188 | p2 | (CT)7 | CAAGTGCAAGGCCAATGTTA | AGAGAATGTGCTGACAGGGG | 208 |
| chr5:35439140-35441188 | p5 | (GATTT)3 | CAAGTGCAAGGCCAATGTTA | AGAGAATGTGCTGACAGGGG | 208 |
| chr5:35439140-35441188 | p2 | (AC)6 | CAAGTGCAAGGCCAATGTTA | AGAGAATGTGCTGACAGGGG | 208 |
| chr5:35439208-35441228 | p2 | (CT)7 | CAAGTGCAAGGCCAATGTTA | AGAGAATGTGCTGACAGGGG | 140 |
| chr5:35439208-35441228 | p5 | (GATTT)3 | CAAGTGCAAGGCCAATGTTA | AGAGAATGTGCTGACAGGGG | 140 |
| chr5:35439208-35441228 | p2 | (AC)6 | CAAGTGCAAGGCCAATGTTA | AGAGAATGTGCTGACAGGGG | 140 |
| chr5:35562209-35564224 | p5 | (AAATG)3 | GCCTCACATACCTTGGTCGT | TTAGCGACGCGTTGTTATTG | 259 |
| chr5:35626119-35628177 | p2 | (AT)29 | CCGGAGTCTATAGGATGCCA | AACCGGATGAAGAACCACAG | 154 |
| chr5:35643881-35645939 | p2 | (AT)29 | CCGGAGTCTATAGGATGCCA | AACCGGATGAAGAACCACAG | 154 |
| chr5:35727433-35729493 | p6 | (ATAAAA)3 | CCCCATATTAAACACCGTGC | TTTCCCGTGAGTTTGGATGT | 245 |
| chr5:35727433-35729493 | p2 | (TA)30 | CCCCATATTAAACACCGTGC | TTTCCCGTGAGTTTGGATGT | 245 |
| chr5:35739669-35741695 | p2 | (AT)13 | GCCAGCGTTGATCACTTTTT | GAGACTTACGCCTGACCGAG | 232 |
| chr5:35739669-35741695 | p2 | (TA)6 | GCCAGCGTTGATCACTTTTT | GAGACTTACGCCTGACCGAG | 232 |
| chr5:35808033-35810057 | p2 | (TA)12 | GCTAATGCTGGGAGTCCAAC | CCGGGAAATTAGCAGTCAAA | 173 |
| chr5:35808033-35810057 | p3 | (GAA)6 | GCTAATGCTGGGAGTCCAAC | CCGGGAAATTAGCAGTCAAA | 173 |
| chr5:35813144-35815158 | p2 | (AT)7 | CCTTGAGGAACGTAATCGGA | ACTTTGGCCATTCCTTTTGA | 138 |
| chr5:35813144-35815158 | p2 | (CT)7 | CCTTGAGGAACGTAATCGGA | ACTTTGGCCATTCCTTTTGA | 138 |
| chr5:35813144-35815158 | p5 | (TTTAT)3 | CCTTGAGGAACGTAATCGGA | ACTTTGGCCATTCCTTTTGA | 138 |
| chr5:35880152-35882167 | p3 | (CTC)5 | TCACAATGCCACTGAACCAT | AACTTGATGAGCGAAATCCG | 222 |
| chr5:36026066-36028080 | p2 | (TA)7 | TACTGAAGGGGTAATTGGCG | CAACATGCATTCCCTGTCAC | 291 |
| chr5:36104056-36106078 | p2 | (AT)11 | AGCAGGGATCATTAGCCAGA | GGTGGCCACTAATCTTTGGA | 232 |
| chr5:36200873-36202891 | p2 | (AT)7 | TTGGACGTCTTAGCATGGTG | GCCTTCCTCCGCTTCTAAGT | 225 |
| chr5:36200873-36202891 | p5 | (CCGGT)3 | TTGGACGTCTTAGCATGGTG | GCCTTCCTCCGCTTCTAAGT | 225 |
| chr5:36226079-36228094 | p5 | (TGCAT)3 | AGAAGGGGAAAGCGAAGAAG | GCATCAAGCCCAAAATCAAT | 207 |
| chr5:36228621-36230641 | p2 | (GA)8 | AATGCCATTTCAGCCTCAAG | AAGAGGGTGAAAACGGAGGT | 236 |
| chr5:36228621-36230641 | p5 | (AAGAA)4 | AATGCCATTTCAGCCTCAAG | AAGAGGGTGAAAACGGAGGT | 236 |
| chr5:36339418-36341433 | p5 | (ATATG)3 | AGAGAAATGGTGGATCGTGG | CCTCACATACCTTTGGCGTT | 249 |
| chr5:36341072-36343090 | p6 | (AAGTGA)3 | TTCCAACCATCCTCGATCTC | CGCCTACTAGCCAATAACGC | 142 |
| chr5:36375973-36377988 | p5 | (GTTTT)3 | ATGCTGGACAAGGCAAAACT | AACGGTCTTTTAGTGCCCCT | 201 |
| chr5:36379878-36381916 | p2 | (TA)19 | TTCGCCTTAGTTTTTGGAGC | GCTATTAATCCATTCGCCCA | 206 |
| chr5:36391228-36393240 | p2 | (AT)6 | TCATCTAACTTACGCGGGCT | TCGAAGGTTTCCAAGGTTTG | 248 |
| chr5:36407466-36409481 | p5 | (GTTTT)3 | ATGCTGGACAAGGCAAAACT | AACGGTCTTTTAGTGCCCCT | 201 |
| chr5:36411371-36413409 | p2 | (TA)19 | TTCGCCTTAGTTTTTGGAGC | GCTATTAATCCATTCGCCCA | 206 |
| chr5:36422721-36424733 | p2 | (AT)6 | TCATCTAACTTACGCGGGCT | TCGAAGGTTTCCAAGGTTTG | 248 |
| chr5:36488051-36490069 | p6 | (TTTTCC)3 | AGGGAAGCATACACAATGCC | ATGTTTTGTCTGCCTGGTCC | 156 |

| Marker ID | SSR type | SSR motif | FORWARD PRIMER (5'-3') | REVERSE PRIMER (5'-3') | size (bp) |
|---|---|---|---|---|---|
| chr5:36642442-36644456 | p2 | (CT)7 | GCAGACTTGTGGCAAACTCA | CCAGACCAGATCCCTTTGAA | 137 |
| chr5:36723260-36725272 | p2 | (AT)6 | GGAGGAGGCCTTCCTGATAG | CTTACGCAGTACGTGGGGAT | 183 |
| chr5:36793671-36795689 | p6 | (TTTTTA)3 | CAGCCACAAAAAGATGAGCA | TCCCATAAGCACGACATGAA | 155 |
| chr5:36793671-36795689 | p2 | (TA)6 | CAGCCACAAAAAGATGAGCA | TCCCATAAGCACGACATGAA | 155 |
| chr5:36823076-36825096 | p2 | (AT)10 | TTGTCTTCGGATGCTGTTTG | GCCCCTGCTTCACAAATCTA | 281 |
| chr5:36869455-36871470 | p5 | (TATAG)3 | CCTCCCCACCAGATCCTAAT | AGCTAGGGTTGGAGCACAGA | 164 |
| chr5:36869455-36871470 | p5 | (AATGC)3 | CCTCCCCACCAGATCCTAAT | AGCTAGGGTTGGAGCACAGA | 164 |
| chr5:36892030-36894045 | p2 | (TC)12 | TTTAAACCATACCCGACCCA | GGGGAGAGTGGGGTAGAGAG | 156 |
| chr5:36892030-36894045 | p5 | (TTTAG)3 | TTTAAACCATACCCGACCCA | GGGGAGAGTGGGGTAGAGAG | 156 |
| chr5:36931885-36933905 | p2 | (TA)11 | CAGGTAGGGGTGGGGTTAAT | GAGTCTTGCCCAAGATCAGG | 203 |
| chr5:36931885-36933905 | p5 | (TTTCT)4 | CAGGTAGGGGTGGGGTTAAT | GAGTCTTGCCCAAGATCAGG | 203 |
| chr5:36992445-36994459 | p2 | (TA)7 | CCAAAATCAAATGCCCTCAT | CAACATGGATATCTTGGGGG | 292 |
| chr5:36996976-36998994 | p3 | (TGT)6 | CCGTCCCTCAAAACTCTTCA | GCCATAAATTTTGCGACCAT | 209 |
| chr5:36996976-36998994 | p3 | (ATG)13 | CCGTCCCTCAAAACTCTTCA | GCCATAAATTTTGCGACCAT | 209 |
| chr5:37084674-37086692 | p6 | (TGAAAC)3 | GGCCGATAAGCAGACTTTTG | ATGCACATAAGCCCTTCGTC | 113 |
| chr5:37162403-37164423 | p5 | (CAAAC)4 | TAGTGGCTGCACCATTTTTG | ATGCAAGTACTGCCCCAAAG | 114 |
| chr5:37178227-37180255 | p2 | (AT)14 | TGAATCCGCCTCTGGATAAG | ATGACCGATCTGCATCACAA | 209 |
| chr5:37262230-37264242 | p2 | (TA)6 | AGCGGTCGATTTTGAACAAC | AAATTAACATGGGGGCACAA | 179 |
| chr5:37394229-37396249 | p5 | (AACCG)4 | TCTGCCAAACCAACTAACCC | GCTTGGACTGCCATTTGATT | 138 |
| chr5:37394229-37396249 | p5 | (AAAAG)3 | TCTGCCAAACCAACTAACCC | GCTTGGACTGCCATTTGATT | 138 |
| chr5:37396703-37398723 | p5 | (TTTTA)4 | GTCCCAGCTTATGCGACACT | AAAGGTGTTTGACTGGGCAC | 196 |
| chr5:37417485-37419497 | p2 | (TA)7 | TACACGGAAACCAGGGAGTC | TTTGGAACTGTTTCATGCCA | 228 |
| chr5:37417485-37419497 | p2 | (TA)6 | TACACGGAAACCAGGGAGTC | TTTGGAACTGTTTCATGCCA | 228 |
| chr5:37500998-37503016 | p2 | (AT)9 | CTGAGGCCTAACTTTGGCTG | CCCTTATTTTGGGGTGGTCT | 119 |
| chr5:37518886-37520901 | p5 | (TTCTT)3 | AGGGAAGCATACACAATGCC | ATGTTTTGTCTGCCTGGTCC | 249 |
| chr5:37530958-37532980 | p2 | (TA)11 | TACGACCCGAGAGGAGAGAA | TCTTCGTATTGCCGCTCTTT | 268 |
| chr5:37530958-37532980 | p2 | (AT)8 | TACGACCCGAGAGGAGAGAA | TCTTCGTATTGCCGCTCTTT | 268 |
| chr5:37839331-37841346 | p5 | (TGCAA)3 | CAATGAAAAAGGCGAAGAGC | TCATTTGTATCGGTTCGCAA | 165 |
| chr5:37920599-37922653 | p2 | (TA)27 | ACGCACGAGTCAGAAGGATT | CATGTGGTTCACTCGGTCAC | 183 |
| chr5:37968489-37970507 | p3 | (GTG)6 | AAACAGCCATAAAACACCGC | AACTTCCACCCCCATTCTTT | 126 |
| chr5:37997714-37999728 | p2 | (GT)7 | TTTCGTGCCGATAGTTAGGG | TTGAAAAATCCGAACCAACC | 178 |
| chr5:38000227-38002259 | p2 | (TA)16 | ACGGAGTGCTTTCTGCTGAT | ACCACCCAAGAAGATGATGC | 114 |
| chr5:38000227-38002259 | p2 | (TA)6 | ACGGAGTGCTTTCTGCTGAT | ACCACCCAAGAAGATGATGC | 114 |
| chr5:38003760-38005804 | p2 | (AT)22 | ATGATAATGGCTGGGAATGC | TTATAAGCCGATCCAAACGG | 125 |
| chr5:38042585-38044606 | p3 | (TGT)7 | AAATCCTGCTGATTGGGATG | TGTATCTGGTGGCGTGTCAT | 102 |
| chr5:38065865-38067880 | p3 | (GTA)5 | ATCCCCATTATGCGAGTCTG | CCCTTTGAAGAACCAGTGGA | 261 |

| Marker ID | SSR type | SSR motif | FORWARD PRIMER (5'-3') | REVERSE PRIMER (5'-3') | size (bp) |
|---|---|---|---|---|---|
| chr5:38088390-38090414 | p2 | (AT)12 | GCCACATTTCTTGCCTCATT | TCCTGTCTCCTCCAGCAGTT | 144 |
| chr5:38113405-38115420 | p5 | (TTTAT)3 | TAGTTTGTTGGGGGCTCTTG | CTTCCGCATGTCCAGGTAAT | 193 |
| chr5:38121901-38123929 | p3 | (GTA)5 | AGCAGCAGGAGGTCTGGTAA | GAACGGACGACGATCTTGTT | 295 |
| chr5:38121901-38123929 | p2 | (TA)14 | AGCAGCAGGAGGTCTGGTAA | GAACGGACGACGATCTTGTT | 295 |
| chr5:38165725-38167740 | p5 | (TTTTA)3 | CATCCCGAGTCCCTGTTAAA | CCTCCACAGTAGCTCCAAGC | 155 |
| chr5:38165725-38167740 | p5 | (TAAAT)3 | CATCCCGAGTCCCTGTTAAA | CCTCCACAGTAGCTCCAAGC | 155 |
| chr5:38214061-38216076 | p5 | (TTTTA)3 | CATCCCGAGTCCCTGTTAAA | CCTCCACAGTAGCTCCAAGC | 155 |
| chr5:38214061-38216076 | p5 | (TAAAT)3 | CATCCCGAGTCCCTGTTAAA | CCTCCACAGTAGCTCCAAGC | 155 |
| chr5:38251131-38253146 | p3 | (GTT)5 | TACCCAGCCATCCAAGAAAG | TTGCCCTTTAGTTTTGGTGG | 100 |
| chr5:38254177-38256191 | p2 | (AG)7 | GAGCCATTGGACATTGAGGT | GTTCCAGAGGTCACTTGGGA | 254 |
| chr5:38263573-38265617 | p2 | (AT)22 | ATTGGAGCCACATTTCTTGC | TCCTGTCTCCTCCAGCAGTT | 137 |
| chr5:38392250-38394278 | p2 | (TA)6 | GCCAGAAAAATGATTCCCAA | TACCATTTCACACGGCGTAA | 215 |
| chr5:38392250-38394278 | p2 | (TA)10 | GCCAGAAAAATGATTCCCAA | TACCATTTCACACGGCGTAA | 215 |
| chr5:38392279-38394311 | p2 | (TA)6 | GCCAGAAAAATGATTCCCAA | TACCATTTCACACGGCGTAA | 186 |
| chr5:38392279-38394311 | p2 | (TA)10 | GCCAGAAAAATGATTCCCAA | TACCATTTCACACGGCGTAA | 186 |
| chr5:38393154-38395174 | p2 | (TA)6 | TGATGTGGGATCGAATGAGA | TTGCTAACCCCCTTTTGATG | 185 |
| chr5:38393154-38395174 | p2 | (TA)10 | TGATGTGGGATCGAATGAGA | TTGCTAACCCCCTTTTGATG | 185 |
| chr5:38508875-38510905 | p2 | (AT)15 | AATCAGAGCTCGTGAATGGC | GCGTGGAAAATCAAAAGTGG | 111 |
| chr5:38582589-38584604 | p3 | (TTC)5 | CCTTTATCCCCTTCAGCTCC | AGGAAACCCACGAAACCTCT | 142 |
| chr5:38582589-38584604 | p3 | (CAA)5 | CCTTTATCCCCTTCAGCTCC | AGGAAACCCACGAAACCTCT | 142 |
| chr5:38695534-38697546 | p2 | (CA)6 | CCCTCGGAATCGACAGAATA | GTTCCTGAGGTAATCGAGCG | 165 |
| chr5:38708739-38710751 | p2 | (CA)6 | TGTATCGGACCATTCAAGCA | TGGTAAACACCTTCCGCTTT | 114 |
| chr5:38844248-38846284 | p3 | (AAT)12 | ATTTGGAGGTGGTATCGTCG | CAAAAGGGTCGTTTTGGGTA | 215 |
| chr5:38870403-38872418 | p5 | (AATAA)3 | GCGTCTTCATGAGCGTGTTA | GGAATGAGAAGTCGATGGGA | 143 |
| chr5:38870403-38872418 | p5 | (AATAA)3 | GCGTCTTCATGAGCGTGTTA | GGAATGAGAAGTCGATGGGA | 143 |
| chr5:38870403-38872418 | p5 | (AATAA)3 | GCGTCTTCATGAGCGTGTTA | GGAATGAGAAGTCGATGGGA | 143 |
| chr5:38870403-38872418 | p3 | (TGG)5 | GCGTCTTCATGAGCGTGTTA | GGAATGAGAAGTCGATGGGA | 143 |
| chr5:38879896-38881930 | p2 | (AT)8 | TGCAATGCATGATACAACAGA | CTGGGTCTGGTTTCCACATT | 165 |
| chr5:38879896-38881930 | p2 | (AT)17 | TGCAATGCATGATACAACAGA | CTGGGTCTGGTTTCCACATT | 165 |
| chr5:38915443-38917458 | p5 | (GTTCG)3 | CCAGGTTGGATCTCTTGCAT | AAATCGAACCCGAAATAGCC | 205 |
| chr5:38915443-38917458 | p5 | (GTTCG)3 | CCAGGTTGGATCTCTTGCAT | AAATCGAACCCGAAATAGCC | 205 |
| chr5:38929068-38931090 | p2 | (TA)11 | CCTCGAAGAGATCGAACCTG | TCAGAAATTGGGAAGGCAAC | 144 |
| chr5:38929068-38931090 | p2 | (AT)17 | CCTCGAAGAGATCGAACCTG | TCAGAAATTGGGAAGGCAAC | 144 |
| chr5:39063197-39065212 | p5 | (GTTTT)3 | AGACAGCGAGTGGGTGAGAT | GGCTCAGATCACAAGCCTTC | 138 |
| chr5:39183310-39185326 | p2 | (CA)8 | ACAACTGATATGCGTGGCAA | GGCGAGTCTGATTTTGCTTC | 203 |
| chr5:39227379-39229425 | p2 | (AT)23 | TGGCATGACATGGGTAGAAG | ACCTTGTTGCACCCTTTTTG | 293 |

| Marker ID | SSR type | SSR motif | FORWARD PRIMER (5'-3') | REVERSE PRIMER (5'-3') | size (bp) |
|---|---|---|---|---|---|
| chr5:39247003-39249017 | p2 | (AT)7 | TGTCCCGACCCTTTTATGAG | ATCTGTGTAGAGGGATGGCG | 228 |
| chr5:39253495-39255507 | p2 | (CT)6 | TCCTTTAATCTCAATGCGGC | ACGGTCTTCAAATGCAATCC | 201 |
| chr5:39342919-39344931 | p2 | (TA)6 | CGGACCGCATTTCAGTTTAT | CGCCCATAATATACCGTTGG | 156 |
| chr5:39363034-39365048 | p2 | (TA)7 | ATTGGGGGTAATCATCAGCA | AGTGGGGCACTGGTTAAGTG | 113 |
| chr5:39400240-39402254 | p2 | (AT)7 | CATGAGGGAGGATTACCCAA | ACGGTGAAGAGTGACATCCC | 188 |
| chr5:39412382-39414397 | p5 | (GCATT)3 | CTGTTGCGTGATTGAGGCTA | TGGGTACGCCAACAATATCA | 145 |
| chr5:39412382-39414397 | p4 | (TTAT)5 | CTGTTGCGTGATTGAGGCTA | TGGGTACGCCAACAATATCA | 145 |
| chr5:39512214-39514230 | p2 | (GA)8 | TAATGTGTTTAGCCCCTCGG | TAGGCCCGAATGGTGATTTA | 137 |
| chr5:39780311-39782341 | p2 | (AT)6 | AATGCAAGTTTGGATTTGGC | TGTAACAGAATGGCGACTGC | 237 |
| chr5:39816337-39818355 | p6 | (TTCTTT)3 | AGTACCCCCTCACCCAAAAC | AGACACTCCTCAGCCTTCCA | 271 |
| chr5:39827184-39829198 | p2 | (TA)7 | TTTAAGTCCCCACAAGTCCG | TTTGGTAATGTCCGCCTCTT | 119 |
| chr5:39854209-39856224 | p3 | (AAT)5 | ATTTGATTGTCGGCCTTGAG | GGGGGTAGACCAACAAAGGT | 181 |
| chr5:39854209-39856224 | p5 | (AAAAT)3 | ATTTGATTGTCGGCCTTGAG | GGGGGTAGACCAACAAAGGT | 181 |
| chr5:39884645-39886657 | p2 | (CT)6 | TTCAACCCTGGGTCAAAAAG | CCGTTTCGAATTGCCTACAT | 233 |
| chr5:39924108-39926126 | p5 | (AAACC)3 | TGAGAAGAACGCCTCCAAGT | GTCGAGAGCCAGAGAGGTTG | 241 |
| chr5:39924156-39926171 | p5 | (AAACC)3 | TGAGAAGAACGCCTCCAAGT | GTCGAGAGCCAGAGAGGTTG | 193 |
| chr5:39924704-39926719 | p5 | (AAACC)3 | CTCTCCATCCACCTTTGAGC | AAAACCGACTCAAACCGATG | 251 |
| chr5:40093658-40095676 | p2 | (TA)9 | CGCTTGGACAATAGCAGTGA | TGCGTACGTTATACCCTCCC | 244 |
| chr5:40199611-40201681 | p2 | (AT)9 | GGATTCATTGTCACATGCGT | GACATGGCGGAATAGCAAAT | 215 |
| chr5:40272011-40274026 | p5 | (ATAAA)3 | TCGACTGTCTGAGCGAAAAA | AGGCCAAATTTAATGGGAGG | 276 |
| chr5:40288152-40290196 | p5 | (TCGGT)3 | ATTTTCACGCAATGGGAGAG | AAAATGAAGACCGCCACATC | 220 |
| chr5:40288152-40290196 | p5 | (AACCG)4 | ATTTTCACGCAATGGGAGAG | AAAATGAAGACCGCCACATC | 220 |
| chr5:40371073-40373091 | p3 | (GTT)5 | ACCTGAAGCCCATGAAAATG | TGCAAAACCATTCTCACCAA | 188 |
| chr5:40371073-40373091 | p2 | (TA)9 | ACCTGAAGCCCATGAAAATG | TGCAAAACCATTCTCACCAA | 188 |
| chr5:40371073-40373091 | p2 | (TA)14 | ACCTGAAGCCCATGAAAATG | TGCAAAACCATTCTCACCAA | 188 |
| chr5:40371073-40373091 | p2 | (AT)10 | ACCTGAAGCCCATGAAAATG | TGCAAAACCATTCTCACCAA | 188 |
| chr5:40437460-40439478 | p6 | (GTGGAT)3 | CAATTAACATGGATGTGGCG | AGAGTGGAACAACGTCCACC | 192 |
| chr5:40544332-40546348 | p2 | (CA)8 | ACCGTATGGTGCCTAATCCA | GGAATGCCAAATAGAAGCCA | 171 |
| chr5:40600117-40602138 | p3 | (ACA)7 | TGACTCTCCGGAAAAACACC | GCTTTCTACCGCATTGAAGC | 231 |
| chr5:40600117-40602138 | p2 | (AG)8 | TGACTCTCCGGAAAAACACC | GCTTTCTACCGCATTGAAGC | 231 |
| chr5:40782596-40784666 | p3 | (TTC)7 | GACCATAGCGGATCCTGAAA | CTTTTGGGGTGCTCAGTTGT | 263 |
| chr5:40782704-40784724 | p3 | (TTC)7 | GACCATAGCGGATCCTGAAA | CTTTTGGGGTGCTCAGTTGT | 155 |
| chr5:40983817-40985832 | p5 | (TTTCT)3 | ATGCAAGCCTCAGCACTTTT | AAAGGGTTGAAACAACACGG | 183 |
| chr5:41084437-41086452 | p5 | (CTTTT)3 | TTCTGCGTTTGTGGAAACAG | TATGTTTTGTGCGCCTATGG | 234 |
| chr5:41096676-41098704 | p5 | (CTAAA)3 | TGCTCCTCACACGTCAAAAG | AACCCCACTCTCGACACAAC | 184 |
| chr5:41096676-41098704 | p2 | (AG)14 | TGCTCCTCACACGTCAAAAG | AACCCCACTCTCGACACAAC | 184 |

| Marker ID | SSR type | SSR motif | FORWARD PRIMER (5'-3') | REVERSE PRIMER (5'-3') | size (bp) |
|---|---|---|---|---|---|
| chr5:41096676-41098704 | p2 | (GT)7 | TGCTCCTCACACGTCAAAAG | AACCCCACTCTCGACACAAC | 184 |
| chr5:41097441-41099455 | p2 | (AG)14 | GGTGACGGGTGTTAATACGG | CCACGGATAGTTGAGAGGGA | 271 |
| chr5:41097441-41099455 | p2 | (GT)7 | GGTGACGGGTGTTAATACGG | CCACGGATAGTTGAGAGGGA | 271 |
| chr5:41109361-41111383 | p2 | (TA)11 | TTTATGAATCCTCCCCCACA | GTCCTTCTGAGCTGAAACCG | 276 |
| chr5:41182576-41184591 | p5 | (TTTTC)3 | TTTCACGAGTGGGGTCTAGG | ATCGTGGCGTCTTCACTCTT | 226 |
| chr5:41236557-41238569 | p2 | (CA)6 | TATATCCAACCCCTCCCTCC | CCTCTTGGTGTGATGGGACT | 255 |
| chr5:41368246-41370258 | p2 | (AG)6 | TGTCCGTTGGTCAGGTACAA | ACCTGCAATCCAAACCAGAG | 105 |
| chr5:41398128-41400143 | p5 | (GGGGT)3 | GCTTGATGCTGATTCCGTTT | GAGCTTTCCCCTTTGGAGTT | 156 |
| chr5:41465393-41467408 | p5 | (AAACC)3 | ATACCTGGCTAAAGAGCGCA | ACCGAACCGATTACAACCAA | 291 |
| chr5:41465393-41467408 | p5 | (AAACC)3 | ATACCTGGCTAAAGAGCGCA | ACCGAACCGATTACAACCAA | 291 |
| chr5:41500836-41502851 | p5 | (ATTTT)3 | GCTTGTCCGATTGAACACCT | AGAGGTTGGTGGGAAGTGTG | 233 |
| chr5:41535539-41537569 | p2 | (TC)7 | CAGTGTCAGGTTCCGGTTTT | CATTTGTCTGGGGAAAGCAT | 243 |
| chr5:41545118-41547132 | p2 | (TG)7 | TGCAGCAGCAATCCAGATAC | CGGCACCCTGTGAACTTTAT | 246 |
| chr5:41564773-41566788 | p5 | (AATAA)3 | TGATATTTCGGGCCAGAGTC | TCGAAGTGTTCGGGAATAGG | 192 |
| chr5:41564773-41566788 | p2 | (TA)7 | TGATATTTCGGGCCAGAGTC | TCGAAGTGTTCGGGAATAGG | 192 |
| chr5:41576135-41578157 | p4 | (GGGT)5 | CAACCTCAACAACTCAGCCA | TCCCACATCGGTTGTTGTTA | 119 |
| chr5:41576135-41578157 | p2 | (TA)11 | CAACCTCAACAACTCAGCCA | TCCCACATCGGTTGTTGTTA | 119 |
| chr5:41576135-41578157 | p2 | (CT)22 | CAACCTCAACAACTCAGCCA | TCCCACATCGGTTGTTGTTA | 119 |
| chr5:41642534-41644570 | p2 | (TA)18 | GAAGCCTGCTAGGGTTTGTG | TCTTCCTTTACCCACATGGC | 146 |
| chr5:41661046-41663064 | p2 | (AT)9 | AAAATCACGCCACATCAACA | GCATGCCGGTAGTTCAGAAT | 278 |
| chr5:41703044-41705058 | p2 | (TC)7 | CCCAATTTGTGCTGCTTTTT | ACCCCTCAAACGAGCTTTCT | 215 |
| chr5:41713927-41715942 | p5 | (TAAAT)3 | CCCCTTACTCCATCCCATTT | TTTTTAGGTCGGGCACACTC | 161 |
| chr5:41747840-41749854 | p5 | (GGGCC)3 | GTGATTGGAAGGGCCAGTAG | CGATATCCAACTTCCACGGT | 256 |
| chr5:41835627-41837642 | p5 | (TTTAT)3 | CTTTTCCCGAAACGATGTGT | TACGGATCTGAAATGCACCA | 217 |
| chr5:42047800-42049812 | p2 | (AT)6 | CGTTAAGGTTCGCTAGGTGC | CCTTCCGTGACCTATTGCAT | 242 |
| chr5:42084220-42086241 | p3 | (ACA)7 | AGGGACTGTCCTCTGCTCTG | AAGGGAAGGGAAATCTTGGA | 196 |
| chr5:42101706-42103721 | p5 | (AAACC)3 | CATTGTGCTAACGTCCCCTT | TAAACCCCGAATGAGATTGG | 118 |
| chr5:42130546-42132558 | p2 | (TA)6 | AAAAGAGCAGTACAGGCCGA | AGGACCTTTCCCAGCTTTGT | 281 |
| chr5:42130700-42132714 | p2 | (TA)6 | AAAAGAGCAGTACAGGCCGA | AGGACCTTTCCCAGCTTTGT | 127 |
| chr5:42151263-42153277 | p2 | (TA)7 | GGACCGGAAAAGGGTATGTT | AAATCCGTCCGAAACTTCCT | 256 |
| chr5:42256158-42258170 | p2 | (AT)8 | TCATATAGGCGGGACAGGAC | CAGGGAATACATAGCGGAGC | 163 |
| chr5:42281259-42283271 | p2 | (CA)6 | TGAAGCATGTATCAACCGGA | GCCCATGACTTCATTGCTTT | 117 |
| chr5:42294367-42296387 | p2 | (AT)10 | GATTGTTCATGGCTCCCACT | CGAAGGCCTAGCACACTTTC | 110 |
| chr5:42294367-42296387 | p3 | (ATC)5 | GATTGTTCATGGCTCCCACT | CGAAGGCCTAGCACACTTTC | 110 |
| chr5:42294975-42296990 | p2 | (AT)10 | GGCACTCATGCTAATGCTCA | ATCACGTATGTGTGCGGTGT | 257 |
| chr5:42294975-42296990 | p3 | (ATC)5 | GGCACTCATGCTAATGCTCA | ATCACGTATGTGTGCGGTGT | 257 |

| Marker ID | SSR type | SSR motif | FORWARD PRIMER (5'-3') | REVERSE PRIMER (5'-3') | size (bp) |
|---|---|---|---|---|---|
| chr5:42338268-42340283 | p2 | (AT)6 | GGGACCCACATGTCAGAGTT | TTGGTGCTTTTCCAAATTCC | 205 |
| chr5:42338268-42340283 | p3 | (TTG)5 | GGGACCCACATGTCAGAGTT | TTGGTGCTTTTCCAAATTCC | 205 |
| chr5:42354318-42356360 | p2 | (AT)21 | TAGATATGGGCCAGGGTGAG | GTACTGGGTACCCGTCATGG | 196 |
| chr5:42432414-42434434 | p2 | (AT)10 | ATTTTGCTCGTGCTTACCGT | AATATAAAGTGTCCCGGCCC | 206 |
| chr5:42481547-42483562 | p3 | (TTG)5 | ATGGTGCCCAATAGAACGAG | GAAGCTGTCAAGGAGATGCC | 191 |
| chr5:42609528-42611543 | p5 | (TTTTC)3 | TCACGAAGGGGATAGTCGTC | GGCCAGGACAGTAGTGGTGT | 109 |
| chr5:42857601-42859615 | p2 | (TA)7 | GAGGAGATCCCCAGTGCATA | TGGCCTCAACACGACAAATA | 211 |
| chr5:42971075-42973087 | p2 | (AG)6 | TCGAAAATTTCAATCGGTCA | CGAGTTGCAGAGTTTCACCA | 211 |
| chr5:42971075-42973087 | p2 | (AT)10 | TCGAAAATTTCAATCGGTCA | CGAGTTGCAGAGTTTCACCA | 211 |
| chr5:42971075-42973087 | p2 | (AG)6 | TCGAAAATTTCAATCGGTCA | CGAGTTGCAGAGTTTCACCA | 211 |
| chr5:42971075-42973087 | p2 | (AT)10 | TCGAAAATTTCAATCGGTCA | CGAGTTGCAGAGTTTCACCA | 211 |
| chr5:42986730-42988752 | p2 | (CT)11 | AGCAACCCTTGAGGGAATTT | CATTAAAGACGCCATCAGCA | 200 |
| chr5:42998623-43000638 | p5 | (TTATT)3 | GTGTGCACCTTCCCAATCTT | CATCATGCTCCATCATTTGC | 286 |
| chr5:42998707-43000731 | p5 | (TTATT)3 | GTGTGCACCTTCCCAATCTT | CATCATGCTCCATCATTTGC | 202 |
| chr5:43150017-43152041 | p3 | (CTT)5 | AACGGCGATGACAAGGATAG | GACCCGGCCCATTATAAACT | 205 |
| chr5:43248826-43250842 | p4 | (TATG)4 | AGGAATTGGACAAACATCGC | ATTCGGGTAGCACCTCCTCT | 124 |
| chr5:43430731-43432746 | p2 | (AG)7 | ATCTAACTTCGGGAGGCCAT | ATACAGGCCCACGACCTATG | 181 |
| chr5:43430731-43432746 | p3 | (TTG)5 | ATCTAACTTCGGGAGGCCAT | ATACAGGCCCACGACCTATG | 181 |
| chr5:43430731-43432746 | p2 | (AT)18 | ATCTAACTTCGGGAGGCCAT | ATACAGGCCCACGACCTATG | 181 |
| chr5:43464653-43466665 | p2 | (TG)6 | GCGGAAACAGTAACCGAAAA | TGGGCAGGAAAGTAGGAATG | 271 |
| chr5:43464653-43466665 | p5 | (TGAAT)3 | GCGGAAACAGTAACCGAAAA | TGGGCAGGAAAGTAGGAATG | 271 |
| chr5:43536691-43538706 | p5 | (AAAGA)3 | GGAAGCGAGCGAGACAATAG | TCGGAAACCACTACTCCTGG | 116 |
| chr5:43715499-43717511 | p2 | (AG)6 | TCCACCCTCTTTTCAGTTGG | GAAGGGACGAAACACGGATA | 172 |
| chr5:43790467-43792481 | p2 | (AT)7 | TCTTTGCCATGGTTCTTTCC | ACCAGTCTCCCCGCTAAAAT | 270 |
| chr5:43797342-43799356 | p2 | (TA)7 | CCATGTGCCAAAATACATGC | CCAACTGTTGTGATGGTTCG | 177 |
| chr5:44000829-44002849 | p4 | (TCTT)5 | ACGATTCCGGTAACAACGTC | TAAGCTGGGCTCAGTTGGAT | 205 |
| chr5:44079548-44081570 | p2 | (AT)11 | AGAAAGGCCAGGTTGAGGTT | CCGAAATTCAAAGTCCCAAA | 261 |
| chr5:44092960-44094981 | p3 | (CAA)7 | GCGAACCATGAACACATCAG | CAGATTGACCCGGAACTTGT | 109 |
| chr5:44284475-44286489 | p2 | (TA)7 | AAAATTTATTGGGGTTGGGG | TGAGATTGGTCGGTTTCACA | 112 |
| chr5:44392406-44394418 | p2 | (TA)6 | TCCCTTCTTTCTCTTTGGCA | TTCTAGGCCTTTCGGAGTGA | 217 |
| chr5:44460885-44462899 | p2 | (TG)7 | TATAGCAGGATCAGGCCCAC | CTAGGTGCATGTGGGGAACT | 168 |
| chr5:44460885-44462899 | p2 | (TA)47 | TATAGCAGGATCAGGCCCAC | CTAGGTGCATGTGGGGAACT | 168 |
| chr5:44493262-44495280 | p6 | (GAAAAA)3 | AATTACCCTTTTGGGATGGC | ATATGTTGTGTGTCGGGCAA | 178 |
| chr5:44513195-44515210 | p5 | (ATGCA)3 | CGTGCAACACATCTACACCC | TTGGATGAAGGAAGGGAATG | 235 |
| chr5:44613707-44615723 | p2 | (TG)8 | ATCAGGCTGTTGGTTTTTGG | AACCACGAAGTGACCGTACC | 243 |
| chr5:44620741-44622774 | p2 | (AT)18 | CGGCAACAAAGTGAGTTGAA | ATGCCTTACCTCGCGTAAAA | 173 |

| Marker ID | SSR type | SSR motif | FORWARD PRIMER (5'-3') | REVERSE PRIMER (5'-3') | size (bp) |
|---|---|---|---|---|---|
| chr5:44674046-44676062 | p2 | (TG)8 | ATCAGGCTGTTGGTTTTTGG | AACCACGAAGTGACCGTACC | 243 |
| chr5:44681080-44683113 | p2 | (AT)18 | CGGCAACAAAGTGAGTTGAA | ATGCCTTACCTCGCGTAAAA | 173 |
| chr5:44707511-44709527 | p5 | (TAAAA)3 | AGCTCCCCAATATGCTCCTT | TTGAATGTCTTTTCTCCCCG | 239 |
| chr5:44707511-44709527 | p2 | (AT)8 | AGCTCCCCAATATGCTCCTT | TTGAATGTCTTTTCTCCCCG | 239 |
| chr5:44707511-44709527 | p3 | (CTT)5 | AGCTCCCCAATATGCTCCTT | TTGAATGTCTTTTCTCCCCG | 239 |
| chr5:44800618-44802633 | p5 | (AATGC)3 | AGTAGCATGGCGGAAAGAAA | TGGGTCGTGACAAGTTGGTA | 191 |
| chr5:44845202-44847226 | p3 | (ACA)6 | CCGAACCAACCACCCTACTA | CGTACACTCTACCCTCCCCA | 137 |
| chr5:44845202-44847226 | p2 | (TA)12 | CCGAACCAACCACCCTACTA | CGTACACTCTACCCTCCCCA | 137 |
| chr5:44848036-44850060 | p2 | (TA)12 | CATTGTAATTCCTTCCCCCA | GATGCCCCACCATTTGTAGT | 123 |
| chr5:44875792-44877830 | p2 | (TA)19 | TTGTTGTGGTTGATGTTGGG | GGATCATGCCAAAGGAAAGA | 174 |
| chr5:44875792-44877830 | p2 | (TA)15 | TTGTTGTGGTTGATGTTGGG | GGATCATGCCAAAGGAAAGA | 174 |
| chr5:44907392-44909410 | p5 | (CAAAC)3 | GTTCATGGTCCGGTTTGACT | GATCGCGTCAAAGCCATATT | 266 |
| chr5:44995076-44997091 | p5 | (ATTGC)3 | CGTCCTGATTTGGACAAGGT | AATCAAGTACCAACAGCGGG | 189 |
| chr5:45014778-45016800 | p2 | (TC)11 | ATGCTTTGCTTCAACGTCCT | GTCACCGGTTTGTTTTTGCT | 228 |
| chr5:45026692-45028704 | p2 | (AT)6 | CCGAACATGAAATGATGACG | AAGGCAATTTTGTCCCTCCT | 147 |
| chr5:45195821-45197865 | p2 | (TA)22 | TAAACGATGGACTTGGAGGC | TTGTTCAAGAGAGGGGGTTG | 275 |
| chr5:45197121-45199137 | p4 | (GTAT)4 | GGATAATCCCTGAGCAACGA | TCAAAACCCAAGAGTCGGTC | 286 |
| chr5:45197121-45199137 | p4 | (GTAT)4 | GGATAATCCCTGAGCAACGA | TCAAAACCCAAGAGTCGGTC | 286 |
| chr5:45425600-45427618 | p6 | (AGGTAG)3 | GTTTTGGCCACCACTAGGAA | TCGGAGTTCTGATGGCTTCT | 141 |
| chr5:45431732-45433794 | p2 | (AG)31 | GGTTTGGGTCGTGACAACTT | AGTGTGGGTTTTGGAAGCAC | 174 |
| chr5:45431732-45433794 | p3 | (ATT)5 | GGTTTGGGTCGTGACAACTT | AGTGTGGGTTTTGGAAGCAC | 174 |
| chr5:45432569-45434584 | p2 | (AG)31 | GGCAAAGGAAAAGGGTCATT | CAACAAAGGGGCCAGTCTTA | 228 |
| chr5:45432569-45434584 | p3 | (ATT)5 | GGCAAAGGAAAAGGGTCATT | CAACAAAGGGGCCAGTCTTA | 228 |
| chr5:45504670-45506682 | p4 | (GAAA)4 | GAAGATCTGGGTGTGGCATT | CGGAATCATGCTTTCATGTG | 105 |
| chr5:45504670-45506682 | p2 | (AT)6 | GAAGATCTGGGTGTGGCATT | CGGAATCATGCTTTCATGTG | 105 |
| chr5:45547822-45549837 | p5 | (ATAAA)3 | TTCGGATTTTGGGGTTGTAA | TGAACCATCCCCTTAACCAA | 118 |
| chr5:45547822-45549837 | p2 | (TA)10 | TTCGGATTTTGGGGTTGTAA | TGAACCATCCCCTTAACCAA | 118 |
| chr5:45682417-45684471 | p2 | (AT)7 | ACATCGACCAACATCGACAA | GCTGGAAGCTTTCCTTTGTG | 128 |
| chr5:45682417-45684471 | p2 | (TA)27 | ACATCGACCAACATCGACAA | GCTGGAAGCTTTCCTTTGTG | 128 |
| chr5:45791245-45793260 | p3 | (TTA)5 | TGTGGGTTTACACCAGGGAT | TGATGCTGACCAACCCTGTA | 245 |
| chr5:45791245-45793260 | p2 | (TA)6 | TGTGGGTTTACACCAGGGAT | TGATGCTGACCAACCCTGTA | 245 |
| chr5:45813874-45815889 | p5 | (AATGC)3 | GGGCCATTTGAGATTCTTGA | CAAAACCGAGTCCCAAAGAA | 107 |
| chr5:45821607-45823659 | p4 | (TTAT)13 | GGAACGATGGAGTGTTCGAT | TACAACATGTCTCGGACGGA | 221 |
| chr5:45884059-45886077 | p6 | (CATATA)3 | TCCCAAAGTATCCCCATCAA | CCATCACACCCTAGCGAGAT | 123 |
| chr5:45946812-45948842 | p2 | (TA)15 | CAAAGATCCGCTTGTCATCA | TGTGTGCCAGAGACTTCGTC | 239 |
| chr5:45995995-45998007 | p2 | (AT)6 | TTATTTTGATGGCCGAGAGG | GCCACGAATAATCACGGAGT | 216 |

| Marker ID | SSR type | SSR motif | FORWARD PRIMER (5'-3') | REVERSE PRIMER (5'-3') | size (bp) |
|---|---|---|---|---|---|
| chr5:46005460-46007492 | p2 | (AT)16 | GTATGGCCTCGGTCACTTGT | CCTCCGGTGAATGTAGAGGA | 293 |
| chr5:46012525-46014539 | p2 | (AT)7 | GCAAAAGATTCAGCGTCCTC | GTGACCGAGGCCATACACTT | 111 |
| chr5:46084212-46086242 | p2 | (AT)15 | GGTAGGTGGACCCTTTCCAT | TAGTGGCTCTTGTACGGCCT | 157 |
| chr5:46211459-46213499 | p2 | (TA)20 | TGGGACTATCATCGTGAGCA | AGGTTGGAAAGTCGGGAAGT | 221 |
| chr5:46255572-46257622 | p2 | (TA)25 | CAGGTGGGTTCACTTGGAGT | TGACAACGTCTGCTTTACGC | 282 |
| chr5:46305327-46307343 | p2 | (AT)25 | GAGCTGTCTGCAAAGCCTCT | ATGCCCGCAGGTGTAGATAG | 287 |
| chr5:46305327-46307343 | p4 | (CATA)4 | GAGCTGTCTGCAAAGCCTCT | ATGCCCGCAGGTGTAGATAG | 287 |
| chr5:46305327-46307343 | p3 | (CAA)7 | GAGCTGTCTGCAAAGCCTCT | ATGCCCGCAGGTGTAGATAG | 287 |
| chr5:46305327-46307343 | p2 | (CT)7 | GAGCTGTCTGCAAAGCCTCT | ATGCCCGCAGGTGTAGATAG | 287 |
| chr5:46357069-46359085 | p4 | (CATA)4 | TCGTCACACTATGACCGAGC | ATTGACGTCCCTTCTTGTGG | 269 |
| chr5:46365301-46367323 | p2 | (TA)11 | GCATGCAATACTCCAGGGAT | GCGAGCTCTTTTGCTGAGAT | 161 |
| chr5:46382735-46384809 | p5 | (AATAA)3 | CATGCTGGACTTTCCCCTAA | AGAGAGAGCGTGGCAAAAAC | 200 |
| chr5:46382735-46384809 | p2 | (AT)37 | CATGCTGGACTTTCCCCTAA | AGAGAGAGCGTGGCAAAAAC | 200 |
| chr5:46382735-46384809 | p2 | (TA)6 | CATGCTGGACTTTCCCCTAA | AGAGAGAGCGTGGCAAAAAC | 200 |
| chr5:46384715-46386735 | p2 | (AT)9 | TAGCACCTCCGGAATAATGG | TCGCACTGACGTCTGTTTTC | 267 |
| chr5:46384744-46386764 | p2 | (AT)9 | TAGCACCTCCGGAATAATGG | TCGCACTGACGTCTGTTTTC | 238 |
| chr5:46416572-46418590 | p6 | (ATTATG)3 | CGAGGACAACTTCGGAAAAA | GGTTCACCATCAGGCAAAAT | 168 |
| chr5:46444687-46446705 | p2 | (AT)9 | AGCGGGGCCATCTAACTATT | GGTTCAGGGCATGACATCTT | 191 |
| chr5:46477312-46479330 | p6 | (ATACAT)3 | AGAGGTCCGGAAGGTCATTT | GACCGAGTCAACTCCCGATA | 257 |
| chr5:46477312-46479330 | p3 | (TCA)6 | AGAGGTCCGGAAGGTCATTT | GACCGAGTCAACTCCCGATA | 257 |
| chr5:46548624-46550646 | p2 | (TG)11 | GATGGGGGAGGGATAACAAT | CCACGTGCCAAGTTTATGTG | 142 |
| chr5:46552983-46554998 | p5 | (ACAAA)3 | TATGTCTGCCTTTGCACTCG | GGCTTGTCAAGATGGCCTTA | 255 |
| chr5:46620152-46622164 | p2 | (CA)6 | AGTAGCCCGCTGACTCATGT | GGTTGACGACCCTTCCAGTA | 176 |
| chr5:46656515-46658533 | p2 | (TA)9 | TCCCTATATGCATTGCCACA | GCTCAAGCTAAGGCCTGCTA | 151 |
| chr5:46656515-46658533 | p2 | (TC)9 | TCCCTATATGCATTGCCACA | GCTCAAGCTAAGGCCTGCTA | 151 |
| chr5:46660302-46662317 | p5 | (ACAAC)3 | TAAGCCCACGACTCTTTCGT | CCGTTGGTCTTACGGATGTT | 104 |
| chr5:46697320-46699383 | p3 | (AAT)21 | CTGCAAACATAAAAGCGCAA | CTCGGGGTTCAACATGAAGT | 265 |
| chr5:46722224-46724239 | p5 | (TTTTC)3 | GGCACCAGACACCAGAAACT | GAATTGAGGTATCGGGGGTT | 170 |
| chr5:46788995-46791007 | p2 | (CT)6 | CAAACGGTAGCCGGTAAAGA | AGCTCCTTGCGTCAATGTCT | 151 |
| chr5:46788995-46791007 | p6 | (AACTGA)3 | CAAACGGTAGCCGGTAAAGA | AGCTCCTTGCGTCAATGTCT | 151 |
| chr5:46922226-46924320 | p2 | (TA)47 | GGTGGATGATTGACCGTCTT | GCAAGGAGTTTTCCACGGTA | 274 |
| chr5:46940350-46942362 | p2 | (TA)6 | GATTTGTGGCACATTCAACG | CTCCGTCCTTTGTCGTCTTC | 234 |
| chr5:47044850-47046865 | p3 | (TTC)5 | CGACCCATTCGTACCAAGAT | ACGGCTTTTGAGGTCTCTGA | 154 |
| chr5:47044850-47046865 | p6 | (TTAAAA)3 | CGACCCATTCGTACCAAGAT | ACGGCTTTTGAGGTCTCTGA | 154 |
| chr5:47089367-47091382 | p5 | (TGAAT)3 | AGAAGCTTGGAGAAGGGGAC | ATTGCTCCAACAACCACTCC | 277 |
| chr5:47100790-47102805 | p2 | (TA)6 | TGTGGCATATGTTCATTGGG | CATGTCCCCTTACAAGCGTT | 111 |

| Marker ID | SSR type | SSR motif | FORWARD PRIMER (5'-3') | REVERSE PRIMER (5'-3') | size (bp) |
|---|---|---|---|---|---|
| chr5:47100790-47102805 | p5 | (TGAAT)3 | TGTGGCATATGTTCATTGGG | CATGTCCCCTTACAAGCGTT | 111 |
| chr5:47101809-47103825 | p2 | (TA)8 | TTTGGCCGAGTTGAATTCTC | CTTTGTATTGGCGCTCGTTT | 103 |
| chr5:47124557-47126569 | p2 | (TA)11 | CATATGCAACACATGCCACA | ATGATTTGAAGAAGCCGTGG | 140 |
| chr5:47124557-47126569 | p2 | (AT)6 | CATATGCAACACATGCCACA | ATGATTTGAAGAAGCCGTGG | 140 |
| chr5:47134373-47136393 | p2 | (AT)10 | TCACCCATAAAAGGTGGGAA | CCGAAAACACGGAGATCATT | 269 |
| chr5:47134373-47136393 | p3 | (CAT)5 | TCACCCATAAAAGGTGGGAA | CCGAAAACACGGAGATCATT | 269 |
| chr5:47141979-47144021 | p2 | (AT)21 | TATAAGTGTGGCGCGTTTCA | GTACTGGGTACCCGTCATGG | 190 |
| chr5:47141979-47144021 | p2 | (TA)28 | TATAAGTGTGGCGCGTTTCA | GTACTGGGTACCCGTCATGG | 190 |
| chr5:47412112-47414130 | p6 | (CATATA)3 | GCACCTCTTATCTCGCGTTC | TGTACATATTTGGGCACGGA | 203 |
| chr5:47457152-47459167 | p5 | (AAACT)3 | TGGCACAAGCCATTTTGATA | TCCCTGTACCTCACATGCCT | 173 |
| chr5:47526005-47528023 | p3 | (AGA)6 | CCAATTTTTGTCAAACCGCT | CTCAGAAGGCGATATCCAGC | 242 |
| chr5:47533135-47535155 | p5 | (TTCAT)4 | CTTCCCATGCTATCTCCCAA | GATCCCAGCAAGCTCAAGAC | 110 |
| chr5:47540868-47542883 | p5 | (TTCAT)3 | TATGCATTTGGGTCGTGTGT | CCACGTATACTTCGTGCCCT | 214 |
| chr5:47593211-47595223 | p2 | (TA)6 | TGAGGGATCCGATAACTTGC | ACCGGGCTGTCCTTTAATTT | 191 |
| chr5:47607593-47609619 | p2 | (AT)13 | ACTTTTGTCACGACCCAACC | TGTCGGCTTGCATATGATGT | 190 |
| chr5:47653111-47655129 | p6 | (TTTTTG)3 | CCCACCCTTACCCTACCACT | GGGTGGGGTGAGTTATTGTG | 221 |
| chr5:47653111-47655129 | p2 | (TG)7 | CCCACCCTTACCCTACCACT | GGGTGGGGTGAGTTATTGTG | 221 |
| chr5:47653111-47655129 | p2 | (GA)7 | CCCACCCTTACCCTACCACT | GGGTGGGGTGAGTTATTGTG | 221 |
| chr5:47668032-47670044 | p2 | (TA)6 | ATGTGCCTTATGGTGCATGA | TATGGCCAAACACAACTCCA | 163 |
| chr5:47688967-47690982 | p3 | (CAA)5 | CCGCCTTAAAACGTCCATAA | ATTCGGGTGATCTGACTTCG | 178 |
| chr5:47761788-47763812 | p6 | (TGTAAG)4 | TTGGTAGGGCCAGGTAGATG | TGCAGGATCGGTGTTTCATA | 163 |
| chr5:47924786-47926828 | p2 | (TA)21 | AAGGCTGCCACATACGAAAC | TTTGTCTAGGGGTACGACGG | 276 |
| chr5:47957511-47959523 | p2 | (AT)6 | TTTTTCGCTGAGGTCGCTAT | CGAGAGATGAAGAGGCCAAC | 227 |
| chr5:48041199-48043223 | p6 | (TGTTTG)4 | CAGCGTTGAGAGAACATGGA | CGATTGACCCGAAATTTGTT | 141 |
| chr5:48205084-48207096 | p2 | (AG)6 | CAACCAAGGCTTCAGGGTTA | AAAGGTGCCTACGGGCTATT | 178 |
| chr5:48212035-48214050 | p5 | (TTCAT)3 | ACGATGTGCAGGTGACAGAG | TTCATCTTGAATCCCTTCCG | 109 |
| chr5:48252015-48254029 | p2 | (TA)7 | CCATGACTAGTGCCCGAGAT | GTCGGCTTGCGTATGGTATT | 215 |
| chr5:48318527-48320545 | p6 | (CTCTAC)3 | CGGGAAAAGGAGTGAAAACA | CCGTTGACTTGGGTTCAGTT | 261 |
| chr5:48336996-48339014 | p6 | (CTCTAC)3 | CGGGAAAAGGAGTGAAAACA | CCGTTGACTTGGGTTCAGTT | 261 |
| chr5:48396508-48398526 | p2 | (AT)10 | CGGCCCAAAGCATTATTAAA | GGGTCGTGACAGGTCGTTAT | 253 |
| chr5:48396508-48398526 | p3 | (ATC)6 | CGGCCCAAAGCATTATTAAA | GGGTCGTGACAGGTCGTTAT | 253 |
| chr5:48599030-48601045 | p3 | (CAA)5 | ACATCACAACAAAGGCCACA | AGGACAGTTGACAGGGGTTG | 121 |
| chr5:48609329-48611347 | p3 | (CAA)6 | ACATCACAACAAAGGCCACA | TAAGGCAAAGTTGACAGGGG | 300 |
| chr5:48708960-48710975 | p5 | (AAAAG)3 | TCTCCGAACCAGATCCAAAC | GGTTTTTGGGACTCAGGTCA | 176 |
| chr5:48740170-48742188 | p2 | (AT)9 | CCAACCGGTATATGTGTCCC | GCTTCACTCACCCTCGTTTC | 113 |
| chr5:48740170-48742188 | p6 | (AGTGGA)3 | CCAACCGGTATATGTGTCCC | GCTTCACTCACCCTCGTTTC | 113 |

| Marker ID | SSR type | SSR motif | FORWARD PRIMER (5'-3') | REVERSE PRIMER (5'-3') | size (bp) |
|---|---|---|---|---|---|
| chr5:48745717-48747749 | p3 | (TTG)6 | TATAAACGGGGGAGATGCTG | GTCCGAGCTCGCAAGTCTAC | 103 |
| chr5:48745717-48747749 | p2 | (TA)16 | TATAAACGGGGGAGATGCTG | GTCCGAGCTCGCAAGTCTAC | 103 |
| chr5:48867148-48869160 | p6 | (ATCCGT)3 | AAGCCGACAAGGCTAACAAA | CATAGCGGGGCTCAATTTTA | 251 |
| chr5:48867148-48869160 | p2 | (TA)6 | AAGCCGACAAGGCTAACAAA | CATAGCGGGGCTCAATTTTA | 251 |
| chr5:48935319-48937331 | p2 | (CT)6 | CACTTCCCCTTTATCGCTCA | TGAGGCCATTGTTGTGTTGT | 125 |
| chr5:48935319-48937331 | p2 | (AT)9 | CACTTCCCCTTTATCGCTCA | TGAGGCCATTGTTGTGTTGT | 125 |
| chr5:49012431-49014445 | p2 | (TG)7 | AAAGGTGCCTACGGGCTATT | TGCATACCCGACCTCTTAGG | 224 |
| chr5:49026030-49028042 | p3 | (AAT)5 | TGTGGTGGATGCCTTGAGTA | AAACAGACGGGCCCTAATCT | 217 |
| chr5:49026030-49028042 | p2 | (AT)6 | TGTGGTGGATGCCTTGAGTA | AAACAGACGGGCCCTAATCT | 217 |
| chr5:49041805-49043859 | p2 | (AT)27 | TCTATGAAACCTTGCGGGTC | AATACTTCTTCCACCCCGCT | 257 |
| chr5:49106271-49108283 | p2 | (AT)6 | CTCCACTTGAACTAAGGGCG | TTAGTTTTCCTGCCGTGGTT | 133 |
| chr5:49120422-49122447 | p5 | (CATAT)5 | CCTAATTCCCAAATGGAGCA | GGTATTGGGTACGTTGGGTG | 111 |
| chr5:49274368-49276380 | p2 | (CT)6 | GAGAAATGACTTGGGAAGCG | CCCCATAACATGGCATTACC | 146 |
| chr5:49299358-49301380 | p2 | (TA)11 | CGTATGCGAAACGTATGTGG | TTACGCCAACCATACAACCA | 284 |
| chr5:49306357-49308377 | p2 | (AT)10 | CGTATGCGAAACGTATGTGG | TTACGCCAACCATACAACCA | 249 |
| chr5:49326406-49328424 | p2 | (AC)9 | CTTCGCAGGGTTGTTCGTAT | CTGCAGAACGAGGGTCAAAT | 207 |
| chr5:49329422-49331450 | p2 | (AT)10 | TTACCCAGGCACCTTTCATC | TTTGTTAGCCTTGTCGGCTT | 143 |
| chr5:49329456-49331468 | p2 | (AT)10 | TTACCCAGGCACCTTTCATC | TTTGTTAGCCTTGTCGGCTT | 109 |
| chr5:49333810-49335828 | p6 | (CATATA)3 | TATCAGGTGGAGGAACTGGG | TGCACTCTTGTTTGTCCTCG | 158 |
| chr5:49338147-49340162 | p5 | (AAATG)3 | ATCGCTCACTTGTTCTCCGT | GAAGCGTTGTCCGACTTCTC | 290 |
| chr5:49338147-49340162 | p2 | (AT)7 | ATCGCTCACTTGTTCTCCGT | GAAGCGTTGTCCGACTTCTC | 290 |
| chr5:49350656-49352671 | p5 | (TATCA)3 | AAATGTTCCCATGCTTCGTC | GGCTTCGCCAGTCAATATGT | 251 |
| chr5:49354475-49356487 | p2 | (TA)8 | CGTCCGTTGTCCCTTAAAAA | CCCATGAGTTTGGAGATGGT | 263 |
| chr5:49354475-49356487 | p2 | (AT)7 | CGTCCGTTGTCCCTTAAAAA | CCCATGAGTTTGGAGATGGT | 263 |
| chr5:49354475-49356487 | p2 | (TA)6 | CGTCCGTTGTCCCTTAAAAA | CCCATGAGTTTGGAGATGGT | 263 |
| chr5:49354475-49356487 | p2 | (AG)6 | CGTCCGTTGTCCCTTAAAAA | CCCATGAGTTTGGAGATGGT | 263 |
| chr5:49372587-49374599 | p2 | (CA)6 | TTGATATGGAAAAGCCCTCG | AGGGCCCACTGATTAGCTTT | 223 |
| chr5:49382591-49384603 | p2 | (CT)6 | GCGGTGACATGGACTTTCTT | AAAGACCATCCCTCATCGTG | 299 |
| chr5:49406988-49409003 | p2 | (AT)17 | GTCGGCGCTTATTTTGTTGT | AAACACCATGCTGGGAACTC | 243 |
| chr5:49406988-49409003 | p3 | (GTT)5 | GTCGGCGCTTATTTTGTTGT | AAACACCATGCTGGGAACTC | 243 |
| chr5:49533750-49535762 | p2 | (GT)6 | TTTCAAAACCACTCCGAAGG | CGCCTTTCTCCATATTTCCA | 142 |
| chr5:49611451-49613466 | p5 | (CATTG)3 | GAAAAGCGAGGGAGTGATTG | AACACAATAGTGGCTTGCCC | 266 |
| chr5:49617753-49619777 | p6 | (TATGTG)4 | AGCCCCAGGTATATTGGTCC | GCACCTTCGCAGAAGAAATC | 190 |
| chr5:49646091-49648106 | p5 | (AAGGA)3 | GGTAACCTTCTCAGCCCCTC | CCCTAGGACGAACTGCTCTG | 283 |
| chr5:49765312-49767327 | p3 | (GTG)5 | TGGTGGTAATTGGTGGGTTT | TTACCACTGGCTCCATCTCC | 218 |
| chr5:49770968-49772988 | p2 | (AT)10 | CGTGTTGTCCCTCTATCGGT | TGCAGCCTGAGGACCTTATT | 134 |

| Marker ID | SSR type | SSR motif | FORWARD PRIMER (5'-3') | REVERSE PRIMER (5'-3') | size (bp) |
|---|---|---|---|---|---|
| chr5:49785329-49787343 | p2 | (AT)7 | TGATAAGGGGTGCTCGGTAG | ATTTCTGCATGAACCCCAAG | 285 |
| chr5:49792644-49794656 | p2 | (AT)6 | AACCTGGGGTCCCAACTTAC | CCTTCTGCCTTAGATCGTGC | 236 |
| chr5:49826578-49828594 | p2 | (TA)8 | CCAAAACCCTTCTCCAAACA | CCACAATGCCCAAGTTTTCT | 244 |
| chr5:49879584-49881599 | p5 | (ACACA)3 | CACATGGTTGACATTCCGAG | AATAGCCCGTAGGCACCTTT | 224 |
| chr5:49879584-49881599 | p2 | (AT)15 | CACATGGTTGACATTCCGAG | AATAGCCCGTAGGCACCTTT | 224 |
| chr5:49885566-49887578 | p2 | (AT)14 | CTCAGCCACACCAGACTTGA | CCCCCACCATAAAAAGAGGT | 189 |
| chr5:49885580-49887592 | p2 | (AT)14 | CTCAGCCACACCAGACTTGA | CCCCCACCATAAAAAGAGGT | 175 |
| chr5:49885594-49887608 | p2 | (AT)14 | CTCAGCCACACCAGACTTGA | CCCCCACCATAAAAAGAGGT | 161 |
| chr5:49886653-49888669 | p2 | (AT)14 | ATGACATAAAATGGCCGGAA | AGAGTCGTGGCACTCCACTT | 239 |
| chr5:49886653-49888669 | p4 | (ATAC)4 | ATGACATAAAATGGCCGGAA | AGAGTCGTGGCACTCCACTT | 239 |
| chr5:49886653-49888669 | p4 | (ATAC)4 | ATGACATAAAATGGCCGGAA | AGAGTCGTGGCACTCCACTT | 239 |
| chr5:49886776-49888792 | p2 | (AT)14 | ATGACATAAAATGGCCGGAA | AGAGTCGTGGCACTCCACTT | 116 |
| chr5:49886776-49888792 | p4 | (ATAC)4 | ATGACATAAAATGGCCGGAA | AGAGTCGTGGCACTCCACTT | 116 |
| chr5:49886776-49888792 | p4 | (ATAC)4 | ATGACATAAAATGGCCGGAA | AGAGTCGTGGCACTCCACTT | 116 |
| chr5:49889998-49892042 | p2 | (TA)22 | AATTGAGTTACGGACCGTGC | GCTTAAGTTGGGGGTGTTGA | 182 |
| chr5:49902530-49904542 | p2 | (AT)6 | TCTTGCAATGGACCGTAACA | GGACGGTCCGTAACTCATGT | 160 |
| chr5:49946822-49948846 | p5 | (ATATC)3 | ATCATAGCCACACAGGGCTC | AGAAGAAAGCGACGCATGAT | 210 |
| chr5:49946822-49948846 | p2 | (AT)7 | ATCATAGCCACACAGGGCTC | AGAAGAAAGCGACGCATGAT | 210 |
| chr5:49946822-49948846 | p2 | (TA)12 | ATCATAGCCACACAGGGCTC | AGAAGAAAGCGACGCATGAT | 210 |
| chr5:50012474-50014492 | p2 | (AT)6 | GCGAGTATCGTGCATGCTTA | TAACACCGAGTCCCAAAAGG | 138 |
| chr5:50012474-50014492 | p4 | (CATA)4 | GCGAGTATCGTGCATGCTTA | TAACACCGAGTCCCAAAAGG | 138 |
| chr5:50012474-50014492 | p3 | (CAA)6 | GCGAGTATCGTGCATGCTTA | TAACACCGAGTCCCAAAAGG | 138 |
| chr5:50012474-50014492 | p2 | (CT)6 | GCGAGTATCGTGCATGCTTA | TAACACCGAGTCCCAAAAGG | 138 |
| chr5:50013632-50015686 | p2 | (CT)6 | GGCCAAAATCTGGTGAAAAA | AGTTTCCAACGCAACAAACC | 174 |
| chr5:50013632-50015686 | p4 | (TTTG)5 | GGCCAAAATCTGGTGAAAAA | AGTTTCCAACGCAACAAACC | 174 |
| chr5:50024893-50026908 | p3 | (ATC)5 | ATGCTCCAGGAAGAAGTGGA | CGTTCAGACTGGCAGGTGTA | 110 |
| chr5:50024893-50026908 | p3 | (AAC)7 | ATGCTCCAGGAAGAAGTGGA | CGTTCAGACTGGCAGGTGTA | 110 |
| chr5:50161050-50163065 | p3 | (ACA)5 | TTGAGGATCATCGTCTTCCC | AACAATGGAGACCAAGCACC | 176 |
| chr5:50161050-50163065 | p2 | (CT)7 | TTGAGGATCATCGTCTTCCC | AACAATGGAGACCAAGCACC | 176 |
| chr5:50207574-50209586 | p2 | (AT)6 | TGCTCCACGCCCTACTACTT | CATTTGGCTGCATCGAGTAA | 136 |
| chr5:50207574-50209586 | p2 | (AT)8 | TGCTCCACGCCCTACTACTT | CATTTGGCTGCATCGAGTAA | 136 |
| chr5:50207574-50209586 | p2 | (TA)16 | TGCTCCACGCCCTACTACTT | CATTTGGCTGCATCGAGTAA | 136 |
| chr5:50207736-50209752 | p2 | (AT)6 | AAGTATCAACCGTGGCTTCG | GTCGCTCCGATCTAATCCAA | 219 |
| chr5:50207736-50209752 | p2 | (AT)8 | AAGTATCAACCGTGGCTTCG | GTCGCTCCGATCTAATCCAA | 219 |
| chr5:50207736-50209752 | p2 | (TA)16 | AAGTATCAACCGTGGCTTCG | GTCGCTCCGATCTAATCCAA | 219 |
| chr5:50272164-50274179 | p2 | (AT)8 | GCAGGTATTGTGTTTGGCCT | CGAGCGAACCAACTGTGATA | 220 |

| Marker ID | SSR type | SSR motif | FORWARD PRIMER (5'-3') | REVERSE PRIMER (5'-3') | size (bp) |
|---|---|---|---|---|---|
| chr5:50272164-50274179 | p3 | (ATT)5 | GCAGGTATTGTGTTTGGCCT | CGAGCGAACCAACTGTGATA | 220 |
| chr5:50290544-50292559 | p2 | (AT)8 | TCCACCAGTCTAGGACACCC | AGTAGGCACACACACCCACA | 207 |
| chr5:50290544-50292559 | p5 | (ATTGC)3 | TCCACCAGTCTAGGACACCC | AGTAGGCACACACACCCACA | 207 |
| chr5:50450493-50452505 | p2 | (TA)6 | TGAAGGCGAAGCTGAAAACT | GCTTATCGACGCTTCAAAGG | 185 |
| chr5:50513468-50515552 | p3 | (AAT)28 | GTTGCGGGTGTACCTGAGAT | GCCTCACAGCTTCTGAATCC | 208 |
| chr5:50591599-50593613 | p2 | (TA)7 | GCAGCATTTGTGAAGAAGCA | CGGGGAACCCAAAAACTATT | 184 |
| chr5:50613012-50615027 | p3 | (TGT)5 | ATGAATTTGGACACACGCAA | CCCCTCATAAGTTCGTCAGC | 250 |
| chr5:50686185-50688200 | p5 | (AGAGA)3 | AGCTCTATGGACTTCCCCGT | GTTGGTACCTCCGGTTGCTA | 282 |
| chr5:50832774-50834789 | p5 | (TTGGT)3 | AACCTGGTCCATGTACACCC | GGGTAGGTTTGGTTCGGTTT | 164 |
| chr5:50877068-50879082 | p2 | (TG)7 | CCGAGTCCAAGACGGAGTAA | TTTCGAGTTAGATGGAGCGG | 212 |
| chr5:50894861-50896875 | p2 | (AT)7 | TGTGGGAGGAGGAATTTTTG | GGGTTCAGAATACGTTTGCC | 280 |
| chr5:50913956-50915970 | p2 | (CA)7 | GGTCATGCCAAAAGGAAGAA | GTTGATGAATTTGGGCGAGT | 185 |
| chr5:50920505-50922519 | p2 | (TC)7 | ATTTGAAGCTTGTGGTTCGG | TTGTTGTGTGTGTGGTGTGG | 237 |
| chr5:50999597-51001619 | p2 | (TA)11 | CGGTTCGTATCCTTGGTTGT | TCGAAGCAGAAAGGTTCGTT | 133 |
| chr5:50999597-51001619 | p3 | (CCT)5 | CGGTTCGTATCCTTGGTTGT | TCGAAGCAGAAAGGTTCGTT | 133 |
| chr5:50999867-51001882 | p3 | (CCT)5 | ATCCAACCTCGTGTGACCTC | TGGGCAAGTTGGAATATGGT | 163 |
| chr5:51082998-51085022 | p2 | (CT)12 | ACTCAACTTGCTTGGGGTTG | CCTGTCCGAGGGTTACAAGA | 256 |
| chr5:51175856-51177874 | p2 | (GA)6 | CGGAGCACCCAAAAATTCTA | GGAAATTTTTAGGCAAGGGC | 287 |
| chr5:51175885-51177899 | p2 | (GA)6 | CGGAGCACCCAAAAATTCTA | GGAAATTTTTAGGCAAGGGC | 258 |
| chr5:51202804-51204819 | p3 | (GAT)5 | ACTGCCCATGAAGGAACAAC | TGGCATTTACCTTGCTTTCC | 220 |
| chr5:51205858-51207874 | p2 | (TA)15 | CAGTTGATGTCCCCATACCC | CCCCCTGTCAAGTGTGTCTT | 164 |
| chr5:51205858-51207874 | p2 | (TA)8 | CAGTTGATGTCCCCATACCC | CCCCCTGTCAAGTGTGTCTT | 164 |
| chr5:51208528-51210546 | p6 | (AAAAAT)3 | AGGTTCGGCTATTTCGGTTT | GAAAAATCGGACCGAACTCA | 159 |
| chr5:51250118-51252132 | p2 | (AT)7 | TGTTGTGAATGTGGGCTGTT | GCGCTTATTTTCGTCACACA | 285 |
| chr5:51275853-51277871 | p6 | (GAATTT)3 | AGGATTCACCTTTGCGATTG | CCAGTGCCTTTGAAGGGTAA | 169 |
| chr5:51339425-51341445 | p4 | (TTTA)5 | TAGATCCATGAGGACAGCCC | GATAACAGCGAACCGCTCTC | 283 |
| chr5:51365917-51367931 | p2 | (GA)7 | GAGCATTCTCAGAACCTCGG | TAAAACGACATCGGAGGAGG | 199 |
| chr5:51416832-51418847 | p5 | (CAAAA)3 | GCGTGAGCACCTATCTTTCC | AGACCAGGTCATGGGAAGTG | 205 |
| chr5:51463128-51465142 | p2 | (TC)7 | ATGATCCGCTGGAAGATACG | GGGCCAAAAATCTTGACAAA | 171 |
| chr5:51516639-51518657 | p6 | (GGAACT)3 | AATCAGCGATGGTGGATCTC | CCAGTCGGTCCTATTTGCAT | 127 |
| chr5:51516639-51518657 | p3 | (ATA)19 | AATCAGCGATGGTGGATCTC | CCAGTCGGTCCTATTTGCAT | 127 |
| chr5:51607538-51609564 | p2 | (AT)13 | TGTTCCCACGACTCATACCA | TCACAGGGACTGTTAAGGGG | 106 |
| chr5:51671905-51673921 | p2 | (AG)8 | GGTCAAGGTCGTGGAGTGAT | ATCGATACCCGCATAAGTGC | 244 |
| chr5:51723807-51725822 | p3 | (GAA)5 | GGGGTAAGGTTTGCGTACAA | GTCTTGTTCCTTTCCACCCA | 128 |
| chr5:51766458-51768473 | p3 | (TTA)5 | TTACCTCCCACGTCCTCATC | TTTATGCTTTGCCCTGATCC | 191 |
| chr5:51766458-51768473 | p5 | (TTATT)3 | TTACCTCCCACGTCCTCATC | TTTATGCTTTGCCCTGATCC | 191 |

| Marker ID | SSR type | SSR motif | FORWARD PRIMER (5'-3') | REVERSE PRIMER (5'-3') | size (bp) |
|---|---|---|---|---|---|
| chr5:51766458-51768473 | p3 | (TTA)6 | TTACCTCCCACGTCCTCATC | TTTATGCTTTGCCCTGATCC | 191 |
| chr5:51802114-51804129 | p3 | (TTA)5 | TTACCTCCCACGTCCTCATC | TTTATGCTTTGCCCTGATCC | 191 |
| chr5:51802114-51804129 | p5 | (TTATT)3 | TTACCTCCCACGTCCTCATC | TTTATGCTTTGCCCTGATCC | 191 |
| chr5:51802114-51804129 | p3 | (TTA)6 | TTACCTCCCACGTCCTCATC | TTTATGCTTTGCCCTGATCC | 191 |
| chr5:51819962-51821978 | p4 | (AAAT)4 | GTCTGTTCCGGAGATGGAAA | TTCTTCCACCTCGTCTGCTT | 279 |
| chr5:51819962-51821978 | p2 | (AT)11 | GTCTGTTCCGGAGATGGAAA | TTCTTCCACCTCGTCTGCTT | 279 |
| chr5:51966350-51968365 | p3 | (ACA)5 | CATGCCAAGAGAAGAGAGGG | AGTTAAATTCTCGGGGCGTT | 195 |
| chr5:51971403-51973437 | p2 | (TA)17 | AATCGGAGGTTCTCGACCTT | ATTAGCCCGCGTTTAGGACT | 289 |
| chr5:52003735-52005761 | p2 | (AT)13 | ACTCTGTCACGACCCAAACC | AGTTGTAATGGGCCATCAGG | 243 |
| chr5:52004078-52006108 | p2 | (AT)13 | CATACCAAAAAGGGGCTGAA | TTGCTTGTGGACGTTGTGTT | 294 |
| chr5:52004110-52006140 | p2 | (AT)13 | CATACCAAAAAGGGGCTGAA | TTGCTTGTGGACGTTGTGTT | 262 |
| chr5:52004142-52006172 | p2 | (AT)13 | CATACCAAAAAGGGGCTGAA | TTGCTTGTGGACGTTGTGTT | 230 |
| chr5:52004174-52006204 | p2 | (AT)13 | CATACCAAAAAGGGGCTGAA | TTGCTTGTGGACGTTGTGTT | 198 |
| chr5:52004206-52006236 | p2 | (AT)13 | CATACCAAAAAGGGGCTGAA | TTGCTTGTGGACGTTGTGTT | 166 |
| chr5:52004238-52006268 | p2 | (AT)13 | CATACCAAAAAGGGGCTGAA | TTGCTTGTGGACGTTGTGTT | 134 |
| chr5:52004270-52006300 | p2 | (AT)13 | CATACCAAAAAGGGGCTGAA | TTGCTTGTGGACGTTGTGTT | 102 |
| chr5:52004398-52006428 | p2 | (AT)13 | CACGCATGCTTAGCTTTTCA | ATGGGGTTATGGGAAAAAGG | 259 |
| chr5:52004430-52006460 | p2 | (AT)13 | CACGCATGCTTAGCTTTTCA | ATGGGGTTATGGGAAAAAGG | 227 |
| chr5:52004462-52006492 | p2 | (AT)13 | CACGCATGCTTAGCTTTTCA | ATGGGGTTATGGGAAAAAGG | 195 |
| chr5:52004494-52006524 | p2 | (AT)13 | CACGCATGCTTAGCTTTTCA | ATGGGGTTATGGGAAAAAGG | 163 |
| chr5:52004526-52006556 | p2 | (AT)13 | CACGCATGCTTAGCTTTTCA | ATGGGGTTATGGGAAAAAGG | 131 |
| chr5:52009278-52011293 | p3 | (CAA)5 | CAAGATCTGCAGGCATGAAA | GCCCATAGAGACTTGACGGA | 291 |
| chr5:52017304-52019340 | p2 | (AT)18 | GCCTTTCTTGGGTTTCTTCC | AGAAATGAAACGGATTTGCG | 277 |
| chr5:52047585-52049597 | p2 | (AT)7 | GGATAACACGAGTCCCGAAA | GTCAAACCCCATTCCTGATG | 142 |
| chr5:52047585-52049597 | p2 | (AT)6 | GGATAACACGAGTCCCGAAA | GTCAAACCCCATTCCTGATG | 142 |
| chr5:52114558-52116570 | p2 | (TA)7 | CATTGTTCACCGAGTCCCTT | ACCGTACTCGGCCCTCTAGT | 139 |
| chr5:52114558-52116570 | p2 | (AT)6 | CATTGTTCACCGAGTCCCTT | ACCGTACTCGGCCCTCTAGT | 139 |
| chr5:52137708-52139746 | p2 | (AT)10 | CCAGTAATTGACGGGTTGCT | ATCGGTTTGGGTCGTAACAG | 281 |
| chr5:52137708-52139746 | p2 | (AT)19 | CCAGTAATTGACGGGTTGCT | ATCGGTTTGGGTCGTAACAG | 281 |
| chr5:52142586-52144642 | p2 | (AT)28 | CGGAAAAAGGTGCAAGGTAA | GGTTCTCGGACCTCACACAT | 179 |
| chr5:52165069-52167081 | p2 | (TA)6 | TTTTGAACCTCGGAGCATCT | TTTGGAGGGTTTCCACTTTG | 151 |
| chr5:52172196-52174208 | p2 | (TA)6 | TTTTGAACCTCGGAGCATCT | TTTGGAGGGTTTCCACTTTG | 151 |
| chr5:52198810-52200825 | p5 | (TTTTC)3 | GATAGCTGAAGCAAGGGCAC | ATACTCCATGGACCCACTGC | 194 |
| chr5:52198810-52200825 | p2 | (AT)28 | GATAGCTGAAGCAAGGGCAC | ATACTCCATGGACCCACTGC | 194 |
| chr5:52241536-52243548 | p2 | (AT)6 | GGTGGGATACTTGGGGCTAT | ATGTTGCACAACCAATCGAA | 183 |
| chr5:52245314-52247398 | p2 | (CT)6 | TTCATCAGCGCAGTTTTACG | CATCATCACAAGCCGACAAG | 279 |

| Marker ID | SSR type | SSR motif | FORWARD PRIMER (5'-3') | REVERSE PRIMER (5'-3') | size (bp) |
|---|---|---|---|---|---|
| chr5:52294672-52296694 | p2 | (AG)11 | ACTGCAGGATGTTGCTCCTT | CCAATGCTTGAACACACGAC | 112 |
| chr5:52335459-52337477 | p6 | (CCCCCA)3 | GTGCATCTCCACCCTTGAGT | CTCTCAACTCAGCAGGTCCC | 202 |
| chr5:52335459-52337477 | p2 | (AT)29 | GTGCATCTCCACCCTTGAGT | CTCTCAACTCAGCAGGTCCC | 202 |
| chr5:52342705-52344723 | p3 | (CAA)5 | TTCCTCCACCCAATTAGCAC | TGTGGCAAGTGGAAAATTGA | 252 |
| chr5:52342705-52344723 | p2 | (CA)6 | TTCCTCCACCCAATTAGCAC | TGTGGCAAGTGGAAAATTGA | 252 |
| chr5:52342705-52344723 | p3 | (TTG)6 | TTCCTCCACCCAATTAGCAC | TGTGGCAAGTGGAAAATTGA | 252 |
| chr5:52386303-52388319 | p4 | (ATAC)4 | GACCAAAATGCTATGGGGAA | GGCAGTTTCAAATAAGGGCA | 279 |
| chr5:52390143-52392167 | p2 | (TA)12 | AAAGCCGACGAAGCTATCAA | ATATAGGTATGGCGGGGTCC | 263 |
| chr5:52390143-52392167 | p2 | (AT)8 | AAAGCCGACGAAGCTATCAA | ATATAGGTATGGCGGGGTCC | 263 |
| chr5:52565183-52567201 | p2 | (TA)9 | TCTTAACGGAAATGCCCAAC | CTTAACGGAAAACCCCAACA | 165 |
| chr5:52603174-52605201 | p3 | (AAT)9 | GGGTTTCAAACTTGGGGAAT | CACCCATATCACTGTCACGC | 108 |
| chr5:52613539-52615554 | p3 | (CAA)5 | TCAAATGGAGACTTTTGGGC | TCCTTCTTTTGGCATGATCC | 229 |
| chr5:52613539-52615554 | p5 | (GTTTT)3 | TCAAATGGAGACTTTTGGGC | TCCTTCTTTTGGCATGATCC | 229 |
| chr5:52706770-52708785 | p5 | (AATGC)3 | CGCAGGCACCTATCATTTCT | TGTGCGAGTCAGACCAGTTC | 152 |
| chr5:52723195-52725213 | p2 | (AT)9 | GTGGTTGTCGCCTTCAATTT | GATGCTGACGAACCAGGAGT | 108 |
| chr5:52753582-52755597 | p3 | (TGT)5 | GATGTCACGCATCGTATTGG | CTTGTCTGCCACAGAGGTCA | 243 |
| chr5:52890577-52892631 | p2 | (AT)27 | AATGAGCTTGGGTCTCCCTT | TGATCCCCAGCATACCTTTC | 299 |
| chr5:52996138-52998156 | p3 | (GTT)6 | TACAACATGGCCGAATCTGA | GCACCACTTGTTCTTCCACA | 105 |
| chr5:53041330-53043362 | p5 | (GTCCC)3 | TATGGAACCCCGAGCTTATG | TCATACCTGGCCCTACCAAG | 126 |
| chr5:53041330-53043362 | p2 | (AT)16 | TATGGAACCCCGAGCTTATG | TCATACCTGGCCCTACCAAG | 126 |
| chr5:53041330-53043362 | p2 | (AT)20 | TATGGAACCCCGAGCTTATG | TCATACCTGGCCCTACCAAG | 126 |
| chr5:53160721-53162737 | p3 | (GGA)6 | CTGCTCCATCCAAAACCCTA | TTTGTGGAGCAAAGAAGGCT | 221 |
| chr5:53160739-53162751 | p3 | (GGA)6 | CTGCTCCATCCAAAACCCTA | TTTGTGGAGCAAAGAAGGCT | 203 |
| chr5:53178056-53180080 | p2 | (AT)12 | CCCATCCAAACAGGCTCTTA | ATGGAAAACCACCATCCAAA | 261 |
| chr5:53178056-53180080 | p5 | (TACAA)3 | CCCATCCAAACAGGCTCTTA | ATGGAAAACCACCATCCAAA | 261 |
| chr5:53275327-53277343 | p2 | (GT)8 | ATAAAGAACCTCCCCTCCGA | TGCCCTTAGCTTTTTCCTCA | 241 |
| chr5:53469779-53471795 | p2 | (AT)8 | TCTCCAAACAGCTGAGCAGA | TAATGGGCCAGGTGTGGTAT | 211 |
| chr5:53538017-53540035 | p6 | (AGAGAA)3 | TGGGGCAAGCAGGATATTAC | GGAGCCGTAGTCAGCTTGTC | 141 |
| chr5:53542039-53544055 | p4 | (TCTG)4 | GCCTTCAAAGCGAACTTTTG | TTATACAGAAGGCATCGGGC | 206 |
| chr5:53647749-53649761 | p2 | (AG)6 | GACGGAAGAGCTGTCCTACG | TGGGGAAGTTTCCTTCATTG | 241 |
| chr5:53649187-53651203 | p2 | (TA)8 | TTGTGATGTGTGGGCTGATT | GTGGCATTTCCCCTACATTG | 276 |
| chr5:53736684-53738696 | p2 | (TG)6 | AAAGGTGCCTACGGGCTATT | TCGATACCTGGCCTACCAAG | 127 |
| chr5:53832589-53834604 | p3 | (TGT)5 | TGGCAGTCTGTGCAATAAGC | AGTCATCCAGGAGCAAATGG | 161 |
| chr5:53845995-53848009 | p3 | (TGT)5 | TATGCCACGTGTGGACTGAT | CACCTCCCCTATATCGCTCA | 254 |
| chr5:53845995-53848009 | p2 | (TA)7 | TATGCCACGTGTGGACTGAT | CACCTCCCCTATATCGCTCA | 254 |
| chr5:53855355-53857370 | p5 | (TATCA)3 | GTGCTTGCTTTCATTTCCGT | GGATTGATTGCGAACAGGAT | 164 |

| Marker ID | SSR type | SSR motif | FORWARD PRIMER (5'-3') | REVERSE PRIMER (5'-3') | size (bp) |
|---|---|---|---|---|---|
| chr5:53914624-53916662 | p2 | (TA)19 | TAATGTGTGGGCTGATTGGA | CACCTCCCCTATATCGCTCA | 280 |
| chr5:53914624-53916662 | p5 | (CGTAT)3 | TAATGTGTGGGCTGATTGGA | CACCTCCCCTATATCGCTCA | 280 |
| chr5:53949231-53951246 | p5 | (AAATG)3 | ATACGAGTGTCCAGTTCGGG | GCATGGTTCCTTGGCTGTAT | 146 |
| chr5:54022838-54024852 | p2 | (AT)7 | CCCGAACTGGACACTCGTAT | TGTTATGCTGTGGTTAGCCG | 285 |
| chr5:54122551-54124566 | p3 | (TCA)5 | GCAATTCTGTGGTGGTGTTG | AGTGTCCCACATGGAAAAGG | 266 |
| chr5:54139585-54141597 | p2 | (TA)6 | ATGGGCCTCAGAGAACCTTT | AGAAACAGTGGGACGTGAGG | 146 |
| chr5:54212595-54214615 | p2 | (AT)10 | TTATTTTATGCCTGAGCGGG | ACTTGCCTCATCGGAGCTTA | 285 |
| chr5:54273037-54275057 | p5 | (AATTC)4 | GGTTCAGCGAACCAAACAAT | GAGTCTGCTTCCTCACCGAG | 164 |
| chr5:54316297-54318343 | p2 | (AT)23 | GGGCAGCTGTAGCTTTCTTG | AGGGTTGTCAAAAGACCGTG | 125 |
| chr5:54316297-54318343 | p2 | (AT)7 | GGGCAGCTGTAGCTTTCTTG | AGGGTTGTCAAAAGACCGTG | 125 |
| chr5:54359661-54361676 | p5 | (AATAA)3 | TACCACCATTTTGCCCCTTA | GCACCAGTGAATCAAGAGCA | 280 |
| chr5:54394873-54396893 | p2 | (TA)6 | CATTTCCCTTCCCTCCAAAT | GTGTGTGTAAGTGATGGCCG | 283 |
| chr5:54394873-54396893 | p2 | (AT)10 | CATTTCCCTTCCCTCCAAAT | GTGTGTGTAAGTGATGGCCG | 283 |
| chr5:54411985-54414005 | p2 | (TA)6 | CATTTCCCTTCCCTCCAAAT | GTGTGTGTAAGTGATGGCCG | 283 |
| chr5:54411985-54414005 | p2 | (AT)10 | CATTTCCCTTCCCTCCAAAT | GTGTGTGTAAGTGATGGCCG | 283 |
| chr5:54478521-54480539 | p6 | (TGATTA)3 | AGATTCGCAAAGACATTCGG | TCCGTGTAACCCGTCTTTTC | 125 |
| chr5:54531623-54533641 | p3 | (CAA)6 | GGTCGTGACACATAGCATGG | TATGAGAGACTGGATCGGGC | 177 |
| chr5:54608892-54610907 | p5 | (CTCAA)3 | TTTTCTTTAGCGGGAACGTG | TTTAGTTGCGGTGCTCAGTG | 181 |
| chr5:54622651-54624671 | p5 | (GGGCC)3 | CTCTCCGCAGTGATTCACAA | CCTGGCTTCCACTTCAGTTC | 255 |
| chr5:54622651-54624671 | p2 | (TC)10 | CTCTCCGCAGTGATTCACAA | CCTGGCTTCCACTTCAGTTC | 255 |
| chr5:54622651-54624671 | p5 | (AGAAA)3 | CTCTCCGCAGTGATTCACAA | CCTGGCTTCCACTTCAGTTC | 255 |
| chr5:54637536-54639554 | p3 | (GTT)5 | ACCAACGATGGAAACTCTGG | CGATGGAAAATTCGACTCGT | 282 |
| chr5:54637536-54639554 | p2 | (AT)9 | ACCAACGATGGAAACTCTGG | CGATGGAAAATTCGACTCGT | 282 |
| chr5:54941086-54943102 | p2 | (AT)8 | TGGCCTTTAAAGTGTGGAGG | GCGTCCCTTCAAGACAACAT | 248 |
| chr5:54998561-55000579 | p6 | (TATTCA)3 | TTCATACCTAAAATGCCCGC | GGTTGGGTCGTGACAAAAGT | 173 |
| chr5:55038917-55040949 | p2 | (TA)16 | CTTGGATGCCTTCTTGCTTC | GGGGAGGTTGGAACACTTTT | 203 |
| chr5:55058578-55060593 | p5 | (TTTTA)3 | AAAATTCCAACCCAAAACCC | CGAAATGGTGTTGTCAATGC | 214 |
| chr5:55096727-55098739 | p2 | (TA)6 | ACGAACTCGTTGGTGCTCTT | CGAAAGATTTTTCACGGCAT | 158 |
| chr5:55157634-55159655 | p3 | (GTA)7 | CTCGTTTCAGGCTGTGTTCA | CATACCCGGCTCGATTTAGA | 157 |
| chr5:55268021-55270042 | p3 | (ACA)7 | GCTCGAGAAAATAGGCGTTG | ATCGAGTCACGGATTAAGCG | 132 |
| chr5:55393872-55395894 | p2 | (AC)11 | GTCCCCCTCACTACTCCCTC | TTTGATACGTGCAGAGCCAG | 212 |
| chr5:55466574-55468608 | p2 | (AT)17 | CCTGCCAGTCTAGTAAGCGG | ACCCGCTGGTTCTCAGTATG | 211 |
| chr5:55466574-55468608 | p2 | (AT)16 | CCTGCCAGTCTAGTAAGCGG | ACCCGCTGGTTCTCAGTATG | 211 |
| chr5:55530855-55532883 | p2 | (AT)14 | CAAGGGCGCAACTGTAATTT | AAGCATGAATTGCCTGTTCC | 193 |
| chr5:55576423-55578444 | p3 | (GAT)7 | TACGGATTAATGTAGCCCGC | TCCTGATGCTCGCTTCTTTT | 102 |
| chr5:55727634-55729649 | p3 | (ATG)5 | ATGGTCCGGACAGCGTATAG | TCCCACACGATGTTTTTCAA | 275 |

| Marker ID | SSR type | SSR motif | FORWARD PRIMER (5'-3') | REVERSE PRIMER (5'-3') | size (bp) |
|---|---|---|---|---|---|
| chr5:55743473-55745493 | p4 | (ATAA)5 | TTCCGCAAAAGCTTCCTTTA | CAAGGCGTTATGATGGTGTG | 190 |
| chr5:55750028-55752049 | p2 | (TA)19 | CTTGCTAGGGCTTGTGGAAC | CTTCATGCTTGACCCCATTT | 189 |
| chr5:55750088-55752132 | p2 | (TA)19 | CTTGCTAGGGCTTGTGGAAC | CTTCATGCTTGACCCCATTT | 129 |
| chr5:55791360-55793375 | p3 | (ATG)5 | ATGGTCCGGACAGCGTATAG | TCCCACACGATGTTTTTCAA | 275 |
| chr5:55807199-55809219 | p4 | (ATAA)5 | TTCCGCAAAAGCTTCCTTTA | CAAGGCGTTATGATGGTGTG | 190 |
| chr5:55813754-55815775 | p2 | (TA)19 | CTTGCTAGGGCTTGTGGAAC | CTTCATGCTTGACCCCATTT | 189 |
| chr5:55813814-55815858 | p2 | (TA)19 | CTTGCTAGGGCTTGTGGAAC | CTTCATGCTTGACCCCATTT | 129 |
| chr5:55912648-55914660 | p2 | (AC)6 | CAAGCTCACTTCCCTTTTCG | GGTTAAGTTCTCGGGGGTTC | 161 |
| chr5:56010872-56012886 | p2 | (TA)7 | GCCCAACCGTACCTAAAACA | GGTTGTGATTCCGACAGCTT | 127 |
| chr5:56063200-56065218 | p5 | (TGAAT)3 | TTGTTGGCATGATTGTTGGT | AACATACCTGCTCCACGTCC | 198 |
| chr5:56063200-56065218 | p6 | (TATATG)3 | TTGTTGGCATGATTGTTGGT | AACATACCTGCTCCACGTCC | 198 |
| chr5:56067932-56069947 | p2 | (TA)6 | AAGCTTGGAGAAGGGGACAT | TTTTTGTACCATGGCCGAAT | 214 |
| chr5:56067932-56069947 | p5 | (TGAAT)3 | AAGCTTGGAGAAGGGGACAT | TTTTTGTACCATGGCCGAAT | 214 |
| chr5:56067932-56069947 | p6 | (TATATG)3 | AAGCTTGGAGAAGGGGACAT | TTTTTGTACCATGGCCGAAT | 214 |
| chr5:56151770-56153785 | p5 | (AAACC)3 | GACCCCGAGGTGACAACTTA | CAACCGTTTGGAATGGTCTT | 156 |
| chr5:56151810-56153825 | p5 | (AAACC)3 | GACCCCGAGGTGACAACTTA | CAACCGTTTGGAATGGTCTT | 116 |
| chr5:56172354-56174366 | p2 | (AT)6 | ACCATACCTCAACAGACGCC | AAATAAGGAGAGGGGCAGGA | 145 |
| chr5:56293430-56295450 | p5 | (GCATT)4 | GTCGCCGTTATTGCGATATT | ACCGCAATCATAGGTTCCAC | 250 |
| chr5:56628253-56630277 | p2 | (TA)12 | CGTTGCATCTGTCTTGTGCT | GACCACTAGCTGAAGCCTGG | 146 |
| chr5:56691797-56693812 | p3 | (TGA)5 | ACCCCTCATTTGCACTCATC | ATCCGAACCTAAATCCCGAG | 248 |
| chr5:56691797-56693812 | p2 | (TA)8 | ACCCCTCATTTGCACTCATC | ATCCGAACCTAAATCCCGAG | 248 |
| chr5:56750354-56752374 | p5 | (CATAT)4 | TTGCATTGTTCCAGTTCTGC | TGGTTTGGGATTATCGGGTA | 203 |
| chr5:56750354-56752374 | p3 | (CAA)8 | TTGCATTGTTCCAGTTCTGC | TGGTTTGGGATTATCGGGTA | 203 |
| chr5:56852862-56854882 | p2 | (AC)29 | AGGGTTTGCACAAAAATGGA | TTCCTTTCCGTAATGCTTCG | 197 |
| chr5:56852862-56854882 | p2 | (AC)14 | AGGGTTTGCACAAAAATGGA | TTCCTTTCCGTAATGCTTCG | 197 |
| chr5:56853102-56855160 | p2 | (AC)29 | GAAAAGCTGTCGCGGAATAG | GCACCATGCCACAAGTCATA | 220 |
| chr5:56858719-56860753 | p3 | (GTT)5 | CTTGTTTTGCGGTGGATTTT | ACGTTGTTTTCCAACTTCCG | 152 |
| chr5:56858719-56860753 | p2 | (AT)17 | CTTGTTTTGCGGTGGATTTT | ACGTTGTTTTCCAACTTCCG | 152 |
| chr5:57057430-57059445 | p5 | (TGAAA)3 | CGTTCGCTTGTTCAACTTCA | CAAAGGAAATGATGCCCAAT | 298 |
| chr5:57100593-57102605 | p2 | (AC)6 | TCGAACAACAATCGGAAACA | TGGCATGATTTTTGGAATGA | 110 |
| chr5:57100593-57102605 | p3 | (TGA)5 | TCGAACAACAATCGGAAACA | TGGCATGATTTTTGGAATGA | 110 |
| chr5:57107297-57109311 | p2 | (AT)7 | CGCATTTAGCCAAGGAAGAC | TTTCGTTAATTTCCCCCACA | 136 |
| chr5:57107297-57109311 | p2 | (AT)7 | CGCATTTAGCCAAGGAAGAC | TTTCGTTAATTTCCCCCACA | 136 |
| chr5:57202412-57204426 | p2 | (CA)7 | CAAGGCCATCCAAATAGCAT | GGCTACAGCTACAGCAAGGG | 298 |
| chr5:57303113-57305145 | p6 | (ATACAT)3 | AACCAGTCAGCTCTAGCCCA | CTTCGGAATTCTTTGGGTCA | 134 |
| chr5:57303113-57305145 | p4 | (CATA)5 | AACCAGTCAGCTCTAGCCCA | CTTCGGAATTCTTTGGGTCA | 134 |

| Marker ID | SSR type | SSR motif | FORWARD PRIMER (5'-3') | REVERSE PRIMER (5'-3') | size (bp) |
|---|---|---|---|---|---|
| chr5:57344320-57346352 | p2 | (AT)16 | AAGGAGTGGAATAACGGGCT | CCAATGGGACTAGCGTGATT | 223 |
| chr5:57437301-57439333 | p2 | (AT)16 | CTTCTCCTGGCTCGTTTCAC | ACCGAGTGGCTGGTTAGATG | 169 |
| chr5:57512847-57514862 | p5 | (ATTGC)3 | TGCTTCTTCGCTTTGGTTTT | TGTATTGATGACTCCGCCAA | 292 |
| chr5:57543359-57545383 | p3 | (AAC)8 | GCCCCATAAGGTCAGTTGAA | TCCAGGAGCAATTGGAGTTC | 236 |
| chr5:57543359-57545383 | p2 | (AT)10 | GCCCCATAAGGTCAGTTGAA | TCCAGGAGCAATTGGAGTTC | 236 |
| chr5:57564557-57566579 | p2 | (TA)11 | CGCAAACTTGAGATGCGATA | ACATTCGGCCAAGAATGAAG | 288 |
| chr5:57567689-57569707 | p6 | (TTAATA)3 | TTTTGCTTTACTTGGGCCTG | AAAGGGGAAATACGGTTTGG | 118 |
| chr5:57609501-57611525 | p2 | (AT)6 | ACAACGTCCAAAGAAAACGG | TGAAAGGCATCCACCCTTAG | 200 |
| chr5:57609501-57611525 | p2 | (TA)12 | ACAACGTCCAAAGAAAACGG | TGAAAGGCATCCACCCTTAG | 200 |
| chr5:57689464-57691479 | p5 | (ATTTC)3 | ATTATCCAGTGGCAGCCAAG | AACCGATCTGCCAATCTTTG | 182 |
| chr5:57720488-57722503 | p3 | (ATC)5 | ACATGCACATGAGCCTACGA | CGCAGGTAGATAGGGAGACG | 146 |
| chr5:57723261-57725273 | p2 | (TA)6 | CTGGTGCTGTGAGGAGCATA | CGGAACGTCGATTCTTCATT | 131 |
| chr5:57741983-57743998 | p3 | (ATC)5 | AGTATAACAATGCAGGGCCG | AGTAGGACTTCCCCTCAGCC | 104 |
| chr5:57854087-57856099 | p2 | (AG)6 | ACATCCGTTTTCTTTGGACG | TCCTGAATAATGGGGTGCTC | 252 |
| chr5:57877778-57879790 | p2 | (TA)6 | AGTCGACGAAGCAAGTCGAT | CCTTGCACTTGATTCTGCTG | 107 |
| chr5:58084230-58086246 | p2 | (AT)7 | TCGAGTGCACGAATGAGTTC | GCTGAAGTGGTCACCAGACA | 106 |
| chr5:58084230-58086246 | p2 | (AT)8 | TCGAGTGCACGAATGAGTTC | GCTGAAGTGGTCACCAGACA | 106 |
| chr5:58085801-58087813 | p3 | (CAA)5 | GGTCATGCCAAAAAGGAAGA | CGGGGATGTTGAATTACAGG | 236 |
| chr5:58085801-58087813 | p2 | (CA)6 | GGTCATGCCAAAAAGGAAGA | CGGGGATGTTGAATTACAGG | 236 |
| chr5:58109152-58111184 | p2 | (AT)16 | TAAGCCGACGAGGCTAAGAA | GGGACTACATAGCGGAGCTG | 251 |
| chr5:58209073-58211091 | p6 | (TTGTTA)3 | ATTCGCAAGGTGGTCTATGG | TTGGCCCTTTGTTTCAAGAC | 203 |
| chr5:58222467-58224485 | p6 | (TTGTTA)3 | ATTCGCAAGGTGGTCTATGG | TTGGCCCTTTGTTTCAAGAC | 203 |
| chr5:58235063-58237078 | p3 | (TTC)5 | AGCTCCTTCATCCCCTTCAT | AGAGGAAACCCACGAAACCT | 288 |
| chr5:58235063-58237078 | p3 | (CAA)5 | AGCTCCTTCATCCCCTTCAT | AGAGGAAACCCACGAAACCT | 288 |
| chr5:58236488-58238503 | p3 | (ATT)5 | GCAATGGAGAGAGAGGAACG | TTGGGGTGTTATGGATGGTT | 296 |
| chr5:58279721-58281733 | p2 | (GA)6 | GAAGTCTTCGGGTTCCTTCC | CAAATGGCCGATGAAGAAAT | 134 |
| chr5:58279721-58281733 | p2 | (TG)6 | GAAGTCTTCGGGTTCCTTCC | CAAATGGCCGATGAAGAAAT | 134 |
| chr5:58393342-58395382 | p2 | (AT)20 | AATTAGGTTCCGAGGGATGG | TGGGTCGTGACATCAACAGT | 210 |
| chr5:58393342-58395382 | p5 | (ATATA)3 | AATTAGGTTCCGAGGGATGG | TGGGTCGTGACATCAACAGT | 210 |
| chr5:58548872-58550888 | p2 | (AT)8 | ATAAATGCAAGCCGACAAGG | CCGGAGTCCAGACTTTGGTA | 278 |
| chr5:58608793-58610808 | p5 | (AAGGC)3 | CTTATTGCTGCCTTTGCTCC | TTACCCCTCCAAAGCTGATG | 193 |
| chr5:58656399-58658417 | p3 | (ATG)6 | GGGCTACACGTTACGCAGAT | TTTCCCTGTCTACCTGTGGC | 267 |
| chr5:58739498-58741518 | p4 | (AAAT)5 | GTCAACACCACATCAGTGCC | TCGCACTTGGGATTTCTTTC | 181 |
| chr5:58739498-58741518 | p5 | (ATCAC)3 | GTCAACACCACATCAGTGCC | TCGCACTTGGGATTTCTTTC | 181 |
| chr5:58860871-58862898 | p3 | (TAT)9 | TTATTTTGATGGCCGAGAGG | CATCCACTGAATCCGATCCT | 106 |
| chr5:58884259-58886274 | p5 | (ATTTC)3 | AAGCATGCGATCCAGATACC | ATAACAATGCCTCACAGGGC | 184 |

| Marker ID | SSR type | SSR motif | FORWARD PRIMER (5'-3') | REVERSE PRIMER (5'-3') | size (bp) |
|---|---|---|---|---|---|
| chr5:58902196-58904214 | p2 | (AT)7 | CCTTACGCTCGTCCAAGTTC | TGGGGGCTGTTTTGATATGT | 142 |
| chr5:58917244-58919278 | p2 | (AT)17 | GGGGGTAGCATTGTCTTTCA | CCTAGGAGGAGTTGAAGGGG | 154 |
| chr5:58958307-58960325 | p2 | (AT)11 | TCCTCCAATTTCTTGGCAAC | TGTGGGGAGGTACAAAAAGC | 230 |
| chr5:58958326-58960350 | p2 | (AT)11 | TCCTCCAATTTCTTGGCAAC | TGTGGGGAGGTACAAAAAGC | 211 |
| chr5:59033462-59035477 | p2 | (TA)6 | AGTGAGTCCCCCAACACTTG | ATTTCCCAATTTCCCCAAAC | 209 |
| chr5:59033511-59035559 | p2 | (TA)6 | AGTGAGTCCCCCAACACTTG | ATTTCCCAATTTCCCCAAAC | 160 |
| chr5:59040036-59042058 | p2 | (TA)6 | AGTCGAGGGTCTTTCGGAAC | CCTCATTGAGTGTCCCGATT | 273 |
| chr5:59040036-59042058 | p6 | (CCGGTT)3 | AGTCGAGGGTCTTTCGGAAC | CCTCATTGAGTGTCCCGATT | 273 |
| chr5:59040129-59042141 | p2 | (TA)6 | AGTCGAGGGTCTTTCGGAAC | CCTCATTGAGTGTCCCGATT | 180 |
| chr5:59040129-59042141 | p6 | (CCGGTT)3 | AGTCGAGGGTCTTTCGGAAC | CCTCATTGAGTGTCCCGATT | 180 |
| chr5:59040378-59042390 | p2 | (TA)6 | CTGGCTTATGACCGCAACAT | TAATCACTCCAAACCGGACC | 210 |
| chr5:59040378-59042390 | p6 | (CCGGTT)3 | CTGGCTTATGACCGCAACAT | TAATCACTCCAAACCGGACC | 210 |
| chr5:59040481-59042505 | p2 | (TA)6 | CTGGCTTATGACCGCAACAT | TAATCACTCCAAACCGGACC | 107 |
| chr5:59040481-59042505 | p6 | (CCGGTT)3 | CTGGCTTATGACCGCAACAT | TAATCACTCCAAACCGGACC | 107 |
| chr5:59058418-59060438 | p4 | (TGTC)4 | CCCGCAAAGATTAGGTGTGT | TGGAAAACACAGGCACAAAA | 254 |
| chr5:59058418-59060438 | p2 | (AG)10 | CCCGCAAAGATTAGGTGTGT | TGGAAAACACAGGCACAAAA | 254 |
| chr5:59058418-59060438 | p3 | (CAC)5 | CCCGCAAAGATTAGGTGTGT | TGGAAAACACAGGCACAAAA | 254 |
| chr5:59058418-59060438 | p3 | (CAC)5 | CCCGCAAAGATTAGGTGTGT | TGGAAAACACAGGCACAAAA | 254 |
| chr5:59067844-59069874 | p3 | (ACA)10 | AAAGCTCTCCGCATCAAGAA | TGGCTGAGAGCCGTATCTTT | 193 |
| chr5:59115004-59117019 | p3 | (ACA)5 | TAAACTGACGCCAGAAGGCT | TGACCCAAGGATGTGTTTGA | 284 |
| chr5:59115004-59117019 | p2 | (AC)6 | TAAACTGACGCCAGAAGGCT | TGACCCAAGGATGTGTTTGA | 284 |
| chr5:59197442-59199454 | p2 | (TA)6 | GTGCTCGGCAAGTAAGGTTC | ACTCAGCTCTATCGCACGGT | 154 |
| chr5:59197442-59199454 | p2 | (AC)7 | GTGCTCGGCAAGTAAGGTTC | ACTCAGCTCTATCGCACGGT | 154 |
| chr5:59197442-59199454 | p6 | (AATAAA)3 | GTGCTCGGCAAGTAAGGTTC | ACTCAGCTCTATCGCACGGT | 154 |
| chr5:59197739-59199753 | p2 | (TA)6 | TGAGGCCCAGATTTAGATGC | AGGTGGCGACTCTTTTTCAA | 204 |
| chr5:59197739-59199753 | p2 | (AC)7 | TGAGGCCCAGATTTAGATGC | AGGTGGCGACTCTTTTTCAA | 204 |
| chr5:59197739-59199753 | p6 | (AATAAA)3 | TGAGGCCCAGATTTAGATGC | AGGTGGCGACTCTTTTTCAA | 204 |
| chr5:59228733-59230779 | p2 | (TA)23 | ACCAATCCTGCCTCATAACG | TGCATATCATAACTCCGCCA | 258 |
| chr5:59336821-59338851 | p2 | (AT)7 | CATAAGCCCTTCGGCTATCA | CCATTTTCTTTGGACGCTGT | 219 |
| chr5:59336821-59338851 | p2 | (AT)15 | CATAAGCCCTTCGGCTATCA | CCATTTTCTTTGGACGCTGT | 219 |
| chr5:59369112-59371137 | p5 | (CATAT)5 | TCAGGACTGCCTCTCACCTT | GCATAGGCCCTACACCAAAA | 147 |
| chr5:59369112-59371137 | p3 | (CAA)5 | TCAGGACTGCCTCTCACCTT | GCATAGGCCCTACACCAAAA | 147 |
| chr5:59406297-59408315 | p6 | (AGAGGC)3 | CGAGGTCCGTCGAGTTAGAG | CCACCATAGTTGGGGCATAC | 291 |
| chr5:59454580-59456595 | p5 | (GAAAG)3 | ATGAGATGGATGAGTTCCGC | CTCCATGGGGTGAGACACTT | 184 |
| chr5:59547337-59549353 | p2 | (AT)8 | ATATATCGTACCCGGCCCTC | ATCGTGCACAGTTTCACTGC | 126 |
| chr5:59547337-59549353 | p5 | (TATTT)3 | ATATATCGTACCCGGCCCTC | ATCGTGCACAGTTTCACTGC | 126 |

| Marker ID | SSR type | SSR motif | FORWARD PRIMER (5'-3') | REVERSE PRIMER (5'-3') | size (bp) |
|---|---|---|---|---|---|
| chr5:59704987-59707007 | p2 | (AT)10 | ATACCAAGTTTGTCACGCCC | TACAGAGTGTGGCCTTGTCG | 132 |
| chr5:59704987-59707007 | p2 | (AT)10 | ATACCAAGTTTGTCACGCCC | TACAGAGTGTGGCCTTGTCG | 132 |
| chr5:59704987-59707007 | p3 | (CAA)5 | ATACCAAGTTTGTCACGCCC | TACAGAGTGTGGCCTTGTCG | 132 |
| chr5:59708577-59710613 | p2 | (AT)8 | TCATGCCTGAGTGGACCATA | GGGTCCTGTCCCATCCTAAT | 279 |
| chr5:59708616-59710674 | p2 | (AT)8 | TCATGCCTGAGTGGACCATA | GGGTCCTGTCCCATCCTAAT | 240 |
| chr5:59829044-59831056 | p2 | (AT)6 | ACACTCCGAGGATGAACTGC | ATCAGTTGTGACTCCCCAGG | 128 |
| chr5:59842813-59844905 | p3 | (CAT)7 | CGTCATCAAGCTCACCTCAA | GGTTGACGGTTCTCCCAGTA | 287 |
| chr5:59887553-59889569 | p4 | (AATT)4 | AGCCTAACATAGGCGGGTTT | GCCATCTGGTGACAGACTGA | 191 |
| chr5:59901112-59903124 | p2 | (CA)6 | AATGGCTCATGGAGGTCAAG | ATTAGCATAGAAAGCGGGCA | 136 |
| chr5:59909666-59911678 | p2 | (CA)6 | ATAGAGTTGTCCCCGAGGCT | CGGATTTGCGACTTGAATTT | 158 |
| chr5:59911735-59913789 | p5 | (GAAAT)3 | TATCCCGACCAGTTCTACGG | CGAAAATGAACTTCGGGAAA | 230 |
| chr5:59911735-59913789 | p3 | (AAC)7 | TATCCCGACCAGTTCTACGG | CGAAAATGAACTTCGGGAAA | 230 |
| chr5:59912298-59914319 | p3 | (AAC)7 | ATGGGCCGATACATATCCAA | TGATGATTATGATGCCCGAA | 271 |
| chr5:60050898-60052914 | p2 | (TA)8 | CTTTGATCGGATTCGTGGAT | CAACAACCGGATCATAGCCT | 280 |
| chr5:60071047-60073067 | p5 | (TCATA)4 | ATGATGAGAGATTGCCCGTC | TGCCTCTACCTTGACCTGCT | 267 |
| chr5:60071047-60073067 | p6 | (TACATA)3 | ATGATGAGAGATTGCCCGTC | TGCCTCTACCTTGACCTGCT | 267 |
| chr5:60101613-60103631 | p5 | (TGAAT)3 | AGCGCCTGTGAAGGAAATAA | GTATCAGCACTCTGGCGTGA | 190 |
| chr5:60139541-60141556 | p3 | (TTG)5 | ATGGAGGAAGGCTAAGGGAA | GGCCTTTGTTTGTTGCTCTC | 220 |
| chr5:60154748-60156776 | p2 | (AT)14 | TTCACAAAGGACTGGGAACC | TGCATATCATAACTCCGCCA | 182 |
| chr5:60187454-60189479 | p5 | (CATAT)5 | GATGATCCGGAGTGAAAGGA | TTGCCTTATCCTTTGTTGGG | 113 |
| chr5:60251929-60253944 | p5 | (TATCA)3 | CGTCACTGACCAAACACGAC | CGGAGCCCATACTTTTGGTA | 247 |
| chr5:60387130-60389144 | p2 | (AT)7 | CCCGCTTCGTTTCATGTATT | ATTGGGCGAGTTTGAACAAC | 233 |
| chr5:60412271-60414285 | p2 | (AT)7 | CCCGCTTCGTTTCATGTATT | ATTGGGCGAGTTTGAACAAC | 233 |
| chr5:60422189-60424207 | p6 | (CATATA)3 | ATGACACATTGGCTGATGGA | TTCCATACTGACGCCCTTTC | 293 |
| chr5:60615798-60617812 | p2 | (AG)7 | GAACCCTCCCGAAATGGTAT | CGGGATACTTAAGGCAACCA | 158 |
| chr5:60628287-60630307 | p2 | (AT)10 | CCAACTTGAAGCACTCGTGA | TCCTTGGACTTTCACGAACC | 221 |
| chr5:60628287-60630307 | p5 | (AAAAG)3 | CCAACTTGAAGCACTCGTGA | TCCTTGGACTTTCACGAACC | 221 |
| chr5:60628287-60630307 | p5 | (TGAAT)3 | CCAACTTGAAGCACTCGTGA | TCCTTGGACTTTCACGAACC | 221 |
| chr5:60632150-60634168 | p2 | (AT)9 | GGCATGGCATGAAGAAAAAT | GGATTTGCACTCACCATCCT | 276 |
| chr5:60635781-60637793 | p2 | (AC)6 | CAAACGAAGAAGTGGGAAGC | AGCTAAGTCTCGGGGGTGTT | 254 |
| chr5:60760771-60762783 | p2 | (AT)6 | GCGCAGGGTTATTTTGGATA | AACAGAAGCCGACAAGGCTA | 130 |
| chr5:60760771-60762783 | p6 | (AGCACC)3 | GCGCAGGGTTATTTTGGATA | AACAGAAGCCGACAAGGCTA | 130 |
| chr5:60761740-60763758 | p2 | (AT)6 | GGGGTCGAGGTAGGAGAAAC | CCTCTGATAAGGGCGGTGTA | 162 |
| chr5:60761740-60763758 | p6 | (AGCACC)3 | GGGGTCGAGGTAGGAGAAAC | CCTCTGATAAGGGCGGTGTA | 162 |
| chr5:60765965-60767980 | p5 | (TGATC)3 | GATTTTGGACGAACTTGGGA | TTTGGAAGTGTTCCGACTCC | 198 |
| chr5:60765965-60767980 | p2 | (AT)9 | GATTTTGGACGAACTTGGGA | TTTGGAAGTGTTCCGACTCC | 198 |

| Marker ID | SSR type | SSR motif | FORWARD PRIMER (5'-3') | REVERSE PRIMER (5'-3') | size (bp) |
|---|---|---|---|---|---|
| chr5:60766830-60768848 | p5 | (TGATC)3 | CGCGTCCTGTACTAAGAGGG | TGCATGCAAGAGGTAGTTGC | 167 |
| chr5:60766830-60768848 | p2 | (AT)9 | CGCGTCCTGTACTAAGAGGG | TGCATGCAAGAGGTAGTTGC | 167 |
| chr5:60772827-60774847 | p2 | (AT)10 | CAGTACTGTGGCATGCGTTT | ATGGAGTGGTTCGACTTTGG | 245 |
| chr5:60793615-60795627 | p2 | (AT)6 | ACATGAACCATCGACGTGAA | TTGTTACCGTTTCATCGCAA | 126 |
| chr5:60793615-60795627 | p5 | (TGAAT)3 | ACATGAACCATCGACGTGAA | TTGTTACCGTTTCATCGCAA | 126 |
| chr5:60847806-60849860 | p2 | (AT)27 | TGGGACCCCTTTAGTTACCC | GGTTTGGGTCGTGACAAGTT | 110 |
| chr5:60866987-60869005 | p6 | (TACATA)3 | GACCCTTACTTAGGGCCGAG | TCCATCAGCCAATGTGTCAT | 173 |
| chr5:60885035-60887049 | p2 | (TA)7 | CATCCGACACCTTCCAACTT | TGATTGTTGTGTTGCGGATT | 175 |
| chr5:60887045-60889057 | p2 | (AT)6 | TTCCTATCTGCCTCCCCTTT | CGACCTTGACAGAAGGAAGC | 181 |
| chr5:60887045-60889057 | p2 | (AT)6 | TTCCTATCTGCCTCCCCTTT | CGACCTTGACAGAAGGAAGC | 181 |
| chr5:60887045-60889057 | p2 | (AT)6 | TTCCTATCTGCCTCCCCTTT | CGACCTTGACAGAAGGAAGC | 181 |
| chr5:61008141-61010161 | p2 | (TA)10 | ATCCTTCAAGGTGCGATTTG | AGTGGTTCCACGATAGGTGC | 222 |
| chr5:61027944-61029962 | p2 | (TA)9 | CCCACTTTGCAAACTTCCAT | GATTGGAGTGGGGAGCATTA | 222 |
| chr5:61052015-61054030 | p2 | (AT)7 | CCAACCAACAAGCTCCATTT | ATATGAGTGTCCAGCTCGGG | 118 |
| chr5:61052025-61054037 | p2 | (AT)7 | CCAACCAACAAGCTCCATTT | ATATGAGTGTCCAGCTCGGG | 108 |
| chr5:61179971-61181989 | p2 | (AT)9 | AGACCAATCATAAGGGCACG | CGAGGGAAATTTGGAGCATA | 225 |
| chr5:61179971-61181989 | p3 | (AAT)7 | AGACCAATCATAAGGGCACG | CGAGGGAAATTTGGAGCATA | 225 |
| chr5:61226068-61228084 | p2 | (CT)8 | ACACATCAACCCCGAGAATC | GCCGGTTGTTGAAATGTTCT | 192 |
| chr5:61448382-61450396 | p2 | (TA)7 | CGTTTCTGAGCCTTCAGAGG | GCGATTCCATTTGCTATGGT | 113 |
| chr5:61540900-61542932 | p2 | (AT)16 | CACAATACCGCCACATCTTG | CGAGTTCAAGTGATCAGGCA | 144 |
| chr5:61706473-61708491 | p2 | (AT)9 | CCGTGCACAACATATCGAAC | GGACGCTCGAAAAATATGGA | 199 |
| chr5:61710065-61712077 | p5 | (ATTCA)3 | TAACTCGGCCCAAATAATCG | GGAAGGTGTCGGGTGAAGTA | 111 |
| chr5:61710065-61712077 | p2 | (TA)6 | TAACTCGGCCCAAATAATCG | GGAAGGTGTCGGGTGAAGTA | 111 |
| chr5:61794407-61796431 | p3 | (GTT)8 | CCACCAAAACTTAAGGGCAA | CTGGGGCATCTCTGTCTCTC | 266 |
| chr5:61806146-61808161 | p3 | (ATG)5 | GCGTAGCTAATAAGGCGTGG | CGTTCGGAAGCATATCCAAT | 142 |
| chr5:61806146-61808161 | p6 | (TATATG)3 | GCGTAGCTAATAAGGCGTGG | CGTTCGGAAGCATATCCAAT | 142 |
| chr5:61838492-61840504 | p2 | (GA)7 | ATATTAAGCGGACACGGTCG | CCCAATATGCAAAATACGGG | 149 |
| chr5:61838492-61840504 | p2 | (TA)6 | ATATTAAGCGGACACGGTCG | CCCAATATGCAAAATACGGG | 149 |
| chr5:61872066-61874078 | p2 | (AT)6 | TCACTTCTTGTGGATGCTGC | GGCGAATACATCGGCTTAAA | 234 |
| chr5:61882331-61884347 | p2 | (AT)8 | TCGTGCATGCCTCAATATGT | ATATATACCGTACCCGGCCC | 140 |
| chr5:61882331-61884347 | p5 | (TATGA)3 | TCGTGCATGCCTCAATATGT | ATATATACCGTACCCGGCCC | 140 |
| chr5:62039817-62041831 | p2 | (AC)7 | ATGTGATGCACCAAACCTCA | TATGGAGGCCACTCCAATTC | 223 |
| chr5:62040538-62042612 | p2 | (AC)7 | TATGGGCCCACAGGTAACTC | TGTACAGGTCAAGTGGCAGC | 202 |
| chr5:62048361-62050375 | p2 | (TA)7 | TGATAATAGTGCAGCCGACG | AGTGCGTCCAACTCCATAGG | 247 |
| chr5:62048361-62050375 | p2 | (AT)7 | TGATAATAGTGCAGCCGACG | AGTGCGTCCAACTCCATAGG | 247 |
| chr5:62106064-62108079 | p5 | (CATAT)3 | GCGCACAACACACCATAAAC | AATTTGTGACGGCCTTATCG | 269 |

| Marker ID | SSR type | SSR motif | FORWARD PRIMER (5'-3') | REVERSE PRIMER (5'-3') | size (bp) |
|---|---|---|---|---|---|
| chr5:62143330-62145350 | p2 | (TA)10 | GTAGCTAATTGGATGCGGGA | GAGCTTCGGAATCACGGATA | 161 |
| chr5:62193061-62195073 | p2 | (TA)6 | CGCACTGGATCTATTGTGGA | AGGGTGGATACGCAAGTACG | 280 |
| chr5:62344367-62346385 | p2 | (AT)9 | CCCAGGAACCAGGGATATTT | GACCCGGCTTATTGTACCCT | 123 |
| chr5:62351398-62353422 | p2 | (AT)10 | AACCCTCCTAAATGCCCTGT | GAGTGGTGCTCGGAGGTTAG | 234 |
| chr5:62351424-62353478 | p2 | (AT)10 | AACCCTCCTAAATGCCCTGT | GAGTGGTGCTCGGAGGTTAG | 208 |
| chr5:62355069-62357081 | p2 | (GT)6 | GACAAAATTGGGCAGCATCT | GAATACGCGAAAATACGGGA | 116 |
| chr5:62380681-62382697 | p2 | (GA)6 | TAAATTTCGGCCTTCCTCCT | GAAGACAACGCAGCGTACAA | 218 |
| chr5:62380681-62382697 | p4 | (CTTT)4 | TAAATTTCGGCCTTCCTCCT | GAAGACAACGCAGCGTACAA | 218 |
| chr5:62399409-62401424 | p5 | (TCATT)3 | GGGCGTGACAGTATCAACCT | ACATAAGCCGATATCGCACC | 151 |
| chr5:62429285-62431305 | p4 | (TTTC)5 | AACTCGGCCAATTCAACAAC | TGGAGATTTCGGGTAAGGTG | 166 |
| chr5:62573215-62575230 | p5 | (TGATA)3 | GCCTACAACGCCTACAGAGG | ATGCAACCTAGTCCGACACC | 169 |
| chr5:62573215-62575230 | p2 | (AT)16 | GCCTACAACGCCTACAGAGG | ATGCAACCTAGTCCGACACC | 169 |
| chr5:62573673-62575705 | p5 | (TGATA)3 | AATTTACGCCAAAGACGGTG | TCTCCCTGCTTTCTGGTTGT | 171 |
| chr5:62573673-62575705 | p2 | (AT)16 | AATTTACGCCAAAGACGGTG | TCTCCCTGCTTTCTGGTTGT | 171 |
| chr5:62661894-62663915 | p3 | (TTG)7 | TGGAATTTGGAAACACACCA | TGTTTGGGGTTGTGTTGAGA | 267 |
| chr5:62730978-62732992 | p2 | (AT)7 | CGGTTTGGCTGTCTTCTAGG | ACTGGTGTACTCAATCGCCC | 168 |
| chr5:62784446-62786464 | p6 | (GGTAGA)3 | CCGGTATGGCAAGAGTTTGT | AGCACTCATGTTACCCTGCC | 241 |
| chr5:63037573-63039603 | p2 | (AT)15 | ACAGGTCGTGGAGTTATCGC | CCATGACTCATAGGCCAGGT | 144 |
| chr5:63037573-63039603 | p2 | (AT)7 | ACAGGTCGTGGAGTTATCGC | CCATGACTCATAGGCCAGGT | 144 |
| chr5:63050501-63052516 | p5 | (AATTG)3 | TCGGAAAAAGGTGAAAGGTG | TTAAAGGGCCGGAATCTTCT | 247 |
| chr5:63072595-63074619 | p6 | (CATATA)4 | TATCACACATAAGCCGGCAG | GGGCGTACCCACTCGTAGTA | 161 |
| chr5:63089485-63091503 | p6 | (CATATA)3 | TCTCCGAAACGAGCGTCTAT | CCTATAATGGTGTGGGGGTG | 159 |
| chr5:63089485-63091503 | p2 | (TA)9 | TCTCCGAAACGAGCGTCTAT | CCTATAATGGTGTGGGGGTG | 159 |
| chr5:63155629-63157643 | p2 | (AT)7 | TTCACGACCTAACCCCGTAG | CCTTGTCGGCTTGCATTTAT | 132 |
| chr5:63198091-63200103 | p2 | (AT)6 | TGAACTGCATCTCTCATGGC | AAGCGACAGCTGCCTAGAAG | 223 |
| chr5:63203624-63205636 | p2 | (TC)6 | AAAACTCGCCTAACCCGAAT | GTTCGTTAGAATTTCGGGCA | 278 |
| chr5:63275539-63277557 | p6 | (ATGTAT)3 | CGACTCAGAATACCGACCGT | TTCTGGTTTACGATCCGTCC | 260 |
| chr5:63275539-63277557 | p2 | (TA)23 | CGACTCAGAATACCGACCGT | TTCTGGTTTACGATCCGTCC | 260 |
| chr5:63289117-63291143 | p2 | (AT)13 | ATAAGCCGACGAGGCTAACA | GTCCTGTCCCGCCTATATGA | 296 |
| chr5:63289117-63291143 | p5 | (GAAAT)3 | ATAAGCCGACGAGGCTAACA | GTCCTGTCCCGCCTATATGA | 296 |
| chr5:63294587-63296602 | p5 | (ACCCG)3 | GAGAGTACTCGCAGGATGGC | TCTGTGGCTGGAGCTCTTTT | 101 |
| chr5:63302761-63304827 | p2 | (AT)33 | CCATTCTTCTGCTGCAATCA | GAGCGGTGCTCGGTAGATAG | 173 |
| chr5:63320606-63322634 | p2 | (AT)14 | TTGCCTCCGTAGCTTTCAAT | CCCTAGTCGGGTCGTGTAAA | 117 |
| chr5:63369426-63371440 | p2 | (AG)7 | ATCCGAAGGTAGGCCAAGTT | AGCCTTCCCAGTCATTTCCT | 115 |
| chr5:63386727-63388769 | p2 | (TA)21 | TCTGTAGGCGATACCCGTTC | TTATAAACAGGAGGCTGCGG | 111 |
| chr5:63386727-63388769 | p3 | (CAT)8 | TCTGTAGGCGATACCCGTTC | TTATAAACAGGAGGCTGCGG | 111 |

| Marker ID | SSR type | SSR motif | FORWARD PRIMER (5'-3') | REVERSE PRIMER (5'-3') | size (bp) |
|---|---|---|---|---|---|
| chr5:63413767-63415779 | p2 | (AT)6 | GCCGGGATCTGTATGTCTGT | AAATACAGATGGCACACGCA | 157 |
| chr5:63417832-63419847 | p2 | (TA)37 | CAAGCCAACGAGGTCGTAAT | CGACGGGTCAGTGAGAATTT | 124 |
| chr5:63417832-63419847 | p3 | (CAA)5 | CAAGCCAACGAGGTCGTAAT | CGACGGGTCAGTGAGAATTT | 124 |
| chr5:63550506-63552526 | p5 | (ATATG)4 | CCTCGGGGGAAACTCTATTC | GCAATCCGATCATGTTTGTG | 271 |
| chr5:63575979-63577999 | p4 | (TGTA)5 | ATGGAAGTTGGAAAACAGCG | CGCAATTCAACAACCAACAC | 253 |
| chr5:63656069-63658085 | p4 | (ACAT)4 | AGGCAGCTAGCTCGACAGAC | GACTTCCTTGCCCTAAACCC | 148 |
| chr5:63696272-63698287 | p3 | (TTA)5 | CTATTTCTTCGGCAGCGAAC | TCAGGCCGTGTGTAGTTCTG | 214 |
| chr5:63699742-63701757 | p3 | (TTA)5 | CTATTTCTTCGGCAGCGAAC | TCAGGCCGTGTGTAGTTCTG | 214 |
| chr5:63707738-63709756 | p2 | (AG)9 | GTGGGAGGGATTTGAAGACA | AAACAGTTTTTCATTGCCCG | 122 |
| chr5:63760434-63762449 | p3 | (TTG)5 | CTCGTAAGTCGTAACCGGGA | TTTTCTTCGCTTTGCTTGGT | 175 |
| chr5:63760434-63762449 | p2 | (TA)25 | CTCGTAAGTCGTAACCGGGA | TTTTCTTCGCTTTGCTTGGT | 175 |
| chr5:63845761-63847773 | p2 | (TA)6 | TCAAATCAGCTCCCCAAATC | TCCTTTTATGACTGGGCTGC | 243 |
| chr5:63927107-63929122 | p5 | (ATTCA)3 | CGACACATGAAGGCTCAGAA | CATTGATGGTGACGAGGATG | 216 |
| chr5:63927107-63929122 | p2 | (TA)6 | CGACACATGAAGGCTCAGAA | CATTGATGGTGACGAGGATG | 216 |
| chr5:63998236-64000260 | p3 | (TGG)8 | TTGGAGAAAAGATGGGCAAG | AGGCTTTTGTTTCGCTTCAA | 258 |
| chr5:64059763-64061788 | p5 | (GATAT)5 | TCGTGTGAAATGTTGGGAAA | GTCAAACGGAGCAATAGGGA | 122 |
| chr5:64073814-64075829 | p3 | (ATA)5 | CTGGGTACTCCGAAGGATCA | GGTCGTACAGAAATGGCGTT | 227 |
| chr5:64107041-64109059 | p2 | (CA)9 | GCCCATTGTTTTAAAAGGCA | GTTGGGCGTTAGGTGTGTTT | 258 |
| chr5:64136319-64138337 | p2 | (CA)9 | GCCCATTGTTTTAAAAGGCA | GTTGGGCGTTAGGTGTGTTT | 258 |
| chr5:64287721-64289736 | p5 | (ATTTC)3 | CGATACTCCTTGAGCTTGGC | TCAGACCATACGATGACGGA | 266 |
| chr5:64287721-64289736 | p2 | (AT)10 | CGATACTCCTTGAGCTTGGC | TCAGACCATACGATGACGGA | 266 |
| chr5:64357871-64359915 | p5 | (ATTGC)3 | AATCCTGGTTGTGCGGTATC | AATCCCAAGTGTGCCAACTC | 271 |
| chr5:64357871-64359915 | p2 | (AT)22 | AATCCTGGTTGTGCGGTATC | AATCCCAAGTGTGCCAACTC | 271 |
| chr5:64385267-64387317 | p2 | (TA)32 | CATCCAGGACCTTTTGCATT | TCCAAAATTGGGCTTGAGTC | 110 |
| chr5:64404431-64406446 | p5 | (AAAGG)3 | TGGGGATTTGGGTCTTCATA | TCCTTTCCTTTGAGCCTTGA | 141 |
| chr5:64404431-64406446 | p5 | (ATTGC)3 | TGGGGATTTGGGTCTTCATA | TCCTTTCCTTTGAGCCTTGA | 141 |
| chr5:64467765-64469780 | p5 | (ATTGC)3 | CACCTCTAGCTGGCTGGAAC | TCCGGGTCTTCAGTTTATGC | 149 |
| chr5:64474275-64476343 | p2 | (AT)34 | GCCACACAGACTGGAAAACA | ATCGATGCTTGGAAGGTTTG | 173 |
| chr5:64489532-64491556 | p3 | (TTG)8 | ACGAATAGTGAATGCCCCAG | GAGATGCCCAGTTTTTCCAA | 213 |
| chr5:64560739-64562757 | p3 | (ACA)6 | TAAGGACGGGATAGGAACCC | GTCCAGATCCATAGGAGCCA | 212 |
| chr5:64602034-64604064 | p5 | (TCATA)6 | CCTTAGACCGACCTGCTCTG | GGAGTCAGTCTGTTAGCGGC | 180 |
| chr5:64602034-64604064 | p6 | (TACATA)5 | CCTTAGACCGACCTGCTCTG | GGAGTCAGTCTGTTAGCGGC | 180 |
| chr5:64602034-64604064 | p5 | (ATTCA)3 | CCTTAGACCGACCTGCTCTG | GGAGTCAGTCTGTTAGCGGC | 180 |
| chr5:64614251-64616297 | p2 | (TA)23 | ATGTCCGTTTTCTTTGGACG | CACATCCGGAACAAAGGAGT | 110 |
| chr5:64614251-64616297 | p2 | (TA)23 | ATGTCCGTTTTCTTTGGACG | CACATCCGGAACAAAGGAGT | 110 |
| chr5:64744842-64746860 | p2 | (TA)9 | ATTCCATTTATTCCGGAGGG | CCCACACAAACGTATCAACG | 274 |

| Marker ID | SSR type | SSR motif | FORWARD PRIMER (5'-3') | REVERSE PRIMER (5'-3') | size (bp) |
|---|---|---|---|---|---|
| chr5:64977420-64979444 | p4 | (CATA)6 | TGCCTAAACGGAAATTCAGG | GCCAAAATTCCTTCCAAACA | 218 |
| chr5:64989564-64991579 | p3 | (GTT)5 | CTTGAGCCCAATCCAAGTGT | TATCACATACAGGCCCACGA | 135 |
| chr5:64989564-64991579 | p2 | (TA)6 | CTTGAGCCCAATCCAAGTGT | TATCACATACAGGCCCACGA | 135 |
| chr5:64994092-64996106 | p2 | (GA)7 | GCGACGGGAGTTAAACACAT | GGTAATCGCATTCGTCCTGT | 262 |
| chr5:65046067-65048085 | p6 | (TATGAC)3 | TGTATGTGGCTGGTTTGCAT | CATGCCAAAAGAAAGAAGGG | 146 |
| chr5:65073497-65075515 | p6 | (TATGAC)3 | TGTATGTGGCTGGTTTGCAT | CATGCCAAAAGAAAGAAGGG | 146 |
| chr5:65159897-65161912 | p3 | (ATC)5 | CCGTCGTGCCCAAAATATAC | ATGCCCACAAGTAGACAGGG | 243 |
| chr5:65162309-65164345 | p2 | (AT)18 | CGTTGTGGCTGATGCTCTTA | AGGCTTGCTAACTGCTGGAG | 192 |
| chr5:65300186-65302201 | p2 | (AT)24 | ATCATGGCTAGGATTTTGCG | TCGATCCCCAACAGTACCTC | 196 |
| chr5:65300186-65302201 | p2 | (AT)6 | ATCATGGCTAGGATTTTGCG | TCGATCCCCAACAGTACCTC | 196 |
| chr5:65300186-65302201 | p5 | (CATTG)3 | ATCATGGCTAGGATTTTGCG | TCGATCCCCAACAGTACCTC | 196 |
| chr5:65414489-65416504 | p3 | (TCT)5 | TAACCCCACTGGGATCCATA | TCGAACGTGAATTACCCCAT | 271 |
| chr5:65426278-65428292 | p2 | (CA)7 | CCCCAAGGTTCAAACTCAAA | GAGTTTGGACGAGCGTAAGG | 217 |
| chr5:65450094-65452109 | p3 | (TCT)5 | TAACCCCACTGGGATCCATA | TCGAACGTGAATTACCCCAT | 271 |
| chr5:65461883-65463897 | p2 | (CA)7 | CCCCAAGGTTCAAACTCAAA | GAGTTTGGACGAGCGTAAGG | 217 |
| chr5:65510919-65512945 | p2 | (AT)13 | CCGGAGTCTATAGGATGCCA | CAGCAAGTACAAAGGCCCAT | 146 |
| chr5:65522189-65524203 | p2 | (TA)7 | ATTGTAGTGTGTGGTCGGCA | TGGCGATCATCTTCCTTACC | 118 |
| chr5:65534129-65536147 | p6 | (TATATG)3 | GTTGATGATTTGGCCGAGTT | TACCCCAAGATGGGTGTGAT | 274 |
| chr5:65534248-65536260 | p6 | (TATATG)3 | GTTGATGATTTGGCCGAGTT | TACCCCAAGATGGGTGTGAT | 155 |
| chr5:65552973-65554985 | p2 | (AT)6 | TTATTGACCCGAAGTAGCCG | AAGGCGGGTAGAAAGAGAGC | 292 |
| chr5:65621663-65623705 | p2 | (TA)8 | GATGGTGGCTAGGATGGCTA | TCTAGTGAGGGGCTCGGTAA | 244 |
| chr5:65621663-65623705 | p2 | (TA)21 | GATGGTGGCTAGGATGGCTA | TCTAGTGAGGGGCTCGGTAA | 244 |
| chr5:65700696-65702710 | p2 | (AT)7 | GGATCCTCATCCAGCTTTGA | GTTGCTTCGGGTCAACAAAT | 265 |
| chr5:65728399-65730413 | p2 | (AG)7 | TGAAATAGGGGAAATGCTGC | AATGAAAGAAGGGTTCGCCT | 102 |
| chr5:65728399-65730413 | p3 | (TTG)6 | TGAAATAGGGGAAATGCTGC | AATGAAAGAAGGGTTCGCCT | 102 |
| chr5:65808042-65810062 | p2 | (AG)10 | AGGTTTCAGCAAAAAGGCAA | CAAAACTAGGGTTTGGGGGT | 109 |
| chr5:65940512-65942527 | p3 | (GTT)5 | TTCGGGTGATCTAACTTCGG | TACGGCCTATGGTTGGAAAG | 223 |
| chr5:66028065-66030119 | p2 | (TC)6 | AGATGAATGGCCTCGATCAC | ACCGCCAATGGATGTAGAAG | 257 |
| chr5:66039196-66041208 | p2 | (TA)6 | CAGGATATGGCCCTGGACTA | TTCAGGAGCATACATCTGCG | 267 |
| chr5:66049157-66051189 | p2 | (AT)16 | GGGCGGTTTCTATTGCATAA | CATAGCTGGGCCTTCAAAAC | 135 |
| chr5:66107498-66109564 | p2 | (TA)33 | GGCGGCATATGATGGTTACT | ACATCCAAAGAAAACGGACG | 205 |
| chr5:66156149-66158161 | p2 | (TA)6 | CCTAGGACCGAACTGCTCTG | TGGGGTTTTGGCATGTTAAT | 155 |
| chr5:66172505-66174537 | p2 | (TA)8 | TACCACTTTAGCCACGACCC | TGTGTATTCCGCAAGACTGC | 125 |
| chr5:66206761-66208776 | p5 | (AATGC)3 | GGACTGTCACGACCCATTTT | TCTTTTCCGCTTGCTTGATT | 296 |
| chr5:66242425-66244440 | p3 | (ATG)5 | GGTTGGTGCAAACCAACTCT | TCAATATCCGCCTTGAATCC | 218 |
| chr5:66242425-66244440 | p2 | (TA)9 | GGTTGGTGCAAACCAACTCT | TCAATATCCGCCTTGAATCC | 218 |

| Marker ID | SSR type | SSR motif | FORWARD PRIMER (5'-3') | REVERSE PRIMER (5'-3') | size (bp) |
|---|---|---|---|---|---|
| chr5:66242425-66244440 | p2 | (TA)7 | GGTTGGTGCAAACCAACTCT | TCAATATCCGCCTTGAATCC | 218 |
| chr5:66255785-66257800 | p5 | (TGATA)3 | GTTTCATCGATTTCGCACCT | ATAGAGTTGTCCCCGAGGCT | 125 |
| chr5:66283757-66285772 | p5 | (ATCAT)3 | GGAATTGCACTGACGGATTT | CTAGTTGGGTCGTGACAGCA | 240 |
| chr5:66353883-66355895 | p2 | (AT)6 | TTAGCCTCGTCGGCTTATGT | GGGTCGTGACAGTAGGCATT | 189 |
| chr5:66431545-66433577 | p2 | (AT)16 | CCTGTAAGGCATCTGTGGGT | TTACCATGTCACCTCGGGAT | 139 |
| chr5:66467084-66469099 | p2 | (AT)7 | CGGGGTCTTAACAACTTCCA | TTTGTCGGACTCCAGTTTCC | 229 |
| chr5:66467084-66469099 | p5 | (TATCA)3 | CGGGGTCTTAACAACTTCCA | TTTGTCGGACTCCAGTTTCC | 229 |
| chr5:66533692-66535708 | p2 | (AT)8 | CAACCCTGTAGGCCATGACT | TGTGTAGTGGCAGCCTTGTC | 223 |
| chr5:66621369-66623381 | p2 | (TA)6 | CAATTTCCGTTTCCTCCAAA | GTTGACATTTTGGCCGAGTT | 199 |
| chr5:66662135-66664150 | p3 | (GAA)5 | CAAGGTGATGTACTGGGCCT | ATGAACTTCCAGGCCTTCCT | 151 |
| chr5:66666383-66668401 | p2 | (CT)9 | CTGTCCACGGTTTTGGACTT | GAAATGGTGGAAGGGTGTTG | 268 |
| chr5:66750532-66752548 | p4 | (AAAG)4 | GTGTCCCGACCCTCTAGTGA | TTCTCGTGCGTACTCCACTG | 197 |
| chr5:66771570-66773584 | p2 | (TA)7 | TTATTTGCTGAGGTGGGACC | AGTCAGATCACCCGAAAACG | 182 |
| chr5:66881973-66883995 | p4 | (CATA)4 | CGGTAGTGCTCCAGGAAGAG | GCCCGCAGGTGTAGATAGAA | 143 |
| chr5:66881973-66883995 | p2 | (AT)11 | CGGTAGTGCTCCAGGAAGAG | GCCCGCAGGTGTAGATAGAA | 143 |
| chr5:66896763-66898801 | p2 | (AT)19 | GAATGAAGGCATCGACGAAT | GTGACCGAGGTCATCCACTT | 145 |
| chr5:66960275-66962293 | p6 | (CAGCTC)3 | CCATTATTCCGGAGGTGCTA | AGCCCTTCGGCTATCAAAAT | 265 |
| chr5:67042764-67044779 | p3 | (TGT)5 | AAGAAAGGCCAGGGGTAAGA | ACATGGCCCCTTTAATTTCC | 216 |
| chr5:67051630-67053650 | p2 | (AT)10 | CGGGGTAAGCCTGTAATGAA | GGCAGGATTACGCTACCAAA | 175 |
| chr5:67075109-67077127 | p6 | (TTCATA)3 | ATGGCTCTATCGCACGATCT | TGTGTCTACGAGCCGTGTTC | 100 |
| chr5:67138297-67140309 | p2 | (AG)6 | AAGCTGCGGACATTTTAGGA | AGCTCTCTCACGCTCCTCAG | 140 |
| chr5:67163546-67165558 | p2 | (TG)6 | AAAGGTGCCTACGGGCTATT | CTCATACCTGGCCCTACCAA | 224 |
| chr5:67170159-67172171 | p2 | (TG)6 | AAAGGTGCCTACGGGCTATT | CTCATACCTGGCCCTACCAA | 224 |
| chr5:67170159-67172171 | p5 | (TCATT)3 | AAAGGTGCCTACGGGCTATT | CTCATACCTGGCCCTACCAA | 224 |
| chr5:67179543-67181558 | p5 | (CAAAA)3 | TGCCTAACCCGAATCATCTC | CATTCAGGCTCATGCTCAGA | 217 |
| chr5:67179543-67181558 | p2 | (GT)6 | TGCCTAACCCGAATCATCTC | CATTCAGGCTCATGCTCAGA | 217 |
| chr5:67204410-67206468 | p2 | (AT)9 | ACCTATCTCGCGAGGAGTCA | TGATATGTTGTGGCAGCGTT | 228 |
| chr5:67204410-67206468 | p2 | (AT)29 | ACCTATCTCGCGAGGAGTCA | TGATATGTTGTGGCAGCGTT | 228 |
| chr5:67225861-67227901 | p2 | (TA)21 | GTCACCGACCTTATGCCACT | ATCATCATTAGCCAGCGTCC | 181 |
| chr5:67225861-67227901 | p2 | (AT)20 | GTCACCGACCTTATGCCACT | ATCATCATTAGCCAGCGTCC | 181 |
| chr5:67233151-67235193 | p2 | (AT)21 | GGTGCATTTTCCTTCCTCAA | CGAGAATTTTTCGGGTCAAA | 149 |
| chr5:67457391-67459406 | p5 | (ACCGA)3 | GACCAGTAGGACCTCCCCTC | CCCTGCCATACCCACATATC | 236 |
| chr5:67500971-67502991 | p4 | (AAAT)5 | CGACCACCCTTTGTGTCTTT | TACCGTGGAAAACTCCTTGC | 299 |
| chr5:67523110-67525126 | p4 | (TTTA)4 | TTGGCCGAACGTAGATAACC | GCAAATTGGTCTCCGTCTGT | 139 |
| chr5:67568600-67570618 | p6 | (CATATA)3 | ATGCCATTACATAGAGGGCG | GATTGTTTTGTGGGCTGCTT | 116 |
| chr5:67613328-67615440 | p2 | (AT)56 | GACTCTGGGGTGACCTTGAA | CGATAACAGGGGTTGGAGAA | 135 |

| Marker ID | SSR type | SSR motif | FORWARD PRIMER (5'-3') | REVERSE PRIMER (5'-3') | size (bp) |
|---|---|---|---|---|---|
| chr5:67620364-67622386 | p2 | (AT)11 | CCGTAGATCGATGGCTGAAT | AGGGCAAGACTACGACGAAA | 252 |
| chr5:67620364-67622386 | p2 | (CA)6 | CCGTAGATCGATGGCTGAAT | AGGGCAAGACTACGACGAAA | 252 |
| chr5:67642117-67644129 | p2 | (AT)6 | AGCAACGACTTTACCGTGCT | CTGGGATTGGCAGGGTACTA | 286 |
| chr5:67779011-67781023 | p2 | (AG)6 | TTGAGCCGTTGCACAATAAG | TTTTGGAGGAAGTTGGGATG | 213 |
| chr5:67779011-67781023 | p2 | (AT)6 | TTGAGCCGTTGCACAATAAG | TTTTGGAGGAAGTTGGGATG | 213 |
| chr5:67829765-67831779 | p4 | (ACAT)4 | AAATGAGAACAGCCATTCCG | TGCGTACACGTTTTTGCTTC | 299 |
| chr5:67851057-67853069 | p2 | (AT)6 | TCCGACAAATCCCAGAAAAC | TAATCGGGAATTCTCATCGC | 186 |
| chr5:67851057-67853069 | p2 | (AT)9 | TCCGACAAATCCCAGAAAAC | TAATCGGGAATTCTCATCGC | 186 |
| chr5:67868661-67870679 | p6 | (GTATAT)3 | GAATTCCCGGATTGTTGTTG | GGGATCATGCCTTGAAAGAA | 195 |
| chr5:67875411-67877431 | p5 | (TTTTC)4 | TATTCGTGAATCCTTTCGCC | GCAATTGGAAATTCGGGATA | 155 |
| chr5:67875411-67877431 | p2 | (CT)6 | TATTCGTGAATCCTTTCGCC | GCAATTGGAAATTCGGGATA | 155 |
| chr5:67923333-67925348 | p3 | (GAA)5 | CAAGGTGATGTACTGGGCCT | TGTCCCGGTGGTTAATGAAT | 154 |
| chr5:67970246-67972296 | p2 | (AT)25 | CACGACTCAGAGACATCCGA | GTTATGGCGACCTTGTCGAT | 249 |
| chr5:68014226-68016260 | p2 | (TA)29 | ACGACCTTATGTCACCTCGG | ATGTATAACCTGCCCGTCCA | 130 |
| chr5:68014226-68016260 | p2 | (TA)17 | ACGACCTTATGTCACCTCGG | ATGTATAACCTGCCCGTCCA | 130 |
| chr5:68088435-68090451 | p4 | (TTTC)4 | GTGTATCCGGAACGAGTGGT | AATCAGCGGAAAACCACAAC | 116 |
| chr5:68151776-68153791 | p5 | (ATTTC)3 | GCCCAGTGAGGCATTGTTAT | GGCCATCAAATGTCATTCCT | 238 |
| chr5:68180754-68182784 | p2 | (AT)12 | CTACCGAGCACCCCTTATCA | GTGGGTCATGGGTGGTTATC | 250 |
| chr5:68180754-68182784 | p2 | (AT)15 | CTACCGAGCACCCCTTATCA | GTGGGTCATGGGTGGTTATC | 250 |
| chr5:68180754-68182784 | p4 | (AAAG)4 | CTACCGAGCACCCCTTATCA | GTGGGTCATGGGTGGTTATC | 250 |
| chr5:68482461-68484473 | p2 | (AT)7 | TACCAAAAGTATGGGCTCCG | GTTTTCTGCATTCCATGCCT | 266 |
| chr5:68482461-68484473 | p2 | (AT)6 | TACCAAAAGTATGGGCTCCG | GTTTTCTGCATTCCATGCCT | 266 |
| chr5:68505410-68507424 | p2 | (AT)7 | AGGCCATCTGCTCACATACC | TGCAGTCAACGAGACCTTTG | 182 |
| chr5:68530204-68532220 | p2 | (TA)8 | AACTGAGGCCCAACCTTCTT | GGGCCTACCTAAGCCCACTA | 200 |
| chr5:68597204-68599236 | p2 | (AT)38 | AAATATGCAAGCCGACAAGG | AGTTTGGTAATCCTCCGGCT | 100 |
| chr5:68597204-68599236 | p2 | (AT)16 | AAATATGCAAGCCGACAAGG | AGTTTGGTAATCCTCCGGCT | 100 |
| chr5:68597204-68599236 | p3 | (CAA)6 | AAATATGCAAGCCGACAAGG | AGTTTGGTAATCCTCCGGCT | 100 |
| chr5:68636694-68638709 | p3 | (TCT)5 | TTGCCAATACAGGCACAAAA | GTTCGCTACCTGGGGTGTTA | 217 |
| chr5:68699605-68701617 | p2 | (AT)6 | CAATGGTGCCTCTTACGGAT | CGTGCAGGTTTTGGGATAGT | 142 |
| chr5:68700297-68702311 | p2 | (AT)6 | ACTTTTGTCACGACCCAACC | GTCGGCTTGCGTATCATTTT | 167 |
| chr5:68788204-68790224 | p2 | (TA)10 | GTGATTTGGCCGAGTTGAAT | TACATACCTTGCCCGCTTTC | 241 |
| chr5:68788204-68790224 | p2 | (AT)6 | GTGATTTGGCCGAGTTGAAT | TACATACCTTGCCCGCTTTC | 241 |
| chr5:68812464-68814492 | p4 | (AATA)7 | AGCGCCTTCGAATACTCTCA | TTTTGTTTCGACGTTAGGGG | 275 |
| chr5:68821647-68823662 | p5 | (CCTTT)3 | ACCTGCAAAGTCCGTGTACC | TTGGACGAGAGTGACGTGAG | 269 |
| chr5:68922302-68924317 | p5 | (GTTTT)3 | GCCAATCTTGGCTAATGGAA | TATCTGCTCCACATGCTTGC | 277 |
| chr5:68922302-68924317 | p6 | (ATATAC)3 | GCCAATCTTGGCTAATGGAA | TATCTGCTCCACATGCTTGC | 277 |

| Marker ID | SSR type | SSR motif | FORWARD PRIMER (5'-3') | REVERSE PRIMER (5'-3') | size (bp) |
|---|---|---|---|---|---|
| chr5:68949510-68951528 | p2 | (TA)11 | TTGCCCTTAGCCTTTCTCAA | GGCCCTATGTTATAGCCCGT | 291 |
| chr5:68949510-68951528 | p2 | (AT)9 | TTGCCCTTAGCCTTTCTCAA | GGCCCTATGTTATAGCCCGT | 291 |
| chr5:68994315-68996333 | p3 | (TTG)6 | GTGGATTCAAACGGAAGCAT | CAAAAGTGCTCTCTCTGCCC | 142 |
| chr5:69008744-69010776 | p2 | (TA)16 | TTAAGGGCGACAGGTCAATC | TGCTGCAAACAAAAACCTTG | 247 |
| chr5:69055680-69057737 | p2 | (AT)13 | TCCTTTTGCCTCTGATTGCT | GCAACCCTGCACATTACCTT | 199 |
| chr5:69055680-69057737 | p3 | (TAT)19 | TCCTTTTGCCTCTGATTGCT | GCAACCCTGCACATTACCTT | 199 |
| chr5:69275762-69277784 | p2 | (AT)11 | GGAGGTGGATCAAACGAAAA | CTGGATTTTTCTTGGGCAAA | 176 |
| chr5:69275762-69277784 | p6 | (TATGTA)3 | GGAGGTGGATCAAACGAAAA | CTGGATTTTTCTTGGGCAAA | 176 |
| chr5:69440057-69442069 | p2 | (TA)6 | ATAAGCCGACGAGGCTAACA | TAGCGGAGCTCCATTTCATT | 243 |
| chr5:69460610-69462664 | p2 | (TA)9 | ACTTTTGTCACGACCCAACC | TTAGCCTCGTCGGCTTATGT | 293 |
| chr5:69506242-69508274 | p2 | (TA)25 | TACATAGGGGAAATGCTGCC | CGTAAATCCGTATCCCATCG | 107 |
| chr5:69589728-69591740 | p2 | (GA)6 | CACCTCTGGCCAAGATTCAT | AACGTGTGGGGAGAGAGAGA | 111 |
| chr5:69695097-69697112 | p3 | (ATG)5 | TTCATCCAAACACCCCCTAA | CCTAGAGCCGCAATGTCTTC | 272 |
| chr5:69739781-69741806 | p3 | (AAC)9 | CACTCAGTCCTCCTTCCAGC | GTCTACGAGCCGTGTCCAAT | 137 |
| chr5:69740674-69742701 | p3 | (AAC)9 | AATCATCTCAAGCTGCCCAC | ACCGACCTTATGTCACTCCG | 233 |
| chr5:69775688-69777702 | p2 | (AT)6 | GGGATGTCACAAAAATTGGG | CTAACTTGCTGCCTGAAGGG | 109 |
| chr5:69775688-69777702 | p2 | (TA)22 | GGGATGTCACAAAAATTGGG | CTAACTTGCTGCCTGAAGGG | 109 |
| chr5:69775688-69777702 | p2 | (AT)7 | GGGATGTCACAAAAATTGGG | CTAACTTGCTGCCTGAAGGG | 109 |
| chr5:69775688-69777702 | p2 | (TA)8 | GGGATGTCACAAAAATTGGG | CTAACTTGCTGCCTGAAGGG | 109 |
| chr5:69790088-69792102 | p2 | (AT)7 | TTATGGGTCTTCAAAAGCCG | CAAGAGGAGCTTCGCTCAGT | 102 |
| chr5:69837410-69839432 | p2 | (GA)11 | TCCTTTAACGATCCGTCCTG | GGTAAGGAATGGTGCGAAAA | 229 |
| chr5:69865404-69867416 | p2 | (TC)7 | ATGTGAATGTTGGATCCGGT | TAGGCCTCATTTGTTGGAGG | 176 |
| chr5:69865404-69867416 | p2 | (TA)6 | ATGTGAATGTTGGATCCGGT | TAGGCCTCATTTGTTGGAGG | 176 |
| chr5:69871111-69873123 | p5 | (ATTCA)3 | CCCCCATTAACCCACTCTTT | GAGTCCAAAGCTCCACCAAA | 206 |
| chr5:69871111-69873123 | p2 | (CT)6 | CCCCCATTAACCCACTCTTT | GAGTCCAAAGCTCCACCAAA | 206 |
| chr5:69953049-69955064 | p3 | (ATT)16 | CGTCATAGGCCAACCAAGAT | GCATTGACATGGTAAAGGGG | 173 |
| chr5:69953049-69955064 | p5 | (ACCCG)3 | CGTCATAGGCCAACCAAGAT | GCATTGACATGGTAAAGGGG | 173 |
| chr5:70034828-70036843 | p5 | (GTTTA)3 | CCTCGTGGTCGTCTTTAAGC | AAGGCTTCAACTTTCTGCCA | 219 |
| chr5:70046495-70048510 | p5 | (GTTTA)3 | CCTCGTGGTCGTCTTTAAGC | AAGGCTTCAACTTTCTGCCA | 219 |
| chr5:70073910-70075942 | p2 | (TA)16 | AGGATGTCGATGATTGCTCC | TCATCTCATGCCACCCACTA | 228 |
| chr5:70118516-70120530 | p2 | (AT)7 | TCCAACTCAAATCCCGTTTC | GATGCTGCCCAAATTTCTGT | 124 |
| chr5:70186515-70188531 | p2 | (AT)8 | AGCTGGACACTCGTATCCGT | GTCGGCTTGCGTACCATATT | 189 |
| chr5:70186515-70188531 | p3 | (ACA)5 | AGCTGGACACTCGTATCCGT | GTCGGCTTGCGTACCATATT | 189 |
| chr5:70186549-70188561 | p2 | (AT)8 | AGCTGGACACTCGTATCCGT | GTCGGCTTGCGTACCATATT | 155 |
| chr5:70186549-70188561 | p3 | (ACA)5 | AGCTGGACACTCGTATCCGT | GTCGGCTTGCGTACCATATT | 155 |
| chr5:70194126-70196138 | p2 | (TA)23 | ACACTCCGAGGATGAACTGC | GTCGGCTTGCGTACCATATT | 110 |

| Marker ID | SSR type | SSR motif | FORWARD PRIMER (5'-3') | REVERSE PRIMER (5'-3') | size (bp) |
|---|---|---|---|---|---|
| chr5:70194126-70196138 | p2 | (AT)6 | ACACTCCGAGGATGAACTGC | GTCGGCTTGCGTACCATATT | 110 |
| chr5:70194126-70196138 | p3 | (ACA)6 | ACACTCCGAGGATGAACTGC | GTCGGCTTGCGTACCATATT | 110 |
| chr5:70330519-70332537 | p6 | (CATATA)3 | ACTGAGTGGGCCGTATGAAC | TACATATTCGAGCACGGCAG | 294 |
| chr5:70402278-70404299 | p3 | (TTC)7 | TCGGACATTTTCCTACAGCC | TTACGCGGCACATCAACTAC | 157 |
| chr5:70478608-70480620 | p2 | (AT)6 | ATCTCGCTAGGGTGTGATGG | CAGAGATTCCCAACCCTTGA | 267 |
| chr5:70509478-70511493 | p5 | (AAACT)3 | CGCGGGCCAAAGTTATACTA | TGGTCGAAGACGTGAGACTG | 162 |
| chr5:70537304-70539322 | p6 | (TATATG)3 | TTGGCCGAGTTAAATTCTCG | GGGATCATGCCTTGAAAGAA | 117 |
| chr5:70537304-70539322 | p2 | (AT)14 | TTGGCCGAGTTAAATTCTCG | GGGATCATGCCTTGAAAGAA | 117 |
| chr5:70557820-70559838 | p3 | (CAA)6 | CGTACCCGACCCTCTAGTCA | GTTATGGCCTCATTTCACCG | 157 |
| chr5:70629928-70631956 | p2 | (TA)14 | CTTTGATCCCGATTCCGTTA | GCATTGTTCCGGTCGTACTT | 168 |
| chr5:70638311-70640323 | p2 | (CA)6 | GGTGGGTTCGTTGTTAATGG | CATCAGGACCCACTCCATCT | 248 |
| chr5:70639523-70641567 | p2 | (TA)22 | AACTGTTGTTGGGACAAGGC | TTGGTGAGCCCGACTTACTC | 184 |
| chr5:70653581-70655596 | p5 | (GTTGG)3 | GGCTCCTGCAGAATTGAAAG | CTGATGATAGCCTGAACGCA | 124 |
| chr5:70748934-70750949 | p2 | (CA)6 | AACGCATCTACCCACACACA | TCGGAACCATATCTCTTGGG | 282 |
| chr5:70748946-70750958 | p2 | (CA)6 | AACGCATCTACCCACACACA | TCGGAACCATATCTCTTGGG | 270 |
| chr5:70753557-70755577 | p5 | (ATTCA)4 | ACATATGTCCCTTCCCCTCC | ACCGATCTTACGTCACTCCG | 100 |
| chr5:70878674-70880692 | p5 | (TGAAT)3 | TTAGATTCTCGGGGATGGTG | AACATACCTGCTCCACGTCC | 179 |
| chr5:70878674-70880692 | p6 | (TATATG)3 | TTAGATTCTCGGGGATGGTG | AACATACCTGCTCCACGTCC | 179 |
| chr5:70878674-70880692 | p2 | (AT)9 | TTAGATTCTCGGGGATGGTG | AACATACCTGCTCCACGTCC | 179 |
| chr5:70908369-70910384 | p5 | (ACAAA)3 | CTTCCTCCTTAAGCGCCTTT | GTTTCCTCGGACATGGAGAA | 120 |
| chr5:70937611-70939626 | p5 | (CCTTT)3 | CAGGAGGGCAGATGATGATT | AGTGAGGGACTCGGTGAATG | 143 |
| chr5:70999914-71001932 | p5 | (TGAAT)3 | TTAGATTCTCGGGGATGGTG | AACATACCTGCTCCACGTCC | 179 |
| chr5:70999914-71001932 | p6 | (TATATG)3 | TTAGATTCTCGGGGATGGTG | AACATACCTGCTCCACGTCC | 179 |
| chr5:70999914-71001932 | p2 | (AT)9 | TTAGATTCTCGGGGATGGTG | AACATACCTGCTCCACGTCC | 179 |
| chr5:71086248-71088272 | p6 | (CATATA)4 | ATAATGGTACGTCGGCTTGC | TTTCATGCCACGAAGGTGTA | 286 |
| chr5:71086248-71088272 | p6 | (CATATA)4 | ATAATGGTACGTCGGCTTGC | TTTCATGCCACGAAGGTGTA | 286 |
| chr5:71086248-71088272 | p2 | (TG)7 | ATAATGGTACGTCGGCTTGC | TTTCATGCCACGAAGGTGTA | 286 |
| chr5:71119998-71122018 | p5 | (TGAAT)3 | ACACATTGACAATCCTGCCA | TTGGTAGCCGCATTCATACA | 217 |
| chr5:71168316-71170338 | p2 | (AT)34 | TTTCGCGATCCTATTGCTTC | GGTGTTTCACATACCCGACC | 203 |
| chr5:71168316-71170338 | p2 | (AT)27 | TTTCGCGATCCTATTGCTTC | GGTGTTTCACATACCCGACC | 203 |
| chr5:71168316-71170338 | p2 | (TA)11 | TTTCGCGATCCTATTGCTTC | GGTGTTTCACATACCCGACC | 203 |
| chr5:71210993-71213013 | p2 | (TA)10 | TTCCACCCTACCAGAGTTGG | GACATTGGGCCTCTTGGTTA | 125 |
| chr5:71336807-71338823 | p4 | (ACAT)4 | TTCCTCTTTACCGGCATCAC | CGGGGGCATTTAGGAATAAT | 209 |
| chr5:71392640-71394656 | p4 | (ACAT)4 | TTCCTCTTTACCGGCATCAC | CGGGGGCATTTAGGAATAAT | 209 |
| chr5:71435942-71437960 | p3 | (ATT)6 | TGGGAACAAATGCAAATTCA | TTCCCGATGAGCCTGTTATC | 196 |
| chr5:71468539-71470557 | p3 | (ATT)6 | TGGGAACAAATGCAAATTCA | TTCCCGATGAGCCTGTTATC | 196 |

| Marker ID | SSR type | SSR motif | FORWARD PRIMER (5'-3') | REVERSE PRIMER (5'-3') | size (bp) |
|---|---|---|---|---|---|
| chr5:71486494-71488514 | p5 | (ATTCA)4 | GATATAACCGGCGACTTGGA | TCTAAGGGGAAATCGGGACT | 130 |
| chr5:71515479-71517494 | p3 | (ACA)5 | TCGGAAACCGTAACACATCA | TGCATATGGTTTCGGCACTA | 101 |
| chr5:71585669-71587684 | p3 | (AGA)5 | ATTCCGTCTTCTGCCCCTAT | TGACACAAAGCCAGAGTTGC | 210 |
| chr5:71774128-71776146 | p2 | (AT)9 | AATGCCCTGCAGTCTCTTGT | CGGTCTGGGTCGTGACTAAT | 147 |
| chr5:71810927-71812949 | p2 | (TC)11 | CCTAAAGCAATTCCAACCCA | GCGGTGCCTATCAACAATTT | 109 |
| chr5:71948556-71950570 | p2 | (TA)7 | AAAGGAAGACCGAGTGGGAT | CAAGGAGCGAAAGAACCAAG | 194 |
| chr5:72030675-72032693 | p3 | (AAC)6 | CCCTTTTCCCATATCCCAGT | AAAGCCACGGCATGATATTC | 149 |
| chr5:72030675-72032693 | p2 | (CT)8 | CCCTTTTCCCATATCCCAGT | AAAGCCACGGCATGATATTC | 149 |
| chr5:72058549-72060561 | p2 | (TG)6 | AAAGGACATGACTTGCACCC | AGATTTTGTGGTTTGCTGGC | 123 |
| chr5:72058549-72060561 | p2 | (AG)11 | AAAGGACATGACTTGCACCC | AGATTTTGTGGTTTGCTGGC | 123 |
| chr5:72173930-72175942 | p2 | (AT)20 | TCGGGAGACCATACTACCCA | ACACATATGTTTGGGCACGA | 209 |
| chr5:72173930-72175942 | p2 | (TA)6 | TCGGGAGACCATACTACCCA | ACACATATGTTTGGGCACGA | 209 |
| chr5:72183905-72185925 | p2 | (TA)10 | CATATGGAAGTTGAGGCCGT | AAATATGCACCCAACAAGCC | 160 |
| chr5:72201565-72203581 | p3 | (TTG)5 | ATGACTAGTGCCCGAGATGG | TGTCATTGTAGCAGCCTTGC | 201 |
| chr5:72201603-72203618 | p3 | (TTG)5 | ATGACTAGTGCCCGAGATGG | TGTCATTGTAGCAGCCTTGC | 163 |
| chr5:72226051-72228065 | p2 | (AT)10 | TACGGAGGATTTCCAAGGTG | TTGGGTCGTGTGTAGCCATA | 292 |
| chr5:72226051-72228065 | p2 | (AT)8 | TACGGAGGATTTCCAAGGTG | TTGGGTCGTGTGTAGCCATA | 292 |
| chr5:72226067-72228079 | p2 | (AT)10 | TACGGAGGATTTCCAAGGTG | TTGGGTCGTGTGTAGCCATA | 276 |
| chr5:72226067-72228079 | p2 | (AT)8 | TACGGAGGATTTCCAAGGTG | TTGGGTCGTGTGTAGCCATA | 276 |
| chr5:72226239-72228259 | p2 | (AT)10 | TACGGAGGATTTCCAAGGTG | TTGGGTCGTGTGTAGCCATA | 104 |
| chr5:72226239-72228259 | p2 | (AT)8 | TACGGAGGATTTCCAAGGTG | TTGGGTCGTGTGTAGCCATA | 104 |
| chr5:72236370-72238382 | p2 | (AT)9 | ATCAAAAATTACGCAAGCCG | TCCGATGTGGTCAGTTGTGT | 287 |
| chr5:72236370-72238382 | p2 | (AT)6 | ATCAAAAATTACGCAAGCCG | TCCGATGTGGTCAGTTGTGT | 287 |
| chr5:72236370-72238382 | p2 | (AT)6 | ATCAAAAATTACGCAAGCCG | TCCGATGTGGTCAGTTGTGT | 287 |
| chr5:72236757-72238769 | p2 | (AT)9 | GCCTCGTCGTACCCTTAACA | AATTTTCCGTACTCACCCCC | 126 |
| chr5:72236757-72238769 | p2 | (AT)6 | GCCTCGTCGTACCCTTAACA | AATTTTCCGTACTCACCCCC | 126 |
| chr5:72236757-72238769 | p2 | (AT)6 | GCCTCGTCGTACCCTTAACA | AATTTTCCGTACTCACCCCC | 126 |
| chr5:72295563-72297583 | p5 | (AATGA)4 | TGAAGGAAGGACGAGCCTTA | CTTCGCTAACGACGACATGA | 192 |
| chr5:72298558-72300573 | p3 | (TTG)5 | GCCCGATCCAATAAAGCATA | AAGCCCAATTTCTTTTGGGT | 107 |
| chr5:72315160-72317178 | p2 | (AT)13 | GCTGCCACTACAGACGAACA | GGTGACCCTTCAGCATAGGA | 115 |
| chr5:72315160-72317178 | p2 | (AT)9 | GCTGCCACTACAGACGAACA | GGTGACCCTTCAGCATAGGA | 115 |
| chr5:72315160-72317178 | p3 | (ACA)8 | GCTGCCACTACAGACGAACA | GGTGACCCTTCAGCATAGGA | 115 |
| chr5:72315590-72317614 | p2 | (AT)13 | TTGGCGCCATAAACACATAA | CGTGCATGCCTCAACATATC | 193 |
| chr5:72315590-72317614 | p2 | (AT)9 | TTGGCGCCATAAACACATAA | CGTGCATGCCTCAACATATC | 193 |
| chr5:72315590-72317614 | p3 | (ACA)8 | TTGGCGCCATAAACACATAA | CGTGCATGCCTCAACATATC | 193 |
| chr5:72425150-72427184 | p2 | (TA)17 | CCCTAGGACGAACTGCTCTG | TTTAGTAGCCCTGACGGTGG | 153 |

| Marker ID | SSR type | SSR motif | FORWARD PRIMER (5'-3') | REVERSE PRIMER (5'-3') | size (bp) |
|---|---|---|---|---|---|
| chr5:72439271-72441285 | p2 | (AT)7 | TGCTAAAGGGTCACCAAACC | GACCTTATCACCGAGTCCCA | 260 |
| chr5:72439271-72441285 | p2 | (TA)7 | TGCTAAAGGGTCACCAAACC | GACCTTATCACCGAGTCCCA | 260 |
| chr5:72439271-72441285 | p3 | (CAA)5 | TGCTAAAGGGTCACCAAACC | GACCTTATCACCGAGTCCCA | 260 |
| chr5:72483593-72485611 | p6 | (TACGGA)3 | CTTTGCACCCAAAAACCCTA | AAGGGACCCAAGCTTCATTT | 214 |
| chr5:72575870-72577908 | p4 | (TGTA)6 | GTGCGGTAAAGAGGCTAACG | AAGAGGGTGGAAGCAAGGAT | 280 |
| chr5:72575906-72577934 | p4 | (TGTA)6 | GTGCGGTAAAGAGGCTAACG | AAGAGGGTGGAAGCAAGGAT | 244 |
| chr5:72586758-72588800 | p2 | (AT)21 | TGATAAAGGTCATGCGGTGA | CCTCCTTTCCGGTCTAATCC | 143 |
| chr5:72591026-72593076 | p4 | (TATG)4 | CTCGGACGAATAAGCGTAGC | TAAGAGGGTGGAAGCAAGGA | 256 |
| chr5:72631826-72633846 | p2 | (AT)10 | TCAGACATTTCAGGTGGCAG | GGGAGGGTTTTGGAGAGTTC | 150 |
| chr5:72684257-72686272 | p5 | (ATTTC)3 | GGCAAAAGGGGGAAACTTAG | CTTGCACATCGCTGAGGTAA | 156 |
| chr5:72748106-72750122 | p2 | (TA)8 | TCAACCATCAAAAGCACAGC | ACAAGTCGTTGTCCTCCTGG | 255 |
| chr5:72755979-72757994 | p3 | (ATG)5 | TAAGGGCGAGGGATTATGTG | TGCTCGATCCCAGAACTTTT | 163 |
| chr5:72755979-72757994 | p2 | (AT)22 | TAAGGGCGAGGGATTATGTG | TGCTCGATCCCAGAACTTTT | 163 |
| chr5:72772244-72774260 | p4 | (TACA)4 | CGGGATTTGGAGTTTGTTGT | ATCCGTGAACTCAAATTGCC | 159 |
| chr5:72868110-72870150 | p2 | (TA)17 | TCCTACTTCGGCTTCCTCAA | ACCTCGGAAAATGTCGTGTC | 266 |
| chr5:72868110-72870150 | p2 | (AT)20 | TCCTACTTCGGCTTCCTCAA | ACCTCGGAAAATGTCGTGTC | 266 |
| chr5:72868191-72870269 | p2 | (TA)17 | TCCTACTTCGGCTTCCTCAA | ACCTCGGAAAATGTCGTGTC | 185 |
| chr5:72868191-72870269 | p2 | (AT)20 | TCCTACTTCGGCTTCCTCAA | ACCTCGGAAAATGTCGTGTC | 185 |
| chr5:73077549-73079565 | p6 | (AACCGG)3 | CCTTCCACCCAAAACATCTC | CGTGAAAGGATTCATGGGTT | 113 |
| chr5:73077549-73079565 | p2 | (TA)14 | CCTTCCACCCAAAACATCTC | CGTGAAAGGATTCATGGGTT | 113 |
| chr5:73077549-73079565 | p2 | (TA)8 | CCTTCCACCCAAAACATCTC | CGTGAAAGGATTCATGGGTT | 113 |
| chr5:73115349-73117361 | p2 | (CA)6 | AAGCATGTATCAACCGGAGC | CTTCGGGCCAAGATGTAAAA | 109 |
| chr5:73234729-73236744 | p5 | (CTAAA)3 | GTTTCGTCGGAAAATCGAAA | CCTCAGTTCCACGGTCACTT | 205 |
| chr5:73234729-73236744 | p2 | (TG)6 | GTTTCGTCGGAAAATCGAAA | CCTCAGTTCCACGGTCACTT | 205 |
| chr5:73235671-73237685 | p5 | (CTAAA)3 | TCCTTGCACTCTTCAACCCT | AACTTGGGATTTTTGGAGCC | 205 |
| chr5:73235671-73237685 | p2 | (TG)6 | TCCTTGCACTCTTCAACCCT | AACTTGGGATTTTTGGAGCC | 205 |
| chr5:73235702-73237728 | p5 | (CTAAA)3 | TCCTTGCACTCTTCAACCCT | AACTTGGGATTTTTGGAGCC | 174 |
| chr5:73235702-73237728 | p2 | (TG)6 | TCCTTGCACTCTTCAACCCT | AACTTGGGATTTTTGGAGCC | 174 |
| chr5:73271675-73273695 | p2 | (TA)19 | ACTAGAGGGCCATGAAGGGT | CGCCATGTTGTTTCACAAAG | 292 |
| chr5:73271675-73273695 | p6 | (GTATCC)4 | ACTAGAGGGCCATGAAGGGT | CGCCATGTTGTTTCACAAAG | 292 |
| chr5:73271740-73273752 | p2 | (TA)19 | ACTAGAGGGCCATGAAGGGT | CGCCATGTTGTTTCACAAAG | 227 |
| chr5:73271740-73273752 | p6 | (GTATCC)4 | ACTAGAGGGCCATGAAGGGT | CGCCATGTTGTTTCACAAAG | 227 |
| chr5:73339536-73341580 | p2 | (TA)22 | GTGATTTGGCCGAGTTGAAT | TTACATACCTTGCCCGCTTC | 232 |
| chr5:73339536-73341580 | p2 | (TA)6 | GTGATTTGGCCGAGTTGAAT | TTACATACCTTGCCCGCTTC | 232 |
| chr5:73339536-73341580 | p2 | (TA)7 | GTGATTTGGCCGAGTTGAAT | TTACATACCTTGCCCGCTTC | 232 |
| chr5:73471106-73473121 | p5 | (GAAAG)3 | CGTGCCTATTTCTGAGGAGC | CCCTATCACCTTGCCCTGTA | 182 |

| Marker ID | SSR type | SSR motif | FORWARD PRIMER (5'-3') | REVERSE PRIMER (5'-3') | size (bp) |
|---|---|---|---|---|---|
| chr5:73480453-73482468 | p5 | (GAAAG)3 | TGGTAAGGAACCGAGGAGTG | CTTTTCGCTCTTTCACGGAC | 276 |
| chr5:73540593-73542608 | p5 | (CATTG)3 | TATTTTTCCCCCGAATTTCC | ATTCACTTTCCCTTTCCGCT | 135 |
| chr5:73580794-73582808 | p2 | (AG)7 | CTCTCTTTGGCCTAAACCCC | TGAGCCCAAAGCCTAAGTGT | 237 |
| chr5:73641396-73643411 | p3 | (ACA)5 | GAGGTGTGGAGAAGGTTGGA | GACGGAGAAGATTGACGCTC | 107 |
| chr5:73641396-73643411 | p2 | (AT)17 | GAGGTGTGGAGAAGGTTGGA | GACGGAGAAGATTGACGCTC | 107 |
| chr5:73650209-73652221 | p2 | (TA)6 | ACTTCACCCCACATCCACTC | ATGGGAGCTTTTTATTGGGG | 152 |
| chr5:73666976-73668996 | p2 | (TA)9 | TTATCCAGGGGTACGACGAG | ATGTCATATGCAAGCCGACA | 125 |
| chr5:73666976-73668996 | p5 | (TATGA)4 | TTATCCAGGGGTACGACGAG | ATGTCATATGCAAGCCGACA | 125 |
| chr5:73761239-73763251 | p2 | (AT)7 | CACGATGTGGGATTTCACTG | CCTTAAGGAAAAGTTCCGGC | 176 |
| chr5:73761239-73763251 | p2 | (TA)6 | CACGATGTGGGATTTCACTG | CCTTAAGGAAAAGTTCCGGC | 176 |
| chr5:73779817-73781829 | p2 | (TC)6 | CACTGGACAACTCTACGCGA | CAACTCCAACGCAAGACAAA | 278 |
| chr5:73779817-73781829 | p2 | (TA)7 | CACTGGACAACTCTACGCGA | CAACTCCAACGCAAGACAAA | 278 |
| chr5:73856569-73858593 | p2 | (TA)12 | GGGCCCTTTACTCTCGAATC | CACACTCGTTGGAGTAGCCA | 249 |
| chr5:73919168-73921188 | p2 | (AT)10 | TCTGGTTGGATCGTGACAAA | CAATTACCTCGTAGCGGCAT | 192 |
| chr5:73929180-73931192 | p2 | (AT)6 | AGAATGTCACACCCCAGTCC | CTCTGCAAGCCGATGTATCA | 116 |
| chr5:73929180-73931192 | p3 | (CAA)5 | AGAATGTCACACCCCAGTCC | CTCTGCAAGCCGATGTATCA | 116 |
| chr5:73971724-73973736 | p2 | (CA)6 | TGCTACCATTAGGAATCCCG | ATTGCCGCATGCTCTTTTAT | 193 |
| chr5:74016405-74018420 | p5 | (GGACT)3 | CCACGTATACTTGCGTGTGC | CATTCGGAGTTGGGCTAAAA | 124 |
| chr5:74016405-74018420 | p3 | (TGT)5 | CCACGTATACTTGCGTGTGC | CATTCGGAGTTGGGCTAAAA | 124 |
| chr5:74032558-74034573 | p2 | (TA)6 | GCAGGAGCACAGGAAGAAAG | ATATACAATGCCGACGAGGC | 140 |
| chr5:74032558-74034573 | p3 | (CAT)5 | GCAGGAGCACAGGAAGAAAG | ATATACAATGCCGACGAGGC | 140 |
| chr5:74062166-74064180 | p2 | (AT)7 | CCACCCGAGCACTCTCTTAG | TTGATGTTGCATGAATGCCT | 129 |
| chr5:74114775-74116805 | p2 | (AT)15 | ACTCGAGCCTGAGTTTGGAA | CCGTATGCCGATAATCCATC | 293 |
| chr5:74114775-74116805 | p2 | (TA)17 | ACTCGAGCCTGAGTTTGGAA | CCGTATGCCGATAATCCATC | 293 |
| chr5:74144366-74146384 | p2 | (CT)9 | CTGTCCACGGTTTTGGACTT | GAAGGGTGTTGTAGCCGTGT | 268 |
| chr5:74229084-74231104 | p2 | (AC)10 | CTTTCACCCGAAAAACTCCA | AGCCCTAGCTCCTATGCCTC | 112 |
| chr5:74246788-74248803 | p5 | (TTCAT)3 | TTGTTGTACGCGAACTCTGC | AATGTACGATTGCATGGGGT | 161 |
| chr5:74299669-74301684 | p3 | (TGA)5 | TTTGTGCCTACTTCTGCACG | AGAAGCCGACAAGGCTATCA | 202 |
| chr5:74308052-74310064 | p2 | (GT)6 | ACTTTATCCCACCGAGTCCC | ACATGAGCATGGTAATGGCA | 289 |
| chr5:74362643-74364661 | p6 | (CATATA)3 | CAGGTAGTGCAGGGATTGGT | CCGCTTAAGATGACGGTTGT | 102 |
| chr5:74362643-74364661 | p2 | (AT)9 | CAGGTAGTGCAGGGATTGGT | CCGCTTAAGATGACGGTTGT | 102 |
| chr5:74397001-74399016 | p5 | (GAAAT)3 | TTGTTATGATGGCCCCAAAT | AGTGTTGTTGTCACGACCCA | 156 |
| chr5:74407968-74409983 | p5 | (GAAAT)3 | TTGTTATGATGGCCCCAAAT | AGTGTTGTTGTCACGACCCA | 156 |
| chr5:74438969-74440985 | p2 | (AT)8 | ATATCATTTTGTCGGCCTCG | TGTCCCATCTTGGTCCTCTC | 283 |
| chr5:74448564-74450576 | p2 | (TG)6 | AAAGGTGCCTTTGGGCTATT | CTGCCAAGGATTCAGGGTTA | 241 |
| chr5:74456380-74458394 | p2 | (TG)7 | AAAGGTGCCTACGGGCTATT | TCATACCTGGCCCTACCAAG | 259 |

| Marker ID | SSR type | SSR motif | FORWARD PRIMER (5'-3') | REVERSE PRIMER (5'-3') | size (bp) |
|---|---|---|---|---|---|
| chr5:74562489-74564503 | p2 | (GT)7 | GCCTAAAATGAAGCTCGTCG | GCGTTTCTGCTCTATCAGCC | 296 |
| chr5:74572058-74574073 | p2 | (AT)8 | TAGAAATGAGGTTGGGGTCG | CGGAGATGTAAGGGCAATGT | 156 |
| chr5:74572058-74574073 | p3 | (TCT)5 | TAGAAATGAGGTTGGGGTCG | CGGAGATGTAAGGGCAATGT | 156 |
| chr5:74572058-74574073 | p2 | (AC)7 | TAGAAATGAGGTTGGGGTCG | CGGAGATGTAAGGGCAATGT | 156 |
| chr5:74572058-74574073 | p3 | (GAA)5 | TAGAAATGAGGTTGGGGTCG | CGGAGATGTAAGGGCAATGT | 156 |
| chr5:74745318-74747330 | p2 | (CT)6 | TTCTAAATTCTGGGCATGGG | AGTGCATCGCTTTGTGAGTG | 265 |
| chr5:74746128-74748140 | p2 | (CT)6 | ATGGAGCCTCATTTGGTGAC | AAACAAGAGGCTACAGGGCA | 144 |
| chr5:74824895-74826915 | p2 | (AT)6 | TGTGGTTGAGATTGGATGGA | GGATCAAATGCCAAAAAGGA | 186 |
| chr5:74824945-74826957 | p2 | (AT)6 | TGTGGTTGAGATTGGATGGA | GGATCAAATGCCAAAAAGGA | 136 |
| chr5:74824963-74826975 | p2 | (AT)6 | TGTGGTTGAGATTGGATGGA | GGATCAAATGCCAAAAAGGA | 118 |
| chr5:74909520-74911538 | p6 | (ATACAT)3 | CGCTGCATCTGTCGAATAAA | GGGCTAGGGGGTAATGATGT | 169 |
| chr5:74909520-74911538 | p2 | (TA)16 | CGCTGCATCTGTCGAATAAA | GGGCTAGGGGGTAATGATGT | 169 |
| chr5:74925508-74927524 | p2 | (TA)8 | TAATGGAATCAGGGGGTTCA | GTGGTGTCTTCCCACGCTAT | 278 |
| chr5:75170149-75172167 | p6 | (TATGGA)3 | GGTAGAGACGTGAGCTTCGG | CACAATCGTGGCGTATCAAG | 272 |
| chr5:75243067-75245083 | p2 | (AT)8 | TTCCGAGGATGAACTGCTCT | TGTATGTGGGTCATGGGTTG | 233 |
| chr5:75297412-75299427 | p5 | (TTTCA)3 | ACTCCACTATGTTCGGCCAC | GGAGGAGTTGTTGGAGCAAG | 235 |
| chr5:75304703-75306715 | p2 | (TA)6 | TCTGCCCAATATCCGTTAGC | GCCTCACATACCTTTTTGCC | 274 |
| chr5:75372596-75374680 | p2 | (AT)42 | TTTATGGTGCAGCTCAAACG | TCTTGAAGGATTGGAGTGCC | 148 |
| chr5:75385187-75387205 | p2 | (AC)9 | GTGCACTCGAAAATCCCAAT | TCGGGGGTATTTGTGTCATT | 299 |
| chr5:75385187-75387205 | p5 | (TCATG)3 | GTGCACTCGAAAATCCCAAT | TCGGGGGTATTTGTGTCATT | 299 |
| chr5:75405262-75407292 | p2 | (AT)15 | TGCGTAATCGGGTTCCTATC | CGGAATTGCGCGTATAAAGT | 153 |
| chr5:75433844-75435904 | p2 | (AT)30 | TCGCTTGCTTGTTCTCCTTT | ATAGTGAACCATAACCGCCG | 187 |
| chr5:75561543-75563577 | p2 | (AT)17 | TCCAATCCATTCTGTCACGA | CGACGTGCCATTATTTTGTG | 296 |
| chr5:75561543-75563577 | p3 | (ATC)5 | TCCAATCCATTCTGTCACGA | CGACGTGCCATTATTTTGTG | 296 |
| chr5:75576466-75578490 | p2 | (TC)12 | TATTCCTTGTTGGCTTTCCG | GAAAGAACGCTGCTCCTACG | 193 |
| chr5:75596248-75598274 | p2 | (AT)13 | ACTTTTCCCTTCATGGGCTT | TTAGGCCTTCCCTCCTCATT | 224 |
| chr5:75623580-75625606 | p2 | (AT)13 | ACTTTTCCCTTCATGGGCTT | TTAGGCCTTCCCTCCTCATT | 224 |
| chr5:75777134-75779149 | p3 | (ATG)5 | CAAATGAAGCTCGGGTTTGT | GGAATGGTTTGGAGTGTTGG | 163 |
| chr5:75881066-75883078 | p2 | (TA)6 | CAATGGGTAAACCTCCAACG | GCTGAAGACTGGGAAGATGC | 138 |
| chr5:75881066-75883078 | p3 | (CAA)5 | CAATGGGTAAACCTCCAACG | GCTGAAGACTGGGAAGATGC | 138 |
| chr5:75982163-75984193 | p2 | (AT)15 | TCCAGGTAACGGAGGAGTTG | TACCTGCAGGCATAGGGAAC | 105 |
| chr5:75994355-75996370 | p5 | (AACAA)3 | CACAAGGATAACCAGCCGAT | AGTGGCTCTTCAGCCAGGTA | 238 |
| chr5:75998122-76000138 | p2 | (AT)8 | TCTCATGGCCCTATCCTTTG | ACTCCTGTCGGCGAGTTCTA | 131 |
| chr5:75998122-76000138 | p2 | (AT)6 | TCTCATGGCCCTATCCTTTG | ACTCCTGTCGGCGAGTTCTA | 131 |
| chr5:75998522-76000534 | p2 | (AT)8 | CTATTGTCACGACCCAACCC | GTTGCGAACCCTTTGTTTGT | 194 |
| chr5:75998522-76000534 | p2 | (AT)6 | CTATTGTCACGACCCAACCC | GTTGCGAACCCTTTGTTTGT | 194 |

| Marker ID | SSR type | SSR motif | FORWARD PRIMER (5'-3') | REVERSE PRIMER (5'-3') | size (bp) |
|---|---|---|---|---|---|
| chr5:76047838-76049850 | p2 | (CT)6 | TGGGTTTCATGGTCTTGTCA | TTGTCGCGTTTTATCTGTGC | 180 |
| chr5:76103408-76105423 | p5 | (ATATA)3 | CCATGACTAGTGCCCGAGAT | CCTCGTCGGCTTGTGTATTT | 103 |
| chr5:76143806-76145840 | p2 | (AT)17 | TACATGAGGCCGAAGATTCC | CTCTAACGCGTAGACGTCCC | 122 |
| chr5:76167897-76169909 | p2 | (AT)6 | TGAACAAAATCTTCCCCGAG | AGGGGAAAGGCTGATGAAAT | 206 |
| chr5:76214383-76216401 | p2 | (TA)9 | CAGCAGCAGTCAGTACGCTC | GCTATGGAAGGGACAGGACA | 117 |
| chr5:76282744-76284762 | p2 | (TA)9 | GCGTGTGTCTGCATTCTTGT | ATTCCAAATCATGCCCACAT | 176 |
| chr5:76314325-76316343 | p2 | (TA)9 | TTGGTCTCACCCACATCAAA | TTGTGGGCCTGTGTTATTGA | 293 |
| chr5:76361954-76363966 | p2 | (AT)6 | AAGTCAAGCTTTGGGGAACA | ATACGAGTGTCCAGTTCGGG | 134 |
| chr5:76418199-76420211 | p2 | (AT)6 | AGATCTTGCGAAGCCAAAGA | TGCCATTCGGAAACATATCA | 201 |
| chr5:76418199-76420211 | p6 | (TATATG)3 | AGATCTTGCGAAGCCAAAGA | TGCCATTCGGAAACATATCA | 201 |
| chr5:76512199-76514228 | p6 | (AAAAGG)3 | CAGAATTCACGTTTCGAGCA | ACGCGTACATAACACCGACA | 240 |
| chr5:76801975-76804027 | p2 | (TA)6 | ATGGCAACCAATTGTGTCAA | ACTACGGTAAAGAGGCGGGT | 243 |
| chr5:76818632-76820648 | p3 | (GTT)5 | ATTGAAAAACGTCGTGAGGG | CCATCCTATATCCTGCCGAA | 256 |
| chr5:76818632-76820648 | p4 | (TGTA)4 | ATTGAAAAACGTCGTGAGGG | CCATCCTATATCCTGCCGAA | 256 |
| chr5:76818632-76820648 | p2 | (AT)20 | ATTGAAAAACGTCGTGAGGG | CCATCCTATATCCTGCCGAA | 256 |
| chr5:76839795-76841807 | p2 | (AT)6 | CCAAATGGCACTCGAAAAAT | TTCGGGTCAATCCGACTTAG | 284 |
| chr5:76839795-76841807 | p2 | (TA)8 | CCAAATGGCACTCGAAAAAT | TTCGGGTCAATCCGACTTAG | 284 |
| chr5:76839795-76841807 | p2 | (TA)10 | CCAAATGGCACTCGAAAAAT | TTCGGGTCAATCCGACTTAG | 284 |
| chr5:76846686-76848698 | p2 | (CA)6 | AATGCCATTCGGAAACAGAC | GAAGCGGGCAAGGTATGTAA | 111 |
| chr5:76855414-76857432 | p6 | (TATATG)3 | CCGAAACTTGTTTCTGAGCC | CACAATGATATAACCGGCCC | 279 |
| chr5:76915194-76917219 | p5 | (CATAT)5 | TACCACAGCTCTATCGCACG | TGGTTTGGGGTTATTGGGTA | 129 |
| chr5:77004851-77006867 | p2 | (AC)8 | TCATCGCGATATGCAATCAT | CCAATCCGGTTGTCATTTTC | 151 |
| chr5:77014767-77016785 | p2 | (AT)9 | GACCCGTCTCTCTGTCTTCG | TGACCCGAGCAACATTCATA | 265 |
| chr5:77028331-77030346 | p3 | (TTG)6 | GAGGAACAGTTGAGGGGTGA | TGATACTACGATCACGGCCA | 187 |
| chr5:77028331-77030346 | p3 | (ATG)5 | GAGGAACAGTTGAGGGGTGA | TGATACTACGATCACGGCCA | 187 |
| chr5:77157227-77159239 | p2 | (AT)6 | AGCGGAATATACATCGCCAG | TGTGGGTAATGGGTCATGTG | 276 |
| chr5:77174211-77176223 | p2 | (AT)6 | AGCGGAATATACATCGCCAG | TGTGGGTAATGGGTCATGTG | 276 |
| chr5:77248582-77250597 | p5 | (CTTTT)3 | TAGCGATATATGGGAACCCG | GCTATGGGAACCGAACTTGA | 102 |
| chr5:77278796-77280811 | p5 | (ATGAT)3 | ACGGGATTTGAATTTGGACA | CTGAATGAAAAAGGGGCAAA | 208 |
| chr5:77278796-77280811 | p5 | (TATTT)3 | ACGGGATTTGAATTTGGACA | CTGAATGAAAAAGGGGCAAA | 208 |
| chr5:77369835-77371850 | p5 | (AAACC)3 | CAGACGATTCGCAGCAGTAG | CGAAAACCCGAGAAAACAGA | 181 |
| chr5:77372184-77374199 | p3 | (ACA)5 | ATTTGCAATCATCGTCGTCA | GGCCGTAGGATTTGGTCATA | 110 |
| chr5:77879810-77881824 | p2 | (TA)7 | GAAGCGGGCAAGGTATGTAA | CGAACGTGCTAGCATTTGAA | 142 |
| chr5:77896399-77898414 | p5 | (TTCGG)3 | GTGTTCTATTCCGCCTGCAT | CCGAACCGAAATTTGAAAAA | 100 |
| chr5:77920502-77922522 | p2 | (CT)9 | CCGTTAGTCCGTTTGATCGT | TGATATGAGGAAGCCAACCC | 138 |
| chr5:77920527-77922557 | p2 | (CT)9 | CCGTTAGTCCGTTTGATCGT | TGATATGAGGAAGCCAACCC | 113 |

| Marker ID | SSR type | SSR motif | FORWARD PRIMER (5'-3') | REVERSE PRIMER (5'-3') | size (bp) |
|---|---|---|---|---|---|
| chr5:78049857-78051872 | p5 | (ATCAT)3 | TTGACGACGTAGGAAAAGGG | TTATTCTCCGATCACTCGGC | 263 |
| chr5:78082155-78084167 | p2 | (GT)6 | TGTCGCAAAACGAGTGAGTC | CCCGAAGTCTCCACATCAAT | 277 |
| chr5:78099197-78101249 | p2 | (TA)26 | AGATAATGCCTTTGGATGCG | CCCATGTCAAAGCACAACAC | 162 |
| chr5:78125804-78127819 | p3 | (TTC)5 | CTTTGCTTTAATTCGCGAGG | GGCCTTATTCGTCACAGCAT | 271 |
| chr5:78255746-78257776 | p5 | (TTTAA)3 | AAGAAGAACCCTTGAGGGGA | CCTGTCATGACCCTCTGGTT | 103 |
| chr5:78261085-78263107 | p2 | (TA)11 | GAACCGCGTTCTTGAATGAT | TTAGGTTCCAAGCAAGGTGG | 146 |
| chr5:78277989-78280001 | p2 | (CA)6 | CATGCATTTCCATTCATTCG | ATGTGTGTGTGTGGGCAGAT | 186 |
| chr5:78277989-78280001 | p2 | (AT)6 | CATGCATTTCCATTCATTCG | ATGTGTGTGTGTGGGCAGAT | 186 |
| chr5:78290309-78292323 | p2 | (TA)7 | GGGGCAGTTTTTGACACCTA | TGATGCCATGAAAATCCCTT | 109 |
| chr5:78290309-78292323 | p5 | (TTTAT)3 | GGGGCAGTTTTTGACACCTA | TGATGCCATGAAAATCCCTT | 109 |
| chr5:78370529-78372569 | p6 | (ATACAT)3 | ATATCACGGTCGGATGGTGT | TGATAGCCTTGTGTGCTTGC | 245 |
| chr5:78370529-78372569 | p2 | (AT)20 | ATATCACGGTCGGATGGTGT | TGATAGCCTTGTGTGCTTGC | 245 |
| chr5:78582630-78584645 | p5 | (AAAAG)3 | ATATTTCGGTGGTCTGCCTG | TTCTATGGCCAAACGCCTAC | 161 |
| chr5:78619016-78621032 | p5 | (GTGTT)4 | AAATGAGATGGATGCGAAGC | TAGTATCTGTGGGGCCTTGG | 166 |
| chr5:78654258-78656302 | p2 | (AT)22 | AAACATGGCGCAACCTTAAC | CCTTGGAATTTTGTGTCGCT | 202 |
| chr5:78654258-78656302 | p2 | (TA)6 | AAACATGGCGCAACCTTAAC | CCTTGGAATTTTGTGTCGCT | 202 |
| chr5:78699763-78701779 | p2 | (AT)8 | CACTCTGCCACATCGACACT | AATAGATCCCCACCCTCACC | 190 |
| chr5:78802409-78804421 | p2 | (AG)6 | TTTGCCGAATCTCATTACCC | GGCCCAGATTTAATTCAGCA | 270 |
| chr5:78851053-78853068 | p3 | (CAA)5 | GCACAATATGCCTGCTCTCA | CGTACGTTCCTCGGCACTAT | 222 |
| chr5:78855231-78857246 | p5 | (AAGAA)3 | TGGAAACATGGAGGAAGGTC | AAGTTTTCCACATGGGCTTG | 242 |
| chr5:78855231-78857246 | p3 | (TTG)5 | TGGAAACATGGAGGAAGGTC | AAGTTTTCCACATGGGCTTG | 242 |
| chr5:78921876-78923892 | p4 | (TATG)4 | TGTCTCGCCATCTGTCTTTG | CAAACATAAGCAGGGCCATT | 179 |
| chr5:78921876-78923892 | p2 | (AT)6 | TGTCTCGCCATCTGTCTTTG | CAAACATAAGCAGGGCCATT | 179 |
| chr5:78946890-78948914 | p3 | (TTA)8 | ATTCATGCCCTTCCAACAAG | GAAAGTTGGGTGAAGGTCCA | 271 |
| chr5:78988727-78990745 | p6 | (TATTCG)3 | CACGGTTAAGCCTTGCTGTT | TGTGCTAAAATGGGTGTGGA | 286 |
| chr5:79015467-79017485 | p2 | (AT)16 | ACAACCCCTAGCACGAATTG | AGTGTGATAAGGGGTGCTCG | 184 |
| chr5:79015467-79017485 | p6 | (CATATA)3 | ACAACCCCTAGCACGAATTG | AGTGTGATAAGGGGTGCTCG | 184 |
| chr5:79041333-79043347 | p2 | (AT)7 | GCAAATTTGAGTAGAGCGGC | ACCCGCTTGTTCTGATTGTC | 250 |
| chr5:79207570-79209616 | p2 | (TA)23 | CCCCAAAAAGGATGTAGCAA | TGGCTCGTGTTATCGTGGTA | 225 |
| chr5:79316692-79318725 | p2 | (TA)22 | CGATGTGTTCATGCCCATAG | AATCATCATAGGCACCTCCG | 170 |
| chr5:79316692-79318725 | p3 | (ACA)11 | CGATGTGTTCATGCCCATAG | AATCATCATAGGCACCTCCG | 170 |
| chr5:79371769-79373781 | p2 | (TA)6 | GTCAATCCAAGCCCTTTCAA | TAAAAGAAACGAGGCATGGG | 180 |
| chr5:79402152-79404164 | p2 | (TA)6 | GTCAATCCAAGCCCTTTCAA | TAAAAGAAACGAGGCATGGG | 180 |
| chr5:79511597-79513617 | p2 | (AT)10 | CCATGGAGGCTGAACTGATT | TTTCGGATTTTGGGGTTGTA | 118 |
| chr5:79533995-79536007 | p2 | (TG)6 | AATTGTGTGCGTAGAAGGGG | CGCCCTACAACTCAGGGTTA | 231 |
| chr5:79537100-79539115 | p5 | (TCCTT)3 | GGAATCTTCGGCCTCATGTA | GACAATCAGAACAAGCGGGT | 180 |

| Marker ID | SSR type | SSR motif | FORWARD PRIMER (5'-3') | REVERSE PRIMER (5'-3') | size (bp) |
|---|---|---|---|---|---|
| chr5:79660456-79662468 | p2 | (AT)6 | ACTAGTGCGCCTCCACAGTT | CATGCAAAGCAAACCTCTGA | 133 |
| chr5:79665399-79667415 | p2 | (TA)8 | CCGCGGTATGTCGAACTTAT | GGAGAAATCAACACGGCAAT | 253 |
| chr5:79671856-79673871 | p5 | (ACACA)3 | ACACGGCCAACTTAGACCAC | TCATGGAGTGGGTGTGTTGT | 204 |
| chr5:79908031-79910049 | p6 | (CATATA)3 | TTACCGATGTCACACCCTGA | CAGACAGAGTCGTGGGCATA | 152 |
| chr5:79928983-79930998 | p3 | (TTG)5 | TGTCCTCGCTGTCATTTGAG | CTTGATGTTTCCCTCCCTGA | 177 |
| chr5:79930139-79932154 | p5 | (TTTGT)3 | GACTTGATGGGCCACTCCTA | TGAAAACAACGTGGAAACCA | 140 |
| chr5:80125099-80127115 | p4 | (GAAA)4 | GGAAGAAAAGGAAATTGGGC | TTTCTGCCACCAAGGCTAGT | 167 |
| chr5:80317033-80319093 | p2 | (AT)30 | ATAAGATGCCAACTCCACGG | GCCTAAATCCTCCGGAGTTC | 168 |
| chr5:80332560-80334575 | p5 | (TTCAT)3 | ACGTACACGAATCACGCAAA | TTTCCTTCCATCTGACCCAG | 249 |
| chr5:80444178-80446190 | p2 | (CT)6 | TTTTGGACTGCCCTTACCAC | GTTAGGCTGTTTTGGACCGA | 266 |
| chr5:80482960-80484975 | p5 | (AAAGG)3 | TTAAAGGACAAGGATTGCGG | GCCACATTCCATTTATTCCG | 115 |
| chr5:80580235-80582247 | p2 | (TA)6 | TCGTATCTGCTCCTCGACCT | TATGGAGTTGGGGGAAATGA | 274 |
| chr5:80586347-80588359 | p2 | (AT)6 | TTGAGCTTCCCTCTGCAGTT | CACCTGAGCATCAATAGCGA | 224 |
| chr5:80592990-80595004 | p2 | (AT)6 | ATTGGAAATTCGGGTTAGGC | GCCAAAAGAAATGAGGGACA | 228 |
| chr5:80592990-80595004 | p2 | (TA)7 | ATTGGAAATTCGGGTTAGGC | GCCAAAAGAAATGAGGGACA | 228 |
| chr5:80653114-80655172 | p2 | (AT)29 | GTTCGAACGGTCCTCAAAAA | ACTGGCTCACCAACTAGCGT | 123 |
| chr5:80703895-80705911 | p2 | (TA)6 | GTCTCGAAGGGGCATCAATA | TCCCCTTGTCAAAAATGGAG | 247 |
| chr5:80703895-80705911 | p2 | (AT)8 | GTCTCGAAGGGGCATCAATA | TCCCCTTGTCAAAAATGGAG | 247 |
| chr5:80905323-80907341 | p2 | (TA)9 | AACGTCGGATTATTTGCTGG | GATTTGGCCCAAGGGTACTT | 162 |
| chr5:81005744-81007762 | p5 | (TTCAT)3 | TAGGCATTATACGGCCCAAC | TCTTGTTTCTACGGGCCAAC | 225 |
| chr5:81005744-81007762 | p2 | (AG)9 | TAGGCATTATACGGCCCAAC | TCTTGTTTCTACGGGCCAAC | 225 |
| chr5:81005744-81007762 | p3 | (TTG)6 | TAGGCATTATACGGCCCAAC | TCTTGTTTCTACGGGCCAAC | 225 |
| chr5:81071522-81073537 | p5 | (AAATT)3 | TGGCTTGAACTTGAGAGGCT | TTGGGGGTAGAATGGACAAA | 207 |
| chr5:81166706-81168722 | p2 | (AT)8 | TCCTCCTTTTTCCCCTTTGT | TAAATTGTTGGTTGGTGGCA | 298 |
| chr5:81215424-81217439 | p5 | (ATCAT)3 | AATGTCTGCAGCCTCCTGTT | AGGGCAGTTTGTCCTAGGGT | 139 |
| chr5:81271658-81273673 | p5 | (ATCAT)3 | AATGTCTGCAGCCTCCTGTT | AGGGCAGTTTGTCCTAGGGT | 139 |
| chr5:81388408-81390434 | p2 | (AT)7 | ACGGCTATGGTATTTTCCCC | ACGACGGGATTCTCAAACAG | 140 |
| chr5:81388408-81390434 | p2 | (TA)13 | ACGGCTATGGTATTTTCCCC | ACGACGGGATTCTCAAACAG | 140 |
| chr5:81388408-81390434 | p2 | (AT)11 | ACGGCTATGGTATTTTCCCC | ACGACGGGATTCTCAAACAG | 140 |
| chr5:81447831-81449925 | p3 | (ATA)5 | AAGCCAGCTAGGTCGCTACA | CAGGTGCCCAGCTTAGTAGG | 216 |
| chr5:81447831-81449925 | p2 | (AT)47 | AAGCCAGCTAGGTCGCTACA | CAGGTGCCCAGCTTAGTAGG | 216 |
| chr5:81447831-81449925 | p2 | (AT)44 | AAGCCAGCTAGGTCGCTACA | CAGGTGCCCAGCTTAGTAGG | 216 |
| chr5:81477661-81479693 | p2 | (AT)16 | AAAAAGCTGGGCTTGGTGTA | TTAAAAATTTTCCCCGACCC | 123 |
| chr5:81489182-81491220 | p2 | (AT)19 | GCAGTGTTAGGACTCGCCTC | GTTAGGTGTGTTTTGGGCGT | 259 |
| chr5:81507679-81509694 | p5 | (TATAT)3 | CGTTGTTGCATGGTTGATTC | TGTGAAGTCTGAGTGCCACC | 188 |
| chr5:81545881-81547896 | p5 | (ATTAT)3 | CGTAGAGTGGCAACAGGACA | GTAGGGGAGATCCTTGAGGC | 228 |

| Marker ID | SSR type | SSR motif | FORWARD PRIMER (5'-3') | REVERSE PRIMER (5'-3') | size (bp) |
|---|---|---|---|---|---|
| chr5:81782294-81784309 | p5 | (ATGAT)3 | GATGCTGCCCAAATTTCTGT | ACAATTTCAAACGCGGATTC | 114 |
| chr5:82009347-82011362 | p3 | (ATG)5 | GCAAACCGACCCTAGAACAA | GCCTCCTTTGAGTCGATGAG | 212 |
| chr5:82009347-82011362 | p2 | (TA)19 | GCAAACCGACCCTAGAACAA | GCCTCCTTTGAGTCGATGAG | 212 |
| chr5:82058551-82060569 | p6 | (TATATG)3 | AGGGGAAATGCTGCCTAAAT | AAGTTACCCCAAGATGCGTG | 252 |
| chr5:82058551-82060569 | p2 | (TA)29 | AGGGGAAATGCTGCCTAAAT | AAGTTACCCCAAGATGCGTG | 252 |
| chr5:82065803-82067823 | p2 | (AG)10 | GTTGATGATTTGGCCGAGTT | CGGGATACTTAAGGCAACCA | 160 |
| chr5:82071231-82073247 | p2 | (AG)8 | GGTAAAGAAGCCATGTCGGA | CGGGATACTTAAGGCAACCA | 112 |
| chr5:82158283-82160349 | p2 | (AT)7 | ACATCAAAAAGGCACAAGGG | GCCAAGAGTGTTCGGGTAAA | 292 |
| chr5:82158283-82160349 | p3 | (TAT)22 | ACATCAAAAAGGCACAAGGG | GCCAAGAGTGTTCGGGTAAA | 292 |
| chr5:82217990-82220005 | p3 | (TTG)5 | CAAGAAGGTTTGGGGACTCA | AAGCCTCATACGGGGCTATT | 101 |
| chr5:82243823-82245841 | p2 | (AT)9 | ATGTTCCACTGCATCCATGA | TTAGCCTCGTCGGCTTATGT | 103 |
| chr5:82310983-82312998 | p5 | (TTCAT)3 | TATGCATTTGGGTCGTGTGT | ATTTGGCCAGAGTTTTCCCT | 292 |
| chr5:82310983-82312998 | p6 | (GAATAC)3 | TATGCATTTGGGTCGTGTGT | ATTTGGCCAGAGTTTTCCCT | 292 |
| chr5:82380542-82382560 | p3 | (ATA)6 | CCACTGTTCACCCAACACAG | CCACTTTGAGCCAAACCCTA | 140 |
| chr5:82380542-82382560 | p2 | (TA)6 | CCACTGTTCACCCAACACAG | CCACTTTGAGCCAAACCCTA | 140 |
| chr5:82633002-82635028 | p2 | (TA)13 | AGGTTCAGCCAATTCGTGTC | AATACGAGAGGAATGCGTGG | 264 |
| chr5:82810947-82812962 | p5 | (TATCG)3 | AGTTCCACCTACCAAACCCC | GGTCGTGACACATGGACTTG | 281 |
| chr5:82810947-82812962 | p5 | (CATAG)3 | AGTTCCACCTACCAAACCCC | GGTCGTGACACATGGACTTG | 281 |
| chr5:82810947-82812962 | p2 | (TA)6 | AGTTCCACCTACCAAACCCC | GGTCGTGACACATGGACTTG | 281 |
| chr5:82950241-82952256 | p3 | (GTT)5 | CCACCAAAACTTAAGGGCAA | TCCTTCTCATCTCGGCAACT | 180 |
| chr5:82973001-82975016 | p3 | (CAA)5 | ATGTACGCGTATACCTGCCC | AATGTTTCCCACACTGAGCC | 134 |
| chr5:83051219-83053253 | p2 | (TG)17 | GTTACACCCCTCGTGTTCGT | CTGGTGTCCTGCAGGTTTTT | 205 |
| chr5:83089966-83091978 | p2 | (TC)6 | AAAGAGGACTGCAAGCTGGA | TCCCCCTAAGGACAAGATCC | 267 |
| chr5:83104283-83106295 | p2 | (AG)6 | CTCATACCTGGCCCTACCAA | GGCCAAACTCCTAATCCCTC | 146 |
| chr5:83178480-83180500 | p2 | (TA)10 | GTACCGCTTGGTATGGGCTA | CCCGTCTTTGGATTTCATGT | 237 |
| chr5:83222421-83224443 | p2 | (AT)11 | GCCCAAGACCTTACAAGCAA | TCTTGTTGAATTCGGAAGGC | 234 |
| chr5:83232006-83234042 | p3 | (TGT)12 | GGAGCCTAGGACAAGTGCAG | ATAAGCTTCCTCCCCACGTT | 258 |
| chr5:83232006-83234042 | p2 | (AT)21 | GGAGCCTAGGACAAGTGCAG | ATAAGCTTCCTCCCCACGTT | 258 |
| chr5:83268839-83270861 | p2 | (AT)11 | TCTGATACACCCCAGAAGGC | ATTTGTTCAACAGCCACCTTG | 129 |
| chr5:83284635-83286653 | p6 | (ATCGTC)3 | AGCCGACAAGGCTAACAAAA | ACTCAATCGCAAGCTCCACT | 161 |
| chr5:83284635-83286653 | p5 | (GAAAT)3 | AGCCGACAAGGCTAACAAAA | ACTCAATCGCAAGCTCCACT | 161 |
| chr5:83358535-83360551 | p2 | (AT)8 | CATGGTAAGCGAGACTGCAA | CCTTCCCAGGTCATCTCTCA | 292 |
| chr5:83360358-83362376 | p6 | (TATATG)3 | TGATAATGTGGCCGAGTTGA | AACATACCTGCTCCACGTCC | 125 |
| chr5:83368252-83370268 | p4 | (ATAA)4 | TGTGATGCTTCTCGTTCAGG | GCAAACACCACAGCTGCTAA | 294 |
| chr5:83393181-83395199 | p6 | (AACAAT)3 | CTGATCAGTGGGATGCAATG | TTGCAATGGTTGATCCTTCA | 295 |
| chr5:83420164-83422189 | p5 | (AAATA)5 | ACAATTCAAAGTCAACCCGC | CAGTAGCAAAACGTGAGGCA | 147 |

| Marker ID | SSR type | SSR motif | FORWARD PRIMER (5'-3') | REVERSE PRIMER (5'-3') | size (bp) |
|---|---|---|---|---|---|
| chr5:83453784-83455802 | p2 | (AT)9 | AGCCGATCCATTCTTTTGTG | CCGTTCTTCAACCACAGGAT | 270 |
| chr5:83493834-83495854 | p5 | (TTTAT)4 | TGTCTACCTGTGGGCATGAA | TACCTATATGGTCGGGCAGC | 197 |
| chr5:83507113-83509164 | p2 | (AT)6 | TCAGAGGCTTGGCAACAAAT | GCCCACTGCAGAATAACCAT | 201 |
| chr5:83507113-83509164 | p5 | (AAATC)3 | TCAGAGGCTTGGCAACAAAT | GCCCACTGCAGAATAACCAT | 201 |
| chr5:83507166-83509220 | p2 | (AT)6 | TCAGAGGCTTGGCAACAAAT | GCCCACTGCAGAATAACCAT | 148 |
| chr5:83507166-83509220 | p5 | (AAATC)3 | TCAGAGGCTTGGCAACAAAT | GCCCACTGCAGAATAACCAT | 148 |
| chr5:83519112-83521127 | p2 | (AT)7 | ACGTTTTCGGATGACCTGAC | TGTTGGCAGCTTAGGGTCTT | 224 |
| chr5:83519112-83521127 | p3 | (TTG)5 | ACGTTTTCGGATGACCTGAC | TGTTGGCAGCTTAGGGTCTT | 224 |
| chr5:83519112-83521127 | p4 | (TATG)6 | ACGTTTTCGGATGACCTGAC | TGTTGGCAGCTTAGGGTCTT | 224 |
| chr5:83691001-83693019 | p6 | (TACTCG)3 | TCGAAACGATGATGTACCCA | CAAAGGAGGACAAGGACCAA | 113 |
| chr5:83790260-83792290 | p3 | (AAT)10 | GTGAATCCAAAAATCCCCCT | CTCTCCCACACCTCCATCAT | 168 |
| chr5:83897807-83899839 | p2 | (AT)16 | GAACCCGTTGGTTCTCGTTA | GCAAGCCGATGTACCATTTT | 284 |
| chr5:83904930-83907024 | p2 | (AT)47 | CCCGACCCTTACTTAGAGCC | AAGCCGATGTACCATTTTGC | 228 |
| chr5:83932556-83934581 | p5 | (CTTTT)5 | AGGGAAGCATACACAATGCC | ATGTTTTGTCTGCCTGGTCC | 267 |
| chr5:83932556-83934581 | p6 | (TTTTCC)3 | AGGGAAGCATACACAATGCC | ATGTTTTGTCTGCCTGGTCC | 267 |
| chr5:83932556-83934581 | p5 | (TTGAT)3 | AGGGAAGCATACACAATGCC | ATGTTTTGTCTGCCTGGTCC | 267 |
| chr5:83932696-83934714 | p5 | (CTTTT)5 | AGGGAAGCATACACAATGCC | ATGTTTTGTCTGCCTGGTCC | 127 |
| chr5:83932696-83934714 | p6 | (TTTTCC)3 | AGGGAAGCATACACAATGCC | ATGTTTTGTCTGCCTGGTCC | 127 |
| chr5:83932696-83934714 | p5 | (TTGAT)3 | AGGGAAGCATACACAATGCC | ATGTTTTGTCTGCCTGGTCC | 127 |
| chr5:84076273-84078297 | p2 | (TA)12 | CTTCCGGAGTGGGGATTATT | CCACTTCGAAAGACGGGATA | 194 |
| chr5:84123417-84125483 | p3 | (TAT)23 | GTGCTCTTCATACGGGCTTG | TGAACAAGCTACTTGGCAACA | 287 |
| chr5:84123417-84125483 | p3 | (ATT)14 | GTGCTCTTCATACGGGCTTG | TGAACAAGCTACTTGGCAACA | 287 |
| chr5:84135177-84137192 | p5 | (ATAAT)3 | TTGGAAGCCACAATACGACA | TTTCCGCGCAGTTAAGAAAT | 218 |
| chr5:84135177-84137192 | p2 | (AT)10 | TTGGAAGCCACAATACGACA | TTTCCGCGCAGTTAAGAAAT | 218 |
| chr5:84135212-84137227 | p5 | (ATAAT)3 | TTGGAAGCCACAATACGACA | TTTCCGCGCAGTTAAGAAAT | 183 |
| chr5:84135212-84137227 | p2 | (AT)10 | TTGGAAGCCACAATACGACA | TTTCCGCGCAGTTAAGAAAT | 183 |
| chr5:84187693-84189719 | p2 | (AC)13 | CACAACAAATTTCAATGCCG | TGTGGGTGTGTGTGAGTGTG | 135 |
| chr5:84187693-84189719 | p2 | (AT)13 | CACAACAAATTTCAATGCCG | TGTGGGTGTGTGTGAGTGTG | 135 |
| chr5:84197672-84199687 | p3 | (TGT)5 | GCCCATTTCCCTTGCTATTT | GCAGCAAACACGTCAACACT | 253 |
| chr5:84241057-84243069 | p2 | (TA)6 | TTGCGAAAGATGAAGAGGCT | TAACGTGGTGTCCTGCCATA | 291 |
| chr5:84272768-84274802 | p2 | (AT)17 | TTAGCAACCCTGCCCATATC | TAGCGGAGCTCCATTTCATT | 244 |
| chr5:84384170-84386182 | p2 | (AT)6 | GTCCTGTCCCGTCCTTATGA | GGGACAAGGTGGCTAACTCA | 110 |
| chr5:84384170-84386182 | p2 | (TA)6 | GTCCTGTCCCGTCCTTATGA | GGGACAAGGTGGCTAACTCA | 110 |
| chr5:84396132-84398144 | p2 | (AT)6 | AGGTCAGAGCATATGGGCAC | TGATTCCTCATTCCGTGTCA | 171 |
| chr5:84396132-84398144 | p3 | (TTG)5 | AGGTCAGAGCATATGGGCAC | TGATTCCTCATTCCGTGTCA | 171 |
| chr5:84640360-84642380 | p4 | (AAAG)5 | ACCGTACCCAGCCCTCTAGT | TATGTTATCGTGCGCACTCC | 272 |

| Marker ID | SSR type | SSR motif | FORWARD PRIMER (5'-3') | REVERSE PRIMER (5'-3') | size (bp) |
|---|---|---|---|---|---|
| chr5:84656425-84658443 | p6 | (TTTCAG)3 | GGTTTTCAAGAAAGCCCCTC | GGTTACAGACGGATTCCCAA | 128 |
| chr5:84777176-84779188 | p2 | (AT)6 | TCGTGTATTCAGGCCCTTTC | AATAACATGCCAAGCCCAAG | 178 |
| chr5:84902915-84904947 | p2 | (AT)16 | TCCTTGGGTTTCTTCCTCCT | AAATTAACACGGTTGCCTCG | 300 |
| chr5:84902915-84904947 | p2 | (AT)9 | TCCTTGGGTTTCTTCCTCCT | AAATTAACACGGTTGCCTCG | 300 |
| chr5:84921141-84923165 | p2 | (TA)12 | TGGGTACGGAGAACCTTGAG | CATATTTTCCGGGCTTTTCA | 250 |
| chr5:85059017-85061032 | p5 | (AAATG)3 | AGTGAAGGGATCGACCAATG | CCAACGTACCAAAATACGGG | 123 |
| chr5:85059017-85061032 | p3 | (TTG)5 | AGTGAAGGGATCGACCAATG | CCAACGTACCAAAATACGGG | 123 |
| chr5:85060716-85062740 | p2 | (TA)28 | CGTTCCACTGCATTGTTCAC | TCCTTGGCACCATAATCACA | 284 |
| chr5:85060716-85062740 | p2 | (AT)12 | CGTTCCACTGCATTGTTCAC | TCCTTGGCACCATAATCACA | 284 |
| chr5:85115419-85117435 | p2 | (TA)8 | TCGAACCATAGTCCGGAAAC | AAGGCGGAAGAACAGATTCA | 104 |
| chr5:85144308-85146320 | p2 | (CA)6 | CCGGTTCCGTTTGTGTAAGT | AACCCTTCCTTCTTTGGCAT | 108 |
| chr5:85144308-85146320 | p2 | (AT)6 | CCGGTTCCGTTTGTGTAAGT | AACCCTTCCTTCTTTGGCAT | 108 |
| chr5:85251998-85254012 | p2 | (AT)7 | CCCGAACTGGACACTCGTAT | CAGCTTGCGTACGGTATTGA | 196 |
| chr5:85338643-85340658 | p5 | (ATATT)3 | ATGATGGAGGTGTAAAGCCG | GATTCCCAATGGTAGCGAAA | 228 |
| chr5:85380146-85382170 | p2 | (AT)12 | TTGGCCGTCCTAGTGTCTCT | TGAATGCTAGAGGGTTCGCT | 206 |
| chr5:85445091-85447109 | p6 | (CATATA)3 | AGCACACCCAGCTGACTCTT | ATTTGCAGGCAAGAGATCGT | 216 |
| chr5:85456256-85458268 | p2 | (TA)6 | ATACGCAAGCTGACAAGGCT | TATGTTTGGGGCATACCCAT | 227 |
| chr5:85485243-85487261 | p6 | (TATATG)3 | GATGCGTACGTGTGATGTCC | CAAACGCGCTATCATTTGAA | 231 |
| chr5:85485243-85487261 | p5 | (TCATA)3 | GATGCGTACGTGTGATGTCC | CAAACGCGCTATCATTTGAA | 231 |
| chr5:85492533-85494565 | p6 | (CATATA)3 | TCAAATTGCCAAGGTCTTCC | CGAATCTGGATTTGCCATCT | 151 |
| chr5:85492576-85494596 | p6 | (CATATA)3 | TCAAATTGCCAAGGTCTTCC | CGAATCTGGATTTGCCATCT | 108 |
| chr5:85492831-85494849 | p6 | (CATATA)3 | GCCACCAAGGCTATCGTAAA | ATATGTGGGTCATGGGTCGT | 175 |
| chr5:85590603-85592633 | p2 | (AT)15 | CCAACCAAGGATCAAAGGAA | TTGCTGATTTACGGTGACCA | 125 |
| chr5:85600743-85602758 | p5 | (AAGAA)3 | CTCCAATCTACGGTTGGGAA | CCTACGTTGAGTGGGTCCAT | 230 |
| chr5:85630827-85632851 | p6 | (CATATA)4 | ACAAACTGACACATCGGCTG | ACATTGCTCGTACTGACCCC | 189 |
| chr5:85630827-85632851 | p6 | (CATATA)4 | ACAAACTGACACATCGGCTG | ACATTGCTCGTACTGACCCC | 189 |
| chr5:85630827-85632851 | p5 | (ATTCA)3 | ACAAACTGACACATCGGCTG | ACATTGCTCGTACTGACCCC | 189 |
| chr5:85640797-85642817 | p2 | (AT)10 | GCCTTTGATGTTCAGCCTTC | TCCAGCTCAGGCACTAGTCA | 105 |
| chr5:85661095-85663110 | p5 | (TCATA)3 | CCCCGGATTTTCCTTATTGT | GCTAATTTGGGGTATCGGGT | 256 |
| chr5:85775414-85777429 | p5 | (TTGTT)3 | TCGACATTCGTTTTGGATGA | TACGTCCTTATTCTTGCGGG | 281 |
| chr5:85777903-85779921 | p2 | (AT)15 | ATGTGAATGCCAGAGGGTTC | AACCCCTAGGACGAACTGCT | 283 |
| chr5:85777903-85779921 | p2 | (AT)9 | ATGTGAATGCCAGAGGGTTC | AACCCCTAGGACGAACTGCT | 283 |
| chr5:85804091-85806106 | p3 | (ACA)6 | ATTTTGATGCGGCCTAAATG | CCCAAACGGACAAACATACC | 234 |
| chr5:85804091-85806106 | p3 | (CAA)5 | ATTTTGATGCGGCCTAAATG | CCCAAACGGACAAACATACC | 234 |
| chr5:85865396-85867412 | p4 | (CAAG)4 | CACGCTTTACGTGGTGAAAA | CCGATGACTGACCCTGAAAT | 178 |
| chr5:85895746-85897758 | p2 | (CT)6 | AGCACCAGACCACCAAAAAC | ACTCTTTCCCAAAATGCCCT | 229 |

| Marker ID | SSR type | SSR motif | FORWARD PRIMER (5'-3') | REVERSE PRIMER (5'-3') | size (bp) |
|---|---|---|---|---|---|
| chr5:85916622-85918634 | p2 | (TA)6 | GATGCCCCGATTTCATTCTA | TCGTGATCGCTTTGTCTCAC | 294 |
| chr5:86020126-86022141 | p3 | (CTG)5 | CCTGGAAAACTGAAAACGGA | TATATTTTCCCGCTCGTTGC | 199 |
| chr5:86020126-86022141 | p3 | (ATC)5 | CCTGGAAAACTGAAAACGGA | TATATTTTCCCGCTCGTTGC | 199 |
| chr5:86122580-86124595 | p3 | (TTG)5 | CCAAGAAGGTTTAGGGGCTC | ACACACATGCCCGTAGCATA | 101 |
| chr5:86238782-86240800 | p2 | (AT)12 | GGTCGTGACACACACACACA | CGACCTTACGTCACTCCGAT | 170 |
| chr5:86238782-86240800 | p3 | (CAA)6 | GGTCGTGACACACACACACA | CGACCTTACGTCACTCCGAT | 170 |
| chr5:86305842-86307854 | p2 | (GT)6 | TGCCGGTATGAATTGTGTGT | TTAGCCTTGTTCGGCACTTT | 205 |
| chr5:86600133-86602148 | p3 | (TTG)5 | CAAGAAGGTTTGGGGAATCA | CTGACACACATGCCCGTAAC | 102 |
| chr5:86621460-86623475 | p5 | (CAAAT)3 | ATGCCTAAACGATCAAACGG | GCAGGACAACCACTAGGCTC | 186 |
| chr5:86621460-86623475 | p2 | (AT)8 | ATGCCTAAACGATCAAACGG | GCAGGACAACCACTAGGCTC | 186 |
| chr5:86695191-86697227 | p2 | (AT)17 | TGATTCAGACGCCTTCTCCT | TATCTTGAAGTGGGGTTGGG | 141 |
| chr5:86695191-86697227 | p2 | (AT)18 | TGATTCAGACGCCTTCTCCT | TATCTTGAAGTGGGGTTGGG | 141 |
| chr5:86755489-86757504 | p5 | (TTCAT)3 | AGGGCATCCTTACTCCGTCT | GACCATGCCAGATTGTTCCT | 215 |
| chr5:86758014-86760062 | p2 | (AT)24 | TGAACCATAGTAGGGCCAGG | TGGAAAACCAAGGTTTCAGG | 278 |
| chr5:86778694-86780721 | p3 | (AAT)9 | AGTGAGGCTTTGGTCCCTTT | CCGTAAACCTCGGTCGTAAA | 168 |
| chr5:86778694-86780721 | p5 | (TGTGT)3 | AGTGAGGCTTTGGTCCCTTT | CCGTAAACCTCGGTCGTAAA | 168 |
| chr5:86780815-86782835 | p2 | (AT)10 | AAGGGCGGAAAGAGGTATGT | TTCCGGAGAATGGAGTTGTC | 104 |
| chr5:86780815-86782835 | p5 | (ATATG)3 | AAGGGCGGAAAGAGGTATGT | TTCCGGAGAATGGAGTTGTC | 104 |
| chr5:86786080-86788096 | p5 | (ATATG)3 | AAGGGCGGAAAGAGGTATGT | TTCCGGAGAATGGAGTTGTC | 104 |
| chr5:87030423-87032437 | p2 | (TA)7 | CAACACGCAACATTCCAAAC | CGGAGCCTAACTTGTGGTGT | 241 |
| chr5:87090841-87092856 | p5 | (ATATA)3 | CAAATCCTGATCAGCAGCAA | TTCACAGACTAGGGTTGGGG | 155 |
| chr5:87174281-87176296 | p5 | (ATATA)3 | CAAATCCTGATCAGCAGCAA | TTCACAGACTAGGGTTGGGG | 155 |
| chr5:87313551-87315566 | p5 | (TTTTA)4 | TGGGCTTGGCTTTGTATTTC | TAGACCTCACCCCAAATTGC | 203 |
| chr5:87313551-87315566 | p5 | (TTTTA)3 | TGGGCTTGGCTTTGTATTTC | TAGACCTCACCCCAAATTGC | 203 |
| chr5:87364955-87366969 | p2 | (AT)7 | TTTCGATTCGTTTGACCTCC | AATTTGAGCCATGTTTTGGG | 163 |
| chr5:87374115-87376129 | p2 | (AT)7 | TTTCGATTCGTTTGACCTCC | AATTTGAGCCATGTTTTGGG | 163 |
| chr5:87473542-87475558 | p2 | (AT)8 | GCGCACAACACACCATAAAC | AGTATGGTCTTGGGTGCCAG | 158 |
| chr5:87473542-87475558 | p3 | (CAA)5 | GCGCACAACACACCATAAAC | AGTATGGTCTTGGGTGCCAG | 158 |
| chr5:87531260-87533272 | p2 | (TA)6 | GAATTTCTAAAGGGGGCGAG | TTGGGAGGGTATCCACTTTG | 110 |
| chr5:87562337-87564351 | p2 | (TG)7 | AAGGAGTCCTCCCCGAAGTA | ATTTTCCGAACGCACAAAAC | 239 |
| chr5:87719255-87721270 | p5 | (TATGA)3 | GATAGCCTTGTCGGCTTCTG | ACATACAGGCCATTTAGGCG | 173 |
| chr5:87765682-87767694 | p3 | (CAA)7 | AGCTGAGGCCATAGCACACT | GAAGCGAGCTTCCAGGTATG | 221 |
| chr5:87765682-87767694 | p2 | (CT)6 | AGCTGAGGCCATAGCACACT | GAAGCGAGCTTCCAGGTATG | 221 |
| chr5:87770726-87772738 | p2 | (CT)6 | AGCTGAGGCCATAGCACACT | GAAGCGAGCTTCCAGGTATG | 249 |
| chr5:87836382-87838397 | p5 | (TTTCA)3 | TTCTTGAATCTCCCCACCAG | TTCACAGACTAGGGTTGGGG | 270 |
| chr5:87894299-87896321 | p2 | (AG)11 | ATGTCACGACCTATTTCGGC | CGGGATACTTAAGGCAACCA | 161 |

| Marker ID | SSR type | SSR motif | FORWARD PRIMER (5'-3') | REVERSE PRIMER (5'-3') | size (bp) |
|---|---|---|---|---|---|
| chr5:87920440-87922455 | p5 | (TTCAT)3 | ATCATCGGACCACTGAAAGG | TTGGGCCATGTTGTGTCTAA | 111 |
| chr5:87937257-87939272 | p5 | (TTCAT)3 | ACTCGAATCCCACAAGAACG | GCATACGTCGTACCCTTCGT | 255 |
| chr5:87967998-87970062 | p2 | (AT)32 | TTAGCAAGATCCGAGCCTGT | AATAGCCCTCAAGCCCAACT | 282 |
| chr5:87998589-88000604 | p5 | (TTCAT)3 | GAAAGGGAACATCCGTCTGA | TGGGCCATGTTGTGTCTAAA | 125 |
| chr5:88123264-88125282 | p3 | (TTA)6 | ACAGTTGTTGGACGTGGACA | ATAGCTGGCCCGTATCACAC | 156 |
| chr5:88139014-88141032 | p3 | (TTA)6 | ACAGTTGTTGGACGTGGACA | ATAGCTGGCCCGTATCACAC | 156 |
| chr5:88158429-88160453 | p2 | (AC)12 | CCGTCTCGACTATCTCTGCC | TGTTCGTTCAATCAAGCAGC | 278 |
| chr5:88183558-88185570 | p2 | (TG)6 | ATGACTCGTTCGTCGTACCC | TAAAGGCTAACCGGCCCTAT | 174 |
| chr5:88264010-88266025 | p5 | (TCATT)3 | CCAGGAGAGATCCCAGATGA | TTTTGGGAACATTCGTCCTC | 280 |
| chr5:88418146-88420158 | p2 | (GT)6 | GAACCACACAAACATGTCGC | CCCGAGGTCTTAGCATCAAA | 276 |
| chr5:88520468-88522483 | p5 | (AATAA)3 | TCATCATCTATAAGCCGCCC | CGTACCTATAAGGCCAGGCA | 132 |
| chr5:88521912-88523954 | p2 | (TA)21 | TCGTGTTTCCGTTGATGTGT | AACTCGCCTAACCCGAACTT | 104 |
| chr5:88549628-88551643 | p5 | (AATAA)3 | TCATCATCTATAAGCCGCCC | CGTACCTATAAGGCCAGGCA | 132 |
| chr5:88555689-88557705 | p4 | (AAAT)4 | ATTCTCTTCTCGCAAACCGA | CCGCCACTACAAAGACCAAT | 126 |
| chr5:88555883-88557921 | p4 | (AAAT)4 | AGTGGCGGTGTTAGTGATCC | ATTCGTAATTCATGCCTCGC | 175 |
| chr5:88562057-88564072 | p5 | (TCTTT)3 | TCCTAACTGGACCAACCTGG | GCTCCATGCTGATGGTTGTA | 176 |
| chr5:88597462-88599478 | p4 | (AAAT)4 | ATTCTCTTCTCGCAAACCGA | CCGCCACTACAAAGACCAAT | 126 |
| chr5:88597656-88599694 | p4 | (AAAT)4 | AGTGGCGGTGTTAGTGATCC | ATTCGTAATTCATGCCTCGC | 175 |
| chr5:88603830-88605845 | p5 | (TCTTT)3 | TCCTAACTGGACCAACCTGG | GCTCCATGCTGATGGTTGTA | 176 |
| chr5:88869722-88871750 | p3 | (CCT)6 | GACAAAATCCCCGAGTGCTA | ACAAGTTTGACCTTGGCTGG | 275 |
| chr5:89047925-89049943 | p2 | (AT)9 | CGTTTCCTTATCACTTGGGG | AATGCCCATCAAAGTTTCCA | 144 |
| chr5:89077735-89079771 | p2 | (TA)18 | GGGAACCGGACACATCATAC | CGTCGAGTCTTGGGATTCAT | 103 |
| chr5:89094032-89096062 | p2 | (TA)15 | CGTCCGGGGTAACTATCTGA | CTCAGCCGAGTAGGGTCTTG | 275 |
| chr5:89094032-89096062 | p5 | (TTCAT)3 | CGTCCGGGGTAACTATCTGA | CTCAGCCGAGTAGGGTCTTG | 275 |
| chr5:89102870-89104888 | p3 | (TTG)6 | GACCCTTGAGCCAAATCAAA | ACCTTCCTGCCTTAAACCGT | 158 |
| chr5:89145057-89147072 | p5 | (TTAAG)3 | ATTCTGGGCAGAACATACGG | CAACGAGTCGACGAGAATGA | 261 |
| chr5:89172514-89174526 | p2 | (AC)6 | TACATACCTTGCCCGCTTTC | TTTGGCCGAGTTGAATTCTC | 268 |
| chr5:89250150-89252172 | p3 | (CAA)5 | CCAGCCATGTGGAATAGCTT | AGCGAGCGATTGAGGTATGT | 143 |
| chr5:89250150-89252172 | p2 | (AC)11 | CCAGCCATGTGGAATAGCTT | AGCGAGCGATTGAGGTATGT | 143 |
| chr5:89250150-89252172 | p2 | (TC)7 | CCAGCCATGTGGAATAGCTT | AGCGAGCGATTGAGGTATGT | 143 |
| chr5:89309245-89311267 | p2 | (AT)11 | GGTTTGTTGCGTTGGAAACT | TCAAACTTGGGGGACGAATA | 243 |
| chr5:89377454-89379466 | p2 | (AT)6 | AGACACTGCCGAGGATGAAC | GAGCCCTTGCAGTTCTTTTG | 110 |
| chr5:89388809-89390821 | p2 | (AT)6 | TAAGAATGGTGCCCCTTACG | GGTATTTCCTGCGTGCAGAT | 139 |
| chr5:89389569-89391603 | p2 | (AT)6 | TTTCCAGATCAACGGTCTCC | TTGTCGGCTTGCATACGATA | 251 |
| chr5:89427527-89429539 | p2 | (AT)6 | TCACATAGGCGGGATAGGAC | TCCTTTTGTTTGAGGACGCT | 209 |
| chr5:89487481-89489493 | p2 | (AT)6 | ATCGTGCGATAGAGCCAAGT | CGTCCGAGCTCTTACTCACC | 198 |

| Marker ID | SSR type | SSR motif | FORWARD PRIMER (5'-3') | REVERSE PRIMER (5'-3') | size (bp) |
|---|---|---|---|---|---|
| chr5:89558749-89560761 | p3 | (GAT)6 | CCAAGGTGAGATATGGCGTT | ATTTAGGCACGGTGGAATTG | 173 |
| chr5:89558749-89560761 | p2 | (TA)6 | CCAAGGTGAGATATGGCGTT | ATTTAGGCACGGTGGAATTG | 173 |
| chr5:89732705-89734720 | p5 | (ATTTC)3 | CGAGACAACAACCGACAAGA | ATTTTGCCAGAGTTTCCCCT | 162 |
| chr5:89758750-89760765 | p5 | (ATTTC)3 | CGAGACAACAACCGACAAGA | ATTTTGCCAGAGTTTCCCCT | 162 |
| chr5:89918340-89920352 | p2 | (TA)7 | CTTTCCCTTCCCCATGAAAT | TGACCTCCTATGTCCCGAAG | 219 |
| chr5:89918356-89920371 | p2 | (TA)7 | CTTTCCCTTCCCCATGAAAT | TGACCTCCTATGTCCCGAAG | 203 |
| chr5:90028398-90030413 | p3 | (TTG)5 | GCATGGGGATAGCCTGAGTA | GCATGTTGGCCTCGTAAAGT | 130 |
| chr5:90136345-90138372 | p3 | (CAA)9 | CACACACGCCAGGATCATAC | AATTACGGGGACAGAAACCC | 115 |
| chr5:90212020-90214032 | p2 | (AC)6 | TTCTTGAATCGTCCCTTTGG | TTTGGAAGCCACATGGTGTA | 181 |
| chr5:90223682-90225694 | p2 | (AC)6 | TTCTTGAATCGTCCCTTTGG | TTTGGAAGCCACATGGTGTA | 181 |
| chr5:90250820-90252832 | p2 | (GT)6 | AAGGAGACCTCCCCGAAGTA | GCGAAAAATGCGGGATATAA | 115 |
| chr5:90261945-90263957 | p6 | (TATATG)3 | TTCTGAGCCTTCATGTGTCG | CCATCGGAGTGACGTAAGGT | 130 |
| chr5:90261945-90263957 | p2 | (AT)6 | TTCTGAGCCTTCATGTGTCG | CCATCGGAGTGACGTAAGGT | 130 |
| chr5:90261945-90263957 | p2 | (AT)7 | TTCTGAGCCTTCATGTGTCG | CCATCGGAGTGACGTAAGGT | 130 |
| chr5:90281915-90283930 | p5 | (TTTCA)3 | AAGAAATGACATGGGAAGCG | ATAACAATGCCTCACAGGGC | 193 |
| chr5:90284345-90286366 | p3 | (ATC)7 | CAGGGACTACATAGCGGAGC | ATAAGCCGACGAGGCTAACA | 206 |
| chr5:90323381-90325399 | p6 | (ACATAT)3 | ATTGCTGGTCAAGCACTCCT | TGTGTTGATTGTTCGGGAGA | 201 |
| chr5:90329545-90331563 | p6 | (CATATA)3 | GCAACAAGTACCCGACCATT | ACGTTGACCGTTAGTTTCCG | 169 |
| chr5:90360166-90362184 | p6 | (ACATAT)3 | ATTGCTGGTCAAGCACTCCT | TGTGTTGATTGTTCGGGAGA | 201 |
| chr5:90366330-90368348 | p6 | (CATATA)3 | GCAACAAGTACCCGACCATT | ACGTTGACCGTTAGTTTCCG | 169 |
| chr5:90373829-90375855 | p5 | (CATTT)3 | TCGGCTCATCATCCTTAACC | TCCTGAAAGGATGACTTGGG | 276 |
| chr5:90373829-90375855 | p2 | (AT)13 | TCGGCTCATCATCCTTAACC | TCCTGAAAGGATGACTTGGG | 276 |
| chr5:90446118-90448136 | p6 | (AAAAAT)3 | TCCGTTTCGGCATAGTTACC | TGTGGGTTTGGAAAATCGTT | 278 |
| chr5:90487370-90489390 | p5 | (TTCAT)4 | AGGCTCGGTGTTGTCCATAC | CGGGGGTTTACCGACTTTAT | 163 |
| chr5:90495047-90497062 | p5 | (TTCAT)3 | AAAAGAAGTCGCTTCGGTCA | TTGAATCTCTTCTGGGGGTG | 175 |
| chr5:90507425-90509449 | p2 | (AT)12 | ATTTTCCAAACACCGACTGC | TGGCATGCATAATTCGTGTT | 183 |
| chr5:90594905-90596923 | p2 | (AG)9 | GATGATATAACACCGCGCCT | TCGGGGTACTATCTCATGCC | 156 |
| chr5:90607482-90609514 | p2 | (AT)7 | CGCTCGGCTCTAAGTAAGGA | CGCCCCAAGTAAGACACAAT | 217 |
| chr5:90607574-90609600 | p2 | (AT)7 | CGCTCGGCTCTAAGTAAGGA | CGCCCCAAGTAAGACACAAT | 125 |
| chr5:90690224-90692246 | p2 | (TG)11 | TGACAAGGACCCGTTTAAGG | CATTCTTGTCTTCCCACCGT | 123 |
| chr5:90895892-90897904 | p2 | (TA)6 | TTGTTGCATTGGGTCCTACA | GCCTCGTCGTACCCTGAATA | 289 |
| chr5:90979693-90981707 | p2 | (AT)7 | GACAGCGCTCCAGTTATTCC | TCAACATTAGCATGCCCAAA | 210 |
| chr5:90993570-90995598 | p2 | (AT)9 | CTTTTGTCATGACCCAACCC | TGTTCGTCTGTAGTGGCAGC | 163 |
| chr5:90993570-90995598 | p2 | (AT)14 | CTTTTGTCATGACCCAACCC | TGTTCGTCTGTAGTGGCAGC | 163 |
| chr5:91013065-91015077 | p2 | (AT)6 | CGCAGCATCTGTCGAATAAA | CAGTTTGAGGGTCCGTGATT | 147 |
| chr5:91055325-91057351 | p2 | (TA)13 | ATAAGATGCCAACTCCACGG | GCAGCAAGTACTAAAGCCCG | 166 |

| Marker ID | SSR type | SSR motif | FORWARD PRIMER (5'-3') | REVERSE PRIMER (5'-3') | size (bp) |
|---|---|---|---|---|---|
| chr5:91085557-91087572 | p5 | (ATTTA)3 | GTTGGCGTCATGGGTCTATT | CCAATTTATCACACCCCACC | 291 |
| chr5:91265628-91267643 | p5 | (AATGC)3 | GCAACAACCATGCACAGACT | CGCACAGTCCTAAAATGGGT | 101 |
| chr5:91481190-91483205 | p5 | (ATATA)3 | GAAAGCTGTCGGATTAAGCG | ATCGGTGTTGGGTCGTACAT | 160 |
| chr5:91481190-91483205 | p2 | (TA)6 | GAAAGCTGTCGGATTAAGCG | ATCGGTGTTGGGTCGTACAT | 160 |
| chr5:91483191-91485209 | p3 | (TTG)6 | GTGCATAAGCCCTCCAAAAA | ACCACGACCTATGGTTGGAA | 183 |
| chr5:91548965-91550977 | p2 | (AT)9 | CCCTGACACCCAATTACACC | GAAAGATGAGACGAGGACCG | 229 |
| chr5:91548978-91550994 | p2 | (AT)9 | CCCTGACACCCAATTACACC | GAAAGATGAGACGAGGACCG | 216 |
| chr5:91634377-91636395 | p6 | (CATATA)3 | CTACCGAGCACCCCTTATCA | ATGGGTGGTTGTTGTCCAGT | 119 |
| chr5:91634377-91636395 | p6 | (CATATA)3 | CTACCGAGCACCCCTTATCA | ATGGGTGGTTGTTGTCCAGT | 119 |
| chr5:91634496-91636514 | p6 | (CATATA)3 | ACTGGACAACAACCACCCAT | CGTCAGCCTAGGACATCCAT | 213 |
| chr5:91634496-91636514 | p6 | (CATATA)3 | ACTGGACAACAACCACCCAT | CGTCAGCCTAGGACATCCAT | 213 |
| chr5:91761388-91763404 | p2 | (TA)8 | TTGTACGAAAACACGGGACA | AACAATCCACAAAACAGCCC | 140 |
| chr5:91800446-91802494 | p6 | (TGAAGT)3 | CCGAGTTTGGAGGTTGAAAA | GACATGTCGATGGTCTGTCG | 289 |
| chr5:91800496-91802524 | p6 | (TGAAGT)3 | CCGAGTTTGGAGGTTGAAAA | GACATGTCGATGGTCTGTCG | 239 |
| chr5:91800526-91802554 | p6 | (TGAAGT)3 | CCGAGTTTGGAGGTTGAAAA | GACATGTCGATGGTCTGTCG | 209 |
| chr5:91800556-91802584 | p6 | (TGAAGT)3 | CCGAGTTTGGAGGTTGAAAA | GACATGTCGATGGTCTGTCG | 179 |
| chr5:91800586-91802614 | p6 | (TGAAGT)3 | CCGAGTTTGGAGGTTGAAAA | TACGGGATCCTATCGACACC | 149 |
| chr5:91829399-91831417 | p2 | (AT)22 | GAACTGTCACGACCCAACCT | TGGCACTTCTCTGCATTACG | 154 |
| chr5:91829399-91831417 | p6 | (CATATA)3 | GAACTGTCACGACCCAACCT | TGGCACTTCTCTGCATTACG | 154 |
| chr5:91901582-91903612 | p2 | (TA)15 | GGTCACCACATAAGTGGGCT | TTTTGGGTCAACTTTGGAGG | 280 |
| chr5:91920163-91922181 | p6 | (TATATG)3 | GATGTACTCGGCGTCTGTGA | CTGGGGCAGATGTAACCTGT | 162 |
| chr5:91932414-91934429 | p3 | (CAA)5 | GGGGTGTAACATCCTTCCCT | TGGCCCAATTCCAATAAGTC | 191 |
| chr5:91974787-91976802 | p3 | (TGT)5 | GGACAGCAATTCTGGGTCAT | GAAGCCATTTGGCAGCTTAG | 283 |
| chr5:91974787-91976802 | p2 | (TA)32 | GGACAGCAATTCTGGGTCAT | GAAGCCATTTGGCAGCTTAG | 283 |
| chr5:92002941-92004956 | p3 | (CTG)5 | GTCCCACTCCGATTCAAAGA | TAGTCGACAGTACCCGGAGG | 292 |
| chr5:92106375-92108390 | p2 | (AT)7 | CCTGATAATCCGCCGATAGA | TGCATGACATTCGCCTTAAA | 222 |
| chr5:92106375-92108390 | p4 | (CATA)4 | CCTGATAATCCGCCGATAGA | TGCATGACATTCGCCTTAAA | 222 |
| chr5:92106375-92108390 | p3 | (ATC)5 | CCTGATAATCCGCCGATAGA | TGCATGACATTCGCCTTAAA | 222 |
| chr5:92106375-92108390 | p3 | (CAA)5 | CCTGATAATCCGCCGATAGA | TGCATGACATTCGCCTTAAA | 222 |
| chr5:92140209-92142224 | p5 | (TATAT)3 | TTCCACCATATCGCAAAACA | GCAAAGGGTTGGTGTGAAGT | 189 |
| chr5:92206860-92208884 | p2 | (AT)12 | TTATCCTCAACGGCCTCAAC | GTTAGTGCAAGAACACGCGA | 190 |
| chr5:92245692-92247716 | p2 | (AT)12 | TTATCCTCAACGGCCTCAAC | GTTAGTGCAAGAACACGCGA | 190 |
| chr5:92394577-92396613 | p2 | (TA)18 | ATCGTGAGACCATGCTACCC | TGTTTTCTTTGGACGCTGTG | 193 |
| chr5:92417205-92419219 | p2 | (TA)7 | AGCCCTTCGGCTATCAAAAT | CCCACAGGTAGACAGGGAGA | 133 |
| chr5:92435356-92437371 | p2 | (GT)7 | GTCATGTGCTTTCCATGTGC | TGGGGTTCAAGAACTCAAGG | 115 |
| chr5:92563419-92565435 | p2 | (AT)6 | TACAGCAACGAGGGCTTTTT | AATCATTGTTACACCCCGGA | 130 |

| Marker ID | SSR type | SSR motif | FORWARD PRIMER (5'-3') | REVERSE PRIMER (5'-3') | size (bp) |
|---|---|---|---|---|---|
| chr5:92563419-92565435 | p2 | (TA)8 | TACAGCAACGAGGGCTTTTT | AATCATTGTTACACCCCGGA | 130 |
| chr5:92572910-92574956 | p2 | (AT)23 | TAAATTGAGCCGTGTAGGGG | ATCAACATGCCCACAAAACA | 117 |
| chr5:92794464-92796482 | p6 | (ATATGT)3 | ATGTTGGATATGAGCCTCGG | GTCGGGTCCTATTTATGCGA | 226 |
| chr5:92830535-92832553 | p6 | (TATATG)3 | TTAGATTCTCGGGGATGGTG | GCGAGCCTTAACATACCTGC | 153 |
| chr5:92851585-92853603 | p6 | (TATATG)3 | GGGCGTACCCACTCGTAGTA | CAAGCCAACAAGGCTACCAT | 186 |
| chr5:92851585-92853603 | p2 | (TA)8 | GGGCGTACCCACTCGTAGTA | CAAGCCAACAAGGCTACCAT | 186 |
| chr5:92899256-92901276 | p2 | (AT)10 | TACCGCCATATCTCGGAGTC | TTGACACAGTTAGTTGCCGC | 147 |
| chr5:92946790-92948805 | p2 | (GT)6 | TTGGTGAAGTGCTCAAGACG | CACACCATGCACAAACACAA | 292 |
| chr5:92946790-92948805 | p3 | (GTT)5 | TTGGTGAAGTGCTCAAGACG | CACACCATGCACAAACACAA | 292 |
| chr5:93012272-93014286 | p2 | (TA)7 | GAGATCGGAACCTTGTTGGA | CATCTTCGATGTTGGTGGTG | 102 |
| chr5:93044949-93046963 | p2 | (TA)7 | GAGATCGGAACCTTGTTGGA | CATCTTCGATGTTGGTGGTG | 102 |
| chr5:93365912-93367926 | p2 | (TG)7 | TTCGAAGGGACGAATGTTTC | AATTTGTCCTTGGCCTTCCT | 295 |
| chr5:93374113-93376128 | p5 | (AACCC)3 | CATCAATGCTTGCCCTTTTT | GCACGAGCCCAACAATTTAT | 227 |
| chr5:93397950-93399962 | p2 | (AT)6 | GGGGGTCCATATTTTGCTTT | GTACCTGCCCCAGTTCTTCA | 284 |
| chr5:93397950-93399962 | p3 | (CAT)5 | GGGGGTCCATATTTTGCTTT | GTACCTGCCCCAGTTCTTCA | 284 |
| chr5:93398572-93400587 | p2 | (AT)6 | AACCCTAGGACGGAACTGCT | TTCATGAGGTCATCCCTTCC | 178 |
| chr5:93398572-93400587 | p3 | (CAT)5 | AACCCTAGGACGGAACTGCT | TTCATGAGGTCATCCCTTCC | 178 |
| chr5:93454168-93456248 | p2 | (AT)40 | GTTGCCCGATTTTCGTTAAA | TATTCGTGAATCCTTTCGCC | 226 |
| chr5:93506322-93508338 | p2 | (AT)6 | AGGCCTTGACTAGTGCCTGA | GTAGTAGCAGCCTTGTCGGC | 208 |
| chr5:93506322-93508338 | p2 | (AT)8 | AGGCCTTGACTAGTGCCTGA | GTAGTAGCAGCCTTGTCGGC | 208 |
| chr5:93570208-93572220 | p2 | (TG)6 | GAAGTCACGGGCTCTCAAAC | CAGAGGAACCTAGGACAGCG | 225 |
| chr5:93570208-93572220 | p4 | (CTTT)4 | GAAGTCACGGGCTCTCAAAC | CAGAGGAACCTAGGACAGCG | 225 |
| chr5:93594667-93596679 | p2 | (TG)6 | AATGCAAGGTCGGAATGAAG | CATACTTGCCTACCCCGAAA | 118 |
| chr5:93647960-93649974 | p2 | (AT)7 | CGTCCTCAGCATCTCATCAA | AGGGGTTTCAGTTCCAGCTT | 199 |
| chr5:93647960-93649974 | p3 | (TTA)5 | CGTCCTCAGCATCTCATCAA | AGGGGTTTCAGTTCCAGCTT | 199 |
| chr5:93898455-93900467 | p2 | (AT)6 | TGGGCATGTAAACCCAAAAT | GCTTAGAACGAGGCTTGTGG | 121 |
| chr5:94049161-94051209 | p5 | (TTTTG)3 | GCCCCTTTTGCTTAAAGACC | TCACGAGGTGGTTCATGGTA | 264 |
| chr5:94049161-94051209 | p2 | (TA)24 | GCCCCTTTTGCTTAAAGACC | TCACGAGGTGGTTCATGGTA | 264 |
| chr5:94184036-94186064 | p4 | (ACAT)7 | ATGGGGAAAAACAGTGCAAG | AGTATGCTCCATTCGATCCG | 165 |
| chr5:94288410-94290425 | p5 | (ATATG)3 | TCCGAGCGTAGTTATGACCC | TGGCGTACGTGAATTATGGA | 282 |
| chr5:94292339-94294351 | p2 | (TA)6 | AGAAAAATAGGGGGCGAAGA | CCTTTTTATCTGGCGATCCA | 248 |
| chr5:94370292-94372330 | p2 | (AT)19 | ATTGTCCGTACTGACGTCCC | GCATACCAAAAAGGGCTGAA | 197 |
| chr5:94444650-94446665 | p5 | (AAACC)3 | GGCCTTGCTATAAGGGCTTC | ATTTTCGCCAAAACATGAGC | 103 |
| chr5:94454407-94456441 | p2 | (AT)17 | TTCCATTGGGCAACAATACA | CAGGGACTACATAGCGGAGC | 246 |
| chr5:94580315-94582345 | p2 | (AT)15 | AGACAGCCTTGTTGGCTCAT | GACAAGGCTCGTAGAAACGC | 253 |
| chr5:94683926-94686000 | p6 | (CAACTC)3 | ATTCGTAGCGTCCTCAGCAT | TAGCAGACGACCGAGGCTAT | 287 |

| Marker ID | SSR type | SSR motif | FORWARD PRIMER (5'-3') | REVERSE PRIMER (5'-3') | size (bp) |
|---|---|---|---|---|---|
| chr5:94684840-94686858 | p6 | (CAACTC)3 | CCTTCCCTCTTTTCCCTCAC | TTTGATGCCCCTATACCTCG | 292 |
| chr5:94811792-94813804 | p5 | (TGAAA)4 | CCAGTGATCCCGAAGTTTGT | GATCTGCCAAGAAAGGCAAG | 297 |
| chr5:94811792-94813804 | p2 | (TA)6 | CCAGTGATCCCGAAGTTTGT | GATCTGCCAAGAAAGGCAAG | 297 |
| chr5:94852349-94854367 | p2 | (TA)9 | ATTCTGATGGTCCTCGTTGC | AGTGCTAAATGGGCCGTATG | 163 |
| chr5:94884441-94886457 | p4 | (TTTC)4 | TCTTTCCGGCATAACGAAAC | AAGTTGTGGGCGAAATATGC | 178 |
| chr5:94902906-94904918 | p2 | (TA)6 | CCGTATTTTTGAAGCTCGGA | TTTGGAGGGTCTCCACTTTG | 145 |
| chr5:94932445-94934459 | p2 | (AT)7 | TTCGACTAGTGGCCTCACCT | TTAGGCGTTCGCTTTCTGAT | 190 |
| chr5:94981703-94983718 | p3 | (TTG)5 | AGGGGCCAAAACGCTATTAT | TTTGGCAGCTTGGAGACTTT | 242 |
| chr5:95010050-95012068 | p6 | (CATATA)3 | GACGTGGCCGACAAAATAAT | GGGAGACAGACGTGATCCAT | 300 |
| chr5:95032484-95034499 | p5 | (TGATC)3 | ACAGCCTACAGGTCGCAGAT | GACGGCCACAAGAAAAACAT | 244 |
| chr5:95032484-95034499 | p2 | (TA)12 | ACAGCCTACAGGTCGCAGAT | GACGGCCACAAGAAAAACAT | 244 |
| chr5:95039338-95041353 | p5 | (TGATC)3 | ACAGCCTACAGGTCGCAGAT | GACGGCCACAAGAAAAACAT | 244 |
| chr5:95039338-95041353 | p2 | (TA)12 | ACAGCCTACAGGTCGCAGAT | GACGGCCACAAGAAAAACAT | 244 |
| chr5:95275481-95277493 | p2 | (AT)6 | TGACTAAATGTTCCCCTCCG | ATGGTTGTGGATGACGTTGA | 109 |
| chr5:95314538-95316550 | p2 | (AT)6 | TGACTAAATGTTCCCCTCCG | ATGGTTGTGGATGACGTTGA | 109 |
| chr5:95460760-95462775 | p5 | (TCCTT)3 | CTGCGGTCCGTCAAACTAAT | CAACAAACCGTTCGTCAATG | 234 |
| chr5:95605713-95607731 | p6 | (TTTGAA)3 | ACGTTTGATGACCTTAGGCG | GCAATTCTGCAACTTCCCAT | 185 |
| chr5:95609820-95611832 | p2 | (AC)6 | TTTGGGACAATTGCATACGA | AACATAATAGGTCGTCGCGG | 167 |
| chr5:95768207-95770222 | p5 | (ATTTC)3 | TCGTGTGTGGGCAAATTTTA | ATTTTGCCAGAGTTTCCCCT | 282 |
| chr5:95815485-95817503 | p3 | (ATT)6 | ATTCCAGGCATGTTGAGACC | TGGCACCAGTACCCAATACA | 156 |
| chr5:95872578-95874612 | p2 | (TA)17 | TGGTGTTTCTAATGCTCCCC | TTGGTCCAAGCCCATTTTAG | 171 |
| chr5:95872578-95874612 | p2 | (TA)11 | TGGTGTTTCTAATGCTCCCC | TTGGTCCAAGCCCATTTTAG | 171 |
| chr5:95964433-95966451 | p2 | (TA)9 | TTGGAATAGCAAAAGGACCG | AAACTTTAGGTGGCCGGATT | 134 |
| chr5:95964433-95966451 | p5 | (GCTAT)4 | TTGGAATAGCAAAAGGACCG | AAACTTTAGGTGGCCGGATT | 134 |
| chr5:96071358-96073374 | p2 | (TA)8 | GCCCGAATTTCTGTAGGTGA | TAGAGTTACCCCAAGGTGCG | 139 |
| chr5:96106808-96108820 | p3 | (CAA)5 | GAAACGGCTTATGGAGGTCA | GGGATTTGATTAGCGAACGA | 149 |
| chr5:96153578-96155593 | p5 | (CTTGA)3 | GGTAGGGGCCGGATTTAATA | TAAGATCCTAAGCGGACGGA | 154 |
| chr5:96220350-96222365 | p5 | (ATTCA)3 | CGGAGTGACATAAGGTCGGT | AACGTTCGTGGAAGGTTCTG | 150 |
| chr5:96243281-96245299 | p2 | (AT)9 | GGGCATGATGGAGTCTTGTT | GCCTCTTGGCCATCAAAATA | 164 |
| chr5:96255905-96257923 | p2 | (CT)9 | AATTTCGGTAGCATTTCCCC | GCGGGTTGTTGAAATGTTCT | 167 |
| chr5:96554600-96556612 | p2 | (CT)6 | ACATGCCAAAGAACAAAGGG | AAGCTGCCTTTTACCCGAAT | 103 |
| chr5:96851671-96853685 | p5 | (TTTAA)3 | CAGTTCGTCCTTGGGAATGT | TGATTTTGCTCTATGGCACG | 213 |
| chr5:96851671-96853685 | p2 | (TA)7 | CAGTTCGTCCTTGGGAATGT | TGATTTTGCTCTATGGCACG | 213 |
| chr5:97091611-97093637 | p2 | (TA)13 | TCAAGAAAACCCATCCCAAG | TATGAAGCGTGGGGAAGAAC | 254 |
| chr5:97129982-97131997 | p5 | (TTTCA)3 | AAATGGGAAGCGGAAGAAAT | ATAACAATGCCTCACAGGGC | 158 |
| chr5:97177607-97179623 | p2 | (AT)8 | AGGTCATGGAAGGATTGCAG | GTTTGCGCAGGGTAAATCAT | 271 |

| Marker ID | SSR type | SSR motif | FORWARD PRIMER (5'-3') | REVERSE PRIMER (5'-3') | size (bp) |
|---|---|---|---|---|---|
| chr5:97191283-97193328 | p3 | (TAT)15 | TTGAAGGTAGAAACGTGGGC | GGAAAAATCATGCCTTTGGA | 246 |
| chr5:97191283-97193328 | p2 | (TA)8 | TTGAAGGTAGAAACGTGGGC | GGAAAAATCATGCCTTTGGA | 246 |
| chr5:97191624-97193640 | p3 | (TAT)15 | CACGACCTTATGCCACCTCT | CCATATAGGCGCGGTGTAAT | 275 |
| chr5:97191624-97193640 | p2 | (TA)8 | CACGACCTTATGCCACCTCT | CCATATAGGCGCGGTGTAAT | 275 |
| chr5:97256847-97258867 | p2 | (AT)10 | CATACCATATGGCTGCGATG | CTCTGGACCCTCCATTTTCA | 234 |
| chr5:97256847-97258867 | p3 | (ATC)5 | CATACCATATGGCTGCGATG | CTCTGGACCCTCCATTTTCA | 234 |
| chr5:97282822-97284840 | p6 | (TATATG)3 | GGTAATTTGGCCGACTTGAA | TTACATACCTTGCCCGCTTC | 149 |
| chr5:97319161-97321176 | p5 | (AAGAA)3 | TATGGAAAGGTGGTGGAAGG | TAGCCATGTTCGAAACCTCC | 257 |
| chr5:97319161-97321176 | p2 | (TA)6 | TATGGAAAGGTGGTGGAAGG | TAGCCATGTTCGAAACCTCC | 257 |
| chr5:97319161-97321176 | p5 | (TGTTG)3 | TATGGAAAGGTGGTGGAAGG | TAGCCATGTTCGAAACCTCC | 257 |
| chr5:97319161-97321176 | p5 | (TGAAT)3 | TATGGAAAGGTGGTGGAAGG | TAGCCATGTTCGAAACCTCC | 257 |
| chr5:97332439-97334451 | p2 | (GT)6 | GAGCTGCAGTCCCTTTATGG | ATATTCCCTTGGCCCAATTC | 293 |
| chr5:97382719-97384737 | p3 | (CAA)6 | CCTTGCGGACTCGATGTTAT | GTAAGGGCCAAGTACGGACA | 185 |
| chr5:97397550-97399564 | p2 | (AT)11 | TGAAGATGCCGAAGTCCTCT | GTCGGCTTGCGTACCATATT | 113 |
| chr5:97397550-97399564 | p2 | (AT)7 | TGAAGATGCCGAAGTCCTCT | GTCGGCTTGCGTACCATATT | 113 |
| chr5:97441025-97443037 | p2 | (GT)6 | CACCAGCCTTCACTCCAAAT | CAAACTTGCATTGGCTTCAA | 276 |
| chr5:97485968-97487983 | p3 | (GTT)5 | ACGTTTTCGGGTGATCTGAC | CATACAGGCCCACGACCTAT | 174 |
| chr5:97557617-97559645 | p4 | (CTTT)7 | CTTAACGAAGGTCCAGGCAG | GTCTTGATCGGAATTTGGGA | 298 |
| chr5:97557617-97559645 | p2 | (TA)27 | CTTAACGAAGGTCCAGGCAG | GTCTTGATCGGAATTTGGGA | 298 |
| chr5:97558913-97560927 | p2 | (TA)27 | TTCTCGTGCGTACTCCACTG | AGTGAGGGACTCGGTGAATG | 170 |
| chr5:97558913-97560927 | p2 | (TA)7 | TTCTCGTGCGTACTCCACTG | AGTGAGGGACTCGGTGAATG | 170 |
| chr5:97579716-97581731 | p3 | (GTT)5 | CGTTTTCGGGTGATCTGACT | TGCCTTAAAACGCCCATAAC | 175 |
| chr5:97651895-97653937 | p2 | (AT)21 | GATCCGGAGTCTCGTTACCA | AATGCATGAAAACACGGGAT | 181 |
| chr5:97813976-97815988 | p2 | (TA)6 | TTGTATAAAAAGGCAGGGCG | AGCCTTGTCAGCTTGCGTAT | 137 |
| chr5:97818887-97820903 | p4 | (AAAG)4 | GAACAGCCTGTGGCCATATT | TGCGGCCCATTATTTATCTC | 256 |
| chr5:97828484-97830522 | p2 | (AT)19 | CGGTGTCCCAGACGAATACT | GCTTCAGACAGTAGGGGTGC | 266 |
| chr5:97828484-97830522 | p4 | (CATA)4 | CGGTGTCCCAGACGAATACT | GCTTCAGACAGTAGGGGTGC | 266 |
| chr5:97836499-97838514 | p3 | (TTG)5 | TGAACGCACCATTCAAACAT | AACAATCGGTCTCGGATGAG | 165 |
| chr5:97903475-97905489 | p2 | (TA)7 | TGACTTCGGCATTGATGTGT | TCGATGCACATTCGTAGGAG | 120 |
| chr5:97903475-97905489 | p5 | (TTGAA)3 | TGACTTCGGCATTGATGTGT | TCGATGCACATTCGTAGGAG | 120 |
| chr5:97909291-97911306 | p5 | (ATTTT)3 | TGAAACGCCGGAAAAATAAG | ATTTCGCCAGAGTTTCCCTT | 262 |
| chr5:97949658-97951678 | p2 | (TA)10 | TACTGATCCAGGTCTTGGGG | ATGCTTCGGACTATGATGGG | 163 |
| chr5:98026587-98028603 | p2 | (AT)8 | GCGTGCCAAAAAGAAAGAAG | CCCGATTTCCGGTAAGTTCT | 140 |
| chr5:98059302-98061320 | p3 | (CAA)6 | CCAGGTGGCTATGCGTCTAT | CTTTCTTTGGGCTCTCATGC | 124 |
| chr5:98072849-98074883 | p2 | (AT)17 | ATGGATTGGTTGGCTTCTTG | CGTGATCATTTTCCTTGCCT | 236 |
| chr5:98129821-98131839 | p2 | (AG)9 | CTCTTTCAATTCTGCGGGAG | GTAGAAGCTGAATCGCCGAC | 286 |

| Marker ID | SSR type | SSR motif | FORWARD PRIMER (5'-3') | REVERSE PRIMER (5'-3') | size (bp) |
|---|---|---|---|---|---|
| chr5:98157262-98159278 | p2 | (AT)8 | GACGTTGAATCGGTGGAGTT | TTCTGGGGGCAAGTAATGTC | 239 |
| chr5:98157262-98159278 | p2 | (TA)8 | GACGTTGAATCGGTGGAGTT | TTCTGGGGGCAAGTAATGTC | 239 |
| chr5:98157381-98159397 | p2 | (AT)8 | GACGTTGAATCGGTGGAGTT | TTCTGGGGGCAAGTAATGTC | 120 |
| chr5:98157381-98159397 | p2 | (TA)8 | GACGTTGAATCGGTGGAGTT | TTCTGGGGGCAAGTAATGTC | 120 |
| chr5:98157381-98159397 | p5 | (TGAAT)3 | GACGTTGAATCGGTGGAGTT | TTCTGGGGGCAAGTAATGTC | 120 |
| chr5:98157591-98159617 | p2 | (AT)8 | GCATTCAAGCCTACACCCAT | CCGGATGTACAAAATACGGG | 276 |
| chr5:98157591-98159617 | p2 | (TA)8 | GCATTCAAGCCTACACCCAT | CCGGATGTACAAAATACGGG | 276 |
| chr5:98157591-98159617 | p5 | (TGAAT)3 | GCATTCAAGCCTACACCCAT | CCGGATGTACAAAATACGGG | 276 |
| chr5:98170764-98172779 | p2 | (AT)16 | AGGGGCAACTCATCATTTTG | CATCCATGACACATGGGAAA | 297 |
| chr5:98170764-98172779 | p2 | (AT)26 | AGGGGCAACTCATCATTTTG | CATCCATGACACATGGGAAA | 297 |
| chr5:98170764-98172779 | p5 | (TGAAT)3 | AGGGGCAACTCATCATTTTG | CATCCATGACACATGGGAAA | 297 |
| chr5:98234516-98236534 | p6 | (TATATG)3 | GTTGATGATTTGGCCGAGTT | TTACATACCTTGCCCGCTTC | 274 |
| chr5:98234516-98236534 | p6 | (TATATG)3 | GTTGATGATTTGGCCGAGTT | TTACATACCTTGCCCGCTTC | 274 |
| chr5:98234516-98236534 | p2 | (TA)6 | GTTGATGATTTGGCCGAGTT | TTACATACCTTGCCCGCTTC | 274 |
| chr5:98341763-98343781 | p2 | (AT)9 | GGGGATCCTTATACGCCAAT | TCAGTCGAAGGAGGAGGAAA | 121 |
| chr5:98357253-98359279 | p2 | (TA)13 | AAAGGGAGCCTGGACAGTTT | AGGGATTGAGGCTTGCTTTT | 135 |
| chr5:98411816-98413832 | p4 | (TTTC)4 | CTCGCTTGCCTCCTAAGCTA | GCCGAAATTTCCGTAGACAA | 138 |
| chr5:98431815-98433829 | p2 | (TA)7 | TTCGATTATTCCATCGAGCC | CAGGAGTCATTTTTGGGGAA | 254 |
| chr5:98431815-98433829 | p6 | (GTATAT)3 | TTCGATTATTCCATCGAGCC | CAGGAGTCATTTTTGGGGAA | 254 |
| chr5:98542915-98544965 | p2 | (AT)6 | CGTTTTCGGGTGATCAGACT | GGTTGGGCTCATTTCACTGT | 135 |
| chr5:98630755-98632770 | p3 | (TGT)5 | CATCGCACACTCATGCTCTT | ATTAACCCTTTGCGAACACG | 162 |
| chr5:98654677-98656699 | p2 | (TA)22 | CATCCAGGAGGCACAACTTT | CTGTGGATAGTGCGTCGAGA | 183 |
| chr5:98654677-98656699 | p2 | (TA)11 | CATCCAGGAGGCACAACTTT | CTGTGGATAGTGCGTCGAGA | 183 |
| chr5:98671014-98673032 | p3 | (CAA)6 | CATATGGTGGTTACCCGGAC | CTCGCCTCCCTGTCTACTTG | 173 |
| chr5:98671014-98673032 | p2 | (AT)20 | CATATGGTGGTTACCCGGAC | CTCGCCTCCCTGTCTACTTG | 173 |
| chr5:98682136-98684151 | p3 | (ATC)5 | CTATATGCGCAAGCCAACAA | GCTTCATCAGCGTTTGAACA | 281 |
| chr5:98709457-98711471 | p2 | (TA)7 | TGCATCATGACCATCCAACT | CGTGCCCTATGATTGGTCTT | 101 |
| chr5:98740887-98742902 | p5 | (CGTTT)3 | ATATAAAGTGTCCCGGCCCT | TAGAGAAACGGAGACGGTGG | 150 |
| chr5:98750400-98752424 | p3 | (GTT)8 | TCAAGGCATCTGAGCATGAG | ACGAAGAGGAGAGGAAAGCC | 134 |
| chr5:98762600-98764615 | p5 | (TGCAA)3 | TGCTGATTTTTCTGGTGCTG | CCACTTGCGACTACTTGGGT | 220 |
| chr5:98762600-98764615 | p2 | (TA)7 | TGCTGATTTTTCTGGTGCTG | CCACTTGCGACTACTTGGGT | 220 |
| chr5:98762600-98764615 | p2 | (AT)19 | TGCTGATTTTTCTGGTGCTG | CCACTTGCGACTACTTGGGT | 220 |
| chr5:98926739-98928755 | p2 | (AT)8 | CCTTGGGTTTCTTCCTCCTC | TTTGAAAGTGGGCATGTGAA | 100 |
| chr5:98926739-98928755 | p2 | (AT)12 | CCTTGGGTTTCTTCCTCCTC | TTTGAAAGTGGGCATGTGAA | 100 |
| chr5:98927723-98929747 | p2 | (AT)8 | TCGATACTGCGCTTCTGCTA | CTGTTTATGGGTGTGTTGCG | 135 |
| chr5:98927723-98929747 | p2 | (AT)12 | TCGATACTGCGCTTCTGCTA | CTGTTTATGGGTGTGTTGCG | 135 |

| Marker ID | SSR type | SSR motif | FORWARD PRIMER (5'-3') | REVERSE PRIMER (5'-3') | size (bp) |
|---|---|---|---|---|---|
| chr5:98927723-98929747 | p2 | (AT)8 | TCGATACTGCGCTTCTGCTA | CTGTTTATGGGTGTGTTGCG | 135 |
| chr5:98927935-98929951 | p2 | (AT)12 | CCCTAGGACGAACTGCTCTG | AACGACGAAAATGGGTTTTG | 116 |
| chr5:98927935-98929951 | p2 | (AT)8 | CCCTAGGACGAACTGCTCTG | AACGACGAAAATGGGTTTTG | 116 |
| chr5:98940784-98942800 | p4 | (TATG)4 | TGAGGTGAACTTGAAAGGGG | TCTCCTCCCAAGCTCTCAAA | 231 |
| chr5:98998801-99000816 | p5 | (AATGC)3 | ATAGCCTCTCCTGGATGGGT | TGGGTCGTGACAAGTTGGTA | 272 |
| chr5:99020939-99022951 | p2 | (AT)6 | ATATAGGGGAAATGCTGCCC | GTCGCCCCTTTTAGCCTATT | 252 |
| chr5:99029629-99031641 | p2 | (CT)6 | GGATCGCTTTCCACCAATAA | TGAAACCGTGTGTGGACTGT | 187 |
| chr5:99029629-99031641 | p2 | (AT)6 | GGATCGCTTTCCACCAATAA | TGAAACCGTGTGTGGACTGT | 187 |
| chr5:99130498-99132510 | p2 | (AT)14 | CGGATAGACACAGCTGGACA | TCGGTTTGGTAACCCTTCAG | 131 |
| chr5:99130498-99132510 | p2 | (TA)6 | CGGATAGACACAGCTGGACA | TCGGTTTGGTAACCCTTCAG | 131 |
| chr5:99183395-99185410 | p5 | (TCATA)3 | TCTGCAAACATAGCCACTCG | GCTAATTTGGGGTATCGGGT | 219 |
| chr5:99190018-99192033 | p5 | (TCATA)3 | TCTGCAAACATAGCCACTCG | GCTAATTTGGGGTATCGGGT | 219 |
| chr5:99298419-99300435 | p2 | (TA)8 | TTTTGTTATTCCTTTCGCCG | CCGTATCCCCGTCACATATC | 171 |
| chr5:99316483-99318498 | p3 | (ATC)5 | AGATTGCAGCTCCGAGGTAA | GTACATCGCTCGTACTGGCA | 159 |
| chr5:99316483-99318498 | p3 | (CAA)5 | AGATTGCAGCTCCGAGGTAA | GTACATCGCTCGTACTGGCA | 159 |
| chr5:99316844-99318859 | p3 | (ATC)5 | CGGTGTTATCCCAAGCTGAT | CCTCATGTCGTTCGTTCCTT | 292 |
| chr5:99316844-99318859 | p3 | (CAA)5 | CGGTGTTATCCCAAGCTGAT | CCTCATGTCGTTCGTTCCTT | 292 |
| chr5:99379061-99381076 | p5 | (AATAT)3 | CATGCATGAGAGCCAAAGAA | CTAAAACGGATTGACCCGAA | 113 |
| chr5:99379061-99381076 | p5 | (ATTCA)3 | CATGCATGAGAGCCAAAGAA | CTAAAACGGATTGACCCGAA | 113 |
| chr5:99382173-99384191 | p6 | (TCATAT)3 | TTGGCACCTATACCACGTCA | TTCGGAGCCATAGTTTTTGG | 156 |
| chr5:99432297-99434312 | p3 | (ATC)5 | GACATGGCCACTACATGTGC | CCGCAGGTTCAGGTAGTCAT | 116 |
| chr5:99474631-99476646 | p5 | (TGAAA)3 | ATCATTATTGGGCTCGCAAC | TATGGAATGGATCCGATGGT | 125 |
| chr5:99510964-99512976 | p2 | (TG)6 | GGGCTATAATGATACCGCGA | AGTGTGCCCGACCATTTAAG | 283 |
| chr5:99545431-99547457 | p2 | (AT)13 | GGCCAAGAACTCACGAGAAG | TCCCAAGTGGGATTACAAGG | 261 |
| chr5:99545431-99547457 | p2 | (TA)7 | GGCCAAGAACTCACGAGAAG | TCCCAAGTGGGATTACAAGG | 261 |
| chr5:99575113-99577127 | p2 | (TA)7 | ATAAACTTCCCCTCCCCCTT | AAGAACAGCCGTGGATTGTC | 139 |
| chr5:99600952-99602967 | p5 | (ACACA)3 | TTACATACCTTGCCCGCTTC | GATAACCTGGCCGAGTTGAA | 266 |
| chr5:99663471-99665501 | p2 | (AT)15 | CCGACCTCTTCTGATCAAGC | GGTTAAAGTTCTTGGCGCAG | 116 |
| chr5:99663471-99665501 | p2 | (TA)6 | CCGACCTCTTCTGATCAAGC | GGTTAAAGTTCTTGGCGCAG | 116 |
| chr5:99740070-99742086 | p2 | (AC)8 | AACGATGGAAGCCACAAAAG | CGAATTTGTACCCCTAGCCA | 231 |
| chr5:99806847-99808862 | p2 | (AT)6 | AACATCTTCCGTTGGTACGC | TTGGAAGCCAAAACATGACA | 137 |
| chr5:99836857-99838869 | p2 | (TC)6 | CACCTAAACGAAGAGCAGGC | TGGTAGTTGGGCCGATGTAT | 219 |
| chr5:99859627-99861675 | p2 | (TA)24 | AATTCAGGGGACATCCTTCC | AGCAGGTTCGTATACGGGTG | 263 |
| chr5:99914540-99916555 | p3 | (GTT)5 | GGGCCTCATGATCGTCTTTA | AGTACATGGAGGCCGATGAG | 150 |
| chr5:99924421-99926436 | p5 | (ACACA)3 | AGTTACCCCGAGGCTCGTAT | GTTGTAGATCTTGCGGAGCC | 246 |
| chr5:99930178-99932190 | p6 | (TTTCAG)3 | GAAGGATTCCTATCCGCTCC | CCAACTTGCCAGGAAACTGT | 223 |

| Marker ID | SSR type | SSR motif | FORWARD PRIMER (5'-3') | REVERSE PRIMER (5'-3') | size (bp) |
|---|---|---|---|---|---|
| chr5:99930178-99932190 | p2 | (TA)6 | GAAGGATTCCTATCCGCTCC | CCAACTTGCCAGGAAACTGT | 223 |
| chr5:99940269-99942284 | p5 | (TGTTT)3 | TCATGCCAAAGGAAGAAAGG | TGCCCGATTTTCGTTAAGTC | 183 |
| chr5:99998614-100000628 | p2 | (AG)7 | ATCAGAAAGCGAACGCCTAA | AAGTGTTCAGGGGTGTTTGG | 233 |
| chr5:99998614-100000628 | p3 | (TTG)6 | ATCAGAAAGCGAACGCCTAA | AAGTGTTCAGGGGTGTTTGG | 233 |
| chr5:100006671-100008689 | p3 | (TTG)6 | CAGGATTGGCAGACAAGTGA | GTTCGTGCAGAGCATTTTCA | 263 |
| chr5:100071711-100073729 | p2 | (TA)32 | TGCAAACAGAGTCATGCACA | GTGCTCGGTATTTAGCTCCG | 185 |
| chr5:100071711-100073729 | p2 | (AT)9 | TGCAAACAGAGTCATGCACA | GTGCTCGGTATTTAGCTCCG | 185 |
| chr5:100098486-100100502 | p2 | (TA)8 | ACTCGGCTGAGTGCCATACT | CTGTGGATAGTGCGTCGAGA | 227 |
| chr5:100161398-100163410 | p2 | (CT)6 | AATTGTCCGTACCGCAAAAC | ACTCTTTCCCAAAATGCCCT | 268 |
| chr5:100275846-100277864 | p5 | (CGGTT)3 | CTAGATGCCCAGCAGGCTAC | CCGAAAACCAAACCGAATTA | 208 |
| chr5:100275846-100277864 | p6 | (AATCCC)3 | CTAGATGCCCAGCAGGCTAC | CCGAAAACCAAACCGAATTA | 208 |
| chr5:100275846-100277864 | p6 | (TTCGGT)3 | CTAGATGCCCAGCAGGCTAC | CCGAAAACCAAACCGAATTA | 208 |
| chr5:100302809-100304824 | p5 | (ATTGC)3 | AAGGCAAGGCAAAAGAGTGA | CGTTTACATGCTGTCTCCGA | 178 |
| chr5:100368650-100370665 | p5 | (GAAAT)3 | CCGACCATGTTATACGGACC | AACGAAAACGGACTCACTGG | 101 |
| chr5:100407108-100409123 | p5 | (TGAAT)3 | CTTGGCATTGGAAGGAGCTA | GAATTGCCCAAATGAATGCT | 249 |
| chr5:100653169-100655183 | p2 | (TA)7 | AAATTTCAATGGAGCGGATG | AGTCATGGCCCAAGAGATTG | 162 |
| chr5:100668371-100670386 | p5 | (TCATA)3 | AATCGGTGATCTGCACCTTC | AGATGACCGCTTGCCTCTTA | 121 |
| chr5:100668371-100670386 | p5 | (TCATA)3 | AATCGGTGATCTGCACCTTC | AGATGACCGCTTGCCTCTTA | 121 |
| chr5:100669204-100671219 | p5 | (TCATA)3 | TCATATCATAGCCGCACAGG | GGAAGTGGTGGCAGACAGTT | 172 |
| chr5:100669204-100671219 | p5 | (TCATA)3 | TCATATCATAGCCGCACAGG | GGAAGTGGTGGCAGACAGTT | 172 |
| chr5:100683515-100685530 | p5 | (AAATG)3 | CCTCACATACCTTGCCGTTT | TCCTAGAACAAGCGATCCGT | 262 |
| chr5:100695192-100697204 | p2 | (CT)6 | GGGTGGTTTGAAACAGGAGA | CTTCATTGGCGACACTTTCA | 141 |
| chr5:100713964-100715978 | p2 | (AT)7 | CAATTCCACCGTGCCTAGAT | GTAACCATCATACGCCACCC | 234 |
| chr5:100760589-100762603 | p2 | (TA)7 | TTCGGAGGCATGAATAATCC | AGTTCCCCCACCAGCTACTT | 258 |
| chr5:100820695-100822711 | p6 | (TCTTCA)3 | TCAGGTTCACGTCTCCTTCC | TCCGTCAACCAAAGATGTGA | 142 |
| chr5:100820695-100822711 | p2 | (TA)8 | TCAGGTTCACGTCTCCTTCC | TCCGTCAACCAAAGATGTGA | 142 |
| chr5:100832449-100834501 | p2 | (AT)26 | TTTGAATCCACAAATGGCAA | TCCAACTAGCCAAACAAGGC | 180 |
| chr5:100832449-100834501 | p2 | (TA)6 | TTTGAATCCACAAATGGCAA | TCCAACTAGCCAAACAAGGC | 180 |
| chr5:100832449-100834501 | p3 | (CAA)5 | TTTGAATCCACAAATGGCAA | TCCAACTAGCCAAACAAGGC | 180 |
| chr5:100832657-100834669 | p2 | (AT)26 | AAGGGCCTTGTTTGGCTAGT | AGGCCTTTGGAGGCTCTTAC | 216 |
| chr5:100832657-100834669 | p2 | (TA)6 | AAGGGCCTTGTTTGGCTAGT | AGGCCTTTGGAGGCTCTTAC | 216 |
| chr5:100832657-100834669 | p3 | (CAA)5 | AAGGGCCTTGTTTGGCTAGT | AGGCCTTTGGAGGCTCTTAC | 216 |
| chr5:100864908-100866930 | p2 | (TA)9 | CGGCTTGCCTATTTTGTCAT | ATGGAAATTGACACACGCAA | 258 |
| chr5:100864908-100866930 | p2 | (AT)8 | CGGCTTGCCTATTTTGTCAT | ATGGAAATTGACACACGCAA | 258 |
| chr5:100864908-100866930 | p2 | (AT)7 | CGGCTTGCCTATTTTGTCAT | ATGGAAATTGACACACGCAA | 258 |
| chr5:100864908-100866930 | p2 | (TA)11 | CGGCTTGCCTATTTTGTCAT | ATGGAAATTGACACACGCAA | 258 |

| Marker ID | SSR type | SSR motif | FORWARD PRIMER (5'-3') | REVERSE PRIMER (5'-3') | size (bp) |
|---|---|---|---|---|---|
| chr5:100864908-100866930 | p3 | (CAA)5 | CGGCTTGCCTATTTTGTCAT | ATGGAAATTGACACACGCAA | 258 |
| chr5:101029963-101031978 | p5 | (TAAAA)3 | CTCACCGCTTCCATCAAAAT | TTTCGGTCGTGCATTTTACA | 250 |
| chr5:101059280-101061292 | p2 | (TA)6 | GGTGGCCCATTTCTAACTCA | AGGGCCTTGTCCTGTCTTTT | 192 |
| chr5:101113908-101115920 | p2 | (AG)6 | GGGCAGAGGTCGTTTGTTTA | ACCGGTCAAACTGTTTCTGC | 106 |
| chr5:101113908-101115920 | p2 | (GT)11 | GGGCAGAGGTCGTTTGTTTA | ACCGGTCAAACTGTTTCTGC | 106 |
| chr5:101149406-101151421 | p5 | (CATAT)3 | AGCAAGAATGGCTCAAAGGA | TTATGCATGACTTGGGGGAT | 141 |
| chr5:101203794-101205814 | p2 | (AT)10 | CGAAGCATTACACTCGACCA | CAGCTTCGGCCTATTTTGAG | 104 |
| chr5:101203794-101205814 | p2 | (AT)8 | CGAAGCATTACACTCGACCA | CAGCTTCGGCCTATTTTGAG | 104 |
| chr5:101248995-101251007 | p2 | (GT)6 | CGAAGTACCCCTCATTTGGA | CACAACGGAACTGGTACACG | 193 |
| chr5:101327441-101329453 | p2 | (GA)6 | AGAATCCATAAGGATGGGGC | GTTGTTGGACGTGTGCATTC | 295 |
| chr5:101335840-101337858 | p6 | (CAACAC)3 | TATACAAGAAAGGCTGCCCG | CGACCTTACGTCACTCCGAT | 216 |
| chr5:101451694-101453706 | p2 | (AT)6 | GGGCCATGAAACAAACTCAT | ATTTCAGCGTTGGTACAGGG | 239 |
| chr5:101451728-101453744 | p2 | (AT)6 | GGGCCATGAAACAAACTCAT | ATTTCAGCGTTGGTACAGGG | 205 |
| chr5:101451745-101453777 | p2 | (AT)6 | GGGCCATGAAACAAACTCAT | ATTTCAGCGTTGGTACAGGG | 188 |
| chr5:101451803-101453829 | p2 | (AT)6 | GGGCCATGAAACAAACTCAT | ATTTCAGCGTTGGTACAGGG | 130 |
| chr5:101488813-101490827 | p2 | (AG)7 | CTGAGCGGCTACTAACTGGG | CCCACTCCAGAGAAGCGTAG | 117 |
| chr5:101488813-101490827 | p2 | (AT)6 | CTGAGCGGCTACTAACTGGG | CCCACTCCAGAGAAGCGTAG | 117 |
| chr5:101523224-101525236 | p2 | (TC)6 | CTCCAGATTGAGGGATGGAA | TTAGGAGCCCAGGGTAGCTT | 124 |
| chr5:101553525-101555549 | p3 | (TTG)8 | AATATGGGCATAAGCCCTCC | TTCGCGCAGGTCTAAATCTT | 293 |
| chr5:101603918-101605933 | p3 | (AAG)5 | TATTGGGGTGAAACCTGAGC | TCTTCCGGAAATTGAAATCG | 200 |
| chr5:101655495-101657515 | p2 | (AT)10 | GACATAGCCATGGTTGCCTT | ATCTTAGCGTGGTGGAATCG | 155 |
| chr5:101807528-101809543 | p5 | (AATGC)3 | CCAATCTGAACAATCTCGCA | ACCCATCCTCGAGAGGCTAT | 184 |
| chr5:101834752-101836818 | p2 | (TA)33 | CGAAAGGGAAACCTCTCTCC | CCGATAACCACGACCAACAT | 166 |
| chr5:101916023-101918039 | p4 | (AATA)4 | CGTAGGTTCAGGTAGGCAGG | AACTGTATGATCCGCCAAGG | 219 |
| chr5:101961951-101964013 | p2 | (AT)31 | CCTCTCAACGGACCTGTGTT | CTCGTTGGGAATTTGAGGAA | 214 |
| chr5:102038936-102041011 | p3 | (ATT)25 | GGGGGTGAATTTGGAAGTTT | AGGTAAGAGGAGAACCCCGA | 278 |
| chr5:102128720-102130738 | p3 | (TTG)6 | GCGACAGAAGGATTCGAAAG | CACGTTCGTGCAGAACACTT | 246 |
| chr5:102259944-102261992 | p2 | (AT)24 | TATCCGCAAGGCAACACATA | CCCAGATTGTGACGACACTG | 205 |
| chr5:102264852-102266867 | p3 | (TGA)5 | AATGCAACAAACCGTGCATA | TTTGGAGAGTTTTGGGGATG | 110 |
| chr5:102321001-102323016 | p3 | (TGT)5 | GGTCATGGAAGGATTGGAGA | TAAATCCCTGCTCTGGCAAC | 215 |
| chr5:102321001-102323016 | p3 | (ATG)5 | GGTCATGGAAGGATTGGAGA | TAAATCCCTGCTCTGGCAAC | 215 |
| chr5:102371916-102373956 | p2 | (AT)20 | GCCACCAGGTTATTCAGCAT | AGGGTCCTTGTACGGTGATG | 245 |
| chr5:102377763-102379779 | p4 | (GAAA)4 | GGAAACGGGACTCGTAAACA | GTTCTTGGGTGATGCCAAGT | 166 |
| chr5:102377763-102379779 | p2 | (AT)13 | GGAAACGGGACTCGTAAACA | GTTCTTGGGTGATGCCAAGT | 166 |
| chr5:102378650-102380676 | p4 | (GAAA)4 | AACCTGCCATGACTAGCACC | TCGGATTTTGGGGTTTGTAA | 269 |
| chr5:102378650-102380676 | p2 | (AT)13 | AACCTGCCATGACTAGCACC | TCGGATTTTGGGGTTTGTAA | 269 |

| Marker ID | SSR type | SSR motif | FORWARD PRIMER (5'-3') | REVERSE PRIMER (5'-3') | size (bp) |
|---|---|---|---|---|---|
| chr5:102471376-102473391 | p5 | (TTCAT)3 | TCTCCGAGCTTTTGAGGTGT | GTCAATCTGCCCGTCAAGTT | 213 |
| chr5:102478731-102480746 | p5 | (TTCAT)3 | TCTGCACTTATTTCAGCCCC | GCCTCACAGGGACACAGAAT | 225 |
| chr5:102482008-102484040 | p6 | (ATCCGT)4 | CATAAGCCGACGAGGCTAAC | TAGCGGAGCTCCATCTCATT | 185 |
| chr5:102482008-102484040 | p2 | (AT)16 | CATAAGCCGACGAGGCTAAC | TAGCGGAGCTCCATCTCATT | 185 |
| chr5:102490033-102492048 | p2 | (AG)6 | ACATAAGGCGACGAGGCTAA | TAGCGGAGCTCCATCTCATT | 199 |
| chr5:102490050-102492064 | p2 | (AG)6 | ACATAAGGCGACGAGGCTAA | TAGCGGAGCTCCATCTCATT | 182 |
| chr5:102490065-102492085 | p2 | (AG)6 | ACATAAGGCGACGAGGCTAA | TAGCGGAGCTCCATCTCATT | 167 |
| chr5:102567031-102569046 | p3 | (GTT)5 | CGAGGACTGGTAATCAGGGA | AATTTCCTTTTCGTTGCCCT | 269 |
| chr5:102587357-102589389 | p3 | (ATA)5 | TGTAAAAAGGACGGGACAGG | CGTACTGACTCCCCATTGCT | 280 |
| chr5:102587357-102589389 | p2 | (AT)16 | TGTAAAAAGGACGGGACAGG | CGTACTGACTCCCCATTGCT | 280 |
| chr5:102653474-102655550 | p2 | (TA)38 | TTATAAGTGTGGCGGGCTTC | CAGCCGAGTAGGGTCTTGAG | 213 |
| chr5:102752955-102755030 | p5 | (TATAT)15 | TGGTTATATTCGGGCATCGT | ACGGAAATATCGACGTCTCG | 165 |
| chr5:102904238-102906253 | p2 | (AT)6 | ACAACCGCACAACAATGAAA | GGAATGTGACAAGCGAGACA | 214 |
| chr5:102904238-102906253 | p5 | (TTATT)3 | ACAACCGCACAACAATGAAA | GGAATGTGACAAGCGAGACA | 214 |
| chr5:102904238-102906253 | p5 | (TTTTC)3 | ACAACCGCACAACAATGAAA | GGAATGTGACAAGCGAGACA | 214 |
| chr5:102913351-102915371 | p4 | (TGAT)5 | TCTCACAAAACCCCAACACA | AGGGGAAGGGAACGATAGAA | 273 |
| chr5:102946232-102948252 | p2 | (AT)10 | GAAACAATCGGCGTTAGCTC | GGGGGAAAGGAAACCAAATA | 294 |
| chr5:103026878-103028914 | p2 | (AT)12 | TACATGCCAACCCAAAGTGA | CCTTGCGAATCCAAACATCT | 272 |
| chr5:103026878-103028914 | p2 | (AT)18 | TACATGCCAACCCAAAGTGA | CCTTGCGAATCCAAACATCT | 272 |
| chr5:103160426-103162441 | p3 | (CAA)5 | ACATCACAACAAAGGCCACA | CACAGTTGACATGGGGTGAC | 188 |
| chr5:103166728-103168743 | p5 | (TTTTA)3 | GGAAAACATTTGAGGCGTGT | TGGCCTTTGTTGTGATGTGT | 169 |
| chr5:103166728-103168743 | p4 | (ATTT)4 | GGAAAACATTTGAGGCGTGT | TGGCCTTTGTTGTGATGTGT | 169 |
| chr5:103258928-103260940 | p2 | (TG)6 | CCCCCACTATGCGAGTAAGA | TCATACCTGGCCCTACCAAG | 148 |
| chr5:103268132-103270144 | p2 | (AT)6 | TGGTCGATATCCGATGTTCA | ATGTCTCGGTAGCACAACCC | 191 |
| chr5:103278294-103280309 | p5 | (GCATT)3 | CCTTCGGGTATGAAGTTGGA | CAGTGGGTATGCCAACAATG | 219 |
| chr5:103304817-103306842 | p5 | (CATTG)5 | TGTTCGGCTATCCAGGTAGG | AGGGCCCAACCTAAGACAAT | 241 |
| chr5:103328697-103330709 | p2 | (TG)6 | AAAGGTGCCTACGGGCTATT | CTCATACCTGGCCCTACCAA | 224 |
| chr5:103337901-103339913 | p2 | (AT)6 | TGGTCGATATCCGATGTTCA | ATGTCTCGGTAGCACAACCC | 191 |
| chr5:103348063-103350078 | p5 | (GCATT)3 | CCTTCGGGTATGAAGTTGGA | CAGTGGGTATGCCAACAATG | 219 |
| chr5:103374586-103376611 | p5 | (CATTG)5 | TGTTCGGCTATCCAGGTAGG | AGGGCCCAACCTAAGACAAT | 241 |
| chr5:103400514-103402534 | p5 | (CTTTT)4 | CTCTTTTATGGCGTCGGTTC | ACAAAGGCACAACCCAAGTC | 221 |
| chr5:103404724-103406744 | p2 | (AT)10 | ATGTGACCTCTCCCATGCTC | CGCGGTCTTCATTTTCTTTC | 263 |
| chr5:103465815-103467833 | p4 | (TTAT)4 | GTTCAGAAGATGAAAGCCGC | CTCCCACCTTGTCTCGGATA | 292 |
| chr5:103465815-103467833 | p3 | (TTG)6 | GTTCAGAAGATGAAAGCCGC | CTCCCACCTTGTCTCGGATA | 292 |
| chr5:103550754-103552772 | p6 | (CTTCTC)3 | TTTGGTTCAAGGCTAGTGGC | TCAGGTATGCACCTCCTTCC | 171 |
| chr5:103584476-103586494 | p3 | (TGT)6 | GCAGGACGGATGGGTATATG | CACGGGGCAAGACACTTTAT | 122 |

| Marker ID | SSR type | SSR motif | FORWARD PRIMER (5'-3') | REVERSE PRIMER (5'-3') | size (bp) |
|---|---|---|---|---|---|
| chr5:103747796-103749836 | p4 | (CATA)4 | GACTATGACGCATTGCCCTT | AATGCAAACCGTAGCACACA | 170 |
| chr5:103747796-103749836 | p2 | (TA)12 | GACTATGACGCATTGCCCTT | AATGCAAACCGTAGCACACA | 170 |
| chr5:103747838-103749850 | p4 | (CATA)4 | GACTATGACGCATTGCCCTT | AATGCAAACCGTAGCACACA | 128 |
| chr5:103747838-103749850 | p2 | (TA)12 | GACTATGACGCATTGCCCTT | AATGCAAACCGTAGCACACA | 128 |
| chr5:103881513-103883541 | p2 | (TA)14 | CACCACCAAAATCACCACAC | CTTCTGAGCAGCGACTGTTG | 299 |
| chr5:103988135-103990169 | p2 | (TA)17 | AACCAAATTGAGAGCATGGC | TTCGAACATGACCCTTGTGA | 109 |
| chr5:104216343-104218358 | p6 | (AAAATA)3 | ATTTTGCCCTCAATTTTCCC | CTGCAACTCAAGCACATGGT | 298 |
| chr5:104216343-104218358 | p5 | (CAAAC)3 | ATTTTGCCCTCAATTTTCCC | CTGCAACTCAAGCACATGGT | 298 |
| chr5:104227191-104229206 | p6 | (AAAATA)3 | ATTTTGCCCTCAATTTTCCC | CTGCAACTCAAGCACATGGT | 298 |
| chr5:104227191-104229206 | p5 | (CAAAC)3 | ATTTTGCCCTCAATTTTCCC | CTGCAACTCAAGCACATGGT | 298 |
| chr5:104350980-104352992 | p2 | (CA)6 | CGAAACCATATGAAACGGCT | TTGGTGCGCATTAATGTTGT | 116 |
| chr5:104354030-104356045 | p5 | (GGAAG)3 | GGTGTATTGGGCCCTTTTCT | GAGAAACTGCTCCTCCATGC | 121 |
| chr5:104440898-104442913 | p2 | (AT)6 | CATGACTAGTGCCCGTGCTA | GTTGCGAACCCTTCATTGTT | 243 |
| chr5:104440917-104442929 | p2 | (AT)6 | CATGACTAGTGCCCGTGCTA | GTTGCGAACCCTTCATTGTT | 224 |
| chr5:104450536-104452586 | p2 | (AT)25 | AGGCAGTTTAAGTGGGAGCA | GTTCCTAATGCCGCACAAGT | 238 |
| chr5:104450536-104452586 | p2 | (AT)10 | AGGCAGTTTAAGTGGGAGCA | GTTCCTAATGCCGCACAAGT | 238 |
| chr5:104464908-104466950 | p3 | (TAT)14 | TACCCGCAGAAGTGGAAATC | CATACCGCCACCATTCTCTT | 131 |
| chr5:104488129-104490199 | p2 | (AT)19 | TCCATGGCTATGTTGGTTGA | CCGAACCCCAAACTTATGTG | 214 |
| chr5:104488206-104490264 | p2 | (AT)19 | GAGAGAGGCAGTTCAGGTGG | TGTTCCTGATGCCGTACAAG | 155 |
| chr5:104728192-104730210 | p6 | (AACACA)3 | GGCTCATGTGGAGCTCTAGG | CCGCTCAAGAAGGAGAACTG | 224 |
| chr5:104728192-104730210 | p5 | (CTTTT)3 | GGCTCATGTGGAGCTCTAGG | CCGCTCAAGAAGGAGAACTG | 224 |
| chr5:104732191-104734212 | p3 | (TGT)7 | ATGTGACAAGAAAGGGCCAC | CCACACATCCACCACAACAT | 229 |
| chr5:104921435-104923449 | p2 | (AT)7 | CCATATCTGAGCGGCATTTT | TCCTGACCTAGATCCCAACG | 215 |
| chr5:104921435-104923449 | p3 | (TCT)7 | CCATATCTGAGCGGCATTTT | TCCTGACCTAGATCCCAACG | 215 |
| chr5:104951598-104953613 | p5 | (GATAT)3 | ACCCGCAAGACAAAATTCAG | GGACCGCCTCAGAACAAATA | 208 |
| chr5:105057485-105059500 | p3 | (TGA)5 | TGGTGTAGGGAGTATTCGGC | TTGGTGTTTTGTCCTCCCTC | 123 |
| chr5:105081526-105083544 | p2 | (TA)9 | TCGTTCCTTCGAGGTGAGAT | GCCATATAGGCGCGGTATTA | 162 |
| chr5:105087994-105090008 | p2 | (AG)7 | GATGGCACCGAAGGAGTAAA | GCCGTGCCATAACTATCCAT | 125 |
| chr5:105087994-105090008 | p3 | (TTG)6 | GATGGCACCGAAGGAGTAAA | GCCGTGCCATAACTATCCAT | 125 |
| chr5:105138072-105140087 | p5 | (CGGTT)3 | GGGTGTCAATGGTACGGTTC | AAAACCGAGATGATTGGACG | 183 |
| chr5:105138072-105140087 | p2 | (AT)8 | GGGTGTCAATGGTACGGTTC | AAAACCGAGATGATTGGACG | 183 |
| chr5:105208660-105210682 | p2 | (AT)11 | AGGCTTCACCCTCCCTTCTA | TGGGGTGGCGTATGTTATTT | 168 |
| chr5:105208660-105210682 | p5 | (GCTAT)3 | AGGCTTCACCCTCCCTTCTA | TGGGGTGGCGTATGTTATTT | 168 |
| chr5:105221095-105223121 | p2 | (AT)10 | CGAACTCCGAAGGTCCATAA | TGTGTAGTGGCAGCCTTGTC | 210 |
| chr5:105221095-105223121 | p2 | (AT)9 | CGAACTCCGAAGGTCCATAA | TGTGTAGTGGCAGCCTTGTC | 210 |
| chr5:105260416-105262438 | p2 | (TA)6 | AGAAGTATTGGGGAGGGGTG | TGCGACCATCGAAGTACAAG | 124 |

| Marker ID | SSR type | SSR motif | FORWARD PRIMER (5'-3') | REVERSE PRIMER (5'-3') | size (bp) |
|---|---|---|---|---|---|
| chr5:105263613-105265634 | p3 | (AAC)7 | GCAGCTCTCCAAGCTTGTTT | ACTTTGATTGAGGGCCTCCT | 235 |
| chr5:105270088-105272103 | p5 | (TATTT)3 | CTTTGGGTGCTAGCAAAAGC | CTCCGGCTATTCAAGAAACG | 189 |
| chr5:105270088-105272103 | p5 | (CAAAA)3 | CTTTGGGTGCTAGCAAAAGC | CTCCGGCTATTCAAGAAACG | 189 |
| chr5:105324518-105326562 | p2 | (AT)17 | TGCAGGTTATATCTTCCGCC | TGTCTACCTGTGGGCATGAA | 156 |
| chr5:105324518-105326562 | p6 | (TGTATA)3 | TGCAGGTTATATCTTCCGCC | TGTCTACCTGTGGGCATGAA | 156 |
| chr5:105324518-105326562 | p2 | (AT)22 | TGCAGGTTATATCTTCCGCC | TGTCTACCTGTGGGCATGAA | 156 |
| chr5:105374682-105376706 | p3 | (GTT)8 | GATTGCCGATGATGTCACAC | TTGCCCTTAAGTTTTGGTGG | 162 |
| chr5:105374682-105376706 | p5 | (ATTTT)6 | GATTGCCGATGATGTCACAC | TTGCCCTTAAGTTTTGGTGG | 162 |
| chr5:105374891-105376921 | p3 | (GTT)8 | CAGTGGAAGACAAGATGCGA | TACTTCTTCGGCCTACCCCT | 237 |
| chr5:105374891-105376921 | p5 | (ATTTT)6 | CAGTGGAAGACAAGATGCGA | TACTTCTTCGGCCTACCCCT | 237 |
| chr5:105670718-105672751 | p4 | (CAGA)4 | GAACGGGGTCAGTACGAAAA | ATCCGTCCGTCTGGACTATG | 256 |
| chr5:105670718-105672751 | p3 | (ATT)11 | GAACGGGGTCAGTACGAAAA | ATCCGTCCGTCTGGACTATG | 256 |
| chr5:105720626-105722640 | p2 | (TA)7 | TATATGTCCGTCCCCGGTTA | AATTCTTCTCCGGTCTCCGT | 194 |
| chr5:105720626-105722640 | p2 | (AT)8 | TATATGTCCGTCCCCGGTTA | AATTCTTCTCCGGTCTCCGT | 194 |
| chr5:105764145-105766157 | p2 | (TA)6 | CAAGGCTTCGAATGAACCAT | ACAGGCGACCATAGTCCAAC | 267 |
| chr5:105766278-105768304 | p2 | (AT)7 | CCCGAGATGGACACTCGTAT | CCTTGTCGACTTGCGTATCA | 201 |
| chr5:105766278-105768304 | p5 | (ATACG)3 | CCCGAGATGGACACTCGTAT | CCTTGTCGACTTGCGTATCA | 201 |
| chr5:105766278-105768304 | p2 | (AT)13 | CCCGAGATGGACACTCGTAT | CCTTGTCGACTTGCGTATCA | 201 |
| chr5:105771328-105773343 | p5 | (TTGGA)3 | GGAAAGTTGCGGTGGTTAAA | ACCTTACCAACCCTCAACCC | 245 |
| chr5:106026011-106028023 | p2 | (TA)6 | GGAGTCGGAACACTTCCAAA | TTATACAGAAGGCATCGGGC | 105 |
| chr5:106113626-106115702 | p3 | (ATG)5 | TGTTGTCCGATTTTCGTTGA | TTCCAACTTGCAAATCCACA | 240 |
| chr5:106113626-106115702 | p2 | (TA)6 | TGTTGTCCGATTTTCGTTGA | TTCCAACTTGCAAATCCACA | 240 |
| chr5:106153744-106155766 | p2 | (AT)11 | TTGCACCGCTTCTTCTTTCT | TGGATCACATTTTGCCTTCA | 175 |
| chr5:106237340-106239354 | p2 | (AT)7 | TTTGGAAAAAGCTCGAAGGA | AAAAATCTTGACGTGTGGGC | 252 |
| chr5:106453465-106455480 | p3 | (GTT)5 | TCCTCTTTTCAGAAGCCCAA | TTGCTTGGCAGATTAAGCCT | 239 |
| chr5:106530982-106532998 | p2 | (GA)8 | AAGAGTGGACCTCAGGGGTT | TTTTCGGCTTGCTCAGTTTT | 251 |
| chr5:106530982-106532998 | p2 | (AT)9 | AAGAGTGGACCTCAGGGGTT | TTTTCGGCTTGCTCAGTTTT | 251 |
| chr5:106560393-106562411 | p3 | (CAA)6 | TCCCATATGTTGGAGGGTGT | TTGACCCATGACTAGGAGGC | 299 |
| chr5:106625283-106627301 | p3 | (AAC)6 | ATCTGGGATGCACGAGATTC | TGATCGGGGTTCGTAAGTTC | 168 |
| chr5:106637566-106639582 | p2 | (AT)8 | GCAGCTGGTTCTTCCTCATC | GCTCAGGTTTCACCCCAATA | 199 |
| chr5:106702782-106704798 | p2 | (TA)8 | TTCAGGGCTGCTGTAGCTTT | GGCAGTTCGTATCCTGGTGT | 178 |
| chr5:106721767-106723803 | p2 | (TA)18 | CACACAATTCAACGGCATTC | TTGAATTTGGAAGGTTTCGG | 146 |
| chr5:106721767-106723803 | p2 | (AT)10 | CACACAATTCAACGGCATTC | TTGAATTTGGAAGGTTTCGG | 146 |
| chr5:106843892-106845914 | p2 | (AT)11 | CTAACCCCGCCCACTAAAAT | TCAGGCACTCATCACGACTC | 187 |
| chr5:106847685-106849721 | p2 | (AT)18 | CACTAGCCGACGAGGCTATC | GGAGCTACAGCTTTTGGGTG | 107 |
| chr5:106849198-106851219 | p3 | (ACA)7 | TTGTGGAAGGTTTTTGGAGC | TGGGTGATCCCTGGTATGAT | 260 |

| Marker ID | SSR type | SSR motif | FORWARD PRIMER (5'-3') | REVERSE PRIMER (5'-3') | size (bp) |
|---|---|---|---|---|---|
| chr5:107096187-107098202 | p5 | (CATAT)3 | TGAGGCATATCGGATGACAA | ACTATTCCCGCTTTGCCTTT | 160 |
| chr5:107124399-107126411 | p2 | (CA)6 | TTGCGCTGAATGAATAGTGC | AGGACGCAATTCCAGGTATG | 107 |
| chr5:107166869-107168889 | p5 | (AATGA)4 | CTCAGGTGGGCAATACCTGT | ACAGCAATGGAAGAACGGAC | 145 |
| chr5:107182598-107184616 | p3 | (TGT)6 | GAAATGAAGCTTGGAACCCA | TGCCTTTAAGTTTTGGTGGC | 107 |
| chr5:107336434-107338449 | p5 | (ATGAT)3 | CCCGAATTTCTTCTATGGCA | GCCACGAGCACTGTACTTCA | 252 |
| chr5:107385741-107387756 | p5 | (AATAA)3 | ACATGGGCACAGTTTGATGA | ACTTGAGTGGGCCATAATGC | 200 |
| chr5:107386999-107389014 | p5 | (TATAT)3 | ATGGGAAATTGTGGGTTGAA | GTCCATAAATCCGTAGCCCA | 205 |
| chr5:107401317-107403333 | p2 | (AT)8 | ATGGATTGGTTGGCTTCTTG | CCTGAACCCGGACTAAATGA | 259 |
| chr5:107401317-107403333 | p4 | (CATA)4 | ATGGATTGGTTGGCTTCTTG | CCTGAACCCGGACTAAATGA | 259 |
| chr5:107583298-107585313 | p2 | (AT)23 | ACCGTCACCCTGTCTACCTG | ACTTACGCATATCATGGCCC | 125 |
| chr5:107583298-107585313 | p3 | (CAA)5 | ACCGTCACCCTGTCTACCTG | ACTTACGCATATCATGGCCC | 125 |
| chr5:107589979-107592011 | p2 | (GA)16 | TAAAACGGTTGGAAAGACGC | CACCTCGGCCAAATAAGTGT | 240 |
| chr5:107589979-107592011 | p2 | (TA)11 | TAAAACGGTTGGAAAGACGC | CACCTCGGCCAAATAAGTGT | 240 |
| chr5:107596466-107598486 | p2 | (TA)10 | CTTGGTTTCCTTGATTCCGA | ATTCCCTATGGGATTCGACC | 238 |
| chr5:107667486-107669498 | p2 | (AT)6 | TTCCTGGAGTGGAAAATTGG | ACCATGGAAGTGCAGGAAAG | 285 |
| chr5:107698984-107701000 | p2 | (AT)8 | ATGGGTTGTGCTACCGAGAC | GAGTTTAGGTGCGCTTGGTC | 167 |
| chr5:107730812-107732826 | p2 | (AT)7 | TCGAGGTAAGGGTTCTGGTG | CGGAAAGTTGCAAGACCATT | 241 |
| chr5:107838366-107840384 | p6 | (AGAAAA)3 | TTGAATCTACGGCAGAGGCT | GTCATGTCAGGCGAAATGTG | 259 |
| chr5:107838366-107840384 | p2 | (AT)6 | TTGAATCTACGGCAGAGGCT | GTCATGTCAGGCGAAATGTG | 259 |
| chr5:107934781-107936799 | p6 | (TCAGTT)3 | TATGGGTGATGGGGAATGAT | ATTCCAGCCTTTGAGCGTAA | 168 |
| chr5:108047800-108049815 | p5 | (TGATC)3 | AATTGGAATTGAAGCCATGC | TGTTCCGACTCCGTTTAAGG | 216 |
| chr5:108292028-108294046 | p2 | (AT)39 | CTGTCTTCCTGCCTACCTGC | TAAGAATGCATGACATCCGC | 272 |
| chr5:108292028-108294046 | p3 | (ATC)6 | CTGTCTTCCTGCCTACCTGC | TAAGAATGCATGACATCCGC | 272 |
| chr5:108328990-108331006 | p4 | (CATT)4 | CAGGACTGCGAAAACCTAGC | TCCCATTTGCTTGTGGTGTA | 212 |
| chr5:108364333-108366348 | p5 | (GTTCG)3 | TTTAGTTTGCAGGTGCGATG | ACACACGACCACATCCAGAA | 157 |
| chr5:108379620-108381744 | p2 | (AT)62 | GTTGTCGTTGTGTTGGTTGC | TCGCATGCGTATCTCAAGTC | 267 |
| chr5:108402974-108404988 | p2 | (TA)8 | GCGTGCATAACCCTCAAAGT | CACAGGGCCAGGATTACTGT | 258 |
| chr5:108402990-108405004 | p2 | (TA)8 | GCGTGCATAACCCTCAAAGT | CACAGGGCCAGGATTACTGT | 242 |
| chr5:108403007-108405023 | p2 | (TA)8 | GCGTGCATAACCCTCAAAGT | CACAGGGCCAGGATTACTGT | 225 |
| chr5:108442511-108444526 | p3 | (TAT)5 | GTCCGTATAATGGACCGCAG | TCAGTAACCATCCCTTTCCG | 246 |
| chr5:108450708-108452723 | p3 | (TAT)5 | GGTCCCATGAGGATTTGAGA | CGATTCTGCGGTCCATTATT | 130 |
| chr5:108540948-108542963 | p3 | (TGT)5 | TGGTTCATGATAAGGCCACA | CCATGGCCGAATACTTCTGT | 287 |
| chr5:108607804-108609825 | p3 | (ATA)7 | TGCAAATTCGGGTACTAGGG | CTGGTTGTCCGCATGTTATG | 172 |
| chr5:108636254-108638276 | p2 | (AC)6 | ACCACCAATGATCCCACATT | TTGGAAGATTTCGGGTGAAG | 251 |
| chr5:108636254-108638276 | p2 | (TA)11 | ACCACCAATGATCCCACATT | TTGGAAGATTTCGGGTGAAG | 251 |
| chr5:108636254-108638276 | p2 | (AT)9 | ACCACCAATGATCCCACATT | TTGGAAGATTTCGGGTGAAG | 251 |

| Marker ID | SSR type | SSR motif | FORWARD PRIMER (5'-3') | REVERSE PRIMER (5'-3') | size (bp) |
|---|---|---|---|---|---|
| chr5:108636383-108638401 | p2 | (AC)6 | ACCACCAATGATCCCACATT | TTGGAAGATTTCGGGTGAAG | 122 |
| chr5:108636383-108638401 | p2 | (TA)11 | ACCACCAATGATCCCACATT | TTGGAAGATTTCGGGTGAAG | 122 |
| chr5:108636383-108638401 | p2 | (AT)9 | ACCACCAATGATCCCACATT | TTGGAAGATTTCGGGTGAAG | 122 |
| chr5:108826776-108828818 | p2 | (AT)21 | CTTTCGCCACAACCCTAGAA | TGCAGCTTCCCGATTAGACT | 158 |
| chr5:109011779-109013794 | p3 | (ATA)5 | CCATGAATTGTGATGGGTCA | AGCCTATCCAAACAGGCTCA | 148 |
| chr5:109192234-109194248 | p2 | (CA)7 | ACCTTACATACCTGTGCCGC | TTACAGGGGAGATACTGCCG | 194 |
| chr5:109257623-109259643 | p2 | (TC)10 | TTCCTGTCCTGGATGCTACC | ATGCTGAGGACGCTGAAACT | 168 |
| chr5:109299475-109301487 | p2 | (TA)6 | TGTCGCGCATGTGATTTATT | CTAAGCCAACCTTGGTGGAA | 252 |
| chr5:109336386-109338398 | p2 | (TA)6 | TGTCGCGCATGTGATTTATT | CTAAGCCAACCTTGGTGGAA | 252 |
| chr5:109425388-109427406 | p6 | (CCTTTT)3 | AGGGAAGCATACACAATGCC | GTTTTGTCTGCCTGGTCCAT | 142 |
| chr5:109561745-109563760 | p2 | (TA)6 | GCCTATGGTGCAGGGTAGAG | ACATACTTGCCAACCAAGGC | 101 |
| chr5:109561745-109563760 | p5 | (TTATG)3 | GCCTATGGTGCAGGGTAGAG | ACATACTTGCCAACCAAGGC | 101 |
| chr5:109568161-109570175 | p2 | (AT)7 | ACCACGGCTAGAACATCCAC | CTGTGACAGACAGCGTCCAT | 220 |
| chr5:109576435-109578453 | p6 | (AAGTGA)3 | TTCCAACCATCCTCGATCTC | CCAATAACGCAAAAAGGGAA | 165 |
| chr5:109585665-109587677 | p2 | (AT)8 | GGTTTTTCTAAGGGCGTTCC | CATTGAAGCGGTTTTACCGT | 281 |
| chr5:109585784-109587800 | p2 | (AT)8 | GGTTTTTCTAAGGGCGTTCC | CATTGAAGCGGTTTTACCGT | 162 |
| chr5:109590307-109592328 | p3 | (TAT)26 | ACAGAACTAAGCCCCACACG | CCTTTTGGTAACGACGAGGA | 229 |
| chr5:109590307-109592328 | p3 | (TAT)7 | ACAGAACTAAGCCCCACACG | CCTTTTGGTAACGACGAGGA | 229 |
| chr5:109590307-109592328 | p2 | (TA)17 | ACAGAACTAAGCCCCACACG | CCTTTTGGTAACGACGAGGA | 229 |
| chr5:109743926-109745944 | p2 | (TA)9 | TTTCGAATGACTTTGGACCC | AAGCCCAAAACACCCTTTCT | 188 |
| chr5:109744845-109746881 | p2 | (TA)9 | CGCAGTATGTTTCCCCTTGT | TTAAGGACTGTTTGGGCTGC | 262 |
| chr5:109786120-109788135 | p5 | (AATGC)3 | CTTTTCCACCTCAGTAGGCG | GTCTAGACCCAGTCGTGGGA | 197 |
| chr5:109835445-109837461 | p4 | (CACG)4 | CCAAGAAGGTTCCACAAGGA | ATGAAAGTAAAGCTGCCCGA | 140 |
| chr5:109835445-109837461 | p2 | (AT)6 | CCAAGAAGGTTCCACAAGGA | ATGAAAGTAAAGCTGCCCGA | 140 |
| chr5:109835445-109837461 | p2 | (AT)13 | CCAAGAAGGTTCCACAAGGA | ATGAAAGTAAAGCTGCCCGA | 140 |
| chr5:110071403-110073449 | p2 | (TA)23 | GGTGGCATCGGATAAAGAGA | TTCGGAAGACCCACAAAAAC | 264 |
| chr5:110071403-110073449 | p2 | (AT)6 | GGTGGCATCGGATAAAGAGA | TTCGGAAGACCCACAAAAAC | 264 |
| chr5:110082018-110084040 | p2 | (AT)9 | ACTTTTGTCACGACCCAACC | CAGTATTGTCGGCTTGCGTA | 142 |
| chr5:110082018-110084040 | p2 | (AT)11 | ACTTTTGTCACGACCCAACC | CAGTATTGTCGGCTTGCGTA | 142 |
| chr5:110095381-110097396 | p2 | (AT)8 | GACGCCTTCCTGTCGTAGTC | AAGAAGCTGCTGGGCATTTA | 133 |
| chr5:110115943-110117958 | p5 | (AAAGG)3 | CCCCAAATTCTAGGGTTGGT | CCCAACTTCATACCCGAAGA | 200 |
| chr5:110115943-110117958 | p5 | (TGCAT)3 | CCCCAAATTCTAGGGTTGGT | CCCAACTTCATACCCGAAGA | 200 |
| chr5:110120918-110122930 | p2 | (CA)6 | TGAGGGTTTTAGGGTGATCG | TACCCTTCAGAATCCGCATC | 201 |
| chr5:110165526-110167542 | p4 | (ACAT)4 | CCGACCCCTTACTTAGAGCC | CTCTGCAAGCCGATGTATCA | 108 |
| chr5:110165526-110167542 | p3 | (ATC)5 | CCGACCCCTTACTTAGAGCC | CTCTGCAAGCCGATGTATCA | 108 |
| chr5:110295513-110297529 | p2 | (AT)8 | AGAGTGCTCCTCGTCATCGT | TGATATACAGGGAGGCGGTC | 182 |

| Marker ID | SSR type | SSR motif | FORWARD PRIMER (5'-3') | REVERSE PRIMER (5'-3') | size (bp) |
|---|---|---|---|---|---|
| chr5:110295513-110297529 | p2 | (AT)6 | AGAGTGCTCCTCGTCATCGT | TGATATACAGGGAGGCGGTC | 182 |
| chr5:110397343-110399357 | p2 | (AG)7 | TCCTCGAGTCCATCAATTCC | GAGGTATTCCGTCCGTAGCA | 178 |
| chr5:110397343-110399357 | p3 | (TTG)7 | TCCTCGAGTCCATCAATTCC | GAGGTATTCCGTCCGTAGCA | 178 |
| chr5:110478249-110480267 | p2 | (AT)6 | TGCTCCTATGCCTCAATCCT | GGCCATGTGGCTTACCTAGA | 165 |
| chr5:110478249-110480267 | p6 | (ATATAC)3 | TGCTCCTATGCCTCAATCCT | GGCCATGTGGCTTACCTAGA | 165 |
| chr5:110483710-110485736 | p4 | (ATGA)4 | TCAGCCGCGTAAGAAAAAGT | ATACTTCGGAGAGGGCCATT | 175 |
| chr5:110483710-110485736 | p2 | (TA)13 | TCAGCCGCGTAAGAAAAAGT | ATACTTCGGAGAGGGCCATT | 175 |
| chr5:110507200-110509215 | p3 | (TCA)5 | AGGCACCTTCAGAACAATGG | GTCCGTTTTCATTGGATGCT | 214 |
| chr5:110517442-110519457 | p5 | (GATTG)3 | TGCGTTGTTAGCGTACCTTG | ACATCGGCCTTCTCACAATC | 291 |
| chr5:110721963-110723975 | p2 | (TA)6 | TAGACTGAGGGGATTGCACC | TTGGAGAAACCGGACTAACG | 117 |
| chr5:110900131-110902147 | p2 | (GA)8 | TGAGTTTGCAGTGAAGCCAG | TTCGAAGTGGGTTGGGTTAG | 298 |
| chr5:110900131-110902147 | p2 | (TA)7 | TGAGTTTGCAGTGAAGCCAG | TTCGAAGTGGGTTGGGTTAG | 298 |
| chr5:110900798-110902812 | p2 | (GA)8 | CAAAAACCGGTTCGGACTTA | CTCCTCCAGTCCGAAGATCA | 149 |
| chr5:110900798-110902812 | p2 | (TA)7 | CAAAAACCGGTTCGGACTTA | CTCCTCCAGTCCGAAGATCA | 149 |
| chr5:111045303-111047318 | p3 | (ATA)5 | CACCATGGGTGATGGTATGA | ACTTTCGGAGGGGTCAGTTT | 213 |
| chr5:111090221-111092241 | p2 | (AT)10 | TCTGAAGGAAACTGGGATGG | CGCAATTTTCATTCGTTGTG | 133 |
| chr5:111204578-111206593 | p5 | (ATGCA)3 | GCCTGAATCATCAGTGCAGA | CGAGTATCCAGGGTGAGCAT | 283 |
| chr5:111305254-111307266 | p2 | (AT)6 | TTATGCCCTCACCAAAGACC | CCGTCCCTGAGCCTACATTA | 182 |
| chr5:111326101-111328121 | p4 | (CAAG)5 | AACTAAGAGGGCATAGCGCA | TTCCCTTCTTTCTTCCCCAT | 201 |
| chr5:111494916-111496931 | p2 | (AT)10 | TTGCATAGGTTACTTCCCCG | GCAATATGGCTTGCACTGAA | 129 |
| chr5:111494916-111496931 | p5 | (AAAAT)3 | TTGCATAGGTTACTTCCCCG | GCAATATGGCTTGCACTGAA | 129 |
| chr5:111501031-111503073 | p2 | (TA)21 | CGTGTTGCCATGGATAAGAA | TTGCCCACATACGCTATCCT | 107 |
| chr5:111534729-111536744 | p3 | (AAT)5 | CAACATCCTTGTTGTCACCG | GGAAGGTTCAACCCATGCTA | 225 |
| chr5:111534729-111536744 | p2 | (GT)6 | CAACATCCTTGTTGTCACCG | GGAAGGTTCAACCCATGCTA | 225 |
| chr5:111701336-111703351 | p4 | (AAGA)6 | AGCCCAGCCCCATATAAATC | GGATTCAAAATCTGGGGGAT | 189 |
| chr5:111701336-111703351 | p3 | (AAG)5 | AGCCCAGCCCCATATAAATC | GGATTCAAAATCTGGGGGAT | 189 |
| chr5:111818898-111820912 | p3 | (TGT)5 | CCAATGGGACTAGCGTGATT | TTCACTTGCTGACAATTCGG | 300 |
| chr5:111855587-111857602 | p5 | (CATTG)3 | TATTATGGCCTTGTGGGCTC | GTCTCACCAAGGGTAAGCCA | 107 |
| chr5:111947673-111949685 | p2 | (CA)6 | TTTTGACCCAAAGAATTCCG | CAGGATGTGAAAGAGGCCAT | 182 |
| chr5:112072207-112074225 | p6 | (CATATA)3 | ATGAGCCGTGACAAGTACCC | TGTATGTGTTGGCCGTGAGT | 173 |
| chr5:112114003-112116021 | p6 | (GGTAGA)3 | GATGCCTTCCTCCGACATTA | GTTGTTGGCCTTCTTCCAAA | 265 |
| chr5:112232090-112234106 | p4 | (GAAA)4 | CGAAAACGCCTGATCTTCTC | TAATGGCAAGTTGTGGTGGA | 130 |
| chr5:112323074-112325089 | p3 | (TAT)5 | TGAATCCGTGCTTCCAAAA | TGCTAGCACAGGCCCAATA | 178 |
| chr5:112323074-112325089 | p5 | (ATATG)3 | TGAATCCGTGCTTCCAAAA | TGCTAGCACAGGCCCAATA | 178 |
| chr5:112382050-112384064 | p2 | (AT)7 | GGTTGAGCTGGAATCCCATA | TTTCTTGGTCGGATGGACTC | 262 |
| chr5:112447422-112449464 | p3 | (TTG)6 | GGAGAGTCGGGTTTGAATGA | AACCTTCAGAAAATTGGGCA | 139 |

| Marker ID | SSR type | SSR motif | FORWARD PRIMER (5'-3') | REVERSE PRIMER (5'-3') | size (bp) |
|---|---|---|---|---|---|
| chr5:112447422-112449464 | p3 | (ATG)6 | GGAGAGTCGGGTTTGAATGA | AACCTTCAGAAAATTGGGCA | 139 |
| chr5:112447422-112449464 | p2 | (TA)21 | GGAGAGTCGGGTTTGAATGA | AACCTTCAGAAAATTGGGCA | 139 |
| chr5:112472257-112474279 | p3 | (TAT)5 | GGGGAGGATAATAAGAGGCG | CAGTTGATCAGGTGGTGGTG | 176 |
| chr5:112472257-112474279 | p2 | (TA)11 | GGGGAGGATAATAAGAGGCG | CAGTTGATCAGGTGGTGGTG | 176 |
| chr5:112485659-112487677 | p6 | (TGTTAA)3 | TCATCTGGACCCTTCTGGAC | GCAAAGAAGTCGGTCCTCAG | 179 |
| chr5:112485659-112487677 | p2 | (TA)16 | TCATCTGGACCCTTCTGGAC | GCAAAGAAGTCGGTCCTCAG | 179 |
| chr5:112521565-112523609 | p2 | (AT)22 | GCCTCTATGCGAGGTTTGAG | GGTCATGGCCAAAACCTTTA | 287 |
| chr5:112651065-112653077 | p2 | (AG)10 | GGCCACATCTCGCAATCTAT | AGAGTGTGGGTCCAGGTGAC | 218 |
| chr5:112651065-112653077 | p2 | (AT)6 | GGCCACATCTCGCAATCTAT | AGAGTGTGGGTCCAGGTGAC | 218 |
| chr5:112726293-112728313 | p2 | (AT)27 | CACAACCCATTTCCAGCTTT | TCCTCCGTGATTTCAAAAGG | 256 |
| chr5:112726293-112728313 | p2 | (AT)10 | CACAACCCATTTCCAGCTTT | TCCTCCGTGATTTCAAAAGG | 256 |
| chr5:112793322-112795366 | p2 | (AT)22 | ACTCACAGCCCATTTCCATC | TCCTCCGTGATTTCAAAAGG | 257 |
| chr5:112890927-112892957 | p2 | (TA)8 | CCGTGAGTTTGGGACTGATT | ATTTTGACGCTCACAATCCC | 189 |
| chr5:112890966-112892988 | p2 | (TA)8 | CCGTGAGTTTGGGACTGATT | ATTTTGACGCTCACAATCCC | 150 |
| chr5:112896240-112898264 | p2 | (TA)12 | CCGTGAGTTTGGGACTGATT | ACTTGCTGCCCAAAAACACT | 204 |
| chr5:112945967-112947979 | p2 | (TC)6 | AGGGACTGCCTCTACCCATT | TTCAAGGTGATCGTACTGCG | 117 |
| chr5:113000112-113002142 | p2 | (AT)15 | TAAATTGAGCCGTGTAGGGG | ACAACATGCCCACAAAACAA | 118 |
| chr5:113000112-113002142 | p2 | (TA)6 | TAAATTGAGCCGTGTAGGGG | ACAACATGCCCACAAAACAA | 118 |
| chr5:113002492-113004526 | p2 | (AG)17 | TAAACCGGCTTCAGAGTGCT | GCAGCTGGGGCATTAGTAAG | 175 |
| chr5:113146401-113148419 | p2 | (TA)8 | CATCACAACCTACCTGGCCT | TCGGAGGTTACGACCTTACG | 210 |
| chr5:113146401-113148419 | p3 | (CAA)6 | CATCACAACCTACCTGGCCT | TCGGAGGTTACGACCTTACG | 210 |
| chr5:113157622-113159640 | p2 | (TA)8 | CATCACAACCTACCTGGCCT | TCGGAGGTTACGACCTTACG | 210 |
| chr5:113157622-113159640 | p3 | (CAA)6 | CATCACAACCTACCTGGCCT | TCGGAGGTTACGACCTTACG | 210 |
| chr5:113188444-113190464 | p2 | (AT)17 | GGGTCGGTGAGACAGAATGT | GGCCAAAGAATGTCCTGAAA | 276 |
| chr5:113311522-113313540 | p2 | (AT)9 | TGCGCTAGTTGGTTGTCTTG | AACTTGCTGCCCAAAAACAC | 264 |
| chr5:113375290-113377302 | p2 | (TC)6 | TGTGTGGCCAAAAGTTTTCA | ACTCTTTCCCAAAATGCCCT | 244 |
| chr5:113475354-113477368 | p2 | (GA)7 | ACGACCACTGTGCACTTCAG | CTCTGAAAATGCGGGGTTTA | 259 |
| chr5:113478574-113480586 | p2 | (GA)6 | GCCGTAGAAAATAAACGCCA | TGTCAATGTCTGAGGAAGCG | 281 |
| chr5:113482004-113484026 | p2 | (TA)11 | AGGAGACCCGGTCAGAATTT | TCCAAATCCTCCAACAGACC | 156 |
| chr5:113544554-113546570 | p2 | (AT)12 | CCCAAATAATGTCACGACCC | TGTTCTGTCGTGGTAGCAGC | 261 |
| chr5:113544554-113546570 | p2 | (AT)8 | CCCAAATAATGTCACGACCC | TGTTCTGTCGTGGTAGCAGC | 261 |
| chr5:113554048-113556106 | p2 | (AT)29 | CGCAGCATCTGTCGAATAAA | GTTCCAGCACCTGAGTCTCC | 101 |
| chr5:113554048-113556106 | p2 | (AT)30 | CGCAGCATCTGTCGAATAAA | GTTCCAGCACCTGAGTCTCC | 101 |
| chr5:113554254-113556314 | p2 | (AT)29 | CTGCTGCACTAGGCTCCTCT | ATGTTTACAGCGATGCCTCC | 174 |
| chr5:113554254-113556314 | p2 | (AT)30 | CTGCTGCACTAGGCTCCTCT | ATGTTTACAGCGATGCCTCC | 174 |
| chr5:113606109-113608121 | p2 | (AT)6 | AGCTACATTCTCTTGCCCGA | GGGACCCGATTACCCTAAGA | 200 |

| Marker ID | SSR type | SSR motif | FORWARD PRIMER (5'-3') | REVERSE PRIMER (5'-3') | size (bp) |
|---|---|---|---|---|---|
| chr5:113807097-113809112 | p2 | (TA)10 | TGTGAATCCTGGCGATACAA | TGCGATGTCGGTTTATGTGT | 159 |
| chr5:113807097-113809112 | p5 | (TTAAA)3 | TGTGAATCCTGGCGATACAA | TGCGATGTCGGTTTATGTGT | 159 |
| chr5:113914158-113916172 | p5 | (ATTCA)3 | GGCCCAAATCATCAATCAAC | GGAAGGTGTCGGGTCAAGTA | 106 |
| chr5:113914158-113916172 | p2 | (AT)7 | GGCCCAAATCATCAATCAAC | GGAAGGTGTCGGGTCAAGTA | 106 |
| chr5:113914158-113916172 | p2 | (AT)11 | GGCCCAAATCATCAATCAAC | GGAAGGTGTCGGGTCAAGTA | 106 |
| chr5:113966617-113968632 | p3 | (GTT)5 | GGTCATGTGCTTTCCATGTG | TGGGGTTCAAGAACTCAAGG | 113 |
| chr5:114081374-114083394 | p4 | (AATA)5 | ATGCCCTAATTTTGCATTGG | AACACCAAACGTGCATGAAA | 248 |
| chr5:114093473-114095521 | p4 | (TTTA)12 | CGTGAAAGGTTCCACATCCT | AGGCCGTACAAGAGCCACTA | 126 |
| chr5:114093473-114095521 | p2 | (TA)6 | CGTGAAAGGTTCCACATCCT | AGGCCGTACAAGAGCCACTA | 126 |
| chr5:114254565-114256583 | p6 | (TGGGTG)3 | GTAACACCATTGCGCCTTTT | TTTCACTCTTTGCTTGGGCT | 236 |
| chr5:114254565-114256583 | p6 | (CACCCG)3 | GTAACACCATTGCGCCTTTT | TTTCACTCTTTGCTTGGGCT | 236 |
| chr5:114397854-114399888 | p2 | (TA)17 | GGATGATGATGATTGGGTCC | ATGGAGTTCCGCTGTATTGC | 289 |
| chr5:114397854-114399888 | p5 | (AATGC)3 | GGATGATGATGATTGGGTCC | ATGGAGTTCCGCTGTATTGC | 289 |
| chr5:114428265-114430285 | p5 | (AATAT)4 | TTATCAGATTGGTCGTGGCA | AGAGACAGTCGCATTGGCTT | 100 |
| chr5:114497597-114499621 | p6 | (ATCATT)4 | CACTGACATCGTCATGGGAC | TAGACAGGGAGGTGACCCAG | 157 |
| chr5:114533350-114535365 | p5 | (AAACA)3 | AATTAACGGAAGCCACAACG | GGTTCGTTCCAAAATCGAGA | 247 |
| chr5:114615956-114617972 | p3 | (TTA)5 | TCAGTTCGTCATCCTCACCA | GTGGTGGGTGTGGGTTAAAG | 283 |
| chr5:114615956-114617972 | p2 | (AT)8 | TCAGTTCGTCATCCTCACCA | GTGGTGGGTGTGGGTTAAAG | 283 |
| chr5:114782155-114784173 | p3 | (CAA)6 | ACATCACAACAAAGGCCACA | GTTGCCGACCTTACGTCACT | 149 |
| chr5:114794581-114796596 | p5 | (CATCT)3 | GGAGAGTCAGCACGAGATCC | TTCCACCAATACAGTTGCCA | 130 |
| chr5:114794581-114796596 | p5 | (TACCA)3 | GGAGAGTCAGCACGAGATCC | TTCCACCAATACAGTTGCCA | 130 |
| chr5:114820586-114822602 | p4 | (AATA)4 | ACGGGAAAGTTGCTCTGAAA | TTTTCTGCTTCCCCTCTTTG | 279 |
| chr5:114877903-114879929 | p2 | (AT)13 | GTAGCTCCGGATGAAATGGA | CGTCATTTTGTTTGTGGACG | 264 |
| chr5:114938043-114940058 | p5 | (TATGA)3 | GCATTCTGGAATTTGGCATT | TCCGTTTGACCCCCATAATA | 178 |
| chr5:115075140-115077164 | p2 | (TA)12 | GGGGCAATGGTATGATGAAC | ATGGGGTTGTATTTGGTGGA | 170 |
| chr5:115134393-115136405 | p5 | (ATGAG)3 | GGGTGGATTTGCAGTGAAGT | AAATGCCACAACATTGGGTT | 179 |
| chr5:115134410-115136450 | p5 | (ATGAG)3 | GGGTGGATTTGCAGTGAAGT | AAATGCCACAACATTGGGTT | 162 |
| chr5:115135680-115137712 | p5 | (ATGAG)3 | CTAGTGGCGGATAGGTTGGA | AACTCATCCTCCCTCCGTTT | 165 |
| chr5:115135680-115137712 | p2 | (TA)16 | CTAGTGGCGGATAGGTTGGA | AACTCATCCTCCCTCCGTTT | 165 |
| chr5:115141509-115143557 | p2 | (AG)6 | TCTGAACGTTGGAGAGCCTT | GTCGATTTGGGAAGGGAAAT | 185 |
| chr5:115141564-115143609 | p2 | (AG)6 | TCTGAACGTTGGAGAGCCTT | GTCGATTTGGGAAGGGAAAT | 130 |
| chr5:115153636-115155648 | p2 | (GA)6 | TCTTCTCCCCTTCTCCACCT | CACAATCAATGTGGAGTGCC | 215 |
| chr5:115166440-115168456 | p4 | (TATT)4 | ACCGTTTTGGGAGAACCTTT | TTGGGACGTTCCTTCAACAT | 166 |
| chr5:115185556-115187607 | p3 | (AAT)17 | CCAAAACTGAGCTCGAAAGG | TCGGAAATACGAAGTCCCTG | 235 |
| chr5:115198362-115200392 | p3 | (TTA)10 | TTTGGAGGGTTTCCACTTTG | CGATAACGGGAACCAAAGAA | 211 |
| chr5:115202410-115204430 | p2 | (TA)10 | TCTTGGGATCAAACCTGAGC | GCAGCTTAACGCCTATGGAG | 216 |

| Marker ID | SSR type | SSR motif | FORWARD PRIMER (5'-3') | REVERSE PRIMER (5'-3') | size (bp) |
|---|---|---|---|---|---|
| chr5:115222935-115224950 | p5 | (GTGGG)3 | CTCCGCCACCATCAAGTTAT | TCATGAGCAAATGGGTTCAA | 270 |
| chr5:115320340-115322358 | p2 | (AT)9 | TGGAAATTATTCAGCCCGTC | GGTTCGCATAAGGGTTCGTA | 235 |
| chr5:115320340-115322358 | p6 | (TTCAGT)3 | TGGAAATTATTCAGCCCGTC | GGTTCGCATAAGGGTTCGTA | 235 |
| chr5:115331380-115333396 | p5 | (CCGGT)3 | AAGTATCAACCGTGGCTTCG | CCTCCGCTTCTGAGTTTTTG | 269 |
| chr5:115331508-115333520 | p5 | (CCGGT)3 | AAGTATCAACCGTGGCTTCG | CCTCCGCTTCTGAGTTTTTG | 141 |
| chr5:115331521-115333569 | p5 | (CCGGT)3 | AAGTATCAACCGTGGCTTCG | CCTCCGCTTCTGAGTTTTTG | 128 |
| chr5:115331678-115333693 | p5 | (CCGGT)3 | ACCAAAAACTCAGAAGCGGA | CATTTCGTTAATTTCCCCCA | 106 |
| chr5:115362708-115364723 | p3 | (GTT)5 | TTCCTGTGACCCAAACACAA | GCTAGAAAAGACCCCAACCC | 118 |
| chr5:115490755-115492775 | p5 | (TGCAA)4 | CGATTCTCCAAGACGCTTTC | GTAGTTGCATGCGCGACTTA | 240 |
| chr5:115490755-115492775 | p5 | (CCTTT)3 | CGATTCTCCAAGACGCTTTC | GTAGTTGCATGCGCGACTTA | 240 |
| chr5:115566950-115568965 | p2 | (CT)12 | TGCAGTCATGCAACAACATC | TTTGTTGTGTCCATGTGCCT | 241 |
| chr5:115566950-115568965 | p3 | (TTG)5 | TGCAGTCATGCAACAACATC | TTTGTTGTGTCCATGTGCCT | 241 |
| chr5:115883656-115885668 | p2 | (TG)6 | ACCTGATCCAGACGAATTGC | TCTGTGGGAATCAGACCACA | 169 |
| chr5:115895028-115897084 | p2 | (TA)28 | CAACGTGAAGGAAATGGCTT | GGCATGACCACCTTTGACTT | 111 |
| chr5:115931310-115933325 | p5 | (TTGAG)3 | GCCTCGTTGTGACCCACTAT | GCTTCAACTTGCATCCAACA | 136 |
| chr5:115931310-115933325 | p5 | (TTGAG)3 | GCCTCGTTGTGACCCACTAT | GCTTCAACTTGCATCCAACA | 136 |
| chr5:115975311-115977335 | p4 | (ATAA)6 | AGAAAAGCTGGGCTAGGGAG | CCACCATTCTCCGAATCACT | 148 |
| chr5:116029998-116032013 | p5 | (ATTTC)3 | GGCGAAATAACAACCGAGAA | ATTTTGCCAGAGTTTCCCCT | 156 |
| chr5:116097091-116099121 | p2 | (AT)29 | ATGTACAAAAGCCGACGAGG | CAGGGACTACATAGCGGAGC | 267 |
| chr5:116097091-116099121 | p6 | (CATATT)3 | ATGTACAAAAGCCGACGAGG | CAGGGACTACATAGCGGAGC | 267 |
| chr5:116097091-116099121 | p2 | (TA)15 | ATGTACAAAAGCCGACGAGG | CAGGGACTACATAGCGGAGC | 267 |
| chr5:116218839-116220854 | p5 | (AAAAT)3 | CGTGCCCAGTCAAATAGGTT | GAATGCCGTGAGATGGAAAT | 279 |
| chr5:116237730-116239745 | p2 | (TA)6 | AAACAGGCTCTTGCTCCAAA | CAAGTCTCTTTCCAGCCTCG | 256 |
| chr5:116237730-116239745 | p5 | (CATTG)3 | AAACAGGCTCTTGCTCCAAA | CAAGTCTCTTTCCAGCCTCG | 256 |
| chr5:116352424-116354439 | p5 | (TTTCC)3 | AGCGAAGACGATGCGTAAGT | AGCAGTAGAAGCGAGAAGCG | 108 |
| chr5:116595285-116597303 | p2 | (AT)9 | GCGTCTCTCATAGCCCTGTC | GACTCCAGTCAGCGATCTCC | 203 |
| chr5:116598668-116600686 | p2 | (AT)9 | GCGTCTCTCATAGCCCTGTC | GACTCCAGTCAGCGATCTCC | 203 |
| chr5:116662612-116664626 | p2 | (AT)7 | ATCGGGTAAGAGTGGCCTTT | TATGCCCAGGGCTGATAGAC | 174 |
| chr5:116674099-116676117 | p2 | (AT)6 | CATCATTTGCCCCTTGAACT | CGGGCCCACCTTTTTATATT | 152 |
| chr5:116674099-116676117 | p6 | (ATCTAT)3 | CATCATTTGCCCCTTGAACT | CGGGCCCACCTTTTTATATT | 152 |
| chr5:116691372-116693426 | p3 | (CTT)18 | GAATGGTGCTGCCCATAAGT | TGTTGGACATGTTCCCGATA | 133 |
| chr5:116691372-116693426 | p3 | (CTT)6 | GAATGGTGCTGCCCATAAGT | TGTTGGACATGTTCCCGATA | 133 |
| chr5:116756113-116758129 | p2 | (AG)8 | ACGGTTTGGCTTGAAAGAAA | AGCCAGTGCGATAAAGCCTA | 115 |
| chr5:116920313-116922337 | p2 | (AG)12 | TTCATGTGCCATGTGTGATG | ACATGGTTGCCGAAGAAAAG | 237 |
| chr5:117006350-117008368 | p2 | (CT)9 | TAGTGATACTGCTGTCGCGG | GGGCACATTCGTAGCTTGTT | 237 |
| chr5:117040920-117042936 | p2 | (AT)8 | CCTGGTCTTGGGGACTGTAA | ATTTGGCCCTTCCAGTTTCT | 128 |

| Marker ID | SSR type | SSR motif | FORWARD PRIMER (5'-3') | REVERSE PRIMER (5'-3') | size (bp) |
|---|---|---|---|---|---|
| chr5:117040920-117042936 | p3 | (TTC)22 | CCTGGTCTTGGGGACTGTAA | ATTTGGCCCTTCCAGTTTCT | 128 |
| chr5:117150830-117152858 | p2 | (AT)14 | GTTGATGATTTGGCCGAGTT | GAAGTCCCATCCTCAACCAA | 176 |
| chr5:117150830-117152858 | p2 | (TG)6 | GTTGATGATTTGGCCGAGTT | GAAGTCCCATCCTCAACCAA | 176 |
| chr5:117165145-117167160 | p5 | (CATTT)3 | CGAAACAATAACCGGGAAGA | ACAGCCACACATGACCAAAA | 142 |
| chr5:117181024-117183052 | p2 | (TA)14 | ATGCGTTTGCTCCTCAAACT | GGTTTGGGGATTTTTGGAGT | 292 |
| chr5:117253868-117255880 | p2 | (TA)6 | CCCGTCATTGAGAATGGAGT | GTGAGGGATTGGGGATTTTT | 135 |
| chr5:117275202-117277222 | p4 | (ATTA)5 | GGTTGACCGCTGGTGTAACT | GACTGGCAGTATAGCGGAGG | 298 |
| chr5:117502123-117504141 | p6 | (AAACTC)3 | TTATCGCTTGTACGTTCCCC | ACGTTATGGAAGGGATGCTG | 178 |
| chr5:117530982-117532997 | p2 | (AT)22 | GCTGATGAAGGTCCAAAGGA | CTGCACTTTCCGCAACACTA | 201 |
| chr5:117530982-117532997 | p3 | (ATC)5 | GCTGATGAAGGTCCAAAGGA | CTGCACTTTCCGCAACACTA | 201 |
| chr5:117589198-117591218 | p5 | (ATTTT)4 | TATTCGGTACGCAGGCTTTT | TGGGCACATTCCAAACACTA | 170 |
| chr5:117614005-117616065 | p2 | (TA)34 | ATATAGGCCACGCTAAGGGG | CCTTGAGCACAATTTTGGGT | 134 |
| chr5:117644567-117646581 | p2 | (TA)7 | ACTTCTATGCCCATCCATCG | GGTGGAGGCCTCTTATGTGA | 276 |
| chr5:117746583-117748647 | p2 | (TA)17 | GAGGAGCGTTTCATTGAAGG | GCCCATACCCCCATTACTTT | 180 |
| chr5:117746583-117748647 | p4 | (TCTA)4 | GAGGAGCGTTTCATTGAAGG | GCCCATACCCCCATTACTTT | 180 |
| chr5:117766094-117768116 | p2 | (AT)11 | CGACACACAACATCCCAAAC | CCGGATCCTAGCTTGTGGTA | 282 |
| chr5:117879830-117881846 | p2 | (TG)8 | GCAACGTCTCTGTATTGCCA | TGGACTCAAGCCTCTCCACT | 253 |
| chr5:117887219-117889285 | p5 | (TTTAT)5 | TTGGCTTTGGTCAACCCTAC | TCGTTTTGGCTTTTTGACCT | 155 |
| chr5:117936332-117938344 | p6 | (AACATG)3 | ACCACTGGACACGGTTTCAT | CGCATTTAGTTCATGCCCTT | 295 |
| chr5:117936346-117938358 | p6 | (AACATG)3 | ACCACTGGACACGGTTTCAT | CGCATTTAGTTCATGCCCTT | 281 |
| chr5:117936360-117938374 | p6 | (AACATG)3 | ACCACTGGACACGGTTTCAT | CGCATTTAGTTCATGCCCTT | 267 |
| chr5:117991600-117993612 | p6 | (CGGTGT)3 | ACTTTTTCAACCCATTCCCC | GATCCACACCCATATCGGAC | 212 |
| chr5:117991600-117993612 | p2 | (TG)6 | ACTTTTTCAACCCATTCCCC | GATCCACACCCATATCGGAC | 212 |
| chr5:118012672-118014684 | p2 | (TG)6 | ATGGAGGATCGACACAGAGG | GAGTCAGCTTGCGAATTTCC | 279 |
| chr5:118067660-118069674 | p2 | (TC)7 | CCGGTCTATCGACCAACCTA | TGCACTGTGCTTATTCCAGC | 240 |
| chr5:118091603-118093615 | p2 | (AT)6 | TGATGCAATTTGCGGAATAA | GCAGATACCGGTTTTCAGGA | 112 |
| chr5:118118204-118120219 | p5 | (CAAGA)3 | ATTGCAGACACAATTGAGCG | TTCCCAATTGCAGTGACAAA | 123 |
| chr5:118201690-118203741 | p3 | (ATA)17 | ACCCCGTTAATGGATGATGA | TCCCTTTCCCTTTACCCCTA | 211 |
| chr5:118390385-118392401 | p2 | (TA)8 | TTCCGACTATCACCCGAGAC | TGTAAACGGACCTTTTTCGG | 278 |
| chr5:118407882-118409942 | p3 | (AAT)20 | ATTTTTGCAGCTGAACCACC | CGTCTCTCCCTCTTTCATCG | 285 |
| chr5:118453202-118455217 | p5 | (TTGAT)3 | TTGCAAACCATTGTCATCGT | TGGTTGTGGCTAGTGGTGAA | 173 |
| chr5:118536130-118538168 | p2 | (TA)19 | CTTCTTTAACCACACCCCGA | CGAAAGGGAAACCTCTCTCC | 221 |
| chr5:118765513-118767527 | p2 | (TA)7 | GGAAAGAAACGAGGGTAGGG | TTGACGGTTGAGTTGGAACA | 256 |
| chr5:118857675-118859690 | p5 | (TATTG)3 | ATCGCCAGTTTGGAATTGAG | TGCTCAGTCCTCATCGACAC | 289 |
| chr5:118857675-118859690 | p5 | (ACAAT)3 | ATCGCCAGTTTGGAATTGAG | TGCTCAGTCCTCATCGACAC | 289 |
| chr5:118908349-118910401 | p2 | (AT)26 | AGTGAACAAACCCCAACCAC | CCTTGTCGATAAACGGCTTC | 165 |

| Marker ID | SSR type | SSR motif | FORWARD PRIMER (5'-3') | REVERSE PRIMER (5'-3') | size (bp) |
|---|---|---|---|---|---|
| chr5:118961470-118963508 | p2 | (AT)19 | GGCTTTCTATCACGACCCAA | ATCATTTTGGCAGCCTTGTC | 185 |
| chr5:118972043-118974061 | p6 | (GGAGAA)3 | AGTGTGCAAAGAGCCACCTT | CAATGGCTAAACGCTGAACA | 253 |
| chr5:118972043-118974061 | p6 | (CTCTAC)5 | AGTGTGCAAAGAGCCACCTT | CAATGGCTAAACGCTGAACA | 253 |
| chr5:119118228-119120240 | p2 | (TA)6 | GAGTTGACTAGGGGCCATGA | TTCTATGGCAACCCATTTCC | 152 |
| chr5:119155358-119157373 | p3 | (TGA)5 | GATAGCCTTGTCGGCTTCTG | ACTTCTGTCACGACCCAACC | 300 |
| chr5:119240713-119242728 | p5 | (ATGCA)3 | TGAATGAAGTCGCACGAGAG | GAAATCGGGTCGTGACAAGT | 181 |
| chr5:119286070-119288082 | p2 | (TA)6 | TCCTCTCGGCAGAAGGATAA | GGCCATTGCACGTTTACTTT | 219 |
| chr5:119286070-119288082 | p2 | (AT)9 | TCCTCTCGGCAGAAGGATAA | GGCCATTGCACGTTTACTTT | 219 |
| chr5:119349714-119351742 | p2 | (TA)14 | AGAACGATGAGGGATGGTTG | TGAAGCTTTCTAACACGCGA | 139 |
| chr5:119380426-119382454 | p2 | (AT)14 | TGCATGAGATCCCAATCAAA | CTTGTTCCTTCAAGGTGGGA | 186 |
| chr5:119390874-119392889 | p3 | (TGT)5 | CCTTGAAATTGCCAGGAAAA | TTCTGCCTAACAGGCGTTCT | 155 |
| chr5:119442203-119444218 | p5 | (GTGTC)3 | ACATGTTCCCCAGTCCAGAG | AAATGCACCAAAAAGCCAAG | 273 |
| chr5:119492169-119494181 | p2 | (AT)6 | GGACCGTCATTTAGCTCCAA | TGCATGAGGACAAGCAAAAG | 181 |
| chr5:119657868-119659883 | p3 | (ATC)5 | CGGCTATATAGGCACGGTGT | CACTACAGTGGGCGGCTTAT | 196 |
| chr5:119657868-119659883 | p3 | (ACA)5 | CGGCTATATAGGCACGGTGT | CACTACAGTGGGCGGCTTAT | 196 |
| chr5:119732838-119734856 | p6 | (TTTTTC)3 | GGAATGAAAGGAGACACGGA | GCATTGTTTCGGCTCAACTT | 111 |
| chr5:119741478-119743493 | p5 | (AATAT)3 | TTCGCAAACCAAACGTCATA | TGGACAAGAGGCGAGCTATT | 101 |
| chr5:119853459-119855474 | p5 | (TACAG)3 | GGCTTCGCAACTTTCTTTTG | TTAGTGCGTTGGTGCTGAAC | 148 |
| chr5:120028388-120030400 | p2 | (AT)6 | AGCGTAGCCAAGGATAGCAA | AGGAATCCACCTTGCTGAGA | 276 |
| chr5:120053711-120055729 | p2 | (TG)9 | TGATTGTGCTGCATGTCAAA | CTGCTTTCCTGCTTTCATCC | 180 |
| chr5:120056769-120058785 | p2 | (AT)8 | CAAGCTCGGAGAGATTACGG | GATGAGCGGTTTATTGCGTT | 290 |
| chr5:120128974-120130992 | p6 | (TATATG)3 | GCATTCTCGGGTTTGTTGTT | TTCACAAGGATCATGCCAAA | 129 |
| chr5:120135369-120137387 | p6 | (TATATG)3 | GCATTCTCGGGTTTGTTGTT | TTCACAAGGATCATGCCAAA | 151 |
| chr5:120152217-120154229 | p2 | (TA)6 | TGGAGGAGCACGAAAAGAAT | AAGGGAATGAGTGGTGCTTG | 117 |
| chr5:120172184-120174223 | p3 | (AAT)13 | AAGGGATGATTCGATGCTTG | AATCGCAGAAATCCCCTTTT | 108 |
| chr5:120199472-120201488 | p4 | (AATT)4 | ATAACACTTGAACCCGCCAG | ATTAACTCTCGGGCACGTTG | 265 |
| chr5:120226547-120228563 | p2 | (AT)8 | AGCGAAGCGCTTATTATGGA | TGAAGCATCCAACAAACGAG | 157 |
| chr5:120242930-120244946 | p4 | (ATTT)4 | GGAACTTCTGCCCTTGAACA | TGGGTTCCTTTACTTGGTCG | 185 |
| chr5:120326467-120328481 | p6 | (ACATAT)4 | GTGATGCTCGTTGGTCACAC | AAGCCGTCGTACCATTATGC | 289 |
| chr5:120326493-120328539 | p6 | (ACATAT)4 | GTGATGCTCGTTGGTCACAC | AAGCCGTCGTACCATTATGC | 263 |
| chr5:120342188-120344203 | p5 | (GAAAG)3 | ATCGAACATCAAAAGCCAGG | GAATCATACTTGCCTCGGGA | 248 |
| chr5:120372163-120374187 | p3 | (TGT)8 | CTTGGAACCAAGGGAAATGA | GGTTTGTTGCGTTGGAAACT | 123 |
| chr5:120372163-120374187 | p3 | (ATG)5 | CTTGGAACCAAGGGAAATGA | GGTTTGTTGCGTTGGAAACT | 123 |
| chr5:120412947-120414962 | p3 | (TGT)5 | GGTCGCTATTGTTCGGTTGT | TCATATAGAAGCCGCCAAGG | 242 |
| chr5:120425521-120427536 | p3 | (TTG)5 | TGGAAGATTCCAGGACCAAG | TCCGATGTCGTATCAACGAA | 164 |
| chr5:120425521-120427536 | p3 | (ATG)5 | TGGAAGATTCCAGGACCAAG | TCCGATGTCGTATCAACGAA | 164 |

| Marker ID | SSR type | SSR motif | FORWARD PRIMER (5'-3') | REVERSE PRIMER (5'-3') | size (bp) |
|---|---|---|---|---|---|
| chr5:120501013-120503051 | p2 | (AT)20 | TAACTAGAGGGCCATGACGG | TTTTTACAGCCTTGTCGGCT | 183 |
| chr5:120501013-120503051 | p2 | (AT)19 | TAACTAGAGGGCCATGACGG | TTTTTACAGCCTTGTCGGCT | 183 |
| chr5:120507443-120509463 | p2 | (AT)10 | GCAGAAAAGGCCCAATCATA | CTTGTAGCGACCTGGTAGGC | 225 |
| chr5:120619611-120621631 | p5 | (TATGA)4 | AGCCTTGTCAGCTTGCGTAT | TCCAAATAACCGGTCTCCTG | 272 |
| chr5:120649919-120651945 | p2 | (AT)13 | GTAACAACGCAAGCCGATTT | TTCAGCCCTTTTTGGTATGC | 252 |
| chr5:120690305-120692320 | p5 | (ATGCA)3 | TGGTAGGCAAACCCTCAATC | TAGACCAGGTCGTGGGAGTC | 170 |
| chr5:120710814-120712834 | p5 | (ATGCA)4 | TCAATAGGCGAATTAACGCC | AGGTTGGCGGAATTCTTTTT | 176 |
| chr5:120720366-120722381 | p5 | (TGCAA)3 | AAATTATCGATCGTGCAGGC | TGACTTTCAAATTTTGGGGG | 199 |
| chr5:120775258-120777292 | p2 | (AT)21 | AGACATATGGACGGGACAGG | TAGACAGGGAGGTGGTCCAG | 197 |
| chr5:120775258-120777292 | p2 | (AT)17 | AGACATATGGACGGGACAGG | TAGACAGGGAGGTGGTCCAG | 197 |
| chr5:120821279-120823297 | p2 | (TA)9 | TGGGGTCACACTGTGTAGGA | CAACGCTATGCATCTGACCA | 133 |
| chr5:120841000-120843018 | p2 | (AT)9 | ATGAGCATTGAGGGGACAAG | AAAGGGAGTAGGGCTGTGGT | 121 |
| chr5:120928861-120930887 | p2 | (TA)13 | AGAGATCCTGAACCCTCGGT | GGATCGGAGAAAGAAAAGGG | 155 |
| chr5:120934607-120936619 | p2 | (TA)6 | ACTGGGCCAACTTTCAAATG | CGCCAGAAACCTCATGGTAT | 196 |
| chr5:120953646-120955661 | p5 | (AATGC)3 | CCCATCCAAACAGGCTCTAA | CATGGGAAGTTGGATCGAGT | 175 |
| chr5:120986658-120988670 | p2 | (AG)6 | CGATCTCATCGTCGATCTCA | ACGGACCGTCTTATTCAACG | 183 |
| chr5:121053438-121055466 | p2 | (GA)14 | ACCATCCTTTCTGATGCAGG | GTGAGGGTTGCATAAGCGTT | 197 |
| chr5:121112076-121114088 | p2 | (TC)6 | CCGAAATCTTCCAAGTCCAA | TCATGTGGGCTGTTTTGGTA | 111 |
| chr5:121112076-121114088 | p2 | (AT)6 | CCGAAATCTTCCAAGTCCAA | TCATGTGGGCTGTTTTGGTA | 111 |
| chr5:121175459-121177489 | p2 | (TA)8 | AACGGGCTTACGGATCTTAAA | TCCGCTTGTGTGTTGAATGT | 114 |
| chr5:121196628-121198646 | p6 | (TTATTT)3 | AACTCTGTCCTGCGCATTTT | AATGGCCCAATTCACAACAT | 194 |
| chr5:121220646-121222664 | p6 | (TTATTT)3 | AACTCTGTCCTGCGCATTTT | AATGGCCCAATTCACAACAT | 194 |
| chr5:121359061-121361073 | p2 | (AT)6 | TCCCTTTACCCTGAGCCTTT | GAAGAAAACAGAACCTGCGG | 158 |
| chr5:121369127-121371143 | p2 | (AT)8 | CGATTCGATTCGCTCAACTT | CTAGTCGGGTATGCCCTTGA | 181 |
| chr5:121542482-121544497 | p5 | (TGATA)3 | CATATCTGTCGGTCGAGGGT | TCCAAATCTGATAGGTCGGC | 153 |
| chr5:121690481-121692519 | p2 | (AT)10 | TGGGTGGCTGTAGCTTTCTT | CCGGTTTGGGTCTTAACAGA | 229 |
| chr5:121690481-121692519 | p2 | (TA)19 | TGGGTGGCTGTAGCTTTCTT | CCGGTTTGGGTCTTAACAGA | 229 |
| chr5:121768888-121770930 | p2 | (AT)21 | TCAGCAGCAGTCAGTACGCT | GCTGACGAAGATGCGTAACA | 240 |
| chr5:121792025-121794039 | p2 | (AG)7 | ACTGAAGGGATCGACCAATG | TCTCCTCCTCGAGCTCACAT | 248 |
| chr5:121792025-121794039 | p3 | (GTT)5 | ACTGAAGGGATCGACCAATG | TCTCCTCCTCGAGCTCACAT | 248 |
| chr5:121819897-121821915 | p3 | (TTA)6 | GCAAAAAGTCGATTGGAGGA | TGGGCATTTGAACAAAATCA | 210 |
| chr5:121819983-121822016 | p3 | (TTA)6 | GCAAAAAGTCGATTGGAGGA | TGGGCATTTGAACAAAATCA | 124 |
| chr5:121833057-121835072 | p3 | (TTA)5 | GAGCTATTTCTTCGGCATCG | GAATCAGGCCGTGTGTAGGT | 132 |
| chr5:121833057-121835072 | p3 | (TTA)5 | GAGCTATTTCTTCGGCATCG | GAATCAGGCCGTGTGTAGGT | 132 |
| chr5:121853995-121856019 | p2 | (AT)12 | TTTACCCTTCGCCAAATTCA | CTCCCAAAAGCTTGGTCAAA | 114 |
| chr5:121853995-121856019 | p2 | (AT)11 | TTTACCCTTCGCCAAATTCA | CTCCCAAAAGCTTGGTCAAA | 114 |

| Marker ID | SSR type | SSR motif | FORWARD PRIMER (5'-3') | REVERSE PRIMER (5'-3') | size (bp) |
|---|---|---|---|---|---|
| chr5:121869067-121871091 | p2 | (AT)12 | TGTCGGAACCATAGCAACAA | GGTCACACCTGAGACCGTTT | 248 |
| chr5:121869067-121871091 | p2 | (AT)6 | TGTCGGAACCATAGCAACAA | GGTCACACCTGAGACCGTTT | 248 |
| chr5:121869203-121871215 | p2 | (AT)12 | TGTCGGAACCATAGCAACAA | GGTCACACCTGAGACCGTTT | 112 |
| chr5:121869203-121871215 | p2 | (AT)6 | TGTCGGAACCATAGCAACAA | GGTCACACCTGAGACCGTTT | 112 |
| chr5:121942894-121944909 | p2 | (TA)7 | CTCAGCCACACACGAAGAAA | GCCTTTGCACAAAAGAGAGG | 227 |
| chr5:121949197-121951213 | p2 | (AT)8 | CGATTATTTTGGCTCTCCCA | TAGCATTGAGCATGACCAGC | 275 |
| chr5:121949197-121951213 | p2 | (AT)12 | CGATTATTTTGGCTCTCCCA | TAGCATTGAGCATGACCAGC | 275 |
| chr5:122063290-122065312 | p3 | (ACA)6 | GTCAATGTTCCAAATGCCCT | GTTGGGTCGTGGACAAAAGT | 228 |
| chr5:122165692-122167707 | p5 | (TGATA)3 | GATGCTGCCCAAATTTCTGT | CGGCTCTCCAGACTCAATTC | 167 |
| chr5:122171996-122174011 | p5 | (TGATA)3 | GAATTGAGTCTGGAGAGGCG | CTGAATGAAAAAGGGGCAAA | 189 |
| chr5:122196311-122198343 | p2 | (TA)16 | CTCCGACCATGGTAGCAAAT | TGCGACTTCTGTCCTCCTCT | 223 |
| chr5:122205413-122207431 | p6 | (CTTACA)3 | TCGGAGTTTGTTGTGTCTGC | CTACAGCAGTCAAATCGCCA | 129 |
| chr5:122218673-122220787 | p2 | (AT)22 | TGTTTTCTTTGGACGCTGTG | AATAATCAATGGCACCTCCG | 178 |
| chr5:122218673-122220787 | p2 | (AT)57 | TGTTTTCTTTGGACGCTGTG | AATAATCAATGGCACCTCCG | 178 |
| chr5:122241114-122243128 | p2 | (TA)7 | GTCTCTAGCCCTATCGCACG | AACGCAAGCATAGGAGTGCT | 101 |
| chr5:122304819-122306839 | p2 | (AT)10 | ATTTCATCGATGGGCTTGAG | GGATTCGACCCCAACCTAGT | 254 |
| chr5:122384847-122386863 | p4 | (TTTC)7 | TCTCGATCCGAGGGAGAGTA | CTGCCAAACCCAAAAACCTA | 100 |
| chr5:122384847-122386863 | p2 | (AC)8 | TCTCGATCCGAGGGAGAGTA | CTGCCAAACCCAAAAACCTA | 100 |
| chr5:122389551-122391571 | p2 | (AT)12 | CCATTTGCATAATGAGGGGA | TACTGCCTCTTGCAAACCAA | 273 |
| chr5:122389612-122391686 | p2 | (AT)12 | CCATTTGCATAATGAGGGGA | TACTGCCTCTTGCAAACCAA | 212 |
| chr5:122511371-122513383 | p2 | (TA)6 | AAGTGTTGGGGGTGTTGTGT | AAGCCAACAATCTCCATTGC | 136 |
| chr5:122543597-122545625 | p4 | (AAGA)7 | TGACCGTCCCTTTTCTTCAG | TTGCCCTGCATGTTATTGAA | 284 |
| chr5:122573815-122575879 | p2 | (AT)32 | CACCTTGTCCCGAATCACTT | GAGGGTGTTGGGCTACAAGA | 295 |
| chr5:122634453-122636473 | p2 | (AT)10 | GACCTACATCTCCGCCCATA | GTAGGTGCTCGCAAGTTTCC | 299 |
| chr5:122695649-122697663 | p2 | (AT)7 | CTCCCTATCGGAGGGAACTC | AAAGGATTGGGTCGTGACAG | 208 |
| chr5:122695649-122697663 | p3 | (CAA)5 | CTCCCTATCGGAGGGAACTC | AAAGGATTGGGTCGTGACAG | 208 |
| chr5:122740829-122742844 | p5 | (AAATA)3 | TCCCCAGCGACTTAGTATGG | CATTTTGAGGTCTCCTCCCA | 277 |
| chr5:122740829-122742844 | p5 | (AAATA)3 | TCCCCAGCGACTTAGTATGG | CATTTTGAGGTCTCCTCCCA | 277 |
| chr5:122740829-122742844 | p5 | (AATAA)3 | TCCCCAGCGACTTAGTATGG | CATTTTGAGGTCTCCTCCCA | 277 |
| chr5:122740829-122742844 | p5 | (TTTAT)4 | TCCCCAGCGACTTAGTATGG | CATTTTGAGGTCTCCTCCCA | 277 |
| chr5:122763196-122765211 | p5 | (AACCT)3 | AACACACAAGAGCAACGCAC | TCCCTTCACCTCTCCATTTG | 237 |
| chr5:122787247-122789262 | p5 | (AAATA)3 | TAACATGGCCATCACATGCT | TTTGTTCAGTCATGGGGGTT | 202 |
| chr5:122794201-122796235 | p2 | (TC)17 | GCCATTCCAGACAGAGGTGT | CCAAAAATGCCTTTCTTCCA | 250 |
| chr5:122839214-122841226 | p2 | (TA)6 | TTCATGCCATTTTGTGCCTA | TTCGTCAAGGAAAGGTGGTT | 234 |
| chr5:122889507-122891545 | p2 | (TA)19 | CTGTGCACCCTATCTCCCAT | GTAGATGCCTTTATCCGCCA | 252 |
| chr5:122897156-122899174 | p2 | (TA)9 | AAACAGGAGGAGGCGAATTT | ATGCTGAGGACGCTGAAACT | 153 |

| Marker ID | SSR type | SSR motif | FORWARD PRIMER (5'-3') | REVERSE PRIMER (5'-3') | size (bp) |
|---|---|---|---|---|---|
| chr5:122906046-122908061 | p5 | (CTAAG)3 | GTAGCAGCGAACAGAGGCTT | TGCTTAACTTGGTGCGACTG | 132 |
| chr5:122906046-122908061 | p2 | (AG)7 | GTAGCAGCGAACAGAGGCTT | TGCTTAACTTGGTGCGACTG | 132 |
| chr5:122937629-122939644 | p5 | (TGGGG)3 | ATTCATTTGGGATGTTGGGA | AGGTTCTCCAGCGGGTATCT | 127 |
| chr5:122990123-122992144 | p3 | (ACA)7 | GAAAAAGGACAAAGGTGGCA | CCCCGAGTTTACCCTTCATT | 247 |
| chr5:123065536-123067557 | p3 | (TGG)7 | TACGGGTTCAAGGTTTCAGG | TTGTTGAAGATCCCCGCTAC | 190 |
| chr5:123078561-123080573 | p5 | (TGAAT)3 | GGAGGACAAGATGCGAGAAG | CCCTACCTCGTCTGAAACCA | 269 |
| chr5:123085554-123087566 | p2 | (GT)6 | TTATACCCCGCGTTTTATGC | ATTTTGCCCTTCGTCTTCCT | 146 |
| chr5:123215063-123217081 | p6 | (TTGTTT)3 | CCTACAGATCCACTCCCGAA | CCCACTTCCTATGCCTTCAA | 192 |
| chr5:123222074-123224089 | p3 | (CAA)5 | CCCCCAAACAATAAGGGAGT | CGTACGTTCCTCGGCACTAT | 133 |
| chr5:123345436-123347451 | p5 | (TTGGT)3 | CGAGTTCAAGCATTCAAGCA | TTCAAGGGATTCAGCAGCTT | 257 |
| chr5:123345436-123347451 | p5 | (TTGGT)3 | CGAGTTCAAGCATTCAAGCA | TTCAAGGGATTCAGCAGCTT | 257 |
| chr5:123451052-123453067 | p5 | (ATAAA)3 | ATGGGCAGATTCAAGAAACG | TTCAGTTGCGTGTGGTGTTT | 172 |
| chr5:123451052-123453067 | p4 | (AAAT)4 | ATGGGCAGATTCAAGAAACG | TTCAGTTGCGTGTGGTGTTT | 172 |
| chr5:123553486-123555501 | p2 | (AT)21 | TCCACAAGGCAACACAAAAA | CTTCTCGCTTATGTGGAGGC | 219 |
| chr5:123553486-123555501 | p3 | (TAT)5 | TCCACAAGGCAACACAAAAA | CTTCTCGCTTATGTGGAGGC | 219 |
| chr5:123553486-123555501 | p5 | (ACCCG)3 | TCCACAAGGCAACACAAAAA | CTTCTCGCTTATGTGGAGGC | 219 |
| chr5:123605509-123607524 | p5 | (TTTTA)3 | GGGTCAGATGATCGGCTAAA | CCAGCTTCCAGATGTCCAGT | 114 |
| chr5:123664546-123666561 | p3 | (AAT)5 | TTGAATTTGGAAGGTTTCGG | CACCATCCCCGAGAATCTAA | 282 |
| chr5:123664546-123666561 | p6 | (TATATG)3 | TTGAATTTGGAAGGTTTCGG | CACCATCCCCGAGAATCTAA | 282 |
| chr5:123676693-123678773 | p4 | (GAAC)4 | CCTAAGGCTCTCCAAAACCC | AACCTTAGCTCCAAGCGTGA | 119 |
| chr5:123676693-123678773 | p2 | (TA)40 | CCTAAGGCTCTCCAAAACCC | AACCTTAGCTCCAAGCGTGA | 119 |
| chr5:123722022-123724066 | p2 | (TA)22 | TGTTGGTGTTGGATTGCTGT | CTGCACTGCCAAGTTGTTGT | 281 |
| chr5:123780803-123782818 | p5 | (TTTTA)3 | AACTTTCCAACCCAAATCCC | ACTTTTGGGTAAACCGACCC | 275 |
| chr5:123784024-123786039 | p5 | (TTTAT)3 | CCTCACGTCTTGTCCTGGTT | TTTAGGGGGAGGGTTGAGTT | 183 |
| chr5:123786349-123788364 | p5 | (TTTAT)3 | CCTCACGTCTTGTCCTGGTT | TTTAGGGGGAGGGTTGAGTT | 183 |
| chr5:123801555-123803575 | p2 | (TA)10 | AATTGTTGGGCTGGTCTGTC | TGAATTCCCATGCAAACAAA | 143 |
| chr5:123849372-123851396 | p3 | (ATC)8 | CATAATGCAACATCGGCTTG | GGCCATTAGCTGGATTGGTA | 130 |
| chr5:123944783-123946798 | p5 | (TTTCA)3 | GAATACGGGTACGGATGTGG | TCATTGGCGACGCATAAATA | 243 |
| chr5:123977151-123979169 | p3 | (TAA)6 | GACAACGAAAGTGGAGGGAA | CCATTTCCCTTTACCCCCTA | 106 |
| chr5:123977151-123979169 | p3 | (ATA)5 | GACAACGAAAGTGGAGGGAA | CCATTTCCCTTTACCCCCTA | 106 |
| chr5:123977284-123979299 | p3 | (TAA)6 | TAGGGGGTAAAGGGAAATGG | TGATCCGACCCGTCTTTATC | 120 |
| chr5:123977284-123979299 | p3 | (ATA)5 | TAGGGGGTAAAGGGAAATGG | TGATCCGACCCGTCTTTATC | 120 |
| chr5:124030229-124032247 | p6 | (CGTATT)3 | ATCATGTCTGTCACGACCCA | TGTATTGTTGCCCAATGGAA | 268 |
| chr5:124069892-124071906 | p2 | (AT)7 | TCAGCAAGCCTTGGAAGAAT | CAGCATCCCCCAAAGTTCTA | 145 |
| chr5:124076859-124078887 | p2 | (TA)6 | GTTGCGGTGACAGAGAGACA | AAGCACCTTTTGGCCTTTTT | 246 |
| chr5:124076859-124078887 | p2 | (AT)14 | GTTGCGGTGACAGAGAGACA | AAGCACCTTTTGGCCTTTTT | 246 |

| Marker ID | SSR type | SSR motif | FORWARD PRIMER (5'-3') | REVERSE PRIMER (5'-3') | size (bp) |
|---|---|---|---|---|---|
| chr5:124077192-124079246 | p2 | (TA)6 | ACGTGGGATATCGTGAGACC | CCGTTTTATTTGGACGCTGT | 253 |
| chr5:124077192-124079246 | p2 | (AT)14 | ACGTGGGATATCGTGAGACC | CCGTTTTATTTGGACGCTGT | 253 |
| chr5:124077248-124079298 | p2 | (TA)6 | ACGTGGGATATCGTGAGACC | CCGTTTTATTTGGACGCTGT | 197 |
| chr5:124077248-124079298 | p2 | (AT)14 | ACGTGGGATATCGTGAGACC | CCGTTTTATTTGGACGCTGT | 197 |
| chr5:124295371-124297386 | p5 | (AAATA)3 | CTTTTGGGAAGTCCCCTCTC | AAAGTTCATGGTGGAGGTGG | 242 |
| chr5:124295371-124297386 | p2 | (TA)7 | CTTTTGGGAAGTCCCCTCTC | AAAGTTCATGGTGGAGGTGG | 242 |
| chr5:124453362-124455378 | p2 | (AT)8 | ATGCCAAACAAAAGAATGGC | TTACAGGGGAAATGCTGACC | 180 |
| chr5:124522919-124524939 | p2 | (TA)10 | AGAAATGACGGGAGAGAGCA | ATAACAGGGCAACAAGCCAC | 171 |
| chr5:124555219-124557239 | p2 | (TA)10 | ACCTTTTGTTGTGCAGCCTT | GGGTTTGAGATTTGGGGATT | 133 |
| chr5:124568946-124570958 | p2 | (AT)6 | CTTTCTGTCGTAGCCTTGCC | TTATGGGTGTAAAGCGCTCC | 203 |
| chr5:124651833-124653845 | p2 | (TA)6 | GGGAAATGTCAGTTGTGCCT | CTAAGTCCAATCCCAGCCAA | 177 |
| chr5:124756374-124758388 | p2 | (AG)7 | TTTTTCACACGGGTTTAGGG | CCTCGAGCTCACATTAAGCC | 167 |
| chr5:124765133-124767149 | p2 | (TA)8 | ATTGGGCTGTTTTGGAAGTG | ATTCGTGCAGCATTTCTCCT | 299 |
| chr5:124765133-124767149 | p2 | (AT)13 | ATTGGGCTGTTTTGGAAGTG | ATTCGTGCAGCATTTCTCCT | 299 |
| chr5:124765133-124767149 | p2 | (AT)11 | ATTGGGCTGTTTTGGAAGTG | ATTCGTGCAGCATTTCTCCT | 299 |
| chr5:124794130-124796157 | p2 | (TA)9 | AATCAAATGCACATCCGTCA | TGGATGGCATCAAAAGTGAA | 212 |
| chr5:124807838-124809853 | p3 | (TGA)5 | AACCCCAATTAACTCGACCC | TTGCAATGATCGGTTTGTGT | 111 |
| chr5:124819017-124821035 | p2 | (TA)6 | AAAATGGAATTGCCATGACC | TCAACAAAGAGGAACCCCAG | 233 |
| chr5:124819017-124821035 | p2 | (AT)9 | AAAATGGAATTGCCATGACC | TCAACAAAGAGGAACCCCAG | 233 |
| chr5:124860838-124862871 | p3 | (CTT)11 | TTTCCGATCCTTCCCTCTTT | AGTAGGAAGGTGGAGCAGCA | 266 |
| chr5:124920607-124922625 | p2 | (AT)9 | CTTCTTGAACCAAACGCCAT | TCCGTTTCTTGAATTCTGCC | 202 |
| chr5:125019378-125021394 | p3 | (TCA)5 | CGTAGGGTGCATGTAGCTGA | GATGATGAATCACAACCCCC | 218 |
| chr5:125019378-125021394 | p2 | (AT)8 | CGTAGGGTGCATGTAGCTGA | GATGATGAATCACAACCCCC | 218 |
| chr5:125019378-125021394 | p2 | (GT)7 | CGTAGGGTGCATGTAGCTGA | GATGATGAATCACAACCCCC | 218 |
| chr5:125027896-125029914 | p6 | (ATAAAT)3 | AATATGGATTGGGAGGAGGC | TCAATTGCTTCATTGGGTGA | 143 |
| chr5:125027896-125029914 | p5 | (GGGCG)4 | AATATGGATTGGGAGGAGGC | TCAATTGCTTCATTGGGTGA | 143 |
| chr5:125041996-125044030 | p5 | (ACCCC)3 | GGGCTTGATCTTGTTTCGAG | CGTGAACAGATTGTCGTGCT | 144 |
| chr5:125092765-125094795 | p2 | (AG)13 | GCACGAAGCATTTTGCATTA | GCCACCTGTTGCAATGTATG | 134 |
| chr5:125092765-125094795 | p2 | (AT)7 | GCACGAAGCATTTTGCATTA | GCCACCTGTTGCAATGTATG | 134 |
| chr5:125092765-125094795 | p3 | (AAC)5 | GCACGAAGCATTTTGCATTA | GCCACCTGTTGCAATGTATG | 134 |
| chr5:125111382-125113400 | p6 | (TGAACA)3 | CCTGAATGGATATGGCTCGT | ATCATTTCCGGATGAGCAAG | 244 |
| chr5:125111382-125113400 | p2 | (TA)9 | CCTGAATGGATATGGCTCGT | ATCATTTCCGGATGAGCAAG | 244 |
| chr5:125170228-125172243 | p5 | (TACAA)3 | TGGTGATTTCTCGTGGTTCA | TGGTGGAGGATGGGGTATAA | 226 |
| chr5:125217693-125219708 | p3 | (TGA)5 | CTTGCTACATCTTGGCGTGA | TCGGACATAAGCTAAACCCG | 167 |
| chr5:125263644-125265662 | p3 | (TTA)6 | GGAGCAAAACGAAGCAGTTC | ATGCCTGATTGCAGAACCTC | 180 |
| chr5:125314966-125317004 | p2 | (TA)19 | GTGCTGCATGTTCTCAAGGA | AAATGGTCCACATGGATGGT | 197 |

| Marker ID | SSR type | SSR motif | FORWARD PRIMER (5'-3') | REVERSE PRIMER (5'-3') | size (bp) |
|---|---|---|---|---|---|
| chr5:125354005-125356020 | p5 | (TTGCA)3 | CACCGATTAGGCAGACCATT | AATTTGCATCTTCGAATGCC | 237 |
| chr5:125397777-125399792 | p5 | (GACCC)3 | TGCCATCGACGATATACGAA | ACCATCAACTCAATCCCTCG | 210 |
| chr5:125562384-125564402 | p3 | (AAC)13 | AACTGTGTCCAACCTCTGGG | GTACTTCTCAGGCGAAACGG | 278 |
| chr5:125562384-125564402 | p6 | (AAAGTA)3 | AACTGTGTCCAACCTCTGGG | GTACTTCTCAGGCGAAACGG | 278 |
| chr5:125563474-125565486 | p2 | (TC)6 | ATGCCCAAACCATCTCAGTC | CTGCCTCAACTTCCCCATAA | 221 |
| chr5:125630364-125632379 | p5 | (GTGAT)3 | TCCGATGTCGTATCAACGAA | TTTGGTCCAACTTGAGGGAG | 192 |
| chr5:125677599-125679614 | p5 | (AAAAG)3 | CCTATGTGCACCCGAAGAAT | GATCAACTGTCAGCCCCAAT | 232 |
| chr5:125678676-125680688 | p2 | (TA)6 | TTGACCTCCTTTTTCGTCGT | TTTGGATGTGGTGAGTTCCA | 159 |
| chr5:125705088-125707106 | p6 | (GTGGAC)3 | GTTCCCCAACAAGCACAAGT | TTCATGTCCACTGTTGCGAT | 195 |
| chr5:125788342-125790363 | p3 | (ACA)7 | ACCCAAAATGATGGGTGCTA | AAACGAGGGTTAGCGTCTCA | 104 |
| chr5:125815944-125817964 | p2 | (AT)10 | ATACAATTTAAAGGGGCGGG | ATCACGCCCTTGTCTGTTTC | 145 |
| chr5:125846189-125848207 | p2 | (TA)9 | AATGCTTCATCGTCCGTTGT | TAACCCACAAAGGGTCAAGC | 290 |
| chr5:125965713-125967725 | p2 | (AT)6 | AATTTCCAAATGAGATGGCG | CCGATCTTTTGTTCATCGGT | 272 |
| chr5:125977882-125979908 | p2 | (AT)13 | TCACCGATCATAACAACCCA | CAGGTACCGACGACAGGTTT | 247 |
| chr5:125986012-125988026 | p2 | (CT)7 | GGAACAACAATCGGGAAGAA | AATATCATCGCCTCACAGGG | 224 |
| chr5:125988104-125990118 | p2 | (CT)7 | GGAACAACAATCGGGAAGAA | TTTCCCCTGTAATCTGGCAC | 224 |
| chr5:126106237-126108271 | p2 | (AT)7 | CCTGCTCCTGAGGATTGAAC | ATTTTGTTTGGCGCTACCAC | 184 |
| chr5:126216734-126218776 | p6 | (CTTTTT)4 | CCCCATTTGCATTAGCCTTA | TTCGCTGCCGAAGAAATAGT | 225 |
| chr5:126313128-126315143 | p3 | (TAA)5 | TGGTTGCGAAATTGAGAAGA | ACCCGAAAGCGAATATGATG | 153 |
| chr5:126313128-126315143 | p3 | (ATA)19 | TGGTTGCGAAATTGAGAAGA | ACCCGAAAGCGAATATGATG | 153 |
| chr5:126322321-126324341 | p2 | (AT)10 | GCACCGATTCTGCTAGATCC | AAGATTTGCCATATCACGCC | 253 |
| chr5:126347991-126350009 | p4 | (CATA)4 | CGGGACTCGGTGGTATCTTA | GATTTCATCTTGCGACGGTT | 253 |
| chr5:126347991-126350009 | p3 | (CAA)6 | CGGGACTCGGTGGTATCTTA | GATTTCATCTTGCGACGGTT | 253 |
| chr5:126347991-126350009 | p2 | (CT)8 | CGGGACTCGGTGGTATCTTA | GATTTCATCTTGCGACGGTT | 253 |
| chr5:126530830-126532842 | p2 | (AT)6 | CAGATGCCACAGCAGTCAGT | TGTGGGTGTGTGTGCCTACT | 213 |
| chr5:126557444-126559480 | p3 | (CTT)12 | CAAAAGGTTAGCTCAAGCGG | TCAGCCCTGATGTTGCATAG | 208 |
| chr5:126557444-126559480 | p2 | (AC)8 | CAAAAGGTTAGCTCAAGCGG | TCAGCCCTGATGTTGCATAG | 208 |
| chr5:126590737-126592751 | p2 | (TA)7 | GAAGCCGGCAAGGTATGTAA | ACAATCCGAGATCGCCTATG | 298 |
| chr5:126628281-126630314 | p3 | (TTA)11 | CTTCGCATGGGACCTTACAT | GGCCAATTAGTTTGGCATTG | 113 |
| chr5:126774465-126776503 | p5 | (GTTTT)3 | CCAACCGCTAAAATTTGGAA | CTGGTTCGTGGGTCTATCGT | 220 |
| chr5:126927056-126929076 | p5 | (CTTTT)4 | GGGATTCCTGTTTGTTTGGA | ACCAACATGGTTTCCCACAT | 289 |
| chr5:127018903-127020918 | p5 | (TAAAA)3 | CTCATTTTGTCACGCTTGGA | CAAGCTTGGGCTTTCTTGAC | 153 |
| chr5:127130328-127132344 | p2 | (AG)8 | GCCTTAAATCAGCCCACAAA | AGCGAGAGTCTTTTGTCCCA | 219 |
| chr5:127134533-127136549 | p2 | (AG)8 | GCCTTAAATCAGCCCACAAA | AGCGAGAGTCTTTTGTCCCA | 219 |
| chr5:127142785-127144799 | p2 | (TG)7 | ATCTTTTCACATCTTGCGGG | GAGTTCAAGCCTCACTTGCC | 152 |
| chr5:127142785-127144799 | p6 | (CCCACC)3 | ATCTTTTCACATCTTGCGGG | GAGTTCAAGCCTCACTTGCC | 152 |

| Marker ID | SSR type | SSR motif | FORWARD PRIMER (5'-3') | REVERSE PRIMER (5'-3') | size (bp) |
|---|---|---|---|---|---|
| chr5:127155451-127157465 | p2 | (TG)7 | ATCTTTTCACATCTTGCGGG | GAGTTCAAGCCTCACTTGCC | 152 |
| chr5:127155451-127157465 | p6 | (CCCACC)3 | ATCTTTTCACATCTTGCGGG | GAGTTCAAGCCTCACTTGCC | 152 |
| chr5:127365720-127367753 | p5 | (AAAAT)3 | AAAGTGGAACCCACCACATC | ATATGGGGCCCACAATTTTT | 102 |
| chr5:127365720-127367753 | p2 | (AT)6 | AAAGTGGAACCCACCACATC | ATATGGGGCCCACAATTTTT | 102 |
| chr5:127365720-127367753 | p2 | (TA)7 | AAAGTGGAACCCACCACATC | ATATGGGGCCCACAATTTTT | 102 |
| chr5:127370989-127373004 | p5 | (TATGA)3 | AGTCAAGTGGTTCGCTTGGT | GGCTCTAAGCTTCATGGCAC | 247 |
| chr5:127383093-127385113 | p4 | (AAAT)5 | CGACATAGTTTTGCAAGCCA | ACTCATGAATTTCGGGTTCG | 152 |
| chr5:127429974-127431998 | p2 | (AT)12 | TTCTGTCATTGTGCCTGAGC | ATCCTCCCCTAAAAGTGCGT | 253 |
| chr5:127651503-127653521 | p5 | (TGAAT)3 | TTAGATTCTCGGGGATGGTG | TTCACAAGGATCATGCCAAA | 201 |
| chr5:127651503-127653521 | p6 | (TATATG)3 | TTAGATTCTCGGGGATGGTG | TTCACAAGGATCATGCCAAA | 201 |
| chr5:127670431-127672446 | p5 | (TGAAA)3 | GCAGAGCCCATTTAATTCCA | TTTTTCCCCACATTTTCACC | 138 |
| chr5:127804911-127806932 | p3 | (TTC)7 | AAGGCCCTTCTCACTTTGGT | TTTTTGGGCTTGAAAAATGG | 114 |
| chr5:127804911-127806932 | p3 | (CTC)7 | AAGGCCCTTCTCACTTTGGT | TTTTTGGGCTTGAAAAATGG | 114 |
| chr5:127844919-127846939 | p2 | (AT)10 | TTGCTCCACGCCCTATTACT | TATCTCGGGCCATTCTTGAG | 237 |
| chr5:127844919-127846939 | p2 | (AT)11 | TTGCTCCACGCCCTATTACT | TATCTCGGGCCATTCTTGAG | 237 |
| chr5:127844919-127846939 | p2 | (TA)16 | TTGCTCCACGCCCTATTACT | TATCTCGGGCCATTCTTGAG | 237 |
| chr5:127869461-127871481 | p3 | (CCA)5 | TAGTTGATTGGGGGCAAAAG | GGAATGAGGGTGGAGGAAAT | 157 |
| chr5:127869461-127871481 | p5 | (AAAAG)4 | TAGTTGATTGGGGGCAAAAG | GGAATGAGGGTGGAGGAAAT | 157 |
| chr5:127869461-127871481 | p2 | (AT)6 | TAGTTGATTGGGGGCAAAAG | GGAATGAGGGTGGAGGAAAT | 157 |
| chr5:128080882-128082897 | p5 | (AACAA)3 | CTGCTTCTTCTTTCCCATGC | AAGGCTCATTAGCCCAAGGT | 199 |
| chr5:128095767-128097817 | p2 | (TA)25 | TTCCCAAAGTATCCCCATCA | AAATGACATGGCGGAATAGC | 175 |
| chr5:128190016-128192032 | p4 | (AAAT)4 | GAGGTGCAGCTCCCTCTATG | TTGGGCGAAAGAAAAAGAGA | 197 |
| chr5:128190016-128192032 | p5 | (GGGGT)3 | GAGGTGCAGCTCCCTCTATG | TTGGGCGAAAGAAAAAGAGA | 197 |
| chr5:128260924-128262939 | p2 | (GA)6 | CTTCGTGTTTTTCTCCTCCG | CCTTTTCCCCCACTTCTCTC | 119 |
| chr5:128260924-128262939 | p3 | (TAA)5 | CTTCGTGTTTTTCTCCTCCG | CCTTTTCCCCCACTTCTCTC | 119 |
| chr5:128260924-128262939 | p3 | (ATA)5 | CTTCGTGTTTTTCTCCTCCG | CCTTTTCCCCCACTTCTCTC | 119 |
| chr5:128408675-128410691 | p4 | (TTTA)4 | ATGTGCACGTTTTCAATCCA | CCTCCTTCCTTGGCTTTCTT | 109 |
| chr5:128440627-128442659 | p5 | (TAAAA)4 | AAAAATAGGGGTGTGGGGAG | CCTTCTGCCCTTGATGAAAA | 152 |
| chr5:128475868-128477896 | p3 | (ACA)11 | GACTATGTTGTATGGGGCGG | ATCAATGGCCAACCTCTCAC | 295 |
| chr5:128475868-128477896 | p5 | (AAGAC)3 | GACTATGTTGTATGGGGCGG | ATCAATGGCCAACCTCTCAC | 295 |
| chr5:128539434-128541452 | p2 | (AG)6 | TCACCATTTCCAGCCTTAGC | AATGACCAGGCAGGAAACAC | 142 |
| chr5:128539434-128541452 | p3 | (TTC)6 | TCACCATTTCCAGCCTTAGC | AATGACCAGGCAGGAAACAC | 142 |
| chr5:128556316-128558332 | p4 | (TCAA)4 | ATTTCGTGCTAGAAAGGCCA | AATCCAAACGGGCTCTAGGT | 145 |
| chr5:128605582-128607594 | p6 | (CACCCA)3 | GCAAGACTCTGGGTGGTGTT | GTGTGTGCGTGTAAGGGATG | 294 |
| chr5:128605582-128607594 | p2 | (TA)6 | GCAAGACTCTGGGTGGTGTT | GTGTGTGCGTGTAAGGGATG | 294 |
| chr5:128619809-128621842 | p5 | (TTTTC)4 | GAACTCCTGGCTCACGTCAT | TGTATCGATTCCACGCAAAA | 147 |

| Marker ID | SSR type | SSR motif | FORWARD PRIMER (5'-3') | REVERSE PRIMER (5'-3') | size (bp) |
|---|---|---|---|---|---|
| chr5:128747714-128749729 | p5 | (AAAAT)3 | ACAGGGTAAACCCAGCTCCT | TTGTTGGTTGGTTGTGCTGT | 128 |
| chr5:128748881-128750911 | p3 | (CTT)10 | CTTCAGTAGAAAGCACGGGG | GTAATACTTTCCCGCACCGA | 106 |
| chr5:128879983-128881997 | p2 | (AT)7 | CCCTTCCAATCAATTTGGTG | GCCAATCCGAATGGTCTTAAT | 144 |
| chr5:128913851-128915901 | p2 | (AT)25 | TTTTGTGCATCCCAAGACAA | TTACCCTTTTCGTTAGGGGG | 103 |
| chr5:128991700-128993718 | p2 | (CA)8 | CCCACAAGTTTTTCACGGTT | TATGTGACATGTGACCCCGT | 138 |
| chr5:128991700-128993718 | p6 | (AAAAAT)3 | CCCACAAGTTTTTCACGGTT | TATGTGACATGTGACCCCGT | 138 |
| chr5:129108164-129110176 | p2 | (TA)6 | AAGCCGACGAAGCTAACAAA | GTCCTGTCCCGCCTATATGA | 260 |
| chr5:129115875-129117889 | p2 | (TA)7 | AAGCCGACGAAGCTAACAAA | TCGGAGCTACAGCTTCTGGT | 268 |
| chr5:129379668-129381694 | p2 | (AT)13 | TGTTAAAAGGAGGTCGGGTG | GGCAGCATTTCCCCTACATA | 248 |
| chr5:129402827-129404845 | p2 | (TA)11 | AATTAATTGATGGGTCCCCC | GGGCATAAGCCATCATGTTT | 221 |
| chr5:129402827-129404845 | p2 | (AT)12 | AATTAATTGATGGGTCCCCC | GGGCATAAGCCATCATGTTT | 221 |
| chr5:129402827-129404845 | p2 | (TA)9 | AATTAATTGATGGGTCCCCC | GGGCATAAGCCATCATGTTT | 221 |
| chr5:129402827-129404845 | p2 | (TA)7 | AATTAATTGATGGGTCCCCC | GGGCATAAGCCATCATGTTT | 221 |
| chr5:129402846-129404860 | p2 | (TA)11 | AATTAATTGATGGGTCCCCC | GGGCATAAGCCATCATGTTT | 202 |
| chr5:129402846-129404860 | p2 | (AT)12 | AATTAATTGATGGGTCCCCC | GGGCATAAGCCATCATGTTT | 202 |
| chr5:129402846-129404860 | p2 | (TA)9 | AATTAATTGATGGGTCCCCC | GGGCATAAGCCATCATGTTT | 202 |
| chr5:129402846-129404860 | p2 | (TA)7 | AATTAATTGATGGGTCCCCC | GGGCATAAGCCATCATGTTT | 202 |
| chr5:129436320-129438332 | p2 | (AG)9 | TGATCCGTGTGGTTGGAATA | GAGTCTCGCAACTTTTTCCG | 161 |
| chr5:129436320-129438332 | p2 | (AT)6 | TGATCCGTGTGGTTGGAATA | GAGTCTCGCAACTTTTTCCG | 161 |
| chr5:129513798-129515814 | p4 | (TTTC)4 | TATCAGTGCGTGGACATGGT | GTCATCCGTGGTCTCCTCAT | 110 |
| chr5:129528350-129530366 | p4 | (ATTT)4 | CCTTCCATCCTTCTCTGGTG | TCCAGTACCGGAATTTGACC | 196 |
| chr5:129644012-129646032 | p4 | (TTCT)5 | TGTGGCTGTGGAGGTTACTG | CTTTTGGAACTCGTCTTCGC | 106 |
| chr5:129788834-129790852 | p6 | (GTATAT)3 | CTGGTTATCAGGGGAGTGGA | ACACAAAATTCCCCAACCAA | 258 |
| chr5:129805367-129807385 | p3 | (ATT)6 | TCACATGACTTGGCACCATT | TCAAACACACATTTTCCCCA | 252 |
| chr5:129931139-129933154 | p5 | (TTTTG)3 | TCGTCCTTAATAACCACGGG | CCGGAAGTAGCCCAGATACA | 248 |
| chr5:129935615-129937633 | p3 | (TGC)6 | TTAGCCTCCGTTGCTTCCTA | TTACCATGCCAAGCATTTCA | 120 |
| chr5:129996272-129998314 | p3 | (AAG)9 | GATGCTCAACCAAACGTTCA | CCTCGATGTGGGTTGCTTAT | 131 |
| chr5:130014867-130016882 | p5 | (ACCCC)3 | CTTGATGATGAATGTTGCCG | ATTCCTACTTCCCCCGATTG | 197 |
| chr5:130056196-130058216 | p5 | (TTGTT)4 | TTCCCCCAAAAACACACAAT | TCTTGGTGGTAAGTGTGCCA | 272 |
| chr5:130206414-130208500 | p2 | (AT)43 | ATCGGTGTGGTTTATGCGAT | GCAGCCCACAATTATTCGAT | 129 |
| chr5:130365333-130367349 | p2 | (AT)8 | GTGAGGCTTCTCCAGCAAAG | AGACATCATATTCCACGGGC | 201 |
| chr5:130365333-130367349 | p2 | (AT)7 | GTGAGGCTTCTCCAGCAAAG | AGACATCATATTCCACGGGC | 201 |
| chr5:130365333-130367349 | p2 | (TA)6 | GTGAGGCTTCTCCAGCAAAG | AGACATCATATTCCACGGGC | 201 |
| chr5:130379561-130381615 | p2 | (CT)6 | AACTATTTCTTCGGCAGCGA | TCAGGTCGTGCGTAGTTCTG | 276 |
| chr5:130379618-130381681 | p2 | (CT)6 | AACTATTTCTTCGGCAGCGA | TCAGGTCGTGCGTAGTTCTG | 219 |
| chr5:130429470-130431484 | p2 | (AT)7 | GTGGTGCAGATGATGATTGG | ACTACTTCGTCCCTCGCAAA | 166 |

| Marker ID | SSR type | SSR motif | FORWARD PRIMER (5'-3') | REVERSE PRIMER (5'-3') | size (bp) |
|---|---|---|---|---|---|
| chr5:130481910-130483922 | p2 | (AT)6 | GGCTAATCTGGGGAAAGGAG | GAAATTGGGTAGTCCATGCC | 134 |
| chr5:130606886-130608904 | p6 | (TCCACT)3 | CGTATGAGAGAGAGGAGCGG | GCGGCCCACTACAATTAAAA | 260 |
| chr5:130606886-130608904 | p6 | (TCCACG)3 | CGTATGAGAGAGAGGAGCGG | GCGGCCCACTACAATTAAAA | 260 |
| chr5:130635720-130637758 | p5 | (TTGTA)3 | GATGTGGGGAACAAAACGAT | CGACGGGAAACGATACAATC | 145 |
| chr5:130665239-130667251 | p2 | (AG)6 | ATCAGAGATCGGAGCAAGGA | TCCGTTTGACCCCCATAATA | 289 |
| chr5:130701558-130703572 | p2 | (AT)7 | CTCAGCCAACACCAGACTCA | ACCCTCACCATGAAAACAGG | 200 |
| chr5:130702085-130704183 | p2 | (AT)7 | AGAGAGAGAAAGGGGAACGG | CAGCTCAAAATAGGCCAGGA | 299 |
| chr5:130756389-130758407 | p3 | (TTA)6 | TTATGTTCCACTGCCACCAA | GAACAGCTGGGCATTTTTGT | 273 |
| chr5:130782508-130784522 | p2 | (TA)7 | AGGTATGGACGTACCGTTGC | AGTTGTTGATCGAATTGGGC | 197 |
| chr5:130820683-130822697 | p2 | (AT)7 | GAGCTTTTGAAGTGCGGAAC | GATGGCCGATGTCTTTGATT | 265 |
| chr5:130820683-130822697 | p2 | (AT)6 | GAGCTTTTGAAGTGCGGAAC | GATGGCCGATGTCTTTGATT | 265 |
| chr5:130834952-130836972 | p2 | (TA)10 | CCTGGGTCGATTTGCTTAAA | TAAAGTGCACCAATTCGCAA | 188 |
| chr5:130897452-130899470 | p3 | (TAT)6 | TTTTCATGATTGCTTGCCTG | GAAGACGAACTCCAAGACGC | 192 |
| chr5:130974086-130976118 | p2 | (AT)16 | AAGGGCCATTTGAAACACAG | GGTGGAGGGTGGGGTATAAT | 298 |
| chr5:130974086-130976118 | p5 | (TACAA)3 | AAGGGCCATTTGAAACACAG | GGTGGAGGGTGGGGTATAAT | 298 |
| chr5:130974264-130976279 | p2 | (AT)16 | AAGGGCCATTTGAAACACAG | GGTGGAGGGTGGGGTATAAT | 120 |
| chr5:130974264-130976279 | p5 | (TACAA)3 | AAGGGCCATTTGAAACACAG | GGTGGAGGGTGGGGTATAAT | 120 |
| chr5:130978984-130981076 | p2 | (AT)46 | CCAGGAACCACCTGACTCAT | AGCTCCACCGGTTCATATTG | 159 |
| chr5:130978984-130981076 | p3 | (CAA)7 | CCAGGAACCACCTGACTCAT | AGCTCCACCGGTTCATATTG | 159 |
| chr5:131046614-131048626 | p2 | (AC)6 | ATCATTTCACAACACGTCCG | TTCGGTTTTCGGCAATAAAA | 263 |
| chr5:131266593-131268615 | p5 | (AAAGT)3 | ATCCCCGCCATAGCTAAAAT | TACTCCCAAAACAGCCCAAC | 162 |
| chr5:131266593-131268615 | p2 | (TA)11 | ATCCCCGCCATAGCTAAAAT | TACTCCCAAAACAGCCCAAC | 162 |
| chr5:131266593-131268615 | p2 | (TA)47 | ATCCCCGCCATAGCTAAAAT | TACTCCCAAAACAGCCCAAC | 162 |
| chr5:131275150-131277238 | p6 | (CCGGTT)3 | CAAACCGGTTACCACCCTTA | GGATGACGCTTTAGATCCCA | 296 |
| chr5:131379193-131381211 | p2 | (CT)9 | TAGTGCCGAGGAAAGTACGG | TACATAGGGGACATGGGGAA | 277 |
| chr5:131386195-131388213 | p2 | (AG)9 | TTTCCACTCCCAAAAACCAC | TTGCTCGTTGTTTCACTTCG | 282 |
| chr5:131406693-131408717 | p2 | (TA)12 | TTCGAGACGAGCGTATGATG | TCATGCCTTTGAAAGAAGGG | 266 |
| chr5:131416617-131418631 | p2 | (AT)7 | CCGAATTCAAGAGGTTTGGA | TCCTTCATCACCCACATGAA | 262 |
| chr5:131457767-131459779 | p2 | (AT)6 | GACCGCGTCACATTATCCTT | AAGTGCAGTGAGTGCAATGG | 138 |
| chr5:131568222-131570237 | p5 | (ACCAA)3 | ATAGGGGTGTACATGGACCG | CCGAGGTTGAAAAACCTGAC | 145 |
| chr5:131568222-131570237 | p5 | (AAAAT)3 | ATAGGGGTGTACATGGACCG | CCGAGGTTGAAAAACCTGAC | 145 |
| chr5:131568222-131570237 | p5 | (AAACC)3 | ATAGGGGTGTACATGGACCG | CCGAGGTTGAAAAACCTGAC | 145 |
| chr5:131568222-131570237 | p4 | (AATA)6 | ATAGGGGTGTACATGGACCG | CCGAGGTTGAAAAACCTGAC | 145 |
| chr5:131570337-131572375 | p2 | (AT)19 | TGGGGAATTACCTACCGTTG | AGGACGTTTTTGTCGAATGG | 214 |
| chr5:131570337-131572375 | p2 | (TA)42 | TGGGGAATTACCTACCGTTG | AGGACGTTTTTGTCGAATGG | 214 |
| chr5:131617991-131620006 | p3 | (TTC)5 | CATCAGGAGAAGCGTGAACA | GCAAATCCTGGAGATGGAGA | 226 |

| Marker ID | SSR type | SSR motif | FORWARD PRIMER (5'-3') | REVERSE PRIMER (5'-3') | size (bp) |
|---|---|---|---|---|---|
| chr5:131638486-131640500 | p2 | (AT)7 | GGGCGAAATACGATTTTTGA | GGGTGAAGGTGGAAGGATCT | 268 |
| chr5:131638486-131640500 | p2 | (AT)10 | GGGCGAAATACGATTTTTGA | GGGTGAAGGTGGAAGGATCT | 268 |
| chr5:131915330-131917396 | p2 | (TA)33 | AACAACCACCCCTTTCTGTG | AATCATCACAGTGAAGGCCC | 216 |
| chr5:131922674-131924689 | p5 | (GGGGT)3 | ATTCAGGGACGAAACTGTGG | TGGGTTGGAGTTAGTGAGGG | 282 |
| chr5:131922736-131924768 | p5 | (GGGGT)3 | ATTCAGGGACGAAACTGTGG | TGGGTTGGAGTTAGTGAGGG | 220 |
| chr5:131959791-131961803 | p2 | (GA)6 | GTTTTCTAGCCCAAAAGGGG | GCGTCTAGCAGATGATGCAA | 295 |
| chr5:131978554-131980596 | p2 | (TA)21 | GGCTCAGCAGGAGTAGCATC | CCCACCCCCAACTAAAGAAT | 118 |
| chr5:131980923-131982937 | p2 | (CT)7 | CGGAACATTGGACAGCTTTT | TGGTTTGTGGGCTGTTGTAA | 148 |
| chr5:131987047-131989093 | p2 | (AT)23 | TGCCCTAGGCTCCTCTGATA | GAAGGGCAAGTCGGTAGTTG | 117 |
| chr5:131987047-131989093 | p2 | (AT)6 | TGCCCTAGGCTCCTCTGATA | GAAGGGCAAGTCGGTAGTTG | 117 |
| chr5:131994991-131997007 | p2 | (TA)8 | CAGTCCTAGGAGGGACCACA | CTGTTGTGGTGAACCAGGTG | 109 |
| chr5:132028384-132030422 | p2 | (TA)36 | TGATGGATTCATTGCCACAT | AAATGACATGGCGGAATAGC | 291 |
| chr5:132028384-132030422 | p2 | (TA)19 | TGATGGATTCATTGCCACAT | AAATGACATGGCGGAATAGC | 291 |
| chr5:132054366-132056384 | p6 | (AAAAAT)3 | TCAGGCAAGGCTTCAAGAAT | GCAAAAATATCCCGGACTCA | 149 |
| chr5:132149357-132151373 | p4 | (TTTA)4 | TGTGTTAAATATGGGCGGGT | CCACAGCAACTAAGCCACAA | 204 |
| chr5:132149357-132151373 | p3 | (TTC)6 | TGTGTTAAATATGGGCGGGT | CCACAGCAACTAAGCCACAA | 204 |
| chr5:132166524-132168540 | p4 | (TTTA)4 | TGTGTTAAATATGGGCGGGT | CCACAGCAACTAAGCCACAA | 204 |
| chr5:132166524-132168540 | p3 | (TTC)6 | TGTGTTAAATATGGGCGGGT | CCACAGCAACTAAGCCACAA | 204 |
| chr5:132210195-132212210 | p3 | (CTT)5 | TTGTGACAAGTTTAGGGGGC | GCATAGCAAGATCGTGAGCA | 195 |
| chr5:132210195-132212210 | p2 | (TA)33 | TTGTGACAAGTTTAGGGGGC | GCATAGCAAGATCGTGAGCA | 195 |
| chr5:132210195-132212210 | p3 | (AGA)5 | TTGTGACAAGTTTAGGGGGC | GCATAGCAAGATCGTGAGCA | 195 |
| chr5:132232083-132234095 | p2 | (AG)6 | TGGTTGCCTTAAGAATTCCG | GCTTCGACGAAACTCCAATC | 241 |
| chr5:132242873-132244888 | p5 | (TATGA)3 | TTTGGAGCCGCTACCATATC | TGTCGTAGGCACTGTTGCTC | 192 |
| chr5:132313304-132315319 | p5 | (TTGTG)3 | AAGGGGTGAGGGAGAGAGAG | TTCCATTTAGGCACCTCCAC | 217 |
| chr5:132326086-132328101 | p5 | (ATATC)3 | GGAACCTGGGTAAAAGGGAA | GGTGGAGTTATTGCCCAGAA | 112 |
| chr5:132486826-132488841 | p5 | (TTCTT)3 | TGATCCCCAAAATTCACTCC | GTACCGCGTCTGTGGAAAAT | 132 |
| chr5:132610773-132612788 | p5 | (TTTTA)3 | ATTGCTCTGCTTCGCAATTT | GCTCCACAAACAAGCCATTT | 181 |
| chr5:132795752-132797773 | p2 | (GA)6 | CTATGCTCGAACACGAACGA | TGTCCCCTGCTCATCTCTCT | 287 |
| chr5:132795752-132797773 | p3 | (TTA)7 | CTATGCTCGAACACGAACGA | TGTCCCCTGCTCATCTCTCT | 287 |
| chr5:132855767-132857833 | p2 | (AT)33 | ACGAGGGTTGCAATAAGTGG | ACCTTTGCATATTGCCAGGA | 246 |
| chr5:132860704-132862722 | p3 | (TTA)6 | TTGGCTGATTTGGGATTCTC | AATCCCCTTCCTCTACCCCT | 226 |
| chr5:132860704-132862722 | p6 | (TTTTAA)3 | TTGGCTGATTTGGGATTCTC | AATCCCCTTCCTCTACCCCT | 226 |
| chr5:132930012-132932024 | p2 | (AT)7 | TGCATATGGTTTCCGCACTA | TCCGTACAACCCTGAACACA | 282 |
| chr5:132930012-132932024 | p2 | (TG)6 | TGCATATGGTTTCCGCACTA | TCCGTACAACCCTGAACACA | 282 |
| chr5:132953339-132955354 | p5 | (TTTTA)3 | ATGTCATTGGCCCTATCGAG | AATCTCCCGCATCAGATCAC | 155 |
| chr5:132984278-132986293 | p3 | (TTG)5 | TTTGTGCATTCGGAAAAACA | GGCCCTATATTCCTTGCCAT | 145 |

| Marker ID | SSR type | SSR motif | FORWARD PRIMER (5'-3') | REVERSE PRIMER (5'-3') | size (bp) |
|---|---|---|---|---|---|
| chr5:133047446-133049462 | p2 | (TA)8 | TTGATGGAAATGCGATGAAA | AAGAGGGAAACGGACACCTT | 243 |
| chr5:133123987-133126005 | p2 | (GT)9 | TGTGGCTTCAACTTCTGCAC | AAGGTCAAAAAGCCCCAACT | 129 |
| chr5:133155803-133157821 | p2 | (TA)9 | AATTGGTTGGACGGTGGTAG | TTGGCTCGTTACATGAGCAG | 119 |
| chr5:133175890-133177906 | p2 | (AT)8 | ACTTTTCCCGCGTCCTTAAT | GCTTTTCTTCCCGTGATTGA | 103 |
| chr5:133175890-133177906 | p5 | (TAAAA)3 | ACTTTTCCCGCGTCCTTAAT | GCTTTTCTTCCCGTGATTGA | 103 |
| chr5:133191194-133193209 | p5 | (CAAAC)3 | GGGGCAATACTTGCATTTTG | CTGCAACTCAAGCACATGGT | 281 |
| chr5:133198717-133200732 | p5 | (TTTTA)3 | CGAATCAATCCGATCCAGTT | ACCAGCAATTGCCTCAAATC | 237 |
| chr5:133317199-133319215 | p2 | (TG)8 | GAAAAGGGTGCAAGTTGGAA | AGCCCATTGGGTTCTCTTTT | 268 |
| chr5:133317199-133319215 | p2 | (AT)17 | GAAAAGGGTGCAAGTTGGAA | AGCCCATTGGGTTCTCTTTT | 268 |
| chr5:133405145-133407157 | p2 | (TA)11 | TTCAAGAAGAGGGCAGTCGT | GGACGACCAGAAAGTTGAGC | 158 |
| chr5:133405145-133407157 | p2 | (AT)6 | TTCAAGAAGAGGGCAGTCGT | GGACGACCAGAAAGTTGAGC | 158 |
| chr5:133405145-133407157 | p5 | (AATTT)3 | TTCAAGAAGAGGGCAGTCGT | GGACGACCAGAAAGTTGAGC | 158 |
| chr5:133425362-133427377 | p3 | (CTG)5 | ACAGGAAGCTAGCCAGTCCA | TCCCCTCTCGTCGCTAGTAA | 247 |
| chr5:133553535-133555604 | p5 | (CGGGT)3 | ACCGGAGATTTTGACTCCCT | CTAGGGTTGCGTCGTCTCTC | 281 |
| chr5:133560817-133562831 | p2 | (AT)7 | AAGGAAGCACCCAAGACAGA | ACTCTCCGGGGTTAAAATGG | 113 |
| chr5:133640421-133642436 | p5 | (TATTG)3 | AGTGTAATCGTGGCCTCACC | GAGCGTGGACTCTTTTACGC | 288 |
| chr5:133649753-133651768 | p3 | (CAC)5 | TATGAAACATGAGTGCCCCA | TGAGATAGAGATCGGGGGTG | 229 |
| chr5:133668236-133670250 | p2 | (TG)7 | AAAGGTGCCTACGGGCTATT | CGACCTCATAGGATCGGTGT | 241 |
| chr5:133683981-133685993 | p2 | (GT)6 | GGATTCTTTGCCTTCGATGA | GCGAATTTCCCTTTTTACCC | 223 |
| chr5:133765596-133767611 | p5 | (ACCAA)3 | CCACCTTATACCGGAAACCA | AGCATAGCCATGTTGAAGGG | 252 |
| chr5:133767355-133769370 | p2 | (AT)6 | TGTTTGGCATGAATGTCGAT | CGGAAACTCTTTCCATCTGC | 254 |
| chr5:133767355-133769370 | p5 | (ATTTT)3 | TGTTTGGCATGAATGTCGAT | CGGAAACTCTTTCCATCTGC | 254 |
| chr5:133799602-133801656 | p6 | (TATATG)9 | CATCCGCCATCTAAGGTGAT | TCCCACATTTATGACCCGAT | 111 |
| chr5:133799602-133801656 | p5 | (TAAAA)3 | CATCCGCCATCTAAGGTGAT | TCCCACATTTATGACCCGAT | 111 |
| chr5:133800631-133802646 | p6 | (GTATAT)5 | TCGCCACCTAGCATTTAAGG | GGATGCCTAACACCTTCCCT | 135 |
| chr5:133800631-133802646 | p5 | (TAAAA)3 | TCGCCACCTAGCATTTAAGG | GGATGCCTAACACCTTCCCT | 135 |
| chr5:133800631-133802646 | p3 | (TAG)8 | TCGCCACCTAGCATTTAAGG | GGATGCCTAACACCTTCCCT | 135 |
| chr5:133860464-133862484 | p2 | (AT)10 | GACGTAGTGTGGTTGCCCTT | ACATAACAGCCCGCATTAGC | 234 |
| chr5:133984106-133986124 | p6 | (ATCAGA)3 | TCAACATCGAGTTGGAGCTG | CCATCTCTGCCCAGGAAGTA | 125 |
| chr5:134074492-134076512 | p5 | (TTGGT)3 | GTTTGGGTCGTGTCACCTCT | TGGTGCAGTTTGTCGGTTTA | 100 |
| chr5:134074492-134076512 | p5 | (TTGGT)4 | GTTTGGGTCGTGTCACCTCT | TGGTGCAGTTTGTCGGTTTA | 100 |
| chr5:134150546-134152561 | p5 | (GAAAA)3 | GCAAGCCTTGGAAGAAGTTG | TGAACAGCAGACAATCTCGG | 202 |
| chr5:134150546-134152561 | p5 | (AAGAA)3 | GCAAGCCTTGGAAGAAGTTG | TGAACAGCAGACAATCTCGG | 202 |
| chr5:134168601-134170616 | p5 | (GAAAA)3 | GCAAGCCTTGGAAGAAGTTG | TGAACAGCAGACAATCTCGG | 202 |
| chr5:134168601-134170616 | p5 | (AAGAA)3 | GCAAGCCTTGGAAGAAGTTG | TGAACAGCAGACAATCTCGG | 202 |
| chr5:134189186-134191198 | p2 | (TA)6 | CTTGATTTAACGTCGCAGCA | GCTCGTGGTCACAAACTTGA | 126 |

| Marker ID | SSR type | SSR motif | FORWARD PRIMER (5'-3') | REVERSE PRIMER (5'-3') | size (bp) |
|---|---|---|---|---|---|
| chr5:134272773-134274785 | p2 | (TA)6 | TGCATTCTCAACAATGGAGC | GCACTTATGTTGGGGCTTGT | 254 |
| chr5:134272773-134274785 | p6 | (AACCGG)3 | TGCATTCTCAACAATGGAGC | GCACTTATGTTGGGGCTTGT | 254 |
| chr5:134272773-134274785 | p2 | (AT)6 | TGCATTCTCAACAATGGAGC | GCACTTATGTTGGGGCTTGT | 254 |
| chr5:134298000-134300015 | p3 | (TAT)21 | TTGCATCCGTCTCTGCATAC | AGGTGATGAGCCCTTGATTG | 105 |
| chr5:134298000-134300015 | p5 | (AGAAA)3 | TTGCATCCGTCTCTGCATAC | AGGTGATGAGCCCTTGATTG | 105 |
| chr5:134319455-134321481 | p5 | (TTGTC)3 | TCAGTGGAAGACAAGATGCG | AAGGGAGGATAAACGCGACT | 249 |
| chr5:134319455-134321481 | p2 | (TA)13 | TCAGTGGAAGACAAGATGCG | AAGGGAGGATAAACGCGACT | 249 |
| chr5:134319455-134321481 | p5 | (TTTAC)4 | TCAGTGGAAGACAAGATGCG | AAGGGAGGATAAACGCGACT | 249 |
| chr5:134319455-134321481 | p2 | (AC)7 | TCAGTGGAAGACAAGATGCG | AAGGGAGGATAAACGCGACT | 249 |
| chr5:134387313-134389325 | p2 | (AT)6 | CACGCCAACTTTTTCTTCGT | GCCTGACAAGTGACAATGGA | 289 |
| chr5:134436124-134438136 | p2 | (CA)7 | ACCTTCAAGGATGATTCCCC | TGGGACCAGAATCCCATTTA | 220 |
| chr5:134436124-134438136 | p2 | (TC)6 | ACCTTCAAGGATGATTCCCC | TGGGACCAGAATCCCATTTA | 220 |
| chr5:134475330-134477348 | p6 | (TTTTAT)3 | GGTTTTTGATTTTGCGGGTA | TCCCCTTCCCTACCTTCATT | 107 |
| chr5:134515111-134517129 | p3 | (ATG)6 | TCAGTAAAACCGAACCGGAC | GGCTCCGTGGATTAAAATGA | 231 |
| chr5:134682826-134684854 | p2 | (AT)14 | GAAAATGCTATCACCCCCAA | CATTTTCAGCAAAGTGGGGT | 294 |
| chr5:134686556-134688572 | p2 | (AT)8 | GCTCCACGCCGTACTACTTC | CGAAGCCACGGTTGATACTT | 118 |
| chr5:134686556-134688572 | p2 | (AT)6 | GCTCCACGCCGTACTACTTC | CGAAGCCACGGTTGATACTT | 118 |
| chr5:134686677-134688689 | p2 | (AT)8 | AAGTATCAACCGTGGCTTCG | TTTCGTTAATTTCCCCCACA | 232 |
| chr5:134686677-134688689 | p2 | (AT)6 | AAGTATCAACCGTGGCTTCG | TTTCGTTAATTTCCCCCACA | 232 |
| chr5:134720171-134722186 | p5 | (TTTTC)3 | AGCCTAATGGCAGAACCAAG | AACCCCTTCCTTATGCAACC | 288 |
| chr5:134733647-134735665 | p6 | (TTCAGT)3 | CTGGGAATTGGGTGAAGAAA | ATGAGGCGTAGGGAGAATCA | 293 |
| chr5:134792306-134794370 | p3 | (TGT)5 | GGAAATGAGAATGCTGCGAT | ACATGCCCAAACCGTCTTAG | 187 |
| chr5:134792306-134794370 | p2 | (AT)32 | GGAAATGAGAATGCTGCGAT | ACATGCCCAAACCGTCTTAG | 187 |
| chr5:134892351-134894366 | p5 | (TATGA)3 | TTGGAGACGTGTATGAAGCG | TTTAAGTGCAGATTTCGGGG | 150 |
| chr5:134921024-134923038 | p2 | (AT)7 | GGCAGCATTACCCAGTTGTT | AAAGTTGCACCGTGTAAGGC | 141 |
| chr5:134935209-134937224 | p3 | (TGT)5 | AAACGGACAGGGTCATTACG | TGGGCCTACCTTTTCATCTG | 102 |
| chr5:134982018-134984072 | p3 | (TGT)7 | CTGGAAGGCCAAGAATTGAG | ACAATGTGAGGTCTGGGGAG | 254 |
| chr5:134990582-134992597 | p5 | (CATTA)3 | GTCCGGTGTTCAAATTGTCC | TGCTGAGCATGGTCTGAAAC | 204 |
| chr5:134990582-134992597 | p5 | (AAATT)3 | GTCCGGTGTTCAAATTGTCC | TGCTGAGCATGGTCTGAAAC | 204 |
| chr5:135048515-135050535 | p2 | (AT)10 | CTACTCGCAAAAGAGGGTCG | AAGTCATGCCTCGGGTACAG | 297 |
| chr5:135062019-135064033 | p2 | (TA)7 | TCCCAAATTGGTCGAATGAT | TCAATTATCATGGAACGCGA | 281 |
| chr5:135062019-135064033 | p2 | (AT)19 | TCCCAAATTGGTCGAATGAT | TCAATTATCATGGAACGCGA | 281 |
| chr5:135063092-135065136 | p2 | (AT)19 | ACTGAGCCGTGTTGTAGGCT | TCTGTGGTGTGGGCTCATTA | 152 |
| chr5:135063092-135065136 | p2 | (AT)22 | ACTGAGCCGTGTTGTAGGCT | TCTGTGGTGTGGGCTCATTA | 152 |
| chr5:135347435-135349450 | p5 | (AAAAT)3 | GTGGTTGAGGGATTTGGAGA | CCTAGCAGAGGCAGATCCAG | 276 |
| chr5:135414322-135416342 | p2 | (AT)10 | GAGGTTCAATGCAGAGCACA | GCCACATTTAGCCAACCAAG | 119 |

| Marker ID | SSR type | SSR motif | FORWARD PRIMER (5'-3') | REVERSE PRIMER (5'-3') | size (bp) |
|---|---|---|---|---|---|
| chr5:135522021-135524036 | p5 | (ACAAA)3 | AAAAACGACCAGCTCCATGT | CCAAGAACGTGACGGTGATA | 298 |
| chr5:135522021-135524036 | p3 | (ATT)9 | AAAAACGACCAGCTCCATGT | CCAAGAACGTGACGGTGATA | 298 |
| chr5:135555806-135557824 | p6 | (TTTTTG)3 | TGAACTGTACCATTGACGGC | GCGAGGGTATTTTGACCTGA | 178 |
| chr5:135653590-135655602 | p2 | (TA)6 | ATCCATTTTCCATGGACTGC | CTCGGACTCATCCTATTCCG | 139 |
| chr5:135653590-135655602 | p2 | (AG)6 | ATCCATTTTCCATGGACTGC | CTCGGACTCATCCTATTCCG | 139 |
| chr5:135684497-135686512 | p5 | (ATTTT)3 | CTGGTTGAGCACTTGATGGA | AGCAAGCCTGCCACAGTAAT | 287 |
| chr5:135696789-135698804 | p3 | (ATA)5 | GGAGGCGGTGGAGTATATGA | ATTTTGACTTACCGGACCCC | 231 |
| chr5:135696919-135698943 | p3 | (ATA)5 | GGAGGCGGTGGAGTATATGA | ATTTTGACTTACCGGACCCC | 101 |
| chr5:135812462-135814477 | p2 | (GA)7 | GCTCAGATTTGAGGCAGAGG | CCCCTCCGTTTCTCTCTCTT | 235 |
| chr5:136018374-136020392 | p6 | (TTTCTT)3 | TCTTGACCCATTTGAAGGGA | CTGGGCTTGATTCTCGGATA | 170 |
| chr5:136027139-136029199 | p2 | (AT)20 | GTACGCCCTAAACACGCCTA | ATTCACCCTCACGACGTTCT | 284 |
| chr5:136047996-136050012 | p5 | (TCTTT)3 | TTGCTTCGCGGTTTTAAAGT | GGCTCCGGTAAATCATGAAA | 195 |
| chr5:136047996-136050012 | p2 | (AT)8 | TTGCTTCGCGGTTTTAAAGT | GGCTCCGGTAAATCATGAAA | 195 |
| chr5:136057632-136059647 | p5 | (TCTTT)3 | ACGCGAGGAATTGTTTTACG | TTGTGGACCATTGAAAGCAA | 173 |
| chr5:136057632-136059647 | p2 | (AT)9 | ACGCGAGGAATTGTTTTACG | TTGTGGACCATTGAAAGCAA | 173 |
| chr5:136058212-136060230 | p5 | (TCTTT)3 | TGCTTCGCGGTTTTAAAGTT | GGCTCCGGTAAATCATGAAA | 196 |
| chr5:136058212-136060230 | p2 | (AT)9 | TGCTTCGCGGTTTTAAAGTT | GGCTCCGGTAAATCATGAAA | 196 |
| chr5:136058212-136060230 | p5 | (GTTCG)4 | TGCTTCGCGGTTTTAAAGTT | GGCTCCGGTAAATCATGAAA | 196 |
| chr5:136095952-136097964 | p5 | (TATTT)3 | GGATTTAGGTGGGCTTAGGG | TGGATTAAAGCAGTGGTGGA | 239 |
| chr5:136095952-136097964 | p2 | (TG)6 | GGATTTAGGTGGGCTTAGGG | TGGATTAAAGCAGTGGTGGA | 239 |
| chr5:136111619-136113634 | p5 | (CCCCA)3 | GAACCCCCTCGGTTACACTT | AAAAATGAAAGACCACCCCC | 252 |
| chr5:136158704-136160719 | p5 | (TTTTA)3 | AAGACACGATTGAGATGCCA | ATGGGACGAAGGGAGTAACA | 151 |
| chr5:136158704-136160719 | p2 | (AT)9 | AAGACACGATTGAGATGCCA | ATGGGACGAAGGGAGTAACA | 151 |
| chr5:136241556-136243581 | p5 | (AAAAT)5 | GCATCCGAGTCCTTTGGTAA | TGTTGTTTCTGGTGAGCTGC | 159 |
| chr5:136295378-136297442 | p6 | (GGAGGT)3 | GGTGGAGGTGGAGGTAGTGA | ACTGTTCGAGAACCACTGGC | 154 |
| chr5:136295378-136297442 | p2 | (AT)32 | GGTGGAGGTGGAGGTAGTGA | ACTGTTCGAGAACCACTGGC | 154 |
| chr5:136362165-136364186 | p2 | (TA)10 | GACCCCCAAGTATCCGAAAT | TCGGGTCGAAATCCTAAATG | 119 |
| chr5:136362165-136364186 | p5 | (ACCCG)3 | GACCCCCAAGTATCCGAAAT | TCGGGTCGAAATCCTAAATG | 119 |
| chr5:136369283-136371298 | p5 | (AAAAG)3 | TTTTGCCCAATAGGATGAGC | TGTGACAAAATTCAGCCCAA | 101 |
| chr5:136394213-136396255 | p2 | (TA)21 | GTGAATTCATTGCCACATGC | CATGGCGGAATAGCGAATAC | 267 |
| chr5:136397254-136399269 | p5 | (GTTCG)3 | AACTTTGCTTTGCGATGCTT | GGAATGCCCTTATTTTGGGT | 285 |
| chr5:136397767-136399782 | p5 | (GTTCG)3 | TTTCCACTCGTCAACCTTCC | CGGTATTTGGTGTCCACCTT | 176 |
| chr5:136397794-136399809 | p5 | (GTTCG)3 | TTTCCACTCGTCAACCTTCC | CGGTATTTGGTGTCCACCTT | 149 |
| chr5:136492180-136494192 | p2 | (TG)6 | TGCAATATCCATACCGCTCA | GAAAAGTGCCCATCTTTCCA | 140 |
| chr5:136497064-136499076 | p2 | (TA)6 | ATGGGCTTGTCCTCTCCTTT | TTTGGCCAGAGTACTGGCTT | 159 |
| chr5:136500896-136502912 | p3 | (TGT)7 | ATCTCTCTCACATCCGTGGG | AAACTGAGGGCAAAACGAGA | 218 |

| Marker ID | SSR type | SSR motif | FORWARD PRIMER (5'-3') | REVERSE PRIMER (5'-3') | size (bp) |
|---|---|---|---|---|---|
| chr5:136500896-136502912 | p4 | (AATT)4 | ATCTCTCTCACATCCGTGGG | AAACTGAGGGCAAAACGAGA | 218 |
| chr5:136504037-136506057 | p5 | (TGGGG)4 | CGGCGAACCATTGTAAATCT | ATCCCGACATCAATTCTTGG | 256 |
| chr5:136553429-136555441 | p2 | (AT)18 | CTGAAGCCGAACTCGCTATT | CGACAAGCCCTTTTTGCTAC | 273 |
| chr5:136553429-136555441 | p2 | (AT)8 | CTGAAGCCGAACTCGCTATT | CGACAAGCCCTTTTTGCTAC | 273 |
| chr5:136553443-136555455 | p2 | (AT)18 | CTGAAGCCGAACTCGCTATT | CGACAAGCCCTTTTTGCTAC | 259 |
| chr5:136553443-136555455 | p2 | (AT)8 | CTGAAGCCGAACTCGCTATT | CGACAAGCCCTTTTTGCTAC | 259 |
| chr5:136560082-136562127 | p3 | (ATT)15 | ATGCATGCGTTGGTTGAATA | GGGGAGAATCATAACGCAAA | 267 |
| chr5:136563374-136565386 | p2 | (TA)6 | CGGGGAGATACACTCGAAAA | GAGGTACTTCCACCGCAAGA | 261 |
| chr5:136563374-136565386 | p2 | (AT)6 | CGGGGAGATACACTCGAAAA | GAGGTACTTCCACCGCAAGA | 261 |
| chr5:136593196-136595256 | p3 | (ATT)20 | TGCGGCATCATATTAGGTGA | ACCCAACCGGTACAACGTAA | 282 |
| chr5:136602714-136604735 | p3 | (ATT)7 | CTCCGAATGCGGGTATCTTA | CGCTGCCAAAGAAAAAGTTC | 258 |
| chr5:136681837-136683931 | p2 | (TA)6 | CTCGGGAATTGGAGTTGTGT | ATTTGGCCATGCATGTTTTT | 177 |
| chr5:136681837-136683931 | p2 | (TA)47 | CTCGGGAATTGGAGTTGTGT | ATTTGGCCATGCATGTTTTT | 177 |
| chr5:136730286-136732304 | p3 | (CAA)6 | ACGAGCGGAGTAGTGAGGAA | TATGTTTTTCATAGGCGGGC | 282 |
| chr5:136742024-136744039 | p3 | (CAA)5 | ACTACCTCCCTCGGGAAAAA | AGGTCTGCCTACACACCACC | 202 |
| chr5:136747620-136749635 | p3 | (ACA)5 | CCAAACTATCTCAGCCTCGC | CTGATTTTGAGCCTCGCTTC | 289 |
| chr5:136747620-136749635 | p3 | (ATA)6 | CCAAACTATCTCAGCCTCGC | CTGATTTTGAGCCTCGCTTC | 289 |
| chr5:136748208-136750226 | p3 | (ACA)5 | ACGTGAACCCATAATCAGGC | ATATGGGATTGGATGAAGCG | 274 |
| chr5:136748208-136750226 | p3 | (ATA)6 | ACGTGAACCCATAATCAGGC | ATATGGGATTGGATGAAGCG | 274 |
| chr5:136775074-136777086 | p2 | (CT)6 | CTCTATCCACCCCACCAAAA | GGAAGGATCCAGAATCCAGA | 179 |
| chr5:136802613-136804625 | p2 | (TA)6 | TTTCGGACACGTGTATTCCA | TTGAGCATGGATTCTTCCCT | 146 |
| chr5:136840903-136842915 | p2 | (TA)6 | TTTCGGACACGTGTATTCCA | TTGAGCATGGATTCTTCCCT | 146 |
| chr5:136899326-136901341 | p3 | (TAA)6 | AAGGTTCAAAGTGGGTGCAG | AAAAACATGTCAGGGGCAAA | 152 |
| chr5:136899326-136901341 | p3 | (AAG)5 | AAGGTTCAAAGTGGGTGCAG | AAAAACATGTCAGGGGCAAA | 152 |
| chr5:136917314-136919329 | p5 | (AAAAT)3 | GCATCAAAACAAGCACCAGA | GACGGTTTTGAAGAGTCCGA | 288 |
| chr5:136917314-136919329 | p3 | (AAT)5 | GCATCAAAACAAGCACCAGA | GACGGTTTTGAAGAGTCCGA | 288 |
| chr5:136971818-136973836 | p2 | (AT)9 | GAGCATTGCCAGTTTCCTTC | CCTGTTCCTTCCAACTCTCG | 265 |
| chr5:136996761-136998776 | p5 | (ATATA)3 | GGCCAAACACTAATTGCTGC | GTTTGGTCACAGCCCAGACT | 224 |
| chr5:137024884-137026902 | p3 | (ACA)6 | CACACCAGGCGAGATAGGAT | CTCCTCCTCACATTCCCAAA | 157 |
| chr5:137027939-137029960 | p3 | (AGC)7 | AGGGAGCCTGGCTTAGTTTC | CGTTGTTACTCAACGGCAGA | 135 |
| chr5:137051420-137053438 | p3 | (TGT)6 | GCCACCAAAACTTAAAGGCA | CATCTGTGCCTCTCCTCCTC | 256 |
| chr5:137051420-137053438 | p3 | (ACA)6 | GCCACCAAAACTTAAAGGCA | CATCTGTGCCTCTCCTCCTC | 256 |
| chr5:137105400-137107415 | p2 | (TA)6 | GTCGGTTCACAAGACGGATT | TTCCTCGATCCCAATGTAGG | 293 |
| chr5:137129166-137131181 | p3 | (TGA)5 | GGTGGGTTCCTGAACTTGAA | GCCTGAAGAATGGGAAACAA | 126 |
| chr5:137131073-137133133 | p3 | (CTT)20 | CAAGTTTAGGGGGTCGCATA | TCCTGAGTGAAAATCCTGGC | 149 |
| chr5:137136160-137138175 | p3 | (TCT)5 | CAAAGGAGGACAAACAGGGA | CAGCATCACTCTAGTGCCCA | 226 |

| Marker ID | SSR type | SSR motif | FORWARD PRIMER (5'-3') | REVERSE PRIMER (5'-3') | size (bp) |
|---|---|---|---|---|---|
| chr5:137136564-137138578 | p3 | (TCT)5 | GGTTAAGGTGCTGGTGCAAT | TATCAAACCCGGCTAGCAAC | 165 |
| chr5:137136601-137138615 | p3 | (TCT)5 | GGTTAAGGTGCTGGTGCAAT | TATCAAACCCGGCTAGCAAC | 128 |
| chr5:137192519-137194534 | p2 | (TA)9 | AACATCACACGGTGGACTCA | CAAGCACAATGAAGAACCGA | 269 |
| chr5:137192535-137194553 | p2 | (TA)9 | AACATCACACGGTGGACTCA | CAAGCACAATGAAGAACCGA | 253 |
| chr5:137196967-137198985 | p2 | (TA)11 | TTGGGTGCCAAACACAACTA | TTTCCGTATCCGAATCTTGC | 270 |
| chr5:137196967-137198985 | p6 | (TTTCTC)3 | TTGGGTGCCAAACACAACTA | TTTCCGTATCCGAATCTTGC | 270 |
| chr5:137201390-137203456 | p3 | (AAT)22 | TGCGTACACTCTACCCTCCC | CGAAGCTTGGCCAAATAATC | 232 |
| chr5:137212086-137214122 | p2 | (AT)18 | CCACTTGATCGGCAGTTTTT | TGTAGTGGGACCGATTTTCC | 218 |
| chr5:137263054-137265075 | p3 | (TGT)6 | ACCTCCATTATGAAAGCCCC | GCAACAACGAAGTAGTGCGA | 156 |
| chr5:137263054-137265075 | p3 | (GAA)7 | ACCTCCATTATGAAAGCCCC | GCAACAACGAAGTAGTGCGA | 156 |
| chr5:137285635-137287649 | p2 | (AT)10 | AAAAGAGTGGCCCCCAATAC | CGGAGATGGTGAGTTCCACT | 148 |
| chr5:137285654-137287668 | p2 | (AT)10 | AAAAGAGTGGCCCCCAATAC | CGGAGATGGTGAGTTCCACT | 129 |
| chr5:137584210-137586228 | p2 | (CA)6 | TTGCAGGGAAAGTAAATGCC | CCCCACTAACCTGACATGCT | 106 |
| chr5:137584210-137586228 | p6 | (AATTCA)3 | TTGCAGGGAAAGTAAATGCC | CCCCACTAACCTGACATGCT | 106 |
| chr5:137598194-137600206 | p2 | (AG)6 | GGAACCTGGCTTTAAAAGGG | GTCTGCAATAAGCCGTCTCC | 230 |
| chr5:137851209-137853224 | p3 | (TCA)5 | TTCCATCTCTGCTGCACAAC | AGAAGTCGTAACAAGCGGGA | 132 |
| chr5:137856793-137858817 | p6 | (TCTATA)4 | GGCCAAACACAAACTGCTTT | CATGACGTGGGAAGAGAGGT | 285 |
| chr5:137939560-137941574 | p2 | (TA)9 | ATCCCGTTTGTCCCAATTTT | ATACGGCATGTGGCAATACA | 184 |
| chr5:137939592-137941607 | p2 | (TA)9 | ATCCCGTTTGTCCCAATTTT | ATACGGCATGTGGCAATACA | 152 |
| chr5:137940379-137942397 | p2 | (TA)9 | AAACAAACACCAAGCCCAAT | TTTTTAGGGACAAACGGACG | 269 |
| chr5:137940379-137942397 | p2 | (AT)8 | AAACAAACACCAAGCCCAAT | TTTTTAGGGACAAACGGACG | 269 |
| chr5:137955980-137958034 | p2 | (AT)27 | CCGACCTATCAAATGGCACT | AATTTCAGCGACCCAGAATG | 252 |
| chr5:137960582-137962663 | p3 | (TTA)27 | CACCCGCATCGCTTTATTAT | AGCCGAGTGTTCGAGGTAGA | 255 |
| chr5:138156627-138158645 | p2 | (CT)6 | TTCACGACTTCAGCAACCAC | TCGAAATCTGAATTGTGCCA | 112 |
| chr5:138156627-138158645 | p6 | (AAAAAT)3 | TTCACGACTTCAGCAACCAC | TCGAAATCTGAATTGTGCCA | 112 |
| chr5:138164908-138166947 | p3 | (AAT)13 | TGCAAATGGACAAAACCTCA | GTTTTGGGGAGGGGAAATTA | 224 |
| chr5:138211337-138213445 | p5 | (AAAAT)3 | CTTACCACGGAGCATCCATT | ATATTCGGCGAAGTTTGTGG | 199 |
| chr5:138211337-138213445 | p3 | (ATA)36 | CTTACCACGGAGCATCCATT | ATATTCGGCGAAGTTTGTGG | 199 |
| chr5:138241186-138243226 | p3 | (TTA)6 | CATTCTGCCTGGATCCTCAT | CGGAGCAAGTGGGATTTAGA | 200 |
| chr5:138241726-138243744 | p3 | (TTA)6 | TTAGTCGGCCCACATTTCTC | GAGTAACACGCAATCGCAAA | 110 |
| chr5:138300492-138302506 | p2 | (TA)7 | TGGTACACACCGAGTCCTGA | CTACCTGAACCTGTGGGCAT | 122 |
| chr5:138300492-138302506 | p2 | (TA)7 | TGGTACACACCGAGTCCTGA | CTACCTGAACCTGTGGGCAT | 122 |
| chr5:138401601-138403622 | p3 | (ACA)5 | TGTGGAGCCAGAATTTGACA | CATATCCATGTTCGAACCCC | 286 |
| chr5:138401601-138403622 | p3 | (ACA)7 | TGTGGAGCCAGAATTTGACA | CATATCCATGTTCGAACCCC | 286 |
| chr5:138404941-138406985 | p2 | (TA)22 | CACGTGCGTATTCGTAGCAA | TGCTTGATTTTGTCAATATGGG | 270 |
| chr5:138456390-138458402 | p4 | (TTTA)4 | TCGTTAACGGGAACACATCA | GAGAAATTGCCCAAAAGCAA | 184 |

| Marker ID | SSR type | SSR motif | FORWARD PRIMER (5'-3') | REVERSE PRIMER (5'-3') | size (bp) |
|---|---|---|---|---|---|
| chr5:138458821-138460836 | p6 | (TTTTTA)3 | ACCCGTCTTCCCAAGACTTT | TGTCACGCCAATATTCAGGA | 272 |
| chr5:138458821-138460836 | p5 | (TTTTG)3 | ACCCGTCTTCCCAAGACTTT | TGTCACGCCAATATTCAGGA | 272 |
| chr5:138462771-138464795 | p4 | (TTTA)6 | CTTAGGGGTCGTTTGGTTGA | CTGAAGAATGTGCAGAGCCA | 203 |
| chr5:138489957-138491975 | p2 | (TG)6 | TAACGCAAAATTTTCCAGGG | CAGTCCATTTGAAGGGCAGT | 208 |
| chr5:138489957-138491975 | p3 | (GAA)6 | TAACGCAAAATTTTCCAGGG | CAGTCCATTTGAAGGGCAGT | 208 |
| chr5:138514565-138516593 | p2 | (AT)14 | CAAACGCCTACTTTTTCCCA | TGGCTGCAAAAATAACCCAT | 149 |
| chr5:138514565-138516593 | p2 | (AT)8 | CAAACGCCTACTTTTTCCCA | TGGCTGCAAAAATAACCCAT | 149 |
| chr5:138540094-138542109 | p5 | (AATAA)3 | TGAAGAAACCCAGATCCCAC | GTGCTTATAAGTGGGGGCAA | 282 |
| chr5:138565185-138567197 | p2 | (AG)6 | CACCTACGGCGGAAATTAAA | AACATATGTCGGCTTCCAGC | 234 |
| chr5:138635339-138637354 | p3 | (TTC)5 | ATTTCCAATGCCCAAAACAA | GAAACGGACGAAGAAAGGTG | 233 |
| chr5:138735039-138737057 | p6 | (AAAAAT)3 | GGAGGCGGATGTAGACTCTG | TTACACACAGAGGCAGACGC | 145 |
| chr5:138847760-138849780 | p4 | (AACA)5 | TCGGTGGATTCACAATTTCA | AAATCATGCACACCTGACGA | 108 |
| chr5:138876626-138878648 | p2 | (AT)20 | GGAGTCCAGAGCCAAAATGA | GCCTTGTGTGGGTATTTCGT | 160 |
| chr5:138876626-138878648 | p2 | (AT)9 | GGAGTCCAGAGCCAAAATGA | GCCTTGTGTGGGTATTTCGT | 160 |
| chr5:138876626-138878648 | p2 | (AT)11 | GGAGTCCAGAGCCAAAATGA | GCCTTGTGTGGGTATTTCGT | 160 |
| chr5:138876626-138878648 | p2 | (AT)12 | GGAGTCCAGAGCCAAAATGA | GCCTTGTGTGGGTATTTCGT | 160 |
| chr5:138896051-138898065 | p2 | (TG)7 | AAGCAACAAATGGAACTGGC | CAACCAAACACCATCAGCAC | 178 |
| chr5:138928142-138930156 | p2 | (AG)7 | TCTATCTCTTCAGGGGCGAA | ATCATGGCACTGCAAATTCA | 146 |
| chr5:138938093-138940108 | p5 | (AACCG)3 | TGGAGGGTTGCTTCAGTCTT | ACCCCATGTCCCTCCTAAAC | 125 |
| chr5:138946909-138948925 | p4 | (TTTG)4 | GGGGAGGCTACAATGACAAA | TCCTAATCATCCAAGCTGCC | 127 |
| chr5:139046270-139048282 | p2 | (AT)6 | CCTCGTGGTCGTCTTTAAGC | TGCCAGCTGAGGAGAGGTAT | 277 |
| chr5:139072182-139074198 | p2 | (TA)8 | ATTAAACGGGTCAGCAGGTG | GGCCAAAAATATCCTCAACG | 233 |
| chr5:139085445-139087457 | p5 | (ACCAA)3 | GGAATTTCTTCATCCCCGTT | ATAGGAACTAAGCCCCGCAT | 152 |
| chr5:139085458-139087586 | p5 | (ACCAA)3 | GGAATTTCTTCATCCCCGTT | ATAGGAACTAAGCCCCGCAT | 139 |
| chr5:139113391-139115406 | p2 | (TA)23 | ACAAGAGAATTTGCGACGGT | TAAATTCGTCTCGAATCCCG | 278 |
| chr5:139115401-139117413 | p2 | (TA)6 | CCACGAGGACCTCCACATAC | TGACGTCATGTTCTCCCTCA | 131 |
| chr5:139277154-139279170 | p4 | (CATT)4 | ATGACCAGCTGCAAATGATG | AACACATCCTTAAGGCCACG | 174 |
| chr5:139278960-139280992 | p2 | (AT)16 | TATGCCACCTCAACATCGAA | TATGCAATTGCCACACCACT | 269 |
| chr5:139278960-139280992 | p5 | (ATAAT)3 | TATGCCACCTCAACATCGAA | TATGCAATTGCCACACCACT | 269 |
| chr5:139332034-139334052 | p6 | (TTGGAG)3 | GCCAAATCAACATCAGCCTT | ACTGGTGGACACAGGACACA | 296 |
| chr5:139332034-139334052 | p2 | (AT)6 | GCCAAATCAACATCAGCCTT | ACTGGTGGACACAGGACACA | 296 |
| chr5:139332857-139334869 | p6 | (TTGGAG)3 | GTAGGCGTTTGGCCATAGAA | GGCCATGAAAATTCCAAAGA | 118 |
| chr5:139332857-139334869 | p2 | (AT)6 | GTAGGCGTTTGGCCATAGAA | GGCCATGAAAATTCCAAAGA | 118 |
| chr5:139341710-139343725 | p5 | (ATTTT)3 | TCAACGTAAAAATCCCGGAG | TTCGGATCGTCATTTTAGGG | 209 |
| chr5:139369845-139371877 | p2 | (AT)16 | TCAACTTGGATGGATCAGCA | CTGTGCATGTGCAAGTTCCT | 220 |
| chr5:139397071-139399091 | p3 | (AAT)6 | ATGTCATTTCTCACCCACCC | GGTTGTGCTGAATTCCAGGT | 193 |

| Marker ID | SSR type | SSR motif | FORWARD PRIMER (5'-3') | REVERSE PRIMER (5'-3') | size (bp) |
|---|---|---|---|---|---|
| chr5:139397071-139399091 | p2 | (TA)10 | ATGTCATTTCTCACCCACCC | GGTTGTGCTGAATTCCAGGT | 193 |
| chr5:139447292-139449312 | p2 | (AT)10 | TCATTTGGGCTGCTATCACA | AGGGTTAAGCTGGTGAGCCT | 120 |
| chr5:139479418-139481448 | p2 | (TA)15 | CGGTTGGGATGACTTCACTT | AAGCTCCCGCTAGCATGTAA | 220 |
| chr5:139479418-139481448 | p2 | (TA)6 | CGGTTGGGATGACTTCACTT | AAGCTCCCGCTAGCATGTAA | 220 |
| chr5:139534343-139536355 | p2 | (AT)6 | TTAAGTCCCCACAGGTCCAC | ACGGACGACGAGAAAGTGAG | 129 |
| chr5:139534343-139536355 | p2 | (AT)18 | TTAAGTCCCCACAGGTCCAC | ACGGACGACGAGAAAGTGAG | 129 |
| chr5:139534343-139536355 | p2 | (AT)12 | TTAAGTCCCCACAGGTCCAC | ACGGACGACGAGAAAGTGAG | 129 |
| chr5:139535988-139538006 | p2 | (AT)12 | ACCCACCATCTCCAAACTCA | ATATGGGGCCCACAGTTTTT | 224 |
| chr5:139535988-139538006 | p2 | (TA)9 | ACCCACCATCTCCAAACTCA | ATATGGGGCCCACAGTTTTT | 224 |
| chr5:139535988-139538006 | p2 | (AT)9 | ACCCACCATCTCCAAACTCA | ATATGGGGCCCACAGTTTTT | 224 |
| chr5:139548398-139550414 | p2 | (AT)8 | CAAATACAGCGCGTTTCTGA | TTTGTCGGTAGCGGAAGACT | 226 |
| chr5:139579911-139581923 | p2 | (AT)6 | TCGCCTCTAAGCTCTTTCCA | TTCACGTTGGGGCTTTTTAG | 262 |
| chr5:139596902-139598920 | p6 | (AAAAAG)3 | CCACTGCCGGTAATGAAGTT | TTTGTCACCCAGTAGTGGCA | 103 |
| chr5:139630817-139632831 | p2 | (AT)7 | AGCAGGAAAACACAAGGGAA | GGAAATAAATAGCCCGCACA | 201 |
| chr5:139683708-139685720 | p2 | (CT)6 | GTCGAGTGAGGCCAAAGTTC | TGGTCTTTAATTTTTGCCCG | 250 |
| chr5:139713187-139715233 | p2 | (TA)23 | TAGGAGGAAACCCATACCCC | GGGAGATCCTTGGTGCATAA | 112 |
| chr5:139824099-139826114 | p3 | (GGA)5 | AGAGGCCAAAGGAATCCAAT | CATTCATGCAGCTTACGCAT | 145 |
| chr5:139879088-139881106 | p6 | (GAGAAA)3 | CACATTGCGGCGTGTATATC | TGTAGTTTCGTTGTCGTCGC | 198 |
| chr5:139909634-139911658 | p2 | (AG)12 | CCACATTGGATACCCTCACC | GCTGGGGACATTGTGAAGAT | 128 |
| chr5:139928956-139930976 | p5 | (TGTGT)4 | AGGACAATAGCACGGTGAGG | GGTCAAAAATCCCGACTTCA | 111 |
| chr5:140033069-140035093 | p5 | (CGGGT)3 | TCGTGAACCCGAAAAGAATC | CCATTTCCCCTCCTCTCTTC | 286 |
| chr5:140033069-140035093 | p3 | (ATA)8 | TCGTGAACCCGAAAAGAATC | CCATTTCCCCTCCTCTCTTC | 286 |
| chr5:140034226-140036247 | p3 | (GTT)7 | ATTCGAACTCGGGTCACAAC | CACCTTGAAGAGTGGGGGTA | 101 |
| chr5:140085260-140087275 | p3 | (AAC)5 | TGATTTGCACCACAAATCGT | AGGGAAACACGTCAGATTCG | 149 |
| chr5:140087522-140089538 | p2 | (TA)10 | CCAATGGAGGATTGTTGCTT | TGTGTGGGGAATAGTGGGTT | 113 |
| chr5:140087522-140089538 | p2 | (TA)8 | CCAATGGAGGATTGTTGCTT | TGTGTGGGGAATAGTGGGTT | 113 |
| chr5:140087522-140089538 | p2 | (AT)14 | CCAATGGAGGATTGTTGCTT | TGTGTGGGGAATAGTGGGTT | 113 |
| chr5:140267803-140269819 | p4 | (GTAT)4 | TTTATGCCTCAACTCCCCTG | TGTTGGGTTGATGCTGAAAA | 141 |
| chr5:140315598-140317614 | p6 | (CAACTC)3 | TTTGTTGACCACATAGCCCA | CCAAACGGGGGATTACTCTAA | 136 |
| chr5:140315598-140317614 | p4 | (TTAT)4 | TTTGTTGACCACATAGCCCA | CCAAACGGGGGATTACTCTAA | 136 |
| chr5:140315598-140317614 | p5 | (TTTAT)3 | TTTGTTGACCACATAGCCCA | CCAAACGGGGGATTACTCTAA | 136 |
| chr5:140341018-140343030 | p6 | (AAAAAC)3 | TCCACATGTGCGGTTACACT | AAATAGCGGGTCGCATAAAA | 173 |
| chr5:140341018-140343030 | p2 | (TA)6 | TCCACATGTGCGGTTACACT | AAATAGCGGGTCGCATAAAA | 173 |
| chr5:140341018-140343030 | p2 | (TG)7 | TCCACATGTGCGGTTACACT | AAATAGCGGGTCGCATAAAA | 173 |
| chr5:140341018-140343030 | p2 | (TG)10 | TCCACATGTGCGGTTACACT | AAATAGCGGGTCGCATAAAA | 173 |
| chr5:140390421-140392485 | p2 | (TA)32 | TAGGGCACGGTGTAGAGCTT | AACCGATTTGGTGTAATCGC | 236 |

| Marker ID | SSR type | SSR motif | FORWARD PRIMER (5'-3') | REVERSE PRIMER (5'-3') | size (bp) |
|---|---|---|---|---|---|
| chr5:140462976-140464994 | p2 | (TA)9 | CCATGTGTGTTGTTTAGCCG | ATTTTGACGCTCACAAACCC | 140 |
| chr5:140503818-140505830 | p2 | (AC)6 | CTTCACCGTTTTCACCACCT | GAAAGCAGCAATTTCGGTTC | 239 |
| chr5:140591731-140593746 | p5 | (TATGA)3 | GTTTAGCCATGTTGGCCACT | ATACCAAGCTTTGTCACGCC | 108 |
| chr5:140743627-140745645 | p3 | (ATA)6 | TCCACTGGAAACCATCCTTC | GTCCCACCGTCATGTAATCC | 171 |
| chr5:140811038-140813054 | p2 | (CA)8 | TGAATAGTTTTGGGGGCAAG | GCAGGGCAAAAGATGGAATA | 297 |
| chr5:140815708-140817724 | p4 | (TTTC)4 | CACCAAACCTCATTTGCCTT | GTGATGCCCCTGGTCTCTTA | 275 |
| chr5:140856166-140858184 | p2 | (AT)9 | ATGTTTTGACCTTCGCTTGG | GTCTGCCCCATAAAGGCATA | 112 |
| chr5:140856166-140858184 | p5 | (ATCAT)3 | ATGTTTTGACCTTCGCTTGG | GTCTGCCCCATAAAGGCATA | 112 |
| chr5:140899965-140901983 | p6 | (AGAAAA)3 | TCAACAAAAGCATCGACACC | AACCATTGCGCTTTTCTCAT | 120 |
| chr5:140984109-140986121 | p2 | (TA)6 | TGTGGATGTAGGCCAGTTCA | TTTAGAGGGTGCCGAACATC | 216 |
| chr5:141057627-141059641 | p2 | (GA)7 | TTGACTGATCGGGAGAAACC | GGCCAAGAAAGAAGCTGTTG | 214 |
| chr5:141132474-141134498 | p3 | (TTG)8 | AGATGAAAGTGGCGGAAATG | CGCATCTTGTCTTCCACTGA | 289 |
| chr5:141132474-141134498 | p2 | (AT)7 | AGATGAAAGTGGCGGAAATG | CGCATCTTGTCTTCCACTGA | 289 |
| chr5:141228627-141230642 | p3 | (ACT)5 | GAGAAATCCCGAGTGTTCCA | CTGTTACGGGCAGGTGAAAT | 122 |
| chr5:141237710-141239724 | p2 | (AT)7 | ACCAAAGGATGTGGTGAAGC | ATATCTGGATTAGCGCCGTG | 117 |
| chr5:141554809-141556824 | p5 | (ACCAA)3 | CTAGGGGTGTACATGGACCG | CGAGAAAGCCGAATAACCTG | 201 |
| chr5:141554809-141556824 | p5 | (AAACC)3 | CTAGGGGTGTACATGGACCG | CGAGAAAGCCGAATAACCTG | 201 |
| chr5:141715486-141717501 | p3 | (AAT)5 | GCAAACAACAACCCAGTGTG | CATAGTGTTCGGGCCAACTT | 184 |
| chr5:141793749-141795765 | p2 | (AT)8 | CAGCCAAATGTGGACATCAC | GTCGCTCCGATCTAATCCAA | 223 |
| chr5:141813474-141815489 | p5 | (AATAA)3 | AGTTTGGCGTGCAATTCTTC | GAGAAGCAACCCCAAATCAA | 284 |
| chr5:141828833-141830851 | p6 | (TATTTT)3 | GGGGTTAAAATAGGGCCAAA | GGGAAAAGGGCCTGATTTAC | 133 |
| chr5:141830520-141832612 | p2 | (AT)46 | TTGGCAGCCACAAGTATGAC | ATTGTGTTCGTCCCTCTTGG | 278 |
| chr5:141875441-141877456 | p3 | (GTT)5 | AATCCCAACACACCGAACTC | TCTGCATCGTGAAGAAAACG | 289 |
| chr5:141909965-141911980 | p3 | (AAT)5 | AGTCAAGCCCTTGGAATCCT | CTGGCTGCTACAGGAAAAGG | 114 |
| chr5:141909965-141911980 | p5 | (CATGT)3 | AGTCAAGCCCTTGGAATCCT | CTGGCTGCTACAGGAAAAGG | 114 |
| chr5:141943159-141945173 | p3 | (TAT)19 | GGTTTGATACTTTCCCCCGT | AGACAAAATTGGGCGTCAAC | 138 |
| chr5:141943159-141945173 | p2 | (CT)7 | GGTTTGATACTTTCCCCCGT | AGACAAAATTGGGCGTCAAC | 138 |
| chr5:142111108-142113123 | p3 | (TCA)5 | TTCAACCGACAAAGTGGACA | CACTGTATTTTCGGCTGGGT | 127 |
| chr5:142125458-142127474 | p2 | (AC)7 | ATCAGGATTCCAACGAGCAG | GATGGGGATCTTTTGGTGAA | 183 |
| chr5:142125458-142127474 | p2 | (CA)8 | ATCAGGATTCCAACGAGCAG | GATGGGGATCTTTTGGTGAA | 183 |
| chr5:142125458-142127474 | p2 | (AC)8 | ATCAGGATTCCAACGAGCAG | GATGGGGATCTTTTGGTGAA | 183 |
| chr5:142161219-142163249 | p5 | (TTTAA)3 | TTCTTCACGTAGACCCCACC | GGGGCTCACTCTTCTGTGAT | 118 |
| chr5:142161219-142163249 | p3 | (ATT)10 | TTCTTCACGTAGACCCCACC | GGGGCTCACTCTTCTGTGAT | 118 |
| chr5:142208564-142210576 | p2 | (AT)6 | CTAGGGAGGTGAGGTGCTTG | GTGTTTGACCACGACTTCCA | 104 |
| chr5:142258117-142260129 | p2 | (AC)6 | TCCACTGCATTTGCTTGTTC | GTGGGCTTTGAACCTGTGAT | 109 |
| chr5:142307539-142309554 | p5 | (TTTTG)3 | TTACAAAGCTGACACCACGG | ATCCCAGGGAGGATTTTGAC | 157 |

| Marker ID | SSR type | SSR motif | FORWARD PRIMER (5'-3') | REVERSE PRIMER (5'-3') | size (bp) |
|---|---|---|---|---|---|
| chr5:142386127-142388145 | p6 | (TTTTTA)3 | GGCATAGTTTCCCTGCAAAA | GCCTAAAACCTCAGGGTTCC | 184 |
| chr5:142400298-142402312 | p2 | (AT)7 | TAACGTGTTTTCACTCCCCC | TGTCCAACAAATCCGTTCAA | 216 |
| chr5:142460505-142462517 | p2 | (TA)6 | CCTATCGTTTGGCTTTAGCG | TTTTTAAGGGACAACGGACG | 123 |
| chr5:142460505-142462517 | p2 | (AT)10 | CCTATCGTTTGGCTTTAGCG | TTTTTAAGGGACAACGGACG | 123 |
| chr5:142461356-142463376 | p2 | (TA)6 | ATGATGGATTCATTGCCACA | AAATGACATCGCGGAATAGC | 253 |
| chr5:142461356-142463376 | p2 | (AT)10 | ATGATGGATTCATTGCCACA | AAATGACATCGCGGAATAGC | 253 |
| chr5:142489983-142491999 | p2 | (AC)8 | CGAAGAGTGCAAACCTGTCA | CAAAGGCTGGGGTAATTGTG | 227 |
| chr5:142493091-142495145 | p2 | (TA)18 | CACTACAAGACCAAGGGCGT | TCAATTGCACCGCTAAGTTG | 175 |
| chr5:142501286-142503328 | p3 | (ACA)5 | CAGTGAAATCCCACAACGTG | TGTTTTTATTGAGCCGAGGG | 262 |
| chr5:142501286-142503328 | p2 | (TA)6 | CAGTGAAATCCCACAACGTG | TGTTTTTATTGAGCCGAGGG | 262 |
| chr5:142501286-142503328 | p2 | (AT)21 | CAGTGAAATCCCACAACGTG | TGTTTTTATTGAGCCGAGGG | 262 |
| chr5:142509123-142511135 | p2 | (TG)6 | TGGTAACCTCACTGCCACAA | CATCCCAAGGCACCTAGAAA | 183 |
| chr5:142509123-142511135 | p2 | (AT)6 | TGGTAACCTCACTGCCACAA | CATCCCAAGGCACCTAGAAA | 183 |
| chr5:142611665-142613717 | p2 | (TA)12 | TTTTGTGACAGCCAAGCAAC | TGGTCAAGACATCTGCAAGC | 178 |
| chr5:142611665-142613717 | p2 | (AT)26 | TTTTGTGACAGCCAAGCAAC | TGGTCAAGACATCTGCAAGC | 178 |
| chr5:142611665-142613717 | p2 | (AT)10 | TTTTGTGACAGCCAAGCAAC | TGGTCAAGACATCTGCAAGC | 178 |
| chr5:142671154-142673169 | p3 | (CTT)5 | TTTAGATGGCAGTTTGTGCG | TTCATGGTTTCATGCCTTCA | 161 |
| chr5:142694098-142696113 | p5 | (AATTA)3 | CGTGCCACGTCATTTGATAC | TTAAGTCCAATTTCACCGGG | 129 |
| chr5:142736097-142738115 | p6 | (TAAAAA)3 | TGGCTTCTGGTTTAGTTGGG | AGCTGGCCAGTCAAACACTC | 178 |
| chr5:142807945-142809960 | p5 | (ATTTG)3 | CCCAGAGCCAGAAAAGTTTG | GCATATTCTTTTCCAGGGCA | 266 |
| chr5:142867869-142869885 | p4 | (TATT)4 | TAGTTCCTGCTTGGGATTGG | ACTCCAAACCGCATTTTCTG | 175 |
| chr5:142882125-142884146 | p3 | (TTC)7 | GTGGGTGTGGATCCAGTTCT | TATGGTGCCCAAAAAGGAAG | 121 |
| chr5:142899178-142901202 | p2 | (TA)12 | TTCGGACCCCATTTTTATCA | ACGAGGGTTCCAGTAGACGA | 161 |
| chr5:143036054-143038070 | p4 | (TATT)4 | TAGTTCCTGCTTGGGATTGG | ACTCCAAACCGCATTTTCTG | 173 |
| chr5:143043929-143045945 | p2 | (AG)8 | AGTTGGGACCGGGAAATAAC | ATAAGCCAATCGAAACACCC | 203 |
| chr5:143066612-143068630 | p3 | (ATT)5 | CATATGTGGCGGTGTTTGAG | GCCAAGAAATCAGATCCCAA | 114 |
| chr5:143066612-143068630 | p3 | (ATT)5 | CATATGTGGCGGTGTTTGAG | GCCAAGAAATCAGATCCCAA | 114 |
| chr5:143066612-143068630 | p3 | (TTA)6 | CATATGTGGCGGTGTTTGAG | GCCAAGAAATCAGATCCCAA | 114 |
| chr5:143066612-143068630 | p3 | (ATT)5 | CATATGTGGCGGTGTTTGAG | GCCAAGAAATCAGATCCCAA | 114 |
| chr5:143122783-143124801 | p3 | (AAT)6 | AATCGGCTCTTTGGGAATTT | TTATCATCTGACGCATCCCA | 112 |
| chr5:143195533-143197548 | p3 | (ACA)5 | TCTTCACCTTGAGGTGCCTT | TTTTGGCAGCAATGTGGTAA | 224 |
| chr5:143236758-143238773 | p3 | (ACA)5 | TCTTCACCTTGAGGTGCCTT | TTTTGGCAGCAATGTGGTAA | 224 |
| chr5:143239566-143241581 | p5 | (TTGGT)3 | TGACTTGGTTTGGTGTTGGA | ACATGTAATGTCGGGCTGGT | 204 |
| chr5:143239566-143241581 | p5 | (TTGGT)3 | TGACTTGGTTTGGTGTTGGA | ACATGTAATGTCGGGCTGGT | 204 |
| chr5:143251062-143253076 | p2 | (TA)7 | TCAGCCTCAAGAGCCAAAAT | AAAACGGACGACGAAAACAC | 166 |
| chr5:143251080-143253092 | p2 | (TA)7 | TCAGCCTCAAGAGCCAAAAT | AAAACGGACGACGAAAACAC | 148 |

| Marker ID | SSR type | SSR motif | FORWARD PRIMER (5'-3') | REVERSE PRIMER (5'-3') | size (bp) |
|---|---|---|---|---|---|
| chr5:143251917-143253931 | p2 | (TA)7 | GTGGGCCATATAGCCTGATG | TTGCTCCCTCTAATGGGTTG | 206 |
| chr5:143251917-143253931 | p2 | (TA)7 | GTGGGCCATATAGCCTGATG | TTGCTCCCTCTAATGGGTTG | 206 |
| chr5:143252349-143254363 | p2 | (TA)7 | TTCTCCAACTCACGGGAAAA | AATGGTACTTTGGTGCTCCG | 141 |
| chr5:143252349-143254363 | p2 | (TA)7 | TTCTCCAACTCACGGGAAAA | AATGGTACTTTGGTGCTCCG | 141 |
| chr5:143265239-143267255 | p2 | (AG)8 | TCCTTTGATCATCACGTCCA | ATAGCAGGGGACATTGATCG | 177 |
| chr5:143341587-143343632 | p3 | (ATT)15 | CGACGTCGTTTGATTCCTTT | CAAACGCTTGAAAACAGGCT | 136 |
| chr5:143470158-143472228 | p3 | (CAA)6 | TAAAATTTTCGACGAAGCCG | CCTCACTTGCCATTTCCACT | 181 |
| chr5:143470158-143472228 | p2 | (TA)35 | TAAAATTTTCGACGAAGCCG | CCTCACTTGCCATTTCCACT | 181 |
| chr5:143474161-143476176 | p5 | (AAAAT)3 | TTATTTTCGCCCACTCCAAG | GATTGGTTGCAATCACCTCA | 135 |
| chr5:143514592-143516610 | p6 | (ATATCT)3 | GTAGACGGCGCAAAGGATTA | CAGGCTGTGGACGTGATAAA | 127 |
| chr5:143536513-143538527 | p2 | (AT)7 | TGGCTGACAGACCAACAGAC | GCAATTTGGACCTTGGAAGA | 237 |
| chr5:143536645-143538657 | p2 | (AT)7 | TGGCTGACAGACCAACAGAC | GCAATTTGGACCTTGGAAGA | 105 |
| chr5:143585664-143587680 | p4 | (GGGT)4 | ATCGTAGTCTCGCAGGTGCT | CCCGACGTAGAACGATGAGT | 110 |
| chr5:143623549-143625570 | p3 | (CAA)7 | TCGCGCAAAAGTGTAGAGTG | ACTGAGCACAGGAGCTGACA | 189 |
| chr5:143623549-143625570 | p2 | (AT)27 | TCGCGCAAAAGTGTAGAGTG | ACTGAGCACAGGAGCTGACA | 189 |
| chr5:143629166-143631181 | p5 | (TTGGT)3 | GGATCTGAAAAGGGACGACA | TGTGCTAAGAGAGCCGTGAA | 241 |
| chr5:143629166-143631181 | p5 | (ATTTT)3 | GGATCTGAAAAGGGACGACA | TGTGCTAAGAGAGCCGTGAA | 241 |
| chr5:143672133-143674163 | p2 | (AT)6 | TTATGCTGAGATGTGAGGCG | CCATCTGTAGGGGGCAGATA | 131 |
| chr5:143692676-143694696 | p2 | (TC)10 | CCAAAATGGGAACTATCGGA | TCTCGATCCAAACATACCCC | 169 |
| chr5:143693869-143695885 | p2 | (AT)8 | CCCCTTTTCCTTTCAGTCAA | GACTGCGTAACGTGCAAGAA | 183 |
| chr5:143742459-143744471 | p2 | (AT)6 | CCTGACCCTTGTTTGGACAT | GTGAATCGGGAAGAGCAAAA | 145 |
| chr5:143746263-143748303 | p2 | (AT)10 | CAGGTGAGTGTTCCATGCAG | TGTAAAACAGAACGCCCACA | 263 |
| chr5:143814288-143816303 | p5 | (TTGTT)3 | AAAAGAACAAATGGTTGCGG | CCAAGTCCATAGCAGGGTGT | 218 |
| chr5:143814288-143816303 | p2 | (AT)16 | AAAAGAACAAATGGTTGCGG | CCAAGTCCATAGCAGGGTGT | 218 |
| chr5:143898768-143900783 | p5 | (TTATT)3 | AGGGTAAACCCAGCTCCTGT | TTGCCTACGGCAAGAGATTT | 214 |
| chr5:143934746-143936761 | p3 | (ACA)5 | TTTGAAAAAGCATCCATCCC | TTCAATCCACAGTTCCCCTC | 146 |
| chr5:143934746-143936761 | p5 | (CCCCA)4 | TTTGAAAAAGCATCCATCCC | TTCAATCCACAGTTCCCCTC | 146 |
| chr5:143935548-143937568 | p3 | (ACA)5 | ATCCCACAAGTGGAGTCTGG | GGGCTAGCCTTTCCATCTCT | 224 |
| chr5:143935548-143937568 | p5 | (CCCCA)4 | ATCCCACAAGTGGAGTCTGG | GGGCTAGCCTTTCCATCTCT | 224 |
| chr5:143945096-143947123 | p3 | (CAA)9 | GAGCAGGGGTGTGCATTATT | AAAAACCCCTTGATTGTCCC | 205 |
| chr5:143965612-143967642 | p2 | (TG)6 | ATTCGCGCTAATACAATGGG | CAGCGCACTTATGAACAACG | 128 |
| chr5:143965612-143967642 | p2 | (AT)15 | ATTCGCGCTAATACAATGGG | CAGCGCACTTATGAACAACG | 128 |
| chr5:143966113-143968127 | p2 | (AT)15 | TGCAAAGCGCAAATAAAGTG | GTTGAGTGGTTCGTCTCGCT | 108 |
| chr5:143988296-143990317 | p3 | (GTT)7 | ACCTCCCCTTCTCAACCACT | TCTCCTCCACTGCCCTACAT | 236 |
| chr5:144008706-144010721 | p6 | (TTTTTC)3 | GCAATGAGGATGGATTTGGT | GCTACAAAAGGGTGCATGGT | 236 |
| chr5:144008706-144010721 | p5 | (TTTTA)3 | GCAATGAGGATGGATTTGGT | GCTACAAAAGGGTGCATGGT | 236 |

| Marker ID | SSR type | SSR motif | FORWARD PRIMER (5'-3') | REVERSE PRIMER (5'-3') | size (bp) |
|---|---|---|---|---|---|
| chr5:144057378-144059414 | p2 | (TA)18 | GAAGGGAAAATGAGGAAGGG | ATTCTCAGACCAAACGCTGC | 260 |
| chr5:144187556-144189572 | p4 | (TTGA)4 | CGAGGAAGAACTCCCTTGTG | TGGGAAACAACCAAACATCA | 146 |
| chr5:144194966-144196981 | p5 | (ATGTT)3 | GACTGGATAAAAGTCCGCCA | CTTGCAATTATCTCCCCGAA | 116 |
| chr5:144322748-144324763 | p5 | (TCATG)3 | AGTTATTGCACCGCCTTGAC | GCGATGACCTAGCAAAGGAG | 105 |
| chr5:144337850-144339862 | p2 | (AT)6 | AATATCCACAAGCACCTCCG | CCGTTTTGTTCTTGGACGTT | 293 |
| chr5:144350908-144352946 | p2 | (CT)8 | ACACCCATCTTTTGCTTTGC | GACCAACTTGGACCCGTAGA | 131 |
| chr5:144423348-144425374 | p3 | (ATG)6 | GCCTGAAGAAGATCAGTGGG | TGGGCTAAAAACATCCCAAC | 166 |
| chr5:144423348-144425374 | p2 | (AT)13 | GCCTGAAGAAGATCAGTGGG | TGGGCTAAAAACATCCCAAC | 166 |
| chr5:144458097-144460109 | p2 | (AT)6 | TGACGTCATTTCCACACGTT | AGATCTTCGAGCTTACGCCA | 248 |
| chr5:144510342-144512357 | p5 | (AAATT)3 | CTTGTCCTCTTCGAGCCCTA | CCTCGGTCGCAATTTATGTT | 181 |
| chr5:144524230-144526245 | p3 | (CTT)5 | TTCTTTTCCATTTTCGGCAG | AGCGACAAGAGGGAACAATG | 129 |
| chr5:144564998-144567010 | p2 | (AT)6 | ATGGAATCCCGTTGAAGATG | AAATCAGGGGACGAACTCCT | 260 |
| chr5:144590420-144592432 | p2 | (AT)6 | ATGGAATCCCGTTGAAGATG | AAATCAGGGGACGAACTCCT | 260 |
| chr5:144596626-144598641 | p5 | (TTAAA)3 | ATATACAAACGCACGGAGGC | GTCAATCTTTTTGGGACGGA | 217 |
| chr5:144603864-144605879 | p5 | (ATTTA)3 | ACGAACCTCAAATTGTTGGC | ATTGCCCTTATGTGTGGGAG | 112 |
| chr5:144612288-144614302 | p2 | (GA)7 | CTCATTGACACGCTGGAAAA | ACATGGTTTGGATCAGGAGC | 161 |
| chr5:144631491-144633506 | p5 | (TATTA)3 | CCCTTCCAAATGCAAATCAC | ACAATATGCAGCGAATGCAG | 254 |
| chr5:144659853-144661874 | p3 | (TGA)7 | AGACCTTCGACATGATTCCG | GGCTGCTAAGAATGACAGGG | 186 |
| chr5:144760529-144762545 | p2 | (AT)8 | AATACAAGCGGCATGAAACC | TGAAGATCCGCAACCACATA | 300 |
| chr5:144760529-144762545 | p3 | (TGT)5 | AATACAAGCGGCATGAAACC | TGAAGATCCGCAACCACATA | 300 |
| chr5:144796344-144798386 | p3 | (CTT)14 | TCTAGCGGAGAGACACGGAT | ATCCTGCTGGGAAGGAAAAT | 112 |
| chr5:144796344-144798386 | p3 | (GAA)12 | TCTAGCGGAGAGACACGGAT | ATCCTGCTGGGAAGGAAAAT | 112 |
| chr5:144797039-144799075 | p3 | (CTT)14 | AAAATTTGATGCCCAGTTGC | TGGAGGAAGAAAATGGGTTG | 125 |
| chr5:144797039-144799075 | p3 | (GAA)12 | AAAATTTGATGCCCAGTTGC | TGGAGGAAGAAAATGGGTTG | 125 |
| chr5:144799302-144801317 | p5 | (ATTTT)3 | TCTCTTTCGCCATTACCACC | CAATTACGGTGATACGGCCT | 165 |
| chr5:144838114-144840135 | p3 | (TTA)7 | CACAGCCATTGAGGAAGGAT | TCCCATTACTCCCTTTTCCC | 262 |
| chr5:144930154-144932169 | p5 | (AGTGG)3 | TACAGTCTTGGGCTTTGGCT | TACCCGTCCCTTTAGCCTTT | 202 |
| chr5:144952160-144954172 | p2 | (TA)6 | CTTTCAAGCAGGTCCTGGAG | CAGGCTCATCATGCATTCAC | 293 |
| chr5:145153564-145155578 | p2 | (TA)7 | AAGACGAACTTTCCTTGCGA | TCTTTTTAAGCGAAGGGCAA | 245 |
| chr6:122044-124108 | p2 | (TA)32 | AAGGGTAGTAGCTGCGGGTT | AATGCGGGTTTCAATTTGTC | 106 |
| chr6:151143-153158 | p3 | (ACA)5 | TAAGAGCCTGTTTGGATGGG | AGTGTTTGGCTGGTCAGCTT | 204 |
| chr6:185196-187260 | p2 | (TA)32 | AAGGGTAGTAGCTGCGGGTT | AATGCGGGTTTCAATTTGTC | 106 |
| chr6:214295-216310 | p3 | (ACA)5 | TAAGAGCCTGTTTGGATGGG | AGTGTTTGGCTGGTCAGCTT | 204 |
| chr6:254221-256236 | p5 | (TTTAT)3 | CTTATCACGATGGGCGAACT | TCGAGTGCAAGAGTGTGGTC | 213 |
| chr6:284634-286650 | p4 | (CTTT)4 | CAAATTGTGCGGGTACACAG | CTGCTGAGGTTGCTGCAATA | 261 |
| chr6:288716-290734 | p6 | (AAAAAC)3 | CGCATCTTGTCTTCCACTGA | GATAAGAAGGTGCCACCGAA | 150 |

| Marker ID | SSR type | SSR motif | FORWARD PRIMER (5'-3') | REVERSE PRIMER (5'-3') | size (bp) |
|---|---|---|---|---|---|
| chr6:379637-381662 | p5 | (TCATA)5 | GATTGTTTATGCGGGCAAGT | CGTATATGTCTCCAGCCCGT | 202 |
| chr6:379637-381662 | p6 | (CATATA)3 | GATTGTTTATGCGGGCAAGT | CGTATATGTCTCCAGCCCGT | 202 |
| chr6:394191-396203 | p2 | (AC)6 | CCCTAGGGATATTGGGGTGT | TGTTGGCCAGACCATTTGTA | 103 |
| chr6:499255-501267 | p2 | (TA)6 | AGCCCTTTAGTGGCGAATTT | GGCCAAAAATAATGCCAAAA | 286 |
| chr6:500990-503005 | p5 | (TATTT)3 | AACGGAAAACGCAAAATACG | AGAGGTGCCAAATCACCTTG | 147 |
| chr6:553561-555579 | p6 | (ATCTTT)3 | ACCATCAACTTGTTGTGCCA | ACGTAGCAGCGTTGCCTTAT | 244 |
| chr6:565477-567505 | p2 | (AT)14 | CGAACTCGAAACCCATGAAT | ACAACACACCCCTTACCCAA | 178 |
| chr6:654919-656933 | p2 | (AT)7 | CGTTGAGGCGTATTCCGTAT | GAACCAAAATGGCTCCAAAA | 122 |
| chr6:800195-802207 | p2 | (AT)6 | TGTACCTAAATTGCCCCGAG | AGGACAACATACGGATTGGG | 130 |
| chr6:843497-845511 | p2 | (TC)7 | TCGGCTTAAGGAAAGGTGAA | AGTTACCATGGATTCGCACC | 179 |
| chr6:853361-855463 | p2 | (TA)51 | CGATGAGCTGTGACGAGTGT | TGTGTTGATCATTCGGGAGA | 177 |
| chr6:853361-855463 | p3 | (AAC)6 | CGATGAGCTGTGACGAGTGT | TGTGTTGATCATTCGGGAGA | 177 |
| chr6:878219-880277 | p2 | (AG)7 | TGATAGGCATACCACCCCAT | TTCACAACCTCTCCTCCACC | 281 |
| chr6:885651-887753 | p2 | (TA)51 | CGATGAGCTGTGACGAGTGT | TGTGTTGATCATTCGGGAGA | 177 |
| chr6:885651-887753 | p3 | (AAC)6 | CGATGAGCTGTGACGAGTGT | TGTGTTGATCATTCGGGAGA | 177 |
| chr6:910535-912561 | p2 | (AG)6 | CAATGGTGCTTTGATGATGG | CTCAAAATGGCAATCACCCT | 150 |
| chr6:910575-912593 | p2 | (AG)6 | CAATGGTGCTTTGATGATGG | CTCAAAATGGCAATCACCCT | 110 |
| chr6:913685-915699 | p2 | (AG)8 | CAATGGTGCTTTGATGATGG | CTCAAAATGGCAATCACCCT | 155 |
| chr6:913685-915699 | p2 | (TA)7 | CAATGGTGCTTTGATGATGG | CTCAAAATGGCAATCACCCT | 155 |
| chr6:994456-996468 | p2 | (TA)6 | GGAGGGGGTGGTGTAAAAAT | TCGTGTTCGTTTGGTTGGTA | 130 |
| chr6:1151112-1153137 | p5 | (GGGGA)5 | GCCGGTTGAAAAGAATGTGT | AATGATGATCCCACCCAAGA | 177 |
| chr6:1155604-1157620 | p4 | (ATTT)4 | TCTGGTGCACAAGGTATCCA | ATAATCGAGTCACGGGTTCG | 248 |
| chr6:1155604-1157620 | p2 | (AC)8 | TCTGGTGCACAAGGTATCCA | ATAATCGAGTCACGGGTTCG | 248 |
| chr6:1203204-1205222 | p2 | (TA)9 | GGATTTACCTTTTCCGAGGG | CAGACCAGCCGAGCTAAAAC | 207 |
| chr6:1203204-1205222 | p2 | (TA)7 | GGATTTACCTTTTCCGAGGG | CAGACCAGCCGAGCTAAAAC | 207 |
| chr6:1280350-1282365 | p5 | (AAAAT)3 | GATCCATCAGCAAAAGCCAT | TGAAGAGGGGCAATTCAGTC | 276 |
| chr6:1327940-1329982 | p2 | (AT)21 | TGCACGTGATTCGGAAATTA | TTTAGGTGCAGATTCCAGGG | 197 |
| chr6:1395179-1397193 | p3 | (TTG)6 | ACACTATTTCCCCAGACCCC | TTGAAAACGTCATGGACGAA | 187 |
| chr6:1395179-1397193 | p2 | (TG)7 | ACACTATTTCCCCAGACCCC | TTGAAAACGTCATGGACGAA | 187 |
| chr6:1401078-1403100 | p2 | (AT)11 | AAACTAAAACTCGGTCCGGC | TTTAAACTGGGCTACACGGG | 109 |
| chr6:1401078-1403100 | p2 | (AT)11 | AAACTAAAACTCGGTCCGGC | TTTAAACTGGGCTACACGGG | 109 |
| chr6:1426813-1428831 | p6 | (ATTTTT)3 | AGCAGGTTCCCAGGAAAGAT | CAACGTGGACGTAGCTCTCA | 260 |
| chr6:1496156-1498171 | p5 | (CAAAC)3 | CTTTGGGGCAATACTTGCAT | TCCGTGGGTATCCTCCTAAA | 293 |
| chr6:1523836-1525858 | p2 | (AT)11 | ACCCACGAGCCTCTTATTCA | TCAGCTTGACGTTGTGGAAG | 116 |
| chr6:1528075-1530091 | p4 | (TTTA)4 | AAAGGGCTACGAATTGGGTT | TCGATTCAAACCGGAAAAAC | 184 |
| chr6:1529468-1531486 | p2 | (TA)9 | TGATGCCATGTCCTCAAGAA | AAATCGCATGTCCAAACTCC | 233 |

| Marker ID | SSR type | SSR motif | FORWARD PRIMER (5'-3') | REVERSE PRIMER (5'-3') | size (bp) |
|---|---|---|---|---|---|
| chr6:1544787-1546803 | p4 | (TTTC)4 | CCCATCATGGAATCTTGGTC | AAAGAGACCAAAAGGGCACA | 234 |
| chr6:1544787-1546803 | p4 | (TATT)6 | CCCATCATGGAATCTTGGTC | AAAGAGACCAAAAGGGCACA | 234 |
| chr6:1606500-1608516 | p2 | (AT)8 | ACAAAATATGAGCCGCCAAG | CAGGGACTGCATAGAGGAGC | 143 |
| chr6:1654059-1656075 | p2 | (AT)8 | ACAAAATATGAGCCGCCAAG | CAGGGACTGCATAGAGGAGC | 143 |
| chr6:1778575-1780591 | p4 | (TATG)4 | AAATGTCGGGTCAAAAGTCG | ATTTGAAGGGTTTTGGGGAC | 215 |
| chr6:1778575-1780591 | p6 | (CTCATG)3 | AAATGTCGGGTCAAAAGTCG | ATTTGAAGGGTTTTGGGGAC | 215 |
| chr6:1781056-1783100 | p2 | (AT)22 | CCTTACGTCACCCCGATAGA | TAATATAAGCCCGCGTCCAG | 203 |
| chr6:1880517-1882533 | p2 | (AT)8 | CCGATAAGCACTTTCGGAAC | TCCGAGGATAACCCGTTAAA | 194 |
| chr6:1948063-1950078 | p5 | (TAATA)3 | CCGTCAGAAATTGGCTTGTT | AGGGACACATATCCAGTGGC | 275 |
| chr6:1974528-1976544 | p4 | (AAAT)4 | ATGTCTGGTTGACCCCCATA | CACTCTACGGGGCATGTTTT | 129 |
| chr6:2022156-2024171 | p2 | (TA)16 | AAACCCGACACACCAAAATC | TATACCCTTTTCGCTGACGG | 160 |
| chr6:2022156-2024171 | p5 | (AATCA)3 | AAACCCGACACACCAAAATC | TATACCCTTTTCGCTGACGG | 160 |
| chr6:2022156-2024171 | p5 | (TTTAA)3 | AAACCCGACACACCAAAATC | TATACCCTTTTCGCTGACGG | 160 |
| chr6:2068000-2070018 | p6 | (TTTTAA)3 | CGCGGAGTCCAAAATACACT | CCATTTGGCAACAATCCTTT | 271 |
| chr6:2072537-2074555 | p6 | (CCCCCA)3 | TACAGCCTACCAAACGACCC | AAGCTTGCCACGTAAATTGC | 114 |
| chr6:2074616-2076655 | p3 | (GTT)7 | CCGATATTTTCCGGTGAGAA | ATTGGCAGAGTGGACACACA | 295 |
| chr6:2074616-2076655 | p2 | (AC)8 | CCGATATTTTCCGGTGAGAA | ATTGGCAGAGTGGACACACA | 295 |
| chr6:2074616-2076655 | p2 | (AT)11 | CCGATATTTTCCGGTGAGAA | ATTGGCAGAGTGGACACACA | 295 |
| chr6:2074677-2076695 | p3 | (GTT)7 | CCGATATTTTCCGGTGAGAA | ATTGGCAGAGTGGACACACA | 234 |
| chr6:2074677-2076695 | p2 | (AC)8 | CCGATATTTTCCGGTGAGAA | ATTGGCAGAGTGGACACACA | 234 |
| chr6:2074677-2076695 | p2 | (AT)11 | CCGATATTTTCCGGTGAGAA | ATTGGCAGAGTGGACACACA | 234 |
| chr6:2206548-2208606 | p2 | (AC)8 | GTCATCATCGTCATCGTCGT | GATCTTGGCCCTCTAGTCCC | 143 |
| chr6:2206548-2208606 | p2 | (AT)29 | GTCATCATCGTCATCGTCGT | GATCTTGGCCCTCTAGTCCC | 143 |
| chr6:2326057-2328072 | p5 | (CAATG)3 | AAGTCCAGGGATGTCACCAG | CGTGACAAAAGCTCATTCCA | 241 |
| chr6:2349751-2351769 | p6 | (TTTCAG)3 | GGGAATTTGTGAAGGAGCAA | TTGGACTGGAAAAGGGATTG | 167 |
| chr6:2400360-2402378 | p2 | (AC)9 | GCTCTTGTCACGACCCAAAT | CTTCTGCACCTCGACATTGA | 260 |
| chr6:2428862-2430874 | p2 | (AT)6 | AAACCGAGGTCGATACCCTT | GACCGAAATATCCGTCCTCA | 233 |
| chr6:2484222-2486284 | p2 | (AT)31 | GATCATTGCACTCAAACCCA | TCAAATTCCACTACCGCTCC | 171 |
| chr6:2486540-2488555 | p2 | (GA)6 | TTTGTTATAATCGGTGGGGC | TCCGCCAAAAACTAAGAGGA | 160 |
| chr6:2486540-2488555 | p5 | (TTTAT)3 | TTTGTTATAATCGGTGGGGC | TCCGCCAAAAACTAAGAGGA | 160 |
| chr6:2527269-2529284 | p5 | (TTGGT)3 | AAATGGGATATCGCAAAGCA | GGTTGGTTTGATTCGGTTTG | 123 |
| chr6:2577475-2579489 | p5 | (ATAAA)3 | ATCGTGCCCCCATTGTATTA | TGACATGGCGAAATAGCAAA | 161 |
| chr6:2577475-2579489 | p2 | (TA)7 | ATCGTGCCCCCATTGTATTA | TGACATGGCGAAATAGCAAA | 161 |
| chr6:2577475-2579489 | p6 | (GAGTTG)3 | ATCGTGCCCCCATTGTATTA | TGACATGGCGAAATAGCAAA | 161 |
| chr6:2673717-2675741 | p4 | (TTTG)6 | CATCGAGTCCCTACCCAAAA | AAGGGACTCACTCACATGGC | 112 |
| chr6:2673717-2675741 | p2 | (CT)23 | CATCGAGTCCCTACCCAAAA | AAGGGACTCACTCACATGGC | 112 |

| Marker ID | SSR type | SSR motif | FORWARD PRIMER (5'-3') | REVERSE PRIMER (5'-3') | size (bp) |
|---|---|---|---|---|---|
| chr6:2673717-2675741 | p3 | (ATT)5 | CATCGAGTCCCTACCCAAAA | AAGGGACTCACTCACATGGC | 112 |
| chr6:2687476-2689491 | p5 | (TTATT)3 | TCCATCAATTTAACGCGACA | CAACAGCTCCAGCAACACAT | 277 |
| chr6:2687476-2689491 | p2 | (CA)6 | TCCATCAATTTAACGCGACA | CAACAGCTCCAGCAACACAT | 277 |
| chr6:2687647-2689659 | p5 | (TTATT)3 | TCCATCAATTTAACGCGACA | TAATGCCATTGCAACAGCTC | 106 |
| chr6:2687647-2689659 | p2 | (CA)6 | TCCATCAATTTAACGCGACA | TAATGCCATTGCAACAGCTC | 106 |
| chr6:2687947-2690035 | p5 | (TTATT)3 | TTTCCTGGTCATTTTGGCTC | AGCATGCATGGTCAATGAGA | 188 |
| chr6:2687947-2690035 | p2 | (CA)6 | TTTCCTGGTCATTTTGGCTC | AGCATGCATGGTCAATGAGA | 188 |
| chr6:2708726-2710742 | p4 | (GTGG)4 | CGAAGTTATTGGGGCAAAGA | ATCACACCTCCTTATTGGCG | 280 |
| chr6:2708726-2710742 | p3 | (GTA)5 | CGAAGTTATTGGGGCAAAGA | ATCACACCTCCTTATTGGCG | 280 |
| chr6:2955247-2957265 | p6 | (AAAAAT)3 | ACTACGCCTCGGTCACAAAC | TCCGCCTCTGTTCAATATCC | 223 |
| chr6:2984941-2986961 | p5 | (AAATG)4 | TTGGAGCTCCTCTGCAAAAC | GAAATTAAACACGGGCGCTA | 239 |
| chr6:3220325-3222369 | p2 | (TA)22 | CATTTTGTTTGTGCGGATTG | TGCAGGTAGACTTGGCATTG | 248 |
| chr6:3239304-3241322 | p3 | (AAC)6 | AACCGTCCTCTGGACTAGCA | TGGTTTGTGTTTGGGCTGTA | 234 |
| chr6:3269451-3271466 | p3 | (TGT)5 | AAAGGAGGGACAAAATTGGG | GTGTGCCTACACATCCATCG | 252 |
| chr6:3298350-3300368 | p3 | (CAA)6 | CCGTCACACATCACAACACA | TTCCTCACTACTCCGCTCGT | 106 |
| chr6:3336441-3338453 | p2 | (TA)6 | TCCCAAAGTATCCCCATCAA | AAATGACATGGCGGAATAGC | 231 |
| chr6:3421075-3423091 | p4 | (ATAA)4 | AAAGCCACATAGGCGCTAAA | TTTGTCGATTTCAACCGGAG | 222 |
| chr6:3501107-3503122 | p3 | (GTT)5 | CACATCGTATTGGGGTAGGG | CCTTGTCCCGAATATCCTCA | 212 |
| chr6:3508905-3510923 | p6 | (AACTGA)3 | GTTCCCCGTAACGTATGGAA | ACGAAATCCTCGTCCCTCTT | 145 |
| chr6:3565216-3567231 | p5 | (TAAAA)3 | CATCAATTGAACGGAGGGAG | GCTTGTTTGGCCATGAAAAT | 248 |
| chr6:3612788-3614813 | p5 | (TGATA)5 | GATGCTGCCCAAATTTCTGT | CTGAATGAAAAAGGGGCAAA | 116 |
| chr6:3612788-3614813 | p2 | (TA)17 | GATGCTGCCCAAATTTCTGT | CTGAATGAAAAAGGGGCAAA | 116 |
| chr6:3682859-3684883 | p3 | (TAT)8 | AACCTCATCGCCCCTTATCT | GCCTGATTCATCGTCCCTAA | 135 |
| chr6:3757707-3759719 | p2 | (AT)6 | TGATTGTGTCACGGCAAGAT | CGCTAACGTCTTGCAAACAA | 160 |
| chr6:3767753-3769783 | p2 | (AT)7 | TTGTGGGCCCATATAACCTG | GTTGATACGCATGTGGCAAT | 113 |
| chr6:3767753-3769783 | p2 | (TA)6 | TTGTGGGCCCATATAACCTG | GTTGATACGCATGTGGCAAT | 113 |
| chr6:3840451-3842466 | p5 | (CAAAC)3 | GTGTGTCGGTTCTGCACAGT | CCATTCTTGATGAATTCGGG | 123 |
| chr6:3846266-3848281 | p5 | (CCCAC)3 | CTCCACCAAAAAGGCGATAA | GTTGAAGGGCTGCGTACAAT | 134 |
| chr6:3846266-3848281 | p5 | (GAAAA)3 | CTCCACCAAAAAGGCGATAA | GTTGAAGGGCTGCGTACAAT | 134 |
| chr6:3861611-3863627 | p2 | (AT)8 | ACAAGCCAAATCCAAACAGG | AGTTCGCCATATTAGCACCG | 191 |
| chr6:3927241-3929259 | p6 | (TAAAAA)3 | GGTTTGTCGCCTATACCGAA | TGAAGCGAAAAGCAAGGAAT | 299 |
| chr6:4036788-4038806 | p6 | (GAACAA)3 | GAAGGGAAGAAAAGGGGTTG | CACACAGTGGTGGATTCAGG | 280 |
| chr6:4081901-4083916 | p3 | (AAC)5 | CCTGGTCCTTGGAGATTTGA | TGCAGTTTGTCAGCACCTTC | 103 |
| chr6:4120730-4122745 | p5 | (AATTA)3 | GCCAAACAAGTGGGACAGTT | ATTCTTGGTAGCGGTCATGC | 284 |
| chr6:4133093-4135105 | p2 | (TG)6 | CCGCTTTGTTATACCGCATT | TGCGAATTTTCCTTTATGCC | 271 |
| chr6:4151445-4153460 | p5 | (CTTTT)3 | TCCAGGCAATGGATAGGTTC | GCACCAACACGCTAGGAAAT | 143 |

| Marker ID | SSR type | SSR motif | FORWARD PRIMER (5'-3') | REVERSE PRIMER (5'-3') | size (bp) |
|---|---|---|---|---|---|
| chr6:4228531-4230547 | p4 | (TTTA)4 | TCACTTCACCACCCCTAACC | GGCCAAATATACCCCTCGTT | 282 |
| chr6:4281342-4283358 | p4 | (TTTA)4 | CTCGACGGGGGAACTATACA | ACTTTTCATCCCCACTGCAC | 297 |
| chr6:4281342-4283358 | p2 | (AT)6 | CTCGACGGGGGAACTATACA | ACTTTTCATCCCCACTGCAC | 297 |
| chr6:4281492-4283504 | p4 | (TTTA)4 | CTCGACGGGGGAACTATACA | ACTTTTCATCCCCACTGCAC | 147 |
| chr6:4281492-4283504 | p2 | (AT)6 | CTCGACGGGGGAACTATACA | ACTTTTCATCCCCACTGCAC | 147 |
| chr6:4378763-4380778 | p3 | (TAG)5 | AGCCAACACCATCAAAGAGG | CCCTAGCTAATCTCCCAGCC | 248 |
| chr6:4401521-4403536 | p3 | (TTC)5 | ACTTAGGCCTAGTGGGGCAT | CCTCTCACTTTTGGGCTGAG | 118 |
| chr6:4487082-4489094 | p2 | (AT)6 | GTTGGCCTCTTTTTGCTGAG | CGTAGCCTTCAATGGCAAAT | 163 |
| chr6:4518371-4520389 | p3 | (ATT)6 | TCATCAAGTGGTTGCCTCAA | TGGAAGAAGCCTCAAATGCT | 256 |
| chr6:4552062-4554074 | p2 | (GA)6 | ATCATCGAAAACCCCTTTCC | TAGGCTTCAGCCGAAGTGAT | 256 |
| chr6:4595628-4597640 | p2 | (TA)6 | CAAGTTTTGATGGGCATCCT | TGGTCTGTGGGAAATCACAA | 229 |
| chr6:4620811-4622826 | p2 | (GA)7 | AAGTGTTTTTGGTTCCGCAG | TAATGCCCTCTCACACCTCC | 254 |
| chr6:4620811-4622826 | p3 | (TAG)5 | AAGTGTTTTTGGTTCCGCAG | TAATGCCCTCTCACACCTCC | 254 |
| chr6:4641453-4643468 | p3 | (ATA)5 | ATAAAAGTGGGGGACAAGGG | TCATTTCCTCCATACCTCCG | 107 |
| chr6:4641575-4643599 | p3 | (ATA)5 | TATTTTGGACGAGAATGGGC | CCCAAATCAAGGTCCGATTA | 128 |
| chr6:4646601-4648616 | p3 | (ATA)5 | ATAAAAGTGGGGGACAAGGG | TCATTTCCTCCATACCTCCG | 107 |
| chr6:4646723-4648747 | p3 | (ATA)5 | TATTTTGGACGAGAATGGGC | CCCAAATCAAGGTCCGATTA | 128 |
| chr6:4666920-4668934 | p2 | (AT)7 | TGGTTGAGGTGAGTGAGTGC | GATTAGTGGGCTTGGTGGAA | 246 |
| chr6:4697636-4699652 | p4 | (TTTC)4 | ATAGGGCACAATTTGCCAAC | TGCCTCCAAATCGTGATGTA | 138 |
| chr6:4773440-4775452 | p2 | (TC)8 | TACCGCCACTTCATCACGTA | TTCGTTGTTGATCTCGATGC | 150 |
| chr6:4773440-4775452 | p2 | (AC)10 | TACCGCCACTTCATCACGTA | TTCGTTGTTGATCTCGATGC | 150 |
| chr6:4773440-4775452 | p2 | (CT)6 | TACCGCCACTTCATCACGTA | TTCGTTGTTGATCTCGATGC | 150 |
| chr6:4806993-4809019 | p2 | (TC)13 | TTTCTCCCCCTTATTTTCCG | GCTATGGGACTGGGGGTAAT | 223 |
| chr6:4825001-4827023 | p2 | (GA)8 | TACAGTTTCTCGACGGAGGG | GTCGGCCCAATTCATTTCTA | 201 |
| chr6:4825001-4827023 | p2 | (TA)11 | TACAGTTTCTCGACGGAGGG | GTCGGCCCAATTCATTTCTA | 201 |
| chr6:4842657-4844675 | p6 | (TTTTTC)3 | ATGGTGCGTCAAGAAAAAGG | CCCATATGTCCCACCTTCAC | 253 |
| chr6:4842657-4844675 | p3 | (TTA)6 | ATGGTGCGTCAAGAAAAAGG | CCCATATGTCCCACCTTCAC | 253 |
| chr6:4864893-4866907 | p2 | (TA)7 | TCCTGGACCTGTACCCACTC | TGAAGGTGGGACATATGGGT | 297 |
| chr6:4864893-4866907 | p2 | (TA)9 | TCCTGGACCTGTACCCACTC | TGAAGGTGGGACATATGGGT | 297 |
| chr6:4865064-4867078 | p2 | (TA)7 | ACCCATATGTCCCACCTTCA | CGGTTAAACAGTGCCAGGTT | 288 |
| chr6:4865064-4867078 | p2 | (TA)9 | ACCCATATGTCCCACCTTCA | CGGTTAAACAGTGCCAGGTT | 288 |
| chr6:4865079-4867091 | p2 | (TA)7 | ACCCATATGTCCCACCTTCA | CGGTTAAACAGTGCCAGGTT | 273 |
| chr6:4865079-4867091 | p2 | (TA)9 | ACCCATATGTCCCACCTTCA | CGGTTAAACAGTGCCAGGTT | 273 |
| chr6:4865098-4867118 | p2 | (TA)7 | ACCCATATGTCCCACCTTCA | CGGTTAAACAGTGCCAGGTT | 254 |
| chr6:4865098-4867118 | p2 | (TA)9 | ACCCATATGTCCCACCTTCA | CGGTTAAACAGTGCCAGGTT | 254 |
| chr6:4865152-4867168 | p2 | (TA)7 | ACCCATATGTCCCACCTTCA | CGGTTAAACAGTGCCAGGTT | 200 |

| Marker ID | SSR type | SSR motif | FORWARD PRIMER (5'-3') | REVERSE PRIMER (5'-3') | size (bp) |
|---|---|---|---|---|---|
| chr6:4865152-4867168 | p2 | (TA)9 | ACCCATATGTCCCACCTTCA | CGGTTAAACAGTGCCAGGTT | 200 |
| chr6:4892192-4894210 | p3 | (TTG)6 | GTGCCCATGCAAATTTTACC | CAACCGTTGTCCGGTACTCT | 199 |
| chr6:4894708-4896720 | p2 | (AC)6 | CCATTCCCCTAAAAATGCCT | TTTGGGGGCAGATATTGAAG | 190 |
| chr6:4901698-4903713 | p5 | (TCGGT)3 | CAACGCTAATGCAAAGAGCA | CCGAAATCGCAAAACTGAAT | 104 |
| chr6:4901698-4903713 | p5 | (TTTTG)4 | CAACGCTAATGCAAAGAGCA | CCGAAATCGCAAAACTGAAT | 104 |
| chr6:4901698-4903713 | p5 | (TTCGG)3 | CAACGCTAATGCAAAGAGCA | CCGAAATCGCAAAACTGAAT | 104 |
| chr6:4901698-4903713 | p5 | (AAAAC)3 | CAACGCTAATGCAAAGAGCA | CCGAAATCGCAAAACTGAAT | 104 |
| chr6:4902516-4904531 | p5 | (TTCGG)3 | CCGAACCGAATTGTTGAAAT | CAAAACTAGGGCTGGGCATA | 112 |
| chr6:4902516-4904531 | p5 | (AAAAC)3 | CCGAACCGAATTGTTGAAAT | CAAAACTAGGGCTGGGCATA | 112 |
| chr6:4910642-4912656 | p2 | (AC)7 | TGAATGGAAACGATAACCCC | GATCTTGGCCCTCTAGTCCC | 187 |
| chr6:4910642-4912656 | p2 | (AT)7 | TGAATGGAAACGATAACCCC | GATCTTGGCCCTCTAGTCCC | 187 |
| chr6:4928045-4930057 | p2 | (AT)10 | TGAAAGATCGGTTCCATCAA | CGTCGAAGCTGATTAGGGTG | 195 |
| chr6:4928045-4930057 | p2 | (TA)6 | TGAAAGATCGGTTCCATCAA | CGTCGAAGCTGATTAGGGTG | 195 |
| chr6:4928045-4930057 | p3 | (AGT)7 | TGAAAGATCGGTTCCATCAA | CGTCGAAGCTGATTAGGGTG | 195 |
| chr6:4928335-4930356 | p2 | (AT)10 | TCAGCTTCGACGTGTGTTTT | TGAGAGTGATGAGCGATCAAA | 148 |
| chr6:4928335-4930356 | p2 | (TA)6 | TCAGCTTCGACGTGTGTTTT | TGAGAGTGATGAGCGATCAAA | 148 |
| chr6:4928335-4930356 | p3 | (AGT)7 | TCAGCTTCGACGTGTGTTTT | TGAGAGTGATGAGCGATCAAA | 148 |
| chr6:5089744-5091756 | p2 | (TA)6 | ACATCCATCCGGATCTCTTG | GAGTGCAAGCAAAAGGGAAG | 167 |
| chr6:5151354-5153372 | p6 | (CTCCAC)3 | GCTCACTCCTCCTGTGAAGG | CCACCGGTGAAGGAGATAGA | 228 |
| chr6:5151354-5153372 | p6 | (CCGCCC)3 | GCTCACTCCTCCTGTGAAGG | CCACCGGTGAAGGAGATAGA | 228 |
| chr6:5151354-5153372 | p3 | (CCT)7 | GCTCACTCCTCCTGTGAAGG | CCACCGGTGAAGGAGATAGA | 228 |
| chr6:5178878-5180900 | p2 | (AT)11 | ACAAATTCGCCCAGTTTTTG | AAAAGGGGAACCCAAATGTC | 195 |
| chr6:5284517-5286537 | p5 | (AGAAA)4 | GGCCTGGACTCCTTAAAACC | TTTTAGGATGCGGGCATAAC | 200 |
| chr6:5303791-5305813 | p2 | (AT)11 | AACGCTTTATTGGGCTTCAA | AATTGGGTTGTTGGACTTGC | 237 |
| chr6:5350630-5352666 | p2 | (AT)18 | TTTGCAATTCCACGAGAATG | CAAACGCTAATTGCTGCTCA | 201 |
| chr6:5350630-5352666 | p2 | (TA)7 | TTTGCAATTCCACGAGAATG | CAAACGCTAATTGCTGCTCA | 201 |
| chr6:5353265-5355280 | p5 | (ATTTT)3 | CCGTGAGTTTGGATGTGATG | ATCCCATATTTTTGGGCTCC | 242 |
| chr6:5353265-5355280 | p5 | (ATTTT)3 | CCGTGAGTTTGGATGTGATG | ATCCCATATTTTTGGGCTCC | 242 |
| chr6:5423372-5425405 | p3 | (AAT)11 | CACCAAGGCAAACGAAGAAT | CCTTTGTCCACGCTCTTCTC | 202 |
| chr6:5423372-5425405 | p2 | (AT)15 | CACCAAGGCAAACGAAGAAT | CCTTTGTCCACGCTCTTCTC | 202 |
| chr6:5430545-5432566 | p2 | (GA)6 | GGAGAAGATGAGGGAGGAGG | TTGATTTACCGGATCCCAAA | 286 |
| chr6:5430545-5432566 | p3 | (TAA)7 | GGAGAAGATGAGGGAGGAGG | TTGATTTACCGGATCCCAAA | 286 |
| chr6:5430545-5432566 | p3 | (ATA)5 | GGAGAAGATGAGGGAGGAGG | TTGATTTACCGGATCCCAAA | 286 |
| chr6:5442817-5444859 | p5 | (GAAAT)3 | CTGATCAAGGGGGAAATGAA | ATCAGGTTCGCAATTCCATC | 126 |
| chr6:5442817-5444859 | p2 | (GA)21 | CTGATCAAGGGGGAAATGAA | ATCAGGTTCGCAATTCCATC | 126 |
| chr6:5541446-5543461 | p5 | (ATTTC)3 | CAGTCAAAGTCTTGTGGCGA | ATAACATTGCCTCACAGGGC | 127 |

| Marker ID | SSR type | SSR motif | FORWARD PRIMER (5'-3') | REVERSE PRIMER (5'-3') | size (bp) |
|---|---|---|---|---|---|
| chr6:5680471-5682489 | p3 | (ACA)6 | GTAGGATGGCTGTTTCCGAG | TTTCTAAGCGCACACCATTG | 291 |
| chr6:5680471-5682489 | p4 | (TTAT)4 | GTAGGATGGCTGTTTCCGAG | TTTCTAAGCGCACACCATTG | 291 |
| chr6:5691844-5693880 | p2 | (AT)18 | AGGACTTGTCGGGTTTGTTG | GGCTAAATGCACTCTCCGAA | 110 |
| chr6:5736747-5738767 | p2 | (AT)10 | GAATCGTCTCATTCGCACCT | CCACGTGCTGTTCACTGTTT | 157 |
| chr6:5736747-5738767 | p2 | (AT)9 | GAATCGTCTCATTCGCACCT | CCACGTGCTGTTCACTGTTT | 157 |
| chr6:5783968-5785983 | p3 | (ACA)5 | TGAATGGGGTTAAGGAGCAG | CACAAAATTCGTGCGTGACT | 154 |
| chr6:5783968-5785983 | p3 | (ATA)7 | TGAATGGGGTTAAGGAGCAG | CACAAAATTCGTGCGTGACT | 154 |
| chr6:5829674-5831689 | p3 | (ACA)5 | TGAATGGGGTTAAGGAGCAG | CACAAAATTCGTGCGTGACT | 154 |
| chr6:5829674-5831689 | p3 | (ATA)7 | TGAATGGGGTTAAGGAGCAG | CACAAAATTCGTGCGTGACT | 154 |
| chr6:5898683-5900707 | p2 | (AG)6 | AGAGGCATTGGGGAGTCTTT | TTTTCCCAAGTTTAAGGGGG | 118 |
| chr6:5920595-5922611 | p2 | (AT)12 | AGGTAGGGTCCTGCCAAACT | GGCCCTAACATGGGTCTGTA | 132 |
| chr6:5920595-5922611 | p2 | (AT)6 | AGGTAGGGTCCTGCCAAACT | GGCCCTAACATGGGTCTGTA | 132 |
| chr6:5920595-5922611 | p2 | (AT)8 | AGGTAGGGTCCTGCCAAACT | GGCCCTAACATGGGTCTGTA | 132 |
| chr6:6018700-6020712 | p2 | (TA)6 | GTCCCACATGCTGTTCACTG | GGCAACCACCCTTGTAAAGA | 185 |
| chr6:6020875-6022890 | p5 | (CGAAC)3 | CAACCCATTAGAGGGAGCAA | GGTTCGGTTCGATTTTTCAA | 143 |
| chr6:6020875-6022890 | p5 | (AAAAT)3 | CAACCCATTAGAGGGAGCAA | GGTTCGGTTCGATTTTTCAA | 143 |
| chr6:6020875-6022890 | p2 | (TA)7 | CAACCCATTAGAGGGAGCAA | GGTTCGGTTCGATTTTTCAA | 143 |
| chr6:6036980-6038995 | p6 | (AAAAAT)3 | AATTTGGTTCACATGCTCCC | GATCCGGCTACAGGACACAT | 154 |
| chr6:6036980-6038995 | p5 | (TTTTG)3 | AATTTGGTTCACATGCTCCC | GATCCGGCTACAGGACACAT | 154 |
| chr6:6092439-6094463 | p6 | (ATATAG)4 | GCCCCAAAGACTTGACAAAA | GGCTGCAGGACTTGGTGTAT | 205 |
| chr6:6127929-6129944 | p5 | (AAAAT)3 | GCACACCTCTTCCTGGTCAT | CAGCTTCCCAATGCTCTTTC | 255 |
| chr6:6141206-6143221 | p5 | (TAACT)3 | TGGAATTCTGCCTCGAGAGT | ACCATGTCAAGTTCCGAAGG | 197 |
| chr6:6549132-6551154 | p4 | (CTTT)5 | TTGTTTTGTTCCTGCACTGG | TCTTCGTCAATCGTCGTCTG | 224 |
| chr6:6549211-6551231 | p4 | (CTTT)5 | TTGTTTTGTTCCTGCACTGG | TCTTCGTCAATCGTCGTCTG | 145 |
| chr6:6658863-6660878 | p5 | (TTTAA)3 | AGACCGTGGACATGGAAAAG | CACTTGAGAGCCCAGGTAGC | 146 |
| chr6:6708761-6710773 | p2 | (AC)7 | ACCTCTGGGACAAAAATGGA | TGCATGGTTCATGTTTCGAT | 130 |
| chr6:6880529-6882547 | p6 | (ATTTGT)3 | TTTTGCCTCCCAATTTGTTC | AAATGCCTGCATCAAAAACC | 291 |
| chr6:6919608-6921626 | p2 | (AT)9 | AGAGGCGAGAAACTCCAACA | TTTGTTTGTGACGGAGACCA | 184 |
| chr6:6966887-6968911 | p6 | (AAATTT)4 | CAGCTCCTCCTCCAGCTATG | CTGGTTCAGGGACAGTGGAT | 173 |
| chr6:7018695-7020707 | p2 | (AT)6 | AGAGGCGAGAAACTCCAACA | TGTGCTGGAACACCTTCTTG | 178 |
| chr6:7026001-7028016 | p3 | (CAA)5 | GAATACTGAGGTTTGCCCCA | CTTTTTGAGCCAAAGCGAAC | 288 |
| chr6:7056635-7058647 | p2 | (AT)6 | AGAGGCGAGAAACTCCAACA | TGTGCTGGAACACCTTCTTG | 178 |
| chr6:7063941-7065956 | p3 | (CAA)5 | GAATACTGAGGTTTGCCCCA | CTTTTTGAGCCAAAGCGAAC | 288 |
| chr6:7196582-7198598 | p2 | (AT)8 | GTGCGTGTCAATCCATGAAA | GACATGGCGGAATAGCAAAT | 244 |
| chr6:7216333-7218403 | p2 | (TA)7 | TTTGCTAAGCGAGGCAGAAT | AAAACAGGGTTTTGTTTGCG | 293 |
| chr6:7216419-7218471 | p2 | (TA)7 | TTTGCTAAGCGAGGCAGAAT | AAAACAGGGTTTTGTTTGCG | 207 |

| Marker ID | SSR type | SSR motif | FORWARD PRIMER (5'-3') | REVERSE PRIMER (5'-3') | size (bp) |
|---|---|---|---|---|---|
| chr6:7291785-7293800 | p5 | (ATAAA)3 | TTTCAGCATTCACTCGTTCG | CGGAACAAAATGGTATGCCT | 219 |
| chr6:7297908-7299924 | p4 | (TTAT)4 | TCTGTATTGATGGCAGAGCG | CGAGTGCGCACACATTATTT | 107 |
| chr6:7297908-7299924 | p2 | (TG)8 | TCTGTATTGATGGCAGAGCG | CGAGTGCGCACACATTATTT | 107 |
| chr6:7427684-7429702 | p2 | (AT)9 | AATCTCCGAACCACCTTTCA | ATACCCTGCTCCATCTGTGG | 192 |
| chr6:7427684-7429702 | p2 | (AT)11 | AATCTCCGAACCACCTTTCA | ATACCCTGCTCCATCTGTGG | 192 |
| chr6:7428435-7430457 | p2 | (AT)9 | ACGGGCCCAACACTTTACAT | AATTTGCCGATTTTGTATTCG | 195 |
| chr6:7428435-7430457 | p2 | (AT)11 | ACGGGCCCAACACTTTACAT | AATTTGCCGATTTTGTATTCG | 195 |
| chr6:7428435-7430457 | p2 | (AT)9 | ACGGGCCCAACACTTTACAT | AATTTGCCGATTTTGTATTCG | 195 |
| chr6:7429678-7431700 | p2 | (AT)9 | ACGGGCCCAACACTTTACAT | AAATTTGCCGATTTTGTATTCG | 195 |
| chr6:7429678-7431700 | p2 | (AT)11 | ACGGGCCCAACACTTTACAT | AAATTTGCCGATTTTGTATTCG | 195 |
| chr6:7429678-7431700 | p2 | (AT)8 | ACGGGCCCAACACTTTACAT | AAATTTGCCGATTTTGTATTCG | 195 |
| chr6:7487127-7489142 | p5 | (ACCGA)3 | AGTGAGTGTGTGTGCGAAGG | CAGCCGTACACTTTTCAGCA | 151 |
| chr6:7490204-7492218 | p2 | (TA)7 | CAGCACCTTTTTGGGGAGTA | GGACGTTAGGCGTGGTTAAG | 201 |
| chr6:7497440-7499456 | p4 | (TACA)4 | ACTGGTATGGAAAGCAACCG | ACCAGACTCCACGTACCCAC | 149 |
| chr6:7570259-7572271 | p2 | (AG)6 | TTCTGATGTTGTGCCAGAGC | GATCAGAGCATGTTGGCCTT | 232 |
| chr6:7633703-7635721 | p2 | (AC)9 | CGTTGATGTGAGGGGCTATT | CGCTTGCTTTGAAAATGACA | 150 |
| chr6:7633703-7635721 | p3 | (TGT)6 | CGTTGATGTGAGGGGCTATT | CGCTTGCTTTGAAAATGACA | 150 |
| chr6:7662224-7664242 | p2 | (AC)9 | CGTTGATGTGAGGGGCTATT | CGCTTGCTTTGAAAATGACA | 150 |
| chr6:7662224-7664242 | p3 | (TGT)6 | CGTTGATGTGAGGGGCTATT | CGCTTGCTTTGAAAATGACA | 150 |
| chr6:7717346-7719366 | p5 | (TTTGG)4 | CGTTGGCCCTAGCCATATTA | ACCATGATTTGAAACCCCAA | 235 |
| chr6:7810644-7812662 | p5 | (TGCAT)3 | CTTGTGGTCCGAGGTTCATT | TGGTGTCCATGTACCAATCG | 133 |
| chr6:7810644-7812662 | p3 | (TAT)6 | CTTGTGGTCCGAGGTTCATT | TGGTGTCCATGTACCAATCG | 133 |
| chr6:7878368-7880383 | p2 | (AT)7 | CGACAAGGACTGCCAAAAAT | TTCTCAGGAACGCTCCACTT | 129 |
| chr6:7878368-7880383 | p2 | (AT)11 | CGACAAGGACTGCCAAAAAT | TTCTCAGGAACGCTCCACTT | 129 |
| chr6:7878368-7880383 | p3 | (CAT)5 | CGACAAGGACTGCCAAAAAT | TTCTCAGGAACGCTCCACTT | 129 |
| chr6:8052024-8054062 | p2 | (AT)19 | CAAGTTTTGATGGGCATCCT | GTTGCTTCGGGGAATTGTTA | 198 |
| chr6:8074341-8076363 | p2 | (TA)11 | TATAAATAAGGGGGAGCGGC | TTCCTTTTTGGTGGTTTTGC | 283 |
| chr6:8081632-8083653 | p3 | (TTG)7 | CCGGAATGTTGTGTGACAAG | CATTTCCGCCACTTTCATCT | 175 |
| chr6:8088614-8090626 | p2 | (TA)6 | GATCCCCTCATTCTCCCAAT | TTTAAACCCGACCCAGCTAA | 164 |
| chr6:8090869-8092881 | p2 | (TA)6 | TTCATAGTGCGACAAATGCC | GCAAACAAAATGGTGGACAA | 137 |
| chr6:8101284-8103296 | p2 | (AT)8 | TAGCTGCGGTCGGATATAGG | GTTGAAACCGCTAATTCCCA | 105 |
| chr6:8170969-8173013 | p3 | (CTA)8 | GACGGCACATCTGCTCAGTA | ACATACTCGCTGGGACAAGG | 263 |
| chr6:8213483-8215779 | p2 | (AT)15 | CGGGACCCCAATTTCTTATT | AGCACGGCATAGACCATGTA | 114 |
| chr6:8262461-8264473 | p2 | (AT)6 | TAACAGCAAACCCCAAAAGC | TGATCCCTGTTGCACTTCTG | 194 |
| chr6:8305411-8307426 | p5 | (AATAA)3 | GTGTTTGCCAAGCGTGTAGA | TTTGGGGGACGTGATATGTT | 146 |
| chr6:8324850-8326864 | p2 | (AT)21 | TGCGATGAATATCGTAAATGGA | AGCACGGGTATAGGCCATCT | 191 |

| Marker ID | SSR type | SSR motif | FORWARD PRIMER (5'-3') | REVERSE PRIMER (5'-3') | size (bp) |
|---|---|---|---|---|---|
| chr6:8324850-8326864 | p2 | (TA)7 | TGCGATGAATATCGTAAATGGA | AGCACGGGTATAGGCCATCT | 191 |
| chr6:8474507-8476551 | p2 | (AT)22 | GGCACTAGGTAAGGCTGGTG | AAGGGTGACCGAGTTTGTTG | 162 |
| chr6:8515887-8517905 | p2 | (TA)9 | ATCTCAATGCCCCATCACTC | CGGGTGTTGTGAGTAGCCTT | 148 |
| chr6:8520720-8522736 | p2 | (AT)8 | CTTGGACATGCGGTTTGTAA | GGTTGGTAAATGGTTGGTGG | 241 |
| chr6:8556842-8558856 | p2 | (TA)7 | TTAGGGGTGCCAAGCTAAAA | ATTTTGGCGGCTAATGTACG | 148 |
| chr6:8579207-8581231 | p5 | (AGAAA)3 | CCAAAGCATAGTCTCGGAGC | TTGTGATAGCAGCCTTGTCG | 197 |
| chr6:8579207-8581231 | p2 | (TA)8 | CCAAAGCATAGTCTCGGAGC | TTGTGATAGCAGCCTTGTCG | 197 |
| chr6:8579207-8581231 | p2 | (AT)12 | CCAAAGCATAGTCTCGGAGC | TTGTGATAGCAGCCTTGTCG | 197 |
| chr6:8703958-8705970 | p2 | (TA)6 | TACTAAGAGGGCCGGGTTCT | AGAAGCAGAAGCCATACGGA | 240 |
| chr6:8709607-8711623 | p4 | (TGCG)4 | CTCATTTGACCCCGAAAAGA | GGTCTTAATCAAGTCGCCCA | 297 |
| chr6:8780396-8782411 | p5 | (ATTTT)3 | GAGAAAAGAAAAGGGGGTGG | ACCATTTGGTCTTGTGCCTC | 281 |
| chr6:8780396-8782411 | p3 | (TTA)5 | GAGAAAAGAAAAGGGGGTGG | ACCATTTGGTCTTGTGCCTC | 281 |
| chr6:8809259-8811287 | p2 | (AT)14 | TCCCAATGCCTTAATTCCAA | TCCAGAATTCCGTCAACTCC | 129 |
| chr6:8949438-8951450 | p2 | (CT)7 | TTCAGAAGCATTGCGTTGTC | TTTCCCTCATTGTTCCCATC | 146 |
| chr6:8949438-8951450 | p2 | (AT)6 | TTCAGAAGCATTGCGTTGTC | TTTCCCTCATTGTTCCCATC | 146 |
| chr6:8980513-8982529 | p2 | (TA)8 | GCAATACCCAAGATGCCTGT | GGACTGGCAAAGGTGATGTT | 197 |
| chr6:8980513-8982529 | p3 | (ATG)7 | GCAATACCCAAGATGCCTGT | GGACTGGCAAAGGTGATGTT | 197 |
| chr6:9025041-9027059 | p3 | (TGT)5 | TCAATCCCATCACGATTCAA | GCTTCTTCCGATCCCTCTTT | 175 |
| chr6:9135997-9138015 | p3 | (TGA)6 | TTGGAGGTTTTCTCTTTGCC | TCTCAGTCAATTCGTCACCG | 247 |
| chr6:9172157-9174172 | p5 | (AATAC)3 | GAATCCCTGCATTTTGGCTA | TGAACCACGAAAAACCATCA | 182 |
| chr6:9172157-9174172 | p2 | (TC)8 | GAATCCCTGCATTTTGGCTA | TGAACCACGAAAAACCATCA | 182 |
| chr6:9271797-9273825 | p2 | (AT)7 | ACCTGTCACACCCCTACGAG | CCTTGTCGGCCTTATTTTGA | 271 |
| chr6:9271968-9273982 | p2 | (AT)7 | ACCTGTCACACCCCTACGAG | CCTTGTCGGCCTTATTTTGA | 100 |
| chr6:9297906-9299933 | p3 | (TCT)9 | GGGTGAAGATGAAGTGCGTT | AGGGTGTGTTTGGTACGGAG | 287 |
| chr6:9297906-9299933 | p2 | (AC)7 | GGGTGAAGATGAAGTGCGTT | AGGGTGTGTTTGGTACGGAG | 287 |
| chr6:9454036-9456078 | p2 | (TA)21 | GACGACACATCTGCTCGGTA | GGCCCCCAACATACCTTAGT | 200 |
| chr6:9473463-9475475 | p2 | (GA)6 | TCAGGAAAGTCCACCAATCC | AAGGATCTCGTCCCTCCACT | 118 |
| chr6:9486399-9488414 | p5 | (AAACC)3 | CCAGCACGCCTAAAATGAAT | GACAATAACCAAACCGGTGG | 256 |
| chr6:9622987-9625002 | p5 | (AATAA)3 | CATCAGAAGGGGATCCTCAA | CCAAATTGAACTGGCTTGCT | 149 |
| chr6:9716423-9718435 | p2 | (TA)8 | GATGTCCGATTTACGCGATT | CCGCAGAAGTGAAATTGACA | 116 |
| chr6:9716423-9718435 | p2 | (TG)7 | GATGTCCGATTTACGCGATT | CCGCAGAAGTGAAATTGACA | 116 |
| chr6:9977023-9979071 | p5 | (CGACA)3 | TCTTTTTGGAAATGCAAGGG | TGGTCATGAATTTCGGTCAA | 232 |
| chr6:9977023-9979071 | p3 | (AAC)8 | TCTTTTTGGAAATGCAAGGG | TGGTCATGAATTTCGGTCAA | 232 |
| chr6:10009310-10011324 | p2 | (TA)7 | CGTGCACTGAACTTCCCTTT | AGTCCATGTGTATGGGGGAA | 218 |
| chr6:10110695-10112709 | p2 | (AG)7 | CACCGCTTCGCTTCTCTTAC | ACCCGAAAGCAACAATCAAC | 245 |
| chr6:10131929-10133944 | p5 | (AAACC)3 | CCTCTTTAAGTGGCTTTGCG | GACTTTAAGCGGAAAGGGCT | 222 |

| Marker ID | SSR type | SSR motif | FORWARD PRIMER (5'-3') | REVERSE PRIMER (5'-3') | size (bp) |
|---|---|---|---|---|---|
| chr6:10131929-10133944 | p5 | (CAAAC)4 | CCTCTTTAAGTGGCTTTGCG | GACTTTAAGCGGAAAGGGCT | 222 |
| chr6:10227896-10229911 | p5 | (AAAAT)3 | GTCTCAAGCCATCCCAATGT | CTTGGTTGGGATGTTGCTAAA | 268 |
| chr6:10227896-10229911 | p3 | (AAT)11 | GTCTCAAGCCATCCCAATGT | CTTGGTTGGGATGTTGCTAAA | 268 |
| chr6:10227896-10229911 | p3 | (AAT)17 | GTCTCAAGCCATCCCAATGT | CTTGGTTGGGATGTTGCTAAA | 268 |
| chr6:10313576-10315594 | p3 | (TTG)6 | TTTACCATTGGTGGAGGAGC | AGAGGTTTGTGCCAAAATGG | 256 |
| chr6:10374252-10376266 | p2 | (TG)7 | GCTAAGGATGACTTGCCTGG | CGGACACGGAATTTAAGGAA | 280 |
| chr6:10482897-10484921 | p2 | (TA)12 | TGGATTGGTGGAGGAACAAT | GAACATTCATTGCGTTGCTG | 129 |
| chr6:10560100-10562116 | p6 | (CATGAG)3 | TAAAAAGGGGACGTCAGCAC | GATCAGACTCCACGCTCCTC | 216 |
| chr6:10560100-10562116 | p4 | (TACA)4 | TAAAAAGGGGACGTCAGCAC | GATCAGACTCCACGCTCCTC | 216 |
| chr6:10715107-10717127 | p2 | (AT)10 | GAACCACCTGACTCATCCGT | CCTTGTCGGCCTTATTTTGA | 213 |
| chr6:10856152-10858167 | p3 | (TAT)5 | GGCCTTTTTGTTTGGTTTCA | CTGAAACGGAACGGAGGTAG | 278 |
| chr6:11011795-11013811 | p4 | (TTTC)4 | ATGGAAAACAGAGGCATTGG | TGCGAAGGAGAAGGAGAGAG | 230 |
| chr6:11063288-11065304 | p2 | (CT)8 | ATCTGTCGATCTTTCCGTCG | TCCCTGCTTCTTTTTCCTCA | 178 |
| chr6:11150572-11152587 | p3 | (CTT)5 | CAACTCGCGGGATATTTTTG | AAGGGGTCCATGTTGTTTGA | 148 |
| chr6:11188136-11190151 | p3 | (TGT)5 | CATTCCAATTGAACATTGCG | TTCAGCTTGCCTCGTTAGGT | 104 |
| chr6:11220286-11222301 | p3 | (GCC)5 | ACCTGTGTGGCTGTGTCAAA | ACACCTTGGACTGATCCTCG | 288 |
| chr6:11375941-11377985 | p2 | (TA)22 | AATGATGCAAGACCAAAGGC | TATTTGCTGGCAAGTGCATC | 213 |
| chr6:11375941-11377985 | p2 | (AT)8 | AATGATGCAAGACCAAAGGC | TATTTGCTGGCAAGTGCATC | 213 |
| chr6:11383259-11385371 | p2 | (AT)56 | TCCGAGAGGTCAGTCTCGAT | CTTCAGACAGTGGGGGTGAT | 262 |
| chr6:11383259-11385371 | p2 | (TA)8 | TCCGAGAGGTCAGTCTCGAT | CTTCAGACAGTGGGGGTGAT | 262 |
| chr6:11413818-11415830 | p2 | (AT)6 | GGTGATGACGGATACTTGGG | GCAGGTGCAGGTCAGTACAA | 113 |
| chr6:11499114-11501126 | p2 | (TA)6 | AGACCAGGTCCCTCTCCATT | TTCGGGAGGAACACGATAAC | 255 |
| chr6:11501982-11504000 | p2 | (CA)9 | TCCCAGAACAAGTTCCCCTA | CTTTCCTGAAGCACTGGTCC | 153 |
| chr6:11513288-11515303 | p5 | (CATTG)3 | GTCCCGTTTTCCCAAATTCT | CGCTACACGAAGTCTCACCA | 120 |
| chr6:11520695-11522717 | p2 | (GA)9 | GAAGGAAATGTTTACCCGCA | GAACACCCCTATCCGTGCTA | 218 |
| chr6:11520695-11522717 | p2 | (TA)13 | GAAGGAAATGTTTACCCGCA | GAACACCCCTATCCGTGCTA | 218 |
| chr6:11520721-11522739 | p2 | (GA)9 | GAAGGAAATGTTTACCCGCA | GAACACCCCTATCCGTGCTA | 192 |
| chr6:11520721-11522739 | p2 | (TA)13 | GAAGGAAATGTTTACCCGCA | GAACACCCCTATCCGTGCTA | 192 |
| chr6:11578617-11580631 | p2 | (TA)14 | TTGGATTAGATCGGAGCGAC | TTTCGTTAATTTCCCCCACA | 114 |
| chr6:11586441-11588459 | p2 | (TA)7 | CGCATTTAGCCAAGGAAGAC | TTTCGTTAATTTCCCCCACA | 234 |
| chr6:11586462-11588476 | p2 | (TA)7 | CGCATTTAGCCAAGGAAGAC | TTTCGTTAATTTCCCCCACA | 213 |
| chr6:11598792-11600810 | p6 | (TCAAGT)3 | TACCCCGTCTTTTGTTTTCG | TAGCCCAGTTGTTACACCCC | 267 |
| chr6:11598792-11600810 | p6 | (TCAAGT)3 | TACCCCGTCTTTTGTTTTCG | TAGCCCAGTTGTTACACCCC | 267 |
| chr6:11660527-11662539 | p2 | (AT)6 | CCAACTGGATGAGGTGGACT | GTTTCTTGGCTAATGCGCTC | 166 |
| chr6:11660527-11662539 | p2 | (AT)9 | CCAACTGGATGAGGTGGACT | GTTTCTTGGCTAATGCGCTC | 166 |
| chr6:11660527-11662539 | p2 | (TA)6 | CCAACTGGATGAGGTGGACT | GTTTCTTGGCTAATGCGCTC | 166 |

| Marker ID | SSR type | SSR motif | FORWARD PRIMER (5'-3') | REVERSE PRIMER (5'-3') | size (bp) |
|---|---|---|---|---|---|
| chr6:11679720-11681756 | p2 | (AT)18 | GATTGTCCGAGCACATCAAA | CTTTGCCTAAGGTGGCTCTG | 134 |
| chr6:11715372-11717387 | p3 | (ACA)5 | ACGTAACAATGGAGGCAAGG | ATGCTATTGTGCATTTGGCA | 223 |
| chr6:11794382-11796402 | p5 | (CCACC)4 | TATTCAATGCTCACCCACCA | ATCAGGGGCAATTTTGTCAG | 109 |
| chr6:11794382-11796402 | p5 | (TACAA)3 | TATTCAATGCTCACCCACCA | ATCAGGGGCAATTTTGTCAG | 109 |
| chr6:11795004-11797019 | p5 | (CCACC)4 | CCTGTTTGGATGGTGGTTTC | TGAACCACGGGTTTACAACA | 155 |
| chr6:11795004-11797019 | p5 | (TACAA)3 | CCTGTTTGGATGGTGGTTTC | TGAACCACGGGTTTACAACA | 155 |
| chr6:11810291-11812315 | p2 | (TA)12 | CATATGCAATAGTGGCGGTG | GAAGTCGATTTACCGTCCCA | 103 |
| chr6:11826859-11828874 | p5 | (TTGGG)3 | AATGACCTATCGCTTGTCGG | GCCAAACCCATATTGACAGC | 198 |
| chr6:11833729-11835744 | p5 | (TAAAA)3 | TTACTCTTACCGCGGTGCTT | CTACATATCAACCGTGGCCC | 166 |
| chr6:11891933-11893945 | p3 | (CAT)6 | TTACGTCGCTGAACTTGCAC | ACTTCTCAGTCGTGATGGGG | 300 |
| chr6:11891933-11893945 | p2 | (AG)6 | TTACGTCGCTGAACTTGCAC | ACTTCTCAGTCGTGATGGGG | 300 |
| chr6:11941006-11943022 | p4 | (TATT)4 | CGATCTCCAAGAGCAACCAT | CGCGTTTGGGTAAAAGGATA | 119 |
| chr6:11960282-11962300 | p6 | (TTTTAT)3 | ATCCGTTCCAAGAAATGACG | TCAAACAACCACAACCCAAA | 205 |
| chr6:11999963-12001978 | p5 | (TTCGG)3 | GACTCCACTGTGGCATCTGA | GAAATTGATGGCATTGGCTT | 227 |
| chr6:11999963-12001978 | p5 | (TCGGT)3 | GACTCCACTGTGGCATCTGA | GAAATTGATGGCATTGGCTT | 227 |
| chr6:12004262-12006284 | p2 | (TA)11 | CTGTTCCGTGCCGAATAAAT | TGCACACACAGTCAGCAAAA | 274 |
| chr6:12027075-12029090 | p5 | (ATTTT)3 | GTGAAGTTGTCTGCGCTTCA | GTCATGTTTCATGCCTGCTG | 164 |
| chr6:12086290-12088310 | p5 | (CATTC)4 | ACTGGAGCGGAGACTGTGTT | TACATGTCAATGAGCCCCAA | 184 |
| chr6:12129272-12131297 | p5 | (TTTGG)5 | TGCAGTCGTGTACTCTTCGG | CGATTGTCGTGAACCTCTGA | 265 |
| chr6:12173943-12175961 | p6 | (CCCCAC)3 | TCCAAGAAAGGGTGGAGATG | ATGACCAAAAAGCCAAGGTG | 289 |
| chr6:12173943-12175961 | p5 | (CTCTT)4 | TCCAAGAAAGGGTGGAGATG | ATGACCAAAAAGCCAAGGTG | 289 |
| chr6:12177138-12179156 | p2 | (CA)6 | ACTCGGCCAAATTCATCAAC | CGGCTTGAAAATTCGTGATT | 229 |
| chr6:12177186-12179200 | p2 | (CA)6 | ACTCGGCCAAATTCATCAAC | CGGCTTGAAAATTCGTGATT | 181 |
| chr6:12211393-12213409 | p2 | (AT)9 | CCATTGCCCTCTGATGTTTT | GCATGGTTGCTTTTCCATTT | 116 |
| chr6:12211393-12213409 | p4 | (TTTG)4 | CCATTGCCCTCTGATGTTTT | GCATGGTTGCTTTTCCATTT | 116 |
| chr6:12224789-12226807 | p6 | (AAATAA)3 | GATAAGTCAGGGGGAAAGCC | ATGTGGATTCAGATGGGCTC | 112 |
| chr6:12239640-12241656 | p2 | (TA)8 | GGCGAGACACTGACAAAACA | TTGAGGTACGCCCAATCTTC | 288 |
| chr6:12314350-12316362 | p2 | (TA)6 | GCGCAAAAGAGAAACCAAAG | ATGTTACAGTGGTTGCGTCG | 290 |
| chr6:12314350-12316362 | p2 | (AT)15 | GCGCAAAAGAGAAACCAAAG | ATGTTACAGTGGTTGCGTCG | 290 |
| chr6:12346137-12348151 | p2 | (TA)7 | ATGAAAACACGAAGCGCAGT | TCAAACCGGTAAACCGAAAC | 206 |
| chr6:12346137-12348151 | p2 | (AT)7 | ATGAAAACACGAAGCGCAGT | TCAAACCGGTAAACCGAAAC | 206 |
| chr6:12346152-12348168 | p2 | (TA)7 | ATGAAAACACGAAGCGCAGT | TCAAACCGGTAAACCGAAAC | 191 |
| chr6:12346152-12348168 | p2 | (AT)7 | ATGAAAACACGAAGCGCAGT | TCAAACCGGTAAACCGAAAC | 191 |
| chr6:12649645-12651661 | p4 | (CTTT)4 | ACGGAAGAAGATGGTTGTGG | TTGCATAAATTGGGACGGTT | 130 |
| chr6:12727930-12729948 | p6 | (GGGGGT)3 | TCCAATAGGCGGTGAAAAAG | TTTCATAAGCGACACCCACA | 237 |
| chr6:12758052-12760067 | p6 | (AGTTGG)3 | CTCTCGGATCCCAATTCAAA | GGATGGTCAATGTTTGGAGG | 212 |

| Marker ID | SSR type | SSR motif | FORWARD PRIMER (5'-3') | REVERSE PRIMER (5'-3') | size (bp) |
|---|---|---|---|---|---|
| chr6:12758052-12760067 | p5 | (AAAAC)3 | CTCTCGGATCCCAATTCAAA | GGATGGTCAATGTTTGGAGG | 212 |
| chr6:12848493-12850508 | p3 | (GTT)5 | CCGTCCTTACTATCCGTCCA | GTATGGCAAGTCTGCCCAGT | 292 |
| chr6:12848493-12850508 | p2 | (TA)6 | CCGTCCTTACTATCCGTCCA | GTATGGCAAGTCTGCCCAGT | 292 |
| chr6:12848493-12850508 | p2 | (AT)6 | CCGTCCTTACTATCCGTCCA | GTATGGCAAGTCTGCCCAGT | 292 |
| chr6:12862764-12864779 | p3 | (TAT)5 | ACATGGCCAAAAAGGTTCAG | AGTCAGGGAACATTCAACGG | 227 |
| chr6:12895726-12897742 | p4 | (TCAT)4 | ATCGCAATGAGGGAAAAGTG | GCCCACCGTACTCTTGTCAT | 148 |
| chr6:12939828-12941840 | p2 | (AT)6 | CGTACGCCCTAAACACACCT | TGAATATCAACGTCGCCTGA | 236 |
| chr6:12960972-12963022 | p4 | (GTGG)4 | TTTGCTGCTCGAAGGAGTTT | TCGACCACTTATCGCATCAA | 101 |
| chr6:13067716-13069731 | p3 | (ATA)5 | AAAAGGAGGAGGCGACTAGG | TCCCTTCATTCCCTTCACAC | 101 |
| chr6:13182141-13184156 | p5 | (AAAAT)3 | AATGGCCACACCACACTTTT | AGATAGGGGGTGGGAAGTTG | 180 |
| chr6:13381390-13383416 | p2 | (AT)6 | GGAATGAGCCCAAGAAGACA | TCAGCTGACATTGGAAGCAC | 262 |
| chr6:13404770-13406795 | p5 | (AATAT)3 | CACCCCAGGGAGGTTAAAAT | TGGAATGCTAACACCACGAC | 216 |
| chr6:13405321-13407336 | p5 | (AATAT)3 | CCACTAGAGCGCCAATCCTA | GAAATGTTGCCCGTTTCACT | 131 |
| chr6:13438991-13441006 | p5 | (AATAT)3 | ATCACCACTAGAGCGCCAAT | GAAATGTTGCCCGTTTCACT | 150 |
| chr6:13556437-13558457 | p2 | (AT)10 | CGCAAAAGGGACATACGATT | TCCAAGGCCACCTGTAGTTC | 150 |
| chr6:13774971-13776986 | p5 | (CAAAT)3 | AGTTTCAAAGCCCCTGAACA | GCTCTCACCGGTCCAAAATA | 299 |
| chr6:13807167-13809179 | p2 | (AT)6 | GGGAGGTGGCAACTATCTCA | AAATGCCGACGTGACACATA | 134 |
| chr6:13939937-13941951 | p2 | (GT)7 | GCAACGAGATTTTTCACGGT | ATTTGGTGGTGTTGCCATTT | 264 |
| chr6:13970055-13972070 | p5 | (AAAAT)3 | TTACATTCATCGTGGCGGTA | TTCAAGGTGGGCACCTAATC | 220 |
| chr6:13970055-13972070 | p6 | (ATTTTT)4 | TTACATTCATCGTGGCGGTA | TTCAAGGTGGGCACCTAATC | 220 |
| chr6:13973376-13975426 | p2 | (CA)7 | ACTCGGCCAAATTCATCAAC | TGTCGAAGGTTGGAGGTTTC | 205 |
| chr6:13973456-13975506 | p2 | (CA)7 | ACTCGGCCAAATTCATCAAC | TGTCGAAGGTTGGAGGTTTC | 125 |
| chr6:13975544-13977559 | p5 | (AATGA)3 | TGTGGCAGCCAACGATATTA | TTGAGGGGAAAAATTGATCG | 245 |
| chr6:13975544-13977559 | p3 | (TTC)5 | TGTGGCAGCCAACGATATTA | TTGAGGGGAAAAATTGATCG | 245 |
| chr6:14027303-14029315 | p2 | (CA)6 | AGAATTTTCCGTTTTGCCCT | GAACAACCACACCTTTGCCT | 177 |
| chr6:14047330-14049342 | p2 | (AC)6 | ACTTGTGCGTACCTTCGCTT | CTGGGCACGGAGTTTAAGAA | 293 |
| chr6:14047330-14049342 | p3 | (TTA)5 | ACTTGTGCGTACCTTCGCTT | CTGGGCACGGAGTTTAAGAA | 293 |
| chr6:14192472-14194492 | p5 | (ATTTA)4 | GAGGAGGCTCAAAACTGCTG | CTCCAAAGCTTGGCTTGAAC | 132 |
| chr6:14250707-14252759 | p4 | (GAAA)13 | TCGTACCCAGCAACAGACAG | CCGTTGACTTGGGTTCAGTT | 150 |
| chr6:14254007-14256033 | p2 | (AC)10 | TTCCGTAACAACTGGAAGGG | TGTATTGGGCCTTTTTCTGC | 244 |
| chr6:14254007-14256033 | p2 | (AT)9 | TTCCGTAACAACTGGAAGGG | TGTATTGGGCCTTTTTCTGC | 244 |
| chr6:14254007-14256033 | p2 | (TA)13 | TTCCGTAACAACTGGAAGGG | TGTATTGGGCCTTTTTCTGC | 244 |
| chr6:14383101-14385115 | p2 | (TA)7 | TGAGAATCGAGTGGATGACG | CTGCCACGTTATTGGGAAGT | 226 |
| chr6:14407979-14410009 | p3 | (AAT)10 | GAAAAGGCAAAAAGGAAGGG | AAAGGGGTGGGATTATACCG | 174 |
| chr6:14424764-14426836 | p3 | (GAA)24 | AGACATTCCCAAGCACCATC | TGATTTCCCAGCATTCCTTC | 167 |
| chr6:14427585-14429600 | p5 | (AATAA)3 | TATAAGCCAATCCAAACGGG | CGAGTTTGGGGTTTGACATT | 110 |

| Marker ID | SSR type | SSR motif | FORWARD PRIMER (5'-3') | REVERSE PRIMER (5'-3') | size (bp) |
|---|---|---|---|---|---|
| chr6:14427585-14429600 | p5 | (AAATA)5 | TATAAGCCAATCCAAACGGG | CGAGTTTGGGGTTTGACATT | 110 |
| chr6:14427585-14429600 | p5 | (AAATA)3 | TATAAGCCAATCCAAACGGG | CGAGTTTGGGGTTTGACATT | 110 |
| chr6:14427585-14429600 | p5 | (AAATA)3 | TATAAGCCAATCCAAACGGG | CGAGTTTGGGGTTTGACATT | 110 |
| chr6:14427585-14429600 | p5 | (AAATA)3 | TATAAGCCAATCCAAACGGG | CGAGTTTGGGGTTTGACATT | 110 |
| chr6:14427748-14429763 | p5 | (AATAA)3 | TCAAACCCCAAACTCGAAAA | TGGGGACTCCCATATGATGT | 162 |
| chr6:14427748-14429763 | p5 | (AAATA)5 | TCAAACCCCAAACTCGAAAA | TGGGGACTCCCATATGATGT | 162 |
| chr6:14427748-14429763 | p5 | (AAATA)3 | TCAAACCCCAAACTCGAAAA | TGGGGACTCCCATATGATGT | 162 |
| chr6:14427748-14429763 | p5 | (AAATA)3 | TCAAACCCCAAACTCGAAAA | TGGGGACTCCCATATGATGT | 162 |
| chr6:14427748-14429763 | p5 | (AAATA)3 | TCAAACCCCAAACTCGAAAA | TGGGGACTCCCATATGATGT | 162 |
| chr6:14427748-14429763 | p5 | (TTTAT)3 | TCAAACCCCAAACTCGAAAA | TGGGGACTCCCATATGATGT | 162 |
| chr6:14636629-14638695 | p2 | (TA)33 | CTTTCCTCACTCGCTCATCC | GTGTGGAAGCAGAGAAAGGC | 132 |
| chr6:14652408-14654460 | p2 | (AT)11 | ACGAGGGTTGGAATCACAAG | TGAATTTTCTCCCACCGAAG | 101 |
| chr6:14652408-14654460 | p2 | (TA)26 | ACGAGGGTTGGAATCACAAG | TGAATTTTCTCCCACCGAAG | 101 |
| chr6:14688989-14691041 | p2 | (TA)26 | GAAGGTACAAGCCATGGGAA | TTGCTTGACCTCGAAACCTT | 177 |
| chr6:14715123-14717138 | p5 | (AAAGG)3 | GGCGATTTGGGTCTTGAATA | CTGGAAAGCCGAACAACAAT | 113 |
| chr6:14715123-14717138 | p4 | (TTAT)7 | GGCGATTTGGGTCTTGAATA | CTGGAAAGCCGAACAACAAT | 113 |
| chr6:14722154-14724169 | p5 | (AAAGG)3 | GCAGATGGCCTTGGGTATTA | GGTATGCCCACAAATGCTCT | 184 |
| chr6:14745922-14747948 | p2 | (AG)14 | TGGGAAAACAATCGCATGTA | TGGGAGGATAGGATGTACGC | 161 |
| chr6:14752371-14754383 | p2 | (CT)6 | GTGATATCCTCCCATTCCCC | CAAGGGGCGAAAATTTTGTA | 253 |
| chr6:14757785-14759801 | p2 | (AT)8 | TTGGCCCTTGATTACTGTCC | GGATCATGAGGCAATGGAGT | 207 |
| chr6:14823389-14825407 | p6 | (ATCTTC)3 | ATATCGGATGGCGTTGAGAG | AGTCATGTCCCACCCAGAAC | 154 |
| chr6:14864394-14866466 | p2 | (AT)36 | TCCCAGACCTGCTTAGCACT | CAATCGTCTGATATGCCAGG | 234 |
| chr6:14891991-14894006 | p3 | (TCA)5 | GAAAATTGTACCGCCCTGAA | GCGTGCGTTTGAGATGATTA | 143 |
| chr6:14941435-14943507 | p2 | (AT)36 | TCCCAGACCTGCTTAGCACT | CAATCGTCTGATATGCCAGG | 234 |
| chr6:14969032-14971047 | p3 | (TCA)5 | GAAAATTGTACCGCCCTGAA | GCGTGCGTTTGAGATGATTA | 143 |
| chr6:15012908-15014922 | p2 | (TA)7 | AGCAGCAGAGTTTGGGGTAA | AGATCTGCAACGTGATTCCC | 237 |
| chr6:15012908-15014922 | p2 | (TA)9 | AGCAGCAGAGTTTGGGGTAA | AGATCTGCAACGTGATTCCC | 237 |
| chr6:15048373-15050387 | p2 | (AG)8 | TGAGCCTTGACGAAATGATG | TCGAAAATGCAACAGACTCG | 151 |
| chr6:15048391-15050413 | p2 | (AG)8 | TGAGCCTTGACGAAATGATG | TCGAAAATGCAACAGACTCG | 133 |
| chr6:15092411-15094426 | p3 | (ACA)5 | GGCTAACTTGCGGAACAAAA | TGGTGAACAAGTGCCAATGT | 122 |
| chr6:15175838-15177854 | p4 | (AAAG)4 | AACGCATGGAAAAGATTGCT | CTGCCTTTTAACGACGAAGG | 293 |
| chr6:15175838-15177854 | p2 | (TC)6 | AACGCATGGAAAAGATTGCT | CTGCCTTTTAACGACGAAGG | 293 |
| chr6:15216315-15218335 | p5 | (GAATT)4 | CCAATTGGAAGGCCCTTATT | ACACCCACGGCTAACTATGC | 252 |
| chr6:15219442-15221468 | p2 | (AT)13 | CGCTTGGAAGATTCGAGAAG | GACCCTTACTTAGGGCCCAG | 263 |
| chr6:15307121-15309139 | p3 | (CTT)9 | AAAATCTGGCCATAATCCCC | ATAACCGTTTCTGGCACCAC | 258 |
| chr6:15307121-15309139 | p6 | (TATTCA)3 | AAAATCTGGCCATAATCCCC | ATAACCGTTTCTGGCACCAC | 258 |

| Marker ID | SSR type | SSR motif | FORWARD PRIMER (5'-3') | REVERSE PRIMER (5'-3') | size (bp) |
|---|---|---|---|---|---|
| chr6:15307121-15309139 | p2 | (AT)6 | AAAATCTGGCCATAATCCCC | ATAACCGTTTCTGGCACCAC | 258 |
| chr6:15347362-15349378 | p4 | (AGAA)4 | CGGATGTTATCCCCTCGTTA | GGACATGGATTCGGTCCTTA | 209 |
| chr6:15408487-15410609 | p2 | (AT)10 | TCTGTTAGCCTTGTCGGCTT | AACACATTTCAACGCAACCA | 270 |
| chr6:15408487-15410609 | p2 | (AG)7 | TCTGTTAGCCTTGTCGGCTT | AACACATTTCAACGCAACCA | 270 |
| chr6:15422427-15424445 | p3 | (TTG)6 | ATAGATGTCGAGTGCCGCTT | AAATGAGCCCTGCTGCTTTA | 107 |
| chr6:15434004-15436019 | p5 | (AAAAT)3 | GGGTTCAAATATTCGGCAAA | TTTCAGTAAAAGGCTTCGCA | 160 |
| chr6:15449404-15451419 | p2 | (AT)7 | AACGATCAATCCAACGCTTC | AAAAGAAGGAGAGGTTCGGC | 231 |
| chr6:15449404-15451419 | p3 | (ATC)5 | AACGATCAATCCAACGCTTC | AAAAGAAGGAGAGGTTCGGC | 231 |
| chr6:15494668-15496686 | p2 | (TA)7 | TTGCAATGATCGTGAAGAGC | GTCTATGGTAACGGTGCGGT | 159 |
| chr6:15494668-15496686 | p6 | (ATGTTT)3 | TTGCAATGATCGTGAAGAGC | GTCTATGGTAACGGTGCGGT | 159 |
| chr6:15516628-15518644 | p2 | (TC)8 | CGAAAAGCCTACCTTATGCG | TTAATTTTTGAAGGTGGCCG | 164 |
| chr6:15562496-15564514 | p3 | (ACA)6 | GGTTTTGTTTGCTGAGGGAA | GAGAACCACCAGAACGGAAA | 193 |
| chr6:15562496-15564514 | p3 | (ACA)5 | GGTTTTGTTTGCTGAGGGAA | GAGAACCACCAGAACGGAAA | 193 |
| chr6:15632808-15634824 | p2 | (TA)8 | TGCATGTCTGTCTACGAGCC | GCTCAGCAGTCAGAGTCGTG | 124 |
| chr6:15651083-15653101 | p2 | (TA)9 | GCTAGGTCTTGGCAGTGTCC | CCCCTATTTCTCAAGCCCTC | 258 |
| chr6:15675835-15677861 | p2 | (TA)13 | ACAATTCACGTGTTTCCGGT | ACATCATGAGTTTGGGCTCG | 128 |
| chr6:15711699-15713739 | p2 | (AG)20 | GAGGTCTTCGATTCACGCTC | GGAGGGTTTTTAGTCGCACA | 215 |
| chr6:15712360-15714477 | p2 | (AG)20 | GAGTATGGGGAGGAGAAGGG | CGCCCCAGTACATCTCATTT | 166 |
| chr6:15923419-15925443 | p2 | (TA)12 | GCCTGCAGGTAGAGTTTTGC | CCCTTCGCCTAATACCATGA | 160 |
| chr6:15923419-15925443 | p2 | (AT)37 | GCCTGCAGGTAGAGTTTTGC | CCCTTCGCCTAATACCATGA | 160 |
| chr6:15937759-15939771 | p2 | (AT)6 | CTTTGCCAACCCTATGGAAA | TCTTCCAGCCGTAAGCAAGT | 135 |
| chr6:16050881-16052896 | p5 | (AAAGG)3 | GAGGCCACCAGTTTGTTCAT | CTAATACCGCTTCTGCAGCC | 109 |
| chr6:16100695-16102719 | p2 | (TA)12 | TTCGACACCGGAATCTAACC | TTTGCACCATTTCTGTTGGA | 279 |
| chr6:16158805-16160821 | p2 | (AG)8 | TTACACCCCGTAAGAGTCCG | CGGGATACTTAAGGCAACCA | 238 |
| chr6:16226549-16228563 | p2 | (CA)7 | GTCATGCTATAATAAATGCGGCT | TCAACTCGATTCTCGTTCTCTTC | 111 |
| chr6:16251393-16253408 | p3 | (GAA)5 | TGTTTGTGGTCCAATGGAGA | TAGCTTTCAACCCATTTGCC | 104 |
| chr6:16363901-16365913 | p2 | (AT)6 | ATCTCACCTGATACCGCCAC | AAGGAACTCCATTCGGGTTT | 121 |
| chr6:16512045-16514095 | p5 | (TTATA)10 | CTGGTATCGTTGTTGCATGG | GGCTAATTGCTCACAGAGGG | 154 |
| chr6:16521911-16523926 | p5 | (AACCG)3 | GTTTGAGGGGGAATTTTCGT | CCAGTTGGCAGAGTGCAGTA | 133 |
| chr6:16526775-16528790 | p5 | (CATTG)3 | CGACTCGTCCGATCGTTATT | ACCAAAAACGCGGTCACTAC | 219 |
| chr6:16604179-16606195 | p2 | (TA)8 | CTTTCCAACCCAGGTCTCAA | TGACCCGAGGAGATAGTTGG | 282 |
| chr6:16698138-16700154 | p2 | (TC)6 | CAAACGAACTGCCACATGAC | TCAGTGGCAGCGATTTACTG | 212 |
| chr6:16698138-16700154 | p2 | (AG)8 | CAAACGAACTGCCACATGAC | TCAGTGGCAGCGATTTACTG | 212 |
| chr6:16698138-16700154 | p4 | (AATT)4 | CAAACGAACTGCCACATGAC | TCAGTGGCAGCGATTTACTG | 212 |
| chr6:16714724-16716742 | p6 | (CCAATG)3 | AGTGACCGTGATTCCTTTGG | AATGGACCATGCAGCACATA | 180 |
| chr6:16738321-16740355 | p2 | (AT)17 | CCCTTCTCCTCGGTTTTACC | GAATTGACATCGGGCAGAGT | 147 |

| Marker ID | SSR type | SSR motif | FORWARD PRIMER (5'-3') | REVERSE PRIMER (5'-3') | size (bp) |
|---|---|---|---|---|---|
| chr6:16852391-16854407 | p2 | (AC)8 | GTCTTATGCTCACCCCCAAA | TGACGTTTGGGTTGTTGTGT | 297 |
| chr6:16854690-16856705 | p5 | (AATAA)3 | GGCCACAATCCACATACACA | GGAGTGGTTGTTGGAGCAAT | 133 |
| chr6:16854690-16856705 | p5 | (TTTAT)3 | GGCCACAATCCACATACACA | GGAGTGGTTGTTGGAGCAAT | 133 |
| chr6:16859373-16861391 | p2 | (AT)9 | AAACATCCAGCATTTCCAGC | TAATAAATGGGGGTGGGGTT | 281 |
| chr6:16860705-16862717 | p2 | (TA)6 | ATCTGTCATTGCCGATTTCC | CCCACCGTCACTTTCTTCAT | 288 |
| chr6:16902275-16904290 | p5 | (GGGGT)3 | TAGACATGGATGCCTGGTCA | GAACCCCTCCAGGTACCCTA | 225 |
| chr6:16991152-16993164 | p2 | (AT)6 | TTTTGAGCATCTTGTGGCTG | TAAATCAGGCAGCATTTCCC | 181 |
| chr6:17032332-17034350 | p6 | (GGGTGG)3 | TGCCTGAACTGGTTGAGTTG | GCACCAGTGCCAAATCCTAT | 272 |
| chr6:17063053-17065068 | p5 | (TATTT)3 | CCGCGTTATAATTCCTGCAT | TTATTTTATGCCTGAGCGGG | 241 |
| chr6:17063053-17065068 | p5 | (TTTTA)3 | CCGCGTTATAATTCCTGCAT | TTATTTTATGCCTGAGCGGG | 241 |
| chr6:17152526-17154544 | p3 | (TGT)6 | ATTTTGGCATCTGGGTGTTT | CATCACCCCTTGTACCCATC | 167 |
| chr6:17172020-17174040 | p2 | (AT)10 | TCATTGTTTGGCACGTGTTT | ATTGATGTCAATCGCAACCA | 142 |
| chr6:17177349-17179367 | p3 | (TTG)29 | TCGCACACTCGAATCAACTC | CAGGTGCATCAAATGGAGTG | 103 |
| chr6:17177349-17179367 | p6 | (CGTCCC)3 | TCGCACACTCGAATCAACTC | CAGGTGCATCAAATGGAGTG | 103 |
| chr6:17194130-17196148 | p2 | (AT)9 | TCATTGTTTGGCACGTGTTT | ATTGATGTCAATCGCAACCA | 142 |
| chr6:17222044-17224059 | p5 | (AAAAT)3 | CCGGACCTTTCAAAATACGA | ACCATTTTCCTTCTCCACCC | 163 |
| chr6:17258557-17260573 | p6 | (AGGAAG)3 | GGTTTCAGGCTTGTTGGCTA | AGGTTAGGTTTGATTGCCCC | 154 |
| chr6:17258557-17260573 | p5 | (AACCT)3 | GGTTTCAGGCTTGTTGGCTA | AGGTTAGGTTTGATTGCCCC | 154 |
| chr6:17258567-17260582 | p6 | (AGGAAG)3 | GGTTTCAGGCTTGTTGGCTA | AGGTTAGGTTTGATTGCCCC | 144 |
| chr6:17258567-17260582 | p5 | (AACCT)3 | GGTTTCAGGCTTGTTGGCTA | AGGTTAGGTTTGATTGCCCC | 144 |
| chr6:17258828-17260846 | p5 | (AACCT)3 | CATTGCAGCAGCTTGTATGG | AGGTTAGGTTTGATTGCCCC | 254 |
| chr6:17386707-17388722 | p3 | (ACA)5 | TTTGGGCCATCTTACACACA | GCCGAAGGCTTGTCTTAATG | 182 |
| chr6:17421565-17423580 | p5 | (AAACC)3 | CTGTTTGTTCGCTTGCAAAA | CGAAAAATCCGAGAAAACCA | 137 |
| chr6:17456818-17458846 | p2 | (AT)14 | CCTGCATCTGAGGGAGATGT | GTGGCATGCTCATGCTCTAA | 275 |
| chr6:17507889-17509917 | p4 | (AAAG)7 | CAGGGGGAGCTATACCATGA | CTAATTTGAGTGCATGCCGA | 154 |
| chr6:17551027-17553059 | p2 | (TA)16 | AACTGGAACTGTCACACCCC | TTGCAGACAGTGTTGTGGGT | 149 |
| chr6:17661411-17663425 | p2 | (AT)7 | TGAAATCACCTACCCCCAAA | CACCGATTCTTTGGTCGATT | 180 |
| chr6:17716818-17718833 | p5 | (AATAA)3 | ACGGATGATTGAGCTTACCG | AAGTCGACGACCGAGAAGAA | 245 |
| chr6:17716818-17718833 | p5 | (AATAA)3 | ACGGATGATTGAGCTTACCG | AAGTCGACGACCGAGAAGAA | 245 |
| chr6:17725239-17727254 | p3 | (TTG)5 | CAAAAGAACCAAGAGGGCAA | AACGTGGAGAATTGGGACAC | 218 |
| chr6:17732925-17734941 | p4 | (TCAA)4 | GCAGAAGCATATCAAGGGGA | CGTTGCCTGAAAAATCCCTA | 245 |
| chr6:17733538-17735568 | p4 | (TCAA)4 | TCCAGTGTTCACCAGTTCCA | GGGAGTCCTAATGCCACAAA | 209 |
| chr6:17789045-17791057 | p2 | (AT)7 | GGGAGACAGCAAATTGGAGA | TCCGCTTCTGAGTTTTGGTT | 248 |
| chr6:17789045-17791057 | p2 | (TA)6 | GGGAGACAGCAAATTGGAGA | TCCGCTTCTGAGTTTTGGTT | 248 |
| chr6:17789045-17791057 | p2 | (TA)9 | GGGAGACAGCAAATTGGAGA | TCCGCTTCTGAGTTTTGGTT | 248 |
| chr6:17789170-17791190 | p2 | (AT)7 | GGGAGACAGCAAATTGGAGA | TCCGCTTCTGAGTTTTGGTT | 123 |

| Marker ID | SSR type | SSR motif | FORWARD PRIMER (5'-3') | REVERSE PRIMER (5'-3') | size (bp) |
|---|---|---|---|---|---|
| chr6:17789170-17791190 | p2 | (TA)6 | GGGAGACAGCAAATTGGAGA | TCCGCTTCTGAGTTTTGGTT | 123 |
| chr6:17789170-17791190 | p2 | (TA)9 | GGGAGACAGCAAATTGGAGA | TCCGCTTCTGAGTTTTGGTT | 123 |
| chr6:17797928-17799948 | p5 | (ATAAT)4 | GATGAGAAACGAGTGGGGTG | GTGGGAGTTTGCAGCATTTT | 199 |
| chr6:17824372-17826387 | p5 | (TGCCC)3 | TGTTATCTCCTTGGCGAACC | TAATGCACGAATGGTCAGGA | 150 |
| chr6:17892520-17894576 | p2 | (AT)28 | GATGAATTCGGCGGTAGAAA | CCGGAGTGATCGTGATTCTT | 205 |
| chr6:18089188-18091203 | p5 | (ATCAG)3 | CCGGAACCCACACATCTATC | TCTAGGGCACAGGTTCTGCT | 260 |
| chr6:18151327-18153343 | p2 | (AC)8 | CGACCCAAAAGAATTCTGGA | TGTTGAAGAGGGTGAGTCCC | 295 |
| chr6:18151327-18153343 | p6 | (TGTATA)4 | CGACCCAAAAGAATTCTGGA | TGTTGAAGAGGGTGAGTCCC | 295 |
| chr6:18153987-18156001 | p2 | (AT)7 | GTATGCGGAAAACGAGGTGT | TTTCAAAGGTCGTGCACTTG | 253 |
| chr6:18196398-18198410 | p2 | (CA)6 | TTTCGACCCAAAGAATTTCG | GCCCAGACTCAGAGACCAAG | 179 |
| chr6:18289243-18291270 | p3 | (AAT)9 | TCTCCTACGAATCGGTTTGG | GAGACATTTTGGGGTGAGGA | 119 |
| chr6:18301032-18303074 | p2 | (AG)8 | TAGGCGGCTCAAAAACTTGT | TTCGTCCGTTTGTGAATCAA | 116 |
| chr6:18475211-18477226 | p5 | (TAAAC)3 | TGATATCGGGCATGCATAGA | GCTTGTTGTTTGGTTGCTGA | 141 |
| chr6:18475211-18477226 | p2 | (GA)7 | TGATATCGGGCATGCATAGA | GCTTGTTGTTTGGTTGCTGA | 141 |
| chr6:18484698-18486713 | p5 | (TAAAC)3 | TGATATCGGGCATGCATAGA | GCTTGTTGTTTGGTTGCTGA | 135 |
| chr6:18484698-18486713 | p2 | (GA)7 | TGATATCGGGCATGCATAGA | GCTTGTTGTTTGGTTGCTGA | 135 |
| chr6:18492989-18495049 | p2 | (AT)30 | TGCTACTCGCGTAGGTCCTT | GTGGGATAGCGACGAGAGAG | 255 |
| chr6:18492989-18495049 | p5 | (TGAAT)3 | TGCTACTCGCGTAGGTCCTT | GTGGGATAGCGACGAGAGAG | 255 |
| chr6:18520612-18522624 | p2 | (GA)12 | CCCTAATCTTTCCACCACCA | GACACATATCTGGCCGTGTG | 165 |
| chr6:18520612-18522624 | p2 | (AC)6 | CCCTAATCTTTCCACCACCA | GACACATATCTGGCCGTGTG | 165 |
| chr6:18583910-18585925 | p5 | (TCTTC)3 | CGTGGTCCATCGGTTTTATT | AATGCAACATTCTGCTGCTG | 279 |
| chr6:18705198-18707213 | p5 | (CATTG)3 | AAATTTTAGGTGCCCACACG | TTCCATTCCGAAGTGCCTAC | 131 |
| chr6:18741727-18743742 | p5 | (AATGC)3 | CATGTTCCATAATCAGCCCC | GATGATATTGGTGCGCTGTG | 270 |
| chr6:18741727-18743742 | p5 | (CCTTT)3 | CATGTTCCATAATCAGCCCC | GATGATATTGGTGCGCTGTG | 270 |
| chr6:18747158-18749240 | p2 | (AT)9 | CTTCTGGGCCACTGATGATT | CCTATTATGGCCCTCCGATT | 120 |
| chr6:18792347-18794362 | p5 | (ATTTT)3 | TATAGGCAGCCCACATAGGG | TATCCCAGCAAGGGAATCAG | 213 |
| chr6:18808249-18810261 | p3 | (ATC)5 | TTTCAAAACTCACTTCGGGG | CTTCGGTCTTTCACCCAAGA | 173 |
| chr6:18808258-18810274 | p3 | (ATC)5 | TTTCAAAACTCACTTCGGGG | CTTCGGTCTTTCACCCAAGA | 164 |
| chr6:18808320-18810338 | p3 | (ATC)5 | TTTCAAAACTCACTTCGGGG | CTTCGGTCTTTCACCCAAGA | 102 |
| chr6:18885762-18887777 | p3 | (GGT)5 | TGATAATGAAAAGCCGAGGG | CTGATGAACACAGCGCCTAA | 155 |
| chr6:19034624-19036636 | p2 | (CA)6 | CCTACAATTTGAGGGGCAAA | ACATTGCCTTTTCGAGCATC | 270 |
| chr6:19034624-19036636 | p2 | (AT)8 | CCTACAATTTGAGGGGCAAA | ACATTGCCTTTTCGAGCATC | 270 |
| chr6:19066623-19068638 | p5 | (AATAT)3 | CAAGAGCTTCAAATGCACCA | TAATTGGGAGGCTACAACGC | 211 |
| chr6:19066623-19068638 | p5 | (AATGC)3 | CAAGAGCTTCAAATGCACCA | TAATTGGGAGGCTACAACGC | 211 |
| chr6:19121425-19123445 | p4 | (TGTA)5 | GGTTATAGGCCCTCACACGA | GGTGTGAGAAAGTTGGGGAA | 239 |
| chr6:19140335-19142399 | p2 | (TA)32 | ATAAATTCCCGGCACATCAA | AGGGGTCCTTGGAAGAAAAA | 260 |

| Marker ID | SSR type | SSR motif | FORWARD PRIMER (5'-3') | REVERSE PRIMER (5'-3') | size (bp) |
|---|---|---|---|---|---|
| chr6:19181613-19183631 | p6 | (TATTCA)3 | TACCACTTTTTCCACGACCC | TGATGTCTTGCAGCCTTGTC | 284 |
| chr6:19182763-19184779 | p4 | (ATAC)4 | CATTACCATCATCGTTTGCG | TTAAGGCATGGTTCCTTTGC | 101 |
| chr6:19247289-19249305 | p4 | (TTTG)4 | TTGTTTGCATCCGTTTTGAA | TGCTACCTTTGGCGGTTATT | 157 |
| chr6:19325951-19327965 | p2 | (TA)7 | TCAAACCGAGCTATGTGCAG | CCTAATCATGGGATCTGCGT | 204 |
| chr6:19390654-19392669 | p5 | (TTCTT)3 | AGAGGAGATGAGTCGGCAAA | GAAGGTTTCGATTTGTGGGA | 123 |
| chr6:19391850-19393870 | p2 | (TA)10 | CTAAGAATCCAGGCACGGAC | CATCATGGACAGCCGTTTTA | 175 |
| chr6:19500456-19502504 | p2 | (AT)24 | GCGAGTACGTAGAAGGGCAC | ACGGGGTATAACATCCTCCC | 184 |
| chr6:19502015-19504079 | p2 | (TA)32 | CTTGAGCAAATAGGACCCGA | CGGAATATTCGTAGCGGAAA | 240 |
| chr6:19502015-19504079 | p5 | (TGTTG)3 | CTTGAGCAAATAGGACCCGA | CGGAATATTCGTAGCGGAAA | 240 |
| chr6:19515808-19517820 | p2 | (TA)6 | AAGAAGAAGCAAGGGAAGCC | GCTGAACCGGTTTTGCATAC | 195 |
| chr6:19515808-19517820 | p5 | (AATAA)3 | AAGAAGAAGCAAGGGAAGCC | GCTGAACCGGTTTTGCATAC | 195 |
| chr6:19515964-19517979 | p2 | (TA)6 | AGCTATTGGCCCCTGTTTTT | ACTTGAGGAGTGATGGCTGC | 204 |
| chr6:19515964-19517979 | p5 | (AATAA)3 | AGCTATTGGCCCCTGTTTTT | ACTTGAGGAGTGATGGCTGC | 204 |
| chr6:19589718-19591736 | p3 | (GTT)6 | ATTTTGTTGGGTTTCCCTCC | GCCAATTTTCTTTGCCTCAA | 107 |
| chr6:19636627-19638661 | p2 | (AT)17 | TCATGCTTTCCCCCTAAATG | ACGTTGGCTTAAGGTTCGTG | 216 |
| chr6:19644098-19646112 | p2 | (TA)7 | GGGTCTTCATCCGTCTTTGA | TATGCCTCCAGTTCCTCCAC | 159 |
| chr6:19694078-19696092 | p5 | (TTGCA)3 | GTCCGAGGTTCAATTCCAGA | TGGCTAATTGCTCATAGGGG | 108 |
| chr6:19694078-19696092 | p2 | (TA)7 | GTCCGAGGTTCAATTCCAGA | TGGCTAATTGCTCATAGGGG | 108 |
| chr6:19694078-19696092 | p6 | (ACACTC)3 | GTCCGAGGTTCAATTCCAGA | TGGCTAATTGCTCATAGGGG | 108 |
| chr6:19843593-19845619 | p2 | (TG)13 | TTCTGATTCGAATGGGGAAG | AGCCGATCGATCCTTAACCT | 226 |
| chr6:19891553-19893573 | p3 | (ATC)6 | GCTTGACCGGCGTTATGTAT | CCTATCAAGCCTCTGTGGGA | 220 |
| chr6:19891553-19893573 | p2 | (GT)10 | GCTTGACCGGCGTTATGTAT | CCTATCAAGCCTCTGTGGGA | 220 |
| chr6:19974357-19976391 | p2 | (TA)6 | CCCCAAACTACACCAGCCTA | AGGGAGATGAGGCACGACTA | 168 |
| chr6:20067562-20069595 | p2 | (AG)6 | AATGGTTTCAAATTGCCAGC | TAAACCCTACACAGACCCGC | 294 |
| chr6:20067602-20069674 | p2 | (AG)6 | AATGGTTTCAAATTGCCAGC | TAAACCCTACACAGACCCGC | 254 |
| chr6:20096839-20098853 | p2 | (TA)7 | ACATGAACACGAGCATTCCA | ATATCGCGGTGCAATACACA | 109 |
| chr6:20114256-20116270 | p2 | (TA)7 | GCCGACAAGGTCGAATACAT | TTTATGGGCATAGCTGGCTC | 272 |
| chr6:20125403-20127433 | p2 | (AT)15 | GCCGACAAGGTCGAATACAT | CAGGGACTACATAGCGGAGC | 277 |
| chr6:20157252-20159282 | p2 | (AT)9 | GCAGAAAAGGCCCAACAATA | ACTTTCCCGACCAGTTGATG | 192 |
| chr6:20157252-20159282 | p2 | (AT)15 | GCAGAAAAGGCCCAACAATA | ACTTTCCCGACCAGTTGATG | 192 |
| chr6:20162013-20164028 | p5 | (AATGC)3 | CGAAAGAAAACAGGCTCGTC | TCATTGGGTTGAGGGTTCTC | 245 |
| chr6:20186400-20188418 | p2 | (AG)9 | GATTTTTGAAATTGCGCGTT | ATAAATACGCACTCCACCGC | 282 |
| chr6:20353557-20355577 | p2 | (TA)10 | TACTCTCGACTACCCGCGTT | AGAGCATAGCCATAGCCGAA | 293 |
| chr6:20379210-20381222 | p2 | (AT)6 | GGGTGTCGTGATCAGCCTAT | TCCCCCTCTTATCTCCAACC | 163 |
| chr6:20416012-20418038 | p2 | (TA)13 | TGTGTCTTCCTGCCCTTTTC | TTGACACCGACCACCTTACA | 261 |
| chr6:20437017-20439047 | p2 | (AT)15 | GGATGCCAACTCCACAGATT | AATTGCTTCGGGTCAACAAG | 165 |

| Marker ID | SSR type | SSR motif | FORWARD PRIMER (5'-3') | REVERSE PRIMER (5'-3') | size (bp) |
|---|---|---|---|---|---|
| chr6:20437017-20439047 | p3 | (CAT)5 | GGATGCCAACTCCACAGATT | AATTGCTTCGGGTCAACAAG | 165 |
| chr6:20540195-20542227 | p2 | (AT)16 | GTTGTTACACCCCGCACTTT | GCCATGTTCTCGTGTTTCAA | 115 |
| chr6:20715205-20717217 | p2 | (TA)6 | TTCAAGGCATGTCATTTGGA | TAAGGGTTTGAATCCCCACA | 164 |
| chr6:20753202-20755230 | p2 | (TA)14 | TTTCGAATTGACCTTGGACC | ATCGCTGTCTTGCCTTCACT | 180 |
| chr6:20770898-20772913 | p5 | (CCCGA)4 | GACATGCCATTTAGCCAGGT | GCACTTAAATAGCCGTTGGG | 130 |
| chr6:20770898-20772913 | p5 | (CCGGC)3 | GACATGCCATTTAGCCAGGT | GCACTTAAATAGCCGTTGGG | 130 |
| chr6:20770898-20772913 | p2 | (AT)12 | GACATGCCATTTAGCCAGGT | GCACTTAAATAGCCGTTGGG | 130 |
| chr6:20802337-20804377 | p2 | (AG)20 | AAAACAACAACGGAACAGGC | TGTGGCATCAGGAACAAGAG | 205 |
| chr6:20825751-20827763 | p2 | (TA)6 | TTCAAGGCATGTCATTTGGA | TCCCCTTTCCCTACCTCCTA | 176 |
| chr6:20844126-20846150 | p2 | (AT)12 | TATCTTGTGCAAGGGAAGGG | ACGGTGACTTTCGGATTTTG | 108 |
| chr6:20866812-20868827 | p5 | (AATGC)3 | ACGACGGAAAGAAAGAGGGT | AGAGCCTTAGGTTAAGCGGC | 205 |
| chr6:20926724-20928739 | p4 | (ATAC)7 | GACGTCCTTTTGCTTTTGGA | GAGCGCTCCTGAGAAATCAC | 275 |
| chr6:20926724-20928739 | p5 | (ATATA)3 | GACGTCCTTTTGCTTTTGGA | GAGCGCTCCTGAGAAATCAC | 275 |
| chr6:21029580-21031607 | p3 | (AAT)9 | TTCGGCTAAAGGGACTGAAA | TCTCTCCCGTGAACGACTCT | 119 |
| chr6:21029580-21031607 | p2 | (AT)16 | TTCGGCTAAAGGGACTGAAA | TCTCTCCCGTGAACGACTCT | 119 |
| chr6:21046145-21048160 | p3 | (TCT)5 | CTACTTGATGCCGCAGTTGA | TGCTGCAACAACTTCCTCAC | 137 |
| chr6:21084433-21086453 | p2 | (AT)10 | TAAATTGAGATCAACGCCCC | GCTATCTCAACCGATGGGAC | 123 |
| chr6:21084433-21086453 | p2 | (AG)10 | TAAATTGAGATCAACGCCCC | GCTATCTCAACCGATGGGAC | 123 |
| chr6:21190575-21192629 | p3 | (ATA)18 | GGAGCATCCATTCACAACCT | ATCGCCCAGATGTCTTTGTC | 119 |
| chr6:21237128-21239143 | p5 | (CCACA)3 | AGCTGGGTCAGCAACTGAAT | AGCTCCCCCTCAGACTTCTC | 257 |
| chr6:21237128-21239143 | p2 | (CA)6 | AGCTGGGTCAGCAACTGAAT | AGCTCCCCCTCAGACTTCTC | 257 |
| chr6:21247281-21249302 | p3 | (ACA)7 | TTAGAGCCTCCACCCAACAC | ATCGGTCACTTGATTCGGTC | 194 |
| chr6:21310734-21312778 | p2 | (TA)22 | GGTCCTCTCCTCATCTTCCC | ATCATGATGAACGAGCATGG | 172 |
| chr6:21408466-21410500 | p2 | (AG)17 | AAATGCTTCAGCCAACTGCT | TCACCTCCTATCATCCAGGG | 204 |
| chr6:21467693-21469720 | p2 | (TC)6 | TTGGGTATGGCAGGTTCCTA | TATGGACCGAGAGGCCATAG | 208 |
| chr6:21602839-21604853 | p2 | (TA)7 | TGGCATGCATGTTTCCTTTA | TGACCGCCTAAAGTAGCGTT | 299 |
| chr6:21611741-21613759 | p6 | (CCTTGG)3 | TCATGGTGCCCAATAGAACA | CTGTCAAGGAAATGTCGGGT | 115 |
| chr6:21635588-21637600 | p2 | (AT)6 | AACTAGCTCGTGGCCTCAAA | TTTGGTTCTCAATGCACTCG | 252 |
| chr6:21656126-21658142 | p2 | (AT)8 | CATAAGCCGACGAGGCTAAC | GTCCTGTCCCGCCTATATGA | 276 |
| chr6:21679015-21681027 | p2 | (TA)6 | CTCGGGGTTGGTTGAACTTA | TTTCACAAGGATCATGCCAA | 295 |
| chr6:21690973-21692991 | p3 | (ATC)6 | TCATAAGGACGGGACAGGAC | CAGGTAGACAGGGAGACGGA | 209 |
| chr6:21708864-21710884 | p5 | (GGTTT)4 | CATCACCACCTCCCAAAGTT | TCTTTGAACGCAAGTTGTGC | 297 |
| chr6:21830242-21832305 | p3 | (GAA)7 | CTACACGAGGGCCAAAACAT | CCATGAATGATGGCTTGTTG | 129 |
| chr6:21894799-21896814 | p3 | (ACA)5 | GACGAACCCAGGTCCAACTA | GCACGTAAAGGACGTGTTGA | 214 |
| chr6:21945256-21947276 | p2 | (AT)14 | TCTCGACCTTATGTCACCCC | GACATGGTGTGTATGCCTGC | 265 |
| chr6:21945256-21947276 | p2 | (TA)10 | TCTCGACCTTATGTCACCCC | GACATGGTGTGTATGCCTGC | 265 |

| Marker ID | SSR type | SSR motif | FORWARD PRIMER (5'-3') | REVERSE PRIMER (5'-3') | size (bp) |
|---|---|---|---|---|---|
| chr6:21974951-21976966 | p5 | (ATCCA)3 | CATCGACCATCTCACGTCAC | GCACTGCATGCACTCTCATT | 118 |
| chr6:22027230-22029245 | p3 | (AAG)16 | CATGGCCGCTAGACTTCTCT | CCATTATGGCAACTGCACTG | 152 |
| chr6:22027230-22029245 | p5 | (AAAAG)3 | CATGGCCGCTAGACTTCTCT | CCATTATGGCAACTGCACTG | 152 |
| chr6:22050504-22052520 | p2 | (TA)8 | TGTTTATGGGTGTGAAGCGA | CAGGGGCATTCATCATCTCT | 252 |
| chr6:22192248-22194269 | p3 | (ATG)7 | AGGGCTAACTCAGCGTGAAA | GCCTTGAATCCAAGCAAAAA | 178 |
| chr6:22192248-22194269 | p2 | (TA)29 | AGGGCTAACTCAGCGTGAAA | GCCTTGAATCCAAGCAAAAA | 178 |
| chr6:22204273-22206288 | p5 | (AAAAG)3 | CTGAGCCTCCATTTTGTGGT | AGGCTGATGGCTCTCATTGT | 224 |
| chr6:22204273-22206288 | p5 | (AAAAG)3 | CTGAGCCTCCATTTTGTGGT | AGGCTGATGGCTCTCATTGT | 224 |
| chr6:22320747-22322762 | p5 | (AAAGG)3 | TGTGTACAGGGTTCGAGCAG | TCATAACTGTGGCGCACTTC | 194 |
| chr6:22321496-22323511 | p5 | (AAAGG)3 | TGAGGCTCAAAGGAAAGGAA | TTTGCATGAATTCTCCGACA | 243 |
| chr6:22321526-22323541 | p5 | (AAAGG)3 | TGAGGCTCAAAGGAAAGGAA | TTTGCATGAATTCTCCGACA | 213 |
| chr6:22339563-22341583 | p2 | (AT)7 | AAGAGTTGTCACAGGCCCC | GCGACGATTTAACCAGGAAA | 203 |
| chr6:22339563-22341583 | p2 | (TA)10 | AAGAGTTGTCACAGGCCCC | GCGACGATTTAACCAGGAAA | 203 |
| chr6:22445039-22447057 | p2 | (TA)7 | TGATGTCCAAGTTCGGTTGA | GCCGTTTTTGGAAGATCAAA | 156 |
| chr6:22445039-22447057 | p6 | (TATATG)3 | TGATGTCCAAGTTCGGTTGA | GCCGTTTTTGGAAGATCAAA | 156 |
| chr6:22517765-22519780 | p2 | (AC)6 | CAACCCATTTTCTTCCTCCA | TGATGCAAACCAAAGCAAAA | 188 |
| chr6:22517765-22519780 | p3 | (GAA)8 | CAACCCATTTTCTTCCTCCA | TGATGCAAACCAAAGCAAAA | 188 |
| chr6:22517765-22519780 | p3 | (GAA)5 | CAACCCATTTTCTTCCTCCA | TGATGCAAACCAAAGCAAAA | 188 |
| chr6:22576543-22578576 | p3 | (TTA)11 | ATTGCGTTTTTGGGAAACAG | GGAAGGAACTTTGGAGGGAC | 218 |
| chr6:22576543-22578576 | p3 | (TTA)5 | ATTGCGTTTTTGGGAAACAG | GGAAGGAACTTTGGAGGGAC | 218 |
| chr6:22576543-22578576 | p3 | (TAT)7 | ATTGCGTTTTTGGGAAACAG | GGAAGGAACTTTGGAGGGAC | 218 |
| chr6:22633110-22635125 | p5 | (AACAA)3 | ATGCAGAAATTGCTTTGCCT | ATAAGCCAACATGAAACGGC | 150 |
| chr6:22756119-22758135 | p4 | (AATT)4 | GCAGGACCTCTTTATGCCAG | TGTCGTGGAGCATCCAATTA | 211 |
| chr6:22760895-22762928 | p3 | (ATA)11 | GTGAGGCTTTCAAATCTCCG | AAAACCCCGTTTATTCTCCG | 192 |
| chr6:22767022-22769040 | p6 | (ATCATT)3 | ATCAACCTCGACGCAAGACT | TCGCCGGAGCACTCTACTAT | 110 |
| chr6:22801776-22803794 | p2 | (TA)9 | GAAGATGCAAGGTGCACAAA | TGTGTTGTTGCCCAATGAGT | 278 |
| chr6:22876030-22878048 | p6 | (CATAAT)3 | GCCAACTAGGCACGATGAAT | TAGCCATGGTTACCTCCCAG | 233 |
| chr6:22935578-22937600 | p2 | (TA)11 | TTCAACCCCTAATGCTCCTG | GTGCATAGCGCTGAATCAAA | 184 |
| chr6:22965625-22967653 | p4 | (AAAC)7 | ACCCGGACAAGCAATAGATG | CGGGTAAGATGGGGGTAAAT | 156 |
| chr6:22973363-22975393 | p3 | (TGT)10 | GAAGGTGCCACCGAAACTTA | CGCATCTTGTCTTCCACTGA | 270 |
| chr6:22973363-22975393 | p2 | (AT)28 | GAAGGTGCCACCGAAACTTA | CGCATCTTGTCTTCCACTGA | 270 |
| chr6:23114126-23116153 | p2 | (AT)19 | TAAGTTGGGGGTGATAAGCG | CCCCTGTGAGGACGACTAAA | 103 |
| chr6:23114126-23116153 | p3 | (ATT)9 | TAAGTTGGGGGTGATAAGCG | CCCCTGTGAGGACGACTAAA | 103 |
| chr6:23187951-23189965 | p2 | (AT)7 | GTGGGGTTAGTTCCCCAAAT | CTTCCTCTTTTCGTGCCTTG | 105 |
| chr6:23355238-23357256 | p2 | (AT)9 | TCCCGATAACCTGTCTCCTG | TGTTCGTCTGTAGTGGCAGC | 253 |
| chr6:23414506-23416521 | p5 | (AATGC)3 | GTAGGCGAACCCTCAATCAA | TCATTCCGGGATAGAGGTTG | 157 |

| Marker ID | SSR type | SSR motif | FORWARD PRIMER (5'-3') | REVERSE PRIMER (5'-3') | size (bp) |
|---|---|---|---|---|---|
| chr6:23541768-23543788 | p5 | (CCCCA)4 | AATTTGCTTCCACCTTCCCT | TGGGATAGGACGGCTAGTTG | 224 |
| chr6:23564871-23566895 | p2 | (AT)12 | TGTCCCTTTTATTGCTTGGG | CTAGATAGCTGACGACGCCC | 238 |
| chr6:23856575-23858590 | p5 | (ATTGC)3 | TGGTGAGACTTCGTGTAGCG | ACGTGGCTAGTCCCAAACAC | 272 |
| chr6:23871176-23873188 | p2 | (AT)6 | GACTTCAGACGACTCTGGGC | TGGGCAGAAGTGTAGATCCC | 199 |
| chr6:23961603-23963618 | p3 | (ATG)5 | TTTAAACCCCAAAGAACCCC | TAATGGCGTGGAAGCTTTTC | 236 |
| chr6:23979469-23981484 | p3 | (ACA)5 | TGAGCAAGCAAATGCAAATC | TTTGGAACATCCTCCTACGG | 256 |
| chr6:24077949-24079971 | p2 | (AG)11 | TCGATTTGGGTCGTGACATA | AATTTGGTAAAGTCGTGCGG | 112 |
| chr6:24149299-24151353 | p2 | (AT)27 | TTTCCATGGCTTCGAGTACC | ATCCCCACCTTTAATCCCAG | 220 |
| chr6:24211163-24213181 | p6 | (AGGTAG)5 | TCCATATCCTTTGCGTAGGG | CTTGTTTCAGGTCCATCGGT | 156 |
| chr6:24211163-24213181 | p6 | (CACCTG)3 | TCCATATCCTTTGCGTAGGG | CTTGTTTCAGGTCCATCGGT | 156 |
| chr6:24374027-24376045 | p6 | (ACCAAA)3 | GTTGGTTCTGACCCCTCTGA | GGTGTTTAGTCGCCACAGGT | 278 |
| chr6:24421499-24423519 | p5 | (TGTGG)4 | CTGCTGGAAGATGACAAGCA | TCGCCGAATTCCAAATTTAC | 181 |
| chr6:24539753-24541769 | p2 | (AT)7 | TTTATTCTGGACTGGGCGAG | AGTGCCAATCATGCCTTACC | 106 |
| chr6:24539753-24541769 | p4 | (CATA)4 | TTTATTCTGGACTGGGCGAG | AGTGCCAATCATGCCTTACC | 106 |
| chr6:24642130-24644160 | p2 | (AT)15 | TCACAAGGATCATGCCAAAA | CTCGGGGTTGGTTGAACTTA | 121 |
| chr6:24644384-24646420 | p3 | (TGT)8 | AGATGAAAGTGGCGGAAATG | AGGCTACCCCAACCTTGTCT | 129 |
| chr6:24644384-24646420 | p2 | (AT)18 | AGATGAAAGTGGCGGAAATG | AGGCTACCCCAACCTTGTCT | 129 |
| chr6:24646361-24648375 | p2 | (TA)7 | CCTTGTTGGTAGCCCGTAGA | TCAACTCAACCTCCTTTCCG | 210 |
| chr6:24718361-24720379 | p3 | (ATG)6 | TTTTGGGTCAACTTTGGAGG | TGGATTTTCTGGATTTTCGC | 171 |
| chr6:24718361-24720379 | p2 | (TA)6 | TTTTGGGTCAACTTTGGAGG | TGGATTTTCTGGATTTTCGC | 171 |
| chr6:24722180-24724195 | p3 | (CAA)5 | ATAACGCTTTGGCCCTTTTT | AATCGGAGGTTACCGACCTT | 144 |
| chr6:24757772-24759784 | p2 | (TA)6 | GCATCGCTGAAATCAGAACA | AGCAGTTCCGTCCTAGGGTT | 245 |
| chr6:24757772-24759784 | p3 | (CAA)5 | GCATCGCTGAAATCAGAACA | AGCAGTTCCGTCCTAGGGTT | 245 |
| chr6:24796598-24798654 | p2 | (AT)28 | TGAAATAGGGGAAATGCTGC | CGTTCACTGCCAAAACTTGA | 247 |
| chr6:24837626-24839640 | p2 | (AT)7 | TTTTGATTTACGGTCCGTCC | TCCAGTGGTCGTATATGGCA | 185 |
| chr6:24975675-24977687 | p2 | (TA)6 | TGTGCATGGGATCTTTACCA | CTCCTCAAGATTTCCCTCCC | 152 |
| chr6:25087541-25089583 | p2 | (TA)31 | ACCTTCCCCAAAAGTTACGG | TTAGACGCTCGCATATTCCC | 255 |
| chr6:25119807-25121822 | p5 | (TTATT)3 | CACAAGGAGAGTGGGGGATA | TCCTTCCCTCATTACATGCC | 252 |
| chr6:25173465-25175523 | p2 | (TA)29 | CTCCCCATAGGGTAAGCCAT | CGTATCGGTATCGGACCTGT | 145 |
| chr6:25185486-25187516 | p2 | (AC)24 | GGAGTAGCAAGGAGACGACG | ATCGTCTGGCATCATCATCA | 238 |
| chr6:25185486-25187516 | p2 | (TA)15 | GGAGTAGCAAGGAGACGACG | ATCGTCTGGCATCATCATCA | 238 |
| chr6:25188489-25190507 | p3 | (CAA)6 | GTAACATCTGCCCGACTGGT | GGTTAACTGATGACCACGGG | 268 |
| chr6:25433516-25435588 | p3 | (CAT)15 | TCAAGCTTTCTGTGGTGGTG | AGTGGACGATTTCCCCTTCT | 202 |
| chr6:25496362-25498413 | p5 | (ATAAA)3 | AGGCCCATAGGGTCATTTTC | TGTGCGACATCAGGAAGAAG | 109 |
| chr6:25555411-25557426 | p5 | (ATTGC)3 | CGAATGTCGCTTGTTTTCCT | CACAAGTACCTCAACGGGGT | 211 |
| chr6:25689477-25691489 | p2 | (CA)6 | TCCATGAAATACTCGCCCTC | CCAAAATGCACGTTTTTCCT | 238 |

| Marker ID | SSR type | SSR motif | FORWARD PRIMER (5'-3') | REVERSE PRIMER (5'-3') | size (bp) |
|---|---|---|---|---|---|
| chr6:25951182-25953228 | p2 | (TA)23 | CAAGTCCCCAATCCCTTTTT | CTTGGGATGGGTTTTCTTGA | 150 |
| chr6:25951182-25953228 | p3 | (ATA)14 | CAAGTCCCCAATCCCTTTTT | CTTGGGATGGGTTTTCTTGA | 150 |
| chr6:26004218-26006239 | p3 | (CAA)7 | ACCGCTGAGTGGAAGTCCTA | ACTCCGCAGAGGCACTTTTA | 189 |
| chr6:26031779-26033797 | p2 | (TA)7 | CCTGCTCAACGTATCTGCAA | GACCCCTCATCACGACTCAT | 249 |
| chr6:26031779-26033797 | p6 | (CATAAG)3 | CCTGCTCAACGTATCTGCAA | GACCCCTCATCACGACTCAT | 249 |
| chr6:26031779-26033797 | p2 | (AT)14 | CCTGCTCAACGTATCTGCAA | GACCCCTCATCACGACTCAT | 249 |
| chr6:26031934-26033962 | p2 | (TA)7 | CTTGCCTTCCCAGGATGATA | GACCCCTCATCACGACTCAT | 119 |
| chr6:26031934-26033962 | p6 | (CATAAG)3 | CTTGCCTTCCCAGGATGATA | GACCCCTCATCACGACTCAT | 119 |
| chr6:26031934-26033962 | p2 | (AT)14 | CTTGCCTTCCCAGGATGATA | GACCCCTCATCACGACTCAT | 119 |
| chr6:26031934-26033962 | p6 | (TCATCT)3 | CTTGCCTTCCCAGGATGATA | GACCCCTCATCACGACTCAT | 119 |
| chr6:26058189-26060204 | p2 | (TG)6 | ACGGGTTCACAACCATGAAT | CACATTCCGCAACACTGACT | 189 |
| chr6:26058189-26060204 | p2 | (TG)10 | ACGGGTTCACAACCATGAAT | CACATTCCGCAACACTGACT | 189 |
| chr6:26058189-26060204 | p5 | (ACTTT)3 | ACGGGTTCACAACCATGAAT | CACATTCCGCAACACTGACT | 189 |
| chr6:26299975-26301989 | p2 | (TA)7 | CCCGAGCAAAGATAGACTCG | CGTCCTGAATCTCCCTACCA | 289 |
| chr6:26337678-26339696 | p6 | (ATAACC)3 | GAACGATGTGCGTGCTAAAA | TTCCAACTCATTCGCCTACC | 162 |
| chr6:26451682-26453697 | p3 | (GTT)5 | CAAAGAGATAGTGCGGCTCC | TATCCTCGAGGGTCTGGATG | 234 |
| chr6:26454889-26456904 | p5 | (TCTTA)3 | ACATTTGCCTCTCGGTCATC | TGAGAGCACTCCATTGTTCG | 147 |
| chr6:26520412-26522430 | p2 | (TA)9 | GTGGCCAGTTATGGAGGAAA | TCCCACACGATGTTTTTCAA | 248 |
| chr6:26633477-26635492 | p5 | (TTTAT)3 | AAAAGGCATCGCGTATTGAC | GGATCAAGGCTGGAGATCAA | 200 |
| chr6:26641298-26643330 | p3 | (ATC)5 | GCCCGATCTAGAACAACTCG | GTCGGCTTGCATAATTGGTT | 212 |
| chr6:26641331-26643343 | p3 | (ATC)5 | GCCCGATCTAGAACAACTCG | GTCGGCTTGCATAATTGGTT | 179 |
| chr6:26687001-26689022 | p3 | (ACA)7 | GGTAGCGTTGGTATTTGCGT | AGTTCCATCCACCTCCACTG | 141 |
| chr6:26687001-26689022 | p2 | (TG)6 | GGTAGCGTTGGTATTTGCGT | AGTTCCATCCACCTCCACTG | 141 |
| chr6:26832370-26834385 | p5 | (TGAAA)3 | CTCGTCATCCGAGGGATTTA | GGTAGAAGCGCTAGCATTGG | 127 |
| chr6:26841973-26843991 | p6 | (CATAGA)3 | GCATCGTTGTATCATGTCCG | CAACAGACAAACGGGAACCT | 283 |
| chr6:26841973-26843991 | p2 | (AG)6 | GCATCGTTGTATCATGTCCG | CAACAGACAAACGGGAACCT | 283 |
| chr6:26885011-26887023 | p2 | (AG)6 | CAAAACTGAGAGCAATGGCA | TTTTATCCACTAGCCGGTCG | 183 |
| chr6:26924466-26926481 | p2 | (AT)28 | CCTGTAACCAACACTGCGAA | TTTCGGTATTTATGCCCAGC | 251 |
| chr6:26924466-26926481 | p3 | (CAA)5 | CCTGTAACCAACACTGCGAA | TTTCGGTATTTATGCCCAGC | 251 |
| chr6:27051608-27053620 | p2 | (GT)6 | TTTGGAGAAAGAGCGACGTT | CATGCACAAACACAACACCA | 125 |
| chr6:27146080-27148095 | p3 | (TGT)5 | GGACCCTTGAAGCAAATCAA | CACCTGTAGACCCACGACCT | 154 |
| chr6:27182645-27184660 | p3 | (TGT)5 | GGAGCAGTCCTCCCATACAA | CTGTGGAGTGCAGCAAAAGA | 119 |
| chr6:27316155-27318185 | p2 | (TC)16 | TACTCGTATGGTCCAAGCCC | AGGCACAAATGGAAGACCAC | 126 |
| chr6:27713817-27715832 | p5 | (GTTCG)3 | GACCAAAAGGAACCTGACCA | ACCACTACAGCGAGGCTTGT | 107 |
| chr6:27713817-27715832 | p5 | (TTCGG)3 | GACCAAAAGGAACCTGACCA | ACCACTACAGCGAGGCTTGT | 107 |
| chr6:27751563-27753621 | p2 | (AT)29 | GTGGTTCGAAGTGGGAGAAA | AGTTTCTGGTGCCCGTAATG | 231 |

| Marker ID | SSR type | SSR motif | FORWARD PRIMER (5'-3') | REVERSE PRIMER (5'-3') | size (bp) |
|---|---|---|---|---|---|
| chr6:27853058-27855072 | p2 | (AT)7 | AATACCCGCAGCCTCCTATT | GTAGCAGAGCAGTTCCGTCC | 293 |
| chr6:27853238-27855252 | p2 | (AT)7 | AAATACCCGCAGCCTCCTAT | GTAGCAGAGCAGTTCCGTCC | 112 |
| chr6:27901817-27903832 | p5 | (TTTTA)3 | TTGCTAACAATCGTGGCAAT | CGCTAGCTTCTATCGTGTGG | 186 |
| chr6:27910379-27912451 | p2 | (AT)6 | TCTTTGGCACCCATTAAAGC | AAGGTCCCCAGCTTATCCAT | 195 |
| chr6:27927780-27929804 | p3 | (TGT)8 | AAATCTGGGACCTCGTGTTG | TGGTCCATCAGACAGCTGAG | 118 |
| chr6:27953462-27955483 | p3 | (CAA)7 | AAGTGTGTCCGGTATGGAGG | GATGAGCTGTGTTGGAGGGT | 166 |
| chr6:28014182-28016197 | p3 | (TGT)5 | TTGGAGAAAAGATGGGCAAG | TTCGCGCAGAGTTAATCCTT | 289 |
| chr6:28145048-28147063 | p5 | (TATTT)3 | TGCCAAGAATGTGCAGAGAG | CATCGCAATCTTTGATGGTG | 115 |
| chr6:28175293-28177313 | p5 | (GAAAT)4 | CATCGGTATCGTCATGTTCG | GCATTGCAGGAGAACAATCA | 115 |
| chr6:28268040-28270055 | p5 | (TTGTT)3 | AGCGATGCTGGTTCTCACTT | CCCACTCGTCTGAGGATGAT | 152 |
| chr6:28319887-28321903 | p2 | (AG)8 | AAATCCTCGAGCGGGTTATT | AAGGTGGGCTCATGACATTC | 136 |
| chr6:28430236-28432248 | p2 | (GA)6 | TGCGCTTCTAGTTCCCTTGT | ACGCCTTCCCCTATGAAGAT | 165 |
| chr6:28439487-28441499 | p2 | (GA)6 | TGCGCTTCTAGTTCCCTTGT | ACGCCTTCCCCTATGAAGAT | 165 |
| chr6:28459913-28461931 | p6 | (TTTTTG)3 | ACGAACTGCTGCTCTCCATT | AAACACACACACCCCAACAA | 198 |
| chr6:28459913-28461931 | p3 | (AAG)20 | ACGAACTGCTGCTCTCCATT | AAACACACACACCCCAACAA | 198 |
| chr6:28494043-28496112 | p3 | (TTA)23 | TTCTGCGAGTGAAGGAGGAT | GCTAAGGCCGATCTTGTCAG | 231 |
| chr6:28497543-28499567 | p2 | (AT)12 | TGCCTTGATTATTTCAGCCC | GGATGGGTCGTGACAGAACT | 107 |
| chr6:28615827-28617839 | p2 | (TA)6 | ATGAAGAGGAGCGGGAAAAT | AAGAAAAGCCGAGGAGGTTC | 123 |
| chr6:28702911-28704965 | p2 | (TA)27 | TTCGTTAGCTTTTGGTCGCT | GTTCTCCTTCCAAGCTCACG | 299 |
| chr6:28745971-28747991 | p2 | (AT)10 | TGCCAAACTTGGTCATGAAA | GCACTGAATAAGGGCTCTGC | 251 |
| chr6:28848272-28850287 | p2 | (GA)6 | TGCCTCTTCAACGTTAACCC | TCCGACCTGCCTTATTCAAC | 292 |
| chr6:28848272-28850287 | p3 | (TAA)5 | TGCCTCTTCAACGTTAACCC | TCCGACCTGCCTTATTCAAC | 292 |
| chr6:29069942-29071957 | p3 | (TGT)5 | ACACGAAGCCTTGGAATGAC | GACTGGCTCTGTCCTTCTGC | 295 |
| chr6:29117460-29119476 | p4 | (TTGA)4 | CGATGCCAACCGACTCTAAT | TTTGAATCCAAGCCAACTCC | 230 |
| chr6:29117460-29119476 | p2 | (AT)6 | CGATGCCAACCGACTCTAAT | TTTGAATCCAAGCCAACTCC | 230 |
| chr6:29247329-29249395 | p2 | (TA)33 | CTTGCCAAACCCATGAAACT | ATTTCAGCTGGCAGTATCCG | 214 |
| chr6:29294601-29296613 | p2 | (TA)6 | GGACTTTTCCATGCTGGTGT | TCTTGGATTGGGGTGGAATA | 118 |
| chr6:29294601-29296613 | p6 | (TATGAA)3 | GGACTTTTCCATGCTGGTGT | TCTTGGATTGGGGTGGAATA | 118 |
| chr6:29312718-29314762 | p2 | (AT)22 | TCCCAAAAGAAGGCAGAGAA | GAGGCCCTTGTTGTAGGTGA | 141 |
| chr6:29312718-29314762 | p2 | (TA)15 | TCCCAAAAGAAGGCAGAGAA | GAGGCCCTTGTTGTAGGTGA | 141 |
| chr6:29317102-29319114 | p2 | (AT)6 | CAAACAAGAGGGCCATGACT | CGAAAGCGCCATTTCATAAT | 300 |
| chr6:29399153-29401173 | p2 | (TA)24 | TAGACCCGGCCTAATTTGTG | ACTTTGCCAAAATGGCTTTG | 275 |
| chr6:29399153-29401173 | p2 | (TA)34 | TAGACCCGGCCTAATTTGTG | ACTTTGCCAAAATGGCTTTG | 275 |
| chr6:29399153-29401173 | p2 | (TA)10 | TAGACCCGGCCTAATTTGTG | ACTTTGCCAAAATGGCTTTG | 275 |
| chr6:29399153-29401173 | p3 | (TAT)5 | TAGACCCGGCCTAATTTGTG | ACTTTGCCAAAATGGCTTTG | 275 |
| chr6:29479429-29481444 | p5 | (ATTTT)3 | AGGACTCCATTCAAACGTGC | GTTCCACGGGCTGAAATAAA | 182 |

| Marker ID | SSR type | SSR motif | FORWARD PRIMER (5'-3') | REVERSE PRIMER (5'-3') | size (bp) |
|---|---|---|---|---|---|
| chr6:29592178-29594196 | p3 | (AAT)6 | CAGTGGCAAATAGCTGAGCA | TATACCCATCAAGGTCGGGA | 178 |
| chr6:29611540-29613602 | p2 | (TA)31 | CTGGCCAAGCACACCTTAAT | CAACACATGTAGTGGCCCTG | 156 |
| chr6:29611540-29613602 | p3 | (CAA)5 | CTGGCCAAGCACACCTTAAT | CAACACATGTAGTGGCCCTG | 156 |
| chr6:29656864-29658878 | p2 | (AT)7 | AAGATCGATTTGCGTCCAAC | CGAAGATTCAGAGCATGCAA | 192 |
| chr6:29666304-29668322 | p3 | (ATC)6 | GAGTGCTCCTGAGATTTCGC | ATTGACGTCCCTTCTTGTGG | 140 |
| chr6:29914226-29916241 | p5 | (TTTCA)3 | ATTCCAAACCTTTGGGCTCT | GCTTTGCTGGTTCCATTGTT | 142 |
| chr6:29945299-29947314 | p3 | (ATA)5 | TGCGTGTGAGTGGAGTTTTC | CAACAACATTTCCGTGTTGC | 276 |
| chr6:29945299-29947314 | p5 | (TCATA)3 | TGCGTGTGAGTGGAGTTTTC | CAACAACATTTCCGTGTTGC | 276 |
| chr6:30049335-30051347 | p2 | (AT)6 | GGCCTCAGAAATCAGTCTCG | TTTTGATAGCTTCGTCGGCT | 273 |
| chr6:30056319-30058334 | p3 | (ATC)5 | GTGTAAGACGACTAGGCCGC | ATGCCCACAGGTTCAGGTAG | 131 |
| chr6:30207762-30209777 | p5 | (TTTTA)3 | CGGCTTGGAACAAAAACAAT | TGCAATCCGAAACGGTATTT | 237 |
| chr6:30593256-30595268 | p5 | (ATAAT)4 | TGGAAAGCAGTGGAAAAACC | CTTTTCCAAAACCACTCCGA | 120 |
| chr6:30593256-30595268 | p2 | (TA)6 | TGGAAAGCAGTGGAAAAACC | CTTTTCCAAAACCACTCCGA | 120 |
| chr6:30593256-30595268 | p4 | (TTTC)4 | TGGAAAGCAGTGGAAAAACC | CTTTTCCAAAACCACTCCGA | 120 |
| chr6:30626823-30628837 | p2 | (TA)7 | AATCATGAAATGACGCCTCC | GTTGAGTAAAGCGGCCATGT | 286 |
| chr6:30630718-30632733 | p2 | (AT)13 | TGCACCAAAGCTTGCATAAG | CTTGATCAATTTCCATGGGG | 107 |
| chr6:30630718-30632733 | p5 | (TGAAT)3 | TGCACCAAAGCTTGCATAAG | CTTGATCAATTTCCATGGGG | 107 |
| chr6:30630718-30632733 | p6 | (TATATG)3 | TGCACCAAAGCTTGCATAAG | CTTGATCAATTTCCATGGGG | 107 |
| chr6:30661381-30663419 | p2 | (AT)19 | AGTCATGGCCTACGGAGTTG | TATCTTTATGGTCCGCCCAG | 293 |
| chr6:30665893-30667911 | p6 | (TTAGCT)3 | GAATTGGTGGCAAAACGAGT | ACTTATGCGCCGTATGTGTG | 151 |
| chr6:30736330-30738350 | p2 | (AT)10 | CATTATGGAGGGTTGGATGG | GCACGAGTGCAGTAGTGAGC | 170 |
| chr6:30769210-30771258 | p2 | (AT)24 | AGGTCCCTGCTTATGGGTCT | TGCAGCTCCAGTACGAACAC | 259 |
| chr6:30888837-30890853 | p2 | (TA)6 | GGATGTCCGGATTGGTATTG | ATTTCACAACGATCATGCCA | 249 |
| chr6:30946560-30948602 | p2 | (AT)21 | TTAGCATGCATGACTCAGCC | CTATGGGCATGAACACAACG | 235 |
| chr6:30965313-30967328 | p5 | (TTCAT)3 | CCCAAGGCCCATCTAGGTAT | TTTCGAGCTGTAACACACGC | 270 |
| chr6:31086748-31088766 | p6 | (GAAAAA)3 | TACAGAACCAGGGGCAGAAC | GATGCCAGAACCCCAGTTTA | 229 |
| chr6:31096693-31098708 | p5 | (ATGAT)3 | AACTTGAGTTTGGATTGGCG | AGGGTCATAATCATGCTCGG | 117 |
| chr6:31157984-31160004 | p2 | (TA)10 | CATCTACACCCCGAAAAGGA | ACATGGCAATGGTAAAGGGA | 152 |
| chr6:31185291-31187318 | p3 | (AAT)9 | GGAAAATGGAGGGAAGGAAA | AGACAAACGGTCGAGAATGG | 196 |
| chr6:31198843-31200855 | p2 | (AT)10 | GTCGAATCATCTCTCCCCAA | AGACTCCCGTCACAGCTCAT | 285 |
| chr6:31198843-31200855 | p2 | (AT)6 | GTCGAATCATCTCTCCCCAA | AGACTCCCGTCACAGCTCAT | 285 |
| chr6:31222385-31224407 | p2 | (AT)11 | GAAGCATGACAGGGTGAGGT | TGGTTGCACTGGGTATGAAG | 119 |
| chr6:31439262-31441283 | p3 | (AAG)7 | TTCTATGACCGAAACAGGGG | CTATGACCGAGGCCTATGGA | 274 |
| chr6:31476166-31478256 | p3 | (TTG)6 | ATAGAGGAGATGCTGCCGAA | CTGGAAACTCGCTTCTCGAC | 291 |
| chr6:31476166-31478256 | p2 | (TA)45 | ATAGAGGAGATGCTGCCGAA | CTGGAAACTCGCTTCTCGAC | 291 |
| chr6:31543178-31545268 | p3 | (TTG)6 | ATAGAGGAGATGCTGCCGAA | CTGGAAACTCGCTTCTCGAC | 291 |

| Marker ID | SSR type | SSR motif | FORWARD PRIMER (5'-3') | REVERSE PRIMER (5'-3') | size (bp) |
|---|---|---|---|---|---|
| chr6:31543178-31545268 | p2 | (TA)45 | ATAGAGGAGATGCTGCCGAA | CTGGAAACTCGCTTCTCGAC | 291 |
| chr6:31642116-31644132 | p4 | (CATA)4 | CACACCATGCACTATTTGGC | GTTGTTTCCCGTTCTACCGA | 158 |
| chr6:31655086-31657101 | p3 | (CAT)5 | ACCATGCATCAATAACGCAA | TTTGATAGCCTTGTCGGCTT | 244 |
| chr6:31704942-31706960 | p6 | (TTTCTT)3 | CGGCGTCCTAGACAAGATTC | TTGGCTTGTAGGCCAATTTC | 279 |
| chr6:31724104-31726148 | p2 | (TA)22 | TCTTTCGTTTTCCTGGATGG | TTCCATAACGATCAGGAGGG | 127 |
| chr6:31729899-31731963 | p2 | (AT)32 | TCCTCTCGCTTGTCTTTGGT | TTCGAGGAGGAAGGAACTCA | 177 |
| chr6:31764308-31766329 | p3 | (TTA)7 | ATTCGCCTAACCCGAATTTT | GGAAAAAGGGATTGTTGGGT | 233 |
| chr6:31816982-31819000 | p6 | (TCATAT)3 | CCAAAAATCACGCCAAAGTT | CGCTGGAAGATACGGATTGT | 168 |
| chr6:31821535-31823571 | p5 | (TTCTC)3 | GCCATCTTTCTTTGGGACAA | CAGACACCAAGCCCATTTCT | 261 |
| chr6:31821574-31823589 | p5 | (TTCTC)3 | GCCATCTTTCTTTGGGACAA | CAGACACCAAGCCCATTTCT | 222 |
| chr6:31863595-31865641 | p2 | (AT)23 | TTGGTAGTGGTGCCATCTGA | AAAAATTCACCCCCTTCACC | 155 |
| chr6:31863595-31865641 | p3 | (ATT)5 | TTGGTAGTGGTGCCATCTGA | AAAAATTCACCCCCTTCACC | 155 |
| chr6:31875250-31877270 | p4 | (ATAA)5 | CACACCATCATAATGCCTCG | GGAGTTCGTGAGAAATGGGA | 263 |
| chr6:31969773-31971793 | p2 | (AT)10 | TCCACCTCATCCTTGCTTTC | GTGACATGGTCGAGATCCCT | 243 |
| chr6:31972311-31974329 | p2 | (AT)9 | TGGGCAAAATAAATGCATCA | GTGGCCTTTGTTGTGATGTG | 259 |
| chr6:32122913-32124935 | p2 | (AT)11 | TTTTCGAAATCGCCTGAAAG | TGCGCTATTGCAACTTCAAC | 214 |
| chr6:32171087-32173102 | p3 | (TTG)5 | GGGCTCAGGGGAGAGTAAAG | ACTGCCAGTTTGACCATTCC | 277 |
| chr6:32260259-32262285 | p5 | (ACTGC)3 | AGACCCATGCCAAATCAGTC | GCAGCAGCACACATCATTCT | 272 |
| chr6:32335970-32337982 | p2 | (AT)16 | CAGATAAGGGGTGCTCGGTA | TTGCCAACTGCATGGTGTAT | 207 |
| chr6:32335970-32337982 | p2 | (TA)6 | CAGATAAGGGGTGCTCGGTA | TTGCCAACTGCATGGTGTAT | 207 |
| chr6:32350085-32352103 | p3 | (TTG)6 | GACCCTTGAGCCAAATCAAA | TAACCCACTGACCTTCCTGC | 162 |
| chr6:32356934-32359000 | p2 | (TA)33 | GTCGCGCAACTCAATCACTA | CTTTCCCAGCCACCATTTTA | 141 |
| chr6:32447466-32449534 | p5 | (ATGCA)3 | AACCCTAGGAGTGCCCATTT | GGAGCATTAAGAACACCCGA | 280 |
| chr6:32462306-32464321 | p5 | (TGTGT)3 | AATGTAAAAAGCCGTGTGGC | TTTCCTTTCATTTTCCGTCG | 140 |
| chr6:32580494-32582512 | p3 | (TTG)9 | AGAATGGAAGGGTGATGCAG | CTTGAGAGTGGGGGTCTGAG | 113 |
| chr6:32580494-32582512 | p6 | (TATATG)3 | AGAATGGAAGGGTGATGCAG | CTTGAGAGTGGGGGTCTGAG | 113 |
| chr6:32831546-32833564 | p3 | (TTG)6 | AAAAGGACCCTTAAGCCGAA | TTTGGCAGCTTGGAGTCTTT | 160 |
| chr6:32836013-32838069 | p2 | (AT)12 | ATAATTGTACAGCGTCCCGC | TGTTCTGATTTCAGCGATGC | 280 |
| chr6:32836013-32838069 | p2 | (TA)28 | ATAATTGTACAGCGTCCCGC | TGTTCTGATTTCAGCGATGC | 280 |
| chr6:32843548-32845566 | p3 | (TTG)6 | AAAAGGACCCTTAAGCCGAA | TTTGGCAGCTTGGAGTCTTT | 160 |
| chr6:32848015-32850069 | p2 | (AT)12 | ATAATTGTACAGCGTCCCGC | TGTTCTGATTTCAGCGATGC | 280 |
| chr6:32848088-32850116 | p2 | (AT)12 | ATAATTGTACAGCGTCCCGC | TGTTCTGATTTCAGCGATGC | 207 |
| chr6:32851328-32853340 | p2 | (TA)6 | ATGTATGCATTGCACTCCCA | GCCATATAGGCGCGGTATTA | 136 |
| chr6:32955385-32957401 | p4 | (AATA)4 | TAGGCCAAAAGCCTAGTGGA | TTTGAGCCCTATTGGCAACT | 229 |
| chr6:32956502-32958520 | p6 | (TTGCAT)3 | CTTCAAAATTGGAGGCCCTT | TGCAATGGATCGCAATAGAG | 137 |
| chr6:33032703-33034723 | p2 | (AT)10 | AATCCCTAGGACGACCTGCT | AATGAGATCCAGTGGGTTCG | 169 |

| Marker ID | SSR type | SSR motif | FORWARD PRIMER (5'-3') | REVERSE PRIMER (5'-3') | size (bp) |
|---|---|---|---|---|---|
| chr6:33074340-33076356 | p3 | (ATC)6 | AAGGCTGTCACGATCCAATC | TAGTTTTGGGCTTGTGTCCC | 171 |
| chr6:33074358-33076380 | p3 | (ATC)6 | AAGGCTGTCACGATCCAATC | TAGTTTTGGGCTTGTGTCCC | 153 |
| chr6:33142994-33145008 | p2 | (AC)7 | AGCTTGGCCTTTGAGTTTCA | TACTGAGCTGGACCGCCTAT | 131 |
| chr6:33147330-33149366 | p2 | (TG)10 | TTCGTGGCTCCTGTTCTTCT | CCTAGGCAATGTCCAACCAT | 184 |
| chr6:33214084-33216099 | p3 | (TTG)5 | GGTGCCCAACAGAATAAGGA | CGTTCAACATGAAAACCGTG | 179 |
| chr6:33387441-33389455 | p2 | (TA)7 | TTCTTTGCAGCCCACTTTTT | AGGTCATAGTTCGGGGTGTG | 142 |
| chr6:33432763-33434778 | p5 | (TTCAT)3 | AAAGAAGTCGCTTCGGTCAA | GCCACACACGACTAAAAGCA | 174 |
| chr6:33517400-33519426 | p2 | (AT)13 | TTGGTATTTGGAGGAAACGG | CAAGGAACGAGAGAGGAACG | 249 |
| chr6:33547317-33549333 | p2 | (AT)8 | AGACGGCGGAATATACATCG | TGGGTCATGTGCGTATTTGT | 152 |
| chr6:33547317-33549333 | p4 | (AAAG)4 | AGACGGCGGAATATACATCG | TGGGTCATGTGCGTATTTGT | 152 |
| chr6:33700286-33702301 | p5 | (AACAC)3 | CTTACACACCTCTTTCCGCC | ATTGGCCAGTTGAAGTCTGG | 279 |
| chr6:33805318-33807336 | p2 | (AT)9 | GGGAGGAAGGGTAGCCATTA | GGGTTGGGTCGTGACACTAT | 128 |
| chr6:33812876-33814891 | p2 | (AT)7 | GCTCCAGAAAAATGGAGCAC | ATTGCCATACTTGCCTCCTG | 140 |
| chr6:33812876-33814891 | p5 | (AGAAA)3 | GCTCCAGAAAAATGGAGCAC | ATTGCCATACTTGCCTCCTG | 140 |
| chr6:33812876-33814891 | p2 | (GT)6 | GCTCCAGAAAAATGGAGCAC | ATTGCCATACTTGCCTCCTG | 140 |
| chr6:34011530-34013576 | p2 | (AT)23 | AGTTTCCCCTTCTTCCCAAA | TTGTAAACAGCGCAAAGTCG | 105 |
| chr6:34095563-34097575 | p2 | (AT)6 | CGTCCTCAGCATCTCATCAA | CTCGCCCAATGTAGAAGAGC | 221 |
| chr6:34167118-34169133 | p5 | (TGCAT)3 | CGGGACTTCCATTTCTCGTA | TGAACCGTCCAACATTAGCA | 221 |
| chr6:34188023-34190053 | p5 | (TGATA)6 | ATGCTGCCCAAATTTCTGTC | TATTCGCTCCTTACGCGTCT | 166 |
| chr6:34241969-34243987 | p2 | (TA)6 | TTGTCTAAGGGTACGACGGG | ATAAATGCAAGCCGACAAGG | 295 |
| chr6:34241969-34243987 | p6 | (TATATG)3 | TTGTCTAAGGGTACGACGGG | ATAAATGCAAGCCGACAAGG | 295 |
| chr6:34247371-34249389 | p3 | (TGT)5 | GTTGATGATTTGGCCGAGTT | TTGTCCGCTTCCTAGGCTAA | 208 |
| chr6:34247371-34249389 | p6 | (TATATG)3 | GTTGATGATTTGGCCGAGTT | TTGTCCGCTTCCTAGGCTAA | 208 |
| chr6:34352802-34354820 | p2 | (AG)9 | ATAAAGGTAATGCGGGGGAG | CGGGATACTTAAGGCAACCA | 184 |
| chr6:34454444-34456472 | p2 | (TA)14 | GATGCTTCCTCCAAAACCAA | GGGCAAGGTGTTTCCTGATA | 159 |
| chr6:34474423-34476457 | p2 | (AT)17 | ATTTGTTTAGACGCGTTCGG | AGGGCAATTTTAGCCCAACT | 243 |
| chr6:34501447-34503462 | p5 | (ATGCA)3 | GGTGCGTGGTCAATAGAGGT | CATTTTTGGGACTTGGGTTG | 183 |
| chr6:34518462-34520476 | p2 | (AT)7 | ACAACTTGGCTGTCAGGACC | CCTTGTCGGCTTGCATTTAT | 127 |
| chr6:34518462-34520476 | p2 | (TA)10 | ACAACTTGGCTGTCAGGACC | CCTTGTCGGCTTGCATTTAT | 127 |
| chr6:34537538-34539572 | p2 | (AT)17 | ACCAAAGAACTGCCACATCC | CGAGCCCGAGATTCTATGAG | 200 |
| chr6:34604258-34606270 | p2 | (AT)6 | GGATGGCTTAGGTGTGTCGT | TTTCCACGCTTCCTTCTGTT | 100 |
| chr6:34617739-34619767 | p2 | (AT)14 | AGGCGCATTTCTTTTCTTGA | GACCCGCAAGGTTTTATTGA | 102 |
| chr6:34633554-34635569 | p3 | (TAT)5 | ATCATCTTGGCGCACTTCTT | TCGACTCAAACCCTGAGCTT | 130 |
| chr6:34753862-34755900 | p2 | (AT)6 | AGCCCTTCGGCTATCAAAAT | TGTTCATGCCCACAGGTAGA | 168 |
| chr6:34753862-34755900 | p2 | (TA)19 | AGCCCTTCGGCTATCAAAAT | TGTTCATGCCCACAGGTAGA | 168 |
| chr6:34813393-34815405 | p2 | (TA)6 | GTAACAAGTGCCCGACCAAT | GTGAGTTCCCGGTATGGTGT | 111 |

| Marker ID | SSR type | SSR motif | FORWARD PRIMER (5'-3') | REVERSE PRIMER (5'-3') | size (bp) |
|---|---|---|---|---|---|
| chr6:34884457-34886484 | p3 | (AAG)9 | GATCGCAGTTGGTCGTTTTT | GACGAATACCAAACCCGAGA | 117 |
| chr6:35029396-35031420 | p3 | (TAT)8 | TAATCACGTTCGCTTTGTCG | TGGAAATTTGGGTTAGGCAA | 189 |
| chr6:35029396-35031420 | p3 | (CAT)5 | TAATCACGTTCGCTTTGTCG | TGGAAATTTGGGTTAGGCAA | 189 |
| chr6:35102478-35104493 | p3 | (ATG)5 | TCCGATAACGTTCCGAAATC | CGTGTGCCTTTAGGAAGGAG | 135 |
| chr6:35123811-35125826 | p3 | (ATG)5 | TCCGATAACGTTCCGAAATC | CGTGTGCCTTTAGGAAGGAG | 135 |
| chr6:35210363-35212435 | p2 | (TA)36 | TCGCCTAACCCGAACTTCTA | CGTGTTGTCGTTGATATGGC | 151 |
| chr6:35223157-35225181 | p2 | (AT)26 | CCCGAGTTGGACAGTCGTAT | CTTGTCGGCTTGCGTATTTT | 128 |
| chr6:35223157-35225181 | p2 | (AT)12 | CCCGAGTTGGACAGTCGTAT | CTTGTCGGCTTGCGTATTTT | 128 |
| chr6:35340978-35342992 | p2 | (TA)7 | ATACGACTGTCCAACTCGGG | AGTTTGAGACAGGTTGCGCT | 195 |
| chr6:35352919-35354949 | p2 | (TA)7 | CGTGCATGCCTCAATATGTC | CACATGCAATACTTGGTGCC | 146 |
| chr6:35352919-35354949 | p2 | (AT)6 | CGTGCATGCCTCAATATGTC | CACATGCAATACTTGGTGCC | 146 |
| chr6:35352919-35354949 | p2 | (AT)15 | CGTGCATGCCTCAATATGTC | CACATGCAATACTTGGTGCC | 146 |
| chr6:35425790-35427814 | p2 | (AT)7 | TGCTAAAGCAGACAGGGGTT | GTCTGAGAGCTCCGTTGTCC | 226 |
| chr6:35425790-35427814 | p2 | (AT)12 | TGCTAAAGCAGACAGGGGTT | GTCTGAGAGCTCCGTTGTCC | 226 |
| chr6:35425790-35427814 | p2 | (TA)6 | TGCTAAAGCAGACAGGGGTT | GTCTGAGAGCTCCGTTGTCC | 226 |
| chr6:35449308-35451320 | p2 | (TA)6 | GAAGAGCATCCCTACCCCTC | GCAGCGTACCTGAAATGTGA | 137 |
| chr6:35467463-35469529 | p2 | (AT)33 | ATGTCATATGCAAGCCGACA | TCTTGGAGAGCTCCGTTGTT | 248 |
| chr6:35467463-35469529 | p2 | (TA)6 | ATGTCATATGCAAGCCGACA | TCTTGGAGAGCTCCGTTGTT | 248 |
| chr6:35468338-35470350 | p2 | (AT)33 | AAGGGACTCGGTGAACAATG | GGTTGCTCGCTACTCTGCTC | 212 |
| chr6:35468338-35470350 | p2 | (TA)6 | AAGGGACTCGGTGAACAATG | GGTTGCTCGCTACTCTGCTC | 212 |
| chr6:35491013-35493025 | p2 | (TA)6 | GAAGAGCATCCCTACCCCTC | GCAGCGTACCTGAAATGTGA | 137 |
| chr6:35562056-35564076 | p2 | (TA)10 | CCGAAATCTTCCAACTTCCA | GGGCTGTTTTTGGAAGACAA | 194 |
| chr6:35659665-35661679 | p2 | (AT)7 | AACATCTGGACCGGACTGAC | TTTCACACCGTGCCTACTTG | 130 |
| chr6:35678206-35680221 | p3 | (CAA)5 | ACATCACAACAAAGGCCACA | CCAGTTGATAGGGGGTGATG | 148 |
| chr6:35685641-35687659 | p6 | (ATATTC)3 | GCATCAACACACAGCCAATC | AGTCGGCAAAATTTCAATGG | 100 |
| chr6:35727729-35729744 | p6 | (CATATA)3 | GCATGAGAGCCCAAAGAAAG | AGCCTTCAGGGTCGGTTTAT | 103 |
| chr6:35727729-35729744 | p5 | (ATTCA)3 | GCATGAGAGCCCAAAGAAAG | AGCCTTCAGGGTCGGTTTAT | 103 |
| chr6:35781972-35783990 | p2 | (GT)9 | TTGTTTTGAGACTTCGTGCG | ATTCCAAATTTACCCTCGGC | 225 |
| chr6:35781972-35783990 | p3 | (TTG)5 | TTGTTTTGAGACTTCGTGCG | ATTCCAAATTTACCCTCGGC | 225 |
| chr6:35836383-35838398 | p3 | (TGT)5 | GCCGAGAAGTCGATTGAAAG | TCGGCCACATATCCAAAAAT | 155 |
| chr6:35836402-35838417 | p3 | (TGT)5 | GCCGAGAAGTCGATTGAAAG | TCGGCCACATATCCAAAAAT | 136 |
| chr6:35841973-35843988 | p3 | (TGT)5 | TTCTAGCCGAGAAGGGGAAT | ATACCACAACCACACAGCCA | 163 |
| chr6:35841992-35844007 | p3 | (TGT)5 | TTCTAGCCGAGAAGGGGAAT | ATACCACAACCACACAGCCA | 144 |
| chr6:35863465-35865480 | p2 | (TA)9 | GTCCTCAACCCGTCCATCTA | ATGGCCATTAGTGGCAGTTC | 138 |
| chr6:35863465-35865480 | p5 | (AAGAG)3 | GTCCTCAACCCGTCCATCTA | ATGGCCATTAGTGGCAGTTC | 138 |
| chr6:35863465-35865480 | p2 | (TA)16 | GTCCTCAACCCGTCCATCTA | ATGGCCATTAGTGGCAGTTC | 138 |

| Marker ID | SSR type | SSR motif | FORWARD PRIMER (5'-3') | REVERSE PRIMER (5'-3') | size (bp) |
|---|---|---|---|---|---|
| chr6:35924286-35926301 | p5 | (TCATA)3 | TTCCGAGGACAAACTGCTCT | AACGGCCCACATGTTTGTAT | 135 |
| chr6:36051681-36053697 | p2 | (AG)8 | GCTTTGCGTAATCTTCAGCC | CCCCGTTCTCGACGTAAATA | 116 |
| chr6:36061532-36063568 | p6 | (TATATG)6 | TTGATGTTGTTGGCACGATT | GCTTAATTGTCCAAGCTCGC | 157 |
| chr6:36073106-36075126 | p2 | (TA)10 | CGTGTGCAAGAATGGATTTG | ATAGAACCCCAACCCTCACC | 186 |
| chr6:36095331-36097346 | p5 | (TCATT)3 | GAATGCTCTGATGGCAGTGA | CCCAAATATGATGGGACCAC | 134 |
| chr6:36097736-36099754 | p2 | (AT)9 | CCGTTTTCTTTGGATGCTGT | ACAATGGGGTGCTTTCAGTC | 195 |
| chr6:36183547-36185635 | p5 | (TGATG)3 | TCGGGGGCTAAAGACCTTAT | AGGCACGGTGGAATCATATC | 154 |
| chr6:36187168-36189182 | p2 | (TA)7 | TGGTTATTTGACGTCGTGGA | ATATGTTGTGCATTTGGCGA | 254 |
| chr6:36206493-36208505 | p2 | (AT)6 | CAGACAACCCCTAGGACGAA | ATGTGAATGCCAGAGGGTTC | 230 |
| chr6:36293956-36295972 | p2 | (AT)8 | ACACTCCGAGGATGAACTGC | AGCCTTGTCGACTTGCGTAT | 141 |
| chr6:36321918-36323938 | p2 | (AT)6 | AAACGGAAACTGTCACGACC | GTCGGCTTGCGTATCATTTT | 114 |
| chr6:36321918-36323938 | p2 | (AT)10 | AAACGGAAACTGTCACGACC | GTCGGCTTGCGTATCATTTT | 114 |
| chr6:36581962-36583988 | p2 | (TA)13 | GGATGGTGGATGGATATTGC | AATGTGGGGCCAGATGTTAG | 156 |
| chr6:36603397-36605424 | p3 | (ATT)9 | TATATGGTCTCGGGCAATCC | ATAGGGATGGGTCTGGCTCT | 280 |
| chr6:36603429-36605450 | p3 | (ATT)9 | TATATGGTCTCGGGCAATCC | ATAGGGATGGGTCTGGCTCT | 248 |
| chr6:36792154-36794169 | p2 | (TA)11 | CAAAACCCCCTTTTCACAGA | ACATTTGCTTGCTCGTTGTG | 173 |
| chr6:36792154-36794169 | p5 | (TTAAA)3 | CAAAACCCCCTTTTCACAGA | ACATTTGCTTGCTCGTTGTG | 173 |
| chr6:36792154-36794169 | p4 | (TTAA)4 | CAAAACCCCCTTTTCACAGA | ACATTTGCTTGCTCGTTGTG | 173 |
| chr6:36806319-36808409 | p2 | (AC)6 | AGACATATGGACGGGACAGG | CCGTTTTCTTTGGACGTTGT | 253 |
| chr6:36806319-36808409 | p2 | (AT)45 | AGACATATGGACGGGACAGG | CCGTTTTCTTTGGACGTTGT | 253 |
| chr6:36838259-36840271 | p2 | (TG)6 | AATCATCATCGTAGCTGCCC | ACCTTGCGAATCTTCCCTTT | 241 |
| chr6:36886019-36888031 | p2 | (AT)15 | ATCGTGAGACCATGCTACCC | TGTTCATGCCCACAGGTAGA | 193 |
| chr6:36886019-36888031 | p2 | (TA)6 | ATCGTGAGACCATGCTACCC | TGTTCATGCCCACAGGTAGA | 193 |
| chr6:36970972-36972996 | p6 | (CATATA)4 | CCCTAGGACGAACTGCTCTG | AGTCTTGTAAGCGGCTCTGC | 208 |
| chr6:37072275-37074360 | p2 | (TA)30 | TTTCTTGTCTTCCCTGCGTT | CATGGTCAACACACAGAGGC | 115 |
| chr6:37179745-37181787 | p2 | (AT)21 | CGGATTTGCACCTTGAATTT | ATAGAGTTGTCCCCGAGGCT | 212 |
| chr6:37179745-37181787 | p2 | (TA)6 | CGGATTTGCACCTTGAATTT | ATAGAGTTGTCCCCGAGGCT | 212 |
| chr6:37210456-37212498 | p2 | (AT)21 | CGGATTTGCACCTTGAATTT | ATAGAGTTGTCCCCGAGGCT | 212 |
| chr6:37372564-37374582 | p6 | (CATATA)3 | GAGCGAACCCCTTTAGCTCT | CAGCCGATGTGTCATTATGC | 220 |
| chr6:37393133-37395151 | p2 | (TA)9 | TGTGGTTGTCGTGTTTCCAT | CCACAGCAATACCAACAACG | 134 |
| chr6:37426981-37429027 | p2 | (AT)23 | GAGAACTCTCGCCACAAAGC | AAAGCAAAATATGGACCCCC | 130 |
| chr6:37426981-37429027 | p6 | (ACATAT)3 | GAGAACTCTCGCCACAAAGC | AAAGCAAAATATGGACCCCC | 130 |
| chr6:37437164-37439226 | p2 | (AT)31 | TCTTCCGCAGTAGGCTGTTT | GGAACTGTTTGCAAAGCCTC | 126 |
| chr6:37468677-37470697 | p4 | (CATA)5 | CACCCTTGGTAAGGCATGAT | GAGGCTTTGCAGACAGTTCC | 127 |
| chr6:37513860-37515875 | p5 | (ACACA)3 | ACACGGCCAACTTAGACCAC | TCATGGAGTGGGTGTGTTGT | 194 |
| chr6:37519461-37521479 | p2 | (AT)9 | TAAAACGCAACAACGAGCAG | GAGATAAGAGGGGGTTTCGG | 254 |

| Marker ID | SSR type | SSR motif | FORWARD PRIMER (5'-3') | REVERSE PRIMER (5'-3') | size (bp) |
|---|---|---|---|---|---|
| chr6:37626777-37628789 | p2 | (AT)6 | TGGTGATTGGGTGTTCTTGA | CTGGAAGGGTCGCCTATACA | 168 |
| chr6:37628573-37630591 | p2 | (TA)9 | CAGTGAGCAAGCAAGTTCCA | CTGAACCCCTCTGGTGACAT | 248 |
| chr6:37725667-37727709 | p6 | (GATAAT)3 | CAGTCTCGTAAGGGGCTCTG | TTACAGCAACGAGGGCTTTT | 155 |
| chr6:37725710-37727734 | p6 | (GATAAT)3 | CAGTCTCGTAAGGGGCTCTG | TTACAGCAACGAGGGCTTTT | 112 |
| chr6:37793613-37795628 | p3 | (CAA)5 | ACATCACAACAAAGGCCACA | AGGCACAGTTGACAGAGGGT | 125 |
| chr6:37799389-37801419 | p5 | (TATAT)6 | AATTAGCCAAGGTCCCGTCT | ACGGAAATATCGACGTCTCG | 276 |
| chr6:37815275-37817290 | p3 | (CAA)5 | ACATCACAACAAAGGCCACA | AGGCACAGTTGACAGAGGGT | 125 |
| chr6:37821051-37823081 | p5 | (TATAT)6 | AATTAGCCAAGGTCCCGTCT | ACGGAAATATCGACGTCTCG | 276 |
| chr6:37840083-37842095 | p2 | (TA)6 | ACAACTTTTGTCACGACCCC | ATATAGGATCCCCTCGTCGG | 299 |
| chr6:38066575-38068591 | p2 | (AT)10 | TCTGGAGACACTCCGAGGAT | GTCGGCTTGCGTATCGTATT | 115 |
| chr6:38066575-38068591 | p2 | (AT)8 | TCTGGAGACACTCCGAGGAT | GTCGGCTTGCGTATCGTATT | 115 |
| chr6:38081059-38083071 | p2 | (AT)6 | ACTTTTGTCACGACCCAACC | AGCCTTGTCAGCTTGCGTAT | 244 |
| chr6:38104250-38106274 | p2 | (TA)12 | CCCATGGAGAAGCACAAAAT | TTTTGCATCTATGGCCCTTC | 227 |
| chr6:38243667-38245681 | p2 | (AT)7 | CTCATTAATTGCCCATGGCT | TGCGAACGAGTGAGAGAATG | 175 |
| chr6:38281750-38283765 | p2 | (TA)8 | AAGCCCTCCAAAAGGATGAT | GTATGAAGGCCCACGACCTA | 202 |
| chr6:38281750-38283765 | p3 | (TTG)5 | AAGCCCTCCAAAAGGATGAT | GTATGAAGGCCCACGACCTA | 202 |
| chr6:38281750-38283765 | p2 | (TA)15 | AAGCCCTCCAAAAGGATGAT | GTATGAAGGCCCACGACCTA | 202 |
| chr6:38408035-38410067 | p2 | (AT)16 | AGACTCTGGACCTTGGGGAT | AGCTCCATAGCCCACTCAGA | 226 |
| chr6:38497796-38499828 | p3 | (ATG)5 | TTTGGCATGGTCTTTGTGAA | ACATCATGGAGACGCATCAA | 165 |
| chr6:38497796-38499828 | p2 | (TA)16 | TTTGGCATGGTCTTTGTGAA | ACATCATGGAGACGCATCAA | 165 |
| chr6:38504239-38506254 | p3 | (ATG)5 | TTAGATTCTCGGGGATGGTG | CTTACATACCTGTCCCGCGT | 153 |
| chr6:38603486-38605520 | p2 | (TA)17 | CGCATTCCAACATCAACAAC | TTGAATTTGGAAGGTTTCGG | 115 |
| chr6:38700475-38702493 | p2 | (AC)6 | CAGCTGGGGTTAAGGATTGA | ATCACCATGCAACCCTCTTC | 109 |
| chr6:38700475-38702493 | p2 | (TG)9 | CAGCTGGGGTTAAGGATTGA | ATCACCATGCAACCCTCTTC | 109 |
| chr6:38746799-38748831 | p5 | (TTTCA)3 | GGAATTCTCGGAGATGACCA | ACTTCAATGTCGACTCGGCT | 177 |
| chr6:38777059-38779073 | p2 | (AC)7 | TCGGCAGCATCTCCTCTATT | TTTGTGAAGGGAGAAATGGG | 101 |
| chr6:38787230-38789260 | p2 | (TA)15 | TGGACGTGTCATAATCGCAT | ATACAATCCGAACGCCTCAC | 235 |
| chr6:38927830-38929842 | p2 | (GA)6 | GGAGTGGCCATAGTCTTGGA | GCCACCTTGTATGGAATGCT | 204 |
| chr6:39026085-39028101 | p2 | (TA)31 | CCCCTAGTTGGGTCTTGACA | CACTGCATGTGTAGCGACCT | 264 |
| chr6:39026085-39028101 | p2 | (TA)8 | CCCCTAGTTGGGTCTTGACA | CACTGCATGTGTAGCGACCT | 264 |
| chr6:39027253-39029267 | p2 | (AT)7 | ACATCGAAAGCCCGTAAATG | GGCTTGTACTGTTCTTCCGC | 199 |
| chr6:39033447-39035465 | p3 | (TAT)6 | TTGTCGCAATTCTCCTTCCT | TTGTTGATGGAATGCTTGGA | 225 |
| chr6:39198162-39200178 | p4 | (ATAC)4 | ATATACGTAATGGGCCTGCG | TGCCAAGTAGACGAGACGTG | 163 |
| chr6:39214071-39216085 | p2 | (TA)7 | AAATCAACCATAGATCGCCG | TTCCCCCTCAATCAGCTATG | 114 |
| chr6:39274941-39276955 | p2 | (TG)7 | CGACGGTATTTGGGAAAGAA | TGAATCGGCCTCATTGTGTA | 101 |
| chr6:39304306-39306322 | p2 | (AT)8 | ACGACCAAACACGAAGGAAC | CACCCCTCCCTTTCTTCTTC | 189 |

| Marker ID | SSR type | SSR motif | FORWARD PRIMER (5'-3') | REVERSE PRIMER (5'-3') | size (bp) |
|---|---|---|---|---|---|
| chr6:39315284-39317358 | p2 | (TA)37 | GCCTAACAAGAGGAAGCGTG | GGTTGTCGACTTTCCCTGAA | 242 |
| chr6:39365690-39367764 | p2 | (TA)37 | GCCTAACAAGAGGAAGCGTG | GGTTGTCGACTTTCCCTGAA | 242 |
| chr6:39389909-39391947 | p2 | (AT)22 | TCCAATGACATTCCTGGTCA | TCAGGTTGAGTTGGGTGACA | 192 |
| chr6:39389948-39391964 | p2 | (AT)22 | TCCAATGACATTCCTGGTCA | TCAGGTTGAGTTGGGTGACA | 153 |
| chr6:39407245-39409283 | p2 | (AT)22 | TCCAATGACATTCCTGGTCA | TCAGGTTGAGTTGGGTGACA | 192 |
| chr6:39407284-39409300 | p2 | (AT)22 | TCCAATGACATTCCTGGTCA | TCAGGTTGAGTTGGGTGACA | 153 |
| chr6:39443270-39445290 | p2 | (TC)19 | TAGTGATTGACGGGGAAAGC | ATGGCGATAAGCCCACATAA | 145 |
| chr6:39443270-39445290 | p5 | (CATTG)4 | TAGTGATTGACGGGGAAAGC | ATGGCGATAAGCCCACATAA | 145 |
| chr6:39459793-39461809 | p4 | (TATG)4 | GGTTCGATGATTATGTGGGG | GGCGTCATACTACTTCGGGA | 209 |
| chr6:39470817-39472832 | p5 | (TCATA)3 | AGCTCAATCGCACGATTCTT | GTAACCCAAGAGGGTGCTCA | 114 |
| chr6:39514595-39516625 | p2 | (TA)15 | TCACCGGAAAACACTGTTCA | TTTAGTCTGCATCTCGGGCT | 296 |
| chr6:39627109-39629135 | p2 | (TA)13 | CAAATCCAATCTCCACCGTT | GTTTCATTCCAACCAGCCAT | 194 |
| chr6:39821312-39823327 | p5 | (ATATC)3 | CCCATGTTTGGCATCTTTTT | GTGGTGGTTGGAGAATGCTT | 181 |
| chr6:39821312-39823327 | p2 | (AT)7 | CCCATGTTTGGCATCTTTTT | GTGGTGGTTGGAGAATGCTT | 181 |
| chr6:39935406-39937418 | p2 | (GT)6 | CGGAGGACGAGATGAAGAAG | ACGCGGGACATAACATTCTC | 247 |
| chr6:40067041-40069079 | p2 | (TA)21 | ATATATGGGGGATGGGGAAA | CGGGTGTAATGGAATAACGG | 211 |
| chr6:40067041-40069079 | p2 | (TA)19 | ATATATGGGGGATGGGGAAA | CGGGTGTAATGGAATAACGG | 211 |
| chr6:40133991-40136003 | p2 | (AT)6 | AATAAGCGGATAGCCTCGGT | GCAAGACTACGGCAAAGAGG | 247 |
| chr6:40182579-40184605 | p2 | (TA)14 | CTAGTCCTGTTCCTGCTCGG | AGTGCTTCGACGATGGAGTT | 213 |
| chr6:40182579-40184605 | p2 | (AT)13 | CTAGTCCTGTTCCTGCTCGG | AGTGCTTCGACGATGGAGTT | 213 |
| chr6:40209272-40211284 | p2 | (AT)6 | CCATAACCTTGTCGGCTTGT | CCGAGCGCCTCTTAACATAC | 140 |
| chr6:40236632-40238650 | p6 | (TATATG)3 | TGCTTCGGTTTTCCAAAAAG | ACAAGTATGCTCACCGACCC | 294 |
| chr6:40421979-40423991 | p2 | (GT)6 | ATTTTGAAGCGGTACATCCG | TACCTGGCGGTCCAAGATAG | 286 |
| chr6:40436446-40438458 | p6 | (TATATG)3 | CACATGGTTGACATTCCGAG | TCTTACCCGGCATCATAAGC | 106 |
| chr6:40436446-40438458 | p2 | (TG)6 | CACATGGTTGACATTCCGAG | TCTTACCCGGCATCATAAGC | 106 |
| chr6:40515601-40517616 | p5 | (TATCG)3 | CACTCTCTTGGCTTTCTCGG | TTGCATTCTAAGATCGTGCG | 170 |
| chr6:40596451-40598479 | p2 | (TA)14 | GTCGTTGCTGTGTTGTTGCT | GGATACTCCAAAACAGCCCA | 155 |
| chr6:40666890-40668908 | p2 | (AT)9 | CTTTTTACCCCTGGCCTTTC | TGCTAAAGTGATCGCAGGTG | 131 |
| chr6:40669690-40671706 | p4 | (ATAC)4 | TTCCTCTTCACCGGCATTAC | GTTGCAGGGCATTTAGGAAA | 169 |
| chr6:40690658-40692672 | p2 | (AT)7 | GACGTCGATTTTTGGCCTTA | GGCTCTAAGTAAGGGTCGGG | 167 |
| chr6:40690658-40692672 | p6 | (CATATA)4 | GACGTCGATTTTTGGCCTTA | GGCTCTAAGTAAGGGTCGGG | 167 |
| chr6:40696432-40698444 | p2 | (AT)6 | CTGCGCATTTGCTAAAATCA | AGCAGGTTGTCCTAGGGGTT | 162 |
| chr6:40696432-40698444 | p6 | (CATATA)4 | CTGCGCATTTGCTAAAATCA | AGCAGGTTGTCCTAGGGGTT | 162 |
| chr6:40886983-40889001 | p2 | (TA)9 | AATGCGTGTGGATGATTTGA | GTGCTCCCCTCCAATACTCA | 119 |
| chr6:40915267-40917287 | p2 | (TA)10 | AATGCGTGTGGATGATTTGA | GTGCTCCCCTCCAATACTCA | 119 |
| chr6:40915267-40917287 | p2 | (TG)16 | AATGCGTGTGGATGATTTGA | GTGCTCCCCTCCAATACTCA | 119 |

| Marker ID | SSR type | SSR motif | FORWARD PRIMER (5'-3') | REVERSE PRIMER (5'-3') | size (bp) |
|---|---|---|---|---|---|
| chr6:40956384-40958404 | p2 | (AT)10 | GGTGACAACTATGGGGCCTA | ATAGCGGTCTGGCCCTATTT | 194 |
| chr6:40956384-40958404 | p2 | (AT)11 | GGTGACAACTATGGGGCCTA | ATAGCGGTCTGGCCCTATTT | 194 |
| chr6:41004414-41006440 | p2 | (AT)13 | TGGACAGAATATTCCCCGAG | TTTGGCCGAGTTGAATTCTC | 222 |
| chr6:41030599-41032613 | p2 | (TA)11 | CCTTCAAGGTGAGGACCAAA | TGCCGGCCATAATCTATTTC | 264 |
| chr6:41030616-41032648 | p2 | (TA)11 | CCTTCAAGGTGAGGACCAAA | TGCCGGCCATAATCTATTTC | 247 |
| chr6:41030658-41032670 | p2 | (TA)11 | CCTTCAAGGTGAGGACCAAA | TGCCGGCCATAATCTATTTC | 205 |
| chr6:41031588-41033610 | p2 | (TA)11 | TGAGTGGTGCTCGGTGTTAG | ACCAGAGCAATCCAGCAAGT | 212 |
| chr6:41074906-41076921 | p5 | (ATTGC)3 | CTTGCAATTATCCAGGCCAT | GCAGTTCTGAGGACCTTTCG | 145 |
| chr6:41089106-41091142 | p2 | (AT)10 | CAAAGGGATAGAAGCCGACA | AGCTGCAGTTTGCTTTTGGT | 271 |
| chr6:41089106-41091142 | p2 | (TA)21 | CAAAGGGATAGAAGCCGACA | AGCTGCAGTTTGCTTTTGGT | 271 |
| chr6:41089106-41091142 | p3 | (CAT)5 | CAAAGGGATAGAAGCCGACA | AGCTGCAGTTTGCTTTTGGT | 271 |
| chr6:41089106-41091142 | p3 | (ACA)14 | CAAAGGGATAGAAGCCGACA | AGCTGCAGTTTGCTTTTGGT | 271 |
| chr6:41100807-41102825 | p3 | (CAA)6 | ATGGGCACGAGCAGAGTAGT | CCGGACTAGTTTGCCTAACG | 235 |
| chr6:41112868-41114880 | p2 | (AT)6 | CTCCGGAGGATTCAAAGTCA | ATTGACCCTGTTAGTTGCCG | 282 |
| chr6:41112868-41114880 | p2 | (AG)7 | CTCCGGAGGATTCAAAGTCA | ATTGACCCTGTTAGTTGCCG | 282 |
| chr6:41181826-41183838 | p2 | (AT)6 | CCGGTGTCCCAAATGAATAC | AAGAGATCAGCCTCGGACAA | 274 |
| chr6:41203562-41205580 | p2 | (AT)9 | CCTTTAAGCTTGATGGCAGC | TGATTCTGATGGAAAAGCCC | 158 |
| chr6:41247911-41249923 | p2 | (AG)6 | CCGTTTTCTTTGGACGTTGT | CACGAAATTTTCTGGGGTGT | 191 |
| chr6:41279738-41281753 | p3 | (TGA)5 | ATAGCCTTGTCGGCTTCTGA | ACTTCTGTCACGACCCAACC | 296 |
| chr6:41420520-41422535 | p5 | (TTCAT)3 | GTTGCCTCCCAACAAACACT | TGAAATTTGCCACACCAAAA | 194 |
| chr6:41428725-41430737 | p2 | (AT)6 | CCTCTGCACTACCCTTCTCG | TCCACCTCCACTTGGATTTC | 228 |
| chr6:41708593-41710611 | p2 | (AC)9 | ATGCCAAATGAAAGAGTGGG | TTTATAGGGAAATGCTGCCG | 251 |
| chr6:41780990-41783010 | p5 | (TATAT)4 | GTGAGCAAGTAGCCAGGGTC | ATCCTGGCACAATCAACACA | 294 |
| chr6:41925477-41927509 | p2 | (TA)16 | TCCTGTCCTTTCTGCAGCTT | GACGAGGCTGCTATGTAGGC | 164 |
| chr6:41925477-41927509 | p2 | (AT)9 | TCCTGTCCTTTCTGCAGCTT | GACGAGGCTGCTATGTAGGC | 164 |
| chr6:42015485-42017497 | p2 | (TG)6 | AAGGTGCCTCGGGCTATTAT | GGTCCTACCAAGGCTTAGGG | 227 |
| chr6:42034959-42036973 | p2 | (TA)7 | GCGTTTTCCCATATTCCAGA | AAACACAACCTGTGAAGGGC | 299 |
| chr6:42235918-42237936 | p6 | (GAAAAA)3 | CGCTCTGTCTAAATCGGGAA | CGAGCTTCAAAAACAAAGCC | 265 |
| chr6:42401209-42403229 | p5 | (ATCAT)4 | CCTAGGACGGAACTGCTCTG | GGGCCCTTGATGTTCAGTAA | 218 |
| chr6:42421361-42423381 | p5 | (ATCAT)4 | CCTAGGACGGAACTGCTCTG | GGGCCCTTGATGTTCAGTAA | 218 |
| chr6:42567850-42569866 | p2 | (AT)8 | AACTCGGCCAAATCATCAAC | GGAAGATTGCGGATTGTGTT | 224 |
| chr6:42607431-42609505 | p2 | (CT)37 | AGATGATTGGGCAGCATTTC | TGGGGTATCTTCGGATCTTG | 206 |
| chr6:42624265-42626301 | p2 | (CA)18 | TGTTATCTCGCTCGCTTCCT | ATGTGAAATGGAAGGCAAGG | 118 |
| chr6:42679841-42681859 | p6 | (CATATA)3 | GGGTTGATCGATTCTTGCAT | GGGTCTAGCCCCATCACATA | 210 |
| chr6:42691949-42693969 | p2 | (AT)10 | GCCTCCAGAGATGAGGTGAG | TACCTGGCTGAAGAGCCACT | 106 |
| chr6:42707283-42709395 | p2 | (TA)14 | AGCCCTTTTGGCTACCACTT | ACGACTTAGTTGGGGACGTG | 236 |

| Marker ID | SSR type | SSR motif | FORWARD PRIMER (5'-3') | REVERSE PRIMER (5'-3') | size (bp) |
|---|---|---|---|---|---|
| chr6:42707283-42709395 | p2 | (TA)56 | AGCCCTTTTGGCTACCACTT | ACGACTTAGTTGGGGACGTG | 236 |
| chr6:42709511-42711527 | p4 | (CATA)4 | ACCTTGGTAAGGCGTGATTG | AGTAGATATGTGGACGCCGC | 109 |
| chr6:42853089-42855104 | p2 | (TA)10 | CTGTCCGAAAAGTTTGCCAT | GGGGTTTAAGCCCATGAAAT | 111 |
| chr6:42853089-42855104 | p5 | (GTCCT)3 | CTGTCCGAAAAGTTTGCCAT | GGGGTTTAAGCCCATGAAAT | 111 |
| chr6:42940718-42942730 | p2 | (TG)6 | TCGGCACTACTCTGTTCGTG | TCGGAAACCGTAACACATCA | 169 |
| chr6:42988678-42990692 | p2 | (TA)7 | TTTGATTTTCCAAAAACGGC | CGACAACTGAAGGCTCACAA | 290 |
| chr6:43193161-43195176 | p3 | (TTG)5 | TTGGCCCTGTAGCACTTTCT | CACGCCTTGGCTCTAATCTC | 128 |
| chr6:43317797-43319811 | p2 | (CA)7 | TGAGATCCTTGTGTGTCCCA | CGGCATGAAGTGGAGGTAGT | 160 |
| chr6:43362658-43364674 | p2 | (AT)8 | GTTTCATGCACGCAGGTAGA | GCCGTACCCCTGAACAAATA | 231 |
| chr6:43379825-43381889 | p2 | (AT)13 | TGCCTAGGGTTTTGAAGAGG | CCCAAATGATGATTTCCACC | 257 |
| chr6:43379914-43381956 | p2 | (AT)13 | TGCCTAGGGTTTTGAAGAGG | CCCAAATGATGATTTCCACC | 168 |
| chr6:43475634-43477662 | p2 | (AT)14 | CCTAGACGGGTCGTGACAAT | CCTCGCACGAATTTTCTCTC | 290 |
| chr6:43535365-43537393 | p2 | (AT)14 | CCTAGACGGGTCGTGACAAT | CCTCGCACGAATTTTCTCTC | 290 |
| chr6:43696572-43698593 | p3 | (ACA)7 | GGCATGCCCGAGTAGAGTAG | GTCCCGGGATAATCCGTAAT | 163 |
| chr6:43696572-43698593 | p2 | (CT)7 | GGCATGCCCGAGTAGAGTAG | GTCCCGGGATAATCCGTAAT | 163 |
| chr6:43737469-43739484 | p3 | (ACA)5 | CTTGCATGGTTCTTTCAGCA | ACGATCCAAACCGATGAGTC | 104 |
| chr6:43768112-43770126 | p2 | (AT)7 | TCCCATGGGGATACTCAAAA | TTGAGGCCAAAGGTAGGATG | 282 |
| chr6:43834442-43836457 | p5 | (TATGA)3 | GCAAGGTACTCCCAGACGAG | CTTACCCTTGAGCTTGGCAG | 270 |
| chr6:43989469-43991507 | p2 | (AT)18 | TTTTGTCACGACCTAACCCC | CCAGGAGACCGGTTATTTGA | 147 |
| chr6:43989469-43991507 | p2 | (TA)19 | TTTTGTCACGACCTAACCCC | CCAGGAGACCGGTTATTTGA | 147 |
| chr6:44040829-44042844 | p3 | (GAA)5 | AAGGTGATGTACTGGGCCTG | TGTCCCGGTGGTTAATGAAT | 156 |
| chr6:44053935-44055947 | p2 | (AT)6 | ACTTCCTTCAAGGTGAGGCA | TGTGTATCTGCCCGTCCATA | 283 |
| chr6:44135929-44137943 | p2 | (AT)8 | ACTATTGTCACGACCCAGCC | ACGTTGTGAACCTTTGGGTC | 125 |
| chr6:44201748-44203766 | p6 | (TCACAT)3 | AACCGGCGTGACATATTAGG | AGTAGAGCCTGTGGAGCCAA | 131 |
| chr6:44244460-44246478 | p3 | (ATG)6 | TTAGATTCTCGGGGATGACG | TATCCCACGACCAACTAGCC | 153 |
| chr6:44244460-44246478 | p6 | (TATATG)3 | TTAGATTCTCGGGGATGACG | TATCCCACGACCAACTAGCC | 153 |
| chr6:44293398-44295428 | p3 | (AAT)23 | CCAACTCCACGGATTCAGTT | GGTCAACGAGTGATCGAGGT | 175 |
| chr6:44293438-44295464 | p3 | (AAT)23 | CCAACTCCACGGATTCAGTT | GGTCAACGAGTGATCGAGGT | 135 |
| chr6:44322193-44324209 | p2 | (AT)8 | CCATGACTAGTGCCCGAGAT | TGTTATGACAGCCTTGTCGC | 173 |
| chr6:44378895-44380910 | p2 | (AT)8 | ATACGGGGGATTTTTCCAAG | TGCTGGATTGAATGGGAAAT | 244 |
| chr6:44473616-44475631 | p3 | (TTG)5 | AGTTTGGACGTTGAATTCCG | TGCCTCAAAACACTCCCTTC | 127 |
| chr6:44473616-44475631 | p3 | (AAC)5 | AGTTTGGACGTTGAATTCCG | TGCCTCAAAACACTCCCTTC | 127 |
| chr6:44517707-44519719 | p2 | (AT)6 | AATTCCATTGGGCAACCATA | GGGTCCTGTCCCACCTTTAT | 187 |
| chr6:44821812-44823864 | p2 | (AT)26 | CGACCTTACGTCACCCTGAT | TTTATCCGTATAGCTGCCCG | 294 |
| chr6:45141857-45143869 | p2 | (TA)6 | GAAACTGGCGTACAGGCTTC | AACTCAGTTTGTGCATCCCC | 101 |
| chr6:45160701-45162743 | p3 | (CAA)5 | ACATGTAAATCAGCCCAGCC | TGTGGTAAGTTTGGCGTTGA | 212 |

| Marker ID | SSR type | SSR motif | FORWARD PRIMER (5'-3') | REVERSE PRIMER (5'-3') | size (bp) |
|---|---|---|---|---|---|
| chr6:45160701-45162743 | p2 | (AT)21 | ACATGTAAATCAGCCCAGCC | TGTGGTAAGTTTGGCGTTGA | 212 |
| chr6:45225516-45227540 | p2 | (AT)12 | AGCAAGCGAAAGGACGAATA | CATCTCTCTTTCCCGCTCTG | 173 |
| chr6:45287390-45289406 | p2 | (AC)8 | CAGTCTCAGAAGCCCTGACC | GGTGATGATGGTGACAGACG | 147 |
| chr6:45329227-45331243 | p4 | (CATA)4 | CAATTTATCACACCCCACCC | TTCATGCCCACAGGTGTAGA | 140 |
| chr6:45377108-45379122 | p2 | (TA)7 | TCGAGACTCCTGTGACATCG | CTATCTTACCCGGCCCTTTC | 149 |
| chr6:45390671-45392689 | p6 | (TATATG)3 | ACCTCCAGGTTGGTGTTGAG | ATGCGTCCGAGCTCATAAGT | 137 |
| chr6:45392184-45394202 | p6 | (CATATA)3 | GAGCGCTTGGCCAATAGTAG | TGTGGTATGCCATTCCGTTA | 297 |
| chr6:45395214-45397234 | p2 | (AT)10 | TTACGCGAACTCGGACTCTT | CAATTGTGCAACATTCGAGG | 103 |
| chr6:45399790-45401808 | p2 | (AT)9 | TTACGCGAACTCGGACTCTT | CAATTGTGCAACATTCGAGG | 101 |
| chr6:45435282-45437300 | p6 | (CATATA)3 | GGCTATCGTAATGCAGGGAA | GTACCAACGCTCGTTACGGT | 267 |
| chr6:45529871-45531891 | p2 | (AT)10 | CAAGCTGCCATACAAGTCCA | TGAATGTGCTCCTTTGTTCG | 238 |
| chr6:45545319-45547335 | p2 | (TA)8 | AAATACACCATCCCCGTGAA | AGGGAGACACGTGTTGGAAG | 151 |
| chr6:45545319-45547335 | p2 | (TA)6 | AAATACACCATCCCCGTGAA | AGGGAGACACGTGTTGGAAG | 151 |
| chr6:45549054-45551072 | p2 | (TA)9 | TGACCATGGAATAAGTGCGA | ATCGTCGTTAGCGAAGCAAT | 154 |
| chr6:45549054-45551072 | p2 | (AT)15 | TGACCATGGAATAAGTGCGA | ATCGTCGTTAGCGAAGCAAT | 154 |
| chr6:45549962-45551992 | p2 | (TA)9 | CGCGGGATATATGGGTTATG | TTTCCTGTCTACCTGTGGGC | 201 |
| chr6:45549962-45551992 | p2 | (AT)15 | CGCGGGATATATGGGTTATG | TTTCCTGTCTACCTGTGGGC | 201 |
| chr6:45699284-45701300 | p4 | (AAGA)4 | GCAGACTCCCCAGACTTCAG | TTTATTGGCTCGGGTTTCAC | 168 |
| chr6:45887647-45889659 | p2 | (TA)6 | GTTGTGTGCAGGTTGGATTG | CCACTATGAACACGAGGGGT | 134 |
| chr6:45946559-45948579 | p2 | (AT)10 | TTGTGGGCTCATCACCATTA | ACAATAACAGGGCAACCAGC | 196 |
| chr6:46008017-46010033 | p2 | (TA)8 | AAATCTTATGCGGACAACCG | AGTTCTGGAGGCAAGTCCAA | 165 |
| chr6:46060054-46062069 | p3 | (GAG)5 | CATTGTAGGCAGTTCCCGTT | TATATCGGGTTCAGGACGGA | 120 |
| chr6:46063786-46065801 | p5 | (AAACA)3 | CAATTTTGGCGACTTCCCTA | CCCAACCATGGGTAAAAATG | 155 |
| chr6:46083505-46085520 | p3 | (GAG)5 | CATTGTAGGCAGTTCCCGTT | TATATCGGGTTCAGGACGGA | 108 |
| chr6:46087260-46089275 | p5 | (AAACA)3 | TGGCGACTTCCCTAAATCAC | TGATCGAGTCCGCATGAATA | 159 |
| chr6:46151343-46153361 | p2 | (AT)9 | CCGATTATGGTGGCTGAGAT | TGACTGACTTGAGTGCCTGG | 176 |
| chr6:46153506-46155518 | p2 | (AT)6 | GGACCCTCAAACTGTCGTGT | GTGCCTGGGACTAGGATTCA | 241 |
| chr6:46179964-46182002 | p2 | (AT)19 | TATGAGGAAATCCCAGTGGC | GAACCTCCTGAGATGACGGA | 291 |
| chr6:46226639-46228657 | p6 | (ATATGA)3 | ATGGATATGGGTGATTGGGA | CCGAAGCTCACGTTTCTACC | 184 |
| chr6:46273172-46275190 | p2 | (CT)9 | TATTGTTCACTTCCCTCCCG | ATGTTATGCGGGCTATCTGG | 261 |
| chr6:46273172-46275190 | p5 | (ATCAT)3 | TATTGTTCACTTCCCTCCCG | ATGTTATGCGGGCTATCTGG | 261 |
| chr6:46364049-46366065 | p2 | (AG)6 | AAAAATTCCTGAGTGCCACG | AATCTCCACATCTTGTCCCG | 100 |
| chr6:46364049-46366065 | p4 | (TTAT)4 | AAAAATTCCTGAGTGCCACG | AATCTCCACATCTTGTCCCG | 100 |
| chr6:46364049-46366065 | p2 | (AT)20 | AAAAATTCCTGAGTGCCACG | AATCTCCACATCTTGTCCCG | 100 |
| chr6:46364049-46366065 | p5 | (TTTAT)4 | AAAAATTCCTGAGTGCCACG | AATCTCCACATCTTGTCCCG | 100 |
| chr6:46411757-46413777 | p5 | (TTCAT)4 | GTCCAAATACCCGCACTTGT | TTAGCCACACATGACCCAAA | 197 |

| Marker ID | SSR type | SSR motif | FORWARD PRIMER (5'-3') | REVERSE PRIMER (5'-3') | size (bp) |
|---|---|---|---|---|---|
| chr6:46442006-46444018 | p2 | (GT)6 | ATTGCGGGAGATTAAGAGCA | GCCTAGCCCAACAATATGGA | 200 |
| chr6:46512755-46514771 | p4 | (ACAT)4 | CGGCAACATTTACCTCCTGT | TATGCCGGAAAGATGGGTAG | 180 |
| chr6:46618073-46620089 | p2 | (TA)10 | TTGTCTAAGGGTACGACGGG | AATATCTGCAAACCGGCAAG | 186 |
| chr6:46618073-46620089 | p2 | (AT)8 | TTGTCTAAGGGTACGACGGG | AATATCTGCAAACCGGCAAG | 186 |
| chr6:46659311-46661325 | p2 | (AT)6 | GGCTCTACCGCACGACTTAG | ACAGGTTCGTATACGGGTGC | 128 |
| chr6:46659311-46661325 | p2 | (TA)7 | GGCTCTACCGCACGACTTAG | ACAGGTTCGTATACGGGTGC | 128 |
| chr6:46733118-46735133 | p5 | (ATATG)3 | TGTTGGAGCAAGCAAAACAC | TAGCCTCCAAATCAACCCAC | 116 |
| chr6:46906157-46908172 | p5 | (TCATA)3 | GGAAGGGGTCATGCTGATAA | GTGGATATCAAGCGGTTCGT | 201 |
| chr6:46940162-46942180 | p6 | (CATATA)3 | CTACCGAGCACCCCTTATCA | TTCCCTGCATTACGATAGCC | 126 |
| chr6:46947007-46949025 | p6 | (ATACAT)3 | TGAGTCACATGATGGCCCTA | AATGTCTTGCAGCCTCCTGT | 136 |
| chr6:47053503-47055518 | p3 | (ACA)5 | CTAAACTGGAGGACCACCCA | CGATGTCGGTGTGTTTGTTC | 297 |
| chr6:47053503-47055518 | p2 | (AC)6 | CTAAACTGGAGGACCACCCA | CGATGTCGGTGTGTTTGTTC | 297 |
| chr6:47132977-47134991 | p2 | (CA)7 | AAATGTACAAGGCCGACGAG | TGCACAGGGAGACAGACTTG | 262 |
| chr6:47164413-47166428 | p5 | (ATTTC)3 | TTCCTTTTTACGCATTTGGG | GGGTCACCAAATGTCATTCC | 300 |
| chr6:47226490-47228505 | p3 | (ATG)5 | AGACGAATGGACGAAGATGG | TGAGTTCGGAATGTCAACCA | 213 |
| chr6:47240742-47242754 | p3 | (AGA)5 | GAACCCCAAGACATGGAAGA | GAAGGCTATGCTCCACGAAG | 218 |
| chr6:47240742-47242754 | p2 | (AT)6 | GAACCCCAAGACATGGAAGA | GAAGGCTATGCTCCACGAAG | 218 |
| chr6:47310805-47312825 | p5 | (CCGGT)4 | AAGTATCAACCGTGGCTTCG | TCTTCCATTTGGTCTGGTCC | 236 |
| chr6:47408849-47410864 | p5 | (CATTG)3 | TGCATATACATTTTCGGCCA | TTTTATCGCCAAGTGAAGGG | 129 |
| chr6:47408849-47410864 | p2 | (TA)20 | TGCATATACATTTTCGGCCA | TTTTATCGCCAAGTGAAGGG | 129 |
| chr6:47434731-47436743 | p2 | (CA)6 | CCGATACCCTAAAACACCCA | CGGTATTAAGGATCCGCTCA | 238 |
| chr6:47499507-47501519 | p2 | (AT)6 | TCCAATAAGTTCTCCGGGTG | TATACCGATTCGATCCGAGC | 231 |
| chr6:47536449-47538463 | p2 | (AT)7 | GATTGCACAGAAGCACTCCA | AAACCTTATATGGGCCACCC | 222 |
| chr6:47555103-47557118 | p5 | (TTAAA)3 | ATCAGCCCCTTTTTCGTTCT | GCGGTTTTGATGGCAATAGT | 168 |
| chr6:47578653-47580671 | p6 | (AACTGA)3 | GGACCCAGGAAACAAGCATA | ATCCGGCCCAAGACTCTTAT | 254 |
| chr6:47655650-47657682 | p2 | (AC)7 | TAAGCCCACCCGAGTTTAGA | TCAAAAATGATTGTGGACCG | 167 |
| chr6:47655864-47657878 | p2 | (AC)7 | GGCAATTTTTAAGGGTTCCA | GGTCTTTCGATGCCCACTAA | 283 |
| chr6:47747510-47749526 | p5 | (TTTTC)3 | GTACGACTGGAGGAAGCAGC | CAATTTCCTGCATTCGGTTT | 225 |
| chr6:47747510-47749526 | p4 | (ATCA)4 | GTACGACTGGAGGAAGCAGC | CAATTTCCTGCATTCGGTTT | 225 |
| chr6:47778590-47780605 | p5 | (TGTGT)3 | CGGGTGTTGGACTAGGTTGT | GCAAGCCGACGTACCATTAT | 293 |
| chr6:47778641-47780656 | p5 | (TGTGT)3 | CGGGTGTTGGACTAGGTTGT | GCAAGCCGACGTACCATTAT | 242 |
| chr6:47842895-47844910 | p5 | (TCATT)3 | AGTGGTTGACCACAATGCAA | AAGTGATCAACGACGGGAAG | 125 |
| chr6:47853370-47855385 | p5 | (GAAAT)3 | TTACCCGGCATCATAAGCTC | CACATGGTTGACATTCCGAG | 117 |
| chr6:47950544-47952578 | p2 | (AT)17 | TTCCAGAGATGGGAAGAACG | CTTTATCGGGCGATGTTGAT | 299 |
| chr6:47950544-47952578 | p2 | (AT)32 | TTCCAGAGATGGGAAGAACG | CTTTATCGGGCGATGTTGAT | 299 |
| chr6:48240112-48242130 | p6 | (TATTCA)3 | TTGAAGGTAGAAACGTGGGC | ATGGGACTTGCATGAATGGT | 170 |

| Marker ID | SSR type | SSR motif | FORWARD PRIMER (5'-3') | REVERSE PRIMER (5'-3') | size (bp) |
|---|---|---|---|---|---|
| chr6:48261284-48263296 | p2 | (AT)6 | CATCCATCTCCCGATGAGTT | GGGTCGTTTACCCAGTCGTA | 166 |
| chr6:48294676-48296694 | p5 | (TTGTC)3 | AGCTGGAGGTTTGGAGACAA | CAAGCTTCACTTCCACGTCA | 114 |
| chr6:48294676-48296694 | p3 | (TTG)6 | AGCTGGAGGTTTGGAGACAA | CAAGCTTCACTTCCACGTCA | 114 |
| chr6:48319655-48321670 | p5 | (AAGCC)3 | GGGCACATTTTGTCACTCCT | GTAGCCTACGGGCAAATCAA | 122 |
| chr6:48328264-48330279 | p5 | (TATTT)3 | CAAAACCGGATAAAGCCAAA | TTTGGACGAGCTTGTAGCCT | 263 |
| chr6:48494648-48496663 | p5 | (AAATG)3 | CCATCGCTTGGTTTTTCTTC | CAAATGCTGCCCAATTTTCT | 299 |
| chr6:48502418-48504430 | p2 | (TG)6 | AAGGGTGTCCCACAGACAAG | GCCTCCCTCCATAATTCCAT | 256 |
| chr6:48631758-48633770 | p2 | (AC)6 | AGTTGTGCCTTATGCTCGCT | TTACAGGGGAAATGCTGACC | 184 |
| chr6:48673500-48675518 | p2 | (TA)9 | TATTGCCCCTTCAAGGTGAG | TGTGTATATCTGCCCGACCA | 217 |
| chr6:48673500-48675518 | p3 | (CAA)6 | TATTGCCCCTTCAAGGTGAG | TGTGTATATCTGCCCGACCA | 217 |
| chr6:48699215-48701245 | p2 | (AT)15 | GGCCATCCAATTGTTGTACC | TTGATGAGATGCTGAGGACG | 146 |
| chr6:48755065-48757081 | p4 | (TTAA)4 | GGGATAATCGTGCAAGTGGT | AGTCCAGGGTATCGCACAAC | 291 |
| chr6:48766307-48768329 | p2 | (AC)11 | TCTCCGTTGGGAATTTCAAG | ATCGACACTCGAAACCATCC | 193 |
| chr6:48840440-48842458 | p2 | (AG)7 | CGTTTTCGGGTGATCTGACT | TGTCACATATAGGCCCACGA | 186 |
| chr6:48840440-48842458 | p3 | (GTT)6 | CGTTTTCGGGTGATCTGACT | TGTCACATATAGGCCCACGA | 186 |
| chr6:48852709-48854727 | p2 | (AG)7 | ACGTTTTCGGGTGATCTGAC | CCCAAGACCTATGGTTGGAA | 170 |
| chr6:48852709-48854727 | p3 | (GTT)6 | ACGTTTTCGGGTGATCTGAC | CCCAAGACCTATGGTTGGAA | 170 |
| chr6:48852709-48854727 | p2 | (TA)8 | ACGTTTTCGGGTGATCTGAC | CCCAAGACCTATGGTTGGAA | 170 |
| chr6:48951423-48953441 | p6 | (TATATG)3 | TGCGTACATACGGTTCCAAA | CCTCAGGAAAAGTAAGCCCC | 221 |
| chr6:48972470-48974496 | p2 | (AT)13 | TACCTTACCCGACATGCCTC | AGAGCCCAGCCATAGGAAAT | 284 |
| chr6:48972470-48974496 | p2 | (AT)11 | TACCTTACCCGACATGCCTC | AGAGCCCAGCCATAGGAAAT | 284 |
| chr6:48980946-48982961 | p3 | (AAT)5 | CCGGAAGTTGGTCGAATAGA | TATTGCCGACACCAAACGTA | 184 |
| chr6:49132476-49134488 | p2 | (GA)6 | GCGTTTGGGAAATGTTTTGT | TCTCTCCATTCCCGAACATC | 114 |
| chr6:49132476-49134488 | p3 | (TTG)5 | GCGTTTGGGAAATGTTTTGT | TCTCTCCATTCCCGAACATC | 114 |
| chr6:49197601-49199635 | p2 | (AT)17 | CTGATATCCAGGACGACGGT | AGGATAGGGGGCTGGAGTAA | 189 |
| chr6:49254940-49256968 | p5 | (AATGC)3 | TTGTAGCTAGGCCTCGGAAA | AGGTGCTTCACCAAAGGCTA | 297 |
| chr6:49254972-49256984 | p5 | (AATGC)3 | TTGTAGCTAGGCCTCGGAAA | AGGTGCTTCACCAAAGGCTA | 265 |
| chr6:49301162-49303176 | p2 | (TA)7 | GTTTAGGGGGTGAGAGAGGG | TGGATGAACAGTAATGCGGA | 279 |
| chr6:49384735-49386759 | p2 | (AT)12 | AAACGGCTCAGTGCTGCTAT | GCCCATCATACCCTAAGCAA | 111 |
| chr6:49392640-49394665 | p5 | (ATATC)5 | GTCATCGCGGAAAACCTTTA | TGACACGGTGGATACCTCAA | 141 |
| chr6:49392640-49394665 | p4 | (AAAG)4 | GTCATCGCGGAAAACCTTTA | TGACACGGTGGATACCTCAA | 141 |
| chr6:49498238-49500253 | p5 | (TGCAA)3 | CAATCTCGGAGACCCATTGT | GTTGAGGGTTCACCTACCGA | 181 |
| chr6:49498238-49500253 | p5 | (ATCAT)3 | CAATCTCGGAGACCCATTGT | GTTGAGGGTTCACCTACCGA | 181 |
| chr6:49580513-49582537 | p3 | (TGG)5 | ATACATCTTCCGCTGCTGCT | CATATCCACAACCACCACCA | 121 |
| chr6:49580513-49582537 | p4 | (CATA)6 | ATACATCTTCCGCTGCTGCT | CATATCCACAACCACCACCA | 121 |
| chr6:49580513-49582537 | p3 | (ATC)5 | ATACATCTTCCGCTGCTGCT | CATATCCACAACCACCACCA | 121 |

| Marker ID | SSR type | SSR motif | FORWARD PRIMER (5'-3') | REVERSE PRIMER (5'-3') | size (bp) |
|---|---|---|---|---|---|
| chr6:49580976-49582991 | p3 | (TGG)5 | GCGGCGTAGTGGTTGATAAT | AATGCCACATCATTTCCACA | 187 |
| chr6:49580976-49582991 | p4 | (CATA)6 | GCGGCGTAGTGGTTGATAAT | AATGCCACATCATTTCCACA | 187 |
| chr6:49580976-49582991 | p3 | (ATC)5 | GCGGCGTAGTGGTTGATAAT | AATGCCACATCATTTCCACA | 187 |
| chr6:49618937-49620952 | p3 | (TTG)5 | AGTGCTCCGGTGTATCTGCT | CCTTCCTCAATATTTCCGCA | 112 |
| chr6:49621182-49623204 | p2 | (AT)11 | ACTGTTCGTTCTGGTGACCC | GACCGCTTAAGGCACAAGTC | 252 |
| chr6:49647353-49649365 | p2 | (AT)6 | ATCATGGAAAGCGTTATCGG | GATTTCTCAAGCCTTGGCAG | 130 |
| chr6:49772382-49774402 | p2 | (AT)10 | TCTTGAAATCAGTCCCGACC | AGGAAACGAAAGGTTGCTCA | 140 |
| chr6:49920950-49922964 | p2 | (AG)7 | TTTGGAAAACATTCACGGGT | GCGACGCGTTCTTATAGTCC | 176 |
| chr6:49944598-49946613 | p5 | (ATTTT)3 | TGTGTGTGGCAAAAACCCTA | TGAATGCAGGTTGGCAGTTA | 220 |
| chr6:50004185-50006212 | p3 | (TTC)9 | AGTAGGAGCATGTGGGATGG | CACCTTGCACTGTCGGTAGA | 119 |
| chr6:50088326-50090341 | p3 | (ATG)5 | AAACCCCCTTCATTTTCACC | ATTCAAGCTGAAATCCACCG | 256 |
| chr6:50088326-50090341 | p2 | (TA)20 | AAACCCCCTTCATTTTCACC | ATTCAAGCTGAAATCCACCG | 256 |
| chr6:50092209-50094236 | p3 | (TTG)9 | GTTTGTTGGGCTTTGGAGAA | ACACGGTTTGTTGCATTGAA | 283 |
| chr6:50092209-50094236 | p3 | (ATG)8 | GTTTGTTGGGCTTTGGAGAA | ACACGGTTTGTTGCATTGAA | 283 |
| chr6:50167824-50169838 | p2 | (AT)7 | TCAAACACATTGCCCAAAGA | CATGGGTCGTTATTGTGCAG | 289 |
| chr6:50232187-50234203 | p2 | (TA)8 | ATCGTATACGCCGTGTCCTC | AGGGGAAACTCTTGCGAAAT | 216 |
| chr6:50351675-50353690 | p5 | (TTCAT)3 | CCTAGCTGGGTCGTGACATT | CGCCTCACAGGTCTACATCA | 226 |
| chr6:50355445-50357460 | p5 | (TTCAT)3 | TCAAAAACTGTCGCAACGTC | AACAGCCTCTTCCAAAAGCA | 127 |
| chr6:50433249-50435261 | p2 | (GT)6 | CGTATTTTCGAACGTCGGAT | AGTCAAATCACCCGAAAACG | 275 |
| chr6:50479499-50481514 | p2 | (GT)6 | AAGACGCTCTCTGACGTGGT | ACACACACACACCCACGACT | 298 |
| chr6:50479499-50481514 | p3 | (TGT)5 | AAGACGCTCTCTGACGTGGT | ACACACACACACCCACGACT | 298 |
| chr6:50714102-50716142 | p2 | (TA)20 | GGTGGGAAGACAAGGATGAA | CCCCTTTAGGATTTTCGCTC | 252 |
| chr6:50813323-50815338 | p5 | (TTCAT)3 | CCTAGCTGGGTCGTGACATT | CGCCTCACAGGTCTACATCA | 226 |
| chr6:50817093-50819108 | p5 | (TTCAT)3 | TCAAAAACTGTCGCAACGTC | AACAGCCTCTTCCAAAAGCA | 127 |
| chr6:50848293-50850308 | p5 | (AAAAT)3 | GCCGTCGTATATAGGGCTCA | AGCAGGTTTGTCCTAGGGGT | 229 |
| chr6:50888669-50890684 | p5 | (TATCA)3 | CCACCGAGTCCCTTAACAAA | ATGATGCATGCCAAGTCGTA | 220 |
| chr6:51003708-51005720 | p5 | (ACACA)3 | CAATTCGATGGCATTCCTTT | TTGAATTTGGAAGGTTTCGG | 143 |
| chr6:51003708-51005720 | p2 | (TA)6 | CAATTCGATGGCATTCCTTT | TTGAATTTGGAAGGTTTCGG | 143 |
| chr6:51039287-51041299 | p2 | (AT)6 | CGACGGGCTGAGTTATTTTC | ATGAGCCCTTGATTTCCCTT | 163 |
| chr6:51039287-51041299 | p6 | (AATATG)3 | CGACGGGCTGAGTTATTTTC | ATGAGCCCTTGATTTCCCTT | 163 |
| chr6:51146437-51148449 | p3 | (CAA)5 | TATGAAACGGCTTACGGAGG | GGACTTGAGTCGGGATTTGA | 147 |
| chr6:51146437-51148449 | p2 | (AC)6 | TATGAAACGGCTTACGGAGG | GGACTTGAGTCGGGATTTGA | 147 |
| chr6:51152517-51154531 | p2 | (AC)7 | TGAAACGGTTTACGGAGGTC | TCGGGATTTGATTAGCGAAC | 149 |
| chr6:51237136-51239154 | p2 | (GT)9 | TGAGAGTGACATGAGGACCG | ATCAATCCTCACAATTCCGC | 105 |
| chr6:51310249-51312265 | p2 | (AT)8 | TCAAGCTCACCTCCCTGTCT | TACCTATATGGACGGGCAGC | 189 |
| chr6:51334202-51336217 | p3 | (TAA)5 | TAACTGCTCCTGGCAGGTTT | TTCGGGTCAAACACAATGAA | 185 |

| Marker ID | SSR type | SSR motif | FORWARD PRIMER (5'-3') | REVERSE PRIMER (5'-3') | size (bp) |
|---|---|---|---|---|---|
| chr6:51380438-51382462 | p2 | (AT)12 | TTACAGCAACGAGGGCTTTT | GGTCATGCGAGGATTAAGGA | 229 |
| chr6:51653356-51655368 | p2 | (AT)6 | TTGAAGGCAAAATCTGACCC | GAGAGTCAGTGGGTTCGCTC | 178 |
| chr6:51684211-51686229 | p2 | (AT)9 | TGGCAAACTCCATTTTCCTC | CCACAGGAAACTGTCTGCAA | 216 |
| chr6:51745990-51748002 | p2 | (AT)6 | ATCCTCCTCATCGTCCTCCT | TAGTTATGAGACGGACCCGC | 143 |
| chr6:51790278-51792293 | p5 | (ATTTT)3 | AGGTGCACAGGGAGACAGAC | GACGAGGCCAACAAAATGAT | 267 |
| chr6:51790278-51792293 | p5 | (TTATT)3 | AGGTGCACAGGGAGACAGAC | GACGAGGCCAACAAAATGAT | 267 |
| chr6:51845019-51847031 | p2 | (AT)9 | GCGGCATTTTCTGAAATCAT | GCGCCAAGCTAATAAACAGG | 133 |
| chr6:51955177-51957189 | p2 | (GT)6 | AACTCGCCTGCTGTCTGTTT | GGGCCTTCTTCCGTATTTTC | 209 |
| chr6:51959294-51961318 | p6 | (ATATGT)4 | TCGGGGATAATCCTACCCTC | GCGTCAATTGTGAGGGTTTT | 244 |
| chr6:51963445-51965457 | p3 | (GTT)5 | AAAAAGAAACTCCCCCGAAA | TTTCCGCATCCGTATTTCTC | 164 |
| chr6:51963462-51965474 | p3 | (GTT)5 | AAAAAGAAACTCCCCCGAAA | TTTCCGCATCCGTATTTCTC | 147 |
| chr6:52066684-52068718 | p2 | (AT)8 | TCCCCACGTTCTCAAGGTAG | CAGTTTGTGGGGGAAGAAAA | 246 |
| chr6:52066684-52068718 | p2 | (TA)11 | TCCCCACGTTCTCAAGGTAG | CAGTTTGTGGGGGAAGAAAA | 246 |
| chr6:52074694-52076728 | p2 | (TA)7 | CTATTTGCTTCCCCCACAAA | TATATGGGTCTCTCCACGCC | 144 |
| chr6:52111617-52113629 | p2 | (AC)6 | GCTTTCGTGCATCCAATTTT | GCTTGTTTGTTGGTTGGGTT | 176 |
| chr6:52111617-52113629 | p2 | (AT)6 | GCTTTCGTGCATCCAATTTT | GCTTGTTTGTTGGTTGGGTT | 176 |
| chr6:52139040-52141060 | p2 | (TA)10 | TCACCCCCTACTTCAACCAC | AGTGAAGGCTAGCCGATTGA | 231 |
| chr6:52278931-52280951 | p6 | (TACATA)3 | AGGACTTCTGGCGACTTTGA | AGTAGCGATGGAAGAAGGCA | 116 |
| chr6:52412934-52414952 | p3 | (TTG)6 | TTGGAAGCCACATGGTGTAA | TTCTTGAATCGTCCCTTTGG | 103 |
| chr6:52563471-52565558 | p3 | (AAT)29 | ACCACAGAGGAACCGTTGAC | CCATGTTTTTAGCGTTGGGT | 263 |
| chr6:52563471-52565558 | p3 | (TAA)7 | ACCACAGAGGAACCGTTGAC | CCATGTTTTTAGCGTTGGGT | 263 |
| chr6:52619394-52621420 | p2 | (TA)20 | ATTTGGGGATTGGTTGTGAA | AGTCTCCACGGCTCGAAGTA | 201 |
| chr6:52619394-52621420 | p2 | (AT)13 | ATTTGGGGATTGGTTGTGAA | AGTCTCCACGGCTCGAAGTA | 201 |
| chr6:52890903-52892979 | p5 | (CACAA)3 | GCTCAACGGCTCAAGAACTC | CGCTCGTCGTGAAATGTTTA | 128 |
| chr6:52890903-52892979 | p2 | (TA)38 | GCTCAACGGCTCAAGAACTC | CGCTCGTCGTGAAATGTTTA | 128 |
| chr6:52930154-52932166 | p5 | (CATAG)3 | AACCCACATCGCTTCAAATC | ACTGACGTCCCTTCTTGTGG | 174 |
| chr6:52930154-52932166 | p2 | (TA)6 | AACCCACATCGCTTCAAATC | ACTGACGTCCCTTCTTGTGG | 174 |
| chr6:52965235-52967257 | p2 | (TA)11 | GCCCTAGGAAGTAAGTGGGG | TGTGCTTTTATTCGACGCAG | 125 |
| chr6:53000147-53002171 | p3 | (AAC)8 | CGAAAAATGAAGACCGGGTA | GTACGAGCAAACCAAGGCTC | 292 |
| chr6:53041127-53043145 | p3 | (ACA)6 | TCCTGCAGCCTCCTGTTTAT | CCCTTTAGTTTGGGTCGTGA | 119 |
| chr6:53191926-53193951 | p3 | (CAA)5 | TCAGCAAACATGTCATCGGT | CTGTTCGTTCTGTTTCCGGT | 204 |
| chr6:53226847-53228865 | p6 | (CATATA)3 | CCGCCATGTTTACATCACAC | TGATAAGGGGTGCTCGGTAG | 140 |
| chr6:53226847-53228865 | p3 | (CAA)5 | CCGCCATGTTTACATCACAC | TGATAAGGGGTGCTCGGTAG | 140 |
| chr6:53274455-53276493 | p2 | (TA)19 | TATGTCGGTTTTGCCTCACA | TGACTGTCTATGACCGAGCG | 111 |
| chr6:53311116-53313134 | p5 | (TCATA)3 | ACGTTTCCCTTCCCCATATC | CCCTCAAGGTGAGACACGAT | 176 |
| chr6:53311116-53313134 | p3 | (CAA)6 | ACGTTTCCCTTCCCCATATC | CCCTCAAGGTGAGACACGAT | 176 |

| Marker ID | SSR type | SSR motif | FORWARD PRIMER (5'-3') | REVERSE PRIMER (5'-3') | size (bp) |
|---|---|---|---|---|---|
| chr6:53317814-53319829 | p5 | (TCATA)3 | CAGTTCACGCGATACCTCAA | GTTGGAGAATGCCTCATGGT | 242 |
| chr6:53317814-53319829 | p2 | (AT)17 | CAGTTCACGCGATACCTCAA | GTTGGAGAATGCCTCATGGT | 242 |
| chr6:53360722-53362736 | p2 | (AT)7 | CTGTTGATGGCTAATCGCAA | ACCGTACCTGGCCCTCTAGT | 230 |
| chr6:53631639-53633657 | p6 | (ATATGT)3 | ATGTTGGATATGAGCCTCGG | TCGGGTCCTATTTATGCGAG | 166 |
| chr6:53636049-53638067 | p6 | (ATATGT)3 | ATGTTGGATATGAGCCTCGG | TCGGGTCCTATTTATGCGAG | 165 |
| chr6:53682555-53684615 | p3 | (TAT)20 | TGCAAATTCGGGTACTAGGG | TATGGGGATGGACTGGTTGT | 204 |
| chr6:53911838-53913856 | p6 | (TTTGTT)3 | ACTGAGAGGCGACGATGTTT | TAGCGAGGGTCTGCAGATTT | 194 |
| chr6:53926725-53928741 | p2 | (AT)8 | AATTGGGTCTAGATGGTGCG | TGGCCTTAGCAACCGTAAAC | 257 |
| chr6:53960664-53962676 | p3 | (TTG)5 | GTTGATTGTTTTGGCCGAGT | ATATGCGTCTTACGTTCGCC | 261 |
| chr6:53960664-53962676 | p2 | (AT)6 | GTTGATTGTTTTGGCCGAGT | ATATGCGTCTTACGTTCGCC | 261 |
| chr6:54013817-54015847 | p2 | (AT)15 | GATGGCAACCGTCAGTTTTT | ACCACACCCAAAACAACCAT | 197 |
| chr6:54122378-54124390 | p2 | (AT)6 | TTGGTGGAAGTTTCCAGAGG | CGCGAACACACAGAGGACTA | 276 |
| chr6:54126844-54128856 | p2 | (TA)6 | TGCTTGAATTGGGGAATCTC | TTCGTCTTGCAACTCGCTTA | 246 |
| chr6:54193823-54195843 | p4 | (AAAT)5 | ACTGCCAGGAAGTAGCAGGA | GCTCAATCCCATGTAGGGAA | 100 |
| chr6:54245006-54247021 | p5 | (ATTTT)3 | GATTGAGATGGTCTGGGCAT | ATATTTCTTTGGCCAACCCC | 200 |
| chr6:54245006-54247021 | p2 | (TA)6 | GATTGAGATGGTCTGGGCAT | ATATTTCTTTGGCCAACCCC | 200 |
| chr6:54245367-54247379 | p5 | (ATTTT)3 | GTGTCGCTTGCGTTACTTCA | AACAACCCAGTGAAATCCCA | 170 |
| chr6:54245367-54247379 | p2 | (TA)6 | GTGTCGCTTGCGTTACTTCA | AACAACCCAGTGAAATCCCA | 170 |
| chr6:54356080-54358098 | p3 | (TTG)6 | GTGAAAAAGGATGCCTTGGA | TTTAGCCATTTTGCGCTTCT | 259 |
| chr6:54356080-54358098 | p6 | (CATATA)3 | GTGAAAAAGGATGCCTTGGA | TTTAGCCATTTTGCGCTTCT | 259 |
| chr6:54401537-54403552 | p5 | (ATATG)3 | TGCATTATTGACGTTTGGGA | GGCCAAAGAAACATGCCTTA | 237 |
| chr6:54463841-54465856 | p5 | (ATATG)3 | TGCATTATTGACGTTTGGGA | GGCCAAAGAAACATGCCTTA | 237 |
| chr6:54556402-54558417 | p5 | (CATAT)3 | CCCCGACCTTACTAGGGTGT | GATAAGGGGTGCTCGGTACA | 134 |
| chr6:54616982-54618998 | p2 | (TA)8 | CAACACGATTCTTTCGCTGA | TCAGAGCAGTTCATCATCGG | 189 |
| chr6:54639002-54641018 | p2 | (TA)8 | CAACACGATTCTTTCGCTGA | TCAGAGCAGTTCATCATCGG | 189 |
| chr6:54726495-54728511 | p3 | (TGT)5 | GTATTGCGGTTCTAGCGGAA | AACAACAACATCAACGCCAA | 137 |
| chr6:54726495-54728511 | p2 | (TA)8 | GTATTGCGGTTCTAGCGGAA | AACAACAACATCAACGCCAA | 137 |
| chr6:54732301-54734319 | p6 | (ATTAAT)3 | AAAGGCACCCGACTACATTG | ACTTCCCGAGGTCTTTGGAT | 192 |
| chr6:54732301-54734319 | p3 | (TTC)7 | AAAGGCACCCGACTACATTG | ACTTCCCGAGGTCTTTGGAT | 192 |
| chr6:54741167-54743183 | p2 | (AT)8 | CCAAGCCTGCTGGTTTACAT | CGTTGATAGATTGGCCGAGT | 193 |
| chr6:54760308-54762326 | p6 | (CAATAA)3 | TCAACCATGGAAGCTTAGGG | TTTGAGTTTGAACCTTGGGG | 299 |
| chr6:54765979-54767994 | p3 | (TCA)5 | GCAGGAAAAACAAGCTCCTG | ATAGTGCAGGACCATCTGGG | 138 |
| chr6:54768303-54770321 | p6 | (CAATAA)3 | TCAACCATGGAAGCTTAGGG | TTTGAGTTTGAACCTTGGGG | 299 |
| chr6:54773974-54775989 | p3 | (TCA)5 | GCAGGAAAAACAAGCTCCTG | ATAGTGCAGGACCATCTGGG | 138 |
| chr6:54802536-54804566 | p6 | (CATATA)5 | CTTTGTCACACCCCGATCTT | AGATGTTCTGGGAGCCCTTT | 101 |
| chr6:54878724-54880738 | p2 | (TA)7 | TGCCATTCTTCCCTGAAATC | GTCCTAGGGGTTGTCTGCAA | 163 |

| Marker ID | SSR type | SSR motif | FORWARD PRIMER (5'-3') | REVERSE PRIMER (5'-3') | size (bp) |
|---|---|---|---|---|---|
| chr6:54937483-54939499 | p2 | (TA)8 | TTATCCACCCCCGAATATCA | CGTGGGGTGTGTGTTTTACA | 152 |
| chr6:54941738-54943752 | p2 | (AT)7 | CCGCTGATAAACTCCTACGG | TACGATAGTTACCCGGACGC | 275 |
| chr6:54951236-54953248 | p2 | (AT)7 | TAACAGGAGGTCGTGAACCC | TTGTGACTCCTCATGGGACA | 114 |
| chr6:54951236-54953248 | p2 | (AT)6 | TAACAGGAGGTCGTGAACCC | TTGTGACTCCTCATGGGACA | 114 |
| chr6:54987734-54989746 | p2 | (AT)6 | GTAAAGCGGACGGGTCAATA | CGTGCAACAGCCAAAAGTAA | 126 |
| chr6:54998413-55000425 | p2 | (TA)6 | ATGATGCTGAGGATTCCAGG | GACCTTTCGCAGGTTCTACG | 183 |
| chr6:55032993-55035007 | p2 | (TA)12 | GTCCGGGTTGGTATTGAATG | CTTTAGCGCTTATTCGCCAC | 202 |
| chr6:55032993-55035007 | p2 | (AT)7 | GTCCGGGTTGGTATTGAATG | CTTTAGCGCTTATTCGCCAC | 202 |
| chr6:55075497-55077512 | p3 | (GCA)5 | CAAGGGCCTGAGATAGCTTG | ATGTTTTCGGATACTCCCCC | 255 |
| chr6:55136212-55138230 | p3 | (TTG)6 | CGTTTTCGGGTGATCTGACT | TGGGTGTTGTGTCCTGAAAA | 218 |
| chr6:55145976-55148072 | p2 | (TA)48 | GATAAGTGGGATGCGTTGCT | CCAGCTTATTCGGATTTCCA | 244 |
| chr6:55145976-55148072 | p2 | (TA)8 | GATAAGTGGGATGCGTTGCT | CCAGCTTATTCGGATTTCCA | 244 |
| chr6:55244318-55246338 | p5 | (GTTTA)4 | ATACATGTTGGTTGGGCCAT | CCTTTTAAACCCGTATGCGA | 233 |
| chr6:55251204-55253224 | p2 | (GA)10 | AGCAGGGACCAACAAACAAC | CGATGTCACCTCGGGATACT | 144 |
| chr6:55300889-55302907 | p2 | (AT)8 | CCTCAGAATGTGCCGGTATT | CAGTTCGCTGCCGAATAAAT | 169 |
| chr6:55300910-55302925 | p2 | (AT)8 | CCTCAGAATGTGCCGGTATT | CAGTTCGCTGCCGAATAAAT | 148 |
| chr6:55450903-55452927 | p5 | (CAGGT)3 | GGTCTTGCTCTGACCGACTC | TAAGGGTCGGCACAAGAATC | 249 |
| chr6:55450903-55452927 | p6 | (GGGGGT)4 | GGTCTTGCTCTGACCGACTC | TAAGGGTCGGCACAAGAATC | 249 |
| chr6:55648694-55650706 | p2 | (TA)6 | AGGCAGGACGTCGATAATTG | ATTTGGCCATGGTGTATCGT | 232 |
| chr6:56027958-56029970 | p6 | (TCCGTA)3 | TGTATTGACGTCACCTGGGA | AAGTACCCTTGGGCCAAATC | 228 |
| chr6:56027958-56029970 | p2 | (AT)6 | TGTATTGACGTCACCTGGGA | AAGTACCCTTGGGCCAAATC | 228 |
| chr6:56091693-56093705 | p2 | (TA)6 | TTGCCATCTTATTTGGAGGC | CGGCTCTCAGTCATTTCCTC | 254 |
| chr6:56234275-56236291 | p4 | (AAAT)4 | TGGTGATTGGGTGTTCTTGA | TAGGAAACCCTGCCTACCCT | 156 |
| chr6:56234275-56236291 | p2 | (AG)10 | TGGTGATTGGGTGTTCTTGA | TAGGAAACCCTGCCTACCCT | 156 |
| chr6:56281149-56283167 | p5 | (TGAAT)3 | TTAGATTCTCGGGGATGGTG | AACATACCTGCTCCACGTCC | 219 |
| chr6:56281149-56283167 | p6 | (TATATG)3 | TTAGATTCTCGGGGATGGTG | AACATACCTGCTCCACGTCC | 219 |
| chr6:56312465-56314480 | p2 | (GA)7 | TTCGGAAATGTTGAGGAAGG | ACCTTGGGCCTCTTCATTTT | 211 |
| chr6:56312465-56314480 | p3 | (TGT)5 | TTCGGAAATGTTGAGGAAGG | ACCTTGGGCCTCTTCATTTT | 211 |
| chr6:56324069-56326084 | p3 | (ATA)5 | GAAACGACGGTTGATGACCT | GCAATTCTGCAACTTCCCAT | 167 |
| chr6:56354103-56356121 | p6 | (TACGAA)3 | TGGTGCTAAAATGGGTGTGA | ACCGTCGTAGCTTGAACCAC | 254 |
| chr6:56381331-56383369 | p2 | (TA)19 | ATGTCCGTGTTATGTCCCGT | CTTGGCCCAATTCGTTAAAA | 154 |
| chr6:56384601-56386613 | p2 | (TC)6 | GGAACAGCCTGTGGCATATT | ATCGATCATGTACCCCCAGA | 291 |
| chr6:56438549-56440563 | p2 | (TA)7 | GTGGTTGGTATTTCCTGCGT | CCCTCCTGAAATGATCTCCA | 183 |
| chr6:56453885-56455913 | p2 | (AT)14 | TGTACCATAACACCGCCTCA | CAACAAGTATCCGAGGCCAT | 162 |
| chr6:56472525-56474537 | p2 | (TG)6 | TCGGTTCGTAAGAGGATTGG | GCCTGCCCCCTTCTATACTC | 280 |
| chr6:56498242-56500300 | p2 | (AT)29 | TACCAATCGGGTGGTTTGAT | TACTTCGGGGAGGTCTCCTT | 155 |

| Marker ID | SSR type | SSR motif | FORWARD PRIMER (5'-3') | REVERSE PRIMER (5'-3') | size (bp) |
|---|---|---|---|---|---|
| chr6:56561258-56563274 | p2 | (TA)18 | GTTGCGGACTCAAAATACCC | TGGTGTCCGTACCACTAGCA | 117 |
| chr6:56561258-56563274 | p2 | (TA)8 | GTTGCGGACTCAAAATACCC | TGGTGTCCGTACCACTAGCA | 117 |
| chr6:56598270-56600288 | p6 | (CATATA)3 | CAAGAGGGCCATGAAAGGTA | TAGCGACCATTTCACCACAA | 226 |
| chr6:56610943-56613009 | p5 | (AATGC)3 | GGGCCTATTCTCATCACCAA | ATCTTCTGATCAGGCCATGC | 190 |
| chr6:56828211-56830223 | p2 | (AT)6 | AAACTCATGAAATCACGGGC | TATGCCTCCAGTTCCTCCAC | 165 |
| chr6:56828211-56830223 | p2 | (AT)8 | AAACTCATGAAATCACGGGC | TATGCCTCCAGTTCCTCCAC | 165 |
| chr6:56923329-56925347 | p3 | (TTG)6 | AATATGGGCATAAGCCCTCC | TTTGGCAGCTTGGAGTCTTT | 174 |
| chr6:56923329-56925347 | p2 | (TA)26 | AATATGGGCATAAGCCCTCC | TTTGGCAGCTTGGAGTCTTT | 174 |
| chr6:56962905-56964920 | p5 | (ATGAT)3 | ACGTTCGAATTTTCCCCTTT | AAAAATCCGGGAATAGTGGG | 167 |
| chr6:57066686-57068698 | p2 | (CA)6 | TGCCAAAGAAAGAAAGGGAA | GCCATAGCTTGGTTGTTGGT | 167 |
| chr6:57169947-57171963 | p2 | (GT)6 | TGTGTGTGTTGTGGTGTTGG | TACCTTCCAGTTCAGCCCAC | 221 |
| chr6:57169947-57171963 | p4 | (CTTT)4 | TGTGTGTGTTGTGGTGTTGG | TACCTTCCAGTTCAGCCCAC | 221 |
| chr6:57272247-57274259 | p2 | (TA)6 | AAATTGCCTTCTGATGTGCC | CTTTGCTTTTTGCTTTTGGG | 285 |
| chr6:57342460-57344475 | p5 | (ATTTC)3 | CAATTGGGGATTCTTGAGGA | TTCCGAGGTGTAATAACGGC | 238 |
| chr6:57406991-57409003 | p2 | (AT)6 | GGGTTAAGTGATGACCACGG | TAGAGCGCAACTACCCGACT | 127 |
| chr6:57412605-57414627 | p2 | (AT)11 | CCAGGAGCGACAAGATTAGG | CCTCGTCTGAAACCATCCAT | 142 |
| chr6:57430499-57432513 | p2 | (AT)7 | TGAACGCATATCCACCTGAA | CATTGCCTGCTTAGCTTTCC | 124 |
| chr6:57430499-57432513 | p2 | (AT)9 | TGAACGCATATCCACCTGAA | CATTGCCTGCTTAGCTTTCC | 124 |
| chr6:57431207-57433225 | p2 | (AT)7 | GATAAGGGGTGCTCGGTACA | TTAGCCTCGTCGGCTTATGT | 128 |
| chr6:57431207-57433225 | p2 | (AT)9 | GATAAGGGGTGCTCGGTACA | TTAGCCTCGTCGGCTTATGT | 128 |
| chr6:57702593-57704605 | p2 | (CG)6 | AGCAAGTGCACCGATAGCTT | TTTCCGGAACATTCAACCTC | 133 |
| chr6:57775557-57777581 | p2 | (TA)12 | CCGACAAGGCTACATCACAA | CTCTTCCAACGGTGGTCAAT | 292 |
| chr6:57798059-57800083 | p3 | (AAC)8 | CACACAGCCTCTTGTCTGGA | GCCACCACCTCCTTTAACAA | 283 |
| chr6:57845432-57847447 | p3 | (ATG)5 | TCAGTTGATGACCACCAGGA | AAATTCTTTATGCGGGGACC | 210 |
| chr6:57882312-57884332 | p5 | (TTCAT)4 | AGTGCGGAGGCTTAGAAACA | ACGACCCCAATGCATAAAAA | 101 |
| chr6:57892228-57894246 | p6 | (ACTGAA)3 | AGGGCTTCATGCTATTGTGG | CGACCCATTTCGTCGTTACT | 230 |
| chr6:57905515-57907529 | p2 | (TA)7 | GGGTTGTGCTACCGAGACAT | AACCAAGTATCCGTCATCGC | 239 |
| chr6:57967077-57969091 | p2 | (TA)7 | ACAACAACCTGTCTCCTGGG | ATGGCTTATGTTTTGGGACG | 259 |
| chr6:57996147-57998165 | p6 | (CAAAAA)3 | GGCATGCATAATTCGGTCTT | ACCTGTCTGTCAACCCGAAC | 233 |
| chr6:58095456-58097648 | p2 | (AT)96 | CAGACCCCTCCCAGTCATAA | GCCCCAAAGAACCCTAAGAG | 171 |
| chr6:58180059-58182077 | p6 | (ATATGT)3 | ATGTTGGATATGAGCCTCGG | TCGGGTCCTATTTATGCGAG | 125 |
| chr6:58199753-58201797 | p2 | (AT)22 | ATGGAAACGTGCAAAAGACC | AGTCTTGTCGGCTTGCGTAT | 246 |
| chr6:58234325-58236345 | p2 | (AT)10 | CCTGTAGCCCGTCCTGTTTA | TTTCGGTTCCAAATAGTGGC | 175 |
| chr6:58245931-58247955 | p2 | (TA)12 | TGCCCGATTTTCGTTAAGTC | TTTGTAAATCCGTTCGCCTC | 276 |
| chr6:58299249-58301261 | p2 | (TA)6 | GCTCAGAACAATTCCCGTGT | AGGGGAATTCAATACGTCCC | 103 |
| chr6:58485848-58487880 | p2 | (AT)16 | GCGGCAACTAACTGTGTCAA | TTCTCCTCCCTAACCGTGTG | 238 |

| Marker ID | SSR type | SSR motif | FORWARD PRIMER (5'-3') | REVERSE PRIMER (5'-3') | size (bp) |
|---|---|---|---|---|---|
| chr6:58502868-58504886 | p6 | (ATATAA)3 | GGATTGACACCGCTTTGATT | GCCATTAGTGTTTGTCGCCT | 300 |
| chr6:58502868-58504886 | p2 | (TA)6 | GGATTGACACCGCTTTGATT | GCCATTAGTGTTTGTCGCCT | 300 |
| chr6:58547971-58549986 | p5 | (ATTTC)3 | GCCCTGTGAGGCAATGTTAT | TGGTGGGAAATTTTCTGAGG | 240 |
| chr6:58575211-58577231 | p5 | (CATAT)4 | ATTTCTGAGTGGGCCGTATG | TTGCAGTGCTCCAGTTGTTC | 213 |
| chr6:58594629-58596644 | p5 | (GAAAA)3 | CGTGACATTGCTTCCTCAGA | CGACTAATCGACGTGCAAAA | 109 |
| chr6:58618232-58620246 | p2 | (AT)6 | TCGATGCTTACCTTGAGCCT | ACATAGTGCGACCGAACACA | 299 |
| chr6:58618232-58620246 | p2 | (TA)7 | TCGATGCTTACCTTGAGCCT | ACATAGTGCGACCGAACACA | 299 |
| chr6:58618232-58620246 | p2 | (AT)12 | TCGATGCTTACCTTGAGCCT | ACATAGTGCGACCGAACACA | 299 |
| chr6:58619085-58621109 | p2 | (TA)7 | TCCGGAGCTCATATTTTTGG | ATCCTATAGGCTGGCGGAAT | 102 |
| chr6:58619085-58621109 | p2 | (AT)12 | TCCGGAGCTCATATTTTTGG | ATCCTATAGGCTGGCGGAAT | 102 |
| chr6:58654941-58656993 | p2 | (TA)8 | CGTATCGTATCGGTCCCTGT | GTATGGGGTATGGGGGAAGT | 183 |
| chr6:58654941-58656993 | p2 | (TA)26 | CGTATCGTATCGGTCCCTGT | GTATGGGGTATGGGGGAAGT | 183 |
| chr6:58655421-58657491 | p2 | (TA)8 | TCATCACTTTCCCCATACCC | GCATATCATTAGCCCTCGGA | 220 |
| chr6:58655421-58657491 | p2 | (TA)26 | TCATCACTTTCCCCATACCC | GCATATCATTAGCCCTCGGA | 220 |
| chr6:58655524-58657539 | p2 | (TA)26 | TCATCACTTTCCCCATACCC | GCATATCATTAGCCCTCGGA | 117 |
| chr6:58672194-58674238 | p2 | (AT)22 | TGCCAACTCCACAGATTCAG | GTTGCTTGGGGTCAACAAGT | 170 |
| chr6:58752788-58754842 | p2 | (AT)27 | GCAGATGGTGGGAAGACAAT | GGTCCCTTTAGGATTTTCGC | 285 |
| chr6:58758429-58760445 | p5 | (TGATA)4 | AATGTTTCCCACACTGAGCC | ACATGGCACAGCATACTCCA | 121 |
| chr6:58795161-58797173 | p3 | (TCA)5 | TCTCTGAAGCCCTGACCAGT | GATGATGATGATGACAGGCG | 190 |
| chr6:58795161-58797173 | p2 | (AT)6 | TCTCTGAAGCCCTGACCAGT | GATGATGATGATGACAGGCG | 190 |
| chr6:58831894-58833909 | p3 | (ATA)5 | GAGCTACTTACCGAGCACCG | TTATTTTGATGGCCGAAAGG | 218 |
| chr6:58881415-58883433 | p3 | (CAA)6 | AGGCACGGTGTTATACTGCC | ATCGACCTTACGTCACTCCG | 112 |
| chr6:58929098-58931113 | p5 | (ATTTC)3 | CAGTCAAAGTCTTGTGGCGA | TCCTGGGGTTGGAATAAGTG | 128 |
| chr6:58929098-58931113 | p2 | (TA)7 | CAGTCAAAGTCTTGTGGCGA | TCCTGGGGTTGGAATAAGTG | 128 |
| chr6:58968528-58970540 | p2 | (AT)6 | AGTGGCGGTGTAGCAGAACT | AAAGCGACAGCTACCCAGAA | 133 |
| chr6:58979537-58981597 | p5 | (ATCAT)12 | AGTGAAATACTGGGCATCGC | GCGGAGCACTCCTCTATCAC | 128 |
| chr6:58979537-58981597 | p6 | (CATATA)3 | AGTGAAATACTGGGCATCGC | GCGGAGCACTCCTCTATCAC | 128 |
| chr6:59042880-59044892 | p2 | (TG)6 | TGCAATCGGATCAATTCAAA | TTCCCTTATGGCCCTTACCT | 164 |
| chr6:59042880-59044892 | p2 | (AT)7 | TGCAATCGGATCAATTCAAA | TTCCCTTATGGCCCTTACCT | 164 |
| chr6:59067092-59069114 | p2 | (AT)11 | TGGCATCTATGCCCATATCA | GTCCTGTCCCGCCTATATGA | 206 |
| chr6:59121918-59123936 | p3 | (TTA)5 | GGCACTGGAAATATCGTCGT | CCATCAACTGCATGTCCAAG | 152 |
| chr6:59121918-59123936 | p3 | (TTA)6 | GGCACTGGAAATATCGTCGT | CCATCAACTGCATGTCCAAG | 152 |
| chr6:59127408-59129453 | p3 | (TTA)5 | CCGCACCTGCTTTGTTTTAT | CCTTGTCGGAAAGTCTGCTC | 254 |
| chr6:59127535-59129553 | p3 | (TTA)5 | CCGCACCTGCTTTGTTTTAT | CCTTGTCGGAAAGTCTGCTC | 127 |
| chr6:59131341-59133361 | p4 | (TAAT)5 | CTATGGGTCATCCATTTCGG | ATTTTTGGTAAGCCACGCAC | 104 |
| chr6:59327289-59329304 | p5 | (AAACA)3 | GGAGGGAGATGCATTGAGAA | CGTTGGAGTGCTAATGGGAT | 232 |

| Marker ID | SSR type | SSR motif | FORWARD PRIMER (5'-3') | REVERSE PRIMER (5'-3') | size (bp) |
|---|---|---|---|---|---|
| chr6:59327289-59329304 | p2 | (TA)6 | GGAGGGAGATGCATTGAGAA | CGTTGGAGTGCTAATGGGAT | 232 |
| chr6:59327289-59329304 | p3 | (TTG)5 | GGAGGGAGATGCATTGAGAA | CGTTGGAGTGCTAATGGGAT | 232 |
| chr6:59526164-59528176 | p3 | (ACA)5 | TCTCAGACGTCGTAAGCCCT | GTTTGTTGATGTGTGGGCTG | 180 |
| chr6:59526164-59528176 | p2 | (AC)6 | TCTCAGACGTCGTAAGCCCT | GTTTGTTGATGTGTGGGCTG | 180 |
| chr6:59551460-59553474 | p2 | (TA)7 | CCTAGGACCGAACTGCTCTG | ACCCGCTGAGAGGTAAAGGT | 130 |
| chr6:59580383-59582399 | p2 | (AT)8 | GCCTATGTTGAAGGGCAAAA | TAAGGCATTCTCCAACCACC | 197 |
| chr6:59580383-59582399 | p2 | (AT)8 | GCCTATGTTGAAGGGCAAAA | TAAGGCATTCTCCAACCACC | 197 |
| chr6:59652965-59654977 | p2 | (AT)6 | GGAGATGTCCCATGATGCTT | CCCCTCCAAACTTGACAAAA | 162 |
| chr6:59757019-59759043 | p2 | (AT)12 | CCCTAGGACGAACTGCTCTG | ACGTATGGGTGCCCAGTTAG | 270 |
| chr6:59810841-59812856 | p5 | (TGTGT)3 | GTGGCCTGGTCTATAACGGA | GTTTTCCCCTAGCCTTCCAC | 113 |
| chr6:59900036-59902054 | p6 | (ATATAG)3 | CACCTTCCCTAAAACCCACA | CGGCAAAAATATCGTCGTTT | 288 |
| chr6:59900036-59902054 | p5 | (TTTAT)3 | CACCTTCCCTAAAACCCACA | CGGCAAAAATATCGTCGTTT | 288 |
| chr6:59914870-59916888 | p6 | (CTTCCA)3 | GTTGCAACCCAAAGTGTCCT | AGGCAGCGACGTATCCTAAA | 205 |
| chr6:59916004-59918030 | p2 | (AT)13 | CTCTCGTTTGTCGTTGGTGA | GGGGAGGAAAGAACTCAAGG | 180 |
| chr6:59927828-59929846 | p3 | (TTG)6 | GCCCTCGTAATCGTAAGCTG | TTTGATTTGGCTGAAGGGTC | 196 |
| chr6:60019173-60021185 | p2 | (TC)6 | ACATCACAACAAAGGCCACA | CAATTAACAGGGGGTGTGCT | 178 |
| chr6:60047525-60049537 | p2 | (TA)6 | AAGTCCTTTGCCTTACCCGT | TGAAGGCATCAATCGAACAG | 152 |
| chr6:60047525-60049537 | p2 | (TA)12 | AAGTCCTTTGCCTTACCCGT | TGAAGGCATCAATCGAACAG | 152 |
| chr6:60049189-60051207 | p2 | (TA)12 | TCGTGTTTTGACACATCGGT | GGAGAATGGTGAGTTTTCCG | 145 |
| chr6:60049189-60051207 | p2 | (AT)19 | TCGTGTTTTGACACATCGGT | GGAGAATGGTGAGTTTTCCG | 145 |
| chr6:60049208-60051236 | p2 | (TA)12 | TCGTGTTTTGACACATCGGT | GGAGAATGGTGAGTTTTCCG | 126 |
| chr6:60049208-60051236 | p2 | (AT)19 | TCGTGTTTTGACACATCGGT | GGAGAATGGTGAGTTTTCCG | 126 |
| chr6:60077032-60079056 | p6 | (ATATGA)4 | ATGTCCTAATGGGGAAAGGG | CATTCTCAAATCCGTTGCCT | 220 |
| chr6:60081275-60083299 | p2 | (AT)12 | ACTTTTGTCACGACCCAACC | AGACGTATTTGGCCTTGTCG | 108 |
| chr6:60115825-60117839 | p2 | (TA)7 | TGTGGTTGGTTTGGCTACAA | CACAACGATCATGCCAAAAG | 264 |
| chr6:60225051-60227066 | p5 | (TGTGT)3 | GGTTGGGCCAGGTATGTATG | TGCTCTTAATCTCCCGCAAT | 184 |
| chr6:60262847-60264863 | p2 | (AG)8 | ATTATATCCCCCGTCTTCGG | GGGGTGTAACAACCATCCTG | 124 |
| chr6:60307679-60309753 | p2 | (TA)37 | GATTCGACGGCATTTCATTT | TTGAATTTGGAAGGTTTCGG | 106 |
| chr6:60313394-60315410 | p2 | (TA)8 | TCGACGGCATTTCATTTACA | TTGAATTTGGAAGGTTTCGG | 168 |
| chr6:60313394-60315410 | p2 | (TA)8 | TCGACGGCATTTCATTTACA | TTGAATTTGGAAGGTTTCGG | 168 |
| chr6:60313394-60315410 | p2 | (TA)9 | TCGACGGCATTTCATTTACA | TTGAATTTGGAAGGTTTCGG | 168 |
| chr6:60322309-60324324 | p3 | (GCT)5 | CTTATCAGGCTTGACGCTCC | CTTATTGCTGAGGCAGAGGG | 279 |
| chr6:60341973-60343988 | p3 | (GTT)5 | AGCGTTGCTATTGCTGGACT | TCATGCCCGAAATCCTAATC | 296 |
| chr6:60495708-60497726 | p6 | (TATGTA)3 | TCATTGGCCTGTGGATATGA | GTGTATTGATGCGGAACGTG | 206 |
| chr6:60591520-60593532 | p2 | (TG)6 | TGATCGGTTATTAGCAGCCC | CCTTGCGCACAGTCTCATAA | 133 |
| chr6:60591520-60593532 | p3 | (TAT)5 | TGATCGGTTATTAGCAGCCC | CCTTGCGCACAGTCTCATAA | 133 |

| Marker ID | SSR type | SSR motif | FORWARD PRIMER (5'-3') | REVERSE PRIMER (5'-3') | size (bp) |
|---|---|---|---|---|---|
| chr6:60607786-60609802 | p2 | (AC)8 | TAACTTTCCTGTGGATCGGG | CGACGAAACGGATTGATTTT | 188 |
| chr6:60698040-60700055 | p5 | (ATCAT)3 | TTCAGGCACATGCTCAGAAC | GGTACGTTTGGGTGTCCAAC | 168 |
| chr6:60698040-60700055 | p3 | (AAC)5 | TTCAGGCACATGCTCAGAAC | GGTACGTTTGGGTGTCCAAC | 168 |
| chr6:60763052-60765080 | p2 | (CA)6 | TGCTCATAAACACGCCGTAG | GACGCCCATGAGACATCTTT | 290 |
| chr6:60763052-60765080 | p3 | (ATC)5 | TGCTCATAAACACGCCGTAG | GACGCCCATGAGACATCTTT | 290 |
| chr6:60763131-60765143 | p2 | (CA)6 | TGCTCATAAACACGCCGTAG | GACGCCCATGAGACATCTTT | 211 |
| chr6:60763131-60765143 | p3 | (ATC)5 | TGCTCATAAACACGCCGTAG | GACGCCCATGAGACATCTTT | 211 |
| chr6:60775091-60777109 | p2 | (TA)9 | AAACTCATGAAATCACGGGC | AGTTCCTCCATCGGCTACCT | 179 |
| chr6:60791161-60793179 | p5 | (AAAAC)3 | CTTTTTCGGAGGCTTCACTG | TGGTATCAGAGCGGTTTTCC | 140 |
| chr6:60791161-60793179 | p6 | (CATATA)3 | CTTTTTCGGAGGCTTCACTG | TGGTATCAGAGCGGTTTTCC | 140 |
| chr6:60795070-60797088 | p2 | (TA)22 | TGGGAGCGAATTGAGAAATC | TTGCACGTTCTCTTTCAACG | 153 |
| chr6:60795070-60797088 | p6 | (ATATGA)3 | TGGGAGCGAATTGAGAAATC | TTGCACGTTCTCTTTCAACG | 153 |
| chr6:60837487-60839505 | p2 | (TA)9 | GGCGAAATAACAACCGAGAA | ATAACATTGCCTCACAGGGC | 292 |
| chr6:60845297-60847331 | p2 | (AT)17 | ATATAAGGGAACATCCCCGC | CAGTCGTGTCTGTGCGATTT | 228 |
| chr6:60856229-60858244 | p5 | (ACACA)3 | ACACACGGCCAACTTAGACC | TCATGGAGTGGGTGTGTTGT | 199 |
| chr6:60970648-60972704 | p2 | (AT)11 | TTTGACCCGAAAAACTCTGG | TATGCCTCCAGTTCCTCCAC | 201 |
| chr6:60970716-60972736 | p2 | (AT)11 | TTTGACCCGAAAAACTCTGG | TATGCCTCCAGTTCCTCCAC | 133 |
| chr6:60993862-60995878 | p4 | (TGTA)4 | GAAAGCCAAACGAGCGTAAG | TTCCATATCGAATTCCGAGG | 154 |
| chr6:61000008-61002026 | p2 | (TA)9 | TTTCTCCAACCCCTGTTGTC | AGGATCAAAGGGGAGCCTTA | 172 |
| chr6:61009547-61011561 | p2 | (AG)7 | GGTTCCGGGGTACTGTGTTA | CTTTGGTGCCACAACACATC | 276 |
| chr6:61009547-61011561 | p2 | (AT)7 | GGTTCCGGGGTACTGTGTTA | CTTTGGTGCCACAACACATC | 276 |
| chr6:61011507-61013523 | p4 | (TGTA)4 | CTCGGAATTCGAGATGGAAA | AAACGGACAGAAAGAAGGCA | 266 |
| chr6:61223190-61225222 | p2 | (AT)16 | GGGGCGCATTATCTTGTCTA | AAGGTCCGCAAGGCTTTATT | 101 |
| chr6:61267493-61269507 | p2 | (AT)7 | AATTGGGAATTCGGGATAGG | CTGTTAACGACGAATCGCAA | 240 |
| chr6:61385758-61387788 | p2 | (TA)15 | ACTCCACGGATTCAGTTTCG | CAAGTACCAAAGCCCGTGAT | 213 |
| chr6:61385758-61387788 | p2 | (AT)7 | ACTCCACGGATTCAGTTTCG | CAAGTACCAAAGCCCGTGAT | 213 |
| chr6:61385758-61387788 | p2 | (TA)7 | ACTCCACGGATTCAGTTTCG | CAAGTACCAAAGCCCGTGAT | 213 |
| chr6:61387254-61389269 | p2 | (TA)7 | TCAGATGTCTTGCAGCCTTG | TGAACCACGTGGCTTACTCA | 275 |
| chr6:61387254-61389269 | p3 | (ATG)5 | TCAGATGTCTTGCAGCCTTG | TGAACCACGTGGCTTACTCA | 275 |
| chr6:61531382-61533397 | p3 | (ACA)5 | TACCTGGGTTGCCTCTCATC | CTGACCTTCGAGTCCCAGAG | 213 |
| chr6:61590168-61592388 | p2 | (TA)15 | GTGATGCCCCTCTTGGATAA | AACGAAATCGGAACAACTGG | 257 |
| chr6:61590168-61592388 | p2 | (AT)6 | GTGATGCCCCTCTTGGATAA | AACGAAATCGGAACAACTGG | 257 |
| chr6:61592323-61594341 | p3 | (CAA)6 | TGCAGCGCATAAGAAAAATG | GTGAGATGGAGCGATGATGA | 222 |
| chr6:61592323-61594341 | p2 | (CA)6 | TGCAGCGCATAAGAAAAATG | GTGAGATGGAGCGATGATGA | 222 |
| chr6:61718125-61720163 | p2 | (AT)19 | GTTGTGAGCGTTTGGGAGTT | CTATTGCTCGCTCGTTCTCC | 170 |
| chr6:61793238-61795262 | p2 | (TA)6 | TTTGACAACGGCCAATTACA | CAAGGGTAAGAGGCAATGGA | 276 |

| Marker ID | SSR type | SSR motif | FORWARD PRIMER (5'-3') | REVERSE PRIMER (5'-3') | size (bp) |
|---|---|---|---|---|---|
| chr6:61793238-61795262 | p2 | (AT)12 | TTTGACAACGGCCAATTACA | CAAGGGTAAGAGGCAATGGA | 276 |
| chr6:61813243-61815255 | p2 | (AT)8 | AAAGAAATAGGGGGCGAAGA | TCATGGGTTTTCTAATCCGC | 177 |
| chr6:61813243-61815255 | p2 | (TA)6 | AAAGAAATAGGGGGCGAAGA | TCATGGGTTTTCTAATCCGC | 177 |
| chr6:61996097-61998115 | p6 | (CATATA)3 | TATCCTTTTCATCGTTCCCG | GCCGGTACATCCAGATGAGT | 227 |
| chr6:62004565-62006581 | p2 | (AT)8 | TGCTCATGCTCTGAAAATGC | CTTCGTTTCCTCCTCGACTG | 234 |
| chr6:62004565-62006581 | p2 | (TG)6 | TGCTCATGCTCTGAAAATGC | CTTCGTTTCCTCCTCGACTG | 234 |
| chr6:62004565-62006581 | p3 | (ATC)5 | TGCTCATGCTCTGAAAATGC | CTTCGTTTCCTCCTCGACTG | 234 |
| chr6:62022294-62024309 | p5 | (ACCGA)3 | GGGTCACGAGGCTTTCATAA | TCAACAATACGCATAACCGC | 282 |
| chr6:62039134-62041152 | p2 | (TA)9 | AACATGGCCCAAAACTCAAG | ATCCTGCATTTTGTCCGTTC | 111 |
| chr6:62065388-62067403 | p5 | (TGATA)3 | CTTTTGATTCGGAGCCCATA | AACAGACGGACCGCATTAAG | 100 |
| chr6:62180913-62182931 | p3 | (TTG)6 | TTGGAGGGTTCATCAAGGAG | TGGACTTGGTTTTTCTTGGG | 189 |
| chr6:62180913-62182931 | p2 | (TA)14 | TTGGAGGGTTCATCAAGGAG | TGGACTTGGTTTTTCTTGGG | 189 |
| chr6:62221750-62223832 | p2 | (AT)10 | ATGACTAGTGCCCGAGTTGG | CGTCGGCTTGCGTATTTATT | 185 |
| chr6:62221750-62223832 | p2 | (TA)41 | ATGACTAGTGCCCGAGTTGG | CGTCGGCTTGCGTATTTATT | 185 |
| chr6:62266402-62268417 | p5 | (TTATT)3 | AGGTTTAATTGGCTCGGGTT | TTTCTGGGCCATGCTATTTC | 230 |
| chr6:62296562-62298594 | p2 | (TA)16 | GGCGTAGCTCATGAACCATT | GCAAAAGGAGTTGAACCTGC | 260 |
| chr6:62345007-62347025 | p6 | (CATATA)3 | CATACAGAAGCCGCTAAGGC | TCATGGGTCGCTATTGTTCA | 290 |
| chr6:62393060-62395078 | p6 | (CATATA)3 | CATACAGAAGCCGCTAAGGC | TCATGGGTCGCTATTGTTCA | 290 |
| chr6:62599888-62601903 | p5 | (AAAAT)3 | CCTCCCAGGATATTCCCAGT | CGCCACAGAACTTTGACAGA | 237 |
| chr6:62599888-62601903 | p2 | (GT)6 | CCTCCCAGGATATTCCCAGT | CGCCACAGAACTTTGACAGA | 237 |
| chr6:62609433-62611445 | p2 | (AT)6 | GGCACCGCTCGTACTATCAT | GAAGAGCACTCCATCTTCCG | 197 |
| chr6:62635246-62637314 | p2 | (AT)34 | CGCCAACTTTCCAATCAGTT | TCCCTTCTCGAGCAAGTCAT | 134 |
| chr6:62635246-62637314 | p5 | (TGAAT)4 | CGCCAACTTTCCAATCAGTT | TCCCTTCTCGAGCAAGTCAT | 134 |
| chr6:62736757-62738771 | p2 | (TA)7 | GGCCGTATAATCCCCAATTT | AGAGAAGGCAAGGCATACGA | 227 |
| chr6:62753448-62755469 | p3 | (GTT)7 | ATTTGGGAATTTCGGAAACC | TGAGGAGGAGAAGGCTGAAA | 216 |
| chr6:62753448-62755469 | p2 | (TA)7 | ATTTGGGAATTTCGGAAACC | TGAGGAGGAGAAGGCTGAAA | 216 |
| chr6:62782511-62784529 | p6 | (AGGTAG)3 | CGTCACTACTTGCCACCTGA | GTTGTCGTTCCTCGAGCTTC | 256 |
| chr6:62785899-62787914 | p3 | (AGA)5 | AACCACATATTCAGCGGAGG | CTTGTCCATCAGTCTGCGAA | 198 |
| chr6:62800603-62802621 | p6 | (TTCATA)3 | TTTTGGACCAACATTCCACA | TGTCCGTGAAACTTTTGCTG | 105 |
| chr6:62811699-62813713 | p2 | (AT)7 | CAAAAGGTAACGGTGGTGCT | AACAAACCTTCAGAGGGGCT | 233 |
| chr6:62811699-62813713 | p2 | (AT)10 | CAAAAGGTAACGGTGGTGCT | AACAAACCTTCAGAGGGGCT | 233 |
| chr6:62812179-62814199 | p2 | (AT)7 | GGTTTTGCCACCTTGTCATT | ACCCTCTGTTGGATCGTGAC | 185 |
| chr6:62812179-62814199 | p2 | (AT)10 | GGTTTTGCCACCTTGTCATT | ACCCTCTGTTGGATCGTGAC | 185 |
| chr6:62815218-62817236 | p6 | (CCAGCA)3 | ATTGCGGTTCCGACAGTTAC | ACGTATCACAATGCAAGCCA | 122 |
| chr6:62845611-62847631 | p2 | (TA)10 | TTACCTCTTAGCAAGGCCGA | TTGTTCCGGAGCCTCATATC | 156 |
| chr6:62852929-62854953 | p2 | (TA)12 | ACGACCAAGGTATGTGAGGC | GGGTGACGTAAGGTCGTGAT | 128 |

| Marker ID | SSR type | SSR motif | FORWARD PRIMER (5'-3') | REVERSE PRIMER (5'-3') | size (bp) |
|---|---|---|---|---|---|
| chr6:62998395-63000407 | p2 | (AT)6 | CAATACCCACGACCCTGTCT | GAAGTTGGCAAGCTCCACTC | 182 |
| chr6:63003432-63005474 | p2 | (TA)6 | GAAAGAAAGAACGCAGTGCC | TGTCCTTCCCAGGTCATTTC | 110 |
| chr6:63003432-63005474 | p2 | (TA)21 | GAAAGAAAGAACGCAGTGCC | TGTCCTTCCCAGGTCATTTC | 110 |
| chr6:63003432-63005474 | p5 | (TGAAT)3 | GAAAGAAAGAACGCAGTGCC | TGTCCTTCCCAGGTCATTTC | 110 |
| chr6:63009195-63011210 | p5 | (TATAT)3 | GAAATTGTGGGTGGAAATGG | ATTTCGCCTAACCCGAATTT | 289 |
| chr6:63009195-63011210 | p2 | (TA)10 | GAAATTGTGGGTGGAAATGG | ATTTCGCCTAACCCGAATTT | 289 |
| chr6:63067126-63069156 | p2 | (GA)13 | ACCTCTTTGAATCCAGCCCT | TCCGGTAAGGTCAAAACAGG | 201 |
| chr6:63113109-63115121 | p2 | (AT)6 | ACGTTTCAGGTCAATCCGAC | CGGAGGTTACTTGGCATGAT | 120 |
| chr6:63319195-63321211 | p2 | (AT)8 | TACGCATTCCCACTCATTCA | TCTGGAATTGATCCTCGGAC | 163 |
| chr6:63327952-63329968 | p2 | (AG)8 | CGCGAAGGTGTTATGAGGTT | TACGTGGAGTGCTCATGCTC | 223 |
| chr6:63327952-63329968 | p3 | (TTG)6 | CGCGAAGGTGTTATGAGGTT | TACGTGGAGTGCTCATGCTC | 223 |
| chr6:63328265-63330283 | p2 | (AG)8 | TCAGAGCATGAGCACTCCAC | TTTTTCGAAGATACGGGGTG | 102 |
| chr6:63328265-63330283 | p3 | (TTG)6 | TCAGAGCATGAGCACTCCAC | TTTTTCGAAGATACGGGGTG | 102 |
| chr6:63344254-63346302 | p2 | (AT)24 | CTCTAACTCGGCAGCACTCC | TTTTTGACCAAAATCCGAGC | 244 |
| chr6:63499586-63501600 | p2 | (CA)7 | CCAAAAGGGAGCTTTTAGGC | TGAATATGAGACCTTCCGGC | 139 |
| chr6:63509359-63511375 | p2 | (AT)8 | ATCGTGCGATAGAGCCAAGT | TCCTAGATTCCCAAATCCCC | 248 |
| chr6:63517294-63519310 | p2 | (AT)8 | ATCGTGCGATAGAGCCAAGT | TCCTAGATTCCCAAATCCCC | 248 |
| chr6:63726184-63728202 | p2 | (TA)9 | AAGTGTGCGACAAAGAGGCT | GAAAACAAGGCGGTTATGGA | 169 |
| chr6:63726184-63728202 | p3 | (CAA)5 | AAGTGTGCGACAAAGAGGCT | GAAAACAAGGCGGTTATGGA | 169 |
| chr6:63726810-63728825 | p2 | (TA)9 | GGATACAACGAGTCCCGAAA | GCCACTGAATGCCCTTTTTA | 250 |
| chr6:63726810-63728825 | p3 | (CAA)5 | GGATACAACGAGTCCCGAAA | GCCACTGAATGCCCTTTTTA | 250 |
| chr6:63795973-63798013 | p2 | (AT)20 | GTTGCAAACAAAAGCCGATT | GAGCAACAACCTGCTTCACA | 101 |
| chr6:63830111-63832126 | p3 | (CAT)5 | TCATAAGGACGGGACAGGAC | CCCACAGGTAGACAGGGAGA | 215 |
| chr6:63856714-63858768 | p3 | (ATG)5 | AAAGGGGATTTTGGGAATTG | GGGCAGCATTTCCTTTGTTA | 192 |
| chr6:63856714-63858768 | p2 | (TA)27 | AAAGGGGATTTTGGGAATTG | GGGCAGCATTTCCTTTGTTA | 192 |
| chr6:63896621-63898639 | p2 | (TA)9 | GGGTGAAATGGTGTTTGTCC | ACCACTACGAAGCGAGGTGT | 190 |
| chr6:64083799-64085814 | p3 | (TCA)5 | CAAAATGTAGGACTTGGCCG | GTCTTATGGCCAGGTTGCAT | 114 |
| chr6:64083799-64085814 | p5 | (ACTCA)3 | CAAAATGTAGGACTTGGCCG | GTCTTATGGCCAGGTTGCAT | 114 |
| chr6:64096610-64098634 | p3 | (ATC)8 | GAGCGGCTGAAAATCATAGG | GTTCATGCACACAGGTGGAC | 260 |
| chr6:64190470-64192482 | p2 | (AT)6 | GCAAGCAGTTGACAAGGTGA | GCCGAATCTCATCACTCCAT | 263 |
| chr6:64190470-64192482 | p2 | (AT)16 | GCAAGCAGTTGACAAGGTGA | GCCGAATCTCATCACTCCAT | 263 |
| chr6:64205298-64207322 | p3 | (ATC)8 | TTTACAAGCAAATCGTGCCA | CTCCACTCCTCCAGAGTTGC | 138 |
| chr6:64218164-64220176 | p4 | (AGAA)4 | AACGGCCTCAACGCTAGATA | CTTGGGCCTAATTTTCTCCC | 171 |
| chr6:64218164-64220176 | p2 | (GT)6 | AACGGCCTCAACGCTAGATA | CTTGGGCCTAATTTTCTCCC | 171 |
| chr6:64464690-64466702 | p2 | (TA)6 | GGTGTTTATGCGAAAAGGGA | GTTTTGCAAGTGGTTGAGCA | 126 |
| chr6:64470003-64472021 | p3 | (TCT)6 | CTAGTGAATTTGTCCGCGCT | AGTCGGGCATTTGTATGCTT | 223 |

| Marker ID | SSR type | SSR motif | FORWARD PRIMER (5'-3') | REVERSE PRIMER (5'-3') | size (bp) |
|---|---|---|---|---|---|
| chr6:64496864-64498876 | p2 | (AT)6 | TGTGGAGTATGAACGGTGGA | ATATGAGTGTCCAGCTCGGG | 199 |
| chr6:64496864-64498876 | p2 | (TA)44 | TGTGGAGTATGAACGGTGGA | ATATGAGTGTCCAGCTCGGG | 199 |
| chr6:64550441-64552459 | p3 | (AAG)5 | CCCAACGGAGGTCAGAATTA | TGACCCTTTTACCCTCGCTA | 251 |
| chr6:64550441-64552459 | p6 | (ATCTAT)3 | CCCAACGGAGGTCAGAATTA | TGACCCTTTTACCCTCGCTA | 251 |
| chr6:64555391-64557403 | p2 | (AG)6 | AGGTGTCGAGATGGAATTGG | GATCTGGTGGAGATGGTGGT | 116 |
| chr6:64555405-64557429 | p2 | (AG)6 | AGGTGTCGAGATGGAATTGG | GATCTGGTGGAGATGGTGGT | 102 |
| chr6:64644333-64646353 | p5 | (ATCAT)4 | ACGTCACTGACCAAACACGA | CGGAGCCCATACTTTTGGTA | 256 |
| chr6:64725036-64727048 | p2 | (TA)6 | TAAACGGACAAAACCTCCCA | AGTTCCCATATTGTCGCCAG | 119 |
| chr6:64811273-64813288 | p5 | (TCATT)3 | GAAGTGCCCATTGCCATACT | GATCGCATACTCTCTTCCGC | 101 |
| chr6:64889537-64891557 | p5 | (TTTAA)4 | AGTTGGCAGGTGATAGTGGG | ATCCTACCGAGCACCTCTGA | 273 |
| chr6:64911932-64913947 | p3 | (AAC)5 | GTTTTTGGAAAATCGAGGCA | CCGATTGGCATTCAAGAGAT | 147 |
| chr6:64965370-64967388 | p3 | (TTG)6 | GGCTGTGTGGCCAGTTTTAT | CATCTCGTCGATGTCTGCAT | 286 |
| chr6:65048794-65050816 | p2 | (TA)11 | ACACTGTCGGGGCATTAGTC | TACAGGGTGCCAAATGGTTT | 167 |
| chr6:65078188-65080206 | p6 | (AAATAA)3 | TTGCCAATAGGGCATAAAGG | TATGGGGTTGATGGGAGTGT | 208 |
| chr6:65090195-65092210 | p5 | (TGATA)3 | GTCAGTTTTTCGCTCTTGGC | GCAGACAACCCCTAGGACAA | 295 |
| chr6:65250626-65252644 | p6 | (ACTGAA)3 | TTGCTCTAAGACATGGCACG | CATGTGGTTCGTTTGGTCAC | 300 |
| chr6:65344319-65346355 | p2 | (AT)18 | AAACCTACTGTCACGACCCG | GCGTGGGTAGTATGCATGTG | 230 |
| chr6:65446266-65448280 | p2 | (TA)7 | ACGGTCCGTATTTCAAGTCG | GAATGACGTTTGACGACCCT | 251 |
| chr6:65446266-65448280 | p2 | (AT)11 | ACGGTCCGTATTTCAAGTCG | GAATGACGTTTGACGACCCT | 251 |
| chr6:65493285-65495303 | p6 | (CAGAGG)3 | CGTCACTACTTGCCACCTGA | ACCATAGTCCGAGCATACCG | 278 |
| chr6:65617682-65619758 | p2 | (AT)6 | ACCGAGCACCTCTGGTCTAA | CCTCGTGGGTTTATGCATCT | 210 |
| chr6:65635893-65637905 | p2 | (TA)6 | CATGCATGCTTAGCTTTCCA | TACACGTTCCGCAGCAATAC | 204 |
| chr6:65635893-65637905 | p2 | (TA)6 | CATGCATGCTTAGCTTTCCA | TACACGTTCCGCAGCAATAC | 204 |
| chr6:65635893-65637905 | p2 | (TA)6 | CATGCATGCTTAGCTTTCCA | TACACGTTCCGCAGCAATAC | 204 |
| chr6:65635893-65637905 | p2 | (TA)6 | CATGCATGCTTAGCTTTCCA | TACACGTTCCGCAGCAATAC | 204 |
| chr6:65635893-65637905 | p2 | (TA)6 | CATGCATGCTTAGCTTTCCA | TACACGTTCCGCAGCAATAC | 204 |
| chr6:65635893-65637905 | p2 | (TA)6 | CATGCATGCTTAGCTTTCCA | TACACGTTCCGCAGCAATAC | 204 |
| chr6:65635893-65637905 | p2 | (TA)7 | CATGCATGCTTAGCTTTCCA | TACACGTTCCGCAGCAATAC | 204 |
| chr6:65659682-65661694 | p2 | (AC)6 | CAAATGCGACAAAAAGGGTT | CGGGAATTGAGATTTGAGGA | 122 |
| chr6:65676220-65678234 | p2 | (AC)7 | ATCAAATCCCGTTACGTCCA | AGTGCTGTCGAAACTTCGGT | 291 |
| chr6:65711175-65713193 | p3 | (CAA)6 | CACATCACAACAAAGGCCAC | CTTCAAGGTGGGATAGAGCG | 279 |
| chr6:65856917-65858933 | p4 | (GAAA)4 | TTTCCTCCGAAAAGTCGCTA | GCTACCTGCGCTCTCCTCTA | 226 |
| chr6:66007194-66009220 | p2 | (AT)13 | TTGACCTGAAGCAACTGGTG | CACCCAAAAGGGCAAGACTA | 268 |
| chr6:66022891-66024903 | p2 | (GT)6 | AGTGCGAGAAGTTAAGGGCA | TTTATTCATTTGGGCCAAGG | 242 |
| chr6:66087061-66089083 | p2 | (AT)9 | CCAAAGGTTCATAACGCCAT | TGTAGTGACAGCCTTGTCGG | 279 |
| chr6:66087061-66089083 | p2 | (AT)11 | CCAAAGGTTCATAACGCCAT | TGTAGTGACAGCCTTGTCGG | 279 |

| Marker ID | SSR type | SSR motif | FORWARD PRIMER (5'-3') | REVERSE PRIMER (5'-3') | size (bp) |
|---|---|---|---|---|---|
| chr6:66201552-66203567 | p3 | (CAA)5 | CGGACGGAATCATAGCAAGT | TTTGCTGACTCCCTCGAGTT | 136 |
| chr6:66277669-66279684 | p5 | (TGTGG)3 | CTCAAGATAATCGCGGGAAG | TTCCTCCATTTTTCCACGAG | 177 |
| chr6:66386059-66388074 | p5 | (AAACC)3 | CGATGGAACGATTGAAAGGT | GAAGAATGGGCTACATGGGA | 137 |
| chr6:66386801-66388816 | p5 | (AAACC)3 | TCCCTACCGTTGAGGGTGTA | CTGAGAAAACCCGAGGTTGA | 206 |
| chr6:66393656-66395678 | p2 | (AT)11 | ATCAGTTCCTCCAATGTCGG | TCTTTCCAAGCCTGCTCAAT | 222 |
| chr6:66404510-66406522 | p2 | (AT)6 | GGTCTTGAAGCGAAAGCAAC | ATAGGTGAAGCCGTGCACTT | 117 |
| chr6:66488662-66490677 | p5 | (ATATA)3 | TGCTCCACGCCCTACTACTT | CCAAAGCCACGGTTGATACT | 293 |
| chr6:66488662-66490677 | p5 | (ATATA)3 | TGCTCCACGCCCTACTACTT | CCAAAGCCACGGTTGATACT | 293 |
| chr6:66750409-66752424 | p5 | (AATGC)3 | GCAACTACCTTTCTGCCTCG | CTTGTTGCGCCCTTAGTTTC | 111 |
| chr6:66754709-66756727 | p3 | (TCG)6 | AACCCCTCGAGGCTACAAAT | CTATCTCGCGCTCCATAAGC | 275 |
| chr6:66806447-66808467 | p2 | (AT)10 | AGTTTCCAACGCAACAAACC | TGGGTCGGGTCTCAACTTAG | 237 |
| chr6:66811737-66813753 | p4 | (AAAT)4 | GGAGAAGGCTCCTCGTTCTT | TCCGTCCTCTCCATGCTATC | 277 |
| chr6:66925483-66927501 | p6 | (TCATTA)3 | TTGGACGGTGACAGAGACTG | TCATTCCTTTCTTTGGACGG | 183 |
| chr6:66931340-66933356 | p4 | (TATT)4 | ATATAGGGGAAATGCTGCCC | TGCTTCACAAAGATCATGCC | 269 |
| chr6:66945764-66947798 | p2 | (AT)17 | CATATGGACAAGGGGGTGTC | AGAAAACCGAACCACAATCG | 195 |
| chr6:66981541-66983555 | p2 | (AT)7 | ACTTTTGTCACGACCCAACC | TGTCTGTAGTGGCAGCCTTG | 173 |
| chr6:67007898-67009916 | p6 | (CATATA)3 | GGCGGCATTACTGAAACAAT | AGCCGGGTCCAAAGTAGAGT | 128 |
| chr6:67130884-67132896 | p2 | (TA)6 | TTTGTTGTGCCGACATGTTT | ATAGGCGCAAAACGAAAGAA | 287 |
| chr6:67149560-67151575 | p5 | (AATTC)3 | GTATCTGTACCTGCGGGCAT | CCTTTTGTGTCCGGTTTGTT | 111 |
| chr6:67202785-67204803 | p6 | (TATATG)3 | TTGAAAAACATCGTGTGGGA | GCTTATTTGTCCAAGCTCGC | 109 |
| chr6:67262757-67264787 | p2 | (TA)15 | TAATCACGCTTTTTCCCGTC | TCCCCAAATTTTTGCTCTTG | 197 |
| chr6:67268360-67270372 | p2 | (TA)6 | AAAGGGCAAGGAGTTCGATT | GGTCTCGCCCTTAACTTTCC | 287 |
| chr6:67317241-67319259 | p3 | (CAT)6 | ACTTTTGTCACGACCCAACC | TATGCGTCGCTTGAGACAAC | 142 |
| chr6:67317257-67319269 | p3 | (CAT)6 | ACTTTTGTCACGACCCAACC | TATGCGTCGCTTGAGACAAC | 126 |
| chr6:67338921-67340937 | p4 | (TTTA)4 | GTCTAGACCAGGTCGTGGGA | CACAACCCAATTTAGGACCG | 188 |
| chr6:67362558-67364612 | p2 | (TA)27 | ACCACCACCCAAGCGTATTA | TCTAGGCTGTGGAAAATGGG | 152 |
| chr6:67362558-67364612 | p2 | (TA)17 | ACCACCACCCAAGCGTATTA | TCTAGGCTGTGGAAAATGGG | 152 |
| chr6:67370166-67372181 | p3 | (CAA)5 | ACAACAAAGGCCACATCACA | CATGCATATGGAAGGCACAG | 232 |
| chr6:67387173-67389185 | p2 | (TA)6 | AGTTATGGGCAGGGCTTTTT | TACACACATGAAGGACCCGA | 255 |
| chr6:67434960-67436975 | p5 | (ATTTC)3 | GCGATCCAGATACCCACACT | CACTCGGGGATTCTACGAAA | 185 |
| chr6:67439864-67441880 | p2 | (AT)8 | GTCAACACCATTCTTCCGCT | AAATCGGGCACTAGTCATGG | 180 |
| chr6:67477795-67479813 | p6 | (CAACAC)3 | CTTTCCCCCTTTCCCATAAA | CTTCTTAAGTCCCGAAGGGG | 230 |
| chr6:67477795-67479813 | p3 | (AAC)6 | CTTTCCCCCTTTCCCATAAA | CTTCTTAAGTCCCGAAGGGG | 230 |
| chr6:67481964-67483986 | p2 | (AT)11 | TTTATTCCCTTTTCCCGACC | ACTACGCACGTGAAAGGACC | 188 |
| chr6:67481964-67483986 | p2 | (TA)12 | TTTATTCCCTTTTCCCGACC | ACTACGCACGTGAAAGGACC | 188 |
| chr6:67488204-67490254 | p2 | (TA)25 | TCCGCTACTGCCTGAAAAGT | GATTGCCTCGGGATTAGACA | 127 |

| Marker ID | SSR type | SSR motif | FORWARD PRIMER (5'-3') | REVERSE PRIMER (5'-3') | size (bp) |
|---|---|---|---|---|---|
| chr6:67498153-67500168 | p3 | (TTA)5 | ATTGGTTTTGCCACTTCTCG | GCGGGAGCCAACATAGAATA | 170 |
| chr6:67523232-67525246 | p3 | (TTG)6 | TGTTGTTGTTGTTGGGCTGT | TAACGCTTATCCTCCCATGC | 162 |
| chr6:67523232-67525246 | p2 | (TA)9 | TGTTGTTGTTGTTGGGCTGT | TAACGCTTATCCTCCCATGC | 162 |
| chr6:67523232-67525246 | p2 | (AT)7 | TGTTGTTGTTGTTGGGCTGT | TAACGCTTATCCTCCCATGC | 162 |
| chr6:67577174-67579189 | p3 | (ACA)5 | ATGGAACGAATCAAACGGAG | TTTATATCCGCCACCTCGAC | 238 |
| chr6:67732093-67734108 | p2 | (TA)9 | CGACCACTATCTTGCCCTTC | TCCATCTTAAGGGATCGTGC | 104 |
| chr6:67732093-67734108 | p3 | (CAT)5 | CGACCACTATCTTGCCCTTC | TCCATCTTAAGGGATCGTGC | 104 |
| chr6:67779477-67781492 | p3 | (ATC)5 | TTTTTGGCTGCTGTAGCCTT | CGGCGGACATTTTAGGAATA | 177 |
| chr6:67786948-67788986 | p2 | (TA)19 | TCTCGACCTTACATCACCCC | ATGTCCAGCCAAATAGGCAC | 102 |
| chr6:67791226-67793240 | p2 | (TA)7 | CTTGTACCGAGCACCCCTTA | TGGTCCGTGTTCTGGTGTAA | 158 |
| chr6:67813666-67815684 | p3 | (TTG)6 | GACCCTTGAGCCAAATCAAA | CCCAGCTTGACCTTTCTGTC | 174 |
| chr6:68096573-68098588 | p2 | (TG)6 | CGGAAATATTGAGGAAGGCA | AAATTAGCACATGGGCTTGG | 260 |
| chr6:68096573-68098588 | p3 | (TTG)5 | CGGAAATATTGAGGAAGGCA | AAATTAGCACATGGGCTTGG | 260 |
| chr6:68245669-68247687 | p6 | (CAAAAT)3 | CTTCCAAGACAACCCTCCAA | TGAAGGAAGGGTGAGCAAGT | 109 |
| chr6:68296124-68298142 | p2 | (AT)6 | CTCCCTGGTATTTGAGGCAA | TGGTTGGGATGTTGAGATGA | 115 |
| chr6:68296124-68298142 | p6 | (TACTCA)3 | CTCCCTGGTATTTGAGGCAA | TGGTTGGGATGTTGAGATGA | 115 |
| chr6:68310645-68312657 | p2 | (CT)6 | AATACCCAGTCGCCAATCTG | CGATGTGGGCGAAACTAAAT | 227 |
| chr6:68312158-68314176 | p2 | (AT)9 | GTCCCTTTCTTCAAATGGCA | ATCCCCCGATCTCTTTATGG | 201 |
| chr6:68394609-68396623 | p2 | (AT)7 | AGAGTCTGAATGGTGCGCTT | GAGGGACTCAGTTCACAGCC | 111 |
| chr6:68486414-68488446 | p2 | (AT)16 | TTGGAGACGCTCCATAAAGG | ACGTGGAAACAGGCCAATAG | 276 |
| chr6:68486414-68488446 | p4 | (GTAT)6 | TTGGAGACGCTCCATAAAGG | ACGTGGAAACAGGCCAATAG | 276 |
| chr6:68500988-68503003 | p3 | (CAT)5 | TTACCGAACAATAGCGACCC | CGTAGGCCATCCATTCTGTT | 232 |
| chr6:68592867-68594895 | p5 | (ATCAT)3 | TAGCATACGGGCCATTTAGG | TGGGTAATGGGTCATGTGTG | 101 |
| chr6:68592867-68594895 | p2 | (AT)14 | TAGCATACGGGCCATTTAGG | TGGGTAATGGGTCATGTGTG | 101 |
| chr6:68723416-68725432 | p2 | (TA)8 | AGACCCAAAGGTTCACAACG | GCCGGCTTGCATATATTGTT | 141 |
| chr6:68827228-68829244 | p2 | (AT)24 | CTCCAGTTAGAGGCTCGTGG | TGACTAGTGGCTTGTCACGG | 221 |
| chr6:68827228-68829244 | p4 | (ATTA)4 | CTCCAGTTAGAGGCTCGTGG | TGACTAGTGGCTTGTCACGG | 221 |
| chr6:68875574-68877589 | p3 | (GAA)5 | CAAGGTGATGTACTGGGCCT | TGTCCCGGTGGTTAATGAAT | 154 |
| chr6:68903788-68905800 | p2 | (AT)6 | GCATGATTGAGCCGGACTAT | CCAAGTCCGAAACGGAAATA | 157 |
| chr6:68991395-68993410 | p3 | (TTG)5 | CCTCAAAATTTTCGCGTCAT | GGAAACTACACCCGACGAGA | 197 |
| chr6:68991655-68993669 | p3 | (TTG)5 | AGTTTCCAACGCAACAAACC | CGGAGAAATTCTGTTCAGGC | 205 |
| chr6:68991694-68993710 | p3 | (TTG)5 | AGTTTCCAACGCAACAAACC | CGGAGAAATTCTGTTCAGGC | 166 |
| chr6:68991711-68993731 | p3 | (TTG)5 | AGTTTCCAACGCAACAAACC | CGGAGAAATTCTGTTCAGGC | 149 |
| chr6:69006208-69008220 | p5 | (ATTCA)3 | TCAATCCCAACAACCCAACT | TGGTGTCGTGGTGGAAGTTA | 269 |
| chr6:69006208-69008220 | p5 | (AGGGG)3 | TCAATCCCAACAACCCAACT | TGGTGTCGTGGTGGAAGTTA | 269 |
| chr6:69045369-69047383 | p2 | (AT)7 | ATACTTGCGTCGAGAATGGG | CAATGCCCTAACTGTCAGCA | 253 |

| Marker ID | SSR type | SSR motif | FORWARD PRIMER (5'-3') | REVERSE PRIMER (5'-3') | size (bp) |
|---|---|---|---|---|---|
| chr6:69045369-69047383 | p2 | (AT)11 | ATACTTGCGTCGAGAATGGG | CAATGCCCTAACTGTCAGCA | 253 |
| chr6:69045834-69047856 | p2 | (AT)7 | CAGTTCCCGAGATGGACACT | TCGTGGGTCGTATTTGTTCA | 253 |
| chr6:69045834-69047856 | p2 | (AT)11 | CAGTTCCCGAGATGGACACT | TCGTGGGTCGTATTTGTTCA | 253 |
| chr6:69148718-69150752 | p2 | (AT)17 | CATTACCAACCACCAGGACC | GGGCTGTTTTTGTAAGCCAA | 171 |
| chr6:69189409-69191424 | p5 | (ATTCA)3 | TTCGGGTCAATCCGACTTAG | CATTGATGGTGACGAGGATG | 244 |
| chr6:69291951-69293969 | p2 | (AG)9 | GGGTGTTTCCTGCAGATGAT | ACAACACTTTAACCGACGCC | 241 |
| chr6:69345510-69347528 | p6 | (ATGTAT)3 | TTGTGAATTTGCCCCTTTTC | GACGTCACCTGTGAGGGTTT | 263 |
| chr6:69361300-69363324 | p2 | (TA)12 | ACTTTTGTCACGACCCAACC | TCTTGATAGCCTTGTCGGCT | 176 |
| chr6:69394040-69396058 | p3 | (CAT)6 | TTACCGAACAATAGCGACCC | GTTGAAGTGCTCCCTCGTTC | 272 |
| chr6:69394040-69396058 | p4 | (AAAG)4 | TTACCGAACAATAGCGACCC | GTTGAAGTGCTCCCTCGTTC | 272 |
| chr6:69407265-69409287 | p2 | (AT)11 | GTCCCTTTCTTCAAATGGCA | ATCCCCCGATCTCTTTATGG | 197 |
| chr6:69407265-69409287 | p2 | (AT)14 | GTCCCTTTCTTCAAATGGCA | ATCCCCCGATCTCTTTATGG | 197 |
| chr6:69412146-69414160 | p2 | (AC)7 | TACAGACCTTGCCCGATTTC | GCATTTACAGGGGTGATGCT | 278 |
| chr6:69421396-69423408 | p2 | (AC)6 | TACATACCTTGCCCGCTTTC | TGAATTCTCGGGGATGTTTC | 284 |
| chr6:69431196-69433211 | p5 | (TTCTT)3 | AGGGAAGCATACACAATGCC | ATGTTTTGTCTGCCTGGTCC | 204 |
| chr6:69487008-69489020 | p2 | (AG)6 | AATATCGATGCCTCACAGGG | GAAGCGGAAGAAAGTATGCG | 224 |
| chr6:69487008-69489020 | p3 | (GTT)5 | AATATCGATGCCTCACAGGG | GAAGCGGAAGAAAGTATGCG | 224 |
| chr6:69657622-69659637 | p3 | (ACA)5 | TCACCCATTGCCTGAGTGTA | ATAAAGGCCTTGTCGGTCCT | 195 |
| chr6:69660388-69662403 | p5 | (AAAAT)3 | CTGTCCCGTCATATTTCGCT | CCATGACTAGTGCCCAACCT | 127 |
| chr6:69660388-69662403 | p2 | (AT)8 | CTGTCCCGTCATATTTCGCT | CCATGACTAGTGCCCAACCT | 127 |
| chr6:69675786-69677802 | p2 | (AT)6 | GCCCGATTTTCTATAAGCCC | TCGCCCCTTTTAGCCTATTT | 263 |
| chr6:69675786-69677802 | p2 | (TA)8 | GCCCGATTTTCTATAAGCCC | TCGCCCCTTTTAGCCTATTT | 263 |
| chr6:69675786-69677802 | p2 | (AT)7 | GCCCGATTTTCTATAAGCCC | TCGCCCCTTTTAGCCTATTT | 263 |
| chr6:69763595-69765655 | p2 | (TA)30 | GCTCCAGTAGGTGCTCTTGG | TGAGAGGAGCAGTAGGGCAT | 300 |
| chr6:69768804-69770819 | p2 | (AT)6 | CCCTTTTAAGGCGAATGTCA | TATTACCGCGCCTTAACGAC | 223 |
| chr6:69768804-69770819 | p3 | (ATC)5 | CCCTTTTAAGGCGAATGTCA | TATTACCGCGCCTTAACGAC | 223 |
| chr6:69860313-69862337 | p2 | (TA)12 | ACCTAAAATGGATGACCCCC | GCAAATTTCCACGTTGTCCT | 287 |
| chr6:69860313-69862337 | p3 | (CAA)6 | ACCTAAAATGGATGACCCCC | GCAAATTTCCACGTTGTCCT | 287 |
| chr6:69936240-69938254 | p2 | (TA)7 | GGCCACATTTATAGGCAGGA | AGGTGAAGTGTGCTCCTCGT | 142 |
| chr6:69957495-69959553 | p2 | (AT)29 | AACAATTGATCTTGACGCCC | GGAGGCATTGGAAGAACAAA | 247 |
| chr6:69991001-69993015 | p2 | (AT)7 | TGCAGCAGTTCTCGTACCAC | GATAAGGGGTGCTCGGTACA | 215 |
| chr6:70088242-70090258 | p2 | (AT)9 | ATGATGCCAAAGGTGCCTAC | TACATACATAGCGTGCCCGA | 147 |
| chr6:70088242-70090258 | p2 | (AT)8 | ATGATGCCAAAGGTGCCTAC | TACATACATAGCGTGCCCGA | 147 |
| chr6:70097555-70099585 | p5 | (TGAAT)6 | GCCCCCTTGAAATCTTTCTC | TTTCCAACTTGCACAACCAA | 276 |
| chr6:70143307-70145322 | p2 | (CA)6 | AACGAACTCACACCCAGACC | TGTGTGAAATTTGGCGATGT | 151 |
| chr6:70143307-70145322 | p3 | (ACA)5 | AACGAACTCACACCCAGACC | TGTGTGAAATTTGGCGATGT | 151 |

| Marker ID | SSR type | SSR motif | FORWARD PRIMER (5'-3') | REVERSE PRIMER (5'-3') | size (bp) |
|---|---|---|---|---|---|
| chr6:70267035-70269053 | p2 | (AT)9 | CAAACCGATGAGTCATGTCG | GAGGCCTAGTCGTCTTGCAC | 107 |
| chr6:70267035-70269053 | p3 | (TCA)5 | CAAACCGATGAGTCATGTCG | GAGGCCTAGTCGTCTTGCAC | 107 |
| chr6:70314469-70316483 | p2 | (CT)7 | TTCCTCACAAAACCACTCCC | CAATTTGGTGGAACCGTAGG | 275 |
| chr6:70435011-70437071 | p2 | (TA)6 | TCCACATTTCACCCCTTCTC | GCTGGGTCGTGACATTTTCT | 127 |
| chr6:70435011-70437071 | p2 | (AT)30 | TCCACATTTCACCCCTTCTC | GCTGGGTCGTGACATTTTCT | 127 |
| chr6:70580303-70582315 | p2 | (TA)6 | CAGTGTCAGTCCGCTTGGTA | TTGGGAACGAGAGCAAGTTT | 232 |
| chr6:70580303-70582315 | p4 | (TTTA)4 | CAGTGTCAGTCCGCTTGGTA | TTGGGAACGAGAGCAAGTTT | 232 |
| chr6:70629970-70631988 | p2 | (TA)9 | TTCTCGACCTTACGTCACCC | TGCCCTTTTGACATGGTGTA | 162 |
| chr6:70667911-70669927 | p2 | (TA)8 | CCCGATTTTCGTTAAACCCT | GCTTTGTCGCACTTTTAGCC | 100 |
| chr6:70667911-70669927 | p2 | (AT)6 | CCCGATTTTCGTTAAACCCT | GCTTTGTCGCACTTTTAGCC | 100 |
| chr6:70674658-70676692 | p2 | (AT)6 | AATGCTGCTCGATTTTCGTT | GTTCGTAAATCGGTAGCCCA | 133 |
| chr6:70780551-70782563 | p2 | (TA)6 | CGTGTTGTTGTTTTGGTTGG | AAACAATTTCACGGCGTTTC | 128 |
| chr6:70780551-70782563 | p2 | (TA)6 | CGTGTTGTTGTTTTGGTTGG | AAACAATTTCACGGCGTTTC | 128 |
| chr6:70824796-70826811 | p5 | (TTCAT)3 | TCCCTTTTCACTCCCCTTCT | CTTGATTTTGCGGGTTTGAT | 143 |
| chr6:70824796-70826811 | p3 | (ATG)5 | TCCCTTTTCACTCCCCTTCT | CTTGATTTTGCGGGTTTGAT | 143 |
| chr6:70824796-70826811 | p2 | (TA)12 | TCCCTTTTCACTCCCCTTCT | CTTGATTTTGCGGGTTTGAT | 143 |
| chr6:70908019-70910034 | p2 | (AT)6 | ATGCTTACTCGTCAATGGGG | AGACGCTAGCAATGGAAGGA | 212 |
| chr6:70908019-70910034 | p3 | (ACA)5 | ATGCTTACTCGTCAATGGGG | AGACGCTAGCAATGGAAGGA | 212 |
| chr6:70943581-70945596 | p5 | (AAAAC)3 | GATGTCTGCGCGTTACAAGA | TACGTCGTTGTGTTTGCTCC | 271 |
| chr6:71016475-71018487 | p2 | (CT)6 | CCTCTGTCCATGACAACCCT | TTTTGTTTATTGCCCGCTTC | 230 |
| chr6:71028694-71030712 | p6 | (ATATGA)3 | GTTATGGCCATGGATATGGG | TCAACTCCCAAACGTCAACA | 162 |
| chr6:71046426-71048441 | p5 | (TTTCA)3 | AAGAAATGACATGGGAAGCG | ATAACAATGCCTCACAGGGC | 165 |
| chr6:71063842-71065854 | p2 | (TG)6 | AAAAAGAAACTCCCCCGAAA | ATTTTGCCCTTGACCTTCCT | 272 |
| chr6:71109794-71111812 | p2 | (AT)9 | CCCGTAAGCGTTATCCAAAA | CCATTCGTCATAACCATCCC | 182 |
| chr6:71130931-71132946 | p5 | (TCTCT)3 | CCATGTCAGTTCAATTCCCC | TAACTCTACCGGCCACCAAG | 213 |
| chr6:71377456-71379470 | p2 | (AT)7 | TACCTGTCGCAGATCAATCG | GGCGGGTTATATAGGGGAAA | 195 |
| chr6:71402216-71404230 | p2 | (AT)7 | TACCTGTCGCAGATCAATCG | GGCGGGTTATATAGGGGAAA | 195 |
| chr6:71507337-71509353 | p2 | (AT)8 | TGTTTTGGCACGTTGTTGTT | GCCCACACCATATTCCAAAC | 145 |
| chr6:71549017-71551035 | p6 | (CATATA)3 | ATACTCAGAGCCGAGCGAAA | CTTTGCAGACAAAGCCATGA | 138 |
| chr6:71628871-71630887 | p4 | (CCCA)4 | CAATCCAAACGGGCTCTAAG | CAAAGCCCAAGGAAAATTCA | 282 |
| chr6:71658091-71660106 | p3 | (CAA)5 | TGCACGATCCGTAATTTTGA | AGGGTCTTGGGTGTCAGTTG | 139 |
| chr6:71658091-71660106 | p5 | (ATATA)3 | TGCACGATCCGTAATTTTGA | AGGGTCTTGGGTGTCAGTTG | 139 |
| chr6:71658091-71660106 | p2 | (AT)13 | TGCACGATCCGTAATTTTGA | AGGGTCTTGGGTGTCAGTTG | 139 |
| chr6:71658091-71660106 | p3 | (ACA)11 | TGCACGATCCGTAATTTTGA | AGGGTCTTGGGTGTCAGTTG | 139 |
| chr6:71683728-71685758 | p2 | (TA)15 | AAGCGGTCCGTTTTGTTATG | GCGGTCGCACAAATAAAAAT | 275 |
| chr6:71740253-71742268 | p5 | (AATGC)3 | CCCTTACCCGTTAACCCATT | TGACTTTCAGATTTTGGGGG | 224 |

| Marker ID | SSR type | SSR motif | FORWARD PRIMER (5'-3') | REVERSE PRIMER (5'-3') | size (bp) |
|---|---|---|---|---|---|
| chr6:71768304-71770316 | p3 | (TGT)5 | CCAGAGCCCATCTTTTGGTA | ACCCTGCACGCAGAAAATAC | 133 |
| chr6:71768304-71770316 | p2 | (TA)6 | CCAGAGCCCATCTTTTGGTA | ACCCTGCACGCAGAAAATAC | 133 |
| chr6:71806546-71808558 | p2 | (CT)6 | ATCCGAGGATACCCCAAAAC | TAGGAGTTGAGGCCATTGCT | 269 |
| chr6:71907887-71909901 | p2 | (TA)7 | TCCACTACATCAACCCACCA | AAGTCCCACATCGGTTGAAG | 214 |
| chr6:72004430-72006450 | p4 | (TATT)5 | GCTAGTGGCAATTACTCCGC | GTGGAATTGACACATGGCAG | 130 |
| chr6:72032516-72034531 | p3 | (CAT)5 | GCTCCTGGGTGACCAGAATA | GGAAATGAGTGGTGCTTGGT | 262 |
| chr6:72121944-72123962 | p5 | (ATACA)3 | TAACTAAAGGGCCATGACGG | ATTGGCTCGTGCCTATGAGT | 175 |
| chr6:72121990-72124044 | p5 | (ATACA)3 | TAACTAAAGGGCCATGACGG | ATTGGCTCGTGCCTATGAGT | 129 |
| chr6:72237965-72239983 | p3 | (CAA)6 | TGAATAGAATTGCCTTCGGG | ACGGGATTTGAATTTTGACG | 220 |
| chr6:72237965-72239983 | p2 | (CA)9 | TGAATAGAATTGCCTTCGGG | ACGGGATTTGAATTTTGACG | 220 |
| chr6:72241907-72243939 | p2 | (AT)16 | TCCTTGGGTTTCTTCCTCCT | AGCCATATTAATGCGGTTGC | 299 |
| chr6:72241907-72243939 | p2 | (AT)6 | TCCTTGGGTTTCTTCCTCCT | AGCCATATTAATGCGGTTGC | 299 |
| chr6:72379067-72381089 | p2 | (TA)11 | CTGGCGGGTTCAAGTGTTAT | ATTAATGTCACGCCCCAAAA | 265 |
| chr6:72379067-72381089 | p2 | (TA)19 | CTGGCGGGTTCAAGTGTTAT | ATTAATGTCACGCCCCAAAA | 265 |
| chr6:72406967-72408982 | p3 | (TTG)5 | TCCTTTGGGGATTCTCCTCT | CGCATCTTGTCTTCCACTGA | 213 |
| chr6:72409966-72411986 | p2 | (AT)10 | CAATGAAACAGACGCCTTCA | TCCGCCTTACGTGAGAGACT | 200 |
| chr6:72415236-72417252 | p2 | (AC)8 | TGCGGTGAAGTACTCAGGTG | TCATATTTTGCAACCCACCA | 179 |
| chr6:72415236-72417252 | p2 | (AT)11 | TGCGGTGAAGTACTCAGGTG | TCATATTTTGCAACCCACCA | 179 |
| chr6:72456713-72458734 | p3 | (ATA)12 | TGTTCTCCCATGTGGACTCA | TTCGCGTTCTAGTTCGGAAT | 273 |
| chr6:72456713-72458734 | p3 | (ATA)5 | TGTTCTCCCATGTGGACTCA | TTCGCGTTCTAGTTCGGAAT | 273 |
| chr6:72456761-72458785 | p3 | (ATA)12 | TGTTCTCCCATGTGGACTCA | TTCGCGTTCTAGTTCGGAAT | 225 |
| chr6:72456794-72458809 | p3 | (ATA)12 | TGTTCTCCCATGTGGACTCA | TTCGCGTTCTAGTTCGGAAT | 192 |
| chr6:72460044-72462059 | p3 | (ATA)5 | TCAAACGGTGCTAAAAGAGC | AAAACTGACTGCCAAGCAGC | 103 |
| chr6:72460044-72462059 | p3 | (TAA)5 | TCAAACGGTGCTAAAAGAGC | AAAACTGACTGCCAAGCAGC | 103 |
| chr6:72462500-72464518 | p6 | (TTTTTC)3 | CACGTGGCATGATCCTATTG | CGTTGGGCTAGTCCATTGAT | 228 |
| chr6:72522814-72524832 | p2 | (TA)9 | CGAGCATCTCTACCCCCATA | CAGAGCTCCATTTCTTTCGG | 251 |
| chr6:72587584-72589602 | p3 | (GTT)6 | ATTGATCCGGTGCTTCAGTC | CAACAAAATACACGCCGATG | 296 |
| chr6:72627996-72630011 | p5 | (AATGC)3 | CAGTAGAACCACCAGCAGCA | ATCATTTGTTCGAGGGTTCG | 273 |
| chr6:72648963-72650978 | p5 | (TTATT)3 | TTGGATTCACCGTCATCAGA | TGCGGTCTGCTAGTGTTGTC | 156 |
| chr6:72739764-72741824 | p2 | (TA)30 | ACGAGCGTATCTGACTGCCT | TTGAATCTCCCACACAACCA | 194 |
| chr6:72745798-72747846 | p2 | (AG)24 | AAGAACATGGCCACACACAA | CATCCGCATTTATTTCCACC | 185 |
| chr6:72769835-72771849 | p2 | (TA)7 | TGCTCTCGGATCTTTCCCTA | TCTCAGGGTTCGTTTTGGTC | 243 |
| chr6:72870199-72872214 | p5 | (ATGAT)3 | TTTGAAGGAAACGGGATTTG | AAAAGTTGCAGGACAATCCG | 200 |
| chr6:72945443-72947473 | p2 | (AT)15 | GTCCACATTCCCCTACCCTT | AAGAATGCTTAGGCTTGCGA | 291 |
| chr6:72955296-72957322 | p2 | (AT)13 | AGGGGAAACTCTGCCAAAAT | TGAGCACGATGAAAAGAACG | 248 |
| chr6:72955296-72957322 | p2 | (AT)7 | AGGGGAAACTCTGCCAAAAT | TGAGCACGATGAAAAGAACG | 248 |

| Marker ID | SSR type | SSR motif | FORWARD PRIMER (5'-3') | REVERSE PRIMER (5'-3') | size (bp) |
|---|---|---|---|---|---|
| chr6:72955423-72957437 | p2 | (AT)13 | AGGGGAAACTCTGCCAAAAT | TGAGCACGATGAAAAGAACG | 121 |
| chr6:72955423-72957437 | p2 | (AT)7 | AGGGGAAACTCTGCCAAAAT | TGAGCACGATGAAAAGAACG | 121 |
| chr6:72965723-72967735 | p2 | (TC)6 | AACACCACCAACAACAACGA | ATTTGAAAATGCGAAGTCGG | 148 |
| chr6:73004071-73006086 | p3 | (GTT)7 | ATTGGGCTGTTTTGGAAGTG | ACATAACGAAATTCGGGCAG | 271 |
| chr6:73004071-73006086 | p3 | (TTG)7 | ATTGGGCTGTTTTGGAAGTG | ACATAACGAAATTCGGGCAG | 271 |
| chr6:73004071-73006086 | p5 | (TATAT)3 | ATTGGGCTGTTTTGGAAGTG | ACATAACGAAATTCGGGCAG | 271 |
| chr6:73004071-73006086 | p2 | (TA)7 | ATTGGGCTGTTTTGGAAGTG | ACATAACGAAATTCGGGCAG | 271 |
| chr6:73010375-73012390 | p2 | (GA)19 | ACGTTTTCGGGTGATCTGAC | CAGGACCACGACCTATGGTT | 157 |
| chr6:73010375-73012390 | p3 | (GTT)5 | ACGTTTTCGGGTGATCTGAC | CAGGACCACGACCTATGGTT | 157 |
| chr6:73035259-73037274 | p5 | (CAAAT)3 | AGAGCCCACTTGGATAGGGT | ACACGGTAAGGGCATGCTAC | 208 |
| chr6:73035259-73037274 | p2 | (AT)6 | AGAGCCCACTTGGATAGGGT | ACACGGTAAGGGCATGCTAC | 208 |
| chr6:73043135-73045153 | p6 | (ATACAT)3 | CACACTACTCCAGGGGGAAA | GGCGGATATCATGCATTCTT | 183 |
| chr6:73182691-73184703 | p2 | (AC)6 | CCCTGAGGTCCAACTTCAAA | GATGCTGCCCAAACTTCTGT | 124 |
| chr6:73318921-73320941 | p2 | (TA)10 | TCGTGCATGCCTCAATGTAT | ACAGTACCCGGCCCTCTAGT | 233 |
| chr6:73333469-73335497 | p2 | (TA)14 | ACGCCTCGGCAATATGATAG | ACATATGATTGCCCTTTGGC | 118 |
| chr6:73400438-73402500 | p2 | (TA)31 | ATACTTATGGGCATGGCGAG | ACCGACAAGGCCAAATACAA | 138 |
| chr6:73425976-73427988 | p2 | (TA)6 | TGGCATATGATCCGATTTGA | AGAAGCAGAAGCCATACGGA | 201 |
| chr6:73483610-73485630 | p2 | (TA)10 | TCACGAACCTGTTTCATCCA | TCAAAGGCAGATGCAGAATG | 276 |
| chr6:73496245-73498260 | p5 | (AATGC)3 | CTCAAGAATGGAGTTGGGGA | TTGCAAATCAGTGAAACCCA | 210 |
| chr6:73573385-73575400 | p5 | (TTGGT)3 | GGACAGTGGCGAAATGCTAT | GTAATGTCGGGTTGGTTTGG | 147 |
| chr6:73583651-73585669 | p3 | (CAA)6 | ATCATCATTAGCCAGCGTCC | AATCGGAGGTTACCGACCTT | 258 |
| chr6:73685515-73687530 | p3 | (TGA)5 | TTTGTTAGCCTTGTCGGCTT | ACGGGCACCTCTCATCATAC | 172 |
| chr6:73685515-73687530 | p2 | (TA)12 | TTTGTTAGCCTTGTCGGCTT | ACGGGCACCTCTCATCATAC | 172 |
| chr6:73685515-73687530 | p2 | (TA)8 | TTTGTTAGCCTTGTCGGCTT | ACGGGCACCTCTCATCATAC | 172 |
| chr6:73833627-73835681 | p2 | (AT)27 | GGGAACGTCCCAGCTATACA | ATTCGTAGTGGCAGCCTTGT | 238 |
| chr6:73851936-73853948 | p2 | (TG)6 | GGCAAAGAATCCGTATCGAA | GCTCGCTTTCGTAATTCCAC | 174 |
| chr6:73922863-73924875 | p2 | (TA)6 | TATGGATAGTCGGGCCACTC | ACAGATTCGTCTGGACACC | 198 |
| chr6:73930049-73932071 | p3 | (TTG)8 | CCCAATTTCGGCAGATTCTA | GGCGAACATCATCAAAACCT | 268 |
| chr6:73930049-73932071 | p3 | (ATG)13 | CCCAATTTCGGCAGATTCTA | GGCGAACATCATCAAAACCT | 268 |
| chr6:73930049-73932071 | p2 | (TA)11 | CCCAATTTCGGCAGATTCTA | GGCGAACATCATCAAAACCT | 268 |
| chr6:73930049-73932071 | p2 | (AT)12 | CCCAATTTCGGCAGATTCTA | GGCGAACATCATCAAAACCT | 268 |
| chr6:73955747-73957765 | p6 | (CATATA)3 | CCCTAGGACGAAGTGCTCTG | TCCACCAGCTGATGTGTCAT | 291 |
| chr6:73957676-73959690 | p2 | (AT)7 | AGCGAACATCATGCTACACG | TGAATTTGGAAAGGTTTCGG | 250 |
| chr6:73957676-73959690 | p2 | (AT)8 | AGCGAACATCATGCTACACG | TGAATTTGGAAAGGTTTCGG | 250 |
| chr6:73957811-73959827 | p2 | (AT)7 | AGCGAACATCATGCTACACG | TGAATTTGGAAAGGTTTCGG | 115 |
| chr6:73957811-73959827 | p2 | (AT)8 | AGCGAACATCATGCTACACG | TGAATTTGGAAAGGTTTCGG | 115 |

| Marker ID | SSR type | SSR motif | FORWARD PRIMER (5'-3') | REVERSE PRIMER (5'-3') | size (bp) |
|---|---|---|---|---|---|
| chr6:73969775-73971787 | p2 | (AC)6 | ACATGCCAAAAGAATGAGGG | TTTATATGGGAGGTGCTGCC | 213 |
| chr6:73971751-73973815 | p2 | (AG)32 | GCTAAAGGGTAACAGCGACG | GCAGCTGGTCGCTAAAAGTC | 293 |
| chr6:74039445-74041460 | p5 | (GAAAT)3 | CCTCGTCCATCTTGGACTGT | CTTGTGTTTTCATGCGTTGG | 197 |
| chr6:74190284-74192296 | p6 | (CATGAG)3 | CTTACTAGGACAAGGCCCCC | CTCCGCTCATTAGGATCTGC | 237 |
| chr6:74190284-74192296 | p2 | (AT)6 | CTTACTAGGACAAGGCCCCC | CTCCGCTCATTAGGATCTGC | 237 |
| chr6:74265807-74267819 | p2 | (GT)6 | CAGGCAACCAGTAGCAACAA | TCCTCCATGAAACCCATCTC | 296 |
| chr6:74417792-74419812 | p2 | (AT)10 | GATGCCTCGGATGTTTCACT | GACTTTCCCGGTACGATTCA | 264 |
| chr6:74433422-74435444 | p2 | (AT)11 | ATCACGACCTAACCGGAGTG | TGCCTTTTCGGCTCATACTT | 273 |
| chr6:74459912-74461930 | p2 | (AT)9 | CGTGTGGAATGTTGGCATAG | ACAACACTCCAAAACAGCCC | 133 |
| chr6:74459912-74461930 | p2 | (AT)12 | CGTGTGGAATGTTGGCATAG | ACAACACTCCAAAACAGCCC | 133 |
| chr6:74470217-74472233 | p4 | (ATTT)4 | AAGTTTCCCGTTTTGCCTTT | AAACACGATGCAATGCCATA | 225 |
| chr6:74606358-74608373 | p3 | (CAT)5 | CTATGCTCTAGCCGGACGAC | GGATCACCAACCGGAGTAGA | 113 |
| chr6:74621834-74623855 | p3 | (TAT)7 | TGTCGTGTCATGAAGGCAAT | GAAGAAACAAGTCGCCCAAA | 104 |
| chr6:74642686-74644706 | p2 | (AT)10 | TCGTGAAACCATGCTAACCA | CCATTTTCTTTGGACGCTGT | 222 |
| chr6:74653572-74655587 | p5 | (ATATG)3 | AAGAGAAATGGTGGAATGCG | TCCTTCGATATCGCGTCTCT | 263 |
| chr6:74653572-74655587 | p2 | (TA)10 | AAGAGAAATGGTGGAATGCG | TCCTTCGATATCGCGTCTCT | 263 |
| chr6:74653572-74655587 | p2 | (AT)8 | AAGAGAAATGGTGGAATGCG | TCCTTCGATATCGCGTCTCT | 263 |
| chr6:74653865-74655885 | p5 | (ATATG)3 | GGAGAACAAGCGAGCGATAG | TCCCCCGATTGTATTATGGA | 193 |
| chr6:74653865-74655885 | p2 | (TA)10 | GGAGAACAAGCGAGCGATAG | TCCCCCGATTGTATTATGGA | 193 |
| chr6:74653865-74655885 | p2 | (AT)8 | GGAGAACAAGCGAGCGATAG | TCCCCCGATTGTATTATGGA | 193 |
| chr6:74729021-74731043 | p2 | (TA)11 | CATCTATCAGGTGTGGCCCT | TTTCTTCGCCATTATTTGCC | 194 |
| chr6:74729021-74731043 | p3 | (TTG)5 | CATCTATCAGGTGTGGCCCT | TTTCTTCGCCATTATTTGCC | 194 |
| chr6:74729560-74731572 | p2 | (TA)11 | ATCTCTCAGACGGTCCCTCA | TCGACCTTCCTCCATATTCG | 252 |
| chr6:74729560-74731572 | p3 | (TTG)5 | ATCTCTCAGACGGTCCCTCA | TCGACCTTCCTCCATATTCG | 252 |
| chr6:74729599-74731619 | p2 | (TA)11 | ATCTCTCAGACGGTCCCTCA | TCGACCTTCCTCCATATTCG | 213 |
| chr6:74729599-74731619 | p3 | (TTG)5 | ATCTCTCAGACGGTCCCTCA | TCGACCTTCCTCCATATTCG | 213 |
| chr6:74729669-74731681 | p2 | (TA)11 | ATCTCTCAGACGGTCCCTCA | TCGACCTTCCTCCATATTCG | 143 |
| chr6:74729669-74731681 | p3 | (TTG)5 | ATCTCTCAGACGGTCCCTCA | TCGACCTTCCTCCATATTCG | 143 |
| chr6:74735178-74737220 | p2 | (AT)21 | GCCCTAGATTACGTGGCAAA | GCCTAGCACGCTTAACTTCG | 202 |
| chr6:74735178-74737220 | p4 | (AAAG)4 | GCCCTAGATTACGTGGCAAA | GCCTAGCACGCTTAACTTCG | 202 |
| chr6:74748442-74750457 | p5 | (GCATT)3 | ACGAATTTCCGCATTTTGAC | AATCACTCATTTGCCTTGCC | 186 |
| chr6:74751364-74753380 | p4 | (CATA)4 | CCGGTAAATACAGGGACGAA | TCATTGCTCTCCAGAAACCC | 138 |
| chr6:74768984-74771050 | p2 | (TA)33 | GTGGTGCTCGGTAGGTTAGC | CTTCAACACTTTTGTCCGCA | 104 |
| chr6:74864651-74866666 | p3 | (ATA)5 | CCCTAATGTCACGACCCAGT | TGGATACGCAGTGTGGGTTA | 172 |
| chr6:74899708-74901730 | p2 | (AT)11 | TCACTTGGACTCGCAATCTG | ATACTTCTCCCACTCCGCCT | 201 |
| chr6:74906554-74908569 | p3 | (CAT)5 | TCGTTGTGATGCTCAACTCC | ATTGTGGTATCCTTGTCGGC | 132 |

| Marker ID | SSR type | SSR motif | FORWARD PRIMER (5'-3') | REVERSE PRIMER (5'-3') | size (bp) |
|---|---|---|---|---|---|
| chr6:74908822-74910884 | p2 | (AT)31 | TGGTTCCAAGGTTGTCCTTC | GGCGACGTTGTCATACCTCT | 278 |
| chr6:74974923-74976941 | p3 | (CAA)6 | ACATCACAACAAAGGCCACA | CAGTTGATAGGGGGTGTGCT | 179 |
| chr6:75006071-75008087 | p2 | (AG)8 | TTGGCAAGAAGGGAAAGCTA | CAATACACATGCGGAGCATC | 156 |
| chr6:75100395-75102407 | p2 | (TA)6 | TGGCGCAATAAACACATCAT | TGCACATTTCCACTGCATTT | 163 |
| chr6:75199688-75201718 | p2 | (TA)15 | GATGATATTACACCGCGCCT | AATCATTAGCCTGCGTCCAG | 131 |
| chr6:75207262-75209282 | p5 | (TTCAT)4 | TCTTCTTCTGTCGCCCAGAT | ATTCACTGCACATCAACGGA | 280 |
| chr6:75219842-75221854 | p2 | (TA)6 | GGATCGAGAACCGAATAGCA | CCTACGGGATTAGGTCGTGA | 251 |
| chr6:75264653-75266681 | p2 | (TA)14 | TTTGAGAGGAAATTGTGGGG | TGGGCAACGTCTCCTTTATC | 278 |
| chr6:75304053-75306081 | p2 | (TA)14 | TTTGAGAGGAAATTGTGGGG | TGGGCAACGTCTCCTTTATC | 278 |
| chr6:75344252-75346264 | p2 | (CA)6 | CAGAGCCATCAGATTCGACA | TGACCCTTCCATCTCCTCAC | 188 |
| chr6:75346469-75348481 | p2 | (AC)6 | AACAAAAGGGTTCAATTGCG | GATTTGAAGAGCGTAAGGCG | 128 |
| chr6:75360029-75362047 | p3 | (TTG)6 | GACCCTTGAGCCAAATCAAA | TGCCTTAAAACGCCCATATC | 167 |
| chr6:75388954-75390969 | p3 | (ATG)5 | TCGGTGGTTACCGACCTTAC | TCATTAGCCCATATCCGAGC | 247 |
| chr6:75388954-75390969 | p3 | (ATC)5 | TCGGTGGTTACCGACCTTAC | TCATTAGCCCATATCCGAGC | 247 |
| chr6:75388954-75390969 | p3 | (ACA)11 | TCGGTGGTTACCGACCTTAC | TCATTAGCCCATATCCGAGC | 247 |
| chr6:75389324-75391357 | p3 | (ATC)5 | GCATGTATGGTATCCGCCTT | TCCGCTGGTAAAGCTCCTAA | 161 |
| chr6:75389324-75391357 | p3 | (ACA)11 | GCATGTATGGTATCCGCCTT | TCCGCTGGTAAAGCTCCTAA | 161 |
| chr6:75444695-75446707 | p2 | (AT)6 | CGTTTTCGTGCGCATATTTA | AGGGGCTCGGTAATAAGGAA | 162 |
| chr6:75444695-75446707 | p2 | (AT)13 | CGTTTTCGTGCGCATATTTA | AGGGGCTCGGTAATAAGGAA | 162 |
| chr6:75473872-75475888 | p2 | (TA)6 | CCCTGGGTTGTTTATGTTGG | TTTGTGACTGAAGCTGCCAC | 229 |
| chr6:75473872-75475888 | p2 | (AT)8 | CCCTGGGTTGTTTATGTTGG | TTTGTGACTGAAGCTGCCAC | 229 |
| chr6:75633077-75635092 | p4 | (ATAA)6 | TGTGGAAGAAGCTTATGGGG | AATATAGTGCCCCCAGCCTT | 121 |
| chr6:75633077-75635092 | p3 | (AAC)5 | TGTGGAAGAAGCTTATGGGG | AATATAGTGCCCCCAGCCTT | 121 |
| chr6:75642046-75644062 | p4 | (AAAT)4 | TTGCGACCATTGTTCTACCA | GGAATGCACACCCTTCTCAT | 215 |
| chr6:75733982-75735997 | p3 | (TTA)11 | CGGGTCTAGGGTAGGTAGGG | GGGTTCATTTCCCATTTCCT | 286 |
| chr6:75733982-75735997 | p3 | (TGT)5 | CGGGTCTAGGGTAGGTAGGG | GGGTTCATTTCCCATTTCCT | 286 |
| chr6:75733982-75735997 | p5 | (GGATG)3 | CGGGTCTAGGGTAGGTAGGG | GGGTTCATTTCCCATTTCCT | 286 |
| chr6:75734127-75736142 | p3 | (TTA)11 | CGGGTCTAGGGTAGGTAGGG | GGGTTCATTTCCCATTTCCT | 141 |
| chr6:75734127-75736142 | p3 | (TGT)5 | CGGGTCTAGGGTAGGTAGGG | GGGTTCATTTCCCATTTCCT | 141 |
| chr6:75734127-75736142 | p5 | (GGATG)3 | CGGGTCTAGGGTAGGTAGGG | GGGTTCATTTCCCATTTCCT | 141 |
| chr6:75886767-75888785 | p3 | (CAA)6 | TCAGCGTGTCGATCAGTACC | ACCCCAGATAAAAGGCAACC | 152 |
| chr6:75972079-75974094 | p3 | (CAT)5 | GCAGGAATTACGGTCCAAGA | AGTTGTGCCCGGATGAATAG | 133 |
| chr6:76003738-76005750 | p2 | (AT)6 | AAAGTATGATGAGCGGTGCC | TGGAAGTCGATGAAAGAGGG | 194 |
| chr6:76003738-76005750 | p2 | (TA)8 | AAAGTATGATGAGCGGTGCC | TGGAAGTCGATGAAAGAGGG | 194 |
| chr6:76005209-76007227 | p2 | (TA)8 | TGGTTCCATCCGGTGATAGT | TCAAAGCAATCCATGAGTGG | 185 |
| chr6:76005209-76007227 | p5 | (ATTAA)3 | TGGTTCCATCCGGTGATAGT | TCAAAGCAATCCATGAGTGG | 185 |

| Marker ID | SSR type | SSR motif | FORWARD PRIMER (5'-3') | REVERSE PRIMER (5'-3') | size (bp) |
|---|---|---|---|---|---|
| chr6:76005209-76007227 | p5 | (ATTTT)3 | TGGTTCCATCCGGTGATAGT | TCAAAGCAATCCATGAGTGG | 185 |
| chr6:76005209-76007227 | p3 | (TTA)6 | TGGTTCCATCCGGTGATAGT | TCAAAGCAATCCATGAGTGG | 185 |
| chr6:76040197-76042212 | p2 | (TA)17 | TATAACGCCTCGCCATTTTC | CGACCTTACGTCACTCCGAT | 207 |
| chr6:76040197-76042212 | p3 | (CAA)5 | TATAACGCCTCGCCATTTTC | CGACCTTACGTCACTCCGAT | 207 |
| chr6:76068759-76070783 | p2 | (AT)7 | CCTCAGAGGCATGGTCAGTT | ACAACATTCCCAAGCAAAGG | 219 |
| chr6:76068759-76070783 | p2 | (TA)12 | CCTCAGAGGCATGGTCAGTT | ACAACATTCCCAAGCAAAGG | 219 |
| chr6:76068759-76070783 | p2 | (AT)9 | CCTCAGAGGCATGGTCAGTT | ACAACATTCCCAAGCAAAGG | 219 |
| chr6:76068970-76070988 | p2 | (AT)7 | CCTTTGCTTGGGAATGTTGT | TAGACTGCAACACCGGATCA | 180 |
| chr6:76068970-76070988 | p2 | (TA)12 | CCTTTGCTTGGGAATGTTGT | TAGACTGCAACACCGGATCA | 180 |
| chr6:76068970-76070988 | p2 | (AT)9 | CCTTTGCTTGGGAATGTTGT | TAGACTGCAACACCGGATCA | 180 |
| chr6:76082113-76084125 | p2 | (AT)6 | CAACTTTGGCTCATCCCATT | AGTGACCCACATCCCCAATA | 262 |
| chr6:76191724-76193749 | p5 | (AATTA)5 | GTTCGGCCAATTGAAAAAGA | AAAATTGCGCATTTGACCTC | 276 |
| chr6:76191724-76193749 | p2 | (AT)10 | GTTCGGCCAATTGAAAAAGA | AAAATTGCGCATTTGACCTC | 276 |
| chr6:76404212-76406227 | p3 | (GCT)5 | AACTTGGACAAACGACGACC | TTGTATGTGGCCGTGACACT | 159 |
| chr6:76408484-76410526 | p3 | (TTA)14 | CATGCACACATGTACACCCA | CCTCACAAATCTCCCTCCAA | 221 |
| chr6:76486884-76488899 | p5 | (ATCAT)3 | AGTTGTCACGTGCTGGACAC | GTAGCGCCATTACAGGGGTA | 287 |
| chr6:76499281-76501299 | p6 | (TGAACA)3 | GGGGACGATCTTATAGGGGA | ATGTGGTTCACTCGGTCACA | 228 |
| chr6:76499281-76501299 | p2 | (TA)13 | GGGGACGATCTTATAGGGGA | ATGTGGTTCACTCGGTCACA | 228 |
| chr6:76518728-76520766 | p2 | (AT)19 | CTCTTTCAATTCTGCGGGAG | AATTTTCGGATGTGTTCCCA | 144 |
| chr6:76552049-76554063 | p2 | (AT)7 | AAATGTTGCCAGGAATTTCG | CCGCTCCTTATGCTATTCCA | 254 |
| chr6:76552049-76554063 | p2 | (AT)7 | AAATGTTGCCAGGAATTTCG | CCGCTCCTTATGCTATTCCA | 254 |
| chr6:76689022-76691098 | p4 | (TTAA)4 | ACTCAACCACGAACCCAAAG | TTCTCAGCCGTTGATGTTTG | 205 |
| chr6:76689112-76691148 | p4 | (TTAA)4 | ACTCAACCACGAACCCAAAG | TTCTCAGCCGTTGATGTTTG | 115 |
| chr6:76689272-76691288 | p4 | (TTAA)4 | AACGGCTGAGAAAAAGCAAA | TTTTCACCAAACTCAACCCC | 209 |
| chr6:76702040-76704088 | p2 | (AT)7 | TTGGCTCTATCGCAGGATCT | TCCTAGGGATGTCCATGAGC | 225 |
| chr6:76702096-76704140 | p2 | (AT)7 | TTGGCTCTATCGCAGGATCT | TCCTAGGGATGTCCATGAGC | 169 |
| chr6:76842850-76844862 | p2 | (AT)6 | ATAAGATGCCAACTCCACGG | GTTGCTTCGGGTCAACAAAT | 162 |
| chr6:76870784-76872802 | p2 | (TA)9 | TCTAAAGACCAAACACCCCG | GGTCGTTGACGGTTCTTGAT | 114 |
| chr6:76939496-76941511 | p5 | (ATCAC)3 | CTGCCTTGGGAAACTCACTC | TGATTTGGTTGTGCGATGTT | 150 |
| chr6:76980912-76982930 | p6 | (ATTATG)3 | TTGGTAAGGCCTTGAGGATG | GTTGTCCTCGTCGTTTCGTT | 213 |
| chr6:77010297-77012315 | p6 | (TATTCG)3 | GTCTCGTAACCCCCTTGACA | GGGTCGTGACACATGACTTG | 130 |
| chr6:77010297-77012315 | p2 | (AT)6 | GTCTCGTAACCCCCTTGACA | GGGTCGTGACACATGACTTG | 130 |
| chr6:77150150-77152174 | p2 | (AT)11 | TCCGGTTTTCCTAATTTCCC | TCGCCATATGTTACGGTTCA | 283 |
| chr6:77150150-77152174 | p2 | (AT)12 | TCCGGTTTTCCTAATTTCCC | TCGCCATATGTTACGGTTCA | 283 |
| chr6:77181752-77183774 | p2 | (TA)11 | TATGAGTCAGGACAAGGCCC | TGTACAGCTAGGATGCGTGG | 192 |
| chr6:77230418-77232436 | p6 | (TTTTTA)3 | ATGGAAGGTGGTGGTGTCAT | TTCACAAACAGAATGCTCGC | 155 |

| Marker ID | SSR type | SSR motif | FORWARD PRIMER (5'-3') | REVERSE PRIMER (5'-3') | size (bp) |
|---|---|---|---|---|---|
| chr6:77478300-77480316 | p2 | (AT)8 | GAAAGAAAGAACGCAGTGCC | GTTGGCTCTCCAGTGTGGAT | 132 |
| chr6:77478300-77480316 | p2 | (TA)8 | GAAAGAAAGAACGCAGTGCC | GTTGGCTCTCCAGTGTGGAT | 132 |
| chr6:77478300-77480316 | p5 | (TGAAT)3 | GAAAGAAAGAACGCAGTGCC | GTTGGCTCTCCAGTGTGGAT | 132 |
| chr6:77483912-77485934 | p2 | (TA)11 | GAAAGAAAGAACGCAGTGCC | TGTCCTTCCCAGGTCATTTC | 114 |
| chr6:77483912-77485934 | p5 | (TGAAT)3 | GAAAGAAAGAACGCAGTGCC | TGTCCTTCCCAGGTCATTTC | 114 |
| chr6:77621444-77623459 | p5 | (TATGA)3 | GAGATAATTGGGAGGCCGTT | CCATCTTCCCTCCCCTTTAC | 156 |
| chr6:77716671-77718699 | p2 | (TA)26 | GGTTGAGAGTTCTCTTGCCG | ATACCTCCACCACGTCTTCG | 122 |
| chr6:77716671-77718699 | p2 | (AT)14 | GGTTGAGAGTTCTCTTGCCG | ATACCTCCACCACGTCTTCG | 122 |
| chr6:77716671-77718699 | p3 | (TCA)6 | GGTTGAGAGTTCTCTTGCCG | ATACCTCCACCACGTCTTCG | 122 |
| chr6:77732691-77734706 | p3 | (TCA)5 | TTTGGGTGTCTTCTTTTGGG | ATTCGAGTGGCATCAAGACC | 148 |
| chr6:77759443-77761458 | p5 | (TTATA)3 | GTGTACATTGCCCGAGGTCT | GATCATGAATGATGGGGGAC | 158 |
| chr6:77975555-77977570 | p3 | (AAC)5 | ATAACAAAGGCCACAGCGTC | CGTCACTCCGGTAAAGTGGT | 103 |
| chr6:77990687-77992702 | p3 | (TGA)5 | TTGGTAATGTCTTGAGGGGC | CGGACCGTATAAGTGGTCGT | 296 |
| chr6:78123084-78125160 | p2 | (AT)6 | CTGCTCACATGGAAACTGGA | CCTTGTCGGCTTGTGGTTAT | 201 |
| chr6:78164416-78166438 | p2 | (AT)11 | TCCCAAGTAAGAATGGTGCC | CGTGCAGGTTTTGGGATAGT | 123 |
| chr6:78195866-78197888 | p2 | (AT)11 | ATCCGGGAACTGGAAACTCT | GTTCACCAAGTGGTGCTCCT | 112 |
| chr6:78218214-78220240 | p2 | (AT)13 | GATCCGAGGTTGTTACCGAA | CCAGGCCTTCTCTCCTTACC | 150 |
| chr6:78260582-78262600 | p6 | (TTGAAC)3 | TTCAGCGATGTGTTCCTGAG | GACATCGAACACCTCCACCT | 297 |
| chr6:78294447-78296459 | p2 | (GA)6 | CGATTGCGACTGCAAAGTAA | GCCTTGAGGAGCTACTGGTG | 188 |
| chr6:78500476-78502548 | p3 | (ATA)24 | TTCGAACTTGCGTATGCTTG | TGGTTGTGAGTGGAATCGAA | 147 |
| chr6:78549743-78551807 | p2 | (TA)32 | TACCAGATATGGGAGACCGC | GCGTGTGCTTGAATACGAGA | 213 |
| chr6:78552914-78554926 | p4 | (TATG)4 | TTGTGATTGATCCCACCTCA | TTTCGGGACTCGGTGTTATC | 261 |
| chr6:78552914-78554926 | p2 | (TA)6 | TTGTGATTGATCCCACCTCA | TTTCGGGACTCGGTGTTATC | 261 |
| chr6:78570489-78572504 | p3 | (TGT)5 | AAGCCAAATGTGGGCATAAG | AATTTTTCTGTCTGCGCGTT | 275 |
| chr6:78733077-78735098 | p3 | (CAA)7 | CAGGGACTACATAGCGGAGC | TCATATAGGCGGGACAGGAC | 251 |
| chr6:78802886-78804902 | p2 | (AT)21 | TCCAAATAACCGGTCTCCTG | CCTTGTCAGCTTGCGTATCA | 218 |
| chr6:78802886-78804902 | p2 | (AT)8 | TCCAAATAACCGGTCTCCTG | CCTTGTCAGCTTGCGTATCA | 218 |
| chr6:78812873-78814885 | p2 | (AT)11 | ATTTCCTATGGCTGGGCTCT | AGGTTGGGTCGTGACAAAAG | 205 |
| chr6:78812873-78814885 | p2 | (AT)6 | ATTTCCTATGGCTGGGCTCT | AGGTTGGGTCGTGACAAAAG | 205 |
| chr6:78917517-78919532 | p3 | (ATC)5 | AAGCCCACGAAGCTACAAAA | CAGCAGACTTGCAGAAGCAC | 144 |
| chr6:79120139-79122169 | p2 | (AT)15 | TTGACCTACCAAGCACCACA | TGTCAGCTTGCATACTTGGG | 267 |
| chr6:79157831-79159849 | p3 | (ATC)6 | TTTATGAGGTCCGGGTCTTG | GTCCGCGACCCTATTTGATA | 239 |
| chr6:79296279-79298293 | p2 | (AT)7 | GGTGTCTTTTGGGCTTACCA | CAGATAAGGGGTGCTCGGTA | 111 |
| chr6:79320507-79322533 | p2 | (AT)13 | TCGGGTCCCACTTAGTCAAG | AAAGGGCGGAAGAGGTATGT | 107 |
| chr6:79375009-79377021 | p2 | (AT)6 | TTCCGATCTACCTCCCCTTT | AGAGCCCAGCCATAGGAAAT | 224 |
| chr6:79375009-79377021 | p2 | (TA)7 | TTCCGATCTACCTCCCCTTT | AGAGCCCAGCCATAGGAAAT | 224 |

| Marker ID | SSR type | SSR motif | FORWARD PRIMER (5'-3') | REVERSE PRIMER (5'-3') | size (bp) |
|---|---|---|---|---|---|
| chr6:79442122-79444140 | p5 | (TTGTG)3 | ATCCACACCGGAGAGTCAAC | TGGGTTTTCTAACCCACTGC | 128 |
| chr6:79475871-79477889 | p3 | (CAT)6 | AATTACCATGCCCCAAATGA | CCTTTAGCCTTGTCGGTTCA | 249 |
| chr6:79573610-79575622 | p2 | (AT)6 | GCTTTCTTATCTACCGCCCC | GGACATAGCGAGAACTGGGA | 231 |
| chr6:79727441-79729481 | p2 | (TC)20 | TGTCAAATTTCATTGCTCGC | CATTGGTGTTTTGCGAAATG | 179 |
| chr6:79753805-79755817 | p2 | (TA)11 | ACACCCTGTCCCCTTCTCTT | TTCGAAAAATGTTTGGGGAC | 217 |
| chr6:79753805-79755817 | p2 | (AC)6 | ACACCCTGTCCCCTTCTCTT | TTCGAAAAATGTTTGGGGAC | 217 |
| chr6:79804168-79806182 | p2 | (AC)7 | AAAGCCATAACGACCAAACG | TGAGCCCAATATGTGTTCCA | 292 |
| chr6:80086926-80088938 | p2 | (TA)6 | TTAAGCCCACCAAGAAGTGG | AGGGAAAATTTGGGATTTGG | 150 |
| chr6:80105802-80107820 | p6 | (TATACA)3 | CCCGTTGGTTCTCGTTATGT | GGGTCTACAAGCCGATGTGT | 225 |
| chr6:80105802-80107820 | p3 | (CAA)5 | CCCGTTGGTTCTCGTTATGT | GGGTCTACAAGCCGATGTGT | 225 |
| chr6:80115294-80117306 | p2 | (AC)6 | ATAAGGCTGCTACCCGGAAT | GCTCCCCCTAGAAAATGAGG | 223 |
| chr6:80154454-80156486 | p2 | (TA)16 | CCGGTATGGCAAGAGTTTGT | CCGAAGCATACCTAGCCAAG | 212 |
| chr6:80154454-80156486 | p2 | (AT)6 | CCGGTATGGCAAGAGTTTGT | CCGAAGCATACCTAGCCAAG | 212 |
| chr6:80237307-80239321 | p2 | (AT)7 | CGCACAGTCCTAAAATGGGT | TGAAGGCACAATCATGGGTA | 252 |
| chr6:80241705-80243725 | p2 | (TA)10 | CAACTCAGCCACACGAAAAA | TGGTTGAACTTTTGGGGTTC | 186 |
| chr6:80294382-80296414 | p2 | (TA)34 | TCGAGGGATCACGACCTTAC | CATGCAATCTAGCCATGGTG | 119 |
| chr6:80294382-80296414 | p2 | (AT)16 | TCGAGGGATCACGACCTTAC | CATGCAATCTAGCCATGGTG | 119 |
| chr6:80307817-80309849 | p2 | (TA)48 | GATAATTGCGGGAAAGCAAA | CGAAGTAGCTTTTCCAAGCG | 227 |
| chr6:80307817-80309849 | p2 | (TA)16 | GATAATTGCGGGAAAGCAAA | CGAAGTAGCTTTTCCAAGCG | 227 |
| chr6:80347998-80350016 | p6 | (CATATA)3 | GGACACGACTTTGCAGACAA | CATCAGCCGATGTGTCATTC | 268 |
| chr6:80378225-80380243 | p6 | (CATATA)3 | ATGAGCCGTGACAAGTACCC | ATGTGTTGGCCTTGAGTTCC | 217 |
| chr6:80382589-80384601 | p2 | (TA)6 | ATTCAGTATCGGAGGCATGG | GAACGGGTCCCAGTGATAGA | 237 |
| chr6:80433427-80435447 | p2 | (TA)10 | TCCAGAGAATGGTGTCCCTC | TGTTGGGTAGGAATCAAGGG | 147 |
| chr6:80526984-80529096 | p2 | (AT)56 | TAAACATAAGCCGACGAGGC | TGCACAGGGAGACAGACTTG | 229 |
| chr6:80526984-80529096 | p2 | (AT)6 | TAAACATAAGCCGACGAGGC | TGCACAGGGAGACAGACTTG | 229 |
| chr6:80549347-80551362 | p5 | (TATAT)3 | GCAGTTCGTCCTAGGGAGTG | TGAGAAGGAAGGGTCAATGG | 197 |
| chr6:80595438-80597453 | p5 | (CATGT)3 | GCTTTTCACCCATAGACGGA | ACACCTTCGCCCATACAAAG | 189 |
| chr6:80608380-80610398 | p6 | (TCTTTT)3 | TCAAGTGTCGCTTAATCCCC | ATCCAAACCAAATTCCGACA | 249 |
| chr6:80639270-80641284 | p2 | (AG)7 | GGGAAGTCATTTTCCCCATT | CACGAAATTTTCTGGGGTGT | 226 |
| chr6:80639270-80641284 | p3 | (GTT)5 | GGGAAGTCATTTTCCCCATT | CACGAAATTTTCTGGGGTGT | 226 |
| chr6:80639794-80641809 | p2 | (AG)7 | ACGTTTTCGGGTGATCTGAC | ATACAGGCCCACGACCTATG | 187 |
| chr6:80639794-80641809 | p3 | (GTT)5 | ACGTTTTCGGGTGATCTGAC | ATACAGGCCCACGACCTATG | 187 |
| chr6:80681888-80683910 | p2 | (AT)11 | CGAATGGCCTTGTGTTTCTT | CCCAAATGGGAGCAAAATAA | 299 |
| chr6:80681888-80683910 | p2 | (AT)14 | CGAATGGCCTTGTGTTTCTT | CCCAAATGGGAGCAAAATAA | 299 |
| chr6:80682379-80684409 | p2 | (AT)11 | GCCGTATCGTGAGGAGAAGA | ATTAAAGGTGGGCCCCATAC | 281 |
| chr6:80682379-80684409 | p2 | (AT)14 | GCCGTATCGTGAGGAGAAGA | ATTAAAGGTGGGCCCCATAC | 281 |

| Marker ID | SSR type | SSR motif | FORWARD PRIMER (5'-3') | REVERSE PRIMER (5'-3') | size (bp) |
|---|---|---|---|---|---|
| chr6:80682411-80684441 | p2 | (AT)11 | GCCGTATCGTGAGGAGAAGA | ATTAAAGGTGGGCCCCATAC | 249 |
| chr6:80682411-80684441 | p2 | (AT)14 | GCCGTATCGTGAGGAGAAGA | ATTAAAGGTGGGCCCCATAC | 249 |
| chr6:80682794-80684822 | p2 | (AT)11 | ACGATGAAGCATGGGAAACT | GGGCGGTTGAAATTGAAATA | 128 |
| chr6:80682794-80684822 | p2 | (AT)14 | ACGATGAAGCATGGGAAACT | GGGCGGTTGAAATTGAAATA | 128 |
| chr6:80714384-80716398 | p2 | (AG)10 | ATCGGCAATAATGGCAACTC | CCTTTATCGTCCATGCCATT | 178 |
| chr6:80795309-80797357 | p2 | (AT)24 | CAGCAACCCACAACAACAAC | TAAAATCATACATCGGCGCA | 115 |
| chr6:80884290-80886305 | p5 | (CATTG)3 | AGGTGGGTTACGGCTTACCT | AAATCACAACACGGCAACAA | 190 |
| chr6:80913207-80915285 | p2 | (AT)39 | AAACAGTTCCCCACAAGCAC | CAATTCACCATTGGAGACCC | 124 |
| chr6:81039453-81041477 | p2 | (TA)12 | GACGACAAGGAGGTGGATGT | AGCCTTGTCGGCTTGTGTAT | 134 |
| chr6:81046641-81048665 | p2 | (AT)18 | TGTCTTACTTCCGGGGTTTG | ACAGTGGTCCACCCTTGTTC | 167 |
| chr6:81046641-81048665 | p2 | (TG)12 | TGTCTTACTTCCGGGGTTTG | ACAGTGGTCCACCCTTGTTC | 167 |
| chr6:81204179-81206199 | p2 | (AT)10 | TCAGTCAAAGTCTTGTGGCG | CGGCCTCCACGTGTTATACT | 229 |
| chr6:81326265-81328283 | p2 | (TA)9 | GGACGTGGCAGTCTTATGGT | TCCCGAGCACTCATATTTCC | 291 |
| chr6:81393054-81395070 | p2 | (TA)8 | CAATCCCATAGGAGGAGCAA | GGGTTGATACGCATGTGAAA | 286 |
| chr6:81457151-81459171 | p2 | (CA)10 | AAGATTTTCACATGGGCCTG | TTAGGCGTGTTTAGGGCGTA | 229 |
| chr6:81483069-81485084 | p5 | (GAAAG)3 | CACAAACTGCTTCCTCCCTC | CCACTCTAACTGGGGAACCA | 108 |
| chr6:81488660-81490675 | p3 | (TGT)5 | AAGTGGTTGTGAGACCGACC | ACTGGAGCATCTGTGCCTCT | 265 |
| chr6:81488660-81490675 | p5 | (AAAAT)3 | AAGTGGTTGTGAGACCGACC | ACTGGAGCATCTGTGCCTCT | 265 |
| chr6:81500020-81502044 | p6 | (AACTTC)4 | GTTGGGGATTTGAGTGGAGA | CGTACAATGAAGTTGGGCCT | 222 |
| chr6:81554252-81556266 | p2 | (AT)7 | TCCCAAAGTATCCCCATCAA | GACATGGCGGAATAGCAAAT | 242 |
| chr6:81554252-81556266 | p2 | (TG)6 | TCCCAAAGTATCCCCATCAA | GACATGGCGGAATAGCAAAT | 242 |
| chr6:81606937-81608952 | p5 | (CAAAG)3 | TCTCCTCCTACCCCGAAACT | GGGTGTGTTGGAGGAAAATG | 101 |
| chr6:81610473-81612535 | p2 | (AT)6 | AAGTTGGCAGCCGTAGACAT | TTCTTGGAATGCTTGACACG | 196 |
| chr6:81610473-81612535 | p2 | (AT)31 | AAGTTGGCAGCCGTAGACAT | TTCTTGGAATGCTTGACACG | 196 |
| chr6:81647176-81649191 | p3 | (TTG)5 | GATGAATGGGCCAAGAGTGT | TCCATGTGGATTTTTCGTCA | 119 |
| chr6:81659510-81661522 | p3 | (TTG)7 | TGTTAGGAAAAATGGCCTGC | TGTGTGGAAGTTGGTGGATG | 134 |
| chr6:81659510-81661522 | p2 | (TA)6 | TGTTAGGAAAAATGGCCTGC | TGTGTGGAAGTTGGTGGATG | 134 |
| chr6:81659510-81661522 | p2 | (AT)39 | TGTTAGGAAAAATGGCCTGC | TGTGTGGAAGTTGGTGGATG | 134 |
| chr6:81659510-81661522 | p2 | (AT)8 | TGTTAGGAAAAATGGCCTGC | TGTGTGGAAGTTGGTGGATG | 134 |
| chr6:81659510-81661522 | p2 | (AT)14 | TGTTAGGAAAAATGGCCTGC | TGTGTGGAAGTTGGTGGATG | 134 |
| chr6:81870061-81872075 | p2 | (AT)7 | CTCGCTTAACCCTTGAGTGC | AGAGCACCTGCTGGAACAGT | 291 |
| chr6:81870061-81872075 | p2 | (AG)7 | CTCGCTTAACCCTTGAGTGC | AGAGCACCTGCTGGAACAGT | 291 |
| chr6:81885476-81887506 | p2 | (TA)15 | GGCCTGAATTTTATGGCAAA | GATATGTATGGGCCACCAGG | 198 |
| chr6:81885476-81887506 | p3 | (ACA)5 | GGCCTGAATTTTATGGCAAA | GATATGTATGGGCCACCAGG | 198 |
| chr6:81898244-81900256 | p2 | (TA)6 | TTCCATTCCCAATGATGGAT | AGTTGGCCGCTTGTTGTAGT | 145 |
| chr6:81913203-81915219 | p2 | (GA)11 | TTGGACATTAAAACGGCCTC | ATACCCCCTGAGTTGTGCAG | 255 |

| Marker ID | SSR type | SSR motif | FORWARD PRIMER (5'-3') | REVERSE PRIMER (5'-3') | size (bp) |
|---|---|---|---|---|---|
| chr6:81919494-81921508 | p2 | (AT)12 | TAATCTCGGAGCTTTCGCAT | TTGTTGTAGCAGCCTTGTCG | 221 |
| chr6:81919494-81921508 | p2 | (AT)7 | TAATCTCGGAGCTTTCGCAT | TTGTTGTAGCAGCCTTGTCG | 221 |
| chr6:82048467-82050483 | p2 | (AT)8 | CGACGACTACTGACCAAGCA | CGAGGTGGAGGTTATTGCAT | 134 |
| chr6:82081776-82083791 | p5 | (TATAT)3 | AGTTGAGACCCGATGCAAAC | TTTGGGAGAGATCATGAGGG | 103 |
| chr6:82106409-82108424 | p5 | (TATAT)3 | AGTTGAGACCCGATGCAAAC | TTTGGGAGAGATCATGAGGG | 103 |
| chr6:82133963-82135989 | p2 | (AT)13 | GGAAGGAACTTCAACCACGA | TTTGGTTTCATCACATCGGA | 212 |
| chr6:82177318-82179336 | p6 | (ACGCAC)3 | TCGCTTGTGTAATCCCATCA | CGAGGAACTAAAGGCACAGC | 198 |
| chr6:82210825-82212847 | p2 | (AT)11 | TGACGATTCGAGACTGTTGC | TAACTTAATGGCGACGAGCC | 150 |
| chr6:82221094-82223121 | p3 | (ATT)9 | AAATGCACAGGCAATGAACA | CTCCCTGTTGTCTCCCATTC | 186 |
| chr6:82338427-82340441 | p2 | (AT)11 | TGTTAATTCGGTCCCCATGT | TCGGGATCCCTGAGTCATAG | 155 |
| chr6:82338427-82340441 | p2 | (TA)7 | TGTTAATTCGGTCCCCATGT | TCGGGATCCCTGAGTCATAG | 155 |
| chr6:82338427-82340441 | p2 | (AT)16 | TGTTAATTCGGTCCCCATGT | TCGGGATCCCTGAGTCATAG | 155 |
| chr6:82407318-82409336 | p2 | (TA)9 | AAGAGGAAAGCCCTCTCTCG | AGCGACTTGAACGCAGAAAT | 234 |
| chr6:82407318-82409336 | p2 | (AT)6 | AAGAGGAAAGCCCTCTCTCG | AGCGACTTGAACGCAGAAAT | 234 |
| chr6:82407318-82409336 | p6 | (CCGGTT)3 | AAGAGGAAAGCCCTCTCTCG | AGCGACTTGAACGCAGAAAT | 234 |
| chr6:82407510-82409522 | p2 | (TA)9 | ACCCTCTTCTCTCTCTCGCC | CAAAAACGTATCGCCCACTT | 157 |
| chr6:82407510-82409522 | p2 | (AT)6 | ACCCTCTTCTCTCTCTCGCC | CAAAAACGTATCGCCCACTT | 157 |
| chr6:82407510-82409522 | p6 | (CCGGTT)3 | ACCCTCTTCTCTCTCTCGCC | CAAAAACGTATCGCCCACTT | 157 |
| chr6:82434019-82436055 | p3 | (CAA)5 | CATTTGGCCCTGAAGAAAGA | GGCCGATAAAGTCGTCAAAA | 116 |
| chr6:82456876-82458933 | p3 | (TAA)19 | CTTACCACGGAGCATCCATT | CAGGGGATGAAGGAGTTCAA | 121 |
| chr6:82462399-82464414 | p5 | (ATTTT)3 | CGAGACAAAAGTCCTCGTCC | CAAATTGGGATGGGACACTC | 247 |
| chr6:82516413-82518461 | p3 | (ATA)16 | ATTTATCCACAATCGGGCAA | CGCATCAGGTCATTTAAGCA | 256 |
| chr6:82549888-82551902 | p5 | (AAAAT)3 | TGTCGGTTTGTCGCATTTTA | TCGGCAGTTCCTTGTTATCC | 150 |
| chr6:82563268-82565308 | p2 | (AC)20 | CGACATGTTTTGATGATGCC | CCTAGTCGGGTCGTGACAAT | 269 |
| chr6:82870263-82872275 | p2 | (TA)6 | TGTGTAGGGTTCGGTGTTGA | TCAGGCACTCGTCACAACTC | 171 |
| chr6:82883193-82885219 | p2 | (TA)10 | CAAAAAGAAAAGTGCACCCC | TGACATGGCGGAATAACAAA | 135 |
| chr6:82883193-82885219 | p2 | (TA)13 | CAAAAAGAAAAGTGCACCCC | TGACATGGCGGAATAACAAA | 135 |
| chr6:82957073-82959089 | p4 | (ATGA)4 | AGCCCGATTCATTCTACCCT | AACACCGGTGATCTCAGGAC | 282 |
| chr6:82957073-82959089 | p4 | (ATGA)4 | AGCCCGATTCATTCTACCCT | AACACCGGTGATCTCAGGAC | 282 |
| chr6:82973432-82975450 | p3 | (AAT)6 | GTTTGACTAGGGGCGTACCA | CAAATGCATAAAACCACCCC | 153 |
| chr6:83035117-83037141 | p3 | (AAG)5 | TTACAGTGGTGGGGGAAGAG | TTTTGCAACCAGACAATGCT | 225 |
| chr6:83035117-83037141 | p2 | (TA)12 | TTACAGTGGTGGGGGAAGAG | TTTTGCAACCAGACAATGCT | 225 |
| chr6:83140102-83142126 | p6 | (TTCAGT)4 | CTGGGAATTGGGTGAAGAAA | TTCCAGCCTTTGAGCGTAAT | 292 |
| chr6:83164540-83166566 | p2 | (AT)13 | TGACCTTCTTACATTCCGGG | AGGTCAAGAACCCCTCGATT | 264 |
| chr6:83281676-83283708 | p6 | (CATATC)4 | TGGCTCCGGATAAAATGAAG | ATCTCCCGATTCTCCCATCT | 252 |
| chr6:83318850-83320866 | p2 | (AT)8 | TCAGAGTGAAGCTTGAGCGA | AACTTCGAATCCTGGCTCTG | 168 |

| Marker ID | SSR type | SSR motif | FORWARD PRIMER (5'-3') | REVERSE PRIMER (5'-3') | size (bp) |
|---|---|---|---|---|---|
| chr6:83401979-83404007 | p2 | (TA)14 | GCCAAAAGGTTTCTCAACCA | GGGTGCAGGTGCTATTTCAT | 243 |
| chr6:83596481-83598495 | p2 | (TA)7 | ACATAGCCATGGTTGCCTTC | CCATGCCCTCTAGACATGGT | 233 |
| chr6:83631199-83633214 | p5 | (AAGAA)3 | CTTGACTTCTTCAGCGGTCC | AGATGGCCTCGGGAATTAGT | 290 |
| chr6:83720021-83722036 | p3 | (TGT)5 | CAAGTCCCAATCACATGCAC | TAGGTGTTTCATGGCTTCCC | 198 |
| chr6:83720021-83722036 | p3 | (TGT)5 | CAAGTCCCAATCACATGCAC | TAGGTGTTTCATGGCTTCCC | 198 |
| chr6:83720021-83722036 | p5 | (TTGAT)3 | CAAGTCCCAATCACATGCAC | TAGGTGTTTCATGGCTTCCC | 198 |
| chr6:83850043-83852055 | p2 | (TG)6 | GATCACCCTTGTGCAAATCC | TTCCCTTTCAAGTTGATGGC | 145 |
| chr6:83898633-83900651 | p6 | (GTATAA)3 | CAAATCCAATATCCAACCCG | TGCAACGTGATAGGCAAAAA | 157 |
| chr6:83942765-83944777 | p2 | (TA)6 | TGTCCAGTTTGGCCTCTTTT | AATTACGCGATATTCGACGC | 233 |
| chr6:83942765-83944777 | p2 | (TA)30 | TGTCCAGTTTGGCCTCTTTT | AATTACGCGATATTCGACGC | 233 |
| chr6:84025219-84027237 | p6 | (AGGTGC)3 | TCTCAGCCTCTGTGGGAAGT | GGAGTGTCGACCAGTTGGAT | 135 |
| chr6:84163647-84165659 | p2 | (TG)6 | AAAGGTGCCTACGGGCTATT | CTCATACCTGGCCCTACCAA | 235 |
| chr6:84167686-84169704 | p6 | (TCCTCA)3 | GGCTCAACATGGGAACAAGT | ATGTGGATCCAGAACTTGCC | 206 |
| chr6:84170627-84172643 | p2 | (GA)8 | TGAAAGACGGTGGGAAAGAC | CCACGGCCAAAGAAGTGTAT | 196 |
| chr6:84343359-84345379 | p2 | (TA)10 | ACAAAACTTCCAAATCCCCC | GCTCAATGGCTTTTGTCCAT | 243 |
| chr6:84438031-84440043 | p2 | (AG)6 | GAAGGGGTAGGATGCAGTGA | GTCGTTGAGGGGAACAAAAA | 230 |
| chr6:84438031-84440043 | p3 | (TTG)5 | GAAGGGGTAGGATGCAGTGA | GTCGTTGAGGGGAACAAAAA | 230 |
| chr6:84445022-84447034 | p2 | (AC)6 | GGGGAATGAGTGTGGATGTT | CATTACCCAGTTCCCCATTG | 229 |
| chr6:84532611-84534627 | p2 | (GT)8 | GTTGGAGGGCTATGTGGTGT | CGTAGGTTCGCTCCTTCTTG | 199 |
| chr6:84532611-84534627 | p2 | (AT)10 | GTTGGAGGGCTATGTGGTGT | CGTAGGTTCGCTCCTTCTTG | 199 |
| chr6:84540979-84542994 | p3 | (GAT)5 | GGTTGGGTCGTGACACTCTT | TAGAGTTTCCAGCGGCACTT | 296 |
| chr6:84564134-84566152 | p3 | (ATG)6 | GGATTAGCTCTACGCGAACG | AGCTAAACTTTTGTGGCGGA | 115 |
| chr6:84573744-84575765 | p3 | (ATG)7 | CTTCCGCCCACAAAAGTTTA | CCTTGTCCCCACTAGAACCA | 296 |
| chr6:84698128-84700230 | p3 | (ATC)5 | CCGACCTTACGTTACTCCGA | ATGCCTAGCGTTCTGGTGTC | 127 |
| chr6:84783747-84785815 | p2 | (TA)28 | AGATCGAAGTTTACCGGCCT | ATTCCCATATATCCCGCGTT | 290 |
| chr6:84794395-84796407 | p2 | (AT)8 | TGGCTCAATTTTGGGAAGAC | AGTTGGGCTTGAACAAGTGG | 294 |
| chr6:84794395-84796407 | p2 | (AT)6 | TGGCTCAATTTTGGGAAGAC | AGTTGGGCTTGAACAAGTGG | 294 |
| chr6:84803832-84805886 | p2 | (AT)27 | GGCTTGGATTCAAGGTGAAA | CAACAAAACACCACCAATGC | 165 |
| chr6:84803832-84805886 | p2 | (TA)7 | GGCTTGGATTCAAGGTGAAA | CAACAAAACACCACCAATGC | 165 |
| chr6:84933490-84935554 | p2 | (AT)32 | TAAGAGAGGGGGTGTTCGTG | ATTGGGAATATGGCCACTCA | 246 |
| chr6:84933490-84935554 | p2 | (TA)39 | TAAGAGAGGGGGTGTTCGTG | ATTGGGAATATGGCCACTCA | 246 |
| chr6:84962026-84964058 | p2 | (AT)16 | CCTGGGTCCAGGAAACTGTA | AGGCCAGGTTTTCAAGGTTT | 184 |
| chr6:85054824-85056856 | p2 | (AT)12 | CCACATAGACAGCTCGTGGA | TGCCCTGTCCTTATGGATTC | 197 |
| chr6:85054873-85056899 | p2 | (AT)12 | CCACATAGACAGCTCGTGGA | TGCCCTGTCCTTATGGATTC | 148 |
| chr6:85054903-85056951 | p2 | (AT)12 | CCACATAGACAGCTCGTGGA | TGCCCTGTCCTTATGGATTC | 118 |
| chr6:85068190-85070202 | p2 | (GT)6 | GACGGGCTGAATAGGTTCAA | TATCATAGCCGGCACATTCA | 241 |

| Marker ID | SSR type | SSR motif | FORWARD PRIMER (5'-3') | REVERSE PRIMER (5'-3') | size (bp) |
|---|---|---|---|---|---|
| chr6:85208461-85210476 | p5 | (TTATT)3 | ACGTCATCTATGCCGGAAAC | GGGGGTGGTAATACCTCGTT | 234 |
| chr6:85208461-85210476 | p5 | (TTATT)3 | ACGTCATCTATGCCGGAAAC | GGGGGTGGTAATACCTCGTT | 234 |
| chr6:85226868-85228880 | p2 | (AG)6 | GTCGCCTACAAATCAGCCTC | AAAAAGATGTTGCAATGGGC | 213 |
| chr6:85245232-85247247 | p5 | (TATTT)3 | CCAAGGTCACTTTCATCCGT | CTGCAAAGCAAGTCAACGAA | 203 |
| chr6:85245232-85247247 | p2 | (AT)24 | CCAAGGTCACTTTCATCCGT | CTGCAAAGCAAGTCAACGAA | 203 |
| chr6:85258161-85260177 | p4 | (TTCT)4 | AATCTCCGGGCTGATTTTCT | ATGAATGGCACACAACCTCA | 142 |
| chr6:85258161-85260177 | p3 | (GAA)8 | AATCTCCGGGCTGATTTTCT | ATGAATGGCACACAACCTCA | 142 |
| chr6:85274840-85276855 | p5 | (AAACC)3 | TCCCAAGTTGGTTCCCAATA | TTGAGGGGTATGTTTGAGCC | 131 |
| chr6:85363095-85365107 | p2 | (AG)6 | AACCCTTTGCAGCCCTAAAT | TACCATTCTGGAGCTTTGGG | 257 |
| chr6:85380815-85382833 | p6 | (ACTGAA)3 | CGAGATTTTGAGTTTGGGGA | CCATCGCTGTGGTACCTCTT | 222 |
| chr6:85464106-85466124 | p3 | (ATC)6 | CAATTGAAAGCACCTGACGA | AGTGAGCCAGACCAGAAGGA | 171 |
| chr6:85623218-85625236 | p6 | (CAAAAA)3 | CGGTTCGGGTTTCAAAGTTA | TTGTTGGGGTTGGGAAATTA | 151 |
| chr6:86025118-86027132 | p2 | (AT)7 | TACGGATTCAGTTTCGGAGG | ACAAGTGACCGAGGCCATAC | 185 |
| chr6:86067701-86069740 | p3 | (AAT)13 | CCCCCTCTTTCGAATCTCTC | CTGACACCCCACACTTTCCT | 213 |
| chr6:86087407-86089421 | p2 | (TA)7 | CAGGTCCATTCCCTGAAGTG | TTGGGAGCAGATTTCAGAGC | 293 |
| chr6:86087407-86089421 | p5 | (TTTAT)4 | CAGGTCCATTCCCTGAAGTG | TTGGGAGCAGATTTCAGAGC | 293 |
| chr6:86181387-86183403 | p2 | (TA)8 | GGAATCGTTAATTGCATGGC | AAGTGTAAGTGCCACCCGTC | 244 |
| chr6:86254735-86256759 | p2 | (TA)12 | GGTTCTAGGTGGGTCGGATT | CATCGTCATGATCGAAATGG | 294 |
| chr6:86254735-86256759 | p2 | (TA)23 | GGTTCTAGGTGGGTCGGATT | CATCGTCATGATCGAAATGG | 294 |
| chr6:86254873-86256913 | p2 | (TA)12 | GGTTCTAGGTGGGTCGGATT | CATCGTCATGATCGAAATGG | 156 |
| chr6:86254873-86256913 | p2 | (TA)23 | GGTTCTAGGTGGGTCGGATT | CATCGTCATGATCGAAATGG | 156 |
| chr6:86254914-86256954 | p2 | (TA)12 | GGTTCTAGGTGGGTCGGATT | CATCGTCATGATCGAAATGG | 115 |
| chr6:86254914-86256954 | p2 | (TA)23 | GGTTCTAGGTGGGTCGGATT | CATCGTCATGATCGAAATGG | 115 |
| chr6:86265552-86267564 | p2 | (TA)6 | CACCTAGGGTCTCATTCGGA | TTTTTGTGGTGCATGCAGAT | 190 |
| chr6:86357832-86359850 | p6 | (CAATCA)3 | TACTTCCAGACGAGATGGGG | CTCGAAGCTTCCAAACGAAC | 241 |
| chr6:86397180-86399192 | p2 | (AG)6 | TTATTCGCGAAGGAATCGAC | GTCATCTTTTGGACATGGCA | 253 |
| chr6:86556472-86558502 | p2 | (AT)15 | CCTGGATGCAGGCTAATGAT | TGTTTACCTGTGGGCATCAA | 227 |
| chr6:86585870-86587882 | p2 | (AT)6 | CACCTCTACGGGAGGTTTGA | GGCCAAAACCATTTTAAGCA | 297 |
| chr6:86625813-86627825 | p2 | (AT)6 | AGGGAAAATTCACCCCAAAC | CCCAGTACAAGCACAAGCAA | 209 |
| chr6:86649866-86651881 | p5 | (AGAAG)3 | ACGGATCGGAAATGAATGAA | CATCCCTGGCTCCTTCATAA | 136 |
| chr6:86727876-86729898 | p2 | (AT)11 | ACAGGACTCTACCGGACACG | GAGAGTCTGTGGGTTCGCTC | 233 |
| chr6:86736105-86738123 | p6 | (TTCAGT)3 | ACTGACCATAATTCAGCCCG | TGATGGTGGAGGAGAAGACC | 134 |
| chr6:86736105-86738123 | p2 | (TA)17 | ACTGACCATAATTCAGCCCG | TGATGGTGGAGGAGAAGACC | 134 |
| chr6:86750157-86752172 | p2 | (AC)6 | CTCGTCAGCTTGCTCAGTTG | ACACCAAGTTTGTCAAGCCC | 212 |
| chr6:86750157-86752172 | p5 | (TTTAA)3 | CTCGTCAGCTTGCTCAGTTG | ACACCAAGTTTGTCAAGCCC | 212 |
| chr6:86750157-86752172 | p6 | (GTACAA)3 | CTCGTCAGCTTGCTCAGTTG | ACACCAAGTTTGTCAAGCCC | 212 |

| Marker ID | SSR type | SSR motif | FORWARD PRIMER (5'-3') | REVERSE PRIMER (5'-3') | size (bp) |
|---|---|---|---|---|---|
| chr6:86772740-86774758 | p3 | (ATC)6 | AGACACAAGCCAACAAGGCT | TTAGAAGCGCTCCATTTGCT | 111 |
| chr6:86774666-86776687 | p3 | (AGA)7 | TTGCTTGATCCCAGAACCTT | GTCGTCTGCGTCGTGAGTAA | 174 |
| chr6:86886983-86888997 | p2 | (AG)9 | CCCAGTCTTGGTTTGGTTGT | TGATGCATTGGAGCGAAATA | 279 |
| chr6:86911939-86913957 | p2 | (AT)9 | TCAACAATGCAACCAGGAAA | TCCAACATCGATTTCAAGCA | 196 |
| chr6:86924121-86926142 | p3 | (AAT)7 | GAATTTCGGTACCAGGCGTA | CAATCTCTGCCCATTCACCT | 194 |
| chr6:86924121-86926142 | p2 | (AT)6 | GAATTTCGGTACCAGGCGTA | CAATCTCTGCCCATTCACCT | 194 |
| chr6:86947453-86949474 | p3 | (TTC)5 | TTATGCAAAAGGGTTGGAGG | GAGGAGGAGGAGGAGAAGGA | 274 |
| chr6:87000033-87002051 | p2 | (TA)9 | AACCACCAACAAGTGCACAA | CAATGAAGCAGACACGCAGT | 106 |
| chr6:87045684-87047704 | p4 | (TATT)5 | CAATTCCCACCTAACTGCGT | TCTTTTAGGGCCTGTTTGGA | 163 |
| chr6:87094516-87096534 | p6 | (AATATA)3 | TCTTCCGAAATCCGACAATC | GAGTTGTTTTGACCCGAGGA | 110 |
| chr6:87148222-87150237 | p5 | (TTTGG)3 | TTTTCCACCCAAACTCATCC | ACCGTGAACGACCCTAGTTG | 220 |
| chr6:87148222-87150237 | p2 | (AG)18 | TTTTCCACCCAAACTCATCC | ACCGTGAACGACCCTAGTTG | 220 |
| chr6:87162421-87164436 | p2 | (AG)15 | GGTGAGCACAGAAGGAGAGG | AACCACCACACCATAACCGT | 223 |
| chr6:87162421-87164436 | p5 | (ATAAA)3 | GGTGAGCACAGAAGGAGAGG | AACCACCACACCATAACCGT | 223 |
| chr6:87189679-87191697 | p2 | (AT)9 | GAACCAATCAGAGGGCCATA | ATCATTTTGGCAGCCTTGTC | 190 |
| chr6:87424229-87426244 | p5 | (CCTTT)3 | GTCTCGGTAGCACAACCCAT | TGAGAGTGATGTGAGGACCG | 248 |
| chr6:87615176-87617224 | p2 | (AT)24 | AGCGGGTATCGTATCAATGC | AGGCAGCAGCTGGTGTTACT | 226 |
| chr6:87656876-87658891 | p3 | (GAT)5 | TCCAACACCTTCCAAACCTC | TTCTCCCGCTAGTACCGTGT | 113 |
| chr6:87666295-87668313 | p3 | (TGA)6 | TCGTATTGCGGTGAGTCTTG | ACAACAGCATCCAACAACCA | 238 |
| chr6:87717432-87719448 | p2 | (CT)6 | ACTTTTGTCACGACCCAACC | ATCGATCTGGTTCGTTTTGG | 246 |
| chr6:87717488-87719522 | p2 | (CT)6 | CCATGAGCTGGACACTCGTA | ATCGATCTGGTTCGTTTTGG | 217 |
| chr6:87755383-87757398 | p3 | (TCA)5 | ATGTAACCATCGGCAACCAT | TACATACTTGGGCACGGTGA | 159 |
| chr6:87755383-87757398 | p3 | (AAC)5 | ATGTAACCATCGGCAACCAT | TACATACTTGGGCACGGTGA | 159 |
| chr6:87796313-87798328 | p5 | (CCTTT)3 | TCCAACAGTAGGGTTACCGC | CGAATTCGTTGGGTGTTTTT | 262 |
| chr6:87819288-87821303 | p5 | (GATGT)3 | ATTCAGCCTGGCTCTCAAAA | TCATTGTTTAGGACGGGGAG | 262 |
| chr6:88036701-88038741 | p2 | (TA)20 | AGACGAAATTAAGGGGAAAACA | GGAGCCACATTATCGTTTTGA | 202 |
| chr6:88121150-88123176 | p2 | (TG)8 | CAAGGTGGTCTGGTTCGTTT | GTGTTTCAACCTCTGCCCAT | 159 |
| chr6:88190129-88192147 | p6 | (GAATAG)3 | AGGCAAAAGATCTTGGCAGA | CGACACTCACAGGAAACGAA | 105 |
| chr6:88274699-88276795 | p2 | (TA)8 | TGGTCATCACCTCCAGTTCA | TGTAGCATCCTAGCTGGCCT | 235 |
| chr6:88287680-88289702 | p2 | (TA)11 | TGCTCAACAAGCATCTGTCC | AAGGAATTAGGGTGGATGCC | 186 |
| chr6:88287680-88289702 | p2 | (TA)10 | TGCTCAACAAGCATCTGTCC | AAGGAATTAGGGTGGATGCC | 186 |
| chr6:88402808-88404824 | p2 | (AT)8 | TCGCTTCCCATAACTTGTCC | GCGGATGCTTAGATGGATGT | 229 |
| chr6:88402808-88404824 | p2 | (AT)10 | TCGCTTCCCATAACTTGTCC | GCGGATGCTTAGATGGATGT | 229 |
| chr6:88405096-88407110 | p2 | (TA)7 | ACGCACCATTTGTGTGTTGT | GACCCCTGAACTTGTCCTGA | 251 |
| chr6:88418727-88420765 | p2 | (TA)19 | GGCACCATCAACGAAGAGTT | AGATTGATACGCATGTGGCA | 203 |
| chr6:88640222-88642264 | p2 | (AT)21 | AGCCCTTCGGCTATCAAAAT | CACAGGTTAGCAGGGAGGAG | 125 |

| Marker ID | SSR type | SSR motif | FORWARD PRIMER (5'-3') | REVERSE PRIMER (5'-3') | size (bp) |
|---|---|---|---|---|---|
| chr6:88698039-88700057 | p3 | (CAT)6 | ACCACAAACCAAAAAGGCAC | GGTGTCCTGTCCCGTCTTTA | 152 |
| chr6:88727813-88729831 | p3 | (GAT)6 | TTTCCACCCAAAGCTTGTTC | CCCGCGTTCTTAACATCATT | 217 |
| chr6:88745584-88747602 | p6 | (CATATA)3 | TAAACTTGGTGGACTTGGGC | GTTGGGGAAGTGTCGAAGAA | 248 |
| chr6:88745584-88747602 | p5 | (ATTCA)3 | TAAACTTGGTGGACTTGGGC | GTTGGGGAAGTGTCGAAGAA | 248 |
| chr6:88750614-88752632 | p6 | (CATATA)3 | CTTTGTCACACCCCGATCTT | ATTGTAATCGGCTCCACCAG | 228 |
| chr6:88776569-88778584 | p5 | (TTTCA)3 | AGATATGACATGGGAAGCGG | AAAATGCCTCACAGGGTCAC | 174 |
| chr6:88776569-88778584 | p3 | (GAT)5 | AGATATGACATGGGAAGCGG | AAAATGCCTCACAGGGTCAC | 174 |
| chr6:88791336-88793351 | p5 | (AAATG)3 | GGTTCTCAACCTCCGATGAA | GTTAACAAAGCACGGCCAAT | 248 |
| chr6:88791336-88793351 | p2 | (AT)6 | GGTTCTCAACCTCCGATGAA | GTTAACAAAGCACGGCCAAT | 248 |
| chr6:88792466-88794478 | p2 | (AT)6 | CTGTCATGCATTTACCACCG | TTACCCGGTGCCTTCATTAG | 262 |
| chr6:88792466-88794478 | p2 | (AT)6 | CTGTCATGCATTTACCACCG | TTACCCGGTGCCTTCATTAG | 262 |
| chr6:88856557-88858569 | p2 | (TA)6 | CTCCGATTAGCACAGCAACA | TGGTACATGGATGGTGGTTG | 161 |
| chr6:88866206-88868224 | p6 | (GGAGAA)3 | AAGAGCCACCTTTCTGGGAT | AAGAAGGGGAGTGCAAGGTT | 287 |
| chr6:88866206-88868224 | p6 | (ACCTCT)3 | AAGAGCCACCTTTCTGGGAT | AAGAAGGGGAGTGCAAGGTT | 287 |
| chr6:88930221-88932313 | p2 | (AT)32 | CAGCCCTGTCACCATAACCT | CTCTCATACTCTGCCCGCTC | 257 |
| chr6:88987289-88989304 | p5 | (TTCTT)3 | TTTGATTTAAGGAGGCGGTG | TTAAATACCCCCTTAGCGGC | 293 |
| chr6:89015766-89017781 | p5 | (ATGCA)3 | CCATGCCTGAAAGAGACACA | CCCTGGGGGTATTTTCCTTA | 258 |
| chr6:89145902-89147922 | p2 | (AT)10 | CTATGGCCAGAAGGATTGGA | CTAATTCTGGCCGATCCGTA | 101 |
| chr6:89145902-89147922 | p3 | (ACA)5 | CTATGGCCAGAAGGATTGGA | CTAATTCTGGCCGATCCGTA | 101 |
| chr6:89150259-89152271 | p2 | (AT)6 | AAGGCCCTACAGCCTCTTGT | AGGGTCTTGGGTGTCAGTTG | 294 |
| chr6:89199414-89201430 | p4 | (CATT)4 | CGTGACAGGGACAAACAATG | TCTTTGCCATTCGCTTCTTT | 265 |
| chr6:89204286-89206301 | p3 | (CTT)5 | CTGGTCAGACCCTGTTTGCT | GGGTCCAAACCATCCCTATT | 232 |
| chr6:89241996-89244014 | p6 | (TATTTT)3 | AAACAATTTTCGTCATGGGC | TGAGGAGGGAGGAGTGAAGA | 213 |
| chr6:89247208-89249222 | p2 | (AC)7 | TAGGAAAACATTTTCGGGGA | GGGTTATCGTTCCCATTCAA | 299 |
| chr6:89247208-89249222 | p2 | (CA)6 | TAGGAAAACATTTTCGGGGA | GGGTTATCGTTCCCATTCAA | 299 |
| chr6:89247208-89249222 | p2 | (AT)6 | TAGGAAAACATTTTCGGGGA | GGGTTATCGTTCCCATTCAA | 299 |
| chr6:89255700-89257715 | p3 | (TTG)5 | ACCATACTCCTCATGACCGC | TCGAGTTTGATGTTCCCCTC | 149 |
| chr6:89387654-89389680 | p2 | (TA)10 | AACAAAAATGTGGGCCAAAG | TTCATGCCCGTAGGTTTAGG | 222 |
| chr6:89387654-89389680 | p2 | (AT)13 | AACAAAAATGTGGGCCAAAG | TTCATGCCCGTAGGTTTAGG | 222 |
| chr6:89489782-89491797 | p5 | (ATATA)3 | TAGGCCGACAAGGCTATCAC | CTGTCCCGTCATATTTCGCT | 254 |
| chr6:89505097-89507115 | p2 | (TA)9 | AATCGGAGGTATCCGACCTT | AGGTACGGTGTTATGCTGCC | 233 |
| chr6:89545339-89547373 | p2 | (TA)9 | GCTTCAGCTCAAGTTTTGGC | AGGAGGAGCACTTTGACGAA | 242 |
| chr6:89545339-89547373 | p2 | (TA)17 | GCTTCAGCTCAAGTTTTGGC | AGGAGGAGCACTTTGACGAA | 242 |
| chr6:89545339-89547373 | p2 | (AT)10 | GCTTCAGCTCAAGTTTTGGC | AGGAGGAGCACTTTGACGAA | 242 |
| chr6:89550495-89552507 | p2 | (TA)6 | GGAGGAATCACAGCAGAAGC | TCAAGGAAACCAAGGTGGAC | 134 |
| chr6:89664094-89666109 | p5 | (ATGCA)3 | ATAAATGACACAGCCTCCGC | TTTGCTACGTCCTTTTTGGG | 113 |

| Marker ID | SSR type | SSR motif | FORWARD PRIMER (5'-3') | REVERSE PRIMER (5'-3') | size (bp) |
|---|---|---|---|---|---|
| chr6:89669426-89671468 | p2 | (TA)21 | CCTTGATTTCGATTGTGCCT | ATTCCCTATGGGATTCGACC | 244 |
| chr6:89671770-89673785 | p5 | (ATGCA)3 | AAATGACACAGCCTCTGCCT | GTGCCCTTTTTGGGTGTAGA | 123 |
| chr6:89675928-89677960 | p2 | (AT)16 | GTCGCTAGGCCAAACATAGC | TCCCTTTGACTACCAAGCGT | 114 |
| chr6:89675928-89677960 | p2 | (AT)16 | GTCGCTAGGCCAAACATAGC | TCCCTTTGACTACCAAGCGT | 114 |
| chr6:89685721-89687766 | p3 | (TAA)15 | GAGTAAGCCACGGACCAAAA | ACGTGAAGGTCGTCAAGTCC | 276 |
| chr6:89700578-89702606 | p2 | (TG)6 | GGGGTAATTTGACGGTTCCT | GCACAGGCTTCTCCAAGTTC | 245 |
| chr6:89700578-89702606 | p2 | (GA)6 | GGGGTAATTTGACGGTTCCT | GCACAGGCTTCTCCAAGTTC | 245 |
| chr6:89714132-89716150 | p6 | (AAGTGA)3 | CATGAATTCCCTTTTTGCGT | CCTCAAATTCGATCCTCCAA | 289 |
| chr6:89727556-89729624 | p2 | (TA)12 | CACTCAGGATGGGGAAAAGA | TCAGGAACAAGCCTCCAATC | 246 |
| chr6:89727556-89729624 | p2 | (AT)34 | CACTCAGGATGGGGAAAAGA | TCAGGAACAAGCCTCCAATC | 246 |
| chr6:89762174-89764189 | p3 | (CAA)5 | CTGTTTCCGGTCAGATTCGT | AAACATGTCATTGTTCGCCA | 211 |
| chr6:89785056-89787093 | p6 | (CAAAAA)3 | ATAAGCTGGGACTGGGGACT | CGGTCGTTCATCCCTTAAAA | 217 |
| chr6:89795108-89797134 | p2 | (AT)13 | ACAAAGAAGCCTAACGGGGT | TGATCGGATCGGAAGAACTC | 185 |
| chr6:89804436-89806476 | p5 | (AAAAT)3 | TGGACGAATATGGGTGGATT | GGTTGTGCAATGTTTGATGG | 235 |
| chr6:89804436-89806476 | p4 | (TTAT)10 | TGGACGAATATGGGTGGATT | GGTTGTGCAATGTTTGATGG | 235 |
| chr6:89859258-89861280 | p2 | (AT)11 | AGCCAAGAACACAAACCCAC | CTGCTGGTCTCAAGGTCCTC | 121 |
| chr6:89859258-89861280 | p2 | (AT)10 | AGCCAAGAACACAAACCCAC | CTGCTGGTCTCAAGGTCCTC | 121 |
| chr6:89859797-89861817 | p2 | (AT)11 | TTGTTGTAAGCTGCCAGTCG | ATCGGCCTAAGTTGAGGGTT | 152 |
| chr6:89859797-89861817 | p2 | (AT)10 | TTGTTGTAAGCTGCCAGTCG | ATCGGCCTAAGTTGAGGGTT | 152 |
| chr6:89905750-89907770 | p3 | (CCT)6 | GGCTATTTTGGTGCTGCAAT | CCACAGGTAAAGGGCAAAAA | 243 |
| chr6:89905750-89907770 | p5 | (AAAAG)3 | GGCTATTTTGGTGCTGCAAT | CCACAGGTAAAGGGCAAAAA | 243 |
| chr6:89907393-89909417 | p5 | (AAAAG)3 | CCTCCTCCTCCTCTTTTGCT | AACCCATGAAGTAATGCCCA | 208 |
| chr6:89907393-89909417 | p2 | (AT)12 | CCTCCTCCTCCTCTTTTGCT | AACCCATGAAGTAATGCCCA | 208 |
| chr6:89931030-89933050 | p2 | (TA)17 | TCCCCCATTCTTTTTATCCC | GCGCGCATTAATTACGGTAT | 162 |
| chr6:89931030-89933050 | p2 | (CA)10 | TCCCCCATTCTTTTTATCCC | GCGCGCATTAATTACGGTAT | 162 |
| chr6:89987533-89989548 | p5 | (CCTTT)3 | CCAAGTGTGCCAAGTCTCAA | GGGTTGTGCTACCGAGATGT | 161 |
| chr6:90026439-90028485 | p2 | (TA)23 | CGAATGTGCCACAATTTCAC | AAAATAGAAAAATGGGGCGG | 255 |
| chr6:90029541-90031573 | p4 | (TTAT)8 | TAGTAATCACCGAGGCCACC | TCAACCGATTCTATTTCGGG | 154 |
| chr6:90137137-90139149 | p2 | (AT)6 | GCGATGCATGATTGATTGAC | TTTCAGCTTCCCTTTTCCAA | 128 |
| chr6:90288273-90290288 | p5 | (ATTGC)3 | TGGCCTATCCTTGGTGAGAC | CACACAACAGGGCCTAACCT | 279 |
| chr6:90349992-90352036 | p2 | (AT)22 | GGCTTTTGAGGATTAAGGGG | GAAAAAGCGAGCCGTAACAG | 186 |
| chr6:90426651-90428715 | p2 | (TA)32 | AATACGTTCCAGCGCATACC | AAGGGGCCAGGCTAAATAAA | 300 |
| chr6:90438279-90440311 | p2 | (AT)16 | AGCCGCCATCATATACGAAC | TTGACTAAGGGTACGACGGG | 292 |
| chr6:90470290-90472305 | p5 | (TTCAT)3 | GAAGAGATTCAAGATGCGGC | TCGAATGTTAAGTCCTCGGG | 228 |
| chr6:90581009-90583021 | p2 | (AT)6 | AGGAAGCGTGGAAAGGAAAT | TCTGGGCTGCTAATCGTCTT | 208 |
| chr6:90663627-90665719 | p2 | (CT)6 | GCAGCTCTAACCAAACGACC | CTAAGCTGACACGACGGACA | 107 |

| Marker ID | SSR type | SSR motif | FORWARD PRIMER (5'-3') | REVERSE PRIMER (5'-3') | size (bp) |
|---|---|---|---|---|---|
| chr6:90663627-90665719 | p2 | (TA)46 | GCAGCTCTAACCAAACGACC | CTAAGCTGACACGACGGACA | 107 |
| chr6:90789160-90791180 | p5 | (CAATG)4 | ATGCATTCTCCCGTCAACTC | GCCCCCAAGAAAAAGAAAAC | 230 |
| chr6:90864802-90866814 | p2 | (TA)6 | AAACGCGGTTGTGACATTTT | CGTGAGGGAAAGGAAGTTTG | 178 |
| chr6:90864802-90866814 | p4 | (TATT)4 | AAACGCGGTTGTGACATTTT | CGTGAGGGAAAGGAAGTTTG | 178 |
| chr6:90870389-90872407 | p3 | (AAT)6 | ATGAGGAGGCAGCTATCGAA | AAACTTGGTGGCCACGTTAC | 282 |
| chr6:90879948-90881972 | p2 | (AT)8 | GAAGAGCGCATTTAGCCAAG | GCCTTCCTCCGCTTCTAAGT | 223 |
| chr6:90879948-90881972 | p2 | (AT)12 | GAAGAGCGCATTTAGCCAAG | GCCTTCCTCCGCTTCTAAGT | 223 |
| chr6:90882318-90884330 | p2 | (AC)6 | CTCTCAATTGCTCTTTGCCC | TCCTTTGGGGTTGACTGTTC | 206 |
| chr6:90884391-90886403 | p2 | (TA)6 | TGTGTGTGTTTGCGTGTGTC | CTGGAAATGACCTGGATGCT | 110 |
| chr6:90956633-90958663 | p5 | (GCATT)3 | AATTCCGAAATCGTTGTTGC | ACGTCTCGGTAGCACAACCT | 204 |
| chr6:90956633-90958663 | p2 | (TA)15 | AATTCCGAAATCGTTGTTGC | ACGTCTCGGTAGCACAACCT | 204 |
| chr6:90956633-90958663 | p2 | (TA)13 | AATTCCGAAATCGTTGTTGC | ACGTCTCGGTAGCACAACCT | 204 |
| chr6:90964438-90966453 | p3 | (GAT)5 | CAGTGGGAGCAACTTAAGGC | AGTGTGCAAAGAGCCACCTT | 144 |
| chr6:90993739-90995754 | p5 | (TTTGG)3 | GCTACGACACACGCGAATAA | CTACCTCAAACTTCGCTCCG | 294 |
| chr6:91011571-91013595 | p6 | (CTGCAC)4 | GTCCCCCATAGGTTGTGTTG | CAAAGCCAATGGTCCAAGTT | 261 |
| chr6:91013513-91015525 | p2 | (TA)6 | CCGAGGGAAGTTGATGATGT | CCAATTTCGCTTTCATGGTT | 239 |
| chr6:91058213-91060225 | p3 | (GCA)5 | CAGACGTTCAATAGGCGGAT | AGGTCAGGGGAGCAATAGGT | 105 |
| chr6:91058373-91060400 | p3 | (GCA)5 | ACCTATTTTGGCTGACGTGG | CACGAACTGGGCTATGTTCA | 214 |
| chr6:91058870-91060885 | p3 | (GCA)5 | CAGCACTCCTCGAGCTTTTT | CAACCCATTTTCTTCCTCCA | 154 |
| chr6:91073663-91075678 | p3 | (GAT)5 | GGGTCTTAAAGTCTTGGGGC | TCCACTTCTCATCCCTGGAC | 281 |
| chr6:91112017-91114032 | p3 | (GAT)5 | GGGTCTTAAAGTCTTGGGGC | TCCACTTCTCATCCCTGGAC | 281 |
| chr6:91168421-91170441 | p2 | (AT)10 | ATCCTTCGTACTGACCCCCT | CACACATGACCCATTACCCA | 145 |
| chr6:91168421-91170441 | p5 | (TATTT)3 | ATCCTTCGTACTGACCCCCT | CACACATGACCCATTACCCA | 145 |
| chr6:91168421-91170441 | p3 | (ACA)6 | ATCCTTCGTACTGACCCCCT | CACACATGACCCATTACCCA | 145 |
| chr6:91229802-91231844 | p2 | (AT)9 | ATGCGTATATAACGCCTCGC | CGTACACTCTACCCTCCCCA | 177 |
| chr6:91229802-91231844 | p3 | (ACA)6 | ATGCGTATATAACGCCTCGC | CGTACACTCTACCCTCCCCA | 177 |
| chr6:91229802-91231844 | p2 | (AT)11 | ATGCGTATATAACGCCTCGC | CGTACACTCTACCCTCCCCA | 177 |
| chr6:91229845-91231869 | p2 | (AT)9 | ATGCGTATATAACGCCTCGC | CGTACACTCTACCCTCCCCA | 134 |
| chr6:91229845-91231869 | p3 | (ACA)6 | ATGCGTATATAACGCCTCGC | CGTACACTCTACCCTCCCCA | 134 |
| chr6:91229845-91231869 | p2 | (AT)11 | ATGCGTATATAACGCCTCGC | CGTACACTCTACCCTCCCCA | 134 |
| chr6:91293580-91295598 | p2 | (TA)9 | TTTTTGAAAACACGTGCTGC | TGCCGTGACTCATCTTCTTG | 249 |
| chr6:91354908-91356952 | p2 | (AT)22 | GTAATAGGACCCCCGCAAAG | GAATTGGTTGGGATTGTGCT | 196 |
| chr6:91361175-91363187 | p2 | (TA)6 | AGCGAAATATGACGGGACAG | CTTGGAGAGCTCCGTTGTTC | 283 |
| chr6:91366530-91368545 | p5 | (ATATC)3 | TCCCTTAGTGTCACAAGCCC | AAGTCCCGTGGTGCTTATTG | 242 |
| chr6:91372913-91374928 | p3 | (CAA)5 | GGGACCACACACATGTACCA | GCGAGCTTGGACAATTAAGC | 262 |
| chr6:91372913-91374928 | p2 | (AT)6 | GGGACCACACACATGTACCA | GCGAGCTTGGACAATTAAGC | 262 |

| Marker ID | SSR type | SSR motif | FORWARD PRIMER (5'-3') | REVERSE PRIMER (5'-3') | size (bp) |
|---|---|---|---|---|---|
| chr6:91547489-91549503 | p2 | (TA)7 | TCAGGCGTTAATTGTAGCCC | TTGAAGCAATCAAGGGGAAC | 252 |
| chr6:91575797-91577811 | p2 | (AT)7 | GTCCTGGGGGTCGATATTTT | CAGGATCAGCATCAGGGATT | 258 |
| chr6:91598597-91600617 | p2 | (TA)10 | TTGCTAGCCTAAAGGGCTCA | GTCCTGTCCCGCCTATATGA | 199 |
| chr6:91624501-91626517 | p4 | (AATT)4 | ATGATGCTGAGGATTCCAGG | GTAACTTGTGGACCCTTCGC | 167 |
| chr6:91693400-91695414 | p2 | (AT)7 | ATCCTTGCAGGTGTCCAAAC | TTCCCTTGCGTCGTTAGAGT | 229 |
| chr6:91712483-91714498 | p5 | (CCTTT)3 | AAGTCCATGTACCTCACGGC | ATATCCGATGGTTGTCTCGG | 278 |
| chr6:91786362-91788392 | p2 | (TA)15 | CCATTGATGCATGGAAGATG | TAAACATGCCGGTTTCAACA | 170 |
| chr6:91886346-91888381 | p2 | (AT)6 | CGTGGATTTTTCCACGAAAT | ACACGAAATGCCAAAACAAA | 267 |
| chr6:91886346-91888381 | p2 | (AT)7 | CGTGGATTTTTCCACGAAAT | ACACGAAATGCCAAAACAAA | 267 |
| chr6:91942047-91944071 | p6 | (AAATCC)4 | GATGCCTCTTAGCCTTGTCG | CAATACCATTTCCATTGGGC | 158 |
| chr6:91959280-91961324 | p2 | (AT)17 | TGTAGAAAGGACGGGACAGG | GGCAGTGGCTAGTATGCTCC | 284 |
| chr6:91959280-91961324 | p5 | (ATTCC)3 | TGTAGAAAGGACGGGACAGG | GGCAGTGGCTAGTATGCTCC | 284 |
| chr6:91983210-91985225 | p5 | (AAAGG)3 | GGAAAATCTGGTGCTGGTGT | AGGCCTTCAAATGAACATGG | 222 |
| chr6:91983210-91985225 | p5 | (ATTGC)3 | GGAAAATCTGGTGCTGGTGT | AGGCCTTCAAATGAACATGG | 222 |
| chr6:92004462-92006477 | p3 | (CAA)5 | CAGATCGTGCAATAGAGCCA | AACCATTGGCGGTACTCTTG | 127 |
| chr6:92060653-92062665 | p2 | (TA)6 | AGGCTTCCATTACACGCAAC | TCTGGAGCCTAGCTTGTGGT | 235 |
| chr6:92092488-92094500 | p2 | (AT)6 | AGGCTTCCATTACACGCAAC | TCTGGAGCCTAGCTTGTGGT | 223 |
| chr6:92100745-92102757 | p2 | (TA)6 | ATTCCGAGGATTTAACCGCT | TTGGAATGTTGCGTGTCTGT | 187 |
| chr6:92100745-92102757 | p2 | (TA)6 | ATTCCGAGGATTTAACCGCT | TTGGAATGTTGCGTGTCTGT | 187 |
| chr6:92118754-92120769 | p5 | (AATAC)3 | TTTGACGCTCAGAATCCCTT | GTTGTTTGGCCGTGAGTTTT | 116 |
| chr6:92118754-92120769 | p3 | (CAA)5 | TTTGACGCTCAGAATCCCTT | GTTGTTTGGCCGTGAGTTTT | 116 |
| chr6:92182625-92184643 | p2 | (AT)6 | CAAACGCTGCATTGGAGTTA | TTGCCACTTTTCAAACAACG | 229 |
| chr6:92182625-92184643 | p2 | (TA)9 | CAAACGCTGCATTGGAGTTA | TTGCCACTTTTCAAACAACG | 229 |
| chr6:92182625-92184643 | p2 | (AT)11 | CAAACGCTGCATTGGAGTTA | TTGCCACTTTTCAAACAACG | 229 |
| chr6:92182652-92184679 | p2 | (AT)6 | CAAACGCTGCATTGGAGTTA | TTGCCACTTTTCAAACAACG | 202 |
| chr6:92182652-92184679 | p2 | (TA)9 | CAAACGCTGCATTGGAGTTA | TTGCCACTTTTCAAACAACG | 202 |
| chr6:92182652-92184679 | p2 | (AT)11 | CAAACGCTGCATTGGAGTTA | TTGCCACTTTTCAAACAACG | 202 |
| chr6:92186642-92188660 | p3 | (AAG)27 | GACCCAAAGGGCAATTATCA | CTGCTCCCCTTCTTCTCCTT | 162 |
| chr6:92186642-92188660 | p3 | (AGA)6 | GACCCAAAGGGCAATTATCA | CTGCTCCCCTTCTTCTCCTT | 162 |
| chr6:92186642-92188660 | p3 | (AAG)6 | GACCCAAAGGGCAATTATCA | CTGCTCCCCTTCTTCTCCTT | 162 |
| chr6:92186672-92188861 | p3 | (AAG)27 | GACCCAAAGGGCAATTATCA | CTGCTCCCCTTCTTCTCCTT | 132 |
| chr6:92186672-92188861 | p3 | (AGA)6 | GACCCAAAGGGCAATTATCA | CTGCTCCCCTTCTTCTCCTT | 132 |
| chr6:92186672-92188861 | p3 | (AAG)6 | GACCCAAAGGGCAATTATCA | CTGCTCCCCTTCTTCTCCTT | 132 |
| chr6:92242323-92244352 | p6 | (CAACTC)3 | AGGAGCTGGAATGGGTAGGT | ACCTGGATCACATTGGGTGT | 117 |
| chr6:92245540-92247558 | p6 | (AAGAAA)3 | TGGACTTGACAGGTGCTGAG | CCGTTCCTCCAACAAAACAT | 281 |
| chr6:92314766-92316798 | p2 | (CT)11 | ACGCTTGTTACTGCCTCGTT | CTACAAAGATGAAGCCCGGA | 282 |

| Marker ID | SSR type | SSR motif | FORWARD PRIMER (5'-3') | REVERSE PRIMER (5'-3') | size (bp) |
|---|---|---|---|---|---|
| chr6:92314812-92316844 | p2 | (CT)11 | ACGCTTGTTACTGCCTCGTT | TACAAAGATGAAGCCCGGAG | 236 |
| chr6:92372887-92374902 | p5 | (ATTGC)3 | GTCGCCTTTTGTCCTGGTTA | TCATCATCGGATATCAGCCA | 254 |
| chr6:92398714-92400729 | p3 | (CTT)5 | TGGAGGAGTTGTTCGCTTCT | GTGAAGCATTATCGGCCATT | 293 |
| chr6:92484680-92486692 | p2 | (CT)6 | ATGCCAAGTCTGCCCATAAG | GAACCCCCACTCAGTCAAAA | 141 |
| chr6:92501914-92503940 | p2 | (AT)13 | TGAATTCCCAACACAATGGA | ATCTCCATAAACCCGCGTAA | 144 |
| chr6:92548139-92550151 | p6 | (GGGTTG)3 | GTTGGGGTTGGAGCTAGTGA | AAGGCCTCATTTGTTTGCAC | 105 |
| chr6:92628428-92630443 | p2 | (TA)13 | GCGAAGTTTTGATTGGTCGT | CAACAAGGAAACAAGCCCAT | 150 |
| chr6:92628428-92630443 | p5 | (CATTG)3 | GCGAAGTTTTGATTGGTCGT | CAACAAGGAAACAAGCCCAT | 150 |
| chr6:92631816-92633858 | p3 | (TGA)5 | CCATTCGACATAAATTGGGG | AGCTCACCGATGAATTTTGG | 156 |
| chr6:92631867-92633896 | p3 | (TGA)5 | CCATTCGACATAAATTGGGG | AGCTCACCGATGAATTTTGG | 105 |
| chr6:92632213-92634233 | p3 | (TGA)5 | CGACATCTCGAGCAATGGTA | TCGTCTCGCCGATAGTCTTT | 296 |
| chr6:92632262-92634287 | p3 | (TGA)5 | CGACATCTCGAGCAATGGTA | TCGTCTCGCCGATAGTCTTT | 247 |
| chr6:92632299-92634341 | p3 | (TGA)5 | CGACATCTCGAGCAATGGTA | TCGTCTCGCCGATAGTCTTT | 210 |
| chr6:92694659-92696674 | p5 | (GGTTT)3 | TGTAGTGCAACTTTGCCAGC | CTGGATGGATGGTCGAGTCT | 206 |
| chr6:92697291-92699303 | p2 | (AT)6 | GCAGCGATGACAAGAACAAA | AAAAGGTCATTTCTTGCGGA | 165 |
| chr6:92697291-92699303 | p2 | (GA)6 | GCAGCGATGACAAGAACAAA | AAAAGGTCATTTCTTGCGGA | 165 |
| chr6:92708481-92710497 | p4 | (TAAT)4 | TCTTCCCCAAACCTTGCTAA | CTCCAGATTTTCACGTGGGT | 158 |
| chr6:92743711-92745727 | p3 | (ATG)5 | TGAGGTGAACTTGAAAGGGG | TCCACCAAAATCCAACACAA | 223 |
| chr6:92743711-92745727 | p4 | (TATG)4 | TGAGGTGAACTTGAAAGGGG | TCCACCAAAATCCAACACAA | 223 |
| chr6:92778565-92780577 | p2 | (GT)6 | CATGGCACGACAAAAGAGAA | TACAACATTGCCAACCCTCA | 181 |
| chr6:92832247-92834262 | p5 | (ATTTT)3 | GCGTGTTGGTATAATTGGGG | GGAACCTCAAAACCTCACCA | 241 |
| chr6:92832247-92834262 | p3 | (ATT)18 | GCGTGTTGGTATAATTGGGG | GGAACCTCAAAACCTCACCA | 241 |
| chr6:92832247-92834262 | p2 | (AT)19 | GCGTGTTGGTATAATTGGGG | GGAACCTCAAAACCTCACCA | 241 |
| chr6:92879928-92881946 | p2 | (AT)6 | GAAACCCACCACATCCAAAC | GGGGCCCACAATTTTATTTT | 243 |
| chr6:92879928-92881946 | p3 | (AAC)6 | GAAACCCACCACATCCAAAC | GGGGCCCACAATTTTATTTT | 243 |
| chr6:92884691-92886711 | p2 | (TA)10 | GGCCCCATATTAAGAACCGT | AGTTTGGATGTGGTGGGTTC | 240 |
| chr6:93025904-93027966 | p2 | (TA)6 | CCCCAACTCTAGTCCGTGAA | CCCCAGAACAGACACAACCT | 262 |
| chr6:93256255-93258273 | p2 | (AT)7 | TGTTATCCTCATTAGCCCGC | CTTGTTCCTTCAAGGTGGGA | 205 |
| chr6:93256255-93258273 | p3 | (CAA)6 | TGTTATCCTCATTAGCCCGC | CTTGTTCCTTCAAGGTGGGA | 205 |
| chr6:93278372-93280387 | p3 | (ATT)5 | TACCTGCGTATGCTCACAGC | TTCGCTGCCGAAGAAATAGT | 262 |
| chr6:93301732-93303800 | p2 | (TA)34 | ACCACTCACAGGTTTGGGTC | TCTAGCGACGAGGATGAGGT | 151 |
| chr6:93309926-93311944 | p2 | (AT)9 | AAACGAGTTGCAGAAACGGT | GGCACAGGCAATGTGATATG | 175 |
| chr6:93432102-93434114 | p3 | (CAA)7 | TCGGAGACATACCCAAGGAC | TGACGATGCGCATAATTGTT | 187 |
| chr6:93432187-93434203 | p3 | (CAA)7 | TCGGAGACATACCCAAGGAC | TGACGATGCGCATAATTGTT | 102 |
| chr6:93475467-93477483 | p2 | (AT)7 | TATAGGAGGGACAGGACCCC | CAGGGACTACATAGCGGAGC | 278 |
| chr6:93570739-93572757 | p6 | (GAAACT)3 | TCCTGGGGTACTTTCTTCCC | GTGTCGTGTTACGTCCATGC | 167 |

| Marker ID | SSR type | SSR motif | FORWARD PRIMER (5'-3') | REVERSE PRIMER (5'-3') | size (bp) |
|---|---|---|---|---|---|
| chr6:93584764-93586782 | p3 | (TCT)6 | CATGGGACGTCTCCGATACT | GGTTTCCCTTCGAACTTTCC | 268 |
| chr6:93603159-93605171 | p3 | (TAT)5 | TCCCGACATCTATCCGAGAC | TGGGTGTCAACATAGGGGTT | 118 |
| chr6:93603159-93605171 | p2 | (TC)6 | TCCCGACATCTATCCGAGAC | TGGGTGTCAACATAGGGGTT | 118 |
| chr6:93728244-93730282 | p2 | (AT)19 | CTCGTGGTCCGAGTTTCATT | AGCGAAAACTACAAGCGGAA | 104 |
| chr6:93762312-93764327 | p2 | (AT)19 | AAAATGAATAGCCCCGAAGG | CCTCATTAGTGGGCTGGGTA | 125 |
| chr6:93776261-93778275 | p2 | (TA)7 | TACGATTTTGGGCTAGGCAT | CATACCTGTTTGGGCGTCTT | 236 |
| chr6:93796384-93798410 | p2 | (AT)10 | TCTGCGGGATCTAAACCAAG | GGCCCATCTCAGATTTTGAA | 114 |
| chr6:93886875-93888937 | p2 | (TA)31 | CATCCCAAATGAATCCAAGG | GCCCACAACTCAACACCTTT | 187 |
| chr6:94011520-94013544 | p6 | (ATCCGT)4 | AGTTCGTTTCGTCGACTCGT | AAGCATCCGATGTTAGACGG | 188 |
| chr6:94011520-94013544 | p2 | (AT)6 | AGTTCGTTTCGTCGACTCGT | AAGCATCCGATGTTAGACGG | 188 |
| chr6:94018433-94020451 | p6 | (TTTTTG)3 | ATCCGAGCTTGAGGTAGCAA | TTCAGCATCGACCTTGACAG | 287 |
| chr6:94031821-94033836 | p5 | (ACCCC)3 | TTTGCACATTGGTCCTTTCA | CATCCTCCAATGCTTCACCT | 146 |
| chr6:94032029-94034054 | p5 | (ACCCC)3 | AGGTGAAGCATTGGAGGATG | GGCCAGCCAAACACTAAAAA | 126 |
| chr6:94032029-94034054 | p5 | (TGGGG)3 | AGGTGAAGCATTGGAGGATG | GGCCAGCCAAACACTAAAAA | 126 |
| chr6:94032514-94034542 | p5 | (ACCCC)3 | TCAGGAAAATACCCACCCAA | GGATGTGGCGGTTTAGAAAA | 268 |
| chr6:94032514-94034542 | p5 | (TGGGG)3 | TCAGGAAAATACCCACCCAA | GGATGTGGCGGTTTAGAAAA | 268 |
| chr6:94032548-94034577 | p5 | (ACCCC)3 | TCAGGAAAATACCCACCCAA | GGATGTGGCGGTTTAGAAAA | 234 |
| chr6:94032548-94034577 | p5 | (TGGGG)3 | TCAGGAAAATACCCACCCAA | GGATGTGGCGGTTTAGAAAA | 234 |
| chr6:94048569-94050595 | p2 | (TA)13 | AGACACATCCCGAAGACGAC | ATTAAATATCCCCGCCCAAC | 142 |
| chr6:94072839-94074851 | p2 | (AG)6 | CGATTCGTTTGACCTCCAAT | TGCAAGTGAGTCCGTTGAAG | 293 |
| chr6:94190771-94192786 | p5 | (TTGGT)3 | TGGCCACAACATTTAGCAAG | CGAATCACCACATCGTCAAG | 200 |
| chr6:94238988-94241006 | p2 | (TA)9 | ACCGAAGCTGGAAACAAATG | CCGTCGCGATATTTTCTTGT | 139 |
| chr6:94256313-94258359 | p2 | (AT)23 | TTTCGCTCTAATAAGGCCCA | TATGCCAGTAGGGGATTTGC | 253 |
| chr6:94274404-94276450 | p2 | (AT)23 | TTTCGCTCTAATAAGGCCCA | TATGCCAGTAGGGGATTTGC | 253 |
| chr6:94372774-94374786 | p2 | (CT)6 | CATCGTTGCATCGGTAGTTG | CACAACCGCCAATATTACCC | 275 |
| chr6:94372774-94374786 | p3 | (TCT)5 | CATCGTTGCATCGGTAGTTG | CACAACCGCCAATATTACCC | 275 |
| chr6:94497365-94499383 | p2 | (TA)9 | GATTTGTTGCACATTCGACG | CTGGCCAATTCTTCTGGTGT | 237 |
| chr6:94665077-94667092 | p5 | (ATAAA)3 | CGATGATCCTTCAGCCATCT | AGATGCTTCGGAAGACCAGA | 176 |
| chr6:94709374-94711388 | p2 | (TA)7 | AAGGAAAAGTTCTGCCCCAT | TGGATACCTGAGGTTCTGGG | 157 |
| chr6:94754995-94757017 | p2 | (AT)11 | CTTATCCTCAACGGGAACCA | ATGCTATATGGGGCGAAGTG | 253 |
| chr6:94769247-94771274 | p3 | (TCT)29 | CCTGTCCCCTTCTTCATTCA | GTCACGCCGGTTAAAACAAT | 174 |
| chr6:94769247-94771274 | p3 | (TCT)35 | CCTGTCCCCTTCTTCATTCA | GTCACGCCGGTTAAAACAAT | 174 |
| chr6:94769307-94771433 | p3 | (TCT)29 | CCTGTCCCCTTCTTCATTCA | GTCACGCCGGTTAAAACAAT | 114 |
| chr6:94769307-94771433 | p3 | (TCT)35 | CCTGTCCCCTTCTTCATTCA | GTCACGCCGGTTAAAACAAT | 114 |
| chr6:94769574-94771661 | p3 | (TCT)29 | CCTCTTTGTTGACGCTTTCC | GAAACTCCGGTGGTCGTAAA | 123 |
| chr6:94769574-94771661 | p3 | (TCT)35 | CCTCTTTGTTGACGCTTTCC | GAAACTCCGGTGGTCGTAAA | 123 |

| Marker ID | SSR type | SSR motif | FORWARD PRIMER (5'-3') | REVERSE PRIMER (5'-3') | size (bp) |
|---|---|---|---|---|---|
| chr6:94769574-94771661 | p2 | (TA)6 | CCTCTTTGTTGACGCTTTCC | GAAACTCCGGTGGTCGTAAA | 123 |
| chr6:94782599-94784651 | p2 | (GA)26 | CAGTCGAGTCTACCCGCTTC | AAAGGTGTTGTCGAACCCAG | 164 |
| chr6:94841121-94843141 | p2 | (TA)10 | CAACGTGACACCATGCTACC | GCCGGATAAGCAGACTTTTG | 226 |
| chr6:94841121-94843141 | p3 | (TCA)5 | CAACGTGACACCATGCTACC | GCCGGATAAGCAGACTTTTG | 226 |
| chr6:94868598-94870614 | p2 | (AT)8 | GAGGCCCCACAGATACAAAA | TTGGGTGAACGAACCTTTTC | 237 |
| chr6:94944531-94946545 | p2 | (AC)7 | TGGTTCACCCTCTCTGGAAC | CTAATCTTGGGTACGGGGGT | 288 |
| chr6:94987524-94989540 | p4 | (TTAA)4 | TGGTTTGGGATTATCGGGTA | TCAGCCATAAACACTGCCTG | 245 |
| chr6:95100315-95102365 | p2 | (AT)25 | GGTCAGGTTGGCATTGTCTT | GTAGGCTGGGAACCACTTGA | 218 |
| chr6:95119707-95121722 | p5 | (TTCTA)3 | GCTTTGCACGCGATATACCT | CCTTTCTTCAGAACGCTTGC | 235 |
| chr6:95183384-95185399 | p5 | (TTTTA)3 | TCGGATTAGCCCAAAATGAC | CAAATGATTAATGGACCGGG | 171 |
| chr6:95253666-95255678 | p2 | (TA)6 | CTTGTGCCCGGGATAAAGTA | GGCCGAAAACGAGAACAATA | 115 |
| chr6:95356728-95358740 | p2 | (AT)6 | GAACCCACCACATCCAAACT | CATGTGGCAATGAGTCCATC | 211 |
| chr6:95362545-95364560 | p5 | (TTGTC)3 | GGAGTGCTGGTGTGACAAGA | CCCACAAACATTTCCAAACC | 285 |
| chr6:95362545-95364560 | p3 | (TTG)14 | GGAGTGCTGGTGTGACAAGA | CCCACAAACATTTCCAAACC | 285 |
| chr6:95362684-95364726 | p5 | (TTGTC)3 | GGAGTGCTGGTGTGACAAGA | CCCACAAACATTTCCAAACC | 146 |
| chr6:95362684-95364726 | p3 | (TTG)14 | GGAGTGCTGGTGTGACAAGA | CCCACAAACATTTCCAAACC | 146 |
| chr6:95489499-95491514 | p5 | (AAAAT)3 | GTCCGCTTATCTCCAACTCG | GGAGCAGTGTTTTCAGAGGC | 123 |
| chr6:95489499-95491514 | p6 | (AAAAAT)3 | GTCCGCTTATCTCCAACTCG | GGAGCAGTGTTTTCAGAGGC | 123 |
| chr6:95501107-95503123 | p5 | (CATTG)3 | TTTGGGAATCGTGAGGTAGC | CAACAGAAGCCAAAGTGCAA | 265 |
| chr6:95501107-95503123 | p4 | (TTAT)4 | TTTGGGAATCGTGAGGTAGC | CAACAGAAGCCAAAGTGCAA | 265 |
| chr6:95568845-95570860 | p5 | (ATTGC)3 | AGTGAGTTGTTGGTGTTGCG | CAAGGCCATACCTAAGGCAA | 258 |
| chr6:95568845-95570860 | p2 | (TA)12 | AGTGAGTTGTTGGTGTTGCG | CAAGGCCATACCTAAGGCAA | 258 |
| chr6:95634775-95636799 | p2 | (AT)12 | ATCACGGGCTTTGGTACTTG | CCCGGTTGAGCTAGGACATA | 298 |
| chr6:95654436-95656451 | p3 | (TGT)5 | AGGCAATGGAAAGATTGACG | CATCACAACATCCGCATCTC | 141 |
| chr6:95692493-95694511 | p3 | (CAA)6 | GCCATAGCACGGTGGTAACT | TAATGGCCTGAGCTTTGCTT | 183 |
| chr6:95730856-95732871 | p3 | (TGT)5 | GGTTGTGAGACCGACCTTGT | CTCGCTTCCCTCATCTTGTC | 270 |
| chr6:95764803-95766821 | p2 | (AT)9 | TCCCGAAGATACAAAGCACC | GTCGGCTTGCATATGTTGTG | 288 |
| chr6:95782560-95784581 | p3 | (ATT)7 | CTGTAAGCCAATCCAAACGG | TTTGCAATTCAGCCCATGTA | 243 |
| chr6:95909188-95911203 | p5 | (AAAAT)3 | GCCCCGATGCAATATCTCTA | ACACCTTCATGGATCCGAAC | 172 |
| chr6:95998390-96000405 | p5 | (AATTA)3 | AGTAAGGGCCTTGTTTGGCT | AAGACGAGTTGCCTCAAGGA | 138 |
| chr6:96043686-96045698 | p2 | (AT)6 | ATATCATCATCAGGAGCGGG | TACTCTCGGTGGGGATTCAC | 224 |
| chr6:96122101-96124113 | p2 | (AT)6 | ATATCATCATCAGGAGCGGG | TACTCTCGGTGGGGATTCAC | 224 |
| chr6:96173808-96175820 | p2 | (AT)7 | AATTTCGCGCGTTGAATTAC | TGCGTGAGTAAATGAGTGGG | 208 |
| chr6:96173858-96175874 | p2 | (AT)7 | AATTTCGCGCGTTGAATTAC | TGCGTGAGTAAATGAGTGGG | 158 |
| chr6:96173888-96175904 | p2 | (AT)7 | AATTTCGCGCGTTGAATTAC | TGCGTGAGTAAATGAGTGGG | 128 |
| chr6:96208788-96210806 | p6 | (ACTACG)3 | ACCACAACAGCGGCTATGAC | CGTCGTTGAAGTAGTCGTCG | 155 |

| Marker ID | SSR type | SSR motif | FORWARD PRIMER (5'-3') | REVERSE PRIMER (5'-3') | size (bp) |
|---|---|---|---|---|---|
| chr6:96208788-96210806 | p6 | (ACTACG)3 | ACCACAACAGCGGCTATGAC | CGTCGTTGAAGTAGTCGTCG | 155 |
| chr6:96208788-96210806 | p3 | (ACT)5 | ACCACAACAGCGGCTATGAC | CGTCGTTGAAGTAGTCGTCG | 155 |
| chr6:96211085-96213118 | p3 | (ACT)8 | CACCACTACGGCCAGTACAA | GGTGGTGGTGGTGGTCAT | 259 |
| chr6:96211147-96213168 | p3 | (ACT)8 | CACCACTACGGCCAGTACAA | GGTGGTGGTGGTGGTCAT | 197 |
| chr6:96211187-96213211 | p3 | (ACT)8 | CACCACTACGGCCAGTACAA | GGTGGTGGTGGTGGTCAT | 157 |
| chr6:96211240-96213267 | p3 | (ACT)8 | CACCACTACGGCCAGTACAA | GGTGGTGGTGGTGGTCAT | 104 |
| chr6:96211495-96213519 | p3 | (ACT)8 | CGACCACCACTACGACCAGTA | AGTCGTCGTAGCAGTAGCAGTA | 160 |
| chr6:96211495-96213519 | p3 | (ACT)84 | CGACCACCACTACGACCAGTA | AGTCGTCGTAGCAGTAGCAGTA | 160 |
| chr6:96228692-96230719 | p6 | (ACTACC)3 | ATGCGATTCAGGGGTATGAC | TGGACACTCAAACGTACCCA | 143 |
| chr6:96228955-96230979 | p6 | (ACTACC)3 | TGGGTACGTTTGAGTGTCCA | TGGTGGTAGTGGTGGTGGTA | 105 |
| chr6:96228955-96230979 | p3 | (ACT)6 | TGGGTACGTTTGAGTGTCCA | TGGTGGTAGTGGTGGTGGTA | 105 |
| chr6:96344789-96346813 | p6 | (TTTATT)4 | GAGTGCAGCAACCATTAGCA | GGCAAATGCTACCTTACGGA | 175 |
| chr6:96344789-96346813 | p2 | (TC)6 | GAGTGCAGCAACCATTAGCA | GGCAAATGCTACCTTACGGA | 175 |
| chr6:96434632-96436647 | p5 | (TTCAT)3 | AAATTGCTAAGCCAAGGGGT | GAATACCAGGAAGATCGCCA | 247 |
| chr6:96520787-96522801 | p2 | (TA)7 | CTTGGTTGTTTGGAGGATGG | AAATTGAGCCGAAAGCAGAA | 243 |
| chr6:96603984-96605999 | p5 | (AAAAT)3 | TGCCCAAAACACTCCTTTTC | TAATTGGGTTAACGCCTTCG | 208 |
| chr6:96656695-96658719 | p4 | (ATAA)6 | GCTATTTGCTTTGCCTTTGC | TCTTTTGTTCCCCCTCTCCT | 185 |
| chr6:96717135-96719147 | p2 | (AT)6 | TTGCACCAAAGAAATGTGGA | TGGTTTCAATTTCACCGACA | 214 |
| chr6:96768284-96770314 | p3 | (TTG)5 | TATGGTGGTAGCATCCGTGA | GGTCCAAGGGGTAATGGACT | 224 |
| chr6:96768284-96770314 | p6 | (GGGGTG)5 | TATGGTGGTAGCATCCGTGA | GGTCCAAGGGGTAATGGACT | 224 |
| chr6:96768284-96770314 | p6 | (GGGGTG)5 | TATGGTGGTAGCATCCGTGA | GGTCCAAGGGGTAATGGACT | 224 |
| chr6:96768284-96770314 | p3 | (TTG)5 | TATGGTGGTAGCATCCGTGA | GGTCCAAGGGGTAATGGACT | 224 |
| chr6:97157733-97159797 | p2 | (AT)32 | TCCGTGTTCGGTCCTACTTT | TTTTCATGAGTTTGGGGAGG | 256 |
| chr6:97157733-97159797 | p2 | (TA)14 | TCCGTGTTCGGTCCTACTTT | TTTTCATGAGTTTGGGGAGG | 256 |
| chr6:97162787-97164801 | p2 | (TA)7 | TAAAGTGGTTGGGTTTTGGG | GGCCCCACCTTAAATTTGTT | 116 |
| chr6:97177528-97179543 | p3 | (AAT)5 | CGATTGATGAAGGGCTGAAT | TCACAAGTCTCCCACATCCA | 262 |
| chr6:97206140-97208155 | p2 | (AG)6 | GCAGTGTGGATGAGGTGATG | CAAATCCGATCCCAATCAAC | 190 |
| chr6:97206140-97208155 | p5 | (TTGTA)3 | GCAGTGTGGATGAGGTGATG | CAAATCCGATCCCAATCAAC | 190 |
| chr6:97206140-97208155 | p5 | (GGGGT)3 | GCAGTGTGGATGAGGTGATG | CAAATCCGATCCCAATCAAC | 190 |
| chr6:97207623-97209638 | p5 | (GGGGT)3 | TGAACCACGGGTTTACAACA | CTGTTTGGATGGTGGTTTCC | 196 |
| chr6:97207623-97209638 | p5 | (TCTAG)3 | TGAACCACGGGTTTACAACA | CTGTTTGGATGGTGGTTTCC | 196 |
| chr6:97282689-97284701 | p2 | (AT)6 | CTTTTTCCCGGTACGATTGA | ATTCGGAGCCACTAGAGCAA | 268 |
| chr6:97339622-97341688 | p2 | (AT)13 | CAATGGCCAAAGGTCAAACT | TGGAGGAAGACCCAAGTGAG | 256 |
| chr6:97339622-97341688 | p2 | (TA)6 | CAATGGCCAAAGGTCAAACT | TGGAGGAAGACCCAAGTGAG | 256 |
| chr6:97340173-97342199 | p2 | (AT)13 | TGGCGCACTACATACCCATA | CTCCGTATGGATCGTGACAA | 114 |
| chr6:97340173-97342199 | p2 | (TA)6 | TGGCGCACTACATACCCATA | CTCCGTATGGATCGTGACAA | 114 |

| Marker ID | SSR type | SSR motif | FORWARD PRIMER (5'-3') | REVERSE PRIMER (5'-3') | size (bp) |
|---|---|---|---|---|---|
| chr6:97403521-97405553 | p2 | (AT)16 | AACCGACGAGGCTAACAAAA | CAGGGACTACATAGCGGAGC | 225 |
| chr6:97408623-97410638 | p6 | (CCCCAC)3 | TGCTCTCTGACAACTCCCCT | GATGTTCATGATTCGGGGAC | 264 |
| chr6:97408623-97410638 | p5 | (AAAAT)3 | TGCTCTCTGACAACTCCCCT | GATGTTCATGATTCGGGGAC | 264 |
| chr6:97505380-97507398 | p2 | (TA)9 | ATGTCCAAAATTCTGCCAGG | AGCCTGAAGAGACCATGGAA | 139 |
| chr6:97541596-97543608 | p2 | (TA)6 | AGCCCTAAAGGGCCATAGAA | AAACTTAAAGGATTGGCCCG | 186 |
| chr6:97541596-97543608 | p2 | (TA)6 | AGCCCTAAAGGGCCATAGAA | AAACTTAAAGGATTGGCCCG | 186 |
| chr6:97541596-97543608 | p3 | (TTA)6 | AGCCCTAAAGGGCCATAGAA | AAACTTAAAGGATTGGCCCG | 186 |
| chr6:97599163-97601179 | p2 | (AT)11 | TGGACAAAAACTGTGCAAGC | GGAGCACTCCTCTTGTTTCG | 231 |
| chr6:97738655-97740667 | p2 | (TA)6 | ACATAAATAGCCGCCAATCG | CCAGGTTCACCTAGTTCCCA | 279 |
| chr6:97781404-97783418 | p2 | (AT)7 | GGGGAACCTGCACAAGTTTA | GTTGCTTCGGGGAATTGTTA | 202 |
| chr6:97820975-97822990 | p5 | (ACCCG)3 | AGCCTCCAATCGTCTCAGAA | TTCCCTACCCGATGAGTCTG | 224 |
| chr6:97872993-97875008 | p5 | (AAAAG)3 | ACACGTGGAAAAGCCAAAAG | ATTAAGGGGGAAGAAATCGC | 174 |
| chr6:97956151-97958165 | p2 | (AT)7 | AGTCGACGAAGAAAAACGGA | TTTGTCCAGTGAGGGTTTCC | 115 |
| chr6:98000211-98002251 | p2 | (TA)20 | CAATCCATGAATTCCCAAGG | AATCGACAAACCGTCGAATC | 201 |
| chr6:98023989-98026001 | p2 | (AT)6 | TGCTCTTTCAGCCCACTTTT | CAGTTCGGTCCTTGGTGTTT | 187 |
| chr6:98023989-98026001 | p3 | (ATT)5 | TGCTCTTTCAGCCCACTTTT | CAGTTCGGTCCTTGGTGTTT | 187 |
| chr6:98057776-98059801 | p5 | (ACAAA)5 | GCTCCAAATTGCGAGAAAAG | CGAGGGGATTCCTAAACACA | 129 |
| chr6:98061177-98063193 | p4 | (CATT)4 | GTCCAAAACGAATTGGGAGA | AGGACGTGCTGAAGCTCAAT | 192 |
| chr6:98089448-98091472 | p2 | (AT)10 | CATGGACTAGTGCCCAACCT | CCTTGGAATTCTTTTTGCGA | 152 |
| chr6:98372419-98374431 | p2 | (TA)6 | GCATGTATGGTATCCGCCTT | TGTATACACCTGCGTGGCAT | 268 |
| chr6:98420759-98422774 | p5 | (TCATA)3 | GAGTTTGTTGGGGTGAGCAT | ATCTCCCCGGTATCTGGTCT | 184 |
| chr6:98420759-98422774 | p2 | (AT)8 | GAGTTTGTTGGGGTGAGCAT | ATCTCCCCGGTATCTGGTCT | 184 |
| chr6:98466527-98468539 | p2 | (AT)6 | CGTTTTCGTGCGCATATTTA | GTGTCCCGACCCTCTAGTGA | 157 |
| chr6:98536739-98538755 | p2 | (AT)8 | CGCTCTAACCTGGGTGTCTC | TGCATGTTTAGAGGTGCTCG | 291 |
| chr6:98692152-98694184 | p2 | (TA)16 | ACGATGGTCGTTGTGTTGAA | CGTTCGAACCATGTGAATTG | 201 |
| chr6:98702690-98704708 | p2 | (GA)9 | GAGTGTCGTCTTCGGGATGT | GAAATTTCCGTCGATTTCCA | 123 |
| chr6:98752983-98754998 | p2 | (AG)6 | CGGGACTCCACTTCGTAAAA | CCCTTTACGTCATAGGCTGC | 152 |
| chr6:98753033-98755057 | p2 | (AG)6 | CGGGACTCCACTTCGTAAAA | CCCTTTACGTCATAGGCTGC | 102 |
| chr6:98819250-98821280 | p6 | (CCCCAC)5 | CATGACACTTGCTTGATGGG | CAATCCCTTCCCAGCACTTA | 185 |
| chr6:98819250-98821280 | p6 | (TTGGAG)4 | CATGACACTTGCTTGATGGG | CAATCCCTTCCCAGCACTTA | 185 |
| chr6:98836555-98838627 | p6 | (TTTCTT)4 | GTCGCACTGGGTAAAGCACT | AATGACAGGAGACGAGTGGG | 132 |
| chr6:98836555-98838627 | p2 | (TA)36 | GTCGCACTGGGTAAAGCACT | AATGACAGGAGACGAGTGGG | 132 |
| chr6:98904753-98906774 | p3 | (GTT)7 | ATATCCGCATTTGAACCCAA | TACAAAAGTCCCCAGGTTCG | 266 |
| chr6:98904753-98906774 | p6 | (AAAAAT)3 | ATATCCGCATTTGAACCCAA | TACAAAAGTCCCCAGGTTCG | 266 |
| chr6:98987815-98989839 | p3 | (ATA)6 | CCCTAACCGGGTATGGACTT | ATCACCACCTTCTTGATCCG | 212 |
| chr6:99021315-99023327 | p2 | (AT)6 | AACGTAGTGGGGAGAACGTG | AAAACCCAATCCCTTCAACC | 250 |

| Marker ID | SSR type | SSR motif | FORWARD PRIMER (5'-3') | REVERSE PRIMER (5'-3') | size (bp) |
|---|---|---|---|---|---|
| chr6:99021482-99023500 | p2 | (AT)6 | GAGAGAGGGAGATGTGCGAC | AAAACCCAATCCCTTCAACC | 132 |
| chr6:99021510-99023532 | p2 | (AT)6 | GAGAGAGGGAGATGTGCGAC | AAAACCCAATCCCTTCAACC | 104 |
| chr6:99059872-99061884 | p2 | (TA)6 | CATTAGGCGGGATTCAGTGT | GCAATTTCCTTGGACCTCAA | 160 |
| chr6:99059872-99061884 | p2 | (TA)7 | CATTAGGCGGGATTCAGTGT | GCAATTTCCTTGGACCTCAA | 160 |
| chr6:99060072-99062086 | p2 | (TA)6 | TTGCGAAAAAGGTATGGGAC | TTAACTGACAACGCCCCTTC | 267 |
| chr6:99060072-99062086 | p2 | (TA)7 | TTGCGAAAAAGGTATGGGAC | TTAACTGACAACGCCCCTTC | 267 |
| chr6:99061114-99063129 | p5 | (TTGCA)3 | ATTCCAATCTTGCCCTTGTG | CATGCTATCCCCGTCACTTT | 232 |
| chr6:99076353-99078367 | p2 | (TA)6 | CCGTTTGGTCGTTTAGCACT | CATTAACTGACAACGCCCCT | 241 |
| chr6:99076353-99078367 | p2 | (TA)7 | CCGTTTGGTCGTTTAGCACT | CATTAACTGACAACGCCCCT | 241 |
| chr6:99098857-99100879 | p2 | (AT)11 | CAATATCTGTTGGGGTTGGG | GCCCTTTAATGGATGGGAGT | 102 |
| chr6:99139120-99141141 | p3 | (ACA)7 | CGGGCAATCCACTACATCTT | TGGATTAGGCTGCAGGAGTT | 184 |
| chr6:99193425-99195440 | p2 | (AT)13 | GCCTCTTTGCCGTAGTCTTG | GTTTATGGTGTGTTGCGTGC | 124 |
| chr6:99193425-99195440 | p2 | (TA)9 | GCCTCTTTGCCGTAGTCTTG | GTTTATGGTGTGTTGCGTGC | 124 |
| chr6:99193425-99195440 | p5 | (ATCAT)3 | GCCTCTTTGCCGTAGTCTTG | GTTTATGGTGTGTTGCGTGC | 124 |
| chr6:99211813-99213829 | p2 | (AT)8 | AACAGCGTCAGTTCGAGGAT | CCAGTACAAGCATCTGGGGT | 227 |
| chr6:99513938-99515956 | p3 | (ATT)10 | AACTCACGGCTTCCAATCAC | CCACTCCAATCTCCCTCAAA | 211 |
| chr6:99517820-99519832 | p2 | (TA)6 | AACTGGAAGCAGTGCCTCAT | TCACCCCTTTCCAGATTCAG | 186 |
| chr6:99624814-99626830 | p2 | (AT)8 | AGTGGGCGGCTATTATGATG | CTGATGGCCTCTTAAGGCTG | 258 |
| chr6:99657558-99659734 | p4 | (GATA)44 | GACAGTGGAGCACCTTGGAT | ATTGCGTTGACCTATCAGGG | 155 |
| chr6:99657558-99659734 | p4 | (GATA)9 | GACAGTGGAGCACCTTGGAT | ATTGCGTTGACCTATCAGGG | 155 |
| chr6:99721953-99723965 | p2 | (TA)6 | AGATTTCCTTCTTGGCCGTT | TGCAGCAAAGTGGATCAGAC | 299 |
| chr6:99866418-99868434 | p2 | (AT)8 | ATATGGGGCCCACAATTTTT | GAAACCCACCACATCCAAAC | 121 |
| chr6:99938097-99940109 | p2 | (TA)6 | CAATGAAATGCAACCGTCAC | AGGTTTCTGCTGGAGAGCAC | 163 |
| chr6:99946888-99948906 | p2 | (AG)9 | TTTTTACGATGATGGTGGCA | GCAAAGAAGAAAAAGCCGAG | 292 |
| chr6:99978498-99980510 | p2 | (TA)6 | CTGCCCCAGCTGTAGTCTTC | TGCAGGACGATGTTGATTGT | 149 |
| chr6:100076049-100078061 | p2 | (AT)9 | TGTAAAGGAGTACGGGGACG | AGGGAATCGCCTCTTACCAT | 200 |
| chr6:100076074-100078088 | p2 | (AT)9 | TGTAAAGGAGTACGGGGACG | AGGGAATCGCCTCTTACCAT | 175 |
| chr6:100113810-100115825 | p5 | (AAAAT)3 | GTTGCTGAAGTTGCGAATGA | TCCTCGACAACAGTCGACAC | 143 |
| chr6:100113810-100115825 | p5 | (AAATA)3 | GTTGCTGAAGTTGCGAATGA | TCCTCGACAACAGTCGACAC | 143 |
| chr6:100113974-100115989 | p5 | (AAAAT)3 | ATTTGGGTGTGGTGGATGAT | ATTCAAATCCACGGAGCAAC | 142 |
| chr6:100113974-100115989 | p5 | (AAATA)3 | ATTTGGGTGTGGTGGATGAT | ATTCAAATCCACGGAGCAAC | 142 |
| chr6:100222099-100224114 | p5 | (TATTA)3 | AAAGGGAACGTTGTTTGACG | AACTGGCCAAAGTGCAAATC | 109 |
| chr6:100290252-100292267 | p3 | (ATG)5 | TAAACCTCTCGGGAGACCCT | CCAACAAGGAAGAGGTTCCA | 138 |
| chr6:100364215-100366236 | p3 | (TTC)7 | TAGCCATATTCAACTCCCCG | TATGTGGACAAAAGTCGGCA | 134 |
| chr6:100394616-100396631 | p5 | (TACAA)3 | TGGATGGTGATTTCCTGTGA | GATAGTGGAGGGTGGAGCAA | 192 |
| chr6:100394616-100396631 | p3 | (TTA)5 | TGGATGGTGATTTCCTGTGA | GATAGTGGAGGGTGGAGCAA | 192 |

| Marker ID | SSR type | SSR motif | FORWARD PRIMER (5'-3') | REVERSE PRIMER (5'-3') | size (bp) |
|---|---|---|---|---|---|
| chr6:100464167-100466188 | p3 | (GTT)7 | TTTGGATGGTCGTTACCCAT | AGGCAACCATCCTTTCAATG | 244 |
| chr6:100464167-100466188 | p2 | (TA)6 | TTTGGATGGTCGTTACCCAT | AGGCAACCATCCTTTCAATG | 244 |
| chr6:100465683-100467711 | p2 | (TA)6 | GCCCTATCGACCAAGTACCA | CGTTGTATTGGGCCTTTTGT | 190 |
| chr6:100465683-100467711 | p2 | (AT)14 | GCCCTATCGACCAAGTACCA | CGTTGTATTGGGCCTTTTGT | 190 |
| chr6:100465683-100467711 | p3 | (CAA)5 | GCCCTATCGACCAAGTACCA | CGTTGTATTGGGCCTTTTGT | 190 |
| chr6:100482182-100484202 | p2 | (AG)21 | GACCACCCCTAAAAGCAACA | GAGCGCATGCCTATAATGGT | 159 |
| chr6:100482182-100484202 | p4 | (TAAT)5 | GACCACCCCTAAAAGCAACA | GAGCGCATGCCTATAATGGT | 159 |
| chr6:100501014-100503026 | p2 | (TA)6 | TGCAGAAAATGGGAGCTTTT | CGTTTGTGGTACAAGCATGG | 271 |
| chr6:100501014-100503026 | p3 | (TCT)5 | TGCAGAAAATGGGAGCTTTT | CGTTTGTGGTACAAGCATGG | 271 |
| chr6:100518774-100520792 | p2 | (AT)9 | AAAAGTGTCGGGTTCTGTGG | CAAAAATGCAGCTTGACCCT | 140 |
| chr6:100518774-100520792 | p2 | (AT)9 | AAAAGTGTCGGGTTCTGTGG | CAAAAATGCAGCTTGACCCT | 140 |
| chr6:100518774-100520792 | p2 | (AT)8 | AAAAGTGTCGGGTTCTGTGG | CAAAAATGCAGCTTGACCCT | 140 |
| chr6:100724238-100726253 | p3 | (GAA)5 | CCCAAATTTCCGAGAGCATA | ACCAATTGTCGCCTAGGATG | 299 |
| chr6:100724238-100726253 | p3 | (TTA)6 | CCCAAATTTCCGAGAGCATA | ACCAATTGTCGCCTAGGATG | 299 |
| chr6:100817897-100819913 | p2 | (AT)8 | CCCGCGTAAAATAGCAACAT | CCTTTTAAGTTCCAGCGACG | 237 |
| chr6:100828197-100830218 | p2 | (TA)24 | CAAGGCCTCATTATGCGATT | AATGCCACCAAAGGTGAAAG | 189 |
| chr6:100828260-100830302 | p2 | (TA)24 | CAAGGCCTCATTATGCGATT | AATGCCACCAAAGGTGAAAG | 126 |
| chr6:100984813-100986829 | p2 | (TA)8 | AAAGACCACGCATTACAGGG | TCAAATATCGGGAGGGACAC | 132 |
| chr6:101018243-101020263 | p2 | (AT)10 | TAGCGCCAAGTACCCAATCT | TCCATACCTTGCCACATTCA | 255 |
| chr6:101067870-101069882 | p2 | (TC)6 | AACTTAGGCTGCTGCATGGT | ACTTTCTGCCAGTCGAGGAA | 127 |
| chr6:101136515-101138529 | p2 | (TA)7 | TGGCCATGAGAATTCCAAAT | CAAAAAGACTTAGGCCCCGT | 155 |
| chr6:101177529-101179541 | p2 | (AT)6 | ACATGTTCGGCGAAACTCTT | TACAATGCTCTTCCGCACAG | 272 |
| chr6:101223639-101225719 | p2 | (TA)40 | TTTCATGATGCAGGTACGGA | TATGAGCCAAACCGGCTAAC | 135 |
| chr6:101236316-101238334 | p6 | (GGGGGT)3 | AGTCCATTCCTGGCAGTTTG | CAGCAATGCCGATATGAAGA | 285 |
| chr6:101268011-101270085 | p2 | (TA)9 | ATGACTACGGTGGTTTTGGC | ATGTGAGGGCGTAAGGATTG | 290 |
| chr6:101268011-101270085 | p2 | (AT)37 | ATGACTACGGTGGTTTTGGC | ATGTGAGGGCGTAAGGATTG | 290 |
| chr6:101311397-101313417 | p2 | (CT)10 | GATCCAAGAACACCTCGGAA | GCTGTTTGGGGCATGTTATT | 236 |
| chr6:101545341-101547359 | p5 | (TCATA)12 | GCGCCTTACCTTTCCCTTAC | TGCATGCTGCTCACATGATA | 181 |
| chr6:101545341-101547359 | p3 | (CAA)6 | GCGCCTTACCTTTCCCTTAC | TGCATGCTGCTCACATGATA | 181 |
| chr6:101607336-101609348 | p2 | (TA)6 | GCCACATGAGGATGAGGTTT | CGTGCAAAAGAGAAGCATGA | 202 |
| chr6:101607336-101609348 | p2 | (AT)6 | GCCACATGAGGATGAGGTTT | CGTGCAAAAGAGAAGCATGA | 202 |
| chr6:101651564-101653582 | p5 | (AACCG)4 | GGGATTAGCCAAGGAAAAGG | GGCTATTTCGGTTTCGGTTT | 144 |
| chr6:101651564-101653582 | p2 | (TA)9 | GGGATTAGCCAAGGAAAAGG | GGCTATTTCGGTTTCGGTTT | 144 |
| chr6:101651564-101653582 | p2 | (TA)27 | GGGATTAGCCAAGGAAAAGG | GGCTATTTCGGTTTCGGTTT | 144 |
| chr6:101667180-101669194 | p5 | (GAAGA)4 | AGGCTGGTTTGGTTTTGATG | ACCCTCCACTTGACACAAGG | 136 |
| chr6:101667180-101669194 | p2 | (TG)7 | AGGCTGGTTTGGTTTTGATG | ACCCTCCACTTGACACAAGG | 136 |

| Marker ID | SSR type | SSR motif | FORWARD PRIMER (5'-3') | REVERSE PRIMER (5'-3') | size (bp) |
|---|---|---|---|---|---|
| chr6:101770868-101772888 | p2 | (TC)10 | ATATGAATCTTGGCTCCCCC | CACTTCAAAGCACTGCCAAA | 109 |
| chr6:101791058-101793070 | p2 | (TA)6 | TCTAACCAATGACCGAAGGC | TGCGTCTATCATTGGAGCTG | 229 |
| chr6:101836073-101838087 | p2 | (AG)7 | ATCCAATGACACACAAGCGA | AAGCACCCATTTCGCTACAC | 174 |
| chr6:101836073-101838087 | p3 | (AAT)15 | ATCCAATGACACACAAGCGA | AAGCACCCATTTCGCTACAC | 174 |
| chr6:102007954-102009966 | p2 | (CA)6 | CCTTCTGAGGGAAGAAGCCT | GGTCAACCAGAAGACAGGGA | 211 |
| chr6:102021886-102023900 | p2 | (TA)7 | TTACCCAATCCCGGTGTTTA | TTTTCACTCAAAAGCGCTCA | 175 |
| chr6:102281037-102283053 | p2 | (AT)6 | AGCATCGTAAAATGGATGCAG | CCAGAATATGCATGGCACAA | 181 |
| chr6:102281037-102283053 | p3 | (CTA)7 | AGCATCGTAAAATGGATGCAG | CCAGAATATGCATGGCACAA | 181 |
| chr6:102281307-102283319 | p2 | (AT)6 | AAAGGAGACGAGACGAGCAA | CGCGATGCGTGTTCATTAT | 276 |
| chr6:102281307-102283319 | p3 | (CTA)7 | AAAGGAGACGAGACGAGCAA | CGCGATGCGTGTTCATTAT | 276 |
| chr6:102281482-102283500 | p2 | (AT)6 | AAAGGAGACGAGACGAGCAA | CGCGATGCGTGTTCATTAT | 101 |
| chr6:102281482-102283500 | p3 | (CTA)7 | AAAGGAGACGAGACGAGCAA | CGCGATGCGTGTTCATTAT | 101 |
| chr6:102305513-102307527 | p2 | (AT)7 | ATAATCAACGATTGCGGAGG | AGCGAGGATGTTAGCCTCAA | 193 |
| chr6:102305513-102307527 | p2 | (AT)7 | ATAATCAACGATTGCGGAGG | AGCGAGGATGTTAGCCTCAA | 193 |
| chr6:102318986-102321004 | p3 | (TTA)6 | CTTGAGCAAAGCCAAGTTCC | CTGATTCAGATATGGCCGGT | 286 |
| chr6:102332189-102334205 | p2 | (AT)8 | TTGAAGCACTTTTCCTTCCC | TTTCGCTCGAAAGTGAGACA | 146 |
| chr6:102486897-102488918 | p2 | (AT)11 | GTTGGAAGTGGTGTGTGTGC | CCGCAACGCTTACTATGTCA | 232 |
| chr6:102486897-102488918 | p3 | (GTT)7 | GTTGGAAGTGGTGTGTGTGC | CCGCAACGCTTACTATGTCA | 232 |
| chr6:102506209-102508230 | p2 | (AT)11 | GTTGGAAGTGGTGTGTGTGC | CCGCAACGCTTACTATGTCA | 232 |
| chr6:102506209-102508230 | p3 | (GTT)7 | GTTGGAAGTGGTGTGTGTGC | CCGCAACGCTTACTATGTCA | 232 |
| chr6:102697730-102699744 | p3 | (TTA)10 | GGAGAGACTGATGGGGTCAA | ATATTTTGGGGAAAGACGGG | 194 |
| chr6:102697750-102699764 | p3 | (TTA)10 | GGAGAGACTGATGGGGTCAA | ATATTTTGGGGAAAGACGGG | 174 |
| chr6:102755624-102757642 | p2 | (AT)9 | GTATAAACCGTGGCTTCGGA | GTCGCTCCGATCTAAACCAA | 181 |
| chr6:102813407-102815421 | p2 | (TG)7 | AAAGGTGCCTACGGGCTATT | CTCCATAGCGATCGGTGTTT | 208 |
| chr6:102830697-102832721 | p2 | (TA)12 | GAAGTCTTCGGGTTCCTTCC | CCCATGTACAAAAGACGGGT | 262 |
| chr6:102847584-102849604 | p5 | (AAAAT)4 | GTCACTTTGCAGGAAAAGGC | GAATGCTCGGAAAGCAAAAA | 121 |
| chr6:102859319-102861334 | p5 | (AAACA)3 | ACCGGTGTAATCCCACACAT | TATGCTTTTCGGTTATCGGG | 152 |
| chr6:102859319-102861334 | p5 | (TTTAT)3 | ACCGGTGTAATCCCACACAT | TATGCTTTTCGGTTATCGGG | 152 |
| chr6:102946679-102948697 | p6 | (TTTTAT)3 | ACACTAAAATTCGCCCGATG | CAGGTGAGAGGGCATTGATT | 138 |
| chr6:102989433-102991451 | p6 | (ACTGAA)3 | TTCAAAGAGCACCTGCACAC | GGCGTGACACTCTTAGAGGC | 250 |
| chr6:102993334-102995370 | p2 | (TA)18 | ACGACGAAGCATGGGTAAAC | TCCTTTGTAGTCCCACCCAC | 237 |
| chr6:103191153-103193169 | p2 | (AT)8 | CAGTCTGCTTACCCCACCAT | TTTTGGGTGTGACGGAAAAT | 241 |
| chr6:103211464-103213482 | p2 | (AT)9 | TTTTTCCACTTTCCCCTTCA | TATCGATGCACTCATGCACA | 268 |
| chr6:103213510-103215562 | p2 | (TA)26 | AATGCCGTTTGGTCGTTTAG | TAACCACCACCTATCCCTGC | 116 |
| chr6:103302314-103304334 | p2 | (TA)10 | GAAGCCGCATGATACCAATTA | AATCGCTTCTCACAATTCGC | 210 |
| chr6:103314288-103316304 | p6 | (TCCACA)3 | CCACGAAGTCATTAGTGCGA | CTGCGTTGTGGATATTGTGG | 269 |

| Marker ID | SSR type | SSR motif | FORWARD PRIMER (5'-3') | REVERSE PRIMER (5'-3') | size (bp) |
|---|---|---|---|---|---|
| chr6:103314288-103316304 | p2 | (AT)8 | CCACGAAGTCATTAGTGCGA | CTGCGTTGTGGATATTGTGG | 269 |
| chr6:103321040-103323062 | p2 | (AT)11 | AACAATGGTGCCTCTTACGG | AGAGAAAACTCCGCCCAAAT | 231 |
| chr6:103342269-103344317 | p2 | (TA)23 | GATCCACTACCACTTCCCCA | TATGTTTCAGCCGAAAAGGG | 159 |
| chr6:103342269-103344317 | p2 | (AT)24 | GATCCACTACCACTTCCCCA | TATGTTTCAGCCGAAAAGGG | 159 |
| chr6:103645769-103647781 | p2 | (AT)27 | AGATCCGCCTCTGAGTTCAA | ATGGGCAAAGAGGTGTTCAG | 208 |
| chr6:103645769-103647781 | p2 | (AT)6 | AGATCCGCCTCTGAGTTCAA | ATGGGCAAAGAGGTGTTCAG | 208 |
| chr6:103645769-103647781 | p6 | (TAAAAC)3 | AGATCCGCCTCTGAGTTCAA | ATGGGCAAAGAGGTGTTCAG | 208 |
| chr6:103724210-103726231 | p3 | (TAT)7 | TTCAACTTTCGAAACAGGGG | TAGTTCGCTGCCGAAGAAAT | 243 |
| chr6:103724210-103726231 | p2 | (CT)9 | TTCAACTTTCGAAACAGGGG | TAGTTCGCTGCCGAAGAAAT | 243 |
| chr6:103734415-103736430 | p5 | (TCGGT)3 | CCGACCAAGAGAGCTTCAAC | TGCCAATTTCATCAGCAAAA | 229 |
| chr6:103799551-103801566 | p3 | (AAT)6 | TGTCAAGCGAGACCCTTTCT | TTTCATCCCACGTGGTTTTT | 219 |
| chr6:103799551-103801566 | p2 | (TA)11 | TGTCAAGCGAGACCCTTTCT | TTTCATCCCACGTGGTTTTT | 219 |
| chr6:103799551-103801566 | p3 | (ATA)5 | TGTCAAGCGAGACCCTTTCT | TTTCATCCCACGTGGTTTTT | 219 |
| chr6:103799551-103801566 | p3 | (TCT)10 | TGTCAAGCGAGACCCTTTCT | TTTCATCCCACGTGGTTTTT | 219 |
| chr6:103835620-103837634 | p2 | (TA)7 | TCTACCATCTCGCGATACCC | CCTCGCTAGGACTCATCCAG | 117 |
| chr6:103838524-103840542 | p6 | (CAACCC)3 | TTCACCTCAACTTGGTGCTG | TGCACGGACCCATTAAAAAT | 165 |
| chr6:103931294-103933312 | p2 | (TC)6 | TTTGCCTTAGGAACCCACAC | TCCCCAAAAAGAAGGGATCT | 202 |
| chr6:103931294-103933312 | p2 | (AT)9 | TTTGCCTTAGGAACCCACAC | TCCCCAAAAAGAAGGGATCT | 202 |
| chr6:103931294-103933312 | p3 | (ACA)6 | TTTGCCTTAGGAACCCACAC | TCCCCAAAAAGAAGGGATCT | 202 |
| chr6:103998543-104000561 | p2 | (AT)9 | GGTTTGGACCCCTACTTGGT | CAGCCAATCTAGCCCTCTTG | 294 |
| chr6:104001857-104003877 | p5 | (TGATA)4 | ATGATATCGTTCACCGGCTC | CCAGAACATCTGGGCTCATT | 156 |
| chr6:104037831-104039846 | p3 | (ATT)5 | TATTCGGCAATCTTCTTCCG | TTCACCTTTTTCCGCAATTC | 225 |
| chr6:104084705-104086729 | p4 | (AAAT)5 | GCCAGCAGAAACTAGCCATC | TCTGGCAACATTTCCCTTTC | 105 |
| chr6:104118569-104120603 | p2 | (TA)17 | TTGGGTCGTAGGCCTAGTTG | TGCATCCATTTTCAGTCCAA | 200 |
| chr6:104209825-104211849 | p6 | (GCAGGT)4 | AGTTGGCATCTGTCCCGTAG | CAATGGCTAAACGCTGAACA | 209 |
| chr6:104209825-104211849 | p3 | (CAA)5 | AGTTGGCATCTGTCCCGTAG | CAATGGCTAAACGCTGAACA | 209 |
| chr6:104323041-104325056 | p5 | (TTTTG)3 | AAGGGTGTCAGGCGTGTATC | TAAATGCACAAATGGCTTGC | 187 |
| chr6:104323041-104325056 | p5 | (TTTAT)3 | AAGGGTGTCAGGCGTGTATC | TAAATGCACAAATGGCTTGC | 187 |
| chr6:104332627-104334642 | p5 | (GTGCA)3 | TCATGAAACGTGATGGCACT | GCCTTACCTTGCATTTCTGC | 107 |
| chr6:104332627-104334642 | p2 | (AT)6 | TCATGAAACGTGATGGCACT | GCCTTACCTTGCATTTCTGC | 107 |
| chr6:104365495-104367510 | p5 | (ATTTT)3 | GCCTAGATCATTCAGCCGTC | TGCTGGTGATCTCAGATTCG | 181 |
| chr6:104365495-104367510 | p4 | (ATTA)4 | GCCTAGATCATTCAGCCGTC | TGCTGGTGATCTCAGATTCG | 181 |
| chr6:104393014-104395032 | p3 | (CTT)6 | TGTGGGTGACGCTATTACCA | GCGGAGAGTGTACTGTGCAA | 112 |
| chr6:104415745-104417761 | p2 | (GA)8 | TCAACCAGAGGATGCATGAG | TCACCCGCCTCTTAAAACAC | 199 |
| chr6:104432763-104434777 | p2 | (TA)7 | AGACGAAAAGAGGTCGGGTT | ACGGTCTGCCACGTTTTATC | 275 |
| chr6:104467436-104469451 | p5 | (TTTAT)3 | GGACCAGCCCAATCCTATTT | GAGAGTGAGGGAAAGCGATG | 105 |

| Marker ID | SSR type | SSR motif | FORWARD PRIMER (5'-3') | REVERSE PRIMER (5'-3') | size (bp) |
|---|---|---|---|---|---|
| chr6:104491923-104493974 | p5 | (AAGAG)3 | ACCTTTGAGCCCCCAAATAC | GGACGAAAGGATGGACAAAA | 159 |
| chr6:104491923-104493974 | p3 | (TAA)17 | ACCTTTGAGCCCCCAAATAC | GGACGAAAGGATGGACAAAA | 159 |
| chr6:104547268-104549280 | p2 | (GA)6 | GCGGAAATACAGACGCTAGG | AATGGGAGGATATTTGCGTG | 152 |
| chr6:104568179-104570193 | p2 | (AT)7 | CGGCAAACCATGAATCCTAT | TGGGATGATAAAGGGGATGA | 182 |
| chr6:104569380-104571394 | p2 | (TA)7 | TTCATCATCATCCTCGTCCA | CCATTGCAGGCTCTTATGCT | 101 |
| chr6:104574550-104576568 | p6 | (ATTTTT)3 | ATAATGCAGTTTCATCCGCC | AGTCGGGCTTCAATGTGAGT | 249 |
| chr6:104616363-104618377 | p2 | (TA)7 | AAAAGCCGGAAAAAGAAGGA | TCCATTGGTAGTTTTCCCCA | 270 |
| chr6:104636765-104638795 | p6 | (TTTCTC)5 | AGCCCCAATGTAGTCCCTCT | TTTTATGCTGCTGCAAATGG | 158 |
| chr6:104661694-104663706 | p2 | (AT)6 | TGCAAGTTCAGTGAAGCACC | CATCGTCAATCTCTCCCGTT | 165 |
| chr6:104761152-104763178 | p2 | (TA)13 | TACCTGGGTTGGAGTTCGTC | AACAGCACATCCTTCATCCC | 269 |
| chr6:104832664-104834682 | p3 | (GTT)6 | CACGAAAATCATGGGCTTCT | ATGTTACATTCGCCTGGACC | 110 |
| chr6:104838490-104840505 | p3 | (GTT)5 | AGGTAGGGGTAGGCCAAAGA | GAAGTAACGCAAGCGACACA | 246 |
| chr6:104838490-104840505 | p5 | (TTTAT)3 | AGGTAGGGGTAGGCCAAAGA | GAAGTAACGCAAGCGACACA | 246 |
| chr6:104911505-104913520 | p5 | (TTTCT)3 | CCCAATCCTACGTGGCTTTA | ACCCAATTAATGGTGGTGGA | 255 |
| chr6:104921659-104923728 | p3 | (AAT)26 | TAGCCAATTTTGTTCCGACC | ACAATTCGAAACGGGTTCAG | 254 |
| chr6:104921659-104923728 | p3 | (TAA)23 | TAGCCAATTTTGTTCCGACC | ACAATTCGAAACGGGTTCAG | 254 |
| chr6:104950074-104952086 | p2 | (TA)6 | CTGGACAGATCGTGCAGAAA | CAAAGGTCTGCTGGTTTCGT | 141 |
| chr6:104950074-104952086 | p2 | (TA)8 | CTGGACAGATCGTGCAGAAA | CAAAGGTCTGCTGGTTTCGT | 141 |
| chr6:104986391-104988409 | p6 | (ATAAAA)3 | AAAATCCCAACAGAATCCCC | TACCGCACCACTCACTCAAC | 112 |
| chr6:104992427-104994442 | p5 | (TCAAG)3 | ATGAATCAATGCCAACGACA | AATGCCACGTAAGATACGGC | 142 |
| chr6:105035750-105037816 | p2 | (TA)33 | TTTGCCCTTCAAAATCGAAC | ATTGCACTGTTTGTCCCTCC | 185 |
| chr6:105154378-105156402 | p3 | (ATT)8 | ACACACGCAAATGTCATCGT | CCCTTCGTCACTTTTGGTTC | 258 |
| chr6:105213081-105215099 | p2 | (AT)9 | ACGAAACTCTTCCTTGCGAA | CGGGCTGGAGAAAGACATAA | 189 |
| chr6:105326465-105328480 | p5 | (GTTCG)3 | GCGCAAAATGCATCAAAATA | TGTTCGGTGTCCATCTTCAA | 186 |
| chr6:105408620-105410635 | p5 | (TCATA)3 | GTTTGATCGTGGCCGTTAGT | GTAGTGCCCTATGGGCTTCA | 182 |
| chr6:105415290-105417305 | p5 | (TATGA)3 | TTGAGGAAACGGGAATTGAG | TGAAAAAGGGGCAAATTCAG | 205 |
| chr6:105415290-105417305 | p5 | (TATGA)3 | TTGAGGAAACGGGAATTGAG | TGAAAAAGGGGCAAATTCAG | 205 |
| chr6:105451482-105453497 | p5 | (AAAAT)3 | ATCTAGCGGCAGTTTGGAGA | TGTCCCAAAAAGATTGTCCC | 128 |
| chr6:105490736-105492760 | p3 | (TGG)8 | AAGCGGTGAAAGGGGTAAAT | TTGTTCATGAATGAGCCCAA | 299 |
| chr6:105490736-105492760 | p2 | (AG)8 | AAGCGGTGAAAGGGGTAAAT | TTGTTCATGAATGAGCCCAA | 299 |
| chr6:105502548-105504563 | p3 | (TCC)5 | TTCCCCATCAATCCCAATTA | TCGCCGCCTTTTATTCTCTA | 130 |
| chr6:105511899-105513913 | p2 | (AT)7 | CCATTTCCCTTTTTCAGCAA | TCACTGGCAATTCCGTTACA | 203 |
| chr6:105511899-105513913 | p3 | (TGG)5 | CCATTTCCCTTTTTCAGCAA | TCACTGGCAATTCCGTTACA | 203 |
| chr6:105531276-105533298 | p2 | (TA)11 | TTGAAAATCCGTTCGAGGAC | CAAGCCCGATGAAGAGAAAG | 253 |
| chr6:105548629-105550703 | p3 | (TTG)5 | AAGATCCGATCCCCAAAAAC | GGCTGGAATACCAAGAACCA | 132 |
| chr6:105564621-105566645 | p6 | (TTGGAG)4 | GAACTAACCATGGTGCCGAT | CAATGCATGGCCCTAGAAAT | 110 |

| Marker ID | SSR type | SSR motif | FORWARD PRIMER (5'-3') | REVERSE PRIMER (5'-3') | size (bp) |
|---|---|---|---|---|---|
| chr6:105676227-105678249 | p2 | (TA)11 | AGCGAATGACAACATCATGC | TGTGTTTGGTACGACGGAAA | 277 |
| chr6:105676227-105678249 | p2 | (AC)6 | AGCGAATGACAACATCATGC | TGTGTTTGGTACGACGGAAA | 277 |
| chr6:105690335-105692355 | p2 | (AT)7 | TTTTTGATGAACCCCTGGAG | TTCATTGCCGGAAGTTCTTT | 242 |
| chr6:105690401-105692421 | p2 | (AT)7 | TTTTTGATGAACCCCTGGAG | TTCATTGCCGGAAGTTCTTT | 176 |
| chr6:105693770-105695782 | p2 | (TA)6 | AGGAAGGGGAAATGAATTGG | CTCAAACCGATAAACCGGAA | 144 |
| chr6:105693770-105695782 | p2 | (AT)11 | AGGAAGGGGAAATGAATTGG | CTCAAACCGATAAACCGGAA | 144 |
| chr6:105693770-105695782 | p2 | (AT)6 | AGGAAGGGGAAATGAATTGG | CTCAAACCGATAAACCGGAA | 144 |
| chr6:105740908-105742933 | p5 | (TTCAT)5 | CCGGAAGAGATCCAAGATGA | ATAACAATGCCTCACAGGGC | 229 |
| chr6:105787544-105789578 | p2 | (TA)17 | GGGCCCCATACTAAACAACA | CCCATGAGTTTGGAGATGGT | 257 |
| chr6:105787544-105789578 | p6 | (AATAAA)3 | GGGCCCCATACTAAACAACA | CCCATGAGTTTGGAGATGGT | 257 |
| chr6:105787544-105789578 | p2 | (AT)17 | GGGCCCCATACTAAACAACA | CCCATGAGTTTGGAGATGGT | 257 |
| chr6:105806632-105808650 | p6 | (TTTAAT)3 | GTGCTCCACGAGATCCATTT | GCGAACAACCCCAATCTAAA | 172 |
| chr6:105924018-105926034 | p2 | (AC)8 | TCATTCGTACGCCATTTTGA | GGGGAGAATCGTATGGACAG | 184 |
| chr6:105924018-105926034 | p2 | (AC)8 | TCATTCGTACGCCATTTTGA | GGGGAGAATCGTATGGACAG | 184 |
| chr6:105966878-105968893 | p5 | (TTCTT)3 | CACGGGCATCCTTGTCTAGT | TACAGCGACGCACCATAGAG | 296 |
| chr6:105978257-105980315 | p2 | (AT)29 | CCACGGATATCTTGGCTGTT | AATTTCCAGGGCCTGTTCTT | 210 |
| chr6:106077446-106079458 | p2 | (AT)6 | TGTGGTTTGGAGTTGTCCAG | TCTTAAGCTCCGTGCCAAGT | 211 |
| chr6:106351847-106353867 | p5 | (CGATC)4 | TCACAAATAACACCGGAGCA | GCGAGTTCGCAGAGGTATTC | 102 |
| chr6:106387102-106389116 | p3 | (AGC)6 | GGGCTAAAATCCCAACACAA | TTTCTTTCGTCATGCTGCTG | 259 |
| chr6:106387102-106389116 | p2 | (AT)7 | GGGCTAAAATCCCAACACAA | TTTCTTTCGTCATGCTGCTG | 259 |
| chr6:106451268-106453283 | p5 | (ATGCA)3 | ACACGCTCTCCTGCGTAGAT | TGGGTCGTGACAAGTTGGTA | 129 |
| chr6:106560950-106562962 | p2 | (AT)6 | CCCTCCGTTTACTCCATCAA | TTCTGGTAAATCGGTCCCTG | 125 |
| chr6:106582924-106584939 | p5 | (AATGC)3 | GGGGATGAGATGAGGATTCA | GCTAATTGGGTCGTGAAGGA | 290 |
| chr6:106582924-106584939 | p2 | (TA)9 | GGGGATGAGATGAGGATTCA | GCTAATTGGGTCGTGAAGGA | 290 |
| chr6:106659628-106661649 | p3 | (TAA)7 | ACCTAGGGCCATGACAAGTG | GTCGGCCATTCTTTTGTTGT | 221 |
| chr6:106738757-106740775 | p2 | (AT)9 | ACAGGTTGGTTGGACTCTGG | ATGGTGCGATTTGGAGTTTC | 275 |
| chr6:106738757-106740775 | p2 | (TA)9 | ACAGGTTGGTTGGACTCTGG | ATGGTGCGATTTGGAGTTTC | 275 |
| chr6:106791722-106793737 | p5 | (ATGCA)3 | TATAGCCTCCTGTGGATGGG | CTTCATTCTGGGATGGAGGA | 249 |
| chr6:106852250-106854264 | p2 | (TA)7 | CAAGTGGAGCAGTACAGCCA | GTTGCGACATGAACACCATC | 190 |
| chr6:106868143-106870158 | p3 | (ATA)5 | AGCCCCCAAATTCTTTCATT | ATCGGGAACATTCAGAGACG | 110 |
| chr6:106907547-106909559 | p2 | (TA)6 | AGTGGGGCATACAGACAAGC | TTTACAGAACCTGGTTGCCC | 167 |
| chr6:106957130-106959146 | p4 | (AATA)4 | TGGACGGTCCTTTAATTTCG | GGAAAGAGGGGTCCAAAAAC | 152 |
| chr6:106971787-106973802 | p3 | (CTT)8 | CGCATTCAACCCATTTTCTT | AAATGGACCGGTATCCCTTC | 226 |
| chr6:106971787-106973802 | p3 | (GAA)5 | CGCATTCAACCCATTTTCTT | AAATGGACCGGTATCCCTTC | 226 |
| chr6:106977650-106979666 | p4 | (ACAA)4 | GTGGAAGATCCCTTGGATGA | AGGGAGGGTATGCTCCATCT | 158 |
| chr6:106994718-106996786 | p2 | (AT)34 | CCAAGGTATTTTTCGGAGCA | TGCAAGCCTGGGTAAAGTCT | 237 |

| Marker ID | SSR type | SSR motif | FORWARD PRIMER (5'-3') | REVERSE PRIMER (5'-3') | size (bp) |
|---|---|---|---|---|---|
| chr6:107013009-107015025 | p2 | (AT)8 | ACCCCTGTCCACAAATACCA | ATTGTGGGGTGTGTGTGCTA | 292 |
| chr6:107124102-107126141 | p2 | (CT)6 | TCATTACCTACCCGCTGAGG | CCACCTTGTAATGAAGCCGT | 210 |
| chr6:107167716-107169752 | p6 | (CCGGTT)3 | TACATTCTGACAGCAAGGCG | GTTGGAGGGGATCGTAGTGA | 101 |
| chr6:107200798-107202813 | p5 | (TTTTG)4 | AGCGGTCTCTCATTGTGCTT | GATGCAAGAACCCAAATGCT | 189 |
| chr6:107200798-107202813 | p5 | (TTTTA)3 | AGCGGTCTCTCATTGTGCTT | GATGCAAGAACCCAAATGCT | 189 |
| chr6:107270701-107272725 | p3 | (TTG)5 | AGGCTATCAAATGGCACGAC | CCGAAAACAGTGGCAAAAAT | 294 |
| chr6:107342071-107344086 | p3 | (ACA)5 | AGAATTATCGGCATGTTGGC | CCTTTTCTCGTGGTCCAAAA | 238 |
| chr6:107401981-107404009 | p2 | (TA)10 | TCCCACGAAAATAGGGATCA | CTACGTCCTGTGCAGTCCAA | 144 |
| chr6:107401981-107404009 | p2 | (AT)14 | TCCCACGAAAATAGGGATCA | CTACGTCCTGTGCAGTCCAA | 144 |
| chr6:107412340-107414370 | p2 | (GA)6 | GTGCAGCTGGTGTGAACCTA | TGGATGGTCTACTGACCCGT | 223 |
| chr6:107412368-107414384 | p2 | (GA)6 | GTGCAGCTGGTGTGAACCTA | TGGATGGTCTACTGACCCGT | 195 |
| chr6:107562608-107564634 | p5 | (CAAAC)3 | ATTTTTCTCGGGTTTTTCGG | CAAACAGTGTTCTGCCTCCA | 186 |
| chr6:107562608-107564634 | p2 | (AT)13 | ATTTTTCTCGGGTTTTTCGG | CAAACAGTGTTCTGCCTCCA | 186 |
| chr6:107562608-107564634 | p3 | (TCT)19 | ATTTTTCTCGGGTTTTTCGG | CAAACAGTGTTCTGCCTCCA | 186 |
| chr6:107586526-107588584 | p2 | (TA)29 | TCTTCCGCTGCCAAATAATC | TCTTTGACGGTCCCAGACTC | 220 |
| chr6:107594635-107596647 | p2 | (AG)6 | TAATTATCTTTGGCCGCCTG | GAAGTCTACGGTGGCAGCAT | 114 |
| chr6:107599545-107601560 | p2 | (AT)8 | TGGTTCCAAGGTTGTCCTTC | ATGTTACAGCCCGCACTTTC | 273 |
| chr6:107599545-107601560 | p5 | (AAAAC)3 | TGGTTCCAAGGTTGTCCTTC | ATGTTACAGCCCGCACTTTC | 273 |
| chr6:107680762-107682777 | p3 | (ATG)5 | GCCCCTACTACCAAACCTCC | TAAAAAGGTGGCAAAATCGC | 100 |
| chr6:107709014-107711078 | p2 | (AT)13 | GAGAGTGCATTTAGCCAGGG | GCCTTCCTCCGCTTCTAAGT | 276 |
| chr6:107709080-107711092 | p2 | (AT)13 | AGTATCAATCGTGGCTTCGG | CTCTTCCTCTTCGCCTTCCT | 191 |
| chr6:107747835-107749853 | p6 | (TCAGTT)3 | GAGTGGGGATTTGGTGAAGA | ATTCCAGCCTTTGAGCGTAA | 234 |
| chr6:107749983-107751999 | p2 | (TA)8 | CATCGGGCAGATTTTTGAGT | ACCAACTTTGTGCAAAACCC | 154 |
| chr6:107941381-107943393 | p2 | (AT)6 | AAGCCCGGATTTGGTAGTCT | GCTAAAGCGCCTAAACATGC | 232 |
| chr6:107993467-107995491 | p2 | (TG)12 | GGGGATCTTTTTGACCCATT | TGGTAAACAAAGGGAGGCTG | 153 |
| chr6:108029147-108031162 | p5 | (TACAA)3 | ATTCCGTACTGACCCCCTTT | GGAATAAATGGAGCTTGCCA | 246 |
| chr6:108114715-108116729 | p2 | (AT)7 | GGCGGATATTAGCCAAATGA | GAAGAAAGAGCAAGGAGCCA | 173 |
| chr6:108130636-108132650 | p2 | (AC)7 | AGGCATGAAGAAGTTGGTCG | TGCTACTCCCCAGAAGCACT | 130 |
| chr6:108152975-108154987 | p2 | (AC)9 | TCGCGAAATTCTGTCTTGTG | CTTCATTTGGCGTGGTCTTT | 296 |
| chr6:108152975-108154987 | p2 | (TA)33 | TCGCGAAATTCTGTCTTGTG | CTTCATTTGGCGTGGTCTTT | 296 |
| chr6:108152975-108154987 | p2 | (AT)16 | TCGCGAAATTCTGTCTTGTG | CTTCATTTGGCGTGGTCTTT | 296 |
| chr6:108153010-108155022 | p2 | (AC)9 | TCGCGAAATTCTGTCTTGTG | CTTCATTTGGCGTGGTCTTT | 261 |
| chr6:108153010-108155022 | p2 | (TA)33 | TCGCGAAATTCTGTCTTGTG | CTTCATTTGGCGTGGTCTTT | 261 |
| chr6:108153010-108155022 | p2 | (AT)16 | TCGCGAAATTCTGTCTTGTG | CTTCATTTGGCGTGGTCTTT | 261 |
| chr6:108195485-108197500 | p5 | (GGCCC)3 | AACGGCTAACATCGATGGAC | CCACCTTCTCCTCAAGTTCG | 259 |
| chr6:108195485-108197500 | p2 | (CT)6 | AACGGCTAACATCGATGGAC | CCACCTTCTCCTCAAGTTCG | 259 |

| Marker ID | SSR type | SSR motif | FORWARD PRIMER (5'-3') | REVERSE PRIMER (5'-3') | size (bp) |
|---|---|---|---|---|---|
| chr6:108253647-108255662 | p3 | (AAT)5 | GAGGGCTACCAGTCCAACAG | CGGTGAAGTAAAGCGTGACA | 124 |
| chr6:108253647-108255662 | p6 | (TAAAAA)3 | GAGGGCTACCAGTCCAACAG | CGGTGAAGTAAAGCGTGACA | 124 |
| chr6:108253647-108255662 | p3 | (CAT)5 | GAGGGCTACCAGTCCAACAG | CGGTGAAGTAAAGCGTGACA | 124 |
| chr6:108301339-108303385 | p2 | (TA)7 | GGGCAGCGTATTCGTATTGT | TGAGTATCAATGGCGACAGC | 164 |
| chr6:108301339-108303385 | p2 | (TA)23 | GGGCAGCGTATTCGTATTGT | TGAGTATCAATGGCGACAGC | 164 |
| chr6:108307389-108309411 | p2 | (TA)11 | GCACTCCAAAGTCCGGTTTA | GAACTCGGTCACAGCTCACA | 281 |
| chr6:108442954-108444969 | p5 | (TAAAA)3 | GTCGGGTCATATAGCGCAGT | CGGAACCAAAATGCTCCTAA | 213 |
| chr6:108442954-108444969 | p4 | (TCTT)4 | GTCGGGTCATATAGCGCAGT | CGGAACCAAAATGCTCCTAA | 213 |
| chr6:108516261-108518279 | p6 | (AAAAAT)3 | GTCCACTCAAACATCCCCAT | AAAGCTTTAATGGCAACCGA | 277 |
| chr6:108594264-108596279 | p5 | (ACCGA)3 | GGGGTGGTGACATCTCATTC | ATGACAACCCCTTTTTGCAG | 111 |
| chr6:108697872-108699887 | p3 | (AGA)5 | GTAACCGCCGCATGTAGATT | ATCCATCATATAGCCGCCAG | 213 |
| chr6:108710033-108712053 | p2 | (TA)10 | TGGATCGATAACCCACCAAT | TAGCGGTGGAACACTTCACA | 120 |
| chr6:108712067-108714082 | p5 | (AAAAG)3 | TTTGTTCGCACACCCAATAA | GGCTTCGGTGACCGTAAATA | 205 |
| chr6:108878783-108880798 | p5 | (TATCG)3 | CACTCTCTTGGCTTTCTCGG | TTGCATTCTAAGATCGTGCG | 170 |
| chr6:108912607-108914622 | p5 | (TTGGT)3 | CTATGGAGAAACCGGACCAA | GACCGAAGGAAGAAAGGGAC | 271 |
| chr6:108961801-108963816 | p5 | (TTTGG)3 | CTGAGCTCGAGGACAAATCC | TGGCTCTCAAAGCACCTTTT | 164 |
| chr6:109102509-109104537 | p3 | (ATC)5 | GCCAGGAACCACCTGACTTA | CCTTGTCGGCCTTATTTTGA | 227 |
| chr6:109102536-109104554 | p3 | (ATC)5 | GCCAGGAACCACCTGACTTA | CCTTGTCGGCCTTATTTTGA | 200 |
| chr6:109102783-109104798 | p3 | (ATC)5 | TCAAAATAAGGCCGACAAGG | GGAGATGGATCAGACCCGTA | 116 |
| chr6:109130576-109132592 | p2 | (AG)8 | TTTCCGGTCTTGTTCTCCAC | TGTGTTTGGAGAAGGGGAAG | 233 |
| chr6:109133051-109135063 | p6 | (ATATGT)3 | TGCATATGGTTTCCGCACTA | GGTGCCACAACACAGCATAC | 145 |
| chr6:109133051-109135063 | p2 | (TA)6 | TGCATATGGTTTCCGCACTA | GGTGCCACAACACAGCATAC | 145 |
| chr6:109133051-109135063 | p2 | (AT)7 | TGCATATGGTTTCCGCACTA | GGTGCCACAACACAGCATAC | 145 |
| chr6:109138243-109140265 | p2 | (AG)11 | CATGCTCCGAAAATGTGTTG | ACTTCTTTGGTGCGCAGTCT | 254 |
| chr6:109210417-109212445 | p2 | (AT)14 | GCCAGGAACCACCTGACTTA | CCTTGTCGGCCTTATTTTGA | 208 |
| chr6:109240704-109242719 | p5 | (AAAAT)3 | ACTGATGTCCCCTCTTGTGG | TTGTGGGAAGGATGTTGTCA | 196 |
| chr6:109240704-109242719 | p2 | (AT)12 | ACTGATGTCCCCTCTTGTGG | TTGTGGGAAGGATGTTGTCA | 196 |
| chr6:109241206-109243230 | p5 | (AAAAT)3 | TGTAAGCTTCCGCATGACAC | GTCTATCTTCGTGCTTCGCC | 148 |
| chr6:109241206-109243230 | p2 | (AT)12 | TGTAAGCTTCCGCATGACAC | GTCTATCTTCGTGCTTCGCC | 148 |
| chr6:109294088-109296103 | p5 | (AAAAT)3 | CATATGTGCTGAATGGTCGG | TTGTGGGAAGGATGTTGTCA | 290 |
| chr6:109294088-109296103 | p2 | (AT)12 | CATATGTGCTGAATGGTCGG | TTGTGGGAAGGATGTTGTCA | 290 |
| chr6:109383480-109385540 | p3 | (AAG)20 | CGATGGCATATTTGAGCCTT | ATCACCCAAATTCATCCCAA | 227 |
| chr6:109434455-109436469 | p2 | (AC)7 | AATTTGGATGCTATGGCGAG | CCCAATAGGTGCCTTTTCAA | 192 |
| chr6:109491732-109493762 | p2 | (AT)15 | TTTTTGTATGAACGCTGCCA | AACGCCATTTTCGTTTGAAG | 120 |
| chr6:109537472-109539487 | p5 | (AAAAT)3 | CAAAGAGAGGCGTTGGAGAC | GAATAGCCTCTTTGCATGGC | 232 |
| chr6:109552527-109554553 | p3 | (TCT)6 | ATCCTCTTCCTCGCACATTG | AAAGCAGAGGTGGAGTTGGA | 223 |

| Marker ID | SSR type | SSR motif | FORWARD PRIMER (5'-3') | REVERSE PRIMER (5'-3') | size (bp) |
|---|---|---|---|---|---|
| chr6:109715036-109717051 | p5 | (CGGGT)3 | CAATCCCAAAAGCAAAGGAA | TTTTTAGCCGCACAAAATCC | 192 |
| chr6:109980964-109983016 | p2 | (AT)26 | CAGTAGTTGGCGTGGATGTG | ATGAGAATGGTGGCAAAAGC | 231 |
| chr6:110095132-110097198 | p3 | (GAA)22 | TAAGGAGTCGGCACTTGTCC | TTGCCTAGCAAAACTTGCCT | 196 |
| chr6:110095132-110097198 | p3 | (AAG)29 | TAAGGAGTCGGCACTTGTCC | TTGCCTAGCAAAACTTGCCT | 196 |
| chr6:110273588-110275612 | p5 | (AAACC)3 | ACACGTTCAAACCTTACCGC | AATCAAGTCAAACCGAACCG | 213 |
| chr6:110273625-110275640 | p5 | (AAACC)3 | ACACGTTCAAACCTTACCGC | AATCAAGTCAAACCGAACCG | 176 |
| chr6:110314573-110316597 | p5 | (AAACC)3 | ACACGTTCAAACCTTACCGC | AATCAAGTCAAACCGAACCG | 213 |
| chr6:110314610-110316625 | p5 | (AAACC)3 | ACACGTTCAAACCTTACCGC | AATCAAGTCAAACCGAACCG | 176 |
| chr6:110367837-110369849 | p2 | (TA)6 | GGAACTCCCCTATAACCCGA | TTGGCATCCCAAGAATTGAT | 275 |
| chr6:110367837-110369849 | p2 | (TA)6 | GGAACTCCCCTATAACCCGA | TTGGCATCCCAAGAATTGAT | 275 |
| chr6:110419493-110421541 | p3 | (ATA)16 | TTTAAATTTAGGGGCAGGGG | TGCTCTGCAAATTGTGTTCC | 177 |
| chr6:110419493-110421541 | p3 | (GTT)7 | TTTAAATTTAGGGGCAGGGG | TGCTCTGCAAATTGTGTTCC | 177 |
| chr6:110592263-110594287 | p3 | (TTA)5 | AATGCCCTACATATGCTCGC | GCTGGAATGTTAACGGGAAA | 291 |
| chr6:110592372-110594390 | p3 | (TTA)5 | AATGCCCTACATATGCTCGC | GCTGGAATGTTAACGGGAAA | 182 |
| chr6:110592391-110594409 | p3 | (TTA)5 | AATGCCCTACATATGCTCGC | GCTGGAATGTTAACGGGAAA | 163 |
| chr6:110594301-110596317 | p2 | (CT)8 | TTCCCCACTCAGCTTTGTCT | TTAGGGAATTTCTGCGGCTA | 105 |
| chr6:110611676-110613733 | p3 | (TAA)19 | ACATGGGCATCAAAACAACA | CGGGTTTCAACATGGAGTCT | 168 |
| chr6:110655552-110657572 | p2 | (AT)10 | ACTGGTGAAATGAACAGGGC | TGCCAATTTTTCTCAGGGAC | 237 |
| chr6:110686405-110688425 | p2 | (AT)10 | ACTGGTGAAATGAACAGGGC | TGCCAATTTTTCTCAGGGAC | 237 |
| chr6:110841408-110843423 | p3 | (GCA)5 | CTTGGGCCGAATTATTCAAA | AAGTACCATGGCATCGAAGG | 257 |
| chr6:110854779-110856797 | p3 | (AGA)6 | GCAAATGCCAGTCAAGGATT | CCTACAATCTCGGCCAATGT | 101 |
| chr6:110871984-110873999 | p2 | (AT)6 | TTTGAGAAGTAGCAGGGCGT | AAGCGACCTCGCTATTGATG | 224 |
| chr6:110872059-110874074 | p2 | (AT)6 | TTTGAGAAGTAGCAGGGCGT | AAGCGACCTCGCTATTGATG | 149 |
| chr6:110879388-110881424 | p2 | (AT)6 | CTCTCCAAGCTCAATCCCAG | GATGCTGCCCAAATTTCTGT | 253 |
| chr6:110890829-110892857 | p2 | (AT)14 | GAACCACCTGACTCATCCGT | CCTTGTCGGCCTTATTTTGA | 131 |
| chr6:110942448-110944463 | p5 | (AATTA)3 | TATCGGGATGGGGTTCAATA | GCAGTTGGAATTGCGAGTTT | 113 |
| chr6:110942448-110944463 | p5 | (AATTA)3 | TATCGGGATGGGGTTCAATA | GCAGTTGGAATTGCGAGTTT | 113 |
| chr6:110942448-110944463 | p5 | (TTTAT)4 | TATCGGGATGGGGTTCAATA | GCAGTTGGAATTGCGAGTTT | 113 |
| chr6:110942448-110944463 | p5 | (TTTAT)3 | TATCGGGATGGGGTTCAATA | GCAGTTGGAATTGCGAGTTT | 113 |
| chr6:110971365-110973380 | p5 | (CAAAC)3 | CCAAATTTTGCCCATCTTGT | TGGAATTCAAGCCTATTGGG | 228 |
| chr6:110982065-110984079 | p2 | (AT)7 | CTAATTGACCAGGGCGGTAA | GGTTGAGATGATTGGTGGCT | 266 |
| chr6:110992792-110994856 | p2 | (TA)32 | ATGAGGAAGCATTGGGTGAC | CCGGGATACTTGAGCAAAAA | 259 |
| chr6:111002204-111004246 | p2 | (TA)21 | CTCCAATGAAAGAGGGACCA | TGATAGGAAATGGGGCAAAG | 239 |
| chr6:111002204-111004246 | p2 | (TA)23 | CTCCAATGAAAGAGGGACCA | TGATAGGAAATGGGGCAAAG | 239 |
| chr6:111038764-111040779 | p5 | (TTGGT)3 | TGTTTTGGTTTGGTGTTGGA | ATGTCGGGTTGGTTTTGTTC | 224 |
| chr6:111131334-111133349 | p5 | (TTTAT)3 | GAAGTGCCAACTGTTTGGGT | TACTGCAATGAATCGTGGCT | 149 |

| Marker ID | SSR type | SSR motif | FORWARD PRIMER (5'-3') | REVERSE PRIMER (5'-3') | size (bp) |
|---|---|---|---|---|---|
| chr6:111183462-111185486 | p2 | (TA)12 | TCTCCAAGCCCTAGAACGAA | CGTGAATAGAAGCCCGACAT | 231 |
| chr6:111243538-111245550 | p2 | (AG)6 | GAGTTACGGGGCGTAAAACA | GCGAACCAAAAGCTATGGAA | 109 |
| chr6:111243538-111245550 | p2 | (GT)6 | GAGTTACGGGGCGTAAAACA | GCGAACCAAAAGCTATGGAA | 109 |
| chr6:111254144-111256172 | p2 | (TA)14 | AACCCATTAGTGGGAGCAAA | TGACATGGCGGAATAACAAA | 153 |
| chr6:111254144-111256172 | p2 | (AT)29 | AACCCATTAGTGGGAGCAAA | TGACATGGCGGAATAACAAA | 153 |
| chr6:111281318-111283333 | p5 | (CATAT)3 | TTGAAGTGATGGGTTGGGTT | ATATGTGGCAGCCTTATCGG | 264 |
| chr6:111310703-111312737 | p2 | (AT)17 | GGGATTCTGGGAGTTGTTCA | GGAATCCCCGAAAGTTGAAT | 212 |
| chr6:111349343-111351361 | p3 | (TTG)6 | TTGCGTTGAATGGTGGAGTA | GCATGTCGTCCACAAAACAC | 179 |
| chr6:111420816-111422846 | p2 | (TA)15 | TCACCGGAAAACACTGTTCA | TTTAGTCTGCATCTCGGGCT | 296 |
| chr6:111479627-111481641 | p2 | (AT)7 | GTCAGACGGGAACATGACCT | CGGAGGCAACACTATCCATT | 238 |
| chr6:111556851-111558866 | p3 | (TAA)5 | TTTCCTTACCACGGAGCATC | GCGAAGTTTGTGGAAAGAGC | 120 |
| chr6:111731850-111733884 | p2 | (AT)17 | CTGCACCGTGGAAAGGTAAT | TGTCCCAATAGATTCCAGCC | 134 |
| chr6:111768762-111770776 | p2 | (AT)7 | GGACCTCTTAGCATGGTGGA | TTCTTCGCCTTTCACAGCTT | 220 |
| chr6:111903999-111906014 | p3 | (GTG)5 | AACTTTTTCGGGGGCTTTTA | TTGTCTGCCAAAATTAGGGG | 136 |
| chr6:111903999-111906014 | p3 | (TTC)6 | AACTTTTTCGGGGGCTTTTA | TTGTCTGCCAAAATTAGGGG | 136 |
| chr6:111990167-111992247 | p2 | (AT)10 | ATGTCACACCCCTGAAAAGG | TGTAGCGACCTTATCGGTCC | 133 |
| chr6:111990167-111992247 | p2 | (AT)40 | ATGTCACACCCCTGAAAAGG | TGTAGCGACCTTATCGGTCC | 133 |
| chr6:112110579-112112625 | p2 | (AT)23 | CACGGCCCAGTAATCTGTCT | GCAGAAAAGGAAAATGAGCG | 204 |
| chr6:112156962-112158974 | p2 | (TA)6 | ATTTGCTTGAGGAGAGCCAA | TCCCATAAGTGGGTTTTGGA | 214 |
| chr6:112163514-112165544 | p5 | (TTTTA)3 | CTTATGCCAAGTCTGCCCAT | ACTCATGGGTTTCAGGTTCG | 185 |
| chr6:112192629-112194644 | p3 | (ATA)5 | AAGTGGGGACAAGGGAGAGT | CCGCCTCTTCTTCTTCCTTT | 241 |
| chr6:112255281-112257297 | p2 | (TA)8 | CTCGCCGTTACACAAAAGGT | GTGCAAATATACCCCGGTTG | 174 |
| chr6:112324311-112326329 | p6 | (TTTTTA)3 | AACATGCCTCCGAATTTTTG | TGGCTTATGCATTTGGTTGA | 218 |
| chr6:112324311-112326329 | p6 | (TTTTTA)3 | AACATGCCTCCGAATTTTTG | TGGCTTATGCATTTGGTTGA | 218 |
| chr6:112324311-112326329 | p6 | (TTTTTA)3 | AACATGCCTCCGAATTTTTG | TGGCTTATGCATTTGGTTGA | 218 |
| chr6:112324352-112326388 | p6 | (TTTTTA)3 | AACATGCCTCCGAATTTTTG | TGGCTTATGCATTTGGTTGA | 177 |
| chr6:112324352-112326388 | p6 | (TTTTTA)3 | AACATGCCTCCGAATTTTTG | TGGCTTATGCATTTGGTTGA | 177 |
| chr6:112324352-112326388 | p6 | (TTTTTA)3 | AACATGCCTCCGAATTTTTG | TGGCTTATGCATTTGGTTGA | 177 |
| chr6:112352870-112354886 | p2 | (TA)8 | AACGGCTGTGACTAATCGCT | TGAATGTGGAATTTGACGGA | 286 |
| chr6:112394589-112396610 | p6 | (TTTTTC)3 | TCCAACCCACCAATTCATTT | CTCCTCTTGTCCAAGGCATC | 104 |
| chr6:112437032-112439044 | p2 | (AT)6 | CTTAACCAGGGGTGGACTCA | GAACCCTCACACTTCCCAAA | 172 |
| chr6:112437032-112439044 | p6 | (TTTTTG)3 | CTTAACCAGGGGTGGACTCA | GAACCCTCACACTTCCCAAA | 172 |
| chr6:112437512-112439536 | p2 | (AT)6 | CACTGCACTTAACGACCGAA | GTAAACAAGGGGCGTTCAGA | 290 |
| chr6:112437512-112439536 | p6 | (TTTTTG)3 | CACTGCACTTAACGACCGAA | GTAAACAAGGGGCGTTCAGA | 290 |
| chr6:112437597-112439619 | p2 | (AT)6 | CACTGCACTTAACGACCGAA | GTAAACAAGGGGCGTTCAGA | 205 |
| chr6:112437597-112439619 | p6 | (TTTTTG)3 | CACTGCACTTAACGACCGAA | GTAAACAAGGGGCGTTCAGA | 205 |

| Marker ID | SSR type | SSR motif | FORWARD PRIMER (5'-3') | REVERSE PRIMER (5'-3') | size (bp) |
|---|---|---|---|---|---|
| chr6:112437680-112439702 | p2 | (AT)6 | CACTGCACTTAACGACCGAA | GTAAACAAGGGGCGTTCAGA | 122 |
| chr6:112437680-112439702 | p6 | (TTTTTG)3 | CACTGCACTTAACGACCGAA | GTAAACAAGGGGCGTTCAGA | 122 |
| chr6:112463277-112465298 | p3 | (TAA)7 | TGAGGTGTCATGATTCAGGC | CGACTAACCCACTTCGCATT | 134 |
| chr6:112463502-112465523 | p3 | (TAA)7 | TGGACAACTGAAATGCGAAG | TGGCTTAACCACATTGCTTG | 153 |
| chr6:112463524-112465539 | p3 | (TAA)7 | TGGACAACTGAAATGCGAAG | TGGCTTAACCACATTGCTTG | 131 |
| chr6:112463542-112465570 | p3 | (TAA)7 | TGGACAACTGAAATGCGAAG | TGGCTTAACCACATTGCTTG | 113 |
| chr6:112482309-112484325 | p2 | (TA)8 | TTGGGGATGTTCATGTGCTA | AAGGCGTACTTTTCGGATCA | 288 |
| chr6:112482309-112484325 | p2 | (AT)20 | TTGGGGATGTTCATGTGCTA | AAGGCGTACTTTTCGGATCA | 288 |
| chr6:112608307-112610325 | p3 | (AAT)6 | TTTTCGCGATAAACGGAAGT | AAGCTTCTCCATCAAAGGCA | 134 |
| chr6:112613299-112615329 | p6 | (AGGTAG)5 | CTTGGACCAGGCAATGAAGT | CCATCAGCATCGCCTTTAAT | 213 |
| chr6:112716192-112718207 | p3 | (AAT)5 | CAGCGTGTCGAGTATTGCAT | CTCACCAGGCTCTCAACACA | 272 |
| chr6:112740586-112742601 | p5 | (TATCA)3 | GACCCTTATTTAGGGCCGAG | TAGTCTGTAAGCGGCTCCGT | 100 |
| chr6:112823456-112825474 | p6 | (AAAAAT)3 | AGAAATTGCACGGTTTGTCC | CAAGCATGATCATTTGCACC | 126 |
| chr6:112823456-112825474 | p2 | (TA)10 | AGAAATTGCACGGTTTGTCC | CAAGCATGATCATTTGCACC | 126 |
| chr6:112824158-112826178 | p6 | (AAAAAT)3 | ACTCAATCACCCATCCCAAA | CAAAAGGGTGTAAGGGCAAA | 281 |
| chr6:112824158-112826178 | p2 | (TA)10 | ACTCAATCACCCATCCCAAA | CAAAAGGGTGTAAGGGCAAA | 281 |
| chr6:112878767-112880779 | p2 | (AT)6 | CCGAGGCTCATATCCAACAT | CGGATTTGCGACTTGAATTT | 213 |
| chr6:112901526-112903544 | p2 | (TA)9 | TCCCACCCTACGACATTTTC | GACACCCTTAGCTGGACTGC | 284 |
| chr6:112946812-112948827 | p5 | (AAAAT)3 | GCTTGCTCACGAAACATGAA | AGGTCAATGTTGGAGGCTTG | 131 |
| chr6:112976682-112978698 | p2 | (TA)6 | TTAGGCCCAATAGGTTGACG | AGATTTGCGAGGTCTGTTGG | 232 |
| chr6:112976682-112978698 | p2 | (AT)8 | TTAGGCCCAATAGGTTGACG | AGATTTGCGAGGTCTGTTGG | 232 |
| chr6:113005668-113007683 | p5 | (AAAAT)3 | GCTTGCTCACGAAACATGAA | AGGTCAATGTTGGAGGCTTG | 131 |
| chr6:113035538-113037554 | p2 | (TA)6 | TTAGGCCCAATAGGTTGACG | AGATTTGCGAGGTCTGTTGG | 232 |
| chr6:113035538-113037554 | p2 | (AT)8 | TTAGGCCCAATAGGTTGACG | AGATTTGCGAGGTCTGTTGG | 232 |
| chr6:113035538-113037554 | p6 | (CCGGTT)3 | TTAGGCCCAATAGGTTGACG | AGATTTGCGAGGTCTGTTGG | 232 |
| chr6:113057349-113059371 | p2 | (GT)10 | ACAGGGGTTAAGGAGCCATT | CCTGATGTCCCTCTTTGCAT | 135 |
| chr6:113057373-113059385 | p2 | (GT)10 | ACAGGGGTTAAGGAGCCATT | CCTGATGTCCCTCTTTGCAT | 111 |
| chr6:113123782-113125798 | p4 | (CTTT)4 | TTGTGGGGTTGTAGGGGTTA | TGTAATGCCTGCAAGAGACG | 281 |
| chr6:113138177-113140191 | p5 | (TAAAA)3 | TACCTGAGGTTCTGGGTTCG | TGCCCCATAAGGCATAGTTC | 139 |
| chr6:113138177-113140191 | p2 | (AT)7 | TACCTGAGGTTCTGGGTTCG | TGCCCCATAAGGCATAGTTC | 139 |
| chr6:113166434-113168473 | p3 | (TAA)13 | AAATTGGGCGGGATATTTTC | TGATTGTACCCCTCCTCGTC | 214 |
| chr6:113280022-113282037 | p3 | (ACT)5 | TTGCTACTTGTGCTGTTCGG | CACGCAGCTCGACTAATTCA | 146 |
| chr6:113294298-113296312 | p5 | (ATGAT)3 | GACTATGCCTCCGTAACCCA | ATACTTGCCCCTTCACATCG | 125 |
| chr6:113294298-113296312 | p2 | (TA)7 | GACTATGCCTCCGTAACCCA | ATACTTGCCCCTTCACATCG | 125 |
| chr6:113295822-113297840 | p6 | (TTGATT)3 | GTCAAATGCCGCAGATCATA | AATGGATATCCCACTGCCTG | 165 |
| chr6:113295822-113297840 | p2 | (AT)8 | GTCAAATGCCGCAGATCATA | AATGGATATCCCACTGCCTG | 165 |

| Marker ID | SSR type | SSR motif | FORWARD PRIMER (5'-3') | REVERSE PRIMER (5'-3') | size (bp) |
|---|---|---|---|---|---|
| chr6:113298795-113300810 | p5 | (AATAA)3 | CCCTTTGCGTAGAGCTTCTG | ATTCCAATCTTGCCCTTGTG | 206 |
| chr6:113363131-113365143 | p2 | (AT)6 | ACTGTGGTGGCTCCCTTATG | TCTCAATATGCTCTGCACGC | 266 |
| chr6:113390639-113392651 | p2 | (AT)6 | ACCGAAGCTGTTGAACTCGT | GTTGGAGGCACGACGTAAAT | 203 |
| chr6:113567450-113569468 | p2 | (TA)9 | CGTGAAGTCTGGAGGGTCAT | GGATACAACCCAAACATGCC | 235 |
| chr6:113732765-113734780 | p3 | (AAT)5 | GCCACGGAAGGAGTATTGAA | TCCTCCGTCATTGGTAAAGC | 110 |
| chr6:113784726-113786752 | p2 | (TA)13 | CAAATCCAATCTCCACCGTT | GTTTCATTCCAACCAGCCAT | 194 |
| chr6:113825030-113827050 | p2 | (TA)17 | TTTGAAAAGAGCCCCAACAG | AAAACCGATAAACCGGAACC | 211 |
| chr6:113825030-113827050 | p2 | (TA)10 | TTTGAAAAGAGCCCCAACAG | AAAACCGATAAACCGGAACC | 211 |
| chr6:113825030-113827050 | p3 | (AGA)5 | TTTGAAAAGAGCCCCAACAG | AAAACCGATAAACCGGAACC | 211 |
| chr6:113837000-113839012 | p2 | (AT)12 | CAAAAATTAGGGGGTGCAAA | TTTGTAAGCACTCGGCAATG | 118 |
| chr6:113837000-113839012 | p2 | (TA)6 | CAAAAATTAGGGGGTGCAAA | TTTGTAAGCACTCGGCAATG | 118 |
| chr6:113840463-113842478 | p3 | (TTG)5 | TCACACATCGAATTAGGGCA | GAAATCTTAACGGGCCAACA | 188 |
| chr6:113850496-113852514 | p6 | (ATGGTG)3 | ACAAAAGCCCATTGTCCTTG | GAGCACGTTTCGCAACATTA | 269 |
| chr6:113925520-113927535 | p5 | (TCGGT)3 | CTTCGGTTCGGTAATCGGTA | CTTGAAAGGTGAATCCCCAA | 227 |
| chr6:114024964-114026982 | p3 | (ACC)6 | TCTTTGTCGGTTGCTGTCAC | TTTTTCGGGAAAATAACCCC | 300 |
| chr6:114029334-114031350 | p2 | (AT)8 | CGCGTTCACTAAATCAGCAA | TGGGGATCTGGATTTCAAAG | 118 |
| chr6:114036696-114038723 | p2 | (AT)7 | GTCGTCCGTTGTCCCTTAAA | AACTTATTGTGCGGGTCCAC | 127 |
| chr6:114036696-114038723 | p3 | (ATT)9 | GTCGTCCGTTGTCCCTTAAA | AACTTATTGTGCGGGTCCAC | 127 |
| chr6:114114198-114116213 | p3 | (GTT)5 | TATTGGTGCAGGATGGTTGA | TTCCGCGACTTTCATCTTCT | 179 |
| chr6:114120787-114122802 | p3 | (GAG)5 | CAGGAGGGGACATTTTGTGT | AGGAGACTGCAAGGTCAGGA | 185 |
| chr6:114288725-114290737 | p3 | (TGT)7 | CTGAGTTGTCCTGCATCGAA | ACCGTTTCCGATAGACCCTT | 188 |
| chr6:114288725-114290737 | p2 | (TA)6 | CTGAGTTGTCCTGCATCGAA | ACCGTTTCCGATAGACCCTT | 188 |
| chr6:114288725-114290737 | p2 | (TC)9 | CTGAGTTGTCCTGCATCGAA | ACCGTTTCCGATAGACCCTT | 188 |
| chr6:114310099-114312124 | p2 | (AT)15 | CTCCCGTGTTCAAAACCAGT | GAAGTCGTAGGCGTTTGAGG | 194 |
| chr6:114310099-114312124 | p5 | (AGCCG)5 | CTCCCGTGTTCAAAACCAGT | GAAGTCGTAGGCGTTTGAGG | 194 |
| chr6:114314752-114316770 | p6 | (CAGTAG)3 | TCTCCGCTGCCTTGTTATCT | TCAAGTTGCGGCAGTGATAG | 280 |
| chr6:114334341-114336357 | p2 | (CT)8 | TCGCTTCTTTCTTAAACGGC | GCTCAAACCCGAGACTTCTG | 223 |
| chr6:114368603-114370621 | p2 | (TA)9 | GAATCAGCATTCGGGTTTGT | TGTTAGGCTCCAAATACGGG | 231 |
| chr6:114486722-114488737 | p3 | (ACA)5 | GAACCGGTGAACGAAGTCAT | TGGACCATTTTGGAATCGTT | 213 |
| chr6:114573964-114575982 | p3 | (CCA)6 | TATAAGCCAATCCAAACGGG | TAACAACCAACCCAAGCCTC | 126 |
| chr6:114796455-114798470 | p5 | (TGCAT)3 | CATGTAAGCAAAGGGGGAAA | TGTGGTATCGCATTGACGTT | 162 |
| chr6:114812447-114814462 | p5 | (TGCAT)3 | CATGTAAGCAAAGGGGGAAA | TGTGGTATCGCATTGACGTT | 162 |
| chr6:114909072-114911092 | p5 | (TAAAA)4 | TTTTGGTGGATTCTTTTCGG | ATTTGTACGCGGTGCTAAGG | 237 |
| chr6:114909072-114911092 | p2 | (AT)8 | TTTTGGTGGATTCTTTTCGG | ATTTGTACGCGGTGCTAAGG | 237 |
| chr6:114955184-114957199 | p5 | (TTGGT)3 | CCGATTAGTGGGTTGGTTTG | GTAATGTCGGGTTGGTTTGG | 242 |
| chr6:114955184-114957199 | p5 | (TTGGT)3 | CCGATTAGTGGGTTGGTTTG | GTAATGTCGGGTTGGTTTGG | 242 |

| Marker ID | SSR type | SSR motif | FORWARD PRIMER (5'-3') | REVERSE PRIMER (5'-3') | size (bp) |
|---|---|---|---|---|---|
| chr6:114955184-114957199 | p2 | (AT)11 | CCGATTAGTGGGTTGGTTTG | GTAATGTCGGGTTGGTTTGG | 242 |
| chr6:114983451-114985469 | p3 | (ACA)6 | GTCTGGTCAAAACGGAGGAA | TGCGTACATTCTACCCTCCC | 241 |
| chr6:114983451-114985469 | p3 | (ACA)6 | GTCTGGTCAAAACGGAGGAA | TGCGTACATTCTACCCTCCC | 241 |
| chr6:115022714-115024729 | p3 | (ATT)5 | ACTCGTCATCTTCCGCATTC | GGCCCAAATTTTCATTTCCT | 181 |
| chr6:115026553-115028571 | p6 | (ACAACC)3 | GTCTGCCCATGTCAAAAGGT | CAAGGTGAGGCAGAATGTCA | 211 |
| chr6:115203177-115205192 | p3 | (AAT)5 | GTGGGACAAAATTGGGTGTC | CCGTGCGATATGTCTGTTTG | 198 |
| chr6:115203177-115205192 | p5 | (CATAT)3 | GTGGGACAAAATTGGGTGTC | CCGTGCGATATGTCTGTTTG | 198 |
| chr6:115262196-115264208 | p2 | (AG)6 | GTTTTGCGAAAATGGCTGAT | TTGGCTCGGTCCTTATATCG | 106 |
| chr6:115344722-115346740 | p3 | (AAG)6 | TATTCAATGCTCACCCACCA | TACAAAGGACGGAGGAATCG | 121 |
| chr6:115367413-115369425 | p2 | (TA)6 | AGCTGTTACTTGGCGGTCAT | GCCACAGGCTCTCTTTTGAG | 194 |
| chr6:115372666-115374694 | p2 | (TA)14 | TAGAGGGTTAGGGGAAGCGT | TCATGATGATTCAGCAAGGC | 178 |
| chr6:115372666-115374694 | p2 | (TA)6 | TAGAGGGTTAGGGGAAGCGT | TCATGATGATTCAGCAAGGC | 178 |
| chr6:115373253-115375265 | p2 | (TA)14 | GTACGGCACGCACTCTTTTT | CATTCAATGCTGCTGACGTT | 168 |
| chr6:115373253-115375265 | p2 | (TA)6 | GTACGGCACGCACTCTTTTT | CATTCAATGCTGCTGACGTT | 168 |
| chr6:115405466-115407481 | p5 | (ATCAT)3 | ATATCATATCCGGCCCCTTC | TGGAGTATGCTGTGCCATGT | 175 |
| chr6:115405466-115407481 | p3 | (CAA)5 | ATATCATATCCGGCCCCTTC | TGGAGTATGCTGTGCCATGT | 175 |
| chr6:115405466-115407481 | p2 | (AT)6 | ATATCATATCCGGCCCCTTC | TGGAGTATGCTGTGCCATGT | 175 |
| chr6:115405959-115407971 | p3 | (CAA)5 | ATGCTCTTTACGCCTTTCCA | GTTGATTGTTTTGGCCGAGT | 299 |
| chr6:115405959-115407971 | p2 | (AT)6 | ATGCTCTTTACGCCTTTCCA | GTTGATTGTTTTGGCCGAGT | 299 |
| chr6:115470918-115472932 | p2 | (AC)7 | TGCCTCCACACACAAGAAAA | TTTTGATCTTTTCCCCATGC | 162 |
| chr6:115470918-115472932 | p2 | (CA)7 | TGCCTCCACACACAAGAAAA | TTTTGATCTTTTCCCCATGC | 162 |
| chr6:115470918-115472932 | p2 | (CA)6 | TGCCTCCACACACAAGAAAA | TTTTGATCTTTTCCCCATGC | 162 |
| chr6:115503853-115505868 | p3 | (ACA)5 | GAGCGCACTGACTACTGCTG | CCTTATTTCTTTGGGGGCTC | 113 |
| chr6:115518776-115520791 | p5 | (TTTTC)3 | GCCTGATCCAGGTGTTGAAT | GCTGGTGTTAGGCACAGACA | 286 |
| chr6:115611087-115613114 | p3 | (TAA)9 | ATTTCGGAATGTCCGCTAAG | GATTGGCCCTACTTCGTTGA | 131 |
| chr6:115765924-115767942 | p3 | (CAA)6 | GGGAAAAGTGGTTTTTGGGT | TGAAGGGAATTTGTAAGGCG | 159 |
| chr6:115912009-115914027 | p6 | (TTAAAA)3 | TCCCAGAAGCAGCAATCTCT | TTTGGGAACAACACAAAGCA | 135 |
| chr6:115912009-115914027 | p6 | (TTTCTT)4 | TCCCAGAAGCAGCAATCTCT | TTTGGGAACAACACAAAGCA | 135 |
| chr6:115935547-115937565 | p6 | (TTAAAA)3 | TCCGTCCCAAAAAGAATGTC | GGGTTTTGGGTACCTCCTGT | 237 |
| chr6:115937534-115939552 | p3 | (AAC)6 | TATCGGAAACAGCCTCCCTA | ATATCGGGTGCTGACACGTT | 114 |
| chr6:115943907-115945922 | p3 | (AAC)5 | TAAGTTCCGCGTACACTCCC | ATATCGGGTGCTGACACGTT | 130 |
| chr6:115955217-115957245 | p2 | (TA)6 | GACCTGCAAGATGAGGGTGT | GCCAACAGAACAAACCAACA | 116 |
| chr6:115955217-115957245 | p2 | (AT)14 | GACCTGCAAGATGAGGGTGT | GCCAACAGAACAAACCAACA | 116 |
| chr6:115955217-115957245 | p2 | (AT)14 | GACCTGCAAGATGAGGGTGT | GCCAACAGAACAAACCAACA | 116 |
| chr6:115955217-115957245 | p2 | (AT)14 | GACCTGCAAGATGAGGGTGT | GCCAACAGAACAAACCAACA | 116 |
| chr6:115955217-115957245 | p6 | (CCGGTT)3 | GACCTGCAAGATGAGGGTGT | GCCAACAGAACAAACCAACA | 116 |

| Marker ID | SSR type | SSR motif | FORWARD PRIMER (5'-3') | REVERSE PRIMER (5'-3') | size (bp) |
|---|---|---|---|---|---|
| chr6:116028940-116030952 | p2 | (AT)8 | TCCATGGGCTTCACTCTTCT | GAACCCACCACATCCAAACT | 103 |
| chr6:116028940-116030952 | p2 | (TA)13 | TCCATGGGCTTCACTCTTCT | GAACCCACCACATCCAAACT | 103 |
| chr6:116028940-116030952 | p2 | (AT)6 | TCCATGGGCTTCACTCTTCT | GAACCCACCACATCCAAACT | 103 |
| chr6:116044937-116046952 | p5 | (AATAA)3 | TGCCGACTTCATGCATTTTA | GGCAATTCGTGCAATTTCTT | 218 |
| chr6:116138314-116140332 | p3 | (ACA)6 | TCGGCTTCTCAGTCCTTGTT | AGACAGGCTCCATCTGCATT | 151 |
| chr6:116256246-116258261 | p5 | (TCGGG)3 | CTCGAAGTATGCCCTGTTCC | GTCTGGGGAGGGTAAAGGAG | 178 |
| chr6:116371170-116373188 | p6 | (ATATAG)3 | CATCTTTCTTCCACCCCAGA | CATAGTGCCTGGCTCGATTT | 195 |
| chr6:116371170-116373188 | p2 | (TA)9 | CATCTTTCTTCCACCCCAGA | CATAGTGCCTGGCTCGATTT | 195 |
| chr6:116385343-116387358 | p5 | (TTTGA)3 | GGGACAACACGAGGTGAGTT | CCAAACGACACCTTTGGATT | 192 |
| chr6:116391607-116393619 | p2 | (TA)6 | TATCGGAGAGAATGGGTTGG | CCGAACTCCCGAACATAAAA | 128 |
| chr6:116428550-116430570 | p4 | (ATTT)5 | ACTTGGCTTGTGCAAGTTCC | CCGGTCCATTATCCATTGAG | 162 |
| chr6:116428550-116430570 | p6 | (TTGGAG)3 | ACTTGGCTTGTGCAAGTTCC | CCGGTCCATTATCCATTGAG | 162 |
| chr6:116505541-116507565 | p4 | (TTTA)6 | CAAATACCGGCACATTTCCT | CCCACTTTCCCAAAATAGCA | 204 |
| chr6:116562900-116564915 | p5 | (CCCGG)3 | GGAGTCTCTTTTTACGGGGC | TTCGAAGCGTCGGGTATTAG | 261 |
| chr6:116562900-116564915 | p3 | (AGG)6 | GGAGTCTCTTTTTACGGGGC | TTCGAAGCGTCGGGTATTAG | 261 |
| chr6:116584541-116586556 | p2 | (CT)6 | ACCTAAGCTCGTTGCAGGAA | GGGTTCAAATTTGCCCTTTT | 279 |
| chr6:116584541-116586556 | p5 | (ATATA)3 | ACCTAAGCTCGTTGCAGGAA | GGGTTCAAATTTGCCCTTTT | 279 |
| chr6:116587268-116589283 | p3 | (ATG)5 | CCCCGAGAAGAGGTTTAAGG | CCATTCCTCCGAGTCTGGTA | 106 |
| chr6:116612778-116614794 | p2 | (CA)8 | GCGAAATGGCAAAAGACTTC | CTGAAGCTTGCCAAAGATCC | 228 |
| chr6:116647737-116649755 | p2 | (TA)10 | TCGGCCCATGTTTTCTTATC | TGGGATTTCAAACCGGTAAA | 257 |
| chr6:116647737-116649755 | p2 | (TA)9 | TCGGCCCATGTTTTCTTATC | TGGGATTTCAAACCGGTAAA | 257 |
| chr6:116647737-116649755 | p6 | (CCGGTT)3 | TCGGCCCATGTTTTCTTATC | TGGGATTTCAAACCGGTAAA | 257 |
| chr6:116675830-116677845 | p5 | (AATTA)3 | GGTATCGTTTCCCATCCCTT | CCGTTTTGGGACGGTAACTA | 197 |
| chr6:116698969-116700990 | p3 | (ATT)7 | CCCCCTCTCAATTGCTGTTA | GCGCCCCCTAAACTTATTCT | 279 |
| chr6:116743152-116745178 | p2 | (AT)13 | GAGCCTTTTTGCTGTTCCAG | GCAACGGGTATGGCTTTCTA | 249 |
| chr6:116808501-116810527 | p2 | (TA)13 | TTATGCCGAGCGAAGTAGGT | CGTGAAGTTTTTCCACGGAT | 138 |
| chr6:116856296-116858347 | p2 | (AG)7 | GGGAAAAATCCCTCCAATGT | TTCATCCTTCCTCCATCAGG | 266 |
| chr6:116856296-116858347 | p3 | (ATA)17 | GGGAAAAATCCCTCCAATGT | TTCATCCTTCCTCCATCAGG | 266 |
| chr6:116921283-116923305 | p3 | (GAA)7 | GGTTGGGAAAAGGAGGAGAA | TTAACCCCTGAACTTGGCAC | 272 |
| chr6:116921283-116923305 | p3 | (TAT)8 | GGTTGGGAAAAGGAGGAGAA | TTAACCCCTGAACTTGGCAC | 272 |
| chr6:116921283-116923305 | p2 | (AT)11 | GGTTGGGAAAAGGAGGAGAA | TTAACCCCTGAACTTGGCAC | 272 |
| chr6:117018908-117020926 | p3 | (AAC)6 | CTAACACTAGCCGTAGCCGC | GAATAGTGTCGCCCATTCGT | 179 |
| chr6:117062929-117064943 | p3 | (AAT)9 | TTTGGCAAGGGTACTTCGAC | CAAATTTTGCAAGCAGGTCA | 145 |
| chr6:117062929-117064943 | p2 | (TA)7 | TTTGGCAAGGGTACTTCGAC | CAAATTTTGCAAGCAGGTCA | 145 |
| chr6:117064875-117066890 | p3 | (AAT)5 | TTCAAAAGGGAAATTCGTGC | CTATCTCTCCCGCTCCTCCT | 221 |
| chr6:117191587-117193602 | p5 | (TATTT)3 | CGTGGAGAACCAAAACCCTA | ACATAGTGCGAGGATCACCC | 237 |

| Marker ID | SSR type | SSR motif | FORWARD PRIMER (5'-3') | REVERSE PRIMER (5'-3') | size (bp) |
|---|---|---|---|---|---|
| chr6:117191637-117193661 | p5 | (TATTT)3 | CGTGGAGAACCAAAACCCTA | ACATAGTGCGAGGATCACCC | 187 |
| chr6:117247581-117249605 | p2 | (TA)12 | TCCGTTTGACCCCCATAATA | TTTACGCCCATGAACAGTGA | 277 |
| chr6:117273724-117275736 | p2 | (TA)6 | GGAGTTGGTCTCGGATGTGT | ATTCAGCTCCCTTTGCTTGA | 267 |
| chr6:117294892-117296913 | p2 | (AT)11 | CAGCCCGTTACAACACCTTT | GCCCAATTCAGTGGCATACT | 255 |
| chr6:117294892-117296913 | p2 | (TC)7 | CAGCCCGTTACAACACCTTT | GCCCAATTCAGTGGCATACT | 255 |
| chr6:117294917-117296932 | p2 | (AT)11 | CAGCCCGTTACAACACCTTT | GCCCAATTCAGTGGCATACT | 230 |
| chr6:117294917-117296932 | p2 | (TC)7 | CAGCCCGTTACAACACCTTT | GCCCAATTCAGTGGCATACT | 230 |
| chr6:117303732-117305747 | p5 | (AAAAG)3 | AAACGGAGTAAGGATGCCCT | AAAATGCGGTCAAAATGAGC | 115 |
| chr6:117384024-117386078 | p2 | (AT)27 | AAAAAGTCAAGGCGAGGTCA | GATTGCCTTGACCAAGGTGT | 238 |
| chr6:117401609-117403633 | p3 | (TTC)29 | TTAGTTTTTGGGTTGCGTCC | AAAAGTGACAAACGCCCAAG | 116 |
| chr6:117498600-117500648 | p2 | (AT)24 | ATTAAAATCTGGCTCCGCCT | AAAAGTGCCCATACACTGGC | 158 |
| chr6:117519211-117521226 | p5 | (CGGTT)3 | GACTTCGCCACTATGCACAA | CCGATACCGAAGAGAAACCA | 100 |
| chr6:117519211-117521226 | p4 | (TTTA)4 | GACTTCGCCACTATGCACAA | CCGATACCGAAGAGAAACCA | 100 |
| chr6:117573123-117575135 | p2 | (AT)6 | TGGTTGTGCCATGAACACTT | GCAAGGTTGGTTGCTAAAGG | 199 |
| chr6:117575418-117577439 | p3 | (CTT)7 | CGACATACGTCGCTTCTTCA | TGGGGAAATAAACCCCAAAT | 150 |
| chr6:117575418-117577439 | p3 | (TTC)7 | CGACATACGTCGCTTCTTCA | TGGGGAAATAAACCCCAAAT | 150 |
| chr6:117585359-117587374 | p3 | (TTC)5 | CTTTGAAAACCCCAGGCATA | GGCAGACAAGAAAGCAAAGG | 219 |
| chr6:117611077-117613098 | p3 | (TAA)7 | GGCTTGACCCTACCTTTTCC | ACGTGTAAAACAAGGGGACG | 120 |
| chr6:117743441-117745457 | p6 | (GCATAT)3 | AAGTTGTGTTGCCCCTGTTC | TGGAGAGGCGGAAGAGAATA | 279 |
| chr6:117743480-117745496 | p6 | (GCATAT)3 | AAGTTGTGTTGCCCCTGTTC | TGGAGAGGCGGAAGAGAATA | 240 |
| chr6:117887133-117889145 | p2 | (TA)6 | TCAAATTTAACGGTGGAGGG | CAACGGCAGCCTCATTTATT | 292 |
| chr6:117893211-117895223 | p2 | (TA)6 | TCGACCATGGATAACGTGAA | GACTTCCAAAAACTTTGGCG | 276 |
| chr6:117893211-117895223 | p3 | (CTA)7 | TCGACCATGGATAACGTGAA | GACTTCCAAAAACTTTGGCG | 276 |
| chr6:117893356-117895386 | p2 | (TA)6 | TCGACCATGGATAACGTGAA | GACTTCCAAAAACTTTGGCG | 131 |
| chr6:117893356-117895386 | p3 | (CTA)7 | TCGACCATGGATAACGTGAA | GACTTCCAAAAACTTTGGCG | 131 |
| chr6:117902499-117904514 | p2 | (AT)10 | GCCTTATGGGGCAGACTTTT | TAAGGCGGAACTTTTCCTCA | 114 |
| chr6:117902499-117904514 | p2 | (AT)12 | GCCTTATGGGGCAGACTTTT | TAAGGCGGAACTTTTCCTCA | 114 |
| chr6:117902499-117904514 | p5 | (AAAAT)3 | GCCTTATGGGGCAGACTTTT | TAAGGCGGAACTTTTCCTCA | 114 |
| chr6:117918788-117920804 | p4 | (TAAT)4 | ACCTGCAGCTGCTTTGTTTT | GGCTGCATGATTTTATCGCT | 282 |
| chr6:117943482-117945500 | p6 | (CAACTC)3 | GTTGCCAACGGTCCATTAAC | AAACCGGAACCGATAGAACC | 240 |
| chr6:117963924-117965940 | p2 | (AT)8 | CTCGACAAGCCAAACGTGTA | AGAGATCCTTTCACGTCGGA | 282 |
| chr6:118014223-118016261 | p2 | (TA)19 | AATCTTTCATGCCCGTTGTC | TCAAATGGTGTGGCATTCAT | 115 |
| chr6:118047876-118049894 | p2 | (AT)9 | TTCCATGCATTCCAAGTCAA | TGCTTTCATGGAAGAACGTG | 240 |
| chr6:118130508-118132523 | p5 | (CCGAC)3 | TCCTCTCCAATGCCAAATTC | AGGATGATGAAGATGCAGGG | 112 |
| chr6:118165015-118167030 | p5 | (GATAT)3 | GATCGAAAGGAACAAAGGCA | AGGCCTTCTCAAGCAGTGAC | 187 |
| chr6:118272577-118274589 | p2 | (AT)6 | CAAGTTGCACAAATGAGCGT | CATGAAGGTTTCAGGGCAAT | 129 |

| Marker ID | SSR type | SSR motif | FORWARD PRIMER (5'-3') | REVERSE PRIMER (5'-3') | size (bp) |
|---|---|---|---|---|---|
| chr6:118294271-118296286 | p3 | (TCA)5 | TTGTTATTAACCCGGAAGCG | GTGGCTTTTCCCCAAATTCT | 202 |
| chr6:118297630-118299644 | p2 | (TA)7 | TGGATTCTCGGGGTTAGTTG | TTACATACCTTGCCCGCTTC | 299 |
| chr6:118322883-118324895 | p2 | (TC)6 | CCATTTTCCTTGATGGGCTA | CAGCATGCACTTCTGGACAT | 198 |
| chr6:118322883-118324895 | p2 | (TC)6 | CCATTTTCCTTGATGGGCTA | CAGCATGCACTTCTGGACAT | 198 |
| chr6:118330404-118332420 | p2 | (AT)11 | CGGGGTCTACGTATCCTTGA | TGAAAAGGTGCCCGAAGTAG | 292 |
| chr6:118330421-118332475 | p2 | (AT)11 | CGGGGTCTACGTATCCTTGA | TGAAAAGGTGCCCGAAGTAG | 275 |
| chr6:118366342-118368358 | p2 | (AT)8 | ATACCCAAAATGGCATCCAA | GAGACGAACGTTTTCTCCCA | 186 |
| chr6:118366342-118368358 | p2 | (AT)7 | ATACCCAAAATGGCATCCAA | GAGACGAACGTTTTCTCCCA | 186 |
| chr6:118367674-118369688 | p2 | (AT)7 | TTTGGATGTGGTGAGTTCCA | TAGACCCCTGCTTCATCGTC | 161 |
| chr6:118367674-118369688 | p2 | (TA)12 | TTTGGATGTGGTGAGTTCCA | TAGACCCCTGCTTCATCGTC | 161 |
| chr6:118377994-118380012 | p2 | (AT)9 | GGCGACATTTGATTGGTTCT | TCGGACTTCACATGATGAGC | 294 |
| chr6:118413438-118415462 | p6 | (CGGTGT)4 | AAAATGGACGAACTGAACCG | TCACTACGGCTGCCACTATG | 152 |
| chr6:118424862-118426877 | p5 | (TAAAT)3 | CCCCAAATGACTTCCAAAGA | AAAGCATTTTGTGTCCAGGG | 241 |
| chr6:118424862-118426877 | p4 | (TATT)4 | CCCCAAATGACTTCCAAAGA | AAAGCATTTTGTGTCCAGGG | 241 |
| chr6:118460366-118462381 | p3 | (CTT)5 | ACCCTGAGCGTCATCATTTC | GACATTGACGGGAAAGTCGT | 269 |
| chr6:118609383-118611398 | p3 | (AAC)5 | ATTCTAACGGGAGGGCAAAT | TGATGGAAACAAATCCACGA | 262 |
| chr6:118703309-118705323 | p2 | (TC)7 | AGCCAATATGCCTCATCACC | CACGAAGGTAAGGGTGGAAA | 122 |
| chr6:118753237-118755251 | p2 | (TC)7 | AGCCAATATGCCTCATCACC | CACGAAGGTAAGGGTGGAAA | 122 |
| chr6:118798122-118800137 | p5 | (TTTTA)3 | ACTCTGAACGGATTTGGTGG | GTGAAGCGAGAAATTGGAGG | 253 |
| chr6:118800618-118802630 | p2 | (TA)6 | TGCAAAATACAAGCCACGAA | ATTGTAACACGGCCTTTTGG | 152 |
| chr6:118800618-118802630 | p2 | (AT)6 | TGCAAAATACAAGCCACGAA | ATTGTAACACGGCCTTTTGG | 152 |
| chr6:118800618-118802630 | p2 | (TA)6 | TGCAAAATACAAGCCACGAA | ATTGTAACACGGCCTTTTGG | 152 |
| chr6:118815637-118817663 | p2 | (AT)13 | AATGTTGTGCCTGCTAAGCC | ACCTCCTGAAGATGGTGTGG | 128 |
| chr6:118815637-118817663 | p5 | (TTTGG)3 | AATGTTGTGCCTGCTAAGCC | ACCTCCTGAAGATGGTGTGG | 128 |
| chr6:118815637-118817663 | p6 | (ATTTTA)3 | AATGTTGTGCCTGCTAAGCC | ACCTCCTGAAGATGGTGTGG | 128 |
| chr6:118854517-118856543 | p2 | (AT)13 | AATGTTGTGCCTGCTAAGCC | ACCTCCTGAAGATGGTGTGG | 128 |
| chr6:118854517-118856543 | p5 | (TTTGG)3 | AATGTTGTGCCTGCTAAGCC | ACCTCCTGAAGATGGTGTGG | 128 |
| chr6:118854517-118856543 | p6 | (ATTTTA)3 | AATGTTGTGCCTGCTAAGCC | ACCTCCTGAAGATGGTGTGG | 128 |
| chr6:118885043-118887065 | p2 | (TA)11 | GCCACATGTGTATCAACCCA | TGACATGGCGAACTAGCAAA | 218 |
| chr6:118959920-118961938 | p3 | (TTG)6 | ACATGCTGGCATTTGCATTA | AGAGTGGGAGGGAATCCAGT | 143 |
| chr6:118966935-118968950 | p3 | (AGA)5 | TGTTTCATCATCGTCGTCGT | AAGACCCGGCTTGGAAGTAT | 259 |
| chr6:119054131-119056149 | p3 | (TTG)6 | ACATGCTGGCATTTGCATTA | AGAGTGGGAGGGAATCCAGT | 143 |
| chr6:119061146-119063161 | p3 | (AGA)5 | TGTTTCATCATCGTCGTCGT | AAGACCCGGCTTGGAAGTAT | 259 |
| chr6:119221171-119223183 | p2 | (TA)6 | TAGCAACAAAACAAACCCCC | GAGAAACAAGCCAAAACCCA | 252 |
| chr6:119357069-119359084 | p3 | (TTC)5 | TTTGGACCTTGGTAGTTCGG | GAGTCCATGGCCAAACTTGT | 278 |
| chr6:119539242-119541264 | p2 | (TA)11 | TCCGAGCACTCTCCAAACTT | ACGCACTAATTGGTGTGCAA | 160 |

| Marker ID | SSR type | SSR motif | FORWARD PRIMER (5'-3') | REVERSE PRIMER (5'-3') | size (bp) |
|---|---|---|---|---|---|
| chr6:119573799-119575817 | p2 | (AT)9 | AAATATCAACCGTGGCTTCG | TCTTCCATTTGGTCCACTCC | 257 |
| chr6:119616866-119618880 | p2 | (AT)7 | ACTTGAACCATGCCTTTTGC | CATGTGACAGCCTACCGAGA | 228 |
| chr6:119730637-119732667 | p3 | (TAA)10 | TCAGATTCTTGGGGTTTTCG | TCCTTTTCATCACCTCGTCC | 131 |
| chr6:119759756-119761836 | p2 | (TC)7 | TGCCAATTCACAGGTTACGA | CTGGCTTTCTCCTCAACACC | 202 |
| chr6:119818192-119820208 | p2 | (TA)8 | CCGAATTCCCCTTTTTCAAT | TCGGGAATATCAACTCAGGC | 167 |
| chr6:119827773-119829788 | p5 | (AACCG)3 | TTAGGGGTGGGCATAAACAC | CGTACCGAAAAACCGAAAAA | 265 |
| chr6:119827773-119829788 | p6 | (AGTTCT)3 | TTAGGGGTGGGCATAAACAC | CGTACCGAAAAACCGAAAAA | 265 |
| chr6:119827773-119829788 | p5 | (AACCG)3 | TTAGGGGTGGGCATAAACAC | CGTACCGAAAAACCGAAAAA | 265 |
| chr6:119834714-119836729 | p5 | (GGTAT)3 | GGAAACAGAGGAGATGGGAA | CACCCTCAATTCCACCAACT | 268 |
| chr6:119869632-119871650 | p2 | (GA)9 | CCGGGTAACTCTATCCAGCA | TTAAAAGGCAAAGGGTGTGG | 205 |
| chr6:119869632-119871650 | p2 | (AG)6 | CCGGGTAACTCTATCCAGCA | TTAAAAGGCAAAGGGTGTGG | 205 |
| chr6:119942512-119944530 | p6 | (AACTGA)3 | AGCAAGAACACATTCACCCC | TAAGAGGCGATTCCCTTGTG | 243 |
| chr6:120142988-120145002 | p2 | (AT)7 | TGCGTATGCCTGTATTCCAA | TTTTTGCCTGGACACCCTTA | 104 |
| chr6:120245215-120247229 | p2 | (GA)7 | ATATAAGAGGGTCCCCCACG | CAACCAACCATGATTGTCCA | 186 |
| chr6:120245215-120247229 | p2 | (TA)33 | ATATAAGAGGGTCCCCCACG | CAACCAACCATGATTGTCCA | 186 |
| chr6:120251713-120253725 | p2 | (TA)6 | AAGTGATGGCTCTGCGAGAT | GCCATATTTGAGCCGACATT | 256 |
| chr6:120278360-120280388 | p2 | (GA)6 | GCAGCCAGGAGAGAGAGAGA | TCAACGTTTCAGTTAGGGGG | 258 |
| chr6:120278360-120280388 | p2 | (AT)14 | GCAGCCAGGAGAGAGAGAGA | TCAACGTTTCAGTTAGGGGG | 258 |
| chr6:120398812-120400830 | p6 | (TCAGTT)3 | CATGGAGCTTTATGCCCACT | ATTCCAGCCTTTGAGCGTAA | 273 |
| chr6:120412763-120414778 | p5 | (TTATG)3 | CATGGCAAAGAAGTGTGCAT | CGGCCCCTTAGGAAATAAAA | 246 |
| chr6:120627106-120629120 | p2 | (TA)7 | TTTCTTTTGGTTTTGGACGG | TGATGCTTGAAATCAGTGGC | 193 |
| chr6:120648007-120650408 | p4 | (AAGA)4 | ACCCACCATCGATCACTACC | AAAATTGTCTCGCGCTCCTA | 218 |
| chr6:120648007-120650408 | p5 | (ATACA)3 | ACCCACCATCGATCACTACC | AAAATTGTCTCGCGCTCCTA | 218 |
| chr6:120657045-120659060 | p3 | (CAA)5 | CAATAACGACCTCATCGGCT | CGAAAAATCCTCGGTTCAGA | 147 |
| chr6:120657045-120659060 | p5 | (CCCAC)3 | CAATAACGACCTCATCGGCT | CGAAAAATCCTCGGTTCAGA | 147 |
| chr6:120657045-120659060 | p2 | (TG)8 | CAATAACGACCTCATCGGCT | CGAAAAATCCTCGGTTCAGA | 147 |
| chr6:120797596-120799611 | p5 | (TTATA)3 | TTGTAATTGCCTCCTCCCAC | CATCACGGTGTATGTCAGGC | 271 |
| chr6:120855585-120857600 | p6 | (CCCACC)3 | CCTTGTTTGGATGGTGGTTT | TGGTGGAGGATGGGGTATAA | 167 |
| chr6:120855585-120857600 | p5 | (TACAA)3 | CCTTGTTTGGATGGTGGTTT | TGGTGGAGGATGGGGTATAA | 167 |
| chr6:120951733-120953775 | p2 | (TC)21 | CAAGGTTGAAGCTCTTTCCG | TTGTCTCCATCAACCATCCA | 250 |
| chr6:120962513-120964531 | p6 | (GAAAAA)3 | AATCACGATCGGTTAGGGTG | AACAATCAAATCCTCCGTCG | 224 |
| chr6:120982737-120984755 | p3 | (CCA)6 | AGTGGTTGAATTTTCCAGCG | TAAGGTGGTGGTGGAGAAGG | 108 |
| chr6:121008722-121010746 | p3 | (CCA)8 | AGTGGTTGAATTTTCCAGCG | GGGTCTGCCGACACTACATT | 108 |
| chr6:121042820-121044832 | p2 | (TA)6 | GGAACCTAACAGGCCTCACA | TGTTTTGTTTGGTTTGGCAG | 174 |
| chr6:121071255-121073273 | p6 | (TATTTT)3 | TCAACCGATCGATCTTTCCT | TTACTGTGCCCAATTAGCCC | 228 |
| chr6:121073724-121075739 | p3 | (AAC)5 | AGCTTCCTAACCCTAGCCG | GAATAGTGTCGCCCATTCGT | 140 |

| Marker ID | SSR type | SSR motif | FORWARD PRIMER (5'-3') | REVERSE PRIMER (5'-3') | size (bp) |
|---|---|---|---|---|---|
| chr6:121097459-121099474 | p5 | (TAATG)3 | TGTTGATGTGGCGTGATTTT | TTCTCCGTGAAACTGCCTCT | 287 |
| chr6:121097459-121099474 | p3 | (ATT)5 | TGTTGATGTGGCGTGATTTT | TTCTCCGTGAAACTGCCTCT | 287 |
| chr6:121152009-121154027 | p6 | (TATTTT)3 | TCAACCGATCGATCTTTCCT | TTACTGTGCCCAATTAGCCC | 226 |
| chr6:121239633-121241659 | p2 | (AT)13 | GATAGTCCGGAGCCAAAATG | AGCCATCAACGAAATACCCA | 232 |
| chr6:121239633-121241659 | p2 | (AT)7 | GATAGTCCGGAGCCAAAATG | AGCCATCAACGAAATACCCA | 232 |
| chr6:121303871-121305883 | p2 | (TA)6 | CAAATTGAAGAGCGCTCAGA | GGGGAACCCGAAAATCTACTT | 160 |
| chr6:121303871-121305883 | p2 | (TG)7 | CAAATTGAAGAGCGCTCAGA | GGGGAACCCGAAAATCTACTT | 160 |
| chr6:121303871-121305883 | p2 | (TA)9 | CAAATTGAAGAGCGCTCAGA | GGGGAACCCGAAAATCTACTT | 160 |
| chr6:121332562-121334577 | p5 | (ATTTC)3 | CCATTTCTTTCGGAGCCATA | TACAAAACATGGGCCAACAA | 178 |
| chr6:121357954-121359969 | p5 | (ACCAA)3 | ACGAATATGTCGGGCAGTTC | CGTTGAAGTGAGGGTGTTCA | 167 |
| chr6:121368870-121370930 | p2 | (TA)30 | TGGTTATGTCCGCCTCTTTT | TTTTAAGTCCCCACGTGTCC | 199 |
| chr6:121398446-121400464 | p3 | (ACA)6 | AAGGCTCGAATCCAAAACCT | ATAATTTCTTGCCGACGGTG | 156 |
| chr6:121502745-121504760 | p5 | (AAAAT)3 | CAATTGCTCCACAAACATGG | TATCACCGCACTCGTGTCAT | 262 |
| chr6:121528226-121530238 | p2 | (AT)6 | TTACGTTCATAGCCGCACAC | TTCGTTGATTGTGTCCGAAA | 285 |
| chr6:121597187-121599202 | p5 | (ACCAA)3 | CACACTTCCCGTCTTGGTTT | CCTCTACCCTCCTCCCAAAG | 134 |
| chr6:121597187-121599202 | p5 | (AAACC)3 | CACACTTCCCGTCTTGGTTT | CCTCTACCCTCCTCCCAAAG | 134 |
| chr6:121705357-121707372 | p5 | (TTGCA)3 | TAAGGTCGTTTTGGGTTTCG | CACGAACGATCAAACCAATG | 270 |
| chr6:121705357-121707372 | p6 | (CCTGCA)3 | TAAGGTCGTTTTGGGTTTCG | CACGAACGATCAAACCAATG | 270 |
| chr6:121713984-121715999 | p5 | (TTGCA)3 | TAAGGTCGTTTTGGGTTTCG | GAACGATCAAACCAATGCCT | 270 |
| chr6:121941426-121943438 | p2 | (AC)6 | GGGGTCGCTAAACTGAAACA | GAGCTCCTGAGAGAGGGGTT | 208 |
| chr6:122001153-122003168 | p3 | (TTG)5 | CTAAGCAGAATGGAGCCAGG | GTTAGTGGCACCGTGTGATG | 205 |
| chr6:122042654-122044670 | p3 | (AAG)5 | CAATGTGGTGGAGAGTGTGC | GCGACCTGTTACACCCTAGC | 118 |
| chr6:122042654-122044670 | p2 | (TA)8 | CAATGTGGTGGAGAGTGTGC | GCGACCTGTTACACCCTAGC | 118 |
| chr6:122042654-122044670 | p5 | (AAATT)3 | CAATGTGGTGGAGAGTGTGC | GCGACCTGTTACACCCTAGC | 118 |
| chr6:122067528-122069543 | p3 | (TTG)5 | CTAAGCAGAATGGAGCCAGG | GTTAGTGGCACCGTGTGATG | 205 |
| chr6:122182778-122184790 | p2 | (TA)6 | AAAAAGTGGAACCCACCACA | CACGGTGTTTAATATGGGGC | 148 |
| chr6:122182778-122184790 | p2 | (TA)7 | AAAAAGTGGAACCCACCACA | CACGGTGTTTAATATGGGGC | 148 |
| chr6:122201657-122203678 | p3 | (AAC)7 | AGGGTCAAATGTGAGTTCGC | GTCGGTTAGCGGTTGGTTTA | 294 |
| chr6:122210730-122212750 | p2 | (AT)7 | TTGCTCCACGCCCTACTACT | CGAAGCCACGGTTGATACTT | 197 |
| chr6:122210781-122212793 | p2 | (AT)7 | TTGCTCCACGCCCTACTACT | CGAAGCCACGGTTGATACTT | 146 |
| chr6:122210781-122212793 | p2 | (TA)6 | TTGCTCCACGCCCTACTACT | CGAAGCCACGGTTGATACTT | 146 |
| chr6:122290823-122292837 | p2 | (AT)7 | TGAGGTACGTGCAAGCTGAC | ATTACCTCCCGTCAGCACAG | 294 |
| chr6:122303221-122305233 | p3 | (TAA)8 | CGGGACAAAATTGGAGATCA | GCACGTGCAAACCTGACTAA | 158 |
| chr6:122303221-122305233 | p2 | (GT)6 | CGGGACAAAATTGGAGATCA | GCACGTGCAAACCTGACTAA | 158 |
| chr6:122303221-122305233 | p2 | (GT)6 | CGGGACAAAATTGGAGATCA | GCACGTGCAAACCTGACTAA | 158 |
| chr6:122303221-122305233 | p2 | (GT)6 | CGGGACAAAATTGGAGATCA | GCACGTGCAAACCTGACTAA | 158 |

| Marker ID | SSR type | SSR motif | FORWARD PRIMER (5'-3') | REVERSE PRIMER (5'-3') | size (bp) |
|---|---|---|---|---|---|
| chr6:122303221-122305233 | p2 | (GT)6 | CGGGACAAAATTGGAGATCA | GCACGTGCAAACCTGACTAA | 158 |
| chr6:122303221-122305233 | p2 | (GT)6 | CGGGACAAAATTGGAGATCA | GCACGTGCAAACCTGACTAA | 158 |
| chr6:122352037-122354052 | p5 | (TGGTT)3 | GGCCACTTAACAAAGGCAAA | CGTAAAATGGAACGGAGGAA | 227 |
| chr6:122352037-122354052 | p5 | (TTGGT)3 | GGCCACTTAACAAAGGCAAA | CGTAAAATGGAACGGAGGAA | 227 |
| chr6:122409453-122411468 | p3 | (TAA)5 | TTTGCGTAGACTCAGATGCG | TTGAGTCGTCCAGAAATCCC | 280 |
| chr6:122471239-122473254 | p3 | (CAA)5 | TCACACAAATGGGACAGAGC | TTATTCTGCATTCTGCACGG | 295 |
| chr6:122492057-122494072 | p3 | (ACA)5 | GGAATAGCTGGCCTCACATC | GTTGAGGACTGCCAAAGAGC | 226 |
| chr6:122540445-122542457 | p2 | (TC)6 | AGCCTGTTTGGATTGGCTTA | CCGTTTGGATGGTCTTATGG | 207 |
| chr6:122540445-122542457 | p5 | (GTGAA)3 | AGCCTGTTTGGATTGGCTTA | CCGTTTGGATGGTCTTATGG | 207 |
| chr6:122544776-122546788 | p2 | (TC)6 | AGCCTGTTTGGATTGGCTTA | CCGTTTGGATGGTCTTATGG | 207 |
| chr6:122544776-122546788 | p5 | (GTGAA)3 | AGCCTGTTTGGATTGGCTTA | CCGTTTGGATGGTCTTATGG | 207 |
| chr6:122616851-122618863 | p2 | (CA)6 | AGGAACTCCACACAGTTGGG | AACTTCCAATCTTGCTCCGA | 202 |
| chr6:122620571-122622586 | p5 | (ATTTA)3 | TATGAAAGGGCCCACTTGAG | CCAGAATCCGAATTGGTGAA | 176 |
| chr6:122620571-122622586 | p2 | (AT)8 | TATGAAAGGGCCCACTTGAG | CCAGAATCCGAATTGGTGAA | 176 |
| chr6:122641356-122643368 | p2 | (AT)6 | CTCTGCAATCCTCGTCGAAT | TGAGCATGAAGCTCCACAAC | 140 |
| chr6:122641356-122643368 | p2 | (TA)14 | CTCTGCAATCCTCGTCGAAT | TGAGCATGAAGCTCCACAAC | 140 |
| chr6:122649966-122651986 | p2 | (AT)10 | TTTTGGGGCTCAAAGTGAGT | TCAGCACCCCTTAAAAATGG | 234 |
| chr6:122657922-122659940 | p3 | (AAG)6 | AATTCGATCAATCTCGCCTG | TTCATCACCCCAAGTGTCAA | 181 |
| chr6:122703205-122705219 | p2 | (AC)7 | TGCACTTCCAAATTTACCCC | TTTGGGGTTGAAAGAATTGG | 173 |
| chr6:122741220-122743253 | p6 | (GGAGAG)3 | GGCTTGTCTCCTCCTCTCCT | TCCCATAAGTGGGGTTTGAG | 121 |
| chr6:122741220-122743253 | p3 | (TAA)11 | GGCTTGTCTCCTCCTCTCCT | TCCCATAAGTGGGGTTTGAG | 121 |
| chr6:122814172-122816196 | p2 | (TA)12 | AATGGGTCTATTGGGGGAAA | GATTGGGCTCCTCTCCATTT | 251 |
| chr6:122853619-122855635 | p4 | (TTCT)4 | TTTTGAAAATGGTCTTGCCC | ACCCAATATTTCCCCCTCAG | 117 |
| chr6:122916992-122919007 | p5 | (TGGCT)3 | AAAGCAACACAAGTTTGGGG | CACAACACCAAGACACCGAC | 185 |
| chr6:123000232-123002258 | p5 | (AAATT)3 | CCGCTAGAAGTCCAAGCAAG | GCTACCTGTGCGGTCTCTTC | 246 |
| chr6:123000232-123002258 | p2 | (TA)13 | CCGCTAGAAGTCCAAGCAAG | GCTACCTGTGCGGTCTCTTC | 246 |
| chr6:123024873-123026897 | p6 | (AACCGG)4 | CAACGTCTTTGCCGAAGATT | TTCGAAGAACAAGCATCACG | 129 |
| chr6:123063283-123065298 | p5 | (TGTAT)3 | TCTCATAGCCAGATGGGTCC | TCGAACACAACCGTTTTACG | 195 |
| chr6:123121506-123123521 | p3 | (TTG)5 | CGCAAGAGACAGATGTGGAA | TGCGACATCATCGTCAATCT | 122 |
| chr6:123124649-123126663 | p2 | (TG)6 | TTGCTTCACCGTTGTTGTTC | TCTGTGCACACAAACGCATA | 247 |
| chr6:123124649-123126663 | p2 | (TG)7 | TTGCTTCACCGTTGTTGTTC | TCTGTGCACACAAACGCATA | 247 |
| chr6:123124673-123126693 | p2 | (TG)6 | TTGCTTCACCGTTGTTGTTC | TCTGTGCACACAAACGCATA | 223 |
| chr6:123124673-123126693 | p2 | (TG)7 | TTGCTTCACCGTTGTTGTTC | TCTGTGCACACAAACGCATA | 223 |
| chr6:123124761-123126861 | p2 | (TG)6 | TTGCTTCACCGTTGTTGTTC | TCTGTGCACACAAACGCATA | 135 |
| chr6:123124761-123126861 | p2 | (TG)7 | TTGCTTCACCGTTGTTGTTC | TCTGTGCACACAAACGCATA | 135 |
| chr6:123124761-123126861 | p3 | (TGT)6 | TTGCTTCACCGTTGTTGTTC | TCTGTGCACACAAACGCATA | 135 |

| Marker ID | SSR type | SSR motif | FORWARD PRIMER (5'-3') | REVERSE PRIMER (5'-3') | size (bp) |
|---|---|---|---|---|---|
| chr6:123125043-123127055 | p2 | (TG)6 | ATGTATGTGCGATCGCTTTG | TCATTTTTGCTCCACACCAA | 119 |
| chr6:123125043-123127055 | p2 | (TG)7 | ATGTATGTGCGATCGCTTTG | TCATTTTTGCTCCACACCAA | 119 |
| chr6:123125043-123127055 | p3 | (TGT)6 | ATGTATGTGCGATCGCTTTG | TCATTTTTGCTCCACACCAA | 119 |
| chr6:123142875-123144893 | p3 | (TAT)6 | CGAAAGGTTTCGAGTTGAGC | CTTTCTTGGGGATGACCTGA | 206 |
| chr6:123176622-123178636 | p2 | (TC)7 | TTTTCCCTCATCGGAACAAG | TTGCAAAACCTGAGCATACG | 131 |
| chr6:123242289-123244304 | p3 | (TTG)5 | CGCAAGAGACAGATGTGGAA | TGCGACATCATCGTCAATCT | 122 |
| chr6:123245432-123247446 | p2 | (TG)6 | TTGCTTCACCGTTGTTGTTC | TCTGTGCACACAAACGCATA | 247 |
| chr6:123245432-123247446 | p2 | (TG)7 | TTGCTTCACCGTTGTTGTTC | TCTGTGCACACAAACGCATA | 247 |
| chr6:123245456-123247476 | p2 | (TG)6 | TTGCTTCACCGTTGTTGTTC | TCTGTGCACACAAACGCATA | 223 |
| chr6:123245456-123247476 | p2 | (TG)7 | TTGCTTCACCGTTGTTGTTC | TCTGTGCACACAAACGCATA | 223 |
| chr6:123245544-123247644 | p2 | (TG)6 | TTGCTTCACCGTTGTTGTTC | TCTGTGCACACAAACGCATA | 135 |
| chr6:123245544-123247644 | p2 | (TG)7 | TTGCTTCACCGTTGTTGTTC | TCTGTGCACACAAACGCATA | 135 |
| chr6:123245544-123247644 | p3 | (TGT)6 | TTGCTTCACCGTTGTTGTTC | TCTGTGCACACAAACGCATA | 135 |
| chr6:123245826-123247838 | p2 | (TG)6 | ATGTATGTGCGATCGCTTTG | TCATTTTTGCTCCACACCAA | 119 |
| chr6:123245826-123247838 | p2 | (TG)7 | ATGTATGTGCGATCGCTTTG | TCATTTTTGCTCCACACCAA | 119 |
| chr6:123245826-123247838 | p3 | (TGT)6 | ATGTATGTGCGATCGCTTTG | TCATTTTTGCTCCACACCAA | 119 |
| chr6:123263658-123265676 | p3 | (TAT)6 | CGAAAGGTTTCGAGTTGAGC | CTTTCTTGGGGATGACCTGA | 206 |
| chr6:123297405-123299419 | p2 | (TC)7 | TTTTCCCTCATCGGAACAAG | TTGCAAAACCTGAGCATACG | 131 |
| chr6:123346535-123348549 | p2 | (AT)7 | CCAAGGCTTAGGCAGATGAG | TCAAAATCCCCAGTTTCCTG | 222 |
| chr6:123356547-123358563 | p2 | (AT)6 | GAAATCTTACAGTTGGCGGC | CCCTCTCTCTTATTTCCCCG | 202 |
| chr6:123356547-123358563 | p2 | (AT)8 | GAAATCTTACAGTTGGCGGC | CCCTCTCTCTTATTTCCCCG | 202 |
| chr6:123440489-123442501 | p2 | (TA)6 | CAAGGAGGAAATGTTCCCAA | CGTTGCTGGTGCTAGTGTGT | 136 |
| chr6:123513328-123515348 | p2 | (TA)10 | CCCTAAAAGCCGATCAACAA | ATGGTACCCCTAGTAGGCCG | 212 |
| chr6:123596804-123598819 | p3 | (AAC)5 | AAAGGTTTCCGAATCTCCGT | ATGATTTACCGAGCCATTCG | 104 |
| chr6:123618865-123620877 | p2 | (AT)6 | CTTTTTGAAAGAATTGCCGC | TTGAGCGGGTTAGAAATTGG | 237 |
| chr6:123650019-123652091 | p2 | (AC)9 | AGGGCGCTACAAAGTGCTTA | TGGGGTCCATTTAGGATTGA | 252 |
| chr6:123650100-123652190 | p2 | (AC)9 | AGGGCGCTACAAAGTGCTTA | TGGGGTCCATTTAGGATTGA | 171 |
| chr6:123749588-123751663 | p3 | (GAG)5 | GTTACAATGGGTGGGCATTC | ATCTGTGTTACCACTCCCGC | 127 |
| chr6:123761550-123763568 | p3 | (CAA)6 | GCCAAACATATGTGGAACCC | TTCCTCACTACTCCGCTCGT | 148 |
| chr6:123897737-123899749 | p2 | (AC)6 | TCGACAAATCATCCGTCAAA | GGAATAAAGAACCGCCATCA | 155 |
| chr6:123946898-123948916 | p6 | (TTTCCT)3 | TGGACTAACGGGTTTATGGC | TCTAGCTTTCCTAGTGCGCC | 282 |
| chr6:123946898-123948916 | p6 | (TTTTTA)4 | TGGACTAACGGGTTTATGGC | TCTAGCTTTCCTAGTGCGCC | 282 |
| chr6:123981302-123983320 | p6 | (TTTTTA)3 | TGGACCCTCGAAGAAAATTG | GTAAGCCATCCCATCGACAT | 280 |
| chr6:124137898-124139918 | p5 | (AATTA)4 | GAAAGCCACAAGGGAATTGA | ACACCCAATTCTCAAACCCA | 101 |
| chr6:124302481-124304496 | p5 | (TTTTC)3 | GGTGGCAGTGTAAGGGTGAT | CCGCACACACAAAGGTTCTA | 264 |
| chr6:124321601-124323625 | p6 | (CAAAGC)4 | TGAAGGATTTACAGACGGGG | CATTTCCCGGATGACCTAGA | 277 |

| Marker ID | SSR type | SSR motif | FORWARD PRIMER (5'-3') | REVERSE PRIMER (5'-3') | size (bp) |
|---|---|---|---|---|---|
| chr6:124359419-124361433 | p2 | (AT)7 | CATAAAATCAGCCCTGCCAT | GAATAGGGACTGGGATGCAA | 153 |
| chr6:124499423-124501439 | p2 | (AT)10 | GAGCCCAACTTTCCAATTCA | GGTAGCTGGGGCTTTTTAGC | 207 |
| chr6:124499423-124501439 | p2 | (AT)10 | GAGCCCAACTTTCCAATTCA | GGTAGCTGGGGCTTTTTAGC | 207 |
| chr6:124499423-124501439 | p2 | (CA)8 | GAGCCCAACTTTCCAATTCA | GGTAGCTGGGGCTTTTTAGC | 207 |
| chr6:124603111-124605123 | p5 | (AAATA)3 | ATAGGCAACTGGAAGCATGG | TGCAAATGCAAAGGTTGAAA | 270 |
| chr6:124603111-124605123 | p2 | (TA)6 | ATAGGCAACTGGAAGCATGG | TGCAAATGCAAAGGTTGAAA | 270 |
| chr6:124640264-124642276 | p5 | (AAATA)3 | ATAGGCAACTGGAAGCATGG | TGCAAATGCAAAGGTTGAAA | 270 |
| chr6:124640264-124642276 | p2 | (TA)6 | ATAGGCAACTGGAAGCATGG | TGCAAATGCAAAGGTTGAAA | 270 |
| chr6:124671007-124673027 | p5 | (TAAAA)4 | TCAAAACGTGCTTGTGGAAG | CACCTTAATGCGATGGGTCT | 105 |
| chr6:124681068-124683094 | p2 | (TC)13 | TCTTGCGTACTTGTTTTGCG | TTGCAGGAAAACTGCAAGTG | 286 |
| chr6:124759863-124761875 | p3 | (GTT)8 | CTATCCAGTGGGGTTTGCAT | GTGTGAAGGCTTGCAGATGA | 256 |
| chr6:124759863-124761875 | p2 | (TC)6 | CTATCCAGTGGGGTTTGCAT | GTGTGAAGGCTTGCAGATGA | 256 |
| chr7:118282-120294 | p2 | (AC)6 | ATGAGCCATTTTGGTAACGG | GGGCTTTTGCACTTCATCAT | 118 |
| chr7:183595-185611 | p2 | (TA)8 | TTCCCAAAGTATCCCCATCA | GACATGGCGGAATAGCAAAT | 156 |
| chr7:202350-204374 | p3 | (AAC)8 | TAACCATGGATTGGCACAGA | TAATGTCCTTTTGGCTTCCG | 246 |
| chr7:231530-233545 | p3 | (TGT)5 | GCTGGTCCCTGAGTTGGTTA | TGTGGACGATTTCGAATCAA | 146 |
| chr7:302062-304077 | p3 | (TTG)5 | TGAACTAAGGGGTCGTTTGG | GGTGCATGGTATAAGCTGGG | 178 |
| chr7:347763-349778 | p5 | (ATCTA)3 | GCATTCCGGAAAAATGAAAA | GTACCTACCGCCCTACCCAT | 227 |
| chr7:349922-351937 | p5 | (ATCTA)3 | GCATTCCGGAAAAATGAAAA | GTACCTACCGCCCTACCCAT | 227 |
| chr7:376321-378346 | p5 | (TCGGG)5 | TCGGACAAGGCAAAAGTTCT | AAATAGTGTTGGCCGAGTGG | 176 |
| chr7:376321-378346 | p2 | (TG)6 | TCGGACAAGGCAAAAGTTCT | AAATAGTGTTGGCCGAGTGG | 176 |
| chr7:428648-430663 | p5 | (ATATT)3 | TAGGTGTGCCGACCACATTA | ATGCATGCAGCTTTGATTTG | 173 |
| chr7:485202-487234 | p2 | (TA)16 | TCCCACTTGTCTTCAAACCC | ACAAAAACGAGAGCGTTGCT | 104 |
| chr7:526139-528159 | p2 | (TA)10 | TTGTCCAAAACCAAGCAACA | TCTACAGCCACAACCTGCTG | 129 |
| chr7:526139-528159 | p2 | (AT)10 | TTGTCCAAAACCAAGCAACA | TCTACAGCCACAACCTGCTG | 129 |
| chr7:526139-528159 | p2 | (TA)8 | TTGTCCAAAACCAAGCAACA | TCTACAGCCACAACCTGCTG | 129 |
| chr7:722337-724351 | p2 | (CT)7 | TCCAGATGGTCAACGTTTCA | GCTCTAACAGTCACCTCGCC | 267 |
| chr7:724662-726682 | p5 | (TCATA)4 | ACAGGTCTTTTACATGCCCG | GGGCTAATTCAATGCCAAGA | 269 |
| chr7:750076-752088 | p2 | (TA)6 | GTAGTCAGATGGGACCCACG | TAAATGGGAGAGCAGGTTGG | 242 |
| chr7:750076-752088 | p2 | (TA)6 | GTAGTCAGATGGGACCCACG | TAAATGGGAGAGCAGGTTGG | 242 |
| chr7:750076-752088 | p2 | (TA)6 | GTAGTCAGATGGGACCCACG | TAAATGGGAGAGCAGGTTGG | 242 |
| chr7:750076-752088 | p2 | (TA)6 | GTAGTCAGATGGGACCCACG | TAAATGGGAGAGCAGGTTGG | 242 |
| chr7:750076-752088 | p2 | (TA)7 | GTAGTCAGATGGGACCCACG | TAAATGGGAGAGCAGGTTGG | 242 |
| chr7:750076-752088 | p2 | (AT)12 | GTAGTCAGATGGGACCCACG | TAAATGGGAGAGCAGGTTGG | 242 |
| chr7:750191-752203 | p2 | (TA)6 | GTAGTCAGATGGGACCCACG | TAAATGGGAGAGCAGGTTGG | 127 |
| chr7:750191-752203 | p2 | (TA)6 | GTAGTCAGATGGGACCCACG | TAAATGGGAGAGCAGGTTGG | 127 |

| Marker ID | SSR type | SSR motif | FORWARD PRIMER (5'-3') | REVERSE PRIMER (5'-3') | size (bp) |
|---|---|---|---|---|---|
| chr7:750191-752203 | p2 | (TA)6 | GTAGTCAGATGGGACCCACG | TAAATGGGAGAGCAGGTTGG | 127 |
| chr7:750191-752203 | p2 | (TA)6 | GTAGTCAGATGGGACCCACG | TAAATGGGAGAGCAGGTTGG | 127 |
| chr7:750191-752203 | p2 | (TA)7 | GTAGTCAGATGGGACCCACG | TAAATGGGAGAGCAGGTTGG | 127 |
| chr7:750191-752203 | p2 | (AT)12 | GTAGTCAGATGGGACCCACG | TAAATGGGAGAGCAGGTTGG | 127 |
| chr7:768705-770717 | p2 | (AC)6 | AGGAAGCAACGACATATGGG | TGAGCTGGTTTGTGATTTGG | 217 |
| chr7:789557-791587 | p3 | (AGA)10 | CGATGGGATTTTCCTGCTAA | CTCCTTCCAGAGCCTTCCTT | 185 |
| chr7:857042-859054 | p2 | (TC)6 | AAGGACGGACCGTAACACAG | AATAGCGCCAAGTTTCTGGA | 180 |
| chr7:857042-859054 | p3 | (ATT)5 | AAGGACGGACCGTAACACAG | AATAGCGCCAAGTTTCTGGA | 180 |
| chr7:858122-860143 | p3 | (ATT)5 | AACCAGGTGAATATGGCAGG | GGAGGAAACACAGCAGAAGC | 260 |
| chr7:858392-860407 | p3 | (ATT)5 | GCTTCTGCTGTGTTTCCTCC | GAAGTGGACAGTCCTCCGAA | 110 |
| chr7:860926-862942 | p2 | (TG)8 | TGTTCCCCATGGTTCTGTTT | TCTGCCAATACAGGGTTTCC | 271 |
| chr7:924675-926695 | p2 | (AT)10 | GCTGTAACCCCAGGCACTAA | GAATTTGACAGGAGCTTGGC | 266 |
| chr7:942754-944766 | p2 | (TC)6 | ACAAGTTTATGCTGGGTCCG | TATGCCTCGTAAGCCGAAAC | 187 |
| chr7:969048-971084 | p2 | (AG)6 | TTCCGAGAGCAGTCTCCATT | TCTTGCCTGATAACGCCTCT | 245 |
| chr7:969048-971084 | p2 | (AC)6 | TTCCGAGAGCAGTCTCCATT | TCTTGCCTGATAACGCCTCT | 245 |
| chr7:969093-971105 | p2 | (AG)6 | TTCCGAGAGCAGTCTCCATT | TCTTGCCTGATAACGCCTCT | 200 |
| chr7:969093-971105 | p2 | (AC)6 | TTCCGAGAGCAGTCTCCATT | TCTTGCCTGATAACGCCTCT | 200 |
| chr7:983766-985792 | p2 | (AT)13 | GATCACGGTCAAAGCCGTAT | TCGGTTTTATCGGTTTGGTT | 252 |
| chr7:1007614-1009674 | p2 | (AT)30 | TCCCATCACCACTCTGTTCA | TGGAGCCTTCCATATGTTCC | 125 |
| chr7:1035994-1038016 | p2 | (CA)11 | CAGCGAGAAGAAAATTTGGC | GAATGGAACGACCAAGGGTA | 257 |
| chr7:1049625-1051640 | p3 | (TGT)5 | CTATAGGGGGTGGGGGTAGA | GACATTCCGCATTGATTGTG | 284 |
| chr7:1192170-1194190 | p2 | (TA)10 | ACAGTGCCATTGGTATGCCT | CCCACTGCTATTCAGTCGCT | 298 |
| chr7:1231516-1233528 | p2 | (TA)6 | GAAGCACAAGCAACAAACGA | GCTGCCCTTTAGTTGACAGG | 163 |
| chr7:1295613-1297631 | p6 | (AATATA)3 | ACGACTGCGGAAAAAGAGAA | CTGAATCGAAGGAACCCAAA | 158 |
| chr7:1385752-1387773 | p3 | (CAA)7 | CAGGGGTCCAAGTTTATCCA | ACTGTTCTTTCCTCCCCCAT | 113 |
| chr7:1406251-1408266 | p2 | (TC)6 | CGATATCGCCAAGGCTAGAG | TTGGAGGATTCTAACCCGTG | 195 |
| chr7:1406251-1408266 | p3 | (CCA)5 | CGATATCGCCAAGGCTAGAG | TTGGAGGATTCTAACCCGTG | 195 |
| chr7:1406251-1408266 | p5 | (TATAT)3 | CGATATCGCCAAGGCTAGAG | TTGGAGGATTCTAACCCGTG | 195 |
| chr7:1406836-1408851 | p2 | (TC)6 | CGCCACGTCAATGTCATATC | GGTTGCGAAAAGTAGGGTGA | 128 |
| chr7:1406836-1408851 | p3 | (CCA)5 | CGCCACGTCAATGTCATATC | GGTTGCGAAAAGTAGGGTGA | 128 |
| chr7:1406836-1408851 | p5 | (TATAT)3 | CGCCACGTCAATGTCATATC | GGTTGCGAAAAGTAGGGTGA | 128 |
| chr7:1406836-1408851 | p3 | (TAT)6 | CGCCACGTCAATGTCATATC | GGTTGCGAAAAGTAGGGTGA | 128 |
| chr7:1445303-1447318 | p3 | (TTG)5 | GAAATGGAGGCTTGCTTCAG | CCGTACAACAAGGTCGGTCT | 126 |
| chr7:1483994-1486009 | p5 | (AAACC)3 | TGAATCGGGTTGGTTCAGAT | TGAAAACCCCGACTTTTGTT | 158 |
| chr7:1483994-1486009 | p5 | (AAACC)3 | TGAATCGGGTTGGTTCAGAT | TGAAAACCCCGACTTTTGTT | 158 |
| chr7:1490601-1492619 | p6 | (TAAAAA)3 | TCGCTTCGCAAATAGGAGTT | TGTTTCGTGTTCATTGGGAA | 277 |

| Marker ID | SSR type | SSR motif | FORWARD PRIMER (5'-3') | REVERSE PRIMER (5'-3') | size (bp) |
|---|---|---|---|---|---|
| chr7:1615528-1617542 | p2 | (TA)7 | TTTTCTTCGGGGACTTGATG | CCGGAGACATCATCCACTTT | 159 |
| chr7:1628212-1630270 | p2 | (AT)29 | CGTTCGAGAATTCAATGGGT | ACATCACCTGGTAAAACGCC | 177 |
| chr7:1773006-1775024 | p6 | (ATTTTG)3 | GTGATGGAGCAGTAGGCACA | GTCGCACTTTGGGAGGTAAA | 262 |
| chr7:1785406-1787424 | p6 | (TTTTTC)3 | ATTTACGTCGTGCCTCCAAG | GGTATCAGGCCACACGTTTT | 162 |
| chr7:1832098-1834114 | p2 | (TA)8 | AGTGTGCACCTGCGTATTCA | CCATCATTAGCTCGCACTCA | 235 |
| chr7:1835315-1837361 | p2 | (AT)6 | AAAAAGTGGAACCCACCACA | CACGGTGTTTAATATGGGGC | 153 |
| chr7:1876847-1878862 | p3 | (AGA)5 | CATCCACCACTATTCCCACC | CACAAACGTCGAGAGTGCAT | 266 |
| chr7:1979139-1981157 | p6 | (TGCTTG)3 | CTTCTGCAGGCTCTTCTGCT | ATGGCAGCTCCTAGAACGAA | 144 |
| chr7:2044741-2046765 | p5 | (AATAC)3 | TCAATGTCGAGGTGCAGAAG | TTGTATTACCTTGCGTTGCG | 103 |
| chr7:2044741-2046765 | p3 | (AAT)8 | TCAATGTCGAGGTGCAGAAG | TTGTATTACCTTGCGTTGCG | 103 |
| chr7:2044741-2046765 | p2 | (AT)12 | TCAATGTCGAGGTGCAGAAG | TTGTATTACCTTGCGTTGCG | 103 |
| chr7:2051148-2053164 | p2 | (AT)8 | CCATCTTCAGGAACCTCCAA | TACGTTGGCCCGTATTCTTC | 199 |
| chr7:2085139-2087163 | p5 | (AATAC)3 | TCAATGTCGAGGTGCAGAAG | TTGTATTACCTTGCGTTGCG | 103 |
| chr7:2085139-2087163 | p3 | (AAT)8 | TCAATGTCGAGGTGCAGAAG | TTGTATTACCTTGCGTTGCG | 103 |
| chr7:2085139-2087163 | p2 | (AT)12 | TCAATGTCGAGGTGCAGAAG | TTGTATTACCTTGCGTTGCG | 103 |
| chr7:2091546-2093562 | p2 | (AT)8 | CCATCTTCAGGAACCTCCAA | TACGTTGGCCCGTATTCTTC | 199 |
| chr7:2114403-2116419 | p2 | (TG)6 | GAGCCAATTTGACACCCCTA | GACACGCGGCAATTTTTATT | 206 |
| chr7:2114421-2116435 | p2 | (TG)6 | GAGCCAATTTGACACCCCTA | GACACGCGGCAATTTTTATT | 188 |
| chr7:2139657-2141691 | p2 | (TA)9 | TACGCATGTGCAATGAATCC | CACCACATCCAAACTCAACG | 272 |
| chr7:2139657-2141691 | p2 | (AT)16 | TACGCATGTGCAATGAATCC | CACCACATCCAAACTCAACG | 272 |
| chr7:2142132-2144147 | p5 | (CCACC)3 | TTTGCTACGAATACGCATGG | TTGGGCCCCATATTAAACAG | 125 |
| chr7:2349442-2351496 | p3 | (TAA)18 | TTCGAGGCAGTTTACAACCC | TTTGGATCGTCGTTTTAGGG | 245 |
| chr7:2440755-2442770 | p5 | (TGGGG)3 | TGGATTAAATGGGCCAAGAG | ATATGGGAAGGCCTTTTGCT | 153 |
| chr7:2482372-2484390 | p3 | (CAA)6 | CATAATACGCACGACAACCG | TATGTTTTTCATAGGCGGGC | 223 |
| chr7:2583496-2585523 | p3 | (ATG)9 | TCCCAATTTAAGCGTCTTGC | ATTGCATACCACTTGCTCCC | 162 |
| chr7:2628674-2630758 | p3 | (GAA)6 | AGCAGATTCGCAGCCATAGT | GCTCTTTCATTTCGAGTCCG | 155 |
| chr7:2628674-2630758 | p2 | (AT)8 | AGCAGATTCGCAGCCATAGT | GCTCTTTCATTTCGAGTCCG | 155 |
| chr7:2709694-2711730 | p2 | (TG)6 | TATGCGGCTCCAAGAACTTT | TTTTCCAGTCAAAAGTGGGG | 215 |
| chr7:2709809-2711824 | p2 | (TG)6 | TATGCGGCTCCAAGAACTTT | TTTTCCAGTCAAAAGTGGGG | 100 |
| chr7:2719319-2721351 | p2 | (TC)16 | GGGACTTCATTTTCCTGCAA | TCAGTTGTTCAATGCCCAAA | 145 |
| chr7:2856230-2858248 | p6 | (TACAAC)3 | GAGGGGAGGAGAAGTTCCAG | TTCAGAATGCGGAGGCTTAT | 185 |
| chr7:2897768-2899788 | p5 | (TCTTT)4 | TAGCTCGGGTGAGGAGAAAA | TGATTCCAAGGAGTGCAGTG | 107 |
| chr7:2898462-2900491 | p5 | (TCTTT)4 | TGGTTGGAAGGTGCTACACA | TGGAGTTCCATGATGGTTGA | 241 |
| chr7:3081423-3083437 | p5 | (AAAGA)3 | GGCTAATGGCTCGTGGAATA | GCAAGCTAAGGATTTGACGC | 181 |
| chr7:3081423-3083437 | p2 | (AT)7 | GGCTAATGGCTCGTGGAATA | GCAAGCTAAGGATTTGACGC | 181 |
| chr7:3081423-3083437 | p6 | (AAAAAT)3 | GGCTAATGGCTCGTGGAATA | GCAAGCTAAGGATTTGACGC | 181 |

| Marker ID | SSR type | SSR motif | FORWARD PRIMER (5'-3') | REVERSE PRIMER (5'-3') | size (bp) |
|---|---|---|---|---|---|
| chr7:3124764-3126780 | p2 | (TA)8 | TAATGGTGGATTCATTGCCA | GACATGGCGGAATAGCAAAT | 296 |
| chr7:3149005-3151039 | p2 | (TA)17 | CATCAAAAGGGGGTTAGCAA | CGGACCCATAAACAGAAGGA | 264 |
| chr7:3157409-3159423 | p2 | (AT)7 | TATCTTTGCCCCTCAAATCG | ACAATTCGCAAGGCTGAGTT | 288 |
| chr7:3183632-3185647 | p2 | (AT)6 | TACCACATAGAGGCGGAACC | ACAACTTCGCGGAATGAGAT | 143 |
| chr7:3183632-3185647 | p5 | (TTTTA)3 | TACCACATAGAGGCGGAACC | ACAACTTCGCGGAATGAGAT | 143 |
| chr7:3200465-3202483 | p2 | (TA)9 | TGCCCACTCCACATGTAAAA | ACCTCACCGTCTGAGGTTTG | 279 |
| chr7:3214136-3216156 | p2 | (TA)10 | TGCAGTGGATGAAACTGCTC | GTTTTATGATTGCGTCGGGT | 272 |
| chr7:3221979-3224021 | p2 | (AT)21 | AACTTCCGGGTTAAAATGGG | TGTGGCGTACATTAGGGTGA | 199 |
| chr7:3224837-3226855 | p2 | (AT)7 | GGCCAAGTAATATGGGACGA | TGTTTTATCGCAAAGGGAGG | 251 |
| chr7:3224837-3226855 | p6 | (TAAAAA)3 | GGCCAAGTAATATGGGACGA | TGTTTTATCGCAAAGGGAGG | 251 |
| chr7:3224837-3226855 | p2 | (TA)16 | GGCCAAGTAATATGGGACGA | TGTTTTATCGCAAAGGGAGG | 251 |
| chr7:3244194-3246212 | p2 | (AT)9 | ATCTTAGCCAACTTCCCCGT | TGGGTACGCAAGTGAATGAA | 114 |
| chr7:3263085-3265103 | p6 | (TTTTTA)3 | AACTTTGTGCTCCGATGTCC | AAAACTTCACGGCAATTTGG | 145 |
| chr7:3340966-3342981 | p3 | (ATA)5 | ATTCAAACTCACAAACGGGC | GACCCGGTCCATTTTCTTTT | 285 |
| chr7:3421691-3423711 | p2 | (AT)10 | AAACATGAGGGAAATGACGG | TTTTGCATGCGAGACAAAAC | 298 |
| chr7:3466386-3468400 | p5 | (TTTTC)4 | CCATGCGTACCCTAGCTGTT | TGAGCATGACAGCAACATCA | 105 |
| chr7:3466386-3468400 | p2 | (CT)7 | CCATGCGTACCCTAGCTGTT | TGAGCATGACAGCAACATCA | 105 |
| chr7:3513989-3516004 | p5 | (TCGGG)3 | TCCTAGCTCTCATTTCGCGT | TAGTTCCTGATGCGAAACCC | 137 |
| chr7:3533491-3535506 | p5 | (AAACC)3 | TCTAATTCCGCACTATCCCG | CGAACAACCCGAGAAAACTC | 109 |
| chr7:3633729-3635741 | p2 | (TG)6 | ACGAAATTAGGGGAGATGGG | ACCGCCCATTCACAATACAC | 234 |
| chr7:3693539-3695577 | p2 | (GA)19 | TGTCTATGGGCTTGAGGACC | TGGGCTGAAAAGGTCTCATC | 202 |
| chr7:3834396-3836414 | p3 | (TAG)6 | GTTCTGCCAGTCGGAACAGT | TAGGGCCGGATGTAATCTTG | 115 |
| chr7:3834396-3836414 | p2 | (AT)11 | GTTCTGCCAGTCGGAACAGT | TAGGGCCGGATGTAATCTTG | 115 |
| chr7:3838893-3840908 | p5 | (AAAAT)3 | GCTAAGGTGCTTCGTGATCC | GCTTTAGTACCCCCGACTCC | 169 |
| chr7:3850139-3852154 | p5 | (TTTTA)3 | TTGATCCGGACAATCACCTT | TGTCGCAGCTTGTTTGTAGC | 125 |
| chr7:3850139-3852154 | p2 | (AT)29 | TTGATCCGGACAATCACCTT | TGTCGCAGCTTGTTTGTAGC | 125 |
| chr7:3886021-3888043 | p2 | (AT)7 | GTGTGTTTAGGGCGTACGGT | TTTGGGACAAACCAAGAAGG | 210 |
| chr7:3886021-3888043 | p2 | (TA)11 | GTGTGTTTAGGGCGTACGGT | TTTGGGACAAACCAAGAAGG | 210 |
| chr7:3947447-3949469 | p2 | (AT)11 | TGAGTCCCTGGTCGCTAAGT | TAGCTCCGTGGCTTCTTCAT | 163 |
| chr7:3947447-3949469 | p2 | (AT)10 | TGAGTCCCTGGTCGCTAAGT | TAGCTCCGTGGCTTCTTCAT | 163 |
| chr7:3948377-3950397 | p2 | (AT)11 | ACAATTAATCTTGGCGGCTG | TTGGCCTCCTAATTCTGCTG | 116 |
| chr7:3948377-3950397 | p2 | (AT)10 | ACAATTAATCTTGGCGGCTG | TTGGCCTCCTAATTCTGCTG | 116 |
| chr7:4122339-4124354 | p5 | (CTTTG)3 | AATGGAGACTGCCTTGGCTA | AACCTGTCACCAGGAACTGC | 296 |
| chr7:4237963-4239975 | p2 | (AG)6 | ACCGCATTCAACGACTCTCT | GGCAGGCTGAAAATTGTGAT | 183 |
| chr7:4318350-4320374 | p3 | (TCA)5 | GTGTGTGATTGTGCAGGACC | TGAGGCAATGCAACAACATT | 121 |
| chr7:4318350-4320374 | p6 | (CCGGTT)4 | GTGTGTGATTGTGCAGGACC | TGAGGCAATGCAACAACATT | 121 |

| Marker ID | SSR type | SSR motif | FORWARD PRIMER (5'-3') | REVERSE PRIMER (5'-3') | size (bp) |
|---|---|---|---|---|---|
| chr7:4318350-4320374 | p6 | (ATAAAA)3 | GTGTGTGATTGTGCAGGACC | TGAGGCAATGCAACAACATT | 121 |
| chr7:4330616-4332631 | p5 | (TTTTA)3 | AATGGCTCCAATCCCTCTTT | AACGGGAACGGAAGTTAGGT | 117 |
| chr7:4402156-4404171 | p3 | (TTG)5 | AAAGGGATTTGCATCAATCG | TTAGTCAATGTTGTGCCCGA | 111 |
| chr7:4408573-4410585 | p2 | (AT)6 | CAGTAACATGACCACCCACG | GCTCCACTTGCTTTGGAGAC | 183 |
| chr7:4491959-4493974 | p5 | (TATTT)3 | AGATTGGGGCTATTTTGGCT | TCCAAACAGGCCCTTAGAGA | 265 |
| chr7:4555201-4557217 | p4 | (AAGA)4 | GCCGAAAAATCATACCGTGT | AATGTATCCTCCCCAGACCC | 149 |
| chr7:4555999-4558035 | p4 | (AAGA)4 | AAATGATTTATGTGCCCCCA | GGAAAAGGGTGAAAAGGGAA | 135 |
| chr7:4555999-4558035 | p3 | (AGA)20 | AAATGATTTATGTGCCCCCA | GGAAAAGGGTGAAAAGGGAA | 135 |
| chr7:4738028-4740040 | p2 | (AT)6 | ATCCTTTAAGCCCTTTCCCA | TTGTCAAAACCACGAACCAA | 251 |
| chr7:4750315-4752333 | p3 | (TAA)6 | AAGCTCCAGGAACTGAGCAA | GTGCAAGGCGGAAACTAGAG | 193 |
| chr7:4750315-4752333 | p6 | (GATGAA)5 | AAGCTCCAGGAACTGAGCAA | GTGCAAGGCGGAAACTAGAG | 193 |
| chr7:4888848-4890863 | p3 | (AAT)5 | ATGGAGGCACAACGAAAGAC | CGGATCCTTCAAAATTGCAT | 126 |
| chr7:4952382-4954403 | p2 | (AG)6 | AAAGGTCACTTTTGGGCCTT | ATTGCCAACCAAAGATCCAC | 115 |
| chr7:4960041-4962056 | p3 | (TGA)5 | CTTGGCCCTATTCACAGAGG | GCCAGTGGTGAGTCAAGGAT | 121 |
| chr7:4960041-4962056 | p3 | (GAA)5 | CTTGGCCCTATTCACAGAGG | GCCAGTGGTGAGTCAAGGAT | 121 |
| chr7:5050628-5052646 | p6 | (AATTGT)3 | TGCAAATGGCTAAGTGGACA | TCGAATTTCATTGTCTCCCC | 282 |
| chr7:5057725-5059743 | p6 | (AATTGT)3 | TGCAAATGGCTAAGTGGACA | TCGAATTTCATTGTCTCCCC | 282 |
| chr7:5134024-5136036 | p2 | (TC)6 | TGATAGAAAGCGTTTTCCCG | AGTCCAAATAGCGGGGACTT | 143 |
| chr7:5244196-5246208 | p2 | (CA)6 | ACATACACTGGCTCATCCCC | ATTCCTGTTGGTGAGGTTGG | 103 |
| chr7:5270370-5272385 | p5 | (ATCAT)3 | AGACAAGTGTCCAACGACCC | TCTGTGACGGCCCTTTAATC | 262 |
| chr7:5288340-5290355 | p5 | (AAAAT)3 | TCGAAAAGGAAAATGCATCA | TCCGGAAGAAGATGAAAAGG | 212 |
| chr7:5288340-5290355 | p5 | (ATTTT)3 | TCGAAAAGGAAAATGCATCA | TCCGGAAGAAGATGAAAAGG | 212 |
| chr7:5288340-5290355 | p2 | (TA)29 | TCGAAAAGGAAAATGCATCA | TCCGGAAGAAGATGAAAAGG | 212 |
| chr7:5400223-5402235 | p2 | (GA)6 | AACCCCCTCGGCTCTATCTA | GTTCAAGCCCAACCCTATCA | 110 |
| chr7:5447934-5449946 | p2 | (TA)8 | AATAATGCCCAAGTTTCCCC | AGCTAGCATGAGAAGAGCCG | 198 |
| chr7:5447934-5449946 | p2 | (TA)6 | AATAATGCCCAAGTTTCCCC | AGCTAGCATGAGAAGAGCCG | 198 |
| chr7:5448029-5450043 | p2 | (TA)8 | AATAATGCCCAAGTTTCCCC | AGCTAGCATGAGAAGAGCCG | 103 |
| chr7:5448029-5450043 | p2 | (TA)6 | AATAATGCCCAAGTTTCCCC | AGCTAGCATGAGAAGAGCCG | 103 |
| chr7:5500057-5502072 | p3 | (AAC)5 | TGATGCTTTCCTTTGGGAAC | CGAACATCAGTTGGGAAACC | 268 |
| chr7:5547588-5549603 | p5 | (AAAAG)3 | CATTTTGCCATTATCACGCA | TGTCGATCATATCCCAGCAA | 278 |
| chr7:5604623-5606639 | p2 | (TC)8 | CACCGCTTATGGATGAACCT | TCACGGCTGATTTTGTGAAA | 195 |
| chr7:5657445-5659459 | p2 | (TA)7 | ACCCGAACTCGCTATTGATG | CGACAAGCCCTTTTTGCTAC | 119 |
| chr7:5657445-5659459 | p2 | (TA)11 | ACCCGAACTCGCTATTGATG | CGACAAGCCCTTTTTGCTAC | 119 |
| chr7:5657735-5659751 | p2 | (TA)7 | GTATAAACCGTGGCTTCGGA | AGTTTTGGTTGTGTCGCTCC | 151 |
| chr7:5657735-5659751 | p2 | (TA)11 | GTATAAACCGTGGCTTCGGA | AGTTTTGGTTGTGTCGCTCC | 151 |
| chr7:5657735-5659751 | p2 | (AT)6 | GTATAAACCGTGGCTTCGGA | AGTTTTGGTTGTGTCGCTCC | 151 |

| Marker ID | SSR type | SSR motif | FORWARD PRIMER (5'-3') | REVERSE PRIMER (5'-3') | size (bp) |
|---|---|---|---|---|---|
| chr7:5743772-5745790 | p6 | (TCTCCT)3 | CCGCAACACACATTTCTCAC | CGTCGACCTACCACAACCTT | 117 |
| chr7:5841168-5843190 | p2 | (AC)11 | ATGTTTAGGGTTTTTGGGGG | CAAATGGCGTTCTTCGAGTT | 198 |
| chr7:5883242-5885270 | p2 | (TC)14 | TCCCATTCAGATCTGTTCCC | AATTTCGTTTCCCGGAGAGT | 218 |
| chr7:5999477-6001492 | p3 | (CCA)5 | GGACGCAAGTACCAGACCAT | TCCTGAAAACCGGTATCTGC | 240 |
| chr7:5999477-6001492 | p3 | (ATT)6 | GGACGCAAGTACCAGACCAT | TCCTGAAAACCGGTATCTGC | 240 |
| chr7:5999477-6001492 | p2 | (TA)7 | GGACGCAAGTACCAGACCAT | TCCTGAAAACCGGTATCTGC | 240 |
| chr7:6013500-6015516 | p2 | (TG)8 | TTCAAGAGGGACATCCCAAC | TCATGGCAAAAATGAACGAA | 285 |
| chr7:6022669-6024689 | p2 | (TA)10 | CTTGTATGGGAAGGAGCAGC | CTTCAAGTGGCTCCAACTCC | 197 |
| chr7:6026862-6028877 | p2 | (AT)8 | TTTTAACTGGAAACTGGCCG | AGGTGGATGACACCTTTTGC | 285 |
| chr7:6026862-6028877 | p2 | (TA)19 | TTTTAACTGGAAACTGGCCG | AGGTGGATGACACCTTTTGC | 285 |
| chr7:6026862-6028877 | p5 | (TTCTC)3 | TTTTAACTGGAAACTGGCCG | AGGTGGATGACACCTTTTGC | 285 |
| chr7:6032891-6034909 | p2 | (TA)9 | CCTGCCTTTTCATTTTCCAA | TGATGTTGAGGCCAACAATG | 219 |
| chr7:6032938-6034964 | p2 | (TA)9 | CCTGCCTTTTCATTTTCCAA | TGATGTTGAGGCCAACAATG | 172 |
| chr7:6097432-6099450 | p2 | (GT)9 | TAATTGGCATCTTTTTGGGG | CGATAACGCTGCCTTCATTT | 249 |
| chr7:6097432-6099450 | p2 | (TA)8 | TAATTGGCATCTTTTTGGGG | CGATAACGCTGCCTTCATTT | 249 |
| chr7:6126494-6128514 | p2 | (TA)10 | AAAAGCCTCTGCGGTTGATA | GGCAGAACCATGACGCTAAT | 124 |
| chr7:6132500-6134515 | p2 | (TA)27 | ACATGAACAATGGGAGAGGC | CCCGAAGTTGAAAAACCTGA | 178 |
| chr7:6132533-6134548 | p2 | (TA)27 | ACATGAACAATGGGAGAGGC | CCCGAAGTTGAAAAACCTGA | 145 |
| chr7:6141915-6143930 | p3 | (TGT)5 | AACATTGCCCATTCAAGAGC | TCCCGTATCTTGTCCTCCAC | 174 |
| chr7:6158673-6160751 | p2 | (AT)39 | AGGCACAGGAGCAGGTAAGA | TCCCTTCCGTCATTCAAGTT | 111 |
| chr7:6158673-6160751 | p2 | (TA)12 | AGGCACAGGAGCAGGTAAGA | TCCCTTCCGTCATTCAAGTT | 111 |
| chr7:6322738-6324762 | p4 | (TTTA)6 | AGCAGGCCTGATTTAGTCCA | AGCAAGATGTATTGGGCCAG | 107 |
| chr7:6526973-6529027 | p2 | (AT)27 | TGACGTTCACATGGCTGTTT | CCACCAGTTAACCACCATCC | 136 |
| chr7:6555103-6557115 | p2 | (AT)6 | CAAAGGGTGGTGTCTCCCTA | TTTGGAGATGGTGGTTCCAT | 206 |
| chr7:6555103-6557115 | p2 | (AT)11 | CAAAGGGTGGTGTCTCCCTA | TTTGGAGATGGTGGTTCCAT | 206 |
| chr7:6556286-6558306 | p2 | (AT)11 | CATGACTCAACGTGATCGCT | GGTATGTTTGGCATTTCCGT | 261 |
| chr7:6556286-6558306 | p2 | (AT)10 | CATGACTCAACGTGATCGCT | GGTATGTTTGGCATTTCCGT | 261 |
| chr7:6556286-6558306 | p2 | (AT)11 | CATGACTCAACGTGATCGCT | GGTATGTTTGGCATTTCCGT | 261 |
| chr7:6556286-6558306 | p2 | (AT)11 | CATGACTCAACGTGATCGCT | GGTATGTTTGGCATTTCCGT | 261 |
| chr7:6639220-6641232 | p5 | (AAAAT)3 | TGGATCGTCAGGTAAGGAGG | GAGGTACCGGAAACAGGTGA | 108 |
| chr7:6662242-6664257 | p5 | (TTTTA)3 | CGGGTCGAGTTTGATAGCTC | TAGCTGCGGCGGTAGTATCT | 150 |
| chr7:6681097-6683109 | p2 | (AC)6 | TTGGCATCCTGAAATCAACA | TAAGCCAGCAACTGCCTCTT | 282 |
| chr7:6726950-6728964 | p2 | (TA)7 | GGCCAAACACAACTCCAACT | AATGAACAAATGGGATTGGC | 231 |
| chr7:6726950-6728964 | p2 | (AT)14 | GGCCAAACACAACTCCAACT | AATGAACAAATGGGATTGGC | 231 |
| chr7:6742459-6744474 | p6 | (CAAAAA)3 | GGTGAGCACGAAAATTGGTT | TTTCTGGGGAAACTCTGGTG | 223 |
| chr7:6742459-6744474 | p3 | (CAA)5 | GGTGAGCACGAAAATTGGTT | TTTCTGGGGAAACTCTGGTG | 223 |

| Marker ID | SSR type | SSR motif | FORWARD PRIMER (5'-3') | REVERSE PRIMER (5'-3') | size (bp) |
|---|---|---|---|---|---|
| chr7:6750511-6752543 | p2 | (TC)16 | TGGGTTTCCATCTGGTTTTC | CCTGAATGTGGAATTGCCTT | 107 |
| chr7:6814239-6816254 | p3 | (AAT)5 | ATGTGAAAATGAAGGCCAGG | CGTCACAGAGGTGGTAGGGT | 279 |
| chr7:6818879-6820909 | p3 | (TAA)5 | GGGGTCCGGTAAGTCAAAAT | AACTTCTGCACACACATCGC | 276 |
| chr7:6818879-6820909 | p6 | (ATAATG)5 | GGGGTCCGGTAAGTCAAAAT | AACTTCTGCACACACATCGC | 276 |
| chr7:6882395-6884410 | p2 | (AG)6 | CGCGTGAAACAAAGCTAACA | TAAATTTTGGCGGTTTTTGC | 293 |
| chr7:6882395-6884410 | p3 | (ATT)5 | CGCGTGAAACAAAGCTAACA | TAAATTTTGGCGGTTTTTGC | 293 |
| chr7:6947905-6949929 | p2 | (AG)12 | AGGTACTGTTGGCGCTGAGT | CATCAAAATGCCCTTGTCCT | 210 |
| chr7:6972868-6974882 | p2 | (TC)7 | CTTTTCTGGCCTCAAAGGCT | AGCTGTCAATATGGGCTTGG | 227 |
| chr7:6974376-6976392 | p2 | (TA)8 | GGCTCCACCAACCAACATAG | CCCACTAAATTGGGCATCAA | 156 |
| chr7:7070627-7072642 | p3 | (TTC)5 | AAACTTCGCTTTTTCAGGCA | TTGGTAGAGGGACAATTCGC | 223 |
| chr7:7145904-7147919 | p5 | (TTTTA)3 | ATTGGAATATGCTTCCACGG | TCACGTGTCAGGTAGCTTCG | 182 |
| chr7:7275222-7277238 | p2 | (TA)8 | ACAACAAAAACCATGGACCC | GACATGGCGGAATAGCAAGT | 232 |
| chr7:7326011-7328026 | p5 | (ATCAT)3 | GTGAAGAAGAAGCAGGGGTG | GGGCATAAAAAGGGGTGTTT | 232 |
| chr7:7346575-7348590 | p5 | (TTGGT)3 | AGTAAACCGACAAACCGCAC | ACCAAACCAATTATGGTCGG | 108 |
| chr7:7379502-7381520 | p3 | (ATT)6 | GCATGGTGGAGGATCTCACT | AAAAAGGGAAAAATGGTCGG | 287 |
| chr7:7379502-7381520 | p6 | (AAATAA)3 | GCATGGTGGAGGATCTCACT | AAAAAGGGAAAAATGGTCGG | 287 |
| chr7:7398121-7400159 | p2 | (TA)19 | GACGAAGGGGGTGATAATGA | CACCTATTGGTGTGAGGGCT | 112 |
| chr7:7479545-7481560 | p3 | (TAA)5 | TGGCAAAAACAGTGGTGTGT | CCCTTTTAGGGTCCCCATTA | 155 |
| chr7:7497779-7499797 | p3 | (ATT)6 | GCAGCCATCTCCTTTTTCAG | AATTGATGAACGGAAGCCAC | 284 |
| chr7:7565363-7567378 | p5 | (AAAAC)3 | ACTAGCCAAATACAACGCCG | GTGGCAACGCCATTTTTAGT | 188 |
| chr7:7572528-7574546 | p6 | (TAAAAA)3 | AATTAAGGATGGGTGGGGAC | ATGGGTCCCACAATTTCAGA | 259 |
| chr7:7580613-7582628 | p5 | (ATATG)3 | TGAATACCGAACCCAAAAGG | TGAGATACGTAGGCAGCCCT | 197 |
| chr7:7583495-7585517 | p2 | (GA)6 | GAAAGGTGTGAGGACAGGGA | GGGACAAAGGGGTATGGTTT | 250 |
| chr7:7583495-7585517 | p2 | (TA)11 | GAAAGGTGTGAGGACAGGGA | GGGACAAAGGGGTATGGTTT | 250 |
| chr7:7583495-7585517 | p2 | (AG)7 | GAAAGGTGTGAGGACAGGGA | GGGACAAAGGGGTATGGTTT | 250 |
| chr7:7584368-7586382 | p2 | (TA)11 | ATGGCATGAGACTCGGTAGG | ATGTCGGACCTGGCATACTC | 252 |
| chr7:7584368-7586382 | p2 | (AG)7 | ATGGCATGAGACTCGGTAGG | ATGTCGGACCTGGCATACTC | 252 |
| chr7:7598201-7600216 | p3 | (ACA)5 | TACGAACTATGGGCGGAAAG | AGGAAAACATTTTCGCGATG | 225 |
| chr7:7640615-7642635 | p2 | (TA)10 | AGCATGGAATGCAACAAACA | TTTTTGAGACGGAGGAAGGA | 178 |
| chr7:7653835-7655856 | p6 | (CAAAAA)3 | AAGCCGTCTGTTCGATTTTC | TCCAACAAATGTTCCACCAA | 213 |
| chr7:7653835-7655856 | p3 | (AAG)7 | AAGCCGTCTGTTCGATTTTC | TCCAACAAATGTTCCACCAA | 213 |
| chr7:7712978-7715028 | p2 | (TA)7 | GCATTTTGATCCCCTTTGAA | GATTGCAAGTGCCTGATGAA | 206 |
| chr7:7774838-7776854 | p3 | (TCT)13 | ACACCCCGATTGGTATCAGA | GAGGCATTTGATGAAAGGGA | 233 |
| chr7:7774838-7776854 | p2 | (AT)8 | ACACCCCGATTGGTATCAGA | GAGGCATTTGATGAAAGGGA | 233 |
| chr7:7841900-7843915 | p5 | (AAAAT)3 | GCCACCAGATCCTTTTTCAA | GATGTGCCACCACTGCTAGA | 230 |
| chr7:7862372-7864393 | p3 | (TAT)7 | CGCGAGTTTATGCTAGCTCC | GCAGCTTATACCATGCACCA | 233 |

| Marker ID | SSR type | SSR motif | FORWARD PRIMER (5'-3') | REVERSE PRIMER (5'-3') | size (bp) |
|---|---|---|---|---|---|
| chr7:7884681-7886697 | p4 | (TAAA)4 | TCTCCAATAACCATCTCGGC | CCAAGCAACAAGCCCATTAT | 149 |
| chr7:7890093-7892138 | p3 | (ATT)15 | CAGGGAGATTGTTTGGGTGT | CCAAAAGGCCTCACATCATT | 159 |
| chr7:8013329-8015345 | p4 | (AATC)4 | AAGAATGTCTTGTGACGCCC | TGCCATGTGAACTTCTGCTC | 119 |
| chr7:8052262-8054277 | p3 | (GCT)5 | GCAGCCCAAGGACATTACAT | AAAATGGGGTGTGACAAGGA | 172 |
| chr7:8162579-8164597 | p3 | (TAT)6 | AAGCAACTACCGGGTGAGAA | ATCCATGTCTTGCTGGTTCC | 187 |
| chr7:8173959-8175971 | p2 | (AT)6 | TTGCTCCACGCCCTACTACT | ACCATGCTAAGACGTCCACC | 113 |
| chr7:8173959-8175971 | p2 | (AT)8 | TTGCTCCACGCCCTACTACT | ACCATGCTAAGACGTCCACC | 113 |
| chr7:8174330-8176346 | p2 | (AT)6 | AACCAAAACTCAGAAGCGGA | TTTCGTTAATTTCCCCCACA | 113 |
| chr7:8174330-8176346 | p2 | (AT)8 | AACCAAAACTCAGAAGCGGA | TTTCGTTAATTTCCCCCACA | 113 |
| chr7:8381062-8383078 | p4 | (AAAT)4 | AACAGCCTCTCTGCCCTACA | CAGCCTCCTAGTTGTGGCTC | 258 |
| chr7:8430671-8432719 | p2 | (TA)24 | CCCGAAGATGAGAACCGTAA | AAGCCTCAAATCCAATCCCT | 177 |
| chr7:8430671-8432719 | p3 | (AAT)5 | CCCGAAGATGAGAACCGTAA | AAGCCTCAAATCCAATCCCT | 177 |
| chr7:8527776-8529794 | p6 | (TCTTTT)3 | CAGAAATCCTGCGTACACCA | CGGTTTGACATTGGGAGAAT | 300 |
| chr7:8607293-8609314 | p2 | (TA)7 | AAGAGAGGTCGCTGACGAAG | AGCATGCATGAACCATCAAA | 262 |
| chr7:8607293-8609314 | p3 | (ATC)7 | AAGAGAGGTCGCTGACGAAG | AGCATGCATGAACCATCAAA | 262 |
| chr7:8618597-8620609 | p2 | (TA)6 | AGGGGTGGCTATGTATGCAG | ATCCATTAGGGTCGATGTGC | 133 |
| chr7:8636072-8638084 | p2 | (AT)6 | TCACATCCAAACTCACGGAA | ATATGGGGCCCACAATTTTT | 220 |
| chr7:8658112-8660139 | p3 | (CAA)9 | TGGGCATCCAACTTGTGTTA | TCCTGGCAAGGTTTTTAACG | 269 |
| chr7:8706919-8708934 | p5 | (GTTTG)3 | AGTTCTCCCCACACCACTTG | TAGAAGGGGGATGGAGGTCT | 228 |
| chr7:8780547-8782563 | p4 | (AAGA)4 | ACTGGGATTTTTGTCGTTGG | AGCCTGATGGGAAAATAGCA | 120 |
| chr7:8817723-8819739 | p4 | (AAGA)4 | ACTGGGATTTTTGTCGTTGG | AGCCTGATGGGAAAATAGCA | 120 |
| chr7:8858645-8860660 | p5 | (AAACC)3 | AGGATGAAGAAGCCAACCCT | CTACCCATCATGCGTGTGTC | 199 |
| chr7:8906511-8908531 | p5 | (AAACA)4 | CTAGGGATTCGCATTACCCA | GCATTTTGAACCAACCAACC | 148 |
| chr7:9115566-9117581 | p5 | (GTCAT)3 | TGAGCAGACGTGTAGTTGGG | GTCGGGGTGTGACAATATCC | 272 |
| chr7:9136583-9138598 | p5 | (ATTTT)3 | TCAGGCAGTTATGGTGTCCA | ACAGTGCGCGTACTTCATTG | 275 |
| chr7:9136583-9138598 | p6 | (CACCCG)3 | TCAGGCAGTTATGGTGTCCA | ACAGTGCGCGTACTTCATTG | 275 |
| chr7:9195242-9197257 | p5 | (GGTTT)3 | TTCATCCCGTTTAGTGGAGG | TAAACCCTGCATTGGGAAAG | 184 |
| chr7:9254141-9256153 | p2 | (TA)6 | GCCTGGTCCAAATCTACGAA | AATTGAAACCCTTTTTGGGG | 198 |
| chr7:9254141-9256153 | p5 | (TTGTT)3 | GCCTGGTCCAAATCTACGAA | AATTGAAACCCTTTTTGGGG | 198 |
| chr7:9262667-9264679 | p2 | (AT)14 | TCCAAATTAACCGCCAGAAG | GACAAACAGCAACAAGGCAA | 218 |
| chr7:9359398-9361414 | p4 | (AAAT)4 | CCGTTTGACGCGATTTTTAT | CTTCATCACCACGGATTTC | 191 |
| chr7:9418649-9420664 | p2 | (TA)9 | CAGGTTTGCATGAACAGGAA | GACAATGCCTGCAATCAAAG | 275 |
| chr7:9418649-9420664 | p2 | (AT)6 | CAGGTTTGCATGAACAGGAA | GACAATGCCTGCAATCAAAG | 275 |
| chr7:9418746-9420758 | p2 | (TA)9 | CAGGTTTGCATGAACAGGAA | GACAATGCCTGCAATCAAAG | 178 |
| chr7:9418746-9420758 | p2 | (AT)6 | CAGGTTTGCATGAACAGGAA | GACAATGCCTGCAATCAAAG | 178 |
| chr7:9565216-9567234 | p3 | (GAA)6 | AAGGCCCGAAGTAAAGCAAT | GAAATTGATTGGGCGCTTTA | 177 |

| Marker ID | SSR type | SSR motif | FORWARD PRIMER (5'-3') | REVERSE PRIMER (5'-3') | size (bp) |
|---|---|---|---|---|---|
| chr7:9594223-9596251 | p4 | (ACAA)7 | CTTCACCTACTGAGGGCAGG | GGCAAACTAGCATTTCTCGC | 209 |
| chr7:9594223-9596251 | p3 | (TAT)5 | CTTCACCTACTGAGGGCAGG | GGCAAACTAGCATTTCTCGC | 209 |
| chr7:9627928-9629940 | p2 | (AT)6 | TTACGGCGATTACCGAAGTC | TACCGGAGTAACGGGAAGTG | 197 |
| chr7:9637361-9639373 | p2 | (TA)6 | ACATTGAGCGGAATACCAGG | AAACCCCATGGAAAAAGGTC | 251 |
| chr7:9718236-9720251 | p3 | (CTT)5 | CCGTCTCTTGTGGAGGGATA | CTGCTCAAGATTTTCGCACA | 102 |
| chr7:9724317-9726335 | p2 | (AT)9 | GCTGAAGTTGTGCCAATCAA | ACTGCCCAAGCTCCTCACTA | 199 |
| chr7:9755200-9757222 | p2 | (AT)11 | ATTGGAAGGAACACCACTGC | GATCATGGACCCGATACAGC | 256 |
| chr7:9755200-9757222 | p2 | (TA)7 | ATTGGAAGGAACACCACTGC | GATCATGGACCCGATACAGC | 256 |
| chr7:9813008-9815020 | p2 | (TA)6 | CAGCATTTGTGGAGAAGCAA | GGAGAGAGGTTTCCCTTTCG | 235 |
| chr7:9890815-9892845 | p5 | (TATTG)6 | GAGAATGTGTGCGGTTGAGA | TGGCAAAGATTTGGTGAACA | 147 |
| chr7:10015868-10017890 | p2 | (AT)8 | GTGCTACACCTGTTGCTCCA | TTTCCCACACAAGAAGGGAG | 182 |
| chr7:10015868-10017890 | p6 | (ATATTC)3 | GTGCTACACCTGTTGCTCCA | TTTCCCACACAAGAAGGGAG | 182 |
| chr7:10015895-10017907 | p2 | (AT)8 | GTGCTACACCTGTTGCTCCA | TTTCCCACACAAGAAGGGAG | 155 |
| chr7:10015895-10017907 | p6 | (ATATTC)3 | GTGCTACACCTGTTGCTCCA | TTTCCCACACAAGAAGGGAG | 155 |
| chr7:10015911-10017929 | p2 | (AT)8 | GTGCTACACCTGTTGCTCCA | TTTCCCACACAAGAAGGGAG | 139 |
| chr7:10015911-10017929 | p6 | (ATATTC)3 | GTGCTACACCTGTTGCTCCA | TTTCCCACACAAGAAGGGAG | 139 |
| chr7:10104614-10106629 | p5 | (TTTTA)3 | GGTTCCACACTTGAGCCAAT | GCTGGAGCAAAGAGTTGAGG | 142 |
| chr7:10108308-10110344 | p2 | (TA)18 | TTTCTTTTTGGTGGTGAGGC | GTTCAATAGGAACGAGGCCA | 216 |
| chr7:10108308-10110344 | p4 | (TTTA)4 | TTTCTTTTTGGTGGTGAGGC | GTTCAATAGGAACGAGGCCA | 216 |
| chr7:10136612-10138627 | p5 | (TTTTA)3 | GGTTCCACACTTGAGCCAAT | GCTGGAGCAAAGAGTTGAGG | 142 |
| chr7:10140306-10142342 | p2 | (TA)18 | TTTCTTTTTGGTGGTGAGGC | GTTCAATAGGAACGAGGCCA | 216 |
| chr7:10140306-10142342 | p4 | (TTTA)4 | TTTCTTTTTGGTGGTGAGGC | GTTCAATAGGAACGAGGCCA | 216 |
| chr7:10515692-10517710 | p6 | (AATTGA)3 | CAGAAGAGGGTGCTTTGAGC | AGCCCTTCACTTCTCTGCAA | 288 |
| chr7:10529122-10531140 | p6 | (TTAATA)3 | TGGCACTGACGTGACACATA | CTTGGTTTGAGCTGGCAAGT | 280 |
| chr7:10583054-10585070 | p5 | (AAACC)3 | GTGTCGGGTTGGTTTGATTC | AAAGCAAACTGACAAACCGC | 239 |
| chr7:10583054-10585070 | p4 | (TTTA)4 | GTGTCGGGTTGGTTTGATTC | AAAGCAAACTGACAAACCGC | 239 |
| chr7:10583054-10585070 | p2 | (AC)9 | GTGTCGGGTTGGTTTGATTC | AAAGCAAACTGACAAACCGC | 239 |
| chr7:10628741-10630757 | p2 | (AC)8 | CCAAGCATGAGAATTTCGCT | AGCATGCACACATTTCGTTC | 300 |
| chr7:10640171-10642183 | p2 | (TA)7 | TGGTATGCTCTGCCCTTTCT | TGGAAGTCATCGTGTTGCTC | 152 |
| chr7:10640171-10642183 | p2 | (AC)6 | TGGTATGCTCTGCCCTTTCT | TGGAAGTCATCGTGTTGCTC | 152 |
| chr7:10728315-10730330 | p3 | (AAT)5 | CAAGCCATTCCAAGCTCTTC | TCGTCATTTCCCGAGGATAG | 298 |
| chr7:10744660-10746675 | p5 | (TTTCA)3 | TCTATGGGCTTACCAGACGG | TGAGCTTTTCCAAATCCCAC | 108 |
| chr7:10779047-10781127 | p6 | (CTCATG)3 | GTGCTGACGTCCCCTTTTTA | GAGATGGGGCTCACCAATAA | 117 |
| chr7:10791157-10793171 | p2 | (AT)10 | ACAACAACCTGTCTCCTGGG | CGATGTTGTTCAGTTGTGCC | 185 |
| chr7:10810198-10812213 | p3 | (TAC)5 | CATTGTAGGACATGGGGGTC | CCGGGCTACTACACCACACT | 242 |
| chr7:10810198-10812213 | p5 | (TTCAT)3 | CATTGTAGGACATGGGGGTC | CCGGGCTACTACACCACACT | 242 |

| Marker ID | SSR type | SSR motif | FORWARD PRIMER (5'-3') | REVERSE PRIMER (5'-3') | size (bp) |
|---|---|---|---|---|---|
| chr7:10820503-10822515 | p2 | (TA)13 | CATGTTGGTCCGTATTGCAG | TTCAAATCAGCCACATTCCA | 282 |
| chr7:10820503-10822515 | p2 | (AT)18 | CATGTTGGTCCGTATTGCAG | TTCAAATCAGCCACATTCCA | 282 |
| chr7:10820559-10822571 | p2 | (TA)13 | CATGTTGGTCCGTATTGCAG | TTCAAATCAGCCACATTCCA | 226 |
| chr7:10820559-10822571 | p2 | (AT)18 | CATGTTGGTCCGTATTGCAG | TTCAAATCAGCCACATTCCA | 226 |
| chr7:10821798-10823834 | p2 | (AT)18 | TATAACGTGTCCCGACCCTC | ACATCCTTCACTGAGTCCCG | 297 |
| chr7:10821798-10823834 | p2 | (AT)18 | TATAACGTGTCCCGACCCTC | ACATCCTTCACTGAGTCCCG | 297 |
| chr7:10829925-10831937 | p5 | (TATGA)3 | GGCCGGAACGTTAATGTCTA | GGTGAGGGACTCGGTGAATA | 270 |
| chr7:10829925-10831937 | p2 | (AT)6 | GGCCGGAACGTTAATGTCTA | GGTGAGGGACTCGGTGAATA | 270 |
| chr7:10829932-10831947 | p5 | (TATGA)3 | GGCCGGAACGTTAATGTCTA | GGTGAGGGACTCGGTGAATA | 263 |
| chr7:10829932-10831947 | p2 | (AT)6 | GGCCGGAACGTTAATGTCTA | GGTGAGGGACTCGGTGAATA | 263 |
| chr7:10829942-10831954 | p5 | (TATGA)3 | GGCCGGAACGTTAATGTCTA | GGTGAGGGACTCGGTGAATA | 253 |
| chr7:10829942-10831954 | p2 | (AT)6 | GGCCGGAACGTTAATGTCTA | GGTGAGGGACTCGGTGAATA | 253 |
| chr7:10829964-10831982 | p5 | (TATGA)3 | GGCCGGAACGTTAATGTCTA | GGTGAGGGACTCGGTGAATA | 231 |
| chr7:10829964-10831982 | p2 | (AT)6 | GGCCGGAACGTTAATGTCTA | GGTGAGGGACTCGGTGAATA | 231 |
| chr7:10829997-10832019 | p5 | (TATGA)3 | GGCCGGAACGTTAATGTCTA | GGTGAGGGACTCGGTGAATA | 198 |
| chr7:10829997-10832019 | p2 | (AT)6 | GGCCGGAACGTTAATGTCTA | GGTGAGGGACTCGGTGAATA | 198 |
| chr7:10831533-10833561 | p2 | (AT)6 | ATACGCAAGCAAACAAGGCT | CATGACTTATGTTTGGGGGC | 202 |
| chr7:10831533-10833561 | p2 | (AT)14 | ATACGCAAGCAAACAAGGCT | CATGACTTATGTTTGGGGGC | 202 |
| chr7:10843354-10845408 | p2 | (TA)27 | TCCTTCCCAATTTCAACCAC | TTAGCACATCCATTTTCGCA | 275 |
| chr7:11068267-11070303 | p2 | (TA)6 | GCGTTATTGGCCTACCTTCA | ATCCACACGAGGTCCAAAAG | 207 |
| chr7:11073163-11075177 | p2 | (GT)7 | AATTCACCCGCATCAGAATC | GAGTAACCACCTTTTGCGGA | 188 |
| chr7:11077545-11079563 | p5 | (TTTGG)3 | CGCGTACGTAGGGAAGTAGG | CCCGTTAGAGATGGCTGAAA | 222 |
| chr7:11077545-11079563 | p6 | (ATTTTA)3 | CGCGTACGTAGGGAAGTAGG | CCCGTTAGAGATGGCTGAAA | 222 |
| chr7:11103448-11105466 | p3 | (TGT)6 | GGCCTAAAATCCTTTGGCTC | AGGCTTATCATGGGTTGGTG | 231 |
| chr7:11127328-11129356 | p2 | (AT)8 | TGCTCCACGCCCTACTACTT | AATGCACTCTCCGAAACCAC | 263 |
| chr7:11127328-11129356 | p3 | (TAT)5 | TGCTCCACGCCCTACTACTT | AATGCACTCTCCGAAACCAC | 263 |
| chr7:11127472-11129488 | p2 | (AT)8 | TGCTCCACGCCCTACTACTT | AATGCACTCTCCGAAACCAC | 119 |
| chr7:11127472-11129488 | p3 | (TAT)5 | TGCTCCACGCCCTACTACTT | AATGCACTCTCCGAAACCAC | 119 |
| chr7:11160528-11162553 | p5 | (TTTGG)5 | CCCAGAACCTCAGGTATCCA | AGGTGGAGGTGCCTTTTCTT | 133 |
| chr7:11200826-11202841 | p3 | (TGA)6 | ACCCTTTCAACCCAATTTCC | CCAGCAACCACAACATCATC | 193 |
| chr7:11200826-11202841 | p5 | (GGGGT)3 | ACCCTTTCAACCCAATTTCC | CCAGCAACCACAACATCATC | 193 |
| chr7:11253763-11255778 | p3 | (AAC)5 | GCACAGACCCACTTGGAAAT | ATTTTGCCCCAATTTTCCTC | 191 |
| chr7:11272201-11274213 | p2 | (TA)6 | CTGGATTAGATCGGAGCGAC | AGTCTTCCTTTTGGTCCGGT | 221 |
| chr7:11272201-11274213 | p2 | (TA)8 | CTGGATTAGATCGGAGCGAC | AGTCTTCCTTTTGGTCCGGT | 221 |
| chr7:11395550-11397565 | p4 | (ATAA)4 | GCTGTTGGAGCTCTCCTTTG | GAGACAACTCCTGCGGCTAC | 153 |
| chr7:11395550-11397565 | p5 | (AATCA)3 | GCTGTTGGAGCTCTCCTTTG | GAGACAACTCCTGCGGCTAC | 153 |

| Marker ID | SSR type | SSR motif | FORWARD PRIMER (5'-3') | REVERSE PRIMER (5'-3') | size (bp) |
|---|---|---|---|---|---|
| chr7:11471422-11473440 | p3 | (ACA)6 | AACCGTCCTTACCCGAGTCT | CACCTTGTGAGGAGGTTGGT | 114 |
| chr7:11514031-11516045 | p2 | (AT)9 | AACCCCGTCTTTGTTCATTG | TTTGAGCTTTGGACTTGGGT | 179 |
| chr7:11514031-11516045 | p2 | (AT)7 | AACCCCGTCTTTGTTCATTG | TTTGAGCTTTGGACTTGGGT | 179 |
| chr7:11514031-11516045 | p2 | (TA)20 | AACCCCGTCTTTGTTCATTG | TTTGAGCTTTGGACTTGGGT | 179 |
| chr7:11590990-11593005 | p3 | (CCT)5 | GGTTAAATTGGGCTTGGGTT | AAGTGCAAATGACCCGTCTC | 269 |
| chr7:11591044-11593062 | p3 | (CCT)5 | GGTTAAATTGGGCTTGGGTT | AAGTGCAAATGACCCGTCTC | 215 |
| chr7:11638792-11640812 | p5 | (CAAAC)4 | GGGGCAGTAATTGCATTTTG | AGACCCTAGCCTTGGGGATA | 255 |
| chr7:11672879-11674894 | p3 | (ACA)5 | AAATGCTACACCCGTTGCTC | CGGATGACTCGCCTTGTATT | 203 |
| chr7:11672879-11674894 | p5 | (CCGGT)3 | AAATGCTACACCCGTTGCTC | CGGATGACTCGCCTTGTATT | 203 |
| chr7:12017462-12019477 | p5 | (ATTTT)3 | GGTAGCCATGGCATGAAACT | ACTTCGTGCTAAATGGGTGC | 156 |
| chr7:12017462-12019477 | p4 | (TTCT)6 | GGTAGCCATGGCATGAAACT | ACTTCGTGCTAAATGGGTGC | 156 |
| chr7:12018390-12020414 | p5 | (ATTTT)3 | CTTTTCAGTGGCGGTGAGAT | CGCATGTGACGTGCTAGAAT | 118 |
| chr7:12018390-12020414 | p4 | (TTCT)6 | CTTTTCAGTGGCGGTGAGAT | CGCATGTGACGTGCTAGAAT | 118 |
| chr7:12081513-12083528 | p5 | (TATTT)3 | ACTCAATTGAAAACAGCGGC | AGATTGTCCGCATCCACTTC | 130 |
| chr7:12144260-12146275 | p5 | (TTGGT)3 | AGTGGGTTGGGTAGAATCCC | TAATGTCGGGTTGGTTTGGT | 138 |
| chr7:12144260-12146275 | p5 | (TTGGT)3 | AGTGGGTTGGGTAGAATCCC | TAATGTCGGGTTGGTTTGGT | 138 |
| chr7:12186122-12188137 | p5 | (GTGTA)3 | CCCAAGTCCCATAATTCCCT | TTCCCAATATCATGCCCATT | 267 |
| chr7:12193781-12195807 | p5 | (TAGAT)3 | ATGGGGTTCACTTTGCTGAG | CGAACCACCTAACCTCCCTT | 279 |
| chr7:12239676-12241690 | p2 | (AT)8 | GTGTCAAAGACGGGGAAAAA | GTTAGTTTGGGTTTTGCCGA | 138 |
| chr7:12239676-12241690 | p2 | (TA)7 | GTGTCAAAGACGGGGAAAAA | GTTAGTTTGGGTTTTGCCGA | 138 |
| chr7:12358658-12360673 | p5 | (AAAAT)3 | AGGAGGAGCAAGACGAGTCA | AGCACGTGTTCAGAAAGGCT | 137 |
| chr7:12358658-12360673 | p5 | (TTTTA)3 | AGGAGGAGCAAGACGAGTCA | AGCACGTGTTCAGAAAGGCT | 137 |
| chr7:12480736-12482752 | p2 | (TA)8 | AAGTATCAACCGTGGCTTCG | TCCGCTTCTGAGTTTTGGTT | 122 |
| chr7:12480736-12482752 | p2 | (AT)6 | AAGTATCAACCGTGGCTTCG | TCCGCTTCTGAGTTTTGGTT | 122 |
| chr7:12503337-12505367 | p2 | (AT)15 | ACAACCTGTCCAATGTGCAA | TGCCCGAATATCCTTACAGC | 131 |
| chr7:12529250-12531265 | p5 | (TTGCA)3 | TTATTGGCATACCCACCGTT | ATCATCGGACATCGGTCTTC | 200 |
| chr7:12529250-12531265 | p3 | (AGA)5 | TTATTGGCATACCCACCGTT | ATCATCGGACATCGGTCTTC | 200 |
| chr7:12534121-12536136 | p5 | (ATTGC)3 | TAAGGCTAGCCAACCCATTG | TGGGTACGCCAACAATATCA | 258 |
| chr7:12553171-12555187 | p2 | (CT)8 | CCTTGAATAACCTGGGAGCA | CTCGAGGAGTAGACGTTGGG | 222 |
| chr7:12608904-12610919 | p5 | (TAAAA)3 | ATCCACTGTAATTGCGGTCC | AGGTCAGAACAGCCGCTAAA | 258 |
| chr7:12665796-12667818 | p2 | (AT)8 | CAGTGGTCTCCCAAGCTCAT | ACTATGGGGCTGGGATAAGG | 108 |
| chr7:12665796-12667818 | p2 | (AT)11 | CAGTGGTCTCCCAAGCTCAT | ACTATGGGGCTGGGATAAGG | 108 |
| chr7:12774642-12776657 | p5 | (AAAAT)3 | GCCAACTCTTCCCACATGAT | TGAAGTCGGAGGGTTGTTTC | 282 |
| chr7:12799887-12801902 | p2 | (TA)6 | CTTATAGAAGGGGGTTCGGC | AAAGGGCATCATAGGCAGTG | 240 |
| chr7:12799887-12801902 | p5 | (TATTG)3 | CTTATAGAAGGGGGTTCGGC | AAAGGGCATCATAGGCAGTG | 240 |
| chr7:12815238-12817254 | p4 | (AATA)4 | CTTTGTCTGCCCCAAACATT | CTATCCGCCATAAATTCCGA | 154 |

| Marker ID | SSR type | SSR motif | FORWARD PRIMER (5'-3') | REVERSE PRIMER (5'-3') | size (bp) |
|---|---|---|---|---|---|
| chr7:12931545-12933561 | p2 | (GA)8 | GATTATTTCGGGCAGCGATA | TTGATCAAGGACGTTTTCCC | 175 |
| chr7:12931545-12933561 | p2 | (AT)6 | GATTATTTCGGGCAGCGATA | TTGATCAAGGACGTTTTCCC | 175 |
| chr7:12987332-12989352 | p5 | (GAATT)4 | GACCTAAGCGTTCAGCCAAG | TGTGCTCGGTTCGACTAAGA | 282 |
| chr7:12992303-12994318 | p5 | (AAAAC)3 | AATAATTCGGCGCAACAATC | ATTTTCGTGAATCCCCATCA | 277 |
| chr7:12992303-12994318 | p5 | (AAATA)3 | AATAATTCGGCGCAACAATC | ATTTTCGTGAATCCCCATCA | 277 |
| chr7:13025931-13027953 | p5 | (AAAAT)3 | AAAGAGTCACCGGGTAGCAA | CAGGCACAATAAATGGCTCA | 133 |
| chr7:13025931-13027953 | p2 | (AT)14 | AAAGAGTCACCGGGTAGCAA | CAGGCACAATAAATGGCTCA | 133 |
| chr7:13025931-13027953 | p2 | (AC)11 | AAAGAGTCACCGGGTAGCAA | CAGGCACAATAAATGGCTCA | 133 |
| chr7:13048366-13050381 | p3 | (TAC)6 | TATGATTTTGGGCGGGAATA | ATGATCATGATGGGGTTGCT | 228 |
| chr7:13048366-13050381 | p5 | (ATTTT)3 | TATGATTTTGGGCGGGAATA | ATGATCATGATGGGGTTGCT | 228 |
| chr7:13087041-13089055 | p2 | (TG)7 | GCCTGTGTCTTTACCCCAAA | TAACGCCAAGCAACAAACTG | 262 |
| chr7:13156962-13158978 | p2 | (TA)8 | GGCTTTTCCGATAGACCCTC | CTTACCCGCGTAAAAACGAA | 177 |
| chr7:13214998-13217010 | p2 | (AT)6 | GCATTCTGGAATTTGGCATT | ACTGCACTGCGTCTCTTCAA | 146 |
| chr7:13279950-13281970 | p2 | (AT)10 | AGCCAAGGAGAAGTGACGAA | CTAATGAATTGCTCCCCCAA | 110 |
| chr7:13288176-13290201 | p5 | (TGATA)5 | TTTGCCCCTTTTTCATTCAG | AATTGAGGTCTATGGTGGCG | 244 |
| chr7:13348892-13350910 | p3 | (GTG)6 | CCCTTCGTTGACCTCACAGT | TTCGATGTAAGGGAGGGATG | 277 |
| chr7:13380158-13382176 | p2 | (AT)9 | CTTCGTCGTCCATGGAAAGT | ATCGCGCGGTATCATAAATC | 256 |
| chr7:13426624-13428642 | p6 | (CAACTC)3 | GAACACATTTGTTGATGGCG | AAGTTCTCTCCGGTTCAGCA | 263 |
| chr7:13428940-13430979 | p3 | (AAT)13 | AAAAGGTACAACTGCCCCCT | GGTCCTTGCTGGACTTGTTC | 247 |
| chr7:13462304-13464316 | p2 | (TA)6 | CTCCAGTTCAGAGGGTGAGC | CTGCCAAAGAAAAGCAGAGG | 119 |
| chr7:13482927-13484945 | p6 | (ATATGT)3 | TAACCCTACCCTCCGGAATC | GCGTCAATTGTGAGGGTTTT | 251 |
| chr7:13493856-13495871 | p5 | (TTGGT)3 | TGAGCAGTGGTTTCAGGATG | TCCATTCTTGGCCTTCAGAC | 144 |
| chr7:13501925-13503937 | p2 | (TA)6 | CAAGGCCCCAGCATACTTTA | CCATGTTGTCTCTTCCGGTT | 208 |
| chr7:13514628-13516640 | p2 | (TG)6 | ATATGCTGGGCTTCTTGTGG | TGACCTTCCTCCATGTTTCC | 255 |
| chr7:13585883-13587901 | p6 | (TTTTTC)3 | AATCAATGCATGTCGCAAAA | TATTGTTTGGTGCCGAATGA | 236 |
| chr7:13608892-13610907 | p5 | (TTATT)3 | CCCTCTCCTACCACCCTTTC | TTAAAGGTTGACCTCACGGG | 145 |
| chr7:13875702-13877716 | p2 | (AT)7 | ACATACCAAAATGGACCCCA | AACGAGAAGCGATTTCCAGA | 162 |
| chr7:13947261-13949273 | p2 | (TA)6 | CGATATACCCTCGGCTCAGA | TGCAACATCCATCAACAGGT | 144 |
| chr7:14015860-14017884 | p2 | (TA)12 | TCCAGTTTGGCCTCTTTTTG | TATGACTGCATGTGCTGCAA | 212 |
| chr7:14015860-14017884 | p2 | (TA)18 | TCCAGTTTGGCCTCTTTTTG | TATGACTGCATGTGCTGCAA | 212 |
| chr7:14015860-14017884 | p3 | (ATT)7 | TCCAGTTTGGCCTCTTTTTG | TATGACTGCATGTGCTGCAA | 212 |
| chr7:14016118-14018133 | p2 | (TA)12 | GCAGTCATAGATGCGTCTCG | AAGAAACGTGACCTGCGAGT | 168 |
| chr7:14016118-14018133 | p2 | (TA)18 | GCAGTCATAGATGCGTCTCG | AAGAAACGTGACCTGCGAGT | 168 |
| chr7:14016118-14018133 | p3 | (ATT)7 | GCAGTCATAGATGCGTCTCG | AAGAAACGTGACCTGCGAGT | 168 |
| chr7:14016118-14018133 | p2 | (TA)7 | GCAGTCATAGATGCGTCTCG | AAGAAACGTGACCTGCGAGT | 168 |
| chr7:14070657-14072677 | p5 | (TATTT)4 | CGAAAAGGGTAAGCTGGTCA | TATCCAACCAGCGCAATGTA | 101 |

| Marker ID | SSR type | SSR motif | FORWARD PRIMER (5'-3') | REVERSE PRIMER (5'-3') | size (bp) |
|---|---|---|---|---|---|
| chr7:14275014-14277029 | p5 | (TTGTA)3 | GTTGATCTTCTCTCAGCCCG | CATGCCTTCGAGGAAATGAT | 160 |
| chr7:14275014-14277029 | p6 | (TGGGGG)3 | GTTGATCTTCTCTCAGCCCG | CATGCCTTCGAGGAAATGAT | 160 |
| chr7:14310087-14312102 | p5 | (AATGC)3 | TGTTAAGAACCAAGGGGTGC | AGGAGGAAATCAATTGCGTG | 143 |
| chr7:14323735-14325751 | p4 | (TCAA)4 | CGCATAGCTTGCTTCATACG | GAAACGCGATCATTCGATTT | 168 |
| chr7:14323735-14325751 | p6 | (TATAGA)3 | CGCATAGCTTGCTTCATACG | GAAACGCGATCATTCGATTT | 168 |
| chr7:14370336-14372351 | p4 | (TAAA)5 | ATTCCACCTTCAAAGCACCA | TGGTTCGGCGGATGTATTAT | 271 |
| chr7:14370336-14372351 | p3 | (GTT)5 | ATTCCACCTTCAAAGCACCA | TGGTTCGGCGGATGTATTAT | 271 |
| chr7:14386872-14388884 | p2 | (AT)6 | CCGATTATGTCCGGATTTGT | AACAGTACCTTGCCGCAAAC | 185 |
| chr7:14404627-14406695 | p2 | (TG)8 | ATGCAAATAGCCAACCGAAC | TTTGCATGCCGATAACAAAA | 158 |
| chr7:14406090-14408108 | p2 | (TG)8 | TACAAAAGTCCCCAGGTTCG | ACCACCTCCACTCATGCTTC | 269 |
| chr7:14410387-14412402 | p5 | (GGTCT)3 | CTAGTTAACTTGACCGCGCC | CAATTTGGCAAGAACTCCGT | 116 |
| chr7:14418403-14420441 | p2 | (TA)19 | TCGGCAAAGTCTCCTTTGTT | TAAATGTTGTGTTTGGGCGA | 143 |
| chr7:14500436-14502456 | p2 | (TA)10 | GCAGAAATCCCCAAGAATCA | ATGCTCGGATTCAAACGAAC | 296 |
| chr7:14591857-14593872 | p3 | (GGT)5 | CCCACTTCATCTGCGTACAA | CCCAAAACTACAAGACCGGA | 259 |
| chr7:14603941-14605955 | p2 | (AT)7 | GGGTTCTGGGACGTCTGTAA | CTTTGTCCGCACTACAAGCA | 169 |
| chr7:14653109-14655139 | p2 | (AT)15 | CTGATCACATGGCAGCAACT | GAGGTGGTGCTCCACATTCT | 278 |
| chr7:14653109-14655139 | p2 | (TC)7 | CTGATCACATGGCAGCAACT | GAGGTGGTGCTCCACATTCT | 278 |
| chr7:14740203-14742219 | p4 | (ATTT)4 | AGTGTTGGCCGATTAAATGC | GTGCTTTGGTTCGGTCACTT | 266 |
| chr7:14881450-14883490 | p2 | (AT)20 | GAATTTTCGCTCCACACCAT | TCCTGAAAGGATGACTTGGG | 214 |
| chr7:14895086-14897150 | p2 | (AT)7 | TGATATTTTGCACCCGTTGA | AAAACCTTCGTCCGTTGAAA | 101 |
| chr7:14895086-14897150 | p2 | (AT)32 | TGATATTTTGCACCCGTTGA | AAAACCTTCGTCCGTTGAAA | 101 |
| chr7:14987258-14989272 | p2 | (TA)7 | ACTTGTGCATGACCCCTTTC | GTGGCCAATGATATGACGTG | 287 |
| chr7:14987258-14989272 | p2 | (AT)8 | ACTTGTGCATGACCCCTTTC | GTGGCCAATGATATGACGTG | 287 |
| chr7:15005442-15007457 | p5 | (TTGTC)3 | ATGCACTGCTGCTGACATTC | CATTTCGGCCACTTTCATCT | 215 |
| chr7:15005442-15007457 | p3 | (TTG)7 | ATGCACTGCTGCTGACATTC | CATTTCGGCCACTTTCATCT | 215 |
| chr7:15005442-15007457 | p5 | (TTGTC)3 | ATGCACTGCTGCTGACATTC | CATTTCGGCCACTTTCATCT | 215 |
| chr7:15005442-15007457 | p3 | (TTG)9 | ATGCACTGCTGCTGACATTC | CATTTCGGCCACTTTCATCT | 215 |
| chr7:15058134-15060149 | p5 | (TTGAT)3 | TTCTTGCTTCTTGCATCACG | GAATGGAACCGAACTCAGGA | 270 |
| chr7:15070329-15072377 | p3 | (AAC)7 | AATGAGGCAACCAAATAGCG | AAAATTGACTTGGTGCCGAA | 223 |
| chr7:15070329-15072377 | p2 | (AT)11 | AATGAGGCAACCAAATAGCG | AAAATTGACTTGGTGCCGAA | 223 |
| chr7:15115313-15117367 | p3 | (AAC)10 | GATGTGGTGGGTTCCATCTT | CGTACACTCTACCCTCCCCA | 179 |
| chr7:15115380-15117432 | p3 | (AAC)10 | GATGTGGTGGGTTCCATCTT | CGTACACTCTACCCTCCCCA | 112 |
| chr7:15161899-15163911 | p2 | (CA)6 | GCCACGCCTTATTAGCTACG | AATTCTCGGGTTTGCTGTTG | 289 |
| chr7:15205563-15207581 | p6 | (AACTGA)3 | CCTCTCCCTGTTGTATCCCA | GGCCCAACTGAGGTTGATTA | 125 |
| chr7:15273746-15275761 | p3 | (TGA)5 | ATAATGCCCACACGTAAGCC | CCGTTGGCCAATAGTTCAGT | 136 |
| chr7:15321986-15324001 | p5 | (TCTTT)3 | TGGTCGGGATATCTTTTTGC | ATTGGCAATTGGAGAAGTGG | 114 |

| Marker ID | SSR type | SSR motif | FORWARD PRIMER (5'-3') | REVERSE PRIMER (5'-3') | size (bp) |
|---|---|---|---|---|---|
| chr7:15417716-15419731 | p5 | (TGGGG)3 | CGTTGCATGGTACGTTTCTG | TCCTCGAGTCCGTCAAGTCT | 162 |
| chr7:15444699-15446717 | p6 | (CGTCCC)3 | GCCCCCATATCCTTCAATTT | CATTCCGAGGGACAACCTTA | 128 |
| chr7:15444699-15446717 | p2 | (TA)6 | GCCCCCATATCCTTCAATTT | CATTCCGAGGGACAACCTTA | 128 |
| chr7:15507128-15509142 | p2 | (AC)7 | TGGGTATTGGTCCGACATTT | CACGACAACAAAACCCAATG | 296 |
| chr7:15530205-15532220 | p5 | (AATAA)3 | TCCGTCTCCCTGTCTACCTG | TACGTCCCGTACTTCTTGGG | 272 |
| chr7:15530205-15532220 | p5 | (TTTAT)3 | TCCGTCTCCCTGTCTACCTG | TACGTCCCGTACTTCTTGGG | 272 |
| chr7:15530205-15532220 | p5 | (TTTAT)3 | TCCGTCTCCCTGTCTACCTG | TACGTCCCGTACTTCTTGGG | 272 |
| chr7:15579751-15581766 | p5 | (GTTCG)3 | AATTCGCCCTTCTACTTGGG | CTCGAAATAGCCAAACCGAA | 210 |
| chr7:15579751-15581766 | p5 | (GTTCG)3 | AATTCGCCCTTCTACTTGGG | CTCGAAATAGCCAAACCGAA | 210 |
| chr7:15609259-15611274 | p2 | (AT)6 | CGACTAGGGGTTTGGTTTGA | GTAATGTCGGGTTGGTTTGG | 242 |
| chr7:15609259-15611274 | p5 | (TTGGT)3 | CGACTAGGGGTTTGGTTTGA | GTAATGTCGGGTTGGTTTGG | 242 |
| chr7:15609299-15611314 | p2 | (AT)6 | CGACTAGGGGTTTGGTTTGA | GTAATGTCGGGTTGGTTTGG | 202 |
| chr7:15609299-15611314 | p5 | (TTGGT)3 | CGACTAGGGGTTTGGTTTGA | GTAATGTCGGGTTGGTTTGG | 202 |
| chr7:15675465-15677514 | p5 | (GTGGT)3 | CGACTTTTACCCCGTCGTTA | ATCCTTGAGGCCAATGTGAG | 120 |
| chr7:15675465-15677514 | p2 | (AT)33 | CGACTTTTACCCCGTCGTTA | ATCCTTGAGGCCAATGTGAG | 120 |
| chr7:15675637-15677652 | p5 | (GTGGT)3 | AACGGGATTGTCATTTCTCG | CAACCAAACCAAATCCAACC | 152 |
| chr7:15675637-15677652 | p2 | (AT)33 | AACGGGATTGTCATTTCTCG | CAACCAAACCAAATCCAACC | 152 |
| chr7:15758066-15760081 | p5 | (TTGGT)3 | TTATTTATGGATCGGGCTGC | GTCCGCAACAAAAAGGACAT | 169 |
| chr7:15758066-15760081 | p6 | (TAAAAT)3 | TTATTTATGGATCGGGCTGC | GTCCGCAACAAAAAGGACAT | 169 |
| chr7:15771934-15773952 | p6 | (TGGATT)3 | CCTGACTGGAAATCCCAAGA | TGCGCTTTATCCTTGTTCCT | 186 |
| chr7:15852663-15854678 | p5 | (AATTA)3 | TTTTGAGGCTTCTTGTTGGG | CTCGTGGATGGTGTGTGTTC | 291 |
| chr7:15867330-15869398 | p2 | (AT)8 | CCAACACCGTGTGAAATGAC | AAATTTTGCACGAGAAGGGA | 283 |
| chr7:15867422-15869480 | p2 | (AT)8 | CCAACACCGTGTGAAATGAC | AAATTTTGCACGAGAAGGGA | 191 |
| chr7:15917875-15919897 | p2 | (AT)11 | GGGACCCCATGTAACACATC | GAAAGAGACAGCAAGCCAGG | 238 |
| chr7:15928057-15930093 | p3 | (ATT)12 | AGGCTTTTGTTTTGCTGGAA | TGTAGGAAGGCTCACGAGGT | 292 |
| chr7:15987401-15989419 | p6 | (GCAGTA)3 | TGTGTCCGATTTTTCCACAA | GAAGGCAAATGGGAGATCAA | 131 |
| chr7:16105650-16107664 | p2 | (AG)7 | TCCCATTCTTTTGTGTGCAA | CTAGCGACCAAACACCCCTA | 233 |
| chr7:16105650-16107664 | p6 | (CAACTC)3 | TCCCATTCTTTTGTGTGCAA | CTAGCGACCAAACACCCCTA | 233 |
| chr7:16130666-16132678 | p2 | (TA)6 | CTTCGTTTTACCTCCCACCA | TCCGTAAAAGATCGTTGGCT | 237 |
| chr7:16140393-16142409 | p4 | (AGTT)4 | TCCCATTCTTTTGTGTGCAA | CGGGCTAACCGTGTAAGAAA | 243 |
| chr7:16140393-16142409 | p2 | (AG)8 | TCCCATTCTTTTGTGTGCAA | CGGGCTAACCGTGTAAGAAA | 243 |
| chr7:16140393-16142409 | p6 | (CAACTC)3 | TCCCATTCTTTTGTGTGCAA | CGGGCTAACCGTGTAAGAAA | 243 |
| chr7:16159212-16161236 | p2 | (AT)12 | TGGGGGCTGTTTTGATATGT | CAACTCAAGTCTTGGGCTCC | 264 |
| chr7:16205610-16207628 | p6 | (AACCGG)3 | GGCAACCCCTTTTGATTTTT | GGAAGAGGCTTGACACGAAA | 160 |
| chr7:16205610-16207628 | p2 | (TA)7 | GGCAACCCCTTTTGATTTTT | GGAAGAGGCTTGACACGAAA | 160 |
| chr7:16254895-16256913 | p6 | (AACCGG)3 | ATGTTTGCCAACACTCCTCC | CGAGACCTCGAACCTGACTC | 273 |

| Marker ID | SSR type | SSR motif | FORWARD PRIMER (5'-3') | REVERSE PRIMER (5'-3') | size (bp) |
|---|---|---|---|---|---|
| chr7:16254895-16256913 | p3 | (GAT)5 | ATGTTTGCCAACACTCCTCC | CGAGACCTCGAACCTGACTC | 273 |
| chr7:16260782-16262800 | p2 | (AT)6 | TTGGATTAGATCGGAGCGAC | TTTCGTTAATTTCCCCCACA | 270 |
| chr7:16410601-16412613 | p2 | (TA)24 | ATGATGGATTCATTGCCACA | AAATGACATGGCGGAATAGC | 176 |
| chr7:16410601-16412613 | p2 | (AT)6 | ATGATGGATTCATTGCCACA | AAATGACATGGCGGAATAGC | 176 |
| chr7:16502694-16504709 | p5 | (TATTG)3 | TGCGTCGCTTTAAACATTTG | AACGGAATCACGAGATGAGG | 172 |
| chr7:16548470-16550482 | p2 | (GA)6 | CTAGCTCGCTCCAGGTATGG | ATGTGTGCTCAACAATCCCA | 263 |
| chr7:16553969-16555987 | p2 | (TA)9 | GCCGGACTAATTTCCAACAA | CCACCCCTATATGAACCCCT | 111 |
| chr7:16826502-16828520 | p3 | (ACA)6 | GCTTCGGGTCAATGTGAAAT | TCAGAAAAGCTGGAAGGGAA | 222 |
| chr7:16826502-16828520 | p5 | (TTGGT)3 | GCTTCGGGTCAATGTGAAAT | TCAGAAAAGCTGGAAGGGAA | 222 |
| chr7:16853329-16855344 | p3 | (TTA)5 | CACTTTGGTGGAGGACTGGT | ACCAATAGGTGGAAGGAGGG | 178 |
| chr7:16866674-16868689 | p2 | (TA)7 | GCCCTTTGAGGTTGAGTTGA | TAGCACGTTGCTCCATCTTG | 254 |
| chr7:16866716-16868737 | p2 | (TA)7 | GCCCTTTGAGGTTGAGTTGA | TAGCACGTTGCTCCATCTTG | 212 |
| chr7:16917320-16919335 | p3 | (TTC)5 | TACATTTTGGTTCAACGCCA | TTCGATGTCATTCTCTCCCC | 267 |
| chr7:16917320-16919335 | p3 | (TTC)10 | TACATTTTGGTTCAACGCCA | TTCGATGTCATTCTCTCCCC | 267 |
| chr7:16982702-16984718 | p4 | (GAAA)4 | TTTCAATGGAAGAGAACGCC | GTCCAACGAGTCATGGAACC | 224 |
| chr7:17121551-17123566 | p3 | (CCG)5 | TTCGTGCCAGCAAGTACAAC | TGGGCCAGCTAGAGTTAGGA | 170 |
| chr7:17249624-17251660 | p2 | (TA)18 | TCATTGACTCTTTGGGGCTT | ATGCAGGGACGTATGTAGGC | 108 |
| chr7:17310255-17312270 | p3 | (ATG)6 | TTTTTGCCTGCTTGCTTTTT | CCGCATCAGCTTCAACATAA | 163 |
| chr7:17426328-17428340 | p2 | (AT)6 | GAACCAAAATTCTGCCCAAA | TCGTAACCGGGAAAGACAAC | 139 |
| chr7:17447567-17449623 | p2 | (TA)28 | AGGTTAATATCGGAACCCCG | CGTTTGGTTGCTTTCTAGCC | 190 |
| chr7:17493083-17495107 | p6 | (CATATA)4 | GGCCGATATGTACAGCCAGT | TACCTCCTAGCAAGCCGTGT | 284 |
| chr7:17572165-17574191 | p2 | (TA)13 | AGGCATCCTCCCTGATTTCT | TTGGTTAGCCCGAGGTGTAG | 243 |
| chr7:17630686-17632701 | p5 | (AATAA)3 | GTAAGGGGTGAGGGGCTAAG | GGGATTAAAAACCACGACCC | 210 |
| chr7:17657530-17659545 | p5 | (TGCAC)3 | CCAGAAAAGACACGCACAAA | GAGTCAGTTGCAATGGGACA | 145 |
| chr7:17657530-17659545 | p3 | (GTT)5 | CCAGAAAAGACACGCACAAA | GAGTCAGTTGCAATGGGACA | 145 |
| chr7:17767931-17769943 | p2 | (AT)6 | GCGCATAGAGGTACGAGGAG | TACACGCACTGCACCTAAGC | 250 |
| chr7:17835625-17837637 | p2 | (AT)6 | TAGGCAATCCACACGTACCA | TGCCGTCGTCGTAATGATAG | 115 |
| chr7:17898811-17900826 | p5 | (TTCGG)3 | GCGAATTAGGTTAGGAGGGG | GAACGTCCAATTGACACCCT | 152 |
| chr7:17981357-17983373 | p2 | (TA)8 | GGAGCAAAGTTTGGAGGTGA | AACTCAGCAGGGAATGATGG | 263 |
| chr7:18140761-18142773 | p2 | (TA)6 | TCTCATTGTTGCCTGCAAAG | AGCGCGTGATACCGTAAGTT | 215 |
| chr7:18145040-18147056 | p4 | (AAAG)4 | AGGGAGTTGAGAAACGCAGA | TTGTTGGCCTTTCACTCACA | 120 |
| chr7:18201134-18203158 | p2 | (AT)7 | TTGGACGTCTTAGCATGGTG | CTCTTCCTCTTCGCCTTCCT | 121 |
| chr7:18201134-18203158 | p2 | (AT)12 | TTGGACGTCTTAGCATGGTG | CTCTTCCTCTTCGCCTTCCT | 121 |
| chr7:18220634-18222650 | p2 | (TA)8 | GTAGCTGCAGATTGAACGCA | CGCAAATAAGAAGCAGGAGG | 162 |
| chr7:18226202-18228244 | p2 | (AT)21 | TCACTGAATGTCGATGGGAA | CTTGAATGAGTTGGGTGGGT | 185 |
| chr7:18276853-18278875 | p2 | (TA)11 | AATTCACGGCAAAGGAAGTG | CCAAACCTTTTCCAGGTTCA | 234 |

| Marker ID | SSR type | SSR motif | FORWARD PRIMER (5'-3') | REVERSE PRIMER (5'-3') | size (bp) |
|---|---|---|---|---|---|
| chr7:18278919-18280945 | p3 | (TTG)6 | GATTGGTTGATGAATTGGGG | TCTTGCGGCTACAACAAGTG | 143 |
| chr7:18528721-18530748 | p3 | (TTA)9 | GCTCTAAGCACCGATGGAAG | GGGTATTAGGGCCTCGACTC | 158 |
| chr7:18538323-18540338 | p3 | (ATT)5 | GAAAATTTCCGCATAAGGCA | TTCGCTGCCGAAGAAATAGT | 221 |
| chr7:18561826-18563846 | p2 | (AT)10 | TTGCTCCACGCCCTACTACT | CGAAGCCACGGTTGATACTT | 185 |
| chr7:18561826-18563846 | p2 | (TA)6 | TTGCTCCACGCCCTACTACT | CGAAGCCACGGTTGATACTT | 185 |
| chr7:18562066-18564084 | p2 | (AT)10 | TTGGACGTCTTAGCATGGTG | GCCTTCCTCCGCTTCTAAGT | 100 |
| chr7:18562066-18564084 | p2 | (TA)6 | TTGGACGTCTTAGCATGGTG | GCCTTCCTCCGCTTCTAAGT | 100 |
| chr7:18563418-18565436 | p2 | (TA)6 | AGTCCACTCTCTCAACCCCA | GTTCGGCTTTGTTGTTGGTT | 297 |
| chr7:18563418-18565436 | p6 | (TAAAAA)3 | AGTCCACTCTCTCAACCCCA | GTTCGGCTTTGTTGTTGGTT | 297 |
| chr7:18578864-18580878 | p2 | (TA)7 | CTAAGCGAGAATTCGAACCG | AAGCCCAACCTAGAAAAGGC | 221 |
| chr7:18694742-18696757 | p3 | (GAA)5 | TCGTCCAGCTTCTTCTCCAT | CAGCATCAAAATGGGGTTCT | 198 |
| chr7:18860978-18862993 | p5 | (TTGCA)3 | GGTCGTTTTTGGGTTTCAGA | AATCCACCTCCACTTCATGC | 294 |
| chr7:18891094-18893112 | p3 | (CAT)6 | CCATATACCCCGACAACGTC | TAAACTTGTGCAGGATCCCC | 204 |
| chr7:18891094-18893112 | p6 | (AATAAA)3 | CCATATACCCCGACAACGTC | TAAACTTGTGCAGGATCCCC | 204 |
| chr7:18915092-18917107 | p3 | (AAC)5 | GCCGTGTTCTTATGCTGTCA | TTTGATTGGGTCTACGGCTC | 245 |
| chr7:18932265-18934285 | p4 | (AATA)5 | TAATACGGGTGGAGCAAAGG | CCGTCGGCTGGTAACTTTTA | 122 |
| chr7:18965723-18967773 | p2 | (GT)6 | GGCTTGCGTCCTTTCAAATA | ACAGGGCATAACAGAATGGC | 293 |
| chr7:19059212-19061227 | p5 | (GTTGG)3 | TAGTTTCGCCCTTGCCTCTA | CGTGCCCTCATTTAATTCGT | 256 |
| chr7:19059212-19061227 | p4 | (AATA)4 | TAGTTTCGCCCTTGCCTCTA | CGTGCCCTCATTTAATTCGT | 256 |
| chr7:19090644-19092700 | p2 | (AT)28 | GACGGACCGTAACTTGGTGT | ACTGAAGCCTATTCGCTGGA | 124 |
| chr7:19115097-19117119 | p2 | (AG)11 | ATCAGAGATCGGAGCAAGGA | TCCGTTTGACCCCCATAATA | 292 |
| chr7:19115097-19117119 | p2 | (AT)6 | ATCAGAGATCGGAGCAAGGA | TCCGTTTGACCCCCATAATA | 292 |
| chr7:19141360-19143384 | p2 | (TA)12 | TCCCTCCACTCACCTTTTTG | GGGGGATGGATCCACTAAAT | 187 |
| chr7:19148725-19150746 | p3 | (ATC)7 | CCGTGATAACGCGGTATTTT | TTGAATATGGAACCAACGCA | 131 |
| chr7:19257396-19259426 | p2 | (TA)15 | TCAAGAAAACCCATCCCAAG | GCGTGGCGAAGAACAATAAT | 252 |
| chr7:19304192-19306207 | p5 | (GGAAA)3 | TTTTGGATGCCCAATAGAGG | TTTGTTCAAATGCAAGACCG | 210 |
| chr7:19323960-19326020 | p2 | (AT)30 | TTTCCAGATGACCGAACCTC | TCTAACCATAGGTCGTGGGC | 151 |
| chr7:19323960-19326020 | p2 | (AT)6 | TTTCCAGATGACCGAACCTC | TCTAACCATAGGTCGTGGGC | 151 |
| chr7:19376878-19378892 | p2 | (TA)6 | CATGCCCACTACCTTGCTTT | CCCTGTGAATTCGTGGAGTT | 224 |
| chr7:19376893-19378905 | p2 | (TA)6 | CATGCCCACTACCTTGCTTT | CCCTGTGAATTCGTGGAGTT | 209 |
| chr7:19389493-19391509 | p2 | (TG)8 | GCCGGACAAGGCATACTTTA | TTTTTCCCTTCGGCACTTTA | 280 |
| chr7:19423371-19425389 | p3 | (CAA)6 | AGAACGAAATCATGGGCATC | TGTGAACATGAGGGCGATAA | 140 |
| chr7:19440894-19442909 | p3 | (CAA)5 | ATATATCGTACCCGGCCCAT | ATGTTCTCGTGCACAGTTCG | 278 |
| chr7:19482312-19484332 | p5 | (CAAAC)4 | TGGCTGCACCATTTTTGTTA | GGGAGAATTGAGGGCAAAAT | 118 |
| chr7:19535435-19537505 | p2 | (TA)35 | GCCCAATTTCCCCTAGAAAA | GCCTTTGAAGGAAAAGGGAC | 245 |
| chr7:19543966-19546000 | p2 | (TA)17 | GAGACCTCAGGACAAGCAGG | CACCCCCAACTAAAGAAGCA | 110 |

| Marker ID | SSR type | SSR motif | FORWARD PRIMER (5'-3') | REVERSE PRIMER (5'-3') | size (bp) |
|---|---|---|---|---|---|
| chr7:19548689-19550704 | p5 | (AAACC)3 | ATGGACCGGGTTGATTTGTA | AATCCGAGAAAACCCGAAGT | 179 |
| chr7:19548689-19550704 | p5 | (AAACC)3 | ATGGACCGGGTTGATTTGTA | AATCCGAGAAAACCCGAAGT | 179 |
| chr7:19606044-19608072 | p2 | (TA)12 | TGCACATATGGCAAAGCAAT | CGGGCATCTTTTCCTACAAA | 202 |
| chr7:19606078-19608090 | p2 | (TA)12 | TGCACATATGGCAAAGCAAT | CGGGCATCTTTTCCTACAAA | 168 |
| chr7:19680893-19682947 | p2 | (AT)27 | GCCGGACAAGGCATACTTTA | ATCGCAAGGCAGGATTTCTA | 284 |
| chr7:19823341-19825389 | p3 | (ATT)16 | ATTTCAGTTCAGGCACCCAC | AGTTCGCTGCCGAAGAAATA | 247 |
| chr7:19829872-19831918 | p2 | (AT)23 | CTGCCTCTTGGTCCTTCTTG | GAATCGAGCATTGTCCCAGT | 122 |
| chr7:19840092-19842107 | p2 | (AT)6 | CTTCGTTTCCCCGCTAAAAT | AAATAAATTGGACGGAGGG | 298 |
| chr7:19848551-19850581 | p3 | (ATG)5 | CGAGGTCACCGACCTTATGT | GGGGTAATCATCTCACACCG | 295 |
| chr7:19848551-19850581 | p6 | (GTATAT)5 | CGAGGTCACCGACCTTATGT | GGGGTAATCATCTCACACCG | 295 |
| chr7:19855010-19857025 | p3 | (TAT)5 | CATCGGGTGCTCCTTATGAT | GACGTTGACCGATGTGTGTC | 100 |
| chr7:19983409-19985439 | p6 | (CATATA)6 | AAACGGCCAATGCAAATATC | TGACAATAAGGCACAACCCA | 230 |
| chr7:19983453-19985468 | p6 | (CATATA)6 | AAACGGCCAATGCAAATATC | TGACAATAAGGCACAACCCA | 186 |
| chr7:19983476-19985491 | p6 | (CATATA)6 | AAACGGCCAATGCAAATATC | TGACAATAAGGCACAACCCA | 163 |
| chr7:19983501-19985521 | p6 | (CATATA)6 | AAACGGCCAATGCAAATATC | TGACAATAAGGCACAACCCA | 138 |
| chr7:19983526-19985541 | p6 | (CATATA)6 | AAACGGCCAATGCAAATATC | TGACAATAAGGCACAACCCA | 113 |
| chr7:19993657-19995682 | p2 | (AT)8 | CACTGCGACCACAAGAGGTA | CACACCCGATGAACACAATC | 183 |
| chr7:20057213-20059225 | p2 | (TC)6 | CAATTAATGTCACCGAGGGG | GACGAGAAAAATGGGGAACA | 116 |
| chr7:20089097-20091112 | p3 | (TAA)5 | CGAGGGTGATTGCTTTCATT | TAATTCAGGCCATCCTGACC | 190 |
| chr7:20100313-20102328 | p5 | (AATCA)3 | AATGTCGAAGCAATCAAGGG | CCGACAAAAATCGAAGCAAT | 249 |
| chr7:20100313-20102328 | p5 | (GATTT)3 | AATGTCGAAGCAATCAAGGG | CCGACAAAAATCGAAGCAAT | 249 |
| chr7:20157708-20159720 | p3 | (TTA)9 | TGTGCTTCCGTGTTTAGCTG | CTAGCTTCTGTGTCCTCGCC | 160 |
| chr7:20157708-20159720 | p2 | (TA)6 | TGTGCTTCCGTGTTTAGCTG | CTAGCTTCTGTGTCCTCGCC | 160 |
| chr7:20197830-20199846 | p4 | (AAAT)4 | GGCAATTGCGTTCACTTTTT | ATTCATAAACGGGCTCGCTA | 198 |
| chr7:20291667-20293682 | p5 | (AAATA)3 | GGGAGGAGTTGGCTAAAAGG | CCGGAAACAGGATCTGAAAA | 283 |
| chr7:20291667-20293682 | p5 | (AAATA)7 | GGGAGGAGTTGGCTAAAAGG | CCGGAAACAGGATCTGAAAA | 283 |
| chr7:20291667-20293682 | p5 | (AATAA)3 | GGGAGGAGTTGGCTAAAAGG | CCGGAAACAGGATCTGAAAA | 283 |
| chr7:20292272-20294307 | p5 | (AAATA)3 | TGGGTGAATGAGAGGAGACC | TTTGGGGTTTGGGGTTTTAT | 110 |
| chr7:20292272-20294307 | p5 | (AAATA)7 | TGGGTGAATGAGAGGAGACC | TTTGGGGTTTGGGGTTTTAT | 110 |
| chr7:20292272-20294307 | p5 | (AATAA)3 | TGGGTGAATGAGAGGAGACC | TTTGGGGTTTGGGGTTTTAT | 110 |
| chr7:20292272-20294307 | p5 | (TTTAT)3 | TGGGTGAATGAGAGGAGACC | TTTGGGGTTTGGGGTTTTAT | 110 |
| chr7:20390665-20392689 | p2 | (TA)12 | TCCGTATGACCCTTCCAGAG | TCAAGGGTTTGGGAGAAGTG | 225 |
| chr7:20453526-20455544 | p2 | (AT)9 | TTGCATCAGGAAGTTTTCCC | TCAGTTTACGCCCATGAACA | 261 |
| chr7:20513743-20515758 | p3 | (ACA)5 | GATCGCAGCAGCACATAAAA | GAAGAAATCCCGGAGTAGGG | 212 |
| chr7:20526749-20528770 | p3 | (ACA)7 | TCCAATTGCCAGCAGTATCA | CGGTTCGATACCCTCCTTTT | 251 |
| chr7:20595236-20597251 | p5 | (AGCCG)3 | TATGAAGGAAATCGATCCCG | GTTAGAGAACAAGGCTGGCG | 126 |

| Marker ID | SSR type | SSR motif | FORWARD PRIMER (5'-3') | REVERSE PRIMER (5'-3') | size (bp) |
|---|---|---|---|---|---|
| chr7:20751731-20753746 | p5 | (TTGGT)3 | AGGACATGCTGAAGAGCGTT | TCAGCAGAGCCAACTCTGAA | 118 |
| chr7:20792065-20794083 | p3 | (CAA)6 | TAACAGAACGCATACGCGAG | TTATCCTCCGTCCGTGAGTC | 225 |
| chr7:20860491-20862511 | p5 | (TGTTT)3 | GGCACATTCGACACCTTTTT | CGGAACGTCGATTCTTCATT | 240 |
| chr7:20860491-20862511 | p2 | (TA)10 | GGCACATTCGACACCTTTTT | CGGAACGTCGATTCTTCATT | 240 |
| chr7:20881477-20883517 | p2 | (AT)20 | TTCCCTCCTTTATTCACCCC | ATTTGGCAATAGGGTTGTGC | 284 |
| chr7:20881477-20883517 | p2 | (TA)9 | TTCCCTCCTTTATTCACCCC | ATTTGGCAATAGGGTTGTGC | 284 |
| chr7:20881477-20883517 | p5 | (TGAAT)3 | TTCCCTCCTTTATTCACCCC | ATTTGGCAATAGGGTTGTGC | 284 |
| chr7:20948793-20950808 | p5 | (ATAAA)3 | TTCGGAACATCAAAGCACTG | TTGGTTTGATCCTTTCCAGC | 120 |
| chr7:20954415-20956430 | p2 | (AT)35 | CGGTACGTTCGATTCGATTT | TTGCTTGAATTGTCGAAACG | 141 |
| chr7:20954415-20956430 | p5 | (CGAAT)3 | CGGTACGTTCGATTCGATTT | TTGCTTGAATTGTCGAAACG | 141 |
| chr7:21006491-21008507 | p2 | (AT)8 | CAACCCAAACAATCATCCCT | ATTTGCGCCACCTAAACTTG | 114 |
| chr7:21042913-21044928 | p5 | (ATTTT)3 | TATCTTGGGACGCCCATAAC | TTTGGTCCAATTTGAGGGAG | 285 |
| chr7:21088604-21090680 | p2 | (CT)27 | GAAGGGAAAAAGCCCAAAAC | TGGTCTCCGGGTATGCTAAC | 195 |
| chr7:21158131-21160159 | p2 | (AG)14 | GGTATCAGAGCAGGCAGAGG | TTGAATTGGGATGGTCCTGT | 266 |
| chr7:21173423-21175438 | p3 | (TTA)5 | TCATGCCCATCCATCCTTAT | TTGGACACTTGAACCCCTTC | 176 |
| chr7:21195817-21197832 | p5 | (CCCAA)3 | TGAGTGCTGTAAGCGTTTGG | GGTGCTTTAATTTGGGGTGA | 177 |
| chr7:21266057-21268081 | p2 | (AC)12 | CCTATTTTGGTCACGCCATT | ACTTTTCCTTGGGAAGCCAT | 254 |
| chr7:21304943-21306957 | p2 | (AG)7 | TACGTAAATGAAGGGGGCAG | CATGATGATTTTGACACGCC | 211 |
| chr7:21304943-21306957 | p2 | (AT)19 | TACGTAAATGAAGGGGGCAG | CATGATGATTTTGACACGCC | 211 |
| chr7:21345912-21347926 | p2 | (TA)7 | CACCCTTCCTTCTATGGCAA | CTTTCCATCATTTGCCCACT | 150 |
| chr7:21397670-21399682 | p2 | (GA)6 | GATTCAGTGCCTTCCTCAGC | TCCAAATTTGTCTCCAAGGC | 236 |
| chr7:21397670-21399682 | p2 | (AC)6 | GATTCAGTGCCTTCCTCAGC | TCCAAATTTGTCTCCAAGGC | 236 |
| chr7:21496196-21498220 | p2 | (AT)12 | GCGATTGTTTTAGCTTTCCG | CATTTGCACTGGTGGTATGC | 249 |
| chr7:21496196-21498220 | p2 | (AT)10 | GCGATTGTTTTAGCTTTCCG | CATTTGCACTGGTGGTATGC | 249 |
| chr7:21555192-21557207 | p3 | (ACA)6 | TTCATTGGTCATTCCTGCAA | CACGTTGTGGGATTTCACTG | 278 |
| chr7:21555192-21557207 | p3 | (ACA)5 | TTCATTGGTCATTCCTGCAA | CACGTTGTGGGATTTCACTG | 278 |
| chr7:21555192-21557207 | p3 | (ACA)5 | TTCATTGGTCATTCCTGCAA | CACGTTGTGGGATTTCACTG | 278 |
| chr7:21555192-21557207 | p3 | (ACA)5 | TTCATTGGTCATTCCTGCAA | CACGTTGTGGGATTTCACTG | 278 |
| chr7:21555192-21557207 | p3 | (ACA)5 | TTCATTGGTCATTCCTGCAA | CACGTTGTGGGATTTCACTG | 278 |
| chr7:21555192-21557207 | p3 | (ACA)5 | TTCATTGGTCATTCCTGCAA | CACGTTGTGGGATTTCACTG | 278 |
| chr7:21555192-21557207 | p3 | (ACA)5 | TTCATTGGTCATTCCTGCAA | CACGTTGTGGGATTTCACTG | 278 |
| chr7:21555192-21557207 | p3 | (ACA)5 | TTCATTGGTCATTCCTGCAA | CACGTTGTGGGATTTCACTG | 278 |
| chr7:21555192-21557207 | p3 | (ACA)5 | TTCATTGGTCATTCCTGCAA | CACGTTGTGGGATTTCACTG | 278 |
| chr7:21555192-21557207 | p3 | (ACA)5 | TTCATTGGTCATTCCTGCAA | CACGTTGTGGGATTTCACTG | 278 |
| chr7:21555192-21557207 | p3 | (ACA)5 | TTCATTGGTCATTCCTGCAA | CACGTTGTGGGATTTCACTG | 278 |
| chr7:21555192-21557207 | p3 | (ACA)5 | TTCATTGGTCATTCCTGCAA | CACGTTGTGGGATTTCACTG | 278 |

| Marker ID | SSR type | SSR motif | FORWARD PRIMER (5'-3') | REVERSE PRIMER (5'-3') | size (bp) |
|---|---|---|---|---|---|
| chr7:21555326-21557341 | p3 | (ACA)6 | TTCATTGGTCATTCCTGCAA | CACGTTGTGGGATTTCACTG | 144 |
| chr7:21555326-21557341 | p3 | (ACA)5 | TTCATTGGTCATTCCTGCAA | CACGTTGTGGGATTTCACTG | 144 |
| chr7:21555326-21557341 | p3 | (ACA)5 | TTCATTGGTCATTCCTGCAA | CACGTTGTGGGATTTCACTG | 144 |
| chr7:21555326-21557341 | p3 | (ACA)5 | TTCATTGGTCATTCCTGCAA | CACGTTGTGGGATTTCACTG | 144 |
| chr7:21555326-21557341 | p3 | (ACA)5 | TTCATTGGTCATTCCTGCAA | CACGTTGTGGGATTTCACTG | 144 |
| chr7:21555326-21557341 | p3 | (ACA)5 | TTCATTGGTCATTCCTGCAA | CACGTTGTGGGATTTCACTG | 144 |
| chr7:21555326-21557341 | p3 | (ACA)5 | TTCATTGGTCATTCCTGCAA | CACGTTGTGGGATTTCACTG | 144 |
| chr7:21555326-21557341 | p3 | (ACA)5 | TTCATTGGTCATTCCTGCAA | CACGTTGTGGGATTTCACTG | 144 |
| chr7:21555326-21557341 | p3 | (ACA)5 | TTCATTGGTCATTCCTGCAA | CACGTTGTGGGATTTCACTG | 144 |
| chr7:21555326-21557341 | p3 | (ACA)5 | TTCATTGGTCATTCCTGCAA | CACGTTGTGGGATTTCACTG | 144 |
| chr7:21555326-21557341 | p3 | (ACA)5 | TTCATTGGTCATTCCTGCAA | CACGTTGTGGGATTTCACTG | 144 |
| chr7:21555326-21557341 | p3 | (ACA)5 | TTCATTGGTCATTCCTGCAA | CACGTTGTGGGATTTCACTG | 144 |
| chr7:21587631-21589643 | p3 | (CAA)7 | GCATGCCAAAAGAAAGAAGG | TCGGGTTGGCCTATTTACAG | 131 |
| chr7:21587631-21589643 | p2 | (AC)6 | GCATGCCAAAAGAAAGAAGG | TCGGGTTGGCCTATTTACAG | 131 |
| chr7:21636995-21639017 | p2 | (AT)11 | CCCGAGCTTCACACAGAAAT | CTTATGCCAATTCTGCCCAT | 282 |
| chr7:21644476-21646494 | p6 | (TAAAAA)3 | TCTAGCCGATCCTTTGCACT | TTTTCTCCCTCACCCCTCTT | 208 |
| chr7:21675044-21677059 | p5 | (TATTT)3 | ATTTCGCCTAACACGTGGAC | CTTCGTTGACCGTGTTGCTA | 277 |
| chr7:21714527-21716545 | p3 | (TTG)6 | AGAGGCCATGGAACCCTTAT | CCTCCAAAGTTGACCCAAAA | 248 |
| chr7:21714527-21716545 | p2 | (AT)16 | AGAGGCCATGGAACCCTTAT | CCTCCAAAGTTGACCCAAAA | 248 |
| chr7:21714527-21716545 | p2 | (AT)9 | AGAGGCCATGGAACCCTTAT | CCTCCAAAGTTGACCCAAAA | 248 |
| chr7:21984659-21986677 | p6 | (AAAAGA)3 | ACCGACTTCAGGAGGGTTTT | GATGGGCCAAGGTAAAGTCA | 227 |
| chr7:22000128-22002146 | p3 | (ATA)6 | AGCTCCACCTCTTCAGTCCA | AAAACCATACCATCACCCCA | 106 |
| chr7:22012311-22014333 | p2 | (AT)11 | AGGAGCCTAGGACATCAGCA | TTGATCCATAGAACTCGGGG | 107 |
| chr7:22038123-22040138 | p3 | (AAT)5 | GCTTGAAGGCACTCCTAACG | ACCGTCAATTCCTTCTCCCT | 107 |
| chr7:22122147-22124165 | p3 | (TAA)13 | AAGAAATCGCAAGGCAGAAA | AACCTCGGCTCAGTCAAAAA | 149 |
| chr7:22137441-22139459 | p3 | (TTG)6 | GGCCAAGCCTAAGAAAAACC | ATCGCTCCAACAACACTTCC | 126 |
| chr7:22159960-22161975 | p5 | (GGGTG)3 | CAAACGTGCATCTAGGGGTT | CCGCGAATTCAAGATTCAGT | 155 |
| chr7:22203243-22205275 | p2 | (AT)16 | CGCATTTAGCCAAGGAAGAC | TCACCTTCCTCCGCTTCTAA | 258 |
| chr7:22203243-22205275 | p2 | (TA)16 | CGCATTTAGCCAAGGAAGAC | TCACCTTCCTCCGCTTCTAA | 258 |
| chr7:22285120-22287134 | p4 | (TTCT)5 | GTGCTCAAATTCTTGTGCCA | TATTGGTGGGGTTCTTTGGA | 292 |
| chr7:22285141-22287165 | p4 | (TTCT)5 | GTGCTCAAATTCTTGTGCCA | GGTGGGGTTCTTTGGAATCT | 271 |
| chr7:22285294-22287314 | p4 | (TTCT)5 | GTGCTCAAATTCTTGTGCCA | GGTGGGGTTCTTTGGAATCT | 118 |
| chr7:22308769-22310785 | p2 | (GT)8 | TTGAGCCATTTAAGTTGGGG | GGATCTGGTAACACGCCAAT | 230 |
| chr7:22365259-22367273 | p2 | (TA)7 | ATTCCCAAAGCATCACCATC | AAATGACATGGCGGAATAGC | 191 |
| chr7:22373745-22375759 | p2 | (TA)7 | AAGAGGGGGTGTATCCTTGG | GCATTTGAGGACAAAGAGCC | 168 |
| chr7:22538773-22540789 | p2 | (TA)7 | AAGGCCTCCTCCTACACGAT | CAGTTCGGTGCAGCTGATTA | 138 |

| Marker ID | SSR type | SSR motif | FORWARD PRIMER (5'-3') | REVERSE PRIMER (5'-3') | size (bp) |
|---|---|---|---|---|---|
| chr7:22538773-22540789 | p4 | (GTTT)4 | AAGGCCTCCTCCTACACGAT | CAGTTCGGTGCAGCTGATTA | 138 |
| chr7:22546381-22548399 | p3 | (ATT)6 | TAACAACCCGCACCCTAAAC | CCACATGGAAGTGGTCTTGA | 130 |
| chr7:22627196-22629214 | p4 | (AAAC)4 | TTAATTAGAGGGCCACCACG | ATGGACGGAGGAAACACTGA | 296 |
| chr7:22627196-22629214 | p3 | (AAT)6 | TTAATTAGAGGGCCACCACG | ATGGACGGAGGAAACACTGA | 296 |
| chr7:22705852-22707884 | p2 | (CA)6 | GCATTCTTTTGCACCCCTTA | GCCCCACCTCAACTTACAAA | 243 |
| chr7:22875457-22877472 | p5 | (TTGTT)3 | TATTGAGCGCAGGATGACTG | ATCCTCCTCCAACTCCTCGT | 283 |
| chr7:22910632-22912644 | p2 | (TA)6 | GACCCTTGAGCCAAATCAAA | TATCTCCTTGTACCGACGGC | 258 |
| chr7:23020558-23022574 | p2 | (TA)8 | GTACACACATGCTATGGCGG | ATGGTTGTGTTGTGGCTTGA | 129 |
| chr7:23092402-23094414 | p2 | (TC)6 | AGGAGGACGGCCTATTGACT | CGTGTTCTCTGGCAGTTTGA | 269 |
| chr7:23149068-23151082 | p2 | (AT)7 | GGGTGGATAGGTATGCTGGA | TGTCTCTGAACCCCACTTCC | 123 |
| chr7:23292423-23294438 | p5 | (TATGA)3 | CTTCCAAACCCTCCGTACAA | TTTGATCCCGTAAAAACCCA | 134 |
| chr7:23292423-23294438 | p2 | (TA)8 | CTTCCAAACCCTCCGTACAA | TTTGATCCCGTAAAAACCCA | 134 |
| chr7:23416473-23418488 | p5 | (CATAT)3 | ACTGGCGAACGTCTCTGACT | CCCTGGGAAGTATGCAAAAA | 254 |
| chr7:23456831-23458846 | p3 | (TGT)5 | TTGGAACACGTGCCAAGTTA | CTACTGTGGACCCACGACCT | 100 |
| chr7:23533325-23535340 | p5 | (TTTTA)3 | AACGGGAACTTTTTGCAATG | CCTCCCTCTTGCCAATAACA | 168 |
| chr7:23646262-23648282 | p5 | (CAACA)4 | AGTGGTGGAAGAACAGTGGC | CTCCCTGCACTAGGCAGTTC | 257 |
| chr7:23661615-23663633 | p6 | (TTATCC)3 | AGGCAAAGACATTGGTGTCC | TCTCTTCCCACTGCGAGTCT | 259 |
| chr7:23665060-23667075 | p3 | (ACA)5 | CCCAAAATTCCCCAAAAGAT | CTCCGATTCACACTCGAACA | 292 |
| chr7:23720743-23722761 | p5 | (TATTT)3 | TGGAGAGCGTCCCAAATATC | GAGGTAGCAACTGACGAGGC | 100 |
| chr7:23725914-23727928 | p2 | (TC)7 | GGTTCGAAACTCAGGAACCA | GGCTAAAATGCGGGTCATAA | 157 |
| chr7:23926876-23928891 | p5 | (TTGGT)3 | CGTAGGACTGGACCTCTGGA | TAATGTCGGGTTGGTTTGGT | 168 |
| chr7:23940851-23942881 | p6 | (AAATTG)5 | AGCACTATCTGTGGCGTGTG | GGTCGCCAGAGGTAACAAGA | 268 |
| chr7:23979159-23981174 | p3 | (ATA)5 | GGACAAGGGAGTGTGGATGT | TTTCCCTTATTCCGCCTTTT | 104 |
| chr7:24125219-24127231 | p2 | (TA)6 | CATTTTCCTCAAAATCGCGT | AATAGCTTGAGTGGCGCATT | 156 |
| chr7:24168225-24170248 | p3 | (ATT)13 | CCACCTCGTTTTCCATCACT | GGCAGTGTAAGTGGGTGGTT | 135 |
| chr7:24168225-24170248 | p2 | (TC)6 | CCACCTCGTTTTCCATCACT | GGCAGTGTAAGTGGGTGGTT | 135 |
| chr7:24199628-24201640 | p2 | (CA)6 | TGGCAATGTCCATGTCTGTT | GCAGATACAGCAGAGGGGTC | 173 |
| chr7:24199628-24201640 | p5 | (TTTTC)3 | TGGCAATGTCCATGTCTGTT | GCAGATACAGCAGAGGGGTC | 173 |
| chr7:24286714-24288744 | p2 | (TA)15 | TCAACGAATTCAAAATGCCA | CTCTTGTGTGGTTTGCAGGA | 216 |
| chr7:24286714-24288744 | p3 | (CAA)5 | TCAACGAATTCAAAATGCCA | CTCTTGTGTGGTTTGCAGGA | 216 |
| chr7:24304572-24306592 | p5 | (TATTC)3 | CAGAGGCCTCGAAAGACATC | AGTTGGTCCTTGTCGCAAAC | 109 |
| chr7:24304572-24306592 | p5 | (TTATT)4 | CAGAGGCCTCGAAAGACATC | AGTTGGTCCTTGTCGCAAAC | 109 |
| chr7:24330608-24332656 | p3 | (AAT)16 | CTTTAAACTCCCTCGTCCCC | AGCAAGAGTCGACACCGATT | 241 |
| chr7:24350742-24352756 | p2 | (CA)7 | GTGAGCATGAAGCTCCACAA | CGGGAAAATACCACCATGAC | 252 |
| chr7:24377735-24379751 | p3 | (TGT)5 | TCCTGTGAACTTCCCAAAGG | AGGTTGGCATAGGTTTTCCC | 130 |
| chr7:24377735-24379751 | p2 | (TC)9 | TCCTGTGAACTTCCCAAAGG | AGGTTGGCATAGGTTTTCCC | 130 |

| Marker ID | SSR type | SSR motif | FORWARD PRIMER (5'-3') | REVERSE PRIMER (5'-3') | size (bp) |
|---|---|---|---|---|---|
| chr7:24377735-24379751 | p4 | (TTTA)4 | TCCTGTGAACTTCCCAAAGG | AGGTTGGCATAGGTTTTCCC | 130 |
| chr7:24534571-24536583 | p2 | (AT)6 | AGGCCTCATTTGTTTGCACT | GGGGTTGGAGCTTGTGATAA | 194 |
| chr7:24612390-24614428 | p2 | (TA)19 | AAAAGCCTGTGTGATTCGCT | ACCAGGTGGTGTTCATGGTT | 203 |
| chr7:24720613-24722631 | p6 | (GAGAAA)3 | TTGCAAACAGACATGCCAAT | CGACGCCTCACTTCACTACA | 188 |
| chr7:25005208-25007259 | p3 | (TAT)17 | ATGGGGCTTAACCCACTTCT | CGTGGTGGTTTTCTTTTGCT | 243 |
| chr7:25005208-25007259 | p5 | (GCATT)4 | ATGGGGCTTAACCCACTTCT | CGTGGTGGTTTTCTTTTGCT | 243 |
| chr7:25055464-25057504 | p2 | (TA)20 | ATTTGCTTGCACTCCCTTGT | GGAATGACAACAGGCAACCT | 242 |
| chr7:25107836-25109904 | p2 | (TA)34 | TCAGTTCCGGAGCGAGTAGT | CACGAAGCCATCAACAAGAA | 129 |
| chr7:25140266-25142278 | p2 | (AG)6 | CGAGCTAGATTTACCAGCGG | ACAACACTTTAACCGACGCC | 163 |
| chr7:25155999-25158017 | p2 | (TA)9 | CGGGCATCTTTTCCTACAAA | GCCCTGATGAGTCGTGAAAT | 130 |
| chr7:25168067-25170082 | p2 | (TA)7 | CCTGGAAGCCCAACACTAAG | GTTCCACGTCCAGATTCCAT | 175 |
| chr7:25168067-25170082 | p3 | (GAA)5 | CCTGGAAGCCCAACACTAAG | GTTCCACGTCCAGATTCCAT | 175 |
| chr7:25177929-25179959 | p3 | (AGA)15 | CGCAAAACAAAACCAATTCA | CCTCCTCATCCTTCTCCTCC | 218 |
| chr7:25177929-25179959 | p3 | (AGA)22 | CGCAAAACAAAACCAATTCA | CCTCCTCATCCTTCTCCTCC | 218 |
| chr7:25205062-25207096 | p2 | (TC)10 | ATCCAAACAACCCAGAACCA | GAACGACACCGTCTTCCTGT | 154 |
| chr7:25206941-25208959 | p6 | (GGAATG)3 | CCAATCGTTTGCAACAACAC | GAAGCTCCATTGACAGCCAT | 116 |
| chr7:25238095-25240109 | p2 | (AT)7 | AAGCACTACAGGACGGAGGA | AATCCGGTTCTCATCGTTTG | 130 |
| chr7:25238095-25240109 | p2 | (TA)6 | AAGCACTACAGGACGGAGGA | AATCCGGTTCTCATCGTTTG | 130 |
| chr7:25299825-25301843 | p6 | (TAAAAT)3 | TGTGTATGCCCAACCAAGAA | CATATTTTGGCAGGGACAGG | 190 |
| chr7:25315494-25317512 | p6 | (TTTTTA)3 | ACCCAACTCGGGAAAATACC | CCCTGATACGGAGTCCAGAA | 200 |
| chr7:25339069-25341084 | p5 | (TTTTA)3 | AAATCCATCGTCACCACCAT | TGAGAGAGGATGTGCAGTGG | 136 |
| chr7:25406207-25408241 | p2 | (AT)17 | CACTCTCACCTGACTGGCAA | CGCCTGGAGTATCAAGAACC | 210 |
| chr7:25411202-25413217 | p5 | (GTTAT)3 | GGTTTGGAGAATTGGGGATT | GTGCAGCCTTGGTTGAGATT | 149 |
| chr7:25423104-25425128 | p6 | (ACTGAA)4 | ACCAGAATCCCCCATTTTTC | GGTTCCCAATGCACTCAGAT | 105 |
| chr7:25585969-25587981 | p2 | (TC)6 | TTACGTCGGGTGAGTTTTCC | CAGCAAGCTGGGGATACAAT | 203 |
| chr7:25723939-25725955 | p4 | (TTTC)4 | GTGGTCCATGTTGTTTGCTG | ACCTTTCGACTCCAGGGTCT | 126 |
| chr7:25833944-25835968 | p3 | (TTC)11 | ATCGGAGGGAAAACCCATTA | TCAAAGACTTCATGTCCCCC | 177 |
| chr7:25833987-25836008 | p3 | (TTC)11 | ATCGGAGGGAAAACCCATTA | TCAAAGACTTCATGTCCCCC | 134 |
| chr7:25833987-25836008 | p5 | (TAATT)3 | ATCGGAGGGAAAACCCATTA | TCAAAGACTTCATGTCCCCC | 134 |
| chr7:25838656-25840678 | p2 | (TA)11 | AGGGCCCAAACACATGTAAA | AATATTCAGCAATACGCCGC | 247 |
| chr7:25838656-25840678 | p2 | (AG)6 | AGGGCCCAAACACATGTAAA | AATATTCAGCAATACGCCGC | 247 |
| chr7:25838656-25840678 | p2 | (TA)8 | AGGGCCCAAACACATGTAAA | AATATTCAGCAATACGCCGC | 247 |
| chr7:25878057-25880077 | p2 | (AT)10 | ACTTGCGTCGATTATGTCCC | GTTCGTCACCTTGGAGCAAT | 219 |
| chr7:25909358-25911373 | p5 | (ACAAC)3 | GGAAAGCATTGACGGGTTTA | GACGGGTGAGCGTAACAAGT | 252 |
| chr7:25909358-25911373 | p2 | (AT)8 | GGAAAGCATTGACGGGTTTA | GACGGGTGAGCGTAACAAGT | 252 |
| chr7:25909358-25911373 | p2 | (CA)7 | GGAAAGCATTGACGGGTTTA | GACGGGTGAGCGTAACAAGT | 252 |

| Marker ID | SSR type | SSR motif | FORWARD PRIMER (5'-3') | REVERSE PRIMER (5'-3') | size (bp) |
|---|---|---|---|---|---|
| chr7:25921257-25923272 | p5 | (AAACC)3 | TTGATCCCGATATGCTGTGA | CATAAGCCATCCCAAAAGGA | 240 |
| chr7:25929309-25931321 | p2 | (AT)6 | ACTTTCACCAAACCAAGCGT | GTGGATGCATGAGCAAAATG | 259 |
| chr7:25959217-25961237 | p5 | (AAAGA)4 | TAACTGCCCTACCAGTTCGG | TTAACAGAGAAGGCCCTGGA | 165 |
| chr7:26155682-26157739 | p2 | (AT)24 | CGAAAGGAAACCACCTTTGA | AAATTGGGATAGAGGGGGTG | 242 |
| chr7:26209848-26211866 | p2 | (AG)7 | ACAGCAAAATGACCTTTGGG | AGATGCCACAATCTGCTGTG | 195 |
| chr7:26209848-26211866 | p3 | (AAC)6 | ACAGCAAAATGACCTTTGGG | AGATGCCACAATCTGCTGTG | 195 |
| chr7:26273110-26275124 | p2 | (AT)7 | TACGTGTGGCAAGGAGAGTG | CAACGACGTGCAACAACTTT | 130 |
| chr7:26273110-26275124 | p2 | (TA)7 | TACGTGTGGCAAGGAGAGTG | CAACGACGTGCAACAACTTT | 130 |
| chr7:26309870-26311885 | p5 | (AAACC)3 | AAAAATTGCATGCCAAAAGG | ACCCGACTTTTGTTGGTTTG | 119 |
| chr7:26428444-26430458 | p2 | (AT)6 | AAGTATCAACCGTGGCTTCG | CTCTTCTCGTTCCTCCCCTT | 213 |
| chr7:26466169-26468184 | p5 | (TTCTT)3 | ACGGTGAAGCGTCAGTTTCT | CAAACCTCAAGCTCAAAGCC | 135 |
| chr7:26509519-26511534 | p5 | (TTCTT)3 | ACGGTGAAGCGTCAGTTTCT | CAAACCTCAAGCTCAAAGCC | 135 |
| chr7:26545127-26547157 | p5 | (ATATT)6 | GATCCAATGCAATCCAGGAC | TTCCCCTACCACACTTCCAG | 192 |
| chr7:26574969-26576984 | p5 | (GATAT)3 | AGGCAGATCCTGTATGCGAC | AGGCCTTCTCAAGCAGTGAC | 182 |
| chr7:26592902-26594940 | p2 | (AT)19 | GGCCAAGTAATATGGGACGA | CTCATGTATACACGCACGCA | 146 |
| chr7:26616688-26618700 | p2 | (TG)6 | CCATTCGCTATGGATTTCGT | TAGACGGTAACCGGCCTATG | 210 |
| chr7:26675131-26677197 | p2 | (TA)21 | AGGAGTTGCACGAATTACCG | GTGGGTACGTGGAGTCTGGT | 281 |
| chr7:26675131-26677197 | p3 | (ATA)22 | AGGAGTTGCACGAATTACCG | GTGGGTACGTGGAGTCTGGT | 281 |
| chr7:26741572-26743587 | p5 | (AAAAG)3 | ATGTCATTTCCAGCCTGTCC | ATTGTGGACAGAACGAAGGC | 161 |
| chr7:26772641-26774704 | p3 | (ATA)21 | TGTTTTCAAAGGCGTTTTCG | GCTTACGCCCCAAATGAATA | 263 |
| chr7:26795683-26797698 | p5 | (ACCCG)3 | CTCTCACAGCTTCGTCCCTC | TTGAGGTGTTCATATGGCGA | 129 |
| chr7:26795683-26797698 | p4 | (AAAG)4 | CTCTCACAGCTTCGTCCCTC | TTGAGGTGTTCATATGGCGA | 129 |
| chr7:26811075-26813093 | p2 | (AT)9 | CAAACCTGCAAAACGACAGA | TTGCTTTACCAAAACACCCC | 227 |
| chr7:26877889-26879905 | p2 | (TA)8 | GACCTCTCCCCAGTTCTTCC | AGCGAGGTTGAAATGGTTTG | 183 |
| chr7:26945147-26947165 | p6 | (ACAGTA)3 | ACCATAGCCACCACCATAGC | GAAGGAATGAATGGCCAGAA | 162 |
| chr7:26974273-26976294 | p3 | (AGA)5 | ACTATTGGCATATGGGCTGC | TTCCAACATGGCACTGATGT | 225 |
| chr7:26974273-26976294 | p5 | (TATTT)3 | ACTATTGGCATATGGGCTGC | TTCCAACATGGCACTGATGT | 225 |
| chr7:26974295-26976313 | p3 | (AGA)5 | ACTATTGGCATATGGGCTGC | TTCCAACATGGCACTGATGT | 203 |
| chr7:26974295-26976313 | p5 | (TATTT)3 | ACTATTGGCATATGGGCTGC | TTCCAACATGGCACTGATGT | 203 |
| chr7:26974808-26976823 | p3 | (AGA)5 | TGACCCTCTCCCATGTTCTC | TTCTATTTTCACCGCCAACC | 173 |
| chr7:26974808-26976823 | p5 | (TATTT)3 | TGACCCTCTCCCATGTTCTC | TTCTATTTTCACCGCCAACC | 173 |
| chr7:27115518-27117532 | p2 | (TA)7 | GTGTAAGCCGCTTCAAGGAG | ACCGTCTATGGTTCTCACGG | 186 |
| chr7:27115518-27117532 | p4 | (AATA)4 | GTGTAAGCCGCTTCAAGGAG | ACCGTCTATGGTTCTCACGG | 186 |
| chr7:27185001-27187021 | p2 | (TA)10 | CAGCAGCCAGTGATGGACTA | AGGTTTCCGGCTACGATTCT | 254 |
| chr7:27189838-27191862 | p2 | (AT)12 | GGCTGCATCAAAGGGATTTA | AAGTGTTGATGAACGGCCTC | 108 |
| chr7:27189838-27191862 | p2 | (TA)7 | GGCTGCATCAAAGGGATTTA | AAGTGTTGATGAACGGCCTC | 108 |

| Marker ID | SSR type | SSR motif | FORWARD PRIMER (5'-3') | REVERSE PRIMER (5'-3') | size (bp) |
|---|---|---|---|---|---|
| chr7:27195621-27197637 | p2 | (TA)8 | CGACCTTACGTCACCTCGAT | CCAACTAGGCTCGGTGTGAT | 227 |
| chr7:27248517-27250561 | p2 | (AT)22 | TAACCTCCCTTCACATGGCT | TTATTGTCTGACGCGGAACA | 212 |
| chr7:27248517-27250561 | p2 | (AT)6 | TAACCTCCCTTCACATGGCT | TTATTGTCTGACGCGGAACA | 212 |
| chr7:27289345-27291389 | p2 | (AT)22 | TAACCTCCCTTCACATGGCT | TTATTGTCTGACGCGGAACA | 212 |
| chr7:27289345-27291389 | p2 | (AT)6 | TAACCTCCCTTCACATGGCT | TTATTGTCTGACGCGGAACA | 212 |
| chr7:27442456-27444483 | p2 | (GA)12 | GCAGGATTGAACCCTTTTCA | CTTTCTTTCTCGCGTGCTTC | 121 |
| chr7:27442456-27444483 | p3 | (TAA)9 | GCAGGATTGAACCCTTTTCA | CTTTCTTTCTCGCGTGCTTC | 121 |
| chr7:27489419-27491457 | p2 | (TA)12 | TAAGAGGTCTTTGCCGGAGA | AGAGGTTGCCTATCGTGGTG | 218 |
| chr7:27511487-27513499 | p2 | (TA)6 | AGGTGTTTGAGGCTTCAGGA | ATCATTTTGGAACACCCTCG | 260 |
| chr7:27575891-27577913 | p2 | (AT)7 | CATAAAGGGTGGACCGTCAT | GGGTGCAGTTTGGAGATGAT | 286 |
| chr7:27575891-27577913 | p2 | (TA)7 | CATAAAGGGTGGACCGTCAT | GGGTGCAGTTTGGAGATGAT | 286 |
| chr7:27576013-27578035 | p2 | (TA)7 | CATAAAGGGTGGACCGTCAT | GGGTGCAGTTTGGAGATGAT | 164 |
| chr7:27697366-27699390 | p2 | (AT)12 | TTTGACACACGATTGCCCTA | GACCCAGAACCAGAGGATGA | 202 |
| chr7:27724761-27726779 | p2 | (AT)9 | GGCGGATTTATGCCTGTAAC | TTCAACAGCGTAACCACCAA | 215 |
| chr7:27761514-27763530 | p4 | (TATT)4 | TGAATCTCCAAGATGCCTCC | CAACTCATCCTGGGGAGAAA | 186 |
| chr7:27768704-27770722 | p2 | (TA)9 | TGGGCCCCATACTAAACAAC | TTTTTAAGGGACAACGGACG | 273 |
| chr7:27768704-27770722 | p2 | (TA)9 | TGGGCCCCATACTAAACAAC | TTTTTAAGGGACAACGGACG | 273 |
| chr7:27788740-27790764 | p4 | (TATT)6 | AGATTTACGCCCTTGCTTCA | CAACTCATCCTGGGGAGAAA | 136 |
| chr7:27860842-27862876 | p2 | (TA)17 | TAGCGGAAATCGATGCCTAT | GATAAAGGAGGAGGGTTGCC | 291 |
| chr7:28025192-28027207 | p2 | (AT)7 | CTTGTGCTTCATTGCCTTCA | TGAGGGTTTGTTTACCGAGG | 274 |
| chr7:28025245-28027260 | p2 | (AT)7 | CTTGTGCTTCATTGCCTTCA | TGAGGGTTTGTTTACCGAGG | 221 |
| chr7:28025271-28027286 | p2 | (AT)7 | CTTGTGCTTCATTGCCTTCA | TGAGGGTTTGTTTACCGAGG | 195 |
| chr7:28038565-28040577 | p2 | (TC)6 | ATCTGACCCGACCTTTCCTT | ATTGCCGATGAAGTTGGTTC | 261 |
| chr7:28166977-28168992 | p5 | (TGATA)3 | AGCCTCGGGGACAACTCTAT | GCAATTGTCTCAACAAGGCA | 267 |
| chr7:28212770-28214788 | p3 | (CAA)6 | CCGACTCGAGGGAATCATTA | ATTCCCGCTTTGCCTTTTAT | 226 |
| chr7:28249370-28251424 | p2 | (AT)27 | GGATGATTGGGAAGGGATTT | GTATACGCAATCGCCCAACT | 248 |
| chr7:28314368-28316434 | p3 | (TTC)5 | ACTGTGGTCTTCACCCGTTC | TTCAGATGGACGTTGCTGAG | 127 |
| chr7:28314368-28316434 | p3 | (TCT)22 | ACTGTGGTCTTCACCCGTTC | TTCAGATGGACGTTGCTGAG | 127 |
| chr7:28315474-28317486 | p2 | (AT)6 | TGTACGAAGTCTTTGCGTCG | TACATGGTGGGTTATGCCCT | 281 |
| chr7:28319718-28321734 | p4 | (ATGT)4 | AGGGGAGATGCTGCCTTAAT | CCTTGCTCCTTAACCTTCCC | 278 |
| chr7:28375363-28377375 | p2 | (TA)6 | CGTGTTGTCGTTGATATGGC | AACTCGCCTAACCCGAACTT | 111 |
| chr7:28399272-28401284 | p2 | (AT)6 | TACGTGTCACGACCCGACTA | GGGTAGCATGGTCTCACGAT | 266 |
| chr7:28464636-28466650 | p2 | (TA)7 | AAGTGTGCGAAGCGTATGTG | CAAGATCCGAGGGTACTCCA | 245 |
| chr7:28514688-28516703 | p3 | (TCA)5 | CTGTCAAATCTCCTAGCGGC | TCGCAGGCTATTCTGGAGTT | 276 |
| chr7:28524852-28526867 | p3 | (TTG)5 | ATCCCAAAGTAATCCCGTCC | GACCAAGTCTGGCCCTATCA | 252 |
| chr7:28541173-28543188 | p3 | (TTG)5 | ATCCCAAAGTAATCCCGTCC | GACCAAGTCTGGCCCTATCA | 252 |

| Marker ID | SSR type | SSR motif | FORWARD PRIMER (5'-3') | REVERSE PRIMER (5'-3') | size (bp) |
|---|---|---|---|---|---|
| chr7:28569657-28571691 | p2 | (TA)17 | TTAATGAAATCGGGGGATCA | GTATGTATGTCTGCCCGCCT | 242 |
| chr7:28575684-28577698 | p2 | (AT)7 | TGTTGTTGATGGTGGCTTGT | CACCAAGGTGCTCCATTTCT | 299 |
| chr7:28589893-28591911 | p6 | (TCAGTT)3 | GGGTGATGGAGAATGCTGTT | AATTCCAGCCTTTGAGCGTA | 173 |
| chr7:28715405-28717420 | p5 | (ATTTC)3 | CAAACTCGCCCTTTAATCCA | GGTAGGGTGTTGGCTTGTGT | 160 |
| chr7:28720883-28722898 | p5 | (ATTTC)3 | TATTTATGCGTCGCCAATGA | GTCGTGCCCTTCGTCAGTAT | 290 |
| chr7:28752607-28754685 | p2 | (TA)39 | TTGCTCCAAAGCTCATTCCT | ACCCCCAAACAACCATAACA | 193 |
| chr7:28757625-28759649 | p3 | (GTT)8 | TGGGAATTTGGGAAAAATCA | TGAGGAGGAGAAGGCTGAAA | 219 |
| chr7:28767790-28769804 | p2 | (AG)7 | GGCAAGGGCAAGAACAAATA | CCTCGAGCTCACATTAAGCC | 262 |
| chr7:28767790-28769804 | p3 | (GTT)5 | GGCAAGGGCAAGAACAAATA | CCTCGAGCTCACATTAAGCC | 262 |
| chr7:28767790-28769804 | p2 | (AT)28 | GGCAAGGGCAAGAACAAATA | CCTCGAGCTCACATTAAGCC | 262 |
| chr7:28845967-28847999 | p2 | (AT)16 | CTCATGACGACATCCAAACG | TCTTCGACCTCCAACAGCTT | 274 |
| chr7:28902572-28904587 | p2 | (TA)14 | ACATCCTTCACTGAGTCCCG | TATAACGTGTCCTGGCCCTC | 130 |
| chr7:28992228-28994243 | p2 | (TA)25 | CTCATACCTGGCCCTACCAA | ACCCCGCGAGTATATGTCAG | 250 |
| chr7:28992228-28994243 | p3 | (TCA)5 | CTCATACCTGGCCCTACCAA | ACCCCGCGAGTATATGTCAG | 250 |
| chr7:28997657-28999677 | p2 | (AT)10 | CTCTGTCACACCCCAACCTT | TAGTTATGTTGGCAGCGTGG | 132 |
| chr7:28997657-28999677 | p3 | (CAA)6 | CTCTGTCACACCCCAACCTT | TAGTTATGTTGGCAGCGTGG | 132 |
| chr7:29205254-29207275 | p3 | (ATT)7 | ATTTTCTCATGCTCGGTTGG | ATATTATGGGGTGTGGGGGT | 212 |
| chr7:29240793-29242808 | p2 | (TA)10 | ATCTCGAAGGATTGGGACCT | AAATCTGCCGAACATTCCAC | 183 |
| chr7:29240917-29242956 | p2 | (TA)10 | TCATAGTGGAATGTTCGGCA | GCTTCCTCTTATCGTCACCG | 266 |
| chr7:29365190-29367208 | p3 | (AGA)6 | TGAATTTCAAACCCTGAGGC | TTGGCAAAATGGAAAACACA | 134 |
| chr7:29485888-29487924 | p2 | (AT)8 | ACACAAGCCCACAAGGCTAC | AGGGAGACGGATCAAATCCT | 253 |
| chr7:29492621-29494633 | p2 | (AT)9 | CCTGTCGTTGTACCTGAGCA | GAATGAGGTTCCATCTCGGA | 299 |
| chr7:29492621-29494633 | p2 | (TA)11 | CCTGTCGTTGTACCTGAGCA | GAATGAGGTTCCATCTCGGA | 299 |
| chr7:29704618-29706630 | p2 | (TC)6 | ATTCTAAATCCGCCTCAGCA | CTGGGTTAGGTCAGATGCGT | 269 |
| chr7:29801164-29803176 | p2 | (AT)6 | TGATGCAGCAAAATGTGGAT | TCTGAGTCTTGGTTGCGTTG | 179 |
| chr7:29801164-29803176 | p2 | (AT)6 | TGATGCAGCAAAATGTGGAT | TCTGAGTCTTGGTTGCGTTG | 179 |
| chr7:29881799-29883814 | p5 | (TACAA)3 | TTACAATGTTGGGCTCGTCA | ATGAACGACGAGAAACCACC | 252 |
| chr7:29923361-29925377 | p5 | (TTTTC)3 | TAAGCGTACGACAACCTCCC | ATGGATTTGAACCTTGAGCG | 212 |
| chr7:29923407-29925425 | p5 | (TTTTC)3 | TAAGCGTACGACAACCTCCC | ATGGATTTGAACCTTGAGCG | 166 |
| chr7:30047964-30049979 | p3 | (TGT)5 | GGCCAAATTTTATTGGCTCA | GCCATGGCTGTTGTAATCCT | 202 |
| chr7:30123995-30126009 | p2 | (AT)7 | TTTTTACGAAGCCCACCATC | GACAGGGCTGTTTTAGAGCG | 162 |
| chr7:30123995-30126009 | p2 | (GT)7 | TTTTTACGAAGCCCACCATC | GACAGGGCTGTTTTAGAGCG | 162 |
| chr7:30183584-30185598 | p2 | (TA)7 | ATGTGGCAATGAATCCATCA | AAAAGTGGAACCCACCACAC | 192 |
| chr7:30279995-30282009 | p2 | (TA)7 | CACAACCAAATGCCAACAAG | CAGTTCCTGGCCATCCTCTA | 179 |
| chr7:30292513-30294535 | p2 | (AT)11 | AAATCACCATGTCCTCAGCC | GGAAACGGAACAGATTTGGA | 109 |
| chr7:30312782-30314798 | p2 | (CA)6 | GTCGACGAGGCCAAATACAT | AGGTGCACAGGGAGACAAAC | 276 |

| Marker ID | SSR type | SSR motif | FORWARD PRIMER (5'-3') | REVERSE PRIMER (5'-3') | size (bp) |
|---|---|---|---|---|---|
| chr7:30312782-30314798 | p2 | (AT)8 | GTCGACGAGGCCAAATACAT | AGGTGCACAGGGAGACAAAC | 276 |
| chr7:30445651-30447679 | p2 | (TA)14 | ACTTGGTGCAAGTCCAGCTT | TTGTACCACGAAATCACCCA | 180 |
| chr7:30471084-30473100 | p2 | (TA)8 | GATTTTCGGAGGCATAACGA | CCTGAGGGGTCATAGGTCTG | 221 |
| chr7:30510529-30512544 | p3 | (ATT)5 | AGCCTCAGCAAAGGCTATCA | TTCGCTGCCGAAGAAATAGT | 236 |
| chr7:30534392-30536407 | p3 | (ATT)5 | AGCCTCAGCAAAGGCTATCA | GTTCGCTGCCGAAGAAATAG | 236 |
| chr7:30552569-30554585 | p3 | (CAA)5 | GGGGTCTGGGGAGAGTAGAG | CCGGACTGTTACTTTCTCGC | 255 |
| chr7:30552569-30554585 | p2 | (GA)8 | GGGGTCTGGGGAGAGTAGAG | CCGGACTGTTACTTTCTCGC | 255 |
| chr7:30630873-30632895 | p2 | (TA)11 | AGTCAACATCCGGCGTAAAC | ACAGAAATACCCCGAGTCCC | 257 |
| chr7:30644405-30646420 | p3 | (TTG)5 | ATATTGGGCGGAGTGTTGTC | CACCCCTACCTTGTCCTGAA | 268 |
| chr7:30783408-30785423 | p5 | (ACCAA)3 | TTTGAGCTTGATTGGTGCAG | TAACGTGGGACGTGAGAACA | 115 |
| chr7:30783408-30785423 | p5 | (AAACC)3 | TTTGAGCTTGATTGGTGCAG | TAACGTGGGACGTGAGAACA | 115 |
| chr7:30813622-30815646 | p2 | (AT)12 | CATGGTCGCTTTACTCAGCA | CCATTTTCTTCCTGGAGCAG | 246 |
| chr7:30898205-30900255 | p2 | (CA)25 | ATTCACCACTGCTGCTCCTT | TCCTACAGGGGTAGGTCGTG | 235 |
| chr7:30931491-30933507 | p4 | (GGGT)4 | GACGCTCATGCTCTCACAAA | AGAACTCACCACACATGGCA | 213 |
| chr7:30931491-30933507 | p5 | (TATTT)3 | GACGCTCATGCTCTCACAAA | AGAACTCACCACACATGGCA | 213 |
| chr7:31032978-31034993 | p5 | (AAGAA)3 | TTGTCCATATTCGGCTTTCC | TGGCGGGATCAGAGTATTTC | 172 |
| chr7:31070327-31072342 | p5 | (TTTAT)3 | TTGCCTTTGCTAGCTTTGCT | TTCACTCGGCTAATGCACAG | 186 |
| chr7:31170774-31172790 | p2 | (TA)8 | GCATTCTGGAATTTGGCAAT | TCCGTTTGACCCCCATAATA | 123 |
| chr7:31203198-31205243 | p3 | (AAT)15 | TGGCGACTCCTCAAAACTCT | GTGGGTTCAACCACGATTTC | 138 |
| chr7:31249221-31251239 | p6 | (GCGGCA)3 | TTGTTTACCTCGCCCTTTTG | TTTTCTCCCCAACAACCAAG | 225 |
| chr7:31259117-31261132 | p3 | (TTC)5 | TGGAATTCAGGTTCTTTGGG | ATCAGGGTTCTTGGTTCGTG | 285 |
| chr7:31323478-31325496 | p6 | (TTTTCT)3 | TTCCCAGCCGAAACAATATC | TGCCTGATGAGCAAAATGAG | 235 |
| chr7:31328893-31330911 | p6 | (TTTTCT)3 | TTCCCAGCCGAAACAATATC | TGCCTGATGAGCAAAATGAG | 235 |
| chr7:31405795-31407810 | p6 | (TCCGGG)3 | CCGATTCACCAGTTGACCTT | GAAGGGCACTCCGTACAAAA | 141 |
| chr7:31405795-31407810 | p5 | (TTAAA)3 | CCGATTCACCAGTTGACCTT | GAAGGGCACTCCGTACAAAA | 141 |
| chr7:31470812-31472864 | p2 | (AT)26 | GCTCAAGGCTCAAGGTTTTG | AACATGGCGTGAGGAAGAAC | 249 |
| chr7:31474666-31476738 | p2 | (TA)36 | AATTTGGGTAGCATTTCCCC | TGTTGTGGCTGTGTTGTTGA | 110 |
| chr7:31474666-31476738 | p2 | (AT)7 | AATTTGGGTAGCATTTCCCC | TGTTGTGGCTGTGTTGTTGA | 110 |
| chr7:31501557-31503602 | p3 | (ATA)15 | GGATTCGAAATTTTTGGGGT | GTTTTTGCTCGAAGACGCTC | 247 |
| chr7:31501557-31503602 | p5 | (GCTTA)3 | GGATTCGAAATTTTTGGGGT | GTTTTTGCTCGAAGACGCTC | 247 |
| chr7:31512212-31514272 | p3 | (ATG)6 | CTCAGGGTTAACGAGCTTGC | AGGGTCATCAAACAACCCAG | 127 |
| chr7:31589009-31591087 | p2 | (TA)39 | ATATGGGACTTAGGGCCTGC | AACAGAGCCCAAGAAAAGCA | 239 |
| chr7:31589009-31591087 | p2 | (TA)17 | ATATGGGACTTAGGGCCTGC | AACAGAGCCCAAGAAAAGCA | 239 |
| chr7:31646187-31648205 | p2 | (TA)12 | TGACATGGCGGAATAACAAA | CATTACCGCGTGCATATCAA | 202 |
| chr7:31646187-31648205 | p2 | (AT)9 | TGACATGGCGGAATAACAAA | CATTACCGCGTGCATATCAA | 202 |
| chr7:31646207-31648219 | p2 | (TA)12 | TGACATGGCGGAATAACAAA | CATTACCGCGTGCATATCAA | 182 |

| Marker ID | SSR type | SSR motif | FORWARD PRIMER (5'-3') | REVERSE PRIMER (5'-3') | size (bp) |
|---|---|---|---|---|---|
| chr7:31646207-31648219 | p2 | (AT)9 | TGACATGGCGGAATAACAAA | CATTACCGCGTGCATATCAA | 182 |
| chr7:31692092-31694112 | p5 | (GTTCG)4 | TCATAACATTGCCACCTGGA | TGGGAACCAAAACCAAGAAG | 117 |
| chr7:31692092-31694112 | p5 | (GTTCG)3 | TCATAACATTGCCACCTGGA | TGGGAACCAAAACCAAGAAG | 117 |
| chr7:31692602-31694617 | p5 | (GTTCG)4 | AATGACCAGCCCATTTGAAG | CCCGAAATAGCCAAACTGAA | 296 |
| chr7:31692602-31694617 | p5 | (GTTCG)3 | AATGACCAGCCCATTTGAAG | CCCGAAATAGCCAAACTGAA | 296 |
| chr7:31727278-31729293 | p5 | (ACCAA)3 | GACCCAGTTTCTTCTTGCCA | GAGATAGCAGGCAGCACTCC | 132 |
| chr7:31757692-31759738 | p2 | (TA)23 | AATCGGAGGTTCTCGACCTT | GGTGTCTGCCCATGCTATCT | 274 |
| chr7:31774920-31776935 | p3 | (ACA)5 | CACAGCATACACGACCCAAC | TGGAAGCATTGTTGGAGTGA | 217 |
| chr7:31774920-31776935 | p2 | (AT)6 | CACAGCATACACGACCCAAC | TGGAAGCATTGTTGGAGTGA | 217 |
| chr7:31774920-31776935 | p2 | (AT)8 | CACAGCATACACGACCCAAC | TGGAAGCATTGTTGGAGTGA | 217 |
| chr7:31783647-31785667 | p2 | (AT)10 | GCTACATTACCCCGGGATTT | TCATAACTTTGCCCTCGGAC | 208 |
| chr7:31885607-31887622 | p5 | (TAAAT)3 | TGGGCAAACATTCGATAACA | TGGCCATTGATGATTTGAGA | 142 |
| chr7:31885607-31887622 | p5 | (TTTAT)3 | TGGGCAAACATTCGATAACA | TGGCCATTGATGATTTGAGA | 142 |
| chr7:31885607-31887622 | p6 | (AAAAAT)3 | TGGGCAAACATTCGATAACA | TGGCCATTGATGATTTGAGA | 142 |
| chr7:32088289-32090307 | p6 | (AACTGA)3 | TGTCTGGGGTTTTGGCTTAG | TTGCACGAACTTCTGAATCG | 126 |
| chr7:32107754-32109776 | p2 | (TA)11 | CAGTTTGAATCTCGGGAAGC | CAGATTAGGAGCCGAGTTGC | 115 |
| chr7:32204011-32206027 | p4 | (AAAG)4 | TAAATGTCGTGCCCAGTCAA | AGTAATTTGCAATTTGCGCC | 107 |
| chr7:32210053-32212069 | p4 | (ACAT)4 | GGCCCAAATTTTATGCCTTT | TTTGGGATGAGCCCAGTAAC | 105 |
| chr7:32226146-32228168 | p2 | (AT)11 | TTCTTGTTGAGGCTCCGACT | CCAGAAGGCATATCCCAAGA | 236 |
| chr7:32226146-32228168 | p5 | (AACCG)3 | TTCTTGTTGAGGCTCCGACT | CCAGAAGGCATATCCCAAGA | 236 |
| chr7:32280035-32282047 | p2 | (AT)17 | TTTGGGAACCAGCTGAAGAC | ATTAAGGTCAGTGGGGGACC | 175 |
| chr7:32280048-32282074 | p2 | (AT)17 | TTTGGGAACCAGCTGAAGAC | ATTAAGGTCAGTGGGGGACC | 162 |
| chr7:32365146-32367161 | p5 | (AAAAT)3 | TTTGCCCTGCTCAAATTACC | TATTTATCACCCCTCGCTGC | 288 |
| chr7:32527593-32529607 | p2 | (CA)7 | TCCTTCTCCAGCTTTTCCAA | ACCATGAGGTTATGGTGGGA | 136 |
| chr7:32527593-32529607 | p2 | (CA)7 | TCCTTCTCCAGCTTTTCCAA | ACCATGAGGTTATGGTGGGA | 136 |
| chr7:32535657-32537697 | p2 | (TA)20 | TCTGCGAAGGAAAAATGACC | AAGGGAAGAGGGAAGGATCA | 204 |
| chr7:32563665-32565677 | p2 | (AT)13 | TGACATAGCTCTACGGCACG | AGCAGTTCGTCCTAGGGGTT | 148 |
| chr7:32563665-32565677 | p2 | (AT)6 | TGACATAGCTCTACGGCACG | AGCAGTTCGTCCTAGGGGTT | 148 |
| chr7:32563665-32565677 | p4 | (CATA)4 | TGACATAGCTCTACGGCACG | AGCAGTTCGTCCTAGGGGTT | 148 |
| chr7:32570451-32572469 | p2 | (TA)9 | ATCATTGTTCTTTCCCACGC | AAGAATGCATGCAAGAACCC | 279 |
| chr7:32578799-32580814 | p5 | (TTTTC)3 | CAATGTGCTAACGTCCCCTT | TACTTGCTGGGACAAGGACC | 217 |
| chr7:32757267-32759279 | p2 | (TA)6 | CCTATGACCCGCTGAACAAT | ATAAGGGAGAGCTAAGGCGG | 159 |
| chr7:32849744-32851785 | p5 | (TAAAC)3 | CCTTTATGCTCCAACTCCCA | TGGGCTGTAGGTTCGAGTTT | 238 |
| chr7:32849744-32851785 | p2 | (TA)6 | CCTTTATGCTCCAACTCCCA | TGGGCTGTAGGTTCGAGTTT | 238 |
| chr7:33082037-33084053 | p4 | (TTTC)4 | TGACAAGCGCTAAGGATGTG | ATCACCAAACACAATCGCAA | 222 |
| chr7:33327875-33329893 | p6 | (TATGTA)3 | GAGATGCGATACGAGCATGA | GGTCGTGAACCACCGACTAT | 133 |

| Marker ID | SSR type | SSR motif | FORWARD PRIMER (5'-3') | REVERSE PRIMER (5'-3') | size (bp) |
|---|---|---|---|---|---|
| chr7:33327875-33329893 | p2 | (TG)14 | GAGATGCGATACGAGCATGA | GGTCGTGAACCACCGACTAT | 133 |
| chr7:33393851-33395869 | p3 | (ACA)6 | GGGGATCATAATCATCGTGG | TTGGAACCAAAAGTTGCTCC | 298 |
| chr7:33395067-33397085 | p3 | (ACA)6 | TAACGCATAATGGCCCTTTC | TTGGAACCAAAAGTTGCTCC | 278 |
| chr7:33468690-33470705 | p5 | (GCATT)3 | AGGCTCGAAGGAAAGGAAAG | CAACAAGGCCCAACCTTAAA | 245 |
| chr7:33531322-33533374 | p4 | (GAAA)5 | ACCACCAACTGCATCAACAA | GAACCCATGGCTTGTCCTAA | 205 |
| chr7:33542365-33544389 | p2 | (AG)12 | AGCATCAAGGGTGTCTTTGG | ACATTACGATGGGTGTGGGT | 210 |
| chr7:33654642-33656654 | p2 | (AT)6 | TCTTTGTGATTCGGAGGGAG | CATTCTCAACCCCTCCTTGA | 274 |
| chr7:33703634-33705646 | p2 | (AT)6 | TACGCATTCCAAGATGGTCA | TCGGGATGGCCTATTTACAG | 206 |
| chr7:33703634-33705646 | p6 | (ATTTTA)3 | TACGCATTCCAAGATGGTCA | TCGGGATGGCCTATTTACAG | 206 |
| chr7:33704442-33706460 | p2 | (AT)6 | AACATTGGGGCCATACTCAG | TGTCGCTTAGCGTCGTAATG | 102 |
| chr7:33704442-33706460 | p6 | (ATTTTA)3 | AACATTGGGGCCATACTCAG | TGTCGCTTAGCGTCGTAATG | 102 |
| chr7:33738785-33740799 | p2 | (AT)7 | TTATTCCTTTTGGATTGGCG | TCTGGTTTACGATGTGCAGC | 276 |
| chr7:33765985-33767999 | p2 | (AT)7 | TTATTCCTTTTGGATTGGCG | TCTGGTTTACGATGTGCAGC | 276 |
| chr7:33814954-33816969 | p5 | (ATAAA)3 | GACAAGTGGCACAGCATCAT | CGGATTACGGATCTTGTTGG | 284 |
| chr7:33860970-33862992 | p2 | (TA)11 | TGTACTAGTAGGGTCGGGCG | CTTGTGGCTGTGGGAAAGTT | 277 |
| chr7:33877789-33879875 | p2 | (TA)43 | TGTGGTGGAATGGTTAGGGT | GAATCTTCATTCCTGGGCAA | 199 |
| chr7:33916018-33918033 | p5 | (TTTGG)3 | AAACGTAATGTTCCGATGCC | CATTTTCATCCAAACACCCC | 284 |
| chr7:33987875-33989890 | p2 | (AT)13 | GCCTACCTGCACACAGACAA | GCAAAATCTCCAACCAAGGA | 162 |
| chr7:33987875-33989890 | p5 | (AGAAA)3 | GCCTACCTGCACACAGACAA | GCAAAATCTCCAACCAAGGA | 162 |
| chr7:34020249-34022293 | p2 | (TA)22 | ATTGGAAATTCGGGTTAGGC | TCGCTTCGTTGCTCCTTTAT | 129 |
| chr7:34039297-34041315 | p3 | (GTT)6 | CGTGTTTTCATGCGTTGAGT | ACGGCCCGTACCTCTATTCT | 257 |
| chr7:34141345-34143367 | p2 | (AT)11 | GATTGGCATGGCTTTGTTTT | ATGCCAATGCGGTATGACTT | 140 |
| chr7:34159109-34161124 | p5 | (AAGCC)3 | ATCCAACCCGGCATATACAA | ACGTCAGGCTGAACACACAG | 101 |
| chr7:34159109-34161124 | p2 | (CA)6 | ATCCAACCCGGCATATACAA | ACGTCAGGCTGAACACACAG | 101 |
| chr7:34239850-34241906 | p2 | (AT)28 | ACCCCCTATGGTGTGATGAA | GGTGCAAAAGAAAGATGGGA | 219 |
| chr7:34301182-34303200 | p6 | (TTTGGA)3 | TCCCCACCAGAAAAATTGAA | TTTGCTGCAAGTGGAGTTTG | 120 |
| chr7:34368269-34370284 | p5 | (CCCGC)3 | GTTCGAACCACCGCTTAAAA | TTTGCAAGGCATAGGTTTCC | 166 |
| chr7:34543470-34545484 | p2 | (AT)11 | GGCGTATCAAACTCACCCTC | TCGTGTTTAGCGTGAGCATC | 191 |
| chr7:34543470-34545484 | p2 | (AT)7 | GGCGTATCAAACTCACCCTC | TCGTGTTTAGCGTGAGCATC | 191 |
| chr7:34543470-34545484 | p2 | (AT)7 | GGCGTATCAAACTCACCCTC | TCGTGTTTAGCGTGAGCATC | 191 |
| chr7:34543710-34545724 | p2 | (AT)11 | GATGCTCACGCTAAACACGA | ATATTCCGGCCATGAACAGT | 131 |
| chr7:34543710-34545724 | p2 | (AT)7 | GATGCTCACGCTAAACACGA | ATATTCCGGCCATGAACAGT | 131 |
| chr7:34543710-34545724 | p2 | (AT)7 | GATGCTCACGCTAAACACGA | ATATTCCGGCCATGAACAGT | 131 |
| chr7:34562674-34564728 | p3 | (AAT)18 | CATCATACCAAACGAGCCCT | TGTGAGATAGGCCCCAAAAC | 212 |
| chr7:34566289-34568311 | p2 | (TA)11 | TGTCAACCTTTGCACTGCTC | ACGATTGAGACACACCCCTC | 101 |
| chr7:34566289-34568311 | p2 | (TA)22 | TGTCAACCTTTGCACTGCTC | ACGATTGAGACACACCCCTC | 101 |

| Marker ID | SSR type | SSR motif | FORWARD PRIMER (5'-3') | REVERSE PRIMER (5'-3') | size (bp) |
|---|---|---|---|---|---|
| chr7:34596810-34598822 | p3 | (ATG)5 | GGGGGTTCTCGACCTTATGT | GTCATAAGCATCCCGCCTTA | 262 |
| chr7:34596810-34598822 | p2 | (TA)6 | GGGGGTTCTCGACCTTATGT | GTCATAAGCATCCCGCCTTA | 262 |
| chr7:34596810-34598822 | p2 | (AT)7 | GGGGGTTCTCGACCTTATGT | GTCATAAGCATCCCGCCTTA | 262 |
| chr7:34596810-34598822 | p2 | (AT)6 | GGGGGTTCTCGACCTTATGT | GTCATAAGCATCCCGCCTTA | 262 |
| chr7:34619050-34621095 | p3 | (GGA)5 | CAGGTCCAAGTGGTGATGTG | TTAGCAGGGGGTGCAGTATC | 201 |
| chr7:34619050-34621095 | p3 | (ATT)15 | CAGGTCCAAGTGGTGATGTG | TTAGCAGGGGGTGCAGTATC | 201 |
| chr7:34619050-34621095 | p2 | (TA)7 | CAGGTCCAAGTGGTGATGTG | TTAGCAGGGGGTGCAGTATC | 201 |
| chr7:34634873-34636885 | p2 | (TA)6 | CCTTTTTGGGCAGATGTGTT | CAGTCAAGAGGGCCATTCAT | 179 |
| chr7:34671781-34673801 | p2 | (TA)9 | TCCCACCTCAGTAGGCAAAC | TGAAGGCCTCCATCATTCTC | 121 |
| chr7:34671781-34673801 | p5 | (CAATG)4 | TCCCACCTCAGTAGGCAAAC | TGAAGGCCTCCATCATTCTC | 121 |
| chr7:34686901-34688925 | p5 | (ATGCA)3 | CCCTTTTGCTAGAGGCACTG | ATGACTCCCACAACCTGGAC | 228 |
| chr7:34729738-34731753 | p3 | (CAT)5 | TCACGACCTAAATCGGAAGG | GGGTCATCCCTTTGTCAAGA | 262 |
| chr7:34729738-34731753 | p2 | (AT)8 | TCACGACCTAAATCGGAAGG | GGGTCATCCCTTTGTCAAGA | 262 |
| chr7:34869745-34871760 | p5 | (TTTTC)3 | CCTTTTGCTAATGGAACCCA | GCTTTGTGTTGCTTTTCGGT | 236 |
| chr7:34869745-34871760 | p2 | (AT)17 | CCTTTTGCTAATGGAACCCA | GCTTTGTGTTGCTTTTCGGT | 236 |
| chr7:34904841-34906856 | p3 | (TTG)5 | CCTTAAGCCAAATATGGGCA | TCGGTTCCACCGACTTATTC | 177 |
| chr7:34904841-34906856 | p4 | (TATG)8 | CCTTAAGCCAAATATGGGCA | TCGGTTCCACCGACTTATTC | 177 |
| chr7:34932114-34934129 | p2 | (AT)21 | ATTTCCTATGGCTGGGCTCT | AGTATCCAGCTCGGGCACTA | 106 |
| chr7:34932114-34934129 | p5 | (CATAT)3 | ATTTCCTATGGCTGGGCTCT | AGTATCCAGCTCGGGCACTA | 106 |
| chr7:34990161-34992207 | p2 | (TA)23 | AATTTTGCTGCCAAGATGCT | GCACGAAAACGGTGACCTAT | 170 |
| chr7:34990161-34992207 | p6 | (CATGAA)3 | AATTTTGCTGCCAAGATGCT | GCACGAAAACGGTGACCTAT | 170 |
| chr7:35066540-35068554 | p2 | (AG)7 | TCCGTTACTCTCGACCATCC | CAATAGCCAGTTGCAAAGCA | 243 |
| chr7:35130182-35132197 | p3 | (ATC)5 | TCATAAGGACGGGACAGGAC | ATCCTTTTGCTTGTGACGCT | 264 |
| chr7:35322469-35324483 | p2 | (AT)7 | GGGTATGAAACTCGCGAAAA | AGGGGTCACTTGCCTTCTTT | 224 |
| chr7:35362145-35364157 | p2 | (GA)6 | TCTGTTCATCACCAGGTCCA | TCCCTGGAAGCACAATTTTC | 164 |
| chr7:35487721-35489739 | p6 | (TATTTT)3 | ATGAGAACTTTTTCGCCGTG | TAGGGGATGTGGTAGGTGGA | 125 |
| chr7:35487721-35489739 | p3 | (AAT)14 | ATGAGAACTTTTTCGCCGTG | TAGGGGATGTGGTAGGTGGA | 125 |
| chr7:35579007-35581035 | p2 | (AT)14 | CACACCACCATCCACAAGAG | TTTTCTCACCACCCTCGAAC | 298 |
| chr7:35638091-35640115 | p3 | (AAT)8 | GTTTGGCCGACAAAGAAAAA | GATAAGTAGACCCCGCCACA | 249 |
| chr7:35812141-35814159 | p3 | (GTT)6 | CAGATTGCCCACAACCTTTT | ACTGGATGGCTTTTCAATGG | 247 |
| chr7:35812141-35814159 | p4 | (GGAA)4 | CAGATTGCCCACAACCTTTT | ACTGGATGGCTTTTCAATGG | 247 |
| chr7:35813996-35816014 | p4 | (GGAA)4 | AAACAATCCAACTTGGCTGC | GCTGGAAGCAGTAGGTGCTC | 129 |
| chr7:35813996-35816014 | p6 | (TTTTAT)3 | AAACAATCCAACTTGGCTGC | GCTGGAAGCAGTAGGTGCTC | 129 |
| chr7:35895540-35897576 | p3 | (TCT)12 | GAAGGAGATGAAACGGTGGA | GAAGAGCGACTGGACCTGAC | 221 |
| chr7:35904429-35906445 | p2 | (TA)8 | TTTTCCCCGATCTTTTGATG | GATGCATCCATGACATCAGC | 245 |
| chr7:36016705-36018741 | p3 | (TCT)12 | GAAGGAGATGAAACGGTGGA | GAAGAGCGACTGGACCTGAC | 221 |

| Marker ID | SSR type | SSR motif | FORWARD PRIMER (5'-3') | REVERSE PRIMER (5'-3') | size (bp) |
|---|---|---|---|---|---|
| chr7:36025594-36027610 | p2 | (TA)8 | TTTTCCCCGATCTTTTGATG | GATGCATCCATGACATCAGC | 245 |
| chr7:36267577-36269589 | p2 | (AC)6 | CAAACATTGCCGACATTGAC | AAGATGAGGTTTGTCCGGTG | 180 |
| chr7:36286546-36288562 | p2 | (TC)8 | ACATCATCCAAAATTCCCGA | CCGTCCTTCATTTCGTCATT | 136 |
| chr7:36286546-36288562 | p2 | (TA)14 | ACATCATCCAAAATTCCCGA | CCGTCCTTCATTTCGTCATT | 136 |
| chr7:36387890-36389908 | p3 | (TTA)5 | CAACTAGCCTCGAAAATGCC | GAGCTATACCAGCCGATCCA | 251 |
| chr7:36387890-36389908 | p3 | (TTA)7 | CAACTAGCCTCGAAAATGCC | GAGCTATACCAGCCGATCCA | 251 |
| chr7:36558113-36560128 | p5 | (TTAAA)3 | CAGCCAAGAGGGTCATTCAT | CGCAAGTTGTCTGGATCTGA | 128 |
| chr7:36570266-36572317 | p2 | (AT)28 | ACCCGAACCTATCCCAAAAC | GGCCCATTTATACTCGACCA | 164 |
| chr7:36677088-36679103 | p5 | (CAACA)3 | TAATCCCAATCTACGCCTCG | CAAGGGAAATTTGCTACCGA | 210 |
| chr7:36713693-36715711 | p2 | (TA)9 | CCCAAAAATGGCTTCTGAAA | ACATAGTGCGGACGATGACA | 229 |
| chr7:36768357-36770379 | p2 | (AT)11 | AATTGGAGTGACGAAGGCAC | GCATATGGGCATTAGCTGGT | 125 |
| chr7:36944652-36946668 | p2 | (TA)8 | TTACATCGAAGCTTGCCCTT | GGGTACTGGGGAAAGGATGT | 287 |
| chr7:36982237-36984252 | p5 | (AAGAA)3 | ATGGTGAGCCGTTTGAAATC | TGCCACCACATTCACCTTTA | 270 |
| chr7:37069145-37071163 | p6 | (TTCTCT)3 | GGAACATCCATTGAACGCTT | GGCCTTCTAGATTTTTGCCC | 252 |
| chr7:37075662-37077674 | p2 | (TA)6 | TTAACGTTCCACGTGTTCCA | ATGGCAAAGAGCATGGTTTC | 206 |
| chr7:37092223-37094239 | p2 | (TA)42 | CAGCAAAGGATGGTTGAGGT | AATGTCCAAAGCCAAACAGG | 155 |
| chr7:37092223-37094239 | p2 | (TA)8 | CAGCAAAGGATGGTTGAGGT | AATGTCCAAAGCCAAACAGG | 155 |
| chr7:37108501-37110519 | p2 | (TA)9 | ACCGAGCACCACATGACATA | ATGGGCACAGTTTGATGACA | 224 |
| chr7:37441928-37443946 | p6 | (TGAAAC)3 | CCGAAATGTTAGGAGGAGCA | CCAGAAAATGGCTCCAAAAA | 100 |
| chr7:37493579-37495594 | p3 | (ATC)5 | CATCTGTCTATGCGCCTCAA | CCCGCAGGTAATTTTCTCAG | 127 |
| chr7:37496389-37498401 | p2 | (AT)6 | TGGTTGGAGGTATTAAGCGG | TTTAGGCGTTCGCTTTTGTT | 161 |
| chr7:37534833-37536854 | p2 | (GT)8 | AACTCATTCACATCCCTCGG | TGGGGAATATGAGTCGAAGC | 181 |
| chr7:37534872-37536905 | p2 | (GT)8 | AACTCATTCACATCCCTCGG | TGGGGAATATGAGTCGAAGC | 142 |
| chr7:37707343-37709363 | p4 | (TTAA)5 | CTCGCCCCTTTAGCCTATTT | CCCGATTTCCGTTAAATCCT | 157 |
| chr7:37760986-37763004 | p6 | (ATATGT)3 | AACGTGAAATTTGGGAGTCG | GTTTAGGCAGTTTCCCGACA | 127 |
| chr7:37781454-37783469 | p5 | (TTGGT)3 | TTGGCAAGAGTTCAAATCCC | ATGTCGGGTTGGTTTGATTC | 106 |
| chr7:37781454-37783469 | p5 | (TTGGT)3 | TTGGCAAGAGTTCAAATCCC | ATGTCGGGTTGGTTTGATTC | 106 |
| chr7:37781454-37783469 | p5 | (TTGTA)3 | TTGGCAAGAGTTCAAATCCC | ATGTCGGGTTGGTTTGATTC | 106 |
| chr7:37841791-37843812 | p3 | (TAC)7 | GGAAAACAAAAGGGACACGA | TCGCTGGGGGTTCTAATATG | 290 |
| chr7:37885297-37887311 | p2 | (TA)7 | GACTGTCACACCCCGACTTT | GCAATGGGTATTGGGTTGTC | 273 |
| chr7:37885297-37887311 | p6 | (CATATA)3 | GACTGTCACACCCCGACTTT | GCAATGGGTATTGGGTTGTC | 273 |
| chr7:37885426-37887444 | p2 | (TA)7 | GACTGTCACACCCCGACTTT | GCAATGGGTATTGGGTTGTC | 144 |
| chr7:37885426-37887444 | p6 | (CATATA)3 | GACTGTCACACCCCGACTTT | GCAATGGGTATTGGGTTGTC | 144 |
| chr7:37931381-37933397 | p4 | (GAAA)4 | TCCACCATAACTTGGCATCA | CCTCGAGAGTTCTTTGGCAC | 236 |
| chr7:37942022-37944034 | p2 | (TA)6 | AAGTATCAACCGTGCTTCGG | TCACCTTCCCTCGCTTCTAA | 192 |
| chr7:37942022-37944034 | p2 | (AT)6 | AAGTATCAACCGTGCTTCGG | TCACCTTCCCTCGCTTCTAA | 192 |

| Marker ID | SSR type | SSR motif | FORWARD PRIMER (5'-3') | REVERSE PRIMER (5'-3') | size (bp) |
|---|---|---|---|---|---|
| chr7:37960848-37962878 | p6 | (CAACAC)5 | GCCAAAGAATGCCCTGAATA | GCCTGGCCTACGAGTTAGTG | 254 |
| chr7:37986915-37988945 | p6 | (CAACAC)5 | GCCAAAGAATGCCCTGAATA | GCCTGGCCTACGAGTTAGTG | 254 |
| chr7:38111415-38113430 | p5 | (ATTTC)3 | CCCACCTTCTAATGCGGATA | GGTGCCTGACCGAAATTAAA | 129 |
| chr7:38209785-38211803 | p2 | (TA)11 | ACCCAGTGCTCCACTCATTC | CGCCTGAACAGTCCATTACA | 152 |
| chr7:38209785-38211803 | p2 | (TA)11 | ACCCAGTGCTCCACTCATTC | CGCCTGAACAGTCCATTACA | 152 |
| chr7:38209785-38211803 | p5 | (TTTAT)3 | ACCCAGTGCTCCACTCATTC | CGCCTGAACAGTCCATTACA | 152 |
| chr7:38209785-38211803 | p6 | (GTTGTA)3 | ACCCAGTGCTCCACTCATTC | CGCCTGAACAGTCCATTACA | 152 |
| chr7:38214429-38216444 | p2 | (AT)6 | TGTCATGTACGCTGTGAGCA | ATAAGCAGGACACAGACCCG | 190 |
| chr7:38214464-38216479 | p2 | (AT)6 | TGTCATGTACGCTGTGAGCA | ATAAGCAGGACACAGACCCG | 155 |
| chr7:38214509-38216533 | p2 | (AT)6 | TGTCATGTACGCTGTGAGCA | ATAAGCAGGACACAGACCCG | 110 |
| chr7:38214608-38216624 | p2 | (AT)6 | CGGGTCTGTGTCCTGCTTAT | TGCCACATCATGGAGAGTGT | 215 |
| chr7:38231470-38233488 | p2 | (TA)9 | GTGTGAGACCTTTTGGGGAA | TCGACGTTTGACCATGAAAA | 168 |
| chr7:38234185-38236203 | p6 | (TTTTTA)3 | TTGTCGAGTTGACCGAATCA | TTCTTCCGCAAATTAACATGG | 121 |
| chr7:38246543-38248558 | p5 | (TTATT)3 | AATTCGTGCATGAAAGAGGG | CCCTAAAACGACGATCCAAA | 199 |
| chr7:38279073-38281087 | p2 | (TA)8 | TGCAGCAGTTGAGAGCATTT | AATGGAGCTAGGACCGGACT | 294 |
| chr7:38279073-38281087 | p2 | (TA)6 | TGCAGCAGTTGAGAGCATTT | AATGGAGCTAGGACCGGACT | 294 |
| chr7:38279073-38281087 | p2 | (AT)6 | TGCAGCAGTTGAGAGCATTT | AATGGAGCTAGGACCGGACT | 294 |
| chr7:38322440-38324458 | p6 | (CTGCTA)3 | GTTGTTCTTTTGCCGGTGTT | ACTCCTTTGAAGAGCCCCAT | 143 |
| chr7:38322440-38324458 | p2 | (AT)8 | GTTGTTCTTTTGCCGGTGTT | ACTCCTTTGAAGAGCCCCAT | 143 |
| chr7:38323561-38325587 | p2 | (AT)8 | GAACTCCGGAATGACCTTGA | TTTTGGATAACCTTGTCGGC | 192 |
| chr7:38323561-38325587 | p2 | (TA)13 | GAACTCCGGAATGACCTTGA | TTTTGGATAACCTTGTCGGC | 192 |
| chr7:38403901-38405935 | p2 | (AT)17 | ACAAAAGAGCTCCAGCCAAA | TCCTGTCACAACAACCCAAA | 179 |
| chr7:38635471-38637515 | p2 | (AT)22 | CCATTGACCCCATCAGTCTT | CGGGTTTATCACACATGCAA | 197 |
| chr7:38770278-38772293 | p5 | (CCCCG)3 | ATGGGAAAGGGAAAGGAAGA | AAGGACACTCCACGATGGTC | 136 |
| chr7:38775457-38777471 | p2 | (TA)7 | ATGCCACGTTCCAAGAAAAC | GTGCATCCTTGGTTGGAGTT | 241 |
| chr7:38933767-38935783 | p2 | (TA)8 | GAACCCGTCATCGTTCTTGT | TGGTCATTCAAACTCCACCA | 291 |
| chr7:38933957-38935973 | p2 | (TA)8 | GAACCCGTCATCGTTCTTGT | TGGTCATTCAAACTCCACCA | 101 |
| chr7:38933957-38935973 | p6 | (CCGGTT)4 | GAACCCGTCATCGTTCTTGT | TGGTCATTCAAACTCCACCA | 101 |
| chr7:38942806-38944821 | p5 | (AAACC)3 | TCCAAGTCGAAGGTTATGGG | CCAAACGGGAGCTCAGAATA | 109 |
| chr7:38942806-38944821 | p5 | (TCGGT)3 | TCCAAGTCGAAGGTTATGGG | CCAAACGGGAGCTCAGAATA | 109 |
| chr7:38959810-38961822 | p2 | (TA)11 | CATAGAAACCCTTTGCCGAA | GCAATTTGATCATTGCCCTT | 251 |
| chr7:38959810-38961822 | p2 | (TA)15 | CATAGAAACCCTTTGCCGAA | GCAATTTGATCATTGCCCTT | 251 |
| chr7:38971295-38973307 | p2 | (TA)6 | CATGATACCTCAGCGTCCCT | CTCTTGCAAATCAGTGCCAA | 104 |
| chr7:39030834-39032849 | p5 | (AATAA)3 | TCCCATCGACTTCTCATTCC | ACCCTGTAGATCACGAACCG | 294 |
| chr7:39030834-39032849 | p5 | (AATAA)3 | TCCCATCGACTTCTCATTCC | ACCCTGTAGATCACGAACCG | 294 |
| chr7:39032526-39034596 | p2 | (AT)35 | GATTTGACAACACACCACGC | TTGTCCTCGACGATACCTCC | 159 |

| Marker ID | SSR type | SSR motif | FORWARD PRIMER (5'-3') | REVERSE PRIMER (5'-3') | size (bp) |
|---|---|---|---|---|---|
| chr7:39110718-39112733 | p2 | (AT)7 | ACGTTTTCGGATGACCTGAC | GGTTTCACCGACTTCCTCAA | 252 |
| chr7:39110718-39112733 | p3 | (TTG)5 | ACGTTTTCGGATGACCTGAC | GGTTTCACCGACTTCCTCAA | 252 |
| chr7:39232463-39234501 | p2 | (AT)19 | TCCGCCTTAAGAGGCAATTA | GTGTCGTCTCCTTGCCTACC | 153 |
| chr7:39314086-39316130 | p2 | (CA)22 | CATTTAGCTACGGATTGGCG | TAGCTACGGCATTTGTCGGT | 152 |
| chr7:39319676-39321704 | p2 | (AT)14 | CGTTGTTTTACCTTGCCGAT | TCACATGCTCGTCTGTTTGA | 197 |
| chr7:39376677-39378692 | p2 | (GT)6 | GCAAGTGGAAAGGAATTGGA | CAACACACAACACACGACCA | 107 |
| chr7:39376677-39378692 | p3 | (TTG)5 | GCAAGTGGAAAGGAATTGGA | CAACACACAACACACGACCA | 107 |
| chr7:39419539-39421554 | p2 | (TA)9 | CTTGTCGGCCTATGATTCGT | TTTGGTAGGTTTTGGGGTGA | 287 |
| chr7:39419539-39421554 | p5 | (TTTTG)3 | CTTGTCGGCCTATGATTCGT | TTTGGTAGGTTTTGGGGTGA | 287 |
| chr7:39435943-39437958 | p5 | (AAAAT)3 | TCCAATTAGGCATGGTGACA | TCATCAAGTGGCACCTTCAA | 132 |
| chr7:39442636-39444651 | p5 | (AATTG)3 | AAGGTAGGGTGTCGTGATGG | CAAAAGGGGAAAAGCACAAA | 262 |
| chr7:39442636-39444651 | p5 | (CTAAT)3 | AAGGTAGGGTGTCGTGATGG | CAAAAGGGGAAAAGCACAAA | 262 |
| chr7:39496070-39498112 | p6 | (ATCTAT)3 | CGGACCCGACATAAACTTGT | GGGTCCGGAATAACTTTGGT | 130 |
| chr7:39496070-39498112 | p2 | (AT)21 | CGGACCCGACATAAACTTGT | GGGTCCGGAATAACTTTGGT | 130 |
| chr7:39554577-39556592 | p2 | (AT)15 | ATGCCAAAAGTTCAATTCCG | GTGACATTACCAGTGCCACG | 111 |
| chr7:39567656-39569668 | p2 | (AT)6 | CCCCGTCTTTCTTCCTCTCT | GAATACCTGACTGGGGGTGA | 130 |
| chr7:39586958-39588976 | p3 | (AAG)6 | CCAGACCCCTGAATTGAAGA | AGCCCTTGAGAATTCAAGCA | 141 |
| chr7:39654478-39656499 | p3 | (AAG)7 | TTCAACAGCCCAACTAACCC | GGACCTGACAATTCATTGGG | 156 |
| chr7:39654478-39656499 | p5 | (CCACC)3 | TTCAACAGCCCAACTAACCC | GGACCTGACAATTCATTGGG | 156 |
| chr7:39674103-39676115 | p2 | (AT)6 | CTGACTCCTTTGACACGCAA | GGGACTCGGTGAAAGACGTA | 159 |
| chr7:39732737-39734752 | p2 | (TA)9 | GAGGCTTTGCAGACAGTTCC | CCACTTTATCACACCCCACC | 210 |
| chr7:39732737-39734752 | p5 | (TTTAA)3 | GAGGCTTTGCAGACAGTTCC | CCACTTTATCACACCCCACC | 210 |
| chr7:39740812-39742827 | p3 | (AAT)5 | ATCCATTCGTGGAGCCATAG | CAATAGGCTACCGCTGGTGT | 124 |
| chr7:39844413-39846479 | p2 | (AT)33 | CAAAACTTACTTCGGGCACC | TTCGGATCGTCATTTTAGGG | 123 |
| chr7:39844413-39846479 | p2 | (TA)8 | CAAAACTTACTTCGGGCACC | TTCGGATCGTCATTTTAGGG | 123 |
| chr7:39844954-39846970 | p2 | (AT)33 | TCCGTCTCCGTGAAGTGTTA | TTTTCACACATCCTTCGCTG | 153 |
| chr7:39844954-39846970 | p2 | (TA)8 | TCCGTCTCCGTGAAGTGTTA | TTTTCACACATCCTTCGCTG | 153 |
| chr7:39879566-39881581 | p3 | (CAA)5 | CTTAAGGCGGATACCATCCA | CGGACGCGAGATATATGGTT | 124 |
| chr7:39969370-39971424 | p2 | (TA)27 | GGTCCTATAGCCGGTGATGA | TTTCATGGGGTTGGATGAAT | 278 |
| chr7:40018524-40020539 | p5 | (AATGC)3 | AGGTCTGGTCAAATCGCAGT | GCAGAGTTTGCTCCAGGTTC | 113 |
| chr7:40156127-40158142 | p5 | (TGCAA)3 | CCCCATTCACCAGACAGAAT | CGACCTACGAGTTGGGTTGT | 272 |
| chr7:40186029-40188061 | p2 | (TA)16 | TGCGTCACTTTTTACCCCTC | CACGTCCCCCAATAATCATC | 170 |
| chr7:40196925-40198943 | p6 | (ATTTTT)3 | TGGGTTCTTGATCCATTGGT | TGAAGGAATGAGGAGGAGGA | 152 |
| chr7:40387906-40389921 | p5 | (TGAAA)3 | GCTATCGTTCGCTTGTCCTC | GTGAGCGTTTGGGAGTTGAT | 284 |
| chr7:40397600-40399618 | p2 | (TA)9 | AAGCCGATGAGGCTAACAAA | GACGCTACGTCATGCCTACA | 246 |
| chr7:40457145-40459160 | p5 | (ATTTC)3 | GGTTTTTGCTTATGCCTCCA | ATTCTAGCCATCGCCTCTGA | 178 |

| Marker ID | SSR type | SSR motif | FORWARD PRIMER (5'-3') | REVERSE PRIMER (5'-3') | size (bp) |
|---|---|---|---|---|---|
| chr7:40463018-40465033 | p5 | (TGCAT)3 | TAGGGAAAGGGATAGGCGTT | GTGCGCATATCATCATCGAG | 206 |
| chr7:40469558-40471573 | p5 | (ATGCA)3 | TTTACTGTTTCGCCCTTTGG | GAATTAGCGTGGTTGCCACT | 261 |
| chr7:40469558-40471573 | p2 | (TA)7 | TTTACTGTTTCGCCCTTTGG | GAATTAGCGTGGTTGCCACT | 261 |
| chr7:40526908-40528941 | p3 | (AAT)11 | CCTCCTTCGACCCCTATCTC | GAAAGGGACAAATGGGGAAT | 242 |
| chr7:40585073-40587101 | p4 | (TATT)7 | GAGTGTTTGTTGCCCAAGGT | GACGGAATGACTCCACGTCT | 182 |
| chr7:40638354-40640369 | p4 | (CATT)4 | GGCCAAATGTGGTCAGAACT | CACCTAAGCCTTTCCCTTCC | 113 |
| chr7:40638354-40640369 | p5 | (CATTG)3 | GGCCAAATGTGGTCAGAACT | CACCTAAGCCTTTCCCTTCC | 113 |
| chr7:40739134-40741146 | p2 | (TA)6 | CAAAAGTCTCATTCCCCTCG | TTGCCAAGCAATGTTAGCTG | 128 |
| chr7:40791236-40793274 | p2 | (AC)19 | TCCATCAGGAACAAACCACA | GGCTAATGGGTTCCTTGACA | 132 |
| chr7:40810532-40812547 | p5 | (ATGCA)3 | AAGTGCGGTGGATTTAGGTG | GTCAGAGAGGTAAGTGCCCG | 169 |
| chr7:40912646-40914670 | p2 | (TA)12 | TGTTCATGCCCACAGGTAGA | CCTCTCGGCCATCAAAATAA | 288 |
| chr7:40944078-40946093 | p2 | (AG)6 | CCTCACGGGCCTTAAATACA | TATGTCAGTTGGTCCGGTCA | 148 |
| chr7:40944078-40946093 | p5 | (GGGCC)3 | CCTCACGGGCCTTAAATACA | TATGTCAGTTGGTCCGGTCA | 148 |
| chr7:40956740-40958755 | p5 | (AACCG)3 | CGAATGCAGAATTCCAGGTT | TTCGGTTTTTCGATTTTTGG | 221 |
| chr7:40977364-40979388 | p3 | (CAT)8 | TTTCAGATCGTGGTGCTCAG | CAATGAGAGGCAAGTGGTCA | 296 |
| chr7:40977572-40979587 | p3 | (CAT)8 | TGACCACTTGCCTCTCATTG | GCCGTTCTTAGTCGCTTTTG | 269 |
| chr7:40977589-40979655 | p3 | (CAT)8 | TGACCACTTGCCTCTCATTG | GCCGTTCTTAGTCGCTTTTG | 252 |
| chr7:41146446-41148512 | p2 | (AT)33 | TGGCAACCCATGAATCCTAT | GTTGTGGAGGACGGGAGATA | 187 |
| chr7:41147788-41149806 | p6 | (CCCCAC)3 | GCAATGGCAGTTTCATGTTG | TTCCGTTTGAATCAGCAATG | 215 |
| chr7:41212169-41214187 | p3 | (TGG)6 | CATCTTGGGTTCCTCCTTGA | TTGTTTGCAATGGGTGAAAA | 183 |
| chr7:41221929-41223944 | p5 | (AATGC)3 | TGAGACACATTTGGACGGAA | GGTGCCTGCACGACTTATTT | 258 |
| chr7:41270640-41272658 | p3 | (TGG)6 | CATCTTGGGTTCCTCCTTGA | TTGTTTGCAATGGGTGAAAA | 183 |
| chr7:41280400-41282415 | p5 | (AATGC)3 | TGAGACACATTTGGACGGAA | GGTGCCTGCACGACTTATTT | 258 |
| chr7:41463807-41465825 | p2 | (AG)9 | AACTTCGGTGGATGAACCAG | AATTTTGCAGGAACAGTGGG | 212 |
| chr7:41519909-41521987 | p2 | (AT)39 | TGTCGGCATGCGTATTGTAT | CCCGAGCTGAACACTCGTAT | 181 |
| chr7:41767779-41769795 | p6 | (TGTATG)3 | CTTTTTCGTCGTCCGTTTGT | GGGGCCCACAATTTATTTTT | 223 |
| chr7:41767779-41769795 | p2 | (AT)10 | CTTTTTCGTCGTCCGTTTGT | GGGGCCCACAATTTATTTTT | 223 |
| chr7:41767779-41769795 | p2 | (TA)8 | CTTTTTCGTCGTCCGTTTGT | GGGGCCCACAATTTATTTTT | 223 |
| chr7:41767779-41769795 | p2 | (TA)28 | CTTTTTCGTCGTCCGTTTGT | GGGGCCCACAATTTATTTTT | 223 |
| chr7:41777030-41779044 | p2 | (TA)7 | TTCGAACAACTGCAAGATGG | TCTTTGAAAGGACGCAAACC | 145 |
| chr7:41956447-41958473 | p2 | (AT)13 | TAGAGGGCCGGATATGACAC | CATGAAAAGAAGCTGGCACA | 281 |
| chr7:41998715-42000733 | p3 | (TAT)6 | AGAGACGGATGTCGCAGAGT | GGGACCATAGACGGAGCATA | 207 |
| chr7:42049720-42051760 | p2 | (AT)20 | AGTTTGCGCGCTTATTTCAT | TCACACCAACCCTTAGCACA | 265 |
| chr7:42143139-42145181 | p3 | (TAT)14 | AATTCACAACGGAGGTCTCG | CGCTTCTCGACATCATCAGA | 300 |
| chr7:42389033-42391045 | p2 | (AT)6 | AAGGCCAAATTCCATTTCCT | ATTTCTGGGGTGCTGTGTTC | 161 |
| chr7:42395254-42397269 | p5 | (CAATG)3 | GATCCCTTGTCCTTGCCATA | CCGCGACCTTCTATCTTTTG | 114 |

| Marker ID | SSR type | SSR motif | FORWARD PRIMER (5'-3') | REVERSE PRIMER (5'-3') | size (bp) |
|---|---|---|---|---|---|
| chr7:42398800-42400818 | p3 | (AGA)6 | CATCACAGGCACCAACAAAC | ATTTGATTTGCCAGCATTCC | 196 |
| chr7:42412846-42414866 | p5 | (AAATA)3 | TTTGTGTCAAACCCTAGCCC | GGGGACTGATTTTTCGATGA | 110 |
| chr7:42412846-42414866 | p5 | (AAATA)4 | TTTGTGTCAAACCCTAGCCC | GGGGACTGATTTTTCGATGA | 110 |
| chr7:42412846-42414866 | p5 | (AAATA)3 | TTTGTGTCAAACCCTAGCCC | GGGGACTGATTTTTCGATGA | 110 |
| chr7:42412846-42414866 | p4 | (TTTA)4 | TTTGTGTCAAACCCTAGCCC | GGGGACTGATTTTTCGATGA | 110 |
| chr7:42412846-42414866 | p5 | (TTTAT)3 | TTTGTGTCAAACCCTAGCCC | GGGGACTGATTTTTCGATGA | 110 |
| chr7:42568472-42570487 | p5 | (GTGCG)3 | TTCGCTGCCGAAGAAATAGT | GCCAAAGGCATCATTCTCAT | 297 |
| chr7:42590217-42592232 | p5 | (CATTG)3 | ACCTCACCCCACTCTGTTTG | GGCGGTCGTATGTCCACTAT | 202 |
| chr7:42701174-42703189 | p3 | (TGT)5 | TACTGGAAAATGGGCCTCTG | AGCCTAACAAACCGTTGCAC | 272 |
| chr7:42788181-42790196 | p3 | (ACA)5 | ACATGGGCAGAGATGGTAGC | ACTGTCAGCTGCATTGTTCG | 273 |
| chr7:42809531-42811546 | p3 | (ACC)5 | ATTTTATGCCAACTACGCGG | AGCTACCCATTTGATCACCG | 204 |
| chr7:42844132-42846152 | p2 | (TA)10 | GTAACAGTTCGCATTTCGCA | CGAAGGATTTTTCACGGTGT | 263 |
| chr7:42974783-42976801 | p6 | (GGTAGA)3 | CCGGTATGGCAAGAGTTTGT | TGGCCAAGGAGTTAAAATGG | 241 |
| chr7:43101782-43103806 | p2 | (AT)12 | TAATTTCCCGACTCTCGTGG | AGGTGGCGACTCTAACCCTT | 275 |
| chr7:43220505-43222525 | p2 | (AG)10 | AGCGGTATCGCAGAAGTGAT | ACCTCAATGAAGAATTGCCG | 199 |
| chr7:43220505-43222525 | p4 | (TAAT)4 | AGCGGTATCGCAGAAGTGAT | ACCTCAATGAAGAATTGCCG | 199 |
| chr7:43409597-43411655 | p2 | (TA)29 | GTCCAGCAAAAGGGAAATCA | CTTGCTAGCAGGAGGAATCG | 106 |
| chr7:43409597-43411655 | p2 | (TA)25 | GTCCAGCAAAAGGGAAATCA | CTTGCTAGCAGGAGGAATCG | 106 |
| chr7:43409597-43411655 | p3 | (ATA)7 | GTCCAGCAAAAGGGAAATCA | CTTGCTAGCAGGAGGAATCG | 106 |
| chr7:43414996-43417038 | p5 | (CTTGA)3 | TCAAGGGCTTTGCTTGTTCT | TCGTAAAGCCAAAAGGGATG | 197 |
| chr7:43414996-43417038 | p2 | (AT)38 | TCAAGGGCTTTGCTTGTTCT | TCGTAAAGCCAAAAGGGATG | 197 |
| chr7:43415040-43417055 | p5 | (CTTGA)3 | TCAAGGGCTTTGCTTGTTCT | TCGTAAAGCCAAAAGGGATG | 153 |
| chr7:43415040-43417055 | p2 | (AT)38 | TCAAGGGCTTTGCTTGTTCT | TCGTAAAGCCAAAAGGGATG | 153 |
| chr7:43421706-43423730 | p5 | (TTTAA)3 | AAAGCGCGATTGCAATTATC | CCTGCAAGCGAAAAAGTGAT | 274 |
| chr7:43421706-43423730 | p2 | (AT)12 | AAAGCGCGATTGCAATTATC | CCTGCAAGCGAAAAAGTGAT | 274 |
| chr7:43430565-43432577 | p2 | (TA)9 | TATTCGCCCTTGGTCTATGC | CCTCTTTCTCTTTTCGGCAA | 128 |
| chr7:43430565-43432577 | p2 | (AT)6 | TATTCGCCCTTGGTCTATGC | CCTCTTTCTCTTTTCGGCAA | 128 |
| chr7:43430565-43432577 | p5 | (AGAAA)3 | TATTCGCCCTTGGTCTATGC | CCTCTTTCTCTTTTCGGCAA | 128 |
| chr7:43496309-43498345 | p2 | (TA)18 | TTCCGGCAGATTCTAAATGG | CCCCTTCGGGACTTAAGAAG | 209 |
| chr7:43566564-43568576 | p2 | (GT)6 | GAGGTATTGCCCCAGACTCA | CACCACCCCTACCCACTATG | 181 |
| chr7:43773740-43775755 | p5 | (ATGCA)3 | GCGCTAGAGGTTGTTTCAGG | CACCCTCTCCCAAACTTTCA | 157 |
| chr7:43796212-43798245 | p3 | (TCT)11 | CAACGGAAGTGCCTATTGGT | CTACCCTGGGGTTTCGTTTT | 192 |
| chr7:43812042-43814062 | p5 | (AACAA)4 | CCGCAGGGAAATTAACTTGA | CGCCTCCAGTCCACTATCTC | 224 |
| chr7:43827433-43829445 | p2 | (AT)6 | CTCAGCTTATTTTCGGCCAC | AATTCAACCCATCTCACCCA | 174 |
| chr7:43844401-43846423 | p2 | (AT)11 | GGGGTTCGAACATGGATATG | TTGGTTTTCTGGTTGGTGCT | 173 |
| chr7:43986486-43988501 | p3 | (GTG)5 | CGATCAAACAGTTAGGGGGA | CGAGGATTTGAGGTGCTTTC | 192 |

| Marker ID | SSR type | SSR motif | FORWARD PRIMER (5'-3') | REVERSE PRIMER (5'-3') | size (bp) |
|---|---|---|---|---|---|
| chr7:43986486-43988501 | p5 | (ACCCG)3 | CGATCAAACAGTTAGGGGGA | CGAGGATTTGAGGTGCTTTC | 192 |
| chr7:44021914-44023932 | p6 | (TTTATT)3 | TGCAGAGTTCCCTTCGTTTT | GGCGAGGACTTTGGTTATGA | 105 |
| chr7:44084787-44086802 | p5 | (ATGCA)3 | TTTCGGGTCCAATCTTTCAC | CGCTGGTTATTGAAAGCACA | 210 |
| chr7:44169929-44171941 | p2 | (AT)6 | CGGGTAAATTTGCTCATGCT | TTCTCATGCGTTGGAAAGTG | 279 |
| chr7:44190392-44192408 | p2 | (TA)8 | TGTGCTAAAATGGGTGTGGA | GCATTTCCCCTATTTCGTCA | 214 |
| chr7:44240647-44242659 | p2 | (TA)6 | GAACGAAGCTACGTGCAACA | CCGGCAGGAATATGTTGTCT | 133 |
| chr7:44247355-44249397 | p2 | (GA)7 | TTCGGGATTAACTTTGGTGC | GCGGGTAGATTCAATGGAGA | 276 |
| chr7:44247413-44249517 | p2 | (GA)7 | TTCGGGATTAACTTTGGTGC | GCGGGTAGATTCAATGGAGA | 218 |
| chr7:44422208-44424241 | p3 | (CTT)11 | CTAGGTCACGTTGCCCACTT | GATGTGCCTCCAGCCATTAT | 179 |
| chr7:44422208-44424241 | p3 | (TTC)7 | CTAGGTCACGTTGCCCACTT | GATGTGCCTCCAGCCATTAT | 179 |
| chr7:44429867-44431879 | p2 | (CA)6 | TATACGACTTACGCTCGCCC | AGATGATTCTCGGGGGCTAT | 190 |
| chr7:44452745-44454760 | p3 | (TTG)5 | TCTCCTGGAAATTGGAATCG | GTACGTTGTTGGTCGCATTG | 121 |
| chr7:44464555-44466577 | p2 | (TA)11 | TCTGGCCAATTTTTCCTTTG | CGCCTTTTCACCATGAATTT | 186 |
| chr7:44464555-44466577 | p3 | (AAT)5 | TCTGGCCAATTTTTCCTTTG | CGCCTTTTCACCATGAATTT | 186 |
| chr7:44548234-44550252 | p3 | (TCT)30 | TATCGGCATGTCGAATCAAA | GTTCTTCGGGTCCCTTTAGC | 292 |
| chr7:44548234-44550252 | p3 | (CTT)5 | TATCGGCATGTCGAATCAAA | GTTCTTCGGGTCCCTTTAGC | 292 |
| chr7:44548336-44550402 | p3 | (TCT)30 | TATCGGCATGTCGAATCAAA | GTTCTTCGGGTCCCTTTAGC | 190 |
| chr7:44548336-44550402 | p3 | (CTT)5 | TATCGGCATGTCGAATCAAA | GTTCTTCGGGTCCCTTTAGC | 190 |
| chr7:44548441-44550459 | p3 | (TCT)30 | AGGTTCTCAATGACCATGCC | GTTCTTCGGGTCCCTTTAGC | 151 |
| chr7:44548441-44550459 | p3 | (CTT)5 | AGGTTCTCAATGACCATGCC | GTTCTTCGGGTCCCTTTAGC | 151 |
| chr7:44548474-44550546 | p3 | (TCT)30 | AGGTTCTCAATGACCATGCC | GTTCTTCGGGTCCCTTTAGC | 118 |
| chr7:44548474-44550546 | p3 | (CTT)5 | AGGTTCTCAATGACCATGCC | GTTCTTCGGGTCCCTTTAGC | 118 |
| chr7:44549546-44551627 | p3 | (TCT)30 | AGCTGGGCCAGTAATCAGAA | TGGGAAGAGACGAAGGAGAA | 221 |
| chr7:44549546-44551627 | p3 | (CTT)5 | AGCTGGGCCAGTAATCAGAA | TGGGAAGAGACGAAGGAGAA | 221 |
| chr7:44549546-44551627 | p3 | (TCT)27 | AGCTGGGCCAGTAATCAGAA | TGGGAAGAGACGAAGGAGAA | 221 |
| chr7:44549546-44551627 | p3 | (TCT)22 | AGCTGGGCCAGTAATCAGAA | TGGGAAGAGACGAAGGAGAA | 221 |
| chr7:44549546-44551627 | p3 | (CTT)20 | AGCTGGGCCAGTAATCAGAA | TGGGAAGAGACGAAGGAGAA | 221 |
| chr7:44550081-44552147 | p3 | (CTT)5 | TGCCTTACCTCCAACGAAAG | GATGGCTACGACGTCCATTT | 116 |
| chr7:44550081-44552147 | p3 | (TCT)27 | TGCCTTACCTCCAACGAAAG | GATGGCTACGACGTCCATTT | 116 |
| chr7:44550081-44552147 | p3 | (TCT)22 | TGCCTTACCTCCAACGAAAG | GATGGCTACGACGTCCATTT | 116 |
| chr7:44550081-44552147 | p3 | (CTT)20 | TGCCTTACCTCCAACGAAAG | GATGGCTACGACGTCCATTT | 116 |
| chr7:44550081-44552147 | p3 | (TCT)35 | TGCCTTACCTCCAACGAAAG | GATGGCTACGACGTCCATTT | 116 |
| chr7:44571897-44573912 | p3 | (ATA)5 | AGGTCATGGCAATCACTTCC | TCTGTAGACAGCTGGCATGG | 125 |
| chr7:44600676-44602704 | p2 | (TA)14 | CCCAACCAAGATTAAGGCAA | TGGGAGCGTCTTTCTTGAGT | 217 |
| chr7:44600676-44602704 | p2 | (TA)6 | CCCAACCAAGATTAAGGCAA | TGGGAGCGTCTTTCTTGAGT | 217 |
| chr7:44652765-44654831 | p2 | (AT)33 | TCAATAAAACCTTGCGGGTC | CTAATGCTTCTTCCACCCCA | 104 |

| Marker ID | SSR type | SSR motif | FORWARD PRIMER (5'-3') | REVERSE PRIMER (5'-3') | size (bp) |
|---|---|---|---|---|---|
| chr7:44670776-44672790 | p2 | (AT)7 | TAACTTTGCTGCCTCGGTCT | GAGCACTCAAGGGTTAAGCG | 153 |
| chr7:44841686-44843701 | p5 | (TTCAT)3 | AGGCACTTCCCTTTTCCACT | AGGGTCTCGTTTTGTGGATG | 223 |
| chr7:45007076-45009092 | p2 | (AT)8 | TGCTCTTTCAGCCCACTTTT | GGGTCTTGGTTATGGGTGTG | 174 |
| chr7:45048277-45050292 | p3 | (ACA)5 | GTCATTTGTTGATTGCACCG | GTTTCGACCTGATTCCCAAA | 162 |
| chr7:45068131-45070143 | p2 | (TA)8 | CTTTCTGAGGGGCCTTATCC | CCAAGTGGCCAGGCTAAATA | 200 |
| chr7:45068131-45070143 | p2 | (TG)6 | CTTTCTGAGGGGCCTTATCC | CCAAGTGGCCAGGCTAAATA | 200 |
| chr7:45068131-45070143 | p2 | (CA)6 | CTTTCTGAGGGGCCTTATCC | CCAAGTGGCCAGGCTAAATA | 200 |
| chr7:45069009-45071021 | p2 | (TG)6 | ACTCCCCGAGAGTCCAGATT | AAAACCAAGTTTGTCACGCC | 236 |
| chr7:45069009-45071021 | p2 | (CA)6 | ACTCCCCGAGAGTCCAGATT | AAAACCAAGTTTGTCACGCC | 236 |
| chr7:45069009-45071021 | p2 | (AT)6 | ACTCCCCGAGAGTCCAGATT | AAAACCAAGTTTGTCACGCC | 236 |
| chr7:45081052-45083076 | p2 | (TA)12 | CTTTCAGAGGGGCCTTATCC | AGGTATGCGTAACTCCGTGG | 167 |
| chr7:45101594-45103614 | p2 | (CA)10 | GCCTTTCGGCTGTCAAAATA | TTGTGTTCATGCCCACAGTT | 173 |
| chr7:45101594-45103614 | p5 | (TATCA)3 | GCCTTTCGGCTGTCAAAATA | TTGTGTTCATGCCCACAGTT | 173 |
| chr7:45151246-45153261 | p3 | (GAA)5 | CAAGGTGATGTACTGGGCCT | TGTCCCGGTGGTTAATGAAT | 154 |
| chr7:45159623-45161638 | p3 | (GAA)5 | CAAGGTGATGTACTGGGCCT | TGTCCCGGTGGTTAATGAAT | 154 |
| chr7:45175498-45177512 | p2 | (TA)7 | GGACTCATCGCTGAAAGAGC | AAATTAGGGCTTTGAGGCGT | 149 |
| chr7:45299423-45301435 | p2 | (TA)6 | GGCTGTTTGGAATATTGCGT | TTCCCAACGTTAGCAACTCC | 277 |
| chr7:45308068-45310083 | p3 | (TTC)5 | ATGATACGCCACCCTGATGT | AACCATTTTCCACGACTTGC | 255 |
| chr7:45448853-45450865 | p2 | (TA)6 | GCTCTTCATTCGGCTTTGAC | CCTCAGCTCCTGCTCCTATG | 235 |
| chr7:45545878-45547892 | p2 | (TA)7 | AGCGAAGCTTGGACGAATAA | TAGAGATACCCCAAGGTGCG | 185 |
| chr7:45545878-45547892 | p2 | (AT)6 | AGCGAAGCTTGGACGAATAA | TAGAGATACCCCAAGGTGCG | 185 |
| chr7:45584415-45586430 | p5 | (TTCAT)3 | CGGGTTGTACGTCCCATAAC | AATTGAATGCATCTGTGGCA | 209 |
| chr7:45716687-45718705 | p6 | (TATGTA)3 | GCCAGCACGTCGATTATTTT | AGTTCCGAGCGTTCGAAGTA | 223 |
| chr7:45767421-45769433 | p2 | (AT)6 | CGAACCCTGGACACTCGTAT | CAGCCTTGTCGACTTGCATA | 169 |
| chr7:45767421-45769433 | p3 | (ACA)5 | CGAACCCTGGACACTCGTAT | CAGCCTTGTCGACTTGCATA | 169 |
| chr7:45834481-45836507 | p2 | (AT)13 | TATGATGCCAACTCCACGAA | CTGGAGTTTTTCGGGTCAAA | 170 |
| chr7:45857618-45859633 | p5 | (AAAGG)3 | TTGCAGACTTGCGAAACAAC | TTTATCCGAAGCCACATTCC | 125 |
| chr7:45902747-45904762 | p5 | (TGATA)3 | TCCCATTAGCCATCCTCAAC | AAGCACACAAGGGAGCCTTA | 161 |
| chr7:45938270-45940284 | p2 | (CA)6 | CCGAAATCTTCCAACTTCCA | CTGGTGTCGTAATTGTTGCG | 103 |
| chr7:45938270-45940284 | p2 | (AT)7 | CCGAAATCTTCCAACTTCCA | CTGGTGTCGTAATTGTTGCG | 103 |
| chr7:45951209-45953224 | p5 | (GATTG)3 | ACTAGCCACCCCATTTTGTG | CCAACAACGTATCGCATCAC | 178 |
| chr7:46020362-46022378 | p2 | (AT)8 | TTCCCTTCATTTCCCACAAG | TAGGGGATAGGCCCAAGAGT | 226 |
| chr7:46056723-46058753 | p2 | (TA)15 | TGCCCGATTTCCATTAAGTC | CGCTTCGTCACTCCTTTAGC | 266 |
| chr7:46255795-46257810 | p5 | (AAAGG)3 | ATGGTTTTTGAGCAGATGGC | GTGACAGTTGGCCCTCAAAT | 149 |
| chr7:46255795-46257810 | p5 | (CATTG)3 | ATGGTTTTTGAGCAGATGGC | GTGACAGTTGGCCCTCAAAT | 149 |
| chr7:46362231-46364243 | p2 | (TA)17 | TTGTTTTGCGATATGGTGGA | ACATCAATGGGATCATGCCT | 260 |

| Marker ID | SSR type | SSR motif | FORWARD PRIMER (5'-3') | REVERSE PRIMER (5'-3') | size (bp) |
|---|---|---|---|---|---|
| chr7:46362231-46364243 | p2 | (TA)15 | TTGTTTTGCGATATGGTGGA | ACATCAATGGGATCATGCCT | 260 |
| chr7:46362300-46364336 | p2 | (TA)17 | TGTTTTGCGATATGGTGGAA | ACATCAATGGGATCATGCCT | 192 |
| chr7:46362300-46364336 | p2 | (TA)15 | TGTTTTGCGATATGGTGGAA | ACATCAATGGGATCATGCCT | 192 |
| chr7:46362300-46364336 | p4 | (TATG)4 | TGTTTTGCGATATGGTGGAA | ACATCAATGGGATCATGCCT | 192 |
| chr7:46505062-46507077 | p2 | (AT)6 | GGGCCATTTGTGAGTTCCTA | TCCTCAGGAGACAGGTTGCT | 260 |
| chr7:46505062-46507077 | p5 | (ATCAT)3 | GGGCCATTTGTGAGTTCCTA | TCCTCAGGAGACAGGTTGCT | 260 |
| chr7:46511929-46513944 | p5 | (TTTTC)3 | ACATTTGTGGCCATGTAGCA | CATCCCCCACTGTCTCTGTT | 199 |
| chr7:46645380-46647412 | p2 | (AT)16 | TACACCCAAAAAGGGCGTAG | GATGAAGGGGAGGGAATTGT | 195 |
| chr7:46682134-46684149 | p2 | (AT)13 | CCTTAAGCCAAATATGGGCA | ATGAGGACCCACGACCTATG | 292 |
| chr7:46682134-46684149 | p3 | (TTG)5 | CCTTAAGCCAAATATGGGCA | ATGAGGACCCACGACCTATG | 292 |
| chr7:46703645-46705663 | p2 | (TA)9 | TCGGGTCAGGCAAAAATTAG | ACGGCAGGTCAAACATTAGG | 135 |
| chr7:46747195-46749207 | p2 | (TA)6 | GCAAGGAGTGCACAGATCAA | TTCTCCTCAAGATGGTCGCT | 283 |
| chr7:46774613-46776637 | p2 | (TA)19 | ATCCTTTTGTTTGTGGACGC | CTCATATAGGCGGGACAGGA | 235 |
| chr7:46774613-46776637 | p3 | (TGT)8 | ATCCTTTTGTTTGTGGACGC | CTCATATAGGCGGGACAGGA | 235 |
| chr7:46826951-46828966 | p3 | (TCA)5 | TGGAGAGCCCTAGGTTCTGA | TTCTGCAGTCCGTCAAAGTG | 280 |
| chr7:46898800-46900815 | p3 | (TCA)5 | TGGAGAGCCCTAGGTTCTGA | TTCTGCAGTCCGTCAAAGTG | 280 |
| chr7:46907709-46909724 | p3 | (CAG)5 | GGTCTGCTACACCCCCACTA | GCCCTGCTACAAGCGTAGTC | 177 |
| chr7:47042464-47044494 | p2 | (TA)8 | GCACCGCTGGAACTTTTTAG | TCTTCTTCACAAGCTCGGGT | 274 |
| chr7:47042496-47044532 | p2 | (TA)8 | GCACCGCTGGAACTTTTTAG | TCTTCTTCACAAGCTCGGGT | 242 |
| chr7:47069387-47071402 | p3 | (ATG)5 | CCGTCCTCCACAGTCGTATT | AACCGTAAAGTGCCATCGTC | 130 |
| chr7:47190334-47192358 | p6 | (ATATAC)4 | CTGAGCAAGTCGAGAAGGCT | GTGAGCTCCCTTTCTGTTCG | 278 |
| chr7:47219750-47221844 | p2 | (TA)47 | TCCAAACTATGGACGAAGGC | TGAGAGTCACTGATTTGGCG | 213 |
| chr7:47270155-47272167 | p2 | (AT)6 | TATCCCCTGCTACCGATTTG | GGTTGGGTCGTGACAAAAGT | 243 |
| chr7:47283728-47285752 | p2 | (TA)12 | TGTGAAATCGTGGGTTGAAA | TCGTAAATCCGTAGCCCATC | 122 |
| chr7:47283728-47285752 | p2 | (AT)8 | TGTGAAATCGTGGGTTGAAA | TCGTAAATCCGTAGCCCATC | 122 |
| chr7:47383929-47385944 | p3 | (ACA)5 | GAACCGGTGAACGAAGTCAT | TTTGGAACATCCTCCTACGG | 215 |
| chr7:47383929-47385944 | p2 | (AC)6 | GAACCGGTGAACGAAGTCAT | TTTGGAACATCCTCCTACGG | 215 |
| chr7:47400232-47402244 | p2 | (AT)6 | AGCTACATTCTCTTGCCCGA | GGGACCCGATTACCCTAAGA | 199 |
| chr7:47435181-47437202 | p3 | (TTG)7 | GATGAAGGCGACCTCTCTTG | AATCTGCCCATTTCAACCAG | 112 |
| chr7:47437338-47439353 | p4 | (ATAC)4 | GCCGTTAAGGCACGGTAATA | ATGGTCGGGCAAGATAACAC | 263 |
| chr7:47437338-47439353 | p3 | (ATC)5 | GCCGTTAAGGCACGGTAATA | ATGGTCGGGCAAGATAACAC | 263 |
| chr7:47495855-47497869 | p2 | (TA)12 | CGTTGAACTCCCGAATCCTA | TCGCTTCCCTTATCATGTCC | 178 |
| chr7:47495855-47497869 | p2 | (AT)6 | CGTTGAACTCCCGAATCCTA | TCGCTTCCCTTATCATGTCC | 178 |
| chr7:47495855-47497869 | p2 | (TA)6 | CGTTGAACTCCCGAATCCTA | TCGCTTCCCTTATCATGTCC | 178 |
| chr7:47495874-47497888 | p2 | (TA)12 | CGTTGAACTCCCGAATCCTA | TCGCTTCCCTTATCATGTCC | 159 |
| chr7:47495874-47497888 | p2 | (AT)6 | CGTTGAACTCCCGAATCCTA | TCGCTTCCCTTATCATGTCC | 159 |

| Marker ID | SSR type | SSR motif | FORWARD PRIMER (5'-3') | REVERSE PRIMER (5'-3') | size (bp) |
|---|---|---|---|---|---|
| chr7:47495874-47497888 | p2 | (TA)6 | CGTTGAACTCCCGAATCCTA | TCGCTTCCCTTATCATGTCC | 159 |
| chr7:47495896-47497910 | p2 | (TA)12 | CGTTGAACTCCCGAATCCTA | TCGCTTCCCTTATCATGTCC | 137 |
| chr7:47495896-47497910 | p2 | (AT)6 | CGTTGAACTCCCGAATCCTA | TCGCTTCCCTTATCATGTCC | 137 |
| chr7:47495896-47497910 | p2 | (TA)6 | CGTTGAACTCCCGAATCCTA | TCGCTTCCCTTATCATGTCC | 137 |
| chr7:47538646-47540658 | p2 | (AT)6 | GACTTAATGGAAATCGGGCA | GTGTGTTGGTTTGTTCGGTG | 271 |
| chr7:47538646-47540658 | p2 | (AT)7 | GACTTAATGGAAATCGGGCA | GTGTGTTGGTTTGTTCGGTG | 271 |
| chr7:47601888-47603954 | p2 | (AT)33 | TTCAAGGTGAGGCCGAATAC | ATCATCGTATGTCTTCCCGC | 206 |
| chr7:47603206-47605228 | p2 | (TG)11 | CCTTACGTCACCCCGATAGA | CATGGTTTATATACCGCCCG | 208 |
| chr7:47756795-47758810 | p5 | (TTATT)3 | TGACATCGAATTGACCTCCA | ATTTGGGTGATTTTCACGGA | 147 |
| chr7:47807536-47809552 | p6 | (AACCGG)3 | CATGCATGACTCAAACCAGC | TCAGTGCATTTCACTCGACG | 253 |
| chr7:47807536-47809552 | p2 | (TA)8 | CATGCATGACTCAAACCAGC | TCAGTGCATTTCACTCGACG | 253 |
| chr7:47899857-47901885 | p2 | (AT)14 | GAGCCACATGGCCTACTCAT | GTCGGCTTGCGTATATGGAT | 238 |
| chr7:47969896-47971911 | p5 | (GAAAT)3 | TTTGAAGGGAATCATAGCCG | TGTTGACGTTTGGGAGTTGA | 208 |
| chr7:48013448-48015460 | p2 | (TA)6 | GTGGCGTGATCCTATTTGCT | AAATGATGAGCCCACTTTGC | 181 |
| chr7:48019359-48021371 | p2 | (AT)8 | AAATTTGCTGATGAGGCAGG | CAACCGATTGGGAACTGATT | 107 |
| chr7:48070951-48072997 | p2 | (GT)6 | CCCAACCATGGGTAAAAATG | CCTACGACCGCTAATTCCAA | 208 |
| chr7:48154417-48156433 | p4 | (TTTA)4 | CCGACAAGGCTATCCCTACA | CAGAGCTGTGTCCGTCGATA | 175 |
| chr7:48173672-48175690 | p3 | (ATG)6 | TGGTTGGGTTGAGGTAAAGG | GCCAAAATCCCCTTTTTGAT | 175 |
| chr7:48207380-48209394 | p4 | (ATTA)5 | GAAGTCGTCATTCGTCCGAT | GAGTCGATTTGGTTGCGAAT | 290 |
| chr7:48207395-48209415 | p4 | (ATTA)5 | GAAGTCGTCATTCGTCCGAT | GAGTCGATTTGGTTGCGAAT | 275 |
| chr7:48207416-48209428 | p4 | (ATTA)5 | GAAGTCGTCATTCGTCCGAT | GAGTCGATTTGGTTGCGAAT | 254 |
| chr7:48244884-48246899 | p5 | (TTTTA)3 | TTTCTCACTGATCGAGCCCT | GGTTAGGTCATGGGTTGGTG | 216 |
| chr7:48293655-48295669 | p2 | (TG)7 | ACATCAGTAGCGGCAAATCC | TTCGAGTTTTCCGTTGGTTC | 277 |
| chr7:48369181-48371201 | p5 | (ATATA)4 | GCTGAGCACCCCTGTTAGAC | TGTTCGTTGTAGCAGCCTTG | 194 |
| chr7:48369181-48371201 | p5 | (TAATA)3 | GCTGAGCACCCCTGTTAGAC | TGTTCGTTGTAGCAGCCTTG | 194 |
| chr7:48380875-48382891 | p4 | (TAAT)4 | CGCTTTGTCGCTCTTTTAGC | ATATAGGGGAAATGCTGCCC | 214 |
| chr7:48401206-48403252 | p2 | (TC)6 | TTTGACCCGAAAAACTCTGG | ACACTTCTTCTACCCCGCCT | 263 |
| chr7:48401315-48403341 | p2 | (TC)6 | TTTGACCCGAAAAACTCTGG | ACACTTCTTCTACCCCGCCT | 154 |
| chr7:48525127-48527145 | p6 | (ATTTTT)3 | CGCTTCCTATGAGAATTGGG | TGTGCTCTGCATTGAACCTC | 114 |
| chr7:48618515-48620527 | p2 | (AT)6 | TGGTCATGAAGAGATTGCCA | TCTGAAAATTTCTGCCCCAG | 118 |
| chr7:48618515-48620527 | p3 | (CAT)5 | TGGTCATGAAGAGATTGCCA | TCTGAAAATTTCTGCCCCAG | 118 |
| chr7:48619181-48621196 | p2 | (AT)6 | GTGCTAGACATAGGGACGGG | TGACGTCCTTTTTCTTTGGG | 286 |
| chr7:48619181-48621196 | p3 | (CAT)5 | GTGCTAGACATAGGGACGGG | TGACGTCCTTTTTCTTTGGG | 286 |
| chr7:48619181-48621196 | p5 | (AAATG)3 | GTGCTAGACATAGGGACGGG | TGACGTCCTTTTTCTTTGGG | 286 |
| chr7:48641696-48643714 | p2 | (AC)9 | CCTAGTCGTCCACCCTGGTA | CCTTACTTCCCCCTCTTTCG | 260 |
| chr7:48674206-48676220 | p2 | (TA)7 | GTGTGTTGTGTGAACAGCCC | GGTTATGTGTCCGACCCAAC | 184 |

| Marker ID | SSR type | SSR motif | FORWARD PRIMER (5'-3') | REVERSE PRIMER (5'-3') | size (bp) |
|---|---|---|---|---|---|
| chr7:48674206-48676220 | p2 | (AT)7 | GTGTGTTGTGTGAACAGCCC | GGTTATGTGTCCGACCCAAC | 184 |
| chr7:48674206-48676220 | p2 | (TA)18 | GTGTGTTGTGTGAACAGCCC | GGTTATGTGTCCGACCCAAC | 184 |
| chr7:48674786-48676822 | p2 | (TA)7 | CGAGGACGAAGAAGAACCAG | CCATTGATGACCAAGGTTCC | 276 |
| chr7:48674786-48676822 | p2 | (AT)7 | CGAGGACGAAGAAGAACCAG | CCATTGATGACCAAGGTTCC | 276 |
| chr7:48674786-48676822 | p2 | (TA)18 | CGAGGACGAAGAAGAACCAG | CCATTGATGACCAAGGTTCC | 276 |
| chr7:48679261-48681311 | p2 | (TA)25 | GAGTAGGCCGAGGAGCCTAT | TATGGAAGGATCTCCGATGC | 113 |
| chr7:48695118-48697154 | p2 | (TA)7 | CGAGGACGAAGAAGAACCAG | CCATTGATGACCAAGGTTCC | 276 |
| chr7:48695118-48697154 | p2 | (AT)7 | CGAGGACGAAGAAGAACCAG | CCATTGATGACCAAGGTTCC | 276 |
| chr7:48695118-48697154 | p2 | (TA)18 | CGAGGACGAAGAAGAACCAG | CCATTGATGACCAAGGTTCC | 276 |
| chr7:48699593-48701643 | p2 | (TA)25 | GAGTAGGCCGAGGAGCCTAT | TATGGAAGGATCTCCGATGC | 113 |
| chr7:48734359-48736374 | p5 | (TCATA)3 | AAATTGTGGGAGACGAATGC | GATTCATGCACTCCTCCGAT | 276 |
| chr7:48773436-48775451 | p5 | (TCATA)3 | GTGGCTGATGACAGACGAGA | ACTTTCCGGACCGAAGTTCT | 234 |
| chr7:48801477-48803495 | p3 | (TGT)6 | AGAGGCTCGCATCTTGTGTT | CAACATTCTCATCTCGGCAA | 178 |
| chr7:48826646-48828712 | p2 | (AT)33 | CCAAACATCCATTGCTCCTT | AGAAGTTCCTCACCGAAGCA | 220 |
| chr7:48877200-48879220 | p2 | (AT)6 | GATGAGCAGATCTCCAAGCC | CTCTTCCAAGGACAAACCCA | 204 |
| chr7:48877224-48879238 | p2 | (AT)6 | GATGAGCAGATCTCCAAGCC | CTCTTCCAAGGACAAACCCA | 180 |
| chr7:48954302-48956314 | p2 | (AG)6 | ACACCTGCTGGAAAATTTGG | AGAATCACGAAAAACACCGC | 124 |
| chr7:48988124-48990140 | p4 | (AAAT)4 | GCTTGGCAATCCGTCTTTAG | GCAAGACAGCAAGACCACAA | 273 |
| chr7:49095042-49097076 | p2 | (AT)17 | AAACCAAAGCTCCAAACCCT | CAACTTGTGCCAAAAACCCT | 234 |
| chr7:49095042-49097076 | p4 | (TATG)4 | AAACCAAAGCTCCAAACCCT | CAACTTGTGCCAAAAACCCT | 234 |
| chr7:49116214-49118234 | p2 | (AT)10 | GTGTGAGGCACCCCAGTTAT | GAAATCCAAAAATGGCCAAA | 248 |
| chr7:49156534-49158549 | p5 | (GCTAT)3 | TATGATGCCCCGATTTCATT | TCGTGATCGCTTTGTCTCAC | 279 |
| chr7:49156534-49158549 | p2 | (AT)7 | TATGATGCCCCGATTTCATT | TCGTGATCGCTTTGTCTCAC | 279 |
| chr7:49217413-49219428 | p3 | (ATC)5 | CTTGTTTTGGTCGTGTGTGG | TGGAATACTTGGTGCCCTTC | 124 |
| chr7:49243828-49245846 | p6 | (TTTCTT)3 | CCAAAGCAGTCGAATCACAA | TACCGCCATAATGGGATGAT | 169 |
| chr7:49279608-49281628 | p2 | (AT)10 | TCGACGTGGAAAATAAAGGG | ACATGTGACACACGCCAAAT | 127 |
| chr7:49279608-49281628 | p5 | (TGAAT)3 | TCGACGTGGAAAATAAAGGG | ACATGTGACACACGCCAAAT | 127 |
| chr7:49319332-49321350 | p6 | (ACAATT)3 | AGGGGTTGAAAAGTGTCACG | TACTCGTCCGGGCAAATAAG | 205 |
| chr7:49341515-49343527 | p2 | (AG)6 | AAATGGTTTGGCGTTTCTTG | CCACAAAAAGGCAAGAGGAG | 270 |
| chr7:49372424-49374436 | p2 | (CT)6 | TTCTTGCTGCCCTCTCAAAT | ACTTCCACTCAGGGCTACGA | 173 |
| chr7:49391206-49393220 | p2 | (TC)7 | TAATCCCAACTGGTGCTTCC | AGCTTGGACTTCCGTTCGTA | 120 |
| chr7:49395612-49397630 | p6 | (CATATA)3 | ATGTGTCACACCCCAACCTT | GTCTTGAAAGCGGCTTCATC | 216 |
| chr7:49395612-49397630 | p5 | (ATTCA)3 | ATGTGTCACACCCCAACCTT | GTCTTGAAAGCGGCTTCATC | 216 |
| chr7:49458410-49460424 | p2 | (TA)7 | GGGGGATTTGTCCAGAATTT | AGCAGTTCCGTCCTAGGGTT | 154 |
| chr7:49458410-49460424 | p2 | (TA)7 | GGGGGATTTGTCCAGAATTT | AGCAGTTCCGTCCTAGGGTT | 154 |
| chr7:49458410-49460424 | p2 | (TA)19 | GGGGGATTTGTCCAGAATTT | AGCAGTTCCGTCCTAGGGTT | 154 |

| Marker ID | SSR type | SSR motif | FORWARD PRIMER (5'-3') | REVERSE PRIMER (5'-3') | size (bp) |
|---|---|---|---|---|---|
| chr7:49517787-49519809 | p6 | (CCCCAC)3 | TTATGGACCACCTCCACCAT | AAAAACCACCAAATCCGTGA | 269 |
| chr7:49517787-49519809 | p5 | (TACAA)3 | TTATGGACCACCTCCACCAT | AAAAACCACCAAATCCGTGA | 269 |
| chr7:49517810-49519842 | p6 | (CCCCAC)3 | TTATGGACCACCTCCACCAT | AAAAACCACCAAATCCGTGA | 246 |
| chr7:49517810-49519842 | p5 | (TACAA)3 | TTATGGACCACCTCCACCAT | AAAAACCACCAAATCCGTGA | 246 |
| chr7:49571904-49573922 | p6 | (CATATA)3 | AGCTACCGAGCACCCCTTAT | GTCCCGTCCTGATGTTCAGT | 290 |
| chr7:49571904-49573922 | p3 | (ATC)7 | AGCTACCGAGCACCCCTTAT | GTCCCGTCCTGATGTTCAGT | 290 |
| chr7:49571904-49573922 | p3 | (CAA)6 | AGCTACCGAGCACCCCTTAT | GTCCCGTCCTGATGTTCAGT | 290 |
| chr7:49574446-49576461 | p5 | (TTGGT)3 | GGGCTTTCATCAAGCAATGT | CGGGTTACGGATGAAGGTTA | 101 |
| chr7:49574446-49576461 | p5 | (TTGGT)3 | GGGCTTTCATCAAGCAATGT | CGGGTTACGGATGAAGGTTA | 101 |
| chr7:49658608-49660620 | p2 | (AT)6 | AGCCTTATATGGGCGGAGAT | CCACTATACCGAGGGCTCAA | 117 |
| chr7:49769060-49771081 | p2 | (AG)6 | AAAAGAAAGAGGGCAGGAGC | TTCCTTTGTTCCCTTTCCCT | 278 |
| chr7:49769087-49771114 | p2 | (AG)6 | AAAAGAAAGAGGGCAGGAGC | TTCCTTTGTTCCCTTTCCCT | 251 |
| chr7:49795354-49797369 | p5 | (GAAAT)3 | CGATCGTAGCGTATCACGAA | CTAGCATTGGAGGAGAACGC | 152 |
| chr7:49968321-49970339 | p3 | (TTG)6 | TCGTGTCATGTGCGTTATGA | CCTCCCAAGTTGATCCAAAA | 212 |
| chr7:49977072-49979088 | p2 | (TA)8 | TATATGGGTAGGGAAGGGGC | TAAACCGGGATACAAACCGA | 232 |
| chr7:50080297-50082315 | p3 | (ACA)6 | TTGCTTCTCCCTCTCTGCAT | TTTACCTCATTCGGTCGGTC | 258 |
| chr7:50228334-50230349 | p5 | (GAAAT)3 | GAGAATTGGCCAAAGAACCA | ACGCTAGCATCGGAGGAGTA | 227 |
| chr7:50242090-50244154 | p2 | (AT)32 | GTTTTGCCTTGTGTGGGAAT | AGGCCCAATCACAAGATCAC | 113 |
| chr7:50269046-50271062 | p2 | (AT)8 | ATACGGGGCGTATTCTTTCC | ATACGTCCACAAGGTACGCC | 128 |
| chr7:50318765-50320780 | p5 | (GAAAT)3 | GAGAATTGGCCAAAGAACCA | ACGCTAGCATCGGAGGAGTA | 227 |
| chr7:50320471-50322483 | p2 | (AT)6 | ATAATTTTCCCCTGATGGGC | ATATGACGGTTCGTGAAGGG | 257 |
| chr7:50353728-50355744 | p4 | (AATT)4 | ACAGCGGAGTAAAGCAAGGA | TTGTAACACGCAGACATCCC | 167 |
| chr7:50353788-50355848 | p4 | (AATT)4 | ACAGCGGAGTAAAGCAAGGA | TTGTAACACGCAGACATCCC | 107 |
| chr7:50364949-50367015 | p2 | (AT)8 | TCTCGACCTTATGTCACCCC | TCATCACTACCATCCTCGCA | 197 |
| chr7:50364949-50367015 | p2 | (AT)33 | TCTCGACCTTATGTCACCCC | TCATCACTACCATCCTCGCA | 197 |
| chr7:50364949-50367015 | p2 | (TA)6 | TCTCGACCTTATGTCACCCC | TCATCACTACCATCCTCGCA | 197 |
| chr7:50415450-50417462 | p2 | (AT)6 | TCCCCATTTGATGTTTCCAT | ATGCCATCCCACCAATACAT | 293 |
| chr7:50415450-50417462 | p3 | (GTT)6 | TCCCCATTTGATGTTTCCAT | ATGCCATCCCACCAATACAT | 293 |
| chr7:50442868-50444880 | p2 | (AT)6 | TCCCCATTTGATGTTTCCAT | ATGCCATCCCACCAATACAT | 293 |
| chr7:50442868-50444880 | p3 | (GTT)6 | TCCCCATTTGATGTTTCCAT | ATGCCATCCCACCAATACAT | 293 |
| chr7:50524179-50526191 | p2 | (TA)6 | ATGGTGATGCTAAAGGTGCC | TCATACCTGGCCCTACCAAG | 192 |
| chr7:50524179-50526191 | p2 | (TG)6 | ATGGTGATGCTAAAGGTGCC | TCATACCTGGCCCTACCAAG | 192 |
| chr7:50538436-50540448 | p2 | (TA)6 | ATGGTGATGCTAAAGGTGCC | TCATACCTGGCCCTACCAAG | 192 |
| chr7:50538436-50540448 | p2 | (TG)6 | ATGGTGATGCTAAAGGTGCC | TCATACCTGGCCCTACCAAG | 192 |
| chr7:50577868-50579883 | p5 | (AAATG)3 | GGGTCGTGACAGAAGGGTTA | TGAAATAGGGGAAATGCTGC | 242 |
| chr7:50599218-50601246 | p2 | (TA)14 | GTGCATACCACGTCTGTTGG | TTTCCACTCGAATTCATCCC | 277 |

| Marker ID | SSR type | SSR motif | FORWARD PRIMER (5'-3') | REVERSE PRIMER (5'-3') | size (bp) |
|---|---|---|---|---|---|
| chr7:50599218-50601246 | p3 | (CAA)6 | GTGCATACCACGTCTGTTGG | TTTCCACTCGAATTCATCCC | 277 |
| chr7:50599352-50601370 | p2 | (TA)14 | GTGCATACCACGTCTGTTGG | TTTCCACTCGAATTCATCCC | 143 |
| chr7:50599352-50601370 | p3 | (CAA)6 | GTGCATACCACGTCTGTTGG | TTTCCACTCGAATTCATCCC | 143 |
| chr7:50654178-50656193 | p5 | (CAAAA)3 | GTGTCGCCTATGAAGGGTGT | GCGACCTCGTCTATCTTTCG | 107 |
| chr7:50654178-50656193 | p2 | (GT)9 | GTGTCGCCTATGAAGGGTGT | GCGACCTCGTCTATCTTTCG | 107 |
| chr7:50654178-50656193 | p3 | (GTT)5 | GTGTCGCCTATGAAGGGTGT | GCGACCTCGTCTATCTTTCG | 107 |
| chr7:50710609-50712621 | p2 | (AT)6 | TGGAGTGGCATTTTCTACCC | AAAGCAAAATATGGACCCCC | 195 |
| chr7:50777854-50779878 | p2 | (AT)12 | GCTGGCATTTGTCTGCAATA | AATGCTCATGAAGAATGCCC | 156 |
| chr7:50785932-50787948 | p2 | (AT)8 | TCAAGAAATCACGGACCCTC | GGCTATGAGAGACGCAGTCC | 273 |
| chr7:50803930-50805944 | p2 | (TA)7 | ATGGATCAAACGGGTCCATA | TCCAGGCCAAATGAGTTTTC | 118 |
| chr7:50850970-50853022 | p2 | (AT)26 | TACACAGCCACATCCCGTAA | CTAAAACAACGGAGGACCCA | 185 |
| chr7:50858748-50860760 | p2 | (AG)6 | GAAGAGTAACACCCCCAGCA | GCAAGTTCCGGCATAAACAT | 124 |
| chr7:50858748-50860760 | p2 | (GT)6 | GAAGAGTAACACCCCCAGCA | GCAAGTTCCGGCATAAACAT | 124 |
| chr7:50930599-50932611 | p2 | (TA)6 | CAGCCACACAGACTCGAAAA | GAACCCCAATCCTCTCACAA | 181 |
| chr7:50941020-50943035 | p5 | (TATAT)3 | ATTAGGGGCTGTTTTGGGAC | ATCAACCAAACGACTACGGC | 295 |
| chr7:51010102-51012122 | p2 | (TA)44 | GCTATGAGCCCAACAACGAT | GGCCTTATAGGCACGGTGTA | 121 |
| chr7:51010102-51012122 | p2 | (TA)10 | GCTATGAGCCCAACAACGAT | GGCCTTATAGGCACGGTGTA | 121 |
| chr7:51049073-51051103 | p2 | (TA)15 | TAATCAAGGTCCGGATAGCG | TTCAATCCACAAGCCAACAA | 232 |
| chr7:51096530-51098548 | p6 | (TGTAAG)3 | TGAGGCCGGGTATGATATGT | TGCAGGATCGGTGTTTCATA | 123 |
| chr7:51180579-51182594 | p3 | (CAA)5 | CACATCTACGTGCCATGACC | CGACCTTACGTCACTCCGAT | 254 |
| chr7:51209573-51211591 | p6 | (TGTGTA)3 | TATTTGTGTTGGCCCTGTGA | GCTACAGACAACCCACGGAT | 230 |
| chr7:51305593-51307607 | p3 | (AAC)5 | GATTAAGCTGGTGGCCCATA | ATATGTAGTGGCCCAGTCGG | 154 |
| chr7:51471148-51473188 | p2 | (TG)20 | GCTGCCAAGAGTGTCATTCA | TTTTTGGGGCGAATACTACG | 296 |
| chr7:51528993-51531029 | p2 | (AT)18 | TTTACCGTTGGCTCCTGTTC | GCGGGGCATTTAGGAATAAT | 116 |
| chr7:51566880-51568898 | p2 | (TA)6 | TAGCCCTTTTCTCTTTCCCC | CTCCGAAGCAGTCTTTTTGG | 297 |
| chr7:51566880-51568898 | p3 | (CAA)6 | TAGCCCTTTTCTCTTTCCCC | CTCCGAAGCAGTCTTTTTGG | 297 |
| chr7:51632051-51634063 | p2 | (AT)6 | TCTTTGCCCACTTTATTCCG | CAACGTCGTACCCAACTCCT | 252 |
| chr7:51633937-51635949 | p2 | (AT)6 | CTAGGGCATGCCAAAAGAAG | GTTGATTGTTTTGGCCGAGT | 112 |
| chr7:51659253-51661268 | p5 | (ATATA)3 | CAAAAGTCCTCGACGTCACA | ATATAATGGCAGCCTCGTCG | 155 |
| chr7:51680579-51682594 | p5 | (AACCG)4 | GACGTCATGAACAACATCGG | TTTGGTATCTATGCCCAGCC | 275 |
| chr7:51680579-51682594 | p5 | (AACCG)3 | GACGTCATGAACAACATCGG | TTTGGTATCTATGCCCAGCC | 275 |
| chr7:51792545-51794563 | p2 | (TA)6 | GTCGATCGATGATGCTGCTA | ATGGATCAACCGATGGCTTA | 168 |
| chr7:51792564-51794578 | p2 | (TA)6 | GTCGATCGATGATGCTGCTA | ATGGATCAACCGATGGCTTA | 149 |
| chr7:51805570-51807590 | p2 | (TA)7 | CAAGGCTGCCATTACAGACA | TCTTGGAGAGCTCCGTTGTT | 198 |
| chr7:51805570-51807590 | p2 | (AT)10 | CAAGGCTGCCATTACAGACA | TCTTGGAGAGCTCCGTTGTT | 198 |
| chr7:51805570-51807590 | p2 | (AT)10 | CAAGGCTGCCATTACAGACA | TCTTGGAGAGCTCCGTTGTT | 198 |

| Marker ID | SSR type | SSR motif | FORWARD PRIMER (5'-3') | REVERSE PRIMER (5'-3') | size (bp) |
|---|---|---|---|---|---|
| chr7:51818903-51820923 | p2 | (TA)10 | TCTGTGGGGATGAGGAAATC | ATCACCTAACCACTCGCACC | 113 |
| chr7:51861224-51863236 | p2 | (AT)6 | GAAGCAAAAACGCGAAGAAC | CCAACCCATAAGCCAACAAC | 287 |
| chr7:51906647-51908663 | p2 | (AT)8 | GTCTGTGTGGCTTGCGTCTA | GGCATACAAATGGCAAATCC | 158 |
| chr7:52056626-52058656 | p3 | (TAT)7 | TTTTGTCCATCCTTTCGTCC | ATGACAGAAGTTTGGACCCG | 227 |
| chr7:52056626-52058656 | p5 | (AAGAG)3 | TTTTGTCCATCCTTTCGTCC | ATGACAGAAGTTTGGACCCG | 227 |
| chr7:52056695-52058710 | p3 | (TAT)7 | TTTTGTCCATCCTTTCGTCC | ATGACAGAAGTTTGGACCCG | 158 |
| chr7:52056695-52058710 | p5 | (AAGAG)3 | TTTTGTCCATCCTTTCGTCC | ATGACAGAAGTTTGGACCCG | 158 |
| chr7:52083751-52085773 | p2 | (AT)11 | CCCAATCCTATTTTCCCGTT | CTGCCTACCGTCTACCTTCG | 277 |
| chr7:52107043-52109058 | p3 | (TGT)5 | TAAATGTTCCGGGATCAAGC | TACCCCTCAAAATTGACCCA | 188 |
| chr7:52107043-52109058 | p2 | (TA)7 | TAAATGTTCCGGGATCAAGC | TACCCCTCAAAATTGACCCA | 188 |
| chr7:52122848-52124866 | p6 | (CTGAAA)3 | CAAAAATGACCGCACATGAC | TCTGGAGGAAATGACTTGGC | 117 |
| chr7:52502948-52504966 | p6 | (TAAAAA)3 | GTCTCCTTGTCTACCTGCGG | CCCTGTCTACCTGTGGCATT | 143 |
| chr7:52616941-52618957 | p2 | (AT)8 | GCTTTGTTTCTTTTCCGTGC | TGCTTTTGCTTGTCACCTTG | 284 |
| chr7:52616941-52618957 | p2 | (AT)9 | GCTTTGTTTCTTTTCCGTGC | TGCTTTTGCTTGTCACCTTG | 284 |
| chr7:52617087-52619105 | p2 | (AT)8 | GCTTTGTTTCTTTTCCGTGC | TGCTTTTGCTTGTCACCTTG | 138 |
| chr7:52617087-52619105 | p2 | (AT)9 | GCTTTGTTTCTTTTCCGTGC | TGCTTTTGCTTGTCACCTTG | 138 |
| chr7:52749733-52751751 | p6 | (CTTACA)3 | ACGGTCCGTAGATTGGACTG | AACCCTCCGAGATAAGGGAA | 241 |
| chr7:52875619-52877685 | p2 | (TA)9 | CTCGTGCATGCCTCAATATG | ACTGTACCCGGCCCTCTAGT | 140 |
| chr7:52875619-52877685 | p2 | (AT)6 | CTCGTGCATGCCTCAATATG | ACTGTACCCGGCCCTCTAGT | 140 |
| chr7:52875619-52877685 | p2 | (AT)33 | CTCGTGCATGCCTCAATATG | ACTGTACCCGGCCCTCTAGT | 140 |
| chr7:52878161-52880179 | p6 | (GATTCT)3 | CGGAAATTTTGGTAAGGCAA | TTTCAGCCGAGCCTATCACT | 166 |
| chr7:52878161-52880179 | p2 | (CG)6 | CGGAAATTTTGGTAAGGCAA | TTTCAGCCGAGCCTATCACT | 166 |
| chr7:52878739-52880751 | p6 | (GATTCT)3 | GCCAGGATGGCACATTATCT | AGAAGCAGAAGCCATACGGA | 150 |
| chr7:52878739-52880751 | p2 | (CG)6 | GCCAGGATGGCACATTATCT | AGAAGCAGAAGCCATACGGA | 150 |
| chr7:52878739-52880751 | p3 | (GAT)5 | GCCAGGATGGCACATTATCT | AGAAGCAGAAGCCATACGGA | 150 |
| chr7:52879668-52881683 | p2 | (CG)6 | GTTGGGTCCGTGACAGAAGT | TTTGAAAGCCTCGCTGAGAT | 205 |
| chr7:52879668-52881683 | p3 | (GAT)5 | GTTGGGTCCGTGACAGAAGT | TTTGAAAGCCTCGCTGAGAT | 205 |
| chr7:52902533-52904548 | p5 | (AATAA)3 | CCGAGCTTTGGTTATGGAAG | ATGAGGCCTACCCAATGCTA | 118 |
| chr7:52932910-52934925 | p5 | (ATTGC)3 | CCCTTGGTGAGACTTCGTGT | CTAGCTCCAATCACGCAACA | 164 |
| chr7:52968098-52970113 | p3 | (AAC)5 | CCCCAGGGATGTACACAAAC | CTGCTATGCCCTCTCCAGAC | 263 |
| chr7:52998335-53000353 | p6 | (ATATGT)3 | GAGGGCACTTCTTTAGTGCG | GTATTGGTGACCCATCCAGG | 252 |
| chr7:53024129-53026147 | p3 | (CAA)6 | ATCCCACGTCACACTCACAA | ACCTATGATATCGTTCGCCG | 131 |
| chr7:53121296-53123311 | p3 | (ATC)5 | GCTCTCTTTCTACCCGCCTT | GCGACTACAGGGCATTTAGG | 144 |
| chr7:53154126-53156192 | p3 | (ATC)6 | TGAAAGGGACACATGCCATA | CACTCCACTTGTTCCGGAGT | 274 |
| chr7:53182806-53184824 | p6 | (ACAATA)3 | CCCTCGACCCTACTTTGTGA | CCTAAGCTGAGGAGGCCTTT | 300 |
| chr7:53224619-53226639 | p2 | (AT)10 | AAGACGATTAGCAGCCCAGA | TTTTACCCTTCACCTCCGTG | 294 |

| Marker ID | SSR type | SSR motif | FORWARD PRIMER (5'-3') | REVERSE PRIMER (5'-3') | size (bp) |
|---|---|---|---|---|---|
| chr7:53241975-53243990 | p5 | (AATGC)3 | TGAGTCCCCAAACCTTCTTG | ATATGGATTTCATCGCAGGC | 107 |
| chr7:53288105-53290120 | p5 | (TTTCA)3 | ACCCTAGCGAGTGAACAGGA | GCAACTCCCAAAAACAGGAA | 128 |
| chr7:53358173-53360221 | p2 | (TA)24 | CCTAGGACGGAACTGCTCTG | ATGTCATCCTTTCGTGAGCC | 115 |
| chr7:53358173-53360221 | p3 | (ATC)5 | CCTAGGACGGAACTGCTCTG | ATGTCATCCTTTCGTGAGCC | 115 |
| chr7:53471027-53473039 | p2 | (TC)10 | AAATTTCATCCCCCAAGACC | AAAATTCACTCGAGGGCTGA | 196 |
| chr7:53471027-53473039 | p2 | (AT)6 | AAATTTCATCCCCCAAGACC | AAAATTCACTCGAGGGCTGA | 196 |
| chr7:53556730-53558745 | p3 | (TTG)5 | AGTTTCCAACGCAACAAACC | TAGAAGTGTGGGGATGAGGG | 232 |
| chr7:53590445-53592459 | p2 | (TA)7 | TAGCCCAGGTCGATGATTTC | CTTGCATTTTGGGGTCTGTT | 287 |
| chr7:53623012-53625028 | p4 | (TTAT)4 | AGCAGGGGAAAAATAGGGAA | TAAATACCGCCCATTGAAGC | 118 |
| chr7:53706705-53708725 | p2 | (TC)10 | GGTCCGACAATGGATTTCAC | CCAACATAAACTCCAACCCG | 160 |
| chr7:53792988-53795003 | p5 | (AAAAT)3 | CCCTATCGACCCTCAAATCA | CCGGACTATGAATCGGAGAA | 110 |
| chr7:53792988-53795003 | p4 | (ATTA)6 | CCCTATCGACCCTCAAATCA | CCGGACTATGAATCGGAGAA | 110 |
| chr7:53792988-53795003 | p3 | (TTC)17 | CCCTATCGACCCTCAAATCA | CCGGACTATGAATCGGAGAA | 110 |
| chr7:53895066-53897080 | p2 | (TA)7 | GCCATCCTATCCATTTGAGC | CATGATTCACCCCTACCCAG | 280 |
| chr7:53975149-53977195 | p2 | (TA)23 | GTGGTTATCCATCCCAATCG | TGCAAAAGGAAAGGAAGCAT | 125 |
| chr7:53975149-53977195 | p2 | (AT)6 | GTGGTTATCCATCCCAATCG | TGCAAAAGGAAAGGAAGCAT | 125 |
| chr7:54077875-54079890 | p3 | (TTG)5 | TCAATGGGTCAAAAGCAACA | TGTGCCCGTGAATGTAGGTA | 270 |
| chr7:54114611-54116623 | p2 | (AG)6 | CCGGTATGGCAAGAGTTTGT | TAGCCAAGGAATTGAAACGG | 230 |
| chr7:54148267-54150281 | p2 | (AG)7 | ATAACGTTGTAGCCGATGCC | CCTCATATTGGCGCTCTTTC | 131 |
| chr7:54148267-54150281 | p3 | (TTG)5 | ATAACGTTGTAGCCGATGCC | CCTCATATTGGCGCTCTTTC | 131 |
| chr7:54179830-54181856 | p2 | (AT)13 | CAGGGACTACATAGCGGAGC | TCATATAGGCGGGACAGGAC | 292 |
| chr7:54212074-54214088 | p2 | (TA)7 | AATGTCAATGAATGCCCCAT | GGTGTATATGCTGCCTGCCT | 240 |
| chr7:54420823-54422838 | p3 | (AAT)5 | TGGATTTGGAAGCTATTCGG | AGAATGTGACCGACCCTCTG | 248 |
| chr7:54420823-54422838 | p2 | (TA)42 | TGGATTTGGAAGCTATTCGG | AGAATGTGACCGACCCTCTG | 248 |
| chr7:54464028-54466050 | p2 | (CT)11 | CCTTCACGCAAAACTCAACA | CAGGCCATTGTGAAGGAACT | 215 |
| chr7:54554980-54556996 | p4 | (TCTT)4 | CGTATACTTGCGTGTGCGTT | GCCATTGAGTGGGTGAGAAT | 286 |
| chr7:54554980-54556996 | p2 | (TC)7 | CGTATACTTGCGTGTGCGTT | GCCATTGAGTGGGTGAGAAT | 286 |
| chr7:54618512-54620536 | p2 | (AT)12 | GTGGTTTTGGGAGTTGTGCT | ACCTCGCCGTCTCCTTAAAT | 267 |
| chr7:54777694-54779706 | p2 | (AT)6 | CTCCAAACCCGAGAACTTCA | GCATTTTAGTTGCCGTCGTT | 285 |
| chr7:54787125-54789140 | p3 | (TTG)5 | GAGGCTTGTTTCAGGAGTGC | TCCCTCATCTTGTCCTCCAC | 194 |
| chr7:54818352-54820364 | p2 | (AT)6 | GACGCCGTATTACCTTTCCA | GTAAGGGGTGCTCGGTACAA | 269 |
| chr7:54841653-54843709 | p2 | (TA)28 | TCAGTAAGGCCCATGAAAGG | CATAGGTCACACCTCCACCC | 175 |
| chr7:54877901-54879913 | p2 | (AT)6 | GGCTGCTCAAGTAAAGACCG | CACTATTGGTGCCGAACCTT | 263 |
| chr7:55008163-55010181 | p6 | (GTGCTG)3 | TGCTCGCTACTGTTGATTGG | AGGCCCATTTCCTTCGTAGT | 268 |
| chr7:55015725-55017741 | p4 | (ATTT)4 | TTTAAAAGTGGGGCATTTCG | ACATTTAATGCCTTTTGCCG | 107 |
| chr7:55133307-55135322 | p5 | (ATTGC)3 | CTATTGTGTTGGCATGACGC | AACAATCGGGTACCCAACAA | 291 |

| Marker ID | SSR type | SSR motif | FORWARD PRIMER (5'-3') | REVERSE PRIMER (5'-3') | size (bp) |
|---|---|---|---|---|---|
| chr7:55210730-55212766 | p5 | (GATAT)3 | GCCATAATAGAATCGGGGGT | ATGTCTGGCCATCTAGGCAC | 242 |
| chr7:55210730-55212766 | p2 | (TA)18 | GCCATAATAGAATCGGGGGT | ATGTCTGGCCATCTAGGCAC | 242 |
| chr7:55236828-55238843 | p5 | (ATATA)3 | GCCGTACCTCTGGGTAAACA | GTTTGGTGACCCTTCAGCAT | 263 |
| chr7:55243398-55245410 | p5 | (ATATC)3 | TCCTTTAACGATCCGTCCTG | GCTGAAGAACGACATCGACA | 209 |
| chr7:55243398-55245410 | p2 | (AT)6 | TCCTTTAACGATCCGTCCTG | GCTGAAGAACGACATCGACA | 209 |
| chr7:55320930-55322945 | p5 | (TTGCA)3 | TTGCTTGAGCTTCGGACTTT | GACCCCCTTTAGTTACCCCA | 289 |
| chr7:55337553-55339565 | p2 | (TC)6 | AACATTGAGCGTTACAGGGC | AAGCACAACCCGAATAGTGG | 118 |
| chr7:55448195-55450210 | p5 | (ATCTG)3 | ACCTAGGCATGTCCTGGATG | TCTGGAAGTGTTCCGACTCC | 260 |
| chr7:55586490-55588508 | p2 | (AT)9 | CCTGACTGGTGGAGAGGGTA | AAAAATCTTGACGTGTGGGC | 151 |
| chr7:55798209-55800224 | p5 | (TACAT)3 | CCAATACCGTTCGCTCATTT | ATGCCCTTCTTTTAGGGGAA | 117 |
| chr7:55835149-55837165 | p3 | (TCT)9 | TTGAAAAGTTGGTTACCGCC | CGCCTTTTAAAACACTGGGA | 291 |
| chr7:55898597-55900617 | p5 | (TTCAT)4 | CGTCTGAACGGGAATGAAAT | CCGAGTGGAAGTAGGAGCAG | 132 |
| chr7:56006377-56008389 | p2 | (TA)6 | AATGAAATCGAGGGTTCACG | CGTATAGCTACCCGCTTTCG | 195 |
| chr7:56040688-56042703 | p5 | (GTTCG)3 | GGTGGGGGTGTGACATAAAC | ATCATCAATTCTTGGGAGCG | 101 |
| chr7:56040688-56042703 | p5 | (GTTCG)3 | GGTGGGGGTGTGACATAAAC | ATCATCAATTCTTGGGAGCG | 101 |
| chr7:56154665-56156677 | p2 | (AT)6 | AGAACTCTCGTCACACGGCT | GCGGATGAGGTTGATTCAGT | 200 |
| chr7:56194090-56196110 | p5 | (TATCA)4 | TTTCGGAGAAGAGCGATGAT | CTTTCCGAAGTTCGTTCTCG | 193 |
| chr7:56213099-56215119 | p2 | (AT)10 | CCAGAGCTTCCAGTTTCAGG | TTGTCCTAGGGGTTGTCTGC | 104 |
| chr7:56213099-56215119 | p6 | (CATATA)3 | CCAGAGCTTCCAGTTTCAGG | TTGTCCTAGGGGTTGTCTGC | 104 |
| chr7:56213519-56215537 | p2 | (AT)10 | ATACTCTGACCCGGCAACAC | TTCATGCCTTACATTGCTCG | 278 |
| chr7:56213519-56215537 | p6 | (CATATA)3 | ATACTCTGACCCGGCAACAC | TTCATGCCTTACATTGCTCG | 278 |
| chr7:56223341-56225361 | p5 | (CATAT)4 | ACAGGAATCTACCGGACACG | TTTTGTTCGTTTGGGGTTGT | 201 |
| chr7:56361893-56363905 | p2 | (TA)8 | CATGCGTCAGTACCACATCC | CTTTAGATTTCATCCCGCCA | 251 |
| chr7:56361893-56363905 | p2 | (TA)6 | CATGCGTCAGTACCACATCC | CTTTAGATTTCATCCCGCCA | 251 |
| chr7:56380310-56382360 | p2 | (TA)25 | CACTCATGCTACCCAGCTCA | TGCAGAGGACTTTGAGGGTT | 209 |
| chr7:56380310-56382360 | p3 | (ATC)5 | CACTCATGCTACCCAGCTCA | TGCAGAGGACTTTGAGGGTT | 209 |
| chr7:56462123-56464138 | p5 | (CAATG)3 | GACCGTCCTTTCTGGTCGTA | GTCGTAAACGGGCATGTCTT | 133 |
| chr7:56504215-56506242 | p2 | (GA)6 | ATTCTTCCCGTTCCCATACC | CTTCTGTCTCCCACTGCTCC | 280 |
| chr7:56504215-56506242 | p3 | (AAT)9 | ATTCTTCCCGTTCCCATACC | CTTCTGTCTCCCACTGCTCC | 280 |
| chr7:56551155-56553167 | p2 | (CA)6 | CTTGCACGTTTCCTTCCAAT | AGACGAATGGACGAAGATGG | 154 |
| chr7:56555281-56557299 | p6 | (TACACA)3 | TACTAAATTTTGGCGGACGG | ACCCATCTGATACGCCTTTG | 218 |
| chr7:56708845-56710857 | p2 | (AT)12 | TGCAACCCATGTAGCAACTC | GTCGGCTTGCGTAATCATTT | 208 |
| chr7:56708859-56710897 | p2 | (AT)12 | TGCAACCCATGTAGCAACTC | GTCGGCTTGCGTAATCATTT | 194 |
| chr7:56717121-56719141 | p2 | (AT)21 | TGCATCTGAGGCAGTTTTTG | TGATGACAAGCCCACACAAT | 154 |
| chr7:56717143-56719181 | p2 | (AT)21 | TGCATCTGAGGCAGTTTTTG | TGATGACAAGCCCACACAAT | 132 |
| chr7:56723720-56725736 | p2 | (AT)8 | CACAAAATAATGGTGCGTCG | GCTTCATCAGCGTTGTCTCA | 300 |

| Marker ID | SSR type | SSR motif | FORWARD PRIMER (5'-3') | REVERSE PRIMER (5'-3') | size (bp) |
|---|---|---|---|---|---|
| chr7:56875881-56877899 | p6 | (CAAACA)3 | ACCAGCCAAATCGTCAAATC | CTTAATGCGGTCCGTCTGTT | 274 |
| chr7:56909272-56911287 | p5 | (TTTAT)3 | AGGTAAGCGACGGTACGAAA | TCCTTTAGCGATCTGCCCTA | 214 |
| chr7:56941481-56943505 | p2 | (TA)12 | GCATTTCCAGTGCTAGGAGG | GTGAGCATGGCAAGAAGACA | 138 |
| chr7:56962856-56964874 | p2 | (TA)6 | TGGAGGACAAGGAAATCAGG | GATGTCCGATTGTTGCACTG | 225 |
| chr7:56962856-56964874 | p3 | (TTG)6 | TGGAGGACAAGGAAATCAGG | GATGTCCGATTGTTGCACTG | 225 |
| chr7:57036392-57038407 | p5 | (ATGCA)3 | ACCGCCTCGTTATGATTCAC | TTCTTTTTCGCTTGCTCGAT | 199 |
| chr7:57042239-57044254 | p5 | (CATAG)3 | TCGAGCACCCCTTGTCTATC | GTTTGGGCAATGCTAATGCT | 191 |
| chr7:57042239-57044254 | p2 | (TA)8 | TCGAGCACCCCTTGTCTATC | GTTTGGGCAATGCTAATGCT | 191 |
| chr7:57124844-57126864 | p2 | (AT)10 | CCCAAAATCTGGAGGAACAA | CTGTTATCCCCTATCCGCAA | 154 |
| chr7:57124844-57126864 | p2 | (AT)6 | CCCAAAATCTGGAGGAACAA | CTGTTATCCCCTATCCGCAA | 154 |
| chr7:57280604-57282619 | p5 | (CCACA)3 | GTCAAACGGAGCAATAGGGA | AGCCTCGGGGACAACTCTAT | 139 |
| chr7:57330398-57332410 | p2 | (AT)6 | AATGAGCCTCGGGGAATAAT | GTCGGGTCCTATTTATGCGA | 105 |
| chr7:57335127-57337151 | p6 | (TTTCTC)4 | ATCAACACAGCAACCCAACA | TTGCCATTGATGAAGTTGGA | 220 |
| chr7:57368849-57370891 | p2 | (TA)21 | TTGTGAGTGCGCAAAACTTC | CAGGTGCTTGAACCAGGTCT | 243 |
| chr7:57394963-57396983 | p5 | (TGCAT)3 | CCAAGTACGGGCTGGAAGTA | TTTGGGTCTGTTTTGGAAGG | 227 |
| chr7:57394979-57396994 | p5 | (TGCAT)3 | CCAAGTACGGGCTGGAAGTA | TTTGGGTCTGTTTTGGAAGG | 211 |
| chr7:57394989-57397025 | p5 | (TGCAT)3 | CCAAGTACGGGCTGGAAGTA | TTTGGGTCTGTTTTGGAAGG | 201 |
| chr7:57395034-57397050 | p5 | (TGCAT)3 | CCAAGTACGGGCTGGAAGTA | TTTGGGTCTGTTTTGGAAGG | 156 |
| chr7:57411061-57413073 | p2 | (CA)6 | ATCGCCAAATTTCACACACA | TTTACGATCTTGAGCGGACC | 121 |
| chr7:57411061-57413073 | p2 | (CA)6 | ATCGCCAAATTTCACACACA | TTTACGATCTTGAGCGGACC | 121 |
| chr7:57418846-57420858 | p2 | (AT)6 | TGAATCCCGTTTTCTCCAAA | AATGTGGCCGAGTCTCATTC | 169 |
| chr7:57440281-57442293 | p2 | (TA)6 | TGTAAACAGACCCACACCGA | TTTGACTTAAGTTTGGGGCG | 111 |
| chr7:57444698-57446713 | p3 | (TTG)5 | TTCGAGTGTCTTCGTTGTGC | CAGCCAAACACACCCTTTTT | 297 |
| chr7:57449236-57451251 | p3 | (GAA)5 | CAAGGTGATGTACTGGGCCT | TGTCCCGGTGGTTAATGAAT | 154 |
| chr7:57470322-57472336 | p2 | (GT)7 | GAGCCAATAGCCATTTTGGA | GAATGCGCAAACACGAGATA | 118 |
| chr7:57475788-57477803 | p5 | (ATTGT)3 | CTTACGCTTTTGGCCTTGAG | TTGGGTGGTTTTAACCCAAA | 208 |
| chr7:57484058-57486072 | p2 | (AT)7 | GACGGACCGAAGAAGCATAG | ATTGCGTCCTCGAATGTTTC | 257 |
| chr7:57527885-57529927 | p2 | (AT)21 | TAATCTTGTCCTTCTGGGCG | TGAAGTGTCTACGAGCCGTG | 132 |
| chr7:57606834-57608850 | p2 | (AT)8 | CCCGTGTTTGGGAGTAAAGA | CTAGTCGGGTCGTGACAGGT | 290 |
| chr7:57615137-57617155 | p3 | (TCA)6 | ACAAGTCTGGATCACCACCC | TTCGTGCCTTGCATACTCAG | 237 |
| chr7:57630656-57632680 | p3 | (TAT)8 | TACACAAAATGGGTGGCTCA | ACACCTGAAGAACAGGTGGG | 111 |
| chr7:57630656-57632680 | p3 | (TAT)19 | TACACAAAATGGGTGGCTCA | ACACCTGAAGAACAGGTGGG | 111 |
| chr7:57639601-57641613 | p2 | (TG)6 | ACACTATGCCGAGGGTGTTC | AAGAAGCCTAAGGGGGTTCA | 241 |
| chr7:57639601-57641613 | p2 | (AT)6 | ACACTATGCCGAGGGTGTTC | AAGAAGCCTAAGGGGGTTCA | 241 |
| chr7:57640096-57642108 | p2 | (TG)6 | TTCTACAGCGGGAAATGGTC | CCCTGGCTTTGTGGTTAAAA | 244 |
| chr7:57640096-57642108 | p2 | (AT)6 | TTCTACAGCGGGAAATGGTC | CCCTGGCTTTGTGGTTAAAA | 244 |

| Marker ID | SSR type | SSR motif | FORWARD PRIMER (5'-3') | REVERSE PRIMER (5'-3') | size (bp) |
|---|---|---|---|---|---|
| chr7:57640439-57642485 | p2 | (TG)6 | TCGTTGGTTGTGTGTGTGTG | ATTTTGGGGTTTGAGCTGTG | 154 |
| chr7:57640439-57642485 | p2 | (AT)6 | TCGTTGGTTGTGTGTGTGTG | ATTTTGGGGTTTGAGCTGTG | 154 |
| chr7:57659240-57661268 | p2 | (AT)9 | TGTGCCCACATATAAGCCAA | TTAGGCTGCCATCTAAGCGT | 285 |
| chr7:57659280-57661298 | p2 | (AT)9 | TGTGCCCACATATAAGCCAA | TTAGGCTGCCATCTAAGCGT | 245 |
| chr7:57659300-57661328 | p2 | (AT)9 | TGTGCCCACATATAAGCCAA | TTAGGCTGCCATCTAAGCGT | 225 |
| chr7:57670848-57672864 | p2 | (AT)8 | TGATTTGGATTGGAAAAGGG | CTTAACGGAAATCGAGCAGC | 212 |
| chr7:57670848-57672864 | p2 | (TA)27 | TGATTTGGATTGGAAAAGGG | CTTAACGGAAATCGAGCAGC | 212 |
| chr7:57716630-57718646 | p2 | (AT)8 | CCCGTGTTTGGGAGTAAAGA | CTAGTCGGGTCGTGACAGGT | 290 |
| chr7:57724933-57726951 | p3 | (TCA)6 | ACAAGTCTGGATCACCACCC | TTCGTGCCTTGCATACTCAG | 237 |
| chr7:57740452-57742476 | p3 | (TAT)8 | TACACAAAATGGGTGGCTCA | ACACCTGAAGAACAGGTGGG | 111 |
| chr7:57740452-57742476 | p3 | (TAT)19 | TACACAAAATGGGTGGCTCA | ACACCTGAAGAACAGGTGGG | 111 |
| chr7:57749397-57751409 | p2 | (TG)6 | ACACTATGCCGAGGGTGTTC | AAGAAGCCTAAGGGGGTTCA | 241 |
| chr7:57749397-57751409 | p2 | (AT)6 | ACACTATGCCGAGGGTGTTC | AAGAAGCCTAAGGGGGTTCA | 241 |
| chr7:57749892-57751904 | p2 | (TG)6 | TTCTACAGCGGGAAATGGTC | CCCTGGCTTTGTGGTTAAAA | 244 |
| chr7:57749892-57751904 | p2 | (AT)6 | TTCTACAGCGGGAAATGGTC | CCCTGGCTTTGTGGTTAAAA | 244 |
| chr7:57750235-57752281 | p2 | (TG)6 | TCGTTGGTTGTGTGTGTGTG | ATTTTGGGGTTTGAGCTGTG | 154 |
| chr7:57750235-57752281 | p2 | (AT)6 | TCGTTGGTTGTGTGTGTGTG | ATTTTGGGGTTTGAGCTGTG | 154 |
| chr7:57769036-57771064 | p2 | (AT)9 | TGTGCCCACATATAAGCCAA | TTAGGCTGCCATCTAAGCGT | 285 |
| chr7:57769076-57771094 | p2 | (AT)9 | TGTGCCCACATATAAGCCAA | TTAGGCTGCCATCTAAGCGT | 245 |
| chr7:57769096-57771124 | p2 | (AT)9 | TGTGCCCACATATAAGCCAA | TTAGGCTGCCATCTAAGCGT | 225 |
| chr7:57780644-57782660 | p2 | (AT)8 | TGATTTGGATTGGAAAAGGG | CTTAACGGAAATCGAGCAGC | 212 |
| chr7:57780644-57782660 | p2 | (TA)27 | TGATTTGGATTGGAAAAGGG | CTTAACGGAAATCGAGCAGC | 212 |
| chr7:57917133-57919157 | p6 | (ATACAT)4 | CGTGCAACACACCCATAAAC | ACGAGTAAGCCATGTGGGTC | 209 |
| chr7:57998990-58001002 | p2 | (TA)6 | ATCGTGAGACCATGCTACCC | TCCATTACTCCGGAGGTGTC | 125 |
| chr7:58004055-58006069 | p2 | (AT)7 | ACAGGTGGACAGGGAGACAG | CGAGCCGACAAGGCTATAAG | 148 |
| chr7:58052869-58054884 | p3 | (TTG)5 | CTAGTCACGACCTACGGGGA | GTTTCATGGCCTCTCTGAGC | 106 |
| chr7:58064352-58066370 | p6 | (CGTTAT)3 | TACACAATAGACCTCCGGGC | ATACTGCGGTTACGGTGAGG | 274 |
| chr7:58072675-58074690 | p5 | (CATAT)3 | CGAACTCAACACCCCAGAAT | CGATGTATATTGCGCCTCCT | 176 |
| chr7:58072675-58074690 | p2 | (AT)30 | CGAACTCAACACCCCAGAAT | CGATGTATATTGCGCCTCCT | 176 |
| chr7:58072675-58074690 | p2 | (AT)31 | CGAACTCAACACCCCAGAAT | CGATGTATATTGCGCCTCCT | 176 |
| chr7:58073574-58075636 | p5 | (CATAT)3 | AACAGGAGTAGCTGCCCAGA | AGGCCCAAAGGAAGTGAAGT | 290 |
| chr7:58073574-58075636 | p2 | (AT)30 | AACAGGAGTAGCTGCCCAGA | AGGCCCAAAGGAAGTGAAGT | 290 |
| chr7:58073574-58075636 | p2 | (AT)31 | AACAGGAGTAGCTGCCCAGA | AGGCCCAAAGGAAGTGAAGT | 290 |
| chr7:58095828-58097854 | p2 | (AT)13 | CAAATGGACAAAAATGGCCT | AGCCTTGTGGCTCACCTAGA | 282 |
| chr7:58164010-58166068 | p6 | (TGAACA)3 | AACGGAGCGATCCTTATCCT | GGTTAGCTCAGCCAGCATTC | 239 |
| chr7:58164010-58166068 | p2 | (AT)29 | AACGGAGCGATCCTTATCCT | GGTTAGCTCAGCCAGCATTC | 239 |

| Marker ID | SSR type | SSR motif | FORWARD PRIMER (5'-3') | REVERSE PRIMER (5'-3') | size (bp) |
|---|---|---|---|---|---|
| chr7:58164010-58166068 | p3 | (ATC)5 | AACGGAGCGATCCTTATCCT | GGTTAGCTCAGCCAGCATTC | 239 |
| chr7:58301942-58303980 | p2 | (TG)19 | ATCATGGAACCAAGATGGGA | TTCAATACCCGAAAACTGGC | 215 |
| chr7:58327566-58329590 | p2 | (TA)12 | ATCCCATTGGGCAAGTCATA | CCCTTGTTTGCCAAAACTGT | 250 |
| chr7:58329629-58331659 | p5 | (TGAAT)3 | ATGTACAGGGGAGATGCTGC | GCTTAATTGTCCAAGCTCGC | 231 |
| chr7:58329629-58331659 | p2 | (TA)15 | ATGTACAGGGGAGATGCTGC | GCTTAATTGTCCAAGCTCGC | 231 |
| chr7:58420536-58422554 | p2 | (AT)9 | GACGGACACCCAAGACCTTA | GAGGTCCCTGATTTGACGAA | 285 |
| chr7:58429952-58432062 | p2 | (TA)6 | CCCTAGAACGAATGGCTCTG | CCCTCAGTTTGTCGGTTCAT | 188 |
| chr7:58438072-58440087 | p5 | (ATATG)3 | TCCGGCCATCTTTACAAAAC | CTTCTCCTCCTCCATTTCCC | 162 |
| chr7:58485818-58487868 | p2 | (AT)10 | CGTGGTGGATTAAAGCCAAA | AAAAATTGGGTTCCGTCTCC | 173 |
| chr7:58538573-58540627 | p2 | (AT)27 | CTCATCTCCTGACCTGAGGC | GCGATGCCGGTAAAGATAAA | 228 |
| chr7:58609816-58611828 | p2 | (CA)6 | CCAAACAAACCAAATACGGG | AATCCAAGTGCAAGTTTGGG | 162 |
| chr7:58638846-58640860 | p2 | (AT)7 | AATCAACAACATTAGCCCGC | ATCACGACCTAACGTCACCC | 285 |
| chr7:58644280-58646300 | p2 | (TA)10 | CCCATCGCTTGAACCTACAT | ACAGCAAAGAGGCGGTAGAA | 218 |
| chr7:58806353-58808368 | p4 | (ATAA)4 | GGTGGAAATTGAGACGAGGA | TCATTAGGTTGAGGGTTCGC | 220 |
| chr7:58806353-58808368 | p5 | (AATGC)3 | GGTGGAAATTGAGACGAGGA | TCATTAGGTTGAGGGTTCGC | 220 |
| chr7:58835114-58837129 | p5 | (TTGTC)3 | ACCCTCTTTGCCAATTGTTG | CAAGTGGGAGCCTATGAGGA | 200 |
| chr7:58839605-58841617 | p5 | (TTTAA)3 | AGCCGTGTCCAGTTGAGTCT | CAATTACCTCGTAGCGGCAT | 238 |
| chr7:58839605-58841617 | p2 | (TA)6 | AGCCGTGTCCAGTTGAGTCT | CAATTACCTCGTAGCGGCAT | 238 |
| chr7:58863064-58865080 | p4 | (TATG)4 | AGTTTGGGCTTTGAAGGGTT | AATGACAACAAGCCCACACA | 270 |
| chr7:58940314-58942329 | p4 | (CATA)4 | AAGTGTCATCCGTACCCAGC | TAACAACGAGTCCCGAAAGG | 185 |
| chr7:58940314-58942329 | p3 | (CAA)5 | AAGTGTCATCCGTACCCAGC | TAACAACGAGTCCCGAAAGG | 185 |
| chr7:58973813-58975849 | p2 | (AG)18 | TGAGTTTCTGATCCTGCCCT | ACAGGTGCCACGAACTATCC | 223 |
| chr7:58973813-58975849 | p2 | (AT)20 | TGAGTTTCTGATCCTGCCCT | ACAGGTGCCACGAACTATCC | 223 |
| chr7:59157465-59159479 | p2 | (TA)7 | ACGCCTCCTCAAGAAACAAA | ATTTGTTCGAAGTTGCCAGG | 219 |
| chr7:59235777-59237799 | p2 | (AC)11 | AGAAGAGCTGAGAGGGGGAC | AGTTTTGAATTGCCTGGTGC | 116 |
| chr7:59277650-59279665 | p3 | (ATC)5 | TCATCATCACAAGCCGACAT | TGCTCCAGAGCTGCAGTTTA | 133 |
| chr7:59283584-59285617 | p2 | (TA)6 | ATGCAAGGCAATCACAACAA | TTGGCTTTGCCAACCTACTT | 191 |
| chr7:59302796-59304811 | p5 | (ATATA)3 | AAGATCGGTCGTCCTTGTTG | CGCATCATCACGGTAAAATG | 230 |
| chr7:59399580-59401606 | p2 | (AT)13 | TGTCGTTGCGGATTCTATGA | ATGCATTGCTACCCGAAAAC | 137 |
| chr7:59522066-59524128 | p2 | (TA)31 | GAGTCAAGCTCTCCGACCAG | GATGGGTGGTGATCGGTATC | 285 |
| chr7:59563679-59565691 | p2 | (TG)6 | AAAGGTGCCTACGGGCTATT | CTCATACCTGGCCCTACCAA | 224 |
| chr7:59573459-59575471 | p2 | (AT)6 | TCGCAAACCATAGGAGGAAG | TAAGCGGGCTGCTTTATGAT | 130 |
| chr7:59583874-59585888 | p2 | (TA)7 | AACAAACTCTTGCCAAACGG | TATGCATGCTTCTGAGACCG | 141 |
| chr7:59633401-59635413 | p2 | (AT)6 | ACGTTGCTTTCAGGACCATC | TATCTTTTGGGCTGGTGTCC | 186 |
| chr7:59698483-59700497 | p2 | (AT)7 | GGTCCCAGAGGCATGAATAA | GCTCAGGCACAACAACAGAA | 224 |
| chr7:59725198-59727210 | p2 | (TA)6 | CGAAACCTTTAGACGCTTGC | GCGTCATGTGCGTTATGAGT | 294 |

| Marker ID | SSR type | SSR motif | FORWARD PRIMER (5'-3') | REVERSE PRIMER (5'-3') | size (bp) |
|---|---|---|---|---|---|
| chr7:59784826-59786844 | p3 | (TTG)6 | ATACGAGTGTCCAGCTTGGG | TATCGTACGAAACCTTGCCC | 133 |
| chr7:59963114-59965132 | p3 | (CAA)6 | TCCGATTGCATTATGGAACA | AAGGCTCCAAAGCGTATGAA | 274 |
| chr7:60053089-60055107 | p6 | (CATATA)3 | ACAATCCCAAGGACGAACTG | GGCCCAGGATAGTCAGTGAA | 118 |
| chr7:60073234-60075250 | p2 | (AT)14 | CTACCGAGCACCCCTTATCA | TACAGCGTTTTGTCCTGCAC | 244 |
| chr7:60073234-60075250 | p2 | (AT)8 | CTACCGAGCACCCCTTATCA | TACAGCGTTTTGTCCTGCAC | 244 |
| chr7:60175562-60177577 | p5 | (AATGC)3 | TGAATTTGGAAGCTTTTGGG | TTAATGGTTCGCCTACCGAG | 183 |
| chr7:60290548-60292568 | p4 | (ATAA)5 | TCTTCAATTTCCCACTTGCC | AGAGAAAGGGAGAGGGTTCG | 210 |
| chr7:60369509-60371529 | p5 | (ATGCA)4 | TGTGATGAGGTGTGGGAAGA | CAGACCCTGGGGGTATTTTT | 276 |
| chr7:60478081-60480093 | p2 | (AT)6 | TAAAATTGGGCAGCATTTCC | TGGGCTGTGATGTTTATGGA | 288 |
| chr7:60555365-60557381 | p2 | (AG)8 | TCAGCATCAGACTCACTGGG | TTATTTCCGAATTTGCCGAG | 220 |
| chr7:60555365-60557381 | p2 | (GT)8 | TCAGCATCAGACTCACTGGG | TTATTTCCGAATTTGCCGAG | 220 |
| chr7:60555365-60557381 | p3 | (GTT)5 | TCAGCATCAGACTCACTGGG | TTATTTCCGAATTTGCCGAG | 220 |
| chr7:60585984-60587996 | p2 | (AG)6 | GTCTGACTCCGACCACGAAT | TCGGTCGTAATACCACGTCA | 291 |
| chr7:60585984-60587996 | p2 | (GT)6 | GTCTGACTCCGACCACGAAT | TCGGTCGTAATACCACGTCA | 291 |
| chr7:60692765-60694777 | p2 | (CT)6 | GAGAGGAAATGACGTGGGAA | TAAAATTTGGCCACACACGA | 190 |
| chr7:60694463-60696475 | p2 | (TG)6 | AAAGGTGCCTACGGGCTATT | CCAACCAAGGCTCAAGGTTA | 211 |
| chr7:60795142-60797162 | p4 | (AGAA)5 | CCGTCTTGTGTGCCTCTGTA | ACCAACAGGTTCAGGACTGG | 117 |
| chr7:60834619-60836639 | p2 | (AT)10 | ATAAGATGCCAACTCCACGG | CAAGTACCAAAGCCCGTGAT | 169 |
| chr7:61120149-61122161 | p2 | (TC)6 | TGACAACAGGCAAAAAGCAG | GAATCTGCCCTACAAAGCCA | 252 |
| chr7:61161002-61163014 | p2 | (TG)6 | ACAGGGTAATGACGACGAGC | TTCCCTTAGCCTTCCTCCAT | 239 |
| chr7:61177252-61179272 | p2 | (AT)10 | CCTACCTTTCGGAGATGCTG | CTCAGCTCCAGTTCCTCCAC | 149 |
| chr7:61379867-61381882 | p5 | (AAAAG)3 | GACACCGGTGGTCAGGTAGT | GACATGACCCGGACAAAAGT | 292 |
| chr7:61379867-61381882 | p5 | (AAAAG)3 | GACACCGGTGGTCAGGTAGT | GACATGACCCGGACAAAAGT | 292 |
| chr7:61403531-61405546 | p5 | (CATGT)3 | CAACTTCTCATCCGCAGACA | CTTCGCCCATACAGAGCTTC | 189 |
| chr7:61489260-61491274 | p2 | (AT)7 | GGTCCGGAGAATCACCATAA | TGGTTGCCTTAAGTATCCCG | 269 |
| chr7:61529572-61531596 | p3 | (TGT)8 | AGGGAATGACGAGATTGACG | ATCTCAGCATCCGCATCTCT | 170 |
| chr7:61675917-61677937 | p2 | (TA)10 | TGAAATCACAGACTCTGGCG | GACGGAGCGAATACCACCTA | 285 |
| chr7:61686554-61688568 | p2 | (CT)7 | CATAACGAGGCCCGAAATTA | TGTGGCTGCTGTTGTTTTTC | 285 |
| chr7:61728647-61730661 | p2 | (AT)7 | CGTCATGACTCATCGGTTTG | TACCAGACTCGAGCGGAAGT | 130 |
| chr7:61737063-61739081 | p2 | (AT)9 | TCCCGATAACCTGTCTCCTG | GTCGGCCTGCGTATCATATT | 296 |
| chr7:61826946-61828958 | p3 | (ACA)6 | CCTGATGTCACGACACAACC | TGCGTACATTCTACCCTCCC | 123 |
| chr7:61826946-61828958 | p2 | (AT)6 | CCTGATGTCACGACACAACC | TGCGTACATTCTACCCTCCC | 123 |
| chr7:61826946-61828958 | p2 | (AT)13 | CCTGATGTCACGACACAACC | TGCGTACATTCTACCCTCCC | 123 |
| chr7:61915534-61917546 | p2 | (AT)33 | GTTCCTTCAAGGTGAGACGC | GGGGTGATAAGAGGCGTGTA | 171 |
| chr7:61915534-61917546 | p2 | (TA)6 | GTTCCTTCAAGGTGAGACGC | GGGGTGATAAGAGGCGTGTA | 171 |
| chr7:61915534-61917546 | p2 | (AT)6 | GTTCCTTCAAGGTGAGACGC | GGGGTGATAAGAGGCGTGTA | 171 |

| Marker ID | SSR type | SSR motif | FORWARD PRIMER (5'-3') | REVERSE PRIMER (5'-3') | size (bp) |
|---|---|---|---|---|---|
| chr7:61936293-61938307 | p2 | (TA)7 | GGGACTTCCACGGACACTTA | ATTGGAGATCGAACGAAACG | 189 |
| chr7:61936775-61938799 | p2 | (TA)7 | CTTCGCTTGATCTCGTAGGG | CCTAGCTGGGTCGTGACAAT | 123 |
| chr7:62023712-62025738 | p2 | (AT)13 | GCGTAAGTGAGGCTCCAAAG | TCTGGCCTGACATAATGCTG | 210 |
| chr7:62023712-62025738 | p2 | (AT)21 | GCGTAAGTGAGGCTCCAAAG | TCTGGCCTGACATAATGCTG | 210 |
| chr7:62115333-62117348 | p5 | (GTTAT)3 | AGCTTGTCCCACATCGATTC | ACTTTCCCCCAAAACTCGAA | 104 |
| chr7:62133076-62135091 | p5 | (GTTAT)3 | ACCGTTGATGCTCATTGTGA | CTTTTGGTGCATCCTTGGTT | 104 |
| chr7:62175631-62177647 | p2 | (TA)8 | GGAGCAGGTATCGCATCAAT | AGGCGGCAGTCAGTGTTACT | 228 |
| chr7:62175631-62177647 | p5 | (TGCAA)3 | GGAGCAGGTATCGCATCAAT | AGGCGGCAGTCAGTGTTACT | 228 |
| chr7:62202318-62204336 | p6 | (TATATG)3 | TTAGATTCTCGGGGATGGTG | AACATACCTGCTCCACGTCC | 203 |
| chr7:62385669-62387684 | p5 | (TATCA)3 | GGCGGGCTACATAATTTTCA | GTGCTTTTGATGGTCCTCGT | 287 |
| chr7:62401418-62403430 | p2 | (AT)6 | TTTGAGCCTTCCAATATCCG | CCATGTCACATATAACGCCG | 283 |
| chr7:62542958-62545006 | p2 | (AT)24 | TCTTAAGGTCGGCTTTCCCT | GCTGTGTGCAGTGTTGAGGT | 221 |
| chr7:62549966-62551978 | p2 | (AT)6 | TTTACGCATCGTGCTGTAGG | ATTCGAGCTCCCAAGACTCA | 223 |
| chr7:62650278-62652292 | p2 | (AG)7 | CCTCCCATGCATCTGACTTT | TCGACGTCGTACCAAACAGA | 164 |
| chr7:62650278-62652292 | p2 | (AT)9 | CCTCCCATGCATCTGACTTT | TCGACGTCGTACCAAACAGA | 164 |
| chr7:62735342-62737357 | p5 | (TTAAA)3 | GTGCATGTTGTGGAAGTTGG | GTGCAAACCGACCCTAGAAA | 140 |
| chr7:62872514-62874534 | p4 | (AGAA)5 | CCGTCTTGTGTGCCTCTGTA | ACCAACAGGTTCAGGACTGG | 117 |
| chr7:62941049-62943069 | p2 | (AT)10 | TAATGGGCTGCAATCAACAA | GGGCCTCCAAACAGATGTAA | 112 |
| chr7:62994813-62996831 | p6 | (CCTCTA)3 | CTCTTTCAATTCTGCGGGAG | CAGCGTCTCCAAGAGGTAGG | 279 |
| chr7:63099275-63101287 | p2 | (AC)6 | CTTCGATTTCCAAGAGCAGG | TGACCATTTGCAGCTTTCTG | 177 |
| chr7:63272075-63274099 | p3 | (TGT)8 | GGCTATGTTGTATGGGGTGG | TGGGGCATCTATGTCTCTCC | 246 |
| chr7:63279725-63281743 | p3 | (TGT)6 | TTTCTGACTTGTCGCAAACG | TGGTCATGTGCTTTCCATGT | 172 |
| chr7:63377191-63379209 | p6 | (AACATG)3 | TGGTTTCTGGTATTGTCGCA | GTACGGAAGAAGCCTTGCAC | 191 |
| chr7:63377191-63379209 | p6 | (AACATG)3 | TGGTTTCTGGTATTGTCGCA | GTACGGAAGAAGCCTTGCAC | 191 |
| chr7:63392169-63394197 | p2 | (TA)14 | CAACAACCATGCCAAAACAC | CAGCAGACAGGTGGACTTGA | 173 |
| chr7:63396804-63398838 | p2 | (AT)17 | AACCTCTGGCGGTAGGAAAT | GCATGTTTCGGATACGGAGT | 135 |
| chr7:63396804-63398838 | p5 | (CATAG)3 | AACCTCTGGCGGTAGGAAAT | GCATGTTTCGGATACGGAGT | 135 |
| chr7:63398163-63400179 | p5 | (CATAG)3 | TCAAGTCCACCTGTCTGCTG | GACGCGATATGCTACGACAA | 156 |
| chr7:63398163-63400179 | p2 | (AT)8 | TCAAGTCCACCTGTCTGCTG | GACGCGATATGCTACGACAA | 156 |
| chr7:63440779-63442794 | p5 | (TGTGT)3 | TAAAAACGGCTAAAGGGGGT | ACGAGGGCCTTCCAATTAGT | 290 |
| chr7:63468864-63470884 | p2 | (AT)10 | CGGAGCTAACTACCGAGCAC | GTGTGGGTTATTGGTGTCCC | 251 |
| chr7:63468864-63470884 | p2 | (AT)6 | CGGAGCTAACTACCGAGCAC | GTGTGGGTTATTGGTGTCCC | 251 |
| chr7:63468864-63470884 | p2 | (AT)10 | CGGAGCTAACTACCGAGCAC | GTGTGGGTTATTGGTGTCCC | 251 |
| chr7:63468864-63470884 | p2 | (AT)6 | CGGAGCTAACTACCGAGCAC | GTGTGGGTTATTGGTGTCCC | 251 |
| chr7:63474648-63476660 | p2 | (AC)6 | AGGTTTCGACATTTGGTTGC | ACCATTTGCAGATTTCGGAC | 118 |
| chr7:63484723-63486738 | p3 | (ATG)5 | TCCCATTTGTTAGCGACTCC | CGGAGTGATGTAAGGTCGGT | 254 |

| Marker ID | SSR type | SSR motif | FORWARD PRIMER (5'-3') | REVERSE PRIMER (5'-3') | size (bp) |
|---|---|---|---|---|---|
| chr7:63484723-63486738 | p5 | (GTCCC)3 | TCCCATTTGTTAGCGACTCC | CGGAGTGATGTAAGGTCGGT | 254 |
| chr7:63501796-63503808 | p2 | (AT)6 | CACGCAAGTCAACAAGGCTA | ATCGTTTTGTTGTGGAAGCC | 268 |
| chr7:63507285-63509300 | p5 | (ATCAT)3 | CCGATAAGGCCATCACAGAT | CCAGAGCCCATCTTTTGGTA | 281 |
| chr7:63515896-63517908 | p2 | (TG)6 | CAAAAGGTGCCTACGGGTTA | CGACCATAAGGTTCGGTGTT | 265 |
| chr7:63536190-63538205 | p5 | (ATGAT)3 | ATTCTGGAGGAAACGGGACT | GGCTCAGATTTCTGGGAACA | 198 |
| chr7:63680201-63682217 | p3 | (CTG)5 | GCCACCAGTGTAGTCAGCAA | TATGCCCAATATGTGCCTGA | 263 |
| chr7:63680201-63682217 | p2 | (AT)8 | GCCACCAGTGTAGTCAGCAA | TATGCCCAATATGTGCCTGA | 263 |
| chr7:63710137-63712155 | p2 | (AT)9 | CACCTCACTCCATTCACCCT | GAATCAAAGAAGGGCCATGA | 194 |
| chr7:63763423-63765435 | p2 | (TG)6 | ACGTTTGTACTACGCACCCC | TATCATAGCCCGTAGGCACC | 147 |
| chr7:63871016-63873031 | p5 | (CTCTC)3 | GGCCGGAAATCTCAAACATA | AATAATTGGCGCGAGCTAGA | 221 |
| chr7:63922982-63925000 | p6 | (CATATA)3 | CATTCACTTTCCTCCGTCGT | AGGAGGCTGCAGGTCATCTA | 102 |
| chr7:63930406-63932424 | p6 | (ATACAT)3 | CGGGCACCTCTCATCATACT | TTTGATAGCCTTGCCAGCTT | 132 |
| chr7:63930406-63932424 | p3 | (CAT)6 | CGGGCACCTCTCATCATACT | TTTGATAGCCTTGCCAGCTT | 132 |
| chr7:64028939-64030951 | p2 | (AG)6 | AGTTGGCCCTGCTTCTAACA | AAGGACATGGACAGACAGGC | 264 |
| chr7:64034576-64036588 | p2 | (TA)6 | GTGCTGGCCGTATTTCAGTT | TTGCGAATTTCCCTTTATGC | 261 |
| chr7:64047407-64049433 | p2 | (AT)13 | CTTGCGAACCTGACAATCAA | TATGAACTTGGGTTGCCACA | 132 |
| chr7:64094547-64096559 | p2 | (AT)6 | TACGGGCTGTATTCTGGGTC | AAAAATTCACCCAGTGGTCG | 211 |
| chr7:64160684-64162702 | p6 | (CATATA)3 | CGCCTCTCACTTCTTCCATC | TGACAAAGTGGCATCAGAGC | 126 |
| chr7:64223390-64225442 | p2 | (TA)26 | TTCCCACATTTCTTCCAAGC | CCCTAGCTACAATGCCCAAG | 226 |
| chr7:64223390-64225442 | p4 | (TGTA)4 | TTCCCACATTTCTTCCAAGC | CCCTAGCTACAATGCCCAAG | 226 |
| chr7:64290744-64292760 | p4 | (AATA)4 | AATTCGCCCGACTTACAATG | ATTAAAACGACCACCGCAAC | 271 |
| chr7:64301919-64303939 | p2 | (AT)10 | CCACATCCTCGTTGTCATTG | ATGCACACAACGGAAATCAA | 139 |
| chr7:64315827-64317849 | p2 | (AC)11 | AGAAGAGCTGAGAGGGGGAC | AGTTTTGAATTGCCTGGTGC | 116 |
| chr7:64362605-64364621 | p4 | (AATA)4 | AATTCGCCCGACTTACAATG | ATTAAAACGACCACCGCAAC | 271 |
| chr7:64373780-64375800 | p2 | (AT)10 | CCACATCCTCGTTGTCATTG | ATGCACACAACGGAAATCAA | 139 |
| chr7:64518474-64520498 | p2 | (AT)12 | GTGAAGATTGCGGGTTGTTT | ATTAGACGACGTTCCGTTCG | 145 |
| chr7:64570680-64572696 | p4 | (AAAG)4 | GTGGGATTACGAAAACGTGG | TTATGGGGCGTCATACCATT | 282 |
| chr7:64593546-64595576 | p2 | (AG)6 | TCAAAATTGAGCGGGATTTC | TTTGGCCATTTGTGAGTTGA | 195 |
| chr7:64593577-64595595 | p2 | (AG)6 | TCAAAATTGAGCGGGATTTC | TTTGGCCATTTGTGAGTTGA | 164 |
| chr7:64593601-64595673 | p2 | (AG)6 | TCAAAATTGAGCGGGATTTC | TTTGGCCATTTGTGAGTTGA | 140 |
| chr7:64714385-64716409 | p6 | (ATAATG)4 | TGCAACCCTCTCCATACCTC | TTCTTGATCTCCACAAGCCC | 126 |
| chr7:64714385-64716409 | p2 | (TA)9 | TGCAACCCTCTCCATACCTC | TTCTTGATCTCCACAAGCCC | 126 |
| chr7:64755600-64757615 | p5 | (ATATA)3 | TTGGCAAGAAGGGAAAGCTA | AAGGAATACTGGATGCACGG | 180 |
| chr7:64755600-64757615 | p5 | (ATTTC)3 | TTGGCAAGAAGGGAAAGCTA | AAGGAATACTGGATGCACGG | 180 |
| chr7:64764118-64766133 | p5 | (ATTTC)3 | CGCATTTATTTCAGCCCCTA | TTAAGTCGTCGGGGATTCTG | 205 |
| chr7:64767638-64769650 | p2 | (AT)6 | CCCGAGCCTATTAACTCACG | GCTTCAGTGGTGGTGGTGTA | 297 |

| Marker ID | SSR type | SSR motif | FORWARD PRIMER (5'-3') | REVERSE PRIMER (5'-3') | size (bp) |
|---|---|---|---|---|---|
| chr7:64877245-64879263 | p6 | (CATGTT)3 | CTCTCCTCATCCTTCCCCAT | GAATCAAACGTAGAGCGGGA | 254 |
| chr7:64880748-64882766 | p6 | (CATGTT)3 | CTCTCCTCATCCTTCCCCAT | GAATCAAACGTAGAGCGGGA | 254 |
| chr7:65085313-65087337 | p2 | (TA)12 | GAGATATGCCGTTGCCATTT | AAATGACCGCGAGGTATGAG | 173 |
| chr7:65126191-65128223 | p2 | (TA)16 | GTGCCAAGGAGGTTCCATAA | TGAGTTTGCAGCCAATATGC | 104 |
| chr7:65126191-65128223 | p2 | (TA)32 | GTGCCAAGGAGGTTCCATAA | TGAGTTTGCAGCCAATATGC | 104 |
| chr7:65137545-65139557 | p2 | (AT)6 | GATGCAAACGAATGAGATGG | CAACCTTGGCTCATACCACA | 263 |
| chr7:65200127-65202139 | p2 | (CT)6 | GGATACTCCAAAACAGCCGA | TGAAGCCGTTGTTGTGTCAT | 287 |
| chr7:65304327-65306339 | p2 | (AT)6 | TCTCGACCTTATGTCACCCC | TATCCCATATGGAACCCCCT | 152 |
| chr7:65304340-65306360 | p2 | (AT)6 | TCTCGACCTTATGTCACCCC | TATCCCATATGGAACCCCCT | 139 |
| chr7:65321127-65323167 | p2 | (AT)6 | TGAAAGGCATCCACCCTTAG | GTCGCCTCCCTGTCTACTTG | 235 |
| chr7:65321127-65323167 | p2 | (TA)11 | TGAAAGGCATCCACCCTTAG | GTCGCCTCCCTGTCTACTTG | 235 |
| chr7:65370053-65372095 | p2 | (TA)21 | AAGGACGCCAGCTTGAGTTA | GAGACCAACCCAGACTCAGC | 150 |
| chr7:65371472-65373492 | p2 | (TA)10 | TACACACATGCTAAGGGCCA | GCTTCAGAGGTCTTTGTCGG | 134 |
| chr7:65454180-65456198 | p2 | (TA)9 | GACATAGCCATGGTTGCCTT | CTGGCGCTGTGGAATTTTAT | 223 |
| chr7:65493466-65495482 | p4 | (CATA)4 | GAGATAGGGTGCCCATGAGA | AGTGTGTGCTCGACTGGTTG | 107 |
| chr7:65580932-65582948 | p2 | (AT)7 | TCCTCTCACGGTCTCCAAGT | TGTCAAAGGAGGGAGGAATG | 215 |
| chr7:65581029-65583043 | p2 | (AT)7 | TCCTCTCACGGTCTCCAAGT | TGTCAAAGGAGGGAGGAATG | 118 |
| chr7:65687372-65689388 | p2 | (AT)8 | GGAATACTCGGAGCACCTTG | CTCGAGAGGAATGGAATGGA | 258 |
| chr7:65750604-65752658 | p2 | (AT)27 | AAGCAGATGGTGGGTAAACG | ACTCGGTTATCGGCCTCTTT | 181 |
| chr7:65789423-65791435 | p2 | (TG)6 | AAAGGTGCCTACGGGCTATT | ACTTAGCCAACCAAGGCTCA | 229 |
| chr7:65913488-65915628 | p2 | (AT)70 | ATTGTCTCGGAGCTTAGCCA | ACACATAAGCCGACAAGGCT | 216 |
| chr7:66240695-66242707 | p2 | (AT)13 | GTGCTCGAGTTGGACACTCA | ATTCGTAGTGGCAGCCTTGT | 159 |
| chr7:66240695-66242707 | p2 | (AT)6 | GTGCTCGAGTTGGACACTCA | ATTCGTAGTGGCAGCCTTGT | 159 |
| chr7:66303302-66305314 | p2 | (TG)6 | TTAGAGCCGGCAGAAGATGT | CAACTTGCCCCTCTTACCAA | 211 |
| chr7:66400759-66402781 | p2 | (TA)11 | GGGGACACTGCAAATGTTTT | TTTGTCAAATTTTCGAGGGG | 207 |
| chr7:66465718-66467733 | p5 | (TGCAA)3 | ACCATTTAGGAGGGAATGGG | TTGGAAACGTAAGTACCCGC | 162 |
| chr7:66543396-66545488 | p3 | (TTG)5 | GCGATGCACACAACAATTTC | TGAATTTCAAGGAAACCCGT | 109 |
| chr7:66643376-66645391 | p3 | (TTA)5 | CCATTCGATTTGTCGCTCTT | ATATAGGGGAAATGCTGCCC | 206 |
| chr7:66669193-66671205 | p2 | (TA)6 | CCAAATCTGTTCCGTCACCT | TGTTGATGTGGCGTGATTTT | 185 |
| chr7:66778266-66780284 | p3 | (TTG)6 | ATGGATGAAATGGAGGCTTG | TTTGACTGGCGAACACTCTG | 152 |
| chr7:66784857-66786872 | p3 | (TTG)5 | TTGTGTGACAAGAAGCTGCC | CAGCATTCTCATTTCTGCCA | 180 |
| chr7:66822257-66824273 | p2 | (TA)8 | CGTAGGCACCTGTTAGGCAT | ACTACGCACCCCATTTGAAG | 280 |
| chr7:66822257-66824273 | p2 | (AT)6 | CGTAGGCACCTGTTAGGCAT | ACTACGCACCCCATTTGAAG | 280 |
| chr7:66892481-66894499 | p6 | (TACATA)3 | GGTTATCCCTCGGTTTAGGC | AAATGATACGCCACCCCATA | 129 |
| chr7:66896102-66898117 | p3 | (CAA)5 | GATTGGGTTATGGCCTCTCA | TAACCGTAAGCTGAGGTGGG | 271 |
| chr7:66898445-66900463 | p3 | (ATG)6 | AGCCATCGGTTTGTGTTAGG | TTTCGATGCCAGTCCACATA | 197 |

| Marker ID | SSR type | SSR motif | FORWARD PRIMER (5'-3') | REVERSE PRIMER (5'-3') | size (bp) |
|---|---|---|---|---|---|
| chr7:66988114-66990129 | p3 | (AAT)5 | GAGGTTTCGCCAATCAAAAA | GTTACCCCCACTCCCTCATT | 243 |
| chr7:67045695-67047710 | p3 | (TTC)5 | AAGTGTTGATGTGGCGTGAA | ATCTATGGTGGCTTTGACGG | 260 |
| chr7:67108100-67110115 | p3 | (CAA)5 | TTTCCGAGGTGTAACAAGCC | GCGGGAAGAAGTCTTAAGGG | 247 |
| chr7:67121302-67123340 | p2 | (AT)9 | AGAATAGGCGAAGTGGCAGA | AAGGCTGCAGAGCATCTAGG | 111 |
| chr7:67121302-67123340 | p2 | (AT)19 | AGAATAGGCGAAGTGGCAGA | AAGGCTGCAGAGCATCTAGG | 111 |
| chr7:67141199-67143214 | p5 | (ATTTC)3 | AACGAGAGGAAATGACGTGG | AAGCCCACAATAACAGTGCC | 149 |
| chr7:67151754-67153766 | p2 | (AT)9 | AGAGCAAACGGGTTCAAATG | AAACCCGAAGGAAAGGAAAA | 183 |
| chr7:67151787-67153799 | p2 | (AT)9 | AGAGCAAACGGGTTCAAATG | AAACCCGAAGGAAAGGAAAA | 150 |
| chr7:67211891-67213913 | p2 | (AT)20 | ATCAAACCGTACGTTCCTCG | TACGGCCCGATCCATAAATA | 269 |
| chr7:67211891-67213913 | p2 | (AT)13 | ATCAAACCGTACGTTCCTCG | TACGGCCCGATCCATAAATA | 269 |
| chr7:67211891-67213913 | p2 | (AT)11 | ATCAAACCGTACGTTCCTCG | TACGGCCCGATCCATAAATA | 269 |
| chr7:67211891-67213913 | p2 | (TA)9 | ATCAAACCGTACGTTCCTCG | TACGGCCCGATCCATAAATA | 269 |
| chr7:67213235-67215251 | p4 | (TTAT)4 | GAGCAGCCCAGCATAGTAGG | TGTAGGAAGGACGGGACAAG | 147 |
| chr7:67237340-67239368 | p2 | (TA)14 | CGAGCACCCCTTGACATAAT | ATACCATTTTGTTGGCCTCG | 230 |
| chr7:67310958-67312976 | p2 | (TA)11 | AGCAAATGGAACCGCTTAGA | GGCCATCTATTCTGCTGCTC | 184 |
| chr7:67310958-67312976 | p6 | (CATATA)3 | AGCAAATGGAACCGCTTAGA | GGCCATCTATTCTGCTGCTC | 184 |
| chr7:67366144-67368159 | p3 | (CAA)5 | GTTGCAAAATTTGGGGAAGA | ACCACCAGTCTTGGCATTTC | 294 |
| chr7:67471751-67473779 | p2 | (AT)14 | GTTTCGGAGGCATGAACAAT | AGTGAACGAGGCCATTCATC | 192 |
| chr7:67535763-67537777 | p2 | (AC)7 | TAAAATGTCCGCAGCTTCCT | GCCCTAGTCGGGTAATGACA | 160 |
| chr7:67549897-67551909 | p2 | (TA)9 | CGTGCATGCCTCAATATGTC | ACCGTACCCAGCCCTCTAGT | 133 |
| chr7:67549897-67551909 | p2 | (TA)6 | CGTGCATGCCTCAATATGTC | ACCGTACCCAGCCCTCTAGT | 133 |
| chr7:67669992-67672008 | p2 | (AT)8 | GACTCCGGTTCGGACTATGA | GCAGAACCTGTGCCCTAGAC | 215 |
| chr7:67693464-67695479 | p3 | (ATG)5 | CGCGTTACAGTGAGCAAAAA | TGATTCGTCGAGATTCGTTG | 177 |
| chr7:67711829-67713853 | p2 | (AT)12 | CGTTTGCATGATTTCGTTTG | CTCCGGATCATGGAGGAGTA | 139 |
| chr7:67810428-67812462 | p2 | (AT)17 | TGTATGCAAACTGGGGCATA | TTTCCCTCCACATGGACTTC | 196 |
| chr7:67810428-67812462 | p2 | (AT)10 | TGTATGCAAACTGGGGCATA | TTTCCCTCCACATGGACTTC | 196 |
| chr7:67995034-67997062 | p2 | (AT)8 | GGGTCGGGTATGTATTCGTG | TGGGACTCGGTTCCATAATC | 149 |
| chr7:67995034-67997062 | p2 | (TA)14 | GGGTCGGGTATGTATTCGTG | TGGGACTCGGTTCCATAATC | 149 |
| chr7:68010493-68012569 | p3 | (ATG)6 | GGGTCGTAAACTGGGTCGTA | CACCTCGAAATTTGTTCCGT | 128 |
| chr7:68010493-68012569 | p2 | (TA)6 | GGGTCGTAAACTGGGTCGTA | CACCTCGAAATTTGTTCCGT | 128 |
| chr7:68018515-68020530 | p6 | (TTTATA)3 | GGTAGTGGTGGGTGAAAGGA | GCCAAATATACCCCCTTCGT | 119 |
| chr7:68018515-68020530 | p6 | (TTTCTT)3 | GGTAGTGGTGGGTGAAAGGA | GCCAAATATACCCCCTTCGT | 119 |
| chr7:68018515-68020530 | p5 | (TATTT)3 | GGTAGTGGTGGGTGAAAGGA | GCCAAATATACCCCCTTCGT | 119 |
| chr7:68123662-68125677 | p3 | (TTG)5 | GGCAAGGATGGAAACCTGTA | ACTGGCGCTAACATTGCTCT | 141 |
| chr7:68159771-68161792 | p3 | (GTT)7 | ACGTTTTCGGGTGATCTGAC | CATACAGGCCCACGACCTAT | 143 |
| chr7:68159771-68161792 | p2 | (TA)6 | ACGTTTTCGGGTGATCTGAC | CATACAGGCCCACGACCTAT | 143 |

| Marker ID | SSR type | SSR motif | FORWARD PRIMER (5'-3') | REVERSE PRIMER (5'-3') | size (bp) |
|---|---|---|---|---|---|
| chr7:68217606-68219618 | p2 | (AG)6 | GGGACCAAATTCAACATTCG | CGTTGGTGTTCCTTCTTGGT | 184 |
| chr7:68235415-68237430 | p5 | (TATCA)3 | TACCCGATACCTGAACCGAG | ATGGTGTTTTGGCATGGTTT | 293 |
| chr7:68317300-68319315 | p5 | (ATTTC)3 | TTTTGGGTCAACTTTGGAGG | GCAGCTTGCATGGGATATTT | 218 |
| chr7:68318525-68320537 | p2 | (TA)6 | ATTGCATGTGAATGGACGAA | CCACGCTTGCGTTTATACCT | 190 |
| chr7:68337892-68339907 | p3 | (GAA)5 | TTGTGGAAATCCTTCCAAGC | ATCCCAATTTCCCTCCAATC | 219 |
| chr7:68364573-68366587 | p2 | (AT)6 | AGTAGAAGAAATGGCCCCGT | TGGATGGTAAAAAGGGCTTG | 279 |
| chr7:68364573-68366587 | p2 | (TA)7 | AGTAGAAGAAATGGCCCCGT | TGGATGGTAAAAAGGGCTTG | 279 |
| chr7:68529115-68531127 | p2 | (TA)6 | GATATGGCCGGACCTGTCTA | TGCACAGGGAGACAGACTTG | 298 |
| chr7:68621018-68623040 | p2 | (AT)11 | TATGGCTGGGTACGGAAGAC | GTGAAGGGCGGCATAAAATA | 104 |
| chr7:68621018-68623040 | p2 | (AT)10 | TATGGCTGGGTACGGAAGAC | GTGAAGGGCGGCATAAAATA | 104 |
| chr7:68621018-68623040 | p2 | (AT)10 | TATGGCTGGGTACGGAAGAC | GTGAAGGGCGGCATAAAATA | 104 |
| chr7:68650967-68652982 | p3 | (ATG)5 | GGGCCAAAACCCTCATATTT | TGCCACTATATGTCACCCGA | 248 |
| chr7:68650967-68652982 | p2 | (TA)23 | GGGCCAAAACCCTCATATTT | TGCCACTATATGTCACCCGA | 248 |
| chr7:68735869-68737921 | p2 | (AT)26 | TGAGAGCACTCCATTGTTCG | AATCCAAGATGATTGTCGCC | 184 |
| chr7:68743491-68745507 | p2 | (AT)8 | GCACCCCTTAAGCACACATT | ATCCAAGAATGATTGTCGCC | 299 |
| chr7:68747933-68749945 | p2 | (GT)6 | GTTGCTGACTGTATCGGGGT | TTCCATTCAAGTACGGGCTC | 115 |
| chr7:68747933-68749945 | p5 | (TTGAA)6 | GTTGCTGACTGTATCGGGGT | TTCCATTCAAGTACGGGCTC | 115 |
| chr7:68795538-68797552 | p2 | (AT)7 | TCACAAGATTCATGCCTTGG | TGAGGACGTGAGCATTGAAG | 152 |
| chr7:68812563-68814575 | p2 | (GA)6 | ATGCGAAGTCCTCCTGAAAA | CGCAATTATTCGTATGCCCT | 233 |
| chr7:68841367-68843381 | p2 | (AT)7 | TCACAAGATTCATGCCTTGG | TGAGGACGTGAGCATTGAAG | 152 |
| chr7:68858392-68860404 | p2 | (GA)6 | ATGCGAAGTCCTCCTGAAAA | CGCAATTATTCGTATGCCCT | 233 |
| chr7:68906899-68908925 | p2 | (TA)13 | TATGGCCACACAAGACCAAA | ACGAGAGGAAATGACATGGG | 144 |
| chr7:68946097-68948133 | p6 | (CATATA)3 | AGGGGATACATAGGCGCTTT | GGCTCTAAGTAAGGGTCGGG | 297 |
| chr7:68946138-68948152 | p6 | (CATATA)3 | AGGGGATACATAGGCGCTTT | GGCTCTAAGTAAGGGTCGGG | 256 |
| chr7:68964412-68966432 | p4 | (AAAT)5 | CCTGAATTGAAACCATGCCT | GGTCCAAACAACTAGCCCAA | 201 |
| chr7:69039597-69041613 | p2 | (AT)8 | GTCCGCATTATGTCGTTGTG | AGCCTCAGTCCCATTTCCTT | 214 |
| chr7:69100616-69102631 | p5 | (ATTTC)3 | CCCTGTGAGGCATCGATATT | ACATCATACACTCGCACCCA | 288 |
| chr7:69197417-69199433 | p2 | (AT)8 | GGTTTCGGAGGCATGAATAA | CAACAAGTAACCGAGGCCAT | 186 |
| chr7:69197417-69199433 | p2 | (TA)7 | GGTTTCGGAGGCATGAATAA | CAACAAGTAACCGAGGCCAT | 186 |
| chr7:69204135-69206149 | p2 | (TA)7 | TTGCAGGGCATTTAGGAATC | CGGATCCTGTGGTCTCAAAT | 276 |
| chr7:69320044-69322100 | p2 | (TA)28 | AATGCGAGGCATAAAAATGG | GTATGCAAAGTTGCTTGCCA | 171 |
| chr7:69342518-69344574 | p2 | (TA)28 | AATGCGAGGCATAAAAATGG | GTATGCAAAGTTGCTTGCCA | 171 |
| chr7:69472982-69474997 | p3 | (TTG)5 | ACGTTTTCGGATGACCTGAC | TCGGTTCCACCGACTTACTC | 226 |
| chr7:69472982-69474997 | p4 | (TATG)5 | ACGTTTTCGGATGACCTGAC | TCGGTTCCACCGACTTACTC | 226 |
| chr7:69479556-69481568 | p2 | (AT)6 | ATAAGATGCCAACTCCACGG | CAGCTCTAGTTCCTCCACCG | 180 |
| chr7:69548360-69550375 | p3 | (TCA)5 | TGCTGCACTGTCAAATCTCC | CAGGGCCAGATTATTCTCCA | 260 |

| Marker ID | SSR type | SSR motif | FORWARD PRIMER (5'-3') | REVERSE PRIMER (5'-3') | size (bp) |
|---|---|---|---|---|---|
| chr7:69626842-69628854 | p5 | (TGATC)3 | ATGCATGTGCCATCAGTGTT | TGGTTCAGACAGACAGACGG | 272 |
| chr7:69626842-69628854 | p2 | (TA)6 | ATGCATGTGCCATCAGTGTT | TGGTTCAGACAGACAGACGG | 272 |
| chr7:69633131-69635143 | p2 | (AT)6 | ACTCCACGGATTCAGTTTCG | CGAGTTCAAGTGATCGGACA | 177 |
| chr7:69639391-69641409 | p6 | (AAAAAT)3 | TACAAGCGGCATGGATGATA | TAGCCAATTCCCGATCTGTC | 100 |
| chr7:69639391-69641409 | p2 | (CA)6 | TACAAGCGGCATGGATGATA | TAGCCAATTCCCGATCTGTC | 100 |
| chr7:69676656-69678668 | p2 | (AT)10 | GTCGTCTCAAAACTGTCGCA | ATGAAAGCCTTGTCGGCTTA | 214 |
| chr7:69676656-69678668 | p2 | (TA)6 | GTCGTCTCAAAACTGTCGCA | ATGAAAGCCTTGTCGGCTTA | 214 |
| chr7:69683678-69685698 | p2 | (TC)10 | GTTTCGCCCAAAACCTACAA | TCGTTAGAATTTCGGGCATC | 177 |
| chr7:69736894-69738909 | p5 | (ATATA)3 | AGTGAAATATGACGGGGCAG | TCATGCTCGCAGGTACAGAC | 299 |
| chr7:69736894-69738909 | p3 | (AAC)5 | AGTGAAATATGACGGGGCAG | TCATGCTCGCAGGTACAGAC | 299 |
| chr7:69763664-69765718 | p2 | (AT)8 | TTAATGAGCCACACCGTTCA | GGAAACAAAGAAGGAAGGGC | 125 |
| chr7:69763664-69765718 | p3 | (ATT)18 | TTAATGAGCCACACCGTTCA | GGAAACAAAGAAGGAAGGGC | 125 |
| chr7:69842969-69844984 | p3 | (GAA)5 | CAAGGTGATGTACTGGGCCT | TGTCCCGGTGGTTAATGAAT | 154 |
| chr7:69851346-69853361 | p3 | (GAA)5 | CAAGGTGATGTACTGGGCCT | TGTCCCGGTGGTTAATGAAT | 154 |
| chr7:69884624-69886650 | p2 | (TA)13 | GCTTTGTCAGTTTTGGGGAA | CGAAGGCGTATTAGAACCCA | 242 |
| chr7:69891096-69893108 | p2 | (TG)6 | CCATGTTGCATCAACACACA | TCCTCCATGTTTCCACATCA | 264 |
| chr7:69958174-69960186 | p3 | (CAA)5 | ACATACCTTCCAATGCGCTC | TGCGTTGTTGTATCGTTGGT | 257 |
| chr7:69958174-69960186 | p5 | (ATTTT)3 | ACATACCTTCCAATGCGCTC | TGCGTTGTTGTATCGTTGGT | 257 |
| chr7:69958250-69960264 | p3 | (CAA)5 | ACATACCTTCCAATGCGCTC | TGCGTTGTTGTATCGTTGGT | 181 |
| chr7:69958250-69960264 | p5 | (ATTTT)3 | ACATACCTTCCAATGCGCTC | TGCGTTGTTGTATCGTTGGT | 181 |
| chr7:69958275-69960309 | p3 | (CAA)5 | ACATACCTTCCAATGCGCTC | TGCGTTGTTGTATCGTTGGT | 156 |
| chr7:69958275-69960309 | p5 | (ATTTT)3 | ACATACCTTCCAATGCGCTC | TGCGTTGTTGTATCGTTGGT | 156 |
| chr7:69958310-69960326 | p3 | (CAA)5 | ACATACCTTCCAATGCGCTC | TGCGTTGTTGTATCGTTGGT | 121 |
| chr7:69958310-69960326 | p5 | (ATTTT)3 | ACATACCTTCCAATGCGCTC | TGCGTTGTTGTATCGTTGGT | 121 |
| chr7:69958501-69960516 | p3 | (CAA)5 | CACCAACGATACAACAACGC | TCAAAGGTTGGAGTTTTCGG | 165 |
| chr7:69958501-69960516 | p5 | (ATTTT)3 | CACCAACGATACAACAACGC | TCAAAGGTTGGAGTTTTCGG | 165 |
| chr7:69961101-69963113 | p2 | (GT)6 | GTGCCTCTCCCATGATCTGT | TTCCACCCCAACGTATTCAT | 280 |
| chr7:70025719-70027737 | p2 | (AT)10 | AAAAGGGCAATGTCACAACC | AGGCCGAAGGGCTTATGTAT | 283 |
| chr7:70025719-70027737 | p2 | (AT)9 | AAAAGGGCAATGTCACAACC | AGGCCGAAGGGCTTATGTAT | 283 |
| chr7:70113087-70115102 | p3 | (TCA)5 | CCAGATCGGGCACTAGTCAT | CCAGTTGCAGGTTGTTTCCT | 299 |
| chr7:70123391-70125406 | p5 | (ATCAT)3 | GATCTGAAAGCCAGCCCATA | TTGGGTCGTGACAATAAGCA | 215 |
| chr7:70123391-70125406 | p2 | (TA)10 | GATCTGAAAGCCAGCCCATA | TTGGGTCGTGACAATAAGCA | 215 |
| chr7:70213835-70215847 | p3 | (CAA)5 | AGTACCCCGAGCTTCCATCT | ACCATTTGCAGTTTTCGGAC | 191 |
| chr7:70213835-70215847 | p2 | (AC)6 | AGTACCCCGAGCTTCCATCT | ACCATTTGCAGTTTTCGGAC | 191 |
| chr7:70303729-70305741 | p2 | (AT)6 | TCCCGGAGTCTATACGATGC | GTTGCTTCGGGTCAACAAAT | 153 |
| chr7:70393184-70395196 | p2 | (TA)8 | TGATAGCCGAAGGGCTTATG | AGGGTTATTGCCCCAAATTC | 257 |

| Marker ID | SSR type | SSR motif | FORWARD PRIMER (5'-3') | REVERSE PRIMER (5'-3') | size (bp) |
|---|---|---|---|---|---|
| chr7:70393184-70395196 | p2 | (AT)6 | TGATAGCCGAAGGGCTTATG | AGGGTTATTGCCCCAAATTC | 257 |
| chr7:70405664-70407679 | p5 | (CATAA)3 | TCTGGGCTGCTAATCGTCTT | TCTGCTGCGTACAGATTTGG | 275 |
| chr7:70405664-70407679 | p5 | (TCATA)6 | TCTGGGCTGCTAATCGTCTT | TCTGCTGCGTACAGATTTGG | 275 |
| chr7:70466815-70468835 | p4 | (AAAG)5 | CCGCATATACACAACCCACA | ATGCCCGCAGGTACAGATAC | 214 |
| chr7:70487571-70489589 | p6 | (TATATG)3 | GTGGATTGATAATTTGGCCG | TTACATACCTGCTCCCCGTC | 265 |
| chr7:70494108-70496122 | p2 | (AT)8 | AATCCGACAAGGCTACATCG | ACAGGGAGACAAACAGGTGG | 264 |
| chr7:70494108-70496122 | p2 | (AT)7 | AATCCGACAAGGCTACATCG | ACAGGGAGACAAACAGGTGG | 264 |
| chr7:70569139-70571203 | p2 | (AT)12 | GAGCATGACGATACCGATGA | AGTTTTCCGGAAGGTAGGGA | 282 |
| chr7:70679734-70681746 | p2 | (TA)6 | ACGATCAGCAGAGGAGCATT | CCGACGATTGCCTAATGAAT | 270 |
| chr7:70786789-70788813 | p2 | (AT)12 | CGGGTTCTCGTGGTATCTGT | TCTGGCGCCATAATACATCA | 240 |
| chr7:70799589-70801605 | p2 | (TA)8 | TTGTGATGTGTGGGCTGATT | GCCCGAAGTGTCTCAACCTA | 256 |
| chr7:70799589-70801605 | p5 | (CGTAT)3 | TTGTGATGTGTGGGCTGATT | GCCCGAAGTGTCTCAACCTA | 256 |
| chr7:70843955-70845967 | p2 | (AT)6 | GGGATCTCGAGGATTGTTGA | TATTCGGCGTTTCACCTTTC | 231 |
| chr7:70847719-70849731 | p2 | (GT)6 | CTTTTCTGGCGCGTATTGTT | CAAAATTGCCCTTAGCCTTG | 285 |
| chr7:70935549-70937564 | p5 | (ATTTA)3 | CCAATCGGACAGACAGAGGT | TACTAAGAGGGCCGGGTTCT | 150 |
| chr7:70945646-70947660 | p2 | (AT)13 | AGAACTGTCACGACCCAACC | TGTTCGTCTGTATTGGCAGC | 146 |
| chr7:70945646-70947660 | p2 | (AT)7 | AGAACTGTCACGACCCAACC | TGTTCGTCTGTATTGGCAGC | 146 |
| chr7:70951977-70953989 | p2 | (AT)6 | TCCAAGTAAGAATGGTGCCC | GCCCAAATTTTCCCAGAAAT | 129 |
| chr7:70975920-70977938 | p6 | (CATGAG)3 | GCGTATTCGGTAGAATGGGA | GGGAGTCTGTGAAACCGTGT | 185 |
| chr7:71000106-71002121 | p5 | (ATTTC)3 | ACGGGCATGTATGTTGGATT | TTGGGGATCTACACTTTCGC | 102 |
| chr7:71000106-71002121 | p2 | (AT)15 | ACGGGCATGTATGTTGGATT | TTGGGGATCTACACTTTCGC | 102 |
| chr7:71085247-71087259 | p2 | (TA)6 | TGAGTTCTCAGGGGAAATGC | CACGAGCTCGCTTTATGACA | 253 |
| chr7:71149695-71151707 | p2 | (AC)6 | CATTCAAGCTTCCTTACGCC | TGGTTTGGGTTGTTGTTTGA | 126 |
| chr7:71201092-71203108 | p2 | (AT)8 | TGATCATGCCCACAGGTAGA | AATCGTCATAAGCACCTCCG | 120 |
| chr7:71227565-71229589 | p2 | (TA)12 | TTCAAGGCTGAAAGATGGCT | GCCTTGTTTGGCTTGTCTTC | 282 |
| chr7:71235889-71237903 | p2 | (TA)7 | TATTCGAGCTTGTCGCTCCT | TATAAAACGAAACGGGCTGC | 152 |
| chr7:71255454-71257466 | p2 | (GT)6 | GACTCCTACCCCGAGCTTTC | ACGAAATCATAGCGAATGGC | 120 |
| chr7:71321431-71323447 | p4 | (TGTA)4 | CGAATTTGATTTTGGGCACT | GACAAACCCGAGGAGACGTA | 247 |
| chr7:71381629-71383645 | p2 | (AC)8 | TCTCCCAAAAGTCATGTCCC | GTTTGTTGATGTGTGGGCTG | 300 |
| chr7:71433294-71435309 | p2 | (TA)6 | CCGGAGCATGTACTTTTGGT | ACCCATCCCACACCAATCTA | 148 |
| chr7:71433294-71435309 | p5 | (ATATG)3 | CCGGAGCATGTACTTTTGGT | ACCCATCCCACACCAATCTA | 148 |
| chr7:71459697-71461712 | p3 | (GTT)5 | CATAGGTCGTGGGCCTGTAT | TCTCTTCATGAGCTTCGGGT | 238 |
| chr7:71459697-71461712 | p2 | (TA)15 | CATAGGTCGTGGGCCTGTAT | TCTCTTCATGAGCTTCGGGT | 238 |
| chr7:71554373-71556389 | p2 | (AT)8 | CAAGGCTCATAGACATGCGA | TTCCAGTCGCGTATTCCTCT | 116 |
| chr7:71585481-71587499 | p3 | (ACA)6 | CAATCACATGCTCGTCGTCT | AATGCTGCCCAATTTTCTTG | 153 |
| chr7:71641249-71643265 | p2 | (TA)8 | CGCTTTCTTTGGAGGCATAG | GGTGGGTCGTGACAAAAAGT | 177 |

| Marker ID | SSR type | SSR motif | FORWARD PRIMER (5'-3') | REVERSE PRIMER (5'-3') | size (bp) |
|---|---|---|---|---|---|
| chr7:71645500-71647526 | p2 | (TA)13 | TGTGATTTGTTGTGGCTGGT | CACCTCCCCTATATCGCTCA | 186 |
| chr7:71671129-71673151 | p2 | (AC)11 | ACATACCTCTTTCCGCCCTT | TTCCCGGATTGTTGTTGATT | 283 |
| chr7:71672547-71674563 | p2 | (TA)8 | GTCATATTGCGCATTTCCCT | AAGGAGACCTCCCCGAAGTA | 288 |
| chr7:71822651-71824665 | p2 | (AT)6 | ATATATACCGTACCCGGCCC | ATACGAGTGTCCAGTTCGGG | 116 |
| chr7:71822651-71824665 | p2 | (AT)7 | ATATATACCGTACCCGGCCC | ATACGAGTGTCCAGTTCGGG | 116 |
| chr7:71884271-71886307 | p2 | (AT)18 | GTCGCTTTTCATGGGATCAT | CTAGCTGGGTCGTGACACCT | 256 |
| chr7:71884271-71886307 | p2 | (AT)9 | GTCGCTTTTCATGGGATCAT | CTAGCTGGGTCGTGACACCT | 256 |
| chr7:71884924-71886942 | p2 | (AT)18 | TGTCTACCTGTGGGCATGAA | CGTTCCGCAGCAATATATGA | 103 |
| chr7:71884924-71886942 | p2 | (AT)9 | TGTCTACCTGTGGGCATGAA | CGTTCCGCAGCAATATATGA | 103 |
| chr7:71929814-71931832 | p6 | (TCATAT)3 | ATGCCCCTTCCTGTTCTTCT | TTAGCCTCGTCGGCTTATGT | 203 |
| chr7:71951494-71953544 | p6 | (TCATAT)5 | TTCTAATTCCATTGGGCAGC | GGTCCTGTCCCGCCTATTAT | 243 |
| chr7:71951494-71953544 | p2 | (TA)25 | TTCTAATTCCATTGGGCAGC | GGTCCTGTCCCGCCTATTAT | 243 |
| chr7:72010071-72012089 | p2 | (AT)9 | GCGGGAAGACTACAAGCAAG | CGATTTAATTTCCAAGCCCA | 187 |
| chr7:72013433-72015445 | p2 | (TG)6 | TACTCGCACTGACCCCTCTT | CTATACCCACGATCTGCCGT | 136 |
| chr7:72072183-72074198 | p5 | (TGTTG)3 | AGGGTCTTTTCCCGTCCTTA | CCTCTCGGCCATCAAAATAA | 284 |
| chr7:72073474-72075486 | p2 | (AT)6 | GTTCTTCAAACCGTCCCTCA | CCGAATGATAAGCCGATTGT | 169 |
| chr7:72083899-72085915 | p2 | (AT)8 | TTGTCTGCCCCTTTGATTTC | TTACTCCAGCCCCCTATCCT | 134 |
| chr7:72083899-72085915 | p2 | (TA)7 | TTGTCTGCCCCTTTGATTTC | TTACTCCAGCCCCCTATCCT | 134 |
| chr7:72083899-72085915 | p2 | (AT)14 | TTGTCTGCCCCTTTGATTTC | TTACTCCAGCCCCCTATCCT | 134 |
| chr7:72150824-72152839 | p5 | (ATCAT)3 | GTTTGGAAAGAAGGGGAAGC | CGAGACATGGAAAACTGGGT | 122 |
| chr7:72356911-72358939 | p2 | (AT)14 | AGGTCGCAAATAAGAAGGCA | GTACCCAGCCGGACAAACTA | 278 |
| chr7:72372973-72374989 | p4 | (CAGA)4 | TCCGAACGGACAACCTAATC | TTGGGTCGTGACAGTGGTTA | 178 |
| chr7:72372973-72374989 | p5 | (GATCA)3 | TCCGAACGGACAACCTAATC | TTGGGTCGTGACAGTGGTTA | 178 |
| chr7:72381563-72383578 | p5 | (GATCA)3 | GTATGCTCCGGATCAAAGGA | TGACTCTGTTCGTCCGTCTG | 297 |
| chr7:72387735-72389750 | p5 | (GATCA)3 | GTATGCTCCGGATCAAAGGA | TGACTCTGTTCGTCCGTCTG | 297 |
| chr7:72618071-72620091 | p2 | (TA)10 | TGTCACAACCCTCAACCGTA | TGGTTCGTTGTTGGGGTATT | 126 |
| chr7:72648491-72650505 | p3 | (ACA)5 | GACGTCGTAAACCCTTCCAA | GTTTGTTGATGTGTGGGCTG | 291 |
| chr7:72648491-72650505 | p2 | (CA)7 | GACGTCGTAAACCCTTCCAA | GTTTGTTGATGTGTGGGCTG | 291 |
| chr7:72697739-72699754 | p6 | (TATATG)3 | CTTTCTTTGGGCTCTCATGC | TCATAGCGCTGAACAACGTC | 164 |
| chr7:72697739-72699754 | p5 | (CGGGT)3 | CTTTCTTTGGGCTCTCATGC | TCATAGCGCTGAACAACGTC | 164 |
| chr7:72697739-72699754 | p6 | (CATATA)3 | CTTTCTTTGGGCTCTCATGC | TCATAGCGCTGAACAACGTC | 164 |
| chr7:72732501-72734513 | p2 | (AT)18 | GAACAACGGAGCTCTCCAAG | CGCACTGACCCTCTTTCTTC | 181 |
| chr7:72732501-72734513 | p2 | (GT)6 | GAACAACGGAGCTCTCCAAG | CGCACTGACCCTCTTTCTTC | 181 |
| chr7:72732501-72734513 | p3 | (TGT)5 | GAACAACGGAGCTCTCCAAG | CGCACTGACCCTCTTTCTTC | 181 |
| chr7:72786629-72788641 | p2 | (AT)18 | CCAGTTTTCACCTTCACCGT | GGCATTCCCCTACACTACGA | 115 |
| chr7:72786629-72788641 | p2 | (AT)6 | CCAGTTTTCACCTTCACCGT | GGCATTCCCCTACACTACGA | 115 |

| Marker ID | SSR type | SSR motif | FORWARD PRIMER (5'-3') | REVERSE PRIMER (5'-3') | size (bp) |
|---|---|---|---|---|---|
| chr7:73070955-73072975 | p4 | (ATAA)5 | TGGGCTAAAAATCTGGTTCG | GTTTGCGATTGGAGTTTCGT | 222 |
| chr7:73095871-73097885 | p2 | (AG)7 | GTGGTGTGCACGTTGAAATC | CCTATCGATCGTCGGCTAAG | 263 |
| chr7:73102665-73104691 | p2 | (AG)13 | ATGACGAAGTGCACCATGAA | CGGTTGTCAGTTGTATTGCG | 188 |
| chr7:73115646-73117662 | p2 | (AT)8 | AAATTGGGCAGCATTTTGTC | GGTTAGCCGTGACTCCTCAG | 240 |
| chr7:73149663-73151709 | p2 | (AT)6 | TACGAGCATGACGATACCGA | TTTCCGGAAGGTAGGGAAGT | 164 |
| chr7:73149710-73151732 | p2 | (AT)6 | TACGAGCATGACGATACCGA | TTTCCGGAAGGTAGGGAAGT | 117 |
| chr7:73157413-73159439 | p2 | (AT)13 | CATGGCTCGCTTATTGGTTT | AACAGGCTTGCGAATGAGAT | 254 |
| chr7:73288546-73290560 | p2 | (CA)7 | ATAGAGTTGTCCCCCAGGCT | TGGCTAAGTTGGATTTTGGG | 204 |
| chr7:73292718-73294736 | p2 | (AT)9 | GACAATTTCAAACGCGGATT | GATGCTGCCCAAATTTCTGT | 184 |
| chr7:73351440-73353455 | p3 | (CAG)5 | GACTCCAACTCCTCAGCCAG | GAACTCTGGAGGGCCACATA | 248 |
| chr7:73385170-73387236 | p2 | (AT)33 | ATTCAAGCTTCCCAAACCCT | AAATTGAAAGGGGGAGGAGA | 193 |
| chr7:73385874-73387892 | p2 | (AT)33 | CGCCTTATAATTGGAGCAGC | CCCTTCCTTCACACGTTCAT | 181 |
| chr7:73385906-73387930 | p2 | (AT)33 | CGCCTTATAATTGGAGCAGC | CCCTTCCTTCACACGTTCAT | 149 |
| chr7:73385932-73387947 | p2 | (AT)33 | CGCCTTATAATTGGAGCAGC | CCCTTCCTTCACACGTTCAT | 123 |
| chr7:73754899-73756913 | p2 | (TA)7 | TGCCCTGTAGCTTCCTGTTT | TCACACCCTAGCGGGATTAG | 101 |
| chr7:73794668-73796683 | p5 | (CATAT)3 | TCCGATCTTCGTCACACATC | TATATGTCCGTCCCCGGTTA | 240 |
| chr7:73869130-73871145 | p5 | (TCATA)3 | TGGACATGGCTCGTAAACAA | TAAGGGGTGGTCGGTACAAG | 196 |
| chr7:73929618-73931636 | p3 | (TAA)6 | CGGCGTTGTGAGTAGCTGTA | CGATTGGAGGGTGAAAAAGA | 244 |
| chr7:73929618-73931636 | p3 | (AAT)13 | CGGCGTTGTGAGTAGCTGTA | CGATTGGAGGGTGAAAAAGA | 244 |
| chr7:73929618-73931636 | p4 | (TATG)5 | CGGCGTTGTGAGTAGCTGTA | CGATTGGAGGGTGAAAAAGA | 244 |
| chr7:73929750-73931765 | p3 | (TAA)6 | CGGCGTTGTGAGTAGCTGTA | CGATTGGAGGGTGAAAAAGA | 112 |
| chr7:73929750-73931765 | p3 | (AAT)13 | CGGCGTTGTGAGTAGCTGTA | CGATTGGAGGGTGAAAAAGA | 112 |
| chr7:73929750-73931765 | p4 | (TATG)5 | CGGCGTTGTGAGTAGCTGTA | CGATTGGAGGGTGAAAAAGA | 112 |
| chr7:73929750-73931765 | p2 | (TA)20 | CGGCGTTGTGAGTAGCTGTA | CGATTGGAGGGTGAAAAAGA | 112 |
| chr7:73967047-73969075 | p2 | (AC)6 | GGGAGTCACTAAGCCGTCTG | TCCAAAACAGCAACCAACAA | 164 |
| chr7:73967047-73969075 | p2 | (TA)14 | GGGAGTCACTAAGCCGTCTG | TCCAAAACAGCAACCAACAA | 164 |
| chr7:73992439-73994455 | p2 | (GA)8 | TGGTTCTAGGCAAACCCAAC | GGGATTGATGGGACATTTTG | 283 |
| chr7:73992439-73994455 | p3 | (TTG)5 | TGGTTCTAGGCAAACCCAAC | GGGATTGATGGGACATTTTG | 283 |
| chr7:73996153-73998168 | p3 | (CAG)5 | TGAGAGATGCGGTTCAGTTG | AATATGGTGGGTACAGGGCA | 294 |
| chr7:73998115-74000127 | p2 | (GA)6 | CAGTGTCGCGAGGTATGAAA | TCTCTCCATTCCCGAACATC | 217 |
| chr7:74039720-74041756 | p5 | (TGAAT)3 | CTTATTCTAGGGTTCGCCCC | CCCTCCCTAATTGTGCGTTA | 135 |
| chr7:74039855-74041869 | p5 | (TGAAT)3 | TAACGCACAATTAGGGAGGG | TACCAGTCACGACCCCTAGC | 122 |
| chr7:74127167-74129183 | p4 | (ATAC)4 | GACCCTTACTTAGGGCCGAG | GTAAGCGGCTCTATCATCCG | 163 |
| chr7:74139835-74141847 | p2 | (AT)6 | CTCTGAAGGTTCATAGCGCC | CTGTCCCGTCATATTTCGCT | 209 |
| chr7:74228207-74230222 | p3 | (TCA)5 | GACGCTCTGGAACTCTGACC | GCCTATGCTCTGGGTATGGA | 259 |
| chr7:74228207-74230222 | p2 | (TA)9 | GACGCTCTGGAACTCTGACC | GCCTATGCTCTGGGTATGGA | 259 |

| Marker ID | SSR type | SSR motif | FORWARD PRIMER (5'-3') | REVERSE PRIMER (5'-3') | size (bp) |
|---|---|---|---|---|---|
| chr7:74267087-74269107 | p5 | (TTCAT)4 | AAGGTGGATACCATGCTTGC | GATGATAACACCGCGCCTAT | 164 |
| chr7:74273603-74275635 | p2 | (AT)16 | ACCCAGGAGCAACAAAAATG | CGGGGTAAGCCTGTAATGAA | 203 |
| chr7:74273603-74275635 | p2 | (AT)17 | ACCCAGGAGCAACAAAAATG | CGGGGTAAGCCTGTAATGAA | 203 |
| chr7:74273603-74275635 | p2 | (AT)9 | ACCCAGGAGCAACAAAAATG | CGGGGTAAGCCTGTAATGAA | 203 |
| chr7:74290238-74292250 | p2 | (AC)6 | GGACATGCCAAAAGAACGAT | TGATGTTGGGCTGTTGTTGT | 238 |
| chr7:74309027-74311043 | p2 | (AT)8 | CGAAGGCTATGGGCGTATAA | TTCCACGTCTTTCCGTTACC | 169 |
| chr7:74309027-74311043 | p3 | (TTG)5 | CGAAGGCTATGGGCGTATAA | TTCCACGTCTTTCCGTTACC | 169 |
| chr7:74309982-74311997 | p2 | (AT)8 | TGACTTGGAGGGGACAAAAC | GGTTTCACCGACTTCCTCAA | 255 |
| chr7:74309982-74311997 | p3 | (TTG)5 | TGACTTGGAGGGGACAAAAC | GGTTTCACCGACTTCCTCAA | 255 |
| chr7:74337188-74339208 | p2 | (AT)10 | TCACCTCCAAACTGAGGACC | TTCCAGCTATTCGAGCCAGT | 181 |
| chr7:74350860-74352872 | p2 | (CT)6 | AACCATACAACACAACGGCA | GATTTCCGAGCATCGTTTGT | 169 |
| chr7:74350860-74352872 | p2 | (TA)7 | AACCATACAACACAACGGCA | GATTTCCGAGCATCGTTTGT | 169 |
| chr7:74350860-74352872 | p2 | (AT)10 | AACCATACAACACAACGGCA | GATTTCCGAGCATCGTTTGT | 169 |
| chr7:74369968-74372000 | p2 | (TA)9 | ATTCCCGGATCTGGAAAATC | TTAGCGGGCTTCTGACAGTT | 129 |
| chr7:74369968-74372000 | p2 | (AT)16 | ATTCCCGGATCTGGAAAATC | TTAGCGGGCTTCTGACAGTT | 129 |
| chr7:74418297-74420315 | p6 | (ATACAT)3 | TGTGACTTTCCATATCGCCA | CTAGCGGGATTAGGGTGTGA | 117 |
| chr7:74418297-74420315 | p3 | (CAT)5 | TGTGACTTTCCATATCGCCA | CTAGCGGGATTAGGGTGTGA | 117 |
| chr7:74483087-74485111 | p6 | (ATATGT)4 | ATCAACCCGTCATTCCAAAA | TACACGTACCCGGAGACACA | 118 |
| chr7:74483340-74485386 | p6 | (ATATGT)4 | GAAAATTTCTTCATCGGCCA | CCTAGGGGCAAATCAAGTTTC | 134 |
| chr7:74543044-74545064 | p2 | (AT)6 | TTCGGGTAATGTGGGGATTA | GACAAGTTGCACCTCGATGA | 266 |
| chr7:74645329-74647344 | p3 | (TGA)5 | TAGCCTTGTCGGCTTCTGAT | ACACCGCCTGTAGACATTCC | 209 |
| chr7:74737146-74739162 | p4 | (TGTA)4 | CATTTGCAGTTTTTGAGCGA | GGCCGAGCCTATTACATTCA | 111 |
| chr7:74912399-74914421 | p2 | (AT)11 | CATAAGCCGACGAGGCTAAC | AGCCAGGGAGACAGACTTGA | 292 |
| chr7:74921170-74923182 | p2 | (AC)6 | CAAATGCGACAAAAAGGGTT | CGGTAGCCAAATATAGCCCA | 129 |
| chr7:74923451-74925463 | p2 | (CA)6 | GACTCTATGGTGGCCCAAAA | ATTCGCTTGTCTCAACGCTT | 105 |
| chr7:74975067-74977079 | p2 | (AT)6 | AGTGGCGGTGTAGCAGAACT | AGCTACCCAAAAGGGCAAAT | 212 |
| chr7:74975067-74977079 | p2 | (AT)12 | AGTGGCGGTGTAGCAGAACT | AGCTACCCAAAAGGGCAAAT | 212 |
| chr7:75038233-75040265 | p2 | (AT)16 | TACCACGTTGAAGAGGACCC | CAAGGCCCAAGTGGAATTTA | 224 |
| chr7:75038233-75040265 | p2 | (AT)20 | TACCACGTTGAAGAGGACCC | CAAGGCCCAAGTGGAATTTA | 224 |
| chr7:75145707-75147719 | p5 | (ATATC)3 | TTATGGCTGACGTGGGTGTA | CCGCTTATTTGAAATTGGGA | 282 |
| chr7:75145757-75147769 | p5 | (ATATC)3 | TTATGGCTGACGTGGGTGTA | CCGCTTATTTGAAATTGGGA | 232 |
| chr7:75163256-75165277 | p3 | (ATT)7 | CCGAGATTCACCATGTCCTT | CACAAACGCGCAGTTAAGAA | 164 |
| chr7:75274465-75276485 | p2 | (AT)10 | CCTTGCATCCAATTCCACTT | GGCCAACTTCAATGATGGAT | 191 |
| chr7:75343074-75345086 | p2 | (AT)6 | AAGGGTTTCGGACACAACAG | CATTTGCAGTTTCTGGACGA | 281 |
| chr7:75381105-75383121 | p2 | (TA)8 | GGTATGAATGCAATGCGATG | TGCACAGTATCTGTAGCGGC | 273 |
| chr7:75398514-75400528 | p2 | (TG)7 | ACGATTGTACTACGCACCCC | AATAGCCCGTAGGCACCTTT | 243 |

| Marker ID | SSR type | SSR motif | FORWARD PRIMER (5'-3') | REVERSE PRIMER (5'-3') | size (bp) |
|---|---|---|---|---|---|
| chr7:75409483-75411521 | p2 | (TA)19 | ACATTCTGGGCGTACTTTGG | GTAGGTCTGTCTTCTGGGCG | 162 |
| chr7:75500748-75502792 | p2 | (TA)22 | GGACATGCCAAAAGAAGGAA | GCCGAAATTTCTGTAGGCAA | 110 |
| chr7:75533271-75535286 | p5 | (ATTTC)3 | GGCAGAATCCTTCATGGAAA | TGGGACCTGATGAACACAGA | 225 |
| chr7:75533271-75535286 | p3 | (GAT)6 | GGCAGAATCCTTCATGGAAA | TGGGACCTGATGAACACAGA | 225 |
| chr7:75597929-75599944 | p5 | (TTCAT)3 | GGATAGATGTTTGACCGCGT | TGCTCGTGTTCACTCCTTTG | 135 |
| chr7:75788722-75790734 | p2 | (AT)6 | GTAAGGGTGGACGAAAGGGT | TCTTCTTCACAAGCTCGGGT | 240 |
| chr7:75788722-75790734 | p2 | (AT)6 | GTAAGGGTGGACGAAAGGGT | TCTTCTTCACAAGCTCGGGT | 240 |
| chr7:75828133-75830153 | p2 | (AT)10 | ACTTTTGTCACGACCCAACC | CTTGTCAGCTTGCGTGTCAT | 189 |
| chr7:75876673-75878701 | p2 | (AT)14 | ATGCTCCACCTCTGCCTCTA | CAGATTCTCAGTTTGGGGGA | 243 |
| chr7:75917496-75919511 | p5 | (GCATT)3 | TGGCAAAGCAAATGAGTGAG | CCATCCAAACTTCATACCCG | 167 |
| chr7:75925881-75927896 | p5 | (ATATG)3 | AGCGCTCTTTTGTTCAGAGC | GACGAGGCTGTCACAGATCA | 196 |
| chr7:75992882-75994894 | p2 | (TA)6 | CGGAAAAACCTCCTAGACCC | AGGAAGCGTGCTAGTTGCAT | 145 |
| chr7:76056330-76058345 | p5 | (AATGC)3 | ATCGTACCGATCCACGAGAC | AAGTTTCCACATGACCCAGG | 228 |
| chr7:76116680-76118695 | p2 | (AT)9 | AGGGACTCGGTGAACAAATG | AAAAAGGGTTCTGCAAGGGT | 213 |
| chr7:76116680-76118695 | p3 | (AAC)5 | AGGGACTCGGTGAACAAATG | AAAAAGGGTTCTGCAAGGGT | 213 |
| chr7:76116680-76118695 | p2 | (AT)7 | AGGGACTCGGTGAACAAATG | AAAAAGGGTTCTGCAAGGGT | 213 |
| chr7:76182022-76184040 | p6 | (TATATG)3 | TGATATGTTTCCGAATGGCA | GCCGTTTTTGGAAAATCAAA | 231 |
| chr7:76204215-76206233 | p3 | (TTG)6 | TGGAAGCCACATGGTGTAAA | TTCTTGAATCGTCCCTTTGG | 125 |
| chr7:76271343-76273359 | p2 | (TA)8 | CTTTGGGCGACAATCATCTT | CAGAGCTCCCATTTCTTTCG | 256 |
| chr7:76279798-76281816 | p3 | (CAA)6 | ATTGCCCAACCAACTGTAGC | CCACGGGTTAAGTGATGACC | 175 |
| chr7:76314766-76316794 | p2 | (AT)14 | CCAACTAAGGAAAGCGCAAC | ATGTTGGCCTTGTCGAACTC | 294 |
| chr7:76314766-76316794 | p2 | (AT)13 | CCAACTAAGGAAAGCGCAAC | ATGTTGGCCTTGTCGAACTC | 294 |
| chr7:76314766-76316794 | p2 | (GT)8 | CCAACTAAGGAAAGCGCAAC | ATGTTGGCCTTGTCGAACTC | 294 |
| chr7:76330986-76333014 | p2 | (AT)14 | CCAACTAAGGAAAGCGCAAC | ATGTTGGCCTTGTCGAACTC | 294 |
| chr7:76330986-76333014 | p2 | (AT)13 | CCAACTAAGGAAAGCGCAAC | ATGTTGGCCTTGTCGAACTC | 294 |
| chr7:76330986-76333014 | p2 | (GT)8 | CCAACTAAGGAAAGCGCAAC | ATGTTGGCCTTGTCGAACTC | 294 |
| chr7:76360111-76362126 | p3 | (ATT)5 | TCTTGCAAAGGACCGGATAC | CGAAGAACAGGGGGTCAGTA | 157 |
| chr7:76435755-76437773 | p2 | (TA)7 | TTCCGGAGCTAGCTTGTTGT | ACACGCAACATCCCAACATA | 251 |
| chr7:76435783-76437795 | p2 | (TA)7 | TTCCGGAGCTAGCTTGTTGT | ACACGCAACATCCCAACATA | 223 |
| chr7:76505845-76507860 | p5 | (ATTTC)3 | TGTTCTTCCGCATGTTGTGT | CTAGCGACCAAACACCCCTA | 261 |
| chr7:76510687-76512699 | p2 | (AT)6 | TACCAACCAAGGCTTGTTCC | TCGTGTCGTGACACAATTCA | 110 |
| chr7:76589793-76591808 | p5 | (ATTTC)3 | GAAGCGGAAGAAAGTATGCG | AATATGGCGCTACCATCCTG | 184 |
| chr7:76637966-76639984 | p2 | (GA)9 | TTCACAAGGATCATGCCAAA | ACACGAACTTTTCTGGGGTG | 231 |
| chr7:76637966-76639984 | p3 | (GTT)5 | TTCACAAGGATCATGCCAAA | ACACGAACTTTTCTGGGGTG | 231 |
| chr7:76656506-76658518 | p2 | (AT)6 | TTTATTCCGTAGACACCGGC | TGGATCGTGACACACCAGTT | 190 |
| chr7:76656506-76658518 | p2 | (AT)22 | TTTATTCCGTAGACACCGGC | TGGATCGTGACACACCAGTT | 190 |

| Marker ID | SSR type | SSR motif | FORWARD PRIMER (5'-3') | REVERSE PRIMER (5'-3') | size (bp) |
|---|---|---|---|---|---|
| chr7:76656506-76658518 | p2 | (TA)6 | TTTATTCCGTAGACACCGGC | TGGATCGTGACACACCAGTT | 190 |
| chr7:76704618-76706634 | p2 | (TA)8 | ACCTTGGCAACCTTTGTCAC | GCATAGCGCGTGACGTAGTA | 138 |
| chr7:76704618-76706634 | p4 | (TATG)5 | ACCTTGGCAACCTTTGTCAC | GCATAGCGCGTGACGTAGTA | 138 |
| chr7:76721159-76723174 | p5 | (AAACA)3 | AACGACATCAACCCTTGGAG | TTCATTCCGTCCTCCACTTC | 254 |
| chr7:76721159-76723174 | p2 | (TA)6 | AACGACATCAACCCTTGGAG | TTCATTCCGTCCTCCACTTC | 254 |
| chr7:76721159-76723174 | p3 | (TTG)5 | AACGACATCAACCCTTGGAG | TTCATTCCGTCCTCCACTTC | 254 |
| chr7:76723740-76725755 | p5 | (AAACA)3 | AACGACATCAACCCTTGGAG | TTCATTCCGTCCTCCACTTC | 254 |
| chr7:76723740-76725755 | p2 | (TA)6 | AACGACATCAACCCTTGGAG | TTCATTCCGTCCTCCACTTC | 254 |
| chr7:76723740-76725755 | p3 | (TTG)5 | AACGACATCAACCCTTGGAG | TTCATTCCGTCCTCCACTTC | 254 |
| chr7:76839636-76841678 | p5 | (ACTTA)3 | GATCTGGAGGCCAAAGACAG | CCGACGGGTTAGTGAGACAT | 128 |
| chr7:76864280-76866320 | p2 | (AT)20 | GCGCCACAGAAGAAGAAAAC | ATCCGTTCCATTCACTCAGG | 234 |
| chr7:77054720-77056734 | p2 | (AG)7 | CCACGGAGGACAGCTATGAT | TCCTCCGACGGATCTGTATC | 143 |
| chr7:77151963-77153977 | p2 | (AT)7 | GAGGTGCATTGACTCCCCTA | GTGCAAGAGCCCATGATTTT | 105 |
| chr7:77200061-77202077 | p2 | (AT)8 | ATGGTTACTCACCACTCCGC | AGTGAGTGGCTCGGTGAACT | 226 |
| chr7:77200061-77202077 | p3 | (TGA)5 | ATGGTTACTCACCACTCCGC | AGTGAGTGGCTCGGTGAACT | 226 |
| chr7:77297109-77299127 | p6 | (ATCCGT)3 | TGACGTCCTTTTGTTTGTGG | TCGTATGAGCGGGATAGGAC | 273 |
| chr7:77318060-77320075 | p5 | (ATGAT)3 | TTGGCCCAGTTAAAATCTCG | AGTTGCACCTTACGCGTTCT | 171 |
| chr7:77428268-77430286 | p6 | (AATATG)3 | AATTCCCGGATTGTTGTTGA | AATTCAAGTTGCAAATCCGC | 280 |
| chr7:77609681-77611709 | p3 | (AAG)6 | TCAAATCCTCCAAATCAGGC | TTAGATGGACGGAGGAGGAA | 112 |
| chr7:77645370-77647384 | p5 | (ATTTC)3 | AGGCAAAAGGAGGGGTAAAA | TTTTGGGACTCGATTTTTCG | 205 |
| chr7:77645402-77647466 | p5 | (ATTTC)3 | AGGCAAAAGGAGGGGTAAAA | TTTTGGGACTCGATTTTTCG | 173 |
| chr7:77659678-77661706 | p2 | (AT)14 | GTCCTCCATGCACATTCCTT | CGAACTTCAAAGTCAACGCA | 128 |
| chr7:77663533-77665548 | p5 | (ATTTC)3 | GCCAACATCAATGCTCACAC | TTGAACTTGGAAGGTTTCGG | 116 |
| chr7:77760820-77762835 | p3 | (TTG)5 | ATCCGAGGCTCATGAAGAGA | ATCCAACTAAAACCGCTCCC | 107 |
| chr7:77787209-77789233 | p2 | (AT)12 | ATATAGAGCACCGGCACACC | CTGGGCGGCTGTATGATATT | 134 |
| chr7:77800840-77802868 | p2 | (TA)14 | TTTGGCCGAGTTGAATTCTC | ATTAAGTCTCGGCTTTCGCA | 277 |
| chr7:77818281-77820296 | p2 | (TA)6 | GCCTTTTCAAAGCATTCTCG | AAACTCGTGGGCTGTGTAGG | 223 |
| chr7:77818281-77820296 | p5 | (CATAT)3 | GCCTTTTCAAAGCATTCTCG | AAACTCGTGGGCTGTGTAGG | 223 |
| chr7:77890869-77892895 | p2 | (TA)25 | TTCGTCACCGTGGTTGATTA | AACAATGATGCCAGAGGGAC | 162 |
| chr7:77902174-77904189 | p5 | (GAAAT)3 | ATTCCGTACTGACCCCCTCT | GGAGCCGCTTACAGACTGAC | 165 |
| chr7:77907729-77909741 | p2 | (AT)6 | AGGATAGGGGGCTGGAGTAA | GATATCTAGGGAGGCGGTCC | 132 |
| chr7:77907729-77909741 | p2 | (AT)25 | AGGATAGGGGGCTGGAGTAA | GATATCTAGGGAGGCGGTCC | 132 |
| chr7:77908178-77910228 | p2 | (AT)6 | AGACCCAAAGGTTCCCATTC | GGCCCTGTCACGTCATATTT | 269 |
| chr7:77908178-77910228 | p2 | (AT)25 | AGACCCAAAGGTTCCCATTC | GGCCCTGTCACGTCATATTT | 269 |
| chr7:77914589-77916604 | p5 | (TGTGT)3 | ATGCAGTTCAATTCCTTGGC | CCACCTGTGCCCTCATATCT | 119 |
| chr7:77976319-77978335 | p2 | (AT)8 | CTCCAAAGACCTGGCAACTC | TATTCTGGGCATTCTTTGGC | 170 |

| Marker ID | SSR type | SSR motif | FORWARD PRIMER (5'-3') | REVERSE PRIMER (5'-3') | size (bp) |
|---|---|---|---|---|---|
| chr7:78001655-78003681 | p2 | (TA)13 | ATTGGAAATTCGGGTTAGGC | TTCGCCCCTTTAGCCTATTT | 201 |
| chr7:78178035-78180061 | p2 | (CA)6 | GCAACGACGCAACTAACAAA | ATGCATGGTGTGTGTGTGTG | 168 |
| chr7:78178078-78180098 | p2 | (CA)6 | GCAACGACGCAACTAACAAA | ATGCATGGTGTGTGTGTGTG | 125 |
| chr7:78310891-78312903 | p2 | (AT)6 | ACGAAAGGAATTACCCCACC | ACAACATTTGCTCTCCCTGG | 120 |
| chr7:78322164-78324176 | p2 | (AG)6 | AGGGAGACTGTCCCCAAAAT | ATCTCACGTCGATTCCTTCG | 136 |
| chr7:78340470-78342482 | p2 | (TC)6 | ACAAGAGCAGCCACGAAAAT | AGTGGGCAGATTTGGAGTTG | 139 |
| chr7:78373596-78375611 | p3 | (AAC)5 | GGGTATTCGGTTAGCGTTCA | AAGGCCCCGACACTTTTTAT | 296 |
| chr7:78470059-78472075 | p2 | (AT)9 | GGCTGCCACTATGAAACGAT | TCTTGGAGAGCTCCGTTGTT | 283 |
| chr7:78470059-78472075 | p2 | (TA)8 | GGCTGCCACTATGAAACGAT | TCTTGGAGAGCTCCGTTGTT | 283 |
| chr7:78500263-78502278 | p3 | (GAA)5 | CAAGGTGATGTACTGGGCCT | TGTCCCGGTGGTTAATGAAT | 154 |
| chr7:78560155-78562170 | p3 | (ATG)5 | GCCCCTTTCTTTAGGGACTG | AGGCTCCGAACAATAGAGCA | 179 |
| chr7:78560155-78562170 | p6 | (CTTATG)3 | GCCCCTTTCTTTAGGGACTG | AGGCTCCGAACAATAGAGCA | 179 |
| chr7:78637824-78639860 | p6 | (CATATA)6 | AGCGGTGGAAAAGATCAATG | TATGGGCCCAGGATAGTCAG | 126 |
| chr7:78689912-78691928 | p4 | (AAGA)4 | CGTGCCACAGAGCTAAATCA | AGAAAACACATCCAATGCCC | 264 |
| chr7:78693744-78695762 | p6 | (TATATG)3 | GGCACTTCCACGACGATAAT | TAGAATGAAATCGGGGCAAC | 158 |
| chr7:78706076-78708091 | p5 | (ATATG)3 | GGCTTCCATGCATGCTATTT | ACGCTGAATACATCGGCTCT | 191 |
| chr7:78706076-78708091 | p5 | (AAAAT)3 | GGCTTCCATGCATGCTATTT | ACGCTGAATACATCGGCTCT | 191 |
| chr7:78706076-78708091 | p3 | (ATG)5 | GGCTTCCATGCATGCTATTT | ACGCTGAATACATCGGCTCT | 191 |
| chr7:78778407-78780421 | p2 | (AT)7 | CCAAAAGAAGAATCGGGACA | GTGTTGTTGTTGGGTTGCTG | 123 |
| chr7:78806897-78808911 | p2 | (TA)7 | GAAAGAAAGAACGCAGTGCC | TGTGGGTCCCCTACAAACAT | 283 |
| chr7:78806897-78808911 | p5 | (TGAAT)3 | GAAAGAAAGAACGCAGTGCC | TGTGGGTCCCCTACAAACAT | 283 |
| chr7:78918031-78920061 | p2 | (AT)15 | CAATCGAACCTCTCGGTCAT | GGAAGAGGGGAGCATTTAGG | 270 |
| chr7:78945117-78947131 | p2 | (AT)7 | GCCAAAACAAATGGGCTAGA | AAATGGCCCTTCCAAGTTTT | 287 |
| chr7:78950566-78952581 | p2 | (TA)11 | TCACCTCCTGTCAACTGTGC | TAGCGATGATGGTTGTTCCA | 166 |
| chr7:78950566-78952581 | p3 | (CAA)5 | TCACCTCCTGTCAACTGTGC | TAGCGATGATGGTTGTTCCA | 166 |
| chr7:79028718-79030738 | p2 | (TA)12 | ATCGTGCACAGTTTCACTGC | ACCGTACCCGACCCTCTAGT | 197 |
| chr7:79028718-79030738 | p2 | (AT)10 | ATCGTGCACAGTTTCACTGC | ACCGTACCCGACCCTCTAGT | 197 |
| chr7:79039103-79041118 | p3 | (TTG)5 | TCCTTTCAAGAACGTTTGGG | AGAGCTTTATACGGGGCCAT | 134 |
| chr7:79165518-79167536 | p6 | (TATATG)3 | AATTGGACGTGGGACAGGTA | TCGGACATCAAAACCACGTA | 212 |
| chr7:79172190-79174208 | p6 | (CATATA)3 | GGCAATGTGTTGTCGTTGTC | AGCCTTGTCGGCTTCTGTTA | 145 |
| chr7:79189499-79191525 | p2 | (TA)13 | CGCCTGTGAAGGGAATAAAA | TGTCCTTCCCAGGTCATTTC | 251 |
| chr7:79189499-79191525 | p2 | (TG)18 | CGCCTGTGAAGGGAATAAAA | TGTCCTTCCCAGGTCATTTC | 251 |
| chr7:79189499-79191525 | p5 | (TGTGT)3 | CGCCTGTGAAGGGAATAAAA | TGTCCTTCCCAGGTCATTTC | 251 |
| chr7:79193064-79195088 | p2 | (TA)12 | AAACTGGAAGCACTGGATGG | CCTTTCCACGCTTCTTTCTG | 214 |
| chr7:79208532-79210550 | p6 | (ATATAC)3 | TTCTAGAGCTCCCCACTCCA | CCCTCAAAAATCTTGTCCCA | 119 |
| chr7:79208532-79210550 | p5 | (ATCAT)4 | TTCTAGAGCTCCCCACTCCA | CCCTCAAAAATCTTGTCCCA | 119 |

| Marker ID | SSR type | SSR motif | FORWARD PRIMER (5'-3') | REVERSE PRIMER (5'-3') | size (bp) |
|---|---|---|---|---|---|
| chr7:79225686-79227698 | p2 | (AT)6 | ACGCCCCAAACATAAGTCAG | CCTGTCCCGTCAATGATTCT | 281 |
| chr7:79229824-79231838 | p2 | (AT)7 | CCAAAACCATAATTCACCCG | TTGTGTTGTTGTGGCCTTGT | 130 |
| chr7:79282381-79284393 | p2 | (TG)6 | TCTTTGGGTCAGAGTGGGAC | ATTACTTCCCAGGGGGTCAC | 105 |
| chr7:79285127-79287142 | p3 | (GTT)5 | CATGGGAGGCAACCTAAAAA | TGCACCGCTTCTTCTTTCTT | 223 |
| chr7:79297216-79299238 | p2 | (AT)11 | AGCCGATAATGCTAAGGCAA | ATAACGCGAGGCATGTTACC | 254 |
| chr7:79306068-79308082 | p2 | (AT)7 | GCATGCAATACTCCAGGGAT | TGTTTCGAGCTGTTCTGCTG | 118 |
| chr7:79312750-79314764 | p2 | (AG)7 | GTTTGAGCCAAATTACCGGA | AATCAGCCCGGAGAATACCT | 251 |
| chr7:79376218-79378230 | p2 | (CA)6 | TCAAATTTGGCTCGGATTTC | CTGCCGAAATTTTTGTAGGC | 216 |
| chr7:79416702-79418718 | p4 | (AGAA)4 | TGGTAAGGAACCGAGGAGTG | CTTTTCGCTCTTTCACGGAC | 264 |
| chr7:79578277-79580295 | p2 | (AT)9 | GGAAAGTCCCTACGAAAGGC | CCTCTCTCCCTGGCTACCTT | 158 |
| chr7:79590219-79592263 | p2 | (AT)22 | ACTTTTGTCACGACCCAACC | AGCCTTGTCGGCTTGTGTAT | 134 |
| chr7:79603471-79605489 | p3 | (TGA)6 | GGTCATGGAGGACCTTGAAA | TGGGCAACATCTCCTTTGTT | 282 |
| chr7:79611662-79613676 | p3 | (AAC)5 | TCAATCTCGGATTTCGAAGG | CGGGAATTAAATTTGAGCGA | 195 |
| chr7:79611662-79613676 | p2 | (CA)7 | TCAATCTCGGATTTCGAAGG | CGGGAATTAAATTTGAGCGA | 195 |
| chr7:79625466-79627480 | p2 | (AG)7 | CCATTTGATCTGGAAGCGAT | CTCTGCTCCCCATGTCATTT | 214 |
| chr7:79755601-79757627 | p2 | (AT)13 | CCGGAGTCTATAGGATGCCA | CAAGTACAAAGGCCCGTGAT | 164 |
| chr7:79764653-79766668 | p3 | (ATA)5 | GATAGCCCCCATGACTCGTA | TAGTTCGAGAGCCCGTGAGT | 256 |
| chr7:79784180-79786194 | p2 | (AT)7 | GGCTGATGAATCATTTGGCT | CAGTGGGGAGGTATTGTGCT | 160 |
| chr7:79784180-79786194 | p3 | (ACA)6 | GGCTGATGAATCATTTGGCT | CAGTGGGGAGGTATTGTGCT | 160 |
| chr7:79835671-79837686 | p3 | (ATC)5 | CGAATGAAATAGAGCTCCGC | AAAGTTCCTGCGTAAGGCAA | 272 |
| chr7:79971346-79973361 | p5 | (ATCAT)3 | AAGGCTCTCTTTCTACCCGC | CATGAGGCTACAGGGCATTT | 122 |
| chr7:79973731-79975763 | p2 | (TA)16 | ACCAACTGGATCAAAGTGCC | AACCATGGACATTCTGCCTC | 200 |
| chr7:80014193-80016205 | p2 | (AT)6 | TAGACAATAAGGGGGATGCG | AGGGAAACACACCTTTGTCG | 197 |
| chr7:80048717-80050729 | p6 | (TGATAA)4 | TTGGGTACGGATAACCCTGA | ACGAAGGTTCGGTGTTTCAC | 223 |
| chr7:80048737-80050765 | p6 | (TGATAA)4 | TTGGGTACGGATAACCCTGA | ACGAAGGTTCGGTGTTTCAC | 203 |
| chr7:80065801-80067817 | p2 | (GT)8 | GGAAGAAAAGACCTGGGAGG | TGTAGAGCATAACGGGGGAG | 126 |
| chr7:80096371-80098383 | p2 | (TG)6 | AAAGGTGCCTACGGGCTATT | TCATACCTGGCCCTACCAAG | 223 |
| chr7:80112235-80114281 | p2 | (AT)23 | GACGGGAATTCAAGTGGAGA | GGCCCAAAGTGTATAGGGGT | 251 |
| chr7:80145693-80147723 | p2 | (TA)15 | GAATGGACGAAGTTGGGTGT | AAGCTAGCGAAAATCGGACA | 250 |
| chr7:80230093-80232109 | p4 | (ATTT)4 | ACTAATGTTGGCGTGCTGTG | GGTTTGTGCTTTCTTTCCCA | 100 |
| chr7:80275831-80277846 | p5 | (ATATG)3 | TTGTGGATCGTGGTCATTGT | TTCGAATAAATCCGCGTTTC | 241 |
| chr7:80286851-80288899 | p2 | (TA)6 | ACCACCTGTCGGTCTACCTG | GCATATTATCCGCCCACAGT | 135 |
| chr7:80286851-80288899 | p2 | (TA)24 | ACCACCTGTCGGTCTACCTG | GCATATTATCCGCCCACAGT | 135 |
| chr7:80396074-80398092 | p3 | (AAT)6 | GCTGGTGATTGGGTGTTCTT | CAACCTTGCCTACCCTACGA | 160 |
| chr7:80401686-80403708 | p2 | (TA)11 | ACGCGTATTCGGTAGAATGG | GTGTCCAGTGGGGTCACTTT | 138 |
| chr7:80401686-80403708 | p5 | (ATATA)3 | ACGCGTATTCGGTAGAATGG | GTGTCCAGTGGGGTCACTTT | 138 |

| Marker ID | SSR type | SSR motif | FORWARD PRIMER (5'-3') | REVERSE PRIMER (5'-3') | size (bp) |
|---|---|---|---|---|---|
| chr7:80455885-80457901 | p2 | (CA)8 | TCATCATTATCCTCCTCCGC | GGACCCCTCTACGACTCCTC | 194 |
| chr7:80498205-80500271 | p2 | (AT)33 | CGTGCTCCTCAGAAATAGGC | AAACAGGAGGCTACGGGAAT | 191 |
| chr7:80498205-80500271 | p2 | (AT)9 | CGTGCTCCTCAGAAATAGGC | AAACAGGAGGCTACGGGAAT | 191 |
| chr7:80522582-80524597 | p3 | (AAT)5 | ATGTTGGCTGTGTGGTTCAA | TGCACCAGAAAAAGTGCAAC | 125 |
| chr7:80545123-80547138 | p5 | (ATATG)3 | TCTTGAAAGTGTCGCCAGTG | TTGCACATCATCTATCGGGA | 243 |
| chr7:80649205-80651220 | p5 | (TTTTC)3 | ACAACACACGACTTCCACCA | GCCGAATGAGGTGTTGATTT | 228 |
| chr7:80715809-80717849 | p2 | (AT)20 | CACCTTGTACCTTCGGGCTA | ATATGAGTGTCCAGCTCGGG | 197 |
| chr7:80804958-80806972 | p2 | (TA)7 | TTTTCACTTTGGGAAAACGC | ACCCATAAGCTTGGGAGGAT | 268 |
| chr7:80804958-80806972 | p2 | (AT)7 | TTTTCACTTTGGGAAAACGC | ACCCATAAGCTTGGGAGGAT | 268 |
| chr7:80895794-80897806 | p2 | (TA)6 | CCTAGGACGGAACTGCTCTG | TCATGCCATTTATGGTCCCT | 264 |
| chr7:80933791-80935806 | p3 | (GTT)5 | CATGTGAGGACCGAGGAGTT | CTTCCCGCAATTATCTCGAA | 104 |
| chr7:81117558-81119586 | p2 | (AT)14 | TGCCCTTAACCTACCACGAC | ATCCTTCCTTCCCTTTCCCT | 121 |
| chr7:81125250-81127265 | p5 | (TGCAA)3 | ATTCCTACTCACACGACGGG | ATCGCAGTGGGAATGATAGG | 170 |
| chr7:81136548-81138564 | p2 | (AT)8 | GCGGGATCTAGCCAATAACA | CGTGACAGGCAAGATGCTAA | 105 |
| chr7:81178953-81180975 | p2 | (TA)11 | TATACAAGTCGGGACAGGGC | CGTCCCTTCTTGTGGATGTT | 286 |
| chr7:81178953-81180975 | p4 | (CATA)4 | TATACAAGTCGGGACAGGGC | CGTCCCTTCTTGTGGATGTT | 286 |
| chr7:81200183-81202253 | p2 | (TA)6 | TGAATGGTTTGATTGCTCCA | ACAAAAAGGGGGAACCATTC | 286 |
| chr7:81338503-81340515 | p2 | (GT)6 | CTTCTAAGTTCTGGGCGTGG | CTCCGTATTTCCGCAACAAT | 190 |
| chr7:81353489-81355509 | p5 | (TTCAA)4 | TTGTTCCGGAGCTAGCTTGT | CGCCCCAAACTTAAGTCAAA | 197 |
| chr7:81380692-81382704 | p2 | (AT)6 | ATAAATGCAAGCCGACAAGG | CCGGACTCCAGACTTTGGTA | 278 |
| chr7:81429741-81431753 | p2 | (AT)6 | TAATATGGAAGCTGCCCGTC | GACATGAGCATTTCACACCG | 218 |
| chr7:81432686-81434700 | p2 | (AT)7 | TGTTCAAGTTGACTTCCCCC | AGTGATGCGTGTGAAACCAG | 179 |
| chr7:81469471-81471499 | p2 | (TA)14 | CCATTCCAATCCATTCCAAG | AATCATGGAAGAAAGTGGCG | 156 |
| chr7:81492340-81494380 | p2 | (AT)6 | CAAGGTCACAAAGGTGGGAT | GACCCTTAGTTGGGTCGTGA | 218 |
| chr7:81497369-81499381 | p2 | (AT)6 | ATTAACGGCCCTGTCACTTG | TGGAAGAAAAAGAAGGGGGT | 294 |
| chr7:81497369-81499381 | p2 | (TA)8 | ATTAACGGCCCTGTCACTTG | TGGAAGAAAAAGAAGGGGGT | 294 |
| chr7:81514651-81516669 | p2 | (TA)9 | GAAGGCCAACAATCCGTAGA | TATCAGGAGGTCGGTGAAGG | 193 |
| chr7:81518916-81520930 | p2 | (AT)7 | AGGCGAGCTTGAACCAGATA | GAAAAAGACTCCGTTCGTGC | 163 |
| chr7:81518916-81520930 | p2 | (TA)9 | AGGCGAGCTTGAACCAGATA | GAAAAAGACTCCGTTCGTGC | 163 |
| chr7:81519630-81521648 | p2 | (AT)7 | CAAAGCTCTATGAGGGACGG | CGCTAGAGGTCCGAAGTACG | 259 |
| chr7:81519630-81521648 | p2 | (TA)9 | CAAAGCTCTATGAGGGACGG | CGCTAGAGGTCCGAAGTACG | 259 |
| chr7:81530030-81532044 | p2 | (AT)7 | TCGGTTAGGGACTGTTTTGG | CATGCGAATATGACACGTCC | 281 |
| chr7:81542647-81544662 | p5 | (TGAAA)3 | TAGCTTTTCTCTTTGGCGGA | ATTTTCCCTAACCGGGTCAC | 152 |
| chr7:81682318-81684338 | p5 | (ATTCA)4 | ATGCCAAGAACGGTATCCAG | GATGTGTTACGGTTCGCTGA | 125 |
| chr7:81760740-81762755 | p5 | (GATAT)3 | CTGCACAGCCTCTCTCAGTG | GGAGCAGCTTCACCATTAGC | 289 |
| chr7:81905681-81907699 | p2 | (AT)9 | GAAATTGCCTCCAAACCAGA | ATCATCCTTAAGGTGGTGCG | 251 |

| Marker ID | SSR type | SSR motif | FORWARD PRIMER (5'-3') | REVERSE PRIMER (5'-3') | size (bp) |
|---|---|---|---|---|---|
| chr7:82014152-82016242 | p2 | (AT)8 | AACATAACGCACGGTTAGGG | ATATCATGGTGGGTGGTGCT | 164 |
| chr7:82014152-82016242 | p2 | (TG)11 | AACATAACGCACGGTTAGGG | ATATCATGGTGGGTGGTGCT | 164 |
| chr7:82070569-82072583 | p2 | (AG)7 | ACATCCAACCAAAAGCGAAC | GTGGAAGGTCACAGCCTCAT | 283 |
| chr7:82180776-82182790 | p2 | (AT)7 | AGTCAGATCACCCGAAAACG | TGGCCCTTGTAACCATAAGC | 257 |
| chr7:82272564-82274580 | p2 | (AT)22 | ATTTCTGGGAAAATTTGGGC | CCCTGTGTGGCTGTGATATG | 104 |
| chr7:82272564-82274580 | p2 | (AT)8 | ATTTCTGGGAAAATTTGGGC | CCCTGTGTGGCTGTGATATG | 104 |
| chr7:82301353-82303369 | p2 | (AT)22 | ATTTCTGGGAAAATTTGGGC | CCCTGTGTGGCTGTGATATG | 104 |
| chr7:82301353-82303369 | p2 | (AT)8 | ATTTCTGGGAAAATTTGGGC | CCCTGTGTGGCTGTGATATG | 104 |
| chr7:82339493-82341505 | p2 | (AT)6 | TTCTTCAGGGTCGGAAGTTG | GAACTTGGCTGCCTTCTGAG | 270 |
| chr7:82400597-82402609 | p2 | (AT)6 | TTTTGTAAGCTGAGGTGGGG | ACCCATGACCTATGGTTGGA | 255 |
| chr7:82453823-82455839 | p5 | (AATGC)3 | GCAATGCAATGCAATAATGG | GTCACACTGGGTTGTGTTGC | 159 |
| chr7:82453823-82455839 | p5 | (AATGC)3 | GCAATGCAATGCAATAATGG | GTCACACTGGGTTGTGTTGC | 159 |
| chr7:82453823-82455839 | p2 | (AT)8 | GCAATGCAATGCAATAATGG | GTCACACTGGGTTGTGTTGC | 159 |
| chr7:82468108-82470120 | p2 | (TA)6 | AAAATCACGCCACATCAACA | CTCCGAGTCAAAACCGATGT | 290 |
| chr7:82482841-82484859 | p2 | (GT)7 | ATGTGTGTGAAGCCGTGTGT | GGTCACAACGACGCAACTAA | 179 |
| chr7:82482841-82484859 | p6 | (AATTTG)3 | ATGTGTGTGAAGCCGTGTGT | GGTCACAACGACGCAACTAA | 179 |
| chr7:82712361-82714377 | p4 | (CATA)4 | CTCTTGAAAACCGAGCGAAC | GGCTTTCCAGACAGTTCCTG | 122 |
| chr7:82768784-82770798 | p2 | (AT)7 | CCCAGTTCCTCCACCAGATA | GCACCTAGAGAATTGCTGCC | 147 |
| chr7:82776030-82778064 | p2 | (TA)17 | GTGAAGGTAGCGACGTGACA | TGCCTCACCTTGAAGGAACT | 127 |
| chr7:82783120-82785134 | p4 | (TTAC)4 | ATAAGCCGACAAGGCTTCAA | TGTCCTTTTGTTTGTGGACG | 184 |
| chr7:82783120-82785134 | p2 | (AT)7 | ATAAGCCGACAAGGCTTCAA | TGTCCTTTTGTTTGTGGACG | 184 |
| chr7:82783120-82785134 | p4 | (TTAC)4 | ATAAGCCGACAAGGCTTCAA | TGTCCTTTTGTTTGTGGACG | 184 |
| chr7:82817492-82819536 | p2 | (AT)22 | CATGACACATGATTCCCGAG | TAACGTACCCGTCCCTCAAG | 246 |
| chr7:82817492-82819536 | p6 | (ATTTTT)3 | CATGACACATGATTCCCGAG | TAACGTACCCGTCCCTCAAG | 246 |
| chr7:82818194-82820212 | p2 | (AT)22 | GTCTTTGGGTGCTCCATTGT | TGCTAACACAGCCGAACAAC | 244 |
| chr7:82818194-82820212 | p6 | (ATTTTT)3 | GTCTTTGGGTGCTCCATTGT | TGCTAACACAGCCGAACAAC | 244 |
| chr7:82926735-82928750 | p3 | (AAC)5 | CAACGACAAGGCAACCTACA | TGGTCGGGTATGGTATCGTT | 116 |
| chr7:83118716-83120732 | p3 | (TGT)5 | CCACTGACTTTGACCCCCTA | CGCCTAAAATCCAAATCGAA | 134 |
| chr7:83118716-83120732 | p2 | (TA)8 | CCACTGACTTTGACCCCCTA | CGCCTAAAATCCAAATCGAA | 134 |
| chr7:83172356-83174380 | p3 | (TTG)8 | AAGCCCTCCAAAAGGATGAT | TTTGGCAGCTTGGAGTCTTT | 193 |
| chr7:83290208-83292224 | p4 | (AATA)4 | CTGGTAGGGTCGGATGAAGA | CACAATGAAATCCATGGCAG | 118 |
| chr7:83317494-83319509 | p5 | (AGAGA)3 | CAAGTGAGCCTTAGTTCGGC | GGGCTAAGTCTGGCCCTATC | 187 |
| chr7:83320576-83322590 | p2 | (AC)7 | ACTTACCTTGCCCGCTTTTT | TGGCCGAGTTAAATTCTTGG | 287 |
| chr7:83398990-83401004 | p2 | (TA)7 | AGTTTCGTTTTGCGTTGGAG | TCCATAAGCCAACAACACCA | 268 |
| chr7:83398990-83401004 | p2 | (AT)23 | AGTTTCGTTTTGCGTTGGAG | TCCATAAGCCAACAACACCA | 268 |
| chr7:83466210-83468224 | p2 | (AT)7 | CACGGGTTGATGATCCTTCT | GGGAAGGAATCGATGTGAGA | 228 |

| Marker ID | SSR type | SSR motif | FORWARD PRIMER (5'-3') | REVERSE PRIMER (5'-3') | size (bp) |
|---|---|---|---|---|---|
| chr7:83489358-83491374 | p5 | (ATCAT)3 | TTTTGGCCATAAGAGCAACC | TCCGGAGCTCATATTTTTGG | 197 |
| chr7:83489358-83491374 | p4 | (ATAA)4 | TTTTGGCCATAAGAGCAACC | TCCGGAGCTCATATTTTTGG | 197 |
| chr7:83536604-83538622 | p2 | (TA)13 | CAGTCCCTTCGGGCATAATA | AAAGGGGAAGAAGTTGCCAT | 282 |
| chr7:83536604-83538622 | p3 | (CAA)6 | CAGTCCCTTCGGGCATAATA | AAAGGGGAAGAAGTTGCCAT | 282 |
| chr7:83792171-83794183 | p2 | (AT)6 | ATGAAACGAGCACCAACACA | GGTTGGCCTCATGTTATGCT | 188 |
| chr7:83820679-83822695 | p2 | (AT)6 | CTGAATCCGGGGAATGTAGA | TGGTAAATTCGGGACCTCAG | 172 |
| chr7:83820679-83822695 | p2 | (AT)8 | CTGAATCCGGGGAATGTAGA | TGGTAAATTCGGGACCTCAG | 172 |
| chr7:83820679-83822695 | p5 | (ATCAT)3 | CTGAATCCGGGGAATGTAGA | TGGTAAATTCGGGACCTCAG | 172 |
| chr7:83981356-83983372 | p4 | (GTCT)4 | TTTTTCGTACTGACCCCCTG | GGTCAAAGGAGCACTCCAAA | 118 |
| chr7:83981356-83983372 | p2 | (AT)20 | TTTTTCGTACTGACCCCCTG | GGTCAAAGGAGCACTCCAAA | 118 |
| chr7:84046955-84048971 | p2 | (TA)8 | ATTTTCCAAAAATGGCTCCC | CCTGAAGGCTCAGAACCAAG | 236 |
| chr7:84049704-84051719 | p3 | (ACA)5 | AACCAAATGCGACAAAAAGG | GATTTGAGAAGCGTAAGGCG | 276 |
| chr7:84049704-84051719 | p2 | (AC)6 | AACCAAATGCGACAAAAAGG | GATTTGAGAAGCGTAAGGCG | 276 |
| chr7:84112368-84114388 | p2 | (TA)10 | GCCGACTAGGCTGCTACATC | GGGGTCCTGTATCGTCCTTT | 259 |
| chr7:84112368-84114388 | p2 | (AT)6 | GCCGACTAGGCTGCTACATC | GGGGTCCTGTATCGTCCTTT | 259 |
| chr7:84113111-84115123 | p2 | (TA)10 | GCGCGATGTTAGGAAATAGC | GCCCCTTTTATTCTTTTGGC | 229 |
| chr7:84113111-84115123 | p2 | (AT)6 | GCGCGATGTTAGGAAATAGC | GCCCCTTTTATTCTTTTGGC | 229 |
| chr7:84150150-84152166 | p2 | (TA)8 | GATTTTTCGTGTAAACGGGC | GCAAAATGAACTTGCCGAAT | 126 |
| chr7:84348024-84350038 | p2 | (AT)7 | GAATGGCCTCTCCTTTCCTC | AGCTTCGAATTTGGGGAGAT | 280 |
| chr7:84355017-84357055 | p2 | (TA)12 | GTCCATTAAGGCCCATGAGA | CTTCACCCCCTGTCAAGTGT | 137 |
| chr7:84409837-84411852 | p5 | (ATCAT)3 | GCCGAGTTTTTGATGGATGT | GCACAGACACTAGTCGCAGC | 155 |
| chr7:84435875-84437890 | p3 | (ATT)5 | GATGTCCCAATGTTTCCCAC | ATCAGAATAGGCGGACATGG | 231 |
| chr7:84435875-84437890 | p6 | (ATTATG)3 | GATGTCCCAATGTTTCCCAC | ATCAGAATAGGCGGACATGG | 231 |
| chr7:84436157-84438175 | p3 | (ATT)5 | TGATTCCTCTGTATTGGGGC | CGGACCTGTCGTCTGTACCT | 156 |
| chr7:84436157-84438175 | p6 | (ATTATG)3 | TGATTCCTCTGTATTGGGGC | CGGACCTGTCGTCTGTACCT | 156 |
| chr7:84623862-84625882 | p2 | (AT)10 | CCGTTGGGCAATTCTACACT | CAGACTGCTTGGGCAGTACA | 191 |
| chr7:84640907-84642929 | p2 | (TA)7 | ACATCCTTCACTGAGTCCCG | AATGAACAGATGCCCTCCTG | 188 |
| chr7:84640973-84642985 | p2 | (TA)7 | ACATCCTTCACTGAGTCCCG | AATGAACAGATGCCCTCCTG | 122 |
| chr7:84640973-84642985 | p2 | (TC)7 | ACATCCTTCACTGAGTCCCG | AATGAACAGATGCCCTCCTG | 122 |
| chr7:84660280-84662298 | p3 | (CAA)6 | TGTATACACCTGCGTGGCAT | GCCATCATTATTAAGCCGGA | 161 |
| chr7:84791742-84793757 | p3 | (TTG)5 | CCTCGTTACGGCAAGCTAAG | GGAGGGCTTATGCCCATATT | 227 |
| chr7:84860755-84862770 | p5 | (AATGC)3 | GGCGAACCCTCAATCAATAA | GGAAGTCCTCCTCCCACTTC | 192 |
| chr7:84911522-84913536 | p2 | (GT)7 | ACGATCTTTCCATACTGGCG | TCAGTCTGGCGCTTATTCCT | 114 |
| chr7:84951790-84953810 | p5 | (TTTCA)4 | ACCAGGATCGTGCCAATAAG | TAGGGGCTGAAATAAATGCG | 118 |
| chr7:85064663-85066675 | p2 | (CA)6 | AAGGGCCTAGGTCCACACTT | CTTGTCGAACCCAACGATCT | 278 |
| chr7:85097549-85099569 | p2 | (TA)6 | GGTGATCCGGACTCGTAGAA | TATGGACAGGACAAGACCCC | 278 |

| Marker ID | SSR type | SSR motif | FORWARD PRIMER (5'-3') | REVERSE PRIMER (5'-3') | size (bp) |
|---|---|---|---|---|---|
| chr7:85097549-85099569 | p2 | (TA)9 | GGTGATCCGGACTCGTAGAA | TATGGACAGGACAAGACCCC | 278 |
| chr7:85097549-85099569 | p2 | (AT)10 | GGTGATCCGGACTCGTAGAA | TATGGACAGGACAAGACCCC | 278 |
| chr7:85346893-85348905 | p2 | (CA)6 | AGTTTCCAACGCAACAAACC | TTTTTGGCGAAGAAAAATGG | 292 |
| chr7:85346893-85348905 | p2 | (AT)6 | AGTTTCCAACGCAACAAACC | TTTTTGGCGAAGAAAAATGG | 292 |
| chr7:85347522-85349534 | p2 | (CA)6 | CGTCTGGGTGAGCCATAAAT | AGATGTGTTTGGCCTTGTCC | 191 |
| chr7:85347522-85349534 | p2 | (AT)6 | CGTCTGGGTGAGCCATAAAT | AGATGTGTTTGGCCTTGTCC | 191 |
| chr7:85373733-85375748 | p5 | (GAAAT)3 | TTGAGCATCGGGAGCTTACT | GCGTGGAAGGGAACGTATAA | 170 |
| chr7:85413708-85415723 | p5 | (ATCAT)3 | CGGATAAAGCACAAGGAAGC | TGCTCTATTGTTCGGAGCCT | 290 |
| chr7:85422905-85424917 | p2 | (AT)6 | CCACATCGTGCTTCAGAAAA | CATTCAATTCCCAACTGCCT | 169 |
| chr7:85493211-85495226 | p5 | (ATGAT)3 | GATCGGGACTTGAGTTTGGA | GGGTCGTAACTATGCTCGGA | 164 |
| chr7:85510052-85512064 | p2 | (TA)8 | GGTCGTGTGAGGCCCTAGTA | ATTTTGGCAGAGTTTCCCCT | 129 |
| chr7:85542785-85544807 | p2 | (AT)31 | GGGATGTTCCCTTTCCCTTA | GTCGTCTCAAAACTGTCGCA | 262 |
| chr7:85542785-85544807 | p2 | (TA)11 | GGGATGTTCCCTTTCCCTTA | GTCGTCTCAAAACTGTCGCA | 262 |
| chr7:85542785-85544807 | p2 | (AT)7 | GGGATGTTCCCTTTCCCTTA | GTCGTCTCAAAACTGTCGCA | 262 |
| chr7:85606251-85608263 | p2 | (CA)6 | ATGGCCTAGCAGAGACTGGA | GAACACCACTCATCGACCCT | 158 |
| chr7:85608519-85610531 | p2 | (AC)6 | AACCAAATGCGACAAAAAGG | GTTTCTGGACAAAACGGGAA | 105 |
| chr7:85709548-85711566 | p2 | (AG)7 | GAATGGGAAGTTGCAGGGTA | CCCCATCACCTTCATTCCTA | 221 |
| chr7:85709548-85711566 | p2 | (AT)9 | GAATGGGAAGTTGCAGGGTA | CCCCATCACCTTCATTCCTA | 221 |
| chr7:85709548-85711566 | p2 | (AT)8 | GAATGGGAAGTTGCAGGGTA | CCCCATCACCTTCATTCCTA | 221 |
| chr7:85722736-85724756 | p2 | (AT)46 | ACCAGCCGACAAGGCTACTA | GTTGACGATGTGTTTGTGCC | 189 |
| chr7:85722736-85724756 | p2 | (TA)10 | ACCAGCCGACAAGGCTACTA | GTTGACGATGTGTTTGTGCC | 189 |
| chr7:85722736-85724756 | p3 | (ATC)6 | ACCAGCCGACAAGGCTACTA | GTTGACGATGTGTTTGTGCC | 189 |
| chr7:85730022-85732082 | p2 | (AT)30 | ATTGGACACGGCTCGTAGAC | TGCTCGGTGTGTTAGCTTTG | 273 |
| chr7:85730022-85732082 | p3 | (CAT)5 | ATTGGACACGGCTCGTAGAC | TGCTCGGTGTGTTAGCTTTG | 273 |
| chr7:85800711-85802727 | p2 | (TC)8 | ACAAGCGGTAAATGGTCGTC | AGCCTGAGTACAGCCTTCCA | 293 |
| chr7:85815183-85817198 | p5 | (AAACC)3 | TGATCCTGTGTCAGTTGGGA | AGCTTCTCTGGTGGAGCAAA | 197 |
| chr7:85815183-85817198 | p5 | (AAAAT)3 | TGATCCTGTGTCAGTTGGGA | AGCTTCTCTGGTGGAGCAAA | 197 |
| chr7:85915550-85917565 | p5 | (TTGTG)3 | GCCACTTGGTGGTGAAGAGT | TTCAATGCCAACTCACTTCG | 113 |
| chr7:85915550-85917565 | p5 | (ATTTC)3 | GCCACTTGGTGGTGAAGAGT | TTCAATGCCAACTCACTTCG | 113 |
| chr7:85968604-85970624 | p2 | (TA)6 | TCGTGCATGCCTCAATATGT | CACACCATCACATCGTCACA | 153 |
| chr7:85968604-85970624 | p2 | (AT)10 | TCGTGCATGCCTCAATATGT | CACACCATCACATCGTCACA | 153 |
| chr7:85968604-85970624 | p5 | (TATGA)3 | TCGTGCATGCCTCAATATGT | CACACCATCACATCGTCACA | 153 |
| chr7:85979049-85981064 | p5 | (CATAT)3 | TGGAGTTTGCTAACCTCGCT | TCAGTTACATCTGTGCCCCA | 218 |
| chr7:85981504-85983516 | p2 | (AT)6 | TTTCCGTTTCGTCCAAAATC | TGTTGATGCTGTTGGGTTGT | 175 |
| chr7:85997004-85999019 | p5 | (ATTTC)3 | AAGCATGCGATCCAGATACC | ATAACAATGCCTCACAGGGC | 201 |
| chr7:86132198-86134218 | p5 | (CAATG)4 | ACTGCTCTCGCATGAGTCCT | AAGGCCTTAGGTTCGTCGAT | 156 |

| Marker ID | SSR type | SSR motif | FORWARD PRIMER (5'-3') | REVERSE PRIMER (5'-3') | size (bp) |
|---|---|---|---|---|---|
| chr7:86220169-86222203 | p2 | (AT)17 | AAACTCATGAAATCACGGGC | CTACGACAGAAAAGGAGCCG | 217 |
| chr7:86360469-86362481 | p2 | (TA)6 | CGAAATCGTGGATTGAAGGT | CGGGACCCTAGATCTTTTCC | 203 |
| chr7:86370967-86372983 | p2 | (AT)8 | TTAATTCGCCTTTTGTTGGC | TTACCCCTCAATGAGTTCGC | 229 |
| chr7:86383937-86385955 | p6 | (TATATG)3 | AAACCGCGCCTAAATTTCTT | GCAACTGGGTCAAATGAGGT | 109 |
| chr7:86383937-86385955 | p2 | (TA)6 | AAACCGCGCCTAAATTTCTT | GCAACTGGGTCAAATGAGGT | 109 |
| chr7:86501753-86503771 | p6 | (ATCCAT)3 | TCACCCTTCCCTGTACCTTG | CTCCTCCACCGGATACTTCA | 263 |
| chr7:86539962-86541980 | p2 | (AT)9 | ATGACTAGTGCCCGAAATGG | GTCGGCTTGCGTATCATTTT | 237 |
| chr7:86583246-86585261 | p2 | (AG)6 | ATAGGTCGTGGGCCTGTATG | TGAGGAGGAGAAGGCTGAAA | 278 |
| chr7:86583246-86585261 | p3 | (GTT)5 | ATAGGTCGTGGGCCTGTATG | TGAGGAGGAGAAGGCTGAAA | 278 |
| chr7:86583246-86585261 | p2 | (TA)6 | ATAGGTCGTGGGCCTGTATG | TGAGGAGGAGAAGGCTGAAA | 278 |
| chr7:86681891-86683911 | p2 | (AT)10 | CCTCTGGAAGTGGCTTTGTC | CAGCAGCAGGGGATACTAGC | 286 |
| chr7:86686479-86688515 | p2 | (AC)18 | CTTGACTGAGGAGCTGGAGG | CAGCAGCAGGGGATACTAGC | 197 |
| chr7:86791816-86793830 | p2 | (AT)7 | TGTAGTGTCAGCCTTGTCGG | ACTTTTGTCACGACCCAACC | 148 |
| chr7:86803848-86805863 | p3 | (TGT)5 | AAGAAAAACATCCGTTTGCG | CACAATCATCCACAAGCCAC | 294 |
| chr7:86803848-86805863 | p5 | (TTGAA)3 | AAGAAAAACATCCGTTTGCG | CACAATCATCCACAAGCCAC | 294 |
| chr7:86820960-86822978 | p2 | (AT)9 | CCGGGTAAGTTATCGTGCAT | AGTCATCATTTGCCCTCCTG | 171 |
| chr7:86820960-86822978 | p2 | (TA)9 | CCGGGTAAGTTATCGTGCAT | AGTCATCATTTGCCCTCCTG | 171 |
| chr7:86834450-86836490 | p2 | (TA)20 | AAGCTGCCAACTCCACAGAT | CCAAGTGATCGGGCAACTAT | 207 |
| chr7:86860239-86862251 | p2 | (AT)15 | GCGTCTCTCATAGCCCTGTC | TACAACGATGCCTCCAATGA | 221 |
| chr7:86860888-86862918 | p2 | (AT)15 | TGCATGTCATAGCCACCTGT | GTTTGGGATGGGATGTTGAG | 189 |
| chr7:86898953-86900977 | p2 | (AT)12 | GACAATTTCAAACGCGGATT | GCCCAAATTTCTGTAGGCAA | 183 |
| chr7:87074779-87076885 | p2 | (AT)53 | TTACAATCGTGGCTGCTCTG | TATTGAGACGATTCCCCCAG | 276 |
| chr7:87134537-87136552 | p5 | (TTTGT)3 | TCAATCTCCCCGTTTTCTTG | GTGCAAGTTCAGTGGCGTAA | 220 |
| chr7:87185253-87187265 | p2 | (AC)6 | TATCATGGCTCTATCGCACG | TGGGTCGTGACAAAGTGGTA | 254 |
| chr7:87185253-87187265 | p3 | (ATC)5 | TATCATGGCTCTATCGCACG | TGGGTCGTGACAAAGTGGTA | 254 |
| chr7:87185689-87187704 | p2 | (AC)6 | TCGTAAGGATGGGACAGGAC | TTTGTGGACGCTGTGTTCAT | 233 |
| chr7:87185689-87187704 | p3 | (ATC)5 | TCGTAAGGATGGGACAGGAC | TTTGTGGACGCTGTGTTCAT | 233 |
| chr7:87268049-87270081 | p2 | (AT)7 | TAGTCTTCATCCCTGACCCG | CAGGCTATTTTGGTAGGCCA | 281 |
| chr7:87268049-87270081 | p2 | (AT)8 | TAGTCTTCATCCCTGACCCG | CAGGCTATTTTGGTAGGCCA | 281 |
| chr7:87268049-87270081 | p2 | (AT)16 | TAGTCTTCATCCCTGACCCG | CAGGCTATTTTGGTAGGCCA | 281 |
| chr7:87364989-87367004 | p5 | (AGGAC)3 | CGGGTTGTAGGAGTTGTCGT | ATCCCTGAACATTTCCCACA | 104 |
| chr7:87374056-87376072 | p2 | (AT)8 | CCCATTACACGGGAAGAAGA | GGGTGCCATAATACTTCGGA | 275 |
| chr7:87391329-87393344 | p5 | (CCACA)3 | CAAACGGAGCAATAGGGAAA | ACCCGACTACCTTGAAACCC | 142 |
| chr7:87587896-87589911 | p5 | (ATATG)3 | GATAGCGAACCCACAACCAT | ATTGGGCATCACTTCCTTTG | 191 |
| chr7:87600973-87603013 | p2 | (AG)20 | AATATCCGTTTGGAGGGCTT | GCCCACACGTCAAGATTTTT | 101 |
| chr7:87633321-87635345 | p2 | (TA)12 | TTGCTGAAATGGATGGTCAA | GTTAAAGCGATCGAGCTTGG | 140 |

| Marker ID | SSR type | SSR motif | FORWARD PRIMER (5'-3') | REVERSE PRIMER (5'-3') | size (bp) |
|---|---|---|---|---|---|
| chr7:87787406-87789418 | p2 | (TA)6 | TATGTCACCCCGACATAGCA | TAGGCACGGTGGGATATCAT | 244 |
| chr7:87790194-87792252 | p2 | (AT)29 | TCACCTGCAACATTCTCAGC | TCTAACTTCAGCACCCCCAC | 264 |
| chr7:87790194-87792252 | p2 | (AT)6 | TCACCTGCAACATTCTCAGC | TCTAACTTCAGCACCCCCAC | 264 |
| chr7:87863446-87865460 | p2 | (AT)7 | CCAACCAAGGCTCAAGGTTA | CCTCTGCAGATGGTACGGAT | 176 |
| chr7:87895109-87897169 | p2 | (AT)30 | GGGTCTTCATCCGACTTTGA | TAATGCTTCTTCCACCCACC | 234 |
| chr7:87910903-87912921 | p3 | (TTG)6 | AAATCGTTCGTCAATCCGAC | CGGGTCAAAACATAAGTGGG | 271 |
| chr7:87949960-87951976 | p2 | (AT)8 | ACATGGCTTCCGTTAAGGTG | TCACATGGTTCCTCATTCCA | 217 |
| chr7:87949960-87951976 | p5 | (TGAAT)3 | ACATGGCTTCCGTTAAGGTG | TCACATGGTTCCTCATTCCA | 217 |
| chr7:88032057-88034077 | p2 | (AT)10 | CATAAGCCGACGAGGCTAAC | TAGCGGAGCTCCATCTCATT | 205 |
| chr7:88034519-88036535 | p4 | (GTTC)4 | GGACAGGGATGCGTCATACT | GCGGTTTTCATGCCTTACAT | 230 |
| chr7:88052957-88054987 | p2 | (AT)7 | CCCACACATACGCATCTACG | GTCCGTTTTCATTGGATGCT | 241 |
| chr7:88052957-88054987 | p2 | (AT)15 | CCCACACATACGCATCTACG | GTCCGTTTTCATTGGATGCT | 241 |
| chr7:88056441-88058457 | p3 | (ATT)5 | CCTCCTTCGTAGCCAAAGTG | TTCCCAATTGTTCAAAAGCC | 285 |
| chr7:88056441-88058457 | p3 | (ATT)6 | CCTCCTTCGTAGCCAAAGTG | TTCCCAATTGTTCAAAAGCC | 285 |
| chr7:88056441-88058457 | p3 | (TTA)6 | CCTCCTTCGTAGCCAAAGTG | TTCCCAATTGTTCAAAAGCC | 285 |
| chr7:88056603-88058618 | p3 | (ATT)5 | CCTCCTTCGTAGCCAAAGTG | TTCCCAATTGTTCAAAAGCC | 123 |
| chr7:88056603-88058618 | p3 | (ATT)6 | CCTCCTTCGTAGCCAAAGTG | TTCCCAATTGTTCAAAAGCC | 123 |
| chr7:88056603-88058618 | p3 | (TTA)6 | CCTCCTTCGTAGCCAAAGTG | TTCCCAATTGTTCAAAAGCC | 123 |
| chr7:88174204-88176219 | p3 | (ATG)5 | TAGGTTGGTGCAACCCTAGC | GCTGCATCTTCACTTCCCTC | 227 |
| chr7:88265705-88267720 | p3 | (ATG)5 | TAGGTTGGTGCAACCCTAGC | GCTGCATCTTCACTTCCCTC | 227 |
| chr7:88335290-88337340 | p2 | (AT)7 | TCCGGAATAGACTCGAATGG | CAGGGAAAGGACTAGCGTTG | 149 |
| chr7:88335290-88337340 | p2 | (AT)25 | TCCGGAATAGACTCGAATGG | CAGGGAAAGGACTAGCGTTG | 149 |
| chr7:88342267-88344281 | p2 | (AT)17 | CCAGCGGCTGTAAGACAAAT | CCTGCTTCGTCCTTTGAACT | 284 |
| chr7:88342267-88344281 | p5 | (AAAGA)3 | CCAGCGGCTGTAAGACAAAT | CCTGCTTCGTCCTTTGAACT | 284 |
| chr7:88382669-88384689 | p2 | (TA)10 | TCGATCCGAGGTATTGGAAG | TGCAATGGCCAATTACTCAA | 256 |
| chr7:88494254-88496272 | p6 | (AAAAAT)3 | TGAAGGGTCAAGAATGGGTC | ACTACGAATCGACCAAACGG | 216 |
| chr7:88548939-88550961 | p2 | (TA)11 | AAAATGAGGCTTGGTCCCTT | GACCGTAAACCTCGGTCGTA | 159 |
| chr7:88563861-88565879 | p6 | (ATACAT)3 | AGGCATGAGGGGTGTTAGTG | AGGCGATCCTTCCTATTCGT | 236 |
| chr7:88647918-88649930 | p2 | (CA)6 | TAGTCGGGTCCCATCTATGC | TATTCTCCGGAATCTGCACC | 110 |
| chr7:88650165-88652183 | p2 | (TA)9 | AACGACCATATGCATGAGCA | AGAAACGTTCATGGGGAGTG | 261 |
| chr7:88733629-88735649 | p2 | (TA)10 | GGGGGTCCATATTTTGCTTT | TGAGATTCAGAATGAGGGGC | 300 |
| chr7:88734862-88736874 | p2 | (TA)6 | CATGCTCATGCCTACCAATG | TGACTCGTGACTTGAAAGCG | 292 |
| chr7:88747509-88749521 | p2 | (AT)11 | TTGGACGCGTGTTCATAAAA | AGTCATATGGACGGGACAGG | 269 |
| chr7:88747509-88749521 | p2 | (AT)6 | TTGGACGCGTGTTCATAAAA | AGTCATATGGACGGGACAGG | 269 |
| chr7:88747509-88749521 | p2 | (AT)6 | TTGGACGCGTGTTCATAAAA | AGTCATATGGACGGGACAGG | 269 |
| chr7:88824586-88826601 | p3 | (TGA)5 | ATTTGTTCAGGGGTACAGCG | TCAGAAGCCGACAAGGCTAT | 118 |

| Marker ID | SSR type | SSR motif | FORWARD PRIMER (5'-3') | REVERSE PRIMER (5'-3') | size (bp) |
|---|---|---|---|---|---|
| chr7:88857855-88859867 | p2 | (AT)6 | ATAGGGGTGGTGATGGAACA | CAGTTGCTTGTTCAGCGGTA | 234 |
| chr7:88878941-88880953 | p2 | (AT)6 | ATAGGGGTGGTGATGGAACA | CAGTTGCTTGTTCAGCGGTA | 234 |
| chr7:88944102-88946114 | p2 | (TA)6 | GTAATAGGAGGGCTCCCGAC | AGCTGCCACTGTACGCCTAT | 216 |
| chr7:89007359-89009374 | p5 | (CATTG)3 | TCAGCGAAAAGGAAAGGCTA | CACAAAACAAGGCCCAAACT | 183 |
| chr7:89203052-89205066 | p2 | (AT)7 | ATTCACAAGGGTTGGCACTC | CAAGTACCAATGGGCGTCTT | 234 |
| chr7:89210582-89212626 | p2 | (AT)22 | CTCCGGAGGATTCAAAATGA | AGTAGTACGAGAGCCGCCAA | 279 |
| chr7:89210582-89212626 | p2 | (AT)23 | CTCCGGAGGATTCAAAATGA | AGTAGTACGAGAGCCGCCAA | 279 |
| chr7:89210761-89212807 | p2 | (AT)22 | CTCCGGAGGATTCAAAATGA | AGTAGTACGAGAGCCGCCAA | 100 |
| chr7:89210761-89212807 | p2 | (AT)23 | CTCCGGAGGATTCAAAATGA | AGTAGTACGAGAGCCGCCAA | 100 |
| chr7:89321927-89323939 | p2 | (TA)6 | CAAAACATTGCATACGTCGC | TCTGCAGTCTCGGTAACACG | 207 |
| chr7:89321927-89323939 | p2 | (AG)6 | CAAAACATTGCATACGTCGC | TCTGCAGTCTCGGTAACACG | 207 |
| chr7:89321927-89323939 | p3 | (TTG)5 | CAAAACATTGCATACGTCGC | TCTGCAGTCTCGGTAACACG | 207 |
| chr7:89322142-89324157 | p2 | (TA)6 | GACTGTTACACCCCGGAAAA | ATATGGCCTCCACTGTCGTC | 145 |
| chr7:89322142-89324157 | p2 | (AG)6 | GACTGTTACACCCCGGAAAA | ATATGGCCTCCACTGTCGTC | 145 |
| chr7:89322142-89324157 | p3 | (TTG)5 | GACTGTTACACCCCGGAAAA | ATATGGCCTCCACTGTCGTC | 145 |
| chr7:89449571-89451595 | p2 | (TA)21 | TTGTAGGGGCATCATGTTCA | GCTAAGGTCCGGTGTCATGT | 187 |
| chr7:89449571-89451595 | p2 | (TC)12 | TTGTAGGGGCATCATGTTCA | GCTAAGGTCCGGTGTCATGT | 187 |
| chr7:89495795-89497813 | p5 | (ACAAA)3 | GAAGGACCCTCCCGAAATAG | TTCCCTTAGGCTTCCTCCAT | 163 |
| chr7:89495795-89497813 | p3 | (TTG)6 | GAAGGACCCTCCCGAAATAG | TTCCCTTAGGCTTCCTCCAT | 163 |
| chr7:89695360-89697386 | p2 | (TA)13 | AAGCCAACGAGGCTAACAAA | GTCCTGTCCCGCCTATATGA | 192 |
| chr7:89710881-89712907 | p2 | (TA)13 | AAGCCAACGAGGCTAACAAA | GTCCTGTCCCGCCTATATGA | 192 |
| chr7:89731455-89733467 | p2 | (CA)6 | ATACCTCGATCGCTCGCTAA | GATGCTGCCGAAATTTTTGT | 287 |
| chr7:89748460-89750475 | p5 | (TTTCA)3 | TCAGTCAAAGTCTTGTGGCG | TAATATTGCCTCGCAGGGAC | 147 |
| chr7:89748460-89750475 | p3 | (ATG)5 | TCAGTCAAAGTCTTGTGGCG | TAATATTGCCTCGCAGGGAC | 147 |
| chr7:89817451-89819463 | p2 | (CT)6 | CGCTTCTTTTGCTATTCCGA | TTCGGTAGCCAAATGTTTCC | 176 |
| chr7:89921279-89923297 | p2 | (TA)9 | TCTTCAAGGTGAGGCAGGAT | GGCCTTATAGGCACGGTGTA | 219 |
| chr7:89987906-89989920 | p2 | (AT)7 | CGCCGTACCCAAATAGTCAT | CGTTTTGGTGATCCTCCAGT | 101 |
| chr7:89999535-90001547 | p2 | (AT)6 | ACAGCAGTACCTCCGGTGTC | GTTAAGCCAAACGGGTTCAA | 201 |
| chr7:90061900-90063927 | p3 | (TTG)9 | CTTTCGTTCAGCAATCGACA | TTGCTGGTTGTATTGGGTGA | 145 |
| chr7:90095028-90097042 | p2 | (AT)7 | CTATCCAGGCTCACTCTCCG | GATGATTGGCCGAGTTGAAT | 193 |
| chr7:90158353-90160375 | p2 | (AT)11 | TGAATTGTGCACTAGAGGCG | GCAGAAGCAACACCAACAGA | 158 |
| chr7:90186603-90188618 | p5 | (TTTTA)3 | TTCCAGCATACAGATGACGC | TAAGATTTTTGTCTGGGCCG | 147 |
| chr7:90240881-90242943 | p2 | (TA)21 | AGCCTGAGAGTTGTGTGGAAA | GCGTGTAAGAGAATGCACCA | 298 |
| chr7:90257046-90259066 | p5 | (TCATT)4 | GATTCAAGATGAAGCCCCAA | ATATCATTGCCTCACAGGGC | 235 |
| chr7:90271269-90273289 | p5 | (TTTCA)4 | TCAGTCAAAGTCTTGTGGCG | TCATGAATTTCTTCTGGGGG | 137 |
| chr7:90285460-90287504 | p6 | (TCTACT)3 | TGGCAAATTAGAGGGCAATC | ACGTAGCATCAAAAGCCAGC | 115 |

| Marker ID | SSR type | SSR motif | FORWARD PRIMER (5'-3') | REVERSE PRIMER (5'-3') | size (bp) |
|---|---|---|---|---|---|
| chr7:90285460-90287504 | p2 | (TA)22 | TGGCAAATTAGAGGGCAATC | ACGTAGCATCAAAAGCCAGC | 115 |
| chr7:90285460-90287504 | p2 | (TA)8 | TGGCAAATTAGAGGGCAATC | ACGTAGCATCAAAAGCCAGC | 115 |
| chr7:90293140-90295156 | p2 | (TC)6 | CCTCCGGTTTCCCAAATTAT | TTATTCGTAAGCCGAGGTGG | 298 |
| chr7:90293140-90295156 | p2 | (AT)8 | CCTCCGGTTTCCCAAATTAT | TTATTCGTAAGCCGAGGTGG | 298 |
| chr7:90299370-90301385 | p5 | (CAATG)3 | AGGAAATGGCTGCAAGAAGA | TTAACAATGGTTTGGTCGCA | 152 |
| chr7:90299370-90301385 | p2 | (TA)8 | AGGAAATGGCTGCAAGAAGA | TTAACAATGGTTTGGTCGCA | 152 |
| chr7:90417204-90419220 | p2 | (TA)7 | CTATGGTTCCCGCTGTTGTT | GGGGATGCGAGATTACAAAA | 223 |
| chr7:90417219-90419234 | p2 | (TA)7 | CTATGGTTCCCGCTGTTGTT | GGGGATGCGAGATTACAAAA | 208 |
| chr7:90418351-90420363 | p2 | (TA)7 | TTTGGCCTACCCCTACCTCT | AAGACAAGATGCGGGAAATG | 292 |
| chr7:90418365-90420415 | p2 | (TA)7 | TTTGGCCTACCCCTACCTCT | AAGACAAGATGCGGGAAATG | 278 |
| chr7:90418416-90420430 | p2 | (TA)7 | TTTGGCCTACCCCTACCTCT | AAGACAAGATGCGGGAAATG | 227 |
| chr7:90418432-90420466 | p2 | (TA)7 | TTTGGCCTACCCCTACCTCT | AAGACAAGATGCGGGAAATG | 211 |
| chr7:90418507-90420523 | p2 | (TA)7 | TTTGGCCTACCCCTACCTCT | AAGACAAGATGCGGGAAATG | 136 |
| chr7:90493950-90495970 | p2 | (TA)10 | GAGGCTCGATGCGAACTATC | GATTGCACCGAGCCATCTAT | 269 |
| chr7:90510028-90512056 | p2 | (CT)14 | CGGATACTCGTACTCCTGGC | CCTTGCTTGGAGTCCTCTTG | 273 |
| chr7:90510028-90512056 | p2 | (CT)8 | CGGATACTCGTACTCCTGGC | CCTTGCTTGGAGTCCTCTTG | 273 |
| chr7:90549346-90551358 | p3 | (AAC)9 | TATTCTGTCGAAAATCGGGC | GCATAAGCCTGGGTATTGGA | 250 |
| chr7:90549346-90551358 | p2 | (AT)6 | TATTCTGTCGAAAATCGGGC | GCATAAGCCTGGGTATTGGA | 250 |
| chr7:90583697-90585712 | p5 | (GACCC)3 | CGGAATTCCACAAGGGTCTA | CTATGTGATCGGGCGAGATT | 237 |
| chr7:90676755-90678771 | p4 | (GAAA)4 | TTGAAAACCACTCTACGGGG | GCTCCGTTCAAGACCCATAA | 275 |
| chr7:90808118-90810136 | p6 | (TATACA)3 | CACAACAATGTTACACGGGC | TGCACAGGGAGACAGACTTG | 228 |
| chr7:90868861-90870879 | p6 | (TCTTTT)3 | AGTTGTTTCCATCACCCCAA | TTACCAGGGGTGTCAATGGT | 254 |
| chr7:90870871-90872886 | p5 | (TTAGA)3 | TCATCTCAACCAATGACGGA | AACCCCCTTCAACGAAAAAT | 145 |
| chr7:90896886-90898898 | p2 | (TA)6 | GACTTTGCAGACAACCCCAT | CTTGAAGAGCCCAGTGAAGG | 267 |
| chr7:91034162-91036174 | p2 | (AT)6 | TAGCTCTACGGCCCGATCTA | CCTGTCATGGTCCTCTGGTT | 138 |
| chr7:91052896-91054912 | p2 | (AT)6 | AACATAGTCACGGGGCTGTC | TTGTCGGCTTATGTACTGCG | 204 |
| chr7:91052896-91054912 | p2 | (AT)8 | AACATAGTCACGGGGCTGTC | TTGTCGGCTTATGTACTGCG | 204 |
| chr7:91052896-91054912 | p2 | (AT)6 | AACATAGTCACGGGGCTGTC | TTGTCGGCTTATGTACTGCG | 204 |
| chr7:91103237-91105252 | p5 | (CAATG)3 | GGTTGATCCTGTGTGGTGTG | GTGACGTGGCAGAGGAATTT | 287 |
| chr7:91110534-91112546 | p2 | (TA)6 | AGGCAGCTAACTGCGTCAAT | TGTCTTCGTCCAGAAAGGCT | 220 |
| chr7:91199190-91201210 | p2 | (TA)10 | GACCTTACGTCACCCCGATA | TGGCCATGGTGTATCGTAGA | 290 |
| chr7:91232524-91234540 | p3 | (CAT)5 | TGTAGAAAGGACGGGACCTG | AGCAGTGGTCAATGTCCTCC | 100 |
| chr7:91232524-91234540 | p2 | (AT)8 | TGTAGAAAGGACGGGACCTG | AGCAGTGGTCAATGTCCTCC | 100 |
| chr7:91240226-91242241 | p2 | (AT)7 | CTGACTGAGGAGATGGAGGC | GCGAGTTCTTCGACAAGTCC | 150 |
| chr7:91240253-91242271 | p2 | (AT)7 | CTGACTGAGGAGATGGAGGC | GCGAGTTCTTCGACAAGTCC | 123 |
| chr7:91403160-91405174 | p2 | (TA)7 | TGATATAGGTAGTGCCGGGG | TGCTTTCTCCCCTTCTGCTA | 171 |

| Marker ID | SSR type | SSR motif | FORWARD PRIMER (5'-3') | REVERSE PRIMER (5'-3') | size (bp) |
|---|---|---|---|---|---|
| chr7:91484246-91486260 | p2 | (AT)6 | GACAAGGCTGCCACTACACA | TCAGCTGTGTCTTTCCGATG | 272 |
| chr7:91484246-91486260 | p2 | (TA)7 | GACAAGGCTGCCACTACACA | TCAGCTGTGTCTTTCCGATG | 272 |
| chr7:91487800-91489818 | p2 | (TA)9 | GTAACATATGGCGGGACAGG | TGCAGATACTCGCTTTGGTG | 300 |
| chr7:91487820-91489838 | p2 | (TA)9 | GTAACATATGGCGGGACAGG | TGCAGATACTCGCTTTGGTG | 280 |
| chr7:91487840-91489860 | p2 | (TA)9 | GTAACATATGGCGGGACAGG | TGCAGATACTCGCTTTGGTG | 260 |
| chr7:91558029-91560047 | p6 | (GAAAAC)3 | TGAAGAGCAACAACGTCCAG | ATCCTCCAAAACCACCACAA | 159 |
| chr7:91614381-91616399 | p6 | (ATATGA)3 | TGGGAGCGAATTGAGAAATC | CGTTTGCACGTTCTCTTTCA | 149 |
| chr7:91615215-91617241 | p6 | (ATATGA)3 | AAACCTTCAGGGTCGGGTAT | ATGCACTTGCTTGCCTTTCT | 106 |
| chr7:91615215-91617241 | p2 | (TA)9 | AAACCTTCAGGGTCGGGTAT | ATGCACTTGCTTGCCTTTCT | 106 |
| chr7:91682687-91684703 | p2 | (AT)8 | CCATGCACAAACACAACCTC | GTGCTCTTCGTGGTGGTACA | 139 |
| chr7:91682687-91684703 | p2 | (AT)7 | CCATGCACAAACACAACCTC | GTGCTCTTCGTGGTGGTACA | 139 |
| chr7:91703543-91705555 | p2 | (TG)6 | GTACCCTCACTTGTTCCCGA | ATCAAGGCAGGAGTTTCCCT | 161 |
| chr7:91958373-91960389 | p4 | (TTAA)4 | TTTCGCCTCATCGTTCTTTT | ATATAGGGGAAATGCTGCCC | 185 |
| chr7:92026745-92028757 | p2 | (TA)6 | CCCTGAGACGTCCCTTGTTA | CGGGACTCAGTGAAGGATGT | 152 |
| chr7:92145821-92147836 | p3 | (CAG)5 | GGTCTGCTACACCCCCACTA | GCCTTCCTCAGGGTCCTATC | 162 |
| chr7:92260810-92262822 | p2 | (TA)6 | AATCCCTAGGACAAACCGCT | TGCTTGGAGGGCTCTAGTGT | 286 |
| chr7:92260810-92262822 | p3 | (CAA)5 | AATCCCTAGGACAAACCGCT | TGCTTGGAGGGCTCTAGTGT | 286 |
| chr7:92261486-92263501 | p2 | (TA)6 | ACATCACAACAAAGGCCACA | ACAGTTGATAGGGGGTGACG | 176 |
| chr7:92261486-92263501 | p3 | (CAA)5 | ACATCACAACAAAGGCCACA | ACAGTTGATAGGGGGTGACG | 176 |
| chr7:92293124-92295144 | p5 | (TATCA)4 | GAAGCCTGGCAACATAGCTC | TAGATAAGGGGTGCTCGGTG | 149 |
| chr7:92341030-92343045 | p5 | (TCAAG)3 | TGGGGTTTGGAAGAGAATTG | TAATGTTTTCATGCCAGCCA | 169 |
| chr7:92358864-92360876 | p2 | (AT)12 | AATGCTGAGCGGAGATAGGA | GTTGAGGACGAAGGAAGTCG | 111 |
| chr7:92358864-92360876 | p2 | (AT)6 | AATGCTGAGCGGAGATAGGA | GTTGAGGACGAAGGAAGTCG | 111 |
| chr7:92358864-92360876 | p2 | (AT)12 | AATGCTGAGCGGAGATAGGA | GTTGAGGACGAAGGAAGTCG | 111 |
| chr7:92416033-92418055 | p2 | (AT)11 | GGTTGAAGGTCATGGCAGTT | GAATAGACCAACGAAGGGCA | 121 |
| chr7:92422611-92424625 | p2 | (AT)6 | ATATAGGGGAAATGCTGCCC | TCGTAAATCCGTAGCCCATC | 261 |
| chr7:92422611-92424625 | p2 | (TA)7 | ATATAGGGGAAATGCTGCCC | TCGTAAATCCGTAGCCCATC | 261 |
| chr7:92445053-92447113 | p2 | (AT)8 | CGCATTTATTTCAGCCCCTA | TCCTGTCACAACAGCCCATA | 211 |
| chr7:92445053-92447113 | p3 | (TCA)5 | CGCATTTATTTCAGCCCCTA | TCCTGTCACAACAGCCCATA | 211 |
| chr7:92450365-92452383 | p6 | (CATAAT)3 | CGTTTCGTTTGGTCTCCAAT | AAAAGGCTAGCGAAGGGAAC | 242 |
| chr7:92463370-92465384 | p2 | (TA)7 | TGCGGGTTCAAAAATAGTCC | TTTGAAACCTGCATTGACCA | 132 |
| chr7:92511036-92513050 | p2 | (AT)7 | AGACTTCGACTTTCTGCCCA | GTCCCTCGTGGTCGTCTTTA | 115 |
| chr7:92511036-92513050 | p2 | (TA)25 | AGACTTCGACTTTCTGCCCA | GTCCCTCGTGGTCGTCTTTA | 115 |
| chr7:92511212-92513262 | p2 | (AT)7 | CCACTGTTCACCCAACACAG | TCCACTTTGAGCAAAAACCC | 136 |
| chr7:92511212-92513262 | p2 | (TA)25 | CCACTGTTCACCCAACACAG | TCCACTTTGAGCAAAAACCC | 136 |
| chr7:92547010-92549028 | p3 | (GTT)6 | GCCCTCGTAACCATAAGCAA | TTATTTCTGGGGGTTTTCCC | 217 |

| Marker ID | SSR type | SSR motif | FORWARD PRIMER (5'-3') | REVERSE PRIMER (5'-3') | size (bp) |
|---|---|---|---|---|---|
| chr7:92576267-92578281 | p2 | (AT)7 | AACGGAGCGGAAAGAGGTAT | TGGGTCCCATTTATGCAAGT | 188 |
| chr7:92580916-92582932 | p4 | (GAAA)4 | CTTCGGGCCAAGATGTAAAA | GCATGTATTAACCGGAGCGT | 158 |
| chr7:92580916-92582932 | p5 | (TGAAT)4 | CTTCGGGCCAAGATGTAAAA | GCATGTATTAACCGGAGCGT | 158 |
| chr7:92580934-92582961 | p4 | (GAAA)4 | CTTCGGGCCAAGATGTAAAA | GCATGTATTAACCGGAGCGT | 140 |
| chr7:92580934-92582961 | p5 | (TGAAT)4 | CTTCGGGCCAAGATGTAAAA | GCATGTATTAACCGGAGCGT | 140 |
| chr7:92590380-92592392 | p2 | (TG)6 | AAAGGTGCCTACGGGCTATT | CGACCTCATAGGATCGGTGT | 220 |
| chr7:92597613-92599628 | p3 | (GAA)5 | CAAGGTGATGTACTGGGCCT | TGTCCCGGTGGTTAATGAAT | 154 |
| chr7:92647276-92649288 | p2 | (TA)6 | AGTGGAGGCGCATTCTCTTA | GACCCGCAAGGTTTTGTAGA | 204 |
| chr7:92759719-92761735 | p2 | (AT)7 | GCCGCTCGATATGGTTCTTA | GCCTCATCCTTATACGCTCG | 218 |
| chr7:92759741-92761755 | p2 | (AT)7 | GTTGGATCGTTTGATGACCC | GCCTCATCCTTATACGCTCG | 291 |
| chr7:92759765-92761777 | p2 | (AT)7 | GTTGGATCGTTTGATGACCC | GCCTCATCCTTATACGCTCG | 267 |
| chr7:92803244-92805259 | p5 | (CATAT)3 | TTGGCGGACAAGTTGTATGA | AAGTCTGTGGGTTGGATTGC | 167 |
| chr7:92803244-92805259 | p2 | (AT)8 | TTGGCGGACAAGTTGTATGA | AAGTCTGTGGGTTGGATTGC | 167 |
| chr7:92850608-92852622 | p2 | (TA)7 | ACCAAAAGCTGTAGCTCCGA | TCGGATGACCTTGACCTTTC | 157 |
| chr7:92877137-92879167 | p2 | (TA)15 | GAAACGGTTGAAGCGAGAAG | GGGGGTTTCGATTTCTTGTT | 218 |
| chr7:92913245-92915257 | p2 | (AT)6 | GGCTCACTCGTTGGTTTCAT | GAGAACTGGGATCAGCGAAG | 245 |
| chr7:92924268-92926286 | p2 | (TA)7 | AGAAATGTGGGCATTGAAGG | AGAGCACCATAGGCTCGTGT | 236 |
| chr7:92924268-92926286 | p6 | (ATACGG)3 | AGAAATGTGGGCATTGAAGG | AGAGCACCATAGGCTCGTGT | 236 |
| chr7:93014737-93016753 | p2 | (AT)8 | AAGCCGACAAGGCTAACAAA | GCGGAGCTCTATTTCATTCG | 238 |
| chr7:93038964-93040978 | p2 | (AT)7 | CCGACAAGGCCAAATACATC | CGGAGCTCTACTTCATTCGG | 275 |
| chr7:93087459-93089479 | p5 | (TTGCA)4 | CTAGCCACCCCGTTATGTGT | AAGTACCAAGAATGGGCACG | 124 |
| chr7:93096494-93098509 | p5 | (TGTTT)3 | TTGTGGCAGCACAGAGTAGG | ATGAACGCCTGATAAAACCG | 217 |
| chr7:93106822-93108836 | p2 | (AT)7 | TCACCATCCTCGTTCTTTCC | AAGAGGGGAGCATTTAGGGA | 131 |
| chr7:93124736-93126760 | p6 | (AAAAAG)4 | ACGATGAGGGATGTCTACGC | TTGTCATCGATTGATTCGGA | 291 |
| chr7:93124736-93126760 | p2 | (AT)9 | ACGATGAGGGATGTCTACGC | TTGTCATCGATTGATTCGGA | 291 |
| chr7:93319851-93321866 | p5 | (GTGTT)3 | AAATTCGCAAAAGGGGCTAT | TTTATCTCGGGCCTTTTCCT | 228 |
| chr7:93345798-93347812 | p2 | (TA)7 | GGAGGACGAGCGAGTGATAG | GGGTGACGTAAGGTCGTGAT | 162 |
| chr7:93351940-93353954 | p2 | (TA)7 | GGAGGACGAGCGAGTGATAG | GGGTGACGTAAGGTCGTGAT | 162 |
| chr7:93366542-93368566 | p2 | (AT)12 | ATTATGCAGATAAACGCGCC | TGCGAAGCATCGAAACATAG | 251 |
| chr7:93366542-93368566 | p2 | (AT)8 | ATTATGCAGATAAACGCGCC | TGCGAAGCATCGAAACATAG | 251 |
| chr7:93366844-93368858 | p2 | (AT)12 | CAGGCAAGTTCGAAGACTCC | CATCGTCTCACAGGGCTACA | 278 |
| chr7:93366844-93368858 | p2 | (AT)8 | CAGGCAAGTTCGAAGACTCC | CATCGTCTCACAGGGCTACA | 278 |
| chr7:93366853-93368868 | p2 | (AT)12 | CAGGCAAGTTCGAAGACTCC | CATCGTCTCACAGGGCTACA | 269 |
| chr7:93366853-93368868 | p2 | (AT)8 | CAGGCAAGTTCGAAGACTCC | CATCGTCTCACAGGGCTACA | 269 |
| chr7:93366872-93368886 | p2 | (AT)12 | CAGGCAAGTTCGAAGACTCC | CATCGTCTCACAGGGCTACA | 250 |
| chr7:93366872-93368886 | p2 | (AT)8 | CAGGCAAGTTCGAAGACTCC | CATCGTCTCACAGGGCTACA | 250 |

| Marker ID | SSR type | SSR motif | FORWARD PRIMER (5'-3') | REVERSE PRIMER (5'-3') | size (bp) |
|---|---|---|---|---|---|
| chr7:93366890-93368938 | p2 | (AT)12 | CAGGCAAGTTCGAAGACTCC | CATCGTCTCACAGGGCTACA | 232 |
| chr7:93366890-93368938 | p2 | (AT)8 | CAGGCAAGTTCGAAGACTCC | CATCGTCTCACAGGGCTACA | 232 |
| chr7:93373391-93375405 | p2 | (AT)7 | CTGACGAGGTTGCCACAGTA | TGTTCTGTCGTGGTAGCAGC | 292 |
| chr7:93433008-93435020 | p2 | (CA)6 | ATTTTCCCCCATTCATCCAT | GCCATAGCTTGGTTGTTGGT | 240 |
| chr7:93433008-93435020 | p2 | (AT)8 | ATTTTCCCCCATTCATCCAT | GCCATAGCTTGGTTGTTGGT | 240 |
| chr7:93510672-93512684 | p2 | (TA)6 | CTCCCTGTCTACCTGTGGGA | TGTACCAGCCGTATGTTGGA | 209 |
| chr7:93526965-93528981 | p2 | (AT)7 | CTGGGTCGTGACAGAACCTT | AGTTGGCTTGCACCGTTATC | 271 |
| chr7:93527001-93529021 | p2 | (AT)7 | CTGGGTCGTGACAGAACCTT | AGTTGGCTTGCACCGTTATC | 235 |
| chr7:93617532-93619548 | p2 | (AT)14 | ATTTCATTCCGGAGTGTTGC | GCATAATGGTACGTCGGCTT | 124 |
| chr7:93617532-93619548 | p2 | (AT)8 | ATTTCATTCCGGAGTGTTGC | GCATAATGGTACGTCGGCTT | 124 |
| chr7:93697100-93699112 | p5 | (AGAAA)3 | ATTGCGGGAAGTTAAGAGCA | TTGGCCCAATTTAGTCAAGC | 173 |
| chr7:93697100-93699112 | p2 | (GT)6 | ATTGCGGGAAGTTAAGAGCA | TTGGCCCAATTTAGTCAAGC | 173 |
| chr7:93767709-93769727 | p2 | (AT)9 | GCATCGCTGAAATCAGAACA | CACTCGTCACGATTCATTGG | 238 |
| chr7:93767709-93769727 | p6 | (CATAAG)3 | GCATCGCTGAAATCAGAACA | CACTCGTCACGATTCATTGG | 238 |
| chr7:93888674-93890686 | p2 | (AT)18 | TATCATACGCAAGCCGACAA | TCTTGGAGAGCTCCGTTGTT | 292 |
| chr7:93888674-93890686 | p2 | (AT)6 | TATCATACGCAAGCCGACAA | TCTTGGAGAGCTCCGTTGTT | 292 |
| chr7:93888674-93890686 | p3 | (ACA)5 | TATCATACGCAAGCCGACAA | TCTTGGAGAGCTCCGTTGTT | 292 |
| chr7:93918213-93920225 | p2 | (AT)18 | TATCATACGCAAGCCGACAA | TCTTGGAGAGCTCCGTTGTT | 292 |
| chr7:93918213-93920225 | p2 | (AT)6 | TATCATACGCAAGCCGACAA | TCTTGGAGAGCTCCGTTGTT | 292 |
| chr7:93918213-93920225 | p3 | (ACA)5 | TATCATACGCAAGCCGACAA | TCTTGGAGAGCTCCGTTGTT | 292 |
| chr7:93972826-93974842 | p2 | (AT)17 | GCCTAAATCCTGTCGCTGAG | ATTTAGCTCCAGCATCCCCT | 185 |
| chr7:93972826-93974842 | p2 | (AT)8 | GCCTAAATCCTGTCGCTGAG | ATTTAGCTCCAGCATCCCCT | 185 |
| chr7:94097714-94099726 | p2 | (TA)6 | TACGAGCCTCGAGGACAACT | GGGTCTAGGAGGTTTTTCCG | 249 |
| chr7:94097714-94099726 | p2 | (AT)6 | TACGAGCCTCGAGGACAACT | GGGTCTAGGAGGTTTTTCCG | 249 |
| chr7:94112180-94114194 | p2 | (AT)7 | GCACTCCATTGTACGGAGGT | AAAAGCCTCTCGTGAGACCA | 107 |
| chr7:94122440-94124460 | p2 | (AT)10 | GCGGGTTAACGATGATGATT | ACAACATCCAAAGAAAGCGG | 196 |
| chr7:94149770-94151800 | p2 | (AT)7 | CTTTTGTCACGACCTAGCCC | GATTGGTTGAGACTCCGCAT | 181 |
| chr7:94149770-94151800 | p2 | (TA)15 | CTTTTGTCACGACCTAGCCC | GATTGGTTGAGACTCCGCAT | 181 |
| chr7:94230700-94232733 | p3 | (AAT)11 | CGGGAAATGGCAAAATAATG | CGGCTTTCCTCCAATTGTTA | 205 |
| chr7:94252110-94254136 | p2 | (TA)13 | AAGCGACCATGTAGGCTGAT | CCCTAGGACGAGTTGCTCTG | 133 |
| chr7:94316524-94318540 | p2 | (AT)8 | CAGTCATGGCCTACGAGGTT | ATATGAGTGTCCAGCTCGGG | 106 |
| chr7:94469836-94471881 | p2 | (AT)8 | TTAAAAATGACGGGACAGGG | CGGTGAGGATGACCATTTCT | 161 |
| chr7:94523431-94525443 | p2 | (AT)6 | GATGGGCATCCTGCATAAGT | GAAAGTCTCAAGGCTGTCGG | 205 |
| chr7:94548539-94550555 | p4 | (TTTC)4 | TGTTTCCACAGGTAGGGTCC | ACTTGTGTGTGCATTTGGGA | 196 |
| chr7:94561213-94563229 | p2 | (TA)8 | AGGACTTGCTTGGTTGAGGA | CGTCATGGAACCTCCTGTTT | 300 |
| chr7:94571162-94573176 | p2 | (TA)7 | CAGTGGTTTCGACCTTGGTT | AGCAACAGTGTCCACTCACG | 190 |

| Marker ID | SSR type | SSR motif | FORWARD PRIMER (5'-3') | REVERSE PRIMER (5'-3') | size (bp) |
|---|---|---|---|---|---|
| chr7:94603098-94605128 | p2 | (AT)15 | CCGTACGACGATAAAGCGAT | TCTGTCCCATAGGGATGCTC | 238 |
| chr7:94605985-94607999 | p2 | (AT)7 | GCGGTGGAGTTTCATTTGAT | TTACACCTCGGAACTTTGGG | 132 |
| chr7:94658021-94660037 | p4 | (CATA)4 | CCCCCGATATACCCTCTTGT | ATGCCCGCAGGTGTAGATAG | 250 |
| chr7:94717102-94719118 | p2 | (AT)8 | CCATGACTAGTGCCCGAGAT | GTCGGCTTGCGTGGTATAAT | 222 |
| chr7:94770133-94772145 | p2 | (AG)6 | TCTCCCTTATGTTGCCCATC | CGTCTGCAGTCTCGGTAACA | 267 |
| chr7:94782104-94784119 | p5 | (TTTTC)3 | TACCTTCCAGTTCAGCCCAC | GGCAGATTTGGTGTAGGCAT | 247 |
| chr7:94791170-94793196 | p2 | (TA)13 | GGGAATCAGCATAGACCCAA | TTTCTGTTTGTTGTGGCAGC | 168 |
| chr7:94892635-94894659 | p2 | (TA)12 | CCTTAGACCGACCTGCTCTG | ATTTCGCCCTTGGTCTTCTT | 286 |
| chr7:94892635-94894659 | p6 | (CATATA)3 | CCTTAGACCGACCTGCTCTG | ATTTCGCCCTTGGTCTTCTT | 286 |
| chr7:95069928-95071946 | p6 | (TGTAAG)3 | TTGGTAGGGCCAGGTAGATG | CAGCCAAGGCTCAAGGTTAC | 124 |
| chr7:95147521-95149536 | p3 | (CTT)5 | TTCCTGGCCGGAGTATTATG | GACATTGACGGGAAAGTCGT | 289 |
| chr7:95244556-95246572 | p2 | (AT)8 | GGGTTCGGACCTTACCAAAT | TGACGGACTGGTTGTTGTGT | 195 |
| chr7:95315498-95317522 | p2 | (TA)12 | TCTTTTAAGAAGGCGGCAAA | TCTGTGTCATTTCGGGTCAA | 165 |
| chr7:95351488-95353552 | p2 | (AT)32 | CGCGGGCTAATGTTATTGAT | TGTCTACCTGTGGGCATGAA | 239 |
| chr7:95351488-95353552 | p2 | (AT)15 | CGCGGGCTAATGTTATTGAT | TGTCTACCTGTGGGCATGAA | 239 |
| chr7:95481293-95483311 | p6 | (CTGAAA)3 | CGAATGGCTCTAATTGCACA | TCAGCAAGCTATGTGGTTCG | 267 |
| chr7:95574347-95576367 | p2 | (CA)10 | TCCAATCCCTCCTAAAGCCT | AATGAAGCAGCGCGAGTATT | 215 |
| chr7:95575810-95577825 | p5 | (TTTGC)3 | TCATCATCCCGCATACGTTA | ATAAGGCGCGAGATGAGCTA | 119 |
| chr7:95578725-95580751 | p2 | (TA)13 | AATGCTACCTTCCGTTGTGG | AATGTACCGCTGTAGCGAGG | 266 |
| chr7:95578725-95580751 | p2 | (TA)12 | AATGCTACCTTCCGTTGTGG | AATGTACCGCTGTAGCGAGG | 266 |
| chr7:95584398-95586412 | p2 | (TA)7 | CCCGATCGGAGTAGACACAT | CAGTTTATGACCGGATGGCT | 290 |
| chr7:95584398-95586412 | p2 | (TA)7 | CCCGATCGGAGTAGACACAT | CAGTTTATGACCGGATGGCT | 290 |
| chr7:95590772-95592784 | p2 | (GT)6 | CTCCCAAGTCTGATGCCATT | TTGGCCACCTTTCGTAATTC | 161 |
| chr7:95612444-95614464 | p2 | (AT)10 | CATATCTTGCGGGGAAAGAA | ACCCGAGATGGTGAAGAAGA | 132 |
| chr7:95646973-95648985 | p2 | (AT)6 | CTACCCACGCATCCGTATCT | AGCGGATTCACAGAAGCACT | 224 |
| chr7:95646973-95648985 | p2 | (AT)7 | CTACCCACGCATCCGTATCT | AGCGGATTCACAGAAGCACT | 224 |
| chr7:95646973-95648985 | p5 | (ATCAT)3 | CTACCCACGCATCCGTATCT | AGCGGATTCACAGAAGCACT | 224 |
| chr7:95653686-95655698 | p2 | (AT)6 | ACATCACGCAACCGATATGA | CCCAGAGTTCTACGGGTCAA | 140 |
| chr7:95653686-95655698 | p5 | (ATCAT)3 | ACATCACGCAACCGATATGA | CCCAGAGTTCTACGGGTCAA | 140 |
| chr7:95654345-95656360 | p2 | (AT)6 | ACTTTTGTCACGACCCAACC | TCCTCAGGAGACAGGTTGCT | 194 |
| chr7:95654345-95656360 | p5 | (ATCAT)3 | ACTTTTGTCACGACCCAACC | TCCTCAGGAGACAGGTTGCT | 194 |
| chr7:95659868-95661888 | p2 | (TA)10 | AGACATATGGACGGGACAGG | ATGCCCACAGGTAGACAAGG | 292 |
| chr7:95712488-95714502 | p2 | (AT)7 | TTTTCTTCTCTCCAAGGCCA | GACGTCGGAATAAGGCGATA | 166 |
| chr7:95718633-95720645 | p2 | (AT)6 | AGAAGGAAGGGTGATGCCTT | AATTCCCGGATTGTGATTGA | 256 |
| chr7:95907585-95909607 | p2 | (TG)11 | TAACAAGAGTGGACCTCGGG | TGATGTCCTTGTCAGCTTGC | 208 |
| chr7:95919138-95921153 | p5 | (TTTAA)3 | ACATAGGGGGTACGGAAAGG | TGGCCACATAGGGATCTAGC | 294 |

| Marker ID | SSR type | SSR motif | FORWARD PRIMER (5'-3') | REVERSE PRIMER (5'-3') | size (bp) |
|---|---|---|---|---|---|
| chr7:95929056-95931146 | p2 | (TA)13 | GCGAGCATCAGACACCAGTA | TGGTTGTGAGTTCTCGTTGC | 285 |
| chr7:95929056-95931146 | p2 | (AT)45 | GCGAGCATCAGACACCAGTA | TGGTTGTGAGTTCTCGTTGC | 285 |
| chr7:95933156-95935178 | p2 | (TG)11 | TAACAAGAGTGGACCTCGGG | TGATGTCCTTGTCAGCTTGC | 208 |
| chr7:95944709-95946724 | p5 | (TTTAA)3 | ACATAGGGGGTACGGAAAGG | TGGCCACATAGGGATCTAGC | 294 |
| chr7:96008879-96010891 | p2 | (TA)6 | CCACCACTCTTTCCCACACT | GTAGGATGCACAGGGGAAGA | 138 |
| chr7:96033583-96035601 | p3 | (TGT)6 | TGGAAGGATTGGAGGACAAG | CGCGCAGAGTTATTCCCTAC | 159 |
| chr7:96131022-96133080 | p2 | (TA)29 | AGTCAGGTCATCCGAAAACG | GTCCCACACGTCGGTACTTT | 142 |
| chr7:96212508-96214546 | p5 | (ACCGG)4 | TCGCACTTGGGATTTCTTTC | ATCTAACCCACTCCCGTTCC | 282 |
| chr7:96212571-96214583 | p5 | (ACCGG)4 | TCGCACTTGGGATTTCTTTC | ATCTAACCCACTCCCGTTCC | 219 |
| chr7:96212656-96214694 | p5 | (ACCGG)4 | TCGCACTTGGGATTTCTTTC | ACACCACATCAGTGCCATGT | 134 |
| chr7:96240723-96242743 | p4 | (ATAA)5 | TCGGCCTTGATTAAAGTGCT | TGCACAAGGTCAAGACGAAG | 209 |
| chr7:96289224-96291264 | p2 | (TA)17 | GCCCACAAGCAACTCAATCT | TGCCGTCCTTTCTACAGCTT | 192 |
| chr7:96289224-96291264 | p2 | (AT)20 | GCCCACAAGCAACTCAATCT | TGCCGTCCTTTCTACAGCTT | 192 |
| chr7:96340090-96342102 | p2 | (AT)15 | CCGAAACCTTCCAAATTCAA | AATGAAGGTTTGGTGGCTTG | 269 |
| chr7:96340090-96342102 | p2 | (AT)6 | CCGAAACCTTCCAAATTCAA | AATGAAGGTTTGGTGGCTTG | 269 |
| chr7:96445914-96447926 | p2 | (TA)6 | ACTTGCGGGCAACTCTTAAA | ATGCTTGTGAGCCGTAGCTT | 158 |
| chr7:96447239-96449261 | p2 | (AT)11 | AGGCCCCTAGTTGGTCATTT | AAAACCAATCTGTTGGGCTG | 234 |
| chr7:96466295-96468307 | p3 | (AAT)7 | AGAAGCTGGCGGGTTTTAAT | TTAACCAAAATGGCGACCTC | 292 |
| chr7:96507117-96509129 | p5 | (ATTCA)4 | TCCCCCAAAGTGCATATCAT | CTGGCCGAGTTAGATTCTCG | 200 |
| chr7:96507117-96509129 | p2 | (CA)6 | TCCCCCAAAGTGCATATCAT | CTGGCCGAGTTAGATTCTCG | 200 |
| chr7:96541092-96543154 | p2 | (AT)31 | TGATGCAGCAAAATGTGGAT | CCTCCGCTTCTGAGTTTTTG | 212 |
| chr7:96544740-96546755 | p3 | (ACA)5 | TTCGATTCGGTTTCCAGTTC | GGTCCAAGTAGGCTCCATCA | 225 |
| chr7:96547971-96549993 | p2 | (AT)11 | TGTTAGGTCGTGAGGTTCCC | ATGGAGGCTTCGGGTTTAGT | 223 |
| chr7:96568067-96570082 | p5 | (CACCC)3 | TGCGCTGCATTATCTTCTTG | GCATGGTGGTCTTGTGATTG | 105 |
| chr7:96641279-96643294 | p5 | (ACAAA)3 | AATTTACGGTCTGCAGTGGG | GAGAACAGTCCCCAACCGTA | 169 |
| chr7:96792879-96794909 | p2 | (AT)15 | AGGTAGAGGGAAGAGCGGAG | TAATCAACTCGGGCTCCATC | 228 |
| chr7:96793901-96795916 | p3 | (TTG)5 | AAGCCCTCCAAAAGGATGAT | AAGCCATTTGGCAGCTTAGA | 190 |
| chr7:96793901-96795916 | p4 | (TGTA)4 | AAGCCCTCCAAAAGGATGAT | AAGCCATTTGGCAGCTTAGA | 190 |
| chr7:96808519-96810549 | p2 | (AT)15 | AGGTAGAGGGAAGAGCGGAG | TAATCAACTCGGGCTCCATC | 228 |
| chr7:96809541-96811556 | p3 | (TTG)5 | AAGCCCTCCAAAAGGATGAT | AAGCCATTTGGCAGCTTAGA | 190 |
| chr7:96809541-96811556 | p4 | (TGTA)4 | AAGCCCTCCAAAAGGATGAT | AAGCCATTTGGCAGCTTAGA | 190 |
| chr7:96843700-96845714 | p2 | (TA)7 | CCAATGTGGTGGTGAACTTG | CATGGCCTCTCCATTTTCAT | 124 |
| chr7:96843700-96845714 | p6 | (CTTATG)3 | CCAATGTGGTGGTGAACTTG | CATGGCCTCTCCATTTTCAT | 124 |
| chr7:96889297-96891309 | p2 | (TA)6 | CGTACTTGCCGTATGCTGAA | TGGTCCCATGTCGAACTGTA | 221 |
| chr7:96894633-96896649 | p2 | (AT)7 | ACATAGCAAGGTGTTTGGGG | CAGGTTCGAGGCTAGACAGG | 250 |
| chr7:96894633-96896649 | p2 | (TA)8 | ACATAGCAAGGTGTTTGGGG | CAGGTTCGAGGCTAGACAGG | 250 |

| Marker ID | SSR type | SSR motif | FORWARD PRIMER (5'-3') | REVERSE PRIMER (5'-3') | size (bp) |
|---|---|---|---|---|---|
| chr7:97014946-97016960 | p2 | (AT)7 | TTTCATTCAAATCAACCGCA | GCTTGTGTATTGGATGGGCT | 119 |
| chr7:97014946-97016960 | p3 | (AAC)6 | TTTCATTCAAATCAACCGCA | GCTTGTGTATTGGATGGGCT | 119 |
| chr7:97014946-97016960 | p2 | (GT)6 | TTTCATTCAAATCAACCGCA | GCTTGTGTATTGGATGGGCT | 119 |
| chr7:97015906-97017918 | p2 | (AT)7 | TTTTTGCCCTTAGCCTCTCA | TGGGCTGTTGTTGTTGTTGT | 297 |
| chr7:97015906-97017918 | p3 | (AAC)6 | TTTTTGCCCTTAGCCTCTCA | TGGGCTGTTGTTGTTGTTGT | 297 |
| chr7:97015906-97017918 | p2 | (GT)6 | TTTTTGCCCTTAGCCTCTCA | TGGGCTGTTGTTGTTGTTGT | 297 |
| chr7:97022824-97024839 | p5 | (TTTCA)3 | GGTGAGCTCCAGTTCTTTCG | CAGAAATGCCGAGATACCGT | 226 |
| chr7:97186087-97188105 | p2 | (TA)6 | GGGAGCAAGGAATGTGTTGT | GAGAATTCAACTCGGCCAAA | 123 |
| chr7:97187546-97189596 | p2 | (AT)25 | TGATAAGGGGTGCTCGGTAG | GTCGGCTTGCGTATTTTGTT | 271 |
| chr7:97187546-97189596 | p3 | (CAA)6 | TGATAAGGGGTGCTCGGTAG | GTCGGCTTGCGTATTTTGTT | 271 |
| chr7:97241431-97243447 | p2 | (GT)8 | ATTCAGGTAACGGAGGGCTT | CGTAGGAAACAAGTGGGGAA | 203 |
| chr7:97373544-97375556 | p6 | (ATACGA)5 | CCTCTGCAGATGGTACGGAT | CCAACCAAGGCTCAAGGTTA | 101 |
| chr7:97373544-97375556 | p2 | (GT)6 | CCTCTGCAGATGGTACGGAT | CCAACCAAGGCTCAAGGTTA | 101 |
| chr7:97394575-97396593 | p6 | (AGCAAC)3 | AAGTCGAGGGGGTTCCTAGA | CGCTACCCTGAACTCTTTGC | 139 |
| chr7:97461441-97463453 | p2 | (AG)6 | GTATCGACGGGTTTGACGAT | CGCAGGTATGAACTCCGTTT | 113 |
| chr7:97478620-97480632 | p2 | (AG)6 | GTATCGACGGGTTTGACGAT | CGCAGGTATGAACTCCGTTT | 113 |
| chr7:97490212-97492227 | p5 | (TTTCA)3 | ATCTGGTTCTGGTCAAACGG | TTTTTCGGATTTTCGAGCAC | 111 |
| chr7:97491390-97493405 | p2 | (TA)12 | CTTTAACAGCTGAAACGGGC | TCGGGTGAATTCAAATGGTT | 209 |
| chr7:97491410-97493430 | p2 | (TA)12 | CTTTAACAGCTGAAACGGGC | TCGGGTGAATTCAAATGGTT | 189 |
| chr7:97566063-97568078 | p5 | (CATGT)3 | AGGCAAACCTCAGCACATCT | GTAACCTACACCTTCGCCCA | 170 |
| chr7:97641861-97643913 | p2 | (AT)26 | CCTTCCGGGAATCTAAGAGC | GGGGTGACATAAGGTCGAGA | 260 |
| chr7:97641861-97643913 | p2 | (AT)11 | CCTTCCGGGAATCTAAGAGC | GGGGTGACATAAGGTCGAGA | 260 |
| chr7:97716977-97718995 | p3 | (GTT)6 | CTGGGCACACTCACTCAAAA | CAGCTCAGTAGGCCTTCCAC | 157 |
| chr7:97737625-97739640 | p5 | (TTCAT)3 | TGCGGTAGGGAACTATGACC | CGCCCTTCGTCAGTCTATTC | 299 |
| chr7:97852235-97854255 | p2 | (AT)10 | TTGGAAAAACGGGACTCAAG | CACGGCTCATCGGATTTAGT | 236 |
| chr7:97852235-97854255 | p2 | (AT)9 | TTGGAAAAACGGGACTCAAG | CACGGCTCATCGGATTTAGT | 236 |
| chr7:97863174-97865216 | p2 | (AT)23 | ATGCATGCTCATGCCTATCA | TATATGATGATGGCCGCAGA | 283 |
| chr7:97863174-97865216 | p2 | (AT)6 | ATGCATGCTCATGCCTATCA | TATATGATGATGGCCGCAGA | 283 |
| chr7:97871967-97873979 | p2 | (AG)6 | CCAAAGTCCGTTCTGGAAAA | TCCCTTAGGGTCCGATTACC | 234 |
| chr7:97871967-97873979 | p3 | (GAT)7 | CCAAAGTCCGTTCTGGAAAA | TCCCTTAGGGTCCGATTACC | 234 |
| chr7:97975535-97977585 | p2 | (AT)25 | AGGCGCAGTCACTTGAGAAT | CGAACGGTTTTGCGATTAGT | 186 |
| chr7:98004186-98006198 | p2 | (AT)6 | GTCCCCTATGTAACCTGCGA | GCACTCAGTTCGTTCGACAA | 240 |
| chr7:98007173-98009191 | p3 | (ATC)6 | TTGATCCCGATATGCTGTGA | ATATGGGCATAAGCCATCCA | 122 |
| chr7:98007173-98009191 | p2 | (AT)6 | TTGATCCCGATATGCTGTGA | ATATGGGCATAAGCCATCCA | 122 |
| chr7:98022872-98024892 | p2 | (TA)10 | ATATAACATCCGTCGGGCAG | GAAACAAGCTTTGGAGCAGC | 153 |
| chr7:98044428-98046448 | p2 | (AT)8 | GGGACAAAGGAGTGCTTTCA | TTCGGCAGTCAGCTACAATG | 280 |

| Marker ID | SSR type | SSR motif | FORWARD PRIMER (5'-3') | REVERSE PRIMER (5'-3') | size (bp) |
|---|---|---|---|---|---|
| chr7:98044428-98046448 | p2 | (TA)10 | GGGACAAAGGAGTGCTTTCA | TTCGGCAGTCAGCTACAATG | 280 |
| chr7:98183785-98185868 | p2 | (AT)36 | TTCCTGCTCCCTCGTTCTTA | CGAGGGGGTTCTTAAACACA | 183 |
| chr7:98183785-98185868 | p5 | (GAAAT)3 | TTCCTGCTCCCTCGTTCTTA | CGAGGGGGTTCTTAAACACA | 183 |
| chr7:98234301-98236384 | p2 | (AT)36 | TTCCTGCTCCCTCGTTCTTA | CGAGGGGGTTCTTAAACACA | 183 |
| chr7:98234301-98236384 | p2 | (AT)37 | TTCCTGCTCCCTCGTTCTTA | CGAGGGGGTTCTTAAACACA | 183 |
| chr7:98234475-98236549 | p2 | (AT)36 | TGAAAATTTGGACCCCTCAG | ACTCACACACACATCGGCAT | 265 |
| chr7:98234475-98236549 | p2 | (AT)37 | TGAAAATTTGGACCCCTCAG | ACTCACACACACATCGGCAT | 265 |
| chr7:98241896-98243914 | p2 | (AC)6 | ACGACCACACCGATGGTTAT | AAAGCAAAGTGGGGTGATTG | 143 |
| chr7:98241896-98243914 | p2 | (TG)9 | ACGACCACACCGATGGTTAT | AAAGCAAAGTGGGGTGATTG | 143 |
| chr7:98290007-98292023 | p2 | (AT)8 | GCAGAAGAATCTCTGGGCTG | CCTAGCTGGGTCGTGACATT | 240 |
| chr7:98349789-98351801 | p2 | (AT)6 | CAACACACACACCCATCACA | CGTTGGAGCGATCAACCTAT | 188 |
| chr7:98349818-98351830 | p2 | (AT)6 | CAACACACACACCCATCACA | CGTTGGAGCGATCAACCTAT | 159 |
| chr7:98349832-98351848 | p2 | (AT)6 | CAACACACACACCCATCACA | CGTTGGAGCGATCAACCTAT | 145 |
| chr7:98350317-98352329 | p2 | (AT)6 | ACACAAGGTGTGGGACCATT | TATTCCGGGATTTGAAGTCG | 144 |
| chr7:98495652-98497668 | p2 | (GT)8 | CTTTCGCAATTTGCATCTCA | TTAACCTCCCGCAATTTGTC | 257 |
| chr7:98495652-98497668 | p3 | (TTG)5 | CTTTCGCAATTTGCATCTCA | TTAACCTCCCGCAATTTGTC | 257 |
| chr7:98603624-98605650 | p2 | (AT)13 | CGATGATTGATCGGTTTCCT | AAAGAGGAGAAGGGCTCTCG | 263 |
| chr7:98704086-98706100 | p2 | (TA)7 | AACCTACCAGGGCACCTCTT | TAAATGTGTTGCTGCCCAAA | 264 |
| chr7:98749265-98751289 | p6 | (CATATT)4 | TCCGCATATGTTTGTGCATT | TTCGGCAGAGTTTCCTTTGT | 111 |
| chr7:98763850-98765865 | p3 | (ACA)5 | CCCCTCCTAGTGTAGGGACC | GCTGCTCATGTATGCCTCAA | 228 |
| chr7:98845454-98847469 | p5 | (ATATG)3 | CATGATTTTCTGGCACCCTT | GTGTGACGACGGTGATTGAG | 202 |
| chr7:98953243-98955255 | p2 | (AT)10 | GGGACCCGATTACCCTAAGA | GGTCCCCTTTGAATTCACCT | 187 |
| chr7:98953243-98955255 | p2 | (TA)6 | GGGACCCGATTACCCTAAGA | GGTCCCCTTTGAATTCACCT | 187 |
| chr7:98966627-98968643 | p4 | (AATT)4 | CATAATTCTTCCGCTTCCCA | CCAGGATCGTGCCAATAAGT | 292 |
| chr7:99013295-99015313 | p3 | (CAA)6 | GCACTCAATATAGCCCCGAA | AGTTTCCCTTGTCGGGAGTT | 286 |
| chr7:99090946-99092960 | p2 | (GT)7 | CCTAGGCCTCTTGCTCACAG | CTCCCGCATCAACATTCTTT | 241 |
| chr7:99118324-99120336 | p2 | (AG)6 | ATTGCTCGTACTGACGTCCC | CTCAGACTCGGCAATTCTCC | 284 |
| chr7:99118324-99120336 | p3 | (TTG)6 | ATTGCTCGTACTGACGTCCC | CTCAGACTCGGCAATTCTCC | 284 |
| chr7:99156110-99158128 | p6 | (TATATG)3 | CAAGTTGGATTCTCGGGTGT | CGTCCGCTTTCTACGCTAAC | 229 |
| chr7:99212978-99214992 | p2 | (AT)7 | AACAAGGGGCTACAGGGAAT | GGCTCTTTGCCGTAGTCTTG | 254 |
| chr7:99212978-99214992 | p2 | (AT)40 | AACAAGGGGCTACAGGGAAT | GGCTCTTTGCCGTAGTCTTG | 254 |
| chr7:99212978-99214992 | p5 | (TGTGT)3 | AACAAGGGGCTACAGGGAAT | GGCTCTTTGCCGTAGTCTTG | 254 |
| chr7:99213132-99215212 | p2 | (AT)7 | AACAAGGGGCTACAGGGAAT | CTCTTTGCCGTAGTCTTGCC | 100 |
| chr7:99213132-99215212 | p2 | (AT)40 | AACAAGGGGCTACAGGGAAT | CTCTTTGCCGTAGTCTTGCC | 100 |
| chr7:99213132-99215212 | p5 | (TGTGT)3 | AACAAGGGGCTACAGGGAAT | CTCTTTGCCGTAGTCTTGCC | 100 |
| chr7:99397137-99399153 | p2 | (AT)12 | ACCCGTTGGGCAATTATACA | GAAGACCTTTCCACCCATCA | 232 |

| Marker ID | SSR type | SSR motif | FORWARD PRIMER (5'-3') | REVERSE PRIMER (5'-3') | size (bp) |
|---|---|---|---|---|---|
| chr7:99397137-99399153 | p3 | (ATT)5 | ACCCGTTGGGCAATTATACA | GAAGACCTTTCCACCCATCA | 232 |
| chr7:99397137-99399153 | p3 | (ATT)6 | ACCCGTTGGGCAATTATACA | GAAGACCTTTCCACCCATCA | 232 |
| chr7:99397137-99399153 | p3 | (TTA)6 | ACCCGTTGGGCAATTATACA | GAAGACCTTTCCACCCATCA | 232 |
| chr7:99397193-99399209 | p2 | (AT)12 | ACCCGTTGGGCAATTATACA | GAAGACCTTTCCACCCATCA | 176 |
| chr7:99397193-99399209 | p3 | (ATT)5 | ACCCGTTGGGCAATTATACA | GAAGACCTTTCCACCCATCA | 176 |
| chr7:99397193-99399209 | p3 | (ATT)6 | ACCCGTTGGGCAATTATACA | GAAGACCTTTCCACCCATCA | 176 |
| chr7:99397193-99399209 | p3 | (TTA)6 | ACCCGTTGGGCAATTATACA | GAAGACCTTTCCACCCATCA | 176 |
| chr7:99397501-99399519 | p3 | (ATT)5 | TGTTCTGCTCGAGGAATGTG | CAACAAGGTTGCATAACCGA | 214 |
| chr7:99397501-99399519 | p3 | (ATT)6 | TGTTCTGCTCGAGGAATGTG | CAACAAGGTTGCATAACCGA | 214 |
| chr7:99397501-99399519 | p3 | (TTA)6 | TGTTCTGCTCGAGGAATGTG | CAACAAGGTTGCATAACCGA | 214 |
| chr7:99411331-99413343 | p2 | (AT)6 | ATTAGCCTTGACCCCTTCGT | AGTTCTGTTCGTTCGGTGCT | 277 |
| chr7:99739226-99741241 | p2 | (TA)6 | ACTGACGTCCCTTCTTGTGG | TCCGGAATAACTGGAGCACT | 295 |
| chr7:99739226-99741241 | p5 | (TATAT)3 | ACTGACGTCCCTTCTTGTGG | TCCGGAATAACTGGAGCACT | 295 |
| chr7:99739226-99741241 | p2 | (TA)8 | ACTGACGTCCCTTCTTGTGG | TCCGGAATAACTGGAGCACT | 295 |
| chr7:99740743-99742761 | p2 | (AT)12 | TGTATTTGCTATCCCGCCAT | AGCTCATTTCCCTTGTGCAT | 214 |
| chr7:99740743-99742761 | p6 | (ATTCTA)3 | TGTATTTGCTATCCCGCCAT | AGCTCATTTCCCTTGTGCAT | 214 |
| chr7:99769069-99771131 | p2 | (TA)31 | TCACCAGTGCTATGGGATCA | ACCCGTCATACTCCTCGTTG | 251 |
| chr7:99917493-99919513 | p2 | (AT)6 | TCGTCGTGGAAAGTTAAGGG | TGCAATTTCTCCTTTTTGCC | 230 |
| chr7:99917493-99919513 | p5 | (TGAAT)4 | TCGTCGTGGAAAGTTAAGGG | TGCAATTTCTCCTTTTTGCC | 230 |
| chr7:99920637-99922663 | p2 | (TA)13 | ATCGTGCGATAGAGCCAAGT | ATTCAAAACCTGTATGCGGC | 137 |
| chr7:99922249-99924281 | p2 | (TA)16 | GAACGGGCATAATGGAAAGA | CCAAACGTACGAAATACGGG | 283 |
| chr7:100009195-100011209 | p2 | (AT)7 | TCCATACGGGCATACAGACA | TACTAAGAGGGCCGGGTTCT | 140 |
| chr7:100069499-100071527 | p2 | (AT)14 | AGGATAGGGGGCTGGAGTAA | TTCCTCCGGAGTTTCTGCTA | 114 |
| chr7:100083649-100085695 | p2 | (AT)23 | ATGGCAAGCTCCAATTGTTC | GGAGCCGCTTACAGACTGAC | 296 |
| chr7:100160164-100162176 | p2 | (AG)6 | GGAAGGAATCATGCTTTGGA | GCGACGCGTTCTTATAATCC | 182 |
| chr7:100207977-100209992 | p5 | (CATTG)3 | TGTTGGCATACCCATTGTTG | CATGAATGGATGAGAATGCG | 241 |
| chr7:100255486-100257544 | p2 | (TA)29 | TGCTTTCCTCTCCCTTTTCA | CCCAAGCAATTTGACGATTT | 167 |
| chr7:100320984-100323002 | p2 | (TA)9 | GCGGAATTTTGTGTTGGATT | CAGCAGTAAAAGCACGGACA | 122 |
| chr7:100329806-100331824 | p6 | (TCCTAT)3 | CTTCAGCTAAACAAACCCGC | CTTTTCCAACCATAGCCCAA | 294 |
| chr7:100622816-100624830 | p2 | (AT)7 | CTCCACTTGCCACATTTCCT | GGCAATTCCCTCATCGTTTA | 132 |
| chr7:100704719-100706735 | p2 | (AG)8 | AGGCTACACACGGGGTAGTG | AGGGCGTACTGCAGAAGAAA | 178 |
| chr7:100714765-100716779 | p2 | (AT)6 | TCGGGTATGTCGTGGAATTT | GGGGTCATATCGATTGGTTG | 277 |
| chr7:100714765-100716779 | p2 | (AT)7 | TCGGGTATGTCGTGGAATTT | GGGGTCATATCGATTGGTTG | 277 |
| chr7:100939872-100941900 | p2 | (AT)14 | CGAATTAGTTCGTGGTCCGT | GGTTTACATGCCTTCTCCGA | 292 |
| chr7:100956066-100958081 | p2 | (TA)6 | CCGGGGTAATCATCTCACAC | CGGAGGCTACTGACCTTACG | 272 |
| chr7:100956066-100958081 | p3 | (ATC)5 | CCGGGGTAATCATCTCACAC | CGGAGGCTACTGACCTTACG | 272 |

| Marker ID | SSR type | SSR motif | FORWARD PRIMER (5'-3') | REVERSE PRIMER (5'-3') | size (bp) |
|---|---|---|---|---|---|
| chr7:100956066-100958081 | p3 | (ACA)5 | CCGGGGTAATCATCTCACAC | CGGAGGCTACTGACCTTACG | 272 |
| chr7:101051521-101053536 | p5 | (TATTT)3 | GCCAAACTAGCCCTTAGCAA | ACACTTGTTGTTCGTGCTGC | 131 |
| chr7:101051521-101053536 | p5 | (TCTTT)3 | GCCAAACTAGCCCTTAGCAA | ACACTTGTTGTTCGTGCTGC | 131 |
| chr7:101192886-101194901 | p2 | (TA)15 | GCCCACCAAAACTACGACAT | TGGGGATGGAAATTATGGAA | 219 |
| chr7:101192899-101194913 | p2 | (TA)15 | GCCCACCAAAACTACGACAT | TGGGGATGGAAATTATGGAA | 206 |
| chr7:101422931-101424946 | p3 | (CAT)5 | TCATATCCCATTGGCCTCTC | TTCCTTCTGTTCATGGGTCC | 116 |
| chr7:101422947-101424965 | p3 | (CAT)5 | TCATATCCCATTGGCCTCTC | TTCCTTCTGTTCATGGGTCC | 100 |
| chr7:101424427-101426443 | p2 | (AT)6 | CAAAGGTTGAAATGCCCATC | TAATGGGGCATTAACAGGGA | 169 |
| chr7:101424427-101426443 | p2 | (AT)8 | CAAAGGTTGAAATGCCCATC | TAATGGGGCATTAACAGGGA | 169 |
| chr7:101454773-101456800 | p3 | (CAA)9 | CCCCAAAATTTGACCACATC | TGGTAACACGATGTTTCGGA | 203 |
| chr7:101515735-101517747 | p2 | (AT)6 | TTTATTTGGTCGGTTGCACA | AGCAAAGCGAGTGATAGGGA | 140 |
| chr7:101624629-101626645 | p2 | (TA)7 | TAATGTCCAAAAGGATGGGC | CTTTAAGGGACAACGGACGA | 247 |
| chr7:101624629-101626645 | p2 | (AT)8 | TAATGTCCAAAAGGATGGGC | CTTTAAGGGACAACGGACGA | 247 |
| chr7:101666263-101668279 | p4 | (TTTA)4 | GTGAGAGTCCGATGTCCGAT | ATGTAAGCATGCCAATGCAA | 265 |
| chr7:101832783-101834861 | p2 | (TA)26 | TGGGGTAGAATCTCACAGGG | ATGTCCGAAAGAGCACCAAC | 135 |
| chr7:101852164-101854182 | p3 | (ATA)6 | TTGCGTCGTTGATTATCTGC | AGGATGGTTCGATGTTGGAG | 122 |
| chr7:101882028-101884043 | p5 | (ATTTC)3 | TTTTGGTCATGTGTGGCCTA | CCCCCACCCTTAAGAAACAT | 258 |
| chr7:102040885-102042900 | p3 | (TGT)5 | AAGGGGACAAAAGATCCCAC | ATCTCCATGCCCAACAAAAG | 160 |
| chr7:102067026-102069041 | p3 | (TGC)5 | CACCCGTTCGATTCAAGTTT | GCCAAGGTCACGTCATAGGT | 178 |
| chr7:102077842-102079862 | p4 | (TATC)5 | TCTCATCTTCACCCTCACCC | TTTCGAGTTATTGCCCCAAG | 239 |
| chr7:102263259-102265291 | p2 | (AT)16 | CCACCCAAGGTTCATTCCTA | AATCATGGCTCAATCCAAGC | 181 |
| chr7:102299235-102301250 | p5 | (AATGC)3 | ATGTCATCATTGGGCCATTT | TCTTTTCCGCTTGCTTGATT | 181 |
| chr7:102478304-102480316 | p2 | (AC)6 | ATCACCTCCCCATCTCCTTT | GGGGAAGGGTCTTAGTGGAG | 253 |
| chr7:102478304-102480316 | p3 | (ACC)5 | ATCACCTCCCCATCTCCTTT | GGGGAAGGGTCTTAGTGGAG | 253 |
| chr7:102482928-102484956 | p2 | (TA)14 | GGCCCTATTGGCAGTTTGTA | ACCCACACTACGGACGAAAG | 294 |
| chr7:102721626-102723644 | p6 | (GCCCTC)3 | TCGGTCTAATGAACCCCTTG | GCGTATTTGGCACATGTTTG | 274 |
| chr7:102875957-102877975 | p2 | (TA)9 | GTAAACCTCCCAGCCCATTT | AATCGTTGTCTTTTGTCGGG | 185 |
| chr7:102926062-102928078 | p2 | (AT)8 | ATAAGATGCCAACTCCACGG | AGGCCATCCAGTTATTGACG | 170 |
| chr7:102963262-102965278 | p2 | (TA)8 | GCCTAGGTGCATTAAGCCAA | AGTTGAGGCTACATGGGTGC | 170 |
| chr7:103016167-103018185 | p3 | (TGT)6 | TGCAGGATTGTTGAAATGGA | CACCTTGTCCCTCATGTCCT | 195 |
| chr7:103034983-103036998 | p5 | (TACAA)3 | TCTGATACCGGATACCCTGC | TTTTTCAGGCCTTGGTTGAG | 132 |
| chr7:103066161-103068181 | p2 | (TA)10 | GCATCGCTGAAATCTGAACA | CTCATCCGAGGGGTTGTCTA | 115 |
| chr7:103066161-103068181 | p3 | (ACA)5 | GCATCGCTGAAATCTGAACA | CTCATCCGAGGGGTTGTCTA | 115 |
| chr7:103150054-103152069 | p3 | (TGT)5 | GGGATCAAGCAAATCGAAGA | GGCTTATCTTGAAATGCCCA | 173 |
| chr7:103170841-103172874 | p3 | (TCT)45 | AACAGCAGCATGCTCATCAC | GAGGAAGAGGAGGAGGAGGA | 206 |
| chr7:103277118-103279178 | p2 | (AT)30 | GCAACCGGAGTAAGAATGGA | AAGACAACCTTTGGGGGTTC | 261 |

| Marker ID | SSR type | SSR motif | FORWARD PRIMER (5'-3') | REVERSE PRIMER (5'-3') | size (bp) |
|---|---|---|---|---|---|
| chr7:103277118-103279178 | p2 | (AT)28 | GCAACCGGAGTAAGAATGGA | AAGACAACCTTTGGGGGTTC | 261 |
| chr7:103301603-103303618 | p3 | (CAA)5 | TAACATTGAGGCATCCACGA | CCCCATTGGATTTGTTTGTC | 173 |
| chr7:103341579-103343595 | p2 | (GA)8 | TCTTTTCTCCGGATTTGGTG | AGTAAACAACCAGGTCCCCC | 133 |
| chr7:103459331-103461351 | p5 | (GTTTT)4 | TGTTGAATTCTGCCCATCAA | GAGAGCTCGGTTTTGGATTG | 259 |
| chr7:103490722-103492770 | p3 | (TAT)16 | GAACCGATGCAAATTCAGGT | GTGGGAGGGTGAAATTGCTA | 196 |
| chr7:103503465-103505483 | p5 | (CAGGG)4 | AGGTCCCCCAAGTTCTGACT | GGACCGCAAAAGGTATCGTA | 146 |
| chr7:103503465-103505483 | p2 | (TA)9 | AGGTCCCCCAAGTTCTGACT | GGACCGCAAAAGGTATCGTA | 146 |
| chr7:103591196-103593214 | p6 | (CATATA)3 | ATCTCGCTAGGGTGTGATGG | GAGTCGTGGGCATAGGATGT | 167 |
| chr7:103647243-103649258 | p5 | (TTTTA)3 | GAACATTGCTCTCCAGCTCC | CTCAACCAGGGAAGGATTCA | 260 |
| chr7:103772441-103774515 | p2 | (TA)37 | GCTTGTCTTTTTCGCTGACC | CCCAAGCACAATCCTCAAAT | 177 |
| chr7:103782411-103784431 | p2 | (AT)10 | CCCGTGTTTGGAGGTAGAAA | CGACGATGATTGGTGTTCTG | 291 |
| chr7:103862777-103864795 | p2 | (AT)9 | ATAGATGCCCACTACGCCTG | CTCAGCCGAGTAAGGTCCTG | 160 |
| chr7:103880609-103882643 | p2 | (TA)17 | ATAGCTCGGAACATCGAGGA | GAATCTCAATTCCAAGGGCA | 169 |
| chr7:103894383-103896415 | p2 | (TA)16 | ATGGAAGATCGGAAATGCTG | ATGAGTATCTCGCCCCCTTT | 205 |
| chr7:103915660-103917684 | p2 | (TA)8 | GCCTCATTAGGCCATCAAAA | CATTGTCCGGAGGTGCTTAT | 112 |
| chr7:103915660-103917684 | p2 | (AT)12 | GCCTCATTAGGCCATCAAAA | CATTGTCCGGAGGTGCTTAT | 112 |
| chr7:103921775-103923787 | p2 | (TA)9 | GCCTCATTAGGCCATCAAAA | CATTGTCCGGAGGTGCTTAT | 112 |
| chr7:103921775-103923787 | p2 | (AT)6 | GCCTCATTAGGCCATCAAAA | CATTGTCCGGAGGTGCTTAT | 112 |
| chr7:103969139-103971151 | p2 | (TG)6 | ACTTGCCTCCAAGTCAAGGA | CCTTAAACTTTCCACGCAGC | 190 |
| chr7:104023378-104025399 | p3 | (AAC)7 | GGATGCTATAGCTTGGACCG | TTCTCATGCGTTGGAAAGTG | 286 |
| chr7:104035885-104037897 | p2 | (CA)6 | ATGTGGGGCACGTTGATAGT | TGGTCGCGCAATCAATATAA | 208 |
| chr7:104093585-104095600 | p5 | (GTTCG)3 | AAACGGACCTTTTTCGGAGT | CACCAGTGCCTACTTCCCAT | 137 |
| chr7:104399132-104401162 | p2 | (AT)12 | GATTGCATCGTAGCCCTTGT | CAAGGCTTATCCGTCTTTCG | 235 |
| chr7:104399132-104401162 | p2 | (AT)15 | GATTGCATCGTAGCCCTTGT | CAAGGCTTATCCGTCTTTCG | 235 |
| chr7:104760498-104762513 | p3 | (CAA)5 | CCAAAGGGAAGTCTCCAACA | TCCGTCAAAGTGGACATCAA | 134 |
| chr7:104802464-104804548 | p3 | (CTT)28 | CATCATGACTCGCGAAATTG | CGGATTCCTTTTGTGTCCAT | 280 |
| chr7:104842837-104844857 | p5 | (TTTAT)3 | GCAAGGAAGAGGCCACATAG | TGAAAACACTAATGCCAGCG | 206 |
| chr7:104842837-104844857 | p5 | (TTTAT)4 | GCAAGGAAGAGGCCACATAG | TGAAAACACTAATGCCAGCG | 206 |
| chr7:105012696-105014711 | p5 | (GAAAG)3 | ACATTTGGTTTGGCTTCGTC | ACACCTTGGCCTCAAGACAC | 174 |
| chr7:105054398-105056410 | p2 | (TG)6 | ACCACCAGTTGAGGATCAGG | GGGAATTGGTAAGCCCAAAT | 274 |
| chr7:105086551-105088605 | p3 | (ATT)18 | TAGGAGAGGGGGCAACTTTT | GAGCTATGGGGCAGGACATA | 148 |
| chr7:105086551-105088605 | p2 | (CT)7 | TAGGAGAGGGGGCAACTTTT | GAGCTATGGGGCAGGACATA | 148 |
| chr7:105100689-105102701 | p3 | (ATT)18 | GACACCCAATTTTGTCCCAC | GCCGCAATAGGATAACGAAA | 273 |
| chr7:105100689-105102701 | p2 | (TC)6 | GACACCCAATTTTGTCCCAC | GCCGCAATAGGATAACGAAA | 273 |
| chr7:105108117-105110183 | p2 | (TA)15 | GGGGGTTGGGGTTCTATTTA | TTGATTTCCATTGGTTGGGT | 292 |
| chr7:105180090-105182110 | p4 | (ATTA)5 | TGGCCTCTTCGAGCTTATGT | TCACACCTCCGTACTAGGGC | 281 |

| Marker ID | SSR type | SSR motif | FORWARD PRIMER (5'-3') | REVERSE PRIMER (5'-3') | size (bp) |
|---|---|---|---|---|---|
| chr7:105301068-105303083 | p5 | (GAGTT)3 | CTACCGAGGTGGGAATGGTA | CCGGTACCCACTCAAGTGTT | 271 |
| chr7:105301068-105303083 | p5 | (TGCAT)3 | CTACCGAGGTGGGAATGGTA | CCGGTACCCACTCAAGTGTT | 271 |
| chr7:105519893-105521917 | p2 | (TA)12 | TCCAAGATTACCGGATCGAC | GGTGGGACACTCGTTCAGTT | 286 |
| chr7:105533158-105535174 | p2 | (TA)8 | GTGGGCCCCATACTAAACAA | TCCCGTGATTTTGGAGATGT | 231 |
| chr7:105533158-105535174 | p2 | (AT)11 | GTGGGCCCCATACTAAACAA | TCCCGTGATTTTGGAGATGT | 231 |
| chr7:105645346-105647412 | p5 | (TCGGT)3 | GGGCTCATGTCCTTTACTGC | CCGAAAATCGAAAAATCGAA | 239 |
| chr7:105645346-105647412 | p2 | (AT)33 | GGGCTCATGTCCTTTACTGC | CCGAAAATCGAAAAATCGAA | 239 |
| chr7:105645346-105647412 | p2 | (AT)12 | GGGCTCATGTCCTTTACTGC | CCGAAAATCGAAAAATCGAA | 239 |
| chr7:105800496-105802508 | p2 | (AT)14 | AGTTTATAGAGCGGGGGCAT | TAGTCGGGAAGAGCATGAGG | 147 |
| chr7:105800496-105802508 | p2 | (AT)6 | AGTTTATAGAGCGGGGGCAT | TAGTCGGGAAGAGCATGAGG | 147 |
| chr7:105858349-105860367 | p6 | (GTGTAT)3 | AGGCTACAGGAGGGTACGGT | CCAAATCTCCAACCGGAGTA | 238 |
| chr7:105897328-105899392 | p2 | (TA)32 | CAAAATCATTGTTGCATGGC | ACATCTCGGTAGCACAACCC | 243 |
| chr7:105961306-105963332 | p2 | (AT)13 | TGCATTTCCATTCCTTGTGA | TGGATTTTTGGGTTAAAGCG | 290 |
| chr7:106062908-106064920 | p2 | (TA)6 | AGTAGTGTGGGCTGTGACCC | GACCAAAAGGGTCAGCAAAA | 280 |
| chr7:106086454-106088468 | p2 | (TG)6 | CGGATGATGACGTAGGTGTG | AGCATAGGAGGGGAGACCAT | 200 |
| chr7:106086454-106088468 | p2 | (TA)7 | CGGATGATGACGTAGGTGTG | AGCATAGGAGGGGAGACCAT | 200 |
| chr7:106100424-106102439 | p3 | (TTG)6 | CACAAAGAGGAGAACCCAGC | GAGGGGTGTTTGGTCTGAAA | 156 |
| chr7:106100424-106102439 | p3 | (ATG)5 | CACAAAGAGGAGAACCCAGC | GAGGGGTGTTTGGTCTGAAA | 156 |
| chr7:106100424-106102439 | p2 | (AT)12 | CACAAAGAGGAGAACCCAGC | GAGGGGTGTTTGGTCTGAAA | 156 |
| chr7:106100935-106102959 | p3 | (TTG)6 | TAGGCTCATCAACAGGGGAG | AGCTTTCGGTCTCCTTAGCC | 166 |
| chr7:106100935-106102959 | p3 | (ATG)5 | TAGGCTCATCAACAGGGGAG | AGCTTTCGGTCTCCTTAGCC | 166 |
| chr7:106100935-106102959 | p2 | (AT)12 | TAGGCTCATCAACAGGGGAG | AGCTTTCGGTCTCCTTAGCC | 166 |
| chr7:106100935-106102959 | p2 | (TA)9 | TAGGCTCATCAACAGGGGAG | AGCTTTCGGTCTCCTTAGCC | 166 |
| chr7:106111241-106113257 | p2 | (AT)8 | TAACGTGGTGTCCTGCCATA | TTGCGAAAGATGAAGAGGCT | 128 |
| chr7:106139855-106141915 | p5 | (AAACT)4 | GCACAAGGGATGGATTAGGA | GCAAGTCCCACATCGGTAGT | 261 |
| chr7:106139855-106141915 | p2 | (TA)30 | GCACAAGGGATGGATTAGGA | GCAAGTCCCACATCGGTAGT | 261 |
| chr7:106248817-106250847 | p2 | (AT)15 | CAAGCAATCCCCTTCGTAAA | CCGCGATTTTGGTATTATGG | 129 |
| chr7:106260472-106262487 | p5 | (CATTG)3 | GCTCTGCGAAATGTAAAGGC | ACAACATGGCCCAACCTAAG | 175 |
| chr7:106344070-106346082 | p2 | (TA)6 | GGGCATTTTGGTCATTTCAC | CACGGCTGTTCACTCTTCAG | 278 |
| chr7:106344070-106346082 | p2 | (AT)15 | GGGCATTTTGGTCATTTCAC | CACGGCTGTTCACTCTTCAG | 278 |
| chr7:106344070-106346082 | p2 | (TA)10 | GGGCATTTTGGTCATTTCAC | CACGGCTGTTCACTCTTCAG | 278 |
| chr7:106379094-106381122 | p2 | (TA)14 | CGTTAGAACAACTTGGGGGA | GGAGAACCCCACTTTTGACA | 140 |
| chr7:106379094-106381122 | p2 | (TA)17 | CGTTAGAACAACTTGGGGGA | GGAGAACCCCACTTTTGACA | 140 |
| chr7:106540025-106542039 | p5 | (TTATA)3 | AACGCAACTGAACCTAAGCC | ACCACCCTCTTACCGGTTTC | 247 |
| chr7:106540025-106542039 | p2 | (TA)7 | AACGCAACTGAACCTAAGCC | ACCACCCTCTTACCGGTTTC | 247 |
| chr7:106540025-106542039 | p2 | (AG)11 | AACGCAACTGAACCTAAGCC | ACCACCCTCTTACCGGTTTC | 247 |

| Marker ID | SSR type | SSR motif | FORWARD PRIMER (5'-3') | REVERSE PRIMER (5'-3') | size (bp) |
|---|---|---|---|---|---|
| chr7:106546925-106548949 | p2 | (AT)6 | CGGTAACCGGACTGGTAAAA | ACAGTCCATCCCATTTCTGC | 171 |
| chr7:106546925-106548949 | p3 | (TGA)8 | CGGTAACCGGACTGGTAAAA | ACAGTCCATCCCATTTCTGC | 171 |
| chr7:106584647-106586662 | p5 | (TAAAA)3 | TTGGCTTCCACTTCAAAACC | GCTCCCGCTACACTTAAACG | 247 |
| chr7:106625876-106627938 | p2 | (AT)31 | TTGGCATCTGATGGTGACAT | AACCGAGGCTGAGATTGAGA | 227 |
| chr7:106665119-106667171 | p2 | (TA)26 | CTGCAAATCCAAGGGAAAAA | TTCTGGCTTGACTCCCATTT | 272 |
| chr7:106665119-106667171 | p2 | (TA)24 | CTGCAAATCCAAGGGAAAAA | TTCTGGCTTGACTCCCATTT | 272 |
| chr7:106805306-106807334 | p2 | (TA)6 | CGGAGTTGCGAGTTCTTAGG | CATGGCTGATGTTGAGCAGT | 169 |
| chr7:106805306-106807334 | p2 | (AT)8 | CGGAGTTGCGAGTTCTTAGG | CATGGCTGATGTTGAGCAGT | 169 |
| chr7:106822809-106824827 | p5 | (ATTGT)4 | GGACCCGACAATGCTGTTAT | GATGGGTGAAAACCTTGTGG | 197 |
| chr7:106822809-106824827 | p6 | (TTCTTT)3 | GGACCCGACAATGCTGTTAT | GATGGGTGAAAACCTTGTGG | 197 |
| chr7:106840220-106842232 | p2 | (AT)6 | TTTCTGCTCCCCATAGGTTG | TTGGAATTGGAGGTAGCTGG | 271 |
| chr7:106847320-106849334 | p2 | (AG)7 | GAGTCCATCAAGCCTCCAAA | TCTACCGGAGAGAGCACGAT | 224 |
| chr7:106877433-106879449 | p4 | (GAAA)4 | CGTTCGAGTTCCCCTAATCA | GTTCTTGGGTGATGCCAAGT | 170 |
| chr7:106956028-106958056 | p2 | (AT)17 | ATAACCCTAGGCCCATGTCC | GATGCAAAGCATGGGTTTTT | 296 |
| chr7:106956028-106958056 | p2 | (AT)7 | ATAACCCTAGGCCCATGTCC | GATGCAAAGCATGGGTTTTT | 296 |
| chr7:107038586-107040626 | p2 | (TA)20 | CAAAATCGTTCCCCAGAAGA | TCGTGACTCCAATGGTTCAA | 126 |
| chr7:107052504-107054519 | p2 | (TA)18 | AAGGCAAGGCAAAAGAGTGA | CACATAACGGGGTGGCTAGT | 138 |
| chr7:107052530-107054555 | p2 | (TA)18 | AAGGCAAGGCAAAAGAGTGA | CACATAACGGGGTGGCTAGT | 112 |
| chr7:107064371-107066422 | p2 | (TA)34 | GCTCAGATCTGCCAAAGTCC | AGGAGCAAGGCTTGTTTGAA | 130 |
| chr7:107117644-107119701 | p4 | (TTTC)4 | GCCGCAGTTCATAGAAGAGG | TCAGCAGTTTCAACACTCGG | 153 |
| chr7:107117644-107119701 | p3 | (ATT)19 | GCCGCAGTTCATAGAAGAGG | TCAGCAGTTTCAACACTCGG | 153 |
| chr7:107118979-107120995 | p4 | (TTTA)4 | TCCTCTCTCTCTCCTCTGCG | TTGCAAACGGACAAAAATCA | 294 |
| chr7:107173542-107175560 | p6 | (GAAAAA)3 | ATCGTCTATTTGAATGGCGG | TGCTCAAATTCCCAATCCTC | 224 |
| chr7:107176897-107178909 | p2 | (TC)6 | GCTCCACATTCTCATGCTCA | GCTCGAACGGATAAGCGTAG | 269 |
| chr7:107220308-107222323 | p5 | (AAATA)3 | CACCTTAAAATCGCCTCCAA | CCATGATGGAGCCGTTAAGT | 100 |
| chr7:107220308-107222323 | p5 | (AAATA)3 | CACCTTAAAATCGCCTCCAA | CCATGATGGAGCCGTTAAGT | 100 |
| chr7:107220308-107222323 | p4 | (ATAA)8 | CACCTTAAAATCGCCTCCAA | CCATGATGGAGCCGTTAAGT | 100 |
| chr7:107220308-107222323 | p5 | (AATAA)3 | CACCTTAAAATCGCCTCCAA | CCATGATGGAGCCGTTAAGT | 100 |
| chr7:107220308-107222323 | p5 | (TTTAT)3 | CACCTTAAAATCGCCTCCAA | CCATGATGGAGCCGTTAAGT | 100 |
| chr7:107220462-107222494 | p5 | (AAATA)3 | GCACTTAACGGCTCCATCAT | CATTTTGAGGTCTCCTCCCA | 110 |
| chr7:107220462-107222494 | p5 | (AAATA)3 | GCACTTAACGGCTCCATCAT | CATTTTGAGGTCTCCTCCCA | 110 |
| chr7:107220462-107222494 | p4 | (ATAA)8 | GCACTTAACGGCTCCATCAT | CATTTTGAGGTCTCCTCCCA | 110 |
| chr7:107220462-107222494 | p5 | (AATAA)3 | GCACTTAACGGCTCCATCAT | CATTTTGAGGTCTCCTCCCA | 110 |
| chr7:107220462-107222494 | p5 | (TTTAT)3 | GCACTTAACGGCTCCATCAT | CATTTTGAGGTCTCCTCCCA | 110 |
| chr7:107319376-107321392 | p3 | (TGT)5 | TTGGTTGAAAAATTGGGGAA | CCGTCAAGCCACCACTTAAT | 293 |
| chr7:107319376-107321392 | p2 | (AT)8 | TTGGTTGAAAAATTGGGGAA | CCGTCAAGCCACCACTTAAT | 293 |

| Marker ID | SSR type | SSR motif | FORWARD PRIMER (5'-3') | REVERSE PRIMER (5'-3') | size (bp) |
|---|---|---|---|---|---|
| chr7:107319376-107321392 | p2 | (TA)8 | TTGGTTGAAAAATTGGGGAA | CCGTCAAGCCACCACTTAAT | 293 |
| chr7:107370895-107372915 | p2 | (TA)10 | AGACAGGGAGGTGAGCTTGA | GCGCATCTACGAGCCTCTAC | 296 |
| chr7:107370895-107372915 | p2 | (TA)6 | AGACAGGGAGGTGAGCTTGA | GCGCATCTACGAGCCTCTAC | 296 |
| chr7:107393909-107395921 | p2 | (AT)6 | ACTCATGAAATCACGGGCTC | AATTCCTCCACCGGCTACTT | 273 |
| chr7:107467317-107469332 | p5 | (AAACC)3 | ATTGGCTTGCCATTCTGTTC | AGGGCTAGGCTTTTGCTGAT | 297 |
| chr7:107467317-107469332 | p2 | (AT)8 | ATTGGCTTGCCATTCTGTTC | AGGGCTAGGCTTTTGCTGAT | 297 |
| chr7:107467317-107469332 | p2 | (TA)6 | ATTGGCTTGCCATTCTGTTC | AGGGCTAGGCTTTTGCTGAT | 297 |
| chr7:107727314-107729326 | p2 | (TA)6 | CCATAAGAAGTCTCCTGCGG | GAGCTTATGCTGCTGGGATT | 256 |
| chr7:107776083-107778098 | p5 | (AAACC)3 | ATGGATCGGATTGGTTTGAA | CCGAGAAAATCCGAAGTTGA | 208 |
| chr7:107776083-107778098 | p2 | (AT)32 | ATGGATCGGATTGGTTTGAA | CCGAGAAAATCCGAAGTTGA | 208 |
| chr7:107796528-107798540 | p2 | (TA)6 | GGCAGCTTCCACAAGAAGTC | GAGCTTATGCTGCTGGGATT | 253 |
| chr7:107837180-107839236 | p5 | (TTTGT)3 | TCGGCTTATCCCGATGATAG | TTCGGAGGGGGTACACATTA | 291 |
| chr7:107837180-107839236 | p2 | (TA)28 | TCGGCTTATCCCGATGATAG | TTCGGAGGGGGTACACATTA | 291 |
| chr7:107840622-107842636 | p5 | (TCATT)4 | GGGACAGCTTTCATGGGTTA | GAACAAGCCTCGAATGGAAA | 146 |
| chr7:107871480-107873537 | p3 | (TTA)19 | GGCAGTCACCTTAGCTTTGC | ACCAACTCAAGCGATGGAAC | 149 |
| chr7:107983660-107985675 | p3 | (ATC)5 | AGCTAACAAGGCTGCCAAAA | CTCAGGAGCGCTTTATTTGC | 105 |
| chr7:108013098-108015113 | p3 | (GAA)9 | GTTGCAACAACTCACGCAGT | GTGTTTTCCGCTCATTTGGT | 149 |
| chr7:108013098-108015113 | p3 | (AAG)10 | GTTGCAACAACTCACGCAGT | GTGTTTTCCGCTCATTTGGT | 149 |
| chr7:108013098-108015113 | p5 | (ATTTT)3 | GTTGCAACAACTCACGCAGT | GTGTTTTCCGCTCATTTGGT | 149 |
| chr7:108013098-108015113 | p4 | (AAAG)4 | GTTGCAACAACTCACGCAGT | GTGTTTTCCGCTCATTTGGT | 149 |
| chr7:108098814-108100834 | p5 | (TGTGT)3 | AAAAACCGTTCGTTATTGCG | TCCATTTCCGCACACTACAA | 230 |
| chr7:108098814-108100834 | p5 | (AATTG)4 | AAAAACCGTTCGTTATTGCG | TCCATTTCCGCACACTACAA | 230 |
| chr7:108098814-108100834 | p6 | (ATATGT)3 | AAAAACCGTTCGTTATTGCG | TCCATTTCCGCACACTACAA | 230 |
| chr7:108104272-108106292 | p5 | (TGTGT)3 | GAAGCAGACGAAAACCGTTC | TTCCATCCATTTTTGCCATT | 214 |
| chr7:108104272-108106292 | p5 | (AATTG)4 | GAAGCAGACGAAAACCGTTC | TTCCATCCATTTTTGCCATT | 214 |
| chr7:108104272-108106292 | p6 | (ATGTAT)3 | GAAGCAGACGAAAACCGTTC | TTCCATCCATTTTTGCCATT | 214 |
| chr7:108237956-108239971 | p5 | (TATAT)3 | TTGTTTTGTTTTGCGTTGGA | CCTTCAAACTTCGCATCCAT | 244 |
| chr7:108251507-108253525 | p3 | (TTG)6 | CGGGAAGGACAATCTTGAAA | CCTCCAAAGTTGACCCAAAA | 212 |
| chr7:108251507-108253525 | p2 | (TA)6 | CGGGAAGGACAATCTTGAAA | CCTCCAAAGTTGACCCAAAA | 212 |
| chr7:108287664-108289684 | p2 | (TA)10 | GTTGTTGGTGTGTTGATGGG | GGACAGCATCTCCCATGTTT | 197 |
| chr7:108287664-108289684 | p2 | (AT)14 | GTTGTTGGTGTGTTGATGGG | GGACAGCATCTCCCATGTTT | 197 |
| chr7:108306575-108308590 | p3 | (AAC)5 | CTTGGGTGATGCTGGGTACT | TTTCAGCTCAACCCCTATGC | 241 |
| chr7:108317498-108319513 | p3 | (TTG)5 | CCTTTCCTATTCCTAGGCGG | TAGTTGACGAGGGCCAAAAC | 177 |
| chr7:108317498-108319513 | p3 | (TTG)5 | CCTTTCCTATTCCTAGGCGG | TAGTTGACGAGGGCCAAAAC | 177 |
| chr7:108317498-108319513 | p3 | (TTG)8 | CCTTTCCTATTCCTAGGCGG | TAGTTGACGAGGGCCAAAAC | 177 |
| chr7:108405430-108407504 | p2 | (AC)9 | CAATCAGGCACACGACAAAC | GGTACATGGACTTCGCTGGT | 105 |

| Marker ID | SSR type | SSR motif | FORWARD PRIMER (5'-3') | REVERSE PRIMER (5'-3') | size (bp) |
|---|---|---|---|---|---|
| chr7:108405430-108407504 | p5 | (CTTTC)4 | CAATCAGGCACACGACAAAC | GGTACATGGACTTCGCTGGT | 105 |
| chr7:108504580-108506630 | p2 | (TA)25 | TTTGCAGAATGTGTTCCCAA | CCAGTTTCAAAAACCCCAAA | 277 |
| chr7:108506631-108508655 | p5 | (TTTTA)3 | CCACTCAAAACGCTCAAACA | ACTTGACGAGCTCCTACCCA | 197 |
| chr7:108506654-108508720 | p5 | (TTTTA)3 | CCACTCAAAACGCTCAAACA | ACTTGACGAGCTCCTACCCA | 174 |
| chr7:108510376-108512394 | p6 | (GAAGAT)3 | TTCACATGACGCCCTGATTA | TGCATCAAATCCAATTTCCA | 217 |
| chr7:108547545-108549563 | p2 | (TA)9 | TAATGTCCAAAAAGGTGGGC | TTTTTAAGGGACAACGGACG | 267 |
| chr7:108547545-108549563 | p2 | (AT)29 | TAATGTCCAAAAAGGTGGGC | TTTTTAAGGGACAACGGACG | 267 |
| chr7:108852829-108854843 | p2 | (TC)7 | CCGTCTATCCGAATTTCCAA | TGGCTGCTGTTTTATTTCCC | 294 |
| chr7:108852829-108854843 | p2 | (AG)6 | CCGTCTATCCGAATTTCCAA | TGGCTGCTGTTTTATTTCCC | 294 |
| chr7:108852829-108854843 | p5 | (GAGAA)3 | CCGTCTATCCGAATTTCCAA | TGGCTGCTGTTTTATTTCCC | 294 |
| chr7:108902569-108904593 | p2 | (TA)12 | GAAGGGGTGTTTTGGGTTTT | TGTGCTGGGGATTTCTTAGG | 221 |
| chr7:108940354-108942406 | p5 | (TGCAT)3 | AAACACTGATGAGCTGGGCT | TTCTCACCAGCACCACTCTG | 148 |
| chr7:109008018-109010036 | p6 | (GGGTTG)3 | GTGTTGTTTGGCCTCTCCAT | TTGGGCTGTCCACTTTCTTC | 149 |
| chr7:109012622-109014642 | p5 | (TTTGG)3 | ATTCATAGCCCTGTCAACCG | AATGGCATGTTCCTTCTTGC | 209 |
| chr7:109012622-109014642 | p5 | (TTCTT)4 | ATTCATAGCCCTGTCAACCG | AATGGCATGTTCCTTCTTGC | 209 |
| chr7:109109978-109111994 | p4 | (CATA)4 | GGAACTGTCTGCAAAGCCTC | TCCCTTCTTATGGACGTTGC | 249 |
| chr7:109145037-109147070 | p3 | (TTC)11 | AGGCATAATCAATCCAAGCG | CTCAAAAAGCACAAGGAGGC | 266 |
| chr7:109145037-109147070 | p3 | (CTT)5 | AGGCATAATCAATCCAAGCG | CTCAAAAAGCACAAGGAGGC | 266 |
| chr7:109145186-109147201 | p3 | (TTC)11 | AGGCATAATCAATCCAAGCG | CTCAAAAAGCACAAGGAGGC | 117 |
| chr7:109145186-109147201 | p3 | (CTT)5 | AGGCATAATCAATCCAAGCG | CTCAAAAAGCACAAGGAGGC | 117 |
| chr7:109165013-109167025 | p2 | (AT)6 | TTGTATGTCTGCGCTCCAAC | TTTGGCAAACTGCAACTGTC | 262 |
| chr7:109288571-109290591 | p2 | (TC)10 | TGCCCCTCAAATTGCTAGTC | TTAAGGCATAACTTTGCCCG | 172 |
| chr7:109539703-109541751 | p2 | (TA)33 | AATCCGAACTTGCTGCTTGT | CTGCAATCCCACACATCAAC | 290 |
| chr7:109539755-109541811 | p2 | (TA)33 | AATCCGAACTTGCTGCTTGT | CTGCAATCCCACACATCAAC | 238 |
| chr7:109547800-109549815 | p3 | (GAT)5 | TGGAGAGATGCTATTTGGGG | AACCCCTTTCACCCCTTATG | 217 |
| chr7:109558216-109560250 | p2 | (AT)17 | GCCAGTATGTCTGGCCGTAT | AGATTCTCAAAGGCATCCGA | 300 |
| chr7:109612697-109614709 | p5 | (AATAA)3 | AAAACTGGGTTTTGGAGCCT | TCCACGGAAACAAGACATCA | 134 |
| chr7:109612697-109614709 | p2 | (AT)6 | AAAACTGGGTTTTGGAGCCT | TCCACGGAAACAAGACATCA | 134 |
| chr7:109689049-109691063 | p2 | (TA)7 | TTGGAATGATTTGGGGTGTT | AGAGGACAATTCCACGGTTG | 270 |
| chr7:109706728-109708743 | p5 | (ATGCA)3 | CGGTGTCCACACATAACTCG | TTCCCTTAGCCTTCCTCGAT | 109 |
| chr7:109706728-109708743 | p3 | (TTG)5 | CGGTGTCCACACATAACTCG | TTCCCTTAGCCTTCCTCGAT | 109 |
| chr7:109793760-109795790 | p3 | (ATC)10 | TGTGTGGGAGGTTGAAATGA | GGCAGAAAAATCCCCAAAAT | 168 |
| chr7:109802883-109804895 | p2 | (AT)6 | TTACCTGTTCAGTGGGCTCC | CGCTTCTTATGGGGATTGAA | 160 |
| chr7:109849701-109851715 | p2 | (AT)7 | GATAAACGGATGGAAGCCAA | CCCTGCGATCCCTTATAACA | 126 |
| chr7:109849701-109851715 | p2 | (AT)6 | GATAAACGGATGGAAGCCAA | CCCTGCGATCCCTTATAACA | 126 |
| chr7:109850074-109852086 | p2 | (AT)7 | TAGTGCCCGACCTAGACACC | TGTGTAGTGGCAGCCTTGTC | 245 |

| Marker ID | SSR type | SSR motif | FORWARD PRIMER (5'-3') | REVERSE PRIMER (5'-3') | size (bp) |
|---|---|---|---|---|---|
| chr7:109850074-109852086 | p2 | (AT)6 | TAGTGCCCGACCTAGACACC | TGTGTAGTGGCAGCCTTGTC | 245 |
| chr7:109904692-109906712 | p2 | (TA)10 | TCGATCCGAGGTATTGGAAG | TGCAATGGCCAATTACTCAA | 256 |
| chr7:109939146-109941190 | p2 | (TA)22 | TGTAATACAGCATCGCCCAA | AAGTCAATCTCCCGCTCTGA | 145 |
| chr7:109939146-109941190 | p2 | (AT)7 | TGTAATACAGCATCGCCCAA | AAGTCAATCTCCCGCTCTGA | 145 |
| chr7:109940371-109942421 | p2 | (AT)7 | TCCGGAATAGACTCGAATGG | CAGGGAAAGGACTAGCGTTG | 149 |
| chr7:109940371-109942421 | p2 | (AT)25 | TCCGGAATAGACTCGAATGG | CAGGGAAAGGACTAGCGTTG | 149 |
| chr7:109947348-109949362 | p2 | (AT)17 | CCAGCGGCTGTAAGACAAAT | CCTGCTTCGTCCTTTGAACT | 284 |
| chr7:109947348-109949362 | p5 | (AAAGA)3 | CCAGCGGCTGTAAGACAAAT | CCTGCTTCGTCCTTTGAACT | 284 |
| chr7:110006835-110008855 | p5 | (CATGT)4 | GGGGATCACTTACCCCTCAT | CGCCTCACGTCTGTGTAAGA | 284 |
| chr7:110071899-110073914 | p5 | (GATTT)3 | ACAATTTCCCAGTCACCGAA | TTTTCGGTTGCAAGATACCC | 147 |
| chr7:110156928-110158943 | p3 | (ACA)5 | CACAAAGTTGCCCCAAAACT | TATTGGCTATCGCTTGGGTC | 248 |
| chr7:110167232-110169250 | p3 | (CAT)6 | TCTCCAACACCAACACCAAA | AGTGAACGTAAAATTGCCCG | 179 |
| chr7:110195496-110197512 | p2 | (AT)8 | AACGCTTGGAACGACTCCTA | GACACACCGGGGTACAGAAC | 172 |
| chr7:110206279-110208303 | p4 | (TTAT)6 | AAATTCGTGCGTGAAAGACC | GAACTGTCGGGTTCTGTGGT | 230 |
| chr7:110206279-110208303 | p4 | (TTAT)6 | AAATTCGTGCGTGAAAGACC | GAACTGTCGGGTTCTGTGGT | 230 |
| chr7:110206279-110208303 | p3 | (TGG)5 | AAATTCGTGCGTGAAAGACC | GAACTGTCGGGTTCTGTGGT | 230 |
| chr7:110338268-110340288 | p5 | (AAACC)3 | GGATGGAAAACCCAAATCCT | AAACCGACCCATGAACACTC | 208 |
| chr7:110427786-110429801 | p5 | (TTTTC)3 | GGGAAGTCAAAACGGACAAA | AGCAATTACAGTTCCGGGTG | 231 |
| chr7:110466746-110468768 | p6 | (AAGATG)3 | ATATAAGCGTCATCACCCGC | TTCCCCAAACTTTTCACGAG | 173 |
| chr7:110466746-110468768 | p2 | (TA)11 | ATATAAGCGTCATCACCCGC | TTCCCCAAACTTTTCACGAG | 173 |
| chr7:110466746-110468768 | p2 | (AT)9 | ATATAAGCGTCATCACCCGC | TTCCCCAAACTTTTCACGAG | 173 |
| chr7:110467427-110469445 | p2 | (TA)11 | GCCGGACTAAGACGTACTCG | TTAAGATTGTGGGCCGAAAG | 228 |
| chr7:110467427-110469445 | p2 | (AT)9 | GCCGGACTAAGACGTACTCG | TTAAGATTGTGGGCCGAAAG | 228 |
| chr7:110513309-110515336 | p3 | (TCA)9 | TAATGAAAATCCTCTCGCGG | GGCAAAATTTGAGGGCCTAT | 232 |
| chr7:110556027-110558047 | p2 | (AC)10 | CGAGTCCAAGAGACTCGTCC | TTTTTGCTATATGGGCCTCG | 245 |
| chr7:110564971-110567007 | p2 | (AT)18 | CACAGCCTCCAACATTGCTA | TTTTGCTAGCACGTGTGGTC | 133 |
| chr7:110693936-110695948 | p3 | (GTT)6 | TCGGGCTAGGCATATAAACG | AAAGGGGCAAGCCTTAACAT | 193 |
| chr7:110693949-110695961 | p3 | (GTT)6 | TCGGGCTAGGCATATAAACG | AAAGGGGCAAGCCTTAACAT | 180 |
| chr7:110808297-110810360 | p6 | (GAAAAG)4 | TCACTGGGTACATGGCAAAA | TAATTGGGCTTTGACTTGGG | 268 |
| chr7:110808357-110810375 | p6 | (GAAAAG)4 | TCACTGGGTACATGGCAAAA | TAATTGGGCTTTGACTTGGG | 208 |
| chr7:110808383-110810398 | p6 | (GAAAAG)4 | TCACTGGGTACATGGCAAAA | TAATTGGGCTTTGACTTGGG | 182 |
| chr7:110866148-110868163 | p5 | (ATAAA)3 | TCCATTCAGCACATGATCAAA | TGGATGGGGACTCTCCTATG | 174 |
| chr7:110866148-110868163 | p5 | (TTTTA)3 | TCCATTCAGCACATGATCAAA | TGGATGGGGACTCTCCTATG | 174 |
| chr7:110875374-110877412 | p2 | (AT)9 | TGGACGTCCTAAGGGACAAC | TTGTGGTCAAGTGGCTTGTT | 279 |
| chr7:110875374-110877412 | p2 | (TA)19 | TGGACGTCCTAAGGGACAAC | TTGTGGTCAAGTGGCTTGTT | 279 |
| chr7:110883065-110885083 | p6 | (TCTATA)3 | TCCGGTCGGATTAACCATTA | GTTGTCACACCCCATTTTCA | 130 |

| Marker ID | SSR type | SSR motif | FORWARD PRIMER (5'-3') | REVERSE PRIMER (5'-3') | size (bp) |
|---|---|---|---|---|---|
| chr7:110890033-110892048 | p3 | (AAC)5 | GCAATATGCAAGGCGGTATT | CCCCTATTTGCAGGTTCTGA | 124 |
| chr7:110892970-110895024 | p3 | (TTA)18 | TCTGGCCGAAAATCTCAACT | TAGAGGCCACTTACCCATCG | 264 |
| chr7:110905014-110907029 | p5 | (TTGGT)3 | AAAGGGATGTGACCGATGAG | AACATGCCCAACACTGACAA | 228 |
| chr7:110945036-110947048 | p2 | (TC)6 | TCAGAAGGACGGACCGTAAC | TTGCCTTAAATGCTCCAACC | 188 |
| chr7:110952459-110954483 | p4 | (AAAT)6 | CCACAGCATGGATTGAAGTG | CACAATTTGCCAAGGGATCT | 232 |
| chr7:110952459-110954483 | p5 | (TTTAT)3 | CCACAGCATGGATTGAAGTG | CACAATTTGCCAAGGGATCT | 232 |
| chr7:110952459-110954483 | p5 | (ATTTC)3 | CCACAGCATGGATTGAAGTG | CACAATTTGCCAAGGGATCT | 232 |
| chr7:110992195-110994231 | p2 | (AT)18 | TCTCATTGGGCAACGATACA | GTCCTGTCCCGCCTATATGA | 235 |
| chr7:111200441-111202457 | p2 | (AT)8 | AACCCGCACACATAAAAAGC | TGGCCACTACACGATCAAAA | 111 |
| chr7:111223182-111225197 | p5 | (GGGAG)3 | AATTATTGGGGAATCCCTGC | AACATGGCCTTAGCAAGTGG | 121 |
| chr7:111265977-111267992 | p5 | (ATGCA)3 | TGATGTTGCCAACGGAAATA | TGCCGTTTGTAGTTTCCACA | 179 |
| chr7:111278939-111280954 | p5 | (TCGGT)3 | TTGTTAAGAAGTCCCACGGC | TGTAAATGGGATGGGCCTAA | 241 |
| chr7:111307215-111309233 | p6 | (ACAATT)3 | AAGCTTTTTGCGCATGTTCT | TTTTCGCGTGTTCTCAAATG | 137 |
| chr7:111325666-111327700 | p2 | (TA)17 | TTGGAACAAGAAAAGGGTCG | GCTTCGTTCGAGTCAAGACC | 230 |
| chr7:111362339-111364403 | p2 | (AT)32 | CAGCCTCCATACCCTTGTGT | AGATGTTGCAACGGGCTATC | 239 |
| chr7:111366420-111368471 | p3 | (ATT)17 | TCACCGCTACCGTTTTATCC | AATCCAAGTCGCCATTTTTG | 156 |
| chr7:111385950-111387962 | p2 | (TA)6 | TAACGATCACTCCTGTGCCA | TTTAAGTCATTGGGCTTGGG | 152 |
| chr7:111433735-111435749 | p2 | (TA)7 | GTGCGCAACACACCATAAAC | GGCACTCAGCCAAGTATGGT | 281 |
| chr7:111450472-111452484 | p2 | (AT)6 | AAGCCATGTCCAGCCATATC | TTTTCCCGTACTTTGATGGC | 140 |
| chr7:111458846-111460861 | p5 | (GTTTG)3 | GTCCTGTCCCGCCTATATGA | CATAAGCCGACGAGGCTAAC | 174 |
| chr7:111460498-111462510 | p2 | (AT)6 | ACGGCTTGATGACCCTTATG | TTAGGCGGTCGCTTATGACT | 182 |
| chr7:111476275-111478296 | p3 | (TGT)7 | AAAATTTATGCAGCGGATGG | GCTCGGTATCGGCATTAGAG | 123 |
| chr7:111514544-111516556 | p2 | (TA)6 | TGTGAAAGGCCAAAGGATTC | CACTCCGTACGACCCTTCAT | 117 |
| chr7:111527715-111529731 | p6 | (TTTTCT)3 | CAGCATGATGAGGTCCATTG | ACCAAGCTCCTCGATGAAAA | 194 |
| chr7:111527715-111529731 | p2 | (TA)8 | CAGCATGATGAGGTCCATTG | ACCAAGCTCCTCGATGAAAA | 194 |
| chr7:111537527-111539539 | p2 | (GT)6 | CAGTTTCAGAGGTTCGGAGC | TGGCATAAGGACACAACCAA | 170 |
| chr7:111651619-111653649 | p2 | (CA)15 | ATGGAGATGCTTGGAGGCTA | TCACCAGGGCGAATAGAAAC | 108 |
| chr7:111651619-111653649 | p5 | (ACCAA)3 | ATGGAGATGCTTGGAGGCTA | TCACCAGGGCGAATAGAAAC | 108 |
| chr7:111652817-111654832 | p5 | (ACCAA)3 | CGGAGTTAGCGATAAGCTGG | TTCGAGAAAACCGAACAACC | 110 |
| chr7:111652817-111654832 | p5 | (AAACC)3 | CGGAGTTAGCGATAAGCTGG | TTCGAGAAAACCGAACAACC | 110 |
| chr7:111668199-111670211 | p2 | (GA)6 | CTATCAAGCCAAAGATGCCC | TGGTGGCGGTAGAGATTTTC | 145 |
| chr7:111668199-111670211 | p2 | (CT)8 | CTATCAAGCCAAAGATGCCC | TGGTGGCGGTAGAGATTTTC | 145 |
| chr7:111871701-111873725 | p2 | (TA)12 | GCGTATTCGGTAGAATGGGA | CCGTGTCCAGTAGGGTCACT | 172 |
| chr7:111936552-111938579 | p3 | (AAT)9 | AATTCTCCACACACAACCCC | TGCACGAGTTGGACTTTGTC | 277 |
| chr7:112140507-112142519 | p2 | (AC)6 | TTCGTGTTTGGAAAGGGAAG | AAAAACCAAATTTTGCGACG | 129 |
| chr7:112157919-112159931 | p2 | (TA)6 | ACCGGATTTCCCAGCTTAAC | CAACCAAATTTGACCAACCC | 128 |

| Marker ID | SSR type | SSR motif | FORWARD PRIMER (5'-3') | REVERSE PRIMER (5'-3') | size (bp) |
|---|---|---|---|---|---|
| chr7:112254174-112256192 | p2 | (TA)9 | AGGTAGCCTTTCAGACCCGT | AACGAACTGCCCTTCAAAAA | 112 |
| chr7:112347667-112349682 | p3 | (GAA)5 | AAATAAAGATGGGCCCCAAG | GCAGCTATGGCGATTAGAGG | 155 |
| chr7:112347667-112349682 | p3 | (GAA)8 | AAATAAAGATGGGCCCCAAG | GCAGCTATGGCGATTAGAGG | 155 |
| chr7:112347667-112349682 | p2 | (AT)7 | AAATAAAGATGGGCCCCAAG | GCAGCTATGGCGATTAGAGG | 155 |
| chr7:112347876-112349900 | p3 | (GAA)5 | CCCATCCTCTAATCGCCATA | CGCAAAAATCTTTGACCCAT | 200 |
| chr7:112347876-112349900 | p3 | (GAA)8 | CCCATCCTCTAATCGCCATA | CGCAAAAATCTTTGACCCAT | 200 |
| chr7:112347876-112349900 | p2 | (AT)7 | CCCATCCTCTAATCGCCATA | CGCAAAAATCTTTGACCCAT | 200 |
| chr7:112348321-112350335 | p3 | (GAA)5 | GATGGTGTGTGGTGAGGATG | CTTCTTCTTCGCCCTTTCCT | 164 |
| chr7:112348321-112350335 | p3 | (GAA)8 | GATGGTGTGTGGTGAGGATG | CTTCTTCTTCGCCCTTTCCT | 164 |
| chr7:112348321-112350335 | p2 | (AT)7 | GATGGTGTGTGGTGAGGATG | CTTCTTCTTCGCCCTTTCCT | 164 |
| chr7:112416257-112418272 | p5 | (CCCTC)3 | CTCTTCAGTGGCAGGGCTAC | GGCCACATGTGCATTCATAG | 173 |
| chr7:112431291-112433321 | p2 | (TA)12 | GCTATCGTGCGAGGCTTAGT | CCCTCACGACGTTCTCAAAT | 269 |
| chr7:112578653-112580677 | p6 | (TGTGAT)4 | AGACCACCATAGCGGTCAAG | TCGGGCCACTATTTTCTGTC | 132 |
| chr7:112588997-112591012 | p3 | (TGT)5 | GTTCCGGTATCGAGCAAATC | GGCTTATCTTGAAATGCCCA | 162 |
| chr7:112588997-112591012 | p3 | (ATG)5 | GTTCCGGTATCGAGCAAATC | GGCTTATCTTGAAATGCCCA | 162 |
| chr7:112596713-112598731 | p3 | (ACA)6 | GCCTCATAAGGCGAATACCA | TGATTACCCCAGACGTAGGC | 139 |
| chr7:112749040-112751054 | p2 | (TA)7 | ACGAAAAATTGTGGGTCTGG | TATATGGGGATCCCAGCAAC | 117 |
| chr7:112749040-112751054 | p5 | (TATAT)3 | ACGAAAAATTGTGGGTCTGG | TATATGGGGATCCCAGCAAC | 117 |
| chr7:112749040-112751054 | p2 | (AT)6 | ACGAAAAATTGTGGGTCTGG | TATATGGGGATCCCAGCAAC | 117 |
| chr7:112857429-112859447 | p3 | (CTT)6 | CAGGGATTCGGAATGCTAAA | TGATACCCCAGCAACAACAA | 279 |
| chr7:112857429-112859447 | p3 | (ATG)5 | CAGGGATTCGGAATGCTAAA | TGATACCCCAGCAACAACAA | 279 |
| chr7:112857770-112859785 | p3 | (CTT)6 | GTGCTGAATCTGCAGTGGAA | ATTGTCATTCGTGGCTCTCC | 204 |
| chr7:112857770-112859785 | p3 | (ATG)5 | GTGCTGAATCTGCAGTGGAA | ATTGTCATTCGTGGCTCTCC | 204 |
| chr7:112881737-112883755 | p3 | (CTT)6 | CAGGGATTCGGAATGCTAAA | TGATACCCCAGCAACAACAA | 279 |
| chr7:112881737-112883755 | p3 | (ATG)5 | CAGGGATTCGGAATGCTAAA | TGATACCCCAGCAACAACAA | 279 |
| chr7:112882078-112884093 | p3 | (CTT)6 | GTGCTGAATCTGCAGTGGAA | ATTGTCATTCGTGGCTCTCC | 204 |
| chr7:112882078-112884093 | p3 | (ATG)5 | GTGCTGAATCTGCAGTGGAA | ATTGTCATTCGTGGCTCTCC | 204 |
| chr7:113030101-113032139 | p2 | (AT)16 | ACCGTACCCGACCCTCTAGT | CCCTGAGTTCTGCTAGTGCC | 158 |
| chr7:113030101-113032139 | p2 | (AT)19 | ACCGTACCCGACCCTCTAGT | CCCTGAGTTCTGCTAGTGCC | 158 |
| chr7:113062840-113064868 | p2 | (AT)14 | CGAAATCCTTACCCTCCACA | CAAGGGGTTCATTAGACCGA | 191 |
| chr7:113075586-113077604 | p6 | (TTTTTA)3 | TGAGGCGAGATTCCTGATTT | AGTTTGCAGCCGATGTATCC | 169 |
| chr7:113085501-113087515 | p2 | (AG)6 | TTGAATCTCCTCCGTCCATC | TGTCCGCGATTAAATTGTGA | 138 |
| chr7:113085516-113087528 | p2 | (AG)6 | TTGAATCTCCTCCGTCCATC | TGTCCGCGATTAAATTGTGA | 123 |
| chr7:113085535-113087551 | p2 | (GA)6 | TTGAATCTCCTCCGTCCATC | TGTCCGCGATTAAATTGTGA | 104 |
| chr7:113102246-113104288 | p2 | (AG)21 | TGGAGATGTTTTCAAAGGGG | AGGTCGTCATCATCCTGGAG | 297 |
| chr7:113191596-113193630 | p2 | (AT)17 | TATATGTAAGCCGGCAAGGC | CCCTTTCTGCTGTCTTGGAG | 287 |

| Marker ID | SSR type | SSR motif | FORWARD PRIMER (5'-3') | REVERSE PRIMER (5'-3') | size (bp) |
|---|---|---|---|---|---|
| chr7:113239587-113241602 | p5 | (TATAT)3 | CGGTTTCGTTGTACCGTTTT | TCCCACAATATTCCAGCACA | 150 |
| chr7:113239587-113241602 | p5 | (TATAT)3 | CGGTTTCGTTGTACCGTTTT | TCCCACAATATTCCAGCACA | 150 |
| chr7:113239587-113241602 | p5 | (ATATT)3 | CGGTTTCGTTGTACCGTTTT | TCCCACAATATTCCAGCACA | 150 |
| chr7:113252795-113254813 | p6 | (TGAAAG)3 | AGCCGGAATAATGTCTGTGG | ATAATCTTCCGAGGGAGCGT | 126 |
| chr7:113330370-113332412 | p2 | (AT)21 | ATGGAAACGAAGGAGATGGA | CCTTGGCTCCAAATACCACA | 249 |
| chr7:113378131-113380149 | p2 | (CA)9 | TTCAAACCAAACGACCCCTA | AGACCTCAGATTAACCGCCC | 223 |
| chr7:113395057-113397075 | p6 | (CATATA)3 | AGCCACACACACATATCCGA | TCACTTGAGTGACCCGACAG | 217 |
| chr7:113444164-113446179 | p5 | (CATGT)3 | TTTCAGATCGTGGTGCTCAG | GACCTACACCTTCGCCCATA | 107 |
| chr7:113447123-113449138 | p5 | (AAAAT)3 | TTGTTTTCGGTTCTTTTCCG | TCCGAGCACATAGCATTCAG | 256 |
| chr7:113478934-113480949 | p5 | (TTTTA)3 | CCGGGTAAGATGCAGGAATA | TGCCAAAATTGATGACCGTA | 191 |
| chr7:113534075-113536095 | p3 | (CTT)7 | CAATTACGCCTCAATCCCAT | GGCAGTTCCGTCTTTAACCA | 287 |
| chr7:113534075-113536095 | p5 | (TTTTA)3 | CAATTACGCCTCAATCCCAT | GGCAGTTCCGTCTTTAACCA | 287 |
| chr7:113562996-113565060 | p2 | (TA)32 | TAGCCAATCGAGCTGCTCTT | CTTGATGACGCTGCATCCTA | 168 |
| chr7:113603983-113605998 | p5 | (AAACC)3 | AGATGGAGTCACGGAGGATG | CGAGAAAACCGAATAACCCA | 172 |
| chr7:113607801-113609816 | p3 | (TAT)5 | TGGAGGTCATCTGTGGACAA | TTGCTGACCAGGTGAAAGTG | 186 |
| chr7:113745637-113747664 | p2 | (CT)9 | CACGCCGAGTTCAAGTATGA | AGGCGTGCGACTCTTAAAAA | 215 |
| chr7:113745637-113747664 | p3 | (AGA)9 | CACGCCGAGTTCAAGTATGA | AGGCGTGCGACTCTTAAAAA | 215 |
| chr7:113810788-113812802 | p2 | (AG)7 | TCTCATTCACGTTCCAGTCG | CACCATTTGTGTATCGGCAG | 184 |
| chr7:113883817-113885843 | p2 | (TA)13 | GACAAGATGAGGAAGCGAGG | CCTAAAGCCATCCATAGCCA | 291 |
| chr7:114096848-114098863 | p3 | (ATT)5 | AGCTGGTCAAAGCGAGAAAA | ACACGACCGCAAGTCTTACC | 252 |
| chr7:114101132-114103148 | p6 | (CATGAG)3 | GCAACCGGAAGAAACAACAT | TTAGCTGATCGGGCAGAGAT | 258 |
| chr7:114101132-114103148 | p4 | (TACA)4 | GCAACCGGAAGAAACAACAT | TTAGCTGATCGGGCAGAGAT | 258 |
| chr7:114138700-114140715 | p3 | (AAC)5 | GCGAGTATCCTTGAGCAACC | TGATCATTGGATCAAGGCAA | 180 |
| chr7:114153213-114155231 | p6 | (TTTTTA)3 | AGTGCCCAACAATAGCAACC | CAGGTTGGCCTGGATATGAT | 154 |
| chr7:114182473-114184503 | p6 | (TTTTAT)5 | TTCAGGCCGAAACCATAAAG | CGCTAGTCAAATCAGTGGCA | 291 |
| chr7:114182473-114184503 | p2 | (AT)6 | TTCAGGCCGAAACCATAAAG | CGCTAGTCAAATCAGTGGCA | 291 |
| chr7:114183794-114185808 | p2 | (AT)6 | GCATGCATTCCATGATGTTC | TATATGGGTGGGGCTCACAA | 280 |
| chr7:114183794-114185808 | p2 | (TA)7 | GCATGCATTCCATGATGTTC | TATATGGGTGGGGCTCACAA | 280 |
| chr7:114183794-114185808 | p2 | (AT)7 | GCATGCATTCCATGATGTTC | TATATGGGTGGGGCTCACAA | 280 |
| chr7:114237872-114239908 | p5 | (CACCC)3 | ACCCTCCACTTCAACCACAC | TTGCGGAACTTCCTAATTGC | 263 |
| chr7:114237872-114239908 | p2 | (AT)18 | ACCCTCCACTTCAACCACAC | TTGCGGAACTTCCTAATTGC | 263 |
| chr7:114253329-114255341 | p2 | (AG)6 | CTACCCCGAACACCACAGAT | AGCTACCCAGAAACCCCTGT | 277 |
| chr7:114253329-114255341 | p3 | (AAT)20 | CTACCCCGAACACCACAGAT | AGCTACCCAGAAACCCCTGT | 277 |
| chr7:114301100-114303112 | p2 | (TA)6 | AGCCGATCTCGCTATTGATG | TTGTATTTCTTCCCCCATGC | 146 |
| chr7:114334946-114336970 | p2 | (TA)12 | GTTTCCCTTGGAGCATTTCA | CATCAAAACCAAACCAGCCT | 131 |
| chr7:114356674-114358689 | p2 | (TC)12 | ACGCCTCTTCCTTCTCCTTC | GGGAAAAATCCCTCCAATGT | 271 |

| Marker ID | SSR type | SSR motif | FORWARD PRIMER (5'-3') | REVERSE PRIMER (5'-3') | size (bp) |
|---|---|---|---|---|---|
| chr7:114356674-114358689 | p5 | (ATTTC)3 | ACGCCTCTTCCTTCTCCTTC | GGGAAAAATCCCTCCAATGT | 271 |
| chr7:114488009-114490037 | p3 | (TTG)5 | TTGAAAAACATCGTGTGGGA | GCTACGCTTATTCGTCCGAG | 275 |
| chr7:114488009-114490037 | p4 | (TATG)7 | TTGAAAAACATCGTGTGGGA | GCTACGCTTATTCGTCCGAG | 275 |
| chr7:114488009-114490037 | p2 | (AT)13 | TTGAAAAACATCGTGTGGGA | GCTACGCTTATTCGTCCGAG | 275 |
| chr7:114494454-114496478 | p3 | (TTG)9 | TTGAAAAACATCGTGTGGGA | GCTACGCTTATTCGTCCGAG | 252 |
| chr7:114494454-114496478 | p4 | (TATG)6 | TTGAAAAACATCGTGTGGGA | GCTACGCTTATTCGTCCGAG | 252 |
| chr7:114604383-114606398 | p5 | (AAAAT)3 | GGCCTTTTCATCACTTTGGA | TTGAGCCTGTAGCCTTTCGT | 186 |
| chr7:114604383-114606398 | p5 | (AAAAC)4 | GGCCTTTTCATCACTTTGGA | TTGAGCCTGTAGCCTTTCGT | 186 |
| chr7:114637702-114639717 | p5 | (TGGAA)3 | CGCTTGTTTTTGGGTTTGTT | CTCAAAGACCTCGGGAACTG | 163 |
| chr7:114767688-114769703 | p5 | (AATAA)3 | CTTGGCTGTTGGACCATCTT | CCAATTCGAAGGAAACCTCA | 205 |
| chr7:114908118-114910133 | p2 | (TA)6 | TGAATCACACCCCTGTTGAA | GCCTCCATCTTCCGTATCAA | 219 |
| chr7:114908118-114910133 | p5 | (ATATA)3 | TGAATCACACCCCTGTTGAA | GCCTCCATCTTCCGTATCAA | 219 |
| chr7:114932163-114934193 | p2 | (TA)15 | GTTAGCCTATGAGGAGGCCC | CACTCGGATGGTTCTTCCAT | 277 |
| chr7:115005188-115007203 | p5 | (TTTTA)3 | CGGAAAAATCTGGCTTTTGA | GTGGCAGGTTTGGTTCAGTT | 270 |
| chr7:115023456-115025494 | p5 | (ACCGG)4 | CAAGGTGGAATACTCGGGAA | AATGTATGCCCTCTTTTGCG | 150 |
| chr7:115023456-115025494 | p3 | (TAT)6 | CAAGGTGGAATACTCGGGAA | AATGTATGCCCTCTTTTGCG | 150 |
| chr7:115139733-115141751 | p3 | (TTG)6 | TTTTGGGTCAACTTTGGAGG | CCGGGTCAAAACATAAATGG | 188 |
| chr7:115146960-115148978 | p6 | (AAAAAG)3 | ATGAAGATGCGCCATGTGTA | TGTTTGATCCACGATCTCCA | 138 |
| chr7:115229842-115231854 | p2 | (TC)6 | GGAAGGGTGATTGTTACCGA | TTTGGGTGTCAACATGGCTA | 244 |
| chr7:115232412-115234427 | p5 | (TGATA)3 | ATAGGGGAGATGCTGCTCAA | TGCCAAAAGAATCGGGATAG | 106 |
| chr7:115324592-115326618 | p2 | (TA)13 | GCTATGTTGCTCGGGATGTT | AGGTGCATGTCCAGATCCTC | 244 |
| chr7:115369090-115371105 | p5 | (ACGAT)3 | AAATTAGGACTTGGGGCTGG | ACGTGTAAAGTGGGGATTGG | 119 |
| chr7:115416092-115418107 | p3 | (CTT)5 | CCGGAAATCCTTCACAGAAA | AGCCACATCTCACATGACCA | 147 |
| chr7:115416092-115418107 | p2 | (AC)9 | CCGGAAATCCTTCACAGAAA | AGCCACATCTCACATGACCA | 147 |
| chr7:115424719-115426731 | p2 | (TC)6 | CAAGGTTGAAGCTCTTTCCG | CCCAAGAAGCCCACAATTTA | 282 |
| chr7:115431262-115433274 | p2 | (TA)6 | TCTTTGTGAGGGCGAGTTCT | CTCCCTCTTTCCTCATTCCC | 148 |
| chr7:115449296-115451314 | p2 | (TC)9 | CATCAGTGTTTGTGCAACCC | TCCCTCCTCCCAAGAAGAAT | 281 |
| chr7:115467566-115469584 | p6 | (GGTAGT)3 | TGAAAATGGAGAGGAGTCGG | ACGTTGGAAAAGGTCACTGC | 248 |
| chr7:115467566-115469584 | p2 | (TA)7 | TGAAAATGGAGAGGAGTCGG | ACGTTGGAAAAGGTCACTGC | 248 |
| chr7:115482111-115484126 | p5 | (GGGGT)3 | TCCTAAAGGGGGCAAGTTTT | GATCTTGCTTTTGTTGGGGA | 174 |
| chr7:115482111-115484126 | p3 | (TAT)8 | TCCTAAAGGGGGCAAGTTTT | GATCTTGCTTTTGTTGGGGA | 174 |
| chr7:115622163-115624179 | p2 | (AT)8 | AAGATTTGGATTGCCACCAC | CCTCCGTTCCTTCTTTGTCA | 113 |
| chr7:115622163-115624179 | p2 | (AT)27 | AAGATTTGGATTGCCACCAC | CCTCCGTTCCTTCTTTGTCA | 113 |
| chr7:115697463-115699481 | p6 | (AAAAAG)3 | CAATCCAAAGTGCTGCTCAA | TCTCAGTGTCGTTCGCTTTG | 101 |
| chr7:115821880-115823895 | p6 | (GTGCAG)3 | TCACCATCTGTGGGAGAACA | ACCCATCCTCGAGAGGCTAT | 145 |
| chr7:115821880-115823895 | p5 | (AATGC)3 | TCACCATCTGTGGGAGAACA | ACCCATCCTCGAGAGGCTAT | 145 |

| Marker ID | SSR type | SSR motif | FORWARD PRIMER (5'-3') | REVERSE PRIMER (5'-3') | size (bp) |
|---|---|---|---|---|---|
| chr7:115821880-115823895 | p5 | (AAATA)3 | TCACCATCTGTGGGAGAACA | ACCCATCCTCGAGAGGCTAT | 145 |
| chr7:115969887-115971899 | p2 | (AC)6 | ATATCCCACAAGTTTTGCCG | GTAGTGTTCCGCAACCGAAT | 211 |
| chr7:115969887-115971899 | p2 | (TA)6 | ATATCCCACAAGTTTTGCCG | GTAGTGTTCCGCAACCGAAT | 211 |
| chr7:116004620-116006632 | p2 | (TA)6 | TCGATTGTGGCGTAAAATCA | CCCTGCCATCCTCTGTATGT | 205 |
| chr7:116008152-116010197 | p3 | (ATA)15 | CCATTCGCAGCCTTGATATT | GCGAAGTTTGTGGAAAGAGC | 128 |
| chr7:116027888-116029902 | p2 | (CT)7 | AGGGTGGTTCCAACTCAATG | ATGACGAGCATGACTTTCCC | 181 |
| chr7:116094800-116096815 | p3 | (TTA)5 | CCTGGTTGCCCATCTATTGT | TTCCTGCCGTTCCATTTAAG | 267 |
| chr7:116223749-116225771 | p2 | (TA)11 | GGGAAGGAAAAATGGGAGAG | GCAAGCTAGGGTCAAACAGG | 124 |
| chr7:116223749-116225771 | p2 | (TA)10 | GGGAAGGAAAAATGGGAGAG | GCAAGCTAGGGTCAAACAGG | 124 |
| chr7:116267681-116269696 | p5 | (ATATG)3 | CATTCAAGGCTACAGTGCCA | AAAACGAATGCACTGGGAAG | 271 |
| chr7:116309192-116311292 | p5 | (ATATG)3 | ACTCGTGCCTACGAATTGCT | TCAATTATCATGGAACGCGA | 213 |
| chr7:116309290-116311308 | p5 | (ATATG)3 | ACTCGTGCCTACGAATTGCT | TCAATTATCATGGAACGCGA | 115 |
| chr7:116333895-116335907 | p2 | (TA)6 | TGTTCAGTGCGCCAAGTTAG | TTGTCCTTCGCTACAATCCC | 235 |
| chr7:116333912-116335932 | p2 | (TA)6 | TGTTCAGTGCGCCAAGTTAG | TTGTCCTTCGCTACAATCCC | 218 |
| chr7:116370061-116372117 | p2 | (TC)15 | CCCAAAATAAGGCCATTCCT | CCGGTAATGGAGAGTTGGAA | 121 |
| chr7:116389882-116391898 | p2 | (TA)8 | ATGCTATGGCGGTATTGAGC | GGAAGTGGATTCACTGGCAT | 299 |
| chr7:116437677-116439692 | p3 | (TGT)5 | AACTTATGCTCCGCGAAAAA | TCGGTTATTGAGCACTTCCC | 174 |
| chr7:116553478-116555490 | p2 | (TA)19 | GTCCCTCGTGGTCGTCTTTA | CTGCCCAGTTGAGAAATGGT | 197 |
| chr7:116553478-116555490 | p2 | (AG)6 | GTCCCTCGTGGTCGTCTTTA | CTGCCCAGTTGAGAAATGGT | 197 |
| chr7:116566581-116568601 | p2 | (TA)10 | AATTCCCTCGAGGTCATCCT | GTAAATGTACCCTTCGGCCA | 293 |
| chr7:116566581-116568601 | p2 | (TA)7 | AATTCCCTCGAGGTCATCCT | GTAAATGTACCCTTCGGCCA | 293 |
| chr7:116566994-116569008 | p2 | (TA)10 | TGGAAGGAGAAAAGGACCAA | CAAGGGGAATAAATGAGCCA | 149 |
| chr7:116566994-116569008 | p2 | (TA)7 | TGGAAGGAGAAAAGGACCAA | CAAGGGGAATAAATGAGCCA | 149 |
| chr7:116566994-116569008 | p5 | (AAATC)3 | TGGAAGGAGAAAAGGACCAA | CAAGGGGAATAAATGAGCCA | 149 |
| chr7:116587604-116589622 | p3 | (GAT)5 | GGTATTGTTAGGGGTGGGCT | GGGGGTTGTGTTTCATCATC | 245 |
| chr7:116587604-116589622 | p6 | (CGACAC)3 | GGTATTGTTAGGGGTGGGCT | GGGGGTTGTGTTTCATCATC | 245 |
| chr7:116639622-116641652 | p6 | (CATATA)5 | CTGGTGGAGCCGTTTACAAT | CATTGCTCGTACTGACCCCT | 285 |
| chr7:116639622-116641652 | p5 | (ATTCA)3 | CTGGTGGAGCCGTTTACAAT | CATTGCTCGTACTGACCCCT | 285 |
| chr7:116648415-116650487 | p2 | (TA)36 | ACGTCAAAATTGAACGGGTC | ACGTCCCGTACATCCAAGTC | 222 |
| chr7:116657854-116659869 | p3 | (TAT)5 | GGTAATTGCATTTCGCCCTA | ATCGACGCACTCCTCCTAAA | 298 |
| chr7:116666613-116668629 | p2 | (AC)8 | AATTAACCCTCAGGGGTTGG | AAGTGCACCACGGGTAATTC | 156 |
| chr7:116666613-116668629 | p2 | (AT)7 | AATTAACCCTCAGGGGTTGG | AAGTGCACCACGGGTAATTC | 156 |
| chr7:116719203-116721218 | p5 | (TCTTT)3 | CTTACGGTCAAGCCTTCCAG | TTTGACGATCCACTTCTCCC | 269 |
| chr7:116749083-116751119 | p2 | (AT)18 | ATTCAAGAGCGAGCCCAATA | AACCCGAGAGGTTTTGGAGT | 233 |
| chr7:116776341-116778359 | p6 | (CAACTC)3 | AACAGGCCTGAGTGTTACGG | TGCACTACGCCAACTAGACG | 209 |
| chr7:116842424-116844436 | p2 | (TA)7 | CATGCTTCGTCGTCTGTTGT | TCCCATAAGTTCGGGAGATG | 196 |

| Marker ID | SSR type | SSR motif | FORWARD PRIMER (5'-3') | REVERSE PRIMER (5'-3') | size (bp) |
|---|---|---|---|---|---|
| chr7:116842888-116844902 | p2 | (TA)7 | CCCATAAAGGCTTGACCAAC | TTTGTTAATACGGGGGTCCA | 110 |
| chr7:116842888-116844902 | p5 | (GAAAT)3 | CCCATAAAGGCTTGACCAAC | TTTGTTAATACGGGGGTCCA | 110 |
| chr7:116842888-116844902 | p2 | (AT)31 | CCCATAAAGGCTTGACCAAC | TTTGTTAATACGGGGGTCCA | 110 |
| chr7:116845397-116847419 | p2 | (TA)11 | CTTGCTTGCAAGGCTAAACC | TAGGAACTGATTATGGCCCG | 202 |
| chr7:116904449-116906475 | p2 | (CA)13 | AGCATATTTCCCTCCAACCC | GGAGTGAGATTTGGAGCAGC | 255 |
| chr7:116930870-116932888 | p6 | (ATCATT)3 | AAGAAGCCTAAGGGGGTTCA | CCTCCGATCTCATGGACAGT | 193 |
| chr7:116944182-116946206 | p3 | (ATT)5 | TGGACGATTCTCACCATGAA | ATCAATCCGTGAAATGCCTC | 287 |
| chr7:116944215-116946233 | p3 | (ATT)5 | TGGACGATTCTCACCATGAA | ATCAATCCGTGAAATGCCTC | 254 |
| chr7:116998486-117000500 | p2 | (AT)7 | ATTTCTTCAACTTGGCGTGG | TCGCACAGTGCACACATAAA | 170 |
| chr7:117026752-117028772 | p5 | (AATGC)4 | ACCTCGGTAGACGAACCCTT | ATCAGACTCTGGGTGGATCG | 283 |
| chr7:117065117-117067132 | p5 | (TATAT)3 | GACTATTCTTCCTTCGGGGG | ACAAACATATGGCGCAGACA | 160 |
| chr7:117076971-117078983 | p2 | (TA)6 | TCGATGTTGTTGGCATGATT | TTACATACCTTGCCCGCTTC | 248 |
| chr7:117076971-117078983 | p2 | (AT)19 | TCGATGTTGTTGGCATGATT | TTACATACCTTGCCCGCTTC | 248 |
| chr7:117130766-117132808 | p2 | (AT)21 | TTATAGGCTGCCGTTTACCG | TGCAAATTCAGGGACATCAA | 277 |
| chr7:117130766-117132808 | p3 | (TTC)5 | TTATAGGCTGCCGTTTACCG | TGCAAATTCAGGGACATCAA | 277 |
| chr7:117142212-117144254 | p2 | (AT)21 | TTATAGGCTGCCGTTTACCG | TGCAAATTCAGGGACATCAA | 277 |
| chr7:117142212-117144254 | p3 | (TTC)5 | TTATAGGCTGCCGTTTACCG | TGCAAATTCAGGGACATCAA | 277 |
| chr7:117176435-117178449 | p5 | (CTTTT)3 | CGTCCGATATTTGGTATCCG | AATTGGAGAGGTAGGGGGAA | 230 |
| chr7:117176435-117178449 | p2 | (AT)7 | CGTCCGATATTTGGTATCCG | AATTGGAGAGGTAGGGGGAA | 230 |
| chr7:117270538-117272553 | p5 | (ATAAA)3 | CCAAGTCAACACGAGGGAGT | AAACACGCCCAAAGAGAAGA | 193 |
| chr7:117313528-117315588 | p3 | (AAT)20 | GGGATTTCGGGTTCAAGATT | TCCCTCTCTCTTCTCCCCTC | 220 |
| chr7:117327168-117329183 | p5 | (ATACA)3 | ACAGGACACCTGACGAGCTT | TTTTATGGGGTTTATGGGCA | 114 |
| chr7:117327168-117329183 | p2 | (AT)19 | ACAGGACACCTGACGAGCTT | TTTTATGGGGTTTATGGGCA | 114 |
| chr7:117356875-117358897 | p2 | (TA)11 | CCTCTTTCGCGGACTTGATA | CAAGGGGAATAAATGAGCCA | 266 |
| chr7:117356875-117358897 | p3 | (TAT)10 | CCTCTTTCGCGGACTTGATA | CAAGGGGAATAAATGAGCCA | 266 |
| chr7:117357919-117359934 | p3 | (TAT)10 | CAAGGGGAATAAATGAGCCA | CTTTCCTGAGCCATTAGCCA | 148 |
| chr7:117357919-117359934 | p3 | (TGT)5 | CAAGGGGAATAAATGAGCCA | CTTTCCTGAGCCATTAGCCA | 148 |
| chr7:117388986-117391001 | p5 | (GGTTC)3 | TTCCTTGATTGCAGAAGGCT | CCTTGCTCACCGGATTAAAA | 248 |
| chr7:117388986-117391001 | p5 | (TTCGG)3 | TTCCTTGATTGCAGAAGGCT | CCTTGCTCACCGGATTAAAA | 248 |
| chr7:117440572-117442584 | p2 | (TA)6 | GGCCCATCAATTTGAAAAGA | GCATGCATAGGGGTTTCACT | 257 |
| chr7:117443133-117445145 | p2 | (AT)6 | CCTTCCTTGACCTCACCAAA | GCACCCTAGGGAGATTAGGG | 134 |
| chr7:117443133-117445145 | p2 | (AT)6 | CCTTCCTTGACCTCACCAAA | GCACCCTAGGGAGATTAGGG | 134 |
| chr7:117443133-117445145 | p2 | (AT)7 | CCTTCCTTGACCTCACCAAA | GCACCCTAGGGAGATTAGGG | 134 |
| chr7:117473566-117475586 | p4 | (AAAT)5 | AGGAGCTGTCACGAGCAAAT | AACTGCGATGATCCAGAAGG | 275 |
| chr7:117475936-117477951 | p2 | (AG)7 | CACCAGGAAAGAAAAACCCA | CAACCAATGGTGCAGATGTC | 201 |
| chr7:117475936-117477951 | p3 | (TTC)5 | CACCAGGAAAGAAAAACCCA | CAACCAATGGTGCAGATGTC | 201 |

| Marker ID | SSR type | SSR motif | FORWARD PRIMER (5'-3') | REVERSE PRIMER (5'-3') | size (bp) |
|---|---|---|---|---|---|
| chr7:117482798-117484814 | p2 | (AT)10 | CGACCCTCTTTTGCGAGTAG | CTGTTATTCCTTTCGCCGAG | 243 |
| chr7:117482798-117484814 | p2 | (GT)8 | CGACCCTCTTTTGCGAGTAG | CTGTTATTCCTTTCGCCGAG | 243 |
| chr7:117482798-117484814 | p3 | (GTT)6 | CGACCCTCTTTTGCGAGTAG | CTGTTATTCCTTTCGCCGAG | 243 |
| chr7:117604347-117606387 | p4 | (CTTT)10 | CCGGAGGTCAAATTTACGAA | TGATGGAGGATGTAGGAGGC | 129 |
| chr7:117611984-117613999 | p5 | (ATTTT)3 | TGGCCAGTCAAACACTCAAA | GCATGAGGTAGGGGACAAAA | 182 |
| chr7:117641181-117643217 | p2 | (TA)18 | GCCGAATGACCCCTTATCTT | GGCCCAAGTCCATGTTCTAA | 100 |
| chr7:117674485-117676500 | p2 | (AT)6 | CATAATTTTGCGCTCGGAGT | AATACCCCGGGATTTTTACG | 137 |
| chr7:117674485-117676500 | p2 | (AT)6 | CATAATTTTGCGCTCGGAGT | AATACCCCGGGATTTTTACG | 137 |
| chr7:117674485-117676500 | p5 | (AATAT)3 | CATAATTTTGCGCTCGGAGT | AATACCCCGGGATTTTTACG | 137 |
| chr7:117786752-117788768 | p2 | (AT)8 | ACCATGAACTCAGTGTGCCA | CCTTCCCCTTGCACAAGATA | 157 |
| chr7:117793341-117795353 | p2 | (AT)8 | CGTAAGATTGGGATTGTGGG | CGGACGACGAGAAAGTGAAC | 217 |
| chr7:117793341-117795353 | p2 | (TA)6 | CGTAAGATTGGGATTGTGGG | CGGACGACGAGAAAGTGAAC | 217 |
| chr7:117793341-117795353 | p2 | (TA)28 | CGTAAGATTGGGATTGTGGG | CGGACGACGAGAAAGTGAAC | 217 |
| chr7:117793341-117795353 | p2 | (TA)8 | CGTAAGATTGGGATTGTGGG | CGGACGACGAGAAAGTGAAC | 217 |
| chr7:117849115-117851136 | p3 | (TAT)7 | GTGGAATGAATAAATGGGCG | TGCAAGCTTTCAATAGCCCT | 197 |
| chr7:117939384-117941414 | p3 | (CAA)10 | TCATTAAAGGGGGACGATGA | AGGATATGCCGCAAAGAATG | 127 |
| chr7:117963994-117966016 | p2 | (AC)11 | CCCTCTTTGCCACTGAGCTA | ATACAAGGGCGGATTGACAC | 159 |
| chr7:117963994-117966016 | p3 | (ATT)5 | CCCTCTTTGCCACTGAGCTA | ATACAAGGGCGGATTGACAC | 159 |
| chr7:117986180-117988198 | p3 | (GAA)12 | AGAGGATGGGCTGATGATTG | AAGCTCACACAGGCGAAGTT | 153 |
| chr7:117986180-117988198 | p3 | (ACA)6 | AGAGGATGGGCTGATGATTG | AAGCTCACACAGGCGAAGTT | 153 |
| chr7:118005647-118007681 | p2 | (AT)17 | TCGGCCAAATTATCACAACA | AGAGATGTCGGGTGAAGTGG | 278 |
| chr7:118014848-118016863 | p5 | (AAAAC)3 | GCCACGGCACAAGGTATATT | AATGGGGTACAACAGCTTGC | 215 |
| chr7:118081677-118083707 | p2 | (TA)15 | TATCTTACGGGGGTCAAACG | CCTGAACAGTGCTGGTCGTA | 259 |
| chr7:118136790-118138808 | p3 | (TAA)19 | GGGACCTTTCGAACATGAAA | TGAATAATGCTTTCCCCCAG | 121 |
| chr7:118136790-118138808 | p3 | (TTG)5 | GGGACCTTTCGAACATGAAA | TGAATAATGCTTTCCCCCAG | 121 |
| chr7:118136790-118138808 | p3 | (TGT)6 | GGGACCTTTCGAACATGAAA | TGAATAATGCTTTCCCCCAG | 121 |
| chr7:118161459-118163501 | p2 | (GA)21 | GTACCAAGCCTCACTGCCTC | GTGTATGTTGTTCGCGGATG | 201 |
| chr7:118202937-118204964 | p3 | (GTT)9 | CCACCAAAACTTAAGGGCAA | GGGCAACTGTGTCTCTCCTC | 269 |
| chr7:118294242-118296257 | p5 | (AAGAA)3 | CAACCATCCAAACAGAGGGT | CTATGGAGTGAGGGGTTGGA | 269 |
| chr7:118294319-118296362 | p5 | (AAGAA)3 | CAACCATCCAAACAGAGGGT | CTATGGAGTGAGGGGTTGGA | 192 |
| chr7:118447923-118449939 | p2 | (AT)8 | GAGAAGAGCACTCCAAACCG | TGCACTTATGTCGGCACTTC | 150 |
| chr7:118472895-118474927 | p2 | (AT)24 | CTGAAGAAAACCCAAACCGA | AATCACCCAAAAGCACAAGG | 259 |
| chr7:118575180-118577198 | p3 | (AAC)6 | TCAAAAGGGGTGAAGGAGAA | TCAAATCCTGGATCCGTCTC | 279 |
| chr7:118577852-118579867 | p5 | (ATTTT)3 | TCCAATTCAAGGGGAAGATG | CAACGTGAACAACAATTGCC | 217 |
| chr7:118621787-118623802 | p3 | (GAA)5 | GGCGGGGTCTAATGGTTAAT | GCTCTATCTACGTCCGCCAC | 139 |
| chr7:118636610-118638625 | p2 | (AT)11 | CCTCAACGTCCTCGTTTGAT | ATCCCCAGGTTCGATAGCTT | 192 |

| Marker ID | SSR type | SSR motif | FORWARD PRIMER (5'-3') | REVERSE PRIMER (5'-3') | size (bp) |
|---|---|---|---|---|---|
| chr7:118659749-118661769 | p2 | (AT)16 | CCCCATATTAAACACCGTGC | GATGTGGTGGGTTCCACTTT | 224 |
| chr7:118659749-118661769 | p2 | (TA)10 | CCCCATATTAAACACCGTGC | GATGTGGTGGGTTCCACTTT | 224 |
| chr7:118699872-118701896 | p3 | (GTT)8 | CAATTGGATTGCTTTCGGAT | AGAAGCCCCTTCCAGTCATT | 289 |
| chr7:118768492-118770512 | p2 | (AG)10 | GCGGAGCCACAGAAATTTTA | CCTAAGGGGAGGAAGGATTG | 169 |
| chr7:118790148-118792163 | p5 | (TACAA)3 | TCAGCCTTTAAACCAGTGGC | GGAAACCACCATCCAAACAG | 148 |
| chr7:118834488-118836506 | p6 | (TTTCAT)3 | GTTGATGGCCTTTGCAACTT | GTGCCGGTCAGATTCTTGTT | 144 |
| chr7:118849630-118851645 | p3 | (GTT)5 | AAGCTGCACGGTCTCAGAAT | GGCCTGATTGAACTTTGGAA | 224 |
| chr7:118864336-118866351 | p3 | (GTT)5 | AAGCTGCACGGTCTCAGAAT | GGCCTGATTGAACTTTGGAA | 224 |
| chr7:118991032-118993044 | p2 | (AT)6 | TGGAACGAACCGTAACATGA | GCTTAAGTTGGGGGTGTTGA | 102 |
| chr7:119031568-119033590 | p2 | (AT)11 | TGCAAATGACACAAAGGCAT | GCAACGGCTTCAGTTTCTTC | 203 |
| chr7:119231566-119233582 | p5 | (CCACC)3 | TGAACAGGTGGTTTGTGGAA | CTAGCCAAGGACCATTGGAA | 248 |
| chr7:119231566-119233582 | p4 | (CCCA)4 | TGAACAGGTGGTTTGTGGAA | CTAGCCAAGGACCATTGGAA | 248 |
| chr7:119330564-119332579 | p5 | (GGGTC)3 | CAACTCAAGAGGAAGCTGGG | TGGATGAGGAGTTGAGGGAC | 112 |
| chr7:119373083-119375108 | p2 | (TG)6 | CTTGAGAATGAGGTTGGGGA | TGCCCTGCAGTAAATAAGGC | 285 |
| chr7:119373150-119375175 | p2 | (TG)6 | CTTGAGAATGAGGTTGGGGA | TGCCCTGCAGTAAATAAGGC | 218 |
| chr7:119373217-119375242 | p2 | (TG)6 | CTTGAGAATGAGGTTGGGGA | TGCCCTGCAGTAAATAAGGC | 151 |
| chr7:119629286-119631316 | p2 | (AG)8 | ACACGGCTAGAACACCCATC | GCGGAAGTGAAGAACAAAGG | 204 |
| chr7:119629286-119631316 | p2 | (AG)15 | ACACGGCTAGAACACCCATC | GCGGAAGTGAAGAACAAAGG | 204 |
| chr7:119655527-119657542 | p5 | (TAAAA)3 | CTGGTAAAGCTGCTGCACTG | CCAAGGCATAGAGCATGTGA | 181 |
| chr7:119655527-119657542 | p2 | (AG)7 | CTGGTAAAGCTGCTGCACTG | CCAAGGCATAGAGCATGTGA | 181 |
| chr7:119722033-119724051 | p6 | (AGAAAG)3 | TTTGTAACGGGAAGTCCCAC | GAATCTCCAAAAACGAGGCA | 110 |
| chr7:119738545-119740563 | p6 | (TTTTTA)3 | GCGATAAACTCTTCCGGCTT | CGACAACAACAACAACCCAG | 144 |
| chr7:119766355-119768371 | p2 | (AT)8 | AGCATTGGAGACCCATGAAG | AAAAAGGCTCCTCTTGCACA | 101 |
| chr7:119779800-119781816 | p2 | (TC)9 | AATGGCAGCTAGGTATGCGT | GGCCGGAAATTTCACACTAA | 115 |
| chr7:119779800-119781816 | p2 | (TA)8 | AATGGCAGCTAGGTATGCGT | GGCCGGAAATTTCACACTAA | 115 |
| chr7:119875803-119877818 | p5 | (TTTTA)3 | AAACGGGAGACCCAAATACC | TGTCACCACATGCTACGGTT | 287 |
| chr7:119875803-119877818 | p2 | (TC)17 | AAACGGGAGACCCAAATACC | TGTCACCACATGCTACGGTT | 287 |
| chr7:119891810-119893828 | p6 | (ACCCCC)3 | CAATCCAAACGGGCTCTAAG | CAAAGCCCAAGGAAAATTCA | 283 |
| chr7:119906040-119908058 | p6 | (GGGGGT)3 | CGATACTTCGCAAGTTGGGT | GTGTTTGTGGAACTTGGGCT | 178 |
| chr7:119941739-119943760 | p3 | (TAA)7 | CAAGGGTTTCTTGAACCGAA | GGAGGGGATTTTTGATTGGT | 113 |
| chr7:120188255-120190271 | p4 | (TATG)4 | TCCAACCATAGGTTGTGGGT | GGGTTGACCCAACTTGAAAA | 141 |
| chr7:120188255-120190271 | p6 | (CTCATG)3 | TCCAACCATAGGTTGTGGGT | GGGTTGACCCAACTTGAAAA | 141 |
| chr7:120204675-120206697 | p2 | (TA)9 | TGGCAATGAATCCACCATTA | ATGCTTCGTTGTCCATCTCC | 145 |
| chr7:120204675-120206697 | p2 | (AT)11 | TGGCAATGAATCCACCATTA | ATGCTTCGTTGTCCATCTCC | 145 |
| chr7:120204675-120206697 | p3 | (TGT)5 | TGGCAATGAATCCACCATTA | ATGCTTCGTTGTCCATCTCC | 145 |
| chr7:120204675-120206697 | p2 | (AT)14 | TGGCAATGAATCCACCATTA | ATGCTTCGTTGTCCATCTCC | 145 |

| Marker ID | SSR type | SSR motif | FORWARD PRIMER (5'-3') | REVERSE PRIMER (5'-3') | size (bp) |
|---|---|---|---|---|---|
| chr7:120205172-120207187 | p2 | (AT)11 | TTGAAGATGGTGGGTCCAAT | GCCTCATCGTCCATCTCCTA | 286 |
| chr7:120205172-120207187 | p3 | (TGT)5 | TTGAAGATGGTGGGTCCAAT | GCCTCATCGTCCATCTCCTA | 286 |
| chr7:120205172-120207187 | p2 | (AT)14 | TTGAAGATGGTGGGTCCAAT | GCCTCATCGTCCATCTCCTA | 286 |
| chr7:120260961-120262979 | p6 | (CATGAG)3 | GGGATCAAGATGACGAGGAA | CTTGTCGCATCCCCACTACT | 156 |
| chr7:120448037-120450058 | p3 | (TTA)11 | ATTCAAACTTGCCCGTGAAC | TCCAACTCGCAAAAACACAC | 164 |
| chr7:120462781-120464799 | p2 | (TA)9 | CGGCGTTAATCCGAAGATAA | CGACGGGAAGAGATGAAATC | 113 |
| chr7:120485169-120487187 | p2 | (TA)11 | TTCAGTTTTGTCACACCCCA | GTTGTTGGGTTCTTCTGCGT | 100 |
| chr7:120485169-120487187 | p2 | (AT)9 | TTCAGTTTTGTCACACCCCA | GTTGTTGGGTTCTTCTGCGT | 100 |
| chr7:120530873-120532885 | p2 | (TA)6 | CATCAAAAGCAGCCACGTTA | AGCCATCACGAAGCCTTCTA | 237 |
| chr7:120550130-120552148 | p3 | (TTA)6 | TGGGAACGGAAAAGTTCTTG | AATCGTTGCCGGATTATTCA | 244 |
| chr7:120595913-120597940 | p3 | (GTT)9 | TGTTTGCTCTTTCGCATTTG | CCTCGTCTGAAACCATCCAT | 279 |
| chr7:120630415-120632429 | p2 | (TA)7 | AAATGACATGGCGGAATAGC | GCCACATGTGTATCAACCCA | 222 |
| chr7:120635823-120637838 | p5 | (TTTTA)3 | CGAACCAAACGAGCCTTTTA | GATGCATGACAAGGGATGAG | 288 |
| chr7:120641913-120643925 | p2 | (AT)6 | AATTCGCTCCGGAAGTCTTT | AGCAAACGACCCCTAAGTCA | 184 |
| chr7:120676978-120678994 | p4 | (AACA)4 | AAAGCATAACCAGGGGGTCT | AGCACTTTTGGCCTCTCAAA | 292 |
| chr7:120691742-120693756 | p3 | (CGT)7 | GGGCTTTGATTTTACCGTCA | ATACCGCAACCAGTTTCACC | 172 |
| chr7:120691742-120693756 | p2 | (TG)7 | GGGCTTTGATTTTACCGTCA | ATACCGCAACCAGTTTCACC | 172 |
| chr7:120691742-120693756 | p2 | (GT)6 | GGGCTTTGATTTTACCGTCA | ATACCGCAACCAGTTTCACC | 172 |
| chr7:120691742-120693756 | p2 | (TA)16 | GGGCTTTGATTTTACCGTCA | ATACCGCAACCAGTTTCACC | 172 |
| chr7:120696879-120698894 | p3 | (CTT)5 | AGTTTGAGCGAGCCATCCTA | CCAAGCTGGGGATGATATGT | 180 |
| chr7:120719032-120721044 | p2 | (TA)6 | GGCTTGCTGCAGTTGATTTT | AAATTGATGGCCAGGATTCA | 281 |
| chr7:120865292-120867352 | p2 | (TA)30 | TTTTTCTCGTTTTCCCCCTT | CAAGTCAACAACATGTGGGC | 263 |
| chr7:120871189-120873204 | p2 | (AT)8 | CGGTTTGGTTTGACTTGGTT | GTCGTGTTGGTTTGGTTTGA | 197 |
| chr7:120871189-120873204 | p5 | (TTGGT)3 | CGGTTTGGTTTGACTTGGTT | GTCGTGTTGGTTTGGTTTGA | 197 |
| chr7:120871189-120873204 | p5 | (TTGGT)3 | CGGTTTGGTTTGACTTGGTT | GTCGTGTTGGTTTGGTTTGA | 197 |
| chr7:120899974-120901989 | p3 | (AAT)5 | TTACCTTCCGCAGTTGTTCC | TTACAAAGTTCAGGGGGCTG | 253 |
| chr7:120955658-120957673 | p3 | (AGA)5 | CCTTTTGCCAAGAGATTCCA | CTTGTGAGTGTCCAGCGAAA | 277 |
| chr7:121090335-121092351 | p4 | (CAAT)4 | GACAGGTTCTGCCGCTGTAT | GTTCAGCATCAGCCAACTCA | 121 |
| chr7:121206541-121208559 | p3 | (TAA)6 | CCGCATTTGTCGATTAAACC | TTGGATCTTTGCTTGTGCTG | 272 |
| chr7:121211936-121213951 | p3 | (ACT)5 | CATGCATTTGATGCCTTCAC | AGCACGTGTAGGGAAATTGG | 300 |
| chr7:121332843-121334861 | p6 | (ATAAAA)3 | TGATTATTAACGGCGCATGA | CCTTCAACCAAGTGGACCAT | 239 |
| chr7:121332843-121334861 | p2 | (AG)6 | TGATTATTAACGGCGCATGA | CCTTCAACCAAGTGGACCAT | 239 |
| chr7:121334612-121336627 | p5 | (TTTTA)3 | ATGCACGAATAGACGGAAGG | AATTGAGAGCGGTAAATGCG | 298 |
| chr7:121368972-121370990 | p6 | (TAAAAA)3 | CATTTGATAGGGGGATGTGC | TCCACAAATTTGCTTCCACA | 268 |
| chr7:121612121-121614145 | p3 | (CCA)8 | CAACCTCTGCCGGAAAATTA | GGGGACCCTAAGAGAGTTGC | 206 |
| chr7:121631628-121633648 | p2 | (AT)10 | GCTGTAACCCCAGGCACTAA | GAATTTGACAGGAGCTTGGC | 266 |

| Marker ID | SSR type | SSR motif | FORWARD PRIMER (5'-3') | REVERSE PRIMER (5'-3') | size (bp) |
|---|---|---|---|---|---|
| chr7:121649707-121651719 | p2 | (TC)6 | ACAAGTTTATGCTGGGTCCG | TATGCCTCGTAAGCCGAAAC | 187 |
| chr7:121718192-121720216 | p3 | (CCA)8 | CAACCTCTGCCGGAAAATTA | GGGGACCCTAAGAGAGTTGC | 206 |
| chr7:121739204-121741219 | p5 | (TTGGT)3 | TAAACCGACAAATCGCATCA | TAATGTCGGGTTGGTTTGGT | 135 |
| chr7:121739204-121741219 | p5 | (TTGGT)3 | TAAACCGACAAATCGCATCA | TAATGTCGGGTTGGTTTGGT | 135 |
| chr7:121806571-121808591 | p4 | (TTTA)5 | GAACCCCCACTCAGTCAAAA | AGTTTGCCCATGAGTATGCC | 273 |
| chr7:121826306-121828321 | p3 | (AAT)5 | ATGTGAAAATGAAGGCCAGG | CGTCACAGAGGTGGTAGGGT | 279 |
| chr7:121830946-121832976 | p3 | (TAA)5 | GGGGTCCGGTAAGTCAAAAT | AACTTCTGCACACACATCGC | 276 |
| chr7:121830946-121832976 | p6 | (ATAATG)5 | GGGGTCCGGTAAGTCAAAAT | AACTTCTGCACACACATCGC | 276 |
| chr7:121848999-121851014 | p5 | (TTGGT)3 | TAAACCGACAAATCGCATCA | TAATGTCGGGTTGGTTTGGT | 135 |
| chr7:121848999-121851014 | p5 | (TTGGT)3 | TAAACCGACAAATCGCATCA | TAATGTCGGGTTGGTTTGGT | 135 |
| chr7:121916366-121918386 | p4 | (TTTA)5 | GAACCCCCACTCAGTCAAAA | AGTTTGCCCATGAGTATGCC | 273 |
| chr8:22991-25005 | p2 | (AG)7 | TTCAAAAGCTCCTGGCAAGT | TCCATTGCGTGCACATTAGT | 122 |
| chr8:32996-35012 | p2 | (AT)8 | AGATTGGGATGCATCGTCTC | TCCAATGGGACAACACTTTG | 155 |
| chr8:59035-61049 | p2 | (TA)7 | ACTCATGGTCCTCATCTGCC | TGAAAAAGATTTGCCCAAGC | 156 |
| chr8:63832-65850 | p2 | (TA)9 | ATGCATCCGTCTTTCACACA | ACATGGCAAGACATTCCACA | 165 |
| chr8:113847-115861 | p2 | (AT)7 | TCGTCGAAATCCAATAAGCC | AATGGGGACGGGAAAATATC | 240 |
| chr8:280836-282852 | p2 | (AT)14 | CCGCATACTAAACGACACCA | TTTTTCAAGGGACCAAAACG | 294 |
| chr8:280894-282938 | p2 | (AT)14 | CCGCATACTAAACGACACCA | TTTTTCAAGGGACCAAAACG | 236 |
| chr8:378977-380992 | p5 | (AAATA)3 | CCACTTGCCTTGTGAAAACA | AACAAATCTTCGGTGGCTTG | 108 |
| chr8:378977-380992 | p5 | (AAATA)3 | CCACTTGCCTTGTGAAAACA | AACAAATCTTCGGTGGCTTG | 108 |
| chr8:423841-425863 | p2 | (TA)18 | TTGGCAAAATGAATGAACCA | ATAGCGTTTCAGAGGAGGCA | 261 |
| chr8:423841-425863 | p2 | (AT)11 | TTGGCAAAATGAATGAACCA | ATAGCGTTTCAGAGGAGGCA | 261 |
| chr8:423841-425863 | p2 | (TA)18 | TTGGCAAAATGAATGAACCA | ATAGCGTTTCAGAGGAGGCA | 261 |
| chr8:423841-425863 | p2 | (AT)11 | TTGGCAAAATGAATGAACCA | ATAGCGTTTCAGAGGAGGCA | 261 |
| chr8:549039-551069 | p2 | (TA)15 | GGTCCATGGCAAAAGGTATG | GGCAAAAGGATGGTCACCTA | 151 |
| chr8:549039-551069 | p5 | (AAACC)3 | GGTCCATGGCAAAAGGTATG | GGCAAAAGGATGGTCACCTA | 151 |
| chr8:602347-604365 | p3 | (AAT)6 | GTCCGGCTTTTAATCGAACA | CCCCCTAAACCTCCTACTCG | 202 |
| chr8:644384-646402 | p3 | (AAT)6 | GTCCGGCTTTTAATCGAACA | CCCCCTAAACCTCCTACTCG | 202 |
| chr8:682240-684255 | p5 | (ACCTT)3 | TCATGGAAATTATCCCCCAA | CGGAAATTATCCCCCAAAAT | 247 |
| chr8:726611-728623 | p2 | (TG)6 | TGCCTGCTCATTGCAGTTAC | GGTGACCTCCATCTTGCAGT | 141 |
| chr8:749224-751238 | p6 | (TTTTAT)3 | CATGCAATCGATGTGTGTGA | AAAGGCAGCTCAATACCGAA | 103 |
| chr8:797149-799169 | p5 | (TTTCT)4 | GAAGGACCATCAAGGACGAA | GGAAGGAGCACTGATGGAAA | 300 |
| chr8:923206-925221 | p2 | (TA)8 | AAGCTTTGCGAGAAGGATCA | TCGGCTAACGGAAGTGAAAG | 275 |
| chr8:923206-925221 | p3 | (CTT)5 | AAGCTTTGCGAGAAGGATCA | TCGGCTAACGGAAGTGAAAG | 275 |
| chr8:930371-932383 | p2 | (GT)6 | GCTTTCACCTTGATCCCAAA | CAAAAACAGCTCTCAAGGCA | 205 |
| chr8:934396-936410 | p2 | (TA)7 | GGCCCATTCGTATGAGAGAA | GGAGTCAGAGACGGATCCAA | 206 |

| Marker ID | SSR type | SSR motif | FORWARD PRIMER (5'-3') | REVERSE PRIMER (5'-3') | size (bp) |
|---|---|---|---|---|---|
| chr8:934396-936410 | p2 | (AT)9 | GGCCCATTCGTATGAGAGAA | GGAGTCAGAGACGGATCCAA | 206 |
| chr8:938035-940059 | p6 | (CCGGTT)4 | TCATTAAGCGCACCCCTAAC | TGGTACCTTCGTTCCCACTC | 294 |
| chr8:938035-940059 | p2 | (CT)8 | TCATTAAGCGCACCCCTAAC | TGGTACCTTCGTTCCCACTC | 294 |
| chr8:1067193-1069261 | p5 | (TTTCT)3 | TTCCCAAAGTATCCCCATCA | GACATGGCGGAATAGCAAAT | 140 |
| chr8:1217934-1219946 | p2 | (AC)6 | ACCGCATACTCATTTGAGGG | TTGATTCCAACTCTCCGCTT | 240 |
| chr8:1217934-1219946 | p3 | (TCT)5 | ACCGCATACTCATTTGAGGG | TTGATTCCAACTCTCCGCTT | 240 |
| chr8:1218295-1220310 | p2 | (AC)6 | CATGCCTGTCCTTCAAAACA | TTGACTGGGCACGACATTTA | 243 |
| chr8:1218295-1220310 | p3 | (TCT)5 | CATGCCTGTCCTTCAAAACA | TTGACTGGGCACGACATTTA | 243 |
| chr8:1239361-1241383 | p2 | (TA)6 | CCCATTCGCTATTTTTGGAA | TCAGCCGAGATAATGAGGCT | 297 |
| chr8:1239361-1241383 | p2 | (AT)6 | CCCATTCGCTATTTTTGGAA | TCAGCCGAGATAATGAGGCT | 297 |
| chr8:1239461-1241473 | p2 | (TA)6 | CCCATTCGCTATTTTTGGAA | TCAGCCGAGATAATGAGGCT | 197 |
| chr8:1239461-1241473 | p2 | (AT)6 | CCCATTCGCTATTTTTGGAA | TCAGCCGAGATAATGAGGCT | 197 |
| chr8:1241311-1243326 | p5 | (ACCAA)3 | TGGGTCTGGATCCCAAATAA | TTGTTGTGAACCGAGTTGGA | 234 |
| chr8:1320732-1322748 | p6 | (TCTTTT)4 | CTCAAGCGCTTTCTTTCACC | AAAGCACTGGAGCATTTTCAA | 114 |
| chr8:1320732-1322748 | p4 | (AAAT)4 | CTCAAGCGCTTTCTTTCACC | AAAGCACTGGAGCATTTTCAA | 114 |
| chr8:1424128-1426140 | p2 | (AT)8 | TGCTTCATTGCAATCCATGT | TGCTGTAGAGTGAAGGCGAG | 169 |
| chr8:1424169-1426183 | p2 | (AT)8 | TGCTTCATTGCAATCCATGT | TGCTGTAGAGTGAAGGCGAG | 128 |
| chr8:1456609-1458657 | p2 | (AT)24 | AGCATACTCGCCTTGCAGTT | GGAGCATCCCTATGGAACAA | 124 |
| chr8:1456609-1458657 | p2 | (AT)32 | AGCATACTCGCCTTGCAGTT | GGAGCATCCCTATGGAACAA | 124 |
| chr8:1630091-1632105 | p2 | (TG)7 | GAAGGAACAAGAGTGGCTCG | GCCTTATTCATCACAAGCCC | 122 |
| chr8:1635045-1637063 | p6 | (TGTGTA)3 | TTGCAGGCTCATCTTTTCCT | CATACCTGGGTGTGCACTTG | 234 |
| chr8:1638493-1640511 | p6 | (TGTGTA)3 | TTGCAGGCTCATCTTTTCCT | CATACCTGGGTGTGCACTTG | 234 |
| chr8:1768900-1770915 | p5 | (TTTTA)3 | TTGCTGGACGTACACCTTTG | GGAAGACAAAAGGGTGGTGA | 221 |
| chr8:1842285-1844337 | p3 | (ATT)5 | CTTCATGCAACAAAGGCAAA | ATTCCCTACAAGGACGGCTT | 128 |
| chr8:1993203-1995218 | p3 | (ACA)5 | TGCCATTAAGCCACACCATA | TTCAGGCTCAGGCTTTTCAT | 248 |
| chr8:2110200-2112221 | p3 | (GAA)7 | ATAGGACCCAGGGAAAGGAA | CTTCCCATGAAACAGAGGGA | 184 |
| chr8:2151928-2153943 | p5 | (AAATA)3 | GTTGGGTTTGGCTTGAGGTA | ACTCCGCTACACGAGCAAAT | 129 |
| chr8:2301187-2303221 | p2 | (AT)17 | AAAATCGCAAGGCAGTGTTT | AAATTTGCACACCCTTCGTC | 165 |
| chr8:2478530-2480542 | p2 | (AG)6 | TGTGAAAGGGCACATTGAAG | GGGATGGAAGCTATTGTGGA | 100 |
| chr8:2576462-2578494 | p2 | (AT)16 | GAAGACCCGTAAAGCAGCAG | CCTTCAAGAGCCAATCAAGC | 294 |
| chr8:2613054-2615086 | p2 | (TA)16 | AACGTGTGGGAGGGTGTTAG | CAAAGAAGATTTGCCCCGTA | 111 |
| chr8:2613054-2615086 | p2 | (TA)14 | AACGTGTGGGAGGGTGTTAG | CAAAGAAGATTTGCCCCGTA | 111 |
| chr8:2637040-2639112 | p2 | (AT)36 | AAAAGATCCAAGAACGGCCT | CAGGGAAGCAGATATTCCCA | 146 |
| chr8:2668327-2670341 | p5 | (ATATT)3 | GAGGGATTGGGAGAGTAGCC | CTCCACCAACTCTGACCGTT | 262 |
| chr8:2668327-2670341 | p2 | (TA)7 | GAGGGATTGGGAGAGTAGCC | CTCCACCAACTCTGACCGTT | 262 |
| chr8:2689313-2691331 | p6 | (GCCTTT)3 | TCTCCCTGCTCTCACAAGGT | TGGTCTGAATATGCTTGCCC | 255 |

| Marker ID | SSR type | SSR motif | FORWARD PRIMER (5'-3') | REVERSE PRIMER (5'-3') | size (bp) |
|---|---|---|---|---|---|
| chr8:2753568-2755583 | p5 | (ATGAT)3 | GATGCTGCCCAAATTTCTGT | TCGTTTCGCCCAAAATCTAC | 107 |
| chr8:2779777-2781805 | p5 | (TATAT)3 | ATGCTTTGGTGATGGTCCTC | AAACTTGCCCCCGTTTAACT | 261 |
| chr8:2779853-2781865 | p5 | (TATAT)3 | ATGCTTTGGTGATGGTCCTC | AAACTTGCCCCCGTTTAACT | 185 |
| chr8:2910329-2912351 | p2 | (AT)11 | CCAAAAGGGTGGACTCGATA | CGGAGATGGTGAGTTCCACT | 128 |
| chr8:2931016-2933034 | p6 | (GGAAGG)3 | TATTTGGCTCCTTTTTCGGA | CAAGGAACTCATAGCGCACA | 143 |
| chr8:3008515-3010553 | p2 | (TA)19 | TTCCATCCCTCTGCGATAAC | AAAAATGGCATGAATGAGCC | 152 |
| chr8:3137874-3139894 | p5 | (ATTGT)4 | CCAAGAGAACTGCGAAAAGG | GCAACGTCACCATCTTGAAA | 111 |
| chr8:3375443-3377458 | p3 | (ACA)5 | ACAACCTATGCCGTCTCTGG | TGCCGGTCTTTAATTTTTGC | 212 |
| chr8:3377037-3379049 | p2 | (AT)6 | ACTTTGGCTTTGCTTCGAGA | TTCGATACGAGTGCGACAAC | 205 |
| chr8:3389228-3391243 | p5 | (GAAAA)3 | CAACGAAAGTCACTCGGACA | TGGAAATTTTGAGGACTCCG | 203 |
| chr8:3423430-3425445 | p5 | (AATTA)3 | GGGTGGCGACTCCTTAAAAT | GTACCGAGTTTCCAGGGTGA | 201 |
| chr8:3437189-3439204 | p5 | (AACCG)3 | AAAGACCGCCCCAAAATAAG | CGGTTCGGTTATTTCGGTTA | 203 |
| chr8:3437189-3439204 | p5 | (TTGAT)3 | AAAGACCGCCCCAAAATAAG | CGGTTCGGTTATTTCGGTTA | 203 |
| chr8:3465088-3467103 | p5 | (AAACC)3 | ACATGGCTCACCTTGGAAAC | TAGTGACTGGCGTGGAAGTG | 283 |
| chr8:3532048-3534060 | p2 | (AT)6 | CGGATATTTCGTTCTTCCCA | CATGGGAGTTGAGTTCGGTT | 289 |
| chr8:3538190-3540208 | p6 | (TTATTT)3 | AGGCACTGGTAGGAAAGGGT | GTGGCAGCAAATCACATGAC | 148 |
| chr8:3567527-3569545 | p6 | (TTTTTG)3 | AACCGCCTACGATTTCATTG | TTTTTACGGCCGAGTCATTC | 220 |
| chr8:3606944-3608959 | p5 | (TATTT)3 | AGGAAGGCTGGGAGTTCATT | CCCAGAAGACGCCACATTAT | 125 |
| chr8:3637510-3639525 | p2 | (AT)6 | CCGTCACACATCACAACACA | CTATGATATCGTTCGCCGGT | 117 |
| chr8:3637510-3639525 | p3 | (CAA)5 | CCGTCACACATCACAACACA | CTATGATATCGTTCGCCGGT | 117 |
| chr8:3766612-3768642 | p3 | (AAT)10 | GTGGTGTAGTGGAGGCAGGT | CTTGGGTTTCCCCTCATTTT | 292 |
| chr8:3793360-3795396 | p2 | (TA)7 | CAACATGCGTTCCTTCTCAC | TAGAAATTTTGGGAGCACGG | 120 |
| chr8:3793360-3795396 | p2 | (AT)7 | CAACATGCGTTCCTTCTCAC | TAGAAATTTTGGGAGCACGG | 120 |
| chr8:3992369-3994384 | p5 | (TGATA)3 | GATGCTGCCCAAATTTCTGT | GTGACAGTTTCAGATGCGGA | 118 |
| chr8:4017931-4019955 | p3 | (AAC)5 | GGTCTTGGAGGGTTCAACTG | GGAAGATGAAGGGAAAAGGC | 158 |
| chr8:4033183-4035195 | p2 | (AT)6 | CCCGATGTTCACCTTTCACT | TGCTCTGTAAGGCAATGCTG | 255 |
| chr8:4150342-4152358 | p2 | (TA)8 | CTCTTTCCCTGTTTGGCAAC | TGTCTGGTTTTGAATTGGCA | 134 |
| chr8:4150342-4152358 | p2 | (AG)6 | CTCTTTCCCTGTTTGGCAAC | TGTCTGGTTTTGAATTGGCA | 134 |
| chr8:4305741-4307756 | p5 | (AAAGA)3 | CTTGGGCATGATCCAAGTTT | CCCCTCTCCCAATTTTCATT | 160 |
| chr8:4374134-4376146 | p3 | (ATC)5 | CGCAAATCAGAAAGTCACGA | AGGGAGGCCGTACAGCTATT | 207 |
| chr8:4374134-4376146 | p2 | (AT)6 | CGCAAATCAGAAAGTCACGA | AGGGAGGCCGTACAGCTATT | 207 |
| chr8:4492969-4494985 | p4 | (ATTT)4 | CTATTTTAGGGCAGGGCTCC | AGATGAGGAAGGATGGGCTT | 132 |
| chr8:4500163-4502178 | p2 | (TA)6 | GGGATGGAGCTAGATAGGGC | AACCCATTAGCGGAACCATA | 165 |
| chr8:4500163-4502178 | p3 | (TGA)5 | GGGATGGAGCTAGATAGGGC | AACCCATTAGCGGAACCATA | 165 |
| chr8:4500163-4502178 | p3 | (TAA)5 | GGGATGGAGCTAGATAGGGC | AACCCATTAGCGGAACCATA | 165 |
| chr8:4555757-4557787 | p2 | (TA)6 | CGAACCAAACCTCCACATTT | GATTCCGCATTTCAAGAAGC | 213 |

| Marker ID | SSR type | SSR motif | FORWARD PRIMER (5'-3') | REVERSE PRIMER (5'-3') | size (bp) |
|---|---|---|---|---|---|
| chr8:4555789-4557833 | p2 | (TA)6 | CGAACCAAACCTCCACATTT | GATTCCGCATTTCAAGAAGC | 181 |
| chr8:4565033-4567053 | p4 | (TATT)5 | AAAGGGCCTACGAAGGAAAA | ACAACAACTTATTTGCGGGG | 294 |
| chr8:4607540-4609555 | p2 | (AG)7 | TTTGGCCTTGAAAAAGATGG | TGCCCGAACTTCTCTCTCTC | 108 |
| chr8:4607540-4609555 | p3 | (TTG)5 | TTTGGCCTTGAAAAAGATGG | TGCCCGAACTTCTCTCTCTC | 108 |
| chr8:4622985-4625000 | p3 | (TTG)5 | CAAGATGGTCTGGTCCGTTT | GTGTTTCAACCTCTGCCCAT | 136 |
| chr8:4730310-4732335 | p5 | (TTAGT)5 | AATTAATGCCCGTCCTTTCC | AAAATATTGGGCGGGATTTC | 296 |
| chr8:4733810-4735828 | p6 | (TATCTA)3 | AAAAGGAAATTCACGCCCTT | TTCCGCAACACCTTGTATGA | 160 |
| chr8:4749974-4752000 | p4 | (ATGC)4 | CGCCTAATTCCTTGAACCAA | TCGTGACACCTGCTACAACC | 284 |
| chr8:4749974-4752000 | p2 | (TC)13 | CGCCTAATTCCTTGAACCAA | TCGTGACACCTGCTACAACC | 284 |
| chr8:4756655-4758676 | p3 | (ATT)7 | CACACTTTCGCACTAAGGCA | TGGAGCAGAGTGTTGACCAG | 170 |
| chr8:4829471-4831486 | p5 | (AATAA)3 | CTTTTTCGTCCTTGCCAAAA | ATGAACCTTAACTGGGCACG | 199 |
| chr8:4843741-4845753 | p2 | (TA)6 | GCCCCATTTCCTCCATATTT | AATCAACATCCCCAAGTTCG | 108 |
| chr8:4963959-4966037 | p2 | (CT)7 | TGAATCATCGGCCTAGAACC | CCCAACAAATTGCCTCAGTT | 208 |
| chr8:4964038-4966053 | p2 | (CT)7 | TGAATCATCGGCCTAGAACC | CCCAACAAATTGCCTCAGTT | 129 |
| chr8:4964055-4966070 | p2 | (CT)7 | TGAATCATCGGCCTAGAACC | CCCAACAAATTGCCTCAGTT | 112 |
| chr8:4969802-4971817 | p3 | (AAT)5 | CTTGTTCTCGGGGAATTTCA | CACCCTCACACTACACCACG | 116 |
| chr8:4983420-4985441 | p3 | (AAT)7 | AAGACTGACCGTTCCACCAA | GGTATCGTTTCCCAACCCTT | 120 |
| chr8:4983420-4985441 | p5 | (AAAAT)3 | AAGACTGACCGTTCCACCAA | GGTATCGTTTCCCAACCCTT | 120 |
| chr8:5068925-5070958 | p3 | (TTC)11 | ATCACATCACCATTTTGGCA | CATCTCCCAACACAATTCCC | 183 |
| chr8:5068925-5070958 | p3 | (CTT)6 | ATCACATCACCATTTTGGCA | CATCTCCCAACACAATTCCC | 183 |
| chr8:5195355-5197369 | p2 | (AG)6 | TAATTCTGGCCCAAATGAGG | AGCAAGATCGAGGTGGAAGA | 246 |
| chr8:5195355-5197369 | p2 | (AT)7 | TAATTCTGGCCCAAATGAGG | AGCAAGATCGAGGTGGAAGA | 246 |
| chr8:5249333-5251347 | p2 | (AC)7 | GAAATCCGGTGACACCATCT | CTGTCGCAACTCCACAAGAC | 126 |
| chr8:5352260-5354282 | p3 | (TTA)11 | GCATTGGATTTGGGAGAGAA | TTTTGCAACAACCAATCGAA | 285 |
| chr8:5352260-5354282 | p2 | (TA)11 | GCATTGGATTTGGGAGAGAA | TTTTGCAACAACCAATCGAA | 285 |
| chr8:5352260-5354282 | p2 | (TG)8 | GCATTGGATTTGGGAGAGAA | TTTTGCAACAACCAATCGAA | 285 |
| chr8:5406704-5408720 | p2 | (TA)8 | TTCTTACCGGAGGCAATGAC | TCGTGTTAGAAGCTAGCGCA | 276 |
| chr8:5527783-5529805 | p2 | (TA)11 | ACTTAGAAGCGGAGGAAGGC | TTTCGTTAATTTCCCCCACA | 214 |
| chr8:5527783-5529805 | p2 | (AT)12 | ACTTAGAAGCGGAGGAAGGC | TTTCGTTAATTTCCCCCACA | 214 |
| chr8:5527783-5529805 | p2 | (TA)11 | ACTTAGAAGCGGAGGAAGGC | TTTCGTTAATTTCCCCCACA | 214 |
| chr8:5532684-5534696 | p2 | (TA)6 | ATAAGCCAATCCAGACGGG | TATACTACGTCGTGGCGCTG | 158 |
| chr8:5575564-5577580 | p4 | (GATG)4 | GAGGAAGAGGATAATGGCCC | CCAAAACGGGTAAAGAACGA | 183 |
| chr8:5589280-5591296 | p4 | (GTGG)4 | TATAAGCCAATCAAACGGGC | CCCATAAGGGGGATTTGAGT | 172 |
| chr8:5613523-5615545 | p2 | (AT)7 | AAGTATCAACCGTGGCTTCG | CTCTTCCTCTTCGCCTTCCT | 281 |
| chr8:5613523-5615545 | p2 | (AT)11 | AAGTATCAACCGTGGCTTCG | CTCTTCCTCTTCGCCTTCCT | 281 |
| chr8:5613523-5615545 | p2 | (TA)6 | AAGTATCAACCGTGGCTTCG | CTCTTCCTCTTCGCCTTCCT | 281 |

| Marker ID | SSR type | SSR motif | FORWARD PRIMER (5'-3') | REVERSE PRIMER (5'-3') | size (bp) |
|---|---|---|---|---|---|
| chr8:5613523-5615545 | p2 | (TA)10 | AAGTATCAACCGTGGCTTCG | CTCTTCCTCTTCGCCTTCCT | 281 |
| chr8:5660049-5662064 | p5 | (TGATT)3 | AGGACTGGAGGTCCCAAGAT | GGTCAAACCTGCCAAACCTA | 201 |
| chr8:5678128-5680148 | p4 | (GTTT)5 | AAAAGCAACCACGTCAGACC | TTTGGACCCTCGGTTATTGA | 123 |
| chr8:5678128-5680148 | p6 | (TTTGGT)3 | AAAAGCAACCACGTCAGACC | TTTGGACCCTCGGTTATTGA | 123 |
| chr8:5691889-5693901 | p2 | (TA)6 | ACCTCAACTCGCCTCACTGT | ACACCTCGATTCGTCTCCAC | 186 |
| chr8:5813971-5815989 | p3 | (TTC)6 | CGGTCAGGTTAGGCAAAAGA | AAACTCGTGGCTTCCAAATG | 226 |
| chr8:5841272-5843287 | p5 | (AAAAC)3 | TGGTTTGATCTGTGGGAACA | CTATCGGGTCCGCATAACAT | 248 |
| chr8:5956465-5958477 | p2 | (TA)6 | GCATATCTTCTCCTCGTCGC | ACCAAAACCCCATTTCCTTC | 156 |
| chr8:6128748-6130784 | p4 | (TTTC)9 | GCAGATTTTTGCCCGAATAG | CTGGGTTTGAAGCCTAGCTG | 227 |
| chr8:6128748-6130784 | p6 | (TTTTGT)5 | GCAGATTTTTGCCCGAATAG | CTGGGTTTGAAGCCTAGCTG | 227 |
| chr8:6198219-6200239 | p5 | (TTTTA)4 | GGTTTCCCGTGTAAATGTCG | CCAGACTCCACGTTGTGAGA | 148 |
| chr8:6208892-6210907 | p3 | (GGA)5 | TTGCAAGGATTCAGCAGTTG | GGGGGAATTTCATCTGGAGT | 100 |
| chr8:6231347-6233368 | p3 | (ATA)7 | TGCAAGGGAATTCTTTTTGG | TCTTTCATCGTCATCTCCCC | 162 |
| chr8:6257203-6259229 | p2 | (TA)13 | GCCTTGTCGGTCTCGTGTAT | CCCCATAGGCGTTGAAACTA | 293 |
| chr8:6310203-6312218 | p5 | (AGTAA)3 | AAAAGCAAGTTTGTGTGGGG | TTGCAACTCTGCCTTACGAA | 182 |
| chr8:6317898-6319930 | p2 | (AT)16 | GGGGGCCACTACAACCTTAT | GCGTGTTTAGGGCGTACAAT | 219 |
| chr8:6317898-6319930 | p2 | (AT)6 | GGGGGCCACTACAACCTTAT | GCGTGTTTAGGGCGTACAAT | 219 |
| chr8:6336401-6338415 | p2 | (CT)7 | GCTTTTCCCAGGAAGCTTTT | TAAAACATTGTGTGCCGCAT | 229 |
| chr8:6585083-6587101 | p3 | (TTG)6 | TATTGGTGCAGGATGGATGA | CGCATCTTGTCTTCCACTGA | 220 |
| chr8:6695731-6697752 | p3 | (TAA)7 | GTCGAGGTTTTGGGTTCAGA | ATCGTAATCGTGGTTGGAGC | 233 |
| chr8:6695731-6697752 | p3 | (ATG)5 | GTCGAGGTTTTGGGTTCAGA | ATCGTAATCGTGGTTGGAGC | 233 |
| chr8:6720923-6722935 | p2 | (TG)6 | TCTTTGGAGAAAATGGTGGC | AATTGTGTGTGCGAGGCATA | 293 |
| chr8:6722468-6724484 | p2 | (AT)8 | AAAGCACTTGCCGCAGTTAT | AAACGGACGACGAGAAAGTG | 204 |
| chr8:6722468-6724484 | p2 | (TA)10 | AAAGCACTTGCCGCAGTTAT | AAACGGACGACGAGAAAGTG | 204 |
| chr8:6849154-6851172 | p3 | (TAT)6 | CTTCCATGGCCTCCTCATAA | GTTGATGAAGTTGGTGGGCT | 111 |
| chr8:6849154-6851172 | p2 | (AT)13 | CTTCCATGGCCTCCTCATAA | GTTGATGAAGTTGGTGGGCT | 111 |
| chr8:6954054-6956074 | p2 | (AT)9 | GCTCAAAAGTTGGGGGATTT | AGCGAGCATATTGTTTTGGG | 249 |
| chr8:6954054-6956074 | p5 | (TTCTT)4 | GCTCAAAAGTTGGGGGATTT | AGCGAGCATATTGTTTTGGG | 249 |
| chr8:7066719-7068737 | p6 | (TTCCCC)3 | CAAAGGTCCAGGCAACAGAT | GAATCCCCGACTTGAACAGA | 291 |
| chr8:7077770-7079794 | p3 | (TGT)5 | TCCTTGTCCCTGGTTCTTTG | TGCCGAAAATTTCACAACAA | 134 |
| chr8:7077770-7079794 | p6 | (GGGGGT)4 | TCCTTGTCCCTGGTTCTTTG | TGCCGAAAATTTCACAACAA | 134 |
| chr8:7087300-7089315 | p3 | (CAA)5 | AGGTGGTTTGACTAGGCACG | TATAAGCCAATCCAAACGGG | 254 |
| chr8:7164176-7166196 | p2 | (TA)10 | AGCGCTAAGTGGAGTTTGGA | TCCTAACCTGTTCCCCTCCT | 168 |
| chr8:7205177-7207189 | p3 | (GAA)6 | CAACCCATATGCCTCGAACT | AACGACAACGATCTTGGAGG | 170 |
| chr8:7205177-7207189 | p2 | (TA)6 | CAACCCATATGCCTCGAACT | AACGACAACGATCTTGGAGG | 170 |
| chr8:7316198-7318213 | p5 | (TTTCT)3 | AGCTTCTTGTTCTTGCCGAA | ATTGCCTGCTTCATTGGAAC | 118 |

| Marker ID | SSR type | SSR motif | FORWARD PRIMER (5'-3') | REVERSE PRIMER (5'-3') | size (bp) |
|---|---|---|---|---|---|
| chr8:7358220-7360232 | p2 | (AT)6 | ATTTGTAAGCGTGAGGTGGG | GTCAGGTCACCCGAAATGTT | 251 |
| chr8:7366279-7368313 | p2 | (AT)17 | CCTTTGGTGTTGGGGTATTG | GGTCGTGAAACAGTTGCTGA | 225 |
| chr8:7383002-7385018 | p2 | (CT)8 | GTAGCAGCCCTGTTTTCTGC | TTGAGTCAACTTTGGAGGGG | 215 |
| chr8:7383002-7385018 | p3 | (ACA)6 | GTAGCAGCCCTGTTTTCTGC | TTGAGTCAACTTTGGAGGGG | 215 |
| chr8:7464966-7466984 | p3 | (TTG)6 | TTACACCTTTTAGCCGGTGG | ATATGCGTCCCTCAAGTTGG | 171 |
| chr8:7506489-7508503 | p2 | (AT)7 | TAGCAAAAAGGGTTTGTCGG | TCCTTGGCTAAATGCGTTCT | 210 |
| chr8:7506489-7508503 | p2 | (AT)8 | TAGCAAAAAGGGTTTGTCGG | TCCTTGGCTAAATGCGTTCT | 210 |
| chr8:7506489-7508503 | p2 | (AT)32 | TAGCAAAAAGGGTTTGTCGG | TCCTTGGCTAAATGCGTTCT | 210 |
| chr8:7506733-7508749 | p2 | (AT)7 | AGAACGCATTTAGCCAAGGA | TCCGCTTCTGAGTTTTGGTT | 205 |
| chr8:7506733-7508749 | p2 | (AT)8 | AGAACGCATTTAGCCAAGGA | TCCGCTTCTGAGTTTTGGTT | 205 |
| chr8:7506733-7508749 | p2 | (AT)32 | AGAACGCATTTAGCCAAGGA | TCCGCTTCTGAGTTTTGGTT | 205 |
| chr8:7510624-7512683 | p2 | (AT)23 | AAAACATAAGCCCCACGACT | CATGCAATGTCACTTCACCC | 130 |
| chr8:7621836-7623852 | p2 | (TC)8 | TTCCATACTCAGGTGGGCTC | TGAGCGTGAGTCAGTAGCGT | 163 |
| chr8:7647979-7649999 | p4 | (AAAT)5 | TCCGTCCTGAAAATCCAAAG | TCACAGCTAGGGGTGCTCTT | 254 |
| chr8:7647979-7649999 | p3 | (TTA)5 | TCCGTCCTGAAAATCCAAAG | TCACAGCTAGGGGTGCTCTT | 254 |
| chr8:7737263-7739277 | p5 | (ATTGC)3 | CTTGGTTTCTGGCTCGTGTT | AATTAGCCACGAGAAACGGA | 158 |
| chr8:7827526-7829538 | p2 | (TA)6 | ACTTAGCCATTTGCACACCC | TTGGTAATTGCTCCCTCCAG | 208 |
| chr8:7927798-7929812 | p2 | (AT)7 | CCCCAAATCATGTGTTTTCC | AGGCAAAAGGCAAGAAGGAT | 197 |
| chr8:7977480-7979534 | p2 | (TA)27 | GAGCGGGAAGCAATCACTAC | ATTGCAGGCCGAGCTTAGTA | 173 |
| chr8:8186393-8188408 | p5 | (TTGGT)3 | CGTCATTCCCTACCATTGCT | CGACTAATCTTCCGCTCTGG | 134 |
| chr8:8195062-8197080 | p3 | (CTT)6 | TGCAACAATCTCTTTTCCCC | AACTCCGATTGGCACATCTC | 128 |
| chr8:8204846-8206858 | p2 | (AT)6 | CCCCTTCCTCCCTTTAATCA | TTGACAATGGCAGTGGTTGT | 209 |
| chr8:8337338-8339420 | p2 | (TA)41 | TTTGAACCTGGCCCTGATAC | AAAATTCCGGGCTTATGCTT | 295 |
| chr8:8337338-8339420 | p2 | (AT)10 | TTTGAACCTGGCCCTGATAC | AAAATTCCGGGCTTATGCTT | 295 |
| chr8:8350185-8352203 | p3 | (TTG)6 | GGTGGAATTGGCAAGTGAAT | AACTGGACACGACGAGCTTT | 283 |
| chr8:8350185-8352203 | p3 | (TGA)5 | GGTGGAATTGGCAAGTGAAT | AACTGGACACGACGAGCTTT | 283 |
| chr8:8350185-8352203 | p4 | (TGTA)4 | GGTGGAATTGGCAAGTGAAT | AACTGGACACGACGAGCTTT | 283 |
| chr8:8499209-8501233 | p2 | (AT)11 | TCGACGATATAGGGGAGGTG | TACCTTATCCGTCGAGCCAC | 174 |
| chr8:8499209-8501233 | p2 | (AT)12 | TCGACGATATAGGGGAGGTG | TACCTTATCCGTCGAGCCAC | 174 |
| chr8:8623831-8625846 | p5 | (TACAA)3 | CCCAATTTTTGTAACCACGG | ATGGGATGGACCATAGTGGA | 239 |
| chr8:8626342-8628377 | p6 | (ATATTA)3 | CTTCGTGGAGAACTCGAACC | CAAACGGAATTTGGTGCTTT | 221 |
| chr8:8698455-8700485 | p5 | (CGGGT)3 | CACACACCGAAAAGCAAAGA | GAGATCCGATCCGATTTTGA | 114 |
| chr8:8698455-8700485 | p2 | (TA)15 | CACACACCGAAAAGCAAAGA | GAGATCCGATCCGATTTTGA | 114 |
| chr8:8770090-8772108 | p2 | (AT)9 | ACACATTTCAAACGCAACCA | CCATATGATTCGGTCGGTCT | 151 |
| chr8:8770090-8772108 | p3 | (CAA)5 | ACACATTTCAAACGCAACCA | CCATATGATTCGGTCGGTCT | 151 |
| chr8:8840754-8842769 | p5 | (ATGAT)3 | GATGCTGCCCAAATTTCTGT | CAAATCCCGTTTCCTTCAAA | 108 |

| Marker ID | SSR type | SSR motif | FORWARD PRIMER (5'-3') | REVERSE PRIMER (5'-3') | size (bp) |
|---|---|---|---|---|---|
| chr8:8852858-8854876 | p6 | (ATCCAC)3 | TGAAAGTTGAAATCCGCACA | TGCACAGTGGAAGTGGATGT | 126 |
| chr8:8852858-8854876 | p6 | (ATCCAC)4 | TGAAAGTTGAAATCCGCACA | TGCACAGTGGAAGTGGATGT | 126 |
| chr8:8921725-8923743 | p5 | (TGTTG)3 | AGGTCCCCAAGTTCTGGAGT | CACGGCTCAAGCTCAACATA | 154 |
| chr8:8921725-8923743 | p3 | (TGT)6 | AGGTCCCCAAGTTCTGGAGT | CACGGCTCAAGCTCAACATA | 154 |
| chr8:9127295-9129310 | p5 | (AATCA)3 | TAGTTCGGCTTGAGTCGGTT | TAACCAAAAACCCCGACAAA | 215 |
| chr8:9127295-9129310 | p5 | (GATTT)3 | TAGTTCGGCTTGAGTCGGTT | TAACCAAAAACCCCGACAAA | 215 |
| chr8:9127295-9129310 | p6 | (AAAATC)3 | TAGTTCGGCTTGAGTCGGTT | TAACCAAAAACCCCGACAAA | 215 |
| chr8:9170932-9172944 | p2 | (CT)6 | TTCAACAATGACTTCGCAGC | CGAAAGCAATAATTGAGGGC | 176 |
| chr8:9170932-9172944 | p2 | (TA)7 | TTCAACAATGACTTCGCAGC | CGAAAGCAATAATTGAGGGC | 176 |
| chr8:9170932-9172944 | p3 | (AAT)9 | TTCAACAATGACTTCGCAGC | CGAAAGCAATAATTGAGGGC | 176 |
| chr8:9170932-9172944 | p2 | (AT)6 | TTCAACAATGACTTCGCAGC | CGAAAGCAATAATTGAGGGC | 176 |
| chr8:9170932-9172944 | p3 | (AAT)8 | TTCAACAATGACTTCGCAGC | CGAAAGCAATAATTGAGGGC | 176 |
| chr8:9324234-9326250 | p4 | (AAGA)4 | CTGAGCCATCCCTCACTTGT | GTCACTCCCTTCGGTTCAAA | 251 |
| chr8:9340379-9342415 | p2 | (TG)8 | CCTCTCGGAATTGACGGATA | TTTTGACCATATTGCCCCAT | 247 |
| chr8:9435817-9437832 | p5 | (ATTTT)3 | GTATGCCTGGCTCCACACTT | TAAAGGGCTTCTTGCTTCCA | 282 |
| chr8:9458589-9460610 | p3 | (TTG)7 | TTTGTCTTGATTTGCATCGC | ACATGGGACGTTTGGATGAT | 184 |
| chr8:9622261-9624275 | p2 | (AT)7 | TCCCTTGTCATTTACCCCAA | GCGGCTATGTAAACCGATGT | 196 |
| chr8:9641076-9643186 | p2 | (AT)55 | TTGATGGAAATGCGATGAAA | GCCAAAGAAAAAGGGGAAAC | 202 |
| chr8:9641076-9643186 | p2 | (AT)8 | TTGATGGAAATGCGATGAAA | GCCAAAGAAAAAGGGGAAAC | 202 |
| chr8:9641512-9643528 | p2 | (AT)55 | AACATTGCACTTTCCTTCCG | AAAAATTGTGCAGGATTCCG | 160 |
| chr8:9641512-9643528 | p2 | (AT)8 | AACATTGCACTTTCCTTCCG | AAAAATTGTGCAGGATTCCG | 160 |
| chr8:9656419-9658437 | p6 | (CATATA)3 | GCTCCCCCATAACAGACTCA | GTGTGACAGCTTCAAGGCAA | 160 |
| chr8:9747935-9749953 | p3 | (TTG)6 | AGAAGATGAAAGTCGCGGAA | GCATCTTGTCTTCCACCGAT | 296 |
| chr8:9763485-9765536 | p3 | (AAT)17 | AAGTCCTGAGGAAATCGGGT | GGCATCTTCTTGGGAAATGA | 276 |
| chr8:9765944-9767966 | p2 | (TA)11 | AAATTCTACCCCCACCAACC | CACGCGATAAAAATATGGGG | 177 |
| chr8:9773757-9775772 | p2 | (AT)6 | CCCTTTCTCTCTCCTCCGTT | AAAGAGGCAGGTGATTGTGG | 209 |
| chr8:9773757-9775772 | p5 | (ATAAT)3 | CCCTTTCTCTCTCCTCCGTT | AAAGAGGCAGGTGATTGTGG | 209 |
| chr8:9807948-9809963 | p5 | (AATAA)3 | AGGTCCAGAGTTGGGATGTG | TATCCTAGGCCATCTGGACG | 150 |
| chr8:9832416-9834432 | p4 | (ATTA)4 | CCCTCAAGGGTAATTTGGGT | TGGGCTGAATTCTTTTGTCC | 232 |
| chr8:9844879-9846893 | p2 | (TA)7 | TCAGACCTCGTCTCGGCTAT | TCTGGCCAATTTTTCCTTTG | 205 |
| chr8:9878623-9880661 | p2 | (AC)7 | TGTGCCTGTTAGATTGGCTG | ACCTACCATTGCTTCCGTTG | 278 |
| chr8:9974654-9976670 | p2 | (TG)7 | TGCCCTTGCGGTATTAAGTC | GTCATTTTCAGTCGGCCAAT | 156 |
| chr8:9974654-9976670 | p4 | (GGGA)4 | TGCCCTTGCGGTATTAAGTC | GTCATTTTCAGTCGGCCAAT | 156 |
| chr8:10071905-10073917 | p2 | (TA)6 | CTTGAGGATTTGTCCCTCCA | GGGGTTTTGAAATTCTGGGT | 124 |
| chr8:10220466-10222484 | p6 | (AAATCC)3 | CAACCTTCTCACGCACAAAA | AACTACATTGTCCGGTTCGC | 162 |
| chr8:10230779-10232797 | p6 | (TGTGTA)3 | GTGGCGTTTCAGGTTCTAGC | AACTCCATATATTCGCCCCC | 262 |

| Marker ID | SSR type | SSR motif | FORWARD PRIMER (5'-3') | REVERSE PRIMER (5'-3') | size (bp) |
|---|---|---|---|---|---|
| chr8:10245929-10247941 | p2 | (AT)11 | AAGATAGATGGGCCATGACG | TTGGTTTCAGCTTGACGATG | 219 |
| chr8:10245949-10248007 | p2 | (AT)11 | AAGATAGATGGGCCATGACG | TTGGTTTCAGCTTGACGATG | 199 |
| chr8:10292893-10294913 | p2 | (TA)6 | AGATGAACACCTTTCGTCCG | CGTTGGACCATCACCTTTTT | 145 |
| chr8:10297168-10299183 | p2 | (AT)6 | CTACCGCACGATCTGAGACA | AGATCGGGGTTGTGACAAAG | 237 |
| chr8:10297187-10299212 | p2 | (AT)6 | CTACCGCACGATCTGAGACA | AGATCGGGGTTGTGACAAAG | 218 |
| chr8:10306597-10308611 | p2 | (AT)7 | TTGATGGAAATGCGATGAAA | CATGCCAAAGAAAAAGGGAA | 134 |
| chr8:10371377-10373425 | p2 | (AT)24 | ATGTCAACCGACACCCCTTA | CCCATGATCTTCCGTTGAAT | 252 |
| chr8:10371377-10373425 | p2 | (TA)9 | ATGTCAACCGACACCCCTTA | CCCATGATCTTCCGTTGAAT | 252 |
| chr8:10371377-10373425 | p2 | (TA)23 | ATGTCAACCGACACCCCTTA | CCCATGATCTTCCGTTGAAT | 252 |
| chr8:10386682-10388696 | p2 | (AT)7 | GAATGTATGCGTGTGCCATC | TCCATACGGACATCAGCAAA | 295 |
| chr8:10386682-10388696 | p3 | (TCA)6 | GAATGTATGCGTGTGCCATC | TCCATACGGACATCAGCAAA | 295 |
| chr8:10513623-10515638 | p3 | (AAC)5 | CGATTTTCCCGTCTCCAATA | CGTGTCCCTGAATTGGCTAT | 121 |
| chr8:10582040-10584055 | p5 | (AACAC)3 | CAACTGAAGCAGCCAATGAA | AAGCTGATTGTGTATCCCGC | 173 |
| chr8:10592072-10594088 | p4 | (AGTT)4 | TGAGGCCTTAAACAGGTGCT | GTTGGGCTCTACCCAAGTGA | 136 |
| chr8:10613638-10615650 | p6 | (TATCTA)3 | CTCCCAATTCTCCATCCTCA | GAAATTGGAGACCGTGCAGT | 253 |
| chr8:10613638-10615650 | p2 | (AT)6 | CTCCCAATTCTCCATCCTCA | GAAATTGGAGACCGTGCAGT | 253 |
| chr8:10623348-10625363 | p3 | (AAC)5 | TGCTTAACGATTTTCCCGTC | CGTGTCCCTGAATTGGCTAT | 114 |
| chr8:10659489-10661507 | p6 | (TATCTA)3 | CCTTGGTCTTCGTTGTGGAT | TTGAGGACCTTCGAGGCTTA | 247 |
| chr8:10723731-10725747 | p4 | (TTTA)4 | GCCTCTTTTGCTGATTTTGC | CCAGATTGTCCCGTTTTTGT | 105 |
| chr8:10845847-10847865 | p2 | (AT)9 | GGGTATCGGGTGGTGTTCTA | ATCGCCGATAAGGGTTTTCT | 134 |
| chr8:10958523-10960537 | p2 | (TG)6 | TCGGATCAAAAAGACGGTTC | TTCTGTGCACCATTTTCGAG | 252 |
| chr8:10958523-10960537 | p2 | (AT)7 | TCGGATCAAAAAGACGGTTC | TTCTGTGCACCATTTTCGAG | 252 |
| chr8:10958523-10960537 | p6 | (AAAAAT)3 | TCGGATCAAAAAGACGGTTC | TTCTGTGCACCATTTTCGAG | 252 |
| chr8:10958523-10960537 | p2 | (GT)7 | TCGGATCAAAAAGACGGTTC | TTCTGTGCACCATTTTCGAG | 252 |
| chr8:11087736-11089751 | p5 | (TTATT)3 | TTTCTTGGGGATGACCTGAG | AACCCTGAAGTCGGAAGGTT | 179 |
| chr8:11109548-11111569 | p3 | (AAC)7 | GAAGCAAAGGGGATTGAACA | CAAGTGTGTTACGGTGGGTG | 164 |
| chr8:11146781-11148796 | p5 | (TTGGT)3 | CCGCAAATAAGCAATGAGGT | CCTGGTTGGCCAAATTTTTA | 173 |
| chr8:11233590-11235614 | p2 | (AC)12 | CGTCGGAATCTTTACCCTCA | ACATCCTTACCCCAGGGTTC | 238 |
| chr8:11290235-11292247 | p2 | (AT)6 | TCTTCCCGCATTTCATAACC | CCCAAATAAGGTTGGGAGGT | 295 |
| chr8:11304779-11306831 | p2 | (AT)26 | AAGGTGTGGGCTGTTTTGAC | ATGCTCAATATCAATCCGGG | 133 |
| chr8:11343348-11345366 | p3 | (TGA)6 | AAGGTTTGTCGAATGATGGC | GTTTTGGTCGTGTGTGTTGG | 188 |
| chr8:11362768-11364805 | p3 | (AAC)6 | TCAGCAACTACCTTCTGGGG | GAAGAACAGAGCCAAGAAGCA | 108 |
| chr8:11362768-11364805 | p5 | (GGGGT)3 | TCAGCAACTACCTTCTGGGG | GAAGAACAGAGCCAAGAAGCA | 108 |
| chr8:11484732-11486747 | p3 | (AGA)5 | TCAAGAACAAGCAAAACCCC | ATTTCAAGATTTTGTGCGGG | 200 |
| chr8:11489638-11491650 | p2 | (AT)6 | ACCAGATATCGCCTTCCCTT | ACACATGAGAACGGCAACAA | 272 |
| chr8:11505152-11507164 | p2 | (AT)6 | CACAACAGCGTCTCCAAGAA | GCATAGCACCCCGTCATTAT | 263 |

| Marker ID | SSR type | SSR motif | FORWARD PRIMER (5'-3') | REVERSE PRIMER (5'-3') | size (bp) |
|---|---|---|---|---|---|
| chr8:11516692-11518710 | p6 | (GCTTCT)3 | TTGTCTCTACCGGGATTTGG | TCACACACAACTCGCTCTCC | 254 |
| chr8:11520226-11522242 | p2 | (TA)8 | AACCATCTGCAGCAAAAAGG | TATATGCTAATGGGCCTCGC | 237 |
| chr8:11520226-11522242 | p3 | (CAA)5 | AACCATCTGCAGCAAAAAGG | TATATGCTAATGGGCCTCGC | 237 |
| chr8:11526237-11528252 | p3 | (TGT)5 | CGTATTGGAGCAGGATGGAT | TATAGCCGGTCTAACCACCG | 207 |
| chr8:11526237-11528252 | p2 | (TA)17 | CGTATTGGAGCAGGATGGAT | TATAGCCGGTCTAACCACCG | 207 |
| chr8:11540432-11542444 | p2 | (AT)6 | TTGTGGGCTTTGATTGTTGA | GCCCACATGTTTTGGGTTAC | 185 |
| chr8:11735194-11737212 | p2 | (TA)9 | GAGGGAGGAAGTATAGGCGG | ACTCCGCTGGGACTTTAGGT | 252 |
| chr8:11800916-11802976 | p2 | (TA)30 | ATTTTCAGTTGGTGCAAGGG | CCCTCCATCCTACCCGTAAT | 103 |
| chr8:11821355-11823413 | p2 | (AT)29 | CGCTGGCTTCCTTTCATAAG | TTTAAGGATGCGTTCAAGGG | 141 |
| chr8:11860630-11862646 | p2 | (TA)8 | TTCCTAGGCAGCCTCTTCTG | TTGTGAATCCGTAGCGTCAG | 296 |
| chr8:11877720-11879735 | p2 | (TA)14 | AAGGTCGTATGGCGAGAGAA | TCCTCCAGTAAATTTTGGCG | 245 |
| chr8:11877720-11879735 | p5 | (AAATT)3 | AAGGTCGTATGGCGAGAGAA | TCCTCCAGTAAATTTTGGCG | 245 |
| chr8:11926437-11928455 | p2 | (AT)13 | GCAGTCCTTGAAAGTTTGGG | AGGATCGTCATCCATCCAAG | 192 |
| chr8:11926437-11928455 | p6 | (TTGATT)3 | GCAGTCCTTGAAAGTTTGGG | AGGATCGTCATCCATCCAAG | 192 |
| chr8:11926437-11928455 | p3 | (TGG)5 | GCAGTCCTTGAAAGTTTGGG | AGGATCGTCATCCATCCAAG | 192 |
| chr8:12045897-12047915 | p6 | (AGCTGT)3 | CACCTGTGGCAGATGAGCTA | CTGACTTGGTTTCTTCCCCA | 275 |
| chr8:12075319-12077331 | p5 | (AAAAT)3 | ATCCAGTGGAATGAGGCAAG | AGTGCACTCTCCAGGCAACT | 249 |
| chr8:12075319-12077331 | p2 | (CA)6 | ATCCAGTGGAATGAGGCAAG | AGTGCACTCTCCAGGCAACT | 249 |
| chr8:12330920-12332938 | p2 | (AC)9 | ACAGGTCGTCCATTACTGCC | CGATTCATTTCCATTGCCTT | 271 |
| chr8:12341769-12343793 | p6 | (CATAAT)4 | CGGTGGAATCGTATCCTCAT | GTCAGGGGTTCACGACCTTA | 242 |
| chr8:12355475-12357499 | p2 | (AT)12 | TTTGACCCGAAAAACTCTGG | TCAGCTCCAGTTCCTCCACT | 143 |
| chr8:12355475-12357499 | p2 | (AC)7 | TTTGACCCGAAAAACTCTGG | TCAGCTCCAGTTCCTCCACT | 143 |
| chr8:12355475-12357499 | p2 | (AT)6 | TTTGACCCGAAAAACTCTGG | TCAGCTCCAGTTCCTCCACT | 143 |
| chr8:12376393-12378405 | p2 | (TA)6 | ATTGATCCGTGACCAGAAGG | AACGTACGACCCGATCCATA | 188 |
| chr8:12381008-12383022 | p2 | (AT)7 | TCACTGAATGTCGATGGGAA | AACCAGCTCACTGGGAAAAA | 192 |
| chr8:12459689-12461703 | p2 | (AT)7 | ATTGTCACCCTTGGGATTTG | CCCCTTTGTTGGCACATTAT | 172 |
| chr8:12459689-12461703 | p2 | (AT)14 | ATTGTCACCCTTGGGATTTG | CCCCTTTGTTGGCACATTAT | 172 |
| chr8:12529360-12531372 | p2 | (TA)6 | GAAAATCAAAAAGGCGTGGA | TGAAGTGTTCGCAAAAATGC | 298 |
| chr8:12819691-12821715 | p2 | (TA)12 | CCCCATATTAAACACCGTGC | GGATGTGGTGGGTTTCACTT | 239 |
| chr8:12836773-12838787 | p2 | (TA)7 | TGTTCCGAGAAAGGCAGAGT | CGAAGAAAGGGGTCAGTACG | 115 |
| chr8:12836773-12838787 | p2 | (TA)7 | TGTTCCGAGAAAGGCAGAGT | CGAAGAAAGGGGTCAGTACG | 115 |
| chr8:12856281-12858296 | p5 | (AAACC)3 | TATGCCGCTAACTAGGGGTG | CGAATAACCCGAGGTTGAAA | 203 |
| chr8:12984078-12986116 | p2 | (TA)19 | TGGGTCGTGACAAAGTGGTA | AAGTGTGGCCCGTTTCATAC | 198 |
| chr8:13005828-13007874 | p2 | (AT)23 | AGTCGACGAGGCTAGCAAAA | GGCCAGGGAGACATACTTGA | 227 |
| chr8:13005884-13007928 | p2 | (AT)23 | AGTCGACGAGGCTAGCAAAA | GGCCAGGGAGACATACTTGA | 171 |
| chr8:13104857-13106882 | p5 | (GGGGC)5 | GGACTGCTCCTCCCTTAACC | GTTTCCACCCTGTGTCCAGT | 175 |

| Marker ID | SSR type | SSR motif | FORWARD PRIMER (5'-3') | REVERSE PRIMER (5'-3') | size (bp) |
|---|---|---|---|---|---|
| chr8:13178135-13180185 | p2 | (TA)25 | GAAGTGGACAATCCTCCGAA | TCGGAATCAAGGAAACCAAG | 255 |
| chr8:13199892-13201956 | p2 | (CT)11 | AAACTTGAAGTGGCCGAATG | AAAGGCCCAATCCCATTATC | 119 |
| chr8:13199892-13201956 | p2 | (AT)32 | AAACTTGAAGTGGCCGAATG | AAAGGCCCAATCCCATTATC | 119 |
| chr8:13449920-13451932 | p2 | (AT)6 | TCTCGCCACAACAGTCAGTC | ATCACCGTTGTAGCGGTCTC | 195 |
| chr8:13449920-13451932 | p2 | (TA)9 | TCTCGCCACAACAGTCAGTC | ATCACCGTTGTAGCGGTCTC | 195 |
| chr8:13465754-13467774 | p2 | (AT)10 | TAGCTTTCGTCATGGACGTG | TCTGCTTTGTGAGAACCCCT | 175 |
| chr8:13490602-13492616 | p2 | (AC)7 | CTGTCTTCGAGGGAAACGTC | GGACATGCATCCTTAATGGG | 215 |
| chr8:13635299-13637319 | p5 | (TGATT)4 | AAGAGCACCTGAGCATTGGT | TTAACCATGTTAGCGGGAGG | 224 |
| chr8:13647245-13649260 | p3 | (GAT)5 | CTTTAGACGGCGAAACAAGC | AATCGTTCGAAAATTGACGG | 158 |
| chr8:13671819-13673834 | p3 | (ACT)5 | CAACGCTTGGTGTAGACCCT | TCTCTCTGCGGCTCCTATGT | 223 |
| chr8:13691617-13693665 | p2 | (AT)24 | GTAGCTCCGGATGAAATGGA | GCCCTACTATCTCCCGATCC | 261 |
| chr8:13705489-13707507 | p6 | (TATTCA)3 | CTGGACACGGCTCATAGACA | CCCTCGTCGGCTTATGTAAA | 240 |
| chr8:13705489-13707507 | p2 | (AT)6 | CTGGACACGGCTCATAGACA | CCCTCGTCGGCTTATGTAAA | 240 |
| chr8:13722793-13724813 | p2 | (AT)10 | AGTCAATGCTCCATAACGGG | GTGTCGAGAAGGGGTTACCA | 102 |
| chr8:13793425-13795440 | p5 | (TGATA)3 | GCCCAAATTTCTGTAGGCAA | ATGCTCTTTACGCCTTTCCA | 124 |
| chr8:13864048-13866076 | p2 | (TA)14 | GACACTGGTTGTTGCGTCAC | ATGCAGGAGCCAAGAGCTAA | 106 |
| chr8:13864048-13866076 | p2 | (AT)8 | GACACTGGTTGTTGCGTCAC | ATGCAGGAGCCAAGAGCTAA | 106 |
| chr8:13864048-13866076 | p2 | (TA)8 | GACACTGGTTGTTGCGTCAC | ATGCAGGAGCCAAGAGCTAA | 106 |
| chr8:13869089-13871104 | p3 | (TAT)5 | GGGTGTTCAAACGGACATCT | TGATGGAAGAGGGTAGGTGG | 247 |
| chr8:13918054-13920069 | p3 | (AAT)5 | GCAACAACAGAGCAAACGAA | GAGGTTTTCCGGTGACAAAA | 113 |
| chr8:13951752-13953770 | p2 | (TA)9 | GCGGCAATGCTAAGGACTAC | CCCTCTCTCTTCTCCCTCGT | 235 |
| chr8:13977586-13979608 | p2 | (TA)8 | TCCTCTCACGGTCTCCAAGT | TCAAAGGAGGGAGGAATTGA | 188 |
| chr8:13989204-13991222 | p5 | (GGGTC)3 | AGAAAGTTGCGGCTGTGTCT | AAACCATTTGGGCTAAGGCT | 292 |
| chr8:13989204-13991222 | p6 | (ATCCGT)3 | AGAAAGTTGCGGCTGTGTCT | AAACCATTTGGGCTAAGGCT | 292 |
| chr8:14007986-14010007 | p3 | (AAC)7 | GCGGGGAAGACACATGTAAT | TGTTAAGTTCGGGTCCAAGC | 277 |
| chr8:14034845-14036859 | p2 | (AC)6 | GGAGCCAACATTCAAAGCTC | TGCAAGAGGTTGTTTTCACG | 262 |
| chr8:14034872-14036894 | p2 | (AC)6 | GGAGCCAACATTCAAAGCTC | TGCAAGAGGTTGTTTTCACG | 235 |
| chr8:14142096-14144124 | p3 | (TGT)5 | AAAATAATCAGCCCCAACCC | TAAGACTGGGGCAGAGAGGA | 140 |
| chr8:14161663-14163675 | p2 | (AC)7 | CCAGTACATCCCACCTGCTT | TAACCACCACTCCGAGCTTC | 123 |
| chr8:14161663-14163675 | p2 | (TG)6 | CCAGTACATCCCACCTGCTT | TAACCACCACTCCGAGCTTC | 123 |
| chr8:14275111-14277126 | p3 | (GAA)5 | ATTTTATTCATCCCGCCACA | TACAGAACAGCGGTCACCAG | 206 |
| chr8:14275111-14277126 | p3 | (AAG)6 | ATTTTATTCATCCCGCCACA | TACAGAACAGCGGTCACCAG | 206 |
| chr8:14384280-14386346 | p2 | (TA)33 | CTGAAACTGTAGCTTCGCCC | ATTAGCCGCGTCTCTCAATG | 271 |
| chr8:14392567-14394582 | p5 | (ATAAA)3 | TGGGCATATGGGTAAGGGTA | TCGAAGTTAGGCTTGGCAAT | 105 |
| chr8:14392663-14394678 | p5 | (ATAAA)3 | AGTGGAGGGTCGGAGAAGAT | TCTCACCCAAGGTATGCTCC | 246 |
| chr8:14395974-14397989 | p2 | (AT)7 | TTGCAACAAACGCACAAAAT | CACCATTGCTGTTTGCTACG | 107 |

| Marker ID | SSR type | SSR motif | FORWARD PRIMER (5'-3') | REVERSE PRIMER (5'-3') | size (bp) |
|---|---|---|---|---|---|
| chr8:14395974-14397989 | p3 | (AAT)5 | TTGCAACAAACGCACAAAAT | CACCATTGCTGTTTGCTACG | 107 |
| chr8:14426758-14428773 | p5 | (ACATA)3 | TCCCCAATCCCAACTGAATA | GGCCTACCTCGGAAAGAAAC | 230 |
| chr8:14458717-14460732 | p5 | (TACAA)3 | TTAATCAACCGCTAGGCCAC | ATAGTGGTCGCCAGGAAATG | 243 |
| chr8:14641977-14644011 | p2 | (AT)17 | CACCCGTATACGAACCTGCT | TGTGTAGTGGCAGCCTTGTC | 240 |
| chr8:14641977-14644011 | p2 | (TA)7 | CACCCGTATACGAACCTGCT | TGTGTAGTGGCAGCCTTGTC | 240 |
| chr8:14664158-14666173 | p3 | (TCC)5 | TCCGCATAATTCACCAAACA | CTGGTTTCCATTGCCACTTT | 172 |
| chr8:14692179-14694205 | p2 | (AT)13 | CGAAGCGTATATTGGTCGGT | CGTCTAACCCGAACTTCCAA | 156 |
| chr8:14692179-14694205 | p5 | (ATATG)3 | CGAAGCGTATATTGGTCGGT | CGTCTAACCCGAACTTCCAA | 156 |
| chr8:14720646-14722664 | p6 | (CTTAAG)3 | ATAATGGAGTGCTCCTGCGT | GGACGCTGGTTAACGATGAT | 123 |
| chr8:14788498-14790518 | p5 | (CATGA)4 | TGTTGGTAAAAGTGGCCCTC | GGAGTTGTCAGGAATGGAGC | 198 |
| chr8:14804097-14806112 | p5 | (ATGCA)3 | CCAAGTGGTGGTGGAGAGAT | TGGGTTGAAGGTTCGCTTAC | 133 |
| chr8:14864546-14866558 | p2 | (CA)6 | TCTCGTCCAAACTCAATCCC | TATAGGACGGGAATTGCTGC | 295 |
| chr8:14871738-14873753 | p5 | (AAACC)3 | GCGTCATCTGCTCAAACAAA | CTACCAAATGCGACTCGGTT | 102 |
| chr8:14943915-14945930 | p5 | (AATGC)3 | ATTCCTGCACCAACTTACCG | TGACAAAAATACACGGGCAA | 207 |
| chr8:14972014-14974029 | p2 | (AT)9 | ACCGTACCCGACCCTCTAGT | TCGTGCATGCCTCAATATGT | 237 |
| chr8:14972014-14974029 | p3 | (CAA)5 | ACCGTACCCGACCCTCTAGT | TCGTGCATGCCTCAATATGT | 237 |
| chr8:15010432-15012447 | p3 | (TAA)5 | GAATGACGAAATGAAGGGGA | ATTTTGACTTACCGGACCCC | 185 |
| chr8:15010432-15012447 | p3 | (ATA)5 | GAATGACGAAATGAAGGGGA | ATTTTGACTTACCGGACCCC | 185 |
| chr8:15010663-15012708 | p3 | (TAA)5 | GGGGTCCGGTAAGTCAAAAT | TACGCATCGCATTTACTTCG | 164 |
| chr8:15010663-15012708 | p3 | (ATA)5 | GGGGTCCGGTAAGTCAAAAT | TACGCATCGCATTTACTTCG | 164 |
| chr8:15010716-15012731 | p3 | (TAA)5 | GGGGTCCGGTAAGTCAAAAT | TACGCATCGCATTTACTTCG | 111 |
| chr8:15010716-15012731 | p3 | (ATA)5 | GGGGTCCGGTAAGTCAAAAT | TACGCATCGCATTTACTTCG | 111 |
| chr8:15237891-15239909 | p6 | (TATATG)3 | AACATGTCGGCACATCAAAA | TCATGTTGGCATGAAGCATT | 257 |
| chr8:15273679-15275694 | p5 | (CAAAC)3 | CTTTGGGGCAGTACTTGCAT | AGACCCTAGCCTTGGGGATA | 286 |
| chr8:15273679-15275694 | p6 | (ATTTTT)3 | CTTTGGGGCAGTACTTGCAT | AGACCCTAGCCTTGGGGATA | 286 |
| chr8:15288393-15290405 | p2 | (AT)6 | ATTCGCAAGCAAATCGACTC | TACACGTGGAAAGTGGGTCA | 245 |
| chr8:15288393-15290405 | p2 | (AT)11 | ATTCGCAAGCAAATCGACTC | TACACGTGGAAAGTGGGTCA | 245 |
| chr8:15291074-15293097 | p2 | (TA)8 | CCATGCTTCGTCGTCAGTAA | CCCGTGAGTTTGGAGACAGT | 176 |
| chr8:15291074-15293097 | p2 | (TA)7 | CCATGCTTCGTCGTCAGTAA | CCCGTGAGTTTGGAGACAGT | 176 |
| chr8:15291527-15293541 | p2 | (TA)8 | TGCTTCATGGTCTGTTGTCC | AGTATTGGGCCCACCTTTTT | 256 |
| chr8:15291527-15293541 | p2 | (TA)7 | TGCTTCATGGTCTGTTGTCC | AGTATTGGGCCCACCTTTTT | 256 |
| chr8:15291527-15293541 | p2 | (AT)9 | TGCTTCATGGTCTGTTGTCC | AGTATTGGGCCCACCTTTTT | 256 |
| chr8:15383266-15385281 | p3 | (ACC)5 | TATAGGCTGGGGTGTTTTGC | AGCAGCGGAGATGTCAACTT | 147 |
| chr8:15384897-15386912 | p5 | (AGAAA)3 | GTTGGGGGTTGGGAGTATTT | TCGAGGAGTCCACAATTTCC | 151 |
| chr8:15400485-15402499 | p6 | (AACCGG)3 | CATGTACACCCCTACCTGGG | ACCGGTAAACTAATGCACGG | 113 |
| chr8:15400485-15402499 | p2 | (TA)7 | CATGTACACCCCTACCTGGG | ACCGGTAAACTAATGCACGG | 113 |

| Marker ID | SSR type | SSR motif | FORWARD PRIMER (5'-3') | REVERSE PRIMER (5'-3') | size (bp) |
|---|---|---|---|---|---|
| chr8:15400485-15402499 | p2 | (AT)13 | CATGTACACCCCTACCTGGG | ACCGGTAAACTAATGCACGG | 113 |
| chr8:15499761-15501776 | p3 | (ACA)5 | GCCTCATAGTCGAGCCACTC | CAATATCCTGGAGCTGAGCC | 202 |
| chr8:15532068-15534086 | p2 | (TG)9 | GAGTGTCAAAACCGTACGCA | GGACTCGGTCCTCAATTCAA | 167 |
| chr8:15577085-15579103 | p6 | (GTTGGA)3 | GAGGTGTTGGGTTGCATTTT | CTCACCCGTGACCTTGAAAT | 230 |
| chr8:15618742-15620758 | p4 | (TTAT)4 | AAATGAAAGTCCGTGATCGG | CAAGGACCTCGAGTATCGGA | 181 |
| chr8:15667591-15669607 | p2 | (TA)8 | AAGGAGACCTCCCCGAAATA | TTGATTATCCGAATGCACGA | 276 |
| chr8:15690215-15692230 | p3 | (ACA)5 | TACGCCATTTTCGTCTTTCC | CAGGCCCAAGATCCATTCTA | 216 |
| chr8:15697801-15699813 | p2 | (AT)11 | AAAATGGTCCCACGTGAAAG | TCGGTACGCCTCAATTTTTC | 131 |
| chr8:15697818-15699830 | p2 | (AT)11 | AAAATGGTCCCACGTGAAAG | TCGGTACGCCTCAATTTTTC | 114 |
| chr8:15709140-15711152 | p2 | (TA)6 | GCCGAGACGAGCAATAAGAC | CCCCAAACCTCACCTGTAGA | 296 |
| chr8:15771708-15773723 | p5 | (GAGAG)3 | TTTGTAGGAAAAGATGCCCG | AACCGTAGCATGTGGTGACA | 119 |
| chr8:15771708-15773723 | p3 | (TAA)5 | TTTGTAGGAAAAGATGCCCG | AACCGTAGCATGTGGTGACA | 119 |
| chr8:15801603-15803618 | p5 | (AAAAT)3 | AGGACAATCCAATGCACACA | AATGCACCACAATTCCAACA | 293 |
| chr8:15832065-15834079 | p2 | (AT)21 | CTATCAACACCAAACCCGCT | GTCGAATCCACATGGGAAAA | 118 |
| chr8:15832065-15834079 | p2 | (AT)7 | CTATCAACACCAAACCCGCT | GTCGAATCCACATGGGAAAA | 118 |
| chr8:15832065-15834079 | p2 | (TA)7 | CTATCAACACCAAACCCGCT | GTCGAATCCACATGGGAAAA | 118 |
| chr8:15833563-15835577 | p2 | (TA)6 | AATAATGTCCAAAAGGGTGGC | TTTAAGGGACAACGGACGAC | 212 |
| chr8:15833563-15835577 | p2 | (AT)7 | AATAATGTCCAAAAGGGTGGC | TTTAAGGGACAACGGACGAC | 212 |
| chr8:15844794-15846809 | p3 | (CCT)5 | TTTCACCCGAATTTTGAAGC | TGTATATCCCGCTCAAGGCT | 130 |
| chr8:15927500-15929515 | p3 | (ATG)5 | TGCATTGAATGATCCCTTGA | CGATTCCACCAGTCGTTTCT | 251 |
| chr8:15994968-15996990 | p2 | (AT)11 | AAAGTGGTGCCACATGTCAA | TACACATGGCGCATCTTCAT | 187 |
| chr8:15994968-15996990 | p3 | (TAA)6 | AAAGTGGTGCCACATGTCAA | TACACATGGCGCATCTTCAT | 187 |
| chr8:16044661-16046676 | p3 | (AGA)5 | GGTGGATGCCATCTTTGACT | TTCGCATAACCCTCCTCATC | 268 |
| chr8:16158594-16160609 | p5 | (TTGGT)3 | AAATAGTTGCACGTGGGAGC | AAACCTATGCCCAAAAAGCA | 101 |
| chr8:16404649-16406667 | p6 | (CCCACA)3 | CGGGCAGAATCAGCTCTAAG | AATTGGCTTGAAGATGGTGG | 187 |
| chr8:16434578-16436618 | p2 | (TA)20 | TCCCTTGTCTCCATCGTACC | CAATTATGGCTCAAACGGGT | 182 |
| chr8:16575930-16577956 | p2 | (TA)13 | CCCAATGTACACATCAAGCG | ATGCGTATGTGTGGGTAGCA | 119 |
| chr8:16598497-16600515 | p6 | (TTCCTG)3 | GGTCTAGTCCCATGTGTCCC | ACCTCTTGACCCCCTCCTTA | 242 |
| chr8:16729835-16731850 | p5 | (AAGCC)3 | GAAATGGAAAAGGATGCGAA | ACAGGTAAGTCAACGTCGGG | 238 |
| chr8:16863843-16865855 | p2 | (CT)20 | ACACATGGCCAATGTCAGAA | CACTTTCCGGACCACAGTTT | 196 |
| chr8:16863843-16865855 | p2 | (AT)6 | ACACATGGCCAATGTCAGAA | CACTTTCCGGACCACAGTTT | 196 |
| chr8:17002777-17004847 | p2 | (TA)35 | TCCAAAGTCCAAACCGAATC | TTCAACATCACGAGGTTTGC | 191 |
| chr8:17164570-17166582 | p2 | (GT)6 | GCTCGATTCGATAGTTTCGG | TGAAACATCGCCTTAACCAA | 150 |
| chr8:17227079-17229101 | p2 | (GT)11 | TCGCCATATGTGATCCTTGA | CTGTAGTCGCCACAACGAAA | 121 |
| chr8:17259578-17261592 | p5 | (TTGGA)3 | ACGGGTGGCACATAGACTTC | TCCAATCCAATCCAACATCA | 232 |
| chr8:17267994-17270009 | p5 | (TATGA)3 | GAGGAATTGGCATCCTCAAA | TTATGGCAATGCTCCAAACA | 224 |

| Marker ID | SSR type | SSR motif | FORWARD PRIMER (5'-3') | REVERSE PRIMER (5'-3') | size (bp) |
|---|---|---|---|---|---|
| chr8:17273637-17275651 | p5 | (TTGGA)3 | ACGGGTGGCACATAGACTTC | TCCAATCCAATCCAACATCA | 232 |
| chr8:17316851-17318863 | p2 | (AT)6 | TGATGAACCAAGGAGAAGCC | ATGGTGGGTTCCACTTTTTG | 196 |
| chr8:17328188-17330203 | p5 | (TATTG)3 | TACTAATGCCAGCCGCCTAT | GCACTTCACCCAAAACCAGT | 259 |
| chr8:17365910-17367934 | p6 | (TGGGGG)4 | AGCAGCAGGGAAACAACACT | GACGAATTGCTTGATGCTGA | 127 |
| chr8:17428971-17430983 | p2 | (AT)6 | CCTAGGACGGAATTGCTCTG | TTTGTTAGCCTCGTTGGCTT | 117 |
| chr8:17439616-17441632 | p2 | (TA)8 | AGCCGAAAAGTAAACGAGCA | AAACCATTCAGCCACAAAGG | 207 |
| chr8:17476310-17478328 | p2 | (AT)9 | TAAAATGCCCGATATTCCCA | TGTGTAGTGGCAGCCTTGTC | 115 |
| chr8:17476310-17478328 | p2 | (AT)6 | TAAAATGCCCGATATTCCCA | TGTGTAGTGGCAGCCTTGTC | 115 |
| chr8:17528360-17530372 | p2 | (TA)6 | GTGGATGACCCAATGTTTCC | GTATACCTGCCCGCTCACAT | 252 |
| chr8:17548740-17550755 | p3 | (GTT)5 | AGGCATGTCAAATCTCCACC | GGAAATGGCAAGAAAGGACA | 214 |
| chr8:17668970-17670985 | p5 | (TGGCC)3 | TGGAGATTTCCGTTGCAGTT | TCCCTTCGTAAGGGCATACA | 171 |
| chr8:17778861-17780876 | p5 | (ATTTT)3 | AGTCGCAGTGTAAAATGCCC | TTATACCTCGTTTGGGTCGC | 253 |
| chr8:17778861-17780876 | p5 | (TAGTT)3 | AGTCGCAGTGTAAAATGCCC | TTATACCTCGTTTGGGTCGC | 253 |
| chr8:17871352-17873373 | p3 | (ATT)7 | AACGCGCGTGCTTAAATTAT | TCAAACCCAGGAACTCCAAG | 268 |
| chr8:17941479-17943491 | p3 | (TAT)5 | GTCGAAGTGTGGGAGGGATA | AGACCTCAAAGCCTCGTTCA | 132 |
| chr8:17941493-17943511 | p3 | (TAT)5 | GTCGAAGTGTGGGAGGGATA | AGACCTCAAAGCCTCGTTCA | 118 |
| chr8:18006600-18008636 | p2 | (TA)11 | GGACCGGACCATAGGTTTCT | ATTGTCCTCCTTGCGAATTG | 159 |
| chr8:18006600-18008636 | p2 | (TC)7 | GGACCGGACCATAGGTTTCT | ATTGTCCTCCTTGCGAATTG | 159 |
| chr8:18161739-18163757 | p6 | (TCTGCT)3 | CCGTAAAACTGCCCCAGTAA | ACGGTCCGTATAACCTGTCG | 193 |
| chr8:18194146-18196191 | p3 | (ATC)5 | AGACCAGCAGCCTCTTCAAA | AGGAACCTATTGTGGTCCCC | 252 |
| chr8:18194188-18196212 | p3 | (ATC)5 | AGACCAGCAGCCTCTTCAAA | AGGAACCTATTGTGGTCCCC | 210 |
| chr8:18194252-18196279 | p3 | (ATC)5 | AGACCAGCAGCCTCTTCAAA | AGGAACCTATTGTGGTCCCC | 146 |
| chr8:18194276-18196294 | p3 | (ATC)5 | AGACCAGCAGCCTCTTCAAA | AGGAACCTATTGTGGTCCCC | 122 |
| chr8:18218162-18220184 | p5 | (ATCTC)3 | GCTAGCAGTTTTGCAAAGGG | TGACTATGTTGGCCACTGGA | 198 |
| chr8:18218162-18220184 | p2 | (AT)11 | GCTAGCAGTTTTGCAAAGGG | TGACTATGTTGGCCACTGGA | 198 |
| chr8:18219732-18221756 | p2 | (AT)12 | GCTCCATCATTTAAACCCGA | TCGTTGTGTTGCAGGATGTT | 259 |
| chr8:18307036-18309051 | p3 | (TGG)5 | ATTTTTGCCCGTCTTCACAC | TGTGGATCCAGGTACGATCA | 100 |
| chr8:18344800-18346816 | p2 | (AT)9 | TATGGATGTTTGGGGAGAGC | CAAGCTAAAGCCGGATAACG | 128 |
| chr8:18345012-18347028 | p2 | (AT)9 | GCCGATGTTGTTATAACCCG | GCTCCGACTTTGAGTTTTGG | 151 |
| chr8:18356402-18358417 | p5 | (ATTTC)3 | AATTTCAGTCAGGCACCCAC | TACCCGCATTAGAAGATGGG | 156 |
| chr8:18359209-18361221 | p2 | (AT)6 | AGCTACATTCTCTTGCCCGA | GGGACCCGATTACCCTAAGA | 199 |
| chr8:18361854-18363870 | p6 | (AACCAC)3 | TCATCCTCTCCCCACAAAAC | GATGGGAGGAGAAGGAAAGG | 240 |
| chr8:18361854-18363870 | p4 | (ATTT)4 | TCATCCTCTCCCCACAAAAC | GATGGGAGGAGAAGGAAAGG | 240 |
| chr8:18381171-18383185 | p2 | (AG)7 | TTAAACAAGGTGAGGCAGGC | TTCTCCTTGAACTGCCACCT | 161 |
| chr8:18396116-18398131 | p5 | (GGGTT)3 | ATTGGGGTGTTCCAACAAAA | GAATGGCCCAGACTCCATTA | 140 |
| chr8:18469049-18471069 | p5 | (GTTTA)4 | CCGGTCAGCCCATACTCTAA | AATCTGCCATTGTTGAAGGC | 157 |

| Marker ID | SSR type | SSR motif | FORWARD PRIMER (5'-3') | REVERSE PRIMER (5'-3') | size (bp) |
|---|---|---|---|---|---|
| chr8:18591876-18593891 | p3 | (AGA)5 | GGTTACCAGCACGTTGGTTT | ATTCCCTATGGGATTCGACC | 175 |
| chr8:18596796-18598814 | p6 | (CTGAAA)3 | TTCGTGGAACTCCCAAAAAC | CTTTGCATTCCATGGGTTCT | 227 |
| chr8:18629620-18631636 | p4 | (TCTT)4 | TTAAATGCCCCTGCTCACTT | AATTTCAGACCCAACCATGC | 150 |
| chr8:18657370-18659386 | p2 | (AT)12 | AATGAACCATAACCGCTTCC | AACCGTGTGGGTGTACATGA | 194 |
| chr8:18657405-18659421 | p2 | (AT)12 | AATGAACCATAACCGCTTCC | AACCGTGTGGGTGTACATGA | 159 |
| chr8:18657422-18659434 | p2 | (AT)12 | AATGAACCATAACCGCTTCC | AACCGTGTGGGTGTACATGA | 142 |
| chr8:18657445-18659460 | p2 | (AT)12 | AATGAACCATAACCGCTTCC | AACCGTGTGGGTGTACATGA | 119 |
| chr8:18859535-18861559 | p2 | (AT)17 | GCCACTTGGCAATTTTAGGA | TCAGCCGTGGGTTTGTAGTT | 127 |
| chr8:18859535-18861559 | p2 | (TA)12 | GCCACTTGGCAATTTTAGGA | TCAGCCGTGGGTTTGTAGTT | 127 |
| chr8:18971195-18973210 | p5 | (AAAAT)3 | TACCCCATCTGGCAGACATT | ACCAGCAACTAAATGGCACC | 275 |
| chr8:18992637-18994695 | p2 | (AT)29 | AGCAAACCTTGGTGGCTCTA | TTCACAGACTAGGGTTGGGG | 113 |
| chr8:18992637-18994695 | p5 | (ATCCA)3 | AGCAAACCTTGGTGGCTCTA | TTCACAGACTAGGGTTGGGG | 113 |
| chr8:18993177-18995192 | p2 | (AT)29 | AGCTCATCGTACCGATCCAC | CATTGACTGGGTTGAGGCTT | 234 |
| chr8:18993177-18995192 | p5 | (ATCCA)3 | AGCTCATCGTACCGATCCAC | CATTGACTGGGTTGAGGCTT | 234 |
| chr8:19014894-19016909 | p5 | (AAAAT)3 | TACCCCATCTGGCAGACATT | CCAGCAACTAAATGGCACCT | 276 |
| chr8:19030587-19032602 | p2 | (AT)30 | AGCTCATCGTACCGATCCAC | AAGGTAAGTGCCCACACGAC | 271 |
| chr8:19030587-19032602 | p5 | (ATCCA)3 | AGCTCATCGTACCGATCCAC | AAGGTAAGTGCCCACACGAC | 271 |
| chr8:19057347-19059361 | p3 | (TTA)5 | CTCCCCAAACTCCATGAAAA | CCATGTTTTTGCTGTGCAAC | 125 |
| chr8:19099055-19101067 | p2 | (TA)6 | TGCAGCTTTTGGCTTCTCTT | CGTGCATCTTTTGAACCAGA | 140 |
| chr8:19133582-19135600 | p2 | (TA)9 | TCAACTTATGGCCAACACCA | GTTTTCACCCGGAGGAATTT | 228 |
| chr8:19166056-19168078 | p2 | (AT)11 | ATCCCGCTACAGTTGTCCAG | GGCCACAGGTGAGTTCGTAT | 240 |
| chr8:19192493-19194505 | p2 | (AT)6 | GAAACCATGCTTTTTCCCAA | TGCGACTCTGCTAGGGATTT | 292 |
| chr8:19233275-19235293 | p6 | (CTCGGA)3 | TCGGGTACATCCCGTATCAT | GGGCTCTGTTTCGTCTCTTG | 222 |
| chr8:19251292-19253316 | p4 | (AAAC)6 | CATTTTTCGCAGCAACTGAA | AATCACTGCCCACTTTGAGG | 118 |
| chr8:19272621-19274645 | p2 | (TA)12 | ATGCATGAATTGTGGCTTCA | TGGATTCAGATCATGGTGGA | 116 |
| chr8:19272621-19274645 | p5 | (CCCCA)3 | ATGCATGAATTGTGGCTTCA | TGGATTCAGATCATGGTGGA | 116 |
| chr8:19273422-19275437 | p2 | (TA)12 | CGGGCCTCTCCTATCTTCTT | GCCCAAATCGGTTGTAGAAA | 225 |
| chr8:19273422-19275437 | p5 | (CCCCA)3 | CGGGCCTCTCCTATCTTCTT | GCCCAAATCGGTTGTAGAAA | 225 |
| chr8:19273422-19275437 | p2 | (TA)8 | CGGGCCTCTCCTATCTTCTT | GCCCAAATCGGTTGTAGAAA | 225 |
| chr8:19273422-19275437 | p2 | (TA)28 | CGGGCCTCTCCTATCTTCTT | GCCCAAATCGGTTGTAGAAA | 225 |
| chr8:19274173-19276189 | p5 | (CCCCA)3 | GCGAATCATCCCCTTAATGA | ATCCATGCCATTCCACCTTA | 275 |
| chr8:19274173-19276189 | p2 | (TA)8 | GCGAATCATCCCCTTAATGA | ATCCATGCCATTCCACCTTA | 275 |
| chr8:19274173-19276189 | p2 | (TA)34 | GCGAATCATCCCCTTAATGA | ATCCATGCCATTCCACCTTA | 275 |
| chr8:19478569-19480581 | p2 | (TA)6 | TAATATGATGATGGGGCGGT | GAGGGACGCGGTAAATCATA | 140 |
| chr8:19532513-19534528 | p5 | (TTCGG)3 | TCATGAATTCAAAAGCAGCG | CGGTATTCGGTGTCCACTTT | 175 |
| chr8:19544657-19546729 | p3 | (ATT)24 | GATGCGTGTAAGGAAAGGGA | CAAATACATGAGCATGCGGT | 163 |

| Marker ID | SSR type | SSR motif | FORWARD PRIMER (5'-3') | REVERSE PRIMER (5'-3') | size (bp) |
|---|---|---|---|---|---|
| chr8:19551013-19553029 | p2 | (TA)8 | TCAGTGAGAGCCCCAGAGAT | CCCATTAACCCCCTATGCTT | 203 |
| chr8:19562181-19564197 | p2 | (TA)8 | CTTGGTTTCCTTGATTCCGA | TCCATCTTCCCTGTGGATTC | 215 |
| chr8:19575610-19577625 | p5 | (GGCCC)3 | ATAGGCGACATAGGTGTGGG | TCATTAGCGGGACTAGTGGG | 255 |
| chr8:19575610-19577625 | p2 | (AT)8 | ATAGGCGACATAGGTGTGGG | TCATTAGCGGGACTAGTGGG | 255 |
| chr8:19663777-19665798 | p3 | (TTC)7 | TGCTCTCGAAATCACCAATG | TGCTCAATCAACCACGTCAT | 263 |
| chr8:19667808-19669823 | p3 | (TCT)5 | TGGCGGACATGTTCACTAAA | ATACGCATTGCTAGCTGCCT | 169 |
| chr8:19738934-19740946 | p2 | (CT)6 | TTTCTAATGGCTCGTCCACC | GATACTCCGCCTCTGTGCTC | 206 |
| chr8:19849185-19851203 | p6 | (CATATA)3 | TCCAATACTCAGCCCCAAAC | TCGAATCTGGACCATTGACA | 239 |
| chr8:19883379-19885394 | p5 | (TCACG)3 | GGACTCAAGAGGTGGAGCTG | TTGGCTGATGATGAGGATGA | 183 |
| chr8:19883379-19885394 | p2 | (AT)8 | GGACTCAAGAGGTGGAGCTG | TTGGCTGATGATGAGGATGA | 183 |
| chr8:19907171-19909261 | p3 | (ATA)30 | GCCAAAAATGGAGGAACAAA | TGACTGGTTCCACAACCAAA | 278 |
| chr8:20033380-20035406 | p2 | (AT)13 | CAGGCAGGGTGGTCAATACT | TCCAATGTGAATACCCGACA | 206 |
| chr8:20117429-20119459 | p2 | (TA)15 | GAAGCTCACCCTTGGAACTG | ACCTTGTTGCACCCTTTTTG | 251 |
| chr8:20126599-20128614 | p5 | (GCATT)3 | ATTTTCTGGTGTCTGGTGCC | AACACAACGGGGTGACTAGC | 255 |
| chr8:20168179-20170191 | p2 | (TA)6 | GAGGGAAAGAGGGAGTACGG | AGTATCACCGGTTGGCTTTG | 170 |
| chr8:20168179-20170191 | p2 | (AT)15 | GAGGGAAAGAGGGAGTACGG | AGTATCACCGGTTGGCTTTG | 170 |
| chr8:20168179-20170191 | p2 | (AT)11 | GAGGGAAAGAGGGAGTACGG | AGTATCACCGGTTGGCTTTG | 170 |
| chr8:20231373-20233385 | p2 | (AT)6 | AAATAAACGGTCTCCTGGGG | CCTTGTCTGCTTGCGTATCA | 265 |
| chr8:20231373-20233385 | p2 | (AT)6 | AAATAAACGGTCTCCTGGGG | CCTTGTCTGCTTGCGTATCA | 265 |
| chr8:20231503-20233515 | p2 | (AT)6 | AAATAAACGGTCTCCTGGGG | CCTTGTCTGCTTGCGTATCA | 135 |
| chr8:20231503-20233515 | p2 | (AT)6 | AAATAAACGGTCTCCTGGGG | CCTTGTCTGCTTGCGTATCA | 135 |
| chr8:20231503-20233515 | p3 | (ACA)5 | AAATAAACGGTCTCCTGGGG | CCTTGTCTGCTTGCGTATCA | 135 |
| chr8:20380977-20382992 | p3 | (CTT)7 | CTTATGCCAAGTCTGCCCAT | GCAGAATTGCCCTTCTTTTG | 201 |
| chr8:20380991-20383011 | p3 | (CTT)7 | CTTATGCCAAGTCTGCCCAT | GCAGAATTGCCCTTCTTTTG | 187 |
| chr8:20389468-20391483 | p5 | (CTTTT)3 | GACCTCTCAAAGCAAAACCG | CTCTCTGATTGTGCCGTGAA | 280 |
| chr8:20429703-20431735 | p2 | (TA)16 | CAAACTGGAGGAGGTGCATT | GACCCGCAAGGTTTTATTGA | 145 |
| chr8:20557055-20559067 | p6 | (AAAGAA)3 | GAAGAAGAGGAGGAGAGGGG | ATACACTAGCCCGCCACATC | 134 |
| chr8:20557055-20559067 | p2 | (CT)7 | GAAGAAGAGGAGGAGAGGGG | ATACACTAGCCCGCCACATC | 134 |
| chr8:20557055-20559067 | p2 | (TG)6 | GAAGAAGAGGAGGAGAGGGG | ATACACTAGCCCGCCACATC | 134 |
| chr8:20562866-20564881 | p6 | (TAAAAA)3 | AACACAGCCTTAAACGCCAC | GCTGAGCACAACCTTCATCA | 244 |
| chr8:20562866-20564881 | p3 | (TCT)5 | AACACAGCCTTAAACGCCAC | GCTGAGCACAACCTTCATCA | 244 |
| chr8:20691822-20693840 | p6 | (TCAGTT)3 | GCTTTATGCCCACCAAGTGT | ATTCCAGCCTTTGAGCGTAA | 278 |
| chr8:20747032-20749066 | p2 | (TA)17 | ATGTGAATGCCAGAGGGTTC | GCAGACAACCCCTAAGACGA | 292 |
| chr8:20815701-20817716 | p3 | (ATG)5 | TACACCCCGAGGATTAGTGC | TTAGAGTTTCCAGCGGCAGT | 204 |
| chr8:20882878-20884894 | p2 | (TA)8 | ACTTAGCAGATGGAAGCGGA | TGATGAGGAGTGTTGTCCCA | 273 |
| chr8:20913757-20915775 | p3 | (TTC)6 | TTGCTGCACTATGCAGATCC | GAGGACCCAACATCAAGGAA | 288 |

| Marker ID | SSR type | SSR motif | FORWARD PRIMER (5'-3') | REVERSE PRIMER (5'-3') | size (bp) |
|---|---|---|---|---|---|
| chr8:20930254-20932266 | p2 | (TA)6 | CGACTTTGTTGTACGGAGCA | TCCCGCATCTTATCTTCCAC | 121 |
| chr8:21174059-21176083 | p3 | (TAT)8 | CAGTGCCATGTAGGCTTGAA | CGATGTAGTCCGTCGGTTTT | 298 |
| chr8:21174059-21176083 | p5 | (ATTTT)3 | CAGTGCCATGTAGGCTTGAA | CGATGTAGTCCGTCGGTTTT | 298 |
| chr8:21187268-21189293 | p2 | (TA)14 | TCGGATTGGTTTGGTTTAGG | ATCAATAAAGCCGTGCGAAT | 188 |
| chr8:21187268-21189293 | p4 | (AATA)5 | TCGGATTGGTTTGGTTTAGG | ATCAATAAAGCCGTGCGAAT | 188 |
| chr8:21187522-21189547 | p2 | (TA)14 | GCACGGCTTTATTGATGTGA | GTTTTAGCTGGGATTCTTGCC | 177 |
| chr8:21187522-21189547 | p4 | (AATA)5 | GCACGGCTTTATTGATGTGA | GTTTTAGCTGGGATTCTTGCC | 177 |
| chr8:21210182-21212218 | p2 | (TA)18 | GGACCGACATTTAGCTCCAA | CCACCGTAGGCATTCTTGTT | 175 |
| chr8:21212554-21214582 | p2 | (AT)9 | GGCCTTCAATTTCTGCACAT | GCTGATGACCACCTACGGAT | 189 |
| chr8:21212591-21214607 | p2 | (AT)9 | GGCCTTCAATTTCTGCACAT | GCTGATGACCACCTACGGAT | 152 |
| chr8:21212615-21214629 | p2 | (AT)9 | GGCCTTCAATTTCTGCACAT | GCTGATGACCACCTACGGAT | 128 |
| chr8:21280653-21282674 | p3 | (AAG)7 | ATGTGGGACCTCATACGCTC | GGGGTCCAACTTATTGCCTT | 204 |
| chr8:21280653-21282674 | p2 | (TA)17 | ATGTGGGACCTCATACGCTC | GGGGTCCAACTTATTGCCTT | 204 |
| chr8:21322613-21324633 | p4 | (ACAA)5 | CGAACCTACTCGCACACTCA | TGGAGGTTAAACGAACTCGG | 130 |
| chr8:21366532-21368554 | p2 | (TA)16 | TGATGGATTCATTGCCACAT | GACATGGCGGAATAGCAAAT | 176 |
| chr8:21366532-21368554 | p2 | (AT)11 | TGATGGATTCATTGCCACAT | GACATGGCGGAATAGCAAAT | 176 |
| chr8:21439777-21441789 | p2 | (AT)6 | ATGGCAAGCTCCATTTGTTC | GACCGACAACCAATACCCAC | 130 |
| chr8:21443186-21445201 | p3 | (AAG)5 | CAACATGGGTGCTACACTGG | TGGACCATTGGCTTAGGTTC | 227 |
| chr8:21489704-21491720 | p5 | (ATTGC)3 | GGTTTCTGTCGGAAAGCGTA | GTACCACTCGCTCCGGAATA | 149 |
| chr8:21489704-21491720 | p4 | (TTAT)4 | GGTTTCTGTCGGAAAGCGTA | GTACCACTCGCTCCGGAATA | 149 |
| chr8:21503714-21505729 | p3 | (CAT)5 | TCATAACCTGAAAGGGACGC | GTCCTGTCCCGTTTTTACGA | 183 |
| chr8:21511150-21513168 | p2 | (TA)6 | GAGGTAAGGGGTGGAAGGAG | GGGGAACTTAAACTCCCAGC | 279 |
| chr8:21511150-21513168 | p2 | (TG)11 | GAGGTAAGGGGTGGAAGGAG | GGGGAACTTAAACTCCCAGC | 279 |
| chr8:21511221-21513239 | p2 | (TA)6 | GAGGTAAGGGGTGGAAGGAG | GGGGAACTTAAACTCCCAGC | 208 |
| chr8:21511221-21513239 | p2 | (TG)11 | GAGGTAAGGGGTGGAAGGAG | GGGGAACTTAAACTCCCAGC | 208 |
| chr8:21511525-21513557 | p2 | (TA)6 | AGGAGCCTTTTTCCCTTTCA | TTAACCACTCCAAACCGGAC | 123 |
| chr8:21511525-21513557 | p2 | (TG)11 | AGGAGCCTTTTTCCCTTTCA | TTAACCACTCCAAACCGGAC | 123 |
| chr8:21559129-21561145 | p2 | (CA)8 | AGGAGCCACGAACTACTCCA | TTTTGAGTAGCCTTTCCCGA | 113 |
| chr8:21723625-21725657 | p2 | (TA)16 | AAAGCCGCTAACGTGACACT | AACTCTTGCCAAAGGAGCAA | 163 |
| chr8:21723625-21725657 | p5 | (ATTTT)4 | AAAGCCGCTAACGTGACACT | AACTCTTGCCAAAGGAGCAA | 163 |
| chr8:21780891-21782915 | p6 | (AATAAC)4 | GCTAAATGGCGGGTGTTTTA | ACTTTGGAGAATCCCACACG | 226 |
| chr8:21852516-21854528 | p2 | (CA)6 | TTTGAGACCCGACCGTTAAG | AGCTACAGCCACAGCAGGAT | 182 |
| chr8:21927347-21929369 | p2 | (TA)11 | AAACCTCGCTCGAGTTCAAA | AGGGTCTTGGTTGTCAGTCG | 165 |
| chr8:21961068-21963106 | p2 | (TA)19 | CCATTGCTTCTGTGCGACTA | TATATTGGGCCTTTTCTGCG | 149 |
| chr8:22004318-22006354 | p3 | (TCT)5 | ATAACCTGACTCGTGGTCCG | AAGAGGAACCGGAGGAAAAA | 170 |
| chr8:22004318-22006354 | p3 | (TAA)12 | ATAACCTGACTCGTGGTCCG | AAGAGGAACCGGAGGAAAAA | 170 |

| Marker ID | SSR type | SSR motif | FORWARD PRIMER (5'-3') | REVERSE PRIMER (5'-3') | size (bp) |
|---|---|---|---|---|---|
| chr8:22057335-22059379 | p5 | (AAAAT)3 | AAAGTGGAACCCACCACATC | GAATACGCACGGCGTTTAAT | 148 |
| chr8:22057335-22059379 | p2 | (AT)6 | AAAGTGGAACCCACCACATC | GAATACGCACGGCGTTTAAT | 148 |
| chr8:22156503-22158521 | p6 | (GTGGAT)3 | TTCAAGGACCAAGTTACCCG | ATTCATGTCCACTGTTGCGA | 110 |
| chr8:22315915-22317941 | p2 | (TA)13 | TACGGTACAAGTTACGGCCC | AATTTCGGCACATTTTACGC | 126 |
| chr8:22315915-22317941 | p6 | (AACAAA)4 | TACGGTACAAGTTACGGCCC | AATTTCGGCACATTTTACGC | 126 |
| chr8:22354562-22356577 | p5 | (TTCAT)3 | ATTTGAGCTTCGTCAATGGC | TGCATGCCAATTAGTTCGAG | 149 |
| chr8:22354562-22356577 | p5 | (TAATT)3 | ATTTGAGCTTCGTCAATGGC | TGCATGCCAATTAGTTCGAG | 149 |
| chr8:22493972-22495987 | p5 | (TAAAA)3 | TGGAATTAGCCACCATAGCC | CTTTGGATGCCACGGTACTT | 149 |
| chr8:22495869-22497887 | p5 | (AAAAT)3 | GACCGCATGAGGTTGAGATT | TATCCGGTGGTGGGTGTATT | 286 |
| chr8:22495869-22497887 | p6 | (AAATTG)3 | GACCGCATGAGGTTGAGATT | TATCCGGTGGTGGGTGTATT | 286 |
| chr8:22550222-22552244 | p2 | (TA)6 | CATTGTGGGCCCCATATTTA | GCACGTGGCAATTAATCCAT | 160 |
| chr8:22550222-22552244 | p2 | (TA)11 | CATTGTGGGCCCCATATTTA | GCACGTGGCAATTAATCCAT | 160 |
| chr8:22550222-22552244 | p2 | (TA)18 | CATTGTGGGCCCCATATTTA | GCACGTGGCAATTAATCCAT | 160 |
| chr8:22602209-22604227 | p6 | (TTTATT)3 | TCAGATGAGCAAGTGTTGGC | GTCTTGGAAGCAATCCGGTA | 244 |
| chr8:22614861-22616879 | p2 | (AT)8 | CGGTTAAAAACCGGAAACTG | TTTAACCACTCCGAACCGAC | 126 |
| chr8:22614861-22616879 | p6 | (ATATAA)3 | CGGTTAAAAACCGGAAACTG | TTTAACCACTCCGAACCGAC | 126 |
| chr8:22614861-22616879 | p6 | (CCGGTT)6 | CGGTTAAAAACCGGAAACTG | TTTAACCACTCCGAACCGAC | 126 |
| chr8:22626219-22628231 | p2 | (TA)6 | GAAACAAAGGAGCCATTGGA | CATTCACTATCTTCGCGGGT | 142 |
| chr8:22857776-22859791 | p5 | (AAGTC)3 | TTCTCACCAAACACAACCCA | TGTGGGACAATTCCCAATTT | 236 |
| chr8:22875710-22877722 | p2 | (AT)6 | AACTCCATGTTGAACCCCAA | GTTGTGCCAGGTTCTTCCAT | 272 |
| chr8:22932019-22934034 | p5 | (AAACC)3 | TGAGGAGGAGAAATGCTGGT | GTTGCTAGTCGTGCCAACAA | 278 |
| chr8:22938733-22940754 | p3 | (TCA)7 | AGCTCTTGGAGGTTGCTTGA | CAATGTGAAGGGTGCCTTTT | 209 |
| chr8:22968534-22970549 | p2 | (AT)21 | ACCTCGTCATGCCCATGTAT | ATTTGTGGATGTTGCGTTCA | 296 |
| chr8:23029551-23031569 | p2 | (TA)9 | TCCCTTTTTAGGAGGGTGCT | CCTTGTCGGCTTGCATATTT | 119 |
| chr8:23057542-23059554 | p2 | (AC)6 | TACATACCTTGCCCGCTTTC | TTTGGCCGAGTTGAATTCTC | 290 |
| chr8:23127062-23129077 | p5 | (ATTGC)3 | TTATTGATGTTCGGCCTTCC | CCAACTTCACACCCGAAAGT | 244 |
| chr8:23280879-23282894 | p3 | (CAA)5 | CGGACGGAATCATAGCAAGT | TTGCTGACTCCCTCGAGTTT | 136 |
| chr8:23329197-23331215 | p6 | (CCACCT)3 | CCTATCGGACAATGGAAGGA | AACAAAAGGCAGCTGAGGAA | 104 |
| chr8:23448928-23450943 | p5 | (AACCG)3 | TCACTTGGAAATAGACCGGG | GAAGGTTCGAGCTTTTGACG | 193 |
| chr8:23448928-23450943 | p5 | (GAACC)3 | TCACTTGGAAATAGACCGGG | GAAGGTTCGAGCTTTTGACG | 193 |
| chr8:23463079-23465094 | p5 | (ATTTC)3 | AAGAAATGACATGGGAAGCG | ACAATGCTTCATAGGGCCAC | 161 |
| chr8:23506718-23508799 | p3 | (ATA)27 | CCATTCGCAGCCTTGATATT | AAAATCATTGCAAAGGGCTG | 153 |
| chr8:23519115-23521139 | p5 | (CAATG)3 | ATGCAATGCAACAGGATTGA | TGAGGTCATATACCGGAGGG | 116 |
| chr8:23519115-23521139 | p2 | (TA)12 | ATGCAATGCAACAGGATTGA | TGAGGTCATATACCGGAGGG | 116 |
| chr8:23542765-23544813 | p3 | (AAG)16 | CTTATTCACCACAAAGCGCA | AGCACTACTCCGGAGGTGAA | 217 |
| chr8:23911484-23913502 | p6 | (GGGGTG)3 | TCACCGAATCAGAAAGAGGG | TGGTGCCTTTGTACTCGACA | 166 |

| Marker ID | SSR type | SSR motif | FORWARD PRIMER (5'-3') | REVERSE PRIMER (5'-3') | size (bp) |
|---|---|---|---|---|---|
| chr8:23967976-23970004 | p4 | (TTTA)7 | AGTTTCTCAAGAACGGGCAA | ATAGCAATTGACCCGGACAG | 250 |
| chr8:24083197-24085213 | p2 | (TA)9 | ACTGCATTATCCAACCGACC | GAGGGACCGGGGAATAAATA | 245 |
| chr8:24083197-24085213 | p2 | (AT)8 | ACTGCATTATCCAACCGACC | GAGGGACCGGGGAATAAATA | 245 |
| chr8:24090472-24092492 | p2 | (AT)10 | AAGATCAGGACGGGTTGATG | TCCCCTTGAAGTCGAATGAC | 174 |
| chr8:24090472-24092492 | p3 | (CAT)6 | AAGATCAGGACGGGTTGATG | TCCCCTTGAAGTCGAATGAC | 174 |
| chr8:24102160-24104175 | p5 | (TTTCT)3 | ACATGCCAAAAGAAAGACGG | TTGGAAATTCGAGTTAGGCG | 212 |
| chr8:24154925-24156941 | p2 | (AT)8 | CCTCAGCCGACGTGTTTTAT | GACCCTTACTTAGGGCCGAG | 142 |
| chr8:24178952-24180979 | p3 | (GTT)9 | CCAGTTAGTAGCCGCTCAGG | AAAATATGGGAGCGCACAAG | 150 |
| chr8:24315311-24317351 | p2 | (AT)20 | TGACTACGCCTTTGTGTTGC | TGATGGGTCGACACTGTGAT | 156 |
| chr8:24385563-24387583 | p5 | (CATAT)4 | GTAGACACTCCCAGGACGGA | TGGTTTGGGATTATCGGGTA | 249 |
| chr8:24385563-24387583 | p3 | (CAA)5 | GTAGACACTCCCAGGACGGA | TGGTTTGGGATTATCGGGTA | 249 |
| chr8:24503595-24505607 | p2 | (CA)6 | TTGACTTGGGTGATCCAACA | GTGCACATGGGCTACTCTGA | 300 |
| chr8:24550682-24552697 | p5 | (TTTTC)3 | AAGTGACCGTCAAATCACCC | GATCAAACGCAGTGACGAGA | 267 |
| chr8:24602160-24604172 | p2 | (TA)6 | TGATATGTTTCCGAGTGGCA | TGCAGTGAAACTGTGCATGA | 300 |
| chr8:24607421-24609437 | p4 | (AATT)4 | AGAATACGGACCGTCACAGC | GCATGCCTCTGGTAACCTGT | 154 |
| chr8:24607421-24609437 | p5 | (ATTTC)3 | AGAATACGGACCGTCACAGC | GCATGCCTCTGGTAACCTGT | 154 |
| chr8:24648034-24650058 | p6 | (TATACA)4 | TTCGTAGAATTCCCGAGTGC | TGAGAACGACGAAACGAATG | 114 |
| chr8:24771220-24773232 | p2 | (TG)6 | CGCGGGAAGATAGTAACCAA | TGCAAATTTCCCTTTATGCC | 230 |
| chr8:24803301-24805313 | p2 | (TG)6 | CGCGGGAAGATAGTAACCAA | TGCAAATTTCCCTTTATGCC | 230 |
| chr8:24949778-24951850 | p2 | (AT)36 | CGAGCAAGGGGATGAAATTA | TACGATAAGAGGTGCCCAGG | 201 |
| chr8:25085455-25087473 | p2 | (TA)9 | AAGCTCGAAGGTCAACCGTA | ATTGCCAGGAATACCTGCTG | 260 |
| chr8:25113134-25115146 | p2 | (GT)6 | TTTCCTGCACCCACATGTAA | CATCAATGATTCCACGAACG | 184 |
| chr8:25239385-25241399 | p2 | (AT)7 | AAAGAAGAATCGGGATGGCT | GCCGGCATATTTCTGTAGGA | 255 |
| chr8:25253221-25255235 | p2 | (AT)7 | CCTCTTTCGCACCGTTACTC | GAGCTGCGAGGACATTTAGG | 289 |
| chr8:25253618-25255632 | p2 | (AT)7 | GCACACCCGTATACGAACCT | GGGCTTGCGGTATCATTTTA | 225 |
| chr8:25253634-25255650 | p2 | (AT)7 | GCACACCCGTATACGAACCT | GGGCTTGCGGTATCATTTTA | 209 |
| chr8:25272141-25274156 | p3 | (TTA)5 | TGTTCATGCCCACAGGTAGA | ACCCACACTGCGTATCAACA | 196 |
| chr8:25419369-25421381 | p2 | (AT)6 | CGCCACATCCCAGAGTCTAT | GTTGCTTCGGGTCAACAAAT | 147 |
| chr8:25501416-25503438 | p2 | (TG)11 | GCCAGCATGTATCACTCCCT | TGGAGCAGGATCACCTCTCT | 154 |
| chr8:25561391-25563409 | p2 | (AT)9 | ATCCCATTGGGCAATCAATA | AAGCTCCATTTCATTCGGTG | 214 |
| chr8:25672715-25674745 | p2 | (AT)15 | TTTTTGGAACGTTGGAGGAC | GTCGGCTTGCGTATCGTATT | 233 |
| chr8:25711378-25713390 | p2 | (AC)6 | ATAGGGCCCTTAGGCTTCAA | TGCAGATTTCAGACGAAACG | 146 |
| chr8:25908032-25910047 | p5 | (ATTTC)3 | TGGGAAGCAGAAGAAAGCAT | ATAACAGTGCCTCACAGGGC | 181 |
| chr8:26126869-26128884 | p3 | (CAA)5 | GAGGCATCCACGAACAGATT | TGACATCTGGAACGCTTCTG | 176 |
| chr8:26137592-26139607 | p5 | (TCATT)3 | GACGTGGGAAGTGGAAGAAA | ACAATGCTTTACAGGGCCAC | 170 |
| chr8:26153219-26155234 | p3 | (CAA)5 | GAGGCATCCACGAACAGATT | TGACATCTGGAACGCTTCTG | 176 |

| Marker ID | SSR type | SSR motif | FORWARD PRIMER (5'-3') | REVERSE PRIMER (5'-3') | size (bp) |
|---|---|---|---|---|---|
| chr8:26163942-26165957 | p5 | (TCATT)3 | GACGTGGGAAGTGGAAGAAA | ACAATGCTTTACAGGGCCAC | 170 |
| chr8:26244726-26246746 | p2 | (AC)10 | TATCCATCAAATGTGGCCCT | CCCCGGTATGGAAAGAATTT | 140 |
| chr8:26268820-26270832 | p2 | (AT)6 | CTAGGTCGCATAGTCTCCGC | CCCAACTCCTTTCAAGACCA | 157 |
| chr8:26268820-26270832 | p6 | (CATATA)3 | CTAGGTCGCATAGTCTCCGC | CCCAACTCCTTTCAAGACCA | 157 |
| chr8:26268820-26270832 | p2 | (AT)6 | CTAGGTCGCATAGTCTCCGC | CCCAACTCCTTTCAAGACCA | 157 |
| chr8:26273050-26275065 | p3 | (TGT)5 | AACTTTGGGGAAAAGTTGGG | CCTTGCTCTGGTAGCCTGTC | 264 |
| chr8:26296061-26298076 | p5 | (CATAT)3 | GCAAATCGGGGTCTTAACAA | AGTGCTAAGAGATTCCGCGA | 244 |
| chr8:26351077-26353103 | p2 | (AT)13 | AGCCAACGAGGCTATCAAAA | CTACATATGGGTATGGCGGG | 103 |
| chr8:26361829-26363859 | p2 | (TA)15 | TAGCTCCGGATGAAATGGAG | TGGCCACTAGGGGTACAAAG | 271 |
| chr8:26415732-26417746 | p2 | (TC)7 | TTACATACCTTGCCCGCTTC | TAGATTCTCGGGGTTGGTTG | 139 |
| chr8:26442259-26444395 | p2 | (AT)68 | ATCACAGACAATGGTGCCAA | AAGCTGTTCATGCCAATTCC | 264 |
| chr8:26450465-26452480 | p5 | (TATTT)3 | TTTGACATGGCTTTCGTGAG | ACATAAGCCCGGATACAACG | 237 |
| chr8:26475102-26477114 | p2 | (TA)6 | ATTTTGATTTTTGGCGTTGC | GCCGTACCCCTGAACAAATA | 245 |
| chr8:26503833-26505849 | p2 | (AT)8 | AGCAAACATTGGCTGAGCTT | TGAGAGTCAGAGGGTTCGCT | 194 |
| chr8:26521933-26523945 | p2 | (TG)6 | GGTCTGTAGAATGGACCCGA | CCAAATCCCAAATTTTCCCT | 257 |
| chr8:26529735-26531765 | p2 | (AT)15 | ATGGACAAGAGTGGCCAAAG | TGTCGCAGCAGTAATTCAGG | 286 |
| chr8:26529735-26531765 | p4 | (TCAT)4 | ATGGACAAGAGTGGCCAAAG | TGTCGCAGCAGTAATTCAGG | 286 |
| chr8:26585537-26587553 | p4 | (ATAA)4 | GCTATTTGCTTTGCCTTTGC | ATGCCATTGCAAAAATCTCC | 206 |
| chr8:26625278-26627294 | p4 | (ATAA)4 | TCACCTATGGCTGGTCCTTC | AAACTAACCGGGTCCCATTC | 189 |
| chr8:26642840-26644855 | p2 | (AT)10 | TGGAGCGCTTCTGAGATTTT | GCAGGTTCCGCAACACTATT | 253 |
| chr8:26642840-26644855 | p3 | (ATC)5 | TGGAGCGCTTCTGAGATTTT | GCAGGTTCCGCAACACTATT | 253 |
| chr8:26863752-26865766 | p2 | (AT)7 | AGGCGAACCCTTTCTCAAAT | CCTTGGACCAGATTCCTTGA | 233 |
| chr8:26936733-26938747 | p2 | (AT)7 | AGTGATGCCAAGGAGGTCAC | TTGGCGGCTGCTAAACTACT | 111 |
| chr8:26940983-26942998 | p5 | (TGTTT)3 | CATCGGTATGCATGATTTGC | AACGGGATGTTCCTAAGGCT | 246 |
| chr8:26969689-26971704 | p5 | (ATTGC)3 | TTATTGTTCGGCTTTCCAGG | ACATGTCATCAAGCCACCAA | 294 |
| chr8:27022170-27024216 | p2 | (TA)23 | GTCGGGAACCACCTGACTTA | TTAACCTTGCCGGCTTTATG | 167 |
| chr8:27058310-27060330 | p2 | (AT)10 | AGTCCCGTTGGACAATCAAC | ATATATTTATGGGCACGGCG | 150 |
| chr8:27065862-27067890 | p2 | (AT)8 | GTAGACACCCCTAGGACCCC | AAAGTCTGTCAATGGGTGCC | 154 |
| chr8:27102066-27104082 | p3 | (GTT)6 | GAAACAGCCGTCCTACCTTG | TTCTTTGAAACTTGCGGCTT | 200 |
| chr8:27102066-27104082 | p4 | (CAAA)4 | GAAACAGCCGTCCTACCTTG | TTCTTTGAAACTTGCGGCTT | 200 |
| chr8:27102066-27104082 | p4 | (ATAA)4 | GAAACAGCCGTCCTACCTTG | TTCTTTGAAACTTGCGGCTT | 200 |
| chr8:27102066-27104082 | p4 | (TAAA)4 | GAAACAGCCGTCCTACCTTG | TTCTTTGAAACTTGCGGCTT | 200 |
| chr8:27102066-27104082 | p4 | (ATAA)4 | GAAACAGCCGTCCTACCTTG | TTCTTTGAAACTTGCGGCTT | 200 |
| chr8:27102066-27104082 | p5 | (AATAG)3 | GAAACAGCCGTCCTACCTTG | TTCTTTGAAACTTGCGGCTT | 200 |
| chr8:27102066-27104082 | p5 | (AATAA)3 | GAAACAGCCGTCCTACCTTG | TTCTTTGAAACTTGCGGCTT | 200 |
| chr8:27102066-27104082 | p5 | (AAATA)3 | GAAACAGCCGTCCTACCTTG | TTCTTTGAAACTTGCGGCTT | 200 |

| Marker ID | SSR type | SSR motif | FORWARD PRIMER (5'-3') | REVERSE PRIMER (5'-3') | size (bp) |
|---|---|---|---|---|---|
| chr8:27119877-27121899 | p2 | (TA)11 | AGCGCACAAAACACTCCTCT | AGCGTGTGTTGCACAGATTC | 257 |
| chr8:27119877-27121899 | p4 | (TTTA)5 | AGCGCACAAAACACTCCTCT | AGCGTGTGTTGCACAGATTC | 257 |
| chr8:27132998-27135028 | p2 | (AT)15 | AACTACGCACACACACCCAA | TTGCCAGGTCTTTTGGAATC | 270 |
| chr8:27200448-27202476 | p2 | (TA)6 | CACCTAGCACCCTTTACCCA | GTGTGAGTTGACTGGCTGGA | 110 |
| chr8:27369111-27371126 | p5 | (ACCCG)3 | AGATGTTGCTTTTGGCATCC | AAAAGAAATGACGTGGTCGG | 198 |
| chr8:27388535-27390550 | p3 | (TCT)5 | GCTCAGTTGCAGTCTCACCA | GAGGCTCTTGCTTATGCACC | 151 |
| chr8:27388535-27390550 | p3 | (ACA)10 | GCTCAGTTGCAGTCTCACCA | GAGGCTCTTGCTTATGCACC | 151 |
| chr8:27411362-27413377 | p5 | (GTTCG)3 | TCTTTAATTTGTGCCCCTGC | TTAGCGTTTAGCTTCGGCAT | 148 |
| chr8:27411362-27413377 | p6 | (AAATCA)3 | TCTTTAATTTGTGCCCCTGC | TTAGCGTTTAGCTTCGGCAT | 148 |
| chr8:27421569-27423581 | p2 | (CA)6 | AGCCTTACATACCTGTGCCG | GCCATAGCTTGGTTGTTGGT | 191 |
| chr8:27421569-27423581 | p2 | (AT)6 | AGCCTTACATACCTGTGCCG | GCCATAGCTTGGTTGTTGGT | 191 |
| chr8:27459349-27461369 | p2 | (AT)10 | CGCAGGCCCCATATAAGTAA | AAGCGATAAAGGATTGGCCT | 120 |
| chr8:27462528-27464548 | p2 | (AT)10 | CGCAGGCCCCATATAAGTAA | AAGCGATAAAGGATTGGCCT | 120 |
| chr8:27490060-27492087 | p3 | (GTT)9 | TTTGGATCAACTTTGGAGGG | TGTTGCACGTCGGTTACTGT | 166 |
| chr8:27490060-27492087 | p2 | (TA)20 | TTTGGATCAACTTTGGAGGG | TGTTGCACGTCGGTTACTGT | 166 |
| chr8:27493788-27495812 | p3 | (CAT)8 | TAAGCCATAAAACTTGGGCG | CGGCTCCTGAGCAGGTATAG | 271 |
| chr8:27522346-27524360 | p2 | (AG)7 | CCTACCAGCGGACTTAGCAG | AACACTTTAACCGACGCCAC | 168 |
| chr8:27537562-27539596 | p2 | (AT)17 | TCTGGACGGGTGTGTTTGTA | TCAGAAGCCGACAAGGCTAT | 230 |
| chr8:27581317-27583332 | p5 | (GAAAA)3 | CCCACAGGTAGACAGGGAGA | TATGCATGAGCTCACAAGGC | 157 |
| chr8:27608744-27610760 | p2 | (AC)8 | CCCGATCACATTTCCATAGG | GCAATTCTGCTCCTCGAAAC | 213 |
| chr8:27608744-27610760 | p2 | (TA)6 | CCCGATCACATTTCCATAGG | GCAATTCTGCTCCTCGAAAC | 213 |
| chr8:27641967-27643989 | p2 | (TA)11 | CCGATGAATCGTGATGAGTG | GCTGTGCTTTCAGCATGTGT | 281 |
| chr8:27646894-27648909 | p3 | (TGA)5 | GGTCAGTTTGCGTTGGAAAT | TAACATCTGCCCGTACCTCC | 245 |
| chr8:27662296-27664311 | p5 | (GCATT)3 | GCTGATTTAGGGAAATGGCA | ACTCATGGCATCAACCATCA | 185 |
| chr8:27665561-27667593 | p2 | (AT)16 | GGCTTGTTGGGTGCATATTT | AGACAATTGGGCAGCATTTC | 240 |
| chr8:27672985-27675023 | p2 | (TA)19 | GTATGAGTTGCGCGTGAAGA | GAGGATTCTCTGCTATGCCG | 269 |
| chr8:27690172-27692216 | p2 | (TA)22 | ACGCTGCCCAATTTTCTCTA | CCTCACATACCTTTGCCGTT | 280 |
| chr8:27690172-27692216 | p2 | (TA)33 | ACGCTGCCCAATTTTCTCTA | CCTCACATACCTTTGCCGTT | 280 |
| chr8:27732020-27734032 | p2 | (AT)6 | AATTCGTCGTTGCCTTCATC | GCCACTATAGTCGGGGCATA | 194 |
| chr8:27743360-27745372 | p2 | (TA)6 | GGATGTGATGTTCCCAAACC | CTAAATTGGCGCGGTGTAAT | 190 |
| chr8:27842477-27844489 | p2 | (TA)6 | AGGGGTCAGTACGAGCAATG | ATGGTCGGGCAGTTTACTTG | 177 |
| chr8:27853268-27855286 | p2 | (AT)9 | GCAGTCGCCCTAGAAGTGTC | CAGTAGCTTTTTGCCTTCGG | 118 |
| chr8:27860081-27862096 | p5 | (TATCA)3 | CCCCGACCTTACTAGGGTGT | TGTTCATGTCACGGGTCCTA | 280 |
| chr8:27915539-27917555 | p2 | (AT)6 | AACCATTTCTGGGTTGACCA | TGATGGGCATAATTGTCGAA | 222 |
| chr8:27915539-27917555 | p4 | (CATA)4 | AACCATTTCTGGGTTGACCA | TGATGGGCATAATTGTCGAA | 222 |
| chr8:27938038-27940068 | p2 | (AT)15 | ACTCCTTCCAAACGGTGATG | TTATTTAGCGCAACGGGAAG | 282 |

| Marker ID | SSR type | SSR motif | FORWARD PRIMER (5'-3') | REVERSE PRIMER (5'-3') | size (bp) |
|---|---|---|---|---|---|
| chr8:27938038-27940068 | p3 | (ATC)5 | ACTCCTTCCAAACGGTGATG | TTATTTAGCGCAACGGGAAG | 282 |
| chr8:28027912-28029924 | p2 | (TA)6 | GTAGCTCCGGATGAAATGGA | TACTATCCCCCGATTCTCCC | 268 |
| chr8:28089107-28091125 | p6 | (CATAAT)3 | CGAAGAGGACCTCGTGAGAC | CCATTATTCCGGAGGTGCTA | 213 |
| chr8:28121711-28123727 | p2 | (AT)7 | ACAACTGTCGACGAGCCTCT | ATCCTTTTGTTTGTGGACGC | 163 |
| chr8:28121711-28123727 | p2 | (TA)7 | ACAACTGTCGACGAGCCTCT | ATCCTTTTGTTTGTGGACGC | 163 |
| chr8:28121711-28123727 | p2 | (AT)8 | ACAACTGTCGACGAGCCTCT | ATCCTTTTGTTTGTGGACGC | 163 |
| chr8:28178678-28180734 | p2 | (AT)28 | CCGACAAGGCTACACCATCT | TCGGTACGCTCCATCTATCC | 267 |
| chr8:28182636-28184652 | p2 | (GT)8 | TGCCTCCATATCTTCAAGGC | ATTTCCAAATGCGCAAAAAC | 254 |
| chr8:28216148-28218162 | p2 | (TA)7 | AAATCTCACTGACCCGTTGG | TCTGGTTGGGTCGTGACATA | 121 |
| chr8:28302453-28304469 | p2 | (AT)8 | AGAGGCTACCAGGCCTTTTC | GGGCACTGTTGTTGATGATG | 239 |
| chr8:28302511-28304535 | p2 | (AT)8 | AGAGGCTACCAGGCCTTTTC | GGGCACTGTTGTTGATGATG | 181 |
| chr8:28376668-28378688 | p5 | (ATCAT)4 | AGATCTGTCACGACCCAACC | ATCTGTGATGGCCTTATCGG | 151 |
| chr8:28380873-28382888 | p5 | (TTACT)3 | CTTCTCGGACAAAATCCCAA | ACGCGCTTAGTCGTGAATCT | 257 |
| chr8:28479766-28481778 | p2 | (TA)6 | GGGGCCCTATCCCAATACTA | CCGAAAGAGCACCAACAAGT | 297 |
| chr8:28479766-28481778 | p2 | (TA)6 | GGGGCCCTATCCCAATACTA | CCGAAAGAGCACCAACAAGT | 297 |
| chr8:28490410-28492424 | p2 | (AT)11 | TATACACGCAAGCCGACAAG | TTGGGATTGCTCCTTTGTTC | 289 |
| chr8:28490410-28492424 | p2 | (AT)7 | TATACACGCAAGCCGACAAG | TTGGGATTGCTCCTTTGTTC | 289 |
| chr8:28532846-28534876 | p2 | (AT)8 | ATAAGCCGACGAGGCTAACA | TAGCGGACCTCCATTTCAGT | 293 |
| chr8:28532846-28534876 | p2 | (AT)15 | ATAAGCCGACGAGGCTAACA | TAGCGGACCTCCATTTCAGT | 293 |
| chr8:28534631-28536647 | p4 | (TTAA)4 | GGAGTGAAGGGGACATGAAA | AAGTTTGCCAAGAGAAGGCA | 230 |
| chr8:28613714-28615732 | p2 | (AT)9 | AGCCCTTCGGCTATCAAAAT | ATGCCCACAAGTTAGCAAGG | 225 |
| chr8:28636254-28638278 | p2 | (TA)12 | CGAAATCTTCCGCAACTCTC | CCTCGGAAAAGCAATGGTTA | 199 |
| chr8:28821022-28823034 | p2 | (CA)6 | TGCGTAACTGACAATGCCTC | TACGCACATACATCAGGCGT | 134 |
| chr8:28821022-28823034 | p2 | (AT)9 | TGCGTAACTGACAATGCCTC | TACGCACATACATCAGGCGT | 134 |
| chr8:28916440-28918458 | p2 | (AT)9 | ACGAGCAGTCGGAGTGAAAT | GCGATGTGTACCGCCTATCT | 208 |
| chr8:29003731-29005746 | p3 | (GTT)5 | GATTGGAGAACAAGCGAAGC | TGGCCAAACTTCTAGCATCC | 209 |
| chr8:29019250-29021262 | p2 | (AT)6 | GGGTGTCCTAAACTGGTGGA | AACTTCTGCTGGCTTTGCAT | 107 |
| chr8:29164508-29166536 | p2 | (AT)14 | TCTGAAGGGGGACATGAGAC | TTCATAGTTTCCCTCGGGTG | 251 |
| chr8:29416522-29418536 | p2 | (AT)7 | ACGAGCCTGGAGCTTTCATA | GAGCAGCAGAGCTGCCTTAT | 297 |
| chr8:29435716-29437731 | p3 | (TCT)5 | TCTCCAGCCTCGATGAAGAT | AGATACGTTCCATGCCTTCG | 113 |
| chr8:29458457-29460469 | p2 | (CA)6 | GGCTTTTGGTGACGACTTGT | AAAAGATTTGGGAGGGGCTA | 119 |
| chr8:29477616-29479631 | p3 | (TCT)5 | TCTCCAGCCTCGATGAAGAT | AGATACGTTCCATGCCTTCG | 113 |
| chr8:29507012-29509024 | p2 | (CA)6 | TGACCCAAAGAATTTCGGAG | CTGAGGAGTGTGAGGAAGCC | 191 |
| chr8:29511322-29513334 | p2 | (CA)6 | ACTCCCATAATCCCTCACCC | GGAGGTGCTTGGAATCAGAG | 170 |
| chr8:29511322-29513334 | p6 | (CAAAAA)3 | ACTCCCATAATCCCTCACCC | GGAGGTGCTTGGAATCAGAG | 170 |
| chr8:29683055-29685075 | p2 | (TA)10 | TCGCTCGAATAGGCATAGGT | GCACGATTGTCCATCATACG | 194 |

| Marker ID | SSR type | SSR motif | FORWARD PRIMER (5'-3') | REVERSE PRIMER (5'-3') | size (bp) |
|---|---|---|---|---|---|
| chr8:29685925-29687945 | p5 | (CAAAC)4 | GCCCTCAATTCTCCCATTTT | AGCATTTGCTGCAACTCAAA | 273 |
| chr8:29760252-29762282 | p3 | (CTC)8 | GGATGTGTCTCGAAGGTGGT | GCCAAAATAGAAGGAGCGTG | 216 |
| chr8:29760296-29762467 | p3 | (CTC)8 | GGATGTGTCTCGAAGGTGGT | GCCAAAATAGAAGGAGCGTG | 172 |
| chr8:29760652-29762730 | p3 | (CTC)8 | CGTAATTTCACGGCACATCA | AGAAGAGGAGGACGACGACA | 117 |
| chr8:29760787-29762853 | p3 | (CTC)8 | TTCTTCCTCGTTGTCGTCGT | GGAAGAGGAGGGAGAGAAGG | 136 |
| chr8:29856630-29858642 | p2 | (AT)6 | ATGACGAGGGTGTTTTGAGG | TTGTAACCCCCGGATGAATA | 162 |
| chr8:29900935-29902947 | p2 | (AT)6 | ATGACGAGGGTGTTTTGAGG | TTGTAACCCCCGGATGAATA | 162 |
| chr8:29933201-29935213 | p2 | (AT)40 | GGACATCCCGAGTTTCTTGA | TGGGAACTTCTTCCCCTTTT | 134 |
| chr8:29933201-29935213 | p2 | (AT)6 | GGACATCCCGAGTTTCTTGA | TGGGAACTTCTTCCCCTTTT | 134 |
| chr8:29937855-29939870 | p3 | (TGT)5 | TTTTGGGTCAACTTTGGAGG | TTGGTGAAACAATGGGGTTT | 213 |
| chr8:30033368-30035382 | p2 | (CT)7 | TGAGCGTGCTATGCTTATGG | CATCAATGAAGGTGGCCTTT | 143 |
| chr8:30075340-30077362 | p2 | (AT)11 | ATTCCGGAGAGGTTCCAAGT | AAGCAGCGAGCAGAGAAAAG | 254 |
| chr8:30188685-30190697 | p2 | (CA)6 | AGCATGAGATGGGAGAATCG | GGTCGGGTATGTGAAACCAC | 127 |
| chr8:30203583-30205601 | p2 | (GA)6 | GGGCATTCTCTCCCCTAGAC | ACGCGGGTTAGGACTTTTCT | 143 |
| chr8:30203583-30205601 | p2 | (TA)9 | GGGCATTCTCTCCCCTAGAC | ACGCGGGTTAGGACTTTTCT | 143 |
| chr8:30342636-30344648 | p2 | (AT)6 | CAAGACTCGGACCCCATAAA | TCAAATCCATCCCCAATCAT | 292 |
| chr8:30391428-30393444 | p4 | (AAGG)4 | ATGCATGGTCTTCAACCCTC | TTCTTCTGCCCCTTGGTATG | 123 |
| chr8:30480302-30482317 | p5 | (GTGGT)3 | TGATATGCTGTGAGGGACCA | ACGTTTTCGGATGACCTGAC | 185 |
| chr8:30557172-30559187 | p5 | (CAAGA)3 | AACCCAGTTGAGGTGTCGTC | CGATTGGTTTCGTGAGGAGT | 295 |
| chr8:30571199-30573217 | p6 | (GTTATG)3 | GTTGAGCCTTGCTTAGCACC | TTCCCCAGAGACATCCATTC | 239 |
| chr8:30573410-30575425 | p5 | (TCATA)3 | ACATATGGCCGAAGCAGAGT | TAAGAAATCCCCGATGCAAG | 207 |
| chr8:30638675-30640693 | p2 | (AT)8 | TTCATCTTTGGTACGCCCTC | CAAGCCAATACGTTCAGCAA | 178 |
| chr8:30688973-30690991 | p6 | (CATATA)3 | TGTCTCCCGAACAATCAACA | GCCTCGTCGACGTGTATTTT | 263 |
| chr8:30798782-30800797 | p3 | (TTG)5 | AATATGTGACGGCCTGATCC | CCACACAGGAATGGGAGAAT | 170 |
| chr8:30798782-30800797 | p5 | (GGTTT)3 | AATATGTGACGGCCTGATCC | CCACACAGGAATGGGAGAAT | 170 |
| chr8:30798782-30800797 | p5 | (CAAAC)3 | AATATGTGACGGCCTGATCC | CCACACAGGAATGGGAGAAT | 170 |
| chr8:30861460-30863476 | p2 | (AT)8 | AACAATCCCTAGATGCCACG | GGGTTGTCTGCAAAGTTCGT | 155 |
| chr8:30966994-30969012 | p2 | (TA)9 | ATCAGCACCAACACCAACAA | ATGGAGTTTGGAAGATTGCG | 244 |
| chr8:30974475-30976499 | p2 | (TA)12 | ATCATCAGCACCAACACCAA | ATGGAGTTTGGAAGATTGCG | 240 |
| chr8:31077216-31079228 | p2 | (TA)6 | TGACACAGGCAAATGGAGAA | GCAGCAAAGATGCAAAAACA | 119 |
| chr8:31101938-31103958 | p2 | (TA)10 | TGTTTTTGCATCTTTGCTGC | CCCTCCTCATTGGGTTTGTA | 218 |
| chr8:31127325-31129341 | p2 | (AT)8 | TGCACCACCTGCTACAAGAG | CATCTCCTTCCGAACTGCTC | 204 |
| chr8:31178394-31180408 | p2 | (AG)7 | TATATAAGCCCTCGCGGTTG | ACCTCACGTTGATTCCTTCG | 269 |
| chr8:31194408-31196422 | p2 | (GA)7 | GTGCGTTGTTTGTGTCAACC | CGGCTACCACCTACACGAAT | 226 |
| chr8:31222520-31224535 | p5 | (ATCAT)3 | TTGTGAGGTGTGTGTCACGA | CTTGTCGGCTTGGGTATTGT | 236 |
| chr8:31631757-31633773 | p4 | (CATA)4 | TCTTAATTCTTGCAACCCCG | ATACATTACCCGCAATCCCA | 219 |

| Marker ID | SSR type | SSR motif | FORWARD PRIMER (5'-3') | REVERSE PRIMER (5'-3') | size (bp) |
|---|---|---|---|---|---|
| chr8:31641030-31643045 | p3 | (ATC)5 | ACCAATAAATTGCAGCTCCG | ATGCATGGTATCCGCCTAAG | 238 |
| chr8:31645548-31647563 | p5 | (TTTTA)3 | GTTCGTAAATCCGTAGCCCA | AATTGGAAATTCGGGGTAGG | 282 |
| chr8:31662967-31665015 | p2 | (AT)24 | GCTGAATGGAGGTATTGGGA | TTCTAAAACGAAACGGGCAG | 222 |
| chr8:31662967-31665015 | p5 | (CGTAT)3 | GCTGAATGGAGGTATTGGGA | TTCTAAAACGAAACGGGCAG | 222 |
| chr8:31841708-31843726 | p6 | (ACTGAA)3 | TGGGTTGTCTCCCAATAAGC | CCAAGAAGTGCAAAATGCAA | 140 |
| chr8:31853095-31855116 | p3 | (CAA)7 | CCCTTTTCCCATATCCCCTA | AAATCGGAGGTTACCGACCT | 221 |
| chr8:31968616-31970630 | p2 | (TG)6 | GCAAGACAAGAACAGGAGGC | TGGTCAACTTCGTCACTTGC | 297 |
| chr8:31968616-31970630 | p2 | (AT)7 | GCAAGACAAGAACAGGAGGC | TGGTCAACTTCGTCACTTGC | 297 |
| chr8:32196676-32198696 | p4 | (ATAA)5 | AAACCGTGCATACCGAAGTC | GTTGGTACGCATGGTCTGTG | 288 |
| chr8:32203826-32205840 | p2 | (TA)7 | TCAACGGTGTTGACCCATTA | CGGAATTGCGCGTATAAAAT | 114 |
| chr8:32362550-32364608 | p2 | (AT)7 | ATTGGGCAACGACACATACA | TTTATGGGCATAGCAGGGTC | 232 |
| chr8:32362550-32364608 | p2 | (AT)29 | ATTGGGCAACGACACATACA | TTTATGGGCATAGCAGGGTC | 232 |
| chr8:32362550-32364608 | p2 | (TA)15 | ATTGGGCAACGACACATACA | TTTATGGGCATAGCAGGGTC | 232 |
| chr8:32366253-32368275 | p2 | (AT)11 | AGGAGGCAGATAGTGAGCCA | GTTCCCGTGCTCATGATTTT | 164 |
| chr8:32366253-32368275 | p3 | (ATC)5 | AGGAGGCAGATAGTGAGCCA | GTTCCCGTGCTCATGATTTT | 164 |
| chr8:32371178-32373196 | p6 | (CTTGAA)3 | AGTATTGCACCCTACCGTGG | AGCTTCTGCCCATTTTGAGA | 202 |
| chr8:32414247-32416259 | p5 | (TGAAT)3 | AGCAAGTAGAGGTGCAGGGA | CTCAAAACCTGAAACCGCAT | 252 |
| chr8:32414359-32416377 | p5 | (TGAAT)3 | AGCAAGTAGAGGTGCAGGGA | CTCAAAACCTGAAACCGCAT | 140 |
| chr8:32436558-32438606 | p2 | (AT)7 | CTTGGTGCTCCAATCCTGTT | TCAAGGACAAAAACCAAGGC | 138 |
| chr8:32436558-32438606 | p6 | (TGTATA)8 | CTTGGTGCTCCAATCCTGTT | TCAAGGACAAAAACCAAGGC | 138 |
| chr8:32529890-32531905 | p3 | (TCT)5 | TATTTCAACGATTGCGGACA | TCCAAAATCAAAATGACGCA | 148 |
| chr8:32644371-32646387 | p2 | (AG)7 | GAGAGATGGGGGAAAAGGAG | AGCCCAATTTAATTCACCCC | 298 |
| chr8:32644371-32646387 | p6 | (TGGGTG)3 | GAGAGATGGGGGAAAAGGAG | AGCCCAATTTAATTCACCCC | 298 |
| chr8:32644393-32646407 | p2 | (AG)7 | GAGAGATGGGGGAAAAGGAG | AGCCCAATTTAATTCACCCC | 276 |
| chr8:32644393-32646407 | p6 | (TGGGTG)3 | GAGAGATGGGGGAAAAGGAG | AGCCCAATTTAATTCACCCC | 276 |
| chr8:32661335-32663357 | p2 | (AT)11 | TCCATGATACGCTGCTTGAG | AGAGAGGATGTGCTGTGGCT | 143 |
| chr8:32816022-32818037 | p5 | (ATTTT)3 | ATGTACAACACCTGGCCCAT | CATGTCAGCACAAAAGGGTG | 216 |
| chr8:32862528-32864540 | p2 | (AC)6 | AAACCAGCACAAAATACGGG | ATTGATTTAGATGGTGCCGC | 202 |
| chr8:32870401-32872413 | p6 | (ATAAAA)3 | GTCACATGCGTATCAACCCA | GACATGGCGGAATAGCAAAT | 142 |
| chr8:32870401-32872413 | p2 | (AT)6 | GTCACATGCGTATCAACCCA | GACATGGCGGAATAGCAAAT | 142 |
| chr8:32870401-32872413 | p2 | (TA)11 | GTCACATGCGTATCAACCCA | GACATGGCGGAATAGCAAAT | 142 |
| chr8:32870563-32872585 | p2 | (AT)6 | GCTATTCCGCCATGTCATTT | CATGTTGTCTTTTGCACGGT | 151 |
| chr8:32870563-32872585 | p2 | (TA)11 | GCTATTCCGCCATGTCATTT | CATGTTGTCTTTTGCACGGT | 151 |
| chr8:32876436-32878450 | p2 | (AT)7 | ATCAGCCAGAAGTCTGCCAC | CCAAAATACATTCGCTGGCT | 158 |
| chr8:32876436-32878450 | p2 | (AT)24 | ATCAGCCAGAAGTCTGCCAC | CCAAAATACATTCGCTGGCT | 158 |
| chr8:32929219-32931243 | p6 | (CATATA)4 | ACATCCATTCGGGTGTTGAT | CGGGAACGATGAAAAGGATA | 138 |

| Marker ID | SSR type | SSR motif | FORWARD PRIMER (5'-3') | REVERSE PRIMER (5'-3') | size (bp) |
|---|---|---|---|---|---|
| chr8:32956005-32958017 | p2 | (AT)7 | GAAGAACAGTCCAAGCCAGC | ACAGGTGGTCCAGCTCAACT | 263 |
| chr8:32956019-32958031 | p2 | (AT)7 | GAAGAACAGTCCAAGCCAGC | ACAGGTGGTCCAGCTCAACT | 249 |
| chr8:33002479-33004499 | p2 | (AT)10 | ATAAGAGACACTGTGCCGGG | CTTTGGTATGCGATTCGGTT | 272 |
| chr8:33095337-33097359 | p2 | (TA)11 | CACAGCCATCGCTGATTCTA | ACCATAGGGACAACCTTCCC | 138 |
| chr8:33106902-33108922 | p4 | (AATA)5 | TGGCACTGCGTTAATACGAG | CAGGTTCGAGATTAGGCTGC | 249 |
| chr8:33118443-33120461 | p6 | (CATGAG)3 | CCTTTCTTCGGACGATGTGT | GCCTCTTGGCCATCAAAATA | 166 |
| chr8:33130274-33132289 | p2 | (TA)26 | AACGTCACTCCAGAGTTGGG | TTTGACGAAACCGTGTGGTA | 242 |
| chr8:33130274-33132289 | p5 | (CCTTT)3 | AACGTCACTCCAGAGTTGGG | TTTGACGAAACCGTGTGGTA | 242 |
| chr8:33133106-33135118 | p2 | (AT)12 | TCGAGGGTTCACGACCTTAC | CGTCTGGGGTACCATCTCAT | 139 |
| chr8:33133106-33135118 | p2 | (AT)6 | TCGAGGGTTCACGACCTTAC | CGTCTGGGGTACCATCTCAT | 139 |
| chr8:33180714-33182735 | p3 | (ACA)7 | AGGTTGAAGAGCAGGACGAA | GTCGACCACCAATTTCCACT | 277 |
| chr8:33374534-33376580 | p2 | (TA)23 | AGAGGTAGAGGTTGGGCGAT | CGTATGCACGGAGCTGACTA | 115 |
| chr8:33465698-33467713 | p5 | (CATCA)3 | CAGCACTCCCTCTGTTCTCC | GGGTTTTGCTAACCCACAAA | 212 |
| chr8:33582709-33584725 | p4 | (CATT)4 | TGCGAATCATAACAGACCCA | ATTCAGCGCGATTGAACTCT | 271 |
| chr8:33804314-33806352 | p2 | (TA)11 | GCTGTCCGGGTTAGTATGGA | CTTTAGCGCTTATTCGCCAC | 227 |
| chr8:33804314-33806352 | p2 | (AT)19 | GCTGTCCGGGTTAGTATGGA | CTTTAGCGCTTATTCGCCAC | 227 |
| chr8:33828015-33830033 | p3 | (TTG)6 | AGCCCTCCAAAAGGATGATT | TTTGGCAGCTTGGAGTCTTT | 193 |
| chr8:33828015-33830033 | p2 | (TA)8 | AGCCCTCCAAAAGGATGATT | TTTGGCAGCTTGGAGTCTTT | 193 |
| chr8:33841988-33844003 | p3 | (CAA)5 | CAGGCTCGAGGGAATCATTA | CCCGCTTTGCTCTCTTATTG | 242 |
| chr8:33933819-33935861 | p2 | (TA)21 | ATTGGAAATTCGGGATAGGC | TATTCGTAAATCCGTTCGCC | 198 |
| chr8:33960302-33962314 | p2 | (TG)6 | TATGATATGCCATGAGGCCA | CCTTGTCTTAAACTTGCCGC | 233 |
| chr8:34050940-34052958 | p2 | (AT)9 | AGCGAACACACATAGGACCC | TCATGCCTGCAGGTACAAAG | 247 |
| chr8:34098253-34100265 | p2 | (AT)6 | TCTTTTTCCCTCCTCCCCTA | ATCCTAATTCCACCTCCGCT | 232 |
| chr8:34141144-34143158 | p2 | (TA)7 | TTTTTGGAGCGCTCTTTTGT | GGGCCAAAAAGCAACATTTA | 168 |
| chr8:34185866-34187881 | p3 | (TTG)6 | GGGCAAGACCCTACAACTCA | GGGGTTTGAATGAGGGAAAT | 137 |
| chr8:34185866-34187881 | p3 | (GAT)5 | GGGCAAGACCCTACAACTCA | GGGGTTTGAATGAGGGAAAT | 137 |
| chr8:34220941-34222956 | p3 | (TTG)6 | GGGCAAGACCCTACAACTCA | GGGGTTTGAATGAGGGAAAT | 137 |
| chr8:34220941-34222956 | p3 | (GAT)5 | GGGCAAGACCCTACAACTCA | GGGGTTTGAATGAGGGAAAT | 137 |
| chr8:34338687-34340702 | p5 | (ATGCA)3 | GTAGCGCGAAGGAATGAAAG | GAGCCTTAGGTTACCCGGTC | 134 |
| chr8:34398981-34401003 | p2 | (AT)11 | TGCTGCTGAGTACCTGTTGG | CATGAAACCTCCTAACGGGA | 205 |
| chr8:34525356-34527372 | p2 | (AT)7 | TACACGCAACATCCCAAAAA | TGAAGTGCTCATTTGTTCCG | 251 |
| chr8:34525356-34527372 | p2 | (AT)8 | TACACGCAACATCCCAAAAA | TGAAGTGCTCATTTGTTCCG | 251 |
| chr8:34533614-34535626 | p2 | (AT)6 | ACGCCCCAAACATAAGTCAG | CCTGTCCCGTCAATGATTCT | 281 |
| chr8:34643906-34645928 | p2 | (TA)11 | CTTGGAGAGCTCCGTTGTTC | CATACTACGCAAGCCGACAA | 110 |
| chr8:34704024-34706036 | p2 | (TA)6 | CATATGGATGGGACAGGACC | CAGGGAGGCGATTCAGATTA | 207 |
| chr8:34798631-34800665 | p2 | (AT)17 | CCCGAGATGGACACTCGTAT | TTCTGATAGCCTTGTCGGCT | 220 |

| Marker ID | SSR type | SSR motif | FORWARD PRIMER (5'-3') | REVERSE PRIMER (5'-3') | size (bp) |
|---|---|---|---|---|---|
| chr8:34817448-34819463 | p5 | (ATATG)3 | AGCGCTCTTTTGTTCAGAGC | GACGAGGCTGTCACAGATCA | 196 |
| chr8:34860059-34862071 | p2 | (TA)6 | ACTCCCTCATGCTCCTCAGA | GACGGTAGCAGAAAAGCAGG | 237 |
| chr8:34886554-34888569 | p2 | (TG)6 | TTACGATCTTAAGTGGCCCG | AATCAGAAAGGAGGCGGAAT | 221 |
| chr8:34886554-34888569 | p5 | (ATCAT)3 | TTACGATCTTAAGTGGCCCG | AATCAGAAAGGAGGCGGAAT | 221 |
| chr8:34886554-34888569 | p2 | (TA)6 | TTACGATCTTAAGTGGCCCG | AATCAGAAAGGAGGCGGAAT | 221 |
| chr8:34915606-34917621 | p3 | (TGT)5 | TGAACCACAAAAAGCCACAA | TGCTGGTCATCACCCATCTA | 265 |
| chr8:34935671-34937683 | p2 | (AT)6 | AAGCTGTAGCTCCGGATGAA | ATCTCCCGATTCTCCCATCT | 261 |
| chr8:35084218-35086282 | p2 | (TA)8 | GACTCCCATTGTTGCGACTT | AATAGCATGGACAAGCAGGG | 294 |
| chr8:35089764-35091798 | p2 | (TA)17 | ACTAGGAGCCCGTTTGGATT | GGCTGGCCAACTTAAAGTCA | 280 |
| chr8:35199415-35201437 | p2 | (TA)7 | TTGGACATGTGTCCCACACT | TTCCGGTTTCGTTTCAAGTC | 129 |
| chr8:35199415-35201437 | p2 | (AT)11 | TTGGACATGTGTCCCACACT | TTCCGGTTTCGTTTCAAGTC | 129 |
| chr8:35309564-35311582 | p2 | (AT)9 | ATCACAGTTGTGCCATTCCA | AGCCTTGTCAGCTTGCGTAT | 173 |
| chr8:35335126-35337150 | p3 | (TTG)8 | GACAAACATGGCCCATTACC | TCGCAATATGACTCGTGGAA | 125 |
| chr8:35402592-35404620 | p2 | (TA)14 | AAACAAGTATCCCGTTCCCC | CTTTGCTTTTTGCTTTTGGG | 133 |
| chr8:35425390-35427402 | p2 | (CA)6 | TGGTTATCGGGCAGGTTAAG | AACATGGACACCCCAAACAT | 189 |
| chr8:35563413-35565439 | p2 | (AT)13 | CCCCACCAATAAAAATGTGC | AGTCATTTGATCACCCTCGG | 189 |
| chr8:35624536-35626562 | p2 | (AT)13 | CCCCACCAATAAAAATGTGC | AGTCATTTGATCACCCTCGG | 189 |
| chr8:35681822-35683837 | p2 | (AT)6 | AATCGGTTTGACTCGGTTTG | GAGGGATCGACGAGACACAT | 296 |
| chr8:35681822-35683837 | p5 | (CAAAC)3 | AATCGGTTTGACTCGGTTTG | GAGGGATCGACGAGACACAT | 296 |
| chr8:35681822-35683837 | p2 | (AG)7 | AATCGGTTTGACTCGGTTTG | GAGGGATCGACGAGACACAT | 296 |
| chr8:35681822-35683837 | p5 | (TTTAT)3 | AATCGGTTTGACTCGGTTTG | GAGGGATCGACGAGACACAT | 296 |
| chr8:35731416-35733436 | p2 | (AT)6 | GTTTGGAAAGCCACCTGGTA | CTTGTGCAACCAAACAGCAC | 162 |
| chr8:35731416-35733436 | p5 | (AATAA)4 | GTTTGGAAAGCCACCTGGTA | CTTGTGCAACCAAACAGCAC | 162 |
| chr8:35802234-35804248 | p2 | (GA)7 | GCGATCCTCGAAAATATGGA | TCCTCGGCCTTCGTAAAGTA | 255 |
| chr8:35940476-35942492 | p4 | (AATT)4 | GGGGAACGCATCTTACTTCA | CGGAGTGAACTCCTCGAAAG | 300 |
| chr8:35947714-35949730 | p2 | (AT)8 | ATAGCGATGAACGAAATGGG | GGTCCTTATTCCGTGCCATA | 205 |
| chr8:35980103-35982163 | p4 | (AAAG)5 | TGTTCATGCCCACAGGTAGA | AGCCCTTCGGCTATCAAAAT | 265 |
| chr8:35980219-35982237 | p4 | (AAAG)5 | TGTTCATGCCCACAGGTAGA | AGCCCTTCGGCTATCAAAAT | 149 |
| chr8:36013771-36015783 | p2 | (TG)6 | TCTGCACTACTCCGTTCGTG | TCCGAAACCGTAACACATCA | 205 |
| chr8:36086645-36088657 | p2 | (TA)6 | GCCATGATTGCCTTCAAAGT | TCGGCACGGTGTAATCATAA | 179 |
| chr8:36086645-36088657 | p2 | (TA)6 | GCCATGATTGCCTTCAAAGT | TCGGCACGGTGTAATCATAA | 179 |
| chr8:36136597-36138613 | p2 | (AG)8 | CAGTCGAGTCTACCCGCTTC | TGCTGAACACAGAAAGGACG | 174 |
| chr8:36220698-36222710 | p2 | (GT)6 | ATTGCTCGTACTGACGTCCC | ATCCGAAGGTGCAAAATACG | 234 |
| chr8:36228090-36230102 | p2 | (GT)6 | ATGGTTTTCTACGCCACAGG | TAACAGATATCGCGTTCCCC | 192 |
| chr8:36264410-36266425 | p5 | (TTATT)3 | GAAGCGGAAGAGAATATGCG | ACCCACAATAGCAATGCCTC | 219 |
| chr8:36269205-36271219 | p2 | (TG)7 | AGTCAGGTCATCCGAAAACG | CGACCTCATAGGATCGGTGT | 247 |

| Marker ID | SSR type | SSR motif | FORWARD PRIMER (5'-3') | REVERSE PRIMER (5'-3') | size (bp) |
|---|---|---|---|---|---|
| chr8:36316910-36318922 | p2 | (AT)6 | ATACTTATGGGCATGGCGAG | ATCATAGGTGCAAGGCCAAC | 272 |
| chr8:36520672-36522692 | p4 | (AGAA)5 | TCCCCACAACAGCATTGTAA | GGAGTGCTTGGAATTTGCTC | 165 |
| chr8:36546091-36548105 | p2 | (AT)7 | ATAAGATGCCAACTCCACGG | CGGTAGTTTTTCGGGTCAAA | 162 |
| chr8:36546091-36548105 | p3 | (ATC)5 | ATAAGATGCCAACTCCACGG | CGGTAGTTTTTCGGGTCAAA | 162 |
| chr8:36550529-36552545 | p4 | (TTTA)4 | GTCCGAGGTTCATTCCAAAA | AATGCAATACACTGGCCCAT | 107 |
| chr8:36625170-36627192 | p2 | (AT)11 | ATCGTGCGATAGAGCCAAGT | CACGCTTCTTCCTGTTAGGC | 138 |
| chr8:36625170-36627192 | p5 | (GAAAT)3 | ATCGTGCGATAGAGCCAAGT | CACGCTTCTTCCTGTTAGGC | 138 |
| chr8:36628867-36630885 | p6 | (TATATG)3 | TTTGGCCGAGTTGAATTCTC | GGGATCATGCCTTGAAAGAA | 182 |
| chr8:36646915-36648927 | p2 | (CT)6 | TGACCAGCTATCCACACGAC | GCATGTTGAAACCGTTTGTG | 226 |
| chr8:36682934-36684946 | p2 | (AT)6 | AGTGTGACTCATGTGGCGAG | CCACACACTGAAGGTTGCAC | 229 |
| chr8:36763548-36765560 | p2 | (AT)6 | AACCCTGCGCATTAAAACAC | TCGTTGTGAGAGGTCTGTGG | 206 |
| chr8:36791899-36793917 | p2 | (AG)9 | TTGCTGAAATGGATGGTCAA | GGGAATCAGATCCTTGCGTA | 244 |
| chr8:36860925-36863165 | p2 | (AT)120 | AACAGCGTCGACCCATTATC | TTGGTCGAAGGAAGAGGAGA | 284 |
| chr8:36912870-36914882 | p2 | (TA)6 | GTAGGGGCAGCAGGATAACA | CATATCTTCCGGGCTTTTCA | 297 |
| chr8:36912870-36914882 | p5 | (AAATG)3 | GTAGGGGCAGCAGGATAACA | CATATCTTCCGGGCTTTTCA | 297 |
| chr8:36946622-36948649 | p3 | (AAG)9 | TCCACTGGAGTTTACGGTCC | CGAAGTGATTTACGACCCGT | 160 |
| chr8:37001637-37003652 | p5 | (TCATT)3 | AGATGTATGGCCTCGGTCAC | TATACAACCGCCCAGAAAGG | 170 |
| chr8:37011245-37013261 | p2 | (TA)6 | ATGCCCACAGATAGACAGGG | TACACAATAGGCCTTTCGGC | 194 |
| chr8:37011269-37013337 | p2 | (TA)6 | ATGCCCACAGATAGACAGGG | TACACAATAGGCCTTTCGGC | 170 |
| chr8:37061233-37063247 | p2 | (TG)7 | ACGATTGTACTACGCACCCC | AATAGCCCGTAGGCACCTTT | 128 |
| chr8:37085460-37087478 | p6 | (CATATA)3 | CATGAATTGGCTCTATCGCA | CAGAGCAGGTCGTTCTAGGG | 152 |
| chr8:37174065-37176081 | p4 | (TAGT)4 | GCGGATTTGGCAAAGAATTA | AATCCATCTGGTTCCTCACG | 283 |
| chr8:37190592-37192612 | p2 | (AG)10 | CACCCGAGAGTTATGCGTTT | AAAATGTTGCAGGAACAGGG | 123 |
| chr8:37196574-37198588 | p2 | (TA)6 | GCAGAACGCCATTTTTCATT | CATTCTGGGCCCTATCAAGA | 166 |
| chr8:37196574-37198588 | p2 | (TA)7 | GCAGAACGCCATTTTTCATT | CATTCTGGGCCCTATCAAGA | 166 |
| chr8:37211603-37213617 | p2 | (TA)7 | AATCCCATTGGGCAACAATA | TAGCGGAGCTCCATTTCATT | 246 |
| chr8:37249029-37251041 | p2 | (TA)6 | AGCTCCACCTCTTCAGTCCA | ACATAAGGTTACCCCCAGCC | 101 |
| chr8:37261409-37263424 | p2 | (AT)10 | CCGGAAACGAATCGACTAGA | TTAGCGAGCGATCGAGGTAT | 292 |
| chr8:37261409-37263424 | p3 | (CAA)5 | CCGGAAACGAATCGACTAGA | TTAGCGAGCGATCGAGGTAT | 292 |
| chr8:37266142-37268178 | p2 | (TA)11 | CGGAGGTGCCATTGACTTAT | AGCCCTTCGGCTATCAAAAT | 300 |
| chr8:37266142-37268178 | p2 | (TA)17 | CGGAGGTGCCATTGACTTAT | AGCCCTTCGGCTATCAAAAT | 300 |
| chr8:37266142-37268178 | p2 | (TA)18 | CGGAGGTGCCATTGACTTAT | AGCCCTTCGGCTATCAAAAT | 300 |
| chr8:37287936-37289951 | p2 | (AG)7 | ACGCTTTTGGGTGATCTGAC | ATGTAGGCCCACGACCTATG | 111 |
| chr8:37287936-37289951 | p3 | (GTT)5 | ACGCTTTTGGGTGATCTGAC | ATGTAGGCCCACGACCTATG | 111 |
| chr8:37287936-37289951 | p2 | (TA)13 | ACGCTTTTGGGTGATCTGAC | ATGTAGGCCCACGACCTATG | 111 |
| chr8:37316361-37318373 | p2 | (CT)6 | GACCAGCTATCCACACGACA | GCATGTGGAAACCAAGTGTG | 223 |

| Marker ID | SSR type | SSR motif | FORWARD PRIMER (5'-3') | REVERSE PRIMER (5'-3') | size (bp) |
|---|---|---|---|---|---|
| chr8:37321917-37323929 | p2 | (CT)6 | GACCAGCTATCCACACGACA | GCATGTGGAAACCAAGTGTG | 223 |
| chr8:37461981-37463999 | p6 | (CCAGCA)3 | GGCGGCAAATTTGTATGAGT | ATGCAAGAATCGATCAACCC | 221 |
| chr8:37692838-37694853 | p5 | (ATCAG)3 | AGTAGGCCCATACAGATGCG | GAAATGGTATGTGGGTTCCG | 134 |
| chr8:37711633-37713648 | p5 | (CATAT)3 | CGGGGACGGACATATAACAC | TGATTTCCTTCACCGAGTCC | 141 |
| chr8:37711633-37713648 | p3 | (CAA)8 | CGGGGACGGACATATAACAC | TGATTTCCTTCACCGAGTCC | 141 |
| chr8:37711891-37713915 | p5 | (CATAT)3 | CGTTTCAGAAGCCTTTTTCG | GTATCCGAAGAGGTTCCGGT | 186 |
| chr8:37711891-37713915 | p3 | (CAA)8 | CGTTTCAGAAGCCTTTTTCG | GTATCCGAAGAGGTTCCGGT | 186 |
| chr8:37776532-37778560 | p2 | (AT)14 | AAAAATCTTGACGTGTGGGC | ATTTCCAGACAATTGGGCAG | 171 |
| chr8:37909432-37911446 | p2 | (AT)7 | TTGCTCGACTTCAAGGGACT | GTACGGTGCCATCCCAATAC | 181 |
| chr8:37911359-37913387 | p2 | (AT)14 | TTCTCGTGCGTACTCCACTG | CATCGACGCCATAACACATC | 113 |
| chr8:37965396-37967411 | p2 | (AT)11 | CATTTCACATCATAACGCCG | CATTATTGACATGTTCGCCG | 175 |
| chr8:37965396-37967411 | p5 | (TACAT)3 | CATTTCACATCATAACGCCG | CATTATTGACATGTTCGCCG | 175 |
| chr8:37983253-37985265 | p2 | (TA)6 | ACCCAATTGTGCCAAGTCTC | ATATGAGGCGTATTGGCTGG | 242 |
| chr8:37985322-37987376 | p6 | (TATATG)3 | TGATATGTTTCCGAATGGCA | TTCGGGTCAATCCGACTTAG | 193 |
| chr8:38026281-38028296 | p2 | (AT)11 | CCCTTTTCCCATATCCCCTA | TGGAGGTTACCGACCTTACG | 205 |
| chr8:38026281-38028296 | p3 | (ACA)5 | CCCTTTTCCCATATCCCCTA | TGGAGGTTACCGACCTTACG | 205 |
| chr8:38104807-38106835 | p2 | (TA)14 | TCATGGGTCCTCGACCTTAC | TCTCCCACTAGGGGTGTCTG | 273 |
| chr8:38121341-38123399 | p3 | (ATG)8 | ATGGAAGATGGAAAACAGCG | TTACGTCTCGGCTCTTCCAC | 131 |
| chr8:38121341-38123399 | p2 | (AT)29 | ATGGAAGATGGAAAACAGCG | TTACGTCTCGGCTCTTCCAC | 131 |
| chr8:38134605-38136623 | p2 | (TA)9 | ATGACTCTGTCTGTCCGGCT | TATATGCTCCGGATCGAAGG | 146 |
| chr8:38293249-38295265 | p2 | (TA)8 | CGAGCATGACAATACCGATG | CGTGGAGAGGTTGGGAAATA | 168 |
| chr8:38369203-38371225 | p2 | (AT)11 | CGGAGGTTCCATTTTTGAGA | CCCACACATACGCATCTACG | 224 |
| chr8:38429812-38431826 | p2 | (AT)7 | CCAAGGGAAGAATTGGGTTT | CGTTTGGGTTGTTGTTGATG | 264 |
| chr8:38429812-38431826 | p3 | (GAT)5 | CCAAGGGAAGAATTGGGTTT | CGTTTGGGTTGTTGTTGATG | 264 |
| chr8:38504267-38506282 | p5 | (CATAT)3 | TGCTCTGATCTCCGTCACAC | TGATTTCCTTCACCGAGTCC | 205 |
| chr8:38553522-38555537 | p3 | (TTG)5 | AACAAAAATTGGGCAGCATC | TCGCCCTTAAACTTACGTCG | 142 |
| chr8:38933638-38935668 | p2 | (TA)15 | GGCTATGGACTTGGCATGAT | CATCAGCCTGTTCGGATTTT | 265 |
| chr8:38953075-38955090 | p5 | (TATTT)3 | GAGCGCTAATATGTGTGGCA | AGATGGCAGGACATTCAAGG | 256 |
| chr8:39144110-39146130 | p2 | (TA)10 | GCCCTTTGGCTATCCACATA | CGGAGGTGCCATTGATTTAC | 202 |
| chr8:39186998-39189010 | p2 | (GT)6 | TGGGGAGTAGTTGGATGGAG | TCACGGTCACACACACACAC | 121 |
| chr8:39201778-39203794 | p2 | (AT)6 | GGAACTGTCTGCAAAGCCTC | ATGCCCGCAGGTGTAGATAG | 270 |
| chr8:39201778-39203794 | p4 | (CATA)4 | GGAACTGTCTGCAAAGCCTC | ATGCCCGCAGGTGTAGATAG | 270 |
| chr8:39233421-39235436 | p5 | (TGTGT)3 | GGATGACACTGAGCCTTGGT | GCCTGGTGTTATACGTGCCT | 124 |
| chr8:39268320-39270335 | p3 | (TTG)5 | CCAGCATCTAAATGGCCCTA | TTGCTGCATCAAGTCTGGTC | 287 |
| chr8:39275112-39277192 | p2 | (AT)40 | AGCATTCTTGGGGCATGTAG | TGAATGTACGCCAGGTGGTA | 198 |
| chr8:39288002-39290017 | p3 | (TTG)5 | CCAGCATCTAAATGGCCCTA | TTGCTGCATCAAGTCTGGTC | 287 |

| Marker ID | SSR type | SSR motif | FORWARD PRIMER (5'-3') | REVERSE PRIMER (5'-3') | size (bp) |
|---|---|---|---|---|---|
| chr8:39423689-39425701 | p2 | (TA)6 | TTACGGACAGTCCCTTTTGC | TGTTGATGTGCCGTGAATTT | 150 |
| chr8:39448876-39450892 | p2 | (AT)8 | CATTTGGTATCGAAATCGGG | TATACGCTACCTCATCCGGC | 288 |
| chr8:39448876-39450892 | p2 | (CA)6 | CATTTGGTATCGAAATCGGG | TATACGCTACCTCATCCGGC | 288 |
| chr8:39483873-39485887 | p2 | (AT)7 | CTCCAGAATTTTTCGGGTCA | CGCCACATCCTGGAGTCTAT | 164 |
| chr8:39502335-39504389 | p2 | (TA)27 | CATTACGTGATTCGCCCTCT | ATGATGTCTGAAAATCCGCC | 116 |
| chr8:39546599-39548613 | p2 | (AG)7 | CAAGCTTAGTCCCCGGTACA | GCCCATTTCTTCAAGACCAA | 173 |
| chr8:39601679-39603691 | p2 | (AG)6 | GTTTAGGGGAGGGCAAGAAC | CCTTCCTCCGTATTTCCACA | 255 |
| chr8:39601679-39603691 | p2 | (GT)6 | GTTTAGGGGAGGGCAAGAAC | CCTTCCTCCGTATTTCCACA | 255 |
| chr8:39643507-39645519 | p2 | (AT)11 | ACACCAAAGAGCCAACAAGG | TCCTTGCCCTCATATTCGTC | 200 |
| chr8:39643507-39645519 | p2 | (TA)6 | ACACCAAAGAGCCAACAAGG | TCCTTGCCCTCATATTCGTC | 200 |
| chr8:39731292-39733322 | p5 | (ATTCA)6 | AGGCCATCTGCTCACATACC | TTGGTTTCCAATGTGCTCAA | 290 |
| chr8:39731292-39733322 | p2 | (TA)7 | AGGCCATCTGCTCACATACC | TTGGTTTCCAATGTGCTCAA | 290 |
| chr8:39786728-39788792 | p2 | (TA)32 | CCCCTTCTTCTTTCACCTCC | AGTTGGGAGTCCGATTGTTG | 235 |
| chr8:39791409-39793427 | p3 | (TTG)6 | AACAATTACCCGTCGTTTGC | CCTCCAAAGTTGACCCAAAA | 222 |
| chr8:39989983-39991995 | p2 | (AT)6 | AGTGTACCCGACCTGACACC | TCCGTCTATAGTGGCAGCCT | 174 |
| chr8:40048080-40050092 | p2 | (AT)6 | CTAAATCCGGGAAATGCAGA | TGAAGGCCCCTAGGGTAAGT | 220 |
| chr8:40048080-40050092 | p2 | (AT)10 | CTAAATCCGGGAAATGCAGA | TGAAGGCCCCTAGGGTAAGT | 220 |
| chr8:40103386-40105398 | p2 | (TA)6 | CATGAGTTTCAGCGTCCTCA | GAAGAGCAAGGAGGAGCCTT | 270 |
| chr8:40141576-40143591 | p2 | (AT)8 | TCGATAACATATGCAAGCCG | CTTGGAGAGCTCCGTTGTTC | 245 |
| chr8:40141576-40143591 | p5 | (ATATA)3 | TCGATAACATATGCAAGCCG | CTTGGAGAGCTCCGTTGTTC | 245 |
| chr8:40177425-40179440 | p5 | (CATAG)3 | TGTACCGAGCACCCCTTATC | GTGTTTGGCATGATGTTTGC | 269 |
| chr8:40177425-40179440 | p2 | (TA)7 | TGTACCGAGCACCCCTTATC | GTGTTTGGCATGATGTTTGC | 269 |
| chr8:40182231-40184253 | p2 | (TA)11 | GAAGCCGGCCACATCTATAA | AGGATTTCCTTTCCACGCTT | 160 |
| chr8:40228766-40230781 | p3 | (CAA)5 | AACTCGAGGGAGTCAGCAAA | GGCCCACTATTTCCGATTTT | 232 |
| chr8:40260626-40262638 | p2 | (TG)6 | AAGGAGACCTCCCCGAAGTA | CGTCTTCCTCCATTTTTCCA | 220 |
| chr8:40260626-40262638 | p5 | (TGTGG)4 | AAGGAGACCTCCCCGAAGTA | CGTCTTCCTCCATTTTTCCA | 220 |
| chr8:40298257-40300283 | p2 | (AT)13 | TTATTTTGATGGCCGAAAGG | GAGCTAGTTACCGAGCACCG | 187 |
| chr8:40370851-40372863 | p5 | (AATTG)3 | TTACAGGCGAAGTGATGCTG | TACCTGTGCCTCGCCTTACT | 258 |
| chr8:40370859-40372877 | p5 | (AATTG)3 | TTACAGGCGAAGTGATGCTG | TACCTGTGCCTCGCCTTACT | 250 |
| chr8:40383736-40385764 | p2 | (AC)14 | GTAGAAGGACGGGATAGGGC | TTATCCGTACTGACTCCCCG | 208 |
| chr8:40505693-40507733 | p2 | (AT)20 | GTGCCACCTGTCCTGAATCT | CGTTGGCTTATGCTTGTGAA | 229 |
| chr8:40549003-40551019 | p2 | (TA)7 | CGGTGGAGGAACTAGAGCTG | AGGCGGATAGAAAGAGAGGC | 266 |
| chr8:40549047-40551067 | p2 | (TA)7 | CGGTGGAGGAACTAGAGCTG | AGGCGGATAGAAAGAGAGGC | 222 |
| chr8:40549071-40551083 | p2 | (TA)7 | CGGTGGAGGAACTAGAGCTG | AGGCGGATAGAAAGAGAGGC | 198 |
| chr8:40550121-40552136 | p2 | (TA)7 | AGAAAAACGGACACGACAGC | TACCTATATGGTCGGGCAGC | 125 |
| chr8:40550121-40552136 | p5 | (TCATT)3 | AGAAAAACGGACACGACAGC | TACCTATATGGTCGGGCAGC | 125 |

| Marker ID | SSR type | SSR motif | FORWARD PRIMER (5'-3') | REVERSE PRIMER (5'-3') | size (bp) |
|---|---|---|---|---|---|
| chr8:40565086-40567101 | p5 | (ATATA)3 | GATTTTTCGCTCCACACCAT | TAATGTGGTTGCCTTGGACA | 235 |
| chr8:40579663-40581677 | p2 | (TA)7 | TTCCCTTAGCCTTCCTCCAT | ACTTTTGTCACGACCCGACT | 126 |
| chr8:40689259-40691293 | p2 | (TA)17 | GCGTTCGTAACGGTTTGATT | CATCCGACGTCATGAACAAC | 143 |
| chr8:40720659-40722677 | p3 | (CAA)6 | AAAACCCTTTGAATGTCCCC | GCTCATGTCTGAGCGTGGTA | 218 |
| chr8:40720659-40722677 | p2 | (CT)8 | AAAACCCTTTGAATGTCCCC | GCTCATGTCTGAGCGTGGTA | 218 |
| chr8:40720976-40722992 | p3 | (CAA)6 | AGCTGAGGCCATAGCACAGT | GAAGCGAGCTTCCAGGTATG | 251 |
| chr8:40720976-40722992 | p2 | (CT)8 | AGCTGAGGCCATAGCACAGT | GAAGCGAGCTTCCAGGTATG | 251 |
| chr8:40730876-40732894 | p6 | (CATATA)3 | CCTAAGCTGTCGGGTCACTC | TTTCTGGCTATGATCCCCTG | 224 |
| chr8:40734819-40736837 | p3 | (CAA)6 | TATTTGCCGGGTTCGTTTAG | CTGAGTGTGCTATGGCCTCA | 244 |
| chr8:40734819-40736837 | p2 | (CT)14 | TATTTGCCGGGTTCGTTTAG | CTGAGTGTGCTATGGCCTCA | 244 |
| chr8:40734819-40736837 | p5 | (CAGTA)3 | TATTTGCCGGGTTCGTTTAG | CTGAGTGTGCTATGGCCTCA | 244 |
| chr8:40844885-40846897 | p2 | (AC)6 | AGATTGCCAAACGAAGGATG | GTGTTTGTTGTTGTGTGGGC | 261 |
| chr8:40850941-40852956 | p5 | (CATTT)3 | TATCCCGGACAGGATCAGAG | ATGTAGCTTACCGCATTGGG | 240 |
| chr8:40900377-40902392 | p2 | (AT)17 | AGCACGACGGAAAGAAAGAA | TTATTGGATTGAGGGTTCGC | 225 |
| chr8:40900377-40902392 | p5 | (AATGC)3 | AGCACGACGGAAAGAAAGAA | TTATTGGATTGAGGGTTCGC | 225 |
| chr8:40966135-40968153 | p3 | (CAA)6 | AAACTGAAGTGGGCCAAATG | ATCTTGCGAAATGGTTTTGG | 266 |
| chr8:40973918-40975930 | p2 | (TA)6 | TTTGGTTGCAATGTTGGTGT | CGACGAACGAAGTCATAGCA | 275 |
| chr8:41076932-41078948 | p2 | (AT)8 | TGATTGCCGAAGTGCTTATG | CGAGGCTAACTACCGAGCAC | 237 |
| chr8:41100630-41102642 | p2 | (AT)6 | GCATTTGAAGAAGTGCCCAT | GGCTTGAGAACGACGAAAAG | 176 |
| chr8:41130811-41132826 | p2 | (AT)6 | ACACATCACAACAAAGGCCA | ACAGTTGATAGGGGGTGACG | 172 |
| chr8:41130811-41132826 | p3 | (CAA)5 | ACACATCACAACAAAGGCCA | ACAGTTGATAGGGGGTGACG | 172 |
| chr8:41139448-41141466 | p6 | (CGTATT)3 | CGCAGCCTCCTGATTAAAAG | GTACGATAAAAGGTGCCCGA | 238 |
| chr8:41273756-41275768 | p2 | (AT)6 | ATGACTAGTGCCCGAGTTGG | GTCGGCTTGCGTATGGTATT | 120 |
| chr8:41364324-41366338 | p5 | (GAAAT)3 | CGTATTCTGGGTCGCATTTT | CGGAATTTTATTTCAGCCCA | 286 |
| chr8:41364324-41366338 | p2 | (AT)7 | CGTATTCTGGGTCGCATTTT | CGGAATTTTATTTCAGCCCA | 286 |
| chr8:41410999-41413011 | p2 | (GT)6 | ACGGGACCTCAACTGTCAAC | ACTGGACTAGCCAAAAGGCA | 164 |
| chr8:41410999-41413011 | p3 | (TGT)6 | ACGGGACCTCAACTGTCAAC | ACTGGACTAGCCAAAAGGCA | 164 |
| chr8:41452333-41454345 | p2 | (GT)6 | ACGGGACCTCAACTGTCAAC | ACTGGACTAGCCAAAAGGCA | 170 |
| chr8:41467590-41469605 | p3 | (GTT)5 | AAGAAGGACCTTGAGCCACA | GGTTGGAAGCGGTATTTCAA | 113 |
| chr8:41467590-41469605 | p2 | (TA)16 | AAGAAGGACCTTGAGCCACA | GGTTGGAAGCGGTATTTCAA | 113 |
| chr8:41531601-41533616 | p5 | (GCATT)3 | TGGAATCGTGGATTGAGTGA | ACATCTCGGTAGCACAACCC | 279 |
| chr8:41531601-41533616 | p3 | (TAT)5 | TGGAATCGTGGATTGAGTGA | ACATCTCGGTAGCACAACCC | 279 |
| chr8:41623851-41625869 | p2 | (TA)7 | GGTCGGGCTGTTTTTGAGTA | GGAGTTACCACAAGCTCGGA | 218 |
| chr8:41623851-41625869 | p6 | (TATATG)3 | GGTCGGGCTGTTTTTGAGTA | GGAGTTACCACAAGCTCGGA | 218 |
| chr8:41623851-41625869 | p2 | (AT)9 | GGTCGGGCTGTTTTTGAGTA | GGAGTTACCACAAGCTCGGA | 218 |
| chr8:41702725-41704737 | p2 | (AT)6 | TGCCCGATTTTCGTTAAGTC | TATTCGTGAATCCTTTCGCC | 265 |

| Marker ID | SSR type | SSR motif | FORWARD PRIMER (5'-3') | REVERSE PRIMER (5'-3') | size (bp) |
|---|---|---|---|---|---|
| chr8:41702725-41704737 | p2 | (TA)7 | TGCCCGATTTTCGTTAAGTC | TATTCGTGAATCCTTTCGCC | 265 |
| chr8:41702849-41704863 | p2 | (AT)6 | TGCCCGATTTTCGTTAAGTC | TATTCGTGAATCCTTTCGCC | 141 |
| chr8:41702849-41704863 | p2 | (TA)7 | TGCCCGATTTTCGTTAAGTC | TATTCGTGAATCCTTTCGCC | 141 |
| chr8:41792142-41794154 | p2 | (AC)6 | ATGGAGTCGGACCTAAGGCT | GTTGTGTTGGTTGTTGGCTG | 239 |
| chr8:41806129-41808159 | p2 | (TA)11 | CGTGCATGCCTCAATATGAC | CATACCATCCTTAGCGCCAT | 135 |
| chr8:41806129-41808159 | p2 | (AT)15 | CGTGCATGCCTCAATATGAC | CATACCATCCTTAGCGCCAT | 135 |
| chr8:41844296-41846314 | p5 | (CAATG)3 | ACAACGAAATACCTCCACGC | TGTATTGGCCGTGAGTTTCA | 120 |
| chr8:41844296-41846314 | p2 | (TA)6 | ACAACGAAATACCTCCACGC | TGTATTGGCCGTGAGTTTCA | 120 |
| chr8:41844296-41846314 | p6 | (CATATA)3 | ACAACGAAATACCTCCACGC | TGTATTGGCCGTGAGTTTCA | 120 |
| chr8:42001938-42003953 | p5 | (ATCAT)3 | TACATGGAAGCCGATAAGGC | AACGACAGGTTTGCTGATCC | 297 |
| chr8:42005338-42007350 | p2 | (TA)6 | TGCTAGAATCCTCCCCCTTT | GCGAAAGACATGGTCCAGAT | 136 |
| chr8:42109739-42111754 | p3 | (CAA)5 | ATAGGCACGCGCAGAGTAGT | CTCCGTTTGATTTGCTCCAT | 119 |
| chr8:42302000-42304036 | p2 | (AT)7 | AGCCGGTTACAAAACTGACG | TAGCTCCACTTCCTTTCGGA | 275 |
| chr8:42302000-42304036 | p6 | (CATATA)6 | AGCCGGTTACAAAACTGACG | TAGCTCCACTTCCTTTCGGA | 275 |
| chr8:42362103-42364119 | p2 | (AT)8 | AGCTCCTGCAGTTCACACCT | TGGAAAACTTGCCATCATCA | 143 |
| chr8:42362103-42364119 | p2 | (AT)7 | AGCTCCTGCAGTTCACACCT | TGGAAAACTTGCCATCATCA | 143 |
| chr8:42421614-42423629 | p5 | (ATCAT)3 | GGAGCAAAATATTGGGCAGA | CGGGTTATTGGGTACGTTTG | 128 |
| chr8:42435097-42437112 | p3 | (CTG)5 | TCTGATCATCTTCCTCGCCT | TTGCCAGAAAAAGGTGGTTC | 112 |
| chr8:42450022-42452037 | p3 | (CTG)5 | TCTGATCATCTTCCTCGCCT | TTGCCAGAAAAAGGTGGTTC | 112 |
| chr8:42465901-42467913 | p2 | (AT)6 | GGTACGCTCAGCTGTCCTTC | CAGGGTCCCCACAGCTATTA | 173 |
| chr8:42474711-42476723 | p2 | (TA)6 | AGCACTCCATTATTCCGGTG | ACAACTCTATCGGGGTGACG | 117 |
| chr8:42533849-42535959 | p2 | (AT)9 | CTTGGAGAGCTCCGTTGTTC | AAAATGATACGCAAGCCGAC | 281 |
| chr8:42685824-42687838 | p2 | (AT)7 | CTTCCTGCCAGTAGGTCAGC | AGTTCACACCTTGCGAATCC | 215 |
| chr8:42697454-42699520 | p2 | (AT)9 | TTCGCTTATGCATTGCACTC | TCACAACTTCACCCCCTCTC | 193 |
| chr8:42697454-42699520 | p2 | (TA)33 | TTCGCTTATGCATTGCACTC | TCACAACTTCACCCCCTCTC | 193 |
| chr8:42754048-42756063 | p3 | (TCT)5 | CGTGCCCAGTCAAATGAGTA | GATTGTTGGGATCCACTGCT | 117 |
| chr8:42755311-42757365 | p2 | (TA)27 | ATTCAAGGCACGTTGAGGAT | TGATTGTGGTTGCACGATTT | 176 |
| chr8:42761884-42763899 | p3 | (TCT)5 | CGTGCCCAGTCAAATGAGTA | GATTGTTGGGATCCACTGCT | 117 |
| chr8:42763147-42765201 | p2 | (TA)27 | ATTCAAGGCACGTTGAGGAT | TGATTGTGGTTGCACGATTT | 176 |
| chr8:42813782-42815797 | p5 | (GGGTC)3 | ATACTCCGACTCGGCAACAC | ACTGACGTCCCTTCTTGTGG | 257 |
| chr8:42944394-42946480 | p2 | (TA)16 | AGGCCTCATTTTGTGATTCG | CGGGCATTTAGGAAAAATGA | 176 |
| chr8:42944394-42946480 | p2 | (AT)43 | AGGCCTCATTTTGTGATTCG | CGGGCATTTAGGAAAAATGA | 176 |
| chr8:42952275-42954289 | p2 | (TA)7 | GAACAGTGTCGTCCATCGTG | GTCCGTAAAATGGCAGTCGT | 207 |
| chr8:42952275-42954289 | p5 | (ATATC)3 | GAACAGTGTCGTCCATCGTG | GTCCGTAAAATGGCAGTCGT | 207 |
| chr8:42987770-42989788 | p6 | (TTCAGT)3 | CTCCTCTCTTTCCGGAGCTT | CTGCACGCAGAAAATACCAA | 103 |
| chr8:43021048-43023060 | p2 | (TA)6 | TGCCAAATCCATCAGATTCA | ATCATGAAAAAGACCCCGTG | 181 |

| Marker ID | SSR type | SSR motif | FORWARD PRIMER (5'-3') | REVERSE PRIMER (5'-3') | size (bp) |
|---|---|---|---|---|---|
| chr8:43021048-43023060 | p6 | (TCTTTA)3 | TGCCAAATCCATCAGATTCA | ATCATGAAAAAGACCCCGTG | 181 |
| chr8:43202291-43204303 | p2 | (AT)6 | AAGGGTAATTTGGGAGGTGC | GTCCTGTCGCTAGTCAAGGC | 288 |
| chr8:43202291-43204303 | p3 | (TGT)5 | AAGGGTAATTTGGGAGGTGC | GTCCTGTCGCTAGTCAAGGC | 288 |
| chr8:43246051-43248066 | p5 | (TTTAT)3 | GAATCTGTTTTGGCTTTCCG | GCGGCTCCAAACTCTCTCTA | 180 |
| chr8:43319085-43321103 | p6 | (GTATAT)3 | CTTCTCTCTTGGCTGGTTGG | GGTGTGGGATTACGAAAGGA | 289 |
| chr8:43319085-43321103 | p2 | (AT)7 | CTTCTCTCTTGGCTGGTTGG | GGTGTGGGATTACGAAAGGA | 289 |
| chr8:43334494-43336514 | p2 | (AT)10 | TATAGCGGGCTTGTTATGCC | TATGCGGCCGTAAACTATCC | 192 |
| chr8:43334494-43336514 | p2 | (TA)11 | TATAGCGGGCTTGTTATGCC | TATGCGGCCGTAAACTATCC | 192 |
| chr8:43347631-43349649 | p6 | (ATATGT)3 | TGTAGGATATGAGCCTCGGG | TCGGGTCCTATTTATGCGAG | 227 |
| chr8:43499516-43501531 | p2 | (TA)6 | ATATCTTTATGGGGGCCAGG | ACGCTTGCCTTTTGAAACAT | 204 |
| chr8:43499516-43501531 | p2 | (TA)20 | ATATCTTTATGGGGGCCAGG | ACGCTTGCCTTTTGAAACAT | 204 |
| chr8:43499516-43501531 | p5 | (TATGA)3 | ATATCTTTATGGGGGCCAGG | ACGCTTGCCTTTTGAAACAT | 204 |
| chr8:43511493-43513513 | p5 | (AAATG)4 | CGCTCACTTTTTCTCCTTCG | CAAATGGATGACGTTTGTCG | 288 |
| chr8:43585742-43587756 | p2 | (TA)7 | AATCGGAGGTTACCGACCTT | TGCTCAGCCATATAAGCACG | 177 |
| chr8:43607444-43609459 | p5 | (ATGGA)3 | CATTGGGCCAAAAATAATGG | GACCAAGTAATGGGCTTGGA | 294 |
| chr8:43769960-43772038 | p3 | (TAT)26 | CCCTCTTCCAACTTCCATCA | GTCACTAGTGCTGCCCCATT | 130 |
| chr8:43771194-43773272 | p3 | (TAT)26 | ATTTTTACCTGACCCGACCC | AGTGGGGGAAGAGGAGCTTA | 227 |
| chr8:43783045-43785060 | p5 | (CACCA)3 | GCACTACCATGGTTGCATTG | TTCTCCAACCTTGGCAAAAC | 109 |
| chr8:43814246-43816258 | p2 | (AT)6 | ACCTACGGGGAGCGTATCTT | GCAAGATGACAAGGACAGCA | 153 |
| chr8:44029923-44031938 | p2 | (CA)6 | AGGCCAACAAGTCCAAACAC | GAGAGCTCCATTTCCATTCG | 228 |
| chr8:44029923-44031938 | p3 | (TTA)5 | AGGCCAACAAGTCCAAACAC | GAGAGCTCCATTTCCATTCG | 228 |
| chr8:44070419-44072447 | p2 | (TA)30 | ATCTGGTTGGCACAGGAAAG | GCATGTTCCCCTTCCAGTTA | 238 |
| chr8:44070419-44072447 | p2 | (AT)14 | ATCTGGTTGGCACAGGAAAG | GCATGTTCCCCTTCCAGTTA | 238 |
| chr8:44087365-44089379 | p2 | (AC)7 | TTGTAGATCGGGTTTCGAGG | AGCTAAGTCTCGGGGGTGTT | 175 |
| chr8:44108963-44110975 | p2 | (TA)6 | TGGATGACATTCCGCTGATA | GCCTGGAGACCCAGACATAA | 254 |
| chr8:44281907-44283922 | p5 | (TTTTA)3 | TCTTTCGTTGCGATTGACTG | AGAATTGACCACGACGGAAG | 127 |
| chr8:44294237-44296285 | p4 | (AAGT)4 | ACGGATACCACTGAGCCTTG | TCCATACCATACCCGACCAT | 193 |
| chr8:44384323-44386341 | p6 | (CTGAAA)3 | GTATTCGGCAGAATGGGAAA | GTGTCCAGTGGGGTCACTTT | 183 |
| chr8:44452106-44454124 | p3 | (TTG)6 | TCTTGACCCTGCCCTATGAC | GTGGCAGGGTTTTATAGCCA | 212 |
| chr8:44527824-44529840 | p2 | (AC)8 | GCATCCCAGGTAGGTTACGA | TCCCAACTTCATACCCGAAG | 198 |
| chr8:44527824-44529840 | p5 | (CATTG)3 | GCATCCCAGGTAGGTTACGA | TCCCAACTTCATACCCGAAG | 198 |
| chr8:44547285-44549300 | p5 | (ATCAT)3 | TGGGTGTGTTTTGGTCTGAA | CCATAGGGTGGAGTTCCCTT | 276 |
| chr8:44727061-44729076 | p5 | (ATTGC)3 | CCACCTCGTTGCGTTATTTT | TAAAGGGGGTATGCCAACAA | 111 |
| chr8:44775327-44777339 | p2 | (AT)6 | AGTTAAGCGAGAGAGCGACG | AATCAATTCACCTCCCCTCC | 292 |
| chr8:44775327-44777339 | p2 | (AT)6 | AGTTAAGCGAGAGAGCGACG | AATCAATTCACCTCCCCTCC | 292 |
| chr8:44775327-44777339 | p2 | (AT)6 | AGTTAAGCGAGAGAGCGACG | AATCAATTCACCTCCCCTCC | 292 |

| Marker ID | SSR type | SSR motif | FORWARD PRIMER (5'-3') | REVERSE PRIMER (5'-3') | size (bp) |
|---|---|---|---|---|---|
| chr8:44798177-44800189 | p2 | (AT)6 | AGTTAAGCGAGAGAGCGACG | AATCAATTCACCTCCCCTCC | 292 |
| chr8:44798177-44800189 | p2 | (AT)6 | AGTTAAGCGAGAGAGCGACG | AATCAATTCACCTCCCCTCC | 292 |
| chr8:44798177-44800189 | p2 | (AT)6 | AGTTAAGCGAGAGAGCGACG | AATCAATTCACCTCCCCTCC | 292 |
| chr8:44837165-44839180 | p3 | (ACA)5 | CCCAACTCCTTTCAAGACCA | TATATCCGCCTCCCACACTC | 127 |
| chr8:44922051-44924069 | p6 | (CATATA)3 | CCCGACCCTTACTTAGAGCC | GAATTGGGTGCCAGTTCTGT | 120 |
| chr8:44937395-44939410 | p5 | (TATAT)3 | GTGATTGTGAAGAGGCCGAT | AACAGACAGATGGGACCCTG | 133 |
| chr8:44944160-44946184 | p6 | (AAAAGG)3 | TCCAGAACTTCATTTTGGGC | GTTTGTGGGACTTTGGAGGA | 126 |
| chr8:44987103-44989119 | p2 | (TA)8 | CGGGTAACCATTATATGCCG | GGGCTGCATATGATTCCACT | 104 |
| chr8:44987103-44989119 | p6 | (CATATA)3 | CGGGTAACCATTATATGCCG | GGGCTGCATATGATTCCACT | 104 |
| chr8:45041997-45044035 | p2 | (AT)19 | CCCTAGGACGAACTGCTCTG | GAAGGCGTCATTTCACGATT | 283 |
| chr8:45151936-45153948 | p3 | (TTG)6 | TTGGAAAACAGTGTGTGGGA | TATGCCTAGCCCGAAATCAC | 259 |
| chr8:45151936-45153948 | p2 | (TA)6 | TTGGAAAACAGTGTGTGGGA | TATGCCTAGCCCGAAATCAC | 259 |
| chr8:45227772-45229784 | p2 | (GA)6 | AGCGACTTCTTGTCCCTGAA | GTCCAAATTCACTAGGGGCA | 236 |
| chr8:45227772-45229784 | p4 | (AAAT)4 | AGCGACTTCTTGTCCCTGAA | GTCCAAATTCACTAGGGGCA | 236 |
| chr8:45295478-45297496 | p6 | (TATATG)3 | GTTTTGAATATCGTGCGGGT | CAATCAAGCCAACACCAATG | 188 |
| chr8:45295478-45297496 | p2 | (TA)11 | GTTTTGAATATCGTGCGGGT | CAATCAAGCCAACACCAATG | 188 |
| chr8:45308697-45310711 | p2 | (AT)19 | GAACTGTTCTTCATCCGCCT | TAGAGGTGGGCAGCAACTTT | 106 |
| chr8:45308697-45310711 | p2 | (AT)7 | GAACTGTTCTTCATCCGCCT | TAGAGGTGGGCAGCAACTTT | 106 |
| chr8:45316653-45318693 | p2 | (TA)20 | TCTTTTTATCACAACCCCGC | GGTGTCAACAGCTTGTCCCT | 241 |
| chr8:45328009-45330059 | p2 | (TA)25 | TTCCCATATCCTTCCATCCA | AGTAGCGGAATGCTCGAAAA | 167 |
| chr8:45457962-45459980 | p6 | (AAAACA)3 | TTAAATGAAAGTGGCAGGGC | GGTGTCAACGACTTGCTTCA | 205 |
| chr8:45483606-45485620 | p6 | (ATACAT)3 | TTATAAATGTGGCCCGCTTC | ACAAGAGTGTCAGCCGGAGT | 167 |
| chr8:45483642-45485660 | p6 | (ATACAT)3 | TTATAAATGTGGCCCGCTTC | ACAAGAGTGTCAGCCGGAGT | 131 |
| chr8:45483661-45485685 | p6 | (ATACAT)3 | TTATAAATGTGGCCCGCTTC | ACAAGAGTGTCAGCCGGAGT | 112 |
| chr8:45483813-45485831 | p6 | (ATACAT)3 | ACTCCGGCTGACACTCTTGT | GCTCCAACAGCCGATATGTT | 120 |
| chr8:45629889-45631903 | p2 | (AT)6 | CTCTGCAAGCCGATGTATCA | ATTACTCAGAGCCGAGCGAA | 112 |
| chr8:45629889-45631903 | p2 | (TA)7 | CTCTGCAAGCCGATGTATCA | ATTACTCAGAGCCGAGCGAA | 112 |
| chr8:45632917-45634932 | p3 | (TTG)5 | ACGTTTTCGGATGACCTGAC | TTGATCCCGATATGCTGTGA | 151 |
| chr8:45632917-45634932 | p4 | (TATG)4 | ACGTTTTCGGATGACCTGAC | TTGATCCCGATATGCTGTGA | 151 |
| chr8:45671076-45673130 | p2 | (TA)27 | CACCCAAACCCTAGCTTGAA | GTGGCCGTACTTTCATCGTT | 161 |
| chr8:45671076-45673130 | p2 | (TA)19 | CACCCAAACCCTAGCTTGAA | GTGGCCGTACTTTCATCGTT | 161 |
| chr8:45732873-45734887 | p2 | (AT)7 | TTTTACGGGTCTTCATTCGG | ATACTTCTCCCACTCCGCCT | 240 |
| chr8:45942870-45944888 | p6 | (TATATG)3 | CATTGGTGTTGGCTTGATTG | GAATTGCCCCAAAGCTCATA | 268 |
| chr8:45977906-45979918 | p2 | (AT)6 | GACGGGCACTCAGAGCTAAC | TTCATAACCCTACCGGTCCA | 123 |
| chr8:45977906-45979918 | p2 | (TA)14 | GACGGGCACTCAGAGCTAAC | TTCATAACCCTACCGGTCCA | 123 |
| chr8:46286565-46288577 | p2 | (AT)6 | TCGCCCAAACACAACAAATA | TAACGACGACAGGTTTGCTG | 220 |

| Marker ID | SSR type | SSR motif | FORWARD PRIMER (5'-3') | REVERSE PRIMER (5'-3') | size (bp) |
|---|---|---|---|---|---|
| chr8:46287933-46289945 | p2 | (AC)6 | CAAAGCTCACTTCCCGTTTC | GTTGAATTCTCGGGGTGCTA | 197 |
| chr8:46308569-46310581 | p2 | (GT)6 | TCGTGGACAGATTGACGAAG | TTCCTCGATATTTCCGCATC | 121 |
| chr8:46417764-46419784 | p2 | (TA)10 | TCTCTCTTCCACGTCACACG | TTTGGGTTGGACTATGGAGG | 251 |
| chr8:46494069-46496083 | p2 | (TA)7 | GTACTGTCACGCCCGAAAAT | TCTTGCCTCGTTATACGCCT | 147 |
| chr8:46494084-46496110 | p2 | (TA)7 | GTACTGTCACGCCCGAAAAT | TCTTGCCTCGTTATACGCCT | 132 |
| chr8:46535749-46537781 | p2 | (TA)16 | TAGGGAAAGTTTGGGGAGGT | AAACCGCTCATCGATACGAC | 128 |
| chr8:46539557-46541581 | p4 | (TATC)6 | TGTGTGATATTGGGACCCCT | AGTAGGCGTCTGCAGCATTT | 190 |
| chr8:46539557-46541581 | p2 | (TA)6 | TGTGTGATATTGGGACCCCT | AGTAGGCGTCTGCAGCATTT | 190 |
| chr8:46758844-46760860 | p4 | (CTAA)4 | GTACAGCCTTCCCCCATTTT | TCAGCTGTTCGGTCAATCTG | 290 |
| chr8:46947766-46949794 | p2 | (GA)14 | CCAAGAAAGGATTGCCTTCA | CAAGTTCAGACCATGCTCCA | 153 |
| chr8:46949519-46951547 | p2 | (GA)14 | ACAATGCCTCTTTCACCTGC | CAAGTTCAGACCATGCTCCA | 114 |
| chr8:46951848-46953866 | p6 | (ATTTTA)3 | ACACGTGTACCCAACCACCT | TGTGGTTTCAAGTGGGTTCA | 100 |
| chr8:47094628-47096648 | p4 | (AGAA)5 | CTTGTTGCTGCTACTGCTGC | TATCGGGATCTGCTTCATCC | 111 |
| chr8:47094628-47096648 | p5 | (GAATT)5 | CTTGTTGCTGCTACTGCTGC | TATCGGGATCTGCTTCATCC | 111 |
| chr8:47169613-47171629 | p4 | (AACG)4 | AAACAAATCGACGACGGAAG | AAGTTGCAGCAGCCTCTGAT | 103 |
| chr8:47169613-47171629 | p2 | (TA)10 | AAACAAATCGACGACGGAAG | AAGTTGCAGCAGCCTCTGAT | 103 |
| chr8:47350019-47352039 | p2 | (AT)10 | CTGCAAGAGGCTGTTTCCTC | ATCAATCCCCACCCCTACTC | 271 |
| chr8:47382355-47384370 | p3 | (ATG)5 | AGCTTACTGCCCAACACCAC | TTTTGGGGTTTAAAAAGGGC | 110 |
| chr8:47401552-47403564 | p2 | (TA)6 | CCCAATTCTCATAGGCTGGA | TCCCAACTTCATACCCGAAG | 179 |
| chr8:47442761-47444782 | p3 | (ATA)7 | CAAAGGGAAGTTCCTAGCCC | CTTTTGCCCTCAATTCTGGA | 134 |
| chr8:47446949-47448997 | p3 | (TCT)16 | TATTTTGATGGCCAAGAGGC | CCGAGCACCAATCGTTCTAT | 118 |
| chr8:47533456-47535524 | p2 | (AT)34 | TTCATTACAGGCTTACCCCG | TGTGGATGAAATGTCGTGCT | 140 |
| chr8:47655819-47657834 | p3 | (GTT)5 | TTTGGGTCAATTTTGGAAGG | GGTTTGTTGCGTTGGAAACT | 211 |
| chr8:47655819-47657834 | p2 | (TA)16 | TTTGGGTCAATTTTGGAAGG | GGTTTGTTGCGTTGGAAACT | 211 |
| chr8:47902236-47904248 | p6 | (TTTTAT)3 | ATGGTAGTTGTGCTGCCCTC | TCATCTCGCCAAACTCAGTG | 127 |
| chr8:47902236-47904248 | p2 | (AT)6 | ATGGTAGTTGTGCTGCCCTC | TCATCTCGCCAAACTCAGTG | 127 |
| chr8:47902236-47904248 | p2 | (AT)14 | ATGGTAGTTGTGCTGCCCTC | TCATCTCGCCAAACTCAGTG | 127 |
| chr8:47928131-47930145 | p2 | (TA)7 | GAACCCCTGACGAAGATGAA | TCGTAAGCAACTCATGTGCC | 278 |
| chr8:47974265-47976277 | p2 | (AT)6 | TACCACTTTGTCACACCCGA | TGTCAGTCTGTAAGCAGCCG | 226 |
| chr8:47986163-47988178 | p3 | (TTG)5 | TGAAAGTGTCGCCAATGAAG | TTTGGAAGGATCTCCAATGC | 120 |
| chr8:47986163-47988178 | p5 | (AGGAC)3 | TGAAAGTGTCGCCAATGAAG | TTTGGAAGGATCTCCAATGC | 120 |
| chr8:48019726-48021738 | p2 | (TG)6 | AAAGACGGAAAGGAAGGCAT | TCGGGGTTATGTCACCGTAT | 102 |
| chr8:48041487-48043499 | p2 | (TG)6 | AAAGACGGAAAGGAAGGCAT | TCGGGGTTATGTCACCGTAT | 102 |
| chr8:48218481-48220493 | p3 | (CAA)5 | ATGCTCTTTACGCCTTTCCA | GCCCAAATTTCTGTAGGCAA | 299 |
| chr8:48218481-48220493 | p2 | (AT)6 | ATGCTCTTTACGCCTTTCCA | GCCCAAATTTCTGTAGGCAA | 299 |
| chr8:48298629-48300645 | p2 | (TA)8 | GTGGTACATTCGACGCCTTT | GCCATCACTGGAAAATCCAT | 244 |

| Marker ID | SSR type | SSR motif | FORWARD PRIMER (5'-3') | REVERSE PRIMER (5'-3') | size (bp) |
|---|---|---|---|---|---|
| chr8:48317858-48319873 | p5 | (CAATG)3 | CCTCTCGCAGATAGGTACGG | TGTTCGTGTGATCCGTGATT | 188 |
| chr8:48399192-48401208 | p2 | (TA)8 | TGTTGGCAGTGGATGATTGT | CACATTACGAAAACCCCACC | 284 |
| chr8:48399192-48401208 | p2 | (AT)6 | TGTTGGCAGTGGATGATTGT | CACATTACGAAAACCCCACC | 284 |
| chr8:48403323-48405335 | p3 | (TCA)6 | TAGCGGTTGGCTTACGAGTT | GATGATGAAACACAACCCCC | 181 |
| chr8:48403323-48405335 | p2 | (TA)6 | TAGCGGTTGGCTTACGAGTT | GATGATGAAACACAACCCCC | 181 |
| chr8:48435493-48437511 | p6 | (ATTGAA)3 | ACGTTCGGTGACCTGACTTT | ACGTTACGACCTTTGGTTGG | 152 |
| chr8:48474915-48476943 | p2 | (TA)14 | AAGTGTGTTGTGGGCGTGTA | TATACGATGCCATTGACGGA | 232 |
| chr8:48500840-48502864 | p2 | (TA)12 | GCTTAATACCTTCCCCTCGG | TTACGCCCTATTTTGAACGG | 281 |
| chr8:48500840-48502864 | p2 | (AT)9 | GCTTAATACCTTCCCCTCGG | TTACGCCCTATTTTGAACGG | 281 |
| chr8:48544884-48546916 | p2 | (AT)16 | CCGACTACTACTGGCCAAGC | CATCGGCGTGTATTTTGTTG | 163 |
| chr8:48690437-48692449 | p2 | (AC)6 | CATTGTTCACCGAGTCCCTT | ATCATCATATGCCCTCCTGG | 227 |
| chr8:48880995-48883010 | p5 | (ATGAT)3 | TTCCATAGCGTAATCGGAGG | TTGAAACTGTGTCCCTTCCC | 180 |
| chr8:48935050-48937068 | p2 | (AT)9 | GTGGAGGAACTGGAGGCATA | AAGGCGGGTAGAAAGAGAGC | 299 |
| chr8:49031983-49034001 | p3 | (CAT)6 | ATCGCACGATCTTAGGGATG | TTGGGTCGTGACAAAAGTGA | 221 |
| chr8:49042560-49044576 | p2 | (AT)8 | AGAGGGAGGGTTTTTGGAGA | CTTTCCAGCCCTGGTCATAA | 149 |
| chr8:49046272-49048284 | p3 | (CAA)6 | AGTCCCCGAGGTCCATATTC | GTTGAGTGTTTTGGCCGAGT | 100 |
| chr8:49046272-49048284 | p2 | (CA)6 | AGTCCCCGAGGTCCATATTC | GTTGAGTGTTTTGGCCGAGT | 100 |
| chr8:49070647-49072659 | p2 | (TA)6 | ACCAGTTCCTCCACCAGATG | AACCCCTGTTACTATGGCCC | 115 |
| chr8:49136527-49138545 | p3 | (CAT)6 | ATCGCACGATCTTAGGGATG | TTGGGTCGTGACAAAAGTGA | 221 |
| chr8:49147104-49149120 | p2 | (AT)8 | AGAGGGAGGGTTTTTGGAGA | CTTTCCAGCCCTGGTCATAA | 149 |
| chr8:49150816-49152828 | p3 | (CAA)6 | AGTCCCCGAGGTCCATATTC | GTTGAGTGTTTTGGCCGAGT | 100 |
| chr8:49150816-49152828 | p2 | (CA)6 | AGTCCCCGAGGTCCATATTC | GTTGAGTGTTTTGGCCGAGT | 100 |
| chr8:49175191-49177203 | p2 | (TA)6 | ACCAGTTCCTCCACCAGATG | AACCCCTGTTACTATGGCCC | 115 |
| chr8:49288387-49290405 | p6 | (ATACAT)3 | ATGGGCATTGTTTCCATGTT | GTGTGACAAACGCGTACACC | 221 |
| chr8:49288387-49290405 | p3 | (CAT)6 | ATGGGCATTGTTTCCATGTT | GTGTGACAAACGCGTACACC | 221 |
| chr8:49292631-49294643 | p2 | (AT)6 | AAAGTGTATCGTTGGACCGC | CCCCACCTAGGAGGAGTTTC | 287 |
| chr8:49303808-49305820 | p5 | (ATTCA)3 | AACATACCTGCTCCACGTCC | TTTGGCCGAGTTGAATTCTC | 183 |
| chr8:49303808-49305820 | p2 | (AT)6 | AACATACCTGCTCCACGTCC | TTTGGCCGAGTTGAATTCTC | 183 |
| chr8:49303808-49305820 | p2 | (AT)8 | AACATACCTGCTCCACGTCC | TTTGGCCGAGTTGAATTCTC | 183 |
| chr8:49328047-49330071 | p6 | (TTTTTG)4 | TCATTCCAGCTTGCAGTTTG | GAGGCAAAGGCATCCAAGTA | 147 |
| chr8:49359971-49361985 | p2 | (AG)7 | TTTCATCCCTTCTTGGTTCG | ATGTTGCAGGAACAGGGAAC | 251 |
| chr8:49365148-49367162 | p2 | (AG)7 | TTGGTCATGTGCTTGGTCAT | CAGGAACAGGGACCTCCATA | 228 |
| chr8:49504384-49506410 | p2 | (AT)13 | CCTCTTCGGGATCAATTTCA | CTCGACTGAGAAGAGGGTGG | 170 |
| chr8:49558196-49560211 | p5 | (TTGGT)4 | AAACCGCACCAAACCAATAA | ACCAAACCAAACCAAACCAA | 213 |
| chr8:49558235-49560250 | p5 | (TTGGT)4 | AAACCGCACCAAACCAATAA | ACCAAACCAAACCAAACCAA | 174 |
| chr8:49580114-49582130 | p2 | (TA)8 | TGACTTGGCCCATGTTGTTA | GGGGAAACCTTTTGGGTAGA | 227 |

| Marker ID | SSR type | SSR motif | FORWARD PRIMER (5'-3') | REVERSE PRIMER (5'-3') | size (bp) |
|---|---|---|---|---|---|
| chr8:49582983-49585007 | p2 | (TA)12 | CCCCAATTCTTTTCTTTGGA | ATTCATTGTGTCTTTCCCGC | 300 |
| chr8:49597819-49599837 | p2 | (AT)7 | AATGTCAACAAGCATTGCCA | CGGAGTGACGTAAGGTCGTT | 287 |
| chr8:49597839-49599857 | p2 | (AT)7 | AATGTCAACAAGCATTGCCA | CGGAGTGACGTAAGGTCGTT | 267 |
| chr8:49597859-49599877 | p2 | (AT)7 | AATGTCAACAAGCATTGCCA | CGGAGTGACGTAAGGTCGTT | 247 |
| chr8:49597879-49599897 | p2 | (AT)7 | AATGTCAACAAGCATTGCCA | CGGAGTGACGTAAGGTCGTT | 227 |
| chr8:49597899-49599917 | p2 | (AT)7 | AATGTCAACAAGCATTGCCA | CGGAGTGACGTAAGGTCGTT | 207 |
| chr8:49597919-49599937 | p2 | (AT)7 | AATGTCAACAAGCATTGCCA | CGGAGTGACGTAAGGTCGTT | 187 |
| chr8:49597939-49599957 | p2 | (AT)7 | AATGTCAACAAGCATTGCCA | CGGAGTGACGTAAGGTCGTT | 167 |
| chr8:49597959-49599977 | p2 | (AT)7 | AATGTCAACAAGCATTGCCA | CGGAGTGACGTAAGGTCGTT | 147 |
| chr8:49597979-49599997 | p2 | (AT)7 | AATGTCAACAAGCATTGCCA | CGGAGTGACGTAAGGTCGTT | 127 |
| chr8:49597999-49600017 | p2 | (AT)7 | AATGTCAACAAGCATTGCCA | CGGAGTGACGTAAGGTCGTT | 107 |
| chr8:49677104-49679158 | p3 | (GAA)18 | AGTGCTTTCTCTTCCCCGAT | GCCCTCAAAAACATCCTTCA | 178 |
| chr8:49714034-49716054 | p2 | (TA)10 | GTTCGTGAATGTGACATCGG | ACCGTCAAAATGGTCCGTAG | 172 |
| chr8:49725065-49727083 | p6 | (ATATGG)3 | TGGGTGTGAAGTTGTGGGTA | CCCCGTTGAAGCTTGAATAA | 182 |
| chr8:49922868-49924884 | p4 | (AGAA)4 | GTGCGGTCCAAAACAAGTCT | TATTCCCCCTTCCCAAAAAG | 137 |
| chr8:49923486-49925518 | p4 | (AGAA)4 | ATGCTTGATTCCCCACAGAG | CACGAGGACGATGATGAAGA | 104 |
| chr8:50132617-50134633 | p2 | (AT)8 | AGAACCCGGCCCTTTTAGTA | GCATATTCCTCGGCATTGAT | 182 |
| chr8:50379579-50381603 | p3 | (TGT)8 | TCGAAGGATAGGCTGCTTGT | TTCCATTTCCATCAAAAGGC | 181 |
| chr8:50426498-50428518 | p2 | (AT)10 | CTGTATGCAAACACCCATGC | ATTGGCTTGCGTATCTTTGG | 229 |
| chr8:50426498-50428518 | p2 | (TA)28 | CTGTATGCAAACACCCATGC | ATTGGCTTGCGTATCTTTGG | 229 |
| chr8:50442330-50444342 | p2 | (TA)6 | CTGTATGCAAACACCCATGC | TGCGTATCTTTGGAGAACCC | 133 |
| chr8:50442330-50444342 | p5 | (TGTGT)3 | CTGTATGCAAACACCCATGC | TGCGTATCTTTGGAGAACCC | 133 |
| chr8:50443191-50445206 | p2 | (TA)6 | TAGGGGAGGTGTAGGTGTCG | CTAGGCTTGGACTTCCATCG | 247 |
| chr8:50443191-50445206 | p5 | (TGTGT)3 | TAGGGGAGGTGTAGGTGTCG | CTAGGCTTGGACTTCCATCG | 247 |
| chr8:50485385-50487400 | p2 | (TA)6 | GGATGACCTGACTTGGAGGA | GCCATTTTGGCAGCTTAGAG | 237 |
| chr8:50485385-50487400 | p3 | (TTG)5 | GGATGACCTGACTTGGAGGA | GCCATTTTGGCAGCTTAGAG | 237 |
| chr8:50485385-50487400 | p4 | (TGTA)5 | GGATGACCTGACTTGGAGGA | GCCATTTTGGCAGCTTAGAG | 237 |
| chr8:50608575-50610614 | p2 | (AC)8 | CAACCCATTTTCTTCCTCCA | AAACAATGAAGGGCAACCAG | 171 |
| chr8:50608575-50610614 | p3 | (GAA)13 | CAACCCATTTTCTTCCTCCA | AAACAATGAAGGGCAACCAG | 171 |
| chr8:50608575-50610614 | p2 | (GT)7 | CAACCCATTTTCTTCCTCCA | AAACAATGAAGGGCAACCAG | 171 |
| chr8:50608878-50610892 | p3 | (GAA)13 | CAAATGGACTGATTGCCCTT | CAACCCATTTTCTTCCTCCA | 278 |
| chr8:50608878-50610892 | p2 | (GT)7 | CAAATGGACTGATTGCCCTT | CAACCCATTTTCTTCCTCCA | 278 |
| chr8:50640560-50642572 | p3 | (CAT)5 | AAACAAAGGTGGACCACGAG | GGAACCTATCCCGACCTTGT | 299 |
| chr8:50640560-50642572 | p2 | (TA)6 | AAACAAAGGTGGACCACGAG | GGAACCTATCCCGACCTTGT | 299 |
| chr8:50640560-50642572 | p2 | (TA)22 | AAACAAAGGTGGACCACGAG | GGAACCTATCCCGACCTTGT | 299 |
| chr8:50964842-50966868 | p2 | (AT)13 | ATATTCCGTCCGCATGTGAT | GACTTTTAGCTCTTCGCCCC | 132 |

| Marker ID | SSR type | SSR motif | FORWARD PRIMER (5'-3') | REVERSE PRIMER (5'-3') | size (bp) |
|---|---|---|---|---|---|
| chr8:51016444-51018462 | p3 | (TTG)6 | CCTTAAGCCAAATATGGGCA | GCCCACGACCTATGGTTAGA | 170 |
| chr8:51121453-51123467 | p2 | (TA)7 | GCTGCACTGTTGTCTGCTTC | TGGACTGCCTTAATCGGTTC | 188 |
| chr8:51242444-51244456 | p2 | (AT)6 | GCGTGCATAGGGAATTAGGA | TTAGTTACGGGTGCCTGTCC | 260 |
| chr8:51315389-51317404 | p5 | (TATAT)3 | TGGTTATATTCGGGCATCGT | TGACTCGCGAATTTTGAGTG | 159 |
| chr8:51437476-51439492 | p4 | (TTAT)4 | AGAGTGACGTGAGGACCGAT | CGGCAATCACTACCAATCCT | 168 |
| chr8:51471779-51473794 | p3 | (TTG)5 | AAGGAGGACCCTTAAGCCAA | GCCTACGACCTACGTTTGGA | 171 |
| chr8:51538285-51540347 | p2 | (TC)31 | CCCTTTGGAGTGGTTTTGAA | ATAAGGTGCCAATTTGAGCG | 200 |
| chr8:51538285-51540347 | p2 | (TA)13 | CCCTTTGGAGTGGTTTTGAA | ATAAGGTGCCAATTTGAGCG | 200 |
| chr8:51622644-51624660 | p5 | (AATGC)3 | AATCCCCAAGGAAGTGGTCT | TTCCACTTCCACTTCCCAAG | 101 |
| chr8:51622644-51624660 | p2 | (GT)8 | AATCCCCAAGGAAGTGGTCT | TTCCACTTCCACTTCCCAAG | 101 |
| chr8:51828699-51830715 | p2 | (TA)16 | AGATCATGACGAATCCGACC | GAGACGTGTGCAGAGGTGAA | 260 |
| chr8:51828699-51830715 | p4 | (GTGG)4 | AGATCATGACGAATCCGACC | GAGACGTGTGCAGAGGTGAA | 260 |
| chr8:51834528-51836561 | p3 | (CAA)11 | TGTGGAAACAGCTGAGATGC | GGCTCATTTTCTAAAGCCCC | 220 |
| chr8:51870272-51872330 | p2 | (AT)7 | GCTGGAGGAGTGAGAACGAC | TGACCATCATGTGACAGCCT | 245 |
| chr8:51995407-51997427 | p5 | (CATAT)4 | GCACACGCAGAAGAAAATCA | TTCCCCGAGACTGTGCTATC | 201 |
| chr8:52181998-52184014 | p2 | (AT)8 | CCGGCTTAATTGTTTCGGTA | TTTCACGTTGCAATCTCGTC | 102 |
| chr8:52181998-52184014 | p2 | (TA)6 | CCGGCTTAATTGTTTCGGTA | TTTCACGTTGCAATCTCGTC | 102 |
| chr8:52358967-52360982 | p5 | (TGAAT)3 | CAAGAAGCATGGAGAGCACA | CAACTACTCCACGTCGCAGA | 187 |
| chr8:52398360-52400372 | p2 | (AC)6 | TCCAAGCTCACTCTCCGTTT | AGGTTATGGCGTTATGTCGG | 192 |
| chr8:52451799-52453817 | p3 | (CAA)6 | GTGGTAACTGCCCAACCAGT | TGATGACCACGGGAAGTGTA | 197 |
| chr8:52461321-52463339 | p6 | (ATACAT)3 | CCCTAGGACGAACTGCTCTG | TTTTGATAGCCTTGTCGGCT | 133 |
| chr8:52461321-52463339 | p3 | (CAT)6 | CCCTAGGACGAACTGCTCTG | TTTTGATAGCCTTGTCGGCT | 133 |
| chr8:52547168-52549184 | p2 | (TA)8 | ATTCGTGGATGTCCCAATGT | TATACCTGCCCCCTCACATC | 285 |
| chr8:52631068-52633086 | p6 | (CAACTC)3 | CCTATTCCCTTGCTGCACAT | TCAACGGGAGCAGCTATTCT | 186 |
| chr8:52634926-52636944 | p2 | (AT)9 | GTTGGCCTCAAGCTGACTTC | AAGCATACAAATCCGCCAAC | 214 |
| chr8:52660873-52662885 | p2 | (AT)6 | TCATGGTTGGACTTTCTCCC | TCCTCCTATGCCACCGTTAG | 225 |
| chr8:52660873-52662885 | p3 | (ATC)5 | TCATGGTTGGACTTTCTCCC | TCCTCCTATGCCACCGTTAG | 225 |
| chr8:52660873-52662885 | p2 | (AC)7 | TCATGGTTGGACTTTCTCCC | TCCTCCTATGCCACCGTTAG | 225 |
| chr8:52675076-52677094 | p3 | (AAC)6 | CTCCATTCCCTCATTTTCCA | CCACACACATGAAGAAACCG | 157 |
| chr8:52730130-52732148 | p6 | (ATTTTT)3 | AAGCTCAAAGCGGCAAATAA | TATTCACAGCCACGGGTACA | 279 |
| chr8:52807300-52809315 | p5 | (ATAAA)3 | CCACCAACCAAACAACTCCT | AAATGGATATGTATGGCCGC | 191 |
| chr8:52898285-52900300 | p5 | (ATAAA)3 | TGCTTGATCTCGGAACCTTT | CAAAAATTTTCGCGTTCGTT | 296 |
| chr8:52898285-52900300 | p2 | (AT)11 | TGCTTGATCTCGGAACCTTT | CAAAAATTTTCGCGTTCGTT | 296 |
| chr8:52971264-52973280 | p2 | (AG)8 | ACACACGACCACGCATTAAA | TCTGCTGCATTTTCTGTTGC | 176 |
| chr8:52991645-52993657 | p5 | (CTCTC)3 | TTAGGGTTTGCTCGCACTCT | CTTGAAGGGAGCTGAACCTG | 280 |
| chr8:52991672-52993684 | p5 | (CTCTC)3 | TTAGGGTTTGCTCGCACTCT | CTTGAAGGGAGCTGAACCTG | 253 |

| Marker ID | SSR type | SSR motif | FORWARD PRIMER (5'-3') | REVERSE PRIMER (5'-3') | size (bp) |
|---|---|---|---|---|---|
| chr8:53041785-53043833 | p3 | (AAG)16 | GAGCCCTGGTACTTGCTGAG | AAAAACGCGTATCCCCTTCT | 180 |
| chr8:53086045-53088063 | p6 | (ATTTTT)3 | TAGGTTAGCTCGCCTGTGCT | GATTTTGTCCCATTCGCCTA | 291 |
| chr8:53086045-53088063 | p5 | (AATCT)4 | TAGGTTAGCTCGCCTGTGCT | GATTTTGTCCCATTCGCCTA | 291 |
| chr8:53109055-53111073 | p6 | (CATATA)3 | GATGGAGCCGCTTACAAAAC | ACACCCAGATGAGTGGAAGG | 295 |
| chr8:53255891-53257909 | p6 | (AAAGAG)3 | ATGATGTGGAAGGAACCGAG | CCCATATAATCGCTAGGCCA | 284 |
| chr8:53260050-53262070 | p4 | (AATA)5 | GAGATGCTCGGAAACTCCAG | CGCTAAATCCTTTGGGTGAA | 168 |
| chr8:53276020-53278040 | p4 | (AAAT)5 | GAATTTCGGTACCAGGCGTA | CAATCTCTGCCCATTCACCT | 186 |
| chr8:53302119-53304133 | p2 | (TG)7 | CGGTGTCCTACGGGCTATTA | CGACCTCATAGGATCGGTGT | 228 |
| chr8:53318651-53320715 | p2 | (AT)32 | ATGACTAGTGCCCGAGTTGG | GACGGCTTGCATATTTGGTT | 116 |
| chr8:53318651-53320715 | p3 | (AAC)5 | ATGACTAGTGCCCGAGTTGG | GACGGCTTGCATATTTGGTT | 116 |
| chr8:53410816-53412846 | p2 | (TA)12 | TTGTCGATTGTTGGTCCGTA | TTTTCTTGGGCCTATTCCCT | 144 |
| chr8:53410816-53412846 | p2 | (TA)15 | TTGTCGATTGTTGGTCCGTA | TTTTCTTGGGCCTATTCCCT | 144 |
| chr8:53502415-53504430 | p3 | (CTT)5 | GATTTTCAGCCATTGTGCCT | ATTTAGTGGAACCGCTGTGG | 236 |
| chr8:53505174-53507189 | p5 | (TTGCA)3 | AGGGTGGACGAAATGTACCA | TCATGACATCAAGCCAGCAT | 232 |
| chr8:53511840-53513852 | p2 | (AT)6 | TGTCTACCTGTGGGCATGAA | GACTTATCCGAGGGCACAAA | 193 |
| chr8:53526121-53528133 | p6 | (AATATG)3 | AAAGGTGCCTACGGGCTATT | CCAACCAAGGCTCAAGGTTA | 218 |
| chr8:53526121-53528133 | p2 | (TG)6 | AAAGGTGCCTACGGGCTATT | CCAACCAAGGCTCAAGGTTA | 218 |
| chr8:53537487-53539505 | p5 | (TGAAT)3 | ATAAGATGCCAACTCCACGG | GTTGCTTCGGGTCAACAAAT | 274 |
| chr8:53537550-53539564 | p5 | (TGAAT)3 | ATAAGATGCCAACTCCACGG | GTTGCTTCGGGTCAACAAAT | 211 |
| chr8:53558978-53560993 | p5 | (ATCAT)3 | ACAACAACCATGCCAAGACA | ATTCGGGCACGACAGTATTC | 249 |
| chr8:53613542-53615560 | p6 | (TTTGCT)3 | AAGATTCTGAGCGTGTGCCT | GCATGCATGGTATCCTCCTT | 126 |
| chr8:53700458-53702473 | p3 | (ATG)5 | CCCATGGCTGCAGAATTTAT | TGCATAAACCAAATCCCCAT | 139 |
| chr8:53710742-53712757 | p5 | (AAATG)3 | GATTCAGGCCATTGGAAAGA | ACATTGTGAATGTTGGGGGT | 232 |
| chr8:53724434-53726452 | p2 | (AT)9 | ACTTTGCGGGTATTCATTCG | ATACTTCTCCCACTCCGCCT | 226 |
| chr8:53724434-53726452 | p2 | (AT)6 | ACTTTGCGGGTATTCATTCG | ATACTTCTCCCACTCCGCCT | 226 |
| chr8:53724434-53726452 | p5 | (AATAA)3 | ACTTTGCGGGTATTCATTCG | ATACTTCTCCCACTCCGCCT | 226 |
| chr8:53724555-53726595 | p2 | (AT)9 | ACTTTGCGGGTATTCATTCG | ATACTTCTCCCACTCCGCCT | 105 |
| chr8:53724555-53726595 | p2 | (AT)6 | ACTTTGCGGGTATTCATTCG | ATACTTCTCCCACTCCGCCT | 105 |
| chr8:53724555-53726595 | p5 | (AATAA)3 | ACTTTGCGGGTATTCATTCG | ATACTTCTCCCACTCCGCCT | 105 |
| chr8:53726108-53728123 | p5 | (AATAA)3 | GTCCAAGAAGGCAACGACAT | TCCAATTCAGCCCGACTTAC | 295 |
| chr8:53726108-53728123 | p3 | (GTT)5 | GTCCAAGAAGGCAACGACAT | TCCAATTCAGCCCGACTTAC | 295 |
| chr8:53726108-53728123 | p2 | (TA)7 | GTCCAAGAAGGCAACGACAT | TCCAATTCAGCCCGACTTAC | 295 |
| chr8:53814087-53816102 | p3 | (TTG)5 | GACCCTTGAACCAAATCGAA | CCTTGTACCGAAGTCACCGT | 157 |
| chr8:53822314-53824328 | p2 | (TA)7 | GAGTGCACAGATGAGTCCGA | GGTGTAACATCCTTCCCCCT | 228 |
| chr8:53842937-53844951 | p2 | (TA)7 | GAGTGCACAGATGAGTCCGA | GGTGTAACATCCTTCCCCCT | 228 |
| chr8:53872009-53874024 | p5 | (TGAAT)3 | GTTTGACCCCGAAAAGATCA | CTAAGGGGGCGACAATGTAA | 218 |

| Marker ID | SSR type | SSR motif | FORWARD PRIMER (5'-3') | REVERSE PRIMER (5'-3') | size (bp) |
|---|---|---|---|---|---|
| chr8:53872050-53874074 | p5 | (TGAAT)3 | GTTTGACCCCGAAAAGATCA | AGGCCATGACTTTCATTTGG | 177 |
| chr8:53916530-53918545 | p3 | (GTT)5 | TGGTGTGGACGAATGTGAGT | TACACCCACATCCCTGACCT | 193 |
| chr8:54071824-54073840 | p2 | (TA)8 | AAAATCACGCCACATCAACA | GCATGCCGGTAGTTCAAAGT | 268 |
| chr8:54121926-54123941 | p5 | (TATAT)3 | GATTGTGAGGAGGCCAATGT | TGGCTAATTTCTCACAGGGG | 156 |
| chr8:54224210-54226222 | p2 | (TA)6 | AGGTACCGACGACAGGTTTG | CCGTCGTTACCCACTGAAAT | 195 |
| chr8:54268703-54270717 | p5 | (AAATT)3 | TACAGGCACCAACATGGAGA | CTCTCGCATGGAAGAGGTTC | 257 |
| chr8:54268789-54270847 | p5 | (AAATT)3 | TACAGGCACCAACATGGAGA | CCCTCTATGCCACACTTGCT | 171 |
| chr8:54355380-54357442 | p4 | (ATAC)6 | TCGTGACATGCTGGTCAAAT | TACCGCACATCGAATGAAAA | 116 |
| chr8:54367227-54369242 | p5 | (AGAAA)3 | AGTTTTGTACGTTGCCTCCG | TCCTCCATTCCTTACAACGC | 171 |
| chr8:54421345-54423361 | p2 | (AC)8 | AATCCACAAAGTGGGGTCTG | TGATGTTTTTGCTGTTGGGA | 149 |
| chr8:54594075-54596145 | p2 | (AT)26 | CGGGATTCAAATTTGGAGAA | GTGGATTCCGAAGAGCAGAG | 108 |
| chr8:54631332-54633398 | p2 | (AT)33 | AGAGTGATTCGGGACCAACA | GGTCCCAGATGTGGCTTTAG | 263 |
| chr8:54742659-54744674 | p5 | (AAATA)3 | AAACCCAAAACAAGCAATCG | TGATGATGATGGGGATGATG | 229 |
| chr8:54873850-54875870 | p2 | (AC)6 | AGAGCCTGGCATAATCCTCA | GGTGGACTGTTGCAGGAGAT | 146 |
| chr8:54873850-54875870 | p2 | (AT)10 | AGAGCCTGGCATAATCCTCA | GGTGGACTGTTGCAGGAGAT | 146 |
| chr8:54942075-54944115 | p2 | (TA)20 | CCCTAGGACGAACTGCTCTG | TGATGGGCACAATTGTTGAT | 153 |
| chr8:54942075-54944115 | p3 | (ATC)9 | CCCTAGGACGAACTGCTCTG | TGATGGGCACAATTGTTGAT | 153 |
| chr8:54942395-54944422 | p2 | (TA)20 | AGGAATCCATAAAGACGGGG | GGAGCTTCATCAACGGAGTC | 155 |
| chr8:54942395-54944422 | p3 | (ATC)9 | AGGAATCCATAAAGACGGGG | GGAGCTTCATCAACGGAGTC | 155 |
| chr8:54976397-54978412 | p5 | (TTTTG)3 | CAAACTTCGACGGTGTTTCA | CGGGTGGAGTGAATTGTTCT | 141 |
| chr8:54994193-54996255 | p2 | (TA)31 | GCTCCAGTTCAGAGGGTGAC | CCCCCTCAGGAAAAGAACTC | 148 |
| chr8:55265208-55267224 | p4 | (AAGA)4 | TTAAAGATCCAGCACCAGGG | TTACCTTGGTTTTTCGCGTT | 205 |
| chr8:55268597-55270612 | p5 | (ACTAT)3 | TCAAGGGCTCCTTTTTGAGA | ACCCCACTTGTGGGATTACA | 110 |
| chr8:55312416-55314438 | p2 | (AT)11 | TCCCCTTTCTAATGGGAACC | TGATAGCCAAAGGGCTTACG | 259 |
| chr8:55312416-55314438 | p3 | (TCA)5 | TCCCCTTTCTAATGGGAACC | TGATAGCCAAAGGGCTTACG | 259 |
| chr8:55497663-55499678 | p2 | (AT)9 | CCTCAACAGATGAAACCCGT | CCGGAGCGTATACTTTTGGA | 259 |
| chr8:55497663-55499678 | p5 | (ATCAG)3 | CCTCAACAGATGAAACCCGT | CCGGAGCGTATACTTTTGGA | 259 |
| chr8:55539220-55541235 | p5 | (ATGCA)3 | TCATTCGGAGATTATTCGGC | CCGTGACACTTAATCACCCC | 188 |
| chr8:55647092-55649108 | p4 | (CAGA)4 | ACATGCCAAAGAATGAAGGG | CCGATTCTGTCAGGTAGGGA | 202 |
| chr8:55782589-55784604 | p3 | (CAA)5 | GGTGAACAATGCAATGCAAC | ACGAACATGCATGACTCCAA | 232 |
| chr8:55782589-55784604 | p3 | (CAA)8 | GGTGAACAATGCAATGCAAC | ACGAACATGCATGACTCCAA | 232 |
| chr8:55874694-55876718 | p6 | (GATGGG)4 | ATGGCAACAAGAATGGAAGG | TCGGGTGTTCTTAATGCTCC | 187 |
| chr8:55952129-55954149 | p2 | (AT)10 | GTGTTGTGGCCTTTGTTGTG | GGAAAACATTTGAGGCGTGT | 185 |
| chr8:55954032-55956050 | p6 | (ATATGT)3 | ATGTTGGATATGAGCCTCGG | GTCGGGTCCTATTTATGCGA | 224 |
| chr8:56008734-56010752 | p6 | (ATATGT)3 | CTCGCATAAATAGGACCCGA | CTCGGTGAACGGGATAAGAA | 296 |
| chr8:56028120-56030135 | p3 | (TAA)10 | AATGGGAAATGAGGGAAACC | TCGGATCCCAAATCAAAGTC | 205 |

| Marker ID | SSR type | SSR motif | FORWARD PRIMER (5'-3') | REVERSE PRIMER (5'-3') | size (bp) |
|---|---|---|---|---|---|
| chr8:56028218-56030239 | p3 | (TAA)10 | AATGGGAAATGAGGGAAACC | GATCCCAAATCAAAGTCCGA | 107 |
| chr8:56062230-56064250 | p2 | (AT)10 | GCATTTTCAGAGCATGAGCA | AAAAATCTCAACGTGTGGGC | 128 |
| chr8:56125198-56127210 | p2 | (CT)6 | AATCGTTTTCGCTGTTGACC | ATTTCCGGGGTGTAACACAA | 225 |
| chr8:56163857-56165879 | p2 | (AC)6 | ATAAATGCAAGCCGACAAGG | TTGGAGAGTTCCGTTGTTCC | 258 |
| chr8:56163857-56165879 | p2 | (AT)11 | ATAAATGCAAGCCGACAAGG | TTGGAGAGTTCCGTTGTTCC | 258 |
| chr8:56251265-56253283 | p5 | (AAATA)3 | GGGCCTCAATAGCCCTTTAC | CCTGGTAAATCCTCGTTCCA | 296 |
| chr8:56251265-56253283 | p6 | (CCCCTT)3 | GGGCCTCAATAGCCCTTTAC | CCTGGTAAATCCTCGTTCCA | 296 |
| chr8:56347926-56349941 | p5 | (GGTTG)3 | AAAGAAGGAGAGGTTTCGGC | GGAAACGACGAAACGGATAA | 157 |
| chr8:56448398-56450410 | p2 | (TA)6 | CACGGCTTGTAGACACTCCA | GTCGGCTTGCGTATCATTTT | 128 |
| chr8:56496478-56498520 | p2 | (TA)9 | GCACGGCTTAGGCAAAATAG | TGGGGAGGGTAGAGTGTACG | 120 |
| chr8:56496478-56498520 | p3 | (GTT)5 | GCACGGCTTAGGCAAAATAG | TGGGGAGGGTAGAGTGTACG | 120 |
| chr8:56505728-56507758 | p2 | (AT)11 | GAGAGGAGAGCAAAGGGGTT | TTGTTCCAACAACCACTCCA | 239 |
| chr8:56505728-56507758 | p5 | (TGAAT)6 | GAGAGGAGAGCAAAGGGGTT | TTGTTCCAACAACCACTCCA | 239 |
| chr8:56552741-56554756 | p3 | (CTG)5 | ACAGGAAGCTAGCCAGTCCA | CACCCGAACCTATAGAGCCA | 247 |
| chr8:56554567-56556581 | p2 | (TA)7 | TTCGGAAAACCTCCTTGATG | ACGTCCAGCAAAATACTGGC | 206 |
| chr8:56573176-56575191 | p5 | (CTTAC)3 | CTTAAGCTCCGTGCCAAATC | CCCTCCGTTTCATGTCTGAT | 174 |
| chr8:56573176-56575191 | p5 | (CTTAC)3 | CTTAAGCTCCGTGCCAAATC | CCCTCCGTTTCATGTCTGAT | 174 |
| chr8:56574158-56576173 | p5 | (CTTAC)3 | AGACACATCAGCCTCCCAAC | CTGGAAACGGTCCAAAGGTA | 258 |
| chr8:56574158-56576173 | p5 | (CTTAC)3 | AGACACATCAGCCTCCCAAC | CTGGAAACGGTCCAAAGGTA | 258 |
| chr8:56625120-56627132 | p2 | (TG)6 | GGCGAAACAATAACCGAGAA | TTACAAGCATTTTGGGGGAG | 105 |
| chr8:56638741-56640756 | p5 | (TTTTA)3 | GGTGTCCATAACCATCAGGG | TTTTATGCGTAGCGGGATTC | 217 |
| chr8:56689351-56691366 | p3 | (TTG)5 | TGCCACCAAAACTTAAAGGC | CGCATCTTGTCTTCCACTGA | 277 |
| chr8:56691801-56693871 | p2 | (TA)28 | GGACCGTCATTTAGCTCCAA | GCTTAAGTTGGGGGTGTTGA | 176 |
| chr8:56724581-56726597 | p2 | (TG)8 | GGCCGACAACGATTAGAAAA | CGGTTTGTGGATTGAGGAGT | 212 |
| chr8:56727612-56729657 | p5 | (AATAT)9 | GTTGGTCATTCATCTGCACG | TTTCGAACTCCCAAAACCAC | 256 |
| chr8:56752639-56754653 | p2 | (AT)7 | TACCCGACAATTCCGGTTTA | GACAGTTTGCGGTGGATTTT | 109 |
| chr8:56799906-56801918 | p2 | (TA)6 | CATGTTGCGAATTTCCCTTT | GGGGAAAATGACAAGCAAGA | 287 |
| chr8:56828519-56830531 | p5 | (TGCAT)3 | CGGGTGGGTAGTTACAGGAA | TAATCATAAGCCATGGGGGA | 231 |
| chr8:56828519-56830531 | p2 | (CT)6 | CGGGTGGGTAGTTACAGGAA | TAATCATAAGCCATGGGGGA | 231 |
| chr8:56877219-56879235 | p2 | (AT)8 | ATCTGGGCCAATCAACTCTG | GAGCACAGCTTTCCATCCTC | 173 |
| chr8:56935887-56937899 | p2 | (CT)6 | ACACGAACTCAACTTGGCCT | TGGCACGATTGTTTGTTGTT | 225 |
| chr8:56975354-56977366 | p2 | (AT)6 | AAATGTTCGCAGCTTCCTGT | TATGTTAGAGGGGTGCCTGG | 151 |
| chr8:57058622-57060647 | p2 | (AT)7 | TTGGCCAATTTCAATCAACA | GTAGTTGGAGGCTTGCCTTG | 233 |
| chr8:57093695-57095713 | p6 | (TATATG)3 | GCACCTGCCAAGGGTAAATA | TAGTACCACGAAAGTGGGGC | 102 |
| chr8:57093695-57095713 | p6 | (GCACCA)3 | GCACCTGCCAAGGGTAAATA | TAGTACCACGAAAGTGGGGC | 102 |
| chr8:57100144-57102159 | p5 | (ATCAT)3 | GTCGGGTACGTAAAACACCG | CCACGTGCCATATCTGTGTC | 218 |

| Marker ID | SSR type | SSR motif | FORWARD PRIMER (5'-3') | REVERSE PRIMER (5'-3') | size (bp) |
|---|---|---|---|---|---|
| chr8:57100144-57102159 | p3 | (CAA)5 | GTCGGGTACGTAAAACACCG | CCACGTGCCATATCTGTGTC | 218 |
| chr8:57106327-57108339 | p2 | (TC)6 | GAATCAGGTTTCGCATGGTT | AAATGAAGTACGTGGCCCAG | 203 |
| chr8:57122121-57124136 | p5 | (TATAT)3 | AAGGGTCCTGGAATTTTGCT | AAGAATGACAGCACACCACG | 263 |
| chr8:57221591-57223607 | p2 | (AG)8 | GCAAAACTGTTACACCCCGT | CGATGTCACCTCGGGATACT | 226 |
| chr8:57233575-57235591 | p2 | (AG)8 | GCAAAACTGTTACACCCCGT | CGATGTCACCTCGGGATACT | 226 |
| chr8:57256862-57258880 | p6 | (ATCATT)3 | ACGTGGAAAATCGTGAGACC | ACAAGTAGACAGGGAGGCGA | 145 |
| chr8:57335181-57337193 | p2 | (CT)6 | TCCCAAACAGCCCACATAAT | GGAGTTGAGGCCATTGTTGT | 284 |
| chr8:57341579-57343613 | p2 | (AT)17 | CACTCGTGTGCGTACCTGTT | GTCGGCTTGCGTACCATATT | 217 |
| chr8:57396805-57398821 | p3 | (CAA)5 | GGACATGCCAAAAGAAAGGA | GAAATTCCGGCAGGATACAA | 117 |
| chr8:57396805-57398821 | p2 | (CA)8 | GGACATGCCAAAAGAAAGGA | GAAATTCCGGCAGGATACAA | 117 |
| chr8:57396805-57398821 | p2 | (AT)8 | GGACATGCCAAAAGAAAGGA | GAAATTCCGGCAGGATACAA | 117 |
| chr8:57396805-57398821 | p5 | (CTTTT)3 | GGACATGCCAAAAGAAAGGA | GAAATTCCGGCAGGATACAA | 117 |
| chr8:57402173-57404189 | p3 | (CAA)5 | AGCCTTACATACCTGTGCCG | ATTCCGGCAGGATACAAGTG | 143 |
| chr8:57402173-57404189 | p2 | (CA)8 | AGCCTTACATACCTGTGCCG | ATTCCGGCAGGATACAAGTG | 143 |
| chr8:57402173-57404189 | p5 | (ATTTC)3 | AGCCTTACATACCTGTGCCG | ATTCCGGCAGGATACAAGTG | 143 |
| chr8:57558323-57560335 | p2 | (CA)6 | AGGAGTGGGGAGCCTTACAT | AGCTAAGTCTCGGGGGTGTT | 271 |
| chr8:57653671-57655683 | p2 | (GT)6 | GTGCTCGCCCTGTAAGACTC | CCTTCCCCAATCACTTAGCA | 232 |
| chr8:57689609-57691621 | p2 | (AT)6 | ATCCAGTCTTCCACGTGTCC | TGGGACGTAACCTTCTCACC | 290 |
| chr8:57693841-57695857 | p4 | (GAAA)4 | ACTCATCACACCTTCGGTCC | GGCTTTTGTTTAGTGCAGGC | 140 |
| chr8:57829620-57831638 | p2 | (TA)9 | TTTTCACCCAAGGTTCATCC | ATACAATGCCGAACCCATGT | 141 |
| chr8:57912075-57914093 | p3 | (TGT)6 | ACCAAGCAAAGCGAAGAAAA | CGGATTGACGAACGGTTTAT | 103 |
| chr8:58008924-58010942 | p2 | (TA)9 | TCTTTTCCTGTGCTCGCTTT | TTGATCAGAAGGGTGAAGGG | 136 |
| chr8:58008924-58010942 | p2 | (GT)6 | TCTTTTCCTGTGCTCGCTTT | TTGATCAGAAGGGTGAAGGG | 136 |
| chr8:58102578-58104598 | p2 | (AT)10 | CCTGTTGTATGTTTGGGGCT | ATGAGCCATCTTTGTGGTCC | 244 |
| chr8:58157207-58159225 | p3 | (TTG)6 | AAACTCATCCCATACCGCTG | GTTAGCTTCGTATCGGCTGC | 282 |
| chr8:58167466-58169490 | p2 | (TA)12 | GTGCCTAGAGAATTGCTGCC | TCTCGTACTTCGCCCACTCT | 266 |
| chr8:58167466-58169490 | p5 | (CATAT)3 | GTGCCTAGAGAATTGCTGCC | TCTCGTACTTCGCCCACTCT | 266 |
| chr8:58203523-58205535 | p2 | (CA)6 | CCATTTCGGATTTCAAGGAA | AGTTTTTGGCGAGCGTAAGA | 196 |
| chr8:58217112-58219127 | p5 | (ATCAT)3 | CCGGTGAGGCTGTCATAGAT | GATTTGTTGACCGCGTTCTT | 294 |
| chr8:58223937-58225952 | p5 | (ATACG)3 | AAGAAGTCACGAGCCCTCAA | AGCTCTACTGCCAGCTCCTG | 132 |
| chr8:58223937-58225952 | p2 | (AT)30 | AAGAAGTCACGAGCCCTCAA | AGCTCTACTGCCAGCTCCTG | 132 |
| chr8:58246104-58248122 | p6 | (TATGTA)3 | GCCCATTCAGCCCACTAATA | TCCCCGATTTTAACAAGTGC | 125 |
| chr8:58251081-58253097 | p4 | (ATAC)4 | ATCATACAGGAGCCACCAGG | TTGCGTCTAAGCGGTCCTAT | 234 |
| chr8:58273887-58275921 | p2 | (TA)23 | CGTGGAATAGAGATCCGAGC | CCGTCAACTTGCCCATAGAT | 273 |
| chr8:58273922-58275944 | p2 | (TA)23 | CGTGGAATAGAGATCCGAGC | CCGTCAACTTGCCCATAGAT | 238 |
| chr8:58328835-58330850 | p3 | (ACA)5 | ATGGAACGAATCAAACGGAG | TTTATATCCGCCACCTCGAC | 179 |

| Marker ID | SSR type | SSR motif | FORWARD PRIMER (5'-3') | REVERSE PRIMER (5'-3') | size (bp) |
|---|---|---|---|---|---|
| chr8:58456251-58458269 | p6 | (CATAAT)3 | AAACGTTGCCTTCTCGCTAA | GGCCCAAATCCATGAACTAA | 198 |
| chr8:58565117-58567135 | p6 | (TATATG)3 | GGATTCTCGGGGTTGGTTAT | TTACATACCTTGCCCGCTTC | 186 |
| chr8:58607015-58609029 | p2 | (TA)7 | CTACACGTTCCGCAGCACTA | GGGGTCAGTACGAGCAATGT | 121 |
| chr8:58607015-58609029 | p2 | (TA)6 | CTACACGTTCCGCAGCACTA | GGGGTCAGTACGAGCAATGT | 121 |
| chr8:58671759-58673777 | p6 | (TTCAGT)3 | TGGGGTGAAGAAACACCATT | CAATGACAAGCCCAAAGGAT | 228 |
| chr8:58708458-58710470 | p2 | (AT)6 | TGTACCATGGAACCCGAAAT | GTTTTGGGAAGCCAATTTGA | 108 |
| chr8:58790920-58792935 | p5 | (TTCAT)3 | GCTATAGCGGAGTGATTCGC | ACCCACACAGCCTCTCAATC | 161 |
| chr8:58812840-58814855 | p2 | (TA)6 | TCGGGGTCTTGACAACTTTC | AGTTTTCATCGGCTGATTGG | 165 |
| chr8:58812840-58814855 | p2 | (TA)22 | TCGGGGTCTTGACAACTTTC | AGTTTTCATCGGCTGATTGG | 165 |
| chr8:58812840-58814855 | p5 | (CATAT)3 | TCGGGGTCTTGACAACTTTC | AGTTTTCATCGGCTGATTGG | 165 |
| chr8:58984878-58986890 | p2 | (AT)6 | GACGGACAGGGTTGTTCCTA | TTCAATGCCGCAAGTACAAG | 270 |
| chr8:59106256-59108271 | p5 | (TTCAT)3 | TTAGTCGTGCCTGCATTCTG | TTGAAATGGCTAACAAGGGG | 284 |
| chr8:59319605-59321619 | p2 | (AT)7 | AAGCCGACAAGGCTACAAGA | TTCCGGAGCTACAACTTTGG | 129 |
| chr8:59322458-59324470 | p2 | (AT)6 | CGACCACTACTGACCAAGCA | TGCAAGTTTTCGATGCAAAG | 173 |
| chr8:59322458-59324470 | p3 | (CAT)5 | CGACCACTACTGACCAAGCA | TGCAAGTTTTCGATGCAAAG | 173 |
| chr8:59322702-59324717 | p2 | (AT)6 | CTCCAGCTCCGAACCAAATA | TTCTTGTGGACGCTGTGTTC | 293 |
| chr8:59322702-59324717 | p3 | (CAT)5 | CTCCAGCTCCGAACCAAATA | TTCTTGTGGACGCTGTGTTC | 293 |
| chr8:59395787-59397805 | p2 | (TA)9 | GACGATAAGTGGGATGCGTT | TCGTCGATTTTCCCTGAATC | 217 |
| chr8:59395787-59397805 | p2 | (CA)6 | GACGATAAGTGGGATGCGTT | TCGTCGATTTTCCCTGAATC | 217 |
| chr8:59415535-59417550 | p5 | (ATTTC)3 | CCGGAAGAGATCCAAGATGA | TCCTGTCACAACAGCCCATA | 222 |
| chr8:59420673-59422697 | p2 | (TA)10 | GGCAGAATTTTGGACAGGAA | TTCCGGTGTTGTTCATAGCA | 296 |
| chr8:59420673-59422697 | p2 | (TA)12 | GGCAGAATTTTGGACAGGAA | TTCCGGTGTTGTTCATAGCA | 296 |
| chr8:59424585-59426603 | p2 | (TA)9 | TGTAAAGGAGACCCGGTCAG | TCCTGCTAAGCCTCGAATGT | 141 |
| chr8:59435024-59437042 | p6 | (GGAGCT)3 | AGGAACATTATGGCACTGGC | AGAAGGGAACGTGAGCAAGA | 104 |
| chr8:59436149-59438175 | p2 | (TA)13 | CCTAGGACCGAACTGCTCTG | GGCTCGCCCAGATGTAAGTA | 246 |
| chr8:59740384-59742402 | p6 | (CATATA)3 | AAATGACACACGGCTGATGA | TGCCATTCCTCTTGGGTTAG | 197 |
| chr8:59784218-59786236 | p2 | (AT)13 | TGATCCATCTCCCTGCCTAC | AGCTAAGCATGCACGGTTTT | 299 |
| chr8:59784218-59786236 | p3 | (ATC)6 | TGATCCATCTCCCTGCCTAC | AGCTAAGCATGCACGGTTTT | 299 |
| chr8:59791771-59793803 | p2 | (AT)22 | CAAGGGAAAGGTGTGGTTGT | TCCCAACTTCATACCCGAAG | 161 |
| chr8:59791771-59793803 | p2 | (AT)7 | CAAGGGAAAGGTGTGGTTGT | TCCCAACTTCATACCCGAAG | 161 |
| chr8:59911371-59913389 | p6 | (AACTGA)3 | TCCCAAGAAACTTCGGACAC | TGAAAAGGGAATTGAGGTCG | 110 |
| chr8:59921095-59923110 | p5 | (CCTCT)3 | CGCCATAGGGACTCAGTCAT | GCTGATAGACTGGCTGGGAG | 215 |
| chr8:60001445-60003463 | p2 | (AT)6 | CCTTGATGTCCATAAGCCGT | GCCATTTTTGGAAAATCGAA | 146 |
| chr8:60001445-60003463 | p6 | (TATATG)3 | CCTTGATGTCCATAAGCCGT | GCCATTTTTGGAAAATCGAA | 146 |
| chr8:60001445-60003463 | p2 | (TA)7 | CCTTGATGTCCATAAGCCGT | GCCATTTTTGGAAAATCGAA | 146 |
| chr8:60080435-60082461 | p2 | (TA)13 | GCTCTGTGAGGCATTGTTGA | CACTTGAAGGCAACCCATTT | 102 |

| Marker ID | SSR type | SSR motif | FORWARD PRIMER (5'-3') | REVERSE PRIMER (5'-3') | size (bp) |
|---|---|---|---|---|---|
| chr8:60082054-60084069 | p3 | (TGT)5 | GAAAGGCCAGGGTAAGAAG | TTGGCCCATTTATGTTGGAT | 209 |
| chr8:60091919-60093934 | p3 | (TGT)5 | TTCAAGATAATCGCGGGAAG | GGCCAAATGACCCCTTTAAT | 117 |
| chr8:60166251-60168275 | p6 | (CATATA)4 | CGGCTGATGGAGCTTCTTAC | AGAGTTGCCGGGTCAGAGTA | 255 |
| chr8:60167860-60169872 | p2 | (AC)6 | GAGAATTCAACTGGGCCAAA | GAATTTGGAAGGTTTCGGGT | 219 |
| chr8:60167860-60169872 | p2 | (TA)6 | GAGAATTCAACTGGGCCAAA | GAATTTGGAAGGTTTCGGGT | 219 |
| chr8:60274264-60276276 | p2 | (CA)6 | GGACATGCCAAAAGAAGGAA | TTATAAGGGAAATGCTGCCG | 239 |
| chr8:60306251-60308267 | p5 | (TGATA)3 | ATTTCAGCTGGCCGAGAGTA | GGCTTACCTCATATGCCCAA | 268 |
| chr8:60306251-60308267 | p2 | (AT)8 | ATTTCAGCTGGCCGAGAGTA | GGCTTACCTCATATGCCCAA | 268 |
| chr8:60306251-60308267 | p2 | (AT)6 | ATTTCAGCTGGCCGAGAGTA | GGCTTACCTCATATGCCCAA | 268 |
| chr8:60320942-60322958 | p2 | (AT)9 | CTGGACACTCGTTTTGCGTA | CAGCCTTGTTGACTTGCGTA | 148 |
| chr8:60320942-60322958 | p2 | (AT)15 | CTGGACACTCGTTTTGCGTA | CAGCCTTGTTGACTTGCGTA | 148 |
| chr8:60320942-60322958 | p2 | (AT)8 | CTGGACACTCGTTTTGCGTA | CAGCCTTGTTGACTTGCGTA | 148 |
| chr8:60381766-60383780 | p2 | (TG)7 | AGGTGCCTACGGGCTATTTT | TCATACCTGGCCCTACCAAG | 240 |
| chr8:60469976-60471988 | p2 | (AT)6 | ATCAATCTCCCGTTTTGGTG | CAGCATCTCCGAGAGGTAGG | 160 |
| chr8:60469976-60471988 | p2 | (TA)18 | ATCAATCTCCCGTTTTGGTG | CAGCATCTCCGAGAGGTAGG | 160 |
| chr8:60469976-60471988 | p2 | (TA)15 | ATCAATCTCCCGTTTTGGTG | CAGCATCTCCGAGAGGTAGG | 160 |
| chr8:60474639-60476657 | p2 | (TA)10 | CGACCTTACGTCACCTCGAT | AACTGGCCATGGTGTATCGT | 262 |
| chr8:60474685-60476707 | p2 | (TA)10 | CGACCTTACGTCACCTCGAT | AACTGGCCATGGTGTATCGT | 216 |
| chr8:60484602-60486620 | p6 | (CATATA)3 | TGTCTCCCGAACAATCAACA | TTGATCTGGAGCTACACCCC | 294 |
| chr8:60529219-60531233 | p2 | (AC)7 | TGACCCAAAGAATTTCGGAG | CGGTCAGGAGATGAGAGAGG | 247 |
| chr8:60540537-60542552 | p3 | (TTG)5 | ATCCAGATGGCCCAATATGA | TTCTGGAGACTCTGGGCTGT | 286 |
| chr8:60558781-60560797 | p4 | (ACAG)4 | ACTTCTGTCACGACCCAACC | TGTGTGGGTTACGGGTTTTT | 221 |
| chr8:60624258-60626273 | p3 | (TTG)5 | GTTTGGCCGTGTGTAGTGTG | ATTTTCGGCCATCACCATAA | 170 |
| chr8:60624258-60626273 | p3 | (TTG)5 | GTTTGGCCGTGTGTAGTGTG | ATTTTCGGCCATCACCATAA | 170 |
| chr8:60652022-60654042 | p4 | (AATT)5 | GCTTTGTCGCTCTTTCATCC | ATTGGAAATTCGGGTTAGGC | 194 |
| chr8:60652022-60654042 | p3 | (TTG)5 | GCTTTGTCGCTCTTTCATCC | ATTGGAAATTCGGGTTAGGC | 194 |
| chr8:60652022-60654042 | p3 | (ATG)5 | GCTTTGTCGCTCTTTCATCC | ATTGGAAATTCGGGTTAGGC | 194 |
| chr8:60684270-60686282 | p2 | (AT)17 | ATTGTCACGACCCGACTAGG | ATGGAGGAACAAGCCTTGAA | 170 |
| chr8:60684270-60686282 | p2 | (TA)6 | ATTGTCACGACCCGACTAGG | ATGGAGGAACAAGCCTTGAA | 170 |
| chr8:60690325-60692339 | p2 | (AT)7 | TTCCTCCGACATTATTTGCC | CTCAGTCACCCATATCCGCT | 245 |
| chr8:60781659-60783677 | p6 | (ATATGA)3 | TCAACACTGGGCAGAACTTG | CAGCATTTCCCCTATTGCAT | 141 |
| chr8:60781659-60783677 | p2 | (TA)10 | TCAACACTGGGCAGAACTTG | CAGCATTTCCCCTATTGCAT | 141 |
| chr8:60836340-60838352 | p2 | (AT)6 | TAAAACATCTTTTTGGGCGG | CTTAACGGACGACGAAGCAT | 250 |
| chr8:60836359-60838373 | p2 | (AT)6 | TAAAACATCTTTTTGGGCGG | CTTAACGGACGACGAAGCAT | 231 |
| chr8:60836437-60838455 | p2 | (AT)6 | TAAAACATCTTTTTGGGCGG | CTTAACGGACGACGAAGCAT | 153 |
| chr8:60867804-60869825 | p3 | (TTG)7 | GCCACCAAAATTTAAAGGCA | TGGGGCATCTATGTCTCTCC | 189 |

| Marker ID | SSR type | SSR motif | FORWARD PRIMER (5'-3') | REVERSE PRIMER (5'-3') | size (bp) |
|---|---|---|---|---|---|
| chr8:60911513-60913539 | p2 | (TA)13 | CGTGTCGAACTGTTGAAGGA | TCCAACTCCAAATAGCCCAC | 112 |
| chr8:60968926-60970941 | p3 | (GTT)5 | GGAGACTTGCTTTCGGAGTG | TTATCTTCCACCAAGGCCAC | 207 |
| chr8:60978878-60980902 | p2 | (AT)12 | GGATGCGAGCTCTACTGGTC | CAAATTAAATTGCCCACGCT | 159 |
| chr8:61015630-61017642 | p2 | (AG)6 | TCTACGCGTTCCGTCCTAGT | ACTTTCCGACAGGCTTCTCA | 274 |
| chr8:61032558-61034596 | p2 | (TA)20 | GTTGGGCGATTTGAGATTGT | TCTGGTCACGGGTCAATGTA | 207 |
| chr8:61032558-61034596 | p2 | (TA)19 | GTTGGGCGATTTGAGATTGT | TCTGGTCACGGGTCAATGTA | 207 |
| chr8:61127844-61129862 | p2 | (CA)9 | TTCAAGGAACAGAATTGCCC | GCAGATTCTGAGCGAAAAGG | 183 |
| chr8:61188810-61190830 | p2 | (AT)10 | CAACCAACCCACCGTAAATC | TTTCAGGTCACTGCTCGATG | 128 |
| chr8:61188810-61190830 | p2 | (AT)18 | CAACCAACCCACCGTAAATC | TTTCAGGTCACTGCTCGATG | 128 |
| chr8:61202673-61204687 | p2 | (AT)7 | AAAGCTGTAGCTCCGGATGA | TATTATCCCCCGATTCTCCC | 269 |
| chr8:61206099-61208113 | p2 | (AT)7 | ATACATTTTCGTTGCTCCCG | TGAGGATCCCGTGCTAATTC | 300 |
| chr8:61218161-61220177 | p2 | (AT)8 | AAAGCTGTAGCTCCGGATGA | TGACGTCCTTTTGTTGTGGA | 269 |
| chr8:61223524-61225539 | p5 | (TTCAT)3 | GATGAGGGATTTCTGCGTGT | AAGCTGTTCCGCTGAAGTGT | 183 |
| chr8:61224784-61226800 | p2 | (TA)8 | TATACAGGGGAAATGCTGCC | AAACCCGTAGCCCATCTCTT | 235 |
| chr8:61224784-61226800 | p2 | (AT)8 | TATACAGGGGAAATGCTGCC | AAACCCGTAGCCCATCTCTT | 235 |
| chr8:61344340-61346366 | p3 | (GTT)7 | GCCACTTGGCTTAATGTGGT | GGGAGGGTAGAGTGTACGCA | 276 |
| chr8:61478267-61480291 | p2 | (AT)9 | TGTTCGTCTGTAGTGGCAGC | TCCCGATAACCTGTCTCCTG | 273 |
| chr8:61478267-61480291 | p3 | (AAC)8 | TGTTCGTCTGTAGTGGCAGC | TCCCGATAACCTGTCTCCTG | 273 |
| chr8:61521645-61523661 | p6 | (AACTCC)3 | TCCTTTTCTTTGGACGCTGT | GCGCATCTACGAGCCTTTAC | 189 |
| chr8:61521645-61523661 | p2 | (GT)8 | TCCTTTTCTTTGGACGCTGT | GCGCATCTACGAGCCTTTAC | 189 |
| chr8:61526771-61528796 | p5 | (ACAAT)5 | GATCAGGGCATGGGATAGAA | CACCACACCTGTCACTTTGG | 253 |
| chr8:61539771-61541786 | p5 | (CTTTG)3 | TGTTGCCCAATTACCCCTAC | AAGGTGGTGGAATGTCGAAG | 210 |
| chr8:61588359-61590375 | p6 | (AACTCC)3 | TCCTTTTCTTTGGACGCTGT | GCGCATCTACGAGCCTTTAC | 189 |
| chr8:61588359-61590375 | p2 | (GT)8 | TCCTTTTCTTTGGACGCTGT | GCGCATCTACGAGCCTTTAC | 189 |
| chr8:61593485-61595510 | p5 | (ACAAT)5 | GATCAGGGCATGGGATAGAA | CACCACACCTGTCACTTTGG | 253 |
| chr8:61645024-61647039 | p5 | (TACTT)3 | TAGAAGCTTGGCCAAACAGG | AAGGGAGCACATTTTGATGG | 262 |
| chr8:61670473-61672488 | p5 | (TTTTC)3 | TGGCATGGCAACTAAATCAA | TCTAAGATCCCGATCCAACG | 171 |
| chr8:61670473-61672488 | p5 | (ACCAA)3 | TGGCATGGCAACTAAATCAA | TCTAAGATCCCGATCCAACG | 171 |
| chr8:61670473-61672488 | p5 | (AAACC)3 | TGGCATGGCAACTAAATCAA | TCTAAGATCCCGATCCAACG | 171 |
| chr8:61696761-61698776 | p3 | (AAT)5 | CACTGCCTCCTTCTTTTTCG | GTCATCGCCAGAGTTCCATT | 201 |
| chr8:61714619-61716631 | p2 | (AT)6 | TTTGGCTGATGTTACGGTGA | CGGACCGTAACTGGATCTGT | 100 |
| chr8:62340732-62342747 | p5 | (TTTCT)3 | AGGGAAGCATACACAATGCC | GGCCAGCCGCATATATTTTA | 294 |
| chr8:62442973-62444988 | p5 | (CCTTT)3 | AACACGAGCCAAAAACCAAG | GGGTTGTGCTACCGAGATGT | 105 |
| chr8:62473032-62475062 | p2 | (TA)15 | CCCGTCATCAACTTGGAACT | ATGAAGCATTTTATTGGCGG | 288 |
| chr8:62528886-62530906 | p2 | (TA)10 | AGTCATGTCGTGAACTCCCC | AACGACAAGCGAGGCTACAT | 277 |
| chr8:62658680-62660695 | p3 | (GAT)5 | ATGATGTCAAGCCACCACAA | GACCTTGAATGTTGAGGGGA | 226 |

| Marker ID | SSR type | SSR motif | FORWARD PRIMER (5'-3') | REVERSE PRIMER (5'-3') | size (bp) |
|---|---|---|---|---|---|
| chr8:62684007-62686023 | p2 | (AT)8 | CTTTTGTCATGACCCAACCC | GTCGGCTTGCGTATCATTTT | 123 |
| chr8:62684007-62686023 | p2 | (AT)8 | CTTTTGTCATGACCCAACCC | GTCGGCTTGCGTATCATTTT | 123 |
| chr8:62699893-62701908 | p5 | (TCATT)3 | AGTGCGAAAGCTCCGAAATA | GCTCACAGGGCTACATCACA | 170 |
| chr8:62728546-62730558 | p2 | (TA)6 | TTTAGGGGTTGTCCTTGCTG | TCTAAGAGGTGCTCGGTGGT | 173 |
| chr8:62742259-62744271 | p2 | (TA)7 | CGAAGGTAAGGGAATGACCA | GAATTTCCTTTTTCGCCCTC | 105 |
| chr8:62742259-62744271 | p2 | (GT)6 | CGAAGGTAAGGGAATGACCA | GAATTTCCTTTTTCGCCCTC | 105 |
| chr8:62800811-62802823 | p2 | (AT)6 | AAATAGGCTAAAGGGGCGAA | CTCGTCGTCTGGTTGTTTCA | 137 |
| chr8:62800825-62802837 | p2 | (AT)6 | AAATAGGCTAAAGGGGCGAA | CTCGTCGTCTGGTTGTTTCA | 123 |
| chr8:62800839-62802867 | p2 | (AT)6 | AAATAGGCTAAAGGGGCGAA | CTCGTCGTCTGGTTGTTTCA | 109 |
| chr8:62810280-62812298 | p2 | (AT)9 | CAAAATCAGCCCATTTTCGT | CTGTCCGTCAGTTTGAGGGT | 283 |
| chr8:62810280-62812298 | p2 | (AT)10 | CAAAATCAGCCCATTTTCGT | CTGTCCGTCAGTTTGAGGGT | 283 |
| chr8:62854282-62856297 | p3 | (ATA)5 | TTTCCCCGAGTCATATTTCG | TCATGAACCCCCGATTGTAT | 276 |
| chr8:62952112-62954132 | p2 | (TA)10 | TCCCTTCTTTCTCTTTGGCA | ACCTCCGAACATCGTTATGG | 245 |
| chr8:62967036-62969062 | p2 | (AT)13 | AATGGACCGAAGATGCTGAC | AGCATTGCCGAGTCTGAGTT | 267 |
| chr8:62967036-62969062 | p3 | (CAA)6 | AATGGACCGAAGATGCTGAC | AGCATTGCCGAGTCTGAGTT | 267 |
| chr8:63076577-63078597 | p4 | (ATAC)5 | TGGACACCCATACACGCTTA | CCTGGTGGCTCCTGTATGAT | 111 |
| chr8:63078054-63080066 | p2 | (AC)6 | CCCGCTTTCTACCCTAGTCC | GTTGAATTCTCGGGCATGTT | 294 |
| chr8:63117989-63120001 | p2 | (AT)6 | TCGAACGGTGAATAAGGAGG | ATGGTATTCGCAGAACAGGG | 278 |
| chr8:63117989-63120001 | p2 | (AT)20 | TCGAACGGTGAATAAGGAGG | ATGGTATTCGCAGAACAGGG | 278 |
| chr8:63252040-63254052 | p2 | (AT)6 | CATTCGGGAACGAATGTTCT | TTCGAAATCCAAATTTTGCC | 288 |
| chr8:63321889-63323901 | p2 | (GT)6 | GTGAAGAAAAATCCGAGGCA | CAACCTCCCGCAATTGTATT | 108 |
| chr8:63353384-63355396 | p3 | (GTT)6 | CTCGGGGTAGCCTATTGTCA | GTCAGTCTGTAAGCGGCTCC | 268 |
| chr8:63353384-63355396 | p2 | (TA)6 | CTCGGGGTAGCCTATTGTCA | GTCAGTCTGTAAGCGGCTCC | 268 |
| chr8:63353384-63355396 | p6 | (TACATA)3 | CTCGGGGTAGCCTATTGTCA | GTCAGTCTGTAAGCGGCTCC | 268 |
| chr8:63353518-63355536 | p3 | (GTT)6 | CTCGGGGTAGCCTATTGTCA | GTCAGTCTGTAAGCGGCTCC | 134 |
| chr8:63353518-63355536 | p2 | (TA)6 | CTCGGGGTAGCCTATTGTCA | GTCAGTCTGTAAGCGGCTCC | 134 |
| chr8:63353518-63355536 | p6 | (TACATA)3 | CTCGGGGTAGCCTATTGTCA | GTCAGTCTGTAAGCGGCTCC | 134 |
| chr8:63358740-63360758 | p6 | (CATATA)3 | TGCAGACAACCCCTTAGACC | GTCAGTCTGTAAGCGGCTCC | 128 |
| chr8:63442476-63444491 | p3 | (CAA)5 | CACATCCCCTGTCAACTGTG | GTGTGGCCTTTGTTGTGATG | 134 |
| chr8:63589449-63591464 | p3 | (ACA)5 | GAACCGGTGAACGAAGTCAT | TTTGGAACATCCTCCTACGG | 215 |
| chr8:63617822-63619838 | p4 | (GAAA)4 | GTGGGCAAAAACTGAAGAGC | GCGTAGAGTTTGCGACCTTC | 110 |
| chr8:63705953-63707971 | p3 | (AAC)6 | TGGTTTACCAGACTCGAGGG | AGTTTCCCACTATTCCCGCT | 216 |
| chr8:63735232-63737247 | p3 | (ATG)5 | GTTGAGCATTTCCCTCCAAA | ATCATGCCTCTCCACCACTC | 298 |
| chr8:63740875-63742889 | p2 | (CA)7 | CGTCATTCTGACTGGCAAGA | TTCATTACAGGCTTACCCCG | 183 |
| chr8:63758253-63760268 | p3 | (ATG)5 | GTTGAGCATTTCCCTCCAAA | ATCATGCCTCTCCACCACTC | 298 |
| chr8:63794288-63796306 | p6 | (ATTCAT)3 | ACTTTTGTCACGACCCGACT | GTGTTTGGCCTTGATGACCT | 289 |

| Marker ID | SSR type | SSR motif | FORWARD PRIMER (5'-3') | REVERSE PRIMER (5'-3') | size (bp) |
|---|---|---|---|---|---|
| chr8:63817558-63819570 | p2 | (AT)6 | ATGGGCACGATACTCAGGAC | CGAGAAGTGATCAAGTGGCA | 230 |
| chr8:63842330-63844344 | p5 | (ACTCA)3 | CCCATAAGATGGGGAAGGAT | CTAGGAGCTCCATCAGTGGC | 146 |
| chr8:63842345-63844359 | p5 | (ACTCA)3 | CCCATAAGATGGGGAAGGAT | CTAGGAGCTCCATCAGTGGC | 131 |
| chr8:63893491-63895509 | p6 | (CATAAT)3 | TGCCATCTAGGCATGGTGTA | TAACGTAATCGGGGGTTCAC | 214 |
| chr8:63979501-63981529 | p2 | (TA)14 | CCAAAACAGACCCAAATGCT | GATTCGCAGGAACAATGGAT | 228 |
| chr8:64001894-64003968 | p2 | (AT)37 | AACGTATCAGCTCCACGTCC | TGAATGCTAGAGGGTTCGCT | 257 |
| chr8:64023602-64025614 | p2 | (TA)6 | AATACCATACGCAAGCCGAC | GGCCCTGTTCCCGTTATAGT | 272 |
| chr8:64136549-64138569 | p5 | (ACACA)4 | CATATCATGGGATCATGGCA | TAGATTTCTCGGGGATGGTG | 210 |
| chr8:64136549-64138569 | p2 | (AT)11 | CATATCATGGGATCATGGCA | TAGATTTCTCGGGGATGGTG | 210 |
| chr8:64228758-64230770 | p5 | (GAAAT)3 | CAAAGTGGAAACCCTCCAAA | TGGGCCTGAATTTGTTTTTC | 277 |
| chr8:64228758-64230770 | p2 | (TA)6 | CAAAGTGGAAACCCTCCAAA | TGGGCCTGAATTTGTTTTTC | 277 |
| chr8:64304983-64307007 | p2 | (AT)12 | GGTGTTTCTTTGAAGCCCAA | ACTTCGGTTTGGATCGTGAC | 248 |
| chr8:64333056-64335082 | p2 | (TA)13 | GCGCACAACACACCATAAAC | ATGGTGTTTTCCACCCAAAA | 292 |
| chr8:64338325-64340340 | p2 | (TA)8 | ACGTGAAGTGCATGATGAGC | AGTGATGAATGGGCCTTGTC | 196 |
| chr8:64338325-64340340 | p5 | (CTTGT)3 | ACGTGAAGTGCATGATGAGC | AGTGATGAATGGGCCTTGTC | 196 |
| chr8:64395757-64397769 | p2 | (TG)6 | CTTTAACATCCGAGGACCGA | CAAGTAAGGCCATTGTGGCT | 186 |
| chr8:64469693-64471711 | p2 | (AT)7 | TCAACAATCCCTAGATGCCC | AGCAGGTTTGTCCTAGGGGT | 218 |
| chr8:64469693-64471711 | p3 | (CAT)6 | TCAACAATCCCTAGATGCCC | AGCAGGTTTGTCCTAGGGGT | 218 |
| chr8:64469709-64471749 | p2 | (AT)7 | TCAACAATCCCTAGATGCCC | AGCAGGTTTGTCCTAGGGGT | 202 |
| chr8:64469709-64471749 | p3 | (CAT)6 | TCAACAATCCCTAGATGCCC | AGCAGGTTTGTCCTAGGGGT | 202 |
| chr8:64485837-64487853 | p2 | (AT)8 | ACGAAATGAGCACCAACACA | TTCCGTCTCTACATGGGTCC | 236 |
| chr8:64485837-64487853 | p5 | (TTGAA)4 | ACGAAATGAGCACCAACACA | TTCCGTCTCTACATGGGTCC | 236 |
| chr8:64513268-64515282 | p2 | (TA)7 | CTGCCACTACACGCAACATC | TTGGGATTGCTCCTTTGTTC | 150 |
| chr8:64576933-64578972 | p3 | (TTC)13 | CCCTTGACTCGGAGCATATC | ACAGAATCATATGGCGGGAC | 257 |
| chr8:64683361-64685376 | p5 | (TGTCC)3 | TGTGTGGCCTTTTGTTGGTA | GAGCACCCCTTATTTCCCTC | 222 |
| chr8:64686645-64688657 | p2 | (AT)6 | GGACTCAGGCGTTACAGGAG | ATGTGAATGCCAGAGGGTTC | 116 |
| chr8:64701148-64703164 | p5 | (CATAT)10 | GTATAACAAATCCCGGCCCT | TCTGTTCACCGTGTCCCATA | 265 |
| chr8:64701148-64703164 | p2 | (TA)8 | GTATAACAAATCCCGGCCCT | TCTGTTCACCGTGTCCCATA | 265 |
| chr8:64714671-64716701 | p5 | (CATAT)6 | GATGGCAAGCTCCATTTGTT | GGAGCCGCTTACAGACTGAC | 213 |
| chr8:64714671-64716701 | p2 | (TA)8 | GATGGCAAGCTCCATTTGTT | GGAGCCGCTTACAGACTGAC | 213 |
| chr8:64715087-64717103 | p5 | (CATAT)6 | GTATAACAAATCCCGGCCCT | CACGTTCTGTTCACCGTGTC | 285 |
| chr8:64715087-64717103 | p2 | (TA)8 | GTATAACAAATCCCGGCCCT | CACGTTCTGTTCACCGTGTC | 285 |
| chr8:64716480-64718495 | p5 | (TATCA)3 | AGCACTCCGGAATAGCAGAA | ATTCCGTACTGACCCCCTCT | 294 |
| chr8:64724409-64726437 | p6 | (CATATT)3 | ATAAGCCGACGAGGCTAACA | GTCCTGTCCCGCCTATATGA | 275 |
| chr8:64724409-64726437 | p2 | (AT)14 | ATAAGCCGACGAGGCTAACA | GTCCTGTCCCGCCTATATGA | 275 |
| chr8:64748799-64750820 | p2 | (TA)10 | TCATAAGGACGGGACAGGAC | GTAGACAGGGAGACGGTCCA | 209 |

| Marker ID | SSR type | SSR motif | FORWARD PRIMER (5'-3') | REVERSE PRIMER (5'-3') | size (bp) |
|---|---|---|---|---|---|
| chr8:64748799-64750820 | p3 | (ATC)7 | TCATAAGGACGGGACAGGAC | GTAGACAGGGAGACGGTCCA | 209 |
| chr8:64764120-64766132 | p2 | (AG)6 | GAGCGTAAGCCGCAACTAAG | CATAGACGCTCATAACGCGA | 206 |
| chr8:64802116-64804198 | p2 | (AT)41 | TTTCAGGTGTTGTCTTGCCA | TTGAGTCGATGAACCAGTGC | 242 |
| chr8:64949633-64951645 | p2 | (GT)6 | TTTGGGCAAAGAAAAGTTGG | ATACGGGACACAACACCCAT | 157 |
| chr8:64949633-64951645 | p3 | (TTG)5 | TTTGGGCAAAGAAAAGTTGG | ATACGGGACACAACACCCAT | 157 |
| chr8:65055596-65057608 | p2 | (TA)11 | AAAGGGATCCTCCCGAATTA | GGGGAAATGATTGTTCCCTT | 116 |
| chr8:65055596-65057608 | p2 | (AT)6 | AAAGGGATCCTCCCGAATTA | GGGGAAATGATTGTTCCCTT | 116 |
| chr8:65055596-65057608 | p5 | (TGAAT)3 | AAAGGGATCCTCCCGAATTA | GGGGAAATGATTGTTCCCTT | 116 |
| chr8:65057268-65059280 | p2 | (TA)6 | TTCGGAATCCGCATCTAAAG | AACAATAGGGAAAATCCGGG | 143 |
| chr8:65091510-65093522 | p2 | (AT)6 | TTAATATGTGTCCCACCGCA | ACCCTAGAAAATTCCGCGTT | 219 |
| chr8:65111106-65113132 | p2 | (AG)13 | GATCTCAGTGGCAATTCCGT | ACGATGTCACCTCGGGATAC | 154 |
| chr8:65209945-65211960 | p5 | (AATGC)3 | ATCATATCGATCCGCGAGAC | GTTCAGTTACCGAGGTGGGA | 263 |
| chr8:65226063-65228081 | p2 | (TA)9 | CGGAAGGAGGTAAATGACGA | TTCCACTCACCATCCGTACA | 192 |
| chr8:65406986-65409002 | p4 | (AAAT)4 | CGGCATTTCCAACTTCCTAA | AGGCTGCAGCTGGTGTTACT | 160 |
| chr8:65473823-65475838 | p5 | (TGTGG)3 | CGGATGGAATATGGAATTGG | TTGCCAACTATGCCAAACAA | 252 |
| chr8:65626558-65628572 | p2 | (TA)6 | GGCACTCATTCCACCTTGTT | CGAAGCGTAAAGTGGGAAAG | 125 |
| chr8:65626568-65628583 | p2 | (TA)6 | GGCACTCATTCCACCTTGTT | CGAAGCGTAAAGTGGGAAAG | 115 |
| chr8:65645577-65647595 | p6 | (TATATG)3 | TTAGATTCTCGGGGATGGTG | GCTTAATTGTCCAAGCTCGC | 162 |
| chr8:65650264-65652278 | p2 | (TC)7 | TCGAGAAATGTGAGGGAAGG | CTTCGAAATGCCACACACAC | 195 |
| chr8:65660819-65662833 | p2 | (AT)7 | GATCTTAACTCCAGGCAGCG | ACGAGTAGTGCTCGGTGGTT | 290 |
| chr8:65701737-65703749 | p2 | (AT)7 | GCACTCAGCCAAGTAGGGTC | GTTTCGTGAGGGTTCGTCAT | 174 |
| chr8:65701751-65703763 | p2 | (AT)7 | GCACTCAGCCAAGTAGGGTC | GTTTCGTGAGGGTTCGTCAT | 160 |
| chr8:65710648-65712660 | p2 | (AT)7 | GCACTCAGCCAAGTAGGGTC | GTTTCGTGAGGGTTCGTCAT | 174 |
| chr8:65710662-65712674 | p2 | (AT)7 | GCACTCAGCCAAGTAGGGTC | GTTTCGTGAGGGTTCGTCAT | 160 |
| chr8:65806734-65808746 | p2 | (CA)6 | TTACCCCAAAGTGCACATCA | GTTGAAATTGGCCGAGTTGT | 223 |
| chr8:65814914-65816932 | p3 | (CAA)6 | TGTCTTAGCCCTTTCCCCTT | TGATGATTATGATGCCCGAA | 204 |
| chr8:65818072-65820087 | p5 | (TGCAA)3 | CGGAGCTCTTTGTACCCATC | AATGGGTTAACGGGTAAGGG | 132 |
| chr8:65822738-65824753 | p3 | (TGA)5 | GATATGATGGCCCCAAAATG | TCCCCTACTGGGTAAATCCC | 205 |
| chr8:65948749-65950767 | p2 | (GA)9 | GAGCAGTCGAGGGATCTTTG | CATTATGAAAGCCCCATGCT | 248 |
| chr8:65948749-65950767 | p6 | (ATTTTT)3 | GAGCAGTCGAGGGATCTTTG | CATTATGAAAGCCCCATGCT | 248 |
| chr8:65977076-65979091 | p5 | (ATTTT)3 | TTGGTCTATGAACGGGAAGG | CCCCTGACATTCCAAGAAGA | 186 |
| chr8:66037221-66039241 | p5 | (ATTTT)4 | ACATTGCCAGAAGGATCCAG | TCCAAAGGGATTTCAACGTC | 269 |
| chr8:66084356-66086428 | p2 | (AT)36 | GATGCGTAATCGGGTTCCTA | AGGAGTGAAATGGAATTGCG | 211 |
| chr8:66161588-66163600 | p2 | (CA)6 | TTTGGAGGGAACAGAACTGG | ACGGGATTTGAGTTTTGACG | 195 |
| chr8:66400621-66402637 | p2 | (TA)8 | GGGAGTTATCTCGGGGAGAG | GGAGCATACCTGGCCATAGA | 122 |
| chr8:66406787-66408855 | p2 | (TA)9 | CGAGAAGTAGGAGTCGGACG | GGACCCATGAACAGAAGGAA | 172 |

| Marker ID | SSR type | SSR motif | FORWARD PRIMER (5'-3') | REVERSE PRIMER (5'-3') | size (bp) |
|---|---|---|---|---|---|
| chr8:66406787-66408855 | p2 | (TA)11 | CGAGAAGTAGGAGTCGGACG | GGACCCATGAACAGAAGGAA | 172 |
| chr8:66406787-66408855 | p2 | (TA)34 | CGAGAAGTAGGAGTCGGACG | GGACCCATGAACAGAAGGAA | 172 |
| chr8:66474752-66476770 | p6 | (TATATG)3 | TGTTGGTTGGGTTGTTTGAA | TTCAACTCGGCCAAATTTTC | 189 |
| chr8:66483337-66485349 | p2 | (TA)6 | TCATCTTGGTAAGCGAACCC | GTCCTTAACCTTGCCAACGA | 296 |
| chr8:66483337-66485349 | p5 | (AATTG)4 | TCATCTTGGTAAGCGAACCC | GTCCTTAACCTTGCCAACGA | 296 |
| chr8:66494584-66496604 | p2 | (TA)10 | GCCGAAAGAAGAACTGAACG | CATGCTCCTCTGGTATGGGT | 252 |
| chr8:66503301-66505325 | p3 | (GTT)6 | CGTGTGTTGGTAATTTTGCG | ACAACGCGACTACAACATCG | 100 |
| chr8:66503301-66505325 | p2 | (TA)12 | CGTGTGTTGGTAATTTTGCG | ACAACGCGACTACAACATCG | 100 |
| chr8:66503301-66505325 | p2 | (TA)11 | CGTGTGTTGGTAATTTTGCG | ACAACGCGACTACAACATCG | 100 |
| chr8:66510272-66512284 | p2 | (TA)24 | GCGCAGGACCAGTTTAACAT | ATTTTCCAAGCAAATCGTGC | 105 |
| chr8:66510272-66512284 | p2 | (TA)6 | GCGCAGGACCAGTTTAACAT | ATTTTCCAAGCAAATCGTGC | 105 |
| chr8:66524331-66526345 | p2 | (TG)7 | AAAGGTGCCTACGGGCTATT | CCAACCAAGGCTCAAGGTTA | 234 |
| chr8:66535713-66537727 | p2 | (TA)7 | GAACAGCCTGTGGCCATATT | CAAACTCCTCGGTCTTTTCG | 242 |
| chr8:66536919-66538934 | p5 | (CATAT)3 | GTGCTGGACACTCGAACGTA | TGACCTGTGACAGCCTCATC | 221 |
| chr8:66559126-66561156 | p2 | (TA)33 | GCAATTTCTGAGTCGGTGGT | TTGATGTGTGGAAGTTGGGA | 254 |
| chr8:66608743-66610755 | p2 | (AT)6 | CCATGGTCTCACATTTGTCG | ATGGGTTCTGTTTAGCGGTG | 193 |
| chr8:66650072-66652084 | p2 | (AT)6 | CAACATGGTGAGGTGATTGC | ATCGCCGTTGTCGTAAATTC | 191 |
| chr8:66692870-66694885 | p5 | (ATGAT)3 | TTGGTCGAAACACGAATTGA | ACAATTTCAAACGCGGATTC | 208 |
| chr8:66700321-66702336 | p5 | (TATGA)3 | GATGCTGCCCAAATTTCTGT | ACAATTTCAAACGCGGATTC | 112 |
| chr8:66762708-66764720 | p2 | (TC)6 | TTACACCCAATTCTCAGCCC | TTGGATCGTTTGTTTGACCA | 158 |
| chr8:66824982-66826994 | p2 | (AT)6 | TCACTCCGACGGATACATGA | CATAGCATAGCACCGAACGA | 252 |
| chr8:66865026-66867042 | p4 | (TATG)4 | GATGGCCATGAGGTGAGTTT | AATCATAATCCGGCAACCAA | 167 |
| chr8:66876862-66878877 | p3 | (GAT)5 | ATGAGGTTGATCGGGATCTG | CAAGCCCACTCTGCACTGTA | 155 |
| chr8:66886160-66888180 | p2 | (AT)10 | GGACACATGCCCCTTCTTTA | GGAAGGCTAGGGGAAAATTG | 157 |
| chr8:66886160-66888180 | p2 | (AT)13 | GGACACATGCCCCTTCTTTA | GGAAGGCTAGGGGAAAATTG | 157 |
| chr8:66911811-66913829 | p6 | (ATTATG)3 | AATGACGAGGACAACTTCGG | TCACCACCATGCAAAACATT | 219 |
| chr8:66938201-66940213 | p2 | (TA)6 | GCCCACGATATACCCCTTTT | CAAGCAAGATGGTACGCAGA | 246 |
| chr8:66984288-66986303 | p5 | (TAAAT)3 | ATATGCAGGGCGCATAAGTT | TGTGTCCAGGTTGTTTGCAT | 179 |
| chr8:67174708-67176723 | p5 | (AAATA)3 | TAATGACCTTCTTCGGGTGC | TTTTCAGCCACCAAAATTCC | 238 |
| chr8:67174894-67176909 | p5 | (AAATA)3 | GGAATTTTGGTGGCTGAAAA | GGCTTTCAACGTTCGGATTA | 243 |
| chr8:67221716-67223752 | p2 | (TA)18 | TGCTCCCCAAAAGCTCTAAA | TGAACTGCGACTTTCTGGTG | 211 |
| chr8:67234440-67236452 | p2 | (TG)12 | CCCTACTTACTGGGCACCAA | GTGTGGCCTTGTCGTCCTAT | 217 |
| chr8:67234440-67236452 | p2 | (TG)11 | CCCTACTTACTGGGCACCAA | GTGTGGCCTTGTCGTCCTAT | 217 |
| chr8:67234440-67236452 | p5 | (ACAAC)3 | CCCTACTTACTGGGCACCAA | GTGTGGCCTTGTCGTCCTAT | 217 |
| chr8:67234481-67236496 | p2 | (TG)12 | CCCTACTTACTGGGCACCAA | GTGTGGCCTTGTCGTCCTAT | 176 |
| chr8:67234481-67236496 | p2 | (TG)11 | CCCTACTTACTGGGCACCAA | GTGTGGCCTTGTCGTCCTAT | 176 |

| Marker ID | SSR type | SSR motif | FORWARD PRIMER (5'-3') | REVERSE PRIMER (5'-3') | size (bp) |
|---|---|---|---|---|---|
| chr8:67234481-67236496 | p5 | (ACAAC)3 | CCCTACTTACTGGGCACCAA | GTGTGGCCTTGTCGTCCTAT | 176 |
| chr8:67235093-67237108 | p2 | (TG)11 | CAACCTGCAACTCTGGGTTT | GATGTCCACAGAGGTCCGAT | 259 |
| chr8:67235093-67237108 | p5 | (ACAAC)3 | CAACCTGCAACTCTGGGTTT | GATGTCCACAGAGGTCCGAT | 259 |
| chr8:67235093-67237108 | p3 | (AAT)17 | CAACCTGCAACTCTGGGTTT | GATGTCCACAGAGGTCCGAT | 259 |
| chr8:67237483-67239495 | p2 | (TG)12 | CCCTACTTACTGGGCACCAA | GTGTGGCCTTGTCGTCCTAT | 217 |
| chr8:67237483-67239495 | p2 | (TG)11 | CCCTACTTACTGGGCACCAA | GTGTGGCCTTGTCGTCCTAT | 217 |
| chr8:67237483-67239495 | p5 | (ACAAC)3 | CCCTACTTACTGGGCACCAA | GTGTGGCCTTGTCGTCCTAT | 217 |
| chr8:67237524-67239539 | p2 | (TG)12 | CCCTACTTACTGGGCACCAA | GTGTGGCCTTGTCGTCCTAT | 176 |
| chr8:67237524-67239539 | p2 | (TG)11 | CCCTACTTACTGGGCACCAA | GTGTGGCCTTGTCGTCCTAT | 176 |
| chr8:67237524-67239539 | p5 | (ACAAC)3 | CCCTACTTACTGGGCACCAA | GTGTGGCCTTGTCGTCCTAT | 176 |
| chr8:67306055-67308070 | p3 | (TGA)5 | ATGGTGGCCCATATGTTTGT | CCACGACTGGTGCCTTATTT | 260 |
| chr8:67341556-67343574 | p3 | (TTG)6 | GCTGAGCATACCACCCAGAT | TTTCTCCCACTTCGAACCAC | 169 |
| chr8:67370840-67372858 | p6 | (TATATG)3 | AGACCATTGGACAAAGACCG | TTGTTGTGCATATGGCTGGT | 224 |
| chr8:67370840-67372858 | p6 | (TATATG)3 | AGACCATTGGACAAAGACCG | TTGTTGTGCATATGGCTGGT | 224 |
| chr8:67467602-67469650 | p2 | (TA)12 | TACCATTGATCAGTTGGGCA | AGCATCCACAAACAAAAGGG | 174 |
| chr8:67467602-67469650 | p2 | (TA)6 | TACCATTGATCAGTTGGGCA | AGCATCCACAAACAAAAGGG | 174 |
| chr8:67468240-67470264 | p2 | (TA)12 | TCGGAGCTAGTCTACCGAGC | TATATGGGGCGGGATTGATA | 215 |
| chr8:67468240-67470264 | p2 | (TA)6 | TCGGAGCTAGTCTACCGAGC | TATATGGGGCGGGATTGATA | 215 |
| chr8:67468240-67470264 | p3 | (ACA)6 | TCGGAGCTAGTCTACCGAGC | TATATGGGGCGGGATTGATA | 215 |
| chr8:67469168-67471186 | p2 | (TA)12 | AAGCAGACACGGCTGCTACT | TACAGACGCTCGTTTTGGTG | 296 |
| chr8:67469168-67471186 | p2 | (TA)6 | AAGCAGACACGGCTGCTACT | TACAGACGCTCGTTTTGGTG | 296 |
| chr8:67469168-67471186 | p3 | (ACA)6 | AAGCAGACACGGCTGCTACT | TACAGACGCTCGTTTTGGTG | 296 |
| chr8:67478997-67481021 | p2 | (TA)8 | TGCTGCCCTAGGATCCTCTA | CAGTCAGACCCGGAGGATTA | 277 |
| chr8:67478997-67481021 | p2 | (AT)12 | TGCTGCCCTAGGATCCTCTA | CAGTCAGACCCGGAGGATTA | 277 |
| chr8:67478997-67481021 | p6 | (CATATA)3 | TGCTGCCCTAGGATCCTCTA | CAGTCAGACCCGGAGGATTA | 277 |
| chr8:67479711-67481729 | p2 | (TA)8 | ATATAGGGGAACGTCCCAGC | CCTAGTAGCAGCCTTGTCGG | 162 |
| chr8:67479711-67481729 | p2 | (AT)12 | ATATAGGGGAACGTCCCAGC | CCTAGTAGCAGCCTTGTCGG | 162 |
| chr8:67479711-67481729 | p6 | (CATATA)3 | ATATAGGGGAACGTCCCAGC | CCTAGTAGCAGCCTTGTCGG | 162 |
| chr8:67577555-67579577 | p2 | (AT)11 | CCGGAGTCTATAGGATGCCA | GCTTCGGGTCAACAAATGAT | 148 |
| chr8:67625328-67627340 | p2 | (TG)6 | TGAGTAACAGACGTGCCTGG | CTTTGGACTCCCTGCCATTA | 166 |
| chr8:67697678-67699693 | p5 | (TCATT)3 | ATCCTCTGGTGGGTTTTCCT | TTGCCTTCTCTTGGCAAACT | 185 |
| chr8:67705471-67707505 | p2 | (TA)17 | GGCACAAAAGTTAGGCAAGG | AAAAATCAAACCATGCCGTC | 269 |
| chr8:67719775-67721800 | p6 | (GTTGAT)3 | ATCTCCCTGCTGCTGTGTCT | AGGGATCTCGACTTCAGCAA | 292 |
| chr8:67719775-67721800 | p5 | (AATAA)5 | ATCTCCCTGCTGCTGTGTCT | AGGGATCTCGACTTCAGCAA | 292 |
| chr8:67724242-67726257 | p5 | (AATTA)3 | ATGAAGGTTTCGTGATTGCC | TATCACGGGACAGAAAAGCC | 211 |
| chr8:67724242-67726257 | p6 | (ATTTAA)3 | ATGAAGGTTTCGTGATTGCC | TATCACGGGACAGAAAAGCC | 211 |

| Marker ID | SSR type | SSR motif | FORWARD PRIMER (5'-3') | REVERSE PRIMER (5'-3') | size (bp) |
|---|---|---|---|---|---|
| chr8:67759482-67761500 | p5 | (AATGC)3 | CGCGTTGCTCTAATGTTGAA | TAGTGGAGTCTCACGGGGTC | 226 |
| chr8:67759502-67761520 | p5 | (AATGC)3 | CGCGTTGCTCTAATGTTGAA | TAGTGGAGTCTCACGGGGTC | 206 |
| chr8:67773407-67775425 | p2 | (AT)6 | GGACACTCATGTCCCCTGTT | CCCATTCGTAATGACAGCCT | 127 |
| chr8:67773407-67775425 | p3 | (CAT)6 | GGACACTCATGTCCCCTGTT | CCCATTCGTAATGACAGCCT | 127 |
| chr8:67919795-67921810 | p5 | (ATTGC)3 | TTCCTATTTTGCCCTTGTGG | GCGAACTATCCGACTTGAGC | 147 |
| chr8:67945607-67947622 | p5 | (TTAGT)3 | TATAATCTCCCCCTCCGTCC | CTCGTTTGGCCTGATTTTGT | 213 |
| chr8:68027733-68029749 | p2 | (AT)8 | ACATAGCCATGGTTGCCTTC | ATCTTGGCGTGGTAGAATCG | 166 |
| chr8:68075409-68077421 | p2 | (CA)6 | ATGGACGGTTGCGGATATTA | TACCCGGCCATATAGACTCG | 276 |
| chr8:68083107-68085119 | p2 | (CA)6 | CCAAATAATTCCGGAGGGTT | TTCGGGCCAAGATGTAAAAG | 194 |
| chr8:68193390-68195404 | p2 | (AT)7 | CGAACGAGTAATCAATGCGA | TACTCGGTTGTTAGGGTCGG | 227 |
| chr8:68193390-68195404 | p2 | (AT)6 | CGAACGAGTAATCAATGCGA | TACTCGGTTGTTAGGGTCGG | 227 |
| chr8:68232225-68234243 | p2 | (AT)31 | CAATACACTGCAGCTCCGAA | CCTTCTTATGGACGTTGCGT | 281 |
| chr8:68232225-68234243 | p3 | (CAT)7 | CAATACACTGCAGCTCCGAA | CCTTCTTATGGACGTTGCGT | 281 |
| chr8:68232245-68234263 | p2 | (AT)31 | CAATACACTGCAGCTCCGAA | CCTTCTTATGGACGTTGCGT | 261 |
| chr8:68232245-68234263 | p3 | (CAT)7 | CAATACACTGCAGCTCCGAA | CCTTCTTATGGACGTTGCGT | 261 |
| chr8:68232265-68234335 | p2 | (AT)31 | CAATACACTGCAGCTCCGAA | CCTTCTTATGGACGTTGCGT | 241 |
| chr8:68232265-68234335 | p3 | (CAT)7 | CAATACACTGCAGCTCCGAA | CCTTCTTATGGACGTTGCGT | 241 |
| chr8:68256484-68258595 | p5 | (ATTGC)3 | GGGCTGGAAATACGTTGAGA | GAGGCCATAACTAGGCCACA | 263 |
| chr8:68256484-68258595 | p3 | (TTA)37 | GGGCTGGAAATACGTTGAGA | GAGGCCATAACTAGGCCACA | 263 |
| chr8:68380430-68382457 | p2 | (TA)26 | ATTTGGGGCGTGACAGTAAG | TACGGTTATTCCTTGCCGAG | 200 |
| chr8:68380430-68382457 | p3 | (AAG)9 | ATTTGGGGCGTGACAGTAAG | TACGGTTATTCCTTGCCGAG | 200 |
| chr8:68548204-68550219 | p3 | (TTG)5 | CTGAGGAGGGGACCACATAA | CGACCTATGGTTGGACAGGT | 103 |
| chr8:68567389-68569421 | p2 | (TC)16 | CCTCAAAATGCCGACAAAAT | CCTCAACCGTGTTTCCATCT | 107 |
| chr8:68567389-68569421 | p3 | (GAA)8 | CCTCAAAATGCCGACAAAAT | CCTCAACCGTGTTTCCATCT | 107 |
| chr8:68586194-68588209 | p5 | (CATTT)3 | ATTTCGTTGATGCATGGTGA | TGCCTTCTCTTGGCAAACTT | 298 |
| chr8:68645768-68647782 | p2 | (TA)7 | CCAAAACCTTCCAAGTCCAA | AATGGGCTGATTTGGTGAAG | 260 |
| chr8:68708624-68710636 | p2 | (TA)6 | CCTTGGGTTTCTTCCTCCTC | TCCTGAAAGGATGACTTGGG | 292 |
| chr8:68758786-68760800 | p2 | (TA)7 | ACATTGCTCGTACTGACCCC | ATTGTCAACTCCGCTGGAAC | 296 |
| chr8:68813210-68815224 | p2 | (TA)7 | AACCGCACCCCTAGAAGTTT | GGTGGGAACAAATGTTGAGG | 112 |
| chr8:68892170-68894190 | p4 | (ATCA)5 | TCTCTTTGAGCCCCGACTAA | GCCATCGGACTGGAGTAAAA | 114 |
| chr8:68936215-68938227 | p2 | (TG)6 | AGGTGCCTACGGGCTATTCT | CGACCTCATAGGATCGGTGT | 250 |
| chr8:68947176-68949192 | p4 | (GAGT)4 | TGGCCCTTTTGAGATTGTTC | AGCAACCAGCTTTTCCTCAA | 230 |
| chr8:69014879-69016921 | p2 | (TA)21 | CGTAAAGCCGTATGTTCGGT | CACGCTTTTTCCTGTTAGGC | 101 |
| chr8:69016586-69018618 | p2 | (AT)16 | GGCATGGCATGAAGAAGAAT | TTCGGATTTCCATTCACCAT | 273 |
| chr8:69027654-69029694 | p2 | (TA)20 | GGCATGGCATGAAGAAGAAT | TTCGGATTTCCATTCACCAT | 283 |
| chr8:69049063-69051077 | p2 | (TA)7 | GGAAGGAATCAAAGGAAGGC | TTGACGATCCAATGTGGCTA | 269 |

| Marker ID | SSR type | SSR motif | FORWARD PRIMER (5'-3') | REVERSE PRIMER (5'-3') | size (bp) |
|---|---|---|---|---|---|
| chr8:69118358-69120370 | p2 | (AT)6 | TTGTTAAGACCCCGACTTGC | CCTTGAACCTTTGGTGCATT | 163 |
| chr8:69118358-69120370 | p2 | (AT)7 | TTGTTAAGACCCCGACTTGC | CCTTGAACCTTTGGTGCATT | 163 |
| chr8:69160256-69162271 | p5 | (ATCAT)3 | ATAAAGCACAAGGAAGCCGA | TGCTCTATTGTTCGGAGCCT | 293 |
| chr8:69160256-69162271 | p2 | (TA)22 | ATAAAGCACAAGGAAGCCGA | TGCTCTATTGTTCGGAGCCT | 293 |
| chr8:69174945-69176960 | p5 | (ATATT)3 | ATGGCCGATTGCAATGTTAT | CATGCCATGAAAATGAGGTG | 187 |
| chr8:69199227-69201251 | p2 | (AT)12 | CTACCAGGGCTAGGTTGCTG | GCAGTGACTCCTTCTCCAGG | 292 |
| chr8:69344180-69346198 | p3 | (TTG)6 | GTGAGAAGGGACATGTGGGT | GCCCACGACCTATGGTTAGA | 171 |
| chr8:69350844-69352859 | p3 | (AAC)5 | GGGTATTCGGTTAGCGTTCA | AAGGCCCCGACACTTTTTAT | 296 |
| chr8:69519983-69521997 | p2 | (AT)7 | TGGACGTGTCATAATCGCAT | CATTCCAAAAAGCCCACATT | 126 |
| chr8:69523817-69525831 | p2 | (TA)7 | TTCAGGTAGCGTTACGGCTT | CATCCCAGCTTTATACCCGA | 230 |
| chr8:69892377-69894403 | p2 | (TA)24 | TAAGCCGACGAGGCTAAAAA | ATTATGGGCATAGCGTGGTC | 231 |
| chr8:69892377-69894403 | p2 | (TA)13 | TAAGCCGACGAGGCTAAAAA | ATTATGGGCATAGCGTGGTC | 231 |
| chr8:69900386-69902406 | p2 | (TA)7 | ATAAGCCGACGAGGCTAACA | GTCCTGTCCCGCCTATATGA | 241 |
| chr8:69900407-69902439 | p2 | (TA)7 | ATAAGCCGACGAGGCTAACA | GTCCTGTCCCGCCTATATGA | 220 |
| chr8:69933974-69936008 | p2 | (AT)17 | GCCATCTGTTACGGTCCACT | CCTTCCCTAGGGGATTACCA | 251 |
| chr8:69944792-69946804 | p2 | (TG)6 | TGTACGCCCTTCAGGGTAAG | ATGCGGGATCTAACAACCTG | 110 |
| chr8:70044751-70046765 | p2 | (GT)7 | TTGAATCAAAATGGCACCAA | ATTATGCGAAAACACGGGAC | 204 |
| chr8:70044751-70046765 | p3 | (TTG)5 | TTGAATCAAAATGGCACCAA | ATTATGCGAAAACACGGGAC | 204 |
| chr8:70062092-70064104 | p2 | (AT)6 | CAACAGGTGAAGGCTGAACA | AATCAAACTTGGCCCTTGTG | 105 |
| chr8:70065596-70067608 | p2 | (AT)6 | CAACAGGTGAAGGCTGAACA | AATCAAACTTGGCCCTTGTG | 105 |
| chr8:70112940-70114952 | p2 | (TA)6 | CCATTGACCCTTCCTTCTCA | ACAGCCCAGAAAACACATCC | 291 |
| chr8:70267697-70269709 | p2 | (TA)6 | TCCCATAACCCAATCCGTAA | AAAATGGAATGCGAAAATCG | 154 |
| chr8:70356786-70358811 | p5 | (TGAAT)5 | AGCATGGGAGAAGTTGGTTG | CTACACGGCCATTCCTCATT | 271 |
| chr8:70361695-70363710 | p6 | (AAAAAT)3 | GGAAAGAATTGGGGCAAAAT | CCATGGTAGATTTCATGGGG | 227 |
| chr8:70361695-70363710 | p5 | (TGTGT)3 | GGAAAGAATTGGGGCAAAAT | CCATGGTAGATTTCATGGGG | 227 |
| chr8:70361695-70363710 | p5 | (TGAAT)5 | GGAAAGAATTGGGGCAAAAT | CCATGGTAGATTTCATGGGG | 227 |
| chr8:70369042-70371060 | p6 | (TATATG)3 | GCGATAGAGCCCAGTCGTAG | TCACCTTCACTTTCATCCCC | 279 |
| chr8:70369042-70371060 | p2 | (TA)7 | GCGATAGAGCCCAGTCGTAG | TCACCTTCACTTTCATCCCC | 279 |
| chr8:70443851-70445867 | p2 | (AT)8 | TGGAACGGTGACTCATGAAA | CTGACCTTGTGGGGTCTTGT | 240 |
| chr8:70443851-70445867 | p5 | (TTGAA)5 | TGGAACGGTGACTCATGAAA | CTGACCTTGTGGGGTCTTGT | 240 |
| chr8:70561890-70563905 | p5 | (ATTTC)3 | CAGGCAGCAGTTCCCTAGAC | CCTTGACTGGCCTGAATCAT | 270 |
| chr8:70570084-70572144 | p2 | (TA)14 | AGGTATAAACGCAAGCGTGG | ACATCGAGCATCAGGAGCTT | 298 |
| chr8:70646291-70648307 | p2 | (TA)8 | TGTTCATGCCCACAGGTAGA | TCATAAGGACGGGACAGGAC | 211 |
| chr8:70669134-70671180 | p2 | (TA)23 | CGGCTTTGCAGATAATCCAT | GCACTCAGCCGAGTAGGTTC | 253 |
| chr8:70723875-70725890 | p3 | (GTT)5 | GTCGTGGTAGGATAAGGGCA | CCATAAGCCACTTTGCGAAT | 187 |
| chr8:70840011-70842031 | p2 | (AC)10 | CATGCCAAAAGAAGATCGGT | GGTGCTGCTAAGAATTTCGG | 166 |

| Marker ID | SSR type | SSR motif | FORWARD PRIMER (5'-3') | REVERSE PRIMER (5'-3') | size (bp) |
|---|---|---|---|---|---|
| chr8:70893226-70895238 | p2 | (GT)6 | CCGTATTTTTGAAGGTCGGA | CTTTTATTTGGCCCAAGGGT | 128 |
| chr8:70905380-70907395 | p3 | (GTT)5 | ACATGCCATAGTGGGACACA | GACGTTTAAATCGCCTTGGA | 142 |
| chr8:70924225-70926243 | p6 | (TATATG)3 | GCATTCTCGGGTTTGTTGTT | ACCTGTCCCACGTCCAATTA | 151 |
| chr8:70941781-70943796 | p5 | (GAAAT)3 | TTACCCGGCATCATAAGCTC | CACATGGTTGACATTCCGAG | 109 |
| chr8:70953564-70955604 | p2 | (AT)20 | ATGGCGAAACCTCTATGCTG | AAAGCATAGCGAGGAAACGA | 191 |
| chr8:71068054-71070074 | p2 | (AT)10 | CGTTTGTTGTTGTTGGGTTG | AACGTTCTCGCCATTACCAC | 164 |
| chr8:71261478-71263490 | p2 | (TA)6 | CTTCATTCCAGGCCATCATT | GGTAAATCCCAACAAAGGCA | 122 |
| chr8:71261478-71263490 | p4 | (CTTT)4 | CTTCATTCCAGGCCATCATT | GGTAAATCCCAACAAAGGCA | 122 |
| chr8:71381585-71383603 | p2 | (AG)8 | GAGATCTCATGCACGCTTCA | TTCCAAATTTTTCCGAATGC | 250 |
| chr8:71381585-71383603 | p2 | (GT)9 | GAGATCTCATGCACGCTTCA | TTCCAAATTTTTCCGAATGC | 250 |
| chr8:71473515-71475530 | p5 | (AAGAA)3 | GTGAAGTGTGCTCCTCGTCA | TCGAGGCCGTACTAGTTGCT | 119 |
| chr8:71473515-71475530 | p2 | (TG)6 | GTGAAGTGTGCTCCTCGTCA | TCGAGGCCGTACTAGTTGCT | 119 |
| chr8:71476630-71478645 | p5 | (ATCAT)3 | AATGGAGTTTGCTGACCCTG | TTCGTACTGACCCCCTGTTC | 263 |
| chr8:71489957-71491973 | p2 | (AT)8 | AACGCTCCAAATCAGTCCAC | ATTGTTGGGCGTTGTGGTAT | 294 |
| chr8:71622584-71624606 | p2 | (TA)11 | AGCCCTTGACACCAAGAGAA | ATGCAATCTCCCGTCAACTC | 288 |
| chr8:71633474-71635510 | p2 | (AT)18 | TGGGTGTGGACACTTCAAAA | ATTTATGTACTGGCTGGGCG | 163 |
| chr8:71633474-71635510 | p2 | (TA)8 | TGGGTGTGGACACTTCAAAA | ATTTATGTACTGGCTGGGCG | 163 |
| chr8:71670391-71672409 | p6 | (CTTCAA)3 | TGTGGAGGTTTCCACAGACA | GCGTGGTCCAAAACAAGTCT | 289 |
| chr8:71670391-71672409 | p2 | (TA)17 | TGTGGAGGTTTCCACAGACA | GCGTGGTCCAAAACAAGTCT | 289 |
| chr8:71772870-71774885 | p5 | (CATAT)3 | CAACTAGTGTCCGTGCTGGA | TGTCGGTTCAGTTAATGCCA | 220 |
| chr8:71772870-71774885 | p3 | (CAA)5 | CAACTAGTGTCCGTGCTGGA | TGTCGGTTCAGTTAATGCCA | 220 |
| chr8:71783366-71785381 | p5 | (TGGGT)3 | GTTCTTCCGCATGTTGTGTG | CTGAAGGCTGGGCTATTGAG | 143 |
| chr8:71825781-71827795 | p2 | (AT)7 | CATATGTGTCGCGTACCAGG | CTACCACGCCATAAGCAGGT | 255 |
| chr8:71892723-71894749 | p2 | (TA)13 | AAATGAACGGACGAAGTTGG | GATTCAACTCCCAAGGGTCA | 215 |
| chr8:71923852-71925916 | p2 | (AT)32 | TCCTCATCCGACTTTGATCC | GTGCCTATTTCTGAGGAGCG | 268 |
| chr8:71975111-71977123 | p2 | (TA)6 | GCCCGAATAGACATCCAAAA | CCATTGCCTTACAACTCGGT | 159 |
| chr8:71992102-71994114 | p2 | (GT)6 | TCCGGAACAATGGATCTCTC | CCTTCCTCCGTATTTCCACA | 240 |
| chr8:71992102-71994114 | p3 | (TTG)5 | TCCGGAACAATGGATCTCTC | CCTTCCTCCGTATTTCCACA | 240 |
| chr8:71993579-71995599 | p3 | (TTG)5 | CAAGATGGGATGCCAAAAGT | AGTCCCCGAGGTCCATATTC | 132 |
| chr8:71993579-71995599 | p5 | (ATATG)4 | CAAGATGGGATGCCAAAAGT | AGTCCCCGAGGTCCATATTC | 132 |
| chr8:72065254-72067312 | p2 | (AT)24 | TGTTCGCCTTAAGAGGCATT | GTACAATCGCTGAGAACGCA | 288 |
| chr8:72065254-72067312 | p2 | (AT)29 | TGTTCGCCTTAAGAGGCATT | GTACAATCGCTGAGAACGCA | 288 |
| chr8:72073415-72075427 | p2 | (TA)6 | ATCGCCCTGCTAGAGCAATA | AAACTTCCAAATTCACCCCC | 298 |
| chr8:72078704-72080722 | p3 | (CAA)6 | GTCTCCTTAGCCACTCGACG | TTCGGGCTAGGCATGTAAAC | 189 |
| chr8:72078704-72080722 | p3 | (CAA)6 | GTCTCCTTAGCCACTCGACG | TTCGGGCTAGGCATGTAAAC | 189 |
| chr8:72104152-72106173 | p3 | (GAA)7 | AACATCGAATGTCACCGTCA | GGCTGTTTATTTCATGGCGT | 282 |

| Marker ID | SSR type | SSR motif | FORWARD PRIMER (5'-3') | REVERSE PRIMER (5'-3') | size (bp) |
|---|---|---|---|---|---|
| chr8:72104152-72106173 | p3 | (CTT)5 | AACATCGAATGTCACCGTCA | GGCTGTTTATTTCATGGCGT | 282 |
| chr8:72106099-72108117 | p2 | (AT)9 | CCCCGGGTTTTACAATTCTT | AAATGGGTGTCAATGCCAAT | 140 |
| chr8:72210951-72212966 | p2 | (AT)7 | TAAGGATGAAACAGTCGGGC | CATGGGCAGTTTTTGTGATG | 125 |
| chr8:72210951-72212966 | p5 | (ATACG)3 | TAAGGATGAAACAGTCGGGC | CATGGGCAGTTTTTGTGATG | 125 |
| chr8:72228304-72230322 | p6 | (CTCATG)3 | CATCCGGAAATTCGTTCACT | CATGGATTGGTTGGCTTCTT | 168 |
| chr8:72320982-72322994 | p2 | (TA)6 | GATCTGTGTATGGGGATGGC | GATGGCACACGCATACATTC | 151 |
| chr8:72392007-72394035 | p2 | (AT)14 | CAACTACAGCCTCGCTTTCC | GACCGCTCAGTACTCAAGGC | 126 |
| chr8:72406012-72408030 | p6 | (GCTGGT)3 | TTCCGCTTATTCGCACTTCT | AGATCGACGCGGATTAAATG | 186 |
| chr8:72434539-72436575 | p2 | (TA)18 | TCGGGTGCATGATCTTGTTA | AGATTGCTAACCCCATGTGC | 127 |
| chr8:72517196-72519211 | p5 | (ATTTC)3 | TTCAATAGAACCTGGCCACC | CGAGAGTTATTCAGGCCAGC | 183 |
| chr8:72611337-72613383 | p2 | (AT)23 | ATAATGACATCGGGGGATCA | AGGCGCGGTGTAATATCATC | 229 |
| chr8:72685192-72687204 | p2 | (TA)6 | TCTTCGTCTTCTCCGACGAT | CCCAGCAACATCCAAAGAGT | 100 |
| chr8:72865016-72867032 | p2 | (TA)8 | CTTCTTGTTGTCGCCTCTCC | ATCGGAGTGACGTAAGGTCG | 260 |
| chr8:72873107-72875121 | p2 | (CA)7 | AGGGGAAGGTCCAAGAAAAA | ACATTAGGATGGGGGAAAGG | 147 |
| chr8:72905102-72907120 | p2 | (TA)9 | CCAACGGCGTTAATTTCCTA | GCTGACGTCTCTCATCCCTC | 284 |
| chr8:72905102-72907120 | p2 | (TA)12 | CCAACGGCGTTAATTTCCTA | GCTGACGTCTCTCATCCCTC | 284 |
| chr8:72905560-72907584 | p2 | (TA)9 | ATCCGTTTGGTCGTTTTGAG | TCATGGTCCACCAAGAATGA | 238 |
| chr8:72905560-72907584 | p2 | (TA)12 | ATCCGTTTGGTCGTTTTGAG | TCATGGTCCACCAAGAATGA | 238 |
| chr8:72907645-72909663 | p2 | (TA)9 | CCAACGGCGTTAATTTCCTA | GCTGACGTCTCTCATCCCTC | 284 |
| chr8:72907645-72909663 | p2 | (TA)12 | CCAACGGCGTTAATTTCCTA | GCTGACGTCTCTCATCCCTC | 284 |
| chr8:72908103-72910127 | p2 | (TA)9 | ATCCGTTTGGTCGTTTTGAG | TCATGGTCCACCAAGAATGA | 238 |
| chr8:72908103-72910127 | p2 | (TA)12 | ATCCGTTTGGTCGTTTTGAG | TCATGGTCCACCAAGAATGA | 238 |
| chr8:72976315-72978327 | p2 | (CA)6 | TCAACCACCTATCACACCGA | GGACGGAGAGATGCGAGTAG | 234 |
| chr8:73245886-73247898 | p2 | (AT)6 | TGACCGTGATTATGGACGAA | TATCGTTCTCCTTCGACGCT | 145 |
| chr8:73273681-73275695 | p2 | (AT)7 | ATCACGGGCCTTTGTACTTG | AGGAAGGCGAATCACACAAC | 291 |
| chr8:73348287-73350299 | p2 | (AT)6 | CCCTATCTAACGGGGACACA | TCCCACAGGGAATATGGTGT | 120 |
| chr8:73348287-73350299 | p2 | (AT)6 | CCCTATCTAACGGGGACACA | TCCCACAGGGAATATGGTGT | 120 |
| chr8:73387606-73389618 | p4 | (TATG)4 | ACGAATGTACTACGCACCCC | AATAGCCCGTAGGCACCTTT | 164 |
| chr8:73387606-73389618 | p2 | (TG)6 | ACGAATGTACTACGCACCCC | AATAGCCCGTAGGCACCTTT | 164 |
| chr8:73410678-73412693 | p5 | (TGTTG)3 | ACACGCTACGCTATGACACG | ATCCGGGTTGTTCCAATGTA | 197 |
| chr8:73437812-73439827 | p3 | (CTT)5 | GAGATCCTGCTCAATCTCGG | TGCTATCATCGTAGGGGTCC | 148 |
| chr8:73459377-73461391 | p2 | (TA)7 | ACTACGCGTTTGTGTTGCAG | CCTGGCCGTGATATCAGATT | 254 |
| chr8:73459377-73461391 | p2 | (TA)7 | ACTACGCGTTTGTGTTGCAG | CCTGGCCGTGATATCAGATT | 254 |
| chr8:73459507-73461521 | p2 | (TA)7 | ACTACGCGTTTGTGTTGCAG | CCTGGCCGTGATATCAGATT | 124 |
| chr8:73459507-73461521 | p2 | (TA)7 | ACTACGCGTTTGTGTTGCAG | CCTGGCCGTGATATCAGATT | 124 |
| chr8:73564642-73566654 | p2 | (TA)6 | TGTTGTTGTGGTTGGTTGCT | TACATACCTTTTCGGTCGCC | 254 |

| Marker ID | SSR type | SSR motif | FORWARD PRIMER (5'-3') | REVERSE PRIMER (5'-3') | size (bp) |
|---|---|---|---|---|---|
| chr8:73564699-73566711 | p2 | (TA)6 | TGTTGTTGTGGTTGGTTGCT | TACATACCTTTTCGGTCGCC | 197 |
| chr8:73564757-73566769 | p2 | (TA)6 | TGTTGTTGTGGTTGGTTGCT | TACATACCTTTTCGGTCGCC | 139 |
| chr8:73624843-73626858 | p5 | (CATAT)3 | ATAAGCACCTCCGGACAATG | AATGACGTCCGTTTTCCTTG | 135 |
| chr8:73624843-73626858 | p3 | (CAA)5 | ATAAGCACCTCCGGACAATG | AATGACGTCCGTTTTCCTTG | 135 |
| chr8:73801683-73803743 | p2 | (TA)10 | AGCGGCCCTAACATAATCCT | CTCACCACGGATTTTGGTCT | 272 |
| chr8:73885608-73887626 | p2 | (AT)27 | AGCCCACACAACAACAACAA | TCTACGACGTGGTATGAGCG | 269 |
| chr8:73885608-73887626 | p3 | (TTA)6 | AGCCCACACAACAACAACAA | TCTACGACGTGGTATGAGCG | 269 |
| chr8:73916913-73918928 | p5 | (AATGC)3 | GTCCTGATCAGACGCAGTCA | GCGGAGGTTCACAGACTAGG | 296 |
| chr8:73943829-73945877 | p2 | (TA)24 | ATATTTCCCACACCCCTTCC | CCAAGTCTTTCAACCCCTCA | 264 |
| chr8:74048681-74050699 | p2 | (AT)7 | CGTACCCGGACCTCTAGTGA | TGAGGTGACAGTTGCCTTTG | 173 |
| chr8:74048681-74050699 | p3 | (CAA)6 | CGTACCCGGACCTCTAGTGA | TGAGGTGACAGTTGCCTTTG | 173 |
| chr8:74140973-74142989 | p4 | (TTAT)4 | AAGTAGACGCACGCACACAG | TCTTTCGGAATCCACCTTTG | 225 |
| chr8:74208812-74210826 | p2 | (GT)7 | TCCCCTATAGGTTGTGCTGC | TCCCACACCACTGTTTTTCA | 259 |
| chr8:74208812-74210826 | p3 | (GTT)5 | TCCCCTATAGGTTGTGCTGC | TCCCACACCACTGTTTTTCA | 259 |
| chr8:74265915-74267947 | p2 | (CT)16 | GGGTGTTGGTGGTTACAAGG | CACACCCACTCATTCCACAG | 158 |
| chr8:74276356-74278371 | p5 | (TCATA)3 | GTGAGCTCCACTTCCTCCTG | CAAACGGAAAAATAAGGCCA | 144 |
| chr8:74304170-74306182 | p2 | (GT)6 | GCCTTTCTGGCGTGGTATTA | ACGAAATCATAGCGAATGGC | 119 |
| chr8:74304170-74306182 | p2 | (AT)21 | GCCTTTCTGGCGTGGTATTA | ACGAAATCATAGCGAATGGC | 119 |
| chr8:74319525-74321537 | p2 | (AC)6 | TGACGACGTTAGAAGATGCG | AGAGTCCCAAGGTTCCCAAT | 177 |
| chr8:74515510-74517530 | p5 | (TTATT)4 | CCACGCAGGATACCAAACTT | TGATGCTGGGAACAAGAGTG | 196 |
| chr8:74647156-74649171 | p2 | (AT)11 | AGAACCAAGGTAAGCGACGA | ATATCTCAGCCACACAGGGC | 191 |
| chr8:74647156-74649171 | p5 | (TTTAT)3 | AGAACCAAGGTAAGCGACGA | ATATCTCAGCCACACAGGGC | 191 |
| chr8:74737625-74739639 | p2 | (AT)7 | TACTTTGACCATGGCTTCCC | GACTTCGCCCACAGATTCAT | 206 |
| chr8:74927064-74929079 | p5 | (TTCAT)3 | TATTGCCTACAAGGTTCGGC | CCGAAGCATACCTAGACCCA | 168 |
| chr8:75010371-75012386 | p5 | (ATTTC)3 | TCTGCGTGGACTATCACGAC | TCGTATGCCTTGCCTTCTCT | 250 |
| chr8:75072251-75074263 | p2 | (TA)6 | TTTGAGAGGGATTTCATGCC | TGACCTGAGCAACATCCAAA | 235 |
| chr8:75136687-75138702 | p5 | (TATCA)3 | GATACCCTCCATGTCATCCG | AATTGCTGGACATTTTTGCC | 126 |
| chr8:75136687-75138702 | p5 | (ATATC)3 | GATACCCTCCATGTCATCCG | AATTGCTGGACATTTTTGCC | 126 |
| chr8:75154835-75156853 | p2 | (CT)9 | ACTCGGCCAAATTCATCAAC | GATGGTTTTTGGTGGGATTG | 172 |
| chr8:75173551-75175569 | p6 | (CATATA)3 | TCCGGAATTGTGGATTGATT | GCAAGCCGACGTACCATTAT | 230 |
| chr8:75227482-75229494 | p2 | (AC)6 | GTTTCGGAGGAATAGTGGCA | AGCATTTCTGCTCCTCGAAA | 114 |
| chr8:75313576-75315594 | p2 | (CT)9 | TCGATCCTCAATATGGCCTC | CCATGCAGATCAGTCCTTCA | 160 |
| chr8:75401376-75403391 | p5 | (ATTTT)3 | ACGAAGTGATCAAAATGGGC | TGTGGTAGGATTGCCATGAA | 253 |
| chr8:75458120-75460132 | p2 | (TA)6 | CCCACCATCTCCAAACTCAC | GACATGGCGGAGTAGCAAAT | 284 |
| chr8:75483542-75485566 | p2 | (AT)12 | GTTGCTTCCTTGTCTGCCAT | AGCCTCAGACACCAGCAGTT | 192 |
| chr8:75558216-75560243 | p3 | (ATG)9 | AATTCCCAACGGAAATACCC | GGCCATATGAATCAATTCGG | 248 |

| Marker ID | SSR type | SSR motif | FORWARD PRIMER (5'-3') | REVERSE PRIMER (5'-3') | size (bp) |
|---|---|---|---|---|---|
| chr8:75594261-75596276 | p3 | (GGT)5 | CGAAGTGAACGGTTTTCCAT | GAAGGGGAAGGAATCCTCTG | 188 |
| chr8:75594261-75596276 | p5 | (CAAAC)3 | CGAAGTGAACGGTTTTCCAT | GAAGGGGAAGGAATCCTCTG | 188 |
| chr8:75626070-75628084 | p2 | (TA)7 | TGGTCGTCCATCATGCTTTA | AAGGCTTCAACTTTCTGCCA | 180 |
| chr8:75637969-75639981 | p2 | (TA)6 | TGATTCCGATTCGAACTTCC | CTGCTGGGGAATACGGTTTA | 244 |
| chr8:75655467-75657482 | p5 | (ATTTC)3 | ACAGAGGAAACTCTGGCGAA | TGCCTTCTCTTGGCAAACTT | 232 |
| chr8:75745742-75747757 | p2 | (AT)8 | CTTTTCTTTTGCGAGGGTTG | ATCAATCGATGGAGAAACGC | 281 |
| chr8:75745742-75747757 | p5 | (AGGAC)3 | CTTTTCTTTTGCGAGGGTTG | ATCAATCGATGGAGAAACGC | 281 |
| chr8:75745742-75747757 | p2 | (AT)7 | CTTTTCTTTTGCGAGGGTTG | ATCAATCGATGGAGAAACGC | 281 |
| chr8:75811643-75813655 | p2 | (AT)6 | CATGTTGACTGTCACGACCC | TGGATACCCTTATCGGCTTG | 207 |
| chr8:75854171-75856185 | p2 | (AC)7 | TCGCTCAAAAACTGCAAATG | TTTGGAGCCGAGTTGAATCT | 208 |
| chr8:75854171-75856185 | p2 | (TC)6 | TCGCTCAAAAACTGCAAATG | TTTGGAGCCGAGTTGAATCT | 208 |
| chr8:75908920-75910934 | p2 | (AC)7 | GGACATGCCAAAGAAAGGAA | TTTTTACAGGGGAGGTGCTG | 112 |
| chr8:75908920-75910934 | p2 | (AT)7 | GGACATGCCAAAGAAAGGAA | TTTTTACAGGGGAGGTGCTG | 112 |
| chr8:76030130-76032144 | p2 | (AT)8 | TACAAGGCAAGCTATCCGGT | TCAAAGTCTTCCATTTGGGC | 250 |
| chr8:76030130-76032144 | p2 | (AT)7 | TACAAGGCAAGCTATCCGGT | TCAAAGTCTTCCATTTGGGC | 250 |
| chr8:76030130-76032144 | p2 | (TA)6 | TACAAGGCAAGCTATCCGGT | TCAAAGTCTTCCATTTGGGC | 250 |
| chr8:76139223-76141235 | p2 | (TG)6 | ACGATTGTACTACGCACCCC | AATAGCCCGTAGGCACCTTT | 152 |
| chr8:76259923-76261937 | p2 | (AT)7 | GCCATATCCCCTCCCTATGT | TGTCTCACTGACCAAGTGGC | 238 |
| chr8:76376034-76378050 | p4 | (AAAG)4 | AGCCTACGGCACATTGAATC | TTCTTTTCCGGGTCCTTCTT | 147 |
| chr8:76385739-76387751 | p2 | (AT)6 | AAATAAGTGGTGCTCGGTGG | CCCAAGGCCTAACTCTCCTT | 213 |
| chr8:76404798-76406816 | p6 | (TATATG)3 | GTGTTTCTGGTCGTGTGTGG | TTTCACCAGAGTTTCCCCTG | 177 |
| chr8:76425458-76427472 | p2 | (AT)7 | GCTAAGTGGTGTAGCGGAGG | GGGATCCTTCCCGAATACAT | 289 |
| chr8:76425458-76427472 | p2 | (AT)6 | GCTAAGTGGTGTAGCGGAGG | GGGATCCTTCCCGAATACAT | 289 |
| chr8:76425458-76427472 | p2 | (AT)20 | GCTAAGTGGTGTAGCGGAGG | GGGATCCTTCCCGAATACAT | 289 |
| chr8:76449864-76451882 | p3 | (TTG)6 | GACCCTTGAGCCAAATCAAA | CTTATTCCGATGTCGCCTGT | 158 |
| chr8:76480122-76482134 | p2 | (TA)6 | AGGTTCTGGTCAGCAGCAGT | ACTTCACCTTGATGCCCAAC | 194 |
| chr8:76604390-76606418 | p3 | (ACA)5 | CTTGCAAAAGAACCCCAAAG | GTCGGCTTGCGTATGGTATT | 256 |
| chr8:76653204-76655216 | p2 | (TA)6 | GCCACACACAGCATAACACC | CCACGACGACTTTGGAATTT | 237 |
| chr8:76693824-76695839 | p3 | (AAC)5 | AAGAGGTCGTGCAATTGTGA | GGATGAACCAACTGGCATCT | 261 |
| chr8:76723608-76725626 | p3 | (TTG)6 | AAATTTGGCCAGTGTTACGC | CCCATATCTTCTCATCCCGA | 282 |
| chr8:76743149-76745164 | p3 | (TTG)5 | GGCCACGTCATAAAGCAAAT | CCCAGGTCTCTCGGAATGTA | 261 |
| chr8:76822391-76824409 | p6 | (GAAAAA)3 | GAGCGTACGCAGACCTTACC | AACCGGTCCTAATCACCCTC | 180 |
| chr8:76840432-76842446 | p5 | (TATTT)3 | CGCCACCCAACTCACTTATT | GTCCCTCTTGCTCTTGGTTG | 106 |
| chr8:76840432-76842446 | p3 | (GGA)5 | CGCCACCCAACTCACTTATT | GTCCCTCTTGCTCTTGGTTG | 106 |
| chr8:76841398-76843413 | p5 | (TATTT)3 | GACATATGTAAAACGCCCCG | TCCAGGATTGAAAATGAGCC | 226 |
| chr8:76841398-76843413 | p3 | (GGA)5 | GACATATGTAAAACGCCCCG | TCCAGGATTGAAAATGAGCC | 226 |

| Marker ID | SSR type | SSR motif | FORWARD PRIMER (5'-3') | REVERSE PRIMER (5'-3') | size (bp) |
|---|---|---|---|---|---|
| chr8:76897996-76900010 | p2 | (TA)7 | TGTGACCGAGTGAACCACAT | ATTTCTTCCGCCCTTACGTT | 152 |
| chr8:76900338-76902352 | p2 | (TA)7 | TGTGACCGAGTGAACCACAT | ATTTCTTCCGCCCTTACGTT | 152 |
| chr8:76900338-76902352 | p5 | (TTGGT)3 | TGTGACCGAGTGAACCACAT | ATTTCTTCCGCCCTTACGTT | 152 |
| chr8:76900814-76902829 | p2 | (TA)7 | GGAATGGGCAGCTCAAATTA | GTTTTGGTTTTCGGTTTGGA | 186 |
| chr8:76900814-76902829 | p5 | (TTGGT)3 | GGAATGGGCAGCTCAAATTA | GTTTTGGTTTTCGGTTTGGA | 186 |
| chr8:76995845-76997863 | p3 | (GGC)6 | CTTCGAGTTCCTTCCGAGTG | CAATTGGCCGATGAAGAAAT | 110 |
| chr8:77008294-77010326 | p3 | (AAC)6 | GGAGCCAGAGGAGAGGAAAT | GAGTGGGTTTTGGAGATGGA | 125 |
| chr8:77034280-77036350 | p2 | (AT)35 | TGCTCCACGCCCTACTACTT | CGAAGCCACGGTTGATACTT | 161 |
| chr8:77034280-77036350 | p2 | (AT)8 | TGCTCCACGCCCTACTACTT | CGAAGCCACGGTTGATACTT | 161 |
| chr8:77191021-77193045 | p4 | (TTAT)6 | GAGAATAATGTGAGGGCCGA | TGGAACATCACATCTCGGAA | 168 |
| chr8:77239884-77241899 | p5 | (ATGCA)3 | CGTGTGTCAGTTCACATCCC | CGCCCACTAGGGGAATTAGT | 230 |
| chr8:77247717-77249731 | p5 | (ATGCA)3 | GCGAACCATCCTTTTAACCA | TTTCGGATTTTGGGGTTGTA | 249 |
| chr8:77247717-77249731 | p2 | (TA)7 | GCGAACCATCCTTTTAACCA | TTTCGGATTTTGGGGTTGTA | 249 |
| chr8:77286277-77288295 | p3 | (GTT)6 | AAAGGACCCTTGAGCCAAAT | CACTGCCCAACTTGACCTTT | 118 |
| chr8:77316610-77318628 | p6 | (ATACAT)3 | TATTTTCCGTCGACCGCTAC | AAATAGCCATGTGGCTCACC | 276 |
| chr8:77331152-77333168 | p2 | (TA)8 | ATAACAAGAGAAAGCGGCGA | GGAGCCTAACCACAATCCAA | 153 |
| chr8:77370810-77372828 | p2 | (TA)9 | GACCAGGTCATGGGAAATTG | TAGGCGAACCCTCTATCGAA | 152 |
| chr8:77534630-77536644 | p2 | (AT)7 | TGCAATAATGCCCATGAGTC | TGGTATGTGGTCAAAGGCAA | 209 |
| chr8:77544554-77546569 | p5 | (TTGCT)3 | AAGCCGTTTGAACACCATTC | TGATGGTTCGTCTGGGTACA | 121 |
| chr8:77559802-77561817 | p5 | (TACAA)3 | CTAGCTCCAGACACCCAAGG | AAGCAATTTTGTAGCACGGG | 257 |
| chr8:77565602-77567618 | p2 | (TG)8 | TCCAATCATCAACTCGACCA | CAGTCACCAAAAAGGAAGGC | 271 |
| chr8:77593152-77595167 | p5 | (TTTTA)3 | GCGGGATTGATGGAGAAGTA | GTCCCAATCGGAGTCTTGAA | 198 |
| chr8:77691565-77693583 | p6 | (GTCCAC)6 | GGATCCCTTCCACGTACAAA | GAAAGTGGACGCATGGATTT | 281 |
| chr8:77691565-77693583 | p6 | (ATCCAC)3 | GGATCCCTTCCACGTACAAA | GAAAGTGGACGCATGGATTT | 281 |
| chr8:77691565-77693583 | p6 | (TCCACG)4 | GGATCCCTTCCACGTACAAA | GAAAGTGGACGCATGGATTT | 281 |
| chr8:77755448-77757460 | p2 | (AT)6 | GCCCTACTTGAACACCCAAA | GCATATATTCGGCGAGGTGT | 104 |
| chr8:77775216-77777236 | p5 | (AAATG)4 | CCTCACATACCTTGCCGTTT | CGAAGGAAGTACTGCCCAAA | 263 |
| chr8:77775216-77777236 | p5 | (GAAAT)3 | CCTCACATACCTTGCCGTTT | CGAAGGAAGTACTGCCCAAA | 263 |
| chr8:77861324-77863339 | p3 | (ATA)5 | ACCTCATCCATACGTGGAGC | AATGGATTGACTGAGGCAGG | 246 |
| chr8:77861324-77863339 | p6 | (ATATAC)4 | ACCTCATCCATACGTGGAGC | AATGGATTGACTGAGGCAGG | 246 |
| chr8:77976237-77978293 | p2 | (AT)28 | CGACGTGAAGCGATACTCAA | TGTCCTTCCCAGGTCATTTC | 236 |
| chr8:77976237-77978293 | p2 | (TA)9 | CGACGTGAAGCGATACTCAA | TGTCCTTCCCAGGTCATTTC | 236 |
| chr8:77976237-77978293 | p5 | (TGAAT)3 | CGACGTGAAGCGATACTCAA | TGTCCTTCCCAGGTCATTTC | 236 |
| chr8:78058244-78060264 | p2 | (TA)10 | TTGGGTACGGATAACCCTGA | CAAACCTGACCACGAAGGTT | 251 |
| chr8:78211240-78213258 | p6 | (CAGCTC)3 | ACCTAGGAGCCTTGTCAGCA | TCATAAGGACGGGACAGGAC | 242 |
| chr8:78224480-78226500 | p5 | (AATGC)4 | CCAATCTGAACAATCTCGCA | TGGGTCGTGACAAGTTGGTA | 184 |

| Marker ID | SSR type | SSR motif | FORWARD PRIMER (5'-3') | REVERSE PRIMER (5'-3') | size (bp) |
|---|---|---|---|---|---|
| chr8:78279709-78281721 | p2 | (TA)9 | CTCTACTCGTGCATGCCTCA | TGTATCATATCCGGCCCATT | 201 |
| chr8:78279709-78281721 | p2 | (AT)6 | CTCTACTCGTGCATGCCTCA | TGTATCATATCCGGCCCATT | 201 |
| chr8:78284848-78286866 | p3 | (TTG)6 | GGAATTTGGAGTTGCCTTCA | TTCACCTGCTCTCCAATCCT | 206 |
| chr8:78284848-78286866 | p2 | (AT)6 | GGAATTTGGAGTTGCCTTCA | TTCACCTGCTCTCCAATCCT | 206 |
| chr8:78285605-78287617 | p3 | (TTG)6 | CATCGCGAGGGATGTTTTAT | TGACACTTATCCTCCCGAGC | 182 |
| chr8:78285605-78287617 | p2 | (AT)6 | CATCGCGAGGGATGTTTTAT | TGACACTTATCCTCCCGAGC | 182 |
| chr8:78304443-78306455 | p2 | (AT)7 | GTTCCTTCGGCAGACATGAT | TTTGGTTGATTTGACGTGGA | 112 |
| chr8:78304443-78306455 | p2 | (TC)6 | GTTCCTTCGGCAGACATGAT | TTTGGTTGATTTGACGTGGA | 112 |
| chr8:78430268-78432283 | p5 | (TTGGT)3 | GAGCTTAACATGGCCTGAGC | TTCCACTCATCCACCATCAA | 126 |
| chr8:78503770-78505816 | p2 | (AT)23 | CGGTATTAAGGATCCGCTCA | CCTAGGTGTCCAAGTCCGAA | 111 |
| chr8:78625535-78627550 | p3 | (ATA)5 | GATAGCCCCCATGACTCGTA | TAGTTCGAGAGCCCGTGAGT | 125 |
| chr8:78698104-78700134 | p2 | (GA)15 | GTATCCATGTGTGGCACGAG | ATCCAAGTAAGTGTTGCGGC | 228 |
| chr8:78698104-78700134 | p3 | (GTT)5 | GTATCCATGTGTGGCACGAG | ATCCAAGTAAGTGTTGCGGC | 228 |
| chr8:78698650-78700665 | p2 | (GA)15 | GGCCTGTATGTGACATCGTG | TTAAATGGCAAGGGAAATGG | 277 |
| chr8:78698650-78700665 | p3 | (GTT)5 | GGCCTGTATGTGACATCGTG | TTAAATGGCAAGGGAAATGG | 277 |
| chr8:78698650-78700665 | p2 | (TA)25 | GGCCTGTATGTGACATCGTG | TTAAATGGCAAGGGAAATGG | 277 |
| chr8:78769892-78771914 | p2 | (TA)11 | ATAAGCCGACGAGGCTAACA | GTCCTGTCCCGCCTATATGA | 286 |
| chr8:78836899-78838941 | p2 | (AT)21 | TCGTGTTCACGCCTACACAT | AAAGCCTAGTGCGGAAGTGA | 119 |
| chr8:78977860-78979881 | p3 | (GTT)7 | CCACCAAAACTTAAGGGCAA | ATGCCCAAACCATCTCAATC | 269 |
| chr8:78985160-78987234 | p3 | (AAC)6 | CCAGCGGCTGTAAGACAAAT | CGTTGATGGCTGTCAGTTTG | 140 |
| chr8:79044830-79046842 | p2 | (CA)6 | CTCGATCCCCCTCTATAGCC | TTAAAAGGGCGACAACAACC | 270 |
| chr8:79054579-79056593 | p2 | (AT)7 | TCTATGAAACCTTGCGGGTC | GTTGCTTCGGGTCAACAAAT | 237 |
| chr8:79068204-79070222 | p6 | (TACCTC)3 | CAATACACATCCGCAACGAC | CCAAATGAGCTCCCATGTCT | 161 |
| chr8:79132823-79134839 | p2 | (TA)8 | CATTTTCCACCCACTGGACT | AAAGCCCTCAAGAATTTGGC | 264 |
| chr8:79144712-79146724 | p2 | (TA)6 | TACCCCTTTTGTTGCTGGTC | AGCATAACATGAGGCCAACC | 161 |
| chr8:79158697-79160751 | p3 | (ATA)18 | TTTTCGTGTGGAATCGTCAA | GCGAAGTTTGTGGAAAGAGC | 184 |
| chr8:79207742-79209760 | p6 | (AATATG)3 | TATCAACAAAGGCTCGTCCC | ACACCCAACTTCGTCCATTC | 102 |
| chr8:79214187-79216199 | p2 | (AT)6 | TTTGGAGGGCTTACACTTGG | ATGTGAGCTCGAGGAGGAGA | 148 |
| chr8:79214187-79216199 | p2 | (AT)7 | TTTGGAGGGCTTACACTTGG | ATGTGAGCTCGAGGAGGAGA | 148 |
| chr8:79278616-79280631 | p5 | (AACAA)3 | CCTCCAAATTCGCTCTCAAG | GTCGTGCCTCGGTGATATTT | 269 |
| chr8:79278616-79280631 | p3 | (GAA)5 | CCTCCAAATTCGCTCTCAAG | GTCGTGCCTCGGTGATATTT | 269 |
| chr8:79279103-79281118 | p5 | (AACAA)3 | TGCGGCCATATGTAAAAACA | ATTTTGTCGGTTCCATCAGC | 176 |
| chr8:79279103-79281118 | p3 | (GAA)5 | TGCGGCCATATGTAAAAACA | ATTTTGTCGGTTCCATCAGC | 176 |
| chr8:79293396-79295408 | p2 | (CT)6 | GGATCATGCCAAGGAAGAAA | TGGTCGTTGTTGTTGTTGGT | 283 |
| chr8:79344511-79346525 | p2 | (AT)7 | ACTCATGAAATCACGGGCTC | AGTCCCTCCATCGGCTACTT | 284 |
| chr8:79358025-79360103 | p2 | (TA)39 | TTTCCCATCGAACACATCAA | CGGAGATGCCAGTGATTTTT | 133 |

| Marker ID | SSR type | SSR motif | FORWARD PRIMER (5'-3') | REVERSE PRIMER (5'-3') | size (bp) |
|---|---|---|---|---|---|
| chr8:79481472-79483490 | p2 | (TA)9 | TTGGAAACTTCTCCCCACTG | AGGCCAATTCAAACACCTTG | 174 |
| chr8:79504084-79506132 | p2 | (AT)24 | CTCCCATGCCTGATACCACT | GCAGAGAGTCAGGGAACTGG | 177 |
| chr8:79504084-79506132 | p2 | (AT)13 | CTCCCATGCCTGATACCACT | GCAGAGAGTCAGGGAACTGG | 177 |
| chr8:79673129-79675149 | p2 | (AT)10 | CACCATCCCCGAGAATCTAA | TGCCGTCGAATTGTGTAAAA | 167 |
| chr8:79701031-79703057 | p2 | (AT)13 | CCAAATGGCATTCGAAAGAT | TTCGGGTCAATCCGACTTAG | 178 |
| chr8:79804817-79806829 | p2 | (GA)6 | AGTTTGCATGTGCCATGTGT | CCAAGGAAGGCCTAACATCA | 275 |
| chr8:79866545-79868560 | p5 | (GCATT)3 | TGGTGAGACTTCGTGTAGCG | CAAGTCACAACACAATGGGG | 252 |
| chr8:80050893-80052915 | p2 | (AT)16 | TCCACCCTTAAAAGTCACGC | GTGGACATGAACACAGCGTC | 283 |
| chr8:80050893-80052915 | p2 | (TA)11 | TCCACCCTTAAAAGTCACGC | GTGGACATGAACACAGCGTC | 283 |
| chr8:80175168-80177184 | p2 | (TA)6 | CAAATTCATGACCTTGTCGG | TGCAAAATAGCTTCAGCCAA | 121 |
| chr8:80175168-80177184 | p2 | (AT)8 | CAAATTCATGACCTTGTCGG | TGCAAAATAGCTTCAGCCAA | 121 |
| chr8:80249270-80251290 | p2 | (AT)8 | AGCAGAATGAGGGGAAGTGA | GACGGAAAGTGAGCCAACAT | 177 |
| chr8:80249270-80251290 | p2 | (AT)6 | AGCAGAATGAGGGGAAGTGA | GACGGAAAGTGAGCCAACAT | 177 |
| chr8:80252767-80254782 | p5 | (GGTTT)3 | AACGATCACGAGAAAAGGGA | ATCAAACCAAACCAAACCCA | 247 |
| chr8:80252767-80254782 | p5 | (TTTGG)3 | AACGATCACGAGAAAAGGGA | ATCAAACCAAACCAAACCCA | 247 |
| chr8:80280930-80282958 | p2 | (AT)14 | GACCGTCCCAGAAAAACAAA | TGTATGCCGGAGTGTGAAAA | 172 |
| chr8:80307177-80309207 | p2 | (TA)6 | TCTGTTGCACCTGTTTTTGC | GGCTTAAATGGCCATGATTG | 242 |
| chr8:80390887-80392901 | p2 | (AT)7 | ATTTCTTCGGCCTACCCCTA | ACCAGGAGTGACAGGATTGG | 218 |
| chr8:80390887-80392901 | p2 | (TA)8 | ATTTCTTCGGCCTACCCCTA | ACCAGGAGTGACAGGATTGG | 218 |
| chr8:80396819-80398837 | p6 | (TATATG)3 | GTTGATGATTTGGCCGAGTT | AAGTCTTTGGCTTCGCAAGA | 149 |
| chr8:80396819-80398837 | p2 | (TA)13 | GTTGATGATTTGGCCGAGTT | AAGTCTTTGGCTTCGCAAGA | 149 |
| chr8:80458209-80460223 | p2 | (AT)7 | TACATACGTGAGCGAGCAGG | CCGAGATCAGGTTGGTGAAT | 250 |
| chr8:80462468-80464483 | p3 | (TAT)5 | TTTTAATCAATGCGGCACAA | CGTCATTCCGAGACAATGTG | 147 |
| chr8:80511243-80513261 | p6 | (AAAGGT)3 | TAGGGTGGTCGCGTAGTTTC | TAGACCCTCGGCTCAAAAGA | 236 |
| chr8:80539334-80541352 | p2 | (TA)9 | ACTTTTGTCACGACCCAACC | TTAGCCTCGTCGGCTTATGT | 124 |
| chr8:80566892-80568926 | p2 | (AT)17 | GCGGACAAGCAACTTTTAGC | CGATATTTCCTCCAAAGCCA | 264 |
| chr8:80637955-80639989 | p2 | (AT)17 | AGTGGGAGCATGTGGGATAG | TGCATCCGTGTTTTGATGAT | 190 |
| chr8:80662710-80664728 | p6 | (ATTTAT)3 | GAACGGCGAATTCAGGATTA | TTTCAAAAGTTGCCACGTCA | 124 |
| chr8:80662710-80664728 | p6 | (ATTTAT)3 | GAACGGCGAATTCAGGATTA | TTTCAAAAGTTGCCACGTCA | 124 |
| chr8:80663161-80665179 | p6 | (ATTTAT)3 | GGAAGCGAGATCAACGCTAA | TTAACACCCACCATCAACCA | 175 |
| chr8:80663161-80665179 | p6 | (ATTTAT)3 | GGAAGCGAGATCAACGCTAA | TTAACACCCACCATCAACCA | 175 |
| chr8:80663161-80665179 | p2 | (TA)13 | GGAAGCGAGATCAACGCTAA | TTAACACCCACCATCAACCA | 175 |
| chr8:80702358-80704373 | p5 | (AATAA)3 | ACCAACAAAGGTCCACTTCG | TTGCCATTTACCCCTTTGAA | 252 |
| chr8:80779007-80781019 | p2 | (AT)6 | AACAATGGTGCCTCTTACGG | GAGCCCTGTGTGGCTATGAT | 141 |
| chr8:80779007-80781019 | p2 | (AT)16 | AACAATGGTGCCTCTTACGG | GAGCCCTGTGTGGCTATGAT | 141 |
| chr8:80779007-80781019 | p2 | (AT)9 | AACAATGGTGCCTCTTACGG | GAGCCCTGTGTGGCTATGAT | 141 |

| Marker ID | SSR type | SSR motif | FORWARD PRIMER (5'-3') | REVERSE PRIMER (5'-3') | size (bp) |
|---|---|---|---|---|---|
| chr8:80791259-80793274 | p5 | (TCAAG)3 | AAGTGAGGCCCTATCTCGGT | CGGTCCGTATTTTGCACTTT | 296 |
| chr8:80868802-80870930 | p2 | (TA)18 | AGTGCCCGACCACTACTGAC | GTCATCCATTGCACGTATGC | 295 |
| chr8:80868802-80870930 | p3 | (CAT)8 | AGTGCCCGACCACTACTGAC | GTCATCCATTGCACGTATGC | 295 |
| chr8:80868802-80870930 | p3 | (CAA)6 | AGTGCCCGACCACTACTGAC | GTCATCCATTGCACGTATGC | 295 |
| chr8:80909806-80911821 | p3 | (TGT)5 | CTGCTGCATCTGTGTGTGTG | GTGACACAGAGTGGTGTGGG | 256 |
| chr8:80939583-80941608 | p5 | (TCCCT)5 | CGGACTCGATGAAGAATGGT | TAACTCATTTGGGCCTGGAC | 179 |
| chr8:81145951-81147975 | p2 | (AT)12 | TGTACCAGGTTTGGCTCCTC | CAACCCCTTGCCACCTACTA | 157 |
| chr8:81184638-81186659 | p2 | (AT)28 | CTCACGACGCTCATCTTCAA | TCGATGAGGCAGCACATAAG | 155 |
| chr8:81184638-81186659 | p3 | (AAT)7 | CTCACGACGCTCATCTTCAA | TCGATGAGGCAGCACATAAG | 155 |
| chr8:81220973-81222988 | p5 | (ACAAA)3 | TGTGGCGAGAAATGTGAGAG | CACTTTGTCACACCCCCTTT | 127 |
| chr8:81235143-81237157 | p2 | (CT)7 | CCAGCAGCTTGCACAGAATA | CTGTACACTTGCTGCCCCTT | 136 |
| chr8:81281555-81283582 | p3 | (GAA)9 | GAACCAACGAGTCCGCTAAG | TTCCGAAACTTTTGGGTGAG | 184 |
| chr8:81291877-81293892 | p5 | (CTATA)3 | GGTATAACACTGATGCCGGG | CTTATGCCCACTTCCTTCCA | 115 |
| chr8:81375854-81377876 | p2 | (AT)7 | GAAAGAGATCCCGAACCCTC | AACCGGATACGGGTAAATCC | 296 |
| chr8:81427133-81429161 | p2 | (AT)14 | TGACGGATCCTCCTCCATAG | ATGAAGAGGACGGCTTCTGA | 236 |
| chr8:81523369-81525384 | p3 | (AAG)5 | AGTCGCACCTTAGCCTGAAA | CGGGATATGGGAATTGTCAC | 199 |
| chr8:81530331-81532346 | p5 | (AAAAT)3 | GAACCCCCACTCAGTCAAAA | CTTATGCCAAGTCTGCCCAT | 135 |
| chr8:81530331-81532346 | p2 | (AT)6 | GAACCCCCACTCAGTCAAAA | CTTATGCCAAGTCTGCCCAT | 135 |
| chr8:81555218-81557233 | p5 | (ATTTC)3 | TCTCGGTTAGCTGGGAAGAA | GGGCGCCTGTAAAAATGTAA | 182 |
| chr8:81566632-81568671 | p3 | (AGG)5 | CAGGTACCGACGACAGGTTT | CCGTACCCATGAACGAGAAT | 242 |
| chr8:81601556-81603574 | p6 | (TAATCT)3 | TGGGCCTTGCATACAACATA | TATCTCCGAACCAAACTCCG | 253 |
| chr8:81678829-81680849 | p3 | (AAC)5 | CTTGATTCGCGTTCAACTCA | TCGGCACACGTACTGTGATT | 244 |
| chr8:81682074-81684098 | p3 | (GTT)8 | TTTTCTTGCCTGTGCCTTCT | GTTATCCAAAGGGGCACTGA | 188 |
| chr8:81821956-81823974 | p3 | (TGT)6 | TTCAGCTTCACACTCCGATG | TCGCCATCTTTCTTTTGCTT | 294 |
| chr8:81821956-81823974 | p3 | (TGT)6 | TTCAGCTTCACACTCCGATG | TCGCCATCTTTCTTTTGCTT | 294 |
| chr8:82195271-82197287 | p2 | (CT)6 | AAAATAAATCCCCAGCCCAC | AGGAAGGGAGCGAAAGAGAG | 233 |
| chr8:82195271-82197287 | p2 | (GT)8 | AAAATAAATCCCCAGCCCAC | AGGAAGGGAGCGAAAGAGAG | 233 |
| chr8:82313892-82315910 | p3 | (TTG)6 | TGGAAGGATTGGAGGACAAG | AGGATTTTGCTTTGCTTGGA | 226 |
| chr8:82337687-82339702 | p5 | (ATTTT)3 | AAATGCTTGGCCAAAGAGAA | ATCAATTTTGACCCCAACCA | 246 |
| chr8:82337687-82339702 | p6 | (ACAAAA)4 | AAATGCTTGGCCAAAGAGAA | ATCAATTTTGACCCCAACCA | 246 |
| chr8:82483878-82485893 | p5 | (CGAAC)3 | CATTACGGTTTGGATACGGC | GAAGTCAGCTTGAGGCCAAC | 243 |
| chr8:82489909-82491921 | p2 | (GT)6 | TTTGGGGCCTAAAGTCACTG | ATTTGCCCTTAGCCTTCCTC | 282 |
| chr8:82603227-82605245 | p6 | (CCTCTA)3 | CTCCCCTTCAGTGGGTAACA | CTGAGGGACCGAACATAGGA | 213 |
| chr8:82618393-82620409 | p2 | (TA)8 | ACAGAACTTCAAACGGTGCC | CAGACAGATGGTCAGGCAGA | 126 |
| chr8:82635820-82637834 | p3 | (CCA)5 | ACTAACTGCCTCCAGCCTCA | GCTTTTGGTTGTGGGGAGTA | 231 |
| chr8:82635820-82637834 | p2 | (AT)25 | ACTAACTGCCTCCAGCCTCA | GCTTTTGGTTGTGGGGAGTA | 231 |

| Marker ID | SSR type | SSR motif | FORWARD PRIMER (5'-3') | REVERSE PRIMER (5'-3') | size (bp) |
|---|---|---|---|---|---|
| chr8:82635859-82637913 | p3 | (CCA)5 | ACTAACTGCCTCCAGCCTCA | GCTTTTGGTTGTGGGGAGTA | 192 |
| chr8:82635859-82637913 | p2 | (AT)25 | ACTAACTGCCTCCAGCCTCA | GCTTTTGGTTGTGGGGAGTA | 192 |
| chr8:82636258-82638308 | p2 | (AT)25 | GCCAATTTTCATAAGCCCAA | GCGTATTCCATTAAGCCCAA | 295 |
| chr8:82705136-82707151 | p5 | (ATTTT)3 | ATTTGGTATGCAAGATGGGC | ATGCAGAATTTGCTCTGGCT | 269 |
| chr8:82797659-82799683 | p3 | (ACA)8 | CCTTGCTAAATCCGTTCCAA | TGGCTGAGATGCAACAGTTC | 197 |
| chr8:82802080-82804095 | p5 | (ATTTT)3 | ATTTGGTATGCAAGATGGGC | CAACTGGCACCAGCAATATG | 275 |
| chr8:82802080-82804095 | p5 | (ATTTT)3 | ATTTGGTATGCAAGATGGGC | CAACTGGCACCAGCAATATG | 275 |
| chr8:82846376-82848390 | p2 | (AT)7 | TCCATAACTGTCACGACCCA | AGCCTCGTTGTCTTGCGTAT | 230 |
| chr8:82859777-82861807 | p2 | (AT)15 | ACCGAAAGGCGTTATGTACG | TATCCGAGGAACGTACAGGC | 225 |
| chr8:82943242-82945260 | p5 | (GGGCC)3 | ACCAGTTGTTGGGCTGAAAC | CTTCGGCATGTTGAAGCATA | 115 |
| chr8:82943242-82945260 | p2 | (TC)9 | ACCAGTTGTTGGGCTGAAAC | CTTCGGCATGTTGAAGCATA | 115 |
| chr8:82954264-82956279 | p5 | (TAAGC)3 | AGGGCCTCTAAAGCCAAAAG | TTTATGCCTGCAATTTTCCC | 197 |
| chr8:82967870-82969885 | p5 | (TTTGG)3 | CCAGCTAGTGCCCAAAATGT | GGTGTGATTTGGCTGGACTT | 204 |
| chr8:82967950-82969965 | p5 | (TTTGG)3 | CCAGCTAGTGCCCAAAATGT | GGTGTGATTTGGCTGGACTT | 124 |
| chr8:83042742-83044758 | p3 | (ATT)6 | ACGGGTGCTATTCATATGGG | CACTTGGGATTTCACTGGCT | 252 |
| chr8:83042742-83044758 | p3 | (AGT)5 | ACGGGTGCTATTCATATGGG | CACTTGGGATTTCACTGGCT | 252 |
| chr8:83042761-83044785 | p3 | (ATT)6 | ACGGGTGCTATTCATATGGG | CACTTGGGATTTCACTGGCT | 233 |
| chr8:83042761-83044785 | p3 | (AGT)5 | ACGGGTGCTATTCATATGGG | CACTTGGGATTTCACTGGCT | 233 |
| chr8:83043024-83045066 | p3 | (ATT)6 | GCCAGTGAAATCCCAAGTGT | ATCTTGAGCCGAGGGTCTTC | 100 |
| chr8:83043024-83045066 | p3 | (AGT)5 | GCCAGTGAAATCCCAAGTGT | ATCTTGAGCCGAGGGTCTTC | 100 |
| chr8:83057194-83059230 | p3 | (TAA)12 | AGAGGCTGGGTTGGAAAAGT | CCATACAGTCGCACCATGTC | 295 |
| chr8:83071035-83073050 | p3 | (GAA)5 | ATTTTTGCAGAGAGACGGGA | CGTTGGGAATTCGACAAGTT | 258 |
| chr8:83251505-83253519 | p2 | (GA)7 | GGTTATGGATGCCCTTGAAA | CTAGAGAAAATTGGGCAGCG | 114 |
| chr8:83251505-83253519 | p3 | (GTT)8 | GGTTATGGATGCCCTTGAAA | CTAGAGAAAATTGGGCAGCG | 114 |
| chr8:83315988-83318006 | p2 | (TC)18 | GTGCCAGACACCACACAATC | ATCATCCACCATGAGCAACA | 261 |
| chr8:83315988-83318006 | p3 | (AGT)6 | GTGCCAGACACCACACAATC | ATCATCCACCATGAGCAACA | 261 |
| chr8:83318606-83320618 | p5 | (ATGCA)5 | ATGCACTGGCCAAATACCTC | GACGATACGGCCTTGAGTGT | 205 |
| chr8:83318606-83320618 | p2 | (AT)6 | ATGCACTGGCCAAATACCTC | GACGATACGGCCTTGAGTGT | 205 |
| chr8:83385794-83387830 | p2 | (TA)18 | TCACATTTTGCCGTGTTGTT | CAGGGAACCAACATTGTCCT | 163 |
| chr8:83407442-83409457 | p5 | (AATGC)3 | GGCGAACCCTCAATCAATAA | GTCTAGACCAGGTCGTGGGA | 189 |
| chr8:83457512-83459534 | p2 | (TA)11 | CCATACCAATCCGCAAGACT | GGTTAAGGGGATAGTTCGCC | 122 |
| chr8:83459601-83461615 | p2 | (AG)7 | ACCCAAAATCCCAACATCAA | TTCCATTGGGGGATGACTTA | 119 |
| chr8:83459601-83461615 | p3 | (TAT)19 | ACCCAAAATCCCAACATCAA | TTCCATTGGGGGATGACTTA | 119 |
| chr8:83621151-83623175 | p2 | (AG)12 | CCTAGCCCTTGTGGACAGAC | CTTGGGGCGTATGTAGCATT | 212 |
| chr8:83674417-83676432 | p5 | (GGTTT)3 | GGATGCCTAACCCTTCCTTT | ACCCTATTTTAACCGGGTCG | 197 |
| chr8:83818523-83820615 | p2 | (AT)46 | GGCGGTAAAGTCAGGTGGTA | TGCAATGGCCAATAACTCAA | 296 |

| Marker ID | SSR type | SSR motif | FORWARD PRIMER (5'-3') | REVERSE PRIMER (5'-3') | size (bp) |
|---|---|---|---|---|---|
| chr8:83850531-83852549 | p6 | (TAAAAA)3 | CTTGTTCCACCAACAACACG | AATCCTCCAAGCGGCTATTT | 209 |
| chr8:83979426-83981444 | p2 | (TA)8 | TTTAGCCTTTCCGAATTCCA | TTTGCTTTCTCTGCCCTCAT | 288 |
| chr8:83979426-83981444 | p2 | (AG)9 | TTTAGCCTTTCCGAATTCCA | TTTGCTTTCTCTGCCCTCAT | 288 |
| chr8:83979426-83981444 | p2 | (AT)6 | TTTAGCCTTTCCGAATTCCA | TTTGCTTTCTCTGCCCTCAT | 288 |
| chr8:84237541-84239556 | p5 | (AGTAA)3 | GTTTGAATGTGGTGGGTTCC | AAAATTGGGCCCCATAAAAA | 140 |
| chr8:84250174-84252192 | p5 | (TTGGT)3 | TGGTGTAGGTTCCGAGAAGG | CAGTCGACTTTGACCCCAAT | 281 |
| chr8:84250174-84252192 | p6 | (TGATTT)3 | TGGTGTAGGTTCCGAGAAGG | CAGTCGACTTTGACCCCAAT | 281 |
| chr8:84258700-84260718 | p2 | (AT)9 | TCCAGATGGTCAACGTTTCA | CATGTGGCAATGAGTCCATC | 266 |
| chr8:84278340-84280372 | p2 | (TA)16 | AGCTTAAGCCCTCCCTGGTA | TATACACGCCCATGGAGACA | 297 |
| chr8:84281444-84283456 | p3 | (CTT)9 | CGTTGCGCCTTAGTCTTACC | AGGAGGAGAAGGAGCAGGAG | 290 |
| chr8:84394072-84396090 | p2 | (AT)13 | ATATACGCAAGCCGACAAGG | TGTGGTTGTTCAGGGTACGA | 201 |
| chr8:84394072-84396090 | p6 | (TTTTTA)3 | ATATACGCAAGCCGACAAGG | TGTGGTTGTTCAGGGTACGA | 201 |
| chr8:84476591-84478607 | p2 | (AT)12 | ATGTGGCAATGAGTCCATCA | TGGAACTCACCACATCCAAA | 171 |
| chr8:84476591-84478607 | p2 | (TA)8 | ATGTGGCAATGAGTCCATCA | TGGAACTCACCACATCCAAA | 171 |
| chr8:84644712-84646856 | p6 | (GAAACT)3 | CTACTAACTCTGCTGCCCCG | GAGACAGCCGAGAGTATCCG | 150 |
| chr8:84644712-84646856 | p2 | (AT)6 | CTACTAACTCTGCTGCCCCG | GAGACAGCCGAGAGTATCCG | 150 |
| chr8:84658676-84660748 | p2 | (TA)36 | CCGTGAGTTTGGGACTGATT | ACTTGCTGCCCAAAAACACT | 163 |
| chr8:84740100-84742214 | p3 | (CTT)7 | TCGAGCAAGGAGGTCATCTT | GATGAGGAGGAGGAAGAGGG | 109 |
| chr8:84839059-84841099 | p2 | (AC)8 | TCAGAGTGCGCTGTTAATGC | ATTTCACCCTCAAGTGGTCG | 228 |
| chr8:84868958-84870970 | p2 | (TA)6 | TTTCGAGTTGACCTTGGACC | TCGGTTTTCTTGCCTTCACT | 119 |
| chr8:84917656-84919692 | p2 | (AT)18 | TCGCTATCCGTCTTGATTCC | AGGACGAATGATCACAAGGG | 230 |
| chr8:85000607-85002621 | p2 | (CT)7 | CAGTGGAGACCTTAGCAGGG | AACCCACAGTTTCAGTTGGC | 116 |
| chr8:85027707-85029727 | p5 | (AAAAG)4 | TGCACTGACTTGTTTCGAGG | TTCATCATCATCGTCGTCGT | 283 |
| chr8:85117564-85119596 | p3 | (GTT)6 | GAAAAGAATTTGCACGGCAT | ATGGCCCTCTAGTTGGGTCT | 175 |
| chr8:85150832-85152847 | p5 | (AAATC)3 | TCGTGAAGGCCATTCATACA | GACCCTCTTGGCATGTTTGT | 198 |
| chr8:85207777-85209795 | p6 | (TTTTTG)3 | TTATGGGCCCAGTTAGATGC | TACGGCTTCGAGCTCATTTT | 183 |
| chr8:85216736-85218754 | p6 | (AAAAGA)3 | GCTACCCCTCATCGATCAAA | GGCAAACCACGATCCTCTAA | 135 |
| chr8:85226832-85228847 | p5 | (AAATT)3 | TCGAATTTGACGCAGAGTTG | AGATGTGCGCCTTCAGAGTT | 129 |
| chr8:85301664-85303685 | p5 | (ACCCG)3 | GAACTATTTCTTCGGCAGCG | TATTTGGAATTGGAGCCGAG | 291 |
| chr8:85370447-85372461 | p6 | (CCCACC)3 | ACCCTCGAATGCACTTCATC | AAGTTTGCGCGGTTATATGG | 130 |
| chr8:85370447-85372461 | p6 | (AAAAAT)3 | ACCCTCGAATGCACTTCATC | AAGTTTGCGCGGTTATATGG | 130 |
| chr8:85370447-85372461 | p2 | (TA)7 | ACCCTCGAATGCACTTCATC | AAGTTTGCGCGGTTATATGG | 130 |
| chr8:85421551-85423608 | p2 | (TG)11 | TGGCACTGATGTGACAGTGA | TGACGTGGCACAATCAAAAT | 246 |
| chr8:85485724-85487744 | p2 | (AT)10 | AACCTACCCTGGCACCTCTT | GTTAGCCTCGTCGGCTTATG | 158 |
| chr8:85565042-85567057 | p5 | (CATTG)3 | TTCCGAACATGCAAAATACG | ACAACACAACACGGTGGCTA | 162 |
| chr8:85660787-85662805 | p6 | (TTTTTA)3 | TTCAATTCCCGACCAGTCTC | CTTCCAGCAAGATGGGGTTA | 276 |

| Marker ID | SSR type | SSR motif | FORWARD PRIMER (5'-3') | REVERSE PRIMER (5'-3') | size (bp) |
|---|---|---|---|---|---|
| chr8:85666846-85668861 | p3 | (AAC)5 | CCTTTCGGAGTGGTTTTGAA | ACTCGGATTGTTCATGGAGG | 179 |
| chr8:85666846-85668861 | p3 | (ACA)5 | CCTTTCGGAGTGGTTTTGAA | ACTCGGATTGTTCATGGAGG | 179 |
| chr8:85729318-85731358 | p2 | (AC)20 | ATTCCACCTTCCAATTTCCC | AAAATGAGCCATGAACCGTC | 242 |
| chr8:85808897-85810911 | p2 | (AT)6 | TCCCTCCGTATGTCCATGAT | CGTAGGCCAGGCATATTTGT | 198 |
| chr8:85808897-85810911 | p2 | (TA)7 | TCCCTCCGTATGTCCATGAT | CGTAGGCCAGGCATATTTGT | 198 |
| chr8:85862559-85864617 | p2 | (TA)29 | CCTTACGTCACCCCGATAGA | TATATGTTGCCCGCCTTACC | 145 |
| chr8:85928523-85930543 | p5 | (TTGGT)3 | ACCCGACTAGTGGTTTGGTG | GTAATGTCGGGTTGGTTTGG | 254 |
| chr8:85928523-85930543 | p5 | (TTGGT)4 | ACCCGACTAGTGGTTTGGTG | GTAATGTCGGGTTGGTTTGG | 254 |
| chr8:85934670-85936686 | p2 | (TA)50 | TCAATTTCAATTTTCCCCCA | TAGGTGTTGAAAAGGTGCCC | 267 |
| chr8:85934670-85936686 | p5 | (AATAA)3 | TCAATTTCAATTTTCCCCCA | TAGGTGTTGAAAAGGTGCCC | 267 |
| chr8:85934683-85936705 | p2 | (TA)50 | TCAATTTCAATTTTCCCCCA | TAGGTGTTGAAAAGGTGCCC | 254 |
| chr8:85934683-85936705 | p5 | (AATAA)3 | TCAATTTCAATTTTCCCCCA | TAGGTGTTGAAAAGGTGCCC | 254 |
| chr8:85934817-85936832 | p2 | (TA)50 | TCAATTTCAATTTTCCCCCA | TAGGTGTTGAAAAGGTGCCC | 120 |
| chr8:85934817-85936832 | p5 | (AATAA)3 | TCAATTTCAATTTTCCCCCA | TAGGTGTTGAAAAGGTGCCC | 120 |
| chr8:85958624-85960652 | p2 | (TA)14 | TGTGCACAAGTAGGCACCTC | ACCATTACACTGACGTGGCA | 205 |
| chr8:85978826-85980862 | p2 | (AT)18 | AAGAATGGACTTGGGAGCCT | GGGAGGTCAGACAACAAGGA | 187 |
| chr8:85997536-85999560 | p2 | (TA)12 | TGATTATCACACCGATCCGA | TGTCTACCTGTGGGCATGAA | 278 |
| chr8:86004418-86006433 | p5 | (TTCTT)3 | AACTCGGCCAAATCATCAAC | GAAATGTCGAGCGGATTGTT | 180 |
| chr8:86014757-86016785 | p2 | (AT)14 | GAACCCGTTGGTTCTCGTTA | CCCTGTCCCAACTTAGGGTT | 154 |
| chr8:86014757-86016785 | p3 | (CAA)7 | GAACCCGTTGGTTCTCGTTA | CCCTGTCCCAACTTAGGGTT | 154 |
| chr8:86017878-86019893 | p3 | (TCT)5 | TGCATTCTTCCAAAATTCCC | TGAAGTGCATCAAGGACGAG | 261 |
| chr8:86111248-86113262 | p2 | (AT)7 | GGGCAAAATTAGGGATGGTT | GCTGCGTCGTGACAAAAGTA | 153 |
| chr8:86111248-86113262 | p3 | (ATC)5 | GGGCAAAATTAGGGATGGTT | GCTGCGTCGTGACAAAAGTA | 153 |
| chr8:86181274-86183289 | p5 | (CTTCT)3 | CAAGCTGCTCCAAAGGAATC | TCACCTGCAAAGTCTGCATC | 249 |
| chr8:86270123-86272138 | p5 | (AAAAT)3 | TTGAATTCATGGAAACCCGT | TGACACTTTGCACCCAACTC | 222 |
| chr8:86270123-86272138 | p3 | (TTC)5 | TTGAATTCATGGAAACCCGT | TGACACTTTGCACCCAACTC | 222 |
| chr8:86273142-86275178 | p3 | (TTC)12 | TGAGGGTGGAGGAAATGAAG | ATGTCAACGCCACATCAAAA | 148 |
| chr8:86273142-86275178 | p3 | (TTC)11 | TGAGGGTGGAGGAAATGAAG | ATGTCAACGCCACATCAAAA | 148 |
| chr8:86275456-86277476 | p2 | (CA)10 | TTCGTTTGCGTGCTTATTTG | ATCCACATCCACCCCATCTA | 130 |
| chr8:86287882-86289897 | p2 | (CT)7 | AATGATCCTGCTGCTGCTTT | GGGAGCATAACCCAGTGAAA | 224 |
| chr8:86287882-86289897 | p5 | (AAAAT)3 | AATGATCCTGCTGCTGCTTT | GGGAGCATAACCCAGTGAAA | 224 |
| chr8:86349275-86351293 | p6 | (TACCAT)3 | CTGAATTCCGGTCCTTTGAA | GGGTTCGACCGAATCAGTAA | 146 |
| chr8:86352473-86354485 | p2 | (TC)17 | TATTTTTCCCGTTTGGCAAG | AGAGGCAGCGTGTAGTTGGT | 293 |
| chr8:86397383-86399399 | p2 | (TA)8 | GAGAGTTTAGGCGCATGGAG | ATGGTGTCCTCCACCAAGAG | 282 |
| chr8:86404159-86406173 | p2 | (CA)7 | GGAAAGGATACTGGTGGGGT | TCCCATCTCATCAATCACCA | 219 |
| chr8:86434224-86436236 | p2 | (CA)6 | AGGCTTCAAACAGCCTGAAA | CTTGCCAGGGACATGAGATT | 165 |

| Marker ID | SSR type | SSR motif | FORWARD PRIMER (5'-3') | REVERSE PRIMER (5'-3') | size (bp) |
|---|---|---|---|---|---|
| chr8:86434224-86436236 | p5 | (AGGAG)3 | AGGCTTCAAACAGCCTGAAA | CTTGCCAGGGACATGAGATT | 165 |
| chr8:86464989-86467009 | p5 | (TTTAT)4 | GACATGAGGCTGGTCGTTTT | TTCTAAGCTGCCACCAACCT | 225 |
| chr8:86501132-86503144 | p2 | (TC)6 | AAATTGCAAGGCAAGGTTTG | CAAGTCTGCCCATAAGGCAT | 273 |
| chr8:86527266-86529278 | p2 | (AT)6 | TGTGAGATTTCACTGGGCTG | TTTTATTTTATGCCTGGGCG | 145 |
| chr8:86557496-86559514 | p6 | (AAAAGA)3 | CTTGAAATTCTCAAAGCGCC | TTTCAACGGAGGAGGAATTG | 141 |
| chr8:86578122-86580138 | p2 | (TA)17 | TGGATTGATACGCTCGTGAC | CAACCGGCGCTTCTTAATAG | 107 |
| chr8:86578122-86580138 | p2 | (AT)21 | TGGATTGATACGCTCGTGAC | CAACCGGCGCTTCTTAATAG | 107 |
| chr8:86578122-86580138 | p2 | (AT)8 | TGGATTGATACGCTCGTGAC | CAACCGGCGCTTCTTAATAG | 107 |
| chr8:86578122-86580138 | p2 | (TA)8 | TGGATTGATACGCTCGTGAC | CAACCGGCGCTTCTTAATAG | 107 |
| chr8:86638480-86640510 | p2 | (GA)6 | ATCCACTTTCCTCCCGTTTT | CCTCTGCACCTTTTCTCGTC | 114 |
| chr8:86643151-86645169 | p6 | (AGAAAA)3 | ATCGTCCAAATCCGTTCATC | GGACACAAGTTGGGAGGAAA | 101 |
| chr8:86643151-86645169 | p2 | (CT)8 | ATCGTCCAAATCCGTTCATC | GGACACAAGTTGGGAGGAAA | 101 |
| chr8:86653982-86656006 | p2 | (AT)12 | GGAGCACTCCACTTTCTTCG | TCAAAATAAGGCCGACAAGG | 228 |
| chr8:86653982-86656006 | p2 | (TA)8 | GGAGCACTCCACTTTCTTCG | TCAAAATAAGGCCGACAAGG | 228 |
| chr8:86683678-86685693 | p2 | (AT)6 | GCCGTGTGGCTTCATTTAGT | ACCAATGCAAAGTGGACACA | 112 |
| chr8:86683678-86685693 | p5 | (TGAAT)3 | GCCGTGTGGCTTCATTTAGT | ACCAATGCAAAGTGGACACA | 112 |
| chr8:86683678-86685693 | p6 | (TATATG)4 | GCCGTGTGGCTTCATTTAGT | ACCAATGCAAAGTGGACACA | 112 |
| chr8:86690153-86692177 | p5 | (TGAAT)3 | CCGAAATTACGGCAGATCAT | GCTTAATTGTCCAAGCTCGC | 246 |
| chr8:86690153-86692177 | p6 | (TATATG)4 | CCGAAATTACGGCAGATCAT | GCTTAATTGTCCAAGCTCGC | 246 |
| chr8:86699467-86701482 | p2 | (AT)6 | GCCGTGTGGCTTCATTTAGT | ACCAATGCAAAGTGGACACA | 112 |
| chr8:86699467-86701482 | p5 | (TGAAT)3 | GCCGTGTGGCTTCATTTAGT | ACCAATGCAAAGTGGACACA | 112 |
| chr8:86699467-86701482 | p6 | (TATATG)4 | GCCGTGTGGCTTCATTTAGT | ACCAATGCAAAGTGGACACA | 112 |
| chr8:86705942-86707966 | p5 | (TGAAT)3 | CCGAAATTACGGCAGATCAT | GCTTAATTGTCCAAGCTCGC | 246 |
| chr8:86705942-86707966 | p6 | (TATATG)4 | CCGAAATTACGGCAGATCAT | GCTTAATTGTCCAAGCTCGC | 246 |
| chr8:86739667-86741685 | p5 | (TGCAC)3 | CTTAGGCTATTTCGCATCGC | TATTTGGGGTTGGAGACGAG | 294 |
| chr8:86739667-86741685 | p3 | (TTA)6 | CTTAGGCTATTTCGCATCGC | TATTTGGGGTTGGAGACGAG | 294 |
| chr8:86780782-86782804 | p2 | (TA)11 | CCCTGTTACGGTTCAATGCT | GCCGTGAATTTAGGCACATT | 121 |
| chr8:86996143-86998158 | p5 | (AATGC)3 | AGCTCATCGTACCGATCCAC | AATTGGGATAAGGGTTTGCC | 276 |
| chr8:87142180-87144204 | p2 | (TC)6 | GAGGGACGTGACGCAACTAT | TCCGGACCGTAGTTTTATCG | 187 |
| chr8:87142180-87144204 | p2 | (TA)12 | GAGGGACGTGACGCAACTAT | TCCGGACCGTAGTTTTATCG | 187 |
| chr8:87189786-87191801 | p5 | (ATAAA)3 | TGTATTTCATGCAGGCGAAG | TAGCACCCTCCAAACTGTCC | 263 |
| chr8:87206240-87208252 | p2 | (TA)6 | TGTTGGGCTTCTTCTGTGTG | CAAGATCCAGAGGGTTTCCA | 206 |
| chr8:87210705-87212720 | p3 | (CAA)5 | ACATCACAACAAAGGCCACA | TGACAAGGGGTGATGCTGTA | 176 |
| chr8:87225375-87227399 | p2 | (TC)6 | GAGGGACGTGACGCAACTAT | TCCGGACCGTAGTTTTATCG | 187 |
| chr8:87225375-87227399 | p2 | (TA)12 | GAGGGACGTGACGCAACTAT | TCCGGACCGTAGTTTTATCG | 187 |
| chr8:87272981-87274996 | p5 | (ATAAA)3 | TGTATTTCATGCAGGCGAAG | TAGCACCCTCCAAACTGTCC | 263 |

| Marker ID | SSR type | SSR motif | FORWARD PRIMER (5'-3') | REVERSE PRIMER (5'-3') | size (bp) |
|---|---|---|---|---|---|
| chr8:87289435-87291447 | p2 | (TA)6 | TGTTGGGCTTCTTCTGTGTG | CAAGATCCAGAGGGTTTCCA | 206 |
| chr8:87293900-87295915 | p3 | (CAA)5 | ACATCACAACAAAGGCCACA | TGACAAGGGGTGATGCTGTA | 176 |
| chr8:87323344-87325376 | p2 | (AT)16 | CCAGTATCAAAGGTTGCGGT | GAACAGCTGAAGGGATGGAG | 109 |
| chr8:87332595-87334610 | p5 | (AAAAG)3 | TGCACCACCAAGAGAAAATG | GCTTGCATGTTTGCTTACGA | 119 |
| chr8:87352661-87354676 | p5 | (CGACC)3 | TTCTTCGCTGCCTTGATTTT | GCTGGACCAACAACCAATTT | 117 |
| chr8:87384672-87386704 | p2 | (AT)16 | CCAGTATCAAAGGTTGCGGT | GAACAGCTGAAGGGATGGAG | 109 |
| chr8:87393923-87395938 | p5 | (AAAAG)3 | TGCACCACCAAGAGAAAATG | GCTTGCATGTTTGCTTACGA | 119 |
| chr8:87467025-87469040 | p5 | (GCATT)3 | CCCTTGGTGAGACTTCGTGT | CACACAACAGGGCCTAACCT | 264 |
| chr8:87533736-87535766 | p4 | (TTTA)4 | TTATAAGGCAAGGTCTGCCG | AAATACCCCGAGGTTATGGG | 153 |
| chr8:87533736-87535766 | p2 | (AT)15 | TTATAAGGCAAGGTCTGCCG | AAATACCCCGAGGTTATGGG | 153 |
| chr8:87540184-87542199 | p5 | (CATTT)3 | ACAAGAGGGCAGCTGATGTT | TCCTTTGGCTTTGATGTTCC | 248 |
| chr8:87591433-87593448 | p5 | (CGACC)3 | TTCTTCGCTGCCTTGATTTT | GCTGGACCAACAACCAATTT | 117 |
| chr8:87802301-87804313 | p5 | (TTTCT)3 | ACGCACCTCCCTAAATGTTG | AAGTTCGTAGATGGCGGATG | 225 |
| chr8:87802301-87804313 | p2 | (GA)6 | ACGCACCTCCCTAAATGTTG | AAGTTCGTAGATGGCGGATG | 225 |
| chr8:87813554-87815596 | p2 | (AT)21 | CAGTTTGGTGGGAGGTCATT | GGTACCGCATCACCTTGTTT | 222 |
| chr8:87938181-87940201 | p2 | (TA)10 | GTTCCGAGAACTGGCAAGAA | AAATGCCCATTGTGGGTAAA | 272 |
| chr8:87941054-87943068 | p2 | (TA)7 | CTTTGCCTTTTGGCATGTTT | ACTCAATCCTCGGAGCAGAA | 275 |
| chr8:87970549-87972561 | p2 | (TA)6 | GAATTGGCATAGCCCCTACA | GTATTTTGCGGGAAAACGAA | 185 |
| chr8:88233018-88235032 | p2 | (AT)9 | CCTTCACCATTTTCACCCAT | GGAACCCCCTATTTTGCATT | 194 |
| chr8:88265283-88267298 | p5 | (TTTGG)3 | TAGGAAGGTGAAGGCACCAC | TTGGAGATGCCACTTAACCC | 169 |
| chr8:88307257-88309272 | p5 | (AATAA)3 | CAAATCCCTCAACCAATGCT | AGCCTTAAGCTTCGAGACCC | 220 |
| chr8:88341726-88343744 | p6 | (GGTCAA)3 | AAACGTCAGACCCAAATTGC | TTTCTAGCTTCGGGCAAAAA | 260 |
| chr8:88341726-88343744 | p2 | (AG)9 | AAACGTCAGACCCAAATTGC | TTTCTAGCTTCGGGCAAAAA | 260 |
| chr8:88364686-88366756 | p2 | (AT)35 | TGGTTCTATGGTGTTGCGAA | TTGGCTTCGTTGATGGTACA | 283 |
| chr8:88560921-88562985 | p2 | (TA)32 | AGGGAGAGGACGAAGGAGAC | CGTTGTACCACCGGATCTTC | 130 |
| chr8:88560921-88562985 | p3 | (AAG)7 | AGGGAGAGGACGAAGGAGAC | CGTTGTACCACCGGATCTTC | 130 |
| chr8:88575115-88577130 | p5 | (ACCAA)3 | GCTGCCACTTGAATTTGTGA | TGCTGCTGTGGTTTGCTTAC | 119 |
| chr8:88784777-88786791 | p2 | (TA)7 | TCAACAGTTTGCAAAGCCAG | GAGACCTTCTCACCCCAACA | 217 |
| chr8:88784777-88786791 | p2 | (TA)7 | TCAACAGTTTGCAAAGCCAG | GAGACCTTCTCACCCCAACA | 217 |
| chr8:88993247-88995262 | p2 | (TA)8 | TTTAAGGGACAACGGACGAC | GGGCCCCATACTAAAACACA | 232 |
| chr8:88993247-88995262 | p2 | (AT)8 | TTTAAGGGACAACGGACGAC | GGGCCCCATACTAAAACACA | 232 |
| chr8:88993247-88995262 | p5 | (TTTTA)3 | TTTAAGGGACAACGGACGAC | GGGCCCCATACTAAAACACA | 232 |
| chr8:88993247-88995262 | p2 | (AT)8 | TTTAAGGGACAACGGACGAC | GGGCCCCATACTAAAACACA | 232 |
| chr8:89059583-89061597 | p2 | (TA)7 | TGTGAGTGATGGGGATGCTA | TGCCCTTACAACCCCAAATA | 253 |
| chr8:89127208-89129223 | p3 | (CTT)5 | CAGTGAGAGCTTCCTTTGGC | AGGTTCTTTTGGTCTGGCCT | 235 |
| chr8:89268202-89270216 | p2 | (CT)7 | CTTCTGATGTTGGCGTCTCA | GCAACAAGACCAATTCAGCA | 155 |

| Marker ID | SSR type | SSR motif | FORWARD PRIMER (5'-3') | REVERSE PRIMER (5'-3') | size (bp) |
|---|---|---|---|---|---|
| chr8:89369907-89371923 | p2 | (AT)8 | CGTAAGCCCATCCAAACAGT | GGGGAGAGAGATGGGAATGT | 142 |
| chr8:89429849-89431861 | p2 | (TA)6 | TGTGGGTAGTGTTCACGCAT | ATCCATTTTTGCCAGTGAGG | 120 |
| chr8:89492242-89494260 | p2 | (TA)9 | CAGTGTGGTGTTCCGAGCTA | CATTGCAAGTTGTTGCTGCT | 153 |
| chr8:89492242-89494260 | p2 | (AT)7 | CAGTGTGGTGTTCCGAGCTA | CATTGCAAGTTGTTGCTGCT | 153 |
| chr8:89492242-89494260 | p2 | (TA)9 | CAGTGTGGTGTTCCGAGCTA | CATTGCAAGTTGTTGCTGCT | 153 |
| chr8:89664168-89666226 | p2 | (TG)10 | CCCACTTTCCTTTCCCCTAC | ATGTTTGCCACCATGTACGA | 139 |
| chr8:89665156-89667176 | p2 | (TG)10 | AGTTGCTTTTCTTGAGCCGA | GTCTCACAAGTGGGGTCTGG | 292 |
| chr8:89710470-89712491 | p3 | (TTG)7 | CCAGGAAGGACAACCTTGAA | CCTCCAAAGTTGACCCAAAA | 207 |
| chr8:89727095-89729110 | p5 | (AAATA)3 | TGTTATGTGGGGAATGGGTT | AAAACCGGTGCAAAAATCTG | 111 |
| chr8:89840390-89842406 | p3 | (CAC)5 | CATTATCACCACCACCACCA | GCCTTCTTTTTCGATTTCCC | 294 |
| chr8:89840390-89842406 | p3 | (GAA)9 | CATTATCACCACCACCACCA | GCCTTCTTTTTCGATTTCCC | 294 |
| chr8:89840480-89842495 | p3 | (CAC)5 | CATTATCACCACCACCACCA | GCCTTCTTTTTCGATTTCCC | 204 |
| chr8:89840480-89842495 | p3 | (GAA)9 | CATTATCACCACCACCACCA | GCCTTCTTTTTCGATTTCCC | 204 |
| chr8:89840498-89842525 | p3 | (CAC)5 | CATTATCACCACCACCACCA | GCCTTCTTTTTCGATTTCCC | 186 |
| chr8:89840498-89842525 | p3 | (GAA)9 | CATTATCACCACCACCACCA | GCCTTCTTTTTCGATTTCCC | 186 |
| chr8:89843087-89845102 | p3 | (TTG)5 | ATGTTTTACGGGGCAGAGTG | TTTCCGTCATCTTGTCCTCC | 268 |
| chr8:89843087-89845102 | p6 | (AACAAA)3 | ATGTTTTACGGGGCAGAGTG | TTTCCGTCATCTTGTCCTCC | 268 |
| chr8:89918877-89920923 | p2 | (AT)23 | GAATTGAGACCCGAACCTGA | TCGTTCAACTTCTAACCCCG | 275 |
| chr8:89918877-89920923 | p2 | (TA)16 | GAATTGAGACCCGAACCTGA | TCGTTCAACTTCTAACCCCG | 275 |
| chr8:89918877-89920923 | p2 | (AT)9 | GAATTGAGACCCGAACCTGA | TCGTTCAACTTCTAACCCCG | 275 |
| chr8:89932086-89934110 | p2 | (TA)12 | TTGTGCGACACTGGGAATAA | GTAAGAGAGGAAAACGGGGG | 277 |
| chr8:89987257-89989303 | p2 | (AT)23 | GAATTGAGACCCGAACCTGA | TCGTTCAACTTCTAACCCCG | 275 |
| chr8:89987257-89989303 | p2 | (TA)16 | GAATTGAGACCCGAACCTGA | TCGTTCAACTTCTAACCCCG | 275 |
| chr8:89987257-89989303 | p2 | (AT)9 | GAATTGAGACCCGAACCTGA | TCGTTCAACTTCTAACCCCG | 275 |
| chr8:90000466-90002490 | p2 | (TA)12 | TTGTGCGACACTGGGAATAA | GTAAGAGAGGAAAACGGGGG | 277 |
| chr8:90242298-90244313 | p3 | (TGT)5 | TTCAAGATAATTGCGGGAGG | TACTTAGGCCCAACCCCTTT | 122 |
| chr8:90264714-90266726 | p2 | (AG)6 | CTTGCCTTCTCTTGGCAAAC | CAGGGCGAGACGAATATGAT | 136 |
| chr8:90296430-90298445 | p3 | (TGT)5 | TTCAAGATAATTGCGGGAGG | TACTTAGGCCCAACCCCTTT | 122 |
| chr8:90367411-90369429 | p3 | (GAA)6 | CTAAGTGGCAGGGACACCAT | GAGGTGTGCGAAAGTGTGAA | 128 |
| chr8:90421019-90423059 | p2 | (GA)20 | AGTTGGGGCATTTGTGAAAG | GAAGTCCGCAGTTCAAGTCC | 192 |
| chr8:90454776-90456791 | p5 | (TTAAT)3 | TGCTTCGTCGTCAGTCACTC | TGATGGGGATACTTTGGGAA | 244 |
| chr8:90454776-90456791 | p5 | (CCGGC)3 | TGCTTCGTCGTCAGTCACTC | TGATGGGGATACTTTGGGAA | 244 |
| chr8:90478151-90480166 | p5 | (TTCTT)3 | CAACCAAGCAAGGCTAAAGG | AGCACTATCTGTGGCGTGTG | 244 |
| chr8:90558876-90560894 | p6 | (AAAAAG)3 | GAAGTTAATTGATCCGCCCA | AATCAAGAAGTCGAAGCCGA | 221 |
| chr8:90586216-90588231 | p3 | (ACA)5 | ATGCTTTGAGTTGGGAGGTG | GAAGAAGCACACCTTCCTCG | 133 |
| chr8:90647748-90649816 | p2 | (AT)34 | GGAGCGAGTGGTACATGGAT | CAGCCACAATCATCAACAGG | 250 |

| Marker ID | SSR type | SSR motif | FORWARD PRIMER (5'-3') | REVERSE PRIMER (5'-3') | size (bp) |
|---|---|---|---|---|---|
| chr8:90647748-90649816 | p2 | (TA)12 | GGAGCGAGTGGTACATGGAT | CAGCCACAATCATCAACAGG | 250 |
| chr8:90647748-90649816 | p4 | (CGTT)4 | GGAGCGAGTGGTACATGGAT | CAGCCACAATCATCAACAGG | 250 |
| chr8:90681117-90683131 | p2 | (TA)7 | CGGTTTTTATCGTGCGTTCT | TCTCCGTACAAACCCGAAAC | 248 |
| chr8:90694860-90696878 | p6 | (AGAAGG)3 | GGCCCAAATGCGTTATATTG | GTGGGGTTGTTCTGCATCTT | 123 |
| chr8:90772273-90774291 | p6 | (AACTGA)3 | TTTGCACACCAAGATTAGCG | GAACCCCAGCAAACACTGAT | 179 |
| chr8:90915608-90917620 | p2 | (CT)6 | CTGATCCCAGCCAAAGAGAG | ACTTTCCGGACCGAAGTTTT | 288 |
| chr8:90915608-90917620 | p5 | (CCCTT)3 | CTGATCCCAGCCAAAGAGAG | ACTTTCCGGACCGAAGTTTT | 288 |
| chr8:90986864-90988876 | p2 | (TG)6 | ATGCCTTTGATCCGAACATC | CTAAGGCAAAGGCGTGTCTC | 108 |
| chr8:90996574-90998586 | p2 | (AT)6 | TTCGATGTGGGTGAAACAAA | CCAGTCACCAACCTGGAACT | 199 |
| chr8:90996574-90998586 | p2 | (AG)10 | TTCGATGTGGGTGAAACAAA | CCAGTCACCAACCTGGAACT | 199 |
| chr8:91055437-91057470 | p3 | (TTA)11 | CGATGCCGGAGAATACAGTT | TTGGCCATGCTCACAATAAA | 237 |
| chr8:91096965-91098993 | p2 | (TA)14 | GGGTTGACCTGCATGAATTT | TCCTGAGAGTGCCATGTGAG | 101 |
| chr8:91104240-91106290 | p2 | (TA)25 | ACCAGCCTCAAAACAGTGCT | GCGGCACGGTGTAGTAAAAT | 112 |
| chr8:91175596-91177610 | p2 | (CA)7 | AAGTCATTGGCGTCCTCATC | ACCAACGATAGTGCAGGGAC | 145 |
| chr8:91317133-91319191 | p2 | (TA)29 | GGCCGATGTTCGATGTTAGT | ATGTGACCCTGGATCTTTGC | 225 |
| chr8:91346813-91348829 | p4 | (TTAT)4 | GGCCAGCTTTGTGACAATTT | AGCAAAACCTATGAAGGGCA | 280 |
| chr8:91372389-91374401 | p2 | (TA)6 | GCTTCTGCGGCTTGTTTTAC | CACAAGGGGGCTTAGATGAA | 135 |
| chr8:91372389-91374401 | p2 | (AT)20 | GCTTCTGCGGCTTGTTTTAC | CACAAGGGGGCTTAGATGAA | 135 |
| chr8:91407430-91409458 | p2 | (TA)7 | TCCACTGCATTATTCACCGA | GCCAACATCCCCATATCATC | 235 |
| chr8:91407430-91409458 | p2 | (AT)14 | TCCACTGCATTATTCACCGA | GCCAACATCCCCATATCATC | 235 |
| chr8:91694853-91696867 | p2 | (AG)7 | GGTCCTATTGACCAGCCTGA | CGCTTTTGTGAGTGGTTTCA | 134 |
| chr8:91753881-91755903 | p2 | (TA)11 | CCTCACTCAGGTCTTGGCTC | CACCTTGTGAGGAGGTTGGT | 146 |
| chr8:91779565-91781583 | p6 | (GTGGCT)3 | CTTCAGCCTTGTCTCGTTCC | TGAGGAGATTTGTTGGGGTC | 282 |
| chr8:91779565-91781583 | p4 | (CAAA)4 | CTTCAGCCTTGTCTCGTTCC | TGAGGAGATTTGTTGGGGTC | 282 |
| chr8:91820283-91822298 | p3 | (ACA)5 | ATCATCCAGGAGGCAATCAG | TCACCTTTTGAGCCTTGCTT | 197 |
| chr8:91827551-91829583 | p2 | (AT)16 | ATTGGAGCCACATTTCTTGC | TCCTGTCTCCTCCAGCAGTT | 137 |
| chr8:91827551-91829583 | p4 | (GTTT)4 | ATTGGAGCCACATTTCTTGC | TCCTGTCTCCTCCAGCAGTT | 137 |
| chr8:91848152-91850200 | p2 | (TA)24 | GGTCAACAAACACCATGTGC | TCAAGCTGTTGAGCCATACG | 282 |
| chr8:91947939-91949954 | p5 | (AAACC)4 | ATCAGTTTCGGTTTTGTCGG | GGGGCTGCGTAATATCAAAA | 278 |
| chr8:92036486-92038504 | p3 | (TTA)6 | CTTAGGAGTGTGGAGGCTGC | GGATACTGTCCCCTCGGAAT | 294 |
| chr8:92036486-92038504 | p3 | (TTA)5 | CTTAGGAGTGTGGAGGCTGC | GGATACTGTCCCCTCGGAAT | 294 |
| chr8:92036855-92038870 | p3 | (TTA)6 | GAGGGGACAGTATCCGAACA | AAAAGGACCGATAATTCGGG | 111 |
| chr8:92036855-92038870 | p3 | (TTA)5 | GAGGGGACAGTATCCGAACA | AAAAGGACCGATAATTCGGG | 111 |
| chr8:92057190-92059206 | p2 | (AG)8 | TCAGTCGAGTCTACCCGCTT | AATGTTGCAGGAACAAAGGG | 176 |
| chr8:92071513-92073543 | p2 | (AT)15 | TCAAGAGTCGGGCTTCTGTT | TCATCATCCAATGCACGAAT | 278 |
| chr8:92152485-92154537 | p2 | (AC)6 | CCTTGGCCTTACCATTCAAA | TGGCCAGAGAGGTTCAGACT | 295 |

| Marker ID | SSR type | SSR motif | FORWARD PRIMER (5'-3') | REVERSE PRIMER (5'-3') | size (bp) |
|---|---|---|---|---|---|
| chr8:92157774-92159789 | p5 | (AAACC)3 | ATTTTCCCCATCCCCAAATA | ATTTTTGGGGGAAGCTATGG | 202 |
| chr8:92314062-92316137 | p3 | (AAT)25 | GCTACAAAAGTCGCGGATTC | GACTCCTCACGGAAAAGCAC | 150 |
| chr8:92317356-92319374 | p2 | (TA)9 | TGATGTGGCAAGCGAATTTA | GAATTTCCCTGGATACGGGT | 105 |
| chr8:92329181-92331195 | p2 | (TC)7 | TGTTGACTATCATTCGGGCA | AAAAATTCGCATCCAAGTGC | 263 |
| chr8:92404943-92406958 | p3 | (TCT)5 | ACCTTTGAATGTCACCCTCG | AGGCGTAGCGACTTTGGTTA | 129 |
| chr8:92431904-92433918 | p2 | (TA)10 | AGAGCAGTGCGTTTTCACCT | TTCTACCGATGAACCCCTTG | 268 |
| chr8:92431904-92433918 | p2 | (TA)7 | AGAGCAGTGCGTTTTCACCT | TTCTACCGATGAACCCCTTG | 268 |
| chr8:92431904-92433918 | p5 | (AAACC)4 | AGAGCAGTGCGTTTTCACCT | TTCTACCGATGAACCCCTTG | 268 |
| chr8:92431904-92433918 | p2 | (TA)9 | AGAGCAGTGCGTTTTCACCT | TTCTACCGATGAACCCCTTG | 268 |
| chr8:92432033-92434053 | p2 | (TA)10 | AGAGCAGTGCGTTTTCACCT | TTCTACCGATGAACCCCTTG | 139 |
| chr8:92432033-92434053 | p2 | (TA)7 | AGAGCAGTGCGTTTTCACCT | TTCTACCGATGAACCCCTTG | 139 |
| chr8:92432033-92434053 | p5 | (AAACC)4 | AGAGCAGTGCGTTTTCACCT | TTCTACCGATGAACCCCTTG | 139 |
| chr8:92432033-92434053 | p2 | (TA)9 | AGAGCAGTGCGTTTTCACCT | TTCTACCGATGAACCCCTTG | 139 |
| chr8:92480732-92482802 | p2 | (AT)35 | TCCCCTCAAAGTCAGGTCAC | GCCCACACGTCAAGATTTTT | 213 |
| chr8:92519322-92521352 | p3 | (GTT)8 | ATGTTCGAACCCCTAATCCC | TGTGGAGCCAGAATTTGACA | 183 |
| chr8:92519322-92521352 | p2 | (TA)15 | ATGTTCGAACCCCTAATCCC | TGTGGAGCCAGAATTTGACA | 183 |
| chr8:92716353-92718367 | p2 | (TC)7 | CGCTTTGGTATCCGTCAAAT | TGAGGCCCAACTTCTGCTAT | 253 |
| chr8:92779397-92781415 | p2 | (AG)9 | AAGTGCATTGGGACCAGAAC | GCTCACTTTTGACTCGAGGG | 293 |
| chr8:92828472-92830505 | p3 | (AAT)11 | GACAAAAGGCCCATGACACT | TGTTTTGGGTCCATTCCATT | 162 |
| chr8:92995139-92997154 | p5 | (TTTTC)3 | ATGGATGTGGGGGATTGTTA | GGAGTCCCTCTTCTCAACCC | 232 |
| chr8:93001327-93003342 | p3 | (TGT)5 | GGTCTCTCCGCTTGCATAAG | AGCGCGAGTGTGGTAGTTCT | 222 |
| chr8:93001327-93003342 | p5 | (TAAAA)3 | GGTCTCTCCGCTTGCATAAG | AGCGCGAGTGTGGTAGTTCT | 222 |
| chr8:93007566-93009608 | p2 | (AG)21 | ACGGTGGTGTTGAAGGGTTA | AGACGCTTCAACTAAGGGCA | 218 |
| chr8:93061487-93063502 | p4 | (TGTA)5 | AAGTCTCACCCCCTCGTCTT | ATGGCCAAACACAAACACAA | 161 |
| chr8:93061487-93063502 | p5 | (TTTTC)3 | AAGTCTCACCCCCTCGTCTT | ATGGCCAAACACAAACACAA | 161 |
| chr8:93061487-93063502 | p5 | (GCAAG)3 | AAGTCTCACCCCCTCGTCTT | ATGGCCAAACACAAACACAA | 161 |
| chr8:93087274-93089294 | p4 | (TATT)5 | TGGTTGTGACGAGCAGTAGG | CCCAAAAAGGGATGTAGCAA | 243 |
| chr8:93360553-93362568 | p5 | (AAAAC)3 | ACAATTACCTCCGGCTCCTT | GTGGCCAACTGAGGTTGAAT | 259 |
| chr8:93487856-93489882 | p5 | (AAAAG)3 | GAAAAGGGCAAATGACGAAA | TGCCAAATTCCCATATCTCC | 265 |
| chr8:93487856-93489882 | p2 | (AT)13 | GAAAAGGGCAAATGACGAAA | TGCCAAATTCCCATATCTCC | 265 |
| chr8:93500342-93502366 | p3 | (AAT)20 | ATTTCCTCCCCCAAAATGAC | TTTTAAATCGACCAGCCCAC | 220 |
| chr8:93500375-93502444 | p3 | (AAT)20 | ATTTCCTCCCCCAAAATGAC | TTTTAAATCGACCAGCCCAC | 187 |
| chr8:93681847-93683861 | p2 | (GT)7 | ACCATCACAAATCTCACGCA | CCTTCCTCAATATTTCCGCA | 247 |
| chr8:93701876-93703904 | p2 | (AT)14 | GACTAAGCGCTCTCCTTCCA | TGCCCAATTTTCGTTAGTCC | 263 |
| chr8:93848821-93850836 | p3 | (TTC)5 | CTTTGAAAACCCCAGGCATA | GGCAGACAAGAAAGCAAAGG | 218 |
| chr8:93868117-93870133 | p2 | (AT)8 | TCATGCCTTTGAAAGAAGGG | TGCCCAATTTTCGTTAGTCC | 210 |

| Marker ID | SSR type | SSR motif | FORWARD PRIMER (5'-3') | REVERSE PRIMER (5'-3') | size (bp) |
|---|---|---|---|---|---|
| chr8:93914296-93916314 | p6 | (TTCAGT)3 | TTGGGAGAGTCGTAAATGGG | TCAAACACTTGGTGGGCATA | 138 |
| chr8:94000676-94002691 | p2 | (AT)16 | CACATCTACGTGTCATGCCC | ATAATCGGAGGCTACCGACC | 118 |
| chr8:94000676-94002691 | p3 | (AAC)5 | CACATCTACGTGTCATGCCC | ATAATCGGAGGCTACCGACC | 118 |
| chr8:94017267-94019282 | p2 | (AT)16 | CACATCTACGTGTCATGCCC | ATAATCGGAGGCTACCGACC | 118 |
| chr8:94017267-94019282 | p3 | (AAC)5 | CACATCTACGTGTCATGCCC | ATAATCGGAGGCTACCGACC | 118 |
| chr8:94128257-94130275 | p3 | (ATA)6 | TGTGGAGCCAGAATTTGACA | ATGTTCGAACCCCTAATCCC | 202 |
| chr8:94128257-94130275 | p4 | (CCCA)4 | TGTGGAGCCAGAATTTGACA | ATGTTCGAACCCCTAATCCC | 202 |
| chr8:94152859-94154874 | p5 | (AAACC)3 | ACATTTTAGGTCCTTCGGGG | TGCAATCAACCAATTCAGGA | 144 |
| chr8:94230807-94232873 | p2 | (AT)33 | GCCGATAAGCCTACAAGACG | GAAGAGCTCCACTTCATCCG | 134 |
| chr8:94236346-94238361 | p3 | (AAC)5 | CACATTATCTTACGCCCCGT | AATGAACAGCCAGTGCAGAA | 259 |
| chr8:94236346-94238361 | p2 | (TA)8 | CACATTATCTTACGCCCCGT | AATGAACAGCCAGTGCAGAA | 259 |
| chr8:94250440-94252455 | p5 | (GGGGT)3 | CAGGTTATAGGGGAGGCACA | CCACTTCCATGTCGGAATCT | 155 |
| chr8:94258159-94260175 | p2 | (AT)8 | GCTACTGGCTTCCCCACATA | AAAATGAGAGCGAGGGGATT | 239 |
| chr8:94273354-94275366 | p2 | (GT)6 | GACTCTGGTCTCGAGTTGCC | GACGATCGAGTTTGCACGTA | 114 |
| chr8:94414966-94416978 | p2 | (AT)6 | CTTCACACACGAGCTCCAGA | CAAGGAAAAATCCAGTGGGA | 272 |
| chr8:94433051-94435091 | p2 | (TA)20 | TCTTGTAGCGCCAAGGCTAT | AATTCCCACAAAAAGGAGGG | 293 |
| chr8:94465779-94467794 | p5 | (AAACC)3 | CTAGGGGTGTACATGGACCG | ACCCGACTTTTGTTGGTTTG | 151 |
| chr8:94500413-94502425 | p5 | (ACCGG)3 | GAAAACCGTCCTTGATGGAA | CAAAACGCAGCGGAAATAAT | 148 |
| chr8:94500413-94502425 | p2 | (AT)6 | GAAAACCGTCCTTGATGGAA | CAAAACGCAGCGGAAATAAT | 148 |
| chr8:94505731-94507779 | p2 | (AT)7 | GCCGACTAATTTCCAACGAA | ACCCCTCTCCATTTCCATTT | 270 |
| chr8:94552566-94554581 | p5 | (AGAGA)3 | GGTTTGTTGCGTTGGAAACT | GAGGATCGGAAAACCAGTGA | 183 |
| chr8:94604461-94606476 | p5 | (TTGGT)3 | CCCAAGCTTTTGAACGACAT | AACCCACATCCAGGCAAATA | 173 |
| chr8:94680519-94682537 | p3 | (ACA)6 | TGCTCCACCAACCTCCTAAC | AGCCATCGGATGAAAGTTTG | 204 |
| chr8:94683336-94685348 | p2 | (TG)6 | TGGCCAGGTAAGAATGGAAG | TCTTTTTGCGTGGATTAGGG | 101 |
| chr8:95041979-95043997 | p6 | (TTTTTA)3 | TAGCCAAGCCAAGCTCATCT | AAAACCGAAAAAGGCCAAAT | 210 |
| chr8:95051520-95053534 | p2 | (TA)7 | ACCAGCCCATTTGAAAGACA | TGCTCTTGGTTGCCCTTAGT | 247 |
| chr8:95171388-95173400 | p2 | (TA)18 | ATGGAAATACCCATGCCAAA | AAAGGGAAGGGAATGACACA | 209 |
| chr8:95182867-95184893 | p2 | (TA)13 | TATGCACAAGTGGCATCACA | AACCGACTTGATGGAACGAC | 149 |
| chr8:95233384-95235400 | p2 | (TA)8 | AAAAATCGAGATGCTGGCCT | TCACATCGAGGTTTGACTGC | 197 |
| chr8:95233384-95235400 | p2 | (TA)7 | AAAAATCGAGATGCTGGCCT | TCACATCGAGGTTTGACTGC | 197 |
| chr8:95370704-95372719 | p3 | (CAT)5 | GTTTTTCCCTTGGGTGGAAT | AAGTCGGGTCGTGACAAAAG | 158 |
| chr8:95397934-95399946 | p2 | (CA)6 | TACAACATCCAAGGCGATCA | GAGTTTTGCCGAGAGCTTTG | 143 |
| chr8:95538994-95541044 | p2 | (TA)25 | TTTTCCATTGTCGTTGACCA | ATGCATTTCTCTACCCCGTG | 176 |
| chr8:95653406-95655420 | p5 | (ACCGG)3 | ATCCAGATGTTTGGTGCCAT | GTCGGAGAATGAAGGGTCAA | 216 |
| chr8:95653406-95655420 | p2 | (TA)7 | ATCCAGATGTTTGGTGCCAT | GTCGGAGAATGAAGGGTCAA | 216 |
| chr8:95661212-95663227 | p5 | (TTGGT)3 | ACCCCGATTAGTGGTTTGGT | ATGTCGGGTTGGTTTGATTC | 245 |

| Marker ID | SSR type | SSR motif | FORWARD PRIMER (5'-3') | REVERSE PRIMER (5'-3') | size (bp) |
|---|---|---|---|---|---|
| chr8:95661212-95663227 | p2 | (AG)6 | ACCCCGATTAGTGGTTTGGT | ATGTCGGGTTGGTTTGATTC | 245 |
| chr8:95661252-95663267 | p5 | (TTGGT)3 | ACCCCGATTAGTGGTTTGGT | ATGTCGGGTTGGTTTGATTC | 205 |
| chr8:95661252-95663267 | p2 | (AG)6 | ACCCCGATTAGTGGTTTGGT | ATGTCGGGTTGGTTTGATTC | 205 |
| chr8:95710676-95712691 | p3 | (ATT)5 | TCCTATGGGCTGGCTGTTAG | TTCCGATTTTTGTATTGGGC | 184 |
| chr8:95710676-95712691 | p2 | (AT)6 | TCCTATGGGCTGGCTGTTAG | TTCCGATTTTTGTATTGGGC | 184 |
| chr8:95721012-95723024 | p2 | (AT)6 | CTCACCACATCCAACTCACG | TGTTGCTACGAATACGGACG | 259 |
| chr8:95836645-95838657 | p2 | (TG)6 | TGATGTGTTACGGTTTCCGA | ACACCAACATCGCAAACAAA | 265 |
| chr8:95907703-95909717 | p2 | (TG)7 | CTTGGATGGGCATTGAAGAT | AAGGGTTGATGTCTCGTTCG | 183 |
| chr8:95958929-95960954 | p5 | (AAATT)5 | CTTGGGAGGGTGTTTCTTCA | TACCGCTGCTATTTGCCTCT | 273 |
| chr8:96037551-96039563 | p2 | (TG)6 | ATGCAAAATGGGAAGTCCTG | TTGTCGGAGGATCACTTTCC | 262 |
| chr8:96043004-96045019 | p2 | (TA)10 | TCAAGGCGAAGGAGTGAGTT | ACCCGAAGGTTCAGATGTTG | 148 |
| chr8:96043027-96045042 | p2 | (TA)10 | TCAAGGCGAAGGAGTGAGTT | ACCCGAAGGTTCAGATGTTG | 125 |
| chr8:96044966-96046981 | p2 | (TA)10 | TCAAGGCGAAGGAGTGAGTT | ACCCGAAGGTTCAGATGTTG | 148 |
| chr8:96044989-96047004 | p2 | (TA)10 | TCAAGGCGAAGGAGTGAGTT | ACCCGAAGGTTCAGATGTTG | 125 |
| chr8:96055032-96057046 | p2 | (AT)6 | ACTAGAGGGCCGAGTACGGT | ACATATTCATATCCGGCCCA | 196 |
| chr8:96055032-96057046 | p2 | (AT)7 | ACTAGAGGGCCGAGTACGGT | ACATATTCATATCCGGCCCA | 196 |
| chr8:96079444-96081456 | p2 | (AT)6 | CCATCTCCTTGGTGGGATAA | GTGAAGAATTGACGGCGTTT | 123 |
| chr8:96100573-96102588 | p5 | (TTATT)3 | TAGCCGATGGCCATAGAAAC | TGTTTGGCCATGAAATTGAA | 164 |
| chr8:96152382-96154394 | p2 | (AT)6 | TAAAGAGCCCGTTTGGATTG | CCTGTTTGGATGGGCTTATG | 233 |
| chr8:96176952-96178964 | p2 | (AG)6 | CGAGAAGAATCCTGGAGCAC | TGGGGTGGTAGAGAGTGAGG | 185 |
| chr8:96176952-96178964 | p2 | (AG)7 | CGAGAAGAATCCTGGAGCAC | TGGGGTGGTAGAGAGTGAGG | 185 |
| chr8:96177369-96179383 | p2 | (AG)6 | TTCCGAGAAAGAGTCGGAGA | GGGCTACGGTTGAGTTGTGT | 264 |
| chr8:96177369-96179383 | p2 | (AG)7 | TTCCGAGAAAGAGTCGGAGA | GGGCTACGGTTGAGTTGTGT | 264 |
| chr8:96230288-96232303 | p3 | (AAG)5 | TCCGGATTGCCTCAAACTAC | TCGGGTTTTGAACACATTGA | 292 |
| chr8:96234981-96236996 | p3 | (AAG)5 | TCCGGATTGCCTCAAACTAC | TCGGGTTTTGAACACATTGA | 292 |
| chr8:96269602-96271632 | p5 | (ATACA)6 | CAACAGGCAACCAATGTGAC | ACAGGGGAGCCTTTGAAGAT | 227 |
| chr8:96325224-96327238 | p2 | (GA)7 | TGATTTTGGCACTGTTTGGA | TCCCAACAACATGTTAGCCA | 269 |
| chr8:96427672-96429688 | p2 | (AT)6 | GCCAAATTCCATCAACAACC | GCTTGAAAAATCCGTAAGCG | 168 |
| chr8:96427672-96429688 | p2 | (CT)8 | GCCAAATTCCATCAACAACC | GCTTGAAAAATCCGTAAGCG | 168 |
| chr8:96458594-96460610 | p2 | (AC)6 | TGGCCATGAAAAATCCAAAT | TTTGGCCATGAAAACCAAAT | 226 |
| chr8:96458594-96460610 | p2 | (TA)7 | TGGCCATGAAAAATCCAAAT | TTTGGCCATGAAAACCAAAT | 226 |
| chr8:96458698-96460716 | p2 | (AC)6 | TGGCCATGAAAAATCCAAAT | TTTGGCCATGAAAACCAAAT | 122 |
| chr8:96458698-96460716 | p2 | (TA)7 | TGGCCATGAAAAATCCAAAT | TTTGGCCATGAAAACCAAAT | 122 |
| chr8:96500776-96502791 | p2 | (TA)6 | GGTTTGGTGTCACCTCCAAT | ATGCGACAATGAATTCACCA | 135 |
| chr8:96500776-96502791 | p5 | (TTTCT)3 | GGTTTGGTGTCACCTCCAAT | ATGCGACAATGAATTCACCA | 135 |
| chr8:96576437-96578449 | p2 | (AT)6 | ATGGTGTATAGCTGACCGCC | CCTGTCATGACCCTCTGGTT | 100 |

| Marker ID | SSR type | SSR motif | FORWARD PRIMER (5'-3') | REVERSE PRIMER (5'-3') | size (bp) |
|---|---|---|---|---|---|
| chr8:96576437-96578449 | p2 | (AT)8 | ATGGTGTATAGCTGACCGCC | CCTGTCATGACCCTCTGGTT | 100 |
| chr8:96576437-96578449 | p3 | (AAC)5 | ATGGTGTATAGCTGACCGCC | CCTGTCATGACCCTCTGGTT | 100 |
| chr8:96578024-96580038 | p3 | (AAC)5 | GAATTCAACTCGGACCCAAA | GCCGTCGAATTATGAGGAAA | 189 |
| chr8:96578024-96580038 | p2 | (TA)7 | GAATTCAACTCGGACCCAAA | GCCGTCGAATTATGAGGAAA | 189 |
| chr8:96584343-96586377 | p2 | (AT)8 | CTACCGAGCACCCCTTATCA | GTTTTGGCAATTTCCTGCAT | 275 |
| chr8:96584343-96586377 | p2 | (AT)17 | CTACCGAGCACCCCTTATCA | GTTTTGGCAATTTCCTGCAT | 275 |
| chr8:96595686-96597698 | p2 | (AT)6 | AACAACATTAGACCGCGTCC | CCTTACGTCACCCCGATAGA | 295 |
| chr8:96623643-96625663 | p2 | (AT)10 | GCCCTTGGACTTCTTCAACA | CTCCCGAAGTGCTACGAGTC | 113 |
| chr8:96705553-96707565 | p2 | (AC)6 | TGAACCGCAACTCACCATAA | GGACCAGATCCCAAGACTGA | 269 |
| chr8:96705553-96707565 | p4 | (TTTA)4 | TGAACCGCAACTCACCATAA | GGACCAGATCCCAAGACTGA | 269 |
| chr8:96724508-96726523 | p5 | (AAATA)3 | AAACGCAACGAAAAGGAAGA | TCGCATGTGATTGGTTTGTT | 201 |
| chr8:96724670-96726754 | p5 | (AAATA)3 | GGGAAAAATCCCTCCAATGT | CGCCTCTCTTCTCCCTCTCT | 275 |
| chr8:96725011-96727035 | p5 | (AAATA)3 | AGAGAGGGAGAAGAGAGGCG | TATCCCAAATCCGTTTGAGC | 180 |
| chr8:96912574-96914589 | p5 | (AAATA)3 | CACCGTGTCCCTTTGAGATT | TAGTTACGGAAGACCCGTGC | 240 |
| chr8:96912574-96914589 | p5 | (AAATA)3 | CACCGTGTCCCTTTGAGATT | TAGTTACGGAAGACCCGTGC | 240 |
| chr8:96912574-96914589 | p5 | (AATAA)4 | CACCGTGTCCCTTTGAGATT | TAGTTACGGAAGACCCGTGC | 240 |
| chr8:96912894-96914909 | p5 | (AAATA)3 | GGTGTCGATTGCAGTCATTG | TTTGGGGTGTGTGGATTTTT | 270 |
| chr8:96912894-96914909 | p5 | (AAATA)3 | GGTGTCGATTGCAGTCATTG | TTTGGGGTGTGTGGATTTTT | 270 |
| chr8:96912894-96914909 | p5 | (AATAA)4 | GGTGTCGATTGCAGTCATTG | TTTGGGGTGTGTGGATTTTT | 270 |
| chr8:96912894-96914909 | p5 | (TTTAT)3 | GGTGTCGATTGCAGTCATTG | TTTGGGGTGTGTGGATTTTT | 270 |
| chr8:96928193-96930208 | p3 | (GAG)5 | TCCCCATCATTTTGTACGGT | AACGGTGTTTTGCCCTACTG | 167 |
| chr8:97039892-97041910 | p2 | (GA)6 | AATGCGAGAGAGAGAGAGCG | CACACTTGGTACCCCCAACT | 108 |
| chr8:97039892-97041910 | p3 | (CAC)6 | AATGCGAGAGAGAGAGAGCG | CACACTTGGTACCCCCAACT | 108 |
| chr8:97039892-97041910 | p3 | (CAC)6 | AATGCGAGAGAGAGAGAGCG | CACACTTGGTACCCCCAACT | 108 |
| chr8:97070797-97072815 | p3 | (ACA)6 | GCTTTCACGGGAAGTTTGAG | ATTGAATGCTGAGGTTTGCC | 106 |
| chr8:97082824-97084838 | p2 | (TA)7 | GAGCCACATGGCCTATTCAT | TTTAGCCTTGTCGGCTCCTA | 152 |
| chr8:97158707-97160755 | p2 | (AT)24 | AGGCTTAGTGGAAGGGGAAA | TAGCCCCAATCACACAACAA | 244 |
| chr8:97158707-97160755 | p5 | (ATTGC)3 | AGGCTTAGTGGAAGGGGAAA | TAGCCCCAATCACACAACAA | 244 |
| chr8:97158707-97160755 | p3 | (GTT)6 | AGGCTTAGTGGAAGGGGAAA | TAGCCCCAATCACACAACAA | 244 |
| chr8:97260056-97262072 | p2 | (CT)8 | CCTTCAACCAACATCCATCC | CCCAATGAGCCACAGAGATT | 212 |
| chr8:97307203-97309226 | p6 | (AATAAA)3 | CCTATTTAGCAGCCCGTTGA | ATTTGAGACGAGGGCAAAAA | 221 |
| chr8:97307203-97309226 | p2 | (AT)8 | CCTATTTAGCAGCCCGTTGA | ATTTGAGACGAGGGCAAAAA | 221 |
| chr8:97346547-97348568 | p2 | (AT)7 | GAAGTGAGGATTGCCCAAGA | TATACGTACGGCCAGTGCAA | 156 |
| chr8:97383590-97385605 | p5 | (AAAAT)3 | ATGTCATCGCGGGAATTTAG | GACATCACATGGCTACACGG | 189 |
| chr8:97433709-97435721 | p2 | (TA)6 | TTCTACAATTTCGCCCCATC | ATATTCTCGAGACCGGCAAA | 284 |
| chr8:97458391-97460425 | p2 | (AT)17 | TTGACCAACATGCCTTGAAA | AACTAAGTGTGGCACAGGGG | 153 |

| Marker ID | SSR type | SSR motif | FORWARD PRIMER (5'-3') | REVERSE PRIMER (5'-3') | size (bp) |
|---|---|---|---|---|---|
| chr8:97503801-97505825 | p2 | (AT)12 | TGATTTCAAGCCCAGATTCC | AACTTTCCCCAACTTCCGTT | 160 |
| chr8:97536036-97538051 | p3 | (ACA)5 | CGAGGGTATTTTTGAACCGA | GAGCTGGTTCTCTCGATTGC | 246 |
| chr8:97705714-97707729 | p2 | (AT)11 | GAGGATTTGTGTTGCCCAGT | TGGATAACATTCCCCTTGGA | 164 |
| chr8:97705714-97707729 | p3 | (TTG)5 | GAGGATTTGTGTTGCCCAGT | TGGATAACATTCCCCTTGGA | 164 |
| chr8:97709916-97711934 | p6 | (AAAAAT)3 | TGGTGGAGGAATTCTTGGAA | ACATGGCAAGGTTTGTGTGA | 136 |
| chr8:97801654-97803680 | p2 | (AC)10 | TGCTCAGAATTTTCCAGCCT | TTCTGTAGACAAGGGGTGGG | 264 |
| chr8:97801654-97803680 | p2 | (AG)13 | TGCTCAGAATTTTCCAGCCT | TTCTGTAGACAAGGGGTGGG | 264 |
| chr8:97911441-97913456 | p5 | (GAAAA)3 | GTTGGGGTTGGGGTTAGAGT | AAGGCCTCATTTGTTTGCAC | 175 |
| chr8:97923765-97925777 | p2 | (TC)6 | TGTGGGATATGATGCGAAAA | TCCTGAGGGAATCAGAATGG | 251 |
| chr8:97923765-97925777 | p2 | (TA)7 | TGTGGGATATGATGCGAAAA | TCCTGAGGGAATCAGAATGG | 251 |
| chr8:97955432-97957447 | p5 | (CATAT)3 | ATATTTCCTAAATGCCCCGC | GATAGGGGTGCTCGGTACAA | 134 |
| chr8:97955432-97957447 | p6 | (TACATA)3 | ATATTTCCTAAATGCCCCGC | GATAGGGGTGCTCGGTACAA | 134 |
| chr8:97957278-97959293 | p5 | (AACAC)3 | CCCGAATTCACGTAGCAAAT | GAATTCCCGGATTGTGATTG | 222 |
| chr8:97973014-97975034 | p4 | (ATAC)5 | GGGTTCAGCGGAGATCAATA | AGCTCAAAAGTCAGAAGCGG | 103 |
| chr8:98040040-98042054 | p2 | (AT)7 | TGACCTACGTCCACAAGCAG | AGTGTCGTATTACGCCCAGG | 161 |
| chr8:98104172-98106190 | p6 | (ATTCAA)3 | AAACAAGGTAGTGATGGCCG | ACGAAAAGAGGGGGAGAAAA | 136 |
| chr8:98174677-98176693 | p4 | (GACG)6 | AAGCCAATACAAATACGGCG | CGGGTTGCTACTGTTCAGGT | 139 |
| chr8:98174677-98176693 | p4 | (ATAC)4 | AAGCCAATACAAATACGGCG | CGGGTTGCTACTGTTCAGGT | 139 |
| chr8:98185200-98187220 | p2 | (TA)30 | CTAAGCCCATGTCACCGATT | CCAAGTCCCTTGCTAATCCA | 187 |
| chr8:98185200-98187220 | p2 | (AT)10 | CTAAGCCCATGTCACCGATT | CCAAGTCCCTTGCTAATCCA | 187 |
| chr8:98185200-98187220 | p2 | (TA)8 | CTAAGCCCATGTCACCGATT | CCAAGTCCCTTGCTAATCCA | 187 |
| chr8:98210867-98212879 | p2 | (TA)6 | GTTGGTTGGAGCTGGACCTA | AGCGCGTGTCTGAAATAACA | 176 |
| chr8:98210867-98212879 | p2 | (TA)6 | GTTGGTTGGAGCTGGACCTA | AGCGCGTGTCTGAAATAACA | 176 |
| chr8:98210867-98212879 | p2 | (AT)7 | GTTGGTTGGAGCTGGACCTA | AGCGCGTGTCTGAAATAACA | 176 |
| chr8:98210986-98213000 | p2 | (TA)6 | GCGCTCAACGTTTTCATACA | ATGAACAGTGCCTTGTGTGG | 170 |
| chr8:98210986-98213000 | p2 | (TA)6 | GCGCTCAACGTTTTCATACA | ATGAACAGTGCCTTGTGTGG | 170 |
| chr8:98210986-98213000 | p2 | (AT)7 | GCGCTCAACGTTTTCATACA | ATGAACAGTGCCTTGTGTGG | 170 |
| chr8:98223175-98225190 | p5 | (GATAT)3 | GGACCGTATAACGAGCCGTA | TGTTCTTGGTTGGAAAAGGG | 118 |
| chr8:98223175-98225190 | p2 | (AT)8 | GGACCGTATAACGAGCCGTA | TGTTCTTGGTTGGAAAAGGG | 118 |
| chr8:98231654-98233670 | p2 | (TA)8 | GTCCCTTTCTTCAAATGGCA | GGGTGACGTAAGGTCGTGAT | 229 |
| chr8:98233926-98235941 | p3 | (TGC)5 | TCCTAATGTCTCGGGAATCG | TATGAAACGACCCGTTAGGC | 289 |
| chr8:98235822-98237837 | p5 | (GTTCG)3 | AACTCGAAGGTGCGCTAAAA | AGGGTTTCCCTTTTCCTCAA | 123 |
| chr8:98235822-98237837 | p5 | (GTTCG)4 | AACTCGAAGGTGCGCTAAAA | AGGGTTTCCCTTTTCCTCAA | 123 |
| chr8:98236251-98238271 | p5 | (GTTCG)3 | GGGGTCTGGAGAGGGTAGAG | TATCAAGGTCCAGGGTGAGC | 186 |
| chr8:98236251-98238271 | p5 | (GTTCG)4 | GGGGTCTGGAGAGGGTAGAG | TATCAAGGTCCAGGGTGAGC | 186 |
| chr8:98440829-98442844 | p5 | (AAATT)3 | TGCGTACATTCTACCCTCCC | GCAAGGGTTTGAAAGTTTGG | 167 |

| Marker ID | SSR type | SSR motif | FORWARD PRIMER (5'-3') | REVERSE PRIMER (5'-3') | size (bp) |
|---|---|---|---|---|---|
| chr8:98465011-98467031 | p2 | (TA)8 | TGGTGGTGGTGGAGTAACAA | AGAGGCGTGAAAAACGCTAA | 272 |
| chr8:98465051-98467063 | p2 | (TA)8 | TGGTGGTGGTGGAGTAACAA | AGAGGCGTGAAAAACGCTAA | 232 |
| chr8:98505984-98507999 | p5 | (TACAA)3 | AGTTGGAACTTTGCAAACCG | TTAATGAAGCACGGAAACC | 218 |
| chr8:98537037-98539073 | p3 | (AAT)12 | CATTGCCACAACCTTCACTG | CCGAAAATCCCTCTCCTTTC | 241 |
| chr8:98629344-98631368 | p4 | (TATT)6 | TTCGACTGAGCTCCTGAACA | GCTTTTGCGTGTGTTCTCAA | 275 |
| chr8:98691834-98693848 | p2 | (TA)7 | CAGTTCGCTGTCACGAAGAA | AGCCCACAGAATCAAACGAG | 258 |
| chr8:98708480-98710574 | p3 | (CTG)5 | ACTGCTGCTGAAACTGCTGA | CGGTGTTTTCAATTCAGGCT | 279 |
| chr8:98708480-98710574 | p2 | (AT)30 | ACTGCTGCTGAAACTGCTGA | CGGTGTTTTCAATTCAGGCT | 279 |
| chr8:98708480-98710574 | p2 | (TA)47 | ACTGCTGCTGAAACTGCTGA | CGGTGTTTTCAATTCAGGCT | 279 |
| chr8:98716611-98718626 | p5 | (TTATT)3 | TGGGAGAACAGTTCAGGGAC | CCCTGCTTTTGCAATTCTGT | 207 |
| chr8:98716611-98718626 | p5 | (AGAAA)3 | TGGGAGAACAGTTCAGGGAC | CCCTGCTTTTGCAATTCTGT | 207 |
| chr8:98716611-98718626 | p2 | (AT)14 | TGGGAGAACAGTTCAGGGAC | CCCTGCTTTTGCAATTCTGT | 207 |
| chr8:98759097-98761111 | p2 | (AT)7 | ATTGGGCTAAGGGTTCGACT | CCTTCATCCCAGCCTTCATA | 218 |
| chr8:98770321-98772363 | p3 | (AAT)14 | GGGAAAATCCCTCCAATGTT | GTGCGGTTATTTGAGGGAAA | 244 |
| chr8:98809717-98811732 | p5 | (ATATC)3 | GTACCACAAGCCCTTCCAAA | TGGAGGATGTTTCATGGTGA | 272 |
| chr8:98882045-98884069 | p2 | (TG)12 | TCCCCGCATAGATTTGACTC | AAAGCAGGCAAGACTTCCAA | 177 |
| chr8:98894286-98896319 | p2 | (AG)7 | AAAATTGGGCGGGATTTAAG | CTTCGTCCATCTGTGAAGCA | 197 |
| chr8:98894321-98896363 | p2 | (AG)7 | AAAATTGGGCGGGATTTAAG | CTTCGTCCATCTGTGAAGCA | 162 |
| chr8:98894366-98896381 | p2 | (AG)7 | AAAATTGGGCGGGATTTAAG | CTTCGTCCATCTGTGAAGCA | 117 |
| chr8:98982385-98984406 | p3 | (TGT)7 | GACTTCTTTTGCCAGCAAGG | TGTACCCCAAACTCCGGTAA | 158 |
| chr8:99038547-99040567 | p2 | (TA)10 | AGCCTTGAATCCGTCTCTGA | GGCTTGTGTGGGTATTTCGT | 293 |
| chr8:99038707-99040721 | p2 | (TA)10 | AGCCTTGAATCCGTCTCTGA | TGTGTGGGTATTTCGTTGGA | 133 |
| chr8:99095080-99097094 | p2 | (TA)7 | TAAAAAGCAACCCCCAGATG | ATCCCAAATCCACTCCATCA | 183 |
| chr8:99100463-99102481 | p6 | (ATTTTT)3 | TTAAACCCTTGGAGGCCTTT | TTCCTCAAACCAACTGACCC | 216 |
| chr8:99102817-99104833 | p5 | (GAGAA)3 | GGGAGGCAGAGGCTTCTAGT | AGTAACCCCTCGTCGCTTTT | 228 |
| chr8:99102817-99104833 | p2 | (TA)8 | GGGAGGCAGAGGCTTCTAGT | AGTAACCCCTCGTCGCTTTT | 228 |
| chr8:99159111-99161127 | p2 | (AT)8 | GTTGCCCCTCAAATTGCTTA | AAACTCGGCTGGAGTTTGAA | 196 |
| chr8:99193542-99195558 | p2 | (TA)15 | TCAGCTAGCAAGTACGCCCT | ATATAGCACTGACGTGGCCC | 200 |
| chr8:99193542-99195558 | p2 | (AT)9 | TCAGCTAGCAAGTACGCCCT | ATATAGCACTGACGTGGCCC | 200 |
| chr8:99193593-99195607 | p2 | (TA)15 | TCAGCTAGCAAGTACGCCCT | ATATAGCACTGACGTGGCCC | 149 |
| chr8:99193593-99195607 | p2 | (AT)9 | TCAGCTAGCAAGTACGCCCT | ATATAGCACTGACGTGGCCC | 149 |
| chr8:99193613-99195641 | p2 | (TA)15 | TCAGCTAGCAAGTACGCCCT | ATATAGCACTGACGTGGCCC | 129 |
| chr8:99193613-99195641 | p2 | (AT)9 | TCAGCTAGCAAGTACGCCCT | ATATAGCACTGACGTGGCCC | 129 |
| chr8:99194076-99196114 | p2 | (TA)15 | AACACGTGCAATACACCTCG | CGGTCCGGTTACCTGTTTTA | 195 |
| chr8:99194076-99196114 | p2 | (AT)9 | AACACGTGCAATACACCTCG | CGGTCCGGTTACCTGTTTTA | 195 |
| chr8:99194153-99196169 | p2 | (TA)15 | AACACGTGCAATACACCTCG | CGGTCCGGTTACCTGTTTTA | 118 |

| Marker ID | SSR type | SSR motif | FORWARD PRIMER (5'-3') | REVERSE PRIMER (5'-3') | size (bp) |
|---|---|---|---|---|---|
| chr8:99194153-99196169 | p2 | (AT)9 | AACACGTGCAATACACCTCG | CGGTCCGGTTACCTGTTTTA | 118 |
| chr8:99216825-99218841 | p2 | (TA)8 | TCGATTTCTCTGCAACCCTT | TCACCTGACTTTTGCACCAA | 227 |
| chr8:99365764-99367794 | p2 | (CT)15 | GGGTTTCAAGTGGCAAGTGT | ATCCTCCCAGTTCCCAAATC | 144 |
| chr8:99407337-99409352 | p5 | (AAATA)3 | TAGCTGCCTTTTCCCAAATG | GGGGTAAAACTAGAACCCCG | 206 |
| chr8:99407337-99409352 | p5 | (AATAA)3 | TAGCTGCCTTTTCCCAAATG | GGGGTAAAACTAGAACCCCG | 206 |
| chr8:99421723-99423807 | p3 | (AAC)28 | TGGATCCCAGTTCCCAATTA | TCGTCAGGCAACTCAAACTG | 133 |
| chr8:99456153-99458171 | p5 | (TTTCT)3 | GCTCTGTTATGCTCTTCGGG | ACAATGACACATCAGCCCAA | 229 |
| chr8:99456174-99458192 | p5 | (TTTCT)3 | GCTCTGTTATGCTCTTCGGG | ACAATGACACATCAGCCCAA | 208 |
| chr8:99456193-99458208 | p5 | (TTTCT)3 | GCTCTGTTATGCTCTTCGGG | ACAATGACACATCAGCCCAA | 189 |
| chr8:99456621-99458649 | p5 | (TTTCT)3 | TTTCCTATCCCTATGGCGTG | CGGTGTTGATAAGGGGCTAA | 299 |
| chr8:99515024-99517036 | p2 | (AT)6 | GTATCAACCTTGGCTTCGGA | TCGTTTCCTCCACTTTGCTT | 270 |
| chr8:99515024-99517036 | p5 | (TATAA)3 | GTATCAACCTTGGCTTCGGA | TCGTTTCCTCCACTTTGCTT | 270 |
| chr8:99518201-99520213 | p2 | (TA)6 | TTGCCACTTGCGTATCAATC | AAATGACATGGCGGAATAGC | 250 |
| chr8:99518201-99520213 | p2 | (AT)7 | TTGCCACTTGCGTATCAATC | AAATGACATGGCGGAATAGC | 250 |
| chr8:99518305-99520353 | p2 | (TA)6 | TTGCCACTTGCGTATCAATC | AAATGACATGGCGGAATAGC | 146 |
| chr8:99518305-99520353 | p2 | (AT)7 | TTGCCACTTGCGTATCAATC | AAATGACATGGCGGAATAGC | 146 |
| chr8:99524497-99526509 | p2 | (AT)7 | TTGCCACTTGCGTATCAATC | AAATGACATGGCGGAATAGC | 250 |
| chr8:99524601-99526649 | p2 | (AT)7 | TTGCCACTTGCGTATCAATC | AAATGACATGGCGGAATAGC | 146 |
| chr8:99556008-99558022 | p2 | (AG)7 | CTGACCCGGCATAAACTTGT | CTTATGGGCAGACTTGGCAT | 107 |
| chr8:99599439-99601451 | p2 | (TA)6 | TGTGGGGTTGTAAAAGGGAG | AAGCAGGGACTGATCTGGAA | 161 |
| chr8:99599439-99601451 | p2 | (TA)37 | TGTGGGGTTGTAAAAGGGAG | AAGCAGGGACTGATCTGGAA | 161 |
| chr8:99648321-99650339 | p2 | (TA)9 | AGTTTCCAACGCAACAAACC | TTTCCGGGGAGATAAGTGTG | 138 |
| chr8:99662383-99664397 | p2 | (TA)7 | TAACGCCATTCGTGTTTCAA | CTGCCCATCCTTACCTCTTG | 218 |
| chr8:99662383-99664397 | p2 | (TA)8 | TAACGCCATTCGTGTTTCAA | CTGCCCATCCTTACCTCTTG | 218 |
| chr8:99704864-99706882 | p2 | (TA)9 | TCCCAACGTCAACTTTAGCC | TATCCCCCAGGATTCAACAA | 180 |
| chr8:99717235-99719293 | p2 | (AT)29 | AATCATGGCTTCGGAAAGTG | TCGCTCCGATCTAATCCAGT | 229 |
| chr8:99780328-99782346 | p6 | (ATTTTT)3 | AGAAAGGGGCTGGCAGTAAT | AGATCCGACAACAAACTCGG | 105 |
| chr8:99780328-99782346 | p2 | (TA)9 | AGAAAGGGGCTGGCAGTAAT | AGATCCGACAACAAACTCGG | 105 |
| chr8:99780490-99782508 | p6 | (ATTTTT)3 | TTTGGGGGTTGTTTGATTTC | GAGTGCACATATTCCATGCG | 227 |
| chr8:99780490-99782508 | p2 | (TA)9 | TTTGGGGGTTGTTTGATTTC | GAGTGCACATATTCCATGCG | 227 |
| chr8:99808958-99810998 | p5 | (TAAAA)3 | AGGCCCTAGAGCATTTTCGT | ATGTCCGACAATTTTTCCCA | 195 |
| chr8:99808958-99810998 | p2 | (TA)20 | AGGCCCTAGAGCATTTTCGT | ATGTCCGACAATTTTTCCCA | 195 |
| chr8:99808958-99810998 | p2 | (AG)9 | AGGCCCTAGAGCATTTTCGT | ATGTCCGACAATTTTTCCCA | 195 |
| chr8:99981894-99983915 | p3 | (TTG)7 | CGCGCACTACAGTAATCGAA | AATTTCGTGGGTGGAAGTTG | 210 |
| chr8:99981894-99983915 | p2 | (TA)15 | CGCGCACTACAGTAATCGAA | AATTTCGTGGGTGGAAGTTG | 210 |
| chr8:99986288-99988303 | p5 | (ATATA)3 | CCAACACGCAACAGCTACAT | CCCATTTCTCACTGGCAACT | 222 |

| Marker ID | SSR type | SSR motif | FORWARD PRIMER (5'-3') | REVERSE PRIMER (5'-3') | size (bp) |
|---|---|---|---|---|---|
| chr8:100045847-100047862 | p5 | (TTAAA)3 | AACCTGGTGCCTTTTCAGTG | CATGTCTGCTACTGCTCCCA | 154 |
| chr8:100045847-100047862 | p5 | (TTTTA)3 | AACCTGGTGCCTTTTCAGTG | CATGTCTGCTACTGCTCCCA | 154 |
| chr8:100045847-100047862 | p5 | (ATTTT)3 | AACCTGGTGCCTTTTCAGTG | CATGTCTGCTACTGCTCCCA | 154 |
| chr8:100097023-100099038 | p5 | (AAAAT)3 | AGTTGGGCATCAGACACCTC | GGCTATTGGTTCGAGCTCAC | 276 |
| chr8:100114231-100116249 | p6 | (GTATAT)3 | CATGGAAGTTGACGGGGTAT | GCACTGTGCAGCAGATGTTT | 253 |
| chr8:100114231-100116249 | p3 | (ACA)5 | CATGGAAGTTGACGGGGTAT | GCACTGTGCAGCAGATGTTT | 253 |
| chr8:100164081-100166099 | p3 | (AAC)6 | TCTACCTTTGGCAGGAGCAT | TGGACGCTTCCAATATGTCA | 117 |
| chr8:100185211-100187235 | p6 | (TGTTAA)4 | GCTCGACCAGAGATCCAGAC | GCATACTTCTCTCCAAGCCG | 234 |
| chr8:100309008-100311023 | p5 | (TATGA)3 | ATATGGGTTATGCTCAGCCG | CCCGAACTGGACACTCGTAT | 121 |
| chr8:100380339-100382355 | p2 | (TA)8 | ATTGCTAACTCGGAAAGCCA | TGGTGTGAGGGGGATACAAT | 182 |
| chr8:100451827-100453862 | p2 | (TA)11 | TGCAATTTGAGTTGGTTTCG | CACAAAAATGGGGGAAGAGA | 170 |
| chr8:100451885-100453907 | p2 | (TA)11 | TGCAATTTGAGTTGGTTTCG | CACAAAAATGGGGGAAGAGA | 112 |
| chr8:100463348-100465368 | p2 | (AT)10 | AAGACCAGCGCAAAATAGGA | AAAGATCAAAGTGGCTCCCA | 105 |
| chr8:100473236-100475251 | p5 | (CTTTT)3 | TGGCCACGAATTTTCATTTT | TTTCCTTGTTCACAGGGTCC | 149 |
| chr8:100544007-100546025 | p2 | (AT)9 | TGGAAGATCGTTAGGGTTGG | CGGAGTCTTGCCTCGTTAAG | 137 |
| chr8:100558357-100560372 | p5 | (TGGTT)3 | ACGAGCTAAATCCCCACCTT | TGTGATCATTCTCGGCGTAA | 246 |
| chr8:100647767-100649785 | p3 | (TTC)6 | AACTCGAACCAAAGCACGAT | TTTGGGCAAATTTGGAACTC | 142 |
| chr8:100681474-100683489 | p5 | (TATTT)3 | TGGGCCTTGCATACAACATA | TGGAGATATTGCACTGGCTG | 226 |
| chr8:100681474-100683489 | p5 | (AAATA)3 | TGGGCCTTGCATACAACATA | TGGAGATATTGCACTGGCTG | 226 |
| chr8:100750182-100752200 | p4 | (CGCA)4 | TGGCAATTCACCATGTCACT | CGCAAGGTGATATTCTGCCT | 202 |
| chr8:100750182-100752200 | p3 | (TAT)6 | TGGCAATTCACCATGTCACT | CGCAAGGTGATATTCTGCCT | 202 |
| chr8:100750182-100752200 | p4 | (CGCA)4 | TGGCAATTCACCATGTCACT | CGCAAGGTGATATTCTGCCT | 202 |
| chr8:100750182-100752200 | p3 | (TAT)6 | TGGCAATTCACCATGTCACT | CGCAAGGTGATATTCTGCCT | 202 |
| chr8:100788763-100790817 | p3 | (ATT)18 | CGGTTAAAATGGGGTGTGAC | AGTGGGATGGATCTGTCAGG | 293 |
| chr8:100857622-100859637 | p5 | (TTTTC)3 | CTCGAGCAAACACAAACCAA | TTTTCGGGAAGGTGTCACTC | 263 |
| chr8:100951591-100953611 | p5 | (AAAAT)4 | AAATGCAGGGTAAGACTGCG | GAAAACCCCTAAAAGGGCAG | 256 |
| chr8:100984661-100986681 | p2 | (TA)10 | GCGTCTTTTGCACACCTCTT | AGTAAAAATGGACGGAGGGG | 199 |
| chr8:100984661-100986681 | p3 | (TTA)14 | GCGTCTTTTGCACACCTCTT | AGTAAAAATGGACGGAGGGG | 199 |
| chr8:101016973-101019001 | p2 | (TA)14 | CGGAACGTCGATTCTTCATT | GTATAGATGTGCCGCGGATT | 155 |
| chr8:101031507-101033522 | p3 | (TTA)5 | TCATCGGAGTCTGAATTCCC | TAATACCCACCGAAAGTGCC | 205 |
| chr8:101096795-101098810 | p5 | (AAAAT)3 | CGAGGGACTATCGTGCAAAT | ATTCAAAATTGGTCCTGCGT | 227 |
| chr8:101096795-101098810 | p5 | (TTTCT)4 | CGAGGGACTATCGTGCAAAT | ATTCAAAATTGGTCCTGCGT | 227 |
| chr8:101121541-101123555 | p2 | (GA)7 | TAACCACCCTCTTTGTTCGG | CCTGTTCACTGCACTCAGGA | 156 |
| chr8:101175260-101177286 | p3 | (TGT)5 | ATGGGTCTACAACCGAAACG | AGAGAAAGAGAAGGGGGCTG | 254 |
| chr8:101175323-101177341 | p3 | (TGT)5 | ATGGGTCTACAACCGAAACG | AGAGAAAGAGAAGGGGGCTG | 191 |
| chr8:101185638-101187658 | p5 | (AAGAG)4 | CATCATCCGGTCATTTACCC | TTTGACGGGGAAGTCCATAG | 239 |

| Marker ID | SSR type | SSR motif | FORWARD PRIMER (5'-3') | REVERSE PRIMER (5'-3') | size (bp) |
|---|---|---|---|---|---|
| chr8:101237498-101239513 | p5 | (TTGGT)3 | GAGCAACAGCATGAAGGTCA | GAGCGACTGAATGGGAAAAG | 127 |
| chr8:101237498-101239513 | p5 | (TTGGT)3 | GAGCAACAGCATGAAGGTCA | GAGCGACTGAATGGGAAAAG | 127 |
| chr8:101281467-101283482 | p3 | (AAT)5 | GAACTTGTTTTAGCGAGGCG | CATTTCCTCTCCCCTTCTCC | 166 |
| chr8:101416615-101418630 | p5 | (TAAAA)3 | TGAATTTGTCCAGCTCAACG | AAGAAGGGGCAATTTCGTTT | 119 |
| chr8:101416615-101418630 | p2 | (AG)24 | TGAATTTGTCCAGCTCAACG | AAGAAGGGGCAATTTCGTTT | 119 |
| chr8:101416832-101418880 | p5 | (TAAAA)3 | AAAACGAAATTGCCCCTTCT | ATCAAAAGAACGCCAAGGAA | 113 |
| chr8:101416832-101418880 | p2 | (AG)24 | AAAACGAAATTGCCCCTTCT | ATCAAAAGAACGCCAAGGAA | 113 |
| chr8:101469734-101471752 | p6 | (ATTTTT)3 | TCAATTGCCATAATCAGCCA | AGGCCTTCTTTTCAAGCTCC | 162 |
| chr8:101498634-101500722 | p2 | (AT)44 | GGAAGTTGTGCAGTTGGGTT | TTCGGACATCAATCCAAACA | 169 |
| chr8:101498634-101500722 | p6 | (AAAAAT)3 | GGAAGTTGTGCAGTTGGGTT | TTCGGACATCAATCCAAACA | 169 |
| chr8:101499679-101501697 | p2 | (TA)21 | AATTAAAGGGTTTCGGGTCG | TCGGTTCAAATTTACCTCGC | 254 |
| chr8:101499679-101501697 | p6 | (AAAAAT)3 | AATTAAAGGGTTTCGGGTCG | TCGGTTCAAATTTACCTCGC | 254 |
| chr8:101526690-101528735 | p2 | (GA)6 | TCACAAACGAACGAAATCCA | TAATGCTCGTCCTTTCCCAG | 285 |
| chr8:101526690-101528735 | p2 | (AT)13 | TCACAAACGAACGAAATCCA | TAATGCTCGTCCTTTCCCAG | 285 |
| chr8:101526743-101528773 | p2 | (GA)6 | TCACAAACGAACGAAATCCA | TAATGCTCGTCCTTTCCCAG | 232 |
| chr8:101526743-101528773 | p2 | (AT)13 | TCACAAACGAACGAAATCCA | TAATGCTCGTCCTTTCCCAG | 232 |
| chr8:101527086-101529112 | p2 | (GA)6 | GTTAATCAATGGGCTGGTCG | TTATCGCACGACCATTACGA | 201 |
| chr8:101527086-101529112 | p2 | (AT)13 | GTTAATCAATGGGCTGGTCG | TTATCGCACGACCATTACGA | 201 |
| chr8:101579135-101581153 | p6 | (ATTTTT)3 | TTGCCATAATCAGCCATCAA | GGCCTTCTCTTCAAGCTCCT | 166 |
| chr8:101603031-101605067 | p2 | (TA)18 | CCCATTGCTCCTGAATTTGT | CCTTTAACTGTCGTCCAGCC | 249 |
| chr8:101783528-101785548 | p5 | (GGGTG)4 | TGGAGAAGTCGAACGAAAGG | TGGGCCAACTAATGTGTGAA | 289 |
| chr8:101820495-101822515 | p5 | (GGGTG)4 | TGGAGAAGTCGAACGAAAGG | TGGGCCAACTAATGTGTGAA | 289 |
| chr8:101983193-101985211 | p6 | (TATGTA)3 | ACTTCAAACGCTCGCAACTT | CAAGCGGAATAGTGAGAGGC | 239 |
| chr8:101983193-101985211 | p2 | (TA)9 | ACTTCAAACGCTCGCAACTT | CAAGCGGAATAGTGAGAGGC | 239 |
| chr8:101992000-101994015 | p5 | (TTTTA)3 | AGCACTTCGTGCAAATCCTT | TTCCCATAAAAGGCAACTCG | 245 |
| chr8:101992000-101994015 | p2 | (CT)6 | AGCACTTCGTGCAAATCCTT | TTCCCATAAAAGGCAACTCG | 245 |
| chr8:102004264-102006278 | p2 | (TA)7 | AGCAGAACCAGCAGAAATCG | TGAATCCCTGAACTTGTCCC | 261 |
| chr8:102037124-102039145 | p3 | (TTG)7 | TGAAGACTTGCTTCCGGAGT | CTCATTTCCGCCACTTTCAT | 142 |
| chr8:102199999-102202013 | p2 | (TA)7 | CTCAAGTGGAAGTGACGCAA | TCTCCCGTGACATGTTCAAA | 163 |
| chr8:102204612-102206626 | p2 | (AT)7 | TTGTCATGAATAACGGCACG | TTGTGGCCCCCAAATAAATA | 219 |
| chr8:102204612-102206626 | p2 | (AT)7 | TTGTCATGAATAACGGCACG | TTGTGGCCCCCAAATAAATA | 219 |
| chr8:102204612-102206626 | p2 | (AT)14 | TTGTCATGAATAACGGCACG | TTGTGGCCCCCAAATAAATA | 219 |
| chr8:102243340-102245358 | p6 | (TGAAAT)3 | AAGCAACCCTGGATGAGCTA | AAGGTTCAAGGGCAAGGTTT | 231 |
| chr8:102422908-102424928 | p2 | (AT)10 | TTATGAACTTTTCGGCACCC | CTGTCAACAGGTTTGGAGCA | 131 |
| chr8:102478299-102480319 | p2 | (AT)10 | AAACCTGTCCCCTGAGGAGT | TGTTGTTTTTCGAGCGTGAG | 194 |
| chr8:102589828-102591843 | p5 | (AAAAG)3 | GTATTTTGGCATGGAGGGTG | TCGGATCCAAGAATTGGAAG | 186 |

| Marker ID | SSR type | SSR motif | FORWARD PRIMER (5'-3') | REVERSE PRIMER (5'-3') | size (bp) |
|---|---|---|---|---|---|
| chr8:102592589-102594603 | p2 | (TA)7 | CCGACGAAGTTCCAACAAAT | AGTTGGGAAGGCAATTAGGG | 203 |
| chr8:102592589-102594603 | p2 | (TA)32 | CCGACGAAGTTCCAACAAAT | AGTTGGGAAGGCAATTAGGG | 203 |
| chr8:102592729-102594793 | p2 | (TA)7 | TAAAGGGAAGCATGCACACA | GTTGGGAAGGCAATTAGGGT | 130 |
| chr8:102592729-102594793 | p2 | (TA)32 | TAAAGGGAAGCATGCACACA | GTTGGGAAGGCAATTAGGGT | 130 |
| chr8:102609444-102611458 | p2 | (AC)7 | ACTGGAGCATCTGTGCCTCT | GGACCGTCATTTAGCTCCAA | 206 |
| chr8:102621233-102623247 | p2 | (AC)7 | ACTGGAGCATCTGTGCCTCT | GGACCGTCATTTAGCTCCAA | 206 |
| chr8:102629372-102631384 | p2 | (CT)6 | TTTGTCGACCCAAAATAGGC | GGCATCCCCCAAAACTTAAT | 197 |
| chr8:102716504-102718516 | p2 | (TA)20 | CTTCGGCGAAACGTTTATGT | AGTTGGGATTGACAGTTCCG | 267 |
| chr8:102716504-102718516 | p2 | (TA)6 | CTTCGGCGAAACGTTTATGT | AGTTGGGATTGACAGTTCCG | 267 |
| chr8:102794939-102796957 | p6 | (TTTGAT)3 | CAGTGGAGAAAACCTCTCCG | AAAGTGTGCTCAGACAGCCA | 252 |
| chr8:102794939-102796957 | p6 | (TTTGAT)3 | CAGTGGAGAAAACCTCTCCG | AAAGTGTGCTCAGACAGCCA | 252 |
| chr8:102794939-102796957 | p3 | (TTC)6 | CAGTGGAGAAAACCTCTCCG | AAAGTGTGCTCAGACAGCCA | 252 |
| chr8:102820522-102822537 | p6 | (TTAAAA)4 | GTGGTGGGTTGACAAGTGTG | ATAGGAAACGGGGAGAAACG | 164 |
| chr8:102820522-102822537 | p5 | (AAACC)3 | GTGGTGGGTTGACAAGTGTG | ATAGGAAACGGGGAGAAACG | 164 |
| chr8:102875110-102877122 | p2 | (AT)6 | GGAGGAAATACCACCACACG | GGGGCCTCGAATCTACTCTC | 277 |
| chr8:102885417-102887433 | p2 | (AG)8 | CCCATCTTCTGGATGGGATA | CCCATCACCCAGAATAATGC | 197 |
| chr8:102934239-102936254 | p3 | (CAA)5 | ATCGAGTCCCACAACTCCAC | GTTTTGGTGAGGGTGCTGAT | 260 |
| chr8:103033256-103035272 | p4 | (AATA)4 | CCATTTGGCCATGAATTTTT | TTGGCCATGAAAACTGTTGA | 196 |
| chr8:103047725-103049746 | p3 | (ACC)7 | GCTGAATTCCACCACCACTT | GCCACCTTTGGTTAAGGACA | 141 |
| chr8:103047725-103049746 | p6 | (TTGAAA)4 | GCTGAATTCCACCACCACTT | GCCACCTTTGGTTAAGGACA | 141 |
| chr8:103047725-103049746 | p2 | (TC)8 | GCTGAATTCCACCACCACTT | GCCACCTTTGGTTAAGGACA | 141 |
| chr8:103048214-103050238 | p3 | (ACC)7 | CAGGTACAAATTCGCCTGGT | CCCGTTGTCAATGGAGAAGT | 213 |
| chr8:103048214-103050238 | p6 | (TTGAAA)4 | CAGGTACAAATTCGCCTGGT | CCCGTTGTCAATGGAGAAGT | 213 |
| chr8:103048214-103050238 | p2 | (TC)8 | CAGGTACAAATTCGCCTGGT | CCCGTTGTCAATGGAGAAGT | 213 |
| chr8:103059751-103061769 | p2 | (TA)9 | TGCACTAGCCGATTGACATC | GTCCAACAATGGCTGGTCTT | 180 |
| chr8:103212274-103214286 | p2 | (TA)6 | TCAAGAAGTGGCAGAGGGAT | TGGCAGTAACAAGACCACCA | 242 |
| chr8:103233503-103235521 | p2 | (GA)6 | AAAAGAGAGAGAGGAGCGGG | ATTCCCGTCCAAAATAAGGG | 298 |
| chr8:103233503-103235521 | p3 | (ATA)6 | AAAAGAGAGAGAGGAGCGGG | ATTCCCGTCCAAAATAAGGG | 298 |
| chr8:103390258-103392318 | p2 | (AT)30 | CATGCATGATCCAAAGGTTG | CCTCGGCACTCGACTTAGAC | 120 |
| chr8:103396509-103398523 | p2 | (TA)7 | TCTTTGAAGGGAAATCGTGC | CGCCACAAAGGTCAATTTTT | 279 |
| chr8:103493083-103495101 | p3 | (CAA)6 | CCGTCACACATCACAACACA | CTATGATATCGTTCGCCGGT | 272 |
| chr8:103529344-103531359 | p3 | (TAA)7 | AGCGGCTGAATGCAACTAAT | ACCTCTCCCCTCTCCTCTGA | 183 |
| chr8:103529344-103531359 | p3 | (ATA)5 | AGCGGCTGAATGCAACTAAT | ACCTCTCCCCTCTCCTCTGA | 183 |
| chr8:103529344-103531359 | p6 | (ATAATG)3 | AGCGGCTGAATGCAACTAAT | ACCTCTCCCCTCTCCTCTGA | 183 |
| chr8:103548151-103550209 | p2 | (AT)8 | AATTTGAGCTCGCTTTTGGA | CACACCCGCTTATCAGAACA | 176 |
| chr8:103612770-103614797 | p2 | (AC)8 | CGCACTTCCTTTACCCTTTG | CAGGCTGTTTTGCTTCAACA | 237 |

| Marker ID | SSR type | SSR motif | FORWARD PRIMER (5'-3') | REVERSE PRIMER (5'-3') | size (bp) |
|---|---|---|---|---|---|
| chr8:103712147-103714167 | p2 | (AT)10 | ACTCACCCTGTTGGGAAAAA | TGCAACAATGGAGGAGTTGT | 211 |
| chr8:103712147-103714167 | p2 | (AT)10 | ACTCACCCTGTTGGGAAAAA | TGCAACAATGGAGGAGTTGT | 211 |
| chr8:103740016-103742028 | p2 | (AT)6 | ATATGTGGTTGCAAGGAGGG | ATCATGAAAATCGTAGCGGG | 158 |
| chr8:103755641-103757714 | p3 | (TTC)5 | CAGCACTGCATTGTGAGGAT | CGCTCAAAAATGTGCAAGAA | 102 |
| chr8:103775135-103777160 | p4 | (TTTA)4 | ATGAATGGGGCTGTTCAAAA | AACGATGGTCGTTTACTGGC | 147 |
| chr8:103957244-103959262 | p6 | (AAAAAC)3 | AATCCGATGCAAACTCATCC | CCTCAAATTGGCGGTAAAAA | 162 |
| chr8:104006239-104008251 | p2 | (TG)6 | CTACCTGCTGCATTTGCTGA | CAACAACAGGTGGTCAATGG | 117 |
| chr8:104006808-104008839 | p2 | (TG)6 | TGTGAGTAGGGCTGAACACG | TAAAGTCGGTTTGGTTCGGT | 232 |
| chr8:104033482-104035498 | p4 | (AAAT)4 | TGTCGCAGAATCAGATGGTC | TGTATGTCCAGACCCGTCAA | 195 |
| chr8:104042556-104044580 | p6 | (CATGAG)4 | AGCGCGATTTAAAGTCCAAA | AGCTCATCATGCACTTCACG | 286 |
| chr8:104064189-104066209 | p4 | (TATT)5 | GAATCATTGGCACACTGCAC | GCTTTGGTGACCATTGTGAA | 226 |
| chr8:104066995-104069010 | p5 | (CAAAC)3 | ACTTGCATTTTGCCCTCAAT | AGACCCTAGCCTTGGGGATA | 298 |
| chr8:104066995-104069010 | p2 | (TA)8 | ACTTGCATTTTGCCCTCAAT | AGACCCTAGCCTTGGGGATA | 298 |
| chr8:104117568-104119580 | p2 | (TA)6 | CGCGCCTAGTAAACTTCCAA | CGAGTCCGACACCCCTACTA | 113 |
| chr8:104133667-104135685 | p6 | (GTATAT)3 | TAAACAGGGGATATGCTGCC | GAAAGAAAAGGGCAAAGCCT | 203 |
| chr8:104175680-104177695 | p3 | (CTT)5 | GAGGTGTTCCAGCGTTTTGT | ATTTTATAATCCACCCCCGC | 236 |
| chr8:104219166-104221181 | p4 | (TAAT)6 | ACCGTGAGAACCATAGACGG | CCTTTGAACTTTCCGTAGCG | 218 |
| chr8:104219166-104221181 | p3 | (TTC)5 | ACCGTGAGAACCATAGACGG | CCTTTGAACTTTCCGTAGCG | 218 |
| chr8:104243707-104245723 | p2 | (AT)8 | GGAGCCGTTCTCGTTAAGTG | TTGGCAAAAACATGGTCAAA | 212 |
| chr8:104259420-104261447 | p3 | (AAT)5 | AGGAATCATGGCGTAATTGG | ACTGGTGCACTAAATTCCCG | 254 |
| chr8:104259420-104261447 | p2 | (AT)21 | AGGAATCATGGCGTAATTGG | ACTGGTGCACTAAATTCCCG | 254 |
| chr8:104307810-104309825 | p3 | (TGT)5 | CAGTCGAACCCCACAAAAAT | AAAAATATTGCGTCGTTGCC | 109 |
| chr8:104333301-104335316 | p5 | (AAAAT)3 | TAAAAGTTACCACCCCAGCG | GGAATTCTGAGACCACGAGC | 204 |
| chr8:104333301-104335316 | p2 | (TA)8 | TAAAAGTTACCACCCCAGCG | GGAATTCTGAGACCACGAGC | 204 |
| chr8:104389972-104391994 | p5 | (ATTTT)3 | AGGGTCAGCATGCTTCAAGT | CAAAGCTCGGTAAGCCTGTC | 168 |
| chr8:104508128-104510148 | p2 | (AT)10 | CATTGGGTTGAGGCAAAAGT | CTTTGGGAAATTTGCACGAT | 109 |
| chr8:104508128-104510148 | p2 | (AT)10 | CATTGGGTTGAGGCAAAAGT | CTTTGGGAAATTTGCACGAT | 109 |
| chr8:104508128-104510148 | p2 | (TA)16 | CATTGGGTTGAGGCAAAAGT | CTTTGGGAAATTTGCACGAT | 109 |
| chr8:104508314-104510334 | p2 | (AT)10 | TGGGGAATCAAACTCTCACC | ACAACGGACGCTGAAAAAGT | 133 |
| chr8:104508314-104510334 | p2 | (AT)10 | TGGGGAATCAAACTCTCACC | ACAACGGACGCTGAAAAAGT | 133 |
| chr8:104508314-104510334 | p2 | (TA)16 | TGGGGAATCAAACTCTCACC | ACAACGGACGCTGAAAAAGT | 133 |
| chr8:104508314-104510334 | p2 | (TA)6 | TGGGGAATCAAACTCTCACC | ACAACGGACGCTGAAAAAGT | 133 |
| chr8:104515281-104517296 | p3 | (TTC)5 | ATTGACACCTGAGGACGGAG | ACGCGAGGTAAATAGGAGCA | 102 |
| chr8:104527514-104529538 | p2 | (AT)7 | AACGTATGGGAAATGGCTCA | TGTCCAATCAAAACAAGCCA | 266 |
| chr8:104527596-104529610 | p2 | (AT)7 | AACGTATGGGAAATGGCTCA | TGTCCAATCAAAACAAGCCA | 184 |
| chr8:104531013-104533027 | p2 | (TA)7 | AAGCGTTGGTGGGTCTCTAA | GCAAGGCTTGAAGAGTTTGG | 107 |

| Marker ID | SSR type | SSR motif | FORWARD PRIMER (5'-3') | REVERSE PRIMER (5'-3') | size (bp) |
|---|---|---|---|---|---|
| chr8:104584068-104586080 | p2 | (AT)6 | GATGAAAGGGGGAAGGGATA | GAAAGTACTCGACGGGCAAG | 238 |
| chr8:104655745-104657760 | p3 | (ATG)5 | GGTTCAAACCAGAATCGGAA | AAAGGGGAGTTTGGAGCTGT | 168 |
| chr8:104668632-104670647 | p6 | (TATATG)3 | GCTAATAGGCGGTCGAACAG | TGCCATTCGGAAACATATCA | 246 |
| chr8:104668642-104670658 | p6 | (TATATG)3 | GCTAATAGGCGGTCGAACAG | TGCCATTCGGAAACATATCA | 236 |
| chr8:104709529-104711544 | p5 | (AACAC)3 | GACATGCCGAAGAAGGAAAG | TTCCCGGATTGTTGTTGATT | 260 |
| chr8:104709529-104711544 | p2 | (TC)8 | GACATGCCGAAGAAGGAAAG | TTCCCGGATTGTTGTTGATT | 260 |
| chr8:104713572-104715588 | p5 | (CCGGT)3 | TTGGACGTCTTAGCATGGTG | AATGCACTTTCCAAAGCCAC | 162 |
| chr8:104753427-104755445 | p3 | (CAA)6 | GATAAAGCCAGCTGACCAGG | TCGGAAGTCACCGACCTTAC | 107 |
| chr8:104884973-104886991 | p3 | (TCT)6 | TGTAAGATTGGTCCCGAAGG | CATGAGTCCAAAAGCGTGAA | 194 |
| chr8:104894082-104896097 | p5 | (GAAAA)3 | TCATGGCGACCTCTTCTTCT | TTGCCCATAAACCCCATAAA | 107 |
| chr8:104908434-104910460 | p2 | (AT)13 | CATTCACACACAGGCAAAGG | TGTTTCGAAGGATTTGGGAC | 158 |
| chr8:104963486-104965516 | p6 | (CCCACC)3 | AGCATTCCCCACGTTACAAG | TACTGCGCAGGTGTATGCTC | 203 |
| chr8:104963486-104965516 | p2 | (AT)15 | AGCATTCCCCACGTTACAAG | TACTGCGCAGGTGTATGCTC | 203 |
| chr8:104963486-104965516 | p5 | (TACAA)3 | AGCATTCCCCACGTTACAAG | TACTGCGCAGGTGTATGCTC | 203 |
| chr8:105014085-105016100 | p5 | (TTATA)3 | CAACGAAAGGTGAGTTGGGT | TGAACGCTAGACTGTTCCCC | 265 |
| chr8:105028967-105030993 | p5 | (ATTTT)3 | GAGATGGCCCAGATAGTGGA | ATCATGGACAAGGTGGAAGC | 218 |
| chr8:105036030-105038048 | p2 | (TA)9 | CCCATAAAGCATCCCTCAAA | GTGCGGAAGGTCTGGAATTA | 110 |
| chr8:105050152-105052172 | p6 | (TTCTCC)3 | TGTCGGATTGTATGAGCCAA | CAACTGCCCTAAATGCCCTA | 281 |
| chr8:105050182-105052200 | p6 | (TTCTCC)3 | TTGCTGAGCACTTTGATTCG | TCCGACATCTGGAACAATGA | 116 |
| chr8:105062340-105064354 | p2 | (AT)7 | GGCCGGATTACTCCACTTTT | TTCCACCACATCCAAACTCA | 270 |
| chr8:105069705-105071719 | p2 | (TA)6 | GGCCCCACAATTCTATGGTA | ATATGGGGCCCACAATTCTT | 243 |
| chr8:105069705-105071719 | p2 | (AT)18 | GGCCCCACAATTCTATGGTA | ATATGGGGCCCACAATTCTT | 243 |
| chr8:105069705-105071719 | p2 | (AT)13 | GGCCCCACAATTCTATGGTA | ATATGGGGCCCACAATTCTT | 243 |
| chr8:105078565-105080577 | p2 | (AT)6 | TAAGGGGTGAATTCGGCATA | CACCTTCCCTACCCAGGTTT | 169 |
| chr8:105208081-105210103 | p3 | (TGA)7 | ATCATGAGGCAATGCAACAA | TTCAAATTTTAGCGGTTGGC | 106 |
| chr8:105249569-105251584 | p5 | (TGTTA)3 | AGCAAGCCAAAGTGGAAAGA | CGACACCTTCGAGCCATATT | 224 |
| chr8:105343973-105345993 | p2 | (TG)10 | TCATTTTGGGGTCTGAGGAG | CTTGAGAGAGGACCAGTGCC | 272 |
| chr8:105404714-105406728 | p2 | (AG)7 | AATATCAGATGGCACCAGGG | GCATGGACTGGTCAAAAGGT | 298 |
| chr8:105404745-105406757 | p2 | (AG)7 | AATATCAGATGGCACCAGGG | GCATGGACTGGTCAAAAGGT | 267 |
| chr8:105458994-105461010 | p2 | (AT)8 | GAACACCCTTCAGTGGGAAA | TGGGCATGTATTGTGCAGAT | 231 |
| chr8:105526653-105528665 | p2 | (TA)6 | GGCACATTCGACACCTTTTT | CGTTGATGTCTCCGTCCTTT | 240 |
| chr8:105526653-105528665 | p5 | (AAATA)3 | GGCACATTCGACACCTTTTT | CGTTGATGTCTCCGTCCTTT | 240 |
| chr8:105528104-105530119 | p5 | (AAATA)3 | CGCGATCTATGATGGACCTT | TTTTCAGATTTGGGGTTTGG | 227 |
| chr8:105528104-105530119 | p5 | (ATAAA)3 | CGCGATCTATGATGGACCTT | TTTTCAGATTTGGGGTTTGG | 227 |
| chr8:105528104-105530119 | p5 | (TTTAT)3 | CGCGATCTATGATGGACCTT | TTTTCAGATTTGGGGTTTGG | 227 |
| chr8:105528619-105530689 | p5 | (ATAAA)3 | TCATACGAGAGTCCCCATCC | CATGAGAGTCCCCGTTCATT | 244 |

| Marker ID | SSR type | SSR motif | FORWARD PRIMER (5'-3') | REVERSE PRIMER (5'-3') | size (bp) |
|---|---|---|---|---|---|
| chr8:105528619-105530689 | p5 | (TTTAT)3 | TCATACGAGAGTCCCCATCC | CATGAGAGTCCCCGTTCATT | 244 |
| chr8:105530929-105532953 | p2 | (GT)6 | AATAAATGGTGGCCTTTCCC | TGTTTAACTCCACCTTCGCC | 255 |
| chr8:105530929-105532953 | p2 | (TG)12 | AATAAATGGTGGCCTTTCCC | TGTTTAACTCCACCTTCGCC | 255 |
| chr8:105530929-105532953 | p2 | (TG)8 | AATAAATGGTGGCCTTTCCC | TGTTTAACTCCACCTTCGCC | 255 |
| chr8:105556513-105558529 | p2 | (AT)8 | ACATCCTTCATTGAGTCCCG | GGACTCGGTGCCATAGTCAT | 182 |
| chr8:105625596-105627618 | p2 | (TA)11 | TATGAGAGGGTTCGGAGTGC | TTTTACCGGTCTGGTTACCG | 130 |
| chr8:105625596-105627618 | p2 | (TA)8 | TATGAGAGGGTTCGGAGTGC | TTTTACCGGTCTGGTTACCG | 130 |
| chr8:105625596-105627618 | p2 | (TA)7 | TATGAGAGGGTTCGGAGTGC | TTTTACCGGTCTGGTTACCG | 130 |
| chr8:105660967-105662979 | p2 | (AT)7 | TAGCATACATGTCAAGCGCC | AGGGCAAAGGTTTCCAATTT | 170 |
| chr8:105660982-105662997 | p2 | (AT)7 | TAGCATACATGTCAAGCGCC | AGGGCAAAGGTTTCCAATTT | 155 |
| chr8:105660992-105663008 | p2 | (AT)7 | TAGCATACATGTCAAGCGCC | AGGGCAAAGGTTTCCAATTT | 145 |
| chr8:105683409-105685423 | p5 | (TTCTT)3 | AAATGTCCCAACCCCTTTTC | GAGAGAGGTTTCCCTTTCGG | 204 |
| chr8:105683427-105685447 | p5 | (TTCTT)3 | AAATGTCCCAACCCCTTTTC | GAGAGAGGTTTCCCTTTCGG | 186 |
| chr8:105684782-105686797 | p5 | (TTCTT)3 | AGACTTCTGGGGTGTTCACG | GTATATGCCGGTTTGGTTCG | 131 |
| chr8:105684782-105686797 | p5 | (CCAAT)3 | AGACTTCTGGGGTGTTCACG | GTATATGCCGGTTTGGTTCG | 131 |
| chr8:105695336-105697356 | p2 | (AT)10 | TTCCAAACAGGCCCTAAGTG | CAACGTTGGAGTTTGTGTCG | 254 |
| chr8:105821761-105823779 | p6 | (TTTAAT)3 | TGCATAGACTCTACCCGCCT | TGAACACCCCTAACTACGCC | 219 |
| chr8:105827831-105829865 | p2 | (AT)17 | ACCAAAAGTCTGTCGATGGG | AGCGGACTCCTGAGGGTATT | 119 |
| chr8:105867377-105869393 | p5 | (ATTTG)3 | CTCGTCCCTGGAAACAGAAG | GAATCGCGCCCTTTGTATTA | 289 |
| chr8:105867377-105869393 | p4 | (TAAA)4 | CTCGTCCCTGGAAACAGAAG | GAATCGCGCCCTTTGTATTA | 289 |
| chr8:105867377-105869393 | p4 | (ATTT)4 | CTCGTCCCTGGAAACAGAAG | GAATCGCGCCCTTTGTATTA | 289 |
| chr8:105867642-105869658 | p5 | (ATTTG)3 | TAATACAAAGGGCGCGATTC | GGGGCGAACTAATTGTCAAA | 188 |
| chr8:105867642-105869658 | p4 | (TAAA)4 | TAATACAAAGGGCGCGATTC | GGGGCGAACTAATTGTCAAA | 188 |
| chr8:105867642-105869658 | p4 | (ATTT)4 | TAATACAAAGGGCGCGATTC | GGGGCGAACTAATTGTCAAA | 188 |
| chr8:105927724-105929739 | p2 | (TA)7 | TACGTGTCACCCAGGATCAA | AGTGGCTAAGGCTGTGCAGT | 192 |
| chr8:105927724-105929739 | p5 | (AAACC)3 | TACGTGTCACCCAGGATCAA | AGTGGCTAAGGCTGTGCAGT | 192 |
| chr8:105927724-105929739 | p5 | (AAACC)3 | TACGTGTCACCCAGGATCAA | AGTGGCTAAGGCTGTGCAGT | 192 |
| chr8:105988541-105990556 | p2 | (TA)7 | TACGTGTCACCCAGGATCAA | AGTGGCTAAGGCTGTGCAGT | 192 |
| chr8:105988541-105990556 | p5 | (AAACC)3 | TACGTGTCACCCAGGATCAA | AGTGGCTAAGGCTGTGCAGT | 192 |
| chr8:105988541-105990556 | p5 | (AAACC)3 | TACGTGTCACCCAGGATCAA | AGTGGCTAAGGCTGTGCAGT | 192 |
| chr8:106028751-106030769 | p5 | (AAATA)3 | GCAAGCAATAACTGGGGAAA | ACTGTTGACTAGCGTGCGTG | 104 |
| chr8:106028751-106030769 | p6 | (CCCCCA)3 | GCAAGCAATAACTGGGGAAA | ACTGTTGACTAGCGTGCGTG | 104 |
| chr8:106146784-106148799 | p5 | (CTTCT)4 | TTTCCCGAAGTGAATGAAGAA | CGTGTTGCAATGCCTTAATTT | 159 |
| chr8:106146784-106148799 | p5 | (TTCTC)3 | TTTCCCGAAGTGAATGAAGAA | CGTGTTGCAATGCCTTAATTT | 159 |
| chr8:106146784-106148799 | p5 | (TCTCT)3 | TTTCCCGAAGTGAATGAAGAA | CGTGTTGCAATGCCTTAATTT | 159 |
| chr8:106171883-106173898 | p3 | (TGT)5 | AAGCCCTCCAAAAGGATGAT | GCCCACGACCTATGGTTAGA | 259 |

| Marker ID | SSR type | SSR motif | FORWARD PRIMER (5'-3') | REVERSE PRIMER (5'-3') | size (bp) |
|---|---|---|---|---|---|
| chr8:106200045-106202063 | p6 | (CATATA)3 | CTCACAATATGCCTTCCGGT | CGCAAGTATGTTGGGGACTT | 115 |
| chr8:106207567-106209653 | p2 | (AT)38 | TCCCTTCTTGCTCCCCTAAT | GTTGCAGGGCATTTAGGAAA | 252 |
| chr8:106207567-106209653 | p6 | (CATATA)3 | TCCCTTCTTGCTCCCCTAAT | GTTGCAGGGCATTTAGGAAA | 252 |
| chr8:106215342-106217360 | p3 | (CAA)6 | GGGACTCGGCGAATTATGTA | TATGTTTTCAATAGGCGGGC | 263 |
| chr8:106215342-106217360 | p2 | (AC)6 | GGGACTCGGCGAATTATGTA | TATGTTTTCAATAGGCGGGC | 263 |
| chr8:106231527-106233542 | p5 | (TTTTC)3 | TATGCCCAAACTGAAGACCC | GTAGGCACCCTGCTGTGATT | 154 |
| chr8:106250809-106252823 | p2 | (TA)7 | TTCCTCTCCATCTCGTCCAT | ACACGGATTCCTTGGTCTTG | 185 |
| chr8:106250809-106252823 | p2 | (TA)6 | TTCCTCTCCATCTCGTCCAT | ACACGGATTCCTTGGTCTTG | 185 |
| chr8:106687657-106689669 | p2 | (AT)6 | GTGCGTGTCAATCCATGAAA | ACGGCGGAATAGCAAATACA | 242 |
| chr8:106793539-106795565 | p5 | (CCGGT)3 | TCCGGCTTTAGTTTGCAAGT | TCTTCCATTTGGTCTGGTCC | 246 |
| chr8:106814712-106816730 | p6 | (GACGTG)3 | ACGGTGGATGTTGTTCCATT | TGGGTTTGGAAGAATTACGC | 110 |
| chr8:107070975-107072989 | p2 | (TA)7 | CGAAACTTGAGGGCAAAAAG | TGTGCAATGGACGTTGAAAT | 262 |
| chr8:107206961-107208979 | p2 | (AT)9 | TCACGTCCACGTTCATTCAT | TCTGTGATGTCTTTGCCAGC | 136 |
| chr8:107227172-107229184 | p2 | (AT)10 | GCCATACGATTGGCTGAAAT | GAAAAGAAGAACGGCCGAC | 130 |
| chr8:107227172-107229184 | p2 | (TC)6 | GCCATACGATTGGCTGAAAT | GAAAAGAAGAACGGCCGAC | 130 |
| chr8:107314465-107316480 | p6 | (CAGTTT)3 | CTAGGGGTGTACATGGACCG | GACAACCCGAGGTTGAAAAA | 210 |
| chr8:107314465-107316480 | p5 | (AAACC)3 | CTAGGGGTGTACATGGACCG | GACAACCCGAGGTTGAAAAA | 210 |
| chr8:107371827-107373842 | p3 | (TTC)5 | CATGCATGCCCTAATGATTG | CAGGGATTTTAGGTTTGGCA | 200 |
| chr8:107371827-107373842 | p2 | (TC)6 | CATGCATGCCCTAATGATTG | CAGGGATTTTAGGTTTGGCA | 200 |
| chr8:107391000-107393014 | p2 | (AC)7 | ATTCCGCTCCTCTCCATTTT | TTGCTCGATTTGTGGAGTTG | 276 |
| chr8:107399281-107401293 | p2 | (AT)6 | CGAGTGTCGTGTTACTCCCA | AAAGTGACCCCACTGGACAC | 167 |
| chr8:107399281-107401293 | p5 | (AAACC)3 | CGAGTGTCGTGTTACTCCCA | AAAGTGACCCCACTGGACAC | 167 |
| chr8:107400133-107402148 | p2 | (AT)6 | GGTCCGGTTTGTGGTTAAAA | GATCACGGTCAAAGCCGTAT | 253 |
| chr8:107400133-107402148 | p5 | (AAACC)3 | GGTCCGGTTTGTGGTTAAAA | GATCACGGTCAAAGCCGTAT | 253 |
| chr8:107400133-107402148 | p3 | (GAA)5 | GGTCCGGTTTGTGGTTAAAA | GATCACGGTCAAAGCCGTAT | 253 |
| chr8:107621413-107623425 | p2 | (GA)6 | CTGCTGGTGGTGAAGGGTAT | GGTACTGCCTCATTCCCTGA | 141 |
| chr8:107635064-107637078 | p2 | (TA)8 | CAGCCCTGTTTGGATTCATT | CGGTTTGGGCTTTCAAACTA | 194 |
| chr8:107716938-107718998 | p3 | (CCA)5 | CTTTAACCCACACCCACCAC | TGCTGCTGTTTTTCATCCAG | 142 |
| chr8:107716938-107718998 | p3 | (GAA)5 | CTTTAACCCACACCCACCAC | TGCTGCTGTTTTTCATCCAG | 142 |
| chr8:107716938-107718998 | p2 | (AT)22 | CTTTAACCCACACCCACCAC | TGCTGCTGTTTTTCATCCAG | 142 |
| chr8:107812595-107814613 | p3 | (TTG)5 | TCTATTTCCATGTCCCCTGC | GCAACAACAACAACAATGGC | 152 |
| chr8:107812595-107814613 | p6 | (CTGGTG)4 | TCTATTTCCATGTCCCCTGC | GCAACAACAACAACAATGGC | 152 |
| chr8:107819719-107821763 | p2 | (AT)7 | AATTGCCTCAAGGGAGAGGT | GAAAAAGGGAATTTGCACGA | 247 |
| chr8:107820029-107822043 | p2 | (AT)7 | AAGACCACCCCCAAATAAGG | ACGACGCACTGCTAATCGTA | 131 |
| chr8:107866872-107868908 | p3 | (ATA)12 | CAGCCCTTTGCAATGATTTT | ATATTCGGCGAAGTTTGTGG | 258 |
| chr8:107987522-107989536 | p2 | (CT)7 | TCGTGTTACGGTCAACCAAA | AAAATCTGGGCACTCACTGG | 247 |

| Marker ID | SSR type | SSR motif | FORWARD PRIMER (5'-3') | REVERSE PRIMER (5'-3') | size (bp) |
|---|---|---|---|---|---|
| chr8:108020806-108022821 | p3 | (GTA)6 | GGCACCACAGGAGTAGGTGT | CATTGTAAACCATTTGCCCC | 148 |
| chr8:108020806-108022821 | p5 | (TATTG)3 | GGCACCACAGGAGTAGGTGT | CATTGTAAACCATTTGCCCC | 148 |
| chr8:108020806-108022821 | p3 | (TTC)5 | GGCACCACAGGAGTAGGTGT | CATTGTAAACCATTTGCCCC | 148 |
| chr8:108078573-108080595 | p5 | (ACCGG)3 | GCGGTGAGAAACCACAGTCT | TGTTTTTAAACCCGAAACCG | 259 |
| chr8:108078573-108080595 | p2 | (TA)52 | GCGGTGAGAAACCACAGTCT | TGTTTTTAAACCCGAAACCG | 259 |
| chr8:108105091-108107109 | p6 | (CATGAG)3 | CAGATTCTCTCCGATGCTCC | CCCAGCTCGATCAATGATTT | 288 |
| chr8:108192038-108194053 | p5 | (TTGGT)3 | TGCTGGTTCACAACCACAAT | GTGTTTACCCTTTTGTGGGG | 260 |
| chr8:108192038-108194053 | p5 | (TTGGT)3 | TGCTGGTTCACAACCACAAT | GTGTTTACCCTTTTGTGGGG | 260 |
| chr8:108192749-108194764 | p5 | (TTGGT)3 | TGACTTGGTTTGGCGTAGAA | ACACGTAATGTTGGGTCGGT | 252 |
| chr8:108192749-108194764 | p5 | (TTGGT)3 | TGACTTGGTTTGGCGTAGAA | ACACGTAATGTTGGGTCGGT | 252 |
| chr8:108210878-108212896 | p6 | (TAATGT)3 | ATGCTATTCTCCGTTGGTGG | ATGTTTCCCTTCCCATTTCC | 253 |
| chr8:108301688-108303702 | p2 | (AT)7 | TTGAAGGAAGTTTTGGGACG | AACATACCTTGTCGCTCGCT | 245 |
| chr8:108301688-108303702 | p4 | (ATGT)4 | TTGAAGGAAGTTTTGGGACG | AACATACCTTGTCGCTCGCT | 245 |
| chr8:108335049-108337064 | p5 | (ACCGG)3 | CTTTCCTCCTCCATTCCTCC | GTGGTTATTCCCACCGAATG | 141 |
| chr8:108466924-108468942 | p6 | (AAGCAG)3 | TTCAAAGCACTGTGACACCC | CCTTTTGGTGCATGAATGTG | 116 |
| chr8:108479743-108481761 | p6 | (AAGCAG)3 | TTCAAAGCACTGTGACACCC | CCTTTTGGTGCATGAATGTG | 116 |
| chr8:108486507-108488555 | p3 | (ACA)5 | TCGGTAAAGAATGGTCCCAC | AATCCAGTCGATGAACACCC | 129 |
| chr8:108512322-108514338 | p4 | (AATA)4 | ACAGCCACGAATAAACGGAC | TTGGACCAAGACCAAGGAAG | 114 |
| chr8:108517058-108519076 | p2 | (AT)9 | TCCATCTAAAATGGTCCCCA | GCTGAATTTTGGGTTTTGGA | 106 |
| chr8:108631479-108633519 | p2 | (TA)20 | CTTGCCTAACACGCCTCTTT | AGGCAAGTAGGGAGGGCTAC | 261 |
| chr8:108678289-108680325 | p2 | (TA)18 | ATTACCCCCTGAACTTGGCT | GTGCGTGCACTGAGAATGTT | 242 |
| chr8:109000198-109002216 | p6 | (CTCATG)3 | ATAGCTTAGCTGATCGGGCA | AACAACACGGGACATGGAAT | 253 |
| chr8:109000198-109002216 | p6 | (CTCATG)3 | ATAGCTTAGCTGATCGGGCA | AACAACACGGGACATGGAAT | 253 |
| chr8:109007244-109009271 | p3 | (TTG)9 | TGTTGGAGATGCTTGTGAGG | TTTCTTCTCCTGCAGCCATT | 211 |
| chr8:109007244-109009271 | p3 | (TTA)9 | TGTTGGAGATGCTTGTGAGG | TTTCTTCTCCTGCAGCCATT | 211 |
| chr8:109023732-109025747 | p5 | (TGGGG)3 | AGAAGGATCCCCCGTACATC | ATCCCGAGAGCAAGGAATTT | 100 |
| chr8:109137284-109139298 | p2 | (CT)7 | CCGTCGAATGAGTTACGGTT | TGTACGGGGGATTTTCTGAG | 278 |
| chr8:109196244-109198258 | p2 | (TC)7 | TGTATGCCCTTCCAATCTCC | CTGGCAGGCATTTGTCTTTT | 244 |
| chr8:109222518-109224542 | p6 | (TTTTTG)4 | ACAAGGTTCGAACAATGCCT | CCACATGTTTAGCCGTTCCT | 206 |
| chr8:109367237-109369255 | p3 | (GAA)6 | ACAGTGGCTTCTGGGAGAGA | AGGTTTTCCAGGGCTCTGTT | 259 |
| chr8:109371656-109373668 | p3 | (TAA)9 | ACGATGTTCCAGCTCCATCT | GCAAGTAACATGGCCACAAA | 198 |
| chr8:109371656-109373668 | p3 | (ATA)17 | ACGATGTTCCAGCTCCATCT | GCAAGTAACATGGCCACAAA | 198 |
| chr8:109371728-109373740 | p3 | (TAA)9 | ACGATGTTCCAGCTCCATCT | AACATGGCCACAAAAACCTG | 126 |
| chr8:109371728-109373740 | p3 | (ATA)19 | ACGATGTTCCAGCTCCATCT | AACATGGCCACAAAAACCTG | 126 |
| chr8:109387653-109389667 | p2 | (TA)7 | TCACAAATTCCACAACCCCT | TGTCCTTTCGCCAAACCTAC | 199 |
| chr8:109465367-109467382 | p3 | (TAA)5 | GATAGTGGAGGTGCGGGTAA | GTCGCCTCCTTCTCCTCTTT | 211 |

| Marker ID | SSR type | SSR motif | FORWARD PRIMER (5'-3') | REVERSE PRIMER (5'-3') | size (bp) |
|---|---|---|---|---|---|
| chr8:109465367-109467382 | p5 | (TTATT)4 | GATAGTGGAGGTGCGGGTAA | GTCGCCTCCTTCTCCTCTTT | 211 |
| chr8:109468501-109470519 | p2 | (AT)9 | TCCAGTCCGGTGACATACAA | CCCGGGGTTAATTGATACCT | 240 |
| chr8:109497125-109499137 | p2 | (AT)6 | AATCCGATGTTCCAAATCCA | TAGAGTCGGTAGGGGTGTGC | 272 |
| chr8:109544083-109546095 | p2 | (TC)6 | AAGGACGGACCGTAACACAG | TTGCCTTAAATGCTCCAACC | 268 |
| chr8:109566233-109568345 | p2 | (TA)7 | GCCTGTTTGCCTTTTTGTGT | GTCTCGGGATCATCACCACT | 250 |
| chr8:109615037-109617055 | p3 | (TTG)6 | TGGACTAGTCACGGGGAAAC | CAGACAATTGAGAGTGCGGA | 119 |
| chr8:110094283-110096298 | p5 | (TACTT)3 | TAGAAGCTTGGCCAAACAGG | AAGGGAGCACATTTTGATGG | 262 |
| chr8:110119732-110121747 | p5 | (TTTTC)3 | TGGCATGGCAACTAAATCAA | TCTAAGATCCCGATCCAACG | 171 |
| chr8:110119732-110121747 | p5 | (ACCAA)3 | TGGCATGGCAACTAAATCAA | TCTAAGATCCCGATCCAACG | 171 |
| chr8:110119732-110121747 | p5 | (AAACC)3 | TGGCATGGCAACTAAATCAA | TCTAAGATCCCGATCCAACG | 171 |
| chr8:110146020-110148035 | p3 | (AAT)5 | CACTGCCTCCTTCTTTTTCG | GTCATCGCCAGAGTTCCATT | 201 |
| chr8:110173581-110175593 | p2 | (AT)26 | TTGGGATGTTCCCGTATGAT | GTCTGAGGAAGCACGTGTCA | 290 |
| chr8:110173581-110175593 | p2 | (GA)6 | TTGGGATGTTCCCGTATGAT | GTCTGAGGAAGCACGTGTCA | 290 |
| chr8:110381931-110383946 | p3 | (ATC)5 | ATGCTCCAGGAAGAAGTGGA | CTTGTGGACGCTGTGTTCAT | 132 |
| chr8:110384309-110386324 | p3 | (ATC)5 | TGTCGACCCTTAAATCCTGC | TTTTAGAGTCACCCATGCCC | 166 |
| chr8:110398196-110400228 | p2 | (TA)16 | GGAATTTTAACAGCCCGTCA | ACATGGGCCTCTTGATTCTG | 119 |
| chr8:110627026-110629038 | p2 | (AT)6 | CGGGTTCCATCAAAATATGC | AACATGTGATTGGTTTGGCA | 204 |
| chr8:110627026-110629038 | p2 | (AT)7 | CGGGTTCCATCAAAATATGC | AACATGTGATTGGTTTGGCA | 204 |
| chr8:110627026-110629038 | p2 | (AT)8 | CGGGTTCCATCAAAATATGC | AACATGTGATTGGTTTGGCA | 204 |
| chr8:110627388-110629402 | p2 | (AT)6 | ACAGCAAGGGAGCAGATCAT | GTTAAGCGGTTCCGGTTTTT | 268 |
| chr8:110627388-110629402 | p2 | (AT)7 | ACAGCAAGGGAGCAGATCAT | GTTAAGCGGTTCCGGTTTTT | 268 |
| chr8:110627388-110629402 | p2 | (AT)8 | ACAGCAAGGGAGCAGATCAT | GTTAAGCGGTTCCGGTTTTT | 268 |
| chr8:110627517-110629533 | p2 | (AT)6 | AAGGGAGCAGATCATGCCTA | CCTTAATCGGTTACCGGTCC | 144 |
| chr8:110627517-110629533 | p2 | (AT)7 | AAGGGAGCAGATCATGCCTA | CCTTAATCGGTTACCGGTCC | 144 |
| chr8:110627517-110629533 | p2 | (AT)8 | AAGGGAGCAGATCATGCCTA | CCTTAATCGGTTACCGGTCC | 144 |
| chr8:110732940-110734960 | p2 | (TA)24 | TTTTAAGGGACAAACGGACG | GGTGCCCCCATACTAAACAA | 296 |
| chr8:110732940-110734960 | p5 | (AGTAA)4 | TTTTAAGGGACAAACGGACG | GGTGCCCCCATACTAAACAA | 296 |
| chr8:110772113-110774135 | p2 | (TA)11 | GCGGGTTCTAGGATTTGTGA | CCAACTTAAACCAGCCCAAA | 235 |
| chr8:110772113-110774135 | p2 | (TA)6 | GCGGGTTCTAGGATTTGTGA | CCAACTTAAACCAGCCCAAA | 235 |
| chr8:110772113-110774135 | p2 | (TA)15 | GCGGGTTCTAGGATTTGTGA | CCAACTTAAACCAGCCCAAA | 235 |
| chr8:110863957-110865973 | p2 | (CA)7 | ATGCCTTTGGAATTGTGCTC | ATGACATGGAAGGCCTCTTG | 218 |
| chr8:110863974-110865996 | p2 | (CA)7 | ATGCCTTTGGAATTGTGCTC | ATGACATGGAAGGCCTCTTG | 201 |
| chr8:110965230-110967245 | p5 | (AATAA)3 | TTTGGTTTGGTTTAGGGGGT | TACCTGGTGGCTTTCCAAAC | 112 |
| chr8:110987631-110989646 | p3 | (GAA)5 | TCCCTTTCTTCCACATCCAC | CGCCTCAGCTCACTTCTTCT | 250 |
| chr8:111104030-111106042 | p2 | (AT)7 | AGCACTTTCGTTTCGGCTAA | CCGGAACAGTGTTGATTGTG | 193 |
| chr8:111104030-111106042 | p2 | (AT)6 | AGCACTTTCGTTTCGGCTAA | CCGGAACAGTGTTGATTGTG | 193 |

| Marker ID | SSR type | SSR motif | FORWARD PRIMER (5'-3') | REVERSE PRIMER (5'-3') | size (bp) |
|---|---|---|---|---|---|
| chr8:111129947-111131965 | p6 | (AAAGAA)3 | AACAAATTGGGTATCCGTGC | GCGACTGCAACCTGTCACTA | 207 |
| chr8:111150989-111153043 | p2 | (TA)27 | TCATTTTTAACTGAGCGGGG | AGTGCTGTTGTGGGCTTTCT | 155 |
| chr8:111225978-111227999 | p3 | (TAA)7 | TCTTTGAACTTCCCCATTGC | GTACTGGGCTAATCGTGCGT | 124 |
| chr8:111225978-111227999 | p2 | (TG)7 | TCTTTGAACTTCCCCATTGC | GTACTGGGCTAATCGTGCGT | 124 |
| chr8:111225978-111227999 | p2 | (AT)13 | TCTTTGAACTTCCCCATTGC | GTACTGGGCTAATCGTGCGT | 124 |
| chr8:111226561-111228587 | p3 | (TAA)7 | CATGTATTGAATTGGCACGC | CCCAATTTATGTGGCACCTT | 139 |
| chr8:111226561-111228587 | p2 | (TG)7 | CATGTATTGAATTGGCACGC | CCCAATTTATGTGGCACCTT | 139 |
| chr8:111226561-111228587 | p2 | (AT)13 | CATGTATTGAATTGGCACGC | CCCAATTTATGTGGCACCTT | 139 |
| chr8:111246525-111248549 | p2 | (AT)6 | TACGTAGGGGTGTTCATGGG | GAAAAACTGACCCAAACCGA | 195 |
| chr8:111270085-111272100 | p5 | (CAAAC)3 | TAGTGGCTGCACCATTTTTG | AATTGAGGGCAAAATGCAAG | 126 |
| chr8:111321146-111323161 | p5 | (CCCTT)3 | AAATGGACAATCAGATCCGC | GTCAATCATGGCAGATCCCT | 175 |
| chr8:111454954-111456969 | p5 | (TTCGG)3 | ATTGGTGTCAATGGGACGAT | AAGGGACGGTCTCGAAAAGT | 232 |
| chr8:111510636-111512688 | p5 | (ACCGG)3 | TTCTTCTTTTTCCGTGTGGC | CGGTTGACGGGCTTACTAGA | 300 |
| chr8:111510636-111512688 | p2 | (TA)20 | TTCTTCTTTTTCCGTGTGGC | CGGTTGACGGGCTTACTAGA | 300 |
| chr8:111510636-111512688 | p2 | (AT)26 | TTCTTCTTTTTCCGTGTGGC | CGGTTGACGGGCTTACTAGA | 300 |
| chr8:111570566-111572618 | p5 | (ACCGG)3 | TTCTTCTTTTTCCGTGTGGC | CGGTTGACGGGCTTACTAGA | 300 |
| chr8:111570566-111572618 | p2 | (TA)20 | TTCTTCTTTTTCCGTGTGGC | CGGTTGACGGGCTTACTAGA | 300 |
| chr8:111570566-111572618 | p2 | (AT)26 | TTCTTCTTTTTCCGTGTGGC | CGGTTGACGGGCTTACTAGA | 300 |
| chr8:111616137-111618155 | p6 | (AACAAA)3 | AAAATTGGCCAGGTGATTTG | GTTGAACTTGTTGTGGTGCG | 181 |
| chr8:111616137-111618155 | p2 | (AG)6 | AAAATTGGCCAGGTGATTTG | GTTGAACTTGTTGTGGTGCG | 181 |
| chr8:111619992-111622004 | p2 | (CT)6 | TGGGTTCGGTTAGGATTCAA | TCCTCCTCACTCCAGCTGTT | 164 |
| chr8:111643587-111645605 | p6 | (TTATTT)3 | ATTCTGCCAGGGTTGTTGTC | ACAAGCGATTGGGAGCTAAA | 114 |
| chr8:111646379-111648394 | p3 | (ATA)5 | TGTTTTTGGAGTTGCACGAA | GAGGTGGTAACGTGCCTGAT | 242 |
| chr8:111646379-111648394 | p2 | (TA)6 | TGTTTTTGGAGTTGCACGAA | GAGGTGGTAACGTGCCTGAT | 242 |
| chr8:111654229-111656245 | p2 | (CA)8 | ATTTTCCCGAATATCCGCTT | TCGAATGCTAACTGTGCGTC | 185 |
| chr8:111668216-111670232 | p2 | (CA)8 | ATTTTCCCGAATATCCGCTT | TCGAATGCTAACTGTGCGTC | 185 |
| chr8:111676741-111678766 | p2 | (AT)20 | AACTCACGGTGGCACATACA | TCACTCCCAAACCACCAAAT | 299 |
| chr8:111676741-111678766 | p5 | (AATAA)5 | AACTCACGGTGGCACATACA | TCACTCCCAAACCACCAAAT | 299 |
| chr8:111700235-111702274 | p6 | (CGATGT)4 | ATTGGGCTTGACATCTTTGG | AACAGCGGGGGAGAGTAGTT | 117 |
| chr8:111751654-111753669 | p5 | (TTAGT)3 | CCGTCTCTCTCATACGCTCC | CGGTTAAAGTTCACCGGAAA | 150 |
| chr8:111816079-111818133 | p3 | (TTA)18 | ATTCAAGCAATCAAATCGGC | AGCTGCTGCTTTCCTTATGC | 228 |
| chr8:111848879-111850893 | p2 | (AT)7 | TTCTTAGCTGCACCCTTCGT | TTATGAGGTGGACGCACAAA | 120 |
| chr8:111863984-111866030 | p2 | (TA)9 | CTCCTTTCATCGTCCGTTGT | ATATGGGGCCCACAATTTTT | 139 |
| chr8:111863984-111866030 | p2 | (TA)6 | CTCCTTTCATCGTCCGTTGT | ATATGGGGCCCACAATTTTT | 139 |
| chr8:111863984-111866030 | p2 | (TA)23 | CTCCTTTCATCGTCCGTTGT | ATATGGGGCCCACAATTTTT | 139 |
| chr8:111863984-111866030 | p5 | (TTTGT)3 | CTCCTTTCATCGTCCGTTGT | ATATGGGGCCCACAATTTTT | 139 |

| Marker ID | SSR type | SSR motif | FORWARD PRIMER (5'-3') | REVERSE PRIMER (5'-3') | size (bp) |
|---|---|---|---|---|---|
| chr8:111869957-111871969 | p2 | (AG)6 | AACGCGACGACAAAGAAACT | CGCACATAGGGACTTTTGGT | 258 |
| chr8:111869957-111871969 | p2 | (TA)27 | AACGCGACGACAAAGAAACT | CGCACATAGGGACTTTTGGT | 258 |
| chr8:111869957-111871969 | p4 | (TTTC)4 | AACGCGACGACAAAGAAACT | CGCACATAGGGACTTTTGGT | 258 |
| chr8:111870488-111872542 | p2 | (AG)6 | GATTTTGGGGTTCCAAACCT | CGCTCCCCTCTCTCTTCTCT | 108 |
| chr8:111870488-111872542 | p2 | (TA)27 | GATTTTGGGGTTCCAAACCT | CGCTCCCCTCTCTCTTCTCT | 108 |
| chr8:111870488-111872542 | p4 | (TTTC)4 | GATTTTGGGGTTCCAAACCT | CGCTCCCCTCTCTCTTCTCT | 108 |
| chr8:111870772-111872788 | p2 | (AG)6 | AAGGGGTTAGGTTTAGGGGG | GGACACACATTTTTCGGACC | 233 |
| chr8:111870772-111872788 | p2 | (TA)27 | AAGGGGTTAGGTTTAGGGGG | GGACACACATTTTTCGGACC | 233 |
| chr8:111870772-111872788 | p4 | (TTTC)4 | AAGGGGTTAGGTTTAGGGGG | GGACACACATTTTTCGGACC | 233 |
| chr8:111870772-111872788 | p5 | (AAAAT)4 | AAGGGGTTAGGTTTAGGGGG | GGACACACATTTTTCGGACC | 233 |
| chr8:111890801-111892813 | p2 | (AC)6 | TGGGATTGGTTAGCAAGGTC | GCAGTATGTGAAGCAGCCAA | 177 |
| chr8:111942975-111945013 | p2 | (GA)6 | CGGGATTGCATTCAAGAGAT | GACGAGCAATCAGAACAGCA | 169 |
| chr8:111997063-111999078 | p5 | (TACAT)3 | CTTCGTTGAGGAAAGCCAAG | GCGAGCTTTACGATGAGTCC | 143 |
| chr8:112024712-112026730 | p2 | (TA)9 | AGGTGGAACAATGAGATGCC | GCAACTGCCATGACACAAAC | 217 |
| chr8:112025795-112027825 | p2 | (TA)15 | TATTCGGTACCCATTGGAGC | AAATAAAGTGCAGCGGATGG | 139 |
| chr8:112071823-112073838 | p5 | (TACAT)3 | CTTCGTTGAGGAAAGCCAAG | GCGAGCTTTACGATGAGTCC | 143 |
| chr8:112099472-112101490 | p2 | (TA)9 | AGGTGGAACAATGAGATGCC | GCAACTGCCATGACACAAAC | 217 |
| chr8:112100555-112102585 | p2 | (TA)15 | TATTCGGTACCCATTGGAGC | AAATAAAGTGCAGCGGATGG | 139 |
| chr8:112154177-112156213 | p3 | (TAA)12 | CCCTATCCCCTTTCTCTTCG | ACGCTCCCCTCCACTTCTAT | 197 |
| chr8:112203098-112205122 | p2 | (GA)13 | AGGTCTGCGATTTTGTTGCT | CCAGTCCACTAACCCTTCCA | 213 |
| chr8:112203125-112205176 | p2 | (GA)13 | AGGTCTGCGATTTTGTTGCT | CCAGTCCACTAACCCTTCCA | 186 |
| chr8:112270130-112272148 | p3 | (TGA)6 | TATTAACCGGATTCGCTTGG | GGCCAAAATGACCGTTTTTA | 258 |
| chr8:112301722-112303754 | p3 | (TGA)5 | AGACGTGGTGCCGGTAATAG | TTAGCGGTTGGCTTACGAGT | 239 |
| chr8:112301722-112303754 | p2 | (AT)16 | AGACGTGGTGCCGGTAATAG | TTAGCGGTTGGCTTACGAGT | 239 |
| chr8:112301722-112303754 | p2 | (AT)10 | AGACGTGGTGCCGGTAATAG | TTAGCGGTTGGCTTACGAGT | 239 |
| chr8:112301942-112303962 | p3 | (TGA)5 | ACTCGTAAGCCAACCGCTAA | ACATTTGAACCCGTTTGCTC | 266 |
| chr8:112301942-112303962 | p2 | (AT)16 | ACTCGTAAGCCAACCGCTAA | ACATTTGAACCCGTTTGCTC | 266 |
| chr8:112301942-112303962 | p2 | (AT)10 | ACTCGTAAGCCAACCGCTAA | ACATTTGAACCCGTTTGCTC | 266 |
| chr8:112319057-112321077 | p2 | (AT)10 | GCAGGGAAATGATCAGGAAA | GCAGCAGCTCACACAGGATA | 151 |
| chr8:112348508-112350523 | p5 | (CCGGT)3 | GTCTTAGCATGGCGGAAGAG | TCCAATCACGAATGCACACT | 161 |
| chr8:112348508-112350523 | p5 | (ATTTT)3 | GTCTTAGCATGGCGGAAGAG | TCCAATCACGAATGCACACT | 161 |
| chr8:112409911-112411935 | p2 | (GA)12 | GGAATGAGGGAAAGGGAGAG | TCAGCCAAATTAATGGCACA | 151 |
| chr8:112439313-112441328 | p5 | (TTTTC)3 | CGGCCAAAAATTAAAGACCA | ATTAGGGTCCAGCAGCATTG | 272 |
| chr8:112471245-112473267 | p2 | (TA)11 | ATGGCTGTCCCGTCTTTATG | TAGCATAATCTGCATGGGCA | 271 |
| chr8:112478005-112480032 | p2 | (TA)8 | ACCTTTGAGCCCCCAAATAC | ACAAAATTGGGCGTCAACAT | 281 |
| chr8:112478042-112480066 | p2 | (TA)8 | ACCTTTGAGCCCCCAAATAC | ACAAAATTGGGCGTCAACAT | 244 |

| Marker ID | SSR type | SSR motif | FORWARD PRIMER (5'-3') | REVERSE PRIMER (5'-3') | size (bp) |
|---|---|---|---|---|---|
| chr8:112478042-112480066 | p5 | (TTCTC)3 | ACCTTTGAGCCCCCAAATAC | ACAAAATTGGGCGTCAACAT | 244 |
| chr8:112478063-112480093 | p2 | (TA)8 | ACCTTTGAGCCCCCAAATAC | ACAAAATTGGGCGTCAACAT | 223 |
| chr8:112478063-112480093 | p5 | (TTCTC)3 | ACCTTTGAGCCCCCAAATAC | ACAAAATTGGGCGTCAACAT | 223 |
| chr8:112478096-112480111 | p2 | (TA)8 | ACCTTTGAGCCCCCAAATAC | ACAAAATTGGGCGTCAACAT | 190 |
| chr8:112478096-112480111 | p5 | (TTCTC)3 | ACCTTTGAGCCCCCAAATAC | ACAAAATTGGGCGTCAACAT | 190 |
| chr8:112478108-112480126 | p2 | (TA)8 | ACCTTTGAGCCCCCAAATAC | ACAAAATTGGGCGTCAACAT | 178 |
| chr8:112478108-112480126 | p5 | (TTCTC)3 | ACCTTTGAGCCCCCAAATAC | ACAAAATTGGGCGTCAACAT | 178 |
| chr8:112493677-112495693 | p2 | (AC)8 | TGTTGCCTAATCCATGACCA | TGGGACGTAGAGGAAGGAGA | 124 |
| chr8:112493677-112495693 | p5 | (TGGTT)3 | TGTTGCCTAATCCATGACCA | TGGGACGTAGAGGAAGGAGA | 124 |
| chr8:112577765-112579783 | p2 | (TA)7 | TCCCCATCATCACCAAAACT | TTAAACCGGAACGGTTATGC | 279 |
| chr8:112604454-112606474 | p4 | (ATAA)5 | TCGGGACTAGCAGGATATGG | GCTAGAGTTAGACCGCCGTG | 116 |
| chr8:112614161-112616194 | p3 | (ATT)11 | GGTTGGAACGGTTGCTATGT | ACCACCTCCGTCTCCATACA | 131 |
| chr8:112616018-112618033 | p5 | (AGTTA)3 | TTTTCTACGCTCCCCATCAC | CCAGATTCTTTGGGGGTTTT | 197 |
| chr8:112628560-112630572 | p2 | (TC)6 | TGCCATTGTTTTTGGTCTGA | CTTTTCACCTTGGCTCTTGC | 300 |
| chr8:112644527-112646539 | p2 | (TC)6 | TGCCATTGTTTTTGGTCTGA | CTTTTCACCTTGGCTCTTGC | 300 |
| chr8:112652879-112654937 | p2 | (AT)29 | TTTTCCCAATTCACGACACA | AATCCGCCTACAAACCCTCT | 243 |
| chr8:112667832-112669847 | p5 | (TTATT)3 | AACTTGCCATCCGTTTTGAC | CTTGGCCTCAATCCACCTTA | 207 |
| chr8:112709292-112711306 | p2 | (AT)7 | CCTTAGTTGGCCTTTCCCTC | TCCCCAGCTTCATATTTTGG | 150 |
| chr8:112771990-112774012 | p2 | (TA)11 | TCCCTTTTTCCTTCTGCTCA | CATTGCCACATGCAAGAACT | 130 |
| chr8:112802838-112804862 | p4 | (AAAT)4 | CCTCGTAGTGGTTGGAGGAA | GTTGGTGATGGGTCGGATAC | 274 |
| chr8:112860294-112862306 | p2 | (AG)6 | TATAGGTGTCAAGGGGTCGG | TGCGGCGAGAGAAATAGAAT | 163 |
| chr8:112883378-112885390 | p2 | (CT)11 | AAAGTATTGTGGCCACTCGC | GGGGCAAGTGAGTGAGAGAG | 182 |
| chr8:112883378-112885390 | p2 | (TA)6 | AAAGTATTGTGGCCACTCGC | GGGGCAAGTGAGTGAGAGAG | 182 |
| chr8:112919088-112921112 | p2 | (TC)12 | CGAGTAGCCGCAATTTCATT | CGTAAAACGGAAAGAACCCA | 181 |
| chr8:112920581-112922617 | p2 | (AT)18 | ATTGCCATGGTTTTTCCAAG | AACACCGAAAATTGCTGAGG | 200 |
| chr8:112920581-112922617 | p2 | (AT)6 | ATTGCCATGGTTTTTCCAAG | AACACCGAAAATTGCTGAGG | 200 |
| chr8:112921978-112923996 | p2 | (AT)6 | CCAAAAGAGTGAGCCCCATA | TAAGAGACAACGGACGACGA | 270 |
| chr8:112921978-112923996 | p2 | (TA)9 | CCAAAAGAGTGAGCCCCATA | TAAGAGACAACGGACGACGA | 270 |
| chr8:112961817-112963839 | p2 | (TC)11 | CTCGGAAGGACGAACGATAG | CTATCCCCGCAAATCAAGAA | 158 |
| chr8:112972295-112974309 | p2 | (AT)7 | AACCGTAAATTCCCCAAAGG | TCCAACAACCAAATCGATCA | 116 |
| chr8:112987376-112989404 | p2 | (TA)14 | CGTCCGTTGTCCCTTAAAAA | TTTCCCGTGAGTTTGGATGT | 291 |
| chr8:113031020-113033088 | p2 | (AT)34 | TGGTAGTGGTTGCGTTTTGA | CAGTGCATTTTGCCAATGTC | 298 |
| chr8:113052704-113054719 | p5 | (TTTTA)3 | CTGAAGGGGGCAATCTCATA | TTTCCCAATTGCCCAAGTAA | 221 |
| chr8:113052704-113054719 | p6 | (CCCACC)3 | CTGAAGGGGGCAATCTCATA | TTTCCCAATTGCCCAAGTAA | 221 |
| chr8:113100424-113102439 | p6 | (AAGAGG)3 | CGGGTCAAATGAAATATGGG | TTAAATAGCGGAGGAAGGCA | 235 |
| chr8:113100424-113102439 | p3 | (ACA)5 | CGGGTCAAATGAAATATGGG | TTAAATAGCGGAGGAAGGCA | 235 |

| Marker ID | SSR type | SSR motif | FORWARD PRIMER (5'-3') | REVERSE PRIMER (5'-3') | size (bp) |
|---|---|---|---|---|---|
| chr8:113177125-113179140 | p5 | (AAATC)3 | CAATTTGGCGTCTTGAGCTT | GTTCAACTCCCTGGTTGGAA | 230 |
| chr8:113177125-113179140 | p2 | (AT)6 | CAATTTGGCGTCTTGAGCTT | GTTCAACTCCCTGGTTGGAA | 230 |
| chr8:113177125-113179140 | p3 | (TTA)5 | CAATTTGGCGTCTTGAGCTT | GTTCAACTCCCTGGTTGGAA | 230 |
| chr8:113190427-113192442 | p5 | (AATTT)3 | GCTTGAGCCTGCACATAACA | AGGCCAGTGCCCTTTATCTT | 233 |
| chr8:113209790-113211817 | p2 | (GT)6 | AACTGCGCTGGAAGTAGGAA | ACCACTAGTCCCACAACCCA | 270 |
| chr8:113209790-113211817 | p2 | (GA)6 | AACTGCGCTGGAAGTAGGAA | ACCACTAGTCCCACAACCCA | 270 |
| chr8:113209790-113211817 | p2 | (GA)6 | AACTGCGCTGGAAGTAGGAA | ACCACTAGTCCCACAACCCA | 270 |
| chr8:113209790-113211817 | p2 | (GA)6 | AACTGCGCTGGAAGTAGGAA | ACCACTAGTCCCACAACCCA | 270 |
| chr8:113209790-113211817 | p2 | (GA)6 | AACTGCGCTGGAAGTAGGAA | ACCACTAGTCCCACAACCCA | 270 |
| chr8:113209790-113211817 | p2 | (GA)6 | AACTGCGCTGGAAGTAGGAA | ACCACTAGTCCCACAACCCA | 270 |
| chr8:113209790-113211817 | p2 | (GA)6 | AACTGCGCTGGAAGTAGGAA | ACCACTAGTCCCACAACCCA | 270 |
| chr8:113209790-113211817 | p2 | (GA)6 | AACTGCGCTGGAAGTAGGAA | ACCACTAGTCCCACAACCCA | 270 |
| chr8:113209790-113211817 | p2 | (GA)6 | AACTGCGCTGGAAGTAGGAA | ACCACTAGTCCCACAACCCA | 270 |
| chr8:113209790-113211817 | p2 | (GA)6 | AACTGCGCTGGAAGTAGGAA | ACCACTAGTCCCACAACCCA | 270 |
| chr8:113209904-113211931 | p2 | (GT)6 | AACTGCGCTGGAAGTAGGAA | ACCACTAGTCCCACAACCCA | 156 |
| chr8:113209904-113211931 | p2 | (GA)6 | AACTGCGCTGGAAGTAGGAA | ACCACTAGTCCCACAACCCA | 156 |
| chr8:113209904-113211931 | p2 | (GA)6 | AACTGCGCTGGAAGTAGGAA | ACCACTAGTCCCACAACCCA | 156 |
| chr8:113209904-113211931 | p2 | (GA)6 | AACTGCGCTGGAAGTAGGAA | ACCACTAGTCCCACAACCCA | 156 |
| chr8:113209904-113211931 | p2 | (GA)6 | AACTGCGCTGGAAGTAGGAA | ACCACTAGTCCCACAACCCA | 156 |
| chr8:113209904-113211931 | p2 | (GA)6 | AACTGCGCTGGAAGTAGGAA | ACCACTAGTCCCACAACCCA | 156 |
| chr8:113209904-113211931 | p2 | (GA)6 | AACTGCGCTGGAAGTAGGAA | ACCACTAGTCCCACAACCCA | 156 |
| chr8:113209904-113211931 | p2 | (GA)6 | AACTGCGCTGGAAGTAGGAA | ACCACTAGTCCCACAACCCA | 156 |
| chr8:113209904-113211931 | p2 | (GA)6 | AACTGCGCTGGAAGTAGGAA | ACCACTAGTCCCACAACCCA | 156 |
| chr8:113209904-113211931 | p2 | (GA)6 | AACTGCGCTGGAAGTAGGAA | ACCACTAGTCCCACAACCCA | 156 |
| chr8:113209904-113211931 | p2 | (GA)6 | AACTGCGCTGGAAGTAGGAA | ACCACTAGTCCCACAACCCA | 156 |
| chr8:113210360-113212387 | p2 | (GT)6 | CAAACGAGGCTTTTCCATGT | TCTCTCTCTCCCACACCCAC | 212 |
| chr8:113210360-113212387 | p2 | (GA)6 | CAAACGAGGCTTTTCCATGT | TCTCTCTCTCCCACACCCAC | 212 |
| chr8:113210360-113212387 | p2 | (GA)6 | CAAACGAGGCTTTTCCATGT | TCTCTCTCTCCCACACCCAC | 212 |
| chr8:113210360-113212387 | p2 | (GA)6 | CAAACGAGGCTTTTCCATGT | TCTCTCTCTCCCACACCCAC | 212 |
| chr8:113210360-113212387 | p2 | (GA)6 | CAAACGAGGCTTTTCCATGT | TCTCTCTCTCCCACACCCAC | 212 |
| chr8:113210360-113212387 | p2 | (GA)6 | CAAACGAGGCTTTTCCATGT | TCTCTCTCTCCCACACCCAC | 212 |
| chr8:113210360-113212387 | p2 | (GA)6 | CAAACGAGGCTTTTCCATGT | TCTCTCTCTCCCACACCCAC | 212 |
| chr8:113210360-113212387 | p2 | (GA)6 | CAAACGAGGCTTTTCCATGT | TCTCTCTCTCCCACACCCAC | 212 |
| chr8:113210360-113212387 | p2 | (GA)6 | CAAACGAGGCTTTTCCATGT | TCTCTCTCTCCCACACCCAC | 212 |
| chr8:113210360-113212387 | p2 | (GA)6 | CAAACGAGGCTTTTCCATGT | TCTCTCTCTCCCACACCCAC | 212 |
| chr8:113210360-113212387 | p2 | (GA)6 | CAAACGAGGCTTTTCCATGT | TCTCTCTCTCCCACACCCAC | 212 |

| Marker ID | SSR type | SSR motif | FORWARD PRIMER (5'-3') | REVERSE PRIMER (5'-3') | size (bp) |
|---|---|---|---|---|---|
| chr8:113258578-113260636 | p2 | (AT)29 | ATCATCAGATCACATGCCGA | AGCTGCATTTGTGTTCAACG | 133 |
| chr8:113261362-113263378 | p4 | (AATT)4 | CGATATCCCTTCGTATCCCA | CAGCCGTCCTATCTCGGTAA | 149 |
| chr8:113323159-113325174 | p3 | (ACA)6 | GCTTGTTCATGCCAGCAGTA | AACATCAGTGCCAAGCTCCT | 194 |
| chr8:113348718-113350733 | p3 | (AAT)7 | TTTTGCACAAACTCTTCCCC | ACCCTCGGCTCAAAAGAAAC | 247 |
| chr8:113348718-113350733 | p3 | (TTA)5 | TTTTGCACAAACTCTTCCCC | ACCCTCGGCTCAAAAGAAAC | 247 |
| chr8:113348718-113350733 | p3 | (TTA)17 | TTTTGCACAAACTCTTCCCC | ACCCTCGGCTCAAAAGAAAC | 247 |
| chr8:113348852-113350903 | p3 | (AAT)7 | TTTTGCACAAACTCTTCCCC | ACCCTCGGCTCAAAAGAAAC | 113 |
| chr8:113348852-113350903 | p3 | (TTA)5 | TTTTGCACAAACTCTTCCCC | ACCCTCGGCTCAAAAGAAAC | 113 |
| chr8:113348852-113350903 | p3 | (TTA)17 | TTTTGCACAAACTCTTCCCC | ACCCTCGGCTCAAAAGAAAC | 113 |
| chr8:113405635-113407677 | p3 | (TTA)14 | TTGGTTGTGGTGTACGGAGA | AAAGGGGTTGGGTTGGTAAG | 235 |
| chr8:113481117-113483139 | p2 | (AT)20 | TTCCCAAAGTATCCCCATCA | AAATGACATGGCGGAATAGC | 205 |
| chr8:113481117-113483139 | p2 | (TA)11 | TTCCCAAAGTATCCCCATCA | AAATGACATGGCGGAATAGC | 205 |
| chr8:113519221-113521233 | p2 | (AT)6 | ATCAGGAATGGGCTTGACAC | TGGCTCAAACAACACCAAAA | 146 |
| chr8:113606461-113608476 | p5 | (AAATT)3 | AACAGCATTCCATAGCCTGG | TACAGGTAAGCGGGGATTTG | 232 |
| chr8:113651493-113653533 | p3 | (TGT)6 | AGCTTTGGCGCTTGATAAAA | GGACAATTCTGCCCTCTCAA | 132 |
| chr8:113701220-113703241 | p3 | (TTA)7 | TCTCCCCTCACTTCTCTCCA | AAGACGGTGGTGTAGTTGGC | 210 |
| chr8:113722176-113724200 | p4 | (AAAT)6 | CCTTTTAGGAGTTGGGGGAG | AAGGTTCAGGCATTTTGCAT | 152 |
| chr8:113896038-113898053 | p5 | (TAAAA)3 | GGGCAAGTAGTGAGACCCAA | GCGAAAAGAACAACACCCAT | 155 |
| chr8:113918689-113920709 | p5 | (TTTTC)3 | AGGGTATCATTTCCCAACCC | TGATCTGTTTCCAGAAGGGG | 259 |
| chr8:113918689-113920709 | p5 | (TAAAT)4 | AGGGTATCATTTCCCAACCC | TGATCTGTTTCCAGAAGGGG | 259 |
| chr8:113979869-113981889 | p5 | (TTAGA)4 | CGGTTTCCTCACAAAATCGT | AATTCATGCGCTCTAATCCG | 188 |
| chr8:113979869-113981889 | p3 | (TAT)5 | CGGTTTCCTCACAAAATCGT | AATTCATGCGCTCTAATCCG | 188 |
| chr8:114059137-114061152 | p2 | (AT)7 | TTCCACTCTTCAGCATCGTG | GGGGGCAAAAATTAAAGACC | 194 |
| chr8:114059137-114061152 | p5 | (TTTAC)3 | TTCCACTCTTCAGCATCGTG | GGGGGCAAAAATTAAAGACC | 194 |
| chr8:114245075-114247090 | p3 | (TAT)5 | CCTCAGCTGGTGGGAGATAG | ACTGCATGACAATGGACCAA | 219 |
| chr8:114322954-114324969 | p5 | (AAACC)3 | CTAGGGGTGTACATGGACCG | AAAACCCGAGAAAACCGAAT | 208 |
| chr8:114361936-114363951 | p5 | (AAACC)3 | CTAGGGGTGTACATGGACCG | AAAACCCGAGAAAACCGAAT | 208 |
| chr8:114392991-114395007 | p2 | (AT)8 | GTATAGATGCGGCGAGAGGA | CAAATTGTAAGGGGGCAAGA | 196 |
| chr8:114395324-114397358 | p2 | (TA)17 | TCTCTCTCTTTCTCTGCCGC | CCACCGCTGGTAATTTCATC | 264 |
| chr8:114513895-114515913 | p2 | (AT)9 | TTTAGACCCCTGCTTCATCG | GATTTTGGCCCCTTGATTTT | 284 |
| chr8:114540485-114542499 | p3 | (TGT)5 | ACTCCGCATTCTGCCTTAGA | CCTGGATCCACCTCCTAACA | 109 |
| chr8:114540485-114542499 | p2 | (TA)7 | ACTCCGCATTCTGCCTTAGA | CCTGGATCCACCTCCTAACA | 109 |
| chr8:114619554-114621569 | p3 | (AGA)5 | TGAGCAGCACTTTGGATTTG | GGCAAAAAGAAAGCAACGAC | 151 |
| chr8:114672988-114675003 | p5 | (ATTTT)3 | TGTTTTGCCACGACACAAAT | TGTAACTTCGCCCGAATAGG | 240 |
| chr8:114692311-114694323 | p3 | (GCA)5 | CGGTCGTCATGGACATACTG | TTCGATCCCACTTTCTCACC | 280 |
| chr8:114692311-114694323 | p3 | (TAT)6 | CGGTCGTCATGGACATACTG | TTCGATCCCACTTTCTCACC | 280 |

| Marker ID | SSR type | SSR motif | FORWARD PRIMER (5'-3') | REVERSE PRIMER (5'-3') | size (bp) |
|---|---|---|---|---|---|
| chr8:114692311-114694323 | p2 | (CT)6 | CGGTCGTCATGGACATACTG | TTCGATCCCACTTTCTCACC | 280 |
| chr8:114777189-114779213 | p2 | (AT)12 | GGGCATACTTTTAATCGGCA | ATGGGCAGACTTGGCATAAG | 127 |
| chr8:114777189-114779213 | p6 | (ATTTTT)3 | GGGCATACTTTTAATCGGCA | ATGGGCAGACTTGGCATAAG | 127 |
| chr8:114777485-114779503 | p2 | (AT)12 | TTTGACTGTGTGAGGGTTCG | CTTGAAATCAGGAAACGGGA | 214 |
| chr8:114777485-114779503 | p6 | (ATTTTT)3 | TTTGACTGTGTGAGGGTTCG | CTTGAAATCAGGAAACGGGA | 214 |
| chr8:114830805-114832843 | p2 | (TA)14 | CGCGGGCATCACTATCTAAT | TAAAAGGTGGGCTTTTGCTC | 295 |
| chr8:114830854-114832868 | p2 | (TA)14 | CGCGGGCATCACTATCTAAT | TAAAAGGTGGGCTTTTGCTC | 246 |
| chr8:114847360-114849398 | p2 | (TA)14 | CGCGGGCATCACTATCTAAT | TAAAAGGTGGGCTTTTGCTC | 295 |
| chr8:114847409-114849423 | p2 | (TA)14 | CGCGGGCATCACTATCTAAT | TAAAAGGTGGGCTTTTGCTC | 246 |
| chr8:114849782-114851806 | p2 | (AT)20 | CCGGCATTGTTTAGTATGGG | TAATGTTCAAAAGGGTGGGC | 213 |
| chr8:114849782-114851806 | p2 | (TA)11 | CCGGCATTGTTTAGTATGGG | TAATGTTCAAAAGGGTGGGC | 213 |
| chr8:114849782-114851806 | p2 | (AT)12 | CCGGCATTGTTTAGTATGGG | TAATGTTCAAAAGGGTGGGC | 213 |
| chr8:114868720-114870738 | p2 | (CA)8 | TAACCCGAACCCATACCAAA | TAGAGGGGCTGTCTCGGTTA | 177 |
| chr8:114868720-114870738 | p6 | (CTTTTT)3 | TAACCCGAACCCATACCAAA | TAGAGGGGCTGTCTCGGTTA | 177 |
| chr8:114868720-114870738 | p2 | (AT)15 | TAACCCGAACCCATACCAAA | TAGAGGGGCTGTCTCGGTTA | 177 |
| chr8:114884508-114886532 | p3 | (TTG)8 | TAGGCGTGATATAGTCCCCG | GGTATCAGGCCACACGTTTT | 297 |
| chr8:114892130-114894148 | p6 | (TGCCTT)3 | TTTTTCGTAACGGACCGAAC | ATGGTTAGAGGTGGGCATTG | 267 |
| chr8:114900670-114902697 | p3 | (TTA)6 | AATAGGCCATAACCGAGCCT | CATCTATTCTCGTGGCGGAT | 266 |
| chr8:114900670-114902697 | p3 | (TAT)9 | AATAGGCCATAACCGAGCCT | CATCTATTCTCGTGGCGGAT | 266 |
| chr8:114905064-114907080 | p4 | (TAAT)4 | TTGGCTCCAAATCCAATCTC | TCCGAATCGCCTACATATCC | 220 |
| chr8:114936685-114938705 | p5 | (CCCAA)4 | CTCCACCGTCAACATCCTCT | AGCTAACGGAGTTGCTGGAA | 101 |
| chr8:114977134-114979160 | p2 | (CT)7 | GGTTACATTTTTGCGATGGG | AAGAGCATGTACGGGGTACG | 140 |
| chr8:114995645-114997663 | p2 | (GA)9 | TCTTGCCGTTAAGATCGGAG | GCGTATCTGATCGACGGAGT | 221 |
| chr8:115127479-115129491 | p2 | (AT)6 | TGCGTGATCCCAAAACATAA | GCGCAAGGCTAGAGAAAATG | 187 |
| chr8:115155292-115157320 | p2 | (TA)6 | CTGGAGACACCCTCATGGTT | GTTCATTGGTTGTGACGTGG | 200 |
| chr8:115155322-115157334 | p2 | (TA)6 | CTGGAGACACCCTCATGGTT | GTTCATTGGTTGTGACGTGG | 170 |
| chr8:115208450-115210468 | p6 | (CCACAA)3 | AAGACAGGCACAGCAAGGTT | ACCAACAGTGGGTTTCCAAG | 198 |
| chr8:115208450-115210468 | p3 | (ATA)6 | AAGACAGGCACAGCAAGGTT | ACCAACAGTGGGTTTCCAAG | 198 |
| chr8:115208450-115210468 | p3 | (TCA)6 | AAGACAGGCACAGCAAGGTT | ACCAACAGTGGGTTTCCAAG | 198 |
| chr8:115290320-115292338 | p3 | (AAT)6 | TCAGCACGCTTACGTTCATC | AACATGGTTGTTTTAGCGGG | 300 |
| chr8:115290320-115292338 | p5 | (GACCC)3 | TCAGCACGCTTACGTTCATC | AACATGGTTGTTTTAGCGGG | 300 |
| chr8:115290441-115292456 | p3 | (AAT)6 | TCAGCACGCTTACGTTCATC | AACATGGTTGTTTTAGCGGG | 179 |
| chr8:115290441-115292456 | p5 | (GACCC)3 | TCAGCACGCTTACGTTCATC | AACATGGTTGTTTTAGCGGG | 179 |
| chr8:115306465-115308477 | p2 | (AG)6 | TCGGGTTTTGAACACATTGA | GGACCAGGCAGACAAAACAT | 137 |
| chr8:115423342-115425357 | p3 | (TTG)5 | TGCAAGCAGTCTTTTTGGTG | AAATCGGTTCCGAAAAACCT | 128 |
| chr8:115455929-115457947 | p2 | (TA)9 | AGGGCCTCAAATTCCTCAAT | CCGTTGCAGGTATTTGTCCT | 100 |

| Marker ID | SSR type | SSR motif | FORWARD PRIMER (5'-3') | REVERSE PRIMER (5'-3') | size (bp) |
|---|---|---|---|---|---|
| chr8:115480616-115482634 | p2 | (TA)9 | ACATTGGGGATGAATTGGAA | TTGTAGAACTTGGGCCTTGC | 105 |
| chr8:115505632-115507730 | p6 | (AATTTA)3 | TGGTCCCCTGTTTCAGACTT | TCCGATCCATCTCTTTGGTT | 298 |
| chr8:115522502-115524517 | p5 | (TTTGG)3 | ACTTTGGGTGGTAGCAGCAG | ATACCCAAACCGTGAACACC | 160 |
| chr8:115555051-115557066 | p5 | (TTTGG)3 | ACTTTGGGTGGTAGCAGCAG | ATACCCAAACCGTGAACACC | 160 |
| chr8:115605855-115607870 | p2 | (AT)8 | TGTTTTCTTTTGCCACGTCA | TCAAAGGCGGGTGTTTTAAG | 103 |
| chr8:115605855-115607870 | p5 | (TTTAA)3 | TGTTTTCTTTTGCCACGTCA | TCAAAGGCGGGTGTTTTAAG | 103 |
| chr8:115617138-115619153 | p5 | (AAAAG)3 | GCTCTCGTGTTTCCTTCTGC | TCACTGTCTGTCGCTGGAAC | 148 |
| chr8:115734728-115736740 | p2 | (TA)6 | ATCACCATACCCCCTATCCC | CCCTTCAGAATCTCTTCCCC | 263 |
| chr8:115782958-115784972 | p2 | (TA)7 | TCCCAAGCCTCCACAATAAG | GATTGGTCGAACAGCCATCT | 257 |
| chr8:115823906-115825941 | p2 | (AT)7 | ACGATCCGGAGAAAGTCAAA | AACATTTTCTGTGCTGTGCG | 226 |
| chr8:115823906-115825941 | p6 | (AAAAAT)3 | ACGATCCGGAGAAAGTCAAA | AACATTTTCTGTGCTGTGCG | 226 |
| chr8:115865051-115867067 | p2 | (GT)6 | TGTCCGTCGATGATTACGAG | GTTGGTTGGCGTACAGTGTG | 278 |
| chr8:115865051-115867067 | p2 | (AT)6 | TGTCCGTCGATGATTACGAG | GTTGGTTGGCGTACAGTGTG | 278 |
| chr8:115865051-115867067 | p2 | (AT)8 | TGTCCGTCGATGATTACGAG | GTTGGTTGGCGTACAGTGTG | 278 |
| chr8:115872176-115874198 | p2 | (AT)11 | GAATTGCGCCGAATAGAAAA | CGAGGTCCAAAACAAGCATT | 127 |
| chr8:115878911-115880925 | p2 | (AT)7 | GCCGAAAAAGGACAAACAAA | GCGGAATGCATCTTCAATTT | 116 |
| chr8:115906242-115908256 | p2 | (TA)6 | AAAGTTTCGCCTAATGGGCT | AGCGAGTCGTCTCAAATTGC | 214 |
| chr8:115986003-115988019 | p2 | (AT)8 | AACAGCCATACCTTTGTCGC | GGGCAAGCACAACAAATCTT | 234 |
| chr8:116075611-116077626 | p3 | (GTT)5 | TCCCACACAGCAATCAACAT | GATAGTGACATGGGATGGGG | 236 |
| chr8:116075611-116077626 | p2 | (AC)9 | TCCCACACAGCAATCAACAT | GATAGTGACATGGGATGGGG | 236 |
| chr8:116174495-116176517 | p2 | (TC)11 | GGATCCATATCCGGTCCTTT | GACCAAGGACTCCGCATAAA | 210 |
| chr8:116174495-116176517 | p2 | (AT)8 | GGATCCATATCCGGTCCTTT | GACCAAGGACTCCGCATAAA | 210 |
| chr8:116303487-116305505 | p6 | (GATGAC)3 | ACAAGATCCTGGTCGATTGG | CTAACGAAATTTGGGGAGCA | 278 |
| chr8:116325965-116327980 | p5 | (ATCAT)3 | ACAACAACCATGCCAAGACA | TCCAGCGGTACTATTTTGGG | 294 |
| chr8:116499882-116501898 | p4 | (AACA)4 | ACGAGCTATTGATTGGGTGG | CCACAATTGAGGTGGAAGGT | 114 |
| chr8:116511857-116513875 | p2 | (AG)9 | TTTAATGGCCAAACGCCTAC | CTAGAACGTCCAAACTCGGC | 141 |
| chr8:116511857-116513875 | p3 | (GTT)6 | TTTAATGGCCAAACGCCTAC | CTAGAACGTCCAAACTCGGC | 141 |
| chr8:116512158-116514176 | p2 | (AG)9 | TCAGCGTATATTTTTCCCGC | ATTCACTTGCCTGCCAATTC | 119 |
| chr8:116512158-116514176 | p3 | (GTT)6 | TCAGCGTATATTTTTCCCGC | ATTCACTTGCCTGCCAATTC | 119 |
| chr8:116539053-116541065 | p2 | (TA)6 | TTCTTGCCAATGTTTCCTCC | TGTCGACGCAAAGTAAATGC | 149 |
| chr8:116605090-116607118 | p3 | (ACA)5 | ATATTTTCATGGCCAAACGC | TCTCTCCGACGCGTAAAAAT | 223 |
| chr8:116625102-116627156 | p3 | (TAA)18 | CCGATTCACCCTCGAAACTA | ACCCTAGCTCAAACCCGAAT | 232 |
| chr8:116630887-116632905 | p3 | (ATA)6 | CGGGAGAGAGAGACAAGTGG | AATTTGACCCGATCCACAAA | 124 |
| chr8:116650491-116652509 | p6 | (TTAACT)3 | CAATTAGAGGCGAGGTCAGG | ATTTGAACCGCAAGTTACCG | 272 |
| chr8:116717979-116719991 | p2 | (AT)6 | TGATTGCCACAAAGGACAAA | AGCAACAAATCAACCCGAAG | 245 |
| chr8:116759154-116761170 | p4 | (AAGA)4 | GAAATTGTTTTGCAGGGGAA | GGATGCAGTTGAAGTCGTGA | 130 |

| Marker ID | SSR type | SSR motif | FORWARD PRIMER (5'-3') | REVERSE PRIMER (5'-3') | size (bp) |
|---|---|---|---|---|---|
| chr8:116773022-116775037 | p2 | (AT)6 | TGTGGGTGTGTGTGCCTACT | AACACATTTCAACGCAACCA | 201 |
| chr8:116773041-116775057 | p2 | (AT)6 | TGTGGGTGTGTGTGCCTACT | AACACATTTCAACGCAACCA | 182 |
| chr8:117089888-117091942 | p2 | (AT)8 | CAAAAGCTAACACCATGCCA | CATCCACGCACGATCTATGT | 195 |
| chr8:117103764-117105779 | p5 | (AACCG)3 | CGGATACGGTCAGCTTCAAT | GGCTAACAAGAAGGCCATGA | 284 |
| chr8:117188357-117190369 | p2 | (TA)6 | GACAAGCGTAACTGCGTTGA | CAATTGGTTGAAAATCGCCT | 245 |
| chr8:117198285-117200305 | p2 | (TA)10 | ACTGCGATGGTATCAGGGAG | ATCCGCTCATGTCCTCTCAT | 131 |
| chr8:117356387-117358399 | p2 | (TA)6 | AAATGTTCCCATGCTTCGTC | TGAGTTTGGAGATGGTGGGT | 276 |
| chr8:117356387-117358399 | p2 | (AT)8 | AAATGTTCCCATGCTTCGTC | TGAGTTTGGAGATGGTGGGT | 276 |
| chr8:117368325-117370343 | p2 | (TG)9 | CGAACCCTGGCTCAGTAAAA | ACATTTGGGCCTGAAAACAG | 242 |
| chr8:117385254-117387284 | p2 | (GA)6 | CGATTGGCAGTACAATGTGG | GCTTCTCCATAGCCGAGTTG | 298 |
| chr8:117385254-117387284 | p2 | (AT)7 | CGATTGGCAGTACAATGTGG | GCTTCTCCATAGCCGAGTTG | 298 |
| chr8:117385254-117387284 | p2 | (TA)15 | CGATTGGCAGTACAATGTGG | GCTTCTCCATAGCCGAGTTG | 298 |
| chr8:117385254-117387284 | p2 | (GA)6 | CGATTGGCAGTACAATGTGG | GCTTCTCCATAGCCGAGTTG | 298 |
| chr8:117386558-117388586 | p2 | (GA)6 | AAGGTTGGTCGTACGGTCTG | GCTTCTCCATAGCCGAGTTG | 269 |
| chr8:117386558-117388586 | p2 | (AT)7 | AAGGTTGGTCGTACGGTCTG | GCTTCTCCATAGCCGAGTTG | 269 |
| chr8:117386558-117388586 | p2 | (TA)14 | AAGGTTGGTCGTACGGTCTG | GCTTCTCCATAGCCGAGTTG | 269 |
| chr8:117394586-117396600 | p4 | (AAAT)4 | AGATGCAAAGCTTGAACCGT | GCAGGAGAGTTGCCACTAGG | 211 |
| chr8:117394586-117396600 | p2 | (TC)7 | AGATGCAAAGCTTGAACCGT | GCAGGAGAGTTGCCACTAGG | 211 |
| chr8:117451613-117453625 | p2 | (CA)6 | ATGGCGTGGCGTATATGATT | GAGCCTCTGGGACAACTCTG | 152 |
| chr8:117461592-117463607 | p3 | (AAG)5 | ACTAGCCAATGCCTGTCCAC | GAATTTGGAGAAACGTCCGA | 283 |
| chr8:117493156-117495189 | p3 | (AAT)11 | GTTCTTTTAGGGGGTTTCCG | CGCTCCACTCTCCTTTGTTC | 132 |
| chr8:117551555-117553567 | p2 | (TA)6 | TGGAGCGATCCTCATCCTAC | TGATATACCATGGGCAGCAA | 134 |
| chr8:117723897-117725915 | p6 | (AACCGG)3 | GCTTTCGGCTGCTATTTCAC | AGCAAAACAATGGAAATCCG | 238 |
| chr8:117723897-117725915 | p3 | (GAT)5 | GCTTTCGGCTGCTATTTCAC | AGCAAAACAATGGAAATCCG | 238 |
| chr8:117809905-117811925 | p3 | (TGT)5 | TGCATTCTGCTTTGAACTGC | ACAATGTGAGGTCTGGGGAG | 239 |
| chr8:117809905-117811925 | p4 | (TTAT)5 | TGCATTCTGCTTTGAACTGC | ACAATGTGAGGTCTGGGGAG | 239 |
| chr8:117824427-117826442 | p5 | (TTTTA)3 | ATTCACATGGGTTTGCTTCC | GGTGACCCACTCACCAAAAT | 184 |
| chr8:117824427-117826442 | p2 | (TA)8 | ATTCACATGGGTTTGCTTCC | GGTGACCCACTCACCAAAAT | 184 |
| chr8:117988186-117990204 | p2 | (TC)9 | CATGCGCTTGCACTTTTCTA | GTTTGAACGGGAGCAGAGAG | 125 |
| chr8:117998957-118000989 | p2 | (TA)16 | TCCTGACTATCTCGGGGCTA | GAGAGGTGGTGGAGGTTCAA | 111 |
| chr8:118080592-118082606 | p2 | (TA)7 | TAGGGCGATCGATGTAAACC | TGCTCATCCGTCATCCATTA | 144 |
| chr8:118118991-118121016 | p2 | (AT)6 | ATTAGAAGCACCGGTTGTGG | GTTTGGATGTGGTGGTTTCC | 211 |
| chr8:118149974-118152040 | p2 | (CT)7 | ATTCAAGTTCAGGCACCCAC | ATTTGTTCGAAGCCACCAAG | 108 |
| chr8:118175559-118177593 | p2 | (TA)17 | TGGCCATAAACCACAACTGA | GCCTTAGGGGCTGGACTACT | 196 |
| chr8:118254237-118256252 | p5 | (AAATA)4 | TGGGCTACGTGGGGTAAATA | TATTGGAGGCCATTGAGAGG | 241 |
| chr8:118254237-118256252 | p3 | (GAT)5 | TGGGCTACGTGGGGTAAATA | TATTGGAGGCCATTGAGAGG | 241 |

| Marker ID | SSR type | SSR motif | FORWARD PRIMER (5'-3') | REVERSE PRIMER (5'-3') | size (bp) |
|---|---|---|---|---|---|
| chr8:118254237-118256252 | p2 | (TA)7 | TGGGCTACGTGGGGTAAATA | TATTGGAGGCCATTGAGAGG | 241 |
| chr8:118367039-118369055 | p2 | (AT)8 | CGGATACTATTCGCCCTTGA | CACTCGGGATAACGGTCACT | 177 |
| chr8:118409043-118411061 | p6 | (ATTTTT)3 | AGGCCGCTTCCTATGAAAAT | GAAGGTGTGCTCTGCATTGA | 213 |
| chr8:118505147-118507177 | p2 | (AC)6 | CCCCCACAAACGTATACTCG | CTGTGCAGCTGGACGTTAAA | 217 |
| chr8:118505147-118507177 | p3 | (CTT)10 | CCCCCACAAACGTATACTCG | CTGTGCAGCTGGACGTTAAA | 217 |
| chr8:118595696-118597740 | p4 | (AAAT)11 | GCAGCTAGTGGCATGTAGCA | GTTGGCCCTTACCAAGTGAA | 171 |
| chr8:118614983-118616998 | p3 | (TGG)5 | ATCACTTTTGAATCCGCGAC | TGCATTTTTGCAGTTTGCTC | 118 |
| chr8:118616628-118618658 | p6 | (CATATA)5 | CCCCTTATCTAGGACCGAGC | GAGGATTGCCAAGTCCATGT | 207 |
| chr8:118619672-118621693 | p3 | (TAT)7 | CCATTTTTCCCCTAAGCTCC | GCGTGGTGTGTTGTCAAATC | 133 |
| chr8:118641924-118643936 | p2 | (AT)6 | AAGTATCAACCGTGGCTTCG | GTCGCTCCGATCTAATCCAA | 207 |
| chr8:118681455-118683471 | p4 | (TTTA)4 | CTTGACGGTGGAGGATTGAT | AAATAAAGGGTTGGGTTGGG | 163 |
| chr8:118732901-118734916 | p3 | (CGG)5 | GCGTTCCCGACTAATTTGAA | ACTGCAAGGGCAAATTGACT | 219 |
| chr8:118793700-118795714 | p2 | (TC)7 | TATCATGTTGCCCATCCCTT | ACCACACACCGGCTAAAAAC | 130 |
| chr8:118857474-118859492 | p6 | (AGAAAA)3 | CACCCTCCGAGGATACTTGA | TGCCCGAAAGTTACATAGCC | 101 |
| chr8:118884018-118886044 | p5 | (AAATA)3 | ACATTGGTAGGGCTATTGCG | TGAAAAACAAACCAGCACCA | 232 |
| chr8:118885657-118887672 | p3 | (TCT)5 | TTGGCTATTGCCAGTGAATG | CGCGACACGTGAAGAAATAA | 216 |
| chr8:118913799-118915817 | p2 | (TA)9 | TTTCATCATCCAAGGCACAA | TTAACCGACCAAACCGTACC | 161 |
| chr8:118925397-118927415 | p6 | (GTTTTT)3 | GCGTAGGTTTACTGCCTTGC | AAGAAGCAGACGAAGAAGCG | 108 |
| chr8:118933511-118935525 | p2 | (AT)7 | TGGTAGTCCAGGCAATGTCC | AGTGAACCTCATGGCTTGCT | 253 |
| chr8:118988956-118990974 | p2 | (AT)9 | TGGATTTTGGCTTCACCAAT | TACAGCAGACACCTCTGGCA | 107 |
| chr8:118996485-118998505 | p2 | (AT)10 | TAGGGATGGAGCCGTATGAG | TTATGTGGGAGACTTCCGCT | 284 |
| chr8:119001402-119003414 | p2 | (AT)6 | CAGGGCCATGTCCTGATAGT | GAACCTCATGGCTTGCTTGT | 256 |
| chr8:119056904-119058922 | p2 | (AT)9 | TGGATTTTGGCTTCACCAAT | TACAGCAGACACCTCTGGCA | 107 |
| chr8:119066293-119068308 | p5 | (ACATA)3 | GTCCAAGGGCAAAATTTGAA | CGAGACCCTTGTTCTTGAGC | 103 |
| chr8:119111888-119113903 | p5 | (AAATA)3 | TAAATTCGCCCCGGTACATA | GTCAGGAAAGGGCAATGTGT | 242 |
| chr8:119139819-119141837 | p2 | (AT)9 | TGCAATGATGCAAAGTGACA | TTTCTGTTTCTTCCTGCGGT | 291 |
| chr8:119139819-119141837 | p3 | (AAG)6 | TGCAATGATGCAAAGTGACA | TTTCTGTTTCTTCCTGCGGT | 291 |
| chr8:119212638-119214660 | p2 | (AT)11 | TGGATTTGGGAGAAACAGGA | TGTTGGAGTACGGCAGCATA | 168 |
| chr8:119212638-119214660 | p2 | (AT)8 | TGGATTTGGGAGAAACAGGA | TGTTGGAGTACGGCAGCATA | 168 |
| chr8:119298056-119300068 | p2 | (TC)6 | TTTTGCCAAACTTTTCTCCG | AAAGCGGCTACTTGTGCAAT | 204 |
| chr8:119355040-119357052 | p2 | (AT)6 | TTCACAAAAATTCATGCCCA | TCGACGAGACGAGCTTATGA | 223 |
| chr8:119496926-119498951 | p5 | (GGGTG)5 | CTTCCTCAGGACCACCAAAA | GCGAGGATGATTCCAACAAT | 160 |
| chr8:119502511-119504526 | p5 | (TGCAT)3 | GAAAGCATGCTAGGCCTTTG | CACCCATCAAATTCAAGCCT | 163 |
| chr8:119617493-119619508 | p3 | (TTG)5 | CAACCGAGCAGAATCTAGCC | GAACAAGCCTTCCGCAATAA | 259 |
| chr8:119653712-119655727 | p3 | (AAC)5 | TGCAAACACACCCTATGGAA | TGTTGTTGATCGGGCAATTA | 251 |
| chr8:119685378-119687393 | p3 | (AAC)5 | TGCAAACACACCCTATGGAA | TGTTGTTGATCGGGCAATTA | 251 |

| Marker ID | SSR type | SSR motif | FORWARD PRIMER (5'-3') | REVERSE PRIMER (5'-3') | size (bp) |
|---|---|---|---|---|---|
| chr8:119696904-119698925 | p3 | (TGT)7 | AATTGCAAGGCAAGATTTCG | GATTGTCCCTATTTCGGGGT | 256 |
| chr8:119773057-119775081 | p2 | (TA)10 | TGCCACATGCGTATCAATCT | TGACATGGCGGAATAACAAA | 226 |
| chr8:119803976-119806002 | p2 | (TA)13 | GTGACAACAAACCAACACGG | GATGGGGGTAAAAGCTCTCC | 116 |
| chr8:119968784-119970802 | p2 | (AT)9 | TCGCCTCAACAGACATTCAG | GGCACCTCTCATTCACACCT | 283 |
| chr8:120039510-120041525 | p5 | (TATTT)3 | ATTATAGCACTGCGGCTGGT | TTTTTCTCCCTTTGTGGGTG | 183 |
| chr8:120062768-120064783 | p2 | (TA)6 | TTCAACAGTAGACCCGTCCC | CCCATACCAGATTTTGTCGG | 259 |
| chr8:120062768-120064783 | p3 | (TAC)5 | TTCAACAGTAGACCCGTCCC | CCCATACCAGATTTTGTCGG | 259 |
| chr8:120073061-120075076 | p3 | (ATT)5 | TAGGGTTTGGTTTTGGTTCG | TTCTTTGGACAAAACAGGGG | 251 |
| chr8:120087239-120089277 | p2 | (AT)19 | TTCGTGCAACTCAAACAAGG | TTAAGGTGTGGGGTGATGGT | 218 |
| chr8:120102737-120104775 | p2 | (AT)19 | TTCGTGCAACTCAAACAAGG | TTAAGGTGTGGGGTGATGGT | 218 |
| chr8:120200982-120202994 | p2 | (TA)6 | TTGCTCCACGCCCTACTACT | ACCATGCTAAGACGTCCACC | 134 |
| chr8:120200982-120202994 | p2 | (AT)9 | TTGCTCCACGCCCTACTACT | ACCATGCTAAGACGTCCACC | 134 |
| chr8:120211268-120213283 | p5 | (TATTT)3 | TTTTCGTGCACCATTTTCAA | AGAGGAGCAAGTTGAGCGAG | 134 |
| chr8:120211268-120213283 | p2 | (AT)6 | TTTTCGTGCACCATTTTCAA | AGAGGAGCAAGTTGAGCGAG | 134 |
| chr8:120214915-120216933 | p3 | (TGT)6 | GTGCACTTTTATGCCCCCTA | CGACTATTTCATAGGCCCCA | 101 |
| chr8:120236127-120238145 | p5 | (ATTTT)3 | CCAAATGGGTATTTCGTTGG | TTCGTGAATTGCACAAAGGA | 260 |
| chr8:120236127-120238145 | p2 | (TG)6 | CCAAATGGGTATTTCGTTGG | TTCGTGAATTGCACAAAGGA | 260 |
| chr8:120236188-120238220 | p5 | (ATTTT)3 | CCAAATGGGTATTTCGTTGG | TTCGTGAATTGCACAAAGGA | 199 |
| chr8:120282457-120284475 | p2 | (AT)9 | ATGAGGAGAGCCCTTCTTCC | GGTATTGCTTCAGGCGAAAA | 220 |
| chr8:120286418-120288433 | p3 | (ATG)5 | TCTTGTTAACCCGAACCCTG | TAATCTATAGGGGGTGGGGG | 186 |
| chr8:120411946-120413960 | p2 | (AG)7 | TTTCTAATGGTGCTGGCCTC | GGCAAACAATTGCTGGTTTT | 216 |
| chr8:120499596-120501614 | p6 | (CTCTGC)3 | TCACACCCCTTAGGATGCTC | CCACCATAACGAAACATCCC | 121 |
| chr8:120622926-120624941 | p5 | (TTTTA)3 | GACCATGGTTGATAGGTGGG | AGCTTTCCATTATTCGGCCT | 287 |
| chr8:120722933-120724948 | p5 | (ATTGG)3 | GGCATTCTGTGGTTGGATCT | CATGCATAGCAATCATTGGG | 227 |
| chr8:120797437-120799519 | p2 | (TA)41 | TTTCATTCGTTCATTTGGCA | TTTTGGTTATGTCCGCCTCT | 154 |
| chr8:120828187-120830205 | p3 | (ATT)6 | TCCATGTTCATGTTTTGGGA | TAGCCGACATGTAACCACCA | 247 |
| chr8:120916615-120918633 | p3 | (TGT)6 | GACGTGCAGCTGGATACTCA | CGAGCTTCCATTTCATCCAT | 105 |
| chr8:120986640-120988661 | p3 | (CAA)7 | GACACTGCAGACCTGTGGAC | TATAAACGCCCAAGGTGGAC | 111 |
| chr8:121031022-121033034 | p2 | (TG)6 | GGGGAAGGTGCTCAACTGTA | CTCATCCTTTGAGGGAGCTG | 153 |
| chr8:121031697-121033750 | p2 | (TG)6 | TAGGGCACCTGCTAATCCAG | CCACCACACACACACAATCA | 101 |
| chr8:121180048-121182066 | p6 | (TAAACA)3 | AATCCCATTCCAAGTGCAAG | CCAAGAATCAAAAGGGGTGA | 170 |
| chr8:121255112-121257139 | p3 | (TAT)9 | TTGAAATCACTCCAAGTGCG | TTTTTGGTCATGAAGTGCCA | 157 |
| chr8:121302443-121304461 | p6 | (TTTTAT)3 | GAAAATGCTCGTGCCAAAAT | GCAAGTGTAGAGGTTGGGGA | 295 |
| chr8:121383367-121385383 | p4 | (GTTT)4 | ACACACTCGGCCGGATATAG | TCCAAGTCACGAGATTTCCC | 157 |
| chr8:121385899-121387923 | p3 | (ACA)8 | ATCACCGCCTCAACTTTGTC | TTCACGGAATCGTTTTAGGC | 150 |
| chr8:121405413-121407428 | p3 | (TTA)5 | AGCAAGGAGACTCTGTCCCA | CGTGTCTACTCCCCATTCGT | 271 |

| Marker ID | SSR type | SSR motif | FORWARD PRIMER (5'-3') | REVERSE PRIMER (5'-3') | size (bp) |
|---|---|---|---|---|---|
| chr8:121442594-121444612 | p3 | (AGA)6 | AGGGCATGAAGGTGTGCTAC | ACGGGACTAGTTGCCATACG | 143 |
| chr8:121510942-121512960 | p3 | (AGA)6 | AGGGCATGAAGGTGTGCTAC | ACGGGACTAGTTGCCATACG | 143 |
| chr8:121568428-121570443 | p5 | (CTTTT)3 | CAACCTCCGGCTAATCAAGA | TTCCTTTTACCCGCTACCCT | 295 |
| chr8:121702255-121704271 | p2 | (AT)8 | CTAAGTACCCGAACTGGGCA | AATCTGAGACCATGGGTTGG | 198 |
| chr8:121702255-121704271 | p2 | (AT)8 | CTAAGTACCCGAACTGGGCA | AATCTGAGACCATGGGTTGG | 198 |
| chr8:121702627-121704643 | p2 | (AT)8 | AAGGGAAATCCTAAGCTGGC | CACCAAGTTCCATAATGGGG | 234 |
| chr8:121702627-121704643 | p2 | (AT)8 | AAGGGAAATCCTAAGCTGGC | CACCAAGTTCCATAATGGGG | 234 |
| chr8:121741619-121743649 | p3 | (GTT)10 | TACCCAAGGGTTATGCCAAA | GTCCCCCACATTTGCTCTTA | 146 |
| chr8:121741619-121743649 | p3 | (GTT)6 | TACCCAAGGGTTATGCCAAA | GTCCCCCACATTTGCTCTTA | 146 |
| chr8:121745116-121747134 | p3 | (ACA)6 | AATTGTGTCTTCAGACGGGC | GCAGGCAGCAGATGAATGTA | 207 |
| chr8:121771140-121773154 | p5 | (TAAAA)3 | CGAGAAAATCAAGCCGATGT | CGTCAAGGTAGTCGTGGGTT | 217 |
| chr8:121771140-121773154 | p2 | (AT)7 | CGAGAAAATCAAGCCGATGT | CGTCAAGGTAGTCGTGGGTT | 217 |
| chr8:121844558-121846574 | p2 | (GA)8 | AATCTGCGTGGGTTCAAAAC | TTTTTGTCCCTCAAATTGCC | 157 |
| chr8:121961017-121963035 | p3 | (AAC)6 | CAAATACGTAGCGGTTCGGT | TAATAACATAGCCGCGGTCC | 146 |
| chr8:121962400-121964414 | p2 | (TA)7 | CCACCAACTTTTCATGCCTT | TGCAAGACCCTCTCCAACTT | 288 |
| chr8:121976853-121978871 | p2 | (TG)9 | ATGATGACACTGAAGCGCAC | AGCTGGTATCAAGTTGGCGT | 245 |
| chr9:96494-98506 | p2 | (AT)6 | GCATGTCCAATTCAGCCTTT | CTTTTTGAAGGAGACTGCGG | 113 |
| chr9:192971-194985 | p2 | (CT)9 | AGGGAAATACTTCTCCGCGT | ATGTTGATGGATTCTTCGCC | 292 |
| chr9:192971-194985 | p2 | (AG)7 | AGGGAAATACTTCTCCGCGT | ATGTTGATGGATTCTTCGCC | 292 |
| chr9:192971-194985 | p2 | (TA)11 | AGGGAAATACTTCTCCGCGT | ATGTTGATGGATTCTTCGCC | 292 |
| chr9:193102-195124 | p2 | (CT)9 | AGGGAAATACTTCTCCGCGT | ATGTTGATGGATTCTTCGCC | 161 |
| chr9:193102-195124 | p2 | (AG)7 | AGGGAAATACTTCTCCGCGT | ATGTTGATGGATTCTTCGCC | 161 |
| chr9:193102-195124 | p2 | (TA)11 | AGGGAAATACTTCTCCGCGT | ATGTTGATGGATTCTTCGCC | 161 |
| chr9:224643-226695 | p2 | (TA)26 | CTTGAGAGCCGTGGTAGAGC | TTTGGTGGACGCACTGTTTA | 242 |
| chr9:322739-324754 | p5 | (GTGTA)3 | AGAACATCCCATCCATGAGC | TCTGCTGGCAAGGCTATTTT | 200 |
| chr9:332077-334092 | p5 | (AAATA)3 | ACTGCCAAATCCCTAATCCC | TAAAGGCGGGCACAAAATAG | 105 |
| chr9:355305-357323 | p6 | (TAAAAA)3 | AGCTCGTACGACGAGTCCAT | GGCCATTTTCGTGGAACTAA | 245 |
| chr9:390618-392672 | p2 | (CT)27 | AACCAATCCTGATCACCTCG | AAAGTTGGTGTAACGACCCG | 256 |
| chr9:409635-411650 | p3 | (ATA)5 | GGGGAGAGGAAATGAAGAGC | GACCCGTACCACAAATGGAC | 135 |
| chr9:409838-411865 | p3 | (ATA)5 | GGGTGCCAGAATATCACCTG | CTTGCATTTCCCAACGTTCT | 294 |
| chr9:465060-467076 | p2 | (TA)8 | TTGTCTGCAAAGTCGTGTCC | CGAAGCAATTTCCTCGTAGC | 228 |
| chr9:576073-578089 | p2 | (AT)18 | TTGCTTACAATCACCCACTCC | TTGGGCCTTTTTCTTTTGAA | 116 |
| chr9:783888-785903 | p3 | (ACC)5 | ATTCTCCACCACCGAACAAC | GTGGAGATGGAATAGGGGGT | 168 |
| chr9:799628-801640 | p2 | (AT)7 | TCAAACCGGTAAACCGAAAC | TTGGGTGTGTTTTTGACCCT | 296 |
| chr9:799628-801640 | p2 | (AT)6 | TCAAACCGGTAAACCGAAAC | TTGGGTGTGTTTTTGACCCT | 296 |
| chr9:806507-808523 | p4 | (TACA)4 | TGAGTTCCAGACGCTGAATG | GGGCTCTGTATGATTGGCAT | 274 |

| Marker ID | SSR type | SSR motif | FORWARD PRIMER (5'-3') | REVERSE PRIMER (5'-3') | size (bp) |
|---|---|---|---|---|---|
| chr9:841909-843921 | p2 | (TA)13 | ATTACGATGGCGAATTCAGG | CGGGAAGAATGATTGTCGTT | 205 |
| chr9:841909-843921 | p2 | (AT)6 | ATTACGATGGCGAATTCAGG | CGGGAAGAATGATTGTCGTT | 205 |
| chr9:855396-857412 | p2 | (TC)8 | CCAGAGGCAAGTCAAAAAGG | AAGCTCTAAGCTTTTGGGGC | 281 |
| chr9:858819-860913 | p2 | (AG)7 | GAGAGAGACCGAGCTCGAAA | AAATTGCGTTCCATTTCTCG | 116 |
| chr9:858819-860913 | p2 | (AT)6 | GAGAGAGACCGAGCTCGAAA | AAATTGCGTTCCATTTCTCG | 116 |
| chr9:858819-860913 | p2 | (AT)47 | GAGAGAGACCGAGCTCGAAA | AAATTGCGTTCCATTTCTCG | 116 |
| chr9:885991-888005 | p3 | (AAC)5 | CGAACGTGTTACGTTTCCTG | AACACGCGCAGATACTCCTT | 105 |
| chr9:954899-956915 | p2 | (CT)8 | CGTCGCAGGATCCATAAACT | ATTTGGAACGGATTGGATGA | 124 |
| chr9:1088308-1090323 | p5 | (AATAA)3 | AGGCAGAGTTTTTCAGGCAA | GCCCTTATTTTGGGATGGTT | 292 |
| chr9:1099349-1101363 | p3 | (AAT)5 | CGATTAAGATGGCCTTGCTC | ATGGCTAAAAATTAGGCGCA | 286 |
| chr9:1099349-1101363 | p2 | (TC)7 | CGATTAAGATGGCCTTGCTC | ATGGCTAAAAATTAGGCGCA | 286 |
| chr9:1122637-1124653 | p2 | (AT)7 | GGAAAGTAGCCAATGCCAAA | ACCTATGCATTGAGATCCCG | 264 |
| chr9:1122958-1124972 | p2 | (AT)7 | CGGGATCTCAATGCATAGGT | TTTGGCTCCGGACTCAATAG | 127 |
| chr9:1146793-1148808 | p3 | (TGT)5 | GGTTGTGAGACCGACCTTGT | ATCTGTGCCTCTCCTCCTCA | 265 |
| chr9:1146793-1148808 | p3 | (TTG)7 | GGTTGTGAGACCGACCTTGT | ATCTGTGCCTCTCCTCCTCA | 265 |
| chr9:1168326-1170341 | p3 | (AAT)5 | TTGTAGGTTCAAACCCCAGC | TCGTTCGTATGGGTTGGTTT | 226 |
| chr9:1297427-1299442 | p5 | (CATAT)3 | ATATGGGCACCCAAACACAT | GTATTTTCTGTGCGCAGGGT | 182 |
| chr9:1451304-1453320 | p2 | (AG)9 | TCTTAAAGGGCGTGCAAAGT | TATCGTGCACGGAACGTTTA | 254 |
| chr9:1451304-1453320 | p2 | (TA)8 | TCTTAAAGGGCGTGCAAAGT | TATCGTGCACGGAACGTTTA | 254 |
| chr9:1504019-1506034 | p5 | (CTTTT)3 | TTTGAACGTGACGAAAGCAG | CCCAAGACATAGATTCCCGA | 187 |
| chr9:1509794-1511812 | p3 | (TCA)5 | CGAGTTCTAGGCGGTTTGTC | TCTAGCCGCCTTCACTTTGT | 262 |
| chr9:1509794-1511812 | p6 | (CCGGTT)3 | CGAGTTCTAGGCGGTTTGTC | TCTAGCCGCCTTCACTTTGT | 262 |
| chr9:1781412-1783427 | p5 | (AAACC)4 | ACCAAAACTGTGAACAGCCC | TGAACTGAGCGCACAAACTC | 200 |
| chr9:1781412-1783427 | p3 | (AAT)5 | ACCAAAACTGTGAACAGCCC | TGAACTGAGCGCACAAACTC | 200 |
| chr9:1801775-1803793 | p3 | (GTT)6 | TACAAGCATATGCAGCGGAG | AATGAATGCGTGGATGGTTT | 294 |
| chr9:1814902-1816917 | p5 | (CAAAC)3 | TAATTCATGCCACCGTTCAA | ATTCCCCCTCGGTTGTAATC | 220 |
| chr9:1814902-1816917 | p5 | (TTGAT)3 | TAATTCATGCCACCGTTCAA | ATTCCCCCTCGGTTGTAATC | 220 |
| chr9:1830896-1832912 | p4 | (TTAT)4 | GGTGCCAGAAACGTTGATTT | AAGTTGGGGGTGTTGATGAG | 169 |
| chr9:1843783-1845828 | p3 | (ATT)15 | TTAATCATCGCCAAAAAGGC | GTTCGCTGCCGAAGAAATAG | 219 |
| chr9:1843783-1845828 | p2 | (CT)12 | TTAATCATCGCCAAAAAGGC | GTTCGCTGCCGAAGAAATAG | 219 |
| chr9:1845097-1847112 | p2 | (CT)12 | ACAATAATCGACCCAACCCA | GTAATCGCGAAAGGCTTCTG | 112 |
| chr9:1845097-1847112 | p5 | (TTAGT)3 | ACAATAATCGACCCAACCCA | GTAATCGCGAAAGGCTTCTG | 112 |
| chr9:1881782-1883797 | p5 | (AATAA)3 | GCGAGGGAATTTGCTGTTAG | TCTTGAGATTCTCTGCGGGT | 190 |
| chr9:1881782-1883797 | p5 | (TAAAA)3 | GCGAGGGAATTTGCTGTTAG | TCTTGAGATTCTCTGCGGGT | 190 |
| chr9:1923675-1925689 | p5 | (TAAAA)3 | TCCGTTGTGAGCATTTTGTC | ATATGGGGCCCACAATTTTT | 247 |
| chr9:1923675-1925689 | p2 | (AT)7 | TCCGTTGTGAGCATTTTGTC | ATATGGGGCCCACAATTTTT | 247 |

| Marker ID | SSR type | SSR motif | FORWARD PRIMER (5'-3') | REVERSE PRIMER (5'-3') | size (bp) |
|---|---|---|---|---|---|
| chr9:1923767-1925779 | p5 | (TAAAA)3 | TCCGTTGTGAGCATTTTGTC | ATATGGGGCCCACAATTTTT | 155 |
| chr9:1923767-1925779 | p2 | (AT)7 | TCCGTTGTGAGCATTTTGTC | ATATGGGGCCCACAATTTTT | 155 |
| chr9:1947460-1949488 | p2 | (AT)11 | CCATTGCTCCATGCCTTACT | CGAAGCCACGGTTGATACTT | 192 |
| chr9:1947460-1949488 | p5 | (CCGGT)3 | CCATTGCTCCATGCCTTACT | CGAAGCCACGGTTGATACTT | 192 |
| chr9:1947494-1949508 | p2 | (AT)11 | CCATTGCTCCATGCCTTACT | CGAAGCCACGGTTGATACTT | 158 |
| chr9:1947494-1949508 | p5 | (CCGGT)3 | CCATTGCTCCATGCCTTACT | CGAAGCCACGGTTGATACTT | 158 |
| chr9:1947692-1949752 | p2 | (AT)11 | AAGTATCAACCGTGGCTTCG | GTCGCTCCGATCTAATCCAA | 200 |
| chr9:1947692-1949752 | p5 | (CCGGT)3 | AAGTATCAACCGTGGCTTCG | GTCGCTCCGATCTAATCCAA | 200 |
| chr9:1967928-1969948 | p4 | (AAGA)5 | AAGGCTTTACAACCAAGCCA | TCGTGCGCAATTTTCATTTA | 134 |
| chr9:2011704-2013736 | p4 | (TTTA)8 | TGGGCCTAACTCAACCTCAC | GAGCCTCAGCAACAACAACA | 288 |
| chr9:2011704-2013736 | p4 | (AAAG)4 | TGGGCCTAACTCAACCTCAC | GAGCCTCAGCAACAACAACA | 288 |
| chr9:2048531-2050549 | p2 | (AT)9 | GTCATGTTTCCCCTTTTCCA | TGGAATAAAAAGGGCAGTCG | 204 |
| chr9:2099517-2101556 | p2 | (TC)10 | TCTTTCTCCCTCCGTTCTCA | AAGCGACGTCGGAAAATAGA | 250 |
| chr9:2146083-2148101 | p6 | (TTTATT)3 | TCTCCTTAGCCGTGAGCAAT | AAAAATGTCATTGTGGCCGT | 106 |
| chr9:2176100-2178115 | p2 | (GA)7 | GAGTGGAGGCAGTGGAGAAG | GTCCGATCCTTTTCATCCAA | 223 |
| chr9:2176100-2178115 | p3 | (ATA)5 | GAGTGGAGGCAGTGGAGAAG | GTCCGATCCTTTTCATCCAA | 223 |
| chr9:2176100-2178115 | p3 | (AAT)10 | GAGTGGAGGCAGTGGAGAAG | GTCCGATCCTTTTCATCCAA | 223 |
| chr9:2176131-2178170 | p2 | (GA)7 | GAGTGGAGGCAGTGGAGAAG | GTCCGATCCTTTTCATCCAA | 192 |
| chr9:2176131-2178170 | p3 | (ATA)5 | GAGTGGAGGCAGTGGAGAAG | GTCCGATCCTTTTCATCCAA | 192 |
| chr9:2176131-2178170 | p3 | (AAT)10 | GAGTGGAGGCAGTGGAGAAG | GTCCGATCCTTTTCATCCAA | 192 |
| chr9:2195310-2197325 | p3 | (AAT)14 | GGAAAGGAGAGGAAAGGGAA | ATTTTGACTTACCCGACCCC | 279 |
| chr9:2195354-2197369 | p3 | (AAT)14 | GGAAAGGAGAGGAAAGGGAA | ATTTTGACTTACCCGACCCC | 235 |
| chr9:2195651-2197693 | p3 | (AAT)14 | GGGGTCGGGTAAGTCAAAAT | CACGGTCATTGCAACATCAT | 191 |
| chr9:2195651-2197693 | p3 | (TAT)6 | GGGGTCGGGTAAGTCAAAAT | CACGGTCATTGCAACATCAT | 191 |
| chr9:2241032-2243048 | p2 | (TA)8 | AGGAGTATGACAGGCTCCGA | TTAAGTATGGCCTTGTCGGC | 112 |
| chr9:2261686-2263704 | p5 | (TCATA)6 | GCCATACCTTTCCCCTTACC | TGCATGCTGCTCACATGATA | 184 |
| chr9:2261686-2263704 | p3 | (CAA)6 | GCCATACCTTTCCCCTTACC | TGCATGCTGCTCACATGATA | 184 |
| chr9:2284352-2286388 | p3 | (AAT)12 | AAAGGCAACATCCTTTGTGG | GTCGAATCTCCAGAAACCCA | 117 |
| chr9:2384421-2386436 | p3 | (GTT)6 | CGCTCCAGGAGAACCTAGTG | CAGGATCTTTTCTCCCACCA | 198 |
| chr9:2384421-2386436 | p3 | (TGG)5 | CGCTCCAGGAGAACCTAGTG | CAGGATCTTTTCTCCCACCA | 198 |
| chr9:2384421-2386436 | p5 | (TCCCC)3 | CGCTCCAGGAGAACCTAGTG | CAGGATCTTTTCTCCCACCA | 198 |
| chr9:2385024-2387039 | p3 | (TGG)5 | GAGTTCTAAGGGCATTTGCG | AAGTGGCCTAAACATGTCCG | 110 |
| chr9:2385024-2387039 | p5 | (TCCCC)3 | GAGTTCTAAGGGCATTTGCG | AAGTGGCCTAAACATGTCCG | 110 |
| chr9:2393339-2395354 | p2 | (CT)7 | AGAGGCGAGTTTAACGCTGA | AGTGGTGTCTGTGGCCAACT | 110 |
| chr9:2393339-2395354 | p5 | (AATAC)3 | AGAGGCGAGTTTAACGCTGA | AGTGGTGTCTGTGGCCAACT | 110 |
| chr9:2531441-2533459 | p2 | (AC)9 | ACACTTTGTAGCCGTCCACC | CCCAAATATCAAGCGAAAGC | 186 |

| Marker ID | SSR type | SSR motif | FORWARD PRIMER (5'-3') | REVERSE PRIMER (5'-3') | size (bp) |
|---|---|---|---|---|---|
| chr9:2595020-2597050 | p2 | (AT)15 | TCAGATTCGACACCAATCCA | CAGGTTTGCCATTCAACCTT | 197 |
| chr9:2714423-2716489 | p3 | (ATA)9 | CTTTCTTCATTGGCCGGTTA | ATATTGCATTTGGCACGACA | 117 |
| chr9:2714423-2716489 | p2 | (AT)33 | CTTTCTTCATTGGCCGGTTA | ATATTGCATTTGGCACGACA | 117 |
| chr9:2937066-2939078 | p2 | (CT)6 | GGAGTGGAGACCCAATTTGA | TTGATTGGCCGTTGTTTGTA | 100 |
| chr9:2937200-2939214 | p2 | (CT)6 | TTACAAACAACGGCCAATCA | TTATCTTCAAATTTGGCGGG | 150 |
| chr9:2937219-2939239 | p2 | (CT)6 | TTACAAACAACGGCCAATCA | TTATCTTCAAATTTGGCGGG | 131 |
| chr9:2977961-2980012 | p6 | (GAAGGA)4 | CACAAATCCTCCCCTTTCAA | CACCCCTTTCTCCTTTCTCC | 276 |
| chr9:2977961-2980012 | p3 | (AAT)17 | CACAAATCCTCCCCTTTCAA | CACCCCTTTCTCCTTTCTCC | 276 |
| chr9:3101503-3103518 | p2 | (TA)16 | AGGGCACCTTTGTCATTTTG | AACACTCGGATAGAATGCGG | 161 |
| chr9:3101503-3103518 | p3 | (TGG)5 | AGGGCACCTTTGTCATTTTG | AACACTCGGATAGAATGCGG | 161 |
| chr9:3110340-3112356 | p2 | (AT)8 | TAATGGGGGTTGCACATTTT | AACTTACCTCTTGCAGGCCA | 206 |
| chr9:3117398-3119416 | p6 | (CAACTC)3 | AGCAAAAGTGCCAAGCAAAT | CGGGCCCTAGTAAGTTGACA | 191 |
| chr9:3156295-3158329 | p2 | (AT)7 | TAACGAACAACACCGATCCA | CTCCGATCTAATCCAGTCGC | 188 |
| chr9:3256233-3258320 | p3 | (TAT)10 | TTCGATTTCGCAATTAAGGC | TTTCACGGCTACAAAATCCC | 239 |
| chr9:3298990-3301005 | p5 | (AAAAT)3 | AGGGGGACCCACTAATGTCT | TTATTTCTGGTGTTTCCGCC | 150 |
| chr9:3347769-3349784 | p5 | (ATGAT)3 | AGCCTCGGGGACAACTCTAT | GTCAAACGGAGCAATAGGGA | 257 |
| chr9:3391843-3393863 | p5 | (AAATA)4 | ATGCTCGGATCAGCAGACTT | GGGCCTAGTCCAACCAATTT | 235 |
| chr9:3391843-3393863 | p2 | (AT)13 | ATGCTCGGATCAGCAGACTT | GGGCCTAGTCCAACCAATTT | 235 |
| chr9:3398215-3400230 | p4 | (AAAC)4 | GAGGGGGACCCACTAAAGTC | ATAGCGAAACCAACTCGTGG | 160 |
| chr9:3398215-3400230 | p5 | (AAAAT)3 | GAGGGGGACCCACTAAAGTC | ATAGCGAAACCAACTCGTGG | 160 |
| chr9:3410515-3412531 | p4 | (TTTA)4 | GCAATGTGAAGGGCAAAAAT | CAGATTGCCCTCCTTTTGAG | 288 |
| chr9:3467304-3469319 | p3 | (ATT)5 | TCAGCACCAACGCACTAAAC | TGAATCATCAAAGAGCACGC | 172 |
| chr9:3564348-3566390 | p3 | (ATT)14 | TATGGTGGTCTCAGGAAGGG | TGTAGAAGTGCCAATGCAGC | 167 |
| chr9:3571373-3573405 | p2 | (AT)16 | CAAAGCTCGGGACTTCTGAC | TGCTGGTGGGAGGTAGTAGG | 169 |
| chr9:3575723-3577741 | p6 | (CCCTAA)3 | TTCAGCTGGCAAAACAGTTG | AAATGACGGTTCTGGGTTTG | 188 |
| chr9:3589912-3591924 | p2 | (CA)6 | ACTGCTTCTGCTACTCCCCA | ACTTTTACTGATTCCGCCCC | 249 |
| chr9:3666982-3668997 | p3 | (AAC)5 | TCTCAACCACCTTGGTTTCC | GACAAGTGCGATGATGCTGT | 275 |
| chr9:3770781-3772799 | p6 | (TTTTAT)3 | TCAATTAGGCGGTTACTGGG | AGCTTTTGGAGCGTGATGTT | 289 |
| chr9:3812902-3814920 | p6 | (TTTTAT)3 | TCAATTAGGCGGTTACTGGG | AGCTTTTGGAGCGTGATGTT | 289 |
| chr9:3858803-3860827 | p2 | (AT)12 | GGAGAAATCCTGGTGGACAA | AAGTTCCGAGGACAACATGG | 278 |
| chr9:3885784-3887798 | p2 | (TA)6 | TTTTTGGGAAGCGGTTAGAA | TTAATCACCCAACCGACGTT | 286 |
| chr9:3885784-3887798 | p2 | (TA)7 | TTTTTGGGAAGCGGTTAGAA | TTAATCACCCAACCGACGTT | 286 |
| chr9:3885830-3887842 | p2 | (TA)6 | TTTTTGGGAAGCGGTTAGAA | TTAATCACCCAACCGACGTT | 240 |
| chr9:3885830-3887842 | p2 | (TA)7 | TTTTTGGGAAGCGGTTAGAA | TTAATCACCCAACCGACGTT | 240 |
| chr9:3885848-3887862 | p2 | (TA)6 | TTTTTGGGAAGCGGTTAGAA | TTAATCACCCAACCGACGTT | 222 |
| chr9:3885848-3887862 | p2 | (TA)7 | TTTTTGGGAAGCGGTTAGAA | TTAATCACCCAACCGACGTT | 222 |

| Marker ID | SSR type | SSR motif | FORWARD PRIMER (5'-3') | REVERSE PRIMER (5'-3') | size (bp) |
|---|---|---|---|---|---|
| chr9:3889364-3891385 | p3 | (GAA)7 | GACTACGGGCGGTAATGAAA | AAAAGAGGGCAAGGAAAAGG | 221 |
| chr9:3926667-3928691 | p2 | (TC)12 | GCATTGCTTCCGACCATTAT | CCAAACGGGCTCTATGTGTT | 198 |
| chr9:3952284-3954356 | p2 | (TA)36 | TCTGTGCGTGAAAAGACAGG | GTTGATGGAATCCGCAACTT | 110 |
| chr9:3964573-3966585 | p6 | (AAAAGA)3 | GTTTGGGCTGGAAAACAAAA | GAAAAGGGCCAAAATTACCC | 135 |
| chr9:3964573-3966585 | p2 | (TA)6 | GTTTGGGCTGGAAAACAAAA | GAAAAGGGCCAAAATTACCC | 135 |
| chr9:4168230-4170245 | p5 | (TATTT)3 | GAGCTAAAAATGCACCGAGG | GGCCTTCGCCTTTAGATTTT | 194 |
| chr9:4174191-4176206 | p5 | (AACCC)3 | TTACGAAGCAAAGAGTGGGG | CGAAGAGGTGCAAAGGAGTC | 153 |
| chr9:4203151-4205163 | p2 | (AT)6 | GTGACCAAACGGGCTCATAG | GCACCAAAATCACCCACCTA | 264 |
| chr9:4241302-4243362 | p3 | (AAT)20 | GATTTTGAGGCGATTCCAGA | AGGAGAAATCCCTTGGAGGA | 164 |
| chr9:4241302-4243362 | p5 | (AATTA)3 | GATTTTGAGGCGATTCCAGA | AGGAGAAATCCCTTGGAGGA | 164 |
| chr9:4242199-4244214 | p3 | (AAT)20 | AATGGATTAATGAAACGCCG | GATTCACGCCGGAAAGTTTA | 194 |
| chr9:4242199-4244214 | p5 | (AATTA)3 | AATGGATTAATGAAACGCCG | GATTCACGCCGGAAAGTTTA | 194 |
| chr9:4250974-4252989 | p3 | (AAC)10 | ATGGGTACGCGTTAAAATCG | ATAATACCCGTTCGGAACCC | 206 |
| chr9:4250974-4252989 | p5 | (ATGTT)3 | ATGGGTACGCGTTAAAATCG | ATAATACCCGTTCGGAACCC | 206 |
| chr9:4270490-4272506 | p2 | (CT)8 | AAAAATCGGGTCAATGCAAG | TGGAGAAACCGAACTAACGG | 192 |
| chr9:4270490-4272506 | p2 | (AT)18 | AAAAATCGGGTCAATGCAAG | TGGAGAAACCGAACTAACGG | 192 |
| chr9:4270490-4272506 | p2 | (AG)8 | AAAAATCGGGTCAATGCAAG | TGGAGAAACCGAACTAACGG | 192 |
| chr9:4272882-4274900 | p6 | (TTTTTC)3 | TTGGCCCTTGTTCCATTTAG | CACCTCATTGCACATCCAAC | 269 |
| chr9:4272882-4274900 | p2 | (TA)9 | TTGGCCCTTGTTCCATTTAG | CACCTCATTGCACATCCAAC | 269 |
| chr9:4285510-4287564 | p2 | (AC)27 | GCACACCCAACAATTGTCAC | GCTTGCACAGTTTTTGTCCA | 107 |
| chr9:4391793-4393808 | p5 | (TTTTC)3 | CTTCACATTAAGCTTCGGGC | ATTTTCATTCCAGGTCGTGC | 264 |
| chr9:4423838-4425853 | p5 | (ATATA)3 | GCATTCCGTGAAATGCTTCT | TAACACAAATTGCAACGGGA | 136 |
| chr9:4423838-4425853 | p3 | (ACA)7 | GCATTCCGTGAAATGCTTCT | TAACACAAATTGCAACGGGA | 136 |
| chr9:4501707-4503723 | p2 | (AT)8 | CACAGAAGGCAGTTGCTGAA | TCAAGAGCCACCTTAGCTCC | 201 |
| chr9:4517126-4519180 | p2 | (TA)27 | TCATGAGCTGGCGACTAGTTT | TGGCAAAACCTCCAATTTTC | 162 |
| chr9:4597132-4599147 | p5 | (TATTG)3 | CTCTCTACCCCTTGGCCTTC | GTGGGCAAAAGGGTGTCTAA | 212 |
| chr9:4722858-4724873 | p3 | (ACC)5 | GGGCCAATTAGAGAAAAGGG | TGAATGGTGGCTTTGTGTGT | 294 |
| chr9:4745974-4747998 | p2 | (TA)12 | AATGTCTACGAGCCGTGTCC | GAAGCAATTTCCTCGTAGCG | 229 |
| chr9:4793209-4795225 | p2 | (AT)34 | CTTGTTGCTGCTTCTTTCCC | GTGGGTCATACAGATTCCGC | 183 |
| chr9:4793250-4795266 | p2 | (AT)34 | CTTGTTGCTGCTTCTTTCCC | CTCCATTAAGCCTTGAGCCA | 142 |
| chr9:4829885-4831900 | p3 | (CCA)5 | GTCAATTGGCAGCTGCTGTA | AGAAATTCGTGCGTGAAACC | 154 |
| chr9:4829885-4831900 | p5 | (AATTT)3 | GTCAATTGGCAGCTGCTGTA | AGAAATTCGTGCGTGAAACC | 154 |
| chr9:4847288-4849304 | p2 | (CT)6 | CATCGAATTTTGGGGAAAGA | AGCCCAAAGTAAGCGGAAAT | 240 |
| chr9:4847300-4849315 | p2 | (CT)6 | CATCGAATTTTGGGGAAAGA | AGCCCAAAGTAAGCGGAAAT | 228 |
| chr9:4853844-4855859 | p6 | (CATATA)5 | AGCCCACAAGGCTACAAAAA | TAGGAGCTTGTCAGGGGAGA | 196 |
| chr9:4853844-4855859 | p3 | (ATC)5 | AGCCCACAAGGCTACAAAAA | TAGGAGCTTGTCAGGGGAGA | 196 |

| Marker ID | SSR type | SSR motif | FORWARD PRIMER (5'-3') | REVERSE PRIMER (5'-3') | size (bp) |
|---|---|---|---|---|---|
| chr9:4873551-4875569 | p5 | (GTCCT)3 | TTTTGTAGCCTTGTCGGCTT | TGTACGGGCATGAATTCTCA | 192 |
| chr9:4873551-4875569 | p6 | (AAAAAT)3 | TTTTGTAGCCTTGTCGGCTT | TGTACGGGCATGAATTCTCA | 192 |
| chr9:4884303-4886318 | p5 | (TTTAA)3 | GCTCTTTTTAATCCGGAGGG | ATAGTGGCGTTAGATCCCCC | 126 |
| chr9:4915906-4917932 | p2 | (AT)10 | TTGCTGCCTAACACATTCCA | TTGAGTCGCCCCTAACAATC | 104 |
| chr9:4915906-4917932 | p2 | (AT)13 | TTGCTGCCTAACACATTCCA | TTGAGTCGCCCCTAACAATC | 104 |
| chr9:4954463-4956514 | p3 | (ATT)17 | GTCCGGCAGAAATTTGGTAA | TGCGGGTCTTTAATTTTTGC | 205 |
| chr9:4954463-4956514 | p3 | (CAA)6 | GTCCGGCAGAAATTTGGTAA | TGCGGGTCTTTAATTTTTGC | 205 |
| chr9:4976570-4978588 | p6 | (GAAAAA)3 | ATCTAGTCCGCTGCATGCTT | GGGATGAGGATTAAACGCAG | 228 |
| chr9:4982776-4984794 | p3 | (TAA)6 | TAATTTCCTGTTTACGCCGC | CAGAATGTTGCTGCTTCCAA | 173 |
| chr9:4982776-4984794 | p5 | (TATTA)7 | TAATTTCCTGTTTACGCCGC | CAGAATGTTGCTGCTTCCAA | 173 |
| chr9:4998958-5000976 | p6 | (AACCGG)3 | CAAACCACCTTTGTGTGTCG | GTCCATGACCAGTTTGCCTT | 133 |
| chr9:4999166-5001180 | p6 | (AACCGG)3 | TGCTTTGTCGCAGTGCTTAT | AGCCAGCAAGAGGATTCAGA | 143 |
| chr9:5039821-5041839 | p2 | (AT)9 | TTAATGTCCAAAAGGGTGGG | TAAAGGGACAACGGACGAAG | 279 |
| chr9:5039851-5041877 | p2 | (AT)9 | TTAATGTCCAAAAGGGTGGG | TAAAGGGACAACGGACGAAG | 249 |
| chr9:5104107-5106128 | p3 | (TGT)7 | CACCTCCACCCAGTTGTTCT | CACGTGACGAGGAGGTGTTA | 282 |
| chr9:5118138-5120158 | p5 | (TTGGT)4 | ACTCGCATAGCGTAGGTGCT | TGGGTGAGCCTTCATTTTTC | 187 |
| chr9:5118138-5120158 | p5 | (TTGGT)3 | ACTCGCATAGCGTAGGTGCT | TGGGTGAGCCTTCATTTTTC | 187 |
| chr9:5123736-5125754 | p6 | (CGATCT)3 | AAGTGATAAATTCACGCGGC | GTTGTTGGGTTCGAGCATTT | 150 |
| chr9:5123736-5125754 | p2 | (TG)8 | AAGTGATAAATTCACGCGGC | GTTGTTGGGTTCGAGCATTT | 150 |
| chr9:5124630-5126646 | p6 | (CGATCT)3 | AATATCCACCAAAACGACGC | CTTCTTCCGCGAAGATCAAC | 215 |
| chr9:5124630-5126646 | p2 | (TG)8 | AATATCCACCAAAACGACGC | CTTCTTCCGCGAAGATCAAC | 215 |
| chr9:5132330-5134345 | p5 | (AATAA)3 | GCGCGCGTCTATACCTACTC | ACATCCCAACTCTGGACCTG | 179 |
| chr9:5132330-5134345 | p3 | (TGG)5 | GCGCGCGTCTATACCTACTC | ACATCCCAACTCTGGACCTG | 179 |
| chr9:5203507-5205521 | p2 | (TA)7 | CTCCATGTTTAACCCCGAGA | CTGCACTTCACGATGCAGAT | 274 |
| chr9:5250490-5252508 | p2 | (TG)14 | AGGCAATTGTTCTCTGGTGG | CCCCTCTCTCTCTCTCCGTT | 269 |
| chr9:5250490-5252508 | p3 | (ACA)6 | AGGCAATTGTTCTCTGGTGG | CCCCTCTCTCTCTCTCCGTT | 269 |
| chr9:5319241-5321256 | p5 | (TAAAA)3 | GTTTGGCCTTGCGACTAAAA | ATTAGCCACACTCAAACGGC | 254 |
| chr9:5319241-5321256 | p5 | (TTTAA)3 | GTTTGGCCTTGCGACTAAAA | ATTAGCCACACTCAAACGGC | 254 |
| chr9:5413607-5415621 | p2 | (TA)7 | AGACTTCATGTTGAACCCCG | TGTTGTGCCATGTTTTTCGT | 253 |
| chr9:5419506-5421542 | p3 | (ACA)12 | TTAAATGTCGTGCCCAGTCA | GGATTCTTCGTTGGTGAGGA | 261 |
| chr9:5432555-5434599 | p2 | (CT)22 | CAATCTGCCCAATAACCACC | AATGAACCAGTTGCTCACCC | 222 |
| chr9:5562106-5564127 | p3 | (AAG)7 | CGAACACGAATGAATTGCAG | TTAGGATGACCGAAGAACCG | 225 |
| chr9:5589294-5591310 | p2 | (TG)8 | AGAAGCCACCACCAAAAATG | ACGTAGCTTCTCAAGGCGAA | 178 |
| chr9:5639548-5641563 | p5 | (AACCG)3 | TACGATCCAAAACCAAAGGC | TGACCGTTCATCCTTTGTGA | 211 |
| chr9:5753408-5755424 | p4 | (AAAT)4 | GTCAACCAACCCAGTGTCCT | GAAAGGCCAAATCAAACGAA | 119 |
| chr9:5753408-5755424 | p3 | (AAT)6 | GTCAACCAACCCAGTGTCCT | GAAAGGCCAAATCAAACGAA | 119 |

| Marker ID | SSR type | SSR motif | FORWARD PRIMER (5'-3') | REVERSE PRIMER (5'-3') | size (bp) |
|---|---|---|---|---|---|
| chr9:5772680-5774696 | p4 | (AAAT)4 | GTCAACCAACCCAGTGTCCT | GAAAGGCCAAATCAAACGAA | 119 |
| chr9:5772680-5774696 | p3 | (AAT)6 | GTCAACCAACCCAGTGTCCT | GAAAGGCCAAATCAAACGAA | 119 |
| chr9:5796705-5798717 | p2 | (TA)6 | ACTGACGGACTGCTCTGGAT | CCGCCTAATTATTTCGTGGA | 138 |
| chr9:5949048-5951073 | p2 | (AT)8 | TTTGTTAAAAATCGGCTGGG | AGGTCATAACTTTGCGCTCG | 168 |
| chr9:5959529-5961544 | p3 | (AAT)5 | TGGTCCCTCTTGTGTCCTCT | TCCCCTAATGTGGGACTCTG | 296 |
| chr9:6019667-6021687 | p2 | (TA)10 | TTCCTTGATTTGCTTGGTCC | AGCCAACAACACCAACACAA | 211 |
| chr9:6109200-6111218 | p3 | (ATT)6 | AGCCATAAGGACATTGCACC | CTGGAATGTGAACGGGAGAT | 186 |
| chr9:6109240-6111255 | p3 | (ATT)6 | AGCCATAAGGACATTGCACC | CTGGAATGTGAACGGGAGAT | 146 |
| chr9:6189669-6191684 | p6 | (TATATG)3 | TGTGGGTGTGTGTGCCTACT | AAAGCCGACAAGGCTATCAA | 281 |
| chr9:6189669-6191684 | p5 | (TGCAT)3 | TGTGGGTGTGTGTGCCTACT | AAAGCCGACAAGGCTATCAA | 281 |
| chr9:6522544-6524559 | p3 | (AAG)5 | GTTCGTAAACAAACCGCCAT | CTCCGTATTGTTTACCCCGA | 278 |
| chr9:6606671-6608716 | p3 | (GAA)8 | TCTTTTTCGCGGTATTCTGG | GAGAGGCGGTACATTCCTGA | 102 |
| chr9:6626308-6628320 | p2 | (AC)6 | TCCCGAAGTGGACGTCTTAG | GCTAAATGCACTTTCCGAGG | 171 |
| chr9:6626308-6628320 | p2 | (CT)26 | TCCCGAAGTGGACGTCTTAG | GCTAAATGCACTTTCCGAGG | 171 |
| chr9:6626322-6628340 | p2 | (AC)6 | TCCCGAAGTGGACGTCTTAG | TTGCCTCACCACTAGCCTTT | 157 |
| chr9:6626322-6628340 | p2 | (CT)26 | TCCCGAAGTGGACGTCTTAG | TTGCCTCACCACTAGCCTTT | 157 |
| chr9:6641989-6644007 | p6 | (TCCACG)3 | AGAGGAATATTTGGGGGTGG | CTATTTCTTCGGCAGCGAAC | 142 |
| chr9:6641989-6644007 | p6 | (TCCACG)3 | AGAGGAATATTTGGGGGTGG | CTATTTCTTCGGCAGCGAAC | 142 |
| chr9:6715045-6717063 | p5 | (TATTT)3 | CGAAATGAAAAGCCGGTAAA | CCCTTGAGAATTGAAGAGCG | 223 |
| chr9:6715045-6717063 | p3 | (ACA)6 | CGAAATGAAAAGCCGGTAAA | CCCTTGAGAATTGAAGAGCG | 223 |
| chr9:6765538-6767556 | p2 | (CT)9 | CTACGAAATCGACCCCAAAA | AGCACAACGTTGATGAACCA | 226 |
| chr9:6776762-6778774 | p2 | (GA)6 | TATCTGTGGGGCAAAGATCC | TAGACATGTGCTCCAGACGG | 161 |
| chr9:6811035-6813050 | p5 | (CACAA)3 | TAATCCGGGCTTGAAAACTG | GCTAGATGGGTTTGCACCAT | 178 |
| chr9:6846326-6848348 | p2 | (AT)11 | CATCAGAATGTCTGGGGGAC | GAAGGATTAGGGCCAAGAGC | 206 |
| chr9:7072240-7074256 | p2 | (AT)8 | CCGTCGTAGAACCTGGTGTT | TTGGCACTTGGAATGTCAGA | 145 |
| chr9:7072240-7074256 | p2 | (TA)31 | CCGTCGTAGAACCTGGTGTT | TTGGCACTTGGAATGTCAGA | 145 |
| chr9:7157083-7159095 | p3 | (ACA)6 | TGGAAATGCAAATCACGAAA | AAGAGGTAGGGGTAGGGCAA | 278 |
| chr9:7157083-7159095 | p2 | (TA)6 | TGGAAATGCAAATCACGAAA | AAGAGGTAGGGGTAGGGCAA | 278 |
| chr9:7229989-7232005 | p2 | (TA)8 | TTTTTCTTTGGCACGAATCC | CTTTTGGAGCGGCATATCAT | 299 |
| chr9:7237575-7239603 | p2 | (TC)14 | AATTGTGGACTCCGTTTGGA | GGGGAATCATCTGAACCCTT | 144 |
| chr9:7295127-7297171 | p2 | (AT)22 | TATTTTGATGGCCAAGAGGC | AACTTTTGTCACGACCCAGC | 168 |
| chr9:7306303-7308321 | p6 | (AAAAAT)3 | GTGTCGAAAGCAATGGTTGA | TTTTGTTTTCAGCCTCGACA | 122 |
| chr9:7341825-7343843 | p6 | (ATTTTT)3 | TCCTATGAGAATTGGGCTCG | GAAGGTGTGCTCTGCATTGA | 276 |
| chr9:7361542-7363556 | p2 | (TC)7 | GTATAGAAATTTCGGCGGCA | GTTCGTTAGAATTTCGGGCA | 114 |
| chr9:7414734-7416752 | p3 | (TGT)6 | GAAGGTGCCACCGAAACTTA | CGCATCTTGTCTTCCACTGA | 277 |
| chr9:7474408-7476423 | p3 | (ACA)5 | TTACACGTGGAGGCCACATA | ACCCCTAAGCTTGGCAAAAT | 285 |

| Marker ID | SSR type | SSR motif | FORWARD PRIMER (5'-3') | REVERSE PRIMER (5'-3') | size (bp) |
|---|---|---|---|---|---|
| chr9:7479011-7481029 | p6 | (GGTGGC)3 | ATCTTGGCGACCCATTACAC | GAAGAAGAAAGGGAGGGTGG | 126 |
| chr9:7536305-7538320 | p5 | (CATTT)3 | CGTGGTTTTGGGTTGACTTT | ATCCCAATTTTTCAAGCACG | 136 |
| chr9:7599808-7601858 | p2 | (AT)25 | ATGCCCACCGAAATCTATCA | AAAAATCTTGACGTGTGGGC | 105 |
| chr9:7641229-7643245 | p5 | (TGCAT)3 | TGGAGAAAGCGAAGGAAGAA | ACCGGCCAACTGTCTCATAC | 131 |
| chr9:7641229-7643245 | p4 | (TATC)4 | TGGAGAAAGCGAAGGAAGAA | ACCGGCCAACTGTCTCATAC | 131 |
| chr9:7682385-7684415 | p2 | (TA)6 | GGGTCGTGACAAAAGATGCT | TCTTTTAGTGCGGGTCCAAG | 190 |
| chr9:7682385-7684415 | p6 | (CCGGTT)5 | GGGTCGTGACAAAAGATGCT | TCTTTTAGTGCGGGTCCAAG | 190 |
| chr9:7736669-7738685 | p2 | (TA)8 | CCGCCATTTAGTTGGTTGAT | CTGTAAACTGGCGGAACCAT | 145 |
| chr9:7736669-7738685 | p2 | (CT)7 | CCGCCATTTAGTTGGTTGAT | CTGTAAACTGGCGGAACCAT | 145 |
| chr9:7765496-7767514 | p2 | (AT)12 | GGAACTGTCTGCAAAGCCTC | TCCACTTCTTCCAGGAGCAT | 133 |
| chr9:7765496-7767514 | p3 | (ATC)6 | GGAACTGTCTGCAAAGCCTC | TCCACTTCTTCCAGGAGCAT | 133 |
| chr9:7918101-7920125 | p2 | (TC)12 | ACCACCTCACTGGAAAAACG | TTTGATTGCATCCGTGAGAA | 284 |
| chr9:7996084-7998105 | p3 | (AAG)7 | ATGACACAGCCTGCATGAAC | AATTATCCGCAAGCAAATCG | 267 |
| chr9:7996084-7998105 | p2 | (TA)9 | ATGACACAGCCTGCATGAAC | AATTATCCGCAAGCAAATCG | 267 |
| chr9:7996084-7998105 | p4 | (TTTC)4 | ATGACACAGCCTGCATGAAC | AATTATCCGCAAGCAAATCG | 267 |
| chr9:8002073-8004091 | p6 | (ATAAAA)3 | TTTCCCCATCCAAAATCAAA | ATGCATTCGTTACTCCAGCC | 214 |
| chr9:8032788-8034808 | p2 | (TC)10 | AACCGCGAAATTTGATTCAC | GTTATGCGATTGGGGATTTG | 129 |
| chr9:8128807-8130834 | p3 | (TTG)9 | AATGACTTTGGAGGTGGGTG | CTTTAAACGGCGCTAATCCA | 109 |
| chr9:8179319-8181334 | p5 | (ATTTT)3 | TTACATCCGCCTTGGCTTAC | CCAAAAGGGAGGGAAAAAGA | 208 |
| chr9:8179319-8181334 | p2 | (CT)7 | TTACATCCGCCTTGGCTTAC | CCAAAAGGGAGGGAAAAAGA | 208 |
| chr9:8317047-8319062 | p3 | (ATT)5 | CTTCACGGATAAAAAGGGCA | CTGGCTTCGTCACCGTATCT | 125 |
| chr9:8372425-8374446 | p3 | (TTA)7 | GGTGTTCATGGGAAATCCAG | TTAGGGGTCGTTTGGTATGG | 173 |
| chr9:8377178-8379193 | p5 | (TTAAA)3 | CCTGTTTGGATGGGCTTAAA | TTTGAGTGTTTGGCTGACCA | 127 |
| chr9:8467031-8469095 | p2 | (TA)32 | AAAGGTGACTGCGCCTTAAA | ACAGCAGACCAAGCCCATAC | 186 |
| chr9:8506515-8508527 | p2 | (TA)6 | GGTTTTTGTTGGGGTTGAGA | TAGGGCTACATATCGGTCGG | 272 |
| chr9:8544442-8546457 | p5 | (TATTA)3 | TGTTGCCTCTCAAATTGCTG | TTTTTATGCTTGAGCTGGGG | 102 |
| chr9:8551026-8553042 | p4 | (TTTA)4 | AAGGCTTGTCCGTGAACATC | CGTGCAGTGACATGTTAGCC | 269 |
| chr9:8631735-8633750 | p5 | (TATGA)3 | TCTTTTCGACCCTGAGCCTA | TCTGGGATAGAGACCGGATG | 215 |
| chr9:8643241-8645256 | p5 | (ATATG)3 | GTTGCTGCCCAAATTTCTGT | ATGCTCTTTACGCCTTTCCA | 105 |
| chr9:8647772-8649784 | p2 | (AG)6 | GTCACCATGAAAACTCGCCT | TTTTGACCCATTTACCCCAA | 184 |
| chr9:8671796-8673808 | p2 | (AT)6 | CGTTCCCGCCAAAATACTAA | CGGGTAGGAATGAACAGCAT | 222 |
| chr9:8679448-8681460 | p2 | (TC)6 | GGTTCTGCTCTTTCACTGCC | GGTTAACGCATATGCAACCC | 181 |
| chr9:8783716-8785768 | p2 | (AT)26 | TGTGCTTGATTTCTGGCTTG | AATCTTCAAGACGGTGGACG | 100 |
| chr9:8803924-8805951 | p3 | (TTA)5 | TTATGACTTGAGTCGCACGG | CGTGCCTTTTAATCCTCCAA | 281 |
| chr9:8893440-8895455 | p5 | (TAAAA)3 | CGCTCTGTTACTCGTTGTCG | CCCTCATACATTAAGGGCCA | 135 |
| chr9:8926574-8928598 | p2 | (CA)12 | TAACCGAGAATAAAACGCCG | TGCTCCAACCAATCACGATA | 295 |

| Marker ID | SSR type | SSR motif | FORWARD PRIMER (5'-3') | REVERSE PRIMER (5'-3') | size (bp) |
|---|---|---|---|---|---|
| chr9:9008073-9010088 | p5 | (TTTGG)3 | AGCCATATGGTTCGCTCAGT | ACGCGTATTCTAGTCCCCCT | 222 |
| chr9:9075026-9077038 | p2 | (GT)6 | TTCCACCCAACACATGCTTA | TCAACAAGTCAGTGGCGAAC | 289 |
| chr9:9100044-9102058 | p2 | (GA)7 | ATGCTCGGATTCAAACGAAC | TCTTGAATCGCCAAATTTCC | 266 |
| chr9:9100569-9102614 | p2 | (GA)7 | AAACCTACAAGCGGACGAAA | CGGTCAGTGCTCAACCAGTA | 200 |
| chr9:9100616-9102658 | p2 | (GA)7 | AAACCTACAAGCGGACGAAA | CGGTCAGTGCTCAACCAGTA | 153 |
| chr9:9259317-9261332 | p5 | (AACCG)3 | ACACCTCAAAGGGTTCATGC | GCGTACCGTAGCTAAGTCGG | 119 |
| chr9:9259317-9261332 | p3 | (GCT)7 | ACACCTCAAAGGGTTCATGC | GCGTACCGTAGCTAAGTCGG | 119 |
| chr9:9366004-9368020 | p4 | (GAAA)4 | TGTTCGCCTAAGTGGACCTC | CAATTCCATAAATCCCGTGG | 177 |
| chr9:9366004-9368020 | p2 | (TA)14 | TGTTCGCCTAAGTGGACCTC | CAATTCCATAAATCCCGTGG | 177 |
| chr9:9447642-9449654 | p2 | (AG)6 | GTCCCTCGCGATATGGTAGA | GCATATACACGGGGCTGAGT | 255 |
| chr9:9461720-9463744 | p3 | (TTC)8 | ACAGCGATGAGAAAGGCACT | CCCTTTTCGATTAACTGGCA | 159 |
| chr9:9468707-9470765 | p2 | (AT)7 | CTCTTTTGCTTTGGAGGTCG | CTTCTTTGACCCGTTTGAGC | 264 |
| chr9:9468707-9470765 | p2 | (AT)8 | CTCTTTTGCTTTGGAGGTCG | CTTCTTTGACCCGTTTGAGC | 264 |
| chr9:9606542-9608560 | p3 | (ACA)6 | GCTGAAGCATTGAGTGGTGA | CAATCTCTTGCCGGACAAAT | 203 |
| chr9:9663505-9665520 | p5 | (GAAAA)3 | GCCAAGTTTGAGCCTCTTTG | GCGATTGTGAACTCAGAGCA | 208 |
| chr9:9669337-9671352 | p5 | (ATTTT)3 | AATCTCGGGATTCAAACGTG | GCGCCAAAGATTAAAACCAA | 175 |
| chr9:9669337-9671352 | p2 | (AT)9 | AATCTCGGGATTCAAACGTG | GCGCCAAAGATTAAAACCAA | 175 |
| chr9:9688562-9690577 | p5 | (GAAAA)3 | GCCAAGTTTGAGCCTCTTTG | GCGATTGTGAACTCAGAGCA | 208 |
| chr9:9694394-9696409 | p5 | (ATTTT)3 | AATCTCGGGATTCAAACGTG | GCGCCAAAGATTAAAACCAA | 175 |
| chr9:9694394-9696409 | p2 | (AT)9 | AATCTCGGGATTCAAACGTG | GCGCCAAAGATTAAAACCAA | 175 |
| chr9:9710111-9712129 | p6 | (TTTTGT)3 | TTTCACTTGCAGCCTTCCTT | AAGGAATGAGTGCCAACACC | 135 |
| chr9:9712426-9714441 | p5 | (AAAAT)3 | GAAAAATGTGCCTGGGAGAA | CATCTCCTGGGTCAGAGCAT | 129 |
| chr9:9804666-9806681 | p5 | (AATAA)3 | TCGGTCTCATCTGAAAACCC | ATAATATGAACATCGCCCGC | 231 |
| chr9:9804666-9806681 | p2 | (AT)11 | TCGGTCTCATCTGAAAACCC | ATAATATGAACATCGCCCGC | 231 |
| chr9:9807006-9809024 | p2 | (TA)9 | GGGCTTCTCAAGAGTTGCTG | GGGACTAAAGTGGGCATTGA | 130 |
| chr9:9854105-9856120 | p5 | (TTTAT)3 | TAGACTAACCAAATGGGCCG | ACGATGCCTAGCTCGAAAAA | 222 |
| chr9:9854105-9856120 | p2 | (CT)9 | TAGACTAACCAAATGGGCCG | ACGATGCCTAGCTCGAAAAA | 222 |
| chr9:9854653-9856671 | p5 | (TTTAT)3 | ACGGGCATGCTGAAATAAAC | TTGGGTGTCAACACCGTCTA | 279 |
| chr9:9854653-9856671 | p2 | (CT)9 | ACGGGCATGCTGAAATAAAC | TTGGGTGTCAACACCGTCTA | 279 |
| chr9:9881971-9883991 | p2 | (AT)10 | AAATCACATGCCATTGCTCA | TCTTCCACAGATGCTGATGC | 116 |
| chr9:9900480-9902504 | p6 | (TATATG)4 | TCCCTTCTTATGGATGCTGC | GCTTATTTGTCCAAGCTCGC | 200 |
| chr9:9918194-9920209 | p2 | (AT)15 | CACCCATGGAAAAATTGACC | ACTCCTCAACTTGCCTCGAA | 178 |
| chr9:9918194-9920209 | p5 | (TGAAT)3 | CACCCATGGAAAAATTGACC | ACTCCTCAACTTGCCTCGAA | 178 |
| chr9:9938820-9940884 | p2 | (AT)32 | TTCATCTGGGCAGTGTATGC | ATGATACCAGGCACCCACAT | 243 |
| chr9:9954524-9956540 | p4 | (CTAA)4 | TGTACTGCTGCCAGTTCCAC | AGCAATCGAGGGAATGACAC | 293 |
| chr9:10016827-10018839 | p2 | (TC)6 | CCATTGCATTCGATTCCTCT | AAATGGCCAAGTGTCAATCC | 157 |

| Marker ID | SSR type | SSR motif | FORWARD PRIMER (5'-3') | REVERSE PRIMER (5'-3') | size (bp) |
|---|---|---|---|---|---|
| chr9:10080072-10082140 | p2 | (TA)34 | GGCACACTGGTCTCCAAACT | GAAGGTCAATCCGCAATCAT | 162 |
| chr9:10087282-10089297 | p3 | (TCT)5 | TCCAATTTCTTGGCAACCTC | TGATTTCATGTGGGCAGGTA | 238 |
| chr9:10087282-10089297 | p2 | (AT)6 | TCCAATTTCTTGGCAACCTC | TGATTTCATGTGGGCAGGTA | 238 |
| chr9:10162356-10164368 | p2 | (TA)6 | AATTTGTGAATCCGAACCGA | AATCCCACCAAACAAGCAAC | 215 |
| chr9:10162356-10164368 | p5 | (AATCA)3 | AATTTGTGAATCCGAACCGA | AATCCCACCAAACAAGCAAC | 215 |
| chr9:10208191-10210239 | p3 | (ATA)16 | TCGGATGACAACCAATCAGA | TCATCAGGGTCCTTTCGTTC | 194 |
| chr9:10265752-10267768 | p4 | (TTTA)4 | TAAATGAACCAGACCCGACC | CGGGTACAAAATGATGCAAA | 195 |
| chr9:10265752-10267768 | p6 | (TTTTTG)3 | TAAATGAACCAGACCCGACC | CGGGTACAAAATGATGCAAA | 195 |
| chr9:10283778-10285796 | p6 | (AGGAAA)3 | CTAGGAGTTAGCGTGGGTGC | GAGACATAAGTCCCGAGGCA | 287 |
| chr9:10354509-10356524 | p5 | (TTTAT)3 | GGAACGTAGGCAATCCAAAA | CCTCCAAAGGAATCAGGTCA | 269 |
| chr9:10463160-10465186 | p2 | (AT)13 | ATCGTCGACGGTCATTTGAT | CCGATCGGTTCCAAAGATAA | 115 |
| chr9:10467811-10469823 | p2 | (AT)6 | AGGGCAATGAGACAAACAGG | CCAAAACTCTTTGCCACCTC | 118 |
| chr9:10467811-10469823 | p2 | (TA)7 | AGGGCAATGAGACAAACAGG | CCAAAACTCTTTGCCACCTC | 118 |
| chr9:10508174-10510189 | p5 | (ATTTA)3 | ACGCTTATTTCACCCTTCCC | TCCATTGCAAAATCATTCCA | 261 |
| chr9:10549477-10551495 | p6 | (TGGGTG)3 | TTTGCCCAGAAAAATGAACC | TTCAAATCCTTATGGGCGTC | 298 |
| chr9:10565742-10567775 | p3 | (AAT)11 | CCGAGATACATTGGGCCTTA | CTTGCGAAGCATTGACACAT | 248 |
| chr9:10611596-10613614 | p6 | (ATTATG)3 | GGATTGGAGATCGAACGAAA | CGGGCATTATACGAACCACT | 294 |
| chr9:10657078-10659142 | p2 | (AT)33 | AAACTTCGTGCCCAATCAAC | TGTGCAAATGGCTAAGTGGA | 214 |
| chr9:10657078-10659142 | p2 | (AT)32 | AAACTTCGTGCCCAATCAAC | TGTGCAAATGGCTAAGTGGA | 214 |
| chr9:10667640-10669667 | p3 | (ATT)9 | TCAGAATTGGGAGTTTTGGG | CAGCAAAACTCAAGCATCCA | 285 |
| chr9:10826978-10828999 | p3 | (AAT)7 | ATCCGAACGTTAGGGCTTCT | TGTCACCACGTCTGCTTCTC | 238 |
| chr9:10954298-10956352 | p2 | (TA)27 | ACTGCTTCTACTGCTCCCCA | TACCCGCAAGACACTTAGCC | 271 |
| chr9:11084494-11086509 | p3 | (TTA)5 | TCCGGTACCTCATTGGAGTC | GTACCGGACACGATATGGGA | 182 |
| chr9:11084494-11086509 | p2 | (TA)8 | TCCGGTACCTCATTGGAGTC | GTACCGGACACGATATGGGA | 182 |
| chr9:11090338-11092404 | p2 | (TA)33 | AACATTGAATTCCTTCAATTTAG[illegible] | ACCATGGGGTTGATTTCTTG | 136 |
| chr9:11093069-11095084 | p5 | (ACCGA)3 | ATTTGTTTGCTTTGGGATGC | GTGAGGTTTGACTGGAGGGA | 239 |
| chr9:11104459-11106474 | p5 | (TGGTT)3 | GGATCACGTCAAAGCCGTAT | TGAGGACATGGAGTTAGGGG | 192 |
| chr9:11104459-11106474 | p5 | (TGGTT)3 | GGATCACGTCAAAGCCGTAT | TGAGGACATGGAGTTAGGGG | 192 |
| chr9:11229716-11231732 | p2 | (AT)8 | CAGCCGGTCAGAAAAGAGTC | ACCGAAGTGGTATCAGAGCG | 138 |
| chr9:11229716-11231732 | p2 | (AT)10 | CAGCCGGTCAGAAAAGAGTC | ACCGAAGTGGTATCAGAGCG | 138 |
| chr9:11231457-11233471 | p2 | (AT)10 | GGTCATGCCAAAAAGGAAGA | TGAATTTACAGGGGAGGTGC | 220 |
| chr9:11231457-11233471 | p2 | (CA)7 | GGTCATGCCAAAAAGGAAGA | TGAATTTACAGGGGAGGTGC | 220 |
| chr9:11231457-11233471 | p5 | (ATTTT)3 | GGTCATGCCAAAAAGGAAGA | TGAATTTACAGGGGAGGTGC | 220 |
| chr9:11291606-11293624 | p6 | (ATATTA)3 | GTCTCTTAGCAGTGCCCGAC | CTCTAACCAGGTGCTAGGCG | 248 |
| chr9:11319058-11321124 | p3 | (ATT)22 | TAGCATGGGTGACATAGCCA | GTTATGCCGGAAAAAGCAAA | 265 |
| chr9:11414051-11416073 | p2 | (TC)11 | GCCCTATGAAAGCGACAGAG | CGCGAGGAAATAAATTGGAA | 260 |

| Marker ID | SSR type | SSR motif | FORWARD PRIMER (5'-3') | REVERSE PRIMER (5'-3') | size (bp) |
|---|---|---|---|---|---|
| chr9:11421027-11423093 | p2 | (AT)28 | TTTCAGAACAAACTCACGGC | TTGTATGGGAGTAACGTGCG | 261 |
| chr9:11504216-11506234 | p6 | (TTCAGT)3 | GTTGTTGCTGATTTGGGTCC | CATACCTCGAGCCTCCCATA | 130 |
| chr9:11504216-11506234 | p3 | (TCT)8 | GTTGTTGCTGATTTGGGTCC | CATACCTCGAGCCTCCCATA | 130 |
| chr9:11508282-11510306 | p3 | (AAT)8 | ATTCTGCCACCATCCCTGTA | GCGTTTCCCGTGTAAAATTG | 263 |
| chr9:11538266-11540278 | p2 | (TC)6 | TGATTTCCAATGCAGGTCAA | TTCCAGGCACTACCCTTTTG | 268 |
| chr9:11546866-11548878 | p2 | (AC)6 | ACCCGGTCTATGTAAACCCC | TGAACAGCGCACAGAGATTT | 227 |
| chr9:11546866-11548878 | p3 | (ATT)7 | ACCCGGTCTATGTAAACCCC | TGAACAGCGCACAGAGATTT | 227 |
| chr9:11550883-11552909 | p2 | (AT)13 | TTGGCAACCCCTTTGTACTC | ACGATTGACTTTGGCACACA | 194 |
| chr9:11550883-11552909 | p2 | (AT)14 | TTGGCAACCCCTTTGTACTC | ACGATTGACTTTGGCACACA | 194 |
| chr9:11601317-11603332 | p5 | (AAGAA)3 | TGGCAAACACTTTGGCATTA | GCGAGCACAAACTGACAATG | 219 |
| chr9:11607024-11609036 | p2 | (CT)6 | GTTGTGGGAGTGAGCAGGTT | GGTTGTTGTGTGCCCTTCTT | 238 |
| chr9:11607024-11609036 | p4 | (TTAT)5 | GTTGTGGGAGTGAGCAGGTT | GGTTGTTGTGTGCCCTTCTT | 238 |
| chr9:11624048-11626076 | p5 | (AACCG)3 | CTTCACCCATGGCTTTGTTT | AAGCATACAAGTCCGCCAAC | 159 |
| chr9:11624048-11626076 | p4 | (AATA)7 | CTTCACCCATGGCTTTGTTT | AAGCATACAAGTCCGCCAAC | 159 |
| chr9:11631569-11633584 | p5 | (TAATA)3 | AGACGAAGAAAAGGGGAAGC | AGCCCTTAAACGAGAACATGA | 288 |
| chr9:11631569-11633584 | p5 | (AATTT)3 | AGACGAAGAAAAGGGGAAGC | AGCCCTTAAACGAGAACATGA | 288 |
| chr9:11675627-11677639 | p2 | (AT)6 | ACAGCAAACTCCACAAACCC | TCCGTCCGATTTCATTGATT | 137 |
| chr9:11696397-11698451 | p2 | (CT)27 | CTCTAAACCGACCCGCAATA | CGACCTCAAAAATTGCCCTA | 102 |
| chr9:11722347-11724361 | p2 | (AT)7 | GTCGAAAAATCCGACCTTCA | CGACAGACATCTGTGGTCGT | 295 |
| chr9:11949411-11951426 | p5 | (AATGA)3 | TCGGGATACGGAAAATCAAG | TCAGCCATGCCAAAATACAA | 300 |
| chr9:11949411-11951426 | p3 | (TTC)5 | TCGGGATACGGAAAATCAAG | TCAGCCATGCCAAAATACAA | 300 |
| chr9:12046733-12048745 | p2 | (CT)6 | ATGGCCTCTCCACACATCTC | CAGTCCTCGGCTCTGTTTTC | 179 |
| chr9:12100719-12102734 | p5 | (TTGGT)3 | GGATTTTGCGTGACGGATAG | TAATGTCGGGTTGGTTTGGT | 127 |
| chr9:12100719-12102734 | p5 | (TTGGT)3 | GGATTTTGCGTGACGGATAG | TAATGTCGGGTTGGTTTGGT | 127 |
| chr9:12131563-12133579 | p2 | (TA)11 | AGACAAAGTCGGCTGCTGAT | CGAGCCTTGAGCGTTAGACT | 107 |
| chr9:12131563-12133579 | p2 | (TA)8 | AGACAAAGTCGGCTGCTGAT | CGAGCCTTGAGCGTTAGACT | 107 |
| chr9:12131563-12133579 | p2 | (TA)10 | AGACAAAGTCGGCTGCTGAT | CGAGCCTTGAGCGTTAGACT | 107 |
| chr9:12131563-12133579 | p2 | (AT)9 | AGACAAAGTCGGCTGCTGAT | CGAGCCTTGAGCGTTAGACT | 107 |
| chr9:12134932-12136956 | p3 | (TGA)8 | AGCCGTTGATTAGTGGGTTG | ATATAGGGGTGGGGGATGAG | 300 |
| chr9:12180879-12182893 | p2 | (AT)7 | AAACCCAGAATTTTGTCCCC | AAATGATACCGCTCCTGTGG | 151 |
| chr9:12221657-12223672 | p5 | (AAAAT)3 | TCGCGCTGGTACGAATACTA | TCCGAAGTGGCAGGTTATTC | 171 |
| chr9:12228705-12230723 | p2 | (TA)9 | AAGGCCCAAAAGGTCACTCT | AGTGGCTCTCTTTTCCGTCA | 112 |
| chr9:12287777-12289795 | p2 | (AG)9 | AACAACTGGACCACCCAGTC | AAATCAAAGATGGTGACGCC | 185 |
| chr9:12383502-12385522 | p5 | (AATTA)4 | ATTTTGTGGGCTGAAGGATG | CTGCGTATGCCTGTATTCCA | 198 |
| chr9:12402102-12404132 | p2 | (TA)7 | TTGAACACAACCCAACGCTA | CCACCATGTGGTTCAGTGAG | 102 |
| chr9:12402102-12404132 | p2 | (AT)15 | TTGAACACAACCCAACGCTA | CCACCATGTGGTTCAGTGAG | 102 |

| Marker ID | SSR type | SSR motif | FORWARD PRIMER (5'-3') | REVERSE PRIMER (5'-3') | size (bp) |
|---|---|---|---|---|---|
| chr9:12402102-12404132 | p3 | (ATC)5 | TTGAACACAACCCAACGCTA | CCACCATGTGGTTCAGTGAG | 102 |
| chr9:12443761-12445779 | p3 | (CAA)6 | CATCACAACAAGGCCACAAC | CAGTTGATAGGGGGTGTGCT | 177 |
| chr9:12456497-12458512 | p5 | (TTTAT)3 | CATTGGCGACACTTTCAAGA | CTGAATTGATCCAGCAAGCA | 182 |
| chr9:12456497-12458512 | p5 | (TTTAT)3 | CATTGGCGACACTTTCAAGA | CTGAATTGATCCAGCAAGCA | 182 |
| chr9:12456497-12458512 | p5 | (TTTAT)3 | CATTGGCGACACTTTCAAGA | CTGAATTGATCCAGCAAGCA | 182 |
| chr9:12493955-12495967 | p2 | (TA)6 | TTTTGCCTCCTTTTCCAATG | GCTGATGAGGGTTGGTCAAT | 156 |
| chr9:12493955-12495967 | p6 | (TTTTTC)3 | TTTTGCCTCCTTTTCCAATG | GCTGATGAGGGTTGGTCAAT | 156 |
| chr9:12558463-12560511 | p4 | (CATA)12 | TATGCCCCTCCAAAGTTGAC | GAAGGTACAAGCCATGGGAA | 143 |
| chr9:12588121-12590136 | p3 | (TTG)5 | GATACGACTCGCATGTGGTG | AATTACGGCATATTCGACGC | 173 |
| chr9:12590496-12592522 | p2 | (AC)13 | TGTATGGGCGCTACACAGAG | GCACCGAGTGTTGTGTTACG | 273 |
| chr9:12595510-12597558 | p2 | (AT)24 | CCAAGGCAATGGTCAATCTT | GTAAATCTATCCGCGCTTCG | 200 |
| chr9:12627705-12629720 | p5 | (TGATA)3 | GATTCGAAATCCGAGCATGT | AATGTTGGAACAATCCGAGC | 300 |
| chr9:12627705-12629720 | p2 | (GT)6 | GATTCGAAATCCGAGCATGT | AATGTTGGAACAATCCGAGC | 300 |
| chr9:12632090-12634106 | p2 | (TA)8 | CCCATAAAATGTGCCATTCC | GAAAAGGTCAGGCACTCGTC | 105 |
| chr9:12632090-12634106 | p2 | (AT)8 | CCCATAAAATGTGCCATTCC | GAAAAGGTCAGGCACTCGTC | 105 |
| chr9:12632090-12634106 | p3 | (ATC)5 | CCCATAAAATGTGCCATTCC | GAAAAGGTCAGGCACTCGTC | 105 |
| chr9:12711286-12713304 | p6 | (AGAGAA)3 | CATGAAACATGTTCCCGACA | ATGAGAGTCAAATGGCCGAC | 146 |
| chr9:12711286-12713304 | p3 | (AAG)10 | CATGAAACATGTTCCCGACA | ATGAGAGTCAAATGGCCGAC | 146 |
| chr9:12738995-12741007 | p2 | (TG)6 | TCATGCTCACAGGTAGACCG | GAGGAGAAACCGTGAGACCA | 153 |
| chr9:12738995-12741007 | p2 | (TA)32 | TCATGCTCACAGGTAGACCG | GAGGAGAAACCGTGAGACCA | 153 |
| chr9:12754982-12756997 | p5 | (TTGTA)3 | CGGTGCCTTTACTCAGCTTC | AGTCAAACCGGTGGAACAAG | 166 |
| chr9:12889863-12891901 | p2 | (TC)19 | AAAGTTCCATGAATCGTGCC | CAAACGTCATTCAATGCAGC | 163 |
| chr9:12951646-12953682 | p2 | (AT)18 | TCAAGAAAAACACCCGTTCC | TGTGGGCATCGTAGAGGATT | 168 |
| chr9:12958307-12960339 | p2 | (TA)16 | AGTCCGTGCCAAAAAGAATG | CACTTTTGACCCTCCTCCAA | 186 |
| chr9:12981341-12983356 | p5 | (TTTGT)3 | GCCCTCTTTTTCTTGCCTCT | TAATTTCCCAACCCCAACAA | 265 |
| chr9:13040646-13042662 | p2 | (TC)6 | CTCTCTCCGTTCCCTTCTCC | GATTGCAACAAGTTGGGCTT | 233 |
| chr9:13040646-13042662 | p4 | (TTTC)4 | CTCTCTCCGTTCCCTTCTCC | GATTGCAACAAGTTGGGCTT | 233 |
| chr9:13082466-13084478 | p2 | (AT)6 | GACGGGAACCACCTGACTTA | CCTTGTCGGCCTTATTTTGA | 208 |
| chr9:13099348-13101363 | p5 | (TGAAT)3 | AAGCTTGGAGAAGGGGACAT | CCAAATGAATGCTTTGGCTT | 213 |
| chr9:13176593-13178608 | p5 | (TTGGT)3 | GGTGACAGCGAACTCACTCA | CCAATTATGGTCGGGTTTTG | 139 |
| chr9:13176593-13178608 | p5 | (TTGGT)3 | GGTGACAGCGAACTCACTCA | CCAATTATGGTCGGGTTTTG | 139 |
| chr9:13176593-13178608 | p5 | (AGAAG)3 | GGTGACAGCGAACTCACTCA | CCAATTATGGTCGGGTTTTG | 139 |
| chr9:13176593-13178608 | p6 | (ATCCAA)3 | GGTGACAGCGAACTCACTCA | CCAATTATGGTCGGGTTTTG | 139 |
| chr9:13269190-13271204 | p2 | (TA)7 | TCCAAACTCATGGGGAAAAA | CCCTTTACGGTGTGCACTTT | 173 |
| chr9:13270334-13272346 | p2 | (AT)6 | AACCCACCACATCCAAACTC | ATATGGGGCCCACAATTTTT | 115 |
| chr9:13274686-13276706 | p2 | (TA)10 | TTCCCAAAGTATCCCCATCA | GACATGGCGGAATAGCAAAT | 247 |

| Marker ID | SSR type | SSR motif | FORWARD PRIMER (5'-3') | REVERSE PRIMER (5'-3') | size (bp) |
|---|---|---|---|---|---|
| chr9:13293337-13295355 | p3 | (TGT)6 | ACGTTGAAATCTATTGCCCG | AATGGTAGGCTGGAAAGGCT | 265 |
| chr9:13306201-13308246 | p3 | (GTT)15 | CGGGAAGTGATAATGCCAGT | TGGGGCATCTGTATCTCTCC | 235 |
| chr9:13306201-13308246 | p3 | (AAC)5 | CGGGAAGTGATAATGCCAGT | TGGGGCATCTGTATCTCTCC | 235 |
| chr9:13307047-13309062 | p3 | (GTT)15 | CGTACACTCTACCCTCCCCA | ACCCCACTTGTGGGATTACA | 108 |
| chr9:13307047-13309062 | p3 | (AAC)5 | CGTACACTCTACCCTCCCCA | ACCCCACTTGTGGGATTACA | 108 |
| chr9:13362536-13364551 | p5 | (ATTTC)3 | TAACCCCAACAACATGAGCA | CTTCGCAGATTCAACATGGA | 227 |
| chr9:13373635-13375650 | p5 | (ATTTC)3 | GAGAGGAAATGTCGTGGGAA | ACCCTGTCACAACAACCCAT | 155 |
| chr9:13470075-13472089 | p5 | (ATTTT)3 | AGGGCTCGCCTAACAGTTCT | CCCTCACGACGTTCTCAAAT | 111 |
| chr9:13470075-13472089 | p2 | (TA)7 | AGGGCTCGCCTAACAGTTCT | CCCTCACGACGTTCTCAAAT | 111 |
| chr9:13470075-13472089 | p2 | (TA)8 | AGGGCTCGCCTAACAGTTCT | CCCTCACGACGTTCTCAAAT | 111 |
| chr9:13535846-13537858 | p2 | (AG)6 | TGGTTGATGTGCAGCATTTT | AGTGATACTGGTGGGGGTCA | 133 |
| chr9:13612658-13614673 | p5 | (GCATT)3 | CTCACCTCACCCCACTCTGT | TTTTTCCGCTCTTAACCCCT | 100 |
| chr9:13668425-13670443 | p2 | (AG)9 | TTCCGACAAAAGTTGGTTCC | CTGAAAAAGCTGGCATCACA | 291 |
| chr9:13712612-13714632 | p2 | (AT)6 | CAGATACCTTGGGTTGCGAT | CCATTAGTGGTGGGGAAGAA | 148 |
| chr9:13712634-13714650 | p2 | (AT)6 | CAGATACCTTGGGTTGCGAT | CCATTAGTGGTGGGGAAGAA | 126 |
| chr9:13860915-13862957 | p2 | (TA)6 | AGTTTATGCCATTTTCCCCC | AACCCAAGAGACTCAAGGCA | 198 |
| chr9:13903856-13905870 | p2 | (AT)7 | TCAAAAATGGTTCTGAGCCC | GGGGCTTTTCAACCGTAAAT | 173 |
| chr9:13959742-13961757 | p5 | (ACCCG)3 | TGCACCCAATAAAGCTAGGG | TTTGGCAGTGTCTGAAGTCG | 128 |
| chr9:13959770-13961803 | p5 | (ACCCG)3 | TGCACCCAATAAAGCTAGGG | TTTGGCAGTGTCTGAAGTCG | 100 |
| chr9:13966835-13968853 | p6 | (CATTAT)3 | TGGAACTATTCCTTCGGCAC | TTCTGGGATGACTACGACCC | 294 |
| chr9:13978078-13980132 | p2 | (AG)27 | TATTGTGGCAAGTGATCCCA | ATTCAAACGTTCTCCCATGC | 241 |
| chr9:13994094-13996109 | p5 | (AAAAT)3 | GCAGATGTCGTGGACAAAGA | ACAGCCACAATTCCCTCAAC | 168 |
| chr9:14091572-14093587 | p3 | (TAT)5 | TTTGGCCCATTTGACTTAGC | CTACAGGGAACGCATGACAA | 114 |
| chr9:14104488-14106503 | p3 | (TAT)5 | TTTGGCCCATTTGACTTAGC | CTACAGGGAACGCATGACAA | 114 |
| chr9:14121215-14123230 | p5 | (GGTTT)3 | AAATTCCATACGCATGGCTC | GGTCATGGGCAACAGAGTTT | 186 |
| chr9:14140049-14142069 | p2 | (AT)34 | CGGCTTACAGATGACACCCT | GCGATCACATGAGGAATTGA | 293 |
| chr9:14156498-14158522 | p4 | (AAAT)6 | CAAAATGCCCAGAAGACTCC | CCTCTCCCAGTCCCCTAAAG | 174 |
| chr9:14156498-14158522 | p5 | (TAAAA)3 | CAAAATGCCCAGAAGACTCC | CCTCTCCCAGTCCCCTAAAG | 174 |
| chr9:14227116-14229132 | p2 | (AT)8 | TCACAGTGGCGTCCATAAAA | TGTTGGAAACACCTGGACAA | 128 |
| chr9:14386121-14388139 | p2 | (AT)15 | GGAGGCTTTTTCGCCTAGAT | TTGCCATCTTTTTGGGAAAG | 197 |
| chr9:14386121-14388139 | p6 | (AAAAAT)3 | GGAGGCTTTTTCGCCTAGAT | TTGCCATCTTTTTGGGAAAG | 197 |
| chr9:14436609-14438653 | p2 | (TA)21 | CCTTCATCCTCTTTTGCTGC | AACCCATGAAGTAATGCCCA | 181 |
| chr9:14436667-14438679 | p2 | (TA)21 | CTGGGCATTACTTCATGGGT | TGCAAAAACTACAACAGCCG | 241 |
| chr9:14449164-14451178 | p2 | (AT)7 | TTTTGAGGAAAAGTGTCGGG | TTAATGCCACTGAGGCTTCC | 202 |
| chr9:14541093-14543109 | p4 | (CAGT)4 | GTTAGCTGCTCAATTCCCCA | CCCATCAAACAACCCAAAAG | 126 |
| chr9:14773889-14775904 | p5 | (TGAAG)3 | TCCAAGAACAGTTGCTGCAC | GTGCAACAGCAGGCTAAACA | 215 |

| Marker ID | SSR type | SSR motif | FORWARD PRIMER (5'-3') | REVERSE PRIMER (5'-3') | size (bp) |
|---|---|---|---|---|---|
| chr9:14834096-14836112 | p4 | (TAAA)4 | AATATGTGCGGGGGTTACAA | TCAAGAACAAGCAAAGCCCT | 202 |
| chr9:14860750-14862765 | p3 | (AAC)5 | AGTGAGGGAGGAGGGATGTT | CTTCGGAGATTCGAGACTGG | 199 |
| chr9:14862353-14864383 | p2 | (AT)33 | TGGCACCCTCTTGTCACATA | GCTGGAGGGTTGGAGACATA | 144 |
| chr9:14862353-14864383 | p2 | (AC)8 | TGGCACCCTCTTGTCACATA | GCTGGAGGGTTGGAGACATA | 144 |
| chr9:14862385-14864400 | p2 | (AT)33 | TGGCACCCTCTTGTCACATA | GCTGGAGGGTTGGAGACATA | 112 |
| chr9:14862385-14864400 | p2 | (AC)8 | TGGCACCCTCTTGTCACATA | GCTGGAGGGTTGGAGACATA | 112 |
| chr9:14877205-14879223 | p6 | (ACCTAA)3 | CCCCTCACCGTTTACCACTA | AACCAATTTCCCATCAACCA | 106 |
| chr9:14939046-14941067 | p3 | (AAC)7 | AAGTGAGGGAGGAGGGATGT | TTGGTCTTGTTCGGTAAGGG | 204 |
| chr9:14940654-14942684 | p2 | (AC)8 | GGTACCAAGCCTCACTTCCA | GATGGATAGATTGACGCGGT | 295 |
| chr9:14940686-14942701 | p2 | (AC)8 | GGTACCAAGCCTCACTTCCA | GATGGATAGATTGACGCGGT | 263 |
| chr9:15065539-15067593 | p3 | (TAA)18 | TCATCAGTATTGCCAGTGCC | CTTTTGAGCGGTCCAACAAT | 178 |
| chr9:15066657-15068675 | p2 | (TA)9 | TGTCCCTGGAACAACATCCT | GCGGTGAAAAAGGAGAGTGA | 284 |
| chr9:15122590-15124608 | p6 | (ACCTAA)3 | CCCCTCACCGTTTACCACTA | AACCAATTTCCCATCAACCA | 106 |
| chr9:15245455-15247475 | p2 | (TA)10 | AGCTGGCTTTGTTTGGAAGA | TATGGCTCATTTTGATCGCA | 233 |
| chr9:15253099-15255111 | p2 | (CT)6 | GCGGCTGTAAACAAAAGAGG | TGCTGAACTAAGTCGCATGG | 132 |
| chr9:15291178-15293198 | p2 | (AT)7 | TGGGTTCGGATTGATTGATT | TTATTGCAGCGTTCATTTGC | 278 |
| chr9:15291193-15293205 | p2 | (AT)7 | TGGGTTCGGATTGATTGATT | TTATTGCAGCGTTCATTTGC | 263 |
| chr9:15292041-15294077 | p2 | (AT)7 | TTTTGCAAGTGCAATCAAGG | GGGTTTTGGACTTTTCCGAT | 224 |
| chr9:15324580-15326610 | p2 | (CT)7 | CGTGACAATTCATCCGACAG | ATGGCGGAGCTAGAATTTCA | 157 |
| chr9:15324613-15326695 | p2 | (CT)7 | CGTGACAATTCATCCGACAG | ATGGCGGAGCTAGAATTTCA | 124 |
| chr9:15329467-15331479 | p2 | (TA)6 | CACCTTCCCACAGGGATAAA | ACGATTAAAGCACGTGGAGC | 120 |
| chr9:15329467-15331479 | p3 | (TTG)10 | CACCTTCCCACAGGGATAAA | ACGATTAAAGCACGTGGAGC | 120 |
| chr9:15358898-15360914 | p2 | (TA)8 | CTTTTCTTGCCACCCAACAT | GATCGATTTTTCCCAGCTCA | 182 |
| chr9:15368844-15370878 | p5 | (AATAA)3 | GTGCGCAAAAGTAGGCATTT | CCTCCAACTTTGGATGTGCT | 193 |
| chr9:15397992-15400004 | p2 | (AC)6 | CACCAATTCCGACGAACTTT | ATCTCGTGAGGCCAGAGGTA | 282 |
| chr9:15435976-15437988 | p2 | (AT)6 | GATGGTTGTCGCACCCTATT | CTCCTTCCCGACCCTTAGAG | 236 |
| chr9:15551163-15553181 | p3 | (CAA)6 | TACCTGTGAGCATGAAACGC | ACTAGAGGGCCGGGTACAGT | 118 |
| chr9:15591778-15593808 | p2 | (TA)15 | GTGCAAGCCCAAATTAAGGA | GCCAAGGTGAACTTGTCGAT | 193 |
| chr9:15637902-15639917 | p5 | (TTTCT)3 | GAGTGTTGGTGCAAAGCAAA | GGATCTTGGAAATGGGAGGT | 199 |
| chr9:15637902-15639917 | p5 | (TTTCT)3 | GAGTGTTGGTGCAAAGCAAA | GGATCTTGGAAATGGGAGGT | 199 |
| chr9:15637902-15639917 | p5 | (TTTCT)3 | GAGTGTTGGTGCAAAGCAAA | GGATCTTGGAAATGGGAGGT | 199 |
| chr9:15638447-15640462 | p5 | (TTTCT)3 | TTCGAAGCTGAGGAGAGGAA | TCAATGATGATTTGCCCAGA | 250 |
| chr9:15638447-15640462 | p5 | (TTTCT)3 | TTCGAAGCTGAGGAGAGGAA | TCAATGATGATTTGCCCAGA | 250 |
| chr9:15638447-15640462 | p5 | (TTTCT)3 | TTCGAAGCTGAGGAGAGGAA | TCAATGATGATTTGCCCAGA | 250 |
| chr9:15860265-15862283 | p6 | (GAAAAA)3 | ACATGAGGCAAGGATGCTCT | GCAGATTGAAAAGGCGAGTC | 245 |
| chr9:15869512-15871527 | p3 | (TTC)5 | GCATCGAGAAATTCGGTAGC | ATGCAATTAAGCAATTGGGC | 209 |

| Marker ID | SSR type | SSR motif | FORWARD PRIMER (5'-3') | REVERSE PRIMER (5'-3') | size (bp) |
|---|---|---|---|---|---|
| chr9:15919640-15921655 | p5 | (CCCCA)3 | ATCAACTTTCAGGGGGCTTT | AACTTTGGGGCAAGTCAAAA | 112 |
| chr9:15946753-15948777 | p2 | (TA)12 | GTGTCCTTGGCACCAATTCT | TGCATTTTAAGATCCCCGAC | 298 |
| chr9:15978159-15980185 | p2 | (CT)13 | GGAGTAGCCCTGGTGAATGA | CGTCCATGTCCAACCTCTCT | 120 |
| chr9:16028889-16030913 | p2 | (TA)12 | GTGTCCTTGGCACCAATTCT | TGCATTTTAAGATCCCCGAC | 298 |
| chr9:16060295-16062321 | p2 | (CT)13 | GGAGTAGCCCTGGTGAATGA | CGTCCATGTCCAACCTCTCT | 120 |
| chr9:16243509-16245560 | p3 | (TAT)17 | AAGCTGCCTGCTTGAGCTAC | CAACAGCTTCCTGAAAAGCC | 180 |
| chr9:16284379-16286391 | p2 | (AT)6 | TCCGAATCTGTTTCGTTTCC | ACAATCGATGCCTTTCTGCT | 164 |
| chr9:16284379-16286391 | p2 | (TA)6 | TCCGAATCTGTTTCGTTTCC | ACAATCGATGCCTTTCTGCT | 164 |
| chr9:16352830-16354868 | p2 | (AT)16 | CTCCGTACAAGAGGTTGGGA | CACACGATGACACTATCGGC | 208 |
| chr9:16352830-16354868 | p2 | (TA)19 | CTCCGTACAAGAGGTTGGGA | CACACGATGACACTATCGGC | 208 |
| chr9:16352830-16354868 | p2 | (TA)8 | CTCCGTACAAGAGGTTGGGA | CACACGATGACACTATCGGC | 208 |
| chr9:16403327-16405342 | p5 | (TTCAA)3 | AGACCCTGGGGGTATTTGTT | TGTGTCACACCCATTTTGCT | 262 |
| chr9:16563136-16565154 | p6 | (TCCCAT)3 | CTTTGAGCCTGGGCAGTTAG | CCTCTGCCCATTTACCAAGA | 220 |
| chr9:16568844-16570859 | p5 | (ATTTT)3 | GGGAGCTGCCAAAAATACTG | GATTAAGGGAGCAGGGGAAG | 110 |
| chr9:16572500-16574542 | p3 | (TAA)14 | CAGCAGCCAAAACTTGTTCA | CCATTTCAAGCAAAGGTCGT | 179 |
| chr9:16583767-16585785 | p6 | (GTGGAC)3 | TAGCAAAAGGAGCGGAAAGA | ATAGAGTTGTCCCCGAGGCT | 121 |
| chr9:16583767-16585785 | p6 | (CGTGGA)3 | TAGCAAAAGGAGCGGAAAGA | ATAGAGTTGTCCCCGAGGCT | 121 |
| chr9:16583767-16585785 | p6 | (GGACGT)3 | TAGCAAAAGGAGCGGAAAGA | ATAGAGTTGTCCCCGAGGCT | 121 |
| chr9:16583767-16585785 | p6 | (GTGGAC)3 | TAGCAAAAGGAGCGGAAAGA | ATAGAGTTGTCCCCGAGGCT | 121 |
| chr9:16609440-16611458 | p6 | (TCCACA)3 | GGATGAGTTTCGGGAGTTGA | AGGGGCATATTTGAGCCTTT | 120 |
| chr9:16621954-16623970 | p3 | (TTG)5 | CTATCCGAGACAAGGTGGGA | AAGGTCATGTCCTCGGTGAG | 106 |
| chr9:16621954-16623970 | p4 | (TTGT)4 | CTATCCGAGACAAGGTGGGA | AAGGTCATGTCCTCGGTGAG | 106 |
| chr9:16640434-16642448 | p2 | (CT)7 | TAGCCACGCTTCATCCTTCT | CGGAGTTCATCCTGAAGAGG | 175 |
| chr9:16649897-16651909 | p5 | (AATCA)3 | TGCACTTACGTGAGCCTCTG | TTCTCCACACGAGGGGTATC | 120 |
| chr9:16649897-16651909 | p2 | (TC)6 | TGCACTTACGTGAGCCTCTG | TTCTCCACACGAGGGGTATC | 120 |
| chr9:16662345-16664357 | p5 | (AATCA)3 | TGCACTTACGTGAGCCTCTG | TTCTCCACACGAGGGGTATC | 120 |
| chr9:16662345-16664357 | p2 | (TC)6 | TGCACTTACGTGAGCCTCTG | TTCTCCACACGAGGGGTATC | 120 |
| chr9:16680921-16682945 | p4 | (TATT)6 | TCCCCAGGATGACTTGCTAC | GCACCGTTTTTCCTTTTTCA | 243 |
| chr9:16796381-16798399 | p6 | (TCTTCA)3 | CTTGGAAGGCCTTGAGTGAG | CCCTGCAAGGTGAAATTTGT | 119 |
| chr9:16833469-16835484 | p3 | (TAT)5 | CTATGCACCATGGAGGAGGT | CTTAAACCCCGTTTGTTCCA | 142 |
| chr9:17185733-17187751 | p3 | (ATT)6 | AGGAACCTGAGAACCCCTGT | ACATCAGCACTTGCCTCCTT | 264 |
| chr9:17403461-17405477 | p2 | (AC)8 | CTCCCGTCGAACTTCTTCTG | ACTATGATCCCAGTGGCGAG | 121 |
| chr9:17459540-17461564 | p2 | (TA)12 | ATCATGAGGTTCTGGGTTCG | AAAGTTATGTCGGACACGGC | 199 |
| chr9:17501627-17503645 | p3 | (TTG)6 | GTACCTCGGAGAAGCACTGG | TGGTTCCAAGGTTGTCCTTC | 103 |
| chr9:17564035-17566051 | p4 | (TTAT)4 | CCAGACTGAAAATGGAGGGA | TCCAGGAGCTCTGCAACTTT | 141 |
| chr9:17650282-17652306 | p4 | (AAAT)6 | AGATTCCGCACACGTTTACC | TGAGTGGCACAGGTATCGAG | 218 |

| Marker ID | SSR type | SSR motif | FORWARD PRIMER (5'-3') | REVERSE PRIMER (5'-3') | size (bp) |
|---|---|---|---|---|---|
| chr9:17801414-17803429 | p5 | (AACAT)3 | CCACTAGTAGGCGGCTTGAG | CATCATTGCACACATCGTCA | 177 |
| chr9:18106339-18108354 | p5 | (ATGCA)3 | TGTGCCGAACATGTCAAAAT | ATAACCAAATGGGGTCGTGA | 160 |
| chr9:18238355-18240375 | p5 | (TGGGC)3 | ATGAGGACGTACTGCTTGGG | CTTGTCCCTCGAACCAGAAA | 109 |
| chr9:18238355-18240375 | p2 | (TC)10 | ATGAGGACGTACTGCTTGGG | CTTGTCCCTCGAACCAGAAA | 109 |
| chr9:18438152-18440164 | p2 | (AT)6 | ACAGGCTCCAGTGCTTCAAT | TCATGGGTTCGATCAGTTCA | 281 |
| chr9:18438152-18440164 | p5 | (ATTAA)3 | ACAGGCTCCAGTGCTTCAAT | TCATGGGTTCGATCAGTTCA | 281 |
| chr9:18438461-18440476 | p2 | (AT)6 | ATTTGGGAAAGGAAAGGGTG | AAGTGGGTAAATGGGGAAGG | 279 |
| chr9:18438461-18440476 | p5 | (ATTAA)3 | ATTTGGGAAAGGAAAGGGTG | AAGTGGGTAAATGGGGAAGG | 279 |
| chr9:18528298-18530313 | p5 | (TAATA)3 | TTTGCTTGACATGCCAGTTG | TTTGCCATCGTTATCCCTTT | 210 |
| chr9:18658240-18660262 | p2 | (TA)11 | GCGGACCCATCAACCTACTA | GGGTGTCCAGGTCTCTTTGA | 114 |
| chr9:18658240-18660262 | p3 | (GAA)6 | GCGGACCCATCAACCTACTA | GGGTGTCCAGGTCTCTTTGA | 114 |
| chr9:18686316-18688331 | p5 | (AGTTT)3 | GAATTCAAGTCCCCTTGGGT | CCCTCATTGAAACCCGACTA | 207 |
| chr9:18752541-18754583 | p2 | (AT)21 | TGTTAAGCCCTATGCGAGGT | ACATGGGTGGCATTATGGTT | 184 |
| chr9:18855995-18858010 | p5 | (TTTTA)3 | ATGCATTGCTGGTGCATAAA | TTTGACCCCTCTACTGTGGC | 205 |
| chr9:18855995-18858010 | p4 | (CATA)4 | ATGCATTGCTGGTGCATAAA | TTTGACCCCTCTACTGTGGC | 205 |
| chr9:18924621-18926641 | p5 | (TAATA)4 | TTGCTGCTGTATGCTGCTCT | GTTCGAGCATTGAGGAGGAG | 141 |
| chr9:18924621-18926641 | p4 | (AAGA)5 | TTGCTGCTGTATGCTGCTCT | GTTCGAGCATTGAGGAGGAG | 141 |
| chr9:18927570-18929585 | p3 | (CCA)5 | TCCAGCCTGTCTTGCCTAAT | TGAAATGTTGACGTGCCAAT | 115 |
| chr9:19011106-19013121 | p5 | (ACAAT)3 | TGTCACGACCCAACTTACGA | TTCCGGGAATTGTTTTTCAG | 169 |
| chr9:19065325-19067340 | p5 | (TACAA)3 | CTCTCCCTCGACGTGTCTTC | CACGGAAAAACCACCAAATC | 237 |
| chr9:19154746-19156761 | p3 | (ACA)5 | ACCCGTACGATTGGCAGTAG | ACCGGTGAGACCAAAGACTG | 144 |
| chr9:19188953-19190967 | p2 | (AT)7 | AAAATTCGGCAAAAATTCCC | TCCTGGGTGTGTCCTAGTCC | 131 |
| chr9:19246921-19248936 | p5 | (CTTTT)3 | AATCAAATTTCGTTGCTCGG | CAAGGACCCTGGGTAATCAA | 125 |
| chr9:19292711-19294732 | p3 | (TGT)5 | CTTTCGGGGGATACTGGATT | TAGACCCTCGGCTCAAGAAA | 228 |
| chr9:19292711-19294732 | p3 | (AAC)7 | CTTTCGGGGGATACTGGATT | TAGACCCTCGGCTCAAGAAA | 228 |
| chr9:19353374-19355394 | p4 | (AATT)5 | TACAATTTGGGACAGAGGGG | GACCCAGACGCCATAGTCAT | 200 |
| chr9:19364523-19366547 | p2 | (AT)12 | ATAAAGGGTTGGGTTGGGTC | CCCACTTTCCTTTATGCCAA | 263 |
| chr9:19470945-19472957 | p2 | (AT)6 | AAAGCAATGGACTTGGATGC | CAAGTCCCACATCGAGGTTT | 229 |
| chr9:19476834-19478876 | p2 | (AT)21 | TGTCGTGGAGAGGACATTTG | CTGTCGGTCCTCATCCAAGT | 133 |
| chr9:19476834-19478876 | p2 | (AT)8 | TGTCGTGGAGAGGACATTTG | CTGTCGGTCCTCATCCAAGT | 133 |
| chr9:19476834-19478876 | p5 | (CATAT)3 | TGTCGTGGAGAGGACATTTG | CTGTCGGTCCTCATCCAAGT | 133 |
| chr9:19525480-19527524 | p2 | (TA)22 | GGATATTGTTTGGTGGGTCG | ATCGGTCACACCCCTTACTG | 119 |
| chr9:19549727-19551743 | p2 | (AT)7 | TAGCATACATGTCAAGCGCC | GTGGTCCTGAAATAGGGCAA | 211 |
| chr9:19549828-19551848 | p2 | (AT)7 | TAGCATACATGTCAAGCGCC | GTGGTCCTGAAATAGGGCAA | 110 |
| chr9:19550061-19552075 | p2 | (AT)7 | CCGAACCAGCTTCATACACA | CATTTCGTCGGGACTTAGGA | 149 |
| chr9:19607480-19609494 | p2 | (AT)7 | TTTGCCTGGATCCTTGATTC | TCTTGTTTCTTCTTGCCCGT | 179 |

| Marker ID | SSR type | SSR motif | FORWARD PRIMER (5'-3') | REVERSE PRIMER (5'-3') | size (bp) |
|---|---|---|---|---|---|
| chr9:19680388-19682436 | p2 | (TA)24 | AGGGACCCAAGCGAACTACT | GCTCGTGGTCTAGCCTTCAC | 203 |
| chr9:19704346-19706362 | p2 | (CA)8 | GGACATGCCAAAAGAAGGAA | CTTATAGGGGAAGTGCTGCG | 246 |
| chr9:19781787-19783802 | p5 | (CGCTT)3 | CTTTTTGGCGCAAATTGTTT | AACAACCACCGGAAAGTCAG | 259 |
| chr9:20006450-20008468 | p3 | (GAA)6 | CATGTCGGCTTCATCATCAC | GCAAGCAAGCACGTAATTCA | 291 |
| chr9:20182943-20184955 | p2 | (AT)6 | AGTAGGGTGCTTGAGGCGTA | AAAAGGCAGTTTCAGGGCTT | 283 |
| chr9:20267381-20269397 | p4 | (TATG)4 | AGCGACGCTTGTTCAATTTT | CATCCCACACGATGTTTCTG | 216 |
| chr9:20267381-20269397 | p2 | (AT)28 | AGCGACGCTTGTTCAATTTT | CATCCCACACGATGTTTCTG | 216 |
| chr9:20304711-20306727 | p2 | (AT)8 | GGACCCTTAGGGGAAGTTCA | CATGTTTAGCAAAGGGGCAT | 116 |
| chr9:20315097-20317109 | p2 | (TA)6 | GCTCCGTTTTGGAACTTCAG | TCCATTTTAATGGCGGAAAG | 203 |
| chr9:20315097-20317109 | p2 | (TA)13 | GCTCCGTTTTGGAACTTCAG | TCCATTTTAATGGCGGAAAG | 203 |
| chr9:20315097-20317109 | p2 | (AC)6 | GCTCCGTTTTGGAACTTCAG | TCCATTTTAATGGCGGAAAG | 203 |
| chr9:20406083-20408099 | p2 | (AT)16 | ACAAGTACTGTTCCGGTGCC | TGATTCACGAAATGCAAAGG | 165 |
| chr9:20433380-20435398 | p2 | (AT)19 | TGAACCATATGGGACGGTTT | CCGATTAACGGGCTTACAAC | 115 |
| chr9:20433380-20435398 | p2 | (TA)8 | TGAACCATATGGGACGGTTT | CCGATTAACGGGCTTACAAC | 115 |
| chr9:20433380-20435398 | p2 | (AT)7 | TGAACCATATGGGACGGTTT | CCGATTAACGGGCTTACAAC | 115 |
| chr9:20433380-20435398 | p2 | (AT)7 | TGAACCATATGGGACGGTTT | CCGATTAACGGGCTTACAAC | 115 |
| chr9:20433380-20435398 | p2 | (AT)7 | TGAACCATATGGGACGGTTT | CCGATTAACGGGCTTACAAC | 115 |
| chr9:20433380-20435398 | p2 | (AT)7 | TGAACCATATGGGACGGTTT | CCGATTAACGGGCTTACAAC | 115 |
| chr9:20434049-20436063 | p2 | (AT)7 | ATGCTTCTATTGAGCCGAGG | AACAACCCAGTGAAATCCCA | 222 |
| chr9:20434049-20436063 | p2 | (AT)7 | ATGCTTCTATTGAGCCGAGG | AACAACCCAGTGAAATCCCA | 222 |
| chr9:20434049-20436063 | p2 | (AT)7 | ATGCTTCTATTGAGCCGAGG | AACAACCCAGTGAAATCCCA | 222 |
| chr9:20434049-20436063 | p2 | (AT)7 | ATGCTTCTATTGAGCCGAGG | AACAACCCAGTGAAATCCCA | 222 |
| chr9:20552618-20554636 | p6 | (TTTTTG)3 | GCCAGGGCATTTGAAAATAA | ATTTGGCCCTGTGTATGCTC | 156 |
| chr9:20619519-20621537 | p3 | (TGT)6 | CAAAGGCGGTTGGTCTTTTA | ACGCAACCGCATCTCTAACT | 122 |
| chr9:20656742-20658760 | p6 | (ATCCAC)3 | AAATCCTTGATTCGTGTCGG | AAAGTCGGCATCACACATCA | 216 |
| chr9:20836684-20838704 | p6 | (GGTGCA)4 | CTCTCGAGGATGGGTATGGA | TGGGTCGTGACAATTTGGTA | 291 |
| chr9:20836684-20838704 | p5 | (AATGC)4 | CTCTCGAGGATGGGTATGGA | TGGGTCGTGACAATTTGGTA | 291 |
| chr9:20891992-20894007 | p5 | (TTTTG)3 | TTGCACATCAAGGGCAACTA | CGCCAGTTCCTTTTGGACTA | 248 |
| chr9:20894717-20896737 | p2 | (TA)10 | ACGGTAGCAGTGAATGGGAC | TTAAGATCTGAGCCGAGGGA | 264 |
| chr9:20920656-20922830 | p2 | (TA)53 | TACTTGCGCTCCAAAACAGA | GGTCACCCGCAAAACTAAAA | 146 |
| chr9:20959551-20961563 | p3 | (ATT)16 | TCATTCGCAAGTGTCGTAGC | TGATGAAAGAATTTTCGGGC | 209 |
| chr9:20959551-20961563 | p2 | (CT)6 | TCATTCGCAAGTGTCGTAGC | TGATGAAAGAATTTTCGGGC | 209 |
| chr9:21103342-21105354 | p2 | (AG)6 | AACTTTGGAAGCAACATCCG | CCCGTTTTTGGAACTTGAAA | 242 |
| chr9:21151229-21153251 | p2 | (TA)11 | TGACGATGACAAGTTCTGGC | TATGAGTCATGGCGGTGGTA | 101 |
| chr9:21217612-21219642 | p2 | (AT)15 | AACCGAAATAGGGCCTCACT | CCTTGGAGTTCTTTTGCGAG | 175 |
| chr9:21246925-21248937 | p2 | (TG)6 | AAAGCTAGGGGGAAGGTCAA | TTCCTCGATATTTCCGCATC | 300 |

| Marker ID | SSR type | SSR motif | FORWARD PRIMER (5'-3') | REVERSE PRIMER (5'-3') | size (bp) |
|---|---|---|---|---|---|
| chr9:21246925-21248937 | p3 | (TTG)5 | AAAGCTAGGGGGAAGGTCAA | TTCCTCGATATTTCCGCATC | 300 |
| chr9:21301612-21303650 | p5 | (TGCAA)3 | AGAACATCCAAGCAACGACC | ACCTACGCTTGCTGCATTCT | 204 |
| chr9:21326968-21328983 | p5 | (AATTG)3 | TGACCACCCTGGTTACCATT | AGCTCAGCCTTGTTGGGTTA | 106 |
| chr9:21392865-21394883 | p5 | (ACAAA)3 | CTGACCCAAAAGTGACCGAT | GGTGTGTACGTGTGTCCAGC | 122 |
| chr9:21392865-21394883 | p2 | (AT)6 | CTGACCCAAAAGTGACCGAT | GGTGTGTACGTGTGTCCAGC | 122 |
| chr9:21444580-21446596 | p2 | (TA)8 | TCGTACCGACTCATCATCCA | AACCCAAGGAAGGCTAAGGA | 102 |
| chr9:21446328-21448346 | p6 | (CATATA)3 | CAGACAACCCCTAGGACGAA | AAGCCGACGTACCATTATGC | 271 |
| chr9:21578597-21580609 | p2 | (AT)6 | TGGAGGTTACCGACCTTACG | CATAAGACGCCGACCTTAGC | 256 |
| chr9:21578613-21580665 | p2 | (AT)6 | TGGAGGTTACCGACCTTACG | CATAAGACGCCGACCTTAGC | 240 |
| chr9:21646187-21648201 | p2 | (AT)7 | ACTCTGCTCGTGCTTACCGT | ATGAACATATGCCCTCCTGG | 160 |
| chr9:21676354-21678366 | p2 | (AC)6 | ACTCGGGTTTCGATGAACAG | GAAGCGGGAATTGAATTTGA | 264 |
| chr9:21737188-21739251 | p3 | (ATA)21 | CCAAATGGACTCCAAATGCT | TGCTCCCTTCCTCTTTCTGA | 219 |
| chr9:21784163-21786179 | p2 | (AT)8 | TTAGCCACCACAACAATGGA | AGTTTCCAAAGTGAGCGGAA | 179 |
| chr9:21793266-21795286 | p2 | (CT)16 | GTAGCGAGCGTTTCTGCTCT | TTTGGTCCAACTTGGGAGAC | 201 |
| chr9:21793266-21795286 | p2 | (TA)10 | GTAGCGAGCGTTTCTGCTCT | TTTGGTCCAACTTGGGAGAC | 201 |
| chr9:21862287-21864319 | p2 | (CT)7 | TTCTTCAGACCTGGGGATTG | TGGAGTAACCGGACTAACGG | 289 |
| chr9:21862332-21864380 | p2 | (CT)7 | TTCTTCAGACCTGGGGATTG | TGGAGTAACCGGACTAACGG | 244 |
| chr9:21889909-21891943 | p2 | (AT)17 | CCTTACGCTCGTCCAGATTC | GGAATGCTTCCAAGTTGCAT | 295 |
| chr9:21889909-21891943 | p6 | (TTTCTC)3 | CCTTACGCTCGTCCAGATTC | GGAATGCTTCCAAGTTGCAT | 295 |
| chr9:21889909-21891943 | p6 | (ACATGA)3 | CCTTACGCTCGTCCAGATTC | GGAATGCTTCCAAGTTGCAT | 295 |
| chr9:21916251-21918267 | p2 | (AT)8 | GGATTTGAACCAGCGACCTA | CAAAATCCAGAAACTCCCGA | 252 |
| chr9:22090965-22092979 | p2 | (TG)7 | TCCCTTCTTTCTCTTTGGCA | TTTCGGAGTGACAGAGGGTC | 236 |
| chr9:22142083-22144098 | p5 | (ACCAA)3 | GTCCGATCGATTGTGGATTT | GTGTGCCCTTTTTGCTGATT | 164 |
| chr9:22142083-22144098 | p5 | (AAACC)3 | GTCCGATCGATTGTGGATTT | GTGTGCCCTTTTTGCTGATT | 164 |
| chr9:22142856-22144871 | p5 | (ACCAA)3 | GGGTGTACATGGACCGAGTT | GAGACAACCCGAGGTTGAAA | 189 |
| chr9:22142856-22144871 | p5 | (AAACC)3 | GGGTGTACATGGACCGAGTT | GAGACAACCCGAGGTTGAAA | 189 |
| chr9:22142856-22144871 | p2 | (TA)8 | GGGTGTACATGGACCGAGTT | GAGACAACCCGAGGTTGAAA | 189 |
| chr9:22187541-22189556 | p5 | (TGCAT)3 | CCGCAATTGTTGTTTGTTTG | CCTACCTCAAAGCCGAACTG | 180 |
| chr9:22209353-22211365 | p2 | (AC)6 | TATTCCGCCTTCGAATTGTC | TCTTTGCGCCTCCAAATACT | 155 |
| chr9:22209353-22211365 | p2 | (AC)6 | TATTCCGCCTTCGAATTGTC | TCTTTGCGCCTCCAAATACT | 155 |
| chr9:22445223-22447238 | p5 | (TAAAA)3 | GGGAAAAGGCCTGATTTACC | GGGGTTAAAAATAGGGGCAA | 147 |
| chr9:22445223-22447238 | p5 | (TTGTA)3 | GGGAAAAGGCCTGATTTACC | GGGGTTAAAAATAGGGGCAA | 147 |
| chr9:22463706-22465733 | p3 | (TGT)9 | GCGGAAGAAATGGAGTGGTA | GTCGTATCGATGAGCGGTTT | 282 |
| chr9:22470202-22472226 | p3 | (TGT)8 | GAAACGAGCGAAAGAAATGG | ATCGATGAGCGGTTTATTGC | 295 |
| chr9:22580006-22582021 | p2 | (TA)11 | TCTCGAGCAAATCAACATGC | CCCAATCATCGCTGAAAAAT | 170 |
| chr9:22580006-22582021 | p5 | (TTGTT)3 | TCTCGAGCAAATCAACATGC | CCCAATCATCGCTGAAAAAT | 170 |

| Marker ID | SSR type | SSR motif | FORWARD PRIMER (5'-3') | REVERSE PRIMER (5'-3') | size (bp) |
|---|---|---|---|---|---|
| chr9:22694523-22696543 | p2 | (TA)10 | TCCATAATGATATCGGGGGA | TATCCGTATAGCTGCCCGTC | 270 |
| chr9:22714663-22716679 | p5 | (TGCAT)3 | CTCTGAGCTGTGTGCAGAGG | CAGTACAAACCATGGGGGAC | 194 |
| chr9:22714663-22716679 | p4 | (TTAT)4 | CTCTGAGCTGTGTGCAGAGG | CAGTACAAACCATGGGGGAC | 194 |
| chr9:22774969-22776985 | p2 | (AT)8 | GACAACCAATACCCACGACC | AGTTGGCAAGCTCCACTTGT | 178 |
| chr9:22810421-22812436 | p5 | (AAGAG)3 | AAAAGCGCCTCTTCTCCTTC | GGGTAAATGACCGGATGATG | 145 |
| chr9:22812159-22814171 | p2 | (AC)6 | GGCTCGGTCCCTTATTTGTT | CAGTTTGGGCCATTTGTCTT | 296 |
| chr9:22967869-22969901 | p2 | (AT)16 | ATAAGATGCCAACTCCACGG | GTTGCTTCGGGTCAACAAAT | 174 |
| chr9:22974909-22976921 | p2 | (AT)6 | TTTACGGGTCTTCATCTGGC | GTTGCTTCGGGTCAACAAAT | 244 |
| chr9:23042051-23044066 | p2 | (AT)11 | CAGGCAACTGCAAAATCTCA | GCCACTTGGTCAGTGAGACA | 281 |
| chr9:23042051-23044066 | p3 | (CAA)5 | CAGGCAACTGCAAAATCTCA | GCCACTTGGTCAGTGAGACA | 281 |
| chr9:23042051-23044066 | p3 | (AAT)5 | CAGGCAACTGCAAAATCTCA | GCCACTTGGTCAGTGAGACA | 281 |
| chr9:23086778-23088870 | p2 | (AT)46 | CGCCAATTTGAACTGTTCCT | CCCTAGCTCTGCTCCTGGTA | 188 |
| chr9:23128564-23130588 | p2 | (AG)12 | GGAAATCAAACTTCCCGACA | TCAGGCCGTGTGTAGTTCTG | 232 |
| chr9:23232356-23234376 | p5 | (GGTGC)4 | GCCGTGTGTAGTTCTGGGAT | CTATTTCTTCGGCAGCGAAC | 266 |
| chr9:23293407-23295431 | p2 | (TA)8 | CGCATTTAGCCAAGGAAGAC | TCTTCCATTTGGTCTGGTCC | 212 |
| chr9:23293437-23295451 | p2 | (TA)8 | CGCATTTAGCCAAGGAAGAC | TCTTCCATTTGGTCTGGTCC | 182 |
| chr9:23323336-23325354 | p6 | (AGTTTC)3 | TCTCCTCATCCTCCCATGTC | GCCCCACATACCTGTAATCG | 270 |
| chr9:23363121-23365136 | p2 | (AT)7 | TTTGCCCTTAGCCTCTCTCA | AACCAGCAACCATGGAAGAC | 140 |
| chr9:23363121-23365136 | p5 | (ATTGT)3 | TTTGCCCTTAGCCTCTCTCA | AACCAGCAACCATGGAAGAC | 140 |
| chr9:23371667-23373681 | p2 | (AT)7 | GTAGCTCAGGTGGCACAACA | TGGGTGACCAAATTCCAAAT | 122 |
| chr9:23395316-23397331 | p5 | (ATTGC)3 | GGAAGGGGAAGGCAATAGAG | TGGCTAGTCCCAAACACACA | 215 |
| chr9:23395844-23397870 | p5 | (ATTGC)3 | TTGGGCTCGGTTTTTACTTG | TTGCCCAATAGACAGAAGGG | 196 |
| chr9:23395874-23397900 | p5 | (ATTGC)3 | TTGGGCTCGGTTTTTACTTG | TTGCCCAATAGACAGAAGGG | 166 |
| chr9:23424829-23426845 | p2 | (TA)24 | CACGGCCCAGATTAAAAGAA | GGCTACACCTCCCCACCTAT | 109 |
| chr9:23424829-23426845 | p2 | (TA)7 | CACGGCCCAGATTAAAAGAA | GGCTACACCTCCCCACCTAT | 109 |
| chr9:23579811-23581827 | p2 | (AT)7 | GTGCGCACTCTTCTGCATTA | CTCCATTGAGGGACTCGGTA | 279 |
| chr9:23579811-23581827 | p2 | (AT)7 | GTGCGCACTCTTCTGCATTA | CTCCATTGAGGGACTCGGTA | 279 |
| chr9:23579811-23581827 | p2 | (AT)8 | GTGCGCACTCTTCTGCATTA | CTCCATTGAGGGACTCGGTA | 279 |
| chr9:23579811-23581827 | p2 | (AT)7 | GTGCGCACTCTTCTGCATTA | CTCCATTGAGGGACTCGGTA | 279 |
| chr9:23636339-23638354 | p5 | (TGAAA)3 | GCGCTCTTTAACGAATCAGG | AAAACCCTGTTGACGGACTG | 171 |
| chr9:23825972-23827984 | p2 | (GT)6 | AAAGGACAAAGCGGTGCTAA | TCCTCACTATTTCCGCATCC | 167 |
| chr9:23825972-23827984 | p2 | (TG)6 | AAAGGACAAAGCGGTGCTAA | TCCTCACTATTTCCGCATCC | 167 |
| chr9:23845829-23847847 | p3 | (TGT)6 | TGCGCAATAGAACCAAACTG | CCGGATTATCTTCGGTCCTT | 290 |
| chr9:23849300-23851321 | p2 | (CT)19 | CCACACAACGTTTCACAAGG | GTAGGCTGGACTTGAGGCAC | 117 |
| chr9:23849300-23851321 | p3 | (TCT)5 | CCACACAACGTTTCACAAGG | GTAGGCTGGACTTGAGGCAC | 117 |
| chr9:23849865-23851903 | p2 | (CT)19 | ATGACGACTGTTGACACCCA | TAGGGTGGTGAAACGTGGAT | 263 |

| Marker ID | SSR type | SSR motif | FORWARD PRIMER (5'-3') | REVERSE PRIMER (5'-3') | size (bp) |
|---|---|---|---|---|---|
| chr9:23849865-23851903 | p3 | (TCT)5 | ATGACGACTGTTGACACCCA | TAGGGTGGTGAAACGTGGAT | 263 |
| chr9:23862091-23864107 | p2 | (AT)6 | AGGCTCCACTACACATTGGG | CCATTGTTCCGAAGCCTAGA | 261 |
| chr9:23862091-23864107 | p2 | (GT)7 | AGGCTCCACTACACATTGGG | CCATTGTTCCGAAGCCTAGA | 261 |
| chr9:23862108-23864122 | p2 | (AT)6 | AGGCTCCACTACACATTGGG | CCATTGTTCCGAAGCCTAGA | 244 |
| chr9:23862108-23864122 | p2 | (GT)7 | AGGCTCCACTACACATTGGG | CCATTGTTCCGAAGCCTAGA | 244 |
| chr9:23888333-23890348 | p5 | (TTGTT)3 | TTAGCATCATGTTAGGGGGC | ACAGTGGGCGAGATATGGTC | 147 |
| chr9:23893170-23895182 | p2 | (AT)6 | AAGGCAAGCAGCTCACTGTT | AAAAGAAGACGCAAAACCGA | 124 |
| chr9:24014053-24016067 | p2 | (GA)7 | CTTCCGAGTGTCTTCTTCGG | TAAACCCAAATGGCCGATAA | 129 |
| chr9:24032121-24034136 | p2 | (TA)23 | CATTGTTCCGGAGGTGCTAT | AGCCCTTCGGCTATCAAAAT | 248 |
| chr9:24032121-24034136 | p5 | (TGCAA)3 | CATTGTTCCGGAGGTGCTAT | AGCCCTTCGGCTATCAAAAT | 248 |
| chr9:24032419-24034509 | p2 | (AT)18 | TGATAGCCGAAGGGCTTATG | CGCCCATACCATTATCCATC | 152 |
| chr9:24032419-24034509 | p5 | (TGCAA)3 | TGATAGCCGAAGGGCTTATG | CGCCCATACCATTATCCATC | 152 |
| chr9:24076646-24078664 | p3 | (AAG)6 | CAGGGGATGCTGTGGTACTT | GAATCAAGTCCTCGCGTAGC | 259 |
| chr9:24094501-24096513 | p2 | (TA)6 | TGAGTTTGGGCCTACTGCTT | ACCAATGGGTTTTGGATGAA | 255 |
| chr9:24197160-24199208 | p2 | (AT)14 | TGATAAAATGGGGCCTTGAG | GCCTGTTGTGATCTGCCTTT | 151 |
| chr9:24197160-24199208 | p2 | (AT)24 | TGATAAAATGGGGCCTTGAG | GCCTGTTGTGATCTGCCTTT | 151 |
| chr9:24402142-24404178 | p3 | (AAG)12 | AATGGATGGTCAAGAGTGGC | CGAGCTTAGAAAAGCGCCTA | 276 |
| chr9:24431790-24433830 | p2 | (AT)20 | TTGAAACCTCGTTGAGGGTC | ACACATCCAACAGCCAACAA | 129 |
| chr9:24451263-24453277 | p2 | (TA)6 | TGGCCTTTGTTGTGATGTGT | GGAAAACATTTGAGGCGTGT | 209 |
| chr9:24451263-24453277 | p2 | (TG)7 | TGGCCTTTGTTGTGATGTGT | GGAAAACATTTGAGGCGTGT | 209 |
| chr9:24457687-24459699 | p2 | (AT)6 | AGGTTATCCCTTGGTTTGGG | TTATGCTTCCGAATTTTGGC | 295 |
| chr9:24643869-24645902 | p3 | (ATC)11 | CCCGTTCATGACTTGGAGTT | TTCCGGTGTTGTTCATAGCA | 114 |
| chr9:24688496-24690514 | p2 | (TA)28 | CTCAGCCGAGGTATGGTGTT | CGGGATACTTAAGGCAACCA | 217 |
| chr9:24688496-24690514 | p2 | (AG)9 | CTCAGCCGAGGTATGGTGTT | CGGGATACTTAAGGCAACCA | 217 |
| chr9:24812730-24814748 | p2 | (AT)9 | CTTTGTTCGAGCTAGCGGTC | ATCCAAAGGGGAGAGGAAAA | 167 |
| chr9:24812730-24814748 | p4 | (TTTA)4 | CTTTGTTCGAGCTAGCGGTC | ATCCAAAGGGGAGAGGAAAA | 167 |
| chr9:24837407-24839425 | p6 | (TCTTTT)3 | TGTAGTTGATCCCACGGTCA | AAATGGGTACGGAGTAGGGG | 261 |
| chr9:24963666-24965684 | p2 | (TA)15 | AGGAAACATGGCCAGTGAAG | AGTTAGGGAGGAAAAGGGCA | 193 |
| chr9:24963666-24965684 | p6 | (AAGGGA)3 | AGGAAACATGGCCAGTGAAG | AGTTAGGGAGGAAAAGGGCA | 193 |
| chr9:25131629-25133641 | p2 | (TA)6 | CATCTCGTCCTGTTGGGACT | TTAGTGACGGTGGTGGTCAA | 112 |
| chr9:25186121-25188136 | p3 | (ACC)5 | ACCAACCCTCAAACATGACC | ATTTGCCAACTTGAAATGCC | 219 |
| chr9:25302136-25304158 | p2 | (TA)11 | AATCTGACGGCATTCCATTC | GAATTTGGAAGGTTTCGGGT | 184 |
| chr9:25365531-25367543 | p2 | (AG)6 | CACCCCGTAATATTCCATGC | GCTTCGACGAAACTCCAATC | 231 |
| chr9:25397680-25399704 | p2 | (TA)12 | AAATGCCCTGTAGCATCCTG | GTGGTATCAGAGCGGTTCGT | 115 |
| chr9:25565052-25567064 | p2 | (AT)38 | CCATGATATGGAGATTGCCC | AGGGAAAGGGTTGAAGGCTA | 108 |
| chr9:25565052-25567064 | p2 | (TG)6 | CCATGATATGGAGATTGCCC | AGGGAAAGGGTTGAAGGCTA | 108 |

| Marker ID | SSR type | SSR motif | FORWARD PRIMER (5'-3') | REVERSE PRIMER (5'-3') | size (bp) |
|---|---|---|---|---|---|
| chr9:25581465-25583505 | p2 | (AG)20 | TCTTGCCAAACAAGCTTCAA | TGTTAGAGGGGGACATGGAC | 182 |
| chr9:25615563-25617590 | p5 | (TTACT)3 | GGTATATGTAACGCCCCCAA | TTTACCCGGTTCTACGACCA | 274 |
| chr9:25615563-25617590 | p3 | (ATA)9 | GGTATATGTAACGCCCCCAA | TTTACCCGGTTCTACGACCA | 274 |
| chr9:25677744-25679756 | p2 | (AT)38 | CCATGATATGGAGATTGCCC | AGGGAAAGGGTTGAAGGCTA | 108 |
| chr9:25677744-25679756 | p2 | (TG)6 | CCATGATATGGAGATTGCCC | AGGGAAAGGGTTGAAGGCTA | 108 |
| chr9:25694157-25696197 | p2 | (AG)20 | TCTTGCCAAACAAGCTTCAA | TGTTAGAGGGGGACATGGAC | 182 |
| chr9:25728355-25730382 | p5 | (TTACT)3 | GGTATATGTAACGCCCCCAA | TTTACCCGGTTCTACGACCA | 274 |
| chr9:25728355-25730382 | p3 | (ATA)9 | GGTATATGTAACGCCCCCAA | TTTACCCGGTTCTACGACCA | 274 |
| chr9:25909439-25911473 | p2 | (AT)6 | TGCATGTCATAGCCACCTGT | ATTCGTAGTGGCAGCCTTGT | 148 |
| chr9:25909439-25911473 | p2 | (AT)17 | TGCATGTCATAGCCACCTGT | ATTCGTAGTGGCAGCCTTGT | 148 |
| chr9:25940500-25942512 | p2 | (AT)6 | TCATATAGGCGGGACAGGAC | CAGGGAAACAGGCTTGATGT | 204 |
| chr9:25948115-25950127 | p2 | (AT)6 | AAACCGACAAGGCTTCAAGA | AGGGGAGCTCCATTTCATCT | 135 |
| chr9:26132743-26134779 | p2 | (AT)16 | ATGAGATTTGTCCACCAGGC | TTTGTGGTTCGAAGTGACCA | 155 |
| chr9:26132743-26134779 | p2 | (AT)18 | ATGAGATTTGTCCACCAGGC | TTTGTGGTTCGAAGTGACCA | 155 |
| chr9:26144758-26146778 | p2 | (TA)10 | CTGGAATTTGGCAGTTGGAT | TCCGTTTGACCCCCATAATA | 158 |
| chr9:26300956-26302972 | p4 | (TATT)4 | ATCATCCGCGAAAAGCATAC | GGCGAGTTGGGATGAAATTA | 102 |
| chr9:26303562-26305578 | p4 | (TATT)4 | ATCATCCGCGAAAAGCATAC | GGCGAGTTGGGATGAAATTA | 102 |
| chr9:26334269-26336284 | p5 | (ACCGA)3 | TTTAGAATTGCACGGGAACC | GTCTAGACCAGGTCGTGGGA | 160 |
| chr9:26483923-26485983 | p4 | (CATA)4 | TGGATCCTTGCGAATGTGTA | CCATTTAGCACCCTGCAGTT | 192 |
| chr9:26492651-26494677 | p2 | (AT)13 | TCCAAAAATCCCCATCAAAA | ACATACACACACCCACACGG | 279 |
| chr9:26496152-26498168 | p2 | (AT)8 | CATGCTGTCAAAAAGAGCGA | TCGGATTATGTTTGAAGCCC | 117 |
| chr9:26496152-26498168 | p2 | (AT)18 | CATGCTGTCAAAAAGAGCGA | TCGGATTATGTTTGAAGCCC | 117 |
| chr9:26547343-26549363 | p2 | (AG)10 | TCTCTCACCCGCTTCTCATT | AAGGCAATCGTCATGGAAAG | 163 |
| chr9:26551385-26553397 | p2 | (TA)6 | CATTCACGTGGCTTCAGTGT | TCCTGCTGCGAGTAGGATTT | 147 |
| chr9:26551385-26553397 | p5 | (GGTTA)3 | CATTCACGTGGCTTCAGTGT | TCCTGCTGCGAGTAGGATTT | 147 |
| chr9:26594314-26596328 | p2 | (TG)7 | GTTGGAACCCATCACTTGCT | TTGCGACATTTTCCAATCAA | 273 |
| chr9:26777078-26779092 | p2 | (TC)7 | GTGAGAGGTTGGCCATGAAT | GGGTCCTCCAAACCCTCTTA | 118 |
| chr9:26777078-26779092 | p2 | (TA)7 | GTGAGAGGTTGGCCATGAAT | GGGTCCTCCAAACCCTCTTA | 118 |
| chr9:26779407-26781427 | p2 | (TA)28 | TGGCCACGTGTCACTTTTAC | ACGGGCTCAACACTTTGAAT | 195 |
| chr9:26779407-26781427 | p5 | (AAAAT)4 | TGGCCACGTGTCACTTTTAC | ACGGGCTCAACACTTTGAAT | 195 |
| chr9:26791025-26793045 | p4 | (AAAT)5 | AAGGAGGTGATTGATGGTGC | GCAATGTTTCCCAGAGCAA | 285 |
| chr9:26806728-26808743 | p5 | (CTTTT)3 | CAGATGCACATGGCCATAAC | CGTTGGAGCAAGAAAACCAT | 179 |
| chr9:26806728-26808743 | p2 | (TA)33 | CAGATGCACATGGCCATAAC | CGTTGGAGCAAGAAAACCAT | 179 |
| chr9:26819426-26821441 | p5 | (GGGTG)3 | CCAAGCGCATTACTTTCCAT | CAGGAGTAAGCGGTCGTTGT | 285 |
| chr9:26819426-26821441 | p5 | (TTTGG)4 | CCAAGCGCATTACTTTCCAT | CAGGAGTAAGCGGTCGTTGT | 285 |
| chr9:27103510-27105522 | p2 | (GT)6 | TCCGCGAGGAATTCATTATC | AGAACGGTTCCCACAATGTC | 292 |

| Marker ID | SSR type | SSR motif | FORWARD PRIMER (5'-3') | REVERSE PRIMER (5'-3') | size (bp) |
|---|---|---|---|---|---|
| chr9:27179796-27181818 | p2 | (AT)11 | TCCCACCTCGGTAAGTGAAC | GACCAGACCCTGGGGTATTT | 121 |
| chr9:27180917-27182983 | p2 | (AT)33 | CAGTTCCAGCCCAAAGGTAA | GACCTGGAAGTCCACCAAAA | 102 |
| chr9:27196047-27198065 | p2 | (AT)12 | CGAACCGACTTTAGCACCTC | GCCTTGAATTCCAACCAAGA | 281 |
| chr9:27437015-27439029 | p2 | (GA)7 | CACAGGTTGGTGCATGTTTC | AGCAGCAGATATGGCTCGAT | 264 |
| chr9:27468051-27470072 | p3 | (ATG)7 | CAAGTGAGGGAACCAAAGGA | TGCCTCCAATCTTCCGATAC | 272 |
| chr9:27468051-27470072 | p2 | (TA)9 | CAAGTGAGGGAACCAAAGGA | TGCCTCCAATCTTCCGATAC | 272 |
| chr9:27496078-27498126 | p5 | (AAACT)4 | CCAACTCCCAACTAAAGCCA | AAAGCCATTCTGGGGTTTCT | 295 |
| chr9:27496078-27498126 | p2 | (TA)24 | CCAACTCCCAACTAAAGCCA | AAAGCCATTCTGGGGTTTCT | 295 |
| chr9:27504023-27506045 | p3 | (ATG)6 | AAGGAATTTCTTCCCCCTGA | TCACGTCCTCCTCCAAAGTT | 170 |
| chr9:27504023-27506045 | p2 | (AT)11 | AAGGAATTTCTTCCCCCTGA | TCACGTCCTCCTCCAAAGTT | 170 |
| chr9:27555910-27557924 | p2 | (AT)7 | TTAATTTTGGAATGACGGGC | TTTGCCTAAACGCCAAAAGT | 218 |
| chr9:27563558-27565572 | p2 | (TC)7 | CGGGTACAGTGAGGGGACTA | CCGCTCATTTGTTCTTCACA | 295 |
| chr9:27612801-27614815 | p2 | (TA)7 | TACCATACCAAGCCACGTCA | AGCGCCCTAGGTTACCTGAT | 256 |
| chr9:27628752-27630767 | p5 | (TGCAA)3 | GCGGGAGTGTTTACAACGAT | CAAGATGGAGGTTGGGTCAT | 267 |
| chr9:27898150-27900178 | p5 | (AATGC)3 | TTCAAACCCAAGCCAGAATC | GCATTGCATGTCACTATGGC | 100 |
| chr9:27935421-27937455 | p2 | (TA)17 | CCTTTCTGTCGTAGCCTTGC | TTATGAGTGTGAAGCGCACC | 201 |
| chr9:28076069-28078083 | p2 | (AG)7 | TTAGAGACCCGCTCTGGAAA | TCCCGAAGGAGAAAACAATG | 256 |
| chr9:28134855-28136869 | p2 | (AT)7 | GGCCTTTTATGTTGACCGAA | CACGTTCTCCCCACTACGTT | 142 |
| chr9:28134855-28136869 | p5 | (TTTTC)3 | GGCCTTTTATGTTGACCGAA | CACGTTCTCCCCACTACGTT | 142 |
| chr9:28158707-28160722 | p3 | (TGT)5 | GGCCGAGAATAACCTTCTCC | ATCGGCTAGCCCTTCACTTT | 272 |
| chr9:28214118-28216146 | p2 | (AT)14 | CTAACCCTAGGACTTGCCCA | GATGAAGGTTATGCCCCAAA | 293 |
| chr9:28243648-28245676 | p2 | (AT)14 | GGCGAACGGATTTACGAATA | TGCCCCAAAGTTCATACCAT | 140 |
| chr9:28376640-28378655 | p5 | (ATAAA)3 | GAAGGGGTTAGGCATCCTTT | ACCCTCTTTGATAAACGGCAT | 148 |
| chr9:28393156-28395200 | p2 | (TA)22 | AGTTTTGGGCTTATTTGGGC | CTTTGGATCTCCGTGTCCAT | 177 |
| chr9:28393156-28395200 | p6 | (CTTTTC)3 | AGTTTTGGGCTTATTTGGGC | CTTTGGATCTCCGTGTCCAT | 177 |
| chr9:28451884-28453899 | p3 | (ATT)5 | ACAATTTGGTTTGAGAGGCG | CACGCAAAGCTTGACACCTA | 125 |
| chr9:28508992-28511032 | p2 | (AT)20 | ATATTCGATCCAACCACCCA | CAGACCCCCATTAGGTGAAA | 105 |
| chr9:28563534-28565555 | p3 | (AAT)7 | CAGGTGGATGGCAGAGGTAT | AGGGTTTCCCCATCATTTTC | 278 |
| chr9:28580698-28582778 | p2 | (AT)40 | TGCTCCGGTATTGTTTGTGA | CGCCCTAATCATCAACTGGT | 134 |
| chr9:28657343-28659359 | p4 | (TGGG)4 | TGTTTGGCCATGAAAATCAA | GACCCGTTTGGCTATGAGAA | 270 |
| chr9:28727824-28729839 | p5 | (AAATC)3 | ATTGCTTCCATTGGCTGTTC | GGTCTAACAAAGGGCAACCA | 144 |
| chr9:28730581-28732596 | p5 | (AAACC)3 | CTGGCCCGATTCAACTCTAC | CCCATGAACACCCCTACTTG | 203 |
| chr9:29031855-29033923 | p2 | (AT)34 | ACAAGGCCTCAACCTGTACG | ACGAGCCCTTGCAGTAGAAA | 284 |
| chr9:29052365-29054385 | p5 | (TTTTC)4 | ACCCCAGTTCTATGGGCTCT | TTTGTGTTTTCGCGAGACTG | 136 |
| chr9:29052365-29054385 | p4 | (TGGG)4 | ACCCCAGTTCTATGGGCTCT | TTTGTGTTTTCGCGAGACTG | 136 |
| chr9:29080901-29082919 | p2 | (GA)9 | CGTTGACCAAGTCAATCCCT | ATGCTTTTCAAGGTGTTCGG | 299 |

| Marker ID | SSR type | SSR motif | FORWARD PRIMER (5'-3') | REVERSE PRIMER (5'-3') | size (bp) |
|---|---|---|---|---|---|
| chr9:29159920-29161932 | p2 | (AC)6 | TATGAGGGATCCCAGACTCG | ATCCATCATAAGTCTGCGGC | 258 |
| chr9:29191486-29193498 | p2 | (AT)6 | CATGGGAATTTGAGACCGAT | CGATTCGATTCGTTCCACTT | 280 |
| chr9:29191486-29193498 | p2 | (TA)6 | CATGGGAATTTGAGACCGAT | CGATTCGATTCGTTCCACTT | 280 |
| chr9:29196543-29198559 | p2 | (TA)22 | ATCGGAGGTTTACCGACCTT | ATAGGTGTCGTATCCGGCTG | 170 |
| chr9:29196543-29198559 | p4 | (ATGT)4 | ATCGGAGGTTTACCGACCTT | ATAGGTGTCGTATCCGGCTG | 170 |
| chr9:29258252-29260270 | p2 | (TA)9 | TCTTCATCTTGGGCAAATCC | TCTTTGGGAGGTTGTGGAAG | 233 |
| chr9:29342517-29344532 | p5 | (CTTCT)3 | CAAGCTGCTCCAAAGGAATC | TCACCTGCAAAGTCTGCATC | 256 |
| chr9:29346135-29348150 | p2 | (TA)6 | ACAAGCTTAGTGGTCCGCAT | TTTCGTGATATCTAGGGCCG | 104 |
| chr9:29346135-29348150 | p3 | (ATC)5 | ACAAGCTTAGTGGTCCGCAT | TTTCGTGATATCTAGGGCCG | 104 |
| chr9:29352491-29354506 | p2 | (AT)24 | GAGTGCTCTTGAGAATCCGC | ATTGACGTCCCTTCTTGTGG | 282 |
| chr9:29352491-29354506 | p2 | (TA)6 | GAGTGCTCTTGAGAATCCGC | ATTGACGTCCCTTCTTGTGG | 282 |
| chr9:29352491-29354506 | p3 | (ATC)5 | GAGTGCTCTTGAGAATCCGC | ATTGACGTCCCTTCTTGTGG | 282 |
| chr9:29352491-29354506 | p3 | (ATA)32 | GAGTGCTCTTGAGAATCCGC | ATTGACGTCCCTTCTTGTGG | 282 |
| chr9:29415654-29417670 | p2 | (TA)8 | CCTGGACCTTTGTTTTGGAA | GCGAGCTTGGACAATTAAGC | 193 |
| chr9:29437254-29439266 | p2 | (TA)6 | GTTTGGCTCAAAGTGGGGTA | CTCCAACTTCGTCCATTCGT | 120 |
| chr9:29469516-29471531 | p3 | (CAA)5 | GCAACTTGAGCACAAGCAAA | GTTCCAAGCTGTTGGCTCTC | 232 |
| chr9:29502447-29504475 | p2 | (AT)14 | GTAGCTCCGGATGAAATGGA | ATCTCCCGATTCTCCCATCT | 291 |
| chr9:29542304-29544324 | p3 | (ATG)7 | GCTCACCCTCAAATCTCTCG | GATTGCTGTTTGTTTGGGCT | 213 |
| chr9:29542304-29544324 | p5 | (TTGTA)4 | GCTCACCCTCAAATCTCTCG | GATTGCTGTTTGTTTGGGCT | 213 |
| chr9:29551352-29553374 | p2 | (AT)11 | TTTTCTTTCATTTTGCGCCT | TGTAAGGATTCTGTGCTGCG | 174 |
| chr9:29652838-29654852 | p2 | (AT)7 | CAGCCGAGGAAATGGTTAAA | TCCTCCATGGGCGTATTTAG | 274 |
| chr9:29715531-29717551 | p5 | (TATGA)4 | TGGGTAATGGGTCATGTGTG | GAATCTGGTAGATTGGCGGA | 143 |
| chr9:29715531-29717551 | p2 | (AT)6 | TGGGTAATGGGTCATGTGTG | GAATCTGGTAGATTGGCGGA | 143 |
| chr9:29880020-29882038 | p6 | (CTCAGG)3 | CTGGATGGCCCAATAGAAGA | CCCACGCATTTGGAGTAGTT | 170 |
| chr9:29946414-29948490 | p2 | (TA)38 | TTGATGTTGGGATTTTGGGT | TCAAAGGAGAACCCACCAAG | 169 |
| chr9:29946414-29948490 | p5 | (TTGCA)3 | TTGATGTTGGGATTTTGGGT | TCAAAGGAGAACCCACCAAG | 169 |
| chr9:29976592-29978604 | p2 | (AT)6 | AAGCATGTATCAACCGGAGC | CTCAGAGGCAAGGTTCAAGG | 137 |
| chr9:30030716-30032732 | p4 | (ATAC)4 | CCGCTGGTTCTCGTTATGTT | CTTTGGAGTGTTGCGAGTCA | 264 |
| chr9:30053809-30055824 | p5 | (ATTTC)3 | TACCCGCACTTATTTCAGGC | ATACCTCACAGGGCCACAAC | 165 |
| chr9:30053809-30055824 | p2 | (AT)6 | TACCCGCACTTATTTCAGGC | ATACCTCACAGGGCCACAAC | 165 |
| chr9:30084132-30086154 | p2 | (AT)11 | CCAAAGGTTCATAGCGTCGT | TGTTCGTCTGTAGTGGCAGC | 243 |
| chr9:30090435-30092463 | p2 | (TA)9 | TCCTTTAACGATCCGTCCTG | CTGAAGACCGACATCGACAA | 207 |
| chr9:30090435-30092463 | p2 | (AT)14 | TCCTTTAACGATCCGTCCTG | CTGAAGACCGACATCGACAA | 207 |
| chr9:30090435-30092463 | p2 | (AT)10 | TCCTTTAACGATCCGTCCTG | CTGAAGACCGACATCGACAA | 207 |
| chr9:30090892-30092912 | p2 | (TA)9 | TGCGAAAGAACTCCAAAGGT | CTGTCCCGTCATATTTCGCT | 241 |
| chr9:30090892-30092912 | p2 | (AT)14 | TGCGAAAGAACTCCAAAGGT | CTGTCCCGTCATATTTCGCT | 241 |

| Marker ID | SSR type | SSR motif | FORWARD PRIMER (5'-3') | REVERSE PRIMER (5'-3') | size (bp) |
|---|---|---|---|---|---|
| chr9:30090892-30092912 | p2 | (AT)10 | TGCGAAAGAACTCCAAAGGT | CTGTCCCGTCATATTTCGCT | 241 |
| chr9:30314835-30316877 | p2 | (AT)21 | TTCGGCGCCTTTTTACTATG | GCTTATCGACGCTTCAAAGG | 239 |
| chr9:30355930-30357946 | p2 | (AC)8 | GGGGATCCAGCACAAGTTTA | GTTGCTTCGGGGAATTGTTA | 205 |
| chr9:30364852-30366958 | p2 | (AT)7 | CCTTGGCGTGCAAAGTTTAT | TTGCGGATCTACCCAATAGC | 103 |
| chr9:30383717-30385759 | p3 | (CAT)14 | TAGGAACCCACTCCCACAAG | AATGAAACTGCCGAGTGCTT | 143 |
| chr9:30419391-30421406 | p5 | (AAATG)3 | CTATTCCACCGCTTGATCGT | AGGGAAATGGTGACTTGTGG | 291 |
| chr9:30433058-30435073 | p5 | (TTTAA)3 | CAGCGGATTCTTAGGAGCAC | GGGAATATGCCGCAAGATAA | 187 |
| chr9:30510491-30512506 | p2 | (AG)6 | TGAATTCGGGAGGCAATAAC | GATGTCACATGCAGACCCAC | 189 |
| chr9:30510491-30512506 | p3 | (TTG)5 | TGAATTCGGGAGGCAATAAC | GATGTCACATGCAGACCCAC | 189 |
| chr9:30510491-30512506 | p2 | (TA)20 | TGAATTCGGGAGGCAATAAC | GATGTCACATGCAGACCCAC | 189 |
| chr9:30551468-30553486 | p2 | (TA)24 | CCACCTAAGAAGGTGGGTGA | TGGGCCGTGATAATAACTCC | 102 |
| chr9:30551468-30553486 | p6 | (ATACAT)3 | CCACCTAAGAAGGTGGGTGA | TGGGCCGTGATAATAACTCC | 102 |
| chr9:30551468-30553486 | p2 | (TA)10 | CCACCTAAGAAGGTGGGTGA | TGGGCCGTGATAATAACTCC | 102 |
| chr9:30560878-30562893 | p5 | (ATAAA)3 | CAATTTCAAGGCACAAGCAA | AGGCTTTTCTGCAAATGGAA | 231 |
| chr9:30564127-30566139 | p2 | (GA)6 | TTTCTTGTACGAGGGTTGGG | AGATCTGGGTCGTCGATTTG | 300 |
| chr9:30564127-30566139 | p2 | (AT)6 | TTTCTTGTACGAGGGTTGGG | AGATCTGGGTCGTCGATTTG | 300 |
| chr9:30564127-30566139 | p4 | (ATTA)4 | TTTCTTGTACGAGGGTTGGG | AGATCTGGGTCGTCGATTTG | 300 |
| chr9:30596755-30598767 | p2 | (AG)6 | ATAGTGTACAAGGTGCCGCC | AGTCCGTTGCGCATTTTTAC | 220 |
| chr9:30796888-30798900 | p2 | (CA)6 | ACAACACTCCGAAATCTGCC | GTTGTAATCGTTGCGGGATT | 281 |
| chr9:30819045-30821063 | p6 | (TGTTTG)3 | TCATGAAGCGAGTGATGAGG | ACCGGAGGTAAAGCCAACTT | 296 |
| chr9:30864133-30866153 | p5 | (CATTG)4 | CCAGCTCGGCACTAAGGTAG | TAATGGCGACAAGCCACATA | 202 |
| chr9:30892511-30894529 | p6 | (AGTTTC)3 | GATATATGGTGGAGCCCGAA | AGGTCGTTTTTCGTGTTTGG | 257 |
| chr9:30892511-30894529 | p2 | (TA)6 | GATATATGGTGGAGCCCGAA | AGGTCGTTTTTCGTGTTTGG | 257 |
| chr9:31075903-31077918 | p5 | (ATTGC)3 | TTATTGTTGATCGGCCTTCC | CCCAACTTCATACCCGAAGA | 202 |
| chr9:31077039-31079059 | p3 | (TTA)5 | GACCTGCACTTGCTGCATTA | GTCGGGTCGTGACAAAGACT | 118 |
| chr9:31097361-31099373 | p3 | (TTG)6 | GTCTTGTTTTGCGGTGGATT | ATAGCCCACACGCTGTTTTC | 145 |
| chr9:31097370-31099438 | p3 | (TTG)6 | ATGGAAGTTGGAAAACAGCG | TATATCCTGCCGAAAATCGC | 295 |
| chr9:31097445-31099461 | p3 | (TTG)6 | ATGGAAGTTGGAAAACAGCG | TATATCCTGCCGAAAATCGC | 220 |
| chr9:31097475-31099491 | p3 | (TTG)6 | ATGGAAGTTGGAAAACAGCG | TATATCCTGCCGAAAATCGC | 190 |
| chr9:31097499-31099515 | p3 | (TTG)6 | ATGGAAGTTGGAAAACAGCG | TATATCCTGCCGAAAATCGC | 166 |
| chr9:31203050-31205065 | p3 | (GTT)5 | TGCCCTAGGATCTCCCTTTT | GACCAAGAAGAAGACGGTGC | 160 |
| chr9:31218609-31220627 | p6 | (CATATA)3 | ATGAGCCGTGACAAGTACCC | TGATGTTATGCAGCCTCGTC | 176 |
| chr9:31245353-31247369 | p4 | (TAAA)4 | ACAGTCACAGCGGGGTAAAG | TCATGGGGCACGTAGTCATA | 143 |
| chr9:31333005-31335020 | p5 | (GTATA)3 | CAAGCTGGGCAAAATTTGAT | AGTACCGGATAGCTTGCCCT | 145 |
| chr9:31333005-31335020 | p2 | (AT)6 | CAAGCTGGGCAAAATTTGAT | AGTACCGGATAGCTTGCCCT | 145 |
| chr9:31359611-31361623 | p2 | (AG)6 | ACGAACAAAATGCTCGTTCC | AGCCCCCTAATTGTTTCCAC | 165 |

| Marker ID | SSR type | SSR motif | FORWARD PRIMER (5'-3') | REVERSE PRIMER (5'-3') | size (bp) |
|---|---|---|---|---|---|
| chr9:31391548-31393562 | p2 | (AG)7 | AGGAGTGCTTGATCCGAAGA | GTACAAGGTGGCAGAACGGT | 145 |
| chr9:31444236-31446268 | p2 | (GA)16 | GGTGAACCCTCCAACGACTA | GGTTCTCGAGCAAGTTGAGG | 260 |
| chr9:31510615-31512649 | p2 | (AT)17 | ATCGGAGTGACGTAAGGTCG | GAGTTGCGAAAGACGAAAGG | 299 |
| chr9:31510615-31512649 | p6 | (TATACA)3 | ATCGGAGTGACGTAAGGTCG | GAGTTGCGAAAGACGAAAGG | 299 |
| chr9:31529795-31531807 | p2 | (CA)6 | TGGCCAATTTCAATCAACAA | ATCGGTTGCGATTATTGGAG | 276 |
| chr9:31529795-31531807 | p2 | (AT)6 | TGGCCAATTTCAATCAACAA | ATCGGTTGCGATTATTGGAG | 276 |
| chr9:31544801-31546816 | p5 | (ATTGC)3 | ATCGTTCCAAGGCTCTCTGA | ACACAATAAGGCGACAAGCC | 111 |
| chr9:31738267-31740307 | p2 | (TA)20 | GATTGCACCGAGCCATCTAT | GTGCATCTCTACCCGACCAT | 273 |
| chr9:31738267-31740307 | p5 | (GAAAA)3 | GATTGCACCGAGCCATCTAT | GTGCATCTCTACCCGACCAT | 273 |
| chr9:31774124-31776139 | p3 | (ATG)5 | CGTTTGCATGATTTCGTTTG | CACCAATGTTTGTACCTGCG | 189 |
| chr9:31864275-31866290 | p3 | (ATG)5 | ACAAGTTCGGGTGACAAAGC | TCCACTTGGAATCCACATCA | 190 |
| chr9:31944030-31946042 | p2 | (AT)6 | GTACCGTACCGTGTGCCTCT | CGTGCATGCCTCAATATGTC | 158 |
| chr9:31946278-31948293 | p5 | (TCCTT)3 | CTCAGCATCATGCGTCACTT | GGACCACAGCGCTATTTGAT | 264 |
| chr9:31953392-31955407 | p5 | (TCCTT)3 | GAGTATCGCATGCCTTTGGT | AAGTGACGCATGATGCTGAG | 131 |
| chr9:31970403-31972415 | p2 | (TA)6 | AGCCGACAAGGCTGCTATTA | AGTGTTATTTGGGTACGGCG | 270 |
| chr9:31995693-31997757 | p2 | (TA)7 | AATGGCCTGGTACGAGAATG | CGAGACTCGGTGTAATCCGT | 142 |
| chr9:31995693-31997757 | p2 | (TA)32 | AATGGCCTGGTACGAGAATG | CGAGACTCGGTGTAATCCGT | 142 |
| chr9:32025817-32027835 | p5 | (TGAAT)3 | GTTGATTGATGATTTGGGGC | GCTTAATTGTCCAAGCTCGC | 207 |
| chr9:32025817-32027835 | p6 | (ATGTAT)3 | GTTGATTGATGATTTGGGGC | GCTTAATTGTCCAAGCTCGC | 207 |
| chr9:32055942-32057969 | p3 | (GAA)9 | AGGCTGTTGCCATTACTGCT | GGCTATGGTGGCTAATTCCA | 120 |
| chr9:32139885-32141899 | p2 | (TA)7 | AGAGTTGGGTTTGGATGACG | TTCGAAAAATCGGGTAGCAC | 157 |
| chr9:32139885-32141899 | p2 | (AG)7 | AGAGTTGGGTTTGGATGACG | TTCGAAAAATCGGGTAGCAC | 157 |
| chr9:32154286-32156354 | p2 | (AT)34 | GCTCATACCCAAATGCCCTA | GGAGGGATGGTACAAAGGGT | 114 |
| chr9:32177196-32179242 | p2 | (TA)23 | ATTGCTCCATAATGGGATCG | TGCCAAAAGGTCATGGTGTA | 185 |
| chr9:32345604-32347622 | p2 | (AT)9 | CTCGCGCTGCTTTTATATCC | GCTTTCACCCATTTGGAAAA | 173 |
| chr9:32345604-32347622 | p2 | (AT)11 | CTCGCGCTGCTTTTATATCC | GCTTTCACCCATTTGGAAAA | 173 |
| chr9:32353325-32355339 | p2 | (AT)11 | AAACGATTATCTGCAAGCCG | CTCAGCTGTTTTGGGAGAGC | 162 |
| chr9:32353325-32355339 | p2 | (GA)7 | AAACGATTATCTGCAAGCCG | CTCAGCTGTTTTGGGAGAGC | 162 |
| chr9:32466179-32468195 | p4 | (GAAA)4 | AAGAGCTGGAAGGTCACGAA | GGTAGAGTTCGTGCTCTGGG | 294 |
| chr9:32472071-32474086 | p3 | (ATC)5 | CCACAAGGAAAAGGCGTTAG | GATGCACTTCCGCAACACTA | 208 |
| chr9:32472071-32474086 | p5 | (ACTCA)3 | CCACAAGGAAAAGGCGTTAG | GATGCACTTCCGCAACACTA | 208 |
| chr9:32483372-32485387 | p2 | (TA)8 | CCCTTTTCCCATATCCCCTA | CGGATGTTACCGACCTTACG | 237 |
| chr9:32483372-32485387 | p5 | (ACTCA)3 | CCCTTTTCCCATATCCCCTA | CGGATGTTACCGACCTTACG | 237 |
| chr9:32558410-32560424 | p2 | (AT)7 | ATGGTTCGCTTGTTGGTTTC | GCGAACGAGGTAAAGGAGTG | 250 |
| chr9:32558410-32560424 | p2 | (AT)30 | ATGGTTCGCTTGTTGGTTTC | GCGAACGAGGTAAAGGAGTG | 250 |
| chr9:32560265-32562299 | p2 | (AT)25 | CTGTGGGCATGAACATAACG | GTGGCCGAATACGGATAATG | 262 |

| Marker ID | SSR type | SSR motif | FORWARD PRIMER (5'-3') | REVERSE PRIMER (5'-3') | size (bp) |
|---|---|---|---|---|---|
| chr9:32560265-32562299 | p2 | (TA)9 | CTGTGGGCATGAACATAACG | GTGGCCGAATACGGATAATG | 262 |
| chr9:32560300-32562332 | p2 | (TA)8 | CTGTGGGCATGAACATAACG | GTGGCCGAATACGGATAATG | 227 |
| chr9:32560300-32562332 | p2 | (TA)9 | CTGTGGGCATGAACATAACG | GTGGCCGAATACGGATAATG | 227 |
| chr9:32624311-32626326 | p5 | (ATACA)3 | GTGGCAACGAGCCAATAAGT | TTGGGTCGTGACACCTATGA | 215 |
| chr9:32624311-32626326 | p3 | (ATC)5 | GTGGCAACGAGCCAATAAGT | TTGGGTCGTGACACCTATGA | 215 |
| chr9:32790877-32792891 | p2 | (AT)10 | CGACTCGTTTGACCTCCAAT | AAGGTTGAGCCATGTTTTGG | 217 |
| chr9:32790877-32792891 | p3 | (TCT)5 | CGACTCGTTTGACCTCCAAT | AAGGTTGAGCCATGTTTTGG | 217 |
| chr9:32790877-32792891 | p2 | (AC)7 | CGACTCGTTTGACCTCCAAT | AAGGTTGAGCCATGTTTTGG | 217 |
| chr9:32790877-32792891 | p3 | (GAA)8 | CGACTCGTTTGACCTCCAAT | AAGGTTGAGCCATGTTTTGG | 217 |
| chr9:32790877-32792891 | p6 | (ATATAC)3 | CGACTCGTTTGACCTCCAAT | AAGGTTGAGCCATGTTTTGG | 217 |
| chr9:32791860-32793878 | p2 | (AC)7 | CTTGGATATTTTGGCTCCCA | AAATTCGACGGTTTCACTGG | 164 |
| chr9:32791860-32793878 | p3 | (GAA)8 | CTTGGATATTTTGGCTCCCA | AAATTCGACGGTTTCACTGG | 164 |
| chr9:32791860-32793878 | p6 | (ATATAC)3 | CTTGGATATTTTGGCTCCCA | AAATTCGACGGTTTCACTGG | 164 |
| chr9:32839544-32841559 | p5 | (TTTTA)3 | ACTAATCGCAAAACCGTTCG | TGTCCGATTTTCGTTAAGCC | 121 |
| chr9:32889219-32891234 | p3 | (ATC)5 | CCGGTGGAGCTACAAAACAT | ACTGACGTCCCTTCTTGTGG | 126 |
| chr9:32968912-32970926 | p2 | (AT)7 | CAAGGCTGCCACGATATACA | TTGTTCCGGAGCTAGCTTGT | 275 |
| chr9:33026145-33028159 | p2 | (GT)7 | TCAAATGCTTGCTATGCCTG | AATGAACCAGCCACTGATCC | 212 |
| chr9:33044150-33046171 | p3 | (AAC)7 | GTGGTAACATCTGCCCGACT | ATGGCCTGAGCTGTGCTTAT | 230 |
| chr9:33062520-33064541 | p3 | (AAC)7 | GTGGTAACATCTGCCCGACT | ATGGCCTGAGCTGTGCTTAT | 230 |
| chr9:33139718-33141730 | p2 | (CT)6 | GAAAGAGTTGAATTTGCGGC | GATCTGCGGCTAAGAGCATC | 147 |
| chr9:33147821-33149833 | p2 | (CT)6 | TACCCGCATTTATTTCAGCC | AACATGGCCCTAACAACCTG | 122 |
| chr9:33157195-33159215 | p2 | (AT)10 | AACTGTATGGCCTCGGTCAC | CAGAAGGGCAAAGCTACGAC | 297 |
| chr9:33277815-33279845 | p2 | (AG)6 | AATAATGGCGACGAGCTCAC | CTTGACCACCTGTTCGGTTT | 138 |
| chr9:33277851-33279871 | p2 | (AG)6 | AATAATGGCGACGAGCTCAC | CTTGACCACCTGTTCGGTTT | 102 |
| chr9:33354042-33356054 | p2 | (GT)6 | CGATGTCGGGATCAAAAGTT | CTTCCCGCGATTATCTTGAA | 126 |
| chr9:33365374-33367398 | p6 | (TATATG)4 | TCGTGTGGGATGTTTATGGA | GCTTATTTGTCCAAGCTCGC | 131 |
| chr9:33567062-33569084 | p2 | (GA)11 | ATTGCAGCCAATTCAAGGTC | GGGCTAGGCTGTGTGTTGAT | 258 |
| chr9:33567062-33569084 | p3 | (GTT)5 | ATTGCAGCCAATTCAAGGTC | GGGCTAGGCTGTGTGTTGAT | 258 |
| chr9:33567600-33569615 | p2 | (GA)11 | TTCGGGTGATCTAACTTCGG | ACGAACTGTGGTTGGAAAGC | 172 |
| chr9:33567600-33569615 | p3 | (GTT)5 | TTCGGGTGATCTAACTTCGG | ACGAACTGTGGTTGGAAAGC | 172 |
| chr9:33625195-33627207 | p2 | (CA)6 | TCCATTGCCCATGTTGTAGA | CCAAACGAAGGAAGAGAACG | 299 |
| chr9:33643259-33645277 | p3 | (GTT)6 | ACGTGCTTCTCGAGGTGTCT | CTGTTGCTGCGTTGGAAATA | 268 |
| chr9:33721133-33723153 | p3 | (ACA)7 | GGATACTCCAAAACAGCCCA | GTCGTTGCTGTGTTGTTGCT | 152 |
| chr9:33721133-33723153 | p2 | (CT)10 | GGATACTCCAAAACAGCCCA | GTCGTTGCTGTGTTGTTGCT | 152 |
| chr9:33727544-33729562 | p2 | (AT)9 | AGCGAATCAAACTCGATGCT | TTGGGAGTTGTCTAGGGGTG | 148 |
| chr9:33764815-33766841 | p4 | (TGTA)5 | ATCAGCTGGGGATAACATCG | ATCTTACTCGGCCCTTTCGT | 187 |

| Marker ID | SSR type | SSR motif | FORWARD PRIMER (5'-3') | REVERSE PRIMER (5'-3') | size (bp) |
|---|---|---|---|---|---|
| chr9:33811473-33813488 | p3 | (ACA)5 | ACATTGCTCGTACTGACCCC | CCAACTTCGCTGGAACATCT | 135 |
| chr9:33811473-33813488 | p2 | (CT)17 | ACATTGCTCGTACTGACCCC | CCAACTTCGCTGGAACATCT | 135 |
| chr9:33813370-33815398 | p2 | (AT)14 | TAGGCCCGCAATGTTATTTC | ACTTCTGTCACGACCCAACC | 119 |
| chr9:33813370-33815398 | p2 | (TA)10 | TAGGCCCGCAATGTTATTTC | ACTTCTGTCACGACCCAACC | 119 |
| chr9:33912145-33914159 | p2 | (TA)7 | TGATGTTGGAGTGAATGGGA | TCGTCAGATCATGCCAAAAG | 218 |
| chr9:33949998-33952012 | p2 | (TA)7 | TGATGTTGGAGTGAATGGGA | TCGTCAGATCATGCCAAAAG | 218 |
| chr9:33989509-33991524 | p5 | (TTTAA)3 | TGCCGGTTCATATCGTTGTA | CTCCGTAGGACGAACTGCTC | 197 |
| chr9:34011828-34013846 | p3 | (CAA)6 | ATGACCCAGGTAGTGGCTTG | CGACCTTACGTCACTCCGAT | 114 |
| chr9:34184600-34186618 | p2 | (AT)8 | TAACCTGTTGTCACGACCCA | TTGATAGCCTGAGGGGCTTA | 186 |
| chr9:34184600-34186618 | p6 | (ATACAT)3 | TAACCTGTTGTCACGACCCA | TTGATAGCCTGAGGGGCTTA | 186 |
| chr9:34218996-34221014 | p5 | (ATGCA)3 | AGGAGTACAACGCAAATGGG | GTCACGATCCGAGGTCTGTT | 272 |
| chr9:34218996-34221014 | p2 | (AT)9 | AGGAGTACAACGCAAATGGG | GTCACGATCCGAGGTCTGTT | 272 |
| chr9:34265116-34267131 | p5 | (TATTT)3 | TTTGCCAACCATCAAATCAA | ATTGGCTACCCTTTTTCCGA | 222 |
| chr9:34275191-34277206 | p2 | (AT)36 | AATAGGTTGCCCTTCCCATC | AAAATCTTGGCTCCACCCTT | 252 |
| chr9:34275191-34277206 | p5 | (TTTTA)3 | AATAGGTTGCCCTTCCCATC | AAAATCTTGGCTCCACCCTT | 252 |
| chr9:34275191-34277206 | p2 | (TA)29 | AATAGGTTGCCCTTCCCATC | AAAATCTTGGCTCCACCCTT | 252 |
| chr9:34281909-34283924 | p5 | (ATGTT)3 | CTGATCCAATGAGGTCGGTT | TCGTCAGTTTTGTTCGTTGC | 274 |
| chr9:34292720-34294735 | p2 | (AT)7 | GTGCGTTAAGGCCGGTATAA | TTTAGATAAGGGGCGACACG | 167 |
| chr9:34292720-34294735 | p5 | (TATCA)3 | GTGCGTTAAGGCCGGTATAA | TTTAGATAAGGGGCGACACG | 167 |
| chr9:34302310-34304328 | p6 | (TGCATG)3 | CTAAGCTGATGTGAACGCCA | AACACCACGTGAAATGGACA | 296 |
| chr9:34331226-34333241 | p5 | (ATTGC)3 | TTATTGTTGTTCGGCCTTCC | CCCAACTTCATACCCGAAGA | 178 |
| chr9:34368486-34370506 | p5 | (CAATG)4 | GCTGCAGCTAACGAAACCTT | CTGTTGTTATGTTGGCGTGG | 268 |
| chr9:34391745-34393783 | p2 | (TA)19 | TTGAGCCCAAAAGGCATTAC | TAACCGCACTAACCCAAAGG | 211 |
| chr9:34434831-34436847 | p4 | (AAAG)4 | CACCCGCCATAGTTTAGCAT | GACGAACGCTTTCCAGACAT | 104 |
| chr9:34464433-34466448 | p5 | (ATGAT)3 | GCCCAAATTTCTGTAGGCAA | TGCTCTTTAGGCCTTTCCAA | 120 |
| chr9:34567217-34569238 | p3 | (TTG)7 | GACCCTTGAGCCAAATCAAA | GCCTAGCCCACTTACCTTCC | 158 |
| chr9:34571966-34574017 | p3 | (GAA)17 | TGCTCCTGTTATTGCTGGTG | TGGCGAGCCTGATTTAGAGT | 206 |
| chr9:34589090-34591102 | p2 | (TA)6 | GGGAGAAGTCCCTGACATGA | CACTTCTGTCACGACCCGTA | 208 |
| chr9:34596233-34598245 | p2 | (AG)6 | GCGAATCCGTCTGATATGCT | ACCAACATTGATGGGGGTAA | 280 |
| chr9:34663706-34665721 | p3 | (AAC)5 | CTCGAGGGAGTAAGCAAACG | TATTCCCGCTTTTCCCTCTT | 260 |
| chr9:34711604-34713619 | p5 | (ATTGC)3 | TATCTGTTCGGCTTTTTGGG | GTGGCTAGCCCCAAATCATA | 288 |
| chr9:34722006-34724021 | p3 | (CAG)5 | CGGGCTCGTGTTTATGATTT | GCCTCCCTCAAGGTCCTATC | 127 |
| chr9:34816054-34818069 | p5 | (GATAT)3 | TTTGCTCAGCTGTGGCTATG | CCCGAACTGGACACTCGTAT | 227 |
| chr9:34860901-34862913 | p2 | (TA)6 | CCAAAACCCTTCTCCAAACA | GCAGCTATCAACTTGTGCCA | 244 |
| chr9:34895848-34897872 | p2 | (TA)12 | GGACCACTTATGCAGTCCGT | CGAGGACAACTTCGGAAAAA | 160 |
| chr9:34962845-34964861 | p2 | (GT)8 | CCCTGATCACGGATCTGTCT | CTCCTCGGTTCCTTACCACA | 283 |

| Marker ID | SSR type | SSR motif | FORWARD PRIMER (5'-3') | REVERSE PRIMER (5'-3') | size (bp) |
|---|---|---|---|---|---|
| chr9:34962845-34964861 | p2 | (TA)6 | CCCTGATCACGGATCTGTCT | CTCCTCGGTTCCTTACCACA | 283 |
| chr9:34971666-34973678 | p2 | (AT)6 | ACTTTTGTCATGACCCAGCC | GCCTTGTCGGCTTACGTATC | 175 |
| chr9:34971666-34973678 | p3 | (CAA)5 | ACTTTTGTCATGACCCAGCC | GCCTTGTCGGCTTACGTATC | 175 |
| chr9:34993533-34995545 | p2 | (CA)6 | GGACATGCCAAAAAGAAGGA | TTGTTGATGATTATGGCCGA | 245 |
| chr9:35017771-35019791 | p2 | (TA)10 | ATTGCAATCCTCGAATCACC | TTCGGTATGGTCGTGACAAA | 195 |
| chr9:35074029-35076041 | p2 | (AC)6 | CGAATGATGGTTGTTGTTCG | TGTTCTCCGTCAATGCTCAC | 297 |
| chr9:35075566-35077581 | p5 | (TGCAT)3 | ACCCGGGTAGTGACACTCAG | ACCTCGAAAGTCGAGAAACG | 196 |
| chr9:35081134-35083152 | p6 | (CATATA)3 | TTTTATCGCCCTTCCACATC | CCTTACAGGGTGCTTTTGGA | 185 |
| chr9:35103097-35105117 | p2 | (AT)10 | CGGTAACGCGACCAACTATT | CGTCTGCTTTCATGCTTTCA | 286 |
| chr9:35125484-35127498 | p2 | (AC)7 | GGACATGCCAAAAGAAGGAA | CGGTAGCCAAGTATAGCCCA | 244 |
| chr9:35201410-35203425 | p3 | (ATC)5 | GAAAAACAAGCTCCAGTCGC | AGGTTATTGCCGGGGTATTC | 151 |
| chr9:35282601-35284616 | p5 | (TATGA)3 | CTGTCCCGTCATATTTCGCT | ACCATATCCTATGCAAGCCG | 164 |
| chr9:35306017-35308031 | p2 | (TA)7 | GTAGACAACCCTTGGACCGA | CTCCGGTGTGAAGCGATATT | 184 |
| chr9:35368883-35370901 | p2 | (TA)9 | TCACCTACAGAAATTCGGGC | GGAAGCGAGGAATGTGTTGT | 178 |
| chr9:35377980-35380004 | p3 | (AAT)8 | ACTCCACAGATTCCGGTTTG | CCTCAAGATTTTTCGGGTCA | 184 |
| chr9:35412055-35414067 | p5 | (ATGTT)3 | TTGTTGTTGTTGTGTGGCCT | GTGACAAGCCCGAATACGAT | 221 |
| chr9:35412070-35414090 | p5 | (ATGTT)3 | TTGTTGTTGTTGTGTGGCCT | GTGACAAGCCCGAATACGAT | 206 |
| chr9:35470880-35472908 | p2 | (AT)14 | TTTCATGTGTTGGAATGCGT | TTTCGAACCTCGACGAAACT | 185 |
| chr9:35635630-35637652 | p2 | (TA)11 | CAGCGCGTATTAACTCCCTC | TGGTTGATTTGCTCACCAAG | 104 |
| chr9:35658827-35660842 | p5 | (ATTTC)3 | ATGTATGTGGGTCGTGGGTT | AAAATATCGCACGTAAGCCG | 260 |
| chr9:35698162-35700177 | p5 | (CCCCA)3 | CACTCTGGTCCTTCACAGCA | AGGATGGGACTGTTCAAACG | 182 |
| chr9:35856588-35858603 | p3 | (TTG)5 | CAAGAAGGTTTGGGGACTCA | AGCCTCATACGGGTCATTTG | 100 |
| chr9:35988651-35990666 | p5 | (ATATA)3 | AATCAAATACGCAAGCCGAC | TTATCGATACGCAGTGGCAG | 135 |
| chr9:35998312-36000327 | p5 | (ATATA)3 | AATCAAATACGCAAGCCGAC | TTATCGATACGCAGTGGCAG | 177 |
| chr9:36103887-36105901 | p2 | (AT)12 | CCAACTCCACAGATTCCGTT | GACCGACGCCATTCATCTAT | 111 |
| chr9:36103887-36105901 | p2 | (AT)7 | CCAACTCCACAGATTCCGTT | GACCGACGCCATTCATCTAT | 111 |
| chr9:36270193-36272223 | p2 | (TA)15 | TTCGAATGTTCGGAAGCTCT | TCACCTCCACACCATACCAA | 190 |
| chr9:36277754-36279769 | p5 | (TGAAT)3 | GGCCAATTCTTACAAGGCAA | CCAACTTGCACAACAAATGC | 250 |
| chr9:36293906-36295936 | p2 | (TA)15 | TTCGAATGTTCGGAAGCTCT | TCACCTCCACACCATACCAA | 190 |
| chr9:36301467-36303482 | p5 | (TGAAT)3 | GGCCAATTCTTACAAGGCAA | CCAACTTGCACAACAAATGC | 250 |
| chr9:36325491-36327506 | p3 | (TCA)5 | CTACATGGCAAAAATGGCCT | TCCGCCTTAAGAGGAAATCA | 256 |
| chr9:36344902-36346920 | p3 | (CAA)6 | GCCATTTTAAGGCACGATGT | TTGTAGATTTCGCTTGGGCT | 163 |
| chr9:36476877-36478923 | p2 | (TA)19 | CCTTAGACCGACCTGCTCTG | GCAATGGGTATTGGGTTGTC | 151 |
| chr9:36476877-36478923 | p2 | (AT)23 | CCTTAGACCGACCTGCTCTG | GCAATGGGTATTGGGTTGTC | 151 |
| chr9:36581521-36583539 | p6 | (TATATG)3 | ATCGTGCGATAGAGCCAAGT | AGGGCATACAGTCATACGCC | 250 |
| chr9:36739434-36741472 | p2 | (AT)7 | CCAGCAAATAATCCGACGTT | TCTGGATCAGGATCACCTCC | 122 |

| Marker ID | SSR type | SSR motif | FORWARD PRIMER (5'-3') | REVERSE PRIMER (5'-3') | size (bp) |
|---|---|---|---|---|---|
| chr9:36739434-36741472 | p2 | (AT)6 | CCAGCAAATAATCCGACGTT | TCTGGATCAGGATCACCTCC | 122 |
| chr9:36739629-36741643 | p2 | (AT)7 | CTTGTCCCGTCCTTGTGTCT | CCTCTCGGCCATCAAAATAA | 135 |
| chr9:36739629-36741643 | p2 | (AT)6 | CTTGTCCCGTCCTTGTGTCT | CCTCTCGGCCATCAAAATAA | 135 |
| chr9:36746388-36748402 | p6 | (GAGCGT)3 | TGAGCAGGGCAGTATGAGTG | GTATGGCCCTAGAACAGCCA | 169 |
| chr9:36746388-36748402 | p2 | (AG)7 | TGAGCAGGGCAGTATGAGTG | GTATGGCCCTAGAACAGCCA | 169 |
| chr9:36964174-36966248 | p2 | (AT)13 | TTGGATAAGTGGACCGAAGG | TTGAAGGGATGTAAGGTCGC | 192 |
| chr9:36964174-36966248 | p2 | (AT)37 | TTGGATAAGTGGACCGAAGG | TTGAAGGGATGTAAGGTCGC | 192 |
| chr9:37055037-37057061 | p2 | (TA)6 | ACGTGGGAAATCATGAGACC | ACAGGGAGGCGACCTAGATT | 138 |
| chr9:37055037-37057061 | p2 | (AT)12 | ACGTGGGAAATCATGAGACC | ACAGGGAGGCGACCTAGATT | 138 |
| chr9:37187273-37189331 | p2 | (AG)6 | CCAACATGCAACACATCACA | CAACCCTAAACCAAGTGGGA | 278 |
| chr9:37187273-37189331 | p2 | (AT)29 | CCAACATGCAACACATCACA | CAACCCTAAACCAAGTGGGA | 278 |
| chr9:37187273-37189331 | p3 | (CAA)6 | CCAACATGCAACACATCACA | CAACCCTAAACCAAGTGGGA | 278 |
| chr9:37238233-37240248 | p2 | (TA)7 | TCTTGCAGCCATGAGTTGTC | TCCGAATCACAAATTCCCAT | 222 |
| chr9:37238289-37240304 | p2 | (TA)7 | TCTTGCAGCCATGAGTTGTC | TCCGAATCACAAATTCCCAT | 166 |
| chr9:37266621-37268636 | p2 | (TA)7 | TCTTGCAGCCATGAGTTGTC | TCCGAATCACAAATTCCCAT | 222 |
| chr9:37266677-37268692 | p2 | (TA)7 | TCTTGCAGCCATGAGTTGTC | TCCGAATCACAAATTCCCAT | 166 |
| chr9:37305871-37307883 | p2 | (AT)6 | TAAGGTATCGGGTGCCAGTC | AATGCCTTGCAAGCCTTCTA | 151 |
| chr9:37329702-37331714 | p2 | (AT)6 | CCCTCCCTAATTGTTCGTCA | AGTTGAGGCTCCCCTTTGAT | 116 |
| chr9:37364016-37366031 | p2 | (AT)19 | AAAACTGTTGCAAGTTGGGG | TATCTTCCGACTGCCGATTC | 222 |
| chr9:37364073-37366087 | p2 | (AT)19 | AAAACTGTTGCAAGTTGGGG | TATCTTCCGACTGCCGATTC | 165 |
| chr9:37410797-37412809 | p2 | (CA)6 | CGGCCTCTAGTCTGATCTGG | GTCCGAGTATTCGTTTGGGA | 124 |
| chr9:37532651-37534757 | p2 | (AT)53 | CGTAAATACAACCATCCCCG | GGTAATTTGGCCGAGTTGAA | 216 |
| chr9:37585024-37587081 | p3 | (ATA)19 | CTCCTCTAGGTTCTTGGGGG | GTTCCCGTTCCTCTTTCTCC | 196 |
| chr9:37653384-37655396 | p2 | (TA)6 | TGCGACGTGAATTTGGAATA | GTCCCACTCGTGCAAGTTCT | 125 |
| chr9:37684865-37686883 | p3 | (TCT)6 | GCCCCTTCGGTATGTAGTCA | TTATTGAGACCCCAGCCTTG | 200 |
| chr9:37756009-37758057 | p2 | (TA)24 | GATGGAGCCTCATTATGGGA | ATAGTGAAACGCCATAGCCG | 126 |
| chr9:37757012-37759042 | p2 | (AT)23 | CAACCACCCAATACCTCTGG | TTGGGGTTTTTCTCAAGTGG | 195 |
| chr9:37867039-37869054 | p3 | (TTG)5 | TGGAAGATTCCAGGACCAAG | AGGTTCGTTGCATTGGAAAC | 292 |
| chr9:37873781-37875796 | p3 | (TTG)5 | TTGTGGAAGATTCCAGGACC | GCCGAACTTGTAACGTCCAT | 300 |
| chr9:37935767-37937782 | p3 | (TTG)5 | TGGAAGATTCCAGGACCAAG | AGGTTCGTTGCATTGGAAAC | 292 |
| chr9:37942509-37944524 | p3 | (TTG)5 | TTGTGGAAGATTCCAGGACC | GCCGAACTTGTAACGTCCAT | 300 |
| chr9:38225445-38227459 | p5 | (ATACG)3 | ACTTTTGTCACGACCCAACC | TTGTCGGTTTGCGTATCGTA | 170 |
| chr9:38225445-38227459 | p2 | (AT)7 | ACTTTTGTCACGACCCAACC | TTGTCGGTTTGCGTATCGTA | 170 |
| chr9:38329693-38331745 | p2 | (TA)11 | AGCAACTGAACCGCATCTCT | GCTAGCAGGTGAGCATAGGG | 295 |
| chr9:38329776-38331790 | p2 | (TA)11 | AGCAACTGAACCGCATCTCT | GCTAGCAGGTGAGCATAGGG | 212 |
| chr9:38360141-38362155 | p2 | (CT)7 | GATCCAGGGCTACAGATCCA | CGGCCTATGATGCCTATGAT | 200 |

| Marker ID | SSR type | SSR motif | FORWARD PRIMER (5'-3') | REVERSE PRIMER (5'-3') | size (bp) |
|---|---|---|---|---|---|
| chr9:38374955-38376970 | p3 | (TGT)5 | AGTGAGGTGAAATTGTGCCC | CACTTGGCCGAAACTTCATT | 120 |
| chr9:38374955-38376970 | p2 | (TA)6 | AGTGAGGTGAAATTGTGCCC | CACTTGGCCGAAACTTCATT | 120 |
| chr9:38394171-38396187 | p4 | (CTTT)4 | AAGTGAAATTTCCGCCCTTT | ATGCAACCACCACGTTATCA | 186 |
| chr9:38560793-38562809 | p2 | (GT)8 | GGTCCAAGAGATTGCACGAT | CCTGACACGCAAATTCAATG | 232 |
| chr9:38673749-38675775 | p2 | (TA)13 | CCCCTTGTTCACTTCGACAT | TGACCGCTCTTCTTCCATCT | 296 |
| chr9:38673749-38675775 | p3 | (CAT)8 | CCCCTTGTTCACTTCGACAT | TGACCGCTCTTCTTCCATCT | 296 |
| chr9:38681480-38683494 | p2 | (AT)7 | AACTTGCGAAAGAGCTCCAA | TATTGATGAGCAGTGGCAGC | 154 |
| chr9:38714713-38716725 | p2 | (AT)7 | CTTTTGTCACGACCCAACCT | CGGCCTGCGTATTATTTCAT | 234 |
| chr9:38714713-38716725 | p2 | (AT)6 | CTTTTGTCACGACCCAACCT | CGGCCTGCGTATTATTTCAT | 234 |
| chr9:38817103-38819129 | p2 | (TA)13 | ATAAGCCGACGAGGGTAACA | GGGACTACATAGCGCAGCTC | 229 |
| chr9:38820292-38822304 | p2 | (AC)6 | GGACATGCCAAAAGAAGGAA | TCGTTGTGGGTGTTGTTGTT | 254 |
| chr9:39078421-39080433 | p2 | (TA)6 | GTGACCATGTTCCTGAGCCT | GGCAGAAAATTTCCAAACCA | 199 |
| chr9:39138406-39140456 | p2 | (TG)25 | TCAGAATTGGGTTTTCTCGG | ATCCCCTTCATGGCATCATA | 234 |
| chr9:39160287-39162311 | p2 | (AT)12 | CACTTAGGATGTGGCCGACT | TGAAACCCTTTTGTTTTGGC | 232 |
| chr9:39160287-39162311 | p2 | (TA)17 | CACTTAGGATGTGGCCGACT | TGAAACCCTTTTGTTTTGGC | 232 |
| chr9:39223656-39225671 | p3 | (CAA)5 | GGTGTTATACTGCCCGGCTA | AATCGGAGGTTACCGACCTT | 250 |
| chr9:39254420-39256435 | p5 | (ATTTC)3 | AAGAAATGACATGGGAAGCG | CACACACGACCAAAAGGATG | 256 |
| chr9:39280863-39282875 | p2 | (TA)6 | CGGGTAACCTATGGCTCTGA | ATTTGGAAAAGGGTGCTCCT | 248 |
| chr9:39447043-39449063 | p5 | (TTTTG)4 | ATTTCCATTCTCCACGCAAC | TGAGAGGGTCTACGGATGCT | 229 |
| chr9:39478144-39480168 | p4 | (AAAT)6 | GCTCGTAACCCCGATACTCA | CGCAAACATCCACAAAACAC | 166 |
| chr9:39485180-39487198 | p2 | (AT)9 | GCGGGTGTTTTCGTAGTGTT | CCACCTCGGCAGATACTCAT | 108 |
| chr9:39516112-39518127 | p3 | (AGA)5 | GAAGCGAGCTTCCAGGTATG | GGTCGCCGTACACAACTCTT | 107 |
| chr9:39516112-39518127 | p2 | (AT)16 | GAAGCGAGCTTCCAGGTATG | GGTCGCCGTACACAACTCTT | 107 |
| chr9:39610034-39612128 | p3 | (CAT)5 | GGCCCTACCTACCGAGTACC | CGTGCTCATCCCTTTTCAAT | 206 |
| chr9:39760591-39762603 | p2 | (AT)6 | ACGGAAAAGAAGGCATCAGA | TTGAAAATGTTGCGATTCCA | 132 |
| chr9:39763280-39765298 | p6 | (TATATG)3 | GGAAATGTGGGACAAACACC | GTCCGAGCCTGCAAGTCTAC | 107 |
| chr9:39794764-39796782 | p2 | (TA)9 | TGCCGTTGATGTGGATTCTA | CTTAACGGAAATCGAGCAGC | 123 |
| chr9:39794764-39796782 | p2 | (AT)7 | TGCCGTTGATGTGGATTCTA | CTTAACGGAAATCGAGCAGC | 123 |
| chr9:39794901-39796915 | p2 | (TA)9 | GCTGCTCGATTTCCGTTAAG | TCGTAAATCCGTAGCCCATC | 245 |
| chr9:39794901-39796915 | p2 | (AT)7 | GCTGCTCGATTTCCGTTAAG | TCGTAAATCCGTAGCCCATC | 245 |
| chr9:40054594-40056610 | p2 | (AC)8 | TTCGATTCCAAATTTCGTCC | TGCCGAAAAGATGGCTAGTT | 234 |
| chr9:40152429-40154441 | p2 | (AT)6 | CCGCAAAATAATCACCGTCT | GCTGCATGATATGGTACCCC | 132 |
| chr9:40166908-40168920 | p2 | (TG)6 | CACATGGTTGACATTCCGAG | TTACCCGGCATCATAAGCTC | 113 |
| chr9:40186001-40188013 | p2 | (AT)6 | CCTCCCTCCCTAGAAACAGG | TCCCACAGGGATGAACTAGC | 222 |
| chr9:40291108-40293120 | p2 | (TA)6 | TACTAAGAGGGCCGGGTTCT | AAAAGCAGAAGCCATACGGA | 242 |
| chr9:40291108-40293120 | p2 | (AT)6 | TACTAAGAGGGCCGGGTTCT | AAAAGCAGAAGCCATACGGA | 242 |

| Marker ID | SSR type | SSR motif | FORWARD PRIMER (5'-3') | REVERSE PRIMER (5'-3') | size (bp) |
|---|---|---|---|---|---|
| chr9:40343967-40345987 | p2 | (TA)10 | CATGCATCGGTGTAGGAATG | CAAATAATCCCAACAAGCGG | 108 |
| chr9:40343967-40345987 | p2 | (AT)8 | CATGCATCGGTGTAGGAATG | CAAATAATCCCAACAAGCGG | 108 |
| chr9:40384611-40386631 | p2 | (TA)10 | CATGCATCGGTGTAGGAATG | CAAATAATCCCAACAAGCGG | 108 |
| chr9:40384611-40386631 | p2 | (AT)8 | CATGCATCGGTGTAGGAATG | CAAATAATCCCAACAAGCGG | 108 |
| chr9:40507964-40509982 | p6 | (AAATTA)3 | ACCATTGGGTGTAAGTTCGC | AGAATTTGGAAGCTTGTGGG | 176 |
| chr9:40581735-40583751 | p2 | (TA)8 | CTCAATAACCCGTGCCTCAT | CCCGAAGGACAATGTTGACT | 236 |
| chr9:40604708-40606728 | p4 | (AAAT)5 | GCCAAACCAAAGAGGAATCA | ATTGGTGAGTGGGAGTGGAG | 291 |
| chr9:40604708-40606728 | p2 | (AT)30 | GCCAAACCAAAGAGGAATCA | ATTGGTGAGTGGGAGTGGAG | 291 |
| chr9:40652444-40654466 | p2 | (TA)11 | TTAACCCCGTGGTCAGTAGC | TTGAGCGTATCAACCACCAA | 172 |
| chr9:40654965-40656980 | p4 | (AGAA)4 | TGGGCTTTTTAAGGTTGACG | ATGTGGATCCAGAACTTGCC | 239 |
| chr9:40654965-40656980 | p3 | (TCA)5 | TGGGCTTTTTAAGGTTGACG | ATGTGGATCCAGAACTTGCC | 239 |
| chr9:40747437-40749479 | p2 | (TA)21 | TCATCAATGCACAACCCCTA | GATGATCATGCTTGCCATTG | 297 |
| chr9:40747437-40749479 | p2 | (AT)8 | TCATCAATGCACAACCCCTA | GATGATCATGCTTGCCATTG | 297 |
| chr9:40747437-40749479 | p5 | (TCGGT)3 | TCATCAATGCACAACCCCTA | GATGATCATGCTTGCCATTG | 297 |
| chr9:40747819-40749835 | p2 | (TA)21 | TCAAAAGTTTGCACCCATGA | ATCCTTTCGTGAAGCCAATG | 144 |
| chr9:40747819-40749835 | p2 | (AT)8 | TCAAAAGTTTGCACCCATGA | ATCCTTTCGTGAAGCCAATG | 144 |
| chr9:40747819-40749835 | p5 | (TCGGT)3 | TCAAAAGTTTGCACCCATGA | ATCCTTTCGTGAAGCCAATG | 144 |
| chr9:40747819-40749835 | p5 | (AAACC)3 | TCAAAAGTTTGCACCCATGA | ATCCTTTCGTGAAGCCAATG | 144 |
| chr9:40840614-40842638 | p3 | (GTT)8 | CCCAGGTCATCCAGAAGAAA | TTGCCCTTAAGTTTTGGTGG | 120 |
| chr9:40893417-40895441 | p3 | (GTT)8 | CCCAGGTCATCCAGAAGAAA | TTGCCCTTAAGTTTTGGTGG | 120 |
| chr9:40983629-40985643 | p2 | (AT)7 | ACATACACACAAGCCGACGA | GATCCGGAGTTGCAGTTGTT | 276 |
| chr9:40983629-40985643 | p2 | (GA)7 | ACATACACACAAGCCGACGA | GATCCGGAGTTGCAGTTGTT | 276 |
| chr9:40996561-40998623 | p2 | (AT)31 | ATAGCTACACGGACCAACGG | ACCGGGGTAATGCTTAGGTC | 117 |
| chr9:41021474-41023486 | p2 | (CA)6 | CTTCCCAACACTGAGCCAAT | CACCCCAAGGTGAGACACTT | 246 |
| chr9:41021474-41023486 | p2 | (TA)7 | CTTCCCAACACTGAGCCAAT | CACCCCAAGGTGAGACACTT | 246 |
| chr9:41082226-41084244 | p3 | (TCA)6 | ACAGGAATCCATGAAGACGG | CTTAGGAGCTTCATCAGCGG | 214 |
| chr9:41132364-41134382 | p3 | (ATC)6 | ACCAATAAGCGACAACTCCG | GCATGCATGACTCCACCTTA | 293 |
| chr9:41253114-41255129 | p5 | (TCATT)3 | TGAAGCGTCGACGATGTAAG | ATGTTCGACACTCGGGAATC | 295 |
| chr9:41255333-41257351 | p3 | (TGT)6 | TGAGAGCACTCCATTGTTCG | CCCACACTACGCATCTACGA | 144 |
| chr9:41255333-41257351 | p6 | (AAGGAT)3 | TGAGAGCACTCCATTGTTCG | CCCACACTACGCATCTACGA | 144 |
| chr9:41262618-41264633 | p3 | (TCT)5 | AAACTAGGCGGTGTTCATGG | ACATGAATGGGGAATGGCTA | 210 |
| chr9:41315679-41317694 | p3 | (GTT)5 | AACGCACAAAATTTCCAAGG | ATTTCCAAAGGTTGCGATTG | 191 |
| chr9:41352599-41354614 | p2 | (AT)19 | GACCCGTTAACCTAGGGCTC | CTTATGCCAAAATGGGTCGT | 118 |
| chr9:41352599-41354614 | p5 | (ATGCA)3 | GACCCGTTAACCTAGGGCTC | CTTATGCCAAAATGGGTCGT | 118 |
| chr9:41426973-41428985 | p3 | (TGT)18 | ATAAACAGGGGACATGCTGC | TGCTTAACGTTTGTCCTCCC | 299 |
| chr9:41462797-41464817 | p2 | (TA)10 | GGATCTGCAAGCCGATGTAT | TGCAGATAACCCCTAGGACG | 104 |

| Marker ID | SSR type | SSR motif | FORWARD PRIMER (5'-3') | REVERSE PRIMER (5'-3') | size (bp) |
|---|---|---|---|---|---|
| chr9:41494665-41496699 | p2 | (TA)17 | CAGGGACATAGGGTGCAGAT | AACTTGGCACAAAAGATGCC | 125 |
| chr9:41581608-41583624 | p2 | (AT)8 | AACAACTGTGCCCATCATCA | GAGCCCTCCATTTGTTTCAA | 299 |
| chr9:41725372-41727390 | p2 | (AT)9 | GCTTTGCTGCAAGTTTGTGA | TGGTGGATATGACCAGGGAT | 129 |
| chr9:41725765-41727805 | p2 | (AT)9 | AGCCGACAAGGCTGCTACTA | CTCAGCTGCTTGGAGAGCTT | 161 |
| chr9:41725801-41727816 | p2 | (AT)9 | AGCCGACAAGGCTGCTACTA | CTCAGCTGCTTGGAGAGCTT | 125 |
| chr9:41745922-41747938 | p2 | (TG)8 | CTCCGTGAATTTCCAGCTTC | TCGCTTTTTGACGGAGAGTT | 130 |
| chr9:41745922-41747938 | p2 | (AT)8 | CTCCGTGAATTTCCAGCTTC | TCGCTTTTTGACGGAGAGTT | 130 |
| chr9:41845598-41847618 | p4 | (TCAT)5 | TTTGGCTGACTTTTTGGGTC | CTATCAGGCTTTCCCCATCA | 208 |
| chr9:41867286-41869316 | p2 | (CA)18 | GGTTTGGAGCTGCAAGTTTC | ACTCTGCGCCAAGAACTCAT | 207 |
| chr9:41867286-41869316 | p2 | (TA)15 | GGTTTGGAGCTGCAAGTTTC | ACTCTGCGCCAAGAACTCAT | 207 |
| chr9:41870901-41872913 | p2 | (TA)6 | CGGTCTAGACACCTTGGGAA | TTAGACAAGGGGACGCAATC | 184 |
| chr9:41921320-41923336 | p4 | (ATAG)4 | GACCAAAGAAGAAGGGAGGG | TCAGCCACCCATTAATCACA | 158 |
| chr9:41941293-41943377 | p2 | (AT)42 | CATGATGTTATGCACCCTCG | TCACTGACTATCTTGGCCCC | 242 |
| chr9:41941293-41943377 | p5 | (AGGAC)3 | CATGATGTTATGCACCCTCG | TCACTGACTATCTTGGCCCC | 242 |
| chr9:41941293-41943377 | p3 | (CAT)5 | CATGATGTTATGCACCCTCG | TCACTGACTATCTTGGCCCC | 242 |
| chr9:41941293-41943377 | p2 | (AT)6 | CATGATGTTATGCACCCTCG | TCACTGACTATCTTGGCCCC | 242 |
| chr9:42103416-42105428 | p2 | (AG)6 | GTGACACACCTGCGAAAATG | ACATTGCAGTGTCCTTGCTG | 219 |
| chr9:42103416-42105428 | p2 | (GT)6 | GTGACACACCTGCGAAAATG | ACATTGCAGTGTCCTTGCTG | 219 |
| chr9:42120593-42122605 | p2 | (CA)6 | CAACCAGCAGGGTCCTTAAA | CAGGGCATGGTGATAGGTCT | 100 |
| chr9:42197489-42199501 | p2 | (TA)6 | GCCGGGTCAGTCAAAAATTA | TTATCCGGGGATTCTGAGTG | 109 |
| chr9:42311542-42313558 | p4 | (AAAC)4 | GAGGTGGACCTCTCATTGGA | AATCTCGGGAAGCGAAAAGT | 280 |
| chr9:42368578-42370610 | p2 | (TA)16 | CACCATCCCCGAGAATCTAA | TTGAATTTGGAAGGTTTCGG | 114 |
| chr9:42498545-42500567 | p4 | (AGTT)5 | TCCACACGATGGTATGATGG | TGACCCCGAGTTGTCACATA | 299 |
| chr9:42498545-42500567 | p2 | (CT)11 | TCCACACGATGGTATGATGG | TGACCCCGAGTTGTCACATA | 299 |
| chr9:42515173-42517188 | p2 | (AT)33 | CGTGACATAGCATTGGTTGG | TATCACCACGGCCATTATCA | 256 |
| chr9:42515173-42517188 | p5 | (ATTTT)3 | CGTGACATAGCATTGGTTGG | TATCACCACGGCCATTATCA | 256 |
| chr9:42600819-42602834 | p3 | (CAT)5 | AACGCGGCTCGATAAGTAAA | GCATTCCACAAAATGTCACG | 297 |
| chr9:42600819-42602834 | p2 | (TA)12 | AACGCGGCTCGATAAGTAAA | GCATTCCACAAAATGTCACG | 297 |
| chr9:42600819-42602834 | p3 | (AAC)5 | AACGCGGCTCGATAAGTAAA | GCATTCCACAAAATGTCACG | 297 |
| chr9:42602910-42604925 | p3 | (AAC)5 | ATTGCATTTGGCCTTAGACG | GCACACCCTTTCATGGACTT | 219 |
| chr9:42831170-42833182 | p5 | (ATTGC)3 | TGGGCTGGGTTTTACTTGAG | TTCTCACTGGCACCACTCTG | 245 |
| chr9:42831170-42833182 | p2 | (TA)6 | TGGGCTGGGTTTTACTTGAG | TTCTCACTGGCACCACTCTG | 245 |
| chr9:42831170-42833182 | p2 | (AT)32 | TGGGCTGGGTTTTACTTGAG | TTCTCACTGGCACCACTCTG | 245 |
| chr9:42842805-42844820 | p5 | (ATTGC)3 | TTCTGAATTTCCCGTTTTGC | ATGGCTAGTCCCAACCACAC | 205 |
| chr9:42842805-42844820 | p2 | (TA)6 | TTCTGAATTTCCCGTTTTGC | ATGGCTAGTCCCAACCACAC | 205 |
| chr9:42861160-42863184 | p4 | (AAAT)6 | AACACTCCCTCCCCATCTCT | GCCTTCTTTGCTCCTACACG | 203 |

| Marker ID | SSR type | SSR motif | FORWARD PRIMER (5'-3') | REVERSE PRIMER (5'-3') | size (bp) |
|---|---|---|---|---|---|
| chr9:42881821-42883837 | p2 | (TA)8 | ATTACCGCTTGGACCACTTG | GCTTTCTATCAGCGGACTCG | 138 |
| chr9:42916856-42918871 | p3 | (ATA)5 | GGGTGGACGTGAAGACTGTT | TCTTTCCAGTCGTGTTGACG | 155 |
| chr9:42933356-42935370 | p2 | (TA)7 | ATTTGCTTCCAAACCAAACG | AGGTGGTGGAAAACGAAATG | 187 |
| chr9:42933356-42935370 | p3 | (ACA)5 | ATTTGCTTCCAAACCAAACG | AGGTGGTGGAAAACGAAATG | 187 |
| chr9:42987748-42989763 | p5 | (TTAAA)3 | GTCCCTTATGGAGTGTCCGA | TCAGAATTGGGAACCCAGAG | 213 |
| chr9:42987748-42989763 | p2 | (AT)23 | GTCCCTTATGGAGTGTCCGA | TCAGAATTGGGAACCCAGAG | 213 |
| chr9:43208507-43210601 | p2 | (TA)47 | TTTCAAAACCATTCTTCCGC | GTTTGGGTAGGGTCTTGGGT | 129 |
| chr9:43208860-43210882 | p2 | (TA)47 | CAACGGTCAACATCTCAACG | GCAGGATTGGTGAACTCCAT | 274 |
| chr9:43220662-43222677 | p3 | (GTT)5 | GTTAGGGGCAAAATGGGAAT | AAATGAACACATGGCCCTTC | 251 |
| chr9:43434801-43436819 | p6 | (AATAAA)3 | GGCTTCATGTGTGGTACGTG | TTACTACCAACCCCGAGCAC | 234 |
| chr9:43460108-43462123 | p5 | (AACCG)3 | CAGCGTCTCCAAGAGGTAGG | GCAGGAGCCATTCTGTAAGG | 208 |
| chr9:43460108-43462123 | p5 | (AACCG)3 | CAGCGTCTCCAAGAGGTAGG | GCAGGAGCCATTCTGTAAGG | 208 |
| chr9:43460624-43462639 | p5 | (AACCG)3 | TCGGGTAATGTCATTTGGGT | TTATGCCCAGCCCTAAAATG | 277 |
| chr9:43460624-43462639 | p5 | (AACCG)3 | TCGGGTAATGTCATTTGGGT | TTATGCCCAGCCCTAAAATG | 277 |
| chr9:43601277-43603329 | p2 | (AT)26 | CATGGCATGAAGAACGATTG | CCCTAAATCCCCCAACAGAT | 275 |
| chr9:43642875-43644893 | p6 | (GCCTCT)3 | TGCAGCATAACTTGCCAAAG | ACCAGCCGGATTACCTTCTT | 279 |
| chr9:43655470-43657484 | p2 | (AT)7 | CAATTTCTTCGTAGCGGCAT | AAACAAGAGGCTACAGGGCA | 112 |
| chr9:43655470-43657484 | p2 | (AT)19 | CAATTTCTTCGTAGCGGCAT | AAACAAGAGGCTACAGGGCA | 112 |
| chr9:43728820-43730838 | p6 | (GCCTCT)3 | TGCAGCATAACTTGCCAAAG | ACCAGCCGGATTACCTTCTT | 279 |
| chr9:43741415-43743429 | p2 | (AT)7 | CAATTTCTTCGTAGCGGCAT | AAACAAGAGGCTACAGGGCA | 112 |
| chr9:43741415-43743429 | p2 | (AT)19 | CAATTTCTTCGTAGCGGCAT | AAACAAGAGGCTACAGGGCA | 112 |
| chr9:43822913-43824935 | p2 | (AT)11 | TTCTCCAACCCCTGTTATCG | GCCACTTTCGAGGATCAGAG | 299 |
| chr9:43890694-43892716 | p2 | (AT)11 | TTCTCCAACCCCTGTTATCG | GCCACTTTCGAGGATCAGAG | 299 |
| chr9:43953298-43955333 | p5 | (ATCAT)7 | TAATCTGATTCAGCGCATCG | CTGTTGGATTTGGGAAGCAT | 186 |
| chr9:44248898-44250910 | p2 | (AC)6 | TATCAAACGGGACTTCGGAC | TTTGGCATGATCTTTGTGGA | 292 |
| chr9:44276631-44278706 | p3 | (AGA)25 | GCAAATGCCAGTCAAGGATT | TGCTCATACGATGTACCCCA | 104 |
| chr9:44301962-44303986 | p4 | (TTTA)6 | TGTGACATCTCTGGCGACTC | AGACTTAGAGGGGTGCACGA | 274 |
| chr9:44375737-44377757 | p2 | (TA)10 | GATGTGTCGTGGATTTGTGG | TCTGGCCAATTTTTCCTTTG | 221 |
| chr9:44492260-44494274 | p2 | (CA)7 | AGACGTCTTACGCTTTCCCA | TGATGTGTGGGCTGTTGTTT | 293 |
| chr9:44597419-44599445 | p2 | (TA)13 | TCTTTCGCCACTTCCTCACT | ATTGTGGCGGATAATGAAGG | 297 |
| chr9:44632799-44634815 | p2 | (TA)8 | CTCCAGAGGTGCCATTTGTT | GAGAAAACTGCTACCCACGC | 157 |
| chr9:44719459-44721501 | p2 | (AT)8 | AACTCGGCCAAATTGTCATC | TGAATCTGGAAGAATTCGGG | 142 |
| chr9:44821326-44823350 | p3 | (ATT)8 | TCCGAAACAGGGGTTACAAG | ATTTGGGTTGGAGACGAGTG | 265 |
| chr9:44895520-44897538 | p2 | (TA)6 | AGGGGGCTTACCCCTTTATT | ACCTTACTCCCGCTAGAGCC | 123 |
| chr9:44895520-44897538 | p6 | (ATGATA)3 | AGGGGGCTTACCCCTTTATT | ACCTTACTCCCGCTAGAGCC | 123 |
| chr9:44895520-44897538 | p2 | (TA)23 | AGGGGGCTTACCCCTTTATT | ACCTTACTCCCGCTAGAGCC | 123 |

| Marker ID | SSR type | SSR motif | FORWARD PRIMER (5'-3') | REVERSE PRIMER (5'-3') | size (bp) |
|---|---|---|---|---|---|
| chr9:44895520-44897538 | p2 | (AT)7 | AGGGGGCTTACCCCTTTATT | ACCTTACTCCCGCTAGAGCC | 123 |
| chr9:44980778-44982796 | p6 | (AAAATG)3 | GTCATTTGAACCCGTTTGCT | CCAACCGCTAAAATTTGGAA | 164 |
| chr9:45023535-45025551 | p2 | (GA)8 | CAACCACTACGCCCTTGACT | AATGATCCCAGCTTGTTTGC | 154 |
| chr9:45038298-45040313 | p5 | (ATTTT)3 | GGCAAGGGTTTGTACCGTAA | ATGAACAAGCCGAATTGGAG | 248 |
| chr9:45038351-45040430 | p5 | (ATTTT)3 | GGCAAGGGTTTGTACCGTAA | ATGAACAAGCCGAATTGGAG | 195 |
| chr9:45081863-45083881 | p2 | (TA)9 | GTAGAAGGGGTTTGTTGGCA | ATAAAACCGGGGGCATAAAC | 147 |
| chr9:45116036-45118087 | p3 | (TTC)17 | CCTGGAAATCAAAACCGAGA | TGATGCAATCAACGGAGGTA | 136 |
| chr9:45192896-45194910 | p2 | (AT)7 | TAGGCGGAGTAGAAGAGGCA | ACTACGGTAGAAAGGCGGGT | 116 |
| chr9:45192896-45194910 | p2 | (TA)24 | TAGGCGGAGTAGAAGAGGCA | ACTACGGTAGAAAGGCGGGT | 116 |
| chr9:45193170-45195218 | p2 | (AT)7 | AGCCCTTCGGCTATCAAAAT | TGTTCATGCCCACAGGTAGA | 269 |
| chr9:45193170-45195218 | p2 | (TA)24 | AGCCCTTCGGCTATCAAAAT | TGTTCATGCCCACAGGTAGA | 269 |
| chr9:45276533-45278548 | p5 | (ATCAT)3 | GTAGACACCCCGAGAACGAA | AGTTGGTAACGCCATTCGAC | 239 |
| chr9:45291881-45293896 | p3 | (ACA)5 | TACGTTTCGACCTGATTCCC | TGCATCAACGGAATTTTTGA | 254 |
| chr9:45411575-45413587 | p2 | (AT)6 | AAGTGAAAGAACCCAAGGCA | CAGAGATGGTGTCTGCCTGA | 164 |
| chr9:45411575-45413587 | p2 | (TA)6 | AAGTGAAAGAACCCAAGGCA | CAGAGATGGTGTCTGCCTGA | 164 |
| chr9:45421660-45423678 | p6 | (CATATA)3 | AACCCCTAGGACAACCTGCT | GTCTTGTAAGCGGCTCCATC | 292 |
| chr9:45433397-45435411 | p2 | (AT)7 | GACCCTTATTTAGGGCCGAG | GCAAGCCGATGTACCATTTT | 276 |
| chr9:45473638-45475706 | p2 | (TA)34 | AAGGGATGGAGATGCAACAG | GAGTAGGGTCCTGGGTGTCA | 152 |
| chr9:45654434-45656462 | p5 | (TTAAA)3 | CGACCTTAGCGATCCTTCAA | AATGAATTCCAATGTCCCCA | 129 |
| chr9:45664544-45666564 | p2 | (AT)10 | TTAACCTGCACATCCGACAA | ATTCGCTCAGTCACATGCAG | 189 |
| chr9:45718437-45720449 | p2 | (GT)6 | CACCTTCAGACATTAGCGCA | CTTGGCCCAATTCGTAAAAA | 281 |
| chr9:45718437-45720449 | p3 | (GTT)5 | CACCTTCAGACATTAGCGCA | CTTGGCCCAATTCGTAAAAA | 281 |
| chr9:45872202-45874226 | p6 | (ATATGT)4 | TTCCTTCTTTGGCATGATCC | AAGGCTTGATGGAACAATCG | 184 |
| chr9:45926117-45928151 | p2 | (AT)17 | TAGAGGTCAGTGGGGGACTG | CTCTAACCACTCGCACACCA | 123 |
| chr9:45993549-45995613 | p2 | (TA)30 | CAGAAGAGTCCAGAGTGCCC | TACTGCCCGACCGTATAAGC | 190 |
| chr9:45993549-45995613 | p2 | (AT)32 | CAGAAGAGTCCAGAGTGCCC | TACTGCCCGACCGTATAAGC | 190 |
| chr9:46000301-46002335 | p2 | (TA)11 | TTTATCGGTGGAGAAATCGG | CATGTTCATAGGCCCGTCTT | 254 |
| chr9:46000301-46002335 | p2 | (TA)17 | TTTATCGGTGGAGAAATCGG | CATGTTCATAGGCCCGTCTT | 254 |
| chr9:46030757-46032775 | p3 | (AAG)6 | CAGACAAATTGCGAAGCAAA | GTGGCTAAGCCTGAGATTGG | 178 |
| chr9:46030757-46032775 | p4 | (TTAA)4 | CAGACAAATTGCGAAGCAAA | GTGGCTAAGCCTGAGATTGG | 178 |
| chr9:46030957-46032973 | p3 | (AAG)6 | CTTCCGGAGAAATCCAATCA | CCCGACTCATTTTACCCCTT | 287 |
| chr9:46030957-46032973 | p4 | (TTAA)4 | CTTCCGGAGAAATCCAATCA | CCCGACTCATTTTACCCCTT | 287 |
| chr9:46032568-46034584 | p4 | (CCTT)4 | GCCTTAGATCTGGGTCGTTG | TTTCCTTTGCAAGAGGGTTG | 290 |
| chr9:46032568-46034584 | p5 | (ATAAA)3 | GCCTTAGATCTGGGTCGTTG | TTTCCTTTGCAAGAGGGTTG | 290 |
| chr9:46052421-46054435 | p2 | (TA)7 | ACAATTTCAAACGCGGATTC | GATGCTGCCCAAATTTCTGT | 179 |
| chr9:46065428-46067444 | p2 | (TA)8 | CTTGTACCGAGCACCCCTTA | TTGGGAGCGCTCTAGTTGTT | 171 |

| Marker ID | SSR type | SSR motif | FORWARD PRIMER (5'-3') | REVERSE PRIMER (5'-3') | size (bp) |
|---|---|---|---|---|---|
| chr9:46256854-46258870 | p2 | (TG)8 | GACCGCATTAGGATTTTCCA | TAGGTCGGCAAGACCCATAC | 230 |
| chr9:46256854-46258870 | p2 | (TA)7 | GACCGCATTAGGATTTTCCA | TAGGTCGGCAAGACCCATAC | 230 |
| chr9:46256854-46258870 | p2 | (TA)8 | GACCGCATTAGGATTTTCCA | TAGGTCGGCAAGACCCATAC | 230 |
| chr9:46256854-46258870 | p3 | (ATC)5 | GACCGCATTAGGATTTTCCA | TAGGTCGGCAAGACCCATAC | 230 |
| chr9:46257161-46259176 | p2 | (TA)7 | CTACATGTCAGAGCCCGTCA | TACAAACCGGACCCTAGTGC | 292 |
| chr9:46257161-46259176 | p2 | (TA)8 | CTACATGTCAGAGCCCGTCA | TACAAACCGGACCCTAGTGC | 292 |
| chr9:46257161-46259176 | p3 | (ATC)5 | CTACATGTCAGAGCCCGTCA | TACAAACCGGACCCTAGTGC | 292 |
| chr9:46299754-46301769 | p3 | (TCA)5 | AACCTACCGAGCACCTCTCA | TCCCTCAGTTTTGTCGGTTC | 192 |
| chr9:46332413-46334425 | p2 | (GT)6 | GTTTCCTACGCCCACAGGTA | ATTATGCGAAAACACGGGAC | 140 |
| chr9:46421932-46423950 | p6 | (TATATG)3 | CGAAAGGTTCTTGACGTGGT | CGAGCGGAGTACTGAGAACC | 214 |
| chr9:46542829-46544841 | p2 | (AT)6 | CATGACTCTGCTCGTGCATT | CCGTATCCCCGTCACATATC | 178 |
| chr9:46621069-46623085 | p2 | (AT)8 | GCCAAGGCTACGACAGAAAG | CCTTGTCGACTTGCGTATCA | 130 |
| chr9:46621069-46623085 | p3 | (ACA)5 | GCCAAGGCTACGACAGAAAG | CCTTGTCGACTTGCGTATCA | 130 |
| chr9:46631857-46633873 | p2 | (AT)8 | TGCCCGATTTTGGTTAAGTC | CGACTATTCGCAAACCCATT | 156 |
| chr9:46631857-46633873 | p2 | (AT)6 | TGCCCGATTTTGGTTAAGTC | CGACTATTCGCAAACCCATT | 156 |
| chr9:46633098-46635116 | p2 | (AT)6 | GGTTGGTGACCTTTCAGCAT | CATGTCATATGCAAGCCGAC | 181 |
| chr9:46633098-46635116 | p2 | (AT)9 | GGTTGGTGACCTTTCAGCAT | CATGTCATATGCAAGCCGAC | 181 |
| chr9:46644603-46646659 | p2 | (AT)28 | CTGTCTCATCTCTAGGCCCG | AATAAGGCCCCACCTCAGTT | 257 |
| chr9:46714575-46716601 | p2 | (AT)13 | TCTCCTCTTCCTTCGACCAA | CGCGTTTGAAGGAATGTTTT | 239 |
| chr9:46726963-46728987 | p2 | (AT)10 | AAGCCGACAAGGCTACAAAA | TCAGCGATGGCATAAGACAG | 157 |
| chr9:46726963-46728987 | p2 | (AT)13 | AAGCCGACAAGGCTACAAAA | TCAGCGATGGCATAAGACAG | 157 |
| chr9:46726963-46728987 | p3 | (ATC)8 | AAGCCGACAAGGCTACAAAA | TCAGCGATGGCATAAGACAG | 157 |
| chr9:46839774-46841789 | p3 | (GTT)5 | CCAAGTGTAAGCCCTCCAAA | CATACAGGCCCACGACCTAT | 138 |
| chr9:46839774-46841789 | p2 | (TA)6 | CCAAGTGTAAGCCCTCCAAA | CATACAGGCCCACGACCTAT | 138 |
| chr9:46842556-46844576 | p2 | (AT)12 | TGCGAAAGAACTCCAAAGGT | TGTTCGTCTGTAGTGGCAGC | 175 |
| chr9:46842556-46844576 | p2 | (AT)10 | TGCGAAAGAACTCCAAAGGT | TGTTCGTCTGTAGTGGCAGC | 175 |
| chr9:46849586-46851612 | p3 | (ACA)8 | CTTTTGTCGATGCCGTTCTT | GCTCGGGCACTAGTCATAGC | 230 |
| chr9:46849586-46851612 | p2 | (AT)13 | CTTTTGTCGATGCCGTTCTT | GCTCGGGCACTAGTCATAGC | 230 |
| chr9:46854994-46857012 | p3 | (CAA)6 | CGGTGAACAATGCAATGAAG | GAGCGAACGACGAACATACA | 232 |
| chr9:46912390-46914402 | p2 | (TA)8 | GCCGACAAGGCTACAACAAT | TTCAGCGGATAGTGTGCAAG | 295 |
| chr9:46912465-46914483 | p2 | (TA)8 | GCCGACAAGGCTACAACAAT | TTCAGCGGATAGTGTGCAAG | 220 |
| chr9:46972456-46974482 | p2 | (TG)13 | CCTGGACCGGAATTCAGTAA | GAACAGAAACGCGTGCTACA | 146 |
| chr9:47035064-47037108 | p2 | (AT)22 | ACGGCCCATAAGTTTGTACG | TTGAAAGTGTCACGACCCAA | 119 |
| chr9:47035064-47037108 | p2 | (AT)23 | ACGGCCCATAAGTTTGTACG | TTGAAAGTGTCACGACCCAA | 119 |
| chr9:47050853-47052868 | p3 | (GTT)5 | CGTGTCAATATGATGGGCAG | GTGAGAGCCAAGATGCTTCC | 146 |
| chr9:47050853-47052868 | p2 | (TA)9 | CGTGTCAATATGATGGGCAG | GTGAGAGCCAAGATGCTTCC | 146 |

| Marker ID | SSR type | SSR motif | FORWARD PRIMER (5'-3') | REVERSE PRIMER (5'-3') | size (bp) |
|---|---|---|---|---|---|
| chr9:47070913-47072928 | p3 | (GTT)5 | CGTGTCAATATGATGGGCAG | GTGAGAGCCAAGATGCTTCC | 146 |
| chr9:47070913-47072928 | p2 | (TA)9 | CGTGTCAATATGATGGGCAG | GTGAGAGCCAAGATGCTTCC | 146 |
| chr9:47212003-47214017 | p2 | (TA)7 | AGTTGTTTTGGGCCTCTGTG | AAGAAGGTTGGGCCTCTAGC | 120 |
| chr9:47253653-47255668 | p3 | (TTC)5 | ACGTTGCGGGTGACTAAATC | GGGACAACTCCCAAGACTCA | 157 |
| chr9:47293031-47295046 | p5 | (TCATT)3 | TAACCACATCTGGGGAGGAC | TTTCAGACAGTGTCGCCAAG | 152 |
| chr9:47395518-47397533 | p5 | (AAAAG)3 | ACAGGTGCGACAAGGATTTT | CACGAAAATTTACAGGGCGT | 219 |
| chr9:47430306-47432321 | p5 | (AAACC)3 | CTACTCGCAAGCCATTTTCC | TTTACACGCTCGACAGATGC | 255 |
| chr9:47430344-47432359 | p5 | (AAACC)3 | CTACTCGCAAGCCATTTTCC | TTTACACGCTCGACAGATGC | 217 |
| chr9:47431974-47433992 | p5 | (AAACC)3 | GTGCAGTTTGTCGGTTTCCT | TGGCTTAACTCTATGGCGCT | 251 |
| chr9:47431974-47433992 | p6 | (GTGGAC)3 | GTGCAGTTTGTCGGTTTCCT | TGGCTTAACTCTATGGCGCT | 251 |
| chr9:47455288-47457351 | p5 | (CTTTT)3 | TTTTGTCCCTTGAGAAACGG | CGAGCTAAAACCTGAGTGCC | 138 |
| chr9:47455288-47457351 | p3 | (TTC)21 | TTTTGTCCCTTGAGAAACGG | CGAGCTAAAACCTGAGTGCC | 138 |
| chr9:47455288-47457351 | p2 | (GA)6 | TTTTGTCCCTTGAGAAACGG | CGAGCTAAAACCTGAGTGCC | 138 |
| chr9:47534318-47536336 | p6 | (AAAGAA)3 | AGAGCCTGTTGGATTCATGG | AAGTGCTGCCTTTTTCCTCA | 232 |
| chr9:47534318-47536336 | p3 | (AAG)6 | AGAGCCTGTTGGATTCATGG | AAGTGCTGCCTTTTTCCTCA | 232 |
| chr9:47558884-47560902 | p2 | (GT)9 | CGGCTCAATCTCAGTTCTCC | TTTCCAAATTTTGCACCCTC | 186 |
| chr9:47590682-47592706 | p2 | (AT)12 | TAAAAGCTCGGGGTCTTCAA | ACCCCATCAACCAAAATGAA | 243 |
| chr9:47610663-47612689 | p2 | (TA)13 | GGACGATGGAAGCCTATTGA | ACAATTGGTGTCACAACGGA | 247 |
| chr9:47621266-47623281 | p5 | (ACACA)3 | ACACGGCCAACTTAGACCAC | TCATGGAGTGGGTGTGTTGT | 205 |
| chr9:47647295-47649307 | p2 | (TA)6 | GCCGGGATCTGTATGTCTGT | AGAACCCGGCCCTCTTAGTA | 143 |
| chr9:47663993-47666011 | p6 | (CATATA)3 | AACTAGGGTGTGATGGGCAC | GGAGTTAGTCTGTCAGCGGC | 167 |
| chr9:47663993-47666011 | p5 | (ATTCA)4 | AACTAGGGTGTGATGGGCAC | GGAGTTAGTCTGTCAGCGGC | 167 |
| chr9:47666549-47668567 | p2 | (AT)6 | TGGCCCCATGTGTTTTTATT | CAAAAGGGAAATTCGCAAAA | 194 |
| chr9:47666549-47668567 | p3 | (ATC)6 | TGGCCCCATGTGTTTTTATT | CAAAAGGGAAATTCGCAAAA | 194 |
| chr9:47813602-47815614 | p2 | (TA)6 | CGTGAAGCATAATTGCGAGA | CCGATTGTCGACTTTCCCTA | 286 |
| chr9:48011238-48013260 | p3 | (GCA)5 | GAGACCCTCAGGAAGCATTG | AAAAATTAAGACGGGGTGGG | 287 |
| chr9:48172162-48174174 | p2 | (AT)6 | CGCAGTGCCAAAAACTTGTA | AAATGATAGGCACGAGGGTG | 140 |
| chr9:48251487-48253502 | p3 | (GAA)5 | CAAGGTGATGTACTGGGCCT | TGTCCCGGTGGTTAATGAAT | 154 |
| chr9:48278317-48280332 | p6 | (TGAGTA)3 | TCAACCCCCAAAAGAGATTG | AACATATCTTGGCCACCAGC | 204 |
| chr9:48278317-48280332 | p5 | (TCATT)3 | TCAACCCCCAAAAGAGATTG | AACATATCTTGGCCACCAGC | 204 |
| chr9:48303977-48305995 | p2 | (AT)9 | AAAGGGTGACGTGTTTCCTG | ACACAGGCTGAACAAGACCC | 148 |
| chr9:48330285-48332300 | p5 | (ATGAT)3 | GATGCTGCCCAAATTTCTGT | GACAATTTCAAACGCGGATT | 114 |
| chr9:48401904-48403918 | p2 | (AT)7 | CATGAATACCCGAACATCCC | GATCGGGTTCTGGTTGTGTT | 158 |
| chr9:48465697-48467709 | p3 | (AAT)5 | CCGTTGAGACAAGGCGTATT | GTGTCCAGTGGGGTCACTTT | 126 |
| chr9:48465697-48467709 | p2 | (TA)6 | CCGTTGAGACAAGGCGTATT | GTGTCCAGTGGGGTCACTTT | 126 |
| chr9:48465715-48467739 | p3 | (AAT)5 | CCGTTGAGACAAGGCGTATT | GTGTCCAGTGGGGTCACTTT | 108 |

| Marker ID | SSR type | SSR motif | FORWARD PRIMER (5'-3') | REVERSE PRIMER (5'-3') | size (bp) |
|---|---|---|---|---|---|
| chr9:48465715-48467739 | p2 | (TA)6 | CCGTTGAGACAAGGCGTATT | GTGTCCAGTGGGGTCACTTT | 108 |
| chr9:48466651-48468663 | p3 | (AAT)5 | GGTACTTGCGTCCTACCAGC | CCGAACTACGTAGCCACCAT | 247 |
| chr9:48466651-48468663 | p2 | (TA)6 | GGTACTTGCGTCCTACCAGC | CCGAACTACGTAGCCACCAT | 247 |
| chr9:48511190-48513210 | p2 | (TA)10 | GTCCCAGCTCAAACCAATGT | GTTTCACTGGTGGGCAATCT | 268 |
| chr9:48521114-48523129 | p5 | (TATCA)3 | GTGTGTTAGGGTTGAGCCGT | AGTTCATACGGCCCACTCAG | 155 |
| chr9:48525182-48527200 | p2 | (TA)40 | GACGGTCTCGGATTGCTTTA | CAGTGGAGTACGCACGAGAA | 147 |
| chr9:48525182-48527200 | p6 | (TATGTA)3 | GACGGTCTCGGATTGCTTTA | CAGTGGAGTACGCACGAGAA | 147 |
| chr9:48535498-48537564 | p2 | (AT)33 | AAGCTCTGGGAAACCTCCAT | TCTCACCTTCCCAACTCGTC | 163 |
| chr9:48539088-48541110 | p6 | (CACAAA)3 | GTGCTTGTGGGGAACTGTTT | CCACAGAGGTGGGATGATTT | 172 |
| chr9:48539088-48541110 | p2 | (TG)11 | GTGCTTGTGGGGAACTGTTT | CCACAGAGGTGGGATGATTT | 172 |
| chr9:48609059-48611117 | p5 | (TTCTT)4 | AAGAAATCAGTCCACGTGCC | GCCCGAATTGTCCATGTAAG | 261 |
| chr9:48609119-48611177 | p5 | (TTCTT)4 | AAGAAATCAGTCCACGTGCC | GCCCGAATTGTCCATGTAAG | 201 |
| chr9:48609179-48611237 | p5 | (TTCTT)4 | AAGAAATCAGTCCACGTGCC | GCCCGAATTGTCCATGTAAG | 141 |
| chr9:48680072-48682086 | p2 | (AT)7 | GGCCATGGACTGACTGACTT | TATGACGTCTCCTTTTCGGG | 289 |
| chr9:48706587-48708602 | p5 | (ATTTT)3 | ATTGGGGGAGTGGAAATACC | AATGTTGGCCGAACTACCTG | 107 |
| chr9:48731282-48733375 | p3 | (TCT)10 | GGCGACCTATTACCCCTCTC | GAAGAAGAAGACGACGTGGC | 216 |
| chr9:48750746-48752760 | p2 | (CT)7 | TAATGTGGCTTGTCAGAGCG | CGACCTTGAGAAGGCTTACG | 134 |
| chr9:48778860-48780896 | p2 | (TC)10 | CGTACACTCTACCCTCCCCA | GTTTGACCACTGGCCTTGAT | 176 |
| chr9:48778860-48780896 | p2 | (TG)18 | CGTACACTCTACCCTCCCCA | GTTTGACCACTGGCCTTGAT | 176 |
| chr9:48820592-48822607 | p5 | (GGGCC)3 | AAGGCAAGTCAGGAATGCAC | TCTTGTGCACCATCTTCTCG | 203 |
| chr9:48820592-48822607 | p5 | (GGGCC)3 | AAGGCAAGTCAGGAATGCAC | TCTTGTGCACCATCTTCTCG | 203 |
| chr9:48894450-48896468 | p3 | (ATG)5 | GACGTGAACTTGAAAGGGGA | AGCGAGCTAACGGAATTTGA | 145 |
| chr9:48894450-48896468 | p2 | (AT)9 | GACGTGAACTTGAAAGGGGA | AGCGAGCTAACGGAATTTGA | 145 |
| chr9:48894450-48896468 | p2 | (TA)8 | GACGTGAACTTGAAAGGGGA | AGCGAGCTAACGGAATTTGA | 145 |
| chr9:48964545-48966561 | p4 | (GAAA)4 | TCAGGAGAGGAGAACCCTCA | CCCTGAGGGATGATCTCAAA | 208 |
| chr9:48964545-48966561 | p5 | (TGCAA)3 | TCAGGAGAGGAGAACCCTCA | CCCTGAGGGATGATCTCAAA | 208 |
| chr9:49010079-49012094 | p3 | (AAG)5 | TGGCCAAAAGCGATAAATTC | TTTTGGGTTAAACGACCGAG | 298 |
| chr9:49013686-49015698 | p2 | (AT)6 | TTCCTCTATGGCCATTTTCG | AGCAAAACCGGGGTAAAATC | 242 |
| chr9:49020417-49022441 | p5 | (TGAAT)3 | TTTGGCCGAGTTGAATTCTC | GCTTAATTGTCCAAGCTCGC | 170 |
| chr9:49020417-49022441 | p6 | (TATATG)4 | TTTGGCCGAGTTGAATTCTC | GCTTAATTGTCCAAGCTCGC | 170 |
| chr9:49062792-49064808 | p4 | (CTTT)4 | TTGGCATGGCATGAAGAATA | TGTCGACTTTCCCTGAATCC | 199 |
| chr9:49187613-49189627 | p3 | (CAA)5 | CATGCCATCAAATCAAGCAC | TAGGGGCTTTGGTCCTTCTT | 283 |
| chr9:49187628-49189662 | p3 | (CAA)5 | CATGCCATCAAATCAAGCAC | TAGGGGCTTTGGTCCTTCTT | 268 |
| chr9:49249640-49251658 | p6 | (CCTCTG)3 | ACAGTACTGGTGCTCCCCAC | TCCTCCGTATTGAATGGCTC | 194 |
| chr9:49261737-49263775 | p2 | (TA)19 | GCCCAGAGTCGACAGAAGTC | CCGAATCTCATCACACCCTT | 134 |
| chr9:49388507-49390527 | p2 | (TA)10 | AGGCTTCACCCTCCCTTCTA | TGGGGTGGCGTATATGATTT | 175 |

| Marker ID | SSR type | SSR motif | FORWARD PRIMER (5'-3') | REVERSE PRIMER (5'-3') | size (bp) |
|---|---|---|---|---|---|
| chr9:49427460-49429492 | p2 | (TA)33 | TCAAATGGCCACACAGACAT | ACCAAGTCCAGTCCCACAAG | 186 |
| chr9:49427460-49429492 | p2 | (TA)16 | TCAAATGGCCACACAGACAT | ACCAAGTCCAGTCCCACAAG | 186 |
| chr9:49447496-49449508 | p2 | (TA)6 | GAGTCATAGACCGTCCGAGC | TTTTGGAACCGTTATGAGGC | 293 |
| chr9:49459006-49461018 | p2 | (TA)6 | GAGTCATAGACCGTCCGAGC | TTTTGGAACCGTTATGAGGC | 293 |
| chr9:49504986-49507034 | p5 | (ATATC)3 | AATGGTGCCTCTTACGGATG | AGAGGGAACTCTGCCCAAAT | 113 |
| chr9:49692327-49694341 | p2 | (AT)7 | TGACGCAGTATTCCCTTTCC | ATGGATTGGTTGGCTTCTTG | 252 |
| chr9:49708572-49710584 | p2 | (AT)6 | TCGATAAAGCCTTGTGGGTC | CTCGCCCAATGTAGAAGAGC | 259 |
| chr9:49770180-49772192 | p2 | (AT)6 | CACCACCACCCTTTGAAGTT | TCACCTGGTTGATTGGTTGA | 248 |
| chr9:49788000-49790034 | p2 | (AT)8 | GTGGGAAATCGTGAGACCAT | CCGTTTTATTTGGACGCTGT | 200 |
| chr9:49788000-49790034 | p2 | (TA)17 | GTGGGAAATCGTGAGACCAT | CCGTTTTATTTGGACGCTGT | 200 |
| chr9:49797341-49799365 | p6 | (TATATG)4 | TGCCCGATTTTCGTTAAGTC | CATGCCAAAAGAAAAAGGGA | 241 |
| chr9:49797341-49799365 | p2 | (TA)12 | TGCCCGATTTTCGTTAAGTC | CATGCCAAAAGAAAAAGGGA | 241 |
| chr9:49797466-49799490 | p6 | (TATATG)4 | TGCCCGATTTTCGTTAAGTC | CATGCCAAAAGAAAAAGGGA | 116 |
| chr9:49797466-49799490 | p2 | (TA)12 | TGCCCGATTTTCGTTAAGTC | CATGCCAAAAGAAAAAGGGA | 116 |
| chr9:49817667-49819682 | p3 | (CAG)5 | GCCCGTGCACATATTATTCC | CTCTGAACCTGCCCTGCTAC | 118 |
| chr9:49847756-49849780 | p2 | (TG)12 | GTCAGTGTCACCAAGGCTCA | GGGCCACTTAGAATGGATGA | 152 |
| chr9:49979514-49981558 | p2 | (AT)22 | CAAATTAGTTTCGGCCCTGA | TTCGATATCCGGTTGACACA | 131 |
| chr9:49979514-49981558 | p2 | (TA)8 | CAAATTAGTTTCGGCCCTGA | TTCGATATCCGGTTGACACA | 131 |
| chr9:50006511-50008529 | p2 | (AT)9 | TTAGATTCTCGGGGATGGTG | TCATGCCTTGAAAGAAAGGG | 110 |
| chr9:50094794-50096830 | p2 | (TA)18 | TTGGTTAAGGGCTGTTTTGG | GCCTAACCCGAATTTCCAAT | 220 |
| chr9:50122000-50124036 | p2 | (TA)18 | TTGGTTAAGGGCTGTTTTGG | GCCTAACCCGAATTTCCAAT | 220 |
| chr9:50169677-50171713 | p2 | (TA)18 | CGTATGTGGTTTGCTTGGTG | GCCTAACCCGAATTTCCAAT | 159 |
| chr9:50199640-50201662 | p2 | (GA)11 | GACTGAAAGACGGTGGGAAA | AAAGCATTCGACGCTCAGTT | 203 |
| chr9:50199884-50201944 | p2 | (GA)11 | TAATACCCTCCAAGCCATCG | TTTTTGCTGGGGGTAGTTTG | 268 |
| chr9:50208785-50210813 | p2 | (TA)14 | GTCCCTCGTGGTCGTCTTTA | CCAGCCGAGAAATGGTAAAA | 257 |
| chr9:50250215-50252230 | p5 | (ATTTC)3 | TATTCTGGAGCCTTCGCACT | TCTTACGCCGGATACCTTTG | 281 |
| chr9:50273029-50275044 | p5 | (ATTTC)3 | TTTGGGTTGTGTGTGGAAAA | CTTGCGTACCAACGGAAAAT | 246 |
| chr9:50375888-50377958 | p2 | (TA)35 | ATTATCTCCTGGGGTGGGAC | TCCCCTCAAAGTCAGGTCAC | 126 |
| chr9:50379212-50381227 | p5 | (GATAT)3 | TGCTCTATTGTTCGGAGCCT | CGGATAAAGCACAAGGAAGC | 214 |
| chr9:50385152-50387167 | p5 | (TATGA)3 | GATGCTGCCCAAATTTCTGT | ACATGCCAAAAGAGGAATCG | 110 |
| chr9:50412271-50414287 | p4 | (TTTA)4 | TCGTAAATCCATAGGCCGTC | TGTGAAATCGTGGGTTGAAA | 293 |
| chr9:50424793-50426811 | p2 | (AT)9 | CGCTGCATCTGTCGAATAAA | TAGGGGGTGATGATGTGGAT | 130 |
| chr9:50424793-50426811 | p2 | (AC)6 | CGCTGCATCTGTCGAATAAA | TAGGGGGTGATGATGTGGAT | 130 |
| chr9:50647506-50649526 | p2 | (AT)10 | TTGTTTTTCCACCCCAAAAA | CTCAGCCGAGTAGGGTCTTG | 172 |
| chr9:50647506-50649526 | p2 | (AT)25 | TTGTTTTTCCACCCCAAAAA | CTCAGCCGAGTAGGGTCTTG | 172 |
| chr9:50654060-50656078 | p3 | (TTC)6 | ACGTTGCGGGTGACTAAATC | TCCCATACTTCTGGTGGAGC | 158 |

| Marker ID | SSR type | SSR motif | FORWARD PRIMER (5'-3') | REVERSE PRIMER (5'-3') | size (bp) |
|---|---|---|---|---|---|
| chr9:50734478-50736493 | p5 | (AGGAG)3 | CAGAAAGGGCCTGAACACAT | GTCCCAAAGACAGCCTTCAG | 114 |
| chr9:50772150-50774162 | p2 | (TA)6 | GAAAGCGGGCAAGGTATGTA | TCGGACATTATGGATACGCA | 143 |
| chr9:50775921-50777936 | p5 | (ATTTC)3 | TCAGTCAAAGTTTTGTGGCG | ACCTCTCCTGGGGGTTAGAA | 123 |
| chr9:50803726-50805744 | p2 | (TA)9 | GTCCCCCATTTTGTTGAGAA | ATATCAGGCGAGAATGACCG | 269 |
| chr9:50926045-50928057 | p2 | (TA)6 | CTCTTGCCGACCTCGTTATC | GGGACCGCTATGTTGAGTGT | 298 |
| chr9:50934500-50936514 | p2 | (AT)7 | CTTACCCCCTGTGCGTTAAA | AGGAGAAGAGCGATCGATGA | 120 |
| chr9:51011368-51013384 | p2 | (AT)11 | GACTGTAGCTTTCTTGCGGG | GATGATGCCATTCTTTGCCT | 246 |
| chr9:51011390-51013408 | p2 | (AT)11 | GACTGTAGCTTTCTTGCGGG | GATGATGCCATTCTTTGCCT | 224 |
| chr9:51065192-51067210 | p6 | (CTTACA)3 | GAACCTCGACAGGACCTCAA | TCCTTCTTTGGTTGGGTACG | 270 |
| chr9:51065192-51067210 | p6 | (ATCCGT)3 | GAACCTCGACAGGACCTCAA | TCCTTCTTTGGTTGGGTACG | 270 |
| chr9:51065192-51067210 | p2 | (AT)6 | GAACCTCGACAGGACCTCAA | TCCTTCTTTGGTTGGGTACG | 270 |
| chr9:51141763-51143795 | p2 | (TA)16 | AGAACGTCCATGATTGCTCC | GGTGTCTGCCTATGCCATCT | 227 |
| chr9:51173567-51175579 | p2 | (TA)6 | CATGAAGGAAGGTTTCCCAA | GTGATCCATTATATGGGCCG | 138 |
| chr9:51173567-51175579 | p5 | (TGAAT)3 | CATGAAGGAAGGTTTCCCAA | GTGATCCATTATATGGGCCG | 138 |
| chr9:51264053-51266071 | p3 | (CAA)6 | ACCGGCTCTGATACCAACTG | CTAGGTGAGACGAAGCGACC | 115 |
| chr9:51333536-51335552 | p2 | (TA)8 | CCAAAAACGGCATCTGAAAT | GGCCCTTTAGTGAGGGACTC | 189 |
| chr9:51394957-51396971 | p2 | (TA)7 | GAATAGGCCACAGCCCACTA | AGAAGGGGAAGGTGGAGAAA | 196 |
| chr9:51498469-51500484 | p5 | (TTTTA)3 | AGCTATATGGACGGGCAGTG | CTACCTGCGAGCATGAACAA | 155 |
| chr9:51551224-51553240 | p4 | (CTTT)4 | TCCAAGATTCCCCGATCATA | TGTTCGAAAGTGGGAACACA | 217 |
| chr9:51572852-51574876 | p6 | (ATACAT)4 | TCTTTTTGCTTTGGCGAGTT | TCCACTTGAACAGCTTGGTG | 190 |
| chr9:51572852-51574876 | p3 | (TCA)5 | TCTTTTTGCTTTGGCGAGTT | TCCACTTGAACAGCTTGGTG | 190 |
| chr9:51661027-51663065 | p2 | (AT)19 | TCTGTCATTGAAGAACCCCC | TATGATGAGAGCGTGGCAAG | 201 |
| chr9:51662367-51664397 | p2 | (TA)15 | CCATTCTTCTGCTGCAATCA | CAGGATGCTACAGGGCATTT | 149 |
| chr9:51662367-51664397 | p2 | (AT)16 | CCATTCTTCTGCTGCAATCA | CAGGATGCTACAGGGCATTT | 149 |
| chr9:51703548-51705586 | p2 | (AT)19 | TCTGTCATTGAAGAACCCCC | TATGATGAGAGCGTGGCAAG | 201 |
| chr9:51704888-51706918 | p2 | (TA)15 | CCATTCTTCTGCTGCAATCA | CAGGATGCTACAGGGCATTT | 149 |
| chr9:51704888-51706918 | p2 | (AT)16 | CCATTCTTCTGCTGCAATCA | CAGGATGCTACAGGGCATTT | 149 |
| chr9:51752962-51754976 | p2 | (AT)7 | GTTCTTGCGGTTCTTGAAGG | GCATTCTTGCACGACTCAAA | 269 |
| chr9:51885561-51887575 | p2 | (TA)7 | AAGTATCAACCGTGGCTTCG | TTCTGAGTTTTGTTGCGTCG | 132 |
| chr9:51885561-51887575 | p2 | (TA)8 | AAGTATCAACCGTGGCTTCG | TTCTGAGTTTTGTTGCGTCG | 132 |
| chr9:51885561-51887575 | p2 | (AT)6 | AAGTATCAACCGTGGCTTCG | TTCTGAGTTTTGTTGCGTCG | 132 |
| chr9:51885561-51887575 | p2 | (AT)6 | AAGTATCAACCGTGGCTTCG | TTCTGAGTTTTGTTGCGTCG | 132 |
| chr9:51952245-51954257 | p2 | (TA)6 | AATTGGACGTGGGACAGGTA | AATCAAGAACGTTGCCATCC | 294 |
| chr9:52037748-52039766 | p5 | (TATTG)3 | TACGAAGCGTATGTGGCTTG | AGCACCTCCCCTATATCGCT | 220 |
| chr9:52037748-52039766 | p3 | (ATG)6 | TACGAAGCGTATGTGGCTTG | AGCACCTCCCCTATATCGCT | 220 |
| chr9:52042186-52044201 | p5 | (ATTTC)3 | TTCAGTAGCATTTTTGGGGC | GGAGCCGTAGTCAGCTTGTC | 168 |

| Marker ID | SSR type | SSR motif | FORWARD PRIMER (5'-3') | REVERSE PRIMER (5'-3') | size (bp) |
|---|---|---|---|---|---|
| chr9:52049520-52051532 | p2 | (AT)6 | CGAAGATCGCTTTCAGAACC | TCCACAACTCGCACTTTGAG | 267 |
| chr9:52087056-52089086 | p2 | (AT)15 | GACTTCTCAGGCGCAGTTCT | TCCATTGGGCTCCTCATAAC | 227 |
| chr9:52103833-52105848 | p3 | (CTG)5 | TGCTGTGGATTCCGATGTTA | GAACCTCAGCCTGCAAAGAC | 271 |
| chr9:52103833-52105848 | p3 | (TCT)5 | TGCTGTGGATTCCGATGTTA | GAACCTCAGCCTGCAAAGAC | 271 |
| chr9:52138060-52140090 | p2 | (AT)15 | TTGCAGGGTTAGATCTTGGG | CGCTTTGGAAAAATTTGAGC | 147 |
| chr9:52138060-52140090 | p3 | (TTC)5 | TTGCAGGGTTAGATCTTGGG | CGCTTTGGAAAAATTTGAGC | 147 |
| chr9:52287960-52289975 | p5 | (TTTCT)3 | GAGGAGACGGCAATGGTAAA | GGGTCTTTGATTGCTTGGAA | 299 |
| chr9:52330767-52332785 | p6 | (TATATG)3 | TTATCTCGGGGGATGGTGTA | CCCTTCGAAAGAAAGGGAAG | 213 |
| chr9:52412559-52414571 | p2 | (GT)6 | GTTTGGTGGCAAGTCCAAAT | ATTCATTTCTTCTCCGCCCT | 191 |
| chr9:52412559-52414571 | p2 | (TA)8 | GTTTGGTGGCAAGTCCAAAT | ATTCATTTCTTCTCCGCCCT | 191 |
| chr9:52412572-52414592 | p2 | (GT)6 | GTTTGGTGGCAAGTCCAAAT | ATTCATTTCTTCTCCGCCCT | 178 |
| chr9:52412572-52414592 | p2 | (TA)8 | GTTTGGTGGCAAGTCCAAAT | ATTCATTTCTTCTCCGCCCT | 178 |
| chr9:52589761-52591779 | p6 | (ATGAGC)3 | GATAGCGAACCCAAAAACCA | GCTATCGTTCGCCTGTTCTC | 189 |
| chr9:52765156-52767171 | p3 | (GTT)5 | CGTTTTCGGGTGATCTGACT | GACCCGGATTGGAATAGGAT | 172 |
| chr9:52765156-52767171 | p2 | (TA)6 | CGTTTTCGGGTGATCTGACT | GACCCGGATTGGAATAGGAT | 172 |
| chr9:52957576-52959591 | p3 | (CAA)5 | CAACCTGGGAGTAACCTGGA | AAAGGGCCATCATGACTACG | 209 |
| chr9:52957576-52959591 | p2 | (CT)6 | CAACCTGGGAGTAACCTGGA | AAAGGGCCATCATGACTACG | 209 |
| chr9:53075757-53077772 | p3 | (TAA)5 | CCCTATCATTCAGGGAGCAA | GAGCTGCGGTTTCTAGGATG | 138 |
| chr9:53143270-53145285 | p3 | (TGG)5 | CAGGTCCAGAGTTGGGATGT | GAGCTACATATCCCAGGCCA | 249 |
| chr9:53149467-53151482 | p5 | (TTCAT)3 | AACACCAAACCTTCGTGGTC | CCTTCAATTGACCCGGAGTA | 130 |
| chr9:53179276-53181288 | p2 | (AG)6 | ACCATCTGATTTGGAGGCAG | CACCTGGACATCATCCACTG | 170 |
| chr9:53290859-53292877 | p3 | (CAA)6 | CATGGCACAGCATACTCCAC | AATGTTTCCCACACTGAGCC | 116 |
| chr9:53290859-53292877 | p2 | (CT)7 | CATGGCACAGCATACTCCAC | AATGTTTCCCACACTGAGCC | 116 |
| chr9:53423954-53426026 | p2 | (AT)15 | TCCTTCAAGGTGAGGCAGAT | TCATCATTACCATCCTCGCA | 145 |
| chr9:53423954-53426026 | p2 | (AT)36 | TCCTTCAAGGTGAGGCAGAT | TCATCATTACCATCCTCGCA | 145 |
| chr9:53526018-53528036 | p3 | (CAA)6 | GTCTCATGTCCCCCTTCAGA | GCCAATGATAGGAAAGCTCG | 244 |
| chr9:53572088-53574112 | p2 | (AT)12 | TCTCTGACCCCACACATTCA | GATTCCAAGAGAGACGACGG | 185 |
| chr9:53600646-53602666 | p2 | (TA)10 | AAAACCAAAGTGGAAGCCCT | TAGGAACCCACGACCTATGG | 271 |
| chr9:53639946-53641964 | p6 | (TTTCTC)3 | ACAGCAACCCAACAACAACA | ATGCAAGGTTGGAATGAAGG | 134 |
| chr9:53658932-53660944 | p2 | (AT)6 | CGTCCCATCATTCCTGTTCT | GAGGAAGAAACCCAAGAGGG | 130 |
| chr9:53665061-53667075 | p2 | (TA)7 | CACCATCCCCGAGAATCTAA | TTGGAGGACTTCGGATTTTG | 182 |
| chr9:53665061-53667075 | p2 | (TA)11 | CACCATCCCCGAGAATCTAA | TTGGAGGACTTCGGATTTTG | 182 |
| chr9:53734652-53736678 | p2 | (AT)12 | GCATGCCTCAATGTCTTTCA | ATATATCGTACCCGGCCCTC | 178 |
| chr9:53734652-53736678 | p2 | (TA)16 | GCATGCCTCAATGTCTTTCA | ATATATCGTACCCGGCCCTC | 178 |
| chr9:53734652-53736678 | p2 | (AT)13 | GCATGCCTCAATGTCTTTCA | ATATATCGTACCCGGCCCTC | 178 |
| chr9:53745635-53747653 | p6 | (ATCCAC)3 | GTTTGCTGCCGAAGAAAAAG | AAATCAGTTCAGGCACCCAC | 192 |

| Marker ID | SSR type | SSR motif | FORWARD PRIMER (5'-3') | REVERSE PRIMER (5'-3') | size (bp) |
|---|---|---|---|---|---|
| chr9:53818383-53820398 | p5 | (ATTGC)3 | AAAAGTGATTCATCGGTGGC | CAACAGGGCCAAACCTAAGA | 261 |
| chr9:53890625-53892649 | p2 | (AT)12 | AGCTCTCGGCTAGCTCCTTT | GCAAGCCGATGTACCATTTT | 216 |
| chr9:53922159-53924177 | p3 | (TGT)6 | GGGGGATGGATGTTATACCC | GGATTAAAAGCCCTTGCCTC | 201 |
| chr9:54007571-54009589 | p2 | (AT)9 | CGAAATGCTTGTCAAACCCT | AGCCTTCCTCAACACTTCCA | 257 |
| chr9:54031940-54033958 | p3 | (ATC)6 | CGTTCCTCGATTTGGAATGT | CGGTAGACTAGTACCCGGCA | 275 |
| chr9:54032013-54034031 | p3 | (ATC)6 | CGTTCCTCGATTTGGAATGT | CGGTAGACTAGTACCCGGCA | 202 |
| chr9:54032269-54034283 | p3 | (ATC)6 | TGAATCCATAAGAGGGGCAG | GCTCATGCCCTCAGGTAGAC | 261 |
| chr9:54032269-54034283 | p3 | (AAC)5 | TGAATCCATAAGAGGGGCAG | GCTCATGCCCTCAGGTAGAC | 261 |
| chr9:54032303-54034315 | p3 | (ATC)6 | TGAATCCATAAGAGGGGCAG | GCTCATGCCCTCAGGTAGAC | 227 |
| chr9:54032303-54034315 | p3 | (AAC)5 | TGAATCCATAAGAGGGGCAG | GCTCATGCCCTCAGGTAGAC | 227 |
| chr9:54114633-54116651 | p5 | (CATAT)3 | AGTGTGGGAAACATTGAGGC | ACTTATTGTGCTTCCGTCCG | 176 |
| chr9:54114633-54116651 | p3 | (CAA)6 | AGTGTGGGAAACATTGAGGC | ACTTATTGTGCTTCCGTCCG | 176 |
| chr9:54116856-54118868 | p2 | (AT)6 | AAAAATCTTGACGTGTGGGC | TCCTTCGAGCCAGGAAGTTA | 197 |
| chr9:54218017-54220032 | p5 | (TTTTA)3 | GGCGATATTTATTTCGGCAA | TTCCTCCATATTTCCCCTCC | 175 |
| chr9:54296286-54298320 | p2 | (AT)17 | CACCCAAACCCTAGCTTGAA | ACTTCTCCCACGTCCATCAG | 198 |
| chr9:54296286-54298320 | p2 | (AT)6 | CACCCAAACCCTAGCTTGAA | ACTTCTCCCACGTCCATCAG | 198 |
| chr9:54311582-54313618 | p2 | (TA)18 | ATGATGGATTCATTGCCACA | AAATGACATGGCGGAATAGC | 159 |
| chr9:54328287-54330302 | p5 | (TATTT)3 | ATTGGCGCTAGAAATCGAGA | GATAAGGGGTGCTCGGTACA | 232 |
| chr9:54328287-54330302 | p5 | (TATCG)3 | ATTGGCGCTAGAAATCGAGA | GATAAGGGGTGCTCGGTACA | 232 |
| chr9:54328287-54330302 | p2 | (TA)11 | ATTGGCGCTAGAAATCGAGA | GATAAGGGGTGCTCGGTACA | 232 |
| chr9:54339935-54341950 | p5 | (AATTA)3 | AAGCCGACAAGGCTATCAAA | TCGGGAGCTGCAGTTTAGTT | 189 |
| chr9:54354669-54356683 | p2 | (TA)7 | TATGGTGCTACGACTGAGCG | AGAAACCCGATTACGCATTG | 270 |
| chr9:54401179-54403191 | p5 | (AAAAT)3 | CATGAGCGAACGACGAGTAA | ATATGCAAAAATGCGGGGTA | 272 |
| chr9:54401179-54403191 | p2 | (TG)6 | CATGAGCGAACGACGAGTAA | ATATGCAAAAATGCGGGGTA | 272 |
| chr9:54402755-54404769 | p2 | (TA)7 | TTAAGTCGGGATTTGTTGGC | AAACCACATCCGGAAACGTA | 114 |
| chr9:54404412-54406426 | p2 | (TA)7 | TGATGATTATGATGCCCGAA | CTGAAAATGCTTCTGAGGGC | 258 |
| chr9:54527995-54530063 | p2 | (TA)7 | ACCCATAAAACTCACGCAGG | CTATGGGCATGAACACAACG | 234 |
| chr9:54556328-54558344 | p4 | (TTTA)4 | AAACTGTTCGCTTTGTCGCT | CTGCCCGATTTTCGTTATGT | 243 |
| chr9:54606336-54608350 | p2 | (AG)7 | CGAAAAAGTTCCCATTTCCA | ATATGGCCTCCACTGTCGTC | 295 |
| chr9:54606336-54608350 | p3 | (TTG)5 | CGAAAAAGTTCCCATTTCCA | ATATGGCCTCCACTGTCGTC | 295 |
| chr9:54619158-54621176 | p2 | (AT)9 | CTCCACGCCTTTAATTTCCA | GTCCAAGGAAGAGGAGGACC | 101 |
| chr9:54714732-54716747 | p2 | (TA)12 | AGCCTTGTCAGCTTGCGTAT | ACCGGTCTCCTGAAGAGTCA | 260 |
| chr9:54714732-54716747 | p5 | (TATGA)3 | AGCCTTGTCAGCTTGCGTAT | ACCGGTCTCCTGAAGAGTCA | 260 |
| chr9:54940655-54942670 | p2 | (TC)10 | ATTTCCTATGGCTGGGCTCT | ATATGAGTGTCCAGCTCGGG | 202 |
| chr9:54940655-54942670 | p5 | (TTTTA)3 | ATTTCCTATGGCTGGGCTCT | ATATGAGTGTCCAGCTCGGG | 202 |
| chr9:54940655-54942670 | p2 | (TC)10 | ATTTCCTATGGCTGGGCTCT | ATATGAGTGTCCAGCTCGGG | 202 |

| Marker ID | SSR type | SSR motif | FORWARD PRIMER (5'-3') | REVERSE PRIMER (5'-3') | size (bp) |
|---|---|---|---|---|---|
| chr9:54953063-54955078 | p3 | (ATT)5 | GCAAACTCGCCACTAACACA | GCATTGTCAGCTCAACAGGA | 244 |
| chr9:54962974-54964988 | p2 | (AT)7 | ATGCCTTGAAAATGTGGAGG | CTCTAGCCATCGCCTTTGAC | 201 |
| chr9:55045624-55047636 | p2 | (CA)6 | GGTCATGCCAAAAGAAGGAA | TTGGCCGAGTTAAATTTTCG | 256 |
| chr9:55103054-55105072 | p6 | (AAAATA)3 | AATCGCTTTCGCTACTTCCA | CCAAATGCTCATCCGTACCT | 297 |
| chr9:55164338-55166350 | p2 | (TA)6 | GTTCACCGAGTCCCTCGATA | CGTACCCGACCCTCTAGTGA | 201 |
| chr9:55164338-55166350 | p2 | (TA)6 | GTTCACCGAGTCCCTCGATA | CGTACCCGACCCTCTAGTGA | 201 |
| chr9:55171424-55173466 | p2 | (TA)9 | TCTCACCACTCTGTTCGTGC | GAGGGACTCGGTGAATCAAA | 243 |
| chr9:55187659-55189673 | p2 | (AC)7 | GTGTGTGAAATTTGGCGATG | TCTACTGGGGACACCACTCC | 145 |
| chr9:55199457-55201497 | p2 | (AT)20 | ACATCATCCCCTCGTAGCAC | CAACCAATCAGGAAGCCAAT | 111 |
| chr9:55241656-55243671 | p5 | (TGTCG)3 | TGTCGGATACGGGTATGGAT | GGTGCTTCCACAAAAAGCTC | 124 |
| chr9:55241656-55243671 | p5 | (CGACA)3 | TGTCGGATACGGGTATGGAT | GGTGCTTCCACAAAAAGCTC | 124 |
| chr9:55332709-55334781 | p2 | (AT)36 | TGCGGTGAATCTTGATTTGA | TATGCCTAGCCCGAAATCAC | 165 |
| chr9:55332709-55334781 | p2 | (AT)13 | TGCGGTGAATCTTGATTTGA | TATGCCTAGCCCGAAATCAC | 165 |
| chr9:55354434-55356449 | p5 | (AAGAA)3 | GACTTACCATCCGACCTGGA | CGCCACAGTGCTTTAACAGA | 158 |
| chr9:55391220-55393236 | p2 | (AT)8 | AAAATGGTCGGACTCACTGC | CTTCATCTTGCGGTGGAGAT | 144 |
| chr9:55402705-55404727 | p2 | (AT)11 | AGTGGCGGTGTAGCAGAACT | CTATGCCTCCAAAGAAAGCG | 184 |
| chr9:55602977-55604991 | p2 | (AT)7 | CAATGGCAAAATCTGTCACG | GTCGGCTTGCGTATCATTTT | 229 |
| chr9:55602977-55604991 | p2 | (TA)8 | CAATGGCAAAATCTGTCACG | GTCGGCTTGCGTATCATTTT | 229 |
| chr9:55603095-55605111 | p2 | (AT)7 | CAATGGCAAAATCTGTCACG | GTCGGCTTGCGTATCATTTT | 111 |
| chr9:55603095-55605111 | p2 | (TA)8 | CAATGGCAAAATCTGTCACG | GTCGGCTTGCGTATCATTTT | 111 |
| chr9:55891188-55893200 | p2 | (TG)6 | ACCCCATGTGACGAGTAAGC | TCATACCTGGCCCTACCAAG | 141 |
| chr9:55935421-55937451 | p2 | (AT)15 | ACCGATGAGCAGAGACGAGT | GAGGCGGTGTTTCTTGTGTT | 115 |
| chr9:55935421-55937451 | p3 | (CAA)6 | ACCGATGAGCAGAGACGAGT | GAGGCGGTGTTTCTTGTGTT | 115 |
| chr9:55979402-55981482 | p2 | (TA)26 | CGTCTTTCTTCCCGACTTTG | AAACGGGCGAAAGGACTAAT | 219 |
| chr9:55979578-55981606 | p2 | (TA)26 | ATTAGTCCTTTCGCCCGTTT | CGTGACATCAGAAAGACGGA | 158 |
| chr9:56046268-56048283 | p3 | (ATG)5 | CTTGGATTCAAGGCGAACAT | ATTCGGGTAGCATCTCCCTT | 150 |
| chr9:56063845-56065857 | p2 | (TG)6 | AGTTGAGTGGGTGGTCAAGG | TCGGAAGTGGAATGGAAGAG | 118 |
| chr9:56066861-56068873 | p2 | (AC)6 | TCGCTCAAAAACTGCAAATG | AAGGGAGGTGCTACCCAAAT | 205 |
| chr9:56152448-56154460 | p2 | (TA)6 | GCATTCTGTCGTTGTGCCTA | AGAAGGGCAAAGCTACGACA | 290 |
| chr9:56268880-56270892 | p2 | (CA)6 | TCTTTGGCTTCGCAAAATCT | TTTACAAGGGAAATGCTGCC | 294 |
| chr9:56333853-56335879 | p2 | (TA)13 | TGTGACGGAGATGTTACGGA | ACAATCCATAAGGACAGGCG | 276 |
| chr9:56462169-56464187 | p2 | (AG)6 | CTACGAAAGCTATGGCCTGG | ACCCATCCTTTCTCCGAAGT | 176 |
| chr9:56462169-56464187 | p3 | (TTG)6 | CTACGAAAGCTATGGCCTGG | ACCCATCCTTTCTCCGAAGT | 176 |
| chr9:56534591-56536609 | p2 | (CT)9 | ACAGTCCACATGACATCCCA | AATTAGGGGTTGTTTTGGGC | 145 |
| chr9:56631006-56633018 | p2 | (GT)6 | TTGAGTGGAAGGGTAATGCC | CTTAGCCTTCCTCGACCCTT | 246 |
| chr9:56694988-56697016 | p2 | (AT)14 | CCCGAAAACTTCCAAATTCA | AATCGTGTGGTGTTCGTTGA | 198 |

| Marker ID | SSR type | SSR motif | FORWARD PRIMER (5'-3') | REVERSE PRIMER (5'-3') | size (bp) |
|---|---|---|---|---|---|
| chr9:56707403-56709433 | p6 | (TATATG)5 | TCTCGGATTGTTGGTTGACA | GGGATCATGCCTTGAAAGAA | 215 |
| chr9:56707403-56709433 | p6 | (TATATG)4 | TCTCGGATTGTTGGTTGACA | GGGATCATGCCTTGAAAGAA | 215 |
| chr9:56707403-56709433 | p2 | (TA)8 | TCTCGGATTGTTGGTTGACA | GGGATCATGCCTTGAAAGAA | 215 |
| chr9:56716961-56718991 | p2 | (TA)15 | CGTCCTCAGCATCTCATCAA | CTCGCCCAATGTAGAAGAGC | 275 |
| chr9:56716961-56718991 | p2 | (AT)11 | CGTCCTCAGCATCTCATCAA | CTCGCCCAATGTAGAAGAGC | 275 |
| chr9:56760057-56762077 | p4 | (AGAA)5 | TTCCAGTTTCCCGATGGTAG | GTACGTTCTGTGCAGCCTGA | 146 |
| chr9:56808832-56810847 | p5 | (ACAAA)3 | CCATCGTGGCTTCTATCCAT | GAGGGTGAGAGAAATGCAGG | 267 |
| chr9:56808832-56810847 | p2 | (GT)6 | CCATCGTGGCTTCTATCCAT | GAGGGTGAGAGAAATGCAGG | 267 |
| chr9:57136818-57138833 | p5 | (ACACA)3 | TTCACAAGGATCATGCCAAA | GTTGATGATTTGGCCGAGTT | 293 |
| chr9:57149447-57151463 | p2 | (AT)8 | TAGCAACCCGGGAACTGTAG | TTTTGGGTCAACTTTGGAGG | 211 |
| chr9:57153401-57155419 | p3 | (CAA)6 | TTTTTAGGCGGATACCATGC | AGTATAACACTGCGCCACCC | 117 |
| chr9:57153401-57155419 | p6 | (AAACTG)3 | TTTTTAGGCGGATACCATGC | AGTATAACACTGCGCCACCC | 117 |
| chr9:57203372-57205428 | p2 | (AT)7 | ATCGTGAGACCATGCTACCC | CATCTTTGAGAGCACCCCAT | 231 |
| chr9:57203372-57205428 | p2 | (AT)7 | ATCGTGAGACCATGCTACCC | CATCTTTGAGAGCACCCCAT | 231 |
| chr9:57203372-57205428 | p2 | (AT)28 | ATCGTGAGACCATGCTACCC | CATCTTTGAGAGCACCCCAT | 231 |
| chr9:57206643-57208669 | p2 | (AT)13 | TCGATAAAACTTTGCGGGTC | TCCACCGACTACTTTGGGTC | 112 |
| chr9:57206643-57208669 | p2 | (AT)7 | TCGATAAAACTTTGCGGGTC | TCCACCGACTACTTTGGGTC | 112 |
| chr9:57248173-57250193 | p5 | (CCCCA)4 | TAATTGATCTCCACCCCAGC | CAAATGAGGCTTTACCCCAA | 282 |
| chr9:57331907-57333935 | p2 | (GT)6 | TTTTGCTCTCCGACCATTCT | TATGCCCGAAGTTAATTGCC | 212 |
| chr9:57336470-57338484 | p2 | (TA)7 | TCGGTATGCCCTTGGAATAG | AACCCTGTCGACCACTTCAC | 213 |
| chr9:57336470-57338484 | p2 | (TA)6 | TCGGTATGCCCTTGGAATAG | AACCCTGTCGACCACTTCAC | 213 |
| chr9:57369566-57371581 | p3 | (ATG)5 | AACCCTCTCCTCCACTCCAT | CCAGGTCTTTCAAACCCTCA | 144 |
| chr9:57370629-57372647 | p2 | (AT)9 | AGATGGCTGGAAATGTCACC | AGGCACGGTGGAATCATAAG | 293 |
| chr9:57373650-57375716 | p2 | (AT)6 | TTCTCATTTTCCTCCATGCC | CCAAGAATCGCTTCACCTTC | 131 |
| chr9:57389094-57391106 | p2 | (CA)6 | CTCCATGGGGTGAGACACTT | GGTGATCCTGTCCTGCTGTT | 211 |
| chr9:57461627-57463665 | p2 | (AT)19 | GACCTCCCATGATCTTCCAA | TGTGTAGTGGCAGCCTTGTC | 129 |
| chr9:57463364-57465376 | p2 | (AT)6 | CCAACCAAAACAACAACACG | AAAACAGTCCAAACCGTGGA | 156 |
| chr9:57493290-57495312 | p2 | (AT)11 | GACCCTCCAGACCTCACAAA | AGGTCGGTTGGAGCCTAAAT | 183 |
| chr9:57493290-57495312 | p2 | (TA)7 | GACCCTCCAGACCTCACAAA | AGGTCGGTTGGAGCCTAAAT | 183 |
| chr9:57509765-57511785 | p2 | (TA)10 | GGTTGACTTTTTGGGCTTGA | CGTGACATTGAGCTTCCAAA | 114 |
| chr9:57583548-57585562 | p2 | (AT)7 | TGATCCTAACAGCCCAGGAC | CCAAAGGCTCCCCTAAGAAC | 274 |
| chr9:57616633-57618647 | p2 | (TC)7 | AACGCTTGCTCATGAGGTCT | GTTACGGGAATGGGTTGTTG | 247 |
| chr9:57641590-57643605 | p2 | (AT)16 | GGAGGGCTTCCACTTTGATT | GCCCACACGTCAAGATTTTT | 155 |
| chr9:57727438-57729453 | p5 | (ATTGC)3 | TACCTTGGCCTTATGTTGCC | GAACGCAGTGTCCACAAGAA | 182 |
| chr9:57776213-57778225 | p2 | (TA)6 | CTTCTTAGGGCATGTTCCCA | CGTAGAATGAGGCCAATGGT | 129 |
| chr9:57783104-57785120 | p2 | (TA)8 | CGTTGGCCTCATTCTACGAT | TACCTAACTGCAGTGGTGCG | 239 |

| Marker ID | SSR type | SSR motif | FORWARD PRIMER (5'-3') | REVERSE PRIMER (5'-3') | size (bp) |
|---|---|---|---|---|---|
| chr9:57783104-57785120 | p2 | (TA)8 | CGTTGGCCTCATTCTACGAT | TACCTAACTGCAGTGGTGCG | 239 |
| chr9:57946363-57948381 | p3 | (TTG)6 | GGCCACACGACAAGATTTTT | TATCGTATGAGTGCCCACGA | 160 |
| chr9:58066848-58068862 | p2 | (TA)7 | AAGGCCTAAGCAAAATGGGT | TTTTCACGCAACCATTTCAA | 288 |
| chr9:58191591-58193606 | p5 | (ATTGC)3 | TCGAGGCTCAAAGGAAAAGA | CATCTCCACATGGCATCAAG | 227 |
| chr9:58296458-58298473 | p5 | (TTGGT)4 | ACCGACAAACCGCACTAAAC | AACCAAACCAAACCAAACCA | 124 |
| chr9:58296458-58298473 | p5 | (TTGGT)3 | ACCGACAAACCGCACTAAAC | AACCAAACCAAACCAAACCA | 124 |
| chr9:58355005-58357020 | p5 | (AACAA)3 | CCCCTTATCGACCTAAAGCC | CTGCTGGTATGACTGCCTCA | 161 |
| chr9:58355005-58357020 | p5 | (TTTTA)4 | CCCCTTATCGACCTAAAGCC | CTGCTGGTATGACTGCCTCA | 161 |
| chr9:58452034-58454049 | p5 | (ACACA)3 | ACGGCCAACTTAGACCACAC | GGAGTGGGTGTGTTGTTGTG | 251 |
| chr9:58452034-58454049 | p5 | (TTAAT)3 | ACGGCCAACTTAGACCACAC | GGAGTGGGTGTGTTGTTGTG | 251 |
| chr9:58492453-58494477 | p2 | (AT)12 | AAAGTCATGAAATCACGGGC | AGAAGAGGGCGAGTCACAAA | 294 |
| chr9:58587084-58589099 | p3 | (ATG)5 | GTCCACTCCTCTTCACCCAA | TCAATGGCCTCCTTTGATTC | 270 |
| chr9:58630749-58632765 | p2 | (TA)8 | CACGACCTTACATCACACCG | TGAAATGCCAATGACATGCT | 268 |
| chr9:58630749-58632765 | p2 | (CT)6 | CACGACCTTACATCACACCG | TGAAATGCCAATGACATGCT | 268 |
| chr9:58630902-58632914 | p2 | (TA)8 | CACGACCTTACATCACACCG | TGAAATGCCAATGACATGCT | 115 |
| chr9:58630902-58632914 | p2 | (CT)6 | CACGACCTTACATCACACCG | TGAAATGCCAATGACATGCT | 115 |
| chr9:58704296-58706314 | p6 | (TCATAA)3 | TCAAAAGGTGATGCAACTCG | CCCCTCAAGTGGTTGAAGAA | 298 |
| chr9:58973357-58975372 | p5 | (AATGC)3 | AGAGAGTGAGGATGCCAGGA | TACTATCGTCGGGTTGAGGG | 202 |
| chr9:58996722-58998740 | p6 | (CTATTC)3 | TGTGCTCCCAGAGACACAAG | TGGGGTAGCAGAAAGGAAGA | 261 |
| chr9:59034016-59036034 | p6 | (TATCCG)3 | AGGATTTGTCACGACCCAAC | TGTGAGCAGGTTTTGGTCAG | 275 |
| chr9:59034016-59036034 | p2 | (AT)9 | AGGATTTGTCACGACCCAAC | TGTGAGCAGGTTTTGGTCAG | 275 |
| chr9:59101410-59103425 | p5 | (ATACG)3 | TGCGAATCTGCAACTTTCAC | GTCGGCTTGCGTATCGTATT | 181 |
| chr9:59101428-59103450 | p5 | (ATACG)3 | TGCGAATCTGCAACTTTCAC | GTCGGCTTGCGTATCGTATT | 163 |
| chr9:59146869-59148881 | p2 | (AT)6 | TGACCCTGATAAAACGCTCC | ATACGGCTATACTGCCCACG | 190 |
| chr9:59257669-59259681 | p2 | (AT)27 | CATGGGCAGTAAAAGAAGCG | TGCATGTGAAAGGTTTTTGG | 158 |
| chr9:59257669-59259681 | p2 | (AT)6 | CATGGGCAGTAAAAGAAGCG | TGCATGTGAAAGGTTTTTGG | 158 |
| chr9:59383377-59385389 | p2 | (AT)27 | CATGGGCAGTAAAAGAAGCG | TGCATGTGAAAGGTTTTTGG | 158 |
| chr9:59383377-59385389 | p2 | (AT)6 | CATGGGCAGTAAAAGAAGCG | TGCATGTGAAAGGTTTTTGG | 158 |
| chr9:59512894-59514914 | p5 | (TCATT)4 | GTTCGCTTCAGCGGACTATC | TGGAGATTTGTCACTGCCTG | 226 |
| chr9:59590361-59592393 | p2 | (AG)16 | AAGGGCTGCATTACCATGAG | ATAAGCACCAAAAACCCACG | 102 |
| chr9:59590361-59592393 | p2 | (GA)16 | AAGGGCTGCATTACCATGAG | ATAAGCACCAAAAACCCACG | 102 |
| chr9:59650200-59652214 | p2 | (AG)7 | AGCGTTTGGCCATAGATTTG | TCGCCATAGATTTTGGAAGC | 255 |
| chr9:59837574-59839592 | p6 | (TATATG)3 | GAATTCTCGGGGTTTGTTGA | TTAAGTCTCGGCTTTCCCAA | 295 |
| chr9:59845573-59847585 | p2 | (TA)6 | ATTCTGTGACGTTTCTGGGC | AGCTTCGTCAATAGGGCTGA | 101 |
| chr9:59883223-59885238 | p5 | (TTATT)3 | CGCTTGTGTTACCGGATTTT | AATGACTTTTGCCTTGCACC | 160 |
| chr9:59922382-59924400 | p3 | (TGC)6 | TCGGGAGGGCTACTTGTATG | ACCTCGGGGAGTATAAGCGT | 231 |

| Marker ID | SSR type | SSR motif | FORWARD PRIMER (5'-3') | REVERSE PRIMER (5'-3') | size (bp) |
|---|---|---|---|---|---|
| chr9:60025408-60027420 | p2 | (AG)6 | GGTTTGAGCCAAATTACCGA | TTAGCCCGAGAAATGCCTTA | 130 |
| chr9:60105260-60107272 | p5 | (ATGAC)3 | TAGCCTCGGGGACAACTCTA | TTTCCGTCTGTGAAGGGTTC | 254 |
| chr9:60105288-60107312 | p5 | (ATGAC)3 | TAGCCTCGGGGACAACTCTA | TTTCCGTCTGTGAAGGGTTC | 226 |
| chr9:60111479-60113495 | p2 | (AC)8 | AAACCCGAACTCTCCCTTGT | TAAAATAAGGCCCAAAGGGG | 154 |
| chr9:60215357-60217372 | p5 | (CATTT)3 | CGAAGTCATTCCGAGTGTCA | CGTCTTGTTTGTCCCAAGGT | 199 |
| chr9:60215357-60217372 | p5 | (CATTT)3 | CGAAGTCATTCCGAGTGTCA | CGTCTTGTTTGTCCCAAGGT | 199 |
| chr9:60273873-60275891 | p6 | (ATACAT)3 | TGACTTGGCTCTATTGCACG | GGCTCTAAGTAAGGGTCGGG | 185 |
| chr9:60275243-60277255 | p2 | (CA)6 | GTGCCTTTCGGATTTCGTTA | TTCACATGGTTCTGATGCGT | 112 |
| chr9:60280524-60282542 | p6 | (ATACAT)3 | CCCTGCAGCATCCTGTTTAT | GTGGTATCAGAGCGGGTTGT | 241 |
| chr9:60302484-60304508 | p2 | (TA)23 | CACCATCCCCGAGAATCTAA | CGTGGCTTGTGAAAGTGAAA | 145 |
| chr9:60339919-60341937 | p6 | (ATATGT)3 | ACAGATTTTGGCGAATTTGG | GTCGGGACCTATTTATGCGA | 131 |
| chr9:60339919-60341937 | p3 | (AAC)5 | ACAGATTTTGGCGAATTTGG | GTCGGGACCTATTTATGCGA | 131 |
| chr9:60340660-60342675 | p6 | (ATATGT)3 | CATGACTTTCCTTGGGCTGT | ATATAACAGTGGCCCTCCCC | 206 |
| chr9:60340660-60342675 | p3 | (AAC)5 | CATGACTTTCCTTGGGCTGT | ATATAACAGTGGCCCTCCCC | 206 |
| chr9:60354195-60356207 | p6 | (CATATA)3 | TGGAGCTTATCCAGGGACAG | CGGCGTCTGTGGAAGTAAAT | 167 |
| chr9:60354195-60356207 | p2 | (TA)6 | TGGAGCTTATCCAGGGACAG | CGGCGTCTGTGGAAGTAAAT | 167 |
| chr9:60382973-60384991 | p3 | (TTG)6 | AAGTAGCGTGGAAGCGTTGT | CCATCACCTCAACACCACAC | 169 |
| chr9:60428649-60430661 | p2 | (TA)6 | CCCGAAGGCAATGACTAAAA | CCTTATTAAAGGGCCGGGTA | 250 |
| chr9:60502171-60504189 | p6 | (TTCAGT)3 | CTGGGAATTGGGTGAAGAAA | TTCCAGCCTTTGAGCGTAAT | 293 |
| chr9:60555445-60557507 | p2 | (AT)7 | ATACGAGTGTCCAGTTCGGG | CCCTATCGCTCACAATGGTT | 256 |
| chr9:60555445-60557507 | p2 | (AT)31 | ATACGAGTGTCCAGTTCGGG | CCCTATCGCTCACAATGGTT | 256 |
| chr9:60625717-60627735 | p6 | (TATATG)3 | TTTACAAGGGAAATGCTGCC | CGTCACCTGCTCGTTTCATA | 191 |
| chr9:60667256-60669280 | p2 | (AT)12 | GAGCGAACGACGAACATACA | ATGTGAATGCCAGAGGGTTC | 114 |
| chr9:60733985-60736000 | p5 | (TTTTG)3 | CCCAACAATCCAACCAAAAC | TGGCGGAACCACTTAAATTC | 149 |
| chr9:60813571-60815603 | p2 | (TA)16 | TGTGAGGCATTGTTGGTGTT | ACGCCATACCCAAATCCATA | 114 |
| chr9:60815442-60817454 | p2 | (AT)6 | ATCCACACCGGAGAGTCAAC | CTCTCCATGCCCATTCCTTA | 294 |
| chr9:60875897-60877935 | p2 | (AT)19 | CGGACTGTTTTGACACGTTG | GCCCACACAATATCCCAAAC | 141 |
| chr9:60990374-60992389 | p6 | (ACATAT)3 | CATGATGTCACACCCCCATA | ATTGTAATCGGCTCCACCAG | 294 |
| chr9:60990516-60992534 | p6 | (ACATAT)3 | CATGATGTCACACCCCCATA | ATTGTAATCGGCTCCACCAG | 152 |
| chr9:60998267-61000283 | p2 | (TA)8 | TTGCTTAATTGGAATTCGGG | AACATACTTGTTCGACCGCC | 252 |
| chr9:61046852-61048867 | p6 | (TATACA)3 | GCCCTTTTTCCCATCTTCTC | CGACCTTACGTCACTCCGAT | 149 |
| chr9:61046852-61048867 | p3 | (CAA)5 | GCCCTTTTTCCCATCTTCTC | CGACCTTACGTCACTCCGAT | 149 |
| chr9:61097909-61099924 | p2 | (GT)6 | TTCAACGAACACCCACAAAA | AACGAAAAGAGCACCTCGAA | 178 |
| chr9:61097909-61099924 | p3 | (TGT)5 | TTCAACGAACACCCACAAAA | AACGAAAAGAGCACCTCGAA | 178 |
| chr9:61105931-61107949 | p2 | (AT)9 | CCGAGCACCCCTTTAACATA | TTGTCGGCCCATATTTTTGT | 285 |
| chr9:61221085-61223099 | p2 | (AT)7 | TGTGGCGGAACAATAACAGA | AGCATTTGGGGAAGGTTCTT | 167 |

| Marker ID | SSR type | SSR motif | FORWARD PRIMER (5'-3') | REVERSE PRIMER (5'-3') | size (bp) |
|---|---|---|---|---|---|
| chr9:61288204-61290216 | p2 | (TA)6 | AGAGGGCCGGGTTATGTTAC | AGAAGCAGAAGCCATACGGA | 255 |
| chr9:61331907-61333919 | p2 | (AG)6 | TTATTCGCGAAGGAATCGAC | CGTTGGTGTTCCTTCTTGGT | 234 |
| chr9:61476570-61478582 | p2 | (AT)6 | ATGAGCATGACACGTGCTTC | AGTCAGATCACCCGAAAACG | 146 |
| chr9:61484307-61486322 | p5 | (CATAT)3 | TGACTTGGCTCCTTATCGCT | CAAGGGGTTGTCTGCAAAGT | 101 |
| chr9:61484307-61486322 | p3 | (AAC)5 | TGACTTGGCTCCTTATCGCT | CAAGGGGTTGTCTGCAAAGT | 101 |
| chr9:61588914-61590938 | p2 | (TA)12 | ATCGTGAGACCATGCTACCC | TGTTCATGCCCACAGGTAGA | 270 |
| chr9:61643157-61645183 | p2 | (TA)13 | TCTCGACCTTATGTCACCCC | ATCTAGGCACGGTGGAATTG | 300 |
| chr9:61720984-61722999 | p3 | (GTT)5 | GCTCACAAGGAGACTCTCGG | ATACCGCATTGGAGAGCATC | 199 |
| chr9:61785576-61787591 | p5 | (TCATA)3 | ATTGTCACGACCCGTCTAGG | CGTTGACATGGCATAAGTGC | 273 |
| chr9:61831604-61833620 | p2 | (AT)11 | GGCTATGAATAGTGCCCGAA | GTAATGGCAACCTTGTCGGT | 110 |
| chr9:61831604-61833620 | p2 | (AT)8 | GGCTATGAATAGTGCCCGAA | GTAATGGCAACCTTGTCGGT | 110 |
| chr9:61840563-61842575 | p2 | (AT)6 | CACCCTTCAATGAAAATGGG | TATGGAAGTGTCGAGGAGGG | 263 |
| chr9:61881870-61883882 | p2 | (AT)6 | CACCCTTCAATGAAAATGGG | TATGGAAGTGTCGAGGAGGG | 263 |
| chr9:62164314-62166336 | p2 | (AT)11 | ACTACACGGCCACATCAACA | CCAATTGAAGGGTGCTCATT | 273 |
| chr9:62202148-62204170 | p2 | (AT)11 | ACTACACGGCCACATCAACA | CCAATTGAAGGGTGCTCATT | 273 |
| chr9:62279062-62281074 | p2 | (AG)6 | TAATTCGACAGATGCAACGC | AGGCAAAAAGTGGCTGAGAA | 117 |
| chr9:62300159-62302173 | p2 | (AT)7 | CAGAGGAGATGCTTTACCCG | CTTCCCGTCTTGCAACTCTC | 283 |
| chr9:62350242-62352262 | p2 | (TA)10 | ATGAGGACATCGTGAGACCC | ACTGACGCCCATTTTCTTTG | 260 |
| chr9:62452400-62454426 | p2 | (AT)13 | AAAAGAGAAATGGCTCGCAA | CACTGCTAGCGACGAGACTG | 248 |
| chr9:62459355-62461370 | p5 | (ATTTC)3 | TATGCATTTGGGTGGTGTGT | AATTTCGCCAGAGTTTCCCT | 239 |
| chr9:62624470-62626485 | p3 | (CAA)5 | AAAACCCTTTGAATGTCCCC | TGGCTTGCCTTCAGCTTTAT | 254 |
| chr9:62762401-62764416 | p5 | (ATCAG)3 | AGCCGATACAGATACGGCAC | CCGGAGCATACACTTTTGGT | 149 |
| chr9:62803274-62805322 | p3 | (ATA)16 | GAGTGGAGGTAACGTGGGAA | ACCCCCATCTCCTCACTTCT | 213 |
| chr9:62845415-62847430 | p5 | (TATCG)3 | CACTCTCTTGGCTTTCTCGG | TTGCATTCTAAGATCGTGCG | 170 |
| chr9:63001860-63003887 | p3 | (ATT)9 | ATCGTGGATTCAACTTTCGG | GTTCGCTGCCGAAGAAATAG | 278 |
| chr9:63005406-63007448 | p2 | (TA)21 | TGGACTTTTGGGTTAAAGCG | TGGCCGAAAGCAATAAATTC | 236 |
| chr9:63005406-63007448 | p2 | (AT)7 | TGGACTTTTGGGTTAAAGCG | TGGCCGAAAGCAATAAATTC | 236 |
| chr9:63235203-63237218 | p3 | (TTG)5 | CCTTAGGCGAGATTGCAAAG | ATGCCAATTTTACGACCTGC | 170 |
| chr9:63279278-63281293 | p5 | (TATAT)3 | TATATCCGGTACGGATCCCA | GGCTTAGATTCTTGGCAACG | 210 |
| chr9:63440425-63442459 | p2 | (AT)17 | CAGTCCGGATTGTAGGGTGT | GATGCAGCCACTAGCAATGA | 178 |
| chr9:63440425-63442459 | p3 | (TCA)5 | CAGTCCGGATTGTAGGGTGT | GATGCAGCCACTAGCAATGA | 178 |
| chr9:63494142-63496157 | p5 | (TCATA)3 | TCTGCAAACATAGCCACTCG | GCTAATTTGGGGTATCGGGT | 220 |
| chr9:63663348-63665360 | p2 | (CT)6 | CTCATGGTTGTCTCCGAGGT | ATCCCAAATTTCCGATTTCC | 132 |
| chr9:63749134-63751149 | p5 | (TATCG)3 | CACTCTCTTGGCTTTCTCGG | TTGCATTCTAAGATCGTGCG | 170 |
| chr9:63752583-63754598 | p5 | (TCATA)3 | TCACGAGTCATGTCGTGGAT | GAAACGCCATTTGTCCCTTA | 170 |
| chr9:63814280-63816295 | p3 | (AAC)5 | TTTTCTCTTGGATTGGGGTG | TCCATCGGGCAGATTTTAAG | 212 |

| Marker ID | SSR type | SSR motif | FORWARD PRIMER (5'-3') | REVERSE PRIMER (5'-3') | size (bp) |
|---|---|---|---|---|---|
| chr9:63822316-63824332 | p2 | (AT)8 | GCCGTGTGCTCTGTGAAGTA | ACCTTTCTGAAGCAACGCAT | 300 |
| chr9:63841743-63843758 | p3 | (AAC)5 | TTTTCTCTTGGATTGGGGTG | TCCATCGGGCAGATTTTAAG | 212 |
| chr9:63849779-63851795 | p2 | (AT)8 | GCCGTGTGCTCTGTGAAGTA | ACCTTTCTGAAGCAACGCAT | 300 |
| chr9:63951599-63953614 | p3 | (TGT)5 | ACAATGTTGTTATGCCCCGT | GCCCTTGTCTCATATCCCAA | 204 |
| chr9:63951599-63953614 | p2 | (TA)12 | ACAATGTTGTTATGCCCCGT | GCCCTTGTCTCATATCCCAA | 204 |
| chr9:64001995-64004007 | p2 | (TA)7 | ATCGTGCACAGTTTCACTGC | CATATCGTACCTGGCCCATT | 250 |
| chr9:64001995-64004007 | p2 | (TA)6 | ATCGTGCACAGTTTCACTGC | CATATCGTACCTGGCCCATT | 250 |
| chr9:64003734-64005750 | p2 | (TA)8 | GAAATCGTGGGTTGAAATGG | GACTTAACGAAAATCGGGCA | 229 |
| chr9:64003734-64005750 | p2 | (TA)7 | GAAATCGTGGGTTGAAATGG | GACTTAACGAAAATCGGGCA | 229 |
| chr9:64033496-64035508 | p2 | (AG)6 | GCTTATGATGCCGGGTAAGA | ACTTAGCCAACCAAGGCTCA | 115 |
| chr9:64055706-64057724 | p3 | (AAC)6 | CAGCCTCACTGAATGGGAAT | AGATTTGCACCTGTGGGTTC | 246 |
| chr9:64055706-64057724 | p2 | (CT)7 | CAGCCTCACTGAATGGGAAT | AGATTTGCACCTGTGGGTTC | 246 |
| chr9:64140329-64142347 | p2 | (AT)9 | ACGTTTGATGACCTTAGGCG | CGGTCCGTATTTTGCACTTT | 103 |
| chr9:64538167-64540187 | p2 | (TA)6 | GGGACCCGATTACCCTAAGA | CCCCCTTTTGATTCACTTCA | 206 |
| chr9:64538167-64540187 | p2 | (TA)10 | GGGACCCGATTACCCTAAGA | CCCCCTTTTGATTCACTTCA | 206 |
| chr9:64544057-64546069 | p2 | (TA)6 | AGAATAGGCGGACATTGCAC | GACCCCATTGGATTTGTTTG | 119 |
| chr9:64558656-64560668 | p2 | (AC)6 | CAAACGAAGAAGTGGGAAGC | CGAGCTAAGTCTCTGGGGTG | 247 |
| chr9:64614544-64616568 | p4 | (CATA)6 | ACTCGGCAATGCTCTAGGAA | TGACGTCCCTTCTTTTGGAC | 159 |
| chr9:64649538-64651550 | p3 | (ACA)5 | TTACCCAAATGGGACCAAAA | CATTTGCAGTTTCTGGACGA | 131 |
| chr9:64649538-64651550 | p2 | (AC)6 | TTACCCAAATGGGACCAAAA | CATTTGCAGTTTCTGGACGA | 131 |
| chr9:64753952-64755982 | p2 | (AT)15 | ACGGAGGTCATTTTGACTCG | ATTCGTGGATTGTTCCTTCG | 133 |
| chr9:64753952-64755982 | p2 | (AT)34 | ACGGAGGTCATTTTGACTCG | ATTCGTGGATTGTTCCTTCG | 133 |
| chr9:64754732-64756800 | p2 | (AT)15 | CCGACAAGGCTACCCAATAA | AGCCAGGAAGATAGATCGCA | 282 |
| chr9:64754732-64756800 | p2 | (AT)34 | CCGACAAGGCTACCCAATAA | AGCCAGGAAGATAGATCGCA | 282 |
| chr9:64831283-64833298 | p5 | (AACCG)3 | CGCTGCAACTGAAAGGTACA | TTTCGGTATTTATGCCCAGC | 224 |
| chr9:64844881-64846909 | p2 | (AG)14 | GTCGCAGGTACCATTCCTGT | ATGTTTGCCGCAGTATTTCC | 252 |
| chr9:64848870-64850886 | p2 | (AT)8 | TTGTGTAAGGGTACCAGCCC | TTCGATACCAGCAAACACCA | 128 |
| chr9:64859848-64861862 | p2 | (TA)7 | GTACCCATCCCACCGTATTG | GGCCTAGCACTATTCCCTCC | 224 |
| chr9:64863506-64865518 | p2 | (AT)6 | CCCAACTTCTTTTGGGTTCA | TGAAAATTTTGGATGTGCGA | 204 |
| chr9:64863506-64865518 | p2 | (TA)6 | CCCAACTTCTTTTGGGTTCA | TGAAAATTTTGGATGTGCGA | 204 |
| chr9:64889572-64891587 | p5 | (ATTTC)3 | CTCCCACCACCCTGTTCTTA | GAGCAATGCAATCGAACTGA | 242 |
| chr9:64964334-64966352 | p3 | (GAT)6 | TGTTGGGTTGATGTCGTTGT | TTTACGGAAAATTTGGCAGC | 194 |
| chr9:64964334-64966352 | p2 | (AT)10 | TGTTGGGTTGATGTCGTTGT | TTTACGGAAAATTTGGCAGC | 194 |
| chr9:65010344-65012359 | p3 | (GTT)5 | GACATCGGAGGTAGTTGGGA | CTTCGAACAACCCAACTGGT | 211 |
| chr9:65088743-65090759 | p2 | (AT)8 | GTCCTAGTCGCAGCACAACA | CCCTCTTTGAGCATGTGGAT | 171 |
| chr9:65149396-65151411 | p5 | (ATTAA)3 | ACTTTCCTGTAGCAGCCGAA | TTGTGAAGGTTATGTGGCGA | 264 |

| Marker ID | SSR type | SSR motif | FORWARD PRIMER (5'-3') | REVERSE PRIMER (5'-3') | size (bp) |
|---|---|---|---|---|---|
| chr9:65167900-65169918 | p2 | (TA)9 | TGCCCTTGAAATGGGAGTAG | AATGAAAATTGGGCAGCATC | 122 |
| chr9:65188483-65190498 | p2 | (TA)6 | CGTACATGTCACCACGAAGG | GTAATCATCATATGCCGCCC | 113 |
| chr9:65188483-65190498 | p3 | (ATG)5 | CGTACATGTCACCACGAAGG | GTAATCATCATATGCCGCCC | 113 |
| chr9:65188483-65190498 | p2 | (TA)10 | CGTACATGTCACCACGAAGG | GTAATCATCATATGCCGCCC | 113 |
| chr9:65188784-65190804 | p3 | (ATG)5 | CCCACAGGTAGACAGGGAGA | GGGACACCAAAAATCCACAC | 157 |
| chr9:65188784-65190804 | p2 | (TA)10 | CCCACAGGTAGACAGGGAGA | GGGACACCAAAAATCCACAC | 157 |
| chr9:65202312-65204338 | p2 | (TA)13 | TAATTTCAGCTGGAGGACGG | GGCACAGGTGACATTTTCCT | 288 |
| chr9:65236541-65238561 | p3 | (CAA)5 | TACATATCCCACTGCCCCAT | TTTATTTTGCCTGCTACGGG | 114 |
| chr9:65236541-65238561 | p2 | (TA)10 | TACATATCCCACTGCCCCAT | TTTATTTTGCCTGCTACGGG | 114 |
| chr9:65291572-65293587 | p5 | (TGTTA)3 | GCTTATGATGCCGGGTAAGA | CGACCTCATAGGATCGGTGT | 130 |
| chr9:65471289-65473369 | p2 | (AT)40 | AGAAGCCCTGACCAGTCTCA | ACGTGACAGTTTGAGGGTCC | 292 |
| chr9:65516800-65518812 | p2 | (AT)7 | GACGATCCATTGATGACGTG | GGTGACGTAAGGTCGAGAGC | 205 |
| chr9:65516800-65518812 | p2 | (AT)6 | GACGATCCATTGATGACGTG | GGTGACGTAAGGTCGAGAGC | 205 |
| chr9:65533704-65535718 | p2 | (AT)7 | GCCTAACAGGAAGAAGCGTG | AATAACCGCGAATGCAACTC | 234 |
| chr9:65558655-65560669 | p2 | (AT)7 | GCCTAACAGGAAGAAGCGTG | AATAACCGCGAATGCAACTC | 234 |
| chr9:65684467-65686523 | p3 | (TTC)8 | TCTTCCTCACTTTTGGTGCC | CAAACACAGCACCCACAAAC | 262 |
| chr9:65684467-65686523 | p2 | (TA)28 | TCTTCCTCACTTTTGGTGCC | CAAACACAGCACCCACAAAC | 262 |
| chr9:65696019-65698074 | p3 | (AGT)5 | CCTACCTGCAAGATCCCAAA | CCATCCATCCACAATGATGA | 193 |
| chr9:65711584-65713596 | p2 | (AC)6 | CCAAAATGTGACAAAAGGGG | GGACCATTTCGATTTGAGGA | 133 |
| chr9:65760571-65762587 | p2 | (AT)8 | CCCGAGTTGGACACTCGTAT | TGTTCGTCTGTAGTGGCAGC | 270 |
| chr9:65767675-65769703 | p2 | (AT)14 | TGGAAACTTGCAAAGGAACC | CCTTGTCAGCTTGCGTATCA | 281 |
| chr9:65798113-65800129 | p2 | (AT)8 | CCCGAGTTGGACACTCGTAT | TGTTCGTCTGTAGTGGCAGC | 270 |
| chr9:65805217-65807245 | p2 | (AT)14 | TGGAAACTTGCAAAGGAACC | CCTTGTCAGCTTGCGTATCA | 281 |
| chr9:65860049-65862064 | p5 | (CAATG)3 | TGTGAAAAAATGCGGTCAAAA | CCCAAGTCCAAAACGGAGTA | 217 |
| chr9:65869095-65871111 | p2 | (AT)8 | GGCGCAATATCTGGCATAGT | AGTGATGCCTCCGAAGAAGA | 292 |
| chr9:65988042-65990076 | p2 | (AT)17 | GGACGTCGTGTGTGTTATGG | ACTCCTTCGATGCTCACGTT | 163 |
| chr9:65994433-65996449 | p2 | (AT)8 | TAGGGTGTTCACCTTCACCC | TCCAGGATTTTCACTCAGGG | 212 |
| chr9:66026907-66028923 | p2 | (AT)8 | ATGCATACGAAAGCCGTAGG | TTATCCGGAGCGTATTTTGG | 267 |
| chr9:66124649-66126661 | p2 | (TA)6 | TTGTATAAAAAGGCAGGGCG | AGCCTTGTCAGCTTGCGTAT | 114 |
| chr9:66149583-66151595 | p2 | (TG)6 | TCGCCACTACTCTGTTCGTG | TCGGAAACCGTAACACATCA | 170 |
| chr9:66159013-66161077 | p5 | (GAAAT)3 | CTGGGATTCCTTGTCTTGGA | TGGAGCGGGATAGATTTGAG | 300 |
| chr9:66165071-66167083 | p3 | (ATG)5 | TTGTTTGGAAGATTGGAGGG | TGCCTTGAGAAAGAAAGGGA | 206 |
| chr9:66165071-66167083 | p2 | (TA)6 | TTGTTTGGAAGATTGGAGGG | TGCCTTGAGAAAGAAAGGGA | 206 |
| chr9:66211651-66213667 | p2 | (TG)8 | AAAGGTGCCTACGGGCTATT | CTCATACCTGGCCCTACCAA | 252 |
| chr9:66221217-66223235 | p6 | (TATATG)3 | AGATGCTGTCCGATTTTTGG | GCTTATTTGTCCAAGCTCGC | 233 |
| chr9:66290796-66292808 | p2 | (AT)6 | TGGGAAAGTTAAGCCTTGGA | TTCGGGTTTTCCCTTCTTTT | 268 |

| Marker ID | SSR type | SSR motif | FORWARD PRIMER (5'-3') | REVERSE PRIMER (5'-3') | size (bp) |
|---|---|---|---|---|---|
| chr9:66405369-66407381 | p2 | (AT)6 | AAGTGTCGGATCACCAAAGC | ATAATGGGCCGGACATGATA | 212 |
| chr9:66434597-66436612 | p5 | (TATCG)3 | CATTCTTCCCCACACCAATC | GATAAGGGGTGCTCGGTACA | 289 |
| chr9:66434597-66436612 | p2 | (TA)11 | CATTCTTCCCCACACCAATC | GATAAGGGGTGCTCGGTACA | 289 |
| chr9:66440474-66442489 | p5 | (TATCG)3 | CCACTCTCTTGGCTTTCTCG | TTGCATTCTAAGATCGTGCG | 169 |
| chr9:66440474-66442489 | p2 | (TA)11 | CCACTCTCTTGGCTTTCTCG | TTGCATTCTAAGATCGTGCG | 169 |
| chr9:66479559-66481573 | p6 | (CATATT)3 | ACTAATCCCATTGGGCAACA | TAGCGGAGCTCCATTTTCAT | 205 |
| chr9:66479574-66481588 | p6 | (CATATT)3 | ACTAATCCCATTGGGCAACA | TAGCGGAGCTCCATTTTCAT | 190 |
| chr9:66479589-66481605 | p6 | (CATATT)3 | ACTAATCCCATTGGGCAACA | TAGCGGAGCTCCATTTTCAT | 175 |
| chr9:66479606-66481624 | p6 | (CATATT)3 | ACTAATCCCATTGGGCAACA | TAGCGGAGCTCCATTTTCAT | 158 |
| chr9:66479625-66481637 | p6 | (CATATT)3 | ACTAATCCCATTGGGCAACA | TAGCGGAGCTCCATTTTCAT | 139 |
| chr9:66509011-66511027 | p4 | (CATA)4 | CACCCGACCCTTCACTTAGA | CTCTGCAAGCCGATGTATCA | 113 |
| chr9:66509011-66511027 | p3 | (CAA)5 | CACCCGACCCTTCACTTAGA | CTCTGCAAGCCGATGTATCA | 113 |
| chr9:66564773-66566788 | p3 | (TGT)5 | TGCGGGAAGTTAAGGACAAG | AACTCAAGCCCACCCTTTTT | 119 |
| chr9:66673204-66675218 | p2 | (AT)7 | TGTTCATGCCCACAGGTAGA | TAAGCCCTTTGGTTGTCCAC | 250 |
| chr9:66764251-66766265 | p2 | (AT)7 | GAGACACGTAGCTGATGGCA | TTCATGCTCGTGCTTGTTTC | 230 |
| chr9:66881287-66883299 | p2 | (CA)6 | AGCCCCACTAACAAAGCTGA | TGAATCACTGCAGCCAACTC | 251 |
| chr9:66893198-66895262 | p3 | (GAT)6 | CCTTGTGGGCTGTTTTGAAT | TCGAGCTCGCAAGTCTACAA | 185 |
| chr9:66893198-66895262 | p2 | (AT)32 | CCTTGTGGGCTGTTTTGAAT | TCGAGCTCGCAAGTCTACAA | 185 |
| chr9:67019524-67021538 | p2 | (TA)7 | GCGGGTGAGAGAAATGTAGG | AGCACCCACTTTGTAATGCC | 120 |
| chr9:67128571-67130591 | p2 | (AG)10 | ATGTGCTGGTGACCCTTCTC | TCTACTTTCGGCTTCCCAGA | 236 |
| chr9:67134612-67136626 | p2 | (AG)7 | CTGTGCTTCACGAATGTGCT | TCTACTTTCGGCTTCCCAGA | 223 |
| chr9:67143724-67145752 | p2 | (AT)6 | TAGAGGGGTCATGGACTTGG | CACCTTCGTCGCCATTACTT | 250 |
| chr9:67143724-67145752 | p2 | (AT)14 | TAGAGGGGTCATGGACTTGG | CACCTTCGTCGCCATTACTT | 250 |
| chr9:67150388-67152400 | p2 | (AT)6 | AGCTACCAGGACTGGGGAGT | TGAGTTCCCAAGGTCCAAAC | 275 |
| chr9:67270321-67272339 | p3 | (CAA)6 | GACTAAGCTGGTCGGCACTC | TATGACCCGTGATGAGCAAA | 259 |
| chr9:67314908-67316920 | p2 | (TG)6 | GTGTGATTCCCTGTTGGCTT | AACCCTCAACTTAGGCCGAT | 166 |
| chr9:67319827-67321877 | p2 | (AT)25 | TATGGATTTGGGTATGGCGT | GACAAATTTGCCTTCGTCGT | 288 |
| chr9:67479627-67481651 | p6 | (GTATCC)4 | TTTTGGGCTGCTGTAGCTTT | TGCGCCACACCTATAAACAA | 123 |
| chr9:67652316-67654334 | p6 | (TATATG)3 | GCGCGTTTGATTATTCCATT | CGAACGTTTCAGAAGCCATT | 249 |
| chr9:67754430-67756490 | p2 | (TA)30 | TGCCCTGTAGCCTCTTGTTT | AGCTGGGTCGTGACAGAAGT | 256 |
| chr9:67754430-67756490 | p2 | (TA)11 | TGCCCTGTAGCCTCTTGTTT | AGCTGGGTCGTGACAGAAGT | 256 |
| chr9:67845000-67847016 | p2 | (AT)8 | TGGACTCATTTCACCCACAA | TGGTGGCACAAGAAAAAGTG | 104 |
| chr9:67897695-67899733 | p2 | (AC)19 | CCACAGGTAGACAGGGAGGA | GCCTCTTGGCCATCAAAATA | 196 |
| chr9:68002727-68004751 | p2 | (AT)12 | TTCATTTTCCCCATTTCCAA | TCCAACTCCAATTCCAAAGC | 212 |
| chr9:68089453-68091479 | p2 | (AT)14 | ATGGGTGAAGGTGCATTAGC | GAGGCAAACCCACAGAATGT | 124 |
| chr9:68090528-68092542 | p2 | (AT)14 | AGACCCAAAGGTTCACAACG | TTGTTGTGGTAGCCTTGCAG | 256 |

| Marker ID | SSR type | SSR motif | FORWARD PRIMER (5'-3') | REVERSE PRIMER (5'-3') | size (bp) |
|---|---|---|---|---|---|
| chr9:68090528-68092542 | p2 | (AT)7 | AGACCCAAAGGTTCACAACG | TTGTTGTGGTAGCCTTGCAG | 256 |
| chr9:68098179-68100194 | p5 | (ATTTC)3 | AGGCTTACAAAGTCAGCCCA | GAGTCCCCAATCCATTCTGA | 186 |
| chr9:68159436-68161451 | p3 | (TTG)5 | GCTCCGTGATCTGTTTGACA | CCCAACCATGACCAGGTATC | 154 |
| chr9:68290942-68292960 | p6 | (CATATA)3 | CGAACCCGCTGACTGTTATT | ACGTCAGTTTTGTAACCGGC | 169 |
| chr9:68371004-68373016 | p2 | (AT)6 | GTCCCTTTCTTCAAATGGCA | CAATTTCGGACATTCTGCCT | 197 |
| chr9:68449864-68451882 | p6 | (CCTCTG)3 | CATCAATCCTTGGGAGAGGA | TGACTTCGCCATTGATGTGT | 101 |
| chr9:68462316-68464331 | p3 | (TTG)5 | CCTTAAGCCAAATATGGGCA | GCCATAACTTTTGATCCCGA | 187 |
| chr9:68478358-68480376 | p6 | (TCCTTT)3 | TGCATGGCATTGAACTAGGA | CTCTACCCCACAAAAAGCCA | 291 |
| chr9:68513880-68515898 | p6 | (GGTAGA)3 | CCTCACAATGGCCTCAAACT | CTGCAGCAGCTTTTTCCTTT | 297 |
| chr9:68542592-68544607 | p3 | (GAA)5 | CAATTGGTATCAGAGCGGGT | CATTCTTGCCTTCCACCATT | 277 |
| chr9:68562978-68564994 | p2 | (TA)8 | CGTAGGGAACTCGCAAAAAG | CGTATTCCTCCTCAAACCCA | 185 |
| chr9:68654232-68656252 | p2 | (TA)10 | GGACAACTGGAAATTTGGGA | TATTCGTGAATCCCTTTCGC | 202 |
| chr9:68669009-68671025 | p2 | (AG)8 | TGGTTGCCTTAAGTATCCCG | GCCTCCAACAATATGGGCTA | 275 |
| chr9:68697235-68699299 | p2 | (AT)7 | GGCATTTACAGACATGGGCT | ATGTGTGCAAAGAGTGCGAG | 266 |
| chr9:68697235-68699299 | p2 | (AT)9 | GGCATTTACAGACATGGGCT | ATGTGTGCAAAGAGTGCGAG | 266 |
| chr9:68727430-68729445 | p2 | (AG)9 | ACCCCAACCCCTAAAACTTG | CCACCGTGAAATCACAACAG | 133 |
| chr9:68727430-68729445 | p5 | (GAAAA)4 | ACCCCAACCCCTAAAACTTG | CCACCGTGAAATCACAACAG | 133 |
| chr9:68727430-68729445 | p5 | (TTATT)3 | ACCCCAACCCCTAAAACTTG | CCACCGTGAAATCACAACAG | 133 |
| chr9:68775287-68777305 | p6 | (ATCCGT)3 | ATAAGCCGACGAGGCTAACA | GTCCCGCCTATATGACCTGA | 280 |
| chr9:68775287-68777305 | p2 | (AT)9 | ATAAGCCGACGAGGCTAACA | GTCCCGCCTATATGACCTGA | 280 |
| chr9:68778240-68780252 | p2 | (TG)6 | CTCGACCCCATATGACGAGT | TCATACCTGGCCCTACCAAG | 135 |
| chr9:68830813-68832843 | p2 | (TA)15 | TCGCAGGGGTATCTAACTGG | ACACTCGTCACGACTCATCG | 101 |
| chr9:68975957-68977972 | p3 | (GTT)5 | CGTGGAAGAGTTGTTTGGGT | GCTCAGGCTCCACATACACA | 181 |
| chr9:68997986-69000025 | p3 | (ATT)13 | GGAAAACATTTGAGGCGTGT | TGGCCTTTGTTGTGATGTGT | 249 |
| chr9:69068129-69070144 | p5 | (CATTG)3 | GTAGTGGTCGGGCACTTGTT | CGTACACTCTACCCTCCCCA | 192 |
| chr9:69108592-69110604 | p2 | (TA)6 | ACACCATCCCCAACAATCAT | GGGCAAGTTTTTGAACGTGT | 208 |
| chr9:69108592-69110604 | p2 | (AT)7 | ACACCATCCCCAACAATCAT | GGGCAAGTTTTTGAACGTGT | 208 |
| chr9:69187848-69189866 | p5 | (CCATT)3 | CCCGTTTTGTCCGAAATCTA | AGGGGAGATGCTGCCTAAAT | 225 |
| chr9:69187848-69189866 | p2 | (AC)9 | CCCGTTTTGTCCGAAATCTA | AGGGGAGATGCTGCCTAAAT | 225 |
| chr9:69320022-69322037 | p6 | (ATACAT)3 | AATCTCATCGGATGTAGGCG | GTGGTCCTGTCCCGTCTTTA | 229 |
| chr9:69320022-69322037 | p3 | (CAT)5 | AATCTCATCGGATGTAGGCG | GTGGTCCTGTCCCGTCTTTA | 229 |
| chr9:69345225-69347237 | p2 | (AC)6 | AGGCATGTATCAACCGAAGC | GCCTATTTCTGAGGAGCACG | 121 |
| chr9:69447993-69450005 | p2 | (TC)6 | AACCGGTTGACAAATTCTCG | AAAGGGCAATGATGTCGAAG | 124 |
| chr9:69487177-69489202 | p5 | (TATAT)5 | CAAATCAGAATTTCCGCGAT | AGATGTCCCATAAGCATGGC | 235 |
| chr9:69502200-69504216 | p2 | (GA)8 | AAAGCGCTCGCAATACAACT | TCATTGGTTGAATCCCGTTT | 234 |
| chr9:69529464-69531478 | p3 | (TTG)7 | ACCATCGTGTGGGATGTTTT | TCGTCATGTGCTTCCTCAAG | 294 |

| Marker ID | SSR type | SSR motif | FORWARD PRIMER (5'-3') | REVERSE PRIMER (5'-3') | size (bp) |
|---|---|---|---|---|---|
| chr9:69529480-69531534 | p3 | (TTG)7 | ACCATCGTGTGGATGTTTT | TCGTCATGTGCTTCCTCAAG | 278 |
| chr9:69530649-69532661 | p2 | (TA)6 | AGGAAAACCGTGTCGTCAAC | CCCTTCGGGCATAATACTCA | 125 |
| chr9:69570379-69572393 | p2 | (AG)7 | CAGACCAGCAGTAGCCATCA | ATCTCGTGAGGCCATCAAAC | 288 |
| chr9:69657720-69659734 | p2 | (GT)7 | GTTTGTGGCATTGATTGACG | AAATTCCAAAGTTTCGCCCT | 222 |
| chr9:69780788-69782938 | p2 | (AT)75 | AGCTAACACACCGAGCACCT | ATTTTAGTCGCATTGTCGGC | 274 |
| chr9:69872586-69874600 | p2 | (AT)7 | GGAGAGCTTTTGTGGAGTGC | GTAGACACAAGCTGGAGGGC | 158 |
| chr9:69887574-69889592 | p5 | (AATAT)3 | GACGCGGATCTTGAACTCTC | TCGTGCCATGAAGCATAGAG | 242 |
| chr9:69887574-69889592 | p6 | (CTGAAA)3 | GACGCGGATCTTGAACTCTC | TCGTGCCATGAAGCATAGAG | 242 |
| chr9:69890167-69892179 | p2 | (TA)6 | TTGAGCTTGTCCAAATGTCG | CTACTTCGGGCACCCATAAA | 177 |
| chr9:69975652-69977664 | p2 | (TA)6 | TGTGGGGCTATTTGGAATGT | ACTTGCTGCCCAAAAACACT | 173 |
| chr9:69975652-69977664 | p2 | (AT)7 | TGTGGGGCTATTTGGAATGT | ACTTGCTGCCCAAAAACACT | 173 |
| chr9:70010246-70012260 | p2 | (TA)7 | CCTTCCGGTGTCGGTAAGTA | ATATACCATACCCGGCCCTC | 194 |
| chr9:70071470-70073482 | p2 | (TA)6 | GGGAAGGTCGACAAGAATGA | TACGCTTGAACTACGACCCC | 177 |
| chr9:70151484-70153502 | p6 | (ATTAGG)3 | AAACTTTGGCGTTTGGTACG | GGATGGGAAGGACTCAATCA | 135 |
| chr9:70151484-70153502 | p5 | (GTGGG)3 | AAACTTTGGCGTTTGGTACG | GGATGGGAAGGACTCAATCA | 135 |
| chr9:70345061-70347073 | p2 | (TA)6 | TATGGCGATCAATTTCACGA | GGGCAACATCCTAAGCAAAA | 272 |
| chr9:70389043-70391058 | p3 | (TGT)5 | GCTAGCCGAGAATGGAGTTG | TAGCACACGACCCACTTCAA | 279 |
| chr9:70389043-70391058 | p2 | (TA)6 | GCTAGCCGAGAATGGAGTTG | TAGCACACGACCCACTTCAA | 279 |
| chr9:70473693-70475713 | p5 | (ATTCA)4 | TCATAGCACCGAACAAGTCG | GACACATGTTTCATCGCCAC | 118 |
| chr9:70493750-70495765 | p5 | (GTTCG)3 | TGGTGATCTCCCTTTATCCG | TCTTCAAAAATCCGAAACCG | 259 |
| chr9:70567906-70569924 | p6 | (AAAACA)3 | GATCTGACTTGGAGGGGACA | CGCTGGGTTAGTTTTTCTGC | 238 |
| chr9:70639530-70641566 | p2 | (TA)18 | TGTGGAAAATCGTGAGACCA | TGTTCATGCCCACAGGTAGA | 230 |
| chr9:70743002-70745017 | p5 | (ATCAT)3 | CATGGCACAGCATACTCCAC | GCCTCAATGTTTCCCACACT | 191 |
| chr9:70743002-70745017 | p3 | (CAA)5 | CATGGCACAGCATACTCCAC | GCCTCAATGTTTCCCACACT | 191 |
| chr9:70824886-70826908 | p2 | (TA)11 | GCATACATACGGGGCTGAGT | TTTGCTATGTTTGCGTCCAG | 227 |
| chr9:70824886-70826908 | p2 | (AT)7 | GCATACATACGGGGCTGAGT | TTTGCTATGTTTGCGTCCAG | 227 |
| chr9:71005101-71007116 | p5 | (ATTTC)3 | GAAGCGGAAGAAAGTATGCG | AATATCGATGCCTCACAGGG | 171 |
| chr9:71008289-71010310 | p3 | (ACA)7 | TTCCCAAGAACGTATCGGAG | GAGCATGCCGACGATTTTAT | 264 |
| chr9:71053094-71055106 | p2 | (AG)6 | ACCTATCGGTTGGTTCGATG | CCCTATGTATCTCGGGCTGA | 124 |
| chr9:71115436-71117460 | p2 | (AT)15 | AAGCTGACGAGGCTAACCAA | TAGCGGAGCTCCATTTCATT | 229 |
| chr9:71115436-71117460 | p6 | (CATATT)5 | AAGCTGACGAGGCTAACCAA | TAGCGGAGCTCCATTTCATT | 229 |
| chr9:71115436-71117460 | p2 | (AT)12 | AAGCTGACGAGGCTAACCAA | TAGCGGAGCTCCATTTCATT | 229 |
| chr9:71132961-71135015 | p3 | (AAT)18 | GTTACGGGTGTGCCTGAGAT | ACGAAGCAAGCCTCACAACT | 205 |
| chr9:71217084-71219104 | p5 | (GTTTT)4 | AAAGTTTTCAGGGGCGTTTT | AGGATCGGTCAGAGCCTGTA | 123 |
| chr9:71226098-71228113 | p5 | (TTTTG)3 | TCGACGACGTTTCATTTACG | TTGGAAGGTTTCGGATGAAG | 131 |
| chr9:71238772-71240790 | p2 | (TA)9 | TGTGTTCATGCCCAGAGGTA | TAAAGGAACCTCCCGAACAA | 261 |

| Marker ID | SSR type | SSR motif | FORWARD PRIMER (5'-3') | REVERSE PRIMER (5'-3') | size (bp) |
|---|---|---|---|---|---|
| chr9:71238772-71240790 | p2 | (AT)23 | TGTGTTCATGCCCAGAGGTA | TAAAGGAACCTCCCGAACAA | 261 |
| chr9:71239732-71241778 | p2 | (TA)9 | CACGTGCGTATCAATCCATC | AAATGACATGCGGAATAGCA | 228 |
| chr9:71239732-71241778 | p2 | (AT)23 | CACGTGCGTATCAATCCATC | AAATGACATGCGGAATAGCA | 228 |
| chr9:71542324-71544352 | p2 | (AT)14 | AATTCATTGGAAAGCGAACG | TCATGACTCATCGGTTTGGA | 270 |
| chr9:71542324-71544352 | p2 | (AT)10 | AATTCATTGGAAAGCGAACG | TCATGACTCATCGGTTTGGA | 270 |
| chr9:71545521-71547533 | p2 | (TA)6 | CATTGGATATTGAATCGGGG | CCATTCCAACCCACAATTTC | 289 |
| chr9:71596671-71598695 | p6 | (CATATA)4 | TGATACCACTTTTGCCACGA | CCATGAGTTCTCGGTACGGT | 145 |
| chr9:71666654-71668666 | p2 | (TA)6 | ATTCCCATTAGCCCGTCAAT | ATGCCATAGAAGGAAAGGGG | 237 |
| chr9:71690869-71692884 | p3 | (AAG)5 | AATTGATGGTCAGGAGTGGC | GTCCTTGCGTAGTGCGTACA | 199 |
| chr9:71845478-71847494 | p2 | (AT)8 | CCCGGTAATTTTCGCAACTA | TGGAATATTAGGCCCATCCA | 252 |
| chr9:71845478-71847494 | p2 | (AT)6 | CCCGGTAATTTTCGCAACTA | TGGAATATTAGGCCCATCCA | 252 |
| chr9:71850043-71852067 | p2 | (AT)8 | CCCGATGCCACACTCTAAAT | GTGTGCCAGTGTGAAGCATT | 190 |
| chr9:72021650-72023666 | p2 | (TA)8 | ACTTTTGTCACGACCCAACC | GTCGGCTTGCGTATCATTTT | 101 |
| chr9:72028944-72030966 | p2 | (TA)11 | TCCCTTCTGTTCATGGGTTC | CATCATCAAACAAGCCCCTT | 207 |
| chr9:72047688-72049704 | p2 | (AT)8 | TCCTAGGGTACGTTTGTCGG | TCGAGCGAACCCTTAGACAT | 179 |
| chr9:72047688-72049704 | p3 | (CTT)5 | TCCTAGGGTACGTTTGTCGG | TCGAGCGAACCCTTAGACAT | 179 |
| chr9:72110647-72112671 | p2 | (AT)12 | TGGATTGGCGAGCTCTATTT | GAGCCGCTTACAGGACTGAC | 254 |
| chr9:72110647-72112671 | p2 | (TA)8 | TGGATTGGCGAGCTCTATTT | GAGCCGCTTACAGGACTGAC | 254 |
| chr9:72157183-72159233 | p2 | (AT)25 | TTTCTGCTGGAGCTTCCACT | GGCCTTTTCTGTTTGCATGT | 280 |
| chr9:72168765-72170809 | p2 | (AT)22 | AAATGCCATGTAGCCTCCAG | CGTCTGGGTGTGACAAAGTG | 233 |
| chr9:72222508-72224523 | p3 | (AAG)5 | CTCCATCTGTCCATGGCTTT | AATTTCAGCTGTCACTGGGG | 185 |
| chr9:72351837-72353849 | p2 | (AT)6 | AGCTACATTCTCTTGCCCGA | GGGACCCGATTACCCTAAGA | 199 |
| chr9:72549765-72551777 | p2 | (CA)6 | TTACATACCTTGCCCGCTTC | TTGTTGGCATTGTTGTTGCT | 272 |
| chr9:72791772-72793812 | p2 | (TA)6 | CTAAGGCGGGCAGCTATATG | GTCAGTACGGGCAATGTCCT | 154 |
| chr9:72849476-72851491 | p5 | (AATTA)3 | CTAGCTATGGCACCATCCGT | TGAGTGGGGTAACTCAAGGC | 133 |
| chr9:72849476-72851491 | p2 | (AT)10 | CTAGCTATGGCACCATCCGT | TGAGTGGGGTAACTCAAGGC | 133 |
| chr9:72898054-72900076 | p2 | (TA)11 | CCCGGCTATAGTGTCCGATA | TCCCGTCTTGTAATGAAGCC | 176 |
| chr9:72968857-72970872 | p3 | (CTT)5 | AGGGTTCTTGGGTCCTTGAT | TGTGAGCAAGTCAACCAACC | 131 |
| chr9:73035820-73037866 | p6 | (TATATG)3 | TGCCCCTCAGTACAGGGTAG | CGTCCAGCCAAAGCATAAAT | 257 |
| chr9:73035820-73037866 | p2 | (TA)23 | TGCCCCTCAGTACAGGGTAG | CGTCCAGCCAAAGCATAAAT | 257 |
| chr9:73057811-73059823 | p2 | (CT)6 | TACCCGCATTTATTTCAGCC | CGCCAGTGTTTCCCCTATAA | 124 |
| chr9:73097206-73099218 | p2 | (TA)6 | GCTGTCGAATGCTGCTGATA | CAGGTTACTTTCTTGCCCCA | 258 |
| chr9:73097206-73099218 | p2 | (AT)6 | GCTGTCGAATGCTGCTGATA | CAGGTTACTTTCTTGCCCCA | 258 |
| chr9:73364677-73366719 | p6 | (ATATAC)7 | ACCATGGCTGTGGATGATTT | GATAAGAGGTGCCCGGTACA | 175 |
| chr9:73379323-73381341 | p3 | (CAA)5 | GAGGCCGAAATCACAACCTA | TGTATAGGGGAAATGCTGCC | 271 |
| chr9:73379323-73381341 | p2 | (CT)9 | GAGGCCGAAATCACAACCTA | TGTATAGGGGAAATGCTGCC | 271 |

| Marker ID | SSR type | SSR motif | FORWARD PRIMER (5'-3') | REVERSE PRIMER (5'-3') | size (bp) |
|---|---|---|---|---|---|
| chr9:73417706-73419738 | p2 | (AT)16 | TGTTATGGGAAATCGTGGGT | CTTAACAGAAATCGGGCAGC | 201 |
| chr9:73417706-73419738 | p2 | (TA)7 | TGTTATGGGAAATCGTGGGT | CTTAACAGAAATCGGGCAGC | 201 |
| chr9:73432572-73434604 | p2 | (AT)16 | TGTTATGGGAAATCGTGGGT | CTTAACAGAAATCGGGCAGC | 201 |
| chr9:73432572-73434604 | p2 | (TA)7 | TGTTATGGGAAATCGTGGGT | CTTAACAGAAATCGGGCAGC | 201 |
| chr9:73534807-73536825 | p2 | (CA)6 | CATGCATTTCCATTCATTCG | TTCGGAGCCATATCTTTTGG | 187 |
| chr9:73534807-73536825 | p2 | (AT)9 | CATGCATTTCCATTCATTCG | TTCGGAGCCATATCTTTTGG | 187 |
| chr9:73554646-73556712 | p2 | (TA)33 | TTCTTTCTCACCCGAAATGG | ATGTCAAGGCCCATGGATTA | 232 |
| chr9:73601678-73603699 | p3 | (ATC)7 | TGCTGATACAGCACCTACGG | ATGCATGGTATCCGCCTAAG | 300 |
| chr9:73606269-73608281 | p2 | (CA)6 | CAATTTGGGGGTTTTCCTTT | CTCCTTTGCCTTGCTCATTC | 253 |
| chr9:73655116-73657131 | p2 | (AT)6 | TGCTCTATTGTGTTGGCTCG | TGAAGAGAGATGTGGATGCG | 147 |
| chr9:73655116-73657131 | p5 | (GCATT)3 | TGCTCTATTGTGTTGGCTCG | TGAAGAGAGATGTGGATGCG | 147 |
| chr9:73666493-73668509 | p2 | (TA)8 | TTCCATGAACCACCTTGTGA | ATCTTGATAGGGGCTTCGGT | 137 |
| chr9:73666493-73668509 | p2 | (TA)10 | TTCCATGAACCACCTTGTGA | ATCTTGATAGGGGCTTCGGT | 137 |
| chr9:73691268-73693282 | p2 | (TA)7 | CGTTCCTCCACCAAATGTCT | TGTTGAACGAACTGAGTGCC | 127 |
| chr9:73707757-73709769 | p2 | (AC)6 | AATCCGCCATCTAATTTCCC | CCTTAGCCTTGCCCTCTCTT | 192 |
| chr9:73711698-73713752 | p2 | (TG)6 | TTCCGTAATTGCCTTGAACC | CCTTGGAATATCCGAGCGTA | 287 |
| chr9:73730600-73732615 | p3 | (TGT)5 | TCGAAATATTCCGGGATCAA | CGATGTCGTTTTGATGAACG | 240 |
| chr9:73730600-73732615 | p3 | (ATG)5 | TCGAAATATTCCGGGATCAA | CGATGTCGTTTTGATGAACG | 240 |
| chr9:73817098-73819110 | p2 | (TG)6 | AAAGGTGCCTACGGGCTATT | CCCAACCAAGCTCCAAGTTA | 177 |
| chr9:73840686-73842720 | p2 | (TA)17 | TGCCAACCCCAGAAAACTAC | GTACTTGGGATACTCCGGCA | 148 |
| chr9:73862888-73864930 | p2 | (AT)21 | CATTTCTGAAGAAACCCCCA | TTGTGACGTGTTGTGGTGTG | 111 |
| chr9:73997263-73999278 | p5 | (CAAAA)3 | ATACGGTCCCAACATACGGA | AGAGAAGGCAAGGCATACGA | 298 |
| chr9:74022363-74024375 | p2 | (TA)6 | TCAATCGGCTATCTGCTCCT | TGTTTGCTTCTTTCCTGCCT | 156 |
| chr9:74058233-74060247 | p2 | (AT)7 | TCTAAAGGGCACTCGAAGGA | TTCGGGTCAATCCGACTTAG | 212 |
| chr9:74153800-74155818 | p2 | (AT)7 | GTCTCCCTGCAGAAGTTTCG | GTCGATACGGAAGGACGAGA | 113 |
| chr9:74153800-74155818 | p2 | (AT)9 | GTCTCCCTGCAGAAGTTTCG | GTCGATACGGAAGGACGAGA | 113 |
| chr9:74177855-74179870 | p3 | (GTT)5 | GACCGACGGAACTTGTGAAT | AATCATCCTTTTGGAGGGCT | 105 |
| chr9:74191826-74193844 | p2 | (AT)9 | GATCGGTTTCCTACTGCCAA | TCGAGGAGGAAGGAACTCAA | 154 |
| chr9:74196518-74198534 | p4 | (CATA)4 | TTTTACCATTTCTCGGCTGG | ATTGCCCAGTGGTCGTAAAG | 280 |
| chr9:74216627-74218645 | p6 | (CATATA)3 | ACCTTAGGGGACTGCTGGTT | AGGGGTGCTCGGTAGCTAAT | 169 |
| chr9:74216627-74218645 | p3 | (CAA)6 | ACCTTAGGGGACTGCTGGTT | AGGGGTGCTCGGTAGCTAAT | 169 |
| chr9:74232658-74234694 | p2 | (AT)18 | CATAAGCCGACGAGGCTAAC | GCGGAGCTTCACTTTATTCG | 233 |
| chr9:74244976-74246994 | p6 | (CTTACA)3 | ATACCGCTTTTGTGGTCCTG | GGTATGTGAGCAGATGGCCT | 155 |
| chr9:74260730-74262756 | p2 | (AT)13 | TTGTTCCGCAGATACCATCA | CCCATCAGAGGTTCAGGAAA | 195 |
| chr9:74275709-74277727 | p2 | (AT)9 | GTGATGGGAAGCCTTTACGA | ACGAATTACCAAACCTCCCC | 294 |
| chr9:74275709-74277727 | p2 | (AT)9 | GTGATGGGAAGCCTTTACGA | ACGAATTACCAAACCTCCCC | 294 |

| Marker ID | SSR type | SSR motif | FORWARD PRIMER (5'-3') | REVERSE PRIMER (5'-3') | size (bp) |
|---|---|---|---|---|---|
| chr9:74353065-74355083 | p2 | (AT)9 | TGCCCATAGGATGGTTTGAT | CGCCCATCTTGAACTCTAGC | 124 |
| chr9:74372305-74374317 | p2 | (AT)6 | TCACACTGCCGTTCTTTCAC | CCGATCAATTGGCAGTTTTT | 246 |
| chr9:74372305-74374317 | p2 | (AT)16 | TCACACTGCCGTTCTTTCAC | CCGATCAATTGGCAGTTTTT | 246 |
| chr9:74412717-74414735 | p3 | (TTG)6 | CCAAACCCAAGAAAAACCAA | ATACACGGCTCACCTCCAAC | 299 |
| chr9:74459901-74461916 | p3 | (GTT)5 | CGTCATGCCCTATCAGGACT | CAACACCGGAAGCACCTTAT | 143 |
| chr9:74540795-74542820 | p5 | (TTCAT)5 | TGGAGAGGAAATGACTTGGG | AAGTTTGGCCATACACGACC | 188 |
| chr9:74568968-74570986 | p3 | (ATC)6 | TCAATCCCATGAGGAAAAGG | CGGAATGCAAACAAAAGGTT | 149 |
| chr9:74568968-74570986 | p3 | (AAC)5 | TCAATCCCATGAGGAAAAGG | CGGAATGCAAACAAAAGGTT | 149 |
| chr9:74569337-74571352 | p3 | (ATC)6 | GACGTTTCATGCGGTTTCTT | GTGACAAAAGGAAAAGGGCA | 138 |
| chr9:74569337-74571352 | p3 | (AAC)5 | GACGTTTCATGCGGTTTCTT | GTGACAAAAGGAAAAGGGCA | 138 |
| chr9:74578396-74580414 | p2 | (AT)9 | GGCCGCCATAACATATCATC | CGTGCATGGCTCAATATGTC | 114 |
| chr9:74578396-74580414 | p3 | (AAC)6 | GGCCGCCATAACATATCATC | CGTGCATGGCTCAATATGTC | 114 |
| chr9:74584756-74586788 | p2 | (AT)16 | AGTGGCGGTGTAGCAGAACT | CAAATCGGTAGCAGGGGATA | 229 |
| chr9:74584756-74586788 | p2 | (AT)7 | AGTGGCGGTGTAGCAGAACT | CAAATCGGTAGCAGGGGATA | 229 |
| chr9:74603805-74605817 | p2 | (AT)6 | TGAATGTGTTATAGCCCGCA | CCCCATTTTTGAATTTCCCT | 248 |
| chr9:74608753-74610765 | p2 | (TC)6 | TTTCATTCAAATCAACCGCA | GCTTGTGTATTGGATGGGCT | 157 |
| chr9:74608753-74610765 | p2 | (AT)6 | TTTCATTCAAATCAACCGCA | GCTTGTGTATTGGATGGGCT | 157 |
| chr9:74623769-74625787 | p3 | (ATA)6 | GATCCCACTCCCCCACTATT | TCTGACACCTCCACTACCCC | 129 |
| chr9:74660455-74662469 | p2 | (AT)7 | AGGTGAAGTGTGCTCCTCGT | TGATATCCAGGGAGGCAGTC | 153 |
| chr9:74685298-74687314 | p2 | (TA)8 | GTATGTCCGTATGGATCCCG | ATCATGATCTGGACCGGAAG | 218 |
| chr9:74685298-74687314 | p2 | (TA)37 | GTATGTCCGTATGGATCCCG | ATCATGATCTGGACCGGAAG | 218 |
| chr9:74698351-74700384 | p3 | (ATC)5 | GCATGGATGCCAATGATATG | CGGTGCTACCCCTTATCGTA | 147 |
| chr9:74698351-74700384 | p3 | (AAC)11 | GCATGGATGCCAATGATATG | CGGTGCTACCCCTTATCGTA | 147 |
| chr9:74730926-74732938 | p2 | (GT)6 | GAGTGCCCGGGAGTATAACA | GAATTTGCCCTTAGCCTTCC | 263 |
| chr9:74859954-74861972 | p2 | (AT)9 | TAGCCTTGTCCCACACTTCC | CTGCCATTCTCCCATGTTTT | 290 |
| chr9:74914867-74916879 | p5 | (TGATA)3 | TGCATGCATTGGTTACGTTT | CAGCAAATCGAGCGTCTGTA | 153 |
| chr9:74999970-75001990 | p5 | (AAACC)4 | AGATACGAGCACAAAGCCGT | AATCCCCCACATGATTCAAA | 221 |
| chr9:75029520-75031535 | p5 | (ATATA)3 | ATTGTCACGACCCGACTAGG | TGGCTCTGTTTACATGGCAG | 144 |
| chr9:75243556-75245576 | p5 | (AAATA)4 | TTTGGAAGAAGAAAAGCGGA | TTGGATCTTTGCTTGCTTCA | 200 |
| chr9:75409689-75411701 | p2 | (CA)6 | CCAATGAATTCTGGGTGGTC | CCAGTTAGTAGCCGCTCAGG | 278 |
| chr9:75412668-75414680 | p2 | (AT)7 | AAGGGCTTCATTCCCACTTT | AGGCTATGCATCCACCAAAC | 178 |
| chr9:75412668-75414680 | p2 | (AT)6 | AAGGGCTTCATTCCCACTTT | AGGCTATGCATCCACCAAAC | 178 |
| chr9:75412721-75414735 | p2 | (AT)7 | AAGGGCTTCATTCCCACTTT | AGGCTATGCATCCACCAAAC | 125 |
| chr9:75412721-75414735 | p2 | (AT)6 | AAGGGCTTCATTCCCACTTT | AGGCTATGCATCCACCAAAC | 125 |
| chr9:75412739-75414757 | p2 | (AT)7 | AAGGGCTTCATTCCCACTTT | AGGCTATGCATCCACCAAAC | 107 |
| chr9:75412739-75414757 | p2 | (AT)6 | AAGGGCTTCATTCCCACTTT | AGGCTATGCATCCACCAAAC | 107 |

| Marker ID | SSR type | SSR motif | FORWARD PRIMER (5'-3') | REVERSE PRIMER (5'-3') | size (bp) |
|---|---|---|---|---|---|
| chr9:75439583-75441601 | p6 | (ATACAT)3 | AAACTCATGAAATCACGGGC | GTTGCTTCGGGTCAACAAGT | 155 |
| chr9:75439583-75441601 | p2 | (AT)26 | AAACTCATGAAATCACGGGC | GTTGCTTCGGGTCAACAAGT | 155 |
| chr9:75530712-75532740 | p2 | (TA)14 | CGTCAATTGCACATACCTGG | TGGGGATTATTTCGTTGGTG | 140 |
| chr9:75604383-75606433 | p2 | (AT)25 | CAGGTGCGTGGTATGCTCTA | GCAGCCGTTCATGGATTATT | 151 |
| chr9:75605618-75607633 | p3 | (CAA)5 | GCTCGGTGGTATCTTACCCA | CTCGAGACTCCCGTGACATT | 236 |
| chr9:75674884-75676922 | p2 | (TA)19 | GCAAATACGGATCGGAAAAA | CTCAGCCGAGTAGGGTCTTG | 237 |
| chr9:75680644-75682659 | p5 | (AATGC)3 | CCCTCTCGTCTAAACAACGC | TTTCATTCCGAGATTGGAGG | 156 |
| chr9:75680644-75682659 | p2 | (AT)31 | CCCTCTCGTCTAAACAACGC | TTTCATTCCGAGATTGGAGG | 156 |
| chr9:75724179-75726199 | p6 | (GCTCCA)4 | GGGTCGTGACACCATTTCTT | GCATGCATGAGAGCACAAGT | 117 |
| chr9:75940324-75942339 | p5 | (TTGTA)3 | TACTTGTGAGCGAGACCCCT | CCTAGGCTTTTTACCTCCCG | 219 |
| chr9:76010958-76012973 | p3 | (GTG)5 | ACCATGCCTAGTTGGTCAGG | TCGCAACTTTGTGTTTGAGG | 271 |
| chr9:76145419-76147433 | p2 | (TA)7 | CCTCTCGGCCATCAAAATAA | GTTCAAGCCCACAGGGTAGA | 167 |
| chr9:76173943-76175957 | p2 | (TA)7 | GCAGTAGGTCAGGGGCTACA | AGAAACAGCCTCCAAAAGCA | 172 |
| chr9:76189096-76191108 | p2 | (GA)6 | TGGGGCCTGTTAGTTCTTTG | CCTTCAGTACCAGACCCCAA | 183 |
| chr9:76257341-76259431 | p3 | (AAT)30 | GTCCCGTGAAAGTTAACCGA | CAGGAACAGTAGGAGCAGGC | 109 |
| chr9:76292341-76294357 | p2 | (AG)8 | CCTTCACTGGTTCCGTGTTT | CCCCCTAAACCTCCTACTCG | 251 |
| chr9:76292341-76294357 | p4 | (TAAA)4 | CCTTCACTGGTTCCGTGTTT | CCCCCTAAACCTCCTACTCG | 251 |
| chr9:76295192-76297207 | p5 | (CCACT)3 | ATGCCCAGCCGTACTACATC | CAAAAATTGAACCCCACACC | 213 |
| chr9:76326774-76328789 | p5 | (TGCAT)3 | GCCATGTGTAGCCGGTAGAT | TCGGCCATGCATGATAAATA | 187 |
| chr9:76386616-76388631 | p3 | (ACA)5 | GAGGGGCTGATGAAAAACAA | AAAAACACACCAGCCCACTC | 258 |
| chr9:76579029-76581075 | p2 | (TA)23 | CCCTAGGACGAACTGCTCTG | CACAGCGAGAGCTTAAAGGG | 293 |
| chr9:76592900-76594940 | p2 | (AT)20 | GGGAATCCATCTTTGTGCAT | TCACTTGCATGCTTCAGACC | 218 |
| chr9:76644875-76646893 | p6 | (TTGTTT)3 | TGGTTGATGATTGGCGTTTA | TCCTGTGACAATCCCTTTCC | 224 |
| chr9:76660499-76662513 | p2 | (TG)7 | GAATCTTTACGTCACCCCGA | AGCTGTTCCATGTATTCCCG | 109 |
| chr9:76708903-76710947 | p2 | (AT)22 | GGGGAGGGGTCACTAACCTA | TTGGTGGTTCTCATTGGTGA | 115 |
| chr9:76711402-76713417 | p5 | (AAAAT)3 | GAACGCAAAGGCAGACATTT | ACGCTTGATTTCACTTGCCT | 238 |
| chr9:76711402-76713417 | p3 | (CAG)5 | GAACGCAAAGGCAGACATTT | ACGCTTGATTTCACTTGCCT | 238 |
| chr9:76720708-76722722 | p2 | (AT)7 | TACAACCAAGAGCAGGGACC | ATGGCTATATGCACCCGGTA | 292 |
| chr9:76837003-76839018 | p5 | (GCATT)3 | CCTTGATGAGGAATTTCCGA | CTTCCTTCACCGCCTACAAA | 195 |
| chr9:76846673-76848688 | p3 | (TTG)5 | CCTTAAGCCAAATATGGGCA | CGGTTCCACCAACCTTCTTA | 164 |
| chr9:76846673-76848688 | p3 | (TGA)6 | CCTTAAGCCAAATATGGGCA | CGGTTCCACCAACCTTCTTA | 164 |
| chr9:76846673-76848688 | p4 | (TGTA)5 | CCTTAAGCCAAATATGGGCA | CGGTTCCACCAACCTTCTTA | 164 |
| chr9:76920560-76922575 | p5 | (GAAAT)3 | AGCACATCAAGCATCGTGAG | AGGTATAAACGCAAGCGTGG | 165 |
| chr9:76997129-76999147 | p6 | (CATATA)3 | TGTCTCCCGAACAATCAACA | CATATATGGGTACGGCAGGG | 267 |
| chr9:77193713-77195781 | p2 | (AT)34 | CGAGGTAAGGGTTCTGGTGA | GTGTGTCCTAGTCCGGCAAT | 246 |
| chr9:77212351-77214366 | p2 | (AT)11 | TTGGGTAATCTGACTTCGGG | TGAGGAGGAGAAGGCTGAAA | 177 |

| Marker ID | SSR type | SSR motif | FORWARD PRIMER (5'-3') | REVERSE PRIMER (5'-3') | size (bp) |
|---|---|---|---|---|---|
| chr9:77212351-77214366 | p3 | (GTT)5 | TTGGGTAATCTGACTTCGGG | TGAGGAGGAGAAGGCTGAAA | 177 |
| chr9:77212351-77214366 | p3 | (TTG)5 | TTGGGTAATCTGACTTCGGG | TGAGGAGGAGAAGGCTGAAA | 177 |
| chr9:77212351-77214366 | p5 | (TATAT)3 | TTGGGTAATCTGACTTCGGG | TGAGGAGGAGAAGGCTGAAA | 177 |
| chr9:77247002-77249017 | p5 | (GCATT)3 | CCTGTTGTGTTGTGTTTGCC | ACCACAGTGGGTATGCCAAT | 173 |
| chr9:77365167-77367185 | p5 | (GAAAT)3 | TTTGGCGTTGTTGATCTTGA | TAAACTCGGTTTACCGCCAC | 290 |
| chr9:77365167-77367185 | p3 | (TTG)6 | TTTGGCGTTGTTGATCTTGA | TAAACTCGGTTTACCGCCAC | 290 |
| chr9:77422292-77424318 | p2 | (TA)13 | GTCTCGGGAGGCATGAATAA | TACCCGATCCCGTGATTTTA | 173 |
| chr9:77475882-77477912 | p2 | (AC)15 | CGGGGTCACCTTTTTAGGTT | CAAACTAGCCAAATCCGAGC | 230 |
| chr9:77515906-77517921 | p3 | (TGT)5 | TCAGTATCTAGATGGCCCCG | CGACGGTTGTCTGCATAAGA | 280 |
| chr9:77515906-77517921 | p2 | (TA)6 | TCAGTATCTAGATGGCCCCG | CGACGGTTGTCTGCATAAGA | 280 |
| chr9:77516590-77518602 | p3 | (TGT)5 | GGAGATGACTTGGGAAGCAG | TCCTGTCACAACAACCCAAA | 167 |
| chr9:77516590-77518602 | p2 | (TA)6 | GGAGATGACTTGGGAAGCAG | TCCTGTCACAACAACCCAAA | 167 |
| chr9:77554872-77556886 | p3 | (CAA)5 | AGTCAAAGACATCGGCCATC | TTAGCGAGCGATCGAGGTAT | 167 |
| chr9:77554872-77556886 | p2 | (CA)7 | AGTCAAAGACATCGGCCATC | TTAGCGAGCGATCGAGGTAT | 167 |
| chr9:77562032-77564046 | p3 | (CAA)5 | AGTCAAAGACATCGGCCATC | TTAGCGAGCGATCGAGGTAT | 167 |
| chr9:77562032-77564046 | p2 | (CA)7 | AGTCAAAGACATCGGCCATC | TTAGCGAGCGATCGAGGTAT | 167 |
| chr9:77612915-77614929 | p3 | (CAA)5 | ATACCTCGATCGCTCGCTAA | CTCGTTCTGGGGTGTTGATT | 277 |
| chr9:77612915-77614929 | p2 | (CA)7 | ATACCTCGATCGCTCGCTAA | CTCGTTCTGGGGTGTTGATT | 277 |
| chr9:77612915-77614929 | p5 | (GTTTT)3 | ATACCTCGATCGCTCGCTAA | CTCGTTCTGGGGTGTTGATT | 277 |
| chr9:77686388-77688414 | p2 | (TA)13 | GACCCCACTGGACACAGTTT | ATTGCCTTGTTACGGTCAGG | 285 |
| chr9:77702726-77704750 | p5 | (CATAT)3 | CCAAGCTGGACACTCGTACA | TTGCCACCTTGCACATGTAT | 169 |
| chr9:77848331-77850351 | p2 | (TA)10 | GTCCTTCTTTTCCCACACCA | ACCCTTTTCCCCTTGTCATC | 261 |
| chr9:77859333-77861361 | p2 | (AT)14 | TACCTCTTGCGGATCCATTC | TGTGGAAGCCACAAACAAAA | 106 |
| chr9:77859333-77861361 | p3 | (TTG)7 | TACCTCTTGCGGATCCATTC | TGTGGAAGCCACAAACAAAA | 106 |
| chr9:77898769-77900835 | p2 | (AT)33 | CACGGTGATTTCCATAGGCT | TGAATGGCATTATGGCAAGA | 133 |
| chr9:77898769-77900835 | p5 | (TTAAA)3 | CACGGTGATTTCCATAGGCT | TGAATGGCATTATGGCAAGA | 133 |
| chr9:78083138-78085152 | p2 | (TA)7 | CACCTCAGAAATGGGCGTAT | TTTGAAGGTGGACAAGCACA | 132 |
| chr9:78329121-78331142 | p3 | (ATC)7 | TGATACCATTTTTGCCACGA | GTCGGCTTGTACAATTGGGT | 141 |
| chr9:78352787-78354799 | p2 | (AC)6 | CTCCAATGCCTAAGGAACCA | TTATTGCCGGCTCAGGTATC | 208 |
| chr9:78387377-78389403 | p2 | (TA)13 | AAGGCCTCCATCCTTCTCAT | CTTTTCCCATGACGTGGTCT | 156 |
| chr9:78387377-78389403 | p5 | (TGAAT)3 | AAGGCCTCCATCCTTCTCAT | CTTTTCCCATGACGTGGTCT | 156 |
| chr9:78393291-78395306 | p5 | (TGAAT)3 | ACATGTGGCCACCCTTGTAT | TTGTTCCAACAACCACTCCA | 263 |
| chr9:78460071-78462086 | p2 | (AT)13 | TTTGGGATATGTAGGCGCTC | CTACATTTTCCGTTTCGGGA | 211 |
| chr9:78460071-78462086 | p5 | (CAAAG)3 | TTTGGGATATGTAGGCGCTC | CTACATTTTCCGTTTCGGGA | 211 |
| chr9:78525257-78527272 | p3 | (ATA)5 | GGGACAAGGTGAGTGGAAAA | CACCCCACTACACCCATACC | 292 |
| chr9:78540273-78542289 | p2 | (AT)8 | ATAGGATCTGGCCCACACTG | CCATAAGATGGTACCCGACG | 251 |

| Marker ID | SSR type | SSR motif | FORWARD PRIMER (5'-3') | REVERSE PRIMER (5'-3') | size (bp) |
|---|---|---|---|---|---|
| chr9:78596991-78599007 | p2 | (TC)8 | ATACGCAAAAGTTCGGATGG | TTGGTTGCTTTCCCTTTGAC | 139 |
| chr9:78659028-78661048 | p2 | (TA)8 | GGCATGTTTTTCCAAGAGGT | TCATGAACAAGTCACTGCGT | 132 |
| chr9:78659028-78661048 | p4 | (TTAT)4 | GGCATGTTTTTCCAAGAGGT | TCATGAACAAGTCACTGCGT | 132 |
| chr9:78659028-78661048 | p2 | (TA)14 | GGCATGTTTTTCCAAGAGGT | TCATGAACAAGTCACTGCGT | 132 |
| chr9:78659028-78661048 | p2 | (TA)11 | GGCATGTTTTTCCAAGAGGT | TCATGAACAAGTCACTGCGT | 132 |
| chr9:78659028-78661048 | p2 | (TA)12 | GGCATGTTTTTCCAAGAGGT | TCATGAACAAGTCACTGCGT | 132 |
| chr9:78659045-78661066 | p2 | (TA)8 | GGCATGTTTTTCCAAGAGGT | TCATGAACAAGTCACTGCGT | 115 |
| chr9:78659045-78661066 | p4 | (TTAT)4 | GGCATGTTTTTCCAAGAGGT | TCATGAACAAGTCACTGCGT | 115 |
| chr9:78659045-78661066 | p2 | (TA)14 | GGCATGTTTTTCCAAGAGGT | TCATGAACAAGTCACTGCGT | 115 |
| chr9:78659045-78661066 | p2 | (TA)11 | GGCATGTTTTTCCAAGAGGT | TCATGAACAAGTCACTGCGT | 115 |
| chr9:78659045-78661066 | p2 | (TA)12 | GGCATGTTTTTCCAAGAGGT | TCATGAACAAGTCACTGCGT | 115 |
| chr9:78659045-78661066 | p2 | (AT)8 | GGCATGTTTTTCCAAGAGGT | TCATGAACAAGTCACTGCGT | 115 |
| chr9:78670410-78672425 | p2 | (TA)6 | CGCTTTGTGTACTTCTGCCA | CTTTGTTCTTCGGGAATCCA | 212 |
| chr9:78670410-78672425 | p3 | (TTA)6 | CGCTTTGTGTACTTCTGCCA | CTTTGTTCTTCGGGAATCCA | 212 |
| chr9:78670486-78672504 | p2 | (TA)6 | CGCTTTGTGTACTTCTGCCA | CTTTGTTCTTCGGGAATCCA | 136 |
| chr9:78670486-78672504 | p3 | (TTA)6 | CGCTTTGTGTACTTCTGCCA | CTTTGTTCTTCGGGAATCCA | 136 |
| chr9:78670486-78672504 | p2 | (TA)6 | CGCTTTGTGTACTTCTGCCA | CTTTGTTCTTCGGGAATCCA | 136 |
| chr9:78670664-78672682 | p2 | (TA)6 | TGCACTGCCTGTTTTGAAAG | ACAACTACCGACTATGCCGC | 116 |
| chr9:78670664-78672682 | p3 | (TTA)6 | TGCACTGCCTGTTTTGAAAG | ACAACTACCGACTATGCCGC | 116 |
| chr9:78677624-78679645 | p3 | (ATT)5 | TTTGTTGCAAAGTGGCTTGT | ATGATCTTCTGCTGGCTGCT | 174 |
| chr9:78677624-78679645 | p2 | (TA)12 | TTTGTTGCAAAGTGGCTTGT | ATGATCTTCTGCTGGCTGCT | 174 |
| chr9:78677624-78679645 | p2 | (AT)11 | TTTGTTGCAAAGTGGCTTGT | ATGATCTTCTGCTGGCTGCT | 174 |
| chr9:78677624-78679645 | p2 | (TA)6 | TTTGTTGCAAAGTGGCTTGT | ATGATCTTCTGCTGGCTGCT | 174 |
| chr9:78677624-78679645 | p3 | (TAG)5 | TTTGTTGCAAAGTGGCTTGT | ATGATCTTCTGCTGGCTGCT | 174 |
| chr9:78677624-78679645 | p2 | (TA)9 | TTTGTTGCAAAGTGGCTTGT | ATGATCTTCTGCTGGCTGCT | 174 |
| chr9:78677624-78679645 | p2 | (AT)8 | TTTGTTGCAAAGTGGCTTGT | ATGATCTTCTGCTGGCTGCT | 174 |
| chr9:78677624-78679645 | p2 | (TA)7 | TTTGTTGCAAAGTGGCTTGT | ATGATCTTCTGCTGGCTGCT | 174 |
| chr9:78677647-78679662 | p3 | (ATT)5 | TTTGTTGCAAAGTGGCTTGT | ATGATCTTCTGCTGGCTGCT | 151 |
| chr9:78677647-78679662 | p2 | (TA)12 | TTTGTTGCAAAGTGGCTTGT | ATGATCTTCTGCTGGCTGCT | 151 |
| chr9:78677647-78679662 | p2 | (AT)11 | TTTGTTGCAAAGTGGCTTGT | ATGATCTTCTGCTGGCTGCT | 151 |
| chr9:78677647-78679662 | p2 | (TA)6 | TTTGTTGCAAAGTGGCTTGT | ATGATCTTCTGCTGGCTGCT | 151 |
| chr9:78677647-78679662 | p3 | (TAG)5 | TTTGTTGCAAAGTGGCTTGT | ATGATCTTCTGCTGGCTGCT | 151 |
| chr9:78677647-78679662 | p2 | (TA)9 | TTTGTTGCAAAGTGGCTTGT | ATGATCTTCTGCTGGCTGCT | 151 |
| chr9:78677647-78679662 | p2 | (AT)8 | TTTGTTGCAAAGTGGCTTGT | ATGATCTTCTGCTGGCTGCT | 151 |
| chr9:78677647-78679662 | p2 | (TA)7 | TTTGTTGCAAAGTGGCTTGT | ATGATCTTCTGCTGGCTGCT | 151 |
| chr9:78815301-78817313 | p2 | (CA)6 | CAAAATCCGAAAGTCACCGT | TGATCCATAGACCGCTACCC | 102 |

| Marker ID | SSR type | SSR motif | FORWARD PRIMER (5'-3') | REVERSE PRIMER (5'-3') | size (bp) |
|---|---|---|---|---|---|
| chr9:78863532-78865574 | p5 | (CAAAT)3 | TAAACAAGGGAAATGCTGCC | GGCTTTTGCAGACAAGAAGG | 176 |
| chr9:78863532-78865574 | p3 | (TTG)6 | TAAACAAGGGAAATGCTGCC | GGCTTTTGCAGACAAGAAGG | 176 |
| chr9:78863593-78865611 | p3 | (TTG)6 | TAAACAAGGGAAATGCTGCC | GGCTTTTGCAGACAAGAAGG | 115 |
| chr9:79021121-79023139 | p6 | (ATTGTT)3 | TCGGAGTGAGTTGCTGACAC | TTCCAACTCCTCGTCTTGCT | 279 |
| chr9:79021121-79023139 | p2 | (AT)6 | TCGGAGTGAGTTGCTGACAC | TTCCAACTCCTCGTCTTGCT | 279 |
| chr9:79347289-79349307 | p2 | (AT)9 | CCCAGTTAGTAGCCGCTCAG | CTGGCTTGAATCACCCCTAA | 210 |
| chr9:79365183-79367198 | p5 | (TACAA)3 | ATGTCTCGAAGCCTTCTCCA | GGTTCTTGTGCTCATGGGAT | 262 |
| chr9:79463896-79465911 | p4 | (TATG)13 | AGACGAATGGACGAAGATGG | ACACTTGCACATTCCCTTCC | 144 |
| chr9:79463896-79465911 | p3 | (ATG)5 | AGACGAATGGACGAAGATGG | ACACTTGCACATTCCCTTCC | 144 |
| chr9:79463896-79465911 | p2 | (TG)6 | AGACGAATGGACGAAGATGG | ACACTTGCACATTCCCTTCC | 144 |
| chr9:79537130-79539180 | p2 | (AT)25 | AGACATATGGACGGGACAGG | GGGAGGTGATCCTGATCTGA | 127 |
| chr9:79537130-79539180 | p3 | (TCA)5 | AGACATATGGACGGGACAGG | GGGAGGTGATCCTGATCTGA | 127 |
| chr9:79630463-79632499 | p2 | (CT)20 | GCCAAAGGCATCATTCTCAT | GTTCGCTGCCGAAGAAATAG | 201 |
| chr9:79630501-79632546 | p2 | (CT)20 | GCCAAAGGCATCATTCTCAT | GTTCGCTGCCGAAGAAATAG | 163 |
| chr9:79716928-79718946 | p6 | (GTAGCA)3 | TGTGCAAATGGGTGAAACAT | CTGCAGCCTATCCCTTCAAA | 207 |
| chr9:79811751-79813763 | p2 | (TA)18 | TGTGGTTGGGCTGATTTACA | TCCCACACGATGTTTTTCAA | 233 |
| chr9:79811751-79813763 | p2 | (AT)6 | TGTGGTTGGGCTGATTTACA | TCCCACACGATGTTTTTCAA | 233 |
| chr9:79937963-79939975 | p2 | (TA)6 | TGGAAAATAAGCGTTTTGGC | CAGCACTCCAGAAAGGAAGG | 141 |
| chr9:79937963-79939975 | p5 | (AAGGA)3 | TGGAAAATAAGCGTTTTGGC | CAGCACTCCAGAAAGGAAGG | 141 |
| chr9:80013551-80015585 | p2 | (TA)10 | GGGTGGAAAAGAACCTGTCA | GCCTTGTTTTGGGTTCTTGA | 150 |
| chr9:80013551-80015585 | p2 | (AT)17 | GGGTGGAAAAGAACCTGTCA | GCCTTGTTTTGGGTTCTTGA | 150 |
| chr9:80044762-80046782 | p2 | (TA)10 | AACAGGAAGAAGCGTGGAGA | GCAAAACCGATTGTCGACTT | 198 |
| chr9:80108299-80110317 | p3 | (CAA)6 | TTATCACCCGACGGTAGAGC | AGTCGGGCAAATGTTACCAC | 121 |
| chr9:80118890-80120916 | p2 | (TA)7 | ACTCATAGTGGCAGCCTCGT | ATGGACCCATAGGACCAACA | 160 |
| chr9:80118890-80120916 | p2 | (TA)13 | ACTCATAGTGGCAGCCTCGT | ATGGACCCATAGGACCAACA | 160 |
| chr9:80149960-80152010 | p2 | (TA)25 | TTTGAAACGAGCAGATGACG | TGGAATGGTCAAACGTGCTA | 171 |
| chr9:80238968-80240983 | p5 | (TCATT)3 | ACAGTAACCCCTGCATCACC | TCAAAGGCGATGGCTAAAGT | 119 |
| chr9:80253819-80255879 | p3 | (TTG)5 | TGGTGTGAATGCTGTTCGAT | TTCAACGGCAGGTTTCTTCT | 128 |
| chr9:80424074-80426096 | p2 | (AT)11 | CTTTTCGTAATTGGGTCCGA | CCCAAATGAGGTCTCCTTGA | 110 |
| chr9:80449402-80451420 | p6 | (CATATA)3 | ATAACGTATCCCGACCCTCC | TCTTGTGCGTACTCCACTGC | 144 |
| chr9:80449402-80451420 | p3 | (CAA)6 | ATAACGTATCCCGACCCTCC | TCTTGTGCGTACTCCACTGC | 144 |
| chr9:80521091-80523105 | p2 | (TA)7 | GTTAGTGGCCGCTTTATCCA | TCTCACTAGCGTGCATCACC | 287 |
| chr9:80521091-80523105 | p5 | (TGAAT)3 | GTTAGTGGCCGCTTTATCCA | TCTCACTAGCGTGCATCACC | 287 |
| chr9:80615377-80617399 | p2 | (AT)11 | TTGCTCTGTGGGTATTGCAG | TCTTCGAAATTACATGCCCC | 134 |
| chr9:80650394-80652408 | p2 | (AT)7 | CTTTTCGTGTAGAGAGCCCG | CTGACTGAGCACCTTCCCTC | 211 |
| chr9:80676730-80678750 | p2 | (TA)10 | CATCAAAACTTGTTGGGCCT | CCACCAGTGCCCATTTTTAC | 151 |

| Marker ID | SSR type | SSR motif | FORWARD PRIMER (5'-3') | REVERSE PRIMER (5'-3') | size (bp) |
|---|---|---|---|---|---|
| chr9:80902487-80904502 | p4 | (CATA)5 | CAGGACTCGGTGTTATCCGT | GATTTCATCTCGCGATGGTT | 212 |
| chr9:80902487-80904502 | p3 | (CAA)5 | CAGGACTCGGTGTTATCCGT | GATTTCATCTCGCGATGGTT | 212 |
| chr9:81006053-81008068 | p5 | (GACCC)3 | ATCCTGACATCCACCGAGAC | AATCCGTTGATTTCGTGAGG | 184 |
| chr9:81006053-81008068 | p3 | (TTG)5 | ATCCTGACATCCACCGAGAC | AATCCGTTGATTTCGTGAGG | 184 |
| chr9:81085011-81087026 | p5 | (GACCC)3 | ATCCTGACATCCACCGAGAC | AATCCGTTGATTTCGTGAGG | 184 |
| chr9:81085011-81087026 | p3 | (TTG)5 | ATCCTGACATCCACCGAGAC | AATCCGTTGATTTCGTGAGG | 184 |
| chr9:81219622-81221636 | p3 | (ACA)6 | TACCCTGGGCCATACACAAT | TGATTAAGGGCCTTGTCCTG | 132 |
| chr9:81219648-81221666 | p3 | (ACA)6 | TACCCTGGGCCATACACAAT | TGATTAAGGGCCTTGTCCTG | 106 |
| chr9:81269277-81271289 | p2 | (TA)6 | GTTGATGATTTGGCCGAGTT | TTCACAAGGATCATGCCAAA | 270 |
| chr9:81278323-81280335 | p2 | (TA)6 | GTTGATGATTTGGCCGAGTT | TTCACAAGGATCATGCCAAA | 270 |
| chr9:81278323-81280335 | p2 | (TA)6 | GTTGATGATTTGGCCGAGTT | TTCACAAGGATCATGCCAAA | 270 |
| chr9:81278438-81280450 | p2 | (TA)6 | GTTGATGATTTGGCCGAGTT | TTCACAAGGATCATGCCAAA | 155 |
| chr9:81278438-81280450 | p2 | (TA)6 | GTTGATGATTTGGCCGAGTT | TTCACAAGGATCATGCCAAA | 155 |
| chr9:81445696-81447716 | p4 | (AAAT)5 | AAACTTACCAGCCGCTTTGA | TCAATCCCAAAATGGGAAAA | 120 |
| chr9:81499969-81501993 | p6 | (TTCATA)4 | GAGCCCCCGAGAAATTAAAC | TGGTTGGGTCGTGACAGTAA | 152 |
| chr9:81571084-81573096 | p2 | (TA)6 | TTGTCGGGTTGTCGTTGATA | CGGAAATCGGGAAACATTTA | 135 |
| chr9:81602251-81604285 | p2 | (AT)17 | ATTCAACTTTGGAGGCATGG | TTGACACAGTTAGTTGCCGC | 220 |
| chr9:81620418-81622433 | p3 | (ATG)5 | ATCGTGGGATCGGAATATGA | CGCTAACCTGACTCGCTACC | 129 |
| chr9:81620418-81622433 | p2 | (TA)6 | ATCGTGGGATCGGAATATGA | CGCTAACCTGACTCGCTACC | 129 |
| chr9:81685620-81687642 | p2 | (AT)6 | TTTCCTTCAAGGTCACTGGG | GACATGGTGTATGTGCTGCC | 129 |
| chr9:81702405-81704420 | p5 | (GATAT)3 | GAGTGAGGAAAAGAGCGGTG | CCCTAGGACGAAATGCTCTG | 183 |
| chr9:81745255-81747270 | p5 | (ATTTC)3 | CTTATTTTAGCCCCCGGAAG | TCGAATGTTAAGTCCTCGGG | 211 |
| chr9:81771198-81773213 | p5 | (ATTTC)3 | TGGTCCGTAGAATCGTCACA | AAGTGACCGTGAAATCGTCC | 163 |
| chr9:81884132-81886148 | p2 | (TA)8 | GCCGTCTTTTGAGCATCTTC | TACAGATTCAGTCGGGGGTC | 243 |
| chr9:81956627-81958643 | p2 | (TA)33 | TGTTGAGCGTCGTCATTCTC | CACCCAACAGAGGGGATAGA | 259 |
| chr9:81956627-81958643 | p4 | (ATGA)4 | TGTTGAGCGTCGTCATTCTC | CACCCAACAGAGGGGATAGA | 259 |
| chr9:82015873-82017887 | p2 | (CT)7 | AGGCATGCCAAAAGAGAGAA | CTCGGGATGGCCTATTTACA | 233 |
| chr9:82047112-82049126 | p2 | (AC)7 | AAAAATTTGGGCAGCATCAC | TTCGCGAAAAGAAGGACCTA | 299 |
| chr9:82066728-82068742 | p2 | (TA)7 | GTCCAGAGATCACCCATGCT | CAAACACCAGATGTGGAACG | 200 |
| chr9:82097438-82099453 | p3 | (TGT)5 | AGACCGACCTTGTTGTACGG | TCTCGCTTCCCTCATCTTGT | 272 |
| chr9:82140850-82142870 | p2 | (AT)10 | ACCTTCCTCGGGAAATCACT | ATGTGGTTCACTCGGTCACA | 289 |
| chr9:82518500-82520514 | p2 | (AC)7 | TGGCCCTGGAAATTTATCTG | TTTGTTTCTGCATTTTCCCC | 166 |
| chr9:82555557-82557597 | p2 | (AT)20 | TGAAAGGGGATTTTGGATTG | TCCCAAGCTCGCAAATCTAC | 299 |
| chr9:82600336-82602351 | p5 | (AGAGA)3 | AAATCCTTCCAAAAACGCCT | GACAAAATGGGCAGGACCTA | 124 |
| chr9:82667039-82669091 | p2 | (TA)26 | ATTTGTCGGTGGAGAAATCG | GGCAGCATCTTCCCTGTTTA | 138 |
| chr9:82735443-82737458 | p3 | (ATG)5 | TCAAATTAGGCTCGGGTTTG | TGAAGAACTTCACCGCACAC | 167 |

| Marker ID | SSR type | SSR motif | FORWARD PRIMER (5'-3') | REVERSE PRIMER (5'-3') | size (bp) |
|---|---|---|---|---|---|
| chr9:82815570-82817582 | p2 | (TA)6 | ACTTCTGTGCTTGCCCTTGT | TGGGAACAGATGGTCACTCA | 178 |
| chr9:82966375-82968387 | p2 | (AT)6 | GGCCAAACGGACTTGTTAAA | GGGGCCACGTTTCAATAGTA | 132 |
| chr9:83031978-83034030 | p2 | (AT)26 | CTGGGTGAGTCTTCCCGTTA | CGTTACAGGAACGCGATACA | 164 |
| chr9:83045163-83047178 | p5 | (GCATT)3 | TGCATCCCGACTTACATTGA | TCCAAGCCAACTTCATACCC | 241 |
| chr9:83045163-83047178 | p5 | (GCATT)3 | TGCATCCCGACTTACATTGA | TCCAAGCCAACTTCATACCC | 241 |
| chr9:83052265-83054289 | p2 | (TG)7 | TTATTGGCGTCTATACCGGC | TCCTCAACCACCCCATACTC | 207 |
| chr9:83052341-83054407 | p2 | (TG)7 | TTATTGGCGTCTATACCGGC | TCCTCAACCACCCCATACTC | 131 |
| chr9:83054312-83056330 | p2 | (CA)9 | CACTTTGTCGCTCATTTCCA | TGCTACACTCGTTGCTCCAC | 105 |
| chr9:83148713-83150777 | p2 | (TA)32 | TCACCTAAGATGCCCAATCC | GGCCCAACCATAAGTGAGAA | 129 |
| chr9:83161314-83163329 | p5 | (TTATT)3 | TGTCAAATCGCACCAACATT | TAATGCGGGATTTAAGCTGG | 147 |
| chr9:83247816-83249831 | p2 | (AT)6 | TGTTCGTCAGTAATGGCAGC | CCCTGTAGGTCGTGACTGGT | 260 |
| chr9:83247816-83249831 | p5 | (TATGA)3 | TGTTCGTCAGTAATGGCAGC | CCCTGTAGGTCGTGACTGGT | 260 |
| chr9:83428051-83430066 | p5 | (AATAA)3 | GCACTGTTTCTTCTCCCAGC | GAAGTTGTCCGAGTTCCTGC | 156 |
| chr9:83459675-83461699 | p2 | (AT)12 | GACACGGCTCGTAGATAGCC | GCATTACGACCCACTCAGGT | 175 |
| chr9:83496383-83498398 | p5 | (ATTTC)3 | CAAACTTTGGGTCGTGTGTG | TCGAATGTTAAGTCCTCGGG | 280 |
| chr9:83582184-83584208 | p6 | (TGAATT)3 | TGAGCTCCATTGTGTTTTGC | CAGAACGCCCTCTAGTCGTC | 160 |
| chr9:83582211-83584244 | p6 | (TGAATT)3 | TGAGCTCCATTGTGTTTTGC | CAGAACGCCCTCTAGTCGTC | 133 |
| chr9:83632382-83634400 | p6 | (CTGAAA)3 | CGTGCGTTTACTGCACATTT | TTACTTCCAGGCCATGGTTC | 230 |
| chr9:83685930-83687996 | p2 | (AT)33 | GCCTCCCTTAAAGCGACTCT | GCTCGTGGATAAGCACCACT | 266 |
| chr9:83708583-83710595 | p2 | (AT)12 | GTGACACAGTGGGGGAAAAT | CACAGGAACCTAAGCTGGGA | 164 |
| chr9:83708583-83710595 | p2 | (TA)8 | GTGACACAGTGGGGGAAAAT | CACAGGAACCTAAGCTGGGA | 164 |
| chr9:83708583-83710595 | p2 | (TA)6 | GTGACACAGTGGGGGAAAAT | CACAGGAACCTAAGCTGGGA | 164 |
| chr9:83739347-83741383 | p2 | (TA)18 | CCTTGTCACACAATTCCACG | GAATTTGGAAGGTTTCGGGT | 206 |
| chr9:83935665-83937677 | p2 | (TC)6 | GCCATAGTTTCCCCAGTCAA | TTCCGTCATTAGGAACCCAG | 101 |
| chr9:84150612-84152638 | p6 | (TGATAT)3 | CTGCTCGTGCATGCTGTTAT | TAGTGAGGGATCCGGTGAAC | 111 |
| chr9:84150612-84152638 | p2 | (AT)13 | CTGCTCGTGCATGCTGTTAT | TAGTGAGGGATCCGGTGAAC | 111 |
| chr9:84253459-84255475 | p3 | (GAT)5 | ACATAGCCATGGTTGCCTTC | AGGCACAGCGGAATCATAAG | 209 |
| chr9:84253459-84255475 | p2 | (TA)8 | ACATAGCCATGGTTGCCTTC | AGGCACAGCGGAATCATAAG | 209 |
| chr9:84253459-84255475 | p5 | (ATTGC)3 | ACATAGCCATGGTTGCCTTC | AGGCACAGCGGAATCATAAG | 209 |
| chr9:84288156-84290174 | p6 | (TCCACG)3 | CTTTGCCAAATTGTCCCAGT | CGTGGAACTATTTCTTCGGC | 286 |
| chr9:84288156-84290174 | p6 | (TCCACG)3 | CTTTGCCAAATTGTCCCAGT | CGTGGAACTATTTCTTCGGC | 286 |
| chr9:84303800-84305814 | p5 | (ATTGC)3 | GATGTCCGATGATCCATTCC | AAATATCGACGTCTCGGTGG | 192 |
| chr9:84303800-84305814 | p2 | (TA)7 | GATGTCCGATGATCCATTCC | AAATATCGACGTCTCGGTGG | 192 |
| chr9:84316826-84318896 | p5 | (TTAAA)3 | CTTCTCCCCTCTCTTTGCCT | ACGTAAAGGATGCGAGGTTG | 227 |
| chr9:84316826-84318896 | p2 | (TA)13 | CTTCTCCCCTCTCTTTGCCT | ACGTAAAGGATGCGAGGTTG | 227 |
| chr9:84335252-84337267 | p5 | (GAAAA)3 | CCAAATATACCCCTCCCGTT | TTTCCGTCCAAGGAGAAGAA | 167 |

| Marker ID | SSR type | SSR motif | FORWARD PRIMER (5'-3') | REVERSE PRIMER (5'-3') | size (bp) |
|---|---|---|---|---|---|
| chr9:84447138-84449150 | p2 | (AT)6 | TTGTCGTTGTTTTGGTTGGA | AAACAATTTGACGGCGTTTC | 277 |
| chr9:84477127-84479145 | p6 | (TCTTTT)3 | TTAAACCCGAGACCTGATGG | TCCGCTTATCTCCAACTCGT | 292 |
| chr9:84536921-84538933 | p2 | (AT)6 | GTAGCTCCGGATGAAATGGA | ATCCTTTTGTTTGTGGACGC | 290 |
| chr9:84634927-84636943 | p2 | (AT)8 | CTTTGATTCGACCTGCCATT | GAATTCATACATGGACCCG | 208 |
| chr9:84725274-84727332 | p2 | (AT)29 | GCATTCACCACCATTGTCTG | TGGGCATTTCCTTTGAAGTC | 185 |
| chr9:84824982-84827000 | p3 | (TCT)6 | ACCCCCTATCGAAATCATCC | GAGGACTTTTGCCACCGATA | 139 |
| chr9:84824982-84827000 | p4 | (AATT)4 | ACCCCCTATCGAAATCATCC | GAGGACTTTTGCCACCGATA | 139 |
| chr9:84825206-84827222 | p3 | (TCT)6 | TATCGGTGGCAAAAGTCCTC | TGATTGTAGCTTGGGTGCAG | 101 |
| chr9:84825206-84827222 | p4 | (AATT)4 | TATCGGTGGCAAAAGTCCTC | TGATTGTAGCTTGGGTGCAG | 101 |
| chr9:84846497-84848517 | p2 | (CA)10 | CCAAAAACACCCATCCATTC | GGCACCAAGTTCCAAGTCAT | 255 |
| chr9:84910490-84912508 | p3 | (TCT)6 | ACCCCCTATCGAAATCATCC | GAGGACTTTTGCCACCGATA | 139 |
| chr9:84910490-84912508 | p4 | (AATT)4 | ACCCCCTATCGAAATCATCC | GAGGACTTTTGCCACCGATA | 139 |
| chr9:84910714-84912730 | p3 | (TCT)6 | TATCGGTGGCAAAAGTCCTC | TGATTGTAGCTTGGGTGCAG | 101 |
| chr9:84910714-84912730 | p4 | (AATT)4 | TATCGGTGGCAAAAGTCCTC | TGATTGTAGCTTGGGTGCAG | 101 |
| chr9:84937164-84939192 | p3 | (GTT)6 | ATACCGGGCTGTTTTGTGAC | AATTCCCCCAATATTCCAGC | 250 |
| chr9:84937164-84939192 | p3 | (ATG)6 | ATACCGGGCTGTTTTGTGAC | AATTCCCCCAATATTCCAGC | 250 |
| chr9:85114567-85116637 | p2 | (TA)37 | TAGTTTATTCGGGCCAACCA | GATGCTGTTGCAGGATGAGA | 158 |
| chr9:85114567-85116637 | p2 | (AT)35 | TAGTTTATTCGGGCCAACCA | GATGCTGTTGCAGGATGAGA | 158 |
| chr9:85114567-85116637 | p2 | (TA)8 | TAGTTTATTCGGGCCAACCA | GATGCTGTTGCAGGATGAGA | 158 |
| chr9:85120959-85122971 | p2 | (AT)6 | TTCCCTAGCCAGGTATGCAC | CTGCACTCGAGCAATATCCA | 198 |
| chr9:85131970-85133985 | p3 | (TCA)5 | GGATGTCACGATTTCTGGCT | TTGATGATTGCATGGGAGAA | 147 |
| chr9:85211948-85213960 | p2 | (AT)6 | TGAGACTGATGCAGTGGAGG | CACCCCCAACTAAAGAAGCA | 124 |
| chr9:85227856-85229876 | p2 | (AT)10 | AAGTCCCTACCGCTGATCCT | AGCGGCAGATGAGGCTAATA | 221 |
| chr9:85283702-85285732 | p2 | (TA)15 | GTACGGGCTGTCTGGAGTTC | ACGACGTTCCGTTCGTATTC | 198 |
| chr9:85329812-85331830 | p2 | (TA)8 | ACGCAGGTTGTATCTCCACC | TAGCACCTCCGGAATAATGG | 140 |
| chr9:85329812-85331830 | p2 | (AG)9 | ACGCAGGTTGTATCTCCACC | TAGCACCTCCGGAATAATGG | 140 |
| chr9:85334800-85336815 | p5 | (TATAT)3 | TGTGAGGAGGCTGATGTCTG | CGTCTCGATAGCACAAACCA | 168 |
| chr9:85334800-85336815 | p5 | (AAAAG)3 | TGTGAGGAGGCTGATGTCTG | CGTCTCGATAGCACAAACCA | 168 |
| chr9:85422780-85424804 | p2 | (TA)12 | GTGCTACACCAGTTGCTCCA | CGAAGCCACGGTTGATACTT | 174 |
| chr9:85422780-85424804 | p2 | (AT)6 | GTGCTACACCAGTTGCTCCA | CGAAGCCACGGTTGATACTT | 174 |
| chr9:85514871-85516895 | p4 | (AATA)6 | AGGCGGATGGAGCACTATAA | ACGCAAAGGCAGATCTAGGA | 134 |
| chr9:85541611-85543629 | p6 | (AAATAA)3 | CCATTCAGGGACAAAATTGG | TGGGAAAAGCTAACACCCAC | 296 |
| chr9:85604051-85606066 | p5 | (GAAAG)3 | CACTGCCAACATTGTGTTCC | TGCGAGTCCTGAGGAGAGAT | 114 |
| chr9:85615516-85617536 | p2 | (AT)10 | TGATGCAGCAAAATGTGGAT | CGAAGCCACGGTTGATATTT | 197 |
| chr9:85615516-85617536 | p5 | (CCGGT)3 | TGATGCAGCAAAATGTGGAT | CGAAGCCACGGTTGATATTT | 197 |
| chr9:85624587-85626605 | p6 | (CTGAAA)3 | TACGTGGAAAGCTCCTTGCT | TCGGTCACATGCAGTAAAGC | 234 |

| Marker ID | SSR type | SSR motif | FORWARD PRIMER (5'-3') | REVERSE PRIMER (5'-3') | size (bp) |
|---|---|---|---|---|---|
| chr9:85785849-85787877 | p2 | (AT)14 | ATCACGGGCTTTGGTACTTG | CTCGCCTAATGTGGAAGAGC | 300 |
| chr9:85785849-85787877 | p3 | (TAT)24 | ATCACGGGCTTTGGTACTTG | CTCGCCTAATGTGGAAGAGC | 300 |
| chr9:85786170-85788242 | p2 | (AT)14 | GCTCTTCCACATTAGGCGAG | CAAGACTACGGCAAAGAGGC | 131 |
| chr9:85786170-85788242 | p3 | (TAT)24 | GCTCTTCCACATTAGGCGAG | CAAGACTACGGCAAAGAGGC | 131 |
| chr9:85970032-85972101 | p3 | (TAA)5 | TCCGTTTGTCTCTCCACTTTG | CTTGTGGGAAAGAGGGGAAT | 114 |
| chr9:85970032-85972101 | p3 | (TTA)23 | TCCGTTTGTCTCTCCACTTTG | CTTGTGGGAAAGAGGGGAAT | 114 |
| chr9:86016213-86018234 | p3 | (TGT)7 | GTGGTTGTGAGACCGACCTT | TCCCTCATCTTGTCCTCCAC | 267 |
| chr9:86016213-86018234 | p2 | (AT)69 | GTGGTTGTGAGACCGACCTT | TCCCTCATCTTGTCCTCCAC | 267 |
| chr9:86039245-86041259 | p2 | (TC)7 | TTGTTATGCCAAGTCCACCA | GCTCTGGCACAACATCAGAA | 169 |
| chr9:86082999-86085014 | p5 | (CCCCA)3 | CCTGTTTGGAAAGCCACCTA | AAAGCTTGCGGTCAAAGTGT | 185 |
| chr9:86157785-86159801 | p2 | (TA)8 | AACCCTCGAACCAACCTTCT | GACCGGAACACTGAAGCATT | 177 |
| chr9:86246123-86248159 | p2 | (AT)13 | TCCACGTCCATTTCCCTATC | TGTCCCTAATGTGGAGGACC | 163 |
| chr9:86246123-86248159 | p2 | (AT)45 | TCCACGTCCATTTCCCTATC | TGTCCCTAATGTGGAGGACC | 163 |
| chr9:86246123-86248159 | p2 | (TA)12 | TCCACGTCCATTTCCCTATC | TGTCCCTAATGTGGAGGACC | 163 |
| chr9:86246123-86248159 | p2 | (AT)18 | TCCACGTCCATTTCCCTATC | TGTCCCTAATGTGGAGGACC | 163 |
| chr9:86246123-86248159 | p5 | (CCGGT)3 | TCCACGTCCATTTCCCTATC | TGTCCCTAATGTGGAGGACC | 163 |
| chr9:86260899-86262951 | p3 | (AAT)15 | ACAAGTGGCATCACCACAAA | ATCTGATAGCTTCCTCCCCC | 258 |
| chr9:86260899-86262951 | p3 | (AAC)6 | ACAAGTGGCATCACCACAAA | ATCTGATAGCTTCCTCCCCC | 258 |
| chr9:86260899-86262951 | p2 | (AT)26 | ACAAGTGGCATCACCACAAA | ATCTGATAGCTTCCTCCCCC | 258 |
| chr9:86318182-86320210 | p2 | (TA)14 | GTGCATGGTACGCAGAGAGA | CCGATAACGACCTCGGATAA | 199 |
| chr9:86458767-86460791 | p3 | (TGT)8 | CATCATACTGGTGCAGGGTG | ACCGCAACATTCTCATTTCC | 209 |
| chr9:86466572-86468608 | p3 | (TTC)12 | ATGGCTAGCAGCCGTGTAAT | ACGCGTTTGAGTTTCCAGAC | 209 |
| chr9:86491407-86493423 | p2 | (TA)8 | TGCATTTCACTGCTCCAAAG | TAGTGCATCAGGCAAAGGTG | 159 |
| chr9:86552860-86554878 | p3 | (CAA)6 | ACTTTGTTAACGGCGGACAC | TCCTCCCTGAAAAGAAGCAA | 221 |
| chr9:86622866-86624908 | p2 | (CA)10 | CTCCCGCCATTTTAGTTGAA | GCCATAACATTTGAGCGGAT | 177 |
| chr9:86720527-86722547 | p2 | (TA)7 | TTATAAGTGTGGTGCGCTGC | GTATGTCATGTGGTGCTCGG | 297 |
| chr9:86720527-86722547 | p2 | (CA)10 | TTATAAGTGTGGTGCGCTGC | GTATGTCATGTGGTGCTCGG | 297 |
| chr9:86758118-86760136 | p3 | (GTT)8 | TTGGGTAACATTTTTGGGGA | CACTTTGAAGAAAGCCCGAC | 132 |
| chr9:86758118-86760136 | p2 | (GT)9 | TTGGGTAACATTTTTGGGGA | CACTTTGAAGAAAGCCCGAC | 132 |
| chr9:86776671-86778686 | p5 | (ATTTC)3 | ATTTTCGACGCAACCTGAAG | GTGGCGGTGTAAGACTGGAT | 276 |
| chr9:86784016-86786028 | p2 | (TA)6 | AGGGCCCAACCTAAGACAAT | CTTTGCTTTTTGCTTTTGGG | 170 |
| chr9:86938754-86940769 | p3 | (CAA)5 | TCAATGTTTGGCCATGAAAA | TTTGGCCATGAAAACCAAAT | 194 |
| chr9:86953881-86955896 | p3 | (AAT)5 | CGAATTGCACGAGGAAGATT | TTTGCACCTTTTGTGCTGAC | 212 |
| chr9:86986256-86988270 | p2 | (AT)7 | GCCTACCATACATGCTGCCT | ATTCCGGCTATGAACAGTGC | 256 |
| chr9:86986256-86988270 | p2 | (AT)7 | GCCTACCATACATGCTGCCT | ATTCCGGCTATGAACAGTGC | 256 |
| chr9:86986274-86988290 | p2 | (AT)7 | GCCTACCATACATGCTGCCT | ATTCCGGCTATGAACAGTGC | 238 |

| Marker ID | SSR type | SSR motif | FORWARD PRIMER (5'-3') | REVERSE PRIMER (5'-3') | size (bp) |
|---|---|---|---|---|---|
| chr9:86986274-86988290 | p2 | (AT)7 | GCCTACCATACATGCTGCCT | ATTCCGGCTATGAACAGTGC | 238 |
| chr9:86986300-86988318 | p2 | (AT)7 | GCCTACCATACATGCTGCCT | ATTCCGGCTATGAACAGTGC | 212 |
| chr9:86986300-86988318 | p2 | (AT)7 | GCCTACCATACATGCTGCCT | ATTCCGGCTATGAACAGTGC | 212 |
| chr9:86987588-86989600 | p2 | (TA)6 | ATCTATGCGGACAAACAGCC | TACCCCGGATTTTTACGTTG | 218 |
| chr9:86987588-86989600 | p2 | (AT)7 | ATCTATGCGGACAAACAGCC | TACCCCGGATTTTTACGTTG | 218 |
| chr9:86987588-86989600 | p2 | (AT)18 | ATCTATGCGGACAAACAGCC | TACCCCGGATTTTTACGTTG | 218 |
| chr9:86987602-86989646 | p2 | (TA)6 | ATCTATGCGGACAAACAGCC | TACCCCGGATTTTTACGTTG | 204 |
| chr9:86987602-86989646 | p2 | (AT)7 | ATCTATGCGGACAAACAGCC | TACCCCGGATTTTTACGTTG | 204 |
| chr9:86987602-86989646 | p2 | (AT)18 | ATCTATGCGGACAAACAGCC | TACCCCGGATTTTTACGTTG | 204 |
| chr9:87006754-87008766 | p2 | (CA)6 | CCTTTCGCTCGTCAAATCTC | TTTATAGGGAAATGCTGCCG | 283 |
| chr9:87006754-87008766 | p6 | (TTTTTG)3 | CCTTTCGCTCGTCAAATCTC | TTTATAGGGAAATGCTGCCG | 283 |
| chr9:87036568-87038602 | p5 | (CATAT)3 | GCCATGACTAATGCCCTAGC | CGTTGTGAACCTTTGGGTCT | 186 |
| chr9:87036568-87038602 | p2 | (AT)17 | GCCATGACTAATGCCCTAGC | CGTTGTGAACCTTTGGGTCT | 186 |
| chr9:87144905-87146925 | p5 | (TTGGT)4 | AATCGGCTTGGTTTGACTTG | CTGTATCGGTTGGTTCGGTT | 205 |
| chr9:87269990-87272004 | p2 | (AG)7 | TTCACAAAAGCTCCTTGCCT | TCCTATTTGCCTCGCTCCTA | 111 |
| chr9:87272392-87274416 | p2 | (TA)12 | CACATGGAACAGGCAGAGAA | TTTTGAACACATTGACCGGA | 244 |
| chr9:87306737-87308752 | p5 | (GCATT)3 | CTCGAGGCTCAAATGAAAGG | GGCTAGCCCCAATCACATAA | 160 |
| chr9:87371082-87373102 | p2 | (AT)10 | CCCGAGACGAACTCTCAGAC | GTTTCACTGGTGGGCAATCT | 180 |
| chr9:87387639-87389653 | p2 | (TA)7 | GAGGATGGTTCGACTCCTGA | TGAAGCACTCAATCGACCAC | 225 |
| chr9:87387639-87389653 | p3 | (TTC)17 | GAGGATGGTTCGACTCCTGA | TGAAGCACTCAATCGACCAC | 225 |
| chr9:87488307-87490361 | p2 | (CT)6 | CTTGGTAAAGTTTGGACCGC | GAAATGTATTTGGCCGGAGA | 233 |
| chr9:87488307-87490361 | p2 | (AT)27 | CTTGGTAAAGTTTGGACCGC | GAAATGTATTTGGCCGGAGA | 233 |
| chr9:87488307-87490361 | p2 | (AT)7 | CTTGGTAAAGTTTGGACCGC | GAAATGTATTTGGCCGGAGA | 233 |
| chr9:87488307-87490361 | p2 | (AT)7 | CTTGGTAAAGTTTGGACCGC | GAAATGTATTTGGCCGGAGA | 233 |
| chr9:87488307-87490361 | p2 | (AT)32 | CTTGGTAAAGTTTGGACCGC | GAAATGTATTTGGCCGGAGA | 233 |
| chr9:87495335-87497347 | p2 | (GT)6 | CCATTCGTTCGAAAAGGAGA | AATGCCAAACTCAAGCCAAC | 206 |
| chr9:87606050-87608065 | p3 | (ATG)5 | ACCGTTCAAACCAGTAACCG | GTTTGGCAACAGCCATTTTT | 224 |
| chr9:87658776-87660792 | p2 | (TA)8 | TGTGCAAGCTGTCCAAAGTC | ATTTTGACGCTCACAATCCC | 229 |
| chr9:87673883-87675901 | p2 | (AT)9 | CTAGCCGACGAGGCTATCAC | TTAGCGGGTTGCCAGATAGT | 269 |
| chr9:87820624-87822644 | p5 | (TATAT)4 | GCATAGCCGATTTCGATGTT | ATGCTATGCCATGCAAATGA | 187 |
| chr9:87911866-87913881 | p3 | (AGA)5 | AGGAGGAGGAGGAGAAGCAG | CCCCAACCAGCTTTCTTTTT | 288 |
| chr9:87911866-87913881 | p3 | (GAA)11 | AGGAGGAGGAGGAGAAGCAG | CCCCAACCAGCTTTCTTTTT | 288 |
| chr9:87911896-87913962 | p3 | (AGA)5 | AGGAGGAGGAGGAGAAGCAG | CCCCAACCAGCTTTCTTTTT | 258 |
| chr9:87911896-87913962 | p3 | (GAA)11 | AGGAGGAGGAGGAGAAGCAG | CCCCAACCAGCTTTCTTTTT | 258 |
| chr9:87911965-87913980 | p3 | (AGA)5 | AGGAGGAGGAGGAGAAGCAG | CCCCAACCAGCTTTCTTTTT | 189 |
| chr9:87911965-87913980 | p3 | (GAA)11 | AGGAGGAGGAGGAGAAGCAG | CCCCAACCAGCTTTCTTTTT | 189 |

| Marker ID | SSR type | SSR motif | FORWARD PRIMER (5'-3') | REVERSE PRIMER (5'-3') | size (bp) |
|---|---|---|---|---|---|
| chr9:88052039-88054053 | p3 | (AAC)5 | GCCAAAACAAGGTCCAAGAA | GGGCTGTTTAGGGCATGTTA | 143 |
| chr9:88052039-88054053 | p2 | (AT)7 | GCCAAAACAAGGTCCAAGAA | GGGCTGTTTAGGGCATGTTA | 143 |
| chr9:88062646-88064660 | p2 | (TA)7 | CCCATGAGTTTGGAGATGGT | CGTCCGTTGTCCCTTAAAAA | 252 |
| chr9:88062646-88064660 | p2 | (TA)7 | CCCATGAGTTTGGAGATGGT | CGTCCGTTGTCCCTTAAAAA | 252 |
| chr9:88187904-88189919 | p5 | (ATAAA)3 | TTATCGGGTGACATGGACAA | GGTCTTATGACACCCTCGGA | 203 |
| chr9:88199141-88201153 | p2 | (TA)6 | ATTTTCGCTTGACCTTGGTG | TATCGGTTCTTCGCATTTCC | 178 |
| chr9:88239751-88241763 | p2 | (TA)6 | AGTCGCCTTTTCAGTCAGGA | ATCATCATCTTCTGGGTCGG | 211 |
| chr9:88284690-88286708 | p6 | (AGGTAG)3 | GGTTTGATCCAAGCCAAAGA | TCCCTACGCAAAGGATATGG | 142 |
| chr9:88284690-88286708 | p6 | (TTCTCC)3 | GGTTTGATCCAAGCCAAAGA | TCCCTACGCAAAGGATATGG | 142 |
| chr9:88405944-88407956 | p2 | (TA)6 | TCTTCGCATTTTCTCAAGCA | AAAAGGGGGAATTGATTTGG | 221 |
| chr9:88405944-88407956 | p2 | (TA)6 | TCTTCGCATTTTCTCAAGCA | AAAAGGGGGAATTGATTTGG | 221 |
| chr9:88480727-88482742 | p5 | (TATTT)3 | CCTCGGGCTAAGGTAGATCC | CTTTTGGCGTGGCATAATTT | 231 |
| chr9:88597734-88599749 | p5 | (TTAAT)3 | GCTGGTTGCAACTTCTGTCA | TGCACCACAGCTGTTTCTGT | 168 |
| chr9:88597734-88599749 | p2 | (TA)12 | GCTGGTTGCAACTTCTGTCA | TGCACCACAGCTGTTTCTGT | 168 |
| chr9:88597734-88599749 | p2 | (TA)28 | GCTGGTTGCAACTTCTGTCA | TGCACCACAGCTGTTTCTGT | 168 |
| chr9:88773602-88775644 | p3 | (ATA)14 | AAGAGTGAAGGATGCGGCTA | AAACGGTCACAACATAGGGC | 286 |
| chr9:88867545-88869560 | p5 | (TTGGT)3 | TCAGAAAGAGGAAGGGAGCA | TAGGGTTTCACTTGCCATCC | 271 |
| chr9:88882929-88884962 | p3 | (ATA)11 | CTTCCCCAAATGAACTCCAA | TGCTCCCTTCCTCTTTCTGA | 212 |
| chr9:88984392-88986404 | p2 | (TA)6 | TGGAATCGTTGATAATGGCA | ATGTCTCGGTAGCACAACCC | 292 |
| chr9:89071196-89073242 | p2 | (TA)23 | AGAATAACAGCATCGGGACG | ACTGTTGATTTTCCGCATCC | 284 |
| chr9:89085616-89087628 | p2 | (TC)6 | AATCGGCTGAGTTAACGGTG | CCATAATGCTTCCAACCCAC | 204 |
| chr9:89085616-89087628 | p3 | (AAG)8 | AATCGGCTGAGTTAACGGTG | CCATAATGCTTCCAACCCAC | 204 |
| chr9:89134981-89137029 | p2 | (TA)24 | CCTAGCGATCGTTCAACTCC | ATTCCGGTGTCACCATTGAT | 123 |
| chr9:89134981-89137029 | p5 | (TTAAA)3 | CCTAGCGATCGTTCAACTCC | ATTCCGGTGTCACCATTGAT | 123 |
| chr9:89175336-89177351 | p5 | (ATATG)3 | TGTTGTTACGATGCTTGGGA | CTAAGCGTTGTCCTTCCGAG | 166 |
| chr9:89180632-89182688 | p5 | (ATATG)3 | TCCAACCTGGAAATTGGAAG | GCTCCGCTGAATATGTGGAT | 193 |
| chr9:89180685-89182703 | p5 | (ATATG)3 | TCCAACCTGGAAATTGGAAG | GCTCCGCTGAATATGTGGAT | 140 |
| chr9:89180886-89182901 | p5 | (ATATG)3 | TGGCATTCCTGTATCCATCA | TCCAAATTCCAATGCACAAA | 128 |
| chr9:89318339-89320357 | p6 | (GGGATG)3 | GCTGAAGCATCCTGCTCTTT | TGTGGGGTAGAAGCCATAGG | 179 |
| chr9:89324381-89326405 | p4 | (AAAT)6 | CCTGGATCAATCACTTTGGG | CAACCCAACAATCCTCCAGT | 121 |
| chr9:89391951-89393975 | p6 | (ATAAAA)4 | GGATTCAACCGGACCAGTAA | AGGGTGATTGACAAACCTCG | 298 |
| chr9:89503805-89505817 | p2 | (CA)6 | GGTTGGGATAATGTCTGCGT | CACAGGAAGAAATCCCCTGA | 237 |
| chr9:89640960-89643016 | p2 | (AT)28 | GCGGATCGTATTCACTTGCT | GTTAAACAGCCAGCTCAGGG | 294 |
| chr9:89691848-89693862 | p2 | (TA)7 | CGATTCATGGGGTTCAAATC | GCGAATCCGAAAAACTCAAG | 258 |
| chr9:89691848-89693862 | p2 | (TA)8 | CGATTCATGGGGTTCAAATC | GCGAATCCGAAAAACTCAAG | 258 |
| chr9:89692959-89695004 | p2 | (TA)8 | GCAAAACTGCTCTTTGCCTC | AATTGCAATATCTCTTGCCGA | 117 |

| Marker ID | SSR type | SSR motif | FORWARD PRIMER (5'-3') | REVERSE PRIMER (5'-3') | size (bp) |
|---|---|---|---|---|---|
| chr9:89692959-89695004 | p3 | (ATA)15 | GCAAAACTGCTCTTTGCCTC | AATTGCAATATCTCTTGCCGA | 117 |
| chr9:89804993-89807011 | p6 | (ATCTAT)3 | TCGGCAAAAAGATCAATTCC | AGCCTTTTCCTCCTTGAACC | 275 |
| chr9:89880104-89882116 | p2 | (TC)6 | GTCCGTAACCCTTCACCGTA | TACGGTCCGTCCTTCATTTC | 118 |
| chr9:89890991-89893003 | p2 | (TC)6 | CTTACGCTTCAGAAGGACGG | TTGCCTTAAATGCTCCAACC | 180 |
| chr9:89965538-89967550 | p3 | (ACA)6 | CTGGGCAGATAGAGAGGCTG | GCTTATCGACGCTTCAAAGG | 180 |
| chr9:89965538-89967550 | p2 | (TA)6 | CTGGGCAGATAGAGAGGCTG | GCTTATCGACGCTTCAAAGG | 180 |
| chr9:90105914-90107949 | p2 | (AT)6 | CGAGCACGTTTCAACCACTA | TAAAGCAGTTAACGGGGACG | 200 |
| chr9:90108015-90110030 | p5 | (ACCGA)3 | CTCATTCCCATGCTCGATTT | AGTCTGGATGCAGGTTGGTC | 110 |
| chr9:90212124-90214138 | p2 | (AT)6 | TAACCCGTTAGTGGGAGCAG | AAATGACATGGCGGAATAGC | 201 |
| chr9:90212139-90214167 | p2 | (AT)6 | TAACCCGTTAGTGGGAGCAG | AAATGACATGGCGGAATAGC | 186 |
| chr9:90225290-90227304 | p2 | (AT)7 | ATGAAGATCATCGGCGATTC | ACTGTCGTCGACCTCTTCGT | 212 |
| chr9:90303825-90305837 | p3 | (TGC)5 | GCTGAAACTGTGGTGCTGAA | TAAGTCTGCATGGGTGTCCA | 234 |
| chr9:90303825-90305837 | p2 | (AT)6 | GCTGAAACTGTGGTGCTGAA | TAAGTCTGCATGGGTGTCCA | 234 |
| chr9:90339912-90341928 | p4 | (ACAT)4 | ACATTTACCGATGGCTCCTG | GGTGTGTTTACGAGCCCTGT | 143 |
| chr9:90401509-90403531 | p2 | (AT)11 | GGCATTGTGTGAAAATGTGC | GCATTTCCCCTATTTCGTCA | 251 |
| chr9:90405902-90407970 | p2 | (AT)34 | AAAGATTGAGGCCGTGAAGA | CTTCGTGAGTGGTGCAGAAA | 122 |
| chr9:90459053-90461068 | p5 | (TGCAA)3 | TAGCCTAAATCGGCTCCAGA | ATCGTTGTGGGAACGATAGG | 155 |
| chr9:90503304-90505354 | p2 | (AT)25 | TTTGCCTCTCTCCTTTAGCG | CTTCAATGGCGGAAAGATGT | 253 |
| chr9:90546853-90548873 | p4 | (ATCA)5 | TTCTGATCCCCAAGTGAACC | ATGCCTCGAACTCTGAAGGA | 195 |
| chr9:90575828-90577852 | p2 | (TA)12 | CAAACGATTTCCCATGCTCT | AAAACAGTGGACCGCAAAAG | 179 |
| chr9:90688849-90690894 | p3 | (AAT)15 | GGTGTCAGAGGAGTGGGTGT | TCCCTCTCCCTTTACCCCTA | 211 |
| chr9:90688849-90690894 | p4 | (TTTA)5 | GGTGTCAGAGGAGTGGGTGT | TCCCTCTCCCTTTACCCCTA | 211 |
| chr9:90757440-90759485 | p3 | (AAT)15 | GGTGTCAGAGGAGTGGGTGT | TCCCTCTCCCTTTACCCCTA | 211 |
| chr9:90757440-90759485 | p4 | (TTTA)5 | GGTGTCAGAGGAGTGGGTGT | TCCCTCTCCCTTTACCCCTA | 211 |
| chr9:90832041-90834056 | p5 | (TTTTA)3 | GGCTGCTTTAAGTGCTGGAC | TGCCCTTTTGTATGCTAGGG | 190 |
| chr9:90895789-90897804 | p2 | (GA)8 | AGGAGGGTGATAACGTGGTG | TCACAATACCAACGGACGAA | 293 |
| chr9:90895789-90897804 | p3 | (TTG)5 | AGGAGGGTGATAACGTGGTG | TCACAATACCAACGGACGAA | 293 |
| chr9:91067565-91069625 | p2 | (GA)7 | TCACAAATCGACGAAAACCA | CTAACAGACGCACCCCATTT | 231 |
| chr9:91067565-91069625 | p3 | (TAA)20 | TCACAAATCGACGAAAACCA | CTAACAGACGCACCCCATTT | 231 |
| chr9:91096964-91099003 | p3 | (ACA)13 | TCCCGAGTCCGTCATATCTC | CAACTAAGGCCCAATCCAAA | 229 |
| chr9:91113263-91115299 | p3 | (TTC)12 | TGTTGGCAATCCATTCAAGA | TTGCTGTTGTGGTGGTTGAT | 276 |
| chr9:91153255-91155269 | p2 | (GA)7 | GCAAATTAAAGACGGTGGGA | ACTACGTGAGGGTTTGCACC | 134 |
| chr9:91169863-91171875 | p2 | (AG)6 | GCTGCTTCTGATGACCCATT | TGATATGACCTCCTTTCGGC | 172 |
| chr9:91178308-91180334 | p2 | (AT)6 | GAATTTCGGTACCAGGCGTA | CAATCTCTGCCCATTCACCT | 199 |
| chr9:91260022-91262037 | p5 | (AACCG)4 | TGGTGGGAAGAAATAGGCTG | CGAAAAACCAAAAATCCGAA | 135 |
| chr9:91260022-91262037 | p5 | (AAATA)3 | TGGTGGGAAGAAATAGGCTG | CGAAAAACCAAAAATCCGAA | 135 |

| Marker ID | SSR type | SSR motif | FORWARD PRIMER (5'-3') | REVERSE PRIMER (5'-3') | size (bp) |
|---|---|---|---|---|---|
| chr9:91260022-91262037 | p3 | (TAG)9 | TGGTGGGAAGAAATAGGCTG | CGAAAAACCAAAAATCCGAA | 135 |
| chr9:91281850-91283862 | p2 | (GT)6 | GGATTGCCAATAGCAAGGAA | TGGTCAAAATTCCCGACTTC | 290 |
| chr9:91314076-91316090 | p2 | (AT)7 | TCATTGTATCCTTCCAGCCC | TTCCTATCTTTGGGCGTGTT | 106 |
| chr9:91391216-91393230 | p2 | (AT)7 | TCATTGTATCCTTCCAGCCC | TTCCTATCTTTGGGCGTGTT | 106 |
| chr9:91642497-91644555 | p3 | (TGT)6 | ATGAGGTTGGATTTGCTTGC | GCGTGCCTTTTCAAACTCTC | 137 |
| chr9:91710330-91712345 | p3 | (TCA)5 | TCGGGCAGTCATGCATAATA | ATAGTGCAGGACCATCTGGG | 104 |
| chr9:91824960-91826978 | p6 | (ATTTAT)3 | CAAGGGTATCTCCGGATCAA | CTTCATTTTGGCCCATGTCT | 201 |
| chr9:91889743-91891761 | p6 | (ACTGAA)3 | CAAGCGTTGAAGATGTGGAA | CAGTTCCAACCATTGCCTTT | 249 |
| chr9:92011613-92013629 | p4 | (GAAA)4 | GGGAGTGGAAACGAGATTCA | GTAGCGGGCATATGTGGAGT | 153 |
| chr9:92318673-92320688 | p5 | (TTATT)3 | GGTCCTTTCACGCACAAAAT | CCCTAAAACGACGATCCAAA | 191 |
| chr9:92440626-92442659 | p3 | (CTT)11 | TTGCTGTGAAACGCAAAAAG | ATGTGCCTCGTGGTGAAACT | 267 |
| chr9:92477493-92479505 | p2 | (AT)6 | GAGGAGTGTGGGGTGGTAGA | GTCCCCTCATTTTCCCATTT | 197 |
| chr9:92477493-92479505 | p3 | (ATA)6 | GAGGAGTGTGGGGTGGTAGA | GTCCCCTCATTTTCCCATTT | 197 |
| chr9:92477493-92479505 | p5 | (TATTT)3 | GAGGAGTGTGGGGTGGTAGA | GTCCCCTCATTTTCCCATTT | 197 |
| chr9:92646919-92648933 | p4 | (AACA)4 | TCTGCTTCGTCGTCATCAAC | ATGTGGCAATGAATCCATCA | 129 |
| chr9:92646919-92648933 | p2 | (TA)7 | TCTGCTTCGTCGTCATCAAC | ATGTGGCAATGAATCCATCA | 129 |
| chr9:92649952-92651966 | p2 | (TA)7 | TGTAATTCCCAAAGCATCCC | AAATGACATGGCGGAATAGC | 183 |
| chr9:92712544-92714572 | p2 | (TA)14 | TCACTTGACATTGCCCAAGA | GGGCAAAACACTTCCTGAAA | 142 |
| chr9:92775195-92777231 | p2 | (TA)11 | TTTTGCCCATAAAAGCGAAG | TTGATTCAGGGTGACAATGG | 219 |
| chr9:92775195-92777231 | p3 | (TAA)12 | TTTTGCCCATAAAAGCGAAG | TTGATTCAGGGTGACAATGG | 219 |
| chr9:92806529-92808587 | p2 | (TA)29 | AACCGTAGCATGTGGTGACA | TTTGTAGGAAAAGATGCCCG | 217 |
| chr9:92806588-92808646 | p2 | (TA)29 | AACCGTAGCATGTGGTGACA | TTTGTAGGAAAAGATGCCCG | 158 |
| chr9:92806588-92808646 | p3 | (TTA)9 | AACCGTAGCATGTGGTGACA | TTTGTAGGAAAAGATGCCCG | 158 |
| chr9:92824504-92826516 | p2 | (TG)6 | CCACAGCTCTTCTTGGAAGG | AAGGAGCCAGCTGAAAAACA | 263 |
| chr9:92939038-92941052 | p2 | (TC)7 | GCATCACCGAAGAGGACATT | TTTGTATGACTTCCCGGAGC | 144 |
| chr9:92939284-92941302 | p2 | (TC)7 | GCTCCGGGAAGTCATACAAA | ATTTCATGAATCGTGCCTCC | 136 |
| chr9:92939305-92941329 | p2 | (TC)7 | GCTCCGGGAAGTCATACAAA | ATTTCATGAATCGTGCCTCC | 115 |
| chr9:93044035-93046055 | p5 | (TTTCA)4 | ATGCGTACTCTGGAAGGGTG | ATCCACTGTGCATCAACGAA | 255 |
| chr9:93044035-93046055 | p2 | (TA)48 | ATGCGTACTCTGGAAGGGTG | ATCCACTGTGCATCAACGAA | 255 |
| chr9:93044778-93046874 | p5 | (TTTCA)4 | CCCCTTCACACAGCACTTCT | TGAACCTTGGGTGGAAACTC | 281 |
| chr9:93044778-93046874 | p2 | (TA)48 | CCCCTTCACACAGCACTTCT | TGAACCTTGGGTGGAAACTC | 281 |
| chr9:93088903-93090929 | p2 | (AT)13 | TCAGGTTCGAACTTCAGGCT | GTGGTCAGCAATGTAGCCAA | 102 |
| chr9:93110458-93112473 | p3 | (TGA)5 | ATCCACACACCGTTTCACAA | TTAGAGTTTCCAGCGGCACT | 261 |
| chr9:93264478-93266583 | p3 | (TTC)35 | TTGGCTATGTTGAACTCCCC | ATGAGAGGGCTGGAGTGAGA | 139 |
| chr9:93275787-93277799 | p2 | (AT)6 | TACCAGTCCCACTACAGCCC | AAACAGAGGTTGATGGTGGC | 251 |
| chr9:93328685-93330700 | p5 | (AAATA)3 | CGAGGTAAGGGTTTTGGTGA | TATCCCGAGTGCGTCCTAGT | 138 |

| Marker ID | SSR type | SSR motif | FORWARD PRIMER (5'-3') | REVERSE PRIMER (5'-3') | size (bp) |
|---|---|---|---|---|---|
| chr9:93328685-93330700 | p5 | (AAATA)3 | CGAGGTAAGGGTTTTGGTGA | TATCCCGAGTGCGTCCTAGT | 138 |
| chr9:93328685-93330700 | p5 | (TTTAT)3 | CGAGGTAAGGGTTTTGGTGA | TATCCCGAGTGCGTCCTAGT | 138 |
| chr9:93328685-93330700 | p5 | (TTTAT)3 | CGAGGTAAGGGTTTTGGTGA | TATCCCGAGTGCGTCCTAGT | 138 |
| chr9:93341830-93343848 | p6 | (ATATAG)3 | TACGCTGGAGTGGGGAATTA | TTCATTAAACACGTGGCAGC | 276 |
| chr9:93341830-93343848 | p5 | (TTGGA)3 | TACGCTGGAGTGGGGAATTA | TTCATTAAACACGTGGCAGC | 276 |
| chr9:93346815-93348830 | p3 | (CAT)5 | ATGGGTTGATACGCATGTGA | AAAAAGTGGAACCCACCACA | 242 |
| chr9:93371364-93373382 | p3 | (GTT)6 | ACCGACTATGCTGTATGGGG | TGGGGCATCTGTATCTCTCC | 182 |
| chr9:93378548-93380564 | p2 | (AT)8 | TGAAAATTTGTGAAGCACGC | CAAACCTCCCGTAGAGGTGA | 132 |
| chr9:93448046-93450070 | p2 | (AT)12 | CACACGGGCATAGTTTGCTA | GATAAATGCGCAACAGACGA | 293 |
| chr9:93513962-93515978 | p5 | (TTTAT)3 | GGAGTGGGAGACATGGAGAA | CTTCTTCCGCCTCTTTCCTT | 239 |
| chr9:93514005-93516021 | p5 | (TTTAT)3 | GGAGTGGGAGACATGGAGAA | CTTTCATCCTCACCCACGTT | 196 |
| chr9:93548238-93550250 | p2 | (GA)6 | TGAGACTCCTCCACCATTCC | TTGGAGGATTCACCCTGAAC | 109 |
| chr9:93555263-93557299 | p5 | (TTCTC)3 | GCATTTCTCAAGGGGTGTGT | GGATATCTGGCTTTCACCCA | 284 |
| chr9:93555263-93557299 | p3 | (TAA)10 | GCATTTCTCAAGGGGTGTGT | GGATATCTGGCTTTCACCCA | 284 |
| chr9:93586449-93588503 | p2 | (TA)27 | TTGGTTAAACACGTGTGCGT | GACGGGTGATCTCCTCATGT | 217 |
| chr9:93604555-93606581 | p2 | (AT)13 | GCGAGGAAGTAGCAGAATGC | TGCAGCAAAGGTGTTTGAAG | 156 |
| chr9:93758767-93760797 | p2 | (CA)15 | TCGTCGGATTTCGTATAGCC | ACTGCCTTCCCAGTCCCTAT | 275 |
| chr9:93805503-93807521 | p2 | (AG)9 | TGCGAAACTTCTCCCACTCT | TCGTAGGTCTCCCGTTCATC | 153 |
| chr9:93885972-93887993 | p3 | (ATC)7 | TCAACAACTGTGCCCATCAT | CCCCCTCTTCATGGATTTTT | 214 |
| chr9:93898842-93900884 | p2 | (AT)21 | AATCGCTGATGAAGCAACCT | AAGCATGCGAGATATCCACC | 204 |
| chr9:93974686-93976740 | p3 | (TTA)18 | ACCAAAACACCACTTACCGC | AGAGCCTTGCAGGAATTGAA | 226 |
| chr9:94005082-94007102 | p5 | (ACCCG)4 | TGGGGGCTGAAATAAATGAG | CTTGAATCGGTCCATCCAGT | 163 |
| chr9:94011089-94013104 | p5 | (AAGCC)3 | TGTCACTCCCAATGCACAAT | TCCACACTTTCGCTTCCTCT | 133 |
| chr9:94035789-94037823 | p3 | (TCA)5 | AAACTCATGAAATCACGGGC | TTCAATCCCAGCTCCTATGC | 108 |
| chr9:94036763-94038778 | p3 | (TCA)5 | CTGTCTACCTGTGGGCATGA | GCGGGTTAACGATGATGACT | 257 |
| chr9:94050661-94052676 | p5 | (CATTG)3 | AGGTCTTAGGCATGGTGTGG | AGCCCACAGAACAATATCGC | 180 |
| chr9:94110404-94112430 | p2 | (TA)13 | CCTGGTGGTTTGGCTTAGAA | GACGGACCGTTGATTCTCAT | 194 |
| chr9:94227792-94229810 | p3 | (TTG)29 | TCGCACACTCGAATCAACTC | CAGGTGCATCAAATGGAGTG | 224 |
| chr9:94227792-94229810 | p6 | (CGTCCC)3 | TCGCACACTCGAATCAACTC | CAGGTGCATCAAATGGAGTG | 224 |
| chr9:94413150-94415174 | p6 | (AAAAAG)4 | TTGGGAAGTGGACGAGTTTC | AAACACACCTTCCTTGGCAC | 265 |
| chr9:94481254-94483269 | p3 | (CCA)5 | CAGACATATGGTCCCGCTTT | ATCGAACCGAACAATCCAAC | 260 |
| chr9:94481254-94483269 | p2 | (AT)17 | CAGACATATGGTCCCGCTTT | ATCGAACCGAACAATCCAAC | 260 |
| chr9:94481254-94483269 | p5 | (TTCGG)3 | CAGACATATGGTCCCGCTTT | ATCGAACCGAACAATCCAAC | 260 |
| chr9:94530449-94532465 | p3 | (AAT)5 | CAAGCAAAACACCCTCGATT | GGGAACGTGTTGGATGATCT | 281 |
| chr9:94530449-94532465 | p2 | (TA)8 | CAAGCAAAACACCCTCGATT | GGGAACGTGTTGGATGATCT | 281 |
| chr9:94541614-94543632 | p6 | (TCCCTT)3 | CACCTGCATGAGACAGCAAT | GGGCAAACCAACAGTAGCAT | 248 |

| Marker ID | SSR type | SSR motif | FORWARD PRIMER (5'-3') | REVERSE PRIMER (5'-3') | size (bp) |
|---|---|---|---|---|---|
| chr9:94575950-94577996 | p2 | (TA)23 | CTTGGCGAAGAAACCGTAAG | AGGGGGCTTGAATACTCGAT | 199 |
| chr9:94694633-94696648 | p5 | (TTCTT)3 | GATGCAAAGGTGTGGTGTTG | CTTTCCGGGACAAACTCTCA | 247 |
| chr9:94870670-94872688 | p2 | (AT)8 | TTGGCTACAATTGCAAGGTTC | TTTGGCCATCATTCTGTCAC | 196 |
| chr9:94870670-94872688 | p2 | (AT)9 | TTGGCTACAATTGCAAGGTTC | TTTGGCCATCATTCTGTCAC | 196 |
| chr9:94913392-94915412 | p2 | (TA)10 | CGTTACCAAAGTCGGTCGTT | TACCGGAGTAACGGGAAGTG | 205 |
| chr9:94956781-94958801 | p3 | (TTA)6 | CTCTGCCACTCCACTCCTTC | GTGGGTTTCTTGGGATTTCA | 261 |
| chr9:94956799-94958817 | p3 | (TTA)6 | CTCTGCCACTCCACTCCTTC | GTGGGTTTCTTGGGATTTCA | 243 |
| chr9:95034509-95036533 | p6 | (TAAAAA)4 | TGGCACGGACTAAAAAGGAA | GCGAATTTTACCGCATTGAT | 204 |
| chr9:95098487-95100505 | p5 | (AACAA)3 | CCCAGTCCTAAACCCCAAAT | TGGGAAGAGTTGAGAATGGG | 120 |
| chr9:95098487-95100505 | p6 | (TTTGAT)3 | CCCAGTCCTAAACCCCAAAT | TGGGAAGAGTTGAGAATGGG | 120 |
| chr9:95194461-95196475 | p2 | (AT)6 | GATACATCCAGCATTGTGCG | TTCATGCCAATGACTTTCCA | 217 |
| chr9:95194461-95196475 | p2 | (TA)7 | GATACATCCAGCATTGTGCG | TTCATGCCAATGACTTTCCA | 217 |
| chr9:95194461-95196475 | p2 | (TA)6 | GATACATCCAGCATTGTGCG | TTCATGCCAATGACTTTCCA | 217 |
| chr9:95194461-95196475 | p2 | (AT)6 | GATACATCCAGCATTGTGCG | TTCATGCCAATGACTTTCCA | 217 |
| chr9:95194505-95196517 | p2 | (AT)6 | GATACATCCAGCATTGTGCG | TTCATGCCAATGACTTTCCA | 173 |
| chr9:95194505-95196517 | p2 | (TA)7 | GATACATCCAGCATTGTGCG | TTCATGCCAATGACTTTCCA | 173 |
| chr9:95194505-95196517 | p2 | (TA)6 | GATACATCCAGCATTGTGCG | TTCATGCCAATGACTTTCCA | 173 |
| chr9:95194505-95196517 | p2 | (AT)6 | GATACATCCAGCATTGTGCG | TTCATGCCAATGACTTTCCA | 173 |
| chr9:95196609-95198625 | p2 | (TA)8 | TCAAACCGTAAAACGGAACC | TGAGGCAATGGGATAGGAGT | 131 |
| chr9:95340631-95342647 | p2 | (AT)8 | AAGCGTTAATGGTGGTTTGG | TACTCCCCTCCGCAACATAC | 219 |
| chr9:95340631-95342647 | p2 | (AT)8 | AAGCGTTAATGGTGGTTTGG | TACTCCCCTCCGCAACATAC | 219 |
| chr9:95340631-95342647 | p2 | (AT)9 | AAGCGTTAATGGTGGTTTGG | TACTCCCCTCCGCAACATAC | 219 |
| chr9:95346511-95348529 | p3 | (ATT)10 | GGGATGGATGACAGGTGTTC | CCCTACACGATTTGTCAGCA | 262 |
| chr9:95346511-95348529 | p6 | (AAATCG)3 | GGGATGGATGACAGGTGTTC | CCCTACACGATTTGTCAGCA | 262 |
| chr9:95362133-95364155 | p2 | (AT)6 | TAACACCAGTGGAACCGACA | AACAAAACGACGGATGGTTC | 279 |
| chr9:95465899-95467941 | p5 | (ACCCG)3 | ATCATCGCAGCTTTTATGGG | AGGCGAGTGTTCGAGGTAGA | 259 |
| chr9:95465899-95467941 | p2 | (TC)22 | ATCATCGCAGCTTTTATGGG | AGGCGAGTGTTCGAGGTAGA | 259 |
| chr9:95465956-95467974 | p5 | (ACCCG)3 | ATCATCGCAGCTTTTATGGG | AGGCGAGTGTTCGAGGTAGA | 202 |
| chr9:95465956-95467974 | p2 | (TC)22 | ATCATCGCAGCTTTTATGGG | AGGCGAGTGTTCGAGGTAGA | 202 |
| chr9:95465990-95468020 | p5 | (ACCCG)3 | ATCATCGCAGCTTTTATGGG | AGGCGAGTGTTCGAGGTAGA | 168 |
| chr9:95465990-95468020 | p2 | (TC)22 | ATCATCGCAGCTTTTATGGG | AGGCGAGTGTTCGAGGTAGA | 168 |
| chr9:95504893-95506908 | p5 | (TTGCA)3 | TTAGAGTGGCAAGGGGTGTC | AGTGCACGTCACCTCACAAG | 264 |
| chr9:95504893-95506908 | p5 | (TTTGT)3 | TTAGAGTGGCAAGGGGTGTC | AGTGCACGTCACCTCACAAG | 264 |
| chr9:95539885-95541900 | p5 | (AATAA)3 | CACCCCTCCCTTTCTCTTTC | AACTCGTCCCCAGAGAGGAT | 245 |
| chr9:95541098-95543124 | p2 | (AT)13 | TACATAAATTGAGGCGGGGA | CACCCTAGCGTGAAGCCTAC | 226 |
| chr9:95648315-95650330 | p5 | (TTGGG)3 | CTCCACCTTCATTCCCTCAA | CATTTGCTCAAGTGCCTCAA | 199 |

| Marker ID | SSR type | SSR motif | FORWARD PRIMER (5'-3') | REVERSE PRIMER (5'-3') | size (bp) |
|---|---|---|---|---|---|
| chr9:95652457-95654469 | p2 | (TG)6 | TGAGAAATGCATGCACAACA | GGTGAAGGGCAACGTTTAAG | 125 |
| chr9:95652457-95654469 | p2 | (TG)6 | TGAGAAATGCATGCACAACA | GGTGAAGGGCAACGTTTAAG | 125 |
| chr9:95670421-95672441 | p5 | (TGAAT)4 | GAGATAGGGGAGGTTCAGCC | GTGCCAACGGATGATTTTCT | 201 |
| chr9:95869850-95871862 | p2 | (AT)6 | CGAGCAACACAGCAATCAGT | GAAGCGTATGTGGGCTTGTT | 290 |
| chr9:95927247-95929261 | p2 | (AT)7 | GCCAAGTCACAATCAAGCAA | GCAGTAGATGTGGGGAAGGA | 153 |
| chr9:96033746-96035761 | p3 | (TCC)5 | CTTGGTACATCCCCTCAGGA | AGCTTGGCATTCAAGCTGAT | 150 |
| chr9:96086967-96088987 | p5 | (ACAAA)3 | ACCAACTTCATTCGCCATCT | TTTTAGGGCAACAATGAGCC | 168 |
| chr9:96086967-96088987 | p5 | (AAAAT)4 | ACCAACTTCATTCGCCATCT | TTTTAGGGCAACAATGAGCC | 168 |
| chr9:96124797-96126817 | p5 | (ACAAA)3 | ACCAACTTCATTCGCCATCT | TTTTAGGGCAACAATGAGCC | 168 |
| chr9:96124797-96126817 | p5 | (AAAAT)4 | ACCAACTTCATTCGCCATCT | TTTTAGGGCAACAATGAGCC | 168 |
| chr9:96162615-96164640 | p5 | (CCCAA)5 | AGTTAAACAAAGCATGCCCG | GGTTGTACCCCGAGTAAGCA | 228 |
| chr9:96169690-96171708 | p2 | (TG)9 | CATCGAGGTCTACCCCAGAA | TTGGTCCTTACATCGAAGCC | 200 |
| chr9:96169690-96171708 | p3 | (GTG)5 | CATCGAGGTCTACCCCAGAA | TTGGTCCTTACATCGAAGCC | 200 |
| chr9:96179322-96181336 | p2 | (TG)7 | CTTTTTGGATTGCGGGTAGA | TGACCTTCCTCCATGTTTCC | 229 |
| chr9:96288632-96290644 | p2 | (AT)6 | CAATAACGGTAACACGCACG | CGGTGGAAATAAGTTGCGTC | 125 |
| chr9:96383377-96385392 | p5 | (CCGAC)3 | TTCGCTTCTAAGAAACCCGA | GTCTCGACCTTTTCCCACAA | 251 |
| chr9:96383377-96385392 | p2 | (CT)6 | TTCGCTTCTAAGAAACCCGA | GTCTCGACCTTTTCCCACAA | 251 |
| chr9:96383377-96385392 | p2 | (AT)14 | TTCGCTTCTAAGAAACCCGA | GTCTCGACCTTTTCCCACAA | 251 |
| chr9:96383445-96385463 | p5 | (CCGAC)3 | TTCGCTTCTAAGAAACCCGA | GTCTCGACCTTTTCCCACAA | 183 |
| chr9:96383445-96385463 | p2 | (CT)6 | TTCGCTTCTAAGAAACCCGA | GTCTCGACCTTTTCCCACAA | 183 |
| chr9:96383445-96385463 | p2 | (AT)14 | TTCGCTTCTAAGAAACCCGA | GTCTCGACCTTTTCCCACAA | 183 |
| chr9:96383714-96385729 | p5 | (CCGAC)3 | AGCTGTCTGGAATGGGTGTC | TTACACCATCACTGGGAGCA | 190 |
| chr9:96383714-96385729 | p2 | (CT)6 | AGCTGTCTGGAATGGGTGTC | TTACACCATCACTGGGAGCA | 190 |
| chr9:96383714-96385729 | p2 | (AT)14 | AGCTGTCTGGAATGGGTGTC | TTACACCATCACTGGGAGCA | 190 |
| chr9:96404400-96406422 | p2 | (AT)24 | TGAGTGCGGGAATAACATGA | CAGGATGACACCTCTGCTCA | 231 |
| chr9:96404400-96406422 | p2 | (AT)11 | TGAGTGCGGGAATAACATGA | CAGGATGACACCTCTGCTCA | 231 |
| chr9:96409298-96411316 | p6 | (AACTGA)3 | TCATGAACGTGACTTCGCTC | GGCTGAAACACAAAAAGGGA | 284 |
| chr9:96409298-96411316 | p5 | (ACTGA)3 | TCATGAACGTGACTTCGCTC | GGCTGAAACACAAAAAGGGA | 284 |
| chr9:96420859-96422874 | p3 | (CAA)5 | TGCCTCTTTTGACCTTGCTT | GCAAGATTTCTCCAACGCTC | 225 |
| chr9:96436827-96438845 | p6 | (CGGTTT)3 | GGGGGTTGTGTTTCATCATC | GGTATTGTTAGGGGTGGGCT | 280 |
| chr9:96442222-96444240 | p4 | (TTCT)4 | GACCGAGATATTTGGGACGA | GTTCAGCTCGGCTCGTTTTA | 166 |
| chr9:96442222-96444240 | p6 | (TTGGAG)3 | GACCGAGATATTTGGGACGA | GTTCAGCTCGGCTCGTTTTA | 166 |
| chr9:96533256-96535272 | p4 | (CAAA)4 | GCAGACCTCTTTGAAGGTGC | ATCTCACGCTATGGAGCCAC | 125 |
| chr9:96563643-96565658 | p5 | (CCTTT)3 | AGTACGAGCACCCACGTCTC | GATTGCTCAATTCGTGCGTA | 242 |
| chr9:96567590-96569612 | p2 | (AT)11 | TAGAAGCATGGGTTTGGACC | TGAGTTTGGAGATGGTGGGT | 205 |
| chr9:96613634-96615646 | p2 | (TA)6 | GATTCAATACGGGGGAAGGT | AGTCCCTGTGTTGTTCACCC | 224 |

| Marker ID | SSR type | SSR motif | FORWARD PRIMER (5'-3') | REVERSE PRIMER (5'-3') | size (bp) |
|---|---|---|---|---|---|
| chr9:96613634-96615646 | p2 | (TG)6 | GATTCAATACGGGGGAAGGT | AGTCCCTGTGTTGTTCACCC | 224 |
| chr9:96614331-96616343 | p2 | (TA)6 | GAACGGACACTAGCCCAAAA | GTCCTGTCCCGTCCTTATGA | 279 |
| chr9:96614331-96616343 | p2 | (TG)6 | GAACGGACACTAGCCCAAAA | GTCCTGTCCCGTCCTTATGA | 279 |
| chr9:96636463-96638483 | p5 | (ATTTT)4 | AACACTCGGACCGATACAGG | CAAGTACCTGGGGTCGGTTA | 196 |
| chr9:96671575-96673641 | p2 | (AT)33 | ACAACGCGTATTTCACTCCC | GAACGAAGCACATTTCACCA | 120 |
| chr9:96683586-96685640 | p2 | (CT)6 | TCCTCAGGTTCTAGGGGGTT | GTCTGTCACCCGCTCTTCTC | 277 |
| chr9:96683707-96685721 | p2 | (CT)6 | TCCTCAGGTTCTAGGGGGTT | GTCTGTCACCCGCTCTTCTC | 156 |
| chr9:96879950-96881968 | p6 | (TGCTGG)3 | CCTGCAGCTGATCAAACAAA | TTGGAGAAGGGTTTCATTCG | 236 |
| chr9:96879950-96881968 | p3 | (CTG)5 | CCTGCAGCTGATCAAACAAA | TTGGAGAAGGGTTTCATTCG | 236 |
| chr9:96901113-96903131 | p6 | (TGCTGG)3 | CCTGCAGCTGATCAAACAAA | TTGGAGAAGGGTTTCATTCG | 236 |
| chr9:96901113-96903131 | p3 | (CTG)5 | CCTGCAGCTGATCAAACAAA | TTGGAGAAGGGTTTCATTCG | 236 |
| chr9:97032842-97034860 | p3 | (CAC)6 | AGCCGTCTGCAGCCTTAGTA | TTTTCTCCATTGGATCCTGG | 259 |
| chr9:97070737-97072752 | p3 | (TCT)5 | AAAACATGTCCCAAAGTGGC | GGGTGAAGATGAGGTGCATT | 157 |
| chr9:97073785-97075809 | p2 | (AT)12 | ATGAGGAGCCATTGGAACAC | ACAAATGCTTCTCACCCGAG | 201 |
| chr9:97112629-97114644 | p2 | (GA)9 | ATAACCCAGGCAAACATCCA | GGGCACCTGGAGTTATTTGA | 275 |
| chr9:97112629-97114644 | p5 | (ATAAA)3 | ATAACCCAGGCAAACATCCA | GGGCACCTGGAGTTATTTGA | 275 |
| chr9:97168712-97170730 | p2 | (AT)9 | TTTATGGGGTTTATGGGCAA | GGCACCTGACGAGCTTCTAC | 254 |
| chr9:97168712-97170730 | p2 | (TA)7 | TTTATGGGGTTTATGGGCAA | GGCACCTGACGAGCTTCTAC | 254 |
| chr9:97170903-97172918 | p2 | (AT)6 | CGGACGACGAGAAAGTGAGT | TTGTCCACGTGCATATTCGT | 132 |
| chr9:97170903-97172918 | p2 | (AT)6 | CGGACGACGAGAAAGTGAGT | TTGTCCACGTGCATATTCGT | 132 |
| chr9:97170903-97172918 | p5 | (AAGTA)3 | CGGACGACGAGAAAGTGAGT | TTGTCCACGTGCATATTCGT | 132 |
| chr9:97202522-97204544 | p6 | (CAACTC)3 | GCTTTTTCCATTTGAGTCGC | TATGGCCAAACGCCTAAAAG | 190 |
| chr9:97202522-97204544 | p2 | (AT)11 | GCTTTTTCCATTTGAGTCGC | TATGGCCAAACGCCTAAAAG | 190 |
| chr9:97251918-97253951 | p2 | (TA)10 | CGGCCTTTTGCTCAAAGTAG | GAAAGGTCCATGCCTCCATA | 158 |
| chr9:97251952-97254000 | p2 | (TA)10 | CGGCCTTTTGCTCAAAGTAG | GAAAGGTCCATGCCTCCATA | 124 |
| chr9:97275856-97277871 | p2 | (AC)7 | CCCGCATCACTGTAAGGAGT | GAAGGGTCAAGGCATTTGAA | 135 |
| chr9:97323447-97325459 | p5 | (TCGGG)4 | TTCAGCTTTCCCTTCCAAAA | AAAGTGAGAGGGTGGGTGTG | 204 |
| chr9:97323447-97325459 | p2 | (AT)6 | TTCAGCTTTCCCTTCCAAAA | AAAGTGAGAGGGTGGGTGTG | 204 |
| chr9:97323447-97325459 | p3 | (TGT)5 | TTCAGCTTTCCCTTCCAAAA | AAAGTGAGAGGGTGGGTGTG | 204 |
| chr9:97351483-97353501 | p6 | (AAAAAG)3 | TTGGCTGCGAGTGATATACG | TACATACGCGGCCCTTAAAC | 168 |
| chr9:97378199-97380243 | p5 | (AAGAA)3 | GCTTTTCTTTCTCACACGGG | CAAGGTTGTGGGATCATCAA | 275 |
| chr9:97424239-97426254 | p5 | (TTTAA)3 | TCGCCTTGCTCAGTTGTATG | AGGCCCGAGAGAACCACTAT | 135 |
| chr9:97470418-97472433 | p5 | (TTTAA)3 | TCGCCTTGCTCAGTTGTATG | AGGCCCGAGAGAACCACTAT | 135 |
| chr9:97575646-97577668 | p2 | (TA)11 | CAAAATTCTGTCCATGTGCG | TAGTAGGGGCAATTTGCAGG | 196 |
| chr9:97584536-97586574 | p2 | (GA)6 | GTTGACCAAGGGGAAGTTGA | CCACTTCCATTCCACCACAT | 183 |
| chr9:97584536-97586574 | p2 | (TC)19 | GTTGACCAAGGGGAAGTTGA | CCACTTCCATTCCACCACAT | 183 |

| Marker ID | SSR type | SSR motif | FORWARD PRIMER (5'-3') | REVERSE PRIMER (5'-3') | size (bp) |
|---|---|---|---|---|---|
| chr9:97620158-97622176 | p3 | (ACA)6 | TGATTGTCTCCTTCCCATCC | GAGATGGGGAATGGAGGTTT | 105 |
| chr9:97636353-97638367 | p2 | (AT)7 | GGGCTCCAACATCAGGTAAA | GTCATGCGTTTGTTGGATTG | 270 |
| chr9:97636353-97638367 | p5 | (TAAAA)3 | GGGCTCCAACATCAGGTAAA | GTCATGCGTTTGTTGGATTG | 270 |
| chr9:97636876-97638891 | p2 | (AT)7 | AGGCTAGCGGTTTACCTTGG | CGTGCTGCATAAGCCAACTA | 173 |
| chr9:97636876-97638891 | p5 | (TAAAA)3 | AGGCTAGCGGTTTACCTTGG | CGTGCTGCATAAGCCAACTA | 173 |
| chr9:97770316-97772328 | p2 | (AT)12 | AAGCCTTTGGAGGAGGAAAA | GAAAGACCAAAAAGTGCCCA | 291 |
| chr9:97770420-97772436 | p2 | (AT)12 | AAGCCTTTGGAGGAGGAAAA | GAAAGACCAAAAAGTGCCCA | 187 |
| chr9:97781437-97783470 | p3 | (TAA)5 | CGGACTTTGATTTGTGGTCC | ACGGTCATTGCGACATCATA | 230 |
| chr9:97781437-97783470 | p3 | (ATA)5 | CGGACTTTGATTTGTGGTCC | ACGGTCATTGCGACATCATA | 230 |
| chr9:97781525-97783561 | p3 | (TAA)5 | CGGACTTTGATTTGTGGTCC | ACGGTCATTGCGACATCATA | 142 |
| chr9:97781525-97783561 | p3 | (ATA)5 | CGGACTTTGATTTGTGGTCC | ACGGTCATTGCGACATCATA | 142 |
| chr9:97892681-97894721 | p2 | (TC)6 | GGCAGTTTATTGACCCAGGA | TCCAGTCCCAACTCCACTTC | 276 |
| chr9:97919016-97921030 | p3 | (AGT)5 | TCCCCGTCCACTTTTACTTG | TATCCATGTTTGACTGGGCA | 224 |
| chr9:97919016-97921030 | p2 | (TC)7 | TCCCCGTCCACTTTTACTTG | TATCCATGTTTGACTGGGCA | 224 |
| chr9:97919016-97921030 | p3 | (ACT)5 | TCCCCGTCCACTTTTACTTG | TATCCATGTTTGACTGGGCA | 224 |
| chr9:97919016-97921030 | p3 | (TCC)5 | TCCCCGTCCACTTTTACTTG | TATCCATGTTTGACTGGGCA | 224 |
| chr9:97919782-97921797 | p3 | (AGT)5 | TCTCTCGCAGGAGGACATTT | ATGTTGGTTGGGGTTGAAAA | 242 |
| chr9:97919782-97921797 | p2 | (TC)7 | TCTCTCGCAGGAGGACATTT | ATGTTGGTTGGGGTTGAAAA | 242 |
| chr9:97919782-97921797 | p3 | (ACT)5 | TCTCTCGCAGGAGGACATTT | ATGTTGGTTGGGGTTGAAAA | 242 |
| chr9:97919782-97921797 | p3 | (TCC)5 | TCTCTCGCAGGAGGACATTT | ATGTTGGTTGGGGTTGAAAA | 242 |
| chr9:97998599-98000619 | p5 | (TGATA)4 | TATTGGACCAGACCCGAGAC | TTCATAACGGTCATGCCAAA | 170 |
| chr9:97998599-98000619 | p2 | (AT)7 | TATTGGACCAGACCCGAGAC | TTCATAACGGTCATGCCAAA | 170 |
| chr9:98018654-98020669 | p6 | (AAAAAT)3 | TGGAAATTTCGACCACATGA | AAACGTTTTTCGCCCTTTTT | 167 |
| chr9:98018654-98020669 | p5 | (AATTA)3 | TGGAAATTTCGACCACATGA | AAACGTTTTTCGCCCTTTTT | 167 |
| chr9:98081314-98083329 | p5 | (GAAAG)3 | CACAGCCCGAGGTAAAACAT | TTGTACCCGGAGTAGGCATC | 282 |
| chr9:98089935-98091947 | p2 | (CT)6 | TCTGACCCCATTGACTCCTC | TGTGGCCTCGTATGAAACAA | 118 |
| chr9:98123687-98125702 | p3 | (ATT)5 | GATGCCTTGGCTAAATGGAA | ATGGTTCCACCTGTGGATGT | 227 |
| chr9:98183584-98185599 | p3 | (TTA)5 | CTCTTGGTCACTAGCCGAGC | ATCAACAAGGGGTTGTAGCG | 241 |
| chr9:98237026-98239040 | p2 | (TA)7 | CGGTTCATAGCCTGGTTGAT | AACAAAAATGGGGATAGGGG | 140 |
| chr9:98243371-98245393 | p2 | (TA)11 | CCATAGGCCCAACATCACTT | TAGCATCCACATGGGTTTCA | 140 |
| chr9:98243371-98245393 | p2 | (TA)50 | CCATAGGCCCAACATCACTT | TAGCATCCACATGGGTTTCA | 140 |
| chr9:98256890-98258908 | p6 | (TTGCAC)3 | TTGTAGGCTCGTTCGCTTTT | AAAATGTCCAATTGAAGGCG | 259 |
| chr9:98292960-98294975 | p2 | (CT)7 | CTATTTCTTCGGCAGCGAAC | GGAATCAGGCCGTGTGTAGT | 290 |
| chr9:98292993-98295035 | p2 | (CT)7 | CTATTTCTTCGGCAGCGAAC | GGAATCAGGCCGTGTGTAGT | 257 |
| chr9:98293044-98295059 | p2 | (CT)7 | CTATTTCTTCGGCAGCGAAC | GGAATCAGGCCGTGTGTAGT | 206 |
| chr9:98293105-98295120 | p2 | (CT)7 | CTATTTCTTCGGCAGCGAAC | GGAATCAGGCCGTGTGTAGT | 145 |

| Marker ID | SSR type | SSR motif | FORWARD PRIMER (5'-3') | REVERSE PRIMER (5'-3') | size (bp) |
|---|---|---|---|---|---|
| chr9:98293128-98295143 | p2 | (CT)7 | CTATTTCTTCGGCAGCGAAC | GGAATCAGGCCGTGTGTAGT | 122 |
| chr9:98484377-98486409 | p2 | (TA)16 | GAGAGAGCACCATTGGGTGT | ATCTTCCCCACAACAAGCAC | 265 |
| chr9:98522621-98524633 | p2 | (TA)6 | GGCACATTCGACACCTTTTT | AGGCCATCACTGGAAAATTG | 252 |
| chr9:98530174-98532186 | p2 | (TA)6 | AGGGTGATCCTCCGAGACTT | TCCGGACCGTAGTTTTATCG | 192 |
| chr9:98547330-98549357 | p3 | (TGT)9 | TGGAAGTCAGGCTGGGTATC | TCATTCACCCTGCACCAATA | 152 |
| chr9:98568166-98570184 | p6 | (ATTGTT)3 | ATTTAAAATGGGACGGAGGG | TGTATTTCACTGCCTGCCAA | 195 |
| chr9:98591325-98593355 | p2 | (AT)15 | AAATTTTTGATAGCCCACGC | TGAGCCTGCCATAAGTGACA | 185 |
| chr9:98597348-98599360 | p2 | (GA)6 | TTGCCCAAAACTCAATAGGC | AAATGCCGACGATAAGATGG | 146 |
| chr9:98597348-98599360 | p4 | (TTTA)4 | TTGCCCAAAACTCAATAGGC | AAATGCCGACGATAAGATGG | 146 |
| chr9:98632336-98634350 | p2 | (TA)7 | TATCGCTATGAGGGCTCCAC | CGCCAAAAGGACATTAGGAA | 112 |
| chr9:98632600-98634614 | p2 | (TA)7 | CGGACGTTCGGTAATAATGC | CAAGCTTGGAATGAAAAGCC | 174 |
| chr9:98632629-98634667 | p2 | (TA)7 | CGGACGTTCGGTAATAATGC | CAAGCTTGGAATGAAAAGCC | 145 |
| chr9:98632787-98634799 | p2 | (TA)7 | CACAAGGCTTTTGTGAGCAA | CGGTTTCCTAACCGATTTCC | 235 |
| chr9:98677823-98679837 | p4 | (AAAT)4 | ATGCCTCTCTTTTGGACCCT | CTGGAGGGGACAAGTCAAGA | 228 |
| chr9:98677823-98679837 | p5 | (TTGAA)3 | ATGCCTCTCTTTTGGACCCT | CTGGAGGGGACAAGTCAAGA | 228 |
| chr9:98677823-98679837 | p2 | (AT)7 | ATGCCTCTCTTTTGGACCCT | CTGGAGGGGACAAGTCAAGA | 228 |
| chr9:98712688-98714700 | p2 | (TA)6 | GTTATCATCGTTGACCGCCT | CCGAAAGCCATAACGATCTC | 295 |
| chr9:98804311-98806326 | p5 | (CAAAC)3 | GGGATATTTTCGGCCACTTT | ATTTCAACAAGGCCCATCAG | 124 |
| chr9:98899718-98901732 | p2 | (TA)7 | GGGAAAACTACACGGCCATA | ACGCCTTCTTTTTCCTCCTC | 208 |
| chr9:98924691-98926703 | p2 | (TA)6 | GGGTGCCCATTCACTGATAC | AGAACCAACCATTCCAGTCG | 246 |
| chr9:98924691-98926703 | p3 | (GAA)6 | GGGTGCCCATTCACTGATAC | AGAACCAACCATTCCAGTCG | 246 |
| chr9:99011976-99013994 | p6 | (AGGAGA)3 | ATGCGAGGATGAAAAGGATG | GTATGGAGCTTCCACGGGTA | 139 |
| chr9:99011976-99013994 | p2 | (AT)6 | ATGCGAGGATGAAAAGGATG | GTATGGAGCTTCCACGGGTA | 139 |
| chr9:99012122-99014134 | p6 | (AGGAGA)3 | TGTGATCGATGCTTCTCCTG | TAAGGCGTATGGAAGTTGGC | 101 |
| chr9:99012122-99014134 | p2 | (AT)6 | TGTGATCGATGCTTCTCCTG | TAAGGCGTATGGAAGTTGGC | 101 |
| chr9:99013262-99015277 | p5 | (AAATA)3 | GCTATCCACAAAGCTCCTGG | AAGTTGCAAGCTTGATGGCT | 209 |
| chr9:99110358-99112384 | p2 | (AT)13 | ATTATCGTAAGCGGTCGCAC | TGTGGGTAATGGGTCATGTG | 107 |
| chr9:99128524-99130538 | p5 | (GAACC)3 | ACAGGCCTAGTCAGAGGGGT | GGAATGATTTGGGAGACCCT | 217 |
| chr9:99128524-99130538 | p2 | (CT)7 | ACAGGCCTAGTCAGAGGGGT | GGAATGATTTGGGAGACCCT | 217 |
| chr9:99264706-99266730 | p6 | (AAGTAA)4 | ATTTCGCTATTGCCCCTTTT | TTTATCTGCTTCAACGCCCT | 153 |
| chr9:99332355-99334367 | p2 | (AG)6 | GGATGAATCCAAGAGGGTCA | TGGAAGGAAGAAGGCTCAAA | 267 |
| chr9:99360321-99362336 | p3 | (TAA)12 | CCGACCAAGAGAGCTTCAAC | TGCCAATTTCATCAGCAAAA | 233 |
| chr9:99360401-99362421 | p3 | (TAA)12 | CCGACCAAGAGAGCTTCAAC | TGCCAATTTCATCAGCAAAA | 153 |
| chr9:99381356-99383372 | p4 | (TTTA)4 | GTTCATGGTCCGGTTTGACT | TACAACCCGGAAAAACCAAA | 289 |
| chr9:99404242-99406256 | p3 | (TGT)5 | AGATGTGGGATTTCACTGGG | AATCCGGTGACCTGTTGAAG | 107 |
| chr9:99441657-99443675 | p6 | (TTTACT)3 | TACCCGGTTTTTCATTGGTG | AACCTGTCAAAAGGAGCGAA | 122 |

| Marker ID | SSR type | SSR motif | FORWARD PRIMER (5'-3') | REVERSE PRIMER (5'-3') | size (bp) |
|---|---|---|---|---|---|
| chr9:99526362-99528377 | p3 | (GTT)5 | AATGGAACCGCTGACTTCTG | TTGTCACCTTGGTCCTGTCA | 183 |
| chr9:99526362-99528377 | p2 | (GA)7 | AATGGAACCGCTGACTTCTG | TTGTCACCTTGGTCCTGTCA | 183 |
| chr9:99527032-99529046 | p3 | (GTT)5 | TGGTGAAGGGAAGAAGAGGA | TGGGTCATAAGTGGCAATGA | 254 |
| chr9:99527032-99529046 | p2 | (GA)7 | TGGTGAAGGGAAGAAGAGGA | TGGGTCATAAGTGGCAATGA | 254 |
| chr9:99570189-99572205 | p2 | (AT)8 | GAAATTTCACGGCCTTCTGA | CATAGACCGAAATACGCGGT | 147 |
| chr9:99570189-99572205 | p2 | (AT)6 | GAAATTTCACGGCCTTCTGA | CATAGACCGAAATACGCGGT | 147 |
| chr9:99620430-99622445 | p3 | (ATA)5 | CATTCGTAAGCCGAGCTTTT | TGGAGTTGCATATGTGGTGG | 168 |
| chr9:99620430-99622445 | p3 | (TAT)5 | CATTCGTAAGCCGAGCTTTT | TGGAGTTGCATATGTGGTGG | 168 |
| chr9:99647377-99649392 | p5 | (AAAAT)3 | GGACAGGAGAACGTGAGGAG | ATCACATTTTACCCGGTCCA | 174 |
| chr9:99647377-99649392 | p3 | (CTT)5 | GGACAGGAGAACGTGAGGAG | ATCACATTTTACCCGGTCCA | 174 |
| chr9:99647377-99649392 | p3 | (GGT)5 | GGACAGGAGAACGTGAGGAG | ATCACATTTTACCCGGTCCA | 174 |
| chr9:99648142-99650157 | p5 | (AAAAT)3 | TCTTCTTCACAATGCCCCTC | GAAGAAGAAGAAGGTGGGGG | 286 |
| chr9:99648142-99650157 | p3 | (CTT)5 | TCTTCTTCACAATGCCCCTC | GAAGAAGAAGAAGGTGGGGG | 286 |
| chr9:99648142-99650157 | p3 | (GGT)5 | TCTTCTTCACAATGCCCCTC | GAAGAAGAAGAAGGTGGGGG | 286 |
| chr9:99681821-99683911 | p3 | (TAA)30 | GGTCAAAACCTCCACCTGAA | CCTTTCCCCGTCAGATAACA | 171 |
| chr9:99683987-99686007 | p4 | (AATA)5 | TGTCTCTTCAAGGGGATTGG | TAAGGGGTAGGAGGAGGGAA | 138 |
| chr9:99854981-99857017 | p3 | (ATT)12 | TATTTCAATGCCAAGGGAGG | CAGCCTGGGAGAAAGAAGTG | 141 |
| chr9:99893418-99895432 | p2 | (AT)7 | TACCGTGCTACAGGTGCTTG | TGGCTATCGAAGGAGACGAT | 228 |
| chr9:99971266-99973296 | p2 | (AG)7 | GCAGAAGTTGCGTTTTGTGA | ACGGGTTTTGCAGGATAATG | 231 |
| chr9:99971266-99973296 | p2 | (AT)15 | GCAGAAGTTGCGTTTTGTGA | ACGGGTTTTGCAGGATAATG | 231 |
| chr9:100005980-100008000 | p2 | (TA)10 | TGGTGGAAATGAACGACAGA | CTCCCATCTAATCCAATCGC | 130 |
| chr9:100079949-100081993 | p2 | (TA)22 | TCTTCAAAAACACCCGAACC | CCATCTGGGCAAAGAGTTGT | 184 |
| chr9:100091399-100093417 | p2 | (AT)11 | ACGTTTGGAAATGTTTTCGC | TCTGTCAAATGGTCCGTCAA | 171 |
| chr9:100091438-100093456 | p2 | (AT)11 | ACGTTTGGAAATGTTTTCGC | TCTGTCAAATGGTCCGTCAA | 132 |
| chr9:100220829-100222845 | p2 | (AG)8 | GTTGTGAGAAAATGGGACCG | AAACATCGAAGTCTGGGTCG | 288 |
| chr9:100306885-100308903 | p6 | (TATTTT)3 | CAAAGCCTTAGCCACTGAGC | TTAATAGCGTCGCAGTGGTG | 108 |
| chr9:100323716-100325740 | p2 | (TA)12 | ATTATTTGGCCCTTTTTCCG | ATTCGCTTGAGTGTCGCTTT | 220 |
| chr9:100387342-100389356 | p2 | (TC)7 | TTTTATGTCAGAGGCGGACC | GGGGCGGACCTATGTAGAAT | 151 |
| chr9:100491463-100493484 | p3 | (TGA)7 | CGAGAGGCTCTTCCACTACG | GTTTCTGCCCACTTGGTGAT | 236 |
| chr9:100561392-100563404 | p2 | (TA)6 | GATTTCCCTTCATTTGGGGT | GAGCAATTAGCAGGATTGGC | 229 |
| chr9:100561392-100563404 | p5 | (GTCAA)3 | GATTTCCCTTCATTTGGGGT | GAGCAATTAGCAGGATTGGC | 229 |
| chr9:100579847-100581861 | p2 | (AG)7 | AGTTAACCAGGTGTGGCAGG | CTGCAAAAGCACAACTCCAA | 203 |
| chr9:100601392-100603459 | p2 | (TA)26 | CGTGGCAGATGACAGAAGAA | GAAGGAGAGCCTTCATGCAC | 277 |
| chr9:100628049-100630063 | p2 | (AT)16 | CTCTTTTTGCAACAACCCGT | GCTCAACCAATAGGCCACAT | 128 |
| chr9:100628049-100630063 | p2 | (TA)7 | CTCTTTTTGCAACAACCCGT | GCTCAACCAATAGGCCACAT | 128 |
| chr9:100628049-100630063 | p2 | (TA)26 | CTCTTTTTGCAACAACCCGT | GCTCAACCAATAGGCCACAT | 128 |

| Marker ID | SSR type | SSR motif | FORWARD PRIMER (5'-3') | REVERSE PRIMER (5'-3') | size (bp) |
|---|---|---|---|---|---|
| chr9:100628049-100630063 | p2 | (TA)6 | CTCTTTTTGCAACAACCCGT | GCTCAACCAATAGGCCACAT | 128 |
| chr9:100628415-100630437 | p2 | (AT)16 | TCACGTAAGTAACCGAGCCC | TAACCGGTAAATCGGAATCG | 102 |
| chr9:100628415-100630437 | p2 | (TA)7 | TCACGTAAGTAACCGAGCCC | TAACCGGTAAATCGGAATCG | 102 |
| chr9:100628415-100630437 | p2 | (TA)26 | TCACGTAAGTAACCGAGCCC | TAACCGGTAAATCGGAATCG | 102 |
| chr9:100628415-100630437 | p2 | (AT)7 | TCACGTAAGTAACCGAGCCC | TAACCGGTAAATCGGAATCG | 102 |
| chr9:100629363-100631377 | p2 | (AT)7 | ATTAGCACGTGCGTTTCGTA | ATACGCATGTTGGCATTGAA | 206 |
| chr9:100706448-100708463 | p3 | (ATT)5 | TTTTGTTGATGCCCTGCATA | AGCCGAGTGTTCGAGGTAGA | 252 |
| chr9:100744065-100746077 | p2 | (TG)6 | TTGTACTACGCACCCCATGA | CTCATACCTGGCCCTACCAA | 232 |
| chr9:100816014-100818030 | p2 | (TA)8 | CAAAAGGGTTGTCGGGTTTA | ATGGTCTCCCATTTGCTGTC | 290 |
| chr9:100816316-100818328 | p2 | (TA)8 | AAGTATCAACCGTGGCTTCG | CCGCTTCTAAGTTTTGGTGC | 191 |
| chr9:100816400-100818414 | p2 | (TA)8 | AAGTATCAACCGTGGCTTCG | CCGCTTCTAAGTTTTGGTGC | 107 |
| chr9:100822119-100824151 | p2 | (TA)16 | TAGCGGCACGTATCCTCTCT | TGCCATGAATTTCGACACAT | 123 |
| chr9:100855919-100857941 | p2 | (AT)6 | TTGGACGTCTTAGCATGGTG | GCCTTCCTCCGCTTCTAAGT | 256 |
| chr9:100855942-100857954 | p2 | (AT)6 | TTGGACGTCTTAGCATGGTG | GCCTTCCTCCGCTTCTAAGT | 233 |
| chr9:100982519-100984534 | p3 | (CTG)5 | ACTTACCCCTCGGCGATACT | GCGGATTTGGTGGAGTTAGA | 149 |
| chr9:100987083-100989098 | p3 | (ACA)5 | AGTTATCGCTGCCCGAAGTA | TCAAGCCATGTCCAACCATA | 106 |
| chr9:101002861-101004883 | p2 | (TA)11 | GCGGACCCATCAACCTACTA | GGGTGTCCAGGTCTCTTTGA | 114 |
| chr9:101002861-101004883 | p3 | (GAA)6 | GCGGACCCATCAACCTACTA | GGGTGTCCAGGTCTCTTTGA | 114 |
| chr9:101030937-101032952 | p5 | (AGTTT)3 | GAATTCAAGTCCCCTTGGGT | CCCTCATTGAAACCCGACTA | 207 |
| chr9:101097162-101099204 | p2 | (AT)21 | TGTTAAGCCCTATGCGAGGT | ACATGGGTGGCATTATGGTT | 184 |
| chr9:101184138-101186153 | p5 | (TTGCA)3 | TGGTGTACCCATTGCCACTA | CGAACATCAGCCTTCTCACA | 289 |
| chr9:101197748-101199763 | p5 | (AAGCC)3 | GGGGCAGATAACACATTGCT | GTGTATGAGGGTCGCCTGTT | 262 |
| chr9:101243242-101245257 | p5 | (TTGCA)3 | TGGTGTACCCATTGCCACTA | CGAACATCAGCCTTCTCACA | 289 |
| chr9:101256852-101258867 | p5 | (AAGCC)3 | GGGGCAGATAACACATTGCT | GTGTATGAGGGTCGCCTGTT | 262 |
| chr9:101313121-101315136 | p5 | (TAGTT)3 | ATAAGCTTTTCGCGTCCTCA | TCGAGCAAAAACGGGATAAC | 167 |
| chr9:101327707-101329722 | p5 | (TAGTT)3 | GGTAATGCACGCTCCATTTT | GGAGGCTTCAATTTTCCTCC | 237 |
| chr9:101356703-101358718 | p5 | (TTTTA)3 | ATGCATTGCTGGTGCATAAA | TTTGACCCCTCTACTGTGGC | 205 |
| chr9:101356703-101358718 | p4 | (CATA)4 | ATGCATTGCTGGTGCATAAA | TTTGACCCCTCTACTGTGGC | 205 |
| chr9:101387132-101389147 | p5 | (TAGTT)3 | ATAAGCTTTTCGCGTCCTCA | TCGAGCAAAAACGGGATAAC | 167 |
| chr9:101401718-101403733 | p5 | (TAGTT)3 | GGTAATGCACGCTCCATTTT | GGAGGCTTCAATTTTCCTCC | 237 |
| chr9:101503468-101505489 | p3 | (TTA)7 | TTTAGGCGGGAAAAGAGACA | CGGCTGATGGAAGTTGGTAT | 114 |
| chr9:101536206-101538218 | p2 | (TA)6 | TAGCCTGTTTTGGTATGGGC | ATCTTGGGAGCCCTTTGATT | 119 |
| chr9:101618573-101620585 | p2 | (TA)6 | TCCGGCGTAGATCTGAGTTT | GACCTTTCACCACCGTCACT | 283 |
| chr9:101618573-101620585 | p2 | (CT)6 | TCCGGCGTAGATCTGAGTTT | GACCTTTCACCACCGTCACT | 283 |
| chr9:101732737-101734783 | p2 | (TA)23 | TACATATGCGAGCCGACAAG | AAGGGTTATTGAGGAGCGGT | 298 |
| chr9:101816988-101819000 | p2 | (CT)6 | GGTGAGGAATGAAATCCCCT | AGCCCGTTTTCAATTTGATG | 179 |

| Marker ID | SSR type | SSR motif | FORWARD PRIMER (5'-3') | REVERSE PRIMER (5'-3') | size (bp) |
|---|---|---|---|---|---|
| chr9:101864748-101866763 | p5 | (GAAAA)3 | AGTCATGGGAAGTCGGAATG | GTAGGCGAACCCTCAATCAA | 150 |
| chr9:101864748-101866763 | p5 | (GTTGT)3 | AGTCATGGGAAGTCGGAATG | GTAGGCGAACCCTCAATCAA | 150 |
| chr9:101865938-101867986 | p2 | (AT)24 | AACGTCGTGGTACGACATGA | TACATGTGGGATTGCGAAGA | 281 |
| chr9:101892793-101894808 | p5 | (GAAAA)3 | AGTCATGGGAAGTCGGAATG | GTAGGCGAACCCTCAATCAA | 150 |
| chr9:101892793-101894808 | p5 | (GTTGT)3 | AGTCATGGGAAGTCGGAATG | GTAGGCGAACCCTCAATCAA | 150 |
| chr9:101893983-101896031 | p2 | (AT)24 | AACGTCGTGGTACGACATGA | TACATGTGGGATTGCGAAGA | 281 |
| chr9:101949694-101951709 | p5 | (CATTA)3 | TACATGGACTTCACGGGTCA | GCCGACAACGATTCGATAAT | 237 |
| chr9:102080696-102082714 | p2 | (AT)9 | TCAGTACACTTCCCAAGGGG | GGAGAGGTGCATGCGATTAT | 297 |
| chr9:102179004-102181030 | p2 | (AT)13 | AGAAGGAAAAATGGGGGAGA | AGCATGCACCAATGATTGAA | 216 |
| chr9:102205759-102207785 | p2 | (AT)13 | AGAAGGAAAAATGGGGGAGA | AGCATGCACCAATGATTGAA | 216 |
| chr9:102292673-102294691 | p3 | (AGG)6 | AGCCATTTGTTGAGGAATCG | CCTCTGTTGGTCCCTCATGT | 131 |
| chr9:102304005-102306023 | p3 | (ACA)6 | ATACCTAGCCAAATGTGCGG | TTGTGCAATGTGCGTCACTA | 123 |
| chr9:102331901-102333913 | p5 | (GTTCG)3 | GAACCTAATGAGCAGAGGCG | TGTTTGGTGTCCACCTTCAA | 139 |
| chr9:102331901-102333913 | p2 | (TA)6 | GAACCTAATGAGCAGAGGCG | TGTTTGGTGTCCACCTTCAA | 139 |
| chr9:102343600-102345615 | p5 | (TTTAT)3 | TTCCAGGGAGGTTGCTTAAA | ATTTCCCTTTGGAAACCTGG | 239 |
| chr9:102369510-102371540 | p3 | (ACA)5 | GCCAAACTGGCCGTATACAT | GGAACATGGTTCCACTTGCT | 229 |
| chr9:102369510-102371540 | p2 | (TA)15 | GCCAAACTGGCCGTATACAT | GGAACATGGTTCCACTTGCT | 229 |
| chr9:102395780-102397798 | p6 | (AAAATA)4 | AATTCGCAGTGCAAAAATCC | ATGCAAAGTCTTGCCGAGTT | 110 |
| chr9:102395780-102397798 | p3 | (TCA)5 | AATTCGCAGTGCAAAAATCC | ATGCAAAGTCTTGCCGAGTT | 110 |
| chr9:102395780-102397798 | p2 | (AT)9 | AATTCGCAGTGCAAAAATCC | ATGCAAAGTCTTGCCGAGTT | 110 |
| chr9:102430554-102432572 | p3 | (TTA)6 | CTTGTGGGATCACTCTGGGT | ATCCAGTTCCCTGTCATTGC | 237 |
| chr9:102445627-102447639 | p5 | (CAAAA)3 | TTACAAGTTGGGTTGCCTCC | CCCCAGAAAGCGTGATAGAA | 110 |
| chr9:102445627-102447639 | p2 | (AG)6 | TTACAAGTTGGGTTGCCTCC | CCCCAGAAAGCGTGATAGAA | 110 |
| chr9:102500682-102502700 | p3 | (AAG)6 | TCGAGAAAACAAAAATCGGG | ATAAGAAGGGGTGCGTTTCC | 150 |
| chr9:102529724-102531742 | p6 | (TAAAAA)3 | CATGTGGCATCTCCATCATC | GGGCCAAATATACCCCTTGT | 118 |
| chr9:102529724-102531742 | p5 | (ATTTT)3 | CATGTGGCATCTCCATCATC | GGGCCAAATATACCCCTTGT | 118 |
| chr9:102530690-102532705 | p6 | (TAAAAA)3 | CCCTTCACTCAACCAAAAGC | TACAAGATGGCTCCACCTCA | 291 |
| chr9:102530690-102532705 | p5 | (ATTTT)3 | CCCTTCACTCAACCAAAAGC | TACAAGATGGCTCCACCTCA | 291 |
| chr9:102568030-102570048 | p3 | (AAG)6 | TCGAGAAAACAAAAATCGGG | ATAAGAAGGGGTGCGTTTCC | 150 |
| chr9:102597835-102599850 | p5 | (ATTTT)3 | CCCTTCACTCAACCAAAAGC | TACAAGATGGCTCCACCTCA | 296 |
| chr9:102638791-102640806 | p5 | (TTATT)3 | TTACCCTTTGTGGGGGTGTA | GCTTGGCGACCACAATAAAT | 116 |
| chr9:102638791-102640806 | p4 | (AAGA)4 | TTACCCTTTGTGGGGGTGTA | GCTTGGCGACCACAATAAAT | 116 |
| chr9:102641502-102643517 | p5 | (AAACC)3 | CCCAAAATAAGGGCATTCCT | CAGAGGTTGAAAAACCCGAA | 148 |
| chr9:102672066-102674081 | p3 | (TTC)5 | ACGTATACGCAGTGGGGAAG | GCATTGGTGGAACAAGGATT | 205 |
| chr9:102698525-102700591 | p2 | (AT)26 | TTCTGGAAGGAAGAAGGGGT | TTCCCCAATTTCAAAACAGC | 248 |
| chr9:102698525-102700591 | p2 | (AT)33 | TTCTGGAAGGAAGAAGGGGT | TTCCCCAATTTCAAAACAGC | 248 |

| Marker ID | SSR type | SSR motif | FORWARD PRIMER (5'-3') | REVERSE PRIMER (5'-3') | size (bp) |
|---|---|---|---|---|---|
| chr9:102821249-102823270 | p3 | (AAT)7 | GCCACATCAGCTTTTCCATT | GAAAAGGTAGTCGCCCAGTG | 158 |
| chr9:102821249-102823270 | p2 | (AT)7 | GCCACATCAGCTTTTCCATT | GAAAAGGTAGTCGCCCAGTG | 158 |
| chr9:102821800-102823814 | p3 | (AAT)7 | GTTGGCAGAACGAATTGTGA | CTGATCACTTGAATGGCGAG | 151 |
| chr9:102821800-102823814 | p2 | (AT)7 | GTTGGCAGAACGAATTGTGA | CTGATCACTTGAATGGCGAG | 151 |
| chr9:102821800-102823814 | p4 | (AATA)4 | GTTGGCAGAACGAATTGTGA | CTGATCACTTGAATGGCGAG | 151 |
| chr9:102822964-102824978 | p4 | (AATA)4 | GGTGCCTCATCACCTTTTTC | AGATAAAGGGGGAGAATGGG | 271 |
| chr9:102822964-102824978 | p2 | (AC)7 | GGTGCCTCATCACCTTTTTC | AGATAAAGGGGGAGAATGGG | 271 |
| chr9:102851602-102853617 | p5 | (AAAAT)3 | CCGCAACGCTTACTATGTCA | GTGATGGGTGTTCTAGCCGT | 163 |
| chr9:102991914-102993926 | p2 | (TA)6 | GCCTTAGCATGGTGGAAGAA | GCTCCGCTTCTGAGTATTGG | 203 |
| chr9:102991914-102993926 | p5 | (CCGGT)3 | GCCTTAGCATGGTGGAAGAA | GCTCCGCTTCTGAGTATTGG | 203 |
| chr9:103002590-103004608 | p3 | (ATT)6 | AGACTGCAACTTGCTGGGTT | CAGGATATATTCGGGCTCCA | 291 |
| chr9:103002590-103004608 | p5 | (GTAGA)3 | AGACTGCAACTTGCTGGGTT | CAGGATATATTCGGGCTCCA | 291 |
| chr9:103002623-103004641 | p3 | (ATT)6 | AGACTGCAACTTGCTGGGTT | CAGGATATATTCGGGCTCCA | 258 |
| chr9:103002623-103004641 | p5 | (GTAGA)3 | AGACTGCAACTTGCTGGGTT | CAGGATATATTCGGGCTCCA | 258 |
| chr9:103002744-103004762 | p3 | (ATT)6 | AGACTGCAACTTGCTGGGTT | CAGGATATATTCGGGCTCCA | 137 |
| chr9:103002744-103004762 | p5 | (GTAGA)3 | AGACTGCAACTTGCTGGGTT | CAGGATATATTCGGGCTCCA | 137 |
| chr9:103003588-103005603 | p3 | (ATT)6 | ACCACCCATACACATTCGGT | TGGTTCCCCTCAATTGAAAC | 297 |
| chr9:103003588-103005603 | p5 | (GTAGA)3 | ACCACCCATACACATTCGGT | TGGTTCCCCTCAATTGAAAC | 297 |
| chr9:103171473-103173488 | p5 | (ATCAA)3 | GTACGGCATCTTTCGGATGT | TTTCTCACTGTGTCGTCGGA | 269 |
| chr9:103171473-103173488 | p5 | (ATTAA)3 | GTACGGCATCTTTCGGATGT | TTTCTCACTGTGTCGTCGGA | 269 |
| chr9:103171611-103173626 | p5 | (ATCAA)3 | GTACGGCATCTTTCGGATGT | TTTCTCACTGTGTCGTCGGA | 131 |
| chr9:103171611-103173626 | p5 | (ATTAA)3 | GTACGGCATCTTTCGGATGT | TTTCTCACTGTGTCGTCGGA | 131 |
| chr9:103197616-103199628 | p2 | (CA)6 | ATGCCACGTAACCAACCAAT | ACCCAATTCCCATTTCATCA | 198 |
| chr9:103251654-103253669 | p5 | (GGCGC)3 | TTATTTGGGCTTTGACCCAG | AAGGTAAATTGGAAACCCCG | 186 |
| chr9:103251654-103253669 | p2 | (TA)18 | TTATTTGGGCTTTGACCCAG | AAGGTAAATTGGAAACCCCG | 186 |
| chr9:103252440-103254476 | p5 | (GGCGC)3 | TCGAAGGGTTAATGGACTGG | CCAATTTTGGGCTCATTTGT | 294 |
| chr9:103252440-103254476 | p2 | (TA)18 | TCGAAGGGTTAATGGACTGG | CCAATTTTGGGCTCATTTGT | 294 |
| chr9:103269182-103271194 | p2 | (TA)6 | AGAGAGTCCGGAGCCAAAAT | GCAATTTCGTGGAACAATCC | 200 |
| chr9:103269182-103271194 | p2 | (TA)8 | AGAGAGTCCGGAGCCAAAAT | GCAATTTCGTGGAACAATCC | 200 |
| chr9:103275932-103277970 | p2 | (AG)19 | CCGGGCCTATGTCTCTCTTT | TTAGCGATTGGAGTGCAGTG | 239 |
| chr9:103329724-103331739 | p5 | (TTTTG)3 | TTGTATGAGGCCCAAAAAGG | ACCGAAATCAGGTGCTTGAG | 219 |
| chr9:103346704-103348728 | p2 | (AT)12 | ATGGGAAGAAAGAGGGTTGG | AAATCGAAGCATGCACACTG | 169 |
| chr9:103365847-103367887 | p4 | (TTAT)4 | TGCGTATCAACCCGTTAGTG | AAATGACATGGCGGAATAGC | 142 |
| chr9:103405057-103407078 | p3 | (TTG)7 | GGTCATCGGTTTGTGCTTCT | GTGTATTACCAGCCGCGTTT | 240 |
| chr9:103510983-103513007 | p3 | (TGT)8 | AATGGGATACCAAGAGCGAA | TTGTTTATTTGGGCTACCGC | 163 |
| chr9:103542571-103544589 | p5 | (TCATA)10 | CGCCTTACCTTTCCCCTTAC | TACCGACCTTACGTCATCCC | 200 |

| Marker ID | SSR type | SSR motif | FORWARD PRIMER (5'-3') | REVERSE PRIMER (5'-3') | size (bp) |
|---|---|---|---|---|---|
| chr9:103542571-103544589 | p6 | (CATATA)3 | CGCCTTACCTTTCCCCTTAC | TACCGACCTTACGTCATCCC | 200 |
| chr9:103542571-103544589 | p3 | (CAA)6 | CGCCTTACCTTTCCCCTTAC | TACCGACCTTACGTCATCCC | 200 |
| chr9:103633329-103635365 | p2 | (AT)18 | TACATTCTTCCCCATGCTCC | TCCCAATAACCGAAATGGAA | 181 |
| chr9:103854568-103856583 | p5 | (AAACC)3 | TACGACCTGGGTTTACGGTC | ACAGTGGTCGTAAAGGGCAC | 205 |
| chr9:103896589-103898641 | p2 | (AT)26 | AAATTACAGTGGCTGGGTCG | AAAGGAGGGAGAATGACGGT | 244 |
| chr9:103912363-103914381 | p5 | (AAGAA)3 | TAAATGCAGACATGGGGTGA | ACCACATACTCTTGCCGTCC | 247 |
| chr9:103912363-103914381 | p2 | (TA)9 | TAAATGCAGACATGGGGTGA | ACCACATACTCTTGCCGTCC | 247 |
| chr9:103967179-103969195 | p2 | (AT)8 | AAGTATCAACCGTGGCTTCG | AGTCTTCATTTGGTCCGGTG | 127 |
| chr9:103995986-103997998 | p2 | (TA)6 | GAAGGCAAGTGAGGGTCAAA | CAAGGCGAACAACACAAAGA | 140 |
| chr9:104023607-104025627 | p5 | (GGGTG)4 | ACGCGATTTCTTTCATTTGC | TGAACCACAGGAAACAACCA | 160 |
| chr9:104038636-104040650 | p2 | (TA)7 | GTCTGTGGCCCTCATCAAGT | TGTGTCATTGAAGGGTTGGA | 149 |
| chr9:104038636-104040650 | p6 | (AAGAAA)3 | GTCTGTGGCCCTCATCAAGT | TGTGTCATTGAAGGGTTGGA | 149 |
| chr9:104055382-104057402 | p5 | (GGGTG)4 | ACGCGATTTCTTTCATTTGC | TGAACCACAGGAAACAACCA | 160 |
| chr9:104068901-104070913 | p2 | (TC)6 | GTTTTGCCGTTCAACACCTT | CCGACTGCTTTTTGGACTTC | 200 |
| chr9:104183558-104185576 | p3 | (AAC)6 | GACTCTCCTCAACGACCGAG | GAGGTGAAGGGAACTGTGGA | 251 |
| chr9:104346915-104348983 | p2 | (AT)34 | TTGGTGTGACAAAAAGGTGC | GGCCTCTCCTACCTTGTCCT | 114 |
| chr9:104393607-104395622 | p3 | (TGT)5 | GGTTGTGAGACCGACCTTGT | TCCCTCATCTTGTCCTCCAC | 269 |
| chr9:104393607-104395622 | p2 | (AT)14 | GGTTGTGAGACCGACCTTGT | TCCCTCATCTTGTCCTCCAC | 269 |
| chr9:104526681-104528699 | p3 | (CAA)6 | AGGCTGGCAAGCAACATAGT | ACCTTACGTCACTCCCGATG | 138 |
| chr9:104553084-104555096 | p2 | (TA)6 | CATTTGGTGCGGGTAAGTTT | ATAGTGGCCCGTCTCATCAC | 266 |
| chr9:104589479-104591494 | p3 | (ATC)5 | AATCACAAGAACGGGACAGG | TTATGTTGCCACACATGCCT | 260 |
| chr9:104767443-104769459 | p2 | (TC)8 | AATGGCCCTCCTGTACACAC | GTGATTTTGGGTTTGGATGC | 169 |
| chr9:104769783-104771799 | p2 | (TC)8 | AATGGCCCTCCTGTACACAC | GTGATTTTGGGTTTGGATGC | 169 |
| chr9:104829312-104831332 | p2 | (AT)11 | CCCCGAAAAGCTTATGTTGA | ACCGTAACAACACACAGCGA | 112 |
| chr9:104829312-104831332 | p2 | (AT)8 | CCCCGAAAAGCTTATGTTGA | ACCGTAACAACACACAGCGA | 112 |
| chr9:104829389-104831405 | p2 | (AT)11 | AGTAAGGGGCGGTATCATCC | AACCGGTTAACGGGCTTACT | 109 |
| chr9:104829389-104831405 | p2 | (AT)8 | AGTAAGGGGCGGTATCATCC | AACCGGTTAACGGGCTTACT | 109 |
| chr9:104861147-104863165 | p6 | (TTTTTG)3 | TTCACTTTTGCATAGCGTCG | ACATCGTCGAGTTCAGCCTT | 114 |
| chr9:104894852-104896902 | p3 | (CAA)5 | GCGGACCAAAATTTAAGGGT | GGTCGGGTCTGGTTCATTTA | 238 |
| chr9:104894852-104896902 | p2 | (AT)25 | GCGGACCAAAATTTAAGGGT | GGTCGGGTCTGGTTCATTTA | 238 |
| chr9:104979469-104981484 | p5 | (TTCTT)3 | TCTTTCCGAGGGTTCATCAC | ATTTGGTGGGATTGGACTGA | 145 |
| chr9:105059418-105061434 | p4 | (CAAA)4 | CGACACGGAGCACAGATTTA | CGCTGCATCCATTCTCATAA | 172 |
| chr9:105072727-105074745 | p2 | (AT)9 | GTTGTGAGCTTCCCCAAAAA | AAGATCGGGGTGTGACAAAG | 250 |
| chr9:105072727-105074745 | p6 | (CATATA)4 | GTTGTGAGCTTCCCCAAAAA | AAGATCGGGGTGTGACAAAG | 250 |
| chr9:105201115-105203130 | p3 | (GTT)5 | TGCTTTCGGAGTGCTATGTG | CCTTGTCCCGAATATCCTCA | 224 |
| chr9:105463353-105465365 | p2 | (AT)6 | GGGAGGCGTTAGCTTCTTTT | AACCTCCCGAAGGCTGTATT | 107 |

| Marker ID | SSR type | SSR motif | FORWARD PRIMER (5'-3') | REVERSE PRIMER (5'-3') | size (bp) |
|---|---|---|---|---|---|
| chr9:105510481-105512496 | p3 | (AGC)5 | ACAGGACAGCAGGCTTCAAT | TGGACACCCCTAGTTGCTTC | 156 |
| chr9:105510481-105512496 | p6 | (TAATAC)3 | ACAGGACAGCAGGCTTCAAT | TGGACACCCCTAGTTGCTTC | 156 |
| chr9:105510481-105512496 | p2 | (AT)10 | ACAGGACAGCAGGCTTCAAT | TGGACACCCCTAGTTGCTTC | 156 |
| chr9:105523867-105525933 | p2 | (AT)33 | TAAGTGGACTGATCCGGAGG | TTTTCGATCGGTTGGGTAAG | 114 |
| chr9:105547494-105549548 | p3 | (AAT)18 | TAGAGAGGTGTGGGGGTGTC | TCCCTCTCTCGTCGTCATCT | 225 |
| chr9:105559788-105561804 | p2 | (AT)8 | CGCCGAGTCCATGGTATATT | ACCTCCCCAGCACATGTAAA | 109 |
| chr9:105561722-105563740 | p2 | (TA)9 | CTTCGTCGTCCGTTAAGCAT | GTGAGTTTGGAGATGGTGGG | 287 |
| chr9:105561722-105563740 | p2 | (AT)16 | CTTCGTCGTCCGTTAAGCAT | GTGAGTTTGGAGATGGTGGG | 287 |
| chr9:105579717-105581729 | p6 | (TGTATT)4 | CCACCACCACCATCTTCTCT | ATCTACGGCATCTGTTTCGG | 244 |
| chr9:105579717-105581729 | p2 | (AT)6 | CCACCACCACCATCTTCTCT | ATCTACGGCATCTGTTTCGG | 244 |
| chr9:105677560-105679574 | p2 | (TA)7 | TTCCCAAAGTATCCCCATCA | GGCGGAAGCAAATACAAAAA | 228 |
| chr9:105751963-105753981 | p6 | (AACTGA)3 | AGATGCGTGGTTGGATTCTC | AAAGGTGATTGATGGCTTGG | 152 |
| chr9:105775435-105777449 | p2 | (AG)7 | AGTGGGTTTGGTCGAGATTG | CCTCGCTAGTGGGAGATTTG | 149 |
| chr9:105823306-105825321 | p5 | (TTTCT)3 | ATCCATAAAAAGGGGATGGG | AATCAGATCAATCCCCACCA | 199 |
| chr9:105823306-105825321 | p5 | (TTTTA)3 | ATCCATAAAAAGGGGATGGG | AATCAGATCAATCCCCACCA | 199 |
| chr9:105823306-105825321 | p2 | (TA)7 | ATCCATAAAAAGGGGATGGG | AATCAGATCAATCCCCACCA | 199 |
| chr9:105832047-105834059 | p2 | (AT)6 | TGTCGGGTTTGTTGCATTTA | CGTCGCTTGTATGTCTTCCA | 125 |
| chr9:105832047-105834059 | p2 | (TA)9 | TGTCGGGTTTGTTGCATTTA | CGTCGCTTGTATGTCTTCCA | 125 |
| chr9:105846049-105848064 | p6 | (AAATAA)3 | TCCTCGGTAAACTGGGACTG | TCAGTGGAGAACAAGATGCG | 135 |
| chr9:105846049-105848064 | p5 | (AAAAG)3 | TCCTCGGTAAACTGGGACTG | TCAGTGGAGAACAAGATGCG | 135 |
| chr9:105934130-105936148 | p2 | (TA)9 | TGATTAGCAAAACCAACCCC | CGATAATCAAGGGCTGCAAT | 260 |
| chr9:105937841-105939857 | p4 | (AATC)4 | GGCACAAACAAACACGCTTA | TTTGGCAATGCATTTTCTTG | 288 |
| chr9:105937841-105939857 | p6 | (AAAATC)3 | GGCACAAACAAACACGCTTA | TTTGGCAATGCATTTTCTTG | 288 |
| chr9:105938025-105940043 | p4 | (AATC)4 | CAGAGGCACAAACAAACACG | TTTGGCAATGCATTTTCTTG | 100 |
| chr9:105938025-105940043 | p6 | (AAAATC)3 | CAGAGGCACAAACAAACACG | TTTGGCAATGCATTTTCTTG | 100 |
| chr9:105955702-105957717 | p3 | (GTT)5 | TTGTGTCTATAGGGCCTCGG | GCTATGGCGGACCTCATTTA | 227 |
| chr9:105973387-105975426 | p3 | (AAT)13 | AAAATTAAAAACCGGCCCAT | GGGCCAAATCGGCTATAAAT | 160 |
| chr9:105978279-105980294 | p3 | (TTA)5 | GGGCAAGCCACGTAGACTAA | CAACATGGGACCACATGAAA | 121 |
| chr9:106032212-106034230 | p5 | (TTTAC)3 | GGTGCTATTTCATCGTCCGT | GTTTTGATGTGGTGGGTTCC | 272 |
| chr9:106105750-106107792 | p2 | (TA)21 | TTTTGGGAAGGCTCAAAATG | TTGATTGGTCCCACACATTG | 263 |
| chr9:106105962-106107980 | p2 | (TA)21 | ATATTGCTTTTGGCCCACAC | TATTCCGGGATTTGAAGTCG | 216 |
| chr9:106106020-106108036 | p2 | (TA)21 | GTGTGGGACCAATCAAGGAC | GAAGTCGGGACTTTTGTCCA | 212 |
| chr9:106106045-106108059 | p2 | (TA)21 | GTGTGGGACCAATCAAGGAC | GAAGTCGGGACTTTTGTCCA | 187 |
| chr9:106176778-106178794 | p2 | (TA)7 | AATAATGTCCAAAAGGGTGGG | TTTTTCCCCATGAGTTGGAG | 298 |
| chr9:106176778-106178794 | p2 | (TA)13 | AATAATGTCCAAAAGGGTGGG | TTTTTCCCCATGAGTTGGAG | 298 |
| chr9:106176778-106178794 | p5 | (TAAAA)3 | AATAATGTCCAAAAGGGTGGG | TTTTTCCCCATGAGTTGGAG | 298 |

| Marker ID | SSR type | SSR motif | FORWARD PRIMER (5'-3') | REVERSE PRIMER (5'-3') | size (bp) |
|---|---|---|---|---|---|
| chr9:106176778-106178794 | p2 | (AT)13 | AATAATGTCCAAAAGGGTGGG | TTTTTCCCCATGAGTTGGAG | 298 |
| chr9:106176802-106178814 | p2 | (TA)7 | AATAATGTCCAAAAGGGTGGG | TTTTTCCCCATGAGTTGGAG | 274 |
| chr9:106176802-106178814 | p2 | (TA)13 | AATAATGTCCAAAAGGGTGGG | TTTTTCCCCATGAGTTGGAG | 274 |
| chr9:106176802-106178814 | p5 | (TAAAA)3 | AATAATGTCCAAAAGGGTGGG | TTTTTCCCCATGAGTTGGAG | 274 |
| chr9:106176802-106178814 | p2 | (AT)13 | AATAATGTCCAAAAGGGTGGG | TTTTTCCCCATGAGTTGGAG | 274 |
| chr9:106245169-106247181 | p2 | (AC)6 | TCGAAAACTGCAATTGGTCA | GAGACAGGGTGCCATATGCT | 207 |
| chr9:106287953-106289971 | p3 | (CAA)6 | CGGCGAAGGAAATAACAAAA | AAAGCTCGGATTGCTTTGAA | 273 |
| chr9:106303440-106305452 | p2 | (AT)6 | TGGCCACCAAGTGTTTGATA | GGATCATGGGGAAGAACGTA | 115 |
| chr9:106348343-106350361 | p2 | (CT)9 | CTGCAAGCCAGCAATGAATA | TTTGCCTGAGCCAGAGTTTT | 126 |
| chr9:106348343-106350361 | p2 | (AT)8 | CTGCAAGCCAGCAATGAATA | TTTGCCTGAGCCAGAGTTTT | 126 |
| chr9:106380554-106382568 | p2 | (AC)7 | CTGGCAGACGACAATACCAA | GTGTTGGTGAACACATTCGC | 179 |
| chr9:106380554-106382568 | p5 | (GAAAA)3 | CTGGCAGACGACAATACCAA | GTGTTGGTGAACACATTCGC | 179 |
| chr9:106392923-106394935 | p2 | (TA)6 | AGATAATTTGCCAACGGACG | ACGCATTGCCTTCTTCATCT | 221 |
| chr9:106392923-106394935 | p2 | (AG)6 | AGATAATTTGCCAACGGACG | ACGCATTGCCTTCTTCATCT | 221 |
| chr9:106413903-106415918 | p3 | (TAA)5 | CTTCATCCACTTCCACCCAT | TTTGCTTGTCCAAAAAGGCT | 261 |
| chr9:106428701-106430725 | p3 | (ATG)8 | AGTGATGCAGGAGAGCGATT | TCACAATTTTGGAGGGAAGC | 176 |
| chr9:106489926-106491941 | p3 | (TAA)5 | CTTCATCCACTTCCACCCAT | TTTGCTTGTCCAAAAAGGCT | 261 |
| chr9:106504724-106506748 | p3 | (ATG)8 | AGTGATGCAGGAGAGCGATT | TCACAATTTTGGAGGGAAGC | 176 |
| chr9:106565864-106567880 | p4 | (ATAA)4 | CTTCTCCTTTTTCTTCCGCC | TTGACAGGCAAATTTGTGGA | 182 |
| chr9:106565864-106567880 | p5 | (AATTT)3 | CTTCTCCTTTTTCTTCCGCC | TTGACAGGCAAATTTGTGGA | 182 |
| chr9:106732230-106734246 | p4 | (TATT)5 | TGCCATCCTGTGACATGTTT | AGCATAGCTACGGGCTTGAG | 267 |
| chr9:106732230-106734246 | p4 | (TTAT)4 | TGCCATCCTGTGACATGTTT | AGCATAGCTACGGGCTTGAG | 267 |
| chr9:106771907-106773922 | p5 | (CCCTG)3 | TGTCCTCCACCACTAGGTCC | TGGTGGCCGTACAGTGATTA | 274 |
| chr9:106799566-106801618 | p5 | (TTTTA)3 | AAAGCTTCCCAATCCGAAAT | GGGCGACGCAAAATCTACTA | 247 |
| chr9:106799566-106801618 | p2 | (TA)26 | AAAGCTTCCCAATCCGAAAT | GGGCGACGCAAAATCTACTA | 247 |
| chr9:106900770-106902792 | p2 | (AT)11 | ACAGAGCCAGTCAGCTCCAT | ACCCCCAGCAAAAAGAAGAT | 105 |
| chr9:106925990-106928010 | p2 | (AG)10 | TTACGCGAGAGCGATGTATG | AACTCGACTAGCGCTGGAAA | 266 |
| chr9:106933593-106935608 | p3 | (CTC)5 | TCGATCATTTTGGAGTCACG | TTGGCTCAAACCCTACAAGG | 101 |
| chr9:107017048-107019062 | p2 | (AT)6 | AGGCAGTTGGGAATTGAATG | AAATCCCGACGAACATCAAC | 186 |
| chr9:107017069-107019081 | p2 | (AT)6 | AGGCAGTTGGGAATTGAATG | AAATCCCGACGAACATCAAC | 165 |
| chr9:107052704-107054718 | p2 | (AT)7 | TGAGTTGCGCTCACCAATAG | GGTCGCGAGGTTTCAGTTTA | 269 |
| chr9:107077708-107079726 | p6 | (ACTGAA)3 | GCCACTTTGAAGGGTCCATA | GCACAGCATAGGATGAAGCA | 194 |
| chr9:107103051-107105066 | p5 | (TAAAT)3 | CCGTTTGCTTTTTCGATGAT | GTTGGAACCGTTGAGATCGT | 268 |
| chr9:107437240-107439267 | p2 | (AT)7 | TAGGAAGAGCCAAAGCCTGA | TGACTGCACGAATGTGTTGA | 194 |
| chr9:107462899-107464915 | p2 | (AG)8 | CAACCTAGCCCAGTCCACAT | AATGATTTTCACCCCGCATA | 112 |
| chr9:107481513-107483528 | p3 | (TTC)5 | ATTGGGGATGAGAAGCACAG | TCCAGCTGAGAATATTGGGC | 180 |

| Marker ID | SSR type | SSR motif | FORWARD PRIMER (5'-3') | REVERSE PRIMER (5'-3') | size (bp) |
|---|---|---|---|---|---|
| chr9:107495417-107497431 | p2 | (TA)7 | CCGTTTCACATATTCAGGGG | CAGATCGAAGCCACACCTTT | 287 |
| chr9:107495417-107497431 | p2 | (AC)7 | CCGTTTCACATATTCAGGGG | CAGATCGAAGCCACACCTTT | 287 |
| chr9:107702924-107704938 | p2 | (TA)7 | CTGGTTCCGGAAAAGACAAA | GACATAGAGGGGCAGAACCA | 181 |
| chr9:107711359-107713398 | p3 | (ATA)13 | TGAAGGACCCCAGGTTACTG | CAAAAGTTTGTGGGGGAAAA | 293 |
| chr9:107723162-107725180 | p3 | (AAG)6 | AGCGCCAAACTCATAATTGG | AGCCTCACTCATTTGCTGCT | 156 |
| chr9:107740628-107742644 | p6 | (AAAAAG)3 | CATTTCTGCCGCCTATTTGT | CGAAAATGTTGAATTGGCCT | 269 |
| chr9:107740628-107742644 | p4 | (TTAT)4 | CATTTCTGCCGCCTATTTGT | CGAAAATGTTGAATTGGCCT | 269 |
| chr9:107740628-107742644 | p3 | (AAG)7 | CATTTCTGCCGCCTATTTGT | CGAAAATGTTGAATTGGCCT | 269 |
| chr9:107756046-107758061 | p3 | (TTC)5 | AACCCTCCTTCCACCACTCT | TTTAAAATGGCGAGAATGCC | 286 |
| chr9:107777766-107779778 | p2 | (TA)6 | TCAGTGGCGGAGCTAAGATT | TCTCCCCCACAAACGTAAAC | 222 |
| chr9:107777766-107779778 | p2 | (AT)6 | TCAGTGGCGGAGCTAAGATT | TCTCCCCCACAAACGTAAAC | 222 |
| chr9:107867068-107869143 | p3 | (ATA)25 | TTATTGTCAGGGGATTTCGC | CCCCATCCCTTCTCTTCTTC | 214 |
| chr9:107867068-107869143 | p2 | (TC)6 | TTATTGTCAGGGGATTTCGC | CCCCATCCCTTCTCTTCTTC | 214 |
| chr9:107970572-107972587 | p5 | (ATTTA)3 | CTGCCAACAGCAGAGATTCA | TGAGACTGGCTCCTCATCCT | 167 |
| chr9:108021549-108023564 | p3 | (TGT)5 | CTTGCCCGGTTAGTCTTTCA | GGCTTTTGGCCTCATATTCA | 225 |
| chr9:108054479-108056497 | p6 | (ATAAAA)3 | AGTCCTTGTGGCTCGAAAAA | GCTGTGCCGGAGATAAGAAG | 132 |
| chr9:108218537-108220549 | p2 | (TA)6 | ATGCCTTTGGCATTCGTTAC | CATTCTGGAGCCGATTCATT | 154 |
| chr9:108218537-108220549 | p2 | (AT)7 | ATGCCTTTGGCATTCGTTAC | CATTCTGGAGCCGATTCATT | 154 |
| chr9:108218537-108220549 | p5 | (TGAAT)3 | ATGCCTTTGGCATTCGTTAC | CATTCTGGAGCCGATTCATT | 154 |
| chr9:108242265-108244289 | p6 | (TTTATT)4 | GCCAAAGATCGGTGAGAAAA | AAACCTTCAATGCCAGGCTA | 212 |
| chr9:108272663-108274679 | p2 | (AT)8 | CCGAAACCGCTATTGATGAT | CGACAAGCCCTTTTTGCTAC | 102 |
| chr9:108272663-108274679 | p2 | (AT)8 | CCGAAACCGCTATTGATGAT | CGACAAGCCCTTTTTGCTAC | 102 |
| chr9:108272795-108274861 | p2 | (AT)8 | AAGTATCAACCGTGGCTTCG | CTCTTCCTCTTCGCCTTCCT | 279 |
| chr9:108272795-108274861 | p2 | (AT)8 | AAGTATCAACCGTGGCTTCG | CTCTTCCTCTTCGCCTTCCT | 279 |
| chr9:108276963-108278975 | p2 | (TA)6 | CCAGATGATCGATAGGGTGC | TCTGATAGCTGACCAACCCC | 296 |
| chr9:108276963-108278975 | p2 | (AT)8 | CCAGATGATCGATAGGGTGC | TCTGATAGCTGACCAACCCC | 296 |
| chr9:108276963-108278975 | p2 | (AT)6 | CCAGATGATCGATAGGGTGC | TCTGATAGCTGACCAACCCC | 296 |
| chr9:108277448-108279460 | p2 | (TA)6 | TTGCAACCTGCAAGGATATG | TAATCACTCCAAACCGGACC | 265 |
| chr9:108277448-108279460 | p2 | (AT)8 | TTGCAACCTGCAAGGATATG | TAATCACTCCAAACCGGACC | 265 |
| chr9:108277448-108279460 | p2 | (AT)6 | TTGCAACCTGCAAGGATATG | TAATCACTCCAAACCGGACC | 265 |
| chr9:108277563-108279591 | p2 | (TA)6 | TTGCAACCTGCAAGGATATG | TAATCACTCCAAACCGGACC | 150 |
| chr9:108277563-108279591 | p2 | (AT)8 | TTGCAACCTGCAAGGATATG | TAATCACTCCAAACCGGACC | 150 |
| chr9:108277563-108279591 | p2 | (AT)6 | TTGCAACCTGCAAGGATATG | TAATCACTCCAAACCGGACC | 150 |
| chr9:108305339-108307353 | p2 | (AT)7 | CAGCGTGCAACAAGGTAGAA | TGTAAAAGTCCCAGCCCAAG | 219 |
| chr9:108318549-108320569 | p2 | (CT)10 | GACGTTCCACTGTAGCGGTT | AAAATCGGTGGTTCCACAAG | 300 |
| chr9:108318693-108320723 | p2 | (CT)10 | GACGTTCCACTGTAGCGGTT | AAAATCGGTGGTTCCACAAG | 156 |

| Marker ID | SSR type | SSR motif | FORWARD PRIMER (5'-3') | REVERSE PRIMER (5'-3') | size (bp) |
|---|---|---|---|---|---|
| chr9:108331536-108333551 | p5 | (TTTAT)3 | AGGTGGAAAACACAAGCACC | CTTGAAGGCAAGATTCGAGG | 287 |
| chr9:108331536-108333551 | p5 | (AATTA)3 | AGGTGGAAAACACAAGCACC | CTTGAAGGCAAGATTCGAGG | 287 |
| chr9:108342944-108344965 | p3 | (CAA)7 | TCTCCCCATATCCCCATACA | CGACCTTACGTCACTCCGAT | 227 |
| chr9:108365272-108367287 | p3 | (GAT)5 | GGTCCGGTTCCGATATTTTT | GGGGGATGGGTTAAAGTGTT | 141 |
| chr9:108456963-108459014 | p3 | (TTC)17 | AAATGCGCATGCAAAAACTA | CGCTTTACATTGCTGGATGA | 182 |
| chr9:108646781-108648793 | p2 | (AG)6 | CAGTCCGATCATTATGCGTG | TGAAGATGCCAACCTTTTCC | 293 |
| chr9:108646781-108648793 | p5 | (AATTA)3 | CAGTCCGATCATTATGCGTG | TGAAGATGCCAACCTTTTCC | 293 |
| chr9:108684045-108686059 | p6 | (AACCGG)4 | GGTTAGTCCCAATTTTCCCC | TCCCCCTAACGCCACTACTA | 194 |
| chr9:108684045-108686059 | p2 | (TA)7 | GGTTAGTCCCAATTTTCCCC | TCCCCCTAACGCCACTACTA | 194 |
| chr9:108684045-108686059 | p2 | (TA)7 | GGTTAGTCCCAATTTTCCCC | TCCCCCTAACGCCACTACTA | 194 |
| chr9:108684045-108686059 | p2 | (AT)6 | GGTTAGTCCCAATTTTCCCC | TCCCCCTAACGCCACTACTA | 194 |
| chr9:108722269-108724308 | p2 | (TA)11 | GGGTCGTAGGTGCTAATGGA | AAGGCTCAAACCCGAAAGTT | 140 |
| chr9:108725239-108727253 | p2 | (AT)7 | ATTACTTGTGGCCTTGTGGC | GCCGCTCATAAAATGAAAGC | 258 |
| chr9:108744485-108746503 | p6 | (AAAAAT)3 | GTGTTTTGCACTTGGGGTTT | CTGTGCCATCTGGTGAAAGA | 144 |
| chr9:108824973-108826988 | p3 | (TCT)5 | TACACCACCTCCACTCCTCC | GATGCACAGAGATTTCGGGT | 163 |
| chr9:108869466-108871506 | p5 | (CATTG)4 | AAGAAAGGGCAAGGCTAAGG | GCGACAAGCCATAGTCAACA | 225 |
| chr9:108957334-108959352 | p2 | (AG)6 | TTCCAGCTTTTATTCGTGGG | ATTTGGACCCAGACTTTCCC | 266 |
| chr9:109025781-109027799 | p2 | (AT)34 | CCTTTCGTCGTAGATTCCCA | CAGTTCCGGATGGACAAAAT | 280 |
| chr9:109025815-109027827 | p2 | (AT)35 | CCTTTCGTCGTAGATTCCCA | CAGTTCCGGATGGACAAAAT | 246 |
| chr9:109098004-109100028 | p2 | (TA)12 | CAAGGCCGCTGAATAACAAT | TGTGGCTAAGGGTAAGTCGG | 227 |
| chr9:109171661-109173679 | p6 | (TAAAAA)3 | ACGAGTAATGAAGCGTTGGG | CTCCACTACACATGCCCCTT | 198 |
| chr9:109173753-109175767 | p2 | (TA)7 | CAGGAACCTAACAGGCCTCA | TTGTTTTGTTTGGTTTGGCA | 183 |
| chr9:109272845-109274897 | p2 | (AT)26 | ACAGGAGCGGATTTACGTTG | GGTGCAGTTGTGTTTTGTCG | 124 |
| chr9:109277497-109279512 | p3 | (ACA)5 | GGAAATTTGCAGAGTTGGGA | GCTTGGGAGGATATGCAGAG | 176 |
| chr9:109293237-109295249 | p2 | (TA)9 | CTCTGCGATTTGTAGGCCAT | GGCGTACAGGTAAATTCGGA | 199 |
| chr9:109293294-109295314 | p2 | (TA)9 | CTCTGCGATTTGTAGGCCAT | GGCGTACAGGTAAATTCGGA | 142 |
| chr9:109454866-109456884 | p3 | (AAC)6 | CTGTCTTCAAAGCACCGTGA | GGAGGAGGCAGGAATAGGAG | 112 |
| chr9:109472049-109474077 | p2 | (TA)7 | AAGGGCCAAATAAATCCCTG | TAAACGGTTACGTCCGGTTC | 285 |
| chr9:109472089-109474101 | p2 | (TA)7 | AAGGGCCAAATAAATCCCTG | TAAACGGTTACGTCCGGTTC | 245 |
| chr9:109686175-109688190 | p3 | (GAG)5 | CCTATGACGAGGAGGGATGA | CGCCATAATCGAGGTGTTTT | 245 |
| chr9:109745642-109747657 | p5 | (CTTTT)3 | TTGTCCCATGGTTTTGGAAT | GCTCAGGAGCATGCTTTAGG | 258 |
| chr9:109837237-109839276 | p3 | (AAT)13 | TAGATTGACCAAGCATCCCC | AGGGGAAGGGACAAAAAGAA | 293 |
| chr9:109866847-109868862 | p5 | (CACCC)4 | GCTTCGTTCGTGCCTATCTC | ACAAACACTACAGGGGTGGG | 117 |
| chr9:109866847-109868862 | p5 | (CACCC)3 | GCTTCGTTCGTGCCTATCTC | ACAAACACTACAGGGGTGGG | 117 |
| chr9:109866847-109868862 | p2 | (TA)17 | GCTTCGTTCGTGCCTATCTC | ACAAACACTACAGGGGTGGG | 117 |
| chr9:109897265-109899283 | p6 | (TAAAAA)3 | CACTTCCCGTTACTCCGGTA | TTCCAGGAAGCTCAAAGCAT | 285 |

| Marker ID | SSR type | SSR motif | FORWARD PRIMER (5'-3') | REVERSE PRIMER (5'-3') | size (bp) |
|---|---|---|---|---|---|
| chr9:109897265-109899283 | p3 | (TAA)12 | CACTTCCCGTTACTCCGGTA | TTCCAGGAAGCTCAAAGCAT | 285 |
| chr9:109897821-109899857 | p6 | (TAAAAA)3 | GCATATTGTTACTCGCCGCT | TGCTCATGGGACAACACATT | 230 |
| chr9:109897821-109899857 | p3 | (TAA)12 | GCATATTGTTACTCGCCGCT | TGCTCATGGGACAACACATT | 230 |
| chr9:109911118-109913136 | p2 | (GA)6 | TAGACCGTTACGTTGGGCTC | ACACCGGCATAAACTCCAAC | 243 |
| chr9:109911118-109913136 | p3 | (GTT)6 | TAGACCGTTACGTTGGGCTC | ACACCGGCATAAACTCCAAC | 243 |
| chr9:109911118-109913136 | p2 | (GT)6 | TAGACCGTTACGTTGGGCTC | ACACCGGCATAAACTCCAAC | 243 |
| chr9:109926017-109928033 | p2 | (AT)8 | TTTTAATGCAACCCCCTCAC | TTCGAAACATGTCGTCGTTC | 246 |
| chr9:109987101-109989117 | p2 | (TA)8 | TATTGGGCAATGGGTCAAAT | TTTTGTGGTGTTTGTTGGGA | 224 |
| chr9:109990144-109992180 | p2 | (TA)11 | ATATGGGGCCCACAATTTTT | GAACCCACCACATCCAAACT | 224 |
| chr9:109990144-109992180 | p2 | (AT)18 | ATATGGGGCCCACAATTTTT | GAACCCACCACATCCAAACT | 224 |
| chr9:110160019-110162031 | p2 | (TA)6 | ACTTGCTATGATTCCGTCCG | GGCTCAGTGTGGGAAACATT | 178 |
| chr9:110160019-110162031 | p5 | (GATAT)3 | ACTTGCTATGATTCCGTCCG | GGCTCAGTGTGGGAAACATT | 178 |
| chr9:110183095-110185115 | p4 | (TAAA)5 | TTTATCGTTGGGCAGAGCTT | CAGGCCAAAGGATGAAGTGT | 105 |
| chr9:110186843-110188858 | p5 | (TAAAA)3 | AATGCGGATTTTTGAGTTCG | ATTCACCAGAAACGGTGGAG | 130 |
| chr9:110202060-110204074 | p2 | (TA)7 | TGCAAAGCTTGATTTGTTCG | GCAACGTTTTCACTGCAACT | 106 |
| chr9:110311827-110313841 | p2 | (AT)7 | AAGTTTGGCCAGGCTAGGAT | AGTGGAAAAGAGCCAAACCC | 139 |
| chr9:110314002-110316036 | p2 | (TA)17 | CTTGGCTCTGGTGTAACGGT | AGCATGGCAGGAAAATTGAC | 121 |
| chr9:110447072-110449084 | p5 | (AAAGA)3 | ACTTTCGGCTCATATGGTGG | AAAAGGTTATGCGCGTGTTT | 145 |
| chr9:110475062-110477077 | p2 | (GA)6 | AAAAATAGGGGTGTGGGGAG | TCGGTTTGGTTCGATTTTTC | 207 |
| chr9:110475062-110477077 | p5 | (CCGAA)3 | AAAAATAGGGGTGTGGGGAG | TCGGTTTGGTTCGATTTTTC | 207 |
| chr9:110475062-110477077 | p2 | (AT)33 | AAAAATAGGGGTGTGGGGAG | TCGGTTTGGTTCGATTTTTC | 207 |
| chr9:110533603-110535618 | p5 | (TTTTA)3 | TTGGAGTTCCAATAGCACCC | CCAGCACAGACACTGGGTAA | 264 |
| chr9:110748665-110750680 | p5 | (GGGTG)3 | CGCTATGTCCACGTAGGGTT | AATCTTTTATTCGGGTCGGG | 263 |
| chr9:110760633-110762649 | p2 | (AT)8 | GAATAAGGGCAGTCGTGGAA | AAGGACATTCCTGCGAATTG | 254 |
| chr9:110760633-110762649 | p3 | (CAA)5 | GAATAAGGGCAGTCGTGGAA | AAGGACATTCCTGCGAATTG | 254 |
| chr9:110772872-110774887 | p3 | (AAG)5 | CCGACTTGCTCAGTGAAGTG | ACCGAGAGAGCAAAGTGGAA | 286 |
| chr9:110786478-110788493 | p5 | (CAACC)3 | GATAATTGCGCAGTGTGGTG | CACAAGGCGGATCCTAAAAA | 223 |
| chr9:110802399-110804411 | p2 | (TG)6 | AGGAGCCATGTTATGAACCG | CCCACAAGTGGGAACCTTTA | 261 |
| chr9:110810698-110812713 | p5 | (AACCG)3 | TGGGACAAACTATGGAAGCC | TTTGGCTATTTCGATTTCGG | 299 |
| chr9:110845569-110847581 | p2 | (TC)6 | AGTGAACAGGGTGTGGAAGG | TCGGTGGTTACCAACAACAA | 148 |
| chr9:110845569-110847581 | p3 | (CTT)5 | AGTGAACAGGGTGTGGAAGG | TCGGTGGTTACCAACAACAA | 148 |
| chr9:110905978-110907993 | p2 | (AT)18 | CGGAGAAGAAAGCGAATGAG | TCTTCCTCCATCACCCAAAC | 197 |
| chr9:110905978-110907993 | p5 | (ATTTT)3 | CGGAGAAGAAAGCGAATGAG | TCTTCCTCCATCACCCAAAC | 197 |
| chr9:110911415-110913467 | p2 | (TA)26 | GAGGCATTCAACGCTAAACC | TCAACCCGCAGTTTGACATA | 136 |
| chr9:110911415-110913467 | p2 | (AT)6 | GAGGCATTCAACGCTAAACC | TCAACCCGCAGTTTGACATA | 136 |
| chr9:110953654-110955669 | p3 | (ATA)5 | GCCCCTTCCTGAGTGGTATT | CTTTCTGTGCTAGGCAGGCT | 136 |

| Marker ID | SSR type | SSR motif | FORWARD PRIMER (5'-3') | REVERSE PRIMER (5'-3') | size (bp) |
|---|---|---|---|---|---|
| chr9:111046749-111048792 | p2 | (TC)10 | GAGAAAGACGCACGGAAGAG | TTAAAATAATCGTTGCCGCC | 138 |
| chr9:111068088-111070106 | p6 | (GCATGT)3 | GAAGCTAAAACCCAATGGCA | TTGATTTTGAAGAATGGGGC | 201 |
| chr9:111109316-111111340 | p3 | (ATA)8 | TCTTCTTCCTTGTCCATCCG | GCTTGGAGAAGACTTGCCAC | 200 |
| chr9:111112896-111114920 | p3 | (ATA)8 | TCTTCTTCCTTGTCCATCCG | GCTTGGAGAAGACTTGCCAC | 200 |
| chr9:111170244-111172262 | p6 | (TAATGA)3 | GGCCGGATTAGACAAATGAA | ATCTTTCTATGCTGGGCGTG | 125 |
| chr9:111170244-111172262 | p3 | (GTG)5 | GGCCGGATTAGACAAATGAA | ATCTTTCTATGCTGGGCGTG | 125 |
| chr9:111207506-111209568 | p2 | (TA)31 | TATACCCCCACCGTTAGCAA | TTGCCACATAAGCACCACAT | 244 |
| chr9:111207506-111209568 | p4 | (AAAT)4 | TATACCCCCACCGTTAGCAA | TTGCCACATAAGCACCACAT | 244 |
| chr9:111229434-111231449 | p3 | (TAA)5 | TCAAGTTGCGGCAGTGATAG | TCTCCGCTGCCTTGTTATCT | 223 |
| chr9:111239340-111241355 | p5 | (TCTTT)3 | GAGACCATCCATCTCCTCCA | GGTTGAGTTACGCCCATGAT | 246 |
| chr9:111239340-111241355 | p2 | (AT)9 | GAGACCATCCATCTCCTCCA | GGTTGAGTTACGCCCATGAT | 246 |
| chr9:111279067-111281081 | p2 | (TG)7 | TTTTTCAGCATTTCAAGCCC | GGGTAATAGGGGCCATGTTT | 241 |
| chr9:111317265-111319286 | p3 | (TTG)7 | CAAGGGAAAACCAAGTCCAA | ATCGCTCCAACAACACTTCC | 134 |
| chr9:111330670-111332712 | p2 | (TA)21 | TTGGCCGTTTTCTTCATTTC | TGTTTCCGATAGACCCTTGG | 268 |
| chr9:111338796-111340816 | p4 | (TTAA)5 | TCAACCCATTCTCGTGATGA | GCCTTTACCATCATTCCCCT | 197 |
| chr9:111359219-111361237 | p2 | (AT)9 | ATTCCAGGGCGAATTTTCTT | ATTGCAAGGCTGAAGAAGGA | 169 |
| chr9:111396892-111398906 | p2 | (CT)7 | GTATATCTTGCCCCAAGCGA | GAGCAGCAGTGTACGCAAGA | 269 |
| chr9:111411796-111413824 | p4 | (TGAT)4 | GTACTTAAGCACCGCCTGGA | TTTCCAAACACGGTTGGTTT | 120 |
| chr9:111411796-111413824 | p2 | (AT)14 | GTACTTAAGCACCGCCTGGA | TTTCCAAACACGGTTGGTTT | 120 |
| chr9:111502387-111504469 | p2 | (TA)41 | TGGGTTGATACGCATGTTTG | ACCCGAGACCAGAAGAGGTT | 216 |
| chr9:111511074-111513089 | p5 | (CTATT)3 | AGTTGAGGGGAACTGCATTG | CACACCCTCTGATCCAACCT | 222 |
| chr9:111618590-111620608 | p2 | (TA)7 | CCCATCCAAACAGGCTCTAA | AAACTTCCAAGTGTGCGGAT | 104 |
| chr9:111618590-111620608 | p6 | (AAAAAT)3 | CCCATCCAAACAGGCTCTAA | AAACTTCCAAGTGTGCGGAT | 104 |
| chr9:111618590-111620608 | p2 | (AT)6 | CCCATCCAAACAGGCTCTAA | AAACTTCCAAGTGTGCGGAT | 104 |
| chr9:111624826-111626841 | p5 | (GGGGT)3 | GGTTGCTGCAGTTGTTCAGA | CTTGGACCCTTTGCTGATTT | 295 |
| chr9:111636374-111638389 | p5 | (TTTGG)3 | TGCCCTTTGTTCTCCATTTC | GCCGGATTATTTCTCCCACT | 130 |
| chr9:111744855-111746870 | p5 | (ATATA)3 | AAAGGAAGAGTGGAAGGGGA | ATTTTGACTTACCGGACCCC | 204 |
| chr9:111802553-111804577 | p2 | (TA)12 | CAACCCATTAGAGGGAGCAA | TGGGTTGATACGCATGTGAC | 251 |
| chr9:111951281-111953296 | p2 | (TA)20 | GTCACGTGCGTATCAATCCA | AAATGACATGGCGGAATAGC | 184 |
| chr9:111951281-111953296 | p5 | (AATAA)3 | GTCACGTGCGTATCAATCCA | AAATGACATGGCGGAATAGC | 184 |
| chr9:111951281-111953296 | p5 | (TTTAT)3 | GTCACGTGCGTATCAATCCA | AAATGACATGGCGGAATAGC | 184 |
| chr9:111961694-111963706 | p2 | (TA)6 | CCGAGCCGAATAATCAAAAA | CCTCAACAAACTCGCCAACT | 119 |
| chr9:112005627-112007651 | p2 | (TA)22 | TATTGGCACCTGTACTGCGA | CCGTTTACCTCGTGAGTCGT | 187 |
| chr9:112005627-112007651 | p2 | (AC)6 | TATTGGCACCTGTACTGCGA | CCGTTTACCTCGTGAGTCGT | 187 |
| chr9:112007667-112009681 | p2 | (AT)7 | GGACTTGACTCAAAGCGGAC | AACTCTCCGTCACCCACATC | 215 |
| chr9:112086274-112088295 | p3 | (TTC)7 | GAACATTGGCCCACTTGAAT | TACCATTCCTCCTGCAGACC | 217 |

| Marker ID | SSR type | SSR motif | FORWARD PRIMER (5'-3') | REVERSE PRIMER (5'-3') | size (bp) |
|---|---|---|---|---|---|
| chr9:112086274-112088295 | p6 | (TTTTTA)3 | GAACATTGGCCCACTTGAAT | TACCATTCCTCCTGCAGACC | 217 |
| chr9:112220694-112222709 | p5 | (AAACC)3 | GTAGGGGTGTACATGGACCG | AACCCGAGATTGAAAAACCC | 183 |
| chr9:112229958-112231976 | p3 | (CAA)6 | TCCCGAGGAGGAACTTCTTT | CAAGCCCTGGGTATGAAAAA | 221 |
| chr9:112252593-112254611 | p6 | (AGTCAA)3 | TGGCCTTTACAAAAACCAGG | GATTCTCAAAGGTGTGGGGA | 154 |
| chr9:112278573-112280606 | p3 | (ATA)5 | AAATACGGACCGAGTGCAAG | TTTCGCATACACATCGCATT | 198 |
| chr9:112278573-112280606 | p3 | (ATA)9 | AAATACGGACCGAGTGCAAG | TTTCGCATACACATCGCATT | 198 |
| chr9:112278573-112280606 | p3 | (AAT)11 | AAATACGGACCGAGTGCAAG | TTTCGCATACACATCGCATT | 198 |
| chr9:112438523-112440538 | p5 | (TTTAT)3 | GACGGGTTTAACTTGTCCGA | CCTGTAACTCTGGCCTCCAC | 235 |
| chr9:112590522-112592534 | p2 | (TA)6 | TACACTACCTCATTCGGCCC | TCATTCGGTAAGGAGATCGG | 246 |
| chr9:112641743-112643761 | p3 | (AAC)6 | CGAGCCTGGGAATAAAGTCA | TGTCCGAGCACTTGATGTTC | 101 |
| chr9:112661047-112663062 | p5 | (TTGGT)3 | CGGAGGCCGCAGATATATTA | CGGTGACAAAATGGGAAATC | 274 |
| chr9:112697305-112699320 | p5 | (AACCG)3 | GGTGAAATTGTTGGGCTCAT | TTTTGCACTCTACACTGGCG | 126 |
| chr9:112709416-112711431 | p3 | (ACA)5 | CATGTCCGTCAAATCTGTGG | TGTAGTGCATATGGGGCTGA | 176 |
| chr9:112824869-112826881 | p2 | (TA)6 | GAGGTTCTAGCCAGTGACCG | GGACAGCTTTGGAGCGATAC | 161 |
| chr9:112824869-112826881 | p2 | (AT)6 | GAGGTTCTAGCCAGTGACCG | GGACAGCTTTGGAGCGATAC | 161 |
| chr9:112824869-112826881 | p2 | (CT)6 | GAGGTTCTAGCCAGTGACCG | GGACAGCTTTGGAGCGATAC | 161 |
| chr9:112825043-112827055 | p2 | (TA)6 | GCTCCAAAGCTGTCCCATAA | TTACCGGGGAGTATGCTCTG | 209 |
| chr9:112825043-112827055 | p2 | (AT)6 | GCTCCAAAGCTGTCCCATAA | TTACCGGGGAGTATGCTCTG | 209 |
| chr9:112825043-112827055 | p2 | (CT)6 | GCTCCAAAGCTGTCCCATAA | TTACCGGGGAGTATGCTCTG | 209 |
| chr9:112845996-112848011 | p5 | (CGTGT)3 | GAGGTGAGGCATTTGTTGGT | TTTCCAAATCACCCCGATTA | 259 |
| chr9:112845996-112848011 | p3 | (ACA)7 | GAGGTGAGGCATTTGTTGGT | TTTCCAAATCACCCCGATTA | 259 |
| chr9:112846126-112848147 | p5 | (CGTGT)3 | GAGGTGAGGCATTTGTTGGT | TTTCCAAATCACCCCGATTA | 129 |
| chr9:112846126-112848147 | p3 | (ACA)7 | GAGGTGAGGCATTTGTTGGT | TTTCCAAATCACCCCGATTA | 129 |
| chr9:112881328-112883340 | p2 | (AT)6 | CGTTACCCGGACTTCGATTA | TACAGTGATGCCGACTTTGC | 143 |
| chr9:112911390-112913410 | p5 | (AAATA)4 | TTGAAAATGGGTCTGGAAGG | TGGATGGGCACTACAAATCA | 278 |
| chr9:112911390-112913410 | p2 | (TA)8 | TTGAAAATGGGTCTGGAAGG | TGGATGGGCACTACAAATCA | 278 |
| chr9:112922626-112924638 | p2 | (AC)6 | GTTTTCCGTTTTCCACCAGA | GTATAGGGAAATGCTGCCGA | 185 |
| chr9:112938827-112940841 | p2 | (TA)7 | CCCGCCAAAATACGAAGATA | CGCCTCATATCCGGTTTCTA | 133 |
| chr9:112938827-112940841 | p3 | (TGA)5 | CCCGCCAAAATACGAAGATA | CGCCTCATATCCGGTTTCTA | 133 |
| chr9:112966416-112968431 | p3 | (ATC)5 | TATGCACAAGCTGACAAGGC | AGGTGCTTCATCAGCGATTT | 131 |
| chr9:112966416-112968431 | p2 | (TA)23 | TATGCACAAGCTGACAAGGC | AGGTGCTTCATCAGCGATTT | 131 |
| chr9:113043755-113045767 | p2 | (AT)6 | GCTGCCTATTGCAGGAGTTC | TCCTTGGAAATAAGCCTCCA | 162 |
| chr9:113043755-113045767 | p2 | (TA)6 | GCTGCCTATTGCAGGAGTTC | TCCTTGGAAATAAGCCTCCA | 162 |
| chr9:113043755-113045767 | p2 | (TA)9 | GCTGCCTATTGCAGGAGTTC | TCCTTGGAAATAAGCCTCCA | 162 |
| chr9:113095297-113097312 | p5 | (TTATT)3 | GGGCCAAGAATAGCAGAGTG | TGCAGCGAACACAAAGAATC | 137 |
| chr9:113128101-113130121 | p2 | (AT)10 | TGATCCATTCTCACAACCCA | AAAGACCACCCCGAATAAGG | 177 |

| Marker ID | SSR type | SSR motif | FORWARD PRIMER (5'-3') | REVERSE PRIMER (5'-3') | size (bp) |
|---|---|---|---|---|---|
| chr9:113138529-113140544 | p3 | (TTC)5 | ACGTTATCACGCGTTCTTCC | GGTGGAAGTGGGAATCAGAA | 126 |
| chr9:113183167-113185185 | p3 | (TAT)12 | TGCTCTTCATATGGGCTTGA | AGCCACTTGGCAACAAATTC | 182 |
| chr9:113183167-113185185 | p3 | (ATT)19 | TGCTCTTCATATGGGCTTGA | AGCCACTTGGCAACAAATTC | 182 |
| chr9:113183167-113185185 | p2 | (TA)7 | TGCTCTTCATATGGGCTTGA | AGCCACTTGGCAACAAATTC | 182 |
| chr9:113183187-113185202 | p3 | (TAT)12 | TGCTCTTCATATGGGCTTGA | AGCCACTTGGCAACAAATTC | 162 |
| chr9:113183187-113185202 | p3 | (ATT)19 | TGCTCTTCATATGGGCTTGA | AGCCACTTGGCAACAAATTC | 162 |
| chr9:113183187-113185202 | p2 | (TA)7 | TGCTCTTCATATGGGCTTGA | AGCCACTTGGCAACAAATTC | 162 |
| chr9:113186500-113188518 | p5 | (TCATG)3 | TTCCCCATATATCCCTTCCC | ACCTTACGTCACTCCGATGG | 173 |
| chr9:113186500-113188518 | p3 | (CAA)6 | TTCCCCATATATCCCTTCCC | ACCTTACGTCACTCCGATGG | 173 |
| chr9:113308367-113310395 | p2 | (TA)9 | CTATAGCGCAGCACGGTGTA | TCGTGGAAGGAAAGCAGAGT | 202 |
| chr9:113308367-113310395 | p2 | (AT)14 | CTATAGCGCAGCACGGTGTA | TCGTGGAAGGAAAGCAGAGT | 202 |
| chr9:113444297-113446313 | p4 | (TTTA)4 | TATCCTTTGAACCCCACCAA | GCCATCCCTCACTTGTCCTA | 120 |
| chr9:113636365-113638395 | p3 | (TTA)5 | GGAATGCAGCAAATGGGTAT | TCAATGTGCTTTGTTCTCGC | 230 |
| chr9:113657900-113659928 | p2 | (AT)14 | TCACAACCAACCCCTTCTTC | GTCTTCCTTGGCTAAATGCG | 242 |
| chr9:113700140-113702152 | p2 | (TA)6 | AACAAGCCCTTTGACACCAG | ATAGTATCCCCTGCAGCCCT | 256 |
| chr9:113780407-113782422 | p2 | (TA)7 | ATTAGATGTAGGGGGCGGAT | ATGCACCTAACCAATGCTCC | 156 |
| chr9:113780407-113782422 | p5 | (ATTTT)3 | ATTAGATGTAGGGGGCGGAT | ATGCACCTAACCAATGCTCC | 156 |
| chr9:113785548-113787563 | p2 | (TA)7 | ATTAGATGTAGGGGGCGGAT | ATGCACCTAACCAATGCTCC | 156 |
| chr9:113785548-113787563 | p5 | (ATTTT)3 | ATTAGATGTAGGGGGCGGAT | ATGCACCTAACCAATGCTCC | 156 |
| chr9:113835936-113837951 | p5 | (AATAA)3 | TGTGCTCAAGTACGGAGTGG | TGCTTTAACCGTCACGAATG | 205 |
| chr9:113838873-113840901 | p2 | (AT)14 | TGGAGACCCTAGCAAGCTGT | TGGCCAAGTTTACGAGAAGC | 117 |
| chr9:113838873-113840901 | p2 | (AT)13 | TGGAGACCCTAGCAAGCTGT | TGGCCAAGTTTACGAGAAGC | 117 |
| chr9:113849925-113851940 | p3 | (ACA)5 | TACATGAAGGCAAGCGAATG | ATTTTGAGGTTGTGGCAAGG | 105 |
| chr9:113904067-113906099 | p3 | (ATT)5 | TCATGGGCAACAAAGACAAA | GGGATGTTTAGTGAAGGGCA | 261 |
| chr9:113904067-113906099 | p2 | (CT)16 | TCATGGGCAACAAAGACAAA | GGGATGTTTAGTGAAGGGCA | 261 |
| chr9:113904067-113906099 | p2 | (TA)28 | TCATGGGCAACAAAGACAAA | GGGATGTTTAGTGAAGGGCA | 261 |
| chr9:113919327-113921339 | p2 | (TA)6 | ACCATTGACCACCTTCCTTG | TTGAAATTTGCATTGGAGCA | 143 |
| chr9:113953578-113955593 | p5 | (AAAAG)3 | ACCACCAATCAAACAATCCC | AAGATGTTGCACGTTGTTGC | 286 |
| chr9:113953578-113955593 | p3 | (GAT)5 | ACCACCAATCAAACAATCCC | AAGATGTTGCACGTTGTTGC | 286 |
| chr9:113958672-113960687 | p5 | (ATGTT)3 | GATATATGCTGGGCCTGGAA | CCTAGGTGACCCGAATTCAA | 148 |
| chr9:113975876-113977894 | p6 | (TTGAAG)3 | GCTTGTTCAGACTCACAGCG | CAAACGAATCAGCCAAGTGA | 142 |
| chr9:113978557-113980572 | p5 | (GTATT)3 | TGGCCCAACTCAAAGAGAAC | AAGGGCATTACTGCGAATTG | 279 |
| chr9:113978557-113980572 | p2 | (TG)10 | TGGCCCAACTCAAAGAGAAC | AAGGGCATTACTGCGAATTG | 279 |
| chr9:113996669-113998689 | p2 | (TA)10 | TTGGTCGTATCGGGTTTCTC | GCCTCCAAAAGTCCAGTCAA | 103 |
| chr9:114008890-114010952 | p5 | (GGGGT)3 | TTTTACCGAGGGTGTTCGAG | AAATACCGACCCACTCCCTC | 293 |
| chr9:114127586-114129604 | p6 | (AATAAA)3 | CGACAAGGTCACCACAACAC | TTTTGTAGCTCCACCGGTTC | 122 |

| Marker ID | SSR type | SSR motif | FORWARD PRIMER (5'-3') | REVERSE PRIMER (5'-3') | size (bp) |
|---|---|---|---|---|---|
| chr9:114205022-114207037 | p5 | (TATAT)3 | AAAATTGGAGTTGCGTTTGG | CCATGGATAGCTTGCACCTT | 162 |
| chr9:114248859-114250873 | p2 | (AT)7 | CTTTTTCGCATGGGCATATT | AACAACTGAAACGTCGAGGG | 221 |
| chr9:114259039-114261053 | p2 | (TA)7 | GATTGTTGAACCCCCTTTGA | TGGCTCCGGACTCAAAATAC | 162 |
| chr9:114259039-114261053 | p5 | (AAAAC)3 | GATTGTTGAACCCCCTTTGA | TGGCTCCGGACTCAAAATAC | 162 |
| chr9:114345230-114347245 | p5 | (AAATT)3 | GTCGTGTGTGAAGACTCCGA | GGCCACCTTTTCAAACAAAA | 234 |
| chr9:114379230-114381245 | p4 | (TTTA)4 | GAGGAGTTTCACCCGAACAA | AGGTTCAGATGTAGTGGCGG | 300 |
| chr9:114379230-114381245 | p5 | (GGGCC)3 | GAGGAGTTTCACCCGAACAA | AGGTTCAGATGTAGTGGCGG | 300 |
| chr9:114379230-114381245 | p2 | (TC)11 | GAGGAGTTTCACCCGAACAA | AGGTTCAGATGTAGTGGCGG | 300 |
| chr9:114467206-114469224 | p6 | (ATCTAT)3 | TTCCCATCTCCCTTTGTCAC | CCACCAGAACCAACTGTTCA | 125 |
| chr9:114522163-114524231 | p2 | (TA)6 | TTATGCAACGAATGGTGGAA | AAGAGGCACGACCCTTTTCT | 276 |
| chr9:114522232-114524266 | p2 | (TA)6 | TTATGCAACGAATGGTGGAA | AAGAGGCACGACCCTTTTCT | 207 |
| chr9:114547178-114549208 | p3 | (AAC)7 | CAGTGAAATCCCACAACGTG | CGCTTTGAATCTCTTAGCCG | 218 |
| chr9:114547178-114549208 | p2 | (TA)15 | CAGTGAAATCCCACAACGTG | CGCTTTGAATCTCTTAGCCG | 218 |
| chr9:114547178-114549208 | p2 | (AT)6 | CAGTGAAATCCCACAACGTG | CGCTTTGAATCTCTTAGCCG | 218 |
| chr9:114572574-114574589 | p2 | (TA)6 | CCATGAGATACCTGCCACCT | TAAACAGAAGCCCAAGGGTG | 111 |
| chr9:114572574-114574589 | p3 | (CAA)5 | CCATGAGATACCTGCCACCT | TAAACAGAAGCCCAAGGGTG | 111 |
| chr9:114587988-114590003 | p5 | (TTGTA)3 | TTAGCTGGCAAGTATGCCCT | GTGCACAAGTAGGCACCTCA | 261 |
| chr9:114620281-114622297 | p2 | (TA)8 | ATCCAAAGAATCCACCCCTC | AAACAATCAGGTGTCTCGGG | 280 |
| chr9:114620281-114622297 | p3 | (TGT)6 | ATCCAAAGAATCCACCCCTC | AAACAATCAGGTGTCTCGGG | 280 |
| chr9:114679354-114681372 | p3 | (AAC)6 | CAAGCATCCCTTCCCAATTA | AATGGACAGAAGCCAAATGC | 114 |
| chr9:114679354-114681372 | p3 | (AGA)5 | CAAGCATCCCTTCCCAATTA | AATGGACAGAAGCCAAATGC | 114 |
| chr9:114721292-114723307 | p5 | (TTTTA)3 | AGCTTTTTGGCTCCCTTTTC | TTGCACAAGGAATAGGTCCC | 100 |
| chr9:114731935-114733950 | p3 | (TTA)5 | GGACAAGCTTGCATTGGATT | TTCAAGAATAGGCGCGAAAT | 105 |
| chr9:114787419-114789437 | p3 | (AAT)6 | TGTTTAGGAGGATTTTGCGG | CTCTTTCCTTCCCTGCCTCT | 241 |
| chr9:114855654-114857669 | p5 | (AAAGA)3 | TGTGCCCCTTCTCTGAGTCT | CTTGCCATATGGGGCATACT | 263 |
| chr9:114861573-114863589 | p2 | (AT)8 | TTAGGACAGTGTTCTCCCGC | CTGCCGTACCATTTTTCCAT | 137 |
| chr9:114955399-114957415 | p2 | (TA)8 | TGGGAACTTTATGGACTCGC | CTGCCCCAAACACAACTCTT | 174 |
| chr9:115010017-115012075 | p2 | (TA)29 | GGGCGAAGGATGTTTTCTTT | CGTTGGACGTTAGGCGTATT | 158 |
| chr9:115028379-115030451 | p2 | (TA)36 | TGCCTTTGATGGGGATTAAC | AAAGCCAGAAAGGCATGCTA | 142 |
| chr9:115050487-115052523 | p5 | (CATTG)3 | ATTTTGTTTGCTACGGGTCG | TATTCCGGCATCCTTAGTGC | 201 |
| chr9:115050568-115052589 | p5 | (CATTG)3 | ATTTTGTTTGCTACGGGTCG | TATTCCGGCATCCTTAGTGC | 120 |
| chr9:115050670-115052691 | p5 | (CATTG)3 | GGGGTCCGGTAAGTCAAAAT | TATTCCGGCATCCTTAGTGC | 186 |
| chr9:115050757-115052784 | p5 | (CATTG)3 | AAATACGGACCGAGTGCAAG | CAACTTTCGCATACACACCG | 115 |
| chr9:115059223-115061241 | p3 | (AAT)6 | ATTGACTGGAGGGTGGTCAG | ATTGAGACCCAAGGGGTAGG | 163 |
| chr9:115121995-115124013 | p2 | (TA)13 | GCACTTTTTCGTCGTTCGTT | GAGATGGTGGGTTCCACTGT | 130 |
| chr9:115121995-115124013 | p2 | (TA)9 | GCACTTTTTCGTCGTTCGTT | GAGATGGTGGGTTCCACTGT | 130 |

| Marker ID | SSR type | SSR motif | FORWARD PRIMER (5'-3') | REVERSE PRIMER (5'-3') | size (bp) |
|---|---|---|---|---|---|
| chr9:115180405-115182417 | p2 | (TA)6 | AAACGCCTCCTAATTTCCAGA | GAACGGTTGGACAACAAACA | 176 |
| chr9:115180467-115182527 | p2 | (TA)6 | AAACGCCTCCTAATTTCCAGA | GAACGGTTGGACAACAAACA | 114 |
| chr9:115182723-115184738 | p5 | (TTTTG)3 | GACGACTAAAAAGCACCCCA | GCGGCTGAGATTTTACTTGC | 281 |
| chr9:115246375-115248391 | p2 | (AT)8 | GGGACACGATTGAAAAAGGA | ACTGCTCTTGATGAACGGCT | 185 |
| chr9:115271854-115273869 | p5 | (TTTTC)3 | ATGGAGAGTGGCGAATAACG | TCTTGGGCTAGCAGAAGGTC | 197 |
| chr9:115327088-115329103 | p3 | (GCG)5 | GGCAGGTATCCGTGGAATTA | ATGCAGTGGTCATGTTCCAA | 234 |
| chr9:115327088-115329103 | p2 | (TA)9 | GGCAGGTATCCGTGGAATTA | ATGCAGTGGTCATGTTCCAA | 234 |
| chr9:115356099-115358111 | p2 | (AT)6 | ACCTTCTACCAAACGACCCC | CGCCGCATATACACGACTTA | 167 |
| chr9:115356974-115359000 | p2 | (AT)6 | CCGCAGAAGTGAAATTGACA | TCGAATTCATAAGCAGCGAA | 297 |
| chr9:115356974-115359000 | p2 | (AT)6 | CCGCAGAAGTGAAATTGACA | TCGAATTCATAAGCAGCGAA | 297 |
| chr9:115356974-115359000 | p2 | (TA)6 | CCGCAGAAGTGAAATTGACA | TCGAATTCATAAGCAGCGAA | 297 |
| chr9:115357165-115359177 | p2 | (AT)6 | CCGCAGAAGTGAAATTGACA | TCGAATTCATAAGCAGCGAA | 106 |
| chr9:115357165-115359177 | p2 | (TA)6 | CCGCAGAAGTGAAATTGACA | TCGAATTCATAAGCAGCGAA | 106 |
| chr9:115452158-115454176 | p6 | (AACTCC)3 | GTGGATTGCACAGCGAGTTA | GCACTCCATTGTAGGGCATT | 191 |
| chr9:115452158-115454176 | p6 | (AACTCC)3 | GTGGATTGCACAGCGAGTTA | GCACTCCATTGTAGGGCATT | 191 |
| chr9:115475990-115478002 | p2 | (CT)6 | TACGAGCTCGGATGGCTAGT | ATCCCAACTTTCAACGAACG | 169 |
| chr9:115495991-115498009 | p6 | (AAATAA)3 | GGCGCTTGACATGTATGCTA | GAACGACACCCCTTGAAAAA | 126 |
| chr9:115497011-115499037 | p2 | (AT)13 | TGGATTGGCGTTTTTGTGTA | CAAACTGAATTTTCGTGCCA | 253 |
| chr9:115540902-115542920 | p6 | (TAAAAA)3 | AGCTCCTGAACCAGATTCCA | GGTAGGTGTTGAACCCCCTT | 143 |
| chr9:115559128-115561146 | p6 | (TAAAAA)3 | AGCTCCTGAACCAGATTCCA | GCCAAGTGTAGTTGAGGGGA | 140 |
| chr9:115618654-115620675 | p3 | (TGT)7 | TGCAGCTCAATTGGTGAAAG | TCTTTGGCCTACCCCTACCT | 275 |
| chr9:115777098-115779120 | p2 | (AT)11 | AGTCGGCTTGCATGGTTATC | GGCATAAGCCTCACAGGAAC | 276 |
| chr9:115788278-115790296 | p5 | (TTTTA)3 | TTGCTGAAGCAATTGTGAGG | GTACACGGATGGAGAGGGAA | 283 |
| chr9:115788278-115790296 | p2 | (TC)9 | TTGCTGAAGCAATTGTGAGG | GTACACGGATGGAGAGGGAA | 283 |
| chr9:115883683-115885711 | p2 | (TA)10 | CCGCAGAAGTGAAATTGACA | TATGCACACAAACTGCCACA | 217 |
| chr9:115999003-116001027 | p2 | (TA)7 | GACGAAAGGATGACAAGGGA | TTACGGTGCACGAATTCAAA | 266 |
| chr9:116004151-116006169 | p6 | (TAAAAA)3 | ACGTTGTTCTGAGGTTCGCT | TGTGCCCAAAAGGCAATTAT | 158 |
| chr9:116007527-116009542 | p5 | (TTGGT)3 | AATGGATTTTACCCCTTGCC | CAGGTTGATTTGGTTCGGTT | 142 |
| chr9:116007527-116009542 | p5 | (TTGGT)3 | AATGGATTTTACCCCTTGCC | CAGGTTGATTTGGTTCGGTT | 142 |
| chr9:116075480-116077498 | p6 | (TATATG)3 | TTTGGCCAAGTTGGATTTTC | GCCACGCCTTATTAGCTACG | 154 |
| chr9:116237964-116239978 | p2 | (TA)7 | TAATCCCAATCTACGCCTCG | TCGGGTTGGCCTATTTACAG | 289 |
| chr9:116250084-116252096 | p2 | (AT)6 | TAGCTTCCATGCCTTGAACA | CTTTTTCGCTGCCACAACTT | 238 |
| chr9:116250084-116252096 | p2 | (AT)8 | TAGCTTCCATGCCTTGAACA | CTTTTTCGCTGCCACAACTT | 238 |
| chr9:116250084-116252096 | p2 | (AT)6 | TAGCTTCCATGCCTTGAACA | CTTTTTCGCTGCCACAACTT | 238 |
| chr9:116250202-116252218 | p2 | (AT)6 | TAGCTTCCATGCCTTGAACA | CTTTTTCGCTGCCACAACTT | 120 |
| chr9:116250202-116252218 | p2 | (AT)8 | TAGCTTCCATGCCTTGAACA | CTTTTTCGCTGCCACAACTT | 120 |

| Marker ID | SSR type | SSR motif | FORWARD PRIMER (5'-3') | REVERSE PRIMER (5'-3') | size (bp) |
|---|---|---|---|---|---|
| chr9:116250202-116252218 | p2 | (AT)6 | TAGCTTCCATGCCTTGAACA | CTTTTTCGCTGCCACAACTT | 120 |
| chr9:116286091-116288103 | p2 | (AT)6 | CTTTTAATGCGGGAGGCATA | ACAACCAGCCACGGAGTAAC | 204 |
| chr9:116301100-116303118 | p3 | (ACT)6 | GCTCGTCGATTAACTGCTCC | AGAAAAAGGCCAGAGGTGGT | 101 |
| chr9:116407387-116409405 | p6 | (AAAAAG)3 | AGCCAATCTTCGAGGTCTGA | CCACTCCTGTGATGTGGATG | 189 |
| chr9:116409310-116411322 | p5 | (AATAA)3 | TAGACGTGGGCCCCATAATA | CAAGGACTTGCAGGTTCGAT | 113 |
| chr9:116409310-116411322 | p2 | (TG)6 | TAGACGTGGGCCCCATAATA | CAAGGACTTGCAGGTTCGAT | 113 |
| chr9:116409310-116411322 | p5 | (TTTAT)5 | TAGACGTGGGCCCCATAATA | CAAGGACTTGCAGGTTCGAT | 113 |
| chr9:116409310-116411322 | p2 | (AT)10 | TAGACGTGGGCCCCATAATA | CAAGGACTTGCAGGTTCGAT | 113 |
| chr9:116465578-116467594 | p4 | (CTTT)4 | ATATCCTGATTGGGCTGTGC | ACACCATCATGTGAAGGCAA | 300 |
| chr9:116534994-116537009 | p3 | (GAT)5 | CCTATTACCCCCTGCACAAA | TATGTGGCAGCTGAGTTTGC | 282 |
| chr9:116546293-116548311 | p2 | (CT)9 | GGGGTATGTGGCATGTTTTC | ATTGGCCCAAATTTAAACCC | 113 |
| chr9:116570564-116572579 | p5 | (ACTTC)3 | GTTTGTTGTCCGACCAGGAT | TTTTGTGGATTAAGCAGGGG | 175 |
| chr9:116606365-116608420 | p5 | (TAAAA)3 | TGTTCACGGGGAGAATTAGG | GGTGTCGTTTAACATTGGGG | 281 |
| chr9:116610607-116612622 | p5 | (TTTTA)3 | GAATGGCAGTTTGCAGTTGA | ACAACAGAGTCTGGCTGGGT | 286 |
| chr9:116627098-116629116 | p6 | (TTTTTG)3 | GACCCGGACACCACAGTTAT | CTCGCAAGAAGAAGCAGGTT | 243 |
| chr9:116627098-116629116 | p2 | (AT)6 | GACCCGGACACCACAGTTAT | CTCGCAAGAAGAAGCAGGTT | 243 |
| chr9:116694007-116696022 | p5 | (GTTCG)3 | GGGGCTTATTTCAAGCACAA | TTTGATAAAACCAAACGCCC | 121 |
| chr9:116694007-116696022 | p5 | (TTCGG)3 | GGGGCTTATTTCAAGCACAA | TTTGATAAAACCAAACGCCC | 121 |
| chr9:116714085-116716100 | p5 | (CTCTC)3 | GGACCATGGCCAATTATCAG | CCGGAAACAATCCTCTCAAA | 156 |
| chr9:116724413-116726428 | p5 | (ATCAA)3 | CGTCTTGTGACTGCCTATCG | CTCAAGGTTGAAAAACCCGA | 126 |
| chr9:116724413-116726428 | p5 | (AAACC)3 | CGTCTTGTGACTGCCTATCG | CTCAAGGTTGAAAAACCCGA | 126 |
| chr9:116731458-116733470 | p2 | (AT)20 | ATTTGAATGGGCACGACATT | TGCCAAAATGAATGACCTCTT | 233 |
| chr9:116731458-116733470 | p2 | (TA)6 | ATTTGAATGGGCACGACATT | TGCCAAAATGAATGACCTCTT | 233 |
| chr9:116735548-116737566 | p2 | (TA)9 | TGAACCGATAGGTGCAATATGA | CATAATGGTTGACGAAGGCA | 137 |
| chr9:116779555-116781571 | p2 | (AT)8 | GCCATTTTTGGCTCATTCAT | CTTAAAACTGGCAAGCAGGC | 275 |
| chr9:116786466-116788487 | p3 | (ATA)7 | CTTATGCCAAGTCTGCCCAT | TTAACGGGCAATTCTTTTGG | 186 |
| chr9:116795091-116797105 | p2 | (AT)7 | TAGTAAGAGCCCAGGGGGTT | TATGCCAAGGGGTTCAAAAA | 132 |
| chr9:116795091-116797105 | p5 | (AATAA)3 | TAGTAAGAGCCCAGGGGGTT | TATGCCAAGGGGTTCAAAAA | 132 |
| chr9:116801222-116803240 | p3 | (TGT)6 | AAGTGATGAGTGCAACGCAG | CCAAATCATTCGCCACTTTT | 197 |
| chr9:116803379-116805419 | p2 | (TA)10 | TCTGCAGCACGCACTAAATC | CTTGCCACATAAATCGCCTC | 192 |
| chr9:116803379-116805419 | p2 | (AT)20 | TCTGCAGCACGCACTAAATC | CTTGCCACATAAATCGCCTC | 192 |
| chr9:116809040-116811068 | p2 | (CA)8 | ACGCCAGCTTTTCTAGGTGA | CTTCCATCTCGTTCCACGAT | 155 |
| chr9:116811475-116813503 | p2 | (CA)8 | ACGCCAGCTTTTCTAGGTGA | CTTCCATCTCGTTCCACGAT | 155 |
| chr9:116817766-116819781 | p3 | (CAA)5 | ACTGGGCCTCAACATACTCG | GCTGGATTGGTGAACTCCAT | 127 |
| chr9:116882567-116884595 | p2 | (AT)14 | ATGTGATCAGGAGGTCAGGG | GGCACGTATTCCTCTTAGCG | 180 |
| chr9:116922216-116924231 | p5 | (TTACT)3 | TTAGGCAGTACGCGTGTCAG | ATAACCCCAGTGTGTGCCTC | 223 |

| Marker ID | SSR type | SSR motif | FORWARD PRIMER (5'-3') | REVERSE PRIMER (5'-3') | size (bp) |
|---|---|---|---|---|---|
| chr9:116965426-116967441 | p3 | (TGT)5 | TGCACTTGCAACCTTCAGAC | TGCAAATTATGTTGCTCGGA | 178 |
| chr9:117121719-117123733 | p2 | (CA)7 | GGACATGCCAAAAGAAGGAA | AGCTAAGCCTCGGGAATGTT | 255 |
| chr9:117146042-117148057 | p5 | (AATAA)3 | TGGCCTGGGATATGTAGCTC | GACGGCTACGAACTCCTCTG | 193 |
| chr9:117146042-117148057 | p5 | (AATAA)3 | TGGCCTGGGATATGTAGCTC | GACGGCTACGAACTCCTCTG | 193 |
| chr9:117146042-117148057 | p3 | (TGG)6 | TGGCCTGGGATATGTAGCTC | GACGGCTACGAACTCCTCTG | 193 |
| chr9:117146042-117148057 | p2 | (TA)7 | TGGCCTGGGATATGTAGCTC | GACGGCTACGAACTCCTCTG | 193 |
| chr9:117146953-117148967 | p5 | (AATAA)3 | GATTGACGAGGAGCCATGTT | GGAGCCTTTAGAGTTCGGCT | 199 |
| chr9:117146953-117148967 | p3 | (TGG)6 | GATTGACGAGGAGCCATGTT | GGAGCCTTTAGAGTTCGGCT | 199 |
| chr9:117146953-117148967 | p2 | (TA)7 | GATTGACGAGGAGCCATGTT | GGAGCCTTTAGAGTTCGGCT | 199 |
| chr9:117184348-117186360 | p2 | (AT)6 | AGCTCCAGTTCAGAGGGTGA | CCCCCTCAGGAAAAGAACTC | 156 |
| chr9:117238640-117240656 | p6 | (ACATGA)3 | GCAACCGGAAGAAACAACAT | CGGCTTATGAAATGCTCTCC | 292 |
| chr9:117238640-117240656 | p4 | (ACAT)4 | GCAACCGGAAGAAACAACAT | CGGCTTATGAAATGCTCTCC | 292 |
| chr9:117260584-117262600 | p6 | (ACATGA)3 | AAACCCCGAATGAGATAGGG | ATGTTGTTTCTTCCGGTTGC | 139 |
| chr9:117260584-117262600 | p4 | (ACAT)4 | AAACCCCGAATGAGATAGGG | ATGTTGTTTCTTCCGGTTGC | 139 |
| chr9:117327412-117329442 | p6 | (AAAAAT)3 | GGGCCTCAACTAAGCATCAA | ATGTGGCAATGAATCCATCA | 170 |
| chr9:117327412-117329442 | p2 | (AT)15 | GGGCCTCAACTAAGCATCAA | ATGTGGCAATGAATCCATCA | 170 |
| chr9:117329226-117331242 | p2 | (AT)8 | AATCGTCTGAAATCCGCCTA | TCCGAAAAACTCTCGGCTTA | 150 |
| chr9:117329226-117331242 | p2 | (AT)8 | AATCGTCTGAAATCCGCCTA | TCCGAAAAACTCTCGGCTTA | 150 |
| chr9:117436695-117438707 | p2 | (TG)6 | TTGTTTAAGGCATTGAGGGG | TGCCTTCCTCAATATTTCCG | 239 |
| chr9:117650525-117652543 | p6 | (TATTTT)3 | CCATCCCATTTCACGCTACT | TGAAATTACTTGCGGGGTTC | 154 |
| chr9:117693079-117695093 | p2 | (AC)7 | ATTTCAAGGCAGTGAGTGGG | GACAGTCTTTTTGGGACCGA | 120 |
| chr9:117755228-117757244 | p2 | (TA)6 | AAGTATCAACCGTGGCTTCG | TCCGCTTCTGAGTTTTGGTT | 119 |
| chr9:117755228-117757244 | p2 | (AT)8 | AAGTATCAACCGTGGCTTCG | TCCGCTTCTGAGTTTTGGTT | 119 |
| chr9:117771296-117773311 | p5 | (AAAAG)3 | CCACATTTGCATGGACAAAC | ATGCTTCCATTCACACACCA | 101 |
| chr9:117822651-117824666 | p3 | (TTG)5 | AACGTCGGTGGATCAAAAAG | GGCTTTATGGCCCTTCTTTC | 125 |
| chr9:117832112-117834138 | p2 | (AT)13 | TACCAAGATTGGGAGCAAGG | AAGCTCGCTAAGCTCCCTTT | 217 |
| chr9:117832112-117834138 | p2 | (AT)14 | TACCAAGATTGGGAGCAAGG | AAGCTCGCTAAGCTCCCTTT | 217 |
| chr9:117832112-117834138 | p2 | (AT)14 | TACCAAGATTGGGAGCAAGG | AAGCTCGCTAAGCTCCCTTT | 217 |
| chr9:117943199-117945215 | p3 | (AGA)5 | CTCTGCATTCCGACAGTCAA | TCAAGATGGAGGGGAGTGTC | 249 |
| chr9:117943199-117945215 | p4 | (TATT)4 | CTCTGCATTCCGACAGTCAA | TCAAGATGGAGGGGAGTGTC | 249 |
| chr9:118000973-118002991 | p6 | (TATTTT)3 | ATTGTTCCTGTGGGATCGAC | CTACCCCTTGACCACTCCAA | 175 |
| chr9:118152616-118154631 | p5 | (AAAGA)3 | GGTGAGTCCCAACGAGTCAT | ATAGTCGGACTTCAATGCGG | 261 |
| chr9:118156569-118158593 | p4 | (AAAT)6 | GACATGTTCAGGGGTCCATC | CGGAAATACACACGCTGAGA | 254 |
| chr9:118196770-118198794 | p2 | (TA)7 | CTGAAGCCAAGCAAACAACA | GTTCGTCACGGTCAATTCCT | 281 |
| chr9:118196863-118198937 | p2 | (TA)7 | CTGAAGCCAAGCAAACAACA | GTTCGTCACGGTCAATTCCT | 188 |
| chr9:118265862-118267878 | p4 | (TACG)4 | ACGTGAACGGAGAAAATTCG | GGATTTTGGCCTATTGGGAT | 237 |

| Marker ID | SSR type | SSR motif | FORWARD PRIMER (5'-3') | REVERSE PRIMER (5'-3') | size (bp) |
|---|---|---|---|---|---|
| chr9:118433063-118435078 | p5 | (TATAA)3 | TCTCTCATGGTCCAACCACA | ACTTACCGTTTTTGCCATGC | 286 |
| chr9:118531799-118533814 | p2 | (TA)6 | TGTGGGTGTGTGTGCCTACT | AACGCATTTCAACACAACCA | 148 |
| chr9:118531799-118533814 | p2 | (AT)9 | TGTGGGTGTGTGTGCCTACT | AACGCATTTCAACACAACCA | 148 |
| chr9:118531799-118533814 | p3 | (GAT)5 | TGTGGGTGTGTGTGCCTACT | AACGCATTTCAACACAACCA | 148 |
| chr9:118566995-118569007 | p2 | (AT)6 | GTGCAGCATCCTTGTGAAGA | GGGACTCCTTTTGCATGAGA | 125 |
| chr9:118571951-118573971 | p4 | (TAAT)5 | CGTTCACCAACCGTTAATCC | CTTTGATCAAGCCCCAAAAA | 261 |
| chr9:118571951-118573971 | p4 | (ACCC)5 | CGTTCACCAACCGTTAATCC | CTTTGATCAAGCCCCAAAAA | 261 |
| chr9:118680064-118682100 | p2 | (TA)18 | ATCCACAAGCAAATTCCCAG | TGTGGCCCCTATCATTTCTC | 161 |
| chr9:118719895-118721929 | p2 | (TA)17 | GGGGCCCACAATTTAGTTTT | CCATCACATCCAAACTCACG | 238 |
| chr9:118774350-118776365 | p5 | (AAAAT)3 | ATTTTAGGGTGGACGAACCC | GCATATGCATCATGTCCTCG | 172 |
| chr9:118789891-118791906 | p5 | (TTGTC)3 | CCACCGAAACTTAAAGGCAA | CGCATCTTGTCTTCCACTGA | 196 |
| chr9:118789891-118791906 | p3 | (TTG)5 | CCACCGAAACTTAAAGGCAA | CGCATCTTGTCTTCCACTGA | 196 |
| chr9:118794069-118796084 | p5 | (AAAAT)3 | ATTTTAGGGTGGACGAACCC | GCATATGCATCATGTCCTCG | 172 |
| chr9:118997009-118999061 | p2 | (AT)7 | TGCAGCCTCTTCAACTCCTT | CAGATGCACCAATTTTGTCG | 114 |
| chr9:118997009-118999061 | p2 | (TA)26 | TGCAGCCTCTTCAACTCCTT | CAGATGCACCAATTTTGTCG | 114 |
| chr9:119058091-119060106 | p3 | (GGT)5 | CAAATTGGTGGTGAATGTGC | GCATGCACTCTACTTTCCCC | 105 |
| chr9:119189852-119191864 | p2 | (TA)6 | CCGCACACTTTCTCTCTTCC | TTTTGGGGGTAAAAGGTTGA | 108 |
| chr9:119218598-119220616 | p3 | (AAC)6 | TCGAAGTTCTTGAGCCGATT | CGATCCAACTACCGTCGAGT | 178 |
| chr9:119226403-119228418 | p5 | (TAATT)3 | GGAAAATCGATCGCCTCATA | CTGATGGCACTTGTGCTACG | 164 |
| chr9:119261421-119263454 | p3 | (AAT)11 | AGAGCGAAAAACGGACAGAA | TTCGATTCATGGAATCCTCC | 238 |
| chr9:119275537-119277551 | p2 | (TC)12 | ATAATACGCCCCTCGTCTCA | ATTGGGCAAAATGAGGAAAA | 296 |
| chr9:119275537-119277551 | p2 | (TC)7 | ATAATACGCCCCTCGTCTCA | ATTGGGCAAAATGAGGAAAA | 296 |
| chr9:119315912-119317933 | p3 | (TAT)7 | ACGACCTTATGTCACCTCGG | GTATACGCTGCCCGCTTTAG | 289 |
| chr9:119315912-119317933 | p3 | (TTC)14 | ACGACCTTATGTCACCTCGG | GTATACGCTGCCCGCTTTAG | 289 |
| chr9:119316459-119318501 | p3 | (TAT)7 | CCGTTTTCTTTGGACGTTGT | TAGCACCTCCGGAATAATGG | 198 |
| chr9:119316459-119318501 | p3 | (TTC)14 | CCGTTTTCTTTGGACGTTGT | TAGCACCTCCGGAATAATGG | 198 |
| chr9:119318773-119320805 | p2 | (AT)16 | ATATGCCCCTACTGTTTGCG | AAAGGTTGGTGCAAAGATGC | 193 |
| chr9:119474261-119476300 | p3 | (ATA)17 | AAAAGCTCTTCCAAAAGCCC | TTGCATTGTGTTCTGGGGTA | 161 |
| chr9:119474303-119476347 | p3 | (ATA)17 | AAAAGCTCTTCCAAAAGCCC | TTGCATTGTGTTCTGGGGTA | 119 |
| chr9:119474863-119476917 | p3 | (ATA)17 | GAGCCTGAGAACTCCATTGC | GTGGAAGGGGTTGGTACTGA | 260 |
| chr9:119474914-119476932 | p3 | (ATA)17 | GAGCCTGAGAACTCCATTGC | ATTGGAGAAGTGGAAGGGGT | 209 |
| chr9:119482599-119484614 | p3 | (TGT)5 | GAGAAGATCGCCAGAAATGG | AGCAGATAGGCATTCCTCCA | 121 |
| chr9:119482599-119484614 | p4 | (AAAT)10 | GAGAAGATCGCCAGAAATGG | AGCAGATAGGCATTCCTCCA | 121 |
| chr9:119532467-119534479 | p2 | (AT)6 | TCAAAGCCTGCCTTCTTGTT | TCCGTATTCGTAGACACCCC | 157 |
| chr9:119550729-119552750 | p5 | (TATCA)5 | CGCCTTACCTTTCCCCTTAC | TTACCGACCTTACGTCACCC | 200 |
| chr9:119550729-119552750 | p6 | (CATATA)3 | CGCCTTACCTTTCCCCTTAC | TTACCGACCTTACGTCACCC | 200 |

| Marker ID | SSR type | SSR motif | FORWARD PRIMER (5'-3') | REVERSE PRIMER (5'-3') | size (bp) |
|---|---|---|---|---|---|
| chr9:119550729-119552750 | p3 | (CAA)7 | CGCCTTACCTTTCCCCTTAC | TTACCGACCTTACGTCACCC | 200 |
| chr9:119656726-119658741 | p3 | (TTA)5 | CCAAATCCTTTGCACAACCT | AAAATGAGGCTGGTCTGGTG | 105 |
| chr9:119736426-119738441 | p3 | (GGT)5 | ATTTTCGAACCCAACAATGC | TCACGGCACGATGATTTTTA | 221 |
| chr9:119818284-119820296 | p4 | (TTTA)4 | CGGAGTCAAAATGGCGTATT | ATTCGTGAATTGCACAACGA | 297 |
| chr9:119818284-119820296 | p2 | (AT)6 | CGGAGTCAAAATGGCGTATT | ATTCGTGAATTGCACAACGA | 297 |
| chr9:119818284-119820296 | p2 | (TA)13 | CGGAGTCAAAATGGCGTATT | ATTCGTGAATTGCACAACGA | 297 |
| chr9:119883263-119885281 | p4 | (AAAT)4 | GTATTAGGGCAGGCGGAAAT | TTCTGTTCCAAGTTTTCGGG | 275 |
| chr9:119883263-119885281 | p3 | (TAA)6 | GTATTAGGGCAGGCGGAAAT | TTCTGTTCCAAGTTTTCGGG | 275 |
| chr9:119883263-119885281 | p6 | (TAATAT)3 | GTATTAGGGCAGGCGGAAAT | TTCTGTTCCAAGTTTTCGGG | 275 |
| chr9:119949663-119951681 | p2 | (AT)9 | GGTGAGCTTACCGTGGATGT | GTGGGGGTGGCACTATCTAA | 127 |
| chr9:119969644-119971662 | p6 | (ATAAAT)3 | ATCGACAGCCTTCTCGTGAT | TTGCTACTCAATCTGGCGTG | 112 |
| chr9:119973585-119975600 | p3 | (AAC)5 | GGGAATAAGGATTCCCACAAA | CACCGGGTATGTTGTTGTTG | 201 |
| chr9:119973585-119975600 | p3 | (AAC)5 | GGGAATAAGGATTCCCACAAA | CACCGGGTATGTTGTTGTTG | 201 |
| chr9:119973585-119975600 | p3 | (AAC)5 | GGGAATAAGGATTCCCACAAA | CACCGGGTATGTTGTTGTTG | 201 |
| chr9:119973585-119975600 | p3 | (AAC)5 | GGGAATAAGGATTCCCACAAA | CACCGGGTATGTTGTTGTTG | 201 |
| chr9:120022197-120024215 | p6 | (AAAATA)3 | CGTTAACAGAAGACGGGCTC | AGAGCACATGTTACGGGGTC | 132 |
| chr9:120088301-120090313 | p2 | (TA)10 | TTCCCCTTCTCTTTCAGCAA | GAGAGAGGTTTCCCTTTCGG | 297 |
| chr9:120088332-120090350 | p2 | (TA)10 | TTCCCCTTCTCTTTCAGCAA | GAGAGAGGTTTCCCTTTCGG | 266 |
| chr9:120090018-120092033 | p5 | (GGGCC)3 | CCCGAATAGGCGTGATACAG | CAAAAGCTATGAACCTGGGG | 144 |
| chr9:120142582-120144597 | p5 | (ATTTT)3 | CTCATTCGTCGTTGGAGGAT | GTACTTGAAAGGGCCGACAA | 200 |
| chr9:120148221-120150237 | p2 | (AT)9 | TAGATTCATTGCCAAATGCG | AATGACATGGCGGAAAAGAA | 155 |
| chr9:120148221-120150237 | p2 | (TA)8 | TAGATTCATTGCCAAATGCG | AATGACATGGCGGAAAAGAA | 155 |
| chr9:120148221-120150237 | p5 | (TTATA)3 | TAGATTCATTGCCAAATGCG | AATGACATGGCGGAAAAGAA | 155 |
| chr9:120148221-120150237 | p4 | (TTAT)4 | TAGATTCATTGCCAAATGCG | AATGACATGGCGGAAAAGAA | 155 |
| chr9:120244442-120246457 | p3 | (TTG)5 | ATTGCTCCACCCATGCTTAG | TGGTACTTGCGTGCTACCAG | 133 |
| chr9:120274575-120276595 | p4 | (ATAC)5 | TTTTTGTAATTCGGCCCTTC | AACAACGACCGAAAAATTGG | 268 |
| chr9:120386278-120388296 | p6 | (CCCCCA)3 | CCAACAATAGAAGCCACCGT | GCAATGTGCTGAGCTTCAAA | 295 |
| chr9:120428867-120430887 | p3 | (GTT)6 | CGTACACTCTACCCTCCCCA | GGAGCGGCACAAACTACAAT | 184 |
| chr9:120428867-120430887 | p5 | (AGTAA)4 | CGTACACTCTACCCTCCCCA | GGAGCGGCACAAACTACAAT | 184 |
| chr9:120428867-120430887 | p3 | (TGT)6 | CGTACACTCTACCCTCCCCA | GGAGCGGCACAAACTACAAT | 184 |
| chr9:120562450-120564468 | p2 | (TA)9 | TACCTTCCTTGCCGAGTTGT | AAGACACTCCTTCGACCCCT | 275 |
| chr9:120604864-120606879 | p5 | (TTAAA)3 | TGTCAGTGATGTGGCAGTGA | TTGCGGATGACCTATTACCC | 221 |
| chr9:120657439-120659455 | p3 | (AAG)5 | GCAAGTGGGCATCAATTTTT | CGGCTCCTAACATTTGGCTA | 236 |
| chr9:120657439-120659455 | p2 | (TA)8 | GCAAGTGGGCATCAATTTTT | CGGCTCCTAACATTTGGCTA | 236 |
| chr9:120695183-120697199 | p4 | (AGTT)4 | GGGCTAGAGGCAATGCAATA | CACACTCGCTCGATACTCCA | 115 |
| chr9:121066684-121068700 | p2 | (TG)8 | TGTCATGCATTCCTTTTCCA | CCAGCGAAGACCTCAAAAAC | 165 |

| Marker ID | SSR type | SSR motif | FORWARD PRIMER (5'-3') | REVERSE PRIMER (5'-3') | size (bp) |
|---|---|---|---|---|---|
| chr9:121124781-121126808 | p3 | (AAG)9 | CAGCCACTCTTTTTGAAGGC | AAGGTGCAAGCTAAGCGAAA | 259 |
| chr9:121127865-121129877 | p2 | (AT)6 | GATTGGGGTTGAGAGACCAA | TGGGTAACTGCCCAGAGTTC | 156 |
| chr9:121158294-121160309 | p5 | (TTTTG)3 | CTCTTTTCAGTCAGGGCAGG | TGTAGTCTGAGGGGTCTGGG | 238 |
| chr9:121196712-121198739 | p3 | (AAG)9 | CAGCCACTCTTTTTGAAGGC | AAGGTGCAAGCTAAGCGAAA | 259 |
| chr9:121199796-121201808 | p2 | (AT)6 | GATTGGGGTTGAGAGACCAA | TGGGTAACTGCCCAGAGTTC | 156 |
| chr9:121330688-121332700 | p2 | (AT)6 | AAACTCATGAAATCACGGGC | AAGGCGGGTAGAAAGAGAGC | 164 |
| chr9:121436978-121439020 | p2 | (GA)12 | CGGCCTTGATGAGGTACTGT | CAACGTGTTCGAAGAAACGA | 194 |
| chr9:121436978-121439020 | p2 | (TA)21 | CGGCCTTGATGAGGTACTGT | CAACGTGTTCGAAGAAACGA | 194 |
| chr9:121436978-121439020 | p4 | (AATA)5 | CGGCCTTGATGAGGTACTGT | CAACGTGTTCGAAGAAACGA | 194 |
| chr9:121436978-121439020 | p5 | (AGAAA)3 | CGGCCTTGATGAGGTACTGT | CAACGTGTTCGAAGAAACGA | 194 |
| chr9:121455997-121458013 | p4 | (CCTT)4 | CCGTATGTTGGCTATGCCTT | CCTTCTTTTTCCGACCTTCC | 269 |
| chr9:121529737-121531752 | p5 | (TCCAT)3 | GGTCTTTTAAGGCCGAATCC | GTTGAATTCTCGGGGTGCTA | 251 |
| chr10:215820-217846 | p2 | (AT)13 | GCAGTGGAGTATGCACGAGA | AACGATCTCGGATTGCTTTG | 112 |
| chr10:360480-362492 | p2 | (AT)6 | AGAAGATTCTCGGGGACGTT | GGACATGCCAAAAGAAGGAA | 282 |
| chr10:379580-381594 | p2 | (AT)7 | TGCATATTCCGATGAGCAAA | GATGCTGCCCAAATTTCTGT | 187 |
| chr10:616786-618804 | p6 | (TTTTTA)3 | CAACCAAATCCATTCCCATC | TCTCACCATGCATTTTTCCA | 152 |
| chr10:641245-643260 | p5 | (ATTTT)3 | AATAACGGGCGATTTGACAG | GGATTCGGATTTGCATAGGA | 272 |
| chr10:648602-650617 | p5 | (ATATC)3 | TGCATGGCAGGACCATATAA | TCGTGACAACAAATTTCCCA | 165 |
| chr10:657835-659847 | p2 | (TA)6 | TTGCCAATCGTGTATCCATC | CCTATGGCAGGGAAGTTGAA | 300 |
| chr10:681714-683732 | p3 | (TTG)6 | GCATCGGTGGAAGACAAGAT | AAGGTCATGTCCTCGGTGAG | 190 |
| chr10:783340-785356 | p4 | (ATCT)4 | GATTGCACAAAGGAAAAGCC | CACTCACGGGAGAATTGGTT | 279 |
| chr10:793715-795730 | p3 | (AAC)5 | ATTGGTAGCGCGATTGGTTA | TCTCCATGGGAGGAAGATTG | 163 |
| chr10:853758-855770 | p2 | (AT)6 | AATTCTGCGTGATTTTCGCT | AGTCCATATCGTGTAGGGCG | 122 |
| chr10:902476-904542 | p2 | (TA)33 | TTCAGGAGTCAATTTTGGGC | TGACCAGACACCAAAACGAA | 183 |
| chr10:935254-937272 | p3 | (ATA)6 | ACTTGTCAACATGGGCATCA | AATTTATGGTAGTCCGGGGC | 137 |
| chr10:943318-945338 | p5 | (TTGGT)4 | AACGCGCGTGCTTATAGATT | ATGAGAAAACGGATGGAACG | 128 |
| chr10:990862-992883 | p3 | (GAA)7 | ACCAGGTGAGTGTTTTTGGC | AAGGTTGGGGAATATTGCATC | 170 |
| chr10:991464-993529 | p3 | (GAA)7 | TCAATTCTCGCGCTATTTTG | TCCCCTTTTCCTTTCTCCTC | 205 |
| chr10:991464-993529 | p3 | (GAT)5 | TCAATTCTCGCGCTATTTTG | TCCCCTTTTCCTTTCTCCTC | 205 |
| chr10:1129327-1131343 | p2 | (AT)8 | TCCACTGCGGTGACATTAAG | TGAGGTGACAATGGCAGAAA | 220 |
| chr10:1154697-1156713 | p2 | (AT)8 | TACTTCCTCCATGGTGAGGC | TGTATGTCTGCCCGTCCATA | 244 |
| chr10:1171990-1174008 | p3 | (TTA)6 | TCATTCATTGTCCGAAAGCA | CCTTCTGCAAACTCTACGGG | 267 |
| chr10:1288755-1290773 | p3 | (TTG)6 | CTGACCAACCCCCATTACAC | CCTGAAGCTATAACAGCCGC | 151 |
| chr10:1319998-1322012 | p2 | (GT)7 | TTCACTGTGCAAAGAGGTGG | AGAACTGGAAAATGGAGGGG | 101 |
| chr10:1508801-1510817 | p4 | (GAAA)4 | GCGCGAAAACCAAAATAAAA | TACGAGGATTATTCGGCTGG | 229 |
| chr10:1518085-1520097 | p2 | (AT)6 | CTTTACCGGCTACCGTTTGA | ACAAAAATGGATGAGACGGG | 290 |

| Marker ID | SSR type | SSR motif | FORWARD PRIMER (5'-3') | REVERSE PRIMER (5'-3') | size (bp) |
|---|---|---|---|---|---|
| chr10:1555258-1557270 | p2 | (AT)6 | CTTTACCGGCTACCGTTTGA | ACAAAAATGGATGAGACGGG | 290 |
| chr10:1612589-1614604 | p5 | (CAAAA)3 | TCAGCAACCCTAGCTCATCC | TCCCCTTTTCTAGGGCATCT | 128 |
| chr10:1617638-1619650 | p2 | (TA)6 | AGGATCCCAACAAGTCATGC | CAAGTGGTGGACCAATGAGA | 114 |
| chr10:1659332-1661347 | p2 | (AG)6 | GAAATTCGTGCGATTAGGGA | GCTTATCTTCCCCACCTTCC | 129 |
| chr10:1659332-1661347 | p3 | (AAT)5 | GAAATTCGTGCGATTAGGGA | GCTTATCTTCCCCACCTTCC | 129 |
| chr10:1854203-1856235 | p2 | (TA)16 | AAGCCAAACGGGCTTTAACT | GGACGGTTTTAATAAGGGGC | 195 |
| chr10:1854203-1856235 | p2 | (TA)6 | AAGCCAAACGGGCTTTAACT | GGACGGTTTTAATAAGGGGC | 195 |
| chr10:1878091-1880133 | p3 | (AAT)14 | TTTTCCTGGTGCGAGTTACC | ACTAGCGCCTTTGAAGGACA | 234 |
| chr10:1878091-1880133 | p2 | (AC)10 | TTTTCCTGGTGCGAGTTACC | ACTAGCGCCTTTGAAGGACA | 234 |
| chr10:1879687-1881705 | p2 | (AC)10 | ACAGAGGAAGAAGAAGCCCC | GACAAGTAGCAGAGGCGGAC | 228 |
| chr10:1879687-1881705 | p2 | (TG)9 | ACAGAGGAAGAAGAAGCCCC | GACAAGTAGCAGAGGCGGAC | 228 |
| chr10:1947757-1949772 | p5 | (AATTT)3 | CTTCCGGTGATGCTGCTTAT | TTTTCTGTTGAAATTCCGGC | 265 |
| chr10:1993594-1995608 | p2 | (CT)7 | ATGCACATAAAGGAGGACCG | AGGCGTAATGGATGAAATGG | 130 |
| chr10:2099810-2101831 | p3 | (ACA)7 | TATGGGGTGGCGTATCATTT | GTTTAGCTTTGTCGGCCTTG | 188 |
| chr10:2103943-2105958 | p5 | (ATTTC)3 | TGTGGAGAAACAACAACCGA | TAAAATTTGGCCACACACGA | 138 |
| chr10:2113639-2115657 | p3 | (ACA)6 | CCCACTTGGGTTCCCTTTAT | AGAGGTGGAGGTGTCCAATG | 200 |
| chr10:2156087-2158105 | p2 | (TA)9 | AAACAGCTTTTCCTCTGCCA | ATTGGCTTCAGAATCAACGG | 211 |
| chr10:2214347-2216374 | p3 | (ATA)6 | TCCCCGTTTCTTCTTCTCCT | TCAATCCCATTTTACCCCAC | 130 |
| chr10:2300071-2302085 | p2 | (TA)7 | GGCCGGATTCACTTTACAGA | AACTAAAATAGCGGGCGGAT | 174 |
| chr10:2310091-2312107 | p4 | (TTTC)4 | CGGATATAGTCGGGCTTCAA | GGGCAATTTCCCTCAATTTT | 212 |
| chr10:2534038-2536054 | p4 | (TTTA)4 | TTTGAGTTCGAGCTCTGGGT | TCCGGTATCCATATTGGAGC | 137 |
| chr10:2534038-2536054 | p6 | (GTGTGC)3 | TTTGAGTTCGAGCTCTGGGT | TCCGGTATCCATATTGGAGC | 137 |
| chr10:2535960-2537976 | p4 | (TTTA)4 | TTTGAGTTCGAGCTCTGGGT | TCCGGTATCCATATTGGAGC | 136 |
| chr10:2535960-2537976 | p6 | (GTGTGC)3 | TTTGAGTTCGAGCTCTGGGT | TCCGGTATCCATATTGGAGC | 136 |
| chr10:2543816-2545828 | p2 | (TA)6 | AGCCGAATGAAGGTGTATGC | AAGTGCAGGAAAGGCATGTT | 203 |
| chr10:2648586-2650600 | p2 | (TA)7 | GGCCGTAAACCAAGTCGTAA | AATGGCCATAACTCTTTGCG | 201 |
| chr10:2823587-2825609 | p2 | (AT)11 | TTGTTTCCACGAAAATGCAA | TTGTTCCACGAAATGCAAAA | 249 |
| chr10:2823587-2825609 | p2 | (TA)7 | TTGTTTCCACGAAAATGCAA | TTGTTCCACGAAATGCAAAA | 249 |
| chr10:2823638-2825664 | p2 | (AT)11 | TTGTTTCCACGAAAATGCAA | TTGTTCCACGAAATGCAAAA | 198 |
| chr10:2823638-2825664 | p2 | (TA)7 | TTGTTTCCACGAAAATGCAA | TTGTTCCACGAAATGCAAAA | 198 |
| chr10:2886859-2888879 | p2 | (AT)6 | CATTGTCGCGTATCAATCCA | TAAATGACATGGCGGTAGCA | 104 |
| chr10:2886859-2888879 | p2 | (TA)10 | CATTGTCGCGTATCAATCCA | TAAATGACATGGCGGTAGCA | 104 |
| chr10:2968603-2970629 | p5 | (AAAAG)3 | GGCCCAGTCTGTTTACCTCA | ACACTTGCACAAGTCCCACA | 233 |
| chr10:3026857-3028902 | p3 | (TAA)15 | GGGTGTCGGTGGTATAATGG | CCGCACACAGATCTCACACT | 242 |
| chr10:3038054-3040072 | p6 | (GTTGGA)3 | CTTTTTGGTCCACAGGGAAA | CCCCAGATCAAGAACCTCAA | 204 |
| chr10:3103350-3105368 | p5 | (ATTTT)3 | AGAAATACCACTTGCCCCCT | AAGCACATGCAGTCATCTCG | 249 |

| Marker ID | SSR type | SSR motif | FORWARD PRIMER (5'-3') | REVERSE PRIMER (5'-3') | size (bp) |
|---|---|---|---|---|---|
| chr10:3103350-3105368 | p6 | (AACCGG)3 | AGAAATACCACTTGCCCCCT | AAGCACATGCAGTCATCTCG | 249 |
| chr10:3103350-3105368 | p2 | (AT)6 | AGAAATACCACTTGCCCCCT | AAGCACATGCAGTCATCTCG | 249 |
| chr10:3135242-3137260 | p6 | (TTTTTG)3 | TTCGCAGGTATCATGAGTCG | GCCGAATAGGGGATACACAA | 194 |
| chr10:3210423-3212438 | p2 | (AT)27 | ATATTCGGGATAAGGTGGGG | TCCTGTCCTCGGTAAGCTGT | 256 |
| chr10:3210423-3212438 | p3 | (GTT)7 | ATATTCGGGATAAGGTGGGG | TCCTGTCCTCGGTAAGCTGT | 256 |
| chr10:3210423-3212438 | p5 | (ATTTT)3 | ATATTCGGGATAAGGTGGGG | TCCTGTCCTCGGTAAGCTGT | 256 |
| chr10:3210423-3212438 | p5 | (TAATA)3 | ATATTCGGGATAAGGTGGGG | TCCTGTCCTCGGTAAGCTGT | 256 |
| chr10:3210909-3212924 | p3 | (GTT)7 | TCCTGTGGTATCTGTGTCGC | GGGAGGGTAGAGTGTACGCA | 142 |
| chr10:3210909-3212924 | p5 | (ATTTT)3 | TCCTGTGGTATCTGTGTCGC | GGGAGGGTAGAGTGTACGCA | 142 |
| chr10:3210909-3212924 | p5 | (TAATA)3 | TCCTGTGGTATCTGTGTCGC | GGGAGGGTAGAGTGTACGCA | 142 |
| chr10:3223190-3225208 | p3 | (GTT)5 | CATATCCATGTTCGAACCCC | CAAATCAAATGTGAGGCCAA | 120 |
| chr10:3223190-3225208 | p6 | (AAAAAG)3 | CATATCCATGTTCGAACCCC | CAAATCAAATGTGAGGCCAA | 120 |
| chr10:3285032-3287056 | p2 | (AT)6 | TCACGATTCTGTCCACTCCA | GGCGCTTGACATGTATGCTA | 111 |
| chr10:3288441-3290469 | p2 | (TG)7 | CTGCCCCTGTTCTTCACTTC | TGGGTATATTTCGGCAAAGC | 260 |
| chr10:3288600-3290614 | p2 | (TG)7 | TGCCCCTGTTCTTCACTTCT | TGGGTATATTTCGGCAAAGC | 102 |
| chr10:3303469-3305484 | p5 | (ACAAA)3 | GGGTTTGTGTTCATGAGCCT | GCTACGAGGGAGTTCGTGAG | 177 |
| chr10:3311681-3313707 | p2 | (AT)13 | TTACACCCAATTCTCAGCCC | CATGTGTTGGGCACTTGAAC | 158 |
| chr10:3336254-3338275 | p3 | (AAG)7 | ATTATTCCCGCCGCTAGAGT | CCATTCTCTGAAGCTCGGTC | 195 |
| chr10:3434724-3436742 | p2 | (TA)31 | AATTACGCATAGCGGTTGCT | TTAGTGTCTTTGCTTCCGCC | 278 |
| chr10:3434745-3436763 | p2 | (TA)22 | AATTACGCATAGCGGTTGCT | TTAGTGTCTTTGCTTCCGCC | 257 |
| chr10:3434766-3436784 | p2 | (AT)12 | AATTACGCATAGCGGTTGCT | TTAGTGTCTTTGCTTCCGCC | 236 |
| chr10:3511018-3513034 | p2 | (TA)8 | CCCACCAGATAATAGCCGAA | GTTACACATTTTGGGTCGGG | 144 |
| chr10:3519256-3521271 | p3 | (TTG)5 | GGCTTACAGCCCAATGTTGT | AACTCGATTCCACGCATTTC | 161 |
| chr10:3553491-3555505 | p2 | (AT)7 | GAAGTTGATGATGCCGACCT | GCAACAAGTCGTTGGGAGTT | 104 |
| chr10:3681085-3683100 | p5 | (CCCGC)3 | CATGTGATGCCCTTGTTCAG | GGGTTAATGCAGGGTGAAAA | 282 |
| chr10:3712963-3714975 | p2 | (AT)6 | ATCAATTGAGCTTTGACCCG | TCCGGACGATAACCCAATAA | 188 |
| chr10:3728484-3730502 | p3 | (TTA)6 | CGAAGAATCGGAAACCTTTG | GTCCCTACCCAGCTGAAATG | 128 |
| chr10:3777754-3779769 | p5 | (GAAAA)3 | CCTGCAGCTGTGATTTTTCA | GTAAAGTCGGTTGATGGGGA | 225 |
| chr10:3789062-3791080 | p3 | (TTG)8 | AGGACATCAGGGACAAGGTG | AGCCCTCCACAACCTCCTAT | 295 |
| chr10:3789062-3791080 | p3 | (TGT)6 | AGGACATCAGGGACAAGGTG | AGCCCTCCACAACCTCCTAT | 295 |
| chr10:3815350-3817371 | p2 | (AT)7 | CCCCATATTAAACACCGTGC | GAGTTTGGATGTGGTGGGTT | 228 |
| chr10:3815350-3817371 | p3 | (GTT)7 | CCCCATATTAAACACCGTGC | GAGTTTGGATGTGGTGGGTT | 228 |
| chr10:3851086-3853106 | p4 | (TCTT)5 | ATACCCGGTGTAATCCCACA | GGAGGAAGCTCACTGTCTCG | 215 |
| chr10:4066435-4068456 | p3 | (TGT)7 | GTATGCGGCGGAATGTTAGT | TTTCTCACTGGGGGCTTCTA | 228 |
| chr10:4123351-4125366 | p5 | (GGTTT)3 | AATAAGGCATGTCCCCTTCC | TGGAAACGAGTATCCTTGGC | 251 |
| chr10:4123351-4125366 | p2 | (AT)9 | AATAAGGCATGTCCCCTTCC | TGGAAACGAGTATCCTTGGC | 251 |

| Marker ID | SSR type | SSR motif | FORWARD PRIMER (5'-3') | REVERSE PRIMER (5'-3') | size (bp) |
|---|---|---|---|---|---|
| chr10:4131868-4133886 | p6 | (ATATAG)3 | CCCGGATCTGACTTGAAAAA | CGAATCCAAACATCTGCCTT | 227 |
| chr10:4168156-4170168 | p2 | (TG)6 | TCATTCCTAACCTTCCCACG | TGTGCACAAAATGAGCCAAT | 229 |
| chr10:4193820-4195844 | p4 | (AAAT)6 | GGTCAAAGTCCACCTCCAAA | GCGGATCCAGAATGTTCAAT | 216 |
| chr10:4228050-4230062 | p2 | (AT)6 | TTTGTGGTGGGATGAACAGA | GCTTCCAAGGCAAATTGTGT | 131 |
| chr10:4264013-4266031 | p2 | (GA)7 | ATGTGCATGCTCAAAGTTGG | ATCTCGGAAAGCGAAACGTA | 252 |
| chr10:4459849-4461864 | p3 | (ATA)5 | ATGGGCCGTTAACTCAAACA | GTCGGAATTGACGCAGTTTT | 271 |
| chr10:4486550-4488604 | p3 | (TAT)18 | ATTCGAATCCCCACGTTGTA | ATGCGGGATCTGTCTGGTAG | 235 |
| chr10:4556435-4558449 | p2 | (AC)7 | CAAAGGAAGCGTGTAGGAGC | TTGGGACTTTCTTTGCGATT | 228 |
| chr10:4560442-4562460 | p6 | (CAACTC)3 | CTCCGGATGTTGACCAAAGT | AGGTTGAAAATCATGGCGTC | 208 |
| chr10:4583896-4585914 | p6 | (CTTTTT)3 | TTACAGGCTCAAGGGTGGAC | TCTCCGCCTCATTCCAATAC | 195 |
| chr10:4617624-4619660 | p2 | (AT)18 | ATGAGGGACACTTTGCAACC | GAGCCAACAACTATGCGTGA | 108 |
| chr10:4693356-4695371 | p5 | (AAAAT)3 | AGTGGTAGAATCAGTGGCGG | ATTGAAGTGAGGGTGTTCGC | 203 |
| chr10:4695939-4697954 | p5 | (AAACC)3 | CCCACTCACTCATTTCGGAT | CAAAGAAACCCGAGGTTGAA | 122 |
| chr10:4731735-4733779 | p2 | (TA)22 | TCAGGGGTTGAGCAGAGAAT | AAAGAAAGCCGAGGAGGTTC | 137 |
| chr10:4804576-4806600 | p2 | (TC)12 | TCCTGGGGTAGATGCAATTC | GTTCAGCAGCCTTCAGTTCC | 274 |
| chr10:4914106-4916122 | p4 | (TAGA)4 | TTGACCGCCAACATGTTAAA | GTAAGCTTCATCCGCACCAT | 104 |
| chr10:4921688-4923708 | p2 | (AT)10 | TAGCCGCTTGGAGTGTTTTT | TTATCGGTTAATTCTGCCCG | 164 |
| chr10:5010000-5012028 | p2 | (AG)14 | GGCATGCCAAGGTGTTAAGT | GCAATCGATCAACTTTGCCT | 282 |
| chr10:5080435-5082465 | p2 | (TA)15 | AAGGGGATGCATTGTTTCAG | TTCCCATATTCCCGATGGTA | 246 |
| chr10:5173884-5175908 | p3 | (TTG)8 | CGTGGATCGCACCTTAATTT | GTTGGGGCCCACTTTTCTAT | 173 |
| chr10:5231987-5234017 | p6 | (AAAAGA)5 | GACGCGGAAGAAAACAAAAG | ACATGTCGACGGACACAAAA | 177 |
| chr10:5231987-5234017 | p6 | (AAAAGA)3 | GACGCGGAAGAAAACAAAAG | ACATGTCGACGGACACAAAA | 177 |
| chr10:5425681-5427693 | p2 | (AG)6 | AAGGTACGGATGTCAGACCG | GCCAACCTAAAAAGGAAGGG | 181 |
| chr10:5548397-5550428 | p2 | (CA)6 | TCTGATGCATTACCAGCAGC | GAAGCTGTCCAAAGAGCTGG | 237 |
| chr10:5591746-5593758 | p2 | (AT)6 | TTTTCTGTTGGCTGCAGATG | ATGCCGAATTCTCGATCTTG | 220 |
| chr10:5645990-5648005 | p3 | (TCT)5 | AGGATCGCCATCATTGAAAG | GAGCGTGACGTAGATGACGA | 185 |
| chr10:5650306-5652363 | p3 | (TCT)8 | GGCTTCTTTCGACCAATCAG | AAGTGGTGCCACATGTCAAA | 105 |
| chr10:5650306-5652363 | p2 | (AC)17 | GGCTTCTTTCGACCAATCAG | AAGTGGTGCCACATGTCAAA | 105 |
| chr10:5650306-5652363 | p2 | (AC)6 | GGCTTCTTTCGACCAATCAG | AAGTGGTGCCACATGTCAAA | 105 |
| chr10:5721298-5723322 | p2 | (TA)12 | TCCACAGTGTCCGTGAAAAA | TTTTCCCACTTCATTCCCAC | 202 |
| chr10:5721298-5723322 | p2 | (AT)9 | TCCACAGTGTCCGTGAAAAA | TTTTCCCACTTCATTCCCAC | 202 |
| chr10:5721298-5723322 | p2 | (AT)36 | TCCACAGTGTCCGTGAAAAA | TTTTCCCACTTCATTCCCAC | 202 |
| chr10:5753192-5755207 | p5 | (TTGGT)3 | AATGGCTTGGTTTGACTTGG | TAATGTCGGGTTGGTTTGGT | 184 |
| chr10:5753192-5755207 | p5 | (TTGGT)3 | AATGGCTTGGTTTGACTTGG | TAATGTCGGGTTGGTTTGGT | 184 |
| chr10:5805584-5807596 | p2 | (AT)6 | GATTCATGGTGACACACCCA | TTAGCCCACCCATTGTTGAT | 185 |
| chr10:5848401-5850431 | p2 | (AT)8 | GAATTTTCGGCCACCTTTTT | GGCGCCATGGTTAATTTCTA | 213 |

| Marker ID | SSR type | SSR motif | FORWARD PRIMER (5'-3') | REVERSE PRIMER (5'-3') | size (bp) |
|---|---|---|---|---|---|
| chr10:5848401-5850431 | p2 | (TA)15 | GAATTTTCGGCCACCTTTTT | GGCGCCATGGTTAATTTCTA | 213 |
| chr10:5943597-5945612 | p3 | (CTT)5 | CCATGTTGGCACATCTATCG | TTCCATAGCAAAAGGCCATC | 256 |
| chr10:5943597-5945612 | p5 | (CTCGA)3 | CCATGTTGGCACATCTATCG | TTCCATAGCAAAAGGCCATC | 256 |
| chr10:5943597-5945612 | p5 | (TAGTT)3 | CCATGTTGGCACATCTATCG | TTCCATAGCAAAAGGCCATC | 256 |
| chr10:5943729-5945744 | p3 | (CTT)5 | CCATGTTGGCACATCTATCG | TTCCATAGCAAAAGGCCATC | 124 |
| chr10:5943729-5945744 | p5 | (CTCGA)3 | CCATGTTGGCACATCTATCG | TTCCATAGCAAAAGGCCATC | 124 |
| chr10:5943729-5945744 | p5 | (TAGTT)3 | CCATGTTGGCACATCTATCG | TTCCATAGCAAAAGGCCATC | 124 |
| chr10:6000779-6002799 | p4 | (AATA)5 | GATCATCGGGCATGAAGTTT | AGGGCCAAATGCACAAATAG | 151 |
| chr10:6003085-6005100 | p3 | (CTT)5 | CCATGTTGGCACATCTATCG | TTCCATAGCAAAAGGCCATC | 256 |
| chr10:6003085-6005100 | p5 | (CTCGA)3 | CCATGTTGGCACATCTATCG | TTCCATAGCAAAAGGCCATC | 256 |
| chr10:6003085-6005100 | p5 | (TAGTT)3 | CCATGTTGGCACATCTATCG | TTCCATAGCAAAAGGCCATC | 256 |
| chr10:6003217-6005232 | p3 | (CTT)5 | CCATGTTGGCACATCTATCG | TTCCATAGCAAAAGGCCATC | 124 |
| chr10:6003217-6005232 | p5 | (CTCGA)3 | CCATGTTGGCACATCTATCG | TTCCATAGCAAAAGGCCATC | 124 |
| chr10:6003217-6005232 | p5 | (TAGTT)3 | CCATGTTGGCACATCTATCG | TTCCATAGCAAAAGGCCATC | 124 |
| chr10:6060267-6062287 | p4 | (AATA)5 | GATCATCGGGCATGAAGTTT | AGGGCCAAATGCACAAATAG | 151 |
| chr10:6118234-6120248 | p2 | (TG)7 | ATTTTGAAAGCCCCACATTG | GAAAGGGTAAAATGTCCGCA | 144 |
| chr10:6118234-6120248 | p6 | (ATCCTA)4 | ATTTTGAAAGCCCCACATTG | GAAAGGGTAAAATGTCCGCA | 144 |
| chr10:6161067-6163091 | p6 | (GATTAG)4 | TTTTGGATCAAAGCCTGGAG | AGATCCTCGTACCTCTCGCA | 230 |
| chr10:6182822-6184837 | p3 | (ATG)5 | AAAACTGTCAATCGGTTCGG | GGAAAAATGGGTTTGGGACT | 126 |
| chr10:6182822-6184837 | p2 | (CA)11 | AAAACTGTCAATCGGTTCGG | GGAAAAATGGGTTTGGGACT | 126 |
| chr10:6183814-6185884 | p3 | (ATG)5 | TATGGATGTTTGGGGAGAGC | CAATTGCGAGTTCGACTTCA | 254 |
| chr10:6183814-6185884 | p2 | (AT)16 | TATGGATGTTTGGGGAGAGC | CAATTGCGAGTTCGACTTCA | 254 |
| chr10:6197534-6199546 | p2 | (GT)6 | GTGGATGTGCAGTTCATTGG | CAACCCTGGAATAACTCGGA | 216 |
| chr10:6226367-6228385 | p3 | (TGA)6 | AAACCTGTCAACTGGTTCGG | GTTTGGCAACGACCATTTTT | 188 |
| chr10:6243197-6245212 | p5 | (AAACC)3 | TGCTAGCACGCTCAATTGTT | CCGAGAAAACCCTAGGTTGA | 178 |
| chr10:6261727-6263741 | p6 | (GATTAG)3 | CAACGGCCTCGAGCTTATAG | TTTTCCATGGACGAATGTCA | 168 |
| chr10:6261727-6263741 | p2 | (CA)7 | CAACGGCCTCGAGCTTATAG | TTTTCCATGGACGAATGTCA | 168 |
| chr10:6274510-6276534 | p6 | (ATTAGG)4 | TTTTGGATCAAAGCCTGGAG | AGATCCTCGTACCTCTCGCA | 169 |
| chr10:6280159-6282175 | p2 | (GA)11 | TTCATGAACAAGGCCAATCA | CCCTGCTTACCACAGACACA | 252 |
| chr10:6280159-6282175 | p4 | (TTAT)4 | TTCATGAACAAGGCCAATCA | CCCTGCTTACCACAGACACA | 252 |
| chr10:6280159-6282175 | p4 | (TGTT)5 | TTCATGAACAAGGCCAATCA | CCCTGCTTACCACAGACACA | 252 |
| chr10:6280804-6282824 | p4 | (TTAT)4 | ATGCGACGGAGCAAGTATTC | GCCGAACACAACTCCATCTT | 180 |
| chr10:6280804-6282824 | p4 | (TGTT)5 | ATGCGACGGAGCAAGTATTC | GCCGAACACAACTCCATCTT | 180 |
| chr10:6338589-6340605 | p2 | (TA)8 | GTTCCGGGAGGGTTTGTAAT | TGTAATGCCTCCAATCACCA | 181 |
| chr10:6360125-6362140 | p5 | (TGAAA)3 | AGTCAACCCCTTTTGAGCCT | AACTTTTCCAACCACCATCG | 226 |
| chr10:6406373-6408385 | p2 | (AG)6 | CCGGGATTGACTCTGATGAT | AAGAAAATCCACACATCGGC | 135 |

| Marker ID | SSR type | SSR motif | FORWARD PRIMER (5'-3') | REVERSE PRIMER (5'-3') | size (bp) |
|---|---|---|---|---|---|
| chr10:6542954-6544998 | p2 | (AT)22 | TCGATGTCACACCCCTATCA | CCTTGTCGGCCTTATTTTGA | 161 |
| chr10:6542954-6544998 | p3 | (CAA)6 | TCGATGTCACACCCCTATCA | CCTTGTCGGCCTTATTTTGA | 161 |
| chr10:6793282-6795298 | p2 | (AT)8 | CATGGTGGCTCTGATACCCT | CGATGATGAACGAAACATGG | 163 |
| chr10:6838690-6840708 | p6 | (CCCCAC)3 | ACAAGAGAAATCGTGGCAGC | TTTGATGACTGGTTTTCGCA | 107 |
| chr10:6854709-6856733 | p3 | (TAA)8 | CTCCAACTGGGGTGTGAACT | GCAGTTTTTCCATTCTCCCA | 164 |
| chr10:6866228-6868243 | p5 | (ATTTT)3 | GCCTTTTCCACCAAATTTCA | CCAGGTGCGATAGCATTTTT | 241 |
| chr10:6873157-6875172 | p3 | (GTT)5 | CCAAACGAGGAAGGTGGTAA | AAGAAACCGGGGTTCAACTT | 197 |
| chr10:7086227-7088239 | p2 | (TA)6 | ATTATCGGGGGTCTAAACGG | CGGTAAACTGCCAATACGGT | 215 |
| chr10:7086227-7088239 | p2 | (TA)6 | ATTATCGGGGGTCTAAACGG | CGGTAAACTGCCAATACGGT | 215 |
| chr10:7090534-7092549 | p5 | (TATAT)3 | GGTCCGCATAACTTTCGGTA | TTTTGCCCTTCGCTTAAAAA | 286 |
| chr10:7211624-7213638 | p2 | (AC)7 | CGCTCTCGTCCCTAAATGAC | ATGACCACACCTTGCACCTT | 149 |
| chr10:7211624-7213638 | p2 | (TA)9 | CGCTCTCGTCCCTAAATGAC | ATGACCACACCTTGCACCTT | 149 |
| chr10:7261890-7263906 | p4 | (ATCA)4 | GGGCTGATTTTGTAACCCCT | GACGAACCAGAACCGGATAA | 115 |
| chr10:7370535-7372547 | p2 | (AC)6 | AGGGAAGATTTGGGTTTGCT | TCGTTTGGGTTGTTGTTGAA | 146 |
| chr10:7442101-7444116 | p3 | (CAG)5 | GAGCCATCTTAGGCAACGTC | CTCTCACGGCCAAGAAAAAG | 299 |
| chr10:7478393-7480411 | p6 | (CAACAG)3 | TAACAGCCAAGCACAGCAAC | TTCTGGCCCCTAACATTTTG | 273 |
| chr10:7482546-7484561 | p3 | (TGT)5 | TGGGGAAATGGGGTATATGA | GTATGAGGCCCATAGAGGCA | 157 |
| chr10:7524569-7526587 | p6 | (CAACAG)3 | TAACAGCCAAGCACAGCAAC | TTCTGGCCCCTAACATTTTG | 279 |
| chr10:7553149-7555173 | p6 | (CAACAG)4 | GTTAACAGCCAAGCACAGCA | CAGGAGGAAGTGCAGGAGTC | 253 |
| chr10:7553149-7555173 | p3 | (GCA)5 | GTTAACAGCCAAGCACAGCA | CAGGAGGAAGTGCAGGAGTC | 253 |
| chr10:7553295-7555310 | p6 | (CAACAG)4 | GTTAACAGCCAAGCACAGCA | CAGGAGGAAGTGCAGGAGTC | 107 |
| chr10:7553295-7555310 | p3 | (GCA)5 | GTTAACAGCCAAGCACAGCA | CAGGAGGAAGTGCAGGAGTC | 107 |
| chr10:7578657-7580673 | p2 | (AT)8 | TGTTCGGGTCATTCAACGTA | CCTTCCGCTGTAAAATCTGC | 135 |
| chr10:7598783-7600804 | p3 | (CAA)7 | GCTGGGCTATGGGTTAATCA | CCTTGGTGTCTCCCTGACAT | 164 |
| chr10:7630101-7632116 | p3 | (TTG)5 | AAGCCCTCCAAAAGGATGAT | GAGCCATTTTGGCAGCTTAG | 216 |
| chr10:7630101-7632116 | p6 | (TATATG)3 | AAGCCCTCCAAAAGGATGAT | GAGCCATTTTGGCAGCTTAG | 216 |
| chr10:7638959-7640974 | p3 | (TTG)5 | AAGCCCTCCAAAAGGATGAT | GAGCCATTTTGGCAGCTTAG | 216 |
| chr10:7638959-7640974 | p6 | (TATATG)3 | AAGCCCTCCAAAAGGATGAT | GAGCCATTTTGGCAGCTTAG | 216 |
| chr10:7692504-7694519 | p3 | (ATG)5 | GGGTAATGGTATGAGCGGTG | AGCTCGAATTGGAATCCCTT | 150 |
| chr10:7752265-7754281 | p2 | (TA)8 | GAACAAATCTTCCAGCAGCC | GGGGGACTCCAATGAATTTT | 244 |
| chr10:7791088-7793205 | p3 | (TTA)39 | TCAGACCTGAATTGGTGTGC | AAGGCACTCAAACGAAATGC | 245 |
| chr10:7809723-7811738 | p3 | (TAT)7 | CTTGCGTCGCTGTATTTCAA | GTTCGCTGCCGAAGAAATAG | 294 |
| chr10:7810266-7812290 | p3 | (TAT)7 | GTAGGCGATCCAGAGACCAG | GATGAAAGAATTTTCGGGCA | 123 |
| chr10:7830451-7832487 | p2 | (TA)18 | TAGCCGTATGCTGTGTGAGG | TCCTAAATTCCCAGATCCCC | 286 |
| chr10:7890106-7892118 | p2 | (TA)6 | TCTTTAGCCCGTTTGAGCAT | TATGCCAGCCAATTTCATCA | 271 |
| chr10:7950406-7952424 | p6 | (ATTTTT)3 | AACAAGCTGAAACCCGTGAC | GCGTGTTACGCTGAATTTGA | 198 |

| Marker ID | SSR type | SSR motif | FORWARD PRIMER (5'-3') | REVERSE PRIMER (5'-3') | size (bp) |
|---|---|---|---|---|---|
| chr10:7957424-7959477 | p5 | (ACCAA)3 | TCCCAAGGATGAAAAACAGC | AAGCAAGTGGGGTTTCAAGA | 130 |
| chr10:7978535-7980550 | p5 | (ATATT)3 | TCCACTCGACCAACATTCAA | ATCATTCAAGCTTTGCACCC | 202 |
| chr10:8019223-8021238 | p3 | (TCT)5 | AAAGAAATGGTCCAAACCCC | ACTGTTTGTCTGGCCAGCTT | 273 |
| chr10:8025947-8027961 | p2 | (TC)7 | CTCGCACTACGTGGTTTCAA | TTTTTGTGGTTTCGTCACCC | 160 |
| chr10:8028264-8030279 | p5 | (TTGGT)3 | AGCTCGCAAACTATACGGGA | CCACATGGGAGACCTGAGAT | 262 |
| chr10:8028264-8030279 | p5 | (TTGGT)3 | AGCTCGCAAACTATACGGGA | CCACATGGGAGACCTGAGAT | 262 |
| chr10:8028994-8031009 | p5 | (TTGGT)3 | CCGAGTCAAACCGAGAAAAA | TAATGTCGGGTTGGTTTGGT | 190 |
| chr10:8028994-8031009 | p5 | (TTGGT)3 | CCGAGTCAAACCGAGAAAAA | TAATGTCGGGTTGGTTTGGT | 190 |
| chr10:8058426-8060450 | p6 | (CACCCC)4 | TCATTCTCTTCCTGGGATGG | GCAATCCAGAGCCATTAAGC | 183 |
| chr10:8075479-8077500 | p3 | (AAT)7 | CGGAAAGAGCACCAAGAAAG | AGGAGGCGTGACAGATTGAT | 138 |
| chr10:8202339-8204369 | p2 | (AT)15 | CCCGCTCCCTAGCTAGAACT | CTCAAGGGGGTGTGAGGTAG | 131 |
| chr10:8211983-8213998 | p5 | (AGAAA)3 | CCGAGAGGAGTGCAATTGAT | AAAAGGTGTTGGGAACCTGA | 206 |
| chr10:8261013-8263029 | p5 | (TTTTA)3 | TTGGTTCAGCTTGCTTTTCC | TATTGAGTTCCGTGCTTCCC | 269 |
| chr10:8283669-8285687 | p3 | (TAT)6 | ATCATGGTGATCTGTTGGCA | GCCATCATCAGTTGTCCCTT | 242 |
| chr10:8337899-8339917 | p5 | (AAAAT)3 | GTTCGGCTCGAAACTGGTAG | TCATGATCAGCTTGGCTCAC | 278 |
| chr10:8337899-8339917 | p6 | (AAAAAT)3 | GTTCGGCTCGAAACTGGTAG | TCATGATCAGCTTGGCTCAC | 278 |
| chr10:8343145-8345159 | p2 | (GT)7 | CCTAAAACCGTCCATAGCCA | TAAGGACATTCGGGACAAGG | 206 |
| chr10:8351050-8353068 | p3 | (TTA)6 | CCTGCCGTTGTAAAAATGGT | CTTACCAGACCCGACACACC | 155 |
| chr10:8351077-8353095 | p3 | (TTA)6 | CCTGCCGTTGTAAAAATGGT | CTTACCAGACCCGACACACC | 128 |
| chr10:8351096-8353138 | p3 | (TTA)6 | CCTGCCGTTGTAAAAATGGT | CTTACCAGACCCGACACACC | 109 |
| chr10:8369663-8371681 | p3 | (TCA)5 | GGGGGTTGTGTTTCATCATC | GACGTGGTGCCGGTAGTAGT | 204 |
| chr10:8369663-8371681 | p6 | (CCGGTT)3 | GGGGGTTGTGTTTCATCATC | GACGTGGTGCCGGTAGTAGT | 204 |
| chr10:8369663-8371681 | p5 | (TAAAT)3 | GGGGGTTGTGTTTCATCATC | GACGTGGTGCCGGTAGTAGT | 204 |
| chr10:8406888-8408915 | p3 | (CCA)5 | GTGGGATGTGGGAATGGTAG | GGCAACCCTCCTCCTTTATC | 203 |
| chr10:8406888-8408915 | p6 | (GACCAA)4 | GTGGGATGTGGGAATGGTAG | GGCAACCCTCCTCCTTTATC | 203 |
| chr10:8428495-8430510 | p5 | (AAATA)3 | GCGTCGATTGCAGTCATAGA | TGAGGTCTCCTCCCATTCAC | 250 |
| chr10:8428495-8430510 | p5 | (AAATA)3 | GCGTCGATTGCAGTCATAGA | TGAGGTCTCCTCCCATTCAC | 250 |
| chr10:8428495-8430510 | p5 | (AATAA)3 | GCGTCGATTGCAGTCATAGA | TGAGGTCTCCTCCCATTCAC | 250 |
| chr10:8428495-8430510 | p4 | (TTTA)4 | GCGTCGATTGCAGTCATAGA | TGAGGTCTCCTCCCATTCAC | 250 |
| chr10:8464197-8466211 | p2 | (AT)7 | GAAACAGAGCTCGTTAGCGG | ACGTGAAGAAGGAGCTGGAA | 117 |
| chr10:8546148-8548160 | p2 | (AC)6 | CCGACTGATCTGCCTTCTTC | GTTGACCAGGACAGGACGTT | 251 |
| chr10:8546148-8548160 | p2 | (TA)6 | CCGACTGATCTGCCTTCTTC | GTTGACCAGGACAGGACGTT | 251 |
| chr10:8546673-8548685 | p2 | (AC)6 | GCCAAACACTCACACGCTTA | ACTCACCCAGGTTTTCATGG | 223 |
| chr10:8546673-8548685 | p2 | (TA)6 | GCCAAACACTCACACGCTTA | ACTCACCCAGGTTTTCATGG | 223 |
| chr10:8580387-8582402 | p3 | (TAA)6 | CGTGCTTTTGTTTGAGGTGA | TGCTTTTGGACTTCGGCTAT | 125 |
| chr10:8580387-8582402 | p3 | (AAT)7 | CGTGCTTTTGTTTGAGGTGA | TGCTTTTGGACTTCGGCTAT | 125 |

| Marker ID | SSR type | SSR motif | FORWARD PRIMER (5'-3') | REVERSE PRIMER (5'-3') | size (bp) |
|---|---|---|---|---|---|
| chr10:8581108-8583126 | p3 | (TAA)6 | CAAAGCACAACTTTGCTCCA | TTCGCCCCATACCTATTCAA | 201 |
| chr10:8581108-8583126 | p3 | (AAT)7 | CAAAGCACAACTTTGCTCCA | TTCGCCCCATACCTATTCAA | 201 |
| chr10:8592734-8594761 | p3 | (AAC)5 | TTTCCACCACAAATCCACAA | TTGATGTGCCTTGAAAGCTG | 217 |
| chr10:8601538-8603556 | p2 | (TC)9 | GGGTCAACTAACGGCAAAAA | GTGCCTGAGGCCTAACTTTG | 275 |
| chr10:8638997-8641011 | p2 | (TA)7 | ATCCATGCTTTTGAAGTGCC | ATCTGCAAACTCCACCTGCT | 170 |
| chr10:8699740-8701754 | p2 | (TA)7 | TATACTTTGCGGCCCTTCAA | TGCCTCATCAGTTTGTTTGC | 108 |
| chr10:8714042-8716060 | p3 | (TGT)6 | AAAAGCACTCGCATCCTCAT | GATTGGTGGACATCTTGGCT | 202 |
| chr10:8733713-8735728 | p5 | (TTTAA)3 | TTAAGCCCGTGCCATCTAGT | ATCTGCTGGATTTCCCTCCT | 165 |
| chr10:8895997-8898012 | p5 | (AGTAA)3 | GCACTGACCACTGCTGGATA | GCTCGCAGGTTCAGGTAGAC | 151 |
| chr10:8916499-8918627 | p2 | (AT)64 | TGCTAGCAAGTTTGGCAATG | GCAAATGTCCTCCCAAGTGT | 100 |
| chr10:8996742-8998787 | p3 | (ATA)15 | CAGCCCTTTGCAATGATTTT | ATATTCGGCGAAGTTTGTGG | 271 |
| chr10:9044052-9046084 | p2 | (CT)16 | ATCTCCAAAACCATGCGAAG | CTGGCAAGGGAACTCAGAAG | 237 |
| chr10:9044052-9046084 | p3 | (AAC)6 | ATCTCCAAAACCATGCGAAG | CTGGCAAGGGAACTCAGAAG | 237 |
| chr10:9129285-9131303 | p2 | (AT)7 | CGAAGGGCAAAAATTAACGA | AACACACGTAGGGCTATCGG | 118 |
| chr10:9129285-9131303 | p3 | (TTG)6 | CGAAGGGCAAAAATTAACGA | AACACACGTAGGGCTATCGG | 118 |
| chr10:9189393-9191408 | p5 | (CCCCG)3 | TATTCCTAACGGCACTTCGC | TTCGGAGAGTCGGTTCAGTT | 147 |
| chr10:9213874-9215889 | p5 | (TATTG)3 | AATCTTGATCTGCGCCTTGT | CTGGAATTTGAACCCGAGAA | 181 |
| chr10:9213874-9215889 | p2 | (TA)6 | AATCTTGATCTGCGCCTTGT | CTGGAATTTGAACCCGAGAA | 181 |
| chr10:9337004-9339016 | p5 | (TTTAT)3 | TTCCATGTGAAAGCATCGAC | GTGTTTAGGGCGTACGGTTG | 184 |
| chr10:9337017-9339033 | p5 | (TTTAT)3 | TTCCATGTGAAAGCATCGAC | GTGTTTAGGGCGTACGGTTG | 171 |
| chr10:9337053-9339071 | p5 | (TTTAT)3 | TTCCATGTGAAAGCATCGAC | GTGTTTAGGGCGTACGGTTG | 135 |
| chr10:9397116-9399131 | p5 | (AATCG)4 | AACCGAATCAAACCGACAAC | ACAGCGATGATTGTTGACCA | 199 |
| chr10:9397116-9399131 | p2 | (GA)8 | AACCGAATCAAACCGACAAC | ACAGCGATGATTGTTGACCA | 199 |
| chr10:9397116-9399131 | p5 | (TTGAT)3 | AACCGAATCAAACCGACAAC | ACAGCGATGATTGTTGACCA | 199 |
| chr10:9479081-9481099 | p2 | (TA)9 | TGACTCCCTACCCAAAGTGG | CATGGAGAGACTGATGGGGT | 260 |
| chr10:9502989-9505003 | p2 | (TA)7 | GATTTCAAAGACTTGCCCCA | CGTTAACCACACCTGCACAC | 244 |
| chr10:9502989-9505003 | p2 | (AG)6 | GATTTCAAAGACTTGCCCCA | CGTTAACCACACCTGCACAC | 244 |
| chr10:9522964-9524988 | p5 | (GTTTG)3 | GGGCTGTTCACAGTTTTGGT | TGGCTCGGTTATTTATTCGG | 128 |
| chr10:9522964-9524988 | p6 | (CATGAG)4 | GGGCTGTTCACAGTTTTGGT | TGGCTCGGTTATTTATTCGG | 128 |
| chr10:9527029-9529041 | p2 | (AC)6 | TAAATTGAGATGGCGGGAAC | ATTTTGGGCTATTGCACCTG | 260 |
| chr10:9527029-9529041 | p3 | (TTC)5 | TAAATTGAGATGGCGGGAAC | ATTTTGGGCTATTGCACCTG | 260 |
| chr10:9551942-9553960 | p6 | (ATCTAT)3 | TCTCAACACTTTGTTTGCCG | CTGATTTACACAACCGGCCT | 194 |
| chr10:9565938-9567956 | p2 | (TA)9 | TCCTGATCCTTATCATCCGC | ATGTTTGGCGTGTCGTTACA | 132 |
| chr10:9618350-9620368 | p6 | (AAAAAT)3 | TCGTGGTGTCAAGGTTTGAG | ATTCGAATCCTGACCTCGTG | 255 |
| chr10:9618350-9620368 | p3 | (TGT)5 | TCGTGGTGTCAAGGTTTGAG | ATTCGAATCCTGACCTCGTG | 255 |
| chr10:9710126-9712156 | p5 | (TTTCT)3 | AGCTAAGACCAGGGGCAGAT | TATGCCCGAATTCAATGTCA | 212 |

| Marker ID | SSR type | SSR motif | FORWARD PRIMER (5'-3') | REVERSE PRIMER (5'-3') | size (bp) |
|---|---|---|---|---|---|
| chr10:9710126-9712156 | p3 | (ATT)10 | AGCTAAGACCAGGGGCAGAT | TATGCCCGAATTCAATGTCA | 212 |
| chr10:9731942-9733972 | p5 | (TTTCT)3 | AGCTAAGACCAGGGGCAGAT | TATGCCCGAATTCAATGTCA | 212 |
| chr10:9731942-9733972 | p3 | (ATT)10 | AGCTAAGACCAGGGGCAGAT | TATGCCCGAATTCAATGTCA | 212 |
| chr10:9754419-9756431 | p2 | (TG)6 | GTTGGAGGCTTGTTTTGGAA | CTTGCTTGTAGTCTTCCCGC | 124 |
| chr10:9842451-9844463 | p2 | (AT)6 | TCATCCGATCGATACCACAA | GGACGACGAAAAATCCTCAA | 103 |
| chr10:9850175-9852190 | p5 | (ACTTT)3 | CAAAAACCAGGTGGCAGTTT | CCCATAACTGAGGGACCAAA | 276 |
| chr10:9852010-9854034 | p2 | (TG)7 | AAGCATCTAATTTGCCGTGG | TCAAAATATTTCGGGGGTCA | 133 |
| chr10:9865428-9867446 | p2 | (AT)9 | AATCTGCCCAAATGAAGCAC | CCGGGTAACTCTATCCACGA | 290 |
| chr10:9865428-9867446 | p2 | (AT)9 | AATCTGCCCAAATGAAGCAC | CCGGGTAACTCTATCCACGA | 290 |
| chr10:9865428-9867446 | p2 | (AT)9 | AATCTGCCCAAATGAAGCAC | CCGGGTAACTCTATCCACGA | 290 |
| chr10:9865428-9867446 | p2 | (AT)9 | AATCTGCCCAAATGAAGCAC | CCGGGTAACTCTATCCACGA | 290 |
| chr10:9865428-9867446 | p2 | (AT)9 | AATCTGCCCAAATGAAGCAC | CCGGGTAACTCTATCCACGA | 290 |
| chr10:9866070-9868088 | p2 | (AT)9 | CCACACAGCAAATCATACCC | TCAGTGCTGCTCGTAATTCA | 234 |
| chr10:9866070-9868088 | p2 | (AT)9 | CCACACAGCAAATCATACCC | TCAGTGCTGCTCGTAATTCA | 234 |
| chr10:9866070-9868088 | p2 | (AT)9 | CCACACAGCAAATCATACCC | TCAGTGCTGCTCGTAATTCA | 234 |
| chr10:9866070-9868088 | p2 | (AT)9 | CCACACAGCAAATCATACCC | TCAGTGCTGCTCGTAATTCA | 234 |
| chr10:9866070-9868088 | p2 | (AT)9 | CCACACAGCAAATCATACCC | TCAGTGCTGCTCGTAATTCA | 234 |
| chr10:9866070-9868088 | p2 | (AT)9 | CCACACAGCAAATCATACCC | TCAGTGCTGCTCGTAATTCA | 234 |
| chr10:9866070-9868088 | p2 | (AT)9 | CCACACAGCAAATCATACCC | TCAGTGCTGCTCGTAATTCA | 234 |
| chr10:9866070-9868088 | p2 | (AT)9 | CCACACAGCAAATCATACCC | TCAGTGCTGCTCGTAATTCA | 234 |
| chr10:9972316-9974418 | p6 | (TTTCTT)3 | TCTGTTTGGAAAGCCACCTC | CCCCCATGAAAATTGAAAAA | 174 |
| chr10:9972316-9974418 | p2 | (AT)51 | TCTGTTTGGAAAGCCACCTC | CCCCCATGAAAATTGAAAAA | 174 |
| chr10:9972316-9974418 | p2 | (AT)16 | TCTGTTTGGAAAGCCACCTC | CCCCCATGAAAATTGAAAAA | 174 |
| chr10:10093095-10095131 | p2 | (TA)18 | ACCGCCAAGGCTTTAGAAAT | TATGGTTGTTTGGGGGTTGT | 271 |
| chr10:10167208-10169229 | p3 | (CTT)10 | CTCTTCACCACTAACCCCCA | GGAAGATGCCAACATTGCTT | 196 |
| chr10:10167208-10169229 | p3 | (GAA)7 | CTCTTCACCACTAACCCCCA | GGAAGATGCCAACATTGCTT | 196 |
| chr10:10226973-10228991 | p2 | (AT)9 | CCCTACTTGGACATGGCATT | CAGCCAAGCATGAAGCATTA | 234 |
| chr10:10226973-10228991 | p5 | (TTGTC)3 | CCCTACTTGGACATGGCATT | CAGCCAAGCATGAAGCATTA | 234 |
| chr10:10232234-10234278 | p2 | (TA)22 | GTTGTGAGCCCTCCATTTGT | GCAGGCCTCAAACACTAAGC | 165 |
| chr10:10232234-10234278 | p2 | (AT)9 | GTTGTGAGCCCTCCATTTGT | GCAGGCCTCAAACACTAAGC | 165 |
| chr10:10232405-10234423 | p2 | (TA)22 | AGTGTTTGAGGCCTGCACTT | ATGACCCAAGGAATGCTCAG | 177 |
| chr10:10232405-10234423 | p2 | (AT)9 | AGTGTTTGAGGCCTGCACTT | ATGACCCAAGGAATGCTCAG | 177 |
| chr10:10374886-10376904 | p2 | (AT)9 | GCCCATTTGAAGGACAAAAA | GAATAGGGCCAAAAGTGCAA | 162 |
| chr10:10374886-10376904 | p2 | (TA)12 | GCCCATTTGAAGGACAAAAA | GAATAGGGCCAAAAGTGCAA | 162 |
| chr10:10389689-10391737 | p2 | (TA)24 | TGCATGTCGGTGAAAAAGAC | GCGTGTTTAGGGCGTACAAT | 211 |
| chr10:10466367-10468379 | p2 | (AT)6 | CCATAACCCAATTGTGACCC | TGAGTATGGGCAGGGGTTAG | 117 |

| Marker ID | SSR type | SSR motif | FORWARD PRIMER (5'-3') | REVERSE PRIMER (5'-3') | size (bp) |
|---|---|---|---|---|---|
| chr10:10576598-10578616 | p3 | (ACA)6 | CCATGACTCATAGGCCAGGT | CGGGTCATGGAGTTGTTACC | 266 |
| chr10:10607499-10609529 | p2 | (AT)15 | AGGCAGAGGGGGAATTAAGA | CAAGGCAAGTAAGCCCTGAG | 120 |
| chr10:10611050-10613064 | p2 | (AT)7 | TCCATTTGAAGGGAAAATCG | AGCTCATCATGCACTTCACG | 298 |
| chr10:10703654-10705674 | p5 | (TTTTC)4 | ACAAATCCAGATGGAGTCCG | CCCTCCACGTTATACCCCTT | 172 |
| chr10:10704936-10706956 | p5 | (TTTTC)4 | AAGGGGTATAACGTGGAGGG | ACGTTGGCTTCCAAGAATTG | 127 |
| chr10:10820799-10822827 | p2 | (AT)14 | ACAAACATAGGCCGACAAGG | GGTGCACAGGGAGACAAACT | 238 |
| chr10:10834438-10836458 | p3 | (CAT)5 | GGGGTAGCTCGAAAAGATCC | GGAGGCATTGATGAAACGAT | 131 |
| chr10:10834462-10836478 | p3 | (CAT)5 | GGGGTAGCTCGAAAAGATCC | GGAGGCATTGATGAAACGAT | 107 |
| chr10:10835188-10837203 | p3 | (CAT)5 | TGACCCACATGGCTAACTCA | AGCAGGTATCCCGTTAGGCT | 182 |
| chr10:10845418-10847480 | p2 | (AT)31 | CACATCACCGCAATGTTCTC | CTCAGGCCAAAAGGTGAGAG | 194 |
| chr10:10845418-10847480 | p3 | (ACA)6 | CACATCACCGCAATGTTCTC | CTCAGGCCAAAAGGTGAGAG | 194 |
| chr10:10845628-10847646 | p2 | (AT)31 | GCTCTATCACCATGACCCGT | ATGAAAGAGCCTCAGCTCCA | 103 |
| chr10:10845628-10847646 | p3 | (ACA)6 | GCTCTATCACCATGACCCGT | ATGAAAGAGCCTCAGCTCCA | 103 |
| chr10:10857996-10860018 | p5 | (TTTCT)3 | TAATGGTGGATTCATTGCCA | GACATGGCGGAATAGCAAAT | 119 |
| chr10:10966622-10968646 | p6 | (TTTTTA)4 | AAAAGGGATTGGGTTTTTGG | TGGGGTCCATCTTCATTTGT | 100 |
| chr10:10969175-10971190 | p5 | (TCTTC)3 | TTATGCCACCGTCGTTATCA | GTCGGGTGTATGGCTCAGAT | 216 |
| chr10:11015391-11017406 | p5 | (ATACA)3 | CAATCCAAAGAGGACCTCCA | GCTCTTTTACACGAGCACCC | 256 |
| chr10:11017175-11019217 | p2 | (TA)7 | GGGTATTTTGGTTCCGGACT | TAAGCAAAGGGACGACCAAG | 125 |
| chr10:11062404-11064416 | p2 | (GA)7 | GCGAAATTTCTTTCCCCTTC | AAATGGGGACTCCAACTCCT | 204 |
| chr10:11062404-11064416 | p2 | (TC)6 | GCGAAATTTCTTTCCCCTTC | AAATGGGGACTCCAACTCCT | 204 |
| chr10:11096409-11098431 | p2 | (AT)11 | GCATGGTACCTTGTAGCGGT | GGCGTCATTCAATCTTTCGT | 122 |
| chr10:11141736-11143751 | p5 | (GTTCG)3 | ACCTAAGCAGTTCCGAAGCA | CGACTTACCCTCCACAGGAA | 202 |
| chr10:11160674-11162740 | p2 | (AT)33 | CTATTTGCTTGCCACACGAA | TCCATGGCAACTCGTGTAAA | 159 |
| chr10:11160674-11162740 | p2 | (AG)7 | CTATTTGCTTGCCACACGAA | TCCATGGCAACTCGTGTAAA | 159 |
| chr10:11168239-11170251 | p2 | (AT)14 | GCCTTTGCATACCTAACCCA | CTAGTGCATCTGCCACCTCA | 224 |
| chr10:11168239-11170251 | p3 | (TCT)6 | GCCTTTGCATACCTAACCCA | CTAGTGCATCTGCCACCTCA | 224 |
| chr10:11168262-11170308 | p2 | (AT)14 | GCCTTTGCATACCTAACCCA | CTAGTGCATCTGCCACCTCA | 201 |
| chr10:11168262-11170308 | p3 | (TCT)6 | GCCTTTGCATACCTAACCCA | CTAGTGCATCTGCCACCTCA | 201 |
| chr10:11240726-11242741 | p5 | (TTTTC)3 | TTGAAAAATGGCATGGATGA | CCGTGTGATAGTGCAAATGG | 250 |
| chr10:11278263-11280309 | p2 | (TA)23 | CCGCATAAGGCTGAACATTT | CGAGTCCTGACACTGCCTTT | 149 |
| chr10:11301701-11303719 | p6 | (CATGAG)3 | GCGTATTCGGTAGAATGGGA | AAACAGTTGCCCACCAAGAC | 200 |
| chr10:11331349-11333364 | p3 | (ACA)5 | GGGGGTTTAAAATTTGCCAT | CATGAATACAACGAGCCCCT | 143 |
| chr10:11331375-11333387 | p3 | (ACA)5 | GGGGGTTTAAAATTTGCCAT | CATGAATACAACGAGCCCCT | 117 |
| chr10:11359822-11361836 | p2 | (TG)12 | ATCACACCCCATCCTTGGTA | TTCGCATTTAGCCTCTCGAT | 154 |
| chr10:11359822-11361836 | p3 | (CAA)5 | ATCACACCCCATCCTTGGTA | TTCGCATTTAGCCTCTCGAT | 154 |
| chr10:11359838-11361876 | p2 | (TG)12 | ATCACACCCCATCCTTGGTA | TTCGCATTTAGCCTCTCGAT | 138 |

| Marker ID | SSR type | SSR motif | FORWARD PRIMER (5'-3') | REVERSE PRIMER (5'-3') | size (bp) |
|---|---|---|---|---|---|
| chr10:11359838-11361876 | p3 | (CAA)5 | ATCACACCCCATCCTTGGTA | TTCGCATTTAGCCTCTCGAT | 138 |
| chr10:11374590-11376602 | p2 | (AC)6 | ATTCCCAAATATGCAGCGAC | TTTTATGGGTTATCTGCGGC | 267 |
| chr10:11572724-11574744 | p5 | (CCCCA)3 | AATCCATGAATCCAAACCCA | TTTGGATTAAGTGCGCAACA | 144 |
| chr10:11578449-11580477 | p2 | (TA)14 | GTCTTCCTAAGACCCGGAGG | CAACTTCAGCCACCCTTCAT | 138 |
| chr10:11578449-11580477 | p2 | (TA)8 | GTCTTCCTAAGACCCGGAGG | CAACTTCAGCCACCCTTCAT | 138 |
| chr10:11597841-11599859 | p6 | (AAAAGA)3 | AGAGAGAGCTCGTGGGTCAA | GGCTTTTGAGCTTGGTCTTG | 109 |
| chr10:11630344-11632376 | p2 | (TA)33 | TGTGAGGCGAGTGGTCTATG | GCCGGCTCTATGTCTCAAAC | 295 |
| chr10:11639312-11641324 | p2 | (AT)6 | TCATTTAGGTTCCGGACTCG | ACTCGCACCATATTCCCATC | 159 |
| chr10:11646720-11648738 | p6 | (GGTGCA)3 | AGTTGGCATCTGTCCGGTAG | CCAGAAAGTTGGCTCTTTGC | 229 |
| chr10:11646720-11648738 | p6 | (GGAGAA)3 | AGTTGGCATCTGTCCGGTAG | CCAGAAAGTTGGCTCTTTGC | 229 |
| chr10:11721483-11723501 | p2 | (CT)6 | TCTTCAATCAATGCCCTTCC | AGATCATCAAGGGCAAATCG | 153 |
| chr10:11721483-11723501 | p6 | (TTTTTG)3 | TCTTCAATCAATGCCCTTCC | AGATCATCAAGGGCAAATCG | 153 |
| chr10:11758493-11760511 | p3 | (TCA)6 | TGGATGCAAAGATGGAATGA | ATGTGGATCCAGAACTTGCC | 106 |
| chr10:11787156-11789186 | p3 | (TCT)10 | GTGTCGTGGCAAGTCACTGT | GAGTGCACGTGTTTGGAAGA | 264 |
| chr10:11913835-11915847 | p2 | (TC)6 | AGAACGGACCGTAACACAGG | TTGCCTTAAATGCTCCAACC | 194 |
| chr10:11948668-11950686 | p2 | (TA)9 | TTCAAGCAGTCGGTTCCTCT | GACAAGATCGGCCTTAGCTG | 238 |
| chr10:12137012-12139027 | p5 | (ATAAA)3 | CACATTCAACCTTTACGGGG | GAGTTCAACAATCGGAGGGA | 280 |
| chr10:12179832-12181850 | p3 | (TTA)6 | AGACACCCTGCCCTTTTTCT | TGGTCAAGGTAGGGAGGAGA | 178 |
| chr10:12189860-12191876 | p2 | (AT)8 | GATCTGGAAATGGAGCTGGA | GAAGGTTCGTGGAAGCTCAC | 281 |
| chr10:12193090-12195112 | p2 | (TA)8 | GTGGGCCCCATACTAAACAA | CCCATGAGTTTGGAGATGGT | 258 |
| chr10:12193090-12195112 | p2 | (TA)11 | GTGGGCCCCATACTAAACAA | CCCATGAGTTTGGAGATGGT | 258 |
| chr10:12193090-12195112 | p2 | (AT)9 | GTGGGCCCCATACTAAACAA | CCCATGAGTTTGGAGATGGT | 258 |
| chr10:12241140-12243152 | p2 | (TA)6 | ATTTTGTGAGTTGCCCAAGC | CCGTGCCTAATGAAACCTTC | 122 |
| chr10:12253719-12255735 | p2 | (AT)8 | ACATCACGAACCAGTGGACA | CCGTAACCCACCTTGATGAC | 145 |
| chr10:12347971-12349991 | p4 | (TTAA)5 | TGACGAACAAAAGGTCACCA | TCGTGGTCCAAAACAAGTCA | 247 |
| chr10:12347971-12349991 | p2 | (TA)6 | TGACGAACAAAAGGTCACCA | TCGTGGTCCAAAACAAGTCA | 247 |
| chr10:12349056-12351068 | p2 | (TA)6 | TTTGTCTCCTCCGTGACTCC | GGTTTCCTAACCGGTTCCAT | 255 |
| chr10:12349056-12351068 | p2 | (TA)6 | TTTGTCTCCTCCGTGACTCC | GGTTTCCTAACCGGTTCCAT | 255 |
| chr10:12389070-12391085 | p5 | (ATTTT)3 | TCCCATAATCGAGCCAAAAG | CACGTACAAATACGCGATGC | 224 |
| chr10:12391293-12393305 | p5 | (TAAAA)3 | CATTTCGTCGTCCATTTGTG | TTTTTCCCGTGAGTTTGGAG | 270 |
| chr10:12391293-12393305 | p5 | (TAAAA)3 | CATTTCGTCGTCCATTTGTG | TTTTTCCCGTGAGTTTGGAG | 270 |
| chr10:12391293-12393305 | p2 | (TA)22 | CATTTCGTCGTCCATTTGTG | TTTTTCCCGTGAGTTTGGAG | 270 |
| chr10:12391876-12393920 | p5 | (TAAAA)3 | TTTTGGTCGTCATTTCGTCA | TTGTTAATACGGAGTCCGCC | 103 |
| chr10:12391876-12393920 | p2 | (TA)22 | TTTTGGTCGTCATTTCGTCA | TTGTTAATACGGAGTCCGCC | 103 |
| chr10:12391876-12393920 | p2 | (AT)7 | TTTTGGTCGTCATTTCGTCA | TTGTTAATACGGAGTCCGCC | 103 |
| chr10:12407948-12409960 | p2 | (TA)23 | ATCATTGTTCTTCCCCATGC | AAATCCATGGAATGGAATGC | 249 |

| Marker ID | SSR type | SSR motif | FORWARD PRIMER (5'-3') | REVERSE PRIMER (5'-3') | size (bp) |
|---|---|---|---|---|---|
| chr10:12407948-12409960 | p2 | (AT)25 | ATCATTGTTCTTCCCCATGC | AAATCCATGGAATGGAATGC | 249 |
| chr10:12407948-12409960 | p2 | (TA)6 | ATCATTGTTCTTCCCCATGC | AAATCCATGGAATGGAATGC | 249 |
| chr10:12468088-12470114 | p2 | (AT)6 | TTGCAATGTTCCGTCTTGAG | TAATTGCTAATTGGGCCTGG | 149 |
| chr10:12468116-12470128 | p2 | (AT)6 | TTGCAATGTTCCGTCTTGAG | TAATTGCTAATTGGGCCTGG | 121 |
| chr10:12468131-12470143 | p2 | (AT)6 | TTGCAATGTTCCGTCTTGAG | TAATTGCTAATTGGGCCTGG | 106 |
| chr10:12543606-12545621 | p5 | (TTTTA)3 | ACGTAAGGTTTGTGTTCGGG | TGGGACCTACACCCAATGTT | 255 |
| chr10:12613082-12615108 | p2 | (TA)8 | TTCATTACCACGTGCGAGTC | GACATGGCGGAATAGCAAAT | 215 |
| chr10:12613082-12615108 | p2 | (TA)6 | TTCATTACCACGTGCGAGTC | GACATGGCGGAATAGCAAAT | 215 |
| chr10:12685789-12687816 | p2 | (AT)6 | CAGCAACCAAACGACGTCTA | GAAGTGGGACGAGATAGCCA | 177 |
| chr10:12685789-12687816 | p3 | (TTA)9 | CAGCAACCAAACGACGTCTA | GAAGTGGGACGAGATAGCCA | 177 |
| chr10:12690388-12692403 | p2 | (AT)7 | TTACGCTCTACGATGCCCTT | TGGGGAAAATTGATTCAAGC | 285 |
| chr10:12690388-12692403 | p3 | (GTA)5 | TTACGCTCTACGATGCCCTT | TGGGGAAAATTGATTCAAGC | 285 |
| chr10:12759938-12761956 | p2 | (TA)9 | ATGGTGGGTTAATGATTCGC | TCACTTGCTCATAGCCCAAC | 229 |
| chr10:12862603-12864621 | p6 | (TTCTTT)3 | GCACTGGAACCTTTGGTGAT | CAATTGAGGGTGCCGTAAGT | 298 |
| chr10:12865629-12867644 | p3 | (AAT)5 | TATCATCAATGCTTGCCCAG | TTCGAGTGAAGAAAGAGGGC | 107 |
| chr10:12865629-12867644 | p3 | (ATA)5 | TATCATCAATGCTTGCCCAG | TTCGAGTGAAGAAAGAGGGC | 107 |
| chr10:12869452-12871482 | p2 | (AT)15 | TATTTTCGAACCAATTCGGG | AGCATGTGCATGTGAAACCT | 150 |
| chr10:12927600-12929615 | p3 | (CAA)5 | GCGAAAGTTTCAAAAGCTGG | ACCATCTAACCTCCGGGTTC | 152 |
| chr10:12927600-12929615 | p2 | (CT)6 | GCGAAAGTTTCAAAAGCTGG | ACCATCTAACCTCCGGGTTC | 152 |
| chr10:12987737-12989749 | p2 | (GT)6 | CGGAGGACGAGATGAAGAAG | TTATGCAAAAACACGGGACA | 138 |
| chr10:13173078-13175100 | p2 | (AT)11 | TTTTTCCCTTGGCAAGACAC | GTGAGGCAGACATTGTGGAA | 228 |
| chr10:13207171-13209186 | p6 | (GATGAA)3 | AGGGTTTTAAAGGGTGGTGG | TTAACCCCTGAACTTGGCAC | 225 |
| chr10:13207171-13209186 | p5 | (TTGGT)3 | AGGGTTTTAAAGGGTGGTGG | TTAACCCCTGAACTTGGCAC | 225 |
| chr10:13207171-13209186 | p5 | (TTGGT)3 | AGGGTTTTAAAGGGTGGTGG | TTAACCCCTGAACTTGGCAC | 225 |
| chr10:13301234-13303250 | p2 | (AT)8 | CACGTGGCACAACGACTTAT | GCCGTTAACTTTGGTGGAAA | 158 |
| chr10:13301234-13303250 | p3 | (CTA)5 | CACGTGGCACAACGACTTAT | GCCGTTAACTTTGGTGGAAA | 158 |
| chr10:13302343-13304357 | p3 | (CTA)5 | TGGCGTAATATGAACATGCC | TTTGTTGCCAAGTGGCTTTT | 253 |
| chr10:13302643-13304661 | p3 | (CTA)5 | AGCCACTTGGCAACAAATTC | TGGCTTGTTCATATTACGCC | 184 |
| chr10:13309390-13311405 | p4 | (AAAT)4 | TGAGGGTGTGGGAGTTTGAT | GAGTCTTCCGTTATGCGGTC | 269 |
| chr10:13309408-13311447 | p4 | (AAAT)4 | TGAGGGTGTGGGAGTTTGAT | GAGTCTTCCGTTATGCGGTC | 251 |
| chr10:13312056-13314071 | p5 | (ATTTT)3 | TCAGGGGATAGCAAAAGTGC | GTTCGAGTAAGCAAAAGGCG | 227 |
| chr10:13312056-13314071 | p2 | (AT)19 | TCAGGGGATAGCAAAAGTGC | GTTCGAGTAAGCAAAAGGCG | 227 |
| chr10:13383022-13385042 | p2 | (TG)10 | ATACTCAGAGCAACGCGGAT | CCATCTCACCAACATCATCG | 157 |
| chr10:13455477-13457498 | p3 | (TCT)7 | TTAGAATCAACCGTCGAGGC | TAGCAAAGAATTATGGGGCG | 229 |
| chr10:13665173-13667185 | p2 | (AT)6 | AGGAATGAAACCTCGTCGTG | CAGAACATTTGACAAGGGGG | 199 |
| chr10:13769372-13771396 | p2 | (AT)12 | ACCTCAATGCCGAATAGGTG | CGGTGTAATTGGTGCAAGTG | 272 |

| Marker ID | SSR type | SSR motif | FORWARD PRIMER (5'-3') | REVERSE PRIMER (5'-3') | size (bp) |
|---|---|---|---|---|---|
| chr10:13812576-13814588 | p2 | (TA)6 | GATCCCAAATTGTCCAATGC | TACGGCCTGCTAAGAGGAAA | 275 |
| chr10:13867586-13869604 | p6 | (GAAATG)3 | AAGTGCGAGTACAGGGATGG | GCAAGGCTAATTTCACTGCC | 269 |
| chr10:13877992-13880010 | p6 | (GAAATG)3 | AAGTGCGAGTACAGGGATGG | GCAAGGCTAATTTCACTGCC | 269 |
| chr10:14275269-14277293 | p2 | (AC)9 | CGGGCACCTCTCATCATACT | TCTTAGCCTTGTCGGCTCAT | 182 |
| chr10:14275269-14277293 | p4 | (CATA)6 | CGGGCACCTCTCATCATACT | TCTTAGCCTTGTCGGCTCAT | 182 |
| chr10:14336242-14338270 | p2 | (TA)14 | AGCATACTCGAGGCAGGAAA | CCACAAGCACGTTTTGATTG | 249 |
| chr10:14435898-14437913 | p5 | (TATTT)3 | AAGGATTCAGGACACATGGC | ACAATCCCCCACTTGTTTCA | 212 |
| chr10:14493838-14495860 | p2 | (AG)11 | GCTTGCTTTGGTTGGATTGT | AACTGGTTCCTGGTCCTCCT | 224 |
| chr10:14497638-14499653 | p5 | (AAAAT)3 | TGAATGTCTGCATCCCAGTG | CAGGTGCGGATTTGACTTTT | 123 |
| chr10:14522580-14524596 | p4 | (AATT)4 | CCTAAATTCTTAAGGCGGGG | GCACAGGGATGGTTACTCGT | 212 |
| chr10:14588261-14590276 | p3 | (ACA)5 | ATAGCTGGTGTGGCAGTCCT | AAGTTTGTTAAACCGCCCCT | 241 |
| chr10:14642960-14644972 | p2 | (AT)6 | CATGACCCATGACCCACATA | AGAGGTTCGTGATCCACCAG | 244 |
| chr10:14800996-14803011 | p3 | (CCA)5 | CATTGCGCATTAAAATGGTG | GATTTGACAACACACCACGC | 237 |
| chr10:14819511-14821609 | p6 | (AATAAA)3 | ATGTGGCAATGAGTCCATCA | CTTCATCGTCCGTTGTCCTT | 134 |
| chr10:14900329-14902347 | p5 | (TTGAA)4 | TACCGAAATTTTGGCAGACC | GCCTTTGGAAGAAAAGGGAC | 223 |
| chr10:14900329-14902347 | p6 | (TATATG)3 | TACCGAAATTTTGGCAGACC | GCCTTTGGAAGAAAAGGGAC | 223 |
| chr10:14930355-14932367 | p2 | (AT)6 | TCTATCAACCACCACTGCCA | CAGCTACAACTTGCCCACAA | 249 |
| chr10:14990602-14992616 | p2 | (AT)7 | ATCACGGGCTTTGGTACTTG | TCCCTCGCCTGCTACTTCTA | 298 |
| chr10:14990602-14992616 | p2 | (TA)9 | ATCACGGGCTTTGGTACTTG | TCCCTCGCCTGCTACTTCTA | 298 |
| chr10:15011208-15013226 | p2 | (TA)9 | GGCACGTGAAGAGGAGAGAC | TAAAAAGGGGTCGAATGTGC | 139 |
| chr10:15149299-15151359 | p2 | (AT)15 | CGGGCACCTCTCATCATACT | TTTTAGCCTTGTCGGCTCAT | 151 |
| chr10:15155858-15157918 | p2 | (AT)15 | CGGGCACCTCTCATCATACT | TTTTAGCCTTGTCGGCTCAT | 151 |
| chr10:15187556-15189574 | p3 | (TTC)6 | TCTTACCCGGGCTTATGTTG | GGTTGAGGGGATTTTGAGGT | 253 |
| chr10:15269949-15271973 | p2 | (TA)6 | AGTCATGGGCATGTGTTTGA | CTCTTCTCCGTTGCCTTTTG | 172 |
| chr10:15269949-15271973 | p2 | (TA)12 | AGTCATGGGCATGTGTTTGA | CTCTTCTCCGTTGCCTTTTG | 172 |
| chr10:15295028-15297067 | p3 | (TTA)13 | TCAAGGGAAATCAAGCGAAC | GGAAGGAATCATGCTTTGGA | 178 |
| chr10:15342471-15344507 | p2 | (TA)18 | TTGAAGCCATTCCAAACACA | TCAGAGATGGCAATGACGAC | 235 |
| chr10:15438497-15440515 | p3 | (CAA)6 | CACATCACAACAAAGGCCAC | AATGGCACAGTTGATAGGGG | 123 |
| chr10:15545952-15547967 | p5 | (AAAGG)3 | GTCAGGGATTTGGGTCTTGA | TCCTTTCCTTTGAGCCTTGA | 138 |
| chr10:15545952-15547967 | p5 | (ATTGC)3 | GTCAGGGATTTGGGTCTTGA | TCCTTTCCTTTGAGCCTTGA | 138 |
| chr10:15573029-15575051 | p2 | (TA)11 | ATCGCACCTTGAAACGTACC | AGAGCCCAGCCATAGGAAAT | 293 |
| chr10:15573029-15575051 | p2 | (TA)8 | ATCGCACCTTGAAACGTACC | AGAGCCCAGCCATAGGAAAT | 293 |
| chr10:15584349-15586364 | p5 | (GGGGT)3 | TGTGACGGAATTGTCACCAC | GCAATTTTGTGACGGAATCA | 283 |
| chr10:15587278-15589294 | p4 | (TTCT)4 | CCATCTGCTCCTGCTTCATT | TTGAAATTCCAACTGTGCCA | 200 |
| chr10:15701756-15703771 | p5 | (CATTG)3 | AGTTATTGGTGCTGCGGTTC | ACAGAACACAACGGGGTAGC | 111 |
| chr10:15701756-15703771 | p2 | (TA)6 | AGTTATTGGTGCTGCGGTTC | ACAGAACACAACGGGGTAGC | 111 |

| Marker ID | SSR type | SSR motif | FORWARD PRIMER (5'-3') | REVERSE PRIMER (5'-3') | size (bp) |
|---|---|---|---|---|---|
| chr10:15750200-15752220 | p4 | (TTCA)5 | AGTATAAGCGCGCTCCAAAA | TATCCTTTCCCATTCGCTTG | 239 |
| chr10:15879211-15881249 | p2 | (AT)19 | AATCCACTAGGGGCTCGTTT | TAGTTAGGGCTCGTTTGGGA | 136 |
| chr10:15895913-15897929 | p4 | (TGTA)4 | TTTTGGCTTGAGCAGGAGTT | TGGGGTGGCTTTAATTTTTG | 268 |
| chr10:15921112-15923133 | p3 | (TTG)7 | GGCCAGTCCAAATCTCACAT | TCTGTCTCGCACCTGAGCTA | 299 |
| chr10:16057916-16059948 | p2 | (AT)14 | GGCTGAGGATTCGCTTACTG | TCCTAAGCCGTGTAGGCACT | 262 |
| chr10:16057916-16059948 | p2 | (AT)15 | GGCTGAGGATTCGCTTACTG | TCCTAAGCCGTGTAGGCACT | 262 |
| chr10:16057916-16059948 | p2 | (AT)15 | GGCTGAGGATTCGCTTACTG | TCCTAAGCCGTGTAGGCACT | 262 |
| chr10:16057916-16059948 | p2 | (AT)15 | GGCTGAGGATTCGCTTACTG | TCCTAAGCCGTGTAGGCACT | 262 |
| chr10:16057916-16059948 | p2 | (AT)15 | GGCTGAGGATTCGCTTACTG | TCCTAAGCCGTGTAGGCACT | 262 |
| chr10:16057916-16059948 | p2 | (AT)15 | GGCTGAGGATTCGCTTACTG | TCCTAAGCCGTGTAGGCACT | 262 |
| chr10:16057916-16059948 | p2 | (AT)16 | GGCTGAGGATTCGCTTACTG | TCCTAAGCCGTGTAGGCACT | 262 |
| chr10:16092533-16094548 | p5 | (CAAAA)3 | GTATTCGCCGACTGAAGCTC | ATGCGCAACACAAAATCAAA | 298 |
| chr10:16143884-16145928 | p2 | (GT)6 | CCGTGTGAGAAGCATGAGAA | AAAAGCAAAAGTTGAGGGGG | 145 |
| chr10:16143884-16145928 | p2 | (AT)22 | CCGTGTGAGAAGCATGAGAA | AAAAGCAAAAGTTGAGGGGG | 145 |
| chr10:16150711-16152723 | p2 | (TA)7 | CAGACCACGCATTGAAGCTA | TACTAGATGGCCTATGCCCG | 167 |
| chr10:16150711-16152723 | p2 | (TA)8 | CAGACCACGCATTGAAGCTA | TACTAGATGGCCTATGCCCG | 167 |
| chr10:16150728-16152756 | p2 | (TA)7 | CAGACCACGCATTGAAGCTA | TACTAGATGGCCTATGCCCG | 150 |
| chr10:16150728-16152756 | p2 | (TA)8 | CAGACCACGCATTGAAGCTA | TACTAGATGGCCTATGCCCG | 150 |
| chr10:16150766-16152790 | p2 | (TA)7 | CAGACCACGCATTGAAGCTA | TACTAGATGGCCTATGCCCG | 112 |
| chr10:16150766-16152790 | p2 | (TA)8 | CAGACCACGCATTGAAGCTA | TACTAGATGGCCTATGCCCG | 112 |
| chr10:16151679-16153695 | p2 | (TA)8 | CGGACGCATATAGCCTGAAT | TATGGGGAGCCACACTTTTT | 170 |
| chr10:16199167-16201185 | p6 | (TTTCAG)3 | ATGGTGGATGGGTGAAGAAA | ATTCCAGCCTTTGAGCGTAA | 277 |
| chr10:16240034-16242052 | p3 | (GTT)6 | CATCGTATTGGGGTAGGGTG | GTCGGTCTGACCACCACTTT | 179 |
| chr10:16240034-16242052 | p2 | (AT)14 | CATCGTATTGGGGTAGGGTG | GTCGGTCTGACCACCACTTT | 179 |
| chr10:16240391-16242419 | p3 | (GTT)6 | GGCAAGAAGTATTGGGGTGA | CAAATAGCGAAAGCGACACA | 224 |
| chr10:16240391-16242419 | p2 | (AT)14 | GGCAAGAAGTATTGGGGTGA | CAAATAGCGAAAGCGACACA | 224 |
| chr10:16240625-16242639 | p3 | (GTT)6 | TGTGTCGCTTTCGCTATTTG | CAGACCTTACGCCTACCTCG | 224 |
| chr10:16240625-16242639 | p2 | (AT)14 | TGTGTCGCTTTCGCTATTTG | CAGACCTTACGCCTACCTCG | 224 |
| chr10:16240703-16242715 | p3 | (GTT)6 | TGTGTCGCTTTCGCTATTTG | CAGACCTTACGCCTACCTCG | 146 |
| chr10:16240703-16242715 | p2 | (AT)14 | TGTGTCGCTTTCGCTATTTG | CAGACCTTACGCCTACCTCG | 146 |
| chr10:16292657-16294671 | p2 | (TA)7 | GAGGACGACCTCCTACCCTC | TTTTGAGGCTTGGGAAAGAA | 127 |
| chr10:16403920-16405932 | p5 | (TTTAT)3 | CCTACAATGCATGTACCCCC | TTCCCAGCTTTACGGAATTG | 194 |
| chr10:16403920-16405932 | p2 | (TA)6 | CCTACAATGCATGTACCCCC | TTCCCAGCTTTACGGAATTG | 194 |
| chr10:16420281-16422296 | p5 | (TTTGG)3 | CCTACGTGGCCAACTTCAAT | ATGCAGCTTGAAATCCTTGG | 236 |
| chr10:16460320-16462340 | p2 | (AT)10 | GGGGCAAAAATGAAAGACAA | TGCTTTGTTGGCTGTGACTC | 134 |
| chr10:16581095-16583107 | p5 | (CCGGT)3 | CTTGGCATGGTGGAAGAAAT | CTCTTCCTCTTCGCCTTCCT | 235 |

| Marker ID | SSR type | SSR motif | FORWARD PRIMER (5'-3') | REVERSE PRIMER (5'-3') | size (bp) |
|---|---|---|---|---|---|
| chr10:16600623-16602639 | p2 | (TA)8 | AGGCTTGGCCAAACAGACTA | TATTGGATCCATAGGCTGGC | 180 |
| chr10:16620700-16622718 | p6 | (TCGGCT)3 | AAAGGGAATGGGATTATGGC | CCTTTTGCGCTTTATCCTTG | 145 |
| chr10:16646895-16648915 | p2 | (AT)6 | TAGCATACATGTCAAGCGCC | ACCGGATGATGTCCAAGTTC | 176 |
| chr10:16733023-16735038 | p5 | (ACCGG)3 | CATTGACCGGTATTCTGCCT | GCAGGATGCCCACTAACCTA | 200 |
| chr10:16803732-16805772 | p2 | (AT)10 | TTGCTCCACGCCCTACTACT | CCGAAGCACGGTTGATATTT | 190 |
| chr10:16803780-16805792 | p2 | (AT)10 | TTGCTCCACGCCCTACTACT | CCGAAGCACGGTTGATATTT | 142 |
| chr10:16810304-16812352 | p3 | (TTG)5 | TCAATTCCGCACTGCTACTG | GTTCCGGCGAGTTTATTTCA | 224 |
| chr10:16810408-16812456 | p3 | (TTG)5 | TCAATTCCGCACTGCTACTG | GTTCCGGCGAGTTTATTTCA | 120 |
| chr10:16810720-16812768 | p3 | (TTG)5 | CCAATGCTACGAAGAGGCAT | TTTCCCTTTGCACTTTCTGG | 232 |
| chr10:16814822-16816837 | p5 | (AAAAT)3 | TGTCTTTTTGGTGTGGCAAA | AATGCAGTTTTGGAGTTGGG | 248 |
| chr10:16814822-16816837 | p5 | (AAAAT)3 | TGTCTTTTTGGTGTGGCAAA | AATGCAGTTTTGGAGTTGGG | 248 |
| chr10:16861802-16863868 | p2 | (AT)33 | GTGGTGATGTCATCGTCCTG | TTCGAGATCGAGTTAGCCGT | 121 |
| chr10:16905219-16907237 | p6 | (TTTTTA)3 | TGATTTGTTTCGGAAGAGGG | GCAATGTGGGTTTAGGGATG | 105 |
| chr10:17104756-17106790 | p2 | (AT)7 | CGTAACCTTAACCGCAAGGA | GCTGGAATTGGCAGCTTTAG | 102 |
| chr10:17104756-17106790 | p2 | (AG)17 | CGTAACCTTAACCGCAAGGA | GCTGGAATTGGCAGCTTTAG | 102 |
| chr10:17122214-17124236 | p2 | (AT)11 | ATCATCATCTTCTGGGTCGG | GTCTTCTCAACAAGCCGAGG | 132 |
| chr10:17160267-17162279 | p2 | (AT)6 | TTTGGGGTCCATTTTGGTTA | ACACATCGTTCACCAGGGAT | 106 |
| chr10:17175504-17177546 | p2 | (TA)21 | GAAGAGAAAGCGACGCAAGT | CATGTGGCAATGAGTCCATC | 118 |
| chr10:17175504-17177546 | p2 | (AT)13 | GAAGAGAAAGCGACGCAAGT | CATGTGGCAATGAGTCCATC | 118 |
| chr10:17344403-17346419 | p4 | (AATT)4 | TCAAGCCCCGTAGGTATGTC | TGACCCAGCTCCTTGTATCC | 252 |
| chr10:17344403-17346419 | p2 | (TC)7 | TCAAGCCCCGTAGGTATGTC | TGACCCAGCTCCTTGTATCC | 252 |
| chr10:17352891-17354906 | p5 | (TTCGG)3 | ATCCACCATTGTTGAACCGT | GCTGAATCTGCACCAATCAA | 166 |
| chr10:17360366-17362381 | p3 | (GAA)5 | AAGGGGCCCAAAAGGTAGTA | CCATTAGGCCAATAAGGCTG | 220 |
| chr10:17365285-17367306 | p3 | (TGA)7 | GTAGTCGTTGGGGCAACCTA | GGGGGATGGGTTAAAGTGTT | 286 |
| chr10:17397329-17399351 | p2 | (TA)8 | TTGTTATTGGGAGTGGGCTC | GGTTAATGCCCAATCGAAAA | 124 |
| chr10:17397329-17399351 | p2 | (AT)11 | TTGTTATTGGGAGTGGGCTC | GGTTAATGCCCAATCGAAAA | 124 |
| chr10:17446582-17448597 | p5 | (GGTCG)3 | AACCAAATGCGGAAACAAAG | TTTTCAACCGCAAAATCTCC | 228 |
| chr10:17446582-17448597 | p3 | (AAT)10 | AACCAAATGCGGAAACAAAG | TTTTCAACCGCAAAATCTCC | 228 |
| chr10:17532983-17535017 | p2 | (GA)17 | CCGACCCAGTTGGTTAAGAA | GCATGGTCGAAAATTCGTTT | 200 |
| chr10:17599430-17601445 | p5 | (GGTTT)3 | CCCAAGAAACTCCACGTGTT | TTGCATACATGGAGGCAGAA | 108 |
| chr10:17612247-17614261 | p2 | (GT)7 | ATTCAGCAATTTTGCCCTTG | TCCCGAACGCTCAAAAATAC | 227 |
| chr10:17656172-17658187 | p3 | (GTT)5 | CCACCAAAACTTAAGGGCAA | CTAATCCTGTCACGCCTGGT | 248 |
| chr10:17661863-17663875 | p2 | (AC)6 | GCCAAATGGAAAGGAATGAA | CTTTCTTCTTGATCCGACCG | 162 |
| chr10:17681626-17683641 | p2 | (AT)11 | TAGCGCCTAACACCCTTCAT | AGGATGGCCCTAAAAATGGT | 196 |
| chr10:17681652-17683666 | p2 | (AT)11 | TGTAGCGCCTAACACCCTTC | AGGATGGCCCTAAAAATGGT | 168 |
| chr10:17681673-17683691 | p2 | (AT)11 | TGTAGCGCCTAACACCCTTC | AGGATGGCCCTAAAAATGGT | 147 |

| Marker ID | SSR type | SSR motif | FORWARD PRIMER (5'-3') | REVERSE PRIMER (5'-3') | size (bp) |
|---|---|---|---|---|---|
| chr10:17721839-17723851 | p2 | (TA)6 | TGATGTCACGGCTTCTTACG | GTCACAGGTTACGCCCAGAT | 155 |
| chr10:17728500-17730532 | p2 | (TC)16 | TGTATGCGAATGTGCAAGGT | AAATAAACGCCTCGGTTGTG | 105 |
| chr10:17787706-17789721 | p3 | (TGT)5 | CGAGGATCTCTCGGAAACAG | GGTAACCGAGCCGATAAACA | 164 |
| chr10:17787706-17789721 | p5 | (AACCG)3 | CGAGGATCTCTCGGAAACAG | GGTAACCGAGCCGATAAACA | 164 |
| chr10:17825753-17827768 | p5 | (TTTAA)3 | GATTCATTGCTTGGTGCCTT | TATAAGCCAATCCAAACGGG | 209 |
| chr10:18070142-18072244 | p5 | (TTTTA)3 | TTGGATGGGATTTGGATTGT | AGATTTTGTTTAAGCGGCCA | 151 |
| chr10:18070142-18072244 | p3 | (TTA)34 | TTGGATGGGATTTGGATTGT | AGATTTTGTTTAAGCGGCCA | 151 |
| chr10:18083530-18085554 | p2 | (TC)12 | GGTTCCATCACGGCTTAGAA | GATGAATACGATCCCTGCGT | 279 |
| chr10:18190784-18192822 | p2 | (AT)14 | TCGCCATGCTCTAGAAAGGT | GCGGTCCTCCATTGAAGTAA | 153 |
| chr10:18190784-18192822 | p2 | (AT)19 | TCGCCATGCTCTAGAAAGGT | GCGGTCCTCCATTGAAGTAA | 153 |
| chr10:18378858-18380873 | p3 | (ACC)5 | AGGGTTATCGCACTCCATTG | AGCACTCCGGTAGCTCAAAA | 284 |
| chr10:18378858-18380873 | p3 | (AGC)5 | AGGGTTATCGCACTCCATTG | AGCACTCCGGTAGCTCAAAA | 284 |
| chr10:18378858-18380873 | p3 | (AAT)6 | AGGGTTATCGCACTCCATTG | AGCACTCCGGTAGCTCAAAA | 284 |
| chr10:18385309-18387367 | p2 | (TA)35 | TGAGACACCCTTTGGGAAAC | CGCCAATCAAAAGGACACTT | 257 |
| chr10:18385309-18387367 | p2 | (TA)29 | TGAGACACCCTTTGGGAAAC | CGCCAATCAAAAGGACACTT | 257 |
| chr10:18450288-18452300 | p2 | (TA)6 | TACTGCAAGCAGGAAAGCAA | AACAAGATTCCAAGGGGACC | 189 |
| chr10:18450288-18452300 | p5 | (AATAA)4 | TACTGCAAGCAGGAAAGCAA | AACAAGATTCCAAGGGGACC | 189 |
| chr10:18509588-18511608 | p5 | (TTTGG)3 | AAATCGAAACCAAACCGATG | AGGGGTTATGCTTAGGGGTG | 258 |
| chr10:18509588-18511608 | p4 | (ATTT)5 | AAATCGAAACCAAACCGATG | AGGGGTTATGCTTAGGGGTG | 258 |
| chr10:18509588-18511608 | p2 | (AT)13 | AAATCGAAACCAAACCGATG | AGGGGTTATGCTTAGGGGTG | 258 |
| chr10:18557872-18559896 | p4 | (GTTT)6 | CCATCCATTCCTTTTTCCCT | TTATTTTACGGGTTCGGCAG | 219 |
| chr10:18571969-18573983 | p5 | (AAAAT)3 | GGTGGTGTAGCCGTTTGACT | ACCATCAACATCATCAGCCA | 181 |
| chr10:18571990-18574046 | p5 | (AAAAT)3 | GGTGGTGTAGCCGTTTGACT | ACCATCAACATCATCAGCCA | 160 |
| chr10:18603760-18605778 | p6 | (TTTTTA)3 | CATGATGTTTCCACACGTCC | CAACCTTGGGGACTCAAAAA | 231 |
| chr10:18603760-18605778 | p5 | (AAATA)3 | CATGATGTTTCCACACGTCC | CAACCTTGGGGACTCAAAAA | 231 |
| chr10:18632927-18634979 | p2 | (AT)7 | ATTGGTCGGCTTAAACTCCA | GCAGAGCCACACCCAATAAT | 107 |
| chr10:18635305-18637321 | p4 | (ATTT)4 | GAAAACCAGAACCGAACCAA | GGGGTTATGCTTAGGGGTGT | 226 |
| chr10:18676778-18678794 | p2 | (TA)8 | AATGATTCTCACGAGGTGGC | AAATAGCGTGGGCAAATCAG | 202 |
| chr10:18676778-18678794 | p2 | (TC)6 | AATGATTCTCACGAGGTGGC | AAATAGCGTGGGCAAATCAG | 202 |
| chr10:18731202-18733217 | p3 | (GTT)5 | ACCGTGGAAGAGTTGTTTGG | AACACCCATTTACACGGCTC | 184 |
| chr10:18847106-18849121 | p3 | (CTT)5 | CCTATGGACCCCAATTACCC | GCCTGGCTGATGATCGTAAT | 251 |
| chr10:18980278-18982296 | p3 | (ATC)6 | AAACGCGAGCATAACTTTGG | TGGCGAACAATTGAACACAT | 117 |
| chr10:18998993-19001008 | p3 | (TTA)5 | GTCGAGCTTGAGGGTGAAAG | GCCAAAATATTGGAAGCGAA | 282 |
| chr10:18998993-19001008 | p5 | (ACCCG)3 | GTCGAGCTTGAGGGTGAAAG | GCCAAAATATTGGAAGCGAA | 282 |
| chr10:19009853-19011867 | p2 | (TA)7 | CCCCTGATCCCACACATAAC | GATGAGGGATAACTGCCCAA | 135 |
| chr10:19040199-19042256 | p3 | (AAT)19 | ACTTGTCAACATGGGCATCA | GCCATGTCCTGGCAGTTAAT | 137 |

| Marker ID | SSR type | SSR motif | FORWARD PRIMER (5'-3') | REVERSE PRIMER (5'-3') | size (bp) |
|---|---|---|---|---|---|
| chr10:19068017-19070029 | p2 | (GA)6 | TCCTATTGGTCGAAATTGGC | TGAAGAAGTAGAGGGGGCCT | 223 |
| chr10:19142275-19144289 | p2 | (TA)7 | AGTGGTGGTCTTCCCAGTTG | CCATTCCCTTCCCTAAGCTC | 132 |
| chr10:19171646-19173666 | p2 | (AT)32 | TCTCTTGACCGCTTGTCCTT | CTGCCCAAAATATAAGCCCA | 293 |
| chr10:19171646-19173666 | p4 | (TTAA)5 | TCTCTTGACCGCTTGTCCTT | CTGCCCAAAATATAAGCCCA | 293 |
| chr10:19220983-19222997 | p2 | (TA)7 | ATATAGGGGAAATGCTGCCC | ACTAATCGCAAAACCGTTCG | 255 |
| chr10:19220983-19222997 | p2 | (AT)11 | ATATAGGGGAAATGCTGCCC | ACTAATCGCAAAACCGTTCG | 255 |
| chr10:19408798-19410813 | p5 | (CCCCT)3 | ACTATCAGATCAGGCACCCG | TTGCCAGTAAGCATCCACAG | 119 |
| chr10:19417577-19419601 | p2 | (TA)12 | TTGGATTTTGGAGTTTTCGC | GAAAAACCAGCAAGCTACGG | 287 |
| chr10:19466324-19468336 | p2 | (AT)25 | CCCTAGGACGAACTGCTCTG | ATCATTTTGGCAGCCTTGTC | 189 |
| chr10:19466324-19468336 | p2 | (AT)6 | CCCTAGGACGAACTGCTCTG | ATCATTTTGGCAGCCTTGTC | 189 |
| chr10:19510346-19512361 | p5 | (TTTGC)3 | AACATTCTCATTTCCGCACC | GCTTCAGGAGTGCTGTGTGA | 147 |
| chr10:19538885-19540957 | p3 | (CAA)5 | CAATCAATTTTACAGGCGCA | CTCAGCCGAGTAGGGTCTTG | 134 |
| chr10:19647354-19649366 | p2 | (CA)6 | AAACAACGACGAGAATCGCT | CAAACTCGGGACCTAAGGGT | 144 |
| chr10:19674907-19676963 | p2 | (AT)9 | AACCGTGACTTCGGAGAGTG | AAGTTTTGGTTGCGTTGCTC | 238 |
| chr10:19685339-19687360 | p3 | (GTT)7 | CTCTTTGTAGGTCCCACCCA | AAGCGAGTTGACGAAATGCT | 164 |
| chr10:19685339-19687360 | p5 | (AAAAG)3 | CTCTTTGTAGGTCCCACCCA | AAGCGAGTTGACGAAATGCT | 164 |
| chr10:19712066-19714080 | p2 | (AT)7 | CCAATAGTGGGAGCAAATGG | GTGAGTTTGGAGATGGGTGG | 190 |
| chr10:19712066-19714080 | p2 | (TA)9 | CCAATAGTGGGAGCAAATGG | GTGAGTTTGGAGATGGGTGG | 190 |
| chr10:19733119-19735134 | p5 | (TGCAA)3 | CCGGTAGTTTCCTTGGATCA | GGGGCGGATTTCATTTACTT | 269 |
| chr10:19780254-19782272 | p5 | (CGAAC)3 | GTGAGGCCAATCCAAGTCAT | CAATGAAACCCCCACCTAGA | 241 |
| chr10:19780254-19782272 | p6 | (TTGGAA)3 | GTGAGGCCAATCCAAGTCAT | CAATGAAACCCCCACCTAGA | 241 |
| chr10:19783614-19785626 | p2 | (TA)6 | TGCCCTTTCAAATTTTCACC | TGTGAGCCATGATCGAGAAG | 242 |
| chr10:19788183-19790197 | p3 | (CTT)5 | GAGCTTCGGCCTGATACAAG | GAACAATCCCACTCCACGTT | 172 |
| chr10:19788183-19790197 | p2 | (AT)6 | GAGCTTCGGCCTGATACAAG | GAACAATCCCACTCCACGTT | 172 |
| chr10:19788183-19790197 | p2 | (TA)7 | GAGCTTCGGCCTGATACAAG | GAACAATCCCACTCCACGTT | 172 |
| chr10:19841856-19843871 | p3 | (TGT)5 | GGGTAGCTAGGGTTTGAGGG | ACAGGGTCGGTTTCCTTTTT | 198 |
| chr10:20000937-20002951 | p2 | (AC)7 | TGGTCATGCATAAAACGGAA | CTTCTGTGGGTTGCCTGAAT | 207 |
| chr10:20038465-20040522 | p3 | (TTC)19 | TGCACTTTCTGTTTGGATCG | GCGGAATGGACAGTGATTTT | 179 |
| chr10:20086533-20088597 | p2 | (AT)32 | ATCAGCCAGGGGGTACTCTT | TAAATCCCGAATCCGTCGTA | 104 |
| chr10:20169198-20171210 | p2 | (AG)6 | TTCCAAAATGCACGCAAATA | AATGAGACCCGCAGTACCAC | 123 |
| chr10:20213822-20215840 | p3 | (TAA)6 | TCAGGTTCCAGAGCTCGTTT | CTTTGTCACCCGCTCTTCTC | 238 |
| chr10:20257891-20259943 | p2 | (TA)26 | CCTTATTCGGGGTGGTCTTT | TTTTAATGCTTGGACCAGGG | 298 |
| chr10:20433781-20435869 | p2 | (AT)13 | GACAAGGTTGACTTCGGCTC | GACGTCATGAACAACATCGG | 177 |
| chr10:20433781-20435869 | p2 | (TA)44 | GACAAGGTTGACTTCGGCTC | GACGTCATGAACAACATCGG | 177 |
| chr10:20433781-20435869 | p2 | (TA)7 | GACAAGGTTGACTTCGGCTC | GACGTCATGAACAACATCGG | 177 |
| chr10:20433781-20435869 | p2 | (AT)17 | GACAAGGTTGACTTCGGCTC | GACGTCATGAACAACATCGG | 177 |

| Marker ID | SSR type | SSR motif | FORWARD PRIMER (5'-3') | REVERSE PRIMER (5'-3') | size (bp) |
|---|---|---|---|---|---|
| chr10:20444599-20446613 | p2 | (TA)7 | CTAAGGACCACCATGTGCCT | CTCCCTCACCTCTTTCCCTC | 283 |
| chr10:20479784-20481800 | p4 | (ATAA)4 | CCCACTTTATCAAGCCCAAA | CCTTTGATTTGACTGGGCAT | 275 |
| chr10:20506656-20508674 | p6 | (GGGGGT)3 | GCAGAAAGGGTCAAGAGCAC | TCCCATTGATTTGAGGGTGT | 300 |
| chr10:20533751-20535778 | p2 | (AT)28 | GGCAATGTGAAAAGCAGGTT | ATATATTTGCAAGCGGGCAC | 292 |
| chr10:20533751-20535778 | p2 | (AT)9 | GGCAATGTGAAAAGCAGGTT | ATATATTTGCAAGCGGGCAC | 292 |
| chr10:20533751-20535778 | p3 | (CTA)7 | GGCAATGTGAAAAGCAGGTT | ATATATTTGCAAGCGGGCAC | 292 |
| chr10:20533751-20535778 | p3 | (ATA)9 | GGCAATGTGAAAAGCAGGTT | ATATATTTGCAAGCGGGCAC | 292 |
| chr10:20568338-20570377 | p2 | (TA)22 | CAATGTGAAAAGCAGGTTGC | ATTAGCATGTGGGTTCTGGC | 275 |
| chr10:20568338-20570377 | p2 | (TA)22 | CAATGTGAAAAGCAGGTTGC | ATTAGCATGTGGGTTCTGGC | 275 |
| chr10:20568338-20570377 | p2 | (TC)17 | CAATGTGAAAAGCAGGTTGC | ATTAGCATGTGGGTTCTGGC | 275 |
| chr10:20568338-20570377 | p2 | (TA)10 | CAATGTGAAAAGCAGGTTGC | ATTAGCATGTGGGTTCTGGC | 275 |
| chr10:20568338-20570377 | p3 | (CTA)9 | CAATGTGAAAAGCAGGTTGC | ATTAGCATGTGGGTTCTGGC | 275 |
| chr10:20572014-20574046 | p2 | (TA)16 | TGCCCATTTTTCGTTAGTCC | CTATGTGCTCCCCTCCGATA | 153 |
| chr10:20583643-20585685 | p2 | (AT)21 | GGGAAATCTCCTATCGAGGC | CCACCGTGGAGGAACTAGAA | 216 |
| chr10:20607353-20609368 | p2 | (TA)18 | TAAGAACCTGTTGGATGGGC | TAGAGCCCGTTTGGATTGAC | 213 |
| chr10:20607353-20609368 | p3 | (CTA)5 | TAAGAACCTGTTGGATGGGC | TAGAGCCCGTTTGGATTGAC | 213 |
| chr10:20607577-20609592 | p2 | (TA)18 | GGCTCTAAGTCGCAATGGTC | ATTGCCAGTGGGACCATAAT | 241 |
| chr10:20607577-20609592 | p3 | (CTA)5 | GGCTCTAAGTCGCAATGGTC | ATTGCCAGTGGGACCATAAT | 241 |
| chr10:20607665-20609743 | p2 | (TA)18 | GGCTCTAAGTCGCAATGGTC | ATTGCCAGTGGGACCATAAT | 153 |
| chr10:20607665-20609743 | p3 | (CTA)5 | GGCTCTAAGTCGCAATGGTC | ATTGCCAGTGGGACCATAAT | 153 |
| chr10:20643124-20645160 | p2 | (TA)18 | TATTTTGGGGCTTGTTGAGG | TAGGCGATCGTGTTCCTACC | 100 |
| chr10:20643124-20645160 | p5 | (ATTTT)3 | TATTTTGGGGCTTGTTGAGG | TAGGCGATCGTGTTCCTACC | 100 |
| chr10:20645274-20647288 | p2 | (TA)6 | GCAACAAAGGTGGAATACCC | CATGCAACATGTGCAACAAA | 191 |
| chr10:20645274-20647288 | p2 | (TA)9 | GCAACAAAGGTGGAATACCC | CATGCAACATGTGCAACAAA | 191 |
| chr10:20681915-20683929 | p2 | (AT)7 | AAGAGGCCGATAACTGAGCA | CGAAACGTTCTCATAACGCA | 271 |
| chr10:20685706-20687778 | p2 | (TA)36 | CTGAAGGGAATTTTGGACGA | TCCAATTCAGCAACCAACAA | 156 |
| chr10:20779960-20781976 | p2 | (AT)8 | CAAGACGAAGCTTTCTTCCG | ACAGAAACTTTGGTCGGTGG | 105 |
| chr10:20786035-20788047 | p2 | (AT)6 | TTGCTCCACGCCCTACTACT | CTCCGAAGCCACGGTTAATA | 190 |
| chr10:20786035-20788047 | p2 | (TA)6 | TTGCTCCACGCCCTACTACT | CTCCGAAGCCACGGTTAATA | 190 |
| chr10:20786035-20788047 | p2 | (TA)7 | TTGCTCCACGCCCTACTACT | CTCCGAAGCCACGGTTAATA | 190 |
| chr10:20802524-20804536 | p2 | (AG)6 | ATCCCCCGATCTCTTTATGG | GTCCCTTTCTTCAAATGGCA | 113 |
| chr10:20819730-20821748 | p2 | (AT)6 | ATGGGTAAACATCCATCCCA | AGGTTCGAATGAACCCCTTT | 265 |
| chr10:20819730-20821748 | p3 | (ATA)6 | ATGGGTAAACATCCATCCCA | AGGTTCGAATGAACCCCTTT | 265 |
| chr10:20835804-20837822 | p6 | (CCAGCA)3 | CAGAATTCTCCGGGACAAAA | TAGGCGATACGAAGCCAACT | 106 |
| chr10:20856203-20858218 | p5 | (AATGC)3 | ATTGGTAGGCGAACCCTCTT | GACACAAGGTCACGGAAAT | 148 |
| chr10:20894094-20896109 | p5 | (AAAAT)3 | AATAGGTGCTGACGGGTGAC | ACATGACCGAAAAGGTCGAG | 186 |

| Marker ID | SSR type | SSR motif | FORWARD PRIMER (5'-3') | REVERSE PRIMER (5'-3') | size (bp) |
|---|---|---|---|---|---|
| chr10:21019302-21021317 | p5 | (AATGC)3 | AGTCTTCAAGGCAGCGAAAG | GTCTGCTTCCTTCTTCGTGG | 152 |
| chr10:21019302-21021317 | p5 | (AAAAC)3 | AGTCTTCAAGGCAGCGAAAG | GTCTGCTTCCTTCTTCGTGG | 152 |
| chr10:21064051-21066087 | p5 | (AAAAT)3 | CGGACAATAACTTGACTGCG | GCCTTAAGTCAGCCTTGCAC | 106 |
| chr10:21064051-21066087 | p2 | (TA)18 | CGGACAATAACTTGACTGCG | GCCTTAAGTCAGCCTTGCAC | 106 |
| chr10:21077885-21079900 | p5 | (GTCCC)3 | TCAGGGATTGGATCAACACA | ATTGACATTGCTGCTTGACG | 111 |
| chr10:21077885-21079900 | p3 | (TGG)5 | TCAGGGATTGGATCAACACA | ATTGACATTGCTGCTTGACG | 111 |
| chr10:21153717-21155729 | p2 | (TA)6 | CACTCCCTCCGTCCATTTTA | TACTCCCTCCGTCCATGTTT | 246 |
| chr10:21153717-21155729 | p2 | (AT)18 | CACTCCCTCCGTCCATTTTA | TACTCCCTCCGTCCATGTTT | 246 |
| chr10:21186355-21188379 | p6 | (ACCCCC)4 | TCGGCATCATTCAATTAGCA | AAAGCCCCCTTGGTAGAAAA | 154 |
| chr10:21212942-21214956 | p2 | (AT)7 | GTAGCCTGCATGCCAAAAAT | TGGATTGATGGGAGAAGAGG | 213 |
| chr10:21223846-21225861 | p5 | (TTCGG)3 | GGGACCCACCTCTACTAGCC | GGGTCATCAGTTCCAGAGGA | 123 |
| chr10:21247105-21249119 | p2 | (TA)7 | TTTTTCTTTGGCACGAATCC | CGGAGTGACATAAGGTCGGT | 276 |
| chr10:21250325-21252340 | p5 | (TTTTA)3 | CCCACTTGGATGCGTAGACT | TGTTCATTTCAGCGAAGACG | 116 |
| chr10:21294379-21296447 | p2 | (TA)34 | GAGCACTCGGGCTTTCATAG | TGTAGTTTGTCTGCAACGGC | 236 |
| chr10:21294379-21296447 | p2 | (TA)11 | GAGCACTCGGGCTTTCATAG | TGTAGTTTGTCTGCAACGGC | 236 |
| chr10:21398876-21400891 | p5 | (GAAAG)3 | TCGGTTCATCCCTATTCTGC | GCAAGCCACTAGGTTTCTGG | 100 |
| chr10:21415166-21417188 | p2 | (TC)11 | ATTGCATGAGACCTTCCTGG | AAGATCCATCATCCACCGAG | 202 |
| chr10:21433716-21435731 | p5 | (TTGGT)3 | CCTCGAGGTATTTCAGCCAC | GGATTTCCCTATAAACGCCC | 295 |
| chr10:21433716-21435731 | p5 | (ATTTT)3 | CCTCGAGGTATTTCAGCCAC | GGATTTCCCTATAAACGCCC | 295 |
| chr10:21450452-21452470 | p6 | (AGAAAG)3 | TTCGTGGTGCCATCTGATAA | AACAAGGAAAGCGCAAAAGA | 101 |
| chr10:21606523-21608545 | p2 | (TA)11 | GTGCAGTGGAGCTTTCTTCC | TCCCGAGTACATCCCGATAG | 107 |
| chr10:21660551-21662574 | p5 | (AAAAT)5 | TCACTTTGCCGCTATCATTG | CCTGCATGTTACTGAAGCGA | 298 |
| chr10:21711936-21713952 | p4 | (AAAT)4 | TATCCGCACCCGAATATGAT | GGGCTCTAGTGGCAGCTATG | 127 |
| chr10:21869000-21871020 | p2 | (AT)10 | CGCACATGTGGGTAGTATGG | CAAGCACCACTCATTCCCTT | 110 |
| chr10:21890969-21892984 | p5 | (GCATT)3 | TATTTGTGCTTGCAGTTCGG | CACCCGAAGGTTTACATGCT | 290 |
| chr10:21890969-21892984 | p2 | (TA)6 | TATTTGTGCTTGCAGTTCGG | CACCCGAAGGTTTACATGCT | 290 |
| chr10:21890969-21892984 | p2 | (TA)79 | TATTTGTGCTTGCAGTTCGG | CACCCGAAGGTTTACATGCT | 290 |
| chr10:21899641-21901656 | p5 | (AATAT)3 | TACACTTCGATTTGAGGGGG | ATCTCTTGAGCCCGATAGCA | 167 |
| chr10:22066741-22068755 | p3 | (TTA)5 | GCACTGTTTGGGCTGGTATT | ATGCCAAAACTGAAAACGGA | 116 |
| chr10:22066741-22068755 | p2 | (TA)7 | GCACTGTTTGGGCTGGTATT | ATGCCAAAACTGAAAACGGA | 116 |
| chr10:22066741-22068755 | p2 | (TA)14 | GCACTGTTTGGGCTGGTATT | ATGCCAAAACTGAAAACGGA | 116 |
| chr10:22069463-22071477 | p2 | (TA)6 | TTTGCTCCCACTAATGGCTT | AAACCACCACGTCCAAACTC | 101 |
| chr10:22069463-22071477 | p5 | (ATTTT)3 | TTTGCTCCCACTAATGGCTT | AAACCACCACGTCCAAACTC | 101 |
| chr10:22154836-22156851 | p5 | (AAATC)3 | AGCTTTAGACAAGCGGGACA | TTTTGAGCGCGTCTGTATTG | 245 |
| chr10:22238861-22240879 | p2 | (TC)8 | AACTCCCAAAGCACACAACC | GAGCGATCAAACCATCCATT | 106 |
| chr10:22238861-22240879 | p6 | (TATATG)3 | AACTCCCAAAGCACACAACC | GAGCGATCAAACCATCCATT | 106 |

| Marker ID | SSR type | SSR motif | FORWARD PRIMER (5'-3') | REVERSE PRIMER (5'-3') | size (bp) |
|---|---|---|---|---|---|
| chr10:22313703-22315770 | p3 | (AAC)5 | TCCGCTTTACTGGTTATCCG | TGGAGAATTAGTTCACCCGC | 145 |
| chr10:22313703-22315770 | p5 | (AATAA)3 | TCCGCTTTACTGGTTATCCG | TGGAGAATTAGTTCACCCGC | 145 |
| chr10:22313703-22315770 | p5 | (AAATA)3 | TCCGCTTTACTGGTTATCCG | TGGAGAATTAGTTCACCCGC | 145 |
| chr10:22317467-22319487 | p5 | (ACAAT)4 | CCGTGTAATTCACTGGGGTC | AATAACCGCAGGCTAGGGAT | 193 |
| chr10:22317467-22319487 | p2 | (AT)8 | CCGTGTAATTCACTGGGGTC | AATAACCGCAGGCTAGGGAT | 193 |
| chr10:22361238-22363254 | p4 | (TATG)4 | AGGTTGTGGGTTCACAGGAG | CTTCACGAGCTTTGGGATTC | 153 |
| chr10:22404411-22406439 | p2 | (AT)14 | GAAAAGGGCCAAATACCCAT | AGGGAGCAAAGCTTCCATTT | 221 |
| chr10:22539506-22541695 | p3 | (TTC)63 | TCCTTTCAACACCTTTTGGG | GCAGAAGAGAACGAAGCCAC | 188 |
| chr10:22670736-22672748 | p2 | (TA)6 | TTAACCCCAAACCCAATCAA | TCCAGTGAAAGATGCCACAA | 275 |
| chr10:22670736-22672748 | p6 | (CCGGTT)3 | TTAACCCCAAACCCAATCAA | TCCAGTGAAAGATGCCACAA | 275 |
| chr10:22849091-22851106 | p5 | (AGAAA)3 | ACCTCCACACCCTCCTTTCT | TCATTCCCGATTGTTTTTCC | 213 |
| chr10:22849091-22851106 | p5 | (ATTTT)3 | ACCTCCACACCCTCCTTTCT | TCATTCCCGATTGTTTTTCC | 213 |
| chr10:23001439-23003453 | p2 | (TA)7 | GGATGCTCATACCCACCTGT | GAGTGGCATTTCACCCCTAA | 109 |
| chr10:23001439-23003453 | p2 | (AT)7 | GGATGCTCATACCCACCTGT | GAGTGGCATTTCACCCCTAA | 109 |
| chr10:23001439-23003453 | p2 | (TA)10 | GGATGCTCATACCCACCTGT | GAGTGGCATTTCACCCCTAA | 109 |
| chr10:23151924-23153939 | p5 | (ATTTT)3 | TTTCGTGACTTGGCTCATTG | CCCACTTAGACTTTGCTCGC | 243 |
| chr10:23165777-23167792 | p5 | (GTTAT)3 | GGTTTGGAGAATTGGGGATT | ATTCTTTCCCCCAAAACTCG | 133 |
| chr10:23179978-23181996 | p5 | (ACCGG)3 | TTTTTGAGTGAGATGCCGTG | TATCCCCGAGGCCTTAAAGT | 228 |
| chr10:23179978-23181996 | p2 | (TA)9 | TTTTTGAGTGAGATGCCGTG | TATCCCCGAGGCCTTAAAGT | 228 |
| chr10:23179978-23181996 | p2 | (AT)7 | TTTTTGAGTGAGATGCCGTG | TATCCCCGAGGCCTTAAAGT | 228 |
| chr10:23397733-23399753 | p5 | (TTAAT)4 | ACCAGAGTGACCAAAATGGC | TCCCGGAGACTCTAGCTCAA | 113 |
| chr10:23474655-23476670 | p5 | (ATCAT)3 | GAAGCTCTACGGAATGGCAG | CCCCCTTATTATCCCCGTTA | 130 |
| chr10:23518891-23520911 | p2 | (AG)10 | CATCGATTAGCTAGCCTCGG | CGATGTCACCTCGGGATACT | 286 |
| chr10:23558928-23560943 | p5 | (ATCAT)3 | GAAGCTCTACGGAATGGCAG | CCCCCTTATTATCCCCGTTA | 130 |
| chr10:23603164-23605184 | p2 | (AG)10 | CATCGATTAGCTAGCCTCGG | CGATGTCACCTCGGGATACT | 286 |
| chr10:23643130-23645142 | p2 | (AT)6 | AGAGCTCAAGGATTTTGCGA | AGGCTGTTCCTTGCCCTTAT | 144 |
| chr10:23643130-23645142 | p2 | (TA)6 | AGAGCTCAAGGATTTTGCGA | AGGCTGTTCCTTGCCCTTAT | 144 |
| chr10:23692766-23694781 | p3 | (GAT)6 | GCCCTATGGGCTATTGAACA | TGGTCATGAATTTCGGTCAA | 240 |
| chr10:23692766-23694781 | p5 | (CGACA)3 | GCCCTATGGGCTATTGAACA | TGGTCATGAATTTCGGTCAA | 240 |
| chr10:23692766-23694781 | p2 | (AT)6 | GCCCTATGGGCTATTGAACA | TGGTCATGAATTTCGGTCAA | 240 |
| chr10:23692766-23694781 | p3 | (ACA)5 | GCCCTATGGGCTATTGAACA | TGGTCATGAATTTCGGTCAA | 240 |
| chr10:23693628-23695650 | p2 | (AT)6 | AATGCGGAGCAGAATTTGAC | GATATCCGCATTTAAGCCCA | 273 |
| chr10:23693628-23695650 | p3 | (ACA)5 | AATGCGGAGCAGAATTTGAC | GATATCCGCATTTAAGCCCA | 273 |
| chr10:23693628-23695650 | p2 | (AT)16 | AATGCGGAGCAGAATTTGAC | GATATCCGCATTTAAGCCCA | 273 |
| chr10:23693666-23695678 | p3 | (ACA)5 | AATGCGGAGCAGAATTTGAC | GATATCCGCATTTAAGCCCA | 235 |
| chr10:23693666-23695678 | p2 | (AT)16 | AATGCGGAGCAGAATTTGAC | GATATCCGCATTTAAGCCCA | 235 |

| Marker ID | SSR type | SSR motif | FORWARD PRIMER (5'-3') | REVERSE PRIMER (5'-3') | size (bp) |
|---|---|---|---|---|---|
| chr10:23722915-23724939 | p2 | (TA)12 | ACCATCCTAGGTGTGCCAAG | GGCTCTCATCACGACTCCTC | 180 |
| chr10:23722915-23724939 | p6 | (CAATAC)4 | ACCATCCTAGGTGTGCCAAG | GGCTCTCATCACGACTCCTC | 180 |
| chr10:23738528-23740543 | p5 | (TTTTA)3 | AGATCTGAGTCGGCGTCTTC | CGCGCGTTCAATAAACTGTA | 235 |
| chr10:23831662-23833674 | p2 | (AG)6 | GCAAGTTGCCTTACCTCCTG | TGGGGATTGGTCATTGACTT | 160 |
| chr10:23831662-23833674 | p2 | (AG)6 | GCAAGTTGCCTTACCTCCTG | TGGGGATTGGTCATTGACTT | 160 |
| chr10:23899336-23901350 | p2 | (GA)7 | AGAAGAAGCCCTGATTCCGT | CGTCGTCATACACGAAGTGG | 181 |
| chr10:23971294-23973309 | p5 | (AAAAT)3 | CTTTCCACAATGGCTTGCTT | ACTTGAACCCCAGGACCTCT | 197 |
| chr10:24023310-24025325 | p5 | (TTTTC)3 | GCGGGTCAAACAGAGAAATC | CTCCAGCTTTGGTGGAAAAA | 165 |
| chr10:24023310-24025325 | p3 | (ATA)5 | GCGGGTCAAACAGAGAAATC | CTCCAGCTTTGGTGGAAAAA | 165 |
| chr10:24048075-24050090 | p2 | (AT)6 | CCAAAAGGGTGGACCTCATA | TTTCCCGTGAGTTTGGATGT | 158 |
| chr10:24048075-24050090 | p5 | (AAAGA)3 | CCAAAAGGGTGGACCTCATA | TTTCCCGTGAGTTTGGATGT | 158 |
| chr10:24135501-24137515 | p2 | (GT)7 | GATGCGGAAATATCGAGGAA | GAGCTTGGGCCCTTTTTACT | 282 |
| chr10:24202573-24204588 | p5 | (TTTGC)3 | TGGAAATGGTGGATTTGGTT | AATCTCAGCATGGAACTGGG | 115 |
| chr10:24222196-24224210 | p2 | (AT)7 | TGGGGTTGTTGGAGCTAAAG | GCTAGACACAGCCCAGAAGG | 281 |
| chr10:24234122-24236154 | p2 | (TA)6 | TGTTTTAAGGGCTGTTTGGG | ATCAACACCCACACGTTCAA | 276 |
| chr10:24234122-24236154 | p2 | (TA)16 | TGTTTTAAGGGCTGTTTGGG | ATCAACACCCACACGTTCAA | 276 |
| chr10:24234122-24236154 | p2 | (AT)10 | TGTTTTAAGGGCTGTTTGGG | ATCAACACCCACACGTTCAA | 276 |
| chr10:24275786-24277828 | p3 | (TAC)14 | TTGATCCCATATCTGCCCAT | TAAAGCACTTGTTTGGCGTG | 215 |
| chr10:24331648-24333666 | p6 | (CATATA)3 | CTCTGGTGGTCATTGGGACT | GTACCCCATGGCCTTACCTT | 111 |
| chr10:24331648-24333666 | p6 | (CATATA)3 | CTCTGGTGGTCATTGGGACT | GTACCCCATGGCCTTACCTT | 111 |
| chr10:24358827-24360845 | p6 | (ATACAT)3 | TAGGAAGAAGGGCAAGCTCA | CTGTCATGGCCATTTGATTG | 287 |
| chr10:24358827-24360845 | p2 | (CA)10 | TAGGAAGAAGGGCAAGCTCA | CTGTCATGGCCATTTGATTG | 287 |
| chr10:24358969-24360989 | p6 | (ATACAT)3 | TAGGAAGAAGGGCAAGCTCA | CTGTCATGGCCATTTGATTG | 145 |
| chr10:24358969-24360989 | p2 | (CA)10 | TAGGAAGAAGGGCAAGCTCA | CTGTCATGGCCATTTGATTG | 145 |
| chr10:24361952-24363970 | p6 | (ATACAT)3 | TAGGAAGAAGGGCAAGCTCA | CTGTCATGGCCATTTGATTG | 287 |
| chr10:24361952-24363970 | p2 | (CA)10 | TAGGAAGAAGGGCAAGCTCA | CTGTCATGGCCATTTGATTG | 287 |
| chr10:24362094-24364114 | p6 | (ATACAT)3 | TAGGAAGAAGGGCAAGCTCA | CTGTCATGGCCATTTGATTG | 145 |
| chr10:24362094-24364114 | p2 | (CA)10 | TAGGAAGAAGGGCAAGCTCA | CTGTCATGGCCATTTGATTG | 145 |
| chr10:24370281-24372299 | p3 | (AAC)6 | GGCTACCTGTAGGCATGAGC | AACACCGAGCCTTGTTATGG | 195 |
| chr10:24431156-24433168 | p2 | (TA)6 | TTCCAGATTCTAGGCATGGG | TTTATCGCCAAAGGTTCGAC | 245 |
| chr10:24492905-24494973 | p2 | (AT)34 | TTGTATCCGAGGGGTCTCAG | GACTTGCCTCCACGAAGAAG | 214 |
| chr10:24822372-24824412 | p2 | (TA)20 | GTTCTTCCCCATGCCTCATA | GGACTACGGACGGATTCTCA | 285 |
| chr10:24850038-24852053 | p5 | (GAATT)3 | ATGACACGAGAACCCGTAGC | AAGCTGCAGAATCATGCAAA | 250 |
| chr10:24887072-24889094 | p2 | (TA)11 | TTGATTCCGATTCGGTCTTC | CTCCATATTCCCTGTGCGAT | 246 |
| chr10:24915645-24917657 | p2 | (TA)8 | GCCGAAAAGGACAAACACAT | GAGCTCCATTTTCATTCGGA | 282 |
| chr10:24915645-24917657 | p2 | (TA)6 | GCCGAAAAGGACAAACACAT | GAGCTCCATTTTCATTCGGA | 282 |

| Marker ID | SSR type | SSR motif | FORWARD PRIMER (5'-3') | REVERSE PRIMER (5'-3') | size (bp) |
|---|---|---|---|---|---|
| chr10:25019777-25021792 | p5 | (AATGC)3 | ATCGTACCGATCCACGAGAC | CGCTTATCGAGGTGAAAAGG | 263 |
| chr10:25069134-25071149 | p5 | (CATTG)3 | CAAGTCGCAGGTTGCACTAA | TAACATGTGCTGGAAGCGAC | 173 |
| chr10:25121541-25123556 | p5 | (AATGC)3 | AGGAACCGAATAAATCGCCT | TCGAATTCGTTTATGCCTCC | 114 |
| chr10:25127948-25129966 | p6 | (TTCAAA)3 | AAACCGGGACCAGAAATACC | TAGTGGTGATGTGGAAGGCA | 206 |
| chr10:25129980-25131998 | p6 | (TTCAAA)3 | TCCCTAGAGAAAACCGAGCA | TTGTTGGTGATGTGGAAGGA | 237 |
| chr10:25194429-25196447 | p6 | (AAAATG)3 | ATTCGGCTGACTCCAATGAC | CCGATTACGAGAAGGGACAA | 293 |
| chr10:25194429-25196447 | p3 | (ACC)5 | ATTCGGCTGACTCCAATGAC | CCGATTACGAGAAGGGACAA | 293 |
| chr10:25194587-25196602 | p6 | (AAAATG)3 | ATTCGGCTGACTCCAATGAC | CCGATTACGAGAAGGGACAA | 135 |
| chr10:25194587-25196602 | p3 | (ACC)5 | ATTCGGCTGACTCCAATGAC | CCGATTACGAGAAGGGACAA | 135 |
| chr10:25214919-25216937 | p6 | (TTCAAA)3 | TCGACGGAGGGAATAAACAC | GATATGAAAGCCACGAGGGA | 286 |
| chr10:25219493-25221508 | p5 | (GCATT)3 | AGCCATCCAGGTAGGTCCTT | ACCTCCATATCGCAACCAAG | 284 |
| chr10:25222073-25224091 | p3 | (CAC)6 | TCTGGACACGTTCTTTGCAC | CCGTCATTGCGAAGGTTATT | 115 |
| chr10:25222073-25224091 | p6 | (TTTCAA)3 | TCTGGACACGTTCTTTGCAC | CCGTCATTGCGAAGGTTATT | 115 |
| chr10:25312120-25314135 | p5 | (TTTTA)3 | TGGCATTGCTGCTAGAGTTG | GCCTATCCAAGGTCGCAATA | 189 |
| chr10:25373707-25375725 | p3 | (ATT)6 | ATTCCAGGCATGTTGAGACC | TGGCACCAGTACCCAATACA | 156 |
| chr10:25385958-25387970 | p2 | (TA)6 | GGCTGTCACTTGGTGTGAGA | CACTAGAGAACCCCAGCAGC | 279 |
| chr10:25477423-25479455 | p2 | (TA)33 | GACAAGTTTGAGCTGGAGGC | TGTCTCCTTAATGCCGGTTC | 114 |
| chr10:25552869-25554881 | p2 | (TA)6 | TTGCTCCACGCCCTACTACT | ACCATGCTAAGACGTCCACC | 133 |
| chr10:25552869-25554881 | p2 | (AT)6 | TTGCTCCACGCCCTACTACT | ACCATGCTAAGACGTCCACC | 133 |
| chr10:25553010-25555022 | p2 | (TA)6 | AAGTATCAACCGTGGCTTCG | TCCGCTTCTGAGTTTTGGTT | 229 |
| chr10:25553010-25555022 | p2 | (AT)6 | AAGTATCAACCGTGGCTTCG | TCCGCTTCTGAGTTTTGGTT | 229 |
| chr10:25673866-25675896 | p2 | (TA)15 | GACCTGTCAGGATTGCGTTT | CATTGGATCACGGGAACTCT | 226 |
| chr10:25692942-25694954 | p2 | (TG)6 | GTCCTAAATCGGGCAACGTA | AACCAGATTGCTTGGGATTG | 167 |
| chr10:25801263-25803291 | p2 | (AT)14 | TTTCGAATTGACCTTGGACC | CATGGTTTCAAAGGGGACAC | 180 |
| chr10:25857268-25859298 | p3 | (ACA)6 | TTGATACCCGCATTAGAGCC | GTTGTGGTGCTCGGTAGGTT | 282 |
| chr10:25857268-25859298 | p2 | (AT)15 | TTGATACCCGCATTAGAGCC | GTTGTGGTGCTCGGTAGGTT | 282 |
| chr10:25924417-25926429 | p2 | (AT)6 | ACCCTTCATAGGCGAAACCT | TTTGTATGGGCGCAGATGTA | 226 |
| chr10:25929917-25931975 | p2 | (TA)29 | GAACCCATGGAATGCAAAAG | GATCTTCCTCACGCCATGTT | 154 |
| chr10:25929917-25931975 | p2 | (TA)6 | GAACCCATGGAATGCAAAAG | GATCTTCCTCACGCCATGTT | 154 |
| chr10:26009052-26011067 | p5 | (TTGAG)3 | CTATGGCTGTGACCGACCTT | GCTTCAACTTGCATCCAACA | 153 |
| chr10:26037354-26039370 | p4 | (ACAT)4 | TTGGCTTCGATTCCAATTTC | TGATGTCATGCTGGGTTGTT | 101 |
| chr10:26037354-26039370 | p2 | (CT)7 | TTGGCTTCGATTCCAATTTC | TGATGTCATGCTGGGTTGTT | 101 |
| chr10:26116588-26118610 | p6 | (AATAGT)3 | TTGCTCCACGCCCTACTACT | CGAAGCCACGGTTGATACTT | 116 |
| chr10:26129148-26131168 | p2 | (TC)10 | TACAACCTTCCCAGTTTGCC | TGTTCATATTTCGCCAGCAA | 241 |
| chr10:26142835-26144853 | p2 | (TA)8 | AACCTACTGGCATGAAACGC | ACTTATGCATATCGTGGCCC | 277 |
| chr10:26149743-26151761 | p6 | (TTGAAG)3 | ATTGTTGCTTCTTTGCGCTT | TCCCAAAGAGCAGTAGCGTT | 271 |

| Marker ID | SSR type | SSR motif | FORWARD PRIMER (5'-3') | REVERSE PRIMER (5'-3') | size (bp) |
|---|---|---|---|---|---|
| chr10:26194229-26196244 | p5 | (TAAAT)3 | GGGGTATTTAGGGGCTCAAA | ATATGTTTGCACCTTTGCCC | 274 |
| chr10:26194229-26196244 | p2 | (TA)6 | GGGGTATTTAGGGGCTCAAA | ATATGTTTGCACCTTTGCCC | 274 |
| chr10:26198976-26200991 | p3 | (TGA)5 | GCCTCATTTTTCACCCAAAA | CCTAATCCCCACAACTCCAA | 126 |
| chr10:26294182-26296208 | p2 | (AT)13 | GGCTTCATCAGCCGATATGT | CCAATCGTGAATGCACTGTC | 294 |
| chr10:26346383-26348418 | p5 | (TTTTA)3 | CGACCCCTTACCTCTCTTCC | GCGCATAAGTTGGCTAAAGG | 279 |
| chr10:26346383-26348418 | p2 | (AT)15 | CGACCCCTTACCTCTCTTCC | GCGCATAAGTTGGCTAAAGG | 279 |
| chr10:26346453-26348483 | p5 | (TTTTA)3 | CGACCCCTTACCTCTCTTCC | GCGCATAAGTTGGCTAAAGG | 209 |
| chr10:26346453-26348483 | p2 | (AT)15 | CGACCCCTTACCTCTCTTCC | GCGCATAAGTTGGCTAAAGG | 209 |
| chr10:26390797-26392819 | p2 | (AT)11 | GCGCTTTGCTATGGGTTTAG | AGCAAGGGATACCAAAGGCT | 177 |
| chr10:26393627-26395645 | p6 | (CTTTTT)3 | GGAGGTTAAAACGTTTGCCA | GGAGAATTCGACATGGGTGT | 161 |
| chr10:26393627-26395645 | p6 | (AAAAAT)3 | GGAGGTTAAAACGTTTGCCA | GGAGAATTCGACATGGGTGT | 161 |
| chr10:26432245-26434299 | p2 | (TA)8 | GCATGCAACTTTTGATGGTG | ACAATGCATTCCCTAACCCA | 269 |
| chr10:26432384-26434398 | p2 | (TA)8 | GCATGCAACTTTTGATGGTG | CACAATGCATTCCCTAACCC | 130 |
| chr10:26464455-26466473 | p6 | (ACGGAG)3 | TTTTCTTTTGACGTTGCGTG | CAAGGAAAGGGAATCGATGA | 173 |
| chr10:26495056-26497071 | p5 | (ATTGC)3 | CGAAAGGCAAGGCAATAGAG | ACCCAAAGGTTCAAATGCTG | 189 |
| chr10:26578175-26580189 | p2 | (TA)7 | TTGTCGACCCCTTTGATTTC | ACAGATTACTCCAGCCCCCT | 130 |
| chr10:26578175-26580189 | p2 | (AT)7 | TTGTCGACCCCTTTGATTTC | ACAGATTACTCCAGCCCCCT | 130 |
| chr10:26600586-26602601 | p5 | (AATGC)3 | CACCCACCTCACCAGAAACT | TAGCTAAATTGGGCCGTGAC | 121 |
| chr10:26615774-26617806 | p2 | (TA)6 | TTGGCACGAATCCAACATAA | ATCTGGGCCTTTCAGAGTGA | 116 |
| chr10:26615774-26617806 | p2 | (AT)16 | TTGGCACGAATCCAACATAA | ATCTGGGCCTTTCAGAGTGA | 116 |
| chr10:26624940-26626958 | p2 | (TA)9 | TATACGTGCAGTGCGACCTC | CCAACACAAGGGCAAAGAGT | 110 |
| chr10:26660106-26662122 | p2 | (AT)8 | TTCTCCAACCCCTGTTATCG | GAACCTCAGAATAAGGCCCC | 298 |
| chr10:26674924-26676936 | p2 | (TA)6 | GGAACCCGACACAACTTGAT | AACCGCACTTGTTTTTGTCC | 212 |
| chr10:26744964-26746979 | p5 | (TTGGT)3 | ATTCAAGGAAAGCATGCACC | GTAATGTCGGGTTGGTTTGG | 133 |
| chr10:26752784-26754808 | p2 | (AT)14 | TCAACAGGATAGGAATGGGG | TAGCGACGACTGTGCTTTTG | 170 |
| chr10:26752784-26754808 | p2 | (AT)12 | TCAACAGGATAGGAATGGGG | TAGCGACGACTGTGCTTTTG | 170 |
| chr10:26752784-26754808 | p2 | (AT)7 | TCAACAGGATAGGAATGGGG | TAGCGACGACTGTGCTTTTG | 170 |
| chr10:26779464-26781479 | p2 | (AT)16 | CCATGTTGAAATCGTGCATC | AACAATCAGGTCCCTCATGC | 251 |
| chr10:26779556-26781625 | p2 | (AT)16 | CCATGTTGAAATCGTGCATC | AACAATCAGGTCCCTCATGC | 159 |
| chr10:27027779-27029791 | p3 | (CAA)5 | TCAACTACCACCACCAACCA | GGCGACCGAAAGGTATGTTA | 267 |
| chr10:27027779-27029791 | p2 | (CT)6 | TCAACTACCACCACCAACCA | GGCGACCGAAAGGTATGTTA | 267 |
| chr10:27329449-27331464 | p3 | (AAC)5 | ACCGTACCTGGCCCTCTAGT | CACTACCCTGCTCGTGCATA | 126 |
| chr10:27329449-27331464 | p2 | (TA)7 | ACCGTACCTGGCCCTCTAGT | CACTACCCTGCTCGTGCATA | 126 |
| chr10:27345814-27347829 | p5 | (AAAGG)3 | GGTTGGGGATTTGAGGTCTT | TACTTGGAAGGCCGAACAAC | 178 |
| chr10:27345814-27347829 | p5 | (CATTG)3 | GGTTGGGGATTTGAGGTCTT | TACTTGGAAGGCCGAACAAC | 178 |
| chr10:27345814-27347829 | p4 | (TTTA)4 | GGTTGGGGATTTGAGGTCTT | TACTTGGAAGGCCGAACAAC | 178 |

| Marker ID | SSR type | SSR motif | FORWARD PRIMER (5'-3') | REVERSE PRIMER (5'-3') | size (bp) |
|---|---|---|---|---|---|
| chr10:27567340-27569356 | p2 | (TA)14 | GTGGGGAGTTTGGTCTTGAA | CCAAAACAGCTCCCAAACAT | 212 |
| chr10:27567416-27569442 | p2 | (TA)14 | GTGGGGAGTTTGGTCTTGAA | CCAAAACAGCTCCCAAACAT | 136 |
| chr10:27567697-27569725 | p2 | (TA)14 | GTTTGGGAGCTGTTTTGGAA | GGAATCATGCCTCAGGAAAA | 116 |
| chr10:27707976-27710006 | p2 | (AT)15 | CAGAAAGCTACGAACCTCCG | TCATTTGCGTACTGTCCTCG | 229 |
| chr10:27791156-27793168 | p2 | (TG)6 | CGGAGGACGAGATGAAGAAG | CAAGCATTTCGGGACAGTTT | 178 |
| chr10:27931143-27933155 | p2 | (TA)6 | TCCTTGAACACCACGACATC | TCCTTGGAAGAAAAACACGC | 176 |
| chr10:27931143-27933155 | p2 | (TA)6 | TCCTTGAACACCACGACATC | TCCTTGGAAGAAAAACACGC | 176 |
| chr10:28061012-28063046 | p2 | (TA)22 | GTCAATCGTACCGGGAAAAA | GGAAGGTCAACCGCATTTTA | 205 |
| chr10:28095599-28097613 | p2 | (GA)6 | CTAGAAATGGAGACGAGGCG | TCAGCCAACCCAAAAGAAAC | 101 |
| chr10:28095599-28097613 | p3 | (TTA)5 | CTAGAAATGGAGACGAGGCG | TCAGCCAACCCAAAAGAAAC | 101 |
| chr10:28095599-28097613 | p2 | (TA)7 | CTAGAAATGGAGACGAGGCG | TCAGCCAACCCAAAAGAAAC | 101 |
| chr10:28157621-28159636 | p2 | (AT)16 | TGCCTTCTAATTGCATCACG | TCCAATGGCTCTTTGCCTAT | 111 |
| chr10:28157621-28159636 | p3 | (ACA)5 | TGCCTTCTAATTGCATCACG | TCCAATGGCTCTTTGCCTAT | 111 |
| chr10:28248029-28250047 | p6 | (CCATAA)3 | CGCATCTGGGGTAAACATCT | TCGGGAGTTCACGACCTTAC | 208 |
| chr10:28386560-28388576 | p2 | (TA)8 | CCGAACTGGACCACTCGTAT | GATTTGGGTTTTGCTTTGGA | 267 |
| chr10:28409493-28411505 | p2 | (AG)6 | TGGAAATTTCTCCATCAGCC | TGGGCTGCTCGTTCTAAGTT | 281 |
| chr10:28432464-28434479 | p5 | (AAATC)3 | ATGCCATCCTCACCAGATTC | AGTGGCCAAAAATTGACAGG | 147 |
| chr10:28432464-28434479 | p2 | (AT)6 | ATGCCATCCTCACCAGATTC | AGTGGCCAAAAATTGACAGG | 147 |
| chr10:28433791-28435809 | p3 | (TTG)6 | GAAAAACCCGACAAAAACGA | GGTACATACGGTTCGGTTGG | 287 |
| chr10:28488448-28490460 | p2 | (CT)6 | ATACACCCGGGATTTCTTCC | GGGAAATTCATGTGGACCAG | 123 |
| chr10:28488448-28490460 | p2 | (CT)7 | ATACACCCGGGATTTCTTCC | GGGAAATTCATGTGGACCAG | 123 |
| chr10:28512762-28514774 | p2 | (CT)6 | ATACACCCGGGATTTCTTCC | GGGAAATTCATGTGGACCAG | 123 |
| chr10:28512762-28514774 | p2 | (CT)7 | ATACACCCGGGATTTCTTCC | GGGAAATTCATGTGGACCAG | 123 |
| chr10:28534596-28536608 | p2 | (AT)6 | TTAGACTCCCGCGTCTGAAT | GCAAAGGAACCAAAACATGG | 139 |
| chr10:28534596-28536608 | p2 | (TA)8 | TTAGACTCCCGCGTCTGAAT | GCAAAGGAACCAAAACATGG | 139 |
| chr10:28534596-28536608 | p2 | (AT)6 | TTAGACTCCCGCGTCTGAAT | GCAAAGGAACCAAAACATGG | 139 |
| chr10:28546323-28548335 | p2 | (TA)6 | CCATGATAGTTGATTCCCCG | CGACCATAGAGGTTCGGTGT | 252 |
| chr10:28576884-28578899 | p3 | (TTG)5 | CCTTAAGCCAAATATGGGCA | TATTGTGTGAGGACCCACGA | 170 |
| chr10:28746803-28748827 | p6 | (TGAGCA)4 | CAGCAGCATCTGGTCCATTA | GCCTATACCAGGCAGACCAA | 147 |
| chr10:28859161-28861209 | p3 | (ATT)16 | GCCAAAGGCATCATTCTCAT | CCCTTGACCTTTGTCAGGAA | 211 |
| chr10:28890410-28892422 | p5 | (ATCCA)5 | GGCGGAGTTCAGAGATGAAG | TGACCCAGTCCAAAAAGGAC | 237 |
| chr10:28982406-28984421 | p5 | (AAGCC)3 | GGGCACATTTTGTCACTCCT | CTGTCTTTGCTTTTGCCTCC | 122 |
| chr10:28998010-29000032 | p2 | (AT)11 | AAATCGAACTCAGCCACACC | CAAGGGAGGACTTGTGGAAA | 276 |
| chr10:29129979-29131993 | p6 | (GTGGGG)3 | GGGTGGGGGTTGTAAGAAAT | ACCCTCGAATGCACTTCATC | 293 |
| chr10:29129979-29131993 | p5 | (GGGTG)3 | GGGTGGGGGTTGTAAGAAAT | ACCCTCGAATGCACTTCATC | 293 |
| chr10:29129979-29131993 | p2 | (TA)7 | GGGTGGGGGTTGTAAGAAAT | ACCCTCGAATGCACTTCATC | 293 |

| Marker ID | SSR type | SSR motif | FORWARD PRIMER (5'-3') | REVERSE PRIMER (5'-3') | size (bp) |
|---|---|---|---|---|---|
| chr10:29153183-29155201 | p6 | (CATATA)3 | CCTGAAAGGGAAACAGGACA | AGGACGAGGCCATAGATGTG | 198 |
| chr10:29171273-29173288 | p5 | (ATAAA)3 | TATCTGTGATCCGTTGCCAG | TGGGCTAACTGGTACTTGGC | 160 |
| chr10:29179594-29181612 | p6 | (TTAAAA)3 | ACCGTATGAAAGCGGCATAG | GGCCATTCACGTCTGAGATT | 147 |
| chr10:29207858-29209876 | p6 | (TAAAGC)3 | CGCAATTTCTTCTCCTCCAG | CCAGCCATGGTTAAAGGAAA | 292 |
| chr10:29254774-29256788 | p2 | (TA)7 | ATGGACGGGCTTGTTTGTAG | TCGCCTCTTAAGGTGCTAGG | 173 |
| chr10:29282883-29284897 | p2 | (TA)7 | ATGGACGGGCTTGTTTGTAG | TCGCCTCTTAAGGTGCTAGG | 173 |
| chr10:29344331-29346347 | p2 | (AT)19 | TATGCCTTGGTCATCGTTCA | AATGGCGGCAAAAATATGTC | 181 |
| chr10:29344331-29346347 | p2 | (AT)6 | TATGCCTTGGTCATCGTTCA | AATGGCGGCAAAAATATGTC | 181 |
| chr10:29344367-29346391 | p2 | (AT)19 | TATGCCTTGGTCATCGTTCA | AATGGCGGCAAAAATATGTC | 145 |
| chr10:29344367-29346391 | p2 | (AT)6 | TATGCCTTGGTCATCGTTCA | AATGGCGGCAAAAATATGTC | 145 |
| chr10:29373434-29375446 | p2 | (TA)6 | CCGCGAGAGGTAAACACAAT | ACGTGATTCAAACGCAACAA | 137 |
| chr10:29383509-29385530 | p3 | (CAA)7 | TACGCACCTTGCACACTCTC | CACGCCAGATGATAATGGTG | 156 |
| chr10:29487606-29489634 | p2 | (AT)14 | CGTAACATCAAGCGAGGGTT | TTTGACTCGCCTCCACTTCT | 276 |
| chr10:29533536-29535554 | p2 | (TA)9 | CCAGGTTCAGGATCACGAGT | GGTCAACAAAACATGCCCTC | 247 |
| chr10:29533680-29535695 | p2 | (TA)9 | CCAGGTTCAGGATCACGAGT | GGTCAACAAAACATGCCCTC | 103 |
| chr10:29533680-29535695 | p2 | (AT)15 | CCAGGTTCAGGATCACGAGT | GGTCAACAAAACATGCCCTC | 103 |
| chr10:29611814-29613846 | p2 | (AT)16 | ATTTCATTTTGGTTGCTGGG | GTGGGATTTTCATGGCAGTT | 110 |
| chr10:29837261-29839276 | p5 | (AATGC)3 | CCCAAAAATTCTCCGACGTA | TGGGTTGAGGGTTTGCTTAC | 147 |
| chr10:29903215-29905231 | p2 | (TA)8 | GCAAACTTGCCAAAACCCTA | CGTCAAATCTGACCCTCCAT | 245 |
| chr10:29939713-29941791 | p3 | (TAT)26 | CAAGTTCCAAGGGGTTTTGA | TTCTGTGCTCCCTCTGTCCT | 175 |
| chr10:29939713-29941791 | p2 | (TC)6 | CAAGTTCCAAGGGGTTTTGA | TTCTGTGCTCCCTCTGTCCT | 175 |
| chr10:29976203-29978281 | p3 | (TAT)26 | CAAGTTCCAAGGGGTTTTGA | TTCTGTGCTCCCTCTGTCCT | 175 |
| chr10:29976203-29978281 | p2 | (TC)6 | CAAGTTCCAAGGGGTTTTGA | TTCTGTGCTCCCTCTGTCCT | 175 |
| chr10:30013416-30015431 | p3 | (AAT)5 | TGAACCAATGGCTAAGGACC | GTGCCAAAAATGGCAAAGTT | 189 |
| chr10:30069468-30071482 | p2 | (AG)7 | AGAATGGACTTGGGAGCCTT | GGGGTCACAACAAGGAGAAA | 296 |
| chr10:30104965-30106980 | p5 | (TTGGT)3 | ATTCGTGCAAAGTAGGGGTG | GTAATGTCGGGTTGGTTTGG | 155 |
| chr10:30104965-30106980 | p5 | (TTGGT)3 | ATTCGTGCAAAGTAGGGGTG | GTAATGTCGGGTTGGTTTGG | 155 |
| chr10:30112132-30114165 | p2 | (AC)10 | GGCATGTCCCATCTTCTCTC | AAAAATTGACAGGACCGCAC | 270 |
| chr10:30143078-30145106 | p2 | (TA)8 | TCCAAACGAACCCAACAAAT | CTTCGGAGCAACTTTAACGC | 169 |
| chr10:30143078-30145106 | p2 | (TA)14 | TCCAAACGAACCCAACAAAT | CTTCGGAGCAACTTTAACGC | 169 |
| chr10:30285976-30288004 | p4 | (TTAT)7 | TGAGAGTGACGTGAGGATCG | CTCGGCAATTGCTACCTCAT | 284 |
| chr10:30297514-30299529 | p5 | (ACCAA)3 | GATATGGTTCGGCATCTGCT | TACTTGCTGAGCCCAAGGTT | 192 |
| chr10:30297514-30299529 | p5 | (ACCAA)3 | GATATGGTTCGGCATCTGCT | TACTTGCTGAGCCCAAGGTT | 192 |
| chr10:30297514-30299529 | p5 | (ACCAA)3 | GATATGGTTCGGCATCTGCT | TACTTGCTGAGCCCAAGGTT | 192 |
| chr10:30297514-30299529 | p5 | (ACCAA)3 | GATATGGTTCGGCATCTGCT | TACTTGCTGAGCCCAAGGTT | 192 |
| chr10:30323763-30325777 | p5 | (ATTTT)3 | CATGCAAATCCGCCTCTATT | AAGAAGTTGTTGCTGGTGGG | 260 |

| Marker ID | SSR type | SSR motif | FORWARD PRIMER (5'-3') | REVERSE PRIMER (5'-3') | size (bp) |
|---|---|---|---|---|---|
| chr10:30323763-30325777 | p2 | (TC)7 | CATGCAAATCCGCCTCTATT | AAGAAGTTGTTGCTGGTGGG | 260 |
| chr10:30325158-30327188 | p2 | (AT)15 | AGGGCAAATGGTCAAAAATG | TTGGATCGGGTTTAAATGGA | 191 |
| chr10:30400176-30402190 | p6 | (CAACTC)6 | TGGAGAATGTGGAGCAATGA | CAGCCGACTTTTGTACCCAT | 177 |
| chr10:30400176-30402190 | p2 | (AT)7 | TGGAGAATGTGGAGCAATGA | CAGCCGACTTTTGTACCCAT | 177 |
| chr10:30455827-30457895 | p5 | (GCATT)3 | TACTGGTGAGACTCCCCCTG | TCCAATCATCAAGCATTCCA | 253 |
| chr10:30487866-30489884 | p6 | (CATATA)3 | ACCCTGTAGGTCGTGACTGG | CTGGTCGGCTTGCGTATTAT | 113 |
| chr10:30633997-30636019 | p2 | (TA)11 | AAATTACGCGTTTGGAGTGG | TTGGTTTTGGGAGGTGGTAG | 171 |
| chr10:30683717-30685753 | p2 | (TA)18 | AGCTATTCGGGCCAAGATTT | CCATTTTTACCCATGGTTGG | 121 |
| chr10:30683717-30685753 | p2 | (TA)20 | AGCTATTCGGGCCAAGATTT | CCATTTTTACCCATGGTTGG | 121 |
| chr10:30760195-30762215 | p2 | (AT)10 | TGGGCACACAAGACCAAGTA | AAACTTGCGAGTGCCATCTT | 244 |
| chr10:30772702-30774814 | p6 | (AAAACA)3 | AACCCATAGCACACTTTGCC | AGAGGAAGATTTTGGGGGTG | 260 |
| chr10:30772702-30774814 | p2 | (TA)56 | AACCCATAGCACACTTTGCC | AGAGGAAGATTTTGGGGGTG | 260 |
| chr10:30831793-30833808 | p5 | (ATTTC)3 | GGAAATATGAGTGCCTGGGA | ATCCCATCTTGCAACAAAGC | 229 |
| chr10:30876036-30878052 | p3 | (AGA)5 | TCTTGGTCCGGAGATATTGG | TGTATTGGGCAAAATGCAAA | 261 |
| chr10:30876049-30878064 | p3 | (AGA)5 | TCTTGGTCCGGAGATATTGG | TGTATTGGGCAAAATGCAAA | 248 |
| chr10:31048938-31050978 | p2 | (AT)20 | CTTCCCCTTCATGGTCTCAA | ATGTAGTGGCAGCCTTGTCC | 130 |
| chr10:31065159-31067171 | p2 | (TA)6 | GGATAACGTGCGAGGCTTAG | CCCTCACCACGTTCTCAAAT | 288 |
| chr10:31089293-31091311 | p2 | (AT)9 | ATTTGATCAGAGGTGTGCCC | GAAATCTTGGTTCGTTGGGA | 298 |
| chr10:31096113-31098125 | p2 | (TA)6 | ATAAAAATTGGGGCGGAAAC | CAAAAACCACACGCACACTC | 258 |
| chr10:31096113-31098125 | p2 | (TA)6 | ATAAAAATTGGGGCGGAAAC | CAAAAACCACACGCACACTC | 258 |
| chr10:31124923-31126935 | p2 | (TA)6 | AGCAGTTCGTCCTAGGGGTT | TCAGCAATTGCCTCGTAGTG | 208 |
| chr10:31231784-31233798 | p2 | (TG)7 | AAAGGTGCCTACGGGCTATT | CTCATACCTGGCCCTACCAA | 225 |
| chr10:31296101-31298116 | p5 | (ATTGC)3 | CCCCTGTAAGCAATTAGCCA | GCAATGCAATGCAATAATGG | 268 |
| chr10:31296101-31298116 | p5 | (TATAT)3 | CCCCTGTAAGCAATTAGCCA | GCAATGCAATGCAATAATGG | 268 |
| chr10:31357485-31359500 | p5 | (AAAAC)3 | TGGGCCTTAAGTGAGGAATG | AGTCCTCCACACCGATGAAG | 219 |
| chr10:31381200-31383218 | p3 | (CAA)6 | GCATGTGGACAAAACTGTGG | TGACGTCACAAATAAGCCCA | 192 |
| chr10:31386891-31388903 | p2 | (TG)6 | GGGAGGCAGAGACATACAGC | TTGTTGATTGCCCCTGTGTA | 192 |
| chr10:31393722-31395734 | p2 | (CA)6 | GGTCCGATATGGACTGCCTA | AGATGGATGGACAGGAGTGG | 156 |
| chr10:31426785-31428800 | p5 | (AAAAC)3 | TGGGCCTTAAGTGAGGAATG | AGTCCTCCACACCGATGAAG | 219 |
| chr10:31450500-31452518 | p3 | (CAA)6 | GCATGTGGACAAAACTGTGG | TGACGTCACAAATAAGCCCA | 192 |
| chr10:31456191-31458203 | p2 | (TG)6 | GGGAGGCAGAGACATACAGC | TTGTTGATTGCCCCTGTGTA | 192 |
| chr10:31463022-31465034 | p2 | (CA)6 | GGTCCGATATGGACTGCCTA | AGATGGATGGACAGGAGTGG | 156 |
| chr10:31704162-31706180 | p2 | (AT)9 | CTTGCATAGCAAGTGGCAAA | CGTGGGTACATTGTGGTGAG | 171 |
| chr10:31704395-31706441 | p2 | (AT)9 | CTCACCACAATGTACCCACG | AGTTCTGCGTTTTGGCCTTA | 110 |
| chr10:31859600-31861618 | p5 | (AGGAC)3 | ATTTGCTATTGAGCACCGCT | GTACATATTTTGGGGCACGG | 170 |
| chr10:31859600-31861618 | p6 | (CATATA)3 | ATTTGCTATTGAGCACCGCT | GTACATATTTTGGGGCACGG | 170 |

| Marker ID | SSR type | SSR motif | FORWARD PRIMER (5'-3') | REVERSE PRIMER (5'-3') | size (bp) |
|---|---|---|---|---|---|
| chr10:31870376-31872394 | p6 | (CAACTC)3 | CCGCATAAGAGGGGTCTGTA | TGTTTTCTCTTGGATTGGGG | 189 |
| chr10:31965704-31967716 | p2 | (AT)6 | GCGAGTCTTCCCTCACAAAG | CCACTCTTGTCCGTCCATTT | 162 |
| chr10:32041836-32043856 | p2 | (AT)10 | TACCAGGCCAAAACCAGTTC | CACATGGGACATTTTGCTTG | 152 |
| chr10:32041836-32043856 | p3 | (TCA)5 | TACCAGGCCAAAACCAGTTC | CACATGGGACATTTTGCTTG | 152 |
| chr10:32063327-32065339 | p2 | (TG)6 | GGGCTTAACGATGTCTTGGA | TGTTTTCGTCGACTCGTTTG | 126 |
| chr10:32107472-32109488 | p2 | (AC)8 | ATGCAAGAGTGGGTTTTTCG | TAAACTCCACCACTCCTGGC | 126 |
| chr10:32126619-32128634 | p2 | (AT)9 | TTGTGTGGATATGGTGTGGG | AGAGAAGGTCGATCCCTGGT | 192 |
| chr10:32126619-32128634 | p3 | (ACA)5 | TTGTGTGGATATGGTGTGGG | AGAGAAGGTCGATCCCTGGT | 192 |
| chr10:32261209-32263225 | p2 | (TA)8 | TGGGCTGTTTGAAATGTTGA | TTCCCAACGTTAGCAACTCC | 298 |
| chr10:32261209-32263225 | p2 | (AT)11 | TGGGCTGTTTGAAATGTTGA | TTCCCAACGTTAGCAACTCC | 298 |
| chr10:32261209-32263225 | p2 | (TA)6 | TGGGCTGTTTGAAATGTTGA | TTCCCAACGTTAGCAACTCC | 298 |
| chr10:32261873-32263885 | p2 | (TA)8 | ATCGTGCACAGTTTCACTGC | ATATACCGTAACCGGCCCTC | 278 |
| chr10:32261873-32263885 | p2 | (AT)11 | ATCGTGCACAGTTTCACTGC | ATATACCGTAACCGGCCCTC | 278 |
| chr10:32261873-32263885 | p2 | (TA)6 | ATCGTGCACAGTTTCACTGC | ATATACCGTAACCGGCCCTC | 278 |
| chr10:32303323-32305341 | p6 | (ACTGAA)3 | CGCGCGTACACACATAAAAG | TCATATACAGCCAGGCACCA | 294 |
| chr10:32353250-32355262 | p2 | (TA)6 | TTTTATTCACCGCAGTTCCC | AGAGGGGGTCAGTACGGAGT | 261 |
| chr10:32417792-32419819 | p3 | (AAT)9 | CAAGTGCTCAAATCGCATGT | TTGGGGCTTACACTTTTTGC | 298 |
| chr10:32637715-32639729 | p2 | (AT)7 | ATACAAGCCGACGAGGCTAA | GTCCTGTCCCGCCTATATGA | 187 |
| chr10:32874829-32876867 | p2 | (TA)19 | TTGGCGATGGAAAGGTAGTC | AAGGCAAGACCCTCTGGTTT | 222 |
| chr10:32923560-32925594 | p2 | (AT)17 | TTTCTCCACACCTCTCCCAC | TCCACCTCCACTTGGATTTC | 106 |
| chr10:32977177-32979192 | p3 | (TTC)5 | AAGGCTGGTATCATGTTGGC | GTAATTGCAGGACGGAGGAA | 173 |
| chr10:32981910-32983922 | p2 | (AT)6 | GTCGGGTTATTGCGTGTTTT | AGCCTTAGGGATCGGACAAT | 300 |
| chr10:32997592-32999604 | p2 | (TA)6 | CTTCGAGGTGAGGCAGTACC | TTTGCGGAACATTGGTCATA | 214 |
| chr10:33003057-33005075 | p3 | (TTA)6 | ACTGCCCAACCAACAGTAGC | CGTTCCACAAAACACAATGC | 180 |
| chr10:33156151-33158225 | p2 | (TA)6 | CTTAGGTTGGGCCATGTTGT | GAATCTCAATTCCAAGGGCA | 152 |
| chr10:33158687-33160875 | p5 | (ATTGC)3 | GAGGCCGATATCCGATGTTA | ACATCTCGGTAGCACAACCC | 240 |
| chr10:33158931-33160945 | p5 | (ATTGC)3 | TAGCATTGCATTGCATCCAT | AACTGATAGGCATCATCGGC | 230 |
| chr10:33159022-33161036 | p5 | (ATTGC)3 | TAGCATTGCATTGCATCCAT | AACTGATAGGCATCATCGGC | 139 |
| chr10:33176922-33178937 | p5 | (TTTTG)3 | AAAGTCATAATTCCGGTGCG | TCGGACCTCTTTTGGACAAC | 269 |
| chr10:33218579-33220675 | p2 | (TA)6 | AATGTACGGAGCGTGAAACC | AGAAGAGGATTGGGGGTTGT | 271 |
| chr10:33327472-33329488 | p4 | (CTTT)4 | TTAAGGCTGTTTCCCATTGC | AGGACTGGTTCTGACCATGC | 139 |
| chr10:33366910-33368925 | p5 | (TTGCA)3 | GATATTGTCATGGGTTGGGC | CATCACCAAGTGGCATCAAG | 116 |
| chr10:33413863-33415881 | p2 | (AT)9 | CACAATACCACCACGTCTCG | TCAAAGTCGGATGAAGACCC | 143 |
| chr10:33449473-33451497 | p4 | (GAAA)6 | GCTGTATCATCTCCCCCTCA | CTTGGGTGATGCCAAGCTAT | 101 |
| chr10:33481454-33483466 | p2 | (TG)6 | TCACCACCCCAGGCTATAAG | CCGTAAAATGGGTCTCCTGA | 239 |
| chr10:33509688-33511700 | p2 | (TA)6 | CATTTTCTCCTCCTCCTCCC | ACGTAGCCCCAATTCCTTCT | 180 |

| Marker ID | SSR type | SSR motif | FORWARD PRIMER (5'-3') | REVERSE PRIMER (5'-3') | size (bp) |
|---|---|---|---|---|---|
| chr10:33528800-33530814 | p2 | (AT)7 | TTTGGGATGACCCTTTCTTG | GATGGCCAAAAGAAACCAAA | 165 |
| chr10:33599641-33601653 | p2 | (AC)6 | CTTTCCCTGTTTGCAGTGGT | CAGATGCGACCATTGATTTG | 219 |
| chr10:33601271-33603286 | p5 | (CCCGC)3 | AGTGTTTGGCTGGTCAGCTT | GTTCCATCGCTGATTGCTCT | 114 |
| chr10:33621700-33623712 | p2 | (AT)6 | GAGGTCGTCTTGACCCGATA | GTCAACATCGGGCGTAAAAT | 135 |
| chr10:33626482-33628517 | p2 | (TA)7 | CGTACCTGCCCATTGCTATT | TCGCACACCTAGACAGAACG | 134 |
| chr10:33626482-33628517 | p2 | (TA)13 | CGTACCTGCCCATTGCTATT | TCGCACACCTAGACAGAACG | 134 |
| chr10:33673280-33675294 | p2 | (GT)7 | TCAATCGTACGGGTGTCAGA | TGATATGAGGAAGCCAACCC | 258 |
| chr10:33673280-33675294 | p2 | (TG)6 | TCAATCGTACGGGTGTCAGA | TGATATGAGGAAGCCAACCC | 258 |
| chr10:33673404-33675416 | p2 | (GT)7 | TCAATCGTACGGGTGTCAGA | TGATATGAGGAAGCCAACCC | 134 |
| chr10:33673404-33675416 | p2 | (TG)6 | TCAATCGTACGGGTGTCAGA | TGATATGAGGAAGCCAACCC | 134 |
| chr10:33678503-33680521 | p5 | (TTGTA)3 | AAATATCAACCGTGGCTTCG | GTCGCTCCGATCTAATCCAA | 206 |
| chr10:33678503-33680521 | p6 | (ATAAAA)3 | AAATATCAACCGTGGCTTCG | GTCGCTCCGATCTAATCCAA | 206 |
| chr10:33708364-33710388 | p5 | (TAAAA)3 | AAGGTAGGAACCAACATGCG | GAGAGCTTTGATCACGGAGG | 204 |
| chr10:33708364-33710388 | p6 | (TACATA)4 | AAGGTAGGAACCAACATGCG | GAGAGCTTTGATCACGGAGG | 204 |
| chr10:33728303-33730318 | p2 | (AT)10 | ATGGAGCAAATCAAACGGAG | AGTCGATACGTTTCCGGATG | 188 |
| chr10:33728303-33730318 | p3 | (CAA)5 | ATGGAGCAAATCAAACGGAG | AGTCGATACGTTTCCGGATG | 188 |
| chr10:33790208-33792224 | p4 | (AAAT)4 | CAGACGAACAAGTCGGTCAA | GGAGTTGGCTAGGCTCACTG | 208 |
| chr10:34023765-34025780 | p5 | (AATGT)3 | ACTTGTGTTGTTTGCCCCTC | CGAGGTGGGAAGGTAAGTGA | 276 |
| chr10:34136541-34138556 | p3 | (AAT)5 | GGGCAAGACATGAGGTTGAT | TCATCGATTCGCACTCTCAC | 203 |
| chr10:34148104-34150143 | p3 | (AAT)13 | TCTCTCCTATCTTCCCCCGT | CCTCCATTTCTAAGGGCCTC | 150 |
| chr10:34205736-34207805 | p3 | (TCT)5 | GGGGCATTGTGTATGCTTCT | TAGCAACCAAGGTCCAAACC | 105 |
| chr10:34209171-34211189 | p6 | (CATGAG)3 | TGGGAACCCCTTCATATTCA | TTGGGGACCCAAATGTTAAA | 127 |
| chr10:34265720-34267736 | p4 | (GAAA)4 | AACCCAAGTCAACGAAGGTG | ATGAAGCCTGCGGTTCTCTA | 244 |
| chr10:34296903-34298918 | p5 | (CCTTT)3 | AATCCCAAGTGTGCCAAGTC | CAATATTGCATTGCGTCCAC | 182 |
| chr10:34349540-34351552 | p6 | (TTTAAT)4 | GCAGAACAAACGCACGACTA | AATTTGCTCGGTAAGCGTTG | 297 |
| chr10:34349553-34351575 | p6 | (TTTAAT)4 | GCAGAACAAACGCACGACTA | AATTTGCTCGGTAAGCGTTG | 284 |
| chr10:34349941-34351965 | p6 | (TTTAAT)4 | TCGGGGGTGCTTGATAATAA | ATTTGATTAGGGGAGGGTGC | 271 |
| chr10:34453666-34455684 | p6 | (AAAAAT)3 | CCTACGTTCCTGTCCCTGAA | GAAGCTATGCATCGGAGGAG | 265 |
| chr10:34453666-34455684 | p2 | (TA)15 | CCTACGTTCCTGTCCCTGAA | GAAGCTATGCATCGGAGGAG | 265 |
| chr10:34454224-34456254 | p6 | (AAAAAT)3 | GACTTCCCCGTTAGAGTCCC | TTCCTGCCTGGTCGAGTAGT | 157 |
| chr10:34454224-34456254 | p2 | (TA)15 | GACTTCCCCGTTAGAGTCCC | TTCCTGCCTGGTCGAGTAGT | 157 |
| chr10:34683311-34685331 | p2 | (AT)6 | GCGTACTCGCTGTTGGTACA | TTCGGACATCGACCTTTTTC | 299 |
| chr10:34683311-34685331 | p5 | (ATTGC)4 | GCGTACTCGCTGTTGGTACA | TTCGGACATCGACCTTTTTC | 299 |
| chr10:34708587-34710637 | p2 | (TA)25 | CCTTACGTCACCCCGATAGA | ATGCGTATATAAGCTGCCCG | 299 |
| chr10:34728133-34730154 | p3 | (TTC)7 | CATGAAAGCCCAAGAGAAGC | AAAGCACAAGGAGGCACTGT | 130 |
| chr10:34735573-34737645 | p2 | (AT)36 | TCCTTTGCCATATTCGGAAC | CCAAAGGGTGACGTTTAGGA | 280 |

| Marker ID | SSR type | SSR motif | FORWARD PRIMER (5'-3') | REVERSE PRIMER (5'-3') | size (bp) |
|---|---|---|---|---|---|
| chr10:34811102-34813117 | p5 | (TGCAA)3 | GGCGAGCCATTCCTATATCA | TAGATCCGCAACCCAGAGAC | 280 |
| chr10:34836057-34838072 | p3 | (TAA)5 | TCCTTCCTATGGTTTGTCCG | GGTGGTAAGGCAATGCAGAT | 161 |
| chr10:34836057-34838072 | p2 | (AT)20 | TCCTTCCTATGGTTTGTCCG | GGTGGTAAGGCAATGCAGAT | 161 |
| chr10:34836057-34838072 | p6 | (AAAAAC)3 | TCCTTCCTATGGTTTGTCCG | GGTGGTAAGGCAATGCAGAT | 161 |
| chr10:35059640-35061652 | p2 | (GT)6 | AGGAGGGTGAAAAATTGCCT | GAATTTGCCCTTAGCCTTCC | 207 |
| chr10:35142812-35144826 | p2 | (TA)7 | GACAGCGAGTGGAGTGCATA | ATGAGGTTGCTCTGCTTCGT | 296 |
| chr10:35339063-35341078 | p5 | (GGTGC)3 | TTTTGAGAGGATTATGGCCG | CTACCTCGAACACTCGCCTC | 283 |
| chr10:35476798-35478824 | p2 | (TC)13 | CAATAAAGAGGACGTGGCGT | TGATAAGTAGGGGCGGACAC | 182 |
| chr10:35494157-35496191 | p2 | (AC)17 | CCGGGTAATCCGAAGGTATT | ATCCTGGTTTCTGGGTTTCC | 143 |
| chr10:35559486-35561504 | p2 | (TC)9 | GATGTGCTTTTCGCTGTTCA | AATTGATTTCCTCAATGGCG | 131 |
| chr10:35586991-35589011 | p2 | (TC)10 | TCGAAAGGGCTCTGTGATTT | TTTTAAACTGATCGGCCCAC | 213 |
| chr10:35695132-35697147 | p5 | (GGGTG)3 | TTACCCCTCGCCTAATACCC | ACGGGAAGAGTTCAAAGCAA | 263 |
| chr10:35695132-35697147 | p2 | (TA)8 | TTACCCCTCGCCTAATACCC | ACGGGAAGAGTTCAAAGCAA | 263 |
| chr10:35695304-35697319 | p5 | (GGGTG)3 | TTGCTTTGAACTCTTCCCGT | CAACCATCCAAACAGAGGGT | 262 |
| chr10:35695304-35697319 | p2 | (TA)8 | TTGCTTTGAACTCTTCCCGT | CAACCATCCAAACAGAGGGT | 262 |
| chr10:35782371-35784383 | p2 | (AT)9 | TAAAATGTATGGCCTCGCGT | CGGGCATAGGCCATCTACTA | 234 |
| chr10:35782371-35784383 | p2 | (AT)6 | TAAAATGTATGGCCTCGCGT | CGGGCATAGGCCATCTACTA | 234 |
| chr10:35857385-35859400 | p3 | (ATC)5 | ACTTTTGTCACGACCCAACC | ATCATTTTGGCAGCCTTGTC | 112 |
| chr10:35859872-35861890 | p3 | (CAT)6 | TTACCAGCAACGAGCTTCCT | GACGGAGGCAGTCTTTTGAG | 132 |
| chr10:35931751-35933817 | p2 | (TA)33 | TATGGGACTTTCGGATTTGC | ACGCAACTTTTTCTTGCGTT | 175 |
| chr10:36154265-36156281 | p2 | (TA)8 | ATGGTGAACCGAAGGTTGAG | TGTCAACAGGTGGTACCGAA | 105 |
| chr10:36154265-36156281 | p6 | (CCCCTT)3 | ATGGTGAACCGAAGGTTGAG | TGTCAACAGGTGGTACCGAA | 105 |
| chr10:36203180-36205204 | p2 | (AT)12 | ACCTTGCGGGTTATCATCAG | TTAACACAGTTGGTTGCCGA | 239 |
| chr10:36229678-36231690 | p2 | (AT)6 | CCCGGAAAGAACAAGAATGA | TGCCCATTCCATGTCTGTTA | 188 |
| chr10:36248565-36250589 | p2 | (AT)12 | ACCTTGCGGGTTATCATCAG | TTAACACAGTTGGTTGCCGA | 239 |
| chr10:36389258-36391273 | p5 | (ACACA)3 | TCGGGTCCTATTTATGCGAG | CGCATAAGAAGCGGAAAAAG | 161 |
| chr10:36389258-36391273 | p2 | (AT)7 | TCGGGTCCTATTTATGCGAG | CGCATAAGAAGCGGAAAAAG | 161 |
| chr10:36397060-36399074 | p2 | (AT)7 | CCGGAGCCTACATTTTGGTA | CAGAAGCCGACAAGGCTATC | 129 |
| chr10:36655825-36657843 | p2 | (AT)9 | TGGCATCGCTGAAATCATAA | GGTGAGTGGTGCTCGGTAAT | 180 |
| chr10:36655825-36657843 | p6 | (ATCATT)3 | TGGCATCGCTGAAATCATAA | GGTGAGTGGTGCTCGGTAAT | 180 |
| chr10:36730044-36732058 | p2 | (AT)7 | TATCGTGTTGGTCCGCATAA | TATCAGAGGAGCCTAGGGCA | 246 |
| chr10:36788623-36790635 | p2 | (CT)6 | AAGAGCTCACGGAGCCATAA | GGCGACCGAAAGGTATGTTA | 292 |
| chr10:36799447-36801497 | p2 | (AT)25 | TGCATTTTGCGATTCACATT | GCGAGCACAAGAAAAGAAGG | 114 |
| chr10:36877590-36879604 | p2 | (AC)7 | TAGCAACATGGGGCCTAAAG | CACGATCCTGGGAGAGACAT | 133 |
| chr10:36920068-36922082 | p2 | (TC)6 | CCCGCCAAAATGAGAAGATA | TTCGATCCAAACACTCCTCC | 154 |
| chr10:36920068-36922082 | p2 | (TA)7 | CCCGCCAAAATGAGAAGATA | TTCGATCCAAACACTCCTCC | 154 |

| Marker ID | SSR type | SSR motif | FORWARD PRIMER (5'-3') | REVERSE PRIMER (5'-3') | size (bp) |
|---|---|---|---|---|---|
| chr10:36920068-36922082 | p4 | (TTAA)4 | CCCGCCAAAATGAGAAGATA | TTCGATCCAAACACTCCTCC | 154 |
| chr10:36949954-36951979 | p5 | (TTTAT)5 | ATGTCGCTCACACACACACA | AAATTCCGGCAGAGTTTCCT | 289 |
| chr10:36992832-36994850 | p6 | (TTTCAG)3 | AAATGCAGCCCGTTTTACAC | ACAACTAGCACGCCAATCCT | 129 |
| chr10:37014238-37016254 | p2 | (AT)8 | GTAGCCGCTCAAAGAACTCG | AACATTGGCTCTTCATTGGC | 228 |
| chr10:37025648-37027666 | p6 | (AGCTGG)3 | ATCGAATCACTGGCCTGAAC | TGAACGTGAGGAGGATAGGG | 187 |
| chr10:37025648-37027666 | p2 | (CA)6 | ATCGAATCACTGGCCTGAAC | TGAACGTGAGGAGGATAGGG | 187 |
| chr10:37174549-37176569 | p2 | (TG)10 | CCCCTCATGTTCGTAGTGGT | TACACGGGGTAAGGCACTTC | 233 |
| chr10:37312056-37314068 | p3 | (ATC)5 | TCCCGCCCTACTATTGTGTC | CGGGCTCGTGTTTTATGATT | 250 |
| chr10:37312056-37314068 | p2 | (CA)6 | TCCCGCCCTACTATTGTGTC | CGGGCTCGTGTTTTATGATT | 250 |
| chr10:37383351-37385367 | p3 | (TTG)5 | TTTGAGCATGTTGTGGTCGT | TCCCACACGATGTTTTTCAA | 108 |
| chr10:37383351-37385367 | p4 | (TGTA)4 | TTTGAGCATGTTGTGGTCGT | TCCCACACGATGTTTTTCAA | 108 |
| chr10:37438125-37440139 | p2 | (TA)7 | GGCCCAGATCCTCAGTGTTA | CTTCATGTTGGGCCTCATTT | 295 |
| chr10:37507535-37509555 | p5 | (ATTGC)4 | CACTTTTCAGGGTGCGTTTT | CATGCTATCCCCGTCACTTT | 262 |
| chr10:37555047-37557062 | p5 | (TACAA)3 | CACTTTTGGGCACACTGAGA | GCAACATGAAAGTGGGGAGT | 264 |
| chr10:37556722-37558746 | p6 | (TTCATG)4 | CGTATAACTCCCGACGCAGT | TTGGAAGGGCTTGAAATGAC | 284 |
| chr10:37818225-37820240 | p3 | (TTG)5 | AGGGAATGACGTTTGACGAC | TTTACGGTCCGTTTTCCAAG | 159 |
| chr10:37895378-37897392 | p2 | (AT)7 | AATGTCCGCAGCTTCTTGTT | AGAAATGTTGCCGTCCAAAG | 117 |
| chr10:37987679-37989697 | p2 | (TA)9 | GTCCGGTCCTTCTGAATCAA | CCATGATCCTTAGGGGGAAT | 111 |
| chr10:38039648-38041664 | p2 | (AG)6 | TGTCCTATTAAAGGCGGTCG | TTCACCAACGCACAACATTT | 152 |
| chr10:38039648-38041664 | p2 | (AT)8 | TGTCCTATTAAAGGCGGTCG | TTCACCAACGCACAACATTT | 152 |
| chr10:38281318-38283336 | p2 | (TA)9 | GCCCAGCATCTCCAGTTTAG | CCCCCTCAGGAAAAGAACTC | 154 |
| chr10:38369819-38371835 | p2 | (AT)8 | CCTTACGTCACCCCGATAGA | TGCCAAAAGGTCATGGTGTA | 276 |
| chr10:38369819-38371835 | p2 | (AT)14 | CCTTACGTCACCCCGATAGA | TGCCAAAAGGTCATGGTGTA | 276 |
| chr10:38466841-38468863 | p2 | (AT)11 | GGTGCATTTTCCTTCCTTGA | GCTAGTTGCCACCAAAGCTC | 135 |
| chr10:38466841-38468863 | p2 | (AT)6 | GGTGCATTTTCCTTCCTTGA | GCTAGTTGCCACCAAAGCTC | 135 |
| chr10:38581001-38583015 | p2 | (TA)7 | ACGGCTCGTAGAAGCACACT | GGCCTTGTGGTTCACCTAGA | 149 |
| chr10:38724627-38726653 | p2 | (AT)13 | CTTGGGTAATTGCGTGAGGT | TACAACCCCGAACCTTTTGA | 123 |
| chr10:38754247-38756263 | p2 | (TA)7 | GAATCCTTTTCGTCTCGTCG | ATTGGAAATTCGGGATAGGC | 138 |
| chr10:38754247-38756263 | p2 | (AT)6 | GAATCCTTTTCGTCTCGTCG | ATTGGAAATTCGGGATAGGC | 138 |
| chr10:38754247-38756263 | p4 | (TAAT)4 | GAATCCTTTTCGTCTCGTCG | ATTGGAAATTCGGGATAGGC | 138 |
| chr10:38774981-38777003 | p2 | (AT)11 | TGGTCGGGCAGATATACACA | GGCATCAAATGAGGAGCAAT | 281 |
| chr10:38774981-38777003 | p5 | (ATCAG)3 | TGGTCGGGCAGATATACACA | GGCATCAAATGAGGAGCAAT | 281 |
| chr10:38805921-38807971 | p2 | (TA)6 | AAGCCCACTTAAAAGCCCAT | GTGAGACACCACCCGAAGTT | 214 |
| chr10:38806005-38808031 | p2 | (TA)6 | AAGCCCACTTAAAAGCCCAT | GTGAGACACCACCCGAAGTT | 130 |
| chr10:38999353-39001368 | p3 | (TAT)5 | GGTTCCCAATAAGCAGGTGA | CGTGGGCAGTGCTAATAGGT | 194 |
| chr10:38999353-39001368 | p2 | (AT)10 | GGTTCCCAATAAGCAGGTGA | CGTGGGCAGTGCTAATAGGT | 194 |

| Marker ID | SSR type | SSR motif | FORWARD PRIMER (5'-3') | REVERSE PRIMER (5'-3') | size (bp) |
|---|---|---|---|---|---|
| chr10:39089427-39091499 | p2 | (AT)27 | CACTGCATTGTTCACCGAGT | GCCTGCCTTTTGAACCATAA | 105 |
| chr10:39089427-39091499 | p2 | (AT)36 | CACTGCATTGTTCACCGAGT | GCCTGCCTTTTGAACCATAA | 105 |
| chr10:39089427-39091499 | p2 | (TA)10 | CACTGCATTGTTCACCGAGT | GCCTGCCTTTTGAACCATAA | 105 |
| chr10:39171607-39173621 | p3 | (TGA)5 | AGTTGTTCATCACAAGGGGC | ATCCCACAAGTGGAGTCTGG | 266 |
| chr10:39171607-39173621 | p3 | (CAC)5 | AGTTGTTCATCACAAGGGGC | ATCCCACAAGTGGAGTCTGG | 266 |
| chr10:39171607-39173621 | p2 | (AT)7 | AGTTGTTCATCACAAGGGGC | ATCCCACAAGTGGAGTCTGG | 266 |
| chr10:39277129-39279144 | p5 | (TCATT)3 | ATAGTGGTGGCAATAAGCGG | ACTGCCATGTTGGATTCTCC | 236 |
| chr10:39430160-39432176 | p2 | (AT)14 | GGAACTGTCTGCAAAGCCTC | GCTGCCTTCAGGTACTGGTC | 267 |
| chr10:39430160-39432176 | p4 | (CATA)4 | GGAACTGTCTGCAAAGCCTC | GCTGCCTTCAGGTACTGGTC | 267 |
| chr10:39440641-39442661 | p4 | (ATAA)5 | GCGTCCTCAGAAGCAGTACC | GCAGACCCTTGAGCAGAGAC | 135 |
| chr10:39543523-39545538 | p2 | (AG)6 | CTCCACCCACGATTTGAACT | CACCATCCCTATCTAGCCCA | 167 |
| chr10:39543523-39545538 | p5 | (GAACC)3 | CTCCACCCACGATTTGAACT | CACCATCCCTATCTAGCCCA | 167 |
| chr10:39687895-39689909 | p2 | (AT)7 | TTATTCCAAAGAACCGCCAC | TTTGCAGTCAAAGCTGGATG | 107 |
| chr10:39755607-39757622 | p5 | (TTTAA)3 | CTTGTCGGTTGGTGATGATG | CCCTAGGACGAACTGCTTTG | 232 |
| chr10:39755847-39757891 | p5 | (TTTAA)3 | GGTTGGGTCGTGACAAAAGT | ATTTAGCTCTGCGGCATGAT | 179 |
| chr10:39866730-39868748 | p6 | (ATATGT)3 | TGGAACGTGAATGTGGAGAA | CTTGTGAGGGTTTTCGGGTA | 136 |
| chr10:39876633-39878645 | p2 | (CT)6 | CGAGATTTCAACTCGGCTTC | TGATATGCGTCCGAATGGTA | 174 |
| chr10:40048607-40050625 | p3 | (TTG)6 | CTAAGGGTTTGGGAGCATGA | GCCCAAAGTACGTCCAGAAA | 109 |
| chr10:40120173-40122185 | p5 | (ATTCA)3 | CTTCGACGACACCCTTCATT | GGTGGAAGTCGTGTGTTGTG | 232 |
| chr10:40120173-40122185 | p5 | (ACACA)3 | CTTCGACGACACCCTTCATT | GGTGGAAGTCGTGTGTTGTG | 232 |
| chr10:40120248-40122263 | p5 | (ATTCA)3 | CTTCGACGACACCCTTCATT | GGTGGAAGTCGTGTGTTGTG | 157 |
| chr10:40120248-40122263 | p5 | (ACACA)3 | CTTCGACGACACCCTTCATT | GGTGGAAGTCGTGTGTTGTG | 157 |
| chr10:40153609-40155623 | p2 | (TA)7 | ATTTGTCGGTGGAGAAATCG | ACCAGCAACCAACAACACAA | 289 |
| chr10:40164897-40166912 | p5 | (TTCAT)3 | GACTTACAGGGGCACACCAT | AGCCCTTCGGCTATCAAAAT | 158 |
| chr10:40192248-40194262 | p2 | (TA)7 | ATTTGTCGGTGGAGAAATCG | ACCAGCAACCAACAACACAA | 289 |
| chr10:40203536-40205551 | p5 | (TTCAT)3 | GACTTACAGGGGCACACCAT | AGCCCTTCGGCTATCAAAAT | 158 |
| chr10:40250733-40252753 | p2 | (TA)10 | CAAGGTGAGGCAGAATGTCA | ATTTGGCCATGGTGTATCGT | 210 |
| chr10:40250733-40252753 | p2 | (AT)14 | CAAGGTGAGGCAGAATGTCA | ATTTGGCCATGGTGTATCGT | 210 |
| chr10:40256657-40258675 | p2 | (AT)9 | CGCTTGTTCTGATCGTCTCA | CAAAATTTGAGAAAGGGGCA | 181 |
| chr10:40312110-40314124 | p2 | (TA)7 | TAAAGATCGGACTTGGGGTG | TCAAGTCAGGCAATCCAACA | 202 |
| chr10:40359883-40361898 | p5 | (AGAAG)3 | AGAAAGAACCTGCACGAGGA | CAAAACCAGTTGGCCTGAAT | 232 |
| chr10:40390576-40392591 | p5 | (ACCCG)3 | TGGTTTTCTCTTGGTCCTCG | CAAAAAGGACCACTTCCCAA | 237 |
| chr10:40476430-40478445 | p5 | (TGTGT)3 | CGATTGTTATATCCCGCGTT | AAATATGGCCCAATGACCAA | 274 |
| chr10:40516052-40518066 | p2 | (TA)7 | GCTTGTAACATGGGGAATGC | CCGTAGGATAAGGGTCGTCA | 117 |
| chr10:40516052-40518066 | p3 | (TTG)5 | GCTTGTAACATGGGGAATGC | CCGTAGGATAAGGGTCGTCA | 117 |
| chr10:40556798-40558810 | p2 | (AT)6 | TCCTCGCAAATTTTAATCCG | TTGTTGTTGATTTGGGCTGA | 106 |

| Marker ID | SSR type | SSR motif | FORWARD PRIMER (5'-3') | REVERSE PRIMER (5'-3') | size (bp) |
|---|---|---|---|---|---|
| chr10:40556798-40558810 | p2 | (CA)9 | TCCTCGCAAATTTTAATCCG | TTGTTGTTGATTTGGGCTGA | 106 |
| chr10:40581771-40583813 | p2 | (TA)21 | CTAAATGTCTCGCAGCCTCC | TGTCTCATGACGGACTTTCG | 235 |
| chr10:40630173-40632187 | p2 | (TA)7 | CTTTTCGTCGTAGCTTTGCC | AGCTGGGTCACGACAGAAGT | 112 |
| chr10:40660529-40662544 | p5 | (AACAC)3 | TCTTTCCGCTCCTTTTGCTA | GAATTCCCGGATTGTTGTTG | 294 |
| chr10:40727029-40729047 | p2 | (AT)35 | GTGTTATGCGTACCTGGCCT | ACGGATACCACTGAGCCTTG | 296 |
| chr10:40727029-40729047 | p2 | (AT)9 | GTGTTATGCGTACCTGGCCT | ACGGATACCACTGAGCCTTG | 296 |
| chr10:40727029-40729047 | p2 | (TA)18 | GTGTTATGCGTACCTGGCCT | ACGGATACCACTGAGCCTTG | 296 |
| chr10:40727171-40729207 | p2 | (AT)9 | GTGTTATGCGTACCTGGCCT | ACGGATACCACTGAGCCTTG | 154 |
| chr10:40727171-40729207 | p2 | (TA)18 | GTGTTATGCGTACCTGGCCT | ACGGATACCACTGAGCCTTG | 154 |
| chr10:40773271-40775301 | p2 | (TA)15 | ACGACATTCCGAAGACAACC | TGCATGCTCTGTCTCGATTC | 236 |
| chr10:40835787-40837811 | p2 | (TA)8 | GCCTCATTAGGCCATCAAAA | CATTGTCCGGAGGTGCTTAT | 148 |
| chr10:40835787-40837811 | p2 | (TA)12 | GCCTCATTAGGCCATCAAAA | CATTGTCCGGAGGTGCTTAT | 148 |
| chr10:40855293-40857313 | p2 | (AT)9 | CGGTGATGTGCTAGGTCTGA | TGACGTTATGAACCGTTGGA | 134 |
| chr10:40855293-40857313 | p2 | (TA)10 | CGGTGATGTGCTAGGTCTGA | TGACGTTATGAACCGTTGGA | 134 |
| chr10:40887999-40890014 | p5 | (TTCAT)3 | ACTCCTTTCATTGGCGACAC | AGATCGATTGTTGACAGCCC | 221 |
| chr10:40894653-40896668 | p5 | (TTCAT)3 | ACGTACACGAATCACGCAAG | TTTTTCCTTCAACTGACCCG | 285 |
| chr10:40906829-40908841 | p2 | (AT)6 | TTCTCAAATGATCTGCACGC | CAAGACCCGATACAGGCATT | 121 |
| chr10:40906829-40908841 | p2 | (AT)7 | TTCTCAAATGATCTGCACGC | CAAGACCCGATACAGGCATT | 121 |
| chr10:40919953-40921977 | p2 | (AT)9 | AAATCTGGGGTTCTCGACCT | AAACTTGGGCTCATCATTGC | 220 |
| chr10:40919953-40921977 | p2 | (AT)12 | AAATCTGGGGTTCTCGACCT | AAACTTGGGCTCATCATTGC | 220 |
| chr10:40959228-40961246 | p6 | (ATATGT)3 | ATGTTGGATATGAGCCTCGG | TCGGGTCCTATTTATGCGAG | 165 |
| chr10:41187482-41189497 | p3 | (GTT)5 | CCAAGTGTAAGCCCTCCAAA | TAAACAGCAAGGGAAGTGGG | 170 |
| chr10:41240866-41242884 | p2 | (TG)8 | CTTCGATCGGCCCATTAGTA | GATATGGTTGCCTCGGACAT | 231 |
| chr10:41240895-41242909 | p2 | (TG)8 | CTTCGATCGGCCCATTAGTA | GATATGGTTGCCTCGGACAT | 202 |
| chr10:41256471-41258489 | p6 | (CATATA)3 | GTGAAGTGTGCTCCTCGTCA | ATCCTCGGTGTCATACCAGC | 274 |
| chr10:41256471-41258489 | p2 | (AT)17 | GTGAAGTGTGCTCCTCGTCA | ATCCTCGGTGTCATACCAGC | 274 |
| chr10:41319749-41321767 | p3 | (TGA)6 | CTGGTAAAACTGGAACCGGA | AGGGTGGGATGGGTTTTAAG | 207 |
| chr10:41319749-41321767 | p2 | (TG)8 | CTGGTAAAACTGGAACCGGA | AGGGTGGGATGGGTTTTAAG | 207 |
| chr10:41320033-41322049 | p3 | (TGA)6 | ACTTAAAACCCATCCCACCC | ACAAAGTGAAGGCGGCTAGA | 103 |
| chr10:41320033-41322049 | p2 | (TG)8 | ACTTAAAACCCATCCCACCC | ACAAAGTGAAGGCGGCTAGA | 103 |
| chr10:41335907-41337921 | p2 | (AC)7 | AAGACACGTGGTATGGGAGC | AATGCCTTTAGCATGGGATG | 149 |
| chr10:41347533-41349551 | p2 | (TA)9 | TCCTACGTCCACTTTTTGGG | CCTTATCCAAACGACCCCTT | 202 |
| chr10:41400476-41402494 | p2 | (AT)6 | ATGGCAAAAAGAGAAATGCG | AAGGACATGTCACACGGTCA | 142 |
| chr10:41400476-41402494 | p3 | (GTT)6 | ATGGCAAAAAGAGAAATGCG | AAGGACATGTCACACGGTCA | 142 |
| chr10:41400476-41402494 | p5 | (TGAAT)3 | ATGGCAAAAAGAGAAATGCG | AAGGACATGTCACACGGTCA | 142 |
| chr10:41401116-41403131 | p3 | (GTT)6 | GATGTTATGTGTGTTGGCCG | TGTCATACCAAATCAGCCCA | 151 |

| Marker ID | SSR type | SSR motif | FORWARD PRIMER (5'-3') | REVERSE PRIMER (5'-3') | size (bp) |
|---|---|---|---|---|---|
| chr10:41401116-41403131 | p5 | (TGAAT)3 | GATGTTATGTGTGTTGGCCG | TGTCATACCAAATCAGCCCA | 151 |
| chr10:41437772-41439784 | p2 | (TA)6 | GGACGTCGTGTGTGTTATGG | TTGCACGTTCTCTTTCAACG | 214 |
| chr10:41582546-41584561 | p5 | (ATTTC)3 | GAGTCCCTTACTGGAGGGCT | ATGGGACTCGGTGAATGAAG | 150 |
| chr10:41623230-41625245 | p5 | (CAATT)3 | TCAGAACAAGCGTACATGGC | TGATCCCTTCCGACTTCATC | 212 |
| chr10:41657777-41659797 | p4 | (CATA)5 | GCGATCCTACTAGCCTGCAC | CCTCCAGACTTCTTTGGCAC | 140 |
| chr10:41657777-41659797 | p2 | (AT)40 | GCGATCCTACTAGCCTGCAC | CCTCCAGACTTCTTTGGCAC | 140 |
| chr10:41733463-41735478 | p5 | (TTCAT)3 | AGTCAGATCACCCGAAAACG | TTATTTGCTGAGGTGGGACC | 227 |
| chr10:41765385-41767397 | p2 | (TA)6 | CTGGAATTTGGCAGTTGGAT | TCCGTTTGACCCCCATAATA | 151 |
| chr10:41765385-41767397 | p2 | (TA)22 | CTGGAATTTGGCAGTTGGAT | TCCGTTTGACCCCCATAATA | 151 |
| chr10:41810281-41812293 | p2 | (CA)6 | CATCAGTCGTAGCGTCCTCA | GACTAAGTTGGTTGCGGCTC | 179 |
| chr10:41821682-41823696 | p2 | (GT)7 | CGAGTGTGGAAAAGGGGATA | TCCAAATTTTCCGAATACGC | 202 |
| chr10:41885148-41887163 | p3 | (TTG)5 | TTGGTGCAAAATACCAACCA | TGAATCCCATCATCACCTCA | 159 |
| chr10:41897654-41899680 | p2 | (AT)13 | ATACGAGCCGACAAGGCTAA | TTCAGCCGGAGCTACAACTT | 136 |
| chr10:42019704-42021724 | p2 | (AT)6 | CAGTTCCGAACAGCGTGTAA | AGCCAACAACACCAACACAA | 248 |
| chr10:42207039-42209057 | p2 | (AT)9 | CCTAGGTGGGTCGTGACACT | ACTAATCATCCCCAAGCACG | 119 |
| chr10:42292172-42294187 | p5 | (TTACT)3 | GTCCATGAGGCTTCTGGTGT | AGTCCGTTTGGTTGGTTCTG | 267 |
| chr10:42292172-42294187 | p2 | (TA)6 | GTCCATGAGGCTTCTGGTGT | AGTCCGTTTGGTTGGTTCTG | 267 |
| chr10:42292827-42294839 | p5 | (TTACT)3 | GGACCCCATGTAAGTGATGC | CAAAGAGGGGGAGTGAATGA | 122 |
| chr10:42292827-42294839 | p2 | (TA)6 | GGACCCCATGTAAGTGATGC | CAAAGAGGGGGAGTGAATGA | 122 |
| chr10:42344102-42346114 | p2 | (AT)6 | GATGACCAGTTCGCCATTCT | ATGGTGTCGACAAGAGGAGG | 262 |
| chr10:42344102-42346114 | p5 | (AATAA)3 | GATGACCAGTTCGCCATTCT | ATGGTGTCGACAAGAGGAGG | 262 |
| chr10:42344616-42346631 | p2 | (AT)6 | ATCCAAATAACCCCGGTCTC | GCTTTGGCGTATCATGTTGA | 221 |
| chr10:42344616-42346631 | p5 | (AATAA)3 | ATCCAAATAACCCCGGTCTC | GCTTTGGCGTATCATGTTGA | 221 |
| chr10:42480403-42482437 | p2 | (AT)17 | GGGAGGCAATTTTCTAAGGC | AATTTGACCAATTGGGCTTG | 211 |
| chr10:42480403-42482437 | p6 | (ATATGA)4 | GGGAGGCAATTTTCTAAGGC | AATTTGACCAATTGGGCTTG | 211 |
| chr10:42549664-42551678 | p2 | (AT)7 | GAAAGAGTTGAATTTGCGGC | GATCTGCAGCTAAGGGCATC | 105 |
| chr10:42590000-42592012 | p2 | (AT)6 | CTTTCCCTTCTTCCCAAACC | TGGGTCGTGACATGAGGTTA | 251 |
| chr10:42590000-42592012 | p2 | (AT)12 | CTTTCCCTTCTTCCCAAACC | TGGGTCGTGACATGAGGTTA | 251 |
| chr10:42674299-42676311 | p2 | (AG)6 | GGAGCCTGGATCAATCAAAA | GCCGTCTGTAGCCTCAGTTC | 149 |
| chr10:42674299-42676311 | p3 | (GTT)5 | GGAGCCTGGATCAATCAAAA | GCCGTCTGTAGCCTCAGTTC | 149 |
| chr10:42674833-42676848 | p2 | (AG)6 | CGTTTTCGGGTGATCTGACT | TGAGGAGGAGAAGGCTGAAA | 171 |
| chr10:42674833-42676848 | p3 | (GTT)5 | CGTTTTCGGGTGATCTGACT | TGAGGAGGAGAAGGCTGAAA | 171 |
| chr10:42674833-42676848 | p2 | (TA)8 | CGTTTTCGGGTGATCTGACT | TGAGGAGGAGAAGGCTGAAA | 171 |
| chr10:42682559-42684577 | p3 | (GTT)5 | ATGGTTCAGTGGGATGCAAT | CACTTCCAAAACAGCCCAAT | 173 |
| chr10:42682576-42684591 | p3 | (GTT)5 | ATGGTTCAGTGGGATGCAAT | CACTTCCAAAACAGCCCAAT | 156 |
| chr10:42688771-42690805 | p2 | (AT)17 | TTTGATTTCCCATTGTGGGT | GGACAGGATTTCCCCAATTT | 290 |

| Marker ID | SSR type | SSR motif | FORWARD PRIMER (5'-3') | REVERSE PRIMER (5'-3') | size (bp) |
|---|---|---|---|---|---|
| chr10:42688771-42690805 | p2 | (TA)21 | TTTGATTTCCCATTGTGGGT | GGACAGGATTTCCCCAATTT | 290 |
| chr10:42689232-42691274 | p2 | (AT)17 | TTGAGCGGTACTGTCAAACG | CGGAGTGACATAAGGTCGGT | 108 |
| chr10:42689232-42691274 | p2 | (TA)21 | TTGAGCGGTACTGTCAAACG | CGGAGTGACATAAGGTCGGT | 108 |
| chr10:42795571-42797583 | p2 | (CA)6 | GGACATGCCAAAAGAAGGAA | TTTATAGGGAAATGCTGCCG | 237 |
| chr10:42795571-42797583 | p6 | (TTTTTG)3 | GGACATGCCAAAAGAAGGAA | TTTATAGGGAAATGCTGCCG | 237 |
| chr10:42810072-42812087 | p5 | (AGGAC)3 | CTTGGGGCCATACTCTACCA | GAGCCACCAACGGGTATAGA | 269 |
| chr10:42813767-42815785 | p6 | (TGTAAG)3 | ATGCGGCTGGGTAAGATATG | CCAACCAAGGCTCAAGGTTA | 100 |
| chr10:42828976-42830991 | p5 | (ATTTC)3 | TACCCGCATTTATTTCAGCC | AATACCAATGCCTCACAGGG | 210 |
| chr10:42988971-42990983 | p2 | (TA)6 | CAGCTTCACACCCATAAGCA | GCCACCAGGTTATTCAGCAT | 121 |
| chr10:42992681-42994696 | p3 | (CAA)5 | TCTCGCGTTCCATATGATGA | CGACCTTACGTCACTCCGAT | 234 |
| chr10:43017870-43019888 | p6 | (TGTATG)3 | GCCGGGATCTGTATGTCTGT | ACTTCTGTCACGACCCAACC | 122 |
| chr10:43121639-43123679 | p2 | (AT)20 | ATTGGTTTGGGTCGTGACAT | GAAATAGTGAACAGGCCCGA | 177 |
| chr10:43195566-43197578 | p2 | (AT)6 | GGCCTTATCCGATTATCCGT | CTTCTCCGTCACCTGGACAT | 280 |
| chr10:43202361-43204379 | p2 | (AT)9 | AGGGGAGCCTCAACTTTAGC | TAGATCCATCAAACCTGGGC | 128 |
| chr10:43230181-43232195 | p2 | (AT)7 | GAGGTGTGGAGAATGAGGGA | CCCTGAGTATGGGGTAGGGT | 143 |
| chr10:43251064-43253079 | p5 | (ATATC)3 | CGCATAATCTGCCTCGGTAT | CTGCTTGCTGCAGATGAAAG | 276 |
| chr10:43289611-43291623 | p2 | (TA)6 | CCGTGCATCAATCAGACATC | AGAGGTCAAACCCACGTCAC | 209 |
| chr10:43289611-43291623 | p2 | (GT)6 | CCGTGCATCAATCAGACATC | AGAGGTCAAACCCACGTCAC | 209 |
| chr10:43370021-43372072 | p3 | (AAT)17 | ACGTGACACCAAGTTCCTCC | CCCTGACCTGATCCGAGATA | 128 |
| chr10:43370021-43372072 | p2 | (TA)8 | ACGTGACACCAAGTTCCTCC | CCCTGACCTGATCCGAGATA | 128 |
| chr10:43385950-43388010 | p2 | (AT)30 | CTCTCGGCCATCAAAACAAT | TGTTCATGCCCACAGGTAGA | 300 |
| chr10:43398832-43400852 | p2 | (TA)10 | CACTCCATTGTTCCGGAGAT | CATAATGGCAACCTCGTGTG | 271 |
| chr10:43398832-43400852 | p2 | (AT)29 | CACTCCATTGTTCCGGAGAT | CATAATGGCAACCTCGTGTG | 271 |
| chr10:43436130-43438146 | p2 | (AT)6 | AACGCCGCATATCATAAACC | GTCGGCTTGCGTATGGTATT | 243 |
| chr10:43436130-43438146 | p2 | (AT)8 | AACGCCGCATATCATAAACC | GTCGGCTTGCGTATGGTATT | 243 |
| chr10:43472929-43474941 | p2 | (AG)6 | GGTGACCCTTCCAGGGTATT | TCCTCTCGGTTATTGTTCCG | 245 |
| chr10:43472929-43474941 | p3 | (TGT)6 | GGTGACCCTTCCAGGGTATT | TCCTCTCGGTTATTGTTCCG | 245 |
| chr10:43481237-43483253 | p2 | (TA)8 | GGCTGTTTGGAATATTGCGT | ACATCACAACCAAGCCAACA | 277 |
| chr10:43504875-43506890 | p5 | (GGTCG)3 | TTAAGCAACGAATGGGAACC | TCTTTAGCCGCGAACAGATT | 292 |
| chr10:43504875-43506890 | p5 | (GAAAT)3 | TTAAGCAACGAATGGGAACC | TCTTTAGCCGCGAACAGATT | 292 |
| chr10:43504896-43506914 | p5 | (GGTCG)3 | TTAAGCAACGAATGGGAACC | TCTTTAGCCGCGAACAGATT | 271 |
| chr10:43504896-43506914 | p5 | (GAAAT)3 | TTAAGCAACGAATGGGAACC | TCTTTAGCCGCGAACAGATT | 271 |
| chr10:43504918-43506933 | p5 | (GGTCG)3 | TTAAGCAACGAATGGGAACC | TCTTTAGCCGCGAACAGATT | 249 |
| chr10:43504918-43506933 | p5 | (GAAAT)3 | TTAAGCAACGAATGGGAACC | TCTTTAGCCGCGAACAGATT | 249 |
| chr10:43508438-43510474 | p2 | (AT)18 | TCTCCCTTTAGGGTCCGATT | TTACGACACGGCAAGTCAAG | 242 |
| chr10:43542077-43544089 | p2 | (AT)6 | TCAAATCTCCCGGTCTTCAC | CAAGAGAAGGAAGGTGCTCG | 157 |

| Marker ID | SSR type | SSR motif | FORWARD PRIMER (5'-3') | REVERSE PRIMER (5'-3') | size (bp) |
|---|---|---|---|---|---|
| chr10:43542077-43544089 | p2 | (TA)14 | TCAAATCTCCCGGTCTTCAC | CAAGAGAAGGAAGGTGCTCG | 157 |
| chr10:43561958-43563973 | p5 | (ATTTC)3 | CATTCGCGAGTATGAGTGGA | GACCGAAGCCCATATCAAGA | 135 |
| chr10:43670360-43672414 | p2 | (AT)27 | CATGGATTGGTTGGCTTCTT | ACCACGGGAACTGACTGAAG | 150 |
| chr10:43819621-43821633 | p2 | (TG)6 | AAAGAACCTCCCCCAAAATG | TTTTGGCCTTAGCCTTCTCA | 244 |
| chr10:43821454-43823474 | p2 | (TA)10 | ATTGACCCGATGCTTGTTTC | CAAATCACATCATAACGCCG | 298 |
| chr10:43836210-43838288 | p3 | (AAG)26 | TCGAAGGGTCAAGAAGGCTA | TTTTCCCATGGGGATTACAA | 177 |
| chr10:43843971-43845983 | p2 | (AT)6 | TCGTAGGCGTGACCTAGTCC | TTAATTGGTTCACGTTGGCA | 235 |
| chr10:43914799-43916817 | p2 | (AT)9 | CAGATCAGTGCATCGCCTAA | GATATCAGGAGCACCGCAAT | 128 |
| chr10:44054269-44056289 | p2 | (TA)10 | TTTGTCACAACCCAACCAGA | CTCTAGGTGCCCGTCATAGC | 206 |
| chr10:44064543-44066559 | p2 | (TA)8 | TCCTCTTCATAGGCATCGCT | CCGTCATGGCCCTCTAGTTA | 148 |
| chr10:44086271-44088285 | p2 | (AG)7 | ATTGGAGCGGATGTGCTATC | TTTCTGGAAGGATCACCGAC | 106 |
| chr10:44086271-44088285 | p2 | (GT)6 | ATTGGAGCGGATGTGCTATC | TTTCTGGAAGGATCACCGAC | 106 |
| chr10:44105579-44107597 | p3 | (TTG)6 | TTGGAGAAAATGTGAAGGGC | GGTAGTGGCGTGTTTGTCCT | 279 |
| chr10:44127425-44129440 | p3 | (TGT)5 | GGGAGGTTAAGAGCAAGCCT | AAATGGCCCCTTATGTTTCC | 128 |
| chr10:44223737-44225801 | p2 | (AT)32 | AAGATCCACCAAAGACGCAC | GCCTTCCCAAGAAAACATCA | 175 |
| chr10:44223737-44225801 | p3 | (ATC)5 | AAGATCCACCAAAGACGCAC | GCCTTCCCAAGAAAACATCA | 175 |
| chr10:44284686-44286702 | p3 | (TAA)5 | GGGAGACAAGGAAATGGTCA | TTAACCGGTGCAAAACCTTC | 192 |
| chr10:44284686-44286702 | p2 | (AC)6 | GGGAGACAAGGAAATGGTCA | TTAACCGGTGCAAAACCTTC | 192 |
| chr10:44442820-44444868 | p2 | (TA)24 | CCAAGGTATGTGAGGCCAGT | TCTATCGGGGTGACGTAAGG | 132 |
| chr10:44458424-44460442 | p2 | (AT)9 | GGTGAAAGGCATGGTCCTAA | CCATGGTTCCCATTCAAATC | 206 |
| chr10:44524729-44526741 | p3 | (TTG)7 | GAGCTCATTTTGAAGGCTCG | GGGATCATGCCAAAAAGAAA | 167 |
| chr10:44524729-44526741 | p2 | (AC)6 | GAGCTCATTTTGAAGGCTCG | GGGATCATGCCAAAAAGAAA | 167 |
| chr10:44615346-44617358 | p2 | (AT)6 | AGTTTAGGCCACCACACGAC | CCCAACCTTCTCACACCACT | 294 |
| chr10:44632632-44634656 | p4 | (CATA)6 | CCCGATCCTTTACTTAGGGC | CTCTGCAAGCCGATGTATCA | 185 |
| chr10:44632632-44634656 | p6 | (AACAAT)3 | CCCGATCCTTTACTTAGGGC | CTCTGCAAGCCGATGTATCA | 185 |
| chr10:44654575-44656590 | p5 | (CAATG)3 | AGCTTGGTTGGTCATTTTCG | AGCACTGTGGGGATTGTTTC | 139 |
| chr10:44654575-44656590 | p2 | (TA)11 | AGCTTGGTTGGTCATTTTCG | AGCACTGTGGGGATTGTTTC | 139 |
| chr10:44779106-44781121 | p3 | (TGT)5 | CATTGCACTCCACCTGTTTG | CACCTGCACTATGCTGTGCT | 251 |
| chr10:44893473-44895487 | p2 | (AC)7 | GGATGGCTGTTTTCCTTGAA | ACATGGATCTGGGGTAGTGC | 227 |
| chr10:44893473-44895487 | p2 | (TA)6 | GGATGGCTGTTTTCCTTGAA | ACATGGATCTGGGGTAGTGC | 227 |
| chr10:44893473-44895487 | p2 | (TA)6 | GGATGGCTGTTTTCCTTGAA | ACATGGATCTGGGGTAGTGC | 227 |
| chr10:45102400-45104460 | p2 | (TA)6 | ATGGCGGCTAAAAAGTTCCT | CGTCCATTTTCGGGTAAGAA | 289 |
| chr10:45102472-45104523 | p2 | (TA)6 | ATGGCGGCTAAAAAGTTCCT | CGTCCATTTTCGGGTAAGAA | 217 |
| chr10:45123099-45125113 | p2 | (CA)7 | AGGAGCATGCCAAAGAAAGA | TGGCCGAGTTAAAATTCTCG | 227 |
| chr10:45123099-45125113 | p2 | (CT)11 | AGGAGCATGCCAAAGAAAGA | TGGCCGAGTTAAAATTCTCG | 227 |
| chr10:45123386-45125408 | p2 | (CA)7 | TCGGCCAAAACATACAACAA | TACAATTGAGACGCCGTGAA | 160 |

| Marker ID | SSR type | SSR motif | FORWARD PRIMER (5'-3') | REVERSE PRIMER (5'-3') | size (bp) |
|---|---|---|---|---|---|
| chr10:45123386-45125408 | p2 | (CT)11 | TCGGCCAAAACATACAACAA | TACAATTGAGACGCCGTGAA | 160 |
| chr10:45162546-45164582 | p3 | (TAT)21 | TCAACTCGGTCATGCTGAAG | AACCTGTTTGGTCCGACATC | 265 |
| chr10:45162585-45164600 | p3 | (TAT)21 | TCAACTCGGTCATGCTGAAG | AACCTGTTTGGTCCGACATC | 226 |
| chr10:45162609-45164675 | p3 | (TAT)21 | TCAACTCGGTCATGCTGAAG | AACCTGTTTGGTCCGACATC | 202 |
| chr10:45164097-45166163 | p3 | (GTA)8 | TAAACGGCACAAAGAAAGGG | TGGTTACCCCCAAAACAAAA | 198 |
| chr10:45164097-45166163 | p3 | (TAA)22 | TAAACGGCACAAAGAAAGGG | TGGTTACCCCCAAAACAAAA | 198 |
| chr10:45277112-45279128 | p2 | (AG)8 | ACCCTGAGCCCTGGTAACTT | AGCCGTCTGCAGCCTTAGTA | 269 |
| chr10:45379069-45381087 | p2 | (TA)9 | ATGATACGAATCCTCCGTGC | AAAATAATTGGAGGGGTGGC | 144 |
| chr10:45381139-45383159 | p2 | (CA)10 | TCTGCCTCGTCTCATCATTG | GGCTTTGGTTCACGTTGAGT | 147 |
| chr10:45381139-45383159 | p2 | (AT)7 | TCTGCCTCGTCTCATCATTG | GGCTTTGGTTCACGTTGAGT | 147 |
| chr10:45506730-45508745 | p3 | (GTT)5 | ACTTCCTCGCACTTCTTCCA | GGCACCTATAGACGAGGGTG | 146 |
| chr10:45586379-45588394 | p6 | (TTATCG)3 | AGCTTTGCAACCTTTGGAGA | TTGAAGGAGAACGTGCAATG | 154 |
| chr10:45586379-45588394 | p5 | (GAAAT)3 | AGCTTTGCAACCTTTGGAGA | TTGAAGGAGAACGTGCAATG | 154 |
| chr10:45698955-45700973 | p6 | (CAGCTC)3 | TACTTGGGTATGGCGGAGTC | ATAACACAAGCCAACCAGGC | 246 |
| chr10:45777112-45779136 | p2 | (TA)24 | TGATCACACCGTGCCTGTAT | TACCTGTCGGCATGAACAAA | 257 |
| chr10:45777112-45779136 | p4 | (GTAT)6 | TGATCACACCGTGCCTGTAT | TACCTGTCGGCATGAACAAA | 257 |
| chr10:45965705-45967725 | p5 | (AATGC)4 | TCTGGCTAAGCAAAGTCCGT | AATTGGGGTCGTGACATGAT | 258 |
| chr10:46093495-46095507 | p2 | (GT)6 | AGCAAGTAGCTCACCCTGGA | TATTTCCTCCACGCACACAA | 160 |
| chr10:46126213-46128229 | p2 | (TA)8 | AGGTGTAACAGATGGGCCAC | GTCCTGTCCCGTCCTTATGA | 261 |
| chr10:46147150-46149165 | p5 | (TCATA)3 | TAGCTTGCGGTTTTGGAGTT | GTAACACCCCACTTGCTGGT | 248 |
| chr10:46173706-46175728 | p2 | (AT)11 | GCCCGAGATCAAATTACCAA | TAACAATGCGTTTGGCTCAG | 106 |
| chr10:46285124-46287140 | p2 | (AT)8 | ATCAACTTCGAGGCTGCACT | CCAAAAAGAGGGTCGAAACA | 179 |
| chr10:46368363-46370383 | p5 | (TGTGG)4 | CGGGAAGACTACAAGCAAGG | TTCCACGAATCATATTGCGA | 193 |
| chr10:46690134-46692158 | p2 | (AT)12 | CGAGCCTATATGGACGGGTA | GTCTTCCTGTCTACCTGCGG | 233 |
| chr10:46708957-46710977 | p2 | (AT)10 | ACGAAGCCTGAAAGCCACTA | GGTTCGCGAGTTTTTAGCAC | 183 |
| chr10:46708957-46710977 | p2 | (AT)25 | ACGAAGCCTGAAAGCCACTA | GGTTCGCGAGTTTTTAGCAC | 183 |
| chr10:46735409-46737424 | p5 | (TATCA)3 | GACTCTAACTGGCACGGCTC | GGTCCTCGTCGCTGTGTAAT | 264 |
| chr10:46755119-46757134 | p5 | (TATAT)3 | CTCCTGCCGAATTGAAAGAG | CTCCGAATATAGGGTCGGGT | 121 |
| chr10:46791448-46793466 | p6 | (TTTTCC)3 | AGGGAAGCATACACAATGCC | ATGTTTTGTCTGCCTGGTCC | 171 |
| chr10:46833149-46835179 | p2 | (AT)15 | AACAGCTCCCAATTTCATGG | TTGAAACTTAAACCGCCACC | 271 |
| chr10:46881765-46883783 | p6 | (AAAATA)3 | TGCTAGGGGAGGGGTTATTT | CTAAAATCGGCAGCCTTGAC | 117 |
| chr10:46931791-46933809 | p3 | (TTA)6 | AAATGGTCCAGAAAATCCCC | CCAGTCTTCTTGGGGACAAA | 166 |
| chr10:46979248-46981276 | p2 | (AT)14 | CATACTCCGTCACTCTGCCA | CCTAGTCGGGTCGTGACATT | 220 |
| chr10:46979248-46981276 | p2 | (TA)9 | CATACTCCGTCACTCTGCCA | CCTAGTCGGGTCGTGACATT | 220 |
| chr10:47026752-47028780 | p2 | (AT)14 | CATACTCCGTCACTCTGCCA | CCTAGTCGGGTCGTGACATT | 220 |
| chr10:47026752-47028780 | p2 | (TA)9 | CATACTCCGTCACTCTGCCA | CCTAGTCGGGTCGTGACATT | 220 |

| Marker ID | SSR type | SSR motif | FORWARD PRIMER (5'-3') | REVERSE PRIMER (5'-3') | size (bp) |
|---|---|---|---|---|---|
| chr10:47144545-47146557 | p2 | (AG)6 | TTGGGCCTTACCAGATTGTC | ACTTCTTTGGTGCGCAGTCT | 151 |
| chr10:47357264-47359276 | p2 | (TA)6 | AATCTGGTCGGCAACTGAAC | AGTAACTAGGGCCCATGCCT | 100 |
| chr10:47373757-47375772 | p3 | (GTT)5 | AAAGATTCCCCCGAAGTGTT | GCATTAAGGCAAAAACGGAA | 154 |
| chr10:47373757-47375772 | p3 | (TGT)5 | AAAGATTCCCCCGAAGTGTT | GCATTAAGGCAAAAACGGAA | 154 |
| chr10:47457661-47459677 | p4 | (CATA)4 | AGTGTATGTGCCGGTAAGGC | CTTCGGAGTTTCCCTGTGAG | 217 |
| chr10:47550566-47552584 | p2 | (TA)9 | ACATAATTGTAGGCGGCGTC | TTGGTAGCGGCAGAACTTTT | 175 |
| chr10:47557068-47559106 | p2 | (AT)19 | GTGGCCTGAGACAACACCTT | GACAGCACAAGCTGGCATAA | 158 |
| chr10:47707241-47709253 | p2 | (TG)6 | TGCACTACTCTGTTCGTGCC | TCGGAAACCGTAACACATCA | 215 |
| chr10:47718033-47720053 | p2 | (AT)10 | CACACACATGTCTTCCAGCC | GAGCTCCATCTCATTCGGAG | 271 |
| chr10:47874180-47876196 | p2 | (AT)7 | TCAAGCTCACCTCCCTGTCT | TACCTATATGGACGGGCAGC | 261 |
| chr10:47874180-47876196 | p2 | (AT)8 | TCAAGCTCACCTCCCTGTCT | TACCTATATGGACGGGCAGC | 261 |
| chr10:47881768-47883834 | p2 | (TA)33 | TGTTCACCTGGTTCACCAAA | CATATTCCCTGTGGGATTCG | 269 |
| chr10:47889030-47891062 | p4 | (CATA)8 | GGAACTGTCTGCAAAGCCTC | ATTGCTCGTACTGACGTCCC | 252 |
| chr10:47946175-47948193 | p3 | (TGT)6 | GGTTGTGAGACCGACCTTGT | CTCGCTTCCCTCATCTTGTC | 273 |
| chr10:47965479-47967491 | p2 | (AT)6 | AGTTTTAGTGGAGGGCCGAT | TAAATTCAACCCAGCCAACC | 219 |
| chr10:47965479-47967491 | p2 | (AT)8 | AGTTTTAGTGGAGGGCCGAT | TAAATTCAACCCAGCCAACC | 219 |
| chr10:48096513-48098541 | p2 | (AT)14 | TTGCCATTGTGGATGAGAAA | AGGTCCAGATGATCCACGTC | 198 |
| chr10:48116780-48118828 | p2 | (TC)11 | TTCCTTTTCCTTTTTGGGCT | TCCGCCACTTGTAATAACCC | 211 |
| chr10:48116780-48118828 | p3 | (TTG)16 | TTCCTTTTCCTTTTTGGGCT | TCCGCCACTTGTAATAACCC | 211 |
| chr10:48250081-48252096 | p3 | (ATG)5 | CGTTGGGGATGTTTTATTGG | TTATCCTCCCAAGCTCGAAA | 268 |
| chr10:48250081-48252096 | p5 | (TTGTA)5 | CGTTGGGGATGTTTTATTGG | TTATCCTCCCAAGCTCGAAA | 268 |
| chr10:48270514-48272578 | p3 | (TTC)5 | TCGACCTGATGTACTTCCCC | ATCTCTTGGCTTTTGCGTGT | 127 |
| chr10:48297598-48299616 | p6 | (TCAGTT)3 | TATGCCCACCAAGTGTTTGA | TTGGAAAGGTAGGTGTTCGG | 210 |
| chr10:48300007-48302039 | p2 | (AC)16 | CTAGGAGCCATTTCCTGACG | CCACTGACGTAGCTCATGGA | 258 |
| chr10:48563913-48565929 | p3 | (CAA)5 | TTCGGCATCATTGTCATCAT | TCCAAGGCATGATTTCCTTC | 249 |
| chr10:48563913-48565929 | p2 | (CT)8 | TTCGGCATCATTGTCATCAT | TCCAAGGCATGATTTCCTTC | 249 |
| chr10:48569881-48571899 | p6 | (CATATA)3 | AACTGATGGGCCGTAACAAG | GTATGCGTTGGCCGTTAGTT | 122 |
| chr10:48569881-48571899 | p3 | (CAA)5 | AACTGATGGGCCGTAACAAG | GTATGCGTTGGCCGTTAGTT | 122 |
| chr10:48635036-48637084 | p2 | (AT)24 | GAGATGTGCCAATCGGAGTT | GCATTTTTGCAATCGGATCT | 146 |
| chr10:48665612-48667633 | p3 | (CAA)7 | CCTCTTTCGTCAGATCCTCG | TCCCCAAGTCATCAGTCTCC | 162 |
| chr10:48685274-48687289 | p5 | (TGTTT)3 | ATCGATGAGCGGTTTATTGC | GTCAACTTTGGAGGGGCATA | 237 |
| chr10:48761940-48763952 | p2 | (TA)8 | GTTTGGTCATCTGGGCATTT | AATTGGGCGATACGACAGAG | 232 |
| chr10:48761940-48763952 | p2 | (AT)6 | GTTTGGTCATCTGGGCATTT | AATTGGGCGATACGACAGAG | 232 |
| chr10:48814402-48816418 | p2 | (CT)8 | GAAGTTGGGGAACGAAATGA | ATGAACACCTTTGCCTTTGG | 185 |
| chr10:48849171-48851215 | p2 | (AT)11 | TGTATGATCCCACTCCCCAT | GGCACAAATATCACACGCTG | 275 |
| chr10:48849171-48851215 | p2 | (AT)25 | TGTATGATCCCACTCCCCAT | GGCACAAATATCACACGCTG | 275 |

| Marker ID | SSR type | SSR motif | FORWARD PRIMER (5'-3') | REVERSE PRIMER (5'-3') | size (bp) |
|---|---|---|---|---|---|
| chr10:48849171-48851215 | p2 | (AT)22 | TGTATGATCCCACTCCCCAT | GGCACAAATATCACACGCTG | 275 |
| chr10:48852151-48854173 | p2 | (AT)11 | ATGCTAACTGGGGTCATTGC | AAGGGACGCGAAGGATAGAT | 144 |
| chr10:48906411-48908431 | p5 | (TATTG)4 | AATCGATTGGAAACCCATGA | GCAGAACTGGAACAATGCAA | 179 |
| chr10:48999402-49001417 | p3 | (TGA)5 | TGTCGGAACTTTGGGAAAAC | GGTTTGTTGCGTTGGAAACT | 199 |
| chr10:49051879-49053897 | p3 | (CAA)6 | GACGTGGGATTTGGAGAAGA | CTATGGCCACCTTGGTCACT | 216 |
| chr10:49058710-49060730 | p4 | (ACGA)5 | TGAGTAGTGGCAGCAAGTGG | GTAAATCAACCATGCCACCC | 122 |
| chr10:49058710-49060730 | p2 | (TG)7 | TGAGTAGTGGCAGCAAGTGG | GTAAATCAACCATGCCACCC | 122 |
| chr10:49094759-49096789 | p2 | (TA)6 | CGCGTCCAAGGTAATCTCAT | GAAGCAAGCGAGTGAGTTCC | 107 |
| chr10:49094759-49096789 | p2 | (TA)15 | CGCGTCCAAGGTAATCTCAT | GAAGCAAGCGAGTGAGTTCC | 107 |
| chr10:49109201-49111253 | p2 | (AT)26 | CCCGGGCAACTTACACTTTA | GAATGCGCAAGAGATGAACA | 194 |
| chr10:49205018-49207033 | p5 | (TTTCA)3 | TATGATCACCGACCCTAGCC | ACTTGGTTTCATAGGCGACG | 124 |
| chr10:49220211-49222226 | p5 | (TTACT)3 | TAACCTACTGGCAATTCGGG | TGGTAGGGAGACGACCTTTG | 298 |
| chr10:49452583-49454598 | p5 | (TTTTC)3 | GGGAATTAGGAAAAGGGCTG | CGTACGTCTACGGTGGGAGT | 142 |
| chr10:49617014-49619032 | p6 | (ACATGA)3 | AAATGCGGCCAAGTTATACG | GGCTTGGAGAACGTGAAAAG | 143 |
| chr10:49617014-49619032 | p6 | (ACATGA)3 | AAATGCGGCCAAGTTATACG | GGCTTGGAGAACGTGAAAAG | 143 |
| chr10:49640900-49642915 | p3 | (CTG)5 | GGCCCTAGCTCGCTTAACTT | AAATTGTTAAGACCCCGGCT | 147 |
| chr10:49707167-49709179 | p3 | (GAT)5 | ACCCCGACATAGCCCTAGTT | ACACCCATCGTCACTCAACA | 228 |
| chr10:49707167-49709179 | p2 | (TA)6 | ACCCCGACATAGCCCTAGTT | ACACCCATCGTCACTCAACA | 228 |
| chr10:49833861-49835877 | p2 | (AT)6 | TCCAGGTGAGGCAGAATACC | GGGGTAATGCCATGAACATC | 238 |
| chr10:49833879-49835893 | p2 | (AT)6 | TCCAGGTGAGGCAGAATACC | GGGGTAATGCCATGAACATC | 220 |
| chr10:49833911-49835945 | p2 | (AT)6 | TCCAGGTGAGGCAGAATACC | GGGGTAATGCCATGAACATC | 188 |
| chr10:49920380-49922480 | p2 | (AT)50 | AGTTGAGAGTGCCTTGGGAA | CAGTTCAGATCGGGTCGATT | 110 |
| chr10:49993563-49995575 | p2 | (TA)6 | CCTGCACACACCTGCTTCTA | CGCCATTGGCTCTTAATCTG | 103 |
| chr10:50017520-50019541 | p3 | (TTG)7 | CCGAGATGATGGGACAGTTT | TTCACTTTTGCATGCTTTCG | 214 |
| chr10:50037236-50039251 | p5 | (ACTGT)3 | AAGCTTGACCTTCTCCACGA | CCCAGACTGATGGACAACCT | 181 |
| chr10:50038444-50040462 | p3 | (AAC)6 | TGTATGCAGGTTGTAGGGCA | TAGTAGGGAAACGGTGACGG | 278 |
| chr10:50083613-50085631 | p3 | (CAA)6 | GGTCTAACCACCGGCTTGTA | ACCCGACCATTACGTCACTC | 259 |
| chr10:50237317-50239329 | p2 | (TG)6 | GCTTGGCAGCTTTTTACAGG | ACACATCCTAGCCCAACACC | 148 |
| chr10:50243987-50246002 | p5 | (ATAAT)3 | TGCAAAAGAGCTCCAAAGGT | CTGTCCCGTCATATTTCGCT | 165 |
| chr10:50526239-50528259 | p3 | (TTG)6 | TGGACGCTATAAGTTCGGCT | TATCAATTCCGCAACCAACA | 300 |
| chr10:50526239-50528259 | p2 | (AT)10 | TGGACGCTATAAGTTCGGCT | TATCAATTCCGCAACCAACA | 300 |
| chr10:50526239-50528259 | p2 | (AT)6 | TGGACGCTATAAGTTCGGCT | TATCAATTCCGCAACCAACA | 300 |
| chr10:50526425-50528437 | p3 | (TTG)6 | TGGACGCTATAAGTTCGGCT | TATCAATTCCGCAACCAACA | 114 |
| chr10:50526425-50528437 | p2 | (AT)10 | TGGACGCTATAAGTTCGGCT | TATCAATTCCGCAACCAACA | 114 |
| chr10:50526425-50528437 | p2 | (AT)6 | TGGACGCTATAAGTTCGGCT | TATCAATTCCGCAACCAACA | 114 |
| chr10:50535155-50537171 | p2 | (TA)15 | CACTAGTGGGCAGCATGAGA | ACATTGTCTACCCGGCCATA | 241 |

| Marker ID | SSR type | SSR motif | FORWARD PRIMER (5'-3') | REVERSE PRIMER (5'-3') | size (bp) |
|---|---|---|---|---|---|
| chr10:50535155-50537171 | p2 | (AT)45 | CACTAGTGGGCAGCATGAGA | ACATTGTCTACCCGGCCATA | 241 |
| chr10:50535175-50537207 | p2 | (TA)15 | CACTAGTGGGCAGCATGAGA | ACATTGTCTACCCGGCCATA | 221 |
| chr10:50535175-50537207 | p2 | (AT)45 | CACTAGTGGGCAGCATGAGA | ACATTGTCTACCCGGCCATA | 221 |
| chr10:50597394-50599409 | p3 | (GAG)5 | CTTCTGCTTCCGTCCATCTC | GCTTCCTCTTCCCAATTTCC | 123 |
| chr10:50601299-50603365 | p2 | (TA)7 | TTTTATATCACCCGGGCAAA | CTGGCCAAAAGTCTCTCCTG | 298 |
| chr10:50606533-50608548 | p5 | (CATTG)3 | GGGATGCTTTGTGGGTAGAA | CTAAGCCCACACGGGTAAAA | 142 |
| chr10:50621069-50623083 | p2 | (AT)12 | GCATCCAAGGAAGGCAATTA | CCTGAACTGCGACTCGTACA | 293 |
| chr10:50621069-50623083 | p2 | (TA)10 | GCATCCAAGGAAGGCAATTA | CCTGAACTGCGACTCGTACA | 293 |
| chr10:50621218-50623242 | p2 | (AT)12 | GCATCCAAGGAAGGCAATTA | CCTGAACTGCGACTCGTACA | 144 |
| chr10:50621218-50623242 | p2 | (TA)10 | GCATCCAAGGAAGGCAATTA | CCTGAACTGCGACTCGTACA | 144 |
| chr10:50621538-50623558 | p2 | (AT)12 | CAGATCATGGCAGAAGCTCA | ATTGAGCCACGAACTCTGCT | 170 |
| chr10:50621538-50623558 | p2 | (TA)10 | CAGATCATGGCAGAAGCTCA | ATTGAGCCACGAACTCTGCT | 170 |
| chr10:50637383-50639397 | p2 | (TA)13 | TAAAGTCAAAATGGGGTGGC | AGTCAAGGTGAAATTTGGCG | 279 |
| chr10:50637383-50639397 | p2 | (AG)7 | TAAAGTCAAAATGGGGTGGC | AGTCAAGGTGAAATTTGGCG | 279 |
| chr10:50637383-50639397 | p2 | (GT)6 | TAAAGTCAAAATGGGGTGGC | AGTCAAGGTGAAATTTGGCG | 279 |
| chr10:50732265-50734281 | p2 | (TA)8 | CCCGTCGTACCCTTAAACAA | ATACTCGCTTTGGTGATCCG | 124 |
| chr10:50745825-50747840 | p3 | (AAT)5 | TTCCTTTCGTGGAACTGTCC | TCCTATTGCTCCTGTTTGGG | 278 |
| chr10:50786743-50788771 | p4 | (CATA)4 | GGTCTTCGGACATGGTGTTT | TCACAATAACAGGGCAACGA | 165 |
| chr10:50786743-50788771 | p2 | (TG)14 | GGTCTTCGGACATGGTGTTT | TCACAATAACAGGGCAACGA | 165 |
| chr10:50982409-50984423 | p2 | (AT)7 | AGTGAGGGACTCGGTGAATG | TTGTTGGCATATAGGGAGCC | 223 |
| chr10:51118555-51120570 | p5 | (ATTTC)3 | ATACCTGTGCCACGACCAAT | GTTGATGAGTTGGCCGAGTT | 250 |
| chr10:51143097-51145157 | p2 | (TA)30 | GGGGGTTCTCGACCTTATGT | GTGGTCATGTCTGGCCTTTT | 300 |
| chr10:51182250-51184276 | p2 | (CA)13 | CCGGACGAAATTGAAACAGT | GAAACACTCGAGCCTTTTGG | 234 |
| chr10:51214275-51216290 | p5 | (AATAA)3 | AACCGAAATACACGGTCCAG | CCTGTCTTCGTCGTGTCTCA | 239 |
| chr10:51239100-51241115 | p3 | (ATT)5 | AGACCCTGCAGAAGCACCTA | TGTCCCCTACTTGCAGGTTC | 300 |
| chr10:51413351-51415369 | p6 | (ATGTAT)3 | TTGCATAAAAAGGACCCGAC | CTCGGTGAACGGAATAAGGA | 287 |
| chr10:51599003-51601018 | p3 | (TGC)5 | ATACCCCTCCTCCATTACCG | TACAAGTAGCCCTCCCGATG | 117 |
| chr10:51603900-51605918 | p6 | (CCAATG)3 | TACTTGTTGCACCAATCCCA | TAGCAAATTTGGGATCCTCG | 237 |
| chr10:51730530-51732545 | p3 | (AAG)5 | TGTTCACCTCGATCCATTCA | CTGTTTGCACAAACGCACTT | 173 |
| chr10:51730530-51732545 | p2 | (TA)6 | TGTTCACCTCGATCCATTCA | CTGTTTGCACAAACGCACTT | 173 |
| chr10:51759294-51761309 | p3 | (CAA)5 | ATGTCCCCTTCGGGAATTAG | CTGAGTGTGCTATGGCCTCA | 282 |
| chr10:51768620-51770650 | p2 | (TG)15 | TGTCCTCCATGATGACTCCA | ATCGGAGTGACGTAAGGTCG | 155 |
| chr10:51795025-51797043 | p2 | (AT)9 | CTTAGGAGCTTCATCAGCGG | GGAATCGATAAAGAAGGGGC | 152 |
| chr10:51795025-51797043 | p2 | (AT)14 | CTTAGGAGCTTCATCAGCGG | GGAATCGATAAAGAAGGGGC | 152 |
| chr10:51795406-51797434 | p2 | (AT)9 | TTATAAACAGGAGGCTGCGG | CTATCTCTGCTGGTGGCTCC | 271 |
| chr10:51795406-51797434 | p2 | (AT)14 | TTATAAACAGGAGGCTGCGG | CTATCTCTGCTGGTGGCTCC | 271 |

| Marker ID | SSR type | SSR motif | FORWARD PRIMER (5'-3') | REVERSE PRIMER (5'-3') | size (bp) |
|---|---|---|---|---|---|
| chr10:51840811-51842823 | p2 | (TA)6 | AAGTTCACTCGGATTGTGCC | CAGCTCCAGTTCCTCTACCG | 206 |
| chr10:52025879-52027897 | p3 | (TTG)5 | TTGAAAAACATCGTGTGGGA | ACCGAAATTTCAGCAGCATC | 140 |
| chr10:52025879-52027897 | p6 | (TATATG)3 | TTGAAAAACATCGTGTGGGA | ACCGAAATTTCAGCAGCATC | 140 |
| chr10:52043275-52045291 | p4 | (GAAC)4 | ATTGAGCACGTCACTTTCCC | AGCCCACACAACAACAACAA | 250 |
| chr10:52055868-52057886 | p6 | (CAACAT)3 | TTATCCATAAGCTGCCCACC | AAATCGGAGGTTACCGACCT | 294 |
| chr10:52091654-52093666 | p4 | (CTTT)4 | CGTGTTAGCATTGGTGTTGG | GAGAATTCAACTCGGCCAAA | 144 |
| chr10:52091654-52093666 | p2 | (TA)6 | CGTGTTAGCATTGGTGTTGG | GAGAATTCAACTCGGCCAAA | 144 |
| chr10:52091654-52093666 | p2 | (TA)9 | CGTGTTAGCATTGGTGTTGG | GAGAATTCAACTCGGCCAAA | 144 |
| chr10:52108009-52110027 | p3 | (GTT)6 | ACGTTTTCGGGTGATCTGAC | TTACTGCCCAGCTTGACCTT | 175 |
| chr10:52119775-52121790 | p5 | (TTGCA)3 | TGTTGAACATGTGGTTGGCT | GTACGGGCATCAGACCCTAA | 257 |
| chr10:52153650-52155668 | p3 | (TCA)6 | ATGCATAAGCCTTTCGGCTA | TTCATGCCCACAGGTAAACA | 119 |
| chr10:52177287-52179299 | p2 | (CT)6 | TGAATCGCCATATCCTCTCC | CCTTCCAGTTGTTGGGTGAT | 140 |
| chr10:52221677-52223692 | p5 | (TTTCA)3 | GCAACGGATTTGAGAATGGT | TTCGTCCTCGAATGATTTCC | 223 |
| chr10:52273325-52275357 | p2 | (AT)16 | ATCCACGGCCTCAACACTAC | GAAATTTGTGCCTTGTGGGT | 208 |
| chr10:52301104-52303134 | p2 | (TA)15 | TTTTGTTTTTCCCAATCCCA | TCCATACATTGGGGCATTTT | 235 |
| chr10:52301104-52303134 | p2 | (TA)9 | TTTTGTTTTTCCCAATCCCA | TCCATACATTGGGGCATTTT | 235 |
| chr10:52306359-52308371 | p3 | (GTT)5 | TTAAGGCACTGGATTCCACC | AAATACGGGACGTAACAGCG | 105 |
| chr10:52479462-52481492 | p2 | (AT)15 | GGACACGACTTTGCAGACAA | CAGCCGATGTGTCATTATGC | 183 |
| chr10:52489458-52491470 | p2 | (CA)6 | TCAAATTTGGCTCGGATTTC | TTTACAGGGAAAATGCTGCC | 215 |
| chr10:52524033-52526045 | p2 | (AC)6 | ATTGGACGGATTGAGACTGG | GAAGCGGGAATTGAATTTGA | 174 |
| chr10:52658404-52660436 | p2 | (TA)16 | TTTGACCCTTTTCCGTTTTG | TTCCAACCTTTATGGGATGC | 213 |
| chr10:52658404-52660436 | p2 | (TA)9 | TTTGACCCTTTTCCGTTTTG | TTCCAACCTTTATGGGATGC | 213 |
| chr10:52686026-52688041 | p2 | (GA)6 | GCTTCAACTGGGAAGAGACG | CGATCCAAGCCCATTATGTT | 176 |
| chr10:52686085-52688103 | p2 | (GA)6 | GCTTCAACTGGGAAGAGACG | CGATCCAAGCCCATTATGTT | 117 |
| chr10:52884243-52886261 | p6 | (TGTGTA)3 | CGGGACACACAGTCTGCTAA | GATAGCACTCCCCAGGATGA | 140 |
| chr10:52907241-52909253 | p2 | (TA)23 | CTGCTCGTGCATGACTCAAT | TCCTTGGCACCATAATCACA | 174 |
| chr10:52907241-52909253 | p2 | (AT)6 | CTGCTCGTGCATGACTCAAT | TCCTTGGCACCATAATCACA | 174 |
| chr10:52926691-52928713 | p5 | (ATTCA)3 | GAGAATTCAACTCGGCCAAA | TTGAATTTGGAAGGTCTCGG | 175 |
| chr10:52926740-52928792 | p5 | (ATTCA)3 | GAGAATTCAACTCGGCCAAA | TTGAATTTGGAAGGTCTCGG | 126 |
| chr10:52938646-52940658 | p2 | (AT)8 | AAGTCTGAGCTCCGGAACAA | ATTCCGTACTGACCCCCTTT | 186 |
| chr10:52938646-52940658 | p2 | (TA)6 | AAGTCTGAGCTCCGGAACAA | ATTCCGTACTGACCCCCTTT | 186 |
| chr10:53094663-53096678 | p5 | (ATCAT)3 | GTGTCGCGAATGCAGTAAGA | GTGACTCCGTTTGCCCTTTA | 261 |
| chr10:53123177-53125192 | p3 | (CAA)5 | AAATCAATGAAGTCGGACGG | TGGGTTGCATTTAGTCTCCC | 122 |
| chr10:53179508-53181528 | p2 | (TA)10 | TCCATAATGATATCGGGGGA | TGCCAAAAGGTCATGGTGTA | 250 |
| chr10:53218912-53220924 | p2 | (GT)6 | TGACTCCTACACCGAGCCTT | CGGGTACTGACGAACGAAAT | 126 |
| chr10:53258698-53260714 | p4 | (GAAG)4 | GAAGGGGTTGAGGAAAGAGG | CAACCCTTTTCTTTTTGGCA | 118 |

| Marker ID | SSR type | SSR motif | FORWARD PRIMER (5'-3') | REVERSE PRIMER (5'-3') | size (bp) |
|---|---|---|---|---|---|
| chr10:53350934-53352966 | p2 | (AT)16 | CTTCAGAGCCCATCTTTTGG | AGCAGAAGAGCAGACTTGGC | 177 |
| chr10:53350934-53352966 | p2 | (TA)7 | CTTCAGAGCCCATCTTTTGG | AGCAGAAGAGCAGACTTGGC | 177 |
| chr10:53360109-53362129 | p5 | (TTTCA)4 | GATTCAAGATGAAGCCCCAA | GGCGAATTTTCCAAGGTGTA | 235 |
| chr10:53422864-53424916 | p2 | (AT)26 | GAAATCGCACGTGCATAAGA | GGTGTGACAACTCCGACCTT | 251 |
| chr10:53441911-53443923 | p2 | (AT)6 | TTGTTGTGGGTGTTGTCGTT | GGACATGCCAAAAGAAGGAA | 252 |
| chr10:53466356-53468376 | p4 | (AAGA)5 | GCCCAGGACAATAAGGTCAA | GATCTTTTGGGGGAGTGTGA | 267 |
| chr10:53469881-53471893 | p2 | (TA)6 | GGGGATGTGCCATATAATCG | TTGAAAGATTTCTGACCGGG | 270 |
| chr10:53471639-53473655 | p4 | (AAGA)4 | GTGGCCTAAGAAGACGAACG | CCACCGGAAGGTAGCATAAA | 292 |
| chr10:53482055-53484091 | p2 | (AT)18 | ACGAGGATGGTGTTGAGGAC | AGTAGCATCATACCCGACCG | 265 |
| chr10:53567955-53569973 | p3 | (ATC)6 | CTGCAACGTCTTGTCAGCAT | GCTCCAATCCCAACAAGGTA | 260 |
| chr10:53567955-53569973 | p5 | (CGACC)3 | CTGCAACGTCTTGTCAGCAT | GCTCCAATCCCAACAAGGTA | 260 |
| chr10:53588254-53590268 | p2 | (CT)7 | ATTCGCCTATCCGAATTTCC | TCGTTAGAATTTCGGGCATC | 157 |
| chr10:53588254-53590268 | p5 | (TTTTC)3 | ATTCGCCTATCCGAATTTCC | TCGTTAGAATTTCGGGCATC | 157 |
| chr10:53623097-53625115 | p6 | (TACGAA)3 | TGTGAGCATCCTATGGGACA | AGCACTCCGAACAAGGATGT | 159 |
| chr10:53673616-53675632 | p2 | (AT)8 | TCCTCCAACCCCTGTTACTG | CCCATACTTCGCCCACTTTA | 179 |
| chr10:53706273-53708285 | p2 | (AC)6 | TTGTAGATCGGGTTTCGAGG | AGCTAAGTCTCGGGGGTGTT | 180 |
| chr10:53913784-53915800 | p4 | (TTAT)4 | CAATCCTGCAAATTGTCCCT | AGAAACACAATGGTGAGGGG | 123 |
| chr10:53913784-53915800 | p2 | (AT)8 | CAATCCTGCAAATTGTCCCT | AGAAACACAATGGTGAGGGG | 123 |
| chr10:54001231-54003279 | p2 | (AG)6 | ACCAACGCAGAGTGTGTCAG | TCCGGATGTCACGATTTGTA | 179 |
| chr10:54001231-54003279 | p2 | (CT)24 | ACCAACGCAGAGTGTGTCAG | TCCGGATGTCACGATTTGTA | 179 |
| chr10:54012113-54014125 | p2 | (AG)6 | TTCTCGTAAGCGGAGCTTGT | TGTCTGAGGTTGCAGGTTTG | 207 |
| chr10:54057183-54059195 | p2 | (AT)6 | AGAAGACTGAGATGGGGCCT | AGAGCTTCAGCTGGCAAGAG | 132 |
| chr10:54151912-54153928 | p2 | (AT)10 | ATCGTCTTCGTATCATGGGC | CCCAACCTTACATTCGGAAA | 170 |
| chr10:54229499-54231517 | p6 | (CATATA)3 | TTTTCTAGGGTGTGATGGGC | GTCAGTCTGTAAGCGGCTCC | 171 |
| chr10:54248121-54250141 | p2 | (AC)10 | ACATACCTCTTTCCGCCCTT | TTCCCGGATTGTTGTTGATT | 283 |
| chr10:54354985-54356997 | p2 | (GT)6 | CAAGCGAAAGACGTGGTACA | TAACCTTTCGGGGAGGTCTT | 102 |
| chr10:54354985-54356997 | p3 | (TTG)5 | CAAGCGAAAGACGTGGTACA | TAACCTTTCGGGGAGGTCTT | 102 |
| chr10:54365077-54367101 | p2 | (AT)12 | TCATGACACTGAGGCAAAGC | GCTGGAGCCATTCTGTAAGG | 292 |
| chr10:54548970-54550988 | p6 | (CATATA)3 | TGCCCACATCAAAATTGAAA | CTGAGCTCCATTTCTTTCGG | 211 |
| chr10:54686038-54688056 | p6 | (CTTACA)3 | CACTGGGCAGAATTTCGTTT | CAAGGAGGAACATTGCCCTA | 218 |
| chr10:54704358-54706376 | p3 | (CAA)6 | CGTCACCCCCTGTCAACTAT | TGGCCTTTGTTGTGATGTGT | 150 |
| chr10:54776725-54778755 | p2 | (TA)7 | CCTTACTAGCCGGAACTCCC | GTGGGCAAGGTTTGTGTCTT | 103 |
| chr10:54776725-54778755 | p2 | (TA)15 | CCTTACTAGCCGGAACTCCC | GTGGGCAAGGTTTGTGTCTT | 103 |
| chr10:54879364-54881394 | p2 | (AT)15 | CGGAGGAAAGCATCAACAAT | CTCCGGAATTCTTTGGATCA | 104 |
| chr10:54985007-54987019 | p2 | (TA)6 | GGGTTGAAACACCTTGGAAA | CCACCTAGCCGAAAGGTGTA | 105 |
| chr10:55026040-55028052 | p2 | (TA)6 | ACTGTCACGACCCGATTAGG | TGTTCGAAAAACAAATGGCA | 180 |

| Marker ID | SSR type | SSR motif | FORWARD PRIMER (5'-3') | REVERSE PRIMER (5'-3') | size (bp) |
|---|---|---|---|---|---|
| chr10:55171922-55173956 | p2 | (TA)16 | AAATTGGAGTTTGTGGGCAG | TAACCACTGCCACGTGTTGT | 181 |
| chr10:55171922-55173956 | p2 | (TA)17 | AAATTGGAGTTTGTGGGCAG | TAACCACTGCCACGTGTTGT | 181 |
| chr10:55171922-55173956 | p2 | (AT)14 | AAATTGGAGTTTGTGGGCAG | TAACCACTGCCACGTGTTGT | 181 |
| chr10:55369628-55371646 | p5 | (TGAAT)3 | TGATTGATGATTTGGGCTGA | GCCTTTGAAAGAAAGGGACC | 219 |
| chr10:55369628-55371646 | p6 | (TATATG)3 | TGATTGATGATTTGGGCTGA | GCCTTTGAAAGAAAGGGACC | 219 |
| chr10:55370797-55372812 | p5 | (GACAA)3 | CAGGTTACGCTTTTACCCCA | TTATTTTGATGGCCGAGAGG | 186 |
| chr10:55377674-55379692 | p5 | (TGAAT)3 | TTAGATTCTCGGGGATGGTG | GCTTAATTGTCCAAGCTCGC | 240 |
| chr10:55377674-55379692 | p6 | (TATATG)3 | TTAGATTCTCGGGGATGGTG | GCTTAATTGTCCAAGCTCGC | 240 |
| chr10:55377674-55379692 | p2 | (TA)7 | TTAGATTCTCGGGGATGGTG | GCTTAATTGTCCAAGCTCGC | 240 |
| chr10:55477846-55479858 | p2 | (TA)6 | TTCAAGAACCTGTCCGCTCT | AATTGGATGGCATCAAGAGG | 164 |
| chr10:55477846-55479858 | p3 | (TTG)5 | TTCAAGAACCTGTCCGCTCT | AATTGGATGGCATCAAGAGG | 164 |
| chr10:55524858-55526874 | p4 | (ACAT)4 | TCTCGCCTACGAAAGGTGTT | ATATCCTGTCTGGTACGCCG | 126 |
| chr10:55531132-55533148 | p4 | (ACAT)4 | ACTTCTGTCACGACCCAACC | GTTAGCCTCATCGGCTTACG | 180 |
| chr10:55618023-55620055 | p6 | (TACATA)3 | CACATTTGTCACGACCCAAC | GTTATCCACGTGCCTCACCT | 206 |
| chr10:55618023-55620055 | p2 | (TA)16 | CACATTTGTCACGACCCAAC | GTTATCCACGTGCCTCACCT | 206 |
| chr10:55680339-55682351 | p2 | (GT)6 | CCTGCACCCACAGGTAATTT | TCGCACCAATACTTCCACAA | 183 |
| chr10:55737213-55739228 | p5 | (TGTGT)3 | GCATGTGACATAAATTGGCG | TTGCGAAACGAATGATTCAC | 187 |
| chr10:55737213-55739228 | p5 | (TGAAT)4 | GCATGTGACATAAATTGGCG | TTGCGAAACGAATGATTCAC | 187 |
| chr10:55834150-55836162 | p2 | (TA)6 | CAATCACAGATCGCATGTCC | CGTTCTGTATTCCGTCCGTT | 145 |
| chr10:55899678-55901694 | p2 | (AG)8 | AAGCTGGTGACAATGAACCC | CCTGTCCTCTTCTCGGTCTG | 223 |
| chr10:55944337-55946352 | p5 | (ATTTC)3 | ACACAGCGTCCACAAACAAA | ATGATGAACAACGCTAGGGG | 291 |
| chr10:55982495-55984531 | p2 | (TA)18 | GGACACGACTTTTGCAGACA | GAGAAGGGTCTTGGGTGTCA | 278 |
| chr10:56083508-56085530 | p4 | (TGTA)5 | TCCCGTGACATTATCGGTTT | CTGATCAGTGGGATGCAATG | 287 |
| chr10:56150578-56152596 | p6 | (AATATG)3 | AGGCCACACGTTCTGCTACT | AACTGACGGACCGCACTAAG | 224 |
| chr10:56202259-56204299 | p2 | (AT)20 | GCTCTTCTATGTTGGGCGAG | AGGGAAAAATCATAACCGCC | 113 |
| chr10:56261156-56263171 | p5 | (GTGTT)3 | TGGTGTGCAAATACCGAAGA | TCTCTTGTTGGGGTTAACGG | 116 |
| chr10:56266460-56268474 | p2 | (TG)7 | CCGTATCATTTGCATCATCG | TCTTCCGTATTTGCACACCA | 174 |
| chr10:56266460-56268474 | p3 | (TGT)5 | CCGTATCATTTGCATCATCG | TCTTCCGTATTTGCACACCA | 174 |
| chr10:56327885-56329897 | p2 | (TA)6 | ATTGGAAATTCGGGTTAGGC | CGTAAATCCGTTCACCTCGT | 227 |
| chr10:56358665-56360680 | p5 | (TGTGT)3 | GGTTGGGTGGGTCCTAAGAT | TTGTGAATTCCCCTTTTTGC | 283 |
| chr10:56400280-56402295 | p5 | (AAAAG)3 | ACCCCGGCTATGGTATTTTC | CTTCACCCCCTGTCAAGTGT | 151 |
| chr10:56429497-56431517 | p2 | (AT)10 | CCAACTCCACGGATTCAGTT | CTCCAGTTCCAATTCCTCCA | 173 |
| chr10:56443226-56445238 | p2 | (CA)6 | CCCCGAGACTCGAATCATAA | CGATGGCCACATTTATAGGG | 258 |
| chr10:56476698-56478740 | p2 | (TA)21 | TTCGTTGGCTAGATATGCCC | TCAATCCCCTTCTATGGTCG | 138 |
| chr10:56584690-56586706 | p2 | (TA)8 | TCTCGTCGCTTCGAGATTTT | CACCGGCAATCGTTTCTATT | 183 |
| chr10:56586758-56588776 | p6 | (ATATGT)3 | ATGTTGGATATGAGCCTCGG | GTCGGGTCCTATTTATGCGA | 165 |

| Marker ID | SSR type | SSR motif | FORWARD PRIMER (5'-3') | REVERSE PRIMER (5'-3') | size (bp) |
|---|---|---|---|---|---|
| chr10:56607428-56609448 | p2 | (TA)10 | ATAAGCCGCTGAGGCTAACA | TTATGGGCATAGAAGGGTGC | 235 |
| chr10:56682690-56684702 | p2 | (TA)6 | GGACTCGGAGGCATGAATAA | ATTGACGCTATTGGTAGCCG | 242 |
| chr10:56745810-56747824 | p2 | (AT)7 | TTTCCAATCTGGTCCGGTAG | GGTGATGAACTGGGCCTAAA | 111 |
| chr10:56848629-56850641 | p2 | (AT)6 | ATGCTCTTTACGCCTTTCCA | GTTGATTGTTTTGGCCGAGT | 294 |
| chr10:56856342-56858362 | p2 | (AT)10 | TTTTGTTTCTTCCGGTTTGC | TTGCTTAAAAGGGGTTGACG | 160 |
| chr10:56856342-56858362 | p2 | (TA)35 | TTTTGTTTCTTCCGGTTTGC | TTGCTTAAAAGGGGTTGACG | 160 |
| chr10:56856598-56858668 | p2 | (AT)10 | CGTCAACCCCTTTTAAGCAA | ACTCATTTGTGGTCAAGGGC | 140 |
| chr10:56856598-56858668 | p2 | (TA)35 | CGTCAACCCCTTTTAAGCAA | ACTCATTTGTGGTCAAGGGC | 140 |
| chr10:56868592-56870608 | p2 | (TA)31 | TTGTTTCGGCTGCTTTTTCT | AGGAAAATTTCGTTGGGCTT | 184 |
| chr10:56868592-56870608 | p4 | (TTAA)4 | TTGTTTCGGCTGCTTTTTCT | AGGAAAATTTCGTTGGGCTT | 184 |
| chr10:56879095-56881127 | p2 | (AT)20 | CCAAACGGCTTTATTTCCAA | TGTGGTTTCCACCGAAAAAT | 250 |
| chr10:56879095-56881127 | p2 | (AT)16 | CCAAACGGCTTTATTTCCAA | TGTGGTTTCCACCGAAAAAT | 250 |
| chr10:56887849-56889861 | p2 | (TA)6 | GCCTTGCTATGCTAACAGCC | AAGAAACGAATAGCGGGGTT | 152 |
| chr10:56887849-56889861 | p4 | (TTAA)4 | GCCTTGCTATGCTAACAGCC | AAGAAACGAATAGCGGGGTT | 152 |
| chr10:56888750-56890766 | p2 | (TA)6 | TCGGTTGCTTGCTTTCTTTT | TCGGCCCAGAATCATTTTAC | 176 |
| chr10:56888750-56890766 | p4 | (TTAA)4 | TCGGTTGCTTGCTTTCTTTT | TCGGCCCAGAATCATTTTAC | 176 |
| chr10:56951527-56953542 | p5 | (TATCA)3 | TGCACTGCCTATTTCACAGC | TTGGTCCAGCAGCCTATACC | 227 |
| chr10:57122309-57124324 | p5 | (AAAAT)3 | CAACGCAACAAACTGCTCAT | TTATACGCACCGACTTTCCC | 173 |
| chr10:57122309-57124324 | p2 | (GT)6 | CAACGCAACAAACTGCTCAT | TTATACGCACCGACTTTCCC | 173 |
| chr10:57141325-57143337 | p2 | (GT)6 | GAACAACAACCGGGAAGAAA | CACAATCATTTTTGGGGGAG | 113 |
| chr10:57178963-57180978 | p2 | (TA)8 | CCCGAATAAATCATGCTGCT | ACACACCTCAAATGGGGAAG | 195 |
| chr10:57178963-57180978 | p5 | (ATTTC)3 | CCCGAATAAATCATGCTGCT | ACACACCTCAAATGGGGAAG | 195 |
| chr10:57180895-57182915 | p4 | (ATAC)5 | GACCCTTATTTAGGGCCGAG | GTCAGTCTGTAAGCGGCTCC | 195 |
| chr10:57182426-57184438 | p2 | (CA)6 | TGATCGCAATAGTGTCCCAA | CTGCCGAAATTTGGTAGGAA | 200 |
| chr10:57209816-57211828 | p2 | (AT)6 | CTAAGGCAGAATGGCAGGAG | CATAAGCCTATAACCGGGCA | 183 |
| chr10:57224399-57226414 | p3 | (AAC)5 | GGGCTCAGTGTGGAAACATT | TCGTCTTGTCACTTGGAACG | 171 |
| chr10:57276411-57278435 | p2 | (TA)12 | GGCATTTTACAAGGAGCCAA | TTTGGGTGTCTTCTTTTGGG | 185 |
| chr10:57296204-57298222 | p2 | (TG)9 | TACAATCCGAGTGGCATCAA | GCATGACCATCTTCAACCCT | 127 |
| chr10:57303119-57305131 | p2 | (GT)6 | GTATGCCTTGGGAATCGAGA | ACAGACAGGTTCAACGACCC | 147 |
| chr10:57311254-57313266 | p2 | (GT)6 | GTTTCCTACGCCCACAGGTA | TTTTAGAACATTCGTCCCCG | 164 |
| chr10:57311254-57313266 | p3 | (GTT)5 | GTTTCCTACGCCCACAGGTA | TTTTAGAACATTCGTCCCCG | 164 |
| chr10:57313985-57316005 | p5 | (TGTCC)3 | GTCCAGTAAGGCCCATGAGA | CTTCACCCCCTGTCAAGTGT | 173 |
| chr10:57313985-57316005 | p2 | (TA)10 | GTCCAGTAAGGCCCATGAGA | CTTCACCCCCTGTCAAGTGT | 173 |
| chr10:57334354-57336366 | p2 | (AT)6 | TTTGACCCGAAAAACTCTGG | AGGATAGGGCCGTGAGAGAT | 112 |
| chr10:57447106-57449124 | p3 | (AAG)6 | AACTGCAAGAGAGGCCATGT | TTGTCCTCACAATTTTGCCA | 120 |
| chr10:57512193-57514211 | p2 | (AT)9 | ATCAGCTGCAGGGTGCTACT | CAAGAGTTCGTCCATGCTCA | 124 |

| Marker ID | SSR type | SSR motif | FORWARD PRIMER (5'-3') | REVERSE PRIMER (5'-3') | size (bp) |
|---|---|---|---|---|---|
| chr10:57512193-57514211 | p2 | (AT)7 | ATCAGCTGCAGGGTGCTACT | CAAGAGTTCGTCCATGCTCA | 124 |
| chr10:57512193-57514211 | p2 | (AT)6 | ATCAGCTGCAGGGTGCTACT | CAAGAGTTCGTCCATGCTCA | 124 |
| chr10:57554961-57556981 | p2 | (TA)10 | TGCATTAGGGATCCTTCCAG | CCTACGAACCTGGGCATCTA | 211 |
| chr10:57607539-57609559 | p2 | (TA)8 | GCGGGTTAACGATGATGAAT | CGTCCAAAGAAAACGGATGT | 291 |
| chr10:57607539-57609559 | p2 | (AT)10 | GCGGGTTAACGATGATGAAT | CGTCCAAAGAAAACGGATGT | 291 |
| chr10:57607539-57609559 | p3 | (CAA)6 | GCGGGTTAACGATGATGAAT | CGTCCAAAGAAAACGGATGT | 291 |
| chr10:57623656-57625690 | p2 | (AT)17 | AGGAGATGCTGTGCGAGTTT | AGCTGGGTCGTGACAAAAGT | 148 |
| chr10:57631264-57633296 | p2 | (AT)16 | CCCTTTGGTGGCATTCTAAA | CCCCTAGCTGGGTAGTGACA | 173 |
| chr10:57642724-57644744 | p2 | (TA)8 | GCGGGTTAACGATGATGAAT | CGTCCAAAGAAAACGGATGT | 291 |
| chr10:57642724-57644744 | p2 | (AT)10 | GCGGGTTAACGATGATGAAT | CGTCCAAAGAAAACGGATGT | 291 |
| chr10:57642724-57644744 | p3 | (CAA)6 | GCGGGTTAACGATGATGAAT | CGTCCAAAGAAAACGGATGT | 291 |
| chr10:57658841-57660875 | p2 | (AT)17 | AGGAGATGCTGTGCGAGTTT | AGCTGGGTCGTGACAAAAGT | 148 |
| chr10:57666449-57668481 | p2 | (AT)16 | CCCTTTGGTGGCATTCTAAA | CCCCTAGCTGGGTAGTGACA | 173 |
| chr10:57716107-57718119 | p3 | (CAA)6 | GGCCGAGCCTAATATGTTCA | TTGGGTTGCTGTGTTGATGT | 238 |
| chr10:57716107-57718119 | p2 | (CA)6 | GGCCGAGCCTAATATGTTCA | TTGGGTTGCTGTGTTGATGT | 238 |
| chr10:57716107-57718119 | p2 | (CT)11 | GGCCGAGCCTAATATGTTCA | TTGGGTTGCTGTGTTGATGT | 238 |
| chr10:57731147-57733159 | p5 | (TTCAA)4 | GCCAGGTTCCAACTCAACTC | TGTCGTCGTTGTGGGTTTTA | 215 |
| chr10:57731147-57733159 | p2 | (TA)6 | GCCAGGTTCCAACTCAACTC | TGTCGTCGTTGTGGGTTTTA | 215 |
| chr10:57774947-57776959 | p2 | (TA)6 | ACACATATGGCAACAAGCCA | AGCATGTTACGCCTTCGACT | 207 |
| chr10:57774947-57776959 | p2 | (AT)25 | ACACATATGGCAACAAGCCA | AGCATGTTACGCCTTCGACT | 207 |
| chr10:57926524-57928542 | p6 | (AAAGAG)3 | AAATATGCAAGCCGACAAGG | TGCTCCGGCTAGACTTCTGT | 249 |
| chr10:58107958-58109990 | p2 | (AC)16 | ACTTACCCGAATTCATCCCC | GATCATAATCGTATGGGCCG | 119 |
| chr10:58208935-58210971 | p2 | (AT)18 | CCCGTTTCGTCAAAAACCTA | TGTATTTATAGGGGCGGTGC | 120 |
| chr10:58484304-58486316 | p2 | (TG)6 | AAAGGTGCCTACGGGCTATT | CTCATACCTGGCCCTACCAA | 235 |
| chr10:58519578-58521590 | p2 | (TA)8 | AAGGACAGTGCAAGCCAACT | TTTGGTAAGCCACCCTTTTG | 164 |
| chr10:58519578-58521590 | p2 | (AT)6 | AAGGACAGTGCAAGCCAACT | TTTGGTAAGCCACCCTTTTG | 164 |
| chr10:58646294-58648312 | p6 | (TATCAT)3 | CCTCTCGGCCATCAAAATAA | GCACTCCATTGTCTGGAGGT | 144 |
| chr10:58692736-58694751 | p5 | (TTATC)3 | GGAGCAGGAACAGTCTGAGG | AGGCTTCCATTCCCTTCCTA | 115 |
| chr10:58729631-58731646 | p4 | (AATT)4 | CCGAAAATAATTCCGCCTTT | GAATGTTAAGTCGTCGGGGA | 281 |
| chr10:58729631-58731646 | p5 | (TTTCA)3 | CCGAAAATAATTCCGCCTTT | GAATGTTAAGTCGTCGGGGA | 281 |
| chr10:58765891-58767903 | p2 | (AT)6 | GTTCTGGATGGCTGGAATGT | CCAGATTCCGATGTAGCGAT | 277 |
| chr10:58777687-58779699 | p2 | (TG)6 | ATGTTATGAGCTTCGCCCAC | TCGTCAGCTAGGCTTGGAGT | 229 |
| chr10:58834287-58836302 | p6 | (TTAACA)3 | TTGTGCTGGTGGTGATTGAT | TCATTTACGGCCCACTTAGG | 171 |
| chr10:58834287-58836302 | p5 | (TATCA)3 | TTGTGCTGGTGGTGATTGAT | TCATTTACGGCCCACTTAGG | 171 |
| chr10:58872666-58874680 | p2 | (AT)7 | CTTTTGTCACGACCCCAACT | ATGTTTATGGCGGCTCTGTC | 212 |
| chr10:59065039-59067053 | p2 | (AT)7 | ACGACAAAGCCATAGTTGCC | AGGCATGGTGGAATCATAGC | 194 |

| Marker ID | SSR type | SSR motif | FORWARD PRIMER (5'-3') | REVERSE PRIMER (5'-3') | size (bp) |
|---|---|---|---|---|---|
| chr10:59091464-59093488 | p2 | (AT)12 | ACTGTTAACACCCGCTGTCC | GTGGTGATGAATGATGGCAG | 105 |
| chr10:59212970-59214986 | p2 | (AT)8 | GTGCTGTCCGGATTGGTATT | TTACGGGGAGATTCAAAACG | 283 |
| chr10:59218836-59220848 | p2 | (TA)6 | TCTTGGGAGCATTCCATTTC | CTTCATGGTCGGGTATGCTT | 168 |
| chr10:59236743-59238799 | p2 | (TA)28 | ATTTTTGACGACCTTGTCGG | CCCTAGGACGAATTGCTCTG | 169 |
| chr10:59307394-59309410 | p2 | (AT)8 | CCCTTCTACAGGTTGTCCCA | GCCACGACTAGGCAGGTAAG | 109 |
| chr10:59307394-59309410 | p3 | (CAT)6 | CCCTTCTACAGGTTGTCCCA | GCCACGACTAGGCAGGTAAG | 109 |
| chr10:59340148-59342168 | p2 | (AT)10 | ATAAGGCAACCCGAAACCTT | GTGGGTTGATTTGGAGGCTA | 287 |
| chr10:59340148-59342168 | p5 | (TTTTC)3 | ATAAGGCAACCCGAAACCTT | GTGGGTTGATTTGGAGGCTA | 287 |
| chr10:59340292-59342307 | p2 | (AT)10 | ATAAGGCAACCCGAAACCTT | GTGGGTTGATTTGGAGGCTA | 143 |
| chr10:59340292-59342307 | p5 | (TTTTC)3 | ATAAGGCAACCCGAAACCTT | GTGGGTTGATTTGGAGGCTA | 143 |
| chr10:59387644-59389656 | p2 | (GT)6 | TACACCATGAGGCCAGTTCA | CCTTCCTCAATATTTCCGCA | 259 |
| chr10:59493879-59495891 | p2 | (AT)6 | TTTGACCCGAAGAATTCCAG | CATAACGAGGCCCCATCTAA | 263 |
| chr10:59493879-59495891 | p5 | (ATCAT)3 | TTTGACCCGAAGAATTCCAG | CATAACGAGGCCCCATCTAA | 263 |
| chr10:59497084-59499099 | p5 | (AAATT)3 | CCTTGAATCCAAGCCATGTT | GACCCGAATTTGACCCACTA | 252 |
| chr10:59536616-59538631 | p5 | (AAATT)3 | CCAATTTCGTCCAATCCAAT | GATTTGCAAGCTCAAGGAGG | 242 |
| chr10:59547846-59549871 | p5 | (TCATA)5 | TTGCCCACTAAGGAATGAGG | GATCCTGCCATCCTGTCAAT | 127 |
| chr10:59548995-59551010 | p5 | (AATTC)3 | AGCCCAAGGAAAGTCATGTG | GTTCCTTCAAGGCGTGACAT | 190 |
| chr10:59548995-59551010 | p5 | (ACACA)3 | AGCCCAAGGAAAGTCATGTG | GTTCCTTCAAGGCGTGACAT | 190 |
| chr10:59729754-59731772 | p2 | (TA)9 | AGAAAAATGGTGGATGTGGC | CTAGAGAAAATTGGGCAGCG | 194 |
| chr10:59857227-59859241 | p2 | (AT)7 | GAAAGGTGAGGCGCTATGAG | CCGATCCATCACCCAATATC | 278 |
| chr10:59857227-59859241 | p5 | (ATTCA)3 | GAAAGGTGAGGCGCTATGAG | CCGATCCATCACCCAATATC | 278 |
| chr10:59881986-59884001 | p5 | (TTCAT)3 | CCTTGGGTGGAAAAACTTCA | ATTCACTGTGCATCAACGGA | 272 |
| chr10:59953511-59955529 | p3 | (CAA)6 | GGTCTCGGCGAACGATATAA | CCGAAACTTGTTTCTGAGCC | 217 |
| chr10:59990532-59992560 | p2 | (TA)14 | CTTCACGTTACACCGTCCCT | GCTCGAGTTTACAGGGCTTG | 130 |
| chr10:60064123-60066153 | p2 | (AT)15 | GATCCTCTGCTGGCTTTGAC | GACAGGGCTATGAGAGACGC | 126 |
| chr10:60064123-60066153 | p2 | (AT)15 | GATCCTCTGCTGGCTTTGAC | GACAGGGCTATGAGAGACGC | 126 |
| chr10:60065086-60067116 | p2 | (AT)15 | GCGTCTCTCATAGCCCTGTC | CAGTCAGACCCGGAGGATTA | 274 |
| chr10:60065086-60067116 | p2 | (AT)15 | GCGTCTCTCATAGCCCTGTC | CAGTCAGACCCGGAGGATTA | 274 |
| chr10:60163093-60165119 | p2 | (AT)13 | TGGGAACTTGCAAAAACTCC | GGCTTGCGTATCGTATTGGT | 267 |
| chr10:60232355-60234370 | p5 | (TCATT)3 | AGTTGCCACAAGAGTTGGCT | ACTTTCTGACTCCTCCGCAA | 253 |
| chr10:60287079-60289094 | p5 | (TCATT)3 | AGTTGCCACAAGAGTTGGCT | ACTTTCTGACTCCTCCGCAA | 253 |
| chr10:60339480-60341495 | p3 | (GTT)5 | CACCAAAAGAGAGTTTCGGC | CTTAAGCGTTTCCGACTTGC | 198 |
| chr10:60383704-60385750 | p2 | (TA)23 | TCTTAAAATGCCCCGTTTTG | GCTGAGGTGTTATGTCCCGT | 257 |
| chr10:60457777-60459792 | p5 | (AAATG)3 | TGCCTCACCTTGAAAGGACT | ACGACGTGGTATCGAAGGAG | 212 |
| chr10:60475842-60477880 | p2 | (TA)9 | CAAGGTGAGGCAGATGTTCA | ATCGTTATATGCCCGCTTTG | 183 |
| chr10:60475842-60477880 | p2 | (TA)19 | CAAGGTGAGGCAGATGTTCA | ATCGTTATATGCCCGCTTTG | 183 |

| Marker ID | SSR type | SSR motif | FORWARD PRIMER (5'-3') | REVERSE PRIMER (5'-3') | size (bp) |
|---|---|---|---|---|---|
| chr10:60486304-60488354 | p2 | (AT)25 | TGTAATGACGGCAAATGGAA | TTCTCGTAGCGAGTCCCACT | 277 |
| chr10:60629165-60631197 | p2 | (AT)16 | GAATGGGGAAAATGGGAAGT | TAATACTTCGCGGCCATTTC | 227 |
| chr10:60657568-60659582 | p2 | (AT)7 | ATCTCTTGGCTGAGCATCGT | GAGTTCAAGCGGCAAGTAGG | 275 |
| chr10:60712680-60714695 | p5 | (TTTCC)3 | GGTCCTGAACTGCAACCAAT | TGGACCAACGCATTAGCATA | 105 |
| chr10:60961128-60963140 | p2 | (TG)6 | ACGACATGTTGAGTCCGACA | TCGGAAACCGTAACACATCA | 118 |
| chr10:60962982-60965000 | p5 | (ATGAA)3 | TGGAAGAATGGTGGAGAAGG | GGTGTGACAATCACAGACGG | 108 |
| chr10:60962982-60965000 | p6 | (CATATA)3 | TGGAAGAATGGTGGAGAAGG | GGTGTGACAATCACAGACGG | 108 |
| chr10:60962982-60965000 | p5 | (ATTCA)3 | TGGAAGAATGGTGGAGAAGG | GGTGTGACAATCACAGACGG | 108 |
| chr10:60970963-60973121 | p2 | (AT)7 | CTTCGACCCGAAGAACTCTG | AGTGACCGAGGCCATACATC | 294 |
| chr10:60971133-60973151 | p2 | (AT)7 | CTTCGACCCGAAGAACTCTG | AGTGACCGAGGCCATACATC | 124 |
| chr10:60971563-60973577 | p2 | (AT)7 | TTGACACGGCTCGTAGACAC | GACCAAAAAGGCTTGTGCAT | 161 |
| chr10:60984712-60986724 | p2 | (AT)6 | AGTCCCTTCATCGCTTAGCA | TATTGCAGGGGAAACTCTGG | 132 |
| chr10:61101630-61103672 | p2 | (AT)21 | CACCACTGCAGTTGGGTATG | TGAAGGATCGGTGTTTCACA | 257 |
| chr10:61153656-61155698 | p2 | (AT)21 | CACCACTGCAGTTGGGTATG | TGAAGGATCGGTGTTTCACA | 257 |
| chr10:61375683-61377709 | p2 | (AT)7 | TTTGGGCAAAAAGAGGAATG | AATAACAGTCAGCGGGTTCG | 119 |
| chr10:61375683-61377709 | p4 | (AAAG)4 | TTTGGGCAAAAAGAGGAATG | AATAACAGTCAGCGGGTTCG | 119 |
| chr10:61463423-61465437 | p2 | (GT)7 | GGGGCTGTTTTGATTACGAA | ATACGCGAAAACACGGGATA | 127 |
| chr10:61575893-61577909 | p2 | (AT)8 | CTCCGGAAGTTCATAGCGTC | CTGTCCCGTCATATTTCGCT | 218 |
| chr10:61759869-61761893 | p3 | (TAA)8 | AAAAATGGGCGATGTCGTAG | ATTTGGGGGTGGAGTAAAGG | 289 |
| chr10:61761514-61763529 | p3 | (ATT)5 | TATGCAGTGGCAGGTCTGAG | AAGCAATGTGTTTCCCTTGC | 123 |
| chr10:61807859-61809871 | p2 | (AC)6 | GTTTCGAGGAGCAGAAATGC | AGCTAAGTCTCGGGGGTGTT | 204 |
| chr10:61810185-61812200 | p2 | (AT)9 | GAAAGGGGAAGGCGATAAAG | AAGACAACATCCATCCCAGC | 209 |
| chr10:61810185-61812200 | p5 | (ATTGC)3 | GAAAGGGGAAGGCGATAAAG | AAGACAACATCCATCCCAGC | 209 |
| chr10:61834150-61836162 | p2 | (TG)6 | TCCGTTCGTGCAATTCATTA | CAAACGGAACCCAGCACTAT | 189 |
| chr10:61834150-61836162 | p5 | (GGGTT)4 | TCCGTTCGTGCAATTCATTA | CAAACGGAACCCAGCACTAT | 189 |
| chr10:61859057-61861075 | p2 | (AT)9 | GACCTGCACTTCCTCCCATA | TCCGACAGAGGAGCTCAGTT | 175 |
| chr10:61917150-61919171 | p3 | (AAC)7 | ACAGCGTCCAAAGAAAATGG | TTATTATCACACCGTGCCGA | 103 |
| chr10:61936753-61938795 | p2 | (AT)21 | CGTAGCTTTCCTCTTCACCG | AGCAGTTCGTCCTAGGGGTT | 156 |
| chr10:62054925-62056949 | p6 | (TACATA)4 | CCGAAAATTCCGAGGTGTAA | GCCGGCCTTGTTATGACTTA | 224 |
| chr10:62054925-62056949 | p2 | (AC)12 | CCGAAAATTCCGAGGTGTAA | GCCGGCCTTGTTATGACTTA | 224 |
| chr10:62135505-62137520 | p5 | (ATTTT)3 | GACTGACTTTTTGGTGGGGA | TTTCCCATTTTCCAACGAAG | 225 |
| chr10:62148879-62150935 | p5 | (AGGAC)3 | ATGGGAAGCTTCAGCAAGAA | CTGCTTGTTTAGAGGGGCAC | 132 |
| chr10:62215496-62217508 | p2 | (AT)6 | TTGACTTACGGTCCGTCCTC | TGCAATTGTCCTTCAGCTTG | 284 |
| chr10:62318668-62320712 | p2 | (AT)22 | ACTCATGAAATCACGGGCTC | AGGTCAACAAATGACCGAGG | 296 |
| chr10:62324605-62326623 | p2 | (AT)9 | CCGTCTCAATTTGACGGTTT | ACTCCTCGGGAGACAGGTTT | 164 |
| chr10:62333843-62335855 | p2 | (AT)6 | TGTCACGACCCAACTCCATA | ACTCCTCGGGAGACAGGTTT | 255 |

| Marker ID | SSR type | SSR motif | FORWARD PRIMER (5'-3') | REVERSE PRIMER (5'-3') | size (bp) |
|---|---|---|---|---|---|
| chr10:62443456-62445484 | p2 | (AT)14 | TTCGGCCACAACATCAATAA | AATGACCCCATGGAAATTGA | 177 |
| chr10:62451035-62453085 | p2 | (AT)25 | CTTTTCACCTTCTTCCGTCG | GGGCATCTAGGGATTGTTGA | 157 |
| chr10:62482056-62484071 | p3 | (GTT)5 | AAGACGCTCTTCGACGTGAT | TAAAGGCCATGACGACACAA | 297 |
| chr10:62576726-62578741 | p5 | (TTCAT)3 | AGAATGACTCAGGAAGCGGA | TCACAAGGCCACAACACAAT | 170 |
| chr10:62742535-62744550 | p5 | (TATCA)3 | GAGCACAAACCAAGCAGACA | CGGAGCCCATACTTTTGGTA | 189 |
| chr10:62753322-62755340 | p3 | (TTG)6 | AATGCTTCTGTTACACCCCG | TTCGTCCCTGAAGTTTGACC | 130 |
| chr10:62753322-62755340 | p6 | (TATATG)4 | AATGCTTCTGTTACACCCCG | TTCGTCCCTGAAGTTTGACC | 130 |
| chr10:62790808-62792823 | p5 | (CATAT)3 | ACTTTTGTCACGACCCAACC | CCGTGCGCTACATCTGTTTA | 287 |
| chr10:62813367-62815435 | p2 | (AT)21 | GCTAGCCTACCAAGCACCAC | CTCTCCCAGCCACGTTATGT | 164 |
| chr10:62813367-62815435 | p2 | (AT)34 | GCTAGCCTACCAAGCACCAC | CTCTCCCAGCCACGTTATGT | 164 |
| chr10:62813367-62815435 | p2 | (AT)35 | GCTAGCCTACCAAGCACCAC | CTCTCCCAGCCACGTTATGT | 164 |
| chr10:62813367-62815435 | p2 | (CA)7 | GCTAGCCTACCAAGCACCAC | CTCTCCCAGCCACGTTATGT | 164 |
| chr10:62825403-62827418 | p3 | (CAG)5 | CACTCGAAACCAGCTCATCA | CCGAACTCTCAGGCTTTGTC | 206 |
| chr10:62976262-62978276 | p2 | (CA)7 | ACATGCCAAAGGAAGGAGTG | TTGTGGGCTGTGTTGTTGAT | 249 |
| chr10:62991340-62993388 | p2 | (TA)24 | ACACACGATCAGGAAAAGGG | AGTTCGTCCTAGGGCTGTCA | 161 |
| chr10:63001672-63003688 | p4 | (AAAG)4 | ATGGGATCAACTGCTGGAAC | TCATTCTCTCACAGATGGCG | 164 |
| chr10:63001672-63003688 | p4 | (AAAG)4 | ATGGGATCAACTGCTGGAAC | TCATTCTCTCACAGATGGCG | 164 |
| chr10:63082005-63084043 | p2 | (AT)19 | TGTCTACCTGTGGGCATGAA | TACCTATATGGACGGGCAGC | 195 |
| chr10:63090093-63092105 | p2 | (TA)6 | TCCGGCGAATGTAGATAACC | ACCAGTTGTCCCTATGGCAG | 276 |
| chr10:63090093-63092105 | p3 | (ATC)6 | TCCGGCGAATGTAGATAACC | ACCAGTTGTCCCTATGGCAG | 276 |
| chr10:63163187-63165199 | p2 | (CA)6 | AGGAACAGTCTCCGAAGCAA | TGTTCCAGGTTACTCCCAGG | 242 |
| chr10:63241669-63243684 | p5 | (TTATT)3 | CCCTATCTACCACCGAACGA | CCCAGGCAAGAGGACTATCA | 271 |
| chr10:63292461-63294479 | p6 | (TGTGAT)3 | GGACAGCTATGTACGGCGTT | AGTATCTTGCCCGAGCTTCA | 184 |
| chr10:63375022-63377037 | p3 | (GTA)5 | AGGTGTGGCAAGAGAGAGGA | TCCGTATATCACCCCTCCAA | 214 |
| chr10:63463985-63466000 | p5 | (TTCAT)3 | GAATCATACTTGCCTCGGGA | AATCGAACACCAAAAGCCTG | 171 |
| chr10:63479910-63481922 | p2 | (TA)6 | AAACTCATGAAATCACGGGC | TCCTCCACTGGTTTCTTTGG | 274 |
| chr10:63648471-63650483 | p2 | (GA)6 | TCGGCATATCACTACCCTCC | ACCTGGAATGATGAGAACCG | 148 |
| chr10:63688083-63690098 | p3 | (AAG)5 | ACCTACTCTCCACCCACCCT | GTTGGTTGGTGCTAATGGCT | 129 |
| chr10:63702937-63704952 | p5 | (AGGAA)3 | CTGCCAACCATTTCTTTGGT | TCTATCGACTGACCGCTGTG | 227 |
| chr10:63769385-63771399 | p2 | (AT)7 | AGGCGAGCCACATAGCTAAA | TGTTAGCCTCGTCGGCTTAT | 162 |
| chr10:63792174-63794186 | p3 | (ACA)5 | TCGATTTCCGAGAGCAAGTT | AGCTAAGTCTCGGGGGTGTT | 195 |
| chr10:63792174-63794186 | p2 | (AC)6 | TCGATTTCCGAGAGCAAGTT | AGCTAAGTCTCGGGGGTGTT | 195 |
| chr10:63907342-63909354 | p6 | (AAGTTG)3 | GACAAGATTCGAGCCCCATA | TCAAATCCATTCCCAACCAT | 281 |
| chr10:63907342-63909354 | p2 | (TA)6 | GACAAGATTCGAGCCCCATA | TCAAATCCATTCCCAACCAT | 281 |
| chr10:63907342-63909354 | p2 | (TA)6 | GACAAGATTCGAGCCCCATA | TCAAATCCATTCCCAACCAT | 281 |
| chr10:63907342-63909354 | p6 | (ATGTAT)3 | GACAAGATTCGAGCCCCATA | TCAAATCCATTCCCAACCAT | 281 |

| Marker ID | SSR type | SSR motif | FORWARD PRIMER (5'-3') | REVERSE PRIMER (5'-3') | size (bp) |
|---|---|---|---|---|---|
| chr10:64076825-64078847 | p2 | (TA)8 | CCTATCACTCCCACAATGCC | TAAGGGAGAAAAGGGGAGGA | 227 |
| chr10:64076825-64078847 | p2 | (AT)11 | CCTATCACTCCCACAATGCC | TAAGGGAGAAAAGGGGAGGA | 227 |
| chr10:64125620-64127632 | p2 | (TA)10 | CTTTGTTCACCGAGTCCCAT | GAAGAAAGGCGGTCAGTACG | 130 |
| chr10:64125620-64127632 | p2 | (GT)6 | CTTTGTTCACCGAGTCCCAT | GAAGAAAGGCGGTCAGTACG | 130 |
| chr10:64195160-64197174 | p2 | (TA)7 | ATGGTGTTGGTGGAATGGTT | CAAGCCAATATCAACGCTCA | 132 |
| chr10:64213747-64215761 | p2 | (AC)6 | CACACATGACCCATTACCCA | ATGCCCGAAGGTACAATGAG | 222 |
| chr10:64213747-64215761 | p2 | (AT)7 | CACACATGACCCATTACCCA | ATGCCCGAAGGTACAATGAG | 222 |
| chr10:64221473-64223491 | p2 | (AT)9 | GATCGGAATACACATCGCCT | TGACTATTTGGGGTAACGGC | 125 |
| chr10:64234049-64236101 | p2 | (TA)26 | CGACCTTACGTCACTCCGAT | GTAACATCATTAGCCCGCGT | 269 |
| chr10:64234049-64236101 | p2 | (AT)7 | CGACCTTACGTCACTCCGAT | GTAACATCATTAGCCCGCGT | 269 |
| chr10:64234591-64236605 | p2 | (TA)26 | GTCCTGTCCCGTCCTTATGA | AACCAATTTCCGAATCTCCC | 263 |
| chr10:64234591-64236605 | p2 | (AT)7 | GTCCTGTCCCGTCCTTATGA | AACCAATTTCCGAATCTCCC | 263 |
| chr10:64234591-64236605 | p3 | (TTG)5 | GTCCTGTCCCGTCCTTATGA | AACCAATTTCCGAATCTCCC | 263 |
| chr10:64287307-64289322 | p3 | (ACA)5 | ATTCTGCCTACTCGGGAGGT | TATCTATGCCTGACCCTCGG | 178 |
| chr10:64293207-64295222 | p2 | (TA)33 | CAATTATGGCCGGGTAGAGA | CCCCAGGAAAAGAGACATCA | 284 |
| chr10:64293207-64295222 | p5 | (TAATT)3 | CAATTATGGCCGGGTAGAGA | CCCCAGGAAAAGAGACATCA | 284 |
| chr10:64383948-64385966 | p5 | (TTTTA)3 | TATGGCATGTAGGCAATGGA | CGATGGTCGCTAACCGTACT | 193 |
| chr10:64383978-64385996 | p5 | (TTTTA)3 | TATGGCATGTAGGCAATGGA | CGATGGTCGCTAACCGTACT | 163 |
| chr10:64384098-64386113 | p5 | (TTTTA)3 | AGTACGGTTAGCGACCATCG | AATCAAACCACTTTGCCAGC | 262 |
| chr10:64616948-64618966 | p6 | (CATATA)3 | TCTGATACCAACTAGGGCCG | TCATAACCTTGCTTGTCCCC | 139 |
| chr10:64688346-64690361 | p5 | (TTCAT)3 | TTCATTTTCCTTTTTCCCCC | GGTGTGGGAATAAGCCTTGA | 107 |
| chr10:64688346-64690361 | p3 | (ATG)5 | TTCATTTTCCTTTTTCCCCC | GGTGTGGGAATAAGCCTTGA | 107 |
| chr10:64688346-64690361 | p2 | (AT)16 | TTCATTTTCCTTTTTCCCCC | GGTGTGGGAATAAGCCTTGA | 107 |
| chr10:64688346-64690361 | p2 | (AT)44 | TTCATTTTCCTTTTTCCCCC | GGTGTGGGAATAAGCCTTGA | 107 |
| chr10:64708640-64710664 | p3 | (AAT)8 | GTGGTTTACGGTCGCAGTTT | CCAGTGGGCCGTAAAATAGA | 285 |
| chr10:64755279-64757291 | p2 | (CT)6 | CCCAAACTCAGCAACACAGA | ATCCTGATCGTTGATTTCGC | 167 |
| chr10:64756274-64758286 | p2 | (TG)6 | GCCAATCATGTAAGTGGGCT | CATGGTTAAAGCAATTGGGG | 157 |
| chr10:64756274-64758286 | p2 | (CT)6 | GCCAATCATGTAAGTGGGCT | CATGGTTAAAGCAATTGGGG | 157 |
| chr10:64860303-64862318 | p5 | (CACAA)3 | AGCAGAGTTACCCCCGAGAT | GTTACTGGACGAAACGGGAA | 177 |
| chr10:64913281-64915315 | p2 | (TA)17 | ATAATGCAATTGGGGGATCA | TCTGGCCATGGTGTATCGTA | 279 |
| chr10:65045733-65047745 | p2 | (TA)7 | CCGAAGGCCATAACTAGTGC | GTCGCCTTGCGTATCATTTT | 135 |
| chr10:65045733-65047745 | p2 | (AT)6 | CCGAAGGCCATAACTAGTGC | GTCGCCTTGCGTATCATTTT | 135 |
| chr10:65359716-65361728 | p2 | (AG)6 | TCCACGAGGACTACCAAACC | CCTGAGTCCGGGACAAAGTA | 114 |
| chr10:65460127-65462145 | p6 | (CATATA)3 | ATTTTGTCACGACACGACCA | AGATGTTTATGGCACCCTCG | 203 |
| chr10:65502874-65504886 | p2 | (AT)6 | TTCCAATCCCTCTGGAAGTG | GCCAAGACCCACAACTCCTA | 156 |
| chr10:65545689-65547704 | p5 | (CTTTG)3 | TTTTTGGACGTCGGAAAGTC | TGAGCAACGAAGAAGAACGA | 260 |

| Marker ID | SSR type | SSR motif | FORWARD PRIMER (5'-3') | REVERSE PRIMER (5'-3') | size (bp) |
|---|---|---|---|---|---|
| chr10:65629268-65631280 | p2 | (TA)6 | TAACTTCGTGCTTTGCCCTT | GCTACCGGTTGAGATTGGAA | 257 |
| chr10:65636687-65638703 | p4 | (TTTG)4 | CTGGGGTGTCGAGTTTGTTT | CTCCACCTCCACCCGTACTA | 235 |
| chr10:65659954-65661972 | p6 | (ATACAT)3 | TAGTTCCCAAGCTGGACACC | TGCGTCTAAGCGGTCCTATT | 181 |
| chr10:65659954-65661972 | p3 | (CAT)5 | TAGTTCCCAAGCTGGACACC | TGCGTCTAAGCGGTCCTATT | 181 |
| chr10:65738440-65740458 | p3 | (ATG)6 | TCCAACATCCATCCACAAGA | GGAGTTTCCCCTCTAGCACC | 216 |
| chr10:65838972-65840987 | p5 | (AAAGG)3 | ATATTGTGGCCACTGAAGGC | ATATACCCGCGCGATCAATA | 234 |
| chr10:65923891-65925909 | p2 | (AT)9 | TTTCATTTCCTATGGCTGGG | AGGTACGTATACGGGGGTCC | 137 |
| chr10:66017171-66019183 | p2 | (CT)6 | AAACATTCCAATCAGCCCAG | ACGTTCGTGAGATTATCGGG | 125 |
| chr10:66131131-66133151 | p2 | (AT)10 | ATATAGGGGAAATGCTGCCC | TCGTAAATCCGTAGCCCATC | 254 |
| chr10:66138444-66140460 | p2 | (AT)7 | CATTATGAATGGGTACGCCC | CCGCCAACTTAGGTCATCAT | 290 |
| chr10:66138444-66140460 | p2 | (AT)8 | CATTATGAATGGGTACGCCC | CCGCCAACTTAGGTCATCAT | 290 |
| chr10:66145425-66147445 | p2 | (AT)7 | AATCCTAAGCAAGCCGACAA | TTTGACTTAAGTTTGGGGCG | 172 |
| chr10:66145425-66147445 | p2 | (AT)10 | AATCCTAAGCAAGCCGACAA | TTTGACTTAAGTTTGGGGCG | 172 |
| chr10:66268869-66270881 | p2 | (AG)6 | TGGAAGCAAAGAAAGTCGCT | TTGCTGCTCTCAACTGCCTA | 192 |
| chr10:66307642-66309662 | p2 | (AT)10 | CTGAGGGAGTGTAGCCTTCG | ATGCACTGGAAAACCTTTCG | 145 |
| chr10:66407706-66409722 | p2 | (AT)8 | CTATTTTTGCACCGCTCACA | TTCCAATCCCTCTGGAAGTG | 147 |
| chr10:66411611-66413623 | p2 | (TA)6 | CGGGGATGTGCCATATAATC | TTGAAAGATTTCTGACCGGG | 256 |
| chr10:66501338-66503356 | p2 | (TA)9 | TTCGCCAAAATTGACCTTTC | ACGGGATGATCAATGGATGT | 131 |
| chr10:66501338-66503356 | p2 | (AT)17 | TTCGCCAAAATTGACCTTTC | ACGGGATGATCAATGGATGT | 131 |
| chr10:66505264-66507279 | p4 | (AAAT)4 | GTGAAGGGGGCAAAGTGTAA | AGTTCCTTTTCCCGGCTAGA | 249 |
| chr10:66505264-66507279 | p5 | (AAGAA)3 | GTGAAGGGGGCAAAGTGTAA | AGTTCCTTTTCCCGGCTAGA | 249 |
| chr10:66505264-66507279 | p5 | (TGTGT)3 | GTGAAGGGGGCAAAGTGTAA | AGTTCCTTTTCCCGGCTAGA | 249 |
| chr10:66512957-66514972 | p5 | (TGTGT)3 | TGGAATAAGGTGAAGGGCAG | GGCCATGCTTGGCTTAGTTA | 113 |
| chr10:66512957-66514972 | p5 | (TGAAT)3 | TGGAATAAGGTGAAGGGCAG | GGCCATGCTTGGCTTAGTTA | 113 |
| chr10:66531775-66533791 | p2 | (TA)8 | GAATGGAATAGGAGGTGCGA | ATGATGATTATGACCGGGGA | 143 |
| chr10:66570088-66572116 | p2 | (AC)14 | TTCTCTTGCTTCAAACGCCT | GCGATGAAAAGAACTCTGGC | 195 |
| chr10:66664009-66666069 | p2 | (TA)30 | GTTCGTAAATCCGTAGCCCA | TCGTTGTGATGCGAAATTGT | 270 |
| chr10:66811757-66813772 | p5 | (ATGAA)3 | GGCCAAAGAATCATGCCTTA | GGTAGAAACGTGAGCTTCGG | 241 |
| chr10:66873464-66875506 | p2 | (AT)21 | CGCTTGATGTGTACAATGGG | AAATGCCCTGTAGCCTCCTT | 266 |
| chr10:67090639-67092654 | p5 | (GAATT)3 | AAATAAAGGGCCAATTTCGG | CTCAATTTCCCACCTGGCTA | 167 |
| chr10:67283909-67285921 | p5 | (CATAT)3 | GGAGCCGATTACAATACCGA | GGAATGCTGCCGAGTTAGAG | 293 |
| chr10:67283909-67285921 | p2 | (AT)6 | GGAGCCGATTACAATACCGA | GGAATGCTGCCGAGTTAGAG | 293 |
| chr10:67376606-67378618 | p2 | (AT)7 | ACCACTTTTGTTCACGACCC | TGAATGTAGTGGCAGCCTTG | 130 |
| chr10:67376606-67378618 | p2 | (AT)6 | ACCACTTTTGTTCACGACCC | TGAATGTAGTGGCAGCCTTG | 130 |
| chr10:67458703-67460719 | p2 | (AT)8 | TGTGCAAGTGACAGAAAGGC | GGTTGAAAGAAGTGCGGGTA | 163 |
| chr10:67773699-67775789 | p2 | (AT)15 | TCCCCCAAACTGAGAATCTG | GCCCAGAATTTTGAAGGACA | 110 |

| Marker ID | SSR type | SSR motif | FORWARD PRIMER (5'-3') | REVERSE PRIMER (5'-3') | size (bp) |
|---|---|---|---|---|---|
| chr10:67788408-67790420 | p2 | (CA)6 | TCGAACAACATTCGGAAACA | ATATCAGGGAGCCCGAGACT | 105 |
| chr10:67794045-67796061 | p2 | (AT)8 | GGAGAGACATAGATGCCCCA | TCCCCGGTAGTCAGAAACTG | 255 |
| chr10:67843477-67845489 | p2 | (AT)6 | TATGCCCAACTGTTCGATGA | AAAGTCCCGTATTTTCCGCT | 142 |
| chr10:67950768-67952850 | p3 | (ATC)5 | CCACCTGGCAGACTTACCAT | TCCAGTTAGGGGCTCGTAGA | 264 |
| chr10:67950873-67952935 | p3 | (ATC)5 | CCACCTGGCAGACTTACCAT | TCCAGTTAGGGGCTCGTAGA | 159 |
| chr10:68065036-68067048 | p2 | (AG)6 | ATGAGGTCAAGAGGGCAATG | GTGGGATTTGGAAGCAAAAA | 157 |
| chr10:68065036-68067048 | p2 | (GT)7 | ATGAGGTCAAGAGGGCAATG | GTGGGATTTGGAAGCAAAAA | 157 |
| chr10:68108313-68110329 | p2 | (AT)8 | GAGGGCGCTCACAATATGAT | TATATGGGTAGGGAAGGGGC | 190 |
| chr10:68108313-68110329 | p3 | (CAA)10 | GAGGGCGCTCACAATATGAT | TATATGGGTAGGGAAGGGGC | 190 |
| chr10:68129341-68131355 | p2 | (AT)7 | TGGGTTGCAAGGTTGTTGTA | AAGTTAAAGGGCACGGGATT | 164 |
| chr10:68129341-68131355 | p6 | (TTCAGT)3 | TGGGTTGCAAGGTTGTTGTA | AAGTTAAAGGGCACGGGATT | 164 |
| chr10:68227344-68229365 | p3 | (TGT)7 | TAAGGCAAGGATGGATGACC | CGAAATGATGAGTGGACGTG | 227 |
| chr10:68229944-68231962 | p2 | (AG)9 | CATAACCCCTTTCCCCCTAA | TCCCGTGGAATTAGTCAAGG | 284 |
| chr10:68229944-68231962 | p6 | (TTTTTA)3 | CATAACCCCTTTCCCCCTAA | TCCCGTGGAATTAGTCAAGG | 284 |
| chr10:68251833-68253848 | p3 | (TGT)5 | TTTTAACTTTTCCATGCGGC | ACGTCGTTGGCTAGGAAGAA | 182 |
| chr10:68251833-68253848 | p5 | (TATAT)3 | TTTTAACTTTTCCATGCGGC | ACGTCGTTGGCTAGGAAGAA | 182 |
| chr10:68308079-68310094 | p3 | (ATA)5 | TCGCTACTACGGACAAGGCT | GCAAGTTTTGAGGGCTTCTG | 206 |
| chr10:68318444-68320466 | p2 | (AT)11 | GCATCTGTGAACTTCTGCCA | AACCAGATGAGGACCCACAG | 136 |
| chr10:68466294-68468308 | p2 | (TC)7 | TCACCCTTCTGCTTCGTTCT | CCAGTGAAATGCACGAATTG | 126 |
| chr10:68532599-68534615 | p2 | (TA)8 | ATTTGACCTATAAGGGGCGG | GCGGTCTTCCCAAGTTGTTA | 290 |
| chr10:68556233-68558248 | p3 | (ATT)5 | CTAGTATCCAGACGGCCTCG | CCTTCAAGAATACCCAGCCA | 288 |
| chr10:68590529-68592637 | p2 | (AT)54 | TACGTCAGACCAGCTACCCC | TAGTCTGGCCTTCGTCGTCT | 118 |
| chr10:68603046-68605061 | p5 | (ATTTC)3 | TCACCACATGACCCAAGAAA | GGTTGACTAATGCGCCTGAT | 216 |
| chr10:68624078-68626094 | p2 | (AT)8 | CAAGGGAAACTCTGCCAAAA | CAAGGGCATACCCGACTAGA | 135 |
| chr10:68662244-68664262 | p2 | (AT)9 | AATGGTGCCTCTTACGGATG | TGGAAATTAGGAGCCCTGTG | 145 |
| chr10:68762526-68764541 | p2 | (AT)13 | CTTAAGAAGGACCCTTGGGC | CGACCTGTCGTTGGAAAGAT | 282 |
| chr10:68762526-68764541 | p3 | (TTG)5 | CTTAAGAAGGACCCTTGGGC | CGACCTGTCGTTGGAAAGAT | 282 |
| chr10:68762526-68764541 | p2 | (TA)8 | CTTAAGAAGGACCCTTGGGC | CGACCTGTCGTTGGAAAGAT | 282 |
| chr10:68776970-68778998 | p2 | (AT)14 | TCATAAGTTGAGGCGCACAC | CGTACCGCCTAGTTCTGCTC | 245 |
| chr10:68923413-68925429 | p2 | (AT)9 | ATATAGGGGAAATGCTGCCC | GCCCACAAGTCTATAGCCCA | 253 |
| chr10:69124789-69126805 | p2 | (TA)8 | GTATACCCCGCCAAAATCCT | TTTCGTTCCCATTTAGTCGC | 262 |
| chr10:69124789-69126805 | p2 | (TA)8 | GTATACCCCGCCAAAATCCT | TTTCGTTCCCATTTAGTCGC | 262 |
| chr10:69228811-69230826 | p5 | (CAAAG)3 | TTTGGGATGCTTTAGGGTTG | TGATGCCGTTGAAATGGATA | 103 |
| chr10:69307160-69309178 | p3 | (CAA)6 | CGGAATTCATACATGGGACC | GTGCTTTCCCAAGCAGGATA | 102 |
| chr10:69324353-69326368 | p5 | (TTCAT)3 | CTCGTGGAAAGGAGAAGTCG | TTCATTCACCTGCGTACCAA | 288 |
| chr10:69370970-69372985 | p5 | (AACAC)3 | CCCGAATTCACGTAGCAAAT | AATTCCCGGATTGTGATTGA | 287 |

| Marker ID | SSR type | SSR motif | FORWARD PRIMER (5'-3') | REVERSE PRIMER (5'-3') | size (bp) |
|---|---|---|---|---|---|
| chr10:69410995-69413010 | p2 | (AT)9 | TTATGCTTCGCAGGTCAGTG | GTTGAGGGTTCGCTTACCAA | 123 |
| chr10:69410995-69413010 | p5 | (AATGC)3 | TTATGCTTCGCAGGTCAGTG | GTTGAGGGTTCGCTTACCAA | 123 |
| chr10:69632135-69634150 | p3 | (CAA)5 | ATGGCACAGCATACTCCACA | AATGTTTCCCACACTGAGCC | 107 |
| chr10:69682041-69684053 | p2 | (TA)19 | AAAACGAATCGTCGCATTTC | AGCTGCTGCGGTCAATAGTT | 190 |
| chr10:69682041-69684053 | p2 | (TA)6 | AAAACGAATCGTCGCATTTC | AGCTGCTGCGGTCAATAGTT | 190 |
| chr10:69682041-69684053 | p2 | (AT)10 | AAAACGAATCGTCGCATTTC | AGCTGCTGCGGTCAATAGTT | 190 |
| chr10:69695928-69698004 | p2 | (TA)32 | GCCAAAGAATGCCTGGAATA | CGATGGGTCAGTGAGACAGA | 140 |
| chr10:69695928-69698004 | p3 | (ACA)5 | GCCAAAGAATGCCTGGAATA | CGATGGGTCAGTGAGACAGA | 140 |
| chr10:69781975-69783987 | p2 | (AG)6 | GGTACACGCGAGGTAAGGAA | CTCCGAAAGTACGGGGTGTA | 266 |
| chr10:69848699-69850713 | p2 | (AT)7 | CCGGAGTCTATAGGATGCCA | GCTTGTTCGGGTCAACAAAT | 154 |
| chr10:69921051-69923063 | p2 | (TA)6 | ACCCTTCCAAGGTCACAATG | AGCGAGAATCGCTTGGTAGA | 252 |
| chr10:70248764-70250782 | p6 | (GCTGGA)3 | TGGTAGGGCGATCAAATCTC | CCCAAAACTTGGAGGAACAA | 171 |
| chr10:70267985-70270003 | p6 | (TATACA)3 | AACCCCTAGGACGAACTGCT | CTCTGCAAGCCGATGTATCA | 117 |
| chr10:70267985-70270003 | p3 | (CAA)5 | AACCCCTAGGACGAACTGCT | CTCTGCAAGCCGATGTATCA | 117 |
| chr10:70272438-70274454 | p2 | (TA)6 | TCGAGGGTTCACGACCTTAC | CGTATAGCTACCCGCTTTCG | 220 |
| chr10:70272438-70274454 | p2 | (TA)8 | TCGAGGGTTCACGACCTTAC | CGTATAGCTACCCGCTTTCG | 220 |
| chr10:70402769-70404781 | p2 | (CA)6 | ACGATTGGTGGCATTTTAGC | AGCATTTCTGCTCCTCGAAA | 117 |
| chr10:70485847-70487875 | p2 | (AT)6 | TCTGCCGCTGAGTCTGAGTA | TATATCAGAGGGGCGTGAGG | 248 |
| chr10:70485847-70487875 | p2 | (AT)14 | TCTGCCGCTGAGTCTGAGTA | TATATCAGAGGGGCGTGAGG | 248 |
| chr10:70485847-70487875 | p2 | (TA)6 | TCTGCCGCTGAGTCTGAGTA | TATATCAGAGGGGCGTGAGG | 248 |
| chr10:70498084-70500099 | p3 | (TGT)5 | TAATTCAAGGTAATCGCGGG | TCTTACCCCTGGCCTTTCTT | 110 |
| chr10:70608108-70610120 | p2 | (AT)6 | CCGATACCTTCCAAGTCCAA | GGGCTGTTTTTGGAAGTCAA | 207 |
| chr10:70632660-70634682 | p2 | (AT)11 | ATCACGGGCTTTGGTACTTG | CTACCCCGCCTAATGTGAAA | 296 |
| chr10:70661198-70663304 | p2 | (AT)53 | TTCCAAATTCACGAACCACA | GTTTTGCGAAGAAAAGGTCG | 292 |
| chr10:70672810-70674822 | p2 | (TG)6 | CTCCGTTCGTGCCATTTATT | TCCATAGACGGTAACCAGCC | 223 |
| chr10:70685823-70687841 | p4 | (ATAC)4 | TTTCGGGACTCGGTGTTATC | GATTTCATCTCGCGATGGTT | 166 |
| chr10:70685823-70687841 | p3 | (CAA)6 | TTTCGGGACTCGGTGTTATC | GATTTCATCTCGCGATGGTT | 166 |
| chr10:70707906-70709924 | p6 | (ATATAC)3 | TGCTACACTGTCTGCCGAAC | AGCCTCGTCAGCGTGTATTT | 198 |
| chr10:70707906-70709924 | p3 | (CAA)5 | TGCTACACTGTCTGCCGAAC | AGCCTCGTCAGCGTGTATTT | 198 |
| chr10:70714402-70716420 | p6 | (CATATA)3 | GGGTCTTGTTAGCACCTCCA | GCAAGCCGTCGTACCATTAT | 107 |
| chr10:70716736-70718750 | p2 | (AG)7 | TCCAAGTTTTCCGAATGTCC | ATAAGGAACGTGCATGGGAG | 185 |
| chr10:70720115-70722183 | p2 | (TA)34 | GTACCCAACTAGGGTGCCAA | GCCATATTGATGTGGTTCCC | 186 |
| chr10:70720115-70722183 | p2 | (AT)12 | GTACCCAACTAGGGTGCCAA | GCCATATTGATGTGGTTCCC | 186 |
| chr10:70722554-70724572 | p6 | (CATATA)3 | AACCCCTAGGACGAACTGCT | GCAAGCCGTCGTACCATTAT | 109 |
| chr10:70814317-70816335 | p2 | (AT)9 | AGGGTCGTCAAACGTCATTC | GTAGTGGTCGGGCACTTGTT | 141 |
| chr10:70855397-70857417 | p2 | (AT)10 | GGCGGCTTTAGCTTTTCTCT | GAAGTGTCTACGGGCCATGT | 181 |

| Marker ID | SSR type | SSR motif | FORWARD PRIMER (5'-3') | REVERSE PRIMER (5'-3') | size (bp) |
|---|---|---|---|---|---|
| chr10:70885344-70887358 | p2 | (TA)7 | CGTAGCTAATAAGGCGTGGC | TAGGTACACATCATCGGGCA | 194 |
| chr10:70885344-70887358 | p2 | (AT)7 | CGTAGCTAATAAGGCGTGGC | TAGGTACACATCATCGGGCA | 194 |
| chr10:70885809-70887823 | p2 | (TA)7 | TAACGATATCCGAAGGCCAC | ATAGTAGACGCACGGAACGG | 123 |
| chr10:70885809-70887823 | p2 | (AT)7 | TAACGATATCCGAAGGCCAC | ATAGTAGACGCACGGAACGG | 123 |
| chr10:70900391-70902403 | p2 | (AT)6 | CCCACTTGAGCTTCACCTTC | AATTTGCCCTTCGTCTTCCT | 242 |
| chr10:70920690-70922705 | p5 | (TGATA)3 | TTCCACGAGGATTGGAAGTC | GATGTCATACGAACCCGCTT | 159 |
| chr10:70938579-70940595 | p2 | (TG)8 | CTAAGGCCACCCCTAAGGAC | GGTGGATGCGATCGTTAGTT | 159 |
| chr10:71241172-71243184 | p2 | (AT)6 | TGTCCGCACTGTTTTGTACC | GTCATAAACTGGCGGCGTAT | 158 |
| chr10:71310410-71312432 | p2 | (TG)8 | ATAATTTGGGAAGGCAAGGG | ACAAAGATGGCCCACAAAAG | 268 |
| chr10:71310410-71312432 | p2 | (TA)11 | ATAATTTGGGAAGGCAAGGG | ACAAAGATGGCCCACAAAAG | 268 |
| chr10:71383488-71385500 | p2 | (TA)6 | AACTGAGGACGGTATCGACG | ACTTACCTCACGTGCCCAAC | 287 |
| chr10:71383488-71385500 | p2 | (AT)8 | AACTGAGGACGGTATCGACG | ACTTACCTCACGTGCCCAAC | 287 |
| chr10:71384204-71386220 | p2 | (TA)6 | CCAACCACATGTTGCTTCAG | ATTCAATTCCCAACTGCCTG | 139 |
| chr10:71384204-71386220 | p2 | (AT)8 | CCAACCACATGTTGCTTCAG | ATTCAATTCCCAACTGCCTG | 139 |
| chr10:71451307-71453322 | p5 | (ATCAT)3 | CCACAACTGCCCATACACAC | TCGGGGTTTTAAGGAAAGGT | 195 |
| chr10:71480309-71482327 | p2 | (AT)9 | ATATTTGCAAGCCGACAAGG | CCCGTCCCGTCATATGATAC | 252 |
| chr10:71558896-71560911 | p3 | (TTA)5 | GTCGCTAACGAGCATAAGGG | CGACAAGAACCCAAGGATGT | 120 |
| chr10:71588298-71590310 | p2 | (TA)6 | TGATCAGTTGTGACTCCCCA | GCCTGCAGGTAGACAGGAAG | 124 |
| chr10:71676985-71679013 | p2 | (TA)14 | GCCACAGGGCCTACTCATAA | CGGGGTACTGTCCTGTCTGT | 254 |
| chr10:71676985-71679013 | p5 | (AAATG)3 | GCCACAGGGCCTACTCATAA | CGGGGTACTGTCCTGTCTGT | 254 |
| chr10:71681637-71683649 | p2 | (CA)6 | TCCATTCATCCAAGGGCTAC | TGTGGCAATCCTACCACAAA | 143 |
| chr10:71747700-71749715 | p3 | (CAA)5 | AGTGGTGGAAACATTGAGGC | CGGATCGGTTTGTCTTTGAT | 216 |
| chr10:71761783-71763805 | p2 | (AT)11 | ATAAGCGGACGAGGCTAACA | GCAGGTGGCTAAGGAGACAG | 186 |
| chr10:71761783-71763805 | p5 | (GAAAT)3 | ATAAGCGGACGAGGCTAACA | GCAGGTGGCTAAGGAGACAG | 186 |
| chr10:71767843-71769855 | p2 | (AC)6 | ATGGAGTCGGACCTAAGGCT | GTTGTGTTGGTTGTTGGCTG | 239 |
| chr10:71794424-71796436 | p2 | (AG)6 | CCCGGTAAACTTGCTTCCTT | ACTAGTCGCGTGAAGCCTGT | 243 |
| chr10:71811381-71813397 | p2 | (TA)8 | GAAAAGAAGTTGAGCGTGGG | CAAAGCCATGCCTTCTCAAT | 198 |
| chr10:71811381-71813397 | p5 | (TTTGG)3 | GAAAAGAAGTTGAGCGTGGG | CAAAGCCATGCCTTCTCAAT | 198 |
| chr10:71878024-71880042 | p3 | (TTA)6 | CTGGGTGGCTTTTGAACATT | ATGAATGTGCTGGAATGCAA | 300 |
| chr10:72017921-72019941 | p2 | (AT)24 | TACGACATGCGGAAGAACAG | TGGTTTGGGGTTACTGGGTA | 144 |
| chr10:72017921-72019941 | p5 | (TATCA)4 | TACGACATGCGGAAGAACAG | TGGTTTGGGGTTACTGGGTA | 144 |
| chr10:72133663-72135681 | p3 | (TTG)6 | CAGCCAAGAACACACGAAAA | ACGATTCTGCAGCGAAAACT | 200 |
| chr10:72133663-72135681 | p6 | (ATAATG)3 | CAGCCAAGAACACACGAAAA | ACGATTCTGCAGCGAAAACT | 200 |
| chr10:72177294-72179308 | p2 | (TA)7 | CCTCAACACCATCAAAGGCT | CCTACTGGGGTAGGTCGTGA | 274 |
| chr10:72184704-72186718 | p2 | (TG)8 | AATGAAAGGACGAGTCGTGG | AAATTCTCCCTGCTCTGGGT | 176 |
| chr10:72184704-72186718 | p2 | (TA)7 | AATGAAAGGACGAGTCGTGG | AAATTCTCCCTGCTCTGGGT | 176 |

| Marker ID | SSR type | SSR motif | FORWARD PRIMER (5'-3') | REVERSE PRIMER (5'-3') | size (bp) |
|---|---|---|---|---|---|
| chr10:72184704-72186718 | p2 | (TA)11 | AATGAAAGGACGAGTCGTGG | AAATTCTCCCTGCTCTGGGT | 176 |
| chr10:72196482-72198496 | p2 | (TG)8 | AATGAAAGGACGAGTCGTGG | AAATTCTCCCTGCTCTGGGT | 176 |
| chr10:72196482-72198496 | p2 | (TA)7 | AATGAAAGGACGAGTCGTGG | AAATTCTCCCTGCTCTGGGT | 176 |
| chr10:72196482-72198496 | p2 | (TA)11 | AATGAAAGGACGAGTCGTGG | AAATTCTCCCTGCTCTGGGT | 176 |
| chr10:72223023-72225038 | p5 | (TATGA)3 | TCGGGTTCTTTCTCCCTTTT | TACCAGGTTTTGGGATCGAG | 269 |
| chr10:72254598-72256610 | p2 | (TA)6 | GGGGTAACGTTATGGCCTTT | TACCCCTTATCGACTGTGCC | 293 |
| chr10:72254598-72256610 | p2 | (AT)6 | GGGGTAACGTTATGGCCTTT | TACCCCTTATCGACTGTGCC | 293 |
| chr10:72254598-72256610 | p2 | (AT)6 | GGGGTAACGTTATGGCCTTT | TACCCCTTATCGACTGTGCC | 293 |
| chr10:72307426-72309444 | p3 | (CAA)5 | CGCTTTTCCAAGCTCACTTC | GTTGATTGTTTTGGCCGAGT | 201 |
| chr10:72307457-72309469 | p3 | (CAA)5 | CGCTTTTCCAAGCTCACTTC | GTTGATTGTTTTGGCCGAGT | 170 |
| chr10:72369936-72371951 | p5 | (ACACA)3 | ACACGGCCAACTTAGACCAC | TCATGGAGTGGGTGTGTTGT | 256 |
| chr10:72370000-72372015 | p5 | (ACACA)3 | ACACGGCCAACTTAGACCAC | TCATGGAGTGGGTGTGTTGT | 192 |
| chr10:72386775-72388793 | p6 | (TAAAAA)3 | AAGGGTGACATGAGAATCGG | CTCGTAGGTCAACTCCTCCG | 203 |
| chr10:72531914-72533928 | p2 | (AT)7 | GGCTGAATTCTGTGGCATTT | CTTCTGTAGGCGTCGTAGGC | 296 |
| chr10:72531914-72533928 | p2 | (AT)7 | GGCTGAATTCTGTGGCATTT | CTTCTGTAGGCGTCGTAGGC | 296 |
| chr10:72531914-72533928 | p2 | (TA)8 | GGCTGAATTCTGTGGCATTT | CTTCTGTAGGCGTCGTAGGC | 296 |
| chr10:72683261-72685273 | p2 | (TA)6 | AAACGGCAAGGTATGTGAGG | TCTATCGGGGTGACGTAAGG | 141 |
| chr10:72687273-72689287 | p2 | (AT)7 | GCGGCAACTAACAGTGTCAA | ATGATATCAGAAATGCCGCC | 249 |
| chr10:72687273-72689287 | p2 | (AT)8 | GCGGCAACTAACAGTGTCAA | ATGATATCAGAAATGCCGCC | 249 |
| chr10:72708631-72710646 | p5 | (AAAGA)3 | TCCGATCCCAGTTACGAATC | CTTTCCGACATGGCTTCTTC | 257 |
| chr10:72770370-72772382 | p2 | (AC)6 | TTTCGAGGAGCAGGATTGTT | GCGGAAATTTCGGTAGACAA | 211 |
| chr10:72770370-72772382 | p2 | (TA)6 | TTTCGAGGAGCAGGATTGTT | GCGGAAATTTCGGTAGACAA | 211 |
| chr10:72770370-72772382 | p5 | (ATTTT)3 | TTTCGAGGAGCAGGATTGTT | GCGGAAATTTCGGTAGACAA | 211 |
| chr10:72898533-72900548 | p2 | (TA)15 | TTGTGGTTGTTGGCTATGGA | ACAAATTTGGCAGCATTTCC | 275 |
| chr10:72919837-72921849 | p2 | (AT)6 | CCAACGGTCAGTCCATCTTT | CAGGGGGAAAGTCAGTTGAA | 166 |
| chr10:72919837-72921849 | p3 | (CAT)5 | CCAACGGTCAGTCCATCTTT | CAGGGGGAAAGTCAGTTGAA | 166 |
| chr10:72965346-72967381 | p5 | (CATGT)7 | AACATTCGCAAGCAGCTTTT | GTGACCTAAAACTTCGCCCA | 161 |
| chr10:73040230-73042248 | p6 | (ATATGT)3 | ATGTTGGATATGAGCCTCGG | TCGGGTCCTATTTATGCGAG | 166 |
| chr10:73067443-73069458 | p5 | (TTGCA)3 | TTGTTGGAATACCCACCGTT | ATCGGCCTTCTCACAATCAC | 118 |
| chr10:73094435-73096449 | p3 | (CAA)6 | CAAAAACATGTCCGTCGTTG | AGGCATGATTCCTATGGCAC | 271 |
| chr10:73094435-73096449 | p2 | (CT)7 | CAAAAACATGTCCGTCGTTG | AGGCATGATTCCTATGGCAC | 271 |
| chr10:73143576-73145588 | p2 | (AT)6 | TGAATAGGGTTGTGGAAGCC | CAGTGCCATACACCAACAGG | 160 |
| chr10:73165993-73168011 | p2 | (TA)9 | ACGGATTCGATGGGTTACAG | GGACCTCTCACACCCACAGT | 261 |
| chr10:73165993-73168011 | p4 | (CATA)4 | ACGGATTCGATGGGTTACAG | GGACCTCTCACACCCACAGT | 261 |
| chr10:73170395-73172410 | p5 | (TATCG)3 | TGGAATTGATGGGAGAAAGC | AGCCACAACCAACAACATGA | 116 |
| chr10:73170395-73172410 | p2 | (TA)10 | TGGAATTGATGGGAGAAAGC | AGCCACAACCAACAACATGA | 116 |

| Marker ID | SSR type | SSR motif | FORWARD PRIMER (5'-3') | REVERSE PRIMER (5'-3') | size (bp) |
|---|---|---|---|---|---|
| chr10:73219579-73221593 | p2 | (AT)6 | CCGACATGCCATAGTTTCCT | ACATGTTCGGCCATTTAAGC | 100 |
| chr10:73219579-73221593 | p2 | (TA)7 | CCGACATGCCATAGTTTCCT | ACATGTTCGGCCATTTAAGC | 100 |
| chr10:73245445-73247459 | p2 | (AT)7 | AAACATGGGCAATGAAGGAG | AAGGAAAAATCATGCCATCG | 201 |
| chr10:73245445-73247459 | p2 | (TA)14 | AAACATGGGCAATGAAGGAG | AAGGAAAAATCATGCCATCG | 201 |
| chr10:73253453-73255468 | p3 | (CTT)5 | GCTTCTCTCTCCGTGATTGG | CATCGGTTTGTAAGTGGGCT | 168 |
| chr10:73303202-73305218 | p4 | (CATA)4 | TTCTAACTCGTGCGCATCAC | AAATCCAATGCAAGCTCCTG | 162 |
| chr10:73319616-73321642 | p2 | (TA)13 | TCGAGGGTTCACGACCTTAC | AGGCGCGGTGTAATATCATC | 241 |
| chr10:73361748-73363763 | p6 | (CATATA)4 | GAGGCAGAATCAGCATGTCA | GGCTCCACCAGTCGATATGT | 259 |
| chr10:73402430-73404460 | p2 | (AT)10 | CCTGTACTGTCACGCCTCAA | GTGGGTCATGGGTGGTTATC | 208 |
| chr10:73402430-73404460 | p2 | (AT)15 | CCTGTACTGTCACGCCTCAA | GTGGGTCATGGGTGGTTATC | 208 |
| chr10:73464510-73466534 | p2 | (AT)12 | CGGGTTCTCGTGGTATCTGT | TCTGGCGCCATAATACATCA | 167 |
| chr10:73470456-73472476 | p2 | (AC)6 | TTATAGGCCCGTTTTCCCTT | CGAACAATCACACATTTGCC | 277 |
| chr10:73470558-73472576 | p2 | (AC)6 | TTATAGGCCCGTTTTCCCTT | CGAACAATCACACATTTGCC | 175 |
| chr10:73485910-73487928 | p6 | (TCTGAT)3 | TTTGGGCCCTTTTTAATTCC | GGCATAAAGGAAAATTCGCA | 189 |
| chr10:73589945-73591969 | p3 | (ATC)5 | CACAATACCGCCACATCTTG | GACCCGCAAGGTTTTATTGA | 195 |
| chr10:73589971-73591987 | p3 | (ATC)5 | CACAATACCGCCACATCTTG | GACCCGCAAGGTTTTATTGA | 169 |
| chr10:73590025-73592041 | p3 | (ATC)5 | CACAATACCGCCACATCTTG | GACCCGCAAGGTTTTATTGA | 115 |
| chr10:73659361-73661385 | p4 | (AATT)4 | TAAAGGCGGAGTCATTTTCG | ATTAAAATTTTCGGGTGGGG | 265 |
| chr10:73659361-73661385 | p2 | (TA)13 | TAAAGGCGGAGTCATTTTCG | ATTAAAATTTTCGGGTGGGG | 265 |
| chr10:73828616-73830654 | p4 | (AAAT)4 | GCTTGCATACCCAACATCCT | CGAGCCTTTCGTCGATACTC | 262 |
| chr10:73846713-73848733 | p2 | (TA)10 | TTGTTGTGTTTGGGCATTGT | CGCCTATTCTCCTGATCTCG | 282 |
| chr10:73875170-73877182 | p2 | (AT)6 | CGTAAACTGCCATCGTCCTT | CGTCCTTCACGACCGTATTT | 179 |
| chr10:73989022-73991100 | p2 | (TA)39 | TCGGGGAATCACGACTTTAC | CACTTAGTGGGGTCTGCCAT | 187 |
| chr10:74096355-74098370 | p2 | (TA)7 | TGAGAAGCTTGGAGAAGGGA | CTTCATGTTGTTAGGGGCGT | 206 |
| chr10:74096355-74098370 | p2 | (TA)7 | TGAGAAGCTTGGAGAAGGGA | CTTCATGTTGTTAGGGGCGT | 206 |
| chr10:74096355-74098370 | p5 | (TGAAT)3 | TGAGAAGCTTGGAGAAGGGA | CTTCATGTTGTTAGGGGCGT | 206 |
| chr10:74096355-74098370 | p6 | (TATATG)3 | TGAGAAGCTTGGAGAAGGGA | CTTCATGTTGTTAGGGGCGT | 206 |
| chr10:74113828-74115858 | p2 | (AG)15 | CTTCCATGGAGGAAGATCCA | GGGGTCAACTCTAGTGCAGC | 222 |
| chr10:74157733-74159745 | p2 | (AT)7 | ATTTGGAGCCCAATCAAATG | CGTCCGTTGTCCCTTAAAAA | 218 |
| chr10:74157733-74159745 | p2 | (AT)6 | ATTTGGAGCCCAATCAAATG | CGTCCGTTGTCCCTTAAAAA | 218 |
| chr10:74226015-74228043 | p2 | (TA)14 | AAGTTTCGGGTTAGGCGAAT | ATTCCTCAGCCCTATCGCTT | 231 |
| chr10:74226015-74228043 | p2 | (AT)14 | AAGTTTCGGGTTAGGCGAAT | ATTCCTCAGCCCTATCGCTT | 231 |
| chr10:74231678-74233693 | p3 | (TAA)5 | ATACGGACTGTAACAGCGGG | GATCTTGTAACGCGCAGACA | 159 |
| chr10:74259738-74261788 | p2 | (AT)25 | ATGAATGCTCCCATCTCTGG | ATCTCAATCCGCCCATGTAG | 273 |
| chr10:74335610-74337670 | p3 | (TTC)20 | AATGGTGATATCTGGGCCAC | CCAGCAGAGGTAAAAGACGC | 176 |
| chr10:74335610-74337670 | p5 | (GAGAA)3 | AATGGTGATATCTGGGCCAC | CCAGCAGAGGTAAAAGACGC | 176 |

| Marker ID | SSR type | SSR motif | FORWARD PRIMER (5'-3') | REVERSE PRIMER (5'-3') | size (bp) |
|---|---|---|---|---|---|
| chr10:74389865-74391880 | p5 | (TCATA)3 | CCCATATCGCCATTCACTTT | CAGAGCAGGTCGGTCTAAGG | 221 |
| chr10:74389918-74391933 | p5 | (TCATA)3 | CCCATATCGCCATTCACTTT | CAGAGCAGGTCGGTCTAAGG | 168 |
| chr10:74407650-74409665 | p3 | (TCA)5 | AGGATTTGATCCGTCTCCCT | ACTTATGGATTGGGCTGCAC | 275 |
| chr10:74411748-74413766 | p2 | (AT)9 | AAACTGCGCTCACACACATC | TAGTTTTCTGGGGTTGGCAG | 300 |
| chr10:74424672-74426692 | p2 | (AT)10 | ATAAGATGCCAACTCCACGG | GTTGCTTCGGGTCAACAAAT | 166 |
| chr10:74468312-74470324 | p2 | (TG)6 | AAGAATGGGTGCTGAATTGC | TAGCCTCATTCCTTGCGAGT | 252 |
| chr10:74485856-74487868 | p2 | (GT)6 | CGAGGGCCTATATTTCACGA | CCTTCCCCAATGACTTAGCA | 299 |
| chr10:74578604-74580620 | p2 | (TA)8 | GTTCTTGTCACACCCCGACT | GCAATGGGTAATGGGTTGTC | 235 |
| chr10:74584056-74586088 | p2 | (TA)16 | TCAACAATCCCTAGATGCCC | AAAGTCGGGGTGTGACAAAG | 114 |
| chr10:74606457-74608469 | p2 | (AT)7 | TCTCCAGTTTGCGAGGTTCT | GTTCTTCCTCCGGGTTTCTC | 294 |
| chr10:74606457-74608469 | p2 | (AT)6 | TCTCCAGTTTGCGAGGTTCT | GTTCTTCCTCCGGGTTTCTC | 294 |
| chr10:74606457-74608469 | p2 | (TA)6 | TCTCCAGTTTGCGAGGTTCT | GTTCTTCCTCCGGGTTTCTC | 294 |
| chr10:74658280-74660295 | p3 | (ACA)5 | TGTTTGGAGAAGGGTTTTGG | GATGAGAACGAAAATCGGGA | 279 |
| chr10:74702517-74704543 | p2 | (TA)13 | CAGGGACTACATAGCGGAGC | TGCCCATCACACCGTAGTAA | 197 |
| chr10:74735609-74737627 | p2 | (AT)9 | TTTCGGTTTACAGACCGTCC | CGTAAACTGCCATGCAAAGA | 227 |
| chr10:74735609-74737627 | p2 | (TA)6 | TTTCGGTTTACAGACCGTCC | CGTAAACTGCCATGCAAAGA | 227 |
| chr10:74741395-74743407 | p2 | (CT)6 | GCCCCGACACCAACTAGATA | CTGCTCCGTCTGGAAGAGTC | 160 |
| chr10:74747066-74749080 | p2 | (AT)7 | CTTTAGCGAAGTTCGGGTTG | TTATACGGATCCGATCTGGC | 235 |
| chr10:74747066-74749080 | p5 | (TTGTA)3 | CTTTAGCGAAGTTCGGGTTG | TTATACGGATCCGATCTGGC | 235 |
| chr10:74763244-74765256 | p2 | (AT)6 | GCTGACCCCTGTCATGATCT | AGTGGCTCTTCAGCCAGGTA | 141 |
| chr10:74773989-74776001 | p2 | (TG)6 | ATGCCAAAGGTGCTTACAGG | CGACCATAAGGTTCGGTGTT | 223 |
| chr10:74914239-74916255 | p4 | (TTAT)4 | CTTGCACCTTCTGTTGGACA | TATGGGAGTGAATGTGGGGT | 158 |
| chr10:75007187-75009203 | p2 | (AT)8 | GTCACCATCTTCACCCCCTA | GCCCAATATGTGCCTGAGAT | 246 |
| chr10:75007187-75009203 | p2 | (TA)6 | GTCACCATCTTCACCCCCTA | GCCCAATATGTGCCTGAGAT | 246 |
| chr10:75022490-75024514 | p3 | (TAT)8 | AATGTTTCCACACTGAGCCC | ATCAGAATAGGCGGACATGG | 246 |
| chr10:75032673-75034691 | p2 | (AT)17 | GTGTGAGGGCCCACTACCTA | CCTTAAGCCAAATATGGGCA | 284 |
| chr10:75032673-75034691 | p3 | (TTG)6 | GTGTGAGGGCCCACTACCTA | CCTTAAGCCAAATATGGGCA | 284 |
| chr10:75051957-75053972 | p5 | (ATGAT)3 | TTTGCCCCTTTTTCATTCAG | ACCTTGGGTCAAATGAGGTG | 210 |
| chr10:75078497-75080512 | p3 | (TTG)5 | GGAGGTCTTTGGGAACATGA | AACCATCCACAACACAAGCA | 258 |
| chr10:75148735-75150750 | p3 | (TGC)5 | CGGACCTATGACGTGACCTT | TTCGAGAAAAATGGATTCCG | 281 |
| chr10:75173503-75175518 | p3 | (TGC)5 | CGGACCTATGACGTGACCTT | TTCGAGAAAAATGGATTCCG | 281 |
| chr10:75223754-75225766 | p2 | (TA)6 | AGACTGATGGAGAAGCCGAA | ATCGATCTCGGATCACCTTG | 200 |
| chr10:75226074-75228089 | p5 | (CATAG)3 | CACCACTAGTGAGCGGCATA | TATCGGGTACTCTTCACGGC | 188 |
| chr10:75226074-75228089 | p5 | (TCATA)3 | CACCACTAGTGAGCGGCATA | TATCGGGTACTCTTCACGGC | 188 |
| chr10:75226074-75228089 | p2 | (TA)7 | CACCACTAGTGAGCGGCATA | TATCGGGTACTCTTCACGGC | 188 |
| chr10:75315813-75317845 | p2 | (AT)16 | TCTATGGGTCCTTGTTTGGC | ACCCAATCAGGATATGCGAG | 233 |

| Marker ID | SSR type | SSR motif | FORWARD PRIMER (5'-3') | REVERSE PRIMER (5'-3') | size (bp) |
|---|---|---|---|---|---|
| chr10:75483814-75485829 | p5 | (GAAAT)3 | GATATTTGGCCGAAGAACCA | GAAGGTGGAAACGTGAGCAT | 239 |
| chr10:75489651-75491666 | p3 | (ATG)5 | CAAAGGATATGCTGCCCAAT | CTATCGCCTGCTTGTTCTCC | 130 |
| chr10:75489651-75491666 | p6 | (ATATGT)3 | CAAAGGATATGCTGCCCAAT | CTATCGCCTGCTTGTTCTCC | 130 |
| chr10:75499518-75501533 | p3 | (GAT)5 | TTGAAGGTAAAAACGTGGGC | AAAGTCATGCCATCGGATTC | 209 |
| chr10:75554624-75556645 | p2 | (AT)9 | ACGAGCCTCTAAATGGAGCA | CAGGTTGACAAGGAGACGGT | 126 |
| chr10:75554624-75556645 | p3 | (CAT)7 | ACGAGCCTCTAAATGGAGCA | CAGGTTGACAAGGAGACGGT | 126 |
| chr10:75575203-75577215 | p5 | (CCCCT)3 | GGGGGCCACACACTCTACTA | GCCCCATAGCTCCTGTATGA | 289 |
| chr10:75641849-75643867 | p2 | (AT)9 | TCTTAAATGCCTTGCAGCCT | GAGTTAGAAATGGGCCACCA | 137 |
| chr10:75641849-75643867 | p6 | (TATTCA)3 | TCTTAAATGCCTTGCAGCCT | GAGTTAGAAATGGGCCACCA | 137 |
| chr10:75663348-75665360 | p2 | (AG)6 | AAAAGGGATGGCCTTGAGTT | CGTACTAAGCCCTCGCAGTC | 162 |
| chr10:75687766-75689786 | p5 | (TGATA)4 | ACACGAGTATCCGGGCATAG | AAAACGGATTTCGATCAACG | 237 |
| chr10:75737829-75739841 | p2 | (CT)6 | AGCCACCAGCTTCTTTTCAA | TTTGCTTGGCATAGTTGCAG | 235 |
| chr10:75790595-75792629 | p2 | (TA)17 | ACCCACTGATGACCCGAATA | CCGGACGGAAGATCATCTAA | 171 |
| chr10:75790595-75792629 | p5 | (TGCAA)3 | ACCCACTGATGACCCGAATA | CCGGACGGAAGATCATCTAA | 171 |
| chr10:75842553-75844565 | p2 | (GA)6 | TCGAGACTCTGGCTGTTCCT | GATTCGGATGTCCTAACGGA | 174 |
| chr10:75919534-75921552 | p6 | (CCAACT)3 | CCTTCCCAACATCTTCTCCA | GGGCTTTTCTAAGGGAGCAC | 138 |
| chr10:75932675-75934693 | p3 | (TTA)6 | AACACCCCAGGACACTCATC | GTTACCCCTGCTAGGGAAGG | 146 |
| chr10:75958367-75960393 | p2 | (TA)6 | TGGGCGACATGAGATGTTTA | CTCTTAAGGCGGAGTCATGC | 137 |
| chr10:75958367-75960393 | p2 | (GA)13 | TGGGCGACATGAGATGTTTA | CTCTTAAGGCGGAGTCATGC | 137 |
| chr10:76164978-76166992 | p2 | (TA)7 | TTTTTGAAAATGGTCTCGCC | TTGAAGGTTATGCCCAAAGC | 184 |
| chr10:76175041-76177053 | p2 | (GA)6 | TTTGGACTGGAACAACACCA | GTGCATTGCATAGGGGAACT | 252 |
| chr10:76178275-76180293 | p2 | (TA)9 | CTCTTTTCCCTTTCTCGGCT | ACCCAACCCAAGGTTTAAGG | 239 |
| chr10:76220171-76222187 | p4 | (AAGA)4 | TCAGTAGGTCGTGGTAGGGG | GTGATAACGTCAGGGGAGGA | 170 |
| chr10:76245606-76247618 | p2 | (AT)6 | CCGGAGTCTATAGGATGCCA | CAAGTACAAAGGCCCGTGAT | 150 |
| chr10:76295009-76297024 | p3 | (ACA)5 | GAGGTTCCAAGAGGTGTGGA | GGTCACAGTGAGGGGTCAGT | 223 |
| chr10:76377066-76379084 | p3 | (AGA)6 | GATTGCATGATGGTGTTTGC | TCCTCATTGGCCCACTAATC | 164 |
| chr10:76411081-76413096 | p2 | (TA)17 | CCCTAGACTCCCCACGGTAT | CATCCCTCAAATAGCCCTCA | 165 |
| chr10:76411081-76413096 | p2 | (TA)7 | CCCTAGACTCCCCACGGTAT | CATCCCTCAAATAGCCCTCA | 165 |
| chr10:76411127-76413183 | p2 | (TA)17 | CCCTAGACTCCCCACGGTAT | CATCCCTCAAATAGCCCTCA | 119 |
| chr10:76411127-76413183 | p2 | (TA)7 | CCCTAGACTCCCCACGGTAT | CATCCCTCAAATAGCCCTCA | 119 |
| chr10:76418036-76420048 | p2 | (TG)6 | TTTGAAACCATGTCCAAACG | TTCAAGCGGACCGTTAACTT | 204 |
| chr10:76418036-76420048 | p2 | (AT)33 | TTTGAAACCATGTCCAAACG | TTCAAGCGGACCGTTAACTT | 204 |
| chr10:76486813-76488879 | p2 | (AT)6 | CCACAAGGGAAAAACAGGAA | CCAAAATTGGGGGTTTCACT | 108 |
| chr10:76486813-76488879 | p3 | (ATT)5 | CCACAAGGGAAAAACAGGAA | CCAAAATTGGGGGTTTCACT | 108 |
| chr10:76487183-76489198 | p2 | (AT)6 | CCATGGCGTAAATCACAAAA | CCTAGTCGGGTCGTGACATC | 284 |
| chr10:76487183-76489198 | p3 | (ATT)5 | CCATGGCGTAAATCACAAAA | CCTAGTCGGGTCGTGACATC | 284 |

| Marker ID | SSR type | SSR motif | FORWARD PRIMER (5'-3') | REVERSE PRIMER (5'-3') | size (bp) |
|---|---|---|---|---|---|
| chr10:76552768-76554816 | p3 | (TAT)16 | ATGGGCATACAATGGGAAAA | CGCTTGAGCCCTTGTTGTAT | 199 |
| chr10:76608551-76610587 | p4 | (ATTT)6 | ATGCTCCAATTTGGGATGAG | CTAATGGAGCGGAGCTGAAC | 202 |
| chr10:76608588-76610600 | p4 | (ATTT)6 | ATGCTCCAATTTGGGATGAG | CTAATGGAGCGGAGCTGAAC | 165 |
| chr10:76608615-76610627 | p4 | (ATTT)6 | ATGCTCCAATTTGGGATGAG | CTAATGGAGCGGAGCTGAAC | 138 |
| chr10:76613591-76615606 | p5 | (TTTAG)3 | TTCGTCTCTCCCTTTCCCTT | CGGGGTTCAAGAACTGTGAT | 151 |
| chr10:76676746-76678760 | p2 | (AT)7 | AAACTCGTGAAATCATGGGC | CCACTTCGCCCAATTTAGAA | 163 |
| chr10:76693091-76695109 | p2 | (AT)9 | GGTGGAGCCACACAAAGATT | GTCCTGCCCATTCCTTATGA | 188 |
| chr10:76738894-76740920 | p2 | (TA)13 | TAGCTGGGGCATTTGTAAGG | TATCTTCCACCCTTGTTGGC | 297 |
| chr10:76826358-76828370 | p2 | (TG)6 | AAAGGTGCCTACGGGCTATT | CCAACCAAGGCTCAAGGTTA | 206 |
| chr10:76876706-76878721 | p5 | (ATGCA)3 | CGCAGCTAGGTACGGAACTC | GGGGTGTTTCCCTAATCCAT | 239 |
| chr10:76891220-76893235 | p5 | (AGAAG)3 | TAATGAGCGATTCGGCTTCT | AATGGCTAGGCATGAACCAC | 220 |
| chr10:76990124-76992139 | p3 | (ATC)5 | AGGAATTCATAAGGACGGGG | CCTTAGGAGCTTCATCAGCG | 232 |
| chr10:77085923-77087959 | p2 | (AT)18 | GACCCTTGCAGGATGTGTTT | CTCTCCCTGAACCTCTGCAC | 144 |
| chr10:77129967-77132031 | p2 | (AT)32 | TCAATAAAACCTTGCGGGTC | TAGTTGCCACTCAGGCACAG | 103 |
| chr10:77185240-77187274 | p2 | (AT)15 | ACTGTGTTTTTGCCTTCGCT | TCGGTTAGTTTGGTATCGGG | 165 |
| chr10:77228005-77230023 | p3 | (TTG)6 | AGCTCGAAGAGGTCATGGAA | CCTCCAAAGTTGACCCAAAA | 201 |
| chr10:77228005-77230023 | p2 | (TA)6 | AGCTCGAAGAGGTCATGGAA | CCTCCAAAGTTGACCCAAAA | 201 |
| chr10:77272296-77274308 | p2 | (AG)6 | TAGAATTTGCCTTGCCAACC | CTCTGCCATTCCAGGTTGAT | 126 |
| chr10:77290406-77292418 | p2 | (AT)6 | AACAGCTTCGGAACAAGTGG | GGGAGCGTTTTGTGTTTCAT | 214 |
| chr10:77417416-77419492 | p3 | (AAG)5 | TCCTTGTCCATCCAGTAGCC | ATGGGTGATGGGAATCAAAA | 261 |
| chr10:77494540-77496556 | p2 | (AT)8 | CTTGCATAAAAAGGACCCGA | ACAATAAGGAAAATCCGGGG | 195 |
| chr10:77528636-77530651 | p5 | (AATGC)3 | CGGTTTGGTTCAAACTTCGT | CACCACTCGCTAGGGTTCTC | 118 |
| chr10:77528636-77530651 | p5 | (CCTTT)3 | CGGTTTGGTTCAAACTTCGT | CACCACTCGCTAGGGTTCTC | 118 |
| chr10:77580975-77583003 | p2 | (AT)14 | AGCCAACTTGCGTCAACTCT | TTTCGACGCTCTTTGACCTT | 264 |
| chr10:77583001-77585029 | p2 | (AT)14 | AGCCAACTTGCGTCAACTCT | TTTCGACGCTCTTTGACCTT | 264 |
| chr10:77583001-77585029 | p2 | (AT)8 | AGCCAACTTGCGTCAACTCT | TTTCGACGCTCTTTGACCTT | 264 |
| chr10:77669227-77671245 | p6 | (TTTCAG)3 | GCATTGACTTAGAGCCGAGG | CACATCGTCCCATCCTTCTT | 156 |
| chr10:77787356-77789374 | p2 | (TA)9 | TTGACGATTCTCATCCCACA | CTCCACAATACCAGCCACCT | 166 |
| chr10:77867414-77869458 | p5 | (TGATA)3 | GAGGCTACAAGGCACGTAGG | CACCTGCTACACTTTCCCGT | 175 |
| chr10:77867414-77869458 | p2 | (AT)22 | GAGGCTACAAGGCACGTAGG | CACCTGCTACACTTTCCCGT | 175 |
| chr10:77917227-77919239 | p2 | (AT)6 | CCGTACGACGATAAAGCGAT | TCTCCTTCGGTGCTCAAAGT | 275 |
| chr10:77917227-77919239 | p2 | (TA)9 | CCGTACGACGATAAAGCGAT | TCTCCTTCGGTGCTCAAAGT | 275 |
| chr10:78001080-78003106 | p2 | (AT)13 | GCATGAGGACTATGAGGGGA | GCCTAGAAATCGGATGGTGA | 142 |
| chr10:78086013-78088025 | p2 | (TA)6 | TGTGCATTTGAGTAGCGGAG | ATTATGATGAGAGGTGCCCG | 223 |
| chr10:78207995-78210007 | p2 | (AT)14 | TTTGGCCATGATTCTGTCAA | TTCGAATTTTCTGTGCATTGAT | 253 |
| chr10:78207995-78210007 | p2 | (AC)6 | TTTGGCCATGATTCTGTCAA | TTCGAATTTTCTGTGCATTGAT | 253 |

| Marker ID | SSR type | SSR motif | FORWARD PRIMER (5'-3') | REVERSE PRIMER (5'-3') | size (bp) |
|---|---|---|---|---|---|
| chr10:78220296-78222310 | p2 | (AT)7 | GGTCAGGCAATGAGTGACAA | TTCGGAGTTCTTCAATGGCT | 246 |
| chr10:78223750-78225764 | p2 | (TA)7 | ATATGGGGCCCACAATTTTT | CCACCAAATCCAAACTCACA | 268 |
| chr10:78338275-78340287 | p2 | (AC)6 | CGAAGTTTTCAATTGCGGAT | GAAGTGAGTTTGGAGAGGCG | 106 |
| chr10:78338275-78340287 | p6 | (TATCGC)3 | CGAAGTTTTCAATTGCGGAT | GAAGTGAGTTTGGAGAGGCG | 106 |
| chr10:78408296-78410332 | p2 | (TA)18 | TGAATTATGACGTTTGGGCA | CATCCGCAATTGGTTCTTCT | 283 |
| chr10:78408296-78410332 | p2 | (AT)8 | TGAATTATGACGTTTGGGCA | CATCCGCAATTGGTTCTTCT | 283 |
| chr10:78606542-78608564 | p2 | (TA)11 | AGTTTGGAATCCGGCTTTTT | AATGTCGGAGGATGACTTCG | 152 |
| chr10:78670290-78672302 | p2 | (CA)6 | TTACATACCTTGCCCGCTTC | TGTTGGGATACTTGTTGGCA | 142 |
| chr10:78670290-78672302 | p4 | (TTTC)4 | TTACATACCTTGCCCGCTTC | TGTTGGGATACTTGTTGGCA | 142 |
| chr10:78670560-78672576 | p2 | (CA)6 | ACACTAATCCGCCACCATTC | CTTGTTGCATTGGGAAGGTT | 238 |
| chr10:78670560-78672576 | p4 | (TTTC)4 | ACACTAATCCGCCACCATTC | CTTGTTGCATTGGGAAGGTT | 238 |
| chr10:78819037-78821049 | p2 | (TA)6 | GGTTAGTCCGCACGTTCATT | GGTTCAATATTCGCCTCCAA | 290 |
| chr10:78870729-78872744 | p5 | (ATGAT)3 | GATGCTGCCCAAATTTCTGT | CGCCTCTCCAAACTCAATTC | 107 |
| chr10:78965531-78967549 | p2 | (AT)9 | GGGGGTGGCTATCTGGTATT | CAAAATTTCTGAGGGGTCCA | 237 |
| chr10:78973477-78975492 | p5 | (CCCGC)3 | TCCCGTCATATGGTATCCGT | TATCACACATAAGCCGGCAA | 150 |
| chr10:78994824-78996840 | p4 | (ATAA)4 | ACTTCCCAAACCTCCGAACT | TCGGATTGATGTAGTGCCAG | 170 |
| chr10:79045755-79047770 | p3 | (GTT)5 | ACAAAGGGGCAAGTGTCAAC | CACGTTGCTGCCTTTTCATA | 270 |
| chr10:79058775-79060793 | p2 | (AT)9 | CTAGCGACCAAACACCCCTA | AGCTGGCCTGCACTGTTATT | 182 |
| chr10:79068392-79070404 | p2 | (AT)17 | TGCAGATAACCCTAGGACGG | TGGGCCCTTGAGTATGTTTC | 115 |
| chr10:79068392-79070404 | p2 | (TA)6 | TGCAGATAACCCTAGGACGG | TGGGCCCTTGAGTATGTTTC | 115 |
| chr10:79219600-79221626 | p2 | (TA)13 | TTCGTTGGCTAGATATGCCC | ATATCACGCTCACCCCTTTG | 137 |
| chr10:79233284-79235296 | p2 | (TG)6 | ATGTGTGCCTATGAAAGGGC | CATACCTGGCCCTACCTGAA | 270 |
| chr10:79238923-79240941 | p2 | (AG)9 | TCGGTGATAGCTAGCCGAGT | CAAGAAGCCAACCAATCCAT | 128 |
| chr10:79244492-79246504 | p2 | (CA)6 | TTGTGAAAAACGGGGAAGTC | TTGTTGGTGTGTTGTGGCTT | 217 |
| chr10:79244492-79246504 | p2 | (TG)6 | TTGTGAAAAACGGGGAAGTC | TTGTTGGTGTGTTGTGGCTT | 217 |
| chr10:79273732-79275748 | p2 | (AG)8 | TTGGGAAGCAGAGGAAGAAA | TCCTGTCACAACAACCCAAA | 168 |
| chr10:79669382-79671400 | p2 | (AT)7 | AGTTACGGATGTCACGACCC | GTCGGCTTGCGTATCATTTT | 102 |
| chr10:79774664-79776700 | p2 | (AT)18 | TTCCAAATCAATGGTGCTGA | GTTCGAGAAGGAAGGAACCC | 103 |
| chr10:79805499-79807515 | p2 | (TA)8 | TGCTGCAAGTTTTAGAGCGA | ATTCCCAACAAGCAAAATGG | 296 |
| chr10:80001867-80003883 | p4 | (ATGT)4 | CATTGGGTGTTGTGACGAAG | TCATGCCTTTGAAAGAAGGG | 274 |
| chr10:80006605-80008623 | p6 | (TATATG)3 | TGTATAGCAGCGCGTTATGG | CCGGAATTCCAACTCAAAAA | 137 |
| chr10:80075934-80077948 | p2 | (TG)7 | GAAGAGATGACTTGGGAGCG | TGCAAAAACACGGGACATAA | 127 |
| chr10:80085451-80087469 | p6 | (ATATGA)3 | ATGGCAGCGAAGTGAGAAAT | TTTGTACGACATTCCCCCAT | 152 |
| chr10:80090900-80092915 | p5 | (TTTCA)3 | AAGAAGGCCAAGACGAGACA | CACTCGGGGATTCTACGAAA | 230 |
| chr10:80214462-80216477 | p5 | (AAAGA)3 | TGAGGAATTTGCTGAGTCGTT | AAAATCATGGGAAAAAGCCC | 271 |
| chr10:80216437-80218469 | p2 | (AT)16 | CCTTCTTATTTGCCGAACCA | CAAAGGAGGTTCATGCCACT | 123 |

| Marker ID | SSR type | SSR motif | FORWARD PRIMER (5'-3') | REVERSE PRIMER (5'-3') | size (bp) |
|---|---|---|---|---|---|
| chr10:80225857-80227873 | p5 | (ATTGC)3 | TCCCGAAATCGAGGTATGAG | GAGCATCACATCTTGGCAGA | 245 |
| chr10:80225857-80227873 | p4 | (TTAT)4 | TCCCGAAATCGAGGTATGAG | GAGCATCACATCTTGGCAGA | 245 |
| chr10:80366348-80368363 | p3 | (AGG)5 | GATTGCCCTTGAATCATCGT | CTCCATGCCTCATTCTCCAT | 175 |
| chr10:80404257-80406272 | p5 | (TGTTG)3 | TACAAGATGATGTGGGGCAA | ACCGTGATTTCAAAACCTGC | 282 |
| chr10:80404257-80406272 | p3 | (CAA)6 | TACAAGATGATGTGGGGCAA | ACCGTGATTTCAAAACCTGC | 282 |
| chr10:80424936-80426980 | p2 | (AT)22 | CTACAAGCCTCCCACCACAT | GCATGCATGATATGGCACTC | 237 |
| chr10:80542817-80544831 | p2 | (AT)13 | TTGGGTCGTGACAGAGTCAG | ACACTTGACAGGGGGTGAAG | 115 |
| chr10:80613106-80615138 | p2 | (AT)16 | AACCGAACCAAACACACACA | CCCATTTTCGAGCATTGATT | 201 |
| chr10:80652678-80654702 | p2 | (AT)12 | CACTCATACTGCCCTGCTCA | ATATGGACCCCCAGGATTTC | 176 |
| chr10:80663123-80665179 | p6 | (CATAAG)3 | TCAATGTGTTCGCACTGGAT | CCGCTCGTCGATACGATATT | 151 |
| chr10:80684846-80686864 | p6 | (TTGACT)3 | CTTCTCCCCAAAGTGAACCA | AGGTGTAACGATCCGGTCAG | 108 |
| chr10:80689099-80691129 | p2 | (AT)15 | AATGGGTTGTGGCTTGATTC | TTGCGATGCTCGACAGATAG | 179 |
| chr10:80689099-80691129 | p2 | (TA)7 | AATGGGTTGTGGCTTGATTC | TTGCGATGCTCGACAGATAG | 179 |
| chr10:80689099-80691129 | p3 | (CAA)5 | AATGGGTTGTGGCTTGATTC | TTGCGATGCTCGACAGATAG | 179 |
| chr10:80823797-80825813 | p4 | (ACAT)4 | GGCCGATGAGGCTAACATAA | CCGGAGCATGTACTTTTGGT | 245 |
| chr10:80943400-80945415 | p3 | (CCA)5 | TCCGAGGCTTCTTATGTTGG | TTTATCGTCGTATGGTGGCA | 163 |
| chr10:81022510-81024530 | p2 | (AT)10 | TACCTATATGGTCGGGCAGC | CTCCTATGGATCTGCACCGT | 235 |
| chr10:81074160-81076172 | p2 | (TG)6 | TACTCCGTTCGTGCTGTCTG | TCGGAAACCGTAACACATCA | 180 |
| chr10:81110333-81112348 | p5 | (GCAAT)3 | TGCACCCAAAACCCTAACTC | GTTCCACGGACCGTACAAGT | 139 |
| chr10:81218890-81220905 | p3 | (ACA)5 | GCAAATAGGATTTGGGAGCA | CATTTGCAGATTCTGAGCGA | 170 |
| chr10:81218890-81220905 | p5 | (CACAA)3 | GCAAATAGGATTTGGGAGCA | CATTTGCAGATTCTGAGCGA | 170 |
| chr10:81222195-81224213 | p2 | (TA)6 | GATCCAATTTTGTCGGGCTA | CTCCGTTCGACCAATGAGAT | 191 |
| chr10:81222220-81224232 | p2 | (TA)6 | GATCCAATTTTGTCGGGCTA | CTCCGTTCGACCAATGAGAT | 166 |
| chr10:81299914-81301926 | p2 | (TC)6 | AAATTGGCATTATTGGTCGC | GGACTGCCTAATGTTTCCCA | 117 |
| chr10:81326909-81328924 | p5 | (TTGCA)3 | ACTTTTTGGGTGCGTTTTTG | ATTCACCTCCACTTCATGCC | 264 |
| chr10:81401054-81403112 | p2 | (AT)29 | TCGACCCTTCTTTTGCCTTA | GGCATTGTTCGTCCTTCAAT | 155 |
| chr10:81419117-81421141 | p2 | (AT)12 | AGTCCAGTTGCGTCAATCCT | CAGAACTTTTGCCTTCTCCG | 158 |
| chr10:81490262-81492283 | p6 | (CATATA)3 | GGGATTCGCCTAAGGTAAGC | CAGTTGGTATCGGAGCTGGT | 100 |
| chr10:81490262-81492283 | p3 | (CAA)7 | GGGATTCGCCTAAGGTAAGC | CAGTTGGTATCGGAGCTGGT | 100 |
| chr10:81490262-81492283 | p2 | (TC)6 | GGGATTCGCCTAAGGTAAGC | CAGTTGGTATCGGAGCTGGT | 100 |
| chr10:81501487-81503499 | p2 | (TC)6 | CATGCCTAGGAACAAAGGGA | GAATTGAGCCGAGCAAAGTC | 295 |
| chr10:81695008-81697023 | p5 | (ATTTT)3 | TACAACCCTCCGGTAGCATC | CATCTTGCCAAAGGAAAAGC | 123 |
| chr10:81727647-81729695 | p2 | (CT)24 | GGTGAAAGAGGACTGCAAGC | AATACCTCGTTTGTTCCCCC | 260 |
| chr10:81834245-81836299 | p5 | (CAAAC)4 | AACCAAACCAAACCAAACCA | CCTTTCTTGTGGCAGCTAGG | 195 |
| chr10:81834245-81836299 | p2 | (TA)9 | AACCAAACCAAACCAAACCA | CCTTTCTTGTGGCAGCTAGG | 195 |
| chr10:81871612-81873666 | p2 | (AT)27 | TCTTTGGAACTCGCTTGCTT | CCTAGTCGGGTCGTGACAAT | 173 |

| Marker ID | SSR type | SSR motif | FORWARD PRIMER (5'-3') | REVERSE PRIMER (5'-3') | size (bp) |
|---|---|---|---|---|---|
| chr10:82057664-82059676 | p2 | (AT)6 | CGAATTCAGCCCCTTCAATA | TAGCGATTGCCCGGTTATAG | 260 |
| chr10:82057664-82059676 | p5 | (ATTTT)3 | CGAATTCAGCCCCTTCAATA | TAGCGATTGCCCGGTTATAG | 260 |
| chr10:82058573-82060588 | p2 | (AT)6 | CCGGATCAGAAATTGAAGGA | CAGTGGCGTGCTGTCATAGT | 244 |
| chr10:82058573-82060588 | p5 | (ATTTT)3 | CCGGATCAGAAATTGAAGGA | CAGTGGCGTGCTGTCATAGT | 244 |
| chr10:82124788-82126804 | p4 | (GAAA)4 | CAGCTGGCTTACACTCCACA | TCATCGACTGTCACGAGAGG | 291 |
| chr10:82145872-82147944 | p2 | (AT)36 | AAGCTGCTGATGTTTTGGCT | TGAGTAGAGGGGGACATGGA | 155 |
| chr10:82153945-82156002 | p3 | (ATA)17 | GCACCCGGTTGAATTAAGAA | GGGGAGGGAAAAGATTTCAG | 115 |
| chr10:82153945-82156002 | p3 | (AAG)19 | GCACCCGGTTGAATTAAGAA | GGGGAGGGAAAAGATTTCAG | 115 |
| chr10:82331649-82333699 | p2 | (TA)11 | TTTTAGGTTGTTGCGTGCAG | ACCACTACGAACACGAAGGG | 111 |
| chr10:82532685-82534700 | p5 | (TTACT)3 | AATTCCGCTCCTATGCAATG | AATGGCATACTACCCGCAAG | 272 |
| chr10:82605964-82607985 | p3 | (TGA)7 | AAGTAGCCGTTGGGACATTG | GGGGGATGGGTTAAAGTGTT | 243 |
| chr10:82611541-82613553 | p2 | (TA)6 | AAATATCAACCGTGGCTTCG | CGATCTAATCAATCGCGACA | 164 |
| chr10:82738506-82740521 | p5 | (TTATT)3 | TCTCCCAAAAAGACCTCCCT | TTGATTTCCATTGGTGGGTT | 172 |
| chr10:82802636-82804651 | p5 | (AACAC)3 | TCCTCACATCCTATGGCCTC | AGGTTGGGCATAGTTTCACG | 133 |
| chr10:82915881-82917896 | p3 | (TTG)5 | TGAAGGAAACGGGAATTGAG | GACAATTTCAAACGCGGATT | 227 |
| chr10:82915881-82917896 | p5 | (TATGA)3 | TGAAGGAAACGGGAATTGAG | GACAATTTCAAACGCGGATT | 227 |
| chr10:82934576-82936591 | p3 | (ACA)5 | CATCCTTGGCCCATAATCAC | CACGTTCTCGTGCACAGTTT | 102 |
| chr10:82945021-82947033 | p2 | (AT)6 | TGGAAAATGGCCTTCAAAAC | GCAAACAACTTTTCACGCAA | 142 |
| chr10:82945021-82947033 | p2 | (AT)10 | TGGAAAATGGCCTTCAAAAC | GCAAACAACTTTTCACGCAA | 142 |
| chr10:82945021-82947033 | p2 | (AT)11 | TGGAAAATGGCCTTCAAAAC | GCAAACAACTTTTCACGCAA | 142 |
| chr10:82945249-82947269 | p2 | (AT)6 | TTGTTGTGAGTGCGAATGGT | CTTGCCATCCCATGAAAACT | 200 |
| chr10:82945249-82947269 | p2 | (AT)10 | TTGTTGTGAGTGCGAATGGT | CTTGCCATCCCATGAAAACT | 200 |
| chr10:82945249-82947269 | p2 | (AT)11 | TTGTTGTGAGTGCGAATGGT | CTTGCCATCCCATGAAAACT | 200 |
| chr10:82980739-82982767 | p2 | (AT)20 | ATCACGCTTCATCCAAGTCC | CATGACAAGGTATTGCGTGG | 184 |
| chr10:82980768-82982786 | p2 | (AT)20 | ATCACGCTTCATCCAAGTCC | CATGACAAGGTATTGCGTGG | 155 |
| chr10:83107878-83109894 | p2 | (TC)8 | GCGCAAAACCTACAAATGGT | TGTTCGTTAGAATTTCGGGC | 176 |
| chr10:83107878-83109894 | p2 | (AT)10 | GCGCAAAACCTACAAATGGT | TGTTCGTTAGAATTTCGGGC | 176 |
| chr10:83347707-83349722 | p5 | (AAAAT)3 | CGGAAGCAACACTATCGTCA | GTGACCATTCTTGCGCTGTA | 204 |
| chr10:83426711-83428799 | p2 | (TC)7 | AAGAAAAAGGTCGGGGAGAA | AGGATGTGCAAACTCTCGCT | 128 |
| chr10:83426711-83428799 | p2 | (AT)44 | AAGAAAAAGGTCGGGGAGAA | AGGATGTGCAAACTCTCGCT | 128 |
| chr10:83524523-83526538 | p5 | (TGCAT)3 | CGTCTGAAGTTGGGATGGAT | ATGCCTCTACATGGCACACA | 126 |
| chr10:83587499-83589513 | p2 | (GA)7 | GTTGAGTCTACCCGCTTTGC | CAGGAGGGATGAAACAAGGA | 168 |
| chr10:83626277-83628301 | p6 | (ACACTC)4 | CCGTCAAAATTCCTCCTTCA | GCCATCGTCAAGCAGTAACA | 140 |
| chr10:83665173-83667195 | p2 | (CT)11 | CTTGGAAGAATGGCCAAAGA | ATGTTTGGTTGAGAAAGCGG | 276 |
| chr10:83665173-83667195 | p6 | (ATTTTT)3 | CTTGGAAGAATGGCCAAAGA | ATGTTTGGTTGAGAAAGCGG | 276 |
| chr10:83671124-83673142 | p2 | (CA)9 | TCCCCCAAGTCTTGTTTTTG | ATGTTTGGTTGAGAAAGCGG | 263 |

| Marker ID | SSR type | SSR motif | FORWARD PRIMER (5'-3') | REVERSE PRIMER (5'-3') | size (bp) |
|---|---|---|---|---|---|
| chr10:83671124-83673142 | p4 | (CATT)4 | TCCCCCAAGTCTTGTTTTTG | ATGTTTGGTTGAGAAAGCGG | 263 |
| chr10:83671124-83673142 | p2 | (TA)15 | TCCCCCAAGTCTTGTTTTTG | ATGTTTGGTTGAGAAAGCGG | 263 |
| chr10:83671495-83673511 | p2 | (CA)9 | CCGCTTTCTCAACCAAACAT | CTAAGGTGGGACCCTCTTCC | 115 |
| chr10:83671495-83673511 | p4 | (CATT)4 | CCGCTTTCTCAACCAAACAT | CTAAGGTGGGACCCTCTTCC | 115 |
| chr10:83671495-83673511 | p2 | (TA)15 | CCGCTTTCTCAACCAAACAT | CTAAGGTGGGACCCTCTTCC | 115 |
| chr10:83685796-83687820 | p6 | (ACACTC)4 | CCGTCAAAATTCCTCCTTCA | GCCATCGTCAAGCAGTAACA | 140 |
| chr10:83768844-83770862 | p6 | (TATTTT)3 | CAGGGTAACCTAAGGGGGAA | CCTAGGGTTGTGTCGTGCTT | 175 |
| chr10:83793883-83795898 | p5 | (ATTAT)3 | TGAATTTGAAAGGTTTCGGG | GACGGCGTTTCATTTACGTT | 158 |
| chr10:83793883-83795898 | p6 | (TATATG)3 | TGAATTTGAAAGGTTTCGGG | GACGGCGTTTCATTTACGTT | 158 |
| chr10:83835447-83837477 | p2 | (AT)15 | TGTTGTGGGCTTGTGTGTTT | AGTGAACCAACCAATCGGAG | 162 |
| chr10:84059290-84061302 | p2 | (TA)6 | GACGTGGCAAAGTAGCCAAT | CACTAACCCCTCCCCATTTT | 118 |
| chr10:84059290-84061302 | p5 | (TTGTA)3 | GACGTGGCAAAGTAGCCAAT | CACTAACCCCTCCCCATTTT | 118 |
| chr10:84059290-84061302 | p6 | (TTGGAG)3 | GACGTGGCAAAGTAGCCAAT | CACTAACCCCTCCCCATTTT | 118 |
| chr10:84059899-84061914 | p2 | (TA)6 | GCCAAGTCAGCGAAAAATGT | AGACAGCTCCTTGTGGTCGT | 149 |
| chr10:84059899-84061914 | p5 | (TTGTA)3 | GCCAAGTCAGCGAAAAATGT | AGACAGCTCCTTGTGGTCGT | 149 |
| chr10:84059899-84061914 | p6 | (TTGGAG)3 | GCCAAGTCAGCGAAAAATGT | AGACAGCTCCTTGTGGTCGT | 149 |
| chr10:84059899-84061914 | p3 | (TTC)5 | GCCAAGTCAGCGAAAAATGT | AGACAGCTCCTTGTGGTCGT | 149 |
| chr10:84090181-84092196 | p5 | (AAAAG)3 | GGAGAACAATGAAGGGGTGA | GCTAGGTTCCACTGCACCAT | 189 |
| chr10:84110680-84112696 | p3 | (TCA)5 | CAGAATAACCTGAAGGCCCA | CATCCTTTTGCAACACATGC | 237 |
| chr10:84110680-84112696 | p5 | (AACAA)3 | CAGAATAACCTGAAGGCCCA | CATCCTTTTGCAACACATGC | 237 |
| chr10:84110697-84112723 | p3 | (TCA)5 | CAGAATAACCTGAAGGCCCA | CATCCTTTTGCAACACATGC | 220 |
| chr10:84110697-84112723 | p5 | (AACAA)3 | CAGAATAACCTGAAGGCCCA | CATCCTTTTGCAACACATGC | 220 |
| chr10:84117035-84119067 | p2 | (AT)16 | TCACTGCAGAGCATGGTTTC | CCCCTAGTTCGTGACTTCCA | 220 |
| chr10:84221050-84223064 | p2 | (AT)7 | TCGTGGGATATCATTTGGGT | GACCTTTCGGAATCGACAAA | 211 |
| chr10:84221050-84223064 | p2 | (AT)8 | TCGTGGGATATCATTTGGGT | GACCTTTCGGAATCGACAAA | 211 |
| chr10:84367917-84369977 | p2 | (TA)30 | GCAAAAAGACCAAGACTCGC | GAAAAATGGGGTGGTCATTG | 205 |
| chr10:84414654-84416669 | p3 | (TCA)5 | TAATCAAGCACCTCCGGAAC | TGTTCATGCCCACAGGTAGA | 145 |
| chr10:84457073-84459088 | p5 | (AAAAG)3 | CCACGAATCATTGCCTTTCT | TCTGAAGAGTTGGAGCCCAT | 270 |
| chr10:84492692-84494726 | p2 | (AT)17 | ATCGTGAGACCATGCTACCC | CCGTTTTCTTTGGATGCTGT | 271 |
| chr10:84513813-84515831 | p6 | (AAAAGA)3 | GCTGGGATAACCCTCGTGTA | GATGGGCCAAGGTAAAGTCA | 173 |
| chr10:84736675-84738702 | p3 | (TAT)9 | GAGAAATGTTCCCAGACGGA | ATGCATATGAGATGCCGTGA | 195 |
| chr10:84846966-84848988 | p2 | (TA)11 | CAACTGCGAGATGGCAAGTA | TGAGTCTTTGGCCGTACACA | 216 |
| chr10:84857330-84859396 | p2 | (CT)6 | CAGACAACGACCAATCGAGA | TTATCTTCAAATTTGGCGGG | 184 |
| chr10:84899450-84901466 | p4 | (TTAA)4 | ACCAACGCAACAACTAACCC | TCGACCAATTCGAACCCTAC | 256 |
| chr10:84914123-84916171 | p2 | (AT)24 | CTGTCTTCTGGGCGATTTTC | ACGTGGACCAAAATTTCAGC | 183 |
| chr10:84923707-84925733 | p2 | (AG)16 | GCAAATTCTGTGCAAAAGCA | CCAAAGTGGATTAGACCCGA | 245 |

| Marker ID | SSR type | SSR motif | FORWARD PRIMER (5'-3') | REVERSE PRIMER (5'-3') | size (bp) |
|---|---|---|---|---|---|
| chr10:84923707-84925733 | p2 | (AT)13 | GCAAATTCTGTGCAAAAGCA | CCAAAGTGGATTAGACCCGA | 245 |
| chr10:84923707-84925733 | p2 | (TG)6 | GCAAATTCTGTGCAAAAGCA | CCAAAGTGGATTAGACCCGA | 245 |
| chr10:84993354-84995366 | p2 | (TG)6 | TGATGTGTTACGGTTTCCGA | ACACCAACATCGCAAACAAA | 264 |
| chr10:85140023-85142041 | p3 | (TGT)6 | GATATGATGCGACGAAAGCA | GTATGAGGCCCATAGAGGCA | 111 |
| chr10:85202714-85204730 | p4 | (TATG)4 | CTATCCTGGCCGTCGATAAA | TCCGCTCGTCGCTATTAAGT | 169 |
| chr10:85219416-85221430 | p2 | (AG)7 | GGTAAAGAAGCCATGTCGGA | CGGGATACTTAAGGCAACCA | 112 |
| chr10:85336537-85338561 | p2 | (AT)12 | GTGTGATTTGGGGGCTAGAA | ACGAGAGTCTAACTCGGGCA | 245 |
| chr10:85345750-85347770 | p2 | (AT)10 | AACCCTCAACTTAGGCCGAT | TTCTCTGGAGGCTTGTCGAT | 122 |
| chr10:85360359-85362387 | p2 | (TA)14 | CGTGTAGGGGGCTGTTCTAA | GACTTAACGAAAATCGGGCA | 109 |
| chr10:85384999-85387015 | p2 | (TA)6 | GAACTCTTACGTTGCGGCTC | GCAGTTTCCATCCAAGGTGT | 134 |
| chr10:85384999-85387015 | p2 | (TA)8 | GAACTCTTACGTTGCGGCTC | GCAGTTTCCATCCAAGGTGT | 134 |
| chr10:85399785-85401801 | p2 | (AT)8 | GCATGAAAGTCATCCACCCT | CCAGTAAATTTGCTAGCGGC | 187 |
| chr10:85412678-85414714 | p2 | (AT)18 | GGGGCGGATTTCATTTACTT | GGGAGTGAAGGTGCAAATGT | 259 |
| chr10:85511490-85513502 | p2 | (AT)10 | TACTAAGGCATTCGGCCATC | TGTTTTCTTTGGACGCTGTG | 201 |
| chr10:85511511-85513526 | p2 | (AT)10 | TACTAAGGCATTCGGCCATC | TGTTTTCTTTGGACGCTGTG | 180 |
| chr10:85511521-85513543 | p2 | (AT)10 | TACTAAGGCATTCGGCCATC | TGTTTTCTTTGGACGCTGTG | 170 |
| chr10:85637812-85639892 | p2 | (AT)40 | CCATCGATGTGGGATAAACC | GGGTCAGATCAACCCAAGAA | 105 |
| chr10:85637812-85639892 | p2 | (AT)39 | CCATCGATGTGGGATAAACC | GGGTCAGATCAACCCAAGAA | 105 |
| chr10:85643206-85645244 | p2 | (TA)19 | ACTGCCAACGCCTCCTACTA | GGCAGGCAAGAAAACAAAAG | 144 |
| chr10:85688059-85690071 | p2 | (GA)6 | TGAGGTGACGAGTCGATGAG | AACCTGATGTCCGAGTTTGG | 292 |
| chr10:85736264-85738279 | p3 | (AAT)5 | CCCAAGACCCACAGGAAGTA | CCTGGTGGGTAGATGAGGAA | 123 |
| chr10:85825185-85827233 | p2 | (TA)24 | CAGACAACCCCTAGGACGAA | ATGGCACTCATCCGAGTAGG | 262 |
| chr10:85831809-85833824 | p5 | (AACCC)3 | GCGGATTTTCAAACCAAAAA | ATCCGCAGAATACGGTTGTC | 161 |
| chr10:85865961-85867985 | p2 | (TA)12 | CAGTCCTAGGAGGGACCACA | TTTGGTGAACCAGGTGAACA | 118 |
| chr10:85940918-85942933 | p5 | (TTTGG)3 | AAAGACGGCTGAAGCTCTTG | TGCTAAAAGCAGTGCCTGAA | 128 |
| chr10:85968563-85970578 | p6 | (TTCAGT)3 | TACAGGACGACGGATCAACA | AAGGCCCCCATCATAACTTT | 183 |
| chr10:85968563-85970578 | p5 | (TTTAT)3 | TACAGGACGACGGATCAACA | AAGGCCCCCATCATAACTTT | 183 |
| chr10:85998865-86000880 | p2 | (AT)9 | TGTGATCGGGGTTTCCTTAG | GAAGCTGCCCAATCTCAGAC | 158 |
| chr10:85998865-86000880 | p5 | (AATGC)3 | TGTGATCGGGGTTTCCTTAG | GAAGCTGCCCAATCTCAGAC | 158 |
| chr10:86054802-86056858 | p2 | (AT)28 | TGTCCAACATCCCTAACGGT | TCCGGACACAGGGTAGTCTC | 119 |
| chr10:86073651-86075672 | p3 | (ATC)7 | TTTGAAGACTTGGGGGAATG | CTAAGGGACGGTCCGTAACA | 182 |
| chr10:86073651-86075672 | p3 | (CAA)7 | TTTGAAGACTTGGGGGAATG | CTAAGGGACGGTCCGTAACA | 182 |
| chr10:86126073-86128094 | p3 | (ATC)7 | TTTGAAGACTTGGGGGAATG | CTAAGGGACGGTCCGTAACA | 182 |
| chr10:86126073-86128094 | p3 | (CAA)7 | TTTGAAGACTTGGGGGAATG | CTAAGGGACGGTCCGTAACA | 182 |
| chr10:86184772-86186787 | p5 | (GGGAA)3 | ACGTCGGTTCGCTTTTCTAA | GGATTCCTGCATTTTGGCTA | 146 |
| chr10:86184772-86186787 | p5 | (GGGAA)3 | ACGTCGGTTCGCTTTTCTAA | GGATTCCTGCATTTTGGCTA | 146 |

| Marker ID | SSR type | SSR motif | FORWARD PRIMER (5'-3') | REVERSE PRIMER (5'-3') | size (bp) |
|---|---|---|---|---|---|
| chr10:86184772-86186787 | p5 | (GGGAA)3 | ACGTCGGTTCGCTTTTCTAA | GGATTCCTGCATTTTGGCTA | 146 |
| chr10:86184772-86186787 | p5 | (AGGGA)3 | ACGTCGGTTCGCTTTTCTAA | GGATTCCTGCATTTTGGCTA | 146 |
| chr10:86184772-86186787 | p3 | (ATC)5 | ACGTCGGTTCGCTTTTCTAA | GGATTCCTGCATTTTGGCTA | 146 |
| chr10:86210312-86212327 | p3 | (AAT)5 | CCCTCTTTCGAATCTTTCCC | TCGTATCCTCCATCCCTCTG | 183 |
| chr10:86222343-86224382 | p3 | (TAA)13 | CCCTCTTTCGAATCTTTCCC | TCGTATCCTCCATCCCTCTG | 216 |
| chr10:86233838-86235853 | p5 | (CCACC)3 | ACCCCCAATTACCTCACTCC | AGGGGGATAGTGGGTCGTAG | 147 |
| chr10:86233838-86235853 | p5 | (CCGGG)3 | ACCCCCAATTACCTCACTCC | AGGGGGATAGTGGGTCGTAG | 147 |
| chr10:86247675-86249702 | p3 | (TCT)6 | AAAATTTGATGCCCAGTTGC | TGGAGGAAGAAAGTGGGTTG | 188 |
| chr10:86247675-86249702 | p3 | (AAG)9 | AAAATTTGATGCCCAGTTGC | TGGAGGAAGAAAGTGGGTTG | 188 |
| chr10:86433003-86435018 | p5 | (TTTAG)3 | GCCTTGTCTCGCTCTTTGAC | GTGAAAGCACAGGCTGAACA | 204 |
| chr10:86450791-86452809 | p3 | (AAG)5 | ATGACCAAACGAGCTAACCG | ATAATTTGCCTCCCGAGCTT | 213 |
| chr10:86450791-86452809 | p6 | (AAACAA)3 | ATGACCAAACGAGCTAACCG | ATAATTTGCCTCCCGAGCTT | 213 |
| chr10:86495568-86497582 | p2 | (CA)11 | CTGGTACTCCGGAGCAATGT | GGACCGTGGCAGTTTAAGAA | 193 |
| chr10:86495610-86497625 | p2 | (CA)11 | CTGGTACTCCGGAGCAATGT | GGACCGTGGCAGTTTAAGAA | 151 |
| chr10:86517167-86519182 | p5 | (ATCAT)3 | AAAACCCACAATTCCCCTTC | CCTAGTCGGGTCGTGACATT | 204 |
| chr10:86571913-86573935 | p2 | (TA)11 | CTGGACAACTTGTGGAGCAA | GTCTTTTCGGTGTGGCAAAT | 186 |
| chr10:86698037-86700061 | p3 | (CAT)8 | GAGCCACATGGCCTACTCAT | AGGCTCGTAGACAGTTGCGT | 192 |
| chr10:86730973-86732988 | p5 | (AACAT)3 | ACTCCAACCAGGCTCAAAGA | CAATTTTCCGCATCCACTTT | 226 |
| chr10:86766819-86768833 | p2 | (AT)7 | AGTGGCGGTGTAGCAGAACT | CAAATCGGTAGCAGGGGATA | 227 |
| chr10:86791757-86793769 | p3 | (CAA)5 | CCCATTGCCTTGGAATCTTA | GGCGACCGAAAGGTATGTTA | 242 |
| chr10:86791757-86793769 | p2 | (CT)6 | CCCATTGCCTTGGAATCTTA | GGCGACCGAAAGGTATGTTA | 242 |
| chr10:86803592-86805606 | p2 | (GT)7 | AAAAAGAAACTCCCCCGAAA | TCGACATTTCCGCATCAATA | 260 |
| chr10:86879056-86881086 | p2 | (TA)15 | AAATCACGAGCTCCATCACC | ATGCTCCACCCATTTCAGAG | 298 |
| chr10:86879056-86881086 | p2 | (AT)8 | AAATCACGAGCTCCATCACC | ATGCTCCACCCATTTCAGAG | 298 |
| chr10:86879700-86881716 | p2 | (TA)15 | GGTGGACCACATGGCTAACT | GGGTCCTGTCCCGTCTTTAT | 256 |
| chr10:86879700-86881716 | p2 | (AT)8 | GGTGGACCACATGGCTAACT | GGGTCCTGTCCCGTCTTTAT | 256 |
| chr10:86899888-86901903 | p5 | (AATTT)3 | TACAACTGGCCCGATTTCTC | TTTTCCCTTTGTTTCCATGC | 206 |
| chr10:87262567-87264581 | p2 | (AT)7 | AGGTTGAGTTCGCCTACCCT | CCAAATCGCGAGGAAGTTTA | 207 |
| chr10:87290943-87292958 | p5 | (ACCAA)3 | GAATGCCCTTATTTTGGGGT | GCAAGGGTTCGAACTTGAAA | 154 |
| chr10:87290943-87292958 | p5 | (TTGGT)3 | GAATGCCCTTATTTTGGGGT | GCAAGGGTTCGAACTTGAAA | 154 |
| chr10:87394513-87396528 | p5 | (AAATA)3 | TTTCAATTTTGGTTCCTGCC | TCCCTAAAATTCAGCCCAGA | 234 |
| chr10:87411439-87413454 | p5 | (AACCG)3 | CAAACTAGGGCTGGGCATAA | TGACCTGGAACCAAACATCA | 252 |
| chr10:87439695-87441709 | p2 | (TA)7 | TATGGGTGATCGTGGACAGA | ACTTGGCTGTAAATTGCGCT | 108 |
| chr10:87439695-87441709 | p5 | (TGTGT)3 | TATGGGTGATCGTGGACAGA | ACTTGGCTGTAAATTGCGCT | 108 |
| chr10:87446053-87448119 | p2 | (TA)6 | CATGGTAAGCGAGACTGCAA | GTTTTCTAATCCACTGCGCC | 185 |
| chr10:87446053-87448119 | p2 | (TA)33 | CATGGTAAGCGAGACTGCAA | GTTTTCTAATCCACTGCGCC | 185 |

| Marker ID | SSR type | SSR motif | FORWARD PRIMER (5'-3') | REVERSE PRIMER (5'-3') | size (bp) |
|---|---|---|---|---|---|
| chr10:87446053-87448119 | p5 | (TGTGT)3 | CATGGTAAGCGAGACTGCAA | GTTTTCTAATCCACTGCGCC | 185 |
| chr10:87463160-87465175 | p5 | (ATGTT)3 | CTTTTGAAGGGTAATCCGCA | CTACCGAACCCACCAGTTGT | 197 |
| chr10:87482588-87484626 | p5 | (TATAT)3 | GGTTTTCGTGCAGGCATATT | GTGTCCCGGACCTCTAGTGA | 242 |
| chr10:87482588-87484626 | p2 | (AT)19 | GGTTTTCGTGCAGGCATATT | GTGTCCCGGACCTCTAGTGA | 242 |
| chr10:87482588-87484626 | p2 | (AT)8 | GGTTTTCGTGCAGGCATATT | GTGTCCCGGACCTCTAGTGA | 242 |
| chr10:87533963-87535977 | p2 | (TA)7 | GGTACAGGTTGTGCGGTCTT | AGTAATGCAATGCTCCGACC | 101 |
| chr10:87607280-87609292 | p3 | (ATT)7 | AATTTTGTCCCGCTTTTCCT | GATGCGTGTGCATAAGATGG | 263 |
| chr10:87607280-87609292 | p2 | (TC)6 | AATTTTGTCCCGCTTTTCCT | GATGCGTGTGCATAAGATGG | 263 |
| chr10:87620679-87622694 | p3 | (CTT)5 | CCCAAGTTTCAAAAGGCAAA | TCAAAACCCATAAACCCCAA | 117 |
| chr10:87620679-87622694 | p5 | (AAAAT)3 | CCCAAGTTTCAAAAGGCAAA | TCAAAACCCATAAACCCCAA | 117 |
| chr10:87620991-87623006 | p3 | (CTT)5 | GAAGTACCATTTGCACTGCG | TCGCATTCACACATAAGGGA | 142 |
| chr10:87620991-87623006 | p5 | (AAAAT)3 | GAAGTACCATTTGCACTGCG | TCGCATTCACACATAAGGGA | 142 |
| chr10:87688234-87690252 | p3 | (TAT)6 | TGAATCATCGGCCTAGAACC | GTTCGCTGCGGAAGAAATAG | 218 |
| chr10:87774560-87776580 | p2 | (AT)6 | GCTACATTACCCCGGGATTT | CCGTTTTACGCGTTTGTTTT | 194 |
| chr10:87774610-87776624 | p2 | (AT)6 | GCTACATTACCCCGGGATTT | CCGTTTTACGCGTTTGTTTT | 144 |
| chr10:87774625-87776685 | p2 | (AT)6 | TGGGGGAAAATTGAAATTGA | GTCCGTTTTACGCGTTTGTT | 214 |
| chr10:87785752-87787773 | p3 | (ACA)7 | GGTTGAGGCAAACATCCACT | AGGTGCTGCCTAAGTTGCAT | 293 |
| chr10:87857000-87859018 | p2 | (TA)9 | GGAGACAGAGAGGATGGCAG | TATTTTCAGCCCAAACGGAC | 199 |
| chr10:87909687-87911705 | p6 | (TGAACA)3 | CTGCGTCAGGTTAATTCGGT | TCACTTTGATTCCCTCGACC | 161 |
| chr10:87967401-87969457 | p2 | (TA)28 | AACCATGCAAGCCAACTAGG | CCATAGGAGACTCAGCCAGC | 275 |
| chr10:88021474-88023492 | p6 | (AGCATG)3 | GAAGCTACAGGGCATTTGGA | TCTGGCTAGTTGTGGGCTTT | 168 |
| chr10:88031761-88033773 | p2 | (TA)6 | GCCATGCTCATGATAACCCT | TGGATTACGGACGGATTCTC | 279 |
| chr10:88107920-88109934 | p2 | (AC)6 | CTTGGAGTTTCTGGGTGAGC | GGAAATACCCCCAATGACCT | 295 |
| chr10:88107946-88109961 | p2 | (AC)6 | CTTGGAGTTTCTGGGTGAGC | GGAAATACCCCCAATGACCT | 269 |
| chr10:88108808-88110820 | p2 | (AC)6 | ATTGGCAGAAGCACAACCTT | GTCTGCTGTGCAATGGCTTA | 184 |
| chr10:88118141-88120162 | p3 | (AAC)7 | GCAATTTGATCCTCTGGCAT | AGAGTGCTGCGCTTTTTGAT | 139 |
| chr10:88150668-88152688 | p6 | (CATATA)3 | CTTCCCTTTCCCCGTACTTC | CGACCTTACGTCACTCCGAT | 170 |
| chr10:88150668-88152688 | p5 | (ATTCA)4 | CTTCCCTTTCCCCGTACTTC | CGACCTTACGTCACTCCGAT | 170 |
| chr10:88150668-88152688 | p2 | (AT)7 | CTTCCCTTTCCCCGTACTTC | CGACCTTACGTCACTCCGAT | 170 |
| chr10:88160427-88162442 | p6 | (GTGGGG)3 | ACCTTTACGGGCGACCTATT | TGGGGGTGGGTGAAATATAA | 292 |
| chr10:88160427-88162442 | p5 | (CGTGG)3 | ACCTTTACGGGCGACCTATT | TGGGGGTGGGTGAAATATAA | 292 |
| chr10:88173474-88175489 | p5 | (CAAAC)3 | CTGTCCTCGGATAGGCTCTG | TGGATCATTGTCATGCCCTA | 125 |
| chr10:88263205-88265221 | p2 | (TA)8 | CTGTTTTGGGTGGTGTAGGG | CGTTCGCATTTAACCCAACT | 216 |
| chr10:88267599-88269611 | p2 | (AT)6 | GGTAAGCATTGATGTGCGTG | CGGGCAAAGGCTGTCTAGT | 210 |
| chr10:88316031-88318059 | p2 | (TA)14 | GAGTGATGATTCAGCTGGCA | GGCGAGAACCTATGCTATGC | 261 |
| chr10:88413844-88415860 | p2 | (TG)8 | GGTTCGCTGTGAACTTGTCA | GGCCTCATTGTTGTCCTGTT | 186 |

| Marker ID | SSR type | SSR motif | FORWARD PRIMER (5'-3') | REVERSE PRIMER (5'-3') | size (bp) |
|---|---|---|---|---|---|
| chr10:88470975-88472993 | p2 | (TA)7 | AGTAATCAATGGCACCTCCG | ACTGACGTCCGTTTTCTTGG | 272 |
| chr10:88470994-88473018 | p2 | (TA)7 | AGTAATCAATGGCACCTCCG | ACTGACGTCCGTTTTCTTGG | 253 |
| chr10:88531072-88533084 | p2 | (TA)6 | CATGCCAAAAGAAAGAAGGG | CCCGATTTCCGTTAAGTCCT | 148 |
| chr10:88561581-88563596 | p2 | (AT)8 | ATAAGTCAATGGCACCTCCG | CCGTTTTCTTTGGATGCTGT | 176 |
| chr10:88561581-88563596 | p3 | (TCA)5 | ATAAGTCAATGGCACCTCCG | CCGTTTTCTTTGGATGCTGT | 176 |
| chr10:88573966-88575984 | p6 | (TGGGGG)3 | TTAGCTGGAACCTCGAAGGA | TGTAACTGACAATTCCGCCA | 251 |
| chr10:88618147-88620205 | p2 | (TA)29 | GTCCATTTTTAGGCCCCATT | TTATGTTGCCGATGCAGGTA | 206 |
| chr10:88657342-88659388 | p2 | (AT)23 | ATGTACAAGCCGACAAGGCT | GGGGTCTTATCCCATCCCTA | 259 |
| chr10:88664578-88666602 | p3 | (TTC)8 | CCATGCTTTGGGCATTAGTT | CACTCAGTCCGGAACCATTT | 166 |
| chr10:88766633-88768675 | p5 | (ATTGC)3 | TTGGAATTGGCCTAAACTCG | GAACAACCACACCTTTGCCT | 223 |
| chr10:88766633-88768675 | p5 | (ATTGC)3 | TTGGAATTGGCCTAAACTCG | GAACAACCACACCTTTGCCT | 223 |
| chr10:88766880-88768895 | p5 | (ATTGC)3 | AAGCATGTTACACCTTCGGG | AGCTTTCCCTTCTTTCCCAA | 204 |
| chr10:88766880-88768895 | p5 | (ATTGC)3 | AAGCATGTTACACCTTCGGG | AGCTTTCCCTTCTTTCCCAA | 204 |
| chr10:88796291-88798311 | p4 | (GAAA)5 | TCTTCCTGCCCCTCAGAGTA | GTTCTTGGGTGATGCCAAGT | 210 |
| chr10:88827717-88829732 | p3 | (AAC)5 | CAGTTGGAATCGCAGTCAGA | AGTTAATGTGCTGGGGCATC | 176 |
| chr10:88849794-88851809 | p5 | (TTTCA)3 | AAGTGAGCCTTAGCACGGAA | CCCAATTACCCCCAAAATCT | 233 |
| chr10:89048553-89050569 | p2 | (TA)13 | CTTCGAGGTCAAGGATGAGC | CTCACGTACGCTCCGATGTA | 104 |
| chr10:89048553-89050569 | p2 | (TA)8 | CTTCGAGGTCAAGGATGAGC | CTCACGTACGCTCCGATGTA | 104 |
| chr10:89048664-89050684 | p2 | (TA)13 | TACATCGGAGCGTACGTGAG | ACTGGCTGGTTTGTTAACCG | 251 |
| chr10:89048664-89050684 | p2 | (TA)8 | TACATCGGAGCGTACGTGAG | ACTGGCTGGTTTGTTAACCG | 251 |
| chr10:89126501-89128519 | p3 | (ATA)6 | TGTCTCCCCCACACCTACTC | CACGACTTGTACGCACGTCT | 113 |
| chr10:89188220-89190232 | p2 | (AT)6 | TGCGAGGTGAAAGGTCTTCT | GTAGAAGGGGGCACAACAAA | 220 |
| chr10:89195952-89197967 | p3 | (CCA)5 | GGATCTGGGTTTGAGTTGGA | TTAACGGCGCTCTCTGAAAT | 134 |
| chr10:89195952-89197967 | p3 | (GAA)5 | GGATCTGGGTTTGAGTTGGA | TTAACGGCGCTCTCTGAAAT | 134 |
| chr10:89195952-89197967 | p2 | (AT)10 | GGATCTGGGTTTGAGTTGGA | TTAACGGCGCTCTCTGAAAT | 134 |
| chr10:89198575-89200590 | p5 | (TTTTC)3 | TAGAATTGAGCCCACCCATC | CAAATGAAACGGAGAAGGGA | 106 |
| chr10:89235051-89237063 | p3 | (ACC)6 | CTTACCCTCCCAACTCCCTC | TGTAATGGCTGAGGTGGTCA | 101 |
| chr10:89235051-89237063 | p3 | (CCA)5 | CTTACCCTCCCAACTCCCTC | TGTAATGGCTGAGGTGGTCA | 101 |
| chr10:89235051-89237063 | p2 | (AT)6 | CTTACCCTCCCAACTCCCTC | TGTAATGGCTGAGGTGGTCA | 101 |
| chr10:89351138-89353156 | p3 | (ATC)6 | CATCATCACCAACCGACAAG | GGAGATGGATCAGACCCGTA | 100 |
| chr10:89487275-89489289 | p2 | (TC)7 | GCAGCAACACGCAAGAATAA | TATGGGACACTTGGCTTTCC | 128 |
| chr10:89519983-89521998 | p5 | (TGCAA)3 | AAATATCGACGTCTCGGTGG | CCGGAATAGAGGTTGTGCAT | 178 |
| chr10:89519983-89521998 | p5 | (CCTTT)3 | AAATATCGACGTCTCGGTGG | CCGGAATAGAGGTTGTGCAT | 178 |
| chr10:89539679-89541694 | p3 | (AAT)5 | TGAGGAGCATGAGGAGGAGT | AGAAGTCGCTCCAGTGGTGT | 129 |
| chr10:89539679-89541694 | p2 | (TA)7 | TGAGGAGCATGAGGAGGAGT | AGAAGTCGCTCCAGTGGTGT | 129 |
| chr10:89575518-89577530 | p2 | (TA)6 | GATTTGTGTATGGGGATGGC | AAATACAGATGGCACACGCA | 176 |

| Marker ID | SSR type | SSR motif | FORWARD PRIMER (5'-3') | REVERSE PRIMER (5'-3') | size (bp) |
|---|---|---|---|---|---|
| chr10:89575518-89577530 | p3 | (AAC)6 | GATTTGTGTATGGGGATGGC | AAATACAGATGGCACACGCA | 176 |
| chr10:89672712-89674727 | p3 | (GAT)5 | ATGGACGTTACAAGTTCGGC | TAAGGTTCAAATTCGGGTCG | 145 |
| chr10:89679708-89681732 | p3 | (TTG)7 | CCGCTCATCGATAGGACATT | TGAAAACAATGGGGGTTTGT | 137 |
| chr10:89679708-89681732 | p3 | (TTG)8 | CCGCTCATCGATAGGACATT | TGAAAACAATGGGGGTTTGT | 137 |
| chr10:89731314-89733329 | p5 | (TATTT)3 | TATTACTCCGGCCACAGTCC | TTTGAGACTTCCAACCCCAG | 175 |
| chr10:89817112-89819127 | p3 | (AAT)5 | AAAAACGTTCGGGATGTACG | AGCAGTGTTTTCAGAGGCGT | 192 |
| chr10:89817112-89819127 | p4 | (AAAT)4 | AAAAACGTTCGGGATGTACG | AGCAGTGTTTTCAGAGGCGT | 192 |
| chr10:89864451-89866491 | p2 | (AT)20 | TCCTTCTTAGGCCTTGCTCA | CCAAGTAGCCTCCTCACTCG | 240 |
| chr10:89917157-89919172 | p5 | (AGAAA)3 | CCTCATAAAGGGGCATCTCA | AGGATATGCGGTTTGATTCG | 288 |
| chr10:89940127-89942142 | p5 | (ATAAA)3 | GCTCCAGTTCAGAGGGTGAG | CCCCCTCAGGAAAAGAACTC | 164 |
| chr10:90003105-90005120 | p2 | (TA)8 | GGGTGGGATAAGATGGGATT | CATCCAATCCCTGTAGGGTG | 190 |
| chr10:90003105-90005120 | p5 | (AAAAC)3 | GGGTGGGATAAGATGGGATT | CATCCAATCCCTGTAGGGTG | 190 |
| chr10:90098138-90100153 | p5 | (CGGGT)3 | ACCCCGAGACACTCAATCAC | TTTTTGAATTCTCGCCCAAC | 187 |
| chr10:90337815-90339830 | p3 | (ACA)5 | CCAACCCCATAGACCATGAC | GTCGGCTTGCGTATTTGTTT | 184 |
| chr10:90337847-90339863 | p3 | (ACA)5 | CCAACCCCATAGACCATGAC | GTCGGCTTGCGTATTTGTTT | 152 |
| chr10:90355801-90357816 | p5 | (TATGA)3 | TTACTCCAGCCCCCTATCCT | ACTTAGTTGTCGGCCCCTTT | 291 |
| chr10:90362135-90364151 | p4 | (GAAA)4 | TCTTGGAGAGCTCCGTTGTT | AGTCGGAAAGGCTGCTACAA | 192 |
| chr10:90449396-90451412 | p4 | (CAGA)4 | CTGAATCTGTGGAGTTGCCA | TAGTGGCGGATAAAGGCATC | 117 |
| chr10:90666361-90668394 | p3 | (TTG)11 | CCACCGAAACTTAAAGGCAA | GTGTGCCCACACATCTATCG | 278 |
| chr10:90703865-90705881 | p2 | (GA)8 | TAGAAAACCACTTGGCACCC | CTCCGGCAGAATTGAAAGAG | 190 |
| chr10:90775884-90777898 | p2 | (AT)7 | GCCTAAGGGTCTGGTCCTTC | GTATGGATGGCATTGAGCCT | 193 |
| chr10:90787183-90789203 | p2 | (AT)10 | TCCCGACTTTGGAAGAATTG | GCATGTATGTCCTGTGGTGC | 174 |
| chr10:90787183-90789203 | p2 | (AT)16 | TCCCGACTTTGGAAGAATTG | GCATGTATGTCCTGTGGTGC | 174 |
| chr10:90819831-90821849 | p3 | (ACA)6 | CGGCTAATGCTTCCAAAGAG | TGAGGTACCAATGTGACCGA | 252 |
| chr10:90828412-90830470 | p2 | (GT)29 | TCCACTGATGGACCACAAAA | TGGGCATGTTTGTTGATGTT | 117 |
| chr10:90865912-90867960 | p2 | (AG)9 | CAGCGTTATGGGCTTCACTT | ATCATTGCAAGATTCAGCCA | 190 |
| chr10:90893237-90895255 | p3 | (AGT)6 | AAAGAAAGGGCGGTGGTACT | GCCACGTAATCCGAAGAAAA | 224 |
| chr10:90893237-90895255 | p2 | (GA)9 | AAAGAAAGGGCGGTGGTACT | GCCACGTAATCCGAAGAAAA | 224 |
| chr10:90926734-90928760 | p2 | (AT)13 | ATGGCTTATTTTTGCAGCCA | TGATTCACGAAATGCAAAGG | 222 |
| chr10:90934096-90936116 | p5 | (AAAAT)3 | TCATCAGAATCGTCACCGTC | TTGGGGTGGTGAAGTTGAGT | 257 |
| chr10:90934118-90936132 | p5 | (AAAAT)3 | TCATCAGAATCGTCACCGTC | TTGGGGTGGTGAAGTTGAGT | 235 |
| chr10:90939886-90941901 | p5 | (TATTG)3 | AAGCGTCAATAGCCCTTTCA | CCGTGGAAAAGATTAGGCAA | 291 |
| chr10:91055271-91057286 | p5 | (GTTTG)3 | GAATTATGCTCCTGTGCCGT | ACCAAAACTGTGAACAGCCC | 232 |
| chr10:91055271-91057286 | p5 | (AACCG)3 | GAATTATGCTCCTGTGCCGT | ACCAAAACTGTGAACAGCCC | 232 |
| chr10:91110775-91112787 | p2 | (GT)6 | AAGTGCACACATTTGCGAAG | TAAGGCATTGTTGCTGTTGC | 138 |
| chr10:91117085-91119103 | p2 | (TA)9 | GGCAATGAGCTCTTCAAAGG | GCTGCTCGATTGCACAATAA | 108 |

| Marker ID | SSR type | SSR motif | FORWARD PRIMER (5'-3') | REVERSE PRIMER (5'-3') | size (bp) |
|---|---|---|---|---|---|
| chr10:91117085-91119103 | p2 | (AT)9 | GGCAATGAGCTCTTCAAAGG | GCTGCTCGATTGCACAATAA | 108 |
| chr10:91121464-91123476 | p2 | (TG)6 | GGGAGTTGATTTTGGGGAAT | AACCAAACAAGCCTCAATGG | 196 |
| chr10:91121464-91123476 | p2 | (TA)8 | GGGAGTTGATTTTGGGGAAT | AACCAAACAAGCCTCAATGG | 196 |
| chr10:91121464-91123476 | p6 | (CATATA)3 | GGGAGTTGATTTTGGGGAAT | AACCAAACAAGCCTCAATGG | 196 |
| chr10:91160178-91162190 | p4 | (CCCA)5 | GAATGCTATCTTCCCACCGA | TCACCATTTTTGGCGTTGTA | 282 |
| chr10:91174437-91176452 | p5 | (TTACT)3 | CTAGAGGGAAGCAATCGTGC | TAACTCCTACCCGCTCAACG | 129 |
| chr10:91190583-91192601 | p6 | (AGCCGA)3 | GAAATTGCGTTTTTCAGGGA | GACATCACGATGACCAAACG | 282 |
| chr10:91253330-91255356 | p2 | (AT)13 | AAGTCGCTTTTGCTGCATTT | GCAAGGTAAAATTTTGGGCA | 162 |
| chr10:91309440-91311506 | p2 | (AT)33 | ATCTTGCAGCTGGAAAGCTC | AGGATAAGGACCCGCAGAAT | 211 |
| chr10:91361192-91363248 | p2 | (AT)28 | AGAGAAGAAGAAAAGCGGGG | TCTAAGCCACTCCTCGGCTA | 128 |
| chr10:91405831-91407849 | p3 | (TTG)6 | GGCCAAGCAAATCAAAGAAA | TTTGGCAGCTTGGAGTCTTT | 165 |
| chr10:91445763-91447799 | p2 | (AT)18 | CGAACTTATGCCACTCCGAT | TAGAACGTACAGCCCGATCC | 223 |
| chr10:91557093-91559108 | p5 | (ACAAT)4 | CCGTGGCCAAATAAAGGTTA | TAAAGGGCACCTAAACCGTG | 300 |
| chr10:91557093-91559108 | p5 | (CAGGC)3 | CCGTGGCCAAATAAAGGTTA | TAAAGGGCACCTAAACCGTG | 300 |
| chr10:91557093-91559108 | p2 | (TC)6 | CCGTGGCCAAATAAAGGTTA | TAAAGGGCACCTAAACCGTG | 300 |
| chr10:91685516-91687588 | p2 | (GA)7 | ATGCCCGTTTTAGTGTGAGG | GTTGTTTGGGGGAAGGAAAT | 152 |
| chr10:91853017-91855029 | p2 | (TA)6 | CGCATTTAGCCAAGGAAGAC | TTGCTCCGCTTCTGAGTTTT | 212 |
| chr10:91853017-91855029 | p2 | (AT)6 | CGCATTTAGCCAAGGAAGAC | TTGCTCCGCTTCTGAGTTTT | 212 |
| chr10:91888301-91890316 | p5 | (CGCCC)3 | ATGCCTTACGACCAGTGCTT | AGGGCGTACAGTTGAGTGCT | 151 |
| chr10:91890192-91892208 | p4 | (TAAA)4 | TTTCAGTCGTGTGGTTGGAA | TGGAATTACATCTGGTGCGA | 281 |
| chr10:91953345-91955357 | p2 | (AT)6 | AGAATAAAGGTTGGGTCGGG | AGATTCTGTGCGTGAAAGGG | 136 |
| chr10:91953345-91955357 | p2 | (AT)8 | AGAATAAAGGTTGGGTCGGG | AGATTCTGTGCGTGAAAGGG | 136 |
| chr10:91953345-91955357 | p2 | (AT)6 | AGAATAAAGGTTGGGTCGGG | AGATTCTGTGCGTGAAAGGG | 136 |
| chr10:91969837-91971861 | p2 | (AC)12 | AAGGCAAATGGCTCATATCG | ATCGGGCGAAGATTAGGTTT | 257 |
| chr10:91969837-91971861 | p3 | (ATC)8 | AAGGCAAATGGCTCATATCG | ATCGGGCGAAGATTAGGTTT | 257 |
| chr10:91998045-92000057 | p2 | (GT)6 | TCGCCGATAATTTCATCTCC | ATCGCTGGAAACTTGTTGCT | 174 |
| chr10:91998045-92000057 | p2 | (CT)6 | TCGCCGATAATTTCATCTCC | ATCGCTGGAAACTTGTTGCT | 174 |
| chr10:92013747-92015765 | p3 | (AAT)6 | TCGGTCGATGTGTTTGTGTT | CCCCCTAAACCTCCTACTCG | 173 |
| chr10:92050161-92052176 | p5 | (AAACC)4 | TTTTCGAAGCACACGTCTTG | TTGGTTTGGTTTGGTTTGGT | 194 |
| chr10:92050161-92052176 | p5 | (AAACC)3 | TTTTCGAAGCACACGTCTTG | TTGGTTTGGTTTGGTTTGGT | 194 |
| chr10:92055047-92057065 | p2 | (TA)36 | GCTCCAATTTCTGTTTGGGA | CAGGGAAAACTCCTCGATCA | 186 |
| chr10:92055047-92057065 | p6 | (GAAAAA)3 | GCTCCAATTTCTGTTTGGGA | CAGGGAAAACTCCTCGATCA | 186 |
| chr10:92061475-92063513 | p2 | (TA)19 | AATGATGGGAGAAGGTGACG | TCGTCGTATGTGAGGCAAAC | 110 |
| chr10:92106873-92108937 | p2 | (AT)32 | ACGTGGCATGTCTTTTAGGC | CTATGGCAAACGGGAGCTTA | 171 |
| chr10:92191830-92193848 | p6 | (AATAAC)3 | ATGGATTCTACGAGGCATGG | TTTGTCTCGCATCAAACTGC | 278 |
| chr10:92243818-92245833 | p5 | (TTGGG)3 | TTTCTCTCCAAACCAATGCC | GCCACCTGATTGCATTTTCT | 110 |

| Marker ID | SSR type | SSR motif | FORWARD PRIMER (5'-3') | REVERSE PRIMER (5'-3') | size (bp) |
|---|---|---|---|---|---|
| chr10:92243818-92245833 | p2 | (GT)8 | TTTCTCTCCAAACCAATGCC | GCCACCTGATTGCATTTTCT | 110 |
| chr10:92301260-92303280 | p5 | (TTTTA)3 | GTGAACAAGAACCGCCTGAT | CTGCGGTTACAAAGACCACA | 204 |
| chr10:92301344-92303359 | p5 | (TTTTA)3 | GTGAACAAGAACCGCCTGAT | CTGCGGTTACAAAGACCACA | 120 |
| chr10:92349262-92351274 | p2 | (TA)6 | GCTAGCTGAACGATCCCTTG | CCGAAAGGAGGGTAGAAAGG | 193 |
| chr10:92407639-92409651 | p2 | (AT)6 | TTTTAAAAGGCGGTGTCAGG | GTTGGGTGTTGGGTGTTAGG | 254 |
| chr10:92454430-92456450 | p2 | (TA)10 | CTTCAACTGGCTGTGCAAAA | GTCCTCCTCCCAAGTCAACA | 262 |
| chr10:92508929-92510941 | p2 | (TA)6 | TGCTCACCATCTGAGCAATC | GATGAAGAGGAGGGAGGGAG | 217 |
| chr10:92514393-92516431 | p2 | (TA)19 | GCCCCTGTTATTCACAGCAT | CTCCTCACTCGTTCTGGAGG | 115 |
| chr10:92574727-92576742 | p3 | (GAG)5 | TCCATTTGGCTTTTGAGTCC | TCTGTTCCTCTGCCTGCTTT | 266 |
| chr10:92577521-92579539 | p3 | (ACA)6 | AGAGCAGCAGAGGGTGTGAT | GATGCAGCATTCAGTAGGCA | 121 |
| chr10:92585724-92587742 | p6 | (CGTCCC)3 | GCCCCCATATCCTTCAATTT | CATTCCGAGGGACAACCTTA | 128 |
| chr10:92595442-92597458 | p2 | (TA)8 | GGTTGGTTGAAGAGATGGGA | TACGCTTCCCAAAACCAATC | 189 |
| chr10:92622870-92624882 | p2 | (TG)6 | AAAGGTGCCTACGGGCTATT | TCATACCTGGCCCTACCAAG | 275 |
| chr10:92640227-92642251 | p2 | (AT)12 | CACAAATAATGACATGGCGG | ATAGGGGATTCATTGCCACA | 242 |
| chr10:92696226-92698244 | p2 | (TA)9 | GAAATTTCCGTCGATTTCCA | CTTCAGGTTCCTTCCGAGTG | 151 |
| chr10:92696226-92698244 | p2 | (TC)11 | GAAATTTCCGTCGATTTCCA | CTTCAGGTTCCTTCCGAGTG | 151 |
| chr10:92774878-92776892 | p2 | (AT)7 | TTGCTAACCCACCATTCCTC | ATTCCAACCACCCAGTTCAG | 211 |
| chr10:92774878-92776892 | p3 | (ATG)7 | TTGCTAACCCACCATTCCTC | ATTCCAACCACCCAGTTCAG | 211 |
| chr10:92798423-92800435 | p2 | (AT)7 | TATGTGTTGTGCGGTGTGTG | GCTCGTGTCCTGGTTGATTT | 221 |
| chr10:92798440-92800456 | p2 | (AT)7 | TATGTGTTGTGCGGTGTGTG | GCTCGTGTCCTGGTTGATTT | 204 |
| chr10:92940879-92942917 | p5 | (TTTAA)3 | GCTCTCTGGTTGTGTCGTGA | AGAATCAGGGGCATCACATC | 284 |
| chr10:92942043-92944055 | p2 | (AT)6 | TGCGATGTCGGTTTATGTGT | CCTAACTGTGAATCCCGGTG | 289 |
| chr10:92977898-92979976 | p2 | (AT)7 | ATTCTCGAAAATGACCGTGC | CTTGTTCCGGAGCTGCTATC | 272 |
| chr10:92978000-92980070 | p2 | (AT)7 | ATTCTCGAAAATGACCGTGC | CTTGTTCCGGAGCTGCTATC | 170 |
| chr10:92984460-92986475 | p5 | (AACTG)3 | GCCCTCGAATCTAGCAAGTG | TGTTGTATTGCAGGTGCGTT | 167 |
| chr10:93051051-93053067 | p2 | (AT)8 | CGCCTTCCATGGGTCTAATA | TTCACAGACTAGGGTTGGGG | 231 |
| chr10:93068853-93070893 | p2 | (TA)20 | GTAACTGTTCCCGCCAAAAA | GGAAGAACTGGACCCAATGA | 147 |
| chr10:93132279-93134294 | p5 | (TTGCA)3 | TTATCAGCCACAGGGGACTC | GCTTGGCTAACTTCTCCGTG | 118 |
| chr10:93137546-93139561 | p3 | (TTA)5 | GATTTGTGTCCATGTGTGGC | TGCAACCTGATCGCAGTTAG | 182 |
| chr10:93182476-93184496 | p5 | (GCATT)4 | ATTTACCCCCGAATTTCCAG | AGCCCGATCAAACAAGTCAC | 240 |
| chr10:93369816-93371828 | p2 | (TA)6 | CTGTTGTGTCCCACATCGTC | GAGAGGTGCCCAGTGGATTA | 228 |
| chr10:93369816-93371828 | p2 | (AG)6 | CTGTTGTGTCCCACATCGTC | GAGAGGTGCCCAGTGGATTA | 228 |
| chr10:93401387-93403399 | p2 | (TA)6 | CCCGAATAGGCATAGGTTCA | CAATTTGCACGATTTCCCTT | 272 |
| chr10:93447494-93449506 | p2 | (TA)6 | CTGTTGTGTCCCACATCGTC | GAGAGGTGCCCAGTGGATTA | 228 |
| chr10:93447494-93449506 | p2 | (AG)6 | CTGTTGTGTCCCACATCGTC | GAGAGGTGCCCAGTGGATTA | 228 |
| chr10:93479065-93481077 | p2 | (TA)6 | CCCGAATAGGCATAGGTTCA | CAATTTGCACGATTTCCCTT | 272 |

| Marker ID | SSR type | SSR motif | FORWARD PRIMER (5'-3') | REVERSE PRIMER (5'-3') | size (bp) |
|---|---|---|---|---|---|
| chr10:93624508-93626528 | p2 | (AT)10 | TAACCCAAAACCCCCTTTTC | ATGCGTTGACCTACGAGCTT | 130 |
| chr10:93630927-93632942 | p5 | (AATGC)3 | AAACCCAAGTGTGCCAAGTC | GCAGGTCCCCTATGGGTTAT | 156 |
| chr10:93630927-93632942 | p5 | (CCTTT)3 | AAACCCAAGTGTGCCAAGTC | GCAGGTCCCCTATGGGTTAT | 156 |
| chr10:93701482-93703494 | p2 | (TG)6 | GGACGGTCCGTAAACTGAAA | GGTATTTAATGGCTTGCCCA | 233 |
| chr10:93769653-93771701 | p2 | (AT)24 | GAGACTAGCGCTCAACCACC | CCACCAGTCCGAAGGATAGA | 291 |
| chr10:93781937-93783959 | p2 | (AT)11 | GATGGCAGAGAGACGTGACA | CCACCAGTCCGAAGGATAGA | 230 |
| chr10:93837774-93839786 | p2 | (CA)6 | ACTCCCATGCCTAGGGAACT | CCATCTTTGTAACGACCCGT | 119 |
| chr10:93854751-93856772 | p3 | (ATA)7 | GTGATCCTTCCCCCTAGCTC | TTTCTCAGACCTACGCCGTT | 118 |
| chr10:93869648-93871717 | p4 | (ATTT)4 | GAGACCCAACTCGTGGTGAT | TGTCACTCCAAATCTCACGC | 298 |
| chr10:93869648-93871717 | p3 | (TAT)23 | GAGACCCAACTCGTGGTGAT | TGTCACTCCAAATCTCACGC | 298 |
| chr10:93974006-93976018 | p2 | (CA)6 | ACCAAACAGACAGACGGGAC | GTTAGCCCCGATCGATTGTA | 121 |
| chr10:93974006-93976018 | p2 | (AT)6 | ACCAAACAGACAGACGGGAC | GTTAGCCCCGATCGATTGTA | 121 |
| chr10:93988171-93990186 | p5 | (TTAGT)3 | CGAGCATTTTTCAGAAAGGG | GGTTAAAGTGGCTGCCAGAG | 202 |
| chr10:93995943-93997955 | p2 | (CA)6 | TGAAGCATGTATCAACCGGA | CGGGCCAAGACATAAGAGAG | 116 |
| chr10:93995943-93997955 | p2 | (AT)6 | TGAAGCATGTATCAACCGGA | CGGGCCAAGACATAAGAGAG | 116 |
| chr10:94007268-94009286 | p6 | (ATGTAT)3 | GTGCGGGGCATAAGAAAGTA | TTAGATGCGGATTCTGAGGG | 181 |
| chr10:94049434-94051470 | p3 | (AGA)12 | TGGGCTCTGAATTTGCTACC | TGACATCGTCTGCTTTACGC | 149 |
| chr10:94154970-94156982 | p2 | (AT)6 | TGATGGCAGAGAGACGTGAC | TCCACCAGTCCGAAGGATAG | 241 |
| chr10:94255181-94257193 | p2 | (AT)7 | GGCTTGGTTCTCTTCAGCAG | GTGCCAATCACGCCTTATCT | 158 |
| chr10:94255181-94257193 | p2 | (AT)6 | GGCTTGGTTCTCTTCAGCAG | GTGCCAATCACGCCTTATCT | 158 |
| chr10:94305671-94307686 | p5 | (AAACA)3 | CAATCATGTCCCAAGCTCCT | TTTTGCTGTTTAGGGATGGC | 136 |
| chr10:94327820-94329834 | p2 | (AT)7 | TTCAGAACATCGGTGGATGA | CAGGAACAGAGGACCACCAT | 229 |
| chr10:94342401-94344451 | p2 | (AT)25 | TGCCTAATTAGCCGGGTATG | AGCACGGTGGAATCATAAGC | 144 |
| chr10:94468489-94470507 | p6 | (TTTTAT)3 | CTGATTCCGTCAGTGGGATT | TGTGGTGTGACCGAACCTTA | 178 |
| chr10:94549838-94551852 | p2 | (AT)7 | CAGCAGTGGGCATGAGTTTA | CCGGTCCTCCATATCTGCTA | 123 |
| chr10:94770941-94772956 | p5 | (AAATA)3 | ACCGGACTAACGACCATGAG | TTGGCCTTACGCTCAAGTTT | 270 |
| chr10:94798344-94800356 | p2 | (GA)6 | AGCTAACCCGCCCATTTACT | AGCGCAACCAATTGAATCTC | 180 |
| chr10:94857917-94859941 | p2 | (TA)14 | TTCGCTTGACGAATAACGTG | TATCGAGGCCAAGGAATGAC | 220 |
| chr10:94857917-94859941 | p2 | (TA)12 | TTCGCTTGACGAATAACGTG | TATCGAGGCCAAGGAATGAC | 220 |
| chr10:94990276-94992296 | p2 | (AT)10 | TGCCCGATTTTCGTTAAGTC | TTCGTAAACCGTAGCCCATC | 149 |
| chr10:95022518-95024582 | p2 | (AT)25 | ACCCAAAACTAGAGACGCGA | GTGATGGCCTTACCAGCCTA | 233 |
| chr10:95022518-95024582 | p2 | (AT)32 | ACCCAAAACTAGAGACGCGA | GTGATGGCCTTACCAGCCTA | 233 |
| chr10:95042383-95044429 | p2 | (TA)28 | GTATTGTGCACCGAGTCCCT | TTATCCTTGGCGCCATAATC | 183 |
| chr10:95042383-95044429 | p2 | (AT)23 | GTATTGTGCACCGAGTCCCT | TTATCCTTGGCGCCATAATC | 183 |
| chr10:95042383-95044429 | p5 | (TATGA)3 | GTATTGTGCACCGAGTCCCT | TTATCCTTGGCGCCATAATC | 183 |
| chr10:95043034-95045049 | p2 | (AT)23 | GTCGGCTTGCGTACCATATT | CAATCCGAGGATGAACTGCT | 286 |

| Marker ID | SSR type | SSR motif | FORWARD PRIMER (5'-3') | REVERSE PRIMER (5'-3') | size (bp) |
|---|---|---|---|---|---|
| chr10:95043034-95045049 | p5 | (TATGA)3 | GTCGGCTTGCGTACCATATT | CAATCCGAGGATGAACTGCT | 286 |
| chr10:95105776-95107790 | p2 | (AT)7 | ACCACTTTTGTTCACGACCC | TTGTGACTTCCCAGGAGACC | 207 |
| chr10:95105776-95107790 | p2 | (AT)7 | ACCACTTTTGTTCACGACCC | TTGTGACTTCCCAGGAGACC | 207 |
| chr10:95154810-95156844 | p2 | (TA)17 | TTATGGACGTCAAACGGACA | TTCTTGATGATGGGAGAGGG | 270 |
| chr10:95200519-95202533 | p2 | (TA)7 | TTAAAACGAGGCGTATTCGG | GGGCGTGATAAAGCTTGGTA | 149 |
| chr10:95207197-95209217 | p2 | (TA)10 | ACTATCCATCGAGTGCAGGG | CAGTGGGAAGGCATGTGTTA | 120 |
| chr10:95214795-95216811 | p4 | (TAAT)4 | CTTCGCAACCACGAATTTTT | GAAGGAATGCCTGAAAACCA | 223 |
| chr10:95253090-95255105 | p5 | (ATAAA)3 | GGCAAAATTCTGTTTTGGGA | TTTTGTGCGGATTACCCTTC | 166 |
| chr10:95272943-95274970 | p3 | (ATT)9 | TCAATTTTTGTGCAACCCAA | TGATGAAACGGGAAAGAAGG | 131 |
| chr10:95296725-95298739 | p2 | (TA)7 | TTAAAACGAGGCGTATTCGG | GGGCGTGATAAAGCTTGGTA | 149 |
| chr10:95303403-95305423 | p2 | (TA)10 | ACTATCCATCGAGTGCAGGG | CAGTGGGAAGGCATGTGTTA | 120 |
| chr10:95311001-95313017 | p4 | (TAAT)4 | CTTCGCAACCACGAATTTTT | GAAGGAATGCCTGAAAACCA | 223 |
| chr10:95349296-95351311 | p5 | (ATAAA)3 | GGCAAAATTCTGTTTTGGGA | TTTTGTGCGGATTACCCTTC | 166 |
| chr10:95369049-95371076 | p3 | (ATT)9 | TCAATTTTTGTGCAACCCAA | TGATGAAACGGGAAAGAAGG | 131 |
| chr10:95383241-95385259 | p6 | (AACTGA)3 | TTGGAGGGTTATTTGCAAGC | CCGTGAGACAATTGTGTTGG | 273 |
| chr10:95386862-95388883 | p3 | (ATT)7 | AGGGAAAAAGTGTCCGTGTG | CTAGTTGTACTCCGCGCTCC | 183 |
| chr10:95397092-95399110 | p3 | (AAG)6 | TTTCACCAAAAACCGATTCA | CTCTCCGGGCCTAAGAAAAC | 203 |
| chr10:95575763-95577826 | p3 | (CTT)7 | TTAAGAGCCCGTTTGGATTG | CCTGTTTGGATGGGCTTAAA | 212 |
| chr10:95575867-95577891 | p3 | (CTT)7 | TTAAGAGCCCGTTTGGATTG | CCTGTTTGGATGGGCTTAAA | 108 |
| chr10:95741362-95743388 | p6 | (CTGTGG)3 | ACACCCATTGTTTCCACCAT | CGAGCTGGGTTGAAGAAGAC | 273 |
| chr10:95741459-95743477 | p6 | (CTGTGG)3 | ACACCCATTGTTTCCACCAT | CGAGCTGGGTTGAAGAAGAC | 176 |
| chr10:95772727-95774747 | p2 | (TA)10 | TACATGGCCATTCAAGTTGG | GAATAGCATCGATGGAGGGA | 153 |
| chr10:95841719-95843745 | p2 | (GA)13 | GAGTTTTCATGTGCCGTGTG | CAAAAGCAGTCCAAGGAAGG | 241 |
| chr10:95892743-95894758 | p5 | (CCACA)3 | AGGACGACCTTTCGATGATG | AATATGGGCTCAAATTGCCA | 262 |
| chr10:95892743-95894758 | p2 | (CA)6 | AGGACGACCTTTCGATGATG | AATATGGGCTCAAATTGCCA | 262 |
| chr10:95901561-95903577 | p2 | (TA)11 | TGCCCCATCCGTATTACATT | AGGCATGAAGGAACAACGAC | 138 |
| chr10:95901561-95903577 | p2 | (AT)8 | TGCCCCATCCGTATTACATT | AGGCATGAAGGAACAACGAC | 138 |
| chr10:95901561-95903577 | p2 | (AT)15 | TGCCCCATCCGTATTACATT | AGGCATGAAGGAACAACGAC | 138 |
| chr10:95901561-95903577 | p2 | (AT)8 | TGCCCCATCCGTATTACATT | AGGCATGAAGGAACAACGAC | 138 |
| chr10:95950421-95952433 | p2 | (AT)6 | CGCATCTAGCCAAGGAAGAC | TTCACCTTCCTCCGCTTCTA | 274 |
| chr10:95950421-95952433 | p2 | (TA)8 | CGCATCTAGCCAAGGAAGAC | TTCACCTTCCTCCGCTTCTA | 274 |
| chr10:95952150-95954164 | p2 | (TA)7 | GAGTCAGGATTTCCGTCGAG | GTTGAACCCCCTTAGGCTTC | 145 |
| chr10:95977291-95979303 | p2 | (AT)6 | GTGGCCGTATTGTTACACCC | AACCTTCCGAAACTCTCCGT | 101 |
| chr10:96030562-96032578 | p4 | (TAAT)4 | TCTTGGTACCGGATTTTTCG | AGGGGAAAGAAGAGGCAGAA | 178 |
| chr10:96185506-96187518 | p2 | (TA)6 | AACGAGATAGGGAGTTGGGC | CCAAATTTGTTGGTGGGTCT | 195 |
| chr10:96277782-96279798 | p2 | (AT)10 | CAAGGTACCCACCGAAAGAA | GGATGGAAGGATGAGTTGGA | 100 |

| Marker ID | SSR type | SSR motif | FORWARD PRIMER (5'-3') | REVERSE PRIMER (5'-3') | size (bp) |
|---|---|---|---|---|---|
| chr10:96287264-96289279 | p5 | (CAATG)3 | CCATGGCCTCTCGAAGATAA | CCAATAGGGGATTTGTGTGG | 218 |
| chr10:96287264-96289279 | p2 | (AC)9 | CCATGGCCTCTCGAAGATAA | CCAATAGGGGATTTGTGTGG | 218 |
| chr10:96287264-96289279 | p2 | (TG)13 | CCATGGCCTCTCGAAGATAA | CCAATAGGGGATTTGTGTGG | 218 |
| chr10:96287672-96289698 | p5 | (CAATG)3 | ACTGGCCTCACGCTAGATGT | CTGTTGTGCCAATCTGCCTA | 272 |
| chr10:96287672-96289698 | p2 | (AC)9 | ACTGGCCTCACGCTAGATGT | CTGTTGTGCCAATCTGCCTA | 272 |
| chr10:96287672-96289698 | p2 | (TG)13 | ACTGGCCTCACGCTAGATGT | CTGTTGTGCCAATCTGCCTA | 272 |
| chr10:96381143-96383155 | p2 | (GA)6 | GCCTTTTTGTTTGGTGGAAA | ATTCCTAGCTGGCAAAGCAA | 247 |
| chr10:96462810-96464822 | p2 | (CT)6 | TCTAGCGGATCCAAAGTGCT | ATTTTCATGGCCAAAAGCTG | 130 |
| chr10:96462810-96464822 | p2 | (CT)7 | TCTAGCGGATCCAAAGTGCT | ATTTTCATGGCCAAAAGCTG | 130 |
| chr10:96506720-96508735 | p3 | (ATT)5 | CCCATTTGGACAACAACACA | TTCGGAGCCATATCTTTTGG | 107 |
| chr10:96609263-96611275 | p2 | (AT)10 | AATTTGCATGGAAGTGGGAC | CTCTACCATGAACCGGAGGA | 261 |
| chr10:96609296-96611308 | p2 | (AT)10 | AATTTGCATGGAAGTGGGAC | CTCTACCATGAACCGGAGGA | 228 |
| chr10:96609536-96611556 | p2 | (AT)10 | TCCTCCGGTTCATGGTAGAG | TGTCCGGTTTCGTTTTAACC | 182 |
| chr10:96649297-96651315 | p2 | (AT)9 | GGGGGAATTTTCTACTTGCC | TTTTTCGAACGCGGAAATAC | 153 |
| chr10:96778406-96780444 | p2 | (TA)19 | CCGCCCAAAATAACCCTAAT | GCTTTGGATCTCCGACTCAC | 114 |
| chr10:96911236-96913256 | p5 | (TTTGG)4 | TGTAACCGATGTCACTGGGA | GCCCACACGTCAAGATTTTT | 273 |
| chr10:97089142-97091154 | p5 | (ATTTT)3 | ATACTCACCCAAACCCCACA | GTAATATGGGTCCACACGGG | 243 |
| chr10:97089142-97091154 | p2 | (AT)6 | ATACTCACCCAAACCCCACA | GTAATATGGGTCCACACGGG | 243 |
| chr10:97089142-97091154 | p5 | (AGAAA)3 | ATACTCACCCAAACCCCACA | GTAATATGGGTCCACACGGG | 243 |
| chr10:97090033-97092048 | p2 | (AT)6 | TTTTGCAAGCTTCTTGAGCA | GTGAGTACATGGCGGAGGAT | 151 |
| chr10:97090033-97092048 | p5 | (AGAAA)3 | TTTTGCAAGCTTCTTGAGCA | GTGAGTACATGGCGGAGGAT | 151 |
| chr10:97137068-97139082 | p2 | (TA)7 | GAATCATTGTTGCTGCATGG | GCATGACTCGCAAAGTCAAA | 221 |
| chr10:97167885-97169899 | p2 | (TA)7 | ACGCAAAAACCCAAAATCTG | TAGCTCAGTGTCCGTGTTGG | 109 |
| chr10:97207819-97209831 | p2 | (AT)6 | ACTCTTCCAGAGGACCCGAT | TTCAGAGCATGAATCAACGC | 257 |
| chr10:97253028-97255042 | p2 | (TA)7 | CAAACCACCTCCTTCCTTCA | GCATGGTATAGCTGGGCATT | 252 |
| chr10:97253028-97255042 | p5 | (TTAAA)3 | CAAACCACCTCCTTCCTTCA | GCATGGTATAGCTGGGCATT | 252 |
| chr10:97257074-97259088 | p2 | (TA)7 | TAGGTCGGGAACCACTTGAC | CCTTGTCGGCCTTATTTTGA | 191 |
| chr10:97310818-97312833 | p5 | (ATGCA)3 | GCTGAAGACTCCCTACACGC | TGAAGGGTTCCATCATGACA | 152 |
| chr10:97474282-97476310 | p2 | (AT)14 | GATTATCGCCTGATGGGAGA | ACCAGTCGGGCAAATGTTAC | 162 |
| chr10:97487741-97489759 | p3 | (ATG)6 | TATTGGGAGCGCTCTTTTGT | AATACATGGAGGCCGATGAG | 149 |
| chr10:97524120-97526140 | p5 | (AGTAA)4 | GCACCAAGTAAATCTGGGGA | GAAAGGTGGAAGGCCACATA | 141 |
| chr10:97567509-97569521 | p2 | (TA)6 | CAAGCGGGTACGACACCTAT | ACTCGGTGTGAAACGTACCC | 260 |
| chr10:97567509-97569521 | p2 | (TA)12 | CAAGCGGGTACGACACCTAT | ACTCGGTGTGAAACGTACCC | 260 |
| chr10:97567509-97569521 | p2 | (TA)6 | CAAGCGGGTACGACACCTAT | ACTCGGTGTGAAACGTACCC | 260 |
| chr10:97567509-97569521 | p2 | (TA)12 | CAAGCGGGTACGACACCTAT | ACTCGGTGTGAAACGTACCC | 260 |
| chr10:97567509-97569521 | p2 | (AC)9 | CAAGCGGGTACGACACCTAT | ACTCGGTGTGAAACGTACCC | 260 |

| Marker ID | SSR type | SSR motif | FORWARD PRIMER (5'-3') | REVERSE PRIMER (5'-3') | size (bp) |
|---|---|---|---|---|---|
| chr10:97568229-97570247 | p2 | (TA)6 | ATTTCATTCGCTCCTCATGC | ACTCGGTGTGAAACGTACCC | 258 |
| chr10:97568229-97570247 | p2 | (TA)12 | ATTTCATTCGCTCCTCATGC | ACTCGGTGTGAAACGTACCC | 258 |
| chr10:97568229-97570247 | p2 | (AC)9 | ATTTCATTCGCTCCTCATGC | ACTCGGTGTGAAACGTACCC | 258 |
| chr10:97568229-97570247 | p2 | (TG)13 | ATTTCATTCGCTCCTCATGC | ACTCGGTGTGAAACGTACCC | 258 |
| chr10:97666210-97668222 | p2 | (TA)6 | AGAATGACGCCGCTAATGTT | AATGTCACGACCCGTCTAGG | 282 |
| chr10:97676846-97678910 | p2 | (TA)32 | TATAAGGGAAAGTGCTGCCG | TGAATCCTTTTTGCCTCGTC | 244 |
| chr10:97969467-97971495 | p2 | (TA)13 | AAATATCAACCGTGGCTTCG | CTCTTTCTCTTCGCCTTCCA | 221 |
| chr10:97969467-97971495 | p2 | (AT)16 | AAATATCAACCGTGGCTTCG | CTCTTTCTCTTCGCCTTCCA | 221 |
| chr10:97969496-97971520 | p2 | (TA)13 | AAATATCAACCGTGGCTTCG | TCTTTCTCTTCGCCTTCCAC | 192 |
| chr10:97969496-97971520 | p2 | (AT)16 | AAATATCAACCGTGGCTTCG | TCTTTCTCTTCGCCTTCCAC | 192 |
| chr10:97983955-97985971 | p4 | (GTCT)4 | TGAAGGCACTGGTGTCTCTG | TATGAGGAGGTTCCCATTGC | 217 |
| chr10:97994101-97996117 | p2 | (AT)8 | ATCCCTAGGACGAACCTGCT | AGTGGGTTCGCTCGACTCTA | 159 |
| chr10:98015347-98017429 | p2 | (AT)41 | CCCCGAGTTGAATGGTTTTA | CCATCATGTGCACCTTTGAG | 116 |
| chr10:98057693-98059713 | p5 | (TGCCC)4 | GTATTGTGGGTAGGGATGGC | CTAAACAGAGCAGGGCAAGG | 130 |
| chr10:98192811-98194826 | p5 | (TAATA)3 | AGCCAATGGAATGACAAAGC | ATCAAGATGGCCAGGAGATG | 238 |
| chr10:98282239-98284251 | p2 | (TA)6 | ACGTAGCGCAACACTGACTG | TGATACACACACGCTTCGGT | 115 |
| chr10:98301516-98303528 | p2 | (AT)6 | AACAGTGGAACCCACCACAT | ACGCATGTGACAATGAGTCC | 208 |
| chr10:98301516-98303528 | p5 | (TTTTC)3 | AACAGTGGAACCCACCACAT | ACGCATGTGACAATGAGTCC | 208 |
| chr10:98321092-98323110 | p6 | (ATAAAA)3 | TCGTTGCCTGCATATCTTTG | ACTTGAGGCCAATCCGTACA | 127 |
| chr10:98343020-98345035 | p5 | (ATACC)3 | ACCCCCAAAAACGAGCTACT | ATTCGTTAACCGTTGGATCG | 277 |
| chr10:98355874-98357904 | p2 | (GA)15 | CACCGTCGATGTTGTTTTTG | AACATTCGACCCTCAATTCG | 251 |
| chr10:98375646-98377662 | p4 | (TATT)4 | GGCATTATGATGGTTACCCG | TGGGTCACCTCCCTGTCTAC | 263 |
| chr10:98395220-98397235 | p5 | (TTGGT)3 | ACCGACAAACCGCATTAAAC | CAAACCAATTATGGTCGGGT | 182 |
| chr10:98395220-98397235 | p2 | (AG)6 | ACCGACAAACCGCATTAAAC | CAAACCAATTATGGTCGGGT | 182 |
| chr10:98395260-98397275 | p5 | (TTGGT)3 | ACCGACAAACCGCATTAAAC | CAAACCAATTATGGTCGGGT | 142 |
| chr10:98395260-98397275 | p2 | (AG)6 | ACCGACAAACCGCATTAAAC | CAAACCAATTATGGTCGGGT | 142 |
| chr10:98495348-98497392 | p2 | (AT)22 | GGTTTGTTGCGTTGGAAACT | TTGGTCCAACTTGAGGGAAG | 182 |
| chr10:98495348-98497392 | p5 | (AATGC)3 | GGTTTGTTGCGTTGGAAACT | TTGGTCCAACTTGAGGGAAG | 182 |
| chr10:98495896-98497911 | p2 | (AT)22 | GTCCCATAGCCTCCTGCATA | ACCGAGGTGGGAAGGTAAGT | 206 |
| chr10:98495896-98497911 | p5 | (AATGC)3 | GTCCCATAGCCTCCTGCATA | ACCGAGGTGGGAAGGTAAGT | 206 |
| chr10:98508749-98510777 | p2 | (AT)14 | GAAGCAATTTCCTCGTAGCG | TTCGTCCTAGGGGTTGTCTG | 113 |
| chr10:98544180-98546195 | p3 | (TTA)5 | CTCGCTTCCCTCATCTTGTC | GGTTGTGAGACCGACCTTGT | 193 |
| chr10:98566504-98568519 | p5 | (GGGGT)3 | TACCCGTCCACATCCTTAGC | CTTTGACGGGGAAGTCCATA | 269 |
| chr10:98608028-98610043 | p5 | (ATTTT)3 | GGCGGAGGTTGTGAGATTAG | AGCTCATCCAAGCACCATCT | 155 |
| chr10:98728470-98730485 | p5 | (TTACT)3 | TAGTGGGTCAGGTCCTCCAG | TGGGGTATTTGGGTTCAAAA | 239 |
| chr10:98735648-98737668 | p2 | (AT)10 | CATGTTGTTTTGGGGGTTTC | AAGGACAACGAACGACGAAG | 240 |

| Marker ID | SSR type | SSR motif | FORWARD PRIMER (5'-3') | REVERSE PRIMER (5'-3') | size (bp) |
|---|---|---|---|---|---|
| chr10:98735648-98737668 | p2 | (TA)15 | CATGTTGTTTTGGGGGTTTC | AAGGACAACGAACGACGAAG | 240 |
| chr10:98747574-98749595 | p3 | (AAT)7 | GGGTTCTTCACGATCCAAAA | CTCACTGTCATCCCCCACTT | 253 |
| chr10:98805104-98807130 | p2 | (AT)13 | AATTTTTGATCCGCCCTTTT | GGGGTATTACGGAGGGGTAA | 211 |
| chr10:98805104-98807130 | p5 | (AAAAT)4 | AATTTTTGATCCGCCCTTTT | GGGGTATTACGGAGGGGTAA | 211 |
| chr10:98824816-98826831 | p5 | (AAAAC)3 | AACGGGTTTGAATGGACAAA | TTTTTAGCCTGGGCTTCTCA | 213 |
| chr10:98858922-98860978 | p2 | (AT)28 | GCTAGACAGAGTCGCAGGCT | TAGACGTAACACGGCACTCG | 158 |
| chr10:98858922-98860978 | p2 | (AT)27 | GCTAGACAGAGTCGCAGGCT | TAGACGTAACACGGCACTCG | 158 |
| chr10:98931732-98933747 | p5 | (AACAA)3 | GAAAGCTCAGTCCGCGATAC | ACCAGGTGTACTGCCTCCAG | 128 |
| chr10:98951053-98953074 | p3 | (AAG)7 | AATTCCATTTAACCCTGCCC | ACGAACACAACGCCAATACA | 204 |
| chr10:98954042-98956063 | p3 | (AAG)7 | AATTCCATTTAACCCTGCCC | ACGAACACAACGCCAATACA | 204 |
| chr10:99057202-99059220 | p2 | (TA)9 | CTCTTTCCAACTCTCCACGC | GCCCACTCCCTTTTTCTTCT | 115 |
| chr10:99205343-99207355 | p2 | (AT)6 | CGAATAGCCTCTGTCCTTGC | CAGAGCCCTAGGTTCGAGTG | 104 |
| chr10:99266459-99268479 | p5 | (AACAT)4 | AAACTCAGGCCACCAAACAC | GAGAAGCTTCCTGGGGTTTC | 251 |
| chr10:99266493-99268505 | p5 | (AACAT)4 | AAACTCAGGCCACCAAACAC | GAGAAGCTTCCTGGGGTTTC | 217 |
| chr10:99321721-99323745 | p3 | (ACA)8 | AGGCTGAGAGACCATCGAAA | CTCACTAAGGAGAATCGCGG | 106 |
| chr10:99326757-99328772 | p5 | (ATCAT)3 | TGTCACGACCCAACCTCATA | GCGCAGGGTTATTTTGGATA | 145 |
| chr10:99448182-99450206 | p3 | (ACA)8 | AGGCTGAGAGACCATCGAAA | CTCACTAAGGAGAATCGCGG | 106 |
| chr10:99453218-99455233 | p5 | (ATCAT)3 | TGTCACGACCCAACCTCATA | GCGCAGGGTTATTTTGGATA | 145 |
| chr10:99533234-99535249 | p3 | (GGT)5 | ATTTGGTCTGGCTGAATTGG | GAATGGGGTTACCGATTCCT | 225 |
| chr10:99542020-99544044 | p3 | (CTT)8 | GTCCCCAGCTGCTGTATCAT | GTTCACCAGTGCTGGTCCTT | 195 |
| chr10:99580837-99582851 | p2 | (AT)7 | CGATAAAACACCATACGGCA | GCTTAATTTTTGGCGTGCAT | 220 |
| chr10:99611712-99613724 | p2 | (TA)6 | CAAGAAATCAAGAGTCGGGC | GCGTGTTTAGGGCGTACAGT | 135 |
| chr10:99613864-99615898 | p3 | (ACA)5 | TTCAAGTCCAGGATTTTCCG | AGACTAAACCTCCCCGCATT | 295 |
| chr10:99613864-99615898 | p2 | (AT)17 | TTCAAGTCCAGGATTTTCCG | AGACTAAACCTCCCCGCATT | 295 |
| chr10:99640070-99642082 | p2 | (TC)6 | GGAGAATTTGCACTTTGGGA | CATAGGGCGGTGGTAGAAAA | 189 |
| chr10:99745307-99747334 | p3 | (TGT)9 | GTGGTTGTGAGACCGACCTT | TCCCTCATCTTGTCCTCCAC | 259 |
| chr10:99914563-99916629 | p2 | (TA)33 | TGGGATTCCAATCATGACCT | TAATTTTGCCTATGGCCTGG | 242 |
| chr10:99934411-99936429 | p3 | (GTT)6 | GCCAAGGGGTTCAACATCTA | AATCCTAGCTTCGCCACTGA | 262 |
| chr10:99934411-99936429 | p5 | (AAAAT)3 | GCCAAGGGGTTCAACATCTA | AATCCTAGCTTCGCCACTGA | 262 |
| chr10:99970118-99972136 | p2 | (AT)9 | AATGCACGTCCACGATGTAA | CGAGTGGAAAGTGGGAAAGA | 291 |
| chr10:99994003-99996021 | p6 | (CAAAAA)3 | TTCCAAATCTCCAAAGCCAC | GAGTCTCACAGCCACCCATT | 164 |
| chr10:100002144-10000415 | p3 | (TTG)5 | TTTGGCACGGACTAAAAAGG | ACCTCTGCACTGGAGCATCT | 237 |
| chr10:100012701-10001471 | p5 | (TTACT)3 | TATTAAGAAGCACGGGTGGG | TGGGGTTCACTTTTGTTTCC | 122 |
| chr10:100012701-10001471 | p2 | (AT)6 | TATTAAGAAGCACGGGTGGG | TGGGGTTCACTTTTGTTTCC | 122 |
| chr10:100012701-10001471 | p6 | (AAAAAT)3 | TATTAAGAAGCACGGGTGGG | TGGGGTTCACTTTTGTTTCC | 122 |
| chr10:100062380-10006439 | p3 | (ATA)5 | GGAAGTTAAATGGTCCCCGT | TATGGTCCACGCCGTATTTT | 127 |

| Marker ID | SSR type | SSR motif | FORWARD PRIMER (5'-3') | REVERSE PRIMER (5'-3') | size (bp) |
|---|---|---|---|---|---|
| chr10:100101608-10010362 | p2 | (TA)7 | GGGGCGTTCAATGAATAAAG | GGCTCCCATTTTACTGGTCA | 227 |
| chr10:100112684-10011470 | p4 | (TTAG)4 | CTAGCGCTGCCTTCACTCTT | GCATGCAAAAAGGGGAGATA | 220 |
| chr10:100119804-10012181 | p2 | (GT)6 | CAGACACTCGCATCACGACT | TCATAGCCGGCACATTCATA | 184 |
| chr10:100149493-10015150 | p5 | (TTTGG)3 | TTGGGTTTGACAGTTGGTGA | ATAGTTCTGCAATGCGGCTT | 283 |
| chr10:100191410-10019342 | p3 | (ACA)5 | TACTGTGTTTTGCGCTCGTC | TCGTGCATCAAAATTCCAAA | 249 |
| chr10:100201031-10020304 | p5 | (AGAAA)3 | TGTTGAAGGCTGTGTGGTCT | AGGCTCTTAGGCACCAGATG | 135 |
| chr10:100276564-10027858 | p5 | (TTGGT)4 | TCTGCCCCATAAGGCATAAC | ATGGGCAGACTTGGCATAAG | 195 |
| chr10:100276564-10027858 | p5 | (TTGGT)3 | TCTGCCCCATAAGGCATAAC | ATGGGCAGACTTGGCATAAG | 195 |
| chr10:100305761-10030777 | p5 | (TAACT)3 | GGAGTGTCAGTGCTTGCTCA | ATCAGGAATCCACAACCTGC | 155 |
| chr10:100305761-10030777 | p4 | (TCTT)4 | GGAGTGTCAGTGCTTGCTCA | ATCAGGAATCCACAACCTGC | 155 |
| chr10:100306000-10030801 | p5 | (TAACT)3 | CCAGAGGCGGGATGTTAGTA | TCTTAAGCTAGCCAGAGGCG | 292 |
| chr10:100306000-10030801 | p4 | (TCTT)4 | CCAGAGGCGGGATGTTAGTA | TCTTAAGCTAGCCAGAGGCG | 292 |
| chr10:100388889-10039090 | p3 | (TTC)5 | ATGTAGTTGGTTGGCGACCT | GTGGTGGTTGAGCCAATCTT | 159 |
| chr10:100388889-10039090 | p2 | (TA)7 | ATGTAGTTGGTTGGCGACCT | GTGGTGGTTGAGCCAATCTT | 159 |
| chr10:100388889-10039090 | p2 | (TA)7 | ATGTAGTTGGTTGGCGACCT | GTGGTGGTTGAGCCAATCTT | 159 |
| chr10:100388889-10039090 | p2 | (TA)7 | ATGTAGTTGGTTGGCGACCT | GTGGTGGTTGAGCCAATCTT | 159 |
| chr10:100400050-10040206 | p2 | (GA)6 | GGTGTGCGGTAAGTTGTCCT | AATAACGGACAATGGCAAGC | 102 |
| chr10:100452471-10045449 | p2 | (TA)10 | AATGATGGTGGCGGAGATAC | GAGTTGTCCGGGAGTTCGTA | 300 |
| chr10:100622299-10062431 | p6 | (ATTTTT)3 | ACCGGTTCATCAACATTCGT | ACGTTTCTCAAAACGCCATT | 261 |
| chr10:100657044-10065906 | p3 | (GTT)8 | GGAACAGTCTCCGAAGCAAG | GGTTACTCCCGGGTTGTTTT | 109 |
| chr10:100657044-10065906 | p3 | (AAC)14 | GGAACAGTCTCCGAAGCAAG | GGTTACTCCCGGGTTGTTTT | 109 |
| chr10:100657044-10065906 | p3 | (AAC)9 | GGAACAGTCTCCGAAGCAAG | GGTTACTCCCGGGTTGTTTT | 109 |
| chr10:100821471-10082348 | p3 | (TTG)5 | CCACCGAAACTTAAAGGCAA | CGCATCTTGTCTTCCACTGA | 212 |
| chr10:100895281-10089729 | p3 | (CTA)5 | CTGGGACTCGATCCATGTTT | TGTGGTTTCCATGTCGAAGA | 297 |
| chr10:100895281-10089729 | p2 | (AC)6 | CTGGGACTCGATCCATGTTT | TGTGGTTTCCATGTCGAAGA | 297 |
| chr10:100911095-10091313 | p3 | (AGA)6 | ATGGAACTTCAAAACACCGC | TACCCGATATTTGGGGATCA | 127 |
| chr10:100965960-10096798 | p5 | (CAAAC)4 | CTTTGGGGCAGTACTTGCAT | CTGCAACTCAAGCACATGGT | 222 |
| chr10:100982971-10098498 | p6 | (AGGGAC)6 | GGCTATGTGTTGCTCGGATT | CTAGGAGGTCACGGGTTTGA | 220 |
| chr10:100982971-10098498 | p6 | (AGGGAC)3 | GGCTATGTGTTGCTCGGATT | CTAGGAGGTCACGGGTTTGA | 220 |
| chr10:101138771-10114082 | p2 | (TA)9 | CTTTGGCTTCGTGGATGATT | CGGAGTGACCTATCGTCGTT | 268 |
| chr10:101289239-10129136 | p3 | (AAG)41 | GGGACCAAGCAAATCAAAGA | GGGTTTCGGCTAGGTTTAGG | 118 |
| chr10:101289239-10129136 | p3 | (AAG)16 | GGGACCAAGCAAATCAAAGA | GGGTTTCGGCTAGGTTTAGG | 118 |
| chr10:101308278-10131030 | p2 | (TA)13 | TTTGTGATTGGGACCATGTG | TTAGGATGTACGGTCGGGAG | 237 |
| chr10:101322086-10132410 | p2 | (AC)7 | GTATGGTGCACAACACGCTC | GGTTGGGTCGTGACAGAAGT | 215 |
| chr10:101364518-10136654 | p3 | (TGA)8 | AAACCGGTAAAATCGGAACC | GGGGGATGGGTTAAAGTGTT | 219 |
| chr10:101439246-10144126 | p5 | (TATTT)4 | GTCACTGCACTTTGAGGCAA | CATCAAGGCATCAGTAGGCA | 282 |

| Marker ID | SSR type | SSR motif | FORWARD PRIMER (5'-3') | REVERSE PRIMER (5'-3') | size (bp) |
|---|---|---|---|---|---|
| chr10:101507433-10150945 | p2 | (TA)13 | CATGTCATATGCAAGCCGAC | TTTTGGAGAGCTCCGTTGTT | 298 |
| chr10:101516144-10151816 | p3 | (TTA)5 | TATGCCTAGCCATTTGGGAG | ATCCTCGGTGACCACTCATC | 113 |
| chr10:101516234-10151824 | p3 | (TTA)5 | TTTTATGCCTAGCCATTCGG | ATCCTCGGTGACCACTCATC | 138 |
| chr10:101585405-10158742 | p3 | (GTG)5 | GTGGTGCTTTTTCTGCCATT | TTCTGCGCCCTTTCTTTCTA | 233 |
| chr10:101606076-10160808 | p2 | (AG)6 | TTGTGGCGAAATAACAACCA | AATTTGCCAGAGTTTCCCCT | 191 |
| chr10:101634444-10163645 | p5 | (TTTTC)3 | TTTCCTTTTCATCTCACCCG | GTTGAAAAACTGACGGGGAA | 271 |
| chr10:101649927-10165195 | p3 | (TAT)9 | TGAATCATCGGCCTAGAACC | CCCAACAAATTGCCTCAGTT | 212 |
| chr10:101651195-10165321 | p5 | (TTTTC)3 | CGGACCAGCCTATCATCACT | CGGGTGAGATGAAAAGGAAA | 120 |
| chr10:101683092-10168511 | p2 | (TA)13 | TGTTGTGAATGTGGGCTGTT | TTCCCAACGTTAGCAACTCC | 264 |
| chr10:101719640-10172165 | p6 | (CATATA)3 | ATATCATACAGAAGCCGCCG | ATGTTCAGGAGTACGTCGGG | 114 |
| chr10:101732989-10173500 | p5 | (ATTTC)3 | ATGCTATTGTGTTTTGGCCC | CTTGCGTACCAACGGAAAAT | 272 |
| chr10:101762346-10176436 | p3 | (TGG)6 | TAGAAAGAAAGGGCGCAGAA | GTGAATGGATACGCCGAGAT | 234 |
| chr10:101762346-10176436 | p3 | (GTT)5 | TAGAAAGAAAGGGCGCAGAA | GTGAATGGATACGCCGAGAT | 234 |
| chr10:101814431-10181645 | p5 | (TTAAT)4 | ACCAGAGTGACCAAAATGGC | TCCCGGAGACTCTAGCTCAA | 113 |
| chr10:101915262-10191735 | p3 | (TCT)32 | TCCTCGCTCCATCTTTGACT | GAGCCAGAACTACAAACGGC | 212 |
| chr10:102295006-10229702 | p4 | (AATA)4 | TCGAACCTTGATATGCTCCC | CCTATAGCTCCAGTGCCACAA | 218 |
| chr10:102311204-10231321 | p3 | (AAT)5 | TTTGGATCATCGGGGACTTA | AACTAATGTCACTTGGCCCG | 132 |
| chr10:102311204-10231321 | p2 | (TA)6 | TTTGGATCATCGGGGACTTA | AACTAATGTCACTTGGCCCG | 132 |
| chr10:102320411-10232242 | p2 | (GT)6 | CCTTGTGCTGCCACTGTAAA | CCCATGAATTGGGAAAAATG | 255 |
| chr10:102460161-10246217 | p5 | (AAACA)3 | TTTTAACCCTGCAACTTCGG | TAAAACCCTCTGCGCTTTGT | 119 |
| chr10:102468514-10247052 | p5 | (AAAAT)3 | GTGAAACCCCCATTTCACAC | TGCACCAGAGCAACACAAAT | 108 |
| chr10:102468514-10247052 | p2 | (AT)7 | GTGAAACCCCCATTTCACAC | TGCACCAGAGCAACACAAAT | 108 |
| chr10:102468514-10247052 | p2 | (TA)7 | GTGAAACCCCCATTTCACAC | TGCACCAGAGCAACACAAAT | 108 |
| chr10:102468514-10247052 | p6 | (CCGGTT)4 | GTGAAACCCCCATTTCACAC | TGCACCAGAGCAACACAAAT | 108 |
| chr10:102468514-10247052 | p2 | (TA)28 | GTGAAACCCCCATTTCACAC | TGCACCAGAGCAACACAAAT | 108 |
| chr10:102469034-10247104 | p2 | (AT)7 | GAACCGGCATTTGAGTGATT | TGGTGAAAGTGCGAAAAGC | 144 |
| chr10:102469034-10247104 | p2 | (TA)7 | GAACCGGCATTTGAGTGATT | TGGTGAAAGTGCGAAAAGC | 144 |
| chr10:102469034-10247104 | p6 | (CCGGTT)4 | GAACCGGCATTTGAGTGATT | TGGTGAAAGTGCGAAAAGC | 144 |
| chr10:102469034-10247104 | p2 | (TA)28 | GAACCGGCATTTGAGTGATT | TGGTGAAAGTGCGAAAAGC | 144 |
| chr10:102529979-10253199 | p2 | (AT)6 | TTGTTGGGGTTGGGAAATTA | ATATCGGTCGGTTCAGTTCG | 251 |
| chr10:102561247-10256326 | p5 | (CGTTT)3 | GCAGAAATGTGGAAAGCCAT | GTCGAGCTTGTTCCGTTAGC | 104 |
| chr10:102611827-10261384 | p2 | (GT)7 | GAGCAACCCTGCTCAAATGT | TGGGGAAATAATTTGGGTGA | 128 |
| chr10:102611827-10261384 | p3 | (ATG)5 | GAGCAACCCTGCTCAAATGT | TGGGGAAATAATTTGGGTGA | 128 |
| chr10:102611827-10261384 | p2 | (AT)33 | GAGCAACCCTGCTCAAATGT | TGGGGAAATAATTTGGGTGA | 128 |
| chr10:102625607-10262763 | p3 | (GTT)8 | CGTGGAAGAGTTGTTTGGGT | AACACCCATTTACACGGCTC | 175 |
| chr10:102681598-10268361 | p5 | (TCTAT)3 | GTCAGCCCCATCAGCTACAT | AGAGGGAATTGGAATGGGTC | 198 |

| Marker ID | SSR type | SSR motif | FORWARD PRIMER (5'-3') | REVERSE PRIMER (5'-3') | size (bp) |
|---|---|---|---|---|---|
| chr10:102681598-10268361 | p5 | (TTATT)3 | GTCAGCCCCATCAGCTACAT | AGAGGGAATTGGAATGGGTC | 198 |
| chr10:102689013-10269103 | p2 | (TA)9 | AAGCTAGCATGAGGCCAAAA | TGGACTCTTTTGACGGGAAG | 174 |
| chr10:102701432-10270348 | p2 | (AT)26 | CGTGCCCTTCGTCAGTCTAT | CTTGGATCTTTGTCCTCCCA | 289 |
| chr10:102756690-10275870 | p3 | (ATT)19 | ATTTTGCAAAGATTCGCACC | ATTTGAGAGAATTGTGGCCG | 111 |
| chr10:102756690-10275870 | p2 | (TC)6 | ATTTTGCAAAGATTCGCACC | ATTTGAGAGAATTGTGGCCG | 111 |
| chr10:102768760-10277077 | p2 | (CT)8 | ATTCTCACGAAACCCGTGAC | TAGCCAATTTTGTTCCGACC | 147 |
| chr10:102842506-10284452 | p2 | (TA)7 | GCTAATTTGGGGTATCGGGT | TTTCGTTAATTTCCCCCACA | 198 |
| chr10:102842506-10284452 | p2 | (TA)7 | GCTAATTTGGGGTATCGGGT | TTTCGTTAATTTCCCCCACA | 198 |
| chr10:102842715-10284473 | p2 | (TA)7 | GGATTAGATCGGAGCGACAC | TTTCGTTAATTTCCCCCACA | 108 |
| chr10:102842715-10284473 | p2 | (TA)7 | GGATTAGATCGGAGCGACAC | TTTCGTTAATTTCCCCCACA | 108 |
| chr10:102846110-10284813 | p2 | (TA)11 | ATTCATGGCCTTTTTGTTGC | AGTTTGCATCCTCAACCACC | 101 |
| chr10:102893343-10289535 | p2 | (AT)7 | TTGCTTCCACTTATCCAGGC | GAGCATTTTGGGGAAAGTGA | 273 |
| chr10:102894440-10289646 | p2 | (AT)7 | CCACTTGAGACCCCAGAGAG | CGTGCGATAATTCCGTTCTT | 136 |
| chr10:102894440-10289646 | p3 | (CTT)7 | CCACTTGAGACCCCAGAGAG | CGTGCGATAATTCCGTTCTT | 136 |
| chr10:102907137-10290915 | p2 | (TA)15 | GTTCCTCTACCCCCAGGAAG | TTCGTAATTAGCCATTCCCG | 221 |
| chr10:102907137-10290915 | p2 | (AT)6 | GTTCCTCTACCCCCAGGAAG | TTCGTAATTAGCCATTCCCG | 221 |
| chr10:102907168-10290918 | p2 | (TA)15 | GTTCCTCTACCCCCAGGAAG | TTCGTAATTAGCCATTCCCG | 190 |
| chr10:102907168-10290918 | p2 | (AT)6 | GTTCCTCTACCCCCAGGAAG | TTCGTAATTAGCCATTCCCG | 190 |
| chr10:102914640-10291665 | p2 | (AT)7 | TTGCTTCCACTTATCCAGGC | GAGCATTTTGGGGAAAGTGA | 273 |
| chr10:102915737-10291775 | p2 | (AT)7 | CCACTTGAGACCCCAGAGAG | CGTGCGATAATTCCGTTCTT | 136 |
| chr10:102915737-10291775 | p3 | (CTT)7 | CCACTTGAGACCCCAGAGAG | CGTGCGATAATTCCGTTCTT | 136 |
| chr10:102929860-10293187 | p2 | (TA)8 | TACATTCTGACAGCAAGGCG | TTGGAGGGGATCGTAGTGAC | 222 |
| chr10:102973506-10297552 | p3 | (GTT)5 | CCCGAGTGTTGATCGGTAGT | AATCCGGTAGGACTCCCAAG | 247 |
| chr10:103008389-10301041 | p2 | (AT)14 | TAAAGACCACCCCAGAAGGA | AGCCCACCAAATAGCAAATG | 179 |
| chr10:103034757-10303677 | p6 | (ACCGCC)4 | TTAACCCCTGAACTTGGCAC | AATGACGTGGTGCGAGTGTA | 104 |
| chr10:103034757-10303677 | p3 | (GAA)6 | TTAACCCCTGAACTTGGCAC | AATGACGTGGTGCGAGTGTA | 104 |
| chr10:103034757-10303677 | p2 | (AT)8 | TTAACCCCTGAACTTGGCAC | AATGACGTGGTGCGAGTGTA | 104 |
| chr10:103034997-10303701 | p6 | (ACCGCC)4 | CATCATCTCCCCCAACAAAT | TGGCCAACAATAACAACCAA | 268 |
| chr10:103034997-10303701 | p3 | (GAA)6 | CATCATCTCCCCCAACAAAT | TGGCCAACAATAACAACCAA | 268 |
| chr10:103034997-10303701 | p2 | (AT)8 | CATCATCTCCCCCAACAAAT | TGGCCAACAATAACAACCAA | 268 |
| chr10:103099938-10310195 | p2 | (TG)6 | GTTTGCCACGTTTGTGATTG | ATAACACTTGAACCCGCCAG | 280 |
| chr10:103125323-10312734 | p6 | (CATATA)3 | CATATCATGTCATAGCGCCG | TATAGACGCACAGCCACCTG | 149 |
| chr10:103125323-10312734 | p3 | (CAA)6 | CATATCATGTCATAGCGCCG | TATAGACGCACAGCCACCTG | 149 |
| chr10:103186349-10318843 | p5 | (AATCA)3 | TTGTTGGCTCAAACATAGCG | GCTTTCCTAACACAAACGGC | 167 |
| chr10:103229814-10323183 | p2 | (TC)12 | GTGCCAAATATCACCATCCC | TGTCGTATTAGCCGAAACCC | 254 |
| chr10:103254406-10325642 | p3 | (TGG)5 | GTGGTCGTCGTAGAGGTGGT | GGAATGAGAAGTCGATGGGA | 100 |

| Marker ID | SSR type | SSR motif | FORWARD PRIMER (5'-3') | REVERSE PRIMER (5'-3') | size (bp) |
|---|---|---|---|---|---|
| chr10:103305892-10330790 | p3 | (TGT)5 | TCCTGAGTGTGTTCCTGCTG | AATGAGACGGATGGACAAGC | 252 |
| chr10:103397104-10339915 | p2 | (TA)18 | CAATCCATTAGTGGGAGCAAA | TGACATGGCGGAATAACAAA | 285 |
| chr10:103397104-10339915 | p2 | (AT)23 | CAATCCATTAGTGGGAGCAAA | TGACATGGCGGAATAACAAA | 285 |
| chr10:103483766-10348579 | p2 | (TA)13 | ATGGCCAACTTAGGGAGCTT | GTGGTCGACAAAACCCATTT | 172 |
| chr10:103483766-10348579 | p4 | (ATTT)4 | ATGGCCAACTTAGGGAGCTT | GTGGTCGACAAAACCCATTT | 172 |
| chr10:103593558-10359563 | p3 | (ATT)26 | GTTCTCTGCTCAGGGGTCTG | CGTGAGAGATCTCCGTCTCC | 194 |
| chr10:103704196-10370622 | p2 | (TA)12 | ACATGCACTGCAATTTGAGC | ATTTGCTGGATGCCTTTTTG | 143 |
| chr10:103704196-10370622 | p5 | (TTTTA)3 | ACATGCACTGCAATTTGAGC | ATTTGCTGGATGCCTTTTTG | 143 |
| chr10:103704573-10370658 | p2 | (TA)12 | GAGCTGGAGGGAGTGTATGC | TGCCCCATAAGGCATAGTTC | 256 |
| chr10:103704573-10370658 | p5 | (TTTTA)3 | GAGCTGGAGGGAGTGTATGC | TGCCCCATAAGGCATAGTTC | 256 |
| chr10:103704587-10370663 | p2 | (TA)12 | GAGCTGGAGGGAGTGTATGC | TGCCCCATAAGGCATAGTTC | 242 |
| chr10:103704587-10370663 | p5 | (TTTTA)3 | GAGCTGGAGGGAGTGTATGC | TGCCCCATAAGGCATAGTTC | 242 |
| chr10:103723021-10372504 | p2 | (TA)12 | ACATGCACTGCAATTTGAGC | ATTTGCTGGATGCCTTTTTG | 143 |
| chr10:103723021-10372504 | p5 | (TTTTA)3 | ACATGCACTGCAATTTGAGC | ATTTGCTGGATGCCTTTTTG | 143 |
| chr10:103723398-10372541 | p2 | (TA)12 | GAGCTGGAGGGAGTGTATGC | TGCCCCATAAGGCATAGTTC | 256 |
| chr10:103723398-10372541 | p5 | (TTTTA)3 | GAGCTGGAGGGAGTGTATGC | TGCCCCATAAGGCATAGTTC | 256 |
| chr10:103723412-10372546 | p2 | (TA)12 | GAGCTGGAGGGAGTGTATGC | TGCCCCATAAGGCATAGTTC | 242 |
| chr10:103723412-10372546 | p5 | (TTTTA)3 | GAGCTGGAGGGAGTGTATGC | TGCCCCATAAGGCATAGTTC | 242 |
| chr10:103837536-10383955 | p5 | (TAATA)3 | TGCACGTTGTTTGAGGAAAG | CCAAGCAACTTCCACGTTTT | 288 |
| chr10:103837536-10383955 | p2 | (AT)10 | TGCACGTTGTTTGAGGAAAG | CCAAGCAACTTCCACGTTTT | 288 |
| chr10:103977423-10397943 | p2 | (TA)12 | CGTGGGGAACTTCAACAACT | TGATCGAGGGTGACATGAAA | 159 |
| chr10:103977469-10397948 | p2 | (TA)12 | CGTGGGGAACTTCAACAACT | TGATCGAGGGTGACATGAAA | 113 |
| chr10:103978076-10398010 | p2 | (TA)12 | AAGAAGCCTTTGTTGGCTCA | GTGTAGCCCCTATCCGAGGT | 285 |
| chr10:104118814-10412082 | p5 | (TGGGG)3 | TCTTGAAGGGTTTGGTGGAG | TCCTCATGTTGCACCCAATA | 274 |
| chr10:104131833-10413384 | p5 | (TTACT)4 | GGTTTTGGTGGACTCGGTAA | AACCGTGGCTGTTGTTTTTC | 275 |
| chr10:104137221-10413924 | p3 | (GTT)8 | CCACCAAAACTTAAGGGCAA | CTCCTCGCATCTTGTCTTCC | 269 |
| chr10:104137221-10413924 | p2 | (AT)7 | CCACCAAAACTTAAGGGCAA | CTCCTCGCATCTTGTCTTCC | 269 |
| chr10:104154089-10415610 | p2 | (AT)6 | TCGTCAACAGTGTGGAAAGC | CAGAAAGACGTCGAAAAGGC | 125 |
| chr10:104169440-10417145 | p5 | (TTATT)3 | CACCGTGGACAAAGACAATG | TTAGCAACCCTTTTCATGCC | 279 |
| chr10:104169440-10417145 | p2 | (CT)8 | CACCGTGGACAAAGACAATG | TTAGCAACCCTTTTCATGCC | 279 |
| chr10:104176113-10417814 | p5 | (AATGC)3 | AATGCAACAATCGCCTCTCT | AGAGTGCCAGAGAGACTCCG | 100 |
| chr10:104176113-10417814 | p4 | (TAGA)9 | AATGCAACAATCGCCTCTCT | AGAGTGCCAGAGAGACTCCG | 100 |
| chr10:104214020-10421604 | p2 | (TA)10 | TGCTTCAAAGTAGGTGGCCT | TTGGGGTGAATGTGTTCAAA | 267 |
| chr10:104214020-10421604 | p5 | (ATTTA)3 | TGCTTCAAAGTAGGTGGCCT | TTGGGGTGAATGTGTTCAAA | 267 |
| chr10:104367883-10436990 | p2 | (AT)9 | TGGCCCTCTCTATGACCAAG | ATGAGCGTGAATGGCTTAGG | 258 |
| chr10:104400937-10440295 | p6 | (GAAAAG)3 | TCTCTTGTTGCCATTGTGGA | GCTTAGGCATGGGTGTTGTT | 242 |

| Marker ID | SSR type | SSR motif | FORWARD PRIMER (5'-3') | REVERSE PRIMER (5'-3') | size (bp) |
|---|---|---|---|---|---|
| chr10:104415340-10441741 | p5 | (AAACT)3 | TGATCGCCCTGTTCCTAATC | CTTATCTTCCCCCTTTTGGC | 117 |
| chr10:104587341-10458940 | p3 | (TAT)21 | TCACCAACTGAGACCACTCG | GAAACCCACCGCTTCATTTA | 168 |
| chr10:104612580-10461459 | p3 | (TTC)5 | AACACGAATCAACCTCTGGG | CTCATACTGCTGGTGGAGCA | 129 |
| chr10:104641465-10464347 | p2 | (AT)7 | AATGAAATCGAGCTCCGCTA | GATGGCCACTAGAGGGACAA | 268 |
| chr10:104691475-10469349 | p2 | (TC)9 | TTAACTTTGAATGCATGGCG | GGTGTGATTCTCCAATGGCT | 198 |
| chr10:104808753-10481077 | p2 | (TA)13 | TAACTCAAACCCCGGTGAAC | TGGCACATGCCTTTTCATAA | 167 |
| chr10:104856517-10485853 | p3 | (GTT)7 | ATTCTAGCGCTCAATCGGAA | AGTTTGGGATGTCCTTGTCG | 275 |
| chr10:104856517-10485853 | p5 | (GGAGT)3 | ATTCTAGCGCTCAATCGGAA | AGTTTGGGATGTCCTTGTCG | 275 |
| chr10:104856517-10485853 | p3 | (TTG)5 | ATTCTAGCGCTCAATCGGAA | AGTTTGGGATGTCCTTGTCG | 275 |
| chr10:104860883-10486289 | p5 | (TGGTT)3 | ACGCAACCAAAATTTTCAGG | AAAAGGAGCCATGGATGATG | 291 |
| chr10:104892389-10489444 | p3 | (TTA)19 | GTCAATTCGGAAGTCTCCCA | CCTTCCTTTCACGCACTAGC | 250 |
| chr10:104919707-10492172 | p2 | (TA)6 | ACTGGTTCAAACCGTAACCG | CCATGTCCCAAGGCTACAAT | 274 |
| chr10:104919707-10492172 | p2 | (AT)10 | ACTGGTTCAAACCGTAACCG | CCATGTCCCAAGGCTACAAT | 274 |
| chr10:104919707-10492172 | p5 | (CCGGT)3 | ACTGGTTCAAACCGTAACCG | CCATGTCCCAAGGCTACAAT | 274 |
| chr10:105067888-10506990 | p2 | (TA)8 | CCCATGGAGGTGTTATACGG | TGGGTACCTTGATTCTTGCC | 127 |
| chr10:105124168-10512618 | p4 | (TATG)4 | GCCCTTTGGCTATCCACATA | TGTTCATGCCCACAGGTAGA | 224 |
| chr10:105225457-10522747 | p3 | (TTG)7 | ATGAAAGTGGCGGAAATGAG | CGCATCTTGTCTTCCACTGA | 300 |
| chr10:105425482-10542750 | p2 | (TA)11 | GTGGGCGACCTACCATTAGA | GTAGGTCTGTCTTCTGGGCG | 202 |
| chr10:105437796-10543981 | p3 | (TAC)6 | CATCTCCACAACACCAAACG | ATTGTTGACTGCTCCGGTTC | 108 |
| chr10:105457600-10545961 | p2 | (TA)9 | GGGACAAAAGTCGTCGGTAA | TTTCCCCCAATGAAAAATCA | 249 |
| chr10:105457600-10545961 | p5 | (TCAAA)3 | GGGACAAAAGTCGTCGGTAA | TTTCCCCCAATGAAAAATCA | 249 |
| chr10:105471724-10547374 | p2 | (TA)8 | TATGTGGTGGGTTCCACCTT | CACCATCTCCAAACTCACGA | 170 |
| chr10:105471724-10547374 | p2 | (AT)6 | TATGTGGTGGGTTCCACCTT | CACCATCTCCAAACTCACGA | 170 |
| chr10:105471724-10547374 | p2 | (AT)6 | TATGTGGTGGGTTCCACCTT | CACCATCTCCAAACTCACGA | 170 |
| chr10:105474686-10547670 | p3 | (AAC)6 | CGTGTTTCAAGGAAATGGGT | AGAATCAACAGCGTGGGAAC | 280 |
| chr10:105487828-10548984 | p2 | (AT)10 | GAGGTGCAAAGGGTGAAAGA | GGGCACAAATATACCCCTCC | 194 |
| chr10:105511616-10551363 | p3 | (GAA)6 | GATGTGGGAGGATGAGGAGA | GGCCTCGTTCACTGAGGTAG | 198 |
| chr10:105594475-10559648 | p2 | (AT)7 | CATAAGCTAGCCGACAAGGC | TGACAGGCAGAGCAGAAGAA | 160 |
| chr10:105594475-10559648 | p2 | (AT)18 | CATAAGCTAGCCGACAAGGC | TGACAGGCAGAGCAGAAGAA | 160 |
| chr10:105634601-10563661 | p3 | (CTT)5 | CCTGTTTGGATGGGCTTAAA | GCCAGCTTAAAGCCATTTTG | 250 |
| chr10:105739716-10574173 | p6 | (AGAAGG)3 | TCGGTATCGTGGATTAAGGC | GGGCCAGAAGTGTACGATGT | 181 |
| chr10:105766718-10576878 | p3 | (TAA)21 | GAGTAATAGGGGTGGGCGTT | GAAATCGAACCGAACCAAAC | 182 |
| chr10:105878558-10588057 | p3 | (CAA)5 | CCCATTGATACCCATTTTCG | GATTCTGGCCAAATTCGTGT | 284 |
| chr10:105903388-10590540 | p5 | (AACCG)3 | TTTCATTTGGCCCTCAAGAC | TGTTAGGCCTCATACCCGAC | 103 |
| chr10:105903388-10590540 | p5 | (AACCG)3 | TTTCATTTGGCCCTCAAGAC | TGTTAGGCCTCATACCCGAC | 103 |
| chr10:105955179-10595719 | p5 | (ACACC)3 | CATTGGCAACCAAACAACTG | GGGTGCACCGGTATTATTTG | 261 |

| Marker ID | SSR type | SSR motif | FORWARD PRIMER (5'-3') | REVERSE PRIMER (5'-3') | size (bp) |
|---|---|---|---|---|---|
| chr10:106005258-10600727 | p4 | (ATTT)5 | GAAGAGAGAGCGGTGGTGTC | TGGACTGGAGAGAAGAGGGA | 295 |
| chr10:106121968-10612398 | p2 | (TA)9 | CTTGCCATGCAACGTAAAGA | TTTTGGCTAAGGAGAGCGAA | 216 |
| chr10:106177093-10617913 | p2 | (AT)6 | TTGTGGCCCCCATATTCTAA | ATATGGGGCCGGATATTTTT | 300 |
| chr10:106177197-10617925 | p2 | (AT)6 | TTGTGGCCCCCATATTCTAA | ATATGGGGCCGGATATTTTT | 196 |
| chr10:106177197-10617925 | p2 | (AT)6 | TTGTGGCCCCCATATTCTAA | ATATGGGGCCGGATATTTTT | 196 |
| chr10:106254007-10625602 | p3 | (ACA)6 | CTCTGTCGGGATGTGAACCT | AGCTCGCTACCGTCTTACCA | 205 |
| chr10:106276090-10627810 | p2 | (AT)6 | TGGTGAAACATCGGCTGTAA | TGGGGTGGTGAAGTTGTGTA | 291 |
| chr10:106516100-10651813 | p4 | (GAAA)13 | CCGAAAATAATTCCGCCTTT | TGGGGTTGGGTACCAAAATA | 153 |
| chr10:106516100-10651813 | p2 | (GT)9 | CCGAAAATAATTCCGCCTTT | TGGGGTTGGGTACCAAAATA | 153 |
| chr10:106516100-10651813 | p6 | (AAAAAT)3 | CCGAAAATAATTCCGCCTTT | TGGGGTTGGGTACCAAAATA | 153 |
| chr10:106545955-10654797 | p2 | (AT)14 | GTTTGGCAACGTCCATTTTT | CAGAATCGGAACCGGTAGAA | 257 |
| chr10:106545955-10654797 | p2 | (TA)10 | GTTTGGCAACGTCCATTTTT | CAGAATCGGAACCGGTAGAA | 257 |
| chr10:106545955-10654797 | p2 | (AT)24 | GTTTGGCAACGTCCATTTTT | CAGAATCGGAACCGGTAGAA | 257 |
| chr10:106637610-10663965 | p5 | (ATATG)3 | ACCTCTCGTTGGCACACTCT | CGGAGTGACCTATCGGTGTT | 227 |
| chr10:106637610-10663965 | p2 | (TA)22 | ACCTCTCGTTGGCACACTCT | CGGAGTGACCTATCGGTGTT | 227 |
| chr10:106847161-10684923 | p2 | (AT)38 | GCTCCGGGCTCTATCTTTCT | CTACCATCAACATCGCCAAA | 244 |
| chr10:106859805-10686182 | p5 | (TTTAA)3 | GTTGTGGAGACCTTGTCGGT | TTGGTTACGACCCCCTACAG | 257 |
| chr10:106859805-10686182 | p2 | (AT)15 | GTTGTGGAGACCTTGTCGGT | TTGGTTACGACCCCCTACAG | 257 |
| chr10:106892669-10689468 | p5 | (AAATA)3 | GAGGACCCCGATGACTGATA | TGGGCAACATCAGAACAAGA | 278 |
| chr10:106892669-10689468 | p4 | (TTTC)4 | GAGGACCCCGATGACTGATA | TGGGCAACATCAGAACAAGA | 278 |
| chr10:107066684-10706869 | p5 | (AAATA)3 | TCGGTAGAATGTGATTGCGA | AAAATTTCGACCACACGAGG | 167 |
| chr10:107095928-10709794 | p4 | (AAAT)4 | GGCCAAACACAAACTGCTTT | TACTTCGCTTTTGACCCGTT | 151 |
| chr10:107230046-10723206 | p3 | (TTA)5 | GACTCATAGCTCTCCCGTGC | GCAAGGCAAACACATCTTCA | 279 |
| chr10:107230046-10723206 | p3 | (TTA)5 | GACTCATAGCTCTCCCGTGC | GCAAGGCAAACACATCTTCA | 279 |
| chr10:107230165-10723218 | p3 | (TTA)5 | GACTCATAGCTCTCCCGTGC | CAATGCAAGGCAAACACATC | 160 |
| chr10:107230165-10723218 | p3 | (TTA)5 | GACTCATAGCTCTCCCGTGC | CAATGCAAGGCAAACACATC | 160 |
| chr10:107230187-10723220 | p3 | (TTA)5 | GACTCATAGCTCTCCCGTGC | CAATGCAAGGCAAACACATC | 138 |
| chr10:107230187-10723220 | p3 | (TTA)5 | GACTCATAGCTCTCCCGTGC | CAATGCAAGGCAAACACATC | 138 |
| chr10:107236936-10723895 | p2 | (AT)7 | TGTTCGATTTTTGCCATCAA | AATTTCAGCGACCCAGAATG | 272 |
| chr10:107322760-10732477 | p6 | (TTTATT)3 | TCAATTTTTCCTTTGGCAGC | GCGATAAGGCTTTTCTGCAC | 134 |
| chr10:107322760-10732477 | p5 | (TTTTA)3 | TCAATTTTTCCTTTGGCAGC | GCGATAAGGCTTTTCTGCAC | 134 |
| chr10:107324887-10732689 | p2 | (TA)6 | GGTTGATGGCTGCAGATTTT | TGACGAACAACCTTTCCTCC | 168 |
| chr10:107342090-10734410 | p6 | (TTTATT)3 | TCAATTTTTCCTTTGGCAGC | GCGATAAGGCTTTTCTGCAC | 134 |
| chr10:107342090-10734410 | p5 | (TTTTA)3 | TCAATTTTTCCTTTGGCAGC | GCGATAAGGCTTTTCTGCAC | 134 |
| chr10:107344217-10734622 | p2 | (TA)6 | GGTTGATGGCTGCAGATTTT | TGACGAACAACCTTTCCTCC | 168 |
| chr10:107394079-10739609 | p2 | (TA)7 | CCTCGTTATCCAGGCTGTGT | GGGCAGCTAGAAGATCATGG | 292 |

| Marker ID | SSR type | SSR motif | FORWARD PRIMER (5'-3') | REVERSE PRIMER (5'-3') | size (bp) |
|---|---|---|---|---|---|
| chr10:107406336-10740834 | p2 | (CA)6 | AAGCTGGCCAGTCAAACACT | TCGTATATTTTGGGCAGGGA | 235 |
| chr10:107406336-10740834 | p2 | (TA)16 | AAGCTGGCCAGTCAAACACT | TCGTATATTTTGGGCAGGGA | 235 |
| chr10:107456827-10745885 | p2 | (TA)14 | CAGACATGGCTGAAAAAGCA | GTGAATCCTCTTTCCCCTCC | 104 |
| chr10:107483305-10748532 | p2 | (TA)9 | TTCTTCGGAGAAGCCACTGT | AAACTCAGATCTACGCCGGA | 150 |
| chr10:107483305-10748532 | p5 | (TATAA)3 | TTCTTCGGAGAAGCCACTGT | AAACTCAGATCTACGCCGGA | 150 |
| chr10:107584843-10758686 | p6 | (AAAGGC)3 | TACCCACATCGAGGCTTACC | GAAGGTTCGATTTTGGACGA | 150 |
| chr10:107587862-10758987 | p5 | (TTTTA)3 | ACAAAACCAGCCCTTTTCCT | CGGCAGTGTCAAAATTAGCA | 148 |
| chr10:107601615-10760375 | p2 | (AT)70 | TATAGGCATGGGGGATTGAA | AAATCGGTGGTTTCACAAGC | 147 |
| chr10:107645752-10764778 | p3 | (TTA)10 | GCTCGTAAACGCACAGCATA | ATGAGAATGATGCCTTTGGC | 125 |
| chr10:107705736-10770775 | p2 | (AC)9 | TAACTTTCCTGTGGATCGGG | AAAGCTCCCCACTCTTCGTT | 159 |
| chr10:107729850-10773190 | p2 | (AT)12 | ATGATGGATTCATTGCCACA | GACATGGCGGAATAGCAAAT | 180 |
| chr10:107729850-10773190 | p2 | (AT)17 | ATGATGGATTCATTGCCACA | GACATGGCGGAATAGCAAAT | 180 |
| chr10:107842246-10784425 | p2 | (AT)6 | CATATGATACATGGGCTGCG | TTCGCTGCCGAAGAAATAGT | 239 |
| chr10:107842246-10784425 | p3 | (ATT)5 | CATATGATACATGGGCTGCG | TTCGCTGCCGAAGAAATAGT | 239 |
| chr10:107886688-10788870 | p2 | (CA)6 | TTACCCTCCCCAGACATCAG | GTCACGCTGAAAAAGAAGGC | 278 |
| chr10:107890528-10789254 | p2 | (TA)9 | GTTCGAGCCTGGGTGTAAAA | TTCGGTACTCATTGGAGCCT | 271 |
| chr10:108007377-10800940 | p2 | (AT)16 | CTTCGGAGAGCGCATTTATC | CCGCTTCTAAGTTTGGTTGC | 297 |
| chr10:108021428-10802344 | p2 | (AT)6 | CAGCTCATTCACAAGCTCCA | ACTCTTTTGTGGCTCTCCGA | 292 |
| chr10:108091817-10809390 | p2 | (AT)43 | TAGTCCCCTATACCGCTGGA | AACCGTTTCCCTTTCTTGGT | 169 |
| chr10:108091817-10809390 | p2 | (AT)14 | TAGTCCCCTATACCGCTGGA | AACCGTTTCCCTTTCTTGGT | 169 |
| chr10:108138398-10814041 | p4 | (AATA)4 | TCTTCCGATGTTGTTCGTGA | TTTGGCCGGATTAATCGTAG | 265 |
| chr10:108160540-10816255 | p2 | (TA)8 | TTTGTGCTCTGTTTCGCTTG | CCGGTAGTTCCGAGTCAAAA | 140 |
| chr10:108167062-10816908 | p2 | (TA)10 | ACCAACGACCCAGATACTCG | GAACTGGAGGAACGCTTGTC | 223 |
| chr10:108167062-10816908 | p2 | (TA)11 | ACCAACGACCCAGATACTCG | GAACTGGAGGAACGCTTGTC | 223 |
| chr10:108177840-10817985 | p2 | (AT)7 | TGCTACACTCGTTGCTCCAC | TCGCCTTGTATTCCTTCCAC | 100 |
| chr10:108197012-10819903 | p3 | (TGT)9 | TGGATTGCATCCAGCATAAA | GTTTCTACACCCGCTACCCA | 204 |
| chr10:108197012-10819903 | p6 | (TGGGTG)4 | TGGATTGCATCCAGCATAAA | GTTTCTACACCCGCTACCCA | 204 |
| chr10:108197012-10819903 | p2 | (TA)16 | TGGATTGCATCCAGCATAAA | GTTTCTACACCCGCTACCCA | 204 |
| chr10:108350037-10835205 | p5 | (TCAAT)3 | TTTGGTGTCTCTTGCATTCG | CTAACCATCCCTCCCTCCTC | 268 |
| chr10:108365379-10836739 | p6 | (CTCATG)3 | TGCCCACCAAGTGTTTGATA | TGGGGAAGAACAAGGATGTC | 181 |
| chr10:108492342-10849435 | p2 | (GA)7 | AGTGCTGGTTTCACCTGCTT | ATTTTCTCGAGTCCCTGGGT | 239 |
| chr10:108503718-10850573 | p3 | (TAA)5 | AAAAATGATGGGGAAAACCC | GCCCGATTGACTCCTGATAA | 283 |
| chr10:108504047-10850606 | p3 | (TAA)5 | TCAATCGGGCTTTAATTTGG | AATTATCCGGGTCGTTTTCC | 137 |
| chr10:108540402-10854241 | p4 | (AGAA)4 | GGATTTGGAATTGAGGCGTA | CCCATGGTTGAATGCCTATC | 287 |
| chr10:108614819-10861683 | p5 | (TCTTT)3 | GTGAGCCCATAAGCCATCAT | TCTGGTTCTGTCAAGTCCCC | 215 |
| chr10:108730031-10873204 | p2 | (CT)6 | CCATAATTCAACACTGGGGC | ATATTATGGGGGCAGAAGGG | 123 |

| Marker ID | SSR type | SSR motif | FORWARD PRIMER (5'-3') | REVERSE PRIMER (5'-3') | size (bp) |
|---|---|---|---|---|---|
| chr10:108767911-10876992 | p5 | (AAAAT)3 | AAATGGACGGAGGGAGTACC | AGGGGCGAAGCTAAGGTAAA | 218 |
| chr10:108776554-10877856 | p3 | (GGT)5 | AGCAAATGCAGTAGCCGAAT | GCAACCTTTTCCCGTACAAA | 237 |
| chr10:108776554-10877856 | p3 | (TAC)5 | AGCAAATGCAGTAGCCGAAT | GCAACCTTTTCCCGTACAAA | 237 |
| chr10:108835983-10883801 | p2 | (AG)6 | AATTTTCAAGGGATTTCGGG | AAATGGTCGCGAGTTGAATC | 231 |
| chr10:108835983-10883801 | p3 | (AAT)10 | AATTTTCAAGGGATTTCGGG | AAATGGTCGCGAGTTGAATC | 231 |
| chr10:108876789-10887880 | p3 | (TAT)5 | ATTTCGTCCGACTTCATTGG | GCGAACAATGCAGAGAATCA | 120 |
| chr10:108954140-10895615 | p6 | (AATAAA)3 | GGCAAACCTCTGTCTGAAGG | GCCCTCAAATTGTTGGTTGT | 118 |
| chr10:108976962-10897898 | p2 | (AT)9 | TTGTTCTTGAGCCTCATCCC | AAAAGCAAATGGTTTCGTGC | 168 |
| chr10:109078937-10908094 | p2 | (TA)6 | TCCGGTCGATGTCTTTTCTT | GGGGACAAAGAGTCGTGAGA | 113 |
| chr10:109079340-10908136 | p2 | (TA)6 | AGATGTGGTTGGGCTAGGTG | GGTTCGTTAACCGGTTCCAT | 159 |
| chr10:109079340-10908136 | p2 | (TA)7 | AGATGTGGTTGGGCTAGGTG | GGTTCGTTAACCGGTTCCAT | 159 |
| chr10:109079340-10908136 | p2 | (AT)7 | AGATGTGGTTGGGCTAGGTG | GGTTCGTTAACCGGTTCCAT | 159 |
| chr10:109079390-10908140 | p2 | (TA)6 | AGATGTGGTTGGGCTAGGTG | GGTTCGTTAACCGGTTCCAT | 109 |
| chr10:109079390-10908140 | p2 | (TA)7 | AGATGTGGTTGGGCTAGGTG | GGTTCGTTAACCGGTTCCAT | 109 |
| chr10:109079390-10908140 | p2 | (AT)7 | AGATGTGGTTGGGCTAGGTG | GGTTCGTTAACCGGTTCCAT | 109 |
| chr10:109117935-10911994 | p2 | (TG)6 | TCCGATGTCATATCGACGAA | TGCGGAGCTTTTCTCTCAAT | 216 |
| chr10:109117935-10911994 | p3 | (TTG)5 | TCCGATGTCATATCGACGAA | TGCGGAGCTTTTCTCTCAAT | 216 |
| chr10:109176054-10917807 | p2 | (TA)8 | TAAAATGGGGCTTACCCCTC | AATCGCGGCTATACGTTGTC | 114 |
| chr10:109176054-10917807 | p2 | (TA)17 | TAAAATGGGGCTTACCCCTC | AATCGCGGCTATACGTTGTC | 114 |
| chr10:109266096-10926813 | p2 | (TA)17 | CAAGCCTCAAGGGCTAACAG | GCCTCATAGGCAAACCTTCA | 231 |
| chr10:109279487-10928150 | p2 | (TA)10 | GCAACAAGTATCCTGCCCAT | GGGTCAATTTTTGTGCACCT | 185 |
| chr10:109279487-10928150 | p2 | (TA)9 | GCAACAAGTATCCTGCCCAT | GGGTCAATTTTTGTGCACCT | 185 |
| chr10:109279487-10928150 | p2 | (AT)6 | GCAACAAGTATCCTGCCCAT | GGGTCAATTTTTGTGCACCT | 185 |
| chr10:109279487-10928150 | p2 | (TA)13 | GCAACAAGTATCCTGCCCAT | GGGTCAATTTTTGTGCACCT | 185 |
| chr10:109279487-10928150 | p5 | (ATATT)3 | GCAACAAGTATCCTGCCCAT | GGGTCAATTTTTGTGCACCT | 185 |
| chr10:109279535-10928154 | p2 | (TA)10 | GCAACAAGTATCCTGCCCAT | GGGTCAATTTTTGTGCACCT | 137 |
| chr10:109279535-10928154 | p2 | (TA)9 | GCAACAAGTATCCTGCCCAT | GGGTCAATTTTTGTGCACCT | 137 |
| chr10:109279535-10928154 | p2 | (AT)6 | GCAACAAGTATCCTGCCCAT | GGGTCAATTTTTGTGCACCT | 137 |
| chr10:109279535-10928154 | p2 | (TA)13 | GCAACAAGTATCCTGCCCAT | GGGTCAATTTTTGTGCACCT | 137 |
| chr10:109308956-10931100 | p2 | (TA)26 | GCCCTTTGGCTATCCACATA | CAAGTAGACAGGGAGGCGAG | 266 |
| chr10:109308956-10931100 | p6 | (TTATCA)3 | GCCCTTTGGCTATCCACATA | CAAGTAGACAGGGAGGCGAG | 266 |
| chr10:109325712-10932772 | p3 | (CTT)5 | AAGCTTTATCCGCCCAGATT | AAGCAAGCGCATCGAACTAT | 266 |
| chr10:109485355-10948737 | p5 | (CTCTC)3 | GGGGGCTTAGAGCCTATTTG | TCAACCGGAGAAAACTGCTT | 117 |
| chr10:109500160-10950217 | p2 | (AC)7 | GGGCTCACATCAATTCACCT | ACAAGCAGGCCAAGCTAAAA | 297 |
| chr10:109578645-10958066 | p3 | (TTG)5 | CCCCTCCCAATTACATTCAA | ATTGTCCGAGGCAAATGAAG | 298 |
| chr10:109582133-10958415 | p3 | (GTT)5 | TCACAATTTTCATGGCCAAA | AAACGGGTTCGGGTTAAATC | 269 |

| Marker ID | SSR type | SSR motif | FORWARD PRIMER (5'-3') | REVERSE PRIMER (5'-3') | size (bp) |
|---|---|---|---|---|---|
| chr10:109582133-10958415 | p2 | (AT)10 | TCACAATTTTCATGGCCAAA | AAACGGGTTCGGGTTAAATC | 269 |
| chr10:109582133-10958415 | p4 | (TTTC)4 | TCACAATTTTCATGGCCAAA | AAACGGGTTCGGGTTAAATC | 269 |
| chr10:109662375-10966439 | p2 | (TA)8 | GTTGTTGGTGAGCGAGTGAA | TGGGTTGCAAGGAGGTTATC | 184 |
| chr10:109662375-10966439 | p2 | (TC)6 | GTTGTTGGTGAGCGAGTGAA | TGGGTTGCAAGGAGGTTATC | 184 |
| chr10:109726767-10972881 | p2 | (TC)26 | TATTTGCCACCTTTCCAAGC | GGATTAGCCCTTCTTTTGGG | 268 |
| chr10:109767367-10976938 | p5 | (GTTCG)3 | ATTGTTCCAAGACTGACGGG | GATAACGGAGAAGGCGATGA | 142 |
| chr10:109951716-10995373 | p3 | (TAG)5 | CCAGGGAACGTGAAACAAAT | GAAAAGCATCCTCCCACAAA | 242 |
| chr10:109951716-10995373 | p2 | (TA)8 | CCAGGGAACGTGAAACAAAT | GAAAAGCATCCTCCCACAAA | 242 |
| chr10:109964134-10996614 | p3 | (TTG)5 | TAAGCTTGCCAAAATCACCC | CCCACTTTGCAAACTTCCAT | 112 |
| chr10:109978764-10998077 | p5 | (TTGGT)3 | AGCGTATTATGCGCACACAG | GTAATGTCGGGTTGGTTTGG | 101 |
| chr10:110138815-11014083 | p2 | (TA)8 | CTTATTGGGGCACTGGACAT | GAGCTTTGGAGAAGGCTGAA | 119 |
| chr10:110138815-11014083 | p2 | (TA)6 | CTTATTGGGGCACTGGACAT | GAGCTTTGGAGAAGGCTGAA | 119 |
| chr10:110265950-11026796 | p3 | (ATA)5 | AAAAGAGAGGGGCAGCTAGG | GGGTCCAATTCATTCCTCAA | 147 |
| chr10:110266205-11026822 | p3 | (ATA)5 | GGGGTCCGGTAAGTCAAAAT | TCGCATTTACTTCGCACATT | 178 |
| chr10:110274781-11027680 | p3 | (AAC)8 | AGCCCGAAAATCCGAGTTAT | TTTCGCAACATGTAATCCGA | 167 |
| chr10:110314637-11031665 | p2 | (TA)7 | TCTGAGGCCAACTCCTGTCT | CACAAGCAGCGCAATAATGT | 204 |
| chr10:110331672-11033369 | p2 | (AT)13 | CGTTGACGTTTTCCCAAAAT | ACAAATCCTGTTTCGCCTTG | 269 |
| chr10:110331672-11033369 | p2 | (AT)6 | CGTTGACGTTTTCCCAAAAT | ACAAATCCTGTTTCGCCTTG | 269 |
| chr10:110372150-11037416 | p2 | (TA)6 | TGCTTCCCCTTCTTACCAAA | TTTAGGACGTACAATCGGGC | 246 |
| chr10:110395931-11039794 | p3 | (TAA)23 | GCAGTGGAGGGTAACGTGAT | CCTCATTTCCTCGTCGTCTC | 108 |
| chr10:110395931-11039794 | p5 | (ATCAT)3 | GCAGTGGAGGGTAACGTGAT | CCTCATTTCCTCGTCGTCTC | 108 |
| chr10:110437006-11043902 | p6 | (TGGGGC)3 | CACATCAGGGACACGTCAAC | TGATGGATATGGTGGTGGTG | 238 |
| chr10:110437047-11043906 | p6 | (TGGGGC)3 | CACATCAGGGACACGTCAAC | TGATGGATATGGTGGTGGTG | 197 |
| chr10:110452396-11045440 | p2 | (TA)6 | TGACGGAAAATCCCTCATTC | GAACTGTCACGTGCGAAAAA | 164 |
| chr10:110466546-11046856 | p5 | (TATTT)4 | TGTTCGTACTCCCAAGTCCC | AAAATAATTGAACGCACGCC | 104 |
| chr10:110466546-11046856 | p2 | (TA)14 | TGTTCGTACTCCCAAGTCCC | AAAATAATTGAACGCACGCC | 104 |
| chr10:110467061-11046908 | p5 | (TATTT)4 | GCACTAACTGGCGACTTTCC | AGAGGTCAGCGAAACGAAAA | 234 |
| chr10:110467061-11046908 | p2 | (TA)14 | GCACTAACTGGCGACTTTCC | AGAGGTCAGCGAAACGAAAA | 234 |
| chr10:110528007-11053001 | p2 | (TA)6 | TCGTTTGTGTAGTTTGCCCA | CCCAAAAGAAAAGGCTAGGG | 132 |
| chr10:110528007-11053001 | p6 | (GGAACC)3 | TCGTTTGTGTAGTTTGCCCA | CCCAAAAGAAAAGGCTAGGG | 132 |
| chr10:110532077-11053409 | p3 | (CTT)5 | TGCTCCACGCCCTACTACTT | GACGTCCACCTCATCCAGTT | 128 |
| chr10:110532077-11053409 | p2 | (AT)7 | TGCTCCACGCCCTACTACTT | GACGTCCACCTCATCCAGTT | 128 |
| chr10:110532100-11053411 | p3 | (CTT)5 | TGCTCCACGCCCTACTACTT | GACGTCCACCTCATCCAGTT | 105 |
| chr10:110532100-11053411 | p2 | (AT)7 | TGCTCCACGCCCTACTACTT | GACGTCCACCTCATCCAGTT | 105 |
| chr10:110705671-11070768 | p2 | (TC)9 | GTGCAATCCAGCCCAATACT | CATCAGGTAAGGTCTGGGGA | 170 |
| chr10:110705671-11070768 | p2 | (TC)7 | GTGCAATCCAGCCCAATACT | CATCAGGTAAGGTCTGGGGA | 170 |

| Marker ID | SSR type | SSR motif | FORWARD PRIMER (5'-3') | REVERSE PRIMER (5'-3') | size (bp) |
|---|---|---|---|---|---|
| chr10:110720032-11072204 | p5 | (CTCTC)3 | GCTGGGCAAACTAACCACAT | ATTTTCGATGTCCCAGATGC | 255 |
| chr10:110752546-11075456 | p2 | (AT)10 | CCATTAGTGGGAGCAAATGAA | AAATGACATGGCGGAATAGC | 272 |
| chr10:110874124-11087615 | p2 | (TA)7 | GGGCCCCATACTAAACAACA | TTTTCCCCATGAGTTTGGAG | 250 |
| chr10:110875138-11087715 | p2 | (AT)7 | TCAATCCATTAGTGGGGAGC | TGACATTGCGGAATAGCAAA | 155 |
| chr10:111051279-11105329 | p3 | (CAC)5 | AAGTGCAAATGACCCCTGTC | GGCAGCAAATACACACGAAA | 242 |
| chr10:111089004-11109101 | p2 | (TA)7 | GCATAACAACGTGTTGGTGG | AATTAATTTGGGGAGTGGGG | 118 |
| chr10:111089004-11109101 | p3 | (AAG)7 | GCATAACAACGTGTTGGTGG | AATTAATTTGGGGAGTGGGG | 118 |
| chr10:111122507-11112452 | p3 | (AGA)7 | ACCAGCTTCTTTAGCTGCCA | TTTGCATGTGCGATAAGTCC | 282 |
| chr10:111160219-11116223 | p2 | (TA)9 | AGGCCATCACTGGAAAATTG | GATTTGTGGCACATACGACG | 108 |
| chr10:111163133-11116514 | p2 | (GT)6 | ATCGCAACCCATTCTCTCAC | GATATCTCAGCCCGCACAAT | 133 |
| chr10:111222684-11122471 | p2 | (TA)13 | TCAGGCATTGCCAGTTGTAA | GCCTTGTTTTTGGTGGATTG | 108 |
| chr10:111222684-11122471 | p2 | (AT)14 | TCAGGCATTGCCAGTTGTAA | GCCTTGTTTTTGGTGGATTG | 108 |
| chr10:111268467-11127048 | p3 | (TTG)5 | TTCCACCAACACATTCATGG | CAACCGAGTGTCGTGTGTCT | 256 |
| chr10:111355411-11135742 | p6 | (GGGTTT)3 | CCGGAGGTGTAGATTGGAGA | TTTGAGCCGAGGGTCTATTG | 209 |
| chr10:111366357-11136837 | p3 | (CAA)6 | AGTGGAAGATGAAGGGGCTT | TTCCAACCTCAGAAACCACC | 147 |
| chr10:111387026-11138904 | p6 | (GGGTTT)3 | CCGGAGGTGTAGATTGGAGA | TTTGAGCCGAGGGTCTATTG | 209 |
| chr10:111397961-11139998 | p3 | (CAA)7 | AGTGGAAGATGAAGGGGCTT | TTCCAACCTCAGAAACCACC | 158 |
| chr10:111603173-11160518 | p4 | (AAAG)4 | TGGTCCCAAAGCCACTTTAC | ATGGCTGCAGACACAGATTG | 239 |
| chr10:111666455-11166847 | p2 | (AT)8 | TGCGGAAATAGCATGAGTTG | GTATTTACAGGACGCCGCAT | 151 |
| chr10:111666455-11166847 | p4 | (GTAT)4 | TGCGGAAATAGCATGAGTTG | GTATTTACAGGACGCCGCAT | 151 |
| chr10:111693807-11169582 | p5 | (GGGAG)3 | CGCCCGAGTGGATAAATAGA | GCCAAAGCACAATACTGGGT | 282 |
| chr10:111773525-11177555 | p2 | (TA)8 | TGCTCCACGCCCTACTACTT | CGAAGCCACGGTTGATACTT | 197 |
| chr10:111773525-11177555 | p2 | (AT)25 | TGCTCCACGCCCTACTACTT | CGAAGCCACGGTTGATACTT | 197 |
| chr10:111773598-11177564 | p2 | (TA)8 | TGCTCCACGCCCTACTACTT | CGAAGCCACGGTTGATACTT | 124 |
| chr10:111773598-11177564 | p2 | (AT)25 | TGCTCCACGCCCTACTACTT | CGAAGCCACGGTTGATACTT | 124 |
| chr10:111773735-11177576 | p2 | (TA)8 | AAGTATCAACCGTGGCTTCG | ACCATTTCGTTAATTCCCCC | 223 |
| chr10:111773735-11177576 | p2 | (AT)25 | AAGTATCAACCGTGGCTTCG | ACCATTTCGTTAATTCCCCC | 223 |
| chr10:111845878-11184789 | p3 | (TAA)6 | GATCAAGTGGGGAACAGGAA | GGGGTTTCCCTTCACATTTT | 197 |
| chr10:111845878-11184789 | p2 | (CA)6 | GATCAAGTGGGGAACAGGAA | GGGGTTTCCCTTCACATTTT | 197 |
| chr10:111845898-11184792 | p3 | (TAA)6 | GATCAAGTGGGGAACAGGAA | GGGGTTTCCCTTCACATTTT | 177 |
| chr10:111845898-11184792 | p2 | (CA)6 | GATCAAGTGGGGAACAGGAA | GGGGTTTCCCTTCACATTTT | 177 |
| chr10:111848446-11185045 | p2 | (TG)6 | CCGCGAGAGGTAAACACAAT | CGATACACAGCCCAACAAGA | 169 |
| chr10:111848446-11185045 | p6 | (ATTTTT)3 | CCGCGAGAGGTAAACACAAT | CGATACACAGCCCAACAAGA | 169 |
| chr10:111862658-11186467 | p2 | (AT)9 | AACCAGCTCATTTGACACCC | AAAGCGCCAAGAAACATACC | 262 |
| chr10:111910485-11191250 | p6 | (CTCCCC)3 | TTATGGGGGATCTTCGACTG | ATTCTAGGGAGGGCCAAAAA | 242 |
| chr10:111910485-11191250 | p6 | (GTTTTT)3 | TTATGGGGGATCTTCGACTG | ATTCTAGGGAGGGCCAAAAA | 242 |

| Marker ID | SSR type | SSR motif | FORWARD PRIMER (5'-3') | REVERSE PRIMER (5'-3') | size (bp) |
|---|---|---|---|---|---|
| chr10:111919393-11192140 | p3 | (ACA)5 | ATACTCCTGACGTGTTGCCC | TGGGTGCATGAAACTGAAAA | 211 |
| chr10:112066441-11206845 | p3 | (CAC)5 | TCCTTCATCTTCCCTCCCTT | ATGAAATGGTGTCGGCTAGG | 145 |
| chr10:112066441-11206845 | p2 | (TA)8 | TCCTTCATCTTCCCTCCCTT | ATGAAATGGTGTCGGCTAGG | 145 |
| chr10:112078943-11208100 | p2 | (TA)7 | ATTTTGATGCGGCCTAAATG | AATATCACACGCTGCCACAA | 197 |
| chr10:112085480-11208752 | p2 | (TA)23 | ATGGCATGGTGTTGCTTGTA | ATAAGCAGCAGGGATTTGGA | 141 |
| chr10:112085480-11208752 | p3 | (TAT)12 | ATGGCATGGTGTTGCTTGTA | ATAAGCAGCAGGGATTTGGA | 141 |
| chr10:112100249-11210226 | p2 | (AT)7 | TTCCTCGTCATCTCACCCTT | CCCCATGGCGTAATTCTCTA | 219 |
| chr10:112100249-11210226 | p2 | (TA)7 | TTCCTCGTCATCTCACCCTT | CCCCATGGCGTAATTCTCTA | 219 |
| chr10:112100276-11210229 | p2 | (AT)7 | TTCCTCGTCATCTCACCCTT | CCCCATGGCGTAATTCTCTA | 192 |
| chr10:112100276-11210229 | p2 | (TA)7 | TTCCTCGTCATCTCACCCTT | CCCCATGGCGTAATTCTCTA | 192 |
| chr10:112141016-11214308 | p3 | (TAT)22 | TTCAGGACATGTTCAAAGCG | CCCCTTTTGACCTGGATTTT | 252 |
| chr10:112156999-11215901 | p3 | (TTG)5 | GTTGATTGTTTTGGCCGAGT | TCTTTAGGACCGCTCGAAGA | 183 |
| chr10:112157048-11215906 | p3 | (TTG)5 | GTTGATTGTTTTGGCCGAGT | TCTTTAGGACCGCTCGAAGA | 134 |
| chr10:112244999-11224701 | p5 | (ATATA)4 | TCTTCAAGATTTGCCCAACC | ATTGCACGCTCTTCCACTTT | 146 |
| chr10:112246204-11224821 | p5 | (AAAAT)3 | AGGTGAAAAGGAAGGCCAGT | CTGGCCACGAAATTTAAGGA | 144 |
| chr10:112274247-11227626 | p3 | (ATT)6 | TTATCAGTTGGAAGCCCTGG | CTTTCGGGACCAAGAAAACA | 120 |
| chr10:112321706-11232372 | p6 | (CTTTGC)3 | GAGAGGGTGGAGCAAATTGA | CGGTCAAACTTTGCTGTTGA | 117 |
| chr10:112321706-11232372 | p5 | (AATTA)3 | GAGAGGGTGGAGCAAATTGA | CGGTCAAACTTTGCTGTTGA | 117 |
| chr10:112419289-11242130 | p5 | (TTTTG)3 | ACACATGGCATCGATTTGAA | TGGGTCTATTAGGGGGAAAA | 223 |
| chr10:112446640-11244865 | p5 | (TTTTG)3 | ACACATGGCATCGATTTGAA | TGGGTCTATTAGGGGGAAAA | 223 |
| chr10:112472891-11247492 | p2 | (TA)8 | GGTTGGCTGAGTGGTTTCTC | ATTGGGCAACCTGCTTACTG | 115 |
| chr10:112472891-11247492 | p2 | (TA)17 | GGTTGGCTGAGTGGTTTCTC | ATTGGGCAACCTGCTTACTG | 115 |
| chr10:112509762-11251178 | p2 | (AT)14 | TGGTCACTGATATCCCAGCA | GGTTGGCCCCATACTAAACA | 265 |
| chr10:112509762-11251178 | p2 | (TA)16 | TGGTCACTGATATCCCAGCA | GGTTGGCCCCATACTAAACA | 265 |
| chr10:112509762-11251178 | p2 | (AT)10 | TGGTCACTGATATCCCAGCA | GGTTGGCCCCATACTAAACA | 265 |
| chr10:112559409-11256144 | p2 | (TA)8 | GGTTGGCTGAGTGGTTTCTC | ATTGGGCAACCTGCTTACTG | 115 |
| chr10:112559409-11256144 | p2 | (TA)17 | GGTTGGCTGAGTGGTTTCTC | ATTGGGCAACCTGCTTACTG | 115 |
| chr10:112596280-11259830 | p2 | (AT)14 | TGGTCACTGATATCCCAGCA | GGTTGGCCCCATACTAAACA | 265 |
| chr10:112596280-11259830 | p2 | (TA)16 | TGGTCACTGATATCCCAGCA | GGTTGGCCCCATACTAAACA | 265 |
| chr10:112596280-11259830 | p2 | (AT)10 | TGGTCACTGATATCCCAGCA | GGTTGGCCCCATACTAAACA | 265 |
| chr10:112607698-11260977 | p2 | (TA)8 | TTACAATTTGGGCCTTCTGC | CAAACCGTGCAATCCTTCTT | 199 |
| chr10:112608029-11261004 | p2 | (TA)8 | TTGGTAATTGGCAAGAAGGC | TCAAATTTTGGCTCCGACTC | 185 |
| chr10:112608108-11261012 | p2 | (TA)8 | TTGGTAATTGGCAAGAAGGC | TCAAATTTTGGCTCCGACTC | 106 |
| chr10:112622490-11262454 | p2 | (AG)9 | GCGGGATCTGAAAACAAAAA | ACGCATGTACCTCCACATCA | 137 |
| chr10:112622490-11262454 | p3 | (TAA)19 | GCGGGATCTGAAAACAAAAA | ACGCATGTACCTCCACATCA | 137 |
| chr10:112640356-11264237 | p3 | (GTT)6 | CCCCTGAGAATAGTGGTGGA | AACACATTGTTTGCATCGGA | 132 |

| Marker ID | SSR type | SSR motif | FORWARD PRIMER (5'-3') | REVERSE PRIMER (5'-3') | size (bp) |
|---|---|---|---|---|---|
| chr10:112659536-11266155 | p4 | (AATT)4 | TTGCAAGCCTGTCATACGTC | ACATTGGTCCTCCAAGCATC | 188 |
| chr10:112695149-11269718 | p2 | (AT)18 | TGAACAAAATTGGGCATCAA | AAAACGAAAAGGGGCCTAAA | 247 |
| chr10:112695149-11269718 | p2 | (AT)10 | TGAACAAAATTGGGCATCAA | AAAACGAAAAGGGGCCTAAA | 247 |
| chr10:112718101-11272011 | p5 | (CCCCA)3 | AGAACCTTTTGGTGCCCTT | TTATCTTTGCGGGGTCATTC | 119 |
| chr10:112729361-11273138 | p3 | (TTC)7 | CCGAACTCAAACGAACACCT | GAATGAGCACTGGATTGGGT | 154 |
| chr10:112733034-11273504 | p5 | (TTGTA)3 | CCGCAACATTGCATAATCAC | CCATTACAGCGAAGTGGGAT | 172 |
| chr10:112733034-11273504 | p5 | (TTGTA)3 | CCGCAACATTGCATAATCAC | CCATTACAGCGAAGTGGGAT | 172 |
| chr10:112733034-11273504 | p2 | (AG)7 | CCGCAACATTGCATAATCAC | CCATTACAGCGAAGTGGGAT | 172 |
| chr10:112740979-11274299 | p5 | (AAATT)3 | TCAAGGTTGAAACTTTGGGC | CAAAAGGAAGGGCAAAAGAA | 123 |
| chr10:112802384-11280440 | p2 | (AT)8 | GGCACACGAATTTGTCATTG | AAGTGATCCATGCTTGACCC | 219 |
| chr10:112818379-11282040 | p5 | (AAAAC)3 | CAAGATAGCCAAAGGGGTGA | GGCACGGAATGTCACTTTTT | 258 |
| chr10:112818379-11282040 | p2 | (AG)9 | CAAGATAGCCAAAGGGGTGA | GGCACGGAATGTCACTTTTT | 258 |
| chr10:112818444-11282046 | p5 | (AAAAC)3 | CAAGATAGCCAAAGGGGTGA | GGCACGGAATGTCACTTTTT | 193 |
| chr10:112818444-11282046 | p2 | (AG)9 | CAAGATAGCCAAAGGGGTGA | GGCACGGAATGTCACTTTTT | 193 |
| chr10:112821617-11282363 | p3 | (TTG)6 | CCTGGAAAGACAAGACCCAA | CTTGCATCAACGCTCTCAAA | 211 |
| chr10:112840552-11284256 | p2 | (AT)7 | TGAAATGGGCAGTCAAGATG | CAAAATTTGCGGTGCTACCT | 195 |
| chr10:112840552-11284256 | p2 | (TA)19 | TGAAATGGGCAGTCAAGATG | CAAAATTTGCGGTGCTACCT | 195 |
| chr10:112862561-11286457 | p2 | (AT)6 | ATTGCCATTGACACCCATTT | ACAGCCCAGTTCGTGAAATC | 145 |
| chr10:112862577-11286458 | p2 | (AT)6 | ATTGCCATTGACACCCATTT | ACAGCCCAGTTCGTGAAATC | 129 |
| chr10:112862597-11286461 | p2 | (AT)6 | ATTGCCATTGACACCCATTT | ACAGCCCAGTTCGTGAAATC | 109 |
| chr10:112862982-11286499 | p2 | (AT)6 | GGCCAGGAGGGTAATAGGTC | GGTATCGTTTCCCAACCCTT | 164 |
| chr10:112880006-11288203 | p3 | (TAT)8 | AAATTGCCATTGACACCCAT | CAAATGGTAAGAGGAGCGGA | 108 |
| chr10:112880006-11288203 | p2 | (TC)16 | AAATTGCCATTGACACCCAT | CAAATGGTAAGAGGAGCGGA | 108 |
| chr10:112961197-11296321 | p6 | (GGTTTT)3 | CAAGAGGTACGCACTCCCAT | TGAGGCTTGTCACCAAGATG | 192 |
| chr10:112961197-11296321 | p5 | (GTTTG)3 | CAAGAGGTACGCACTCCCAT | TGAGGCTTGTCACCAAGATG | 192 |
| chr10:112973448-11297546 | p2 | (AT)7 | CAGGGAGGGTAATGGTCTCA | TAGTTCCCCCAACGTGTAGC | 233 |
| chr10:113069872-11307188 | p2 | (AT)8 | ATAGCGTGCCGAAATTGTTC | GGGGCGATAGTGAAAGTCAA | 150 |
| chr10:113082113-11308412 | p2 | (TA)7 | AATTAGGGGTCAAGGGTTCG | TGGTTTTCTATCCAGTGCCC | 133 |
| chr10:113085553-11308756 | p3 | (CAA)5 | CACCATCCTCTCCCAAGAAA | GATTACGGCCAAGCAAGGTA | 297 |
| chr10:113085553-11308756 | p5 | (CCACC)3 | CACCATCCTCTCCCAAGAAA | GATTACGGCCAAGCAAGGTA | 297 |
| chr10:113085553-11308756 | p6 | (ATAAAA)3 | CACCATCCTCTCCCAAGAAA | GATTACGGCCAAGCAAGGTA | 297 |
| chr10:113198604-11320061 | p2 | (TA)6 | ACCAAAAACCAGGTGGAGAA | TATCGAAAATCTGTGCGCTG | 274 |
| chr10:113294675-11329669 | p2 | (GA)10 | CTGGGGAGCTTTTCATACCA | TTCGGAGGATTGTCCACTTC | 293 |
| chr10:113379902-11338196 | p2 | (TA)32 | GACCATTGTCATGCTGGTTG | CACGCTAGTGTATGCGGTGT | 283 |
| chr10:113414637-11341667 | p2 | (AT)9 | TGGGGGAAAATTGAAATTGA | GTCCGTTTTACGCGTTTGTT | 159 |
| chr10:113414637-11341667 | p2 | (AT)7 | TGGGGGAAAATTGAAATTGA | GTCCGTTTTACGCGTTTGTT | 159 |

| Marker ID | SSR type | SSR motif | FORWARD PRIMER (5'-3') | REVERSE PRIMER (5'-3') | size (bp) |
|---|---|---|---|---|---|
| chr10:113414677-11341669 | p2 | (AT)9 | TGGGGGAAAATTGAAATTGA | GTCCGTTTTACGCGTTTGTT | 119 |
| chr10:113414677-11341669 | p2 | (AT)7 | TGGGGGAAAATTGAAATTGA | GTCCGTTTTACGCGTTTGTT | 119 |
| chr10:113456583-11345861 | p2 | (AT)14 | AACAAAATATTCCCCGAGGC | TTGCCCAAATTTCTGTAGGC | 100 |
| chr10:113536963-11353897 | p2 | (AT)11 | GGGCCCATCCCTTTATTTTA | GCAACAAGTGGCATCAAGTG | 102 |
| chr10:113536963-11353897 | p3 | (AGT)5 | GGGCCCATCCCTTTATTTTA | GCAACAAGTGGCATCAAGTG | 102 |
| chr10:113579471-11358148 | p3 | (ATT)5 | GGCCGTAAGCAAACACTAGC | ACTAAGTGCAAAACCGCTCG | 285 |
| chr10:113579471-11358148 | p2 | (AC)6 | GGCCGTAAGCAAACACTAGC | ACTAAGTGCAAAACCGCTCG | 285 |
| chr10:113579471-11358148 | p2 | (AT)17 | GGCCGTAAGCAAACACTAGC | ACTAAGTGCAAAACCGCTCG | 285 |
| chr10:113579623-11358168 | p2 | (AC)6 | GGCCGTAAGCAAACACTAGC | ACTAAGTGCAAAACCGCTCG | 133 |
| chr10:113748338-11375040 | p3 | (CTT)22 | ACAACAAGGCGTGAATTTCC | CACAAGGGCAAAGTTGGAAT | 268 |
| chr10:113764473-11376648 | p2 | (AC)7 | CCCAATCGAAGTGACGTTTT | CAAATCAAGGCTTGGGACAT | 137 |
| chr10:113958060-11396007 | p2 | (TC)7 | AGGTATCGACCCGCTTCTTT | TAGCCAATTTTGTTCCGACC | 118 |
| chr10:113975390-11397740 | p5 | (AACCG)3 | CATCTGCATCCGTATCATGC | GAATTTCGATAACGCACGGT | 199 |
| chr10:114040943-11404296 | p2 | (AG)9 | CATGGTGTACGGACCTGTTG | AGGAATGTGCACCATGTCAA | 159 |
| chr10:114046000-11404801 | p2 | (CT)7 | GGTGATGGAACAATTCCTGG | TCCGAAGGGATCTCAATGTC | 212 |
| chr10:114198864-11420087 | p2 | (AT)6 | AACCGCGAGTTCAAGAAGAA | CTTCGCTCTCTCGTGGAGTT | 113 |
| chr10:114250250-11425226 | p2 | (AG)9 | ATTAGTGTCAGCCAGGGACG | TTTCTGGGAATTGAACCTGC | 220 |
| chr10:114250250-11425226 | p2 | (TA)7 | ATTAGTGTCAGCCAGGGACG | TTTCTGGGAATTGAACCTGC | 220 |
| chr10:114289269-11429128 | p5 | (TCGGT)3 | GGGAAAAACACTCCCTCACA | CCATCAAAAGGTCCATGCTT | 254 |
| chr10:114325720-11432773 | p5 | (TCGGT)3 | GGGAAAAACACTCCCTCACA | CCATCAAAAGGTCCATGCTT | 254 |
| chr10:114384312-11438632 | p2 | (TA)7 | TATATGTCCGTCCCCGGTTA | AACAAAACCACGGTACTCGG | 125 |
| chr10:114419782-11442184 | p3 | (ATT)21 | AGATACTTGTGGACGGCTGG | AAGGCACCAATTGACGTTTC | 121 |
| chr10:114436676-11443869 | p6 | (AAACAG)3 | ACCAACGGAAGCTGATTCAC | CAGCACCCCAAACCTCTTTA | 169 |
| chr10:114436676-11443869 | p2 | (GT)6 | ACCAACGGAAGCTGATTCAC | CAGCACCCCAAACCTCTTTA | 169 |
| chr10:114487172-11448923 | p5 | (TTTTA)3 | GCATCCAAATTCCAAAGTCG | CGATGATCCACGTTGATAGG | 103 |
| chr10:114606563-11460857 | p2 | (AG)7 | GAAAGACCCTCAGCTCAACG | TAAGACTCTGCGCACCAAAA | 111 |
| chr10:114678619-11468063 | p5 | (AATAA)4 | CGTGGACAAGCAAAAGTGAA | TTTTTAAATGCCCAACGGAC | 294 |
| chr10:114678619-11468063 | p2 | (TA)14 | CGTGGACAAGCAAAAGTGAA | TTTTTAAATGCCCAACGGAC | 294 |
| chr10:114708394-11471041 | p6 | (TTTTTC)3 | TCCTAGGCAATTGAAGCACA | TGGCAGCTCCTTATACTGCC | 160 |
| chr10:114756519-11475854 | p6 | (AAGTGA)3 | TGAGACACGTGGCTGAAGTC | ACTTACCCTCGGCTCCAGAT | 206 |
| chr10:114756519-11475854 | p2 | (AT)12 | TGAGACACGTGGCTGAAGTC | ACTTACCCTCGGCTCCAGAT | 206 |
| chr10:114756519-11475854 | p2 | (AT)14 | TGAGACACGTGGCTGAAGTC | ACTTACCCTCGGCTCCAGAT | 206 |
| chr10:114894818-11489683 | p2 | (TA)6 | GAAACCCATCAGCACCTGTT | TGTTCCAGTCCCCTATTTGC | 166 |
| chr10:114928124-11493014 | p3 | (AAC)5 | GCTCCGTTTCCACGTTTTTA | CCTGCTAAGATCCCCGTGTA | 106 |
| chr10:114928124-11493014 | p2 | (AT)6 | GCTCCGTTTCCACGTTTTTA | CCTGCTAAGATCCCCGTGTA | 106 |
| chr10:115025071-11502708 | p6 | (ATAAAA)3 | ATACAAGCAGTGGCGGATTC | TTCAAGTGTTCCAGGGTGTTC | 107 |

| Marker ID | SSR type | SSR motif | FORWARD PRIMER (5'-3') | REVERSE PRIMER (5'-3') | size (bp) |
|---|---|---|---|---|---|
| chr10:115046142-11504815 | p2 | (TC)8 | TTAAGAAAGGATGGCCCCTT | AAAGGGATTGGGAGAGCCTA | 145 |
| chr10:115063151-11506516 | p2 | (AT)44 | ATGCTCTTTACGCCTTTCCA | TGTATAGGGGAAATGCTGCC | 109 |
| chr10:115080256-11508229 | p2 | (AT)18 | TATGTGATGGCAGCCTTGTC | GCCCGACCTAGACACCTGTA | 253 |
| chr10:115094513-11509652 | p2 | (TA)6 | CTTGGAAACGATCCCAGAAA | GGGCTCAGTGTGGAAAACAT | 232 |
| chr10:115105115-11510713 | p2 | (TA)8 | CAATTGGGCTCCAAAAAGAA | GATTTGGAGGGGGTGATTTT | 103 |
| chr10:115130271-11513229 | p3 | (TGT)5 | CCAAGGGTCTATCGGAAACA | ATGCTCAAGGGATGGTCTTG | 251 |
| chr10:115130271-11513229 | p2 | (TA)13 | CCAAGGGTCTATCGGAAACA | ATGCTCAAGGGATGGTCTTG | 251 |
| chr10:115178899-11518093 | p5 | (AAAAT)3 | CCCATTAGTGGGAGCAAATG | AAATGACATGGCGGAATAGC | 176 |
| chr10:115178899-11518093 | p2 | (TA)18 | CCCATTAGTGGGAGCAAATG | AAATGACATGGCGGAATAGC | 176 |
| chr10:115210291-11521232 | p3 | (ATA)11 | CGGTTGTTCTTCAAAGGCAT | TTTTTGTTGATTAAGCGGGG | 278 |
| chr10:115310350-11531236 | p2 | (AT)7 | CCAGCTATGCCAGGTCTCTC | AGCTGAATCAAGCCAAGGAA | 208 |
| chr10:115411206-11541321 | p2 | (TG)6 | TACCCGGGCACAATAGGATA | AACTTGTCGAGTGAGCCGAT | 285 |
| chr10:115544091-11554610 | p2 | (AT)8 | CGCATCTTGTCTTCCACTGA | AAGTGGCGGAAATGAGAATG | 254 |
| chr10:115572916-11557493 | p4 | (AAAG)4 | CTGCTTAACTGATGCCCCTC | CAGCTAAAACAGGGACTGCC | 292 |
| chr10:115592551-11559458 | p2 | (CT)8 | AAGCACACTCCACCTCCAAC | CTGCACACTACCCCAACCTT | 269 |
| chr10:115599601-11560161 | p4 | (ACAA)4 | TGCATCCAAGAGGTGTCAAC | CCCCTCAGCTCATGGTATGT | 157 |
| chr10:115603869-11560590 | p2 | (AT)15 | CCTGCATAATGCGAACTGAA | GGACAGTTGGAGATTTGCGT | 276 |
| chr10:115603869-11560590 | p2 | (AT)20 | CCTGCATAATGCGAACTGAA | GGACAGTTGGAGATTTGCGT | 276 |
| chr10:115603869-11560590 | p3 | (TGT)5 | CCTGCATAATGCGAACTGAA | GGACAGTTGGAGATTTGCGT | 276 |
| chr10:115647373-11564939 | p6 | (CCCCCA)3 | GGTTCGAGGTCCAGTGTTGT | ACTAGCCACCCAAAATGCTG | 221 |
| chr10:115708652-11571066 | p2 | (TA)7 | GGTCTCGGAGGCATGAATAA | CCTATGCCTCCAATTCCTCA | 214 |
| chr10:115754351-11575637 | p6 | (AGCCCC)4 | ACGCCCCAAAAACTCACTTA | GGCTTTTCGTCGTTCCATAA | 265 |
| chr10:115814181-11581619 | p2 | (TA)6 | CGTTAGTCCCGAGATTTCCA | TCTCCGGATTGAATACGACC | 123 |
| chr10:115814181-11581619 | p4 | (TTTA)4 | CGTTAGTCCCGAGATTTCCA | TCTCCGGATTGAATACGACC | 123 |
| chr10:115824181-11582622 | p2 | (AT)8 | TAATTATAGCCTCCACGGCG | TGGATTTTCTTATGGCCTCG | 276 |
| chr10:115824181-11582622 | p3 | (TTA)5 | TAATTATAGCCTCCACGGCG | TGGATTTTCTTATGGCCTCG | 276 |
| chr10:115824181-11582622 | p3 | (TTA)6 | TAATTATAGCCTCCACGGCG | TGGATTTTCTTATGGCCTCG | 276 |
| chr10:115824332-11582634 | p2 | (AT)8 | TAATTATAGCCTCCACGGCG | TGGATTTTCTTATGGCCTCG | 125 |
| chr10:115824332-11582634 | p3 | (TTA)5 | TAATTATAGCCTCCACGGCG | TGGATTTTCTTATGGCCTCG | 125 |
| chr10:115824332-11582634 | p3 | (TTA)6 | TAATTATAGCCTCCACGGCG | TGGATTTTCTTATGGCCTCG | 125 |
| chr10:115926226-11592824 | p6 | (TATTTT)3 | ATCGGGTCAAGTGCACAAAT | TTCTTGCAACTTCAATCCCC | 281 |
| chr10:115941166-11594317 | p2 | (TA)6 | TACCTCCAAGTTGACAGGGC | ATTTGGGAAACGAGAACGTG | 206 |
| chr10:115951877-11595389 | p5 | (TTTTA)3 | AGCAACGGAAAGAATTGCTC | CTATGCGCTCAGACGAATCA | 117 |
| chr10:115951877-11595389 | p6 | (ATTCTA)3 | AGCAACGGAAAGAATTGCTC | CTATGCGCTCAGACGAATCA | 117 |
| chr10:115967578-11596959 | p5 | (ATAAA)3 | CTAGCACAGCCCACAACAAA | AGGCAAAGCTTTTCAAGCAA | 130 |
| chr10:116001068-11600308 | p4 | (TAGT)4 | AATTCAAAGGTGCAGGAGGA | TCTGCATCACAAAAGGCTTG | 161 |

| Marker ID | SSR type | SSR motif | FORWARD PRIMER (5'-3') | REVERSE PRIMER (5'-3') | size (bp) |
|---|---|---|---|---|---|
| chr10:116001068-11600308 | p2 | (TA)9 | AATTCAAAGGTGCAGGAGGA | TCTGCATCACAAAAGGCTTG | 161 |
| chr10:116001493-11600351 | p4 | (TAGT)4 | AATCCGGTATTCCATGAACG | GATAAACATGTTGGGGTGCC | 249 |
| chr10:116001493-11600351 | p2 | (TA)9 | AATCCGGTATTCCATGAACG | GATAAACATGTTGGGGTGCC | 249 |
| chr10:116003936-11600595 | p2 | (TA)7 | GGGACAAACTTGATCGGAGA | ACAAATTGGGACGAGCAGAG | 197 |
| chr10:116003936-11600595 | p5 | (TTGGT)3 | GGGACAAACTTGATCGGAGA | ACAAATTGGGACGAGCAGAG | 197 |
| chr10:116025910-11602793 | p2 | (AT)12 | CAAACCGCCAAAAACAACTT | CCGCAGAAGTGAAATTGACA | 217 |
| chr10:116037543-11603955 | p3 | (CTT)5 | TAGAATTTTTGGGTGCTCGG | TCCAGGAATTTTGAGTTCGG | 140 |
| chr10:116037543-11603955 | p2 | (AG)20 | TAGAATTTTTGGGTGCTCGG | TCCAGGAATTTTGAGTTCGG | 140 |
| chr10:116039714-11604172 | p2 | (GT)19 | AGGAAGGCCAAGGTCAAAAT | ATGGCCTTCTTTTGTGATGC | 244 |
| chr10:116039714-11604172 | p3 | (TTG)5 | AGGAAGGCCAAGGTCAAAAT | ATGGCCTTCTTTTGTGATGC | 244 |
| chr10:116110561-11611257 | p6 | (GGGGAG)3 | ACCGGAAGACATGGAAACAC | ATATCTCGAGGTGCAGTGGG | 184 |
| chr10:116119281-11612129 | p5 | (ATAAA)3 | CATCAGCGCACTAAGCACAT | CAGTCGAAACGGGAGGTTTA | 137 |
| chr10:116136322-11613834 | p6 | (TTTTTC)3 | CTTCCATTTGCCATGAGCTT | CCATCGCTAAGGATAAAGGC | 172 |
| chr10:116271078-11627309 | p5 | (ATTAT)3 | TTGCTCGGTCAGAGAGTGAA | GCACTGACAAGTCCGTGAGA | 279 |
| chr10:116424178-11642619 | p5 | (TATTT)3 | CAATAGAGGGCTGGGATCTG | GTAAGGGTCGCGGTAAACAA | 104 |
| chr10:116493045-11649506 | p5 | (CGACA)3 | TCCAACTTCTGCAGCATCAC | TGATGCCAAACCAACACATT | 102 |
| chr10:116619246-11662126 | p2 | (AT)9 | GGGGACAGCAAGTCAAATGT | CCGCTTCTTCATTTCACCAT | 113 |
| chr10:116630432-11663248 | p2 | (AT)24 | GTATGCTGGAATGGGCTGTT | TTTTAGGCGACGTTTGCTCT | 148 |
| chr10:116639928-11664194 | p2 | (AT)6 | CTTCATCCACCCATTTTTGG | ATTGGCCCGAGAAAGGTACT | 252 |
| chr10:116645057-11664707 | p3 | (GTT)5 | GCTTGAGCAATCCCACAAAT | GTCTGAACACCTGATGCGAA | 141 |
| chr10:116804144-11680615 | p2 | (TG)8 | TGGCTATAGGATACCCACCG | TGAGCTGAAAACCGAAACCT | 219 |
| chr10:116804144-11680615 | p2 | (TG)6 | TGGCTATAGGATACCCACCG | TGAGCTGAAAACCGAAACCT | 219 |
| chr10:116804144-11680615 | p5 | (TTTTG)3 | TGGCTATAGGATACCCACCG | TGAGCTGAAAACCGAAACCT | 219 |
| chr10:116825942-11682795 | p3 | (ATT)5 | AATTTGGCATCTTTGTTGGC | GTCCCGTAGTAGCGTTGGAA | 260 |
| chr10:116825942-11682795 | p2 | (TC)7 | AATTTGGCATCTTTGTTGGC | GTCCCGTAGTAGCGTTGGAA | 260 |
| chr10:116885127-11688717 | p5 | (AACCC)3 | GAGTTCGAACCTGAAACCCA | ATTAGGACACGTGGCAGACC | 179 |
| chr10:116908096-11691011 | p2 | (TA)7 | ACTGTTCACGCCGGAAATAC | TCTCCTTCTAGCTGGTCCGA | 250 |
| chr10:116908096-11691011 | p2 | (TA)6 | ACTGTTCACGCCGGAAATAC | TCTCCTTCTAGCTGGTCCGA | 250 |
| chr10:116908096-11691011 | p2 | (TA)12 | ACTGTTCACGCCGGAAATAC | TCTCCTTCTAGCTGGTCCGA | 250 |
| chr10:117104027-11710604 | p6 | (ATTTTT)3 | CGACTCCCACATAATGGCTT | CCTCAGCCATAGTAGCTGCC | 175 |
| chr10:117116586-11711859 | p2 | (CA)6 | GACGGGATAAAAACGGTGTG | TGTGTCCACTTTGGACCGTA | 178 |
| chr10:117135528-11713754 | p6 | (TCGGCT)3 | AGGTGGTTGAAAGATCACCG | TCATCACGAGAATGGACCAA | 137 |
| chr10:117143174-11714519 | p2 | (AT)12 | TGACCACACGCTCCTCTATG | CAAAAAGGCTAAGCGGACAC | 145 |
| chr10:117164954-11716696 | p3 | (CCA)6 | GGCCAGCTTAAGCACATCTC | GTAGTGGTGCTTTGTGGGGT | 259 |
| chr10:117164954-11716696 | p2 | (AT)7 | GGCCAGCTTAAGCACATCTC | GTAGTGGTGCTTTGTGGGGT | 259 |
| chr10:117201250-11720326 | p3 | (AAC)7 | GGGCGTATCTAGCACGGTAA | CCCACAAAGGTAGAGGCAAA | 170 |

| Marker ID | SSR type | SSR motif | FORWARD PRIMER (5'-3') | REVERSE PRIMER (5'-3') | size (bp) |
|---|---|---|---|---|---|
| chr10:117201250-11720326 | p3 | (TGT)5 | GGGCGTATCTAGCACGGTAA | CCCACAAAGGTAGAGGCAAA | 170 |
| chr10:117239835-11724184 | p2 | (AT)7 | AAGAATGTGGGCAGCCATAC | AACGTTTGGGGCATCTTATG | 139 |
| chr10:117263211-11726522 | p5 | (AAAAT)3 | GTCAAAGAGCTTCGGATTGC | AGTGCGATCTGGAGAGGGTA | 283 |
| chr10:117312576-11731458 | p2 | (TC)6 | TCTACCAACGTCTGGCCTCT | CAACTTGTCCCTCTTCGACC | 265 |
| chr10:117312576-11731458 | p5 | (TTTTG)3 | TCTACCAACGTCTGGCCTCT | CAACTTGTCCCTCTTCGACC | 265 |
| chr10:117405313-11740732 | p5 | (ACCAA)3 | TGGACCAGGTTGGTTCAGAT | CCTGAGGTTGAAAAACCCAA | 235 |
| chr10:117405313-11740732 | p5 | (AAACC)3 | TGGACCAGGTTGGTTCAGAT | CCTGAGGTTGAAAAACCCAA | 235 |
| chr10:117437197-11743921 | p4 | (TTGT)4 | AGCCCTTTGCTCAAGACTGA | TGTCACCGGTTTTGTTCTCA | 278 |
| chr10:117439907-11744194 | p2 | (AT)11 | AACCCAACCGATGTCCATTA | TAGTGCAATTCCCCTCCATC | 178 |
| chr10:117439907-11744194 | p3 | (GAA)6 | AACCCAACCGATGTCCATTA | TAGTGCAATTCCCCTCCATC | 178 |
| chr10:117439907-11744194 | p2 | (AT)9 | AACCCAACCGATGTCCATTA | TAGTGCAATTCCCCTCCATC | 178 |
| chr10:117439907-11744194 | p2 | (TA)20 | AACCCAACCGATGTCCATTA | TAGTGCAATTCCCCTCCATC | 178 |
| chr10:117465988-11746800 | p3 | (GTG)5 | TGATGACGGTGAAAGGATGA | GCCCCATAATGACCAAACAC | 120 |
| chr10:117503878-11750589 | p3 | (GTT)6 | CACGATGTGGGATTTCACTG | CTCGCTTCCCTCATCTTGTC | 246 |
| chr10:117503878-11750589 | p3 | (TGT)5 | CACGATGTGGGATTTCACTG | CTCGCTTCCCTCATCTTGTC | 246 |
| chr10:117558368-11756038 | p5 | (AAACC)3 | TATTTCGCTTCTCTGCGACA | TTTGGGAAACCTGAACCAAC | 147 |
| chr10:117558368-11756038 | p5 | (ATAAA)3 | TATTTCGCTTCTCTGCGACA | TTTGGGAAACCTGAACCAAC | 147 |
| chr10:117570292-11757231 | p6 | (CCAACC)3 | ATTGAATGCCCGTGAGAAAC | GAGCTCTTGTTGGGTCAAGG | 253 |
| chr10:117608170-11761018 | p5 | (AAAAC)3 | ACGTACGCTCAAAGTTGGCT | TGTGGTACGTTGCACGAGTT | 156 |
| chr10:117626109-11762812 | p5 | (TTTTC)3 | GATGAAGGAAGCAGCTCTGG | CTGCAGTATCCCAGGCAAAT | 259 |
| chr10:117658207-11766022 | p4 | (AAAG)4 | CCCGTTCCAATGTGAGTTTT | ACCCATGACAAGAGCCAAAG | 121 |
| chr10:117691407-11769342 | p2 | (AT)7 | CTCCCAAAGTATCCCCATCA | AAATGACATGGCGGAATAGC | 159 |
| chr10:117822352-11782436 | p2 | (TA)8 | CGAAATATCCAACGAGCCAT | TTTCAGTTTTCCCCGATCAC | 113 |
| chr10:117825809-11782782 | p2 | (TA)7 | CTTTTTCGTCGTGTTGGACC | CGTGAGTTTTGGATGTGGTG | 121 |
| chr10:117839928-11784194 | p5 | (AATAA)3 | TCCCAAAAATTACCGACAGC | ATTTGCTTTTCGAGGTGGTG | 300 |
| chr10:117839928-11784194 | p2 | (AT)10 | TCCCAAAAATTACCGACAGC | ATTTGCTTTTCGAGGTGGTG | 300 |
| chr10:117839928-11784194 | p5 | (AATTT)3 | TCCCAAAAATTACCGACAGC | ATTTGCTTTTCGAGGTGGTG | 300 |
| chr10:117911728-11791374 | p6 | (TAAAAA)3 | ACCGAGGAGCTCTTTTCACA | TGCCCTCCACACTTCCTTAC | 283 |
| chr10:117911728-11791374 | p2 | (TC)9 | ACCGAGGAGCTCTTTTCACA | TGCCCTCCACACTTCCTTAC | 283 |
| chr10:117940990-11794300 | p2 | (AG)9 | CAGTTCCAACTCCAACAGCA | CACTGCGACGCTCGTATCTA | 276 |
| chr10:118016747-11801876 | p3 | (TGT)5 | AAACGTTTGGGTCCTTCCTT | GCATGCCTCATTGATCAGAA | 126 |
| chr10:118016747-11801876 | p3 | (GAA)6 | AAACGTTTGGGTCCTTCCTT | GCATGCCTCATTGATCAGAA | 126 |
| chr10:118117420-11811945 | p2 | (GT)6 | TTGCATGCACCTCGACTAAC | ACACCCACACCAGGTGATTT | 189 |
| chr10:118117420-11811945 | p2 | (TA)17 | TTGCATGCACCTCGACTAAC | ACACCCACACCAGGTGATTT | 189 |
| chr10:118202200-11820421 | p3 | (TTG)6 | TACACGTGAGCAAAAACCCA | ATGACCGAAACTATGCGAGG | 256 |
| chr10:118251031-11825305 | p2 | (GT)11 | TTGGTGTGGGACATTTAGCA | TCCAAAATTCCAATTCCTGC | 256 |

| Marker ID | SSR type | SSR motif | FORWARD PRIMER (5'-3') | REVERSE PRIMER (5'-3') | size (bp) |
|---|---|---|---|---|---|
| chr10:118251031-11825305 | p3 | (TTG)6 | TTGGTGTGGGACATTTAGCA | TCCAAAATTCCAATTCCTGC | 256 |
| chr10:118274171-11827619 | p2 | (GT)11 | CCCCCGCAATACTTGTAAGA | CCTTCCTTGATATTTCCGCA | 279 |
| chr10:118274171-11827619 | p3 | (TTG)5 | CCCCCGCAATACTTGTAAGA | CCTTCCTTGATATTTCCGCA | 279 |
| chr10:118279206-11828122 | p5 | (TTATT)3 | TATTTTTGGCCTTTGCGAAC | CCATCCCGGTATGATTTTTG | 176 |
| chr10:118282942-11828495 | p5 | (TTAGA)3 | TTATACTCCGCTTCAACCCG | AAAACACCCCAAACATCCAA | 289 |
| chr10:118311549-11831356 | p5 | (GATTT)3 | TACGAAGCTGCGAAATGATG | ATTGCGTACCATCACTGCAA | 138 |
| chr10:118356158-11835817 | p5 | (AAAGG)3 | AGGGACAGCAGCCAATAGAA | AGGTCTGGATCGAGCTTTCA | 192 |
| chr10:118391109-11839313 | p2 | (AT)7 | TCCTTATCCTGCCAACCAAA | GCGGGCTCTAAATTCAAATG | 131 |
| chr10:118464715-11846673 | p3 | (TAA)5 | TTCACCTGTGGGACAACGTA | AGGGTCCATCACATAGCACC | 158 |
| chr10:118468060-11847008 | p2 | (AT)6 | CTTGTCCGGCTTCAAACAAT | ATGCCTCTCTTACCGCTTGA | 154 |
| chr10:118468060-11847008 | p2 | (AG)11 | CTTGTCCGGCTTCAAACAAT | ATGCCTCTCTTACCGCTTGA | 154 |
| chr10:118472612-11847462 | p5 | (TTTTA)3 | CACACTGATCCCACAACCTG | CCTACCCCTTGGTGGAGTTT | 122 |
| chr10:118477635-11847965 | p5 | (GAAAA)3 | AGGTCAACCGTCCCCTTAAC | ATGTGCATTTTTCACCCCAT | 262 |
| chr10:118486564-11848857 | p5 | (GAAAA)3 | GCCAAGATTTGTCTGTGCAA | CCCGCATTTGCTGTTAGATT | 201 |
| chr10:118510896-11851291 | p2 | (TA)7 | ATAATTCCTGGCAATCGCAC | GGAGCCAATGCCAACATAAG | 278 |
| chr10:118525590-11852760 | p2 | (CA)6 | CCGGCGAGTCTAATAGTTGC | CCCTCAATCTTCCTCCATCA | 112 |
| chr10:118548838-11855085 | p5 | (TTGGT)3 | TTTGTTGGTTCTAGCTGGGG | TGGTTTGGTTCGGTTTGACT | 285 |
| chr10:118548838-11855085 | p3 | (TTG)6 | TTTGTTGGTTCTAGCTGGGG | TGGTTTGGTTCGGTTTGACT | 285 |
| chr10:118548838-11855085 | p5 | (TTGGT)3 | TTTGTTGGTTCTAGCTGGGG | TGGTTTGGTTCGGTTTGACT | 285 |
| chr10:118602081-11860411 | p2 | (TA)7 | TTCGGTTGACACAATTCACG | ATACGAGACGAAAGGGCAAA | 159 |
| chr10:118602081-11860411 | p2 | (TA)6 | TTCGGTTGACACAATTCACG | ATACGAGACGAAAGGGCAAA | 159 |
| chr10:118625599-11862766 | p2 | (AT)35 | GGGTGACAAAACTAGCCCAA | GATTGAGCATTTTCACGGGT | 256 |
| chr10:118639235-11864125 | p3 | (CAT)6 | CTGTGGCAGTACTTGGAGCA | ACCGCTTGTTACCTACCCCT | 113 |
| chr10:118694941-11869695 | p3 | (TTG)6 | TTCGCCATTGATTTTCTTCC | ACTTCATCCCTCCATTTCCC | 143 |
| chr10:118722240-11872425 | p5 | (AAACC)3 | TGACCACTACTCCCACCCTT | TGAGCCTCATGCTGATAACG | 109 |
| chr10:118769774-11877179 | p3 | (CAA)6 | ACGAGCGGAGTAGTGAGGAA | TATGTTTTTCATAGGCGGGC | 284 |
| chr10:118924230-11892624 | p2 | (TA)14 | GGAAGAAGTGCGATGGGATA | CGGTGTTACTGGAAAACGCT | 159 |
| chr10:118924230-11892624 | p5 | (ACCAA)3 | GGAAGAAGTGCGATGGGATA | CGGTGTTACTGGAAAACGCT | 159 |
| chr10:119041332-11904334 | p2 | (TA)8 | AGTCCAGTTGCGTCAATCCT | GTTTTGATGATGTGAGGGGG | 189 |
| chr10:119076382-11907840 | p2 | (CA)6 | AAACAGGGATGCCAATGAAA | TTTGTGTGTGTGTGGGTGTG | 146 |
| chr10:119076382-11907840 | p2 | (AT)10 | AAACAGGGATGCCAATGAAA | TTTGTGTGTGTGTGGGTGTG | 146 |
| chr10:119164901-11916692 | p2 | (AT)10 | CGTATTCGGTAGAACGGGAA | GTGTCCAGTGGGGTCACTTT | 164 |
| chr10:119177774-11917978 | p5 | (TCGGG)3 | AACGAGGAGTGGGCCTTTAT | AACCGAACAAGCAATTTTGG | 104 |
| chr10:119214906-11921692 | p3 | (ACC)5 | CAACCTTTTCCGGTGACCTA | GAAGAGAGGGAGGGAAGGTG | 228 |
| chr10:119214906-11921692 | p4 | (AAAT)4 | CAACCTTTTCCGGTGACCTA | GAAGAGAGGGAGGGAAGGTG | 228 |
| chr10:119214906-11921692 | p2 | (TG)7 | CAACCTTTTCCGGTGACCTA | GAAGAGAGGGAGGGAAGGTG | 228 |

| Marker ID | SSR type | SSR motif | FORWARD PRIMER (5'-3') | REVERSE PRIMER (5'-3') | size (bp) |
|---|---|---|---|---|---|
| chr10:119214906-11921692 | p2 | (TG)6 | CAACCTTTTCCGGTGACCTA | GAAGAGAGGGAGGGAAGGTG | 228 |
| chr10:119214906-11921692 | p2 | (TG)10 | CAACCTTTTCCGGTGACCTA | GAAGAGAGGGAGGGAAGGTG | 228 |
| chr10:119216179-11921822 | p2 | (AG)23 | ATCCCATCATCAATTGCCTC | GGTCCGGTCAAAACTCGTAA | 263 |
| chr10:119249065-11925107 | p2 | (AT)6 | TTTTGTACCCCAAATCCCAA | CGAACCCAGTAGCTTTTTCG | 163 |
| chr10:119358508-11936052 | p5 | (ATAAA)3 | ATTTCGGCAAAAGGAAGGAT | ATCTATTCCAGGGCGAGCTT | 271 |
| chr10:119437227-11943926 | p5 | (TATTG)4 | TGTTTGGTACGCAAATTGGA | AACCTCCCCACCTGTACTCC | 211 |
| chr10:119451612-11945362 | p5 | (ATATA)3 | AACAGTCGCGGAAAAAGAAA | TGTTCATCCTTCACACGGAG | 229 |
| chr10:119614606-11961661 | p2 | (TA)6 | GGTGACCCTTCAGCATAGGA | CCGACAAGGCTGTCACTACA | 150 |
| chr10:119614606-11961661 | p3 | (ATA)5 | GGTGACCCTTCAGCATAGGA | CCGACAAGGCTGTCACTACA | 150 |
| chr10:119719037-11972106 | p3 | (ATT)8 | CATGAATGTGATTGTCCGCT | AAATACCAACTGCACAGGGG | 251 |
| chr10:119774854-11977686 | p2 | (TA)24 | ACCACGTGCGTGTCAATCTA | AAATGACATGGCGGAATAGC | 232 |
| chr10:119881568-11988358 | p2 | (AT)9 | TTGTCCTTTAAATGGGCTGG | TAATTTTTGGCCCTCAATGC | 154 |
| chr10:119942848-11994486 | p5 | (TACAA)3 | ACGCCTTGTTTGGATAGTGG | TGAACCACGGGTTTACAACA | 204 |
| chr10:119972370-11997438 | p2 | (TG)7 | AGCAGAAGCAGGTAGGGACA | AGTCTGCGTATTCCGGCTAA | 115 |
| chr10:119972370-11997438 | p5 | (CCCCA)3 | AGCAGAAGCAGGTAGGGACA | AGTCTGCGTATTCCGGCTAA | 115 |
| chr10:119973543-11997555 | p5 | (CCCCA)3 | CTGTTTGGATGGTGGTTTCC | CCGTGGTTACAAAAATTGGG | 291 |
| chr10:119973543-11997555 | p5 | (TACAA)3 | CTGTTTGGATGGTGGTTTCC | CCGTGGTTACAAAAATTGGG | 291 |
| chr10:119987515-11998953 | p5 | (TACAA)3 | ACGCCTTGTTTGGATAGTGG | TGAACCACGGGTTTACAACA | 204 |
| chr10:120061864-12006388 | p6 | (AATTGT)3 | GAACCCATACCACACCAACC | ACCAACCAGCCTATCACCAC | 163 |
| chr10:120110947-12011296 | p6 | (AATTGT)3 | GAACCCATACCACACCAACC | ACCAACCAGCCTATCACCAC | 163 |
| chr10:120190628-12019264 | p3 | (TGT)6 | GCACCATAGCGACTGACTGA | GGGTTATCTGTTGTGGGTGG | 286 |
| chr10:120286269-12028828 | p3 | (TGT)6 | GCACCATAGCGACTGACTGA | GGGTTATCTGTTGTGGGTGG | 286 |
