## Supplementary figures and images for "Genome-wide identification of microsatellite markers and their application in genetic studies of wolfberry (*Lycium barbarum*)"

### Supplementary Figure S2.docx

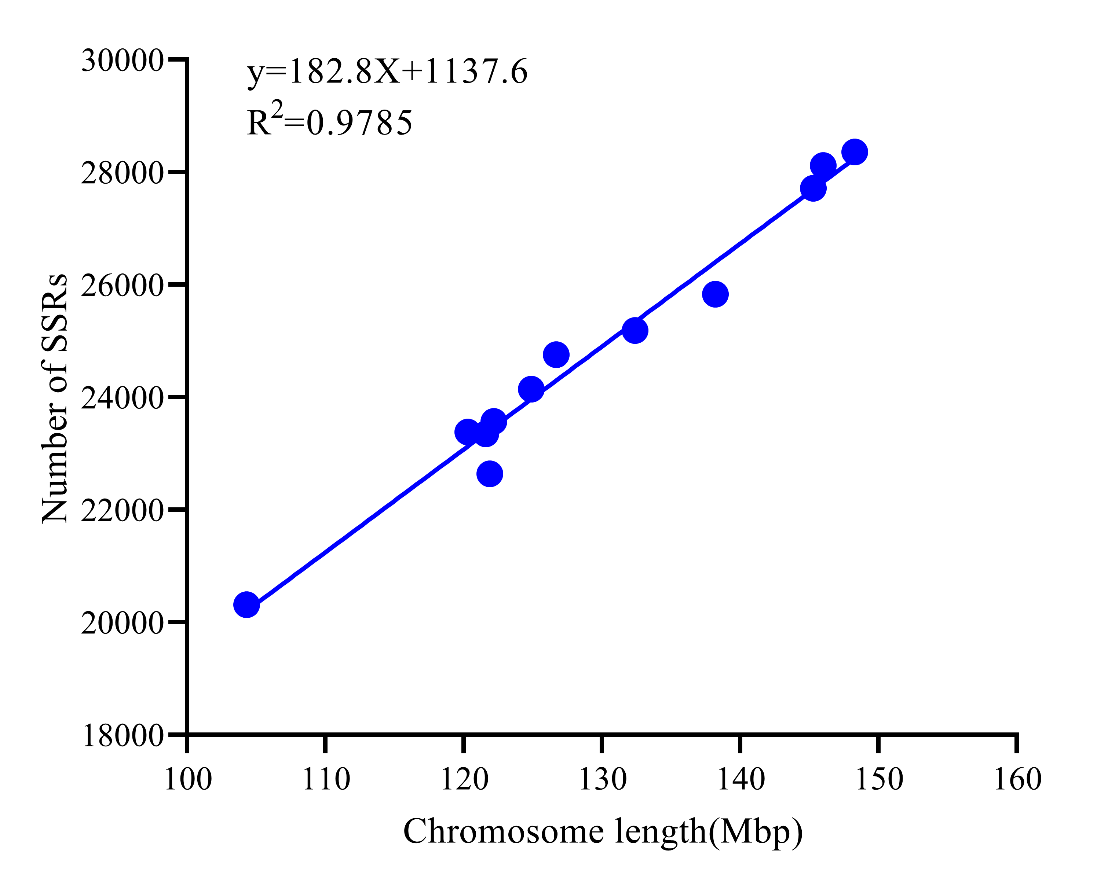

**Figure S2** The relationship between chromosome length and the number of SSRs of each chromosome
